# Methylome-based cell-of-origin modeling (Methyl-COOM) identifies aberrant expression of immune regulatory molecules in CLL

**DOI:** 10.1101/2020.02.04.933937

**Authors:** Justyna A. Wierzbinska, Reka Toth, Naveed Ishaque, Karsten Rippe, Jan-Philipp Mallm, Lara Klett, Daniel Mertens, Thorsten Zenz, Thomas Hielscher, Marc Seifert, Ralf Küppers, Yassen Assenov, Pavlo Lutsik, Stephan Stilgenbauer, Philipp M. Roessner, Martina Seiffert, John Byrd, Christopher C. Oakes, Christoph Plass, Daniel B. Lipka

## Abstract

**Background:** In cancer, normal epigenetic patterns are disturbed and contribute to gene expression changes, disease onset and progression. The cancer epigenome is composed of the epigenetic patterns present in the tumor-initiating cell at the time of transformation, and the tumor-specific epigenetic alterations that are acquired during tumor initiation and progression. The precise dissection of these two components of the tumor epigenome will facilitate a better understanding of the biological mechanisms underlying malignant transformation. Chronic lymphocytic leukemia (CLL) originates from differentiating B cells, which undergo extensive epigenetic programming. This poses the challenge to precisely determine the epigenomic ground-state of the cell-of-origin in order to identify CLL-specific epigenetic aberrations.

**Methods:** We developed a linear regression model, methylome-based cell-of-origin modeling (Methyl-COOM), to map the cell-of-origin for individual CLL patients based on the continuum of epigenomic changes during normal B cell differentiation.

**Results:** Methyl-COOM accurately maps the cell-of-origin of CLL and identifies CLL-specific aberrant DNA methylation events that are not confounded by physiologic epigenetic B cell programming. Furthermore, Methyl-COOM unmasks abnormal action of transcription factors, altered super-enhancer activities, and aberrant transcript expression in CLL. Among the aberrantly regulated transcripts were many genes that have previously been implicated in T cell biology. Flow cytometry analysis of these markers confirmed their aberrant expression on malignant B cells at the protein level.

**Conclusions:** Methyl-COOM analysis of CLL identified disease-specific aberrant gene regulation. The aberrantly expressed genes identified in this study might play a role in immune-evasion in CLL and might serve as novel targets for immunotherapy approaches. In summary, we propose a novel framework for *in silico* modeling of reference DNA methylomes and for the identification of cancer-specific epigenetic changes, a concept that can be broadly applied to other human malignancies.

## BACKGROUND

In cancer, normal epigenetic patterns are disturbed and contribute to gene expression changes, disease onset and progression [1]. This seems to be a universal characteristic of all cancers, including chronic lymphocytic leukemia (CLL). CLL originates from rapidly differentiating B cells. Although several mutations creating a pre-leukemic clone, including variants in *SF3B1*, *NOTCH1* or *TP53*, have been identified in the hematopoietic stem cell (HSC) compartment of CLL patients, additional genetic or epigenetic driver events are required for full transformation[2]. Normal B cells undergo extensive epigenetic programming during differentiation [3, 4]. The epigenetic fingerprint of the B cell that has acquired the transforming hit is ‘frozen’ and stably propagated in the leukemic cells [4]. This demonstrates that two factors contribute to the epigenomic landscape of CLL: first, epigenetic patterns that were present in the tumor-initiating B cell at the time of transformation, and second, CLL-specific epigenetic alterations that are acquired during leukemia initiation and progression. For the purpose of this study, we define the cell-of-origin of CLL as the normal B cell differentiation stage with the highest overlap to the CLL methylome. Consequently, the cell-of-origin of CLL represents the differentiation stage at which the clonal B cells deviate significantly from the normal differentiation trajectory and therefore the cell-of-origin defines the first cell that has acquired sufficient oncogenic hits to initiate leukemic transformation[5].

Numerous publications have reported extensive epigenetic alterations in CLL resulting in deregulation of protein coding genes [6–11] or miRNAs [12–19]. In this context, most studies used the epigenome of CD19^+^ B cells as controls, but such an approach neglects the epigenetic programming occurring during B cell differentiation. As a result, the genes found to be deregulated mainly reflected the changes occurring during normal B cell differentiation rather than CLL-specific pathogenic events. Refined analyses should aim at discriminating between epigenetic changes occurring during normal B cell differentiation and CLL-specific epigenetic aberrations. Here we outline a novel framework for cancer methylome analysis, termed methylome-based cell-of-origin modeling (Methyl-COOM). We show how Methyl-COOM can be applied to epigenomic datasets from CLL patients to identify disease-specific epigenetic events and demonstrate its power to detect epigenetically deregulated transcripts which encode for proteins that are involved in immune regulatory processes.

## METHODS

### Flow cytometry analysis

Patients’ samples were obtained from the Department of Internal Medicine III of Ulm University after approval of the study protocol by the local ethics committee according to the declaration of Helsinki, and after obtaining informed consent of patients. Patients met standard diagnosis criteria for CLL. Patients’ characteristics such as age, gender, mutational state and Binet stage are depicted in Table 1.

**Table 1:** Characteristics of the CLL patients used for flow cytometric analysis.

|  |  |
| --- | --- |
| **# of patients** | 7 |
| **Age [years]** | 57.1 (mean)<br>52 (median) |
| **Sex** | 5/7 male<br>2/7 female |
| **Prior therapies** | 7/7 no prior treatment |
| **Binet stage** | 7/7 A<br>0/7 B<br>0/7 C |
| **IGHV status** | 6/7 mutated<br>1/7 unmutated |
| **Genetics (FISH)** | 1/7 Trisomy 12<br>5/7 del(13q)<br>1/7 no aberration |
| **TP53 mutation status** | 4/7 WT<br>3/7 not tested |

Peripheral blood was drawn using Ethylenediaminetetraacetic acid (EDTA)-coated tubes (Sarstedt, Nümbrecht, Germany). PBMCs were isolated by Ficoll (Biochrom, Berlin, Germany) density gradient centrifugation. PBMCs were viably frozen and, when needed, thawed and further processed.

After blockade of Fc-receptors using Human TruStain FcX™ (BioLegend, London, United Kingdom), 5*10^6^ PBMCs were stained with fluorescently labelled antibodies in phosphate-buffered saline (PBS) with addition Fixable Viability Dye eFluor® (ThermoFisher Scientific, Dreieich, Germany) for 30 min at 4°C. Cells were fixed using eBioscience™ IC Fixation Buffer (ThermoFisher Scientific, Dreieich, Germany) for 30 min at room temperature. The antibodies used are listed in Table 2. If necessary, cells were permeabilized with eBioscience™ Permeabilization Buffer (ThermoFisher Scientific) and stained intracellularly for 30 min at room temperature. CTLA-4 was stained as surface as well as intracellular marker. Samples were stored at 4°C in the dark until acquisition. Data was acquired using a BD LSR Fortessa (BD Biosciences, Heidelberg, Germany) FACS analyzer. Flow cytometric data was analyzed using FlowJo X 10.0.7 software (FlowJo, Ashland, OR, USA). Paired Wilcoxon signed-rank test was used to determine statistical significance of changes between CLL B cells and normal B cells.

**Table 2:**
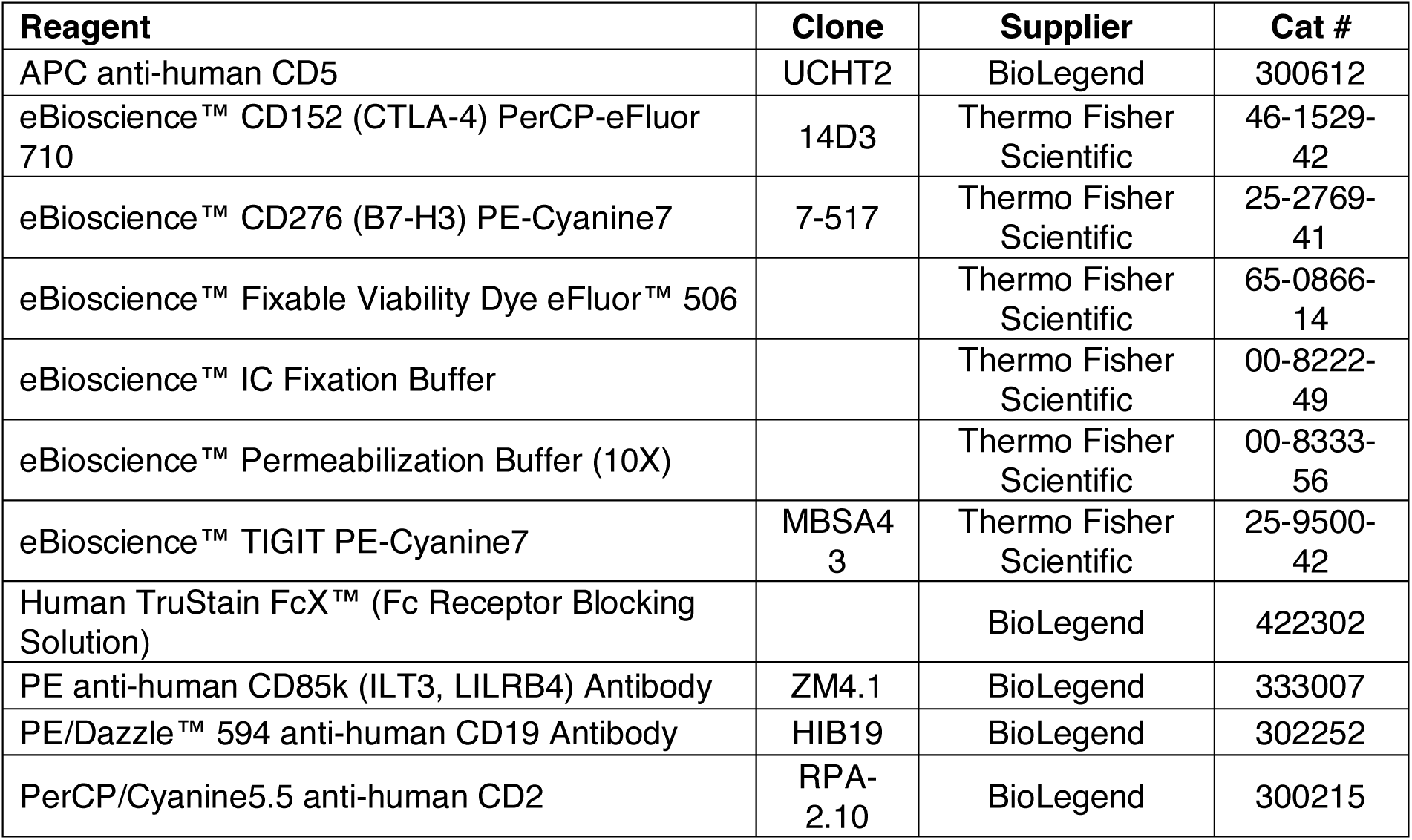
List of FACS antibodies and reagents.

### Analysis of RNA-seq / sncRNA-seq data

Expression data (RNA-Seq) from CLLs were obtained from our previous study [4]. RNA-Seq data from normal B cells was obtained from International Cancer Genome Consortium (ICGC). Reads per kilo base per million mapped reads (RPKM) normalized values were used for the comparison of gene expression levels. sncRNA-seq data from CLLs was obtained from our previous study [20]. Differential miRNA expression was assessed using normalized counts, reads per million (RPM).

### Analysis of 450k methylome array data

450K data from B cells was obtained from Oakes et al. [4]. CLL 450k data for the discovery and validation cohorts were both obtained from previous studies [4, 21]. The analysis of 450K data was performed using RnBeads software [22]. Both datasets (normal B cells and CLLs) were processed simultaneously. Briefly, raw 450K data for both CLL and healthy B cell sample sets were normalized by the BMIQ method [23] without the background subtraction. The probes overlapping SNPs and the X and Y chromosomes were removed and remaining probes (n=464,743 CpGs) were considered for the downstream analysis, for the identification of CLL-specific methylation events (Method Section: ‘Identification of disease-specific methylation events’).

### Inference of the cell-of-origin and identification of disease-specific methylation events

We studied the DNA methylation programming during normal B cell differentiation, using six discrete B cell subpopulations including naïve to mature B cells: referred to as naïve B cells (NBCs), germinal center founder cells (GCFs), low- and intermediate-memory B cells (loMBCs, intMBC), splenic marginal zone B cells (sMGZs), and high maturity memory B cells (hiMBCs). DNA methylomes from 2-4 donors per normal B cell subpopulation. In addition 34 CLL samples were analyzed using Illumina 450k Bead Chip arrays.

### Cell-of-origin based methylome analysis, Methyl-COOM

For analysis, we determined the DNA methylation dynamics during normal B cell differentiation (differentiation axis). Here we assumed that changes in DNA methylation during the cellular differentiation process are reminiscent of the DNA nucleotide changes over the evolutionary time. CpG sites showing a statistically significant gain or loss of methylation of more than 20% during B cell differentiation defined our set of so-called B cell-specific CpGs (n=74,333 CpGs; student’s t-test). A Manhattan distance matrix was calculated and used to build a methylation-based phylogenetic tree of normal B cell differentiation by applying the minimum evolution method (*fastme.bal* function, R package “ape”; Desper and Gascuel, [24]). Each node in the phylogenetic tree corresponds to a certain differentiation stage reached by the B cell. Using this approach, we observed a non-branched differentiation trajectory of normal B cell differentiation. Therefore, we initially used all B cell-specific CpGs to generate a linear regression model of DNA methylation programming during normal B cell differentiation. Linear behavior between the differentiation stage of every B cell subset and the methylation profiles at B cell-specific CpGs were tested at the single CpG level using F-test. The majority of the B cell-specific CpGs (79.8%, n=59,326 CpGs) showed linear methylation dynamics across the six B cell differentiation states. To exclude a potential bias on differentiation stage assignment, we re-created both the phylogeny and the regression model of normal B cell differentiation, this time using the linearly behaving B cell specific CpGs, only. The final regression model was designed to infer DNA methylation levels of all CpGs included in our analysis.

Next, we mapped all CLL samples onto the normal B cell differentiation trajectory in order to infer the closest virtual normal B cell methylome (cell-of-origin) defined as the position of the closest normal B cell node in the phylogenetic tree. Then, we applied the linear regression model to infer the DNA methylation levels for each CpG site in the putative cell-of-origin for every patient, according to the formula:

M = α + β *d.s.

, where

M denotes the calculated beta methylation value for a CpG site of cell-of-origin, d.s. denotes the differentiation stage (defined as the distance between the NBC and the cell-of-origin nodes as determined by the phylogenetic analysis),

β denotes the slope of the regression line,

α denotes the vertical (y-axis) intercept.

To test our cell-of-origin assignment, we applied a cross-validation model on our phylogenetic analysis. The patient cohort was repeatedly divided into two subgroups; 70% and 30% (5000 repetitions). To minimize the likelihood of selecting the same sample multiple times, a random sampling was allowed in the 70%-group, while sample replacement was restricted only to the 30%-group. Using this approach, we observed that our original cell-of-origin is located between interquartile ranges of the cross-validation assignments, confirming the robustness of the cell-of-origin definition (**Supplementary Figure S2 f**).

#### Identification of CLL-specific DNA methylation

Subsequently, the inferred DNA methylome of the cell-of-origin was used as a reference to determine aberrantly methylated CpG sites in each sample. Disease-specific CpGs were defined as sites with significant deviation from the expected methylation levels as compared to the corresponding cell-of-origin.

#### Sites with epigenetic B cell programming

Sites undergoing epigenetic B cell programming (i.e. B cell-specific CpGs) could still show disease-specific methylation events if their actual methylation status massively deviates from what would be expected based on the regression model (sites with “epigenetic B cell programming”). We used a conservative cut-off of more than 20% methylation loss (class A) or gain (class B) relative to the calculated cell-of-origin methylation value (M value) in at least 75% of the CLL patients.

#### Sites without epigenetic B cell programming

Sites with no epigenetic B cell programming (i.e. non-B cell-specific CpGs) were defined to have CLL-specific aberrant DNA methylation if they displayed either methylation loss (class C) or gain (class D) of more than 20% relative to the cell-of-origin in at least 75% of the CLL patients.

### Identification of CLL-specific protein-coding genes

To identify CLL-specific protein-coding genes, disease-specific methylation events were overlapped with promoter regions (−2.5kb, +0.5kb to TSS) of protein-coding genes. Next, correlation between aberrant DNA methylation and gene expression was determined (Pearson correlation test, p-value <0.05; correlation coefficient < −0.7). A full list of identified CLL-specific protein-coding genes is available in **Supplementary Table S1**.

### Identification of CLL-specific SE-associated genes

To identify CLL-specific Super-enhancer (SE)-associated genes, SE data from DKFZ PRECiSe consortium was used [28]. All statistically significant, differential super-enhancers being gained in CLLs (“gained”, p<0.05, FC>0) and consensus super-enhancers shared between normal B cells and CLLs (“stable”) were used for the analysis. Firstly, SEs were associated with the closest gene in the vicinity. CLL-specific methylation events were then overlapped with SE coordinates. Next, correlation between aberrant DNA methylation in SE region and gene expression of the SE-closest gene (Pearson correlation test, p-value <0.05; correlation coefficient < −0.7) was used to identify CLL-specific Super-enhancer (SE)-associated genes. A full list of identified SE-associated genes is available in **Supplementary Table S2**.

### Super-enhancer (SE) enrichment analysis

For the super-enhancer enrichment analysis two sets of super-enhancers were used, SE data from DKFZ PRECiSe consortium [25] and SE data from Ott et al. [26]. From DKFZ PRECiSe consortium all statistically significant, differential super-enhancers being gained in CLLs (“gained”, p<0.05, FC>0) and consensus super-enhancers shared between normal B cells and CLLs (“stable”) were used for the analysis. From Ott et al. paper a unified SE region was created using reduce function in GenomicRanges package, providing a SE data from individual CLL patients (n=18). All CpG probes present on the 450k array were used as a background in the enrichment analysis.

### Identification of micro-RNA promoters

To identify miRNA promoters, the promoter segmentation data from CLLs (DKFZ PRECiSe consortium; promoter segmentation data is deposited under GSE113336; raw ChIP-seq data can be found in the European Genome-phenome Archive under the accession number EGAS00001002518) and normal cell lines (Encyclopedia of DNA Elements – ENCODE; ENCODE Mar 2012 Freeze, UCSC accession numbers: wgEncodeEH000784, wgEncodeEH000785, wgEncodeEH000790, wgEncodeEH000789, wgEncodeEH000788, wgEncodeEH000786, wgEncodeEH000787, wgEncodeEH000791, wgEncodeEH000792) was used. To define constant promoter segments, the *reduce* function from the “GenomicRanges” R package was used to create simplified promoter regions, present in all datasets (CLL and ENCODE segmentation data). Putative promoters of pri-miRNAs were assigned based on their distance to the pri-miRNA TSSs. The genomic coordinates of pri-miRNAs/miRNAs were downloaded from miRBase (version 20; v20). Any promoter located within 100kb upstream of a pri-miRNA TSS was considered as a putative pri-miRNA promoter. The distance of 100 kb was chosen based on similar approaches that have been used in the past by Corcoran *et al.*, Fujita *et al.* and Fukao *et al.* [27–29]. The larger distance of putative promoters to pri-miRNA TSSs is especially important in case of intergenic miRNAs, which are originating from intronic sequences and which are considered to be transcribed together with their host gene.

### Identification of CLL-specific micro-RNAs

To identify CLL-specific microRNAs, disease-specific methylation events were overlapped with potential pri-miRNA promoters. To identify candidate CLL-specific miRNAs, correlation between aberrant DNA methylation and pri-miRNA expression was determined (Spearman correlation test, p-value <0.05; abs(correlation coefficient ρ) ≥ 0.35). Since many mature miRNAs are derived from the same pri-miRNAs, correlations were calculated using pri-miRNA expression levels determined by sncRNA-seq. A full list of identified CLL-specific microRNAs is available in **Supplementary Table S3**.

**Table 3:** List of datasets used in the manuscript.

| Dataset | Source |
| --- | --- |
| Illumina 450 data, normal B cells | Oakes et al. [4] |
| Illumina 450 data, CLLs discovery cohort | Dietrich et al. [21] |
| Illumina 450 data, CLLs validation cohort | Oakes et al. [4] |
| RNAseq, CLLs | Dietrich et al. [21] |
| RNAseq, normal B cells | International Cancer Genome Consortium (ICGC); EGAD00001000258 |
| sncRNAseq, CLL | Blume et al. [20] |
| ENCODE TF ChIP-seq GM12878 | ENCODE project [89] |
| ATAC-seq, normal B cells and CLLs | DKFZ PRECiSE consortium [25] |
| ChIP-seq, normal B cells and CLLs | DKFZ PRECiSE consortium [25]; EGAS00001002518 |
| Promoter segmentation data CLL | DKFZ PRECiSE consortium [25]; GSE113336 |
| ChromHMM GM12878 data | ENCODE ENCSR212BHV [89] |

### Target genes of CLL-specific microRNAs

To link CLL-specific microRNAs with their pathogenetic effects, two databases of experimentally validated microRNA-target gene interactions were used, TarBase v8.0 and miRTarBase. A full list of experimentally validated CLL-specific microRNA targets is included in **Supplementary Table S4**. To find whether CLL-specific microRNAs are targeting epigenetic regulators, the comprehensive list of epigenetic regulators was used (**Supplementary Table S5**). The list of epigenetic regulators was further used as a query for the list of CLL-specific microRNA targets defined above. The epigenetic regulators targeted by CLL-specific microRNAs are included in the **Supplementary Table S6**.

### Transcription factor enrichment analysis

Transcription factor motif analysis in disease-specific methylation events was performed using HOMER software v4.5 [30] using only the results for the ‘known motifs’ analysis. All CpGs present on the 450k array were used as a background and adjustment for GC- and CpG-content was used. Furthermore, enrichment of actual binding events of TFs and other DNA-binding proteins was analyzed using available ChIP-seq data from the tier 1 ENCODE cell line GM12878 (for a complete list of datasets used for this analysis, please refer to **Supplementary Table S7**). The ChIP-seq enrichment analysis was performed using the LOLA tool [31] providing all CpG probes present on the 450k array as the ‘universe’. Unsupervised hierarchical clustering and data visualization were performed using R.

## RESULTS

### Modeling of normal B cell differentiation

CLL epigenomes are shaped by two major components. The first component constitutes signatures that stem from the leukemia-initiating B cell. The second component is formed by epigenetic alterations acquired during leukemogenesis and progression of the disease. To discriminate these components, we developed an *in silico* approach to infer DNA methylation dynamics during normal B cell differentiation and to model the epigenome of the cell-of-origin, utilizing previously published Illumina 450k array DNA methylome data from six distinct B cell subpopulations [4] and from 34 CLL samples [21](**Figure 1a**). Our approach to this was based on classical phylogeny analysis (minimum evolution method, Desper and Gascuel [24]), which is typically used to reconstruct evolutionary processes based on inherent characters. Similarly to copy number or mutational studies [32, 33], phylogeny analysis on DNA methylation has been used successfully to reconstruct the developmental processes occuring during cell proliferation and differentiation [4, 34]. Therefore, to model B cell differentiation, we inferred the hierarchical relationship between normal B cell subsets ranging from naïve to memory B cells based on their DNA methylation patterns. The normal B cell methylomes were used to identify CpG sites that show dynamic DNA methylation during B cell differentiation (B cell-specific CpGs; see also **Methods**). A total of 74,333 B cell-specific CpGs were identified (≥ 20% DNA methylation change between naïve and differentiated memory B cells, Student’s t-test, p-value<0.05 [4, 35]). Pairwise Manhattan distances based on DNA methylation profiles at B cell-specific CpGs for normal B cell subsets were used to build a methylation-based phylogenetic tree revealing a non-branched trajectory of B cell differentiation (**Supplementary Figure S1a)**. This suggested that linear regression might be suitable to model DNA methylation dynamics. The initial linear regression model of B cell differentiation considered all B cell-specific CpGs. Testing the linearity between the differentiation stage of every normal B cell subset and the methylation profiles at B cell-specific CpGs, revealed that the vast majority of the differentiation-specific CpGs (79.8%, n=59,326 CpGs) showed linear behavior across all B cell differentiation states (F-test, p-value <0.05; **Supplementary Figure S1b-g, Supplementary Table S8**). To exclude a potential bias on the model from the non-linear CpG sites, we re-generated both the phylogeny and the regression model of normal B cell differentiation using only the linearly behaving B cell-specific CpGs.

**Figure 1.**
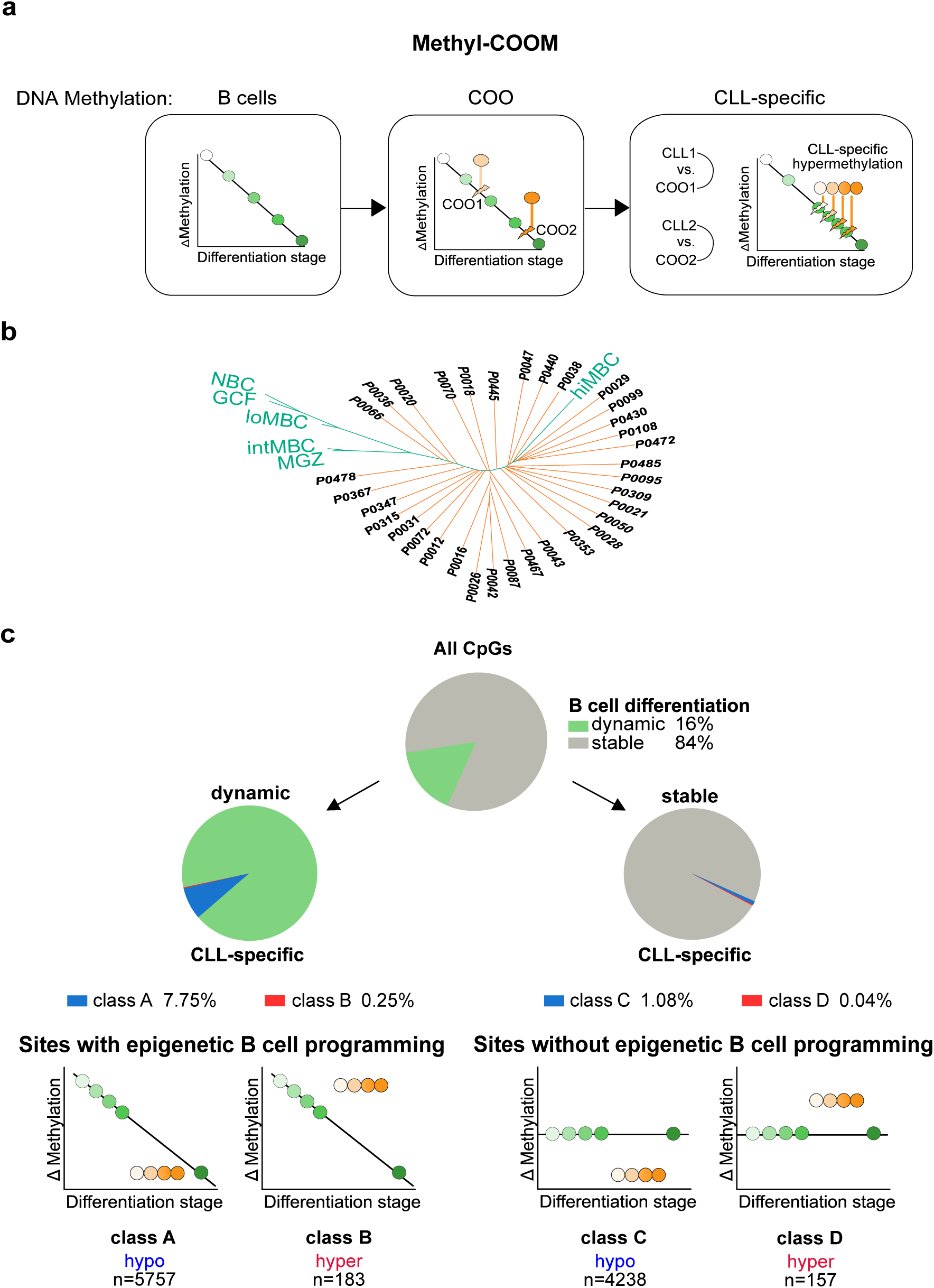
Identification of CLL-specific DNA methylation events using Methyl-COOM. **a)** Schematic outline of the Methyl-COOM pipeline used for the identification of CLL-specific DNA methylation events. Methylome data of six distinct B cell subpopulations, representing different stages of B cell differentiation were used to infer normal B cell differentiation. A linear regression model was applied to model DNA methylation dynamics during normal B cell differentiation (‘DNA methylation: B cells’). DNA methylomes of 34 primary CLL samples were used to identify the closest virtual normal B cell (cell-of-origin; COO) based on phylogeny analysis. The linear regression model was then used to infer the DNA methylome of the COO (‘DNA methylation: COO’). Next, the DNA methylome of each CLL was compared to the DNA methylome of its COO. CLL-specific aberrant DNA methylation was defined as a significant deviation from the inferred COO methylome (‘DNA methylation: CLL-specific’). **b)** Identification of the cell-of-origin in CLL samples using phylogenetic analysis. A phylogenetic tree was generated using a set of linear CpG sites that show dynamic DNA methylation changes during normal B cell differentiation (linear B cell-specific CpGs, 59,326 CpGs). Pairwise Manhattan distances were calculated between DNA methylation profiles of normal B cells and CLL samples at B cell-specific CpGs and were subsequently used to assign the closest normal (virtual) B cell methylome (location of the node on the phylogenetic tree = differentiation stage of the cell-of-origin) to each CLL case. NBCs - naïve B cells; GCFs – germinal center founder B cells; loMBCs – early non class-switched memory B cells; intMBCs – non class-switched memory B cells; sMGZs – splenic marginal zone B cells; hiMBCs – class-switched memory B cells (mature B cells). CLL samples are depicted in orange color. Normal B cells are represented in green. **c)** Summary of CLL-specific DNA methylation events. <u>Top:</u> pie chart displays the frequency of CpGs that are either dynamic (green) or stable (grey) during normal B cell differentiation. <u>Middle:</u> pie charts depict the frequency of CLL-specific DNA methylation events as fractions of the dynamic (class A and B; left), and stable (class C and D; right) sites. <u>Bottom</u>: schematic depicting the classification of CLL-specific DNA methylation events. We identified two groups: ‘sites with epigenetic B cell programming’ and ‘sites without epigenetic B cell programming’. ‘Sites with epigenetic B cell programming’ undergo DNA methylation programming during normal B cell differentiation, encompassing hypomethylation (class A) and hypermethylation events (class B) relative to the DNA methylome of the COO. ‘Sites without epigenetic B cell programming’ are defined as CpG sites without significant DNA methylation changes during normal B cell differentiation and are classified as either hypo- or hypermethylation (class C and D, respectively). Numbers of CLL-specific DNA methylation events (CLL-specific CpGs) resolved by class are indicated at the bottom.

### Identification of disease-specific DNA methylation patterns in CLL

This B cell differentiation model was applied to a CLL patient cohort (n= 34) in order to determine the closest virtual normal B cell methylome (i.e. cell-of-origin or B cell differentiation stage) for each CLL case (**Figure 1b**). As expected, our model confirmed that good-prognosis *IGHV* mutated CLL originates from more mature B cells, as opposed to *IGHV* unmutated CLL, which develops from more immature B cells (**Supplementary Figure S2a-e**). Next, we tested the stability of cell-of-origin assignment using a cross-validation model (5000 repetitions; for details see **Methods** section). Using this approach, we observed that the predicted cell-of-origin is located between interquartile ranges of the cross-validation assignments, confirming the robustness of the cell-of-origin definition (**Supplementary Figure S2f**). The linear regression model was then used to infer DNA methylation levels for all 464,743 CpG sites in the predicted cell-of-origin of every patient. These inferred cell-of-origin methylomes were subsequently used as controls to identify aberrant (i.e. CLL-specific) DNA methylation patterns for each sample individually (see **Figure 1a** for a schematic overview of Methyl-COOM). CLL-specific aberrant DNA methylation was defined as CpG sites with >20% deviation from the expected DNA methylation level of the cell-of-origin, and which were aberrantly methylated in at least 75% of patients. This analysis revealed two categories of CLL-specific DNA methylation events; 1) aberrant DNA methylation occurring at sites undergoing epigenetic programming during B cell differentiation (‘Sites with epigenetic B cell programming’) and 2) aberrant DNA methylation occurring at CpG sites that normally do not change during B cell differentiation (‘Sites with no epigenetic B cell programming’) (see **Figure 1c**). The first category was further subdivided into class A, showing a loss, and class B, showing a gain of DNA methylation relative to the differentiation stage achieved. The second group of CpG sites without DNA methylation programming during normal B cell differentiation was subdivided into class C and class D displaying hypo- and hypermethylation, respectively (**Figure 1c**). Overall, only 2.2% of all CpG-sites (10,335 CpGs) represented on the 450k array were affected by disease-specific methylation programming, the majority of which were ‘sites with epigenetic B cell programming’ (class A & B, 5,940 CpG sites; **Figure 1c, Supplementary Table S9**). The majority of CLL-specific DNA methylation events were characterized by hypomethylation (9,995 hypomethylated CLL-specific CpGs; class A: 5,757 CpGs, class C: 4,238 CpGs), while only a small proportion of CpGs were hypermethylated as compared to their inferred cell-of-origin (340 hypermethylated CLL-specific CpGs; class B: 183 CpGs, class D: 157 CpGs) (**Figure 1c, Supplementary Figure S2g, h**).

### CLL-specific aberrant DNA methylation patterns are independent of the differentiation stage achieved

CLL-specific DNA methylation changes were quantified for each CpG site in each sample as compared to the cell-of-origin and inspected by unsupervised hierarchical clustering. For all classes, consistent patterns of either loss or gain in methylation relative to the cell-of-origin were observed, irrespective of the differentiation stage achieved (**Figure 2a**, **Supplementary Figure S2i**). Hypomethylation at class A sites resulted from an exaggerated loss of DNA methylation at sites which show loss of methylation during normal B cell differentiation (**Figure 2b, c**, **Supplementary Figure S2i**; class A, hypomethylation). Aberrant hypermethylation observed at class B sites results from exaggeration of hypermethylation normally occurring during B cell differentiation, and from failed hypomethylation during normal B cell programming (**Figure 2b, c**, **Supplementary Figure S2i**; class B, hypermethylation). Class C and class D sites do not undergo any significant DNA methylation programming during normal B cell differentiation, highlighting the potential importance of these sites for CLL pathogenesis (**Figure 2a-c Supplementary Figure S2i;** class C, class D). Overall, the observed CLL-specific aberrant methylation patterns are largely independent of the differentiation stage achieved by the CLL cell-of-origin.

**Figure 2.**
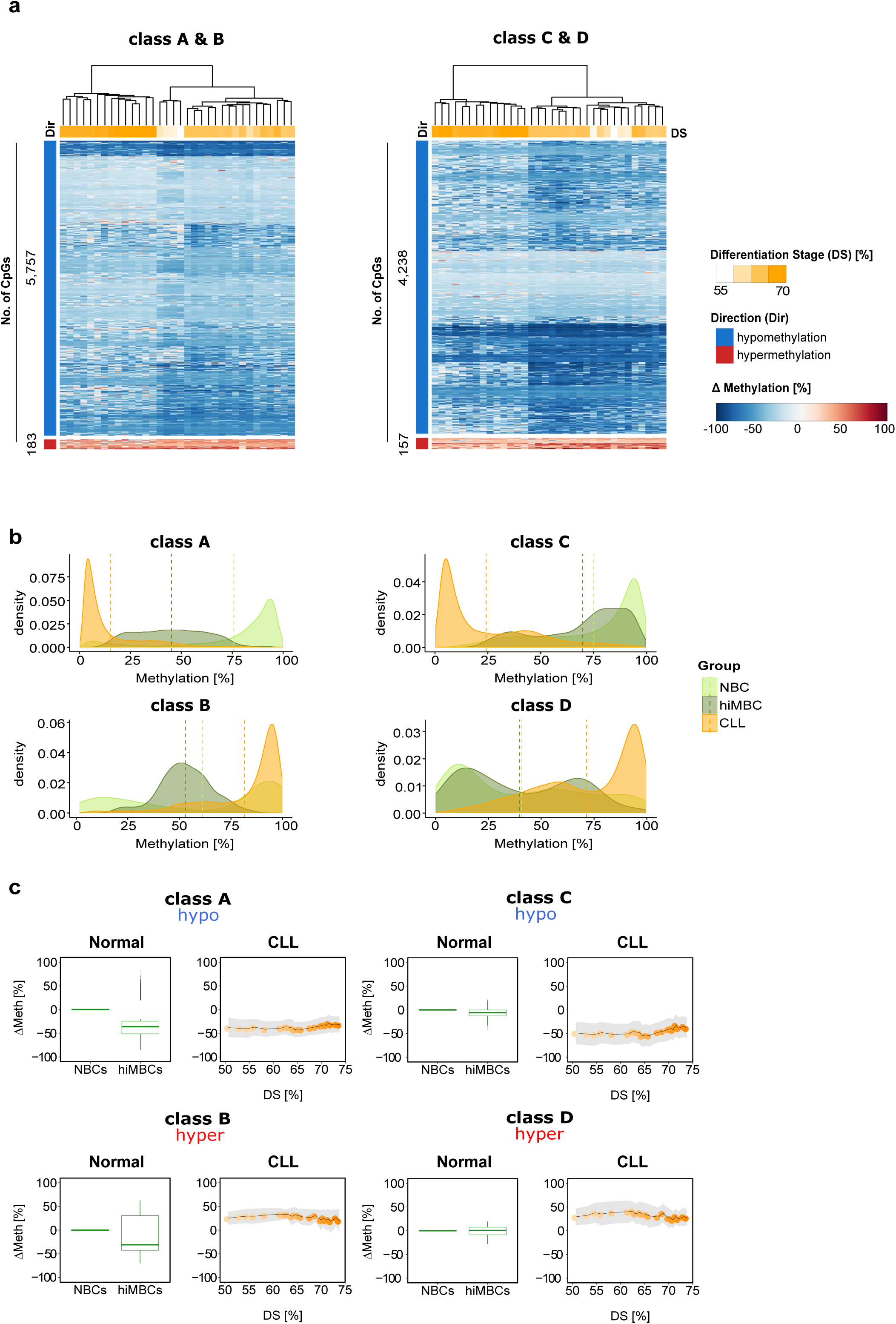
Programming of disease-specific DNA methylation patterns in CLL. **a)** Heatmap depicting DNA methylation changes (ΔMethylation [%]) at CLL-specific CpG sites relative to the samples’ COO. Unsupervised hierarchical clustering of CLL-specific CpGs, class A and B sites (left), class C and D sites (right). The direction of DNA methylation change (Dir [%]) is indicated as blue and red bars for hypo- and hypermethylation, respectively, and the numbers of CpG sites plotted are indicated next to the bars. Differentiation stages (DS) are denoted as a color gradient (white-orange), where CLL samples with immature COO are represented in white and samples with a more mature COO in orange. DS refers to % normal differentiation programming achieved (relative to hiMBCs). **b)** Density plots summarizing the distribution of absolute DNA methylation levels for all CLL-specific CpG sites stratified by class (classes A - D). CLL patients (CLL): orange, naïve B cells (NBC): light green, class-switched memory B cells (hiMBC): dark green. **c)** Box plots and ribbon plots displaying the average DNA methylation change for each class of CLL-specific alterations across normal B cells and CLLs. <u>Left (normal)</u>: average DNA methylation change (ΔMeth) of CLL-specific CpGs during normal B cell differentiation from naïve B cells (NBCs) to class-switched memory B cells (hiMBCs) plotted for all classes (classes A [n=5757 CpG sites], B [n=183 CpG sites], C [n=4238 CpG sites], and D [n=157 CpG sites]). <u>Right (CLL)</u>: ΔMeth for CLL-specific CpGs in CLL. ΔMeth [%] is represented as the mean DNA methylation change relative to the expected DNA methylation level of the COO. Standard deviation is depicted as grey shaded ribbons. DS refers to % normal differentiation programming achieved (relative to hiMBCs).

### CLL-specific DNA methylation affects super-enhancers

To test for functional implications of CLL-specific DNA methylation events, we tested their enrichment in ENCODE ChromHMM genome segments in the GM12878 lymphoblastoid cell line. Aberrantly methylated CpG sites from classes A, B & C were enriched for enhancer elements (**Figure 3a)**. A recent systematic assessment of transcription factor dependencies in CLL has implicated super-enhancer (SE) based transcription factor (TF) rewiring in CLL pathogenesis [26, 36]. In line with this, enrichment of CLL-specific CpGs was detected in SE regions identified in a recently published CLL data set from Ott et al. (**Supplementary Figure S3a**) [26]. Using another SE data set from Rippe and colleagues [25, 37] enabled us to distinguish between SEs that are either present in normal B cells (“stable”) or that have been acquired *de novo* in CLL (“gained”). Enrichment of *de novo* SEs was found in class A and class C sites (**Figure 3b**). *De novo* SEs overlapping with CLL-specific CpG sites harbor many known genes with relevance in CLL biology (e.g. *CD5*, *CLLU1*, *IRF2*; **Supplementary Figure S3b, Supplementary Table S2**).

**Figure 3.**
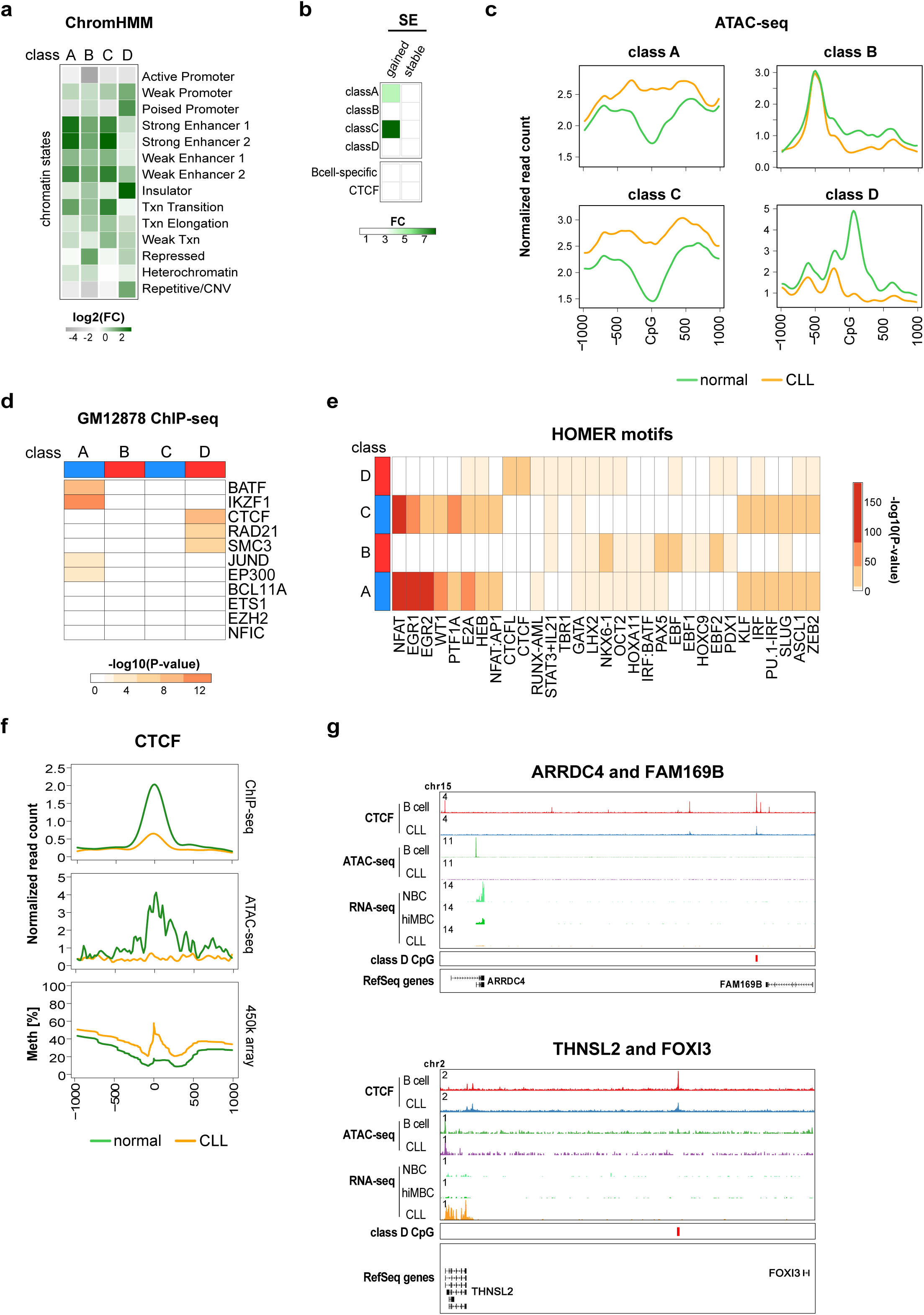
CLL-specific DNA methylation differences result from aberrant transcription factor programming. **a)** Enrichment of chromatin states in sequences representing CLL-specific DNA methylation. Chromatin states were defined using the 15-state ChromHMM model from immortalized B cells [88] for CLL-specific methylation sites of the classes A - D. The enrichment in category ‘Repetitive/CNV’ represents the averaged enrichment value of ChromHMM states called ‘Repetitive/CNV’. Log2 fold change (log2 FC) was calculated using all 450k probes as a background. **b)** Enrichment of super-enhancers (SE) in sequences representing CLL-specific DNA methylation. SE were defined as either being gained in CLLs (gained) or consensus between CLLs and B cells (stable). Fold change (FC) was calculated using all 450k probes as a background. **c)** ATAC-seq read density (normalized read counts *10^-3^) at CLL-specific CpG sites (±1kb) for categories of classes A, B, C and D. CLL samples (n=18) are represented in orange, normal CD19^+^ B cells (n=3) in green. **d)** Transcription factor enrichment analysis using ENCODE ChIP-seq peaks from the B-cell lymphoblastoid cell line, GM12878. Displayed are –log10(p-values) resulting from Fisher’s exact test with false discovery rate correction. **e)** Transcription factor motif enrichment analysis using HOMER. The top 10 most enriched TF motifs for each class are displayed. The colors represent –log10(p-values) derived from a cumulative binomial distribution function as implemented in HOMER. **f)** ATAC-seq & ChIP-seq read density (normalized read counts *10^-3^) and DNA methylation profiles at class D CpGs co-locating with CTCF motifs (23 CpGs) (±1kb). CLL samples (n=7 CTCF ChIP-seq, n=18 ATAC-seq) are represented in orange, normal CD19^+^ B cells (n=4 CTCF ChIP-seq, n=3 ATAC-seq) in green. **g)** Locus plots of exemplary genes associated with CTCF/classD events. Locus plots include data from CTCF ChIP-seq on normal B cells (red) and CLL (blue); ATAC-seq on normal B cells (green) and CLL (purple); RNA-seq on NBC (light green), hiMBC (dark green) and CLL (orange). The class D CpGs are annotated in red.

### CLL-specific DNA methylation differences result from aberrant transcription factor programming

Recent SE perturbation studies implicated rewiring of TF regulatory circuitries in CLL pathogenesis [26]. These findings motivated us to ask whether CLL-specific DNA methylation patterns would be indicative of aberrant TF programming. To address this hypothesis, we used ATAC-seq to test whether CLL-specific DNA methylation patterns were reflected at the level of chromatin accessibility. Indeed, we found that CLL-specific hypo- and hypermethylation events were associated with inverse changes in chromatin accessibility in CLL as compared to normal B cells (**Figure 3c**). These concomitant changes in DNA methylation and chromatin accessibility indicated that CLL-specific DNA methylation patterns reflect global epigenomic changes and further demonstrated that disease-specific DNA methylation changes identify functionally relevant *cis*-regulatory sequences in CLL. In line with this, transcription factor (TF) binding sites enriched in class A (e.g. IKZF1, BATF, NFAT, EGR1/2) and in class C sequences (e.g. NFAT, EGR1/2, E2A) were predominantly associated with B cell biology, e.g. BATF controling the expression of activation-induced cytidine deaminase (AID) and of IH-CH germline transcripts or E2A controlling B cell lineage commitment. This suggested involvement of altered TF binding patterns in CLL pathogenesis: class A CpG sites are characterized by stronger than normal TF binding and class C sites are likely de novo bound by B cell specific TFs (**Figure 3d, e**). Class B sites were enriched in motifs for EBF, NKX6-1 and PAX5, but overall the motif enrichment as well as the associated changes in chromatin accessibility were only moderate (**Figure 3c-e**). Binding of proteins related to genome architecture (CTCF, RAD21, SMC3) was overrepresented in class D sites (**Figure 3d, e**). Aberrant DNA methylation patterns at TF binding sites in CLL might be associated with disturbed TF expression levels. TF expression analysis revealed transcriptional deregulation of MAFB, JUN, KLF14, KLF4, IRF2 and EBF1, none of which showed major changes in their promoter DNA methylation status (**Supplementary Figure S4a, b**). Among the deregulated TFs, EBF1 showed the strongest and most consistent transcriptional deregulation with almost complete loss of expression in CLL samples (log2-FC: −7.98 [CLL - hiMBC]; **Supplementary Figure S4a**). The EBF1 downregulation potentially explains the observed CLL-specific hypermethylation at class B sites, as EBF1 has been shown to possess pioneering activity [38]. Similarly, upregulation of KLF4, JUN and IRF2 (**Supplementary Figure S4a**) could explain hypomethylation programming observed at class A and C CpG sites as all of these TFs have been reported to possess pioneering activity [39–41].

### Class D hypermethylation is associated with reduced CTCF binding and potentially deregulates expression of neighboring genes

The enrichment of CTCF binding sites and motifs as well as the enrichment of ChromHMM insulator regions (**Figure 3a, d, e**) led us to investigate the effects of aberrant CTCF binding in CLL in more detail. We found that class D sites had lower CTCF occupancy and reduced chromatin accessibility in CLL samples as compared to normal B cells (**Figure 3f**) while globally, these patterns were identical (**Supplementary Figure S5a, b**). The differences in CTCF binding were associated with changes in gene expression of neighboring genes (**Figure 3g**). This further highlights the importance of aberrant CTCF binding at class D CpGs and might point towards a novel pathogenetic mechanism in CLL. Unfortunately, the low absolute number of class D sites does not allow a comprehensive analysis of associated gene expression changes and further studies involving whole-genome bisulfite sequencing will be required to systematically address this observation.

### Identification of epigenetically deregulated transcripts in CLL

The promoter DNA methylation status is widely used as a marker for gene regulation and significant correlation of promoter DNA methylation with gene expression has been demonstrated before [12,42–44]. Previous studies in CLL identified many epigenetic events potentially deregulating the expression of protein-coding genes and miRNAs. However, all of the work published so far used CD19^+^ B cells as controls to call aberrant DNA methylation [6,9,11,45–53]. To stress the importance of using appropriate controls to delineate disease-specific DNA methylation events, we compared our cell-of-origin model to the classical approach using bulk CD19^+^ B cells as a reference. We correlated DNA methylation levels of all aberrant promoter CpGs with gene-expression. The classical approach resulted in a ∼1.5-fold overcalling of epigenetically deregulated protein-coding genes (**Supplementary Figure S6a**).

For miRNAs this difference was even more pronounced (about 5- to 7-fold; **Supplementary Figure S6b**). Interestingly, previously identified differentially methylated promoters of *TCL1*, *HOXA4*, *TWIST2* or *DAPK1* did not pass the stringent filtering criteria of our correlation analysis. This suggested that applying Methyl-COOM results in the identification of a more relevant set of epigenetically deregulated candidate genes.

Using the cell-of-origin model, corrleation between promoter DNA methylation and miRNA expression levels identified 8 CLL-specific miRNAs (**Figure 4a, b**). Seven out of these miRNAs have been demonstrated to regulate epigenetic key players, and, even more importantly, they regulate genes that have been shown to be recurrently mutated in CLL, namely *ARID1A*, *ASXL1*, *CHD2*, *SETD1A*, *SETD2* and *KMT2D*. Reasoning that miRNA binding to their target genes results in gene expression changes, we compared expression levels between miRNAs and their target genes in CLL and normal B cells. Indeed, concordant with the pattern of miRNA promoter hypomethylation and subsequent upregulation of miRNA transcript levels, we found that known target genes of CLL-specific miRNAs were significantly downregulated in CLL as compared to normal B cells while non-target genes were unaffected (**Figure 4c**).

**Figure 4.**
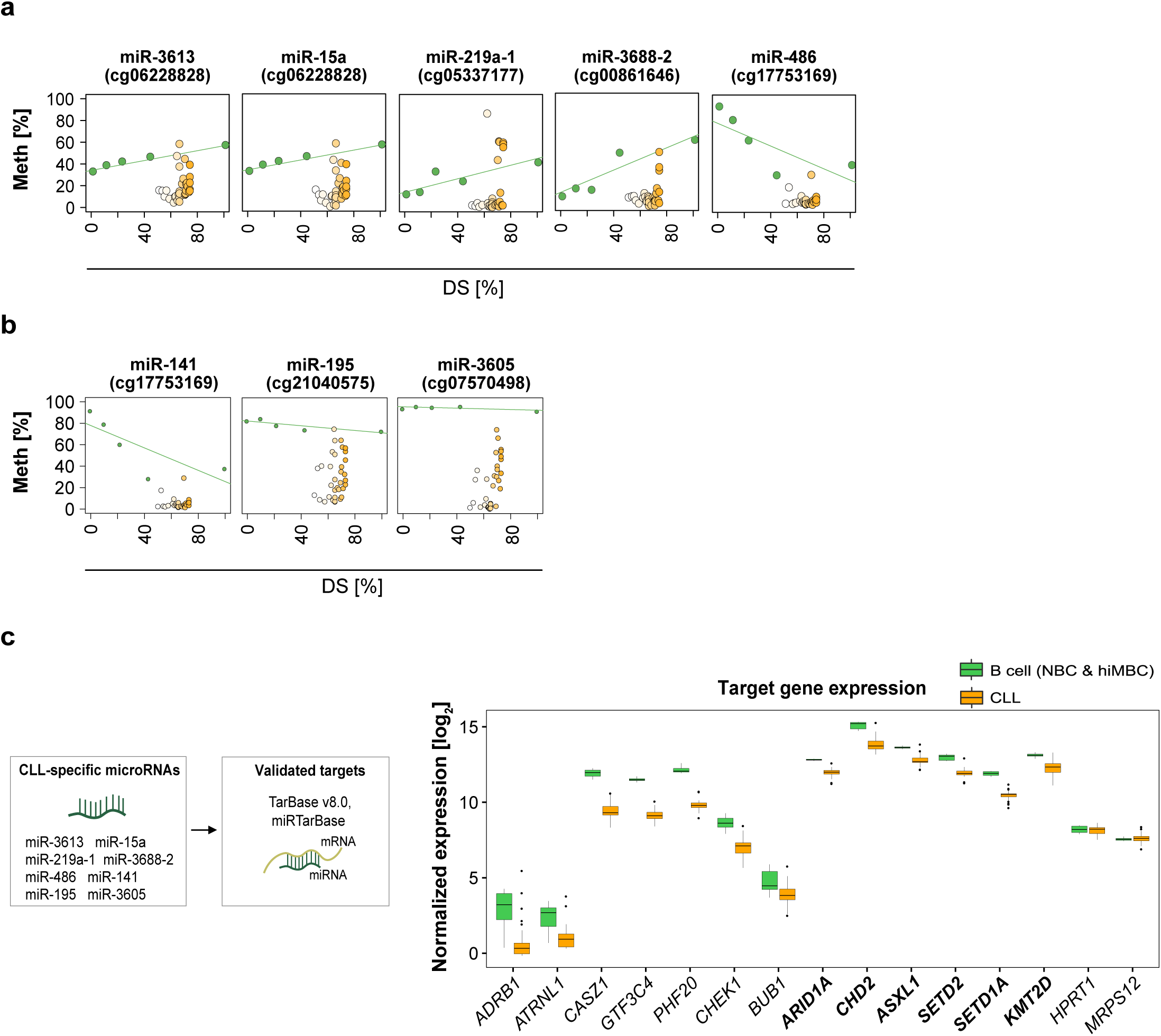
microRNAs associated with CLL-specific DNA methylation. **a)** Candidate CLL-specific microRNAs deregulated by class A events in their promoter regions. Epigenetic programming during normal B cell differentiation is represented as a green line. Average DNA methylation values are represented as dots; normal B cell subpopulations (green dots); CLL samples (white-orange dots). The y-axis represents DNA methylation levels (%), while the x-axis depicts the differentiation stage of normal B cell subpopulations and of CLL samples relative to hiMBCs (DS). **b)** Candidate CLL-specific microRNAs deregulated by class C events in their promoter regions. Epigenetic programming during normal B cell differentiation is represented as a green line. Average DNA methylation values are represented as dots; normal B cell subpopulations (green dots); CLL samples (white-orange dots). The y-axis represents DNA methylation levels (%), while the x-axis depicts the differentiation stage of normal B cell subpopulations and of CLL samples relative to hiMBCs (DS). **c)** CLL-specific microRNAs target epigenetic regulators. <u>Left panel</u>: schematic outline of microRNA-target gene prediction. Two databases of experimentally validated targets of microRNAs, TarBase v8.0 and miRTarBase, were used to define a set of CLL-specific microRNA targets. <u>Right panel</u>: normalized gene expression levels (rlog normalized) of epigenetic regulators being targeted by CLL-specific microRNAs as well as gene expression levels of non-target genes (negative controls; *HPRT1* and *MRPS12*) are shown. Recurrently mutated epigenetic regulators in CLL are presented in bold. Statistical significance of expression change between normal B cells (NBCs, hiMBCs) and CLLs was tested using Wilcoxon rank sum test (p-values: *ARDB1*=0.002; *ATRNL1*=0.0013; *CASZ1*= 0.000014; *GTF3C4*=0.000014; *PHF20*=0.000014; *CHEK1*= 0.000025; *BUB1*= 0.007; *ARID1A*=0.000014; *CHD2*=0.00003; *ASXL1*=0.00005; *SETD2*=0.00002; *SETD1A*=0.000014; *KMT2D*= 0.00007; *HPRT1*=0.43, *MRPS12*=0.45).

A similar correlation analysis on protein-coding genes revealed statistically significant correlations between DNA methylation and gene expression for 491 (class A), 20 (class B), 390 (class C), and 20 (class D) genes. The majority of correlations observed were negative (i.e. a decrease in DNA methylation was associated with an increase in gene expression and vice versa; **Supplementary Figure S6c**), and, as expected, the negative correlation with gene expression was most unambiguous for hypermethylation events (59% class A, 95% class B, 70% class C, 85% class D; **Figure 5a**, **Supplementary Figure S6d**). A detailed analysis of the top correlating genes (Pearson correlation test, p-value <0.05; correlation coefficient < −0.7) encompassing 102 transcripts demonstrated a tight link between CLL-specific aberrant DNA methylation and the expression levels of the corresponding genes (**Figure 5b**; **Supplementary Figure S6a**). Normal B cell differentiation-related epigenetic and transcriptional changes were exaggerated in class A and B whereas the changes detected in class C and D were observed exclusively in CLL. Aberrantly methylated CpGs of class A and C converged in promoters of 12/102 transcripts (*TIGIT*, *SH3D21*, *LAX1*, *LILRB4*, *CD5*, *NOD2*, *POLR3GL*, *IGFBP4*, *ZAP70*, *KSR2*, *XXYLT1*−*AS2*, and *LAG3*), highlighting the potential functional relevance of the associated genes in CLL pathogenesis. In order to validate our findings, we applied Methyl-COOM to 107 CLL samples that have been published previously by Oakes and colleagues (**Supplementary Figure S7a**; [4]). This analysis identified 11,059 CLL-specific CpGs, of which 8,440 (76%) overlapped with the 10,339 CpGs identified in our discovery cohort (**Supplementary Figure S7b**). Furthermore, CLL-specific CpGs identified in our validation cohort recapitulated 92/102 (90%) of the top correlating candidate genes found in the discovery cohort (**Supplementary Figure S7c**.

**Figure 5.**
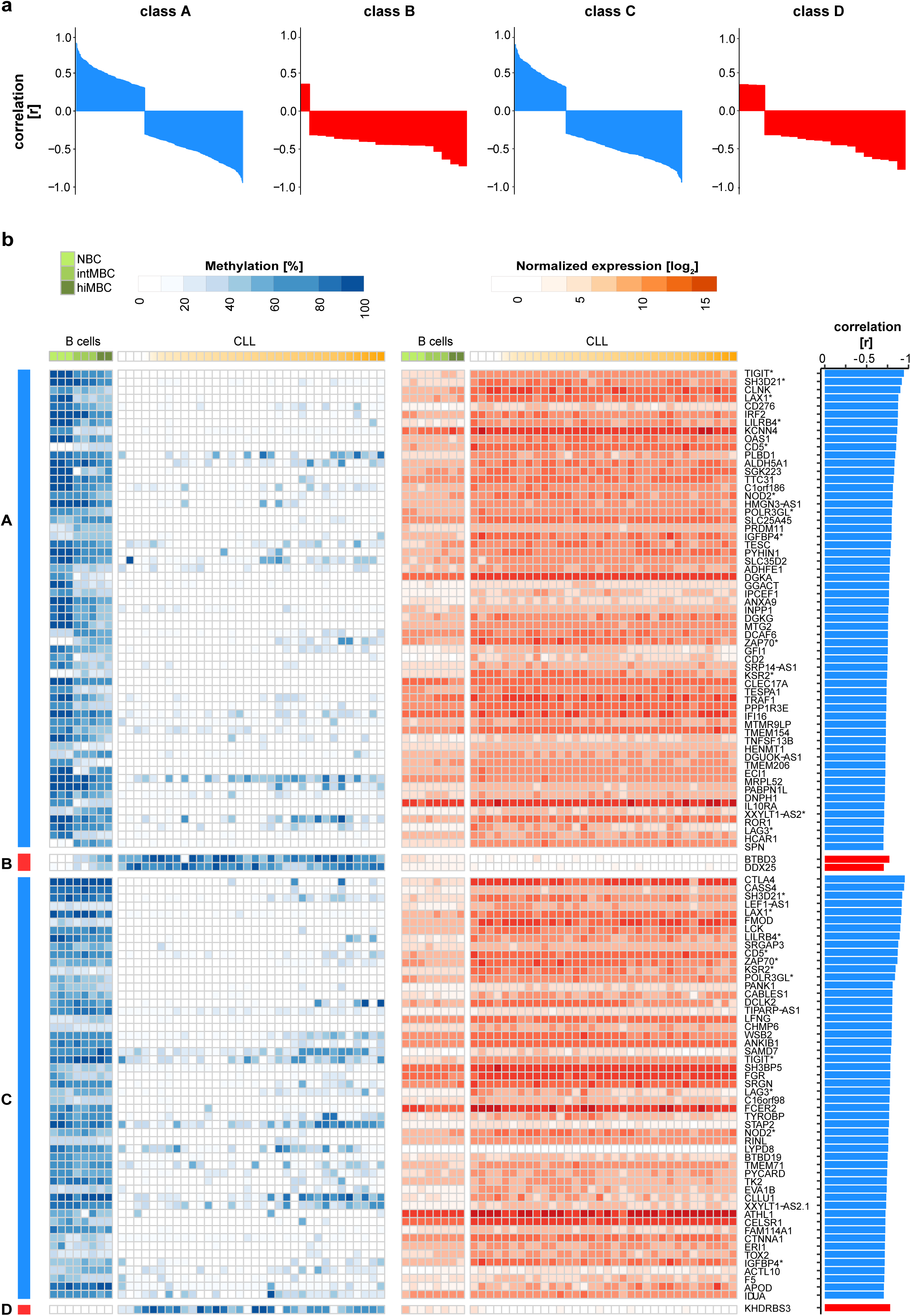
Protein-coding genes associated with CLL-specific aberrant DNA methylation. **a)** Waterfall plots summarizing the correlation coefficients [r] between DNA methylation in the promoters and gene expression profiles of protein-coding genes for each class of CLL-specific alterations (classes A - D). The direction of DNA methylation change is indicated in blue and red for hypo- and hypermethylation, respectively. **b)** CLL-specific epigenetically deregulated transcripts. <u>Left panel</u>: heatmap depicting absolute DNA methylation levels [%] at CLL-specific CpG sites (classes A - D) in the promoter regions of protein-coding genes. Samples were sorted according to the differentiation stage. Differentiation stages are denoted as color gradients, CLLs (white - orange), normal B cells (light - dark green). <u>Middle panel</u>: heatmap depicting normalized gene expression levels (rlog normalization) of protein-coding genes in B cells (light - dark green) and CLLs (white - orange). Transcripts enriched for more than one class of CLL-specific events in their promoter regions are marked with asterisks. <u>Right panel</u>: barplots summarizing correlation coefficients [r] from Pearson correlation analysis between DNA methylation at CLL-specific CpGs in the promoter region and protein-coding gene expression levels. The direction of DNA methylation change is indicated in blue and red for hypo- and hypermethylation, respectively.

### Epigenetically deregulated transcripts are enriched for T cell-related and immune-modulating genes

Some of the top correlating genes have already been implicated to play a role in CLL biology, e.g. *ZAP70*, *CD5*, *LCK*, *LAG3* or *CLLU1* (**Supplementary Figure S8a, b**), while for others their role in CLL pathogenesis is currently unknown. To gain insights into the potential functional role of these epigenetically deregulated genes, we performed enrichment analysis of known biological functions, interactions, or pathways. MSigDB and GO analysis revealed strong enrichment of gene sets related to immune response, immune system processes, hematopoietic stem cells, CLL, and NOTCH signaling (**Supplementary Figure S8a, b**). Ingenunity Pathway Analysis (IPA) and Metascape analysis resulted in enrichment of T-lymphocyte-related processes (*<u>Metascape:</u>* ‘Reguation of T cell activation’, ‘Reguation of T cell receptor signaling pathway’, ‘T cell costimulation’, ‘T cell differentiation’; *<u>IPA</u>*: ‘Cell Proliferation of T Lymphocytes’, ‘T cell homeostasis’, ‘Proliferation of lymphocytes’ (**Supplementary Figure S8a**). These findings are in line with recent reports demonstrating that CD8^+^ T cells from patients with chronic lymphocytic leukemia exhibit features of T cell exhaustion, i.e. lower proliferative and cytotoxic capacity and increased expression of inhibitory receptors (e.g. CTLA-4, TIGIT, Lag3, PD-1), suggesting both CLL and T cell specific changes leading to decreased ability to eliminate malignant cells [54–57].

### Epigenetically deregulated transcripts show aberrant protein expression in CLL

Cancer cells express immune regulatory molecules that might represent potential targets for novel immunotherapies. These proteins modulate the activity of tumor-infiltrating immune cells and mediate immune-escape of tumor cells. Among the epigenetically deregulated genes we identified several with immune regulatory function. Therefore, we aimed to determine whether these are also aberrantly expressed at the protein level in CLL cells. We selected 5 candidates from the list of top correlated genes which are known to be involved in lymphocyte/T-lymphocyte related processes (*TIGIT*, *CTLA-4*, *CD276*, *LILRB4*, and *CD2*; **Figure 6a**). Flow cytometry was utilized for the differential analysis of protein expression in malignant (CD19^+^CD5^+^) and normal (CD19^+^CD5^-^) B cells of 7 CLL patients’ blood samples (gating strategy in **Supplementary Figure S9a**). We found that CTLA-4, TIGIT, LILRB4 and CD276 showed statistically significant increased expression in malignant B cells as compared to normal B cells (CTLA-4, p-val=0.047; TIGIT, p-val=0.016; CD276, p-val=0.016; LILRB4, p-val=0.016 [Wilcoxon paired signed-rank test]), while CD2 surface expression was not detectable neither in normal nor CLL B cells (**Figure 6b**; **Supplementary Figure S9b**). Despite the fact that the functional relevance of some of these aberrantly expressed proteins (TIGIT, CD276 or LILRB4) still remains to be elucidated in the context of CLL, our observation is of particular interest for the development of new therapeutic strategies in CLL. Options to interfere with the signaling of these receptors are currently investigated as potential novel therapeutic strategies in several cancer entities.

**Figure 6.**
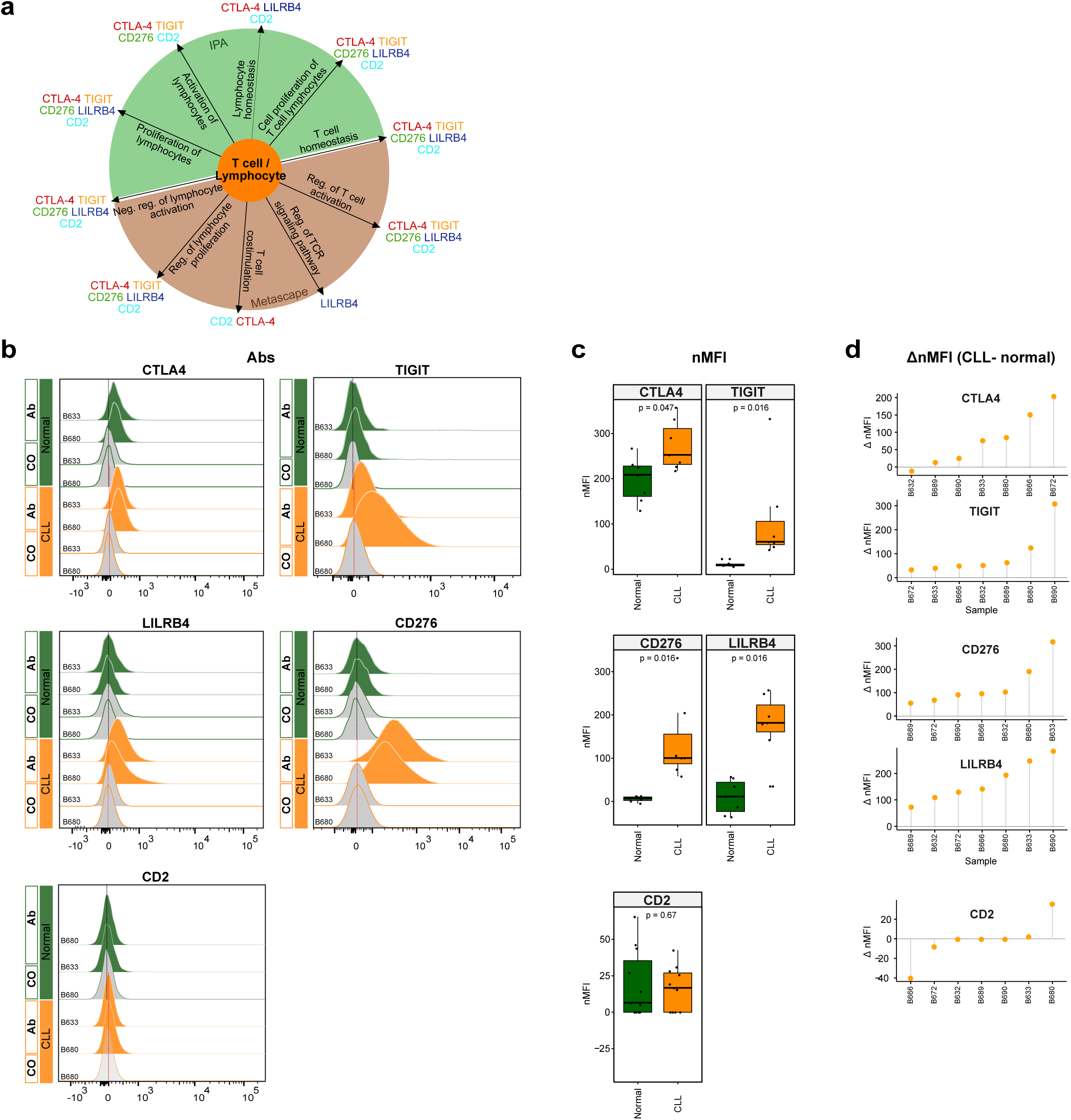
Flow cytometry analysis of T cell-/lymphocyte-specific markers on normal and malignant B cells from CLL patients. **a)** Summary scheme representing functional implications of CLL-specific candidate genes selected for flow cytometric analysis. **b)** Flow cytometric analysis of expression of CTLA-4, TIGIT, CD276, LILRB4, and CD2 on peripheral blood B cells of CLL patients. The expression was determined for non-malignant B cells (‘Normal’; CD19^+^ CD5^-^ B cells, represented in green) and neoplastic B cells (‘CLL’, CD19^+^ CD5^+^ B cells, represented in orange) detected in the same samples. ‘Co’, no antibody staining control; ‘Ab’, staining with the antibody of interest as indicated. **c)** <u>N</u>ormalized median fluorescence intensities (target MFI - MFI of negative control [Co]; nMFI). **d)** Δ normalized median fluorescence intensities between CLL cells and normal B cells (ΔnMFI (CLL-normal)) for each patient tested.

## DISCUSSION

Applying Methyl-COOM analysis to CLL cells, we identified a number of microRNAs and protein-coding genes that are epigenetically deregulated and validated the CLL-specific epigentic deregulation for the vast majority of target genes in an independent patient cohort. These epigentically deregulated transcripts are likely involved in the pathogenesis or maintenance of CLL and are functionally enriched for immune system- and lymphocyte-related processes. The expression levels of these transcripts are very low in normal B cells, which is in stark contrast to the strong overexpression observed in CLL cells. These epigenetically deregulated transcripts are further expressed and detectable on the surface of malignant B cells. CLL patients are known to progressively develop an immunosuppressive state including dysfunctional T cells [57] and our data suggest that CLL cells contribute to the immunosuppressive microenvironment as well as T cell exhaustion by expressing immune regulatory molecules. Immune dysregulation is known to worsen over the course of the disease, e.g. effector T cells are increased in early-stage disease and show progressive accumulation and exhaustion in the late-stage [57, 58]. This, together with the fact that CLL frequently affects older patients with co-morbidities, makes CLL an ideal candidate for the development of effective immunotherapies. CD276, TIGIT and LILRB4 would be of particular interest, since to our knowledge they were not yet considered as immunotherapeutic targets in CLL. TIGIT is a recently identified inhibitory receptor expressed on T cells and natural killer (NK) cells. In T cells, TIGIT expression inhibits cell proliferation, cytokine production, and T cell receptor signaling [59]. In tumors, TIGIT is involved in mediating a T cell exhaustion phenotype, which is manifested by poor effector function of T cells and, consequently, decreased ability to eliminate tumor cells. In non-Hodgkin B cell lymphomas, PD1- and TIGIT-expressing intratumoral T cells were shown to mark dysfunctional or exhausted effector T cells [60]. CLL patients with an advanced disease stage display elevated numbers of TIGIT^+^ CD4^+^ T cells compared to low risk patients [61]. In preclinical models of colorectal and breast carcinoma, TIGIT blockade was shown to reverse the exhaustion phenotype of cytotoxic T cells and to inhibit tumor growth [62]. Another immune inhibitory receptor, LILRB4, was reported as tumor-associated antigen that is highly expressed on monocytic AML cells [63, 64]. It was also reported as a selective marker of neoplastic B cells and HSCs from CLL patients [65]. LILRB4 targeting, either by antibodies or by CAR-T cells, impeded AML development [55, 56]. CD276 overexpression, on the other hand, was linked to anti-apoptosis in colorectal cancer through activation of Jak2-STAT3 signaling pathway, and as a result, increased expression of anti-apoptotic protein Bcl-2 [66]. High CD276 expression levels were already linked to poor prognosis in CLL, prostate and pancreatic cancer [67–70]. Altogether, TIGIT, LILRB4 and CD276 represent attractive therapeutic targets for treatment of CLL.

The present study demonstrates that Methyl-COOM delineates cancer-specific DNA methylation patterns and identifies deregulated pathways involved in the pathogenesis or maintenance of CLL. Our work serves as a proof-of-concept that tracing the cell-of-origin by comparison to normal differentiation trajectories is of great conceptual importance in cancer epigenetics. Identifying the cell-of-origin is not only crucial for the precise analysis of epigenetic data, but it is also important for clinical translation. The cell-of-origin impacts on tumor biology, affects chemo- and radiosensitivity and influences disease outcome. For instance, studies in a murine model of MLL-rearranged AML have shown that the cell-of-origin can influence the phenotype and the aggressiveness of the resulting leukemia [71]. Likewise, glioma subtypes vary in their response to therapy and share molecular signatures with different normal neural lineages, suggesting a difference in their cellular origin [72–76]. So far, the identification of a cancer’s cellular origin is based on genetic lineage-tracing experiments in mice, like the ones from Blanpain and colleagues demonstrating the presence of distinct cells-of-origin for two types of skin cancer [77]. In colorectal cancer the cell-of-origin has been studied intensively, pointing towards three potential cell types as founder cells: intestinal stem cells [78–82], transit amplifying cells [78, 83], and differentiated villus cells [83]. In most instances, however, the precise cell-of-origin, in which transformation occurs, remains undefined.

Methyl-COOM can, in principle, be applicable to any type of DNA methylation data as a source of epigenetic information. In contrast to previous reports in CLL and other malignancies, epigenetic pathomechanisms were investigated using an approach that systematically avoids confounding factors introduced by epigenome dynamics occurring in the context of physiological differentiation processes. It has been demonstrated that similar concepts apply to other lymphatic neoplasms, e.g. T-ALL, DLBCL or MCL [84–87]. However, for other tumors, including myeloid malignancies, the knowledge on the cell-of-origin is still scarce. Therefore, beyond the field of CLL research this study could serve as a template for the analysis of epigenomic data in other cancer entities.

## CONCLUSIONS

Our work describes a new analytical framework, Methyl-COOM, to delineate cancer-specific DNA methylation patterns, a concept that should, in principle, be applicable to all tumor entities. Using Methyl-COOM, we interrogated DNA methylomes of CLL samples in the context of normal B cell differentiation. This enabled us to unmask abnormal transcription factor and super enhancer activities, as well as to identify aberrant transcript expression in CLL. Furthermore, we were able to demonstrate that epigenetically deregulated transcripts are enriched in immune regulatory molecules which are also expressed at the protein level in CLL cells, suggesting that CLL cells contribute to immunosuppression and T cell exhaustion by upregulation of immune regulatory molecules. This finding might serve as a starting point for the development of novel therapeutic strategies to overcome immune evasion of CLL cells.

## DECLARATIONS

### Ethics approval and consent to participate

The study was conducted in accordance with the declaration of Helsinki and was approved by the Ethics Committee Heidelberg (University of Heidelberg, Germany; S-206/2011; S-356/2013) and by the Ethics Committee Ulm (Ulm University; 130/2002). Samples were taken after patients gave their written informed consent.

### Availability of data and materials

The datasets used and analysed in the current study were published previously as indicated in Table 3. The Methyl-COOM framework is accessible via GitHub (https://github.com/justannwska/Methyl-COOM)[88].

## URLs

Bioconductor http://bioconductor.org/ [90]

Human genome (hg19, GRCh37) http://genome.ucsc.edu/downloads.html

LOLA https://bioconductor.org/packages/release/bioc/html/LOLA.html [31]

ENCODE https://www.encodeproject.org/ [89]

HOMER http://homer.ucsd.edu/homer/ [30]

miRTarBase: http://mirtarbase.mbc.nctu.edu.tw/php/index.php [91]

TarBase v8.0 http://carolina.imis.athena-innovation.gr/diana_tools/web/index.php?r=tarbasev8%2Findex [92]

microRNA.org http://www.microrna.org [93]

miRBase v.18.0 http://www.mirbase.org [94]

## Author contributions

J.A. W, C.P. and D.B.L. developed the research concept, designed the analysis workflow and experiments, and collected and interpreted the data. J.A.W., R.T., N.I., T.H., Y.A. and P.L. analyzed data. J.A.W. performed experiments. K.R., J.M., L.K., D.M., T.Z., Marc.S., R.K., S.S., J.B and C.C.O. provided clinical samples or data. P.M.R. and Mart.S. performed flow cytometry experiments and analyzed data. J.A.W, C.P. and D.B.L. prepared the figures and wrote the manuscript. C.P. and D.B.L. jointly supervised the project. All authors contributed to the writing of the manuscript and approved the final version.

## Acknowledgments

We would like to thank Thomas Höfer, Stefan Fröhling, Annika Baude, and Simin Öz for helpful discussions.

