## Supplementary Figures S1-S9 for "Methylome-based cell-of-origin modeling (Methyl-COOM) identifies aberrant expression of immune regulatory molecules in CLL"

#### Supplementary Figure 1

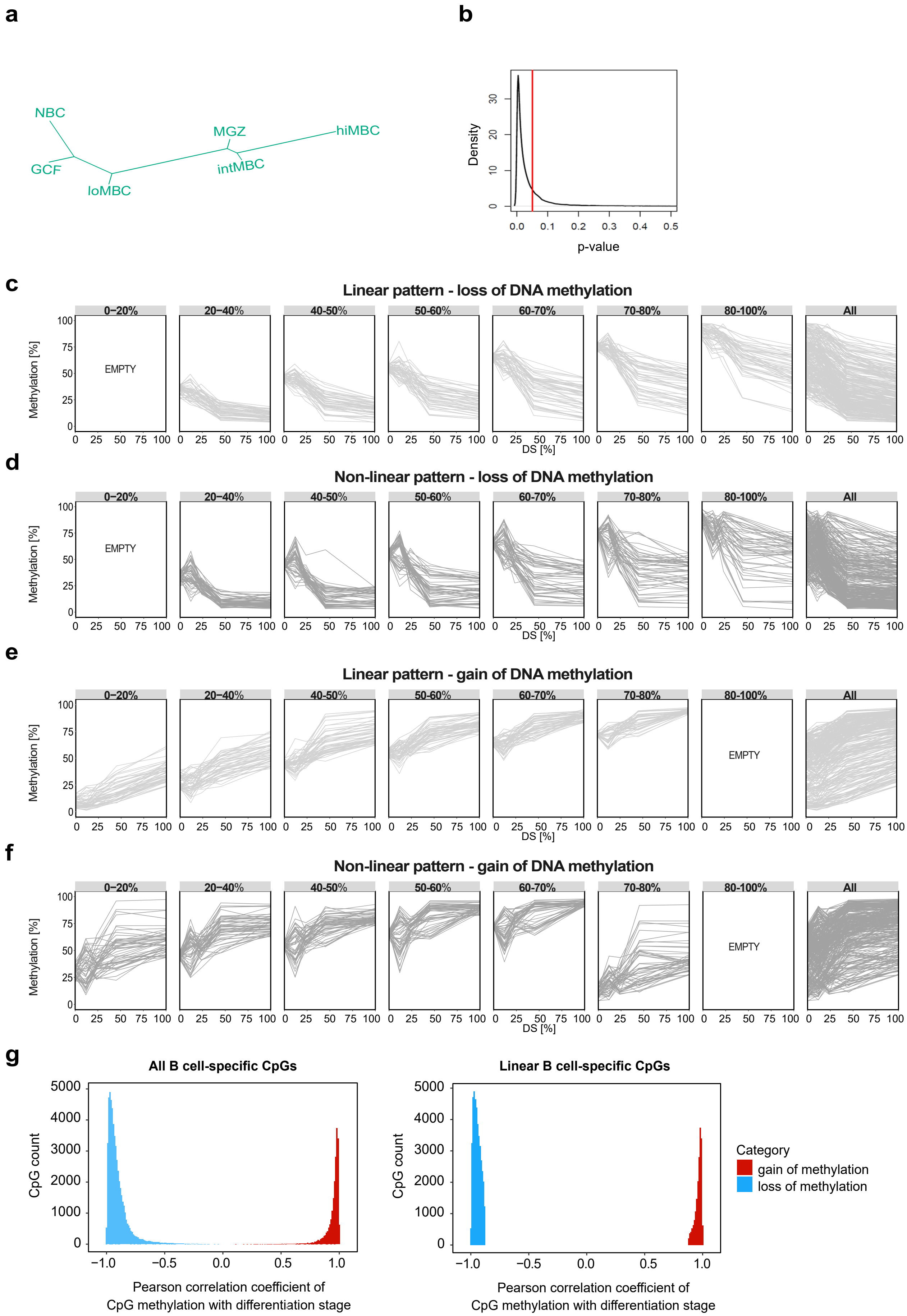

**Supplementary Figure 1. DNA methylation dynamics during normal B cell differentiation can be described by a linear model.**

**a) DNA methylation-based phylogenetic tree of normal B cell development**. The phylogenetic tree was generated using a set of CpG sites (total of 74,333CpGs, >20% DNA methylation change) that show dynamic DNA methylation changes during normal B cell differentiation (B cell-specific CpGs, minimum evolution method, R package ape). Manhattan pairwise distances between DNA methylation profiles of normal B cells at B cell-specific CpGs were used to determine the mode of methylation progression from naïve to memory B cell. Each branch represents a different B-cell subtype. NBC – naïve B cells; GCF –germinal center founder B cells; loMBC – early non-class switched memory B cells; intMBC – non class-switched memory B cells; sMGZ – splenic marginal zone B cells; hiMBC – class-switched memory B cells.

**b) Linear relationship between the differentiation stage of every B cell and the DNA methylation profiles at B cell-specific CpGs.** F-test statistics was used to test for linear relationship between the assigned differentiation stage for every B cell and the DNA methylation values at B cell-specific sites at a single CpG level. The vast majority of the B cell-specific CpGs (79.8%, 59,326 CpGs, p-value <0.05) showed linear DNA methylation dynamics across the six B cell differentiation stages. The y axis represents the density of B cell-specific CpG sites. The x axis represents p-values from F-test. The red line indicates p-value=0.05.

**c-f) Linearity of DNA methylation changes during normal B cell differentiation.** Absolute DNA methylation (Methylation [%]) was categorized into eight bins (0-20%; 20-40%, 40-50%; 50-60%; 60-70%; 70-80%; 80-100%, all events) according to the DNA methylation status of naïve B cells. For each DNA methylation bin, 50 B cell-specific CpGs were randomly selected from a total pool of 41,244 linear **(c)**, or 13,034 non-linear **(d)** B cell-specific CpGs that are losing methylation during B cell differentiation. For each DNA methylation bin with gain in methylation, 50 B-cell specific CpGs were randomly selected from a total pool of 18,102 linear **(e)**, or 1,973 non-linear **(f)** B cell specific CpGs. The y-axes represent absolute DNA methylation levels (%), while the x-axes depict the differentiation stage (DS) of normal B cells relative to hiMBCs.

**g) Histogram of Pearson correlation coefficients for DNA methylation status and differentiation stage of normal B cells**. Left panel: the histogram depicts the distribution of Pearson correlation coefficients for absolute DNA methylation levels with differentiation stage of all B cell-specific CpGs. Right panel: the histogram depicts the distribution of Pearson correlation coefficients for absolute DNA methylation levels with differentiation stage of linear B cell-specific CpG, only. Correlation is depicted depending on the direction of DNA methylation changes during B cell differentiation. DNA methylation loss (blue), DNA methylation gain (red).

#### Supplementary Figure 2

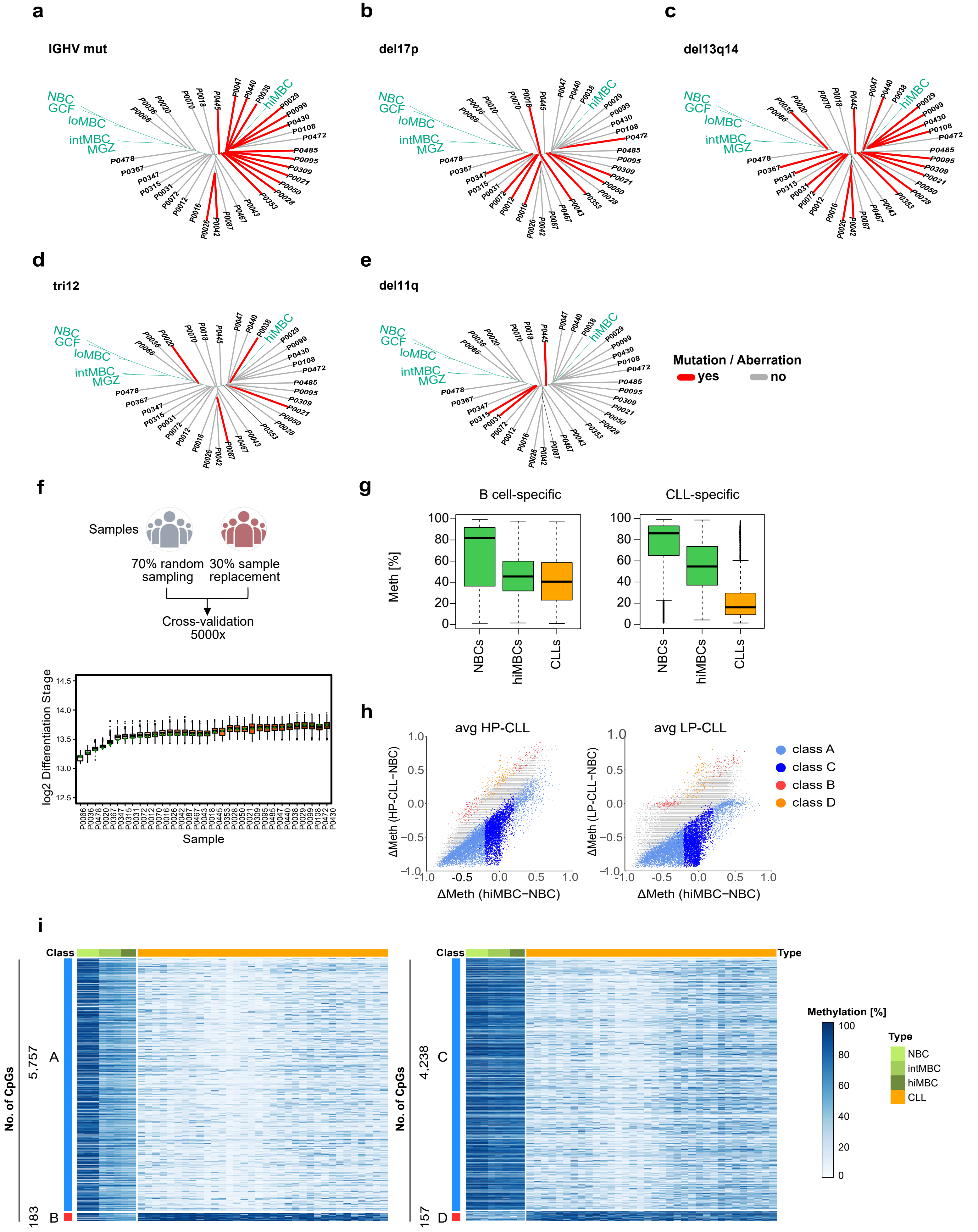

**Supplementary Figure 2. DNA methylation patterns of CLL in relation to normal B cell differentiation.**

**a-e) Identification of the cell-of-origin in CLL samples and their molecular/cytogenetic status**. The presence of IGVH mutation (a), del17p (b), del13q14 (c), tri12 (d), del11q (e) are indicated in red, absence thereof in grey. Normal B cells are represented in green. NBCs - naïve B cells; GCFs – germinal center founder B cells; loMBCs – early non class-switched memory B cells; intMBCs – non class-switched memory B cells; sMGZs – splenic marginal zone B cells; hiMBCs – class-switched memory B cells (mature B cells).

**f) Robustness of the cell-of-origin assignment.** Bootstrapping (5000x) with a random sample replacement was used to infer phylogenetic relationships and the closest normal B cell methylome (cell-of-origin) for every CLL sample. CLL patient cohort was repeatedly divided into two subgroups; 70% and 30% (5000 bootstraps). To minimize the likelihood of selection of the same sample multiple times, that would result in a bias towards few samples in the bootstrap analysis, a random sampling was allowed in the 70%-group, while sample replacement was restricted only to the 30%-group. The x axis represents CLL samples, the y axis denotes log2 distances from the assigned cell-of-origin to NBCs (log2 DS). Green points are representing the assign cell-of-origin using with the full CLL sample set (original assignment is represented in Supplementary Figure 1c).

**g) Net DNA methylation changes at B cell- and CLL-specific CpGs.** Left panel: net DNA methylation change at B cell-specific CpGs for B cells (green) and CLL samples (orange). Right panel: net DNA methylation change at CLL-specific CpGs for B cells (green) and CLL samples (orange). The x-axes represent different cell types; NBCs – naïve B cells; hiMBCs - class-switched memory B cells; CLL – chronic lymphocytic leukemia B cells. The y-axes denote absolute DNA methylation levels (%).

**h) CLL-specific DNA methylation events in the context of the classification by Oakes *et al.*** Differences in DNA methylation were represented from naive B cells to high-mature memory B cells (x-axis) and to CLLs (y-axis). Data for CpGs were averaged for each CLL subtype (average LP-CLL, average HP-CLL). CpGs were categorized as class A (light blue), class B (red), class C (dark blue), and class D (orange).

**i) Heatmap depicting absolute DNA methylation changes (Methylation, [%]) at CLL-specific CpG sites.** Unsupervised hierarchical clustering of CLL-specific CpGs, class A and B sites (left), class C and D sites (right). The direction of DNA methylation change is indicated as blue and red bars for hypo- and hypermethylation, respectively. CLL samples are represented in orange. Normal B cells are represented in green (NBCs – naïve B cells; intMBC – non class-switched memory B cells, hiMBCs - class-switched memory B cells).

### Supplementary Figure 3

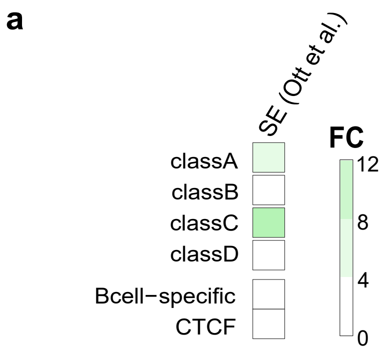


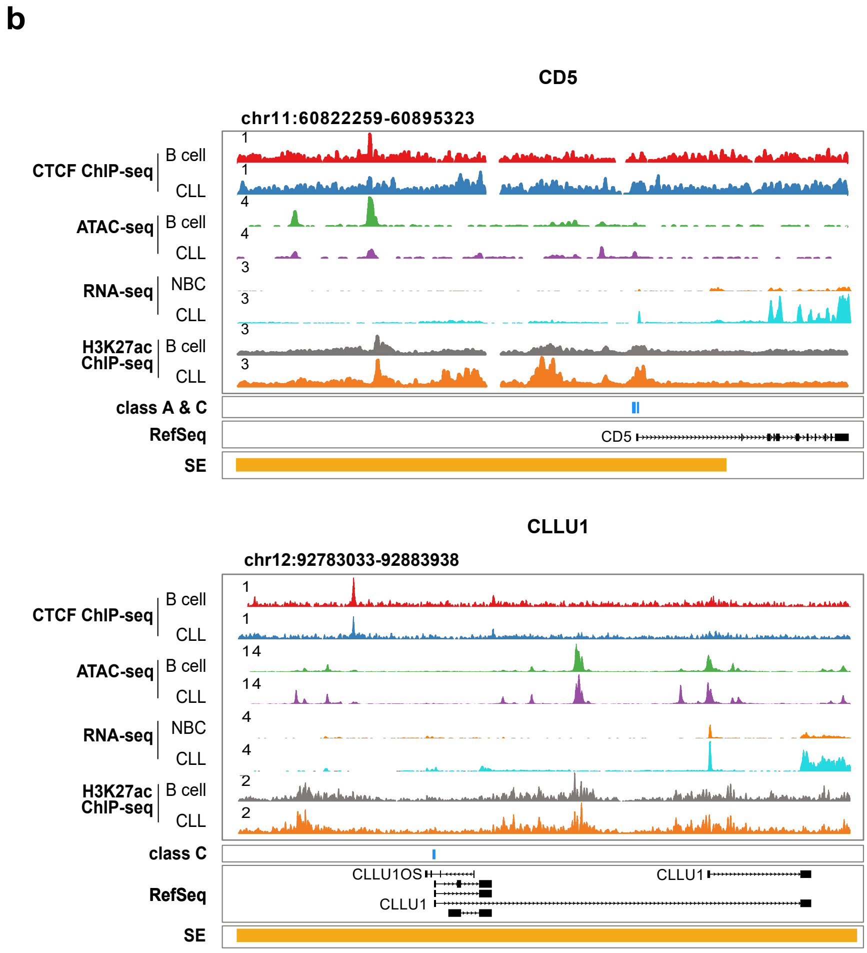

**Supplementary Figure 3. CLL-specific DNA methylation affects super-enhancers.**

**a) Enrichment of unified super-enhancer regions (SE) from Ott et al. in sequences representing CLL-specific methylation** (PMID:30503705). Union of SEs was defined based on individual patient data in Ott et al. (n=18). Fold change (FC) was calculated using all 450k probes as a background. B-cell specific CpGs and CTCF motifs were used as controls.

**b) Locus plots of exemplary CLL-specific SE-associated genes.** CpG sites overlapping SEs were associated with the closest gene and the correlation analysis between DNA methylation and gene expression was used to identify CLL-specific SE-associated genes. Locus plots include data from CTCF ChIP-seq on normal B cell (red) and CLL (blue); ATAC-seq on normal B cells (green) and CLL (purple); RNA-seq on normal B cells (orange) and CLL (cyan); H3K27ac on normal B cell (grey) and CLLs (orange). CLL-specific CpGs are annotated in blue. SE annotations are represented in orange.

#### Supplementary Figure 4

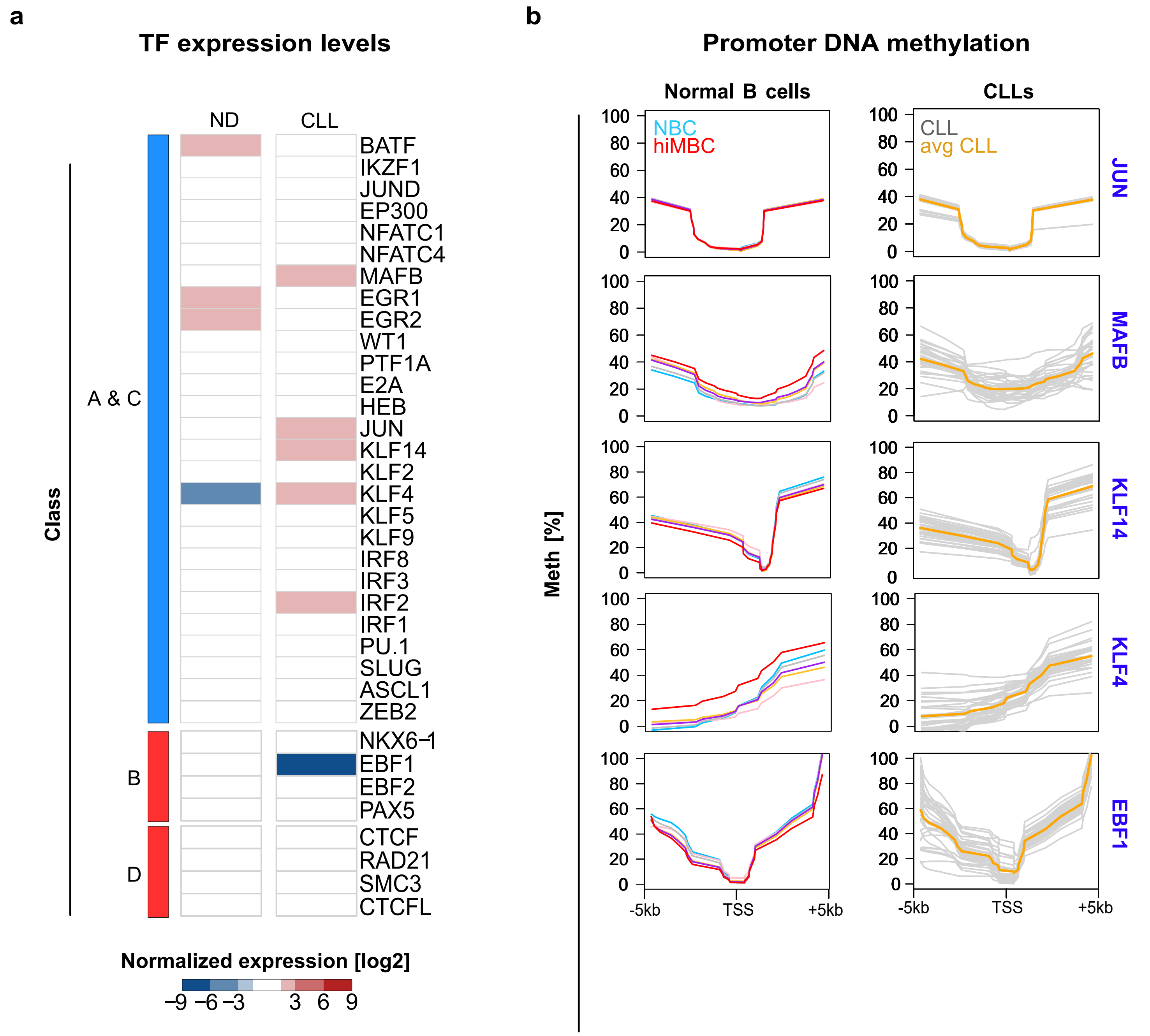

**Supplementary Figure 4. Aberrant TF programming in CLL.**

**a) Comparison of TF expression levels in CLL and in normal B cells**. TFs were selected based on the enrichment analysis presented in Figure 5d and 5e. Two different comparisons of expression patterns were represented, normal B cell differentiation-associated (ND, ΔhiMBC-NBC), and CLL-associated (CLL; ΔCLL-hiMBC). Normalized expression values were used (rlog normalization, log2). The direction of DNA methylation change observed at motif-associated CLL-specific CpG sites is indicated as blue and red bars for hypo- and hypermethylation, respectively.

**b) DNA methylation profiles of promoters of differentially expressed TFs identified at CLL-specific CpG sites**. DNA methylation in the promoter regions is shown for six normal B cell subsets, representing different stages of B cell differentiation (left) and the CLL (right). NBC: naïve B cell; hiMBC – class-switched memory B cell; avg CLL – average DNA methylation change in CLL. The y-axis represents DNA methylation levels (%). The x-axis represents the transcription start site (TSS) +/-5kb.

### Supplementary Figure 5

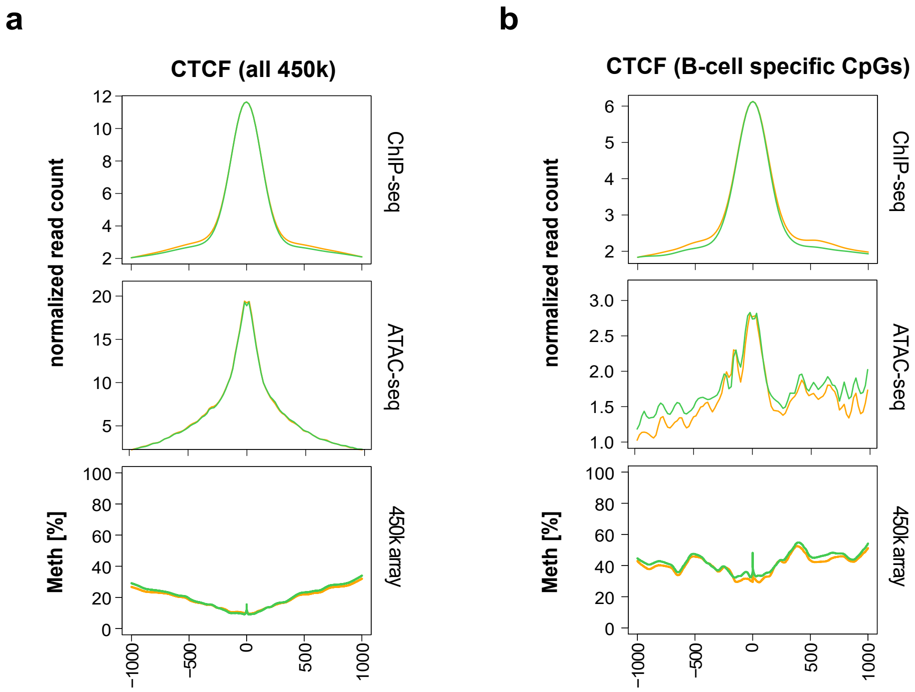

**Supplementary Figure 5. Aberrant CTCF programming in CLL.**

**a)** ATAC-seq and ChIP-seq read density and DNA methylation levels (%) at all 450k CpG probes co-locating with CTCF motifs (n=21,810 CpGs). Depicted are the genomic regions surrounding the CTCF motifs (±400bp). CLL samples (n=7 CTCF ChIP-seq, n=18 ATAC-seq) are represented in orange, normal $CD19^+$ B cells (n=4 CTCF ChIP-seq, n=3 ATAC-seq) are depicted in green.

**b)** ATAC-seq and ChIP-seq read density and DNA methylation levels (%) at B cell-specific CpG sites co-locating with CTCF motifs (n=1,587 CpGs). Depicted are the genomic regions surrounding the CTCF motifs (±400bp). CLLs (n=7 CTCF ChIP-seq, n=18 ATAC-seq) are represented in orange, normal $CD19^+$ B cells (n=4 CTCF ChIP-seq, n=3 ATAC-seq) are depicted in green.

#### Supplementary Figure 6

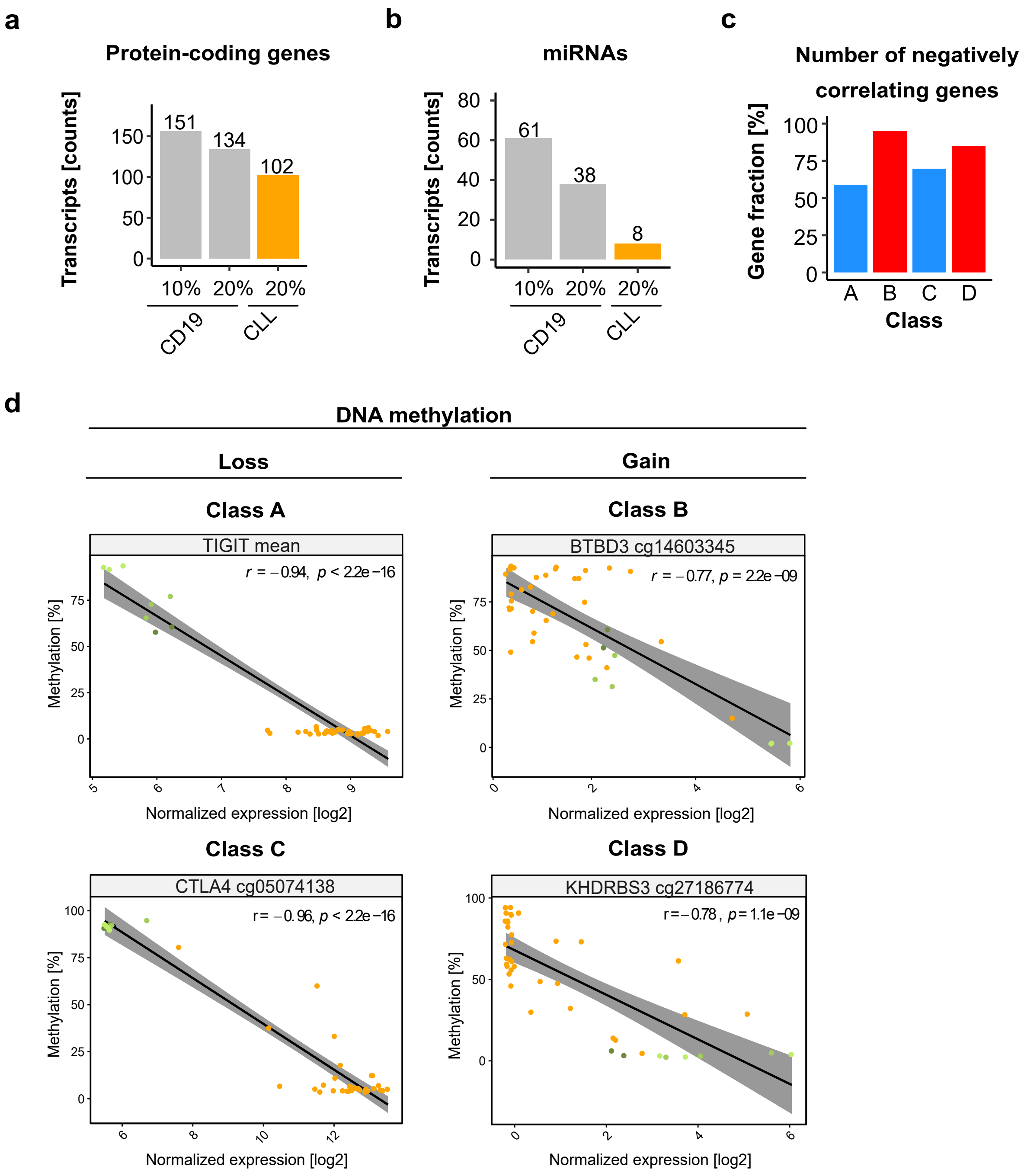

**Supplementary Figure 6. Transcripts associated with CLL-specific aberrant DNA methylation.**

**a) Usage of $CD19^+$ B cells as a reference overestimates the number of CLL-specific protein-coding genes.** Differential DNA methylation between control B cells and CLL samples was calculated using different DNA methylation thresholds (10% or 20%). The bar plot illustrates the proportion of transcripts defined as CLL-specific using different control B cell sources ($CD19^+$ B cells are represented in grey, individual cell-of-origin is represented in orange). For CLL-specific protein coding genes, we used a correlation coefficient cutoff $<-0.7$. The numbers of uniquely identified CLL-specific genes are annotated on the top of the bars.

**b) Usage of $CD19^+$ B cells as a reference overestimates the number of CLL-specific microRNAs.** Differential DNA methylation between control B cells and CLL samples was calculated using different DNA methylation thresholds (10% or 20%). The bar plot illustrates the proportion of microRNAs defined as CLL-specific using different control B cell sources ($CD19^+$ B cells are represented in grey, individual cell-of-origin is represented in orange). For CLL-specific microRNAs we used an correlation coefficient cutoff $\leq-0.35$. The numbers of uniquely identified CLL-specific microRNAs are annotated on the top of the bars.

**c) Fraction of negatively correlating CLL-specific protein-coding genes.** DNA methylation at CLL-specific CpGs in the promoters of protein-coding genes was correlated with gene expression levels (Pearson correlation). Depicted is the percentage of negatively correlating genes (correlation coefficient r, $r<0$).

**d) Exemplary correlation plots between CLL-specific DNA methylation and gene expression.** DNA methylation levels at promoter regions of protein-coding genes were correlated with the expression levels of associated genes. Examples for negative correlations are represented for each class of CLL-specific CpGs. The x-axes depict gene expression levels (log2 normalized counts) and the y-axes represent absolute DNA methylation [%]. Green: normal B cells. Orange: CLL samples.

#### Supplementary Figure 7

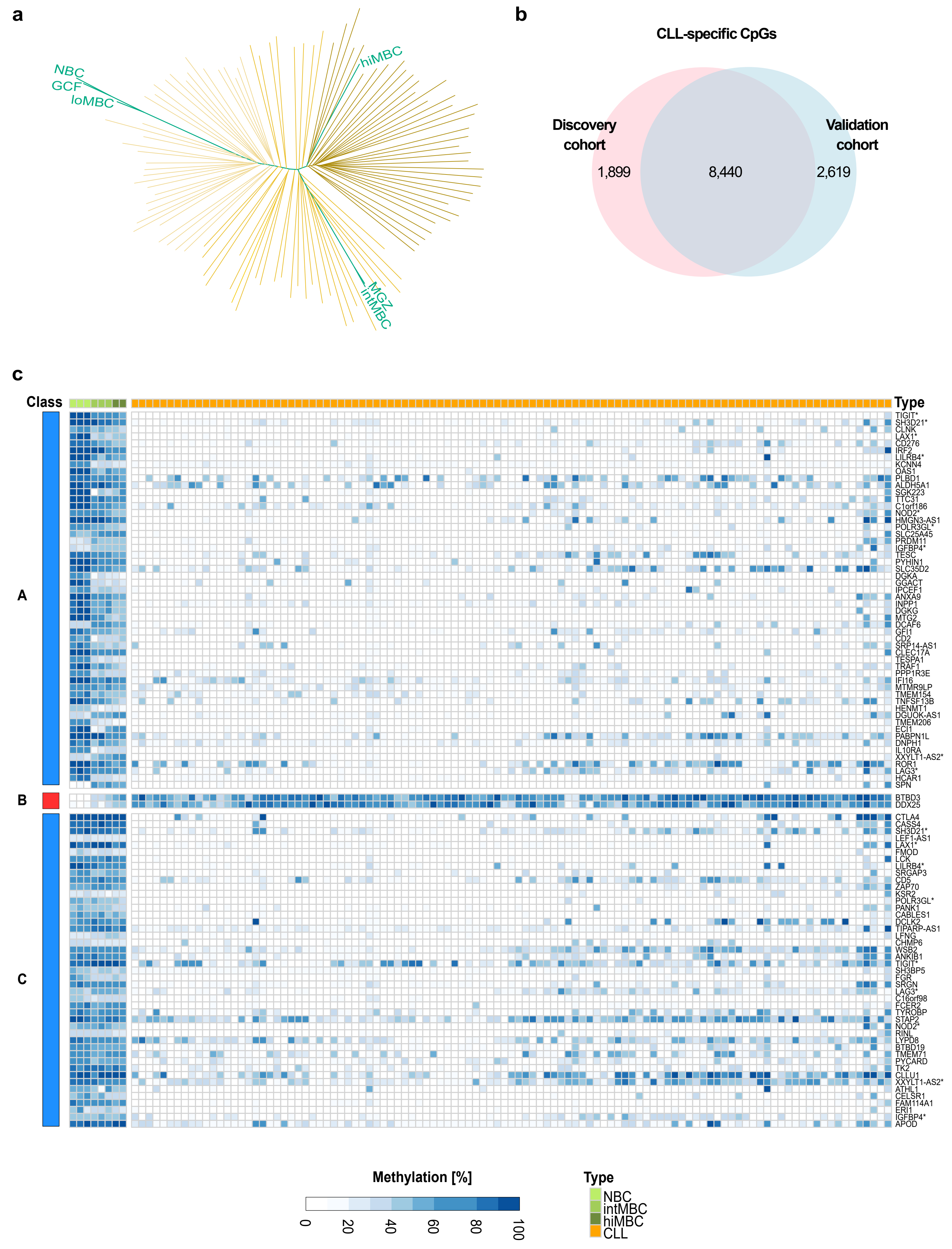

**Supplementary Figure 7. Validation of CLL-specific DNA methylation events in an independent cohort of CLL patients.**

**a) Identification of the cell-of-origin in CLL samples using phylogenetic analysis**. The Methyl-COOM framework was applied to an independent cohort of CLL samples (Oakes et al., Nat Genet 2016). Depicted is the phylogenetic tree that was generated using a set of linear CpG sites that show dynamic methylation changes during normal B cell differentiation as (linear B cell-specific CpGs, 59,326 CpGs). NBCs - naïve B cells; GCFs – germinal center founder B cells; loMBCs – early non class-switched memory B cells; intMBCs – non class-switched memory B cells; sMGZs – splenic marginal zone B cells; hiMBCs – class-switched memory B cells (mature B cells). The gradient color code of CLL samples corresponds to different levels of maturity reached by the cell-of-origin during the transformation event. CLL samples with a relatively immature cell-of-origin that are reprogrammed early during the differentiation process are represented in light orange color. CLLs with a cell-of-origin reprogrammed at a later stage of B cell differentiation are depicted in dark orange color. Normal B cells are represented in green.

**b) High degree of concordance between CLL-specific CpGs identified in the discovery and validation cohorts.** The Venn diagram summarizes the overlap between the CLL-specific CpGs identified in the discovery (n=34 samples; 10,339 CpGs) and in the validation (n=107; 11,029 CpGs) cohorts.

**c) Protein-coding genes associated with CLL-specific aberrant DNA methylation**. Heatmap depicting absolute DNA methylation levels [%] at CLL-specific CpG sites (classes A, B, C, and D) in the promoter regions of protein-coding genes. CLLs are represented in orange, normal B cells in green. Transcripts associated with more than one class of CLL-specific events in their promoter regions are marked with asterisks.

#### Supplementary Figure 8

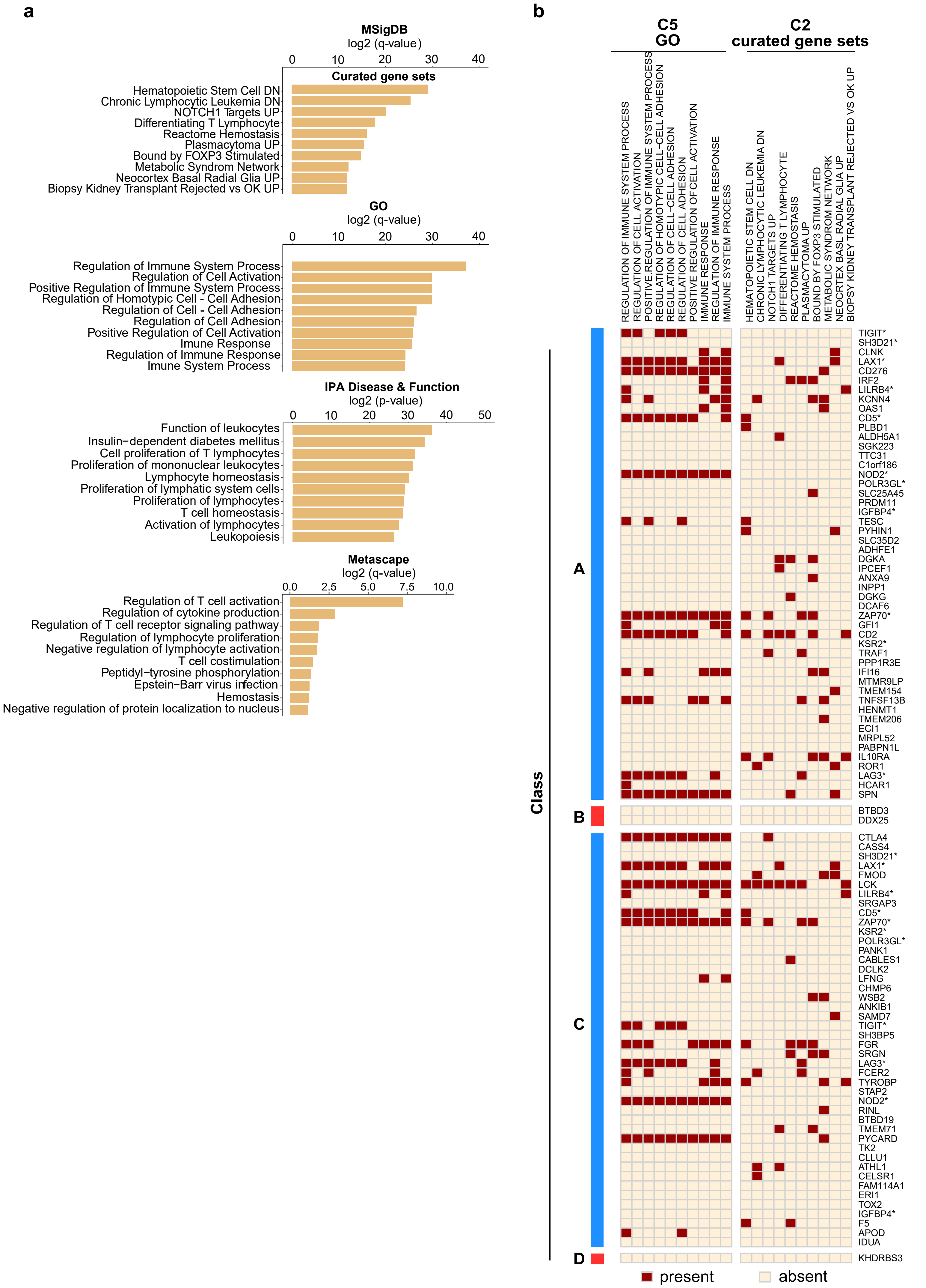

**Supplementary Figure 8. Transcripts associated with CLL-specific aberrant DNA methylation.**

**a) Functional enrichment analysis of CLL-specific protein-coding genes.** First panel: enrichment results from MSigDB 'Curated gene set'. Second panel: gene ontology (GO) enrichment analysis of CLL-specific epigenetically deregulated transcripts. Third panel: ingenuity (IPA) pathway disease & function enrichment. Fourth panel: metascape enrichment analysis of CLL-specific epigenetically deregulated transcripts

**b) Heatmap depicting results from MSigDB functional enrichment analysis of CLL-specific protein-coding genes.** MSigDB 'C2 Curated gene set' and MSigDB 'C5 GO analysis'. The direction of DNA methylation change is indicated on the left side of the heatmap in blue and red for hypo- and hypermethylation, respectively**.**

### Supplementary Figure 9

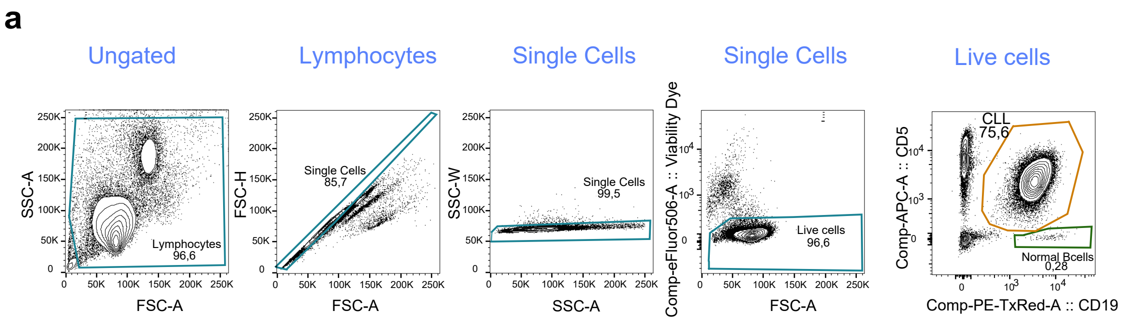


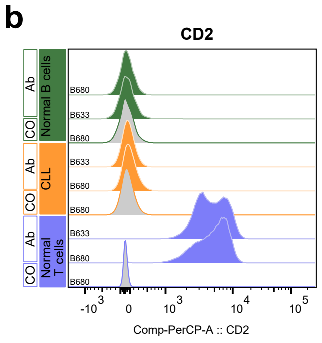

**Supplementary Figure 9. Flow cytometry analysis of B cells from CLL patients.**

**a) Gating strategy for the isolation of normal and malignant B cells from CLL patients.** Normal B cells were identified by the sole expression of $CD19^+$, while neoplastic B cells are positive for both $CD19^+$ and $CD5^+$.

**b) Flow cytometry analysis of CD2.** Normal B cells ('Normal B cells'; $CD19^+$ B cells; green), CLL cells ('CLL', $CD19^+$ $CD5^+$ B cells; orange) and normal T cells ('Normal T cells', $CD5^+$ T cells, blue). 'Co', no antibody staining control; 'Ab', staining with an anti-CD2 antibody.
