## Supplementary Table S1 for "Methylome-based cell-of-origin modeling (Methyl-COOM) identifies aberrant expression of immune regulatory molecules in CLL"

**Supplementary Table S1: List of CLL-specific protein-coding genes**

| Gene | Promoter | CpG | p-value | correlation coefficient r | class |
|---|---|---|---|---|---|
| ADHFE1 | chr8:67342217-67345217 | mean | 1.59e-09 | -0.775837250729239 | classA |
| ALDH5A1 | chr6:24492696-24495696 | cg03967327 | 1.00E-11 | -0.832091356011116 | classA |
| ANXA9 | chr1:150951998-150954998 | cg13320146 | 3.73e-09 | -0.764744929323204 | classA |
| C1orf186 | chr1:206287736-206290736 | cg13685151 | 9.38e-09 | -0.752056172411214 | classA |
| C1orf186 | chr1:206287736-206290736 | cg24056232 | 2.64e-09 | -0.769315732700692 | classA |
| C1orf186 | chr1:206287736-206290736 | mean | 6.00E-11 | -0.813500698809516 | classA |
| CD2 | chr1:117294585-117297585 | cg16719404 | 1.28e-08 | -0.747600522077558 | classA |
| CD276 | chr15:73974121-73977121 | cg04289575 | 0 | -0.874527181733735 | classA |
| CD5 | chr11:60867429-60870429 | cg06417460 | 0 | -0.851852052685724 | classA |
| CD5 | chr11:60867429-60870429 | cg24674703 | 2.2e-10 | -0.799491802538728 | classA |
| CD5 | chr11:60867429-60870429 | mean | 2.00E-11 | -0.822284217248575 | classA |
| CLEC17A | chr19:14691395-14694395 | cg25770495 | 4.14E-05 | -0.729915634966355 | classA |
| CLEC17A | chr19:14691395-14694395 | cg25945273 | 2.44E-05 | -0.738064096698458 | classA |
| CLEC17A | chr19:14691395-14694395 | mean | 3.02E-05 | -0.734820163263315 | classA |
| CLNK | chr4:10685886-10688886 | cg03485217 | 0 | -0.902138940405541 | classA |
| DCAF6 | chr1:167903296-167906296 | cg12749559 | 8.51e-09 | -0.753440217870646 | classA |
| DGKA | chr12:56322445-56325445 | cg08338281 | 5.97e-09 | -0.758366680493838 | classA |
| DGKA | chr12:56322445-56325445 | cg21587469 | 2.62e-09 | -0.769404486622536 | classA |
| DGKA | chr12:56322445-56325445 | mean | 3.5e-09 | -0.765594073561009 | classA |
| DGKG | chr3:186079523-186082523 | cg00434119 | 6.96e-09 | -0.756246419253565 | classA |
| DGUOK-AS1 | chr2:74208066-74211066 | cg05981785 | 5.83E-05 | -0.724477661281641 | classA |
| DNPH1 | chr6:43196711-43199711 | cg00741333 | 8.84E-05 | -0.717685969276767 | classA |
| ECI1 | chr16:2301102-2304102 | cg09970941 | 6.50E-05 | -0.722731689525009 | classA |
| GFI1 | chr1:92951933-92954933 | cg04777348 | 1.98e-08 | -0.741190569026367 | classA |
| GFI1 | chr1:92951933-92954933 | cg24517501 | 4.57E-05 | -0.728356366023144 | classA |
| GFI1 | chr1:92951933-92954933 | mean | 9.45e-09 | -0.751949870523501 | classA |
| GGACT | chr13:101240546-101243546 | cg27248148 | 3.02e-09 | -0.767548032706135 | classA |
| HCAR1 | chr12:123214629-123217629 | cg00357958 | 2.43E-03 | -0.700368020041794 | classA |
| HENMT1 | chr1:109203648-109206648 | cg00328227 | 5.66E-05 | -0.72496185800815 | classA |
| HMGN3-AS1 | chr6:79940890-79943890 | cg16343616 | 1.00E-10 | -0.807533344420226 | classA |
| IFI16 | chr1:158977181-158980181 | cg19939130 | 4.14E-05 | -0.729901659598959 | classA |
| IFI16 | chr1:158977181-158980181 | mean | 1.03E-04 | -0.715083005703993 | classA |
| IGFBP4 | chr17:38597175-38600175 | cg11961138 | 2.6e-10 | -0.797233146952221 | classA |
| IL10RA | chr11:117854605-117857605 | cg01697865 | 1.43E-04 | -0.709611644879929 | classA |
| INPP1 | chr2:191205695-191208695 | cg17516156 | 6.05e-09 | -0.758203706920682 | classA |
| IPCEF1 | chr6:154677400-154680400 | cg19690214 | 3.27e-09 | -0.766487769861641 | classA |
| IRF2 | chr4:185395226-185398226 | cg21960364 | 0 | -0.872806717076645 | classA |
| KCNN4 | chr19:44284909-44287909 | cg04833845 | 2.6e-10 | -0.79739098179341 | classA |
| KCNN4 | chr19:44284909-44287909 | cg18506018 | 9.00E-11 | -0.80921244685823 | classA |
| KCNN4 | chr19:44284909-44287909 | cg24900654 | 0 | -0.86457564495141 | classA |
| KCNN4 | chr19:44284909-44287909 | cg26890181 | 3.6e-10 | -0.793679429228187 | classA |
| KCNN4 | chr19:44284909-44287909 | mean | 2.00E-11 | -0.823838784220103 | classA |
| KSR2 | chr12:118405528-118408528 | cg00791253 | 1.62E-05 | -0.744132060949442 | classA |
| LAG3 | chr12:6879169-6882169 | cg04153135 | 2.35E-03 | -0.700972300571603 | classA |
| LAX1 | chr1:203731783-203734783 | cg00942920 | 2.83e-09 | -0.768391494276638 | classA |
| LAX1 | chr1:203731783-203734783 | cg06958535 | 0 | -0.846292199442119 | classA |
| LAX1 | chr1:203731783-203734783 | cg10117369 | 2.65e-09 | -0.769291392946307 | classA |
| LAX1 | chr1:203731783-203734783 | cg23893629 | 0 | -0.878182011463705 | classA |
| LAX1 | chr1:203731783-203734783 | mean | 1.00E-11 | -0.83110589001876 | classA |
| LILRB4 | chr19:55171770-55174770 | cg05329879 | 0 | -0.869977324233555 | classA |

| | | | | | |
|---|---|---|---|---|---|
| LILRB4 | chr19:55171770-55174770 | cg24140775 | 1.1e-10 | -0.806822986698213 | classA |
| LILRB4 | chr19:55171770-55174770 | mean | 0 | -0.845037733390338 | classA |
| MRPL52 | chr14:23296591-23299591 | cg07436208 | 8.03E-05 | -0.71927874503728 | classA |
| MTG2 | chr20:60755580-60758580 | cg06914270 | 7.66e-09 | -0.754908705435461 | classA |
| MTMR9LP | chr1:32706811-32709811 | cg02830936 | 4.18E-05 | -0.729740099649233 | classA |
| NOD2 | chr16:50728549-50731549 | cg18177814 | 7.00E-11 | -0.811278424325301 | classA |
| NOD2 | chr16:50728549-50731549 | cg26954174 | 6.62e-09 | -0.756941629112072 | classA |
| NOD2 | chr16:50728549-50731549 | mean | 6.4e-10 | -0.787080076741518 | classA |
| OAS1 | chr12:113342238-113345238 | cg09151598 | 0 | -0.842243492128602 | classA |
| OAS1 | chr12:113342238-113345238 | cg17445535 | 4.00E-11 | -0.818274617592851 | classA |
| OAS1 | chr12:113342238-113345238 | cg25668626 | 0 | -0.855295081267569 | classA |
| OAS1 | chr12:113342238-113345238 | mean | 0 | -0.849520868792348 | classA |
| PABPN1L | chr16:88932568-88935568 | cg00056002 | 8.61E-05 | -0.718132257251161 | classA |
| PABPN1L | chr16:88932568-88935568 | mean | 1.45E-03 | -0.709336282013582 | classA |
| PLBD1 | chr12:14720291-14723291 | cg04473902 | 0 | -0.841047701244261 | classA |
| POLR3GL | chr1:145469887-145472887 | cg19913118 | 2.00E-10 | -0.80050561909857 | classA |
| PPP1R3E | chr14:23771557-23774557 | cg13408636 | 3.11e-08 | -0.73435307144483 | classA |
| PPP1R3E | chr14:23771557-23774557 | mean | 2.02E-03 | -0.703586032203606 | classA |
| PRDM11 | chr11:45113063-45116063 | cg26821540 | 2.4e-10 | -0.798525626943073 | classA |
| PRDM11 | chr11:45113063-45116063 | mean | 1.89e-09 | -0.773664105938001 | classA |
| PYHIN1 | chr1:158898836-158901836 | cg15036529 | 8.8e-10 | -0.783214866128537 | classA |
| ROR1 | chr1:64237189-64240189 | cg24502478 | 1.67E-03 | -0.706934283803974 | classA |
| SGK223 | chr8:8238757-8241757 | cg02239258 | 1.00E-11 | -0.829042519609816 | classA |
| SH3D21 | chr1:36769493-36772493 | cg21418854 | 0 | -0.920320523381099 | classA |
| SLC25A45 | chr11:65150672-65153672 | cg24215776 | 2.3e-10 | -0.798885066618466 | classA |
| SLC35D2 | chr9:99145492-99148492 | cg13771161 | 1.55e-09 | -0.77617235803577 | classA |
| SLC35D2 | chr9:99145492-99148492 | mean | 1.41E-05 | -0.746174871537122 | classA |
| SPN | chr16:29671770-29674770 | cg03549705 | 2.44E-03 | -0.700251928294752 | classA |
| SRP14-AS1 | chr15:40329011-40332011 | cg20752233 | 1.32E-05 | -0.747142722045878 | classA |
| TESC | chr12:117536751-117539751 | cg02867305 | 5.7e-10 | -0.788478351126211 | classA |
| TESPA1 | chr12:55378030-55381030 | cg07880109 | 2.55E-05 | -0.737403285905437 | classA |
| TIGIT | chr3:114010332-114013332 | cg13669740 | 0 | -0.941782332028607 | classA |
| TIGIT | chr3:114010332-114013332 | cg19421218 | 0 | -0.943328385018044 | classA |
| TIGIT | chr3:114010332-114013332 | cg19456938 | 0 | -0.942519068577294 | classA |
| TIGIT | chr3:114010332-114013332 | cg20832020 | 0 | -0.94336868518533 | classA |
| TIGIT | chr3:114010332-114013332 | mean | 0 | -0.944244163144143 | classA |
| TMEM154 | chr4:153600817-153603817 | mean | 4.25E-05 | -0.729501104084562 | classA |
| TMEM206 | chr1:212587767-212590767 | cg21400549 | 6.39E-05 | -0.722987229334421 | classA |
| TNFSF13B | chr13:108919476-108922476 | cg09646392 | 4.42E-05 | -0.728871843132472 | classA |
| TRAF1 | chr9:123690951-123693951 | cg14580859 | 3.01E-05 | -0.734837672431118 | classA |
| TTC31 | chr2:74707699-74710699 | cg01584932 | 3.00E-11 | -0.821141954037798 | classA |
| XXYLT1-AS2 | chr3:194866099-194869099 | cg19760965 | 1.44E-03 | -0.709422515837849 | classA |
| ZAP70 | chr2:98327530-98330530 | cg21773162 | 8.88e-09 | -0.752830444305011 | classA |
| ZAP70 | chr2:98327530-98330530 | mean | 1.17E-05 | -0.748935645589701 | classA |
| BTBD3 | chr20:11868976-11871976 | cg12894649 | 3.89e-08 | -0.730882985414123 | classB |
| BTBD3 | chr20:11868976-11871976 | cg14603345 | 2.21e-09 | -0.77159750387462 | classB |
| BTBD3 | chr20:11868976-11871976 | cg23944804 | 2.46e-09 | -0.770243744798231 | classB |
| BTBD3 | chr20:11868976-11871976 | mean | 3.09e-09 | -0.767232236130368 | classB |
| DDX25 | chr11:125771760-125774760 | cg19792599 | 1.93E-03 | -0.704430792155692 | classB |
| ACTL10 | chr20:32251803-32254803 | cg12710480 | 1.20E-03 | -0.712605220341696 | classC |
| ANKIB1 | chr7:91873047-91876047 | cg00035937 | 3.1e-10 | -0.79531822654729 | classC |
| APOD | chr3:195310576-195313576 | cg05624196 | 1.77E-03 | -0.705890728264477 | classC |
| ATHL1 | chr11:286637-289637 | cg08829299 | 5.81E-05 | -0.724523272372594 | classC |

| | | | | | |
|---|---|---|---|---|---|
| ATHL1 | chr11:286637-289637 | cg18727742 | 1.18E-03 | -0.712875470138527 | classC |
| ATHL1 | chr11:286637-289637 | mean | 6.63E-05 | -0.722387394461075 | classC |
| BTBD19 | chr1:45271653-45274653 | cg00043599 | 1.03E-05 | -0.750684621796029 | classC |
| BTBD19 | chr1:45271653-45274653 | cg03434029 | 4.53E-05 | -0.728481793756599 | classC |
| BTBD19 | chr1:45271653-45274653 | cg11448683 | 1.63E-05 | -0.744090460593045 | classC |
| BTBD19 | chr1:45271653-45274653 | cg14565781 | 1.42e-08 | -0.746094104409361 | classC |
| BTBD19 | chr1:45271653-45274653 | mean | 1.21E-05 | -0.748463442745605 | classC |
| C16orf98 | chr16:31210705-31213705 | cg27615603 | 2.14e-09 | -0.772066101044905 | classC |
| CABLES1 | chr18:20712027-20715027 | cg22158648 | 1.00E-10 | -0.807867607930747 | classC |
| CASS4 | chr20:54984667-54987667 | cg00387658 | 0 | -0.891079457653141 | classC |
| CASS4 | chr20:54984667-54987667 | cg23676369 | 0 | -0.889820853353139 | classC |
| CASS4 | chr20:54984667-54987667 | cg24519157 | 0 | -0.947037834873521 | classC |
| CASS4 | chr20:54984667-54987667 | cg26712809 | 0 | -0.896289325413418 | classC |
| CASS4 | chr20:54984667-54987667 | mean | 0 | -0.93982394364249 | classC |
| CD5 | chr11:60867429-60870429 | cg16001613 | 3.32e-09 | -0.766296937980816 | classC |
| CD5 | chr11:60867429-60870429 | cg22202088 | 1.00E-11 | -0.831869672005622 | classC |
| CD5 | chr11:60867429-60870429 | cg27301655 | 0 | -0.870918755345629 | classC |
| CD5 | chr11:60867429-60870429 | mean | 0 | -0.8704877147807 | classC |
| CELSR1 | chr22:46932567-46935567 | cg11533712 | 6.23e-08 | -0.7234032224443 | classC |
| CHMP6 | chr17:78963140-78966140 | cg09164913 | 2.1e-10 | -0.800077183178358 | classC |
| CLLU1 | chr12:92812806-92815806 | cg04845867 | 3.26E-05 | -0.733628428342836 | classC |
| CTLA4 | chr2:204730010-204733010 | cg05074138 | 0 | -0.955258463676382 | classC |
| CTNNA1 | chr5:138086606-138089606 | cg02276665 | 7.72E-05 | -0.719928030684773 | classC |
| DCLK2 | chr4:150996925-150999925 | cg05561193 | 1.2e-10 | -0.806186902731367 | classC |
| ERI1 | chr8:8857813-8860813 | cg18237548 | 8.89E-05 | -0.717599820394129 | classC |
| EVA1B | chr1:36789255-36792255 | cg15095906 | 2.3e-08 | -0.738941390076494 | classC |
| F5 | chr1:169555269-169558269 | cg16054275 | 1.37E-03 | -0.710393216643297 | classC |
| FAM114A1 | chr4:38866853-38869853 | cg16030869 | 1.66E-03 | -0.707060394541062 | classC |
| FAM114A1 | chr4:38866853-38869853 | cg20484352 | 6.92E-05 | -0.721698052776808 | classC |
| FAM114A1 | chr4:38866853-38869853 | mean | 8.60E-05 | -0.718144465611258 | classC |
| FCER2 | chr19:7766532-7769532 | cg05641903 | 2.46e-09 | -0.77026320877382 | classC |
| FGR | chr1:27961227-27964227 | cg09845000 | 9.7e-10 | -0.781999583162846 | classC |
| FGR | chr1:27961227-27964227 | mean | 2.47E-05 | -0.737898480208753 | classC |
| FMOD | chr1:203320057-203323057 | cg03764585 | 0 | -0.88142652812359 | classC |
| FMOD | chr1:203320057-203323057 | cg04704856 | 0 | -0.907789604877199 | classC |
| FMOD | chr1:203320057-203323057 | cg16289210 | 1.00E-11 | -0.836711666434786 | classC |
| FMOD | chr1:203320057-203323057 | cg26987645 | 1.00E-11 | -0.832781309092702 | classC |
| FMOD | chr1:203320057-203323057 | mean | 0 | -0.888163832125579 | classC |
| IDUA | chr4:978284-981284 | mean | 1.98E-03 | -0.704009523613509 | classC |
| IGFBP4 | chr17:38597175-38600175 | cg02627216 | 1.01E-03 | -0.715497356920456 | classC |
| KSR2 | chr12:118405528-118408528 | cg01386425 | 6.1e-10 | -0.787653555752696 | classC |
| KSR2 | chr12:118405528-118408528 | cg16300531 | 0 | -0.845214881030986 | classC |
| KSR2 | chr12:118405528-118408528 | mean | 1.00E-11 | -0.828726719907436 | classC |
| LAG3 | chr12:6879169-6882169 | cg04671742 | 6.51E-05 | -0.722697287349605 | classC |
| LAG3 | chr12:6879169-6882169 | cg22777668 | 8.55e-09 | -0.753379541596865 | classC |
| LAG3 | chr12:6879169-6882169 | mean | 1.41e-09 | -0.777372826901823 | classC |
| LAX1 | chr1:203731783-203734783 | cg25625457 | 0 | -0.914901531537427 | classC |
| LCK | chr1:32714339-32717339 | cg01525376 | 0 | -0.859786478743944 | classC |
| LCK | chr1:32714339-32717339 | cg11683242 | 0 | -0.901245160202916 | classC |
| LCK | chr1:32714339-32717339 | mean | 0 | -0.892795088057934 | classC |
| LEF1-AS1 | chr4:109086180-109089180 | cg08460812 | 0 | -0.901734074946273 | classC |
| LEF1-AS1 | chr4:109086180-109089180 | cg24023289 | 0 | -0.917631346546461 | classC |
| LEF1-AS1 | chr4:109086180-109089180 | mean | 0 | -0.919304862463799 | classC |

| | | | | | |
|---|---|---|---|---|---|
| LFNG | chr7:2549662-2552662 | cg21531604 | 1.6e-10 | -0.802718951049662 | classC |
| LILRB4 | chr19:55171770-55174770 | cg05922591 | 0 | -0.89391550574407 | classC |
| LYPD8 | chr1:248902651-248905651 | cg04024799 | 6.74e-09 | -0.756696609713039 | classC |
| NOD2 | chr16:50728549-50731549 | cg16771652 | 5.92e-09 | -0.758496795248592 | classC |
| NOD2 | chr16:50728549-50731549 | mean | 8.33E-05 | -0.718679694322141 | classC |
| PANK1 | chr10:91404829-91407829 | cg08661112 | 7.10E-05 | -0.721279365653109 | classC |
| PANK1 | chr10:91404829-91407829 | cg25770176 | 8.00E-11 | -0.810111260601639 | classC |
| PANK1 | chr10:91404829-91407829 | mean | 1.83e-09 | -0.774049182050759 | classC |
| POLR3GL | chr1:145469887-145472887 | cg08916374 | 0 | -0.838150538399143 | classC |
| PYCARD | chr16:31213597-31216597 | cg07461837 | 1.64E-05 | -0.744006728610458 | classC |
| RINL | chr19:39368419-39371419 | cg15581429 | 6.00E-09 | -0.758302059024826 | classC |
| SAMD7 | chr3:169626981-169629981 | cg05721199 | 4.9e-10 | -0.790055561159422 | classC |
| SH3BP5 | chr3:15382401-15385401 | cg01788682 | 2.6e-08 | -0.737087998881298 | classC |
| SH3BP5 | chr3:15382401-15385401 | cg05046996 | 9.6e-10 | -0.782126741240425 | classC |
| SH3BP5 | chr3:15382401-15385401 | mean | 6.43e-09 | -0.757355238186218 | classC |
| SH3D21 | chr1:36769493-36772493 | cg02865595 | 2.4e-09 | -0.770549526347736 | classC |
| SH3D21 | chr1:36769493-36772493 | cg10622536 | 0 | -0.917771231016952 | classC |
| SH3D21 | chr1:36769493-36772493 | cg19373545 | 0 | -0.921507136966518 | classC |
| SH3D21 | chr1:36769493-36772493 | mean | 0 | -0.924052799011841 | classC |
| SRGAP3 | chr3:9290869-9293869 | cg26184741 | 0 | -0.876012040719286 | classC |
| SRGN | chr10:70845327-70848327 | cg06486994 | 1.3e-09 | -0.778318826305982 | classC |
| STAP2 | chr19:4338347-4341347 | cg15797834 | 3.3e-09 | -0.766394635853633 | classC |
| STAP2 | chr19:4338347-4341347 | mean | 6.25E-05 | -0.72335657399481 | classC |
| TIGIT | chr3:114010332-114013332 | cg22577252 | 8.3e-10 | -0.783847469307881 | classC |
| TIPARP-AS1 | chr3:156393002-156396002 | cg09379146 | 1.4e-10 | -0.804187702134629 | classC |
| TIPARP-AS1 | chr3:156393002-156396002 | mean | 1.92E-05 | -0.741620449895668 | classC |
| TK2 | chr16:66583815-66586815 | cg09238666 | 2.23E-05 | -0.739389112228126 | classC |
| TK2 | chr16:66583815-66586815 | mean | 2.36E-03 | -0.700908762626889 | classC |
| TMEM71 | chr8:133772414-133775414 | cg05598678 | 1.80E-05 | -0.742630099790969 | classC |
| TMEM71 | chr8:133772414-133775414 | cg10054641 | 1.57E-05 | -0.744619663926485 | classC |
| TMEM71 | chr8:133772414-133775414 | mean | 1.28E-05 | -0.747639118131929 | classC |
| TOX2 | chr20:42540991-42543991 | cg00789528 | 9.52e-08 | -0.716460736074329 | classC |
| TYROBP | chr19:36398711-36401711 | cg21495704 | 3.01e-09 | -0.767613640712531 | classC |
| WSB2 | chr12:118499479-118502479 | cg01932247 | 3.96E-05 | -0.730591551221142 | classC |
| WSB2 | chr12:118499479-118502479 | cg14474811 | 6.6e-10 | -0.786703591721742 | classC |
| WSB2 | chr12:118499479-118502479 | mean | 2.4e-10 | -0.798231282133102 | classC |
| XXYLT1-AS2 | chr3:194866099-194869099 | cg02041946 | 3.44E-05 | -0.732803957918885 | classC |
| XXYLT1-AS2 | chr3:194866099-194869099 | cg07463963 | 6.93E-05 | -0.721686754371287 | classC |
| XXYLT1-AS2 | chr3:194866099-194869099 | mean | 4.43E-05 | -0.72885615074238 | classC |
| ZAP70 | chr2:98327530-98330530 | cg08859278 | 0 | -0.844812783413939 | classC |
| ZAP70 | chr2:98327530-98330530 | cg09006159 | 2.00E-11 | -0.82206607742438 | classC |
| ZAP70 | chr2:98327530-98330530 | cg15933451 | 0 | -0.839690104550259 | classC |
| ZAP70 | chr2:98327530-98330530 | mean | 0 | -0.867532791913483 | classC |
| KHDRBS3 | chr8:136467215-136470215 | cg27186774 | 1.08e-09 | -0.780670565780852 | classD |
