## Supplementary Table S2 for "Methylome-based cell-of-origin modeling (Methyl-COOM) identifies aberrant expression of immune regulatory molecules in CLL"

**Supplementary Table S2: List of super-enhancer associated protein coding genes**

| gene | region | cpg | p | correlation coefficient | class | SE |
|---|---|---|---|---|---|---|
| ABCB10 | chr1:229671416-229695012 | cg00355656 | 1.66E-02 | -0.707024217195626 | classA | consensus |
| ACTG1 | chr17:79467015-79549577 | cg14316944 | 1.78E-02 | -0.705845226357013 | classA | consensus |
| ADRB2 | chr5:148184825-148248568 | cg08370787 | 9.05E-06 | -0.752574523815796 | classA | consensus |
| ADTRP | chr6:11692858-11853115 | cg06193328 | 0 | -0.918711420338961 | classA | consensus |
| ADTRP | chr6:11692858-11853115 | cg16533359 | 0 | -0.935179876128049 | classA | consensus |
| ADTRP | chr6:11692858-11853115 | cg25589070 | 0 | -0.906928339923155 | classA | consensus |
| ADTRP | chr6:11692858-11853115 | mean | 0 | -0.919421215940654 | classA | consensus |
| AIM2 | chr1:158974678-159048845 | cg19939130 | 6.52E-06 | -0.757163008296939 | classA | consensus |
| AIM2 | chr1:158974678-159048845 | cg24539090 | 1.12E-02 | -0.713770403165335 | classA | consensus |
| AIM2 | chr1:158974678-159048845 | mean | 9.05e-09 | -0.75256875555797 | classA | consensus |
| AKAP8 | chr19:15351808-15492926 | cg02681400 | 1.07E-05 | 0.750171439046837 | classA | consensus |
| AKAP8 | chr19:15351808-15492926 | cg10225149 | 1.62E-04 | 0.744188348915589 | classA | consensus |
| AKAP8 | chr19:15351808-15492926 | mean | 1.41E-04 | 0.746233584709234 | classA | consensus |
| ARFRP1 | chr20:62150820-62340527 | cg22298487 | 4.86e-10 | -0.790258810931462 | classA | consensus |
| ARL4C | chr2:235295407-235363318 | cg17342709 | 2.1e-11 | -0.823892330699382 | classA | consensus |
| ARL4C | chr2:235295407-235363318 | cg17342709 | 2.1e-11 | -0.823892330699382 | classA | consensus |
| ATOX1 | chr5:151115934-151160506 | cg01201215 | 1.48E-06 | -0.77672510894645 | classA | consensus |
| ATOX1 | chr5:151115934-151160506 | cg19939793 | 4.00E-04 | -0.730458632556588 | classA | consensus |
| ATOX1 | chr5:151115934-151160506 | mean | 3.27E-06 | -0.766513108657799 | classA | consensus |
| ATP11A | chr13:113318512-113446398 | cg18748064 | 2.06E-04 | -0.740634800018126 | classA | consensus |
| ATP11A | chr13:113318512-113446398 | mean | 5.60E-04 | -0.725127295852944 | classA | consensus |
| ATP6V0D1 | chr16:67465524-67635531 | cg01872947 | 6.03E-04 | 0.723921816089769 | classA | consensus |
| ATXN1 | chr6:16620807-16795056 | cg00464814 | 0 | -0.890921251349027 | classA | consensus |
| ATXN1 | chr6:16620807-16795056 | cg26843612 | 5.00E-11 | -0.815049556277753 | classA | consensus |
| ATXN1 | chr6:16620807-16795056 | mean | 0 | -0.858897843007339 | classA | consensus |
| BICD1 | chr12:32253683-32338900 | cg15061954 | 3.66E-06 | -0.765016457632844 | classA | consensus |
| BIN3 | chr8:22516805-22643451 | cg22631350 | 0 | 0.886986219629904 | classA | consensus |
| BSG | chr19:570931-681129 | cg03233876 | 2.99E-04 | -0.734977551552284 | classA | consensus |
| BSG | chr19:570931-681129 | cg12111149 | 6.86E-04 | -0.721846270181844 | classA | consensus |
| BSG | chr19:570931-681129 | mean | 1.99E-02 | -0.703836514885598 | classA | consensus |
| C8orf46 | chr8:67394992-67465261 | mean | 2.46E-02 | -0.700123234518148 | classA | consensus |
| CABLES1 | chr18:20708779-20896507 | cg26052586 | 1.85E-04 | -0.742196897994612 | classA | consensus |
| CBX7 | chr22:39345734-39549865 | cg08665930 | 1.8e-11 | -0.825004688750587 | classA | consensus |
| CBX7 | chr22:39345734-39549865 | cg18696237 | 3.3e-09 | -0.766384922248524 | classA | consensus |
| CBX7 | chr22:39345734-39549865 | mean | 2.4e-11 | -0.822514955221212 | classA | consensus |
| CD200 | chr3:112050350-112244018 | cg02919122 | 0 | -0.86833919688152 | classA | consensus |
| CD200 | chr3:112050350-112244018 | cg07681748 | 0 | -0.894788143010009 | classA | consensus |
| CD200 | chr3:112050350-112244018 | cg13671536 | 9.71e-10 | -0.781962995200776 | classA | consensus |
| CD200 | chr3:112050350-112244018 | mean | 8.00E-11 | -0.810236646586522 | classA | consensus |
| CDK14 | chr7:90324837-90431656 | cg21700774 | 1.73E-02 | -0.706352516838746 | classA | consensus |
| CHD7 | chr8:61590710-61615719 | cg10232395 | 2.91E-05 | 0.735355304493799 | classA | consensus |
| CHIT1 | chr1:203232464-203504070 | cg06373167 | 8.07e-10 | -0.784213548363276 | classA | consensus |
| CHIT1 | chr1:203232464-203504070 | cg20138067 | 1.96E-02 | -0.704175865626893 | classA | consensus |
| CHIT1 | chr1:203232464-203504070 | mean | 1.17E-04 | -0.748898570492836 | classA | consensus |
| CLIC4 | chr1:25018827-25101619 | cg10546487 | 1.53E-02 | 0.708394898588731 | classA | consensus |
| CLNK | chr4:10575279-10819110 | cg03485217 | 0 | -0.902138940405541 | classA | consensus |
| CLNK | chr4:10575279-10819110 | cg03485217 | 0 | -0.902138940405541 | classA | consensus |
| COL9A2 | chr1:40764961-40783088 | cg10512203 | 2.17E-02 | -0.702344071754918 | classA | consensus |
| OMMD3-BMI1 | chr10:22604623-22630204 | cg10763374 | 1.47E-04 | 0.745564859717203 | classA | consensus |
| CORO6 | chr17:27992137-28117624 | cg23534301 | 4.25E-04 | -0.729506969301656 | classA | consensus |
| CORO6 | chr17:27882320-27952581 | cg00574412 | 2.00E-04 | -0.741062413411059 | classA | consensus |
| CORO6 | chr17:27882320-27952581 | mean | 1.01E-04 | -0.751018170320333 | classA | consensus |
| CORO6 | chr17:27992137-28117624 | cg23534301 | 4.25E-04 | -0.729506969301656 | classA | consensus |
| CORO6 | chr17:27882320-27952581 | cg00574412 | 2.00E-04 | -0.741062413411059 | classA | consensus |
| CORO6 | chr17:27882320-27952581 | mean | 1.01E-04 | -0.751018170320333 | classA | consensus |
| COTL1 | chr16:84578659-84655244 | cg03738384 | 1.04E-03 | -0.714931319361127 | classA | consensus |
| DEGS2 | chr14:100525300-100660987 | cg14245199 | 8.36e-10 | -0.783790922047767 | classA | consensus |
| DEGS2 | chr14:100525300-100660987 | mean | 3.77E-06 | -0.764607357060142 | classA | consensus |
| DGKA | chr12:56308002-56336273 | cg08338281 | 5.97E-06 | -0.758366680493838 | classA | consensus |
| DGKA | chr12:56308002-56336273 | cg21587469 | 2.62E-06 | -0.769404486622536 | classA | consensus |
| DGKA | chr12:56308002-56336273 | mean | 3.50E-06 | -0.765594073561009 | classA | consensus |
| DGKG | chr3:186035120-186101676 | cg00434119 | 6.96E-06 | -0.756246419253565 | classA | consensus |
| DGKG | chr3:186035120-186101676 | cg00434119 | 6.96E-06 | -0.756246419253565 | classA | consensus |
| DNPH1 | chr6:43078670-43199471 | cg00741333 | 8.84E-04 | -0.717685969276767 | classA | consensus |

| | | | | | | |
|---|---|---|---|---|---|---|
| ECI1 | chr16:2289427-2319834 | cg09970941 | 6.49E-04 | -0.722731689525009 | classA | consensus |
| EPS8L2 | chr11:680428-727241 | cg07161179 | 2.18E-03 | -0.702280771697621 | classA | consensus |
| EPS8L2 | chr11:680428-727241 | cg18770763 | 1.60E-02 | -0.707651186157645 | classA | consensus |
| EPS8L2 | chr11:680428-727241 | mean | 2.11E-02 | -0.702830849982584 | classA | consensus |
| ERCC1 | chr19:45942641-45992367 | cg14890509 | 2.71E-05 | -0.736469719673141 | classA | consensus |
| ERCC1 | chr19:45942641-45992367 | mean | 1.95E-05 | -0.741426186286545 | classA | consensus |
| FFAR1 | chr19:35684861-35844023 | cg14554776 | 4.35E-04 | 0.729113537131159 | classA | consensus |
| FMNL1 | chr17:43296471-43432314 | cg08823240 | 2.43E-04 | 0.73811994337214 | classA | consensus |
| FMNL1 | chr17:43296471-43432314 | cg16826777 | 1.36E-04 | 0.746770348343407 | classA | consensus |
| FMNL1 | chr17:43296471-43432314 | mean | 1.60E-06 | 0.77578790626753 | classA | consensus |
| FOXN2 | chr2:48540146-48544409 | cg00032884 | 4.75e-10 | 0.790514366373985 | classA | consensus |
| FOXP1 | chr3:71014005-71643844 | cg19628600 | 4.28E-04 | 0.729387688647067 | classA | consensus |
| GAB1 | chr4:144256345-144366279 | cg06855485 | 0 | -0.87280730644136 | classA | consensus |
| GAB1 | chr4:144256345-144366279 | cg17453767 | 2.1e-11 | -0.823560851267405 | classA | consensus |
| GAB1 | chr4:144256345-144366279 | cg25911551 | 4.3e-11 | -0.816638984533236 | classA | consensus |
| GAB1 | chr4:144256345-144366279 | mean | 0 | -0.909313269127043 | classA | consensus |
| GATAD2A | chr19:19495632-19798225 | cg19379721 | 1.97E-06 | 0.773089821292984 | classA | consensus |
| GATAD2A | chr19:19495632-19798225 | mean | 6.87E-06 | 0.756436906856216 | classA | consensus |
| GRAMD1B | chr11:123320379-123473034 | cg26678144 | 1.2e-11 | -0.828669925803785 | classA | consensus |
| GRAMD1B | chr11:123320379-123473034 | mean | 4.66E-06 | -0.761758727008032 | classA | consensus |
| IL10RA | chr11:117839701-117930886 | cg01697865 | 1.43E-02 | -0.709611644879929 | classA | consensus |
| INO80B | chr2:74682132-74742444 | cg01584932 | 1.83E-04 | -0.742355963285444 | classA | consensus |
| INO80B | chr2:74682132-74742444 | cg08973382 | 1.79E-02 | -0.705774823748899 | classA | consensus |
| INO80B | chr2:74682132-74742444 | mean | 6.63E-06 | -0.756934016938781 | classA | consensus |
| INPP5F | chr10:121438946-121576868 | cg20341089 | 0 | -0.87558006963543 | classA | consensus |
| IRF2 | chr4:185174927-185435847 | cg12453504 | 7.99E-04 | -0.719347564664962 | classA | consensus |
| IRF2 | chr4:185174927-185435847 | cg21960364 | 0 | -0.872806717076645 | classA | consensus |
| IRF2 | chr4:185174927-185435847 | cg25556051 | 0 | -0.86263231313682 | classA | consensus |
| IRF2 | chr4:185174927-185435847 | mean | 8.00E-12 | -0.832679082100789 | classA | consensus |
| IST1 | chr16:71906867-71930934 | cg26585899 | 7.36E-06 | 0.755471672678476 | classA | consensus |
| ITPR1 | chr3:4533777-4594716 | cg24131262 | 3.40E-06 | 0.765994446093569 | classA | consensus |
| JDP2 | chr14:75885576-75947457 | cg04407490 | 1.68E-02 | -0.706836200725772 | classA | consensus |
| KCNN4 | chr19:44223140-44295389 | cg04833845 | 2.61e-10 | -0.79739098179341 | classA | consensus |
| KCNN4 | chr19:44223140-44295389 | cg18506018 | 8.8e-11 | -0.80921244685823 | classA | consensus |
| KCNN4 | chr19:44223140-44295389 | cg24900654 | 0 | -0.86457564495141 | classA | consensus |
| KCNN4 | chr19:44223140-44295389 | cg26890181 | 3.62e-10 | -0.793679429228187 | classA | consensus |
| KCNN4 | chr19:44223140-44295389 | mean | 2.1e-11 | -0.823838784220103 | classA | consensus |
| KCTD18 | chr2:201310653-201333611 | cg04945753 | 4.9e-11 | 0.815264888033454 | classA | consensus |
| KCTD18 | chr2:201310653-201333611 | cg06240652 | 4.24e-10 | 0.791833856406979 | classA | consensus |
| KCTD18 | chr2:201310653-201333611 | mean | 5.4e-11 | 0.814284956829064 | classA | consensus |
| KLHDC9 | chr1:161002074-161072381 | cg02957185 | 1.25E-04 | -0.747942024493383 | classA | consensus |
| KLHDC9 | chr1:161002074-161072381 | cg27582585 | 4.18E-04 | -0.729769383087602 | classA | consensus |
| KLHDC9 | chr1:161002074-161072381 | mean | 3.00E-04 | -0.734880302012131 | classA | consensus |
| KSR1 | chr17:25782569-25881105 | cg08041350 | 2.19E-04 | 0.739654902137596 | classA | consensus |
| LAG3 | chr12:6874667-6887901 | cg04153135 | 2.35E-02 | -0.700972300571603 | classA | consensus |
| LAG3 | chr12:6874667-6887901 | mean | 3.47E-04 | -0.732643196468662 | classA | consensus |
| LFNG | chr7:2410726-2582633 | cg02613295 | 1.30E-06 | -0.778347930183869 | classA | consensus |
| LFNG | chr7:2410726-2582633 | cg23510807 | 4.96e-10 | -0.790003819010922 | classA | consensus |
| LFNG | chr7:2410726-2582633 | cg27186851 | 2.1e-09 | -0.772285715622474 | classA | consensus |
| LFNG | chr7:2410726-2582633 | mean | 8.94e-10 | -0.782968899632789 | classA | consensus |
| LIX1L | chr1:145426814-145479979 | cg19913118 | 1.91E-02 | 0.704624012519546 | classA | consensus |
| MAP3K14 | chr17:43443900-43529580 | cg26980978 | 7.54E-04 | 0.720309781859269 | classA | consensus |
| MAP3K14 | chr17:43443900-43529580 | mean | 1.56E-06 | 0.77609897788944 | classA | consensus |
| MAP3K14 | chr17:43443900-43529580 | cg26980978 | 7.54E-04 | 0.720309781859269 | classA | consensus |
| MAP3K14 | chr17:43443900-43529580 | mean | 1.56E-06 | 0.77609897788944 | classA | consensus |
| MBNL1-AS1 | chr3:151955764-152032762 | cg13702222 | 1.31E-02 | 0.711090339214665 | classA | consensus |
| MIR3128 | chr2:178110476-178130561 | cg27507284 | 6.85E-04 | 0.721871357683247 | classA | consensus |
| MYO1C | chr17:1336059-1402600 | cg05981034 | 1.80E-02 | 0.705635466237926 | classA | consensus |
| MYO1C | chr17:1336059-1402600 | mean | 1.19E-03 | 0.712787171435902 | classA | consensus |
| OGDH | chr7:44599534-44753708 | mean | 2.44E-02 | -0.700278599503742 | classA | consensus |
| PARP12 | chr7:139753823-139763284 | cg03873220 | 4.96E-06 | 0.760905054295963 | classA | consensus |
| PDCD1 | chr2:242782277-242857257 | cg09034753 | 2.2e-10 | -0.79930748723764 | classA | consensus |
| PDCD1 | chr2:242782277-242857257 | cg11654325 | 7.00E-12 | -0.833661468475583 | classA | consensus |
| PDCD1 | chr2:242782277-242857257 | mean | 1.9e-11 | -0.824415684838076 | classA | consensus |
| PLXND1 | chr3:129200451-129353480 | cg09463220 | 1.12E-04 | -0.749590413790649 | classA | consensus |
| PPTC7 | chr12:110964850-111300733 | cg07414574 | 1.77E-04 | 0.742892103126958 | classA | consensus |
| PPTC7 | chr12:110964850-111300733 | cg15698107 | 1.28E-04 | 0.747588144344519 | classA | consensus |

| | | | | | | |
|---|---|---|---|---|---|---|
| PPTC7 | chr12:110964850-111300733 | mean | 1.81E-02 | 0.705548817583568 | classA | consensus |
| PRKCD | chr3:53157353-53233297 | cg07687398 | 4.64E-04 | 0.728108143764366 | classA | consensus |
| PSMD3 | chr17:38062953-38153023 | cg24910161 | 4.29e-10 | 0.791695837351697 | classA | consensus |
| PSMD3 | chr17:38062953-38153023 | cg26162295 | 1.02E-06 | 0.781409227278079 | classA | consensus |
| PSMD3 | chr17:38062953-38153023 | mean | 6.23e-10 | 0.787318039519299 | classA | consensus |
| PTP4A2 | chr1:32392622-32423903 | cg09373727 | 1.87E-02 | 0.705008940253793 | classA | consensus |
| PTP4A2 | chr1:32392622-32423903 | mean | 1.84E-02 | 0.705234635663359 | classA | consensus |
| RASGRF1 | chr15:79199877-79385468 | cg02587988 | 5.80E-06 | -0.758785729218541 | classA | consensus |
| RASGRF1 | chr15:79199877-79385468 | cg21840434 | 0 | -0.888183128646504 | classA | consensus |
| RASGRF1 | chr15:79199877-79385468 | cg22821300 | 1.00E-12 | -0.853301105139947 | classA | consensus |
| RASGRF1 | chr15:79199877-79385468 | mean | 0 | -0.88531604365507 | classA | consensus |
| RELA | chr11:65096483-65434058 | cg02345399 | 6.37E-06 | 0.757487434549028 | classA | consensus |
| RELA | chr11:65096483-65434058 | cg02652361 | 3.32E-05 | 0.733338677728195 | classA | consensus |
| RELA | chr11:65096483-65434058 | cg09801334 | 2.14E-02 | 0.702575655530683 | classA | consensus |
| RELA | chr11:65096483-65434058 | cg11573318 | 1.86E-02 | 0.705057723487242 | classA | consensus |
| RELA | chr11:65096483-65434058 | cg12589431 | 2.10E-02 | 0.70290788528844 | classA | consensus |
| RELA | chr11:65096483-65434058 | cg17451760 | 8.00E-12 | 0.832508733967886 | classA | consensus |
| RELA | chr11:65096483-65434058 | cg24215776 | 0 | 0.858432616304556 | classA | consensus |
| RELA | chr11:65096483-65434058 | mean | 1.55e-10 | 0.803170513903774 | classA | consensus |
| RGS10 | chr10:120966371-121302914 | cg00291478 | 0 | -0.901153077557479 | classA | consensus |
| RGS10 | chr10:120966371-121302914 | cg05241536 | 9.48e-10 | -0.782252463783973 | classA | consensus |
| RGS10 | chr10:120966371-121302914 | cg07551060 | 0 | -0.882301927466377 | classA | consensus |
| RGS10 | chr10:120966371-121302914 | cg15611912 | 0 | -0.876694527042908 | classA | consensus |
| RGS10 | chr10:120966371-121302914 | mean | 0 | -0.885212572718762 | classA | consensus |
| ROR1 | chr1:64237486-64489243 | cg24502478 | 1.67E-02 | -0.706934283803974 | classA | consensus |
| RPUSD2 | chr15:40817601-40866485 | cg17140476 | 1.29E-02 | 0.711344786711647 | classA | consensus |
| RREB1 | chr6:7098272-7209426 | cg20893838 | 5.72E-04 | 0.724773851250192 | classA | consensus |
| RREB1 | chr6:7098272-7209426 | mean | 2.40E-03 | 0.70059017982946 | classA | consensus |
| RSAD1 | chr17:48532628-48565255 | cg09151131 | 1.15E-05 | -0.749187405154563 | classA | consensus |
| SAMD8 | chr10:76771419-76872476 | cg01354426 | 1.10E-02 | 0.714102453807693 | classA | consensus |
| SBNO1 | chr12:123841599-123979974 | cg05861255 | 3.13E-04 | 0.734236685532627 | classA | consensus |
| SBNO1 | chr12:123841599-123979974 | cg06160572 | 9.09E-04 | 0.717234984578664 | classA | consensus |
| SEMA4D | chr9:91962974-92169084 | cg13584258 | 1.23E-02 | -0.712120494161879 | classA | consensus |
| SERINC5 | chr5:79388087-79555168 | cg01267797 | 3.85E-06 | -0.764328420934851 | classA | consensus |
| SERINC5 | chr5:79388087-79555168 | cg14711743 | 1.81e-09 | -0.774185593435448 | classA | consensus |
| SERINC5 | chr5:79388087-79555168 | mean | 7.78e-10 | -0.784657329851218 | classA | consensus |
| SKAP1 | chr17:46484641-46523025 | cg05106502 | 2.3e-11 | -0.822767863840405 | classA | consensus |
| SNX18 | chr5:53804511-53851756 | cg26118943 | 1.73E-02 | -0.706368280131888 | classA | consensus |
| SRGN | chr10:70794826-70862131 | cg24417845 | 7.00E-11 | -0.811601594530483 | classA | consensus |
| STRN4 | chr19:47198585-47255913 | cg00838415 | 1.15E-04 | 0.749138641350944 | classA | consensus |
| STRN4 | chr19:47198585-47255913 | cg26482609 | 0 | 0.878320814741321 | classA | consensus |
| STRN4 | chr19:47198585-47255913 | mean | 1.00E-12 | 0.847768722682479 | classA | consensus |
| TBRG4 | chr7:44992641-45152327 | cg12573289 | 1.30E-02 | 0.711237676504797 | classA | consensus |
| TESC | chr12:117461624-117616567 | cg02867305 | 5.65e-10 | -0.788478351126211 | classA | consensus |
| TESC | chr12:117461624-117616567 | cg06597431 | 7.58E-06 | -0.755061354070138 | classA | consensus |
| TESC | chr12:117461624-117616567 | mean | 1.11e-10 | -0.806799901204625 | classA | consensus |
| TET3 | chr2:74191698-74281436 | cg02251243 | 1.27E-02 | 0.711661428167861 | classA | consensus |
| TET3 | chr2:74191698-74281436 | cg05981785 | 2.59E-06 | 0.769586911051677 | classA | consensus |
| TET3 | chr2:74191698-74281436 | cg19445690 | 6.12E-04 | 0.723696017566352 | classA | consensus |
| TET3 | chr2:74191698-74281436 | mean | 7.73e-10 | 0.784733245767552 | classA | consensus |
| TGFBR3 | chr1:92171519-92352408 | cg15326069 | 1.12e-09 | -0.780211582434955 | classA | consensus |
| TGFBR3 | chr1:92171519-92352408 | cg21116284 | 0 | -0.878833087448292 | classA | consensus |
| TGFBR3 | chr1:92171519-92352408 | mean | 0 | -0.867764718244152 | classA | consensus |
| TGIF1 | chr18:3444750-3480409 | cg26701815 | 2.03E-02 | -0.703573653305932 | classA | consensus |
| TGIF1 | chr18:3444750-3480409 | mean | 8.01E-04 | -0.719304475163224 | classA | consensus |
| TMBIM1 | chr2:219106984-219163678 | cg13835894 | 1.97E-02 | 0.704023506552651 | classA | consensus |
| TMEM154 | chr4:153548459-153630443 | mean | 1.16E-02 | -0.713180290840589 | classA | consensus |
| TPST2 | chr22:26920248-27039444 | cg09856467 | 3.67E-04 | -0.731799999163512 | classA | consensus |
| TPST2 | chr22:26920248-27039444 | cg18895476 | 1.44E-04 | -0.745915419977729 | classA | consensus |
| TPST2 | chr22:26920248-27039444 | mean | 2.8e-11 | -0.820718331069353 | classA | consensus |
| TRAM2 | chr6:52354447-52476581 | cg22388471 | 2.62E-06 | -0.769437179521233 | classA | consensus |
| TRAM2 | chr6:52354447-52476581 | cg26020069 | 2.41E-04 | -0.738242843755085 | classA | consensus |
| TRAM2 | chr6:52354447-52476581 | mean | 1.01e-09 | -0.781476451381996 | classA | consensus |
| TRAPPC10 | chr21:45425337-45801001 | cg04643920 | 6.00E-11 | 0.813250718756081 | classA | consensus |
| TRAPPC10 | chr21:45425337-45801001 | cg08039560 | 7.38e-08 | 0.720654718910295 | classA | consensus |
| TRAPPC10 | chr21:45425337-45801001 | mean | 7.86e-10 | 0.784533428111105 | classA | consensus |
| TRAPPC10 | chr21:45425337-45801001 | cg04643920 | 6.00E-11 | 0.813250718756081 | classA | consensus |

| | | | | | | |
|---|---|---|---|---|---|---|
| TRAPPC10 | chr21:45425337-45801001 | cg08039560 | 7.38e-08 | 0.720654718910295 | classA | consensus |
| TRAPPC10 | chr21:45425337-45801001 | mean | 7.86e-10 | 0.784533428111105 | classA | consensus |
| TSTA3 | chr8:144504468-144700876 | cg11210357 | 1.10E-04 | -0.749783755225239 | classA | consensus |
| TSTA3 | chr8:144504468-144700876 | cg18047920 | 1.35E-02 | -0.710569886201223 | classA | consensus |
| TSTA3 | chr8:144504468-144700876 | mean | 1.03E-02 | -0.715094190030724 | classA | consensus |
| USP7 | chr16:9035541-9058525 | cg07636761 | 2.44E-04 | 0.738056576318808 | classA | consensus |
| USP7 | chr16:9035541-9058525 | cg10359807 | 8.69E-04 | 0.717975560074612 | classA | consensus |
| USP7 | chr16:9035541-9058525 | mean | 1.62E-04 | 0.744149167056782 | classA | consensus |
| WDR11-AS1 | chr10:122544458-122621403 | cg13789938 | 7.08E-06 | -0.756021710807748 | classA | consensus |
| XXYLT1 | chr3:194779122-194992398 | cg05486924 | 3.57E-04 | -0.732239995611384 | classA | consensus |
| XXYLT1 | chr3:194779122-194992398 | mean | 8.01E-04 | -0.719320967228831 | classA | consensus |
| ZAP70 | chr2:98328846-98336068 | cg21773162 | 8.88E-06 | -0.752830444305011 | classA | consensus |
| ZAP70 | chr2:98328846-98336068 | mean | 1.17E-04 | -0.748935645589701 | classA | consensus |
| ZFPM1 | chr16:88441354-88624317 | cg00216061 | 2.44E-04 | -0.73804980353996 | classA | consensus |
| ZFPM1 | chr16:88441354-88624317 | cg00352652 | 1.11E-04 | -0.749662616635929 | classA | consensus |
| ZFPM1 | chr16:88441354-88624317 | cg03080043 | 7.17E-05 | -0.721132434046399 | classA | consensus |
| ZFPM1 | chr16:88441354-88624317 | cg04160201 | 1.46E-06 | -0.776945856011835 | classA | consensus |
| ZFPM1 | chr16:88441354-88624317 | cg05067156 | 1.91E-02 | -0.704576049365894 | classA | consensus |
| ZFPM1 | chr16:88441354-88624317 | cg06928887 | 3.35E-04 | -0.733197115763135 | classA | consensus |
| ZFPM1 | chr16:88441354-88624317 | cg08466082 | 3.60E-06 | -0.7652114649885 | classA | consensus |
| ZFPM1 | chr16:88441354-88624317 | cg26899496 | 6.02E-05 | -0.723948996200522 | classA | consensus |
| ZFPM1 | chr16:88441354-88624317 | mean | 1.40E-06 | -0.777398317203062 | classA | consensus |
| ACSF3 | chr16:89098347-89206120 | cg03406609 | 0 | -0.858326223615184 | classA | diff |
| ACSF3 | chr16:89098347-89206120 | cg04033022 | 2.00E-12 | -0.845266750809226 | classA | diff |
| ACSF3 | chr16:89098347-89206120 | cg05231226 | 1.18E-04 | -0.748758910768003 | classA | diff |
| ACSF3 | chr16:89098347-89206120 | cg09427016 | 0 | -0.876227425688333 | classA | diff |
| ACSF3 | chr16:89098347-89206120 | cg10217183 | 1.00E-12 | -0.853840682137293 | classA | diff |
| ACSF3 | chr16:89098347-89206120 | mean | 0 | -0.860598691175185 | classA | diff |
| CD5 | chr11:60822259-60879990 | cg06417460 | 1.00E-12 | -0.851852052685724 | classA | diff |
| CD5 | chr11:60822259-60879990 | cg24674703 | 2.16e-10 | -0.799491802538728 | classA | diff |
| CD5 | chr11:60822259-60879990 | mean | 2.4e-11 | -0.822284217248575 | classA | diff |
| CHIT1 | chr1:203236286-203353767 | cg06373167 | 8.07e-10 | -0.784213548363276 | classA | diff |
| CHIT1 | chr1:203236286-203353767 | cg20138067 | 1.96E-02 | -0.704175865626893 | classA | diff |
| CHIT1 | chr1:203236286-203353767 | mean | 1.17E-04 | -0.748898570492836 | classA | diff |
| CTSH | chr15:79265754-79325467 | cg21840434 | 8.73E-06 | -0.753072196871874 | classA | diff |
| CTSH | chr15:79265754-79325467 | cg22821300 | 8.83E-06 | -0.752911465847083 | classA | diff |
| CTSH | chr15:79265754-79325467 | mean | 5.21E-06 | -0.760251653670166 | classA | diff |
| DAD1 | chr14:22932348-23038986 | cg01536878 | 1.25E-04 | -0.747996216660009 | classA | diff |
| DAD1 | chr14:22932348-23038986 | cg16302916 | 7.54e-10 | -0.785040502078318 | classA | diff |
| DAD1 | chr14:22932348-23038986 | cg20073893 | 1.03E-04 | -0.750662382628425 | classA | diff |
| DAD1 | chr14:22932348-23038986 | cg26053876 | 1.82e-10 | -0.801403228395917 | classA | diff |
| IL10RA | chr11:117842673-117887699 | cg01697865 | 1.43E-02 | -0.709611644879929 | classA | diff |
| IRF2 | chr4:185306708-185403585 | cg12453504 | 7.99E-04 | -0.719347564664962 | classA | diff |
| IRF2 | chr4:185306708-185403585 | cg21960364 | 0 | -0.872806717076645 | classA | diff |
| IRF2 | chr4:185306708-185403585 | mean | 3.00E-12 | -0.842829974046622 | classA | diff |
| PITPNM2 | chr12:123515700-123637201 | cg01973483 | 0 | -0.881379429123945 | classA | diff |
| PITPNM2 | chr12:123515700-123637201 | cg19515108 | 6.43e-10 | -0.786944193072471 | classA | diff |
| PITPNM2 | chr12:123515700-123637201 | mean | 0 | -0.902394999858563 | classA | diff |
| SELPLG | chr12:109006555-109042818 | cg25165932 | 6.80E-04 | -0.721982020830158 | classA | diff |
| SH3BP5 | chr3:15298050-15404967 | cg04858987 | 7.23e-09 | -0.755722894040086 | classA | diff |
| SH3BP5 | chr3:15298050-15404967 | cg08359336 | 9.54E-04 | -0.716427251100218 | classA | diff |
| SH3BP5 | chr3:15298050-15404967 | mean | 8.36E-04 | -0.718605752848992 | classA | diff |
| SLC25A19 | chr17:73283975-73357209 | cg11818853 | 2.86E-06 | -0.768276946378715 | classA | diff |
| SLC25A19 | chr17:73283975-73357209 | cg14852456 | 4.61E-04 | -0.728208118087003 | classA | diff |
| SLC25A19 | chr17:73283975-73357209 | mean | 1.01E-04 | -0.750958645033277 | classA | diff |
| TESC | chr12:117490227-117600511 | cg02867305 | 5.65e-10 | -0.788478351126211 | classA | diff |
| TESC | chr12:117490227-117600511 | cg06597431 | 7.58E-06 | -0.755061354070138 | classA | diff |
| TESC | chr12:117490227-117600511 | mean | 1.11e-10 | -0.806799901204625 | classA | diff |
| TRAPPC2L | chr16:88919603-89070961 | cg00056002 | 6.05e-09 | -0.758193141436958 | classA | diff |
| TRAPPC2L | chr16:88919603-89070961 | cg01502457 | 5.62E-04 | -0.725062081680588 | classA | diff |
| TRAPPC2L | chr16:88919603-89070961 | cg03353765 | 1.79E-02 | -0.705704977579105 | classA | diff |
| TRAPPC2L | chr16:88919603-89070961 | cg03797660 | 1.25E-02 | -0.711957513904313 | classA | diff |
| TRAPPC2L | chr16:88919603-89070961 | cg05579598 | 1.00E-12 | -0.851011338132066 | classA | diff |
| TRAPPC2L | chr16:88919603-89070961 | cg06204040 | 0 | -0.881385012204044 | classA | diff |
| TRAPPC2L | chr16:88919603-89070961 | cg06326865 | 4.38E-04 | -0.729012747490419 | classA | diff |
| TRAPPC2L | chr16:88919603-89070961 | cg07625774 | 7.92E-04 | -0.719501059022488 | classA | diff |
| TRAPPC2L | chr16:88919603-89070961 | cg08698997 | 8.9e-08 | -0.717576771722375 | classA | diff |

| | | | | | | |
|---|---|---|---|---|---|---|
| TRAPPC2L | chr16:88919603-89070961 | cg26776806 | 5.89e-10 | -0.787988382732691 | classA | diff |
| TRAPPC2L | chr16:88919603-89070961 | cg27061485 | 3.42E-04 | -0.732865238241398 | classA | diff |
| TRAPPC2L | chr16:88919603-89070961 | mean | 0 | -0.88722412947842 | classA | diff |
| XXYLT1-AS2 | chr3:194796146-194885613 | cg12085101 | 3.86E-04 | -0.731009997717873 | classA | diff |
| XXYLT1-AS2 | chr3:194796146-194885613 | cg19760965 | 1.44E-02 | -0.709422515837849 | classA | diff |
| XXYLT1-AS2 | chr3:194796146-194885613 | mean | 1.01E-05 | -0.75099442336873 | classA | diff |
| ZFPM1 | chr16:88509664-88551933 | cg00216061 | 2.44E-04 | -0.73804980353996 | classA | diff |
| ZFPM1 | chr16:88509664-88551933 | cg00352652 | 1.11E-04 | -0.749662616635929 | classA | diff |
| ZFPM1 | chr16:88509664-88551933 | cg03080043 | 7.17E-05 | -0.721132434046399 | classA | diff |
| ZFPM1 | chr16:88509664-88551933 | cg04160201 | 1.46E-06 | -0.776945856011835 | classA | diff |
| ZFPM1 | chr16:88509664-88551933 | cg06928887 | 3.35E-04 | -0.733197115763135 | classA | diff |
| ZFPM1 | chr16:88509664-88551933 | cg08466082 | 3.60E-06 | -0.7652114649885 | classA | diff |
| ZFPM1 | chr16:88509664-88551933 | cg26899496 | 6.02E-05 | -0.723948996200522 | classA | diff |
| ZFPM1 | chr16:88509664-88551933 | mean | 1.26E-06 | -0.778718395385321 | classA | diff |
| ZNF710 | chr15:90541151-90611052 | cg00624799 | 1.19E-02 | -0.712687210896193 | classA | diff |
| ZNF710 | chr15:90541151-90611052 | mean | 1.26E-04 | -0.747848244005227 | classA | diff |
| ACBD3 | chr1:226270365-226383660 | cg15260921 | 1.60E-03 | 0.707675713557636 | classC | consensus |
| ANKRD33B | chr5:10552543-10711025 | cg16362140 | 5.61e-10 | 0.788561292159291 | classC | consensus |
| ARAP1 | chr11:72264596-72510098 | cg02470043 | 1.58E-02 | -0.707875290303629 | classC | consensus |
| ARAP1 | chr11:72264596-72510098 | mean | 1.99E-02 | -0.703895002177188 | classC | consensus |
| ARFRP1 | chr20:62150820-62340527 | cg00121366 | 1.59E-02 | -0.707809190103492 | classC | consensus |
| ARFRP1 | chr20:62150820-62340527 | mean | 2.83E-06 | -0.768393497805217 | classC | consensus |
| ARID1A | chr1:27018553-27133574 | cg24313303 | 6.11E-06 | 0.758064485891578 | classC | consensus |
| ARL4C | chr2:235374001-235413914 | cg05204104 | 0 | -0.865664849223155 | classC | consensus |
| ARL4C | chr2:235374001-235413914 | cg15016771 | 9.5e-11 | -0.80841356191954 | classC | consensus |
| ARL4C | chr2:235374001-235413914 | cg24441922 | 0 | -0.856367296034818 | classC | consensus |
| ARL4C | chr2:235374001-235413914 | mean | 4.00E-12 | -0.840031513123482 | classC | consensus |
| ARL4C | chr2:235374001-235413914 | cg05204104 | 0 | -0.865664849223155 | classC | consensus |
| ARL4C | chr2:235374001-235413914 | cg15016771 | 9.5e-11 | -0.80841356191954 | classC | consensus |
| ARL4C | chr2:235374001-235413914 | cg24441922 | 0 | -0.856367296034818 | classC | consensus |
| ARL4C | chr2:235374001-235413914 | mean | 4.00E-12 | -0.840031513123482 | classC | consensus |
| ATP11A | chr13:113318512-113446398 | cg12917253 | 2.07E-06 | -0.772449259569865 | classC | consensus |
| ATP11A | chr13:113318512-113446398 | cg14864184 | 7.58e-10 | -0.784968904762727 | classC | consensus |
| ATP11A | chr13:113318512-113446398 | cg21513610 | 4.48e-09 | -0.762293813302839 | classC | consensus |
| ATP11A | chr13:113318512-113446398 | cg21897676 | 1.02E-02 | -0.715275800372377 | classC | consensus |
| ATP11A | chr13:113318512-113446398 | mean | 1.19e-10 | -0.806048274664118 | classC | consensus |
| ATP6V0D1 | chr16:67465524-67635531 | cg03737424 | 1.00E-09 | 0.781600382415951 | classC | consensus |
| ATXN1 | chr6:16620807-16795056 | cg10262747 | 1.07e-10 | -0.807145510254306 | classC | consensus |
| ATXN1 | chr6:16620807-16795056 | cg14791639 | 2.81e-09 | -0.768500162615994 | classC | consensus |
| ATXN1 | chr6:16620807-16795056 | mean | 3.13e-10 | -0.79533052109723 | classC | consensus |
| B3GNT3 | chr19:17811190-17914494 | cg06649165 | 3.52E-04 | 0.73243814513164 | classC | consensus |
| BIN3 | chr8:22516805-22643451 | cg03720745 | 5.45E-04 | 0.725560929686346 | classC | consensus |
| BIN3 | chr8:22516805-22643451 | mean | 4.90E-05 | 0.727257012980567 | classC | consensus |
| BLNK | chr10:97967726-98060192 | cg23715056 | 4.22E-06 | -0.763105953024114 | classC | consensus |
| C8orf46 | chr8:67394992-67465261 | cg09622557 | 7.23E-04 | -0.7209804142861 | classC | consensus |
| C8orf46 | chr8:67394992-67465261 | cg15070801 | 2.02E-02 | -0.703608338001738 | classC | consensus |
| C8orf46 | chr8:67394992-67465261 | cg27131563 | 4.83E-06 | -0.761265325172991 | classC | consensus |
| C8orf46 | chr8:67394992-67465261 | mean | 1.73E-04 | -0.743226778897071 | classC | consensus |
| CABLES1 | chr18:20708779-20896507 | cg22158648 | 1.00E-10 | -0.807867607930747 | classC | consensus |
| CACNB2 | chr10:18620712-18659631 | cg05719612 | 6.53E-04 | -0.722644374499814 | classC | consensus |
| CACNB2 | chr10:18620712-18659631 | cg19998294 | 3.00E-04 | -0.734913928803241 | classC | consensus |
| CACNB2 | chr10:18620712-18659631 | cg22858500 | 3.20E-04 | -0.733919164509448 | classC | consensus |
| CACNB2 | chr10:18620712-18659631 | mean | 1.95E-05 | -0.741441211308523 | classC | consensus |
| CACNB2 | chr10:18620712-18659631 | cg05719612 | 6.53E-04 | -0.722644374499814 | classC | consensus |
| CACNB2 | chr10:18620712-18659631 | cg19998294 | 3.00E-04 | -0.734913928803241 | classC | consensus |
| CACNB2 | chr10:18620712-18659631 | cg22858500 | 3.20E-04 | -0.733919164509448 | classC | consensus |
| CACNB2 | chr10:18620712-18659631 | mean | 1.95E-05 | -0.741441211308523 | classC | consensus |
| CAPN12 | chr19:39127220-39236589 | cg25383568 | 1.00E-02 | -0.715560839506439 | classC | consensus |
| CBX7 | chr22:39345734-39549865 | cg10016348 | 8.23E-06 | -0.753916578193137 | classC | consensus |
| CBX7 | chr22:39345734-39549865 | cg13256546 | 6.62E-06 | -0.756959899743302 | classC | consensus |
| CBX7 | chr22:39345734-39549865 | cg24424889 | 2.31E-04 | -0.738888571191293 | classC | consensus |
| CBX7 | chr22:39345734-39549865 | mean | 1.94E-02 | -0.704357420531317 | classC | consensus |
| CCDC88A | chr2:55640643-55653423 | cg01953317 | 0 | -0.920706418227191 | classC | consensus |
| CDIP1 | chr16:4490764-4594619 | cg02642565 | 4.00E-06 | -0.763826418343075 | classC | consensus |
| CDIPT | chr16:29776570-29874841 | cg02876297 | 3.39E-04 | -0.733016355004285 | classC | consensus |
| CDIPT | chr16:29776570-29874841 | mean | 3.71E-05 | -0.731612356367572 | classC | consensus |
| CDK14 | chr7:90324837-90431656 | cg08583610 | 2.31E-06 | -0.771061664312234 | classC | consensus |

| | | | | | | |
|---|---|---|---|---|---|---|
| CDK14 | chr7:90324837-90431656 | cg27403658 | 5.21E-06 | -0.760250885063005 | classC | consensus |
| CDK14 | chr7:90324837-90431656 | mean | 7.00E-11 | -0.811607040197263 | classC | consensus |
| CELSR1 | chr22:46742806-47314341 | cg11533712 | 6.23E-04 | -0.7234032224443 | classC | consensus |
| CHIT1 | chr1:203232464-203504070 | cg03764585 | 1.42E-06 | -0.777246800088499 | classC | consensus |
| CHIT1 | chr1:203232464-203504070 | cg04704856 | 1.82e-10 | -0.80139899446331 | classC | consensus |
| CHIT1 | chr1:203232464-203504070 | cg12067421 | 9.00E-11 | -0.808948942559685 | classC | consensus |
| CHIT1 | chr1:203232464-203504070 | cg16289210 | 7.63E-06 | -0.754972557081319 | classC | consensus |
| CHIT1 | chr1:203232464-203504070 | cg25143508 | 1.08e-09 | -0.780657890148094 | classC | consensus |
| CHIT1 | chr1:203232464-203504070 | cg26987645 | 3.24e-09 | -0.766627597512917 | classC | consensus |
| CHIT1 | chr1:203232464-203504070 | mean | 1.34e-10 | -0.804722665170269 | classC | consensus |
| CHMP4A | chr14:24520383-24685857 | cg13167816 | 2.1e-11 | 0.82362573809388 | classC | consensus |
| CLIC4 | chr1:25018827-25101619 | cg07191152 | 2.97e-10 | 0.795931978425408 | classC | consensus |
| CLLU1 | chr12:92746178-92913479 | cg04845867 | 3.26E-04 | -0.733628428342836 | classC | consensus |
| CLLU1 | chr12:92746178-92913479 | mean | 1.80E-04 | -0.742629901903054 | classC | consensus |
| COQ10A | chr12:56493010-56662406 | cg00668685 | 1.54E-02 | -0.70832321183565 | classC | consensus |
| COQ10A | chr12:56493010-56662406 | mean | 6.26E-04 | -0.723320561441604 | classC | consensus |
| CORO6 | chr17:27882320-27952581 | cg13872326 | 2.73E-06 | -0.768861187911738 | classC | consensus |
| CORO6 | chr17:27882320-27952581 | cg20649847 | 6.1e-11 | -0.813034530249863 | classC | consensus |
| CORO6 | chr17:27882320-27952581 | cg22961402 | 3.03E-06 | -0.767518318444297 | classC | consensus |
| CORO6 | chr17:27882320-27952581 | mean | 3.00E-12 | -0.842595431664262 | classC | consensus |
| CORO6 | chr17:27882320-27952581 | cg13872326 | 2.73E-06 | -0.768861187911738 | classC | consensus |
| CORO6 | chr17:27882320-27952581 | cg20649847 | 6.1e-11 | -0.813034530249863 | classC | consensus |
| CORO6 | chr17:27882320-27952581 | cg22961402 | 3.03E-06 | -0.767518318444297 | classC | consensus |
| CORO6 | chr17:27882320-27952581 | mean | 3.00E-12 | -0.842595431664262 | classC | consensus |
| COX4I1 | chr16:85582796-85843772 | cg08774871 | 1.75E-02 | 0.706158895770373 | classC | consensus |
| CREBBP | chr16:3899488-3931563 | cg08364730 | 1.19e-10 | 0.806064017206093 | classC | consensus |
| CRTC1 | chr19:18793517-18853326 | cg25561382 | 2.07E-04 | -0.740535126526299 | classC | consensus |
| CTLA4 | chr2:204705078-204774245 | cg05074138 | 0 | -0.955258463676382 | classC | consensus |
| CTLA4 | chr2:204705078-204774245 | cg18914852 | 1.81E-02 | -0.705581304747191 | classC | consensus |
| CTLA4 | chr2:204705078-204774245 | cg26091609 | 0 | -0.960482128368801 | classC | consensus |
| CTLA4 | chr2:204705078-204774245 | mean | 0 | -0.936828196298022 | classC | consensus |
| DCLK2 | chr4:150998214-151040659 | cg03779973 | 0 | -0.873002373516405 | classC | consensus |
| DCLK2 | chr4:150998214-151040659 | cg05561193 | 1.17e-10 | -0.806186902731367 | classC | consensus |
| DCLK2 | chr4:150998214-151040659 | cg12115081 | 1.00E-12 | -0.854219822365538 | classC | consensus |
| DCLK2 | chr4:150998214-151040659 | mean | 0 | -0.88003414145288 | classC | consensus |
| DEGS2 | chr14:100525300-100660987 | cg19741660 | 3.23E-06 | -0.766656006968834 | classC | consensus |
| DEGS2 | chr14:100525300-100660987 | mean | 1.69E-02 | -0.706761059061396 | classC | consensus |
| DENND4A | chr15:66055350-66127032 | cg27665985 | 2.72E-04 | 0.736381191871896 | classC | consensus |
| DNMBP | chr10:101714734-101785799 | cg02732134 | 2.27E-02 | -0.701594082487507 | classC | consensus |
| ECHDC3 | chr10:11691053-11758626 | cg00594191 | 3.95E-04 | 0.730630044274807 | classC | consensus |
| EP300 | chr22:41444573-41511966 | cg22822910 | 2.82E-04 | 0.735852830223008 | classC | consensus |
| FAM167A | chr8:11269191-11458958 | cg14702231 | 2.62E-05 | 0.736984500730161 | classC | consensus |
| FAM167A | chr8:11269191-11458958 | cg15685006 | 2.40E-02 | 0.700583964252854 | classC | consensus |
| FAM167A | chr8:11269191-11458958 | cg26161004 | 2.87E-04 | 0.735605515130926 | classC | consensus |
| FAM167A | chr8:11269191-11458958 | mean | 1.43E-04 | 0.745962796803634 | classC | consensus |
| FAM222B | chr17:27166807-27312962 | cg17445097 | 3.03E-04 | 0.734731549370124 | classC | consensus |
| FFAR1 | chr19:35684861-35844023 | mean | 4.56E-04 | 0.728402502477189 | classC | consensus |
| FGD3 | chr9:95709261-95931180 | mean | 3.20E-04 | -0.733907075924499 | classC | consensus |
| FRY | chr13:32591425-32622735 | cg16995086 | 1.6e-11 | -0.826249327455518 | classC | consensus |
| FRY | chr13:32591425-32622735 | mean | 2.3e-11 | -0.822628421237847 | classC | consensus |
| GAB1 | chr4:144256345-144366279 | cg12710519 | 1.47e-10 | -0.803777140488745 | classC | consensus |
| GAB1 | chr4:144256345-144366279 | cg14142817 | 1.00E-12 | -0.854165170652032 | classC | consensus |
| GAB1 | chr4:144256345-144366279 | mean | 3.00E-12 | -0.841600568079304 | classC | consensus |
| GATAD2A | chr19:19495632-19798225 | cg16640973 | 1.87E-02 | 0.704969813305409 | classC | consensus |
| GDF7 | chr2:20767862-20900102 | cg00011239 | 1.23E-02 | -0.712204763103036 | classC | consensus |
| GDF7 | chr2:20767862-20900102 | mean | 1.76E-04 | -0.742945106511503 | classC | consensus |
| GNLY | chr2:85864558-85939217 | cg03380645 | 1.45E-04 | -0.745817687286982 | classC | consensus |
| GNLY | chr2:85864558-85939217 | cg07313956 | 6.36E-04 | -0.723074149230447 | classC | consensus |
| GNLY | chr2:85864558-85939217 | mean | 1.54E-02 | -0.708373360954735 | classC | consensus |
| GPT2 | chr16:46912994-46970598 | cg00270789 | 0 | -0.913999852981742 | classC | consensus |
| GPT2 | chr16:46912994-46970598 | cg00622166 | 0 | -0.944167126424136 | classC | consensus |
| GPT2 | chr16:46912994-46970598 | cg03533472 | 0 | -0.935182426830102 | classC | consensus |
| GPT2 | chr16:46912994-46970598 | cg05380921 | 1.35e-10 | -0.804650076309172 | classC | consensus |
| GPT2 | chr16:46912994-46970598 | cg07710971 | 0 | -0.926745471323912 | classC | consensus |
| GPT2 | chr16:46912994-46970598 | cg23684449 | 0 | -0.940875032752331 | classC | consensus |
| GPT2 | chr16:46912994-46970598 | mean | 0 | -0.942054984232004 | classC | consensus |
| GPX1 | chr3:49371824-49396680 | cg05383531 | 6.60E-04 | -0.722466066243523 | classC | consensus |

| | | | | | | |
|---|---|---|---|---|---|---|
| GPX1 | chr3:49371824-49396680 | mean | 6.64E-04 | -0.722378301108136 | classC | consensus |
| GRAMD1B | chr11:123320379-123473034 | cg25243766 | 1.21E-03 | -0.712430484871225 | classC | consensus |
| GRASP | chr12:52388002-52484654 | cg12541478 | 2.48E-04 | 0.737833066306695 | classC | consensus |
| GRASP | chr12:52388002-52484654 | cg19107511 | 1.07E-02 | 0.714488031055373 | classC | consensus |
| GRASP | chr12:52388002-52484654 | mean | 8.9e-10 | 0.783031045960322 | classC | consensus |
| HEATR6 | chr17:58153617-58185439 | cg03120558 | 2.18E-04 | -0.739751000678711 | classC | consensus |
| HS3ST1 | chr4:11388923-11442762 | mean | 5.77E-04 | -0.72464140354835 | classC | consensus |
| IL17RB | chr3:53737038-53902022 | mean | 1.70E-04 | -0.743494116160621 | classC | consensus |
| IL24 | chr1:207067277-207141157 | cg05721773 | 1.43E-06 | -0.777173520651256 | classC | consensus |
| IL24 | chr1:207067277-207141157 | cg21607172 | 1.00E-12 | -0.850352150655553 | classC | consensus |
| IL24 | chr1:207067277-207141157 | mean | 4.19E-04 | -0.729725987505409 | classC | consensus |
| IL2RA | chr10:6053438-6132784 | mean | 1.89E-02 | -0.704783915939389 | classC | consensus |
| IL6R | chr1:154374725-154443028 | cg17427986 | 2.84E-04 | -0.73572864829091 | classC | consensus |
| IL6R | chr1:154374725-154443028 | mean | 5.87E-04 | -0.72437514334195 | classC | consensus |
| INPP5F | chr10:121438946-121576868 | cg07003611 | 0 | -0.916468675353853 | classC | consensus |
| INPP5F | chr10:121438946-121576868 | cg10053779 | 0 | -0.891650535477444 | classC | consensus |
| INPP5F | chr10:121438946-121576868 | cg11788307 | 0 | -0.930644335523718 | classC | consensus |
| INPP5F | chr10:121438946-121576868 | cg20704028 | 0 | -0.916864125353366 | classC | consensus |
| INPP5F | chr10:121438946-121576868 | mean | 0 | -0.931503852005095 | classC | consensus |
| IRF2 | chr4:185174927-185435847 | cg02354388 | 0 | -0.87700880407083 | classC | consensus |
| IRF2 | chr4:185174927-185435847 | cg11802666 | 3.56e-10 | -0.793863227945134 | classC | consensus |
| IRF2 | chr4:185174927-185435847 | mean | 7.2e-11 | -0.811295460541147 | classC | consensus |
| KAT6B | chr10:76573721-76622870 | cg00432328 | 2.49E-02 | 0.70619704100086 | classC | consensus |
| KMT2E | chr7:104561168-104657460 | cg16314851 | 1.20E-02 | 0.71259565691453 | classC | consensus |
| KSR1 | chr17:25782569-25881105 | cg04561005 | 4.45E-04 | 0.728776062525132 | classC | consensus |
| KSR2 | chr12:118300832-118376402 | cg06711448 | 0 | -0.86563078563151 | classC | consensus |
| LAG3 | chr12:6874667-6887901 | cg04671742 | 6.51E-04 | -0.722697287349605 | classC | consensus |
| LAG3 | chr12:6874667-6887901 | cg20652042 | 7.99E-06 | -0.754332744917271 | classC | consensus |
| LAG3 | chr12:6874667-6887901 | cg22777668 | 8.55E-06 | -0.753379541596865 | classC | consensus |
| LAG3 | chr12:6874667-6887901 | mean | 1.49e-10 | -0.803593362293405 | classC | consensus |
| LEF1-AS1 | chr4:108943438-109094928 | cg08460812 | 0 | -0.901734074946273 | classC | consensus |
| LEF1-AS1 | chr4:108943438-109094928 | cg19296305 | 3.00E-12 | -0.84270825777743 | classC | consensus |
| LEF1-AS1 | chr4:108943438-109094928 | cg24023289 | 0 | -0.917631346546461 | classC | consensus |
| LEF1-AS1 | chr4:108943438-109094928 | mean | 0 | -0.903157386533191 | classC | consensus |
| LFNG | chr7:2410726-2582633 | cg03676485 | 4.69E-06 | -0.761661269854988 | classC | consensus |
| LFNG | chr7:2410726-2582633 | cg09125127 | 3.87e-10 | -0.792913411064934 | classC | consensus |
| LFNG | chr7:2410726-2582633 | cg09723161 | 1.99e-10 | -0.80043520479131 | classC | consensus |
| LFNG | chr7:2410726-2582633 | cg21531604 | 1.62e-10 | -0.802718951049662 | classC | consensus |
| LFNG | chr7:2410726-2582633 | mean | 1.09E-06 | -0.780569410277952 | classC | consensus |
| LIX1L | chr1:145426814-145479979 | cg08916374 | 8.39E-04 | 0.718554014115345 | classC | consensus |
| LIX1L | chr1:145426814-145479979 | mean | 1.34E-02 | 0.710740841767711 | classC | consensus |
| LRRC25 | chr19:18498742-18641172 | cg22161383 | 3.26E-04 | -0.733605250945466 | classC | consensus |
| MCFD2 | chr2:47118860-47326683 | cg17647823 | 4.44E-04 | 0.728815356387401 | classC | consensus |
| MCFD2 | chr2:47118860-47326683 | cg22360765 | 1.49E-02 | 0.70884163888426 | classC | consensus |
| MCFD2 | chr2:47118860-47326683 | mean | 1.63E-04 | 0.744072938947013 | classC | consensus |
| MEF2A | chr15:99947214-99995771 | cg02378208 | 1.4e-11 | 0.827569291720352 | classC | consensus |
| MEF2A | chr15:99947214-99995771 | cg22700246 | 0 | 0.867414367267783 | classC | consensus |
| MEF2A | chr15:99947214-99995771 | mean | 0 | 0.862737971217625 | classC | consensus |
| MEF2A | chr15:99947214-99995771 | cg02378208 | 1.4e-11 | 0.827569291720352 | classC | consensus |
| MEF2A | chr15:99947214-99995771 | cg22700246 | 0 | 0.867414367267783 | classC | consensus |
| MEF2A | chr15:99947214-99995771 | mean | 0 | 0.862737971217625 | classC | consensus |
| MMS19 | chr10:99250107-99310055 | cg07052063 | 2.00E-12 | -0.844863156481634 | classC | consensus |
| MS4A6A | chr11:59949995-59962333 | mean | 7.38E-04 | -0.720665519154648 | classC | consensus |
| MSI2 | chr17:55320808-55784787 | cg05251593 | 1.15E-02 | -0.713346172060826 | classC | consensus |
| MSI2 | chr17:55320808-55784787 | cg06247967 | 1.83E-02 | -0.705298002173515 | classC | consensus |
| MSI2 | chr17:55320808-55784787 | cg16323330 | 1.3e-11 | -0.827998215873414 | classC | consensus |
| MSI2 | chr17:55320808-55784787 | mean | 1.43E-06 | -0.777140751887042 | classC | consensus |
| MYO1C | chr17:1336059-1402600 | cg00946187 | 1.06e-10 | 0.807284972647156 | classC | consensus |
| PDCD1 | chr2:242782277-242857257 | cg07728865 | 0 | -0.881048489695388 | classC | consensus |
| PDCD1 | chr2:242782277-242857257 | cg10057601 | 1.96E-06 | -0.773140210347633 | classC | consensus |
| PDCD1 | chr2:242782277-242857257 | mean | 0 | -0.867228783287569 | classC | consensus |
| PEX26 | chr22:18205753-18575249 | cg00040575 | 1.58E-04 | 0.744565243808952 | classC | consensus |
| PEX26 | chr22:18205753-18575249 | cg03291548 | 7.37E-04 | 0.720684525269443 | classC | consensus |
| PHTF1 | chr1:114292593-114302988 | cg12587818 | 0 | -0.888904876573824 | classC | consensus |
| PHTF1 | chr1:114292593-114302988 | cg22495801 | 1.00E-12 | -0.848230940739632 | classC | consensus |
| PHTF1 | chr1:114292593-114302988 | mean | 0 | -0.882155716743315 | classC | consensus |
| POC1A | chr3:51963239-52189508 | cg05504759 | 5.98E-06 | -0.758356270472029 | classC | consensus |

| | | | | | | |
|---|---|---|---|---|---|---|
| POLDIP3 | chr22:42999942-43060411 | cg02935298 | 1.17E-02 | 0.713019940136935 | classC | consensus |
| PPP6R3 | chr11:68109655-68213779 | mean | 6.66E-06 | 0.756869178769745 | classC | consensus |
| PPTC7 | chr12:110964850-111300733 | cg10453481 | 1.35E-04 | 0.746781505808766 | classC | consensus |
| PPTC7 | chr12:110964850-111300733 | mean | 1.13E-02 | 0.713614857475213 | classC | consensus |
| RAB43 | chr3:128799801-128848089 | cg05859076 | 7.32e-09 | -0.755549788848422 | classC | consensus |
| RAB43 | chr3:128799801-128848089 | mean | 3.28E-06 | -0.766471416124908 | classC | consensus |
| RAI1 | chr17:17565666-17825929 | cg23875758 | 7.88E-04 | -0.719577010440995 | classC | consensus |
| RAI1 | chr17:17565666-17825929 | mean | 1.85E-02 | -0.705118570435423 | classC | consensus |
| RELA | chr11:65096483-65434058 | cg03390187 | 5.80E-06 | 0.758773189553872 | classC | consensus |
| RELA | chr11:65096483-65434058 | cg04238983 | 3.5e-11 | 0.818776575046025 | classC | consensus |
| RELA | chr11:65096483-65434058 | cg04261607 | 1.91E-02 | 0.704628932500398 | classC | consensus |
| RELA | chr11:65096483-65434058 | cg10416861 | 0 | 0.870349426209499 | classC | consensus |
| RELA | chr11:65096483-65434058 | cg13167340 | 2.00E-12 | 0.845073659017576 | classC | consensus |
| RELA | chr11:65096483-65434058 | cg24772412 | 0 | 0.869056836889456 | classC | consensus |
| RELA | chr11:65096483-65434058 | cg26615176 | 0 | 0.874282922268681 | classC | consensus |
| RELA | chr11:65096483-65434058 | mean | 1.2e-11 | 0.82902740363356 | classC | consensus |
| RFC5 | chr12:118454470-118526839 | cg01932247 | 2.72e-10 | -0.796924079705611 | classC | consensus |
| RFC5 | chr12:118454470-118526839 | cg10617224 | 3.00E-11 | -0.820294730048158 | classC | consensus |
| RFC5 | chr12:118454470-118526839 | cg14474811 | 1.35e-10 | -0.804695542427171 | classC | consensus |
| RFC5 | chr12:118454470-118526839 | mean | 1.00E-12 | -0.849186680042747 | classC | consensus |
| RGS10 | chr10:120966371-121302914 | cg04182483 | 0 | -0.877368331595613 | classC | consensus |
| RGS10 | chr10:120966371-121302914 | cg08650639 | 1.55E-04 | -0.744777015465125 | classC | consensus |
| RGS10 | chr10:120966371-121302914 | cg17675386 | 0 | -0.873647948240536 | classC | consensus |
| RGS10 | chr10:120966371-121302914 | mean | 0 | -0.875866622013908 | classC | consensus |
| RLF | chr1:40626505-40627976 | cg18646240 | 2.2e-07 | 0.702116542104918 | classC | consensus |
| RNF214 | chr11:117040826-117105397 | cg11029270 | 2.29E-02 | -0.70140737522914 | classC | consensus |
| RNF4 | chr4:2445790-2515495 | cg07799157 | 5.53E-06 | 0.759426036495073 | classC | consensus |
| ROR1 | chr1:64237486-64489243 | cg03140289 | 0 | -0.900699279907852 | classC | consensus |
| ROR1 | chr1:64237486-64489243 | cg03950655 | 0 | -0.893898218867578 | classC | consensus |
| ROR1 | chr1:64237486-64489243 | cg06743782 | 0 | -0.87782318571676 | classC | consensus |
| ROR1 | chr1:64237486-64489243 | mean | 0 | -0.890012348111721 | classC | consensus |
| RREB1 | chr6:7098272-7209426 | cg09754197 | 2.3e-11 | 0.822710949158672 | classC | consensus |
| RREB1 | chr6:7098272-7209426 | cg26991433 | 1.1e-11 | 0.8299489996466 | classC | consensus |
| RREB1 | chr6:7098272-7209426 | mean | 9.00E-12 | 0.832127165489742 | classC | consensus |
| RUNX3 | chr1:25222331-25435095 | cg08544331 | 1.59E-06 | 0.775868044951279 | classC | consensus |
| RUNX3 | chr1:25222331-25435095 | cg20670361 | 2.41E-04 | 0.738243138684123 | classC | consensus |
| RUNX3 | chr1:25222331-25435095 | mean | 7.14E-06 | 0.75590575449577 | classC | consensus |
| SEMA4A | chr1:156063457-156147595 | cg12593793 | 1.69E-04 | -0.74353548770467 | classC | consensus |
| SEMA4A | chr1:156063457-156147595 | cg26667550 | 1.52E-02 | -0.708512619442562 | classC | consensus |
| SEMA4A | chr1:156063457-156147595 | mean | 6.29E-05 | -0.723241991290081 | classC | consensus |
| SEMA4D | chr9:91962974-92169084 | cg02702041 | 1.59E-06 | -0.775802758156494 | classC | consensus |
| SEMA4D | chr9:91962974-92169084 | cg14541848 | 3.00E-06 | -0.767649292230504 | classC | consensus |
| SEMA4D | chr9:91962974-92169084 | mean | 8.16e-10 | -0.784080988648402 | classC | consensus |
| SERINC5 | chr5:79388087-79555168 | cg10531355 | 1.57e-09 | -0.775991688054534 | classC | consensus |
| SERINC5 | chr5:79388087-79555168 | cg14978242 | 8.02E-06 | -0.754274975366666 | classC | consensus |
| SERINC5 | chr5:79388087-79555168 | mean | 3.37e-10 | -0.794484716231973 | classC | consensus |
| SFMBT1 | chr3:52948330-53124324 | cg04503182 | 9.12E-06 | -0.75245580619779 | classC | consensus |
| SFMBT1 | chr3:52948330-53124324 | cg26850469 | 8.91E-06 | -0.75279462640362 | classC | consensus |
| SFMBT1 | chr3:52948330-53124324 | mean | 5.54E-06 | -0.759417350830363 | classC | consensus |
| SH3BP5-AS1 | chr3:15284041-15451011 | cg01788682 | 2.13E-02 | -0.702706792653964 | classC | consensus |
| SH3BP5-AS1 | chr3:15284041-15451011 | mean | 9.13E-04 | -0.717157693955047 | classC | consensus |
| SLA | chr8:133989865-134141673 | cg11883141 | 1.64E-04 | -0.743987766312096 | classC | consensus |
| SMAD3 | chr15:67325517-67491383 | cg27130493 | 1.15E-02 | 0.713346350953164 | classC | consensus |
| SMCO4 | chr11:93250073-93277017 | cg12895631 | 1.2e-10 | -0.805970662165475 | classC | consensus |
| SNX18 | chr5:53804511-53851756 | cg01582396 | 0 | -0.925819320341889 | classC | consensus |
| SNX18 | chr5:53804511-53851756 | cg08544436 | 0 | -0.924446029025362 | classC | consensus |
| SNX18 | chr5:53804511-53851756 | cg14684434 | 2.00E-12 | -0.846994799832747 | classC | consensus |
| SNX18 | chr5:53804511-53851756 | cg14952488 | 1.99E-02 | -0.703919800410889 | classC | consensus |
| SNX18 | chr5:53804511-53851756 | cg16358034 | 1.00E-12 | -0.849689570071092 | classC | consensus |
| SNX18 | chr5:53804511-53851756 | cg21995347 | 0 | -0.898387637739902 | classC | consensus |
| SNX18 | chr5:53804511-53851756 | cg23001370 | 0 | -0.926986389361029 | classC | consensus |
| SNX18 | chr5:53804511-53851756 | mean | 0 | -0.918305000462116 | classC | consensus |
| SNX9 | chr6:158225529-158251805 | cg04064963 | 8.72E-04 | 0.717915811623504 | classC | consensus |
| SNX9 | chr6:158175110-158200357 | cg17071432 | 1.15E-02 | 0.713228715027926 | classC | consensus |
| SNX9 | chr6:158225529-158251805 | cg04064963 | 8.72E-04 | 0.717915811623504 | classC | consensus |
| SNX9 | chr6:158175110-158200357 | cg17071432 | 1.15E-02 | 0.713228715027926 | classC | consensus |
| SPOCK2 | chr10:73785554-73871197 | cg20571967 | 1.65e-10 | 0.802519905195252 | classC | consensus |

| | | | | | | |
|---|---|---|---|---|---|---|
| SPOCK2 | chr10:73785554-73871197 | cg23723835 | 2.29e-09 | 0.771165788495478 | classC | consensus |
| SPOCK2 | chr10:73785554-73871197 | mean | 6.69e-10 | 0.78647215691992 | classC | consensus |
| SRGN | chr10:70794826-70862131 | cg06486994 | 1.30E-06 | -0.778318826305982 | classC | consensus |
| STAMBPL1 | chr10:90636916-90704080 | cg14785527 | 0 | -0.920426081593454 | classC | consensus |
| STAP2 | chr19:4322691-4410395 | cg15797834 | 3.30E-06 | -0.766394635853633 | classC | consensus |
| STAP2 | chr19:4322691-4410395 | mean | 6.25E-04 | -0.72335657399481 | classC | consensus |
| TGFBI | chr5:135359409-135398951 | cg05838308 | 2.2e-11 | -0.823324370810826 | classC | consensus |
| TGFBI | chr5:135359409-135398951 | cg09096234 | 1.15E-04 | -0.749118068630648 | classC | consensus |
| TGFBI | chr5:135359409-135398951 | cg21583694 | 1.7e-10 | -0.802173626897387 | classC | consensus |
| TGFBI | chr5:135359409-135398951 | mean | 4.00E-12 | -0.838730840261316 | classC | consensus |
| TGFBR3 | chr1:92171519-92352408 | cg00342358 | 1.00E-12 | -0.848469339622927 | classC | consensus |
| TGFBR3 | chr1:92171519-92352408 | cg01336162 | 1.00E-12 | -0.853697907790304 | classC | consensus |
| TGFBR3 | chr1:92171519-92352408 | cg17074704 | 5.6e-11 | -0.813850119992126 | classC | consensus |
| TGFBR3 | chr1:92171519-92352408 | mean | 0 | -0.872899897378816 | classC | consensus |
| TGIF1 | chr18:3444750-3480409 | cg18204321 | 4.76E-04 | -0.72769425662416 | classC | consensus |
| TMEM104 | chr17:72728289-72780881 | cg22312249 | 9.54E-06 | -0.751815599615748 | classC | consensus |
| TMEM222 | chr1:27648154-28000134 | cg04089743 | 1.57E-04 | 0.744601638103384 | classC | consensus |
| TMEM222 | chr1:27648154-28000134 | mean | 6.96E-04 | 0.721605718298935 | classC | consensus |
| TNIP2 | chr4:2730793-2770921 | cg25585279 | 7.01e-09 | 0.756153579285156 | classC | consensus |
| TOX2 | chr20:42522068-42715360 | cg00789528 | 9.52E-04 | -0.716460736074329 | classC | consensus |
| TOX2 | chr20:42522068-42715360 | cg21293933 | 2.09E-06 | -0.772360350644319 | classC | consensus |
| TOX2 | chr20:42522068-42715360 | cg24403644 | 8.71E-04 | -0.717933655063413 | classC | consensus |
| TOX2 | chr20:42522068-42715360 | mean | 1.12e-10 | -0.806669244760544 | classC | consensus |
| TPST2 | chr22:26920248-27039444 | cg02455383 | 2.00E-12 | -0.84310960472665 | classC | consensus |
| TRAK1 | chr3:42041988-42191047 | cg01486558 | 4.3e-11 | 0.816524157391741 | classC | consensus |
| TRAK1 | chr3:42041988-42191047 | cg09654300 | 9.00E-12 | 0.831174698247483 | classC | consensus |
| TRAK1 | chr3:42041988-42191047 | mean | 3.77E-06 | 0.764620451232129 | classC | consensus |
| TRAPPC10 | chr21:45425337-45801001 | cg03738656 | 2.99E-06 | 0.767691051585551 | classC | consensus |
| TRAPPC10 | chr21:45425337-45801001 | cg21705505 | 2.42E-02 | 0.7004642555418 | classC | consensus |
| TRAPPC10 | chr21:45425337-45801001 | mean | 3.66E-06 | 0.765026351613073 | classC | consensus |
| TRAPPC10 | chr21:45336443-45405278 | cg04312999 | 3.20E-06 | 0.766781322093537 | classC | consensus |
| TRAPPC10 | chr21:45425337-45801001 | cg03738656 | 2.99E-06 | 0.767691051585551 | classC | consensus |
| TRAPPC10 | chr21:45425337-45801001 | cg21705505 | 2.42E-02 | 0.7004642555418 | classC | consensus |
| TRAPPC10 | chr21:45425337-45801001 | mean | 3.66E-06 | 0.765026351613073 | classC | consensus |
| TRAPPC10 | chr21:45336443-45405278 | cg04312999 | 3.20E-06 | 0.766781322093537 | classC | consensus |
| TSTA3 | chr8:144504468-144700876 | mean | 7.39E-04 | -0.720633922010797 | classC | consensus |
| UBL7-AS1 | chr15:74664329-74754207 | cg19251514 | 3.03E-06 | -0.767497678108892 | classC | consensus |
| WNT3 | chr17:44788117-44913504 | cg00140914 | 3.77E-06 | -0.764623451597125 | classC | consensus |
| WNT3 | chr17:44788117-44913504 | mean | 4.71E-04 | -0.727887360191764 | classC | consensus |
| WNT3 | chr17:44788117-44913504 | cg00140914 | 3.77E-06 | -0.764623451597125 | classC | consensus |
| WNT3 | chr17:44788117-44913504 | mean | 4.71E-04 | -0.727887360191764 | classC | consensus |
| XXYLT1 | chr3:194779122-194992398 | cg02041946 | 1.70E-06 | -0.774976141044074 | classC | consensus |
| XXYLT1 | chr3:194779122-194992398 | cg03916038 | 2.03e-10 | -0.800210337181105 | classC | consensus |
| XXYLT1 | chr3:194779122-194992398 | cg06704969 | 1.40E-04 | -0.746331143863356 | classC | consensus |
| XXYLT1 | chr3:194779122-194992398 | cg07463963 | 6.75E-06 | -0.756688154878191 | classC | consensus |
| XXYLT1 | chr3:194779122-194992398 | cg15722952 | 1.00E-12 | -0.852586353003877 | classC | consensus |
| XXYLT1 | chr3:194779122-194992398 | cg25921813 | 1.45E-06 | -0.776998077375551 | classC | consensus |
| XXYLT1 | chr3:194779122-194992398 | cg26062560 | 0 | -0.868250180871928 | classC | consensus |
| XXYLT1 | chr3:194779122-194992398 | mean | 0 | -0.873166941498145 | classC | consensus |
| ZAP70 | chr2:98328846-98336068 | cg08859278 | 2.00E-12 | -0.844812783413939 | classC | consensus |
| ZAP70 | chr2:98328846-98336068 | cg09006159 | 2.5e-11 | -0.82206607742438 | classC | consensus |
| ZAP70 | chr2:98328846-98336068 | cg15933451 | 4.00E-12 | -0.839690104550259 | classC | consensus |
| ZAP70 | chr2:98328846-98336068 | mean | 0 | -0.867532791913483 | classC | consensus |
| ZFPM1 | chr16:88441354-88624317 | cg02383666 | 1.42e-10 | -0.804093105399661 | classC | consensus |
| ZFPM1 | chr16:88441354-88624317 | cg04387347 | 3.77E-04 | -0.731365236344687 | classC | consensus |
| ZFPM1 | chr16:88441354-88624317 | cg05183538 | 3.08E-06 | -0.767308000217987 | classC | consensus |
| ZFPM1 | chr16:88441354-88624317 | cg05446615 | 4.60E-04 | -0.728254037660527 | classC | consensus |
| ZFPM1 | chr16:88441354-88624317 | mean | 2.09E-04 | -0.740414029386954 | classC | consensus |
| ZHX2 | chr8:123782932-123897246 | cg03314365 | 2.01e-10 | -0.800298272260102 | classC | consensus |
| ZPBP2 | chr17:37771665-38026697 | cg04587617 | 5.21E-05 | -0.726284019877596 | classC | consensus |
| ACSF3 | chr16:89098347-89206120 | cg01937601 | 2.18e-10 | -0.799393191030196 | classC | diff |
| ACSF3 | chr16:89098347-89206120 | cg03060214 | 1.58E-06 | -0.775919406500386 | classC | diff |
| CD5 | chr11:60822259-60879990 | cg16001613 | 3.32E-06 | -0.766296937980816 | classC | diff |
| CD5 | chr11:60822259-60879990 | cg20418308 | 0 | -0.934550284313043 | classC | diff |
| CD5 | chr11:60822259-60879990 | cg22202088 | 9.00E-12 | -0.831869672005622 | classC | diff |
| CD5 | chr11:60822259-60879990 | cg27301655 | 0 | -0.870918755345629 | classC | diff |
| CD5 | chr11:60822259-60879990 | mean | 0 | -0.914642281883123 | classC | diff |

| | | | | | | |
|---|---|---|---|---|---|---|
| CHIT1 | chr1:203236286-203353767 | cg03764585 | 1.42E-06 | -0.777246800088499 | classC | diff |
| CHIT1 | chr1:203236286-203353767 | cg04704856 | 1.82e-10 | -0.80139899446331 | classC | diff |
| CHIT1 | chr1:203236286-203353767 | cg12067421 | 9.00E-11 | -0.808948942559685 | classC | diff |
| CHIT1 | chr1:203236286-203353767 | cg16289210 | 7.63E-06 | -0.754972557081319 | classC | diff |
| CHIT1 | chr1:203236286-203353767 | cg26987645 | 3.24e-09 | -0.766627597512917 | classC | diff |
| CHIT1 | chr1:203236286-203353767 | mean | 1.17e-10 | -0.806206021996567 | classC | diff |
| CLLU1 | chr12:92783033-92883938 | cg04845867 | 3.26E-04 | -0.733628428342836 | classC | diff |
| CLLU1 | chr12:92783033-92883938 | mean | 1.80E-04 | -0.742629901903054 | classC | diff |
| DAD1 | chr14:22932348-23038986 | cg09953583 | 2.64E-06 | -0.769323487914752 | classC | diff |
| DAD1 | chr14:22932348-23038986 | cg15849446 | 6.04E-04 | -0.72391375882452 | classC | diff |
| DAD1 | chr14:22932348-23038986 | cg18887769 | 4.72e-10 | -0.790583481057587 | classC | diff |
| DAD1 | chr14:22932348-23038986 | cg23062357 | 1.04E-05 | -0.750659831755264 | classC | diff |
| DAD1 | chr14:22932348-23038986 | mean | 5.6e-11 | -0.813993538058702 | classC | diff |
| GTSF1L | chr20:42307202-42383663 | cg15099490 | 3.89e-10 | -0.792843715271909 | classC | diff |
| GTSF1L | chr20:42307202-42383663 | cg20774413 | 2.71E-06 | -0.768988425980799 | classC | diff |
| GTSF1L | chr20:42307202-42383663 | cg21426218 | 2.09E-04 | -0.740401223827821 | classC | diff |
| GTSF1L | chr20:42307202-42383663 | mean | 1.04e-10 | -0.807480928079539 | classC | diff |
| IRF2 | chr4:185306708-185403585 | cg02354388 | 0 | -0.87700880407083 | classC | diff |
| IRF2 | chr4:185306708-185403585 | cg11802666 | 3.56e-10 | -0.793863227945134 | classC | diff |
| IRF2 | chr4:185306708-185403585 | mean | 1.00E-12 | -0.84934476478526 | classC | diff |
| MSI2 | chr17:55509468-55686246 | cg05251593 | 1.15E-02 | -0.713346172060826 | classC | diff |
| MSI2 | chr17:55509468-55686246 | cg06247967 | 1.83E-02 | -0.705298002173515 | classC | diff |
| MSI2 | chr17:55509468-55686246 | mean | 5.02E-04 | -0.726853188099458 | classC | diff |
| PITPNM2 | chr12:123515700-123637201 | cg05384139 | 0 | -0.886618833347691 | classC | diff |
| SELPLG | chr12:109006555-109042818 | cg23429240 | 1.07E-04 | -0.750204777233409 | classC | diff |
| SELPLG | chr12:109006555-109042818 | cg23816140 | 1.00E-12 | -0.849470708302224 | classC | diff |
| SELPLG | chr12:109006555-109042818 | mean | 4.00E-12 | -0.837957483226188 | classC | diff |
| SH3BP5 | chr3:15298050-15404967 | cg01788682 | 2.60E-04 | -0.737087998881298 | classC | diff |
| SH3BP5 | chr3:15298050-15404967 | cg05046996 | 9.58e-10 | -0.782126741240425 | classC | diff |
| SH3BP5 | chr3:15298050-15404967 | cg18444702 | 1.56E-04 | -0.744716660289901 | classC | diff |
| SH3BP5 | chr3:15298050-15404967 | cg27375330 | 2.61e-09 | -0.769467534193077 | classC | diff |
| SH3BP5 | chr3:15298050-15404967 | mean | 2.2e-10 | -0.799329643526301 | classC | diff |
| TNFRSF1B | chr1:12184973-12271597 | cg06740893 | 5.35E-06 | -0.759887313034418 | classC | diff |
| TNFRSF1B | chr1:12184973-12271597 | cg07707505 | 1.60E-03 | -0.707629413967496 | classC | diff |
| TNFRSF1B | chr1:12184973-12271597 | cg11665613 | 1.47E-02 | -0.709158747053645 | classC | diff |
| TNFRSF1B | chr1:12184973-12271597 | cg15622556 | 2.55E-06 | -0.769780423892631 | classC | diff |
| TNFRSF1B | chr1:12184973-12271597 | cg18124517 | 3.11E-04 | -0.734331298859289 | classC | diff |
| TNFRSF1B | chr1:12184973-12271597 | mean | 1.43E-04 | -0.745943462476781 | classC | diff |
| TRAPPC2L | chr16:88919603-89070961 | cg00121135 | 1.3e-11 | -0.828313167727404 | classC | diff |
| TRAPPC2L | chr16:88919603-89070961 | cg01379825 | 1.45E-02 | -0.709306526821463 | classC | diff |
| TRAPPC2L | chr16:88919603-89070961 | cg05531174 | 6.23e-10 | -0.787315680520641 | classC | diff |
| TRAPPC2L | chr16:88919603-89070961 | cg07630078 | 0 | -0.91843657428251 | classC | diff |
| TRAPPC2L | chr16:88919603-89070961 | cg08345246 | 1.44e-10 | -0.803975400727001 | classC | diff |
| TRAPPC2L | chr16:88919603-89070961 | cg08484992 | 3.07E-04 | -0.734542814229392 | classC | diff |
| TRAPPC2L | chr16:88919603-89070961 | cg08843248 | 1.12E-02 | -0.713753598698726 | classC | diff |
| TRAPPC2L | chr16:88919603-89070961 | cg08940410 | 0 | -0.858770248468139 | classC | diff |
| TRAPPC2L | chr16:88919603-89070961 | cg26655856 | 1.00E-12 | -0.850032797078137 | classC | diff |
| TRAPPC2L | chr16:88919603-89070961 | cg27248474 | 0 | -0.903522219276505 | classC | diff |
| TRAPPC2L | chr16:88919603-89070961 | cg27292079 | 2.00E-12 | -0.845638015869319 | classC | diff |
| TRAPPC2L | chr16:88919603-89070961 | mean | 0 | -0.874221854858184 | classC | diff |
| XXYLT1-AS2 | chr3:194796146-194885613 | cg02041946 | 3.44E-04 | -0.732803957918885 | classC | diff |
| XXYLT1-AS2 | chr3:194796146-194885613 | cg07463963 | 6.93E-05 | -0.721686754371287 | classC | diff |
| XXYLT1-AS2 | chr3:194796146-194885613 | mean | 2.25E-06 | -0.7714242619335 | classC | diff |
| ZFPM1 | chr16:88509664-88551933 | cg02383666 | 1.42e-10 | -0.804093105399661 | classC | diff |
| ZFPM1 | chr16:88509664-88551933 | cg04387347 | 3.77E-04 | -0.731365236344687 | classC | diff |
| ZFPM1 | chr16:88509664-88551933 | cg05183538 | 3.08E-06 | -0.767308000217987 | classC | diff |
| ZFPM1 | chr16:88509664-88551933 | cg05446615 | 4.60E-04 | -0.728254037660527 | classC | diff |
| ZFPM1 | chr16:88509664-88551933 | mean | 2.09E-04 | -0.740414029386954 | classC | diff |
| ZNF710 | chr15:90541151-90611052 | cg16558770 | 2.33E-04 | -0.738748147736841 | classC | diff |
| ZNF710 | chr15:90541151-90611052 | mean | 1.58E-04 | -0.744503009355451 | classC | diff |
