## Supplementary Table S3 for "Methylome-based cell-of-origin modeling (Methyl-COOM) identifies aberrant expression of immune regulatory molecules in CLL"

**Supplementary Table S3: List of CLL-specific miRNAs**

| Class | microRNA | promoter | CpG | p-value | Correlation coefficient (rho) |
|---|---|---|---|---|---|
| classA | mir-3613 | chr13:50648999-50659799 | cg06228828 | 0.008261949 | -0.45 |
| classA | mir-15a | chr13:50648999-50659799 | cg06228828 | 0.033058248 | -0.37 |
| classA | mir-3688-2 | chr4:160023200-160029550 | cg00861646 | 0.019395319 | -0.40 |
| classA | mir-219a-1 | chr6:33084200-33086822 | cg05337177 | 0.02407845 | -0.39 |
| classA | mir-486 | chr8:41520600-41524000 | cg17753169 | 0.017795959 | -0.41 |
| classC | mir-3605 | chr1:33811613-33817213 | cg07570498 | 0.011054525 | -0.43 |
| classC | mir-141 | chr12:7022139-7025739 | cg04604946 | 0.047770444 | 0.35 |
| classC | mir-195 | chr17:6921076-6923876 | cg21040575 | 0.034632919 | 0.36 |
