## Supplementary Table S4 for "Methylome-based cell-of-origin modeling (Methyl-COOM) identifies aberrant expression of immune regulatory molecules in CLL"

**Supplementary Table S4: List of candidate target genes of CLL-specific microRNAs**

| miRNA | Target.Gene | Experiments | Database |
|---|---|---|---|
| hsa-miR-195-5p | WEE1 | ELISA//GFP reporter assay//Luciferase reporter assay//Microarray//qRT-PCR//Western blot | miRTarbase |
| hsa-miR-195-5p | WEE1 | Immunoblot//Immunohistochemistry//Luciferase reporter assay//qRT-PCR | miRTarbase |
| hsa-miR-195-5p | WEE1 | PAR-CLIP | miRTarbase |
| hsa-miR-195-5p | WEE1 | PAR-CLIP | miRTarbase |
| hsa-miR-195-5p | E2F3 | Immunohistochemistry//Luciferase reporter assay//Northern blot//qRT-PCR//Western blot//Reporter assay;Western blot;qRT-PCR;Other | miRTarbase |
| hsa-miR-195-5p | E2F3 | Luciferase reporter assay//Western blot | miRTarbase |
| hsa-miR-195-5p | E2F3 | HITS-CLIP | miRTarbase |
| hsa-miR-195-5p | CDK6 | Reporter assay;Western blot;qRT-PCR;Other | miRTarbase |
| hsa-miR-195-5p | CDK6 | Luciferase reporter assay//qRT-PCR//Western blot | miRTarbase |
| hsa-miR-195-5p | CDK6 | PAR-CLIP | miRTarbase |
| hsa-miR-195-5p | CCND1 | Luciferase reporter assay//Western blot | miRTarbase |
| hsa-miR-195-5p | CCND1 | Luciferase reporter assay | miRTarbase |
| hsa-miR-195-5p | CCND1 | Immunohistochemistry//Luciferase reporter assay//qRT-PCR//Western blot | miRTarbase |
| hsa-miR-195-5p | CCND1 | Luciferase reporter assay//qRT-PCR//Western blot | miRTarbase |
| hsa-miR-195-5p | CCND1 | PAR-CLIP | miRTarbase |
| hsa-miR-15a-5p | BMI1 | Luciferase reporter assay//Western blot | miRTarbase |
| hsa-miR-15a-5p | WNT3A | Luciferase reporter assay | miRTarbase |
| hsa-miR-15a-5p | WNT3A | Luciferase reporter assay//qRT-PCR | miRTarbase |
| hsa-miR-15a-5p | MYB | Luciferase reporter assay//Western blot | miRTarbase |
| hsa-miR-15a-5p | MYB | Luciferase reporter assay//Microarray//qRT-PCR | miRTarbase |
| hsa-miR-15a-5p | CDC25A | Luciferase reporter assay | miRTarbase |
| hsa-miR-15a-5p | CDC25A | HITS-CLIP | miRTarbase |
| hsa-miR-15a-5p | CDC25A | PAR-CLIP | miRTarbase |
| hsa-miR-15a-5p | CCND2 | Luciferase reporter assay | miRTarbase |
| hsa-miR-15a-5p | CCND2 | PAR-CLIP | miRTarbase |
| hsa-miR-15a-5p | CCND2 | PAR-CLIP | miRTarbase |
| hsa-miR-15a-5p | CCND2 | CLASH | miRTarbase |
| hsa-miR-141-3p | ZEB2 | Luciferase reporter assay//qRT-PCR//Western blot | miRTarbase |
| hsa-miR-141-3p | ZEB2 | Luciferase reporter assay | miRTarbase |
| hsa-miR-141-3p | ZEB2 | Luciferase reporter assay | miRTarbase |
| hsa-miR-141-3p | ZEB2 | Luciferase reporter assay//qRT-PCR//Western blot | miRTarbase |
| hsa-miR-141-3p | ZEB2 | Luciferase reporter assay//qRT-PCR//Western blot | miRTarbase |
| hsa-miR-141-3p | ZEB2 | qRT-PCR | miRTarbase |
| hsa-miR-141-3p | ZEB2 | TRAP | miRTarbase |
| hsa-miR-141-3p | ZEB2 | Luciferase reporter assay//qRT-PCR | miRTarbase |
| hsa-miR-141-3p | ZEB1 | Luciferase reporter assay | miRTarbase |
| hsa-miR-141-3p | ZEB1 | In situ hybridization//qRT-PCR//Western blot | miRTarbase |
| hsa-miR-141-3p | ZEB1 | Luciferase reporter assay//qRT-PCR//Western blot | miRTarbase |
| hsa-miR-141-3p | ZEB1 | qRT-PCR;Other | miRTarbase |
| hsa-miR-141-3p | ZEB1 | Luciferase reporter assay//qRT-PCR | miRTarbase |
| hsa-miR-141-3p | DLX5 | Luciferase reporter assay | miRTarbase |
| hsa-miR-141-3p | BAP1 | Reporter assay | miRTarbase |
| hsa-miR-141-3p | BAP1 | Microarray | miRTarbase |
| hsa-miR-195-5p | BCL2L11 | Microarray//Northern blot | miRTarbase |
| hsa-miR-195-5p | MECP2 | Microarray//Northern blot | miRTarbase |
| hsa-miR-15a-5p | RAB9B | proteomics analysis | miRTarbase |
| hsa-miR-15a-5p | ACTR1A | proteomics analysis | miRTarbase |
| hsa-miR-15a-5p | TPI1 | proteomics analysis | miRTarbase |
| hsa-miR-15a-5p | PDCD4 | qRT-PCR | miRTarbase |
| hsa-miR-15a-5p | PDCD4 | HITS-CLIP | miRTarbase |
| hsa-miR-15a-5p | RAB21 | qRT-PCR | miRTarbase |
| hsa-miR-15a-5p | BCL2 | Luciferase reporter assay//qRT-PCR//Western blot | miRTarbase |
| hsa-miR-15a-5p | BCL2 | Luciferase reporter assay | miRTarbase |
| hsa-miR-15a-5p | BCL2 | Luciferase reporter assay | miRTarbase |
| hsa-miR-15a-5p | BCL2 | Western blot | miRTarbase |
| hsa-miR-15a-5p | BCL2 | Western blot | miRTarbase |
| hsa-miR-15a-5p | BCL2 | Microarray//qRT-PCR | miRTarbase |
| hsa-miR-15a-5p | BCL2 | qRT-PCR//proteomics analysis | miRTarbase |
| hsa-miR-15a-5p | BCL2 | HITS-CLIP | miRTarbase |
| hsa-miR-15a-5p | WT1 | qRT-PCR//proteomics analysis | miRTarbase |
| hsa-miR-15a-5p | ASXL2 | Microarray | miRTarbase |
| hsa-miR-15a-5p | TMEM251 | Microarray | miRTarbase |
| hsa-miR-15a-5p | CARD8 | Microarray | miRTarbase |
| hsa-miR-15a-5p | CDC14B | Microarray | miRTarbase |
| hsa-miR-15a-5p | CENPJ | Microarray | miRTarbase |
| hsa-miR-15a-5p | CEP63 | Microarray | miRTarbase |
| hsa-miR-15a-5p | CREBL2 | Microarray | miRTarbase |
| hsa-miR-15a-5p | CREBL2 | PAR-CLIP | miRTarbase |
| hsa-miR-15a-5p | CREBL2 | PAR-CLIP | miRTarbase |
| hsa-miR-15a-5p | ECHDC1 | Microarray | miRTarbase |
| hsa-miR-15a-5p | GOLGA5 | Microarray | miRTarbase |
| hsa-miR-15a-5p | GOLPH3L | Microarray | miRTarbase |
| hsa-miR-15a-5p | GTF2H1 | Microarray | miRTarbase |
| hsa-miR-15a-5p | H3F3B | Microarray | miRTarbase |
| hsa-miR-15a-5p | HACE1 | Microarray | miRTarbase |
| hsa-miR-15a-5p | HDHD2 | Microarray | miRTarbase |
| hsa-miR-15a-5p | HERC6 | Microarray | miRTarbase |
| hsa-miR-15a-5p | HRSP12 | Microarray | miRTarbase |
| hsa-miR-15a-5p | HSDL2 | Microarray | miRTarbase |
| hsa-miR-15a-5p | HSPA1A | Microarray | miRTarbase |
| hsa-miR-15a-5p | JUN | Microarray | miRTarbase |
| hsa-miR-15a-5p | MCL1 | Microarray | miRTarbase |
| hsa-miR-15a-5p | MSH2 | Microarray | miRTarbase |
| hsa-miR-15a-5p | OMA1 | Microarray | miRTarbase |
| hsa-miR-15a-5p | OSGEPL1 | Microarray | miRTarbase |
| hsa-miR-15a-5p | PDCD6IP | Microarray | miRTarbase |
| hsa-miR-15a-5p | PHKB | Microarray | miRTarbase |
| hsa-miR-15a-5p | PMS1 | Microarray | miRTarbase |
| hsa-miR-15a-5p | PNN | Microarray | miRTarbase |
| hsa-miR-15a-5p | PRIM1 | Microarray | miRTarbase |
| hsa-miR-15a-5p | RAD51C | Microarray | miRTarbase |
| hsa-miR-15a-5p | RHOT1 | Microarray | miRTarbase |
| hsa-miR-15a-5p | RNASEL | Microarray | miRTarbase |
| hsa-miR-15a-5p | SLC35A1 | Microarray | miRTarbase |
| hsa-miR-15a-5p | SLC35B3 | Microarray | miRTarbase |
| hsa-miR-15a-5p | TIA1 | Microarray | miRTarbase |
| hsa-miR-15a-5p | UGDH | Microarray | miRTarbase |
| hsa-miR-15a-5p | UGP2 | Microarray | miRTarbase |
| hsa-miR-15a-5p | ZNF559 | Microarray | miRTarbase |
| hsa-miR-15a-5p | CCND1 | Luciferase reporter assay | miRTarbase |
| hsa-miR-15a-5p | CCND1 | Luciferase reporter assay | miRTarbase |
| hsa-miR-15a-5p | CCND1 | qRT-PCR//Luciferase reporter assay//Western blot | miRTarbase |
| hsa-miR-15a-5p | CCND1 | CLASH | miRTarbase |
| hsa-miR-15a-5p | CCND1 | PAR-CLIP | miRTarbase |
| hsa-miR-15a-5p | CCNE1 | immunohistochemistry//Luciferase reporter assay//Microarray//qRT-PCR | miRTarbase |
| hsa-miR-15a-5p | CCNE1 | Luciferase reporter assay | miRTarbase |
| hsa-miR-15a-5p | CCNE1 | qRT-PCR//Luciferase reporter assay//Western blot | miRTarbase |
| hsa-miR-15a-5p | CCNE1 | Luciferase reporter assay//qRT-PCR//Western blot | miRTarbase |
| hsa-miR-15a-5p | CCNE1 | PAR-CLIP | miRTarbase |
| hsa-miR-15a-5p | CCNE1 | PAR-CLIP | miRTarbase |
| hsa-miR-15a-5p | CCNE1 | HITS-CLIP | miRTarbase |
| hsa-miR-15a-5p | BACE1 | Luciferase reporter assay//Reporter assay;Other | miRTarbase |
| hsa-miR-141-3p | KLF5 | Luciferase reporter assay//Reporter assay;Other | miRTarbase |
| hsa-miR-141-3p | STK3 | Luciferase reporter assay//Reporter assay;Other | miRTarbase |
| hsa-miR-141-3p | TGFB2 | Immunoblot//Luciferase reporter assay//qRT-PCR | miRTarbase |
| hsa-miR-141-3p | SFPQ | Immunoprecipitaion//Luciferase reporter assay//Western blot | miRTarbase |
| hsa-miR-15a-5p | DMTF1 | Luciferase reporter assay//Reporter assay | miRTarbase |
| hsa-miR-15a-5p | DMTF1 | PAR-CLIP | miRTarbase |
| hsa-miR-15a-5p | DMTF1 | HITS-CLIP | miRTarbase |
| hsa-miR-141-3p | CLOCK | Luciferase reporter assay//Reporter assay | miRTarbase |
| hsa-miR-141-3p | CLOCK | Northern blot//qRT-PCR//Western blot | miRTarbase |
| hsa-miR-141-3p | BRD3 | Luciferase reporter assay//Western blot//Reporter assay;Western blot;Other | miRTarbase |
| hsa-miR-141-3p | BRD3 | PAR-CLIP | miRTarbase |
| hsa-miR-141-3p | UBAP1 | Luciferase reporter assay//Western blot//Reporter assay;Western blot;Other | miRTarbase |
| hsa-miR-141-3p | PTEN | Luciferase reporter assay//Western blot//Reporter assay;Western blot;Other | miRTarbase |
| hsa-miR-141-3p | PTEN | PAR-CLIP | miRTarbase |
| hsa-miR-15a-5p | BRCA1 | Luciferase reporter assay | miRTarbase |
| hsa-miR-15a-5p | AKT3 | Luciferase reporter assay | miRTarbase |
| hsa-miR-15a-5p | AKT3 | PAR-CLIP | miRTarbase |
| hsa-miR-15a-5p | AKT3 | PAR-CLIP | miRTarbase |
| hsa-miR-15a-5p | WIPF1 | Microarray | miRTarbase |
| hsa-miR-15a-5p | VPS45 | Microarray | miRTarbase |
| hsa-miR-15a-5p | HSP90B1 | Microarray | miRTarbase |
| hsa-miR-15a-5p | SKAP2 | Microarray//qRT-PCR | miRTarbase |
| hsa-miR-15a-5p | NT5DC1 | Microarray | miRTarbase |
| hsa-miR-15a-5p | FAM69A | Microarray | miRTarbase |
| hsa-miR-15a-5p | C2orf74 | Microarray | miRTarbase |
| hsa-miR-15a-5p | FAM122C | Microarray | miRTarbase |
| hsa-miR-15a-5p | PWWP2A | Microarray | miRTarbase |
| hsa-miR-15a-5p | C17orf80 | Microarray | miRTarbase |
| hsa-miR-15a-5p | CCDC111 | Microarray | miRTarbase |
| hsa-miR-15a-5p | C2orf43 | Microarray | miRTarbase |
| hsa-miR-15a-5p | C4orf27 | Microarray | miRTarbase |
| hsa-miR-15a-5p | NIPAL2 | Microarray | miRTarbase |
| hsa-miR-15a-5p | TRMT13 | Microarray | miRTarbase |
| hsa-miR-15a-5p | ANAPC16 | Microarray | miRTarbase |

| | | | |
|---|---|---|---|
| hsa-miR-15a-5p | CADM1 | Reporter assay//qRT-PCR | miRTarbase |
| hsa-miR-15a-5p | TMEM184B | Luciferase reporter assay//Reporter assay | miRTarbase |
| hsa-miR-15a-5p | APP | PAR-CLIP | miRTarbase |
| hsa-miR-15a-5p | APP | Luciferase reporter assay | miRTarbase |
| hsa-miR-15a-5p | UCP2 | Luciferase reporter assay//qRT-PCR | miRTarbase |
| hsa-miR-195-5p | VEGFA | ELISA//Luciferase reporter assay | miRTarbase |
| hsa-miR-195-5p | VEGFA | Luciferase reporter assay | miRTarbase |
| hsa-miR-195-5p | VEGFA | PAR-CLIP | miRTarbase |
| hsa-miR-195-5p | VEGFA | PAR-CLIP//HITS-CLIP | miRTarbase |
| hsa-miR-195-5p | VEGFA | PAR-CLIP | miRTarbase |
| hsa-miR-195-5p | VEGFA | HITS-CLIP | miRTarbase |
| hsa-miR-195-5p | VEGFA | PAR-CLIP | miRTarbase |
| hsa-miR-195-5p | VEGFA | PAR-CLIP | miRTarbase |
| hsa-miR-195-5p | VEGFA | PAR-CLIP | miRTarbase |
| hsa-miR-15a-5p | VEGFA | ELISA//Luciferase reporter assay | miRTarbase |
| hsa-miR-15a-5p | VEGFA | Luciferase reporter assay//qRT-PCR//Western blot | miRTarbase |
| hsa-miR-15a-5p | VEGFA | Luciferase reporter assay | miRTarbase |
| hsa-miR-15a-5p | VEGFA | PAR-CLIP | miRTarbase |
| hsa-miR-15a-5p | VEGFA | PAR-CLIP//HITS-CLIP | miRTarbase |
| hsa-miR-15a-5p | VEGFA | PAR-CLIP | miRTarbase |
| hsa-miR-15a-5p | VEGFA | HITS-CLIP | miRTarbase |
| hsa-miR-15a-5p | VEGFA | PAR-CLIP | miRTarbase |
| hsa-miR-15a-5p | VEGFA | PAR-CLIP | miRTarbase |
| hsa-miR-15a-5p | VEGFA | PAR-CLIP | miRTarbase |
| hsa-miR-195-5p | SKI | Microarray//Northern blot | miRTarbase |
| hsa-miR-195-5p | SKI | PAR-CLIP | miRTarbase |
| hsa-miR-195-5p | SKI | HITS-CLIP | miRTarbase |
| hsa-miR-486-5p | CD40 | Microarray//qRT-PCR | miRTarbase |
| hsa-miR-141-3p | ZFPM2 | Luciferase reporter assay//Western blot//Reporter assay | miRTarbase |
| hsa-miR-195-5p | CCL4 | ELISA//Immunoprecipitaion//qRT-PCR//Western blot | miRTarbase |
| hsa-miR-15a-5p | TSPYL2 | Luciferase reporter assay | miRTarbase |
| hsa-miR-15a-5p | NFKB1 | Microarray//qRT-PCR | miRTarbase |
| hsa-miR-141-3p | TRAPPC2P1 | Luciferase reporter assay | miRTarbase |
| hsa-miR-195-5p | KRT7 | Immunoblot//qRT-PCR | miRTarbase |
| hsa-miR-195-5p | BCL2 | Luciferase reporter assay//qRT-PCR//Western blot | miRTarbase |
| hsa-miR-195-5p | BCL2 | Immunohistochemistry//Luciferase reporter assay//qRT-PCR//Western blot | miRTarbase |
| hsa-miR-195-5p | BCL2 | HITS-CLIP | miRTarbase |
| hsa-miR-141-3p | EIF4E | Luciferase reporter assay//Northern blot//qRT-PCR//Western blot | miRTarbase |
| hsa-miR-15a-5p | CHUK | Luciferase reporter assay//qRT-PCR//Western blot | miRTarbase |
| hsa-miR-15a-5p | TP53 | Immunoblot//Luciferase reporter assay | miRTarbase |
| hsa-miR-15a-5p | FGF7 | Luciferase reporter assay//qRT-PCR | miRTarbase |
| hsa-miR-195-5p | RAF1 | Luciferase reporter assay//Northern blot//qRT-PCR//Western blot | miRTarbase |
| hsa-miR-195-5p | RAF1 | Flow//Immunofluorescence//qRT-PCR//Western blot | miRTarbase |
| hsa-miR-195-5p | RUNX2 | Luciferase reporter assay//Northern blot//qRT-PCR//Western blot | miRTarbase |
| hsa-miR-141-3p | CTBP2 | Luciferase reporter assay//Western blot | miRTarbase |
| hsa-miR-141-3p | CDYL | Luciferase reporter assay//Western blot | miRTarbase |
| hsa-miR-15a-5p | CLCN3 | Luciferase reporter assay//Microarray//Northern blot//qRT-PCR//Western blot | miRTarbase |
| hsa-miR-15a-5p | CRKL | Luciferase reporter assay//Microarray//Northern blot//qRT-PCR//Western blot | miRTarbase |
| hsa-miR-15a-5p | CRKL | PAR-CLIP | miRTarbase |
| hsa-miR-15a-5p | MN1 | Luciferase reporter assay//Microarray//Northern blot//qRT-PCR//Western blot | miRTarbase |
| hsa-miR-195-5p | SLC2A3 | Luciferase reporter assay//Western blot | miRTarbase |
| hsa-miR-195-5p | SLC2A3 | PAR-CLIP | miRTarbase |
| hsa-miR-195-5p | SLC2A3 | PAR-CLIP | miRTarbase |
| hsa-miR-195-5p | TBCCD1 | Luciferase reporter assay//Western blot | miRTarbase |
| hsa-miR-195-5p | TBCCD1 | HITS-CLIP | miRTarbase |
| hsa-miR-195-5p | CCND3 | Luciferase reporter assay//Western blot | miRTarbase |
| hsa-miR-195-5p | CCND3 | Luciferase reporter assay//Western blot | miRTarbase |
| hsa-miR-195-5p | CDK4 | Luciferase reporter assay | miRTarbase |
| hsa-miR-195-5p | CDK4 | Luciferase reporter assay//Western blot | miRTarbase |
| hsa-miR-141-3p | ACVR2B | Luciferase reporter assay//Western blot | miRTarbase |
| hsa-miR-141-3p | MAPK14 | Luciferase reporter assay//Microarray//Western blot | miRTarbase |
| hsa-miR-141-3p | MAPK14 | HITS-CLIP | miRTarbase |
| hsa-miR-141-3p | PPARA | Luciferase reporter assay//qRT-PCR//Western blot | miRTarbase |
| hsa-miR-141-3p | NR0B2 | Luciferase reporter assay | miRTarbase |
| hsa-miR-141-3p | YWHAG | Luciferase reporter assay//qRT-PCR//Western blot | miRTarbase |
| hsa-miR-15a-5p | HMGA1 | Luciferase reporter assay//qRT-PCR//Western blot | miRTarbase |
| hsa-miR-15a-5p | HMGA1 | HITS-CLIP | miRTarbase |
| hsa-miR-15a-5p | HMGA2 | Luciferase reporter assay//qRT-PCR//Western blot | miRTarbase |
| hsa-miR-15a-5p | IFNG | Luciferase reporter assay//qRT-PCR | miRTarbase |
| hsa-miR-195-5p | CDC42 | Luciferase reporter assay | miRTarbase |
| hsa-miR-195-5p | CDC42 | Luciferase reporter assay//qRT-PCR//Western blot | miRTarbase |
| hsa-miR-15a-5p | PURA | Luciferase reporter assay | miRTarbase |
| hsa-miR-15a-5p | PURA | HITS-CLIP | miRTarbase |
| hsa-miR-195-5p | CAB39 | Immunocytochemistry//In situ hybridization//Northern blot//qRT-PCR//Western blot | miRTarbase |
| hsa-miR-15a-5p | RECK | GFP reporter assay//qRT-PCR | miRTarbase |
| hsa-miR-15a-5p | RECK | PAR-CLIP | miRTarbase |
| hsa-miR-195-5p | CHUK | Luciferase reporter assay | miRTarbase |
| hsa-miR-195-5p | TAB3 | Luciferase reporter assay | miRTarbase |
| hsa-miR-195-5p | MBD1 | Luciferase reporter assay | miRTarbase |
| hsa-miR-195-5p | CCNE1 | Luciferase reporter assay | miRTarbase |
| hsa-miR-195-5p | CCNE1 | Luciferase reporter assay//Western blot | miRTarbase |
| hsa-miR-195-5p | CCNE1 | PAR-CLIP | miRTarbase |
| hsa-miR-195-5p | CCNE1 | PAR-CLIP | miRTarbase |
| hsa-miR-195-5p | CCNE1 | HITS-CLIP | miRTarbase |
| hsa-miR-195-5p | BCL2L2 | Luciferase reporter assay//Western blot | miRTarbase |
| hsa-miR-141-3p | ELAVL4 | qRT-PCR | miRTarbase |
| hsa-miR-141-3p | MAPK9 | Western blot | miRTarbase |
| hsa-miR-141-3p | TFDP2 | Western blot | miRTarbase |
| hsa-miR-141-3p | E2F3 | Western blot | miRTarbase |
| hsa-miR-141-3p | SHC1 | Reporter assay | miRTarbase |
| hsa-miR-141-3p | VAC14 | Reporter assay | miRTarbase |
| hsa-miR-141-3p | TCF7L1 | Reporter assay | miRTarbase |
| hsa-miR-141-3p | ELMO2 | Reporter assay | miRTarbase |
| hsa-miR-141-3p | RASSF2 | Reporter assay | miRTarbase |
| hsa-miR-141-3p | KLHL20 | Reporter assay | miRTarbase |
| hsa-miR-141-3p | RIN2 | Reporter assay | miRTarbase |
| hsa-miR-141-3p | SEPT7 | Reporter assay | miRTarbase |
| hsa-miR-141-3p | HOXB5 | Reporter assay | miRTarbase |
| hsa-miR-141-3p | ERBB2IP | Reporter assay | miRTarbase |
| hsa-miR-141-3p | KLF11 | Reporter assay | miRTarbase |
| hsa-miR-141-3p | PTPRD | Reporter assay | miRTarbase |
| hsa-miR-141-3p | WDR37 | Reporter assay | miRTarbase |
| hsa-miR-15a-5p | DLK1 | qRT-PCR;Other | miRTarbase |
| hsa-miR-15a-3p | LEPROTL1 | CLASH | miRTarbase |
| hsa-miR-15a-3p | RSPO2 | CLASH | miRTarbase |
| hsa-miR-15a-3p | DHX57 | CLASH | miRTarbase |
| hsa-miR-15a-3p | RASAL2 | CLASH | miRTarbase |
| hsa-miR-15a-3p | ND4L | CLASH | miRTarbase |
| hsa-miR-15a-3p | TFAM | CLASH | miRTarbase |
| hsa-miR-15a-3p | CNOT6 | CLASH | miRTarbase |
| hsa-miR-486-5p | TMED1 | CLASH | miRTarbase |
| hsa-miR-195-5p | JAK2 | CLASH | miRTarbase |
| hsa-miR-195-5p | CAMKV | CLASH | miRTarbase |
| hsa-miR-195-5p | AGER | CLASH | miRTarbase |
| hsa-miR-195-5p | LSM11 | PAR-CLIP | miRTarbase |
| hsa-miR-195-5p | LSM11 | HITS-CLIP | miRTarbase |
| hsa-miR-195-5p | LSM11 | CLASH | miRTarbase |
| hsa-miR-195-5p | ABCB7 | CLASH | miRTarbase |
| hsa-miR-195-5p | ZNF280C | CLASH | miRTarbase |
| hsa-miR-195-5p | SPTBN1 | CLASH | miRTarbase |
| hsa-miR-195-5p | NOLC1 | CLASH | miRTarbase |
| hsa-miR-195-5p | CAND1 | CLASH | miRTarbase |
| hsa-miR-195-5p | COPB1 | CLASH | miRTarbase |
| hsa-miR-195-5p | RPL10 | CLASH | miRTarbase |
| hsa-miR-195-5p | TMC6 | CLASH | miRTarbase |
| hsa-miR-195-5p | TPI1 | CLASH | miRTarbase |
| hsa-miR-195-5p | SH3BGRL2 | CLASH | miRTarbase |
| hsa-miR-195-5p | AGO1 | CLASH | miRTarbase |
| hsa-miR-15a-5p | PLA2G2D | CLASH | miRTarbase |
| hsa-miR-15a-5p | ACVR1B | CLASH | miRTarbase |
| hsa-miR-15a-5p | IKBKG | CLASH | miRTarbase |
| hsa-miR-15a-5p | GCLM | CLASH | miRTarbase |
| hsa-miR-15a-5p | PCF11 | CLASH | miRTarbase |
| hsa-miR-15a-5p | HIST1H2BK | CLASH | miRTarbase |
| hsa-miR-15a-5p | ODC1 | CLASH | miRTarbase |
| hsa-miR-15a-5p | CALD1 | CLASH | miRTarbase |
| hsa-miR-15a-5p | RPP30 | CLASH | miRTarbase |
| hsa-miR-15a-5p | ASNSD1 | CLASH | miRTarbase |
| hsa-miR-15a-5p | CCNYL1 | CLASH | miRTarbase |
| hsa-miR-15a-5p | RGPD5 | CLASH | miRTarbase |
| hsa-miR-15a-5p | PREB | CLASH | miRTarbase |
| hsa-miR-15a-5p | PDHX | CLASH | miRTarbase |
| hsa-miR-15a-5p | SNX6 | CLASH | miRTarbase |
| hsa-miR-15a-5p | CNN3 | CLASH | miRTarbase |
| hsa-miR-15a-5p | KIF1A | CLASH | miRTarbase |
| hsa-miR-15a-5p | NAB1 | CLASH | miRTarbase |
| hsa-miR-15a-5p | CCT6B | CLASH | miRTarbase |
| hsa-miR-15a-5p | CHD4 | CLASH | miRTarbase |
| hsa-miR-15a-5p | CLCC1 | CLASH | miRTarbase |

| | | | |
|---|---|---|---|
| hsa-miR-15a-5p | GDI2 | CLASH | miRTarbase |
| hsa-miR-15a-5p | BRWD1 | CLASH | miRTarbase |
| hsa-miR-15a-5p | MAPK6 | CLASH | miRTarbase |
| hsa-miR-15a-5p | PSMC4 | CLASH | miRTarbase |
| hsa-miR-15a-5p | ATF2 | CLASH | miRTarbase |
| hsa-miR-15a-5p | ATP6AP1 | CLASH | miRTarbase |
| hsa-miR-15a-5p | FBXO3 | CLASH | miRTarbase |
| hsa-miR-15a-5p | PRDX3 | CLASH | miRTarbase |
| hsa-miR-15a-5p | CABIN1 | CLASH | miRTarbase |
| hsa-miR-15a-5p | FASN | PAR-CLIP | miRTarbase |
| hsa-miR-15a-5p | FASN | PAR-CLIP | miRTarbase |
| hsa-miR-15a-5p | FASN | CLASH | miRTarbase |
| hsa-miR-15a-5p | SEC63 | CLASH | miRTarbase |
| hsa-miR-15a-5p | PTAR1 | CLASH | miRTarbase |
| hsa-miR-15a-5p | DSTYK | CLASH | miRTarbase |
| hsa-miR-15a-5p | FOXO1 | Luciferase reporter assay//qRT-PCR//Western blot | miRTarbase |
| hsa-miR-15a-5p | FOXO1 | CLASH | miRTarbase |
| hsa-miR-15a-5p | TMEM214 | CLASH | miRTarbase |
| hsa-miR-15a-5p | TRIM28 | CLASH | miRTarbase |
| hsa-miR-15a-5p | NOP2 | CLASH | miRTarbase |
| hsa-miR-15a-5p | MYBL1 | CLASH | miRTarbase |
| hsa-miR-15a-5p | TTC1 | CLASH | miRTarbase |
| hsa-miR-15a-5p | BTRC | HITS-CLIP | miRTarbase |
| hsa-miR-15a-5p | BTRC | CLASH | miRTarbase |
| hsa-miR-3605-3p | RPLP1 | CLASH | miRTarbase |
| hsa-miR-3605-3p | VIM | CLASH | miRTarbase |
| hsa-miR-141-3p | STAT4 | Luciferase reporter assay//qRT-PCR//Western blot//Immunohistochemistry | miRTarbase |
| hsa-miR-141-3p | YAP1 | Luciferase reporter assay//Western blot | miRTarbase |
| hsa-miR-15a-5p | REPIN1 | Luciferase reporter assay//Western//qRT-PCR | miRTarbase |
| hsa-miR-15a-5p | KLF4 | Western blot | miRTarbase |
| hsa-miR-141-3p | CDC25C | Luciferase reporter assay//Western blot | miRTarbase |
| hsa-miR-195-5p | TAR | qRT-PCR//Western blot | miRTarbase |
| hsa-miR-195-5p | DICER1 | qRT-PCR//Western blot | miRTarbase |
| hsa-miR-195-5p | DICER1 | HITS-CLIP | miRTarbase |
| hsa-miR-195-5p | FASN | Luciferase reporter assay//qRT-PCR//Western blot | miRTarbase |
| hsa-miR-195-5p | FASN | PAR-CLIP | miRTarbase |
| hsa-miR-195-5p | FASN | PAR-CLIP | miRTarbase |
| hsa-miR-486-5p | ARHGAP5 | Immunohistochemistry//Luciferase reporter assay//qRT-PCR//Western blot | miRTarbase |
| hsa-miR-486-3p | BCL11A | Luciferase reporter assay//qRT-PCR//Western blot | miRTarbase |
| hsa-miR-195-5p | ARL2 | Flow//Luciferase reporter assay//Microarray//qRT-PCR//Western blot | miRTarbase |
| hsa-miR-486-5p | OLFM4 | In situ hybridization//Luciferase reporter assay//Microarray//qRT-PCR//Western blot | miRTarbase |
| hsa-miR-141-3p | HDGF | Luciferase reporter assay//qRT-PCR//Western blot | miRTarbase |
| hsa-miR-141-3p | KEAP1 | Luciferase reporter assay//Microarray//qRT-PCR//Western blot | miRTarbase |
| hsa-miR-141-3p | KEAP1 | Luciferase reporter assay//qRT-PCR//Western blot | miRTarbase |
| hsa-miR-486-5p | SERPINE1 | qRT-PCR | miRTarbase |
| hsa-miR-141-3p | TAZ | Luciferase reporter assay//Microarray//qRT-PCR//Western blot | miRTarbase |
| hsa-miR-15a-5p | YAP1 | Flow//Luciferase reporter assay//qRT-PCR//Western blot | miRTarbase |
| hsa-miR-195-5p | BIRC5 | Luciferase reporter assay//qRT-PCR | miRTarbase |
| hsa-miR-195-5p | WNT7A | Luciferase reporter assay//Western blot | miRTarbase |
| hsa-miR-15a-5p | CARM1 | Luciferase reporter assay//qRT-PCR//Western blot | miRTarbase |
| hsa-miR-141-3p | TIAM1 | Luciferase reporter assay//qRT-PCR//Western blot | miRTarbase |
| hsa-miR-141-3p | TM4SF1 | Luciferase reporter assay//qRT-PCR//Western blot | miRTarbase |
| hsa-miR-141-3p | CDC25A | Luciferase reporter assay//Immunoblot//Western blot | miRTarbase |
| hsa-miR-141-3p | PHLPP1 | Luciferase reporter assay//qRT-PCR | miRTarbase |
| hsa-miR-141-3p | PHLPP2 | Luciferase reporter assay//qRT-PCR | miRTarbase |
| hsa-miR-195-5p | MYB | Luciferase reporter assay//qRT-PCR//Western blot | miRTarbase |
| hsa-miR-15a-5p | SOX5 | Luciferase reporter assay//Western blot | miRTarbase |
| hsa-miR-195-5p | ATG14 | Immunofluorescence//Luciferase reporter assay//qRT-PCR | miRTarbase |
| hsa-miR-195-5p | ATG14 | PAR-CLIP | miRTarbase |
| hsa-miR-15a-5p | SHOC2 | PAR-CLIP | miRTarbase |
| hsa-miR-15a-5p | SHOC2 | PAR-CLIP | miRTarbase |
| hsa-miR-15a-5p | SHOC2 | HITS-CLIP | miRTarbase |
| hsa-miR-15a-5p | SHOC2 | PAR-CLIP | miRTarbase |
| hsa-miR-15a-5p | SHOC2 | PAR-CLIP | miRTarbase |
| hsa-miR-15a-5p | SHOC2 | PAR-CLIP | miRTarbase |
| hsa-miR-195-5p | SHOC2 | PAR-CLIP | miRTarbase |
| hsa-miR-195-5p | SHOC2 | PAR-CLIP | miRTarbase |
| hsa-miR-195-5p | SHOC2 | HITS-CLIP | miRTarbase |
| hsa-miR-195-5p | SHOC2 | PAR-CLIP | miRTarbase |
| hsa-miR-195-5p | SHOC2 | PAR-CLIP | miRTarbase |
| hsa-miR-195-5p | SHOC2 | PAR-CLIP | miRTarbase |
| hsa-miR-15a-5p | PLEKHA1 | PAR-CLIP | miRTarbase |
| hsa-miR-195-5p | PLEKHA1 | PAR-CLIP | miRTarbase |
| hsa-miR-3613-3p | OTUD1 | PAR-CLIP | miRTarbase |
| hsa-miR-195-3p | PPIF | PAR-CLIP | miRTarbase |
| hsa-miR-195-3p | PPIF | PAR-CLIP | miRTarbase |
| hsa-miR-15a-5p | CEP55 | PAR-CLIP | miRTarbase |
| hsa-miR-15a-5p | CEP55 | PAR-CLIP | miRTarbase |
| hsa-miR-15a-5p | CEP55 | PAR-CLIP | miRTarbase |
| hsa-miR-15a-5p | CEP55 | PAR-CLIP | miRTarbase |
| hsa-miR-195-5p | CEP55 | PAR-CLIP | miRTarbase |
| hsa-miR-195-5p | CEP55 | PAR-CLIP | miRTarbase |
| hsa-miR-195-5p | CEP55 | PAR-CLIP | miRTarbase |
| hsa-miR-195-5p | CEP55 | PAR-CLIP | miRTarbase |
| hsa-miR-3613-5p | LCOR | PAR-CLIP | miRTarbase |
| hsa-miR-15a-5p | ZDHHC16 | PAR-CLIP | miRTarbase |
| hsa-miR-195-5p | ZDHHC16 | PAR-CLIP | miRTarbase |
| hsa-miR-15a-5p | STXBP3 | HITS-CLIP | miRTarbase |
| hsa-miR-195-5p | STXBP3 | HITS-CLIP | miRTarbase |
| hsa-miR-195-3p | RSBN1 | PAR-CLIP | miRTarbase |
| hsa-miR-195-3p | RSBN1 | PAR-CLIP | miRTarbase |
| hsa-miR-195-3p | RSBN1 | PAR-CLIP | miRTarbase |
| hsa-miR-195-3p | RSBN1 | PAR-CLIP | miRTarbase |
| hsa-miR-15a-5p | C1ORF21 | PAR-CLIP | miRTarbase |
| hsa-miR-15a-5p | C1ORF21 | PAR-CLIP | miRTarbase |
| hsa-miR-15a-5p | C1ORF21 | PAR-CLIP | miRTarbase |
| hsa-miR-195-5p | C1ORF21 | PAR-CLIP | miRTarbase |
| hsa-miR-195-5p | C1ORF21 | PAR-CLIP | miRTarbase |
| hsa-miR-195-5p | C1ORF21 | PAR-CLIP | miRTarbase |
| hsa-miR-15a-5p | AMOTL1 | PAR-CLIP | miRTarbase |
| hsa-miR-15a-5p | AMOTL1 | PAR-CLIP//HITS-CLIP | miRTarbase |
| hsa-miR-15a-5p | AMOTL1 | PAR-CLIP | miRTarbase |
| hsa-miR-15a-5p | AMOTL1 | PAR-CLIP | miRTarbase |
| hsa-miR-15a-5p | AMOTL1 | PAR-CLIP | miRTarbase |
| hsa-miR-15a-5p | AMOTL1 | PAR-CLIP | miRTarbase |
| hsa-miR-195-5p | AMOTL1 | PAR-CLIP | miRTarbase |
| hsa-miR-195-5p | AMOTL1 | PAR-CLIP//HITS-CLIP | miRTarbase |
| hsa-miR-195-5p | AMOTL1 | PAR-CLIP | miRTarbase |
| hsa-miR-195-5p | AMOTL1 | PAR-CLIP | miRTarbase |
| hsa-miR-195-5p | AMOTL1 | PAR-CLIP | miRTarbase |
| hsa-miR-195-5p | AMOTL1 | PAR-CLIP | miRTarbase |
| hsa-miR-15a-5p | BTG2 | HITS-CLIP | miRTarbase |
| hsa-miR-195-5p | BTG2 | HITS-CLIP | miRTarbase |
| hsa-miR-141-3p | PPP1R15B | PAR-CLIP | miRTarbase |
| hsa-miR-3613-3p | ZNF664 | PAR-CLIP | miRTarbase |
| hsa-miR-3613-3p | ZNF664 | PAR-CLIP | miRTarbase |
| hsa-miR-3613-3p | ZNF664 | PAR-CLIP | miRTarbase |
| hsa-miR-3613-3p | ZNF664 | PAR-CLIP | miRTarbase |
| hsa-miR-15a-5p | ETNK1 | HITS-CLIP | miRTarbase |
| hsa-miR-195-5p | ETNK1 | HITS-CLIP | miRTarbase |
| hsa-miR-195-5p | CCND2 | PAR-CLIP | miRTarbase |
| hsa-miR-195-5p | CCND2 | PAR-CLIP | miRTarbase |
| hsa-miR-3613-3p | CCND2 | PAR-CLIP | miRTarbase |
| hsa-miR-3613-3p | CCND2 | PAR-CLIP | miRTarbase |
| hsa-miR-15a-3p | ZBTB18 | PAR-CLIP | miRTarbase |
| hsa-miR-15a-5p | TARBP2 | PAR-CLIP | miRTarbase |
| hsa-miR-15a-5p | TARBP2 | PAR-CLIP | miRTarbase |
| hsa-miR-195-5p | TARBP2 | PAR-CLIP | miRTarbase |
| hsa-miR-195-5p | TARBP2 | PAR-CLIP | miRTarbase |
| hsa-miR-15a-5p | MTFR1L | PAR-CLIP | miRTarbase |
| hsa-miR-195-5p | MTFR1L | PAR-CLIP | miRTarbase |
| hsa-miR-15a-5p | USP15 | PAR-CLIP | miRTarbase |
| hsa-miR-195-5p | USP15 | PAR-CLIP | miRTarbase |
| hsa-miR-15a-5p | AKAP11 | HITS-CLIP | miRTarbase |
| hsa-miR-195-5p | AKAP11 | HITS-CLIP | miRTarbase |
| hsa-miR-219a-1-3p | KPNA3 | PAR-CLIP | miRTarbase |
| hsa-miR-219a-1-3p | KPNA3 | HITS-CLIP | miRTarbase |
| hsa-miR-219a-1-3p | KPNA3 | PAR-CLIP | miRTarbase |
| hsa-miR-3613-3p | SLC39A9 | HITS-CLIP | miRTarbase |
| hsa-miR-15a-5p | FCF1 | PAR-CLIP | miRTarbase |
| hsa-miR-195-5p | FCF1 | PAR-CLIP | miRTarbase |
| hsa-miR-3613-3p | FCF1 | PAR-CLIP | miRTarbase |
| hsa-miR-141-3p | RAB8B | PAR-CLIP | miRTarbase |
| hsa-miR-15a-5p | ARIH1 | PAR-CLIP | miRTarbase |
| hsa-miR-15a-5p | ARIH1 | PAR-CLIP | miRTarbase |
| hsa-miR-15a-5p | ARIH1 | HITS-CLIP | miRTarbase |
| hsa-miR-195-5p | ARIH1 | PAR-CLIP | miRTarbase |
| hsa-miR-195-5p | ARIH1 | PAR-CLIP | miRTarbase |
| hsa-miR-195-5p | ARIH1 | HITS-CLIP | miRTarbase |

| | | | |
|---|---|---|---|
| hsa-miR-195-3p | NFATC2IP | PAR-CLIP | miRTarbase |
| hsa-miR-195-3p | NFATC2IP | PAR-CLIP | miRTarbase |
| hsa-miR-15a-5p | PAGR1 | PAR-CLIP | miRTarbase |
| hsa-miR-15a-5p | PAGR1 | PAR-CLIP | miRTarbase |
| hsa-miR-195-5p | PAGR1 | PAR-CLIP | miRTarbase |
| hsa-miR-195-5p | PAGR1 | PAR-CLIP | miRTarbase |
| hsa-miR-15a-5p | SNTB2 | PAR-CLIP | miRTarbase |
| hsa-miR-15a-5p | SNTB2 | PAR-CLIP | miRTarbase |
| hsa-miR-195-5p | SNTB2 | PAR-CLIP | miRTarbase |
| hsa-miR-195-5p | SNTB2 | PAR-CLIP | miRTarbase |
| hsa-miR-15a-5p | VPS4A | PAR-CLIP | miRTarbase |
| hsa-miR-15a-5p | VPS4A | PAR-CLIP | miRTarbase |
| hsa-miR-15a-5p | VPS4A | PAR-CLIP | miRTarbase |
| hsa-miR-15a-5p | VPS4A | PAR-CLIP | miRTarbase |
| hsa-miR-195-5p | VPS4A | PAR-CLIP | miRTarbase |
| hsa-miR-195-5p | VPS4A | PAR-CLIP | miRTarbase |
| hsa-miR-195-5p | VPS4A | PAR-CLIP | miRTarbase |
| hsa-miR-195-5p | VPS4A | PAR-CLIP | miRTarbase |
| hsa-miR-15a-5p | C16ORF72 | PAR-CLIP//HITS-CLIP | miRTarbase |
| hsa-miR-15a-5p | C16ORF72 | HITS-CLIP | miRTarbase |
| hsa-miR-15a-5p | C16ORF72 | PAR-CLIP | miRTarbase |
| hsa-miR-15a-5p | C16ORF72 | PAR-CLIP | miRTarbase |
| hsa-miR-195-5p | C16ORF72 | PAR-CLIP//HITS-CLIP | miRTarbase |
| hsa-miR-195-5p | C16ORF72 | HITS-CLIP | miRTarbase |
| hsa-miR-195-5p | C16ORF72 | PAR-CLIP | miRTarbase |
| hsa-miR-195-5p | C16ORF72 | PAR-CLIP | miRTarbase |
| hsa-miR-486-3p | PHF12 | PAR-CLIP | miRTarbase |
| hsa-miR-15a-5p | GOSR1 | PAR-CLIP | miRTarbase |
| hsa-miR-195-5p | GOSR1 | PAR-CLIP | miRTarbase |
| hsa-miR-3613-3p | UBE2Z | PAR-CLIP | miRTarbase |
| hsa-miR-15a-5p | MINK1 | HITS-CLIP | miRTarbase |
| hsa-miR-195-5p | MINK1 | HITS-CLIP | miRTarbase |
| hsa-miR-15a-5p | RPS6KB1 | PAR-CLIP | miRTarbase |
| hsa-miR-195-5p | RPS6KB1 | PAR-CLIP | miRTarbase |
| hsa-miR-141-3p | MED13 | PAR-CLIP | miRTarbase |
| hsa-miR-3613-3p | MAP3K3 | PAR-CLIP | miRTarbase |
| hsa-miR-3613-3p | BIRC5 | PAR-CLIP | miRTarbase |
| hsa-miR-3613-3p | BIRC5 | PAR-CLIP | miRTarbase |
| hsa-miR-15a-5p | NAPG | PAR-CLIP | miRTarbase |
| hsa-miR-15a-5p | NAPG | PAR-CLIP//HITS-CLIP | miRTarbase |
| hsa-miR-15a-5p | NAPG | PAR-CLIP | miRTarbase |
| hsa-miR-15a-5p | NAPG | PAR-CLIP | miRTarbase |
| hsa-miR-15a-5p | NAPG | PAR-CLIP | miRTarbase |
| hsa-miR-15a-5p | NAPG | PAR-CLIP | miRTarbase |
| hsa-miR-195-5p | NAPG | PAR-CLIP | miRTarbase |
| hsa-miR-195-5p | NAPG | PAR-CLIP//HITS-CLIP | miRTarbase |
| hsa-miR-195-5p | NAPG | PAR-CLIP | miRTarbase |
| hsa-miR-195-5p | NAPG | PAR-CLIP | miRTarbase |
| hsa-miR-195-5p | NAPG | PAR-CLIP | miRTarbase |
| hsa-miR-195-5p | NAPG | PAR-CLIP | miRTarbase |
| hsa-miR-15a-5p | GALNT1 | PAR-CLIP | miRTarbase |
| hsa-miR-15a-5p | GALNT1 | PAR-CLIP | miRTarbase |
| hsa-miR-195-5p | GALNT1 | PAR-CLIP | miRTarbase |
| hsa-miR-195-5p | GALNT1 | PAR-CLIP | miRTarbase |
| hsa-miR-3613-3p | LDLR | PAR-CLIP | miRTarbase |
| hsa-miR-3613-3p | LDLR | PAR-CLIP | miRTarbase |
| hsa-miR-3613-3p | LDLR | PAR-CLIP | miRTarbase |
| hsa-miR-195-3p | ZNF431 | PAR-CLIP | miRTarbase |
| hsa-miR-195-3p | ZNF431 | PAR-CLIP | miRTarbase |
| hsa-miR-486-3p | HNRNPUL1 | PAR-CLIP | miRTarbase |
| hsa-miR-486-3p | HNRNPUL1 | PAR-CLIP | miRTarbase |
| hsa-miR-15a-5p | PNPLA6 | PAR-CLIP | miRTarbase |
| hsa-miR-195-5p | PNPLA6 | PAR-CLIP | miRTarbase |
| hsa-miR-15a-5p | ZCCHC3 | PAR-CLIP | miRTarbase |
| hsa-miR-15a-5p | ZCCHC3 | PAR-CLIP | miRTarbase |
| hsa-miR-15a-5p | ZCCHC3 | PAR-CLIP | miRTarbase |
| hsa-miR-195-5p | ZCCHC3 | PAR-CLIP | miRTarbase |
| hsa-miR-195-5p | ZCCHC3 | PAR-CLIP | miRTarbase |
| hsa-miR-195-5p | ZCCHC3 | PAR-CLIP | miRTarbase |
| hsa-miR-15a-5p | SOWAHC | PAR-CLIP | miRTarbase |
| hsa-miR-15a-5p | SOWAHC | PAR-CLIP//HITS-CLIP | miRTarbase |
| hsa-miR-195-5p | SOWAHC | PAR-CLIP | miRTarbase |
| hsa-miR-195-5p | SOWAHC | PAR-CLIP//HITS-CLIP | miRTarbase |
| hsa-miR-15a-5p | CCNT2 | HITS-CLIP | miRTarbase |
| hsa-miR-195-5p | CCNT2 | HITS-CLIP | miRTarbase |
| hsa-miR-15a-5p | UBR3 | PAR-CLIP//HITS-CLIP | miRTarbase |
| hsa-miR-195-5p | UBR3 | PAR-CLIP//HITS-CLIP | miRTarbase |
| hsa-miR-15a-5p | ZNRF3 | PAR-CLIP | miRTarbase |
| hsa-miR-195-5p | ZNRF3 | PAR-CLIP | miRTarbase |
| hsa-miR-15a-5p | YWHAH | PAR-CLIP | miRTarbase |
| hsa-miR-195-5p | YWHAH | PAR-CLIP | miRTarbase |
| hsa-miR-3613-3p | PTMA | PAR-CLIP | miRTarbase |
| hsa-miR-3613-3p | PTMA | PAR-CLIP | miRTarbase |
| hsa-miR-15a-5p | SEPT2 | HITS-CLIP | miRTarbase |
| hsa-miR-195-5p | SEPT2 | HITS-CLIP | miRTarbase |
| hsa-miR-3613-3p | FOXN2 | PAR-CLIP | miRTarbase |
| hsa-miR-3613-3p | FOXN2 | PAR-CLIP | miRTarbase |
| hsa-miR-3613-3p | FOXN2 | PAR-CLIP | miRTarbase |
| hsa-miR-3613-3p | FOXN2 | PAR-CLIP | miRTarbase |
| hsa-miR-3613-3p | FOXN2 | PAR-CLIP | miRTarbase |
| hsa-miR-3613-3p | FOXN2 | PAR-CLIP | miRTarbase |
| hsa-miR-15a-5p | B3GNT2 | PAR-CLIP | miRTarbase |
| hsa-miR-15a-5p | B3GNT2 | PAR-CLIP | miRTarbase |
| hsa-miR-195-5p | B3GNT2 | PAR-CLIP | miRTarbase |
| hsa-miR-195-5p | B3GNT2 | PAR-CLIP | miRTarbase |
| hsa-miR-15a-5p | ACTR2 | PAR-CLIP | miRTarbase |
| hsa-miR-15a-5p | ACTR2 | HITS-CLIP | miRTarbase |
| hsa-miR-195-5p | ACTR2 | PAR-CLIP | miRTarbase |
| hsa-miR-195-5p | ACTR2 | HITS-CLIP | miRTarbase |
| hsa-miR-15a-5p | CDV3 | HITS-CLIP | miRTarbase |
| hsa-miR-195-5p | CDV3 | HITS-CLIP | miRTarbase |
| hsa-miR-15a-5p | U2SURP | HITS-CLIP | miRTarbase |
| hsa-miR-195-5p | U2SURP | HITS-CLIP | miRTarbase |
| hsa-miR-15a-5p | RARB | PAR-CLIP | miRTarbase |
| hsa-miR-15a-5p | RARB | PAR-CLIP | miRTarbase |
| hsa-miR-15a-5p | RARB | PAR-CLIP | miRTarbase |
| hsa-miR-195-5p | RARB | PAR-CLIP | miRTarbase |
| hsa-miR-195-5p | RARB | PAR-CLIP | miRTarbase |
| hsa-miR-195-5p | RARB | PAR-CLIP | miRTarbase |
| hsa-miR-15a-5p | ITPR1 | HITS-CLIP | miRTarbase |
| hsa-miR-195-5p | ITPR1 | HITS-CLIP | miRTarbase |
| hsa-miR-15a-5p | BHLHE40 | HITS-CLIP | miRTarbase |
| hsa-miR-195-5p | BHLHE40 | HITS-CLIP | miRTarbase |
| hsa-miR-15a-5p | PI4K2B | PAR-CLIP | miRTarbase |
| hsa-miR-15a-5p | PI4K2B | PAR-CLIP | miRTarbase |
| hsa-miR-15a-5p | PI4K2B | PAR-CLIP | miRTarbase |
| hsa-miR-195-5p | PI4K2B | PAR-CLIP | miRTarbase |
| hsa-miR-195-5p | PI4K2B | PAR-CLIP | miRTarbase |
| hsa-miR-195-5p | PI4K2B | PAR-CLIP | miRTarbase |
| hsa-miR-3613-3p | SH3BP2 | HITS-CLIP | miRTarbase |
| hsa-miR-195-5p | PURA | HITS-CLIP | miRTarbase |
| hsa-miR-15a-5p | CANX | PAR-CLIP | miRTarbase |
| hsa-miR-195-5p | CANX | PAR-CLIP | miRTarbase |
| hsa-miR-3613-3p | CANX | PAR-CLIP | miRTarbase |
| hsa-miR-195-3p | ERBB2IP | PAR-CLIP | miRTarbase |
| hsa-miR-3613-3p | MAP7 | PAR-CLIP | miRTarbase |
| hsa-miR-15a-5p | PCMT1 | HITS-CLIP | miRTarbase |
| hsa-miR-195-5p | PCMT1 | HITS-CLIP | miRTarbase |
| hsa-miR-141-3p | QKI | PAR-CLIP | miRTarbase |
| hsa-miR-486-5p | ID4 | PAR-CLIP | miRTarbase |
| hsa-miR-486-5p | ID4 | PAR-CLIP | miRTarbase |
| hsa-miR-486-5p | ID4 | PAR-CLIP | miRTarbase |
| hsa-miR-15a-5p | E2F3 | HITS-CLIP | miRTarbase |
| hsa-miR-3613-3p | SOX4 | PAR-CLIP | miRTarbase |
| hsa-miR-3613-3p | SOX4 | PAR-CLIP | miRTarbase |
| hsa-miR-3613-3p | SOX4 | PAR-CLIP | miRTarbase |
| hsa-miR-3613-3p | SOX4 | PAR-CLIP | miRTarbase |
| hsa-miR-15a-5p | PPP1R11 | HITS-CLIP | miRTarbase |
| hsa-miR-195-5p | PPP1R11 | HITS-CLIP | miRTarbase |
| hsa-miR-15a-5p | HSPA1B | Luciferase reporter assay | miRTarbase |
| hsa-miR-15a-5p | HSPA1B | PAR-CLIP | miRTarbase |
| hsa-miR-15a-5p | HSPA1B | PAR-CLIP | miRTarbase |
| hsa-miR-15a-5p | HSPA1B | HITS-CLIP | miRTarbase |
| hsa-miR-15a-5p | HSPA1B | PAR-CLIP | miRTarbase |
| hsa-miR-195-5p | HSPA1B | PAR-CLIP | miRTarbase |
| hsa-miR-195-5p | HSPA1B | PAR-CLIP | miRTarbase |
| hsa-miR-195-5p | HSPA1B | HITS-CLIP | miRTarbase |
| hsa-miR-195-5p | HSPA1B | PAR-CLIP | miRTarbase |
| hsa-miR-3613-3p | HSPA1B | PAR-CLIP | miRTarbase |

| | | | |
|---|---|---|---|
| hsa-miR-15a-5p | PIM1 | PAR-CLIP | miRTarbase |
| hsa-miR-195-5p | PIM1 | PAR-CLIP | miRTarbase |
| hsa-miR-15a-5p | CD2AP | PAR-CLIP | miRTarbase |
| hsa-miR-195-5p | CD2AP | PAR-CLIP | miRTarbase |
| hsa-miR-15a-5p | CALU | HITS-CLIP | miRTarbase |
| hsa-miR-15a-5p | CALU | PAR-CLIP | miRTarbase |
| hsa-miR-195-5p | CALU | HITS-CLIP | miRTarbase |
| hsa-miR-195-5p | CALU | PAR-CLIP | miRTarbase |
| hsa-miR-15a-5p | UBN2 | PAR-CLIP | miRTarbase |
| hsa-miR-15a-5p | UBN2 | PAR-CLIP | miRTarbase |
| hsa-miR-15a-5p | UBN2 | PAR-CLIP | miRTarbase |
| hsa-miR-15a-5p | UBN2 | HITS-CLIP | miRTarbase |
| hsa-miR-15a-5p | UBN2 | PAR-CLIP | miRTarbase |
| hsa-miR-15a-5p | UBN2 | PAR-CLIP | miRTarbase |
| hsa-miR-195-5p | UBN2 | PAR-CLIP | miRTarbase |
| hsa-miR-195-5p | UBN2 | PAR-CLIP | miRTarbase |
| hsa-miR-195-5p | UBN2 | PAR-CLIP | miRTarbase |
| hsa-miR-195-5p | UBN2 | HITS-CLIP | miRTarbase |
| hsa-miR-195-5p | UBN2 | PAR-CLIP | miRTarbase |
| hsa-miR-195-5p | UBN2 | PAR-CLIP | miRTarbase |
| hsa-miR-15a-5p | EN2 | PAR-CLIP | miRTarbase |
| hsa-miR-15a-5p | EN2 | PAR-CLIP | miRTarbase |
| hsa-miR-15a-5p | EN2 | PAR-CLIP | miRTarbase |
| hsa-miR-195-5p | EN2 | PAR-CLIP | miRTarbase |
| hsa-miR-195-5p | EN2 | PAR-CLIP | miRTarbase |
| hsa-miR-195-5p | EN2 | PAR-CLIP | miRTarbase |
| hsa-miR-15a-5p | MAFK | PAR-CLIP | miRTarbase |
| hsa-miR-15a-5p | MAFK | PAR-CLIP | miRTarbase |
| hsa-miR-15a-5p | MAFK | HITS-CLIP | miRTarbase |
| hsa-miR-195-5p | MAFK | PAR-CLIP | miRTarbase |
| hsa-miR-195-5p | MAFK | PAR-CLIP | miRTarbase |
| hsa-miR-195-5p | MAFK | HITS-CLIP | miRTarbase |
| hsa-miR-3613-3p | MAFK | PAR-CLIP | miRTarbase |
| hsa-miR-3613-3p | MAFK | PAR-CLIP | miRTarbase |
| hsa-miR-3613-3p | CBX3 | PAR-CLIP | miRTarbase |
| hsa-miR-3613-3p | CBX3 | HITS-CLIP | miRTarbase |
| hsa-miR-15a-5p | FOXK1 | PAR-CLIP | miRTarbase |
| hsa-miR-15a-5p | FOXK1 | HITS-CLIP | miRTarbase |
| hsa-miR-195-5p | FOXK1 | PAR-CLIP | miRTarbase |
| hsa-miR-195-5p | FOXK1 | HITS-CLIP | miRTarbase |
| hsa-miR-15a-5p | USP42 | PAR-CLIP | miRTarbase |
| hsa-miR-15a-5p | USP42 | PAR-CLIP | miRTarbase |
| hsa-miR-15a-5p | USP42 | PAR-CLIP | miRTarbase |
| hsa-miR-195-5p | USP42 | PAR-CLIP | miRTarbase |
| hsa-miR-195-5p | USP42 | PAR-CLIP | miRTarbase |
| hsa-miR-195-5p | USP42 | PAR-CLIP | miRTarbase |
| hsa-miR-195-5p | DMTF1 | PAR-CLIP | miRTarbase |
| hsa-miR-195-5p | DMTF1 | HITS-CLIP | miRTarbase |
| hsa-miR-3605-5p | PEG10 | PAR-CLIP | miRTarbase |
| hsa-miR-195-3p | MYC | PAR-CLIP | miRTarbase |
| hsa-miR-15a-5p | ZFHX4 | PAR-CLIP | miRTarbase |
| hsa-miR-15a-5p | ZFHX4 | PAR-CLIP//HITS-CLIP | miRTarbase |
| hsa-miR-15a-5p | ZFHX4 | PAR-CLIP | miRTarbase |
| hsa-miR-195-5p | ZFHX4 | PAR-CLIP | miRTarbase |
| hsa-miR-195-5p | ZFHX4 | PAR-CLIP//HITS-CLIP | miRTarbase |
| hsa-miR-195-5p | ZFHX4 | PAR-CLIP | miRTarbase |
| hsa-miR-15a-5p | RAD23B | PAR-CLIP | miRTarbase |
| hsa-miR-15a-5p | RAD23B | HITS-CLIP | miRTarbase |
| hsa-miR-195-5p | RAD23B | PAR-CLIP | miRTarbase |
| hsa-miR-195-5p | RAD23B | HITS-CLIP | miRTarbase |
| hsa-miR-15a-5p | ZBTB34 | PAR-CLIP | miRTarbase |
| hsa-miR-195-5p | ZBTB34 | PAR-CLIP | miRTarbase |
| hsa-miR-195-5p | RECK | PAR-CLIP | miRTarbase |
| hsa-miR-195-3p | PTAR1 | PAR-CLIP | miRTarbase |
| hsa-miR-195-3p | PTAR1 | PAR-CLIP | miRTarbase |
| hsa-miR-15a-5p | SLC9A6 | HITS-CLIP | miRTarbase |
| hsa-miR-195-5p | SLC9A6 | HITS-CLIP | miRTarbase |
| hsa-miR-15a-5p | ZNF275 | PAR-CLIP | miRTarbase |
| hsa-miR-195-5p | ZNF275 | PAR-CLIP | miRTarbase |
| hsa-miR-195-3p | KLHL15 | PAR-CLIP | miRTarbase |
| hsa-miR-195-3p | KLHL15 | PAR-CLIP | miRTarbase |
| hsa-miR-195-3p | KLHL15 | PAR-CLIP | miRTarbase |
| hsa-miR-15a-5p | OGT | PAR-CLIP | miRTarbase |
| hsa-miR-15a-5p | OGT | PAR-CLIP | miRTarbase |
| hsa-miR-15a-5p | OGT | HITS-CLIP | miRTarbase |
| hsa-miR-15a-5p | OGT | PAR-CLIP | miRTarbase |
| hsa-miR-195-5p | OGT | PAR-CLIP | miRTarbase |
| hsa-miR-195-5p | OGT | PAR-CLIP | miRTarbase |
| hsa-miR-195-5p | OGT | HITS-CLIP | miRTarbase |
| hsa-miR-195-5p | OGT | PAR-CLIP | miRTarbase |
| hsa-miR-486-3p | OGT | PAR-CLIP | miRTarbase |
| hsa-miR-486-3p | OGT | PAR-CLIP | miRTarbase |
| hsa-miR-486-3p | OGT | PAR-CLIP | miRTarbase |
| hsa-miR-3613-3p | FDX1 | HITS-CLIP | miRTarbase |
| hsa-miR-219a-1-3p | BTG2 | PAR-CLIP | miRTarbase |
| hsa-miR-15a-5p | LUZP1 | PAR-CLIP | miRTarbase |
| hsa-miR-15a-5p | LUZP1 | PAR-CLIP | miRTarbase |
| hsa-miR-15a-5p | LUZP1 | PAR-CLIP | miRTarbase |
| hsa-miR-195-5p | LUZP1 | PAR-CLIP | miRTarbase |
| hsa-miR-195-5p | LUZP1 | PAR-CLIP | miRTarbase |
| hsa-miR-195-5p | LUZP1 | PAR-CLIP | miRTarbase |
| hsa-miR-3613-3p | CFL2 | PAR-CLIP | miRTarbase |
| hsa-miR-141-5p | ZFYVE9 | PAR-CLIP | miRTarbase |
| hsa-miR-15a-5p | CHAC1 | PAR-CLIP | miRTarbase |
| hsa-miR-195-5p | CHAC1 | PAR-CLIP | miRTarbase |
| hsa-miR-141-3p | ADNP2 | HITS-CLIP | miRTarbase |
| hsa-miR-15a-5p | SCAMP4 | PAR-CLIP | miRTarbase |
| hsa-miR-195-5p | SCAMP4 | PAR-CLIP | miRTarbase |
| hsa-miR-3613-3p | BTBD3 | PAR-CLIP | miRTarbase |
| hsa-miR-3613-3p | BTBD3 | PAR-CLIP//HITS-CLIP | miRTarbase |
| hsa-miR-3613-3p | BTBD3 | PAR-CLIP | miRTarbase |
| hsa-miR-3613-3p | BTBD3 | PAR-CLIP | miRTarbase |
| hsa-miR-3613-3p | BTBD3 | PAR-CLIP | miRTarbase |
| hsa-miR-3613-3p | BTBD3 | PAR-CLIP | miRTarbase |
| hsa-miR-3613-3p | MAVS | PAR-CLIP | miRTarbase |
| hsa-miR-3613-3p | MAVS | PAR-CLIP | miRTarbase |
| hsa-miR-219a-1-3p | NCOA3 | PAR-CLIP | miRTarbase |
| hsa-miR-15a-5p | PAK2 | PAR-CLIP | miRTarbase |
| hsa-miR-195-5p | PAK2 | PAR-CLIP | miRTarbase |
| hsa-miR-3605-5p | GINS4 | PAR-CLIP | miRTarbase |
| hsa-miR-3605-5p | GINS4 | PAR-CLIP | miRTarbase |
| hsa-miR-3605-3p | GINS4 | PAR-CLIP | miRTarbase |
| hsa-miR-3605-3p | GINS4 | PAR-CLIP | miRTarbase |
| hsa-miR-195-5p | PDCD4 | HITS-CLIP | miRTarbase |
| hsa-miR-141-3p | ARL5B | PAR-CLIP | miRTarbase |
| hsa-miR-141-3p | ARL5B | PAR-CLIP | miRTarbase |
| hsa-miR-141-3p | ARL5B | PAR-CLIP | miRTarbase |
| hsa-miR-141-3p | ARL5B | PAR-CLIP | miRTarbase |
| hsa-miR-141-3p | ARL5B | PAR-CLIP | miRTarbase |
| hsa-miR-141-3p | TNKS2 | PAR-CLIP | miRTarbase |
| hsa-miR-15a-5p | ENTPD1 | PAR-CLIP | miRTarbase |
| hsa-miR-15a-5p | ENTPD1 | HITS-CLIP | miRTarbase |
| hsa-miR-195-5p | ENTPD1 | PAR-CLIP | miRTarbase |
| hsa-miR-195-5p | ENTPD1 | HITS-CLIP | miRTarbase |
| hsa-miR-15a-5p | UBE4A | HITS-CLIP | miRTarbase |
| hsa-miR-195-5p | UBE4A | HITS-CLIP | miRTarbase |
| hsa-miR-15a-5p | ARCN1 | HITS-CLIP | miRTarbase |
| hsa-miR-195-5p | ARCN1 | HITS-CLIP | miRTarbase |
| hsa-miR-15a-5p | PTPRJ | HITS-CLIP | miRTarbase |
| hsa-miR-195-5p | PTPRJ | HITS-CLIP | miRTarbase |
| hsa-miR-15a-5p | TMEM138 | HITS-CLIP | miRTarbase |
| hsa-miR-195-5p | TMEM138 | HITS-CLIP | miRTarbase |
| hsa-miR-15a-5p | RASSF5 | HITS-CLIP | miRTarbase |
| hsa-miR-195-5p | RASSF5 | HITS-CLIP | miRTarbase |
| hsa-miR-15a-5p | MAPKAPK2 | HITS-CLIP | miRTarbase |
| hsa-miR-195-5p | MAPKAPK2 | HITS-CLIP | miRTarbase |
| hsa-miR-15a-5p | BCL7A | HITS-CLIP | miRTarbase |
| hsa-miR-195-5p | BCL7A | HITS-CLIP | miRTarbase |
| hsa-miR-15a-5p | SKI | PAR-CLIP | miRTarbase |
| hsa-miR-15a-5p | SKI | HITS-CLIP | miRTarbase |
| hsa-miR-141-3p | SPRYD4 | PAR-CLIP | miRTarbase |
| hsa-miR-15a-5p | RCOR1 | HITS-CLIP | miRTarbase |
| hsa-miR-195-5p | RCOR1 | HITS-CLIP | miRTarbase |
| hsa-miR-3613-3p | BTF3L4 | PAR-CLIP | miRTarbase |
| hsa-miR-15a-5p | SPRED1 | HITS-CLIP | miRTarbase |
| hsa-miR-15a-5p | SPRED1 | PAR-CLIP | miRTarbase |
| hsa-miR-195-5p | SPRED1 | HITS-CLIP | miRTarbase |
| hsa-miR-195-5p | SPRED1 | PAR-CLIP | miRTarbase |
| hsa-miR-15a-5p | SMAD3 | HITS-CLIP | miRTarbase |

| | | | |
|---|---|---|---|
| hsa-miR-195-5p | SMAD3 | HITS-CLIP | miRTarbase |
| hsa-miR-15a-5p | SCAMP5 | HITS-CLIP | miRTarbase |
| hsa-miR-195-5p | SCAMP5 | HITS-CLIP | miRTarbase |
| hsa-miR-15a-5p | UBE2Q2 | HITS-CLIP | miRTarbase |
| hsa-miR-195-5p | UBE2Q2 | HITS-CLIP | miRTarbase |
| hsa-miR-15a-5p | DCTN5 | PAR-CLIP | miRTarbase |
| hsa-miR-15a-5p | DCTN5 | PAR-CLIP | miRTarbase |
| hsa-miR-15a-5p | DCTN5 | HITS-CLIP | miRTarbase |
| hsa-miR-15a-5p | DCTN5 | PAR-CLIP | miRTarbase |
| hsa-miR-15a-5p | DCTN5 | PAR-CLIP | miRTarbase |
| hsa-miR-195-5p | DCTN5 | PAR-CLIP | miRTarbase |
| hsa-miR-195-5p | DCTN5 | PAR-CLIP | miRTarbase |
| hsa-miR-195-5p | DCTN5 | HITS-CLIP | miRTarbase |
| hsa-miR-195-5p | DCTN5 | PAR-CLIP | miRTarbase |
| hsa-miR-195-5p | DCTN5 | PAR-CLIP | miRTarbase |
| hsa-miR-15a-5p | PSKH1 | HITS-CLIP | miRTarbase |
| hsa-miR-195-5p | PSKH1 | HITS-CLIP | miRTarbase |
| hsa-miR-15a-5p | ANKRD13B | HITS-CLIP | miRTarbase |
| hsa-miR-195-5p | ANKRD13B | HITS-CLIP | miRTarbase |
| hsa-miR-15a-5p | EZH1 | HITS-CLIP | miRTarbase |
| hsa-miR-195-5p | EZH1 | HITS-CLIP | miRTarbase |
| hsa-miR-3613-3p | GPATCH8 | PAR-CLIP | miRTarbase |
| hsa-miR-15a-5p | PNPO | HITS-CLIP | miRTarbase |
| hsa-miR-195-5p | PNPO | HITS-CLIP | miRTarbase |
| hsa-miR-15a-5p | SNX11 | HITS-CLIP | miRTarbase |
| hsa-miR-195-5p | SNX11 | HITS-CLIP | miRTarbase |
| hsa-miR-3613-3p | FOXK2 | HITS-CLIP | miRTarbase |
| hsa-miR-15a-5p | RNF138 | HITS-CLIP | miRTarbase |
| hsa-miR-195-5p | RNF138 | HITS-CLIP | miRTarbase |
| hsa-miR-15a-5p | IER2 | HITS-CLIP | miRTarbase |
| hsa-miR-195-5p | IER2 | HITS-CLIP | miRTarbase |
| hsa-miR-15a-5p | TNFSF9 | HITS-CLIP | miRTarbase |
| hsa-miR-15a-5p | TNFSF9 | PAR-CLIP | miRTarbase |
| hsa-miR-195-5p | TNFSF9 | HITS-CLIP | miRTarbase |
| hsa-miR-195-5p | TNFSF9 | PAR-CLIP | miRTarbase |
| hsa-miR-15a-5p | ENTPD6 | HITS-CLIP | miRTarbase |
| hsa-miR-195-5p | ENTPD6 | HITS-CLIP | miRTarbase |
| hsa-miR-3613-3p | PLAGL2 | PAR-CLIP | miRTarbase |
| hsa-miR-15a-5p | KIF3B | HITS-CLIP | miRTarbase |
| hsa-miR-195-5p | KIF3B | HITS-CLIP | miRTarbase |
| hsa-miR-15a-5p | NOL4L | HITS-CLIP | miRTarbase |
| hsa-miR-195-5p | NOL4L | HITS-CLIP | miRTarbase |
| hsa-miR-15a-5p | RASSF2 | PAR-CLIP | miRTarbase |
| hsa-miR-195-5p | RASSF2 | PAR-CLIP | miRTarbase |
| hsa-miR-15a-5p | CDS2 | HITS-CLIP | miRTarbase |
| hsa-miR-195-5p | CDS2 | HITS-CLIP | miRTarbase |
| hsa-miR-15a-5p | ATP5G3 | PAR-CLIP | miRTarbase |
| hsa-miR-195-5p | ATP5G3 | PAR-CLIP | miRTarbase |
| hsa-miR-15a-5p | TNRC6B | HITS-CLIP | miRTarbase |
| hsa-miR-15a-5p | TNRC6B | HITS-CLIP | miRTarbase |
| hsa-miR-15a-5p | TNRC6B | HITS-CLIP | miRTarbase |
| hsa-miR-195-5p | TNRC6B | HITS-CLIP | miRTarbase |
| hsa-miR-195-5p | TNRC6B | HITS-CLIP | miRTarbase |
| hsa-miR-195-5p | TNRC6B | HITS-CLIP | miRTarbase |
| hsa-miR-15a-5p | EPT1 | HITS-CLIP | miRTarbase |
| hsa-miR-195-5p | EPT1 | HITS-CLIP | miRTarbase |
| hsa-miR-15a-5p | PEX13 | HITS-CLIP | miRTarbase |
| hsa-miR-195-5p | PEX13 | HITS-CLIP | miRTarbase |
| hsa-miR-15a-5p | TET3 | HITS-CLIP | miRTarbase |
| hsa-miR-195-5p | TET3 | HITS-CLIP | miRTarbase |
| hsa-miR-3613-3p | TET3 | PAR-CLIP | miRTarbase |
| hsa-miR-3613-3p | CDV3 | HITS-CLIP | miRTarbase |
| hsa-miR-3613-3p | CDV3 | PAR-CLIP | miRTarbase |
| hsa-miR-15a-5p | PRKCD | HITS-CLIP | miRTarbase |
| hsa-miR-195-5p | PRKCD | HITS-CLIP | miRTarbase |
| hsa-miR-15a-5p | CPEB2 | HITS-CLIP | miRTarbase |
| hsa-miR-195-5p | CPEB2 | HITS-CLIP | miRTarbase |
| hsa-miR-15a-5p | TADA2B | HITS-CLIP | miRTarbase |
| hsa-miR-195-5p | TADA2B | HITS-CLIP | miRTarbase |
| hsa-miR-15a-5p | GRAMD3 | HITS-CLIP | miRTarbase |
| hsa-miR-15a-5p | GRAMD3 | HITS-CLIP | miRTarbase |
| hsa-miR-141-5p | GRAMD3 | HITS-CLIP | miRTarbase |
| hsa-miR-195-5p | GRAMD3 | HITS-CLIP | miRTarbase |
| hsa-miR-195-5p | GRAMD3 | HITS-CLIP | miRTarbase |
| hsa-miR-15a-5p | CREBRF | HITS-CLIP | miRTarbase |
| hsa-miR-15a-5p | CREBRF | PAR-CLIP | miRTarbase |
| hsa-miR-195-5p | CREBRF | HITS-CLIP | miRTarbase |
| hsa-miR-195-5p | CREBRF | PAR-CLIP | miRTarbase |
| hsa-miR-195-3p | PAPD7 | PAR-CLIP | miRTarbase |
| hsa-miR-195-3p | PAPD7 | PAR-CLIP | miRTarbase |
| hsa-miR-195-3p | PAPD7 | PAR-CLIP | miRTarbase |
| hsa-miR-3613-3p | TNPO1 | HITS-CLIP | miRTarbase |
| hsa-miR-195-5p | HMGA1 | HITS-CLIP | miRTarbase |
| hsa-miR-15a-5p | CDKN1A | HITS-CLIP | miRTarbase |
| hsa-miR-195-5p | CDKN1A | HITS-CLIP | miRTarbase |
| hsa-miR-15a-5p | IRF4 | HITS-CLIP | miRTarbase |
| hsa-miR-195-5p | IRF4 | HITS-CLIP | miRTarbase |
| hsa-miR-141-5p | VEGFA | HITS-CLIP | miRTarbase |
| hsa-miR-15a-5p | KLHDC10 | HITS-CLIP | miRTarbase |
| hsa-miR-195-5p | KLHDC10 | HITS-CLIP | miRTarbase |
| hsa-miR-15a-5p | UBE3C | HITS-CLIP | miRTarbase |
| hsa-miR-195-5p | UBE3C | HITS-CLIP | miRTarbase |
| hsa-miR-195-3p | YWHAG | PAR-CLIP | miRTarbase |
| hsa-miR-15a-5p | SUN1 | HITS-CLIP | miRTarbase |
| hsa-miR-195-5p | SUN1 | HITS-CLIP | miRTarbase |
| hsa-miR-15a-5p | HMBOX1 | HITS-CLIP | miRTarbase |
| hsa-miR-195-5p | HMBOX1 | HITS-CLIP | miRTarbase |
| hsa-miR-15a-5p | RNF38 | HITS-CLIP | miRTarbase |
| hsa-miR-195-5p | RNF38 | HITS-CLIP | miRTarbase |
| hsa-miR-3613-3p | RNF38 | HITS-CLIP | miRTarbase |
| hsa-miR-15a-5p | PSAT1 | PAR-CLIP | miRTarbase |
| hsa-miR-15a-5p | PSAT1 | PAR-CLIP | miRTarbase |
| hsa-miR-15a-5p | PSAT1 | HITS-CLIP | miRTarbase |
| hsa-miR-15a-5p | PSAT1 | PAR-CLIP | miRTarbase |
| hsa-miR-195-5p | PSAT1 | PAR-CLIP | miRTarbase |
| hsa-miR-195-5p | PSAT1 | PAR-CLIP | miRTarbase |
| hsa-miR-195-5p | PSAT1 | HITS-CLIP | miRTarbase |
| hsa-miR-195-5p | PSAT1 | PAR-CLIP | miRTarbase |
| hsa-miR-15a-5p | ZBTB33 | HITS-CLIP | miRTarbase |
| hsa-miR-195-5p | ZBTB33 | HITS-CLIP | miRTarbase |
| hsa-miR-141-3p | OGT | PAR-CLIP | miRTarbase |
| hsa-miR-141-3p | OGT | PAR-CLIP | miRTarbase |
| hsa-miR-141-3p | PGK1 | PAR-CLIP | miRTarbase |
| hsa-miR-3613-5p | SMC3 | PAR-CLIP | miRTarbase |
| hsa-miR-219a-1-3p | UBE2D1 | PAR-CLIP | miRTarbase |
| hsa-miR-219a-1-3p | UBE2D1 | PAR-CLIP | miRTarbase |
| hsa-miR-219a-1-3p | UBE2D1 | PAR-CLIP | miRTarbase |
| hsa-miR-15a-5p | PAFAH1B2 | PAR-CLIP | miRTarbase |
| hsa-miR-195-5p | PAFAH1B2 | PAR-CLIP | miRTarbase |
| hsa-miR-3613-3p | PAFAH1B2 | PAR-CLIP | miRTarbase |
| hsa-miR-3613-3p | PAFAH1B2 | PAR-CLIP | miRTarbase |
| hsa-miR-3613-3p | SIKE1 | PAR-CLIP | miRTarbase |
| hsa-miR-15a-5p | RPRD2 | PAR-CLIP | miRTarbase |
| hsa-miR-15a-5p | RPRD2 | PAR-CLIP | miRTarbase |
| hsa-miR-15a-5p | RPRD2 | PAR-CLIP | miRTarbase |
| hsa-miR-15a-5p | RPRD2 | PAR-CLIP | miRTarbase |
| hsa-miR-195-5p | RPRD2 | PAR-CLIP | miRTarbase |
| hsa-miR-195-5p | RPRD2 | PAR-CLIP | miRTarbase |
| hsa-miR-195-5p | RPRD2 | PAR-CLIP | miRTarbase |
| hsa-miR-195-5p | RPRD2 | PAR-CLIP | miRTarbase |
| hsa-miR-3613-3p | ADIPOR2 | HITS-CLIP | miRTarbase |
| hsa-miR-3613-3p | ZNF678 | PAR-CLIP | miRTarbase |
| hsa-miR-15a-5p | PNRC2 | PAR-CLIP | miRTarbase |
| hsa-miR-195-5p | PNRC2 | PAR-CLIP | miRTarbase |
| hsa-miR-3613-3p | RAP1B | PAR-CLIP | miRTarbase |
| hsa-miR-3605-5p | TXLNA | PAR-CLIP | miRTarbase |
| hsa-miR-3605-5p | TXLNA | PAR-CLIP | miRTarbase |
| hsa-miR-15a-5p | CDADC1 | PAR-CLIP | miRTarbase |
| hsa-miR-195-5p | CDADC1 | PAR-CLIP | miRTarbase |
| hsa-miR-15a-5p | AGO4 | PAR-CLIP | miRTarbase |
| hsa-miR-195-5p | AGO4 | PAR-CLIP | miRTarbase |
| hsa-miR-15a-5p | GPR180 | PAR-CLIP | miRTarbase |
| hsa-miR-15a-5p | GPR180 | PAR-CLIP | miRTarbase |
| hsa-miR-15a-5p | GPR180 | PAR-CLIP | miRTarbase |
| hsa-miR-195-5p | GPR180 | PAR-CLIP | miRTarbase |
| hsa-miR-195-5p | GPR180 | PAR-CLIP | miRTarbase |
| hsa-miR-195-5p | GPR180 | PAR-CLIP | miRTarbase |
| hsa-miR-3613-3p | EXOC5 | HITS-CLIP | miRTarbase |
| hsa-miR-15a-5p | PPM1A | PAR-CLIP | miRTarbase |

| | | | |
|---|---|---|---|
| hsa-miR-195-5p | PPM1A | PAR-CLIP | miRTarbase |
| hsa-miR-15a-5p | SLC39A9 | PAR-CLIP | miRTarbase |
| hsa-miR-15a-5p | SLC39A9 | PAR-CLIP | miRTarbase |
| hsa-miR-15a-5p | SLC39A9 | PAR-CLIP | miRTarbase |
| hsa-miR-195-5p | SLC39A9 | PAR-CLIP | miRTarbase |
| hsa-miR-195-5p | SLC39A9 | PAR-CLIP | miRTarbase |
| hsa-miR-195-5p | SLC39A9 | PAR-CLIP | miRTarbase |
| hsa-miR-3613-3p | ELMSAN1 | HITS-CLIP | miRTarbase |
| hsa-miR-195-3p | B2M | PAR-CLIP | miRTarbase |
| hsa-miR-15a-5p | FAM103A1 | PAR-CLIP | miRTarbase |
| hsa-miR-15a-5p | FAM103A1 | PAR-CLIP | miRTarbase |
| hsa-miR-15a-5p | FAM103A1 | PAR-CLIP | miRTarbase |
| hsa-miR-195-5p | FAM103A1 | PAR-CLIP | miRTarbase |
| hsa-miR-195-5p | FAM103A1 | PAR-CLIP | miRTarbase |
| hsa-miR-195-5p | FAM103A1 | PAR-CLIP | miRTarbase |
| hsa-miR-15a-5p | RBBP6 | PAR-CLIP | miRTarbase |
| hsa-miR-15a-5p | RBBP6 | PAR-CLIP | miRTarbase |
| hsa-miR-15a-5p | RBBP6 | PAR-CLIP | miRTarbase |
| hsa-miR-15a-5p | RBBP6 | PAR-CLIP | miRTarbase |
| hsa-miR-195-5p | RBBP6 | PAR-CLIP | miRTarbase |
| hsa-miR-195-5p | RBBP6 | PAR-CLIP | miRTarbase |
| hsa-miR-195-5p | RBBP6 | PAR-CLIP | miRTarbase |
| hsa-miR-195-5p | RBBP6 | PAR-CLIP | miRTarbase |
| hsa-miR-195-3p | RBBP6 | PAR-CLIP | miRTarbase |
| hsa-miR-195-3p | RBBP6 | PAR-CLIP | miRTarbase |
| hsa-miR-195-3p | RBBP6 | PAR-CLIP | miRTarbase |
| hsa-miR-195-3p | RBBP6 | PAR-CLIP | miRTarbase |
| hsa-miR-3613-3p | GADD45A | PAR-CLIP | miRTarbase |
| hsa-miR-15a-5p | PAFAH1B1 | HITS-CLIP | miRTarbase |
| hsa-miR-195-5p | PAFAH1B1 | HITS-CLIP | miRTarbase |
| hsa-miR-15a-5p | TAOK1 | PAR-CLIP | miRTarbase |
| hsa-miR-195-5p | TAOK1 | PAR-CLIP | miRTarbase |
| hsa-miR-3613-3p | C18ORF25 | PAR-CLIP | miRTarbase |
| hsa-miR-15a-5p | SNRPB2 | PAR-CLIP | miRTarbase |
| hsa-miR-15a-5p | SNRPB2 | PAR-CLIP | miRTarbase |
| hsa-miR-15a-5p | SNRPB2 | PAR-CLIP | miRTarbase |
| hsa-miR-15a-5p | SNRPB2 | PAR-CLIP | miRTarbase |
| hsa-miR-195-5p | SNRPB2 | PAR-CLIP | miRTarbase |
| hsa-miR-195-5p | SNRPB2 | PAR-CLIP | miRTarbase |
| hsa-miR-195-5p | SNRPB2 | PAR-CLIP | miRTarbase |
| hsa-miR-195-5p | SNRPB2 | PAR-CLIP | miRTarbase |
| hsa-miR-15a-3p | ZSWIM1 | PAR-CLIP | miRTarbase |
| hsa-miR-3613-3p | CEBPB | PAR-CLIP | miRTarbase |
| hsa-miR-15a-5p | HSPE1-MOB4 | PAR-CLIP | miRTarbase |
| hsa-miR-15a-5p | HSPE1-MOB4 | PAR-CLIP | miRTarbase |
| hsa-miR-15a-5p | HSPE1-MOB4 | PAR-CLIP | miRTarbase |
| hsa-miR-15a-5p | HSPE1-MOB4 | PAR-CLIP | miRTarbase |
| hsa-miR-195-5p | HSPE1-MOB4 | PAR-CLIP | miRTarbase |
| hsa-miR-195-5p | HSPE1-MOB4 | PAR-CLIP | miRTarbase |
| hsa-miR-195-5p | HSPE1-MOB4 | PAR-CLIP | miRTarbase |
| hsa-miR-195-5p | HSPE1-MOB4 | PAR-CLIP | miRTarbase |
| hsa-miR-15a-5p | MOB4 | PAR-CLIP | miRTarbase |
| hsa-miR-15a-5p | MOB4 | PAR-CLIP | miRTarbase |
| hsa-miR-15a-5p | MOB4 | PAR-CLIP | miRTarbase |
| hsa-miR-15a-5p | MOB4 | PAR-CLIP | miRTarbase |
| hsa-miR-195-5p | MOB4 | PAR-CLIP | miRTarbase |
| hsa-miR-195-5p | MOB4 | PAR-CLIP | miRTarbase |
| hsa-miR-195-5p | MOB4 | PAR-CLIP | miRTarbase |
| hsa-miR-195-5p | MOB4 | PAR-CLIP | miRTarbase |
| hsa-miR-15a-5p | BZW1 | PAR-CLIP | miRTarbase |
| hsa-miR-15a-5p | BZW1 | PAR-CLIP | miRTarbase |
| hsa-miR-15a-5p | BZW1 | PAR-CLIP | miRTarbase |
| hsa-miR-15a-5p | BZW1 | PAR-CLIP | miRTarbase |
| hsa-miR-15a-5p | BZW1 | PAR-CLIP | miRTarbase |
| hsa-miR-15a-5p | BZW1 | PAR-CLIP | miRTarbase |
| hsa-miR-195-5p | BZW1 | PAR-CLIP | miRTarbase |
| hsa-miR-195-5p | BZW1 | PAR-CLIP | miRTarbase |
| hsa-miR-195-5p | BZW1 | PAR-CLIP | miRTarbase |
| hsa-miR-195-5p | BZW1 | PAR-CLIP | miRTarbase |
| hsa-miR-195-5p | BZW1 | PAR-CLIP | miRTarbase |
| hsa-miR-195-5p | BZW1 | PAR-CLIP | miRTarbase |
| hsa-miR-15a-5p | NUP50 | PAR-CLIP | miRTarbase |
| hsa-miR-15a-5p | NUP50 | PAR-CLIP | miRTarbase |
| hsa-miR-15a-5p | NUP50 | HITS-CLIP | miRTarbase |
| hsa-miR-15a-5p | NUP50 | PAR-CLIP | miRTarbase |
| hsa-miR-195-5p | NUP50 | PAR-CLIP | miRTarbase |
| hsa-miR-195-5p | NUP50 | PAR-CLIP | miRTarbase |
| hsa-miR-195-5p | NUP50 | HITS-CLIP | miRTarbase |
| hsa-miR-195-5p | NUP50 | PAR-CLIP | miRTarbase |
| hsa-miR-3613-3p | SKIL | PAR-CLIP | miRTarbase |
| hsa-miR-3613-3p | SKIL | PAR-CLIP | miRTarbase |
| hsa-miR-3613-3p | SKIL | PAR-CLIP | miRTarbase |
| hsa-miR-3613-3p | SKIL | HITS-CLIP | miRTarbase |
| hsa-miR-3613-3p | SKIL | PAR-CLIP | miRTarbase |
| hsa-miR-3613-3p | TBL1XR1 | PAR-CLIP | miRTarbase |
| hsa-miR-3613-3p | ZMAT3 | PAR-CLIP | miRTarbase |
| hsa-miR-3613-3p | ZMAT3 | PAR-CLIP | miRTarbase |
| hsa-miR-3613-3p | ZMAT3 | PAR-CLIP | miRTarbase |
| hsa-miR-3613-3p | ZMAT3 | PAR-CLIP | miRTarbase |
| hsa-miR-15a-5p | FGF2 | PAR-CLIP | miRTarbase |
| hsa-miR-15a-5p | FGF2 | PAR-CLIP | miRTarbase |
| hsa-miR-195-5p | FGF2 | PAR-CLIP | miRTarbase |
| hsa-miR-195-5p | FGF2 | PAR-CLIP | miRTarbase |
| hsa-miR-15a-5p | HSPA4L | PAR-CLIP | miRTarbase |
| hsa-miR-15a-5p | HSPA4L | PAR-CLIP | miRTarbase |
| hsa-miR-195-5p | HSPA4L | PAR-CLIP | miRTarbase |
| hsa-miR-195-5p | HSPA4L | PAR-CLIP | miRTarbase |
| hsa-miR-15a-5p | RBPJ | PAR-CLIP | miRTarbase |
| hsa-miR-15a-5p | RBPJ | PAR-CLIP | miRTarbase |
| hsa-miR-15a-5p | RBPJ | PAR-CLIP | miRTarbase |
| hsa-miR-15a-5p | RBPJ | PAR-CLIP | miRTarbase |
| hsa-miR-195-5p | RBPJ | PAR-CLIP | miRTarbase |
| hsa-miR-195-5p | RBPJ | PAR-CLIP | miRTarbase |
| hsa-miR-195-5p | RBPJ | PAR-CLIP | miRTarbase |
| hsa-miR-195-5p | RBPJ | PAR-CLIP | miRTarbase |
| hsa-miR-3613-3p | DCP2 | PAR-CLIP | miRTarbase |
| hsa-miR-195-3p | SUB1 | PAR-CLIP | miRTarbase |
| hsa-miR-219a-1-3p | C5ORF51 | PAR-CLIP | miRTarbase |
| hsa-miR-219a-1-3p | C5ORF51 | PAR-CLIP | miRTarbase |
| hsa-miR-219a-1-3p | C5ORF51 | PAR-CLIP | miRTarbase |
| hsa-miR-219a-1-3p | C5ORF51 | PAR-CLIP | miRTarbase |
| hsa-miR-219a-1-3p | C5ORF51 | PAR-CLIP | miRTarbase |
| hsa-miR-3613-3p | MTX3 | HITS-CLIP | miRTarbase |
| hsa-miR-15a-5p | TBPL1 | PAR-CLIP | miRTarbase |
| hsa-miR-15a-5p | TBPL1 | HITS-CLIP | miRTarbase |
| hsa-miR-195-5p | TBPL1 | PAR-CLIP | miRTarbase |
| hsa-miR-195-5p | TBPL1 | HITS-CLIP | miRTarbase |
| hsa-miR-219a-1-3p | C6ORF120 | PAR-CLIP | miRTarbase |
| hsa-miR-486-5p | PIM1 | qRT-PCR | miRTarbase |
| hsa-miR-195-3p | ATP6V1C1 | PAR-CLIP | miRTarbase |
| hsa-miR-195-3p | ATP6V1C1 | PAR-CLIP | miRTarbase |
| hsa-miR-15a-5p | FZD6 | PAR-CLIP | miRTarbase |
| hsa-miR-15a-5p | FZD6 | HITS-CLIP | miRTarbase |
| hsa-miR-15a-5p | FZD6 | PAR-CLIP | miRTarbase |
| hsa-miR-195-5p | FZD6 | PAR-CLIP | miRTarbase |
| hsa-miR-195-5p | FZD6 | HITS-CLIP | miRTarbase |
| hsa-miR-195-5p | FZD6 | PAR-CLIP | miRTarbase |
| hsa-miR-15a-5p | BAG4 | PAR-CLIP | miRTarbase |
| hsa-miR-195-5p | BAG4 | PAR-CLIP | miRTarbase |
| hsa-miR-3613-5p | ANP32B | PAR-CLIP | miRTarbase |
| hsa-miR-3613-5p | ANP32B | PAR-CLIP | miRTarbase |
| hsa-miR-141-3p | ZBTB34 | PAR-CLIP | miRTarbase |
| hsa-miR-15a-5p | ZNF449 | PAR-CLIP | miRTarbase |
| hsa-miR-195-5p | ZNF449 | PAR-CLIP | miRTarbase |
| hsa-miR-3613-3p | EIF2S3 | HITS-CLIP | miRTarbase |
| hsa-miR-15a-5p | YIPF6 | PAR-CLIP | miRTarbase |
| hsa-miR-195-5p | YIPF6 | PAR-CLIP | miRTarbase |
| hsa-miR-15a-5p | DDX3Y | HITS-CLIP | miRTarbase |
| hsa-miR-195-5p | DDX3Y | HITS-CLIP | miRTarbase |
| hsa-miR-3613-3p | EIF4G3 | HITS-CLIP | miRTarbase |
| hsa-miR-15a-5p | MSL1 | PAR-CLIP | miRTarbase |
| hsa-miR-15a-5p | MSL1 | PAR-CLIP | miRTarbase |
| hsa-miR-15a-5p | MSL1 | PAR-CLIP | miRTarbase |
| hsa-miR-15a-5p | MSL1 | HITS-CLIP | miRTarbase |
| hsa-miR-195-5p | MSL1 | PAR-CLIP | miRTarbase |
| hsa-miR-195-5p | MSL1 | PAR-CLIP | miRTarbase |
| hsa-miR-195-5p | MSL1 | PAR-CLIP | miRTarbase |
| hsa-miR-195-5p | MSL1 | HITS-CLIP | miRTarbase |
| hsa-miR-195-3p | AMOTL1 | PAR-CLIP | miRTarbase |

| | | | |
|---|---|---|---|
| hsa-miR-195-3p | AMOTL1 | PAR-CLIP | miRTarbase |
| hsa-miR-3613-3p | TMC5 | PAR-CLIP | miRTarbase |
| hsa-miR-3613-3p | TMC5 | HITS-CLIP | miRTarbase |
| hsa-miR-3613-3p | MCM4 | PAR-CLIP | miRTarbase |
| hsa-miR-15a-5p | ENTPD7 | HITS-CLIP | miRTarbase |
| hsa-miR-195-5p | ENTPD7 | HITS-CLIP | miRTarbase |
| hsa-miR-15a-5p | PRRC2C | HITS-CLIP | miRTarbase |
| hsa-miR-195-5p | PRRC2C | HITS-CLIP | miRTarbase |
| hsa-miR-15a-5p | WEE1 | PAR-CLIP | miRTarbase |
| hsa-miR-15a-5p | WEE1 | PAR-CLIP | miRTarbase |
| hsa-miR-15a-5p | ELK4 | PAR-CLIP | miRTarbase |
| hsa-miR-15a-5p | ELK4 | PAR-CLIP | miRTarbase |
| hsa-miR-195-5p | ELK4 | PAR-CLIP | miRTarbase |
| hsa-miR-195-5p | ELK4 | PAR-CLIP | miRTarbase |
| hsa-miR-15a-5p | GABARAPL1 | PAR-CLIP | miRTarbase |
| hsa-miR-15a-5p | GABARAPL1 | PAR-CLIP | miRTarbase |
| hsa-miR-15a-5p | GABARAPL1 | PAR-CLIP | miRTarbase |
| hsa-miR-195-5p | GABARAPL1 | PAR-CLIP | miRTarbase |
| hsa-miR-195-5p | GABARAPL1 | PAR-CLIP | miRTarbase |
| hsa-miR-195-5p | GABARAPL1 | PAR-CLIP | miRTarbase |
| hsa-miR-3613-3p | GABARAPL1 | PAR-CLIP | miRTarbase |
| hsa-miR-3613-3p | GABARAPL1 | PAR-CLIP | miRTarbase |
| hsa-miR-3613-3p | GABARAPL1 | PAR-CLIP | miRTarbase |
| hsa-miR-15a-5p | PDIK1L | HITS-CLIP | miRTarbase |
| hsa-miR-195-5p | PDIK1L | HITS-CLIP | miRTarbase |
| hsa-miR-15a-5p | ATXN7L3B | PAR-CLIP | miRTarbase |
| hsa-miR-15a-5p | ATXN7L3B | PAR-CLIP | miRTarbase |
| hsa-miR-195-5p | ATXN7L3B | PAR-CLIP | miRTarbase |
| hsa-miR-195-5p | ATXN7L3B | PAR-CLIP | miRTarbase |
| hsa-miR-141-5p | BCL2L2-PABPN1 | PAR-CLIP | miRTarbase |
| hsa-miR-15a-5p | ZNF691 | PAR-CLIP | miRTarbase |
| hsa-miR-15a-5p | ZNF691 | PAR-CLIP | miRTarbase |
| hsa-miR-195-5p | ZNF691 | PAR-CLIP | miRTarbase |
| hsa-miR-195-5p | ZNF691 | PAR-CLIP | miRTarbase |
| hsa-miR-15a-5p | DYNLL2 | PAR-CLIP | miRTarbase |
| hsa-miR-15a-5p | DYNLL2 | PAR-CLIP | miRTarbase |
| hsa-miR-195-5p | DYNLL2 | PAR-CLIP | miRTarbase |
| hsa-miR-195-5p | DYNLL2 | PAR-CLIP | miRTarbase |
| hsa-miR-15a-3p | KIAA0195 | PAR-CLIP | miRTarbase |
| hsa-miR-15a-5p | SRPRB | PAR-CLIP | miRTarbase |
| hsa-miR-15a-5p | SRPRB | PAR-CLIP | miRTarbase |
| hsa-miR-15a-5p | SRPRB | HITS-CLIP | miRTarbase |
| hsa-miR-195-5p | SRPRB | PAR-CLIP | miRTarbase |
| hsa-miR-195-5p | SRPRB | PAR-CLIP | miRTarbase |
| hsa-miR-195-5p | SRPRB | HITS-CLIP | miRTarbase |
| hsa-miR-15a-5p | CDC42SE2 | PAR-CLIP | miRTarbase |
| hsa-miR-195-5p | CDC42SE2 | PAR-CLIP | miRTarbase |
| hsa-miR-15a-5p | WIPI2 | PAR-CLIP | miRTarbase |
| hsa-miR-15a-5p | WIPI2 | HITS-CLIP | miRTarbase |
| hsa-miR-195-5p | WIPI2 | PAR-CLIP | miRTarbase |
| hsa-miR-195-5p | WIPI2 | HITS-CLIP | miRTarbase |
| hsa-miR-3613-3p | TCF7L2 | PAR-CLIP | miRTarbase |
| hsa-miR-3613-3p | USP6NL | HITS-CLIP | miRTarbase |
| hsa-miR-3613-3p | C11ORF57 | PAR-CLIP | miRTarbase |
| hsa-miR-15a-5p | TBRG1 | PAR-CLIP | miRTarbase |
| hsa-miR-195-5p | TBRG1 | PAR-CLIP | miRTarbase |
| hsa-miR-15a-5p | CHEK1 | HITS-CLIP | miRTarbase |
| hsa-miR-15a-5p | CHEK1 | HITS-CLIP | miRTarbase |
| hsa-miR-195-5p | CHEK1 | HITS-CLIP | miRTarbase |
| hsa-miR-195-5p | CHEK1 | HITS-CLIP | miRTarbase |
| hsa-miR-15a-5p | TMEM109 | HITS-CLIP | miRTarbase |
| hsa-miR-195-5p | TMEM109 | HITS-CLIP | miRTarbase |
| hsa-miR-15a-5p | C1ORF226 | PAR-CLIP | miRTarbase |
| hsa-miR-195-5p | C1ORF226 | PAR-CLIP | miRTarbase |
| hsa-miR-15a-5p | SIRT4 | HITS-CLIP | miRTarbase |
| hsa-miR-195-5p | SIRT4 | HITS-CLIP | miRTarbase |
| hsa-miR-15a-5p | SETD1B | PAR-CLIP | miRTarbase |
| hsa-miR-195-5p | SETD1B | PAR-CLIP | miRTarbase |
| hsa-miR-15a-5p | HOXC8 | PAR-CLIP | miRTarbase |
| hsa-miR-195-5p | HOXC8 | PAR-CLIP | miRTarbase |
| hsa-miR-15a-5p | RAB3IP | PAR-CLIP | miRTarbase |
| hsa-miR-195-5p | RAB3IP | PAR-CLIP | miRTarbase |
| hsa-miR-3613-3p | AKIRIN1 | PAR-CLIP | miRTarbase |
| hsa-miR-15a-5p | PPP2R5C | PAR-CLIP | miRTarbase |
| hsa-miR-15a-5p | PPP2R5C | PAR-CLIP | miRTarbase |
| hsa-miR-195-5p | PPP2R5C | PAR-CLIP | miRTarbase |
| hsa-miR-195-5p | PPP2R5C | PAR-CLIP | miRTarbase |
| hsa-miR-15a-5p | SLCO3A1 | HITS-CLIP | miRTarbase |
| hsa-miR-195-5p | SLCO3A1 | HITS-CLIP | miRTarbase |
| hsa-miR-3613-3p | IGF1R | PAR-CLIP | miRTarbase |
| hsa-miR-3613-3p | SYNM | PAR-CLIP | miRTarbase |
| hsa-miR-3613-3p | ATF7IP2 | HITS-CLIP | miRTarbase |
| hsa-miR-3605-5p | KCTD5 | PAR-CLIP | miRTarbase |
| hsa-miR-3613-5p | MT2A | PAR-CLIP | miRTarbase |
| hsa-miR-15a-5p | MLLT6 | HITS-CLIP | miRTarbase |
| hsa-miR-195-5p | MLLT6 | HITS-CLIP | miRTarbase |
| hsa-miR-3613-3p | DYNLL2 | PAR-CLIP | miRTarbase |
| hsa-miR-15a-5p | CBX2 | PAR-CLIP | miRTarbase |
| hsa-miR-195-5p | CBX2 | PAR-CLIP | miRTarbase |
| hsa-miR-195-3p | SH3GLB1 | PAR-CLIP | miRTarbase |
| hsa-miR-486-3p | MKNK2 | PAR-CLIP | miRTarbase |
| hsa-miR-486-3p | MKNK2 | PAR-CLIP | miRTarbase |
| hsa-miR-486-3p | MKNK2 | PAR-CLIP | miRTarbase |
| hsa-miR-15a-5p | ZFP28 | PAR-CLIP | miRTarbase |
| hsa-miR-195-5p | ZFP28 | PAR-CLIP | miRTarbase |
| hsa-miR-3613-3p | RPS28 | PAR-CLIP | miRTarbase |
| hsa-miR-15a-5p | CHMP4B | PAR-CLIP | miRTarbase |
| hsa-miR-195-5p | CHMP4B | PAR-CLIP | miRTarbase |
| hsa-miR-15a-5p | GABPA | PAR-CLIP | miRTarbase |
| hsa-miR-15a-5p | GABPA | PAR-CLIP | miRTarbase |
| hsa-miR-195-5p | GABPA | PAR-CLIP | miRTarbase |
| hsa-miR-195-5p | GABPA | PAR-CLIP | miRTarbase |
| hsa-miR-15a-5p | STRADB | PAR-CLIP | miRTarbase |
| hsa-miR-195-5p | STRADB | PAR-CLIP | miRTarbase |
| hsa-miR-195-5p | STRADB | Microarray//qRT-PCR | miRTarbase |
| hsa-miR-15a-5p | MTMR3 | PAR-CLIP | miRTarbase |
| hsa-miR-195-5p | MTMR3 | PAR-CLIP | miRTarbase |
| hsa-miR-15a-5p | CRIM1 | PAR-CLIP | miRTarbase |
| hsa-miR-15a-5p | CRIM1 | PAR-CLIP | miRTarbase |
| hsa-miR-15a-5p | CRIM1 | PAR-CLIP | miRTarbase |
| hsa-miR-195-5p | CRIM1 | PAR-CLIP | miRTarbase |
| hsa-miR-195-5p | CRIM1 | PAR-CLIP | miRTarbase |
| hsa-miR-195-5p | CRIM1 | PAR-CLIP | miRTarbase |
| hsa-miR-15a-5p | SOCS5 | PAR-CLIP | miRTarbase |
| hsa-miR-195-5p | SOCS5 | PAR-CLIP | miRTarbase |
| hsa-miR-3613-3p | CALM2 | PAR-CLIP | miRTarbase |
| hsa-miR-3613-3p | CALM2 | PAR-CLIP | miRTarbase |
| hsa-miR-3613-3p | CALM2 | PAR-CLIP | miRTarbase |
| hsa-miR-15a-5p | CTDSPL | PAR-CLIP | miRTarbase |
| hsa-miR-15a-5p | CTDSPL | PAR-CLIP | miRTarbase |
| hsa-miR-195-5p | CTDSPL | PAR-CLIP | miRTarbase |
| hsa-miR-195-5p | CTDSPL | PAR-CLIP | miRTarbase |
| hsa-miR-3613-3p | RHOA | HITS-CLIP | miRTarbase |
| hsa-miR-3613-3p | ZNF654 | PAR-CLIP | miRTarbase |
| hsa-miR-15a-3p | LMNB1 | PAR-CLIP | miRTarbase |
| hsa-miR-141-3p | HNRNPAB | PAR-CLIP | miRTarbase |
| hsa-miR-15a-5p | ITGA2 | PAR-CLIP | miRTarbase |
| hsa-miR-195-5p | ITGA2 | PAR-CLIP | miRTarbase |
| hsa-miR-15a-5p | PIK3R1 | PAR-CLIP//HITS-CLIP | miRTarbase |
| hsa-miR-15a-5p | PIK3R1 | HITS-CLIP | miRTarbase |
| hsa-miR-15a-5p | PIK3R1 | PAR-CLIP | miRTarbase |
| hsa-miR-15a-5p | PIK3R1 | HITS-CLIP | miRTarbase |
| hsa-miR-195-5p | PIK3R1 | PAR-CLIP//HITS-CLIP | miRTarbase |
| hsa-miR-195-5p | PIK3R1 | HITS-CLIP | miRTarbase |
| hsa-miR-195-5p | PIK3R1 | PAR-CLIP | miRTarbase |
| hsa-miR-195-5p | PIK3R1 | HITS-CLIP | miRTarbase |
| hsa-miR-15a-5p | PPIL1 | PAR-CLIP | miRTarbase |
| hsa-miR-15a-5p | PPIL1 | PAR-CLIP | miRTarbase |
| hsa-miR-15a-5p | PPIL1 | HITS-CLIP | miRTarbase |
| hsa-miR-15a-5p | PPIL1 | PAR-CLIP | miRTarbase |
| hsa-miR-195-5p | PPIL1 | PAR-CLIP | miRTarbase |
| hsa-miR-195-5p | PPIL1 | PAR-CLIP | miRTarbase |
| hsa-miR-195-5p | PPIL1 | HITS-CLIP | miRTarbase |
| hsa-miR-195-5p | PPIL1 | PAR-CLIP | miRTarbase |
| hsa-miR-15a-5p | CAPZA2 | PAR-CLIP | miRTarbase |
| hsa-miR-195-5p | CAPZA2 | PAR-CLIP | miRTarbase |
| hsa-miR-15a-5p | ZNRF2 | PAR-CLIP | miRTarbase |
| hsa-miR-195-5p | ZNRF2 | PAR-CLIP | miRTarbase |
| hsa-miR-486-5p | ZNRF2 | PAR-CLIP | miRTarbase |

| | | | |
|---|---|---|---|
| hsa-miR-141-5p | RBM12B | PAR-CLIP | miRTarbase |
| hsa-miR-3613-3p | AKAP2 | HITS-CLIP | miRTarbase |
| hsa-miR-3613-3p | PALM2-AKAP2 | HITS-CLIP | miRTarbase |
| hsa-miR-15a-5p | IFT74 | HITS-CLIP | miRTarbase |
| hsa-miR-195-5p | IFT74 | HITS-CLIP | miRTarbase |
| hsa-miR-15a-5p | OCRL | HITS-CLIP | miRTarbase |
| hsa-miR-195-5p | OCRL | HITS-CLIP | miRTarbase |
| hsa-miR-15a-5p | CHIC1 | PAR-CLIP | miRTarbase |
| hsa-miR-15a-5p | CHIC1 | PAR-CLIP | miRTarbase |
| hsa-miR-15a-5p | CHIC1 | PAR-CLIP | miRTarbase |
| hsa-miR-195-5p | CHIC1 | PAR-CLIP | miRTarbase |
| hsa-miR-195-5p | CHIC1 | PAR-CLIP | miRTarbase |
| hsa-miR-195-5p | CHIC1 | PAR-CLIP | miRTarbase |
| hsa-miR-486-3p | U2AF2 | PAR-CLIP | miRTarbase |
| hsa-miR-219a-1-3p | ACTR2 | PAR-CLIP | miRTarbase |
| hsa-miR-3613-3p | SGMS2 | PAR-CLIP | miRTarbase |
| hsa-miR-3613-3p | SGMS2 | PAR-CLIP | miRTarbase |
| hsa-miR-3605-5p | GDI1 | PAR-CLIP | miRTarbase |
| hsa-miR-3605-5p | GDI1 | PAR-CLIP | miRTarbase |
| hsa-miR-195-5p | CRKL | PAR-CLIP | miRTarbase |
| hsa-miR-141-3p | IPO5 | PAR-CLIP | miRTarbase |
| hsa-miR-219a-1-3p | NUFIP2 | PAR-CLIP | miRTarbase |
| hsa-miR-486-3p | MBD6 | PAR-CLIP | miRTarbase |
| hsa-miR-3613-3p | TMSB4X | PAR-CLIP | miRTarbase |
| hsa-miR-3613-3p | DFFA | HITS-CLIP | miRTarbase |
| hsa-miR-3613-3p | CELF1 | PAR-CLIP | miRTarbase |
| hsa-miR-3613-3p | PTBP3 | PAR-CLIP | miRTarbase |
| hsa-miR-3613-5p | FLVCR1 | PAR-CLIP | miRTarbase |
| hsa-miR-3613-5p | MT1E | PAR-CLIP | miRTarbase |
| hsa-miR-3613-3p | IGFBP5 | PAR-CLIP | miRTarbase |
| hsa-miR-141-3p | MAP4K4 | Luciferase reporter assay//qRT-PCR//Western blot | miRTarbase |
| hsa-miR-195-5p | ALOX12 | Microarray//qRT-PCR | miRTarbase |
| hsa-miR-195-5p | PHACTR2 | Microarray//qRT-PCR | miRTarbase |
| hsa-miR-195-5p | CGNL1 | Microarray//qRT-PCR | miRTarbase |
| hsa-miR-195-5p | KIF21A | Microarray//qRT-PCR | miRTarbase |
| hsa-miR-195-5p | RASGEF1B | Microarray//qRT-PCR | miRTarbase |
| hsa-miR-195-5p | TUFT1 | Microarray//qRT-PCR | miRTarbase |
| hsa-miR-195-5p | PDIA6 | PAR-CLIP | miRTarbase |
| hsa-miR-195-5p | PDIA6 | Microarray//qRT-PCR | miRTarbase |
| hsa-miR-195-5p | PDIA6 | HITS-CLIP | miRTarbase |
| hsa-miR-195-5p | SNX16 | PAR-CLIP | miRTarbase |
| hsa-miR-195-5p | SNX16 | PAR-CLIP | miRTarbase |
| hsa-miR-195-5p | SNX16 | Microarray//qRT-PCR | miRTarbase |
| hsa-miR-195-5p | SNX16 | PAR-CLIP | miRTarbase |
| hsa-miR-195-5p | TRIP10 | Microarray//qRT-PCR | miRTarbase |
| hsa-miR-195-5p | CDC25A | Luciferase reporter assay//Western blot | miRTarbase |
| hsa-miR-195-5p | CDC25A | HITS-CLIP | miRTarbase |
| hsa-miR-195-5p | CDC25A | PAR-CLIP | miRTarbase |
| hsa-miR-195-5p | BTRC | Luciferase reporter assay//Western blot | miRTarbase |
| hsa-miR-195-5p | BTRC | HITS-CLIP | miRTarbase |
| hsa-miR-141-3p | HNRNPD | Luciferase reporter assay//Western blot | miRTarbase |
| hsa-miR-15a-5p | KLF6 | Microarray//qRT-PCR// | miRTarbase |
| hsa-miR-15a-5p | PHLPP1 | Luciferase reporter assay//Northern blot//qRT-PCR//Western blot | miRTarbase |
| hsa-miR-141-3p | ZMPSTE24 | Luciferase reporter assay | miRTarbase |
| hsa-miR-195-5p | INSR | Luciferase reporter assay//qRT-PCR//Western blot | miRTarbase |
| hsa-miR-195-5p | ELN | Luciferase reporter assay//Western blot//qRT-PCR | miRTarbase |
| hsa-miR-195-5p | RET | Flow//Luciferase reporter assay | miRTarbase |
| hsa-miR-15a-5p | RET | Flow//Luciferase reporter assay | miRTarbase |
| hsa-miR-486-5p | IGF1R | Luciferase reporter assay//qRT-PCR//Western blot | miRTarbase |
| hsa-miR-486-5p | DOCK3 | Immunofluorescence//Immunohistochemistry//Immunoprecipitaion//Luciferase reporter assay//qRT-PCR//Western blot | miRTarbase |
| hsa-miR-486-5p | CADM1 | PAR-CLIP | miRTarbase |
| hsa-miR-486-5p | CADM1 | qRT-PCR | miRTarbase |
| hsa-miR-486-5p | H3F3B | PAR-CLIP | miRTarbase |
| hsa-miR-486-5p | H3F3B | PAR-CLIP | miRTarbase |
| hsa-miR-486-5p | H3F3B | qRT-PCR | miRTarbase |
| hsa-miR-486-5p | H3F3B | PAR-CLIP | miRTarbase |
| hsa-miR-486-5p | H3F3B | PAR-CLIP | miRTarbase |
| hsa-miR-195-5p | Bace1 | Luciferase reporter assay | miRTarbase |
| hsa-miR-3613-3p | PHKG2 | PAR-CLIP | miRTarbase |
| hsa-miR-3613-3p | AK4 | PAR-CLIP | miRTarbase |
| hsa-miR-3613-3p | AK4 | PAR-CLIP | miRTarbase |
| hsa-miR-141-3p | STXBP2 | PAR-CLIP | miRTarbase |
| hsa-miR-3613-3p | CCNB1IP1 | PAR-CLIP | miRTarbase |
| hsa-miR-3613-3p | CCNB1IP1 | PAR-CLIP | miRTarbase |
| hsa-miR-3613-3p | CCNB1IP1 | PAR-CLIP | miRTarbase |
| hsa-miR-141-3p | SEPT8 | PAR-CLIP | miRTarbase |
| hsa-miR-3613-3p | NINJ1 | PAR-CLIP | miRTarbase |
| hsa-miR-15a-3p | C17orf96 | PAR-CLIP | miRTarbase |
| hsa-miR-15a-3p | C17orf96 | PAR-CLIP | miRTarbase |
| hsa-miR-15a-3p | KCNH5 | PAR-CLIP | miRTarbase |
| hsa-miR-141-3p | DPY19L1 | PAR-CLIP | miRTarbase |
| hsa-miR-15a-3p | ZNF514 | PAR-CLIP | miRTarbase |
| hsa-miR-3613-3p | UHRF1BP1L | PAR-CLIP | miRTarbase |
| hsa-miR-3613-3p | SIX1 | PAR-CLIP | miRTarbase |
| hsa-miR-3613-3p | SIX1 | PAR-CLIP | miRTarbase |
| hsa-miR-3613-3p | TMEM158 | PAR-CLIP | miRTarbase |
| hsa-miR-3613-3p | CCBE1 | PAR-CLIP | miRTarbase |
| hsa-miR-3613-3p | CTNNA3 | PAR-CLIP | miRTarbase |
| hsa-miR-3613-3p | CTNNA3 | HITS-CLIP | miRTarbase |
| hsa-miR-15a-3p | CTNNA3 | PAR-CLIP | miRTarbase |
| hsa-miR-15a-3p | ENAM | PAR-CLIP | miRTarbase |
| hsa-miR-15a-3p | ECHDC3 | PAR-CLIP | miRTarbase |
| hsa-miR-3613-3p | FBXO48 | PAR-CLIP | miRTarbase |
| hsa-miR-3613-3p | FBXO48 | PAR-CLIP | miRTarbase |
| hsa-miR-3613-3p | FBXO48 | PAR-CLIP | miRTarbase |
| hsa-miR-219a-1-3p | TAF8 | PAR-CLIP | miRTarbase |
| hsa-miR-219a-1-3p | TAF8 | PAR-CLIP | miRTarbase |
| hsa-miR-141-3p | CHML | PAR-CLIP | miRTarbase |
| hsa-miR-15a-3p | PLSCR1 | PAR-CLIP | miRTarbase |
| hsa-miR-3613-3p | CDR2L | PAR-CLIP | miRTarbase |
| hsa-miR-3613-3p | SOWAHB | PAR-CLIP | miRTarbase |
| hsa-miR-3613-3p | TRAPPC6B | PAR-CLIP | miRTarbase |
| hsa-miR-15a-3p | TRAPPC6B | PAR-CLIP | miRTarbase |
| hsa-miR-3613-3p | SLC14A1 | PAR-CLIP | miRTarbase |
| hsa-miR-15a-5p | SIDT2 | PAR-CLIP | miRTarbase |
| hsa-miR-195-5p | SIDT2 | PAR-CLIP | miRTarbase |
| hsa-miR-3613-3p | ZNF257 | PAR-CLIP | miRTarbase |
| hsa-miR-3613-3p | S1PR2 | PAR-CLIP | miRTarbase |
| hsa-miR-219a-1-3p | FAM9C | PAR-CLIP | miRTarbase |
| hsa-miR-3613-3p | FAM46D | PAR-CLIP | miRTarbase |
| hsa-miR-195-3p | ABCG2 | PAR-CLIP | miRTarbase |
| hsa-miR-3613-3p | GSTCD | PAR-CLIP | miRTarbase |
| hsa-miR-3613-3p | GSTCD | HITS-CLIP | miRTarbase |
| hsa-miR-3613-3p | NCOA4 | PAR-CLIP | miRTarbase |
| hsa-miR-3613-3p | PHF14 | PAR-CLIP | miRTarbase |
| hsa-miR-3613-3p | PHF14 | PAR-CLIP | miRTarbase |
| hsa-miR-3613-3p | PTCHD1 | PAR-CLIP | miRTarbase |
| hsa-miR-3613-3p | PTCHD1 | PAR-CLIP | miRTarbase |
| hsa-miR-3605-5p | KDM6A | PAR-CLIP | miRTarbase |
| hsa-miR-3613-3p | CRNKL1 | PAR-CLIP | miRTarbase |
| hsa-miR-3613-3p | CRNKL1 | PAR-CLIP | miRTarbase |
| hsa-miR-486-5p | PCCA | PAR-CLIP | miRTarbase |
| hsa-miR-3605-5p | ATP5C1 | PAR-CLIP | miRTarbase |
| hsa-miR-3605-5p | SYNCRIP | PAR-CLIP | miRTarbase |
| hsa-miR-219a-1-3p | SNX1 | PAR-CLIP | miRTarbase |
| hsa-miR-195-5p | ASCC1 | PAR-CLIP | miRTarbase |
| hsa-miR-15a-5p | ASCC1 | PAR-CLIP | miRTarbase |
| hsa-miR-3613-3p | FUT10 | PAR-CLIP | miRTarbase |
| hsa-miR-3613-3p | SLC36A4 | PAR-CLIP | miRTarbase |
| hsa-miR-3605-5p | ZNF554 | PAR-CLIP | miRTarbase |
| hsa-miR-3613-3p | MCM9 | PAR-CLIP | miRTarbase |
| hsa-miR-3605-5p | SLCO4C1 | PAR-CLIP | miRTarbase |
| hsa-miR-3613-3p | KCNAB1 | PAR-CLIP | miRTarbase |
| hsa-miR-3613-3p | SLFN5 | PAR-CLIP | miRTarbase |
| hsa-miR-3613-3p | ADM | PAR-CLIP | miRTarbase |
| hsa-miR-15a-3p | TEX261 | PAR-CLIP | miRTarbase |
| hsa-miR-3613-5p | CUL3 | PAR-CLIP | miRTarbase |
| hsa-miR-141-3p | NDST4 | PAR-CLIP | miRTarbase |
| hsa-miR-195-5p | DMRT2 | PAR-CLIP | miRTarbase |
| hsa-miR-15a-5p | DMRT2 | PAR-CLIP | miRTarbase |
| hsa-miR-3605-5p | SLC25A43 | PAR-CLIP | miRTarbase |
| hsa-miR-15a-5p | TLL1 | PAR-CLIP | miRTarbase |
| hsa-miR-195-5p | TLL1 | PAR-CLIP | miRTarbase |
| hsa-miR-3613-3p | CLCN6 | PAR-CLIP | miRTarbase |
| hsa-miR-3605-5p | LUC7L3 | PAR-CLIP | miRTarbase |
| hsa-miR-195-5p | LUC7L3 | PAR-CLIP | miRTarbase |
| hsa-miR-15a-5p | LUC7L3 | PAR-CLIP | miRTarbase |
| hsa-miR-3613-3p | SHOX2 | PAR-CLIP | miRTarbase |
| hsa-miR-3613-3p | HAT1 | PAR-CLIP | miRTarbase |

| | | | |
|---|---|---|---|
| hsa-miR-219a-1-3p | HES2 | PAR-CLIP | miRTarbase |
| hsa-miR-3605-5p | C1orf109 | PAR-CLIP | miRTarbase |
| hsa-miR-3613-3p | ADH5 | PAR-CLIP | miRTarbase |
| hsa-miR-141-5p | TNFAIP3 | PAR-CLIP | miRTarbase |
| hsa-miR-3613-5p | RORA | PAR-CLIP | miRTarbase |
| hsa-miR-486-3p | PIN1 | PAR-CLIP | miRTarbase |
| hsa-miR-3605-5p | PIN1 | PAR-CLIP | miRTarbase |
| hsa-miR-486-3p | CIAPIN1 | PAR-CLIP | miRTarbase |
| hsa-miR-3605-5p | ALDH3B1 | PAR-CLIP | miRTarbase |
| hsa-miR-195-5p | ALDH3B1 | PAR-CLIP | miRTarbase |
| hsa-miR-15a-5p | ALDH3B1 | PAR-CLIP | miRTarbase |
| hsa-miR-3605-5p | SOD2 | PAR-CLIP | miRTarbase |
| hsa-miR-3605-5p | SOD2 | PAR-CLIP | miRTarbase |
| hsa-miR-3605-5p | SOD2 | PAR-CLIP | miRTarbase |
| hsa-miR-3605-5p | SOD2 | PAR-CLIP | miRTarbase |
| hsa-miR-486-3p | TMPRSS5 | PAR-CLIP | miRTarbase |
| hsa-miR-3605-5p | NUCB2 | PAR-CLIP | miRTarbase |
| hsa-miR-141-3p | TRAM1 | PAR-CLIP | miRTarbase |
| hsa-miR-486-3p | EIF5AL1 | PAR-CLIP | miRTarbase |
| hsa-miR-486-3p | LY6E | PAR-CLIP | miRTarbase |
| hsa-miR-486-3p | LY6E | PAR-CLIP | miRTarbase |
| hsa-miR-141-5p | SORCS2 | PAR-CLIP | miRTarbase |
| hsa-miR-486-3p | REPIN1 | PAR-CLIP | miRTarbase |
| hsa-miR-486-3p | TANGO2 | PAR-CLIP | miRTarbase |
| hsa-miR-195-5p | EFTUD2 | PAR-CLIP | miRTarbase |
| hsa-miR-15a-5p | EFTUD2 | PAR-CLIP | miRTarbase |
| hsa-miR-486-3p | CACNA2D2 | PAR-CLIP | miRTarbase |
| hsa-miR-486-3p | SLC4A2 | PAR-CLIP | miRTarbase |
| hsa-miR-195-5p | CSNK1E | PAR-CLIP | miRTarbase |
| hsa-miR-15a-5p | CSNK1E | PAR-CLIP | miRTarbase |
| hsa-miR-3605-5p | DDT | PAR-CLIP | miRTarbase |
| hsa-miR-486-3p | SLC29A1 | PAR-CLIP | miRTarbase |
| hsa-miR-195-5p | TPM2 | PAR-CLIP | miRTarbase |
| hsa-miR-15a-5p | TPM2 | PAR-CLIP | miRTarbase |
| hsa-miR-486-3p | MEN1 | PAR-CLIP | miRTarbase |
| hsa-miR-3605-5p | C6orf106 | PAR-CLIP | miRTarbase |
| hsa-miR-141-3p | YRDC | PAR-CLIP | miRTarbase |
| hsa-miR-141-3p | YRDC | PAR-CLIP//HITS-CLIP | miRTarbase |
| hsa-miR-141-3p | YRDC | PAR-CLIP | miRTarbase |
| hsa-miR-141-3p | YRDC | PAR-CLIP | miRTarbase |
| hsa-miR-141-3p | YRDC | PAR-CLIP | miRTarbase |
| hsa-miR-141-3p | YRDC | PAR-CLIP | miRTarbase |
| hsa-miR-486-3p | HECTD3 | PAR-CLIP | miRTarbase |
| hsa-miR-3613-5p | F11R | PAR-CLIP | miRTarbase |
| hsa-miR-3605-5p | MRPL18 | PAR-CLIP | miRTarbase |
| hsa-miR-3605-5p | TOMM20 | PAR-CLIP | miRTarbase |
| hsa-miR-486-3p | NID1 | PAR-CLIP | miRTarbase |
| hsa-miR-486-3p | MTHFSD | PAR-CLIP | miRTarbase |
| hsa-miR-195-5p | ZNF460 | PAR-CLIP | miRTarbase |
| hsa-miR-195-5p | ZNF460 | PAR-CLIP//HITS-CLIP | miRTarbase |
| hsa-miR-195-5p | ZNF460 | PAR-CLIP | miRTarbase |
| hsa-miR-195-5p | ZNF460 | PAR-CLIP | miRTarbase |
| hsa-miR-195-5p | ZNF460 | PAR-CLIP | miRTarbase |
| hsa-miR-15a-5p | ZNF460 | PAR-CLIP | miRTarbase |
| hsa-miR-15a-5p | ZNF460 | PAR-CLIP//HITS-CLIP | miRTarbase |
| hsa-miR-15a-5p | ZNF460 | PAR-CLIP | miRTarbase |
| hsa-miR-15a-5p | ZNF460 | PAR-CLIP | miRTarbase |
| hsa-miR-15a-5p | ZNF460 | PAR-CLIP | miRTarbase |
| hsa-miR-486-3p | CAMK2A | PAR-CLIP | miRTarbase |
| hsa-miR-486-3p | ZMAT5 | PAR-CLIP | miRTarbase |
| hsa-miR-3605-5p | MRPL12 | PAR-CLIP | miRTarbase |
| hsa-miR-141-5p | FOXL2 | PAR-CLIP | miRTarbase |
| hsa-miR-486-3p | KLHL38 | PAR-CLIP | miRTarbase |
| hsa-miR-486-3p | KLHL38 | HITS-CLIP | miRTarbase |
| hsa-miR-486-3p | KLHL38 | PAR-CLIP | miRTarbase |
| hsa-miR-3605-5p | TNFRSF10D | PAR-CLIP | miRTarbase |
| hsa-miR-219a-1-3p | HSPA6 | PAR-CLIP | miRTarbase |
| hsa-miR-219a-1-3p | HSPA6 | HITS-CLIP | miRTarbase |
| hsa-miR-3605-5p | POTED | PAR-CLIP | miRTarbase |
| hsa-miR-3605-5p | POTED | PAR-CLIP | miRTarbase |
| hsa-miR-3605-5p | POTED | PAR-CLIP | miRTarbase |
| hsa-miR-3605-5p | POTED | PAR-CLIP | miRTarbase |
| hsa-miR-3605-5p | POTED | PAR-CLIP | miRTarbase |
| hsa-miR-3605-5p | POC1A | PAR-CLIP | miRTarbase |
| hsa-miR-195-5p | FGFR4 | PAR-CLIP | miRTarbase |
| hsa-miR-15a-5p | FGFR4 | PAR-CLIP | miRTarbase |
| hsa-miR-486-3p | TXNDC16 | HITS-CLIP | miRTarbase |
| hsa-miR-486-3p | TXNDC16 | PAR-CLIP | miRTarbase |
| hsa-miR-3605-5p | TNFRSF10B | PAR-CLIP | miRTarbase |
| hsa-miR-3605-5p | TNFRSF10B | PAR-CLIP | miRTarbase |
| hsa-miR-195-5p | DOCK11 | PAR-CLIP | miRTarbase |
| hsa-miR-15a-5p | DOCK11 | PAR-CLIP | miRTarbase |
| hsa-miR-3605-5p | METTL1 | PAR-CLIP | miRTarbase |
| hsa-miR-195-5p | ACTR3B | PAR-CLIP | miRTarbase |
| hsa-miR-15a-5p | ACTR3B | PAR-CLIP | miRTarbase |
| hsa-miR-3605-5p | ZNF426 | PAR-CLIP | miRTarbase |
| hsa-miR-3605-5p | MPDU1 | PAR-CLIP | miRTarbase |
| hsa-miR-486-3p | FXR2 | PAR-CLIP | miRTarbase |
| hsa-miR-486-3p | C20orf27 | PAR-CLIP | miRTarbase |
| hsa-miR-486-3p | C20orf27 | PAR-CLIP | miRTarbase |
| hsa-miR-3605-5p | TTLL12 | PAR-CLIP | miRTarbase |
| hsa-miR-486-3p | NTN1 | PAR-CLIP | miRTarbase |
| hsa-miR-486-3p | BCL3 | PAR-CLIP | miRTarbase |
| hsa-miR-3605-5p | CYP51A1 | PAR-CLIP | miRTarbase |
| hsa-miR-195-5p | ZNF367 | PAR-CLIP | miRTarbase |
| hsa-miR-195-5p | ZNF367 | PAR-CLIP//HITS-CLIP | miRTarbase |
| hsa-miR-195-5p | ZNF367 | PAR-CLIP | miRTarbase |
| hsa-miR-195-5p | ZNF367 | PAR-CLIP | miRTarbase |
| hsa-miR-195-5p | ZNF367 | PAR-CLIP | miRTarbase |
| hsa-miR-15a-5p | ZNF367 | PAR-CLIP | miRTarbase |
| hsa-miR-15a-5p | ZNF367 | PAR-CLIP//HITS-CLIP | miRTarbase |
| hsa-miR-15a-5p | ZNF367 | PAR-CLIP | miRTarbase |
| hsa-miR-15a-5p | ZNF367 | PAR-CLIP | miRTarbase |
| hsa-miR-15a-5p | ZNF367 | PAR-CLIP | miRTarbase |
| hsa-miR-195-3p | ZFX | PAR-CLIP | miRTarbase |
| hsa-miR-195-3p | ZFX | PAR-CLIP | miRTarbase |
| hsa-miR-486-3p | ZFHX3 | PAR-CLIP | miRTarbase |
| hsa-miR-486-5p | ZDHHC20 | PAR-CLIP | miRTarbase |
| hsa-miR-486-3p | ZC3HAV1L | PAR-CLIP | miRTarbase |
| hsa-miR-486-3p | WAC | PAR-CLIP | miRTarbase |
| hsa-miR-219a-1-3p | VPS28 | PAR-CLIP | miRTarbase |
| hsa-miR-486-3p | UST | PAR-CLIP | miRTarbase |
| hsa-miR-195-5p | UBE2V1 | PAR-CLIP | miRTarbase |
| hsa-miR-195-5p | UBE2V1 | PAR-CLIP | miRTarbase |
| hsa-miR-195-5p | UBE2V1 | PAR-CLIP | miRTarbase |
| hsa-miR-195-5p | UBE2V1 | PAR-CLIP | miRTarbase |
| hsa-miR-15a-5p | UBE2V1 | PAR-CLIP | miRTarbase |
| hsa-miR-15a-5p | UBE2V1 | PAR-CLIP | miRTarbase |
| hsa-miR-15a-5p | UBE2V1 | PAR-CLIP | miRTarbase |
| hsa-miR-15a-5p | UBE2V1 | PAR-CLIP | miRTarbase |
| hsa-miR-195-5p | UBE2Q1 | HITS-CLIP | miRTarbase |
| hsa-miR-195-5p | UBE2Q1 | PAR-CLIP | miRTarbase |
| hsa-miR-15a-5p | UBE2Q1 | HITS-CLIP | miRTarbase |
| hsa-miR-15a-5p | UBE2Q1 | PAR-CLIP | miRTarbase |
| hsa-miR-486-3p | UBB | PAR-CLIP | miRTarbase |
| hsa-miR-486-3p | UBB | PAR-CLIP | miRTarbase |
| hsa-miR-486-3p | UBB | PAR-CLIP | miRTarbase |
| hsa-miR-486-3p | UBB | PAR-CLIP | miRTarbase |
| hsa-miR-486-3p | TXLNA | PAR-CLIP | miRTarbase |
| hsa-miR-486-3p | TXLNA | PAR-CLIP | miRTarbase |
| hsa-miR-3605-5p | TWIST1 | PAR-CLIP | miRTarbase |
| hsa-miR-486-3p | TUBB2A | PAR-CLIP | miRTarbase |
| hsa-miR-486-3p | TUBB2A | PAR-CLIP | miRTarbase |
| hsa-miR-486-3p | TUBB2A | PAR-CLIP | miRTarbase |
| hsa-miR-486-3p | TUBB2A | PAR-CLIP | miRTarbase |
| hsa-miR-486-3p | TUBB2A | PAR-CLIP | miRTarbase |
| hsa-miR-3605-5p | TTC14 | PAR-CLIP | miRTarbase |
| hsa-miR-3605-5p | TSR1 | PAR-CLIP | miRTarbase |
| hsa-miR-3613-3p | TSC22D3 | PAR-CLIP | miRTarbase |
| hsa-miR-3613-3p | TSC22D3 | PAR-CLIP | miRTarbase |
| hsa-miR-195-5p | TSC22D2 | PAR-CLIP | miRTarbase |
| hsa-miR-15a-5p | TSC22D2 | PAR-CLIP | miRTarbase |
| hsa-miR-219a-1-3p | TOR1B | PAR-CLIP | miRTarbase |
| hsa-miR-3605-5p | TOR1B | PAR-CLIP | miRTarbase |
| hsa-miR-195-5p | TOB2 | PAR-CLIP | miRTarbase |
| hsa-miR-15a-5p | TOB2 | PAR-CLIP | miRTarbase |
| hsa-miR-486-3p | TNRC6B | PAR-CLIP | miRTarbase |
| hsa-miR-15a-3p | TMEM64 | PAR-CLIP | miRTarbase |
| hsa-miR-15a-3p | TMEM64 | PAR-CLIP | miRTarbase |

| | | | |
|---|---|---|---|
| hsa-miR-15a-5p | TMEM189-UBE2V1 | PAR-CLIP | miRTarbase |
| hsa-miR-15a-5p | TMEM189-UBE2V1 | PAR-CLIP | miRTarbase |
| hsa-miR-15a-5p | TMEM189-UBE2V1 | PAR-CLIP | miRTarbase |
| hsa-miR-15a-5p | TMEM189-UBE2V1 | PAR-CLIP | miRTarbase |
| hsa-miR-195-5p | TMEM189-UBE2V1 | PAR-CLIP | miRTarbase |
| hsa-miR-195-5p | TMEM189-UBE2V1 | PAR-CLIP | miRTarbase |
| hsa-miR-195-5p | TMEM189-UBE2V1 | PAR-CLIP | miRTarbase |
| hsa-miR-195-5p | TMEM189-UBE2V1 | PAR-CLIP | miRTarbase |
| hsa-miR-15a-5p | TMEM189 | PAR-CLIP | miRTarbase |
| hsa-miR-15a-5p | TMEM189 | PAR-CLIP | miRTarbase |
| hsa-miR-15a-5p | TMEM189 | PAR-CLIP | miRTarbase |
| hsa-miR-15a-5p | TMEM189 | PAR-CLIP | miRTarbase |
| hsa-miR-195-5p | TMEM189 | PAR-CLIP | miRTarbase |
| hsa-miR-195-5p | TMEM189 | PAR-CLIP | miRTarbase |
| hsa-miR-195-5p | TMEM189 | PAR-CLIP | miRTarbase |
| hsa-miR-195-5p | TMEM189 | PAR-CLIP | miRTarbase |
| hsa-miR-3613-3p | TMEM170A | PAR-CLIP | miRTarbase |
| hsa-miR-3613-3p | TMEM170A | PAR-CLIP | miRTarbase |
| hsa-miR-3613-3p | TMEM170A | PAR-CLIP | miRTarbase |
| hsa-miR-3613-3p | TMEM170A | PAR-CLIP | miRTarbase |
| hsa-miR-3613-3p | TMEM170A | PAR-CLIP | miRTarbase |
| hsa-miR-3613-3p | TMEM170A | HITS-CLIP | miRTarbase |
| hsa-miR-3613-3p | TMEM170A | PAR-CLIP | miRTarbase |
| hsa-miR-195-5p | TM4SF1 | PAR-CLIP | miRTarbase |
| hsa-miR-15a-5p | TM4SF1 | PAR-CLIP | miRTarbase |
| hsa-miR-195-5p | TFAP2A | PAR-CLIP | miRTarbase |
| hsa-miR-195-5p | TFAP2A | PAR-CLIP | miRTarbase |
| hsa-miR-195-5p | TFAP2A | PAR-CLIP | miRTarbase |
| hsa-miR-195-5p | TFAP2A | PAR-CLIP | miRTarbase |
| hsa-miR-15a-5p | TFAP2A | PAR-CLIP | miRTarbase |
| hsa-miR-15a-5p | TFAP2A | PAR-CLIP | miRTarbase |
| hsa-miR-15a-5p | TFAP2A | PAR-CLIP | miRTarbase |
| hsa-miR-15a-5p | TFAP2A | PAR-CLIP | miRTarbase |
| hsa-miR-219a-1-3p | TEAD1 | PAR-CLIP | miRTarbase |
| hsa-miR-15a-5p | STK38 | PAR-CLIP | miRTarbase |
| hsa-miR-15a-5p | STK38 | PAR-CLIP | miRTarbase |
| hsa-miR-15a-5p | STK38 | PAR-CLIP | miRTarbase |
| hsa-miR-15a-5p | STK38 | PAR-CLIP | miRTarbase |
| hsa-miR-15a-5p | STK38 | PAR-CLIP | miRTarbase |
| hsa-miR-195-5p | STK38 | PAR-CLIP | miRTarbase |
| hsa-miR-195-5p | STK38 | PAR-CLIP | miRTarbase |
| hsa-miR-195-5p | STK38 | PAR-CLIP | miRTarbase |
| hsa-miR-195-5p | STK38 | PAR-CLIP | miRTarbase |
| hsa-miR-195-5p | STK38 | PAR-CLIP | miRTarbase |
| hsa-miR-195-5p | SSRP1 | PAR-CLIP | miRTarbase |
| hsa-miR-195-5p | SSRP1 | HITS-CLIP | miRTarbase |
| hsa-miR-195-5p | SSRP1 | PAR-CLIP | miRTarbase |
| hsa-miR-15a-5p | SSRP1 | PAR-CLIP | miRTarbase |
| hsa-miR-15a-5p | SSRP1 | HITS-CLIP | miRTarbase |
| hsa-miR-15a-5p | SSRP1 | PAR-CLIP | miRTarbase |
| hsa-miR-486-3p | SREBF2 | PAR-CLIP | miRTarbase |
| hsa-miR-486-3p | SPRED1 | PAR-CLIP | miRTarbase |
| hsa-miR-219a-1-3p | SLC36A1 | PAR-CLIP | miRTarbase |
| hsa-miR-486-3p | SLC25A34 | PAR-CLIP | miRTarbase |
| hsa-miR-486-3p | SIK1 | PAR-CLIP | miRTarbase |
| hsa-miR-15a-5p | SIK1 | HITS-CLIP | miRTarbase |
| hsa-miR-15a-5p | SIK1 | PAR-CLIP | miRTarbase |
| hsa-miR-195-5p | SIK1 | HITS-CLIP | miRTarbase |
| hsa-miR-195-5p | SIK1 | PAR-CLIP | miRTarbase |
| hsa-miR-486-3p | SH3PXD2A | PAR-CLIP | miRTarbase |
| hsa-miR-195-5p | SH3BP4 | PAR-CLIP | miRTarbase |
| hsa-miR-15a-5p | SH3BP4 | PAR-CLIP | miRTarbase |
| hsa-miR-15a-5p | SEC24A | PAR-CLIP | miRTarbase |
| hsa-miR-15a-5p | SEC24A | PAR-CLIP | miRTarbase |
| hsa-miR-195-5p | SEC24A | PAR-CLIP | miRTarbase |
| hsa-miR-195-5p | SEC24A | PAR-CLIP | miRTarbase |
| hsa-miR-219a-1-3p | RPS14 | PAR-CLIP | miRTarbase |
| hsa-miR-486-3p | RNF41 | PAR-CLIP | miRTarbase |
| hsa-miR-195-5p | RNF168 | PAR-CLIP | miRTarbase |
| hsa-miR-15a-5p | RNF168 | PAR-CLIP | miRTarbase |
| hsa-miR-195-5p | REL | PAR-CLIP | miRTarbase |
| hsa-miR-195-5p | REL | PAR-CLIP | miRTarbase |
| hsa-miR-195-5p | REL | PAR-CLIP | miRTarbase |
| hsa-miR-15a-5p | REL | PAR-CLIP | miRTarbase |
| hsa-miR-15a-5p | REL | PAR-CLIP | miRTarbase |
| hsa-miR-15a-5p | REL | PAR-CLIP | miRTarbase |
| hsa-miR-141-5p | RACGAP1 | PAR-CLIP | miRTarbase |
| hsa-miR-141-5p | RACGAP1 | PAR-CLIP | miRTarbase |
| hsa-miR-141-5p | RACGAP1 | PAR-CLIP | miRTarbase |
| hsa-miR-141-5p | RACGAP1 | PAR-CLIP | miRTarbase |
| hsa-miR-486-3p | PTPRF | PAR-CLIP | miRTarbase |
| hsa-miR-486-3p | PTPRF | PAR-CLIP | miRTarbase |
| hsa-miR-486-3p | PTPRF | HITS-CLIP | miRTarbase |
| hsa-miR-486-3p | PTPRF | PAR-CLIP | miRTarbase |
| hsa-miR-141-3p | PSMD11 | PAR-CLIP | miRTarbase |
| hsa-miR-3613-5p | PRKAA1 | PAR-CLIP | miRTarbase |
| hsa-miR-3605-5p | PPP2R5E | PAR-CLIP | miRTarbase |
| hsa-miR-15a-3p | COASY | PAR-CLIP | miRTarbase |
| hsa-miR-3605-3p | PODXL | PAR-CLIP | miRTarbase |
| hsa-miR-195-5p | PISD | PAR-CLIP | miRTarbase |
| hsa-miR-195-5p | PISD | PAR-CLIP | miRTarbase |
| hsa-miR-195-5p | PISD | PAR-CLIP | miRTarbase |
| hsa-miR-195-5p | PISD | PAR-CLIP | miRTarbase |
| hsa-miR-195-5p | PISD | PAR-CLIP | miRTarbase |
| hsa-miR-15a-5p | PISD | PAR-CLIP | miRTarbase |
| hsa-miR-15a-5p | PISD | PAR-CLIP | miRTarbase |
| hsa-miR-15a-5p | PISD | PAR-CLIP | miRTarbase |
| hsa-miR-15a-5p | PISD | PAR-CLIP | miRTarbase |
| hsa-miR-15a-5p | PISD | PAR-CLIP | miRTarbase |
| hsa-miR-486-3p | PGAM4 | PAR-CLIP | miRTarbase |
| hsa-miR-3605-5p | PDPR | PAR-CLIP | miRTarbase |
| hsa-miR-195-5p | PDE4D | PAR-CLIP | miRTarbase |
| hsa-miR-195-5p | PDE4D | PAR-CLIP | miRTarbase |
| hsa-miR-15a-5p | PDE4D | PAR-CLIP | miRTarbase |
| hsa-miR-15a-5p | PDE4D | PAR-CLIP | miRTarbase |
| hsa-miR-3613-5p | PCGF3 | PAR-CLIP | miRTarbase |
| hsa-miR-195-3p | NUS1 | PAR-CLIP | miRTarbase |
| hsa-miR-195-3p | NUS1 | PAR-CLIP | miRTarbase |
| hsa-miR-195-3p | NUS1 | PAR-CLIP | miRTarbase |
| hsa-miR-195-3p | NUS1 | PAR-CLIP | miRTarbase |
| hsa-miR-3605-5p | NUAK2 | PAR-CLIP | miRTarbase |
| hsa-miR-3613-3p | NRAS | PAR-CLIP | miRTarbase |
| hsa-miR-195-5p | NR6A1 | PAR-CLIP | miRTarbase |
| hsa-miR-15a-5p | NR6A1 | PAR-CLIP | miRTarbase |
| hsa-miR-141-5p | NPEPPS | PAR-CLIP | miRTarbase |
| hsa-miR-15a-5p | NFIC | PAR-CLIP | miRTarbase |
| hsa-miR-195-5p | NFIC | PAR-CLIP | miRTarbase |
| hsa-miR-486-3p | NDRG3 | PAR-CLIP | miRTarbase |
| hsa-miR-486-3p | NDRG3 | HITS-CLIP | miRTarbase |
| hsa-miR-15a-5p | NAA25 | PAR-CLIP | miRTarbase |
| hsa-miR-15a-5p | NAA25 | PAR-CLIP | miRTarbase |
| hsa-miR-195-5p | NAA25 | PAR-CLIP | miRTarbase |
| hsa-miR-195-5p | NAA25 | PAR-CLIP | miRTarbase |
| hsa-miR-486-3p | MTMR12 | PAR-CLIP | miRTarbase |
| hsa-miR-3605-5p | MTMR10 | PAR-CLIP | miRTarbase |
| hsa-miR-3605-5p | MTMR10 | PAR-CLIP | miRTarbase |
| hsa-miR-3605-5p | MRAS | PAR-CLIP | miRTarbase |
| hsa-miR-486-3p | MBD4 | PAR-CLIP | miRTarbase |
| hsa-miR-486-3p | MBD4 | PAR-CLIP | miRTarbase |
| hsa-miR-486-3p | MBD4 | PAR-CLIP | miRTarbase |
| hsa-miR-486-3p | MDM4 | PAR-CLIP | miRTarbase |
| hsa-miR-141-3p | MCL1 | PAR-CLIP | miRTarbase |
| hsa-miR-3605-5p | MAPK1 | PAR-CLIP | miRTarbase |
| hsa-miR-486-3p | LGSN | PAR-CLIP | miRTarbase |
| hsa-miR-195-5p | LAMC1 | PAR-CLIP | miRTarbase |
| hsa-miR-15a-5p | LAMC1 | PAR-CLIP | miRTarbase |
| hsa-miR-486-3p | KMT2D | PAR-CLIP | miRTarbase |
| hsa-miR-3605-5p | KIF5A | PAR-CLIP | miRTarbase |
| hsa-miR-3613-3p | KIF13A | PAR-CLIP | miRTarbase |
| hsa-miR-3613-3p | KIF13A | PAR-CLIP | miRTarbase |
| hsa-miR-3613-3p | KIF13A | PAR-CLIP | miRTarbase |
| hsa-miR-15a-5p | KIAA0226 | PAR-CLIP | miRTarbase |
| hsa-miR-195-5p | KIAA0226 | PAR-CLIP | miRTarbase |
| hsa-miR-3605-5p | KCND3 | PAR-CLIP | miRTarbase |
| hsa-miR-195-5p | IVNS1ABP | PAR-CLIP | miRTarbase |
| hsa-miR-195-5p | IVNS1ABP | PAR-CLIP | miRTarbase |
| hsa-miR-195-5p | IVNS1ABP | PAR-CLIP | miRTarbase |
| hsa-miR-15a-5p | IVNS1ABP | PAR-CLIP | miRTarbase |
| hsa-miR-15a-5p | IVNS1ABP | PAR-CLIP | miRTarbase |

| | | | |
|---|---|---|---|
| hsa-miR-15a-5p | IVNS1ABP | PAR-CLIP | miRTarbase |
| hsa-miR-195-5p | IPPK | PAR-CLIP | miRTarbase |
| hsa-miR-15a-5p | IPPK | PAR-CLIP | miRTarbase |
| hsa-miR-195-5p | HOXA3 | PAR-CLIP | miRTarbase |
| hsa-miR-195-5p | HOXA3 | PAR-CLIP//HITS-CLIP | miRTarbase |
| hsa-miR-195-5p | HOXA3 | PAR-CLIP | miRTarbase |
| hsa-miR-195-5p | HOXA3 | PAR-CLIP | miRTarbase |
| hsa-miR-15a-5p | HOXA3 | PAR-CLIP | miRTarbase |
| hsa-miR-15a-5p | HOXA3 | PAR-CLIP//HITS-CLIP | miRTarbase |
| hsa-miR-15a-5p | HOXA3 | PAR-CLIP | miRTarbase |
| hsa-miR-15a-5p | HOXA3 | PAR-CLIP | miRTarbase |
| hsa-miR-219a-1-3p | HMGB1 | PAR-CLIP | miRTarbase |
| hsa-miR-195-5p | HEYL | PAR-CLIP | miRTarbase |
| hsa-miR-15a-5p | HEYL | PAR-CLIP | miRTarbase |
| hsa-miR-15a-3p | HDGF | PAR-CLIP | miRTarbase |
| hsa-miR-195-5p | HDGF | PAR-CLIP | miRTarbase |
| hsa-miR-195-5p | HDGF | PAR-CLIP | miRTarbase |
| hsa-miR-15a-5p | HDGF | PAR-CLIP | miRTarbase |
| hsa-miR-15a-5p | HDGF | PAR-CLIP | miRTarbase |
| hsa-miR-3613-5p | H3F3C | PAR-CLIP | miRTarbase |
| hsa-miR-3613-5p | H3F3C | PAR-CLIP | miRTarbase |
| hsa-miR-3605-5p | GPR137C | PAR-CLIP | miRTarbase |
| hsa-miR-3605-5p | GPR137C | PAR-CLIP | miRTarbase |
| hsa-miR-3605-5p | GPR137C | PAR-CLIP | miRTarbase |
| hsa-miR-3605-5p | GPR137C | PAR-CLIP | miRTarbase |
| hsa-miR-195-5p | GNB1 | PAR-CLIP | miRTarbase |
| hsa-miR-195-5p | GNB1 | PAR-CLIP | miRTarbase |
| hsa-miR-195-5p | GNB1 | HITS-CLIP | miRTarbase |
| hsa-miR-195-5p | GNB1 | PAR-CLIP | miRTarbase |
| hsa-miR-15a-5p | GNB1 | PAR-CLIP | miRTarbase |
| hsa-miR-15a-5p | GNB1 | PAR-CLIP | miRTarbase |
| hsa-miR-15a-5p | GNB1 | HITS-CLIP | miRTarbase |
| hsa-miR-15a-5p | GNB1 | PAR-CLIP | miRTarbase |
| hsa-miR-195-5p | GNAL | PAR-CLIP | miRTarbase |
| hsa-miR-195-5p | GNAL | PAR-CLIP//HITS-CLIP | miRTarbase |
| hsa-miR-195-5p | GNAL | PAR-CLIP | miRTarbase |
| hsa-miR-15a-5p | GNAL | PAR-CLIP | miRTarbase |
| hsa-miR-15a-5p | GNAL | PAR-CLIP//HITS-CLIP | miRTarbase |
| hsa-miR-15a-5p | GNAL | PAR-CLIP | miRTarbase |
| hsa-miR-195-5p | FURIN | PAR-CLIP | miRTarbase |
| hsa-miR-15a-5p | FURIN | PAR-CLIP | miRTarbase |
| hsa-miR-3605-5p | EPHA2 | PAR-CLIP | miRTarbase |
| hsa-miR-486-3p | ELOVL5 | PAR-CLIP | miRTarbase |
| hsa-miR-195-5p | EIF1AX | PAR-CLIP | miRTarbase |
| hsa-miR-195-5p | EIF1AX | PAR-CLIP | miRTarbase |
| hsa-miR-195-5p | EIF1AX | PAR-CLIP | miRTarbase |
| hsa-miR-195-5p | EIF1AX | PAR-CLIP | miRTarbase |
| hsa-miR-15a-5p | EIF1AX | PAR-CLIP | miRTarbase |
| hsa-miR-15a-5p | EIF1AX | PAR-CLIP | miRTarbase |
| hsa-miR-15a-5p | EIF1AX | PAR-CLIP | miRTarbase |
| hsa-miR-15a-5p | EIF1AX | PAR-CLIP | miRTarbase |
| hsa-miR-3605-5p | DYRK3 | PAR-CLIP | miRTarbase |
| hsa-miR-195-5p | DYRK3 | PAR-CLIP | miRTarbase |
| hsa-miR-15a-5p | DYRK3 | PAR-CLIP | miRTarbase |
| hsa-miR-3605-5p | DNAJC10 | PAR-CLIP | miRTarbase |
| hsa-miR-3605-5p | DNAJC10 | PAR-CLIP | miRTarbase |
| hsa-miR-3605-5p | DNAJC10 | PAR-CLIP | miRTarbase |
| hsa-miR-3605-5p | DNAJC10 | PAR-CLIP | miRTarbase |
| hsa-miR-3605-5p | DNAJC10 | PAR-CLIP | miRTarbase |
| hsa-miR-486-3p | DLGAP4 | PAR-CLIP | miRTarbase |
| hsa-miR-3613-3p | DLG5 | PAR-CLIP | miRTarbase |
| hsa-miR-3613-3p | DLG5 | PAR-CLIP | miRTarbase |
| hsa-miR-3613-3p | DLG5 | PAR-CLIP | miRTarbase |
| hsa-miR-3613-3p | DDX6 | HITS-CLIP | miRTarbase |
| hsa-miR-3613-3p | DDX6 | PAR-CLIP | miRTarbase |
| hsa-miR-3605-5p | DDX3X | PAR-CLIP | miRTarbase |
| hsa-miR-3605-5p | DDX3X | PAR-CLIP | miRTarbase |
| hsa-miR-486-3p | DAZAP2 | PAR-CLIP | miRTarbase |
| hsa-miR-141-3p | CYP1B1 | PAR-CLIP | miRTarbase |
| hsa-miR-219a-1-3p | CTNND1 | PAR-CLIP | miRTarbase |
| hsa-miR-486-3p | CTDNEP1 | PAR-CLIP | miRTarbase |
| hsa-miR-195-5p | CPSF7 | PAR-CLIP | miRTarbase |
| hsa-miR-195-5p | CPSF7 | PAR-CLIP | miRTarbase |
| hsa-miR-195-5p | CPSF7 | PAR-CLIP | miRTarbase |
| hsa-miR-15a-5p | CPSF7 | PAR-CLIP | miRTarbase |
| hsa-miR-15a-5p | CPSF7 | PAR-CLIP | miRTarbase |
| hsa-miR-15a-5p | CPSF7 | PAR-CLIP | miRTarbase |
| hsa-miR-3613-3p | COIL | PAR-CLIP | miRTarbase |
| hsa-miR-3613-3p | COIL | PAR-CLIP | miRTarbase |
| hsa-miR-3613-3p | COIL | PAR-CLIP | miRTarbase |
| hsa-miR-15a-5p | CDK6 | PAR-CLIP | miRTarbase |
| hsa-miR-195-3p | CDC25A | PAR-CLIP | miRTarbase |
| hsa-miR-219a-1-3p | CCNF | PAR-CLIP | miRTarbase |
| hsa-miR-195-5p | CARD10 | PAR-CLIP | miRTarbase |
| hsa-miR-15a-5p | CARD10 | PAR-CLIP | miRTarbase |
| hsa-miR-486-3p | CALR | PAR-CLIP | miRTarbase |
| hsa-miR-486-3p | CAD | PAR-CLIP | miRTarbase |
| hsa-miR-3605-5p | C16orf72 | PAR-CLIP | miRTarbase |
| hsa-miR-3605-5p | C16orf72 | PAR-CLIP | miRTarbase |
| hsa-miR-3605-5p | C16orf72 | PAR-CLIP | miRTarbase |
| hsa-miR-486-3p | BAZ2A | PAR-CLIP | miRTarbase |
| hsa-miR-195-5p | AVL9 | PAR-CLIP | miRTarbase |
| hsa-miR-195-5p | AVL9 | PAR-CLIP | miRTarbase |
| hsa-miR-195-5p | AVL9 | PAR-CLIP | miRTarbase |
| hsa-miR-15a-5p | AVL9 | PAR-CLIP | miRTarbase |
| hsa-miR-15a-5p | AVL9 | PAR-CLIP | miRTarbase |
| hsa-miR-15a-5p | AVL9 | PAR-CLIP | miRTarbase |
| hsa-miR-486-3p | ATXN7L3B | PAR-CLIP | miRTarbase |
| hsa-miR-486-3p | ATXN7L3B | PAR-CLIP | miRTarbase |
| hsa-miR-486-3p | ATXN7L3 | PAR-CLIP | miRTarbase |
| hsa-miR-195-3p | ARL6IP1 | PAR-CLIP | miRTarbase |
| hsa-miR-195-3p | ARL6IP1 | PAR-CLIP | miRTarbase |
| hsa-miR-195-3p | ARL6IP1 | PAR-CLIP | miRTarbase |
| hsa-miR-195-3p | ARL6IP1 | PAR-CLIP | miRTarbase |
| hsa-miR-3613-5p | ARID1A | PAR-CLIP | miRTarbase |
| hsa-miR-3605-5p | ANKRD50 | PAR-CLIP | miRTarbase |
| hsa-miR-195-5p | AKT3 | PAR-CLIP | miRTarbase |
| hsa-miR-195-5p | AKT3 | PAR-CLIP | miRTarbase |
| hsa-miR-15a-5p | AGO2 | PAR-CLIP | miRTarbase |
| hsa-miR-195-5p | AGO2 | PAR-CLIP | miRTarbase |
| hsa-miR-141-5p | ADRB1 | PAR-CLIP | miRTarbase |
| hsa-miR-141-5p | ADRB1 | PAR-CLIP | miRTarbase |
| hsa-miR-141-5p | ADRB1 | PAR-CLIP | miRTarbase |
| hsa-miR-141-5p | ADRB1 | PAR-CLIP | miRTarbase |
| hsa-miR-141-5p | ADRB1 | PAR-CLIP | miRTarbase |
| hsa-miR-3605-5p | ADAR | PAR-CLIP | miRTarbase |
| hsa-miR-3605-5p | ADAR | PAR-CLIP | miRTarbase |
| hsa-miR-3613-3p | ACTB | PAR-CLIP | miRTarbase |
| hsa-miR-3613-3p | ACTB | PAR-CLIP | miRTarbase |
| hsa-miR-195-5p | ABL2 | PAR-CLIP | miRTarbase |
| hsa-miR-195-5p | ABL2 | PAR-CLIP | miRTarbase |
| hsa-miR-195-5p | ABL2 | PAR-CLIP | miRTarbase |
| hsa-miR-195-5p | ABL2 | PAR-CLIP | miRTarbase |
| hsa-miR-195-5p | ABL2 | PAR-CLIP | miRTarbase |
| hsa-miR-15a-5p | ABL2 | PAR-CLIP | miRTarbase |
| hsa-miR-15a-5p | ABL2 | PAR-CLIP | miRTarbase |
| hsa-miR-15a-5p | ABL2 | PAR-CLIP | miRTarbase |
| hsa-miR-15a-5p | ABL2 | PAR-CLIP | miRTarbase |
| hsa-miR-15a-5p | ABL2 | PAR-CLIP | miRTarbase |
| hsa-miR-195-5p | ABHD2 | PAR-CLIP | miRTarbase |
| hsa-miR-15a-5p | ABHD2 | PAR-CLIP | miRTarbase |
| hsa-miR-486-3p | COPZ1 | PAR-CLIP | miRTarbase |
| hsa-miR-195-3p | SH3BP5 | PAR-CLIP | miRTarbase |
| hsa-miR-219a-1-3p | PIP4K2A | PAR-CLIP | miRTarbase |
| hsa-miR-486-3p | HIVEP3 | PAR-CLIP | miRTarbase |
| hsa-miR-486-3p | HIVEP3 | PAR-CLIP | miRTarbase |
| hsa-miR-15a-3p | TAGLN2 | PAR-CLIP | miRTarbase |
| hsa-miR-486-3p | CASKIN1 | PAR-CLIP | miRTarbase |
| hsa-miR-486-3p | CASKIN1 | PAR-CLIP | miRTarbase |
| hsa-miR-3613-3p | TGIF1 | PAR-CLIP | miRTarbase |
| hsa-miR-3613-3p | TGIF1 | HITS-CLIP | miRTarbase |
| hsa-miR-219a-1-3p | SUGT1 | PAR-CLIP | miRTarbase |
| hsa-miR-219a-1-3p | SUGT1 | PAR-CLIP | miRTarbase |
| hsa-miR-195-5p | ABCC6 | PAR-CLIP | miRTarbase |
| hsa-miR-195-5p | ABCC6 | PAR-CLIP | miRTarbase |
| hsa-miR-15a-5p | ABCC6 | PAR-CLIP | miRTarbase |
| hsa-miR-15a-5p | ABCC6 | PAR-CLIP | miRTarbase |
| hsa-miR-219a-1-3p | ZNF652 | PAR-CLIP | miRTarbase |
| hsa-miR-3613-3p | ZFYVE26 | PAR-CLIP | miRTarbase |

| | | | |
|---|---|---|---|
| hsa-miR-3613-3p | ZFYVE26 | PAR-CLIP | miRTarbase |
| hsa-miR-15a-5p | PRKAR2A | PAR-CLIP | miRTarbase |
| hsa-miR-15a-5p | PRKAR2A | PAR-CLIP | miRTarbase |
| hsa-miR-15a-5p | PRKAR2A | PAR-CLIP | miRTarbase |
| hsa-miR-15a-5p | PRKAR2A | PAR-CLIP | miRTarbase |
| hsa-miR-195-5p | PRKAR2A | PAR-CLIP | miRTarbase |
| hsa-miR-195-5p | PRKAR2A | PAR-CLIP | miRTarbase |
| hsa-miR-195-5p | PRKAR2A | PAR-CLIP | miRTarbase |
| hsa-miR-195-5p | PRKAR2A | PAR-CLIP | miRTarbase |
| hsa-miR-3613-3p | DICER1 | PAR-CLIP | miRTarbase |
| hsa-miR-3613-3p | DICER1 | PAR-CLIP | miRTarbase |
| hsa-miR-15a-3p | CASP16 | PAR-CLIP | miRTarbase |
| hsa-miR-15a-3p | CASP16 | PAR-CLIP | miRTarbase |
| hsa-miR-15a-3p | CASP16 | PAR-CLIP | miRTarbase |
| hsa-miR-15a-3p | CASP16 | PAR-CLIP | miRTarbase |
| hsa-miR-15a-3p | CASP16 | PAR-CLIP | miRTarbase |
| hsa-miR-486-3p | PPP1R14A | PAR-CLIP | miRTarbase |
| hsa-miR-3613-5p | UBBP4 | PAR-CLIP | miRTarbase |
| hsa-miR-486-3p | RXRA | PAR-CLIP | miRTarbase |
| hsa-miR-3613-3p | CNOT4 | PAR-CLIP | miRTarbase |
| hsa-miR-3613-3p | CNOT4 | PAR-CLIP | miRTarbase |
| hsa-miR-3613-3p | CNOT4 | PAR-CLIP | miRTarbase |
| hsa-miR-486-3p | SCARF2 | PAR-CLIP | miRTarbase |
| hsa-miR-195-5p | C10orf54 | PAR-CLIP | miRTarbase |
| hsa-miR-15a-5p | C10orf54 | PAR-CLIP | miRTarbase |
| hsa-miR-486-3p | BRSK2 | PAR-CLIP | miRTarbase |
| hsa-miR-486-3p | BRSK2 | PAR-CLIP | miRTarbase |
| hsa-miR-486-3p | CASZ1 | PAR-CLIP | miRTarbase |
| hsa-miR-486-3p | CASZ1 | PAR-CLIP | miRTarbase |
| hsa-miR-486-3p | CST9 | PAR-CLIP | miRTarbase |
| hsa-miR-486-3p | POU3F1 | PAR-CLIP | miRTarbase |
| hsa-miR-486-3p | PAX2 | PAR-CLIP | miRTarbase |
| hsa-miR-3613-5p | VPS13B | PAR-CLIP | miRTarbase |
| hsa-miR-3613-5p | VPS13B | PAR-CLIP | miRTarbase |
| hsa-miR-141-3p | BICD2 | PAR-CLIP | miRTarbase |
| hsa-miR-141-3p | BICD2 | PAR-CLIP | miRTarbase |
| hsa-miR-486-3p | POFUT2 | PAR-CLIP | miRTarbase |
| hsa-miR-486-3p | PTGES | PAR-CLIP | miRTarbase |
| hsa-miR-486-3p | B4GALNT4 | PAR-CLIP | miRTarbase |
| hsa-miR-486-3p | B4GALNT4 | PAR-CLIP | miRTarbase |
| hsa-miR-486-3p | SYNGR1 | PAR-CLIP | miRTarbase |
| hsa-miR-486-3p | SYNGR1 | PAR-CLIP | miRTarbase |
| hsa-miR-486-3p | SHMT1 | PAR-CLIP | miRTarbase |
| hsa-miR-486-3p | SHMT1 | PAR-CLIP | miRTarbase |
| hsa-miR-486-3p | KRT80 | PAR-CLIP | miRTarbase |
| hsa-miR-486-3p | KRT80 | PAR-CLIP | miRTarbase |
| hsa-miR-3605-5p | ANK1 | PAR-CLIP | miRTarbase |
| hsa-miR-3605-5p | ANK1 | PAR-CLIP | miRTarbase |
| hsa-miR-486-3p | SRCIN1 | PAR-CLIP | miRTarbase |
| hsa-miR-486-3p | STRN4 | PAR-CLIP | miRTarbase |
| hsa-miR-486-3p | MRPL34 | PAR-CLIP | miRTarbase |
| hsa-miR-486-3p | GFER | PAR-CLIP | miRTarbase |
| hsa-miR-486-3p | MNT | PAR-CLIP | miRTarbase |
| hsa-miR-486-3p | MNT | PAR-CLIP | miRTarbase |
| hsa-miR-486-5p | UBASH3B | PAR-CLIP | miRTarbase |
| hsa-miR-195-5p | PHYHIP | PAR-CLIP | miRTarbase |
| hsa-miR-15a-5p | PHYHIP | PAR-CLIP | miRTarbase |
| hsa-miR-486-3p | NAV1 | PAR-CLIP | miRTarbase |
| hsa-miR-3605-5p | GRAP2 | PAR-CLIP | miRTarbase |
| hsa-miR-486-3p | ENG | PAR-CLIP | miRTarbase |
| hsa-miR-141-5p | ECT2 | PAR-CLIP | miRTarbase |
| hsa-miR-195-5p | CASKIN1 | PAR-CLIP | miRTarbase |
| hsa-miR-15a-5p | CASKIN1 | PAR-CLIP | miRTarbase |
| hsa-miR-486-3p | SOLH | PAR-CLIP | miRTarbase |
| hsa-miR-486-3p | BCL7A | PAR-CLIP | miRTarbase |
| hsa-miR-486-3p | AP1G1 | PAR-CLIP | miRTarbase |
| hsa-miR-15a-3p | TRADD | PAR-CLIP | miRTarbase |
| hsa-miR-195-5p | ZNRF1 | PAR-CLIP | miRTarbase |
| hsa-miR-15a-5p | ZNRF1 | PAR-CLIP | miRTarbase |
| hsa-miR-15a-3p | LONRF3 | PAR-CLIP | miRTarbase |
| hsa-miR-3605-5p | PLEKHA3 | PAR-CLIP | miRTarbase |
| hsa-miR-3613-3p | SMAD2 | PAR-CLIP | miRTarbase |
| hsa-miR-3613-3p | SMAD2 | HITS-CLIP | miRTarbase |
| hsa-miR-15a-3p | SLC1A5 | PAR-CLIP | miRTarbase |
| hsa-miR-3613-3p | EIF3H | PAR-CLIP | miRTarbase |
| hsa-miR-15a-3p | UQCR10 | PAR-CLIP | miRTarbase |
| hsa-miR-15a-3p | AGBL3 | PAR-CLIP | miRTarbase |
| hsa-miR-15a-3p | CLCN6 | PAR-CLIP | miRTarbase |
| hsa-miR-15a-3p | DPY19L1 | PAR-CLIP | miRTarbase |
| hsa-miR-3613-3p | ZNF641 | PAR-CLIP | miRTarbase |
| hsa-miR-15a-3p | FBXO41 | PAR-CLIP | miRTarbase |
| hsa-miR-195-5p | CD180 | PAR-CLIP | miRTarbase |
| hsa-miR-15a-5p | CD180 | PAR-CLIP | miRTarbase |
| hsa-miR-15a-3p | SLC36A1 | PAR-CLIP | miRTarbase |
| hsa-miR-15a-3p | SH3BGRL | PAR-CLIP | miRTarbase |
| hsa-miR-3613-3p | PARVB | PAR-CLIP | miRTarbase |
| hsa-miR-486-3p | TAGLN | PAR-CLIP | miRTarbase |
| hsa-miR-3613-3p | WNT2B | PAR-CLIP | miRTarbase |
| hsa-miR-3613-3p | NPM3 | PAR-CLIP | miRTarbase |
| hsa-miR-3613-3p | AAED1 | PAR-CLIP | miRTarbase |
| hsa-miR-3613-3p | AAED1 | HITS-CLIP | miRTarbase |
| hsa-miR-486-3p | RALY | PAR-CLIP | miRTarbase |
| hsa-miR-3605-5p | SLC23A1 | PAR-CLIP | miRTarbase |
| hsa-miR-3613-3p | DAZAP1 | PAR-CLIP | miRTarbase |
| hsa-miR-195-5p | KIAA0895 | PAR-CLIP | miRTarbase |
| hsa-miR-15a-5p | KIAA0895 | PAR-CLIP | miRTarbase |
| hsa-miR-141-5p | SNX5 | PAR-CLIP | miRTarbase |
| hsa-miR-141-5p | SNX5 | PAR-CLIP | miRTarbase |
| hsa-miR-141-5p | SNX5 | PAR-CLIP | miRTarbase |
| hsa-miR-486-5p | RCOR3 | PAR-CLIP | miRTarbase |
| hsa-miR-141-3p | PRELID2 | PAR-CLIP | miRTarbase |
| hsa-miR-15a-3p | PRMT3 | PAR-CLIP | miRTarbase |
| hsa-miR-486-3p | KRT8 | PAR-CLIP | miRTarbase |
| hsa-miR-15a-3p | RPL37A | PAR-CLIP | miRTarbase |
| hsa-miR-195-5p | ORC4 | PAR-CLIP | miRTarbase |
| hsa-miR-195-5p | ORC4 | PAR-CLIP | miRTarbase |
| hsa-miR-195-5p | ORC4 | PAR-CLIP | miRTarbase |
| hsa-miR-195-5p | ORC4 | PAR-CLIP | miRTarbase |
| hsa-miR-15a-5p | ORC4 | PAR-CLIP | miRTarbase |
| hsa-miR-15a-5p | ORC4 | PAR-CLIP | miRTarbase |
| hsa-miR-15a-5p | ORC4 | PAR-CLIP | miRTarbase |
| hsa-miR-15a-5p | ORC4 | PAR-CLIP | miRTarbase |
| hsa-miR-486-3p | RFWD3 | PAR-CLIP | miRTarbase |
| hsa-miR-195-5p | ODF2L | PAR-CLIP | miRTarbase |
| hsa-miR-195-5p | ODF2L | PAR-CLIP | miRTarbase |
| hsa-miR-195-5p | ODF2L | PAR-CLIP | miRTarbase |
| hsa-miR-195-5p | ODF2L | PAR-CLIP | miRTarbase |
| hsa-miR-15a-5p | ODF2L | PAR-CLIP | miRTarbase |
| hsa-miR-15a-5p | ODF2L | PAR-CLIP | miRTarbase |
| hsa-miR-15a-5p | ODF2L | PAR-CLIP | miRTarbase |
| hsa-miR-15a-5p | ODF2L | PAR-CLIP | miRTarbase |
| hsa-miR-195-5p | DNAJA1 | PAR-CLIP | miRTarbase |
| hsa-miR-195-5p | DNAJA1 | PAR-CLIP//HITS-CLIP | miRTarbase |
| hsa-miR-195-5p | DNAJA1 | PAR-CLIP | miRTarbase |
| hsa-miR-195-5p | DNAJA1 | PAR-CLIP | miRTarbase |
| hsa-miR-15a-5p | DNAJA1 | PAR-CLIP | miRTarbase |
| hsa-miR-15a-5p | DNAJA1 | PAR-CLIP//HITS-CLIP | miRTarbase |
| hsa-miR-15a-5p | DNAJA1 | PAR-CLIP | miRTarbase |
| hsa-miR-15a-5p | DNAJA1 | PAR-CLIP | miRTarbase |
| hsa-miR-3613-3p | KCTD2 | PAR-CLIP | miRTarbase |
| hsa-miR-3613-3p | KCTD2 | PAR-CLIP | miRTarbase |
| hsa-miR-195-5p | L2HGDH | PAR-CLIP | miRTarbase |
| hsa-miR-195-5p | L2HGDH | PAR-CLIP | miRTarbase |
| hsa-miR-195-5p | L2HGDH | PAR-CLIP | miRTarbase |
| hsa-miR-195-5p | L2HGDH | PAR-CLIP | miRTarbase |
| hsa-miR-15a-5p | L2HGDH | PAR-CLIP | miRTarbase |
| hsa-miR-15a-5p | L2HGDH | PAR-CLIP | miRTarbase |
| hsa-miR-15a-5p | L2HGDH | PAR-CLIP | miRTarbase |
| hsa-miR-15a-5p | L2HGDH | PAR-CLIP | miRTarbase |
| hsa-miR-219a-1-3p | POLD3 | PAR-CLIP | miRTarbase |
| hsa-miR-219a-1-3p | POLD3 | PAR-CLIP//HITS-CLIP | miRTarbase |
| hsa-miR-219a-1-3p | POLD3 | PAR-CLIP | miRTarbase |
| hsa-miR-219a-1-3p | POLD3 | PAR-CLIP | miRTarbase |
| hsa-miR-15a-5p | ZNF622 | PAR-CLIP | miRTarbase |
| hsa-miR-15a-5p | ZNF622 | PAR-CLIP | miRTarbase |
| hsa-miR-15a-5p | ZNF622 | HITS-CLIP | miRTarbase |
| hsa-miR-15a-5p | ZNF622 | PAR-CLIP | miRTarbase |
| hsa-miR-15a-5p | ZNF622 | PAR-CLIP | miRTarbase |
| hsa-miR-195-5p | ZNF622 | PAR-CLIP | miRTarbase |

| | | | |
|---|---|---|---|
| hsa-miR-195-5p | ZNF622 | PAR-CLIP | miRTarbase |
| hsa-miR-195-5p | ZNF622 | HITS-CLIP | miRTarbase |
| hsa-miR-195-5p | ZNF622 | PAR-CLIP | miRTarbase |
| hsa-miR-195-5p | ZNF622 | PAR-CLIP | miRTarbase |
| hsa-miR-195-5p | ZMAT3 | PAR-CLIP | miRTarbase |
| hsa-miR-195-5p | ZMAT3 | PAR-CLIP | miRTarbase |
| hsa-miR-15a-5p | ZMAT3 | PAR-CLIP | miRTarbase |
| hsa-miR-15a-5p | ZMAT3 | PAR-CLIP | miRTarbase |
| hsa-miR-195-5p | USP53 | PAR-CLIP | miRTarbase |
| hsa-miR-15a-5p | USP53 | PAR-CLIP | miRTarbase |
| hsa-miR-141-3p | USP53 | PAR-CLIP | miRTarbase |
| hsa-miR-15a-5p | SYPL1 | PAR-CLIP | miRTarbase |
| hsa-miR-15a-5p | SYPL1 | PAR-CLIP | miRTarbase |
| hsa-miR-15a-5p | SYPL1 | PAR-CLIP | miRTarbase |
| hsa-miR-15a-5p | SYPL1 | PAR-CLIP | miRTarbase |
| hsa-miR-195-5p | SYPL1 | PAR-CLIP | miRTarbase |
| hsa-miR-195-5p | SYPL1 | PAR-CLIP | miRTarbase |
| hsa-miR-195-5p | SYPL1 | PAR-CLIP | miRTarbase |
| hsa-miR-195-5p | SYPL1 | PAR-CLIP | miRTarbase |
| hsa-miR-219a-1-3p | STARD13 | PAR-CLIP | miRTarbase |
| hsa-miR-219a-1-3p | STARD13 | PAR-CLIP | miRTarbase |
| hsa-miR-15a-5p | SRPR | PAR-CLIP | miRTarbase |
| hsa-miR-15a-5p | SRPR | HITS-CLIP | miRTarbase |
| hsa-miR-15a-5p | SRPR | HITS-CLIP | miRTarbase |
| hsa-miR-15a-5p | SRPR | PAR-CLIP | miRTarbase |
| hsa-miR-195-5p | SRPR | PAR-CLIP | miRTarbase |
| hsa-miR-195-5p | SRPR | HITS-CLIP | miRTarbase |
| hsa-miR-195-5p | SRPR | HITS-CLIP | miRTarbase |
| hsa-miR-195-5p | SRPR | PAR-CLIP | miRTarbase |
| hsa-miR-195-5p | SREK1 | PAR-CLIP | miRTarbase |
| hsa-miR-195-5p | SREK1 | PAR-CLIP//HITS-CLIP | miRTarbase |
| hsa-miR-195-5p | SREK1 | PAR-CLIP | miRTarbase |
| hsa-miR-195-5p | SREK1 | PAR-CLIP | miRTarbase |
| hsa-miR-15a-5p | SREK1 | PAR-CLIP | miRTarbase |
| hsa-miR-15a-5p | SREK1 | PAR-CLIP//HITS-CLIP | miRTarbase |
| hsa-miR-15a-5p | SREK1 | PAR-CLIP | miRTarbase |
| hsa-miR-15a-5p | SREK1 | PAR-CLIP | miRTarbase |
| hsa-miR-3613-3p | SPTSSA | PAR-CLIP | miRTarbase |
| hsa-miR-3613-3p | SMEK1 | PAR-CLIP | miRTarbase |
| hsa-miR-195-5p | SMAD7 | PAR-CLIP | miRTarbase |
| hsa-miR-195-5p | SMAD7 | PAR-CLIP | miRTarbase |
| hsa-miR-195-5p | SMAD7 | PAR-CLIP | miRTarbase |
| hsa-miR-195-5p | SMAD7 | PAR-CLIP | miRTarbase |
| hsa-miR-15a-5p | SMAD7 | PAR-CLIP | miRTarbase |
| hsa-miR-15a-5p | SMAD7 | PAR-CLIP | miRTarbase |
| hsa-miR-15a-5p | SMAD7 | PAR-CLIP | miRTarbase |
| hsa-miR-15a-5p | SMAD7 | PAR-CLIP | miRTarbase |
| hsa-miR-195-5p | PRICKLE2 | PAR-CLIP | miRTarbase |
| hsa-miR-15a-5p | PRICKLE2 | PAR-CLIP | miRTarbase |
| hsa-miR-486-3p | NSD1 | PAR-CLIP | miRTarbase |
| hsa-miR-195-5p | LRIG2 | PAR-CLIP | miRTarbase |
| hsa-miR-15a-5p | LRIG2 | PAR-CLIP | miRTarbase |
| hsa-miR-195-5p | KIF5B | PAR-CLIP | miRTarbase |
| hsa-miR-195-5p | KIF5B | PAR-CLIP | miRTarbase |
| hsa-miR-195-5p | KIF5B | HITS-CLIP | miRTarbase |
| hsa-miR-195-5p | KIF5B | PAR-CLIP | miRTarbase |
| hsa-miR-195-5p | KIF5B | PAR-CLIP | miRTarbase |
| hsa-miR-15a-5p | KIF5B | PAR-CLIP | miRTarbase |
| hsa-miR-15a-5p | KIF5B | PAR-CLIP | miRTarbase |
| hsa-miR-15a-5p | KIF5B | HITS-CLIP | miRTarbase |
| hsa-miR-15a-5p | KIF5B | PAR-CLIP | miRTarbase |
| hsa-miR-15a-5p | KIF5B | PAR-CLIP | miRTarbase |
| hsa-miR-141-3p | HNRNPF | PAR-CLIP | miRTarbase |
| hsa-miR-141-3p | HNRNPF | PAR-CLIP | miRTarbase |
| hsa-miR-195-3p | GRPEL2 | PAR-CLIP | miRTarbase |
| hsa-miR-195-3p | GRPEL2 | PAR-CLIP | miRTarbase |
| hsa-miR-195-3p | GRPEL2 | PAR-CLIP | miRTarbase |
| hsa-miR-195-3p | GRPEL2 | PAR-CLIP | miRTarbase |
| hsa-miR-141-3p | GNA13 | PAR-CLIP | miRTarbase |
| hsa-miR-141-3p | FUT11 | PAR-CLIP | miRTarbase |
| hsa-miR-141-3p | FUT11 | PAR-CLIP | miRTarbase |
| hsa-miR-15a-5p | FAM122B | PAR-CLIP | miRTarbase |
| hsa-miR-15a-5p | FAM122B | PAR-CLIP | miRTarbase |
| hsa-miR-15a-5p | FAM122B | PAR-CLIP | miRTarbase |
| hsa-miR-15a-5p | FAM122B | PAR-CLIP | miRTarbase |
| hsa-miR-195-5p | FAM122B | PAR-CLIP | miRTarbase |
| hsa-miR-195-5p | FAM122B | PAR-CLIP | miRTarbase |
| hsa-miR-195-5p | FAM122B | PAR-CLIP | miRTarbase |
| hsa-miR-195-5p | FAM122B | PAR-CLIP | miRTarbase |
| hsa-miR-141-3p | EPHA2 | PAR-CLIP | miRTarbase |
| hsa-miR-141-3p | EPHA2 | PAR-CLIP | miRTarbase |
| hsa-miR-195-5p | E2F7 | PAR-CLIP | miRTarbase |
| hsa-miR-195-5p | E2F7 | PAR-CLIP | miRTarbase |
| hsa-miR-195-5p | E2F7 | PAR-CLIP | miRTarbase |
| hsa-miR-195-5p | E2F7 | HITS-CLIP | miRTarbase |
| hsa-miR-195-5p | E2F7 | PAR-CLIP | miRTarbase |
| hsa-miR-195-5p | E2F7 | PAR-CLIP | miRTarbase |
| hsa-miR-15a-5p | E2F7 | PAR-CLIP | miRTarbase |
| hsa-miR-15a-5p | E2F7 | PAR-CLIP | miRTarbase |
| hsa-miR-15a-5p | E2F7 | PAR-CLIP | miRTarbase |
| hsa-miR-15a-5p | E2F7 | HITS-CLIP | miRTarbase |
| hsa-miR-15a-5p | E2F7 | PAR-CLIP | miRTarbase |
| hsa-miR-15a-5p | E2F7 | PAR-CLIP | miRTarbase |
| hsa-miR-15a-5p | DDX3X | PAR-CLIP | miRTarbase |
| hsa-miR-15a-5p | DDX3X | PAR-CLIP | miRTarbase |
| hsa-miR-15a-5p | DDX3X | PAR-CLIP | miRTarbase |
| hsa-miR-15a-5p | DDX3X | PAR-CLIP | miRTarbase |
| hsa-miR-195-5p | DDX3X | PAR-CLIP | miRTarbase |
| hsa-miR-195-5p | DDX3X | PAR-CLIP | miRTarbase |
| hsa-miR-195-5p | DDX3X | PAR-CLIP | miRTarbase |
| hsa-miR-195-5p | DDX3X | PAR-CLIP | miRTarbase |
| hsa-miR-219a-1-3p | CELSR3 | PAR-CLIP | miRTarbase |
| hsa-miR-219a-1-3p | CELSR3 | PAR-CLIP | miRTarbase |
| hsa-miR-219a-1-3p | CELSR3 | PAR-CLIP | miRTarbase |
| hsa-miR-195-5p | CDCA4 | PAR-CLIP | miRTarbase |
| hsa-miR-195-5p | CDCA4 | PAR-CLIP | miRTarbase |
| hsa-miR-195-5p | CDCA4 | HITS-CLIP | miRTarbase |
| hsa-miR-195-5p | CDCA4 | PAR-CLIP | miRTarbase |
| hsa-miR-195-5p | CDCA4 | PAR-CLIP | miRTarbase |
| hsa-miR-15a-5p | CDCA4 | PAR-CLIP | miRTarbase |
| hsa-miR-15a-5p | CDCA4 | PAR-CLIP | miRTarbase |
| hsa-miR-15a-5p | CDCA4 | HITS-CLIP | miRTarbase |
| hsa-miR-15a-5p | CDCA4 | PAR-CLIP | miRTarbase |
| hsa-miR-15a-5p | CDCA4 | PAR-CLIP | miRTarbase |
| hsa-miR-195-5p | CDC37L1 | PAR-CLIP | miRTarbase |
| hsa-miR-195-5p | CDC37L1 | PAR-CLIP//HITS-CLIP | miRTarbase |
| hsa-miR-195-5p | CDC37L1 | HITS-CLIP | miRTarbase |
| hsa-miR-195-5p | CDC37L1 | PAR-CLIP | miRTarbase |
| hsa-miR-195-5p | CDC37L1 | PAR-CLIP | miRTarbase |
| hsa-miR-15a-5p | CDC37L1 | PAR-CLIP | miRTarbase |
| hsa-miR-15a-5p | CDC37L1 | PAR-CLIP//HITS-CLIP | miRTarbase |
| hsa-miR-15a-5p | CDC37L1 | HITS-CLIP | miRTarbase |
| hsa-miR-15a-5p | CDC37L1 | PAR-CLIP | miRTarbase |
| hsa-miR-15a-5p | CDC37L1 | PAR-CLIP | miRTarbase |
| hsa-miR-3613-3p | CCNL1 | PAR-CLIP | miRTarbase |
| hsa-miR-3613-3p | CCNL1 | PAR-CLIP | miRTarbase |
| hsa-miR-3613-3p | CCNL1 | PAR-CLIP | miRTarbase |
| hsa-miR-3613-3p | CCNL1 | PAR-CLIP | miRTarbase |
| hsa-miR-15a-5p | ATG9A | PAR-CLIP | miRTarbase |
| hsa-miR-15a-5p | ATG9A | PAR-CLIP | miRTarbase |
| hsa-miR-15a-5p | ATG9A | HITS-CLIP | miRTarbase |
| hsa-miR-15a-5p | ATG9A | PAR-CLIP | miRTarbase |
| hsa-miR-195-5p | ATG9A | PAR-CLIP | miRTarbase |
| hsa-miR-195-5p | ATG9A | PAR-CLIP | miRTarbase |
| hsa-miR-195-5p | ATG9A | HITS-CLIP | miRTarbase |
| hsa-miR-195-5p | ATG9A | PAR-CLIP | miRTarbase |
| hsa-miR-3613-3p | AGO2 | PAR-CLIP | miRTarbase |
| hsa-miR-3613-3p | AGO2 | PAR-CLIP | miRTarbase |
| hsa-miR-195-5p | ASGR2 | PAR-CLIP | miRTarbase |
| hsa-miR-195-5p | ASGR2 | PAR-CLIP | miRTarbase |
| hsa-miR-195-5p | ASGR2 | PAR-CLIP | miRTarbase |
| hsa-miR-15a-5p | ASGR2 | PAR-CLIP | miRTarbase |
| hsa-miR-15a-5p | ASGR2 | PAR-CLIP | miRTarbase |
| hsa-miR-15a-5p | ASGR2 | PAR-CLIP | miRTarbase |
| hsa-miR-195-5p | ZNF620 | PAR-CLIP | miRTarbase |
| hsa-miR-195-5p | ZNF620 | PAR-CLIP | miRTarbase |
| hsa-miR-195-5p | ZNF620 | PAR-CLIP | miRTarbase |
| hsa-miR-15a-5p | ZNF620 | PAR-CLIP | miRTarbase |
| hsa-miR-15a-5p | ZNF620 | PAR-CLIP | miRTarbase |

| | | | |
|---|---|---|---|
| hsa-miR-15a-5p | ZNF620 | PAR-CLIP | miRTarbase |
| hsa-miR-3605-5p | ZNF117 | PAR-CLIP | miRTarbase |
| hsa-miR-195-5p | HAUS3 | PAR-CLIP | miRTarbase |
| hsa-miR-195-5p | HAUS3 | PAR-CLIP//HITS-CLIP | miRTarbase |
| hsa-miR-195-5p | HAUS3 | PAR-CLIP | miRTarbase |
| hsa-miR-15a-5p | HAUS3 | PAR-CLIP | miRTarbase |
| hsa-miR-15a-5p | HAUS3 | PAR-CLIP//HITS-CLIP | miRTarbase |
| hsa-miR-15a-5p | HAUS3 | PAR-CLIP | miRTarbase |
| hsa-miR-3613-3p | CD38 | PAR-CLIP | miRTarbase |
| hsa-miR-3613-3p | CD38 | PAR-CLIP | miRTarbase |
| hsa-miR-486-5p | ZNF460 | PAR-CLIP | miRTarbase |
| hsa-miR-486-5p | ZNF460 | PAR-CLIP | miRTarbase |
| hsa-miR-3613-3p | ZNF226 | PAR-CLIP | miRTarbase |
| hsa-miR-3613-3p | ZNF226 | PAR-CLIP | miRTarbase |
| hsa-miR-486-5p | ZIC5 | PAR-CLIP | miRTarbase |
| hsa-miR-486-5p | ZIC5 | PAR-CLIP | miRTarbase |
| hsa-miR-486-5p | ZIC5 | PAR-CLIP | miRTarbase |
| hsa-miR-15a-5p | YTHDC1 | PAR-CLIP | miRTarbase |
| hsa-miR-15a-5p | YTHDC1 | PAR-CLIP | miRTarbase |
| hsa-miR-15a-5p | YTHDC1 | PAR-CLIP | miRTarbase |
| hsa-miR-195-5p | YTHDC1 | PAR-CLIP | miRTarbase |
| hsa-miR-195-5p | YTHDC1 | PAR-CLIP | miRTarbase |
| hsa-miR-195-5p | YTHDC1 | PAR-CLIP | miRTarbase |
| hsa-miR-141-3p | TMOD3 | PAR-CLIP | miRTarbase |
| hsa-miR-141-3p | TMOD3 | PAR-CLIP | miRTarbase |
| hsa-miR-195-5p | TMEM245 | PAR-CLIP | miRTarbase |
| hsa-miR-195-5p | TMEM245 | PAR-CLIP//HITS-CLIP | miRTarbase |
| hsa-miR-195-5p | TMEM245 | PAR-CLIP | miRTarbase |
| hsa-miR-15a-5p | TMEM245 | PAR-CLIP | miRTarbase |
| hsa-miR-15a-5p | TMEM245 | PAR-CLIP//HITS-CLIP | miRTarbase |
| hsa-miR-15a-5p | TMEM245 | PAR-CLIP | miRTarbase |
| hsa-miR-195-5p | TMEM100 | PAR-CLIP | miRTarbase |
| hsa-miR-15a-5p | TMEM100 | PAR-CLIP | miRTarbase |
| hsa-miR-195-5p | SRSF1 | PAR-CLIP | miRTarbase |
| hsa-miR-195-5p | SRSF1 | PAR-CLIP | miRTarbase |
| hsa-miR-195-5p | SRSF1 | PAR-CLIP | miRTarbase |
| hsa-miR-15a-5p | SRSF1 | PAR-CLIP | miRTarbase |
| hsa-miR-15a-5p | SRSF1 | PAR-CLIP | miRTarbase |
| hsa-miR-15a-5p | SRSF1 | PAR-CLIP | miRTarbase |
| hsa-miR-15a-5p | SNX16 | PAR-CLIP | miRTarbase |
| hsa-miR-15a-5p | SNX16 | PAR-CLIP | miRTarbase |
| hsa-miR-15a-5p | SNX16 | PAR-CLIP | miRTarbase |
| hsa-miR-195-5p | SESTD1 | PAR-CLIP | miRTarbase |
| hsa-miR-195-5p | SESTD1 | PAR-CLIP | miRTarbase |
| hsa-miR-195-5p | SESTD1 | PAR-CLIP | miRTarbase |
| hsa-miR-15a-5p | SESTD1 | PAR-CLIP | miRTarbase |
| hsa-miR-15a-5p | SESTD1 | PAR-CLIP | miRTarbase |
| hsa-miR-15a-5p | SESTD1 | PAR-CLIP | miRTarbase |
| hsa-miR-141-5p | RNF219 | PAR-CLIP | miRTarbase |
| hsa-miR-195-5p | RIMS3 | PAR-CLIP | miRTarbase |
| hsa-miR-195-5p | RIMS3 | PAR-CLIP | miRTarbase |
| hsa-miR-195-5p | RIMS3 | PAR-CLIP | miRTarbase |
| hsa-miR-15a-5p | RIMS3 | PAR-CLIP | miRTarbase |
| hsa-miR-15a-5p | RIMS3 | PAR-CLIP | miRTarbase |
| hsa-miR-15a-5p | RIMS3 | PAR-CLIP | miRTarbase |
| hsa-miR-195-5p | RCAN3 | PAR-CLIP | miRTarbase |
| hsa-miR-195-5p | RCAN3 | PAR-CLIP | miRTarbase |
| hsa-miR-15a-5p | RCAN3 | PAR-CLIP | miRTarbase |
| hsa-miR-15a-5p | RCAN3 | PAR-CLIP | miRTarbase |
| hsa-miR-195-5p | PPIG | PAR-CLIP | miRTarbase |
| hsa-miR-195-5p | PPIG | PAR-CLIP | miRTarbase |
| hsa-miR-195-5p | PPIG | PAR-CLIP | miRTarbase |
| hsa-miR-15a-5p | PPIG | PAR-CLIP | miRTarbase |
| hsa-miR-15a-5p | PPIG | PAR-CLIP | miRTarbase |
| hsa-miR-15a-5p | PPIG | PAR-CLIP | miRTarbase |
| hsa-miR-195-5p | PLRG1 | PAR-CLIP | miRTarbase |
| hsa-miR-195-5p | PLRG1 | PAR-CLIP | miRTarbase |
| hsa-miR-15a-5p | PLRG1 | PAR-CLIP | miRTarbase |
| hsa-miR-15a-5p | PLRG1 | PAR-CLIP | miRTarbase |
| hsa-miR-15a-5p | PLAG1 | PAR-CLIP | miRTarbase |
| hsa-miR-15a-5p | PLAG1 | PAR-CLIP | miRTarbase |
| hsa-miR-15a-5p | PLAG1 | PAR-CLIP | miRTarbase |
| hsa-miR-15a-5p | PLAG1 | HITS-CLIP | miRTarbase |
| hsa-miR-15a-5p | PLAG1 | PAR-CLIP | miRTarbase |
| hsa-miR-195-5p | PLAG1 | PAR-CLIP | miRTarbase |
| hsa-miR-195-5p | PLAG1 | PAR-CLIP | miRTarbase |
| hsa-miR-195-5p | PLAG1 | PAR-CLIP | miRTarbase |
| hsa-miR-195-5p | PLAG1 | HITS-CLIP | miRTarbase |
| hsa-miR-195-5p | PLAG1 | PAR-CLIP | miRTarbase |
| hsa-miR-195-5p | PHKA1 | PAR-CLIP | miRTarbase |
| hsa-miR-195-5p | PHKA1 | PAR-CLIP//HITS-CLIP | miRTarbase |
| hsa-miR-195-5p | PHKA1 | PAR-CLIP | miRTarbase |
| hsa-miR-15a-5p | PHKA1 | PAR-CLIP | miRTarbase |
| hsa-miR-15a-5p | PHKA1 | PAR-CLIP//HITS-CLIP | miRTarbase |
| hsa-miR-15a-5p | PHKA1 | PAR-CLIP | miRTarbase |
| hsa-miR-486-3p | NACC2 | PAR-CLIP | miRTarbase |
| hsa-miR-486-3p | NACC2 | PAR-CLIP | miRTarbase |
| hsa-miR-15a-5p | MYO5A | PAR-CLIP | miRTarbase |
| hsa-miR-15a-5p | MYO5A | PAR-CLIP | miRTarbase |
| hsa-miR-15a-5p | MYO5A | HITS-CLIP | miRTarbase |
| hsa-miR-15a-5p | MYO5A | PAR-CLIP | miRTarbase |
| hsa-miR-195-5p | MYO5A | PAR-CLIP | miRTarbase |
| hsa-miR-195-5p | MYO5A | PAR-CLIP | miRTarbase |
| hsa-miR-195-5p | MYO5A | HITS-CLIP | miRTarbase |
| hsa-miR-195-5p | MYO5A | PAR-CLIP | miRTarbase |
| hsa-miR-141-3p | MARCH6 | PAR-CLIP | miRTarbase |
| hsa-miR-141-3p | MARCH6 | PAR-CLIP | miRTarbase |
| hsa-miR-141-3p | MARCH6 | PAR-CLIP | miRTarbase |
| hsa-miR-141-3p | MARCH6 | PAR-CLIP | miRTarbase |
| hsa-miR-15a-5p | KIF23 | PAR-CLIP | miRTarbase |
| hsa-miR-15a-5p | KIF23 | PAR-CLIP | miRTarbase |
| hsa-miR-15a-5p | KIF23 | HITS-CLIP | miRTarbase |
| hsa-miR-15a-5p | KIF23 | PAR-CLIP | miRTarbase |
| hsa-miR-195-5p | KIF23 | PAR-CLIP | miRTarbase |
| hsa-miR-195-5p | KIF23 | PAR-CLIP | miRTarbase |
| hsa-miR-195-5p | KIF23 | HITS-CLIP | miRTarbase |
| hsa-miR-195-5p | KIF23 | PAR-CLIP | miRTarbase |
| hsa-miR-141-3p | KIAA1147 | PAR-CLIP | miRTarbase |
| hsa-miR-3613-3p | HNRNPDL | PAR-CLIP | miRTarbase |
| hsa-miR-3613-3p | HNRNPDL | PAR-CLIP | miRTarbase |
| hsa-miR-3613-3p | HNRNPDL | PAR-CLIP | miRTarbase |
| hsa-miR-195-5p | HNRNPDL | PAR-CLIP | miRTarbase |
| hsa-miR-195-5p | HNRNPDL | PAR-CLIP | miRTarbase |
| hsa-miR-195-5p | HNRNPDL | PAR-CLIP | miRTarbase |
| hsa-miR-15a-5p | HNRNPDL | PAR-CLIP | miRTarbase |
| hsa-miR-15a-5p | HNRNPDL | PAR-CLIP | miRTarbase |
| hsa-miR-15a-5p | HNRNPDL | PAR-CLIP | miRTarbase |
| hsa-miR-219a-1-3p | H2AFX | PAR-CLIP | miRTarbase |
| hsa-miR-219a-1-3p | H2AFX | PAR-CLIP | miRTarbase |
| hsa-miR-3605-5p | CDKN1B | PAR-CLIP | miRTarbase |
| hsa-miR-15a-5p | CDK1 | PAR-CLIP | miRTarbase |
| hsa-miR-15a-5p | CDK1 | PAR-CLIP | miRTarbase |
| hsa-miR-15a-5p | CDK1 | PAR-CLIP | miRTarbase |
| hsa-miR-195-5p | CDK1 | PAR-CLIP | miRTarbase |
| hsa-miR-195-5p | CDK1 | PAR-CLIP | miRTarbase |
| hsa-miR-195-5p | CDK1 | PAR-CLIP | miRTarbase |
| hsa-miR-195-3p | CCNT1 | PAR-CLIP | miRTarbase |
| hsa-miR-195-5p | CCNE2 | PAR-CLIP | miRTarbase |
| hsa-miR-195-5p | CCNE2 | PAR-CLIP | miRTarbase |
| hsa-miR-195-5p | CCNE2 | PAR-CLIP | miRTarbase |
| hsa-miR-15a-5p | CCNE2 | PAR-CLIP | miRTarbase |
| hsa-miR-15a-5p | CCNE2 | PAR-CLIP | miRTarbase |
| hsa-miR-15a-5p | CCNE2 | PAR-CLIP | miRTarbase |
| hsa-miR-15a-5p | CBX6 | PAR-CLIP | miRTarbase |
| hsa-miR-195-5p | CBX6 | PAR-CLIP | miRTarbase |
| hsa-miR-219a-1-3p | BTG3 | PAR-CLIP | miRTarbase |
| hsa-miR-219a-1-3p | BTG3 | PAR-CLIP | miRTarbase |
| hsa-miR-219a-1-3p | BTG3 | PAR-CLIP | miRTarbase |
| hsa-miR-486-5p | BTF3L4 | PAR-CLIP | miRTarbase |
| hsa-miR-486-5p | BTF3L4 | PAR-CLIP | miRTarbase |
| hsa-miR-486-5p | BTF3L4 | PAR-CLIP | miRTarbase |
| hsa-miR-195-5p | AXIN2 | PAR-CLIP | miRTarbase |
| hsa-miR-195-5p | AXIN2 | PAR-CLIP//HITS-CLIP | miRTarbase |
| hsa-miR-195-5p | AXIN2 | PAR-CLIP | miRTarbase |
| hsa-miR-15a-5p | AXIN2 | PAR-CLIP | miRTarbase |
| hsa-miR-15a-5p | AXIN2 | PAR-CLIP//HITS-CLIP | miRTarbase |
| hsa-miR-15a-5p | AXIN2 | PAR-CLIP | miRTarbase |
| hsa-miR-195-3p | MTRNR2L6 | HITS-CLIP | miRTarbase |
| hsa-miR-195-3p | MTRNR2L6 | PAR-CLIP | miRTarbase |

| | | | |
|---|---|---|---|
| hsa-miR-195-5p | CASK | HITS-CLIP | miRTarbase |
| hsa-miR-195-5p | CASK | PAR-CLIP | miRTarbase |
| hsa-miR-15a-5p | CASK | HITS-CLIP | miRTarbase |
| hsa-miR-15a-5p | CASK | PAR-CLIP | miRTarbase |
| hsa-miR-3605-5p | PTPLB | PAR-CLIP | miRTarbase |
| hsa-miR-15a-3p | SNRPB | PAR-CLIP | miRTarbase |
| hsa-miR-15a-3p | SNRPB | PAR-CLIP | miRTarbase |
| hsa-miR-195-5p | DMPK | PAR-CLIP | miRTarbase |
| hsa-miR-195-5p | DMPK | HITS-CLIP | miRTarbase |
| hsa-miR-195-5p | DMPK | PAR-CLIP | miRTarbase |
| hsa-miR-195-5p | DMPK | HITS-CLIP | miRTarbase |
| hsa-miR-195-5p | DMPK | HITS-CLIP | miRTarbase |
| hsa-miR-15a-5p | DMPK | PAR-CLIP | miRTarbase |
| hsa-miR-15a-5p | DMPK | HITS-CLIP | miRTarbase |
| hsa-miR-15a-5p | DMPK | PAR-CLIP | miRTarbase |
| hsa-miR-15a-5p | DMPK | HITS-CLIP | miRTarbase |
| hsa-miR-15a-5p | DMPK | HITS-CLIP | miRTarbase |
| hsa-miR-3613-3p | GALK2 | HITS-CLIP | miRTarbase |
| hsa-miR-3613-3p | GALK2 | PAR-CLIP | miRTarbase |
| hsa-miR-3613-3p | GALK2 | HITS-CLIP | miRTarbase |
| hsa-miR-3613-3p | GALK2 | PAR-CLIP | miRTarbase |
| hsa-miR-195-5p | ATAD5 | PAR-CLIP | miRTarbase |
| hsa-miR-195-5p | ATAD5 | PAR-CLIP | miRTarbase |
| hsa-miR-15a-5p | ATAD5 | PAR-CLIP | miRTarbase |
| hsa-miR-15a-5p | ATAD5 | PAR-CLIP | miRTarbase |
| hsa-miR-3613-3p | NIPAL1 | PAR-CLIP | miRTarbase |
| hsa-miR-195-5p | AKR1B10 | PAR-CLIP | miRTarbase |
| hsa-miR-195-5p | AKR1B10 | PAR-CLIP | miRTarbase |
| hsa-miR-15a-5p | AKR1B10 | PAR-CLIP | miRTarbase |
| hsa-miR-15a-5p | AKR1B10 | PAR-CLIP | miRTarbase |
| hsa-miR-3613-3p | PLEKHG3 | PAR-CLIP | miRTarbase |
| hsa-miR-3613-3p | PLEKHG3 | PAR-CLIP//HITS-CLIP | miRTarbase |
| hsa-miR-3613-3p | PLEKHG3 | PAR-CLIP | miRTarbase |
| hsa-miR-3613-3p | PLEKHG3 | HITS-CLIP | miRTarbase |
| hsa-miR-486-3p | MED28 | PAR-CLIP | miRTarbase |
| hsa-miR-3613-3p | ZNFX1 | PAR-CLIP | miRTarbase |
| hsa-miR-3613-3p | ZNFX1 | PAR-CLIP | miRTarbase |
| hsa-miR-195-3p | UBE2D3 | PAR-CLIP | miRTarbase |
| hsa-miR-195-3p | UBE2D3 | PAR-CLIP | miRTarbase |
| hsa-miR-195-3p | UBE2D3 | PAR-CLIP | miRTarbase |
| hsa-miR-3605-5p | TMED7 | PAR-CLIP | miRTarbase |
| hsa-miR-3605-5p | TMED7 | PAR-CLIP | miRTarbase |
| hsa-miR-3613-3p | RAN | PAR-CLIP | miRTarbase |
| hsa-miR-3613-3p | RAN | PAR-CLIP//HITS-CLIP | miRTarbase |
| hsa-miR-3613-3p | RAN | PAR-CLIP | miRTarbase |
| hsa-miR-3613-3p | RAN | PAR-CLIP | miRTarbase |
| hsa-miR-195-5p | GPATCH8 | PAR-CLIP | miRTarbase |
| hsa-miR-195-5p | GPATCH8 | HITS-CLIP | miRTarbase |
| hsa-miR-195-5p | GPATCH8 | PAR-CLIP | miRTarbase |
| hsa-miR-15a-5p | GPATCH8 | PAR-CLIP | miRTarbase |
| hsa-miR-15a-5p | GPATCH8 | HITS-CLIP | miRTarbase |
| hsa-miR-15a-5p | GPATCH8 | PAR-CLIP | miRTarbase |
| hsa-miR-195-5p | ARHGDIA | PAR-CLIP | miRTarbase |
| hsa-miR-195-5p | ARHGDIA | PAR-CLIP | miRTarbase |
| hsa-miR-195-5p | ARHGDIA | HITS-CLIP | miRTarbase |
| hsa-miR-195-5p | ARHGDIA | PAR-CLIP | miRTarbase |
| hsa-miR-15a-5p | ARHGDIA | PAR-CLIP | miRTarbase |
| hsa-miR-15a-5p | ARHGDIA | PAR-CLIP | miRTarbase |
| hsa-miR-15a-5p | ARHGDIA | HITS-CLIP | miRTarbase |
| hsa-miR-15a-5p | ARHGDIA | PAR-CLIP | miRTarbase |
| hsa-miR-195-5p | CPEB3 | PAR-CLIP | miRTarbase |
| hsa-miR-195-5p | CPEB3 | HITS-CLIP | miRTarbase |
| hsa-miR-195-5p | CPEB3 | PAR-CLIP | miRTarbase |
| hsa-miR-15a-5p | CPEB3 | PAR-CLIP | miRTarbase |
| hsa-miR-15a-5p | CPEB3 | HITS-CLIP | miRTarbase |
| hsa-miR-15a-5p | CPEB3 | PAR-CLIP | miRTarbase |
| hsa-miR-219a-1-3p | GLRX | PAR-CLIP | miRTarbase |
| hsa-miR-3613-3p | PRKCI | PAR-CLIP | miRTarbase |
| hsa-miR-486-3p | EMP1 | PAR-CLIP | miRTarbase |
| hsa-miR-486-3p | EMP1 | PAR-CLIP | miRTarbase |
| hsa-miR-486-3p | EMP1 | HITS-CLIP | miRTarbase |
| hsa-miR-15a-5p | JARID2 | PAR-CLIP | miRTarbase |
| hsa-miR-15a-5p | JARID2 | PAR-CLIP | miRTarbase |
| hsa-miR-15a-5p | JARID2 | PAR-CLIP | miRTarbase |
| hsa-miR-15a-5p | JARID2 | PAR-CLIP | miRTarbase |
| hsa-miR-195-5p | JARID2 | PAR-CLIP | miRTarbase |
| hsa-miR-195-5p | JARID2 | PAR-CLIP | miRTarbase |
| hsa-miR-195-5p | JARID2 | PAR-CLIP | miRTarbase |
| hsa-miR-195-5p | JARID2 | PAR-CLIP | miRTarbase |
| hsa-miR-3613-3p | GRB10 | PAR-CLIP | miRTarbase |
| hsa-miR-3613-3p | GRB10 | PAR-CLIP | miRTarbase |
| hsa-miR-3613-3p | GRB10 | PAR-CLIP | miRTarbase |
| hsa-miR-15a-5p | CAMSAP1 | PAR-CLIP | miRTarbase |
| hsa-miR-15a-5p | CAMSAP1 | HITS-CLIP | miRTarbase |
| hsa-miR-15a-5p | CAMSAP1 | PAR-CLIP | miRTarbase |
| hsa-miR-195-5p | CAMSAP1 | PAR-CLIP | miRTarbase |
| hsa-miR-195-5p | CAMSAP1 | HITS-CLIP | miRTarbase |
| hsa-miR-195-5p | CAMSAP1 | PAR-CLIP | miRTarbase |
| hsa-miR-15a-5p | ATG14 | PAR-CLIP | miRTarbase |
| hsa-miR-486-3p | UBBP4 | PAR-CLIP | miRTarbase |
| hsa-miR-486-3p | UBBP4 | PAR-CLIP | miRTarbase |
| hsa-miR-486-3p | UBBP4 | PAR-CLIP | miRTarbase |
| hsa-miR-3605-5p | SHISA9 | PAR-CLIP | miRTarbase |
| hsa-miR-3605-5p | SHISA9 | PAR-CLIP | miRTarbase |
| hsa-miR-3613-3p | SRPX2 | PAR-CLIP | miRTarbase |
| hsa-miR-3613-3p | SRPX2 | PAR-CLIP | miRTarbase |
| hsa-miR-3605-5p | RAB32 | PAR-CLIP | miRTarbase |
| hsa-miR-3605-5p | RAB32 | PAR-CLIP | miRTarbase |
| hsa-miR-486-3p | SYTL3 | HITS-CLIP | miRTarbase |
| hsa-miR-486-3p | SYTL3 | PAR-CLIP | miRTarbase |
| hsa-miR-486-3p | TUBB8 | PAR-CLIP | miRTarbase |
| hsa-miR-195-3p | RPS4X | PAR-CLIP | miRTarbase |
| hsa-miR-195-3p | RPS4X | PAR-CLIP | miRTarbase |
| hsa-miR-195-5p | TRIM35 | PAR-CLIP | miRTarbase |
| hsa-miR-15a-5p | TRIM35 | PAR-CLIP | miRTarbase |
| hsa-miR-195-5p | FLCN | PAR-CLIP | miRTarbase |
| hsa-miR-195-5p | FLCN | PAR-CLIP | miRTarbase |
| hsa-miR-195-5p | FLCN | PAR-CLIP | miRTarbase |
| hsa-miR-15a-5p | FLCN | PAR-CLIP | miRTarbase |
| hsa-miR-15a-5p | FLCN | PAR-CLIP | miRTarbase |
| hsa-miR-15a-5p | FLCN | PAR-CLIP | miRTarbase |
| hsa-miR-219a-1-3p | ZNF117 | PAR-CLIP | miRTarbase |
| hsa-miR-195-5p | NNT | PAR-CLIP | miRTarbase |
| hsa-miR-195-5p | NNT | PAR-CLIP | miRTarbase |
| hsa-miR-15a-5p | NNT | PAR-CLIP | miRTarbase |
| hsa-miR-15a-5p | NNT | PAR-CLIP | miRTarbase |
| hsa-miR-3605-3p | FKBP14 | PAR-CLIP | miRTarbase |
| hsa-miR-141-3p | ZNF805 | PAR-CLIP | miRTarbase |
| hsa-miR-141-3p | ZNF805 | PAR-CLIP | miRTarbase |
| hsa-miR-141-5p | ZNF354B | PAR-CLIP | miRTarbase |
| hsa-miR-141-5p | ZNF354B | PAR-CLIP | miRTarbase |
| hsa-miR-141-5p | ZNF354B | PAR-CLIP | miRTarbase |
| hsa-miR-141-5p | ZNF354B | HITS-CLIP | miRTarbase |
| hsa-miR-141-3p | ZMAT3 | PAR-CLIP | miRTarbase |
| hsa-miR-141-3p | ZMAT3 | PAR-CLIP | miRTarbase |
| hsa-miR-3613-3p | ZER1 | PAR-CLIP | miRTarbase |
| hsa-miR-3613-3p | TMF1 | PAR-CLIP | miRTarbase |
| hsa-miR-15a-5p | SLC2A3 | PAR-CLIP | miRTarbase |
| hsa-miR-15a-5p | SLC2A3 | PAR-CLIP | miRTarbase |
| hsa-miR-195-5p | SBNO1 | PAR-CLIP | miRTarbase |
| hsa-miR-195-5p | SBNO1 | PAR-CLIP | miRTarbase |
| hsa-miR-195-5p | SBNO1 | PAR-CLIP | miRTarbase |
| hsa-miR-15a-5p | SBNO1 | PAR-CLIP | miRTarbase |
| hsa-miR-15a-5p | SBNO1 | PAR-CLIP | miRTarbase |
| hsa-miR-15a-5p | SBNO1 | PAR-CLIP | miRTarbase |
| hsa-miR-219a-1-3p | RAD51 | PAR-CLIP | miRTarbase |
| hsa-miR-195-3p | PRKAA1 | PAR-CLIP | miRTarbase |
| hsa-miR-195-3p | PRKAA1 | PAR-CLIP//HITS-CLIP | miRTarbase |
| hsa-miR-195-3p | PRKAA1 | PAR-CLIP | miRTarbase |
| hsa-miR-195-3p | PRKAA1 | PAR-CLIP | miRTarbase |
| hsa-miR-195-5p | POM121C | PAR-CLIP | miRTarbase |
| hsa-miR-15a-5p | POM121C | PAR-CLIP | miRTarbase |
| hsa-miR-141-5p | NUP50 | PAR-CLIP | miRTarbase |
| hsa-miR-141-5p | NUP50 | PAR-CLIP | miRTarbase |
| hsa-miR-15a-5p | NUFIP2 | PAR-CLIP | miRTarbase |
| hsa-miR-15a-5p | NUFIP2 | HITS-CLIP | miRTarbase |
| hsa-miR-15a-5p | NUFIP2 | PAR-CLIP | miRTarbase |
| hsa-miR-195-5p | NUFIP2 | PAR-CLIP | miRTarbase |
| hsa-miR-195-5p | NUFIP2 | HITS-CLIP | miRTarbase |

| | | | |
|---|---|---|---|
| hsa-miR-195-5p | NUFIP2 | PAR-CLIP | miRTarbase |
| hsa-miR-195-5p | LAMP2 | PAR-CLIP | miRTarbase |
| hsa-miR-195-5p | LAMP2 | PAR-CLIP | miRTarbase |
| hsa-miR-195-5p | LAMP2 | PAR-CLIP | miRTarbase |
| hsa-miR-15a-5p | LAMP2 | PAR-CLIP | miRTarbase |
| hsa-miR-15a-5p | LAMP2 | PAR-CLIP | miRTarbase |
| hsa-miR-15a-5p | LAMP2 | PAR-CLIP | miRTarbase |
| hsa-miR-3613-3p | INPP4A | PAR-CLIP | miRTarbase |
| hsa-miR-3613-3p | INPP4A | PAR-CLIP | miRTarbase |
| hsa-miR-219a-1-3p | HIST1H2AI | PAR-CLIP | miRTarbase |
| hsa-miR-219a-1-3p | HIST1H2AI | PAR-CLIP | miRTarbase |
| hsa-miR-219a-1-3p | HIST1H2AI | PAR-CLIP | miRTarbase |
| hsa-miR-3613-3p | GUCD1 | PAR-CLIP | miRTarbase |
| hsa-miR-3613-3p | GUCD1 | HITS-CLIP | miRTarbase |
| hsa-miR-3605-5p | C4orf32 | PAR-CLIP | miRTarbase |
| hsa-miR-195-3p | FRK | PAR-CLIP | miRTarbase |
| hsa-miR-3613-3p | HEPH | PAR-CLIP | miRTarbase |
| hsa-miR-3613-3p | C4orf17 | PAR-CLIP | miRTarbase |
| hsa-miR-141-5p | UGT2A1 | PAR-CLIP | miRTarbase |
| hsa-miR-486-5p | CCDC14 | PAR-CLIP | miRTarbase |
| hsa-miR-141-5p | UGT2A2 | PAR-CLIP | miRTarbase |
| hsa-miR-486-5p | CMSS1 | PAR-CLIP | miRTarbase |
| hsa-miR-195-3p | UBE2E3 | PAR-CLIP | miRTarbase |
| hsa-miR-3613-3p | CLASP1 | PAR-CLIP | miRTarbase |
| hsa-miR-3613-3p | TMEM196 | PAR-CLIP | miRTarbase |
| hsa-miR-3613-5p | DDX17 | PAR-CLIP | miRTarbase |
| hsa-miR-3613-3p | ORC4 | PAR-CLIP | miRTarbase |
| hsa-miR-3613-3p | ZNF552 | PAR-CLIP | miRTarbase |
| hsa-miR-3613-3p | TRAF3IP1 | PAR-CLIP | miRTarbase |
| hsa-miR-3613-3p | PSMB2 | PAR-CLIP | miRTarbase |
| hsa-miR-141-5p | TPBG | PAR-CLIP | miRTarbase |
| hsa-miR-3613-3p | ISLR2 | PAR-CLIP | miRTarbase |
| hsa-miR-3613-3p | ISLR2 | PAR-CLIP | miRTarbase |
| hsa-miR-3613-3p | ATRNL1 | PAR-CLIP | miRTarbase |
| hsa-miR-141-3p | MALT1 | PAR-CLIP | miRTarbase |
| hsa-miR-3613-3p | MALT1 | PAR-CLIP | miRTarbase |
| hsa-miR-195-3p | EI24 | PAR-CLIP | miRTarbase |
| hsa-miR-141-5p | TRPV2 | PAR-CLIP | miRTarbase |
| hsa-miR-3613-3p | PSMG1 | PAR-CLIP | miRTarbase |
| hsa-miR-3613-3p | PSMG1 | PAR-CLIP | miRTarbase |
| hsa-miR-3613-3p | HCCS | PAR-CLIP | miRTarbase |
| hsa-miR-3613-3p | PLXDC1 | PAR-CLIP | miRTarbase |
| hsa-miR-15a-3p | TNFRSF10D | PAR-CLIP | miRTarbase |
| hsa-miR-195-3p | SULT1B1 | PAR-CLIP | miRTarbase |
| hsa-miR-3613-3p | ZNF582 | PAR-CLIP | miRTarbase |
| hsa-miR-15a-3p | TRUB1 | PAR-CLIP | miRTarbase |
| hsa-miR-15a-3p | TRUB1 | PAR-CLIP | miRTarbase |
| hsa-miR-3613-3p | GPR85 | PAR-CLIP | miRTarbase |
| hsa-miR-3605-5p | CYGB | PAR-CLIP | miRTarbase |
| hsa-miR-3613-3p | PLA2G4D | PAR-CLIP | miRTarbase |
| hsa-miR-3613-3p | FANCC | PAR-CLIP | miRTarbase |
| hsa-miR-3613-3p | NAA38 | PAR-CLIP | miRTarbase |
| hsa-miR-3613-3p | NAA38 | PAR-CLIP | miRTarbase |
| hsa-miR-3613-3p | HAUS3 | HITS-CLIP | miRTarbase |
| hsa-miR-3613-3p | HAUS3 | PAR-CLIP | miRTarbase |
| hsa-miR-3613-3p | LINC00598 | PAR-CLIP | miRTarbase |
| hsa-miR-3613-3p | WDR13 | HITS-CLIP | miRTarbase |
| hsa-miR-3613-3p | WDR13 | PAR-CLIP | miRTarbase |
| hsa-miR-3613-3p | WASF3 | PAR-CLIP | miRTarbase |
| hsa-miR-3613-3p | WASF3 | PAR-CLIP | miRTarbase |
| hsa-miR-486-3p | UNC119B | PAR-CLIP | miRTarbase |
| hsa-miR-141-5p | UBE2W | PAR-CLIP | miRTarbase |
| hsa-miR-195-3p | TBC1D15 | PAR-CLIP | miRTarbase |
| hsa-miR-15a-3p | TAF1D | PAR-CLIP | miRTarbase |
| hsa-miR-141-5p | SUZ12 | PAR-CLIP | miRTarbase |
| hsa-miR-3613-3p | SPPL3 | PAR-CLIP | miRTarbase |
| hsa-miR-141-5p | SMG1 | PAR-CLIP | miRTarbase |
| hsa-miR-3605-5p | SENP3 | PAR-CLIP | miRTarbase |
| hsa-miR-3613-3p | RNF20 | PAR-CLIP | miRTarbase |
| hsa-miR-3613-3p | RAB18 | PAR-CLIP | miRTarbase |
| hsa-miR-3613-5p | QSER1 | PAR-CLIP | miRTarbase |
| hsa-miR-3613-3p | PLK2 | PAR-CLIP | miRTarbase |
| hsa-miR-3613-5p | MYO10 | PAR-CLIP | miRTarbase |
| hsa-miR-3613-5p | MYO10 | PAR-CLIP | miRTarbase |
| hsa-miR-3605-5p | MED13 | PAR-CLIP | miRTarbase |
| hsa-miR-3613-3p | MAPK8 | PAR-CLIP | miRTarbase |
| hsa-miR-3613-3p | MAPK8 | PAR-CLIP | miRTarbase |
| hsa-miR-3605-3p | MAML2 | PAR-CLIP | miRTarbase |
| hsa-miR-3613-5p | LIMA1 | PAR-CLIP | miRTarbase |
| hsa-miR-3613-3p | LIMA1 | PAR-CLIP | miRTarbase |
| hsa-miR-141-3p | KIAA1549 | PAR-CLIP | miRTarbase |
| hsa-miR-219a-1-3p | KIAA0319L | PAR-CLIP | miRTarbase |
| hsa-miR-195-3p | IPO7 | PAR-CLIP | miRTarbase |
| hsa-miR-3613-3p | HOOK3 | PAR-CLIP | miRTarbase |
| hsa-miR-3613-3p | HOOK3 | PAR-CLIP | miRTarbase |
| hsa-miR-3613-3p | HNRNPD | PAR-CLIP | miRTarbase |
| hsa-miR-3613-3p | HIVEP1 | PAR-CLIP | miRTarbase |
| hsa-miR-3605-5p | GPR180 | PAR-CLIP | miRTarbase |
| hsa-miR-3613-3p | GOLGA3 | PAR-CLIP | miRTarbase |
| hsa-miR-3613-3p | GOLGA3 | PAR-CLIP | miRTarbase |
| hsa-miR-3613-3p | GOLGA3 | HITS-CLIP | miRTarbase |
| hsa-miR-3613-3p | GDE1 | PAR-CLIP | miRTarbase |
| hsa-miR-3613-3p | GDE1 | PAR-CLIP | miRTarbase |
| hsa-miR-3613-3p | FOXP2 | PAR-CLIP | miRTarbase |
| hsa-miR-219a-1-3p | FBXL7 | PAR-CLIP | miRTarbase |
| hsa-miR-3613-3p | ERGIC2 | PAR-CLIP | miRTarbase |
| hsa-miR-195-3p | ERGIC2 | PAR-CLIP | miRTarbase |
| hsa-miR-195-3p | ERGIC2 | PAR-CLIP | miRTarbase |
| hsa-miR-141-3p | ELAVL2 | PAR-CLIP | miRTarbase |
| hsa-miR-15a-5p | EFNB2 | PAR-CLIP | miRTarbase |
| hsa-miR-15a-5p | EFNB2 | PAR-CLIP | miRTarbase |
| hsa-miR-195-5p | EFNB2 | PAR-CLIP | miRTarbase |
| hsa-miR-195-5p | EFNB2 | PAR-CLIP | miRTarbase |
| hsa-miR-3613-3p | DPP8 | PAR-CLIP | miRTarbase |
| hsa-miR-3613-3p | DCDC2 | PAR-CLIP | miRTarbase |
| hsa-miR-3613-3p | CTC1 | PAR-CLIP | miRTarbase |
| hsa-miR-3613-3p | CTC1 | PAR-CLIP | miRTarbase |
| hsa-miR-3605-5p | CSNK2A1 | PAR-CLIP | miRTarbase |
| hsa-miR-195-3p | CLCN3 | PAR-CLIP | miRTarbase |
| hsa-miR-141-5p | CD2AP | PAR-CLIP | miRTarbase |
| hsa-miR-141-5p | CD2AP | PAR-CLIP | miRTarbase |
| hsa-miR-141-3p | ATXN7L1 | PAR-CLIP | miRTarbase |
| hsa-miR-195-3p | ANP32E | PAR-CLIP | miRTarbase |
| hsa-miR-195-3p | ANP32E | PAR-CLIP | miRTarbase |
| hsa-miR-195-3p | ANP32E | PAR-CLIP | miRTarbase |
| hsa-miR-195-3p | AGO2 | PAR-CLIP | miRTarbase |
| hsa-miR-195-3p | AGO2 | PAR-CLIP | miRTarbase |
| hsa-miR-3613-3p | RPL32 | PAR-CLIP | miRTarbase |
| hsa-miR-195-5p | RPL14 | PAR-CLIP | miRTarbase |
| hsa-miR-195-5p | RPL14 | HITS-CLIP | miRTarbase |
| hsa-miR-15a-5p | RPL14 | PAR-CLIP | miRTarbase |
| hsa-miR-15a-5p | RPL14 | HITS-CLIP | miRTarbase |
| hsa-miR-195-3p | CCT4 | PAR-CLIP | miRTarbase |
| hsa-miR-3613-3p | CDC5L | PAR-CLIP | miRTarbase |
| hsa-miR-15a-3p | OCLN | PAR-CLIP | miRTarbase |
| hsa-miR-15a-3p | OCLN | PAR-CLIP | miRTarbase |
| hsa-miR-15a-3p | OCLN | HITS-CLIP | miRTarbase |
| hsa-miR-195-5p | GNAT1 | PAR-CLIP | miRTarbase |
| hsa-miR-195-5p | GNAT1 | HITS-CLIP | miRTarbase |
| hsa-miR-15a-5p | GNAT1 | PAR-CLIP | miRTarbase |
| hsa-miR-15a-5p | GNAT1 | HITS-CLIP | miRTarbase |
| hsa-miR-3613-3p | ZNF136 | PAR-CLIP | miRTarbase |
| hsa-miR-486-3p | RAF1 | PAR-CLIP | miRTarbase |
| hsa-miR-195-3p | HOXA10 | PAR-CLIP | miRTarbase |
| hsa-miR-15a-5p | HOXA10 | PAR-CLIP | miRTarbase |
| hsa-miR-195-5p | HOXA10 | PAR-CLIP | miRTarbase |
| hsa-miR-195-5p | CBX4 | PAR-CLIP | miRTarbase |
| hsa-miR-195-5p | CBX4 | HITS-CLIP | miRTarbase |
| hsa-miR-15a-5p | CBX4 | PAR-CLIP | miRTarbase |
| hsa-miR-15a-5p | CBX4 | HITS-CLIP | miRTarbase |
| hsa-miR-3613-3p | C18orf32 | PAR-CLIP | miRTarbase |
| hsa-miR-3613-3p | C18orf32 | HITS-CLIP | miRTarbase |
| hsa-miR-3613-3p | ASB11 | PAR-CLIP//HITS-CLIP | miRTarbase |
| hsa-miR-219a-1-3p | PLD5 | PAR-CLIP | miRTarbase |
| hsa-miR-219a-1-3p | PLD5 | PAR-CLIP | miRTarbase |
| hsa-miR-3613-3p | QTRTD1 | PAR-CLIP | miRTarbase |
| hsa-miR-3613-3p | SESN3 | PAR-CLIP | miRTarbase |
| hsa-miR-195-5p | PHC3 | PAR-CLIP | miRTarbase |
| hsa-miR-195-5p | PHC3 | HITS-CLIP | miRTarbase |

| | | | |
|---|---|---|---|
| hsa-miR-15a-5p | PHC3 | PAR-CLIP | miRTarbase |
| hsa-miR-15a-5p | PHC3 | HITS-CLIP | miRTarbase |
| hsa-miR-195-5p | PDCD1 | PAR-CLIP | miRTarbase |
| hsa-miR-195-5p | PDCD1 | HITS-CLIP | miRTarbase |
| hsa-miR-195-5p | PDCD1 | HITS-CLIP | miRTarbase |
| hsa-miR-15a-5p | PDCD1 | PAR-CLIP | miRTarbase |
| hsa-miR-15a-5p | PDCD1 | HITS-CLIP | miRTarbase |
| hsa-miR-15a-5p | PDCD1 | HITS-CLIP | miRTarbase |
| hsa-miR-15a-3p | NR6A1 | PAR-CLIP | miRTarbase |
| hsa-miR-3613-3p | DYRK2 | PAR-CLIP | miRTarbase |
| hsa-miR-195-5p | BAZ2A | PAR-CLIP | miRTarbase |
| hsa-miR-15a-5p | BAZ2A | PAR-CLIP | miRTarbase |
| hsa-miR-195-5p | APP | PAR-CLIP | miRTarbase |
| hsa-miR-219a-1-3p | FICD | PAR-CLIP | miRTarbase |
| hsa-miR-195-3p | TMEM117 | PAR-CLIP | miRTarbase |
| hsa-miR-3613-3p | STXBP2 | PAR-CLIP | miRTarbase |
| hsa-miR-195-5p | ZNF585B | PAR-CLIP | miRTarbase |
| hsa-miR-15a-5p | ZNF585B | PAR-CLIP | miRTarbase |
| hsa-miR-195-3p | DROSHA | PAR-CLIP | miRTarbase |
| hsa-miR-195-5p | ANAPC13 | PAR-CLIP | miRTarbase |
| hsa-miR-15a-5p | ANAPC13 | PAR-CLIP | miRTarbase |
| hsa-miR-195-5p | PRSS21 | PAR-CLIP | miRTarbase |
| hsa-miR-15a-5p | PRSS21 | PAR-CLIP | miRTarbase |
| hsa-miR-486-3p | RPL10 | PAR-CLIP | miRTarbase |
| hsa-miR-195-5p | RALGAPB | PAR-CLIP | miRTarbase |
| hsa-miR-195-5p | RALGAPB | PAR-CLIP | miRTarbase |
| hsa-miR-15a-5p | RALGAPB | PAR-CLIP | miRTarbase |
| hsa-miR-15a-5p | RALGAPB | PAR-CLIP | miRTarbase |
| hsa-miR-195-5p | GSG1 | PAR-CLIP | miRTarbase |
| hsa-miR-15a-5p | GSG1 | PAR-CLIP | miRTarbase |
| hsa-miR-3613-3p | ITM2A | PAR-CLIP | miRTarbase |
| hsa-miR-3613-3p | LRIF1 | PAR-CLIP | miRTarbase |
| hsa-miR-3613-3p | ZNF860 | PAR-CLIP | miRTarbase |
| hsa-miR-195-5p | POLDIP3 | PAR-CLIP | miRTarbase |
| hsa-miR-15a-5p | POLDIP3 | PAR-CLIP | miRTarbase |
| hsa-miR-195-5p | AP5Z1 | PAR-CLIP | miRTarbase |
| hsa-miR-195-5p | AP5Z1 | PAR-CLIP | miRTarbase |
| hsa-miR-15a-5p | AP5Z1 | PAR-CLIP | miRTarbase |
| hsa-miR-15a-5p | AP5Z1 | PAR-CLIP | miRTarbase |
| hsa-miR-141-5p | ATXN2L | PAR-CLIP | miRTarbase |
| hsa-miR-3613-3p | INTU | PAR-CLIP | miRTarbase |
| hsa-miR-3605-5p | C8orf33 | PAR-CLIP | miRTarbase |
| hsa-miR-195-5p | CLSPN | PAR-CLIP | miRTarbase |
| hsa-miR-15a-5p | CLSPN | PAR-CLIP | miRTarbase |
| hsa-miR-3613-3p | BVES | PAR-CLIP | miRTarbase |
| hsa-miR-3605-5p | MIS18BP1 | PAR-CLIP//HITS-CLIP | miRTarbase |
| hsa-miR-195-5p | UGT2B4 | PAR-CLIP | miRTarbase |
| hsa-miR-15a-5p | UGT2B4 | PAR-CLIP | miRTarbase |
| hsa-miR-3613-3p | PDZRN4 | PAR-CLIP | miRTarbase |
| hsa-miR-3613-3p | ADAMTS9 | PAR-CLIP | miRTarbase |
| hsa-miR-195-5p | MAP4K2 | PAR-CLIP | miRTarbase |
| hsa-miR-195-5p | MAP4K2 | PAR-CLIP | miRTarbase |
| hsa-miR-15a-5p | MAP4K2 | PAR-CLIP | miRTarbase |
| hsa-miR-15a-5p | MAP4K2 | PAR-CLIP | miRTarbase |
| hsa-miR-195-5p | CCDC83 | PAR-CLIP | miRTarbase |
| hsa-miR-15a-5p | CCDC83 | PAR-CLIP | miRTarbase |
| hsa-miR-195-5p | DECR1 | PAR-CLIP | miRTarbase |
| hsa-miR-15a-5p | DECR1 | PAR-CLIP | miRTarbase |
| hsa-miR-195-5p | ZBTB10 | PAR-CLIP | miRTarbase |
| hsa-miR-15a-5p | ZBTB10 | PAR-CLIP | miRTarbase |
| hsa-miR-15a-5p | YWHAQ | PAR-CLIP | miRTarbase |
| hsa-miR-195-5p | YWHAQ | PAR-CLIP | miRTarbase |
| hsa-miR-3613-5p | VEZF1 | PAR-CLIP | miRTarbase |
| hsa-miR-195-5p | USP48 | PAR-CLIP | miRTarbase |
| hsa-miR-195-5p | USP48 | PAR-CLIP | miRTarbase |
| hsa-miR-15a-5p | USP48 | PAR-CLIP | miRTarbase |
| hsa-miR-15a-5p | USP48 | PAR-CLIP | miRTarbase |
| hsa-miR-3613-3p | UBE3A | PAR-CLIP | miRTarbase |
| hsa-miR-195-5p | SALL1 | PAR-CLIP | miRTarbase |
| hsa-miR-195-5p | SALL1 | PAR-CLIP | miRTarbase |
| hsa-miR-15a-5p | SALL1 | PAR-CLIP | miRTarbase |
| hsa-miR-15a-5p | SALL1 | PAR-CLIP | miRTarbase |
| hsa-miR-195-5p | RUNX1T1 | PAR-CLIP | miRTarbase |
| hsa-miR-15a-5p | RUNX1T1 | PAR-CLIP | miRTarbase |
| hsa-miR-195-5p | RTN4 | PAR-CLIP | miRTarbase |
| hsa-miR-15a-5p | RTN4 | PAR-CLIP | miRTarbase |
| hsa-miR-195-3p | RPS6KA5 | PAR-CLIP | miRTarbase |
| hsa-miR-3613-3p | RNF115 | PAR-CLIP | miRTarbase |
| hsa-miR-195-3p | RAB1A | PAR-CLIP | miRTarbase |
| hsa-miR-3613-3p | POGZ | PAR-CLIP | miRTarbase |
| hsa-miR-195-5p | PNISR | PAR-CLIP | miRTarbase |
| hsa-miR-195-5p | PNISR | HITS-CLIP | miRTarbase |
| hsa-miR-15a-5p | PNISR | PAR-CLIP | miRTarbase |
| hsa-miR-15a-5p | PNISR | HITS-CLIP | miRTarbase |
| hsa-miR-141-3p | PIGW | PAR-CLIP | miRTarbase |
| hsa-miR-195-5p | PHLPP2 | PAR-CLIP | miRTarbase |
| hsa-miR-15a-5p | PHLPP2 | PAR-CLIP | miRTarbase |
| hsa-miR-195-5p | PAG1 | PAR-CLIP | miRTarbase |
| hsa-miR-195-5p | PAG1 | PAR-CLIP | miRTarbase |
| hsa-miR-15a-5p | PAG1 | PAR-CLIP | miRTarbase |
| hsa-miR-15a-5p | PAG1 | PAR-CLIP | miRTarbase |
| hsa-miR-195-5p | NUCKS1 | PAR-CLIP | miRTarbase |
| hsa-miR-195-5p | NUCKS1 | HITS-CLIP | miRTarbase |
| hsa-miR-15a-5p | NUCKS1 | PAR-CLIP | miRTarbase |
| hsa-miR-15a-5p | NUCKS1 | HITS-CLIP | miRTarbase |
| hsa-miR-15a-5p | MKX | PAR-CLIP | miRTarbase |
| hsa-miR-195-5p | MKX | PAR-CLIP | miRTarbase |
| hsa-miR-195-5p | MBD4 | PAR-CLIP | miRTarbase |
| hsa-miR-15a-5p | MBD4 | PAR-CLIP | miRTarbase |
| hsa-miR-3613-3p | MBNL3 | PAR-CLIP | miRTarbase |
| hsa-miR-3613-3p | MBNL3 | PAR-CLIP | miRTarbase |
| hsa-miR-15a-5p | LRRFIP2 | PAR-CLIP | miRTarbase |
| hsa-miR-15a-5p | LRRFIP2 | PAR-CLIP | miRTarbase |
| hsa-miR-195-5p | LRRFIP2 | PAR-CLIP | miRTarbase |
| hsa-miR-195-5p | LRRFIP2 | PAR-CLIP | miRTarbase |
| hsa-miR-15a-5p | KPNA3 | PAR-CLIP//HITS-CLIP | miRTarbase |
| hsa-miR-195-5p | KPNA3 | PAR-CLIP//HITS-CLIP | miRTarbase |
| hsa-miR-195-5p | KPNA1 | PAR-CLIP | miRTarbase |
| hsa-miR-195-5p | KPNA1 | PAR-CLIP | miRTarbase |
| hsa-miR-15a-5p | KPNA1 | PAR-CLIP | miRTarbase |
| hsa-miR-15a-5p | KPNA1 | PAR-CLIP | miRTarbase |
| hsa-miR-15a-5p | HIGD1A | PAR-CLIP | miRTarbase |
| hsa-miR-15a-5p | HIGD1A | PAR-CLIP | miRTarbase |
| hsa-miR-195-5p | HIGD1A | PAR-CLIP | miRTarbase |
| hsa-miR-195-5p | HIGD1A | PAR-CLIP | miRTarbase |
| hsa-miR-195-5p | HCFC2 | PAR-CLIP | miRTarbase |
| hsa-miR-195-5p | HCFC2 | PAR-CLIP | miRTarbase |
| hsa-miR-15a-5p | HCFC2 | PAR-CLIP | miRTarbase |
| hsa-miR-15a-5p | HCFC2 | PAR-CLIP | miRTarbase |
| hsa-miR-195-5p | GRB2 | PAR-CLIP | miRTarbase |
| hsa-miR-195-5p | GRB2 | PAR-CLIP | miRTarbase |
| hsa-miR-15a-5p | GRB2 | PAR-CLIP | miRTarbase |
| hsa-miR-15a-5p | GRB2 | PAR-CLIP | miRTarbase |
| hsa-miR-3613-3p | GIGYF1 | PAR-CLIP | miRTarbase |
| hsa-miR-141-3p | GATA6 | PAR-CLIP | miRTarbase |
| hsa-miR-15a-5p | FKBP1A | PAR-CLIP | miRTarbase |
| hsa-miR-195-5p | FKBP1A | PAR-CLIP | miRTarbase |
| hsa-miR-15a-5p | FBXL20 | PAR-CLIP | miRTarbase |
| hsa-miR-195-5p | FBXL20 | PAR-CLIP | miRTarbase |
| hsa-miR-141-3p | EPHA7 | PAR-CLIP | miRTarbase |
| hsa-miR-15a-5p | CRK | PAR-CLIP | miRTarbase |
| hsa-miR-195-5p | CRK | PAR-CLIP | miRTarbase |
| hsa-miR-219a-1-3p | CREBBP | PAR-CLIP | miRTarbase |
| hsa-miR-195-5p | CLIP4 | PAR-CLIP | miRTarbase |
| hsa-miR-195-5p | CLIP4 | PAR-CLIP | miRTarbase |
| hsa-miR-15a-5p | CLIP4 | PAR-CLIP | miRTarbase |
| hsa-miR-15a-5p | CLIP4 | PAR-CLIP | miRTarbase |
| hsa-miR-3613-3p | CHD1 | PAR-CLIP | miRTarbase |
| hsa-miR-3613-3p | CHD1 | PAR-CLIP | miRTarbase |
| hsa-miR-15a-5p | CDK17 | PAR-CLIP | miRTarbase |
| hsa-miR-15a-5p | CDK17 | HITS-CLIP | miRTarbase |
| hsa-miR-195-5p | CDK17 | PAR-CLIP | miRTarbase |
| hsa-miR-195-5p | CDK17 | HITS-CLIP | miRTarbase |
| hsa-miR-15a-5p | CACUL1 | PAR-CLIP | miRTarbase |
| hsa-miR-195-5p | CACUL1 | PAR-CLIP | miRTarbase |
| hsa-miR-141-3p | C18orf25 | PAR-CLIP | miRTarbase |
| hsa-miR-15a-5p | C11orf24 | PAR-CLIP | miRTarbase |
| hsa-miR-15a-5p | C11orf24 | PAR-CLIP | miRTarbase |
| hsa-miR-195-5p | C11orf24 | PAR-CLIP | miRTarbase |

| | | | |
|---|---|---|---|
| hsa-miR-195-5p | C11orf24 | PAR-CLIP | miRTarbase |
| hsa-miR-3613-3p | BMP3 | PAR-CLIP | miRTarbase |
| hsa-miR-195-5p | ASH1L | PAR-CLIP | miRTarbase |
| hsa-miR-195-5p | ASH1L | HITS-CLIP | miRTarbase |
| hsa-miR-15a-5p | ASH1L | PAR-CLIP | miRTarbase |
| hsa-miR-15a-5p | ASH1L | HITS-CLIP | miRTarbase |
| hsa-miR-3613-3p | ARHGAP12 | PAR-CLIP | miRTarbase |
| hsa-miR-3613-3p | ARHGAP12 | PAR-CLIP | miRTarbase |
| hsa-miR-195-5p | AMOT | PAR-CLIP | miRTarbase |
| hsa-miR-15a-5p | AMOT | PAR-CLIP | miRTarbase |
| hsa-miR-141-5p | TLE3 | PAR-CLIP | miRTarbase |
| hsa-miR-141-5p | TLE3 | PAR-CLIP | miRTarbase |
| hsa-miR-195-3p | MRPS30 | PAR-CLIP//HITS-CLIP | miRTarbase |
| hsa-miR-195-5p | SLC29A1 | PAR-CLIP | miRTarbase |
| hsa-miR-195-5p | SLC29A1 | PAR-CLIP | miRTarbase |
| hsa-miR-15a-5p | SLC29A1 | PAR-CLIP | miRTarbase |
| hsa-miR-15a-5p | SLC29A1 | PAR-CLIP | miRTarbase |
| hsa-miR-195-5p | OSCAR | PAR-CLIP | miRTarbase |
| hsa-miR-195-5p | OSCAR | PAR-CLIP | miRTarbase |
| hsa-miR-15a-5p | OSCAR | PAR-CLIP | miRTarbase |
| hsa-miR-15a-5p | OSCAR | PAR-CLIP | miRTarbase |
| hsa-miR-3613-3p | PLCXD1 | PAR-CLIP//HITS-CLIP | miRTarbase |
| hsa-miR-195-5p | MTHFR | PAR-CLIP | miRTarbase |
| hsa-miR-15a-5p | MTHFR | PAR-CLIP | miRTarbase |
| hsa-miR-195-5p | FAM229B | PAR-CLIP | miRTarbase |
| hsa-miR-15a-5p | FAM229B | PAR-CLIP | miRTarbase |
| hsa-miR-486-5p | TTC8 | PAR-CLIP | miRTarbase |
| hsa-miR-195-5p | EPM2AIP1 | PAR-CLIP | miRTarbase |
| hsa-miR-15a-5p | EPM2AIP1 | PAR-CLIP | miRTarbase |
| hsa-miR-195-5p | ZNF267 | PAR-CLIP | miRTarbase |
| hsa-miR-15a-5p | ZNF267 | PAR-CLIP | miRTarbase |
| hsa-miR-195-5p | SSU72 | PAR-CLIP | miRTarbase |
| hsa-miR-15a-5p | SSU72 | PAR-CLIP | miRTarbase |
| hsa-miR-141-5p | TMEM138 | PAR-CLIP | miRTarbase |
| hsa-miR-3613-5p | MMS22L | PAR-CLIP | miRTarbase |
| hsa-miR-195-5p | DNAJC10 | PAR-CLIP | miRTarbase |
| hsa-miR-15a-5p | DNAJC10 | PAR-CLIP | miRTarbase |
| hsa-miR-3613-5p | MED10 | PAR-CLIP | miRTarbase |
| hsa-miR-195-5p | ZNF704 | PAR-CLIP | miRTarbase |
| hsa-miR-15a-5p | ZNF704 | PAR-CLIP | miRTarbase |
| hsa-miR-219a-1-3p | ZNF460 | PAR-CLIP | miRTarbase |
| hsa-miR-219a-1-3p | YY1 | PAR-CLIP | miRTarbase |
| hsa-miR-15a-5p | YRDC | PAR-CLIP | miRTarbase |
| hsa-miR-195-5p | YRDC | PAR-CLIP | miRTarbase |
| hsa-miR-3613-3p | WNK3 | PAR-CLIP//HITS-CLIP | miRTarbase |
| hsa-miR-195-3p | USP46 | PAR-CLIP | miRTarbase |
| hsa-miR-3613-3p | TUBB2A | PAR-CLIP | miRTarbase |
| hsa-miR-3613-3p | TUBB2A | PAR-CLIP | miRTarbase |
| hsa-miR-195-5p | TPM3 | PAR-CLIP | miRTarbase |
| hsa-miR-15a-5p | TPM3 | PAR-CLIP | miRTarbase |
| hsa-miR-195-5p | TMEM161B | PAR-CLIP | miRTarbase |
| hsa-miR-15a-5p | TMEM161B | PAR-CLIP | miRTarbase |
| hsa-miR-195-5p | TM7SF3 | PAR-CLIP | miRTarbase |
| hsa-miR-15a-5p | TM7SF3 | PAR-CLIP | miRTarbase |
| hsa-miR-195-5p | TAF13 | PAR-CLIP | miRTarbase |
| hsa-miR-195-5p | TAF13 | PAR-CLIP | miRTarbase |
| hsa-miR-15a-5p | TAF13 | PAR-CLIP | miRTarbase |
| hsa-miR-15a-5p | TAF13 | PAR-CLIP | miRTarbase |
| hsa-miR-195-5p | SZRD1 | PAR-CLIP | miRTarbase |
| hsa-miR-195-5p | SZRD1 | PAR-CLIP | miRTarbase |
| hsa-miR-15a-5p | SZRD1 | PAR-CLIP | miRTarbase |
| hsa-miR-15a-5p | SZRD1 | PAR-CLIP | miRTarbase |
| hsa-miR-141-3p | SLC35D1 | PAR-CLIP | miRTarbase |
| hsa-miR-219a-1-3p | SIPA1L2 | PAR-CLIP | miRTarbase |
| hsa-miR-195-5p | RNF149 | PAR-CLIP//HITS-CLIP | miRTarbase |
| hsa-miR-15a-5p | RNF149 | PAR-CLIP//HITS-CLIP | miRTarbase |
| hsa-miR-486-5p | RBM12B | PAR-CLIP | miRTarbase |
| hsa-miR-195-5p | RACGAP1 | PAR-CLIP | miRTarbase |
| hsa-miR-15a-5p | RACGAP1 | PAR-CLIP | miRTarbase |
| hsa-miR-195-5p | RAB23 | PAR-CLIP | miRTarbase |
| hsa-miR-15a-5p | RAB23 | PAR-CLIP | miRTarbase |
| hsa-miR-15a-5p | PTPRD | PAR-CLIP | miRTarbase |
| hsa-miR-195-5p | PTPRD | PAR-CLIP | miRTarbase |
| hsa-miR-195-5p | PRKAA1 | PAR-CLIP | miRTarbase |
| hsa-miR-195-5p | PRKAA1 | PAR-CLIP | miRTarbase |
| hsa-miR-15a-5p | PRKAA1 | PAR-CLIP | miRTarbase |
| hsa-miR-15a-5p | PRKAA1 | PAR-CLIP | miRTarbase |
| hsa-miR-195-3p | PREPL | PAR-CLIP | miRTarbase |
| hsa-miR-195-5p | PRDM4 | PAR-CLIP | miRTarbase |
| hsa-miR-15a-5p | PRDM4 | PAR-CLIP | miRTarbase |
| hsa-miR-15a-5p | PPAP2B | PAR-CLIP | miRTarbase |
| hsa-miR-195-5p | PPAP2B | PAR-CLIP | miRTarbase |
| hsa-miR-141-3p | PGAM4 | PAR-CLIP | miRTarbase |
| hsa-miR-195-3p | NRIP1 | PAR-CLIP//HITS-CLIP | miRTarbase |
| hsa-miR-3613-5p | MTF2 | PAR-CLIP | miRTarbase |
| hsa-miR-3613-5p | MTF2 | PAR-CLIP | miRTarbase |
| hsa-miR-486-5p | MBD4 | PAR-CLIP | miRTarbase |
| hsa-miR-195-5p | LURAP1L | PAR-CLIP | miRTarbase |
| hsa-miR-15a-5p | LURAP1L | PAR-CLIP | miRTarbase |
| hsa-miR-3613-5p | LMNB2 | PAR-CLIP | miRTarbase |
| hsa-miR-3613-5p | LMNB2 | PAR-CLIP | miRTarbase |
| hsa-miR-3613-3p | LMNB2 | PAR-CLIP | miRTarbase |
| hsa-miR-3613-3p | LMNB2 | PAR-CLIP | miRTarbase |
| hsa-miR-195-5p | KANK1 | PAR-CLIP | miRTarbase |
| hsa-miR-195-5p | KANK1 | PAR-CLIP | miRTarbase |
| hsa-miR-15a-5p | KANK1 | PAR-CLIP | miRTarbase |
| hsa-miR-15a-5p | KANK1 | PAR-CLIP | miRTarbase |
| hsa-miR-195-3p | HOXA9 | PAR-CLIP | miRTarbase |
| hsa-miR-3613-3p | HNRNPU | PAR-CLIP | miRTarbase |
| hsa-miR-3613-3p | HNRNPU | HITS-CLIP | miRTarbase |
| hsa-miR-195-5p | HIST2H2BE | PAR-CLIP | miRTarbase |
| hsa-miR-15a-5p | HIST2H2BE | PAR-CLIP | miRTarbase |
| hsa-miR-141-3p | H2AFZ | PAR-CLIP | miRTarbase |
| hsa-miR-141-3p | H2AFZ | PAR-CLIP | miRTarbase |
| hsa-miR-195-5p | GPR27 | PAR-CLIP | miRTarbase |
| hsa-miR-195-5p | GPR27 | PAR-CLIP | miRTarbase |
| hsa-miR-15a-5p | GPR27 | PAR-CLIP | miRTarbase |
| hsa-miR-15a-5p | GPR27 | PAR-CLIP | miRTarbase |
| hsa-miR-486-5p | FAM217B | PAR-CLIP | miRTarbase |
| hsa-miR-195-5p | EXT1 | PAR-CLIP | miRTarbase |
| hsa-miR-15a-5p | EXT1 | PAR-CLIP | miRTarbase |
| hsa-miR-3613-3p | ESCO2 | PAR-CLIP | miRTarbase |
| hsa-miR-3613-3p | DYNC1LI2 | PAR-CLIP | miRTarbase |
| hsa-miR-195-5p | CYP26B1 | PAR-CLIP | miRTarbase |
| hsa-miR-195-5p | CYP26B1 | PAR-CLIP | miRTarbase |
| hsa-miR-15a-5p | CYP26B1 | PAR-CLIP | miRTarbase |
| hsa-miR-15a-5p | CYP26B1 | PAR-CLIP | miRTarbase |
| hsa-miR-195-5p | CREBL2 | PAR-CLIP | miRTarbase |
| hsa-miR-195-5p | CREBL2 | PAR-CLIP | miRTarbase |
| hsa-miR-195-5p | CNKSR3 | PAR-CLIP | miRTarbase |
| hsa-miR-15a-5p | CNKSR3 | PAR-CLIP | miRTarbase |
| hsa-miR-15a-5p | CA8 | PAR-CLIP | miRTarbase |
| hsa-miR-195-5p | CA8 | PAR-CLIP | miRTarbase |
| hsa-miR-15a-5p | BTN3A3 | PAR-CLIP | miRTarbase |
| hsa-miR-195-5p | BTN3A3 | PAR-CLIP | miRTarbase |
| hsa-miR-195-5p | ARHGAP12 | PAR-CLIP | miRTarbase |
| hsa-miR-195-5p | ARHGAP12 | PAR-CLIP | miRTarbase |
| hsa-miR-195-5p | ARHGAP12 | HITS-CLIP | miRTarbase |
| hsa-miR-15a-5p | ARHGAP12 | PAR-CLIP | miRTarbase |
| hsa-miR-15a-5p | ARHGAP12 | PAR-CLIP | miRTarbase |
| hsa-miR-15a-5p | ARHGAP12 | HITS-CLIP | miRTarbase |
| hsa-miR-141-5p | AP1AR | PAR-CLIP//HITS-CLIP | miRTarbase |
| hsa-miR-3613-3p | TRMT112 | PAR-CLIP | miRTarbase |
| hsa-miR-141-3p | TRMT112 | PAR-CLIP | miRTarbase |
| hsa-miR-219a-1-3p | ZWINT | PAR-CLIP | miRTarbase |
| hsa-miR-195-5p | OSBPL3 | PAR-CLIP | miRTarbase |
| hsa-miR-15a-5p | OSBPL3 | PAR-CLIP | miRTarbase |
| hsa-miR-195-5p | KRT33B | PAR-CLIP | miRTarbase |
| hsa-miR-15a-5p | KRT33B | PAR-CLIP | miRTarbase |
| hsa-miR-3613-3p | YY1 | PAR-CLIP | miRTarbase |
| hsa-miR-195-3p | TUBB2A | PAR-CLIP | miRTarbase |
| hsa-miR-195-5p | TUBB2A | PAR-CLIP | miRTarbase |
| hsa-miR-15a-5p | TUBB2A | PAR-CLIP | miRTarbase |
| hsa-miR-486-3p | TMCC1 | PAR-CLIP | miRTarbase |
| hsa-miR-195-3p | RNF219 | PAR-CLIP | miRTarbase |
| hsa-miR-3605-5p | PPP2R2A | PAR-CLIP | miRTarbase |
| hsa-miR-3613-5p | NREP | PAR-CLIP | miRTarbase |
| hsa-miR-219a-1-3p | NREP | PAR-CLIP | miRTarbase |
| hsa-miR-195-5p | MSANTD4 | PAR-CLIP | miRTarbase |

| | | | |
|---|---|---|---|
| hsa-miR-15a-5p | MSANTD4 | PAR-CLIP | miRTarbase |
| hsa-miR-195-5p | LANCL1 | PAR-CLIP | miRTarbase |
| hsa-miR-15a-5p | LANCL1 | PAR-CLIP | miRTarbase |
| hsa-miR-195-5p | HNRNPA2B1 | PAR-CLIP | miRTarbase |
| hsa-miR-15a-5p | HNRNPA2B1 | PAR-CLIP | miRTarbase |
| hsa-miR-3613-5p | CDK6 | PAR-CLIP | miRTarbase |
| hsa-miR-195-5p | KIAA1456 | PAR-CLIP | miRTarbase |
| hsa-miR-15a-5p | KIAA1456 | PAR-CLIP | miRTarbase |
| hsa-miR-3613-5p | GPR141 | PAR-CLIP | miRTarbase |
| hsa-miR-3613-3p | ZNF28 | PAR-CLIP | miRTarbase |
| hsa-miR-3613-3p | ZNF28 | HITS-CLIP | miRTarbase |
| hsa-miR-3613-3p | SLC25A12 | PAR-CLIP | miRTarbase |
| hsa-miR-195-5p | SLC25A12 | PAR-CLIP | miRTarbase |
| hsa-miR-195-5p | SLC25A12 | HITS-CLIP | miRTarbase |
| hsa-miR-15a-5p | SLC25A12 | PAR-CLIP | miRTarbase |
| hsa-miR-15a-5p | SLC25A12 | HITS-CLIP | miRTarbase |
| hsa-miR-195-3p | PABPC4L | PAR-CLIP | miRTarbase |
| hsa-miR-3613-3p | ZWINT | PAR-CLIP | miRTarbase |
| hsa-miR-195-5p | DLGAP3 | PAR-CLIP | miRTarbase |
| hsa-miR-15a-5p | DLGAP3 | PAR-CLIP | miRTarbase |
| hsa-miR-3613-5p | RBM41 | PAR-CLIP | miRTarbase |
| hsa-miR-3613-3p | PFDN2 | PAR-CLIP | miRTarbase |
| hsa-miR-195-5p | THRAP3 | PAR-CLIP | miRTarbase |
| hsa-miR-15a-5p | THRAP3 | PAR-CLIP | miRTarbase |
| hsa-miR-3613-3p | HUWE1 | PAR-CLIP | miRTarbase |
| hsa-miR-195-5p | SMDT1 | PAR-CLIP | miRTarbase |
| hsa-miR-15a-5p | SMDT1 | PAR-CLIP | miRTarbase |
| hsa-miR-195-5p | RAPH1 | PAR-CLIP | miRTarbase |
| hsa-miR-15a-5p | RAPH1 | PAR-CLIP | miRTarbase |
| hsa-miR-3613-3p | STAT2 | PAR-CLIP | miRTarbase |
| hsa-miR-3613-3p | GPBP1L1 | PAR-CLIP | miRTarbase |
| hsa-miR-195-5p | CCNT1 | PAR-CLIP | miRTarbase |
| hsa-miR-15a-5p | CCNT1 | PAR-CLIP | miRTarbase |
| hsa-miR-219a-1-3p | MRPS18B | PAR-CLIP | miRTarbase |
| hsa-miR-195-5p | ZNF391 | PAR-CLIP | miRTarbase |
| hsa-miR-15a-5p | ZNF391 | PAR-CLIP | miRTarbase |
| hsa-miR-195-5p | CCDC80 | PAR-CLIP | miRTarbase |
| hsa-miR-15a-5p | CCDC80 | PAR-CLIP | miRTarbase |
| hsa-miR-15a-3p | ZNF35 | PAR-CLIP | miRTarbase |
| hsa-miR-3605-5p | ZNF322P1 | PAR-CLIP | miRTarbase |
| hsa-miR-3613-3p | ZFC3H1 | PAR-CLIP | miRTarbase |
| hsa-miR-15a-5p | ZBTB16 | PAR-CLIP | miRTarbase |
| hsa-miR-195-5p | ZBTB16 | PAR-CLIP | miRTarbase |
| hsa-miR-141-3p | XPOT | PAR-CLIP | miRTarbase |
| hsa-miR-195-5p | XKR7 | PAR-CLIP | miRTarbase |
| hsa-miR-15a-5p | XKR7 | PAR-CLIP | miRTarbase |
| hsa-miR-15a-5p | WNK3 | PAR-CLIP | miRTarbase |
| hsa-miR-195-5p | WNK3 | PAR-CLIP | miRTarbase |
| hsa-miR-3613-3p | WEE1 | PAR-CLIP | miRTarbase |
| hsa-miR-3613-3p | VIM | PAR-CLIP | miRTarbase |
| hsa-miR-15a-5p | VAV2 | PAR-CLIP | miRTarbase |
| hsa-miR-195-5p | VAV2 | PAR-CLIP | miRTarbase |
| hsa-miR-195-5p | TGFBR3 | PAR-CLIP | miRTarbase |
| hsa-miR-15a-5p | TGFBR3 | PAR-CLIP | miRTarbase |
| hsa-miR-219a-1-3p | SESN3 | PAR-CLIP | miRTarbase |
| hsa-miR-195-3p | RNF44 | PAR-CLIP | miRTarbase |
| hsa-miR-15a-3p | RFX1 | PAR-CLIP | miRTarbase |
| hsa-miR-195-5p | RASEF | PAR-CLIP | miRTarbase |
| hsa-miR-15a-5p | RASEF | PAR-CLIP | miRTarbase |
| hsa-miR-3613-3p | NFYA | PAR-CLIP | miRTarbase |
| hsa-miR-3613-3p | NFYA | HITS-CLIP | miRTarbase |
| hsa-miR-195-5p | NCKAP1 | PAR-CLIP | miRTarbase |
| hsa-miR-15a-5p | NCKAP1 | PAR-CLIP | miRTarbase |
| hsa-miR-195-5p | MAP3K7 | PAR-CLIP | miRTarbase |
| hsa-miR-15a-5p | MAP3K7 | PAR-CLIP | miRTarbase |
| hsa-miR-3613-3p | LMNB1 | PAR-CLIP | miRTarbase |
| hsa-miR-219a-1-3p | KPNA1 | PAR-CLIP | miRTarbase |
| hsa-miR-15a-5p | KLHL15 | PAR-CLIP | miRTarbase |
| hsa-miR-195-5p | KLHL15 | PAR-CLIP | miRTarbase |
| hsa-miR-486-5p | IGSF3 | PAR-CLIP | miRTarbase |
| hsa-miR-141-5p | HSP90AA1 | PAR-CLIP | miRTarbase |
| hsa-miR-15a-3p | HNRNPA0 | PAR-CLIP | miRTarbase |
| hsa-miR-141-5p | H3F3B | PAR-CLIP | miRTarbase |
| hsa-miR-195-5p | GNG12 | PAR-CLIP | miRTarbase |
| hsa-miR-15a-5p | GNG12 | PAR-CLIP | miRTarbase |
| hsa-miR-195-5p | FZD9 | PAR-CLIP | miRTarbase |
| hsa-miR-15a-5p | FZD9 | PAR-CLIP | miRTarbase |
| hsa-miR-3613-3p | FCHSD2 | PAR-CLIP | miRTarbase |
| hsa-miR-3613-3p | FAM60A | PAR-CLIP | miRTarbase |
| hsa-miR-195-5p | CMTM4 | PAR-CLIP | miRTarbase |
| hsa-miR-15a-5p | CMTM4 | PAR-CLIP | miRTarbase |
| hsa-miR-141-3p | CDV3 | PAR-CLIP | miRTarbase |
| hsa-miR-3613-3p | CCNY | PAR-CLIP | miRTarbase |
| hsa-miR-15a-5p | CCDC88C | PAR-CLIP | miRTarbase |
| hsa-miR-195-5p | CCDC88C | PAR-CLIP | miRTarbase |
| hsa-miR-195-5p | ARMC12 | PAR-CLIP | miRTarbase |
| hsa-miR-15a-5p | ARMC12 | PAR-CLIP | miRTarbase |
| hsa-miR-195-5p | AHNAK2 | PAR-CLIP | miRTarbase |
| hsa-miR-15a-5p | AHNAK2 | PAR-CLIP | miRTarbase |
| hsa-miR-15a-5p | ACVR2A | PAR-CLIP | miRTarbase |
| hsa-miR-195-5p | ACVR2A | PAR-CLIP | miRTarbase |
| hsa-miR-3613-3p | MYBL1 | PAR-CLIP | miRTarbase |
| hsa-miR-15a-3p | CADM2 | PAR-CLIP | miRTarbase |
| hsa-miR-3605-3p | TMC5 | PAR-CLIP | miRTarbase |
| hsa-miR-3605-3p | TMC5 | HITS-CLIP | miRTarbase |
| hsa-miR-486-3p | CC2D1B | PAR-CLIP | miRTarbase |
| hsa-miR-3605-5p | KLK10 | PAR-CLIP | miRTarbase |
| hsa-miR-3613-3p | HIPK3 | PAR-CLIP | miRTarbase |
| hsa-miR-15a-3p | VAV3 | PAR-CLIP | miRTarbase |
| hsa-miR-15a-5p | TLK1 | PAR-CLIP | miRTarbase |
| hsa-miR-15a-5p | TLK1 | HITS-CLIP | miRTarbase |
| hsa-miR-195-5p | TLK1 | PAR-CLIP | miRTarbase |
| hsa-miR-195-5p | TLK1 | HITS-CLIP | miRTarbase |
| hsa-miR-3605-5p | GPRC5C | PAR-CLIP | miRTarbase |
| hsa-miR-141-5p | TOR3A | PAR-CLIP | miRTarbase |
| hsa-miR-195-5p | UBE2H | PAR-CLIP | miRTarbase |
| hsa-miR-15a-5p | UBE2H | PAR-CLIP | miRTarbase |
| hsa-miR-3613-3p | OR9Q1 | PAR-CLIP | miRTarbase |
| hsa-miR-195-5p | TTLL5 | PAR-CLIP | miRTarbase |
| hsa-miR-15a-5p | TTLL5 | PAR-CLIP | miRTarbase |
| hsa-miR-3613-3p | TTPAL | PAR-CLIP | miRTarbase |
| hsa-miR-3613-3p | TTPAL | HITS-CLIP | miRTarbase |
| hsa-miR-195-5p | RIF1 | PAR-CLIP | miRTarbase |
| hsa-miR-15a-5p | RIF1 | PAR-CLIP | miRTarbase |
| hsa-miR-486-3p | SKI | PAR-CLIP | miRTarbase |
| hsa-miR-195-5p | SERBP1 | PAR-CLIP | miRTarbase |
| hsa-miR-15a-5p | SERBP1 | PAR-CLIP | miRTarbase |
| hsa-miR-15a-5p | PHF19 | PAR-CLIP | miRTarbase |
| hsa-miR-195-5p | PHF19 | PAR-CLIP | miRTarbase |
| hsa-miR-3613-3p | LSM11 | PAR-CLIP | miRTarbase |
| hsa-miR-15a-5p | LSM11 | PAR-CLIP | miRTarbase |
| hsa-miR-15a-5p | LSM11 | HITS-CLIP | miRTarbase |
| hsa-miR-3605-5p | CALU | PAR-CLIP | miRTarbase |
| hsa-miR-3613-3p | RNF103-CHMP3 | PAR-CLIP | miRTarbase |
| hsa-miR-195-3p | CCDC14 | PAR-CLIP | miRTarbase |
| hsa-miR-141-5p | FAM204A | PAR-CLIP | miRTarbase |
| hsa-miR-3613-3p | CHMP3 | PAR-CLIP | miRTarbase |
| hsa-miR-3613-3p | PDE3A | PAR-CLIP | miRTarbase |
| hsa-miR-141-3p | KIAA1549L | PAR-CLIP | miRTarbase |
| hsa-miR-486-3p | RPL18A | PAR-CLIP | miRTarbase |
| hsa-miR-3613-3p | SLC24A4 | HITS-CLIP | miRTarbase |
| hsa-miR-3613-3p | SLC24A4 | PAR-CLIP | miRTarbase |
| hsa-miR-3613-3p | SLC24A4 | HITS-CLIP | miRTarbase |
| hsa-miR-195-5p | PROSC | PAR-CLIP | miRTarbase |
| hsa-miR-15a-5p | PROSC | PAR-CLIP | miRTarbase |
| hsa-miR-3613-3p | CCDC132 | HITS-CLIP | miRTarbase |
| hsa-miR-3613-3p | CCDC132 | PAR-CLIP | miRTarbase |
| hsa-miR-3613-3p | CCDC132 | HITS-CLIP | miRTarbase |
| hsa-miR-195-5p | CLEC2D | PAR-CLIP | miRTarbase |
| hsa-miR-15a-5p | CLEC2D | PAR-CLIP | miRTarbase |
| hsa-miR-3605-5p | ZBTB37 | PAR-CLIP | miRTarbase |
| hsa-miR-3613-5p | YY1 | PAR-CLIP | miRTarbase |
| hsa-miR-195-3p | TM9SF3 | PAR-CLIP | miRTarbase |
| hsa-miR-3613-3p | RPS16 | PAR-CLIP | miRTarbase |
| hsa-miR-15a-5p | PDIA6 | PAR-CLIP | miRTarbase |
| hsa-miR-15a-5p | PDIA6 | HITS-CLIP | miRTarbase |
| hsa-miR-15a-5p | N4BP1 | PAR-CLIP | miRTarbase |
| hsa-miR-15a-5p | N4BP1 | HITS-CLIP | miRTarbase |
| hsa-miR-195-5p | N4BP1 | PAR-CLIP | miRTarbase |

| | | | |
|---|---|---|---|
| hsa-miR-195-5p | N4BP1 | HITS-CLIP | miRTarbase |
| hsa-miR-3613-3p | HMGB2 | PAR-CLIP | miRTarbase |
| hsa-miR-3613-5p | EIF4A1 | PAR-CLIP | miRTarbase |
| hsa-miR-486-5p | YAE1D1 | HITS-CLIP | miRTarbase |
| hsa-miR-486-5p | YAE1D1 | HITS-CLIP | miRTarbase |
| hsa-miR-486-5p | YAE1D1 | HITS-CLIP | miRTarbase |
| hsa-miR-3613-3p | TANGO2 | HITS-CLIP | miRTarbase |
| hsa-miR-195-3p | VSNL1 | HITS-CLIP | miRTarbase |
| hsa-miR-486-3p | HS6ST3 | HITS-CLIP | miRTarbase |
| hsa-miR-486-3p | HS6ST3 | HITS-CLIP | miRTarbase |
| hsa-miR-486-3p | HS6ST3 | HITS-CLIP | miRTarbase |
| hsa-miR-486-3p | ERBB2 | HITS-CLIP | miRTarbase |
| hsa-miR-3613-3p | PARP15 | HITS-CLIP | miRTarbase |
| hsa-miR-141-5p | SERPINA4 | HITS-CLIP | miRTarbase |
| hsa-miR-3613-5p | MTX3 | HITS-CLIP | miRTarbase |
| hsa-miR-141-5p | TRPC4AP | HITS-CLIP | miRTarbase |
| hsa-miR-3613-5p | NHSL1 | HITS-CLIP | miRTarbase |
| hsa-miR-3613-3p | SMS | HITS-CLIP | miRTarbase |
| hsa-miR-486-5p | SEL1L3 | HITS-CLIP | miRTarbase |
| hsa-miR-486-5p | SEL1L3 | HITS-CLIP | miRTarbase |
| hsa-miR-141-5p | HSPA2 | HITS-CLIP | miRTarbase |
| hsa-miR-3613-3p | TMEM44 | HITS-CLIP | miRTarbase |
| hsa-miR-3613-3p | TMEM44 | HITS-CLIP | miRTarbase |
| hsa-miR-141-5p | BARHL2 | HITS-CLIP | miRTarbase |
| hsa-miR-15a-3p | SPTA1 | HITS-CLIP | miRTarbase |
| hsa-miR-15a-3p | SPTA1 | HITS-CLIP | miRTarbase |
| hsa-miR-15a-3p | SPTA1 | HITS-CLIP | miRTarbase |
| hsa-miR-195-3p | CTGF | HITS-CLIP | miRTarbase |
| hsa-miR-141-5p | AGPAT4 | HITS-CLIP | miRTarbase |
| hsa-miR-3613-3p | SGOL2 | HITS-CLIP | miRTarbase |
| hsa-miR-3613-3p | NUDCD3 | HITS-CLIP | miRTarbase |
| hsa-miR-3613-3p | HINFP | HITS-CLIP | miRTarbase |
| hsa-miR-3613-3p | BSDC1 | HITS-CLIP | miRTarbase |
| hsa-miR-141-5p | RRP1B | HITS-CLIP | miRTarbase |
| hsa-miR-141-5p | ZWILCH | HITS-CLIP | miRTarbase |
| hsa-miR-3613-3p | HSPA4 | HITS-CLIP | miRTarbase |
| hsa-miR-3613-3p | HSPA4 | HITS-CLIP | miRTarbase |
| hsa-miR-3613-3p | GDPD1 | HITS-CLIP | miRTarbase |
| hsa-miR-141-5p | PRSS23 | HITS-CLIP | miRTarbase |
| hsa-miR-3613-3p | UTP20 | HITS-CLIP | miRTarbase |
| hsa-miR-3613-3p | PCNXL2 | HITS-CLIP | miRTarbase |
| hsa-miR-3613-3p | PCNXL2 | HITS-CLIP | miRTarbase |
| hsa-miR-3613-3p | PCNXL2 | HITS-CLIP | miRTarbase |
| hsa-miR-141-5p | MICALL1 | HITS-CLIP | miRTarbase |
| hsa-miR-141-5p | SP1 | HITS-CLIP | miRTarbase |
| hsa-miR-3613-3p | PSPC1 | HITS-CLIP | miRTarbase |
| hsa-miR-3613-3p | GNAQ | HITS-CLIP | miRTarbase |
| hsa-miR-141-5p | GBX2 | HITS-CLIP | miRTarbase |
| hsa-miR-486-5p | FOXP1 | HITS-CLIP | miRTarbase |
| hsa-miR-141-5p | EXTL3 | HITS-CLIP | miRTarbase |
| hsa-miR-141-5p | BACE2 | HITS-CLIP | miRTarbase |
| hsa-miR-141-5p | ABCC12 | HITS-CLIP | miRTarbase |
| hsa-miR-141-5p | TMEM132C | HITS-CLIP | miRTarbase |
| hsa-miR-3613-3p | SCN1A | HITS-CLIP | miRTarbase |
| hsa-miR-3613-5p | NTNG2 | HITS-CLIP | miRTarbase |
| hsa-miR-3613-3p | INTS7 | HITS-CLIP | miRTarbase |
| hsa-miR-141-5p | DNAJB2 | HITS-CLIP | miRTarbase |
| hsa-miR-486-5p | TM4SF20 | HITS-CLIP | miRTarbase |
| hsa-miR-3613-3p | TM4SF20 | HITS-CLIP | miRTarbase |
| hsa-miR-3613-3p | TMTC1 | HITS-CLIP | miRTarbase |
| hsa-miR-141-3p | LHX1 | HITS-CLIP | miRTarbase |
| hsa-miR-15a-5p | TRAK1 | HITS-CLIP | miRTarbase |
| hsa-miR-195-5p | TRAK1 | HITS-CLIP | miRTarbase |
| hsa-miR-3605-5p | PLEKHA6 | HITS-CLIP | miRTarbase |
| hsa-miR-3605-5p | PLEKHA6 | HITS-CLIP | miRTarbase |
| hsa-miR-3613-3p | BRD4 | HITS-CLIP | miRTarbase |
| hsa-miR-3613-3p | FGF5 | HITS-CLIP | miRTarbase |
| hsa-miR-3613-3p | DNTTIP2 | HITS-CLIP | miRTarbase |
| hsa-miR-3613-3p | DLG4 | HITS-CLIP | miRTarbase |
| hsa-miR-3613-3p | CCNF | HITS-CLIP | miRTarbase |
| hsa-miR-3613-3p | SLC35C2 | HITS-CLIP | miRTarbase |
| hsa-miR-141-5p | CYP2A7 | HITS-CLIP | miRTarbase |
| hsa-miR-3613-3p | PSMD11 | HITS-CLIP | miRTarbase |
| hsa-miR-3613-3p | GGCX | HITS-CLIP | miRTarbase |
| hsa-miR-141-5p | HMGB1 | HITS-CLIP | miRTarbase |
| hsa-miR-3613-3p | FAM117B | HITS-CLIP | miRTarbase |
| hsa-miR-3613-3p | ACTR10 | HITS-CLIP | miRTarbase |
| hsa-miR-3613-3p | DPF3 | HITS-CLIP | miRTarbase |
| hsa-miR-3613-3p | FAM126B | HITS-CLIP | miRTarbase |
| hsa-miR-195-5p | ADRA2B | HITS-CLIP | miRTarbase |
| hsa-miR-15a-5p | ADRA2B | HITS-CLIP | miRTarbase |
| hsa-miR-3613-3p | PARD6B | HITS-CLIP | miRTarbase |
| hsa-miR-3613-3p | PARD6B | HITS-CLIP | miRTarbase |
| hsa-miR-141-5p | ZNF629 | HITS-CLIP | miRTarbase |
| hsa-miR-141-5p | ZNF638 | HITS-CLIP | miRTarbase |
| hsa-miR-141-5p | SPIB | HITS-CLIP | miRTarbase |
| hsa-miR-3613-3p | HECTD1 | HITS-CLIP | miRTarbase |
| hsa-miR-3613-3p | SPIC | HITS-CLIP | miRTarbase |
| hsa-miR-3613-3p | MICA | HITS-CLIP | miRTarbase |
| hsa-miR-3613-3p | IGSF11 | HITS-CLIP | miRTarbase |
| hsa-miR-3613-3p | IGSF11 | HITS-CLIP | miRTarbase |
| hsa-miR-3613-3p | ZNF568 | HITS-CLIP | miRTarbase |
| hsa-miR-3613-3p | SPECC1 | HITS-CLIP | miRTarbase |
| hsa-miR-141-5p | HLA-DRB5 | HITS-CLIP | miRTarbase |
| hsa-miR-195-5p | ANKMY1 | HITS-CLIP | miRTarbase |
| hsa-miR-15a-5p | ANKMY1 | HITS-CLIP | miRTarbase |
| hsa-miR-3613-3p | FUNDC2 | HITS-CLIP | miRTarbase |
| hsa-miR-3613-3p | CWF19L1 | HITS-CLIP | miRTarbase |
| hsa-miR-3613-3p | LMBR1 | HITS-CLIP | miRTarbase |
| hsa-miR-195-5p | RNF41 | HITS-CLIP | miRTarbase |
| hsa-miR-3613-3p | TMEM220 | HITS-CLIP | miRTarbase |
| hsa-miR-3613-3p | SRFBP1 | HITS-CLIP | miRTarbase |
| hsa-miR-3613-3p | NRIP2 | HITS-CLIP | miRTarbase |
| hsa-miR-3613-3p | CCDC65 | HITS-CLIP | miRTarbase |
| hsa-miR-3613-3p | RAPH1 | HITS-CLIP | miRTarbase |
| hsa-miR-3613-3p | TLN1 | HITS-CLIP | miRTarbase |
| hsa-miR-141-5p | C10orf10 | HITS-CLIP | miRTarbase |
| hsa-miR-141-5p | C10orf10 | HITS-CLIP | miRTarbase |
| hsa-miR-3613-3p | POLR2D | HITS-CLIP | miRTarbase |
| hsa-miR-3613-3p | TM4SF5 | HITS-CLIP | miRTarbase |
| hsa-miR-3613-3p | RPL3L | HITS-CLIP | miRTarbase |
| hsa-miR-3613-3p | ITGB8 | HITS-CLIP | miRTarbase |
| hsa-miR-195-5p | GPRC5A | HITS-CLIP | miRTarbase |
| hsa-miR-195-5p | GPRC5A | HITS-CLIP | miRTarbase |
| hsa-miR-15a-5p | GPRC5A | HITS-CLIP | miRTarbase |
| hsa-miR-15a-5p | GPRC5A | HITS-CLIP | miRTarbase |
| hsa-miR-3613-3p | ZNF514 | HITS-CLIP | miRTarbase |
| hsa-miR-3613-3p | SV2C | HITS-CLIP | miRTarbase |
| hsa-miR-3613-3p | STK17B | HITS-CLIP | miRTarbase |
| hsa-miR-3613-3p | SEH1L | HITS-CLIP | miRTarbase |
| hsa-miR-3613-3p | RRAGC | HITS-CLIP | miRTarbase |
| hsa-miR-486-3p | PLXNA4 | HITS-CLIP | miRTarbase |
| hsa-miR-486-5p | MARCH4 | HITS-CLIP | miRTarbase |
| hsa-miR-195-3p | LPP | HITS-CLIP | miRTarbase |
| hsa-miR-3605-5p | LHX4 | HITS-CLIP | miRTarbase |
| hsa-miR-3613-3p | HINT1 | HITS-CLIP | miRTarbase |
| hsa-miR-3613-3p | GATSL2 | HITS-CLIP | miRTarbase |
| hsa-miR-141-5p | GALNT1 | HITS-CLIP | miRTarbase |
| hsa-miR-3613-3p | FGF12 | HITS-CLIP | miRTarbase |
| hsa-miR-3613-3p | FBN2 | HITS-CLIP | miRTarbase |
| hsa-miR-3613-3p | FAM104A | HITS-CLIP | miRTarbase |
| hsa-miR-3613-3p | EDEM1 | HITS-CLIP | miRTarbase |
| hsa-miR-3613-3p | CRISPLD2 | HITS-CLIP | miRTarbase |
| hsa-miR-3613-3p | CNOT6L | HITS-CLIP | miRTarbase |
| hsa-miR-141-5p | CISD1 | HITS-CLIP | miRTarbase |
| hsa-miR-3613-3p | C7orf73 | HITS-CLIP | miRTarbase |
| hsa-miR-141-5p | BLMH | HITS-CLIP | miRTarbase |
| hsa-miR-195-3p | AR | HITS-CLIP | miRTarbase |
| hsa-miR-141-5p | MARCH2 | HITS-CLIP | miRTarbase |
| hsa-miR-3613-3p | PAK4 | HITS-CLIP | miRTarbase |
| hsa-miR-3613-3p | OGFOD1 | HITS-CLIP | miRTarbase |
| hsa-miR-3613-3p | CCS | HITS-CLIP | miRTarbase |
| hsa-miR-3613-3p | ZNF619 | HITS-CLIP | miRTarbase |
| hsa-miR-3613-3p | TP73 | HITS-CLIP | miRTarbase |
| hsa-miR-3613-3p | PAQR7 | HITS-CLIP | miRTarbase |
| hsa-miR-486-3p | ARHGAP18 | HITS-CLIP | miRTarbase |
| hsa-miR-3613-3p | PIK3C2B | HITS-CLIP | miRTarbase |
| hsa-miR-141-5p | BID | HITS-CLIP | miRTarbase |
| hsa-miR-3613-3p | ASAP2 | HITS-CLIP | miRTarbase |

| | | | |
|---|---|---|---|
| hsa-miR-3613-3p | SLC19A1 | HITS-CLIP | miRTarbase |
| hsa-miR-3613-3p | ZBTB3 | HITS-CLIP | miRTarbase |
| hsa-miR-141-5p | STRN | HITS-CLIP | miRTarbase |
| hsa-miR-3613-3p | STRIP2 | HITS-CLIP | miRTarbase |
| hsa-miR-486-5p | SP4 | HITS-CLIP | miRTarbase |
| hsa-miR-3613-3p | SNIP1 | HITS-CLIP | miRTarbase |
| hsa-miR-486-5p | SEC23IP | HITS-CLIP | miRTarbase |
| hsa-miR-3613-3p | RFFL | HITS-CLIP | miRTarbase |
| hsa-miR-3613-3p | RAD51L3-RFFL | HITS-CLIP | miRTarbase |
| hsa-miR-486-3p | OLFML2A | HITS-CLIP | miRTarbase |
| hsa-miR-3613-3p | FREM2 | HITS-CLIP | miRTarbase |
| hsa-miR-3613-3p | FKBP9 | HITS-CLIP | miRTarbase |
| hsa-miR-3613-3p | C8orf37 | HITS-CLIP | miRTarbase |
| hsa-miR-3613-3p | ANAPC16 | HITS-CLIP | miRTarbase |
| hsa-miR-486-5p | BAG2 | HITS-CLIP | miRTarbase |
| hsa-miR-486-5p | WBSCR27 | HITS-CLIP | miRTarbase |
| hsa-miR-15a-3p | IGSF6 | HITS-CLIP | miRTarbase |
| hsa-miR-3605-3p | KDM2B | HITS-CLIP | miRTarbase |
| hsa-miR-15a-3p | CLEC17A | HITS-CLIP | miRTarbase |
| hsa-miR-3613-5p | COG6 | HITS-CLIP | miRTarbase |
| hsa-miR-3605-3p | RNASEH2B | HITS-CLIP | miRTarbase |
| hsa-miR-195-3p | ALDH1A2 | HITS-CLIP | miRTarbase |
| hsa-miR-141-3p | YME1L1 | HITS-CLIP | miRTarbase |
| hsa-miR-15a-3p | SRRD | HITS-CLIP | miRTarbase |
| hsa-miR-15a-3p | RBM3 | HITS-CLIP | miRTarbase |
| hsa-miR-486-5p | DENND5B | HITS-CLIP | miRTarbase |
| hsa-miR-15a-3p | DSN1 | HITS-CLIP | miRTarbase |
| hsa-miR-3613-3p | PLEKHA3 | HITS-CLIP | miRTarbase |
| hsa-miR-141-3p | APOOL | HITS-CLIP | miRTarbase |
| hsa-miR-3613-3p | CD59 | HITS-CLIP | miRTarbase |
| hsa-miR-141-5p | FAM179A | HITS-CLIP | miRTarbase |
| hsa-miR-3613-3p | PRRG3 | HITS-CLIP | miRTarbase |
| hsa-miR-3613-3p | PRRG3 | HITS-CLIP | miRTarbase |
| hsa-miR-3613-3p | ATP5A1 | HITS-CLIP | miRTarbase |
| hsa-miR-3613-3p | GLTSCR2 | HITS-CLIP | miRTarbase |
| hsa-miR-3613-5p | LHFP | HITS-CLIP | miRTarbase |
| hsa-miR-3613-5p | LHFP | HITS-CLIP | miRTarbase |
| hsa-miR-3613-5p | AFAP1 | HITS-CLIP | miRTarbase |
| hsa-miR-3613-3p | DEFB105B | HITS-CLIP | miRTarbase |
| hsa-miR-3613-3p | DEFB105A | HITS-CLIP | miRTarbase |
| hsa-miR-3613-3p | B3GNT7 | HITS-CLIP | miRTarbase |
| hsa-miR-486-3p | SC5D | HITS-CLIP | miRTarbase |
| hsa-miR-486-3p | IER5 | HITS-CLIP | miRTarbase |
| hsa-miR-3613-3p | CDK9 | HITS-CLIP | miRTarbase |
| hsa-miR-3613-3p | RNF2 | HITS-CLIP | miRTarbase |
| hsa-miR-141-5p | ZNF429 | HITS-CLIP | miRTarbase |
| hsa-miR-3613-3p | ACER2 | HITS-CLIP | miRTarbase |
| hsa-miR-141-3p | ZNF621 | HITS-CLIP | miRTarbase |
| hsa-miR-141-5p | SLC25A32 | HITS-CLIP | miRTarbase |
| hsa-miR-141-5p | SLC25A32 | HITS-CLIP | miRTarbase |
| hsa-miR-141-5p | AHCY | HITS-CLIP | miRTarbase |
| hsa-miR-141-5p | SLIT1 | HITS-CLIP | miRTarbase |
| hsa-miR-195-5p | C3orf36 | HITS-CLIP | miRTarbase |
| hsa-miR-15a-5p | C3orf36 | HITS-CLIP | miRTarbase |
| hsa-miR-141-5p | TM9SF4 | HITS-CLIP | miRTarbase |
| hsa-miR-141-5p | ZBTB26 | HITS-CLIP | miRTarbase |
| hsa-miR-141-5p | POU2F1 | HITS-CLIP | miRTarbase |
| hsa-miR-3613-3p | ZBTB33 | HITS-CLIP | miRTarbase |
| hsa-miR-141-5p | COL19A1 | HITS-CLIP | miRTarbase |
| hsa-miR-141-5p | SLX4 | HITS-CLIP | miRTarbase |
| hsa-miR-3613-3p | RABGEF1 | HITS-CLIP | miRTarbase |
| hsa-miR-3605-5p | POLA2 | HITS-CLIP | miRTarbase |
| hsa-miR-141-5p | SNW1 | HITS-CLIP | miRTarbase |
| hsa-miR-486-5p | MACROD2 | HITS-CLIP | miRTarbase |
| hsa-miR-141-5p | SMIM17 | HITS-CLIP | miRTarbase |
| hsa-miR-3613-3p | ZNF99 | HITS-CLIP | miRTarbase |
| hsa-miR-141-5p | EPPIN | HITS-CLIP | miRTarbase |
| hsa-miR-141-5p | EPPIN-WFDC6 | HITS-CLIP | miRTarbase |
| hsa-miR-141-5p | KANSL3 | HITS-CLIP | miRTarbase |
| hsa-miR-195-3p | CBS | HITS-CLIP | miRTarbase |
| hsa-miR-141-5p | FRMD6 | HITS-CLIP | miRTarbase |
| hsa-miR-3613-3p | C17orf77 | HITS-CLIP | miRTarbase |
| hsa-miR-3613-3p | SRSF2 | HITS-CLIP | miRTarbase |
| hsa-miR-3613-3p | PGM2 | HITS-CLIP | miRTarbase |
| hsa-miR-3613-3p | STEAP4 | HITS-CLIP | miRTarbase |
| hsa-miR-3613-3p | RPP40 | HITS-CLIP | miRTarbase |
| hsa-miR-141-5p | MAGEB4 | HITS-CLIP | miRTarbase |
| hsa-miR-141-3p | ANGPTL7 | HITS-CLIP | miRTarbase |
| hsa-miR-3613-3p | C9orf64 | HITS-CLIP | miRTarbase |
| hsa-miR-3613-3p | C9orf47 | HITS-CLIP | miRTarbase |
| hsa-miR-141-5p | MKX | HITS-CLIP | miRTarbase |
| hsa-miR-195-5p | BSPRY | HITS-CLIP | miRTarbase |
| hsa-miR-15a-5p | BSPRY | HITS-CLIP | miRTarbase |
| hsa-miR-141-5p | HLA-DRB1 | HITS-CLIP | miRTarbase |
| hsa-miR-195-5p | ANKRD36 | HITS-CLIP | miRTarbase |
| hsa-miR-15a-5p | ANKRD36 | HITS-CLIP | miRTarbase |
| hsa-miR-3613-3p | CDCA2 | HITS-CLIP | miRTarbase |
| hsa-miR-486-3p | ZNF616 | HITS-CLIP | miRTarbase |
| hsa-miR-3605-5p | FAM109B | HITS-CLIP | miRTarbase |
| hsa-miR-141-5p | FAM109B | HITS-CLIP | miRTarbase |
| hsa-miR-3613-3p | IGSF9B | HITS-CLIP | miRTarbase |
| hsa-miR-195-3p | BID | HITS-CLIP | miRTarbase |
| hsa-miR-486-5p | CIRH1A | HITS-CLIP | miRTarbase |
| hsa-miR-3613-3p | ZNF562 | HITS-CLIP | miRTarbase |
| hsa-miR-195-5p | KLHL40 | HITS-CLIP | miRTarbase |
| hsa-miR-15a-5p | KLHL40 | HITS-CLIP | miRTarbase |
| hsa-miR-15a-3p | SLC1A2 | HITS-CLIP | miRTarbase |
| hsa-miR-486-5p | LTBP2 | HITS-CLIP | miRTarbase |
| hsa-miR-3613-3p | LPCAT3 | HITS-CLIP | miRTarbase |
| hsa-miR-141-5p | TRAFD1 | HITS-CLIP | miRTarbase |
| hsa-miR-141-5p | KRBA1 | HITS-CLIP | miRTarbase |
| hsa-miR-141-5p | MSANTD2 | HITS-CLIP | miRTarbase |
| hsa-miR-486-3p | WNT16 | HITS-CLIP | miRTarbase |
| hsa-miR-15a-3p | PGRMC1 | HITS-CLIP | miRTarbase |
| hsa-miR-3613-3p | ZNF518B | HITS-CLIP | miRTarbase |
| hsa-miR-3613-5p | XKR4 | HITS-CLIP | miRTarbase |
| hsa-miR-141-5p | WASF3 | HITS-CLIP | miRTarbase |
| hsa-miR-195-3p | WASF2 | HITS-CLIP | miRTarbase |
| hsa-miR-3613-3p | VPS37A | HITS-CLIP | miRTarbase |
| hsa-miR-3613-3p | TRA2B | HITS-CLIP | miRTarbase |
| hsa-miR-141-5p | TMEM2 | HITS-CLIP | miRTarbase |
| hsa-miR-3613-3p | SNAP29 | HITS-CLIP | miRTarbase |
| hsa-miR-141-5p | SLC38A7 | HITS-CLIP | miRTarbase |
| hsa-miR-141-5p | SKIL | HITS-CLIP | miRTarbase |
| hsa-miR-141-5p | SIRPA | HITS-CLIP | miRTarbase |
| hsa-miR-141-5p | RSPH4A | HITS-CLIP | miRTarbase |
| hsa-miR-3613-3p | RASSF2 | HITS-CLIP | miRTarbase |
| hsa-miR-3605-3p | PSMB5 | HITS-CLIP | miRTarbase |
| hsa-miR-141-3p | PRKAA2 | HITS-CLIP | miRTarbase |
| hsa-miR-3613-3p | PNO1 | HITS-CLIP | miRTarbase |
| hsa-miR-3605-3p | PLXDC2 | HITS-CLIP | miRTarbase |
| hsa-miR-141-5p | PHAX | HITS-CLIP | miRTarbase |
| hsa-miR-3613-3p | PAK3 | HITS-CLIP | miRTarbase |
| hsa-miR-3613-3p | NUDT21 | HITS-CLIP | miRTarbase |
| hsa-miR-195-5p | NOTCH2 | HITS-CLIP | miRTarbase |
| hsa-miR-195-5p | NOTCH2 | HITS-CLIP | miRTarbase |
| hsa-miR-15a-5p | NOTCH2 | HITS-CLIP | miRTarbase |
| hsa-miR-15a-5p | NOTCH2 | HITS-CLIP | miRTarbase |
| hsa-miR-3613-3p | NGDN | HITS-CLIP | miRTarbase |
| hsa-miR-141-5p | MYO5A | HITS-CLIP | miRTarbase |
| hsa-miR-141-5p | MYO5A | HITS-CLIP | miRTarbase |
| hsa-miR-141-5p | MTA3 | HITS-CLIP | miRTarbase |
| hsa-miR-141-5p | MFSD6 | HITS-CLIP | miRTarbase |
| hsa-miR-3613-3p | MBOAT2 | HITS-CLIP | miRTarbase |
| hsa-miR-141-3p | MACC1 | HITS-CLIP | miRTarbase |
| hsa-miR-3613-3p | MACC1 | HITS-CLIP | miRTarbase |
| hsa-miR-3613-3p | KCNN3 | HITS-CLIP | miRTarbase |
| hsa-miR-141-5p | INO80D | HITS-CLIP | miRTarbase |
| hsa-miR-141-5p | HOXB5 | HITS-CLIP | miRTarbase |
| hsa-miR-3613-3p | C21orf33 | HITS-CLIP | miRTarbase |
| hsa-miR-3613-3p | GJD3 | HITS-CLIP | miRTarbase |
| hsa-miR-141-5p | GFPT1 | HITS-CLIP | miRTarbase |
| hsa-miR-141-5p | GAN | HITS-CLIP | miRTarbase |
| hsa-miR-3613-3p | FRMD3 | HITS-CLIP | miRTarbase |
| hsa-miR-486-3p | FAM167B | HITS-CLIP | miRTarbase |
| hsa-miR-141-5p | EPB41L4B | HITS-CLIP | miRTarbase |
| hsa-miR-141-5p | EIF4EBP2 | HITS-CLIP | miRTarbase |
| hsa-miR-195-5p | EIF2B2 | HITS-CLIP | miRTarbase |
| hsa-miR-15a-5p | EIF2B2 | HITS-CLIP | miRTarbase |

| | | | |
|---|---|---|---|
| hsa-miR-486-3p | DNAJC6 | HITS-CLIP | miRTarbase |
| hsa-miR-195-5p | CUL3 | HITS-CLIP | miRTarbase |
| hsa-miR-15a-5p | CUL3 | HITS-CLIP | miRTarbase |
| hsa-miR-3613-3p | CLSPN | HITS-CLIP | miRTarbase |
| hsa-miR-141-5p | CLDN1 | HITS-CLIP | miRTarbase |
| hsa-miR-486-3p | CDC42EP4 | HITS-CLIP | miRTarbase |
| hsa-miR-141-5p | CAPRIN1 | HITS-CLIP | miRTarbase |
| hsa-miR-141-5p | B3GALT5 | HITS-CLIP | miRTarbase |
| hsa-miR-486-5p | ARF6 | HITS-CLIP | miRTarbase |
| hsa-miR-3613-3p | APLF | HITS-CLIP | miRTarbase |
| hsa-miR-3613-3p | ANTXR2 | HITS-CLIP | miRTarbase |
| hsa-miR-3613-3p | AGPS | HITS-CLIP | miRTarbase |
| hsa-miR-141-5p | SERPINB5 | HITS-CLIP | miRTarbase |
| hsa-miR-15a-3p | HM13 | HITS-CLIP | miRTarbase |
| hsa-miR-15a-3p | TMEM216 | HITS-CLIP | miRTarbase |
| hsa-miR-141-3p | CCDC18 | HITS-CLIP | miRTarbase |
| hsa-miR-15a-3p | TRIM72 | HITS-CLIP | miRTarbase |
| hsa-miR-3613-3p | DHODH | HITS-CLIP | miRTarbase |
| hsa-miR-3613-3p | SHMT1 | HITS-CLIP | miRTarbase |
| hsa-miR-15a-3p | RNF157 | HITS-CLIP | miRTarbase |
| hsa-miR-141-5p | MYOCD | HITS-CLIP | miRTarbase |
| hsa-miR-3613-3p | KCNB1 | HITS-CLIP | miRTarbase |
| hsa-miR-3613-3p | HCFC2 | HITS-CLIP | miRTarbase |
| hsa-miR-3613-3p | GINS2 | HITS-CLIP | miRTarbase |
| hsa-miR-141-5p | GABRA1 | HITS-CLIP | miRTarbase |
| hsa-miR-3613-3p | ENAH | HITS-CLIP | miRTarbase |
| hsa-miR-141-5p | CHRFAM7A | HITS-CLIP | miRTarbase |
| hsa-miR-141-5p | C19orf44 | HITS-CLIP | miRTarbase |
| hsa-miR-486-5p | RBM22 | HITS-CLIP | miRTarbase |
| hsa-miR-3613-3p | KLHDC8A | HITS-CLIP | miRTarbase |
| hsa-miR-15a-3p | NDUFA7 | HITS-CLIP | miRTarbase |
| hsa-miR-15a-3p | SLU7 | HITS-CLIP | miRTarbase |
| hsa-miR-486-3p | ITGB3 | HITS-CLIP | miRTarbase |
| hsa-miR-3613-3p | TBPL1 | HITS-CLIP | miRTarbase |
| hsa-miR-195-5p | DCAF17 | HITS-CLIP | miRTarbase |
| hsa-miR-15a-5p | DCAF17 | HITS-CLIP | miRTarbase |
| hsa-miR-219a-1-3p | CIITA | HITS-CLIP | miRTarbase |
| hsa-miR-15a-5p | RS1 | HITS-CLIP | miRTarbase |
| hsa-miR-195-5p | RS1 | HITS-CLIP | miRTarbase |
| hsa-miR-195-5p | GLP2R | HITS-CLIP | miRTarbase |
| hsa-miR-15a-5p | GLP2R | HITS-CLIP | miRTarbase |
| hsa-miR-3605-5p | COX6B1 | HITS-CLIP | miRTarbase |
| hsa-miR-3613-3p | CPM | HITS-CLIP | miRTarbase |
| hsa-miR-141-3p | DNAJC28 | HITS-CLIP | miRTarbase |
| hsa-miR-141-3p | ZNF292 | HITS-CLIP | miRTarbase |
| hsa-miR-141-3p | RPL12 | HITS-CLIP | miRTarbase |
| hsa-miR-141-3p | A1BG | HITS-CLIP | miRTarbase |
| hsa-miR-3613-3p | ZNF581 | HITS-CLIP | miRTarbase |
| hsa-miR-141-5p | MICB | HITS-CLIP | miRTarbase |
| hsa-miR-3613-3p | MCM10 | HITS-CLIP | miRTarbase |
| hsa-miR-3613-3p | IFIT3 | HITS-CLIP | miRTarbase |
| hsa-miR-3613-3p | MCTS1 | HITS-CLIP | miRTarbase |
| hsa-miR-3613-3p | RPL10A | HITS-CLIP | miRTarbase |
| hsa-miR-3613-3p | COX18 | HITS-CLIP | miRTarbase |
| hsa-miR-3613-3p | FBXO25 | HITS-CLIP | miRTarbase |
| hsa-miR-3613-3p | LSG1 | HITS-CLIP | miRTarbase |
| hsa-miR-141-5p | TXK | HITS-CLIP | miRTarbase |
| hsa-miR-3613-3p | TVP23C | HITS-CLIP | miRTarbase |
| hsa-miR-15a-3p | PTGFRN | HITS-CLIP | miRTarbase |
| hsa-miR-141-5p | NSUN4 | HITS-CLIP | miRTarbase |
| hsa-miR-3613-3p | KIAA1467 | HITS-CLIP | miRTarbase |
| hsa-miR-3613-3p | HHIP | HITS-CLIP | miRTarbase |
| hsa-miR-3613-3p | DNA2 | HITS-CLIP | miRTarbase |
| hsa-miR-486-3p | ATP6AP1 | HITS-CLIP | miRTarbase |
| hsa-miR-195-3p | AGMAT | HITS-CLIP | miRTarbase |
| hsa-miR-3613-3p | ZNF574 | HITS-CLIP | miRTarbase |
| hsa-miR-3613-3p | C5AR2 | HITS-CLIP | miRTarbase |
| hsa-miR-3613-3p | NXN | HITS-CLIP | miRTarbase |
| hsa-miR-486-3p | WDR73 | HITS-CLIP | miRTarbase |
| hsa-miR-3613-3p | PLA2G2C | HITS-CLIP | miRTarbase |
| hsa-miR-3613-3p | SGSM2 | HITS-CLIP | miRTarbase |
| hsa-miR-3605-5p | GET4 | HITS-CLIP | miRTarbase |
| hsa-miR-195-5p | FLOT2 | HITS-CLIP | miRTarbase |
| hsa-miR-195-5p | FLOT2 | HITS-CLIP | miRTarbase |
| hsa-miR-15a-5p | FLOT2 | HITS-CLIP | miRTarbase |
| hsa-miR-15a-5p | FLOT2 | HITS-CLIP | miRTarbase |
| hsa-miR-3613-3p | PTCD2 | HITS-CLIP | miRTarbase |
| hsa-miR-3613-3p | RNF207 | HITS-CLIP | miRTarbase |
| hsa-miR-15a-3p | POLR1E | HITS-CLIP | miRTarbase |
| hsa-miR-141-3p | POLR3F | HITS-CLIP | miRTarbase |
| hsa-miR-3613-3p | ORC1 | HITS-CLIP | miRTarbase |
| hsa-miR-3613-3p | MXRA7 | HITS-CLIP | miRTarbase |
| hsa-miR-3613-5p | HNRNPA1L2 | HITS-CLIP | miRTarbase |
| hsa-miR-195-5p | HNRNPA1L2 | HITS-CLIP | miRTarbase |
| hsa-miR-15a-5p | HNRNPA1L2 | HITS-CLIP | miRTarbase |
| hsa-miR-3613-5p | NCR3LG1 | HITS-CLIP | miRTarbase |
| hsa-miR-486-3p | MTA1 | HITS-CLIP | miRTarbase |
| hsa-miR-3613-3p | GSG2 | HITS-CLIP | miRTarbase |
| hsa-miR-3613-3p | LRTOMT | HITS-CLIP | miRTarbase |
| hsa-miR-3613-3p | PHAX | HITS-CLIP | miRTarbase |
| hsa-miR-486-3p | ALPI | HITS-CLIP | miRTarbase |
| hsa-miR-3613-3p | CLPB | HITS-CLIP | miRTarbase |
| hsa-miR-141-5p | WDR35 | HITS-CLIP | miRTarbase |
| hsa-miR-3613-3p | GLP2R | HITS-CLIP | miRTarbase |
| hsa-miR-3613-3p | LYZ | HITS-CLIP | miRTarbase |
| hsa-miR-486-3p | EIF2S3 | HITS-CLIP | miRTarbase |
| hsa-miR-141-3p | COX6B1 | HITS-CLIP | miRTarbase |
| hsa-miR-486-3p | CRIPT | HITS-CLIP | miRTarbase |
| hsa-miR-3613-3p | APOC3 | HITS-CLIP | miRTarbase |
| hsa-miR-141-3p | RBM28 | HITS-CLIP | miRTarbase |
| hsa-miR-3613-3p | SAMD15 | HITS-CLIP | miRTarbase |
| hsa-miR-15a-3p | USP8 | HITS-CLIP | miRTarbase |
| hsa-miR-3605-5p | TPM1 | HITS-CLIP | miRTarbase |
| hsa-miR-195-3p | TNPO1 | HITS-CLIP | miRTarbase |
| hsa-miR-3613-3p | TNFRSF13C | HITS-CLIP | miRTarbase |
| hsa-miR-3613-5p | TMEM66 | HITS-CLIP | miRTarbase |
| hsa-miR-219a-1-3p | TMEM2 | HITS-CLIP | miRTarbase |
| hsa-miR-3613-3p | TMEM127 | HITS-CLIP | miRTarbase |
| hsa-miR-219a-1-3p | TMED2 | HITS-CLIP | miRTarbase |
| hsa-miR-219a-1-3p | TEX261 | HITS-CLIP | miRTarbase |
| hsa-miR-3613-3p | SOS1 | HITS-CLIP | miRTarbase |
| hsa-miR-141-5p | RUNX1 | HITS-CLIP | miRTarbase |
| hsa-miR-195-3p | PTPN14 | HITS-CLIP | miRTarbase |
| hsa-miR-219a-1-3p | ORAI2 | HITS-CLIP | miRTarbase |
| hsa-miR-3613-3p | NT5DC3 | HITS-CLIP | miRTarbase |
| hsa-miR-195-5p | NEGR1 | HITS-CLIP | miRTarbase |
| hsa-miR-15a-5p | NEGR1 | HITS-CLIP | miRTarbase |
| hsa-miR-486-3p | MYADM | HITS-CLIP | miRTarbase |
| hsa-miR-195-5p | MCFD2 | HITS-CLIP | miRTarbase |
| hsa-miR-15a-5p | MCFD2 | HITS-CLIP | miRTarbase |
| hsa-miR-15a-3p | LPP | HITS-CLIP | miRTarbase |
| hsa-miR-141-3p | LPP | HITS-CLIP | miRTarbase |
| hsa-miR-3613-3p | IFNLR1 | HITS-CLIP | miRTarbase |
| hsa-miR-3613-5p | HNRNPA1 | HITS-CLIP | miRTarbase |
| hsa-miR-195-5p | HNRNPA1 | HITS-CLIP | miRTarbase |
| hsa-miR-15a-5p | HNRNPA1 | HITS-CLIP | miRTarbase |
| hsa-miR-3613-3p | FNBP1 | HITS-CLIP | miRTarbase |
| hsa-miR-3613-3p | ERGIC1 | HITS-CLIP | miRTarbase |
| hsa-miR-195-3p | DYRK2 | HITS-CLIP | miRTarbase |
| hsa-miR-3613-3p | DCAF16 | HITS-CLIP | miRTarbase |
| hsa-miR-3613-3p | CLN8 | HITS-CLIP | miRTarbase |
| hsa-miR-3613-3p | CDC73 | HITS-CLIP | miRTarbase |
| hsa-miR-3613-3p | C5orf51 | HITS-CLIP | miRTarbase |
| hsa-miR-195-3p | ACBD5 | HITS-CLIP | miRTarbase |
| hsa-miR-486-3p | PKD1 | HITS-CLIP | miRTarbase |
| hsa-miR-3605-5p | TMEM186 | HITS-CLIP | miRTarbase |
| hsa-miR-3613-3p | TARDBP | HITS-CLIP | miRTarbase |
| hsa-miR-3613-3p | TBX20 | HITS-CLIP | miRTarbase |
| hsa-miR-3613-3p | CERCAM | HITS-CLIP | miRTarbase |
| hsa-miR-3613-3p | SPATS2 | HITS-CLIP | miRTarbase |
| hsa-miR-3613-3p | OLR1 | HITS-CLIP | miRTarbase |
| hsa-miR-141-3p | CCDC71L | HITS-CLIP | miRTarbase |
| hsa-miR-141-3p | TROVE2 | HITS-CLIP | miRTarbase |
| hsa-miR-486-5p | SAPCD2 | HITS-CLIP | miRTarbase |
| hsa-miR-141-5p | SACM1L | HITS-CLIP | miRTarbase |
| hsa-miR-141-5p | FAM46A | HITS-CLIP | miRTarbase |
| hsa-miR-3613-5p | RWDD2A | HITS-CLIP | miRTarbase |
| hsa-miR-486-3p | JMJD4 | HITS-CLIP | miRTarbase |
| hsa-miR-141-5p | METTL15 | HITS-CLIP | miRTarbase |

| | | | |
|---|---|---|---|
| hsa-miR-141-5p | CSTF2T | HITS-CLIP | miRTarbase |
| hsa-miR-3613-3p | PHF7 | HITS-CLIP | miRTarbase |
| hsa-miR-486-5p | BASP1 | HITS-CLIP | miRTarbase |
| hsa-miR-3613-3p | KIAA2018 | HITS-CLIP | miRTarbase |
| hsa-miR-141-3p | GLTSCR1L | HITS-CLIP | miRTarbase |
| hsa-miR-486-3p | CCDC106 | HITS-CLIP | miRTarbase |
| hsa-miR-486-5p | CENPN | HITS-CLIP | miRTarbase |
| hsa-miR-3613-3p | SKA1 | HITS-CLIP | miRTarbase |
| hsa-miR-486-5p | CRIPT | HITS-CLIP | miRTarbase |
| hsa-miR-486-3p | MACROD2 | HITS-CLIP | miRTarbase |
| hsa-miR-486-3p | CYTH2 | HITS-CLIP | miRTarbase |
| hsa-miR-141-5p | RNF103-CHMP3 | HITS-CLIP | miRTarbase |
| hsa-miR-3613-3p | LAMTOR1 | HITS-CLIP | miRTarbase |
| hsa-miR-3613-3p | KCNMB3 | HITS-CLIP | miRTarbase |
| hsa-miR-486-5p | DCTN4 | HITS-CLIP | miRTarbase |
| hsa-miR-3613-3p | RPF2 | HITS-CLIP | miRTarbase |
| hsa-miR-141-3p | ERO1L | HITS-CLIP | miRTarbase |
| hsa-miR-486-5p | SPRTN | HITS-CLIP | miRTarbase |
| hsa-miR-195-5p | SLC35E2B | HITS-CLIP | miRTarbase |
| hsa-miR-15a-5p | SLC35E2B | HITS-CLIP | miRTarbase |
| hsa-miR-3613-3p | KCNK5 | HITS-CLIP | miRTarbase |
| hsa-miR-3613-3p | SYDE2 | HITS-CLIP | miRTarbase |
| hsa-miR-3613-3p | WIPF2 | HITS-CLIP | miRTarbase |
| hsa-miR-3613-3p | PDP2 | HITS-CLIP | miRTarbase |
| hsa-miR-3613-3p | FBXO31 | HITS-CLIP | miRTarbase |
| hsa-miR-195-5p | ARHGAP32 | HITS-CLIP | miRTarbase |
| hsa-miR-15a-5p | ARHGAP32 | HITS-CLIP | miRTarbase |
| hsa-miR-141-3p | HSPA4 | HITS-CLIP | miRTarbase |
| hsa-miR-3613-3p | ZBTB39 | HITS-CLIP | miRTarbase |
| hsa-miR-3613-5p | TNFRSF13C | HITS-CLIP | miRTarbase |
| hsa-miR-3613-3p | CENPH | HITS-CLIP | miRTarbase |
| hsa-miR-3613-3p | ADAMTS17 | HITS-CLIP | miRTarbase |
| hsa-miR-141-5p | CHMP3 | HITS-CLIP | miRTarbase |
| hsa-miR-219a-1-3p | USP6NL | HITS-CLIP | miRTarbase |
| hsa-miR-15a-3p | HEBP2 | HITS-CLIP | miRTarbase |
| hsa-miR-195-3p | COMMD3-BMI1 | HITS-CLIP | miRTarbase |
| hsa-miR-195-5p | RAB15 | HITS-CLIP | miRTarbase |
| hsa-miR-15a-5p | RAB15 | HITS-CLIP | miRTarbase |
| hsa-miR-3613-3p | ZNF394 | HITS-CLIP | miRTarbase |
| hsa-miR-486-5p | EPGN | HITS-CLIP | miRTarbase |
| hsa-miR-486-3p | C1orf229 | HITS-CLIP | miRTarbase |
| hsa-miR-195-3p | BMI1 | HITS-CLIP | miRTarbase |
| hsa-miR-141-5p | RNF8 | HITS-CLIP | miRTarbase |
| hsa-miR-195-5p | ADORA3 | HITS-CLIP | miRTarbase |
| hsa-miR-15a-5p | ADORA3 | HITS-CLIP | miRTarbase |
| hsa-miR-219a-1-3p | MTMR14 | HITS-CLIP | miRTarbase |
| hsa-miR-486-3p | SYNDIG1L | HITS-CLIP | miRTarbase |
| hsa-miR-141-3p | SCD5 | HITS-CLIP | miRTarbase |
| hsa-miR-141-5p | VWA9 | HITS-CLIP | miRTarbase |
| hsa-miR-3613-3p | TRIM66 | HITS-CLIP | miRTarbase |
| hsa-miR-486-5p | HPGD | HITS-CLIP | miRTarbase |
| hsa-miR-3605-5p | CAMK4 | HITS-CLIP | miRTarbase |
| hsa-miR-3613-3p | VGLL4 | HITS-CLIP | miRTarbase |
| hsa-miR-3613-3p | AP5M1 | HITS-CLIP | miRTarbase |
| hsa-miR-3613-3p | JAG1 | HITS-CLIP | miRTarbase |
| hsa-miR-3613-3p | PPIP5K2 | HITS-CLIP | miRTarbase |
| hsa-miR-195-5p | PPIP5K2 | HITS-CLIP | miRTarbase |
| hsa-miR-15a-5p | PPIP5K2 | HITS-CLIP | miRTarbase |
| hsa-miR-486-5p | FAM46A | HITS-CLIP | miRTarbase |
| hsa-miR-3613-3p | ZNF549 | HITS-CLIP | miRTarbase |
| hsa-miR-3613-3p | C11orf54 | HITS-CLIP | miRTarbase |
| hsa-miR-486-3p | FAM83A | HITS-CLIP | miRTarbase |
| hsa-miR-3613-5p | STRBP | HITS-CLIP | miRTarbase |
| hsa-miR-141-5p | PSMC2 | HITS-CLIP | miRTarbase |
| hsa-miR-3613-3p | MYOCD | HITS-CLIP | miRTarbase |
| hsa-miR-141-5p | VTI1A | HITS-CLIP | miRTarbase |
| hsa-miR-141-5p | HEATR5A | HITS-CLIP | miRTarbase |
| hsa-miR-141-5p | DOK6 | HITS-CLIP | miRTarbase |
| hsa-miR-3613-3p | MAT1A | HITS-CLIP | miRTarbase |
| hsa-miR-3613-3p | BUB3 | HITS-CLIP | miRTarbase |
| hsa-miR-141-5p | ARHGAP19 | HITS-CLIP | miRTarbase |
| hsa-miR-3613-3p | PAPD5 | HITS-CLIP | miRTarbase |
| hsa-miR-3613-3p | MPRIP | HITS-CLIP | miRTarbase |
| hsa-miR-141-3p | CLDND1 | HITS-CLIP | miRTarbase |
| hsa-miR-3613-3p | AVL9 | HITS-CLIP | miRTarbase |
| hsa-miR-3613-3p | TMED10 | HITS-CLIP | miRTarbase |
| hsa-miR-141-3p | PEX11A | HITS-CLIP | miRTarbase |
| hsa-miR-3613-3p | KIAA0146 | HITS-CLIP | miRTarbase |
| hsa-miR-486-5p | ABCF2 | HITS-CLIP | miRTarbase |
| hsa-miR-3613-3p | HRH4 | HITS-CLIP | miRTarbase |
| hsa-miR-195-3p | AP3B1 | HITS-CLIP | miRTarbase |
| hsa-miR-195-3p | IGFBP5 | HITS-CLIP | miRTarbase |
| hsa-miR-141-5p | TXNL1 | HITS-CLIP | miRTarbase |
| hsa-miR-195-5p | SYNRG | HITS-CLIP | miRTarbase |
| hsa-miR-15a-5p | SYNRG | HITS-CLIP | miRTarbase |
| hsa-miR-195-3p | LRIG2 | HITS-CLIP | miRTarbase |
| hsa-miR-141-3p | CELF1 | HITS-CLIP | miRTarbase |
| hsa-miR-195-5p | ZNF91 | HITS-CLIP | miRTarbase |
| hsa-miR-15a-5p | ZNF91 | HITS-CLIP | miRTarbase |
| hsa-miR-195-5p | ZBTB5 | HITS-CLIP | miRTarbase |
| hsa-miR-15a-5p | ZBTB5 | HITS-CLIP | miRTarbase |
| hsa-miR-195-5p | VPS33B | HITS-CLIP | miRTarbase |
| hsa-miR-15a-5p | VPS33B | HITS-CLIP | miRTarbase |
| hsa-miR-195-5p | CHMP3 | HITS-CLIP | miRTarbase |
| hsa-miR-15a-5p | CHMP3 | HITS-CLIP | miRTarbase |
| hsa-miR-195-5p | VCL | HITS-CLIP | miRTarbase |
| hsa-miR-15a-5p | VCL | HITS-CLIP | miRTarbase |
| hsa-miR-195-5p | USP3 | HITS-CLIP | miRTarbase |
| hsa-miR-15a-5p | USP3 | HITS-CLIP | miRTarbase |
| hsa-miR-195-5p | USP31 | HITS-CLIP | miRTarbase |
| hsa-miR-15a-5p | USP31 | HITS-CLIP | miRTarbase |
| hsa-miR-195-5p | TUBB | HITS-CLIP | miRTarbase |
| hsa-miR-15a-5p | TUBB | HITS-CLIP | miRTarbase |
| hsa-miR-195-5p | TRAM1 | HITS-CLIP | miRTarbase |
| hsa-miR-15a-5p | TRAM1 | HITS-CLIP | miRTarbase |
| hsa-miR-195-5p | TMEM69 | HITS-CLIP | miRTarbase |
| hsa-miR-15a-5p | TMEM69 | HITS-CLIP | miRTarbase |
| hsa-miR-195-5p | TMEM55B | HITS-CLIP | miRTarbase |
| hsa-miR-15a-5p | TMEM55B | HITS-CLIP | miRTarbase |
| hsa-miR-195-5p | TMEM135 | HITS-CLIP | miRTarbase |
| hsa-miR-15a-5p | TMEM135 | HITS-CLIP | miRTarbase |
| hsa-miR-195-5p | TLE4 | HITS-CLIP | miRTarbase |
| hsa-miR-15a-5p | TLE4 | HITS-CLIP | miRTarbase |
| hsa-miR-195-5p | TKTL1 | HITS-CLIP | miRTarbase |
| hsa-miR-15a-5p | TKTL1 | HITS-CLIP | miRTarbase |
| hsa-miR-195-5p | TIMM13 | HITS-CLIP | miRTarbase |
| hsa-miR-15a-5p | TIMM13 | HITS-CLIP | miRTarbase |
| hsa-miR-195-5p | TFB1M | HITS-CLIP | miRTarbase |
| hsa-miR-15a-5p | TFB1M | HITS-CLIP | miRTarbase |
| hsa-miR-195-5p | TCF3 | HITS-CLIP | miRTarbase |
| hsa-miR-15a-5p | TCF3 | HITS-CLIP | miRTarbase |
| hsa-miR-195-5p | TBL1XR1 | HITS-CLIP | miRTarbase |
| hsa-miR-15a-5p | TBL1XR1 | HITS-CLIP | miRTarbase |
| hsa-miR-15a-5p | TBCCD1 | HITS-CLIP | miRTarbase |
| hsa-miR-195-5p | TBC1D20 | HITS-CLIP | miRTarbase |
| hsa-miR-15a-5p | TBC1D20 | HITS-CLIP | miRTarbase |
| hsa-miR-195-5p | TBC1D14 | HITS-CLIP | miRTarbase |
| hsa-miR-15a-5p | TBC1D14 | HITS-CLIP | miRTarbase |
| hsa-miR-195-5p | TASP1 | HITS-CLIP | miRTarbase |
| hsa-miR-15a-5p | TASP1 | HITS-CLIP | miRTarbase |
| hsa-miR-195-5p | SUPT16H | HITS-CLIP | miRTarbase |
| hsa-miR-15a-5p | SUPT16H | HITS-CLIP | miRTarbase |
| hsa-miR-195-5p | STX17 | HITS-CLIP | miRTarbase |
| hsa-miR-15a-5p | STX17 | HITS-CLIP | miRTarbase |
| hsa-miR-195-5p | SRPK1 | HITS-CLIP | miRTarbase |
| hsa-miR-15a-5p | SRPK1 | HITS-CLIP | miRTarbase |
| hsa-miR-195-5p | SPTLC1 | HITS-CLIP | miRTarbase |
| hsa-miR-15a-5p | SPTLC1 | HITS-CLIP | miRTarbase |
| hsa-miR-195-5p | SMURF1 | HITS-CLIP | miRTarbase |
| hsa-miR-15a-5p | SMURF1 | HITS-CLIP | miRTarbase |
| hsa-miR-195-5p | SLC9A1 | HITS-CLIP | miRTarbase |
| hsa-miR-15a-5p | SLC9A1 | HITS-CLIP | miRTarbase |
| hsa-miR-195-5p | SLC7A5 | HITS-CLIP | miRTarbase |
| hsa-miR-15a-5p | SLC7A5 | HITS-CLIP | miRTarbase |
| hsa-miR-195-5p | SLC25A29 | HITS-CLIP | miRTarbase |
| hsa-miR-15a-5p | SLC25A29 | HITS-CLIP | miRTarbase |
| hsa-miR-195-5p | SLC25A22 | HITS-CLIP | miRTarbase |
| hsa-miR-15a-5p | SLC25A22 | HITS-CLIP | miRTarbase |
| hsa-miR-195-5p | RPS6KA3 | HITS-CLIP | miRTarbase |

| | | | |
|---|---|---|---|
| hsa-miR-15a-5p | RPS6KA3 | HITS-CLIP | miRTarbase |
| hsa-miR-195-5p | RPS5 | HITS-CLIP | miRTarbase |
| hsa-miR-15a-5p | RPS5 | HITS-CLIP | miRTarbase |
| hsa-miR-195-5p | RPL36 | HITS-CLIP | miRTarbase |
| hsa-miR-15a-5p | RPL36 | HITS-CLIP | miRTarbase |
| hsa-miR-195-5p | RNPS1 | HITS-CLIP | miRTarbase |
| hsa-miR-15a-5p | RNPS1 | HITS-CLIP | miRTarbase |
| hsa-miR-195-5p | RNMT | HITS-CLIP | miRTarbase |
| hsa-miR-15a-5p | RNMT | HITS-CLIP | miRTarbase |
| hsa-miR-195-5p | RNH1 | HITS-CLIP | miRTarbase |
| hsa-miR-15a-5p | RNH1 | HITS-CLIP | miRTarbase |
| hsa-miR-195-5p | RFWD2 | HITS-CLIP | miRTarbase |
| hsa-miR-15a-5p | RFWD2 | HITS-CLIP | miRTarbase |
| hsa-miR-195-5p | REXO1 | HITS-CLIP | miRTarbase |
| hsa-miR-15a-5p | REXO1 | HITS-CLIP | miRTarbase |
| hsa-miR-195-5p | RELT | HITS-CLIP | miRTarbase |
| hsa-miR-15a-5p | RELT | HITS-CLIP | miRTarbase |
| hsa-miR-195-5p | RAP2C | HITS-CLIP | miRTarbase |
| hsa-miR-15a-5p | RAP2C | HITS-CLIP | miRTarbase |
| hsa-miR-195-5p | RAB40B | HITS-CLIP | miRTarbase |
| hsa-miR-15a-5p | RAB40B | HITS-CLIP | miRTarbase |
| hsa-miR-195-5p | RAB11FIP2 | HITS-CLIP | miRTarbase |
| hsa-miR-15a-5p | RAB11FIP2 | HITS-CLIP | miRTarbase |
| hsa-miR-195-5p | PSMB5 | HITS-CLIP | miRTarbase |
| hsa-miR-15a-5p | PSMB5 | HITS-CLIP | miRTarbase |
| hsa-miR-195-5p | PPP6C | HITS-CLIP | miRTarbase |
| hsa-miR-15a-5p | PPP6C | HITS-CLIP | miRTarbase |
| hsa-miR-195-5p | POU2AF1 | HITS-CLIP | miRTarbase |
| hsa-miR-15a-5p | POU2AF1 | HITS-CLIP | miRTarbase |
| hsa-miR-195-5p | POLE4 | HITS-CLIP | miRTarbase |
| hsa-miR-15a-5p | POLE4 | HITS-CLIP | miRTarbase |
| hsa-miR-195-5p | PGD | HITS-CLIP | miRTarbase |
| hsa-miR-15a-5p | PGD | HITS-CLIP | miRTarbase |
| hsa-miR-195-5p | PEX12 | HITS-CLIP | miRTarbase |
| hsa-miR-15a-5p | PEX12 | HITS-CLIP | miRTarbase |
| hsa-miR-195-5p | PANK1 | HITS-CLIP | miRTarbase |
| hsa-miR-15a-5p | PANK1 | HITS-CLIP | miRTarbase |
| hsa-miR-195-5p | TM9SF2 | HITS-CLIP | miRTarbase |
| hsa-miR-15a-5p | TM9SF2 | HITS-CLIP | miRTarbase |
| hsa-miR-195-5p | OTUB1 | HITS-CLIP | miRTarbase |
| hsa-miR-15a-5p | OTUB1 | HITS-CLIP | miRTarbase |
| hsa-miR-195-5p | NR2C2 | HITS-CLIP | miRTarbase |
| hsa-miR-15a-5p | NR2C2 | HITS-CLIP | miRTarbase |
| hsa-miR-195-5p | NCOR2 | HITS-CLIP | miRTarbase |
| hsa-miR-15a-5p | NCOR2 | HITS-CLIP | miRTarbase |
| hsa-miR-195-5p | MTMR4 | HITS-CLIP | miRTarbase |
| hsa-miR-15a-5p | MTMR4 | HITS-CLIP | miRTarbase |
| hsa-miR-195-5p | MRPL40 | HITS-CLIP | miRTarbase |
| hsa-miR-15a-5p | MRPL40 | HITS-CLIP | miRTarbase |
| hsa-miR-195-5p | MLXIP | HITS-CLIP | miRTarbase |
| hsa-miR-15a-5p | MLXIP | HITS-CLIP | miRTarbase |
| hsa-miR-195-5p | MIB1 | HITS-CLIP | miRTarbase |
| hsa-miR-15a-5p | MIB1 | HITS-CLIP | miRTarbase |
| hsa-miR-195-5p | MED11 | HITS-CLIP | miRTarbase |
| hsa-miR-15a-5p | MED11 | HITS-CLIP | miRTarbase |
| hsa-miR-195-5p | MCM3AP-AS1 | HITS-CLIP | miRTarbase |
| hsa-miR-15a-5p | MCM3AP-AS1 | HITS-CLIP | miRTarbase |
| hsa-miR-195-5p | LYRM5 | HITS-CLIP | miRTarbase |
| hsa-miR-15a-5p | LYRM5 | HITS-CLIP | miRTarbase |
| hsa-miR-195-5p | LRRC57 | HITS-CLIP | miRTarbase |
| hsa-miR-15a-5p | LRRC57 | HITS-CLIP | miRTarbase |
| hsa-miR-195-5p | LRPPRC | HITS-CLIP | miRTarbase |
| hsa-miR-15a-5p | LRPPRC | HITS-CLIP | miRTarbase |
| hsa-miR-195-5p | LITAF | HITS-CLIP | miRTarbase |
| hsa-miR-15a-5p | LITAF | HITS-CLIP | miRTarbase |
| hsa-miR-195-5p | KLC2 | HITS-CLIP | miRTarbase |
| hsa-miR-15a-5p | KLC2 | HITS-CLIP | miRTarbase |
| hsa-miR-195-5p | TECPR2 | HITS-CLIP | miRTarbase |
| hsa-miR-15a-5p | TECPR2 | HITS-CLIP | miRTarbase |
| hsa-miR-195-5p | KATNAL1 | HITS-CLIP | miRTarbase |
| hsa-miR-15a-5p | KATNAL1 | HITS-CLIP | miRTarbase |
| hsa-miR-195-5p | IRAK1BP1 | HITS-CLIP | miRTarbase |
| hsa-miR-15a-5p | IRAK1BP1 | HITS-CLIP | miRTarbase |
| hsa-miR-195-5p | HYOU1 | HITS-CLIP | miRTarbase |
| hsa-miR-15a-5p | HYOU1 | HITS-CLIP | miRTarbase |
| hsa-miR-195-5p | GSK3B | HITS-CLIP | miRTarbase |
| hsa-miR-15a-5p | GSK3B | HITS-CLIP | miRTarbase |
| hsa-miR-195-5p | GGA3 | HITS-CLIP | miRTarbase |
| hsa-miR-15a-5p | GGA3 | HITS-CLIP | miRTarbase |
| hsa-miR-195-5p | GANAB | HITS-CLIP | miRTarbase |
| hsa-miR-15a-5p | GANAB | HITS-CLIP | miRTarbase |
| hsa-miR-195-5p | GABARAP | HITS-CLIP | miRTarbase |
| hsa-miR-15a-5p | GABARAP | HITS-CLIP | miRTarbase |
| hsa-miR-195-5p | FRYL | HITS-CLIP | miRTarbase |
| hsa-miR-15a-5p | FRYL | HITS-CLIP | miRTarbase |
| hsa-miR-195-5p | FAM73A | HITS-CLIP | miRTarbase |
| hsa-miR-15a-5p | FAM73A | HITS-CLIP | miRTarbase |
| hsa-miR-195-5p | AMER1 | HITS-CLIP | miRTarbase |
| hsa-miR-15a-5p | AMER1 | HITS-CLIP | miRTarbase |
| hsa-miR-195-5p | EDC3 | HITS-CLIP | miRTarbase |
| hsa-miR-15a-5p | EDC3 | HITS-CLIP | miRTarbase |
| hsa-miR-195-5p | DSCR3 | HITS-CLIP | miRTarbase |
| hsa-miR-15a-5p | DSCR3 | HITS-CLIP | miRTarbase |
| hsa-miR-195-5p | DPP8 | HITS-CLIP | miRTarbase |
| hsa-miR-15a-5p | DPP8 | HITS-CLIP | miRTarbase |
| hsa-miR-195-5p | DNAJC9 | HITS-CLIP | miRTarbase |
| hsa-miR-15a-5p | DNAJC9 | HITS-CLIP | miRTarbase |
| hsa-miR-15a-5p | DICER1 | HITS-CLIP | miRTarbase |
| hsa-miR-195-5p | CYLD | HITS-CLIP | miRTarbase |
| hsa-miR-15a-5p | CYLD | HITS-CLIP | miRTarbase |
| hsa-miR-195-5p | CYB561A3 | HITS-CLIP | miRTarbase |
| hsa-miR-15a-5p | CYB561A3 | HITS-CLIP | miRTarbase |
| hsa-miR-195-5p | CUL2 | HITS-CLIP | miRTarbase |
| hsa-miR-15a-5p | CUL2 | HITS-CLIP | miRTarbase |
| hsa-miR-195-5p | CSDE1 | HITS-CLIP | miRTarbase |
| hsa-miR-15a-5p | CSDE1 | HITS-CLIP | miRTarbase |
| hsa-miR-195-5p | CREG1 | HITS-CLIP | miRTarbase |
| hsa-miR-15a-5p | CREG1 | HITS-CLIP | miRTarbase |
| hsa-miR-195-5p | CPNE1 | HITS-CLIP | miRTarbase |
| hsa-miR-15a-5p | CPNE1 | HITS-CLIP | miRTarbase |
| hsa-miR-195-5p | RHOV | HITS-CLIP | miRTarbase |
| hsa-miR-15a-5p | RHOV | HITS-CLIP | miRTarbase |
| hsa-miR-195-5p | CDKN2AIPNL | HITS-CLIP | miRTarbase |
| hsa-miR-15a-5p | CDKN2AIPNL | HITS-CLIP | miRTarbase |
| hsa-miR-195-5p | CDC27 | HITS-CLIP | miRTarbase |
| hsa-miR-15a-5p | CDC27 | HITS-CLIP | miRTarbase |
| hsa-miR-195-5p | CBFA2T3 | HITS-CLIP | miRTarbase |
| hsa-miR-15a-5p | CBFA2T3 | HITS-CLIP | miRTarbase |
| hsa-miR-195-5p | C6orf106 | HITS-CLIP | miRTarbase |
| hsa-miR-15a-5p | C6orf106 | HITS-CLIP | miRTarbase |
| hsa-miR-195-5p | C2orf42 | HITS-CLIP | miRTarbase |
| hsa-miR-15a-5p | C2orf42 | HITS-CLIP | miRTarbase |
| hsa-miR-195-5p | LRIF1 | HITS-CLIP | miRTarbase |
| hsa-miR-15a-5p | LRIF1 | HITS-CLIP | miRTarbase |
| hsa-miR-195-5p | C15orf39 | HITS-CLIP | miRTarbase |
| hsa-miR-15a-5p | C15orf39 | HITS-CLIP | miRTarbase |
| hsa-miR-195-5p | BSG | HITS-CLIP | miRTarbase |
| hsa-miR-15a-5p | BSG | HITS-CLIP | miRTarbase |
| hsa-miR-195-5p | B4GALT1 | HITS-CLIP | miRTarbase |
| hsa-miR-15a-5p | B4GALT1 | HITS-CLIP | miRTarbase |
| hsa-miR-195-5p | ATP13A3 | HITS-CLIP | miRTarbase |
| hsa-miR-15a-5p | ATP13A3 | HITS-CLIP | miRTarbase |
| hsa-miR-195-5p | ASXL1 | HITS-CLIP | miRTarbase |
| hsa-miR-15a-5p | ASXL1 | HITS-CLIP | miRTarbase |
| hsa-miR-195-5p | AP3M1 | HITS-CLIP | miRTarbase |
| hsa-miR-15a-5p | AP3M1 | HITS-CLIP | miRTarbase |
| hsa-miR-195-5p | AFF4 | HITS-CLIP | miRTarbase |
| hsa-miR-15a-5p | AFF4 | HITS-CLIP | miRTarbase |
| hsa-miR-195-5p | ACOX1 | HITS-CLIP | miRTarbase |
| hsa-miR-15a-5p | ACOX1 | HITS-CLIP | miRTarbase |
| hsa-miR-141-3p | DP2(hsa) | Western Blot | Tarbase |
| hsa-miR-15a-3p | DPM1 | HITS-CLIP | Tarbase |
| hsa-miR-3613-5p | DPM1 | HITS-CLIP | Tarbase |
| hsa-miR-195-5p | C1orf112 | Microarrays | Tarbase |
| hsa-miR-15a-5p | FUCA2 | HITS-CLIP | Tarbase |
| hsa-miR-195-5p | FUCA2 | HITS-CLIP | Tarbase |
| hsa-miR-141-3p | GCLC | HITS-CLIP | Tarbase |
| hsa-miR-195-5p | GCLC | Microarrays | Tarbase |
| hsa-miR-3605-5p | LAS1L | HITS-CLIP | Tarbase |

| | | | |
|---|---|---|---|
| hsa-miR-15a-5p | ENPP4 | HITS-CLIP | Tarbase |
| hsa-miR-195-5p | ENPP4 | HITS-CLIP | Tarbase |
| hsa-miR-15a-5p | ANKIB1 | HITS-CLIP | Tarbase |
| hsa-miR-195-5p | ANKIB1 | HITS-CLIP | Tarbase |
| hsa-miR-15a-5p | ANKIB1 | HITS-CLIP | Tarbase |
| hsa-miR-195-5p | ANKIB1 | HITS-CLIP | Tarbase |
| hsa-miR-141-5p | CYP51A1 | HITS-CLIP | Tarbase |
| hsa-miR-15a-5p | CYP51A1 | HITS-CLIP | Tarbase |
| hsa-miR-195-5p | CYP51A1 | HITS-CLIP | Tarbase |
| hsa-miR-15a-5p | CYP51A1 | HITS-CLIP | Tarbase |
| hsa-miR-195-5p | CYP51A1 | HITS-CLIP | Tarbase |
| hsa-miR-141-3p | CYP26B1 | HITS-CLIP | Tarbase |
| hsa-miR-15a-5p | CYP26B1 | PAR-CLIP | Tarbase |
| hsa-miR-15a-5p | CYP26B1 | Chimeric fragments | Tarbase |
| hsa-miR-15a-5p | ALS2 | HITS-CLIP | Tarbase |
| hsa-miR-195-5p | ALS2 | HITS-CLIP | Tarbase |
| hsa-miR-15a-5p | CFLAR | HITS-CLIP | Tarbase |
| hsa-miR-195-5p | CFLAR | HITS-CLIP | Tarbase |
| hsa-miR-195-5p | CFLAR | qPCR | Tarbase |
| hsa-miR-3613-3p | CFLAR | HITS-CLIP | Tarbase |
| hsa-miR-15a-5p | TFPI | HITS-CLIP | Tarbase |
| hsa-miR-195-5p | TFPI | HITS-CLIP | Tarbase |
| hsa-miR-141-3p | SLC7A2 | HITS-CLIP | Tarbase |
| hsa-miR-15a-5p | SLC7A2 | HITS-CLIP | Tarbase |
| hsa-miR-15a-5p | SARM1 | PAR-CLIP | Tarbase |
| hsa-miR-15a-5p | SARM1 | PAR-CLIP | Tarbase |
| hsa-miR-141-3p | AK2 | HITS-CLIP | Tarbase |
| hsa-miR-141-5p | RBM6 | HITS-CLIP | Tarbase |
| hsa-miR-141-5p | RBM6 | HITS-CLIP | Tarbase |
| hsa-miR-15a-5p | RBM6 | PAR-CLIP | Tarbase |
| hsa-miR-195-5p | CAMKK1 | Microarrays | Tarbase |
| hsa-miR-219a-1-3p | NDUFAB1 | PAR-CLIP | Tarbase |
| hsa-miR-15a-5p | PDK4 | Chimeric fragments | Tarbase |
| hsa-miR-141-5p | SLC25A13 | HITS-CLIP | Tarbase |
| hsa-miR-195-5p | SLC25A13 | Microarrays | Tarbase |
| hsa-miR-15a-5p | CDC27 | HITS-CLIP | Tarbase |
| hsa-miR-15a-5p | CDC27 | PAR-CLIP | Tarbase |
| hsa-miR-195-5p | CDC27 | HITS-CLIP | Tarbase |
| hsa-miR-195-5p | HCCS | Microarrays | Tarbase |
| hsa-miR-15a-5p | UPF1 | HITS-CLIP | Tarbase |
| hsa-miR-141-3p | SKAP2 | HITS-CLIP | Tarbase |
| hsa-miR-141-3p | DHX33 | HITS-CLIP | Tarbase |
| hsa-miR-15a-5p | DHX33 | PAR-CLIP | Tarbase |
| hsa-miR-15a-5p | DHX33 | PAR-CLIP | Tarbase |
| hsa-miR-195-5p | DHX33 | PAR-CLIP | Tarbase |
| hsa-miR-15a-5p | DHX33 | Chimeric fragments | Tarbase |
| hsa-miR-195-5p | THSD7A | Microarrays | Tarbase |
| hsa-miR-3613-5p | LIG3 | HITS-CLIP | Tarbase |
| hsa-miR-195-5p | ACSM3 | Microarrays | Tarbase |
| hsa-miR-195-5p | AC004381.6 | Microarrays | Tarbase |
| hsa-miR-15a-5p | SPPL2B | HITS-CLIP | Tarbase |
| hsa-miR-195-5p | SPPL2B | HITS-CLIP | Tarbase |
| hsa-miR-219a-1-3p | MSL3 | PAR-CLIP | Tarbase |
| hsa-miR-3613-3p | MSL3 | PAR-CLIP | Tarbase |
| hsa-miR-141-5p | CREBBP | HITS-CLIP | Tarbase |
| hsa-miR-15a-5p | CREBBP | HITS-CLIP | Tarbase |
| hsa-miR-195-5p | CREBBP | HITS-CLIP | Tarbase |
| hsa-miR-141-5p | ZFX | HITS-CLIP | Tarbase |
| hsa-miR-141-5p | ZFX | HITS-CLIP | Tarbase |
| hsa-miR-15a-5p | LAMP2 | PAR-CLIP | Tarbase |
| hsa-miR-15a-5p | LAMP2 | PAR-CLIP | Tarbase |
| hsa-miR-15a-5p | LAMP2 | PAR-CLIP | Tarbase |
| hsa-miR-15a-5p | LAMP2 | HITS-CLIP | Tarbase |
| hsa-miR-195-5p | LAMP2 | HITS-CLIP | Tarbase |
| hsa-miR-15a-5p | GDE1 | HITS-CLIP | Tarbase |
| hsa-miR-195-5p | GDE1 | HITS-CLIP | Tarbase |
| hsa-miR-15a-5p | OSBPL7 | PAR-CLIP | Tarbase |
| hsa-miR-141-3p | AP2B1 | HITS-CLIP | Tarbase |
| hsa-miR-141-3p | AP2B1 | HITS-CLIP | Tarbase |
| hsa-miR-141-5p | AP2B1 | HITS-CLIP | Tarbase |
| hsa-miR-15a-5p | AP2B1 | PAR-CLIP | Tarbase |
| hsa-miR-15a-5p | AP2B1 | HITS-CLIP | Tarbase |
| hsa-miR-15a-5p | AP2B1 | HITS-CLIP | Tarbase |
| hsa-miR-195-5p | AP2B1 | HITS-CLIP | Tarbase |
| hsa-miR-15a-5p | AP2B1 | HITS-CLIP | Tarbase |
| hsa-miR-195-5p | AP2B1 | HITS-CLIP | Tarbase |
| hsa-miR-15a-5p | TNFRSF12A | HITS-CLIP | Tarbase |
| hsa-miR-195-5p | TNFRSF12A | HITS-CLIP | Tarbase |
| hsa-miR-195-5p | TNFRSF12A | Microarrays | Tarbase |
| hsa-miR-141-5p | MAP3K9 | HITS-CLIP | Tarbase |
| hsa-miR-141-5p | MAP3K9 | HITS-CLIP | Tarbase |
| hsa-miR-141-5p | MAP3K9 | HITS-CLIP | Tarbase |
| hsa-miR-15a-5p | MAP3K9 | HITS-CLIP | Tarbase |
| hsa-miR-195-5p | MAP3K9 | HITS-CLIP | Tarbase |
| hsa-miR-141-5p | RALA | HITS-CLIP | Tarbase |
| hsa-miR-195-5p | ETV1 | Microarrays | Tarbase |
| hsa-miR-15a-5p | AGK | PAR-CLIP | Tarbase |
| hsa-miR-141-3p | PHTF2 | HITS-CLIP | Tarbase |
| hsa-miR-15a-3p | GGCT | PAR-CLIP | Tarbase |
| hsa-miR-141-5p | IFRD1 | HITS-CLIP | Tarbase |
| hsa-miR-15a-5p | PNPLA4 | HITS-CLIP | Tarbase |
| hsa-miR-195-5p | PNPLA4 | HITS-CLIP | Tarbase |
| hsa-miR-15a-3p | ADIPOR2 | PAR-CLIP | Tarbase |
| hsa-miR-486-5p | ADIPOR2 | PAR-CLIP | Tarbase |
| hsa-miR-141-5p | PAFAH1B1 | HITS-CLIP | Tarbase |
| hsa-miR-15a-5p | PAFAH1B1 | PAR-CLIP | Tarbase |
| hsa-miR-15a-5p | PAFAH1B1 | HITS-CLIP | Tarbase |
| hsa-miR-15a-5p | PAFAH1B1 | HITS-CLIP | Tarbase |
| hsa-miR-15a-5p | PAFAH1B1 | HITS-CLIP | Tarbase |
| hsa-miR-15a-5p | PAFAH1B1 | PAR-CLIP | Tarbase |
| hsa-miR-15a-5p | PAFAH1B1 | HITS-CLIP | Tarbase |
| hsa-miR-15a-5p | PAFAH1B1 | HITS-CLIP | Tarbase |
| hsa-miR-195-5p | PAFAH1B1 | PAR-CLIP | Tarbase |
| hsa-miR-195-5p | PAFAH1B1 | HITS-CLIP | Tarbase |
| hsa-miR-195-5p | PAFAH1B1 | HITS-CLIP | Tarbase |
| hsa-miR-195-5p | PAFAH1B1 | HITS-CLIP | Tarbase |
| hsa-miR-195-5p | PAFAH1B1 | HITS-CLIP | Tarbase |
| hsa-miR-3613-3p | PAFAH1B1 | PAR-CLIP | Tarbase |
| hsa-miR-141-3p | KIAA0100 | HITS-CLIP | Tarbase |
| hsa-miR-15a-5p | KIAA0100 | HITS-CLIP | Tarbase |
| hsa-miR-15a-5p | KIAA0100 | PAR-CLIP | Tarbase |
| hsa-miR-195-5p | KIAA0100 | HITS-CLIP | Tarbase |
| hsa-miR-195-5p | KIAA0100 | PAR-CLIP | Tarbase |
| hsa-miR-15a-5p | CRAMP1L | HITS-CLIP | Tarbase |
| hsa-miR-195-5p | CRAMP1L | HITS-CLIP | Tarbase |
| hsa-miR-486-5p | CRAMP1L | HITS-CLIP | Tarbase |
| hsa-miR-141-5p | MYLIP | HITS-CLIP | Tarbase |
| hsa-miR-141-5p | MYLIP | HITS-CLIP | Tarbase |
| hsa-miR-15a-5p | MYLIP | HITS-CLIP | Tarbase |
| hsa-miR-15a-5p | MYLIP | HITS-CLIP | Tarbase |
| hsa-miR-219a-1-3p | MYLIP | PAR-CLIP | Tarbase |
| hsa-miR-195-5p | E2F2 | Microarrays | Tarbase |
| hsa-miR-141-5p | JARID2 | HITS-CLIP | Tarbase |
| hsa-miR-141-5p | JARID2 | HITS-CLIP | Tarbase |
| hsa-miR-15a-3p | JARID2 | HITS-CLIP | Tarbase |
| hsa-miR-15a-5p | JARID2 | PAR-CLIP | Tarbase |
| hsa-miR-15a-5p | JARID2 | PAR-CLIP | Tarbase |
| hsa-miR-15a-5p | JARID2 | PAR-CLIP | Tarbase |
| hsa-miR-15a-5p | JARID2 | HITS-CLIP | Tarbase |
| hsa-miR-3613-5p | JARID2 | HITS-CLIP | Tarbase |
| hsa-miR-15a-5p | JARID2 | Chimeric fragments | Tarbase |
| hsa-miR-195-5p | CYTH3 | Microarrays | Tarbase |
| hsa-miR-15a-5p | SYPL1 | PAR-CLIP | Tarbase |
| hsa-miR-15a-5p | SYPL1 | PAR-CLIP | Tarbase |
| hsa-miR-3613-5p | SYPL1 | PAR-CLIP | Tarbase |
| hsa-miR-486-5p | SYPL1 | PAR-CLIP | Tarbase |
| hsa-miR-15a-5p | SYPL1 | HITS-CLIP | Tarbase |
| hsa-miR-195-5p | SYPL1 | HITS-CLIP | Tarbase |
| hsa-miR-15a-5p | SPAG9 | HITS-CLIP | Tarbase |
| hsa-miR-195-5p | SPAG9 | HITS-CLIP | Tarbase |
| hsa-miR-486-5p | CELSR3 | HITS-CLIP | Tarbase |
| hsa-miR-15a-5p | AASS | HITS-CLIP | Tarbase |
| hsa-miR-141-5p | ST3GAL1 | HITS-CLIP | Tarbase |
| hsa-miR-141-5p | SEC62 | HITS-CLIP | Tarbase |
| hsa-miR-15a-5p | CSDE1 | PAR-CLIP | Tarbase |
| hsa-miR-15a-5p | CSDE1 | HITS-CLIP | Tarbase |
| hsa-miR-15a-5p | CSDE1 | HITS-CLIP | Tarbase |
| hsa-miR-15a-5p | CSDE1 | HITS-CLIP | Tarbase |
| hsa-miR-15a-5p | CSDE1 | HITS-CLIP | Tarbase |

| | | | |
|---|---|---|---|
| hsa-miR-15a-5p | CSDE1 | HITS-CLIP | Tarbase |
| hsa-miR-15a-5p | CSDE1 | HITS-CLIP | Tarbase |
| hsa-miR-195-5p | CSDE1 | HITS-CLIP | Tarbase |
| hsa-miR-195-5p | CSDE1 | HITS-CLIP | Tarbase |
| hsa-miR-195-5p | CSDE1 | HITS-CLIP | Tarbase |
| hsa-miR-195-5p | CSDE1 | HITS-CLIP | Tarbase |
| hsa-miR-3613-5p | CSDE1 | HITS-CLIP | Tarbase |
| hsa-miR-3613-5p | CSDE1 | HITS-CLIP | Tarbase |
| hsa-miR-141-3p | CSDE1 | Chimeric fragments | Tarbase |
| hsa-miR-141-3p | UBE3C | HITS-CLIP | Tarbase |
| hsa-miR-195-5p | UBE3C | HITS-CLIP | Tarbase |
| hsa-miR-219a-1-3p | UBE3C | PAR-CLIP | Tarbase |
| hsa-miR-219a-1-3p | UBE3C | PAR-CLIP | Tarbase |
| hsa-miR-219a-1-3p | UBE3C | PAR-CLIP | Tarbase |
| hsa-miR-3613-5p | UBE3C | HITS-CLIP | Tarbase |
| hsa-miR-3613-5p | UBE3C | HITS-CLIP | Tarbase |
| hsa-miR-3613-5p | UBE3C | PAR-CLIP | Tarbase |
| hsa-miR-3613-5p | UBE3C | PAR-CLIP | Tarbase |
| hsa-miR-3613-5p | UBE3C | PAR-CLIP | Tarbase |
| hsa-miR-15a-3p | REV3L | HITS-CLIP | Tarbase |
| hsa-miR-15a-5p | REV3L | PAR-CLIP | Tarbase |
| hsa-miR-15a-5p | REV3L | PAR-CLIP | Tarbase |
| hsa-miR-15a-5p | REV3L | PAR-CLIP | Tarbase |
| hsa-miR-15a-5p | REV3L | PAR-CLIP | Tarbase |
| hsa-miR-15a-5p | REV3L | PAR-CLIP | Tarbase |
| hsa-miR-195-5p | REV3L | PAR-CLIP | Tarbase |
| hsa-miR-195-5p | REV3L | PAR-CLIP | Tarbase |
| hsa-miR-3613-5p | REV3L | PAR-CLIP | Tarbase |
| hsa-miR-3613-5p | REV3L | PAR-CLIP | Tarbase |
| hsa-miR-3613-5p | POMT2 | HITS-CLIP | Tarbase |
| hsa-miR-141-5p | VTA1 | HITS-CLIP | Tarbase |
| hsa-miR-15a-5p | VTA1 | HITS-CLIP | Tarbase |
| hsa-miR-195-5p | VTA1 | HITS-CLIP | Tarbase |
| hsa-miR-15a-3p | BAZ1B | HITS-CLIP | Tarbase |
| hsa-miR-141-3p | RANBP9 | HITS-CLIP | Tarbase |
| hsa-miR-15a-5p | RANBP9 | HITS-CLIP | Tarbase |
| hsa-miR-195-5p | RANBP9 | HITS-CLIP | Tarbase |
| hsa-miR-15a-5p | SPRTN | HITS-CLIP | Tarbase |
| hsa-miR-15a-5p | SPRTN | HITS-CLIP | Tarbase |
| hsa-miR-15a-5p | SPRTN | PAR-CLIP | Tarbase |
| hsa-miR-195-5p | SPRTN | HITS-CLIP | Tarbase |
| hsa-miR-195-5p | SPRTN | HITS-CLIP | Tarbase |
| hsa-miR-486-5p | SPRTN | HITS-CLIP | Tarbase |
| hsa-miR-15a-5p | ZNF207 | PAR-CLIP | Tarbase |
| hsa-miR-141-5p | NCAPD2 | HITS-CLIP | Tarbase |
| hsa-miR-15a-5p | NCAPD2 | PAR-CLIP | Tarbase |
| hsa-miR-15a-5p | NCAPD2 | PAR-CLIP | Tarbase |
| hsa-miR-195-5p | NCAPD2 | PAR-CLIP | Tarbase |
| hsa-miR-195-5p | NCAPD2 | Microarrays | Tarbase |
| hsa-miR-15a-5p | NCAPD2 | Chimeric fragments | Tarbase |
| hsa-miR-141-5p | IDS | HITS-CLIP | Tarbase |
| hsa-miR-486-5p | CD4 | PAR-CLIP | Tarbase |
| hsa-miR-15a-5p | HIVEP2 | PAR-CLIP | Tarbase |
| hsa-miR-15a-5p | HIVEP2 | PAR-CLIP | Tarbase |
| hsa-miR-195-5p | HIVEP2 | PAR-CLIP | Tarbase |
| hsa-miR-486-3p | HIVEP2 | HITS-CLIP | Tarbase |
| hsa-miR-3613-3p | TCEB3 | PAR-CLIP | Tarbase |
| hsa-miR-15a-5p | LYPLA2 | PAR-CLIP | Tarbase |
| hsa-miR-15a-5p | LYPLA2 | HITS-CLIP | Tarbase |
| hsa-miR-195-5p | LYPLA2 | HITS-CLIP | Tarbase |
| hsa-miR-195-5p | LYPLA2 | Microarrays | Tarbase |
| hsa-miR-15a-5p | BTBD7 | HITS-CLIP | Tarbase |
| hsa-miR-486-5p | BTBD7 | PAR-CLIP | Tarbase |
| hsa-miR-141-5p | KAL1 | HITS-CLIP | Tarbase |
| hsa-miR-195-5p | KAL1 | Microarrays | Tarbase |
| hsa-miR-141-3p | MBTD1 | HITS-CLIP | Tarbase |
| hsa-miR-3605-5p | RNF216 | PAR-CLIP | Tarbase |
| hsa-miR-15a-5p | PTBP1 | HITS-CLIP | Tarbase |
| hsa-miR-486-3p | PTBP1 | HITS-CLIP | Tarbase |
| hsa-miR-195-5p | PLAUR | Microarrays | Tarbase |
| hsa-miR-141-3p | ANLN | HITS-CLIP | Tarbase |
| hsa-miR-141-3p | ANLN | HITS-CLIP | Tarbase |
| hsa-miR-141-5p | ANLN | HITS-CLIP | Tarbase |
| hsa-miR-15a-5p | ANLN | HITS-CLIP | Tarbase |
| hsa-miR-15a-5p | ANLN | PAR-CLIP | Tarbase |
| hsa-miR-15a-5p | ANLN | HITS-CLIP | Tarbase |
| hsa-miR-195-5p | ANLN | HITS-CLIP | Tarbase |
| hsa-miR-195-5p | ANLN | Microarrays | Tarbase |
| hsa-miR-141-3p | RABGAP1 | HITS-CLIP | Tarbase |
| hsa-miR-15a-5p | TMEM159 | HITS-CLIP | Tarbase |
| hsa-miR-15a-5p | TMEM159 | HITS-CLIP | Tarbase |
| hsa-miR-15a-5p | TMEM159 | HITS-CLIP | Tarbase |
| hsa-miR-195-5p | TMEM159 | HITS-CLIP | Tarbase |
| hsa-miR-195-5p | TMEM159 | HITS-CLIP | Tarbase |
| hsa-miR-195-5p | TMEM159 | HITS-CLIP | Tarbase |
| hsa-miR-15a-3p | BRCA1 | HITS-CLIP | Tarbase |
| hsa-miR-195-5p | BRCA1 | Microarrays | Tarbase |
| hsa-miR-3605-3p | SEMA3B | PAR-CLIP | Tarbase |
| hsa-miR-3605-3p | SEMA3B | PAR-CLIP | Tarbase |
| hsa-miR-141-3p | MBTPS2 | PAR-CLIP | Tarbase |
| hsa-miR-141-3p | MBTPS2 | HITS-CLIP | Tarbase |
| hsa-miR-15a-5p | MBTPS2 | PAR-CLIP | Tarbase |
| hsa-miR-195-5p | NR1H4 | Microarrays | Tarbase |
| hsa-miR-15a-5p | ELOVL5 | HITS-CLIP | Tarbase |
| hsa-miR-195-5p | ELOVL5 | HITS-CLIP | Tarbase |
| hsa-miR-3605-5p | ELOVL5 | HITS-CLIP | Tarbase |
| hsa-miR-15a-5p | ELOVL5 | HITS-CLIP | Tarbase |
| hsa-miR-195-5p | ELOVL5 | HITS-CLIP | Tarbase |
| hsa-miR-195-5p | ELOVL5 | Microarrays | Tarbase |
| hsa-miR-195-5p | UBR7 | Microarrays | Tarbase |
| hsa-miR-195-5p | EHD3 | Microarrays | Tarbase |
| hsa-miR-141-5p | MAN2B2 | HITS-CLIP | Tarbase |
| hsa-miR-15a-5p | SLC25A39 | PAR-CLIP | Tarbase |
| hsa-miR-141-5p | PGM3 | HITS-CLIP | Tarbase |
| hsa-miR-15a-3p | PGM3 | PAR-CLIP | Tarbase |
| hsa-miR-195-5p | RWDD2A | Microarrays | Tarbase |
| hsa-miR-195-5p | RNF14 | Microarrays | Tarbase |
| hsa-miR-195-5p | GPRC5A | Microarrays | Tarbase |
| hsa-miR-141-5p | TACC3 | HITS-CLIP | Tarbase |
| hsa-miR-15a-5p | TACC3 | PAR-CLIP | Tarbase |
| hsa-miR-195-5p | TACC3 | Microarrays | Tarbase |
| hsa-miR-141-5p | UFL1 | HITS-CLIP | Tarbase |
| hsa-miR-219a-1-3p | UFL1 | PAR-CLIP | Tarbase |
| hsa-miR-195-5p | POLA2 | Microarrays | Tarbase |
| hsa-miR-15a-5p | MDH1 | PAR-CLIP | Tarbase |
| hsa-miR-15a-5p | MDH1 | HITS-CLIP | Tarbase |
| hsa-miR-195-5p | MDH1 | HITS-CLIP | Tarbase |
| hsa-miR-3613-5p | MDH1 | HITS-CLIP | Tarbase |
| hsa-miR-195-5p | MTMR11 | Microarrays | Tarbase |
| hsa-miR-15a-5p | COX15 | HITS-CLIP | Tarbase |
| hsa-miR-3613-3p | CCDC88C | HITS-CLIP | Tarbase |
| hsa-miR-141-3p | ZMYND11 | HITS-CLIP | Tarbase |
| hsa-miR-15a-5p | ZMYND11 | HITS-CLIP | Tarbase |
| hsa-miR-15a-5p | ZMYND11 | HITS-CLIP | Tarbase |
| hsa-miR-195-5p | ZMYND11 | HITS-CLIP | Tarbase |
| hsa-miR-15a-5p | ZMYND11 | Chimeric fragments | Tarbase |
| hsa-miR-141-3p | MATR3 | HITS-CLIP | Tarbase |
| hsa-miR-15a-5p | MATR3 | HITS-CLIP | Tarbase |
| hsa-miR-15a-5p | MATR3 | HITS-CLIP | Tarbase |
| hsa-miR-15a-5p | MATR3 | HITS-CLIP | Tarbase |
| hsa-miR-195-5p | MATR3 | HITS-CLIP | Tarbase |
| hsa-miR-195-5p | MATR3 | HITS-CLIP | Tarbase |
| hsa-miR-15a-5p | MATR3 | HITS-CLIP | Tarbase |
| hsa-miR-195-5p | MATR3 | HITS-CLIP | Tarbase |
| hsa-miR-15a-5p | MATR3 | Chimeric fragments | Tarbase |
| hsa-miR-141-5p | XYLT2 | HITS-CLIP | Tarbase |
| hsa-miR-195-5p | XYLT2 | Microarrays | Tarbase |
| hsa-miR-15a-3p | ISL1 | HITS-CLIP | Tarbase |
| hsa-miR-15a-5p | CLCA1 | HITS-CLIP | Tarbase |
| hsa-miR-195-5p | CLCA1 | HITS-CLIP | Tarbase |
| hsa-miR-15a-5p | ATP2C1 | HITS-CLIP | Tarbase |
| hsa-miR-195-5p | ATP2C1 | HITS-CLIP | Tarbase |
| hsa-miR-219a-1-3p | IGF1 | PAR-CLIP | Tarbase |
| hsa-miR-141-5p | RUFY3 | HITS-CLIP | Tarbase |
| hsa-miR-141-3p | WWTR1 | Luciferase Reporter Assay | Tarbase |
| hsa-miR-141-3p | WWTR1 | Luciferase Reporter Assay | Tarbase |
| hsa-miR-15a-5p | HGF | PAR-CLIP | Tarbase |
| hsa-miR-195-5p | HGF | PAR-CLIP | Tarbase |
| hsa-miR-195-5p | HGF | Microarrays | Tarbase |

| | | | |
|---|---|---|---|
| hsa-miR-3605-3p | SAMD4A | HITS-CLIP | Tarbase |
| hsa-miR-195-5p | SAMD4A | Microarrays | Tarbase |
| hsa-miR-486-5p | MRE11A | HITS-CLIP | Tarbase |
| hsa-miR-486-5p | SERPINB1 | HITS-CLIP | Tarbase |
| hsa-miR-15a-5p | SPAST | PAR-CLIP | Tarbase |
| hsa-miR-195-5p | SPAST | PAR-CLIP | Tarbase |
| hsa-miR-15a-5p | GABRA1 | HITS-CLIP | Tarbase |
| hsa-miR-195-5p | GABRA1 | HITS-CLIP | Tarbase |
| hsa-miR-195-5p | GABRA1 | Microarrays | Tarbase |
| hsa-miR-15a-3p | GRAMD1B | HITS-CLIP | Tarbase |
| hsa-miR-195-5p | RNH1 | HITS-CLIP | Tarbase |
| hsa-miR-15a-5p | NDUFS1 | HITS-CLIP | Tarbase |
| hsa-miR-195-5p | NDUFS1 | HITS-CLIP | Tarbase |
| hsa-miR-141-3p | RB1CC1 | HITS-CLIP | Tarbase |
| hsa-miR-141-3p | RB1CC1 | HITS-CLIP | Tarbase |
| hsa-miR-141-5p | ALAS1 | HITS-CLIP | Tarbase |
| hsa-miR-15a-5p | AKAP11 | PAR-CLIP | Tarbase |
| hsa-miR-15a-5p | AKAP11 | PAR-CLIP | Tarbase |
| hsa-miR-15a-5p | AKAP11 | HITS-CLIP | Tarbase |
| hsa-miR-195-5p | AKAP11 | PAR-CLIP | Tarbase |
| hsa-miR-195-5p | AKAP11 | HITS-CLIP | Tarbase |
| hsa-miR-219a-1-3p | AKAP11 | PAR-CLIP | Tarbase |
| hsa-miR-15a-5p | AKAP11 | HITS-CLIP | Tarbase |
| hsa-miR-195-5p | AKAP11 | HITS-CLIP | Tarbase |
| hsa-miR-141-3p | STRAP | HITS-CLIP | Tarbase |
| hsa-miR-141-5p | UBR2 | HITS-CLIP | Tarbase |
| hsa-miR-15a-3p | DEPDC1 | HITS-CLIP | Tarbase |
| hsa-miR-195-5p | DEPDC1 | Microarrays | Tarbase |
| hsa-miR-15a-5p | RRAGD | HITS-CLIP | Tarbase |
| hsa-miR-15a-5p | RRAGD | PAR-CLIP | Tarbase |
| hsa-miR-195-5p | RRAGD | HITS-CLIP | Tarbase |
| hsa-miR-141-5p | PHF20 | HITS-CLIP | Tarbase |
| hsa-miR-195-5p | NCAPH2 | Microarrays | Tarbase |
| hsa-miR-195-5p | TOMM34 | Microarrays | Tarbase |
| hsa-miR-486-5p | SEC63 | HITS-CLIP | Tarbase |
| hsa-miR-15a-5p | KPNA6 | HITS-CLIP | Tarbase |
| hsa-miR-15a-5p | KPNA6 | PAR-CLIP | Tarbase |
| hsa-miR-219a-1-3p | KPNA6 | PAR-CLIP | Tarbase |
| hsa-miR-219a-1-3p | KPNA6 | PAR-CLIP | Tarbase |
| hsa-miR-15a-5p | VIM | HITS-CLIP | Tarbase |
| hsa-miR-15a-5p | VIM | HITS-CLIP | Tarbase |
| hsa-miR-15a-5p | VIM | HITS-CLIP | Tarbase |
| hsa-miR-195-5p | VIM | HITS-CLIP | Tarbase |
| hsa-miR-195-5p | VIM | HITS-CLIP | Tarbase |
| hsa-miR-195-5p | VIM | HITS-CLIP | Tarbase |
| hsa-miR-486-5p | VIM | HITS-CLIP | Tarbase |
| hsa-miR-141-5p | CD44 | HITS-CLIP | Tarbase |
| hsa-miR-195-5p | CD44 | qPCR | Tarbase |
| hsa-miR-3605-5p | CD44 | HITS-CLIP | Tarbase |
| hsa-miR-486-3p | SLAMF7 | PAR-CLIP | Tarbase |
| hsa-miR-3605-5p | MIPEP | HITS-CLIP | Tarbase |
| hsa-miR-15a-5p | VEZT | HITS-CLIP | Tarbase |
| hsa-miR-195-5p | VEZT | HITS-CLIP | Tarbase |
| hsa-miR-486-5p | VEZT | HITS-CLIP | Tarbase |
| hsa-miR-486-5p | VEZT | PAR-CLIP | Tarbase |
| hsa-miR-486-3p | POU2F2 | HITS-CLIP | Tarbase |
| hsa-miR-15a-5p | TBPL1 | HITS-CLIP | Tarbase |
| hsa-miR-15a-5p | TBPL1 | HITS-CLIP | Tarbase |
| hsa-miR-15a-5p | TBPL1 | HITS-CLIP | Tarbase |
| hsa-miR-15a-5p | TBPL1 | PAR-CLIP | Tarbase |
| hsa-miR-15a-5p | TBPL1 | PAR-CLIP | Tarbase |
| hsa-miR-15a-5p | TBPL1 | HITS-CLIP | Tarbase |
| hsa-miR-195-5p | TBPL1 | HITS-CLIP | Tarbase |
| hsa-miR-195-5p | TBPL1 | HITS-CLIP | Tarbase |
| hsa-miR-141-3p | ARNTL2 | HITS-CLIP | Tarbase |
| hsa-miR-141-5p | ARNTL2 | HITS-CLIP | Tarbase |
| hsa-miR-141-5p | BCLAF1 | HITS-CLIP | Tarbase |
| hsa-miR-141-3p | SLC39A9 | HITS-CLIP | Tarbase |
| hsa-miR-141-5p | HMGB3 | HITS-CLIP | Tarbase |
| hsa-miR-141-3p | NUP160 | HITS-CLIP | Tarbase |
| hsa-miR-15a-5p | NUP160 | HITS-CLIP | Tarbase |
| hsa-miR-195-5p | NUP160 | HITS-CLIP | Tarbase |
| hsa-miR-3613-3p | NUP160 | PAR-CLIP | Tarbase |
| hsa-miR-3613-3p | NUP160 | PAR-CLIP | Tarbase |
| hsa-miR-3613-3p | NUP160 | PAR-CLIP | Tarbase |
| hsa-miR-3605-5p | IKZF2 | PAR-CLIP | Tarbase |
| hsa-miR-15a-3p | GRN | HITS-CLIP | Tarbase |
| hsa-miR-486-5p | FAM13B | HITS-CLIP | Tarbase |
| hsa-miR-141-5p | CENPQ | HITS-CLIP | Tarbase |
| hsa-miR-141-5p | CENPQ | HITS-CLIP | Tarbase |
| hsa-miR-195-5p | CENPQ | Microarrays | Tarbase |
| hsa-miR-141-5p | ARID4A | HITS-CLIP | Tarbase |
| hsa-miR-15a-5p | PNPLA6 | HITS-CLIP | Tarbase |
| hsa-miR-195-5p | PNPLA6 | HITS-CLIP | Tarbase |
| hsa-miR-195-5p | PNPLA6 | Microarrays | Tarbase |
| hsa-miR-15a-5p | ABCF2 | HITS-CLIP | Tarbase |
| hsa-miR-15a-5p | ABCF2 | PAR-CLIP | Tarbase |
| hsa-miR-15a-5p | ABCF2 | HITS-CLIP | Tarbase |
| hsa-miR-15a-5p | ABCF2 | HITS-CLIP | Tarbase |
| hsa-miR-195-5p | ABCF2 | HITS-CLIP | Tarbase |
| hsa-miR-195-5p | ABCF2 | PAR-CLIP | Tarbase |
| hsa-miR-195-5p | ABCF2 | HITS-CLIP | Tarbase |
| hsa-miR-15a-5p | CHPF2 | HITS-CLIP | Tarbase |
| hsa-miR-195-5p | CHPF2 | HITS-CLIP | Tarbase |
| hsa-miR-195-5p | CHPF2 | Microarrays | Tarbase |
| hsa-miR-15a-5p | LRRC7 | HITS-CLIP | Tarbase |
| hsa-miR-195-5p | LRRC7 | HITS-CLIP | Tarbase |
| hsa-miR-15a-5p | UBA6 | PAR-CLIP | Tarbase |
| hsa-miR-15a-5p | UBA6 | HITS-CLIP | Tarbase |
| hsa-miR-195-5p | UBA6 | PAR-CLIP | Tarbase |
| hsa-miR-195-5p | UBA6 | HITS-CLIP | Tarbase |
| hsa-miR-3605-5p | UBA6 | HITS-CLIP | Tarbase |
| hsa-miR-15a-5p | PIAS1 | HITS-CLIP | Tarbase |
| hsa-miR-195-5p | PIAS1 | HITS-CLIP | Tarbase |
| hsa-miR-195-5p | APBA2 | Microarrays | Tarbase |
| hsa-miR-141-5p | TMSB10 | HITS-CLIP | Tarbase |
| hsa-miR-141-3p | VCL | HITS-CLIP | Tarbase |
| hsa-miR-141-5p | VCL | HITS-CLIP | Tarbase |
| hsa-miR-195-5p | VCL | Microarrays | Tarbase |
| hsa-miR-141-3p | DEPDC1B | HITS-CLIP | Tarbase |
| hsa-miR-15a-5p | DEPDC1B | HITS-CLIP | Tarbase |
| hsa-miR-195-5p | DEPDC1B | Microarrays | Tarbase |
| hsa-miR-15a-5p | ADSS | HITS-CLIP | Tarbase |
| hsa-miR-15a-5p | ADSS | HITS-CLIP | Tarbase |
| hsa-miR-195-5p | ADSS | HITS-CLIP | Tarbase |
| hsa-miR-141-5p | TIMP2 | HITS-CLIP | Tarbase |
| hsa-miR-15a-3p | TIMP2 | HITS-CLIP | Tarbase |
| hsa-miR-15a-3p | TIMP2 | HITS-CLIP | Tarbase |
| hsa-miR-15a-3p | TIMP2 | HITS-CLIP | Tarbase |
| hsa-miR-15a-3p | ZZZ3 | HITS-CLIP | Tarbase |
| hsa-miR-15a-3p | ZZZ3 | HITS-CLIP | Tarbase |
| hsa-miR-15a-5p | NSUN2 | HITS-CLIP | Tarbase |
| hsa-miR-195-5p | NSUN2 | HITS-CLIP | Tarbase |
| hsa-miR-219a-1-3p | MFAP3 | HITS-CLIP | Tarbase |
| hsa-miR-195-5p | MFAP3 | Microarrays | Tarbase |
| hsa-miR-15a-5p | PI4K2B | HITS-CLIP | Tarbase |
| hsa-miR-15a-5p | PI4K2B | HITS-CLIP | Tarbase |
| hsa-miR-195-5p | PI4K2B | HITS-CLIP | Tarbase |
| hsa-miR-141-5p | BOD1L1 | HITS-CLIP | Tarbase |
| hsa-miR-3613-3p | TRIO | HITS-CLIP | Tarbase |
| hsa-miR-195-5p | TRIO | Microarrays | Tarbase |
| hsa-miR-195-5p | VCAN | Microarrays | Tarbase |
| hsa-miR-15a-5p | CDH1 | HITS-CLIP | Tarbase |
| hsa-miR-195-5p | CDH1 | HITS-CLIP | Tarbase |
| hsa-miR-486-5p | FAM65A | PAR-CLIP | Tarbase |
| hsa-miR-195-5p | C6 | Microarrays | Tarbase |
| hsa-miR-15a-5p | RAI14 | HITS-CLIP | Tarbase |
| hsa-miR-195-5p | RAI14 | HITS-CLIP | Tarbase |
| hsa-miR-486-3p | RAI14 | HITS-CLIP | Tarbase |
| hsa-miR-141-3p | PHLPP2 | PAR-CLIP | Tarbase |
| hsa-miR-15a-5p | PHLPP2 | HITS-CLIP | Tarbase |
| hsa-miR-195-5p | PHLPP2 | HITS-CLIP | Tarbase |
| hsa-miR-15a-5p | PHLPP2 | Luciferase Reporter Assay | Tarbase |
| hsa-miR-3605-3p | PHF23 | HITS-CLIP | Tarbase |
| hsa-miR-195-5p | CDH10 | Microarrays | Tarbase |
| hsa-miR-15a-5p | INPP4A | HITS-CLIP | Tarbase |
| hsa-miR-15a-5p | INPP4A | HITS-CLIP | Tarbase |
| hsa-miR-195-5p | INPP4A | HITS-CLIP | Tarbase |
| hsa-miR-195-5p | RAB27B | Microarrays | Tarbase |

| | | | |
|---|---|---|---|
| hsa-miR-141-5p | PSMA4 | HITS-CLIP | Tarbase |
| hsa-miR-15a-5p | LSG1 | HITS-CLIP | Tarbase |
| hsa-miR-195-5p | LSG1 | HITS-CLIP | Tarbase |
| hsa-miR-15a-3p | TNC | HITS-CLIP | Tarbase |
| hsa-miR-15a-3p | TNC | HITS-CLIP | Tarbase |
| hsa-miR-195-5p | FAM65C | Microarrays | Tarbase |
| hsa-miR-15a-5p | MED17 | HITS-CLIP | Tarbase |
| hsa-miR-195-5p | MED17 | HITS-CLIP | Tarbase |
| hsa-miR-195-5p | TG | Microarrays | Tarbase |
| hsa-miR-141-5p | ZIC2 | HITS-CLIP | Tarbase |
| hsa-miR-195-5p | LCP2 | Microarrays | Tarbase |
| hsa-miR-195-5p | TRIT1 | Microarrays | Tarbase |
| hsa-miR-141-3p | ADRB1 | HITS-CLIP | Tarbase |
| hsa-miR-195-5p | CUL7 | Microarrays | Tarbase |
| hsa-miR-141-5p | HSPA5 | HITS-CLIP | Tarbase |
| hsa-miR-15a-3p | HSPA5 | HITS-CLIP | Tarbase |
| hsa-miR-15a-5p | HSPA5 | HITS-CLIP | Tarbase |
| hsa-miR-15a-5p | HSPA5 | HITS-CLIP | Tarbase |
| hsa-miR-195-5p | HSPA5 | HITS-CLIP | Tarbase |
| hsa-miR-195-5p | HSPA5 | HITS-CLIP | Tarbase |
| hsa-miR-141-3p | DSG2 | HITS-CLIP | Tarbase |
| hsa-miR-15a-5p | DSG2 | HITS-CLIP | Tarbase |
| hsa-miR-15a-5p | DSG2 | Chimeric fragments | Tarbase |
| hsa-miR-141-5p | OFD1 | HITS-CLIP | Tarbase |
| hsa-miR-15a-5p | GPM6B | HITS-CLIP | Tarbase |
| hsa-miR-195-5p | GPM6B | HITS-CLIP | Tarbase |
| hsa-miR-141-3p | WDR37 | Luciferase Reporter Assay | Tarbase |
| hsa-miR-141-3p | WDR37 | Luciferase Reporter Assay | Tarbase |
| hsa-miR-141-3p | WDR37 | HITS-CLIP | Tarbase |
| hsa-miR-141-3p | WDR37 | HITS-CLIP | Tarbase |
| hsa-miR-15a-5p | CTPS2 | HITS-CLIP | Tarbase |
| hsa-miR-195-5p | CTPS2 | HITS-CLIP | Tarbase |
| hsa-miR-141-3p | POLR2B | HITS-CLIP | Tarbase |
| hsa-miR-141-3p | POLR2B | HITS-CLIP | Tarbase |
| hsa-miR-15a-5p | POLR2B | HITS-CLIP | Tarbase |
| hsa-miR-15a-5p | POLR2B | HITS-CLIP | Tarbase |
| hsa-miR-15a-5p | FAM214A | HITS-CLIP | Tarbase |
| hsa-miR-15a-5p | FAM214A | HITS-CLIP | Tarbase |
| hsa-miR-195-5p | FAM214A | HITS-CLIP | Tarbase |
| hsa-miR-195-5p | FAM214A | HITS-CLIP | Tarbase |
| hsa-miR-15a-5p | FAM214A | HITS-CLIP | Tarbase |
| hsa-miR-195-5p | FAM214A | HITS-CLIP | Tarbase |
| hsa-miR-15a-5p | ARAP2 | HITS-CLIP | Tarbase |
| hsa-miR-195-5p | ARAP2 | HITS-CLIP | Tarbase |
| hsa-miR-15a-3p | SCML1 | HITS-CLIP | Tarbase |
| hsa-miR-15a-5p | SCML1 | PAR-CLIP | Tarbase |
| hsa-miR-15a-5p | SCML1 | PAR-CLIP | Tarbase |
| hsa-miR-15a-5p | SCML1 | HITS-CLIP | Tarbase |
| hsa-miR-15a-5p | SCML1 | PAR-CLIP | Tarbase |
| hsa-miR-195-5p | SCML1 | HITS-CLIP | Tarbase |
| hsa-miR-15a-5p | MAP4 | HITS-CLIP | Tarbase |
| hsa-miR-195-5p | MAP4 | HITS-CLIP | Tarbase |
| hsa-miR-486-5p | MAP4 | HITS-CLIP | Tarbase |
| hsa-miR-141-3p | GOPC | HITS-CLIP | Tarbase |
| hsa-miR-15a-5p | RRM2B | PAR-CLIP | Tarbase |
| hsa-miR-15a-5p | RRM2B | PAR-CLIP | Tarbase |
| hsa-miR-195-5p | RRM2B | PAR-CLIP | Tarbase |
| hsa-miR-3605-3p | ZNF800 | HITS-CLIP | Tarbase |
| hsa-miR-486-5p | ZNF800 | PAR-CLIP | Tarbase |
| hsa-miR-15a-5p | MRPS10 | HITS-CLIP | Tarbase |
| hsa-miR-3605-5p | RSF1 | HITS-CLIP | Tarbase |
| hsa-miR-15a-5p | FAM120A | PAR-CLIP | Tarbase |
| hsa-miR-15a-5p | FAM120A | HITS-CLIP | Tarbase |
| hsa-miR-195-5p | FAM120A | HITS-CLIP | Tarbase |
| hsa-miR-15a-5p | FAM120A | Chimeric fragments | Tarbase |
| hsa-miR-195-5p | KITLG | Microarrays | Tarbase |
| hsa-miR-195-5p | ADAMTS6 | Microarrays | Tarbase |
| hsa-miR-141-3p | VAMP3 | HITS-CLIP | Tarbase |
| hsa-miR-15a-3p | VAMP3 | HITS-CLIP | Tarbase |
| hsa-miR-195-5p | PER3 | Microarrays | Tarbase |
| hsa-miR-195-5p | TNFRSF9 | Microarrays | Tarbase |
| hsa-miR-15a-5p | LTBP1 | HITS-CLIP | Tarbase |
| hsa-miR-15a-5p | LTBP1 | HITS-CLIP | Tarbase |
| hsa-miR-195-5p | LTBP1 | HITS-CLIP | Tarbase |
| hsa-miR-195-5p | RFC2 | Microarrays | Tarbase |
| hsa-miR-15a-3p | ARID1B | HITS-CLIP | Tarbase |
| hsa-miR-141-3p | JKAMP | HITS-CLIP | Tarbase |
| hsa-miR-141-5p | JKAMP | HITS-CLIP | Tarbase |
| hsa-miR-15a-5p | JKAMP | HITS-CLIP | Tarbase |
| hsa-miR-15a-5p | JKAMP | HITS-CLIP | Tarbase |
| hsa-miR-195-5p | JKAMP | HITS-CLIP | Tarbase |
| hsa-miR-15a-5p | MAPK9 | PAR-CLIP | Tarbase |
| hsa-miR-15a-5p | MAPK9 | HITS-CLIP | Tarbase |
| hsa-miR-15a-5p | MAPK9 | HITS-CLIP | Tarbase |
| hsa-miR-195-5p | MAPK9 | HITS-CLIP | Tarbase |
| hsa-miR-195-5p | BCAR1 | Microarrays | Tarbase |
| hsa-miR-195-5p | RAD51 | Microarrays | Tarbase |
| hsa-miR-195-5p | POLQ | Microarrays | Tarbase |
| hsa-miR-15a-5p | PLEKHA5 | PAR-CLIP | Tarbase |
| hsa-miR-486-3p | SIKE1 | HITS-CLIP | Tarbase |
| hsa-miR-141-3p | FNIP2 | HITS-CLIP | Tarbase |
| hsa-miR-15a-5p | TTC17 | HITS-CLIP | Tarbase |
| hsa-miR-15a-5p | TTC17 | HITS-CLIP | Tarbase |
| hsa-miR-195-5p | TTC17 | HITS-CLIP | Tarbase |
| hsa-miR-219a-1-3p | THRAP3 | HITS-CLIP | Tarbase |
| hsa-miR-141-3p | LY75 | HITS-CLIP | Tarbase |
| hsa-miR-141-5p | ARID4B | HITS-CLIP | Tarbase |
| hsa-miR-141-5p | ARID4B | HITS-CLIP | Tarbase |
| hsa-miR-486-5p | ARID4B | HITS-CLIP | Tarbase |
| hsa-miR-141-3p | KIF1B | HITS-CLIP | Tarbase |
| hsa-miR-15a-5p | KIF1B | HITS-CLIP | Tarbase |
| hsa-miR-195-5p | KIF1B | HITS-CLIP | Tarbase |
| hsa-miR-195-5p | KIF1B | HITS-CLIP | Tarbase |
| hsa-miR-141-3p | FOXC1 | HITS-CLIP | Tarbase |
| hsa-miR-141-3p | SYNE2 | HITS-CLIP | Tarbase |
| hsa-miR-15a-3p | SYNE2 | HITS-CLIP | Tarbase |
| hsa-miR-15a-5p | PLEKHH1 | PAR-CLIP | Tarbase |
| hsa-miR-15a-5p | PLEKHH1 | HITS-CLIP | Tarbase |
| hsa-miR-141-5p | ATP9A | HITS-CLIP | Tarbase |
| hsa-miR-3605-5p | ATP9A | HITS-CLIP | Tarbase |
| hsa-miR-3613-3p | ATP9A | HITS-CLIP | Tarbase |
| hsa-miR-486-5p | RELT | HITS-CLIP | Tarbase |
| hsa-miR-15a-5p | SZRD1 | PAR-CLIP | Tarbase |
| hsa-miR-195-5p | CUL1 | HITS-CLIP | Tarbase |
| hsa-miR-141-3p | FAM114A2 | HITS-CLIP | Tarbase |
| hsa-miR-3605-5p | CYFIP2 | HITS-CLIP | Tarbase |
| hsa-miR-195-5p | CYFIP2 | Microarrays | Tarbase |
| hsa-miR-15a-5p | TAB2 | HITS-CLIP | Tarbase |
| hsa-miR-195-5p | TAB2 | HITS-CLIP | Tarbase |
| hsa-miR-486-5p | TAB2 | PAR-CLIP | Tarbase |
| hsa-miR-141-5p | GINM1 | HITS-CLIP | Tarbase |
| hsa-miR-141-3p | EIF2AK2 | HITS-CLIP | Tarbase |
| hsa-miR-15a-5p | EIF2AK2 | PAR-CLIP | Tarbase |
| hsa-miR-15a-5p | EIF2AK2 | PAR-CLIP | Tarbase |
| hsa-miR-15a-5p | EIF2AK2 | PAR-CLIP | Tarbase |
| hsa-miR-195-5p | EIF2AK2 | PAR-CLIP | Tarbase |
| hsa-miR-3613-3p | USP36 | HITS-CLIP | Tarbase |
| hsa-miR-141-5p | KMT2C | HITS-CLIP | Tarbase |
| hsa-miR-141-5p | KMT2C | HITS-CLIP | Tarbase |
| hsa-miR-15a-5p | KMT2C | HITS-CLIP | Tarbase |
| hsa-miR-15a-5p | KMT2C | HITS-CLIP | Tarbase |
| hsa-miR-15a-5p | KMT2C | HITS-CLIP | Tarbase |
| hsa-miR-195-5p | KMT2C | HITS-CLIP | Tarbase |
| hsa-miR-195-5p | KMT2C | HITS-CLIP | Tarbase |
| hsa-miR-3613-3p | KMT2C | HITS-CLIP | Tarbase |
| hsa-miR-3613-3p | KMT2C | PAR-CLIP | Tarbase |
| hsa-miR-15a-5p | KMT2C | HITS-CLIP | Tarbase |
| hsa-miR-141-5p | PUM2 | HITS-CLIP | Tarbase |
| hsa-miR-15a-5p | PUM2 | HITS-CLIP | Tarbase |
| hsa-miR-15a-5p | PUM2 | PAR-CLIP | Tarbase |
| hsa-miR-15a-5p | PUM2 | PAR-CLIP | Tarbase |
| hsa-miR-15a-5p | PUM2 | PAR-CLIP | Tarbase |
| hsa-miR-15a-5p | PUM2 | PAR-CLIP | Tarbase |
| hsa-miR-15a-5p | PUM2 | PAR-CLIP | Tarbase |
| hsa-miR-15a-5p | PUM2 | PAR-CLIP | Tarbase |
| hsa-miR-15a-5p | PUM2 | PAR-CLIP | Tarbase |
| hsa-miR-15a-5p | PUM2 | PAR-CLIP | Tarbase |
| hsa-miR-195-5p | PUM2 | HITS-CLIP | Tarbase |
| hsa-miR-195-5p | PUM2 | PAR-CLIP | Tarbase |

| | | | |
|---|---|---|---|
| hsa-miR-195-5p | PUM2 | PAR-CLIP | Tarbase |
| hsa-miR-195-5p | PUM2 | PAR-CLIP | Tarbase |
| hsa-miR-195-5p | PUM2 | PAR-CLIP | Tarbase |
| hsa-miR-486-5p | PUM2 | HITS-CLIP | Tarbase |
| hsa-miR-486-5p | PUM2 | HITS-CLIP | Tarbase |
| hsa-miR-486-5p | PUM2 | HITS-CLIP | Tarbase |
| hsa-miR-15a-5p | ZFR | HITS-CLIP | Tarbase |
| hsa-miR-15a-5p | ZFR | HITS-CLIP | Tarbase |
| hsa-miR-195-5p | ZFR | HITS-CLIP | Tarbase |
| hsa-miR-195-5p | ZFR | HITS-CLIP | Tarbase |
| hsa-miR-3613-5p | ZFR | HITS-CLIP | Tarbase |
| hsa-miR-141-3p | RC3H2 | HITS-CLIP | Tarbase |
| hsa-miR-141-5p | RC3H2 | HITS-CLIP | Tarbase |
| hsa-miR-141-5p | RC3H2 | HITS-CLIP | Tarbase |
| hsa-miR-141-5p | RC3H2 | HITS-CLIP | Tarbase |
| hsa-miR-141-5p | RC3H2 | HITS-CLIP | Tarbase |
| hsa-miR-15a-5p | RC3H2 | PAR-CLIP | Tarbase |
| hsa-miR-195-5p | RC3H2 | PAR-CLIP | Tarbase |
| hsa-miR-486-3p | IL17RB | PAR-CLIP | Tarbase |
| hsa-miR-141-3p | TRAF3IP2 | HITS-CLIP | Tarbase |
| hsa-miR-15a-5p | DCBLD2 | HITS-CLIP | Tarbase |
| hsa-miR-15a-5p | DCBLD2 | HITS-CLIP | Tarbase |
| hsa-miR-15a-5p | DCBLD2 | HITS-CLIP | Tarbase |
| hsa-miR-195-5p | DCBLD2 | HITS-CLIP | Tarbase |
| hsa-miR-195-5p | DCBLD2 | HITS-CLIP | Tarbase |
| hsa-miR-15a-5p | GDI2 | HITS-CLIP | Tarbase |
| hsa-miR-15a-5p | GDI2 | HITS-CLIP | Tarbase |
| hsa-miR-195-5p | GDI2 | HITS-CLIP | Tarbase |
| hsa-miR-15a-3p | PRDM1 | HITS-CLIP | Tarbase |
| hsa-miR-15a-5p | PRDM1 | PAR-CLIP | Tarbase |
| hsa-miR-195-5p | PRDM1 | Microarrays | Tarbase |
| hsa-miR-3613-3p | ATP11B | PAR-CLIP | Tarbase |
| hsa-miR-15a-5p | LAMC2 | HITS-CLIP | Tarbase |
| hsa-miR-195-5p | LAMC2 | HITS-CLIP | Tarbase |
| hsa-miR-15a-5p | SEC61A1 | PAR-CLIP | Tarbase |
| hsa-miR-15a-5p | SEC61A1 | HITS-CLIP | Tarbase |
| hsa-miR-15a-5p | SEC61A1 | PAR-CLIP | Tarbase |
| hsa-miR-195-5p | SEC61A1 | HITS-CLIP | Tarbase |
| hsa-miR-219a-1-3p | SEC61A1 | HITS-CLIP | Tarbase |
| hsa-miR-141-5p | PPP1R12A | HITS-CLIP | Tarbase |
| hsa-miR-141-5p | ZC3H11A | HITS-CLIP | Tarbase |
| hsa-miR-15a-5p | ZC3H11A | HITS-CLIP | Tarbase |
| hsa-miR-15a-5p | ZC3H11A | HITS-CLIP | Tarbase |
| hsa-miR-195-5p | ZC3H11A | HITS-CLIP | Tarbase |
| hsa-miR-15a-3p | NDC1 | HITS-CLIP | Tarbase |
| hsa-miR-15a-5p | NDC1 | HITS-CLIP | Tarbase |
| hsa-miR-195-5p | NDC1 | HITS-CLIP | Tarbase |
| hsa-miR-486-3p | NDC1 | HITS-CLIP | Tarbase |
| hsa-miR-195-5p | TBXAS1 | Microarrays | Tarbase |
| hsa-miR-15a-5p | ALDH18A1 | HITS-CLIP | Tarbase |
| hsa-miR-195-5p | ALDH18A1 | HITS-CLIP | Tarbase |
| hsa-miR-15a-5p | CDK17 | PAR-CLIP | Tarbase |
| hsa-miR-15a-5p | CDK17 | PAR-CLIP | Tarbase |
| hsa-miR-486-5p | CDK17 | HITS-CLIP | Tarbase |
| hsa-miR-15a-5p | CDK17 | HITS-CLIP | Tarbase |
| hsa-miR-195-5p | CDK17 | HITS-CLIP | Tarbase |
| hsa-miR-15a-5p | SLC2A3 | HITS-CLIP | Tarbase |
| hsa-miR-15a-5p | SLC2A3 | PAR-CLIP | Tarbase |
| hsa-miR-195-5p | SLC2A3 | HITS-CLIP | Tarbase |
| hsa-miR-15a-5p | SLC2A3 | HITS-CLIP | Tarbase |
| hsa-miR-195-5p | SLC2A3 | HITS-CLIP | Tarbase |
| hsa-miR-15a-5p | PSD | HITS-CLIP | Tarbase |
| hsa-miR-195-5p | PSD | HITS-CLIP | Tarbase |
| hsa-miR-141-3p | WNK1 | HITS-CLIP | Tarbase |
| hsa-miR-141-3p | WNK1 | HITS-CLIP | Tarbase |
| hsa-miR-486-3p | WNK1 | HITS-CLIP | Tarbase |
| hsa-miR-15a-5p | OGFR | HITS-CLIP | Tarbase |
| hsa-miR-195-5p | OGFR | HITS-CLIP | Tarbase |
| hsa-miR-15a-5p | PIGV | HITS-CLIP | Tarbase |
| hsa-miR-195-5p | PIGV | HITS-CLIP | Tarbase |
| hsa-miR-15a-5p | QSER1 | HITS-CLIP | Tarbase |
| hsa-miR-195-5p | QSER1 | HITS-CLIP | Tarbase |
| hsa-miR-195-5p | BCAT1 | Microarrays | Tarbase |
| hsa-miR-195-5p | LZTS1 | Microarrays | Tarbase |
| hsa-miR-3613-5p | NCKAP1 | HITS-CLIP | Tarbase |
| hsa-miR-195-5p | GUCY1B3 | Microarrays | Tarbase |
| hsa-miR-141-3p | SFSWAP | HITS-CLIP | Tarbase |
| hsa-miR-15a-3p | TNK2 | HITS-CLIP | Tarbase |
| hsa-miR-15a-5p | TNK2 | HITS-CLIP | Tarbase |
| hsa-miR-195-5p | TNK2 | HITS-CLIP | Tarbase |
| hsa-miR-3605-3p | TNK2 | HITS-CLIP | Tarbase |
| hsa-miR-141-3p | MON2 | HITS-CLIP | Tarbase |
| hsa-miR-141-3p | MON2 | HITS-CLIP | Tarbase |
| hsa-miR-15a-5p | MON2 | PAR-CLIP | Tarbase |
| hsa-miR-195-5p | MON2 | PAR-CLIP | Tarbase |
| hsa-miR-141-3p | ELMO2 | Luciferase Reporter Assay | Tarbase |
| hsa-miR-141-3p | ELMO2 | Luciferase Reporter Assay | Tarbase |
| hsa-miR-15a-5p | ELMO2 | HITS-CLIP | Tarbase |
| hsa-miR-195-5p | ELMO2 | HITS-CLIP | Tarbase |
| hsa-miR-15a-5p | WAPAL | HITS-CLIP | Tarbase |
| hsa-miR-195-5p | WAPAL | HITS-CLIP | Tarbase |
| hsa-miR-15a-5p | VMP1 | HITS-CLIP | Tarbase |
| hsa-miR-195-5p | VMP1 | HITS-CLIP | Tarbase |
| hsa-miR-141-3p | APPBP2 | HITS-CLIP | Tarbase |
| hsa-miR-141-3p | APPBP2 | HITS-CLIP | Tarbase |
| hsa-miR-3613-3p | APPBP2 | HITS-CLIP | Tarbase |
| hsa-miR-15a-3p | SPHK2 | HITS-CLIP | Tarbase |
| hsa-miR-486-5p | SPHK2 | PAR-CLIP | Tarbase |
| hsa-miR-15a-5p | U2AF2 | HITS-CLIP | Tarbase |
| hsa-miR-195-5p | U2AF2 | HITS-CLIP | Tarbase |
| hsa-miR-141-3p | EPN1 | HITS-CLIP | Tarbase |
| hsa-miR-15a-5p | EPN1 | HITS-CLIP | Tarbase |
| hsa-miR-195-5p | EPN1 | HITS-CLIP | Tarbase |
| hsa-miR-3613-3p | AHRR | PAR-CLIP | Tarbase |
| hsa-miR-15a-5p | ZNF275 | HITS-CLIP | Tarbase |
| hsa-miR-3613-3p | ZNF275 | PAR-CLIP | Tarbase |
| hsa-miR-3613-5p | ZNF275 | HITS-CLIP | Tarbase |
| hsa-miR-486-5p | ZNF275 | HITS-CLIP | Tarbase |
| hsa-miR-15a-5p | ZNF275 | Chimeric fragments | Tarbase |
| hsa-miR-15a-5p | MTMR1 | HITS-CLIP | Tarbase |
| hsa-miR-195-5p | MTMR1 | HITS-CLIP | Tarbase |
| hsa-miR-195-5p | GPC1 | Microarrays | Tarbase |
| hsa-miR-141-3p | RNF4 | HITS-CLIP | Tarbase |
| hsa-miR-15a-5p | RNF4 | HITS-CLIP | Tarbase |
| hsa-miR-15a-3p | LIMCH1 | HITS-CLIP | Tarbase |
| hsa-miR-219a-1-3p | LIMCH1 | HITS-CLIP | Tarbase |
| hsa-miR-141-3p | ST3GAL6 | HITS-CLIP | Tarbase |
| hsa-miR-195-5p | CDON | Microarrays | Tarbase |
| hsa-miR-15a-5p | HIPK2 | HITS-CLIP | Tarbase |
| hsa-miR-15a-5p | HIPK2 | PAR-CLIP | Tarbase |
| hsa-miR-15a-5p | HIPK2 | HITS-CLIP | Tarbase |
| hsa-miR-15a-5p | HIPK2 | HITS-CLIP | Tarbase |
| hsa-miR-195-5p | HIPK2 | HITS-CLIP | Tarbase |
| hsa-miR-195-5p | HIPK2 | HITS-CLIP | Tarbase |
| hsa-miR-3605-5p | HIPK2 | HITS-CLIP | Tarbase |
| hsa-miR-15a-5p | HIPK2 | HITS-CLIP | Tarbase |
| hsa-miR-195-5p | HIPK2 | HITS-CLIP | Tarbase |
| hsa-miR-15a-5p | HIPK2 | Chimeric fragments | Tarbase |
| hsa-miR-15a-5p | TNPO3 | HITS-CLIP | Tarbase |
| hsa-miR-195-5p | TNPO3 | HITS-CLIP | Tarbase |
| hsa-miR-195-5p | TNPO3 | Microarrays | Tarbase |
| hsa-miR-15a-5p | TMEM161A | HITS-CLIP | Tarbase |
| hsa-miR-195-5p | TMEM161A | HITS-CLIP | Tarbase |
| hsa-miR-15a-5p | CTSA | HITS-CLIP | Tarbase |
| hsa-miR-15a-5p | CTSA | HITS-CLIP | Tarbase |
| hsa-miR-3613-3p | SUGP2 | HITS-CLIP | Tarbase |
| hsa-miR-3613-5p | SLC12A2 | HITS-CLIP | Tarbase |
| hsa-miR-15a-5p | SLC12A2 | HITS-CLIP | Tarbase |
| hsa-miR-195-5p | SLC12A2 | HITS-CLIP | Tarbase |
| hsa-miR-195-5p | CNN2 | Microarrays | Tarbase |
| hsa-miR-15a-3p | ABCA7 | HITS-CLIP | Tarbase |
| hsa-miR-141-5p | TAF11 | HITS-CLIP | Tarbase |
| hsa-miR-195-5p | TAF11 | Microarrays | Tarbase |
| hsa-miR-141-5p | AP3D1 | HITS-CLIP | Tarbase |
| hsa-miR-15a-5p | UHRF1BP1 | PAR-CLIP | Tarbase |
| hsa-miR-15a-5p | UHRF1BP1 | PAR-CLIP | Tarbase |
| hsa-miR-195-5p | UHRF1BP1 | PAR-CLIP | Tarbase |
| hsa-miR-15a-5p | GNAI3 | PAR-CLIP | Tarbase |
| hsa-miR-15a-5p | GNAI3 | HITS-CLIP | Tarbase |

| | | | |
|---|---|---|---|
| hsa-miR-195-5p | GNAI3 | HITS-CLIP | Tarbase |
| hsa-miR-219a-1-3p | GNAI3 | PAR-CLIP | Tarbase |
| hsa-miR-141-3p | IPO5 | HITS-CLIP | Tarbase |
| hsa-miR-141-5p | IPO5 | HITS-CLIP | Tarbase |
| hsa-miR-15a-5p | IPO5 | HITS-CLIP | Tarbase |
| hsa-miR-195-5p | OAT | Microarrays | Tarbase |
| hsa-miR-195-3p | PKN2 | HITS-CLIP | Tarbase |
| hsa-miR-3613-3p | TRAM2 | HITS-CLIP | Tarbase |
| hsa-miR-15a-5p | NTN1 | HITS-CLIP | Tarbase |
| hsa-miR-195-5p | NTN1 | HITS-CLIP | Tarbase |
| hsa-miR-195-5p | GLP2R | Microarrays | Tarbase |
| hsa-miR-15a-5p | KARS | HITS-CLIP | Tarbase |
| hsa-miR-15a-5p | KARS | HITS-CLIP | Tarbase |
| hsa-miR-15a-5p | KARS | HITS-CLIP | Tarbase |
| hsa-miR-195-5p | KARS | HITS-CLIP | Tarbase |
| hsa-miR-195-5p | KARS | HITS-CLIP | Tarbase |
| hsa-miR-3613-3p | PDIA5 | HITS-CLIP | Tarbase |
| hsa-miR-141-5p | SPEN | HITS-CLIP | Tarbase |
| hsa-miR-15a-5p | SPEN | HITS-CLIP | Tarbase |
| hsa-miR-15a-5p | SPEN | PAR-CLIP | Tarbase |
| hsa-miR-15a-5p | SPEN | HITS-CLIP | Tarbase |
| hsa-miR-195-5p | SPEN | HITS-CLIP | Tarbase |
| hsa-miR-195-5p | SPEN | HITS-CLIP | Tarbase |
| hsa-miR-486-5p | SPEN | HITS-CLIP | Tarbase |
| hsa-miR-195-5p | MYLK | Microarrays | Tarbase |
| hsa-miR-15a-5p | MAP2K4 | HITS-CLIP | Tarbase |
| hsa-miR-195-5p | MAP2K4 | HITS-CLIP | Tarbase |
| hsa-miR-15a-5p | TMEM206 | PAR-CLIP | Tarbase |
| hsa-miR-195-5p | TMEM206 | PAR-CLIP | Tarbase |
| hsa-miR-3613-3p | SLK | HITS-CLIP | Tarbase |
| hsa-miR-15a-5p | SLK | HITS-CLIP | Tarbase |
| hsa-miR-195-5p | SLK | HITS-CLIP | Tarbase |
| hsa-miR-15a-5p | SEC61A2 | HITS-CLIP | Tarbase |
| hsa-miR-195-5p | SEC61A2 | HITS-CLIP | Tarbase |
| hsa-miR-141-5p | PRKCQ | HITS-CLIP | Tarbase |
| hsa-miR-141-5p | ASB1 | HITS-CLIP | Tarbase |
| hsa-miR-141-5p | ASB1 | HITS-CLIP | Tarbase |
| hsa-miR-195-5p | ASB1 | HITS-CLIP | Tarbase |
| hsa-miR-195-5p | FAM107B | Microarrays | Tarbase |
| hsa-miR-195-5p | ME1 | Microarrays | Tarbase |
| hsa-miR-141-3p | CDK13 | HITS-CLIP | Tarbase |
| hsa-miR-15a-5p | MTHFD2 | Chimeric fragments | Tarbase |
| hsa-miR-141-3p | DIP2B | HITS-CLIP | Tarbase |
| hsa-miR-141-3p | ASPM | HITS-CLIP | Tarbase |
| hsa-miR-15a-5p | ASPM | HITS-CLIP | Tarbase |
| hsa-miR-15a-5p | ASPM | HITS-CLIP | Tarbase |
| hsa-miR-195-5p | ASPM | HITS-CLIP | Tarbase |
| hsa-miR-195-5p | ASPM | Microarrays | Tarbase |
| hsa-miR-195-5p | MPPED2 | Microarrays | Tarbase |
| hsa-miR-141-5p | ZBTB11 | HITS-CLIP | Tarbase |
| hsa-miR-15a-5p | ZBTB11 | PAR-CLIP | Tarbase |
| hsa-miR-141-5p | FGFR2 | HITS-CLIP | Tarbase |
| hsa-miR-15a-5p | ISOC1 | PAR-CLIP | Tarbase |
| hsa-miR-15a-5p | ISOC1 | HITS-CLIP | Tarbase |
| hsa-miR-141-3p | THUMPD1 | HITS-CLIP | Tarbase |
| hsa-miR-15a-5p | THUMPD1 | PAR-CLIP | Tarbase |
| hsa-miR-15a-5p | THUMPD1 | HITS-CLIP | Tarbase |
| hsa-miR-15a-5p | THUMPD1 | HITS-CLIP | Tarbase |
| hsa-miR-15a-3p | KIF26A | HITS-CLIP | Tarbase |
| hsa-miR-141-3p | ATG2B | HITS-CLIP | Tarbase |
| hsa-miR-141-3p | ATG2B | HITS-CLIP | Tarbase |
| hsa-miR-15a-3p | ATG2B | HITS-CLIP | Tarbase |
| hsa-miR-15a-5p | ATG2B | HITS-CLIP | Tarbase |
| hsa-miR-15a-5p | ARFGEF1 | Chimeric fragments | Tarbase |
| hsa-miR-15a-5p | FECH | HITS-CLIP | Tarbase |
| hsa-miR-195-5p | FECH | HITS-CLIP | Tarbase |
| hsa-miR-3613-3p | PFKP | HITS-CLIP | Tarbase |
| hsa-miR-195-5p | PFKP | Microarrays | Tarbase |
| hsa-miR-15a-3p | SP100 | HITS-CLIP | Tarbase |
| hsa-miR-141-3p | KLF6 | HITS-CLIP | Tarbase |
| hsa-miR-141-3p | KLF6 | HITS-CLIP | Tarbase |
| hsa-miR-195-5p | KLF6 | Microarrays | Tarbase |
| hsa-miR-195-5p | PPAP2A | Microarrays | Tarbase |
| hsa-miR-141-3p | TRAM1 | HITS-CLIP | Tarbase |
| hsa-miR-15a-5p | TRAM1 | HITS-CLIP | Tarbase |
| hsa-miR-15a-5p | TRAM1 | HITS-CLIP | Tarbase |
| hsa-miR-15a-5p | TRAM1 | HITS-CLIP | Tarbase |
| hsa-miR-195-5p | TRAM1 | HITS-CLIP | Tarbase |
| hsa-miR-195-5p | TRAM1 | HITS-CLIP | Tarbase |
| hsa-miR-3605-5p | TRAM1 | HITS-CLIP | Tarbase |
| hsa-miR-3613-3p | TRAM1 | PAR-CLIP | Tarbase |
| hsa-miR-15a-5p | PHKA1 | PAR-CLIP | Tarbase |
| hsa-miR-15a-5p | PHKA1 | PAR-CLIP | Tarbase |
| hsa-miR-15a-5p | PHKA1 | PAR-CLIP | Tarbase |
| hsa-miR-195-5p | PHKA1 | PAR-CLIP | Tarbase |
| hsa-miR-141-5p | PKM | HITS-CLIP | Tarbase |
| hsa-miR-141-5p | PKM | HITS-CLIP | Tarbase |
| hsa-miR-15a-5p | PKM | HITS-CLIP | Tarbase |
| hsa-miR-15a-5p | PKM | HITS-CLIP | Tarbase |
| hsa-miR-15a-5p | PKM | HITS-CLIP | Tarbase |
| hsa-miR-15a-5p | PKM | HITS-CLIP | Tarbase |
| hsa-miR-195-5p | PKM | HITS-CLIP | Tarbase |
| hsa-miR-195-5p | PKM | HITS-CLIP | Tarbase |
| hsa-miR-195-5p | PKM | HITS-CLIP | Tarbase |
| hsa-miR-195-5p | PKM | HITS-CLIP | Tarbase |
| hsa-miR-3605-5p | PKM | HITS-CLIP | Tarbase |
| hsa-miR-141-5p | RHOA | HITS-CLIP | Tarbase |
| hsa-miR-141-5p | RHOA | HITS-CLIP | Tarbase |
| hsa-miR-3613-5p | DHX8 | HITS-CLIP | Tarbase |
| hsa-miR-3613-3p | ZFY | HITS-CLIP | Tarbase |
| hsa-miR-141-5p | IARS2 | HITS-CLIP | Tarbase |
| hsa-miR-15a-5p | ROGDI | HITS-CLIP | Tarbase |
| hsa-miR-195-5p | ROGDI | HITS-CLIP | Tarbase |
| hsa-miR-15a-5p | ROGDI | HITS-CLIP | Tarbase |
| hsa-miR-195-5p | ROGDI | HITS-CLIP | Tarbase |
| hsa-miR-15a-5p | CBFB | HITS-CLIP | Tarbase |
| hsa-miR-141-3p | FGFR3 | HITS-CLIP | Tarbase |
| hsa-miR-15a-5p | HEATR6 | HITS-CLIP | Tarbase |
| hsa-miR-195-5p | HEATR6 | HITS-CLIP | Tarbase |
| hsa-miR-15a-5p | COASY | PAR-CLIP | Tarbase |
| hsa-miR-15a-5p | COASY | HITS-CLIP | Tarbase |
| hsa-miR-195-5p | COASY | HITS-CLIP | Tarbase |
| hsa-miR-15a-5p | COASY | Chimeric fragments | Tarbase |
| hsa-miR-141-3p | MEF2A | HITS-CLIP | Tarbase |
| hsa-miR-3613-5p | MEF2A | PAR-CLIP | Tarbase |
| hsa-miR-15a-5p | TBC1D25 | HITS-CLIP | Tarbase |
| hsa-miR-195-5p | TBC1D25 | HITS-CLIP | Tarbase |
| hsa-miR-15a-5p | ACSL4 | PAR-CLIP | Tarbase |
| hsa-miR-195-5p | ACSL4 | PAR-CLIP | Tarbase |
| hsa-miR-15a-5p | ACSL4 | HITS-CLIP | Tarbase |
| hsa-miR-195-5p | ACSL4 | HITS-CLIP | Tarbase |
| hsa-miR-195-5p | FTSJ1 | Microarrays | Tarbase |
| hsa-miR-195-5p | PRR11 | Microarrays | Tarbase |
| hsa-miR-15a-5p | POLR1A | PAR-CLIP | Tarbase |
| hsa-miR-141-5p | LAPTM4A | HITS-CLIP | Tarbase |
| hsa-miR-15a-5p | LAPTM4A | HITS-CLIP | Tarbase |
| hsa-miR-195-5p | LAPTM4A | HITS-CLIP | Tarbase |
| hsa-miR-3613-3p | LAPTM4A | HITS-CLIP | Tarbase |
| hsa-miR-3613-3p | LAPTM4A | PAR-CLIP | Tarbase |
| hsa-miR-15a-5p | LAPTM4A | Chimeric fragments | Tarbase |
| hsa-miR-141-3p | SRBD1 | HITS-CLIP | Tarbase |
| hsa-miR-15a-5p | KIF2A | HITS-CLIP | Tarbase |
| hsa-miR-195-5p | KIF2A | HITS-CLIP | Tarbase |
| hsa-miR-3613-3p | KIF2A | HITS-CLIP | Tarbase |
| hsa-miR-195-5p | PPP2R5B | Microarrays | Tarbase |
| hsa-miR-141-3p | MAST4 | HITS-CLIP | Tarbase |
| hsa-miR-141-3p | MAST4 | HITS-CLIP | Tarbase |
| hsa-miR-15a-5p | MAST4 | HITS-CLIP | Tarbase |
| hsa-miR-15a-5p | MAST4 | HITS-CLIP | Tarbase |
| hsa-miR-195-5p | MAST4 | HITS-CLIP | Tarbase |
| hsa-miR-3613-3p | MAST4 | HITS-CLIP | Tarbase |
| hsa-miR-15a-5p | GPR116 | HITS-CLIP | Tarbase |
| hsa-miR-195-5p | GPR116 | HITS-CLIP | Tarbase |
| hsa-miR-141-5p | NUCKS1 | HITS-CLIP | Tarbase |
| hsa-miR-141-5p | NUCKS1 | HITS-CLIP | Tarbase |
| hsa-miR-141-5p | NUCKS1 | HITS-CLIP | Tarbase |
| hsa-miR-15a-5p | NUCKS1 | HITS-CLIP | Tarbase |
| hsa-miR-15a-5p | NUCKS1 | PAR-CLIP | Tarbase |
| hsa-miR-15a-5p | NUCKS1 | HITS-CLIP | Tarbase |

| | | | |
|---|---|---|---|
| hsa-miR-15a-5p | NUCKS1 | HITS-CLIP | Tarbase |
| hsa-miR-15a-5p | NUCKS1 | HITS-CLIP | Tarbase |
| hsa-miR-15a-5p | NUCKS1 | HITS-CLIP | Tarbase |
| hsa-miR-15a-5p | NUCKS1 | HITS-CLIP | Tarbase |
| hsa-miR-15a-5p | NUCKS1 | HITS-CLIP | Tarbase |
| hsa-miR-15a-5p | NUCKS1 | HITS-CLIP | Tarbase |
| hsa-miR-195-5p | NUCKS1 | HITS-CLIP | Tarbase |
| hsa-miR-195-5p | NUCKS1 | PAR-CLIP | Tarbase |
| hsa-miR-195-5p | NUCKS1 | HITS-CLIP | Tarbase |
| hsa-miR-195-5p | NUCKS1 | HITS-CLIP | Tarbase |
| hsa-miR-195-5p | NUCKS1 | HITS-CLIP | Tarbase |
| hsa-miR-195-5p | NUCKS1 | HITS-CLIP | Tarbase |
| hsa-miR-195-5p | NUCKS1 | HITS-CLIP | Tarbase |
| hsa-miR-3613-5p | NUCKS1 | HITS-CLIP | Tarbase |
| hsa-miR-486-5p | NUCKS1 | PAR-CLIP | Tarbase |
| hsa-miR-15a-5p | NUCKS1 | HITS-CLIP | Tarbase |
| hsa-miR-195-5p | NUCKS1 | HITS-CLIP | Tarbase |
| hsa-miR-141-3p | VPS35 | HITS-CLIP | Tarbase |
| hsa-miR-3613-5p | VPS35 | PAR-CLIP | Tarbase |
| hsa-miR-3613-5p | VPS35 | PAR-CLIP | Tarbase |
| hsa-miR-15a-5p | DNAJA2 | PAR-CLIP | Tarbase |
| hsa-miR-486-5p | KCNAB2 | PAR-CLIP | Tarbase |
| hsa-miR-15a-5p | RORA | HITS-CLIP | Tarbase |
| hsa-miR-195-5p | RORA | HITS-CLIP | Tarbase |
| hsa-miR-195-5p | RORA | Microarrays | Tarbase |
| hsa-miR-141-5p | TGFBR3 | HITS-CLIP | Tarbase |
| hsa-miR-219a-1-3p | TGFBR3 | HITS-CLIP | Tarbase |
| hsa-miR-141-5p | KIAA1107 | HITS-CLIP | Tarbase |
| hsa-miR-141-3p | ATP1B3 | HITS-CLIP | Tarbase |
| hsa-miR-141-3p | ATP1B3 | HITS-CLIP | Tarbase |
| hsa-miR-141-3p | NEDD4 | HITS-CLIP | Tarbase |
| hsa-miR-15a-5p | PIGB | PAR-CLIP | Tarbase |
| hsa-miR-15a-5p | PIGB | PAR-CLIP | Tarbase |
| hsa-miR-15a-5p | PIGB | PAR-CLIP | Tarbase |
| hsa-miR-15a-5p | PIGB | PAR-CLIP | Tarbase |
| hsa-miR-195-5p | PIGB | PAR-CLIP | Tarbase |
| hsa-miR-195-5p | PIGB | PAR-CLIP | Tarbase |
| hsa-miR-141-3p | MAPK6 | HITS-CLIP | Tarbase |
| hsa-miR-15a-5p | MAPK6 | PAR-CLIP | Tarbase |
| hsa-miR-15a-5p | MAPK6 | HITS-CLIP | Tarbase |
| hsa-miR-3613-5p | RAB27A | PAR-CLIP | Tarbase |
| hsa-miR-3613-5p | RAB27A | PAR-CLIP | Tarbase |
| hsa-miR-141-3p | IKBKAP | HITS-CLIP | Tarbase |
| hsa-miR-15a-5p | PFN2 | HITS-CLIP | Tarbase |
| hsa-miR-15a-5p | PFN2 | HITS-CLIP | Tarbase |
| hsa-miR-195-5p | PFN2 | HITS-CLIP | Tarbase |
| hsa-miR-195-5p | PFN2 | HITS-CLIP | Tarbase |
| hsa-miR-15a-5p | PTPN3 | HITS-CLIP | Tarbase |
| hsa-miR-15a-5p | PTPN3 | PAR-CLIP | Tarbase |
| hsa-miR-15a-5p | PTPN3 | HITS-CLIP | Tarbase |
| hsa-miR-195-5p | PTPN3 | HITS-CLIP | Tarbase |
| hsa-miR-15a-5p | SLC44A1 | HITS-CLIP | Tarbase |
| hsa-miR-15a-5p | SLC44A1 | HITS-CLIP | Tarbase |
| hsa-miR-195-5p | SLC44A1 | HITS-CLIP | Tarbase |
| hsa-miR-141-3p | EXOC5 | HITS-CLIP | Tarbase |
| hsa-miR-195-5p | FSTL3 | Microarrays | Tarbase |
| hsa-miR-141-5p | DGCR2 | HITS-CLIP | Tarbase |
| hsa-miR-141-5p | DGCR2 | HITS-CLIP | Tarbase |
| hsa-miR-15a-3p | DGCR2 | HITS-CLIP | Tarbase |
| hsa-miR-15a-5p | MNT | HITS-CLIP | Tarbase |
| hsa-miR-15a-5p | MNT | HITS-CLIP | Tarbase |
| hsa-miR-195-5p | MNT | HITS-CLIP | Tarbase |
| hsa-miR-195-5p | MNT | HITS-CLIP | Tarbase |
| hsa-miR-141-3p | ZXDC | HITS-CLIP | Tarbase |
| hsa-miR-195-5p | JMJD6 | Microarrays | Tarbase |
| hsa-miR-195-5p | WIPI1 | Microarrays | Tarbase |
| hsa-miR-141-3p | NDST1 | HITS-CLIP | Tarbase |
| hsa-miR-15a-5p | NDST1 | HITS-CLIP | Tarbase |
| hsa-miR-195-5p | NDST1 | HITS-CLIP | Tarbase |
| hsa-miR-219a-1-3p | AP3M2 | HITS-CLIP | Tarbase |
| hsa-miR-195-5p | ST6GALNAC2 | HITS-CLIP | Tarbase |
| hsa-miR-15a-3p | PABPC1 | HITS-CLIP | Tarbase |
| hsa-miR-15a-5p | CFAP20 | HITS-CLIP | Tarbase |
| hsa-miR-141-3p | CDC42 | PAR-CLIP | Tarbase |
| hsa-miR-15a-5p | CDC42 | PAR-CLIP | Tarbase |
| hsa-miR-15a-5p | CDC42 | HITS-CLIP | Tarbase |
| hsa-miR-15a-5p | CDC42 | HITS-CLIP | Tarbase |
| hsa-miR-15a-5p | CDC42 | HITS-CLIP | Tarbase |
| hsa-miR-15a-5p | CDC42 | HITS-CLIP | Tarbase |
| hsa-miR-15a-5p | CDC42 | HITS-CLIP | Tarbase |
| hsa-miR-195-5p | CDC42 | HITS-CLIP | Tarbase |
| hsa-miR-195-5p | CDC42 | HITS-CLIP | Tarbase |
| hsa-miR-195-5p | CDC42 | HITS-CLIP | Tarbase |
| hsa-miR-15a-3p | RAD18 | HITS-CLIP | Tarbase |
| hsa-miR-141-5p | ATP2B1 | HITS-CLIP | Tarbase |
| hsa-miR-141-5p | ATP2B1 | HITS-CLIP | Tarbase |
| hsa-miR-141-5p | ATP2B1 | HITS-CLIP | Tarbase |
| hsa-miR-141-5p | ATP2B1 | HITS-CLIP | Tarbase |
| hsa-miR-15a-5p | ATP2B1 | HITS-CLIP | Tarbase |
| hsa-miR-15a-5p | ATP2B1 | HITS-CLIP | Tarbase |
| hsa-miR-15a-5p | ATP2B1 | PAR-CLIP | Tarbase |
| hsa-miR-195-5p | ATP2B1 | HITS-CLIP | Tarbase |
| hsa-miR-195-5p | ATP2B1 | HITS-CLIP | Tarbase |
| hsa-miR-15a-5p | ATP2B1 | HITS-CLIP | Tarbase |
| hsa-miR-195-5p | ATP2B1 | HITS-CLIP | Tarbase |
| hsa-miR-141-3p | MAP4K4 | Luciferase Reporter Assay | Tarbase |
| hsa-miR-141-3p | MAP4K4 | qPCR | Tarbase |
| hsa-miR-141-3p | MAP4K4 | qPCR | Tarbase |
| hsa-miR-141-3p | MAP4K4 | qPCR | Tarbase |
| hsa-miR-141-3p | MAP4K4 | Western Blot | Tarbase |
| hsa-miR-141-3p | MAP4K4 | Immunohistochemistry | Tarbase |
| hsa-miR-141-3p | MAP4K4 | qPCR | Tarbase |
| hsa-miR-141-3p | MAP4K4 | Western Blot | Tarbase |
| hsa-miR-141-3p | MAP4K4 | qPCR | Tarbase |
| hsa-miR-141-5p | MAP4K4 | HITS-CLIP | Tarbase |
| hsa-miR-141-3p | MGAT4A | HITS-CLIP | Tarbase |
| hsa-miR-15a-5p | MGAT4A | PAR-CLIP | Tarbase |
| hsa-miR-15a-5p | MGAT4A | PAR-CLIP | Tarbase |
| hsa-miR-15a-5p | MGAT4A | HITS-CLIP | Tarbase |
| hsa-miR-195-5p | MGAT4A | HITS-CLIP | Tarbase |
| hsa-miR-15a-3p | WDR1 | HITS-CLIP | Tarbase |
| hsa-miR-195-5p | WDR1 | Microarrays | Tarbase |
| hsa-miR-141-5p | ING3 | HITS-CLIP | Tarbase |
| hsa-miR-195-5p | TRIP13 | Microarrays | Tarbase |
| hsa-miR-486-5p | ATP6AP1 | HITS-CLIP | Tarbase |
| hsa-miR-15a-5p | TCF3 | HITS-CLIP | Tarbase |
| hsa-miR-195-5p | TCF3 | HITS-CLIP | Tarbase |
| hsa-miR-195-5p | TRIB2 | Microarrays | Tarbase |
| hsa-miR-15a-3p | MBD3 | HITS-CLIP | Tarbase |
| hsa-miR-15a-3p | MBD3 | HITS-CLIP | Tarbase |
| hsa-miR-15a-5p | MBD3 | HITS-CLIP | Tarbase |
| hsa-miR-195-5p | MBD3 | HITS-CLIP | Tarbase |
| hsa-miR-15a-3p | CYBRD1 | HITS-CLIP | Tarbase |
| hsa-miR-195-5p | CYBRD1 | Microarrays | Tarbase |
| hsa-miR-141-5p | PDCD2 | HITS-CLIP | Tarbase |
| hsa-miR-195-5p | SLC6A15 | Microarrays | Tarbase |
| hsa-miR-15a-5p | RDH11 | PAR-CLIP | Tarbase |
| hsa-miR-15a-5p | RDH11 | PAR-CLIP | Tarbase |
| hsa-miR-195-5p | RDH11 | PAR-CLIP | Tarbase |
| hsa-miR-195-5p | RDH11 | Microarrays | Tarbase |
| hsa-miR-15a-5p | PRKACA | HITS-CLIP | Tarbase |
| hsa-miR-195-5p | PRKACA | HITS-CLIP | Tarbase |
| hsa-miR-195-5p | PRKACA | Microarrays | Tarbase |
| hsa-miR-15a-5p | LPHN1 | PAR-CLIP | Tarbase |
| hsa-miR-195-5p | ACTN1 | Microarrays | Tarbase |
| hsa-miR-141-5p | ZFYVE26 | HITS-CLIP | Tarbase |
| hsa-miR-15a-5p | ALDH3A2 | HITS-CLIP | Tarbase |
| hsa-miR-195-5p | ALDH3A2 | Microarrays | Tarbase |
| hsa-miR-141-3p | TFRC | HITS-CLIP | Tarbase |
| hsa-miR-15a-5p | AFF4 | PAR-CLIP | Tarbase |
| hsa-miR-15a-5p | AFF4 | HITS-CLIP | Tarbase |
| hsa-miR-15a-5p | AFF4 | HITS-CLIP | Tarbase |
| hsa-miR-195-5p | AFF4 | HITS-CLIP | Tarbase |
| hsa-miR-15a-5p | UBE2D1 | HITS-CLIP | Tarbase |
| hsa-miR-195-5p | UBE2D1 | HITS-CLIP | Tarbase |
| hsa-miR-195-5p | RHOBTB1 | Microarrays | Tarbase |
| hsa-miR-3613-5p | SMC1A | PAR-CLIP | Tarbase |
| hsa-miR-195-5p | SMC1A | Microarrays | Tarbase |
| hsa-miR-195-5p | HMMR | Microarrays | Tarbase |

| | | | |
|---|---|---|---|
| hsa-miR-195-5p | TRHDE | Microarrays | Tarbase |
| hsa-miR-15a-5p | NFATC3 | HITS-CLIP | Tarbase |
| hsa-miR-15a-5p | NFATC3 | HITS-CLIP | Tarbase |
| hsa-miR-195-5p | NFATC3 | HITS-CLIP | Tarbase |
| hsa-miR-141-3p | TRNT1 | HITS-CLIP | Tarbase |
| hsa-miR-141-3p | FBXW11 | PAR-CLIP | Tarbase |
| hsa-miR-141-3p | FBXW11 | HITS-CLIP | Tarbase |
| hsa-miR-141-3p | FBXW11 | PAR-CLIP | Tarbase |
| hsa-miR-195-5p | MRVI1 | Microarrays | Tarbase |
| hsa-miR-195-5p | PVR | Microarrays | Tarbase |
| hsa-miR-195-5p | MCM2 | Microarrays | Tarbase |
| hsa-miR-195-5p | MOV10L1 | Microarrays | Tarbase |
| hsa-miR-3613-5p | SELO | HITS-CLIP | Tarbase |
| hsa-miR-15a-5p | CLCN4 | HITS-CLIP | Tarbase |
| hsa-miR-195-5p | CLCN4 | HITS-CLIP | Tarbase |
| hsa-miR-195-5p | CLCN4 | Microarrays | Tarbase |
| hsa-miR-15a-5p | SMARCE1 | HITS-CLIP | Tarbase |
| hsa-miR-3613-5p | SMARCE1 | HITS-CLIP | Tarbase |
| hsa-miR-141-3p | KDM5A | HITS-CLIP | Tarbase |
| hsa-miR-141-3p | PTGS2 | HITS-CLIP | Tarbase |
| hsa-miR-195-5p | PTGS2 | Microarrays | Tarbase |
| hsa-miR-195-5p | FRY | Microarrays | Tarbase |
| hsa-miR-141-3p | PICALM | HITS-CLIP | Tarbase |
| hsa-miR-141-3p | PICALM | HITS-CLIP | Tarbase |
| hsa-miR-15a-5p | NSF | PAR-CLIP | Tarbase |
| hsa-miR-15a-5p | NSF | HITS-CLIP | Tarbase |
| hsa-miR-15a-5p | NSF | HITS-CLIP | Tarbase |
| hsa-miR-195-5p | NSF | HITS-CLIP | Tarbase |
| hsa-miR-15a-5p | GLI2 | HITS-CLIP | Tarbase |
| hsa-miR-195-5p | GLI2 | HITS-CLIP | Tarbase |
| hsa-miR-15a-5p | CLASP1 | HITS-CLIP | Tarbase |
| hsa-miR-15a-5p | CLASP1 | PAR-CLIP | Tarbase |
| hsa-miR-195-5p | CLASP1 | HITS-CLIP | Tarbase |
| hsa-miR-195-5p | CLASP1 | HITS-CLIP | Tarbase |
| hsa-miR-3605-3p | CLASP1 | HITS-CLIP | Tarbase |
| hsa-miR-15a-5p | NOTCH3 | HITS-CLIP | Tarbase |
| hsa-miR-195-5p | NOTCH3 | HITS-CLIP | Tarbase |
| hsa-miR-15a-5p | CLNS1A | HITS-CLIP | Tarbase |
| hsa-miR-15a-5p | CLNS1A | HITS-CLIP | Tarbase |
| hsa-miR-15a-5p | CLNS1A | HITS-CLIP | Tarbase |
| hsa-miR-15a-5p | CLNS1A | PAR-CLIP | Tarbase |
| hsa-miR-195-5p | CLNS1A | HITS-CLIP | Tarbase |
| hsa-miR-195-5p | CLNS1A | HITS-CLIP | Tarbase |
| hsa-miR-195-5p | CLNS1A | HITS-CLIP | Tarbase |
| hsa-miR-3613-3p | EED | HITS-CLIP | Tarbase |
| hsa-miR-3613-5p | CA12 | PAR-CLIP | Tarbase |
| hsa-miR-195-5p | MGLL | Microarrays | Tarbase |
| hsa-miR-15a-3p | NTN4 | HITS-CLIP | Tarbase |
| hsa-miR-15a-3p | BCS1L | HITS-CLIP | Tarbase |
| hsa-miR-3605-5p | NUAK1 | HITS-CLIP | Tarbase |
| hsa-miR-15a-5p | ZNF532 | HITS-CLIP | Tarbase |
| hsa-miR-195-5p | ZNF532 | HITS-CLIP | Tarbase |
| hsa-miR-15a-5p | LMAN1 | PAR-CLIP | Tarbase |
| hsa-miR-15a-5p | LMAN1 | PAR-CLIP | Tarbase |
| hsa-miR-15a-5p | LMAN1 | PAR-CLIP | Tarbase |
| hsa-miR-15a-5p | LMAN1 | HITS-CLIP | Tarbase |
| hsa-miR-15a-5p | LMAN1 | PAR-CLIP | Tarbase |
| hsa-miR-195-5p | LMAN1 | PAR-CLIP | Tarbase |
| hsa-miR-195-5p | LMAN1 | HITS-CLIP | Tarbase |
| hsa-miR-195-5p | LMAN1 | PAR-CLIP | Tarbase |
| hsa-miR-15a-5p | IPCEF1 | HITS-CLIP | Tarbase |
| hsa-miR-195-5p | IPCEF1 | HITS-CLIP | Tarbase |
| hsa-miR-141-5p | EIF4G3 | HITS-CLIP | Tarbase |
| hsa-miR-15a-5p | EIF4G3 | HITS-CLIP | Tarbase |
| hsa-miR-195-5p | EIF4G3 | HITS-CLIP | Tarbase |
| hsa-miR-15a-5p | SEMA3A | PAR-CLIP | Tarbase |
| hsa-miR-195-5p | GTSE1 | Microarrays | Tarbase |
| hsa-miR-15a-5p | SEMA3C | PAR-CLIP | Tarbase |
| hsa-miR-15a-3p | SLC25A40 | HITS-CLIP | Tarbase |
| hsa-miR-195-5p | SLC25A40 | Microarrays | Tarbase |
| hsa-miR-195-5p | ADD2 | Microarrays | Tarbase |
| hsa-miR-141-5p | MARK3 | HITS-CLIP | Tarbase |
| hsa-miR-141-3p | SLC25A3 | HITS-CLIP | Tarbase |
| hsa-miR-141-3p | SLC25A3 | HITS-CLIP | Tarbase |
| hsa-miR-141-3p | SLC25A3 | HITS-CLIP | Tarbase |
| hsa-miR-141-3p | SLC25A3 | HITS-CLIP | Tarbase |
| hsa-miR-141-3p | SLC25A3 | HITS-CLIP | Tarbase |
| hsa-miR-141-5p | SLC25A3 | HITS-CLIP | Tarbase |
| hsa-miR-15a-5p | SLC25A3 | HITS-CLIP | Tarbase |
| hsa-miR-15a-5p | SLC25A3 | PAR-CLIP | Tarbase |
| hsa-miR-195-5p | SLC25A3 | HITS-CLIP | Tarbase |
| hsa-miR-195-5p | SLC25A3 | PAR-CLIP | Tarbase |
| hsa-miR-15a-3p | FNDC3B | HITS-CLIP | Tarbase |
| hsa-miR-15a-5p | FNDC3B | PAR-CLIP | Tarbase |
| hsa-miR-195-5p | FNDC3B | PAR-CLIP | Tarbase |
| hsa-miR-141-3p | FOSL2 | HITS-CLIP | Tarbase |
| hsa-miR-15a-5p | FOSL2 | HITS-CLIP | Tarbase |
| hsa-miR-15a-5p | FOSL2 | HITS-CLIP | Tarbase |
| hsa-miR-15a-5p | FOSL2 | PAR-CLIP | Tarbase |
| hsa-miR-15a-5p | FOSL2 | HITS-CLIP | Tarbase |
| hsa-miR-15a-5p | FOSL2 | HITS-CLIP | Tarbase |
| hsa-miR-15a-5p | FOSL2 | HITS-CLIP | Tarbase |
| hsa-miR-195-5p | FOSL2 | HITS-CLIP | Tarbase |
| hsa-miR-195-5p | FOSL2 | HITS-CLIP | Tarbase |
| hsa-miR-195-5p | FOSL2 | HITS-CLIP | Tarbase |
| hsa-miR-195-5p | CACNG4 | Microarrays | Tarbase |
| hsa-miR-15a-5p | FRYL | PAR-CLIP | Tarbase |
| hsa-miR-15a-5p | FRYL | HITS-CLIP | Tarbase |
| hsa-miR-195-5p | FRYL | PAR-CLIP | Tarbase |
| hsa-miR-3613-5p | FRYL | PAR-CLIP | Tarbase |
| hsa-miR-141-3p | TMEM131 | PAR-CLIP | Tarbase |
| hsa-miR-15a-5p | ACTB | HITS-CLIP | Tarbase |
| hsa-miR-15a-5p | ACTB | HITS-CLIP | Tarbase |
| hsa-miR-15a-5p | ACTB | HITS-CLIP | Tarbase |
| hsa-miR-15a-5p | ACTB | PAR-CLIP | Tarbase |
| hsa-miR-15a-5p | ACTB | PAR-CLIP | Tarbase |
| hsa-miR-15a-5p | ACTB | Western Blot | Tarbase |
| hsa-miR-195-5p | ACTB | HITS-CLIP | Tarbase |
| hsa-miR-195-5p | ACTB | HITS-CLIP | Tarbase |
| hsa-miR-195-5p | ACTB | HITS-CLIP | Tarbase |
| hsa-miR-195-5p | ACTB | Western Blot | Tarbase |
| hsa-miR-195-5p | ACTB | Western Blot | Tarbase |
| hsa-miR-195-5p | ACTB | Western Blot | Tarbase |
| hsa-miR-486-5p | ACTB | HITS-CLIP | Tarbase |
| hsa-miR-486-5p | ACTB | PAR-CLIP | Tarbase |
| hsa-miR-195-5p | WDR62 | Microarrays | Tarbase |
| hsa-miR-15a-5p | BCAP29 | HITS-CLIP | Tarbase |
| hsa-miR-195-5p | BCAP29 | HITS-CLIP | Tarbase |
| hsa-miR-15a-3p | SEC31B | HITS-CLIP | Tarbase |
| hsa-miR-15a-5p | KIFAP3 | PAR-CLIP | Tarbase |
| hsa-miR-15a-5p | KIFAP3 | PAR-CLIP | Tarbase |
| hsa-miR-15a-5p | KIFAP3 | PAR-CLIP | Tarbase |
| hsa-miR-15a-5p | KIFAP3 | PAR-CLIP | Tarbase |
| hsa-miR-195-5p | KIFAP3 | PAR-CLIP | Tarbase |
| hsa-miR-195-5p | KIFAP3 | PAR-CLIP | Tarbase |
| hsa-miR-195-5p | KIFAP3 | PAR-CLIP | Tarbase |
| hsa-miR-195-5p | MCM6 | Microarrays | Tarbase |
| hsa-miR-15a-5p | RBM7 | HITS-CLIP | Tarbase |
| hsa-miR-15a-5p | RBM7 | HITS-CLIP | Tarbase |
| hsa-miR-195-5p | RBM7 | HITS-CLIP | Tarbase |
| hsa-miR-141-3p | RBMS2 | HITS-CLIP | Tarbase |
| hsa-miR-141-5p | BAZ2A | HITS-CLIP | Tarbase |
| hsa-miR-141-5p | BAZ2A | HITS-CLIP | Tarbase |
| hsa-miR-15a-5p | BAZ2A | HITS-CLIP | Tarbase |
| hsa-miR-15a-5p | BAZ2A | HITS-CLIP | Tarbase |
| hsa-miR-195-5p | BAZ2A | HITS-CLIP | Tarbase |
| hsa-miR-141-3p | UNG | PAR-CLIP | Tarbase |
| hsa-miR-15a-5p | UNG | HITS-CLIP | Tarbase |
| hsa-miR-15a-5p | UNG | HITS-CLIP | Tarbase |
| hsa-miR-195-5p | UNG | HITS-CLIP | Tarbase |
| hsa-miR-195-5p | UNG | HITS-CLIP | Tarbase |
| hsa-miR-195-5p | UNG | Microarrays | Tarbase |
| hsa-miR-195-5p | FMO4 | Microarrays | Tarbase |
| hsa-miR-141-3p | KLHL20 | Luciferase Reporter Assay | Tarbase |
| hsa-miR-141-3p | KLHL20 | Luciferase Reporter Assay | Tarbase |
| hsa-miR-486-5p | KLHL20 | HITS-CLIP | Tarbase |
| hsa-miR-15a-5p | PLXNA2 | PAR-CLIP | Tarbase |
| hsa-miR-195-5p | PLXNA2 | Microarrays | Tarbase |
| hsa-miR-195-5p | SPAG5 | Microarrays | Tarbase |

| | | | |
|---|---|---|---|
| hsa-miR-15a-3p | TPD52 | HITS-CLIP | Tarbase |
| hsa-miR-15a-5p | TRAF4 | HITS-CLIP | Tarbase |
| hsa-miR-15a-5p | TRAF4 | HITS-CLIP | Tarbase |
| hsa-miR-195-5p | TRAF4 | HITS-CLIP | Tarbase |
| hsa-miR-15a-5p | PAG1 | PAR-CLIP | Tarbase |
| hsa-miR-15a-5p | PAG1 | PAR-CLIP | Tarbase |
| hsa-miR-15a-5p | PAG1 | PAR-CLIP | Tarbase |
| hsa-miR-15a-5p | PAG1 | PAR-CLIP | Tarbase |
| hsa-miR-15a-5p | PAG1 | PAR-CLIP | Tarbase |
| hsa-miR-15a-5p | PAG1 | PAR-CLIP | Tarbase |
| hsa-miR-15a-5p | PAG1 | PAR-CLIP | Tarbase |
| hsa-miR-15a-5p | PAG1 | PAR-CLIP | Tarbase |
| hsa-miR-195-5p | PAG1 | PAR-CLIP | Tarbase |
| hsa-miR-195-5p | PAG1 | PAR-CLIP | Tarbase |
| hsa-miR-195-5p | PAG1 | PAR-CLIP | Tarbase |
| hsa-miR-15a-5p | GPC4 | HITS-CLIP | Tarbase |
| hsa-miR-195-5p | GPC4 | HITS-CLIP | Tarbase |
| hsa-miR-195-5p | GPC4 | Microarrays | Tarbase |
| hsa-miR-195-5p | RAP1GAP | Microarrays | Tarbase |
| hsa-miR-15a-5p | CTTNBP2 | PAR-CLIP | Tarbase |
| hsa-miR-15a-5p | CTTNBP2 | PAR-CLIP | Tarbase |
| hsa-miR-15a-5p | CTTNBP2 | PAR-CLIP | Tarbase |
| hsa-miR-195-5p | RARB | Microarrays | Tarbase |
| hsa-miR-15a-5p | TM9SF3 | HITS-CLIP | Tarbase |
| hsa-miR-195-5p | TM9SF3 | HITS-CLIP | Tarbase |
| hsa-miR-15a-5p | UBE2T | PAR-CLIP | Tarbase |
| hsa-miR-15a-5p | UBE2T | HITS-CLIP | Tarbase |
| hsa-miR-15a-5p | UBE2T | PAR-CLIP | Tarbase |
| hsa-miR-195-5p | UBE2T | HITS-CLIP | Tarbase |
| hsa-miR-195-5p | UBE2T | PAR-CLIP | Tarbase |
| hsa-miR-195-5p | UBE2T | Microarrays | Tarbase |
| hsa-miR-15a-5p | PPP1R12B | PAR-CLIP | Tarbase |
| hsa-miR-15a-5p | DNAJC10 | PAR-CLIP | Tarbase |
| hsa-miR-141-3p | USP33 | PAR-CLIP | Tarbase |
| hsa-miR-195-5p | CAPN6 | Microarrays | Tarbase |
| hsa-miR-195-5p | DCX | Microarrays | Tarbase |
| hsa-miR-15a-5p | POLD3 | PAR-CLIP | Tarbase |
| hsa-miR-219a-1-3p | POLD3 | PAR-CLIP | Tarbase |
| hsa-miR-3605-5p | CAPZB | HITS-CLIP | Tarbase |
| hsa-miR-141-3p | GPR137B | HITS-CLIP | Tarbase |
| hsa-miR-15a-5p | GPR137B | HITS-CLIP | Tarbase |
| hsa-miR-195-5p | GPR137B | HITS-CLIP | Tarbase |
| hsa-miR-195-5p | GPR137B | Microarrays | Tarbase |
| hsa-miR-141-3p | FGFR1 | HITS-CLIP | Tarbase |
| hsa-miR-15a-5p | FGFR1 | PAR-CLIP | Tarbase |
| hsa-miR-195-5p | FGFR1 | Microarrays | Tarbase |
| hsa-miR-486-3p | FBLN1 | HITS-CLIP | Tarbase |
| hsa-miR-195-5p | CST7 | Microarrays | Tarbase |
| hsa-miR-15a-5p | PIAS2 | HITS-CLIP | Tarbase |
| hsa-miR-195-5p | PIAS2 | HITS-CLIP | Tarbase |
| hsa-miR-3605-3p | PIAS2 | HITS-CLIP | Tarbase |
| hsa-miR-15a-5p | PIAS2 | HITS-CLIP | Tarbase |
| hsa-miR-195-5p | PIAS2 | HITS-CLIP | Tarbase |
| hsa-miR-141-3p | FAP | HITS-CLIP | Tarbase |
| hsa-miR-141-3p | UBE2K | HITS-CLIP | Tarbase |
| hsa-miR-141-3p | UBE2K | HITS-CLIP | Tarbase |
| hsa-miR-141-3p | UBE2K | HITS-CLIP | Tarbase |
| hsa-miR-141-5p | N4BP2 | HITS-CLIP | Tarbase |
| hsa-miR-195-5p | N4BP2 | Microarrays | Tarbase |
| hsa-miR-15a-5p | C12orf5 | PAR-CLIP | Tarbase |
| hsa-miR-195-5p | C12orf5 | PAR-CLIP | Tarbase |
| hsa-miR-195-5p | ADCY2 | Microarrays | Tarbase |
| hsa-miR-141-3p | PPP2R5C | HITS-CLIP | Tarbase |
| hsa-miR-15a-5p | PPP2R5C | HITS-CLIP | Tarbase |
| hsa-miR-15a-5p | PPP2R5C | PAR-CLIP | Tarbase |
| hsa-miR-195-5p | PPP2R5C | HITS-CLIP | Tarbase |
| hsa-miR-141-5p | GNB1 | HITS-CLIP | Tarbase |
| hsa-miR-15a-5p | GNB1 | HITS-CLIP | Tarbase |
| hsa-miR-195-5p | GNB1 | HITS-CLIP | Tarbase |
| hsa-miR-141-5p | HOXA9 | HITS-CLIP | Tarbase |
| hsa-miR-195-5p | EDN1 | Microarrays | Tarbase |
| hsa-miR-141-3p | EDN1 | Luciferase Reporter Assay | Tarbase |
| hsa-miR-141-5p | MLLT10 | HITS-CLIP | Tarbase |
| hsa-miR-3613-5p | MLLT10 | PAR-CLIP | Tarbase |
| hsa-miR-195-5p | ITM2A | Microarrays | Tarbase |
| hsa-miR-15a-3p | NRD1 | HITS-CLIP | Tarbase |
| hsa-miR-15a-5p | VDAC3 | HITS-CLIP | Tarbase |
| hsa-miR-15a-5p | VDAC3 | HITS-CLIP | Tarbase |
| hsa-miR-15a-5p | VDAC3 | HITS-CLIP | Tarbase |
| hsa-miR-15a-5p | VDAC3 | HITS-CLIP | Tarbase |
| hsa-miR-195-5p | VDAC3 | HITS-CLIP | Tarbase |
| hsa-miR-195-5p | VDAC3 | HITS-CLIP | Tarbase |
| hsa-miR-195-5p | VDAC3 | HITS-CLIP | Tarbase |
| hsa-miR-141-3p | PCM1 | HITS-CLIP | Tarbase |
| hsa-miR-141-5p | PCM1 | HITS-CLIP | Tarbase |
| hsa-miR-141-5p | PCM1 | HITS-CLIP | Tarbase |
| hsa-miR-15a-5p | PCM1 | HITS-CLIP | Tarbase |
| hsa-miR-195-5p | PCM1 | HITS-CLIP | Tarbase |
| hsa-miR-141-3p | TNRC6C | HITS-CLIP | Tarbase |
| hsa-miR-15a-5p | TNRC6C | HITS-CLIP | Tarbase |
| hsa-miR-195-5p | TNRC6C | HITS-CLIP | Tarbase |
| hsa-miR-15a-5p | TNRC6C | Chimeric fragments | Tarbase |
| hsa-miR-141-3p | ITCH | HITS-CLIP | Tarbase |
| hsa-miR-15a-5p | RUNX1T1 | Chimeric fragments | Tarbase |
| hsa-miR-195-5p | SLC1A3 | Microarrays | Tarbase |
| hsa-miR-15a-5p | XRCC5 | PAR-CLIP | Tarbase |
| hsa-miR-141-3p | SAR1A | PAR-CLIP | Tarbase |
| hsa-miR-141-3p | SAR1A | HITS-CLIP | Tarbase |
| hsa-miR-141-3p | SAR1A | PAR-CLIP | Tarbase |
| hsa-miR-141-3p | SAR1A | PAR-CLIP | Tarbase |
| hsa-miR-486-3p | SAR1A | HITS-CLIP | Tarbase |
| hsa-miR-486-5p | SAR1A | HITS-CLIP | Tarbase |
| hsa-miR-15a-5p | CDC14A | HITS-CLIP | Tarbase |
| hsa-miR-195-5p | CDC14A | HITS-CLIP | Tarbase |
| hsa-miR-15a-3p | CIC | HITS-CLIP | Tarbase |
| hsa-miR-15a-3p | CIC | HITS-CLIP | Tarbase |
| hsa-miR-15a-5p | CIC | Chimeric fragments | Tarbase |
| hsa-miR-15a-5p | FDFT1 | HITS-CLIP | Tarbase |
| hsa-miR-195-5p | KIF22 | Microarrays | Tarbase |
| hsa-miR-15a-5p | KIF22 | Chimeric fragments | Tarbase |
| hsa-miR-486-5p | LRRC16A | PAR-CLIP | Tarbase |
| hsa-miR-141-3p | DDX1 | HITS-CLIP | Tarbase |
| hsa-miR-15a-5p | DDX1 | HITS-CLIP | Tarbase |
| hsa-miR-15a-5p | DDX1 | HITS-CLIP | Tarbase |
| hsa-miR-15a-5p | DDX1 | PAR-CLIP | Tarbase |
| hsa-miR-15a-5p | DDX1 | HITS-CLIP | Tarbase |
| hsa-miR-195-5p | DDX1 | HITS-CLIP | Tarbase |
| hsa-miR-195-5p | DDX1 | HITS-CLIP | Tarbase |
| hsa-miR-195-5p | DDX1 | HITS-CLIP | Tarbase |
| hsa-miR-15a-3p | DNM2 | HITS-CLIP | Tarbase |
| hsa-miR-195-5p | EPB41L2 | HITS-CLIP | Tarbase |
| hsa-miR-195-5p | EPB41L2 | Microarrays | Tarbase |
| hsa-miR-195-5p | RIMS1 | Microarrays | Tarbase |
| hsa-miR-141-5p | STX7 | HITS-CLIP | Tarbase |
| hsa-miR-141-3p | CRYBG3 | HITS-CLIP | Tarbase |
| hsa-miR-195-5p | EPHA6 | Microarrays | Tarbase |
| hsa-miR-15a-5p | RFX3 | HITS-CLIP | Tarbase |
| hsa-miR-141-5p | RIF1 | HITS-CLIP | Tarbase |
| hsa-miR-15a-5p | RIF1 | HITS-CLIP | Tarbase |
| hsa-miR-15a-5p | RIF1 | HITS-CLIP | Tarbase |
| hsa-miR-195-5p | RIF1 | HITS-CLIP | Tarbase |
| hsa-miR-15a-5p | RIF1 | HITS-CLIP | Tarbase |
| hsa-miR-195-5p | RIF1 | HITS-CLIP | Tarbase |
| hsa-miR-3613-5p | RAB21 | HITS-CLIP | Tarbase |
| hsa-miR-486-5p | RAB21 | HITS-CLIP | Tarbase |
| hsa-miR-3605-5p | SLC4A4 | HITS-CLIP | Tarbase |
| hsa-miR-141-3p | SESN1 | PAR-CLIP | Tarbase |
| hsa-miR-15a-5p | SESN1 | HITS-CLIP | Tarbase |
| hsa-miR-15a-5p | SESN1 | HITS-CLIP | Tarbase |
| hsa-miR-15a-5p | SESN1 | PAR-CLIP | Tarbase |
| hsa-miR-15a-5p | SESN1 | HITS-CLIP | Tarbase |
| hsa-miR-195-5p | SESN1 | HITS-CLIP | Tarbase |
| hsa-miR-195-5p | SESN1 | HITS-CLIP | Tarbase |
| hsa-miR-195-5p | SESN1 | HITS-CLIP | Tarbase |
| hsa-miR-141-3p | SRCAP | HITS-CLIP | Tarbase |
| hsa-miR-141-3p | SRCAP | HITS-CLIP | Tarbase |
| hsa-miR-15a-5p | SRCAP | HITS-CLIP | Tarbase |
| hsa-miR-15a-5p | SRCAP | HITS-CLIP | Tarbase |
| hsa-miR-15a-5p | SRCAP | HITS-CLIP | Tarbase |

| | | | |
|---|---|---|---|
| hsa-miR-195-5p | SRCAP | HITS-CLIP | Tarbase |
| hsa-miR-15a-5p | CNOT4 | PAR-CLIP | Tarbase |
| hsa-miR-3613-5p | CNOT4 | PAR-CLIP | Tarbase |
| hsa-miR-141-3p | PSEN1 | HITS-CLIP | Tarbase |
| hsa-miR-15a-5p | CLDND1 | HITS-CLIP | Tarbase |
| hsa-miR-195-5p | CLDND1 | HITS-CLIP | Tarbase |
| hsa-miR-141-3p | HSP90AA1 | HITS-CLIP | Tarbase |
| hsa-miR-141-5p | HSP90AA1 | HITS-CLIP | Tarbase |
| hsa-miR-15a-3p | HSP90AA1 | HITS-CLIP | Tarbase |
| hsa-miR-15a-5p | HSP90AA1 | HITS-CLIP | Tarbase |
| hsa-miR-15a-5p | HSP90AA1 | HITS-CLIP | Tarbase |
| hsa-miR-15a-5p | HSP90AA1 | HITS-CLIP | Tarbase |
| hsa-miR-15a-5p | HSP90AA1 | HITS-CLIP | Tarbase |
| hsa-miR-15a-5p | HSP90AA1 | PAR-CLIP | Tarbase |
| hsa-miR-15a-5p | HSP90AA1 | HITS-CLIP | Tarbase |
| hsa-miR-195-5p | HSP90AA1 | HITS-CLIP | Tarbase |
| hsa-miR-195-5p | HSP90AA1 | HITS-CLIP | Tarbase |
| hsa-miR-195-5p | HSP90AA1 | HITS-CLIP | Tarbase |
| hsa-miR-195-5p | HSP90AA1 | HITS-CLIP | Tarbase |
| hsa-miR-15a-5p | RBL1 | PAR-CLIP | Tarbase |
| hsa-miR-15a-5p | RBL1 | PAR-CLIP | Tarbase |
| hsa-miR-15a-5p | RBL1 | PAR-CLIP | Tarbase |
| hsa-miR-15a-5p | RBL1 | PAR-CLIP | Tarbase |
| hsa-miR-195-5p | RBL1 | PAR-CLIP | Tarbase |
| hsa-miR-195-5p | RBL1 | PAR-CLIP | Tarbase |
| hsa-miR-15a-3p | DLGAP4 | HITS-CLIP | Tarbase |
| hsa-miR-195-5p | NDC80 | Microarrays | Tarbase |
| hsa-miR-15a-5p | AP4E1 | HITS-CLIP | Tarbase |
| hsa-miR-15a-5p | RSBN1 | PAR-CLIP | Tarbase |
| hsa-miR-15a-5p | RSBN1 | HITS-CLIP | Tarbase |
| hsa-miR-15a-5p | RSBN1 | HITS-CLIP | Tarbase |
| hsa-miR-15a-5p | RSBN1 | PAR-CLIP | Tarbase |
| hsa-miR-195-5p | RSBN1 | HITS-CLIP | Tarbase |
| hsa-miR-141-3p | CXCL2 | Luciferase Reporter Assay | Tarbase |
| hsa-miR-195-5p | CDH7 | Microarrays | Tarbase |
| hsa-miR-195-5p | ARG2 | Microarrays | Tarbase |
| hsa-miR-3613-3p | STK17B | PAR-CLIP | Tarbase |
| hsa-miR-15a-5p | CDC14B | HITS-CLIP | Tarbase |
| hsa-miR-15a-5p | ZNF510 | PAR-CLIP | Tarbase |
| hsa-miR-195-5p | AACS | Microarrays | Tarbase |
| hsa-miR-486-5p | KIAA0141 | PAR-CLIP | Tarbase |
| hsa-miR-141-3p | ATP8B1 | HITS-CLIP | Tarbase |
| hsa-miR-141-5p | BZW1 | HITS-CLIP | Tarbase |
| hsa-miR-15a-5p | BZW1 | PAR-CLIP | Tarbase |
| hsa-miR-15a-5p | BZW1 | PAR-CLIP | Tarbase |
| hsa-miR-15a-5p | BZW1 | PAR-CLIP | Tarbase |
| hsa-miR-15a-5p | BZW1 | PAR-CLIP | Tarbase |
| hsa-miR-195-5p | BZW1 | PAR-CLIP | Tarbase |
| hsa-miR-141-3p | PGR | HITS-CLIP | Tarbase |
| hsa-miR-141-3p | CCNT2 | PAR-CLIP | Tarbase |
| hsa-miR-141-3p | CCNT2 | HITS-CLIP | Tarbase |
| hsa-miR-15a-5p | CCNT2 | PAR-CLIP | Tarbase |
| hsa-miR-15a-5p | CCNT2 | HITS-CLIP | Tarbase |
| hsa-miR-195-5p | CCNT2 | HITS-CLIP | Tarbase |
| hsa-miR-3605-5p | CCNT2 | HITS-CLIP | Tarbase |
| hsa-miR-195-5p | CCNT2 | HITS-CLIP | Tarbase |
| hsa-miR-15a-5p | FAM135A | PAR-CLIP | Tarbase |
| hsa-miR-3613-3p | FAM135A | PAR-CLIP | Tarbase |
| hsa-miR-195-5p | COBLL1 | Microarrays | Tarbase |
| hsa-miR-15a-5p | TRAF5 | PAR-CLIP | Tarbase |
| hsa-miR-15a-5p | TRAF5 | HITS-CLIP | Tarbase |
| hsa-miR-195-5p | TRAF5 | HITS-CLIP | Tarbase |
| hsa-miR-141-5p | NFE2L1 | HITS-CLIP | Tarbase |
| hsa-miR-15a-5p | NFE2L1 | HITS-CLIP | Tarbase |
| hsa-miR-15a-5p | NFE2L1 | PAR-CLIP | Tarbase |
| hsa-miR-15a-5p | NFE2L1 | HITS-CLIP | Tarbase |
| hsa-miR-15a-5p | NFE2L1 | HITS-CLIP | Tarbase |
| hsa-miR-15a-5p | NFE2L1 | PAR-CLIP | Tarbase |
| hsa-miR-15a-5p | NFE2L1 | HITS-CLIP | Tarbase |
| hsa-miR-195-5p | NFE2L1 | HITS-CLIP | Tarbase |
| hsa-miR-195-5p | NFE2L1 | HITS-CLIP | Tarbase |
| hsa-miR-195-5p | NFE2L1 | HITS-CLIP | Tarbase |
| hsa-miR-195-5p | NFE2L1 | HITS-CLIP | Tarbase |
| hsa-miR-195-5p | ITGB5 | Microarrays | Tarbase |
| hsa-miR-141-5p | ERC1 | HITS-CLIP | Tarbase |
| hsa-miR-141-5p | XPO1 | HITS-CLIP | Tarbase |
| hsa-miR-15a-5p | XPO1 | HITS-CLIP | Tarbase |
| hsa-miR-15a-5p | XPO1 | HITS-CLIP | Tarbase |
| hsa-miR-15a-5p | XPO1 | HITS-CLIP | Tarbase |
| hsa-miR-195-3p | XPO1 | HITS-CLIP | Tarbase |
| hsa-miR-195-5p | XPO1 | HITS-CLIP | Tarbase |
| hsa-miR-195-5p | XPO1 | HITS-CLIP | Tarbase |
| hsa-miR-195-5p | XPO1 | HITS-CLIP | Tarbase |
| hsa-miR-3613-5p | XPO1 | HITS-CLIP | Tarbase |
| hsa-miR-486-5p | XPO1 | HITS-CLIP | Tarbase |
| hsa-miR-15a-5p | KAT6A | HITS-CLIP | Tarbase |
| hsa-miR-15a-5p | KAT6A | HITS-CLIP | Tarbase |
| hsa-miR-15a-5p | KAT6A | HITS-CLIP | Tarbase |
| hsa-miR-15a-5p | KAT6A | HITS-CLIP | Tarbase |
| hsa-miR-195-5p | KAT6A | HITS-CLIP | Tarbase |
| hsa-miR-195-5p | KAT6A | HITS-CLIP | Tarbase |
| hsa-miR-141-3p | ULK2 | HITS-CLIP | Tarbase |
| hsa-miR-141-3p | TNPO1 | HITS-CLIP | Tarbase |
| hsa-miR-141-3p | TNPO1 | HITS-CLIP | Tarbase |
| hsa-miR-15a-5p | TNPO1 | HITS-CLIP | Tarbase |
| hsa-miR-195-5p | TNPO1 | HITS-CLIP | Tarbase |
| hsa-miR-3613-3p | TNPO1 | HITS-CLIP | Tarbase |
| hsa-miR-486-5p | DIS3 | PAR-CLIP | Tarbase |
| hsa-miR-141-3p | PDS5B | HITS-CLIP | Tarbase |
| hsa-miR-141-5p | PDS5B | HITS-CLIP | Tarbase |
| hsa-miR-15a-5p | PDS5B | HITS-CLIP | Tarbase |
| hsa-miR-195-5p | PDS5B | HITS-CLIP | Tarbase |
| hsa-miR-15a-5p | PDS5B | HITS-CLIP | Tarbase |
| hsa-miR-195-5p | EPYC | Microarrays | Tarbase |
| hsa-miR-141-3p | CYLD | HITS-CLIP | Tarbase |
| hsa-miR-15a-5p | CYLD | HITS-CLIP | Tarbase |
| hsa-miR-195-5p | CYLD | HITS-CLIP | Tarbase |
| hsa-miR-141-3p | ZNF416 | PAR-CLIP | Tarbase |
| hsa-miR-141-3p | ZNF416 | HITS-CLIP | Tarbase |
| hsa-miR-15a-5p | ZNF264 | HITS-CLIP | Tarbase |
| hsa-miR-15a-5p | ZNF264 | HITS-CLIP | Tarbase |
| hsa-miR-195-5p | ZNF264 | HITS-CLIP | Tarbase |
| hsa-miR-195-5p | ZNF264 | HITS-CLIP | Tarbase |
| hsa-miR-15a-5p | ZNF264 | Chimeric fragments | Tarbase |
| hsa-miR-141-3p | FAT1 | HITS-CLIP | Tarbase |
| hsa-miR-141-3p | FAT1 | HITS-CLIP | Tarbase |
| hsa-miR-15a-5p | FAT1 | HITS-CLIP | Tarbase |
| hsa-miR-15a-5p | FAT1 | PAR-CLIP | Tarbase |
| hsa-miR-15a-5p | FAT1 | HITS-CLIP | Tarbase |
| hsa-miR-195-5p | FAT1 | HITS-CLIP | Tarbase |
| hsa-miR-195-5p | FAT1 | HITS-CLIP | Tarbase |
| hsa-miR-3613-5p | FAT1 | HITS-CLIP | Tarbase |
| hsa-miR-195-5p | FAT1 | Microarrays | Tarbase |
| hsa-miR-141-5p | YTHDC1 | HITS-CLIP | Tarbase |
| hsa-miR-141-5p | YTHDC1 | HITS-CLIP | Tarbase |
| hsa-miR-15a-5p | YTHDC1 | PAR-CLIP | Tarbase |
| hsa-miR-15a-5p | YTHDC1 | HITS-CLIP | Tarbase |
| hsa-miR-15a-5p | YTHDC1 | HITS-CLIP | Tarbase |
| hsa-miR-195-5p | YTHDC1 | HITS-CLIP | Tarbase |
| hsa-miR-141-5p | ZMPSTE24 | HITS-CLIP | Tarbase |
| hsa-miR-195-5p | ZMPSTE24 | HITS-CLIP | Tarbase |
| hsa-miR-3605-5p | ZMPSTE24 | PAR-CLIP | Tarbase |
| hsa-miR-3605-5p | ZMPSTE24 | PAR-CLIP | Tarbase |
| hsa-miR-141-3p | STARD7 | HITS-CLIP | Tarbase |
| hsa-miR-141-3p | REST | HITS-CLIP | Tarbase |
| hsa-miR-141-3p | REST | HITS-CLIP | Tarbase |
| hsa-miR-141-3p | REST | HITS-CLIP | Tarbase |
| hsa-miR-15a-5p | REST | PAR-CLIP | Tarbase |
| hsa-miR-15a-5p | REST | HITS-CLIP | Tarbase |
| hsa-miR-15a-5p | REST | PAR-CLIP | Tarbase |
| hsa-miR-15a-5p | GSTP1 | PAR-CLIP | Tarbase |
| hsa-miR-195-5p | APLP2 | HITS-CLIP | Tarbase |
| hsa-miR-219a-1-3p | APLP2 | HITS-CLIP | Tarbase |
| hsa-miR-15a-5p | KIAA1467 | HITS-CLIP | Tarbase |
| hsa-miR-195-5p | KIAA1467 | HITS-CLIP | Tarbase |
| hsa-miR-195-5p | WBP11 | Microarrays | Tarbase |
| hsa-miR-3613-5p | EIF3I | HITS-CLIP | Tarbase |
| hsa-miR-195-5p | COL16A1 | HITS-CLIP | Tarbase |
| hsa-miR-15a-5p | TXLNA | PAR-CLIP | Tarbase |

| | | | |
|---|---|---|---|
| hsa-miR-15a-5p | TXLNA | PAR-CLIP | Tarbase |
| hsa-miR-15a-5p | TXLNA | PAR-CLIP | Tarbase |
| hsa-miR-195-5p | TXLNA | PAR-CLIP | Tarbase |
| hsa-miR-195-5p | TXLNA | PAR-CLIP | Tarbase |
| hsa-miR-141-5p | NCOA1 | HITS-CLIP | Tarbase |
| hsa-miR-3613-3p | NCOA1 | PAR-CLIP | Tarbase |
| hsa-miR-195-5p | AGBL5 | Microarrays | Tarbase |
| hsa-miR-141-3p | RAB10 | HITS-CLIP | Tarbase |
| hsa-miR-15a-5p | RAB10 | PAR-CLIP | Tarbase |
| hsa-miR-15a-5p | RAB10 | HITS-CLIP | Tarbase |
| hsa-miR-195-5p | RAB10 | HITS-CLIP | Tarbase |
| hsa-miR-15a-5p | RAB10 | HITS-CLIP | Tarbase |
| hsa-miR-195-5p | RAB10 | HITS-CLIP | Tarbase |
| hsa-miR-15a-5p | HADHA | HITS-CLIP | Tarbase |
| hsa-miR-195-5p | HADHA | HITS-CLIP | Tarbase |
| hsa-miR-195-5p | MAPRE3 | HITS-CLIP | Tarbase |
| hsa-miR-15a-3p | CAD | HITS-CLIP | Tarbase |
| hsa-miR-15a-3p | CAD | HITS-CLIP | Tarbase |
| hsa-miR-195-5p | CAD | HITS-CLIP | Tarbase |
| hsa-miR-195-5p | CD59 | Microarrays | Tarbase |
| hsa-miR-15a-3p | BCORL1 | HITS-CLIP | Tarbase |
| hsa-miR-15a-3p | BCORL1 | HITS-CLIP | Tarbase |
| hsa-miR-3613-5p | BCORL1 | HITS-CLIP | Tarbase |
| hsa-miR-15a-3p | PREP | HITS-CLIP | Tarbase |
| hsa-miR-15a-5p | PREP | HITS-CLIP | Tarbase |
| hsa-miR-195-5p | PREP | HITS-CLIP | Tarbase |
| hsa-miR-195-5p | HACE1 | Microarrays | Tarbase |
| hsa-miR-141-3p | SEH1L | HITS-CLIP | Tarbase |
| hsa-miR-141-5p | WDR47 | HITS-CLIP | Tarbase |
| hsa-miR-15a-5p | ABCB1 | HITS-CLIP | Tarbase |
| hsa-miR-195-5p | ABCB1 | HITS-CLIP | Tarbase |
| hsa-miR-15a-5p | ABCB1 | HITS-CLIP | Tarbase |
| hsa-miR-195-5p | ABCB1 | HITS-CLIP | Tarbase |
| hsa-miR-15a-5p | RRN3 | PAR-CLIP | Tarbase |
| hsa-miR-141-3p | CTTN | HITS-CLIP | Tarbase |
| hsa-miR-3613-3p | CTTN | HITS-CLIP | Tarbase |
| hsa-miR-195-5p | DDHD2 | Microarrays | Tarbase |
| hsa-miR-141-5p | EPS15 | HITS-CLIP | Tarbase |
| hsa-miR-15a-5p | EPS15 | PAR-CLIP | Tarbase |
| hsa-miR-15a-5p | EPS15 | PAR-CLIP | Tarbase |
| hsa-miR-15a-5p | EPS15 | HITS-CLIP | Tarbase |
| hsa-miR-195-5p | EPS15 | HITS-CLIP | Tarbase |
| hsa-miR-195-5p | ORC1 | Microarrays | Tarbase |
| hsa-miR-141-3p | ATG16L1 | HITS-CLIP | Tarbase |
| hsa-miR-486-5p | USP40 | HITS-CLIP | Tarbase |
| hsa-miR-15a-3p | RAD54L | HITS-CLIP | Tarbase |
| hsa-miR-195-5p | RAD54L | Microarrays | Tarbase |
| hsa-miR-3605-3p | MAST2 | HITS-CLIP | Tarbase |
| hsa-miR-15a-5p | DNAJA1 | PAR-CLIP | Tarbase |
| hsa-miR-15a-5p | DNAJA1 | PAR-CLIP | Tarbase |
| hsa-miR-15a-5p | DNAJA1 | PAR-CLIP | Tarbase |
| hsa-miR-15a-5p | DNAJA1 | PAR-CLIP | Tarbase |
| hsa-miR-15a-5p | DNAJA1 | HITS-CLIP | Tarbase |
| hsa-miR-15a-5p | DNAJA1 | PAR-CLIP | Tarbase |
| hsa-miR-15a-5p | DNAJA1 | PAR-CLIP | Tarbase |
| hsa-miR-195-5p | DNAJA1 | PAR-CLIP | Tarbase |
| hsa-miR-195-5p | DNAJA1 | PAR-CLIP | Tarbase |
| hsa-miR-195-5p | DNAJA1 | PAR-CLIP | Tarbase |
| hsa-miR-15a-5p | DNAJA1 | Chimeric fragments | Tarbase |
| hsa-miR-15a-3p | B4GALT1 | HITS-CLIP | Tarbase |
| hsa-miR-15a-5p | B4GALT1 | HITS-CLIP | Tarbase |
| hsa-miR-195-5p | B4GALT1 | HITS-CLIP | Tarbase |
| hsa-miR-15a-3p | EIF2AK1 | HITS-CLIP | Tarbase |
| hsa-miR-141-5p | SNX10 | HITS-CLIP | Tarbase |
| hsa-miR-3613-5p | SEPHS1 | PAR-CLIP | Tarbase |
| hsa-miR-15a-5p | RBM22 | HITS-CLIP | Tarbase |
| hsa-miR-195-5p | RBM22 | HITS-CLIP | Tarbase |
| hsa-miR-15a-5p | RBM22 | HITS-CLIP | Tarbase |
| hsa-miR-195-5p | RBM22 | HITS-CLIP | Tarbase |
| hsa-miR-15a-5p | TMED2 | HITS-CLIP | Tarbase |
| hsa-miR-195-5p | TMED2 | HITS-CLIP | Tarbase |
| hsa-miR-3613-3p | TMED2 | HITS-CLIP | Tarbase |
| hsa-miR-15a-5p | TMED2 | Chimeric fragments | Tarbase |
| hsa-miR-3613-3p | ERO1LB | HITS-CLIP | Tarbase |
| hsa-miR-15a-5p | ZFAND6 | PAR-CLIP | Tarbase |
| hsa-miR-15a-5p | ZFAND6 | PAR-CLIP | Tarbase |
| hsa-miR-15a-5p | ZFAND6 | PAR-CLIP | Tarbase |
| hsa-miR-195-5p | ZFAND6 | PAR-CLIP | Tarbase |
| hsa-miR-3613-3p | ZFAND6 | HITS-CLIP | Tarbase |
| hsa-miR-486-5p | ZFAND6 | HITS-CLIP | Tarbase |
| hsa-miR-3605-5p | TXLNG | HITS-CLIP | Tarbase |
| hsa-miR-15a-3p | HUWE1 | HITS-CLIP | Tarbase |
| hsa-miR-15a-5p | HUWE1 | HITS-CLIP | Tarbase |
| hsa-miR-15a-5p | HUWE1 | HITS-CLIP | Tarbase |
| hsa-miR-195-5p | HUWE1 | HITS-CLIP | Tarbase |
| hsa-miR-15a-5p | ZW10 | PAR-CLIP | Tarbase |
| hsa-miR-195-5p | ZW10 | PAR-CLIP | Tarbase |
| hsa-miR-141-3p | ALG9 | HITS-CLIP | Tarbase |
| hsa-miR-15a-5p | ACOX3 | HITS-CLIP | Tarbase |
| hsa-miR-195-5p | ACOX3 | HITS-CLIP | Tarbase |
| hsa-miR-15a-5p | MTMR2 | PAR-CLIP | Tarbase |
| hsa-miR-195-5p | MTMR2 | PAR-CLIP | Tarbase |
| hsa-miR-486-5p | MTMR2 | HITS-CLIP | Tarbase |
| hsa-miR-141-3p | SRRT | HITS-CLIP | Tarbase |
| hsa-miR-15a-5p | SRRT | PAR-CLIP | Tarbase |
| hsa-miR-141-3p | BAX | Western Blot | Tarbase |
| hsa-miR-15a-3p | NLK | HITS-CLIP | Tarbase |
| hsa-miR-15a-5p | PIGS | HITS-CLIP | Tarbase |
| hsa-miR-195-5p | PIGS | HITS-CLIP | Tarbase |
| hsa-miR-3613-5p | PIGS | HITS-CLIP | Tarbase |
| hsa-miR-15a-5p | ATXN7L3 | PAR-CLIP | Tarbase |
| hsa-miR-15a-5p | ATXN7L3 | HITS-CLIP | Tarbase |
| hsa-miR-15a-5p | ATXN7L3 | PAR-CLIP | Tarbase |
| hsa-miR-15a-5p | ATXN7L3 | PAR-CLIP | Tarbase |
| hsa-miR-195-5p | ATXN7L3 | HITS-CLIP | Tarbase |
| hsa-miR-195-5p | ATXN7L3 | PAR-CLIP | Tarbase |
| hsa-miR-195-5p | ATXN7L3 | PAR-CLIP | Tarbase |
| hsa-miR-486-3p | ATXN7L3 | HITS-CLIP | Tarbase |
| hsa-miR-486-5p | ATXN7L3 | HITS-CLIP | Tarbase |
| hsa-miR-15a-5p | ATXN7L3 | HITS-CLIP | Tarbase |
| hsa-miR-195-5p | ATXN7L3 | HITS-CLIP | Tarbase |
| hsa-miR-195-5p | ATXN7L3 | Microarrays | Tarbase |
| hsa-miR-141-3p | MMP2 | Western Blot | Tarbase |
| hsa-miR-195-5p | LPCAT2 | Microarrays | Tarbase |
| hsa-miR-15a-5p | OGFOD1 | HITS-CLIP | Tarbase |
| hsa-miR-15a-5p | OGFOD1 | HITS-CLIP | Tarbase |
| hsa-miR-15a-5p | OGFOD1 | PAR-CLIP | Tarbase |
| hsa-miR-195-5p | OGFOD1 | HITS-CLIP | Tarbase |
| hsa-miR-15a-5p | SH3BP2 | PAR-CLIP | Tarbase |
| hsa-miR-3613-3p | SH3BP2 | HITS-CLIP | Tarbase |
| hsa-miR-195-5p | SH3BP2 | Microarrays | Tarbase |
| hsa-miR-3613-3p | ADD1 | PAR-CLIP | Tarbase |
| hsa-miR-3613-3p | GMCL1 | HITS-CLIP | Tarbase |
| hsa-miR-141-5p | KLHL42 | HITS-CLIP | Tarbase |
| hsa-miR-141-5p | KLHL42 | HITS-CLIP | Tarbase |
| hsa-miR-486-5p | KLHL42 | PAR-CLIP | Tarbase |
| hsa-miR-15a-3p | DNM1L | HITS-CLIP | Tarbase |
| hsa-miR-15a-5p | DNM1L | HITS-CLIP | Tarbase |
| hsa-miR-15a-5p | DNM1L | HITS-CLIP | Tarbase |
| hsa-miR-195-5p | DNM1L | HITS-CLIP | Tarbase |
| hsa-miR-195-5p | DNM1L | HITS-CLIP | Tarbase |
| hsa-miR-141-3p | TFAP2C | HITS-CLIP | Tarbase |
| hsa-miR-15a-5p | TFAP2C | HITS-CLIP | Tarbase |
| hsa-miR-195-5p | TFAP2C | Microarrays | Tarbase |
| hsa-miR-195-5p | AURKA | Microarrays | Tarbase |
| hsa-miR-3605-5p | CNOT3 | HITS-CLIP | Tarbase |
| hsa-miR-15a-5p | PTPN4 | HITS-CLIP | Tarbase |
| hsa-miR-195-5p | PTPN4 | HITS-CLIP | Tarbase |
| hsa-miR-15a-5p | PTPN4 | HITS-CLIP | Tarbase |
| hsa-miR-195-5p | PTPN4 | HITS-CLIP | Tarbase |
| hsa-miR-15a-3p | GNA11 | HITS-CLIP | Tarbase |
| hsa-miR-195-5p | TPX2 | Microarrays | Tarbase |
| hsa-miR-141-3p | ANKRD10 | HITS-CLIP | Tarbase |
| hsa-miR-195-5p | TMEM40 | Microarrays | Tarbase |
| hsa-miR-141-3p | MAVS | HITS-CLIP | Tarbase |
| hsa-miR-15a-3p | MAVS | HITS-CLIP | Tarbase |
| hsa-miR-15a-5p | MAVS | HITS-CLIP | Tarbase |
| hsa-miR-15a-5p | MAVS | HITS-CLIP | Tarbase |
| hsa-miR-195-5p | MAVS | HITS-CLIP | Tarbase |

| | | | |
|---|---|---|---|
| hsa-miR-141-5p | XRN2 | HITS-CLIP | Tarbase |
| hsa-miR-141-5p | XRN2 | HITS-CLIP | Tarbase |
| hsa-miR-141-5p | SNX5 | HITS-CLIP | Tarbase |
| hsa-miR-141-5p | SNX5 | HITS-CLIP | Tarbase |
| hsa-miR-141-5p | GCN1L1 | HITS-CLIP | Tarbase |
| hsa-miR-15a-5p | GCN1L1 | HITS-CLIP | Tarbase |
| hsa-miR-15a-5p | PXN | HITS-CLIP | Tarbase |
| hsa-miR-15a-5p | PXN | HITS-CLIP | Tarbase |
| hsa-miR-15a-5p | KIF16B | PAR-CLIP | Tarbase |
| hsa-miR-141-5p | CHGB | HITS-CLIP | Tarbase |
| hsa-miR-15a-5p | BRAP | HITS-CLIP | Tarbase |
| hsa-miR-141-3p | FUS | HITS-CLIP | Tarbase |
| hsa-miR-141-3p | FUS | HITS-CLIP | Tarbase |
| hsa-miR-15a-3p | FUS | HITS-CLIP | Tarbase |
| hsa-miR-141-5p | GANAB | HITS-CLIP | Tarbase |
| hsa-miR-3613-5p | GANAB | HITS-CLIP | Tarbase |
| hsa-miR-15a-5p | BIRC5 | PAR-CLIP | Tarbase |
| hsa-miR-15a-5p | BIRC5 | PAR-CLIP | Tarbase |
| hsa-miR-15a-5p | BIRC5 | HITS-CLIP | Tarbase |
| hsa-miR-15a-5p | BIRC5 | PAR-CLIP | Tarbase |
| hsa-miR-15a-5p | BIRC5 | PAR-CLIP | Tarbase |
| hsa-miR-15a-5p | BIRC5 | HITS-CLIP | Tarbase |
| hsa-miR-195-5p | BIRC5 | PAR-CLIP | Tarbase |
| hsa-miR-195-5p | BIRC5 | HITS-CLIP | Tarbase |
| hsa-miR-195-5p | BIRC5 | qPCR | Tarbase |
| hsa-miR-195-5p | BIRC5 | Microarrays | Tarbase |
| hsa-miR-195-5p | BIRC5 | Luciferase Reporter Assay | Tarbase |
| hsa-miR-141-3p | NECAP1 | HITS-CLIP | Tarbase |
| hsa-miR-141-5p | NECAP1 | HITS-CLIP | Tarbase |
| hsa-miR-15a-5p | NECAP1 | HITS-CLIP | Tarbase |
| hsa-miR-195-5p | NECAP1 | HITS-CLIP | Tarbase |
| hsa-miR-219a-1-3p | DHX32 | HITS-CLIP | Tarbase |
| hsa-miR-141-3p | RCOR1 | HITS-CLIP | Tarbase |
| hsa-miR-15a-5p | RCOR1 | HITS-CLIP | Tarbase |
| hsa-miR-195-3p | RCOR1 | HITS-CLIP | Tarbase |
| hsa-miR-195-5p | RCOR1 | HITS-CLIP | Tarbase |
| hsa-miR-15a-5p | GPATCH2L | HITS-CLIP | Tarbase |
| hsa-miR-219a-1-3p | LTBP4 | HITS-CLIP | Tarbase |
| hsa-miR-15a-5p | BLVRB | HITS-CLIP | Tarbase |
| hsa-miR-195-5p | BLVRB | HITS-CLIP | Tarbase |
| hsa-miR-195-5p | BLVRB | Microarrays | Tarbase |
| hsa-miR-15a-5p | SLC9A1 | HITS-CLIP | Tarbase |
| hsa-miR-141-3p | PAPOLA | HITS-CLIP | Tarbase |
| hsa-miR-141-3p | PAPOLA | HITS-CLIP | Tarbase |
| hsa-miR-195-5p | ICAM1 | qPCR | Tarbase |
| hsa-miR-195-5p | ICAM1 | Western Blot | Tarbase |
| hsa-miR-486-3p | ICAM1 | HITS-CLIP | Tarbase |
| hsa-miR-3605-3p | STRN4 | HITS-CLIP | Tarbase |
| hsa-miR-15a-5p | TFAP4 | HITS-CLIP | Tarbase |
| hsa-miR-141-5p | DNAJB11 | HITS-CLIP | Tarbase |
| hsa-miR-15a-5p | LEPREL1 | PAR-CLIP | Tarbase |
| hsa-miR-195-5p | FLT3LG | Microarrays | Tarbase |
| hsa-miR-141-5p | RAB11FIP3 | HITS-CLIP | Tarbase |
| hsa-miR-15a-5p | GOLGA3 | HITS-CLIP | Tarbase |
| hsa-miR-195-5p | GOLGA3 | HITS-CLIP | Tarbase |
| hsa-miR-3605-5p | GOLGA3 | HITS-CLIP | Tarbase |
| hsa-miR-141-3p | PABPC4 | HITS-CLIP | Tarbase |
| hsa-miR-15a-3p | PABPC4 | HITS-CLIP | Tarbase |
| hsa-miR-15a-5p | PABPC4 | HITS-CLIP | Tarbase |
| hsa-miR-15a-5p | PABPC4 | HITS-CLIP | Tarbase |
| hsa-miR-195-5p | PABPC4 | HITS-CLIP | Tarbase |
| hsa-miR-195-5p | PABPC4 | HITS-CLIP | Tarbase |
| hsa-miR-486-3p | PABPC4 | HITS-CLIP | Tarbase |
| hsa-miR-486-3p | PABPC4 | HITS-CLIP | Tarbase |
| hsa-miR-195-5p | PABPC4 | Microarrays | Tarbase |
| hsa-miR-141-5p | AARS | HITS-CLIP | Tarbase |
| hsa-miR-15a-3p | AARS | HITS-CLIP | Tarbase |
| hsa-miR-15a-5p | GLG1 | HITS-CLIP | Tarbase |
| hsa-miR-195-5p | GLG1 | HITS-CLIP | Tarbase |
| hsa-miR-15a-5p | KIF4A | HITS-CLIP | Tarbase |
| hsa-miR-195-5p | KIF4A | Microarrays | Tarbase |
| hsa-miR-141-3p | TNRC6A | HITS-CLIP | Tarbase |
| hsa-miR-141-3p | TNRC6A | HITS-CLIP | Tarbase |
| hsa-miR-141-3p | TNRC6A | HITS-CLIP | Tarbase |
| hsa-miR-141-3p | TNRC6A | HITS-CLIP | Tarbase |
| hsa-miR-141-3p | TNRC6A | HITS-CLIP | Tarbase |
| hsa-miR-141-5p | TNRC6A | HITS-CLIP | Tarbase |
| hsa-miR-15a-5p | TNRC6A | PAR-CLIP | Tarbase |
| hsa-miR-15a-5p | TNRC6A | PAR-CLIP | Tarbase |
| hsa-miR-15a-5p | TNRC6A | HITS-CLIP | Tarbase |
| hsa-miR-15a-5p | TNRC6A | HITS-CLIP | Tarbase |
| hsa-miR-15a-5p | TNRC6A | HITS-CLIP | Tarbase |
| hsa-miR-15a-5p | TNRC6A | PAR-CLIP | Tarbase |
| hsa-miR-15a-5p | TNRC6A | PAR-CLIP | Tarbase |
| hsa-miR-15a-5p | TNRC6A | PAR-CLIP | Tarbase |
| hsa-miR-15a-5p | TNRC6A | HITS-CLIP | Tarbase |
| hsa-miR-15a-5p | TNRC6A | PAR-CLIP | Tarbase |
| hsa-miR-15a-5p | TNRC6A | PAR-CLIP | Tarbase |
| hsa-miR-15a-5p | TNRC6A | PAR-CLIP | Tarbase |
| hsa-miR-15a-5p | TNRC6A | PAR-CLIP | Tarbase |
| hsa-miR-15a-5p | TNRC6A | PAR-CLIP | Tarbase |
| hsa-miR-15a-5p | TNRC6A | HITS-CLIP | Tarbase |
| hsa-miR-195-5p | TNRC6A | PAR-CLIP | Tarbase |
| hsa-miR-195-5p | TNRC6A | HITS-CLIP | Tarbase |
| hsa-miR-195-5p | TNRC6A | HITS-CLIP | Tarbase |
| hsa-miR-195-5p | TNRC6A | HITS-CLIP | Tarbase |
| hsa-miR-195-5p | TNRC6A | HITS-CLIP | Tarbase |
| hsa-miR-195-5p | TNRC6A | PAR-CLIP | Tarbase |
| hsa-miR-195-5p | TNRC6A | PAR-CLIP | Tarbase |
| hsa-miR-195-5p | TNRC6A | PAR-CLIP | Tarbase |
| hsa-miR-195-5p | TNRC6A | PAR-CLIP | Tarbase |
| hsa-miR-195-5p | TNRC6A | PAR-CLIP | Tarbase |
| hsa-miR-3613-3p | TNRC6A | HITS-CLIP | Tarbase |
| hsa-miR-15a-3p | PITPNM2 | HITS-CLIP | Tarbase |
| hsa-miR-15a-5p | PITPNM2 | HITS-CLIP | Tarbase |
| hsa-miR-195-5p | PITPNM2 | HITS-CLIP | Tarbase |
| hsa-miR-141-3p | EXOC1 | HITS-CLIP | Tarbase |
| hsa-miR-141-3p | OSBPL8 | HITS-CLIP | Tarbase |
| hsa-miR-15a-5p | DLD | HITS-CLIP | Tarbase |
| hsa-miR-15a-5p | DLD | PAR-CLIP | Tarbase |
| hsa-miR-15a-5p | DLD | HITS-CLIP | Tarbase |
| hsa-miR-195-5p | DLD | HITS-CLIP | Tarbase |
| hsa-miR-195-5p | DLD | HITS-CLIP | Tarbase |
| hsa-miR-15a-5p | DLD | Chimeric fragments | Tarbase |
| hsa-miR-141-3p | CMTM6 | HITS-CLIP | Tarbase |
| hsa-miR-15a-3p | CMTM6 | HITS-CLIP | Tarbase |
| hsa-miR-141-3p | ITGA6 | HITS-CLIP | Tarbase |
| hsa-miR-141-3p | ITGA6 | HITS-CLIP | Tarbase |
| hsa-miR-195-5p | ITGA6 | Microarrays | Tarbase |
| hsa-miR-195-5p | RAPGEF4 | Microarrays | Tarbase |
| hsa-miR-15a-5p | CDV3 | PAR-CLIP | Tarbase |
| hsa-miR-15a-5p | CDV3 | HITS-CLIP | Tarbase |
| hsa-miR-15a-5p | CDV3 | HITS-CLIP | Tarbase |
| hsa-miR-15a-5p | CDV3 | PAR-CLIP | Tarbase |
| hsa-miR-15a-5p | CDV3 | HITS-CLIP | Tarbase |
| hsa-miR-15a-5p | CDV3 | HITS-CLIP | Tarbase |
| hsa-miR-15a-5p | CDV3 | PAR-CLIP | Tarbase |
| hsa-miR-195-5p | CDV3 | HITS-CLIP | Tarbase |
| hsa-miR-195-5p | CDV3 | HITS-CLIP | Tarbase |
| hsa-miR-195-5p | CDV3 | HITS-CLIP | Tarbase |
| hsa-miR-195-5p | CDV3 | PAR-CLIP | Tarbase |
| hsa-miR-15a-5p | CDV3 | HITS-CLIP | Tarbase |
| hsa-miR-195-5p | CDV3 | HITS-CLIP | Tarbase |
| hsa-miR-15a-5p | NLRP1 | PAR-CLIP | Tarbase |
| hsa-miR-15a-5p | NLRP1 | PAR-CLIP | Tarbase |
| hsa-miR-195-5p | NLRP1 | PAR-CLIP | Tarbase |
| hsa-miR-195-5p | SPAG7 | Microarrays | Tarbase |
| hsa-miR-15a-5p | ORC6 | PAR-CLIP | Tarbase |
| hsa-miR-195-5p | ORC6 | Microarrays | Tarbase |
| hsa-miR-15a-5p | ZFHX4 | PAR-CLIP | Tarbase |
| hsa-miR-15a-5p | ZFHX4 | PAR-CLIP | Tarbase |
| hsa-miR-195-5p | ZFHX4 | Microarrays | Tarbase |
| hsa-miR-15a-5p | ZC3HC1 | HITS-CLIP | Tarbase |
| hsa-miR-195-5p | ZC3HC1 | HITS-CLIP | Tarbase |
| hsa-miR-15a-3p | RGS17 | HITS-CLIP | Tarbase |
| hsa-miR-195-5p | SLC7A8 | Microarrays | Tarbase |
| hsa-miR-15a-5p | SCFD1 | PAR-CLIP | Tarbase |
| hsa-miR-15a-5p | SCFD1 | PAR-CLIP | Tarbase |
| hsa-miR-195-5p | SCFD1 | PAR-CLIP | Tarbase |

| | | | |
|---|---|---|---|
| hsa-miR-141-3p | HECTD1 | HITS-CLIP | Tarbase |
| hsa-miR-141-5p | HECTD1 | HITS-CLIP | Tarbase |
| hsa-miR-15a-5p | HECTD1 | HITS-CLIP | Tarbase |
| hsa-miR-15a-5p | HECTD1 | HITS-CLIP | Tarbase |
| hsa-miR-195-5p | HECTD1 | HITS-CLIP | Tarbase |
| hsa-miR-141-3p | HNRNPC | HITS-CLIP | Tarbase |
| hsa-miR-15a-5p | SUPT16H | HITS-CLIP | Tarbase |
| hsa-miR-15a-5p | SUPT16H | HITS-CLIP | Tarbase |
| hsa-miR-195-5p | SUPT16H | HITS-CLIP | Tarbase |
| hsa-miR-486-3p | SUPT16H | HITS-CLIP | Tarbase |
| hsa-miR-141-3p | TOX4 | HITS-CLIP | Tarbase |
| hsa-miR-15a-5p | TOX4 | PAR-CLIP | Tarbase |
| hsa-miR-15a-5p | TOX4 | PAR-CLIP | Tarbase |
| hsa-miR-195-5p | TOX4 | PAR-CLIP | Tarbase |
| hsa-miR-141-3p | SEMA6A | HITS-CLIP | Tarbase |
| hsa-miR-141-3p | SEMA6A | HITS-CLIP | Tarbase |
| hsa-miR-141-5p | TRPM7 | HITS-CLIP | Tarbase |
| hsa-miR-195-5p | TYRO3 | Microarrays | Tarbase |
| hsa-miR-195-5p | WDR76 | Microarrays | Tarbase |
| hsa-miR-141-3p | SNAP23 | HITS-CLIP | Tarbase |
| hsa-miR-141-3p | SNAP23 | HITS-CLIP | Tarbase |
| hsa-miR-486-5p | PHGDH | PAR-CLIP | Tarbase |
| hsa-miR-3613-5p | EZR | HITS-CLIP | Tarbase |
| hsa-miR-141-5p | MYL6 | HITS-CLIP | Tarbase |
| hsa-miR-195-5p | MYL6 | Microarrays | Tarbase |
| hsa-miR-141-3p | AGO1 | HITS-CLIP | Tarbase |
| hsa-miR-15a-3p | AGO1 | HITS-CLIP | Tarbase |
| hsa-miR-15a-5p | AGO1 | HITS-CLIP | Tarbase |
| hsa-miR-15a-5p | AGO1 | HITS-CLIP | Tarbase |
| hsa-miR-15a-5p | AGO1 | HITS-CLIP | Tarbase |
| hsa-miR-15a-5p | AGO1 | HITS-CLIP | Tarbase |
| hsa-miR-15a-5p | AGO1 | PAR-CLIP | Tarbase |
| hsa-miR-15a-5p | AGO1 | HITS-CLIP | Tarbase |
| hsa-miR-195-5p | AGO1 | HITS-CLIP | Tarbase |
| hsa-miR-195-5p | AGO1 | HITS-CLIP | Tarbase |
| hsa-miR-195-5p | AGO1 | HITS-CLIP | Tarbase |
| hsa-miR-195-5p | AGO1 | HITS-CLIP | Tarbase |
| hsa-miR-195-5p | AGO1 | HITS-CLIP | Tarbase |
| hsa-miR-195-5p | TGFB2 | Microarrays | Tarbase |
| hsa-miR-15a-5p | GPATCH2 | PAR-CLIP | Tarbase |
| hsa-miR-15a-5p | GPATCH2 | PAR-CLIP | Tarbase |
| hsa-miR-195-5p | GPATCH2 | PAR-CLIP | Tarbase |
| hsa-miR-15a-5p | NUP50 | PAR-CLIP | Tarbase |
| hsa-miR-195-5p | CDC45 | Microarrays | Tarbase |
| hsa-miR-15a-5p | COMT | HITS-CLIP | Tarbase |
| hsa-miR-195-5p | COMT | HITS-CLIP | Tarbase |
| hsa-miR-141-3p | ECHDC1 | HITS-CLIP | Tarbase |
| hsa-miR-15a-5p | ECHDC1 | HITS-CLIP | Tarbase |
| hsa-miR-15a-5p | ECHDC1 | PAR-CLIP | Tarbase |
| hsa-miR-195-5p | ECHDC1 | HITS-CLIP | Tarbase |
| hsa-miR-15a-5p | SEC22C | HITS-CLIP | Tarbase |
| hsa-miR-195-5p | SEC22C | HITS-CLIP | Tarbase |
| hsa-miR-486-5p | CDC23 | HITS-CLIP | Tarbase |
| hsa-miR-195-5p | AAAS | Microarrays | Tarbase |
| hsa-miR-141-3p | CBX5 | HITS-CLIP | Tarbase |
| hsa-miR-141-3p | CBX5 | HITS-CLIP | Tarbase |
| hsa-miR-141-5p | CBX5 | HITS-CLIP | Tarbase |
| hsa-miR-141-5p | CBX5 | HITS-CLIP | Tarbase |
| hsa-miR-15a-5p | CBX5 | HITS-CLIP | Tarbase |
| hsa-miR-15a-5p | CBX5 | HITS-CLIP | Tarbase |
| hsa-miR-15a-5p | CBX5 | HITS-CLIP | Tarbase |
| hsa-miR-15a-5p | CBX5 | HITS-CLIP | Tarbase |
| hsa-miR-195-5p | CBX5 | HITS-CLIP | Tarbase |
| hsa-miR-195-5p | CBX5 | HITS-CLIP | Tarbase |
| hsa-miR-195-5p | CBX5 | HITS-CLIP | Tarbase |
| hsa-miR-219a-1-3p | CBX5 | HITS-CLIP | Tarbase |
| hsa-miR-195-5p | CBX5 | Microarrays | Tarbase |
| hsa-miR-141-5p | SUCO | HITS-CLIP | Tarbase |
| hsa-miR-141-5p | SUCO | HITS-CLIP | Tarbase |
| hsa-miR-141-5p | SUCO | HITS-CLIP | Tarbase |
| hsa-miR-15a-5p | SUCO | PAR-CLIP | Tarbase |
| hsa-miR-141-3p | MSH2 | PAR-CLIP | Tarbase |
| hsa-miR-141-3p | MSH2 | PAR-CLIP | Tarbase |
| hsa-miR-141-5p | MSH2 | HITS-CLIP | Tarbase |
| hsa-miR-141-3p | MAP3K1 | HITS-CLIP | Tarbase |
| hsa-miR-15a-5p | ARCN1 | HITS-CLIP | Tarbase |
| hsa-miR-15a-5p | ARCN1 | HITS-CLIP | Tarbase |
| hsa-miR-195-5p | ARCN1 | HITS-CLIP | Tarbase |
| hsa-miR-141-5p | EPB41L4B | HITS-CLIP | Tarbase |
| hsa-miR-15a-3p | PSMD5 | HITS-CLIP | Tarbase |
| hsa-miR-15a-5p | PSMD5 | HITS-CLIP | Tarbase |
| hsa-miR-195-5p | PSMD5 | HITS-CLIP | Tarbase |
| hsa-miR-219a-1-3p | NUP188 | HITS-CLIP | Tarbase |
| hsa-miR-3613-5p | NUP188 | PAR-CLIP | Tarbase |
| hsa-miR-141-3p | BTAF1 | HITS-CLIP | Tarbase |
| hsa-miR-15a-5p | BTAF1 | HITS-CLIP | Tarbase |
| hsa-miR-15a-5p | BTAF1 | HITS-CLIP | Tarbase |
| hsa-miR-15a-5p | BTAF1 | HITS-CLIP | Tarbase |
| hsa-miR-15a-5p | BTAF1 | HITS-CLIP | Tarbase |
| hsa-miR-195-5p | BTAF1 | HITS-CLIP | Tarbase |
| hsa-miR-195-5p | BTAF1 | HITS-CLIP | Tarbase |
| hsa-miR-486-5p | BTAF1 | PAR-CLIP | Tarbase |
| hsa-miR-15a-5p | BTAF1 | HITS-CLIP | Tarbase |
| hsa-miR-195-5p | BTAF1 | HITS-CLIP | Tarbase |
| hsa-miR-15a-5p | BTAF1 | Chimeric fragments | Tarbase |
| hsa-miR-141-3p | TDRD1 | HITS-CLIP | Tarbase |
| hsa-miR-15a-5p | BAMBI | HITS-CLIP | Tarbase |
| hsa-miR-195-5p | BAMBI | HITS-CLIP | Tarbase |
| hsa-miR-141-5p | WAC | HITS-CLIP | Tarbase |
| hsa-miR-219a-1-3p | WAC | PAR-CLIP | Tarbase |
| hsa-miR-219a-1-3p | WAC | PAR-CLIP | Tarbase |
| hsa-miR-195-5p | FKBP5 | Microarrays | Tarbase |
| hsa-miR-15a-5p | SRPK1 | PAR-CLIP | Tarbase |
| hsa-miR-15a-5p | BRPF3 | HITS-CLIP | Tarbase |
| hsa-miR-141-3p | HSP90AB1 | HITS-CLIP | Tarbase |
| hsa-miR-141-3p | HSP90AB1 | HITS-CLIP | Tarbase |
| hsa-miR-141-3p | HSP90AB1 | HITS-CLIP | Tarbase |
| hsa-miR-141-5p | HSP90AB1 | HITS-CLIP | Tarbase |
| hsa-miR-141-5p | HSP90AB1 | HITS-CLIP | Tarbase |
| hsa-miR-15a-5p | HSP90AB1 | HITS-CLIP | Tarbase |
| hsa-miR-195-5p | HSP90AB1 | HITS-CLIP | Tarbase |
| hsa-miR-141-5p | CDC5L | HITS-CLIP | Tarbase |
| hsa-miR-141-5p | DSP | HITS-CLIP | Tarbase |
| hsa-miR-486-5p | DSP | HITS-CLIP | Tarbase |
| hsa-miR-486-5p | SIRT1 | HITS-CLIP | Tarbase |
| hsa-miR-141-3p | HNRNPH3 | HITS-CLIP | Tarbase |
| hsa-miR-141-5p | HNRNPH3 | HITS-CLIP | Tarbase |
| hsa-miR-141-5p | HNRNPH3 | HITS-CLIP | Tarbase |
| hsa-miR-15a-5p | IFT74 | HITS-CLIP | Tarbase |
| hsa-miR-195-5p | IFT74 | HITS-CLIP | Tarbase |
| hsa-miR-141-3p | ABL1 | HITS-CLIP | Tarbase |
| hsa-miR-15a-5p | CDC7 | HITS-CLIP | Tarbase |
| hsa-miR-15a-5p | CDC7 | PAR-CLIP | Tarbase |
| hsa-miR-195-5p | CDC7 | HITS-CLIP | Tarbase |
| hsa-miR-195-5p | CDC7 | Microarrays | Tarbase |
| hsa-miR-141-3p | SCD | HITS-CLIP | Tarbase |
| hsa-miR-141-3p | SCD | HITS-CLIP | Tarbase |
| hsa-miR-141-3p | SCD | HITS-CLIP | Tarbase |
| hsa-miR-141-3p | SCD | HITS-CLIP | Tarbase |
| hsa-miR-141-5p | SCD | HITS-CLIP | Tarbase |
| hsa-miR-141-5p | SCD | HITS-CLIP | Tarbase |
| hsa-miR-141-5p | SCD | HITS-CLIP | Tarbase |
| hsa-miR-141-5p | SCD | HITS-CLIP | Tarbase |
| hsa-miR-141-5p | SCD | HITS-CLIP | Tarbase |
| hsa-miR-219a-1-3p | SCD | HITS-CLIP | Tarbase |
| hsa-miR-486-3p | SCD | HITS-CLIP | Tarbase |
| hsa-miR-486-5p | ABLIM1 | PAR-CLIP | Tarbase |
| hsa-miR-15a-5p | ERMP1 | HITS-CLIP | Tarbase |
| hsa-miR-195-5p | ERMP1 | HITS-CLIP | Tarbase |
| hsa-miR-15a-5p | RAB18 | PAR-CLIP | Tarbase |
| hsa-miR-15a-5p | RAB18 | PAR-CLIP | Tarbase |
| hsa-miR-15a-5p | RAB18 | HITS-CLIP | Tarbase |
| hsa-miR-195-5p | RAB18 | HITS-CLIP | Tarbase |
| hsa-miR-195-5p | NRP1 | Microarrays | Tarbase |
| hsa-miR-15a-5p | NRP1 | Chimeric fragments | Tarbase |
| hsa-miR-195-5p | PRTFDC1 | Microarrays | Tarbase |
| hsa-miR-141-5p | PALMD | HITS-CLIP | Tarbase |
| hsa-miR-141-5p | PALMD | HITS-CLIP | Tarbase |
| hsa-miR-141-5p | H2AFY2 | HITS-CLIP | Tarbase |

| | | | |
|---|---|---|---|
| hsa-miR-15a-5p | FBXL19 | HITS-CLIP | Tarbase |
| hsa-miR-195-5p | FBXL19 | HITS-CLIP | Tarbase |
| hsa-miR-195-5p | FBXL19 | HITS-CLIP | Tarbase |
| hsa-miR-3605-3p | FBXL19 | HITS-CLIP | Tarbase |
| hsa-miR-3605-3p | FBXL19 | HITS-CLIP | Tarbase |
| hsa-miR-15a-3p | HSD3B7 | HITS-CLIP | Tarbase |
| hsa-miR-486-5p | SETD1A | PAR-CLIP | Tarbase |
| hsa-miR-15a-3p | CIRBP | HITS-CLIP | Tarbase |
| hsa-miR-15a-3p | CIRBP | HITS-CLIP | Tarbase |
| hsa-miR-15a-5p | CIRBP | HITS-CLIP | Tarbase |
| hsa-miR-15a-5p | CIRBP | HITS-CLIP | Tarbase |
| hsa-miR-195-5p | CIRBP | HITS-CLIP | Tarbase |
| hsa-miR-195-5p | CIRBP | HITS-CLIP | Tarbase |
| hsa-miR-141-5p | HNRNPM | HITS-CLIP | Tarbase |
| hsa-miR-141-5p | HNRNPM | HITS-CLIP | Tarbase |
| hsa-miR-15a-3p | HNRNPM | HITS-CLIP | Tarbase |
| hsa-miR-15a-3p | HNRNPM | HITS-CLIP | Tarbase |
| hsa-miR-15a-3p | HNRNPM | HITS-CLIP | Tarbase |
| hsa-miR-195-5p | MARCH2 | Microarrays | Tarbase |
| hsa-miR-15a-5p | TIMM13 | HITS-CLIP | Tarbase |
| hsa-miR-195-5p | TIMM13 | HITS-CLIP | Tarbase |
| hsa-miR-141-5p | CEP170B | HITS-CLIP | Tarbase |
| hsa-miR-15a-3p | CEP170B | HITS-CLIP | Tarbase |
| hsa-miR-141-5p | MKNK2 | HITS-CLIP | Tarbase |
| hsa-miR-15a-3p | MKNK2 | PAR-CLIP | Tarbase |
| hsa-miR-195-5p | MKNK2 | HITS-CLIP | Tarbase |
| hsa-miR-3605-3p | MKNK2 | HITS-CLIP | Tarbase |
| hsa-miR-3613-3p | MKNK2 | HITS-CLIP | Tarbase |
| hsa-miR-3613-3p | MKNK2 | HITS-CLIP | Tarbase |
| hsa-miR-3613-3p | MKNK2 | PAR-CLIP | Tarbase |
| hsa-miR-486-3p | MKNK2 | PAR-CLIP | Tarbase |
| hsa-miR-486-3p | MKNK2 | PAR-CLIP | Tarbase |
| hsa-miR-486-5p | MKNK2 | PAR-CLIP | Tarbase |
| hsa-miR-195-5p | MKNK2 | Microarrays | Tarbase |
| hsa-miR-141-3p | CRKL | HITS-CLIP | Tarbase |
| hsa-miR-15a-5p | CRKL | HITS-CLIP | Tarbase |
| hsa-miR-15a-5p | CRKL | PAR-CLIP | Tarbase |
| hsa-miR-15a-5p | CRKL | HITS-CLIP | Tarbase |
| hsa-miR-15a-5p | CRKL | HITS-CLIP | Tarbase |
| hsa-miR-195-5p | CRKL | HITS-CLIP | Tarbase |
| hsa-miR-195-5p | CRKL | HITS-CLIP | Tarbase |
| hsa-miR-219a-1-3p | CRKL | HITS-CLIP | Tarbase |
| hsa-miR-15a-5p | CRKL | HITS-CLIP | Tarbase |
| hsa-miR-15a-5p | CRKL | Luciferase Reporter Assay | Tarbase |
| hsa-miR-15a-5p | SPECC1L | HITS-CLIP | Tarbase |
| hsa-miR-15a-5p | SNRPD3 | PAR-CLIP | Tarbase |
| hsa-miR-3613-3p | SLC35E4 | PAR-CLIP | Tarbase |
| hsa-miR-15a-5p | CARD10 | PAR-CLIP | Tarbase |
| hsa-miR-15a-5p | SLC25A1 | HITS-CLIP | Tarbase |
| hsa-miR-15a-5p | SRRD | HITS-CLIP | Tarbase |
| hsa-miR-141-5p | MICALL1 | HITS-CLIP | Tarbase |
| hsa-miR-141-3p | TTC28 | HITS-CLIP | Tarbase |
| hsa-miR-195-5p | CENPM | Microarrays | Tarbase |
| hsa-miR-15a-5p | SEPT3 | HITS-CLIP | Tarbase |
| hsa-miR-195-5p | SEPT3 | HITS-CLIP | Tarbase |
| hsa-miR-15a-3p | KDELR3 | HITS-CLIP | Tarbase |
| hsa-miR-195-5p | KDELR3 | Microarrays | Tarbase |
| hsa-miR-141-5p | DDX17 | HITS-CLIP | Tarbase |
| hsa-miR-141-5p | DDX17 | HITS-CLIP | Tarbase |
| hsa-miR-141-5p | DDX17 | HITS-CLIP | Tarbase |
| hsa-miR-15a-3p | DDX17 | HITS-CLIP | Tarbase |
| hsa-miR-15a-5p | DDX17 | HITS-CLIP | Tarbase |
| hsa-miR-15a-5p | DDX17 | HITS-CLIP | Tarbase |
| hsa-miR-195-5p | DDX17 | HITS-CLIP | Tarbase |
| hsa-miR-141-3p | TCF20 | HITS-CLIP | Tarbase |
| hsa-miR-15a-5p | TCF20 | HITS-CLIP | Tarbase |
| hsa-miR-15a-5p | TCF20 | PAR-CLIP | Tarbase |
| hsa-miR-15a-5p | TCF20 | HITS-CLIP | Tarbase |
| hsa-miR-195-5p | TCF20 | HITS-CLIP | Tarbase |
| hsa-miR-15a-5p | RTCB | HITS-CLIP | Tarbase |
| hsa-miR-195-5p | RTCB | HITS-CLIP | Tarbase |
| hsa-miR-141-3p | JOSD1 | HITS-CLIP | Tarbase |
| hsa-miR-15a-3p | RAB36 | PAR-CLIP | Tarbase |
| hsa-miR-15a-5p | TIMP3 | HITS-CLIP | Tarbase |
| hsa-miR-15a-5p | TIMP3 | HITS-CLIP | Tarbase |
| hsa-miR-195-5p | TIMP3 | HITS-CLIP | Tarbase |
| hsa-miR-15a-3p | SBF1 | HITS-CLIP | Tarbase |
| hsa-miR-195-5p | SUN2 | Microarrays | Tarbase |
| hsa-miR-15a-5p | LMF2 | HITS-CLIP | Tarbase |
| hsa-miR-195-5p | LMF2 | HITS-CLIP | Tarbase |
| hsa-miR-15a-5p | RHBDD3 | HITS-CLIP | Tarbase |
| hsa-miR-15a-5p | RHBDD3 | HITS-CLIP | Tarbase |
| hsa-miR-195-5p | RHBDD3 | HITS-CLIP | Tarbase |
| hsa-miR-195-5p | RHBDD3 | HITS-CLIP | Tarbase |
| hsa-miR-141-5p | AP1B1 | HITS-CLIP | Tarbase |
| hsa-miR-486-5p | AP1B1 | HITS-CLIP | Tarbase |
| hsa-miR-141-3p | HMGXB4 | HITS-CLIP | Tarbase |
| hsa-miR-195-5p | BIK | AGO-IP | Tarbase |
| hsa-miR-195-5p | HMOX1 | Microarrays | Tarbase |
| hsa-miR-141-5p | THOC5 | HITS-CLIP | Tarbase |
| hsa-miR-15a-5p | THOC5 | HITS-CLIP | Tarbase |
| hsa-miR-195-5p | THOC5 | HITS-CLIP | Tarbase |
| hsa-miR-141-5p | MCM5 | HITS-CLIP | Tarbase |
| hsa-miR-195-5p | MCM5 | Microarrays | Tarbase |
| hsa-miR-141-5p | CBX7 | HITS-CLIP | Tarbase |
| hsa-miR-15a-5p | RBFOX2 | HITS-CLIP | Tarbase |
| hsa-miR-195-5p | RBFOX2 | HITS-CLIP | Tarbase |
| hsa-miR-486-3p | RBFOX2 | HITS-CLIP | Tarbase |
| hsa-miR-141-3p | MTMR3 | HITS-CLIP | Tarbase |
| hsa-miR-141-3p | MTMR3 | HITS-CLIP | Tarbase |
| hsa-miR-15a-5p | MTMR3 | HITS-CLIP | Tarbase |
| hsa-miR-195-5p | MTMR3 | HITS-CLIP | Tarbase |
| hsa-miR-195-5p | PNPLA3 | Microarrays | Tarbase |
| hsa-miR-3605-3p | MYH9 | HITS-CLIP | Tarbase |
| hsa-miR-486-3p | MYH9 | HITS-CLIP | Tarbase |
| hsa-miR-486-3p | MYH9 | HITS-CLIP | Tarbase |
| hsa-miR-486-3p | MYH9 | HITS-CLIP | Tarbase |
| hsa-miR-195-5p | MYH9 | Microarrays | Tarbase |
| hsa-miR-195-5p | TXN2 | Microarrays | Tarbase |
| hsa-miR-195-5p | FOXRED2 | Microarrays | Tarbase |
| hsa-miR-141-5p | TNRC6B | HITS-CLIP | Tarbase |
| hsa-miR-15a-5p | TNRC6B | HITS-CLIP | Tarbase |
| hsa-miR-15a-5p | TNRC6B | HITS-CLIP | Tarbase |
| hsa-miR-15a-5p | TNRC6B | PAR-CLIP | Tarbase |
| hsa-miR-15a-5p | TNRC6B | HITS-CLIP | Tarbase |
| hsa-miR-195-5p | TNRC6B | HITS-CLIP | Tarbase |
| hsa-miR-195-5p | TNRC6B | HITS-CLIP | Tarbase |
| hsa-miR-3613-3p | TNRC6B | HITS-CLIP | Tarbase |
| hsa-miR-3613-3p | TNRC6B | PAR-CLIP | Tarbase |
| hsa-miR-15a-5p | FAM118A | HITS-CLIP | Tarbase |
| hsa-miR-195-5p | FAM118A | HITS-CLIP | Tarbase |
| hsa-miR-15a-3p | IL2RB | HITS-CLIP | Tarbase |
| hsa-miR-195-5p | IL2RB | PAR-CLIP | Tarbase |
| hsa-miR-195-5p | IL2RB | PAR-CLIP | Tarbase |
| hsa-miR-195-5p | IL2RB | PAR-CLIP | Tarbase |
| hsa-miR-195-5p | IL2RB | PAR-CLIP | Tarbase |
| hsa-miR-15a-5p | RBX1 | PAR-CLIP | Tarbase |
| hsa-miR-15a-5p | EP300 | HITS-CLIP | Tarbase |
| hsa-miR-15a-5p | EP300 | HITS-CLIP | Tarbase |
| hsa-miR-195-5p | EP300 | HITS-CLIP | Tarbase |
| hsa-miR-3613-3p | EP300 | HITS-CLIP | Tarbase |
| hsa-miR-15a-5p | RANGAP1 | PAR-CLIP | Tarbase |
| hsa-miR-3605-5p | RANGAP1 | HITS-CLIP | Tarbase |
| hsa-miR-486-3p | RANGAP1 | HITS-CLIP | Tarbase |
| hsa-miR-15a-3p | POLR3H | HITS-CLIP | Tarbase |
| hsa-miR-15a-5p | POLR3H | HITS-CLIP | Tarbase |
| hsa-miR-195-5p | POLR3H | HITS-CLIP | Tarbase |
| hsa-miR-141-5p | TRMU | HITS-CLIP | Tarbase |
| hsa-miR-15a-5p | DESI1 | HITS-CLIP | Tarbase |
| hsa-miR-15a-5p | DESI1 | PAR-CLIP | Tarbase |
| hsa-miR-195-5p | DESI1 | HITS-CLIP | Tarbase |
| hsa-miR-486-5p | DESI1 | HITS-CLIP | Tarbase |
| hsa-miR-195-5p | DESI1 | Microarrays | Tarbase |
| hsa-miR-141-3p | KHNYN | HITS-CLIP | Tarbase |
| hsa-miR-15a-5p | KHNYN | HITS-CLIP | Tarbase |
| hsa-miR-195-5p | KHNYN | HITS-CLIP | Tarbase |
| hsa-miR-15a-3p | RBM23 | HITS-CLIP | Tarbase |
| hsa-miR-195-5p | POLE2 | Microarrays | Tarbase |
| hsa-miR-141-5p | NIN | HITS-CLIP | Tarbase |

| | | | |
|---|---|---|---|
| hsa-miR-15a-5p | NIN | PAR-CLIP | Tarbase |
| hsa-miR-15a-5p | NIN | PAR-CLIP | Tarbase |
| hsa-miR-15a-5p | NIN | PAR-CLIP | Tarbase |
| hsa-miR-15a-5p | NIN | PAR-CLIP | Tarbase |
| hsa-miR-15a-5p | NIN | PAR-CLIP | Tarbase |
| hsa-miR-15a-5p | NIN | PAR-CLIP | Tarbase |
| hsa-miR-15a-5p | NIN | PAR-CLIP | Tarbase |
| hsa-miR-15a-5p | NIN | PAR-CLIP | Tarbase |
| hsa-miR-195-5p | NIN | PAR-CLIP | Tarbase |
| hsa-miR-195-5p | NIN | PAR-CLIP | Tarbase |
| hsa-miR-195-5p | NIN | PAR-CLIP | Tarbase |
| hsa-miR-195-5p | NIN | PAR-CLIP | Tarbase |
| hsa-miR-15a-5p | NIN | Chimeric fragments | Tarbase |
| hsa-miR-195-5p | CDKN3 | Microarrays | Tarbase |
| hsa-miR-141-3p | CNIH1 | HITS-CLIP | Tarbase |
| hsa-miR-15a-5p | CNIH1 | HITS-CLIP | Tarbase |
| hsa-miR-195-5p | CNIH1 | HITS-CLIP | Tarbase |
| hsa-miR-195-5p | CNIH1 | HITS-CLIP | Tarbase |
| hsa-miR-195-5p | CNIH1 | HITS-CLIP | Tarbase |
| hsa-miR-486-5p | C14orf105 | HITS-CLIP | Tarbase |
| hsa-miR-141-5p | PSMA3 | HITS-CLIP | Tarbase |
| hsa-miR-15a-5p | VTI1B | PAR-CLIP | Tarbase |
| hsa-miR-15a-5p | VTI1B | HITS-CLIP | Tarbase |
| hsa-miR-195-5p | VTI1B | HITS-CLIP | Tarbase |
| hsa-miR-195-5p | VTI1B | Microarrays | Tarbase |
| hsa-miR-141-5p | TMED8 | HITS-CLIP | Tarbase |
| hsa-miR-15a-5p | TMED8 | HITS-CLIP | Tarbase |
| hsa-miR-3613-3p | TMED8 | PAR-CLIP | Tarbase |
| hsa-miR-195-5p | TMED8 | Microarrays | Tarbase |
| hsa-miR-15a-5p | SPTLC2 | HITS-CLIP | Tarbase |
| hsa-miR-195-5p | SPTLC2 | HITS-CLIP | Tarbase |
| hsa-miR-486-5p | ITPK1 | PAR-CLIP | Tarbase |
| hsa-miR-141-5p | PPM1A | HITS-CLIP | Tarbase |
| hsa-miR-15a-5p | PPM1A | PAR-CLIP | Tarbase |
| hsa-miR-15a-5p | PPM1A | PAR-CLIP | Tarbase |
| hsa-miR-195-3p | PPM1A | HITS-CLIP | Tarbase |
| hsa-miR-219a-1-3p | PPM1A | PAR-CLIP | Tarbase |
| hsa-miR-3613-3p | PPM1A | PAR-CLIP | Tarbase |
| hsa-miR-15a-5p | PPM1A | HITS-CLIP | Tarbase |
| hsa-miR-195-5p | PPM1A | HITS-CLIP | Tarbase |
| hsa-miR-141-3p | SIX4 | HITS-CLIP | Tarbase |
| hsa-miR-195-5p | CEP128 | Microarrays | Tarbase |
| hsa-miR-3613-3p | HIF1A | PAR-CLIP | Tarbase |
| hsa-miR-195-5p | HIF1A | Microarrays | Tarbase |
| hsa-miR-15a-5p | KIAA0247 | HITS-CLIP | Tarbase |
| hsa-miR-195-5p | KIAA0247 | HITS-CLIP | Tarbase |
| hsa-miR-141-3p | EIF5 | HITS-CLIP | Tarbase |
| hsa-miR-141-5p | EIF5 | HITS-CLIP | Tarbase |
| hsa-miR-141-3p | DICER1 | HITS-CLIP | Tarbase |
| hsa-miR-141-5p | DICER1 | HITS-CLIP | Tarbase |
| hsa-miR-141-5p | DICER1 | HITS-CLIP | Tarbase |
| hsa-miR-15a-5p | DICER1 | HITS-CLIP | Tarbase |
| hsa-miR-15a-5p | DICER1 | HITS-CLIP | Tarbase |
| hsa-miR-15a-5p | DICER1 | HITS-CLIP | Tarbase |
| hsa-miR-195-5p | DICER1 | HITS-CLIP | Tarbase |
| hsa-miR-195-5p | DICER1 | HITS-CLIP | Tarbase |
| hsa-miR-15a-5p | ZFYVE21 | PAR-CLIP | Tarbase |
| hsa-miR-141-3p | ZC3H14 | HITS-CLIP | Tarbase |
| hsa-miR-15a-5p | ZC3H14 | PAR-CLIP | Tarbase |
| hsa-miR-486-5p | ZC3H14 | PAR-CLIP | Tarbase |
| hsa-miR-486-5p | ZC3H14 | PAR-CLIP | Tarbase |
| hsa-miR-15a-5p | RPS6KA5 | HITS-CLIP | Tarbase |
| hsa-miR-195-5p | RPS6KA5 | HITS-CLIP | Tarbase |
| hsa-miR-15a-5p | SMEK1 | HITS-CLIP | Tarbase |
| hsa-miR-15a-5p | SMEK1 | PAR-CLIP | Tarbase |
| hsa-miR-195-5p | SMEK1 | HITS-CLIP | Tarbase |
| hsa-miR-15a-5p | SMEK1 | Chimeric fragments | Tarbase |
| hsa-miR-486-3p | PSMB5 | HITS-CLIP | Tarbase |
| hsa-miR-15a-5p | TRIP11 | PAR-CLIP | Tarbase |
| hsa-miR-15a-5p | TRIP11 | Chimeric fragments | Tarbase |
| hsa-miR-141-5p | PABPN1 | HITS-CLIP | Tarbase |
| hsa-miR-3613-3p | ARHGAP5 | HITS-CLIP | Tarbase |
| hsa-miR-15a-5p | ARHGAP5 | HITS-CLIP | Tarbase |
| hsa-miR-195-5p | ARHGAP5 | HITS-CLIP | Tarbase |
| hsa-miR-15a-5p | CINP | HITS-CLIP | Tarbase |
| hsa-miR-15a-5p | DHRS2 | HITS-CLIP | Tarbase |
| hsa-miR-195-5p | DHRS2 | HITS-CLIP | Tarbase |
| hsa-miR-141-3p | CHD8 | HITS-CLIP | Tarbase |
| hsa-miR-15a-5p | CHD8 | HITS-CLIP | Tarbase |
| hsa-miR-486-3p | PCK2 | HITS-CLIP | Tarbase |
| hsa-miR-15a-5p | KIAA0391 | HITS-CLIP | Tarbase |
| hsa-miR-195-5p | KIAA0391 | HITS-CLIP | Tarbase |
| hsa-miR-141-3p | NFKBIA | HITS-CLIP | Tarbase |
| hsa-miR-141-3p | NFKBIA | HITS-CLIP | Tarbase |
| hsa-miR-141-3p | BRMS1L | HITS-CLIP | Tarbase |
| hsa-miR-141-3p | SEC23A | HITS-CLIP | Tarbase |
| hsa-miR-141-3p | SEC23A | HITS-CLIP | Tarbase |
| hsa-miR-141-5p | SEC23A | HITS-CLIP | Tarbase |
| hsa-miR-15a-5p | SEC23A | HITS-CLIP | Tarbase |
| hsa-miR-15a-5p | SEC23A | HITS-CLIP | Tarbase |
| hsa-miR-15a-5p | SEC23A | HITS-CLIP | Tarbase |
| hsa-miR-15a-5p | SEC23A | HITS-CLIP | Tarbase |
| hsa-miR-15a-5p | SEC23A | HITS-CLIP | Tarbase |
| hsa-miR-195-5p | SEC23A | HITS-CLIP | Tarbase |
| hsa-miR-195-5p | SEC23A | HITS-CLIP | Tarbase |
| hsa-miR-195-5p | SEC23A | HITS-CLIP | Tarbase |
| hsa-miR-195-5p | SEC23A | HITS-CLIP | Tarbase |
| hsa-miR-3613-5p | SEC23A | HITS-CLIP | Tarbase |
| hsa-miR-141-5p | PNN | HITS-CLIP | Tarbase |
| hsa-miR-15a-5p | PNN | HITS-CLIP | Tarbase |
| hsa-miR-195-5p | PNN | HITS-CLIP | Tarbase |
| hsa-miR-15a-3p | GSS | HITS-CLIP | Tarbase |
| hsa-miR-141-3p | MMP9 | Western Blot | Tarbase |
| hsa-miR-195-5p | MMP9 | qPCR | Tarbase |
| hsa-miR-486-3p | TRPC4AP | PAR-CLIP | Tarbase |
| hsa-miR-15a-3p | GINS1 | HITS-CLIP | Tarbase |
| hsa-miR-195-5p | GINS1 | Microarrays | Tarbase |
| hsa-miR-15a-5p | GINS1 | Chimeric fragments | Tarbase |
| hsa-miR-15a-5p | ZMYND8 | HITS-CLIP | Tarbase |
| hsa-miR-15a-5p | IFT52 | HITS-CLIP | Tarbase |
| hsa-miR-195-5p | MYBL2 | Microarrays | Tarbase |
| hsa-miR-486-5p | C20orf24 | PAR-CLIP | Tarbase |
| hsa-miR-141-3p | STK4 | HITS-CLIP | Tarbase |
| hsa-miR-15a-5p | STK4 | HITS-CLIP | Tarbase |
| hsa-miR-141-3p | ADNP | HITS-CLIP | Tarbase |
| hsa-miR-141-3p | ADNP | HITS-CLIP | Tarbase |
| hsa-miR-141-3p | ADNP | HITS-CLIP | Tarbase |
| hsa-miR-141-3p | ADNP | HITS-CLIP | Tarbase |
| hsa-miR-141-5p | ADNP | HITS-CLIP | Tarbase |
| hsa-miR-15a-3p | ADNP | HITS-CLIP | Tarbase |
| hsa-miR-15a-5p | ADNP | Chimeric fragments | Tarbase |
| hsa-miR-141-3p | CSTF1 | HITS-CLIP | Tarbase |
| hsa-miR-141-3p | RAE1 | HITS-CLIP | Tarbase |
| hsa-miR-141-5p | DNAJC5 | HITS-CLIP | Tarbase |
| hsa-miR-15a-5p | DNAJC5 | HITS-CLIP | Tarbase |
| hsa-miR-15a-5p | DNAJC5 | HITS-CLIP | Tarbase |
| hsa-miR-15a-5p | DNAJC5 | PAR-CLIP | Tarbase |
| hsa-miR-15a-5p | DNAJC5 | HITS-CLIP | Tarbase |
| hsa-miR-15a-5p | DNAJC5 | HITS-CLIP | Tarbase |
| hsa-miR-195-5p | DNAJC5 | HITS-CLIP | Tarbase |
| hsa-miR-195-5p | DNAJC5 | HITS-CLIP | Tarbase |
| hsa-miR-195-5p | DNAJC5 | HITS-CLIP | Tarbase |
| hsa-miR-141-3p | SLMO2 | HITS-CLIP | Tarbase |
| hsa-miR-141-3p | MRGBP | HITS-CLIP | Tarbase |
| hsa-miR-15a-5p | TCFL5 | HITS-CLIP | Tarbase |
| hsa-miR-195-5p | TCFL5 | HITS-CLIP | Tarbase |
| hsa-miR-141-3p | DIDO1 | HITS-CLIP | Tarbase |
| hsa-miR-3605-5p | GID8 | HITS-CLIP | Tarbase |
| hsa-miR-486-5p | GID8 | PAR-CLIP | Tarbase |
| hsa-miR-195-5p | ARFGAP1 | Microarrays | Tarbase |
| hsa-miR-15a-5p | C20orf27 | HITS-CLIP | Tarbase |
| hsa-miR-141-3p | CDC25B | PAR-CLIP | Tarbase |
| hsa-miR-141-3p | CDC25B | HITS-CLIP | Tarbase |
| hsa-miR-141-3p | CDC25B | HITS-CLIP | Tarbase |
| hsa-miR-141-3p | CDC25B | Western Blot | Tarbase |
| hsa-miR-141-3p | CDC25B | Luciferase Reporter Assay | Tarbase |
| hsa-miR-15a-5p | CDC25B | HITS-CLIP | Tarbase |
| hsa-miR-486-3p | CDC25B | HITS-CLIP | Tarbase |
| hsa-miR-486-3p | CDC25B | HITS-CLIP | Tarbase |

| | | | |
|---|---|---|---|
| hsa-miR-486-3p | CDC25B | HITS-CLIP | Tarbase |
| hsa-miR-195-5p | CDC25B | Microarrays | Tarbase |
| hsa-miR-141-3p | RASSF2 | Luciferase Reporter Assay | Tarbase |
| hsa-miR-141-3p | RASSF2 | Luciferase Reporter Assay | Tarbase |
| hsa-miR-195-5p | RASSF2 | Microarrays | Tarbase |
| hsa-miR-141-3p | CSNK2A1 | HITS-CLIP | Tarbase |
| hsa-miR-141-3p | CSNK2A1 | HITS-CLIP | Tarbase |
| hsa-miR-15a-5p | CSNK2A1 | HITS-CLIP | Tarbase |
| hsa-miR-15a-5p | CSNK2A1 | HITS-CLIP | Tarbase |
| hsa-miR-141-5p | CDS2 | HITS-CLIP | Tarbase |
| hsa-miR-15a-5p | CDS2 | HITS-CLIP | Tarbase |
| hsa-miR-195-5p | CDS2 | HITS-CLIP | Tarbase |
| hsa-miR-15a-5p | CDS2 | HITS-CLIP | Tarbase |
| hsa-miR-195-5p | CDS2 | HITS-CLIP | Tarbase |
| hsa-miR-195-5p | CDS2 | Microarrays | Tarbase |
| hsa-miR-141-5p | SEC23B | HITS-CLIP | Tarbase |
| hsa-miR-195-5p | SEC23B | Microarrays | Tarbase |
| hsa-miR-15a-5p | POFUT1 | PAR-CLIP | Tarbase |
| hsa-miR-15a-3p | SAMHD1 | HITS-CLIP | Tarbase |
| hsa-miR-15a-5p | MAPRE1 | HITS-CLIP | Tarbase |
| hsa-miR-15a-5p | CDK5RAP1 | HITS-CLIP | Tarbase |
| hsa-miR-3605-3p | SNTA1 | HITS-CLIP | Tarbase |
| hsa-miR-15a-3p | E2F1 | HITS-CLIP | Tarbase |
| hsa-miR-195-5p | E2F1 | Microarrays | Tarbase |
| hsa-miR-486-3p | CST3 | HITS-CLIP | Tarbase |
| hsa-miR-195-5p | FAM83D | Microarrays | Tarbase |
| hsa-miR-15a-5p | ADNP2 | HITS-CLIP | Tarbase |
| hsa-miR-15a-5p | ADNP2 | HITS-CLIP | Tarbase |
| hsa-miR-195-5p | ADNP2 | HITS-CLIP | Tarbase |
| hsa-miR-15a-5p | USP14 | HITS-CLIP | Tarbase |
| hsa-miR-15a-5p | USP14 | PAR-CLIP | Tarbase |
| hsa-miR-15a-5p | USP14 | HITS-CLIP | Tarbase |
| hsa-miR-195-5p | USP14 | HITS-CLIP | Tarbase |
| hsa-miR-195-5p | USP14 | HITS-CLIP | Tarbase |
| hsa-miR-15a-5p | USP14 | Chimeric fragments | Tarbase |
| hsa-miR-141-3p | VAPA | HITS-CLIP | Tarbase |
| hsa-miR-15a-5p | VAPA | HITS-CLIP | Tarbase |
| hsa-miR-15a-5p | VAPA | HITS-CLIP | Tarbase |
| hsa-miR-195-5p | VAPA | HITS-CLIP | Tarbase |
| hsa-miR-15a-5p | METTL4 | PAR-CLIP | Tarbase |
| hsa-miR-15a-5p | METTL4 | PAR-CLIP | Tarbase |
| hsa-miR-15a-5p | METTL4 | PAR-CLIP | Tarbase |
| hsa-miR-195-5p | METTL4 | PAR-CLIP | Tarbase |
| hsa-miR-195-5p | METTL4 | PAR-CLIP | Tarbase |
| hsa-miR-141-3p | LPIN2 | HITS-CLIP | Tarbase |
| hsa-miR-15a-5p | MYL12A | HITS-CLIP | Tarbase |
| hsa-miR-195-5p | MYL12A | HITS-CLIP | Tarbase |
| hsa-miR-195-5p | MYL12A | Microarrays | Tarbase |
| hsa-miR-15a-5p | RNMT | PAR-CLIP | Tarbase |
| hsa-miR-15a-5p | RNMT | HITS-CLIP | Tarbase |
| hsa-miR-15a-5p | SMAD7 | PAR-CLIP | Tarbase |
| hsa-miR-15a-5p | SMAD7 | PAR-CLIP | Tarbase |
| hsa-miR-195-5p | SMAD7 | PAR-CLIP | Tarbase |
| hsa-miR-15a-5p | ANKRD12 | PAR-CLIP | Tarbase |
| hsa-miR-15a-5p | ANKRD12 | HITS-CLIP | Tarbase |
| hsa-miR-15a-5p | ANKRD12 | PAR-CLIP | Tarbase |
| hsa-miR-195-5p | ANKRD12 | PAR-CLIP | Tarbase |
| hsa-miR-195-5p | ANKRD12 | HITS-CLIP | Tarbase |
| hsa-miR-15a-5p | ANKRD12 | HITS-CLIP | Tarbase |
| hsa-miR-195-5p | ANKRD12 | HITS-CLIP | Tarbase |
| hsa-miR-15a-5p | MIB1 | HITS-CLIP | Tarbase |
| hsa-miR-15a-5p | MIB1 | HITS-CLIP | Tarbase |
| hsa-miR-15a-5p | MIB1 | PAR-CLIP | Tarbase |
| hsa-miR-15a-5p | MIB1 | HITS-CLIP | Tarbase |
| hsa-miR-195-5p | MIB1 | HITS-CLIP | Tarbase |
| hsa-miR-195-5p | MIB1 | HITS-CLIP | Tarbase |
| hsa-miR-3613-5p | MIB1 | HITS-CLIP | Tarbase |
| hsa-miR-141-5p | PGRMC1 | HITS-CLIP | Tarbase |
| hsa-miR-15a-5p | PGRMC1 | HITS-CLIP | Tarbase |
| hsa-miR-195-5p | PGRMC1 | HITS-CLIP | Tarbase |
| hsa-miR-15a-5p | NKAP | PAR-CLIP | Tarbase |
| hsa-miR-195-5p | NKAP | PAR-CLIP | Tarbase |
| hsa-miR-15a-3p | AMMECR1 | HITS-CLIP | Tarbase |
| hsa-miR-15a-3p | AMMECR1 | PAR-CLIP | Tarbase |
| hsa-miR-486-3p | SUV39H1 | HITS-CLIP | Tarbase |
| hsa-miR-195-5p | SUV39H1 | Microarrays | Tarbase |
| hsa-miR-141-5p | XIAP | HITS-CLIP | Tarbase |
| hsa-miR-15a-5p | XIAP | HITS-CLIP | Tarbase |
| hsa-miR-15a-5p | XIAP | HITS-CLIP | Tarbase |
| hsa-miR-195-5p | XIAP | qPCR | Tarbase |
| hsa-miR-195-5p | XIAP | Western Blot | Tarbase |
| hsa-miR-195-5p | XIAP | HITS-CLIP | Tarbase |
| hsa-miR-15a-5p | XIAP | HITS-CLIP | Tarbase |
| hsa-miR-195-5p | XIAP | HITS-CLIP | Tarbase |
| hsa-miR-3613-3p | STAG2 | HITS-CLIP | Tarbase |
| hsa-miR-141-5p | PLS3 | HITS-CLIP | Tarbase |
| hsa-miR-15a-5p | SMARCA1 | PAR-CLIP | Tarbase |
| hsa-miR-15a-5p | SMARCA1 | PAR-CLIP | Tarbase |
| hsa-miR-195-5p | ASB9 | Microarrays | Tarbase |
| hsa-miR-141-5p | RBBP7 | HITS-CLIP | Tarbase |
| hsa-miR-141-5p | RBBP7 | HITS-CLIP | Tarbase |
| hsa-miR-15a-3p | RBBP7 | HITS-CLIP | Tarbase |
| hsa-miR-141-5p | FMR1 | HITS-CLIP | Tarbase |
| hsa-miR-15a-5p | PIM2 | HITS-CLIP | Tarbase |
| hsa-miR-15a-5p | PIM2 | PAR-CLIP | Tarbase |
| hsa-miR-195-5p | PIM2 | HITS-CLIP | Tarbase |
| hsa-miR-15a-5p | SCML2 | PAR-CLIP | Tarbase |
| hsa-miR-15a-3p | PCSK1N | HITS-CLIP | Tarbase |
| hsa-miR-15a-5p | EMD | HITS-CLIP | Tarbase |
| hsa-miR-15a-5p | EMD | HITS-CLIP | Tarbase |
| hsa-miR-195-5p | EMD | HITS-CLIP | Tarbase |
| hsa-miR-195-5p | EMD | HITS-CLIP | Tarbase |
| hsa-miR-141-3p | PGK1 | HITS-CLIP | Tarbase |
| hsa-miR-141-5p | PGK1 | HITS-CLIP | Tarbase |
| hsa-miR-141-3p | UBL4A | HITS-CLIP | Tarbase |
| hsa-miR-141-3p | UBL4A | HITS-CLIP | Tarbase |
| hsa-miR-486-5p | EEA1 | PAR-CLIP | Tarbase |
| hsa-miR-15a-3p | CDK16 | HITS-CLIP | Tarbase |
| hsa-miR-15a-5p | USP11 | PAR-CLIP | Tarbase |
| hsa-miR-3605-5p | MAGED2 | PAR-CLIP | Tarbase |
| hsa-miR-141-5p | SYTL4 | HITS-CLIP | Tarbase |
| hsa-miR-195-5p | CENPI | Microarrays | Tarbase |
| hsa-miR-15a-5p | GLA | HITS-CLIP | Tarbase |
| hsa-miR-195-5p | ARMCX3 | Microarrays | Tarbase |
| hsa-miR-141-3p | FNDC3A | HITS-CLIP | Tarbase |
| hsa-miR-15a-5p | FNDC3A | PAR-CLIP | Tarbase |
| hsa-miR-15a-5p | FNDC3A | PAR-CLIP | Tarbase |
| hsa-miR-15a-5p | FNDC3A | PAR-CLIP | Tarbase |
| hsa-miR-15a-5p | FNDC3A | PAR-CLIP | Tarbase |
| hsa-miR-15a-5p | FNDC3A | PAR-CLIP | Tarbase |
| hsa-miR-15a-5p | FNDC3A | PAR-CLIP | Tarbase |
| hsa-miR-15a-5p | FNDC3A | PAR-CLIP | Tarbase |
| hsa-miR-15a-5p | FNDC3A | PAR-CLIP | Tarbase |
| hsa-miR-195-5p | FNDC3A | PAR-CLIP | Tarbase |
| hsa-miR-195-5p | FNDC3A | PAR-CLIP | Tarbase |
| hsa-miR-195-5p | FNDC3A | PAR-CLIP | Tarbase |
| hsa-miR-195-5p | FNDC3A | PAR-CLIP | Tarbase |
| hsa-miR-141-3p | KLF5 | Luciferase Reporter Assay | Tarbase |
| hsa-miR-141-3p | KLF5 | Luciferase Reporter Assay | Tarbase |
| hsa-miR-15a-5p | KLF5 | PAR-CLIP | Tarbase |
| hsa-miR-15a-5p | KLF5 | PAR-CLIP | Tarbase |
| hsa-miR-15a-5p | KLF5 | HITS-CLIP | Tarbase |
| hsa-miR-15a-5p | KLF5 | PAR-CLIP | Tarbase |
| hsa-miR-195-5p | KLF5 | PAR-CLIP | Tarbase |
| hsa-miR-195-5p | KLF5 | HITS-CLIP | Tarbase |
| hsa-miR-195-5p | KLF5 | PAR-CLIP | Tarbase |
| hsa-miR-219a-1-3p | ARHGEF7 | PAR-CLIP | Tarbase |
| hsa-miR-15a-5p | FGF9 | PAR-CLIP | Tarbase |
| hsa-miR-141-3p | MRPS31 | PAR-CLIP | Tarbase |
| hsa-miR-141-3p | MRPS31 | PAR-CLIP | Tarbase |
| hsa-miR-141-3p | MRPS31 | PAR-CLIP | Tarbase |
| hsa-miR-141-3p | MRPS31 | PAR-CLIP | Tarbase |
| hsa-miR-141-3p | MRPS31 | PAR-CLIP | Tarbase |
| hsa-miR-141-3p | MRPS31 | PAR-CLIP | Tarbase |
| hsa-miR-195-5p | RGCC | Microarrays | Tarbase |
| hsa-miR-141-5p | DGKH | HITS-CLIP | Tarbase |
| hsa-miR-15a-5p | KATNAL1 | PAR-CLIP | Tarbase |
| hsa-miR-15a-5p | KATNAL1 | PAR-CLIP | Tarbase |
| hsa-miR-195-5p | KATNAL1 | PAR-CLIP | Tarbase |

| | | | |
|---|---|---|---|
| hsa-miR-15a-5p | KATNAL1 | HITS-CLIP | Tarbase |
| hsa-miR-195-5p | KATNAL1 | HITS-CLIP | Tarbase |
| hsa-miR-15a-5p | TSC22D1 | HITS-CLIP | Tarbase |
| hsa-miR-195-5p | TSC22D1 | HITS-CLIP | Tarbase |
| hsa-miR-195-5p | MAPK3 | Microarrays | Tarbase |
| hsa-miR-15a-5p | NFAT5 | HITS-CLIP | Tarbase |
| hsa-miR-3613-5p | NFAT5 | PAR-CLIP | Tarbase |
| hsa-miR-15a-5p | N4BP1 | PAR-CLIP | Tarbase |
| hsa-miR-15a-5p | N4BP1 | PAR-CLIP | Tarbase |
| hsa-miR-15a-5p | N4BP1 | PAR-CLIP | Tarbase |
| hsa-miR-15a-5p | N4BP1 | PAR-CLIP | Tarbase |
| hsa-miR-15a-5p | N4BP1 | HITS-CLIP | Tarbase |
| hsa-miR-15a-5p | N4BP1 | PAR-CLIP | Tarbase |
| hsa-miR-15a-5p | N4BP1 | PAR-CLIP | Tarbase |
| hsa-miR-15a-5p | N4BP1 | PAR-CLIP | Tarbase |
| hsa-miR-15a-5p | N4BP1 | PAR-CLIP | Tarbase |
| hsa-miR-15a-5p | N4BP1 | PAR-CLIP | Tarbase |
| hsa-miR-15a-5p | N4BP1 | PAR-CLIP | Tarbase |
| hsa-miR-15a-5p | N4BP1 | PAR-CLIP | Tarbase |
| hsa-miR-195-5p | N4BP1 | PAR-CLIP | Tarbase |
| hsa-miR-195-5p | N4BP1 | HITS-CLIP | Tarbase |
| hsa-miR-195-5p | N4BP1 | PAR-CLIP | Tarbase |
| hsa-miR-195-5p | N4BP1 | PAR-CLIP | Tarbase |
| hsa-miR-195-5p | N4BP1 | PAR-CLIP | Tarbase |
| hsa-miR-195-5p | N4BP1 | PAR-CLIP | Tarbase |
| hsa-miR-195-5p | N4BP1 | PAR-CLIP | Tarbase |
| hsa-miR-15a-3p | CCL22 | PAR-CLIP | Tarbase |
| hsa-miR-15a-3p | CCL22 | PAR-CLIP | Tarbase |
| hsa-miR-15a-5p | CTCF | HITS-CLIP | Tarbase |
| hsa-miR-3605-3p | POLR2C | PAR-CLIP | Tarbase |
| hsa-miR-3605-5p | CYB5B | HITS-CLIP | Tarbase |
| hsa-miR-195-5p | CCDC113 | Microarrays | Tarbase |
| hsa-miR-15a-5p | NME3 | HITS-CLIP | Tarbase |
| hsa-miR-195-5p | NME3 | HITS-CLIP | Tarbase |
| hsa-miR-486-5p | NME3 | HITS-CLIP | Tarbase |
| hsa-miR-195-5p | NDRG4 | Microarrays | Tarbase |
| hsa-miR-15a-5p | PSMD7 | PAR-CLIP | Tarbase |
| hsa-miR-15a-5p | PSMD7 | PAR-CLIP | Tarbase |
| hsa-miR-195-5p | PSMD7 | PAR-CLIP | Tarbase |
| hsa-miR-141-3p | VAC14 | Luciferase Reporter Assay | Tarbase |
| hsa-miR-141-3p | VAC14 | Luciferase Reporter Assay | Tarbase |
| hsa-miR-15a-5p | SLC7A6 | HITS-CLIP | Tarbase |
| hsa-miR-15a-5p | SLC7A6 | HITS-CLIP | Tarbase |
| hsa-miR-486-5p | SLC7A6 | HITS-CLIP | Tarbase |
| hsa-miR-15a-5p | PLA2G15 | PAR-CLIP | Tarbase |
| hsa-miR-15a-5p | PLA2G15 | HITS-CLIP | Tarbase |
| hsa-miR-195-5p | PLA2G15 | HITS-CLIP | Tarbase |
| hsa-miR-3613-5p | PLA2G15 | PAR-CLIP | Tarbase |
| hsa-miR-195-5p | PLA2G15 | Microarrays | Tarbase |
| hsa-miR-15a-5p | MON1B | HITS-CLIP | Tarbase |
| hsa-miR-195-5p | MON1B | HITS-CLIP | Tarbase |
| hsa-miR-195-3p | USP10 | HITS-CLIP | Tarbase |
| hsa-miR-15a-3p | NME4 | HITS-CLIP | Tarbase |
| hsa-miR-15a-5p | NME4 | HITS-CLIP | Tarbase |
| hsa-miR-195-5p | NME4 | HITS-CLIP | Tarbase |
| hsa-miR-3605-3p | NME4 | HITS-CLIP | Tarbase |
| hsa-miR-141-3p | ABCC1 | HITS-CLIP | Tarbase |
| hsa-miR-195-5p | FOXF1 | Microarrays | Tarbase |
| hsa-miR-141-3p | SLC7A5 | PAR-CLIP | Tarbase |
| hsa-miR-15a-5p | SLC7A5 | HITS-CLIP | Tarbase |
| hsa-miR-195-5p | SLC7A5 | HITS-CLIP | Tarbase |
| hsa-miR-15a-3p | FBXO31 | HITS-CLIP | Tarbase |
| hsa-miR-15a-5p | UBE2I | HITS-CLIP | Tarbase |
| hsa-miR-15a-3p | CAPN15 | HITS-CLIP | Tarbase |
| hsa-miR-141-5p | PIEZO1 | HITS-CLIP | Tarbase |
| hsa-miR-195-5p | PIEZO1 | Microarrays | Tarbase |
| hsa-miR-15a-3p | GSPT1 | HITS-CLIP | Tarbase |
| hsa-miR-15a-5p | GSPT1 | PAR-CLIP | Tarbase |
| hsa-miR-15a-5p | UBFD1 | HITS-CLIP | Tarbase |
| hsa-miR-195-5p | UBFD1 | HITS-CLIP | Tarbase |
| hsa-miR-141-3p | GGA2 | HITS-CLIP | Tarbase |
| hsa-miR-141-5p | GGA2 | HITS-CLIP | Tarbase |
| hsa-miR-15a-5p | GGA2 | PAR-CLIP | Tarbase |
| hsa-miR-141-3p | USP31 | HITS-CLIP | Tarbase |
| hsa-miR-15a-5p | USP31 | PAR-CLIP | Tarbase |
| hsa-miR-15a-5p | USP31 | PAR-CLIP | Tarbase |
| hsa-miR-195-5p | USP31 | PAR-CLIP | Tarbase |
| hsa-miR-15a-5p | USP31 | Chimeric fragments | Tarbase |
| hsa-miR-15a-5p | DNAJA3 | HITS-CLIP | Tarbase |
| hsa-miR-15a-5p | DNAJA3 | HITS-CLIP | Tarbase |
| hsa-miR-195-5p | DNAJA3 | HITS-CLIP | Tarbase |
| hsa-miR-15a-5p | BFAR | HITS-CLIP | Tarbase |
| hsa-miR-195-5p | BFAR | HITS-CLIP | Tarbase |
| hsa-miR-3605-5p | BFAR | HITS-CLIP | Tarbase |
| hsa-miR-15a-5p | SALL1 | HITS-CLIP | Tarbase |
| hsa-miR-15a-5p | SALL1 | HITS-CLIP | Tarbase |
| hsa-miR-15a-5p | SALL1 | HITS-CLIP | Tarbase |
| hsa-miR-15a-5p | SALL1 | PAR-CLIP | Tarbase |
| hsa-miR-195-5p | SALL1 | HITS-CLIP | Tarbase |
| hsa-miR-195-5p | SALL1 | HITS-CLIP | Tarbase |
| hsa-miR-195-5p | SALL1 | HITS-CLIP | Tarbase |
| hsa-miR-195-5p | SALL1 | Microarrays | Tarbase |
| hsa-miR-15a-5p | SALL1 | Chimeric fragments | Tarbase |
| hsa-miR-15a-5p | RBL2 | PAR-CLIP | Tarbase |
| hsa-miR-195-5p | RBL2 | PAR-CLIP | Tarbase |
| hsa-miR-15a-5p | QPRT | HITS-CLIP | Tarbase |
| hsa-miR-15a-5p | MAZ | HITS-CLIP | Tarbase |
| hsa-miR-15a-5p | CDIPT | PAR-CLIP | Tarbase |
| hsa-miR-15a-5p | CDIPT | PAR-CLIP | Tarbase |
| hsa-miR-195-5p | CDIPT | PAR-CLIP | Tarbase |
| hsa-miR-141-3p | TMC5 | HITS-CLIP | Tarbase |
| hsa-miR-15a-5p | HERC1 | HITS-CLIP | Tarbase |
| hsa-miR-141-3p | RAB11A | PAR-CLIP | Tarbase |
| hsa-miR-141-3p | RAB11A | HITS-CLIP | Tarbase |
| hsa-miR-15a-5p | RAB11A | PAR-CLIP | Tarbase |
| hsa-miR-3613-3p | CTSH | HITS-CLIP | Tarbase |
| hsa-miR-141-5p | CD276 | HITS-CLIP | Tarbase |
| hsa-miR-141-5p | CD276 | HITS-CLIP | Tarbase |
| hsa-miR-195-5p | CEMIP | Microarrays | Tarbase |
| hsa-miR-195-5p | HOMER2 | Microarrays | Tarbase |
| hsa-miR-15a-5p | TMEM87A | PAR-CLIP | Tarbase |
| hsa-miR-15a-5p | ZNF106 | HITS-CLIP | Tarbase |
| hsa-miR-141-3p | GABPB1 | HITS-CLIP | Tarbase |
| hsa-miR-15a-5p | DMXL2 | PAR-CLIP | Tarbase |
| hsa-miR-15a-5p | DMXL2 | PAR-CLIP | Tarbase |
| hsa-miR-195-5p | DMXL2 | PAR-CLIP | Tarbase |
| hsa-miR-141-5p | SPG11 | HITS-CLIP | Tarbase |
| hsa-miR-15a-5p | RHOV | HITS-CLIP | Tarbase |
| hsa-miR-195-5p | RHOV | HITS-CLIP | Tarbase |
| hsa-miR-195-5p | OIP5 | Microarrays | Tarbase |
| hsa-miR-141-3p | SGK3 | HITS-CLIP | Tarbase |
| hsa-miR-195-5p | PDGFRL | Microarrays | Tarbase |
| hsa-miR-15a-5p | TRIM35 | PAR-CLIP | Tarbase |
| hsa-miR-15a-5p | TRIM35 | PAR-CLIP | Tarbase |
| hsa-miR-15a-5p | TRIM35 | PAR-CLIP | Tarbase |
| hsa-miR-15a-5p | TRIM35 | PAR-CLIP | Tarbase |
| hsa-miR-195-5p | TRIM35 | PAR-CLIP | Tarbase |
| hsa-miR-195-5p | TRIM35 | PAR-CLIP | Tarbase |
| hsa-miR-15a-5p | TRIM35 | Chimeric fragments | Tarbase |
| hsa-miR-15a-5p | CA2 | HITS-CLIP | Tarbase |
| hsa-miR-195-5p | CA2 | AGO-IP | Tarbase |
| hsa-miR-141-5p | FZD3 | HITS-CLIP | Tarbase |
| hsa-miR-15a-3p | FZD3 | HITS-CLIP | Tarbase |
| hsa-miR-3613-5p | EYA1 | HITS-CLIP | Tarbase |
| hsa-miR-195-5p | EYA1 | Microarrays | Tarbase |
| hsa-miR-141-3p | IMPAD1 | HITS-CLIP | Tarbase |
| hsa-miR-141-3p | IMPAD1 | HITS-CLIP | Tarbase |
| hsa-miR-141-3p | IMPAD1 | HITS-CLIP | Tarbase |
| hsa-miR-195-5p | SFRP1 | Microarrays | Tarbase |
| hsa-miR-141-3p | LAPTM4B | HITS-CLIP | Tarbase |
| hsa-miR-15a-5p | LAPTM4B | HITS-CLIP | Tarbase |
| hsa-miR-3605-5p | UBE2W | HITS-CLIP | Tarbase |
| hsa-miR-15a-5p | IKBKB | PAR-CLIP | Tarbase |
| hsa-miR-15a-5p | IKBKB | PAR-CLIP | Tarbase |
| hsa-miR-15a-5p | IKBKB | PAR-CLIP | Tarbase |
| hsa-miR-195-5p | IKBKB | PAR-CLIP | Tarbase |
| hsa-miR-195-5p | IKBKB | PAR-CLIP | Tarbase |
| hsa-miR-195-5p | PLAT | Microarrays | Tarbase |
| hsa-miR-195-5p | JPH1 | Microarrays | Tarbase |

| | | | |
|---|---|---|---|
| hsa-miR-141-3p | STK3 | Luciferase Reporter Assay | Tarbase |
| hsa-miR-141-3p | STK3 | Luciferase Reporter Assay | Tarbase |
| hsa-miR-141-3p | STK3 | HITS-CLIP | Tarbase |
| hsa-miR-195-5p | GDAP1 | Microarrays | Tarbase |
| hsa-miR-15a-3p | EMC2 | PAR-CLIP | Tarbase |
| hsa-miR-15a-5p | EMC2 | PAR-CLIP | Tarbase |
| hsa-miR-15a-5p | EMC2 | PAR-CLIP | Tarbase |
| hsa-miR-3605-3p | ESRP1 | HITS-CLIP | Tarbase |
| hsa-miR-195-5p | WISP1 | Microarrays | Tarbase |
| hsa-miR-3605-5p | NDRG1 | HITS-CLIP | Tarbase |
| hsa-miR-15a-5p | ARMC1 | HITS-CLIP | Tarbase |
| hsa-miR-15a-5p | ARMC1 | PAR-CLIP | Tarbase |
| hsa-miR-195-5p | ARMC1 | HITS-CLIP | Tarbase |
| hsa-miR-141-3p | TRPS1 | HITS-CLIP | Tarbase |
| hsa-miR-15a-5p | TRPS1 | HITS-CLIP | Tarbase |
| hsa-miR-195-5p | TRPS1 | HITS-CLIP | Tarbase |
| hsa-miR-3613-3p | TRPS1 | PAR-CLIP | Tarbase |
| hsa-miR-486-5p | NCALD | HITS-CLIP | Tarbase |
| hsa-miR-15a-5p | SNX16 | PAR-CLIP | Tarbase |
| hsa-miR-15a-5p | SNX16 | HITS-CLIP | Tarbase |
| hsa-miR-195-5p | SNX16 | HITS-CLIP | Tarbase |
| hsa-miR-141-3p | EEF1D | HITS-CLIP | Tarbase |
| hsa-miR-141-5p | EEF1D | HITS-CLIP | Tarbase |
| hsa-miR-15a-3p | EEF1D | HITS-CLIP | Tarbase |
| hsa-miR-15a-3p | EEF1D | HITS-CLIP | Tarbase |
| hsa-miR-15a-3p | EEF1D | HITS-CLIP | Tarbase |
| hsa-miR-15a-5p | EEF1D | HITS-CLIP | Tarbase |
| hsa-miR-15a-5p | EEF1D | HITS-CLIP | Tarbase |
| hsa-miR-195-5p | EEF1D | HITS-CLIP | Tarbase |
| hsa-miR-195-5p | EEF1D | HITS-CLIP | Tarbase |
| hsa-miR-15a-5p | SQLE | HITS-CLIP | Tarbase |
| hsa-miR-195-5p | SQLE | HITS-CLIP | Tarbase |
| hsa-miR-141-5p | SLC39A14 | HITS-CLIP | Tarbase |
| hsa-miR-15a-5p | SLC39A14 | HITS-CLIP | Tarbase |
| hsa-miR-15a-5p | SLC39A14 | PAR-CLIP | Tarbase |
| hsa-miR-15a-5p | SLC39A14 | PAR-CLIP | Tarbase |
| hsa-miR-15a-5p | SLC39A14 | PAR-CLIP | Tarbase |
| hsa-miR-195-5p | SLC39A14 | HITS-CLIP | Tarbase |
| hsa-miR-486-5p | SLC39A14 | HITS-CLIP | Tarbase |
| hsa-miR-141-3p | MTMR9 | HITS-CLIP | Tarbase |
| hsa-miR-195-5p | GSR | Microarrays | Tarbase |
| hsa-miR-15a-5p | PPP2CB | HITS-CLIP | Tarbase |
| hsa-miR-195-5p | PPP2CB | HITS-CLIP | Tarbase |
| hsa-miR-15a-5p | ERICH1 | HITS-CLIP | Tarbase |
| hsa-miR-195-5p | ERICH1 | HITS-CLIP | Tarbase |
| hsa-miR-141-3p | TUSC3 | HITS-CLIP | Tarbase |
| hsa-miR-3605-3p | TUSC3 | HITS-CLIP | Tarbase |
| hsa-miR-15a-5p | TUSC3 | HITS-CLIP | Tarbase |
| hsa-miR-195-5p | MCM4 | Microarrays | Tarbase |
| hsa-miR-486-5p | BNIP3L | PAR-CLIP | Tarbase |
| hsa-miR-15a-5p | KCNN4 | PAR-CLIP | Tarbase |
| hsa-miR-195-5p | KCNN4 | PAR-CLIP | Tarbase |
| hsa-miR-195-5p | GYS1 | Microarrays | Tarbase |
| hsa-miR-15a-3p | MAP4K1 | PAR-CLIP | Tarbase |
| hsa-miR-141-5p | HNRNPL | HITS-CLIP | Tarbase |
| hsa-miR-141-5p | HNRNPL | HITS-CLIP | Tarbase |
| hsa-miR-195-5p | TUBB4A | Microarrays | Tarbase |
| hsa-miR-3605-3p | SNRNP70 | HITS-CLIP | Tarbase |
| hsa-miR-486-5p | CLPTM1 | HITS-CLIP | Tarbase |
| hsa-miR-15a-3p | PPP1R37 | HITS-CLIP | Tarbase |
| hsa-miR-15a-5p | ARHGEF18 | PAR-CLIP | Tarbase |
| hsa-miR-195-5p | ARHGEF18 | PAR-CLIP | Tarbase |
| hsa-miR-141-5p | ERCC2 | HITS-CLIP | Tarbase |
| hsa-miR-195-5p | RNASEH2A | Microarrays | Tarbase |
| hsa-miR-15a-5p | SF3A2 | PAR-CLIP | Tarbase |
| hsa-miR-15a-5p | TBC1D17 | HITS-CLIP | Tarbase |
| hsa-miR-195-5p | TBC1D17 | HITS-CLIP | Tarbase |
| hsa-miR-195-5p | PTOV1 | HITS-CLIP | Tarbase |
| hsa-miR-141-5p | AES | HITS-CLIP | Tarbase |
| hsa-miR-195-5p | NOVA2 | Microarrays | Tarbase |
| hsa-miR-3605-5p | CCDC61 | PAR-CLIP | Tarbase |
| hsa-miR-141-5p | ASF1B | HITS-CLIP | Tarbase |
| hsa-miR-15a-5p | ASF1B | PAR-CLIP | Tarbase |
| hsa-miR-195-5p | ASF1B | Microarrays | Tarbase |
| hsa-miR-195-5p | SYDE1 | Microarrays | Tarbase |
| hsa-miR-141-3p | CCNE1 | HITS-CLIP | Tarbase |
| hsa-miR-15a-5p | CCNE1 | HITS-CLIP | Tarbase |
| hsa-miR-15a-5p | CCNE1 | HITS-CLIP | Tarbase |
| hsa-miR-15a-5p | CCNE1 | HITS-CLIP | Tarbase |
| hsa-miR-15a-5p | CCNE1 | PAR-CLIP | Tarbase |
| hsa-miR-15a-5p | CCNE1 | HITS-CLIP | Tarbase |
| hsa-miR-15a-5p | CCNE1 | HITS-CLIP | Tarbase |
| hsa-miR-195-5p | CCNE1 | HITS-CLIP | Tarbase |
| hsa-miR-195-5p | CCNE1 | HITS-CLIP | Tarbase |
| hsa-miR-195-5p | CCNE1 | AGO-IP | Tarbase |
| hsa-miR-195-5p | CCNE1 | Luciferase Reporter Assay | Tarbase |
| hsa-miR-195-5p | CCNE1 | Western Blot | Tarbase |
| hsa-miR-195-5p | CCNE1 | Western Blot | Tarbase |
| hsa-miR-195-5p | CCNE1 | HITS-CLIP | Tarbase |
| hsa-miR-195-5p | CCNE1 | HITS-CLIP | Tarbase |
| hsa-miR-141-5p | PDCD5 | HITS-CLIP | Tarbase |
| hsa-miR-141-5p | FBL | HITS-CLIP | Tarbase |
| hsa-miR-15a-5p | DYRK1B | HITS-CLIP | Tarbase |
| hsa-miR-15a-5p | DYRK1B | HITS-CLIP | Tarbase |
| hsa-miR-15a-5p | DYRK1B | HITS-CLIP | Tarbase |
| hsa-miR-15a-5p | DYRK1B | PAR-CLIP | Tarbase |
| hsa-miR-195-5p | DYRK1B | HITS-CLIP | Tarbase |
| hsa-miR-195-5p | DYRK1B | HITS-CLIP | Tarbase |
| hsa-miR-195-5p | DYRK1B | HITS-CLIP | Tarbase |
| hsa-miR-141-3p | GPI | HITS-CLIP | Tarbase |
| hsa-miR-141-3p | GPI | HITS-CLIP | Tarbase |
| hsa-miR-15a-3p | AKT2 | HITS-CLIP | Tarbase |
| hsa-miR-141-5p | FSD1 | HITS-CLIP | Tarbase |
| hsa-miR-3613-5p | CLIP3 | PAR-CLIP | Tarbase |
| hsa-miR-15a-5p | SLC1A5 | HITS-CLIP | Tarbase |
| hsa-miR-15a-5p | SLC1A5 | HITS-CLIP | Tarbase |
| hsa-miR-15a-5p | SLC1A5 | HITS-CLIP | Tarbase |
| hsa-miR-15a-5p | SLC1A5 | HITS-CLIP | Tarbase |
| hsa-miR-195-5p | SLC1A5 | HITS-CLIP | Tarbase |
| hsa-miR-195-5p | SLC1A5 | HITS-CLIP | Tarbase |
| hsa-miR-195-5p | SLC1A5 | HITS-CLIP | Tarbase |
| hsa-miR-195-5p | SLC1A5 | HITS-CLIP | Tarbase |
| hsa-miR-15a-3p | PRKD2 | HITS-CLIP | Tarbase |
| hsa-miR-15a-5p | APLP1 | HITS-CLIP | Tarbase |
| hsa-miR-195-5p | APLP1 | HITS-CLIP | Tarbase |
| hsa-miR-141-5p | HNRNPUL1 | HITS-CLIP | Tarbase |
| hsa-miR-15a-5p | BBC3 | Western Blot | Tarbase |
| hsa-miR-15a-5p | BBC3 | Western Blot | Tarbase |
| hsa-miR-195-5p | RABAC1 | AGO-IP | Tarbase |
| hsa-miR-195-5p | ATP1A3 | Microarrays | Tarbase |
| hsa-miR-3605-3p | MEGF8 | HITS-CLIP | Tarbase |
| hsa-miR-15a-5p | KDELR1 | PAR-CLIP | Tarbase |
| hsa-miR-3613-5p | KDELR1 | PAR-CLIP | Tarbase |
| hsa-miR-3613-5p | KDELR1 | PAR-CLIP | Tarbase |
| hsa-miR-3613-5p | KDELR1 | PAR-CLIP | Tarbase |
| hsa-miR-15a-3p | CYTH2 | HITS-CLIP | Tarbase |
| hsa-miR-15a-3p | GRWD1 | HITS-CLIP | Tarbase |
| hsa-miR-195-5p | LIG1 | Microarrays | Tarbase |
| hsa-miR-141-3p | RAB3D | HITS-CLIP | Tarbase |
| hsa-miR-141-3p | RAB3D | HITS-CLIP | Tarbase |
| hsa-miR-195-5p | TMEM205 | Microarrays | Tarbase |
| hsa-miR-3613-3p | BCAT2 | HITS-CLIP | Tarbase |
| hsa-miR-141-5p | PPP2R1A | HITS-CLIP | Tarbase |
| hsa-miR-15a-5p | PPP2R1A | HITS-CLIP | Tarbase |
| hsa-miR-195-5p | PPP2R1A | HITS-CLIP | Tarbase |
| hsa-miR-15a-3p | TNPO2 | HITS-CLIP | Tarbase |
| hsa-miR-15a-5p | TNPO2 | HITS-CLIP | Tarbase |
| hsa-miR-195-5p | TNPO2 | HITS-CLIP | Tarbase |
| hsa-miR-3605-3p | TNPO2 | HITS-CLIP | Tarbase |
| hsa-miR-15a-5p | WDR83OS | HITS-CLIP | Tarbase |
| hsa-miR-15a-3p | CACNG7 | HITS-CLIP | Tarbase |
| hsa-miR-486-3p | PRPF31 | HITS-CLIP | Tarbase |
| hsa-miR-486-3p | PRPF31 | HITS-CLIP | Tarbase |
| hsa-miR-195-5p | RAB3A | Microarrays | Tarbase |
| hsa-miR-15a-5p | ISYNA1 | HITS-CLIP | Tarbase |
| hsa-miR-15a-5p | ISYNA1 | HITS-CLIP | Tarbase |
| hsa-miR-15a-5p | ISYNA1 | HITS-CLIP | Tarbase |
| hsa-miR-15a-5p | ISYNA1 | HITS-CLIP | Tarbase |
| hsa-miR-15a-5p | ISYNA1 | PAR-CLIP | Tarbase |

| | | | |
|---|---|---|---|
| hsa-miR-195-5p | ISYNA1 | HITS-CLIP | Tarbase |
| hsa-miR-195-5p | ISYNA1 | HITS-CLIP | Tarbase |
| hsa-miR-195-5p | ISYNA1 | HITS-CLIP | Tarbase |
| hsa-miR-195-5p | ISYNA1 | HITS-CLIP | Tarbase |
| hsa-miR-15a-5p | ELL | PAR-CLIP | Tarbase |
| hsa-miR-15a-5p | ELL | PAR-CLIP | Tarbase |
| hsa-miR-3613-3p | USF2 | Chimeric fragments | Tarbase |
| hsa-miR-15a-5p | LSR | PAR-CLIP | Tarbase |
| hsa-miR-15a-3p | FKBP8 | HITS-CLIP | Tarbase |
| hsa-miR-15a-3p | ERF | HITS-CLIP | Tarbase |
| hsa-miR-195-5p | SIPA1L3 | HITS-CLIP | Tarbase |
| hsa-miR-195-5p | ZNF85 | Microarrays | Tarbase |
| hsa-miR-15a-5p | AVL9 | HITS-CLIP | Tarbase |
| hsa-miR-15a-5p | AVL9 | HITS-CLIP | Tarbase |
| hsa-miR-195-5p | AVL9 | Microarrays | Tarbase |
| hsa-miR-15a-5p | CDK6 | HITS-CLIP | Tarbase |
| hsa-miR-15a-5p | CDK6 | HITS-CLIP | Tarbase |
| hsa-miR-15a-5p | CDK6 | HITS-CLIP | Tarbase |
| hsa-miR-195-5p | CDK6 | HITS-CLIP | Tarbase |
| hsa-miR-195-5p | CDK6 | Luciferase Reporter Assay | Tarbase |
| hsa-miR-195-5p | CDK6 | Western Blot | Tarbase |
| hsa-miR-195-5p | CDK6 | Western Blot | Tarbase |
| hsa-miR-195-5p | CDK6 | HITS-CLIP | Tarbase |
| hsa-miR-195-5p | CDK6 | Luciferase Reporter Assay | Tarbase |
| hsa-miR-195-5p | CDK6 | qPCR | Tarbase |
| hsa-miR-195-5p | CDK6 | Western Blot | Tarbase |
| hsa-miR-3613-5p | CDK6 | PAR-CLIP | Tarbase |
| hsa-miR-195-5p | CDK6 | Luciferase Reporter Assay | Tarbase |
| hsa-miR-141-5p | DNAJC2 | HITS-CLIP | Tarbase |
| hsa-miR-195-5p | TFPI2 | Microarrays | Tarbase |
| hsa-miR-15a-5p | TWISTNB | PAR-CLIP | Tarbase |
| hsa-miR-195-5p | TWISTNB | PAR-CLIP | Tarbase |
| hsa-miR-141-3p | ITGB8 | HITS-CLIP | Tarbase |
| hsa-miR-486-5p | ITGB8 | PAR-CLIP | Tarbase |
| hsa-miR-486-5p | ITGB8 | PAR-CLIP | Tarbase |
| hsa-miR-15a-5p | HBP1 | HITS-CLIP | Tarbase |
| hsa-miR-15a-5p | HBP1 | HITS-CLIP | Tarbase |
| hsa-miR-195-5p | HBP1 | HITS-CLIP | Tarbase |
| hsa-miR-486-5p | HBP1 | HITS-CLIP | Tarbase |
| hsa-miR-3613-3p | CBLL1 | PAR-CLIP | Tarbase |
| hsa-miR-3613-3p | DFNA5 | PAR-CLIP | Tarbase |
| hsa-miR-141-3p | H2AFV | HITS-CLIP | Tarbase |
| hsa-miR-141-3p | H2AFV | HITS-CLIP | Tarbase |
| hsa-miR-15a-5p | H2AFV | HITS-CLIP | Tarbase |
| hsa-miR-195-5p | H2AFV | HITS-CLIP | Tarbase |
| hsa-miR-141-3p | CAV2 | HITS-CLIP | Tarbase |
| hsa-miR-195-5p | CAV2 | Microarrays | Tarbase |
| hsa-miR-141-3p | MET | HITS-CLIP | Tarbase |
| hsa-miR-486-5p | MET | HITS-CLIP | Tarbase |
| hsa-miR-141-3p | LMBR1 | HITS-CLIP | Tarbase |
| hsa-miR-15a-5p | HOXA3 | Chimeric fragments | Tarbase |
| hsa-miR-141-3p | LFNG | HITS-CLIP | Tarbase |
| hsa-miR-15a-5p | LFNG | HITS-CLIP | Tarbase |
| hsa-miR-15a-5p | LFNG | HITS-CLIP | Tarbase |
| hsa-miR-15a-5p | LFNG | HITS-CLIP | Tarbase |
| hsa-miR-195-5p | LFNG | HITS-CLIP | Tarbase |
| hsa-miR-195-5p | HIBADH | Microarrays | Tarbase |
| hsa-miR-141-5p | TAX1BP1 | HITS-CLIP | Tarbase |
| hsa-miR-141-5p | TAX1BP1 | HITS-CLIP | Tarbase |
| hsa-miR-3605-3p | CHN2 | HITS-CLIP | Tarbase |
| hsa-miR-195-5p | CHN2 | Microarrays | Tarbase |
| hsa-miR-219a-1-3p | GRB10 | HITS-CLIP | Tarbase |
| hsa-miR-141-3p | PLEKHA8 | HITS-CLIP | Tarbase |
| hsa-miR-141-3p | GARS | HITS-CLIP | Tarbase |
| hsa-miR-141-5p | GARS | HITS-CLIP | Tarbase |
| hsa-miR-141-5p | GARS | HITS-CLIP | Tarbase |
| hsa-miR-15a-5p | GARS | PAR-CLIP | Tarbase |
| hsa-miR-15a-5p | GARS | HITS-CLIP | Tarbase |
| hsa-miR-195-5p | GARS | HITS-CLIP | Tarbase |
| hsa-miR-195-5p | CASP2 | Microarrays | Tarbase |
| hsa-miR-15a-5p | NPTX2 | HITS-CLIP | Tarbase |
| hsa-miR-15a-5p | ZKSCAN1 | HITS-CLIP | Tarbase |
| hsa-miR-3613-5p | ZKSCAN1 | PAR-CLIP | Tarbase |
| hsa-miR-141-5p | EIF3B | HITS-CLIP | Tarbase |
| hsa-miR-3605-3p | SNX8 | HITS-CLIP | Tarbase |
| hsa-miR-15a-5p | USP42 | PAR-CLIP | Tarbase |
| hsa-miR-15a-5p | USP42 | HITS-CLIP | Tarbase |
| hsa-miR-15a-5p | USP42 | PAR-CLIP | Tarbase |
| hsa-miR-195-5p | USP42 | HITS-CLIP | Tarbase |
| hsa-miR-195-5p | USP42 | PAR-CLIP | Tarbase |
| hsa-miR-195-5p | SERPINE1 | Microarrays | Tarbase |
| hsa-miR-141-3p | C1GALT1 | HITS-CLIP | Tarbase |
| hsa-miR-15a-3p | PLOD3 | HITS-CLIP | Tarbase |
| hsa-miR-141-3p | GLCCI1 | HITS-CLIP | Tarbase |
| hsa-miR-195-5p | EZH2 | Microarrays | Tarbase |
| hsa-miR-195-5p | ANKMY2 | Microarrays | Tarbase |
| hsa-miR-195-5p | TSPAN13 | Microarrays | Tarbase |
| hsa-miR-195-5p | GLI3 | Microarrays | Tarbase |
| hsa-miR-3605-3p | URGCP | HITS-CLIP | Tarbase |
| hsa-miR-15a-5p | TMEM248 | HITS-CLIP | Tarbase |
| hsa-miR-141-3p | RHEB | HITS-CLIP | Tarbase |
| hsa-miR-141-3p | RHEB | HITS-CLIP | Tarbase |
| hsa-miR-141-3p | RHEB | HITS-CLIP | Tarbase |
| hsa-miR-195-5p | SLC1A1 | Microarrays | Tarbase |
| hsa-miR-15a-5p | FKTN | HITS-CLIP | Tarbase |
| hsa-miR-141-5p | SPIN1 | HITS-CLIP | Tarbase |
| hsa-miR-15a-5p | SPIN1 | HITS-CLIP | Tarbase |
| hsa-miR-15a-5p | TMEM245 | HITS-CLIP | Tarbase |
| hsa-miR-15a-5p | TMEM245 | HITS-CLIP | Tarbase |
| hsa-miR-15a-5p | TMEM245 | HITS-CLIP | Tarbase |
| hsa-miR-195-5p | TMEM245 | HITS-CLIP | Tarbase |
| hsa-miR-195-5p | TMEM245 | HITS-CLIP | Tarbase |
| hsa-miR-195-5p | TMEM245 | HITS-CLIP | Tarbase |
| hsa-miR-15a-5p | TMEM245 | HITS-CLIP | Tarbase |
| hsa-miR-195-5p | TMEM245 | HITS-CLIP | Tarbase |
| hsa-miR-195-5p | TMEM245 | Microarrays | Tarbase |
| hsa-miR-141-5p | PRUNE2 | HITS-CLIP | Tarbase |
| hsa-miR-15a-5p | MEGF9 | HITS-CLIP | Tarbase |
| hsa-miR-15a-5p | MEGF9 | HITS-CLIP | Tarbase |
| hsa-miR-195-5p | MEGF9 | HITS-CLIP | Tarbase |
| hsa-miR-195-5p | TGFBR1 | Microarrays | Tarbase |
| hsa-miR-195-5p | C5 | Microarrays | Tarbase |
| hsa-miR-15a-5p | TLE4 | HITS-CLIP | Tarbase |
| hsa-miR-15a-5p | TLE4 | PAR-CLIP | Tarbase |
| hsa-miR-195-5p | TLE4 | HITS-CLIP | Tarbase |
| hsa-miR-15a-5p | TLE4 | HITS-CLIP | Tarbase |
| hsa-miR-195-5p | TLE4 | HITS-CLIP | Tarbase |
| hsa-miR-15a-3p | LHX6 | HITS-CLIP | Tarbase |
| hsa-miR-15a-5p | DNM1 | HITS-CLIP | Tarbase |
| hsa-miR-195-5p | DNM1 | HITS-CLIP | Tarbase |
| hsa-miR-15a-5p | CDC37L1 | PAR-CLIP | Tarbase |
| hsa-miR-15a-5p | CDC37L1 | PAR-CLIP | Tarbase |
| hsa-miR-195-5p | CDC37L1 | PAR-CLIP | Tarbase |
| hsa-miR-15a-5p | TBC1D13 | HITS-CLIP | Tarbase |
| hsa-miR-15a-5p | TBC1D13 | PAR-CLIP | Tarbase |
| hsa-miR-15a-5p | TBC1D13 | HITS-CLIP | Tarbase |
| hsa-miR-195-5p | TBC1D13 | HITS-CLIP | Tarbase |
| hsa-miR-195-5p | TBC1D13 | HITS-CLIP | Tarbase |
| hsa-miR-195-5p | TBC1D13 | Microarrays | Tarbase |
| hsa-miR-141-3p | RIC1 | HITS-CLIP | Tarbase |
| hsa-miR-15a-5p | RIC1 | PAR-CLIP | Tarbase |
| hsa-miR-15a-5p | RIC1 | HITS-CLIP | Tarbase |
| hsa-miR-15a-5p | RIC1 | HITS-CLIP | Tarbase |
| hsa-miR-15a-5p | RIC1 | PAR-CLIP | Tarbase |
| hsa-miR-195-5p | RIC1 | PAR-CLIP | Tarbase |
| hsa-miR-195-5p | RIC1 | HITS-CLIP | Tarbase |
| hsa-miR-195-5p | RIC1 | HITS-CLIP | Tarbase |
| hsa-miR-3613-3p | RIC1 | PAR-CLIP | Tarbase |
| hsa-miR-15a-5p | KANK1 | PAR-CLIP | Tarbase |
| hsa-miR-3605-5p | KANK1 | HITS-CLIP | Tarbase |
| hsa-miR-15a-5p | KANK1 | HITS-CLIP | Tarbase |
| hsa-miR-195-5p | KANK1 | HITS-CLIP | Tarbase |
| hsa-miR-15a-5p | ELAVL2 | HITS-CLIP | Tarbase |
| hsa-miR-195-5p | ELAVL2 | HITS-CLIP | Tarbase |
| hsa-miR-15a-5p | CA9 | HITS-CLIP | Tarbase |
| hsa-miR-195-5p | CA9 | HITS-CLIP | Tarbase |
| hsa-miR-195-5p | CA9 | Microarrays | Tarbase |
| hsa-miR-141-5p | EDF1 | HITS-CLIP | Tarbase |
| hsa-miR-195-5p | GLIS3 | Microarrays | Tarbase |

| | | | |
|---|---|---|---|
| hsa-miR-195-5p | BAG1 | Microarrays | Tarbase |
| hsa-miR-15a-3p | RAPGEF1 | HITS-CLIP | Tarbase |
| hsa-miR-15a-5p | RAPGEF1 | HITS-CLIP | Tarbase |
| hsa-miR-195-5p | RAPGEF1 | HITS-CLIP | Tarbase |
| hsa-miR-15a-3p | NPDC1 | HITS-CLIP | Tarbase |
| hsa-miR-3605-3p | NPDC1 | HITS-CLIP | Tarbase |
| hsa-miR-141-3p | SETX | HITS-CLIP | Tarbase |
| hsa-miR-141-3p | SETX | HITS-CLIP | Tarbase |
| hsa-miR-15a-5p | SETX | HITS-CLIP | Tarbase |
| hsa-miR-195-5p | SETX | HITS-CLIP | Tarbase |
| hsa-miR-15a-5p | SETX | Chimeric fragments | Tarbase |
| hsa-miR-195-5p | SHB | Microarrays | Tarbase |
| hsa-miR-15a-5p | UBE2R2 | HITS-CLIP | Tarbase |
| hsa-miR-195-5p | UBE2R2 | HITS-CLIP | Tarbase |
| hsa-miR-15a-5p | ABHD17B | HITS-CLIP | Tarbase |
| hsa-miR-15a-5p | ABHD17B | HITS-CLIP | Tarbase |
| hsa-miR-15a-5p | ABHD17B | HITS-CLIP | Tarbase |
| hsa-miR-195-5p | ABHD17B | HITS-CLIP | Tarbase |
| hsa-miR-195-5p | ABHD17B | HITS-CLIP | Tarbase |
| hsa-miR-195-5p | ABHD17B | HITS-CLIP | Tarbase |
| hsa-miR-141-3p | ZFAND5 | HITS-CLIP | Tarbase |
| hsa-miR-141-5p | CCNJ | HITS-CLIP | Tarbase |
| hsa-miR-15a-5p | CCNJ | PAR-CLIP | Tarbase |
| hsa-miR-141-3p | GATA3 | HITS-CLIP | Tarbase |
| hsa-miR-141-3p | GATA3 | HITS-CLIP | Tarbase |
| hsa-miR-15a-5p | GATA3 | PAR-CLIP | Tarbase |
| hsa-miR-195-5p | GATA3 | PAR-CLIP | Tarbase |
| hsa-miR-195-5p | ATRNL1 | Microarrays | Tarbase |
| hsa-miR-15a-5p | DNMBP | HITS-CLIP | Tarbase |
| hsa-miR-486-3p | DNMBP | PAR-CLIP | Tarbase |
| hsa-miR-15a-5p | RAB11FIP2 | HITS-CLIP | Tarbase |
| hsa-miR-3613-5p | RAB11FIP2 | PAR-CLIP | Tarbase |
| hsa-miR-141-3p | CXCL12 | HITS-CLIP | Tarbase |
| hsa-miR-15a-3p | ERLIN1 | HITS-CLIP | Tarbase |
| hsa-miR-15a-5p | ERLIN1 | PAR-CLIP | Tarbase |
| hsa-miR-195-5p | ERLIN1 | PAR-CLIP | Tarbase |
| hsa-miR-195-5p | ERLIN1 | Microarrays | Tarbase |
| hsa-miR-15a-5p | EIF3A | HITS-CLIP | Tarbase |
| hsa-miR-195-5p | EIF3A | HITS-CLIP | Tarbase |
| hsa-miR-15a-5p | MAPK8 | HITS-CLIP | Tarbase |
| hsa-miR-195-5p | MAPK8 | HITS-CLIP | Tarbase |
| hsa-miR-141-5p | SEC23IP | HITS-CLIP | Tarbase |
| hsa-miR-141-5p | SEC23IP | HITS-CLIP | Tarbase |
| hsa-miR-15a-5p | SEC23IP | HITS-CLIP | Tarbase |
| hsa-miR-15a-5p | SEC23IP | HITS-CLIP | Tarbase |
| hsa-miR-195-5p | SEC23IP | HITS-CLIP | Tarbase |
| hsa-miR-3613-3p | ATE1 | PAR-CLIP | Tarbase |
| hsa-miR-15a-5p | PLEKHA1 | HITS-CLIP | Tarbase |
| hsa-miR-15a-5p | PLEKHA1 | PAR-CLIP | Tarbase |
| hsa-miR-195-5p | PLEKHA1 | HITS-CLIP | Tarbase |
| hsa-miR-15a-5p | UNC5B | PAR-CLIP | Tarbase |
| hsa-miR-141-5p | PPP3CB | HITS-CLIP | Tarbase |
| hsa-miR-141-3p | CCSER2 | HITS-CLIP | Tarbase |
| hsa-miR-486-3p | ACTA2 | HITS-CLIP | Tarbase |
| hsa-miR-141-5p | LIPA | HITS-CLIP | Tarbase |
| hsa-miR-195-5p | SFXN3 | Microarrays | Tarbase |
| hsa-miR-141-3p | TNKS2 | HITS-CLIP | Tarbase |
| hsa-miR-141-3p | TNKS2 | HITS-CLIP | Tarbase |
| hsa-miR-15a-5p | TNKS2 | HITS-CLIP | Tarbase |
| hsa-miR-195-5p | TNKS2 | HITS-CLIP | Tarbase |
| hsa-miR-219a-1-3p | TNKS2 | HITS-CLIP | Tarbase |
| hsa-miR-15a-5p | GBF1 | HITS-CLIP | Tarbase |
| hsa-miR-15a-5p | GBF1 | HITS-CLIP | Tarbase |
| hsa-miR-15a-5p | GBF1 | HITS-CLIP | Tarbase |
| hsa-miR-15a-5p | GBF1 | HITS-CLIP | Tarbase |
| hsa-miR-15a-5p | GBF1 | HITS-CLIP | Tarbase |
| hsa-miR-195-5p | GBF1 | HITS-CLIP | Tarbase |
| hsa-miR-195-5p | GBF1 | HITS-CLIP | Tarbase |
| hsa-miR-195-5p | GBF1 | HITS-CLIP | Tarbase |
| hsa-miR-195-5p | GBF1 | HITS-CLIP | Tarbase |
| hsa-miR-15a-5p | GBF1 | HITS-CLIP | Tarbase |
| hsa-miR-195-5p | GBF1 | HITS-CLIP | Tarbase |
| hsa-miR-15a-3p | ARHGAP21 | HITS-CLIP | Tarbase |
| hsa-miR-141-3p | CPEB3 | HITS-CLIP | Tarbase |
| hsa-miR-15a-5p | CPEB3 | PAR-CLIP | Tarbase |
| hsa-miR-15a-5p | CPEB3 | PAR-CLIP | Tarbase |
| hsa-miR-195-5p | CPEB3 | PAR-CLIP | Tarbase |
| hsa-miR-3605-3p | CPEB3 | HITS-CLIP | Tarbase |
| hsa-miR-15a-5p | CPEB3 | Chimeric fragments | Tarbase |
| hsa-miR-15a-5p | ANKRD26 | HITS-CLIP | Tarbase |
| hsa-miR-195-5p | ANKRD26 | HITS-CLIP | Tarbase |
| hsa-miR-15a-5p | ACBD5 | HITS-CLIP | Tarbase |
| hsa-miR-15a-5p | ACBD5 | HITS-CLIP | Tarbase |
| hsa-miR-15a-5p | ACBD5 | HITS-CLIP | Tarbase |
| hsa-miR-195-5p | ACBD5 | HITS-CLIP | Tarbase |
| hsa-miR-195-5p | ACBD5 | HITS-CLIP | Tarbase |
| hsa-miR-3613-5p | ACBD5 | PAR-CLIP | Tarbase |
| hsa-miR-15a-5p | LARP4B | HITS-CLIP | Tarbase |
| hsa-miR-195-5p | LARP4B | HITS-CLIP | Tarbase |
| hsa-miR-3613-3p | LARP4B | PAR-CLIP | Tarbase |
| hsa-miR-141-5p | EDRF1 | HITS-CLIP | Tarbase |
| hsa-miR-141-3p | MTPAP | PAR-CLIP | Tarbase |
| hsa-miR-195-5p | DKK1 | Microarrays | Tarbase |
| hsa-miR-486-3p | GLRX3 | HITS-CLIP | Tarbase |
| hsa-miR-141-3p | FAM208B | HITS-CLIP | Tarbase |
| hsa-miR-15a-5p | FAM208B | PAR-CLIP | Tarbase |
| hsa-miR-15a-5p | FAM208B | HITS-CLIP | Tarbase |
| hsa-miR-486-5p | FAM208B | HITS-CLIP | Tarbase |
| hsa-miR-141-5p | SMC3 | HITS-CLIP | Tarbase |
| hsa-miR-15a-5p | SMC3 | PAR-CLIP | Tarbase |
| hsa-miR-195-5p | SMC3 | PAR-CLIP | Tarbase |
| hsa-miR-15a-5p | SHOC2 | Chimeric fragments | Tarbase |
| hsa-miR-486-5p | TFAM | PAR-CLIP | Tarbase |
| hsa-miR-3613-5p | CUL2 | HITS-CLIP | Tarbase |
| hsa-miR-141-5p | CCNY | HITS-CLIP | Tarbase |
| hsa-miR-15a-5p | CCNY | Chimeric fragments | Tarbase |
| hsa-miR-195-5p | UBE2S | Microarrays | Tarbase |
| hsa-miR-3613-5p | RPL28 | HITS-CLIP | Tarbase |
| hsa-miR-486-5p | RPL28 | PAR-CLIP | Tarbase |
| hsa-miR-15a-5p | PPIF | HITS-CLIP | Tarbase |
| hsa-miR-15a-5p | PPIF | HITS-CLIP | Tarbase |
| hsa-miR-195-5p | PPIF | Microarrays | Tarbase |
| hsa-miR-141-3p | TSPAN14 | HITS-CLIP | Tarbase |
| hsa-miR-15a-5p | TSPAN14 | HITS-CLIP | Tarbase |
| hsa-miR-141-5p | NUFIP2 | HITS-CLIP | Tarbase |
| hsa-miR-141-5p | NUFIP2 | HITS-CLIP | Tarbase |
| hsa-miR-141-5p | NUFIP2 | HITS-CLIP | Tarbase |
| hsa-miR-15a-5p | NUFIP2 | HITS-CLIP | Tarbase |
| hsa-miR-15a-5p | NUFIP2 | HITS-CLIP | Tarbase |
| hsa-miR-15a-5p | NUFIP2 | PAR-CLIP | Tarbase |
| hsa-miR-15a-5p | NUFIP2 | HITS-CLIP | Tarbase |
| hsa-miR-15a-5p | NUFIP2 | PAR-CLIP | Tarbase |
| hsa-miR-15a-5p | NUFIP2 | HITS-CLIP | Tarbase |
| hsa-miR-15a-5p | NUFIP2 | PAR-CLIP | Tarbase |
| hsa-miR-15a-5p | NUFIP2 | HITS-CLIP | Tarbase |
| hsa-miR-15a-5p | NUFIP2 | HITS-CLIP | Tarbase |
| hsa-miR-15a-5p | NUFIP2 | HITS-CLIP | Tarbase |
| hsa-miR-195-5p | NUFIP2 | HITS-CLIP | Tarbase |
| hsa-miR-195-5p | NUFIP2 | HITS-CLIP | Tarbase |
| hsa-miR-195-5p | NUFIP2 | HITS-CLIP | Tarbase |
| hsa-miR-195-5p | NUFIP2 | HITS-CLIP | Tarbase |
| hsa-miR-195-5p | NUFIP2 | HITS-CLIP | Tarbase |
| hsa-miR-3613-5p | NUFIP2 | PAR-CLIP | Tarbase |
| hsa-miR-15a-5p | NUFIP2 | HITS-CLIP | Tarbase |
| hsa-miR-195-5p | NUFIP2 | HITS-CLIP | Tarbase |
| hsa-miR-15a-5p | NUFIP2 | Chimeric fragments | Tarbase |
| hsa-miR-15a-3p | GIT1 | HITS-CLIP | Tarbase |
| hsa-miR-15a-3p | GIT1 | HITS-CLIP | Tarbase |
| hsa-miR-141-5p | RPL19 | HITS-CLIP | Tarbase |
| hsa-miR-15a-5p | RPL19 | HITS-CLIP | Tarbase |
| hsa-miR-195-5p | RPL19 | HITS-CLIP | Tarbase |
| hsa-miR-141-3p | CSF3 | HITS-CLIP | Tarbase |
| hsa-miR-141-5p | PSMD3 | HITS-CLIP | Tarbase |
| hsa-miR-15a-5p | RAPGEFL1 | HITS-CLIP | Tarbase |
| hsa-miR-195-5p | RGS9 | Microarrays | Tarbase |
| hsa-miR-15a-5p | RNF43 | PAR-CLIP | Tarbase |
| hsa-miR-15a-5p | RNF43 | HITS-CLIP | Tarbase |
| hsa-miR-195-5p | RNF43 | HITS-CLIP | Tarbase |
| hsa-miR-15a-5p | RAD51C | HITS-CLIP | Tarbase |

| | | | |
|---|---|---|---|
| hsa-miR-195-5p | RAD51C | HITS-CLIP | Tarbase |
| hsa-miR-195-5p | SEPT4 | Microarrays | Tarbase |
| hsa-miR-15a-5p | TRIM37 | HITS-CLIP | Tarbase |
| hsa-miR-15a-5p | TRIM37 | HITS-CLIP | Tarbase |
| hsa-miR-195-5p | TRIM37 | HITS-CLIP | Tarbase |
| hsa-miR-15a-5p | TRIM37 | HITS-CLIP | Tarbase |
| hsa-miR-195-5p | TRIM37 | HITS-CLIP | Tarbase |
| hsa-miR-141-3p | DHX40 | HITS-CLIP | Tarbase |
| hsa-miR-141-5p | KPNB1 | HITS-CLIP | Tarbase |
| hsa-miR-15a-3p | KPNB1 | HITS-CLIP | Tarbase |
| hsa-miR-15a-5p | KPNB1 | HITS-CLIP | Tarbase |
| hsa-miR-15a-5p | KPNB1 | HITS-CLIP | Tarbase |
| hsa-miR-15a-5p | KPNB1 | HITS-CLIP | Tarbase |
| hsa-miR-15a-5p | KPNB1 | HITS-CLIP | Tarbase |
| hsa-miR-15a-5p | KPNB1 | HITS-CLIP | Tarbase |
| hsa-miR-15a-5p | KPNB1 | PAR-CLIP | Tarbase |
| hsa-miR-15a-5p | KPNB1 | HITS-CLIP | Tarbase |
| hsa-miR-195-5p | KPNB1 | HITS-CLIP | Tarbase |
| hsa-miR-195-5p | KPNB1 | HITS-CLIP | Tarbase |
| hsa-miR-195-5p | KPNB1 | HITS-CLIP | Tarbase |
| hsa-miR-195-5p | KPNB1 | HITS-CLIP | Tarbase |
| hsa-miR-195-5p | KPNB1 | HITS-CLIP | Tarbase |
| hsa-miR-195-5p | KPNB1 | HITS-CLIP | Tarbase |
| hsa-miR-486-5p | RPS6KB1 | PAR-CLIP | Tarbase |
| hsa-miR-15a-5p | CAMTA2 | HITS-CLIP | Tarbase |
| hsa-miR-195-5p | CAMTA2 | HITS-CLIP | Tarbase |
| hsa-miR-141-3p | MED13 | PAR-CLIP | Tarbase |
| hsa-miR-15a-5p | MED13 | PAR-CLIP | Tarbase |
| hsa-miR-15a-5p | MED13 | PAR-CLIP | Tarbase |
| hsa-miR-3613-3p | PFN1 | HITS-CLIP | Tarbase |
| hsa-miR-141-3p | RNF167 | Chimeric fragments | Tarbase |
| hsa-miR-15a-5p | RAI1 | HITS-CLIP | Tarbase |
| hsa-miR-15a-5p | RAI1 | HITS-CLIP | Tarbase |
| hsa-miR-195-5p | RAI1 | HITS-CLIP | Tarbase |
| hsa-miR-15a-5p | C1QBP | PAR-CLIP | Tarbase |
| hsa-miR-195-5p | BLMH | Microarrays | Tarbase |
| hsa-miR-15a-5p | CPD | HITS-CLIP | Tarbase |
| hsa-miR-195-5p | CPD | HITS-CLIP | Tarbase |
| hsa-miR-141-3p | GOSR1 | HITS-CLIP | Tarbase |
| hsa-miR-15a-5p | GOSR1 | HITS-CLIP | Tarbase |
| hsa-miR-195-5p | GOSR1 | HITS-CLIP | Tarbase |
| hsa-miR-15a-5p | CCDC47 | HITS-CLIP | Tarbase |
| hsa-miR-195-5p | CCDC47 | HITS-CLIP | Tarbase |
| hsa-miR-15a-5p | AKAP10 | PAR-CLIP | Tarbase |
| hsa-miR-15a-5p | AKAP10 | PAR-CLIP | Tarbase |
| hsa-miR-15a-5p | AKAP10 | PAR-CLIP | Tarbase |
| hsa-miR-15a-5p | AKAP10 | HITS-CLIP | Tarbase |
| hsa-miR-15a-5p | AKAP10 | PAR-CLIP | Tarbase |
| hsa-miR-15a-5p | AKAP10 | HITS-CLIP | Tarbase |
| hsa-miR-195-5p | AKAP10 | PAR-CLIP | Tarbase |
| hsa-miR-195-5p | AKAP10 | HITS-CLIP | Tarbase |
| hsa-miR-195-5p | AKAP10 | HITS-CLIP | Tarbase |
| hsa-miR-195-5p | SYNGR2 | Microarrays | Tarbase |
| hsa-miR-195-5p | UTP6 | Microarrays | Tarbase |
| hsa-miR-141-5p | DDX5 | HITS-CLIP | Tarbase |
| hsa-miR-15a-3p | DDX5 | HITS-CLIP | Tarbase |
| hsa-miR-15a-3p | DDX5 | HITS-CLIP | Tarbase |
| hsa-miR-15a-5p | DDX5 | HITS-CLIP | Tarbase |
| hsa-miR-195-5p | DDX5 | HITS-CLIP | Tarbase |
| hsa-miR-15a-5p | DDX5 | HITS-CLIP | Tarbase |
| hsa-miR-195-5p | DDX5 | HITS-CLIP | Tarbase |
| hsa-miR-195-5p | CCL2 | Microarrays | Tarbase |
| hsa-miR-15a-5p | CNTNAP1 | HITS-CLIP | Tarbase |
| hsa-miR-15a-5p | EZH1 | HITS-CLIP | Tarbase |
| hsa-miR-195-5p | EZH1 | HITS-CLIP | Tarbase |
| hsa-miR-15a-5p | LRRC59 | HITS-CLIP | Tarbase |
| hsa-miR-15a-5p | LRRC59 | HITS-CLIP | Tarbase |
| hsa-miR-15a-5p | LRRC59 | PAR-CLIP | Tarbase |
| hsa-miR-15a-5p | LRRC59 | HITS-CLIP | Tarbase |
| hsa-miR-195-5p | LRRC59 | HITS-CLIP | Tarbase |
| hsa-miR-195-5p | LRRC59 | HITS-CLIP | Tarbase |
| hsa-miR-15a-5p | ABCC3 | HITS-CLIP | Tarbase |
| hsa-miR-195-5p | ABCC3 | HITS-CLIP | Tarbase |
| hsa-miR-141-3p | LUC7L3 | HITS-CLIP | Tarbase |
| hsa-miR-15a-3p | LUC7L3 | HITS-CLIP | Tarbase |
| hsa-miR-15a-5p | MPP2 | PAR-CLIP | Tarbase |
| hsa-miR-195-5p | MPP2 | Microarrays | Tarbase |
| hsa-miR-15a-5p | SMURF2 | PAR-CLIP | Tarbase |
| hsa-miR-15a-5p | SMURF2 | HITS-CLIP | Tarbase |
| hsa-miR-15a-5p | SMURF2 | HITS-CLIP | Tarbase |
| hsa-miR-15a-5p | SMURF2 | PAR-CLIP | Tarbase |
| hsa-miR-195-5p | SMURF2 | HITS-CLIP | Tarbase |
| hsa-miR-195-5p | SMURF2 | HITS-CLIP | Tarbase |
| hsa-miR-141-3p | DUSP3 | HITS-CLIP | Tarbase |
| hsa-miR-141-3p | DUSP3 | HITS-CLIP | Tarbase |
| hsa-miR-195-5p | HLF | Microarrays | Tarbase |
| hsa-miR-141-3p | SLC16A6 | PAR-CLIP | Tarbase |
| hsa-miR-195-5p | SLC16A6 | Microarrays | Tarbase |
| hsa-miR-141-3p | PRKAR1A | HITS-CLIP | Tarbase |
| hsa-miR-141-3p | PRKAR1A | HITS-CLIP | Tarbase |
| hsa-miR-141-3p | PRKAR1A | HITS-CLIP | Tarbase |
| hsa-miR-15a-5p | PRKAR1A | HITS-CLIP | Tarbase |
| hsa-miR-15a-5p | PRKAR1A | HITS-CLIP | Tarbase |
| hsa-miR-15a-5p | PRKAR1A | HITS-CLIP | Tarbase |
| hsa-miR-15a-5p | PRKAR1A | HITS-CLIP | Tarbase |
| hsa-miR-15a-5p | PRKAR1A | HITS-CLIP | Tarbase |
| hsa-miR-15a-5p | PRKAR1A | PAR-CLIP | Tarbase |
| hsa-miR-15a-5p | PRKAR1A | HITS-CLIP | Tarbase |
| hsa-miR-15a-5p | PRKAR1A | HITS-CLIP | Tarbase |
| hsa-miR-195-5p | PRKAR1A | HITS-CLIP | Tarbase |
| hsa-miR-195-5p | PRKAR1A | HITS-CLIP | Tarbase |
| hsa-miR-195-5p | PRKAR1A | HITS-CLIP | Tarbase |
| hsa-miR-195-5p | PRKAR1A | HITS-CLIP | Tarbase |
| hsa-miR-195-5p | PRKAR1A | HITS-CLIP | Tarbase |
| hsa-miR-195-5p | PRKAR1A | HITS-CLIP | Tarbase |
| hsa-miR-3613-3p | PRKAR1A | PAR-CLIP | Tarbase |
| hsa-miR-3605-5p | FAM20A | HITS-CLIP | Tarbase |
| hsa-miR-15a-5p | YWHAE | PAR-CLIP | Tarbase |
| hsa-miR-15a-5p | YWHAE | HITS-CLIP | Tarbase |
| hsa-miR-15a-5p | MMD | PAR-CLIP | Tarbase |
| hsa-miR-15a-5p | MMD | HITS-CLIP | Tarbase |
| hsa-miR-195-5p | MMD | HITS-CLIP | Tarbase |
| hsa-miR-195-5p | MMD | Microarrays | Tarbase |
| hsa-miR-15a-5p | MMD | Chimeric fragments | Tarbase |
| hsa-miR-15a-5p | WSB1 | HITS-CLIP | Tarbase |
| hsa-miR-15a-5p | WSB1 | HITS-CLIP | Tarbase |
| hsa-miR-15a-5p | WSB1 | HITS-CLIP | Tarbase |
| hsa-miR-195-5p | WSB1 | HITS-CLIP | Tarbase |
| hsa-miR-195-5p | WSB1 | HITS-CLIP | Tarbase |
| hsa-miR-15a-5p | WSB1 | HITS-CLIP | Tarbase |
| hsa-miR-195-5p | WSB1 | HITS-CLIP | Tarbase |
| hsa-miR-15a-5p | TNFAIP1 | HITS-CLIP | Tarbase |
| hsa-miR-15a-5p | TNFAIP1 | HITS-CLIP | Tarbase |
| hsa-miR-195-5p | TNFAIP1 | HITS-CLIP | Tarbase |
| hsa-miR-195-5p | TMEM97 | Microarrays | Tarbase |
| hsa-miR-195-5p | CDR2L | Microarrays | Tarbase |
| hsa-miR-15a-5p | PMP22 | HITS-CLIP | Tarbase |
| hsa-miR-195-5p | PMP22 | HITS-CLIP | Tarbase |
| hsa-miR-195-5p | PMP22 | Microarrays | Tarbase |
| hsa-miR-141-5p | UNC119 | HITS-CLIP | Tarbase |
| hsa-miR-195-5p | ALDOC | Microarrays | Tarbase |
| hsa-miR-141-5p | SUPT6H | HITS-CLIP | Tarbase |
| hsa-miR-15a-5p | PHF12 | HITS-CLIP | Tarbase |
| hsa-miR-15a-5p | PHF12 | HITS-CLIP | Tarbase |
| hsa-miR-15a-5p | PHF12 | HITS-CLIP | Tarbase |
| hsa-miR-195-5p | PHF12 | HITS-CLIP | Tarbase |
| hsa-miR-195-5p | PHF12 | HITS-CLIP | Tarbase |
| hsa-miR-3613-3p | TMEM33 | PAR-CLIP | Tarbase |
| hsa-miR-486-5p | SLAIN2 | PAR-CLIP | Tarbase |
| hsa-miR-195-5p | SULT1E1 | Microarrays | Tarbase |
| hsa-miR-15a-5p | LAMTOR3 | HITS-CLIP | Tarbase |
| hsa-miR-195-5p | LAMTOR3 | HITS-CLIP | Tarbase |
| hsa-miR-15a-5p | UBE2D3 | PAR-CLIP | Tarbase |
| hsa-miR-3613-3p | UBE2D3 | PAR-CLIP | Tarbase |
| hsa-miR-195-5p | MAPK10 | Microarrays | Tarbase |
| hsa-miR-15a-5p | TBC1D9 | HITS-CLIP | Tarbase |
| hsa-miR-195-5p | TBC1D9 | HITS-CLIP | Tarbase |
| hsa-miR-3613-3p | INPP4B | HITS-CLIP | Tarbase |
| hsa-miR-15a-5p | KLHL2 | HITS-CLIP | Tarbase |

| | | | |
|---|---|---|---|
| hsa-miR-195-5p | KLHL2 | HITS-CLIP | Tarbase |
| hsa-miR-195-5p | KLHL2 | Microarrays | Tarbase |
| hsa-miR-195-5p | WFS1 | Microarrays | Tarbase |
| hsa-miR-141-5p | FRG1 | HITS-CLIP | Tarbase |
| hsa-miR-15a-5p | CLCN3 | PAR-CLIP | Tarbase |
| hsa-miR-15a-5p | CLCN3 | PAR-CLIP | Tarbase |
| hsa-miR-15a-5p | CLCN3 | PAR-CLIP | Tarbase |
| hsa-miR-195-5p | CLCN3 | PAR-CLIP | Tarbase |
| hsa-miR-15a-5p | CLCN3 | HITS-CLIP | Tarbase |
| hsa-miR-195-5p | CLCN3 | HITS-CLIP | Tarbase |
| hsa-miR-195-5p | CLCN3 | Microarrays | Tarbase |
| hsa-miR-15a-5p | CLCN3 | Luciferase Reporter Assay | Tarbase |
| hsa-miR-195-5p | AADAT | Microarrays | Tarbase |
| hsa-miR-141-3p | GALNT7 | HITS-CLIP | Tarbase |
| hsa-miR-15a-5p | GALNT7 | HITS-CLIP | Tarbase |
| hsa-miR-195-5p | GALNT7 | HITS-CLIP | Tarbase |
| hsa-miR-141-3p | DHX15 | HITS-CLIP | Tarbase |
| hsa-miR-15a-3p | DHX15 | HITS-CLIP | Tarbase |
| hsa-miR-15a-5p | DHX15 | PAR-CLIP | Tarbase |
| hsa-miR-15a-5p | DHX15 | PAR-CLIP | Tarbase |
| hsa-miR-195-5p | DHX15 | PAR-CLIP | Tarbase |
| hsa-miR-15a-5p | TRIM2 | HITS-CLIP | Tarbase |
| hsa-miR-195-5p | TRIM2 | HITS-CLIP | Tarbase |
| hsa-miR-195-5p | TRIM2 | Microarrays | Tarbase |
| hsa-miR-141-3p | FBXW7 | HITS-CLIP | Tarbase |
| hsa-miR-141-3p | FBXW7 | HITS-CLIP | Tarbase |
| hsa-miR-15a-5p | FBXW7 | PAR-CLIP | Tarbase |
| hsa-miR-15a-5p | FBXW7 | PAR-CLIP | Tarbase |
| hsa-miR-15a-5p | FBXW7 | PAR-CLIP | Tarbase |
| hsa-miR-15a-5p | FBXW7 | HITS-CLIP | Tarbase |
| hsa-miR-15a-5p | FBXW7 | HITS-CLIP | Tarbase |
| hsa-miR-15a-5p | FBXW7 | PAR-CLIP | Tarbase |
| hsa-miR-15a-5p | FBXW7 | PAR-CLIP | Tarbase |
| hsa-miR-15a-5p | FBXW7 | PAR-CLIP | Tarbase |
| hsa-miR-15a-5p | FBXW7 | HITS-CLIP | Tarbase |
| hsa-miR-15a-5p | FBXW7 | HITS-CLIP | Tarbase |
| hsa-miR-15a-5p | FBXW7 | PAR-CLIP | Tarbase |
| hsa-miR-15a-5p | FBXW7 | PAR-CLIP | Tarbase |
| hsa-miR-15a-5p | FBXW7 | PAR-CLIP | Tarbase |
| hsa-miR-15a-5p | FBXW7 | PAR-CLIP | Tarbase |
| hsa-miR-15a-5p | FBXW7 | PAR-CLIP | Tarbase |
| hsa-miR-195-5p | FBXW7 | PAR-CLIP | Tarbase |
| hsa-miR-195-5p | FBXW7 | HITS-CLIP | Tarbase |
| hsa-miR-195-5p | FBXW7 | HITS-CLIP | Tarbase |
| hsa-miR-195-5p | FBXW7 | HITS-CLIP | Tarbase |
| hsa-miR-195-5p | FBXW7 | PAR-CLIP | Tarbase |
| hsa-miR-195-5p | FBXW7 | PAR-CLIP | Tarbase |
| hsa-miR-195-5p | FBXW7 | PAR-CLIP | Tarbase |
| hsa-miR-195-5p | FBXW7 | PAR-CLIP | Tarbase |
| hsa-miR-195-5p | FBXW7 | PAR-CLIP | Tarbase |
| hsa-miR-15a-5p | FBXW7 | HITS-CLIP | Tarbase |
| hsa-miR-195-5p | FBXW7 | HITS-CLIP | Tarbase |
| hsa-miR-15a-5p | FBXW7 | Chimeric fragments | Tarbase |
| hsa-miR-195-5p | NEIL3 | Microarrays | Tarbase |
| hsa-miR-486-3p | WHSC1 | HITS-CLIP | Tarbase |
| hsa-miR-195-5p | WHSC1 | Microarrays | Tarbase |
| hsa-miR-141-3p | SH3D19 | HITS-CLIP | Tarbase |
| hsa-miR-15a-5p | SH3D19 | HITS-CLIP | Tarbase |
| hsa-miR-195-5p | SH3D19 | HITS-CLIP | Tarbase |
| hsa-miR-15a-5p | RAPGEF2 | HITS-CLIP | Tarbase |
| hsa-miR-195-5p | RAPGEF2 | HITS-CLIP | Tarbase |
| hsa-miR-141-3p | KLF3 | HITS-CLIP | Tarbase |
| hsa-miR-141-5p | KLHL5 | HITS-CLIP | Tarbase |
| hsa-miR-195-5p | NCAPG | Microarrays | Tarbase |
| hsa-miR-195-5p | PPARGC1A | Microarrays | Tarbase |
| hsa-miR-195-5p | CRYAB | Microarrays | Tarbase |
| hsa-miR-15a-5p | FNBP4 | PAR-CLIP | Tarbase |
| hsa-miR-15a-5p | FNBP4 | PAR-CLIP | Tarbase |
| hsa-miR-15a-5p | FNBP4 | PAR-CLIP | Tarbase |
| hsa-miR-15a-5p | FNBP4 | PAR-CLIP | Tarbase |
| hsa-miR-195-5p | FNBP4 | PAR-CLIP | Tarbase |
| hsa-miR-195-5p | FNBP4 | PAR-CLIP | Tarbase |
| hsa-miR-195-5p | FNBP4 | PAR-CLIP | Tarbase |
| hsa-miR-15a-5p | TECTA | HITS-CLIP | Tarbase |
| hsa-miR-195-5p | TECTA | HITS-CLIP | Tarbase |
| hsa-miR-141-5p | HSPA8 | HITS-CLIP | Tarbase |
| hsa-miR-141-5p | HSPA8 | HITS-CLIP | Tarbase |
| hsa-miR-15a-5p | HSPA8 | PAR-CLIP | Tarbase |
| hsa-miR-15a-5p | HSPA8 | PAR-CLIP | Tarbase |
| hsa-miR-15a-5p | HSPA8 | HITS-CLIP | Tarbase |
| hsa-miR-195-5p | SIAE | Microarrays | Tarbase |
| hsa-miR-195-5p | LPXN | Microarrays | Tarbase |
| hsa-miR-15a-3p | ATG2A | HITS-CLIP | Tarbase |
| hsa-miR-486-3p | ATG2A | HITS-CLIP | Tarbase |
| hsa-miR-141-3p | OSBP | HITS-CLIP | Tarbase |
| hsa-miR-15a-5p | SUV420H1 | HITS-CLIP | Tarbase |
| hsa-miR-141-5p | PPP6R3 | HITS-CLIP | Tarbase |
| hsa-miR-15a-5p | NRXN2 | PAR-CLIP | Tarbase |
| hsa-miR-195-5p | ST3GAL4 | Microarrays | Tarbase |
| hsa-miR-15a-5p | CPT1A | HITS-CLIP | Tarbase |
| hsa-miR-195-5p | CPT1A | HITS-CLIP | Tarbase |
| hsa-miR-141-3p | CCND1 | Western Blot | Tarbase |
| hsa-miR-141-3p | CCND1 | Western Blot | Tarbase |
| hsa-miR-141-5p | CCND1 | HITS-CLIP | Tarbase |
| hsa-miR-141-5p | CCND1 | HITS-CLIP | Tarbase |
| hsa-miR-141-5p | CCND1 | HITS-CLIP | Tarbase |
| hsa-miR-141-5p | CCND1 | HITS-CLIP | Tarbase |
| hsa-miR-15a-5p | CCND1 | HITS-CLIP | Tarbase |
| hsa-miR-15a-5p | CCND1 | HITS-CLIP | Tarbase |
| hsa-miR-15a-5p | CCND1 | PAR-CLIP | Tarbase |
| hsa-miR-15a-5p | CCND1 | PAR-CLIP | Tarbase |
| hsa-miR-15a-5p | CCND1 | HITS-CLIP | Tarbase |
| hsa-miR-15a-5p | CCND1 | HITS-CLIP | Tarbase |
| hsa-miR-15a-5p | CCND1 | HITS-CLIP | Tarbase |
| hsa-miR-15a-5p | CCND1 | PAR-CLIP | Tarbase |
| hsa-miR-15a-5p | CCND1 | HITS-CLIP | Tarbase |
| hsa-miR-15a-5p | CCND1 | HITS-CLIP | Tarbase |
| hsa-miR-15a-5p | CCND1 | HITS-CLIP | Tarbase |
| hsa-miR-195-5p | CCND1 | HITS-CLIP | Tarbase |
| hsa-miR-195-5p | CCND1 | HITS-CLIP | Tarbase |
| hsa-miR-195-5p | CCND1 | Luciferase Reporter Assay | Tarbase |
| hsa-miR-195-5p | CCND1 | Western Blot | Tarbase |
| hsa-miR-195-5p | CCND1 | Western Blot | Tarbase |
| hsa-miR-195-5p | CCND1 | qPCR | Tarbase |
| hsa-miR-195-5p | CCND1 | Western Blot | Tarbase |
| hsa-miR-195-5p | CCND1 | HITS-CLIP | Tarbase |
| hsa-miR-195-5p | CCND1 | HITS-CLIP | Tarbase |
| hsa-miR-195-5p | CCND1 | HITS-CLIP | Tarbase |
| hsa-miR-219a-1-3p | CCND1 | HITS-CLIP | Tarbase |
| hsa-miR-486-3p | CCND1 | HITS-CLIP | Tarbase |
| hsa-miR-195-5p | CCND1 | Luciferase Reporter Assay | Tarbase |
| hsa-miR-195-5p | CCND1 | Luciferase Reporter Assay | Tarbase |
| hsa-miR-3613-3p | PRPF19 | HITS-CLIP | Tarbase |
| hsa-miR-15a-3p | TMEM109 | HITS-CLIP | Tarbase |
| hsa-miR-15a-3p | TMEM109 | HITS-CLIP | Tarbase |
| hsa-miR-15a-5p | TMEM109 | HITS-CLIP | Tarbase |
| hsa-miR-195-5p | TMEM109 | Microarrays | Tarbase |
| hsa-miR-15a-5p | CHORDC1 | HITS-CLIP | Tarbase |
| hsa-miR-195-5p | CHORDC1 | HITS-CLIP | Tarbase |
| hsa-miR-195-5p | ANAPC15 | Microarrays | Tarbase |
| hsa-miR-141-5p | EIF4G2 | HITS-CLIP | Tarbase |
| hsa-miR-15a-5p | EIF4G2 | HITS-CLIP | Tarbase |
| hsa-miR-15a-5p | EIF4G2 | HITS-CLIP | Tarbase |
| hsa-miR-15a-5p | EIF4G2 | HITS-CLIP | Tarbase |
| hsa-miR-195-5p | EIF4G2 | HITS-CLIP | Tarbase |
| hsa-miR-195-5p | EIF4G2 | HITS-CLIP | Tarbase |
| hsa-miR-219a-1-3p | EIF4G2 | HITS-CLIP | Tarbase |
| hsa-miR-219a-1-3p | EIF4G2 | HITS-CLIP | Tarbase |
| hsa-miR-141-5p | BIRC2 | HITS-CLIP | Tarbase |
| hsa-miR-15a-5p | UBE4A | HITS-CLIP | Tarbase |
| hsa-miR-195-5p | UBE4A | HITS-CLIP | Tarbase |
| hsa-miR-141-3p | DDX6 | HITS-CLIP | Tarbase |
| hsa-miR-15a-5p | DDX6 | HITS-CLIP | Tarbase |
| hsa-miR-15a-5p | DDX6 | HITS-CLIP | Tarbase |
| hsa-miR-195-5p | DDX6 | HITS-CLIP | Tarbase |
| hsa-miR-141-3p | CBL | HITS-CLIP | Tarbase |
| hsa-miR-15a-3p | CBL | HITS-CLIP | Tarbase |
| hsa-miR-3605-5p | CBL | PAR-CLIP | Tarbase |
| hsa-miR-141-3p | PVRL1 | HITS-CLIP | Tarbase |

| | | | |
|---|---|---|---|
| hsa-miR-15a-5p | PVRL1 | HITS-CLIP | Tarbase |
| hsa-miR-195-5p | PVRL1 | HITS-CLIP | Tarbase |
| hsa-miR-195-5p | PVRL1 | Microarrays | Tarbase |
| hsa-miR-15a-5p | HIPK3 | PAR-CLIP | Tarbase |
| hsa-miR-195-5p | HIPK3 | Microarrays | Tarbase |
| hsa-miR-15a-5p | KIAA1549L | HITS-CLIP | Tarbase |
| hsa-miR-195-5p | KIAA1549L | HITS-CLIP | Tarbase |
| hsa-miR-15a-3p | FBXO3 | HITS-CLIP | Tarbase |
| hsa-miR-15a-3p | FBXO3 | HITS-CLIP | Tarbase |
| hsa-miR-15a-5p | FBXO3 | CLASH | Tarbase |
| hsa-miR-15a-5p | FBXO3 | HITS-CLIP | Tarbase |
| hsa-miR-15a-5p | FBXO3 | PAR-CLIP | Tarbase |
| hsa-miR-195-5p | FBXO3 | HITS-CLIP | Tarbase |
| hsa-miR-15a-5p | PDHX | PAR-CLIP | Tarbase |
| hsa-miR-195-5p | PDHX | PAR-CLIP | Tarbase |
| hsa-miR-141-5p | MDK | HITS-CLIP | Tarbase |
| hsa-miR-195-5p | SLC22A18 | Microarrays | Tarbase |
| hsa-miR-15a-5p | CD81 | PAR-CLIP | Tarbase |
| hsa-miR-15a-5p | CD81 | PAR-CLIP | Tarbase |
| hsa-miR-3613-3p | SOX6 | HITS-CLIP | Tarbase |
| hsa-miR-15a-5p | C11orf58 | HITS-CLIP | Tarbase |
| hsa-miR-195-5p | C11orf58 | HITS-CLIP | Tarbase |
| hsa-miR-15a-5p | AIP | HITS-CLIP | Tarbase |
| hsa-miR-195-5p | AIP | HITS-CLIP | Tarbase |
| hsa-miR-3605-5p | AIP | HITS-CLIP | Tarbase |
| hsa-miR-141-3p | NUP98 | HITS-CLIP | Tarbase |
| hsa-miR-15a-5p | NUP98 | HITS-CLIP | Tarbase |
| hsa-miR-195-5p | NUP98 | HITS-CLIP | Tarbase |
| hsa-miR-141-3p | EXPH5 | HITS-CLIP | Tarbase |
| hsa-miR-195-5p | EXPH5 | Microarrays | Tarbase |
| hsa-miR-141-3p | HPS5 | HITS-CLIP | Tarbase |
| hsa-miR-141-3p | HPS5 | HITS-CLIP | Tarbase |
| hsa-miR-15a-5p | GTF2H1 | PAR-CLIP | Tarbase |
| hsa-miR-141-5p | VWF | HITS-CLIP | Tarbase |
| hsa-miR-141-5p | PPFIBP1 | HITS-CLIP | Tarbase |
| hsa-miR-15a-5p | PRDM4 | HITS-CLIP | Tarbase |
| hsa-miR-15a-5p | PRDM4 | HITS-CLIP | Tarbase |
| hsa-miR-15a-5p | PRDM4 | PAR-CLIP | Tarbase |
| hsa-miR-15a-5p | PRDM4 | HITS-CLIP | Tarbase |
| hsa-miR-195-5p | PRDM4 | HITS-CLIP | Tarbase |
| hsa-miR-195-5p | PRDM4 | HITS-CLIP | Tarbase |
| hsa-miR-195-5p | PRDM4 | HITS-CLIP | Tarbase |
| hsa-miR-15a-5p | COQ5 | HITS-CLIP | Tarbase |
| hsa-miR-195-5p | COQ5 | HITS-CLIP | Tarbase |
| hsa-miR-3613-5p | CORO1C | HITS-CLIP | Tarbase |
| hsa-miR-486-3p | CORO1C | HITS-CLIP | Tarbase |
| hsa-miR-15a-5p | KCTD10 | HITS-CLIP | Tarbase |
| hsa-miR-195-5p | KCTD10 | HITS-CLIP | Tarbase |
| hsa-miR-195-5p | KCTD10 | Microarrays | Tarbase |
| hsa-miR-141-3p | SLC11A2 | HITS-CLIP | Tarbase |
| hsa-miR-15a-5p | SLC11A2 | PAR-CLIP | Tarbase |
| hsa-miR-15a-5p | SLC11A2 | HITS-CLIP | Tarbase |
| hsa-miR-15a-5p | SLC11A2 | HITS-CLIP | Tarbase |
| hsa-miR-195-5p | SLC11A2 | PAR-CLIP | Tarbase |
| hsa-miR-141-5p | MLEC | HITS-CLIP | Tarbase |
| hsa-miR-15a-5p | MLEC | HITS-CLIP | Tarbase |
| hsa-miR-15a-5p | MLEC | HITS-CLIP | Tarbase |
| hsa-miR-195-5p | MLEC | HITS-CLIP | Tarbase |
| hsa-miR-3613-5p | MLEC | HITS-CLIP | Tarbase |
| hsa-miR-15a-3p | CAMKK2 | HITS-CLIP | Tarbase |
| hsa-miR-15a-5p | ATP5B | PAR-CLIP | Tarbase |
| hsa-miR-141-3p | PTGES3 | HITS-CLIP | Tarbase |
| hsa-miR-3613-5p | PTGES3 | Chimeric fragments | Tarbase |
| hsa-miR-15a-5p | BCL7A | PAR-CLIP | Tarbase |
| hsa-miR-486-5p | RSRC2 | PAR-CLIP | Tarbase |
| hsa-miR-486-5p | RSRC2 | PAR-CLIP | Tarbase |
| hsa-miR-195-5p | LIN7A | Microarrays | Tarbase |
| hsa-miR-15a-5p | KRT18 | PAR-CLIP | Tarbase |
| hsa-miR-15a-5p | KRT18 | HITS-CLIP | Tarbase |
| hsa-miR-15a-5p | KRT18 | HITS-CLIP | Tarbase |
| hsa-miR-195-5p | KRT18 | HITS-CLIP | Tarbase |
| hsa-miR-195-5p | KRT18 | HITS-CLIP | Tarbase |
| hsa-miR-15a-5p | TENC1 | HITS-CLIP | Tarbase |
| hsa-miR-195-5p | TENC1 | HITS-CLIP | Tarbase |
| hsa-miR-195-5p | GLI1 | Microarrays | Tarbase |
| hsa-miR-141-5p | METAP2 | HITS-CLIP | Tarbase |
| hsa-miR-15a-5p | METAP2 | HITS-CLIP | Tarbase |
| hsa-miR-15a-5p | METAP2 | Chimeric fragments | Tarbase |
| hsa-miR-3613-5p | LTA4H | HITS-CLIP | Tarbase |
| hsa-miR-15a-3p | ELK3 | HITS-CLIP | Tarbase |
| hsa-miR-141-5p | FOXM1 | HITS-CLIP | Tarbase |
| hsa-miR-195-5p | FOXM1 | Microarrays | Tarbase |
| hsa-miR-195-5p | RAD51AP1 | Microarrays | Tarbase |
| hsa-miR-3613-3p | SH2B3 | PAR-CLIP | Tarbase |
| hsa-miR-141-5p | DUSP16 | HITS-CLIP | Tarbase |
| hsa-miR-219a-1-3p | CREBL2 | HITS-CLIP | Tarbase |
| hsa-miR-486-5p | CREBL2 | HITS-CLIP | Tarbase |
| hsa-miR-15a-5p | ALDH2 | PAR-CLIP | Tarbase |
| hsa-miR-141-5p | CDKN1B | HITS-CLIP | Tarbase |
| hsa-miR-141-5p | CDKN1B | HITS-CLIP | Tarbase |
| hsa-miR-141-5p | CDKN1B | HITS-CLIP | Tarbase |
| hsa-miR-3613-5p | CDKN1B | HITS-CLIP | Tarbase |
| hsa-miR-3613-5p | CDKN1B | HITS-CLIP | Tarbase |
| hsa-miR-15a-5p | NAA25 | PAR-CLIP | Tarbase |
| hsa-miR-15a-5p | LTBR | HITS-CLIP | Tarbase |
| hsa-miR-3613-5p | OGFOD2 | HITS-CLIP | Tarbase |
| hsa-miR-195-5p | CDK2AP1 | HITS-CLIP | Tarbase |
| hsa-miR-219a-1-3p | OAS2 | PAR-CLIP | Tarbase |
| hsa-miR-15a-5p | MGP | HITS-CLIP | Tarbase |
| hsa-miR-195-5p | MGP | HITS-CLIP | Tarbase |
| hsa-miR-141-5p | ARHGDIB | HITS-CLIP | Tarbase |
| hsa-miR-15a-5p | SLC38A1 | HITS-CLIP | Tarbase |
| hsa-miR-15a-5p | SLC38A1 | HITS-CLIP | Tarbase |
| hsa-miR-195-5p | SLC38A1 | HITS-CLIP | Tarbase |
| hsa-miR-195-5p | C12orf49 | Microarrays | Tarbase |
| hsa-miR-195-5p | RFC5 | Microarrays | Tarbase |
| hsa-miR-141-3p | COPZ1 | HITS-CLIP | Tarbase |
| hsa-miR-141-5p | CAND1 | HITS-CLIP | Tarbase |
| hsa-miR-141-5p | CAND1 | HITS-CLIP | Tarbase |
| hsa-miR-141-5p | CAND1 | HITS-CLIP | Tarbase |
| hsa-miR-141-5p | CAND1 | HITS-CLIP | Tarbase |
| hsa-miR-141-5p | CAND1 | HITS-CLIP | Tarbase |
| hsa-miR-15a-5p | CAND1 | HITS-CLIP | Tarbase |
| hsa-miR-195-5p | CAND1 | HITS-CLIP | Tarbase |
| hsa-miR-141-5p | RAB5B | HITS-CLIP | Tarbase |
| hsa-miR-195-5p | TIMELESS | Microarrays | Tarbase |
| hsa-miR-141-3p | CPSF6 | HITS-CLIP | Tarbase |
| hsa-miR-15a-5p | CPSF6 | PAR-CLIP | Tarbase |
| hsa-miR-15a-5p | CPSF6 | HITS-CLIP | Tarbase |
| hsa-miR-195-5p | CPSF6 | HITS-CLIP | Tarbase |
| hsa-miR-15a-5p | CPSF6 | HITS-CLIP | Tarbase |
| hsa-miR-195-5p | CPSF6 | HITS-CLIP | Tarbase |
| hsa-miR-141-3p | GAPDH | Western Blot | Tarbase |
| hsa-miR-141-3p | GAPDH | Western Blot | Tarbase |
| hsa-miR-141-5p | NOP2 | HITS-CLIP | Tarbase |
| hsa-miR-15a-3p | CHD4 | HITS-CLIP | Tarbase |
| hsa-miR-15a-3p | CHD4 | HITS-CLIP | Tarbase |
| hsa-miR-15a-5p | CHD4 | HITS-CLIP | Tarbase |
| hsa-miR-15a-5p | CHD4 | HITS-CLIP | Tarbase |
| hsa-miR-15a-5p | CHD4 | HITS-CLIP | Tarbase |
| hsa-miR-195-5p | CHD4 | HITS-CLIP | Tarbase |
| hsa-miR-195-5p | CHD4 | HITS-CLIP | Tarbase |
| hsa-miR-195-5p | CHD4 | HITS-CLIP | Tarbase |
| hsa-miR-3613-5p | CHD4 | HITS-CLIP | Tarbase |
| hsa-miR-15a-5p | COPS7A | HITS-CLIP | Tarbase |
| hsa-miR-15a-5p | COPS7A | PAR-CLIP | Tarbase |
| hsa-miR-195-5p | COPS7A | HITS-CLIP | Tarbase |
| hsa-miR-195-5p | CDCA3 | Microarrays | Tarbase |
| hsa-miR-15a-5p | CHPT1 | PAR-CLIP | Tarbase |
| hsa-miR-15a-5p | CHPT1 | HITS-CLIP | Tarbase |
| hsa-miR-195-5p | CHPT1 | HITS-CLIP | Tarbase |
| hsa-miR-195-5p | ENO2 | Microarrays | Tarbase |
| hsa-miR-486-5p | NT5DC3 | HITS-CLIP | Tarbase |
| hsa-miR-15a-5p | GOLT1B | PAR-CLIP | Tarbase |
| hsa-miR-141-5p | LDHB | HITS-CLIP | Tarbase |
| hsa-miR-141-5p | CMAS | HITS-CLIP | Tarbase |
| hsa-miR-195-5p | ST8SIA1 | Microarrays | Tarbase |
| hsa-miR-3613-5p | C2CD5 | HITS-CLIP | Tarbase |
| hsa-miR-486-5p | C2CD5 | PAR-CLIP | Tarbase |

| | | | |
|---|---|---|---|
| hsa-miR-15a-5p | RAB35 | PAR-CLIP | Tarbase |
| hsa-miR-15a-5p | COL12A1 | HITS-CLIP | Tarbase |
| hsa-miR-15a-5p | COL12A1 | HITS-CLIP | Tarbase |
| hsa-miR-195-5p | COL12A1 | HITS-CLIP | Tarbase |
| hsa-miR-195-5p | COL12A1 | HITS-CLIP | Tarbase |
| hsa-miR-195-5p | COL12A1 | Microarrays | Tarbase |
| hsa-miR-486-5p | BTN3A3 | PAR-CLIP | Tarbase |
| hsa-miR-195-3p | TDP2 | HITS-CLIP | Tarbase |
| hsa-miR-15a-5p | TDP2 | HITS-CLIP | Tarbase |
| hsa-miR-195-5p | TDP2 | HITS-CLIP | Tarbase |
| hsa-miR-15a-5p | FRK | HITS-CLIP | Tarbase |
| hsa-miR-195-5p | FRK | HITS-CLIP | Tarbase |
| hsa-miR-15a-5p | DSE | PAR-CLIP | Tarbase |
| hsa-miR-195-5p | DSE | PAR-CLIP | Tarbase |
| hsa-miR-195-5p | GCNT2 | Microarrays | Tarbase |
| hsa-miR-141-5p | SMIM8 | HITS-CLIP | Tarbase |
| hsa-miR-195-5p | NEDD9 | Microarrays | Tarbase |
| hsa-miR-15a-5p | MAN1A1 | PAR-CLIP | Tarbase |
| hsa-miR-195-5p | MAN1A1 | PAR-CLIP | Tarbase |
| hsa-miR-141-3p | SERINC1 | HITS-CLIP | Tarbase |
| hsa-miR-195-5p | TPD52L1 | Microarrays | Tarbase |
| hsa-miR-141-3p | NCOA7 | HITS-CLIP | Tarbase |
| hsa-miR-141-3p | NCOA7 | HITS-CLIP | Tarbase |
| hsa-miR-141-5p | NCOA7 | HITS-CLIP | Tarbase |
| hsa-miR-15a-5p | UST | HITS-CLIP | Tarbase |
| hsa-miR-15a-5p | UST | HITS-CLIP | Tarbase |
| hsa-miR-195-5p | UST | HITS-CLIP | Tarbase |
| hsa-miR-195-5p | UST | HITS-CLIP | Tarbase |
| hsa-miR-141-5p | FBXO5 | HITS-CLIP | Tarbase |
| hsa-miR-141-5p | FBXO5 | HITS-CLIP | Tarbase |
| hsa-miR-195-5p | FANCE | Microarrays | Tarbase |
| hsa-miR-141-3p | MAPK14 | HITS-CLIP | Tarbase |
| hsa-miR-141-3p | MAPK14 | HITS-CLIP | Tarbase |
| hsa-miR-15a-5p | MAPK14 | HITS-CLIP | Tarbase |
| hsa-miR-15a-5p | MAPK14 | HITS-CLIP | Tarbase |
| hsa-miR-195-5p | MAPK14 | HITS-CLIP | Tarbase |
| hsa-miR-486-5p | MAPK14 | PAR-CLIP | Tarbase |
| hsa-miR-15a-5p | KCTD20 | HITS-CLIP | Tarbase |
| hsa-miR-195-5p | KCTD20 | HITS-CLIP | Tarbase |
| hsa-miR-141-3p | STK38 | HITS-CLIP | Tarbase |
| hsa-miR-141-3p | STK38 | HITS-CLIP | Tarbase |
| hsa-miR-15a-5p | STK38 | PAR-CLIP | Tarbase |
| hsa-miR-15a-5p | STK38 | PAR-CLIP | Tarbase |
| hsa-miR-15a-5p | STK38 | PAR-CLIP | Tarbase |
| hsa-miR-15a-5p | STK38 | PAR-CLIP | Tarbase |
| hsa-miR-15a-5p | STK38 | PAR-CLIP | Tarbase |
| hsa-miR-195-5p | STK38 | PAR-CLIP | Tarbase |
| hsa-miR-195-5p | STK38 | PAR-CLIP | Tarbase |
| hsa-miR-486-5p | SRSF3 | HITS-CLIP | Tarbase |
| hsa-miR-3613-3p | SOD2 | HITS-CLIP | Tarbase |
| hsa-miR-195-5p | MCM3 | Microarrays | Tarbase |
| hsa-miR-15a-5p | FBXO9 | PAR-CLIP | Tarbase |
| hsa-miR-15a-5p | MDN1 | HITS-CLIP | Tarbase |
| hsa-miR-195-5p | MDN1 | HITS-CLIP | Tarbase |
| hsa-miR-486-5p | MDN1 | HITS-CLIP | Tarbase |
| hsa-miR-195-5p | BMP5 | Microarrays | Tarbase |
| hsa-miR-195-5p | BACH2 | Microarrays | Tarbase |
| hsa-miR-15a-5p | ZNF451 | HITS-CLIP | Tarbase |
| hsa-miR-3613-3p | ZNF451 | PAR-CLIP | Tarbase |
| hsa-miR-486-5p | ZNF451 | HITS-CLIP | Tarbase |
| hsa-miR-15a-5p | RAB23 | Chimeric fragments | Tarbase |
| hsa-miR-15a-5p | GPR63 | PAR-CLIP | Tarbase |
| hsa-miR-195-5p | GPR63 | Microarrays | Tarbase |
| hsa-miR-141-5p | CCNC | HITS-CLIP | Tarbase |
| hsa-miR-15a-5p | CCNC | HITS-CLIP | Tarbase |
| hsa-miR-195-5p | CCNC | HITS-CLIP | Tarbase |
| hsa-miR-141-3p | E2F3 | HITS-CLIP | Tarbase |
| hsa-miR-141-3p | E2F3 | Western Blot | Tarbase |
| hsa-miR-141-3p | E2F3 | HITS-CLIP | Tarbase |
| hsa-miR-141-3p | E2F3 | Western Blot | Tarbase |
| hsa-miR-15a-5p | E2F3 | HITS-CLIP | Tarbase |
| hsa-miR-15a-5p | E2F3 | HITS-CLIP | Tarbase |
| hsa-miR-15a-5p | E2F3 | PAR-CLIP | Tarbase |
| hsa-miR-15a-5p | E2F3 | HITS-CLIP | Tarbase |
| hsa-miR-195-5p | E2F3 | Luciferase Reporter Assay | Tarbase |
| hsa-miR-195-5p | E2F3 | Western Blot | Tarbase |
| hsa-miR-195-5p | E2F3 | Western Blot | Tarbase |
| hsa-miR-195-5p | E2F3 | HITS-CLIP | Tarbase |
| hsa-miR-195-5p | E2F3 | HITS-CLIP | Tarbase |
| hsa-miR-195-5p | E2F3 | HITS-CLIP | Tarbase |
| hsa-miR-195-5p | E2F3 | Luciferase Reporter Assay | Tarbase |
| hsa-miR-195-5p | E2F3 | qPCR | Tarbase |
| hsa-miR-195-5p | E2F3 | Western Blot | Tarbase |
| hsa-miR-195-5p | E2F3 | Luciferase Reporter Assay | Tarbase |
| hsa-miR-15a-5p | E2F3 | Chimeric fragments | Tarbase |
| hsa-miR-141-3p | PTP4A1 | PAR-CLIP | Tarbase |
| hsa-miR-141-3p | PTP4A1 | PAR-CLIP | Tarbase |
| hsa-miR-141-3p | PTP4A1 | HITS-CLIP | Tarbase |
| hsa-miR-141-3p | PTP4A1 | HITS-CLIP | Tarbase |
| hsa-miR-141-3p | PTP4A1 | Chimeric fragments | Tarbase |
| hsa-miR-141-3p | ASCC3 | HITS-CLIP | Tarbase |
| hsa-miR-15a-5p | ASCC3 | HITS-CLIP | Tarbase |
| hsa-miR-195-5p | ASCC3 | HITS-CLIP | Tarbase |
| hsa-miR-15a-5p | BVES | HITS-CLIP | Tarbase |
| hsa-miR-195-5p | BVES | HITS-CLIP | Tarbase |
| hsa-miR-15a-3p | ALDH5A1 | HITS-CLIP | Tarbase |
| hsa-miR-15a-5p | AIM1 | PAR-CLIP | Tarbase |
| hsa-miR-15a-5p | AIM1 | PAR-CLIP | Tarbase |
| hsa-miR-195-5p | AIM1 | PAR-CLIP | Tarbase |
| hsa-miR-195-5p | AIM1 | PAR-CLIP | Tarbase |
| hsa-miR-195-5p | AIM1 | Microarrays | Tarbase |
| hsa-miR-15a-3p | RPS12 | HITS-CLIP | Tarbase |
| hsa-miR-15a-3p | C6orf62 | PAR-CLIP | Tarbase |
| hsa-miR-486-5p | C6orf62 | HITS-CLIP | Tarbase |
| hsa-miR-141-5p | GMNN | HITS-CLIP | Tarbase |
| hsa-miR-195-5p | GMNN | Microarrays | Tarbase |
| hsa-miR-15a-3p | SOBP | HITS-CLIP | Tarbase |
| hsa-miR-15a-5p | SOBP | HITS-CLIP | Tarbase |
| hsa-miR-15a-5p | SOBP | HITS-CLIP | Tarbase |
| hsa-miR-195-5p | SOBP | HITS-CLIP | Tarbase |
| hsa-miR-195-5p | SOBP | HITS-CLIP | Tarbase |
| hsa-miR-141-5p | SNX3 | HITS-CLIP | Tarbase |
| hsa-miR-141-5p | HBS1L | HITS-CLIP | Tarbase |
| hsa-miR-15a-5p | PERP | HITS-CLIP | Tarbase |
| hsa-miR-3613-3p | KIAA1244 | HITS-CLIP | Tarbase |
| hsa-miR-195-5p | KIAA1244 | Microarrays | Tarbase |
| hsa-miR-141-3p | GPR126 | HITS-CLIP | Tarbase |
| hsa-miR-15a-5p | PHACTR2 | PAR-CLIP | Tarbase |
| hsa-miR-195-5p | PHACTR2 | Microarrays | Tarbase |
| hsa-miR-3605-3p | SLC39A7 | HITS-CLIP | Tarbase |
| hsa-miR-141-3p | QKI | HITS-CLIP | Tarbase |
| hsa-miR-15a-5p | QKI | HITS-CLIP | Tarbase |
| hsa-miR-195-5p | QKI | HITS-CLIP | Tarbase |
| hsa-miR-15a-5p | QKI | HITS-CLIP | Tarbase |
| hsa-miR-195-5p | QKI | HITS-CLIP | Tarbase |
| hsa-miR-15a-5p | CCND3 | PAR-CLIP | Tarbase |
| hsa-miR-15a-5p | CCND3 | HITS-CLIP | Tarbase |
| hsa-miR-195-5p | CCND3 | Luciferase Reporter Assay | Tarbase |
| hsa-miR-195-5p | CCND3 | Western Blot | Tarbase |
| hsa-miR-195-5p | CCND3 | Western Blot | Tarbase |
| hsa-miR-195-5p | CCND3 | Microarrays | Tarbase |
| hsa-miR-195-5p | CCND3 | Luciferase Reporter Assay | Tarbase |
| hsa-miR-15a-5p | BYSL | HITS-CLIP | Tarbase |
| hsa-miR-195-5p | BYSL | HITS-CLIP | Tarbase |
| hsa-miR-15a-3p | FAM120B | HITS-CLIP | Tarbase |
| hsa-miR-15a-3p | TBP | HITS-CLIP | Tarbase |
| hsa-miR-15a-5p | TBP | HITS-CLIP | Tarbase |
| hsa-miR-15a-5p | TBP | PAR-CLIP | Tarbase |
| hsa-miR-15a-5p | TBP | HITS-CLIP | Tarbase |
| hsa-miR-195-5p | TBP | HITS-CLIP | Tarbase |
| hsa-miR-195-5p | TBP | HITS-CLIP | Tarbase |
| hsa-miR-15a-5p | TMEM30A | PAR-CLIP | Tarbase |
| hsa-miR-15a-3p | SENP6 | HITS-CLIP | Tarbase |
| hsa-miR-141-3p | VEGFA | Luciferase Reporter Assay | Tarbase |
| hsa-miR-141-3p | VEGFA | qPCR | Tarbase |
| hsa-miR-141-5p | VEGFA | HITS-CLIP | Tarbase |
| hsa-miR-141-5p | VEGFA | HITS-CLIP | Tarbase |
| hsa-miR-141-5p | VEGFA | HITS-CLIP | Tarbase |
| hsa-miR-15a-5p | VEGFA | Luciferase Reporter Assay | Tarbase |

| | | | |
|---|---|---|---|
| hsa-miR-15a-5p | VEGFA | Western Blot | Tarbase |
| hsa-miR-15a-5p | VEGFA | Luciferase Reporter Assay | Tarbase |
| hsa-miR-15a-5p | VEGFA | Luciferase Reporter Assay | Tarbase |
| hsa-miR-15a-5p | VEGFA | qPCR | Tarbase |
| hsa-miR-15a-5p | VEGFA | Western Blot | Tarbase |
| hsa-miR-15a-5p | VEGFA | HITS-CLIP | Tarbase |
| hsa-miR-15a-5p | VEGFA | HITS-CLIP | Tarbase |
| hsa-miR-15a-5p | VEGFA | PAR-CLIP | Tarbase |
| hsa-miR-195-5p | VEGFA | HITS-CLIP | Tarbase |
| hsa-miR-195-5p | VEGFA | HITS-CLIP | Tarbase |
| hsa-miR-15a-5p | VEGFA | Chimeric fragments | Tarbase |
| hsa-miR-141-3p | PRPF4B | HITS-CLIP | Tarbase |
| hsa-miR-15a-5p | TTK | PAR-CLIP | Tarbase |
| hsa-miR-195-5p | TTK | PAR-CLIP | Tarbase |
| hsa-miR-486-5p | TTK | PAR-CLIP | Tarbase |
| hsa-miR-195-5p | TTK | Microarrays | Tarbase |
| hsa-miR-195-5p | LAMA4 | Microarrays | Tarbase |
| hsa-miR-15a-5p | FAM46A | HITS-CLIP | Tarbase |
| hsa-miR-195-5p | FAM46A | HITS-CLIP | Tarbase |
| hsa-miR-141-3p | ENPP5 | HITS-CLIP | Tarbase |
| hsa-miR-141-3p | ERBB2IP | Luciferase Reporter Assay | Tarbase |
| hsa-miR-141-3p | ERBB2IP | Luciferase Reporter Assay | Tarbase |
| hsa-miR-195-5p | CEP72 | Microarrays | Tarbase |
| hsa-miR-15a-3p | PAPD7 | HITS-CLIP | Tarbase |
| hsa-miR-15a-5p | PAPD7 | Chimeric fragments | Tarbase |
| hsa-miR-141-3p | HMGCS1 | HITS-CLIP | Tarbase |
| hsa-miR-141-5p | HMGCS1 | HITS-CLIP | Tarbase |
| hsa-miR-15a-5p | HMGCS1 | HITS-CLIP | Tarbase |
| hsa-miR-15a-5p | HMGCS1 | PAR-CLIP | Tarbase |
| hsa-miR-15a-5p | HMGCS1 | PAR-CLIP | Tarbase |
| hsa-miR-15a-5p | HMGCS1 | HITS-CLIP | Tarbase |
| hsa-miR-3613-3p | HMGCS1 | HITS-CLIP | Tarbase |
| hsa-miR-3613-5p | HMGCS1 | HITS-CLIP | Tarbase |
| hsa-miR-15a-5p | HMGCS1 | HITS-CLIP | Tarbase |
| hsa-miR-195-5p | KIF20A | Microarrays | Tarbase |
| hsa-miR-3605-5p | MRPS30 | HITS-CLIP | Tarbase |
| hsa-miR-3605-5p | MRPS30 | HITS-CLIP | Tarbase |
| hsa-miR-15a-5p | HSPA9 | HITS-CLIP | Tarbase |
| hsa-miR-15a-5p | HSPA9 | HITS-CLIP | Tarbase |
| hsa-miR-15a-5p | HSPA9 | HITS-CLIP | Tarbase |
| hsa-miR-195-5p | HSPA9 | HITS-CLIP | Tarbase |
| hsa-miR-195-5p | HSPA9 | HITS-CLIP | Tarbase |
| hsa-miR-195-5p | HSPA9 | HITS-CLIP | Tarbase |
| hsa-miR-195-5p | HSPA9 | HITS-CLIP | Tarbase |
| hsa-miR-3605-5p | MRPS27 | PAR-CLIP | Tarbase |
| hsa-miR-486-5p | MRPS27 | PAR-CLIP | Tarbase |
| hsa-miR-3613-5p | LOX | HITS-CLIP | Tarbase |
| hsa-miR-195-5p | LOX | Microarrays | Tarbase |
| hsa-miR-195-5p | CDH9 | Microarrays | Tarbase |
| hsa-miR-141-5p | APBB3 | HITS-CLIP | Tarbase |
| hsa-miR-15a-5p | SPARC | HITS-CLIP | Tarbase |
| hsa-miR-15a-5p | SPARC | HITS-CLIP | Tarbase |
| hsa-miR-195-5p | SPARC | HITS-CLIP | Tarbase |
| hsa-miR-195-5p | SPARC | HITS-CLIP | Tarbase |
| hsa-miR-141-5p | HMGCR | HITS-CLIP | Tarbase |
| hsa-miR-15a-5p | HMGCR | PAR-CLIP | Tarbase |
| hsa-miR-15a-5p | HMGCR | HITS-CLIP | Tarbase |
| hsa-miR-141-3p | FAF2 | HITS-CLIP | Tarbase |
| hsa-miR-486-5p | FAF2 | PAR-CLIP | Tarbase |
| hsa-miR-195-5p | PCDHB3 | Microarrays | Tarbase |
| hsa-miR-141-5p | CLINT1 | HITS-CLIP | Tarbase |
| hsa-miR-15a-5p | CNOT6 | HITS-CLIP | Tarbase |
| hsa-miR-195-5p | CNOT6 | HITS-CLIP | Tarbase |
| hsa-miR-3605-3p | CNOT6 | HITS-CLIP | Tarbase |
| hsa-miR-486-5p | CNOT6 | Chimeric fragments | Tarbase |
| hsa-miR-141-5p | MSH3 | HITS-CLIP | Tarbase |
| hsa-miR-195-5p | RASGRF2 | Microarrays | Tarbase |
| hsa-miR-195-5p | GABRG2 | Microarrays | Tarbase |
| hsa-miR-141-3p | CCNG1 | HITS-CLIP | Tarbase |
| hsa-miR-141-3p | CCNG1 | HITS-CLIP | Tarbase |
| hsa-miR-141-3p | CCNG1 | HITS-CLIP | Tarbase |
| hsa-miR-141-3p | CCNG1 | HITS-CLIP | Tarbase |
| hsa-miR-141-5p | DROSHA | HITS-CLIP | Tarbase |
| hsa-miR-195-5p | CDH6 | Microarrays | Tarbase |
| hsa-miR-15a-3p | LMNB1 | PAR-CLIP | Tarbase |
| hsa-miR-15a-3p | LMNB1 | PAR-CLIP | Tarbase |
| hsa-miR-15a-5p | LMNB1 | PAR-CLIP | Tarbase |
| hsa-miR-195-5p | LMNB1 | Microarrays | Tarbase |
| hsa-miR-141-3p | ARRDC3 | HITS-CLIP | Tarbase |
| hsa-miR-3613-3p | ARRDC3 | HITS-CLIP | Tarbase |
| hsa-miR-141-3p | SUB1 | HITS-CLIP | Tarbase |
| hsa-miR-141-5p | SUB1 | HITS-CLIP | Tarbase |
| hsa-miR-3605-5p | TARS | HITS-CLIP | Tarbase |
| hsa-miR-15a-3p | LNPEP | HITS-CLIP | Tarbase |
| hsa-miR-15a-3p | LNPEP | HITS-CLIP | Tarbase |
| hsa-miR-15a-3p | LNPEP | HITS-CLIP | Tarbase |
| hsa-miR-3605-5p | LNPEP | HITS-CLIP | Tarbase |
| hsa-miR-15a-5p | PDE4D | PAR-CLIP | Tarbase |
| hsa-miR-141-5p | RAD1 | HITS-CLIP | Tarbase |
| hsa-miR-141-5p | BRIX1 | HITS-CLIP | Tarbase |
| hsa-miR-141-3p | PRLR | HITS-CLIP | Tarbase |
| hsa-miR-141-3p | PRLR | HITS-CLIP | Tarbase |
| hsa-miR-15a-5p | PRLR | HITS-CLIP | Tarbase |
| hsa-miR-195-5p | PRLR | Microarrays | Tarbase |
| hsa-miR-195-5p | SLC12A7 | HITS-CLIP | Tarbase |
| hsa-miR-195-5p | ST8SIA4 | Microarrays | Tarbase |
| hsa-miR-141-3p | NUP155 | HITS-CLIP | Tarbase |
| hsa-miR-141-3p | NUP155 | HITS-CLIP | Tarbase |
| hsa-miR-15a-3p | PPP2CA | HITS-CLIP | Tarbase |
| hsa-miR-15a-5p | PPP2CA | HITS-CLIP | Tarbase |
| hsa-miR-195-5p | PPP2CA | HITS-CLIP | Tarbase |
| hsa-miR-15a-5p | PPP2CA | HITS-CLIP | Tarbase |
| hsa-miR-195-5p | PPP2CA | HITS-CLIP | Tarbase |
| hsa-miR-195-5p | FGF1 | Microarrays | Tarbase |
| hsa-miR-141-3p | NR3C1 | HITS-CLIP | Tarbase |
| hsa-miR-15a-5p | NR3C1 | HITS-CLIP | Tarbase |
| hsa-miR-195-5p | NR3C1 | HITS-CLIP | Tarbase |
| hsa-miR-3613-5p | NR3C1 | PAR-CLIP | Tarbase |
| hsa-miR-3613-5p | NR3C1 | PAR-CLIP | Tarbase |
| hsa-miR-3613-5p | NR3C1 | PAR-CLIP | Tarbase |
| hsa-miR-486-5p | NR3C1 | HITS-CLIP | Tarbase |
| hsa-miR-219a-1-3p | C5orf15 | HITS-CLIP | Tarbase |
| hsa-miR-141-3p | LIFR | HITS-CLIP | Tarbase |
| hsa-miR-141-3p | LIFR | HITS-CLIP | Tarbase |
| hsa-miR-141-5p | LIFR | HITS-CLIP | Tarbase |
| hsa-miR-195-5p | LIFR | Microarrays | Tarbase |
| hsa-miR-141-3p | SEC24A | HITS-CLIP | Tarbase |
| hsa-miR-141-3p | SEC24A | HITS-CLIP | Tarbase |
| hsa-miR-195-5p | SEC24A | Microarrays | Tarbase |
| hsa-miR-15a-5p | RARS | HITS-CLIP | Tarbase |
| hsa-miR-15a-5p | RARS | HITS-CLIP | Tarbase |
| hsa-miR-15a-5p | RARS | HITS-CLIP | Tarbase |
| hsa-miR-15a-5p | RARS | PAR-CLIP | Tarbase |
| hsa-miR-15a-5p | RARS | HITS-CLIP | Tarbase |
| hsa-miR-15a-5p | RARS | HITS-CLIP | Tarbase |
| hsa-miR-195-5p | RARS | HITS-CLIP | Tarbase |
| hsa-miR-195-5p | RARS | HITS-CLIP | Tarbase |
| hsa-miR-195-5p | RARS | HITS-CLIP | Tarbase |
| hsa-miR-195-5p | RARS | HITS-CLIP | Tarbase |
| hsa-miR-195-5p | WWC1 | Microarrays | Tarbase |
| hsa-miR-141-3p | TCERG1 | HITS-CLIP | Tarbase |
| hsa-miR-141-3p | TCERG1 | HITS-CLIP | Tarbase |
| hsa-miR-141-5p | TCERG1 | HITS-CLIP | Tarbase |
| hsa-miR-219a-1-3p | SMAD5 | HITS-CLIP | Tarbase |
| hsa-miR-15a-5p | CSNK1A1 | HITS-CLIP | Tarbase |
| hsa-miR-15a-5p | CSNK1A1 | HITS-CLIP | Tarbase |
| hsa-miR-195-5p | CSNK1A1 | HITS-CLIP | Tarbase |
| hsa-miR-195-5p | CSNK1A1 | HITS-CLIP | Tarbase |
| hsa-miR-3613-3p | CSNK1A1 | HITS-CLIP | Tarbase |
| hsa-miR-3613-5p | CSNK1A1 | HITS-CLIP | Tarbase |
| hsa-miR-141-3p | ATP6V0E1 | HITS-CLIP | Tarbase |
| hsa-miR-15a-5p | ATP6V0E1 | HITS-CLIP | Tarbase |
| hsa-miR-141-5p | CPEB4 | HITS-CLIP | Tarbase |
| hsa-miR-15a-5p | CPEB4 | HITS-CLIP | Tarbase |
| hsa-miR-15a-5p | CPEB4 | HITS-CLIP | Tarbase |
| hsa-miR-195-5p | CPEB4 | HITS-CLIP | Tarbase |
| hsa-miR-195-5p | CPEB4 | HITS-CLIP | Tarbase |
| hsa-miR-15a-3p | ZNF346 | HITS-CLIP | Tarbase |
| hsa-miR-15a-5p | ZNF346 | PAR-CLIP | Tarbase |

| | | | |
|---|---|---|---|
| hsa-miR-195-5p | UNC5A | Microarrays | Tarbase |
| hsa-miR-141-3p | SMC4 | HITS-CLIP | Tarbase |
| hsa-miR-195-5p | SMC4 | Microarrays | Tarbase |
| hsa-miR-15a-5p | TBCCD1 | PAR-CLIP | Tarbase |
| hsa-miR-195-5p | AMOTL2 | Microarrays | Tarbase |
| hsa-miR-195-5p | FAM162A | Microarrays | Tarbase |
| hsa-miR-15a-5p | KPNA1 | PAR-CLIP | Tarbase |
| hsa-miR-15a-5p | KPNA1 | PAR-CLIP | Tarbase |
| hsa-miR-15a-5p | KPNA1 | PAR-CLIP | Tarbase |
| hsa-miR-15a-5p | KPNA1 | HITS-CLIP | Tarbase |
| hsa-miR-195-5p | KPNA1 | PAR-CLIP | Tarbase |
| hsa-miR-486-3p | KPNA1 | HITS-CLIP | Tarbase |
| hsa-miR-141-5p | UBE3A | HITS-CLIP | Tarbase |
| hsa-miR-15a-5p | UBE3A | HITS-CLIP | Tarbase |
| hsa-miR-15a-5p | UBE3A | HITS-CLIP | Tarbase |
| hsa-miR-15a-5p | UBE3A | HITS-CLIP | Tarbase |
| hsa-miR-195-5p | UBE3A | HITS-CLIP | Tarbase |
| hsa-miR-195-5p | UBE3A | HITS-CLIP | Tarbase |
| hsa-miR-15a-5p | UBE3A | HITS-CLIP | Tarbase |
| hsa-miR-195-5p | UBE3A | HITS-CLIP | Tarbase |
| hsa-miR-141-3p | SLC25A36 | PAR-CLIP | Tarbase |
| hsa-miR-15a-5p | SLC25A36 | HITS-CLIP | Tarbase |
| hsa-miR-195-5p | SLC25A36 | Microarrays | Tarbase |
| hsa-miR-15a-5p | SLC25A36 | Chimeric fragments | Tarbase |
| hsa-miR-141-3p | RNF7 | HITS-CLIP | Tarbase |
| hsa-miR-141-3p | XRN1 | HITS-CLIP | Tarbase |
| hsa-miR-15a-5p | XRN1 | HITS-CLIP | Tarbase |
| hsa-miR-15a-5p | XRN1 | HITS-CLIP | Tarbase |
| hsa-miR-3613-3p | XRN1 | HITS-CLIP | Tarbase |
| hsa-miR-3613-3p | XRN1 | HITS-CLIP | Tarbase |
| hsa-miR-141-5p | WNT5A | HITS-CLIP | Tarbase |
| hsa-miR-195-5p | WNT5A | Microarrays | Tarbase |
| hsa-miR-15a-5p | PRKAR2A | PAR-CLIP | Tarbase |
| hsa-miR-15a-5p | PRKAR2A | PAR-CLIP | Tarbase |
| hsa-miR-15a-5p | PRKAR2A | HITS-CLIP | Tarbase |
| hsa-miR-15a-5p | PRKAR2A | PAR-CLIP | Tarbase |
| hsa-miR-195-5p | PRKAR2A | PAR-CLIP | Tarbase |
| hsa-miR-195-5p | PRKAR2A | HITS-CLIP | Tarbase |
| hsa-miR-15a-5p | HES1 | HITS-CLIP | Tarbase |
| hsa-miR-15a-5p | HES1 | HITS-CLIP | Tarbase |
| hsa-miR-195-5p | HES1 | HITS-CLIP | Tarbase |
| hsa-miR-195-5p | HES1 | Microarrays | Tarbase |
| hsa-miR-141-5p | TFG | HITS-CLIP | Tarbase |
| hsa-miR-15a-5p | TFG | HITS-CLIP | Tarbase |
| hsa-miR-15a-5p | TFG | PAR-CLIP | Tarbase |
| hsa-miR-15a-5p | TFG | PAR-CLIP | Tarbase |
| hsa-miR-15a-5p | TFG | PAR-CLIP | Tarbase |
| hsa-miR-15a-5p | TFG | PAR-CLIP | Tarbase |
| hsa-miR-195-5p | TFG | HITS-CLIP | Tarbase |
| hsa-miR-195-5p | TFG | PAR-CLIP | Tarbase |
| hsa-miR-195-5p | TFG | PAR-CLIP | Tarbase |
| hsa-miR-15a-5p | TUSC2 | HITS-CLIP | Tarbase |
| hsa-miR-15a-5p | TUSC2 | HITS-CLIP | Tarbase |
| hsa-miR-195-5p | TUSC2 | HITS-CLIP | Tarbase |
| hsa-miR-195-5p | TUSC2 | HITS-CLIP | Tarbase |
| hsa-miR-141-3p | C3orf14 | HITS-CLIP | Tarbase |
| hsa-miR-195-5p | GBE1 | Microarrays | Tarbase |
| hsa-miR-486-3p | SNX4 | HITS-CLIP | Tarbase |
| hsa-miR-15a-5p | FRMD4B | PAR-CLIP | Tarbase |
| hsa-miR-15a-5p | PLXNA1 | HITS-CLIP | Tarbase |
| hsa-miR-195-5p | PLXNA1 | HITS-CLIP | Tarbase |
| hsa-miR-3613-5p | PLXNA1 | HITS-CLIP | Tarbase |
| hsa-miR-15a-3p | ATP6V1A | HITS-CLIP | Tarbase |
| hsa-miR-3605-5p | ATP6V1A | HITS-CLIP | Tarbase |
| hsa-miR-15a-5p | ATP6V1A | HITS-CLIP | Tarbase |
| hsa-miR-195-5p | ATP6V1A | HITS-CLIP | Tarbase |
| hsa-miR-195-5p | ATP6V1A | Microarrays | Tarbase |
| hsa-miR-15a-3p | KLHL18 | PAR-CLIP | Tarbase |
| hsa-miR-15a-5p | SCAP | HITS-CLIP | Tarbase |
| hsa-miR-15a-5p | PLSCR4 | PAR-CLIP | Tarbase |
| hsa-miR-15a-5p | PLSCR4 | PAR-CLIP | Tarbase |
| hsa-miR-195-5p | PLSCR4 | Microarrays | Tarbase |
| hsa-miR-3605-3p | HEMK1 | HITS-CLIP | Tarbase |
| hsa-miR-15a-5p | ACVR2B | HITS-CLIP | Tarbase |
| hsa-miR-141-3p | ACVR2B | Luciferase Reporter Assay | Tarbase |
| hsa-miR-15a-5p | ABCC5 | HITS-CLIP | Tarbase |
| hsa-miR-195-5p | ABCC5 | HITS-CLIP | Tarbase |
| hsa-miR-195-5p | KLHL24 | Microarrays | Tarbase |
| hsa-miR-15a-3p | SSR3 | HITS-CLIP | Tarbase |
| hsa-miR-15a-3p | SSR3 | HITS-CLIP | Tarbase |
| hsa-miR-15a-5p | ZBTB47 | HITS-CLIP | Tarbase |
| hsa-miR-195-5p | ZBTB47 | HITS-CLIP | Tarbase |
| hsa-miR-141-5p | NKTR | HITS-CLIP | Tarbase |
| hsa-miR-3605-5p | NKTR | HITS-CLIP | Tarbase |
| hsa-miR-141-3p | FOXP1 | HITS-CLIP | Tarbase |
| hsa-miR-141-5p | EIF4G1 | HITS-CLIP | Tarbase |
| hsa-miR-141-5p | EIF4G1 | HITS-CLIP | Tarbase |
| hsa-miR-141-5p | EIF4G1 | HITS-CLIP | Tarbase |
| hsa-miR-15a-3p | EIF4G1 | HITS-CLIP | Tarbase |
| hsa-miR-15a-5p | EIF4G1 | HITS-CLIP | Tarbase |
| hsa-miR-195-5p | EIF4G1 | HITS-CLIP | Tarbase |
| hsa-miR-3605-3p | EIF4G1 | HITS-CLIP | Tarbase |
| hsa-miR-141-3p | INO80D | HITS-CLIP | Tarbase |
| hsa-miR-141-5p | INO80D | HITS-CLIP | Tarbase |
| hsa-miR-15a-5p | ADAM23 | HITS-CLIP | Tarbase |
| hsa-miR-195-5p | ADAM23 | HITS-CLIP | Tarbase |
| hsa-miR-3613-3p | MOB1A | HITS-CLIP | Tarbase |
| hsa-miR-195-5p | KANSL3 | Microarrays | Tarbase |
| hsa-miR-15a-5p | LMAN2L | PAR-CLIP | Tarbase |
| hsa-miR-15a-5p | TTL | HITS-CLIP | Tarbase |
| hsa-miR-219a-1-3p | TTL | PAR-CLIP | Tarbase |
| hsa-miR-141-3p | PIKFYVE | HITS-CLIP | Tarbase |
| hsa-miR-141-5p | NCL | HITS-CLIP | Tarbase |
| hsa-miR-141-5p | NCL | HITS-CLIP | Tarbase |
| hsa-miR-15a-5p | SLC35F5 | PAR-CLIP | Tarbase |
| hsa-miR-15a-5p | SLC35F5 | PAR-CLIP | Tarbase |
| hsa-miR-195-5p | SLC35F5 | PAR-CLIP | Tarbase |
| hsa-miR-3613-5p | SLC35F5 | HITS-CLIP | Tarbase |
| hsa-miR-15a-5p | ACTR3 | HITS-CLIP | Tarbase |
| hsa-miR-15a-5p | ACTR3 | HITS-CLIP | Tarbase |
| hsa-miR-195-5p | ACTR3 | HITS-CLIP | Tarbase |
| hsa-miR-3605-3p | ACTR3 | HITS-CLIP | Tarbase |
| hsa-miR-141-5p | GPD2 | HITS-CLIP | Tarbase |
| hsa-miR-141-5p | TANC1 | HITS-CLIP | Tarbase |
| hsa-miR-486-5p | TANC1 | HITS-CLIP | Tarbase |
| hsa-miR-141-3p | GTF3C2 | HITS-CLIP | Tarbase |
| hsa-miR-15a-5p | GTF3C2 | PAR-CLIP | Tarbase |
| hsa-miR-15a-5p | GTF3C2 | PAR-CLIP | Tarbase |
| hsa-miR-15a-5p | NRBP1 | HITS-CLIP | Tarbase |
| hsa-miR-195-5p | NRBP1 | HITS-CLIP | Tarbase |
| hsa-miR-195-5p | NRBP1 | Microarrays | Tarbase |
| hsa-miR-15a-5p | GPR75-ASB3 | PAR-CLIP | Tarbase |
| hsa-miR-141-5p | NDUFS7 | HITS-CLIP | Tarbase |
| hsa-miR-141-3p | SPTBN1 | HITS-CLIP | Tarbase |
| hsa-miR-141-5p | SPTBN1 | HITS-CLIP | Tarbase |
| hsa-miR-141-5p | SPTBN1 | HITS-CLIP | Tarbase |
| hsa-miR-3605-5p | SPTBN1 | HITS-CLIP | Tarbase |
| hsa-miR-3613-5p | SPTBN1 | HITS-CLIP | Tarbase |
| hsa-miR-15a-5p | AUP1 | HITS-CLIP | Tarbase |
| hsa-miR-15a-5p | AUP1 | PAR-CLIP | Tarbase |
| hsa-miR-195-5p | AUP1 | HITS-CLIP | Tarbase |
| hsa-miR-141-3p | RTN4 | HITS-CLIP | Tarbase |
| hsa-miR-141-3p | RTN4 | HITS-CLIP | Tarbase |
| hsa-miR-15a-5p | RTN4 | HITS-CLIP | Tarbase |
| hsa-miR-15a-5p | RTN4 | HITS-CLIP | Tarbase |
| hsa-miR-15a-5p | RTN4 | HITS-CLIP | Tarbase |
| hsa-miR-15a-5p | RTN4 | HITS-CLIP | Tarbase |
| hsa-miR-15a-5p | RTN4 | PAR-CLIP | Tarbase |
| hsa-miR-15a-5p | RTN4 | HITS-CLIP | Tarbase |
| hsa-miR-15a-5p | RTN4 | PAR-CLIP | Tarbase |
| hsa-miR-195-5p | RTN4 | HITS-CLIP | Tarbase |
| hsa-miR-15a-5p | RTN4 | HITS-CLIP | Tarbase |
| hsa-miR-195-5p | RTN4 | HITS-CLIP | Tarbase |
| hsa-miR-15a-3p | DOK1 | HITS-CLIP | Tarbase |
| hsa-miR-141-3p | GALNT3 | HITS-CLIP | Tarbase |
| hsa-miR-15a-5p | GALNT3 | HITS-CLIP | Tarbase |
| hsa-miR-195-5p | GALNT3 | HITS-CLIP | Tarbase |
| hsa-miR-15a-5p | POLE4 | HITS-CLIP | Tarbase |
| hsa-miR-15a-5p | CCDC88A | HITS-CLIP | Tarbase |

| | | | |
|---|---|---|---|
| hsa-miR-195-5p | CCDC88A | HITS-CLIP | Tarbase |
| hsa-miR-195-5p | EVA1A | Microarrays | Tarbase |
| hsa-miR-3613-5p | LANCL1 | HITS-CLIP | Tarbase |
| hsa-miR-141-5p | WDR75 | HITS-CLIP | Tarbase |
| hsa-miR-15a-5p | EFEMP1 | HITS-CLIP | Tarbase |
| hsa-miR-195-5p | EFEMP1 | HITS-CLIP | Tarbase |
| hsa-miR-3613-5p | FN1 | HITS-CLIP | Tarbase |
| hsa-miR-141-3p | GLS | HITS-CLIP | Tarbase |
| hsa-miR-141-5p | GLS | HITS-CLIP | Tarbase |
| hsa-miR-141-5p | GLS | HITS-CLIP | Tarbase |
| hsa-miR-15a-5p | GLS | HITS-CLIP | Tarbase |
| hsa-miR-195-5p | ELMOD3 | Microarrays | Tarbase |
| hsa-miR-15a-5p | IGFBP5 | HITS-CLIP | Tarbase |
| hsa-miR-195-5p | IGFBP5 | HITS-CLIP | Tarbase |
| hsa-miR-195-5p | IGFBP5 | Microarrays | Tarbase |
| hsa-miR-141-5p | CCT4 | HITS-CLIP | Tarbase |
| hsa-miR-15a-3p | CCT4 | HITS-CLIP | Tarbase |
| hsa-miR-15a-3p | CCT4 | HITS-CLIP | Tarbase |
| hsa-miR-15a-5p | CCT4 | HITS-CLIP | Tarbase |
| hsa-miR-15a-5p | CCT4 | HITS-CLIP | Tarbase |
| hsa-miR-195-5p | CCT4 | HITS-CLIP | Tarbase |
| hsa-miR-195-5p | CCT4 | HITS-CLIP | Tarbase |
| hsa-miR-3613-3p | EHBP1 | HITS-CLIP | Tarbase |
| hsa-miR-141-5p | SF3B1 | HITS-CLIP | Tarbase |
| hsa-miR-141-5p | SF3B1 | HITS-CLIP | Tarbase |
| hsa-miR-141-5p | SF3B1 | HITS-CLIP | Tarbase |
| hsa-miR-15a-5p | SF3B1 | HITS-CLIP | Tarbase |
| hsa-miR-195-5p | SF3B1 | HITS-CLIP | Tarbase |
| hsa-miR-195-5p | SF3B1 | HITS-CLIP | Tarbase |
| hsa-miR-195-5p | SF3B1 | HITS-CLIP | Tarbase |
| hsa-miR-195-5p | SF3B1 | HITS-CLIP | Tarbase |
| hsa-miR-3613-5p | SF3B1 | PAR-CLIP | Tarbase |
| hsa-miR-15a-5p | ST3GAL5 | HITS-CLIP | Tarbase |
| hsa-miR-195-5p | ST3GAL5 | HITS-CLIP | Tarbase |
| hsa-miR-195-5p | ST3GAL5 | Microarrays | Tarbase |
| hsa-miR-15a-5p | MOB4 | HITS-CLIP | Tarbase |
| hsa-miR-195-5p | MOB4 | HITS-CLIP | Tarbase |
| hsa-miR-15a-5p | HSPE1 | HITS-CLIP | Tarbase |
| hsa-miR-195-5p | HSPE1 | HITS-CLIP | Tarbase |
| hsa-miR-3605-5p | KDM3A | HITS-CLIP | Tarbase |
| hsa-miR-15a-5p | CHMP3 | PAR-CLIP | Tarbase |
| hsa-miR-15a-5p | CHMP3 | HITS-CLIP | Tarbase |
| hsa-miR-195-5p | CHMP3 | HITS-CLIP | Tarbase |
| hsa-miR-15a-5p | ZNF142 | Chimeric fragments | Tarbase |
| hsa-miR-195-5p | IL1R2 | Microarrays | Tarbase |
| hsa-miR-195-5p | IL1R1 | Microarrays | Tarbase |
| hsa-miR-195-5p | IL18R1 | Microarrays | Tarbase |
| hsa-miR-15a-5p | SLC9A2 | HITS-CLIP | Tarbase |
| hsa-miR-195-5p | SLC9A2 | HITS-CLIP | Tarbase |
| hsa-miR-15a-5p | FHL2 | HITS-CLIP | Tarbase |
| hsa-miR-486-3p | FHL2 | HITS-CLIP | Tarbase |
| hsa-miR-195-5p | FHL2 | Microarrays | Tarbase |
| hsa-miR-195-5p | MLPH | Microarrays | Tarbase |
| hsa-miR-15a-3p | STK25 | HITS-CLIP | Tarbase |
| hsa-miR-15a-5p | ID2 | PAR-CLIP | Tarbase |
| hsa-miR-141-3p | TAF1B | PAR-CLIP | Tarbase |
| hsa-miR-15a-5p | ODC1 | HITS-CLIP | Tarbase |
| hsa-miR-15a-5p | ODC1 | HITS-CLIP | Tarbase |
| hsa-miR-15a-5p | ODC1 | PAR-CLIP | Tarbase |
| hsa-miR-15a-5p | ODC1 | HITS-CLIP | Tarbase |
| hsa-miR-15a-5p | ODC1 | PAR-CLIP | Tarbase |
| hsa-miR-15a-5p | ODC1 | PAR-CLIP | Tarbase |
| hsa-miR-15a-5p | ODC1 | HITS-CLIP | Tarbase |
| hsa-miR-195-5p | ODC1 | HITS-CLIP | Tarbase |
| hsa-miR-195-5p | ODC1 | HITS-CLIP | Tarbase |
| hsa-miR-195-5p | ODC1 | PAR-CLIP | Tarbase |
| hsa-miR-195-5p | ODC1 | HITS-CLIP | Tarbase |
| hsa-miR-195-5p | ODC1 | Microarrays | Tarbase |
| hsa-miR-141-3p | BIRC6 | HITS-CLIP | Tarbase |
| hsa-miR-15a-5p | BIRC6 | HITS-CLIP | Tarbase |
| hsa-miR-486-5p | BIRC6 | PAR-CLIP | Tarbase |
| hsa-miR-486-5p | BIRC6 | PAR-CLIP | Tarbase |
| hsa-miR-141-3p | PLEKHB2 | HITS-CLIP | Tarbase |
| hsa-miR-15a-5p | PLEKHB2 | HITS-CLIP | Tarbase |
| hsa-miR-15a-5p | PLEKHB2 | HITS-CLIP | Tarbase |
| hsa-miR-195-5p | PLEKHB2 | HITS-CLIP | Tarbase |
| hsa-miR-15a-5p | GORASP2 | HITS-CLIP | Tarbase |
| hsa-miR-141-3p | STRN | HITS-CLIP | Tarbase |
| hsa-miR-141-3p | STRN | HITS-CLIP | Tarbase |
| hsa-miR-141-3p | STRN | HITS-CLIP | Tarbase |
| hsa-miR-15a-5p | PRKD3 | Chimeric fragments | Tarbase |
| hsa-miR-15a-5p | DCAF17 | HITS-CLIP | Tarbase |
| hsa-miR-15a-5p | QPCT | PAR-CLIP | Tarbase |
| hsa-miR-3605-5p | RAB3GAP1 | PAR-CLIP | Tarbase |
| hsa-miR-15a-5p | SLC25A12 | HITS-CLIP | Tarbase |
| hsa-miR-195-5p | SLC25A12 | HITS-CLIP | Tarbase |
| hsa-miR-195-5p | DLX2 | Microarrays | Tarbase |
| hsa-miR-15a-3p | DARS | HITS-CLIP | Tarbase |
| hsa-miR-15a-5p | DARS | HITS-CLIP | Tarbase |
| hsa-miR-195-5p | DARS | HITS-CLIP | Tarbase |
| hsa-miR-15a-5p | SLC1A4 | HITS-CLIP | Tarbase |
| hsa-miR-195-5p | SLC1A4 | HITS-CLIP | Tarbase |
| hsa-miR-141-5p | SOS1 | HITS-CLIP | Tarbase |
| hsa-miR-141-3p | ORC2 | PAR-CLIP | Tarbase |
| hsa-miR-15a-5p | ATF2 | PAR-CLIP | Tarbase |
| hsa-miR-15a-5p | ATF2 | HITS-CLIP | Tarbase |
| hsa-miR-195-5p | ATF2 | PAR-CLIP | Tarbase |
| hsa-miR-195-5p | ATF2 | HITS-CLIP | Tarbase |
| hsa-miR-141-3p | EPAS1 | HITS-CLIP | Tarbase |
| hsa-miR-141-3p | EPAS1 | HITS-CLIP | Tarbase |
| hsa-miR-15a-3p | EPAS1 | HITS-CLIP | Tarbase |
| hsa-miR-3613-3p | ARID3A | PAR-CLIP | Tarbase |
| hsa-miR-3613-3p | ARID3A | PAR-CLIP | Tarbase |
| hsa-miR-141-3p | SUMO1 | HITS-CLIP | Tarbase |
| hsa-miR-141-5p | NFE2L2 | HITS-CLIP | Tarbase |
| hsa-miR-15a-3p | MSH6 | HITS-CLIP | Tarbase |
| hsa-miR-15a-3p | MSH6 | HITS-CLIP | Tarbase |
| hsa-miR-15a-3p | MSH6 | HITS-CLIP | Tarbase |
| hsa-miR-486-5p | MSH6 | PAR-CLIP | Tarbase |
| hsa-miR-486-5p | MSH6 | HITS-CLIP | Tarbase |
| hsa-miR-486-5p | MSH6 | HITS-CLIP | Tarbase |
| hsa-miR-15a-5p | PLEKHA3 | HITS-CLIP | Tarbase |
| hsa-miR-195-5p | PLEKHA3 | HITS-CLIP | Tarbase |
| hsa-miR-195-5p | EPHA4 | Microarrays | Tarbase |
| hsa-miR-195-5p | BCL9 | Microarrays | Tarbase |
| hsa-miR-219a-1-3p | PRRX1 | HITS-CLIP | Tarbase |
| hsa-miR-15a-3p | DHCR24 | HITS-CLIP | Tarbase |
| hsa-miR-15a-3p | DHCR24 | HITS-CLIP | Tarbase |
| hsa-miR-15a-5p | DNAJC16 | HITS-CLIP | Tarbase |
| hsa-miR-195-5p | DNAJC16 | HITS-CLIP | Tarbase |
| hsa-miR-15a-5p | MARK1 | HITS-CLIP | Tarbase |
| hsa-miR-195-5p | MARK1 | HITS-CLIP | Tarbase |
| hsa-miR-195-5p | PAPPA2 | Microarrays | Tarbase |
| hsa-miR-15a-5p | TMEM59 | HITS-CLIP | Tarbase |
| hsa-miR-195-5p | TMEM59 | HITS-CLIP | Tarbase |
| hsa-miR-219a-1-3p | TMEM59 | HITS-CLIP | Tarbase |
| hsa-miR-15a-5p | TMEM59 | HITS-CLIP | Tarbase |
| hsa-miR-195-5p | TMEM59 | HITS-CLIP | Tarbase |
| hsa-miR-15a-5p | ICMT | PAR-CLIP | Tarbase |
| hsa-miR-141-5p | QSOX1 | HITS-CLIP | Tarbase |
| hsa-miR-15a-5p | QSOX1 | HITS-CLIP | Tarbase |
| hsa-miR-195-5p | QSOX1 | HITS-CLIP | Tarbase |
| hsa-miR-15a-5p | QSOX1 | HITS-CLIP | Tarbase |
| hsa-miR-195-5p | QSOX1 | HITS-CLIP | Tarbase |
| hsa-miR-15a-5p | STXBP3 | PAR-CLIP | Tarbase |
| hsa-miR-15a-5p | STXBP3 | HITS-CLIP | Tarbase |
| hsa-miR-195-5p | STXBP3 | HITS-CLIP | Tarbase |
| hsa-miR-15a-5p | STXBP3 | HITS-CLIP | Tarbase |
| hsa-miR-195-5p | STXBP3 | HITS-CLIP | Tarbase |
| hsa-miR-141-3p | PHF13 | HITS-CLIP | Tarbase |
| hsa-miR-141-3p | ERRFI1 | HITS-CLIP | Tarbase |
| hsa-miR-141-3p | ERRFI1 | HITS-CLIP | Tarbase |
| hsa-miR-219a-1-3p | RAP1A | PAR-CLIP | Tarbase |
| hsa-miR-15a-5p | HDAC1 | Chimeric fragments | Tarbase |
| hsa-miR-15a-5p | CAPZA1 | PAR-CLIP | Tarbase |
| hsa-miR-15a-5p | CAPZA1 | PAR-CLIP | Tarbase |
| hsa-miR-141-3p | S100PBP | HITS-CLIP | Tarbase |
| hsa-miR-141-3p | S100PBP | HITS-CLIP | Tarbase |
| hsa-miR-3613-3p | S100PBP | HITS-CLIP | Tarbase |

| | | | |
|---|---|---|---|
| hsa-miR-15a-5p | SCAMP3 | HITS-CLIP | Tarbase |
| hsa-miR-195-5p | SCAMP3 | HITS-CLIP | Tarbase |
| hsa-miR-195-5p | TRIM62 | Microarrays | Tarbase |
| hsa-miR-141-3p | ASH1L | HITS-CLIP | Tarbase |
| hsa-miR-141-3p | ASH1L | PAR-CLIP | Tarbase |
| hsa-miR-15a-3p | ASH1L | HITS-CLIP | Tarbase |
| hsa-miR-15a-5p | ASH1L | PAR-CLIP | Tarbase |
| hsa-miR-15a-5p | ASH1L | PAR-CLIP | Tarbase |
| hsa-miR-15a-5p | ASH1L | HITS-CLIP | Tarbase |
| hsa-miR-15a-5p | ASH1L | PAR-CLIP | Tarbase |
| hsa-miR-15a-5p | ASH1L | HITS-CLIP | Tarbase |
| hsa-miR-15a-5p | ASH1L | HITS-CLIP | Tarbase |
| hsa-miR-15a-5p | ASH1L | PAR-CLIP | Tarbase |
| hsa-miR-15a-5p | ASH1L | HITS-CLIP | Tarbase |
| hsa-miR-15a-5p | ASH1L | PAR-CLIP | Tarbase |
| hsa-miR-195-5p | ASH1L | PAR-CLIP | Tarbase |
| hsa-miR-195-5p | ASH1L | HITS-CLIP | Tarbase |
| hsa-miR-195-5p | ASH1L | HITS-CLIP | Tarbase |
| hsa-miR-195-5p | ASH1L | PAR-CLIP | Tarbase |
| hsa-miR-141-3p | SFPQ | HITS-CLIP | Tarbase |
| hsa-miR-141-5p | SFPQ | HITS-CLIP | Tarbase |
| hsa-miR-141-5p | SFPQ | HITS-CLIP | Tarbase |
| hsa-miR-141-5p | SFPQ | HITS-CLIP | Tarbase |
| hsa-miR-141-5p | SFPQ | HITS-CLIP | Tarbase |
| hsa-miR-15a-3p | SFPQ | HITS-CLIP | Tarbase |
| hsa-miR-15a-5p | SFPQ | HITS-CLIP | Tarbase |
| hsa-miR-195-5p | SFPQ | HITS-CLIP | Tarbase |
| hsa-miR-3605-3p | SFPQ | HITS-CLIP | Tarbase |
| hsa-miR-141-3p | SRM | HITS-CLIP | Tarbase |
| hsa-miR-141-3p | SRM | HITS-CLIP | Tarbase |
| hsa-miR-15a-3p | SRM | HITS-CLIP | Tarbase |
| hsa-miR-15a-3p | SRM | HITS-CLIP | Tarbase |
| hsa-miR-15a-3p | SRM | HITS-CLIP | Tarbase |
| hsa-miR-15a-3p | SRM | HITS-CLIP | Tarbase |
| hsa-miR-3605-5p | IVNS1ABP | HITS-CLIP | Tarbase |
| hsa-miR-141-5p | MFN2 | HITS-CLIP | Tarbase |
| hsa-miR-15a-5p | MFN2 | HITS-CLIP | Tarbase |
| hsa-miR-219a-1-3p | SMG7 | PAR-CLIP | Tarbase |
| hsa-miR-195-5p | NCF2 | Microarrays | Tarbase |
| hsa-miR-3605-3p | SLC35D1 | HITS-CLIP | Tarbase |
| hsa-miR-3613-3p | SLC35D1 | HITS-CLIP | Tarbase |
| hsa-miR-195-5p | PLA2G4A | Microarrays | Tarbase |
| hsa-miR-141-5p | TROVE2 | HITS-CLIP | Tarbase |
| hsa-miR-141-3p | SRSF11 | HITS-CLIP | Tarbase |
| hsa-miR-15a-5p | PLEKHM2 | HITS-CLIP | Tarbase |
| hsa-miR-195-5p | PLEKHM2 | HITS-CLIP | Tarbase |
| hsa-miR-141-3p | CRYZ | HITS-CLIP | Tarbase |
| hsa-miR-3613-5p | CD58 | PAR-CLIP | Tarbase |
| hsa-miR-3613-5p | CD58 | PAR-CLIP | Tarbase |
| hsa-miR-15a-3p | KIF21B | HITS-CLIP | Tarbase |
| hsa-miR-141-3p | MAP7D1 | HITS-CLIP | Tarbase |
| hsa-miR-141-3p | MAP7D1 | HITS-CLIP | Tarbase |
| hsa-miR-3605-3p | MAP7D1 | HITS-CLIP | Tarbase |
| hsa-miR-219a-1-3p | EXOC8 | PAR-CLIP | Tarbase |
| hsa-miR-141-5p | MTR | HITS-CLIP | Tarbase |
| hsa-miR-486-5p | MTR | HITS-CLIP | Tarbase |
| hsa-miR-15a-5p | SIPA1L2 | PAR-CLIP | Tarbase |
| hsa-miR-195-5p | SIPA1L2 | Microarrays | Tarbase |
| hsa-miR-3613-5p | RLF | PAR-CLIP | Tarbase |
| hsa-miR-15a-5p | AKT3 | HITS-CLIP | Tarbase |
| hsa-miR-15a-5p | AKT3 | Luciferase Reporter Assay | Tarbase |
| hsa-miR-15a-5p | AKT3 | Western Blot | Tarbase |
| hsa-miR-15a-5p | AKT3 | PAR-CLIP | Tarbase |
| hsa-miR-15a-5p | AKT3 | PAR-CLIP | Tarbase |
| hsa-miR-15a-5p | AKT3 | PAR-CLIP | Tarbase |
| hsa-miR-15a-5p | AKT3 | PAR-CLIP | Tarbase |
| hsa-miR-15a-5p | AKT3 | PAR-CLIP | Tarbase |
| hsa-miR-15a-5p | AKT3 | PAR-CLIP | Tarbase |
| hsa-miR-15a-5p | AKT3 | PAR-CLIP | Tarbase |
| hsa-miR-195-5p | AKT3 | HITS-CLIP | Tarbase |
| hsa-miR-195-5p | AKT3 | PAR-CLIP | Tarbase |
| hsa-miR-195-5p | AKT3 | PAR-CLIP | Tarbase |
| hsa-miR-195-5p | AKT3 | PAR-CLIP | Tarbase |
| hsa-miR-195-5p | AKT3 | PAR-CLIP | Tarbase |
| hsa-miR-15a-5p | AKT3 | HITS-CLIP | Tarbase |
| hsa-miR-195-5p | AKT3 | HITS-CLIP | Tarbase |
| hsa-miR-141-3p | ETV3 | HITS-CLIP | Tarbase |
| hsa-miR-3605-5p | CD48 | HITS-CLIP | Tarbase |
| hsa-miR-3613-5p | CD48 | PAR-CLIP | Tarbase |
| hsa-miR-15a-5p | MFAP2 | HITS-CLIP | Tarbase |
| hsa-miR-195-5p | MFAP2 | HITS-CLIP | Tarbase |
| hsa-miR-141-5p | KDM5B | HITS-CLIP | Tarbase |
| hsa-miR-141-3p | KIF17 | HITS-CLIP | Tarbase |
| hsa-miR-141-5p | RAB29 | HITS-CLIP | Tarbase |
| hsa-miR-15a-5p | RAB29 | HITS-CLIP | Tarbase |
| hsa-miR-15a-5p | RAB29 | HITS-CLIP | Tarbase |
| hsa-miR-195-5p | RAB29 | HITS-CLIP | Tarbase |
| hsa-miR-486-5p | ECE1 | HITS-CLIP | Tarbase |
| hsa-miR-15a-5p | ID3 | PAR-CLIP | Tarbase |
| hsa-miR-15a-5p | ID3 | HITS-CLIP | Tarbase |
| hsa-miR-141-5p | CD46 | HITS-CLIP | Tarbase |
| hsa-miR-141-5p | CD46 | HITS-CLIP | Tarbase |
| hsa-miR-141-5p | CD46 | HITS-CLIP | Tarbase |
| hsa-miR-141-5p | CD46 | HITS-CLIP | Tarbase |
| hsa-miR-141-5p | CD46 | HITS-CLIP | Tarbase |
| hsa-miR-15a-5p | CD46 | HITS-CLIP | Tarbase |
| hsa-miR-195-5p | CD46 | HITS-CLIP | Tarbase |
| hsa-miR-15a-5p | APH1A | PAR-CLIP | Tarbase |
| hsa-miR-3613-3p | APH1A | HITS-CLIP | Tarbase |
| hsa-miR-486-5p | APH1A | HITS-CLIP | Tarbase |
| hsa-miR-195-5p | CDC20 | Microarrays | Tarbase |
| hsa-miR-15a-5p | IPO13 | PAR-CLIP | Tarbase |
| hsa-miR-15a-5p | ATP6V0B | HITS-CLIP | Tarbase |
| hsa-miR-195-5p | ATP6V0B | HITS-CLIP | Tarbase |
| hsa-miR-195-5p | PTCH2 | AGO-IP | Tarbase |
| hsa-miR-15a-5p | AKR1A1 | HITS-CLIP | Tarbase |
| hsa-miR-195-5p | AKR1A1 | HITS-CLIP | Tarbase |
| hsa-miR-141-3p | SLC19A2 | HITS-CLIP | Tarbase |
| hsa-miR-15a-5p | SLC19A2 | PAR-CLIP | Tarbase |
| hsa-miR-15a-5p | SLC19A2 | HITS-CLIP | Tarbase |
| hsa-miR-15a-5p | DR1 | PAR-CLIP | Tarbase |
| hsa-miR-15a-5p | CNN3 | CLASH | Tarbase |
| hsa-miR-195-5p | CNN3 | Microarrays | Tarbase |
| hsa-miR-141-5p | PRRC2C | HITS-CLIP | Tarbase |
| hsa-miR-15a-5p | PRRC2C | HITS-CLIP | Tarbase |
| hsa-miR-15a-5p | PRRC2C | HITS-CLIP | Tarbase |
| hsa-miR-15a-5p | PRRC2C | PAR-CLIP | Tarbase |
| hsa-miR-195-5p | PRRC2C | HITS-CLIP | Tarbase |
| hsa-miR-195-5p | PRRC2C | HITS-CLIP | Tarbase |
| hsa-miR-3605-3p | PRRC2C | HITS-CLIP | Tarbase |
| hsa-miR-486-5p | PRRC2C | PAR-CLIP | Tarbase |
| hsa-miR-195-5p | PTBP2 | Microarrays | Tarbase |
| hsa-miR-15a-5p | PTBP2 | Chimeric fragments | Tarbase |
| hsa-miR-141-3p | IRF6 | PAR-CLIP | Tarbase |
| hsa-miR-15a-5p | DIEXF | PAR-CLIP | Tarbase |
| hsa-miR-15a-5p | DIEXF | PAR-CLIP | Tarbase |
| hsa-miR-195-5p | DIEXF | PAR-CLIP | Tarbase |
| hsa-miR-195-5p | LPPR5 | Microarrays | Tarbase |
| hsa-miR-195-5p | LPPR4 | Microarrays | Tarbase |
| hsa-miR-15a-5p | RCAN3 | HITS-CLIP | Tarbase |
| hsa-miR-195-5p | RCAN3 | HITS-CLIP | Tarbase |
| hsa-miR-3613-3p | RCAN3 | HITS-CLIP | Tarbase |
| hsa-miR-486-5p | SLC35A3 | HITS-CLIP | Tarbase |
| hsa-miR-141-5p | RCOR3 | HITS-CLIP | Tarbase |
| hsa-miR-15a-5p | MTFR1L | PAR-CLIP | Tarbase |
| hsa-miR-141-3p | MAN1C1 | HITS-CLIP | Tarbase |
| hsa-miR-195-5p | NEK2 | Microarrays | Tarbase |
| hsa-miR-195-5p | RPS6KA1 | Microarrays | Tarbase |
| hsa-miR-15a-5p | ARID1A | HITS-CLIP | Tarbase |
| hsa-miR-486-5p | ARID1A | HITS-CLIP | Tarbase |
| hsa-miR-141-3p | CENPF | HITS-CLIP | Tarbase |
| hsa-miR-141-5p | CENPF | HITS-CLIP | Tarbase |
| hsa-miR-15a-3p | CENPF | HITS-CLIP | Tarbase |
| hsa-miR-219a-1-3p | CENPF | PAR-CLIP | Tarbase |
| hsa-miR-195-5p | CENPF | Microarrays | Tarbase |
| hsa-miR-141-5p | RPA2 | HITS-CLIP | Tarbase |
| hsa-miR-15a-5p | RPA2 | HITS-CLIP | Tarbase |
| hsa-miR-15a-5p | RPA2 | PAR-CLIP | Tarbase |
| hsa-miR-15a-5p | RPA2 | PAR-CLIP | Tarbase |

| | | | |
|---|---|---|---|
| hsa-miR-15a-5p | RPA2 | HITS-CLIP | Tarbase |
| hsa-miR-195-5p | RPA2 | HITS-CLIP | Tarbase |
| hsa-miR-195-5p | OSBPL9 | Microarrays | Tarbase |
| hsa-miR-15a-5p | MESDC2 | PAR-CLIP | Tarbase |
| hsa-miR-15a-5p | MESDC2 | HITS-CLIP | Tarbase |
| hsa-miR-195-5p | MESDC2 | HITS-CLIP | Tarbase |
| hsa-miR-141-5p | RCN2 | HITS-CLIP | Tarbase |
| hsa-miR-141-5p | RCN2 | HITS-CLIP | Tarbase |
| hsa-miR-141-5p | RCN2 | HITS-CLIP | Tarbase |
| hsa-miR-3613-5p | RCN2 | HITS-CLIP | Tarbase |
| hsa-miR-15a-5p | STK11 | HITS-CLIP | Tarbase |
| hsa-miR-195-5p | STK11 | HITS-CLIP | Tarbase |
| hsa-miR-141-3p | KMT2A | HITS-CLIP | Tarbase |
| hsa-miR-141-5p | KMT2A | HITS-CLIP | Tarbase |
| hsa-miR-15a-5p | KMT2A | PAR-CLIP | Tarbase |
| hsa-miR-15a-5p | KMT2A | HITS-CLIP | Tarbase |
| hsa-miR-3613-5p | KMT2A | HITS-CLIP | Tarbase |
| hsa-miR-486-3p | KMT2A | HITS-CLIP | Tarbase |
| hsa-miR-486-5p | KMT2A | HITS-CLIP | Tarbase |
| hsa-miR-15a-5p | RPS25 | HITS-CLIP | Tarbase |
| hsa-miR-15a-5p | RPS25 | HITS-CLIP | Tarbase |
| hsa-miR-15a-5p | RPS25 | HITS-CLIP | Tarbase |
| hsa-miR-15a-5p | RPS25 | HITS-CLIP | Tarbase |
| hsa-miR-15a-5p | RPS25 | HITS-CLIP | Tarbase |
| hsa-miR-15a-5p | RPS25 | PAR-CLIP | Tarbase |
| hsa-miR-195-5p | RPS25 | HITS-CLIP | Tarbase |
| hsa-miR-195-5p | RPS25 | HITS-CLIP | Tarbase |
| hsa-miR-195-5p | RPS25 | HITS-CLIP | Tarbase |
| hsa-miR-195-5p | RPS25 | HITS-CLIP | Tarbase |
| hsa-miR-195-5p | RPS25 | HITS-CLIP | Tarbase |
| hsa-miR-15a-5p | KIF14 | PAR-CLIP | Tarbase |
| hsa-miR-195-5p | KIF14 | PAR-CLIP | Tarbase |
| hsa-miR-195-5p | KIF14 | Microarrays | Tarbase |
| hsa-miR-141-3p | CAMSAP2 | HITS-CLIP | Tarbase |
| hsa-miR-141-3p | CAMSAP2 | HITS-CLIP | Tarbase |
| hsa-miR-141-5p | CAMSAP2 | HITS-CLIP | Tarbase |
| hsa-miR-15a-5p | CAMSAP2 | HITS-CLIP | Tarbase |
| hsa-miR-195-5p | CAMSAP2 | HITS-CLIP | Tarbase |
| hsa-miR-15a-5p | CAMSAP2 | HITS-CLIP | Tarbase |
| hsa-miR-195-5p | CAMSAP2 | HITS-CLIP | Tarbase |
| hsa-miR-195-5p | ATF6 | Microarrays | Tarbase |
| hsa-miR-195-5p | NRP2 | Microarrays | Tarbase |
| hsa-miR-15a-5p | CREB1 | HITS-CLIP | Tarbase |
| hsa-miR-195-5p | CREB1 | HITS-CLIP | Tarbase |
| hsa-miR-15a-5p | KLF7 | HITS-CLIP | Tarbase |
| hsa-miR-195-5p | KLF7 | HITS-CLIP | Tarbase |
| hsa-miR-3613-3p | KLF7 | PAR-CLIP | Tarbase |
| hsa-miR-195-5p | KLF7 | Microarrays | Tarbase |
| hsa-miR-141-3p | B4GALT6 | HITS-CLIP | Tarbase |
| hsa-miR-141-5p | B4GALT6 | HITS-CLIP | Tarbase |
| hsa-miR-3613-3p | B4GALT6 | PAR-CLIP | Tarbase |
| hsa-miR-195-5p | ATP10B | Microarrays | Tarbase |
| hsa-miR-15a-3p | CASP8AP2 | HITS-CLIP | Tarbase |
| hsa-miR-15a-3p | CASP8AP2 | HITS-CLIP | Tarbase |
| hsa-miR-195-5p | SGIP1 | Microarrays | Tarbase |
| hsa-miR-141-5p | PHF3 | HITS-CLIP | Tarbase |
| hsa-miR-15a-5p | PHF3 | HITS-CLIP | Tarbase |
| hsa-miR-15a-5p | PHF3 | HITS-CLIP | Tarbase |
| hsa-miR-195-5p | PHF3 | HITS-CLIP | Tarbase |
| hsa-miR-486-5p | PHF3 | HITS-CLIP | Tarbase |
| hsa-miR-195-5p | PLAGL1 | Microarrays | Tarbase |
| hsa-miR-15a-3p | TNFAIP3 | HITS-CLIP | Tarbase |
| hsa-miR-15a-3p | TNFAIP3 | HITS-CLIP | Tarbase |
| hsa-miR-15a-3p | TNFAIP3 | HITS-CLIP | Tarbase |
| hsa-miR-219a-1-3p | TNFAIP3 | PAR-CLIP | Tarbase |
| hsa-miR-141-3p | RAB32 | HITS-CLIP | Tarbase |
| hsa-miR-15a-3p | MYB | HITS-CLIP | Tarbase |
| hsa-miR-15a-5p | MYB | HITS-CLIP | Tarbase |
| hsa-miR-15a-5p | MYB | HITS-CLIP | Tarbase |
| hsa-miR-195-5p | MYB | HITS-CLIP | Tarbase |
| hsa-miR-15a-5p | SGK1 | HITS-CLIP | Tarbase |
| hsa-miR-15a-5p | SGK1 | HITS-CLIP | Tarbase |
| hsa-miR-15a-5p | SGK1 | PAR-CLIP | Tarbase |
| hsa-miR-195-5p | SGK1 | HITS-CLIP | Tarbase |
| hsa-miR-195-5p | SGK1 | HITS-CLIP | Tarbase |
| hsa-miR-195-5p | SGK1 | Microarrays | Tarbase |
| hsa-miR-15a-3p | CTGF | HITS-CLIP | Tarbase |
| hsa-miR-3613-5p | CTGF | HITS-CLIP | Tarbase |
| hsa-miR-195-5p | CTGF | Microarrays | Tarbase |
| hsa-miR-141-3p | FBXL5 | HITS-CLIP | Tarbase |
| hsa-miR-15a-3p | FBXL5 | HITS-CLIP | Tarbase |
| hsa-miR-3613-3p | MED28 | HITS-CLIP | Tarbase |
| hsa-miR-15a-5p | VAMP8 | PAR-CLIP | Tarbase |
| hsa-miR-195-5p | VAMP8 | Microarrays | Tarbase |
| hsa-miR-195-5p | DCLRE1B | Microarrays | Tarbase |
| hsa-miR-141-3p | FOXO3 | HITS-CLIP | Tarbase |
| hsa-miR-141-3p | FOXO3 | HITS-CLIP | Tarbase |
| hsa-miR-15a-3p | RPN2 | HITS-CLIP | Tarbase |
| hsa-miR-141-3p | TGIF2 | HITS-CLIP | Tarbase |
| hsa-miR-15a-5p | TGIF2 | HITS-CLIP | Tarbase |
| hsa-miR-195-5p | TGIF2 | HITS-CLIP | Tarbase |
| hsa-miR-3613-3p | TGIF2 | PAR-CLIP | Tarbase |
| hsa-miR-195-5p | SPP1 | Microarrays | Tarbase |
| hsa-miR-3605-3p | FAM47E-STBD1 | HITS-CLIP | Tarbase |
| hsa-miR-3613-5p | CCNI | HITS-CLIP | Tarbase |
| hsa-miR-486-3p | CCNI | HITS-CLIP | Tarbase |
| hsa-miR-3613-3p | RAB3GAP2 | HITS-CLIP | Tarbase |
| hsa-miR-15a-5p | PPL | HITS-CLIP | Tarbase |
| hsa-miR-141-3p | CCND2 | HITS-CLIP | Tarbase |
| hsa-miR-141-5p | CCND2 | HITS-CLIP | Tarbase |
| hsa-miR-141-5p | CCND2 | HITS-CLIP | Tarbase |
| hsa-miR-141-5p | CCND2 | HITS-CLIP | Tarbase |
| hsa-miR-15a-3p | CCND2 | HITS-CLIP | Tarbase |
| hsa-miR-15a-5p | CCND2 | HITS-CLIP | Tarbase |
| hsa-miR-15a-5p | CCND2 | PAR-CLIP | Tarbase |
| hsa-miR-15a-5p | CCND2 | PAR-CLIP | Tarbase |
| hsa-miR-15a-5p | CCND2 | HITS-CLIP | Tarbase |
| hsa-miR-15a-5p | CCND2 | HITS-CLIP | Tarbase |
| hsa-miR-15a-5p | CCND2 | CLASH | Tarbase |
| hsa-miR-15a-5p | CCND2 | HITS-CLIP | Tarbase |
| hsa-miR-15a-5p | CCND2 | HITS-CLIP | Tarbase |
| hsa-miR-15a-5p | CCND2 | PAR-CLIP | Tarbase |
| hsa-miR-15a-5p | CCND2 | PAR-CLIP | Tarbase |
| hsa-miR-195-5p | CCND2 | HITS-CLIP | Tarbase |
| hsa-miR-195-5p | CCND2 | PAR-CLIP | Tarbase |
| hsa-miR-195-5p | CCND2 | HITS-CLIP | Tarbase |
| hsa-miR-195-5p | CCND2 | HITS-CLIP | Tarbase |
| hsa-miR-195-5p | CCND2 | HITS-CLIP | Tarbase |
| hsa-miR-195-5p | CCND2 | HITS-CLIP | Tarbase |
| hsa-miR-195-5p | CCND2 | PAR-CLIP | Tarbase |
| hsa-miR-3605-5p | CCND2 | HITS-CLIP | Tarbase |
| hsa-miR-3613-3p | CCND2 | HITS-CLIP | Tarbase |
| hsa-miR-3613-3p | CCND2 | PAR-CLIP | Tarbase |
| hsa-miR-15a-5p | ELL2 | HITS-CLIP | Tarbase |
| hsa-miR-195-5p | ELL2 | HITS-CLIP | Tarbase |
| hsa-miR-219a-1-3p | ELL2 | PAR-CLIP | Tarbase |
| hsa-miR-219a-1-3p | ELL2 | PAR-CLIP | Tarbase |
| hsa-miR-219a-1-3p | ELL2 | PAR-CLIP | Tarbase |
| hsa-miR-486-5p | GDA | PAR-CLIP | Tarbase |
| hsa-miR-195-5p | KLF9 | Microarrays | Tarbase |
| hsa-miR-15a-5p | SENP5 | HITS-CLIP | Tarbase |
| hsa-miR-195-5p | SENP5 | HITS-CLIP | Tarbase |
| hsa-miR-3613-3p | SENP5 | PAR-CLIP | Tarbase |
| hsa-miR-3613-3p | SENP5 | PAR-CLIP | Tarbase |
| hsa-miR-195-5p | C1orf198 | Microarrays | Tarbase |
| hsa-miR-141-5p | HEATR1 | HITS-CLIP | Tarbase |
| hsa-miR-15a-5p | HEATR1 | PAR-CLIP | Tarbase |
| hsa-miR-195-5p | HEATR1 | PAR-CLIP | Tarbase |
| hsa-miR-195-5p | PTBP3 | HITS-CLIP | Tarbase |
| hsa-miR-219a-1-3p | PTBP3 | PAR-CLIP | Tarbase |
| hsa-miR-15a-5p | RAD23B | HITS-CLIP | Tarbase |
| hsa-miR-195-5p | RAD23B | HITS-CLIP | Tarbase |
| hsa-miR-15a-5p | RAD23B | HITS-CLIP | Tarbase |
| hsa-miR-195-5p | RAD23B | HITS-CLIP | Tarbase |
| hsa-miR-15a-5p | RAD23B | Chimeric fragments | Tarbase |
| hsa-miR-195-5p | WDR34 | Microarrays | Tarbase |
| hsa-miR-141-5p | SET | HITS-CLIP | Tarbase |
| hsa-miR-141-5p | SET | HITS-CLIP | Tarbase |
| hsa-miR-15a-5p | PPP2R4 | PAR-CLIP | Tarbase |

| | | | |
|---|---|---|---|
| hsa-miR-141-3p | RAB14 | HITS-CLIP | Tarbase |
| hsa-miR-15a-5p | RAB14 | HITS-CLIP | Tarbase |
| hsa-miR-15a-5p | RAB14 | PAR-CLIP | Tarbase |
| hsa-miR-15a-5p | RAB14 | HITS-CLIP | Tarbase |
| hsa-miR-15a-5p | RAB14 | HITS-CLIP | Tarbase |
| hsa-miR-195-5p | RAB14 | HITS-CLIP | Tarbase |
| hsa-miR-195-5p | RAB14 | HITS-CLIP | Tarbase |
| hsa-miR-3613-5p | RAB14 | HITS-CLIP | Tarbase |
| hsa-miR-195-5p | CNTRL | Microarrays | Tarbase |
| hsa-miR-15a-3p | TRIM32 | PAR-CLIP | Tarbase |
| hsa-miR-141-3p | FBXW2 | HITS-CLIP | Tarbase |
| hsa-miR-15a-5p | FBXW2 | HITS-CLIP | Tarbase |
| hsa-miR-195-5p | FBXW2 | HITS-CLIP | Tarbase |
| hsa-miR-3605-5p | FBXW2 | HITS-CLIP | Tarbase |
| hsa-miR-486-5p | FBXW2 | PAR-CLIP | Tarbase |
| hsa-miR-141-5p | PHF19 | HITS-CLIP | Tarbase |
| hsa-miR-141-5p | PHF19 | HITS-CLIP | Tarbase |
| hsa-miR-141-5p | PHF19 | HITS-CLIP | Tarbase |
| hsa-miR-15a-3p | PHF19 | HITS-CLIP | Tarbase |
| hsa-miR-15a-3p | PHF19 | HITS-CLIP | Tarbase |
| hsa-miR-195-5p | PHF19 | Microarrays | Tarbase |
| hsa-miR-141-3p | NEK6 | HITS-CLIP | Tarbase |
| hsa-miR-141-3p | NEK6 | HITS-CLIP | Tarbase |
| hsa-miR-195-5p | NEK6 | Microarrays | Tarbase |
| hsa-miR-141-3p | PPP6C | HITS-CLIP | Tarbase |
| hsa-miR-15a-3p | PPP6C | PAR-CLIP | Tarbase |
| hsa-miR-15a-3p | PPP6C | HITS-CLIP | Tarbase |
| hsa-miR-15a-3p | PPP6C | PAR-CLIP | Tarbase |
| hsa-miR-15a-5p | PPP6C | PAR-CLIP | Tarbase |
| hsa-miR-15a-5p | HSDL2 | HITS-CLIP | Tarbase |
| hsa-miR-195-5p | HSDL2 | HITS-CLIP | Tarbase |
| hsa-miR-15a-5p | MAPKAP1 | HITS-CLIP | Tarbase |
| hsa-miR-195-5p | MAPKAP1 | HITS-CLIP | Tarbase |
| hsa-miR-195-5p | GALNT12 | Microarrays | Tarbase |
| hsa-miR-141-3p | ALG2 | HITS-CLIP | Tarbase |
| hsa-miR-15a-5p | ALG2 | HITS-CLIP | Tarbase |
| hsa-miR-195-5p | ALG2 | Microarrays | Tarbase |
| hsa-miR-15a-5p | KDSR | HITS-CLIP | Tarbase |
| hsa-miR-15a-5p | KDSR | PAR-CLIP | Tarbase |
| hsa-miR-15a-5p | KDSR | HITS-CLIP | Tarbase |
| hsa-miR-195-5p | KDSR | HITS-CLIP | Tarbase |
| hsa-miR-195-5p | KDSR | HITS-CLIP | Tarbase |
| hsa-miR-486-5p | VPS4B | HITS-CLIP | Tarbase |
| hsa-miR-15a-5p | ONECUT2 | PAR-CLIP | Tarbase |
| hsa-miR-15a-5p | ZBTB45 | HITS-CLIP | Tarbase |
| hsa-miR-195-5p | ZBTB45 | HITS-CLIP | Tarbase |
| hsa-miR-141-3p | DCAF4 | HITS-CLIP | Tarbase |
| hsa-miR-15a-5p | FCF1 | PAR-CLIP | Tarbase |
| hsa-miR-141-3p | NPC2 | HITS-CLIP | Tarbase |
| hsa-miR-15a-5p | IRF2BPL | HITS-CLIP | Tarbase |
| hsa-miR-15a-5p | IRF2BPL | HITS-CLIP | Tarbase |
| hsa-miR-195-5p | IRF2BPL | HITS-CLIP | Tarbase |
| hsa-miR-195-5p | LTBP2 | Microarrays | Tarbase |
| hsa-miR-15a-5p | AREL1 | PAR-CLIP | Tarbase |
| hsa-miR-15a-5p | AREL1 | PAR-CLIP | Tarbase |
| hsa-miR-195-5p | AREL1 | PAR-CLIP | Tarbase |
| hsa-miR-3613-3p | AREL1 | HITS-CLIP | Tarbase |
| hsa-miR-195-5p | AREL1 | Microarrays | Tarbase |
| hsa-miR-15a-5p | FLVCR2 | PAR-CLIP | Tarbase |
| hsa-miR-195-5p | FLVCR2 | PAR-CLIP | Tarbase |
| hsa-miR-195-5p | PPP4R4 | Microarrays | Tarbase |
| hsa-miR-486-5p | SLIRP | PAR-CLIP | Tarbase |
| hsa-miR-15a-5p | RBM25 | PAR-CLIP | Tarbase |
| hsa-miR-15a-5p | RBM25 | HITS-CLIP | Tarbase |
| hsa-miR-15a-5p | RBM25 | HITS-CLIP | Tarbase |
| hsa-miR-195-5p | RBM25 | HITS-CLIP | Tarbase |
| hsa-miR-195-5p | RBM25 | HITS-CLIP | Tarbase |
| hsa-miR-195-5p | TMEM214 | HITS-CLIP | Tarbase |
| hsa-miR-141-3p | ATL2 | HITS-CLIP | Tarbase |
| hsa-miR-15a-3p | ATL2 | HITS-CLIP | Tarbase |
| hsa-miR-15a-3p | ATL2 | PAR-CLIP | Tarbase |
| hsa-miR-15a-5p | ATL2 | Chimeric fragments | Tarbase |
| hsa-miR-141-3p | YPEL5 | HITS-CLIP | Tarbase |
| hsa-miR-141-5p | FAM98A | HITS-CLIP | Tarbase |
| hsa-miR-15a-5p | FAM98A | HITS-CLIP | Tarbase |
| hsa-miR-15a-5p | FAM98A | HITS-CLIP | Tarbase |
| hsa-miR-195-5p | FAM98A | HITS-CLIP | Tarbase |
| hsa-miR-195-5p | FAM98A | HITS-CLIP | Tarbase |
| hsa-miR-15a-5p | SLC17A5 | PAR-CLIP | Tarbase |
| hsa-miR-15a-5p | SLC17A5 | HITS-CLIP | Tarbase |
| hsa-miR-195-5p | SLC17A5 | HITS-CLIP | Tarbase |
| hsa-miR-195-5p | SLC17A5 | Microarrays | Tarbase |
| hsa-miR-15a-5p | OGFRL1 | PAR-CLIP | Tarbase |
| hsa-miR-15a-3p | IDE | HITS-CLIP | Tarbase |
| hsa-miR-15a-5p | IFIT3 | HITS-CLIP | Tarbase |
| hsa-miR-195-5p | IFIT3 | HITS-CLIP | Tarbase |
| hsa-miR-15a-5p | GPAM | PAR-CLIP | Tarbase |
| hsa-miR-141-5p | SMNDC1 | HITS-CLIP | Tarbase |
| hsa-miR-15a-5p | HELLS | HITS-CLIP | Tarbase |
| hsa-miR-195-5p | HELLS | HITS-CLIP | Tarbase |
| hsa-miR-141-3p | KCNIP2 | HITS-CLIP | Tarbase |
| hsa-miR-141-3p | GNA13 | PAR-CLIP | Tarbase |
| hsa-miR-141-3p | GNA13 | HITS-CLIP | Tarbase |
| hsa-miR-15a-5p | GNA13 | PAR-CLIP | Tarbase |
| hsa-miR-15a-5p | GNA13 | PAR-CLIP | Tarbase |
| hsa-miR-15a-5p | GNA13 | PAR-CLIP | Tarbase |
| hsa-miR-15a-5p | GNA13 | PAR-CLIP | Tarbase |
| hsa-miR-15a-5p | GNA13 | PAR-CLIP | Tarbase |
| hsa-miR-195-5p | GNA13 | PAR-CLIP | Tarbase |
| hsa-miR-195-5p | GNA13 | PAR-CLIP | Tarbase |
| hsa-miR-141-3p | HOXB5 | Luciferase Reporter Assay | Tarbase |
| hsa-miR-141-3p | HOXB5 | Luciferase Reporter Assay | Tarbase |
| hsa-miR-15a-3p | HOXB3 | HITS-CLIP | Tarbase |
| hsa-miR-3613-3p | DUSP1 | HITS-CLIP | Tarbase |
| hsa-miR-195-5p | DUSP1 | Microarrays | Tarbase |
| hsa-miR-141-3p | PANK3 | HITS-CLIP | Tarbase |
| hsa-miR-195-5p | MSX2 | Microarrays | Tarbase |
| hsa-miR-141-5p | CAAP1 | HITS-CLIP | Tarbase |
| hsa-miR-141-5p | CAAP1 | HITS-CLIP | Tarbase |
| hsa-miR-15a-5p | CAAP1 | PAR-CLIP | Tarbase |
| hsa-miR-141-5p | MOB3B | HITS-CLIP | Tarbase |
| hsa-miR-15a-5p | MOB3B | PAR-CLIP | Tarbase |
| hsa-miR-15a-5p | MOB3B | HITS-CLIP | Tarbase |
| hsa-miR-15a-5p | MOB3B | HITS-CLIP | Tarbase |
| hsa-miR-195-5p | MOB3B | HITS-CLIP | Tarbase |
| hsa-miR-15a-5p | PCMT1 | PAR-CLIP | Tarbase |
| hsa-miR-195-5p | PCMT1 | Luciferase Reporter Assay | Tarbase |
| hsa-miR-15a-5p | PCMT1 | HITS-CLIP | Tarbase |
| hsa-miR-195-5p | PCMT1 | HITS-CLIP | Tarbase |
| hsa-miR-195-5p | PCMT1 | Microarrays | Tarbase |
| hsa-miR-195-5p | PCDHB14 | Microarrays | Tarbase |
| hsa-miR-195-5p | CENPL | Microarrays | Tarbase |
| hsa-miR-15a-5p | TCP1 | HITS-CLIP | Tarbase |
| hsa-miR-15a-5p | TCP1 | HITS-CLIP | Tarbase |
| hsa-miR-15a-5p | TCP1 | HITS-CLIP | Tarbase |
| hsa-miR-15a-5p | TCP1 | HITS-CLIP | Tarbase |
| hsa-miR-15a-5p | TCP1 | HITS-CLIP | Tarbase |
| hsa-miR-195-5p | TCP1 | HITS-CLIP | Tarbase |
| hsa-miR-195-5p | TCP1 | HITS-CLIP | Tarbase |
| hsa-miR-195-5p | TCP1 | HITS-CLIP | Tarbase |
| hsa-miR-195-5p | TCP1 | HITS-CLIP | Tarbase |
| hsa-miR-3613-5p | SNX19 | HITS-CLIP | Tarbase |
| hsa-miR-15a-5p | SLC10A7 | PAR-CLIP | Tarbase |
| hsa-miR-15a-5p | NUDCD1 | HITS-CLIP | Tarbase |
| hsa-miR-15a-5p | NUDCD1 | HITS-CLIP | Tarbase |
| hsa-miR-15a-5p | NUDCD1 | HITS-CLIP | Tarbase |
| hsa-miR-15a-5p | NUDCD1 | HITS-CLIP | Tarbase |
| hsa-miR-195-5p | NUDCD1 | HITS-CLIP | Tarbase |
| hsa-miR-195-5p | NUDCD1 | HITS-CLIP | Tarbase |
| hsa-miR-195-5p | NUDCD1 | HITS-CLIP | Tarbase |
| hsa-miR-195-5p | MASTL | Microarrays | Tarbase |
| hsa-miR-15a-5p | PLXDC2 | HITS-CLIP | Tarbase |
| hsa-miR-15a-5p | EPC1 | PAR-CLIP | Tarbase |
| hsa-miR-15a-5p | EPC1 | HITS-CLIP | Tarbase |
| hsa-miR-15a-5p | EPC1 | PAR-CLIP | Tarbase |
| hsa-miR-195-5p | EPC1 | PAR-CLIP | Tarbase |
| hsa-miR-195-5p | CCDC77 | Microarrays | Tarbase |
| hsa-miR-15a-5p | PROSER1 | HITS-CLIP | Tarbase |
| hsa-miR-15a-5p | HSPH1 | PAR-CLIP | Tarbase |

| | | | |
|---|---|---|---|
| hsa-miR-195-5p | EXOSC8 | Microarrays | Tarbase |
| hsa-miR-486-5p | ETF1 | HITS-CLIP | Tarbase |
| hsa-miR-141-3p | PAIP2 | HITS-CLIP | Tarbase |
| hsa-miR-3613-3p | EGR1 | HITS-CLIP | Tarbase |
| hsa-miR-15a-5p | SERP1 | HITS-CLIP | Tarbase |
| hsa-miR-141-3p | UTP20 | HITS-CLIP | Tarbase |
| hsa-miR-15a-3p | UTP20 | HITS-CLIP | Tarbase |
| hsa-miR-195-5p | TMPO | Microarrays | Tarbase |
| hsa-miR-195-5p | DUSP4 | Microarrays | Tarbase |
| hsa-miR-141-5p | TNFRSF10B | HITS-CLIP | Tarbase |
| hsa-miR-141-5p | TNFRSF10B | HITS-CLIP | Tarbase |
| hsa-miR-15a-5p | TNFRSF10B | HITS-CLIP | Tarbase |
| hsa-miR-195-5p | TNFRSF10B | HITS-CLIP | Tarbase |
| hsa-miR-3613-3p | TNFRSF10B | PAR-CLIP | Tarbase |
| hsa-miR-486-5p | TNFRSF10B | HITS-CLIP | Tarbase |
| hsa-miR-15a-5p | SORBS3 | HITS-CLIP | Tarbase |
| hsa-miR-195-5p | SORBS3 | HITS-CLIP | Tarbase |
| hsa-miR-141-3p | RNF170 | PAR-CLIP | Tarbase |
| hsa-miR-141-3p | TARDBP | HITS-CLIP | Tarbase |
| hsa-miR-195-5p | CRISPLD1 | Microarrays | Tarbase |
| hsa-miR-141-3p | AKAP1 | HITS-CLIP | Tarbase |
| hsa-miR-3613-3p | AKAP1 | HITS-CLIP | Tarbase |
| hsa-miR-141-3p | TRIM25 | HITS-CLIP | Tarbase |
| hsa-miR-141-5p | SPOP | HITS-CLIP | Tarbase |
| hsa-miR-15a-5p | TBX2 | HITS-CLIP | Tarbase |
| hsa-miR-195-5p | NCAPH | Microarrays | Tarbase |
| hsa-miR-195-5p | MND1 | Microarrays | Tarbase |
| hsa-miR-15a-3p | LHX4 | HITS-CLIP | Tarbase |
| hsa-miR-3613-5p | TRMT1L | PAR-CLIP | Tarbase |
| hsa-miR-195-5p | CSTA | Microarrays | Tarbase |
| hsa-miR-141-3p | NAA50 | HITS-CLIP | Tarbase |
| hsa-miR-15a-5p | NAA50 | HITS-CLIP | Tarbase |
| hsa-miR-195-5p | KIF18A | Microarrays | Tarbase |
| hsa-miR-141-3p | DESI2 | HITS-CLIP | Tarbase |
| hsa-miR-3613-5p | CAT | HITS-CLIP | Tarbase |
| hsa-miR-15a-3p | PILRB | HITS-CLIP | Tarbase |
| hsa-miR-15a-5p | PILRB | HITS-CLIP | Tarbase |
| hsa-miR-15a-5p | PILRB | HITS-CLIP | Tarbase |
| hsa-miR-195-5p | PILRB | HITS-CLIP | Tarbase |
| hsa-miR-195-5p | PILRB | HITS-CLIP | Tarbase |
| hsa-miR-15a-3p | ZMYM2 | HITS-CLIP | Tarbase |
| hsa-miR-15a-5p | GJB6 | HITS-CLIP | Tarbase |
| hsa-miR-195-5p | GJB6 | HITS-CLIP | Tarbase |
| hsa-miR-15a-5p | BAI2 | HITS-CLIP | Tarbase |
| hsa-miR-195-5p | BAI2 | HITS-CLIP | Tarbase |
| hsa-miR-15a-5p | PIK3CA | HITS-CLIP | Tarbase |
| hsa-miR-15a-5p | PIK3CA | HITS-CLIP | Tarbase |
| hsa-miR-195-5p | PIK3CA | HITS-CLIP | Tarbase |
| hsa-miR-195-5p | PIK3CA | HITS-CLIP | Tarbase |
| hsa-miR-15a-5p | PDS5A | HITS-CLIP | Tarbase |
| hsa-miR-15a-5p | PDS5A | HITS-CLIP | Tarbase |
| hsa-miR-195-5p | PDS5A | HITS-CLIP | Tarbase |
| hsa-miR-195-5p | LIAS | Microarrays | Tarbase |
| hsa-miR-195-5p | CPXM2 | HITS-CLIP | Tarbase |
| hsa-miR-486-5p | CLCC1 | HITS-CLIP | Tarbase |
| hsa-miR-486-5p | CLCC1 | HITS-CLIP | Tarbase |
| hsa-miR-195-5p | GPSM2 | Microarrays | Tarbase |
| hsa-miR-3605-5p | CXCR4 | HITS-CLIP | Tarbase |
| hsa-miR-15a-5p | ACVR2A | HITS-CLIP | Tarbase |
| hsa-miR-195-5p | ACVR2A | HITS-CLIP | Tarbase |
| hsa-miR-3613-3p | POLK | PAR-CLIP | Tarbase |
| hsa-miR-141-3p | FYTTD1 | HITS-CLIP | Tarbase |
| hsa-miR-141-3p | FYTTD1 | HITS-CLIP | Tarbase |
| hsa-miR-195-5p | OCRL | Microarrays | Tarbase |
| hsa-miR-15a-5p | MRPS2 | HITS-CLIP | Tarbase |
| hsa-miR-195-5p | MRPS2 | HITS-CLIP | Tarbase |
| hsa-miR-15a-3p | COPA | HITS-CLIP | Tarbase |
| hsa-miR-15a-3p | COPA | HITS-CLIP | Tarbase |
| hsa-miR-15a-5p | LY9 | PAR-CLIP | Tarbase |
| hsa-miR-15a-5p | LY9 | PAR-CLIP | Tarbase |
| hsa-miR-195-5p | LY9 | PAR-CLIP | Tarbase |
| hsa-miR-141-3p | RBBP6 | HITS-CLIP | Tarbase |
| hsa-miR-141-5p | RBBP6 | HITS-CLIP | Tarbase |
| hsa-miR-15a-5p | RBBP6 | PAR-CLIP | Tarbase |
| hsa-miR-15a-5p | RBBP6 | HITS-CLIP | Tarbase |
| hsa-miR-15a-5p | RBBP6 | HITS-CLIP | Tarbase |
| hsa-miR-195-5p | RBBP6 | HITS-CLIP | Tarbase |
| hsa-miR-3605-3p | RBBP6 | HITS-CLIP | Tarbase |
| hsa-miR-15a-5p | RBBP6 | Chimeric fragments | Tarbase |
| hsa-miR-15a-5p | ANXA11 | HITS-CLIP | Tarbase |
| hsa-miR-15a-5p | ANXA11 | HITS-CLIP | Tarbase |
| hsa-miR-195-5p | ANXA11 | HITS-CLIP | Tarbase |
| hsa-miR-195-5p | ANXA11 | HITS-CLIP | Tarbase |
| hsa-miR-486-3p | NAA60 | HITS-CLIP | Tarbase |
| hsa-miR-141-5p | RPL5 | HITS-CLIP | Tarbase |
| hsa-miR-3613-5p | RPL5 | HITS-CLIP | Tarbase |
| hsa-miR-141-5p | ZNF644 | HITS-CLIP | Tarbase |
| hsa-miR-3605-3p | ZNF644 | HITS-CLIP | Tarbase |
| hsa-miR-486-5p | ZNF644 | HITS-CLIP | Tarbase |
| hsa-miR-141-3p | SEPT7 | Luciferase Reporter Assay | Tarbase |
| hsa-miR-141-3p | SEPT7 | Luciferase Reporter Assay | Tarbase |
| hsa-miR-141-3p | SEPT7 | HITS-CLIP | Tarbase |
| hsa-miR-486-5p | SEPT7 | HITS-CLIP | Tarbase |
| hsa-miR-195-5p | KIAA0087 | Microarrays | Tarbase |
| hsa-miR-15a-3p | CBX3 | HITS-CLIP | Tarbase |
| hsa-miR-15a-5p | CBX3 | HITS-CLIP | Tarbase |
| hsa-miR-15a-5p | CBX3 | HITS-CLIP | Tarbase |
| hsa-miR-141-5p | HNRNPA2B1 | HITS-CLIP | Tarbase |
| hsa-miR-15a-5p | HNRNPA2B1 | HITS-CLIP | Tarbase |
| hsa-miR-195-5p | INHBA | Microarrays | Tarbase |
| hsa-miR-3613-3p | FKBP9 | HITS-CLIP | Tarbase |
| hsa-miR-141-5p | FTSJ2 | HITS-CLIP | Tarbase |
| hsa-miR-141-5p | CLTA | HITS-CLIP | Tarbase |
| hsa-miR-15a-5p | RECK | PAR-CLIP | Tarbase |
| hsa-miR-15a-5p | RECK | HITS-CLIP | Tarbase |
| hsa-miR-15a-5p | RECK | PAR-CLIP | Tarbase |
| hsa-miR-15a-5p | RECK | PAR-CLIP | Tarbase |
| hsa-miR-15a-5p | RECK | PAR-CLIP | Tarbase |
| hsa-miR-15a-5p | RECK | HITS-CLIP | Tarbase |
| hsa-miR-195-5p | RECK | PAR-CLIP | Tarbase |
| hsa-miR-195-5p | RECK | HITS-CLIP | Tarbase |
| hsa-miR-195-5p | RECK | PAR-CLIP | Tarbase |
| hsa-miR-15a-5p | ACO1 | HITS-CLIP | Tarbase |
| hsa-miR-195-5p | ACO1 | HITS-CLIP | Tarbase |
| hsa-miR-15a-5p | DCAF10 | PAR-CLIP | Tarbase |
| hsa-miR-195-5p | C7orf49 | HITS-CLIP | Tarbase |
| hsa-miR-195-5p | PLAU | qPCR | Tarbase |
| hsa-miR-195-5p | PLAU | Microarrays | Tarbase |
| hsa-miR-195-5p | BICC1 | Microarrays | Tarbase |
| hsa-miR-195-5p | ZWINT | Microarrays | Tarbase |
| hsa-miR-15a-5p | RBM19 | HITS-CLIP | Tarbase |
| hsa-miR-195-5p | RBM19 | HITS-CLIP | Tarbase |
| hsa-miR-141-3p | MED13L | HITS-CLIP | Tarbase |
| hsa-miR-141-3p | MED13L | HITS-CLIP | Tarbase |
| hsa-miR-141-3p | MED13L | HITS-CLIP | Tarbase |
| hsa-miR-195-5p | CDKN2C | Microarrays | Tarbase |
| hsa-miR-141-3p | RNF11 | HITS-CLIP | Tarbase |
| hsa-miR-141-3p | RNF11 | HITS-CLIP | Tarbase |
| hsa-miR-141-5p | RNF11 | HITS-CLIP | Tarbase |
| hsa-miR-141-5p | RNF11 | HITS-CLIP | Tarbase |
| hsa-miR-15a-5p | RASSF8 | PAR-CLIP | Tarbase |
| hsa-miR-195-3p | RASSF8 | HITS-CLIP | Tarbase |
| hsa-miR-15a-5p | ITPR2 | HITS-CLIP | Tarbase |
| hsa-miR-195-5p | ITPR2 | HITS-CLIP | Tarbase |
| hsa-miR-15a-5p | NECAB1 | HITS-CLIP | Tarbase |
| hsa-miR-195-5p | NECAB1 | HITS-CLIP | Tarbase |
| hsa-miR-195-5p | ACOT9 | Microarrays | Tarbase |
| hsa-miR-195-5p | DDX39A | Microarrays | Tarbase |
| hsa-miR-195-5p | CENPK | Microarrays | Tarbase |
| hsa-miR-3613-5p | ATF1 | HITS-CLIP | Tarbase |
| hsa-miR-486-5p | NR4A1 | PAR-CLIP | Tarbase |
| hsa-miR-15a-5p | NR4A1 | HITS-CLIP | Tarbase |
| hsa-miR-195-5p | NR4A1 | HITS-CLIP | Tarbase |
| hsa-miR-141-3p | HOXC13 | PAR-CLIP | Tarbase |
| hsa-miR-141-3p | CDK2 | Western Blot | Tarbase |
| hsa-miR-195-5p | CDK2 | Microarrays | Tarbase |
| hsa-miR-141-3p | SMUG1 | HITS-CLIP | Tarbase |
| hsa-miR-195-5p | STIL | Microarrays | Tarbase |
| hsa-miR-195-5p | HJURP | Microarrays | Tarbase |

| | | | |
|---|---|---|---|
| hsa-miR-141-3p | AMD1 | PAR-CLIP | Tarbase |
| hsa-miR-141-5p | AMD1 | HITS-CLIP | Tarbase |
| hsa-miR-141-3p | MORF4L2 | HITS-CLIP | Tarbase |
| hsa-miR-141-5p | MORF4L2 | HITS-CLIP | Tarbase |
| hsa-miR-141-5p | MORF4L2 | HITS-CLIP | Tarbase |
| hsa-miR-141-5p | MORF4L2 | HITS-CLIP | Tarbase |
| hsa-miR-15a-3p | MORF4L2 | HITS-CLIP | Tarbase |
| hsa-miR-3605-5p | MORF4L2 | HITS-CLIP | Tarbase |
| hsa-miR-141-3p | FAM199X | HITS-CLIP | Tarbase |
| hsa-miR-15a-5p | FAM199X | HITS-CLIP | Tarbase |
| hsa-miR-195-5p | FAM199X | HITS-CLIP | Tarbase |
| hsa-miR-141-3p | BAZ2B | HITS-CLIP | Tarbase |
| hsa-miR-141-3p | LPGAT1 | HITS-CLIP | Tarbase |
| hsa-miR-141-5p | LPGAT1 | HITS-CLIP | Tarbase |
| hsa-miR-141-5p | LPGAT1 | HITS-CLIP | Tarbase |
| hsa-miR-3613-3p | LPGAT1 | PAR-CLIP | Tarbase |
| hsa-miR-141-3p | RAP2C | HITS-CLIP | Tarbase |
| hsa-miR-15a-3p | RAP2C | HITS-CLIP | Tarbase |
| hsa-miR-15a-5p | PLA2G12A | HITS-CLIP | Tarbase |
| hsa-miR-195-5p | PLA2G12A | HITS-CLIP | Tarbase |
| hsa-miR-141-5p | AGO2 | HITS-CLIP | Tarbase |
| hsa-miR-141-5p | AGO2 | HITS-CLIP | Tarbase |
| hsa-miR-141-3p | MXD4 | HITS-CLIP | Tarbase |
| hsa-miR-3613-3p | CKS2 | PAR-CLIP | Tarbase |
| hsa-miR-195-5p | CKS2 | Microarrays | Tarbase |
| hsa-miR-141-3p | ACSL3 | HITS-CLIP | Tarbase |
| hsa-miR-141-5p | ACSL3 | HITS-CLIP | Tarbase |
| hsa-miR-141-5p | ACSL3 | HITS-CLIP | Tarbase |
| hsa-miR-15a-3p | ACSL3 | HITS-CLIP | Tarbase |
| hsa-miR-15a-5p | ACSL3 | PAR-CLIP | Tarbase |
| hsa-miR-15a-5p | ACSL3 | HITS-CLIP | Tarbase |
| hsa-miR-15a-5p | ACSL3 | PAR-CLIP | Tarbase |
| hsa-miR-195-5p | ACSL3 | HITS-CLIP | Tarbase |
| hsa-miR-15a-5p | CHPF | HITS-CLIP | Tarbase |
| hsa-miR-15a-5p | CHPF | HITS-CLIP | Tarbase |
| hsa-miR-195-5p | CHPF | HITS-CLIP | Tarbase |
| hsa-miR-15a-5p | OBSL1 | HITS-CLIP | Tarbase |
| hsa-miR-195-5p | OBSL1 | HITS-CLIP | Tarbase |
| hsa-miR-15a-5p | SLC12A4 | HITS-CLIP | Tarbase |
| hsa-miR-195-5p | SLC12A4 | Microarrays | Tarbase |
| hsa-miR-141-3p | TTPAL | HITS-CLIP | Tarbase |
| hsa-miR-15a-5p | TTPAL | HITS-CLIP | Tarbase |
| hsa-miR-195-5p | TTPAL | HITS-CLIP | Tarbase |
| hsa-miR-15a-3p | PREX1 | HITS-CLIP | Tarbase |
| hsa-miR-15a-5p | SLC12A5 | HITS-CLIP | Tarbase |
| hsa-miR-195-5p | SLC12A5 | HITS-CLIP | Tarbase |
| hsa-miR-15a-5p | SDC4 | HITS-CLIP | Tarbase |
| hsa-miR-15a-5p | SDC4 | HITS-CLIP | Tarbase |
| hsa-miR-195-5p | SDC4 | HITS-CLIP | Tarbase |
| hsa-miR-141-3p | NCOA3 | HITS-CLIP | Tarbase |
| hsa-miR-141-3p | NCOA3 | HITS-CLIP | Tarbase |
| hsa-miR-141-5p | PIGT | HITS-CLIP | Tarbase |
| hsa-miR-15a-5p | VAPB | PAR-CLIP | Tarbase |
| hsa-miR-219a-1-3p | VAPB | HITS-CLIP | Tarbase |
| hsa-miR-141-3p | PARD6B | HITS-CLIP | Tarbase |
| hsa-miR-15a-5p | PARD6B | PAR-CLIP | Tarbase |
| hsa-miR-15a-5p | PARD6B | HITS-CLIP | Tarbase |
| hsa-miR-15a-5p | PARD6B | HITS-CLIP | Tarbase |
| hsa-miR-15a-5p | PARD6B | HITS-CLIP | Tarbase |
| hsa-miR-195-5p | PARD6B | HITS-CLIP | Tarbase |
| hsa-miR-195-5p | PARD6B | HITS-CLIP | Tarbase |
| hsa-miR-3613-5p | PARD6B | PAR-CLIP | Tarbase |
| hsa-miR-15a-5p | CHD6 | HITS-CLIP | Tarbase |
| hsa-miR-195-5p | CHD6 | HITS-CLIP | Tarbase |
| hsa-miR-486-3p | CHD6 | HITS-CLIP | Tarbase |
| hsa-miR-486-5p | PLCG1 | HITS-CLIP | Tarbase |
| hsa-miR-3613-5p | SRSF6 | HITS-CLIP | Tarbase |
| hsa-miR-15a-5p | ARFGEF2 | PAR-CLIP | Tarbase |
| hsa-miR-15a-5p | ARFGEF2 | PAR-CLIP | Tarbase |
| hsa-miR-15a-5p | ARFGEF2 | HITS-CLIP | Tarbase |
| hsa-miR-15a-5p | ARFGEF2 | HITS-CLIP | Tarbase |
| hsa-miR-15a-5p | ARFGEF2 | HITS-CLIP | Tarbase |
| hsa-miR-195-5p | ARFGEF2 | HITS-CLIP | Tarbase |
| hsa-miR-3613-3p | ARFGEF2 | HITS-CLIP | Tarbase |
| hsa-miR-141-5p | ZNFX1 | HITS-CLIP | Tarbase |
| hsa-miR-15a-5p | ZNFX1 | HITS-CLIP | Tarbase |
| hsa-miR-15a-5p | ZNFX1 | HITS-CLIP | Tarbase |
| hsa-miR-195-5p | ZNFX1 | HITS-CLIP | Tarbase |
| hsa-miR-195-5p | ZNFX1 | HITS-CLIP | Tarbase |
| hsa-miR-141-3p | TMEM189-UBE2V1 | HITS-CLIP | Tarbase |
| hsa-miR-141-3p | TMEM189-UBE2V1 | HITS-CLIP | Tarbase |
| hsa-miR-15a-3p | TMEM189-UBE2V1 | HITS-CLIP | Tarbase |
| hsa-miR-15a-5p | TMEM189-UBE2V1 | HITS-CLIP | Tarbase |
| hsa-miR-15a-5p | TMEM189-UBE2V1 | HITS-CLIP | Tarbase |
| hsa-miR-195-5p | TMEM189-UBE2V1 | HITS-CLIP | Tarbase |
| hsa-miR-195-5p | TMEM189-UBE2V1 | HITS-CLIP | Tarbase |
| hsa-miR-15a-5p | TMEM189-UBE2V1 | HITS-CLIP | Tarbase |
| hsa-miR-195-5p | TMEM189-UBE2V1 | HITS-CLIP | Tarbase |
| hsa-miR-195-5p | TMEM189-UBE2V1 | Microarrays | Tarbase |
| hsa-miR-141-3p | RAB22A | HITS-CLIP | Tarbase |
| hsa-miR-15a-5p | RAB22A | PAR-CLIP | Tarbase |
| hsa-miR-15a-5p | STAU1 | PAR-CLIP | Tarbase |
| hsa-miR-15a-5p | STX16 | HITS-CLIP | Tarbase |
| hsa-miR-15a-5p | STX16 | HITS-CLIP | Tarbase |
| hsa-miR-195-5p | STX16 | HITS-CLIP | Tarbase |
| hsa-miR-486-5p | STX16 | HITS-CLIP | Tarbase |
| hsa-miR-141-3p | PMEPA1 | HITS-CLIP | Tarbase |
| hsa-miR-195-5p | PMEPA1 | Microarrays | Tarbase |
| hsa-miR-15a-5p | PEPD | HITS-CLIP | Tarbase |
| hsa-miR-195-5p | PEPD | HITS-CLIP | Tarbase |
| hsa-miR-195-5p | PEPD | Microarrays | Tarbase |
| hsa-miR-15a-5p | VAMP7 | HITS-CLIP | Tarbase |
| hsa-miR-195-5p | VAMP7 | HITS-CLIP | Tarbase |
| hsa-miR-195-5p | ATP8A1 | Microarrays | Tarbase |
| hsa-miR-141-3p | USP22 | HITS-CLIP | Tarbase |
| hsa-miR-15a-3p | USP22 | HITS-CLIP | Tarbase |
| hsa-miR-15a-3p | USP22 | HITS-CLIP | Tarbase |
| hsa-miR-15a-5p | USP22 | HITS-CLIP | Tarbase |
| hsa-miR-195-5p | POF1B | AGO-IP | Tarbase |
| hsa-miR-141-3p | USP9X | HITS-CLIP | Tarbase |
| hsa-miR-141-3p | USP9X | HITS-CLIP | Tarbase |
| hsa-miR-15a-3p | USP9X | HITS-CLIP | Tarbase |
| hsa-miR-195-5p | F13A1 | Microarrays | Tarbase |
| hsa-miR-15a-5p | SNRPC | Chimeric fragments | Tarbase |
| hsa-miR-15a-5p | ABCC10 | HITS-CLIP | Tarbase |
| hsa-miR-195-5p | ABCC10 | HITS-CLIP | Tarbase |
| hsa-miR-195-5p | HIST1H1D | Microarrays | Tarbase |
| hsa-miR-195-5p | NQO2 | Microarrays | Tarbase |
| hsa-miR-3613-5p | HIST1H2BJ | HITS-CLIP | Tarbase |
| hsa-miR-195-5p | HIST1H2BJ | Microarrays | Tarbase |
| hsa-miR-15a-5p | KLHDC3 | HITS-CLIP | Tarbase |
| hsa-miR-195-5p | KLHDC3 | HITS-CLIP | Tarbase |
| hsa-miR-141-3p | CDKN1A | PAR-CLIP | Tarbase |
| hsa-miR-141-3p | CDKN1A | HITS-CLIP | Tarbase |
| hsa-miR-141-3p | CDKN1A | HITS-CLIP | Tarbase |
| hsa-miR-15a-3p | CDKN1A | PAR-CLIP | Tarbase |
| hsa-miR-15a-3p | CDKN1A | HITS-CLIP | Tarbase |
| hsa-miR-15a-5p | CDKN1A | Western Blot | Tarbase |
| hsa-miR-15a-5p | CDKN1A | Western Blot | Tarbase |
| hsa-miR-15a-5p | CDKN1A | HITS-CLIP | Tarbase |
| hsa-miR-15a-5p | CDKN1A | PAR-CLIP | Tarbase |
| hsa-miR-195-5p | CDKN1A | HITS-CLIP | Tarbase |
| hsa-miR-15a-5p | RREB1 | PAR-CLIP | Tarbase |
| hsa-miR-15a-5p | RREB1 | PAR-CLIP | Tarbase |
| hsa-miR-15a-5p | RREB1 | PAR-CLIP | Tarbase |
| hsa-miR-195-5p | RREB1 | PAR-CLIP | Tarbase |
| hsa-miR-141-3p | SSR1 | HITS-CLIP | Tarbase |
| hsa-miR-141-5p | SSR1 | HITS-CLIP | Tarbase |
| hsa-miR-15a-3p | SSR1 | HITS-CLIP | Tarbase |
| hsa-miR-3613-5p | SSR1 | HITS-CLIP | Tarbase |
| hsa-miR-141-5p | RIOK1 | HITS-CLIP | Tarbase |
| hsa-miR-219a-1-3p | ATXN1 | PAR-CLIP | Tarbase |
| hsa-miR-3613-5p | ATXN1 | PAR-CLIP | Tarbase |
| hsa-miR-141-3p | NUP153 | HITS-CLIP | Tarbase |
| hsa-miR-141-3p | NUP153 | HITS-CLIP | Tarbase |
| hsa-miR-15a-5p | NUP153 | HITS-CLIP | Tarbase |
| hsa-miR-15a-5p | NUP153 | HITS-CLIP | Tarbase |
| hsa-miR-195-5p | NUP153 | HITS-CLIP | Tarbase |
| hsa-miR-141-3p | DEK | HITS-CLIP | Tarbase |

| | | | |
|---|---|---|---|
| hsa-miR-141-3p | DEK | HITS-CLIP | Tarbase |
| hsa-miR-195-5p | RUNX2 | Microarrays | Tarbase |
| hsa-miR-219a-1-3p | LRRFIP1 | PAR-CLIP | Tarbase |
| hsa-miR-3605-3p | LRRFIP1 | HITS-CLIP | Tarbase |
| hsa-miR-486-5p | LRRFIP1 | HITS-CLIP | Tarbase |
| hsa-miR-141-3p | AHNAK | HITS-CLIP | Tarbase |
| hsa-miR-141-5p | AHNAK | HITS-CLIP | Tarbase |
| hsa-miR-141-5p | AHNAK | HITS-CLIP | Tarbase |
| hsa-miR-3605-5p | AHNAK | HITS-CLIP | Tarbase |
| hsa-miR-3605-5p | AHNAK | HITS-CLIP | Tarbase |
| hsa-miR-486-3p | AHNAK | HITS-CLIP | Tarbase |
| hsa-miR-15a-3p | CNOT1 | HITS-CLIP | Tarbase |
| hsa-miR-15a-5p | CNOT1 | PAR-CLIP | Tarbase |
| hsa-miR-15a-5p | CNOT1 | HITS-CLIP | Tarbase |
| hsa-miR-15a-5p | CNOT1 | PAR-CLIP | Tarbase |
| hsa-miR-195-5p | CNOT1 | PAR-CLIP | Tarbase |
| hsa-miR-195-5p | CNOT1 | HITS-CLIP | Tarbase |
| hsa-miR-195-5p | CNOT1 | PAR-CLIP | Tarbase |
| hsa-miR-3613-3p | MT1G | HITS-CLIP | Tarbase |
| hsa-miR-15a-3p | C16orf70 | HITS-CLIP | Tarbase |
| hsa-miR-15a-5p | TMTC4 | HITS-CLIP | Tarbase |
| hsa-miR-195-5p | TMTC4 | HITS-CLIP | Tarbase |
| hsa-miR-3613-5p | TMTC4 | PAR-CLIP | Tarbase |
| hsa-miR-15a-5p | RAP2A | HITS-CLIP | Tarbase |
| hsa-miR-195-5p | RAP2A | HITS-CLIP | Tarbase |
| hsa-miR-195-5p | ABCC4 | Microarrays | Tarbase |
| hsa-miR-141-5p | EFNB2 | HITS-CLIP | Tarbase |
| hsa-miR-15a-5p | TM9SF2 | HITS-CLIP | Tarbase |
| hsa-miR-15a-5p | TM9SF2 | HITS-CLIP | Tarbase |
| hsa-miR-15a-5p | TM9SF2 | HITS-CLIP | Tarbase |
| hsa-miR-195-5p | TM9SF2 | HITS-CLIP | Tarbase |
| hsa-miR-195-5p | TM9SF2 | HITS-CLIP | Tarbase |
| hsa-miR-195-5p | C17orf53 | Microarrays | Tarbase |
| hsa-miR-141-5p | IRF1 | HITS-CLIP | Tarbase |
| hsa-miR-15a-5p | IRF1 | PAR-CLIP | Tarbase |
| hsa-miR-195-5p | IRF1 | PAR-CLIP | Tarbase |
| hsa-miR-15a-5p | RNF113A | HITS-CLIP | Tarbase |
| hsa-miR-141-5p | SEPT6 | HITS-CLIP | Tarbase |
| hsa-miR-195-5p | TMEM255A | Microarrays | Tarbase |
| hsa-miR-195-5p | BMP4 | Microarrays | Tarbase |
| hsa-miR-3613-3p | FAM193A | PAR-CLIP | Tarbase |
| hsa-miR-195-5p | HS3ST3B1 | HITS-CLIP | Tarbase |
| hsa-miR-141-5p | MRPS7 | HITS-CLIP | Tarbase |
| hsa-miR-141-5p | MRPS7 | HITS-CLIP | Tarbase |
| hsa-miR-3605-5p | GGA3 | HITS-CLIP | Tarbase |
| hsa-miR-195-5p | NUP85 | Microarrays | Tarbase |
| hsa-miR-15a-5p | GTF3C4 | HITS-CLIP | Tarbase |
| hsa-miR-195-5p | GTF3C4 | HITS-CLIP | Tarbase |
| hsa-miR-219a-1-3p | GTF3C4 | HITS-CLIP | Tarbase |
| hsa-miR-3605-3p | DDX31 | HITS-CLIP | Tarbase |
| hsa-miR-3613-3p | PPP1R12C | HITS-CLIP | Tarbase |
| hsa-miR-15a-5p | MBOAT7 | HITS-CLIP | Tarbase |
| hsa-miR-15a-5p | MBOAT7 | HITS-CLIP | Tarbase |
| hsa-miR-195-5p | MBOAT7 | HITS-CLIP | Tarbase |
| hsa-miR-195-5p | SLC2A4RG | Microarrays | Tarbase |
| hsa-miR-15a-5p | INSIG2 | HITS-CLIP | Tarbase |
| hsa-miR-195-5p | INSIG2 | HITS-CLIP | Tarbase |
| hsa-miR-141-5p | SLC25A23 | HITS-CLIP | Tarbase |
| hsa-miR-195-5p | TNFSF9 | AGO-IP | Tarbase |
| hsa-miR-3613-3p | THOC2 | HITS-CLIP | Tarbase |
| hsa-miR-141-5p | MED1 | HITS-CLIP | Tarbase |
| hsa-miR-15a-5p | MED1 | HITS-CLIP | Tarbase |
| hsa-miR-195-5p | MED1 | HITS-CLIP | Tarbase |
| hsa-miR-219a-1-3p | MED1 | HITS-CLIP | Tarbase |
| hsa-miR-141-3p | RPL23 | HITS-CLIP | Tarbase |
| hsa-miR-141-5p | TRIP10 | HITS-CLIP | Tarbase |
| hsa-miR-195-5p | TRIP10 | Microarrays | Tarbase |
| hsa-miR-3613-3p | FOSB | HITS-CLIP | Tarbase |
| hsa-miR-15a-5p | SYMPK | HITS-CLIP | Tarbase |
| hsa-miR-195-5p | SYMPK | HITS-CLIP | Tarbase |
| hsa-miR-141-3p | PANK2 | HITS-CLIP | Tarbase |
| hsa-miR-15a-5p | GZF1 | Chimeric fragments | Tarbase |
| hsa-miR-141-5p | CENPB | HITS-CLIP | Tarbase |
| hsa-miR-3613-5p | PSMF1 | HITS-CLIP | Tarbase |
| hsa-miR-141-3p | TMX4 | HITS-CLIP | Tarbase |
| hsa-miR-141-3p | TMX4 | HITS-CLIP | Tarbase |
| hsa-miR-15a-5p | STK35 | HITS-CLIP | Tarbase |
| hsa-miR-15a-5p | STK35 | HITS-CLIP | Tarbase |
| hsa-miR-15a-5p | STK35 | PAR-CLIP | Tarbase |
| hsa-miR-15a-5p | STK35 | HITS-CLIP | Tarbase |
| hsa-miR-195-5p | STK35 | HITS-CLIP | Tarbase |
| hsa-miR-195-5p | STK35 | HITS-CLIP | Tarbase |
| hsa-miR-195-5p | STK35 | HITS-CLIP | Tarbase |
| hsa-miR-195-5p | STK35 | Microarrays | Tarbase |
| hsa-miR-15a-5p | SNRPB | HITS-CLIP | Tarbase |
| hsa-miR-195-5p | SNRPB | HITS-CLIP | Tarbase |
| hsa-miR-195-5p | AP5S1 | Microarrays | Tarbase |
| hsa-miR-141-3p | RRBP1 | HITS-CLIP | Tarbase |
| hsa-miR-141-3p | RRBP1 | HITS-CLIP | Tarbase |
| hsa-miR-3613-5p | RRBP1 | HITS-CLIP | Tarbase |
| hsa-miR-195-5p | FLRT3 | Microarrays | Tarbase |
| hsa-miR-15a-5p | SNRPB2 | PAR-CLIP | Tarbase |
| hsa-miR-15a-5p | TBC1D20 | HITS-CLIP | Tarbase |
| hsa-miR-15a-5p | TBC1D20 | HITS-CLIP | Tarbase |
| hsa-miR-195-5p | TBC1D20 | HITS-CLIP | Tarbase |
| hsa-miR-195-5p | TBC1D20 | HITS-CLIP | Tarbase |
| hsa-miR-15a-5p | MCM8 | HITS-CLIP | Tarbase |
| hsa-miR-195-5p | MCM8 | Microarrays | Tarbase |
| hsa-miR-195-5p | FAM110A | Microarrays | Tarbase |
| hsa-miR-15a-5p | NCLN | HITS-CLIP | Tarbase |
| hsa-miR-195-5p | NCLN | HITS-CLIP | Tarbase |
| hsa-miR-15a-5p | HNRNPR | PAR-CLIP | Tarbase |
| hsa-miR-3613-3p | HNRNPR | HITS-CLIP | Tarbase |
| hsa-miR-15a-5p | MAX | PAR-CLIP | Tarbase |
| hsa-miR-15a-5p | ID1 | HITS-CLIP | Tarbase |
| hsa-miR-195-5p | ID1 | HITS-CLIP | Tarbase |
| hsa-miR-141-3p | ERGIC3 | HITS-CLIP | Tarbase |
| hsa-miR-15a-5p | ROMO1 | HITS-CLIP | Tarbase |
| hsa-miR-195-5p | ROMO1 | HITS-CLIP | Tarbase |
| hsa-miR-195-5p | ROMO1 | HITS-CLIP | Tarbase |
| hsa-miR-141-5p | PLAGL2 | HITS-CLIP | Tarbase |
| hsa-miR-15a-3p | PLAGL2 | HITS-CLIP | Tarbase |
| hsa-miR-15a-5p | PLAGL2 | HITS-CLIP | Tarbase |
| hsa-miR-15a-5p | PLAGL2 | HITS-CLIP | Tarbase |
| hsa-miR-15a-5p | PLAGL2 | HITS-CLIP | Tarbase |
| hsa-miR-195-5p | PLAGL2 | HITS-CLIP | Tarbase |
| hsa-miR-195-5p | PLAGL2 | HITS-CLIP | Tarbase |
| hsa-miR-486-5p | PLAGL2 | PAR-CLIP | Tarbase |
| hsa-miR-3613-5p | KDM5C | PAR-CLIP | Tarbase |
| hsa-miR-15a-5p | AMOT | HITS-CLIP | Tarbase |
| hsa-miR-15a-5p | AMOT | HITS-CLIP | Tarbase |
| hsa-miR-195-5p | AMOT | HITS-CLIP | Tarbase |
| hsa-miR-195-5p | AMOT | HITS-CLIP | Tarbase |
| hsa-miR-3605-3p | AMOT | HITS-CLIP | Tarbase |
| hsa-miR-15a-5p | AMOT | HITS-CLIP | Tarbase |
| hsa-miR-195-5p | AMOT | HITS-CLIP | Tarbase |
| hsa-miR-15a-5p | TMEM115 | HITS-CLIP | Tarbase |
| hsa-miR-15a-5p | TMEM115 | HITS-CLIP | Tarbase |
| hsa-miR-195-5p | TMEM115 | HITS-CLIP | Tarbase |
| hsa-miR-3605-5p | PSMB2 | HITS-CLIP | Tarbase |
| hsa-miR-15a-5p | AGO3 | PAR-CLIP | Tarbase |
| hsa-miR-15a-5p | AGO3 | PAR-CLIP | Tarbase |
| hsa-miR-15a-5p | AGO3 | HITS-CLIP | Tarbase |
| hsa-miR-195-5p | AGO3 | HITS-CLIP | Tarbase |
| hsa-miR-15a-5p | KLC1 | HITS-CLIP | Tarbase |
| hsa-miR-195-5p | KLC1 | HITS-CLIP | Tarbase |
| hsa-miR-195-5p | XRCC3 | Microarrays | Tarbase |
| hsa-miR-141-5p | TUBGCP3 | HITS-CLIP | Tarbase |
| hsa-miR-15a-3p | CAPNS1 | HITS-CLIP | Tarbase |
| hsa-miR-15a-5p | RBM42 | Chimeric fragments | Tarbase |
| hsa-miR-15a-5p | PRDX5 | HITS-CLIP | Tarbase |
| hsa-miR-195-5p | PRDX5 | HITS-CLIP | Tarbase |
| hsa-miR-15a-5p | BCL2L12 | HITS-CLIP | Tarbase |
| hsa-miR-15a-5p | BCL2L12 | HITS-CLIP | Tarbase |
| hsa-miR-15a-5p | BCL2L12 | PAR-CLIP | Tarbase |
| hsa-miR-195-5p | BCL2L12 | HITS-CLIP | Tarbase |
| hsa-miR-195-5p | BCL2L12 | HITS-CLIP | Tarbase |
| hsa-miR-195-5p | BCL2L12 | Microarrays | Tarbase |
| hsa-miR-15a-3p | IRF3 | HITS-CLIP | Tarbase |

| | | | |
|---|---|---|---|
| hsa-miR-3605-5p | PRMT1 | HITS-CLIP | Tarbase |
| hsa-miR-15a-5p | PRR12 | HITS-CLIP | Tarbase |
| hsa-miR-195-5p | STATH | Microarrays | Tarbase |
| hsa-miR-15a-5p | BECN1 | HITS-CLIP | Tarbase |
| hsa-miR-15a-5p | BECN1 | HITS-CLIP | Tarbase |
| hsa-miR-195-5p | BECN1 | HITS-CLIP | Tarbase |
| hsa-miR-15a-3p | TRAP1 | HITS-CLIP | Tarbase |
| hsa-miR-15a-5p | TRAP1 | HITS-CLIP | Tarbase |
| hsa-miR-195-5p | TRAP1 | HITS-CLIP | Tarbase |
| hsa-miR-141-5p | NSRP1 | HITS-CLIP | Tarbase |
| hsa-miR-15a-5p | NSRP1 | HITS-CLIP | Tarbase |
| hsa-miR-195-5p | NSRP1 | HITS-CLIP | Tarbase |
| hsa-miR-15a-5p | AHDC1 | HITS-CLIP | Tarbase |
| hsa-miR-3605-5p | ZNF384 | PAR-CLIP | Tarbase |
| hsa-miR-15a-3p | CFP | HITS-CLIP | Tarbase |
| hsa-miR-486-5p | PCNXL4 | HITS-CLIP | Tarbase |
| hsa-miR-15a-5p | ATG14 | HITS-CLIP | Tarbase |
| hsa-miR-195-5p | ATG14 | Microarrays | Tarbase |
| hsa-miR-15a-5p | ATG14 | Chimeric fragments | Tarbase |
| hsa-miR-141-3p | KTN1 | HITS-CLIP | Tarbase |
| hsa-miR-15a-5p | KTN1 | HITS-CLIP | Tarbase |
| hsa-miR-15a-5p | KTN1 | HITS-CLIP | Tarbase |
| hsa-miR-15a-5p | KTN1 | HITS-CLIP | Tarbase |
| hsa-miR-195-5p | KTN1 | HITS-CLIP | Tarbase |
| hsa-miR-195-5p | KTN1 | HITS-CLIP | Tarbase |
| hsa-miR-195-5p | KTN1 | HITS-CLIP | Tarbase |
| hsa-miR-195-5p | KTN1 | HITS-CLIP | Tarbase |
| hsa-miR-195-5p | KTN1 | HITS-CLIP | Tarbase |
| hsa-miR-195-5p | RHOJ | Microarrays | Tarbase |
| hsa-miR-195-5p | DLGAP5 | Microarrays | Tarbase |
| hsa-miR-195-5p | L3HYPDH | Microarrays | Tarbase |
| hsa-miR-15a-5p | HSPA2 | HITS-CLIP | Tarbase |
| hsa-miR-15a-5p | OMG | HITS-CLIP | Tarbase |
| hsa-miR-195-5p | OMG | HITS-CLIP | Tarbase |
| hsa-miR-141-3p | HNRNPH2 | HITS-CLIP | Tarbase |
| hsa-miR-141-3p | HNRNPH2 | HITS-CLIP | Tarbase |
| hsa-miR-15a-3p | ARMCX1 | HITS-CLIP | Tarbase |
| hsa-miR-15a-5p | ARMCX1 | HITS-CLIP | Tarbase |
| hsa-miR-195-5p | ARMCX1 | HITS-CLIP | Tarbase |
| hsa-miR-15a-5p | ARMCX1 | HITS-CLIP | Tarbase |
| hsa-miR-195-5p | ARMCX1 | HITS-CLIP | Tarbase |
| hsa-miR-141-3p | CANX | HITS-CLIP | Tarbase |
| hsa-miR-141-5p | CANX | HITS-CLIP | Tarbase |
| hsa-miR-15a-3p | CANX | HITS-CLIP | Tarbase |
| hsa-miR-15a-5p | CANX | HITS-CLIP | Tarbase |
| hsa-miR-15a-5p | CANX | HITS-CLIP | Tarbase |
| hsa-miR-15a-5p | CANX | HITS-CLIP | Tarbase |
| hsa-miR-15a-5p | CANX | HITS-CLIP | Tarbase |
| hsa-miR-195-5p | CANX | HITS-CLIP | Tarbase |
| hsa-miR-15a-5p | CANX | HITS-CLIP | Tarbase |
| hsa-miR-195-5p | CANX | HITS-CLIP | Tarbase |
| hsa-miR-15a-5p | IPPK | PAR-CLIP | Tarbase |
| hsa-miR-3613-3p | ZNF484 | HITS-CLIP | Tarbase |
| hsa-miR-195-5p | TSPAN8 | Microarrays | Tarbase |
| hsa-miR-141-3p | DYRK2 | HITS-CLIP | Tarbase |
| hsa-miR-15a-5p | DYRK2 | PAR-CLIP | Tarbase |
| hsa-miR-15a-5p | DYRK2 | HITS-CLIP | Tarbase |
| hsa-miR-15a-5p | DYRK2 | HITS-CLIP | Tarbase |
| hsa-miR-195-5p | DYRK2 | HITS-CLIP | Tarbase |
| hsa-miR-195-5p | DYRK2 | HITS-CLIP | Tarbase |
| hsa-miR-195-5p | DYRK2 | Microarrays | Tarbase |
| hsa-miR-141-5p | YEATS4 | HITS-CLIP | Tarbase |
| hsa-miR-141-5p | YEATS4 | HITS-CLIP | Tarbase |
| hsa-miR-141-5p | YEATS4 | HITS-CLIP | Tarbase |
| hsa-miR-141-5p | YEATS4 | HITS-CLIP | Tarbase |
| hsa-miR-141-5p | YEATS4 | HITS-CLIP | Tarbase |
| hsa-miR-195-5p | AUNIP | Microarrays | Tarbase |
| hsa-miR-141-5p | PIN1 | HITS-CLIP | Tarbase |
| hsa-miR-141-5p | EMC1 | HITS-CLIP | Tarbase |
| hsa-miR-195-5p | PLA2G5 | Microarrays | Tarbase |
| hsa-miR-15a-3p | UBR4 | HITS-CLIP | Tarbase |
| hsa-miR-15a-3p | UBR4 | HITS-CLIP | Tarbase |
| hsa-miR-15a-3p | UBR4 | HITS-CLIP | Tarbase |
| hsa-miR-15a-5p | UBR4 | HITS-CLIP | Tarbase |
| hsa-miR-195-5p | UBR4 | HITS-CLIP | Tarbase |
| hsa-miR-3613-5p | UBR4 | HITS-CLIP | Tarbase |
| hsa-miR-195-5p | EMR2 | Microarrays | Tarbase |
| hsa-miR-3613-5p | SLC35E1 | HITS-CLIP | Tarbase |
| hsa-miR-195-5p | PKMYT1 | Microarrays | Tarbase |
| hsa-miR-141-3p | FBXL16 | HITS-CLIP | Tarbase |
| hsa-miR-141-5p | FBXL16 | HITS-CLIP | Tarbase |
| hsa-miR-15a-3p | CHTF18 | HITS-CLIP | Tarbase |
| hsa-miR-15a-3p | CHTF18 | HITS-CLIP | Tarbase |
| hsa-miR-195-5p | CHTF18 | Microarrays | Tarbase |
| hsa-miR-15a-5p | MACF1 | HITS-CLIP | Tarbase |
| hsa-miR-15a-5p | MACF1 | HITS-CLIP | Tarbase |
| hsa-miR-195-5p | MACF1 | HITS-CLIP | Tarbase |
| hsa-miR-195-5p | MACF1 | HITS-CLIP | Tarbase |
| hsa-miR-141-5p | SMARCA4 | HITS-CLIP | Tarbase |
| hsa-miR-141-5p | SMARCA4 | HITS-CLIP | Tarbase |
| hsa-miR-15a-5p | SMARCA4 | HITS-CLIP | Tarbase |
| hsa-miR-15a-5p | SMARCA4 | PAR-CLIP | Tarbase |
| hsa-miR-15a-5p | SMARCA4 | HITS-CLIP | Tarbase |
| hsa-miR-15a-5p | SMARCA4 | HITS-CLIP | Tarbase |
| hsa-miR-195-5p | SMARCA4 | HITS-CLIP | Tarbase |
| hsa-miR-3613-5p | KDM4B | HITS-CLIP | Tarbase |
| hsa-miR-141-3p | METTL25 | HITS-CLIP | Tarbase |
| hsa-miR-141-5p | METTL25 | HITS-CLIP | Tarbase |
| hsa-miR-15a-5p | EMC6 | HITS-CLIP | Tarbase |
| hsa-miR-195-5p | TUBA4A | Microarrays | Tarbase |
| hsa-miR-15a-5p | AAMP | HITS-CLIP | Tarbase |
| hsa-miR-15a-5p | AKAP9 | HITS-CLIP | Tarbase |
| hsa-miR-141-5p | PTPN12 | HITS-CLIP | Tarbase |
| hsa-miR-15a-5p | PTPN12 | HITS-CLIP | Tarbase |
| hsa-miR-15a-5p | PTPN12 | PAR-CLIP | Tarbase |
| hsa-miR-15a-5p | PTPN12 | HITS-CLIP | Tarbase |
| hsa-miR-195-5p | PTPN12 | HITS-CLIP | Tarbase |
| hsa-miR-195-5p | PTPN12 | HITS-CLIP | Tarbase |
| hsa-miR-141-5p | STYXL1 | HITS-CLIP | Tarbase |
| hsa-miR-195-5p | STEAP4 | Microarrays | Tarbase |
| hsa-miR-141-5p | PEX1 | HITS-CLIP | Tarbase |
| hsa-miR-195-5p | LRFN1 | Microarrays | Tarbase |
| hsa-miR-486-3p | ZFP36 | HITS-CLIP | Tarbase |
| hsa-miR-486-3p | ZFP36 | HITS-CLIP | Tarbase |
| hsa-miR-486-3p | ZFP36 | HITS-CLIP | Tarbase |
| hsa-miR-141-5p | DGCR8 | HITS-CLIP | Tarbase |
| hsa-miR-141-5p | YWHAH | HITS-CLIP | Tarbase |
| hsa-miR-15a-5p | YWHAH | Chimeric fragments | Tarbase |
| hsa-miR-3613-3p | MGAT3 | PAR-CLIP | Tarbase |
| hsa-miR-3613-3p | MGAT3 | PAR-CLIP | Tarbase |
| hsa-miR-486-3p | CDC42EP1 | HITS-CLIP | Tarbase |
| hsa-miR-195-5p | LIF | Microarrays | Tarbase |
| hsa-miR-15a-5p | EMC4 | HITS-CLIP | Tarbase |
| hsa-miR-15a-5p | EMC4 | HITS-CLIP | Tarbase |
| hsa-miR-195-5p | EMC4 | HITS-CLIP | Tarbase |
| hsa-miR-141-5p | SPECC1 | HITS-CLIP | Tarbase |
| hsa-miR-195-5p | CPA4 | Microarrays | Tarbase |
| hsa-miR-195-5p | DOCK4 | Microarrays | Tarbase |
| hsa-miR-3605-3p | VGF | HITS-CLIP | Tarbase |
| hsa-miR-141-5p | FOXP2 | HITS-CLIP | Tarbase |
| hsa-miR-3613-5p | DNAJB9 | HITS-CLIP | Tarbase |
| hsa-miR-141-3p | CALU | HITS-CLIP | Tarbase |
| hsa-miR-15a-5p | CALU | HITS-CLIP | Tarbase |
| hsa-miR-195-5p | CALU | HITS-CLIP | Tarbase |
| hsa-miR-3613-3p | CALU | HITS-CLIP | Tarbase |
| hsa-miR-486-3p | CALU | HITS-CLIP | Tarbase |
| hsa-miR-141-3p | SMO | HITS-CLIP | Tarbase |
| hsa-miR-141-3p | KLHDC10 | HITS-CLIP | Tarbase |
| hsa-miR-486-5p | KLHDC10 | HITS-CLIP | Tarbase |
| hsa-miR-486-5p | MYO1B | HITS-CLIP | Tarbase |
| hsa-miR-15a-5p | HOXD10 | PAR-CLIP | Tarbase |
| hsa-miR-15a-5p | HOXD10 | Chimeric fragments | Tarbase |
| hsa-miR-486-3p | HERC2 | HITS-CLIP | Tarbase |
| hsa-miR-219a-1-3p | TWSG1 | PAR-CLIP | Tarbase |
| hsa-miR-141-3p | EIF2AK4 | HITS-CLIP | Tarbase |
| hsa-miR-15a-5p | MYO5C | HITS-CLIP | Tarbase |
| hsa-miR-15a-5p | MYO5C | HITS-CLIP | Tarbase |
| hsa-miR-195-5p | MYO5C | HITS-CLIP | Tarbase |
| hsa-miR-15a-5p | TTBK2 | PAR-CLIP | Tarbase |

| | | | |
|---|---|---|---|
| hsa-miR-15a-5p | TTBK2 | PAR-CLIP | Tarbase |
| hsa-miR-195-5p | TTBK2 | PAR-CLIP | Tarbase |
| hsa-miR-15a-5p | ICE2 | HITS-CLIP | Tarbase |
| hsa-miR-195-5p | ICE2 | HITS-CLIP | Tarbase |
| hsa-miR-141-3p | FAM63B | PAR-CLIP | Tarbase |
| hsa-miR-3613-5p | FAM63B | PAR-CLIP | Tarbase |
| hsa-miR-3613-5p | FAM63B | PAR-CLIP | Tarbase |
| hsa-miR-141-3p | IVD | HITS-CLIP | Tarbase |
| hsa-miR-15a-5p | IVD | HITS-CLIP | Tarbase |
| hsa-miR-141-3p | KNSTRN | HITS-CLIP | Tarbase |
| hsa-miR-195-5p | KNSTRN | Microarrays | Tarbase |
| hsa-miR-15a-5p | CHAC1 | HITS-CLIP | Tarbase |
| hsa-miR-15a-5p | CHAC1 | HITS-CLIP | Tarbase |
| hsa-miR-15a-5p | CHAC1 | PAR-CLIP | Tarbase |
| hsa-miR-195-5p | CHAC1 | HITS-CLIP | Tarbase |
| hsa-miR-195-5p | CHAC1 | HITS-CLIP | Tarbase |
| hsa-miR-486-5p | CHAC1 | PAR-CLIP | Tarbase |
| hsa-miR-486-5p | CHAC1 | PAR-CLIP | Tarbase |
| hsa-miR-15a-3p | ARPP19 | PAR-CLIP | Tarbase |
| hsa-miR-15a-3p | ARPP19 | PAR-CLIP | Tarbase |
| hsa-miR-15a-5p | ARPP19 | HITS-CLIP | Tarbase |
| hsa-miR-15a-5p | ARPP19 | HITS-CLIP | Tarbase |
| hsa-miR-15a-5p | ARPP19 | PAR-CLIP | Tarbase |
| hsa-miR-15a-5p | ARPP19 | HITS-CLIP | Tarbase |
| hsa-miR-15a-5p | ARPP19 | HITS-CLIP | Tarbase |
| hsa-miR-195-5p | ARPP19 | HITS-CLIP | Tarbase |
| hsa-miR-195-5p | ARPP19 | HITS-CLIP | Tarbase |
| hsa-miR-195-5p | ARPP19 | HITS-CLIP | Tarbase |
| hsa-miR-15a-5p | ARPP19 | HITS-CLIP | Tarbase |
| hsa-miR-195-5p | ARPP19 | HITS-CLIP | Tarbase |
| hsa-miR-141-5p | VPS13C | HITS-CLIP | Tarbase |
| hsa-miR-195-5p | VPS13C | Microarrays | Tarbase |
| hsa-miR-195-5p | CALML4 | Microarrays | Tarbase |
| hsa-miR-15a-5p | ANAPC13 | PAR-CLIP | Tarbase |
| hsa-miR-15a-5p | ANAPC13 | PAR-CLIP | Tarbase |
| hsa-miR-15a-5p | ANAPC13 | PAR-CLIP | Tarbase |
| hsa-miR-15a-5p | ANAPC13 | PAR-CLIP | Tarbase |
| hsa-miR-15a-5p | ANAPC13 | PAR-CLIP | Tarbase |
| hsa-miR-15a-5p | ANAPC13 | PAR-CLIP | Tarbase |
| hsa-miR-15a-5p | ANAPC13 | PAR-CLIP | Tarbase |
| hsa-miR-15a-5p | ANAPC13 | HITS-CLIP | Tarbase |
| hsa-miR-195-5p | ANAPC13 | PAR-CLIP | Tarbase |
| hsa-miR-195-5p | ANAPC13 | PAR-CLIP | Tarbase |
| hsa-miR-195-5p | ANAPC13 | PAR-CLIP | Tarbase |
| hsa-miR-195-5p | ANAPC13 | Microarrays | Tarbase |
| hsa-miR-15a-5p | COPB1 | PAR-CLIP | Tarbase |
| hsa-miR-195-5p | COPB1 | PAR-CLIP | Tarbase |
| hsa-miR-141-5p | PSMA1 | HITS-CLIP | Tarbase |
| hsa-miR-141-5p | SERGEF | HITS-CLIP | Tarbase |
| hsa-miR-15a-3p | SERGEF | HITS-CLIP | Tarbase |
| hsa-miR-195-5p | E2F8 | Microarrays | Tarbase |
| hsa-miR-195-5p | FAM64A | Microarrays | Tarbase |
| hsa-miR-141-3p | KIF1C | HITS-CLIP | Tarbase |
| hsa-miR-141-3p | KIF1C | HITS-CLIP | Tarbase |
| hsa-miR-15a-5p | KIF1C | PAR-CLIP | Tarbase |
| hsa-miR-15a-5p | KIF1C | PAR-CLIP | Tarbase |
| hsa-miR-141-3p | CCNT1 | HITS-CLIP | Tarbase |
| hsa-miR-141-3p | CCNT1 | HITS-CLIP | Tarbase |
| hsa-miR-15a-5p | CCNT1 | PAR-CLIP | Tarbase |
| hsa-miR-15a-5p | CCNT1 | HITS-CLIP | Tarbase |
| hsa-miR-15a-5p | CCNT1 | PAR-CLIP | Tarbase |
| hsa-miR-15a-5p | CCNT1 | HITS-CLIP | Tarbase |
| hsa-miR-15a-5p | CCNT1 | HITS-CLIP | Tarbase |
| hsa-miR-195-5p | CCNT1 | HITS-CLIP | Tarbase |
| hsa-miR-195-5p | CCNT1 | HITS-CLIP | Tarbase |
| hsa-miR-141-3p | ILF3 | PAR-CLIP | Tarbase |
| hsa-miR-15a-5p | ILF3 | HITS-CLIP | Tarbase |
| hsa-miR-15a-5p | ILF3 | PAR-CLIP | Tarbase |
| hsa-miR-15a-5p | ILF3 | HITS-CLIP | Tarbase |
| hsa-miR-195-5p | ILF3 | HITS-CLIP | Tarbase |
| hsa-miR-195-5p | ILF3 | HITS-CLIP | Tarbase |
| hsa-miR-3613-5p | ILF3 | HITS-CLIP | Tarbase |
| hsa-miR-15a-5p | ILF3 | HITS-CLIP | Tarbase |
| hsa-miR-195-5p | ILF3 | HITS-CLIP | Tarbase |
| hsa-miR-15a-3p | BCL2L2 | HITS-CLIP | Tarbase |
| hsa-miR-15a-5p | BCL2L2 | HITS-CLIP | Tarbase |
| hsa-miR-15a-5p | BCL2L2 | HITS-CLIP | Tarbase |
| hsa-miR-195-5p | BCL2L2 | HITS-CLIP | Tarbase |
| hsa-miR-195-5p | BCL2L2 | HITS-CLIP | Tarbase |
| hsa-miR-195-5p | BCL2L2 | HITS-CLIP | Tarbase |
| hsa-miR-486-3p | BCL2L2 | HITS-CLIP | Tarbase |
| hsa-miR-141-3p | AJUBA | HITS-CLIP | Tarbase |
| hsa-miR-15a-5p | AJUBA | HITS-CLIP | Tarbase |
| hsa-miR-195-5p | PARP2 | Microarrays | Tarbase |
| hsa-miR-15a-5p | HEATR5A | HITS-CLIP | Tarbase |
| hsa-miR-195-5p | HEATR5A | HITS-CLIP | Tarbase |
| hsa-miR-141-3p | FOXA1 | HITS-CLIP | Tarbase |
| hsa-miR-141-3p | FOXA1 | HITS-CLIP | Tarbase |
| hsa-miR-15a-5p | FOXA1 | HITS-CLIP | Tarbase |
| hsa-miR-141-3p | SNX6 | HITS-CLIP | Tarbase |
| hsa-miR-141-3p | SNX6 | HITS-CLIP | Tarbase |
| hsa-miR-141-5p | EAPP | HITS-CLIP | Tarbase |
| hsa-miR-141-5p | EAPP | HITS-CLIP | Tarbase |
| hsa-miR-195-5p | RNASE1 | Microarrays | Tarbase |
| hsa-miR-15a-5p | EPB41L4A | HITS-CLIP | Tarbase |
| hsa-miR-195-5p | EPB41L4A | HITS-CLIP | Tarbase |
| hsa-miR-141-5p | REEP5 | HITS-CLIP | Tarbase |
| hsa-miR-141-5p | REEP5 | HITS-CLIP | Tarbase |
| hsa-miR-141-5p | REEP5 | HITS-CLIP | Tarbase |
| hsa-miR-15a-5p | REEP5 | HITS-CLIP | Tarbase |
| hsa-miR-195-5p | REEP5 | HITS-CLIP | Tarbase |
| hsa-miR-15a-5p | REEP5 | HITS-CLIP | Tarbase |
| hsa-miR-195-5p | REEP5 | HITS-CLIP | Tarbase |
| hsa-miR-3605-5p | ITFG1 | HITS-CLIP | Tarbase |
| hsa-miR-15a-5p | CHRNA10 | PAR-CLIP | Tarbase |
| hsa-miR-195-5p | SGOL1 | Microarrays | Tarbase |
| hsa-miR-195-5p | KLF16 | Microarrays | Tarbase |
| hsa-miR-15a-5p | TMEM8A | HITS-CLIP | Tarbase |
| hsa-miR-15a-5p | TMEM8A | HITS-CLIP | Tarbase |
| hsa-miR-195-5p | TMEM8A | HITS-CLIP | Tarbase |
| hsa-miR-486-3p | TMEM8A | HITS-CLIP | Tarbase |
| hsa-miR-195-5p | LBP | Microarrays | Tarbase |
| hsa-miR-15a-5p | CBFA2T3 | HITS-CLIP | Tarbase |
| hsa-miR-3605-5p | HDHD1 | HITS-CLIP | Tarbase |
| hsa-miR-195-5p | PHF10 | Microarrays | Tarbase |
| hsa-miR-195-5p | STARD8 | Microarrays | Tarbase |
| hsa-miR-15a-5p | SAT1 | PAR-CLIP | Tarbase |
| hsa-miR-141-3p | SH3BP4 | HITS-CLIP | Tarbase |
| hsa-miR-195-5p | SH3BP4 | Microarrays | Tarbase |
| hsa-miR-15a-5p | MOSPD2 | PAR-CLIP | Tarbase |
| hsa-miR-195-5p | MOSPD2 | PAR-CLIP | Tarbase |
| hsa-miR-15a-5p | DOCK6 | HITS-CLIP | Tarbase |
| hsa-miR-195-5p | DOCK6 | HITS-CLIP | Tarbase |
| hsa-miR-141-5p | LDLR | HITS-CLIP | Tarbase |
| hsa-miR-15a-3p | LDLR | HITS-CLIP | Tarbase |
| hsa-miR-486-5p | LDLR | HITS-CLIP | Tarbase |
| hsa-miR-15a-5p | PRKCSH | HITS-CLIP | Tarbase |
| hsa-miR-15a-5p | PVRL2 | HITS-CLIP | Tarbase |
| hsa-miR-141-5p | APOE | HITS-CLIP | Tarbase |
| hsa-miR-15a-5p | XPO7 | PAR-CLIP | Tarbase |
| hsa-miR-15a-5p | XPO7 | HITS-CLIP | Tarbase |
| hsa-miR-15a-5p | XPO7 | PAR-CLIP | Tarbase |
| hsa-miR-15a-5p | XPO7 | HITS-CLIP | Tarbase |
| hsa-miR-15a-5p | XPO7 | HITS-CLIP | Tarbase |
| hsa-miR-195-5p | XPO7 | PAR-CLIP | Tarbase |
| hsa-miR-195-5p | XPO7 | HITS-CLIP | Tarbase |
| hsa-miR-195-5p | XPO7 | HITS-CLIP | Tarbase |
| hsa-miR-195-5p | KIF1A | Microarrays | Tarbase |
| hsa-miR-141-3p | GTPBP3 | HITS-CLIP | Tarbase |
| hsa-miR-15a-5p | COLGALT1 | PAR-CLIP | Tarbase |
| hsa-miR-486-5p | COLGALT1 | PAR-CLIP | Tarbase |
| hsa-miR-486-5p | COLGALT1 | PAR-CLIP | Tarbase |
| hsa-miR-486-5p | COLGALT1 | PAR-CLIP | Tarbase |
| hsa-miR-219a-1-3p | TULP4 | HITS-CLIP | Tarbase |
| hsa-miR-3613-3p | TULP4 | PAR-CLIP | Tarbase |
| hsa-miR-141-3p | MLLT4 | HITS-CLIP | Tarbase |
| hsa-miR-15a-5p | MLLT4 | HITS-CLIP | Tarbase |
| hsa-miR-195-5p | MLLT4 | HITS-CLIP | Tarbase |
| hsa-miR-219a-1-3p | ACTN4 | HITS-CLIP | Tarbase |

| | | | |
|---|---|---|---|
| hsa-miR-3605-5p | ACTN4 | HITS-CLIP | Tarbase |
| hsa-miR-15a-5p | STK33 | HITS-CLIP | Tarbase |
| hsa-miR-195-5p | STK33 | HITS-CLIP | Tarbase |
| hsa-miR-15a-5p | ARPC1B | HITS-CLIP | Tarbase |
| hsa-miR-195-5p | ARPC1B | HITS-CLIP | Tarbase |
| hsa-miR-15a-5p | ZSWIM6 | HITS-CLIP | Tarbase |
| hsa-miR-15a-5p | ZSWIM6 | HITS-CLIP | Tarbase |
| hsa-miR-15a-5p | ZSWIM6 | HITS-CLIP | Tarbase |
| hsa-miR-195-5p | ZSWIM6 | HITS-CLIP | Tarbase |
| hsa-miR-195-5p | ZSWIM6 | HITS-CLIP | Tarbase |
| hsa-miR-195-5p | ZSWIM6 | HITS-CLIP | Tarbase |
| hsa-miR-15a-5p | UNC13A | HITS-CLIP | Tarbase |
| hsa-miR-195-5p | UNC13A | HITS-CLIP | Tarbase |
| hsa-miR-195-5p | PXDN | Microarrays | Tarbase |
| hsa-miR-195-5p | SSBP4 | Microarrays | Tarbase |
| hsa-miR-3613-5p | PGPEP1 | PAR-CLIP | Tarbase |
| hsa-miR-3613-5p | PGPEP1 | PAR-CLIP | Tarbase |
| hsa-miR-195-5p | LSM4 | Microarrays | Tarbase |
| hsa-miR-141-3p | JUND | HITS-CLIP | Tarbase |
| hsa-miR-3605-3p | JUND | HITS-CLIP | Tarbase |
| hsa-miR-486-3p | JUND | HITS-CLIP | Tarbase |
| hsa-miR-141-3p | ZNF557 | HITS-CLIP | Tarbase |
| hsa-miR-15a-5p | CAMSAP1 | PAR-CLIP | Tarbase |
| hsa-miR-195-5p | UBAC1 | Microarrays | Tarbase |
| hsa-miR-15a-5p | ZBTB46 | HITS-CLIP | Tarbase |
| hsa-miR-195-5p | ZBTB46 | HITS-CLIP | Tarbase |
| hsa-miR-195-5p | COL5A1 | HITS-CLIP | Tarbase |
| hsa-miR-195-5p | COL5A1 | Microarrays | Tarbase |
| hsa-miR-141-3p | MNX1 | HITS-CLIP | Tarbase |
| hsa-miR-141-3p | MNX1 | HITS-CLIP | Tarbase |
| hsa-miR-195-5p | CEP85 | Microarrays | Tarbase |
| hsa-miR-15a-3p | PRRC2B | HITS-CLIP | Tarbase |
| hsa-miR-15a-5p | PRRC2B | HITS-CLIP | Tarbase |
| hsa-miR-15a-5p | PRRC2B | HITS-CLIP | Tarbase |
| hsa-miR-15a-5p | PRRC2B | HITS-CLIP | Tarbase |
| hsa-miR-15a-5p | PRRC2B | HITS-CLIP | Tarbase |
| hsa-miR-15a-5p | PRRC2B | HITS-CLIP | Tarbase |
| hsa-miR-195-5p | PRRC2B | HITS-CLIP | Tarbase |
| hsa-miR-195-5p | PRRC2B | HITS-CLIP | Tarbase |
| hsa-miR-195-5p | PRRC2B | HITS-CLIP | Tarbase |
| hsa-miR-195-5p | PRRC2B | HITS-CLIP | Tarbase |
| hsa-miR-141-5p | CHMP2A | HITS-CLIP | Tarbase |
| hsa-miR-141-5p | TRIM28 | HITS-CLIP | Tarbase |
| hsa-miR-15a-3p | TRIM28 | HITS-CLIP | Tarbase |
| hsa-miR-15a-3p | TRIM28 | HITS-CLIP | Tarbase |
| hsa-miR-15a-3p | TRIM28 | HITS-CLIP | Tarbase |
| hsa-miR-3605-3p | TRIM28 | HITS-CLIP | Tarbase |
| hsa-miR-141-5p | C16orf13 | HITS-CLIP | Tarbase |
| hsa-miR-3605-3p | C16orf13 | HITS-CLIP | Tarbase |
| hsa-miR-15a-5p | YIPF2 | HITS-CLIP | Tarbase |
| hsa-miR-195-5p | YIPF2 | HITS-CLIP | Tarbase |
| hsa-miR-15a-5p | EIF2S3 | HITS-CLIP | Tarbase |
| hsa-miR-15a-5p | EIF2S3 | HITS-CLIP | Tarbase |
| hsa-miR-195-5p | EIF2S3 | HITS-CLIP | Tarbase |
| hsa-miR-195-5p | EIF2S3 | HITS-CLIP | Tarbase |
| hsa-miR-15a-5p | LRRC47 | HITS-CLIP | Tarbase |
| hsa-miR-15a-5p | LRRC47 | HITS-CLIP | Tarbase |
| hsa-miR-15a-5p | LRRC47 | HITS-CLIP | Tarbase |
| hsa-miR-195-5p | LRRC47 | HITS-CLIP | Tarbase |
| hsa-miR-195-5p | LRRC47 | HITS-CLIP | Tarbase |
| hsa-miR-195-5p | LRRC47 | HITS-CLIP | Tarbase |
| hsa-miR-15a-5p | MED18 | HITS-CLIP | Tarbase |
| hsa-miR-486-5p | CLIP1 | HITS-CLIP | Tarbase |
| hsa-miR-195-5p | DNMT1 | Microarrays | Tarbase |
| hsa-miR-15a-5p | SLC6A8 | HITS-CLIP | Tarbase |
| hsa-miR-195-5p | SLC6A8 | HITS-CLIP | Tarbase |
| hsa-miR-486-3p | SLC6A8 | HITS-CLIP | Tarbase |
| hsa-miR-141-3p | DKC1 | HITS-CLIP | Tarbase |
| hsa-miR-141-5p | DKC1 | HITS-CLIP | Tarbase |
| hsa-miR-15a-5p | DKC1 | PAR-CLIP | Tarbase |
| hsa-miR-15a-5p | ZNF236 | PAR-CLIP | Tarbase |
| hsa-miR-3613-3p | ZNF236 | PAR-CLIP | Tarbase |
| hsa-miR-3613-3p | ZNF236 | PAR-CLIP | Tarbase |
| hsa-miR-3613-3p | ZNF236 | PAR-CLIP | Tarbase |
| hsa-miR-15a-5p | LRP3 | HITS-CLIP | Tarbase |
| hsa-miR-141-3p | NOL11 | HITS-CLIP | Tarbase |
| hsa-miR-141-3p | NOL11 | HITS-CLIP | Tarbase |
| hsa-miR-15a-5p | NOL11 | HITS-CLIP | Tarbase |
| hsa-miR-15a-5p | UBE4B | PAR-CLIP | Tarbase |
| hsa-miR-15a-5p | UBE4B | PAR-CLIP | Tarbase |
| hsa-miR-15a-5p | UBE4B | HITS-CLIP | Tarbase |
| hsa-miR-195-5p | UBE4B | PAR-CLIP | Tarbase |
| hsa-miR-195-5p | UBE4B | Microarrays | Tarbase |
| hsa-miR-219a-1-3p | PRRG1 | HITS-CLIP | Tarbase |
| hsa-miR-195-5p | ULBP2 | Microarrays | Tarbase |
| hsa-miR-141-3p | AKAP12 | HITS-CLIP | Tarbase |
| hsa-miR-141-5p | AKAP12 | HITS-CLIP | Tarbase |
| hsa-miR-141-3p | SYNE1 | HITS-CLIP | Tarbase |
| hsa-miR-141-5p | SYNE1 | HITS-CLIP | Tarbase |
| hsa-miR-195-5p | ULBP3 | Microarrays | Tarbase |
| hsa-miR-141-3p | LATS1 | HITS-CLIP | Tarbase |
| hsa-miR-141-5p | LATS1 | HITS-CLIP | Tarbase |
| hsa-miR-15a-5p | LATS1 | PAR-CLIP | Tarbase |
| hsa-miR-15a-5p | LATS1 | PAR-CLIP | Tarbase |
| hsa-miR-15a-5p | LATS1 | PAR-CLIP | Tarbase |
| hsa-miR-15a-5p | LATS1 | PAR-CLIP | Tarbase |
| hsa-miR-15a-5p | LATS1 | PAR-CLIP | Tarbase |
| hsa-miR-15a-5p | LATS1 | PAR-CLIP | Tarbase |
| hsa-miR-15a-5p | LATS1 | PAR-CLIP | Tarbase |
| hsa-miR-15a-5p | LATS1 | PAR-CLIP | Tarbase |
| hsa-miR-15a-5p | LATS1 | HITS-CLIP | Tarbase |
| hsa-miR-195-5p | LATS1 | PAR-CLIP | Tarbase |
| hsa-miR-195-5p | LATS1 | PAR-CLIP | Tarbase |
| hsa-miR-195-5p | LATS1 | PAR-CLIP | Tarbase |
| hsa-miR-195-5p | LATS1 | PAR-CLIP | Tarbase |
| hsa-miR-141-5p | ACSS2 | HITS-CLIP | Tarbase |
| hsa-miR-15a-5p | GFAP | HITS-CLIP | Tarbase |
| hsa-miR-195-5p | GFAP | HITS-CLIP | Tarbase |
| hsa-miR-15a-5p | ZNF428 | HITS-CLIP | Tarbase |
| hsa-miR-195-5p | ZNF428 | HITS-CLIP | Tarbase |
| hsa-miR-219a-1-3p | COX4I1 | HITS-CLIP | Tarbase |
| hsa-miR-15a-5p | GSE1 | HITS-CLIP | Tarbase |
| hsa-miR-3605-3p | GSE1 | HITS-CLIP | Tarbase |
| hsa-miR-195-5p | GINS2 | Microarrays | Tarbase |
| hsa-miR-15a-5p | PPT1 | HITS-CLIP | Tarbase |
| hsa-miR-15a-5p | PPT1 | HITS-CLIP | Tarbase |
| hsa-miR-15a-5p | PPT1 | HITS-CLIP | Tarbase |
| hsa-miR-195-5p | PPT1 | HITS-CLIP | Tarbase |
| hsa-miR-195-5p | PPT1 | HITS-CLIP | Tarbase |
| hsa-miR-486-5p | PPT1 | HITS-CLIP | Tarbase |
| hsa-miR-195-5p | PPT1 | Microarrays | Tarbase |
| hsa-miR-15a-3p | RLIM | HITS-CLIP | Tarbase |
| hsa-miR-15a-5p | TRAF3 | PAR-CLIP | Tarbase |
| hsa-miR-195-5p | HAUS8 | Microarrays | Tarbase |
| hsa-miR-15a-5p | MRPS25 | HITS-CLIP | Tarbase |
| hsa-miR-195-5p | MRPS25 | HITS-CLIP | Tarbase |
| hsa-miR-15a-5p | ZFYVE20 | PAR-CLIP | Tarbase |
| hsa-miR-15a-5p | ZFYVE20 | HITS-CLIP | Tarbase |
| hsa-miR-15a-5p | ZFYVE20 | HITS-CLIP | Tarbase |
| hsa-miR-195-5p | ZFYVE20 | HITS-CLIP | Tarbase |
| hsa-miR-195-5p | GFPT2 | Microarrays | Tarbase |
| hsa-miR-141-5p | PSME3 | HITS-CLIP | Tarbase |
| hsa-miR-141-5p | ACLY | HITS-CLIP | Tarbase |
| hsa-miR-15a-5p | ACLY | HITS-CLIP | Tarbase |
| hsa-miR-195-5p | ACLY | HITS-CLIP | Tarbase |
| hsa-miR-195-5p | ACLY | Microarrays | Tarbase |
| hsa-miR-15a-5p | DIAPH1 | PAR-CLIP | Tarbase |
| hsa-miR-141-3p | PPFIA1 | HITS-CLIP | Tarbase |
| hsa-miR-15a-5p | MAP1B | PAR-CLIP | Tarbase |
| hsa-miR-15a-5p | MAP1B | PAR-CLIP | Tarbase |
| hsa-miR-486-5p | MAP1B | HITS-CLIP | Tarbase |
| hsa-miR-15a-5p | IL13RA1 | HITS-CLIP | Tarbase |
| hsa-miR-141-3p | TOP2A | HITS-CLIP | Tarbase |
| hsa-miR-141-5p | TOP2A | HITS-CLIP | Tarbase |
| hsa-miR-195-5p | TOP2A | Microarrays | Tarbase |
| hsa-miR-15a-5p | PRKAB2 | HITS-CLIP | Tarbase |
| hsa-miR-15a-5p | PRKAB2 | HITS-CLIP | Tarbase |
| hsa-miR-195-5p | PRKAB2 | HITS-CLIP | Tarbase |
| hsa-miR-141-3p | PDHA1 | HITS-CLIP | Tarbase |

| | | | |
|---|---|---|---|
| hsa-miR-141-3p | PDHA1 | HITS-CLIP | Tarbase |
| hsa-miR-15a-5p | MCCC2 | HITS-CLIP | Tarbase |
| hsa-miR-15a-5p | VIMP | PAR-CLIP | Tarbase |
| hsa-miR-141-5p | C19orf12 | HITS-CLIP | Tarbase |
| hsa-miR-195-5p | C19orf40 | Microarrays | Tarbase |
| hsa-miR-15a-3p | DNAJB1 | HITS-CLIP | Tarbase |
| hsa-miR-195-5p | ZSWIM4 | Microarrays | Tarbase |
| hsa-miR-141-3p | HGNC:9982 | HITS-CLIP | Tarbase |
| hsa-miR-15a-3p | CC2D1A | HITS-CLIP | Tarbase |
| hsa-miR-141-5p | SPATA6 | HITS-CLIP | Tarbase |
| hsa-miR-15a-5p | LRRC41 | HITS-CLIP | Tarbase |
| hsa-miR-15a-5p | LRRC41 | HITS-CLIP | Tarbase |
| hsa-miR-195-5p | LRRC41 | HITS-CLIP | Tarbase |
| hsa-miR-195-5p | LRRC41 | HITS-CLIP | Tarbase |
| hsa-miR-486-3p | LRRC41 | HITS-CLIP | Tarbase |
| hsa-miR-195-5p | RAF1 | Luciferase Reporter Assay | Tarbase |
| hsa-miR-15a-5p | NUP210 | HITS-CLIP | Tarbase |
| hsa-miR-15a-5p | NUP210 | HITS-CLIP | Tarbase |
| hsa-miR-15a-5p | NUP210 | HITS-CLIP | Tarbase |
| hsa-miR-15a-5p | NUP210 | PAR-CLIP | Tarbase |
| hsa-miR-195-5p | NUP210 | HITS-CLIP | Tarbase |
| hsa-miR-195-5p | NUP210 | HITS-CLIP | Tarbase |
| hsa-miR-195-5p | NUP210 | PAR-CLIP | Tarbase |
| hsa-miR-195-5p | NUP210 | Microarrays | Tarbase |
| hsa-miR-15a-5p | TIMM10B | PAR-CLIP | Tarbase |
| hsa-miR-141-5p | PTCD3 | HITS-CLIP | Tarbase |
| hsa-miR-141-5p | IMMT | HITS-CLIP | Tarbase |
| hsa-miR-15a-5p | PER2 | HITS-CLIP | Tarbase |
| hsa-miR-195-5p | PER2 | HITS-CLIP | Tarbase |
| hsa-miR-3605-5p | PTPRE | HITS-CLIP | Tarbase |
| hsa-miR-195-5p | PTPRE | Microarrays | Tarbase |
| hsa-miR-15a-5p | RAN | HITS-CLIP | Tarbase |
| hsa-miR-195-5p | RAN | HITS-CLIP | Tarbase |
| hsa-miR-141-3p | PRKAA1 | HITS-CLIP | Tarbase |
| hsa-miR-141-5p | PRKAA1 | HITS-CLIP | Tarbase |
| hsa-miR-141-5p | PRKAA1 | HITS-CLIP | Tarbase |
| hsa-miR-15a-5p | CLUH | HITS-CLIP | Tarbase |
| hsa-miR-195-5p | CLUH | HITS-CLIP | Tarbase |
| hsa-miR-195-5p | RPA1 | Microarrays | Tarbase |
| hsa-miR-15a-5p | PNISR | HITS-CLIP | Tarbase |
| hsa-miR-195-5p | PNISR | HITS-CLIP | Tarbase |
| hsa-miR-15a-5p | PNISR | Chimeric fragments | Tarbase |
| hsa-miR-15a-5p | GRSF1 | HITS-CLIP | Tarbase |
| hsa-miR-15a-5p | GRSF1 | HITS-CLIP | Tarbase |
| hsa-miR-195-5p | GRSF1 | HITS-CLIP | Tarbase |
| hsa-miR-141-3p | ANKRD17 | HITS-CLIP | Tarbase |
| hsa-miR-141-3p | ANKRD17 | HITS-CLIP | Tarbase |
| hsa-miR-141-3p | ANKRD17 | HITS-CLIP | Tarbase |
| hsa-miR-141-5p | ANKRD17 | HITS-CLIP | Tarbase |
| hsa-miR-15a-5p | ANKRD17 | HITS-CLIP | Tarbase |
| hsa-miR-15a-5p | ANKRD17 | HITS-CLIP | Tarbase |
| hsa-miR-15a-5p | ANKRD17 | HITS-CLIP | Tarbase |
| hsa-miR-195-5p | ANKRD17 | HITS-CLIP | Tarbase |
| hsa-miR-195-5p | ANKRD17 | HITS-CLIP | Tarbase |
| hsa-miR-195-5p | ANKRD17 | HITS-CLIP | Tarbase |
| hsa-miR-141-3p | H3F3B | HITS-CLIP | Tarbase |
| hsa-miR-3605-5p | H3F3B | HITS-CLIP | Tarbase |
| hsa-miR-486-5p | H3F3B | HITS-CLIP | Tarbase |
| hsa-miR-486-5p | H3F3B | HITS-CLIP | Tarbase |
| hsa-miR-486-5p | H3F3B | HITS-CLIP | Tarbase |
| hsa-miR-15a-5p | EIF5A | HITS-CLIP | Tarbase |
| hsa-miR-195-5p | EIF5A | HITS-CLIP | Tarbase |
| hsa-miR-141-3p | VPS13B | HITS-CLIP | Tarbase |
| hsa-miR-15a-5p | VPS13B | HITS-CLIP | Tarbase |
| hsa-miR-195-5p | VPS13B | HITS-CLIP | Tarbase |
| hsa-miR-141-5p | FLOT2 | HITS-CLIP | Tarbase |
| hsa-miR-15a-5p | FLOT2 | HITS-CLIP | Tarbase |
| hsa-miR-15a-5p | FLOT2 | HITS-CLIP | Tarbase |
| hsa-miR-486-3p | FLOT2 | HITS-CLIP | Tarbase |
| hsa-miR-141-5p | TERF2 | HITS-CLIP | Tarbase |
| hsa-miR-486-5p | TERF2 | PAR-CLIP | Tarbase |
| hsa-miR-486-5p | TERF2 | PAR-CLIP | Tarbase |
| hsa-miR-195-5p | VPS4A | Microarrays | Tarbase |
| hsa-miR-15a-3p | MTSS1L | HITS-CLIP | Tarbase |
| hsa-miR-15a-5p | MTSS1L | HITS-CLIP | Tarbase |
| hsa-miR-15a-5p | MTSS1L | HITS-CLIP | Tarbase |
| hsa-miR-195-5p | MTSS1L | HITS-CLIP | Tarbase |
| hsa-miR-15a-5p | ANKEF1 | PAR-CLIP | Tarbase |
| hsa-miR-15a-5p | ANKEF1 | HITS-CLIP | Tarbase |
| hsa-miR-195-5p | ANKEF1 | HITS-CLIP | Tarbase |
| hsa-miR-141-5p | PCED1A | HITS-CLIP | Tarbase |
| hsa-miR-15a-5p | BTBD3 | HITS-CLIP | Tarbase |
| hsa-miR-195-5p | BTBD3 | HITS-CLIP | Tarbase |
| hsa-miR-15a-5p | BTBD3 | Chimeric fragments | Tarbase |
| hsa-miR-15a-5p | POLR3F | PAR-CLIP | Tarbase |
| hsa-miR-15a-5p | POLR3F | HITS-CLIP | Tarbase |
| hsa-miR-195-5p | POLR3F | HITS-CLIP | Tarbase |
| hsa-miR-141-3p | RIN2 | Luciferase Reporter Assay | Tarbase |
| hsa-miR-141-3p | RIN2 | Luciferase Reporter Assay | Tarbase |
| hsa-miR-195-5p | RIN2 | Microarrays | Tarbase |
| hsa-miR-15a-5p | DAP3 | HITS-CLIP | Tarbase |
| hsa-miR-15a-5p | KIAA0907 | PAR-CLIP | Tarbase |
| hsa-miR-15a-5p | ARHGEF11 | HITS-CLIP | Tarbase |
| hsa-miR-195-5p | ARHGEF11 | HITS-CLIP | Tarbase |
| hsa-miR-15a-5p | DCAF8 | PAR-CLIP | Tarbase |
| hsa-miR-195-5p | SYT11 | Microarrays | Tarbase |
| hsa-miR-15a-3p | MMACHC | PAR-CLIP | Tarbase |
| hsa-miR-195-5p | NASP | Microarrays | Tarbase |
| hsa-miR-141-3p | RBM38 | HITS-CLIP | Tarbase |
| hsa-miR-15a-5p | SERINC3 | HITS-CLIP | Tarbase |
| hsa-miR-195-5p | SERINC3 | HITS-CLIP | Tarbase |
| hsa-miR-15a-5p | AP3B1 | HITS-CLIP | Tarbase |
| hsa-miR-3605-5p | INADL | HITS-CLIP | Tarbase |
| hsa-miR-195-5p | INADL | Microarrays | Tarbase |
| hsa-miR-15a-5p | SYT4 | HITS-CLIP | Tarbase |
| hsa-miR-195-5p | SYT4 | HITS-CLIP | Tarbase |
| hsa-miR-195-5p | SYT4 | Microarrays | Tarbase |
| hsa-miR-486-5p | CASP9 | HITS-CLIP | Tarbase |
| hsa-miR-15a-5p | DCTN4 | HITS-CLIP | Tarbase |
| hsa-miR-195-5p | DCTN4 | HITS-CLIP | Tarbase |
| hsa-miR-3613-3p | ATP8A2 | PAR-CLIP | Tarbase |
| hsa-miR-15a-3p | ZMYM5 | PAR-CLIP | Tarbase |
| hsa-miR-15a-5p | XPO4 | HITS-CLIP | Tarbase |
| hsa-miR-15a-5p | CHRM3 | HITS-CLIP | Tarbase |
| hsa-miR-195-5p | CHRM3 | HITS-CLIP | Tarbase |
| hsa-miR-195-5p | CHRM3 | Microarrays | Tarbase |
| hsa-miR-141-3p | MYH10 | HITS-CLIP | Tarbase |
| hsa-miR-141-3p | MYH10 | HITS-CLIP | Tarbase |
| hsa-miR-141-3p | MPRIP | HITS-CLIP | Tarbase |
| hsa-miR-141-3p | DSTYK | HITS-CLIP | Tarbase |
| hsa-miR-3605-3p | DSTYK | HITS-CLIP | Tarbase |
| hsa-miR-195-5p | SLC41A1 | Microarrays | Tarbase |
| hsa-miR-15a-5p | DCLK1 | HITS-CLIP | Tarbase |
| hsa-miR-195-5p | DCLK1 | HITS-CLIP | Tarbase |
| hsa-miR-195-5p | CCNA1 | Microarrays | Tarbase |
| hsa-miR-195-5p | EPSTI1 | Microarrays | Tarbase |
| hsa-miR-195-5p | TRPC4 | Microarrays | Tarbase |
| hsa-miR-3613-5p | POSTN | HITS-CLIP | Tarbase |
| hsa-miR-3605-5p | KL | HITS-CLIP | Tarbase |
| hsa-miR-195-5p | RFC3 | Microarrays | Tarbase |
| hsa-miR-15a-5p | STARD13 | HITS-CLIP | Tarbase |
| hsa-miR-15a-5p | STARD13 | HITS-CLIP | Tarbase |
| hsa-miR-195-5p | STARD13 | HITS-CLIP | Tarbase |
| hsa-miR-195-5p | STARD13 | HITS-CLIP | Tarbase |
| hsa-miR-141-5p | IRS4 | HITS-CLIP | Tarbase |
| hsa-miR-141-5p | IRS4 | HITS-CLIP | Tarbase |
| hsa-miR-15a-5p | IRS4 | PAR-CLIP | Tarbase |
| hsa-miR-15a-5p | IRS4 | HITS-CLIP | Tarbase |
| hsa-miR-195-5p | IRS4 | HITS-CLIP | Tarbase |
| hsa-miR-141-5p | BEX2 | HITS-CLIP | Tarbase |
| hsa-miR-195-5p | EPHB2 | Microarrays | Tarbase |
| hsa-miR-15a-5p | SRRM1 | HITS-CLIP | Tarbase |
| hsa-miR-195-5p | SRRM1 | HITS-CLIP | Tarbase |
| hsa-miR-15a-5p | RTN3 | HITS-CLIP | Tarbase |
| hsa-miR-195-5p | RTN3 | HITS-CLIP | Tarbase |
| hsa-miR-195-5p | RARRES3 | Microarrays | Tarbase |
| hsa-miR-15a-5p | PDZD2 | HITS-CLIP | Tarbase |
| hsa-miR-195-5p | PDZD2 | HITS-CLIP | Tarbase |
| hsa-miR-195-5p | PDZD2 | Microarrays | Tarbase |

| | | | |
|---|---|---|---|
| hsa-miR-141-5p | MKRN1 | HITS-CLIP | Tarbase |
| hsa-miR-15a-5p | AGAP3 | HITS-CLIP | Tarbase |
| hsa-miR-195-5p | AGAP3 | HITS-CLIP | Tarbase |
| hsa-miR-141-5p | ATP13A3 | HITS-CLIP | Tarbase |
| hsa-miR-15a-5p | ATP13A3 | HITS-CLIP | Tarbase |
| hsa-miR-15a-5p | ATP13A3 | HITS-CLIP | Tarbase |
| hsa-miR-15a-5p | ATP13A3 | HITS-CLIP | Tarbase |
| hsa-miR-15a-5p | ATP13A3 | PAR-CLIP | Tarbase |
| hsa-miR-195-5p | ATP13A3 | HITS-CLIP | Tarbase |
| hsa-miR-195-5p | ATP13A3 | HITS-CLIP | Tarbase |
| hsa-miR-195-5p | ATP13A3 | HITS-CLIP | Tarbase |
| hsa-miR-3613-3p | ATP13A3 | HITS-CLIP | Tarbase |
| hsa-miR-15a-5p | TMTC1 | Chimeric fragments | Tarbase |
| hsa-miR-15a-5p | KRAS | PAR-CLIP | Tarbase |
| hsa-miR-15a-5p | IPO8 | HITS-CLIP | Tarbase |
| hsa-miR-15a-5p | IPO8 | HITS-CLIP | Tarbase |
| hsa-miR-15a-5p | IPO8 | PAR-CLIP | Tarbase |
| hsa-miR-15a-5p | IPO8 | HITS-CLIP | Tarbase |
| hsa-miR-15a-5p | IPO8 | HITS-CLIP | Tarbase |
| hsa-miR-195-5p | IPO8 | HITS-CLIP | Tarbase |
| hsa-miR-195-3p | LARS | HITS-CLIP | Tarbase |
| hsa-miR-15a-5p | LARS | Chimeric fragments | Tarbase |
| hsa-miR-15a-5p | E2F5 | PAR-CLIP | Tarbase |
| hsa-miR-15a-5p | E2F5 | PAR-CLIP | Tarbase |
| hsa-miR-15a-5p | CCDC59 | HITS-CLIP | Tarbase |
| hsa-miR-15a-5p | CCDC59 | HITS-CLIP | Tarbase |
| hsa-miR-15a-5p | CCDC59 | HITS-CLIP | Tarbase |
| hsa-miR-195-5p | CCDC59 | HITS-CLIP | Tarbase |
| hsa-miR-195-5p | CCDC59 | HITS-CLIP | Tarbase |
| hsa-miR-195-5p | CCDC59 | HITS-CLIP | Tarbase |
| hsa-miR-195-5p | SWAP70 | Microarrays | Tarbase |
| hsa-miR-15a-5p | SBF2 | HITS-CLIP | Tarbase |
| hsa-miR-195-5p | SBF2 | HITS-CLIP | Tarbase |
| hsa-miR-195-5p | MICAL2 | Microarrays | Tarbase |
| hsa-miR-15a-5p | RRAS2 | PAR-CLIP | Tarbase |
| hsa-miR-195-5p | RRAS2 | PAR-CLIP | Tarbase |
| hsa-miR-15a-5p | HSD17B4 | HITS-CLIP | Tarbase |
| hsa-miR-15a-5p | HSD17B4 | HITS-CLIP | Tarbase |
| hsa-miR-195-5p | HSD17B4 | HITS-CLIP | Tarbase |
| hsa-miR-195-5p | HSD17B4 | HITS-CLIP | Tarbase |
| hsa-miR-3613-5p | TEX15 | HITS-CLIP | Tarbase |
| hsa-miR-141-3p | SARAF | HITS-CLIP | Tarbase |
| hsa-miR-15a-3p | RNF122 | HITS-CLIP | Tarbase |
| hsa-miR-15a-5p | NUMB | PAR-CLIP | Tarbase |
| hsa-miR-219a-1-3p | NUMB | HITS-CLIP | Tarbase |
| hsa-miR-219a-1-3p | NUMB | HITS-CLIP | Tarbase |
| hsa-miR-486-3p | NUMB | HITS-CLIP | Tarbase |
| hsa-miR-15a-5p | VRTN | PAR-CLIP | Tarbase |
| hsa-miR-486-5p | COX16 | PAR-CLIP | Tarbase |
| hsa-miR-3613-5p | ADAM20 | PAR-CLIP | Tarbase |
| hsa-miR-195-5p | LOXL2 | Microarrays | Tarbase |
| hsa-miR-141-3p | IER3IP1 | HITS-CLIP | Tarbase |
| hsa-miR-15a-3p | IER3IP1 | HITS-CLIP | Tarbase |
| hsa-miR-195-5p | CCNB1 | Microarrays | Tarbase |
| hsa-miR-219a-1-3p | THUMPD3 | PAR-CLIP | Tarbase |
| hsa-miR-219a-1-3p | THUMPD3 | PAR-CLIP | Tarbase |
| hsa-miR-15a-5p | VHL | HITS-CLIP | Tarbase |
| hsa-miR-195-5p | VHL | HITS-CLIP | Tarbase |
| hsa-miR-15a-5p | BHLHE40 | HITS-CLIP | Tarbase |
| hsa-miR-486-3p | BHLHE40 | HITS-CLIP | Tarbase |
| hsa-miR-15a-5p | ARL8B | HITS-CLIP | Tarbase |
| hsa-miR-15a-5p | ARL8B | PAR-CLIP | Tarbase |
| hsa-miR-195-5p | ARL8B | HITS-CLIP | Tarbase |
| hsa-miR-141-3p | EDEM1 | HITS-CLIP | Tarbase |
| hsa-miR-15a-5p | EDEM1 | Chimeric fragments | Tarbase |
| hsa-miR-15a-5p | CHL1 | HITS-CLIP | Tarbase |
| hsa-miR-195-5p | CHL1 | HITS-CLIP | Tarbase |
| hsa-miR-15a-5p | KATNBL1 | HITS-CLIP | Tarbase |
| hsa-miR-141-5p | EMC7 | HITS-CLIP | Tarbase |
| hsa-miR-15a-5p | EMC7 | HITS-CLIP | Tarbase |
| hsa-miR-195-5p | EMC7 | HITS-CLIP | Tarbase |
| hsa-miR-15a-3p | PRPF38B | HITS-CLIP | Tarbase |
| hsa-miR-141-3p | VAV3 | HITS-CLIP | Tarbase |
| hsa-miR-195-5p | VAV3 | Microarrays | Tarbase |
| hsa-miR-195-5p | HMGCS2 | Microarrays | Tarbase |
| hsa-miR-195-5p | SORT1 | Microarrays | Tarbase |
| hsa-miR-15a-3p | LAMTOR5 | HITS-CLIP | Tarbase |
| hsa-miR-15a-5p | NOTCH2 | HITS-CLIP | Tarbase |
| hsa-miR-15a-5p | NOTCH2 | HITS-CLIP | Tarbase |
| hsa-miR-15a-5p | NOTCH2 | HITS-CLIP | Tarbase |
| hsa-miR-195-5p | NOTCH2 | HITS-CLIP | Tarbase |
| hsa-miR-15a-5p | VTCN1 | HITS-CLIP | Tarbase |
| hsa-miR-195-5p | VTCN1 | HITS-CLIP | Tarbase |
| hsa-miR-15a-5p | NAPG | PAR-CLIP | Tarbase |
| hsa-miR-15a-5p | NAPG | PAR-CLIP | Tarbase |
| hsa-miR-15a-5p | NAPG | PAR-CLIP | Tarbase |
| hsa-miR-15a-5p | NAPG | HITS-CLIP | Tarbase |
| hsa-miR-15a-5p | NAPG | PAR-CLIP | Tarbase |
| hsa-miR-15a-5p | NAPG | PAR-CLIP | Tarbase |
| hsa-miR-195-5p | NAPG | PAR-CLIP | Tarbase |
| hsa-miR-195-5p | NAPG | HITS-CLIP | Tarbase |
| hsa-miR-195-5p | NAPG | PAR-CLIP | Tarbase |
| hsa-miR-15a-5p | NAPG | HITS-CLIP | Tarbase |
| hsa-miR-195-5p | NAPG | HITS-CLIP | Tarbase |
| hsa-miR-15a-5p | NAPG | Chimeric fragments | Tarbase |
| hsa-miR-141-3p | PPHLN1 | HITS-CLIP | Tarbase |
| hsa-miR-141-5p | PPHLN1 | HITS-CLIP | Tarbase |
| hsa-miR-15a-5p | ARF3 | HITS-CLIP | Tarbase |
| hsa-miR-195-5p | ARF3 | HITS-CLIP | Tarbase |
| hsa-miR-195-5p | TMEM106C | Microarrays | Tarbase |
| hsa-miR-141-5p | SLC38A2 | HITS-CLIP | Tarbase |
| hsa-miR-141-5p | SLC38A2 | HITS-CLIP | Tarbase |
| hsa-miR-219a-1-3p | SLC38A2 | HITS-CLIP | Tarbase |
| hsa-miR-3613-3p | SLC38A2 | HITS-CLIP | Tarbase |
| hsa-miR-195-5p | SLC38A2 | Microarrays | Tarbase |
| hsa-miR-15a-5p | YWHAQ | PAR-CLIP | Tarbase |
| hsa-miR-141-3p | GRHL1 | HITS-CLIP | Tarbase |
| hsa-miR-141-3p | LDHA | HITS-CLIP | Tarbase |
| hsa-miR-141-3p | LDHA | HITS-CLIP | Tarbase |
| hsa-miR-141-3p | LDHA | HITS-CLIP | Tarbase |
| hsa-miR-3613-5p | LDHA | HITS-CLIP | Tarbase |
| hsa-miR-3613-5p | LDHA | HITS-CLIP | Tarbase |
| hsa-miR-486-5p | LDHA | HITS-CLIP | Tarbase |
| hsa-miR-141-3p | IL6ST | HITS-CLIP | Tarbase |
| hsa-miR-15a-5p | IL6ST | HITS-CLIP | Tarbase |
| hsa-miR-195-5p | IL6ST | HITS-CLIP | Tarbase |
| hsa-miR-141-3p | FST | HITS-CLIP | Tarbase |
| hsa-miR-195-5p | FST | Microarrays | Tarbase |
| hsa-miR-15a-5p | NAV1 | HITS-CLIP | Tarbase |
| hsa-miR-195-5p | NAV1 | HITS-CLIP | Tarbase |
| hsa-miR-486-5p | NAV1 | PAR-CLIP | Tarbase |
| hsa-miR-15a-5p | CDC73 | PAR-CLIP | Tarbase |
| hsa-miR-15a-5p | CDC73 | PAR-CLIP | Tarbase |
| hsa-miR-195-5p | CDC73 | PAR-CLIP | Tarbase |
| hsa-miR-141-5p | TIMM17A | HITS-CLIP | Tarbase |
| hsa-miR-195-5p | ERN2 | AGO-IP | Tarbase |
| hsa-miR-15a-5p | NARS | PAR-CLIP | Tarbase |
| hsa-miR-15a-5p | NARS | PAR-CLIP | Tarbase |
| hsa-miR-195-5p | NARS | HITS-CLIP | Tarbase |
| hsa-miR-15a-5p | KIAA1468 | HITS-CLIP | Tarbase |
| hsa-miR-195-5p | KIAA1468 | HITS-CLIP | Tarbase |
| hsa-miR-3613-3p | KIAA1468 | PAR-CLIP | Tarbase |
| hsa-miR-195-5p | CABLES1 | Microarrays | Tarbase |
| hsa-miR-486-5p | EMP1 | HITS-CLIP | Tarbase |
| hsa-miR-195-5p | EMP1 | Microarrays | Tarbase |
| hsa-miR-15a-5p | SOX5 | PAR-CLIP | Tarbase |
| hsa-miR-15a-5p | SOX5 | PAR-CLIP | Tarbase |
| hsa-miR-195-5p | SOX5 | PAR-CLIP | Tarbase |
| hsa-miR-195-5p | SOX5 | PAR-CLIP | Tarbase |
| hsa-miR-141-3p | ACP2 | HITS-CLIP | Tarbase |
| hsa-miR-15a-5p | ACP2 | PAR-CLIP | Tarbase |
| hsa-miR-15a-5p | ACP2 | PAR-CLIP | Tarbase |
| hsa-miR-195-5p | ACP2 | PAR-CLIP | Tarbase |
| hsa-miR-15a-5p | FAM127A | HITS-CLIP | Tarbase |
| hsa-miR-141-5p | PUM1 | HITS-CLIP | Tarbase |
| hsa-miR-141-5p | PUM1 | HITS-CLIP | Tarbase |
| hsa-miR-15a-5p | PUM1 | PAR-CLIP | Tarbase |
| hsa-miR-195-5p | PUM1 | PAR-CLIP | Tarbase |
| hsa-miR-219a-1-3p | PUM1 | HITS-CLIP | Tarbase |

| | | | |
|---|---|---|---|
| hsa-miR-195-5p | SPOCD1 | Microarrays | Tarbase |
| hsa-miR-141-5p | YARS | HITS-CLIP | Tarbase |
| hsa-miR-486-5p | CDCA8 | PAR-CLIP | Tarbase |
| hsa-miR-195-5p | CDCA8 | Microarrays | Tarbase |
| hsa-miR-15a-5p | AGO4 | PAR-CLIP | Tarbase |
| hsa-miR-15a-5p | AGO4 | PAR-CLIP | Tarbase |
| hsa-miR-15a-5p | AGO4 | PAR-CLIP | Tarbase |
| hsa-miR-15a-5p | AGO4 | PAR-CLIP | Tarbase |
| hsa-miR-195-5p | AGO4 | PAR-CLIP | Tarbase |
| hsa-miR-195-5p | AGO4 | PAR-CLIP | Tarbase |
| hsa-miR-195-5p | AGO4 | PAR-CLIP | Tarbase |
| hsa-miR-3613-3p | BTF3L4 | PAR-CLIP | Tarbase |
| hsa-miR-15a-5p | ZCCHC11 | HITS-CLIP | Tarbase |
| hsa-miR-15a-5p | ZCCHC11 | PAR-CLIP | Tarbase |
| hsa-miR-195-5p | ZCCHC11 | HITS-CLIP | Tarbase |
| hsa-miR-195-5p | ZCCHC11 | PAR-CLIP | Tarbase |
| hsa-miR-3605-3p | DSC2 | HITS-CLIP | Tarbase |
| hsa-miR-15a-3p | RNF138 | HITS-CLIP | Tarbase |
| hsa-miR-15a-5p | RNF138 | HITS-CLIP | Tarbase |
| hsa-miR-15a-5p | RNF138 | PAR-CLIP | Tarbase |
| hsa-miR-195-5p | RNF138 | HITS-CLIP | Tarbase |
| hsa-miR-15a-3p | ELP2 | HITS-CLIP | Tarbase |
| hsa-miR-15a-5p | DTNA | HITS-CLIP | Tarbase |
| hsa-miR-195-5p | DTNA | HITS-CLIP | Tarbase |
| hsa-miR-195-5p | FHOD3 | Microarrays | Tarbase |
| hsa-miR-15a-3p | SLC43A3 | HITS-CLIP | Tarbase |
| hsa-miR-15a-5p | FADS2 | HITS-CLIP | Tarbase |
| hsa-miR-195-5p | FADS2 | HITS-CLIP | Tarbase |
| hsa-miR-15a-3p | TMEM258 | HITS-CLIP | Tarbase |
| hsa-miR-15a-5p | TMEM258 | HITS-CLIP | Tarbase |
| hsa-miR-15a-5p | TMEM258 | HITS-CLIP | Tarbase |
| hsa-miR-195-5p | TMEM258 | HITS-CLIP | Tarbase |
| hsa-miR-195-5p | TMEM258 | HITS-CLIP | Tarbase |
| hsa-miR-195-5p | TMEM258 | Microarrays | Tarbase |
| hsa-miR-141-3p | CLOCK | Luciferase Reporter Assay | Tarbase |
| hsa-miR-141-3p | CLOCK | Luciferase Reporter Assay | Tarbase |
| hsa-miR-141-5p | CLOCK | HITS-CLIP | Tarbase |
| hsa-miR-3613-5p | CLOCK | HITS-CLIP | Tarbase |
| hsa-miR-195-5p | COL4A2 | Microarrays | Tarbase |
| hsa-miR-15a-5p | DZIP1 | HITS-CLIP | Tarbase |
| hsa-miR-195-5p | DZIP1 | HITS-CLIP | Tarbase |
| hsa-miR-15a-5p | ARGLU1 | PAR-CLIP | Tarbase |
| hsa-miR-486-5p | ARGLU1 | HITS-CLIP | Tarbase |
| hsa-miR-141-5p | ERCC5 | HITS-CLIP | Tarbase |
| hsa-miR-15a-5p | ARHGAP32 | PAR-CLIP | Tarbase |
| hsa-miR-15a-5p | ARHGAP32 | HITS-CLIP | Tarbase |
| hsa-miR-15a-5p | ARHGAP32 | HITS-CLIP | Tarbase |
| hsa-miR-195-5p | ARHGAP32 | HITS-CLIP | Tarbase |
| hsa-miR-15a-5p | ARHGAP32 | HITS-CLIP | Tarbase |
| hsa-miR-195-5p | ARHGAP32 | HITS-CLIP | Tarbase |
| hsa-miR-195-5p | ARHGAP32 | Microarrays | Tarbase |
| hsa-miR-15a-5p | STT3A | HITS-CLIP | Tarbase |
| hsa-miR-15a-5p | STT3A | HITS-CLIP | Tarbase |
| hsa-miR-195-5p | STT3A | HITS-CLIP | Tarbase |
| hsa-miR-219a-1-3p | ETS1 | PAR-CLIP | Tarbase |
| hsa-miR-195-5p | TMED7 | HITS-CLIP | Tarbase |
| hsa-miR-15a-5p | APC | HITS-CLIP | Tarbase |
| hsa-miR-195-5p | APC | HITS-CLIP | Tarbase |
| hsa-miR-15a-5p | RFK | HITS-CLIP | Tarbase |
| hsa-miR-15a-5p | RFK | PAR-CLIP | Tarbase |
| hsa-miR-15a-5p | RFK | PAR-CLIP | Tarbase |
| hsa-miR-15a-5p | RFK | HITS-CLIP | Tarbase |
| hsa-miR-195-5p | RFK | HITS-CLIP | Tarbase |
| hsa-miR-15a-5p | RFK | HITS-CLIP | Tarbase |
| hsa-miR-195-5p | RFK | HITS-CLIP | Tarbase |
| hsa-miR-15a-5p | RFK | Chimeric fragments | Tarbase |
| hsa-miR-15a-3p | CTSL | HITS-CLIP | Tarbase |
| hsa-miR-15a-5p | CTSL | PAR-CLIP | Tarbase |
| hsa-miR-15a-5p | CTSL | HITS-CLIP | Tarbase |
| hsa-miR-141-5p | TMEM2 | HITS-CLIP | Tarbase |
| hsa-miR-15a-5p | TMEM2 | HITS-CLIP | Tarbase |
| hsa-miR-15a-5p | TMEM2 | PAR-CLIP | Tarbase |
| hsa-miR-15a-5p | TMEM2 | HITS-CLIP | Tarbase |
| hsa-miR-15a-5p | TMEM2 | HITS-CLIP | Tarbase |
| hsa-miR-195-5p | TMEM2 | HITS-CLIP | Tarbase |
| hsa-miR-195-5p | TMEM2 | HITS-CLIP | Tarbase |
| hsa-miR-195-5p | TMEM2 | Microarrays | Tarbase |
| hsa-miR-486-5p | AGTPBP1 | HITS-CLIP | Tarbase |
| hsa-miR-15a-5p | PSAT1 | PAR-CLIP | Tarbase |
| hsa-miR-15a-5p | PSAT1 | HITS-CLIP | Tarbase |
| hsa-miR-15a-5p | PSAT1 | PAR-CLIP | Tarbase |
| hsa-miR-15a-5p | PSAT1 | HITS-CLIP | Tarbase |
| hsa-miR-15a-5p | PSAT1 | PAR-CLIP | Tarbase |
| hsa-miR-195-5p | PSAT1 | HITS-CLIP | Tarbase |
| hsa-miR-195-5p | PSAT1 | PAR-CLIP | Tarbase |
| hsa-miR-195-5p | PSAT1 | HITS-CLIP | Tarbase |
| hsa-miR-195-5p | ADAM19 | Microarrays | Tarbase |
| hsa-miR-15a-5p | FBXO21 | PAR-CLIP | Tarbase |
| hsa-miR-15a-3p | TBX3 | HITS-CLIP | Tarbase |
| hsa-miR-15a-3p | HRK | HITS-CLIP | Tarbase |
| hsa-miR-141-5p | DMTF1 | HITS-CLIP | Tarbase |
| hsa-miR-141-5p | DMTF1 | HITS-CLIP | Tarbase |
| hsa-miR-15a-5p | DMTF1 | Luciferase Reporter Assay | Tarbase |
| hsa-miR-15a-5p | DMTF1 | Luciferase Reporter Assay | Tarbase |
| hsa-miR-15a-5p | DMTF1 | PAR-CLIP | Tarbase |
| hsa-miR-15a-5p | DMTF1 | PAR-CLIP | Tarbase |
| hsa-miR-195-5p | DMTF1 | PAR-CLIP | Tarbase |
| hsa-miR-195-5p | TMEM243 | Microarrays | Tarbase |
| hsa-miR-141-3p | TMEM60 | HITS-CLIP | Tarbase |
| hsa-miR-195-5p | CD36 | Microarrays | Tarbase |
| hsa-miR-15a-5p | KHDC1 | PAR-CLIP | Tarbase |
| hsa-miR-3605-5p | SYNCRIP | HITS-CLIP | Tarbase |
| hsa-miR-195-5p | NT5E | Microarrays | Tarbase |
| hsa-miR-195-5p | EPHA7 | Microarrays | Tarbase |
| hsa-miR-15a-5p | MAP3K7 | PAR-CLIP | Tarbase |
| hsa-miR-195-5p | PRR5L | Microarrays | Tarbase |
| hsa-miR-141-3p | EHF | HITS-CLIP | Tarbase |
| hsa-miR-3605-5p | PRRG4 | PAR-CLIP | Tarbase |
| hsa-miR-15a-5p | CAPRIN1 | HITS-CLIP | Tarbase |
| hsa-miR-15a-5p | CAPRIN1 | HITS-CLIP | Tarbase |
| hsa-miR-195-5p | CAPRIN1 | HITS-CLIP | Tarbase |
| hsa-miR-15a-3p | CD63 | HITS-CLIP | Tarbase |
| hsa-miR-195-5p | CDK4 | Luciferase Reporter Assay | Tarbase |
| hsa-miR-195-5p | CDK4 | Western Blot | Tarbase |
| hsa-miR-195-5p | CDK4 | Western Blot | Tarbase |
| hsa-miR-195-5p | TROAP | Microarrays | Tarbase |
| hsa-miR-195-5p | B4GALNT1 | Microarrays | Tarbase |
| hsa-miR-15a-5p | TFCP2 | HITS-CLIP | Tarbase |
| hsa-miR-15a-5p | TFCP2 | HITS-CLIP | Tarbase |
| hsa-miR-15a-5p | TFCP2 | PAR-CLIP | Tarbase |
| hsa-miR-195-5p | TFCP2 | HITS-CLIP | Tarbase |
| hsa-miR-195-5p | TFCP2 | PAR-CLIP | Tarbase |
| hsa-miR-15a-3p | ESPL1 | HITS-CLIP | Tarbase |
| hsa-miR-195-5p | ESPL1 | Microarrays | Tarbase |
| hsa-miR-141-3p | HNRNPA1 | HITS-CLIP | Tarbase |
| hsa-miR-141-3p | ACVR1B | HITS-CLIP | Tarbase |
| hsa-miR-141-3p | ACVR1B | HITS-CLIP | Tarbase |
| hsa-miR-141-5p | ACVR1B | HITS-CLIP | Tarbase |
| hsa-miR-15a-5p | ACVR1B | PAR-CLIP | Tarbase |
| hsa-miR-195-5p | ACVR1B | Microarrays | Tarbase |
| hsa-miR-141-5p | LTV1 | HITS-CLIP | Tarbase |
| hsa-miR-141-3p | MAP7 | HITS-CLIP | Tarbase |
| hsa-miR-15a-5p | MAP7 | HITS-CLIP | Tarbase |
| hsa-miR-195-5p | MAP7 | HITS-CLIP | Tarbase |
| hsa-miR-141-5p | CD164 | HITS-CLIP | Tarbase |
| hsa-miR-141-5p | CD164 | HITS-CLIP | Tarbase |
| hsa-miR-15a-5p | CD164 | HITS-CLIP | Tarbase |
| hsa-miR-15a-5p | CD164 | HITS-CLIP | Tarbase |
| hsa-miR-15a-5p | CD164 | HITS-CLIP | Tarbase |
| hsa-miR-195-5p | CD164 | HITS-CLIP | Tarbase |
| hsa-miR-195-5p | CD164 | HITS-CLIP | Tarbase |
| hsa-miR-195-3p | AHI1 | HITS-CLIP | Tarbase |
| hsa-miR-15a-5p | REPS1 | HITS-CLIP | Tarbase |
| hsa-miR-195-5p | REPS1 | HITS-CLIP | Tarbase |
| hsa-miR-15a-3p | PRADC1 | HITS-CLIP | Tarbase |
| hsa-miR-15a-5p | RAB11FIP5 | PAR-CLIP | Tarbase |
| hsa-miR-15a-5p | SMYD5 | HITS-CLIP | Tarbase |
| hsa-miR-195-5p | SMYD5 | HITS-CLIP | Tarbase |
| hsa-miR-195-5p | SMYD5 | Microarrays | Tarbase |

| | | | |
|---|---|---|---|
| hsa-miR-15a-5p | USP15 | PAR-CLIP | Tarbase |
| hsa-miR-15a-5p | GNS | PAR-CLIP | Tarbase |
| hsa-miR-195-5p | CPM | Microarrays | Tarbase |
| hsa-miR-15a-3p | MDM2 | HITS-CLIP | Tarbase |
| hsa-miR-15a-3p | MDM2 | PAR-CLIP | Tarbase |
| hsa-miR-15a-5p | MDM2 | HITS-CLIP | Tarbase |
| hsa-miR-15a-5p | MDM2 | PAR-CLIP | Tarbase |
| hsa-miR-195-5p | MDM2 | HITS-CLIP | Tarbase |
| hsa-miR-3613-3p | MDM2 | PAR-CLIP | Tarbase |
| hsa-miR-3613-3p | MDM2 | Chimeric fragments | Tarbase |
| hsa-miR-486-5p | BCO1 | HITS-CLIP | Tarbase |
| hsa-miR-15a-5p | DYNC1LI2 | HITS-CLIP | Tarbase |
| hsa-miR-15a-5p | DYNC1LI2 | HITS-CLIP | Tarbase |
| hsa-miR-195-5p | DYNC1LI2 | HITS-CLIP | Tarbase |
| hsa-miR-15a-5p | AGT | HITS-CLIP | Tarbase |
| hsa-miR-195-5p | AGT | HITS-CLIP | Tarbase |
| hsa-miR-141-5p | URB2 | HITS-CLIP | Tarbase |
| hsa-miR-15a-5p | URB2 | HITS-CLIP | Tarbase |
| hsa-miR-15a-5p | EGLN1 | HITS-CLIP | Tarbase |
| hsa-miR-195-5p | EGLN1 | HITS-CLIP | Tarbase |
| hsa-miR-15a-5p | TAF5L | HITS-CLIP | Tarbase |
| hsa-miR-195-5p | TAF5L | Microarrays | Tarbase |
| hsa-miR-141-5p | DHX9 | HITS-CLIP | Tarbase |
| hsa-miR-15a-5p | DHX9 | PAR-CLIP | Tarbase |
| hsa-miR-15a-5p | DHX9 | HITS-CLIP | Tarbase |
| hsa-miR-141-5p | CEP350 | HITS-CLIP | Tarbase |
| hsa-miR-141-5p | CEP350 | HITS-CLIP | Tarbase |
| hsa-miR-15a-5p | CEP350 | PAR-CLIP | Tarbase |
| hsa-miR-195-5p | FAM129A | Microarrays | Tarbase |
| hsa-miR-141-5p | MRPL44 | HITS-CLIP | Tarbase |
| hsa-miR-15a-5p | MRPL44 | PAR-CLIP | Tarbase |
| hsa-miR-141-3p | ITM2C | HITS-CLIP | Tarbase |
| hsa-miR-15a-5p | ITM2C | PAR-CLIP | Tarbase |
| hsa-miR-195-5p | SERPINE2 | Microarrays | Tarbase |
| hsa-miR-141-5p | CAB39 | HITS-CLIP | Tarbase |
| hsa-miR-15a-5p | CAB39 | HITS-CLIP | Tarbase |
| hsa-miR-15a-5p | CAB39 | HITS-CLIP | Tarbase |
| hsa-miR-195-5p | CAB39 | HITS-CLIP | Tarbase |
| hsa-miR-195-5p | CAB39 | HITS-CLIP | Tarbase |
| hsa-miR-3613-3p | CAB39 | PAR-CLIP | Tarbase |
| hsa-miR-15a-5p | CAB39 | HITS-CLIP | Tarbase |
| hsa-miR-195-5p | CAB39 | HITS-CLIP | Tarbase |
| hsa-miR-486-3p | COX5B | HITS-CLIP | Tarbase |
| hsa-miR-141-5p | TSGA10 | HITS-CLIP | Tarbase |
| hsa-miR-3613-3p | MFSD9 | PAR-CLIP | Tarbase |
| hsa-miR-15a-5p | TMEM127 | HITS-CLIP | Tarbase |
| hsa-miR-15a-5p | TMEM127 | HITS-CLIP | Tarbase |
| hsa-miR-195-5p | TMEM127 | HITS-CLIP | Tarbase |
| hsa-miR-486-3p | TMEM127 | HITS-CLIP | Tarbase |
| hsa-miR-3605-3p | MRPS9 | HITS-CLIP | Tarbase |
| hsa-miR-15a-5p | ANKRD36 | HITS-CLIP | Tarbase |
| hsa-miR-195-5p | ANKRD36 | HITS-CLIP | Tarbase |
| hsa-miR-141-5p | EPC2 | HITS-CLIP | Tarbase |
| hsa-miR-3613-5p | EPC2 | PAR-CLIP | Tarbase |
| hsa-miR-15a-5p | USP44 | PAR-CLIP | Tarbase |
| hsa-miR-195-3p | SCYL2 | HITS-CLIP | Tarbase |
| hsa-miR-195-5p | CKAP4 | HITS-CLIP | Tarbase |
| hsa-miR-3613-3p | PLXNC1 | HITS-CLIP | Tarbase |
| hsa-miR-3613-3p | APPL2 | HITS-CLIP | Tarbase |
| hsa-miR-195-5p | DRAM1 | Microarrays | Tarbase |
| hsa-miR-141-5p | KIAA1033 | HITS-CLIP | Tarbase |
| hsa-miR-195-3p | KIAA1033 | HITS-CLIP | Tarbase |
| hsa-miR-15a-5p | SLC41A2 | PAR-CLIP | Tarbase |
| hsa-miR-195-5p | SLC41A2 | PAR-CLIP | Tarbase |
| hsa-miR-195-5p | SLC41A2 | Microarrays | Tarbase |
| hsa-miR-141-5p | FLNB | HITS-CLIP | Tarbase |
| hsa-miR-141-5p | FLNB | HITS-CLIP | Tarbase |
| hsa-miR-15a-5p | FLNB | HITS-CLIP | Tarbase |
| hsa-miR-15a-5p | VPS36 | HITS-CLIP | Tarbase |
| hsa-miR-195-5p | VPS36 | HITS-CLIP | Tarbase |
| hsa-miR-15a-3p | CKAP2 | HITS-CLIP | Tarbase |
| hsa-miR-141-3p | TBC1D4 | HITS-CLIP | Tarbase |
| hsa-miR-195-5p | TBC1D4 | Microarrays | Tarbase |
| hsa-miR-195-5p | BORA | Microarrays | Tarbase |
| hsa-miR-3613-5p | LRCH1 | Chimeric fragments | Tarbase |
| hsa-miR-15a-5p | SUCLA2 | HITS-CLIP | Tarbase |
| hsa-miR-195-5p | SUCLA2 | HITS-CLIP | Tarbase |
| hsa-miR-3613-5p | RCBTB1 | HITS-CLIP | Tarbase |
| hsa-miR-141-3p | COG3 | HITS-CLIP | Tarbase |
| hsa-miR-15a-5p | COG3 | HITS-CLIP | Tarbase |
| hsa-miR-195-5p | COG3 | HITS-CLIP | Tarbase |
| hsa-miR-15a-5p | LMO7 | HITS-CLIP | Tarbase |
| hsa-miR-15a-5p | LMO7 | HITS-CLIP | Tarbase |
| hsa-miR-195-5p | LMO7 | HITS-CLIP | Tarbase |
| hsa-miR-3613-3p | EDNRB | HITS-CLIP | Tarbase |
| hsa-miR-3613-3p | LCP1 | PAR-CLIP | Tarbase |
| hsa-miR-141-3p | SCRN1 | HITS-CLIP | Tarbase |
| hsa-miR-15a-5p | SCRN1 | HITS-CLIP | Tarbase |
| hsa-miR-15a-5p | SCRN1 | HITS-CLIP | Tarbase |
| hsa-miR-195-5p | SCRN1 | HITS-CLIP | Tarbase |
| hsa-miR-15a-5p | SCRN1 | HITS-CLIP | Tarbase |
| hsa-miR-195-5p | SCRN1 | HITS-CLIP | Tarbase |
| hsa-miR-141-3p | TNS3 | HITS-CLIP | Tarbase |
| hsa-miR-15a-5p | TNS3 | PAR-CLIP | Tarbase |
| hsa-miR-195-5p | TNS3 | Microarrays | Tarbase |
| hsa-miR-486-3p | IGF2BP3 | HITS-CLIP | Tarbase |
| hsa-miR-195-5p | GPNMB | Microarrays | Tarbase |
| hsa-miR-141-3p | RAPGEF5 | HITS-CLIP | Tarbase |
| hsa-miR-141-3p | RAC1 | HITS-CLIP | Tarbase |
| hsa-miR-15a-5p | RAC1 | PAR-CLIP | Tarbase |
| hsa-miR-3613-5p | RAC1 | PAR-CLIP | Tarbase |
| hsa-miR-141-3p | KDELR2 | PAR-CLIP | Tarbase |
| hsa-miR-141-5p | KDELR2 | HITS-CLIP | Tarbase |
| hsa-miR-141-5p | KDELR2 | HITS-CLIP | Tarbase |
| hsa-miR-15a-5p | KDELR2 | HITS-CLIP | Tarbase |
| hsa-miR-195-5p | KDELR2 | HITS-CLIP | Tarbase |
| hsa-miR-195-5p | IL6 | Microarrays | Tarbase |
| hsa-miR-15a-5p | ZDHHC4 | PAR-CLIP | Tarbase |
| hsa-miR-195-5p | AOAH | Microarrays | Tarbase |
| hsa-miR-15a-5p | BZW2 | HITS-CLIP | Tarbase |
| hsa-miR-15a-5p | TBRG4 | PAR-CLIP | Tarbase |
| hsa-miR-15a-5p | MYO1G | PAR-CLIP | Tarbase |
| hsa-miR-195-5p | MYO1G | PAR-CLIP | Tarbase |
| hsa-miR-15a-3p | TTYH3 | HITS-CLIP | Tarbase |
| hsa-miR-15a-5p | TTYH3 | HITS-CLIP | Tarbase |
| hsa-miR-15a-5p | TTYH3 | HITS-CLIP | Tarbase |
| hsa-miR-195-5p | TTYH3 | HITS-CLIP | Tarbase |
| hsa-miR-195-5p | TTYH3 | HITS-CLIP | Tarbase |
| hsa-miR-15a-5p | CIDEB | PAR-CLIP | Tarbase |
| hsa-miR-195-5p | CIDEB | PAR-CLIP | Tarbase |
| hsa-miR-3613-5p | ZFHX2 | HITS-CLIP | Tarbase |
| hsa-miR-15a-5p | ABHD17C | HITS-CLIP | Tarbase |
| hsa-miR-141-3p | IREB2 | HITS-CLIP | Tarbase |
| hsa-miR-141-5p | IREB2 | HITS-CLIP | Tarbase |
| hsa-miR-15a-5p | IREB2 | HITS-CLIP | Tarbase |
| hsa-miR-195-5p | IREB2 | HITS-CLIP | Tarbase |
| hsa-miR-195-5p | CIB2 | Microarrays | Tarbase |
| hsa-miR-3613-5p | NMT1 | PAR-CLIP | Tarbase |
| hsa-miR-141-3p | SRSF1 | HITS-CLIP | Tarbase |
| hsa-miR-141-5p | SRSF1 | HITS-CLIP | Tarbase |
| hsa-miR-15a-5p | SRSF1 | Chimeric fragments | Tarbase |
| hsa-miR-15a-5p | VEZF1 | HITS-CLIP | Tarbase |
| hsa-miR-3605-5p | VEZF1 | HITS-CLIP | Tarbase |
| hsa-miR-15a-5p | TEX2 | HITS-CLIP | Tarbase |
| hsa-miR-141-3p | DCAF7 | HITS-CLIP | Tarbase |
| hsa-miR-141-5p | DCAF7 | HITS-CLIP | Tarbase |
| hsa-miR-15a-5p | DCAF7 | HITS-CLIP | Tarbase |
| hsa-miR-15a-5p | DCAF7 | HITS-CLIP | Tarbase |
| hsa-miR-195-5p | DCAF7 | HITS-CLIP | Tarbase |
| hsa-miR-3613-5p | DCAF7 | HITS-CLIP | Tarbase |
| hsa-miR-486-3p | DCAF7 | HITS-CLIP | Tarbase |
| hsa-miR-15a-5p | BRIP1 | HITS-CLIP | Tarbase |
| hsa-miR-219a-1-3p | NDUFB5 | HITS-CLIP | Tarbase |
| hsa-miR-141-3p | TRA2B | HITS-CLIP | Tarbase |
| hsa-miR-141-3p | MARCH7 | HITS-CLIP | Tarbase |
| hsa-miR-141-3p | MARCH7 | HITS-CLIP | Tarbase |
| hsa-miR-15a-5p | MARCH7 | HITS-CLIP | Tarbase |
| hsa-miR-15a-5p | MARCH7 | HITS-CLIP | Tarbase |
| hsa-miR-195-5p | MARCH7 | HITS-CLIP | Tarbase |

| | | | |
|---|---|---|---|
| hsa-miR-141-5p | TANK | HITS-CLIP | Tarbase |
| hsa-miR-15a-5p | SKIL | HITS-CLIP | Tarbase |
| hsa-miR-15a-5p | SKIL | HITS-CLIP | Tarbase |
| hsa-miR-195-5p | SKIL | HITS-CLIP | Tarbase |
| hsa-miR-15a-5p | SKIL | HITS-CLIP | Tarbase |
| hsa-miR-195-5p | SKIL | HITS-CLIP | Tarbase |
| hsa-miR-141-3p | EPRS | HITS-CLIP | Tarbase |
| hsa-miR-195-5p | SMPD4 | Microarrays | Tarbase |
| hsa-miR-3613-3p | WDR33 | HITS-CLIP | Tarbase |
| hsa-miR-195-5p | BIN1 | Microarrays | Tarbase |
| hsa-miR-15a-5p | STAM | HITS-CLIP | Tarbase |
| hsa-miR-141-5p | DNAJC1 | HITS-CLIP | Tarbase |
| hsa-miR-486-3p | LRRC8A | HITS-CLIP | Tarbase |
| hsa-miR-15a-5p | KIAA0368 | HITS-CLIP | Tarbase |
| hsa-miR-195-5p | KIAA0368 | HITS-CLIP | Tarbase |
| hsa-miR-3613-3p | KIAA0368 | PAR-CLIP | Tarbase |
| hsa-miR-15a-3p | SMC2 | HITS-CLIP | Tarbase |
| hsa-miR-219a-1-3p | SMC2 | HITS-CLIP | Tarbase |
| hsa-miR-486-5p | SMC2 | PAR-CLIP | Tarbase |
| hsa-miR-195-5p | SMC2 | Microarrays | Tarbase |
| hsa-miR-486-5p | RALGPS1 | HITS-CLIP | Tarbase |
| hsa-miR-195-5p | CDK5RAP2 | Microarrays | Tarbase |
| hsa-miR-15a-5p | SLC31A1 | HITS-CLIP | Tarbase |
| hsa-miR-195-5p | SLC31A1 | Microarrays | Tarbase |
| hsa-miR-15a-5p | STX17 | HITS-CLIP | Tarbase |
| hsa-miR-141-5p | PRPF4 | HITS-CLIP | Tarbase |
| hsa-miR-141-5p | PRPF4 | HITS-CLIP | Tarbase |
| hsa-miR-15a-5p | ATP6V1G1 | HITS-CLIP | Tarbase |
| hsa-miR-195-5p | ATP6V1G1 | HITS-CLIP | Tarbase |
| hsa-miR-15a-5p | TEX10 | Chimeric fragments | Tarbase |
| hsa-miR-15a-5p | GABBR2 | HITS-CLIP | Tarbase |
| hsa-miR-195-5p | GABBR2 | HITS-CLIP | Tarbase |
| hsa-miR-15a-5p | GOLGA1 | PAR-CLIP | Tarbase |
| hsa-miR-15a-5p | GOLGA1 | HITS-CLIP | Tarbase |
| hsa-miR-195-5p | GOLGA1 | PAR-CLIP | Tarbase |
| hsa-miR-195-5p | GOLGA1 | HITS-CLIP | Tarbase |
| hsa-miR-486-5p | NCBP1 | HITS-CLIP | Tarbase |
| hsa-miR-141-5p | ANP32B | HITS-CLIP | Tarbase |
| hsa-miR-486-3p | ANP32B | HITS-CLIP | Tarbase |
| hsa-miR-486-3p | ANP32B | HITS-CLIP | Tarbase |
| hsa-miR-195-5p | ANP32B | Microarrays | Tarbase |
| hsa-miR-195-5p | CTSV | Microarrays | Tarbase |
| hsa-miR-15a-5p | ENPP2 | HITS-CLIP | Tarbase |
| hsa-miR-195-5p | ENPP2 | HITS-CLIP | Tarbase |
| hsa-miR-195-5p | DSCC1 | Microarrays | Tarbase |
| hsa-miR-15a-3p | MYC | HITS-CLIP | Tarbase |
| hsa-miR-195-5p | MYC | qPCR | Tarbase |
| hsa-miR-195-5p | MYC | Western Blot | Tarbase |
| hsa-miR-3605-3p | MYC | HITS-CLIP | Tarbase |
| hsa-miR-141-3p | RANBP6 | HITS-CLIP | Tarbase |
| hsa-miR-141-3p | RANBP6 | HITS-CLIP | Tarbase |
| hsa-miR-141-3p | RANBP6 | HITS-CLIP | Tarbase |
| hsa-miR-15a-5p | RANBP6 | HITS-CLIP | Tarbase |
| hsa-miR-195-5p | RANBP6 | HITS-CLIP | Tarbase |
| hsa-miR-15a-5p | PLAA | HITS-CLIP | Tarbase |
| hsa-miR-15a-5p | PLAA | HITS-CLIP | Tarbase |
| hsa-miR-195-5p | PLAA | HITS-CLIP | Tarbase |
| hsa-miR-195-5p | PLAA | HITS-CLIP | Tarbase |
| hsa-miR-15a-5p | UBAP2 | HITS-CLIP | Tarbase |
| hsa-miR-195-5p | UBAP2 | HITS-CLIP | Tarbase |
| hsa-miR-141-3p | RNF38 | PAR-CLIP | Tarbase |
| hsa-miR-141-3p | RNF38 | HITS-CLIP | Tarbase |
| hsa-miR-141-5p | RNF38 | HITS-CLIP | Tarbase |
| hsa-miR-15a-5p | RNF38 | HITS-CLIP | Tarbase |
| hsa-miR-141-5p | TLN1 | HITS-CLIP | Tarbase |
| hsa-miR-15a-5p | TLN1 | HITS-CLIP | Tarbase |
| hsa-miR-195-5p | TLN1 | HITS-CLIP | Tarbase |
| hsa-miR-195-5p | TLN1 | Microarrays | Tarbase |
| hsa-miR-195-5p | DNAJB5 | Microarrays | Tarbase |
| hsa-miR-141-3p | DCTN3 | HITS-CLIP | Tarbase |
| hsa-miR-141-3p | DCTN3 | HITS-CLIP | Tarbase |
| hsa-miR-195-5p | ALDH1B1 | Microarrays | Tarbase |
| hsa-miR-195-5p | ARHGEF39 | Microarrays | Tarbase |
| hsa-miR-141-3p | DENND4C | PAR-CLIP | Tarbase |
| hsa-miR-15a-5p | PPIL1 | PAR-CLIP | Tarbase |
| hsa-miR-219a-1-3p | PIM1 | PAR-CLIP | Tarbase |
| hsa-miR-195-5p | GMPR | Microarrays | Tarbase |
| hsa-miR-3605-5p | CMTR1 | HITS-CLIP | Tarbase |
| hsa-miR-3613-5p | CMTR1 | HITS-CLIP | Tarbase |
| hsa-miR-15a-5p | TFAP2A | PAR-CLIP | Tarbase |
| hsa-miR-15a-5p | TFAP2A | HITS-CLIP | Tarbase |
| hsa-miR-15a-5p | TFAP2A | HITS-CLIP | Tarbase |
| hsa-miR-15a-5p | YIPF3 | HITS-CLIP | Tarbase |
| hsa-miR-15a-5p | YIPF3 | HITS-CLIP | Tarbase |
| hsa-miR-195-5p | YIPF3 | HITS-CLIP | Tarbase |
| hsa-miR-15a-5p | TMEM63B | HITS-CLIP | Tarbase |
| hsa-miR-486-3p | TJAP1 | PAR-CLIP | Tarbase |
| hsa-miR-15a-5p | LRRC1 | HITS-CLIP | Tarbase |
| hsa-miR-15a-5p | LRRC1 | HITS-CLIP | Tarbase |
| hsa-miR-195-5p | LRRC1 | HITS-CLIP | Tarbase |
| hsa-miR-195-5p | FOXF2 | Microarrays | Tarbase |
| hsa-miR-15a-5p | HMGA1 | HITS-CLIP | Tarbase |
| hsa-miR-15a-5p | HMGA1 | HITS-CLIP | Tarbase |
| hsa-miR-15a-5p | HMGA1 | PAR-CLIP | Tarbase |
| hsa-miR-15a-5p | HMGA1 | HITS-CLIP | Tarbase |
| hsa-miR-15a-5p | HMGA1 | HITS-CLIP | Tarbase |
| hsa-miR-195-5p | HMGA1 | HITS-CLIP | Tarbase |
| hsa-miR-195-5p | HMGA1 | HITS-CLIP | Tarbase |
| hsa-miR-195-5p | HMGA1 | HITS-CLIP | Tarbase |
| hsa-miR-15a-5p | HMGA1 | Luciferase Reporter Assay | Tarbase |
| hsa-miR-195-5p | TCF19 | Microarrays | Tarbase |
| hsa-miR-195-5p | IER3 | Microarrays | Tarbase |
| hsa-miR-195-5p | MDC1 | Microarrays | Tarbase |
| hsa-miR-15a-3p | PGBD1 | HITS-CLIP | Tarbase |
| hsa-miR-141-3p | RNF144B | HITS-CLIP | Tarbase |
| hsa-miR-15a-5p | RNF144B | PAR-CLIP | Tarbase |
| hsa-miR-15a-5p | RNF144B | PAR-CLIP | Tarbase |
| hsa-miR-15a-5p | RNF144B | HITS-CLIP | Tarbase |
| hsa-miR-195-5p | RNF144B | Microarrays | Tarbase |
| hsa-miR-195-5p | NRM | Microarrays | Tarbase |
| hsa-miR-15a-3p | TAF8 | PAR-CLIP | Tarbase |
| hsa-miR-3613-5p | FAM8A1 | PAR-CLIP | Tarbase |
| hsa-miR-15a-5p | CPEB2 | HITS-CLIP | Tarbase |
| hsa-miR-15a-5p | CPEB2 | HITS-CLIP | Tarbase |
| hsa-miR-195-5p | CPEB2 | HITS-CLIP | Tarbase |
| hsa-miR-195-5p | CPEB2 | HITS-CLIP | Tarbase |
| hsa-miR-3613-3p | CPEB2 | HITS-CLIP | Tarbase |
| hsa-miR-3613-3p | CPEB2 | PAR-CLIP | Tarbase |
| hsa-miR-15a-5p | CPEB2 | HITS-CLIP | Tarbase |
| hsa-miR-195-5p | CPEB2 | HITS-CLIP | Tarbase |
| hsa-miR-15a-5p | CPEB2 | Chimeric fragments | Tarbase |
| hsa-miR-15a-5p | FHDC1 | HITS-CLIP | Tarbase |
| hsa-miR-15a-5p | FHDC1 | HITS-CLIP | Tarbase |
| hsa-miR-195-5p | FHDC1 | HITS-CLIP | Tarbase |
| hsa-miR-195-5p | SLCO2B1 | Microarrays | Tarbase |
| hsa-miR-195-5p | ANKRD42 | Microarrays | Tarbase |
| hsa-miR-15a-3p | IL18BP | HITS-CLIP | Tarbase |
| hsa-miR-195-5p | SYTL2 | Microarrays | Tarbase |
| hsa-miR-15a-5p | CREBZF | PAR-CLIP | Tarbase |
| hsa-miR-15a-5p | CREBZF | HITS-CLIP | Tarbase |
| hsa-miR-15a-5p | PRCP | PAR-CLIP | Tarbase |
| hsa-miR-195-5p | SULF1 | Microarrays | Tarbase |
| hsa-miR-15a-5p | SDCBP | HITS-CLIP | Tarbase |
| hsa-miR-195-5p | SDCBP | HITS-CLIP | Tarbase |
| hsa-miR-15a-5p | YAP1 | HITS-CLIP | Tarbase |
| hsa-miR-15a-5p | YAP1 | HITS-CLIP | Tarbase |
| hsa-miR-15a-5p | YAP1 | PAR-CLIP | Tarbase |
| hsa-miR-15a-5p | YAP1 | HITS-CLIP | Tarbase |
| hsa-miR-195-5p | YAP1 | HITS-CLIP | Tarbase |
| hsa-miR-195-5p | YAP1 | HITS-CLIP | Tarbase |
| hsa-miR-141-5p | PPP2R1B | HITS-CLIP | Tarbase |
| hsa-miR-15a-5p | PPP2R1B | HITS-CLIP | Tarbase |
| hsa-miR-15a-5p | PPP2R1B | HITS-CLIP | Tarbase |
| hsa-miR-15a-5p | PPP2R1B | HITS-CLIP | Tarbase |
| hsa-miR-15a-5p | PPP2R1B | HITS-CLIP | Tarbase |
| hsa-miR-195-5p | PPP2R1B | HITS-CLIP | Tarbase |
| hsa-miR-195-5p | PPP2R1B | HITS-CLIP | Tarbase |
| hsa-miR-195-5p | PPP2R1B | HITS-CLIP | Tarbase |
| hsa-miR-15a-5p | UNC13C | HITS-CLIP | Tarbase |

| | | | |
|---|---|---|---|
| hsa-miR-195-5p | UNC13C | HITS-CLIP | Tarbase |
| hsa-miR-141-5p | CTDSPL2 | HITS-CLIP | Tarbase |
| hsa-miR-141-3p | THBS1 | HITS-CLIP | Tarbase |
| hsa-miR-141-3p | THBS1 | HITS-CLIP | Tarbase |
| hsa-miR-195-5p | THBS1 | Microarrays | Tarbase |
| hsa-miR-219a-1-3p | NUSAP1 | PAR-CLIP | Tarbase |
| hsa-miR-219a-1-3p | NUSAP1 | PAR-CLIP | Tarbase |
| hsa-miR-195-5p | NUSAP1 | Microarrays | Tarbase |
| hsa-miR-141-5p | KIF23 | HITS-CLIP | Tarbase |
| hsa-miR-15a-5p | KIF23 | PAR-CLIP | Tarbase |
| hsa-miR-15a-5p | KIF23 | PAR-CLIP | Tarbase |
| hsa-miR-195-5p | KIF23 | PAR-CLIP | Tarbase |
| hsa-miR-195-5p | KIF23 | Microarrays | Tarbase |
| hsa-miR-15a-5p | KIF23 | Chimeric fragments | Tarbase |
| hsa-miR-195-5p | CASC5 | Microarrays | Tarbase |
| hsa-miR-15a-5p | HAUS2 | HITS-CLIP | Tarbase |
| hsa-miR-15a-3p | RPLP1 | HITS-CLIP | Tarbase |
| hsa-miR-15a-3p | RPLP1 | HITS-CLIP | Tarbase |
| hsa-miR-15a-3p | RPLP1 | HITS-CLIP | Tarbase |
| hsa-miR-15a-3p | RPLP1 | HITS-CLIP | Tarbase |
| hsa-miR-15a-3p | RPLP1 | HITS-CLIP | Tarbase |
| hsa-miR-15a-3p | RPLP1 | HITS-CLIP | Tarbase |
| hsa-miR-15a-3p | RPLP1 | HITS-CLIP | Tarbase |
| hsa-miR-15a-3p | RPLP1 | HITS-CLIP | Tarbase |
| hsa-miR-15a-5p | RPLP1 | HITS-CLIP | Tarbase |
| hsa-miR-486-3p | RPLP1 | HITS-CLIP | Tarbase |
| hsa-miR-195-5p | PAQR5 | Microarrays | Tarbase |
| hsa-miR-195-5p | TUBGCP4 | Microarrays | Tarbase |
| hsa-miR-141-5p | ADAM10 | HITS-CLIP | Tarbase |
| hsa-miR-15a-5p | ADAM10 | PAR-CLIP | Tarbase |
| hsa-miR-15a-5p | ADAM10 | PAR-CLIP | Tarbase |
| hsa-miR-15a-5p | ADAM10 | PAR-CLIP | Tarbase |
| hsa-miR-15a-5p | ADAM10 | HITS-CLIP | Tarbase |
| hsa-miR-195-5p | ADAM10 | PAR-CLIP | Tarbase |
| hsa-miR-195-5p | ADAM10 | HITS-CLIP | Tarbase |
| hsa-miR-195-5p | ADAM10 | PAR-CLIP | Tarbase |
| hsa-miR-195-5p | ADAM10 | PAR-CLIP | Tarbase |
| hsa-miR-195-5p | ADAM10 | PAR-CLIP | Tarbase |
| hsa-miR-195-5p | ADAM10 | PAR-CLIP | Tarbase |
| hsa-miR-195-5p | ADAM10 | PAR-CLIP | Tarbase |
| hsa-miR-3605-5p | ADAM10 | HITS-CLIP | Tarbase |
| hsa-miR-3613-5p | ZNF280D | PAR-CLIP | Tarbase |
| hsa-miR-195-5p | SEMA6D | Microarrays | Tarbase |
| hsa-miR-195-5p | BCAR3 | Microarrays | Tarbase |
| hsa-miR-141-3p | FNBP1L | HITS-CLIP | Tarbase |
| hsa-miR-3613-3p | IFI44L | PAR-CLIP | Tarbase |
| hsa-miR-3613-5p | SLC44A5 | HITS-CLIP | Tarbase |
| hsa-miR-15a-5p | RTCA | Chimeric fragments | Tarbase |
| hsa-miR-141-3p | EPT1 | HITS-CLIP | Tarbase |
| hsa-miR-141-3p | HADHB | HITS-CLIP | Tarbase |
| hsa-miR-15a-5p | PPM1B | HITS-CLIP | Tarbase |
| hsa-miR-195-5p | PPM1B | HITS-CLIP | Tarbase |
| hsa-miR-141-3p | CYP1B1 | PAR-CLIP | Tarbase |
| hsa-miR-141-5p | RAB1A | HITS-CLIP | Tarbase |
| hsa-miR-141-5p | RAB1A | HITS-CLIP | Tarbase |
| hsa-miR-15a-5p | RAB1A | PAR-CLIP | Tarbase |
| hsa-miR-3613-5p | RAB1A | HITS-CLIP | Tarbase |
| hsa-miR-15a-5p | ACTR2 | PAR-CLIP | Tarbase |
| hsa-miR-15a-5p | ACTR2 | HITS-CLIP | Tarbase |
| hsa-miR-195-5p | ACTR2 | HITS-CLIP | Tarbase |
| hsa-miR-219a-1-3p | ACTR2 | HITS-CLIP | Tarbase |
| hsa-miR-15a-5p | ACTR2 | HITS-CLIP | Tarbase |
| hsa-miR-195-5p | ACTR2 | HITS-CLIP | Tarbase |
| hsa-miR-15a-5p | SLC5A6 | HITS-CLIP | Tarbase |
| hsa-miR-141-3p | PREPL | HITS-CLIP | Tarbase |
| hsa-miR-141-3p | PREPL | HITS-CLIP | Tarbase |
| hsa-miR-141-3p | PREPL | HITS-CLIP | Tarbase |
| hsa-miR-141-5p | PREPL | HITS-CLIP | Tarbase |
| hsa-miR-141-5p | FBXO11 | HITS-CLIP | Tarbase |
| hsa-miR-3613-3p | FBXO11 | PAR-CLIP | Tarbase |
| hsa-miR-195-5p | CENPO | Microarrays | Tarbase |
| hsa-miR-15a-5p | LRPPRC | HITS-CLIP | Tarbase |
| hsa-miR-15a-5p | LRPPRC | HITS-CLIP | Tarbase |
| hsa-miR-195-5p | LRPPRC | HITS-CLIP | Tarbase |
| hsa-miR-15a-5p | ACTR1A | HITS-CLIP | Tarbase |
| hsa-miR-195-5p | TMEM180 | Microarrays | Tarbase |
| hsa-miR-195-5p | MYOF | Microarrays | Tarbase |
| hsa-miR-141-3p | DUSP5 | HITS-CLIP | Tarbase |
| hsa-miR-486-3p | DUSP5 | HITS-CLIP | Tarbase |
| hsa-miR-195-5p | DUSP5 | Microarrays | Tarbase |
| hsa-miR-15a-5p | CEP55 | PAR-CLIP | Tarbase |
| hsa-miR-195-5p | CEP55 | Microarrays | Tarbase |
| hsa-miR-15a-5p | CEP55 | Chimeric fragments | Tarbase |
| hsa-miR-195-5p | KIF20B | Microarrays | Tarbase |
| hsa-miR-141-5p | DNAJC13 | HITS-CLIP | Tarbase |
| hsa-miR-141-5p | ANXA7 | HITS-CLIP | Tarbase |
| hsa-miR-195-5p | ADAMTS14 | Microarrays | Tarbase |
| hsa-miR-195-5p | DNA2 | Microarrays | Tarbase |
| hsa-miR-15a-5p | BARD1 | HITS-CLIP | Tarbase |
| hsa-miR-195-5p | BARD1 | HITS-CLIP | Tarbase |
| hsa-miR-195-5p | BARD1 | Microarrays | Tarbase |
| hsa-miR-15a-5p | ASNSD1 | HITS-CLIP | Tarbase |
| hsa-miR-15a-5p | ASNSD1 | HITS-CLIP | Tarbase |
| hsa-miR-195-5p | ASNSD1 | HITS-CLIP | Tarbase |
| hsa-miR-15a-5p | ASNSD1 | Chimeric fragments | Tarbase |
| hsa-miR-15a-5p | NAB1 | HITS-CLIP | Tarbase |
| hsa-miR-195-5p | NAB1 | HITS-CLIP | Tarbase |
| hsa-miR-15a-5p | PPIG | HITS-CLIP | Tarbase |
| hsa-miR-195-5p | PPIG | HITS-CLIP | Tarbase |
| hsa-miR-15a-5p | PPIG | HITS-CLIP | Tarbase |
| hsa-miR-195-5p | PPIG | HITS-CLIP | Tarbase |
| hsa-miR-141-5p | IDH1 | HITS-CLIP | Tarbase |
| hsa-miR-219a-1-3p | SSFA2 | PAR-CLIP | Tarbase |
| hsa-miR-195-5p | CHRNA1 | Microarrays | Tarbase |
| hsa-miR-15a-5p | ABI2 | HITS-CLIP | Tarbase |
| hsa-miR-15a-5p | ABI2 | HITS-CLIP | Tarbase |
| hsa-miR-195-5p | ABI2 | HITS-CLIP | Tarbase |
| hsa-miR-195-5p | ABI2 | HITS-CLIP | Tarbase |
| hsa-miR-3605-3p | ABI2 | HITS-CLIP | Tarbase |
| hsa-miR-15a-5p | ABI2 | HITS-CLIP | Tarbase |
| hsa-miR-195-5p | ABI2 | HITS-CLIP | Tarbase |
| hsa-miR-195-5p | ABI2 | Microarrays | Tarbase |
| hsa-miR-141-5p | ITGAV | HITS-CLIP | Tarbase |
| hsa-miR-141-5p | ITGAV | HITS-CLIP | Tarbase |
| hsa-miR-141-5p | SECISBP2L | HITS-CLIP | Tarbase |
| hsa-miR-15a-5p | SECISBP2L | PAR-CLIP | Tarbase |
| hsa-miR-15a-5p | SECISBP2L | HITS-CLIP | Tarbase |
| hsa-miR-195-5p | SECISBP2L | HITS-CLIP | Tarbase |
| hsa-miR-141-3p | TMOD3 | HITS-CLIP | Tarbase |
| hsa-miR-15a-5p | TMOD3 | PAR-CLIP | Tarbase |
| hsa-miR-195-5p | TMOD3 | PAR-CLIP | Tarbase |
| hsa-miR-141-3p | GLCE | HITS-CLIP | Tarbase |
| hsa-miR-15a-5p | GLCE | HITS-CLIP | Tarbase |
| hsa-miR-15a-5p | GLCE | PAR-CLIP | Tarbase |
| hsa-miR-15a-5p | GLCE | PAR-CLIP | Tarbase |
| hsa-miR-195-5p | GLCE | HITS-CLIP | Tarbase |
| hsa-miR-486-5p | GLCE | HITS-CLIP | Tarbase |
| hsa-miR-195-5p | GLCE | Microarrays | Tarbase |
| hsa-miR-141-3p | APH1B | PAR-CLIP | Tarbase |
| hsa-miR-15a-5p | PARP16 | PAR-CLIP | Tarbase |
| hsa-miR-15a-5p | PARP16 | HITS-CLIP | Tarbase |
| hsa-miR-15a-5p | PARP16 | HITS-CLIP | Tarbase |
| hsa-miR-15a-5p | PARP16 | PAR-CLIP | Tarbase |
| hsa-miR-15a-5p | PARP16 | HITS-CLIP | Tarbase |
| hsa-miR-195-5p | PARP16 | HITS-CLIP | Tarbase |
| hsa-miR-195-5p | PARP16 | PAR-CLIP | Tarbase |
| hsa-miR-195-5p | PARP16 | Microarrays | Tarbase |
| hsa-miR-195-5p | PCDH10 | Microarrays | Tarbase |
| hsa-miR-195-5p | NDST4 | Microarrays | Tarbase |
| hsa-miR-195-5p | ZGRF1 | Microarrays | Tarbase |
| hsa-miR-141-3p | HNRNPD | HITS-CLIP | Tarbase |
| hsa-miR-15a-5p | HNRNPD | HITS-CLIP | Tarbase |
| hsa-miR-15a-5p | RASGEF1B | PAR-CLIP | Tarbase |
| hsa-miR-195-5p | RASGEF1B | Microarrays | Tarbase |
| hsa-miR-195-5p | AGPAT9 | Microarrays | Tarbase |
| hsa-miR-15a-5p | FGF2 | HITS-CLIP | Tarbase |
| hsa-miR-15a-5p | FGF2 | PAR-CLIP | Tarbase |
| hsa-miR-15a-5p | FGF2 | PAR-CLIP | Tarbase |
| hsa-miR-15a-5p | FGF2 | PAR-CLIP | Tarbase |

| | | | |
|---|---|---|---|
| hsa-miR-195-5p | FGF2 | HITS-CLIP | Tarbase |
| hsa-miR-195-5p | FGF2 | PAR-CLIP | Tarbase |
| hsa-miR-15a-5p | FGF2 | Chimeric fragments | Tarbase |
| hsa-miR-15a-5p | BBS7 | HITS-CLIP | Tarbase |
| hsa-miR-15a-3p | NUP54 | HITS-CLIP | Tarbase |
| hsa-miR-141-3p | G3BP2 | HITS-CLIP | Tarbase |
| hsa-miR-15a-5p | G3BP2 | HITS-CLIP | Tarbase |
| hsa-miR-486-5p | G3BP2 | HITS-CLIP | Tarbase |
| hsa-miR-15a-5p | SCARB2 | HITS-CLIP | Tarbase |
| hsa-miR-195-5p | SCARB2 | HITS-CLIP | Tarbase |
| hsa-miR-15a-5p | CCNG2 | HITS-CLIP | Tarbase |
| hsa-miR-195-5p | CCNG2 | HITS-CLIP | Tarbase |
| hsa-miR-15a-5p | CNOT6L | PAR-CLIP | Tarbase |
| hsa-miR-195-5p | CNOT6L | Microarrays | Tarbase |
| hsa-miR-15a-5p | USO1 | PAR-CLIP | Tarbase |
| hsa-miR-195-5p | USO1 | PAR-CLIP | Tarbase |
| hsa-miR-195-5p | CENPE | Microarrays | Tarbase |
| hsa-miR-15a-5p | GSTCD | PAR-CLIP | Tarbase |
| hsa-miR-195-5p | GSTCD | Microarrays | Tarbase |
| hsa-miR-195-5p | HADH | Microarrays | Tarbase |
| hsa-miR-195-5p | EGF | Microarrays | Tarbase |
| hsa-miR-3613-5p | PAPSS1 | HITS-CLIP | Tarbase |
| hsa-miR-15a-3p | FBN2 | HITS-CLIP | Tarbase |
| hsa-miR-15a-5p | FBN2 | HITS-CLIP | Tarbase |
| hsa-miR-195-5p | FBN2 | HITS-CLIP | Tarbase |
| hsa-miR-3605-3p | MAPK8IP3 | HITS-CLIP | Tarbase |
| hsa-miR-15a-5p | GUCD1 | HITS-CLIP | Tarbase |
| hsa-miR-195-5p | GUCD1 | HITS-CLIP | Tarbase |
| hsa-miR-141-3p | RNF185 | HITS-CLIP | Tarbase |
| hsa-miR-15a-5p | B4GALNT3 | HITS-CLIP | Tarbase |
| hsa-miR-195-5p | B4GALNT3 | HITS-CLIP | Tarbase |
| hsa-miR-15a-5p | GABARAPL1 | PAR-CLIP | Tarbase |
| hsa-miR-15a-5p | GABARAPL1 | Chimeric fragments | Tarbase |
| hsa-miR-15a-5p | KIF21A | PAR-CLIP | Tarbase |
| hsa-miR-15a-5p | KIF21A | PAR-CLIP | Tarbase |
| hsa-miR-15a-5p | KIF21A | PAR-CLIP | Tarbase |
| hsa-miR-15a-5p | KIF21A | PAR-CLIP | Tarbase |
| hsa-miR-195-5p | KIF21A | PAR-CLIP | Tarbase |
| hsa-miR-195-5p | KIF21A | PAR-CLIP | Tarbase |
| hsa-miR-195-5p | KIF21A | PAR-CLIP | Tarbase |
| hsa-miR-15a-5p | KIF21A | HITS-CLIP | Tarbase |
| hsa-miR-195-5p | KIF21A | HITS-CLIP | Tarbase |
| hsa-miR-141-3p | FGD4 | HITS-CLIP | Tarbase |
| hsa-miR-15a-5p | FGD4 | HITS-CLIP | Tarbase |
| hsa-miR-195-5p | FGD4 | HITS-CLIP | Tarbase |
| hsa-miR-15a-5p | FAM60A | HITS-CLIP | Tarbase |
| hsa-miR-15a-5p | AEBP2 | PAR-CLIP | Tarbase |
| hsa-miR-15a-5p | AEBP2 | HITS-CLIP | Tarbase |
| hsa-miR-15a-5p | AEBP2 | PAR-CLIP | Tarbase |
| hsa-miR-195-5p | AEBP2 | HITS-CLIP | Tarbase |
| hsa-miR-486-5p | METTL20 | PAR-CLIP | Tarbase |
| hsa-miR-141-5p | ETNK1 | HITS-CLIP | Tarbase |
| hsa-miR-15a-5p | ETNK1 | HITS-CLIP | Tarbase |
| hsa-miR-195-5p | ETNK1 | HITS-CLIP | Tarbase |
| hsa-miR-15a-5p | ZCRB1 | HITS-CLIP | Tarbase |
| hsa-miR-195-5p | ZCRB1 | HITS-CLIP | Tarbase |
| hsa-miR-219a-1-3p | TMEM117 | PAR-CLIP | Tarbase |
| hsa-miR-141-3p | AMIGO2 | HITS-CLIP | Tarbase |
| hsa-miR-195-5p | AMIGO2 | Microarrays | Tarbase |
| hsa-miR-141-3p | SCAF11 | HITS-CLIP | Tarbase |
| hsa-miR-15a-5p | SCAF11 | HITS-CLIP | Tarbase |
| hsa-miR-486-3p | SCAF11 | HITS-CLIP | Tarbase |
| hsa-miR-141-5p | LRIG3 | HITS-CLIP | Tarbase |
| hsa-miR-486-5p | GLIPR1 | HITS-CLIP | Tarbase |
| hsa-miR-195-5p | GLIPR1 | Microarrays | Tarbase |
| hsa-miR-15a-5p | TPH2 | HITS-CLIP | Tarbase |
| hsa-miR-195-5p | TPH2 | HITS-CLIP | Tarbase |
| hsa-miR-195-5p | PHLDA1 | Microarrays | Tarbase |
| hsa-miR-195-5p | DUSP6 | Microarrays | Tarbase |
| hsa-miR-141-3p | TMTC3 | HITS-CLIP | Tarbase |
| hsa-miR-141-3p | TMTC3 | HITS-CLIP | Tarbase |
| hsa-miR-15a-5p | TMTC3 | PAR-CLIP | Tarbase |
| hsa-miR-15a-5p | TMTC3 | HITS-CLIP | Tarbase |
| hsa-miR-195-5p | TMTC3 | PAR-CLIP | Tarbase |
| hsa-miR-3613-5p | LUM | HITS-CLIP | Tarbase |
| hsa-miR-195-5p | LUM | Microarrays | Tarbase |
| hsa-miR-195-5p | ASCL1 | Microarrays | Tarbase |
| hsa-miR-195-5p | GAS2L3 | Microarrays | Tarbase |
| hsa-miR-15a-5p | SLC15A4 | PAR-CLIP | Tarbase |
| hsa-miR-15a-5p | SLC15A4 | PAR-CLIP | Tarbase |
| hsa-miR-15a-5p | SLC15A4 | PAR-CLIP | Tarbase |
| hsa-miR-15a-5p | SLC15A4 | PAR-CLIP | Tarbase |
| hsa-miR-15a-5p | SLC15A4 | PAR-CLIP | Tarbase |
| hsa-miR-15a-5p | SLC15A4 | PAR-CLIP | Tarbase |
| hsa-miR-15a-5p | SLC15A4 | PAR-CLIP | Tarbase |
| hsa-miR-15a-5p | SLC15A4 | PAR-CLIP | Tarbase |
| hsa-miR-195-5p | SLC15A4 | PAR-CLIP | Tarbase |
| hsa-miR-195-5p | SLC15A4 | PAR-CLIP | Tarbase |
| hsa-miR-195-5p | SLC15A4 | PAR-CLIP | Tarbase |
| hsa-miR-141-3p | NUPL1 | HITS-CLIP | Tarbase |
| hsa-miR-141-3p | NUPL1 | HITS-CLIP | Tarbase |
| hsa-miR-141-3p | NUPL1 | HITS-CLIP | Tarbase |
| hsa-miR-141-3p | NUPL1 | HITS-CLIP | Tarbase |
| hsa-miR-15a-3p | NUPL1 | HITS-CLIP | Tarbase |
| hsa-miR-15a-5p | NUPL1 | HITS-CLIP | Tarbase |
| hsa-miR-15a-5p | NUPL1 | PAR-CLIP | Tarbase |
| hsa-miR-195-5p | NUPL1 | HITS-CLIP | Tarbase |
| hsa-miR-15a-3p | SLC7A1 | HITS-CLIP | Tarbase |
| hsa-miR-15a-3p | SUOX | HITS-CLIP | Tarbase |
| hsa-miR-15a-5p | TARBP2 | PAR-CLIP | Tarbase |
| hsa-miR-15a-5p | TARBP2 | PAR-CLIP | Tarbase |
| hsa-miR-195-5p | TARBP2 | Microarrays | Tarbase |
| hsa-miR-15a-5p | SMARCC2 | PAR-CLIP | Tarbase |
| hsa-miR-195-5p | BRCA2 | Microarrays | Tarbase |
| hsa-miR-15a-5p | KANSL2 | HITS-CLIP | Tarbase |
| hsa-miR-195-5p | KANSL2 | HITS-CLIP | Tarbase |
| hsa-miR-195-5p | LMBR1L | Microarrays | Tarbase |
| hsa-miR-15a-3p | TMBIM6 | HITS-CLIP | Tarbase |
| hsa-miR-15a-3p | TMBIM6 | HITS-CLIP | Tarbase |
| hsa-miR-15a-3p | TMBIM6 | HITS-CLIP | Tarbase |
| hsa-miR-15a-3p | TMBIM6 | HITS-CLIP | Tarbase |
| hsa-miR-15a-5p | TMBIM6 | HITS-CLIP | Tarbase |
| hsa-miR-15a-5p | TMBIM6 | HITS-CLIP | Tarbase |
| hsa-miR-195-5p | TMBIM6 | HITS-CLIP | Tarbase |
| hsa-miR-195-5p | TMBIM6 | HITS-CLIP | Tarbase |
| hsa-miR-141-3p | ANKRD52 | HITS-CLIP | Tarbase |
| hsa-miR-141-3p | ANKRD52 | HITS-CLIP | Tarbase |
| hsa-miR-141-3p | ANKRD52 | HITS-CLIP | Tarbase |
| hsa-miR-15a-5p | ANKRD52 | HITS-CLIP | Tarbase |
| hsa-miR-15a-5p | ANKRD52 | HITS-CLIP | Tarbase |
| hsa-miR-15a-5p | ANKRD52 | HITS-CLIP | Tarbase |
| hsa-miR-15a-5p | ANKRD52 | PAR-CLIP | Tarbase |
| hsa-miR-195-5p | ANKRD52 | HITS-CLIP | Tarbase |
| hsa-miR-195-5p | ANKRD52 | HITS-CLIP | Tarbase |
| hsa-miR-3605-3p | ANKRD52 | HITS-CLIP | Tarbase |
| hsa-miR-3605-5p | ANKRD52 | HITS-CLIP | Tarbase |
| hsa-miR-3605-5p | ANKRD52 | HITS-CLIP | Tarbase |
| hsa-miR-195-5p | ANKRD52 | Microarrays | Tarbase |
| hsa-miR-141-5p | ZNF740 | HITS-CLIP | Tarbase |
| hsa-miR-195-5p | LPAR6 | Microarrays | Tarbase |
| hsa-miR-141-3p | SBNO1 | HITS-CLIP | Tarbase |
| hsa-miR-15a-3p | SBNO1 | HITS-CLIP | Tarbase |
| hsa-miR-15a-5p | SBNO1 | HITS-CLIP | Tarbase |
| hsa-miR-219a-1-3p | SBNO1 | PAR-CLIP | Tarbase |
| hsa-miR-219a-1-3p | SBNO1 | PAR-CLIP | Tarbase |
| hsa-miR-141-3p | SETD1B | HITS-CLIP | Tarbase |
| hsa-miR-141-3p | SETD1B | HITS-CLIP | Tarbase |
| hsa-miR-141-5p | SETD1B | HITS-CLIP | Tarbase |
| hsa-miR-15a-5p | SETD1B | PAR-CLIP | Tarbase |
| hsa-miR-15a-5p | SETD1B | HITS-CLIP | Tarbase |
| hsa-miR-15a-5p | SETD1B | HITS-CLIP | Tarbase |
| hsa-miR-195-5p | SETD1B | HITS-CLIP | Tarbase |
| hsa-miR-141-3p | DENR | HITS-CLIP | Tarbase |
| hsa-miR-15a-5p | DENR | PAR-CLIP | Tarbase |
| hsa-miR-15a-5p | DENR | HITS-CLIP | Tarbase |
| hsa-miR-195-5p | DENR | HITS-CLIP | Tarbase |
| hsa-miR-141-5p | DIAPH3 | HITS-CLIP | Tarbase |
| hsa-miR-195-5p | DIAPH3 | Microarrays | Tarbase |
| hsa-miR-141-3p | SLAIN1 | HITS-CLIP | Tarbase |

| | | | |
|---|---|---|---|
| hsa-miR-141-5p | RBM26 | HITS-CLIP | Tarbase |
| hsa-miR-141-5p | RBM26 | HITS-CLIP | Tarbase |
| hsa-miR-15a-5p | RBM26 | PAR-CLIP | Tarbase |
| hsa-miR-15a-5p | RBM26 | PAR-CLIP | Tarbase |
| hsa-miR-15a-5p | RBM26 | PAR-CLIP | Tarbase |
| hsa-miR-195-5p | RBM26 | PAR-CLIP | Tarbase |
| hsa-miR-195-5p | RBM26 | PAR-CLIP | Tarbase |
| hsa-miR-3613-3p | RBM26 | HITS-CLIP | Tarbase |
| hsa-miR-15a-5p | MBNL2 | PAR-CLIP | Tarbase |
| hsa-miR-15a-5p | MBNL2 | PAR-CLIP | Tarbase |
| hsa-miR-195-5p | MBNL2 | PAR-CLIP | Tarbase |
| hsa-miR-219a-1-3p | MBNL2 | HITS-CLIP | Tarbase |
| hsa-miR-15a-5p | ZIC5 | HITS-CLIP | Tarbase |
| hsa-miR-195-5p | ZIC5 | HITS-CLIP | Tarbase |
| hsa-miR-3613-3p | ABHD13 | PAR-CLIP | Tarbase |
| hsa-miR-15a-5p | ABHD13 | HITS-CLIP | Tarbase |
| hsa-miR-195-5p | ABHD13 | HITS-CLIP | Tarbase |
| hsa-miR-195-5p | RAB20 | Microarrays | Tarbase |
| hsa-miR-141-5p | CUL4A | HITS-CLIP | Tarbase |
| hsa-miR-195-5p | NOVA1 | Microarrays | Tarbase |
| hsa-miR-15a-5p | FITM1 | HITS-CLIP | Tarbase |
| hsa-miR-15a-5p | FITM1 | HITS-CLIP | Tarbase |
| hsa-miR-15a-5p | FITM1 | HITS-CLIP | Tarbase |
| hsa-miR-195-5p | FITM1 | HITS-CLIP | Tarbase |
| hsa-miR-195-5p | FITM1 | HITS-CLIP | Tarbase |
| hsa-miR-195-5p | FITM1 | HITS-CLIP | Tarbase |
| hsa-miR-195-5p | MDGA2 | Microarrays | Tarbase |
| hsa-miR-3613-5p | FRMD6 | HITS-CLIP | Tarbase |
| hsa-miR-15a-5p | PELI2 | PAR-CLIP | Tarbase |
| hsa-miR-3613-5p | PELI2 | HITS-CLIP | Tarbase |
| hsa-miR-486-5p | SYT16 | HITS-CLIP | Tarbase |
| hsa-miR-141-3p | NAA30 | HITS-CLIP | Tarbase |
| hsa-miR-141-5p | NAA30 | HITS-CLIP | Tarbase |
| hsa-miR-15a-5p | NAA30 | PAR-CLIP | Tarbase |
| hsa-miR-195-5p | RDH12 | Microarrays | Tarbase |
| hsa-miR-15a-5p | RAB15 | PAR-CLIP | Tarbase |
| hsa-miR-15a-5p | NIPA2 | HITS-CLIP | Tarbase |
| hsa-miR-141-5p | MFAP1 | HITS-CLIP | Tarbase |
| hsa-miR-15a-5p | MFAP1 | PAR-CLIP | Tarbase |
| hsa-miR-195-5p | MFAP1 | PAR-CLIP | Tarbase |
| hsa-miR-15a-5p | TCF12 | PAR-CLIP | Tarbase |
| hsa-miR-195-5p | TCF12 | PAR-CLIP | Tarbase |
| hsa-miR-195-5p | SORD | Microarrays | Tarbase |
| hsa-miR-141-3p | SERF2 | HITS-CLIP | Tarbase |
| hsa-miR-15a-5p | FGF7 | Luciferase Reporter Assay | Tarbase |
| hsa-miR-141-5p | CDAN1 | HITS-CLIP | Tarbase |
| hsa-miR-141-5p | ANP32A | HITS-CLIP | Tarbase |
| hsa-miR-3605-3p | ANP32A | HITS-CLIP | Tarbase |
| hsa-miR-141-3p | UBE2Q2 | HITS-CLIP | Tarbase |
| hsa-miR-141-3p | HMG20A | HITS-CLIP | Tarbase |
| hsa-miR-15a-5p | TSPAN3 | PAR-CLIP | Tarbase |
| hsa-miR-15a-5p | TSPAN3 | PAR-CLIP | Tarbase |
| hsa-miR-195-5p | TSPAN3 | PAR-CLIP | Tarbase |
| hsa-miR-219a-1-3p | TSPAN3 | HITS-CLIP | Tarbase |
| hsa-miR-141-3p | NCOA2 | HITS-CLIP | Tarbase |
| hsa-miR-141-3p | NCOA2 | HITS-CLIP | Tarbase |
| hsa-miR-15a-5p | NCOA2 | HITS-CLIP | Tarbase |
| hsa-miR-486-5p | NCOA2 | HITS-CLIP | Tarbase |
| hsa-miR-141-3p | IGF1R | HITS-CLIP | Tarbase |
| hsa-miR-141-5p | IGF1R | HITS-CLIP | Tarbase |
| hsa-miR-15a-5p | IGF1R | HITS-CLIP | Tarbase |
| hsa-miR-15a-5p | IGF1R | HITS-CLIP | Tarbase |
| hsa-miR-15a-5p | IGF1R | PAR-CLIP | Tarbase |
| hsa-miR-15a-5p | IGF1R | HITS-CLIP | Tarbase |
| hsa-miR-195-5p | IGF1R | HITS-CLIP | Tarbase |
| hsa-miR-195-5p | IGF1R | HITS-CLIP | Tarbase |
| hsa-miR-3613-5p | IGF1R | HITS-CLIP | Tarbase |
| hsa-miR-141-3p | ARRDC4 | HITS-CLIP | Tarbase |
| hsa-miR-15a-5p | ARRDC4 | HITS-CLIP | Tarbase |
| hsa-miR-15a-5p | ARRDC4 | HITS-CLIP | Tarbase |
| hsa-miR-15a-5p | ARRDC4 | PAR-CLIP | Tarbase |
| hsa-miR-15a-5p | ARRDC4 | HITS-CLIP | Tarbase |
| hsa-miR-195-5p | ARRDC4 | HITS-CLIP | Tarbase |
| hsa-miR-195-5p | ARRDC4 | HITS-CLIP | Tarbase |
| hsa-miR-15a-5p | PIF1 | PAR-CLIP | Tarbase |
| hsa-miR-15a-5p | PIF1 | PAR-CLIP | Tarbase |
| hsa-miR-15a-5p | PML | HITS-CLIP | Tarbase |
| hsa-miR-195-5p | PML | HITS-CLIP | Tarbase |
| hsa-miR-15a-3p | ADAMTS17 | HITS-CLIP | Tarbase |
| hsa-miR-15a-5p | SCAMP2 | PAR-CLIP | Tarbase |
| hsa-miR-15a-5p | SCAMP2 | PAR-CLIP | Tarbase |
| hsa-miR-195-5p | SCAMP2 | PAR-CLIP | Tarbase |
| hsa-miR-195-5p | SCAMP2 | PAR-CLIP | Tarbase |
| hsa-miR-15a-5p | FANCI | HITS-CLIP | Tarbase |
| hsa-miR-195-5p | FANCI | Microarrays | Tarbase |
| hsa-miR-15a-3p | ABHD2 | HITS-CLIP | Tarbase |
| hsa-miR-15a-5p | ABHD2 | HITS-CLIP | Tarbase |
| hsa-miR-3605-5p | ABHD2 | HITS-CLIP | Tarbase |
| hsa-miR-195-5p | TICRR | Microarrays | Tarbase |
| hsa-miR-195-5p | NTRK3 | Microarrays | Tarbase |
| hsa-miR-15a-5p | DET1 | HITS-CLIP | Tarbase |
| hsa-miR-15a-5p | UNC45A | HITS-CLIP | Tarbase |
| hsa-miR-195-5p | UNC45A | HITS-CLIP | Tarbase |
| hsa-miR-195-5p | MCTP2 | Microarrays | Tarbase |
| hsa-miR-15a-5p | FURIN | PAR-CLIP | Tarbase |
| hsa-miR-195-5p | FURIN | Microarrays | Tarbase |
| hsa-miR-141-5p | IQGAP1 | HITS-CLIP | Tarbase |
| hsa-miR-15a-5p | IQGAP1 | HITS-CLIP | Tarbase |
| hsa-miR-141-5p | CRTC3 | HITS-CLIP | Tarbase |
| hsa-miR-15a-5p | CRTC3 | HITS-CLIP | Tarbase |
| hsa-miR-141-3p | SEC11A | HITS-CLIP | Tarbase |
| hsa-miR-195-5p | PMM2 | Microarrays | Tarbase |
| hsa-miR-195-5p | ITGAX | Microarrays | Tarbase |
| hsa-miR-195-5p | TGFB1I1 | Microarrays | Tarbase |
| hsa-miR-195-5p | NKD1 | Microarrays | Tarbase |
| hsa-miR-141-3p | ZFHX3 | HITS-CLIP | Tarbase |
| hsa-miR-141-3p | ZFHX3 | HITS-CLIP | Tarbase |
| hsa-miR-15a-5p | ZFHX3 | HITS-CLIP | Tarbase |
| hsa-miR-15a-5p | ZFHX3 | HITS-CLIP | Tarbase |
| hsa-miR-195-5p | ZFHX3 | HITS-CLIP | Tarbase |
| hsa-miR-195-5p | ZFHX3 | HITS-CLIP | Tarbase |
| hsa-miR-486-5p | KIFC3 | HITS-CLIP | Tarbase |
| hsa-miR-195-5p | KIFC3 | Microarrays | Tarbase |
| hsa-miR-15a-5p | CMTM3 | PAR-CLIP | Tarbase |
| hsa-miR-195-5p | CDH11 | Microarrays | Tarbase |
| hsa-miR-141-5p | MBTPS1 | HITS-CLIP | Tarbase |
| hsa-miR-141-3p | ZCCHC14 | HITS-CLIP | Tarbase |
| hsa-miR-3605-3p | IRF8 | PAR-CLIP | Tarbase |
| hsa-miR-15a-5p | GID4 | Chimeric fragments | Tarbase |
| hsa-miR-141-3p | KSR1 | HITS-CLIP | Tarbase |
| hsa-miR-3613-3p | RANBP10 | HITS-CLIP | Tarbase |
| hsa-miR-141-3p | NOB1 | HITS-CLIP | Tarbase |
| hsa-miR-15a-5p | NOB1 | Chimeric fragments | Tarbase |
| hsa-miR-15a-5p | UNC45B | PAR-CLIP | Tarbase |
| hsa-miR-195-5p | PCTP | Microarrays | Tarbase |
| hsa-miR-219a-1-3p | C17orf80 | PAR-CLIP | Tarbase |
| hsa-miR-219a-1-3p | C17orf80 | PAR-CLIP | Tarbase |
| hsa-miR-219a-1-3p | C17orf80 | PAR-CLIP | Tarbase |
| hsa-miR-15a-5p | NPEPPS | HITS-CLIP | Tarbase |
| hsa-miR-15a-5p | NPEPPS | HITS-CLIP | Tarbase |
| hsa-miR-15a-5p | NPEPPS | HITS-CLIP | Tarbase |
| hsa-miR-195-5p | NPEPPS | HITS-CLIP | Tarbase |
| hsa-miR-195-5p | NPEPPS | HITS-CLIP | Tarbase |
| hsa-miR-195-5p | ARSG | Microarrays | Tarbase |
| hsa-miR-15a-3p | G6PC3 | HITS-CLIP | Tarbase |
| hsa-miR-15a-3p | CLTC | HITS-CLIP | Tarbase |
| hsa-miR-15a-5p | CLTC | HITS-CLIP | Tarbase |
| hsa-miR-15a-5p | CLTC | HITS-CLIP | Tarbase |
| hsa-miR-15a-5p | CLTC | PAR-CLIP | Tarbase |
| hsa-miR-15a-5p | CLTC | HITS-CLIP | Tarbase |
| hsa-miR-195-5p | CLTC | HITS-CLIP | Tarbase |
| hsa-miR-195-5p | CLTC | HITS-CLIP | Tarbase |
| hsa-miR-219a-1-3p | CLTC | HITS-CLIP | Tarbase |
| hsa-miR-3613-5p | CLTC | HITS-CLIP | Tarbase |
| hsa-miR-219a-1-3p | SS18 | HITS-CLIP | Tarbase |
| hsa-miR-15a-5p | AFG3L2 | HITS-CLIP | Tarbase |
| hsa-miR-141-5p | SLC39A6 | HITS-CLIP | Tarbase |
| hsa-miR-15a-5p | SLC39A6 | HITS-CLIP | Tarbase |
| hsa-miR-141-3p | RPRD1A | HITS-CLIP | Tarbase |

| | | | |
|---|---|---|---|
| hsa-miR-15a-5p | GALNT1 | HITS-CLIP | Tarbase |
| hsa-miR-195-5p | GALNT1 | HITS-CLIP | Tarbase |
| hsa-miR-15a-5p | GALNT1 | HITS-CLIP | Tarbase |
| hsa-miR-195-5p | GALNT1 | HITS-CLIP | Tarbase |
| hsa-miR-141-5p | ASXL3 | HITS-CLIP | Tarbase |
| hsa-miR-195-5p | ADCYAP1 | Microarrays | Tarbase |
| hsa-miR-15a-5p | GAREM | PAR-CLIP | Tarbase |
| hsa-miR-15a-5p | GAREM | HITS-CLIP | Tarbase |
| hsa-miR-195-5p | GAREM | Microarrays | Tarbase |
| hsa-miR-141-3p | GATA6 | HITS-CLIP | Tarbase |
| hsa-miR-141-3p | GREB1L | HITS-CLIP | Tarbase |
| hsa-miR-141-5p | PELP1 | HITS-CLIP | Tarbase |
| hsa-miR-15a-5p | PELP1 | HITS-CLIP | Tarbase |
| hsa-miR-3613-3p | NPC1 | HITS-CLIP | Tarbase |
| hsa-miR-486-5p | NPC1 | HITS-CLIP | Tarbase |
| hsa-miR-195-5p | NPC1 | Microarrays | Tarbase |
| hsa-miR-195-5p | SLC14A1 | Microarrays | Tarbase |
| hsa-miR-15a-3p | MINK1 | PAR-CLIP | Tarbase |
| hsa-miR-15a-5p | MINK1 | HITS-CLIP | Tarbase |
| hsa-miR-15a-5p | ASGR1 | PAR-CLIP | Tarbase |
| hsa-miR-15a-5p | ASGR1 | PAR-CLIP | Tarbase |
| hsa-miR-15a-5p | TP53 | Luciferase Reporter Assay | Tarbase |
| hsa-miR-15a-5p | TP53 | Western Blot | Tarbase |
| hsa-miR-15a-3p | ARHGDIA | HITS-CLIP | Tarbase |
| hsa-miR-15a-5p | ARHGDIA | PAR-CLIP | Tarbase |
| hsa-miR-15a-5p | ARHGDIA | HITS-CLIP | Tarbase |
| hsa-miR-15a-5p | ARHGDIA | HITS-CLIP | Tarbase |
| hsa-miR-15a-5p | ARHGDIA | HITS-CLIP | Tarbase |
| hsa-miR-195-5p | ARHGDIA | HITS-CLIP | Tarbase |
| hsa-miR-195-5p | ARHGDIA | HITS-CLIP | Tarbase |
| hsa-miR-486-3p | ARHGDIA | HITS-CLIP | Tarbase |
| hsa-miR-486-3p | ARHGDIA | HITS-CLIP | Tarbase |
| hsa-miR-486-3p | ARHGDIA | PAR-CLIP | Tarbase |
| hsa-miR-195-5p | ARHGDIA | Microarrays | Tarbase |
| hsa-miR-15a-3p | TMC6 | HITS-CLIP | Tarbase |
| hsa-miR-141-5p | SLC16A3 | HITS-CLIP | Tarbase |
| hsa-miR-141-5p | SLC16A3 | HITS-CLIP | Tarbase |
| hsa-miR-15a-3p | EIF4A3 | HITS-CLIP | Tarbase |
| hsa-miR-141-5p | CSNK1D | HITS-CLIP | Tarbase |
| hsa-miR-195-5p | ANAPC11 | Microarrays | Tarbase |
| hsa-miR-486-5p | TBCD | PAR-CLIP | Tarbase |
| hsa-miR-195-5p | FN3KRP | Microarrays | Tarbase |
| hsa-miR-15a-5p | NARF | PAR-CLIP | Tarbase |
| hsa-miR-15a-5p | NARF | HITS-CLIP | Tarbase |
| hsa-miR-15a-5p | NARF | HITS-CLIP | Tarbase |
| hsa-miR-15a-5p | NARF | PAR-CLIP | Tarbase |
| hsa-miR-15a-5p | NARF | PAR-CLIP | Tarbase |
| hsa-miR-15a-5p | NARF | PAR-CLIP | Tarbase |
| hsa-miR-195-5p | NARF | PAR-CLIP | Tarbase |
| hsa-miR-195-5p | NARF | HITS-CLIP | Tarbase |
| hsa-miR-195-5p | NARF | HITS-CLIP | Tarbase |
| hsa-miR-195-5p | NARF | PAR-CLIP | Tarbase |
| hsa-miR-15a-5p | NARF | Chimeric fragments | Tarbase |
| hsa-miR-15a-3p | TRIM65 | HITS-CLIP | Tarbase |
| hsa-miR-15a-3p | TRIM65 | HITS-CLIP | Tarbase |
| hsa-miR-3605-5p | RNF157 | HITS-CLIP | Tarbase |
| hsa-miR-15a-3p | WDR45B | HITS-CLIP | Tarbase |
| hsa-miR-219a-1-3p | WDR45B | PAR-CLIP | Tarbase |
| hsa-miR-219a-1-3p | WDR45B | PAR-CLIP | Tarbase |
| hsa-miR-15a-5p | CBX4 | HITS-CLIP | Tarbase |
| hsa-miR-15a-5p | CBX4 | HITS-CLIP | Tarbase |
| hsa-miR-15a-5p | CBX4 | PAR-CLIP | Tarbase |
| hsa-miR-15a-5p | CBX4 | HITS-CLIP | Tarbase |
| hsa-miR-15a-5p | CBX4 | HITS-CLIP | Tarbase |
| hsa-miR-15a-5p | CBX4 | HITS-CLIP | Tarbase |
| hsa-miR-195-5p | CBX4 | HITS-CLIP | Tarbase |
| hsa-miR-195-5p | CBX4 | HITS-CLIP | Tarbase |
| hsa-miR-195-5p | CBX4 | HITS-CLIP | Tarbase |
| hsa-miR-486-5p | SMAD4 | HITS-CLIP | Tarbase |
| hsa-miR-141-5p | ZCCHC2 | HITS-CLIP | Tarbase |
| hsa-miR-15a-5p | ZCCHC2 | PAR-CLIP | Tarbase |
| hsa-miR-195-5p | ZCCHC2 | PAR-CLIP | Tarbase |
| hsa-miR-15a-5p | CBLN2 | HITS-CLIP | Tarbase |
| hsa-miR-195-5p | CBLN2 | HITS-CLIP | Tarbase |
| hsa-miR-195-5p | CBLN2 | Microarrays | Tarbase |
| hsa-miR-195-5p | PMAIP1 | Microarrays | Tarbase |
| hsa-miR-486-3p | ERBB2 | HITS-CLIP | Tarbase |
| hsa-miR-486-3p | ERBB2 | HITS-CLIP | Tarbase |
| hsa-miR-486-3p | ERBB2 | Luciferase Reporter Assay | Tarbase |
| hsa-miR-141-5p | FKBP10 | HITS-CLIP | Tarbase |
| hsa-miR-15a-3p | FKBP10 | HITS-CLIP | Tarbase |
| hsa-miR-141-5p | SAMD1 | HITS-CLIP | Tarbase |
| hsa-miR-486-3p | SAMD1 | HITS-CLIP | Tarbase |
| hsa-miR-486-5p | SAMD1 | PAR-CLIP | Tarbase |
| hsa-miR-141-3p | BRD4 | HITS-CLIP | Tarbase |
| hsa-miR-141-3p | BRD4 | HITS-CLIP | Tarbase |
| hsa-miR-15a-3p | BRD4 | HITS-CLIP | Tarbase |
| hsa-miR-486-5p | BRD4 | PAR-CLIP | Tarbase |
| hsa-miR-15a-3p | NFIC | HITS-CLIP | Tarbase |
| hsa-miR-15a-3p | PFKL | HITS-CLIP | Tarbase |
| hsa-miR-15a-3p | PFKL | HITS-CLIP | Tarbase |
| hsa-miR-15a-3p | SH3GL1 | HITS-CLIP | Tarbase |
| hsa-miR-15a-3p | SH3GL1 | HITS-CLIP | Tarbase |
| hsa-miR-486-5p | DPP9 | HITS-CLIP | Tarbase |
| hsa-miR-15a-5p | CCDC97 | Chimeric fragments | Tarbase |
| hsa-miR-141-3p | IFNAR1 | HITS-CLIP | Tarbase |
| hsa-miR-15a-5p | SIK1 | HITS-CLIP | Tarbase |
| hsa-miR-15a-5p | SIK1 | HITS-CLIP | Tarbase |
| hsa-miR-15a-5p | SIK1 | HITS-CLIP | Tarbase |
| hsa-miR-195-5p | SIK1 | HITS-CLIP | Tarbase |
| hsa-miR-195-5p | SIK1 | HITS-CLIP | Tarbase |
| hsa-miR-486-3p | SIK1 | HITS-CLIP | Tarbase |
| hsa-miR-195-5p | TMEM50B | Microarrays | Tarbase |
| hsa-miR-141-5p | APP | HITS-CLIP | Tarbase |
| hsa-miR-15a-5p | APP | PAR-CLIP | Tarbase |
| hsa-miR-15a-5p | APP | HITS-CLIP | Tarbase |
| hsa-miR-15a-5p | APP | PAR-CLIP | Tarbase |
| hsa-miR-15a-5p | APP | HITS-CLIP | Tarbase |
| hsa-miR-15a-5p | APP | HITS-CLIP | Tarbase |
| hsa-miR-195-5p | APP | HITS-CLIP | Tarbase |
| hsa-miR-195-5p | APP | HITS-CLIP | Tarbase |
| hsa-miR-15a-5p | APP | Luciferase Reporter Assay | Tarbase |
| hsa-miR-15a-5p | DOPEY2 | HITS-CLIP | Tarbase |
| hsa-miR-195-5p | DOPEY2 | HITS-CLIP | Tarbase |
| hsa-miR-141-5p | URB1 | HITS-CLIP | Tarbase |
| hsa-miR-15a-3p | AKT1 | HITS-CLIP | Tarbase |
| hsa-miR-15a-5p | SAE1 | HITS-CLIP | Tarbase |
| hsa-miR-195-5p | CBLC | AGO-IP | Tarbase |
| hsa-miR-15a-3p | ADAMTS10 | HITS-CLIP | Tarbase |
| hsa-miR-3613-5p | CARM1 | HITS-CLIP | Tarbase |
| hsa-miR-195-5p | CARM1 | Microarrays | Tarbase |
| hsa-miR-3613-5p | RPS11 | HITS-CLIP | Tarbase |
| hsa-miR-3613-5p | RPS11 | HITS-CLIP | Tarbase |
| hsa-miR-195-5p | IGLON5 | Microarrays | Tarbase |
| hsa-miR-195-5p | SLC2A5 | Microarrays | Tarbase |
| hsa-miR-141-3p | RERE | HITS-CLIP | Tarbase |
| hsa-miR-141-3p | RERE | HITS-CLIP | Tarbase |
| hsa-miR-15a-3p | RERE | HITS-CLIP | Tarbase |
| hsa-miR-3613-3p | RERE | HITS-CLIP | Tarbase |
| hsa-miR-3613-5p | RERE | PAR-CLIP | Tarbase |
| hsa-miR-141-3p | EPHA2 | PAR-CLIP | Tarbase |
| hsa-miR-141-3p | EPHA2 | HITS-CLIP | Tarbase |
| hsa-miR-195-5p | EPHA2 | Microarrays | Tarbase |
| hsa-miR-141-5p | PEX14 | HITS-CLIP | Tarbase |
| hsa-miR-141-5p | PEX14 | HITS-CLIP | Tarbase |
| hsa-miR-15a-5p | PGD | PAR-CLIP | Tarbase |
| hsa-miR-15a-5p | PGD | HITS-CLIP | Tarbase |
| hsa-miR-195-5p | PGD | Microarrays | Tarbase |
| hsa-miR-15a-5p | KIAA0319L | HITS-CLIP | Tarbase |
| hsa-miR-195-5p | PLK4 | Microarrays | Tarbase |
| hsa-miR-195-5p | WDTC1 | Microarrays | Tarbase |
| hsa-miR-15a-3p | HSPG2 | HITS-CLIP | Tarbase |
| hsa-miR-15a-5p | HSPG2 | HITS-CLIP | Tarbase |
| hsa-miR-15a-5p | HSPG2 | HITS-CLIP | Tarbase |
| hsa-miR-195-5p | HSPG2 | HITS-CLIP | Tarbase |
| hsa-miR-195-5p | HSPG2 | HITS-CLIP | Tarbase |
| hsa-miR-3613-5p | CYR61 | HITS-CLIP | Tarbase |
| hsa-miR-195-5p | CYR61 | Microarrays | Tarbase |
| hsa-miR-15a-5p | KIF2C | PAR-CLIP | Tarbase |

| | | | |
|---|---|---|---|
| hsa-miR-195-5p | KIF2C | Microarrays | Tarbase |
| hsa-miR-15a-5p | PTPRF | HITS-CLIP | Tarbase |
| hsa-miR-195-5p | PTPRF | HITS-CLIP | Tarbase |
| hsa-miR-141-3p | LMO4 | HITS-CLIP | Tarbase |
| hsa-miR-15a-5p | LMO4 | HITS-CLIP | Tarbase |
| hsa-miR-195-5p | LMO4 | HITS-CLIP | Tarbase |
| hsa-miR-15a-5p | LMO4 | Chimeric fragments | Tarbase |
| hsa-miR-141-3p | MTF2 | HITS-CLIP | Tarbase |
| hsa-miR-141-3p | MTF2 | HITS-CLIP | Tarbase |
| hsa-miR-141-3p | MTF2 | HITS-CLIP | Tarbase |
| hsa-miR-15a-5p | MTF2 | HITS-CLIP | Tarbase |
| hsa-miR-3613-5p | MTF2 | HITS-CLIP | Tarbase |
| hsa-miR-486-5p | MTF2 | HITS-CLIP | Tarbase |
| hsa-miR-195-5p | SLC44A3 | Microarrays | Tarbase |
| hsa-miR-141-3p | ZNF697 | HITS-CLIP | Tarbase |
| hsa-miR-15a-5p | ZNF697 | HITS-CLIP | Tarbase |
| hsa-miR-195-5p | ZNF697 | HITS-CLIP | Tarbase |
| hsa-miR-219a-1-3p | CTTNBP2NL | PAR-CLIP | Tarbase |
| hsa-miR-15a-5p | STRIP1 | HITS-CLIP | Tarbase |
| hsa-miR-195-5p | STRIP1 | HITS-CLIP | Tarbase |
| hsa-miR-141-5p | PSMA5 | HITS-CLIP | Tarbase |
| hsa-miR-3613-5p | FNDC7 | HITS-CLIP | Tarbase |
| hsa-miR-15a-5p | CELSR2 | HITS-CLIP | Tarbase |
| hsa-miR-195-5p | CELSR2 | HITS-CLIP | Tarbase |
| hsa-miR-3605-5p | ALDH9A1 | PAR-CLIP | Tarbase |
| hsa-miR-141-3p | ATP1B1 | HITS-CLIP | Tarbase |
| hsa-miR-15a-5p | ATP1B1 | HITS-CLIP | Tarbase |
| hsa-miR-3613-3p | TIPRL | HITS-CLIP | Tarbase |
| hsa-miR-141-5p | CREG1 | HITS-CLIP | Tarbase |
| hsa-miR-15a-5p | CREG1 | PAR-CLIP | Tarbase |
| hsa-miR-15a-5p | CREG1 | HITS-CLIP | Tarbase |
| hsa-miR-195-5p | CREG1 | PAR-CLIP | Tarbase |
| hsa-miR-195-5p | CREG1 | HITS-CLIP | Tarbase |
| hsa-miR-3605-5p | CREG1 | HITS-CLIP | Tarbase |
| hsa-miR-141-5p | DCAF6 | HITS-CLIP | Tarbase |
| hsa-miR-195-5p | GPA33 | Microarrays | Tarbase |
| hsa-miR-141-5p | POU2F1 | HITS-CLIP | Tarbase |
| hsa-miR-15a-5p | POU2F1 | HITS-CLIP | Tarbase |
| hsa-miR-195-5p | POU2F1 | HITS-CLIP | Tarbase |
| hsa-miR-15a-3p | ILDR2 | HITS-CLIP | Tarbase |
| hsa-miR-15a-5p | UFC1 | PAR-CLIP | Tarbase |
| hsa-miR-486-3p | NUF2 | HITS-CLIP | Tarbase |
| hsa-miR-195-5p | NUF2 | Microarrays | Tarbase |
| hsa-miR-15a-5p | RGS5 | HITS-CLIP | Tarbase |
| hsa-miR-15a-5p | RGS5 | HITS-CLIP | Tarbase |
| hsa-miR-195-5p | RGS5 | HITS-CLIP | Tarbase |
| hsa-miR-141-5p | SDHC | HITS-CLIP | Tarbase |
| hsa-miR-15a-5p | PFDN2 | PAR-CLIP | Tarbase |
| hsa-miR-486-5p | PRCC | PAR-CLIP | Tarbase |
| hsa-miR-15a-5p | ISG20L2 | HITS-CLIP | Tarbase |
| hsa-miR-15a-5p | ISG20L2 | HITS-CLIP | Tarbase |
| hsa-miR-15a-5p | ISG20L2 | PAR-CLIP | Tarbase |
| hsa-miR-195-5p | ISG20L2 | HITS-CLIP | Tarbase |
| hsa-miR-195-5p | ISG20L2 | HITS-CLIP | Tarbase |
| hsa-miR-195-5p | CRABP2 | Microarrays | Tarbase |
| hsa-miR-15a-3p | HDGF | HITS-CLIP | Tarbase |
| hsa-miR-15a-5p | HDGF | HITS-CLIP | Tarbase |
| hsa-miR-15a-5p | HDGF | HITS-CLIP | Tarbase |
| hsa-miR-195-5p | HDGF | HITS-CLIP | Tarbase |
| hsa-miR-195-5p | HDGF | HITS-CLIP | Tarbase |
| hsa-miR-141-3p | ABL2 | HITS-CLIP | Tarbase |
| hsa-miR-141-3p | ABL2 | HITS-CLIP | Tarbase |
| hsa-miR-15a-5p | ABL2 | HITS-CLIP | Tarbase |
| hsa-miR-15a-5p | ABL2 | HITS-CLIP | Tarbase |
| hsa-miR-195-5p | ABL2 | HITS-CLIP | Tarbase |
| hsa-miR-15a-5p | ABL2 | Chimeric fragments | Tarbase |
| hsa-miR-3613-5p | XPR1 | PAR-CLIP | Tarbase |
| hsa-miR-15a-5p | HMCN1 | HITS-CLIP | Tarbase |
| hsa-miR-195-5p | HMCN1 | Microarrays | Tarbase |
| hsa-miR-15a-3p | RGL1 | HITS-CLIP | Tarbase |
| hsa-miR-3613-5p | RGL1 | HITS-CLIP | Tarbase |
| hsa-miR-15a-5p | LYPLAL1 | Chimeric fragments | Tarbase |
| hsa-miR-195-5p | RORC | Microarrays | Tarbase |
| hsa-miR-141-5p | CGN | HITS-CLIP | Tarbase |
| hsa-miR-141-3p | SNX27 | HITS-CLIP | Tarbase |
| hsa-miR-141-3p | SETDB1 | HITS-CLIP | Tarbase |
| hsa-miR-141-3p | MCL1 | PAR-CLIP | Tarbase |
| hsa-miR-141-3p | MCL1 | HITS-CLIP | Tarbase |
| hsa-miR-15a-3p | MCL1 | HITS-CLIP | Tarbase |
| hsa-miR-15a-5p | MCL1 | HITS-CLIP | Tarbase |
| hsa-miR-15a-5p | MCL1 | HITS-CLIP | Tarbase |
| hsa-miR-15a-5p | MCL1 | HITS-CLIP | Tarbase |
| hsa-miR-195-5p | MCL1 | HITS-CLIP | Tarbase |
| hsa-miR-195-5p | MCL1 | HITS-CLIP | Tarbase |
| hsa-miR-195-5p | MCL1 | HITS-CLIP | Tarbase |
| hsa-miR-195-5p | MCL1 | HITS-CLIP | Tarbase |
| hsa-miR-141-5p | RFX5 | HITS-CLIP | Tarbase |
| hsa-miR-141-5p | RFX5 | HITS-CLIP | Tarbase |
| hsa-miR-141-3p | ANP32E | HITS-CLIP | Tarbase |
| hsa-miR-3605-5p | CERS2 | HITS-CLIP | Tarbase |
| hsa-miR-141-5p | ENSA | HITS-CLIP | Tarbase |
| hsa-miR-15a-3p | ENSA | HITS-CLIP | Tarbase |
| hsa-miR-15a-5p | ENSA | PAR-CLIP | Tarbase |
| hsa-miR-486-5p | ENSA | PAR-CLIP | Tarbase |
| hsa-miR-15a-5p | SEMA6C | HITS-CLIP | Tarbase |
| hsa-miR-195-5p | SEMA6C | HITS-CLIP | Tarbase |
| hsa-miR-141-3p | ARNT | HITS-CLIP | Tarbase |
| hsa-miR-141-5p | ARNT | HITS-CLIP | Tarbase |
| hsa-miR-141-5p | ARNT | HITS-CLIP | Tarbase |
| hsa-miR-15a-3p | ARNT | HITS-CLIP | Tarbase |
| hsa-miR-219a-1-3p | POGZ | HITS-CLIP | Tarbase |
| hsa-miR-3613-3p | GABPB2 | PAR-CLIP | Tarbase |
| hsa-miR-3613-5p | DTL | HITS-CLIP | Tarbase |
| hsa-miR-3613-5p | DTL | HITS-CLIP | Tarbase |
| hsa-miR-3613-5p | DTL | HITS-CLIP | Tarbase |
| hsa-miR-195-5p | DTL | Microarrays | Tarbase |
| hsa-miR-3613-3p | INTS7 | PAR-CLIP | Tarbase |
| hsa-miR-195-5p | INTS7 | Microarrays | Tarbase |
| hsa-miR-486-5p | VASH2 | PAR-CLIP | Tarbase |
| hsa-miR-15a-5p | SUSD4 | HITS-CLIP | Tarbase |
| hsa-miR-195-5p | DUSP10 | Microarrays | Tarbase |
| hsa-miR-195-5p | ATP8B2 | HITS-CLIP | Tarbase |
| hsa-miR-15a-3p | SLC39A1 | HITS-CLIP | Tarbase |
| hsa-miR-15a-5p | SLC39A1 | HITS-CLIP | Tarbase |
| hsa-miR-195-5p | SLC39A1 | HITS-CLIP | Tarbase |
| hsa-miR-15a-3p | C1orf43 | HITS-CLIP | Tarbase |
| hsa-miR-141-3p | GATAD2B | HITS-CLIP | Tarbase |
| hsa-miR-141-5p | GATAD2B | HITS-CLIP | Tarbase |
| hsa-miR-15a-5p | GATAD2B | HITS-CLIP | Tarbase |
| hsa-miR-15a-5p | GATAD2B | HITS-CLIP | Tarbase |
| hsa-miR-195-5p | GATAD2B | HITS-CLIP | Tarbase |
| hsa-miR-195-5p | RIT1 | Microarrays | Tarbase |
| hsa-miR-15a-5p | GALNT2 | HITS-CLIP | Tarbase |
| hsa-miR-195-5p | GALNT2 | HITS-CLIP | Tarbase |
| hsa-miR-141-5p | LYST | HITS-CLIP | Tarbase |
| hsa-miR-15a-5p | MLK4 | HITS-CLIP | Tarbase |
| hsa-miR-195-5p | MLK4 | HITS-CLIP | Tarbase |
| hsa-miR-486-5p | MLK4 | HITS-CLIP | Tarbase |
| hsa-miR-15a-5p | CEP170 | HITS-CLIP | Tarbase |
| hsa-miR-195-5p | CEP170 | HITS-CLIP | Tarbase |
| hsa-miR-486-3p | CEP170 | HITS-CLIP | Tarbase |
| hsa-miR-486-3p | CEP170 | HITS-CLIP | Tarbase |
| hsa-miR-15a-5p | ACP1 | HITS-CLIP | Tarbase |
| hsa-miR-15a-5p | ACP1 | HITS-CLIP | Tarbase |
| hsa-miR-195-5p | ACP1 | HITS-CLIP | Tarbase |
| hsa-miR-195-5p | DEGS1 | Microarrays | Tarbase |
| hsa-miR-15a-5p | CNIH4 | PAR-CLIP | Tarbase |
| hsa-miR-141-5p | PARP1 | HITS-CLIP | Tarbase |
| hsa-miR-141-5p | PYCR2 | HITS-CLIP | Tarbase |
| hsa-miR-141-5p | PYCR2 | HITS-CLIP | Tarbase |
| hsa-miR-141-5p | PYCR2 | HITS-CLIP | Tarbase |
| hsa-miR-486-5p | PYCR2 | HITS-CLIP | Tarbase |
| hsa-miR-486-5p | PYCR2 | PAR-CLIP | Tarbase |
| hsa-miR-141-3p | LBR | HITS-CLIP | Tarbase |
| hsa-miR-141-5p | LBR | HITS-CLIP | Tarbase |
| hsa-miR-15a-5p | LBR | HITS-CLIP | Tarbase |
| hsa-miR-15a-5p | LBR | HITS-CLIP | Tarbase |
| hsa-miR-15a-5p | LBR | HITS-CLIP | Tarbase |
| hsa-miR-15a-5p | LBR | HITS-CLIP | Tarbase |

| | | | |
|---|---|---|---|
| hsa-miR-15a-5p | LBR | HITS-CLIP | Tarbase |
| hsa-miR-195-5p | LBR | HITS-CLIP | Tarbase |
| hsa-miR-195-5p | LBR | HITS-CLIP | Tarbase |
| hsa-miR-195-5p | LBR | HITS-CLIP | Tarbase |
| hsa-miR-195-5p | LBR | HITS-CLIP | Tarbase |
| hsa-miR-15a-5p | WNT9A | HITS-CLIP | Tarbase |
| hsa-miR-195-5p | WNT9A | HITS-CLIP | Tarbase |
| hsa-miR-141-5p | SOX13 | HITS-CLIP | Tarbase |
| hsa-miR-195-5p | PLEKHA6 | Microarrays | Tarbase |
| hsa-miR-15a-5p | ARL8A | PAR-CLIP | Tarbase |
| hsa-miR-195-5p | OSR1 | Microarrays | Tarbase |
| hsa-miR-15a-5p | PDIA6 | HITS-CLIP | Tarbase |
| hsa-miR-15a-5p | PDIA6 | HITS-CLIP | Tarbase |
| hsa-miR-15a-5p | PDIA6 | HITS-CLIP | Tarbase |
| hsa-miR-15a-5p | PDIA6 | HITS-CLIP | Tarbase |
| hsa-miR-15a-5p | PDIA6 | HITS-CLIP | Tarbase |
| hsa-miR-15a-5p | PDIA6 | HITS-CLIP | Tarbase |
| hsa-miR-15a-5p | PDIA6 | PAR-CLIP | Tarbase |
| hsa-miR-15a-5p | PDIA6 | HITS-CLIP | Tarbase |
| hsa-miR-15a-5p | PDIA6 | HITS-CLIP | Tarbase |
| hsa-miR-195-5p | PDIA6 | HITS-CLIP | Tarbase |
| hsa-miR-195-5p | PDIA6 | HITS-CLIP | Tarbase |
| hsa-miR-195-5p | PDIA6 | HITS-CLIP | Tarbase |
| hsa-miR-195-5p | PDIA6 | HITS-CLIP | Tarbase |
| hsa-miR-195-5p | PDIA6 | HITS-CLIP | Tarbase |
| hsa-miR-195-5p | PDIA6 | HITS-CLIP | Tarbase |
| hsa-miR-195-5p | PDIA6 | HITS-CLIP | Tarbase |
| hsa-miR-141-3p | RHOB | HITS-CLIP | Tarbase |
| hsa-miR-195-5p | RHOB | Microarrays | Tarbase |
| hsa-miR-15a-5p | HNRNPLL | HITS-CLIP | Tarbase |
| hsa-miR-195-5p | HNRNPLL | HITS-CLIP | Tarbase |
| hsa-miR-15a-5p | EML4 | PAR-CLIP | Tarbase |
| hsa-miR-195-5p | EML4 | PAR-CLIP | Tarbase |
| hsa-miR-15a-5p | EML4 | Chimeric fragments | Tarbase |
| hsa-miR-141-3p | CALM2 | HITS-CLIP | Tarbase |
| hsa-miR-219a-1-3p | CALM2 | HITS-CLIP | Tarbase |
| hsa-miR-15a-5p | ASXL2 | HITS-CLIP | Tarbase |
| hsa-miR-195-5p | ASXL2 | HITS-CLIP | Tarbase |
| hsa-miR-15a-5p | MEIS1 | PAR-CLIP | Tarbase |
| hsa-miR-15a-5p | SNRNP200 | HITS-CLIP | Tarbase |
| hsa-miR-195-5p | SNRNP200 | HITS-CLIP | Tarbase |
| hsa-miR-3613-5p | SNRNP200 | HITS-CLIP | Tarbase |
| hsa-miR-3613-5p | SNRNP200 | HITS-CLIP | Tarbase |
| hsa-miR-3613-5p | SNRNP200 | HITS-CLIP | Tarbase |
| hsa-miR-3613-5p | SNRNP200 | PAR-CLIP | Tarbase |
| hsa-miR-15a-5p | TEX261 | PAR-CLIP | Tarbase |
| hsa-miR-195-5p | TEX261 | PAR-CLIP | Tarbase |
| hsa-miR-15a-5p | TEX261 | Chimeric fragments | Tarbase |
| hsa-miR-15a-3p | DUSP11 | HITS-CLIP | Tarbase |
| hsa-miR-141-3p | SLC20A1 | HITS-CLIP | Tarbase |
| hsa-miR-141-3p | SLC20A1 | HITS-CLIP | Tarbase |
| hsa-miR-195-5p | SLC20A1 | HITS-CLIP | Tarbase |
| hsa-miR-3605-5p | SLC20A1 | HITS-CLIP | Tarbase |
| hsa-miR-3613-5p | SLC20A1 | HITS-CLIP | Tarbase |
| hsa-miR-195-5p | SLC20A1 | Microarrays | Tarbase |
| hsa-miR-15a-5p | LIPT1 | HITS-CLIP | Tarbase |
| hsa-miR-195-5p | LIPT1 | HITS-CLIP | Tarbase |
| hsa-miR-195-5p | AFF3 | Microarrays | Tarbase |
| hsa-miR-141-3p | UBXN4 | HITS-CLIP | Tarbase |
| hsa-miR-141-5p | NXPH2 | HITS-CLIP | Tarbase |
| hsa-miR-141-3p | SPOPL | PAR-CLIP | Tarbase |
| hsa-miR-195-5p | SPOPL | Microarrays | Tarbase |
| hsa-miR-15a-5p | POLR2D | PAR-CLIP | Tarbase |
| hsa-miR-15a-5p | POLR2D | HITS-CLIP | Tarbase |
| hsa-miR-15a-5p | POLR2D | PAR-CLIP | Tarbase |
| hsa-miR-15a-5p | POLR2D | PAR-CLIP | Tarbase |
| hsa-miR-15a-5p | POLR2D | PAR-CLIP | Tarbase |
| hsa-miR-195-5p | POLR2D | PAR-CLIP | Tarbase |
| hsa-miR-195-5p | POLR2D | HITS-CLIP | Tarbase |
| hsa-miR-195-5p | POLR2D | PAR-CLIP | Tarbase |
| hsa-miR-15a-5p | AMMECR1L | PAR-CLIP | Tarbase |
| hsa-miR-15a-5p | AMMECR1L | HITS-CLIP | Tarbase |
| hsa-miR-195-5p | AMMECR1L | HITS-CLIP | Tarbase |
| hsa-miR-195-5p | TMEFF2 | Microarrays | Tarbase |
| hsa-miR-195-5p | CDCA7 | Microarrays | Tarbase |
| hsa-miR-15a-5p | UBR3 | PAR-CLIP | Tarbase |
| hsa-miR-15a-5p | UBR3 | PAR-CLIP | Tarbase |
| hsa-miR-15a-5p | UBR3 | HITS-CLIP | Tarbase |
| hsa-miR-15a-5p | UBR3 | HITS-CLIP | Tarbase |
| hsa-miR-15a-5p | UBR3 | PAR-CLIP | Tarbase |
| hsa-miR-15a-5p | UBR3 | HITS-CLIP | Tarbase |
| hsa-miR-195-5p | UBR3 | PAR-CLIP | Tarbase |
| hsa-miR-195-5p | UBR3 | HITS-CLIP | Tarbase |
| hsa-miR-195-5p | UBR3 | HITS-CLIP | Tarbase |
| hsa-miR-195-5p | UBR3 | HITS-CLIP | Tarbase |
| hsa-miR-15a-5p | UBR3 | HITS-CLIP | Tarbase |
| hsa-miR-195-5p | UBR3 | HITS-CLIP | Tarbase |
| hsa-miR-15a-5p | GULP1 | PAR-CLIP | Tarbase |
| hsa-miR-15a-5p | FAM171B | HITS-CLIP | Tarbase |
| hsa-miR-195-5p | FAM171B | HITS-CLIP | Tarbase |
| hsa-miR-15a-5p | FAM171B | Chimeric fragments | Tarbase |
| hsa-miR-195-5p | FAM171B | Chimeric fragments | Tarbase |
| hsa-miR-141-5p | HSPD1 | HITS-CLIP | Tarbase |
| hsa-miR-3605-5p | UNC80 | HITS-CLIP | Tarbase |
| hsa-miR-15a-5p | UNC80 | HITS-CLIP | Tarbase |
| hsa-miR-195-5p | UNC80 | HITS-CLIP | Tarbase |
| hsa-miR-3613-5p | ABCA12 | PAR-CLIP | Tarbase |
| hsa-miR-3613-5p | ABCA12 | PAR-CLIP | Tarbase |
| hsa-miR-3613-5p | ABCA12 | PAR-CLIP | Tarbase |
| hsa-miR-15a-5p | RHBDD1 | PAR-CLIP | Tarbase |
| hsa-miR-15a-5p | RHBDD1 | PAR-CLIP | Tarbase |
| hsa-miR-15a-5p | RHBDD1 | PAR-CLIP | Tarbase |
| hsa-miR-195-5p | RHBDD1 | PAR-CLIP | Tarbase |
| hsa-miR-195-5p | RHBDD1 | PAR-CLIP | Tarbase |
| hsa-miR-15a-5p | ACKR3 | HITS-CLIP | Tarbase |
| hsa-miR-15a-5p | COPS7B | HITS-CLIP | Tarbase |
| hsa-miR-195-5p | COPS7B | HITS-CLIP | Tarbase |
| hsa-miR-195-5p | COPS7B | Microarrays | Tarbase |
| hsa-miR-195-5p | FANCD2 | Microarrays | Tarbase |
| hsa-miR-141-5p | VGLL4 | HITS-CLIP | Tarbase |
| hsa-miR-15a-5p | VGLL4 | HITS-CLIP | Tarbase |
| hsa-miR-195-5p | VGLL4 | HITS-CLIP | Tarbase |
| hsa-miR-3613-3p | VGLL4 | HITS-CLIP | Tarbase |
| hsa-miR-141-3p | RAB5A | HITS-CLIP | Tarbase |
| hsa-miR-486-5p | RAB5A | PAR-CLIP | Tarbase |
| hsa-miR-15a-5p | FAM134A | PAR-CLIP | Tarbase |
| hsa-miR-15a-5p | CTDSP1 | PAR-CLIP | Tarbase |
| hsa-miR-15a-5p | CTDSP1 | HITS-CLIP | Tarbase |
| hsa-miR-195-5p | CTDSP1 | Microarrays | Tarbase |
| hsa-miR-3613-3p | RQCD1 | PAR-CLIP | Tarbase |
| hsa-miR-3605-5p | EAF1 | HITS-CLIP | Tarbase |
| hsa-miR-15a-3p | FAM198A | HITS-CLIP | Tarbase |
| hsa-miR-141-5p | GOLGA4 | HITS-CLIP | Tarbase |
| hsa-miR-141-3p | IQSEC1 | HITS-CLIP | Tarbase |
| hsa-miR-141-3p | PTPRG | HITS-CLIP | Tarbase |
| hsa-miR-195-5p | IL17RD | Microarrays | Tarbase |
| hsa-miR-141-5p | TMF1 | HITS-CLIP | Tarbase |
| hsa-miR-219a-1-3p | TMF1 | PAR-CLIP | Tarbase |
| hsa-miR-486-5p | TMF1 | HITS-CLIP | Tarbase |
| hsa-miR-15a-5p | LRIG1 | HITS-CLIP | Tarbase |
| hsa-miR-195-5p | LRIG1 | HITS-CLIP | Tarbase |
| hsa-miR-141-3p | NFKBIZ | HITS-CLIP | Tarbase |
| hsa-miR-195-5p | COL8A1 | Microarrays | Tarbase |
| hsa-miR-195-5p | PHLDB2 | Microarrays | Tarbase |
| hsa-miR-141-3p | ABHD10 | HITS-CLIP | Tarbase |
| hsa-miR-486-3p | ABHD10 | HITS-CLIP | Tarbase |
| hsa-miR-195-5p | PLA1A | Microarrays | Tarbase |
| hsa-miR-15a-5p | SRPRB | PAR-CLIP | Tarbase |
| hsa-miR-195-5p | MED12L | Microarrays | Tarbase |
| hsa-miR-141-3p | NCEH1 | HITS-CLIP | Tarbase |
| hsa-miR-486-5p | NCEH1 | HITS-CLIP | Tarbase |
| hsa-miR-141-3p | LPP | HITS-CLIP | Tarbase |
| hsa-miR-141-3p | LPP | HITS-CLIP | Tarbase |
| hsa-miR-15a-5p | VPRBP | HITS-CLIP | Tarbase |
| hsa-miR-195-5p | MANF | Microarrays | Tarbase |
| hsa-miR-141-5p | SLIT2 | HITS-CLIP | Tarbase |
| hsa-miR-141-5p | SLIT2 | HITS-CLIP | Tarbase |
| hsa-miR-15a-5p | SLIT2 | HITS-CLIP | Tarbase |
| hsa-miR-15a-5p | SLIT2 | HITS-CLIP | Tarbase |

| | | | |
|---|---|---|---|
| hsa-miR-15a-5p | SLIT2 | HITS-CLIP | Tarbase |
| hsa-miR-15a-5p | SLIT2 | PAR-CLIP | Tarbase |
| hsa-miR-195-5p | SLIT2 | HITS-CLIP | Tarbase |
| hsa-miR-195-5p | SLIT2 | HITS-CLIP | Tarbase |
| hsa-miR-195-5p | SLIT2 | HITS-CLIP | Tarbase |
| hsa-miR-195-5p | SLIT2 | Microarrays | Tarbase |
| hsa-miR-15a-5p | EIF2B5 | HITS-CLIP | Tarbase |
| hsa-miR-195-5p | EIF2B5 | HITS-CLIP | Tarbase |
| hsa-miR-195-5p | ECE2 | Microarrays | Tarbase |
| hsa-miR-195-5p | EPHA5 | Microarrays | Tarbase |
| hsa-miR-195-5p | SLC10A6 | Microarrays | Tarbase |
| hsa-miR-195-5p | SNCA | Microarrays | Tarbase |
| hsa-miR-15a-5p | ANK2 | HITS-CLIP | Tarbase |
| hsa-miR-195-5p | ANK2 | HITS-CLIP | Tarbase |
| hsa-miR-141-3p | CCNA2 | HITS-CLIP | Tarbase |
| hsa-miR-141-3p | CCNA2 | HITS-CLIP | Tarbase |
| hsa-miR-15a-5p | CCNA2 | HITS-CLIP | Tarbase |
| hsa-miR-195-5p | CCNA2 | HITS-CLIP | Tarbase |
| hsa-miR-195-5p | CCNA2 | Microarrays | Tarbase |
| hsa-miR-141-3p | USP53 | PAR-CLIP | Tarbase |
| hsa-miR-141-3p | SETD7 | HITS-CLIP | Tarbase |
| hsa-miR-15a-5p | PDGFC | HITS-CLIP | Tarbase |
| hsa-miR-195-3p | PDGFC | HITS-CLIP | Tarbase |
| hsa-miR-195-5p | PDGFC | HITS-CLIP | Tarbase |
| hsa-miR-15a-5p | MARCH6 | PAR-CLIP | Tarbase |
| hsa-miR-195-5p | CDH18 | Microarrays | Tarbase |
| hsa-miR-219a-1-3p | MYO10 | HITS-CLIP | Tarbase |
| hsa-miR-195-5p | MYO10 | Microarrays | Tarbase |
| hsa-miR-195-5p | OSMR | Microarrays | Tarbase |
| hsa-miR-195-5p | PLK2 | Microarrays | Tarbase |
| hsa-miR-141-3p | PIK3R1 | HITS-CLIP | Tarbase |
| hsa-miR-141-5p | PIK3R1 | HITS-CLIP | Tarbase |
| hsa-miR-195-5p | LHFPL2 | Microarrays | Tarbase |
| hsa-miR-15a-3p | RASA1 | HITS-CLIP | Tarbase |
| hsa-miR-15a-5p | RASA1 | HITS-CLIP | Tarbase |
| hsa-miR-15a-5p | RASA1 | PAR-CLIP | Tarbase |
| hsa-miR-15a-5p | RASA1 | PAR-CLIP | Tarbase |
| hsa-miR-195-5p | RASA1 | HITS-CLIP | Tarbase |
| hsa-miR-195-5p | RASA1 | PAR-CLIP | Tarbase |
| hsa-miR-195-5p | RASA1 | PAR-CLIP | Tarbase |
| hsa-miR-15a-5p | PAM | HITS-CLIP | Tarbase |
| hsa-miR-15a-5p | PAM | PAR-CLIP | Tarbase |
| hsa-miR-195-5p | PAM | HITS-CLIP | Tarbase |
| hsa-miR-15a-5p | PAM | HITS-CLIP | Tarbase |
| hsa-miR-195-5p | PAM | HITS-CLIP | Tarbase |
| hsa-miR-3613-3p | BDP1 | HITS-CLIP | Tarbase |
| hsa-miR-15a-5p | BDP1 | HITS-CLIP | Tarbase |
| hsa-miR-195-5p | BDP1 | HITS-CLIP | Tarbase |
| hsa-miR-141-3p | FBXL17 | HITS-CLIP | Tarbase |
| hsa-miR-3613-3p | TNFAIP8 | HITS-CLIP | Tarbase |
| hsa-miR-15a-5p | FEM1C | Chimeric fragments | Tarbase |
| hsa-miR-195-5p | COMMD10 | Microarrays | Tarbase |
| hsa-miR-15a-5p | YIPF5 | PAR-CLIP | Tarbase |
| hsa-miR-195-5p | CXCL14 | Microarrays | Tarbase |
| hsa-miR-15a-5p | DDX46 | HITS-CLIP | Tarbase |
| hsa-miR-195-5p | DDX46 | HITS-CLIP | Tarbase |
| hsa-miR-141-5p | RNF145 | HITS-CLIP | Tarbase |
| hsa-miR-141-3p | G3BP1 | HITS-CLIP | Tarbase |
| hsa-miR-15a-3p | G3BP1 | HITS-CLIP | Tarbase |
| hsa-miR-15a-5p | G3BP1 | PAR-CLIP | Tarbase |
| hsa-miR-195-5p | G3BP1 | PAR-CLIP | Tarbase |
| hsa-miR-3613-3p | G3BP1 | PAR-CLIP | Tarbase |
| hsa-miR-486-5p | G3BP1 | HITS-CLIP | Tarbase |
| hsa-miR-486-5p | G3BP1 | HITS-CLIP | Tarbase |
| hsa-miR-195-5p | KCNMB1 | Microarrays | Tarbase |
| hsa-miR-141-5p | TBC1D7 | HITS-CLIP | Tarbase |
| hsa-miR-15a-5p | LRRTM2 | HITS-CLIP | Tarbase |
| hsa-miR-195-5p | LRRTM2 | HITS-CLIP | Tarbase |
| hsa-miR-486-5p | HIGD2A | HITS-CLIP | Tarbase |
| hsa-miR-486-5p | HIGD2A | HITS-CLIP | Tarbase |
| hsa-miR-486-5p | HIGD2A | HITS-CLIP | Tarbase |
| hsa-miR-486-5p | HIGD2A | HITS-CLIP | Tarbase |
| hsa-miR-141-3p | RNF44 | HITS-CLIP | Tarbase |
| hsa-miR-141-5p | RNF44 | HITS-CLIP | Tarbase |
| hsa-miR-195-5p | MUT | HITS-CLIP | Tarbase |
| hsa-miR-15a-5p | PPP1R18 | PAR-CLIP | Tarbase |
| hsa-miR-486-3p | PPP1R18 | HITS-CLIP | Tarbase |
| hsa-miR-195-5p | DAAM2 | Microarrays | Tarbase |
| hsa-miR-141-3p | TPBG | HITS-CLIP | Tarbase |
| hsa-miR-141-3p | TPBG | HITS-CLIP | Tarbase |
| hsa-miR-141-3p | IRAK1BP1 | PAR-CLIP | Tarbase |
| hsa-miR-141-3p | PHIP | HITS-CLIP | Tarbase |
| hsa-miR-141-3p | PHIP | HITS-CLIP | Tarbase |
| hsa-miR-195-5p | PHIP | HITS-CLIP | Tarbase |
| hsa-miR-486-5p | PHIP | HITS-CLIP | Tarbase |
| hsa-miR-195-5p | PHIP | Microarrays | Tarbase |
| hsa-miR-195-5p | MMS22L | Microarrays | Tarbase |
| hsa-miR-3613-3p | PNRC1 | HITS-CLIP | Tarbase |
| hsa-miR-141-5p | RNF217 | HITS-CLIP | Tarbase |
| hsa-miR-141-5p | RNF217 | HITS-CLIP | Tarbase |
| hsa-miR-15a-5p | RNF217 | PAR-CLIP | Tarbase |
| hsa-miR-15a-5p | RNF217 | PAR-CLIP | Tarbase |
| hsa-miR-15a-5p | RNF217 | PAR-CLIP | Tarbase |
| hsa-miR-15a-5p | RNF217 | HITS-CLIP | Tarbase |
| hsa-miR-15a-5p | RNF217 | HITS-CLIP | Tarbase |
| hsa-miR-195-5p | RNF217 | HITS-CLIP | Tarbase |
| hsa-miR-15a-5p | ARHGAP18 | HITS-CLIP | Tarbase |
| hsa-miR-141-5p | ABRACL | HITS-CLIP | Tarbase |
| hsa-miR-195-5p | MTFR2 | Microarrays | Tarbase |
| hsa-miR-15a-5p | SHPRH | HITS-CLIP | Tarbase |
| hsa-miR-195-5p | SHPRH | HITS-CLIP | Tarbase |
| hsa-miR-15a-5p | AIG1 | HITS-CLIP | Tarbase |
| hsa-miR-195-5p | AIG1 | HITS-CLIP | Tarbase |
| hsa-miR-3605-5p | TIAM2 | PAR-CLIP | Tarbase |
| hsa-miR-141-5p | TMEM181 | HITS-CLIP | Tarbase |
| hsa-miR-15a-5p | TMEM181 | HITS-CLIP | Tarbase |
| hsa-miR-195-5p | TMEM181 | HITS-CLIP | Tarbase |
| hsa-miR-3613-5p | TMEM181 | HITS-CLIP | Tarbase |
| hsa-miR-195-5p | TMEM181 | Microarrays | Tarbase |
| hsa-miR-141-3p | WTAP | HITS-CLIP | Tarbase |
| hsa-miR-15a-5p | WTAP | HITS-CLIP | Tarbase |
| hsa-miR-195-5p | WTAP | HITS-CLIP | Tarbase |
| hsa-miR-15a-5p | WTAP | HITS-CLIP | Tarbase |
| hsa-miR-195-5p | WTAP | HITS-CLIP | Tarbase |
| hsa-miR-141-3p | ZMYM4 | HITS-CLIP | Tarbase |
| hsa-miR-141-3p | ZMYM4 | HITS-CLIP | Tarbase |
| hsa-miR-141-3p | ZMYM4 | HITS-CLIP | Tarbase |
| hsa-miR-141-3p | GNA12 | HITS-CLIP | Tarbase |
| hsa-miR-15a-3p | GNA12 | HITS-CLIP | Tarbase |
| hsa-miR-15a-5p | GNA12 | HITS-CLIP | Tarbase |
| hsa-miR-195-5p | GNA12 | HITS-CLIP | Tarbase |
| hsa-miR-15a-5p | CDCA5 | PAR-CLIP | Tarbase |
| hsa-miR-195-5p | CDCA5 | Microarrays | Tarbase |
| hsa-miR-15a-5p | IGFBP3 | HITS-CLIP | Tarbase |
| hsa-miR-195-5p | IGFBP3 | HITS-CLIP | Tarbase |
| hsa-miR-195-5p | IGFBP3 | Microarrays | Tarbase |
| hsa-miR-15a-5p | MDH2 | PAR-CLIP | Tarbase |
| hsa-miR-195-5p | MDH2 | PAR-CLIP | Tarbase |
| hsa-miR-141-5p | ZNF92 | HITS-CLIP | Tarbase |
| hsa-miR-141-3p | GIGYF1 | PAR-CLIP | Tarbase |
| hsa-miR-141-3p | GIGYF1 | HITS-CLIP | Tarbase |
| hsa-miR-141-5p | GIGYF1 | HITS-CLIP | Tarbase |
| hsa-miR-141-5p | GIGYF1 | HITS-CLIP | Tarbase |
| hsa-miR-15a-3p | GIGYF1 | HITS-CLIP | Tarbase |
| hsa-miR-15a-3p | GIGYF1 | HITS-CLIP | Tarbase |
| hsa-miR-15a-3p | GIGYF1 | HITS-CLIP | Tarbase |
| hsa-miR-15a-5p | GIGYF1 | HITS-CLIP | Tarbase |
| hsa-miR-15a-5p | GIGYF1 | HITS-CLIP | Tarbase |
| hsa-miR-15a-5p | GIGYF1 | HITS-CLIP | Tarbase |
| hsa-miR-15a-5p | GIGYF1 | PAR-CLIP | Tarbase |
| hsa-miR-15a-5p | GIGYF1 | HITS-CLIP | Tarbase |
| hsa-miR-15a-5p | GIGYF1 | HITS-CLIP | Tarbase |
| hsa-miR-195-5p | GIGYF1 | HITS-CLIP | Tarbase |
| hsa-miR-195-5p | GIGYF1 | HITS-CLIP | Tarbase |
| hsa-miR-195-5p | GIGYF1 | HITS-CLIP | Tarbase |
| hsa-miR-195-5p | GIGYF1 | Microarrays | Tarbase |
| hsa-miR-195-5p | ZC3HAV1L | Microarrays | Tarbase |
| hsa-miR-195-5p | NCAPG2 | Microarrays | Tarbase |
| hsa-miR-141-5p | C7orf55-LUC7L2 | HITS-CLIP | Tarbase |

| | | | |
|---|---|---|---|
| hsa-miR-141-5p | C7orf55-LUC7L2 | HITS-CLIP | Tarbase |
| hsa-miR-195-5p | DENND2A | Microarrays | Tarbase |
| hsa-miR-15a-5p | TMEM47 | HITS-CLIP | Tarbase |
| hsa-miR-195-5p | TMEM47 | HITS-CLIP | Tarbase |
| hsa-miR-195-5p | SYTL5 | Microarrays | Tarbase |
| hsa-miR-141-5p | KDM6A | HITS-CLIP | Tarbase |
| hsa-miR-15a-5p | KDM6A | PAR-CLIP | Tarbase |
| hsa-miR-15a-5p | KDM6A | HITS-CLIP | Tarbase |
| hsa-miR-195-5p | KDM6A | HITS-CLIP | Tarbase |
| hsa-miR-15a-5p | MSN | HITS-CLIP | Tarbase |
| hsa-miR-195-5p | MSN | HITS-CLIP | Tarbase |
| hsa-miR-141-5p | TAF1 | HITS-CLIP | Tarbase |
| hsa-miR-486-5p | TAF1 | HITS-CLIP | Tarbase |
| hsa-miR-195-5p | GPR174 | Microarrays | Tarbase |
| hsa-miR-141-3p | NONO | HITS-CLIP | Tarbase |
| hsa-miR-15a-5p | NONO | HITS-CLIP | Tarbase |
| hsa-miR-195-5p | NONO | HITS-CLIP | Tarbase |
| hsa-miR-141-3p | OGT | HITS-CLIP | Tarbase |
| hsa-miR-15a-5p | OGT | PAR-CLIP | Tarbase |
| hsa-miR-15a-5p | OGT | HITS-CLIP | Tarbase |
| hsa-miR-15a-5p | OGT | HITS-CLIP | Tarbase |
| hsa-miR-195-5p | OGT | HITS-CLIP | Tarbase |
| hsa-miR-219a-1-3p | OGT | HITS-CLIP | Tarbase |
| hsa-miR-15a-5p | OGT | HITS-CLIP | Tarbase |
| hsa-miR-195-5p | OGT | HITS-CLIP | Tarbase |
| hsa-miR-15a-5p | OGT | Chimeric fragments | Tarbase |
| hsa-miR-141-5p | ZNF711 | HITS-CLIP | Tarbase |
| hsa-miR-141-5p | ZNF711 | HITS-CLIP | Tarbase |
| hsa-miR-141-5p | ZNF711 | HITS-CLIP | Tarbase |
| hsa-miR-3613-5p | ZNF711 | HITS-CLIP | Tarbase |
| hsa-miR-486-5p | ZNF711 | HITS-CLIP | Tarbase |
| hsa-miR-3613-5p | ZNF711 | Chimeric fragments | Tarbase |
| hsa-miR-195-5p | CPXCR1 | Microarrays | Tarbase |
| hsa-miR-15a-5p | FRMPD3 | HITS-CLIP | Tarbase |
| hsa-miR-195-5p | FRMPD3 | HITS-CLIP | Tarbase |
| hsa-miR-15a-5p | GPC3 | PAR-CLIP | Tarbase |
| hsa-miR-15a-5p | MFHAS1 | HITS-CLIP | Tarbase |
| hsa-miR-195-5p | CSGALNACT1 | Microarrays | Tarbase |
| hsa-miR-15a-5p | ATP6V1B2 | HITS-CLIP | Tarbase |
| hsa-miR-141-5p | HMBOX1 | HITS-CLIP | Tarbase |
| hsa-miR-3605-5p | HMBOX1 | PAR-CLIP | Tarbase |
| hsa-miR-195-5p | HMBOX1 | Microarrays | Tarbase |
| hsa-miR-3613-3p | SLC25A37 | HITS-CLIP | Tarbase |
| hsa-miR-15a-3p | DOCK5 | HITS-CLIP | Tarbase |
| hsa-miR-15a-5p | PROSC | PAR-CLIP | Tarbase |
| hsa-miR-15a-5p | ERLIN2 | HITS-CLIP | Tarbase |
| hsa-miR-15a-5p | ERLIN2 | HITS-CLIP | Tarbase |
| hsa-miR-15a-5p | ERLIN2 | PAR-CLIP | Tarbase |
| hsa-miR-195-5p | ERLIN2 | HITS-CLIP | Tarbase |
| hsa-miR-15a-3p | TACC1 | HITS-CLIP | Tarbase |
| hsa-miR-15a-5p | TACC1 | HITS-CLIP | Tarbase |
| hsa-miR-195-5p | TACC1 | HITS-CLIP | Tarbase |
| hsa-miR-3605-5p | TACC1 | HITS-CLIP | Tarbase |
| hsa-miR-3613-3p | TACC1 | PAR-CLIP | Tarbase |
| hsa-miR-15a-5p | TACC1 | HITS-CLIP | Tarbase |
| hsa-miR-195-5p | TACC1 | HITS-CLIP | Tarbase |
| hsa-miR-195-5p | GINS4 | Microarrays | Tarbase |
| hsa-miR-15a-5p | WHSC1L1 | HITS-CLIP | Tarbase |
| hsa-miR-195-5p | WHSC1L1 | HITS-CLIP | Tarbase |
| hsa-miR-15a-5p | PMP2 | HITS-CLIP | Tarbase |
| hsa-miR-195-5p | PMP2 | HITS-CLIP | Tarbase |
| hsa-miR-195-5p | TERF1 | Microarrays | Tarbase |
| hsa-miR-195-5p | ATP6V0D2 | Microarrays | Tarbase |
| hsa-miR-195-5p | SYBU | Microarrays | Tarbase |
| hsa-miR-219a-1-3p | MTDH | PAR-CLIP | Tarbase |
| hsa-miR-15a-5p | LRP12 | PAR-CLIP | Tarbase |
| hsa-miR-15a-3p | FAM83A | HITS-CLIP | Tarbase |
| hsa-miR-141-3p | NFIB | HITS-CLIP | Tarbase |
| hsa-miR-141-3p | NFIB | HITS-CLIP | Tarbase |
| hsa-miR-15a-5p | NFIB | HITS-CLIP | Tarbase |
| hsa-miR-195-5p | NFIB | HITS-CLIP | Tarbase |
| hsa-miR-15a-5p | NFIB | CLEAR-CLIP | Tarbase |
| hsa-miR-15a-5p | FBXO10 | PAR-CLIP | Tarbase |
| hsa-miR-15a-5p | NTRK2 | HITS-CLIP | Tarbase |
| hsa-miR-195-5p | NTRK2 | HITS-CLIP | Tarbase |
| hsa-miR-15a-3p | HIATL1 | HITS-CLIP | Tarbase |
| hsa-miR-15a-5p | HIATL1 | HITS-CLIP | Tarbase |
| hsa-miR-15a-5p | HIATL1 | HITS-CLIP | Tarbase |
| hsa-miR-195-5p | HIATL1 | HITS-CLIP | Tarbase |
| hsa-miR-195-5p | INIP | Microarrays | Tarbase |
| hsa-miR-3613-3p | UGCG | HITS-CLIP | Tarbase |
| hsa-miR-195-5p | UGCG | Microarrays | Tarbase |
| hsa-miR-15a-5p | SNX30 | PAR-CLIP | Tarbase |
| hsa-miR-195-5p | SNX30 | PAR-CLIP | Tarbase |
| hsa-miR-15a-5p | STOM | HITS-CLIP | Tarbase |
| hsa-miR-195-5p | STOM | HITS-CLIP | Tarbase |
| hsa-miR-141-3p | NR6A1 | HITS-CLIP | Tarbase |
| hsa-miR-141-5p | NR6A1 | HITS-CLIP | Tarbase |
| hsa-miR-15a-3p | NR6A1 | HITS-CLIP | Tarbase |
| hsa-miR-195-5p | ALAD | HITS-CLIP | Tarbase |
| hsa-miR-141-5p | WDR31 | HITS-CLIP | Tarbase |
| hsa-miR-141-5p | POLE3 | HITS-CLIP | Tarbase |
| hsa-miR-15a-5p | SURF4 | HITS-CLIP | Tarbase |
| hsa-miR-195-5p | SURF4 | HITS-CLIP | Tarbase |
| hsa-miR-3605-3p | MED22 | HITS-CLIP | Tarbase |
| hsa-miR-486-5p | RPL7A | PAR-CLIP | Tarbase |
| hsa-miR-15a-5p | NTMT1 | HITS-CLIP | Tarbase |
| hsa-miR-195-5p | NTMT1 | HITS-CLIP | Tarbase |
| hsa-miR-15a-5p | INPP5E | HITS-CLIP | Tarbase |
| hsa-miR-15a-5p | INPP5E | HITS-CLIP | Tarbase |
| hsa-miR-195-5p | INPP5E | HITS-CLIP | Tarbase |
| hsa-miR-195-5p | INPP5E | HITS-CLIP | Tarbase |
| hsa-miR-3605-5p | SEC16A | HITS-CLIP | Tarbase |
| hsa-miR-486-5p | SEC16A | PAR-CLIP | Tarbase |
| hsa-miR-15a-5p | NACC2 | PAR-CLIP | Tarbase |
| hsa-miR-15a-5p | NACC2 | HITS-CLIP | Tarbase |
| hsa-miR-195-5p | NACC2 | HITS-CLIP | Tarbase |
| hsa-miR-141-3p | USP6NL | HITS-CLIP | Tarbase |
| hsa-miR-15a-5p | FAM171A1 | PAR-CLIP | Tarbase |
| hsa-miR-195-5p | FAM171A1 | Microarrays | Tarbase |
| hsa-miR-195-5p | RSU1 | Microarrays | Tarbase |
| hsa-miR-141-3p | ZEB1 | Other | Tarbase |
| hsa-miR-141-3p | ZEB1 | qPCR | Tarbase |
| hsa-miR-141-3p | ZEB1 | HITS-CLIP | Tarbase |
| hsa-miR-141-3p | ZEB1 | Luciferase Reporter Assay | Tarbase |
| hsa-miR-141-3p | ZEB1 | Luciferase Reporter Assay | Tarbase |
| hsa-miR-15a-5p | POLR3A | PAR-CLIP | Tarbase |
| hsa-miR-195-5p | ANKRD1 | Microarrays | Tarbase |
| hsa-miR-195-5p | HTR7 | Microarrays | Tarbase |
| hsa-miR-141-5p | EIF4EBP2 | HITS-CLIP | Tarbase |
| hsa-miR-195-5p | PLEKHS1 | Microarrays | Tarbase |
| hsa-miR-141-3p | MKI67 | HITS-CLIP | Tarbase |
| hsa-miR-195-5p | MKI67 | Microarrays | Tarbase |
| hsa-miR-195-5p | TAF5 | Microarrays | Tarbase |
| hsa-miR-15a-5p | PPRC1 | HITS-CLIP | Tarbase |
| hsa-miR-15a-5p | PPRC1 | PAR-CLIP | Tarbase |
| hsa-miR-195-5p | PPRC1 | HITS-CLIP | Tarbase |
| hsa-miR-3605-5p | PPRC1 | HITS-CLIP | Tarbase |
| hsa-miR-15a-5p | CNNM2 | Chimeric fragments | Tarbase |
| hsa-miR-195-5p | ADAM12 | Microarrays | Tarbase |
| hsa-miR-141-5p | ADM | HITS-CLIP | Tarbase |
| hsa-miR-141-5p | ADM | HITS-CLIP | Tarbase |
| hsa-miR-195-5p | ADM | Microarrays | Tarbase |
| hsa-miR-3613-3p | LIN7C | HITS-CLIP | Tarbase |
| hsa-miR-195-5p | LRRC4C | Microarrays | Tarbase |
| hsa-miR-15a-5p | TUT1 | HITS-CLIP | Tarbase |
| hsa-miR-15a-5p | TUT1 | PAR-CLIP | Tarbase |
| hsa-miR-195-5p | TUT1 | HITS-CLIP | Tarbase |
| hsa-miR-15a-5p | TNKS1BP1 | HITS-CLIP | Tarbase |
| hsa-miR-195-5p | TNKS1BP1 | HITS-CLIP | Tarbase |
| hsa-miR-3605-5p | SSRP1 | PAR-CLIP | Tarbase |
| hsa-miR-15a-5p | C11orf49 | HITS-CLIP | Tarbase |
| hsa-miR-195-5p | C11orf49 | HITS-CLIP | Tarbase |
| hsa-miR-15a-5p | ARFGAP2 | HITS-CLIP | Tarbase |
| hsa-miR-141-3p | CELF1 | HITS-CLIP | Tarbase |
| hsa-miR-15a-5p | CELF1 | HITS-CLIP | Tarbase |
| hsa-miR-15a-5p | CELF1 | HITS-CLIP | Tarbase |
| hsa-miR-195-5p | CELF1 | HITS-CLIP | Tarbase |
| hsa-miR-195-5p | CELF1 | HITS-CLIP | Tarbase |

| | | | |
|---|---|---|---|
| hsa-miR-219a-1-3p | CELF1 | HITS-CLIP | Tarbase |
| hsa-miR-3605-5p | CELF1 | HITS-CLIP | Tarbase |
| hsa-miR-3613-5p | CELF1 | HITS-CLIP | Tarbase |
| hsa-miR-15a-5p | SESN3 | HITS-CLIP | Tarbase |
| hsa-miR-15a-5p | SESN3 | HITS-CLIP | Tarbase |
| hsa-miR-15a-5p | SESN3 | PAR-CLIP | Tarbase |
| hsa-miR-195-5p | SESN3 | HITS-CLIP | Tarbase |
| hsa-miR-195-5p | SESN3 | HITS-CLIP | Tarbase |
| hsa-miR-15a-5p | SESN3 | HITS-CLIP | Tarbase |
| hsa-miR-195-5p | SESN3 | HITS-CLIP | Tarbase |
| hsa-miR-141-3p | ENDOD1 | HITS-CLIP | Tarbase |
| hsa-miR-486-5p | ENDOD1 | PAR-CLIP | Tarbase |
| hsa-miR-15a-5p | CCDC82 | PAR-CLIP | Tarbase |
| hsa-miR-15a-5p | SERPINH1 | HITS-CLIP | Tarbase |
| hsa-miR-3613-5p | ZC3H12C | HITS-CLIP | Tarbase |
| hsa-miR-195-5p | NCAM1 | Microarrays | Tarbase |
| hsa-miR-141-3p | SLX4IP | HITS-CLIP | Tarbase |
| hsa-miR-15a-5p | LAMTOR1 | HITS-CLIP | Tarbase |
| hsa-miR-15a-5p | LAMTOR1 | HITS-CLIP | Tarbase |
| hsa-miR-195-5p | LAMTOR1 | HITS-CLIP | Tarbase |
| hsa-miR-195-5p | LAMTOR1 | HITS-CLIP | Tarbase |
| hsa-miR-141-5p | HYOU1 | HITS-CLIP | Tarbase |
| hsa-miR-15a-5p | CSRP2BP | HITS-CLIP | Tarbase |
| hsa-miR-195-5p | CSRP2BP | HITS-CLIP | Tarbase |
| hsa-miR-15a-5p | TMEM138 | Chimeric fragments | Tarbase |
| hsa-miR-15a-3p | PLCH2 | HITS-CLIP | Tarbase |
| hsa-miR-3605-5p | PLCH2 | HITS-CLIP | Tarbase |
| hsa-miR-15a-3p | CPSF7 | HITS-CLIP | Tarbase |
| hsa-miR-15a-5p | CPSF7 | PAR-CLIP | Tarbase |
| hsa-miR-15a-5p | CPSF7 | PAR-CLIP | Tarbase |
| hsa-miR-486-5p | CPSF7 | PAR-CLIP | Tarbase |
| hsa-miR-141-3p | EI24 | HITS-CLIP | Tarbase |
| hsa-miR-3613-5p | EI24 | HITS-CLIP | Tarbase |
| hsa-miR-195-5p | CCDC15 | Microarrays | Tarbase |
| hsa-miR-15a-5p | CHEK1 | PAR-CLIP | Tarbase |
| hsa-miR-219a-1-3p | CHEK1 | HITS-CLIP | Tarbase |
| hsa-miR-141-5p | FEZ1 | HITS-CLIP | Tarbase |
| hsa-miR-195-5p | MPZL2 | Microarrays | Tarbase |
| hsa-miR-195-5p | TAGLN | Microarrays | Tarbase |
| hsa-miR-195-5p | DSN1 | Microarrays | Tarbase |
| hsa-miR-141-3p | SOGA1 | HITS-CLIP | Tarbase |
| hsa-miR-141-3p | SOGA1 | HITS-CLIP | Tarbase |
| hsa-miR-15a-5p | SOGA1 | HITS-CLIP | Tarbase |
| hsa-miR-195-5p | SOGA1 | HITS-CLIP | Tarbase |
| hsa-miR-3613-5p | SOGA1 | PAR-CLIP | Tarbase |
| hsa-miR-15a-5p | SOGA1 | HITS-CLIP | Tarbase |
| hsa-miR-195-5p | SOGA1 | HITS-CLIP | Tarbase |
| hsa-miR-141-3p | LSM14B | HITS-CLIP | Tarbase |
| hsa-miR-15a-5p | LSM14B | HITS-CLIP | Tarbase |
| hsa-miR-195-5p | LSM14B | HITS-CLIP | Tarbase |
| hsa-miR-15a-5p | YTHDF1 | HITS-CLIP | Tarbase |
| hsa-miR-15a-5p | YTHDF1 | HITS-CLIP | Tarbase |
| hsa-miR-195-5p | YTHDF1 | HITS-CLIP | Tarbase |
| hsa-miR-3613-5p | YTHDF1 | HITS-CLIP | Tarbase |
| hsa-miR-15a-5p | YTHDF1 | HITS-CLIP | Tarbase |
| hsa-miR-195-5p | YTHDF1 | HITS-CLIP | Tarbase |
| hsa-miR-195-5p | CABLES2 | Microarrays | Tarbase |
| hsa-miR-15a-5p | TM7SF2 | PAR-CLIP | Tarbase |
| hsa-miR-15a-5p | TM7SF2 | HITS-CLIP | Tarbase |
| hsa-miR-195-5p | TM7SF2 | HITS-CLIP | Tarbase |
| hsa-miR-15a-5p | PPP4C | Chimeric fragments | Tarbase |
| hsa-miR-195-5p | ALDOA | HITS-CLIP | Tarbase |
| hsa-miR-486-3p | ALDOA | PAR-CLIP | Tarbase |
| hsa-miR-15a-3p | TAOK2 | HITS-CLIP | Tarbase |
| hsa-miR-15a-5p | TAOK2 | HITS-CLIP | Tarbase |
| hsa-miR-15a-5p | TAOK2 | HITS-CLIP | Tarbase |
| hsa-miR-195-5p | TAOK2 | HITS-CLIP | Tarbase |
| hsa-miR-195-5p | TAOK2 | HITS-CLIP | Tarbase |
| hsa-miR-15a-3p | TMEM219 | HITS-CLIP | Tarbase |
| hsa-miR-15a-5p | HMGA2 | PAR-CLIP | Tarbase |
| hsa-miR-15a-5p | HMGA2 | Luciferase Reporter Assay | Tarbase |
| hsa-miR-195-5p | MMP3 | Microarrays | Tarbase |
| hsa-miR-195-5p | CLEC1A | Microarrays | Tarbase |
| hsa-miR-195-5p | PCDH15 | Microarrays | Tarbase |
| hsa-miR-141-3p | ARID5B | HITS-CLIP | Tarbase |
| hsa-miR-15a-5p | ARID5B | HITS-CLIP | Tarbase |
| hsa-miR-15a-5p | ARID5B | PAR-CLIP | Tarbase |
| hsa-miR-15a-5p | ARID5B | PAR-CLIP | Tarbase |
| hsa-miR-195-5p | ARID5B | PAR-CLIP | Tarbase |
| hsa-miR-219a-1-3p | ARID5B | HITS-CLIP | Tarbase |
| hsa-miR-3605-5p | ARID5B | HITS-CLIP | Tarbase |
| hsa-miR-195-5p | ARID5B | Microarrays | Tarbase |
| hsa-miR-486-5p | DCUN1D2 | HITS-CLIP | Tarbase |
| hsa-miR-195-5p | DCUN1D2 | Microarrays | Tarbase |
| hsa-miR-15a-5p | LATS2 | HITS-CLIP | Tarbase |
| hsa-miR-15a-5p | LATS2 | PAR-CLIP | Tarbase |
| hsa-miR-15a-5p | LATS2 | PAR-CLIP | Tarbase |
| hsa-miR-15a-5p | LATS2 | HITS-CLIP | Tarbase |
| hsa-miR-15a-5p | KIAA1328 | HITS-CLIP | Tarbase |
| hsa-miR-195-5p | KIAA1328 | HITS-CLIP | Tarbase |
| hsa-miR-195-5p | LYPD1 | Microarrays | Tarbase |
| hsa-miR-195-5p | LYPD6B | Microarrays | Tarbase |
| hsa-miR-15a-5p | PDCD4 | HITS-CLIP | Tarbase |
| hsa-miR-15a-5p | PDCD4 | HITS-CLIP | Tarbase |
| hsa-miR-15a-5p | PDCD4 | HITS-CLIP | Tarbase |
| hsa-miR-15a-5p | PDCD4 | PAR-CLIP | Tarbase |
| hsa-miR-15a-5p | PDCD4 | HITS-CLIP | Tarbase |
| hsa-miR-15a-5p | PDCD4 | HITS-CLIP | Tarbase |
| hsa-miR-195-5p | PDCD4 | HITS-CLIP | Tarbase |
| hsa-miR-195-5p | PDCD4 | HITS-CLIP | Tarbase |
| hsa-miR-195-5p | PDCD4 | HITS-CLIP | Tarbase |
| hsa-miR-195-5p | PDCD4 | HITS-CLIP | Tarbase |
| hsa-miR-15a-5p | PDCD4 | HITS-CLIP | Tarbase |
| hsa-miR-195-5p | PDCD4 | HITS-CLIP | Tarbase |
| hsa-miR-15a-5p | CCDC102B | PAR-CLIP | Tarbase |
| hsa-miR-195-5p | CCDC102B | PAR-CLIP | Tarbase |
| hsa-miR-195-5p | RGS18 | Microarrays | Tarbase |
| hsa-miR-195-5p | PRSS23 | Microarrays | Tarbase |
| hsa-miR-141-3p | MTMR12 | HITS-CLIP | Tarbase |
| hsa-miR-195-5p | PPP1R1C | Microarrays | Tarbase |
| hsa-miR-15a-5p | CCT5 | Chimeric fragments | Tarbase |
| hsa-miR-195-5p | DOCK1 | Microarrays | Tarbase |
| hsa-miR-15a-5p | DIXDC1 | PAR-CLIP | Tarbase |
| hsa-miR-195-5p | FREM2 | HITS-CLIP | Tarbase |
| hsa-miR-141-3p | FOXO1 | HITS-CLIP | Tarbase |
| hsa-miR-15a-5p | FOXO1 | Chimeric fragments | Tarbase |
| hsa-miR-15a-3p | CRIM1 | HITS-CLIP | Tarbase |
| hsa-miR-3605-5p | SEC24D | HITS-CLIP | Tarbase |
| hsa-miR-195-5p | ABCB9 | HITS-CLIP | Tarbase |
| hsa-miR-141-5p | UBC | HITS-CLIP | Tarbase |
| hsa-miR-15a-5p | ITPR1 | HITS-CLIP | Tarbase |
| hsa-miR-195-5p | ITPR1 | HITS-CLIP | Tarbase |
| hsa-miR-141-5p | CCRN4L | HITS-CLIP | Tarbase |
| hsa-miR-141-5p | UEVLD | HITS-CLIP | Tarbase |
| hsa-miR-15a-5p | TMEM263 | HITS-CLIP | Tarbase |
| hsa-miR-15a-3p | UBE3B | HITS-CLIP | Tarbase |
| hsa-miR-219a-1-3p | UBE3B | HITS-CLIP | Tarbase |
| hsa-miR-486-5p | UBE3B | HITS-CLIP | Tarbase |
| hsa-miR-15a-5p | ANK3 | HITS-CLIP | Tarbase |
| hsa-miR-15a-5p | ANK3 | HITS-CLIP | Tarbase |
| hsa-miR-195-5p | ANK3 | HITS-CLIP | Tarbase |
| hsa-miR-141-5p | IPMK | HITS-CLIP | Tarbase |
| hsa-miR-3613-3p | IPMK | HITS-CLIP | Tarbase |
| hsa-miR-195-5p | PLBD2 | Microarrays | Tarbase |
| hsa-miR-486-5p | GXYLT1 | HITS-CLIP | Tarbase |
| hsa-miR-486-5p | GXYLT1 | HITS-CLIP | Tarbase |
| hsa-miR-486-5p | GXYLT1 | HITS-CLIP | Tarbase |
| hsa-miR-486-5p | GXYLT1 | HITS-CLIP | Tarbase |
| hsa-miR-195-5p | AKAP6 | Microarrays | Tarbase |
| hsa-miR-15a-5p | MBIP | HITS-CLIP | Tarbase |
| hsa-miR-195-5p | MBIP | HITS-CLIP | Tarbase |
| hsa-miR-15a-5p | MIPOL1 | HITS-CLIP | Tarbase |
| hsa-miR-195-5p | MIPOL1 | HITS-CLIP | Tarbase |
| hsa-miR-15a-5p | EXT2 | PAR-CLIP | Tarbase |
| hsa-miR-15a-3p | NDUFC2 | HITS-CLIP | Tarbase |
| hsa-miR-141-3p | ANKRD50 | HITS-CLIP | Tarbase |
| hsa-miR-3613-3p | ANKRD50 | PAR-CLIP | Tarbase |
| hsa-miR-141-5p | UPF2 | HITS-CLIP | Tarbase |
| hsa-miR-486-5p | CDC123 | HITS-CLIP | Tarbase |

| | | | |
|---|---|---|---|
| hsa-miR-195-5p | SCLT1 | Microarrays | Tarbase |
| hsa-miR-195-5p | CCDC3 | Microarrays | Tarbase |
| hsa-miR-3613-5p | EPS8 | HITS-CLIP | Tarbase |
| hsa-miR-195-5p | NCAPD3 | Microarrays | Tarbase |
| hsa-miR-3613-3p | VTI1A | PAR-CLIP | Tarbase |
| hsa-miR-141-3p | FAM160B1 | HITS-CLIP | Tarbase |
| hsa-miR-141-5p | FAM160B1 | HITS-CLIP | Tarbase |
| hsa-miR-141-5p | FAM160B1 | HITS-CLIP | Tarbase |
| hsa-miR-141-5p | FAM160B1 | HITS-CLIP | Tarbase |
| hsa-miR-141-5p | FAM160B1 | HITS-CLIP | Tarbase |
| hsa-miR-15a-5p | FAM160B1 | HITS-CLIP | Tarbase |
| hsa-miR-15a-5p | FAM160B1 | PAR-CLIP | Tarbase |
| hsa-miR-195-5p | FAM160B1 | HITS-CLIP | Tarbase |
| hsa-miR-3613-5p | FAM160B1 | PAR-CLIP | Tarbase |
| hsa-miR-195-5p | ANO4 | Microarrays | Tarbase |
| hsa-miR-15a-5p | QTRTD1 | PAR-CLIP | Tarbase |
| hsa-miR-15a-5p | QTRTD1 | PAR-CLIP | Tarbase |
| hsa-miR-15a-5p | QTRTD1 | PAR-CLIP | Tarbase |
| hsa-miR-195-5p | QTRTD1 | PAR-CLIP | Tarbase |
| hsa-miR-15a-5p | ZNF827 | PAR-CLIP | Tarbase |
| hsa-miR-15a-5p | ZNF827 | HITS-CLIP | Tarbase |
| hsa-miR-195-5p | ZNF827 | HITS-CLIP | Tarbase |
| hsa-miR-195-5p | AKR1C2 | Microarrays | Tarbase |
| hsa-miR-486-5p | ASAP2 | HITS-CLIP | Tarbase |
| hsa-miR-141-5p | ADAM17 | HITS-CLIP | Tarbase |
| hsa-miR-15a-5p | CENPU | HITS-CLIP | Tarbase |
| hsa-miR-195-5p | CENPU | HITS-CLIP | Tarbase |
| hsa-miR-195-5p | CENPU | Microarrays | Tarbase |
| hsa-miR-195-5p | SLC25A4 | Microarrays | Tarbase |
| hsa-miR-15a-5p | AMN1 | HITS-CLIP | Tarbase |
| hsa-miR-195-5p | AMN1 | HITS-CLIP | Tarbase |
| hsa-miR-141-3p | ZNF385D | HITS-CLIP | Tarbase |
| hsa-miR-195-5p | TDO2 | Microarrays | Tarbase |
| hsa-miR-195-5p | GABRA2 | Microarrays | Tarbase |
| hsa-miR-15a-5p | SACS | HITS-CLIP | Tarbase |
| hsa-miR-15a-5p | SACS | HITS-CLIP | Tarbase |
| hsa-miR-195-5p | SACS | HITS-CLIP | Tarbase |
| hsa-miR-15a-5p | SACS | HITS-CLIP | Tarbase |
| hsa-miR-195-5p | SACS | HITS-CLIP | Tarbase |
| hsa-miR-15a-5p | GFRA1 | PAR-CLIP | Tarbase |
| hsa-miR-15a-5p | GFRA1 | PAR-CLIP | Tarbase |
| hsa-miR-15a-5p | GFRA1 | PAR-CLIP | Tarbase |
| hsa-miR-15a-5p | CACUL1 | HITS-CLIP | Tarbase |
| hsa-miR-15a-5p | CACUL1 | HITS-CLIP | Tarbase |
| hsa-miR-15a-5p | CACUL1 | HITS-CLIP | Tarbase |
| hsa-miR-15a-5p | CACUL1 | PAR-CLIP | Tarbase |
| hsa-miR-15a-5p | CACUL1 | PAR-CLIP | Tarbase |
| hsa-miR-195-5p | CACUL1 | HITS-CLIP | Tarbase |
| hsa-miR-195-5p | CACUL1 | HITS-CLIP | Tarbase |
| hsa-miR-195-5p | CACUL1 | HITS-CLIP | Tarbase |
| hsa-miR-195-5p | CACUL1 | PAR-CLIP | Tarbase |
| hsa-miR-486-5p | CACUL1 | PAR-CLIP | Tarbase |
| hsa-miR-141-5p | DST | HITS-CLIP | Tarbase |
| hsa-miR-15a-3p | DST | HITS-CLIP | Tarbase |
| hsa-miR-15a-5p | DST | HITS-CLIP | Tarbase |
| hsa-miR-195-5p | DST | HITS-CLIP | Tarbase |
| hsa-miR-3613-5p | TMEM132D | HITS-CLIP | Tarbase |
| hsa-miR-141-5p | TMEM56 | HITS-CLIP | Tarbase |
| hsa-miR-141-3p | FAM168B | HITS-CLIP | Tarbase |
| hsa-miR-195-5p | MGAT5 | Microarrays | Tarbase |
| hsa-miR-141-3p | POU4F1 | HITS-CLIP | Tarbase |
| hsa-miR-195-5p | GRID2 | Microarrays | Tarbase |
| hsa-miR-195-5p | RIT2 | Microarrays | Tarbase |
| hsa-miR-141-5p | ARL14EP | HITS-CLIP | Tarbase |
| hsa-miR-15a-5p | EPG5 | HITS-CLIP | Tarbase |
| hsa-miR-195-5p | EPG5 | HITS-CLIP | Tarbase |
| hsa-miR-195-5p | SPC25 | Microarrays | Tarbase |
| hsa-miR-195-5p | PDK1 | Microarrays | Tarbase |
| hsa-miR-195-5p | PDE3B | Microarrays | Tarbase |
| hsa-miR-141-3p | TCF7L1 | Luciferase Reporter Assay | Tarbase |
| hsa-miR-141-3p | TCF7L1 | Luciferase Reporter Assay | Tarbase |
| hsa-miR-15a-5p | TGOLN2 | HITS-CLIP | Tarbase |
| hsa-miR-195-5p | TGOLN2 | HITS-CLIP | Tarbase |
| hsa-miR-3613-3p | TGOLN2 | HITS-CLIP | Tarbase |
| hsa-miR-15a-5p | TGOLN2 | Chimeric fragments | Tarbase |
| hsa-miR-486-5p | SH2D6 | HITS-CLIP | Tarbase |
| hsa-miR-141-3p | UHMK1 | HITS-CLIP | Tarbase |
| hsa-miR-15a-3p | UHMK1 | HITS-CLIP | Tarbase |
| hsa-miR-15a-5p | UHMK1 | PAR-CLIP | Tarbase |
| hsa-miR-15a-5p | UHMK1 | HITS-CLIP | Tarbase |
| hsa-miR-15a-5p | UHMK1 | HITS-CLIP | Tarbase |
| hsa-miR-15a-5p | UHMK1 | HITS-CLIP | Tarbase |
| hsa-miR-15a-5p | UHMK1 | PAR-CLIP | Tarbase |
| hsa-miR-15a-5p | UHMK1 | HITS-CLIP | Tarbase |
| hsa-miR-15a-5p | UHMK1 | HITS-CLIP | Tarbase |
| hsa-miR-195-5p | UHMK1 | PAR-CLIP | Tarbase |
| hsa-miR-195-5p | UHMK1 | HITS-CLIP | Tarbase |
| hsa-miR-195-5p | UHMK1 | HITS-CLIP | Tarbase |
| hsa-miR-195-5p | UHMK1 | HITS-CLIP | Tarbase |
| hsa-miR-195-5p | UHMK1 | HITS-CLIP | Tarbase |
| hsa-miR-486-5p | UHMK1 | PAR-CLIP | Tarbase |
| hsa-miR-486-5p | UHMK1 | PAR-CLIP | Tarbase |
| hsa-miR-15a-5p | UHMK1 | HITS-CLIP | Tarbase |
| hsa-miR-195-5p | UHMK1 | HITS-CLIP | Tarbase |
| hsa-miR-15a-5p | GUCY1A2 | HITS-CLIP | Tarbase |
| hsa-miR-195-5p | GUCY1A2 | HITS-CLIP | Tarbase |
| hsa-miR-3613-5p | NMT2 | PAR-CLIP | Tarbase |
| hsa-miR-15a-5p | TRIM36 | HITS-CLIP | Tarbase |
| hsa-miR-141-3p | PFKM | HITS-CLIP | Tarbase |
| hsa-miR-15a-5p | TMEM123 | PAR-CLIP | Tarbase |
| hsa-miR-15a-5p | TMEM123 | HITS-CLIP | Tarbase |
| hsa-miR-195-5p | TMEM123 | HITS-CLIP | Tarbase |
| hsa-miR-141-3p | MBNL1 | PAR-CLIP | Tarbase |
| hsa-miR-141-3p | MBNL1 | HITS-CLIP | Tarbase |
| hsa-miR-141-3p | MBNL1 | HITS-CLIP | Tarbase |
| hsa-miR-141-3p | MBNL1 | HITS-CLIP | Tarbase |
| hsa-miR-141-5p | MBNL1 | HITS-CLIP | Tarbase |
| hsa-miR-15a-5p | MBNL1 | HITS-CLIP | Tarbase |
| hsa-miR-195-5p | MBNL1 | HITS-CLIP | Tarbase |
| hsa-miR-15a-5p | MBNL1 | Chimeric fragments | Tarbase |
| hsa-miR-141-3p | SLC30A6 | HITS-CLIP | Tarbase |
| hsa-miR-195-5p | SLC30A6 | Microarrays | Tarbase |
| hsa-miR-195-5p | FARP1 | Microarrays | Tarbase |
| hsa-miR-15a-5p | PANK1 | HITS-CLIP | Tarbase |
| hsa-miR-141-3p | HNRNPDL | HITS-CLIP | Tarbase |
| hsa-miR-15a-5p | HNRNPDL | HITS-CLIP | Tarbase |
| hsa-miR-15a-5p | HNRNPDL | HITS-CLIP | Tarbase |
| hsa-miR-15a-5p | HNRNPDL | PAR-CLIP | Tarbase |
| hsa-miR-195-5p | HNRNPDL | HITS-CLIP | Tarbase |
| hsa-miR-195-5p | HNRNPDL | HITS-CLIP | Tarbase |
| hsa-miR-195-5p | HNRNPDL | HITS-CLIP | Tarbase |
| hsa-miR-3605-5p | GGPS1 | HITS-CLIP | Tarbase |
| hsa-miR-15a-5p | CNTNAP4 | HITS-CLIP | Tarbase |
| hsa-miR-195-5p | PLOD2 | Microarrays | Tarbase |
| hsa-miR-195-3p | CPB1 | HITS-CLIP | Tarbase |
| hsa-miR-3613-3p | SRP19 | PAR-CLIP | Tarbase |
| hsa-miR-195-5p | CENPH | Microarrays | Tarbase |
| hsa-miR-3613-5p | DAB2 | HITS-CLIP | Tarbase |
| hsa-miR-15a-5p | BCL2L11 | PAR-CLIP | Tarbase |
| hsa-miR-195-5p | BCL2L11 | qPCR | Tarbase |
| hsa-miR-3605-5p | BCL2L11 | HITS-CLIP | Tarbase |
| hsa-miR-15a-5p | CAST | PAR-CLIP | Tarbase |
| hsa-miR-141-3p | SCOC | HITS-CLIP | Tarbase |
| hsa-miR-141-5p | SMARCA5 | HITS-CLIP | Tarbase |
| hsa-miR-15a-5p | SMARCA5 | PAR-CLIP | Tarbase |
| hsa-miR-3613-3p | SMARCA5 | HITS-CLIP | Tarbase |
| hsa-miR-141-5p | HNRNPU | HITS-CLIP | Tarbase |
| hsa-miR-141-5p | HNRNPU | HITS-CLIP | Tarbase |
| hsa-miR-15a-5p | HNRNPU | HITS-CLIP | Tarbase |
| hsa-miR-195-5p | HNRNPU | HITS-CLIP | Tarbase |
| hsa-miR-15a-5p | RANBP2 | PAR-CLIP | Tarbase |
| hsa-miR-15a-5p | AHCTF1 | HITS-CLIP | Tarbase |
| hsa-miR-195-5p | AHCTF1 | HITS-CLIP | Tarbase |
| hsa-miR-15a-5p | TMEM87B | HITS-CLIP | Tarbase |
| hsa-miR-195-5p | TMEM87B | HITS-CLIP | Tarbase |
| hsa-miR-15a-5p | RBMS1 | HITS-CLIP | Tarbase |
| hsa-miR-15a-5p | RBMS1 | HITS-CLIP | Tarbase |
| hsa-miR-15a-5p | RBMS1 | HITS-CLIP | Tarbase |
| hsa-miR-195-5p | RBMS1 | HITS-CLIP | Tarbase |

| | | | |
|---|---|---|---|
| hsa-miR-195-5p | RBMS1 | HITS-CLIP | Tarbase |
| hsa-miR-195-5p | CD96 | Microarrays | Tarbase |
| hsa-miR-141-3p | ASAP1 | HITS-CLIP | Tarbase |
| hsa-miR-15a-5p | ASAP1 | HITS-CLIP | Tarbase |
| hsa-miR-195-5p | ASAP1 | HITS-CLIP | Tarbase |
| hsa-miR-15a-5p | INO80C | Chimeric fragments | Tarbase |
| hsa-miR-141-3p | LPCAT1 | HITS-CLIP | Tarbase |
| hsa-miR-3605-5p | LPCAT1 | HITS-CLIP | Tarbase |
| hsa-miR-15a-5p | PLEKHG4B | HITS-CLIP | Tarbase |
| hsa-miR-195-5p | PLEKHG4B | HITS-CLIP | Tarbase |
| hsa-miR-195-5p | UBALD1 | Microarrays | Tarbase |
| hsa-miR-141-3p | RMND5A | HITS-CLIP | Tarbase |
| hsa-miR-15a-5p | RMND5A | HITS-CLIP | Tarbase |
| hsa-miR-195-5p | RMND5A | HITS-CLIP | Tarbase |
| hsa-miR-141-3p | PTPRD | Luciferase Reporter Assay | Tarbase |
| hsa-miR-141-3p | PTPRD | Luciferase Reporter Assay | Tarbase |
| hsa-miR-15a-5p | LURAP1L | HITS-CLIP | Tarbase |
| hsa-miR-141-5p | CNKSR3 | HITS-CLIP | Tarbase |
| hsa-miR-15a-5p | CNKSR3 | HITS-CLIP | Tarbase |
| hsa-miR-195-5p | CNKSR3 | HITS-CLIP | Tarbase |
| hsa-miR-141-3p | JAZF1 | HITS-CLIP | Tarbase |
| hsa-miR-15a-5p | TRIP12 | HITS-CLIP | Tarbase |
| hsa-miR-15a-5p | DDAH1 | Chimeric fragments | Tarbase |
| hsa-miR-15a-5p | SREK1 | PAR-CLIP | Tarbase |
| hsa-miR-15a-5p | SREK1 | PAR-CLIP | Tarbase |
| hsa-miR-15a-5p | SREK1 | PAR-CLIP | Tarbase |
| hsa-miR-195-5p | SREK1 | PAR-CLIP | Tarbase |
| hsa-miR-486-5p | CHD1 | PAR-CLIP | Tarbase |
| hsa-miR-141-5p | MSI2 | HITS-CLIP | Tarbase |
| hsa-miR-15a-5p | MSI2 | HITS-CLIP | Tarbase |
| hsa-miR-195-5p | MSI2 | HITS-CLIP | Tarbase |
| hsa-miR-195-5p | MSI2 | Microarrays | Tarbase |
| hsa-miR-15a-5p | CACNA2D1 | HITS-CLIP | Tarbase |
| hsa-miR-15a-5p | CACNA2D1 | PAR-CLIP | Tarbase |
| hsa-miR-195-5p | CACNA2D1 | HITS-CLIP | Tarbase |
| hsa-miR-195-5p | CACNA2D1 | Microarrays | Tarbase |
| hsa-miR-195-5p | SEMA3D | Microarrays | Tarbase |
| hsa-miR-195-5p | AK5 | Microarrays | Tarbase |
| hsa-miR-195-5p | ANKRD29 | Microarrays | Tarbase |
| hsa-miR-3605-5p | CHST9 | HITS-CLIP | Tarbase |
| hsa-miR-15a-5p | TBCEL | HITS-CLIP | Tarbase |
| hsa-miR-15a-5p | ANKH | HITS-CLIP | Tarbase |
| hsa-miR-195-5p | ANKH | HITS-CLIP | Tarbase |
| hsa-miR-195-5p | UBASH3B | Microarrays | Tarbase |
| hsa-miR-195-5p | NRGN | Microarrays | Tarbase |
| hsa-miR-195-5p | CDH12 | Microarrays | Tarbase |
| hsa-miR-15a-3p | TOMM70A | HITS-CLIP | Tarbase |
| hsa-miR-195-5p | PITPNC1 | Microarrays | Tarbase |
| hsa-miR-15a-5p | CC2D1B | HITS-CLIP | Tarbase |
| hsa-miR-15a-5p | CC2D1B | PAR-CLIP | Tarbase |
| hsa-miR-15a-5p | CC2D1B | HITS-CLIP | Tarbase |
| hsa-miR-195-5p | CC2D1B | Microarrays | Tarbase |
| hsa-miR-3605-3p | PRKCA | HITS-CLIP | Tarbase |
| hsa-miR-15a-5p | LRRK1 | HITS-CLIP | Tarbase |
| hsa-miR-15a-5p | LRRK1 | PAR-CLIP | Tarbase |
| hsa-miR-195-5p | LRRK1 | HITS-CLIP | Tarbase |
| hsa-miR-195-5p | LRRK1 | PAR-CLIP | Tarbase |
| hsa-miR-15a-5p | MIA3 | HITS-CLIP | Tarbase |
| hsa-miR-195-5p | MIA3 | HITS-CLIP | Tarbase |
| hsa-miR-3605-5p | MIA3 | PAR-CLIP | Tarbase |
| hsa-miR-195-5p | DISP1 | Microarrays | Tarbase |
| hsa-miR-15a-5p | WNT3A | HITS-CLIP | Tarbase |
| hsa-miR-195-5p | WNT3A | HITS-CLIP | Tarbase |
| hsa-miR-15a-5p | WNT3A | Luciferase Reporter Assay | Tarbase |
| hsa-miR-141-5p | TRIM11 | HITS-CLIP | Tarbase |
| hsa-miR-3605-3p | TRIM11 | HITS-CLIP | Tarbase |
| hsa-miR-15a-5p | ENAH | HITS-CLIP | Tarbase |
| hsa-miR-15a-5p | ENAH | PAR-CLIP | Tarbase |
| hsa-miR-15a-5p | ENAH | HITS-CLIP | Tarbase |
| hsa-miR-15a-5p | ENAH | HITS-CLIP | Tarbase |
| hsa-miR-195-5p | ENAH | HITS-CLIP | Tarbase |
| hsa-miR-195-5p | SH3RF1 | Microarrays | Tarbase |
| hsa-miR-15a-5p | BUB3 | PAR-CLIP | Tarbase |
| hsa-miR-3613-5p | BUB3 | PAR-CLIP | Tarbase |
| hsa-miR-141-3p | ATP5G3 | HITS-CLIP | Tarbase |
| hsa-miR-15a-5p | ATP5G3 | HITS-CLIP | Tarbase |
| hsa-miR-15a-5p | ATP5G3 | PAR-CLIP | Tarbase |
| hsa-miR-195-5p | ATP5G3 | HITS-CLIP | Tarbase |
| hsa-miR-141-3p | SORBS2 | HITS-CLIP | Tarbase |
| hsa-miR-3605-5p | SORBS2 | HITS-CLIP | Tarbase |
| hsa-miR-195-5p | LY96 | Microarrays | Tarbase |
| hsa-miR-219a-1-3p | BTG3 | HITS-CLIP | Tarbase |
| hsa-miR-219a-1-3p | BTG3 | HITS-CLIP | Tarbase |
| hsa-miR-219a-1-3p | BTG3 | HITS-CLIP | Tarbase |
| hsa-miR-15a-3p | JAM2 | HITS-CLIP | Tarbase |
| hsa-miR-486-3p | ADAMTS1 | HITS-CLIP | Tarbase |
| hsa-miR-141-5p | ADAMTS5 | HITS-CLIP | Tarbase |
| hsa-miR-15a-5p | ADAMTS5 | HITS-CLIP | Tarbase |
| hsa-miR-195-5p | ADAMTS5 | Microarrays | Tarbase |
| hsa-miR-195-5p | WNT7A | Luciferase Reporter Assay | Tarbase |
| hsa-miR-195-5p | XPC | Microarrays | Tarbase |
| hsa-miR-141-3p | PLCL2 | HITS-CLIP | Tarbase |
| hsa-miR-3613-5p | PLCL2 | HITS-CLIP | Tarbase |
| hsa-miR-15a-5p | PLCL2 | Chimeric fragments | Tarbase |
| hsa-miR-15a-5p | CXXC1 | HITS-CLIP | Tarbase |
| hsa-miR-195-5p | CXXC1 | HITS-CLIP | Tarbase |
| hsa-miR-195-5p | SKA1 | Microarrays | Tarbase |
| hsa-miR-141-3p | PPP4R1 | HITS-CLIP | Tarbase |
| hsa-miR-141-5p | PPP4R1 | HITS-CLIP | Tarbase |
| hsa-miR-195-5p | APCDD1 | Microarrays | Tarbase |
| hsa-miR-195-5p | PIEZO2 | Microarrays | Tarbase |
| hsa-miR-195-5p | EME1 | Microarrays | Tarbase |
| hsa-miR-15a-5p | ANKRD40 | PAR-CLIP | Tarbase |
| hsa-miR-15a-5p | ANKRD40 | HITS-CLIP | Tarbase |
| hsa-miR-195-5p | ANKRD40 | HITS-CLIP | Tarbase |
| hsa-miR-15a-5p | APOOL | HITS-CLIP | Tarbase |
| hsa-miR-195-5p | DKK2 | Microarrays | Tarbase |
| hsa-miR-141-5p | KLF10 | HITS-CLIP | Tarbase |
| hsa-miR-141-5p | KLF10 | HITS-CLIP | Tarbase |
| hsa-miR-15a-5p | KLF10 | Chimeric fragments | Tarbase |
| hsa-miR-15a-5p | PTPRN2 | HITS-CLIP | Tarbase |
| hsa-miR-195-5p | PTPRN2 | HITS-CLIP | Tarbase |
| hsa-miR-141-5p | AZIN1 | HITS-CLIP | Tarbase |
| hsa-miR-141-5p | AZIN1 | HITS-CLIP | Tarbase |
| hsa-miR-15a-5p | AZIN1 | HITS-CLIP | Tarbase |
| hsa-miR-195-5p | AZIN1 | HITS-CLIP | Tarbase |
| hsa-miR-486-5p | ATP6V1C1 | HITS-CLIP | Tarbase |
| hsa-miR-15a-5p | TMEM55A | PAR-CLIP | Tarbase |
| hsa-miR-15a-5p | TMEM55A | PAR-CLIP | Tarbase |
| hsa-miR-195-5p | TMEM55A | PAR-CLIP | Tarbase |
| hsa-miR-15a-5p | TMEM55A | HITS-CLIP | Tarbase |
| hsa-miR-195-5p | TMEM55A | HITS-CLIP | Tarbase |
| hsa-miR-15a-5p | TTC39B | HITS-CLIP | Tarbase |
| hsa-miR-195-5p | TTC39B | HITS-CLIP | Tarbase |
| hsa-miR-141-3p | AGPAT5 | HITS-CLIP | Tarbase |
| hsa-miR-141-3p | AGPAT5 | HITS-CLIP | Tarbase |
| hsa-miR-195-5p | MMS19 | Microarrays | Tarbase |
| hsa-miR-486-5p | SLC25A28 | PAR-CLIP | Tarbase |
| hsa-miR-141-3p | HSPA13 | HITS-CLIP | Tarbase |
| hsa-miR-15a-5p | USP25 | PAR-CLIP | Tarbase |
| hsa-miR-15a-5p | GRAMD3 | HITS-CLIP | Tarbase |
| hsa-miR-195-5p | GRAMD3 | HITS-CLIP | Tarbase |
| hsa-miR-141-3p | C16orf87 | HITS-CLIP | Tarbase |
| hsa-miR-195-5p | MOV10 | Microarrays | Tarbase |
| hsa-miR-219a-1-3p | SLC16A1 | HITS-CLIP | Tarbase |
| hsa-miR-141-5p | HEATR3 | HITS-CLIP | Tarbase |
| hsa-miR-15a-3p | LARP1 | HITS-CLIP | Tarbase |
| hsa-miR-15a-3p | LARP1 | PAR-CLIP | Tarbase |
| hsa-miR-15a-5p | LARP1 | HITS-CLIP | Tarbase |
| hsa-miR-15a-5p | LARP1 | HITS-CLIP | Tarbase |
| hsa-miR-195-5p | LARP1 | HITS-CLIP | Tarbase |
| hsa-miR-195-5p | LARP1 | HITS-CLIP | Tarbase |
| hsa-miR-3605-3p | LARP1 | HITS-CLIP | Tarbase |
| hsa-miR-15a-5p | CNOT8 | PAR-CLIP | Tarbase |
| hsa-miR-15a-5p | CNOT8 | HITS-CLIP | Tarbase |
| hsa-miR-195-5p | CNOT8 | HITS-CLIP | Tarbase |
| hsa-miR-141-5p | NUP205 | HITS-CLIP | Tarbase |
| hsa-miR-15a-5p | ZKSCAN2 | PAR-CLIP | Tarbase |
| hsa-miR-141-5p | TTN | HITS-CLIP | Tarbase |

| | | | |
|---|---|---|---|
| hsa-miR-15a-5p | PDIA4 | PAR-CLIP | Tarbase |
| hsa-miR-15a-5p | FAM126B | PAR-CLIP | Tarbase |
| hsa-miR-15a-5p | FAM126B | PAR-CLIP | Tarbase |
| hsa-miR-15a-5p | FAM126B | PAR-CLIP | Tarbase |
| hsa-miR-195-5p | FAM126B | PAR-CLIP | Tarbase |
| hsa-miR-195-5p | TMEM237 | Microarrays | Tarbase |
| hsa-miR-141-3p | FMN2 | HITS-CLIP | Tarbase |
| hsa-miR-15a-3p | FMN2 | HITS-CLIP | Tarbase |
| hsa-miR-141-3p | RNF20 | HITS-CLIP | Tarbase |
| hsa-miR-15a-5p | SLC26A2 | HITS-CLIP | Tarbase |
| hsa-miR-15a-5p | LSM11 | PAR-CLIP | Tarbase |
| hsa-miR-15a-5p | LSM11 | HITS-CLIP | Tarbase |
| hsa-miR-195-5p | LSM11 | Microarrays | Tarbase |
| hsa-miR-15a-5p | LSM11 | Chimeric fragments | Tarbase |
| hsa-miR-15a-5p | RRAGA | HITS-CLIP | Tarbase |
| hsa-miR-195-5p | RRAGA | HITS-CLIP | Tarbase |
| hsa-miR-15a-5p | RRAGA | HITS-CLIP | Tarbase |
| hsa-miR-195-5p | RRAGA | HITS-CLIP | Tarbase |
| hsa-miR-15a-5p | SLC24A2 | HITS-CLIP | Tarbase |
| hsa-miR-195-5p | SLC24A2 | HITS-CLIP | Tarbase |
| hsa-miR-141-5p | PXYLP1 | HITS-CLIP | Tarbase |
| hsa-miR-15a-3p | PXYLP1 | HITS-CLIP | Tarbase |
| hsa-miR-15a-5p | PXYLP1 | HITS-CLIP | Tarbase |
| hsa-miR-195-5p | PXYLP1 | HITS-CLIP | Tarbase |
| hsa-miR-141-3p | RASA2 | HITS-CLIP | Tarbase |
| hsa-miR-141-5p | KIF5A | HITS-CLIP | Tarbase |
| hsa-miR-15a-5p | KIF5A | PAR-CLIP | Tarbase |
| hsa-miR-219a-1-3p | PSD3 | HITS-CLIP | Tarbase |
| hsa-miR-15a-5p | PSD3 | HITS-CLIP | Tarbase |
| hsa-miR-195-5p | PSD3 | HITS-CLIP | Tarbase |
| hsa-miR-15a-3p | C9orf41 | HITS-CLIP | Tarbase |
| hsa-miR-195-5p | MCU | Microarrays | Tarbase |
| hsa-miR-15a-5p | ELMSAN1 | HITS-CLIP | Tarbase |
| hsa-miR-15a-5p | ELMSAN1 | HITS-CLIP | Tarbase |
| hsa-miR-195-5p | ELMSAN1 | HITS-CLIP | Tarbase |
| hsa-miR-3605-5p | ELMSAN1 | PAR-CLIP | Tarbase |
| hsa-miR-486-5p | ELMSAN1 | HITS-CLIP | Tarbase |
| hsa-miR-141-3p | GNAQ | HITS-CLIP | Tarbase |
| hsa-miR-141-3p | GNAQ | HITS-CLIP | Tarbase |
| hsa-miR-15a-5p | GNAQ | PAR-CLIP | Tarbase |
| hsa-miR-15a-5p | GNAQ | PAR-CLIP | Tarbase |
| hsa-miR-15a-5p | GNAQ | PAR-CLIP | Tarbase |
| hsa-miR-15a-5p | GNAQ | HITS-CLIP | Tarbase |
| hsa-miR-195-5p | GNAQ | PAR-CLIP | Tarbase |
| hsa-miR-195-5p | GNAQ | HITS-CLIP | Tarbase |
| hsa-miR-15a-5p | GNAQ | HITS-CLIP | Tarbase |
| hsa-miR-195-5p | GNAQ | HITS-CLIP | Tarbase |
| hsa-miR-3605-5p | KCNMA1 | HITS-CLIP | Tarbase |
| hsa-miR-195-5p | KCNMA1 | Microarrays | Tarbase |
| hsa-miR-141-5p | NDUFAF6 | HITS-CLIP | Tarbase |
| hsa-miR-195-5p | MAP3K7CL | Microarrays | Tarbase |
| hsa-miR-15a-5p | BACH1 | HITS-CLIP | Tarbase |
| hsa-miR-195-5p | BACH1 | HITS-CLIP | Tarbase |
| hsa-miR-195-5p | TSPAN7 | Microarrays | Tarbase |
| hsa-miR-486-5p | TIAM1 | HITS-CLIP | Tarbase |
| hsa-miR-15a-5p | SCAF4 | HITS-CLIP | Tarbase |
| hsa-miR-15a-5p | SCAF4 | PAR-CLIP | Tarbase |
| hsa-miR-195-5p | SCAF4 | HITS-CLIP | Tarbase |
| hsa-miR-15a-5p | SCAF4 | Chimeric fragments | Tarbase |
| hsa-miR-15a-5p | C14orf2 | PAR-CLIP | Tarbase |
| hsa-miR-195-5p | FBXO43 | Microarrays | Tarbase |
| hsa-miR-15a-3p | HK1 | HITS-CLIP | Tarbase |
| hsa-miR-195-5p | HK1 | HITS-CLIP | Tarbase |
| hsa-miR-141-5p | PHF6 | HITS-CLIP | Tarbase |
| hsa-miR-15a-5p | MED19 | Chimeric fragments | Tarbase |
| hsa-miR-219a-1-3p | ZFAND3 | HITS-CLIP | Tarbase |
| hsa-miR-15a-5p | NPTN | HITS-CLIP | Tarbase |
| hsa-miR-195-5p | NPTN | HITS-CLIP | Tarbase |
| hsa-miR-15a-5p | KAT6B | HITS-CLIP | Tarbase |
| hsa-miR-141-3p | SAMD8 | HITS-CLIP | Tarbase |
| hsa-miR-15a-5p | SAMD8 | PAR-CLIP | Tarbase |
| hsa-miR-3613-3p | SAMD8 | HITS-CLIP | Tarbase |
| hsa-miR-15a-5p | SAMD8 | HITS-CLIP | Tarbase |
| hsa-miR-195-5p | SAMD8 | HITS-CLIP | Tarbase |
| hsa-miR-15a-5p | RAB11FIP1 | HITS-CLIP | Tarbase |
| hsa-miR-195-5p | RAB11FIP1 | HITS-CLIP | Tarbase |
| hsa-miR-3613-5p | RAB11FIP1 | HITS-CLIP | Tarbase |
| hsa-miR-195-5p | RAB11FIP1 | Microarrays | Tarbase |
| hsa-miR-15a-5p | BAG4 | PAR-CLIP | Tarbase |
| hsa-miR-141-3p | ATAD2 | HITS-CLIP | Tarbase |
| hsa-miR-195-5p | FBXO32 | Microarrays | Tarbase |
| hsa-miR-141-5p | ZNF689 | HITS-CLIP | Tarbase |
| hsa-miR-15a-3p | FBRS | HITS-CLIP | Tarbase |
| hsa-miR-486-3p | FBRS | PAR-CLIP | Tarbase |
| hsa-miR-15a-5p | PHKG2 | HITS-CLIP | Tarbase |
| hsa-miR-141-5p | HIAT1 | HITS-CLIP | Tarbase |
| hsa-miR-15a-3p | HIAT1 | HITS-CLIP | Tarbase |
| hsa-miR-15a-5p | HIAT1 | HITS-CLIP | Tarbase |
| hsa-miR-15a-5p | HIAT1 | PAR-CLIP | Tarbase |
| hsa-miR-15a-5p | HIAT1 | HITS-CLIP | Tarbase |
| hsa-miR-15a-5p | HIAT1 | HITS-CLIP | Tarbase |
| hsa-miR-15a-5p | HIAT1 | PAR-CLIP | Tarbase |
| hsa-miR-195-5p | HIAT1 | HITS-CLIP | Tarbase |
| hsa-miR-195-5p | HIAT1 | HITS-CLIP | Tarbase |
| hsa-miR-195-5p | HIAT1 | HITS-CLIP | Tarbase |
| hsa-miR-219a-1-3p | HIAT1 | HITS-CLIP | Tarbase |
| hsa-miR-195-5p | BUB1B | Microarrays | Tarbase |
| hsa-miR-15a-5p | EIF4A2 | HITS-CLIP | Tarbase |
| hsa-miR-15a-5p | EIF4A2 | HITS-CLIP | Tarbase |
| hsa-miR-15a-5p | EIF4A2 | HITS-CLIP | Tarbase |
| hsa-miR-195-5p | EIF4A2 | HITS-CLIP | Tarbase |
| hsa-miR-195-5p | EIF4A2 | HITS-CLIP | Tarbase |
| hsa-miR-141-5p | BRPF1 | HITS-CLIP | Tarbase |
| hsa-miR-15a-5p | TATDN2 | HITS-CLIP | Tarbase |
| hsa-miR-195-5p | TATDN2 | HITS-CLIP | Tarbase |
| hsa-miR-15a-5p | NMNAT2 | HITS-CLIP | Tarbase |
| hsa-miR-195-5p | NMNAT2 | HITS-CLIP | Tarbase |
| hsa-miR-141-5p | ZFYVE9 | HITS-CLIP | Tarbase |
| hsa-miR-141-5p | ZFYVE9 | HITS-CLIP | Tarbase |
| hsa-miR-141-5p | ZFYVE9 | HITS-CLIP | Tarbase |
| hsa-miR-141-5p | ZFYVE9 | HITS-CLIP | Tarbase |
| hsa-miR-141-3p | SMG1 | HITS-CLIP | Tarbase |
| hsa-miR-3613-3p | SMG1 | HITS-CLIP | Tarbase |
| hsa-miR-3613-3p | SMG1 | HITS-CLIP | Tarbase |
| hsa-miR-195-5p | RBPMS | Microarrays | Tarbase |
| hsa-miR-195-5p | TMEM171 | Microarrays | Tarbase |
| hsa-miR-141-5p | STEAP2 | HITS-CLIP | Tarbase |
| hsa-miR-15a-3p | STEAP2 | HITS-CLIP | Tarbase |
| hsa-miR-3605-3p | STEAP2 | HITS-CLIP | Tarbase |
| hsa-miR-15a-5p | CLDN12 | HITS-CLIP | Tarbase |
| hsa-miR-15a-5p | CLDN12 | HITS-CLIP | Tarbase |
| hsa-miR-195-5p | CLDN12 | HITS-CLIP | Tarbase |
| hsa-miR-195-5p | CLDN12 | HITS-CLIP | Tarbase |
| hsa-miR-3613-3p | CLDN12 | HITS-CLIP | Tarbase |
| hsa-miR-15a-5p | CACNA2D3 | PAR-CLIP | Tarbase |
| hsa-miR-141-3p | RNF111 | HITS-CLIP | Tarbase |
| hsa-miR-141-3p | RNF111 | HITS-CLIP | Tarbase |
| hsa-miR-141-5p | RNF111 | HITS-CLIP | Tarbase |
| hsa-miR-15a-5p | RNF111 | HITS-CLIP | Tarbase |
| hsa-miR-195-5p | CCNB2 | Microarrays | Tarbase |
| hsa-miR-195-5p | MYO1E | Microarrays | Tarbase |
| hsa-miR-15a-5p | APPL1 | PAR-CLIP | Tarbase |
| hsa-miR-15a-5p | TSC22D3 | HITS-CLIP | Tarbase |
| hsa-miR-141-3p | DSCR3 | HITS-CLIP | Tarbase |
| hsa-miR-141-3p | DSCR3 | HITS-CLIP | Tarbase |
| hsa-miR-219a-1-3p | DSCR3 | PAR-CLIP | Tarbase |
| hsa-miR-15a-5p | DYRK1A | PAR-CLIP | Tarbase |
| hsa-miR-15a-5p | DYRK1A | PAR-CLIP | Tarbase |
| hsa-miR-15a-5p | DYRK1A | PAR-CLIP | Tarbase |
| hsa-miR-15a-5p | DYRK1A | PAR-CLIP | Tarbase |
| hsa-miR-15a-5p | DYRK1A | PAR-CLIP | Tarbase |
| hsa-miR-15a-5p | DYRK1A | HITS-CLIP | Tarbase |
| hsa-miR-195-5p | DYRK1A | HITS-CLIP | Tarbase |
| hsa-miR-3613-3p | DYRK1A | PAR-CLIP | Tarbase |
| hsa-miR-486-5p | DYRK1A | PAR-CLIP | Tarbase |
| hsa-miR-195-5p | KCNJ6 | Microarrays | Tarbase |
| hsa-miR-141-3p | C2CD2 | PAR-CLIP | Tarbase |
| hsa-miR-15a-5p | TAB3 | Luciferase Reporter Assay | Tarbase |
| hsa-miR-15a-5p | TAB3 | qPCR | Tarbase |

| | | | |
|---|---|---|---|
| hsa-miR-15a-5p | TAB3 | Western Blot | Tarbase |
| hsa-miR-195-5p | TAB3 | qPCR | Tarbase |
| hsa-miR-195-5p | TAB3 | Western Blot | Tarbase |
| hsa-miR-15a-3p | PALM2-AKAP2 | PAR-CLIP | Tarbase |
| hsa-miR-15a-5p | PALM2-AKAP2 | HITS-CLIP | Tarbase |
| hsa-miR-15a-5p | PALM2-AKAP2 | PAR-CLIP | Tarbase |
| hsa-miR-195-5p | PALM2-AKAP2 | HITS-CLIP | Tarbase |
| hsa-miR-195-5p | PALM2-AKAP2 | PAR-CLIP | Tarbase |
| hsa-miR-15a-5p | ZNF618 | HITS-CLIP | Tarbase |
| hsa-miR-3613-5p | ZNF618 | HITS-CLIP | Tarbase |
| hsa-miR-195-5p | C9orf91 | HITS-CLIP | Tarbase |
| hsa-miR-15a-5p | C9orf91 | Chimeric fragments | Tarbase |
| hsa-miR-141-3p | UBN2 | HITS-CLIP | Tarbase |
| hsa-miR-15a-5p | UBN2 | PAR-CLIP | Tarbase |
| hsa-miR-15a-5p | UBN2 | PAR-CLIP | Tarbase |
| hsa-miR-15a-5p | UBN2 | PAR-CLIP | Tarbase |
| hsa-miR-15a-5p | UBN2 | PAR-CLIP | Tarbase |
| hsa-miR-195-5p | UBN2 | PAR-CLIP | Tarbase |
| hsa-miR-15a-5p | UBN2 | Chimeric fragments | Tarbase |
| hsa-miR-3613-3p | UBN2 | Chimeric fragments | Tarbase |
| hsa-miR-141-5p | BRAF | HITS-CLIP | Tarbase |
| hsa-miR-141-5p | BRAF | HITS-CLIP | Tarbase |
| hsa-miR-15a-3p | BRAF | HITS-CLIP | Tarbase |
| hsa-miR-195-5p | FMNL2 | Microarrays | Tarbase |
| hsa-miR-15a-5p | SPPL3 | HITS-CLIP | Tarbase |
| hsa-miR-141-5p | DPYSL5 | HITS-CLIP | Tarbase |
| hsa-miR-141-5p | DPYSL5 | HITS-CLIP | Tarbase |
| hsa-miR-141-5p | DPYSL5 | HITS-CLIP | Tarbase |
| hsa-miR-141-5p | DPYSL5 | HITS-CLIP | Tarbase |
| hsa-miR-486-5p | RER1 | HITS-CLIP | Tarbase |
| hsa-miR-141-5p | SKI | HITS-CLIP | Tarbase |
| hsa-miR-15a-5p | SKI | HITS-CLIP | Tarbase |
| hsa-miR-15a-5p | SKI | HITS-CLIP | Tarbase |
| hsa-miR-15a-5p | SKI | PAR-CLIP | Tarbase |
| hsa-miR-15a-5p | SKI | HITS-CLIP | Tarbase |
| hsa-miR-15a-5p | SKI | HITS-CLIP | Tarbase |
| hsa-miR-15a-5p | SKI | HITS-CLIP | Tarbase |
| hsa-miR-15a-5p | SKI | HITS-CLIP | Tarbase |
| hsa-miR-195-5p | SKI | HITS-CLIP | Tarbase |
| hsa-miR-195-5p | SKI | HITS-CLIP | Tarbase |
| hsa-miR-195-5p | SKI | HITS-CLIP | Tarbase |
| hsa-miR-486-3p | SKI | HITS-CLIP | Tarbase |
| hsa-miR-15a-5p | SKI | HITS-CLIP | Tarbase |
| hsa-miR-195-5p | SKI | HITS-CLIP | Tarbase |
| hsa-miR-15a-5p | WIPI2 | HITS-CLIP | Tarbase |
| hsa-miR-15a-5p | WIPI2 | HITS-CLIP | Tarbase |
| hsa-miR-15a-5p | WIPI2 | PAR-CLIP | Tarbase |
| hsa-miR-195-5p | WIPI2 | HITS-CLIP | Tarbase |
| hsa-miR-195-5p | WIPI2 | HITS-CLIP | Tarbase |
| hsa-miR-15a-3p | AGAP1 | HITS-CLIP | Tarbase |
| hsa-miR-15a-5p | PAFAH2 | HITS-CLIP | Tarbase |
| hsa-miR-195-5p | PAFAH2 | HITS-CLIP | Tarbase |
| hsa-miR-195-5p | WDR66 | Microarrays | Tarbase |
| hsa-miR-15a-5p | DUSP2 | PAR-CLIP | Tarbase |
| hsa-miR-195-5p | DUSP2 | PAR-CLIP | Tarbase |
| hsa-miR-141-5p | NCK1 | HITS-CLIP | Tarbase |
| hsa-miR-15a-5p | TPRG1L | HITS-CLIP | Tarbase |
| hsa-miR-195-5p | TPRG1L | HITS-CLIP | Tarbase |
| hsa-miR-3605-3p | TPRG1L | HITS-CLIP | Tarbase |
| hsa-miR-195-5p | TPRG1L | Microarrays | Tarbase |
| hsa-miR-15a-3p | XKR8 | HITS-CLIP | Tarbase |
| hsa-miR-195-5p | XKR8 | AGO-IP | Tarbase |
| hsa-miR-141-5p | FANCC | HITS-CLIP | Tarbase |
| hsa-miR-195-5p | FANCC | Microarrays | Tarbase |
| hsa-miR-195-5p | MRAS | Microarrays | Tarbase |
| hsa-miR-195-5p | CLSTN2 | Microarrays | Tarbase |
| hsa-miR-195-5p | COLEC12 | Microarrays | Tarbase |
| hsa-miR-15a-5p | CUL4B | PAR-CLIP | Tarbase |
| hsa-miR-15a-3p | AUTS2 | HITS-CLIP | Tarbase |
| hsa-miR-141-3p | HIST1H2BD | HITS-CLIP | Tarbase |
| hsa-miR-15a-5p | HIST1H2BD | PAR-CLIP | Tarbase |
| hsa-miR-486-3p | HIST1H2BD | HITS-CLIP | Tarbase |
| hsa-miR-195-5p | HIST1H2BD | Microarrays | Tarbase |
| hsa-miR-195-5p | CDC25C | Microarrays | Tarbase |
| hsa-miR-15a-3p | HIST1H4H | HITS-CLIP | Tarbase |
| hsa-miR-15a-5p | EIF5B | HITS-CLIP | Tarbase |
| hsa-miR-15a-5p | EIF5B | PAR-CLIP | Tarbase |
| hsa-miR-195-5p | EIF5B | HITS-CLIP | Tarbase |
| hsa-miR-3613-5p | TSPAN33 | PAR-CLIP | Tarbase |
| hsa-miR-141-5p | B4GALT5 | HITS-CLIP | Tarbase |
| hsa-miR-3613-5p | B4GALT5 | HITS-CLIP | Tarbase |
| hsa-miR-141-3p | SPATA2 | HITS-CLIP | Tarbase |
| hsa-miR-141-5p | SPATA2 | HITS-CLIP | Tarbase |
| hsa-miR-15a-5p | SPATA2 | PAR-CLIP | Tarbase |
| hsa-miR-195-5p | SPATA2 | PAR-CLIP | Tarbase |
| hsa-miR-195-5p | SPATA2 | PAR-CLIP | Tarbase |
| hsa-miR-15a-5p | GDPD5 | HITS-CLIP | Tarbase |
| hsa-miR-195-5p | GDPD5 | HITS-CLIP | Tarbase |
| hsa-miR-3613-3p | DYNC1I1 | HITS-CLIP | Tarbase |
| hsa-miR-141-3p | TMED4 | HITS-CLIP | Tarbase |
| hsa-miR-219a-1-3p | TMED4 | PAR-CLIP | Tarbase |
| hsa-miR-141-5p | PPP1R15B | HITS-CLIP | Tarbase |
| hsa-miR-219a-1-3p | PPP1R15B | HITS-CLIP | Tarbase |
| hsa-miR-141-5p | C11orf30 | HITS-CLIP | Tarbase |
| hsa-miR-15a-5p | C11orf30 | PAR-CLIP | Tarbase |
| hsa-miR-141-3p | AGPAT6 | HITS-CLIP | Tarbase |
| hsa-miR-15a-5p | AGPAT6 | HITS-CLIP | Tarbase |
| hsa-miR-15a-5p | ELK4 | PAR-CLIP | Tarbase |
| hsa-miR-15a-5p | ELK4 | HITS-CLIP | Tarbase |
| hsa-miR-15a-5p | ELK4 | Chimeric fragments | Tarbase |
| hsa-miR-15a-3p | SLAMF8 | HITS-CLIP | Tarbase |
| hsa-miR-195-5p | DUSP23 | Microarrays | Tarbase |
| hsa-miR-15a-3p | F11R | HITS-CLIP | Tarbase |
| hsa-miR-141-5p | USF1 | HITS-CLIP | Tarbase |
| hsa-miR-195-5p | DEDD | Microarrays | Tarbase |
| hsa-miR-15a-5p | EDA | PAR-CLIP | Tarbase |
| hsa-miR-15a-5p | PINK1 | HITS-CLIP | Tarbase |
| hsa-miR-15a-5p | PINK1 | HITS-CLIP | Tarbase |
| hsa-miR-195-5p | PINK1 | HITS-CLIP | Tarbase |
| hsa-miR-195-5p | PINK1 | HITS-CLIP | Tarbase |
| hsa-miR-15a-3p | CDC42SE2 | PAR-CLIP | Tarbase |
| hsa-miR-15a-5p | CDC42SE2 | HITS-CLIP | Tarbase |
| hsa-miR-15a-5p | CDC42SE2 | HITS-CLIP | Tarbase |
| hsa-miR-15a-5p | CDC42SE2 | HITS-CLIP | Tarbase |
| hsa-miR-15a-5p | CDC42SE2 | PAR-CLIP | Tarbase |
| hsa-miR-15a-5p | CDC42SE2 | HITS-CLIP | Tarbase |
| hsa-miR-195-5p | CDC42SE2 | HITS-CLIP | Tarbase |
| hsa-miR-195-5p | CDC42SE2 | HITS-CLIP | Tarbase |
| hsa-miR-195-5p | CDC42SE2 | HITS-CLIP | Tarbase |
| hsa-miR-141-3p | RAPGEF6 | HITS-CLIP | Tarbase |
| hsa-miR-195-5p | MIS18A | Microarrays | Tarbase |
| hsa-miR-15a-5p | SYNJ1 | PAR-CLIP | Tarbase |
| hsa-miR-15a-5p | IFNGR2 | HITS-CLIP | Tarbase |
| hsa-miR-195-5p | IFNGR2 | HITS-CLIP | Tarbase |
| hsa-miR-15a-5p | GART | HITS-CLIP | Tarbase |
| hsa-miR-195-5p | GART | HITS-CLIP | Tarbase |
| hsa-miR-141-3p | SON | HITS-CLIP | Tarbase |
| hsa-miR-15a-5p | SON | PAR-CLIP | Tarbase |
| hsa-miR-15a-5p | SON | PAR-CLIP | Tarbase |
| hsa-miR-15a-5p | SON | HITS-CLIP | Tarbase |
| hsa-miR-15a-5p | SON | HITS-CLIP | Tarbase |
| hsa-miR-15a-5p | SON | PAR-CLIP | Tarbase |
| hsa-miR-15a-5p | SON | HITS-CLIP | Tarbase |
| hsa-miR-195-5p | SON | HITS-CLIP | Tarbase |
| hsa-miR-195-5p | SON | HITS-CLIP | Tarbase |
| hsa-miR-195-5p | SON | HITS-CLIP | Tarbase |
| hsa-miR-15a-5p | SON | HITS-CLIP | Tarbase |
| hsa-miR-195-5p | SON | HITS-CLIP | Tarbase |
| hsa-miR-195-5p | STC1 | Microarrays | Tarbase |
| hsa-miR-15a-5p | C1QC | HITS-CLIP | Tarbase |
| hsa-miR-195-5p | C1QC | HITS-CLIP | Tarbase |
| hsa-miR-195-5p | RCAN1 | Microarrays | Tarbase |
| hsa-miR-141-3p | UBE2Z | HITS-CLIP | Tarbase |
| hsa-miR-141-5p | UBE2Z | HITS-CLIP | Tarbase |
| hsa-miR-15a-5p | UBE2Z | HITS-CLIP | Tarbase |
| hsa-miR-141-5p | IGF2BP1 | HITS-CLIP | Tarbase |
| hsa-miR-195-5p | CHAF1B | Microarrays | Tarbase |
| hsa-miR-486-5p | HLCS | Chimeric fragments | Tarbase |
| hsa-miR-141-3p | ADPGK | HITS-CLIP | Tarbase |

| | | | |
|---|---|---|---|
| hsa-miR-15a-5p | ADIPOR1 | HITS-CLIP | Tarbase |
| hsa-miR-15a-5p | ADIPOR1 | HITS-CLIP | Tarbase |
| hsa-miR-15a-5p | ADIPOR1 | HITS-CLIP | Tarbase |
| hsa-miR-195-5p | ADIPOR1 | HITS-CLIP | Tarbase |
| hsa-miR-195-5p | ADIPOR1 | HITS-CLIP | Tarbase |
| hsa-miR-195-5p | ADIPOR1 | HITS-CLIP | Tarbase |
| hsa-miR-3613-5p | PSMB4 | PAR-CLIP | Tarbase |
| hsa-miR-141-5p | BTG2 | HITS-CLIP | Tarbase |
| hsa-miR-15a-3p | BTG2 | HITS-CLIP | Tarbase |
| hsa-miR-15a-5p | BTG2 | HITS-CLIP | Tarbase |
| hsa-miR-15a-5p | BTG2 | PAR-CLIP | Tarbase |
| hsa-miR-15a-5p | BTG2 | HITS-CLIP | Tarbase |
| hsa-miR-15a-5p | BTG2 | HITS-CLIP | Tarbase |
| hsa-miR-15a-5p | BTG2 | HITS-CLIP | Tarbase |
| hsa-miR-15a-5p | BTG2 | HITS-CLIP | Tarbase |
| hsa-miR-15a-5p | BTG2 | PAR-CLIP | Tarbase |
| hsa-miR-195-5p | BTG2 | HITS-CLIP | Tarbase |
| hsa-miR-195-5p | BTG2 | PAR-CLIP | Tarbase |
| hsa-miR-195-5p | BTG2 | HITS-CLIP | Tarbase |
| hsa-miR-195-5p | BTG2 | HITS-CLIP | Tarbase |
| hsa-miR-195-5p | BTG2 | HITS-CLIP | Tarbase |
| hsa-miR-195-5p | BTG2 | HITS-CLIP | Tarbase |
| hsa-miR-195-5p | BTG2 | PAR-CLIP | Tarbase |
| hsa-miR-3613-3p | BTG2 | PAR-CLIP | Tarbase |
| hsa-miR-3613-5p | BTG2 | HITS-CLIP | Tarbase |
| hsa-miR-3613-5p | BTG2 | HITS-CLIP | Tarbase |
| hsa-miR-3613-5p | BTG2 | HITS-CLIP | Tarbase |
| hsa-miR-195-5p | BTG2 | Microarrays | Tarbase |
| hsa-miR-3613-3p | HK2 | HITS-CLIP | Tarbase |
| hsa-miR-195-5p | HK2 | Microarrays | Tarbase |
| hsa-miR-15a-5p | AMFR | HITS-CLIP | Tarbase |
| hsa-miR-195-5p | AMFR | HITS-CLIP | Tarbase |
| hsa-miR-15a-5p | RGL4 | HITS-CLIP | Tarbase |
| hsa-miR-195-5p | RGL4 | HITS-CLIP | Tarbase |
| hsa-miR-15a-5p | GPBP1L1 | PAR-CLIP | Tarbase |
| hsa-miR-15a-5p | GPBP1L1 | PAR-CLIP | Tarbase |
| hsa-miR-15a-5p | GPBP1L1 | PAR-CLIP | Tarbase |
| hsa-miR-15a-5p | GPBP1L1 | PAR-CLIP | Tarbase |
| hsa-miR-15a-5p | GPBP1L1 | PAR-CLIP | Tarbase |
| hsa-miR-15a-5p | GPBP1L1 | PAR-CLIP | Tarbase |
| hsa-miR-15a-5p | GPBP1L1 | PAR-CLIP | Tarbase |
| hsa-miR-15a-5p | GPBP1L1 | PAR-CLIP | Tarbase |
| hsa-miR-15a-5p | GPBP1L1 | PAR-CLIP | Tarbase |
| hsa-miR-195-5p | GPBP1L1 | PAR-CLIP | Tarbase |
| hsa-miR-195-5p | GPBP1L1 | PAR-CLIP | Tarbase |
| hsa-miR-195-5p | GPBP1L1 | PAR-CLIP | Tarbase |
| hsa-miR-195-5p | GPBP1L1 | PAR-CLIP | Tarbase |
| hsa-miR-486-5p | GPBP1L1 | HITS-CLIP | Tarbase |
| hsa-miR-15a-5p | NAE1 | HITS-CLIP | Tarbase |
| hsa-miR-195-5p | NAE1 | HITS-CLIP | Tarbase |
| hsa-miR-195-5p | EFCAB14 | Microarrays | Tarbase |
| hsa-miR-15a-3p | ZFYVE28 | HITS-CLIP | Tarbase |
| hsa-miR-15a-5p | PSKH1 | HITS-CLIP | Tarbase |
| hsa-miR-15a-5p | PSKH1 | HITS-CLIP | Tarbase |
| hsa-miR-15a-5p | PSKH1 | HITS-CLIP | Tarbase |
| hsa-miR-195-5p | PSKH1 | HITS-CLIP | Tarbase |
| hsa-miR-195-5p | PSKH1 | HITS-CLIP | Tarbase |
| hsa-miR-195-5p | PSKH1 | Microarrays | Tarbase |
| hsa-miR-141-3p | CCDC117 | HITS-CLIP | Tarbase |
| hsa-miR-141-5p | CCDC117 | HITS-CLIP | Tarbase |
| hsa-miR-15a-5p | CCDC117 | PAR-CLIP | Tarbase |
| hsa-miR-486-5p | CCDC117 | PAR-CLIP | Tarbase |
| hsa-miR-486-5p | CCDC117 | PAR-CLIP | Tarbase |
| hsa-miR-195-5p | CCDC117 | Microarrays | Tarbase |
| hsa-miR-486-5p | ZNF233 | HITS-CLIP | Tarbase |
| hsa-miR-219a-1-3p | GNE | HITS-CLIP | Tarbase |
| hsa-miR-15a-5p | ARHGAP35 | PAR-CLIP | Tarbase |
| hsa-miR-15a-5p | ARHGAP35 | HITS-CLIP | Tarbase |
| hsa-miR-219a-1-3p | ARHGAP35 | HITS-CLIP | Tarbase |
| hsa-miR-3605-3p | ARHGAP35 | HITS-CLIP | Tarbase |
| hsa-miR-15a-3p | CALM3 | PAR-CLIP | Tarbase |
| hsa-miR-3613-3p | DFFA | PAR-CLIP | Tarbase |
| hsa-miR-15a-5p | BSDC1 | HITS-CLIP | Tarbase |
| hsa-miR-15a-5p | BSDC1 | HITS-CLIP | Tarbase |
| hsa-miR-195-5p | BSDC1 | HITS-CLIP | Tarbase |
| hsa-miR-195-5p | BSDC1 | HITS-CLIP | Tarbase |
| hsa-miR-15a-5p | SSU72 | PAR-CLIP | Tarbase |
| hsa-miR-141-3p | VMA21 | HITS-CLIP | Tarbase |
| hsa-miR-15a-5p | VMA21 | HITS-CLIP | Tarbase |
| hsa-miR-195-5p | VMA21 | HITS-CLIP | Tarbase |
| hsa-miR-15a-5p | TFF3 | HITS-CLIP | Tarbase |
| hsa-miR-141-3p | U2AF1 | HITS-CLIP | Tarbase |
| hsa-miR-141-5p | U2AF1 | HITS-CLIP | Tarbase |
| hsa-miR-141-5p | RRP1B | HITS-CLIP | Tarbase |
| hsa-miR-141-5p | RRP1B | HITS-CLIP | Tarbase |
| hsa-miR-15a-3p | PDXK | HITS-CLIP | Tarbase |
| hsa-miR-15a-5p | PDXK | Chimeric fragments | Tarbase |
| hsa-miR-195-5p | G6PD | Microarrays | Tarbase |
| hsa-miR-15a-3p | AGPAT3 | HITS-CLIP | Tarbase |
| hsa-miR-15a-5p | AGPAT3 | HITS-CLIP | Tarbase |
| hsa-miR-195-5p | AGPAT3 | HITS-CLIP | Tarbase |
| hsa-miR-486-3p | AGPAT3 | HITS-CLIP | Tarbase |
| hsa-miR-141-3p | TRAPPC10 | HITS-CLIP | Tarbase |
| hsa-miR-3613-3p | TRAPPC10 | HITS-CLIP | Tarbase |
| hsa-miR-15a-5p | ICOSLG | HITS-CLIP | Tarbase |
| hsa-miR-15a-3p | RALGDS | HITS-CLIP | Tarbase |
| hsa-miR-195-5p | FTCD | HITS-CLIP | Tarbase |
| hsa-miR-15a-5p | SPATC1L | HITS-CLIP | Tarbase |
| hsa-miR-141-5p | MCM3AP | HITS-CLIP | Tarbase |
| hsa-miR-195-5p | C21orf58 | Microarrays | Tarbase |
| hsa-miR-141-5p | PCNT | HITS-CLIP | Tarbase |
| hsa-miR-3613-5p | ADAMTS13 | HITS-CLIP | Tarbase |
| hsa-miR-195-5p | SLC2A6 | Microarrays | Tarbase |
| hsa-miR-15a-3p | GPSM1 | HITS-CLIP | Tarbase |
| hsa-miR-15a-3p | GPSM1 | HITS-CLIP | Tarbase |
| hsa-miR-486-5p | GPSM1 | HITS-CLIP | Tarbase |
| hsa-miR-486-5p | ST6GALNAC6 | HITS-CLIP | Tarbase |
| hsa-miR-15a-5p | ZER1 | HITS-CLIP | Tarbase |
| hsa-miR-195-5p | ZER1 | HITS-CLIP | Tarbase |
| hsa-miR-15a-5p | ZDHHC12 | HITS-CLIP | Tarbase |
| hsa-miR-195-5p | ZDHHC12 | HITS-CLIP | Tarbase |
| hsa-miR-195-5p | ZDHHC12 | Microarrays | Tarbase |
| hsa-miR-195-5p | PKN3 | Microarrays | Tarbase |
| hsa-miR-3605-5p | BRSK1 | HITS-CLIP | Tarbase |
| hsa-miR-141-3p | TAOK1 | HITS-CLIP | Tarbase |
| hsa-miR-15a-5p | TAOK1 | HITS-CLIP | Tarbase |
| hsa-miR-15a-5p | TAOK1 | HITS-CLIP | Tarbase |
| hsa-miR-15a-5p | TAOK1 | PAR-CLIP | Tarbase |
| hsa-miR-15a-5p | TAOK1 | HITS-CLIP | Tarbase |
| hsa-miR-15a-5p | TAOK1 | HITS-CLIP | Tarbase |
| hsa-miR-195-5p | TAOK1 | HITS-CLIP | Tarbase |
| hsa-miR-195-5p | TAOK1 | HITS-CLIP | Tarbase |
| hsa-miR-195-5p | TAOK1 | HITS-CLIP | Tarbase |
| hsa-miR-3613-3p | TAOK1 | PAR-CLIP | Tarbase |
| hsa-miR-15a-5p | TAOK1 | HITS-CLIP | Tarbase |
| hsa-miR-195-5p | TAOK1 | HITS-CLIP | Tarbase |
| hsa-miR-15a-5p | SIK3 | PAR-CLIP | Tarbase |
| hsa-miR-15a-5p | MPZL3 | HITS-CLIP | Tarbase |
| hsa-miR-195-5p | MPZL3 | Microarrays | Tarbase |
| hsa-miR-15a-5p | TLCD1 | HITS-CLIP | Tarbase |
| hsa-miR-486-3p | ZBTB7B | HITS-CLIP | Tarbase |
| hsa-miR-141-3p | SHC1 | Luciferase Reporter Assay | Tarbase |
| hsa-miR-141-3p | SHC1 | Luciferase Reporter Assay | Tarbase |
| hsa-miR-15a-3p | SHC1 | HITS-CLIP | Tarbase |
| hsa-miR-15a-5p | NLRX1 | PAR-CLIP | Tarbase |
| hsa-miR-195-5p | NLRX1 | Microarrays | Tarbase |
| hsa-miR-141-3p | ADAR | HITS-CLIP | Tarbase |
| hsa-miR-141-3p | ADAR | HITS-CLIP | Tarbase |
| hsa-miR-15a-3p | ADAR | HITS-CLIP | Tarbase |
| hsa-miR-15a-5p | ADAR | HITS-CLIP | Tarbase |
| hsa-miR-141-3p | IL6R | HITS-CLIP | Tarbase |
| hsa-miR-3613-5p | IL6R | HITS-CLIP | Tarbase |
| hsa-miR-15a-5p | UBE2Q1 | PAR-CLIP | Tarbase |
| hsa-miR-3605-5p | CRTC2 | PAR-CLIP | Tarbase |
| hsa-miR-141-3p | FDPS | HITS-CLIP | Tarbase |
| hsa-miR-141-3p | FDPS | HITS-CLIP | Tarbase |
| hsa-miR-141-5p | FDPS | HITS-CLIP | Tarbase |
| hsa-miR-15a-3p | FDPS | HITS-CLIP | Tarbase |
| hsa-miR-15a-5p | RUSC1 | HITS-CLIP | Tarbase |

| | | | |
|---|---|---|---|
| hsa-miR-195-5p | RUSC1 | HITS-CLIP | Tarbase |
| hsa-miR-15a-3p | FAM189B | HITS-CLIP | Tarbase |
| hsa-miR-15a-5p | FAM189B | HITS-CLIP | Tarbase |
| hsa-miR-15a-5p | FAM189B | HITS-CLIP | Tarbase |
| hsa-miR-15a-5p | FAM189B | HITS-CLIP | Tarbase |
| hsa-miR-15a-5p | FAM189B | HITS-CLIP | Tarbase |
| hsa-miR-15a-5p | FAM189B | PAR-CLIP | Tarbase |
| hsa-miR-15a-5p | FAM189B | HITS-CLIP | Tarbase |
| hsa-miR-195-5p | FAM189B | HITS-CLIP | Tarbase |
| hsa-miR-195-5p | FAM189B | HITS-CLIP | Tarbase |
| hsa-miR-195-5p | FAM189B | HITS-CLIP | Tarbase |
| hsa-miR-195-5p | FAM189B | HITS-CLIP | Tarbase |
| hsa-miR-195-5p | FAM189B | Microarrays | Tarbase |
| hsa-miR-15a-5p | FAM189B | Chimeric fragments | Tarbase |
| hsa-miR-15a-5p | PAQR6 | HITS-CLIP | Tarbase |
| hsa-miR-195-5p | PAQR6 | HITS-CLIP | Tarbase |
| hsa-miR-195-5p | SLC25A44 | Microarrays | Tarbase |
| hsa-miR-195-5p | FGFR4 | Microarrays | Tarbase |
| hsa-miR-15a-3p | NACC1 | HITS-CLIP | Tarbase |
| hsa-miR-15a-3p | NACC1 | HITS-CLIP | Tarbase |
| hsa-miR-15a-3p | NACC1 | HITS-CLIP | Tarbase |
| hsa-miR-15a-3p | NACC1 | HITS-CLIP | Tarbase |
| hsa-miR-3613-3p | NACC1 | HITS-CLIP | Tarbase |
| hsa-miR-141-3p | LY6K | HITS-CLIP | Tarbase |
| hsa-miR-219a-1-3p | LY6K | HITS-CLIP | Tarbase |
| hsa-miR-15a-5p | LY6E | HITS-CLIP | Tarbase |
| hsa-miR-15a-5p | LY6E | HITS-CLIP | Tarbase |
| hsa-miR-15a-5p | LY6E | HITS-CLIP | Tarbase |
| hsa-miR-195-5p | LY6E | HITS-CLIP | Tarbase |
| hsa-miR-195-5p | LY6E | HITS-CLIP | Tarbase |
| hsa-miR-486-3p | LY6E | HITS-CLIP | Tarbase |
| hsa-miR-15a-5p | SQSTM1 | PAR-CLIP | Tarbase |
| hsa-miR-15a-5p | SQSTM1 | HITS-CLIP | Tarbase |
| hsa-miR-15a-5p | SQSTM1 | HITS-CLIP | Tarbase |
| hsa-miR-195-5p | SQSTM1 | HITS-CLIP | Tarbase |
| hsa-miR-3613-3p | SQSTM1 | PAR-CLIP | Tarbase |
| hsa-miR-15a-5p | MGAT4B | HITS-CLIP | Tarbase |
| hsa-miR-195-5p | MGAT4B | HITS-CLIP | Tarbase |
| hsa-miR-486-5p | RPL8 | PAR-CLIP | Tarbase |
| hsa-miR-15a-3p | MAML1 | HITS-CLIP | Tarbase |
| hsa-miR-15a-5p | PGLYRP2 | HITS-CLIP | Tarbase |
| hsa-miR-195-5p | PGLYRP2 | HITS-CLIP | Tarbase |
| hsa-miR-15a-5p | PSMC2 | HITS-CLIP | Tarbase |
| hsa-miR-15a-5p | PSMC2 | HITS-CLIP | Tarbase |
| hsa-miR-195-5p | PSMC2 | HITS-CLIP | Tarbase |
| hsa-miR-15a-5p | YDJC | Chimeric fragments | Tarbase |
| hsa-miR-15a-3p | CCDC116 | HITS-CLIP | Tarbase |
| hsa-miR-15a-5p | DVL3 | HITS-CLIP | Tarbase |
| hsa-miR-15a-5p | DVL3 | HITS-CLIP | Tarbase |
| hsa-miR-195-5p | DVL3 | HITS-CLIP | Tarbase |
| hsa-miR-486-5p | DVL3 | PAR-CLIP | Tarbase |
| hsa-miR-15a-5p | ABCF3 | PAR-CLIP | Tarbase |
| hsa-miR-195-5p | ABCF3 | Microarrays | Tarbase |
| hsa-miR-15a-5p | FDXR | HITS-CLIP | Tarbase |
| hsa-miR-141-5p | SAP30BP | HITS-CLIP | Tarbase |
| hsa-miR-15a-3p | SAP30BP | HITS-CLIP | Tarbase |
| hsa-miR-15a-5p | ACOX1 | HITS-CLIP | Tarbase |
| hsa-miR-195-5p | ACOX1 | HITS-CLIP | Tarbase |
| hsa-miR-15a-5p | SRSF2 | HITS-CLIP | Tarbase |
| hsa-miR-195-5p | SRSF2 | HITS-CLIP | Tarbase |
| hsa-miR-486-3p | ITGA5 | HITS-CLIP | Tarbase |
| hsa-miR-219a-1-3p | ZNF385A | HITS-CLIP | Tarbase |
| hsa-miR-15a-3p | EMC10 | HITS-CLIP | Tarbase |
| hsa-miR-15a-5p | JOSD2 | HITS-CLIP | Tarbase |
| hsa-miR-195-5p | JOSD2 | HITS-CLIP | Tarbase |
| hsa-miR-195-5p | FAM171A2 | Microarrays | Tarbase |
| hsa-miR-195-5p | DBF4B | Microarrays | Tarbase |
| hsa-miR-141-5p | PLCD3 | HITS-CLIP | Tarbase |
| hsa-miR-486-3p | FMNL3 | PAR-CLIP | Tarbase |
| hsa-miR-15a-5p | RACGAP1 | PAR-CLIP | Tarbase |
| hsa-miR-15a-5p | RACGAP1 | PAR-CLIP | Tarbase |
| hsa-miR-15a-5p | RACGAP1 | PAR-CLIP | Tarbase |
| hsa-miR-15a-5p | RACGAP1 | PAR-CLIP | Tarbase |
| hsa-miR-15a-5p | RACGAP1 | PAR-CLIP | Tarbase |
| hsa-miR-15a-5p | RACGAP1 | PAR-CLIP | Tarbase |
| hsa-miR-195-5p | RACGAP1 | PAR-CLIP | Tarbase |
| hsa-miR-195-5p | RACGAP1 | PAR-CLIP | Tarbase |
| hsa-miR-195-5p | RACGAP1 | PAR-CLIP | Tarbase |
| hsa-miR-195-5p | RACGAP1 | PAR-CLIP | Tarbase |
| hsa-miR-195-5p | RACGAP1 | Microarrays | Tarbase |
| hsa-miR-15a-5p | LARP4 | PAR-CLIP | Tarbase |
| hsa-miR-15a-5p | LARP4 | PAR-CLIP | Tarbase |
| hsa-miR-15a-5p | LARP4 | PAR-CLIP | Tarbase |
| hsa-miR-15a-5p | LARP4 | PAR-CLIP | Tarbase |
| hsa-miR-15a-5p | LARP4 | HITS-CLIP | Tarbase |
| hsa-miR-195-5p | LARP4 | PAR-CLIP | Tarbase |
| hsa-miR-195-5p | SPC24 | Microarrays | Tarbase |
| hsa-miR-15a-3p | EIF4A1 | HITS-CLIP | Tarbase |
| hsa-miR-15a-3p | EIF4A1 | HITS-CLIP | Tarbase |
| hsa-miR-141-5p | RPL26 | HITS-CLIP | Tarbase |
| hsa-miR-141-5p | RPL26 | HITS-CLIP | Tarbase |
| hsa-miR-15a-5p | POLR3K | HITS-CLIP | Tarbase |
| hsa-miR-195-5p | POLR3K | HITS-CLIP | Tarbase |
| hsa-miR-195-5p | CCNF | Microarrays | Tarbase |
| hsa-miR-15a-3p | AMDHD2 | HITS-CLIP | Tarbase |
| hsa-miR-141-5p | ADCY9 | HITS-CLIP | Tarbase |
| hsa-miR-141-5p | ADCY9 | HITS-CLIP | Tarbase |
| hsa-miR-15a-5p | SHANK2 | HITS-CLIP | Tarbase |
| hsa-miR-195-5p | SHANK2 | HITS-CLIP | Tarbase |
| hsa-miR-15a-5p | CLPB | Chimeric fragments | Tarbase |
| hsa-miR-195-5p | CYB561A3 | Microarrays | Tarbase |
| hsa-miR-15a-5p | LRP5 | HITS-CLIP | Tarbase |
| hsa-miR-141-3p | CMPK1 | HITS-CLIP | Tarbase |
| hsa-miR-3613-3p | CMPK1 | HITS-CLIP | Tarbase |
| hsa-miR-141-3p | COA7 | HITS-CLIP | Tarbase |
| hsa-miR-141-5p | COA7 | HITS-CLIP | Tarbase |
| hsa-miR-15a-5p | ZYG11B | HITS-CLIP | Tarbase |
| hsa-miR-195-5p | SLC1A7 | Microarrays | Tarbase |
| hsa-miR-15a-5p | USP24 | HITS-CLIP | Tarbase |
| hsa-miR-15a-5p | PPAP2B | PAR-CLIP | Tarbase |
| hsa-miR-15a-5p | PPAP2B | PAR-CLIP | Tarbase |
| hsa-miR-3605-5p | PPAP2B | HITS-CLIP | Tarbase |
| hsa-miR-15a-3p | SEPN1 | HITS-CLIP | Tarbase |
| hsa-miR-195-5p | PDPN | Microarrays | Tarbase |
| hsa-miR-15a-5p | SDC3 | HITS-CLIP | Tarbase |
| hsa-miR-195-5p | SDC3 | HITS-CLIP | Tarbase |
| hsa-miR-195-5p | SDC3 | Microarrays | Tarbase |
| hsa-miR-15a-5p | KIAA1522 | HITS-CLIP | Tarbase |
| hsa-miR-15a-5p | KIAA1522 | HITS-CLIP | Tarbase |
| hsa-miR-195-5p | KIAA1522 | HITS-CLIP | Tarbase |
| hsa-miR-195-5p | CAMK2N1 | Microarrays | Tarbase |
| hsa-miR-141-3p | NFIA | HITS-CLIP | Tarbase |
| hsa-miR-486-5p | NFIA | HITS-CLIP | Tarbase |
| hsa-miR-15a-5p | OMA1 | HITS-CLIP | Tarbase |
| hsa-miR-141-5p | MYSM1 | HITS-CLIP | Tarbase |
| hsa-miR-15a-5p | MYSM1 | HITS-CLIP | Tarbase |
| hsa-miR-195-5p | MYSM1 | HITS-CLIP | Tarbase |
| hsa-miR-195-5p | USP1 | Microarrays | Tarbase |
| hsa-miR-141-3p | FUBP1 | HITS-CLIP | Tarbase |
| hsa-miR-3613-3p | FUBP1 | PAR-CLIP | Tarbase |
| hsa-miR-15a-5p | DNAJB4 | HITS-CLIP | Tarbase |
| hsa-miR-15a-5p | DNAJB4 | HITS-CLIP | Tarbase |
| hsa-miR-15a-5p | DNAJB4 | PAR-CLIP | Tarbase |
| hsa-miR-195-5p | DNAJB4 | HITS-CLIP | Tarbase |
| hsa-miR-195-5p | DNAJB4 | HITS-CLIP | Tarbase |
| hsa-miR-15a-5p | DNAJB4 | HITS-CLIP | Tarbase |
| hsa-miR-195-5p | DNAJB4 | HITS-CLIP | Tarbase |
| hsa-miR-195-5p | B3GALT2 | Microarrays | Tarbase |
| hsa-miR-195-5p | NTNG1 | Microarrays | Tarbase |
| hsa-miR-195-5p | C1orf52 | PAR-CLIP | Tarbase |
| hsa-miR-486-5p | ATXN7L2 | PAR-CLIP | Tarbase |
| hsa-miR-141-5p | ZNF326 | HITS-CLIP | Tarbase |
| hsa-miR-15a-5p | ZNF326 | PAR-CLIP | Tarbase |
| hsa-miR-15a-5p | ZNF326 | PAR-CLIP | Tarbase |
| hsa-miR-195-5p | ZNF326 | PAR-CLIP | Tarbase |
| hsa-miR-486-5p | ZNF326 | HITS-CLIP | Tarbase |
| hsa-miR-141-3p | AGL | HITS-CLIP | Tarbase |
| hsa-miR-195-5p | VCAM1 | qPCR | Tarbase |
| hsa-miR-195-5p | VCAM1 | Western Blot | Tarbase |

| | | | |
|---|---|---|---|
| hsa-miR-195-5p | VCAM1 | Microarrays | Tarbase |
| hsa-miR-141-3p | EXTL2 | HITS-CLIP | Tarbase |
| hsa-miR-195-5p | EXTL2 | Microarrays | Tarbase |
| hsa-miR-141-5p | SLC30A7 | HITS-CLIP | Tarbase |
| hsa-miR-15a-5p | SLC30A7 | PAR-CLIP | Tarbase |
| hsa-miR-15a-5p | SLC30A7 | PAR-CLIP | Tarbase |
| hsa-miR-195-5p | SLC30A7 | PAR-CLIP | Tarbase |
| hsa-miR-141-3p | ZNF281 | HITS-CLIP | Tarbase |
| hsa-miR-15a-5p | ZNF281 | HITS-CLIP | Tarbase |
| hsa-miR-15a-5p | ZNF281 | HITS-CLIP | Tarbase |
| hsa-miR-15a-5p | ZNF281 | PAR-CLIP | Tarbase |
| hsa-miR-195-5p | ZNF281 | HITS-CLIP | Tarbase |
| hsa-miR-195-5p | ZNF281 | HITS-CLIP | Tarbase |
| hsa-miR-195-5p | ZNF281 | PAR-CLIP | Tarbase |
| hsa-miR-219a-1-3p | ZNF281 | HITS-CLIP | Tarbase |
| hsa-miR-141-3p | ARPC5 | HITS-CLIP | Tarbase |
| hsa-miR-141-3p | ARPC5 | HITS-CLIP | Tarbase |
| hsa-miR-15a-5p | PEX19 | PAR-CLIP | Tarbase |
| hsa-miR-15a-5p | PEX19 | PAR-CLIP | Tarbase |
| hsa-miR-195-5p | PEX19 | PAR-CLIP | Tarbase |
| hsa-miR-486-5p | PEX19 | PAR-CLIP | Tarbase |
| hsa-miR-15a-5p | RBM15 | HITS-CLIP | Tarbase |
| hsa-miR-15a-5p | BPNT1 | HITS-CLIP | Tarbase |
| hsa-miR-195-5p | BPNT1 | HITS-CLIP | Tarbase |
| hsa-miR-15a-5p | CNST | HITS-CLIP | Tarbase |
| hsa-miR-195-5p | CNST | HITS-CLIP | Tarbase |
| hsa-miR-141-3p | PPP1R21 | HITS-CLIP | Tarbase |
| hsa-miR-15a-5p | PPP1R21 | HITS-CLIP | Tarbase |
| hsa-miR-15a-5p | PPP1R21 | HITS-CLIP | Tarbase |
| hsa-miR-195-5p | PPP1R21 | HITS-CLIP | Tarbase |
| hsa-miR-141-5p | B3GALNT2 | HITS-CLIP | Tarbase |
| hsa-miR-195-5p | IL24 | Microarrays | Tarbase |
| hsa-miR-195-5p | C1orf145 | Microarrays | Tarbase |
| hsa-miR-15a-5p | WDR26 | PAR-CLIP | Tarbase |
| hsa-miR-141-3p | REL | HITS-CLIP | Tarbase |
| hsa-miR-3613-3p | REL | PAR-CLIP | Tarbase |
| hsa-miR-15a-5p | C2orf47 | HITS-CLIP | Tarbase |
| hsa-miR-195-5p | C2orf47 | HITS-CLIP | Tarbase |
| hsa-miR-141-5p | ARL5A | HITS-CLIP | Tarbase |
| hsa-miR-195-5p | FAM84A | Microarrays | Tarbase |
| hsa-miR-15a-5p | CLHC1 | CLEAR-CLIP | Tarbase |
| hsa-miR-141-5p | NUP35 | HITS-CLIP | Tarbase |
| hsa-miR-141-3p | CCDC138 | HITS-CLIP | Tarbase |
| hsa-miR-195-5p | FBXO41 | HITS-CLIP | Tarbase |
| hsa-miR-195-5p | ACTG2 | Microarrays | Tarbase |
| hsa-miR-15a-5p | SLC16A14 | HITS-CLIP | Tarbase |
| hsa-miR-195-5p | SLC16A14 | HITS-CLIP | Tarbase |
| hsa-miR-15a-5p | SGPP2 | PAR-CLIP | Tarbase |
| hsa-miR-15a-5p | SGPP2 | PAR-CLIP | Tarbase |
| hsa-miR-195-5p | SGPP2 | PAR-CLIP | Tarbase |
| hsa-miR-15a-5p | PDLIM5 | HITS-CLIP | Tarbase |
| hsa-miR-141-3p | STPG2 | HITS-CLIP | Tarbase |
| hsa-miR-141-3p | RPRD2 | PAR-CLIP | Tarbase |
| hsa-miR-15a-5p | RPRD2 | PAR-CLIP | Tarbase |
| hsa-miR-195-5p | RPRD2 | PAR-CLIP | Tarbase |
| hsa-miR-219a-1-3p | RPRD2 | PAR-CLIP | Tarbase |
| hsa-miR-15a-5p | ANKRD23 | HITS-CLIP | Tarbase |
| hsa-miR-195-5p | ANKRD23 | HITS-CLIP | Tarbase |
| hsa-miR-15a-5p | VPS72 | PAR-CLIP | Tarbase |
| hsa-miR-141-5p | ERCC3 | HITS-CLIP | Tarbase |
| hsa-miR-141-3p | RNF149 | HITS-CLIP | Tarbase |
| hsa-miR-15a-5p | RNF149 | PAR-CLIP | Tarbase |
| hsa-miR-195-5p | RNF149 | PAR-CLIP | Tarbase |
| hsa-miR-15a-5p | BOLA3 | HITS-CLIP | Tarbase |
| hsa-miR-195-5p | CDC42EP3 | Microarrays | Tarbase |
| hsa-miR-219a-1-3p | S100A11 | HITS-CLIP | Tarbase |
| hsa-miR-141-5p | DHX57 | HITS-CLIP | Tarbase |
| hsa-miR-195-5p | FZD5 | Microarrays | Tarbase |
| hsa-miR-195-5p | NPPC | HITS-CLIP | Tarbase |
| hsa-miR-195-5p | GABRG1 | Microarrays | Tarbase |
| hsa-miR-195-5p | GABRB1 | Microarrays | Tarbase |
| hsa-miR-15a-5p | PAQR3 | HITS-CLIP | Tarbase |
| hsa-miR-195-5p | ANTXR2 | Microarrays | Tarbase |
| hsa-miR-141-3p | CGGBP1 | HITS-CLIP | Tarbase |
| hsa-miR-15a-5p | CGGBP1 | HITS-CLIP | Tarbase |
| hsa-miR-15a-5p | CGGBP1 | HITS-CLIP | Tarbase |
| hsa-miR-195-5p | CGGBP1 | HITS-CLIP | Tarbase |
| hsa-miR-15a-5p | GPR155 | HITS-CLIP | Tarbase |
| hsa-miR-195-5p | GPR155 | Microarrays | Tarbase |
| hsa-miR-15a-5p | PBXIP1 | HITS-CLIP | Tarbase |
| hsa-miR-195-5p | PBXIP1 | HITS-CLIP | Tarbase |
| hsa-miR-195-5p | PBXIP1 | HITS-CLIP | Tarbase |
| hsa-miR-3605-5p | PBXIP1 | HITS-CLIP | Tarbase |
| hsa-miR-195-5p | CLDN1 | Microarrays | Tarbase |
| hsa-miR-141-3p | HIPK1 | HITS-CLIP | Tarbase |
| hsa-miR-141-3p | HIPK1 | HITS-CLIP | Tarbase |
| hsa-miR-141-3p | HIPK1 | HITS-CLIP | Tarbase |
| hsa-miR-15a-5p | HIPK1 | PAR-CLIP | Tarbase |
| hsa-miR-15a-5p | HIPK1 | PAR-CLIP | Tarbase |
| hsa-miR-15a-5p | HIPK1 | HITS-CLIP | Tarbase |
| hsa-miR-3605-5p | HIPK1 | HITS-CLIP | Tarbase |
| hsa-miR-3613-3p | HIPK1 | PAR-CLIP | Tarbase |
| hsa-miR-486-5p | HIPK1 | PAR-CLIP | Tarbase |
| hsa-miR-195-5p | COL6A3 | Microarrays | Tarbase |
| hsa-miR-15a-3p | C1orf106 | HITS-CLIP | Tarbase |
| hsa-miR-141-5p | YY1AP1 | HITS-CLIP | Tarbase |
| hsa-miR-141-3p | EOGT | HITS-CLIP | Tarbase |
| hsa-miR-141-5p | ATP1A1 | HITS-CLIP | Tarbase |
| hsa-miR-219a-1-3p | ATP1A1 | HITS-CLIP | Tarbase |
| hsa-miR-219a-1-3p | ATP1A1 | HITS-CLIP | Tarbase |
| hsa-miR-486-5p | ATP1A1 | PAR-CLIP | Tarbase |
| hsa-miR-141-3p | LRRC58 | HITS-CLIP | Tarbase |
| hsa-miR-141-5p | LRRC58 | HITS-CLIP | Tarbase |
| hsa-miR-15a-5p | LRRC58 | HITS-CLIP | Tarbase |
| hsa-miR-15a-5p | LRRC58 | PAR-CLIP | Tarbase |
| hsa-miR-141-5p | FSTL1 | HITS-CLIP | Tarbase |
| hsa-miR-15a-5p | FSTL1 | HITS-CLIP | Tarbase |
| hsa-miR-15a-5p | FSTL1 | HITS-CLIP | Tarbase |
| hsa-miR-195-5p | FSTL1 | AGO-IP | Tarbase |
| hsa-miR-195-5p | FSTL1 | Microarrays | Tarbase |
| hsa-miR-195-5p | ELF3 | Microarrays | Tarbase |
| hsa-miR-15a-5p | TMEM183A | PAR-CLIP | Tarbase |
| hsa-miR-15a-5p | TMEM183A | HITS-CLIP | Tarbase |
| hsa-miR-195-5p | TMEM183A | HITS-CLIP | Tarbase |
| hsa-miR-486-3p | IGFBP7 | HITS-CLIP | Tarbase |
| hsa-miR-195-5p | IGFBP7 | Microarrays | Tarbase |
| hsa-miR-15a-5p | CCT3 | Chimeric fragments | Tarbase |
| hsa-miR-195-5p | IHH | AGO-IP | Tarbase |
| hsa-miR-195-5p | KIAA1524 | Microarrays | Tarbase |
| hsa-miR-141-5p | STT3B | HITS-CLIP | Tarbase |
| hsa-miR-3605-3p | NFASC | HITS-CLIP | Tarbase |
| hsa-miR-195-5p | SGOL2 | Microarrays | Tarbase |
| hsa-miR-141-3p | CLASP2 | PAR-CLIP | Tarbase |
| hsa-miR-141-3p | CLASP2 | HITS-CLIP | Tarbase |
| hsa-miR-15a-5p | CLASP2 | HITS-CLIP | Tarbase |
| hsa-miR-15a-5p | CLASP2 | HITS-CLIP | Tarbase |
| hsa-miR-195-5p | CLASP2 | HITS-CLIP | Tarbase |
| hsa-miR-15a-5p | SUCLG1 | HITS-CLIP | Tarbase |
| hsa-miR-15a-5p | SUCLG1 | HITS-CLIP | Tarbase |
| hsa-miR-195-5p | SUCLG1 | HITS-CLIP | Tarbase |
| hsa-miR-195-5p | SUCLG1 | HITS-CLIP | Tarbase |
| hsa-miR-195-5p | RPL22L1 | Microarrays | Tarbase |
| hsa-miR-15a-5p | RYBP | HITS-CLIP | Tarbase |
| hsa-miR-15a-5p | PPP4R2 | PAR-CLIP | Tarbase |
| hsa-miR-15a-5p | PPP4R2 | HITS-CLIP | Tarbase |
| hsa-miR-195-5p | PPP4R2 | HITS-CLIP | Tarbase |
| hsa-miR-141-3p | C3orf17 | HITS-CLIP | Tarbase |
| hsa-miR-15a-5p | CDS1 | HITS-CLIP | Tarbase |
| hsa-miR-195-5p | CDS1 | HITS-CLIP | Tarbase |
| hsa-miR-141-3p | WDFY3 | HITS-CLIP | Tarbase |
| hsa-miR-15a-5p | WDFY3 | PAR-CLIP | Tarbase |
| hsa-miR-195-5p | PTPN13 | Microarrays | Tarbase |
| hsa-miR-15a-5p | ATXN7 | HITS-CLIP | Tarbase |
| hsa-miR-195-5p | PRICKLE2 | Microarrays | Tarbase |
| hsa-miR-195-5p | ADAMTS9 | Microarrays | Tarbase |
| hsa-miR-15a-5p | PPM1K | PAR-CLIP | Tarbase |
| hsa-miR-15a-5p | PPM1K | HITS-CLIP | Tarbase |
| hsa-miR-195-5p | PPM1K | PAR-CLIP | Tarbase |
| hsa-miR-195-5p | PPM1K | HITS-CLIP | Tarbase |

| | | | |
|---|---|---|---|
| hsa-miR-15a-5p | PPM1K | HITS-CLIP | Tarbase |
| hsa-miR-195-5p | PPM1K | HITS-CLIP | Tarbase |
| hsa-miR-141-5p | SLMAP | HITS-CLIP | Tarbase |
| hsa-miR-141-3p | SMIM14 | HITS-CLIP | Tarbase |
| hsa-miR-15a-5p | C3orf67 | HITS-CLIP | Tarbase |
| hsa-miR-195-5p | C3orf67 | HITS-CLIP | Tarbase |
| hsa-miR-195-5p | APBB2 | Microarrays | Tarbase |
| hsa-miR-141-3p | U2SURP | HITS-CLIP | Tarbase |
| hsa-miR-141-3p | U2SURP | HITS-CLIP | Tarbase |
| hsa-miR-141-3p | U2SURP | HITS-CLIP | Tarbase |
| hsa-miR-141-5p | U2SURP | HITS-CLIP | Tarbase |
| hsa-miR-141-5p | U2SURP | HITS-CLIP | Tarbase |
| hsa-miR-15a-5p | U2SURP | HITS-CLIP | Tarbase |
| hsa-miR-15a-5p | U2SURP | PAR-CLIP | Tarbase |
| hsa-miR-15a-5p | U2SURP | HITS-CLIP | Tarbase |
| hsa-miR-195-5p | U2SURP | HITS-CLIP | Tarbase |
| hsa-miR-3605-3p | MTMR14 | HITS-CLIP | Tarbase |
| hsa-miR-15a-5p | TTC14 | HITS-CLIP | Tarbase |
| hsa-miR-195-5p | TTC14 | HITS-CLIP | Tarbase |
| hsa-miR-15a-5p | CXCL5 | HITS-CLIP | Tarbase |
| hsa-miR-195-5p | CXCL5 | HITS-CLIP | Tarbase |
| hsa-miR-15a-5p | HPS3 | HITS-CLIP | Tarbase |
| hsa-miR-195-5p | HPS3 | HITS-CLIP | Tarbase |
| hsa-miR-486-3p | TOPBP1 | HITS-CLIP | Tarbase |
| hsa-miR-141-5p | SNRK | HITS-CLIP | Tarbase |
| hsa-miR-15a-5p | SNRK | HITS-CLIP | Tarbase |
| hsa-miR-15a-5p | SNRK | HITS-CLIP | Tarbase |
| hsa-miR-195-5p | SNRK | HITS-CLIP | Tarbase |
| hsa-miR-195-5p | SNRK | HITS-CLIP | Tarbase |
| hsa-miR-195-5p | KIF15 | Microarrays | Tarbase |
| hsa-miR-141-5p | WDR43 | HITS-CLIP | Tarbase |
| hsa-miR-15a-5p | WDR43 | PAR-CLIP | Tarbase |
| hsa-miR-15a-5p | WDR43 | PAR-CLIP | Tarbase |
| hsa-miR-15a-3p | ZDHHC3 | HITS-CLIP | Tarbase |
| hsa-miR-195-5p | CDCP1 | Microarrays | Tarbase |
| hsa-miR-15a-5p | FYCO1 | PAR-CLIP | Tarbase |
| hsa-miR-15a-5p | FYCO1 | PAR-CLIP | Tarbase |
| hsa-miR-195-5p | FYCO1 | PAR-CLIP | Tarbase |
| hsa-miR-3605-3p | FYCO1 | HITS-CLIP | Tarbase |
| hsa-miR-486-5p | FYCO1 | HITS-CLIP | Tarbase |
| hsa-miR-195-5p | FYCO1 | Microarrays | Tarbase |
| hsa-miR-195-5p | CCR1 | Microarrays | Tarbase |
| hsa-miR-141-5p | ELP6 | HITS-CLIP | Tarbase |
| hsa-miR-15a-3p | ELP6 | HITS-CLIP | Tarbase |
| hsa-miR-15a-3p | ELP6 | HITS-CLIP | Tarbase |
| hsa-miR-195-5p | DTX3L | Microarrays | Tarbase |
| hsa-miR-141-3p | ZNF148 | HITS-CLIP | Tarbase |
| hsa-miR-141-3p | ZNF148 | HITS-CLIP | Tarbase |
| hsa-miR-3605-5p | YEATS2 | HITS-CLIP | Tarbase |
| hsa-miR-15a-3p | ZC3H12A | HITS-CLIP | Tarbase |
| hsa-miR-141-3p | SNIP1 | HITS-CLIP | Tarbase |
| hsa-miR-15a-5p | POLR2H | HITS-CLIP | Tarbase |
| hsa-miR-195-5p | POLR2H | HITS-CLIP | Tarbase |
| hsa-miR-195-5p | POLR2H | HITS-CLIP | Tarbase |
| hsa-miR-15a-3p | TMEM41A | HITS-CLIP | Tarbase |
| hsa-miR-195-5p | RFC4 | Microarrays | Tarbase |
| hsa-miR-195-5p | RPL39L | Microarrays | Tarbase |
| hsa-miR-141-3p | BAP1 | Luciferase Reporter Assay | Tarbase |
| hsa-miR-141-3p | BAP1 | Luciferase Reporter Assay | Tarbase |
| hsa-miR-195-5p | TKT | Microarrays | Tarbase |
| hsa-miR-15a-5p | PRKCD | HITS-CLIP | Tarbase |
| hsa-miR-195-5p | PRKCD | HITS-CLIP | Tarbase |
| hsa-miR-195-5p | PRKCD | Microarrays | Tarbase |
| hsa-miR-15a-5p | RFT1 | HITS-CLIP | Tarbase |
| hsa-miR-195-5p | SFMBT1 | Microarrays | Tarbase |
| hsa-miR-15a-5p | UVSSA | HITS-CLIP | Tarbase |
| hsa-miR-195-5p | UVSSA | HITS-CLIP | Tarbase |
| hsa-miR-15a-5p | FAM208A | HITS-CLIP | Tarbase |
| hsa-miR-195-5p | FAM208A | HITS-CLIP | Tarbase |
| hsa-miR-3613-3p | SLBP | PAR-CLIP | Tarbase |
| hsa-miR-141-3p | UBXN7 | HITS-CLIP | Tarbase |
| hsa-miR-141-5p | UBXN7 | HITS-CLIP | Tarbase |
| hsa-miR-3613-3p | UBXN7 | PAR-CLIP | Tarbase |
| hsa-miR-3613-3p | UBXN7 | PAR-CLIP | Tarbase |
| hsa-miR-15a-5p | RNF168 | PAR-CLIP | Tarbase |
| hsa-miR-195-5p | RNF168 | PAR-CLIP | Tarbase |
| hsa-miR-15a-5p | RNF168 | Chimeric fragments | Tarbase |
| hsa-miR-195-5p | S100P | Microarrays | Tarbase |
| hsa-miR-15a-3p | ZNF691 | HITS-CLIP | Tarbase |
| hsa-miR-15a-5p | DNAJB14 | HITS-CLIP | Tarbase |
| hsa-miR-195-5p | DNAJB14 | HITS-CLIP | Tarbase |
| hsa-miR-141-3p | H2AFZ | HITS-CLIP | Tarbase |
| hsa-miR-195-5p | H2AFZ | Microarrays | Tarbase |
| hsa-miR-141-3p | CDC25A | HITS-CLIP | Tarbase |
| hsa-miR-141-3p | CDC25A | Luciferase Reporter Assay | Tarbase |
| hsa-miR-141-3p | CDC25A | Western Blot | Tarbase |
| hsa-miR-15a-5p | CDC25A | PAR-CLIP | Tarbase |
| hsa-miR-195-5p | CDC25A | Luciferase Reporter Assay | Tarbase |
| hsa-miR-195-5p | CDC25A | Western Blot | Tarbase |
| hsa-miR-195-5p | CDC25A | Western Blot | Tarbase |
| hsa-miR-195-5p | CDC25A | Microarrays | Tarbase |
| hsa-miR-15a-5p | SHISA5 | HITS-CLIP | Tarbase |
| hsa-miR-195-5p | SHISA5 | HITS-CLIP | Tarbase |
| hsa-miR-15a-5p | SPRY1 | PAR-CLIP | Tarbase |
| hsa-miR-195-5p | SPRY1 | Microarrays | Tarbase |
| hsa-miR-195-3p | INTU | HITS-CLIP | Tarbase |
| hsa-miR-141-5p | MFSD8 | HITS-CLIP | Tarbase |
| hsa-miR-141-5p | RAD54L2 | HITS-CLIP | Tarbase |
| hsa-miR-141-3p | WDR82 | HITS-CLIP | Tarbase |
| hsa-miR-15a-5p | WDR82 | HITS-CLIP | Tarbase |
| hsa-miR-15a-5p | WDR82 | PAR-CLIP | Tarbase |
| hsa-miR-15a-5p | WDR82 | HITS-CLIP | Tarbase |
| hsa-miR-195-5p | WDR82 | HITS-CLIP | Tarbase |
| hsa-miR-195-5p | WDR82 | HITS-CLIP | Tarbase |
| hsa-miR-195-5p | PITX2 | Microarrays | Tarbase |
| hsa-miR-195-5p | PRSS12 | Microarrays | Tarbase |
| hsa-miR-141-5p | HMGB2 | HITS-CLIP | Tarbase |
| hsa-miR-15a-5p | HMGB2 | PAR-CLIP | Tarbase |
| hsa-miR-195-5p | HMGB2 | Microarrays | Tarbase |
| hsa-miR-141-5p | ANXA5 | HITS-CLIP | Tarbase |
| hsa-miR-15a-5p | ANXA5 | HITS-CLIP | Tarbase |
| hsa-miR-195-5p | ANXA5 | HITS-CLIP | Tarbase |
| hsa-miR-195-5p | GUCY1A3 | Microarrays | Tarbase |
| hsa-miR-195-5p | HPGD | Microarrays | Tarbase |
| hsa-miR-15a-5p | TMEM144 | HITS-CLIP | Tarbase |
| hsa-miR-195-5p | FAM198B | Microarrays | Tarbase |
| hsa-miR-141-3p | NPY1R | HITS-CLIP | Tarbase |
| hsa-miR-141-3p | NAA15 | HITS-CLIP | Tarbase |
| hsa-miR-15a-3p | FAM160A1 | HITS-CLIP | Tarbase |
| hsa-miR-15a-5p | FAM160A1 | HITS-CLIP | Tarbase |
| hsa-miR-195-5p | FAM160A1 | HITS-CLIP | Tarbase |
| hsa-miR-195-5p | HHIP | Microarrays | Tarbase |
| hsa-miR-141-3p | OTUD4 | HITS-CLIP | Tarbase |
| hsa-miR-15a-5p | OTUD4 | HITS-CLIP | Tarbase |
| hsa-miR-15a-5p | OTUD4 | HITS-CLIP | Tarbase |
| hsa-miR-15a-5p | OTUD4 | HITS-CLIP | Tarbase |
| hsa-miR-195-5p | OTUD4 | HITS-CLIP | Tarbase |
| hsa-miR-195-5p | OTUD4 | HITS-CLIP | Tarbase |
| hsa-miR-219a-1-3p | OTUD4 | PAR-CLIP | Tarbase |
| hsa-miR-219a-1-3p | OTUD4 | HITS-CLIP | Tarbase |
| hsa-miR-15a-5p | ITGA2 | PAR-CLIP | Tarbase |
| hsa-miR-195-5p | ITGA2 | Microarrays | Tarbase |
| hsa-miR-15a-5p | ELOVL7 | PAR-CLIP | Tarbase |
| hsa-miR-15a-5p | ELOVL7 | HITS-CLIP | Tarbase |
| hsa-miR-15a-5p | ELOVL7 | HITS-CLIP | Tarbase |
| hsa-miR-195-5p | ELOVL7 | HITS-CLIP | Tarbase |
| hsa-miR-486-5p | LMBRD2 | HITS-CLIP | Tarbase |
| hsa-miR-141-5p | NIPBL | HITS-CLIP | Tarbase |
| hsa-miR-15a-5p | NIPBL | PAR-CLIP | Tarbase |
| hsa-miR-3613-5p | NIPBL | PAR-CLIP | Tarbase |
| hsa-miR-141-5p | RNF180 | HITS-CLIP | Tarbase |
| hsa-miR-195-5p | GPR98 | Microarrays | Tarbase |
| hsa-miR-141-3p | SLC25A46 | PAR-CLIP | Tarbase |
| hsa-miR-15a-3p | STARD4 | HITS-CLIP | Tarbase |
| hsa-miR-3613-3p | STARD4 | PAR-CLIP | Tarbase |
| hsa-miR-3613-3p | STARD4 | HITS-CLIP | Tarbase |
| hsa-miR-195-5p | F2RL2 | Microarrays | Tarbase |
| hsa-miR-141-3p | CMBL | HITS-CLIP | Tarbase |
| hsa-miR-195-5p | CMBL | Microarrays | Tarbase |

| | | | |
|---|---|---|---|
| hsa-miR-15a-5p | PRRC1 | HITS-CLIP | Tarbase |
| hsa-miR-195-5p | PRRC1 | HITS-CLIP | Tarbase |
| hsa-miR-195-5p | F2RL1 | Microarrays | Tarbase |
| hsa-miR-141-3p | RHOBTB3 | HITS-CLIP | Tarbase |
| hsa-miR-15a-5p | RHOBTB3 | HITS-CLIP | Tarbase |
| hsa-miR-15a-5p | RHOBTB3 | HITS-CLIP | Tarbase |
| hsa-miR-195-5p | RHOBTB3 | HITS-CLIP | Tarbase |
| hsa-miR-3605-5p | ERAP1 | HITS-CLIP | Tarbase |
| hsa-miR-141-3p | CFAP97 | HITS-CLIP | Tarbase |
| hsa-miR-141-3p | RICTOR | HITS-CLIP | Tarbase |
| hsa-miR-15a-5p | RICTOR | PAR-CLIP | Tarbase |
| hsa-miR-195-5p | RICTOR | PAR-CLIP | Tarbase |
| hsa-miR-15a-5p | UBLCP1 | HITS-CLIP | Tarbase |
| hsa-miR-195-5p | UBLCP1 | HITS-CLIP | Tarbase |
| hsa-miR-15a-5p | UTP15 | HITS-CLIP | Tarbase |
| hsa-miR-195-5p | UTP15 | HITS-CLIP | Tarbase |
| hsa-miR-15a-5p | CCDC127 | HITS-CLIP | Tarbase |
| hsa-miR-195-5p | CCDC127 | HITS-CLIP | Tarbase |
| hsa-miR-15a-5p | ACSL6 | HITS-CLIP | Tarbase |
| hsa-miR-195-5p | ACSL6 | HITS-CLIP | Tarbase |
| hsa-miR-141-3p | SEPT8 | HITS-CLIP | Tarbase |
| hsa-miR-141-3p | SEPT8 | HITS-CLIP | Tarbase |
| hsa-miR-141-5p | SHROOM1 | HITS-CLIP | Tarbase |
| hsa-miR-141-5p | CITED2 | HITS-CLIP | Tarbase |
| hsa-miR-15a-5p | CREBRF | HITS-CLIP | Tarbase |
| hsa-miR-15a-5p | CREBRF | PAR-CLIP | Tarbase |
| hsa-miR-15a-5p | CREBRF | PAR-CLIP | Tarbase |
| hsa-miR-15a-5p | CREBRF | HITS-CLIP | Tarbase |
| hsa-miR-15a-5p | CREBRF | PAR-CLIP | Tarbase |
| hsa-miR-15a-5p | CREBRF | PAR-CLIP | Tarbase |
| hsa-miR-15a-5p | CREBRF | PAR-CLIP | Tarbase |
| hsa-miR-195-5p | CREBRF | HITS-CLIP | Tarbase |
| hsa-miR-195-5p | CREBRF | PAR-CLIP | Tarbase |
| hsa-miR-195-5p | CREBRF | HITS-CLIP | Tarbase |
| hsa-miR-195-5p | CREBRF | PAR-CLIP | Tarbase |
| hsa-miR-195-5p | CREBRF | PAR-CLIP | Tarbase |
| hsa-miR-219a-1-3p | CREBRF | PAR-CLIP | Tarbase |
| hsa-miR-141-3p | DCBLD1 | HITS-CLIP | Tarbase |
| hsa-miR-15a-5p | DCBLD1 | HITS-CLIP | Tarbase |
| hsa-miR-195-5p | DCBLD1 | HITS-CLIP | Tarbase |
| hsa-miR-141-3p | SFXN1 | HITS-CLIP | Tarbase |
| hsa-miR-15a-5p | SFXN1 | PAR-CLIP | Tarbase |
| hsa-miR-195-5p | KIAA0895 | Microarrays | Tarbase |
| hsa-miR-195-5p | GALNT10 | Microarrays | Tarbase |
| hsa-miR-15a-5p | RPS14 | HITS-CLIP | Tarbase |
| hsa-miR-15a-5p | RPS14 | PAR-CLIP | Tarbase |
| hsa-miR-15a-5p | RPS14 | HITS-CLIP | Tarbase |
| hsa-miR-15a-5p | RPS14 | HITS-CLIP | Tarbase |
| hsa-miR-15a-5p | RPS14 | HITS-CLIP | Tarbase |
| hsa-miR-15a-5p | RPS14 | HITS-CLIP | Tarbase |
| hsa-miR-15a-5p | RPS14 | PAR-CLIP | Tarbase |
| hsa-miR-195-5p | RPS14 | HITS-CLIP | Tarbase |
| hsa-miR-195-5p | RPS14 | AGO-IP | Tarbase |
| hsa-miR-195-5p | RPS14 | HITS-CLIP | Tarbase |
| hsa-miR-195-5p | RPS14 | HITS-CLIP | Tarbase |
| hsa-miR-195-5p | RPS14 | HITS-CLIP | Tarbase |
| hsa-miR-195-5p | RPS14 | HITS-CLIP | Tarbase |
| hsa-miR-486-3p | RPS14 | HITS-CLIP | Tarbase |
| hsa-miR-15a-5p | RPS14 | Chimeric fragments | Tarbase |
| hsa-miR-141-3p | COG5 | HITS-CLIP | Tarbase |
| hsa-miR-195-5p | PTTG1 | Microarrays | Tarbase |
| hsa-miR-15a-5p | KCNK5 | PAR-CLIP | Tarbase |
| hsa-miR-3605-5p | KCNK5 | HITS-CLIP | Tarbase |
| hsa-miR-195-5p | KIF6 | Microarrays | Tarbase |
| hsa-miR-141-3p | ZNF12 | HITS-CLIP | Tarbase |
| hsa-miR-141-5p | ZNF12 | HITS-CLIP | Tarbase |
| hsa-miR-3613-5p | ZNF12 | HITS-CLIP | Tarbase |
| hsa-miR-195-5p | CDCA7L | Microarrays | Tarbase |
| hsa-miR-15a-5p | ZNF704 | PAR-CLIP | Tarbase |
| hsa-miR-141-3p | LMTK2 | HITS-CLIP | Tarbase |
| hsa-miR-141-3p | LMTK2 | HITS-CLIP | Tarbase |
| hsa-miR-141-5p | LMTK2 | HITS-CLIP | Tarbase |
| hsa-miR-15a-3p | CTSB | HITS-CLIP | Tarbase |
| hsa-miR-486-3p | CTSB | HITS-CLIP | Tarbase |
| hsa-miR-141-3p | DLC1 | PAR-CLIP | Tarbase |
| hsa-miR-15a-5p | DLC1 | PAR-CLIP | Tarbase |
| hsa-miR-141-3p | RAD21 | HITS-CLIP | Tarbase |
| hsa-miR-141-5p | RAD21 | HITS-CLIP | Tarbase |
| hsa-miR-195-5p | RAD21 | Microarrays | Tarbase |
| hsa-miR-15a-5p | SLC30A8 | HITS-CLIP | Tarbase |
| hsa-miR-195-5p | SLC30A8 | HITS-CLIP | Tarbase |
| hsa-miR-219a-1-3p | TNFRSF11B | HITS-CLIP | Tarbase |
| hsa-miR-15a-5p | EN2 | PAR-CLIP | Tarbase |
| hsa-miR-141-3p | HEATR2 | HITS-CLIP | Tarbase |
| hsa-miR-15a-5p | HEATR2 | HITS-CLIP | Tarbase |
| hsa-miR-195-5p | HEATR2 | HITS-CLIP | Tarbase |
| hsa-miR-141-3p | OSGIN2 | HITS-CLIP | Tarbase |
| hsa-miR-141-3p | SUN1 | HITS-CLIP | Tarbase |
| hsa-miR-3613-3p | SUN1 | PAR-CLIP | Tarbase |
| hsa-miR-15a-5p | OXR1 | HITS-CLIP | Tarbase |
| hsa-miR-195-5p | CDK5 | Microarrays | Tarbase |
| hsa-miR-15a-5p | SLC4A2 | PAR-CLIP | Tarbase |
| hsa-miR-15a-5p | SLC4A2 | HITS-CLIP | Tarbase |
| hsa-miR-195-5p | SLC4A2 | HITS-CLIP | Tarbase |
| hsa-miR-486-3p | SLC4A2 | HITS-CLIP | Tarbase |
| hsa-miR-15a-3p | FASTK | HITS-CLIP | Tarbase |
| hsa-miR-15a-5p | FASTK | HITS-CLIP | Tarbase |
| hsa-miR-15a-5p | FASTK | HITS-CLIP | Tarbase |
| hsa-miR-15a-5p | FASTK | HITS-CLIP | Tarbase |
| hsa-miR-15a-5p | FASTK | PAR-CLIP | Tarbase |
| hsa-miR-195-5p | FASTK | HITS-CLIP | Tarbase |
| hsa-miR-195-5p | FASTK | HITS-CLIP | Tarbase |
| hsa-miR-195-5p | FASTK | HITS-CLIP | Tarbase |
| hsa-miR-15a-5p | TMUB1 | HITS-CLIP | Tarbase |
| hsa-miR-195-5p | TMUB1 | HITS-CLIP | Tarbase |
| hsa-miR-15a-5p | TMUB1 | HITS-CLIP | Tarbase |
| hsa-miR-195-5p | TMUB1 | HITS-CLIP | Tarbase |
| hsa-miR-195-5p | TMUB1 | Microarrays | Tarbase |
| hsa-miR-15a-5p | PHAX | HITS-CLIP | Tarbase |
| hsa-miR-195-5p | PHAX | HITS-CLIP | Tarbase |
| hsa-miR-15a-5p | FOXK1 | HITS-CLIP | Tarbase |
| hsa-miR-15a-5p | FOXK1 | PAR-CLIP | Tarbase |
| hsa-miR-15a-5p | FOXK1 | PAR-CLIP | Tarbase |
| hsa-miR-15a-5p | FOXK1 | HITS-CLIP | Tarbase |
| hsa-miR-15a-5p | FOXK1 | HITS-CLIP | Tarbase |
| hsa-miR-195-5p | FOXK1 | HITS-CLIP | Tarbase |
| hsa-miR-195-5p | FOXK1 | PAR-CLIP | Tarbase |
| hsa-miR-195-5p | FOXK1 | HITS-CLIP | Tarbase |
| hsa-miR-15a-3p | FZD6 | HITS-CLIP | Tarbase |
| hsa-miR-15a-5p | FZD6 | HITS-CLIP | Tarbase |
| hsa-miR-15a-5p | FZD6 | HITS-CLIP | Tarbase |
| hsa-miR-195-5p | FZD6 | HITS-CLIP | Tarbase |
| hsa-miR-195-5p | FZD6 | Microarrays | Tarbase |
| hsa-miR-195-5p | CTHRC1 | Microarrays | Tarbase |
| hsa-miR-195-5p | SLC25A32 | Microarrays | Tarbase |
| hsa-miR-141-3p | DCAF13 | HITS-CLIP | Tarbase |
| hsa-miR-141-5p | TP53INP1 | HITS-CLIP | Tarbase |
| hsa-miR-15a-5p | TP53INP1 | PAR-CLIP | Tarbase |
| hsa-miR-195-5p | TP53INP1 | PAR-CLIP | Tarbase |
| hsa-miR-3613-3p | TP53INP1 | HITS-CLIP | Tarbase |
| hsa-miR-195-5p | TP53INP1 | Microarrays | Tarbase |
| hsa-miR-15a-5p | PDP1 | HITS-CLIP | Tarbase |
| hsa-miR-195-5p | PDP1 | Microarrays | Tarbase |
| hsa-miR-15a-5p | KIAA1161 | HITS-CLIP | Tarbase |
| hsa-miR-15a-5p | KIAA1161 | HITS-CLIP | Tarbase |
| hsa-miR-195-5p | KIAA1161 | HITS-CLIP | Tarbase |
| hsa-miR-141-3p | UBAP1 | Luciferase Reporter Assay | Tarbase |
| hsa-miR-141-3p | UBAP1 | Western Blot | Tarbase |
| hsa-miR-15a-5p | UBAP1 | HITS-CLIP | Tarbase |
| hsa-miR-15a-5p | UBAP1 | PAR-CLIP | Tarbase |
| hsa-miR-195-5p | UBAP1 | HITS-CLIP | Tarbase |
| hsa-miR-15a-5p | DIRAS2 | HITS-CLIP | Tarbase |
| hsa-miR-195-5p | ABCA1 | Microarrays | Tarbase |
| hsa-miR-195-5p | ALDH1A1 | Microarrays | Tarbase |
| hsa-miR-3613-3p | KDM1B | PAR-CLIP | Tarbase |
| hsa-miR-15a-5p | HGSNAT | HITS-CLIP | Tarbase |
| hsa-miR-195-5p | HGSNAT | HITS-CLIP | Tarbase |
| hsa-miR-15a-5p | RASEF | PAR-CLIP | Tarbase |
| hsa-miR-15a-5p | RASEF | PAR-CLIP | Tarbase |

| | | | |
|---|---|---|---|
| hsa-miR-15a-5p | ANKS6 | HITS-CLIP | Tarbase |
| hsa-miR-195-5p | ANKS6 | HITS-CLIP | Tarbase |
| hsa-miR-15a-5p | ZHX1 | HITS-CLIP | Tarbase |
| hsa-miR-15a-5p | ZHX1 | HITS-CLIP | Tarbase |
| hsa-miR-195-5p | ZHX1 | HITS-CLIP | Tarbase |
| hsa-miR-15a-5p | ZHX1 | HITS-CLIP | Tarbase |
| hsa-miR-195-5p | ZHX1 | HITS-CLIP | Tarbase |
| hsa-miR-15a-5p | ZHX1 | Chimeric fragments | Tarbase |
| hsa-miR-15a-3p | DYNLT3 | HITS-CLIP | Tarbase |
| hsa-miR-15a-5p | DYNLT3 | HITS-CLIP | Tarbase |
| hsa-miR-195-5p | DYNLT3 | HITS-CLIP | Tarbase |
| hsa-miR-195-5p | PTCHD1 | Microarrays | Tarbase |
| hsa-miR-195-5p | GAPVD1 | Microarrays | Tarbase |
| hsa-miR-141-5p | C9orf89 | HITS-CLIP | Tarbase |
| hsa-miR-15a-3p | WNK2 | HITS-CLIP | Tarbase |
| hsa-miR-141-5p | ATP7A | HITS-CLIP | Tarbase |
| hsa-miR-141-5p | ZNF367 | HITS-CLIP | Tarbase |
| hsa-miR-141-5p | ZNF367 | HITS-CLIP | Tarbase |
| hsa-miR-15a-5p | ZNF367 | PAR-CLIP | Tarbase |
| hsa-miR-15a-5p | ZNF367 | PAR-CLIP | Tarbase |
| hsa-miR-15a-5p | ZNF367 | HITS-CLIP | Tarbase |
| hsa-miR-15a-5p | ZNF367 | HITS-CLIP | Tarbase |
| hsa-miR-15a-5p | ZNF367 | HITS-CLIP | Tarbase |
| hsa-miR-15a-5p | ZNF367 | PAR-CLIP | Tarbase |
| hsa-miR-15a-5p | ZNF367 | HITS-CLIP | Tarbase |
| hsa-miR-15a-5p | ZNF367 | HITS-CLIP | Tarbase |
| hsa-miR-195-5p | ZNF367 | PAR-CLIP | Tarbase |
| hsa-miR-195-5p | ZNF367 | HITS-CLIP | Tarbase |
| hsa-miR-195-5p | ZNF367 | HITS-CLIP | Tarbase |
| hsa-miR-195-5p | ZNF367 | HITS-CLIP | Tarbase |
| hsa-miR-486-5p | ZNF367 | HITS-CLIP | Tarbase |
| hsa-miR-486-5p | ZNF367 | HITS-CLIP | Tarbase |
| hsa-miR-486-5p | ZNF367 | HITS-CLIP | Tarbase |
| hsa-miR-486-5p | ZNF367 | HITS-CLIP | Tarbase |
| hsa-miR-486-5p | ZNF367 | HITS-CLIP | Tarbase |
| hsa-miR-15a-5p | ZNF367 | Chimeric fragments | Tarbase |
| hsa-miR-195-5p | NOL6 | Microarrays | Tarbase |
| hsa-miR-15a-3p | VCP | HITS-CLIP | Tarbase |
| hsa-miR-15a-3p | VCP | PAR-CLIP | Tarbase |
| hsa-miR-15a-5p | VCP | HITS-CLIP | Tarbase |
| hsa-miR-195-5p | VCP | HITS-CLIP | Tarbase |
| hsa-miR-15a-5p | PIGO | HITS-CLIP | Tarbase |
| hsa-miR-15a-5p | PIGO | HITS-CLIP | Tarbase |
| hsa-miR-195-5p | PIGO | HITS-CLIP | Tarbase |
| hsa-miR-195-5p | PIGO | HITS-CLIP | Tarbase |
| hsa-miR-195-5p | MELK | Microarrays | Tarbase |
| hsa-miR-141-5p | ARHGAP12 | HITS-CLIP | Tarbase |
| hsa-miR-15a-5p | ARHGAP12 | HITS-CLIP | Tarbase |
| hsa-miR-15a-5p | ARHGAP12 | HITS-CLIP | Tarbase |
| hsa-miR-195-5p | ARHGAP12 | HITS-CLIP | Tarbase |
| hsa-miR-486-5p | ARHGAP12 | HITS-CLIP | Tarbase |
| hsa-miR-195-5p | LRFN5 | Microarrays | Tarbase |
| hsa-miR-141-5p | SPTSSA | HITS-CLIP | Tarbase |
| hsa-miR-141-5p | WRN | HITS-CLIP | Tarbase |
| hsa-miR-141-3p | CFL2 | HITS-CLIP | Tarbase |
| hsa-miR-15a-5p | SUGT1 | HITS-CLIP | Tarbase |
| hsa-miR-195-5p | SUGT1 | HITS-CLIP | Tarbase |
| hsa-miR-141-5p | GTF2A1 | HITS-CLIP | Tarbase |
| hsa-miR-15a-5p | PGM2L1 | HITS-CLIP | Tarbase |
| hsa-miR-195-5p | PGM2L1 | HITS-CLIP | Tarbase |
| hsa-miR-15a-5p | SLC16A9 | HITS-CLIP | Tarbase |
| hsa-miR-15a-3p | INPPL1 | HITS-CLIP | Tarbase |
| hsa-miR-141-3p | REEP3 | HITS-CLIP | Tarbase |
| hsa-miR-15a-5p | REEP3 | HITS-CLIP | Tarbase |
| hsa-miR-195-5p | SKA3 | Microarrays | Tarbase |
| hsa-miR-195-5p | DDIAS | Microarrays | Tarbase |
| hsa-miR-195-5p | PKNOX2 | Microarrays | Tarbase |
| hsa-miR-141-3p | RPL36AL | HITS-CLIP | Tarbase |
| hsa-miR-141-3p | RPL36AL | HITS-CLIP | Tarbase |
| hsa-miR-141-3p | KLHDC2 | HITS-CLIP | Tarbase |
| hsa-miR-141-3p | KLHDC2 | HITS-CLIP | Tarbase |
| hsa-miR-15a-5p | NEMF | HITS-CLIP | Tarbase |
| hsa-miR-15a-5p | NEMF | HITS-CLIP | Tarbase |
| hsa-miR-15a-5p | NEMF | PAR-CLIP | Tarbase |
| hsa-miR-195-5p | NEMF | HITS-CLIP | Tarbase |
| hsa-miR-195-5p | NEMF | HITS-CLIP | Tarbase |
| hsa-miR-15a-5p | RPUSD4 | PAR-CLIP | Tarbase |
| hsa-miR-195-5p | RPUSD4 | Microarrays | Tarbase |
| hsa-miR-486-3p | ARF6 | HITS-CLIP | Tarbase |
| hsa-miR-195-5p | ARF6 | Chimeric fragments | Tarbase |
| hsa-miR-141-3p | KBTBD6 | PAR-CLIP | Tarbase |
| hsa-miR-15a-5p | SLC18A2 | HITS-CLIP | Tarbase |
| hsa-miR-195-5p | SLC18A2 | HITS-CLIP | Tarbase |
| hsa-miR-3605-5p | SLC18A2 | HITS-CLIP | Tarbase |
| hsa-miR-141-3p | PDZD8 | HITS-CLIP | Tarbase |
| hsa-miR-15a-3p | PDZD8 | HITS-CLIP | Tarbase |
| hsa-miR-15a-5p | PDZD8 | PAR-CLIP | Tarbase |
| hsa-miR-15a-5p | PDZD8 | HITS-CLIP | Tarbase |
| hsa-miR-141-3p | ZNF503 | HITS-CLIP | Tarbase |
| hsa-miR-141-3p | ZNF503 | HITS-CLIP | Tarbase |
| hsa-miR-15a-5p | ZNF503 | PAR-CLIP | Tarbase |
| hsa-miR-15a-5p | NSD1 | PAR-CLIP | Tarbase |
| hsa-miR-15a-5p | NSD1 | HITS-CLIP | Tarbase |
| hsa-miR-15a-5p | NSD1 | PAR-CLIP | Tarbase |
| hsa-miR-15a-5p | NSD1 | HITS-CLIP | Tarbase |
| hsa-miR-15a-5p | NSD1 | HITS-CLIP | Tarbase |
| hsa-miR-195-5p | NSD1 | PAR-CLIP | Tarbase |
| hsa-miR-195-5p | NSD1 | HITS-CLIP | Tarbase |
| hsa-miR-195-5p | NSD1 | HITS-CLIP | Tarbase |
| hsa-miR-15a-5p | PMPCA | HITS-CLIP | Tarbase |
| hsa-miR-195-5p | PMPCA | HITS-CLIP | Tarbase |
| hsa-miR-15a-3p | TSC1 | HITS-CLIP | Tarbase |
| hsa-miR-141-5p | FAM69B | HITS-CLIP | Tarbase |
| hsa-miR-141-5p | FAM69B | HITS-CLIP | Tarbase |
| hsa-miR-141-5p | FAM69B | HITS-CLIP | Tarbase |
| hsa-miR-15a-5p | RET | HITS-CLIP | Tarbase |
| hsa-miR-15a-5p | RET | Luciferase Reporter Assay | Tarbase |
| hsa-miR-195-5p | RET | Luciferase Reporter Assay | Tarbase |
| hsa-miR-15a-3p | DDX21 | HITS-CLIP | Tarbase |
| hsa-miR-15a-5p | DDX21 | HITS-CLIP | Tarbase |
| hsa-miR-195-5p | DDX21 | HITS-CLIP | Tarbase |
| hsa-miR-486-5p | DDX21 | PAR-CLIP | Tarbase |
| hsa-miR-195-5p | KIAA1462 | Microarrays | Tarbase |
| hsa-miR-15a-5p | TMEM55B | PAR-CLIP | Tarbase |
| hsa-miR-15a-5p | TMEM55B | HITS-CLIP | Tarbase |
| hsa-miR-15a-5p | TMEM55B | HITS-CLIP | Tarbase |
| hsa-miR-195-5p | TMEM55B | HITS-CLIP | Tarbase |
| hsa-miR-15a-3p | NSMF | HITS-CLIP | Tarbase |
| hsa-miR-15a-3p | NSMF | HITS-CLIP | Tarbase |
| hsa-miR-15a-3p | TRUB1 | PAR-CLIP | Tarbase |
| hsa-miR-15a-5p | TRUB1 | HITS-CLIP | Tarbase |
| hsa-miR-195-5p | TRUB1 | HITS-CLIP | Tarbase |
| hsa-miR-15a-5p | ZFYVE1 | PAR-CLIP | Tarbase |
| hsa-miR-15a-5p | ZFYVE1 | PAR-CLIP | Tarbase |
| hsa-miR-15a-5p | ZFYVE1 | PAR-CLIP | Tarbase |
| hsa-miR-15a-5p | ZFYVE1 | PAR-CLIP | Tarbase |
| hsa-miR-15a-5p | ZFYVE1 | PAR-CLIP | Tarbase |
| hsa-miR-15a-5p | ZFYVE1 | PAR-CLIP | Tarbase |
| hsa-miR-195-5p | ZFYVE1 | PAR-CLIP | Tarbase |
| hsa-miR-195-5p | ZFYVE1 | PAR-CLIP | Tarbase |
| hsa-miR-15a-5p | E2F7 | PAR-CLIP | Tarbase |
| hsa-miR-486-5p | E2F7 | PAR-CLIP | Tarbase |
| hsa-miR-486-5p | E2F7 | PAR-CLIP | Tarbase |
| hsa-miR-486-5p | E2F7 | PAR-CLIP | Tarbase |
| hsa-miR-15a-5p | GYLTL1B | HITS-CLIP | Tarbase |
| hsa-miR-195-5p | GYLTL1B | HITS-CLIP | Tarbase |
| hsa-miR-195-5p | SLC39A13 | Microarrays | Tarbase |
| hsa-miR-486-5p | CPSF2 | HITS-CLIP | Tarbase |
| hsa-miR-15a-5p | CPSF2 | Chimeric fragments | Tarbase |
| hsa-miR-15a-5p | MOAP1 | PAR-CLIP | Tarbase |
| hsa-miR-3605-3p | MOAP1 | HITS-CLIP | Tarbase |
| hsa-miR-15a-5p | CLMN | HITS-CLIP | Tarbase |
| hsa-miR-195-5p | CLMN | HITS-CLIP | Tarbase |
| hsa-miR-3613-3p | CLMN | HITS-CLIP | Tarbase |
| hsa-miR-15a-5p | PTER | HITS-CLIP | Tarbase |
| hsa-miR-15a-5p | C1QL3 | HITS-CLIP | Tarbase |
| hsa-miR-195-5p | C1QL3 | HITS-CLIP | Tarbase |
| hsa-miR-195-5p | CACNB2 | Microarrays | Tarbase |
| hsa-miR-141-3p | ARL5B | HITS-CLIP | Tarbase |

| | | | |
|---|---|---|---|
| hsa-miR-15a-5p | ARL5B | PAR-CLIP | Tarbase |
| hsa-miR-15a-5p | ARL5B | HITS-CLIP | Tarbase |
| hsa-miR-15a-5p | ARL5B | PAR-CLIP | Tarbase |
| hsa-miR-195-5p | ARL5B | PAR-CLIP | Tarbase |
| hsa-miR-195-5p | ARL5B | HITS-CLIP | Tarbase |
| hsa-miR-15a-5p | ARL5B | HITS-CLIP | Tarbase |
| hsa-miR-195-5p | ARL5B | HITS-CLIP | Tarbase |
| hsa-miR-15a-5p | ARL5B | Chimeric fragments | Tarbase |
| hsa-miR-15a-5p | KCNC2 | HITS-CLIP | Tarbase |
| hsa-miR-195-5p | KCNC2 | HITS-CLIP | Tarbase |
| hsa-miR-15a-5p | AMOTL1 | HITS-CLIP | Tarbase |
| hsa-miR-195-5p | AMOTL1 | HITS-CLIP | Tarbase |
| hsa-miR-15a-5p | HTRA1 | HITS-CLIP | Tarbase |
| hsa-miR-195-5p | HTRA1 | HITS-CLIP | Tarbase |
| hsa-miR-15a-5p | SPRED1 | PAR-CLIP | Tarbase |
| hsa-miR-15a-5p | SPRED1 | PAR-CLIP | Tarbase |
| hsa-miR-15a-5p | SPRED1 | HITS-CLIP | Tarbase |
| hsa-miR-15a-5p | SPRED1 | PAR-CLIP | Tarbase |
| hsa-miR-195-5p | SPRED1 | PAR-CLIP | Tarbase |
| hsa-miR-195-5p | SPRED1 | HITS-CLIP | Tarbase |
| hsa-miR-486-3p | SPRED1 | HITS-CLIP | Tarbase |
| hsa-miR-15a-5p | SPRED1 | HITS-CLIP | Tarbase |
| hsa-miR-195-5p | SPRED1 | HITS-CLIP | Tarbase |
| hsa-miR-15a-5p | SPRED1 | Chimeric fragments | Tarbase |
| hsa-miR-141-3p | RAB8B | PAR-CLIP | Tarbase |
| hsa-miR-141-5p | HIF1AN | HITS-CLIP | Tarbase |
| hsa-miR-3613-5p | FBN1 | HITS-CLIP | Tarbase |
| hsa-miR-3613-5p | FBN1 | HITS-CLIP | Tarbase |
| hsa-miR-486-3p | FBN1 | HITS-CLIP | Tarbase |
| hsa-miR-486-5p | FBN1 | HITS-CLIP | Tarbase |
| hsa-miR-15a-5p | LRTM2 | HITS-CLIP | Tarbase |
| hsa-miR-195-5p | LRTM2 | HITS-CLIP | Tarbase |
| hsa-miR-15a-3p | CKB | HITS-CLIP | Tarbase |
| hsa-miR-15a-3p | CKB | HITS-CLIP | Tarbase |
| hsa-miR-15a-5p | CKB | HITS-CLIP | Tarbase |
| hsa-miR-15a-5p | CKB | PAR-CLIP | Tarbase |
| hsa-miR-15a-5p | CKB | PAR-CLIP | Tarbase |
| hsa-miR-195-5p | CKB | HITS-CLIP | Tarbase |
| hsa-miR-3605-3p | CKB | HITS-CLIP | Tarbase |
| hsa-miR-141-3p | BTRC | HITS-CLIP | Tarbase |
| hsa-miR-15a-5p | BTRC | PAR-CLIP | Tarbase |
| hsa-miR-15a-5p | BTRC | HITS-CLIP | Tarbase |
| hsa-miR-195-5p | BTRC | HITS-CLIP | Tarbase |
| hsa-miR-195-5p | BTRC | Luciferase Reporter Assay | Tarbase |
| hsa-miR-195-5p | BTRC | Western Blot | Tarbase |
| hsa-miR-195-5p | BTRC | Western Blot | Tarbase |
| hsa-miR-15a-5p | BTRC | HITS-CLIP | Tarbase |
| hsa-miR-195-5p | BTRC | HITS-CLIP | Tarbase |
| hsa-miR-15a-5p | BTRC | Chimeric fragments | Tarbase |
| hsa-miR-15a-5p | BAG5 | HITS-CLIP | Tarbase |
| hsa-miR-195-5p | BAG5 | HITS-CLIP | Tarbase |
| hsa-miR-141-3p | API5 | HITS-CLIP | Tarbase |
| hsa-miR-141-5p | COPS2 | HITS-CLIP | Tarbase |
| hsa-miR-486-5p | GABRB3 | HITS-CLIP | Tarbase |
| hsa-miR-141-5p | SGPL1 | HITS-CLIP | Tarbase |
| hsa-miR-15a-5p | FRS2 | HITS-CLIP | Tarbase |
| hsa-miR-15a-5p | FRS2 | PAR-CLIP | Tarbase |
| hsa-miR-15a-5p | FRS2 | HITS-CLIP | Tarbase |
| hsa-miR-195-5p | FRS2 | HITS-CLIP | Tarbase |
| hsa-miR-195-5p | FRS2 | HITS-CLIP | Tarbase |
| hsa-miR-15a-5p | FRS2 | Chimeric fragments | Tarbase |
| hsa-miR-15a-5p | CCT2 | HITS-CLIP | Tarbase |
| hsa-miR-3613-3p | ARIH1 | PAR-CLIP | Tarbase |
| hsa-miR-15a-5p | ARIH1 | Chimeric fragments | Tarbase |
| hsa-miR-15a-5p | ZNF202 | HITS-CLIP | Tarbase |
| hsa-miR-195-5p | WBP1L | Microarrays | Tarbase |
| hsa-miR-15a-5p | TMEM100 | PAR-CLIP | Tarbase |
| hsa-miR-15a-3p | SMPD1 | HITS-CLIP | Tarbase |
| hsa-miR-195-5p | SMPD1 | Microarrays | Tarbase |
| hsa-miR-195-5p | APBB1 | HITS-CLIP | Tarbase |
| hsa-miR-15a-5p | TRIM44 | HITS-CLIP | Tarbase |
| hsa-miR-15a-5p | TRIM44 | PAR-CLIP | Tarbase |
| hsa-miR-15a-5p | TRIM44 | PAR-CLIP | Tarbase |
| hsa-miR-195-5p | TRIM44 | HITS-CLIP | Tarbase |
| hsa-miR-195-5p | TRIM44 | PAR-CLIP | Tarbase |
| hsa-miR-195-5p | TRIM44 | Microarrays | Tarbase |
| hsa-miR-15a-5p | ILK | HITS-CLIP | Tarbase |
| hsa-miR-195-3p | USP54 | HITS-CLIP | Tarbase |
| hsa-miR-15a-5p | PPFIBP2 | HITS-CLIP | Tarbase |
| hsa-miR-195-5p | PPFIBP2 | HITS-CLIP | Tarbase |
| hsa-miR-219a-1-3p | SERPINB7 | HITS-CLIP | Tarbase |
| hsa-miR-3605-3p | KIAA0355 | HITS-CLIP | Tarbase |
| hsa-miR-15a-5p | KIAA0355 | Chimeric fragments | Tarbase |
| hsa-miR-15a-3p | RPL27A | HITS-CLIP | Tarbase |
| hsa-miR-15a-5p | RPL27A | HITS-CLIP | Tarbase |
| hsa-miR-15a-5p | RPL27A | PAR-CLIP | Tarbase |
| hsa-miR-195-5p | RPL27A | HITS-CLIP | Tarbase |
| hsa-miR-195-5p | RPL27A | Microarrays | Tarbase |
| hsa-miR-15a-5p | RPL27A | Chimeric fragments | Tarbase |
| hsa-miR-195-5p | ST5 | Microarrays | Tarbase |
| hsa-miR-486-5p | TMEM130 | HITS-CLIP | Tarbase |
| hsa-miR-15a-5p | TMEM130 | HITS-CLIP | Tarbase |
| hsa-miR-195-5p | TMEM130 | HITS-CLIP | Tarbase |
| hsa-miR-15a-5p | CENPN | HITS-CLIP | Tarbase |
| hsa-miR-195-5p | CENPN | HITS-CLIP | Tarbase |
| hsa-miR-195-5p | CENPN | Microarrays | Tarbase |
| hsa-miR-141-3p | ATMIN | HITS-CLIP | Tarbase |
| hsa-miR-141-3p | TMEM41B | HITS-CLIP | Tarbase |
| hsa-miR-195-5p | TMX3 | HITS-CLIP | Tarbase |
| hsa-miR-195-5p | MFAP4 | Microarrays | Tarbase |
| hsa-miR-141-5p | WEE1 | HITS-CLIP | Tarbase |
| hsa-miR-141-5p | WEE1 | HITS-CLIP | Tarbase |
| hsa-miR-15a-5p | WEE1 | HITS-CLIP | Tarbase |
| hsa-miR-15a-5p | WEE1 | HITS-CLIP | Tarbase |
| hsa-miR-15a-5p | WEE1 | HITS-CLIP | Tarbase |
| hsa-miR-15a-5p | WEE1 | HITS-CLIP | Tarbase |
| hsa-miR-15a-5p | WEE1 | PAR-CLIP | Tarbase |
| hsa-miR-15a-5p | WEE1 | HITS-CLIP | Tarbase |
| hsa-miR-15a-5p | WEE1 | HITS-CLIP | Tarbase |
| hsa-miR-15a-5p | WEE1 | HITS-CLIP | Tarbase |
| hsa-miR-15a-5p | WEE1 | HITS-CLIP | Tarbase |
| hsa-miR-15a-5p | WEE1 | HITS-CLIP | Tarbase |
| hsa-miR-15a-5p | WEE1 | PAR-CLIP | Tarbase |
| hsa-miR-15a-5p | WEE1 | HITS-CLIP | Tarbase |
| hsa-miR-15a-5p | WEE1 | HITS-CLIP | Tarbase |
| hsa-miR-195-5p | WEE1 | HITS-CLIP | Tarbase |
| hsa-miR-195-5p | WEE1 | HITS-CLIP | Tarbase |
| hsa-miR-195-5p | WEE1 | HITS-CLIP | Tarbase |
| hsa-miR-195-5p | WEE1 | HITS-CLIP | Tarbase |
| hsa-miR-195-5p | WEE1 | HITS-CLIP | Tarbase |
| hsa-miR-195-5p | WEE1 | HITS-CLIP | Tarbase |
| hsa-miR-195-5p | WEE1 | HITS-CLIP | Tarbase |
| hsa-miR-195-5p | WEE1 | HITS-CLIP | Tarbase |
| hsa-miR-195-5p | WEE1 | HITS-CLIP | Tarbase |
| hsa-miR-195-5p | WEE1 | HITS-CLIP | Tarbase |
| hsa-miR-15a-5p | WEE1 | Chimeric fragments | Tarbase |
| hsa-miR-15a-5p | HDGFRP3 | HITS-CLIP | Tarbase |
| hsa-miR-15a-5p | HDGFRP3 | HITS-CLIP | Tarbase |
| hsa-miR-195-5p | HDGFRP3 | HITS-CLIP | Tarbase |
| hsa-miR-141-5p | MCM7 | HITS-CLIP | Tarbase |
| hsa-miR-15a-5p | MCM7 | PAR-CLIP | Tarbase |
| hsa-miR-195-5p | MCM7 | Microarrays | Tarbase |
| hsa-miR-486-5p | ZSCAN21 | PAR-CLIP | Tarbase |
| hsa-miR-15a-5p | RIMKLB | HITS-CLIP | Tarbase |
| hsa-miR-195-5p | RIMKLB | HITS-CLIP | Tarbase |
| hsa-miR-15a-5p | TMED3 | HITS-CLIP | Tarbase |
| hsa-miR-195-5p | TMED3 | HITS-CLIP | Tarbase |
| hsa-miR-141-5p | SEC11C | HITS-CLIP | Tarbase |
| hsa-miR-15a-5p | TMEM135 | Chimeric fragments | Tarbase |
| hsa-miR-486-3p | NDEL1 | HITS-CLIP | Tarbase |
| hsa-miR-141-5p | HSP90B1 | HITS-CLIP | Tarbase |
| hsa-miR-15a-5p | BLCAP | HITS-CLIP | Tarbase |
| hsa-miR-3613-5p | COG1 | HITS-CLIP | Tarbase |
| hsa-miR-141-3p | ZNF606 | HITS-CLIP | Tarbase |
| hsa-miR-141-3p | ZNF592 | HITS-CLIP | Tarbase |
| hsa-miR-15a-5p | ZNF592 | PAR-CLIP | Tarbase |
| hsa-miR-141-3p | CASC4 | HITS-CLIP | Tarbase |
| hsa-miR-15a-3p | CASC4 | HITS-CLIP | Tarbase |
| hsa-miR-3613-3p | CASC4 | HITS-CLIP | Tarbase |

| | | | |
|---|---|---|---|
| hsa-miR-486-5p | KIAA0430 | HITS-CLIP | Tarbase |
| hsa-miR-3613-3p | PPIB | HITS-CLIP | Tarbase |
| hsa-miR-15a-5p | FAM111A | HITS-CLIP | Tarbase |
| hsa-miR-195-5p | FAM111A | HITS-CLIP | Tarbase |
| hsa-miR-195-5p | KIAA0101 | Microarrays | Tarbase |
| hsa-miR-15a-5p | TMEM170A | Chimeric fragments | Tarbase |
| hsa-miR-141-5p | ANPEP | HITS-CLIP | Tarbase |
| hsa-miR-15a-3p | ANPEP | HITS-CLIP | Tarbase |
| hsa-miR-141-3p | DCTN5 | HITS-CLIP | Tarbase |
| hsa-miR-15a-5p | DCTN5 | HITS-CLIP | Tarbase |
| hsa-miR-15a-5p | DCTN5 | PAR-CLIP | Tarbase |
| hsa-miR-15a-5p | DCTN5 | HITS-CLIP | Tarbase |
| hsa-miR-195-5p | DCTN5 | HITS-CLIP | Tarbase |
| hsa-miR-195-5p | DCTN5 | HITS-CLIP | Tarbase |
| hsa-miR-15a-5p | DCTN5 | HITS-CLIP | Tarbase |
| hsa-miR-195-5p | DCTN5 | HITS-CLIP | Tarbase |
| hsa-miR-195-5p | PLK1 | Microarrays | Tarbase |
| hsa-miR-15a-5p | CLPX | HITS-CLIP | Tarbase |
| hsa-miR-15a-5p | CLPX | HITS-CLIP | Tarbase |
| hsa-miR-15a-5p | CLPX | HITS-CLIP | Tarbase |
| hsa-miR-195-5p | CLPX | HITS-CLIP | Tarbase |
| hsa-miR-195-5p | CLPX | HITS-CLIP | Tarbase |
| hsa-miR-15a-5p | ZBTB39 | HITS-CLIP | Tarbase |
| hsa-miR-15a-5p | ZBTB39 | PAR-CLIP | Tarbase |
| hsa-miR-195-5p | ZBTB39 | HITS-CLIP | Tarbase |
| hsa-miR-195-5p | TMEM194A | Microarrays | Tarbase |
| hsa-miR-141-5p | PATL1 | HITS-CLIP | Tarbase |
| hsa-miR-3605-5p | PATL1 | HITS-CLIP | Tarbase |
| hsa-miR-486-5p | STX3 | PAR-CLIP | Tarbase |
| hsa-miR-141-3p | MTMR10 | HITS-CLIP | Tarbase |
| hsa-miR-15a-5p | MTMR10 | PAR-CLIP | Tarbase |
| hsa-miR-486-5p | MTMR10 | HITS-CLIP | Tarbase |
| hsa-miR-15a-5p | YWHAB | HITS-CLIP | Tarbase |
| hsa-miR-195-5p | YWHAB | HITS-CLIP | Tarbase |
| hsa-miR-141-5p | SCG5 | HITS-CLIP | Tarbase |
| hsa-miR-3613-5p | DIS3L | HITS-CLIP | Tarbase |
| hsa-miR-3613-5p | DIS3L | PAR-CLIP | Tarbase |
| hsa-miR-3613-5p | DIS3L | PAR-CLIP | Tarbase |
| hsa-miR-15a-3p | SMAD3 | HITS-CLIP | Tarbase |
| hsa-miR-15a-5p | MAP1A | HITS-CLIP | Tarbase |
| hsa-miR-195-5p | MAP1A | HITS-CLIP | Tarbase |
| hsa-miR-141-5p | AKTIP | HITS-CLIP | Tarbase |
| hsa-miR-15a-5p | AKTIP | PAR-CLIP | Tarbase |
| hsa-miR-15a-5p | MAPRE2 | PAR-CLIP | Tarbase |
| hsa-miR-141-5p | MARS | HITS-CLIP | Tarbase |
| hsa-miR-15a-5p | MARS | HITS-CLIP | Tarbase |
| hsa-miR-15a-5p | MARS | PAR-CLIP | Tarbase |
| hsa-miR-15a-5p | MARS | HITS-CLIP | Tarbase |
| hsa-miR-15a-5p | NKX3-1 | HITS-CLIP | Tarbase |
| hsa-miR-195-5p | NKX3-1 | HITS-CLIP | Tarbase |
| hsa-miR-15a-5p | PBX3 | HITS-CLIP | Tarbase |
| hsa-miR-195-5p | PBX3 | HITS-CLIP | Tarbase |
| hsa-miR-195-5p | PBX3 | Microarrays | Tarbase |
| hsa-miR-15a-3p | PHB | HITS-CLIP | Tarbase |
| hsa-miR-141-3p | SNRPD1 | HITS-CLIP | Tarbase |
| hsa-miR-141-3p | FAM102A | HITS-CLIP | Tarbase |
| hsa-miR-3613-5p | GOLGA2 | HITS-CLIP | Tarbase |
| hsa-miR-15a-5p | CERCAM | HITS-CLIP | Tarbase |
| hsa-miR-195-5p | CERCAM | HITS-CLIP | Tarbase |
| hsa-miR-486-3p | CERCAM | HITS-CLIP | Tarbase |
| hsa-miR-486-5p | C15orf39 | HITS-CLIP | Tarbase |
| hsa-miR-195-5p | SP2 | Microarrays | Tarbase |
| hsa-miR-15a-5p | PRR15L | HITS-CLIP | Tarbase |
| hsa-miR-195-5p | PRR15L | HITS-CLIP | Tarbase |
| hsa-miR-195-5p | GPRC5B | Microarrays | Tarbase |
| hsa-miR-141-3p | CRK | PAR-CLIP | Tarbase |
| hsa-miR-15a-5p | CRK | PAR-CLIP | Tarbase |
| hsa-miR-15a-5p | CRK | PAR-CLIP | Tarbase |
| hsa-miR-15a-5p | CRK | HITS-CLIP | Tarbase |
| hsa-miR-15a-5p | CRK | HITS-CLIP | Tarbase |
| hsa-miR-195-5p | CRK | HITS-CLIP | Tarbase |
| hsa-miR-15a-5p | HDHD2 | HITS-CLIP | Tarbase |
| hsa-miR-195-5p | HDHD2 | HITS-CLIP | Tarbase |
| hsa-miR-219a-1-3p | ZNF91 | HITS-CLIP | Tarbase |
| hsa-miR-15a-5p | ZNF91 | HITS-CLIP | Tarbase |
| hsa-miR-195-5p | ZNF91 | HITS-CLIP | Tarbase |
| hsa-miR-3613-5p | IGF2 | HITS-CLIP | Tarbase |
| hsa-miR-141-3p | CDK12 | HITS-CLIP | Tarbase |
| hsa-miR-15a-3p | CDK12 | HITS-CLIP | Tarbase |
| hsa-miR-15a-5p | CDK12 | PAR-CLIP | Tarbase |
| hsa-miR-15a-5p | CDK12 | Chimeric fragments | Tarbase |
| hsa-miR-15a-5p | ENGASE | HITS-CLIP | Tarbase |
| hsa-miR-195-5p | ENGASE | HITS-CLIP | Tarbase |
| hsa-miR-15a-5p | ATP5L | HITS-CLIP | Tarbase |
| hsa-miR-15a-3p | TBC1D16 | HITS-CLIP | Tarbase |
| hsa-miR-15a-5p | MYO5B | HITS-CLIP | Tarbase |
| hsa-miR-15a-3p | STIM1 | HITS-CLIP | Tarbase |
| hsa-miR-15a-5p | RRM1 | HITS-CLIP | Tarbase |
| hsa-miR-195-5p | RRM1 | Microarrays | Tarbase |
| hsa-miR-15a-5p | FN3K | HITS-CLIP | Tarbase |
| hsa-miR-195-5p | FN3K | HITS-CLIP | Tarbase |
| hsa-miR-15a-5p | FN3K | HITS-CLIP | Tarbase |
| hsa-miR-195-5p | FN3K | HITS-CLIP | Tarbase |
| hsa-miR-15a-5p | IRGQ | PAR-CLIP | Tarbase |
| hsa-miR-141-5p | PPP2R3B | HITS-CLIP | Tarbase |
| hsa-miR-141-3p | ZNF668 | HITS-CLIP | Tarbase |
| hsa-miR-3605-5p | SMG8 | HITS-CLIP | Tarbase |
| hsa-miR-219a-1-3p | TPM4 | HITS-CLIP | Tarbase |
| hsa-miR-219a-1-3p | TPM4 | HITS-CLIP | Tarbase |
| hsa-miR-15a-3p | GPX4 | HITS-CLIP | Tarbase |
| hsa-miR-3613-5p | GPX4 | HITS-CLIP | Tarbase |
| hsa-miR-15a-5p | MIDN | HITS-CLIP | Tarbase |
| hsa-miR-15a-5p | MIDN | HITS-CLIP | Tarbase |
| hsa-miR-15a-5p | MIDN | HITS-CLIP | Tarbase |
| hsa-miR-195-5p | MIDN | HITS-CLIP | Tarbase |
| hsa-miR-195-5p | MIDN | HITS-CLIP | Tarbase |
| hsa-miR-195-5p | MIDN | HITS-CLIP | Tarbase |
| hsa-miR-3605-3p | MIDN | HITS-CLIP | Tarbase |
| hsa-miR-3613-3p | MIDN | PAR-CLIP | Tarbase |
| hsa-miR-15a-5p | KLHL26 | HITS-CLIP | Tarbase |
| hsa-miR-195-5p | KLHL26 | HITS-CLIP | Tarbase |
| hsa-miR-15a-5p | GATAD2A | HITS-CLIP | Tarbase |
| hsa-miR-195-5p | GATAD2A | HITS-CLIP | Tarbase |
| hsa-miR-15a-5p | GATAD2A | Chimeric fragments | Tarbase |
| hsa-miR-195-5p | CDT1 | Microarrays | Tarbase |
| hsa-miR-15a-5p | DHRS13 | HITS-CLIP | Tarbase |
| hsa-miR-195-5p | DHRS13 | HITS-CLIP | Tarbase |
| hsa-miR-141-3p | KMT2D | HITS-CLIP | Tarbase |
| hsa-miR-141-5p | KMT2D | HITS-CLIP | Tarbase |
| hsa-miR-15a-3p | KMT2D | HITS-CLIP | Tarbase |
| hsa-miR-15a-5p | KMT2D | HITS-CLIP | Tarbase |
| hsa-miR-15a-5p | KMT2D | PAR-CLIP | Tarbase |
| hsa-miR-15a-5p | KMT2D | PAR-CLIP | Tarbase |
| hsa-miR-15a-5p | KMT2D | HITS-CLIP | Tarbase |
| hsa-miR-15a-5p | KMT2D | HITS-CLIP | Tarbase |
| hsa-miR-195-5p | KMT2D | HITS-CLIP | Tarbase |
| hsa-miR-195-5p | KMT2D | HITS-CLIP | Tarbase |
| hsa-miR-486-3p | KMT2D | HITS-CLIP | Tarbase |
| hsa-miR-15a-5p | TUBA1A | HITS-CLIP | Tarbase |
| hsa-miR-195-5p | TUBA1A | HITS-CLIP | Tarbase |
| hsa-miR-195-5p | TUBA1A | AGO-IP | Tarbase |
| hsa-miR-486-3p | TUBA1C | HITS-CLIP | Tarbase |
| hsa-miR-486-3p | TUBA1C | HITS-CLIP | Tarbase |
| hsa-miR-15a-5p | ZNF701 | Chimeric fragments | Tarbase |
| hsa-miR-15a-5p | SERTAD3 | PAR-CLIP | Tarbase |
| hsa-miR-195-5p | SERTAD3 | PAR-CLIP | Tarbase |
| hsa-miR-195-5p | PROSER3 | Microarrays | Tarbase |
| hsa-miR-195-5p | NFKBID | Microarrays | Tarbase |
| hsa-miR-141-3p | LENG8 | HITS-CLIP | Tarbase |
| hsa-miR-141-5p | LENG8 | HITS-CLIP | Tarbase |
| hsa-miR-15a-5p | LENG8 | HITS-CLIP | Tarbase |
| hsa-miR-15a-5p | LENG8 | HITS-CLIP | Tarbase |
| hsa-miR-195-5p | LENG8 | HITS-CLIP | Tarbase |
| hsa-miR-195-5p | LENG8 | HITS-CLIP | Tarbase |
| hsa-miR-3613-3p | LENG8 | HITS-CLIP | Tarbase |
| hsa-miR-141-5p | ZNF526 | HITS-CLIP | Tarbase |
| hsa-miR-486-5p | TRAPPC9 | HITS-CLIP | Tarbase |
| hsa-miR-15a-5p | ZNF146 | HITS-CLIP | Tarbase |
| hsa-miR-195-5p | ZNF146 | HITS-CLIP | Tarbase |

| | | | |
|---|---|---|---|
| hsa-miR-15a-5p | SPINT2 | PAR-CLIP | Tarbase |
| hsa-miR-15a-5p | YIF1B | HITS-CLIP | Tarbase |
| hsa-miR-15a-3p | EEF2 | HITS-CLIP | Tarbase |
| hsa-miR-15a-3p | EEF2 | HITS-CLIP | Tarbase |
| hsa-miR-195-5p | CHAF1A | Microarrays | Tarbase |
| hsa-miR-195-5p | PLIN4 | Microarrays | Tarbase |
| hsa-miR-195-5p | NXN | Microarrays | Tarbase |
| hsa-miR-15a-5p | TSR1 | Chimeric fragments | Tarbase |
| hsa-miR-15a-3p | C19orf48 | HITS-CLIP | Tarbase |
| hsa-miR-15a-5p | C19orf48 | HITS-CLIP | Tarbase |
| hsa-miR-15a-5p | C19orf48 | HITS-CLIP | Tarbase |
| hsa-miR-195-5p | C19orf48 | HITS-CLIP | Tarbase |
| hsa-miR-195-5p | KRT80 | Microarrays | Tarbase |
| hsa-miR-15a-5p | OTUB1 | HITS-CLIP | Tarbase |
| hsa-miR-195-5p | OTUB1 | HITS-CLIP | Tarbase |
| hsa-miR-141-3p | SPRYD3 | HITS-CLIP | Tarbase |
| hsa-miR-141-3p | SPRYD3 | HITS-CLIP | Tarbase |
| hsa-miR-15a-5p | SPRYD3 | PAR-CLIP | Tarbase |
| hsa-miR-15a-5p | SPRYD3 | HITS-CLIP | Tarbase |
| hsa-miR-15a-5p | SPRYD3 | HITS-CLIP | Tarbase |
| hsa-miR-195-5p | SPRYD3 | HITS-CLIP | Tarbase |
| hsa-miR-195-5p | SPRYD3 | Microarrays | Tarbase |
| hsa-miR-15a-5p | CD300A | PAR-CLIP | Tarbase |
| hsa-miR-195-5p | CD300A | PAR-CLIP | Tarbase |
| hsa-miR-15a-5p | HID1 | HITS-CLIP | Tarbase |
| hsa-miR-15a-5p | TMC8 | HITS-CLIP | Tarbase |
| hsa-miR-195-5p | TMC8 | HITS-CLIP | Tarbase |
| hsa-miR-195-5p | TK1 | Microarrays | Tarbase |
| hsa-miR-3605-3p | GSDMA | HITS-CLIP | Tarbase |
| hsa-miR-141-5p | SRRM2 | HITS-CLIP | Tarbase |
| hsa-miR-15a-5p | SRRM2 | PAR-CLIP | Tarbase |
| hsa-miR-15a-5p | SRRM2 | HITS-CLIP | Tarbase |
| hsa-miR-15a-5p | SRRM2 | PAR-CLIP | Tarbase |
| hsa-miR-195-5p | SRRM2 | HITS-CLIP | Tarbase |
| hsa-miR-219a-1-3p | SRRM2 | HITS-CLIP | Tarbase |
| hsa-miR-3605-3p | SRRM2 | HITS-CLIP | Tarbase |
| hsa-miR-486-3p | SRRM2 | HITS-CLIP | Tarbase |
| hsa-miR-486-5p | SRRM2 | HITS-CLIP | Tarbase |
| hsa-miR-15a-5p | DDB1 | HITS-CLIP | Tarbase |
| hsa-miR-15a-5p | DDB1 | HITS-CLIP | Tarbase |
| hsa-miR-15a-5p | DDB1 | HITS-CLIP | Tarbase |
| hsa-miR-195-5p | DDB1 | HITS-CLIP | Tarbase |
| hsa-miR-195-5p | DDB1 | HITS-CLIP | Tarbase |
| hsa-miR-195-5p | DDB1 | HITS-CLIP | Tarbase |
| hsa-miR-15a-5p | VPS37C | HITS-CLIP | Tarbase |
| hsa-miR-195-5p | VPS37C | HITS-CLIP | Tarbase |
| hsa-miR-141-3p | FTH1 | HITS-CLIP | Tarbase |
| hsa-miR-486-5p | SLC3A2 | PAR-CLIP | Tarbase |
| hsa-miR-15a-5p | C2CD3 | HITS-CLIP | Tarbase |
| hsa-miR-141-3p | RPSA | HITS-CLIP | Tarbase |
| hsa-miR-141-3p | RPSA | HITS-CLIP | Tarbase |
| hsa-miR-141-3p | CTNNB1 | HITS-CLIP | Tarbase |
| hsa-miR-15a-3p | CTNNB1 | HITS-CLIP | Tarbase |
| hsa-miR-15a-5p | CTNNB1 | HITS-CLIP | Tarbase |
| hsa-miR-195-5p | CTNNB1 | HITS-CLIP | Tarbase |
| hsa-miR-195-5p | CTNNB1 | Microarrays | Tarbase |
| hsa-miR-195-5p | SCARA3 | Microarrays | Tarbase |
| hsa-miR-195-5p | PBK | Microarrays | Tarbase |
| hsa-miR-141-3p | PAFAH1B2 | HITS-CLIP | Tarbase |
| hsa-miR-141-5p | PAFAH1B2 | HITS-CLIP | Tarbase |
| hsa-miR-141-5p | PAFAH1B2 | HITS-CLIP | Tarbase |
| hsa-miR-141-5p | PAFAH1B2 | HITS-CLIP | Tarbase |
| hsa-miR-15a-5p | PAFAH1B2 | PAR-CLIP | Tarbase |
| hsa-miR-15a-5p | PAFAH1B2 | PAR-CLIP | Tarbase |
| hsa-miR-15a-5p | PAFAH1B2 | PAR-CLIP | Tarbase |
| hsa-miR-15a-5p | PAFAH1B2 | HITS-CLIP | Tarbase |
| hsa-miR-15a-5p | PAFAH1B2 | PAR-CLIP | Tarbase |
| hsa-miR-15a-5p | PAFAH1B2 | HITS-CLIP | Tarbase |
| hsa-miR-195-5p | PAFAH1B2 | PAR-CLIP | Tarbase |
| hsa-miR-195-5p | PAFAH1B2 | HITS-CLIP | Tarbase |
| hsa-miR-15a-3p | SETD5 | HITS-CLIP | Tarbase |
| hsa-miR-15a-3p | SETD5 | HITS-CLIP | Tarbase |
| hsa-miR-15a-5p | SETD5 | PAR-CLIP | Tarbase |
| hsa-miR-15a-5p | SETD5 | HITS-CLIP | Tarbase |
| hsa-miR-195-5p | SETD5 | HITS-CLIP | Tarbase |
| hsa-miR-3613-5p | SETD5 | HITS-CLIP | Tarbase |
| hsa-miR-15a-5p | RNF187 | HITS-CLIP | Tarbase |
| hsa-miR-195-5p | RNF187 | HITS-CLIP | Tarbase |
| hsa-miR-486-5p | RNF187 | PAR-CLIP | Tarbase |
| hsa-miR-195-5p | RNF187 | HITS-CLIP | Tarbase |
| hsa-miR-15a-5p | MAPK1IP1L | HITS-CLIP | Tarbase |
| hsa-miR-141-3p | DDIT4 | HITS-CLIP | Tarbase |
| hsa-miR-141-3p | DDIT4 | HITS-CLIP | Tarbase |
| hsa-miR-15a-5p | DDIT4 | HITS-CLIP | Tarbase |
| hsa-miR-141-3p | RBPJ | HITS-CLIP | Tarbase |
| hsa-miR-15a-3p | RBPJ | PAR-CLIP | Tarbase |
| hsa-miR-15a-5p | RBPJ | PAR-CLIP | Tarbase |
| hsa-miR-15a-5p | PTGDR | PAR-CLIP | Tarbase |
| hsa-miR-15a-5p | IRF2BP2 | PAR-CLIP | Tarbase |
| hsa-miR-15a-5p | IRF2BP2 | HITS-CLIP | Tarbase |
| hsa-miR-195-5p | IRF2BP2 | HITS-CLIP | Tarbase |
| hsa-miR-15a-5p | KIF5C | HITS-CLIP | Tarbase |
| hsa-miR-195-5p | KIF5C | HITS-CLIP | Tarbase |
| hsa-miR-486-5p | KIF5C | HITS-CLIP | Tarbase |
| hsa-miR-15a-5p | KIF5C | HITS-CLIP | Tarbase |
| hsa-miR-195-5p | KIF5C | HITS-CLIP | Tarbase |
| hsa-miR-15a-5p | BMI1 | HITS-CLIP | Tarbase |
| hsa-miR-15a-5p | HIST1H1E | PAR-CLIP | Tarbase |
| hsa-miR-195-5p | HIST1H1E | Microarrays | Tarbase |
| hsa-miR-15a-5p | IRF2 | PAR-CLIP | Tarbase |
| hsa-miR-15a-5p | IRF2 | PAR-CLIP | Tarbase |
| hsa-miR-15a-5p | IRF2 | PAR-CLIP | Tarbase |
| hsa-miR-195-5p | IRF2 | PAR-CLIP | Tarbase |
| hsa-miR-195-5p | IRF2 | PAR-CLIP | Tarbase |
| hsa-miR-15a-5p | SEPT2 | PAR-CLIP | Tarbase |
| hsa-miR-15a-5p | SEPT2 | PAR-CLIP | Tarbase |
| hsa-miR-15a-5p | SEPT2 | HITS-CLIP | Tarbase |
| hsa-miR-15a-5p | SEPT2 | HITS-CLIP | Tarbase |
| hsa-miR-15a-5p | SEPT2 | HITS-CLIP | Tarbase |
| hsa-miR-15a-5p | SEPT2 | PAR-CLIP | Tarbase |
| hsa-miR-15a-5p | SEPT2 | PAR-CLIP | Tarbase |
| hsa-miR-15a-5p | SEPT2 | PAR-CLIP | Tarbase |
| hsa-miR-15a-5p | SEPT2 | HITS-CLIP | Tarbase |
| hsa-miR-15a-5p | SEPT2 | HITS-CLIP | Tarbase |
| hsa-miR-15a-5p | SEPT2 | PAR-CLIP | Tarbase |
| hsa-miR-15a-5p | SEPT2 | PAR-CLIP | Tarbase |
| hsa-miR-15a-5p | SEPT2 | PAR-CLIP | Tarbase |
| hsa-miR-15a-5p | SEPT2 | PAR-CLIP | Tarbase |
| hsa-miR-15a-5p | SEPT2 | PAR-CLIP | Tarbase |
| hsa-miR-15a-5p | SEPT2 | HITS-CLIP | Tarbase |
| hsa-miR-195-5p | SEPT2 | PAR-CLIP | Tarbase |
| hsa-miR-195-5p | SEPT2 | HITS-CLIP | Tarbase |
| hsa-miR-195-5p | SEPT2 | HITS-CLIP | Tarbase |
| hsa-miR-195-5p | SEPT2 | HITS-CLIP | Tarbase |
| hsa-miR-195-5p | SEPT2 | HITS-CLIP | Tarbase |
| hsa-miR-195-5p | SEPT2 | PAR-CLIP | Tarbase |
| hsa-miR-195-5p | SEPT2 | PAR-CLIP | Tarbase |
| hsa-miR-195-5p | SEPT2 | PAR-CLIP | Tarbase |
| hsa-miR-195-5p | SEPT2 | PAR-CLIP | Tarbase |
| hsa-miR-195-5p | SEPT2 | PAR-CLIP | Tarbase |
| hsa-miR-15a-5p | SEPT2 | HITS-CLIP | Tarbase |
| hsa-miR-195-5p | SEPT2 | HITS-CLIP | Tarbase |
| hsa-miR-195-5p | FILIP1L | Microarrays | Tarbase |
| hsa-miR-195-5p | MFSD2A | Microarrays | Tarbase |
| hsa-miR-195-5p | DTYMK | Microarrays | Tarbase |
| hsa-miR-15a-5p | BDKRB2 | HITS-CLIP | Tarbase |
| hsa-miR-195-5p | BDKRB2 | HITS-CLIP | Tarbase |
| hsa-miR-15a-5p | RFWD3 | HITS-CLIP | Tarbase |
| hsa-miR-195-5p | RFWD3 | HITS-CLIP | Tarbase |
| hsa-miR-486-3p | RFWD3 | HITS-CLIP | Tarbase |
| hsa-miR-195-5p | RHOH | Microarrays | Tarbase |
| hsa-miR-141-3p | CDC40 | HITS-CLIP | Tarbase |
| hsa-miR-141-5p | CDC40 | HITS-CLIP | Tarbase |
| hsa-miR-15a-5p | STIP1 | HITS-CLIP | Tarbase |
| hsa-miR-195-5p | HR | Microarrays | Tarbase |
| hsa-miR-195-5p | REEP4 | Microarrays | Tarbase |
| hsa-miR-15a-3p | ATXN2L | HITS-CLIP | Tarbase |
| hsa-miR-195-5p | FEN1 | Microarrays | Tarbase |
| hsa-miR-15a-5p | CDKN2AIP | PAR-CLIP | Tarbase |

| | | | |
|---|---|---|---|
| hsa-miR-15a-5p | SNRNP48 | PAR-CLIP | Tarbase |
| hsa-miR-15a-5p | SLC20A2 | HITS-CLIP | Tarbase |
| hsa-miR-195-5p | SLC20A2 | HITS-CLIP | Tarbase |
| hsa-miR-3613-5p | SLC20A2 | HITS-CLIP | Tarbase |
| hsa-miR-195-5p | SLC20A2 | Microarrays | Tarbase |
| hsa-miR-15a-5p | STAT3 | HITS-CLIP | Tarbase |
| hsa-miR-195-5p | STAT3 | HITS-CLIP | Tarbase |
| hsa-miR-15a-5p | ZSWIM1 | PAR-CLIP | Tarbase |
| hsa-miR-141-3p | ADAM9 | HITS-CLIP | Tarbase |
| hsa-miR-15a-5p | AXIN2 | PAR-CLIP | Tarbase |
| hsa-miR-15a-5p | AXIN2 | Chimeric fragments | Tarbase |
| hsa-miR-3613-3p | LDLRAD4 | HITS-CLIP | Tarbase |
| hsa-miR-195-5p | SLC16A4 | Microarrays | Tarbase |
| hsa-miR-141-5p | NPNT | HITS-CLIP | Tarbase |
| hsa-miR-15a-5p | SEMA4C | HITS-CLIP | Tarbase |
| hsa-miR-195-5p | SEMA4C | HITS-CLIP | Tarbase |
| hsa-miR-15a-5p | SEMA4C | HITS-CLIP | Tarbase |
| hsa-miR-3613-3p | TET2 | PAR-CLIP | Tarbase |
| hsa-miR-15a-5p | ZBTB5 | HITS-CLIP | Tarbase |
| hsa-miR-195-5p | ZBTB5 | HITS-CLIP | Tarbase |
| hsa-miR-219a-1-3p | CHTF8 | PAR-CLIP | Tarbase |
| hsa-miR-15a-5p | SNTB2 | PAR-CLIP | Tarbase |
| hsa-miR-15a-5p | SNTB2 | HITS-CLIP | Tarbase |
| hsa-miR-15a-5p | SNTB2 | HITS-CLIP | Tarbase |
| hsa-miR-195-5p | SNTB2 | HITS-CLIP | Tarbase |
| hsa-miR-486-3p | SNTB2 | PAR-CLIP | Tarbase |
| hsa-miR-15a-5p | MAT2A | HITS-CLIP | Tarbase |
| hsa-miR-15a-5p | MAT2A | PAR-CLIP | Tarbase |
| hsa-miR-195-5p | MAT2A | HITS-CLIP | Tarbase |
| hsa-miR-195-5p | MAT2A | PAR-CLIP | Tarbase |
| hsa-miR-3605-3p | SPRY3 | HITS-CLIP | Tarbase |
| hsa-miR-195-5p | SPRY3 | Microarrays | Tarbase |
| hsa-miR-141-3p | FEM1B | HITS-CLIP | Tarbase |
| hsa-miR-15a-5p | FEM1B | PAR-CLIP | Tarbase |
| hsa-miR-15a-5p | FEM1B | HITS-CLIP | Tarbase |
| hsa-miR-15a-5p | FEM1B | HITS-CLIP | Tarbase |
| hsa-miR-195-5p | FEM1B | HITS-CLIP | Tarbase |
| hsa-miR-486-5p | FEM1B | HITS-CLIP | Tarbase |
| hsa-miR-15a-5p | FEM1B | HITS-CLIP | Tarbase |
| hsa-miR-195-5p | FEM1B | HITS-CLIP | Tarbase |
| hsa-miR-15a-5p | MAP2K1 | PAR-CLIP | Tarbase |
| hsa-miR-195-5p | MAP2K1 | Microarrays | Tarbase |
| hsa-miR-486-5p | HNRNPH1 | HITS-CLIP | Tarbase |
| hsa-miR-141-3p | IRS1 | HITS-CLIP | Tarbase |
| hsa-miR-195-5p | IRS1 | Microarrays | Tarbase |
| hsa-miR-141-3p | MECP2 | HITS-CLIP | Tarbase |
| hsa-miR-15a-5p | HSPBAP1 | HITS-CLIP | Tarbase |
| hsa-miR-195-5p | HSPBAP1 | HITS-CLIP | Tarbase |
| hsa-miR-3613-5p | PARM1 | PAR-CLIP | Tarbase |
| hsa-miR-15a-5p | CSNK1G1 | HITS-CLIP | Tarbase |
| hsa-miR-15a-5p | FAM110B | HITS-CLIP | Tarbase |
| hsa-miR-195-5p | FAM110B | HITS-CLIP | Tarbase |
| hsa-miR-141-5p | AFAP1L2 | HITS-CLIP | Tarbase |
| hsa-miR-15a-5p | PCSK9 | HITS-CLIP | Tarbase |
| hsa-miR-141-3p | XPO6 | HITS-CLIP | Tarbase |
| hsa-miR-15a-5p | XPO6 | HITS-CLIP | Tarbase |
| hsa-miR-195-5p | XPO6 | HITS-CLIP | Tarbase |
| hsa-miR-15a-5p | MN1 | Luciferase Reporter Assay | Tarbase |
| hsa-miR-141-3p | RAB3B | HITS-CLIP | Tarbase |
| hsa-miR-195-5p | CA5B | Microarrays | Tarbase |
| hsa-miR-15a-5p | EFNA1 | HITS-CLIP | Tarbase |
| hsa-miR-195-5p | EFNA1 | HITS-CLIP | Tarbase |
| hsa-miR-195-5p | EFNA1 | Microarrays | Tarbase |
| hsa-miR-141-5p | MRPL1 | HITS-CLIP | Tarbase |
| hsa-miR-15a-5p | MRPL1 | HITS-CLIP | Tarbase |
| hsa-miR-195-5p | MRPL1 | HITS-CLIP | Tarbase |
| hsa-miR-3605-5p | CRADD | HITS-CLIP | Tarbase |
| hsa-miR-141-5p | SIN3A | HITS-CLIP | Tarbase |
| hsa-miR-486-5p | SIN3A | HITS-CLIP | Tarbase |
| hsa-miR-195-5p | CXCL8 | qPCR | Tarbase |
| hsa-miR-195-5p | CXCL8 | Microarrays | Tarbase |
| hsa-miR-195-5p | SCN9A | Microarrays | Tarbase |
| hsa-miR-15a-5p | SDC2 | PAR-CLIP | Tarbase |
| hsa-miR-3613-5p | SDC2 | HITS-CLIP | Tarbase |
| hsa-miR-15a-5p | SDC2 | HITS-CLIP | Tarbase |
| hsa-miR-195-5p | SDC2 | HITS-CLIP | Tarbase |
| hsa-miR-195-5p | CD52 | Microarrays | Tarbase |
| hsa-miR-3613-3p | MMGT1 | PAR-CLIP | Tarbase |
| hsa-miR-141-3p | ZEB2 | TRAP | Tarbase |
| hsa-miR-3613-5p | ZEB2 | HITS-CLIP | Tarbase |
| hsa-miR-141-3p | ZEB2 | Luciferase Reporter Assay | Tarbase |
| hsa-miR-141-3p | ZEB2 | Luciferase Reporter Assay | Tarbase |
| hsa-miR-15a-5p | HINT1 | HITS-CLIP | Tarbase |
| hsa-miR-15a-5p | HINT1 | HITS-CLIP | Tarbase |
| hsa-miR-15a-5p | HINT1 | PAR-CLIP | Tarbase |
| hsa-miR-195-5p | HINT1 | HITS-CLIP | Tarbase |
| hsa-miR-195-5p | HINT1 | HITS-CLIP | Tarbase |
| hsa-miR-195-5p | DFFB | Microarrays | Tarbase |
| hsa-miR-141-5p | ANTXR1 | HITS-CLIP | Tarbase |
| hsa-miR-195-5p | ANTXR1 | Microarrays | Tarbase |
| hsa-miR-195-5p | CKAP2L | Microarrays | Tarbase |
| hsa-miR-15a-5p | LUZP1 | PAR-CLIP | Tarbase |
| hsa-miR-195-5p | BUB1 | Microarrays | Tarbase |
| hsa-miR-15a-5p | CHRNA5 | PAR-CLIP | Tarbase |
| hsa-miR-141-5p | STRA13 | HITS-CLIP | Tarbase |
| hsa-miR-15a-3p | AGPAT2 | HITS-CLIP | Tarbase |
| hsa-miR-15a-5p | AGPAT2 | HITS-CLIP | Tarbase |
| hsa-miR-195-5p | AGPAT2 | HITS-CLIP | Tarbase |
| hsa-miR-15a-3p | FASN | HITS-CLIP | Tarbase |
| hsa-miR-15a-5p | FASN | PAR-CLIP | Tarbase |
| hsa-miR-15a-5p | FASN | HITS-CLIP | Tarbase |
| hsa-miR-15a-5p | CNBP | HITS-CLIP | Tarbase |
| hsa-miR-15a-5p | CNBP | HITS-CLIP | Tarbase |
| hsa-miR-15a-5p | CNBP | HITS-CLIP | Tarbase |
| hsa-miR-195-5p | CNBP | HITS-CLIP | Tarbase |
| hsa-miR-195-5p | CNBP | HITS-CLIP | Tarbase |
| hsa-miR-195-5p | CNBP | HITS-CLIP | Tarbase |
| hsa-miR-15a-3p | DUS1L | HITS-CLIP | Tarbase |
| hsa-miR-15a-5p | DUS1L | HITS-CLIP | Tarbase |
| hsa-miR-15a-5p | DUS1L | PAR-CLIP | Tarbase |
| hsa-miR-195-5p | DUS1L | HITS-CLIP | Tarbase |
| hsa-miR-195-5p | NRG4 | Microarrays | Tarbase |
| hsa-miR-15a-3p | LIMS1 | HITS-CLIP | Tarbase |
| hsa-miR-15a-5p | TAPT1 | HITS-CLIP | Tarbase |
| hsa-miR-195-5p | TAPT1 | HITS-CLIP | Tarbase |
| hsa-miR-141-3p | HNRNPF | PAR-CLIP | Tarbase |
| hsa-miR-141-3p | HNRNPF | HITS-CLIP | Tarbase |
| hsa-miR-141-3p | HNRNPF | HITS-CLIP | Tarbase |
| hsa-miR-486-3p | PCDH7 | HITS-CLIP | Tarbase |
| hsa-miR-486-5p | ROBO1 | HITS-CLIP | Tarbase |
| hsa-miR-15a-5p | REPS2 | HITS-CLIP | Tarbase |
| hsa-miR-15a-5p | SYAP1 | HITS-CLIP | Tarbase |
| hsa-miR-195-5p | SYAP1 | HITS-CLIP | Tarbase |
| hsa-miR-3613-3p | SYAP1 | HITS-CLIP | Tarbase |
| hsa-miR-141-5p | TOR1AIP2 | HITS-CLIP | Tarbase |
| hsa-miR-141-5p | TOR1AIP2 | HITS-CLIP | Tarbase |
| hsa-miR-141-5p | TOR1AIP2 | HITS-CLIP | Tarbase |
| hsa-miR-141-5p | TOR1AIP2 | HITS-CLIP | Tarbase |
| hsa-miR-141-5p | TOR1AIP2 | HITS-CLIP | Tarbase |
| hsa-miR-141-5p | TOR1AIP2 | HITS-CLIP | Tarbase |
| hsa-miR-15a-5p | TM4SF1 | HITS-CLIP | Tarbase |
| hsa-miR-141-3p | OTUD3 | HITS-CLIP | Tarbase |
| hsa-miR-141-5p | OTUD3 | HITS-CLIP | Tarbase |
| hsa-miR-141-5p | OTUD3 | HITS-CLIP | Tarbase |
| hsa-miR-15a-3p | OTUD3 | HITS-CLIP | Tarbase |
| hsa-miR-15a-5p | OTUD3 | HITS-CLIP | Tarbase |
| hsa-miR-195-5p | OTUD3 | HITS-CLIP | Tarbase |
| hsa-miR-15a-5p | GUSB | HITS-CLIP | Tarbase |
| hsa-miR-195-5p | GUSB | HITS-CLIP | Tarbase |
| hsa-miR-141-3p | BRD3 | PAR-CLIP | Tarbase |
| hsa-miR-141-3p | BRD3 | HITS-CLIP | Tarbase |
| hsa-miR-141-3p | BRD3 | Luciferase Reporter Assay | Tarbase |
| hsa-miR-141-3p | BRD3 | Western Blot | Tarbase |
| hsa-miR-15a-5p | BRD3 | HITS-CLIP | Tarbase |
| hsa-miR-195-5p | BRD3 | HITS-CLIP | Tarbase |
| hsa-miR-141-5p | ZFPM2 | HITS-CLIP | Tarbase |
| hsa-miR-15a-3p | ZNF747 | HITS-CLIP | Tarbase |
| hsa-miR-15a-5p | NLGN2 | HITS-CLIP | Tarbase |
| hsa-miR-195-5p | NLGN2 | HITS-CLIP | Tarbase |

| | | | |
|---|---|---|---|
| hsa-miR-141-5p | CHD3 | HITS-CLIP | Tarbase |
| hsa-miR-15a-5p | CHD3 | PAR-CLIP | Tarbase |
| hsa-miR-15a-5p | CHD3 | PAR-CLIP | Tarbase |
| hsa-miR-15a-5p | CHD3 | HITS-CLIP | Tarbase |
| hsa-miR-3605-3p | CHD3 | HITS-CLIP | Tarbase |
| hsa-miR-141-3p | TMEM154 | PAR-CLIP | Tarbase |
| hsa-miR-3613-3p | TMEM154 | PAR-CLIP | Tarbase |
| hsa-miR-15a-5p | ALCAM | HITS-CLIP | Tarbase |
| hsa-miR-195-5p | ALCAM | HITS-CLIP | Tarbase |
| hsa-miR-15a-5p | ALCAM | HITS-CLIP | Tarbase |
| hsa-miR-195-5p | ALCAM | HITS-CLIP | Tarbase |
| hsa-miR-141-3p | YWHAG | HITS-CLIP | Tarbase |
| hsa-miR-141-3p | YWHAG | HITS-CLIP | Tarbase |
| hsa-miR-141-5p | YWHAG | HITS-CLIP | Tarbase |
| hsa-miR-15a-5p | YWHAG | HITS-CLIP | Tarbase |
| hsa-miR-15a-5p | YWHAG | HITS-CLIP | Tarbase |
| hsa-miR-15a-5p | YWHAG | HITS-CLIP | Tarbase |
| hsa-miR-15a-5p | YWHAG | HITS-CLIP | Tarbase |
| hsa-miR-195-5p | YWHAG | HITS-CLIP | Tarbase |
| hsa-miR-195-5p | YWHAG | HITS-CLIP | Tarbase |
| hsa-miR-3605-5p | YWHAG | PAR-CLIP | Tarbase |
| hsa-miR-195-5p | YWHAG | Microarrays | Tarbase |
| hsa-miR-141-3p | TMEM192 | HITS-CLIP | Tarbase |
| hsa-miR-195-5p | TMEM192 | Microarrays | Tarbase |
| hsa-miR-3613-5p | NIPA1 | HITS-CLIP | Tarbase |
| hsa-miR-15a-5p | GPR25 | HITS-CLIP | Tarbase |
| hsa-miR-195-5p | GPR25 | HITS-CLIP | Tarbase |
| hsa-miR-3613-3p | UBE2E1 | Chimeric fragments | Tarbase |
| hsa-miR-141-5p | RNF150 | HITS-CLIP | Tarbase |
| hsa-miR-15a-5p | USP38 | PAR-CLIP | Tarbase |
| hsa-miR-15a-5p | USP38 | Chimeric fragments | Tarbase |
| hsa-miR-141-3p | PWWP2A | HITS-CLIP | Tarbase |
| hsa-miR-486-5p | PWWP2A | HITS-CLIP | Tarbase |
| hsa-miR-141-5p | ZNF212 | HITS-CLIP | Tarbase |
| hsa-miR-3613-3p | ZNF282 | HITS-CLIP | Tarbase |
| hsa-miR-141-5p | CRTAP | HITS-CLIP | Tarbase |
| hsa-miR-15a-5p | ELP5 | PAR-CLIP | Tarbase |
| hsa-miR-141-5p | CDK1 | HITS-CLIP | Tarbase |
| hsa-miR-195-5p | CDK1 | Microarrays | Tarbase |
| hsa-miR-141-3p | NFRKB | HITS-CLIP | Tarbase |
| hsa-miR-219a-1-3p | NFRKB | HITS-CLIP | Tarbase |
| hsa-miR-141-3p | B3GNT2 | HITS-CLIP | Tarbase |
| hsa-miR-15a-5p | B3GNT2 | PAR-CLIP | Tarbase |
| hsa-miR-15a-5p | B3GNT2 | PAR-CLIP | Tarbase |
| hsa-miR-15a-5p | B3GNT2 | Chimeric fragments | Tarbase |
| hsa-miR-15a-5p | TMED10 | HITS-CLIP | Tarbase |
| hsa-miR-15a-5p | TMED10 | HITS-CLIP | Tarbase |
| hsa-miR-195-5p | TMED10 | HITS-CLIP | Tarbase |
| hsa-miR-195-5p | TMED10 | HITS-CLIP | Tarbase |
| hsa-miR-219a-1-3p | TMED10 | HITS-CLIP | Tarbase |
| hsa-miR-195-5p | SEMA3E | Microarrays | Tarbase |
| hsa-miR-3613-3p | SLC30A1 | PAR-CLIP | Tarbase |
| hsa-miR-195-5p | DCLK2 | Microarrays | Tarbase |
| hsa-miR-15a-3p | ADORA2B | HITS-CLIP | Tarbase |
| hsa-miR-15a-5p | ADORA2B | HITS-CLIP | Tarbase |
| hsa-miR-195-5p | ADORA2B | HITS-CLIP | Tarbase |
| hsa-miR-15a-5p | WIBG | HITS-CLIP | Tarbase |
| hsa-miR-15a-5p | WIBG | HITS-CLIP | Tarbase |
| hsa-miR-195-5p | WIBG | HITS-CLIP | Tarbase |
| hsa-miR-195-5p | NPAS2 | Microarrays | Tarbase |
| hsa-miR-15a-3p | LONRF2 | HITS-CLIP | Tarbase |
| hsa-miR-195-5p | HSD17B13 | Microarrays | Tarbase |
| hsa-miR-15a-3p | PA2G4 | HITS-CLIP | Tarbase |
| hsa-miR-15a-3p | PFKFB3 | HITS-CLIP | Tarbase |
| hsa-miR-15a-5p | ARL6IP1 | PAR-CLIP | Tarbase |
| hsa-miR-486-5p | ARL6IP1 | HITS-CLIP | Tarbase |
| hsa-miR-15a-5p | SERPINB9 | HITS-CLIP | Tarbase |
| hsa-miR-195-5p | SERPINB9 | HITS-CLIP | Tarbase |
| hsa-miR-141-3p | CDH2 | HITS-CLIP | Tarbase |
| hsa-miR-141-5p | EMB | HITS-CLIP | Tarbase |
| hsa-miR-15a-5p | EMB | PAR-CLIP | Tarbase |
| hsa-miR-195-5p | STAT2 | Microarrays | Tarbase |
| hsa-miR-3605-5p | NUDCD2 | HITS-CLIP | Tarbase |
| hsa-miR-15a-5p | IRF2BP1 | HITS-CLIP | Tarbase |
| hsa-miR-15a-5p | IRF2BP1 | PAR-CLIP | Tarbase |
| hsa-miR-195-5p | IRF2BP1 | HITS-CLIP | Tarbase |
| hsa-miR-141-3p | HSPA4 | HITS-CLIP | Tarbase |
| hsa-miR-195-5p | HSPA4 | HITS-CLIP | Tarbase |
| hsa-miR-15a-5p | RNF34 | HITS-CLIP | Tarbase |
| hsa-miR-195-5p | RNF34 | HITS-CLIP | Tarbase |
| hsa-miR-15a-5p | ATF7 | PAR-CLIP | Tarbase |
| hsa-miR-15a-5p | ATF7 | PAR-CLIP | Tarbase |
| hsa-miR-15a-5p | ATF7 | HITS-CLIP | Tarbase |
| hsa-miR-195-5p | ATF7 | PAR-CLIP | Tarbase |
| hsa-miR-195-5p | ATF7 | HITS-CLIP | Tarbase |
| hsa-miR-15a-5p | SOCS6 | HITS-CLIP | Tarbase |
| hsa-miR-15a-5p | SOCS6 | PAR-CLIP | Tarbase |
| hsa-miR-15a-5p | SOCS6 | HITS-CLIP | Tarbase |
| hsa-miR-195-5p | SOCS6 | HITS-CLIP | Tarbase |
| hsa-miR-195-5p | SOCS6 | HITS-CLIP | Tarbase |
| hsa-miR-141-5p | HOXB9 | HITS-CLIP | Tarbase |
| hsa-miR-15a-3p | HOXB9 | HITS-CLIP | Tarbase |
| hsa-miR-141-3p | KIF5B | HITS-CLIP | Tarbase |
| hsa-miR-15a-5p | KIF5B | PAR-CLIP | Tarbase |
| hsa-miR-15a-5p | KIF5B | PAR-CLIP | Tarbase |
| hsa-miR-15a-5p | KIF5B | PAR-CLIP | Tarbase |
| hsa-miR-15a-5p | KIF5B | HITS-CLIP | Tarbase |
| hsa-miR-15a-5p | KIF5B | PAR-CLIP | Tarbase |
| hsa-miR-15a-5p | KIF5B | HITS-CLIP | Tarbase |
| hsa-miR-15a-5p | KIF5B | PAR-CLIP | Tarbase |
| hsa-miR-195-5p | KIF5B | PAR-CLIP | Tarbase |
| hsa-miR-195-5p | KIF5B | HITS-CLIP | Tarbase |
| hsa-miR-195-5p | KIF5B | HITS-CLIP | Tarbase |
| hsa-miR-195-5p | KIF5B | PAR-CLIP | Tarbase |
| hsa-miR-486-5p | KIF5B | PAR-CLIP | Tarbase |
| hsa-miR-15a-5p | KIF5B | HITS-CLIP | Tarbase |
| hsa-miR-195-5p | KIF5B | HITS-CLIP | Tarbase |
| hsa-miR-15a-5p | KIF5B | Chimeric fragments | Tarbase |
| hsa-miR-15a-3p | AKAP13 | HITS-CLIP | Tarbase |
| hsa-miR-219a-1-3p | AKAP13 | PAR-CLIP | Tarbase |
| hsa-miR-219a-1-3p | AKAP13 | PAR-CLIP | Tarbase |
| hsa-miR-219a-1-3p | AKAP13 | PAR-CLIP | Tarbase |
| hsa-miR-15a-5p | CDCA4 | PAR-CLIP | Tarbase |
| hsa-miR-15a-5p | CDCA4 | PAR-CLIP | Tarbase |
| hsa-miR-15a-5p | CDCA4 | HITS-CLIP | Tarbase |
| hsa-miR-15a-5p | CDCA4 | PAR-CLIP | Tarbase |
| hsa-miR-15a-5p | CDCA4 | PAR-CLIP | Tarbase |
| hsa-miR-15a-5p | CDCA4 | PAR-CLIP | Tarbase |
| hsa-miR-15a-5p | CDCA4 | HITS-CLIP | Tarbase |
| hsa-miR-15a-5p | CDCA4 | PAR-CLIP | Tarbase |
| hsa-miR-15a-5p | CDCA4 | PAR-CLIP | Tarbase |
| hsa-miR-15a-5p | CDCA4 | PAR-CLIP | Tarbase |
| hsa-miR-15a-5p | CDCA4 | PAR-CLIP | Tarbase |
| hsa-miR-15a-5p | CDCA4 | PAR-CLIP | Tarbase |
| hsa-miR-195-5p | CDCA4 | HITS-CLIP | Tarbase |
| hsa-miR-195-5p | CDCA4 | PAR-CLIP | Tarbase |
| hsa-miR-195-5p | CDCA4 | PAR-CLIP | Tarbase |
| hsa-miR-195-5p | CDCA4 | PAR-CLIP | Tarbase |
| hsa-miR-195-5p | CDCA4 | PAR-CLIP | Tarbase |
| hsa-miR-195-5p | CDCA4 | PAR-CLIP | Tarbase |
| hsa-miR-195-5p | CDCA4 | Microarrays | Tarbase |
| hsa-miR-15a-5p | CDCA4 | Chimeric fragments | Tarbase |
| hsa-miR-15a-3p | PPM1D | HITS-CLIP | Tarbase |
| hsa-miR-15a-5p | PPM1D | PAR-CLIP | Tarbase |
| hsa-miR-15a-5p | PPM1D | PAR-CLIP | Tarbase |
| hsa-miR-141-3p | KBTBD2 | HITS-CLIP | Tarbase |
| hsa-miR-141-5p | KBTBD2 | HITS-CLIP | Tarbase |
| hsa-miR-15a-5p | KBTBD2 | HITS-CLIP | Tarbase |
| hsa-miR-15a-5p | KBTBD2 | HITS-CLIP | Tarbase |
| hsa-miR-195-5p | KBTBD2 | HITS-CLIP | Tarbase |
| hsa-miR-15a-5p | TRIAP1 | Chimeric fragments | Tarbase |
| hsa-miR-141-5p | KIAA0232 | HITS-CLIP | Tarbase |
| hsa-miR-15a-5p | KIAA0232 | HITS-CLIP | Tarbase |
| hsa-miR-195-5p | KIAA0232 | HITS-CLIP | Tarbase |
| hsa-miR-15a-5p | TMEM43 | HITS-CLIP | Tarbase |
| hsa-miR-15a-5p | TMEM43 | PAR-CLIP | Tarbase |
| hsa-miR-15a-5p | TMEM43 | HITS-CLIP | Tarbase |
| hsa-miR-195-5p | TMEM43 | HITS-CLIP | Tarbase |
| hsa-miR-195-5p | TMEM43 | HITS-CLIP | Tarbase |

| | | | |
|---|---|---|---|
| hsa-miR-141-3p | RNF139 | HITS-CLIP | Tarbase |
| hsa-miR-15a-5p | RNF139 | HITS-CLIP | Tarbase |
| hsa-miR-195-5p | CYTL1 | Microarrays | Tarbase |
| hsa-miR-3613-5p | MSANTD4 | PAR-CLIP | Tarbase |
| hsa-miR-195-5p | OSCAR | AGO-IP | Tarbase |
| hsa-miR-15a-5p | TANC2 | HITS-CLIP | Tarbase |
| hsa-miR-15a-5p | TANC2 | HITS-CLIP | Tarbase |
| hsa-miR-195-5p | TANC2 | HITS-CLIP | Tarbase |
| hsa-miR-15a-5p | DNAJC24 | HITS-CLIP | Tarbase |
| hsa-miR-15a-5p | DNAJC24 | HITS-CLIP | Tarbase |
| hsa-miR-195-5p | DNAJC24 | HITS-CLIP | Tarbase |
| hsa-miR-195-5p | DNAJC24 | HITS-CLIP | Tarbase |
| hsa-miR-195-5p | PDGFD | Microarrays | Tarbase |
| hsa-miR-195-5p | PLAC1 | Microarrays | Tarbase |
| hsa-miR-15a-5p | S1PR1 | PAR-CLIP | Tarbase |
| hsa-miR-195-5p | S1PR1 | PAR-CLIP | Tarbase |
| hsa-miR-15a-5p | S1PR1 | HITS-CLIP | Tarbase |
| hsa-miR-195-5p | S1PR1 | HITS-CLIP | Tarbase |
| hsa-miR-195-5p | HS6ST2 | Microarrays | Tarbase |
| hsa-miR-195-5p | PKIA | Microarrays | Tarbase |
| hsa-miR-3613-5p | ALK | PAR-CLIP | Tarbase |
| hsa-miR-195-5p | INSR | Microarrays | Tarbase |
| hsa-miR-141-3p | PRKCE | HITS-CLIP | Tarbase |
| hsa-miR-3605-3p | ZNF692 | HITS-CLIP | Tarbase |
| hsa-miR-3613-3p | JUNB | HITS-CLIP | Tarbase |
| hsa-miR-195-5p | TMEM37 | Microarrays | Tarbase |
| hsa-miR-195-5p | SHCBP1 | Microarrays | Tarbase |
| hsa-miR-141-3p | FAM98B | Chimeric fragments | Tarbase |
| hsa-miR-15a-3p | GAA | HITS-CLIP | Tarbase |
| hsa-miR-15a-5p | CANT1 | HITS-CLIP | Tarbase |
| hsa-miR-3613-3p | CANT1 | HITS-CLIP | Tarbase |
| hsa-miR-15a-5p | ZDHHC16 | Chimeric fragments | Tarbase |
| hsa-miR-15a-5p | CHD7 | PAR-CLIP | Tarbase |
| hsa-miR-15a-5p | CHD7 | HITS-CLIP | Tarbase |
| hsa-miR-195-5p | CHD7 | HITS-CLIP | Tarbase |
| hsa-miR-141-5p | ESCO2 | HITS-CLIP | Tarbase |
| hsa-miR-15a-5p | CLCN5 | PAR-CLIP | Tarbase |
| hsa-miR-3605-5p | KSR2 | HITS-CLIP | Tarbase |
| hsa-miR-15a-5p | POLR1C | HITS-CLIP | Tarbase |
| hsa-miR-15a-5p | POLR1C | HITS-CLIP | Tarbase |
| hsa-miR-15a-5p | POLR1C | PAR-CLIP | Tarbase |
| hsa-miR-15a-5p | POLR1C | HITS-CLIP | Tarbase |
| hsa-miR-15a-5p | POLR1C | HITS-CLIP | Tarbase |
| hsa-miR-195-5p | POLR1C | HITS-CLIP | Tarbase |
| hsa-miR-195-5p | POLR1C | HITS-CLIP | Tarbase |
| hsa-miR-195-5p | POLR1C | HITS-CLIP | Tarbase |
| hsa-miR-141-3p | ASXL1 | HITS-CLIP | Tarbase |
| hsa-miR-15a-5p | ASXL1 | PAR-CLIP | Tarbase |
| hsa-miR-15a-5p | ASXL1 | PAR-CLIP | Tarbase |
| hsa-miR-195-5p | ASXL1 | PAR-CLIP | Tarbase |
| hsa-miR-15a-5p | ASXL1 | Chimeric fragments | Tarbase |
| hsa-miR-3613-3p | ZNF562 | HITS-CLIP | Tarbase |
| hsa-miR-141-3p | ZNF318 | HITS-CLIP | Tarbase |
| hsa-miR-141-5p | ZNF318 | HITS-CLIP | Tarbase |
| hsa-miR-141-3p | OR1L3 | PAR-CLIP | Tarbase |
| hsa-miR-141-3p | OR1L3 | PAR-CLIP | Tarbase |
| hsa-miR-141-3p | OR1L3 | PAR-CLIP | Tarbase |
| hsa-miR-141-5p | LRRC8D | HITS-CLIP | Tarbase |
| hsa-miR-141-5p | PPID | HITS-CLIP | Tarbase |
| hsa-miR-15a-5p | LPAR3 | PAR-CLIP | Tarbase |
| hsa-miR-195-5p | PTGER4 | Microarrays | Tarbase |
| hsa-miR-141-5p | MAP6 | HITS-CLIP | Tarbase |
| hsa-miR-195-5p | FGG | Microarrays | Tarbase |
| hsa-miR-195-5p | FGB | Microarrays | Tarbase |
| hsa-miR-195-5p | CXXC5 | Microarrays | Tarbase |
| hsa-miR-195-5p | PIK3CD | Microarrays | Tarbase |
| hsa-miR-15a-5p | ENC1 | HITS-CLIP | Tarbase |
| hsa-miR-15a-5p | ENC1 | HITS-CLIP | Tarbase |
| hsa-miR-195-5p | ENC1 | HITS-CLIP | Tarbase |
| hsa-miR-195-5p | ENC1 | Microarrays | Tarbase |
| hsa-miR-141-5p | BPTF | HITS-CLIP | Tarbase |
| hsa-miR-195-5p | ZIK1 | Microarrays | Tarbase |
| hsa-miR-141-5p | TCEA2 | HITS-CLIP | Tarbase |
| hsa-miR-141-5p | CAMTA1 | HITS-CLIP | Tarbase |
| hsa-miR-195-5p | CAMTA1 | HITS-CLIP | Tarbase |
| hsa-miR-141-5p | SPATA5L1 | HITS-CLIP | Tarbase |
| hsa-miR-141-3p | BCL2 | Western Blot | Tarbase |
| hsa-miR-15a-5p | BCL2 | Western Blot | Tarbase |
| hsa-miR-15a-5p | BCL2 | Western Blot | Tarbase |
| hsa-miR-15a-5p | BCL2 | Western Blot | Tarbase |
| hsa-miR-15a-5p | BCL2 | PAR-CLIP | Tarbase |
| hsa-miR-15a-5p | BCL2 | PAR-CLIP | Tarbase |
| hsa-miR-15a-5p | BCL2 | HITS-CLIP | Tarbase |
| hsa-miR-195-5p | BCL2 | PAR-CLIP | Tarbase |
| hsa-miR-486-5p | BCL2 | HITS-CLIP | Tarbase |
| hsa-miR-15a-5p | BCL2 | Luciferase Reporter Assay | Tarbase |
| hsa-miR-195-5p | RHNO1 | Microarrays | Tarbase |
| hsa-miR-141-5p | CTPS1 | HITS-CLIP | Tarbase |
| hsa-miR-141-3p | WDR87 | HITS-CLIP | Tarbase |
| hsa-miR-15a-5p | PWWP2B | HITS-CLIP | Tarbase |
| hsa-miR-15a-5p | PWWP2B | HITS-CLIP | Tarbase |
| hsa-miR-195-5p | PWWP2B | HITS-CLIP | Tarbase |
| hsa-miR-141-3p | RRM2 | HITS-CLIP | Tarbase |
| hsa-miR-195-5p | RRM2 | Microarrays | Tarbase |
| hsa-miR-486-5p | RPS21 | HITS-CLIP | Tarbase |
| hsa-miR-486-5p | RPS21 | HITS-CLIP | Tarbase |
| hsa-miR-195-5p | C3AR1 | Microarrays | Tarbase |
| hsa-miR-141-3p | PTEN | HITS-CLIP | Tarbase |
| hsa-miR-141-3p | PTEN | Luciferase Reporter Assay | Tarbase |
| hsa-miR-141-3p | PTEN | Western Blot | Tarbase |
| hsa-miR-141-3p | PTEN | Western Blot | Tarbase |
| hsa-miR-141-3p | PTEN | Luciferase Reporter Assay | Tarbase |
| hsa-miR-141-3p | PTEN | Luciferase Reporter Assay | Tarbase |
| hsa-miR-141-3p | PTEN | Western Blot | Tarbase |
| hsa-miR-486-5p | PTEN | HITS-CLIP | Tarbase |
| hsa-miR-195-5p | FRMD5 | Microarrays | Tarbase |
| hsa-miR-195-5p | CYP4F11 | Microarrays | Tarbase |
| hsa-miR-141-3p | ZNF217 | HITS-CLIP | Tarbase |
| hsa-miR-141-3p | ZNF217 | HITS-CLIP | Tarbase |
| hsa-miR-219a-1-3p | ZNF217 | PAR-CLIP | Tarbase |
| hsa-miR-141-5p | SCG2 | HITS-CLIP | Tarbase |
| hsa-miR-15a-5p | PPIH | HITS-CLIP | Tarbase |
| hsa-miR-195-5p | PPIH | Microarrays | Tarbase |
| hsa-miR-141-5p | JMJD1C | HITS-CLIP | Tarbase |
| hsa-miR-141-5p | JMJD1C | HITS-CLIP | Tarbase |
| hsa-miR-15a-5p | JMJD1C | HITS-CLIP | Tarbase |
| hsa-miR-195-5p | JMJD1C | HITS-CLIP | Tarbase |
| hsa-miR-486-5p | JMJD1C | HITS-CLIP | Tarbase |
| hsa-miR-141-3p | ORMDL3 | HITS-CLIP | Tarbase |
| hsa-miR-15a-5p | ORMDL3 | HITS-CLIP | Tarbase |
| hsa-miR-195-5p | ORMDL3 | HITS-CLIP | Tarbase |
| hsa-miR-3613-5p | ORMDL3 | PAR-CLIP | Tarbase |
| hsa-miR-141-3p | KLF11 | Luciferase Reporter Assay | Tarbase |
| hsa-miR-141-3p | KLF11 | Luciferase Reporter Assay | Tarbase |
| hsa-miR-15a-3p | KLF11 | HITS-CLIP | Tarbase |
| hsa-miR-15a-3p | LRRC15 | HITS-CLIP | Tarbase |
| hsa-miR-195-5p | CALB2 | Microarrays | Tarbase |
| hsa-miR-486-3p | SNTB1 | HITS-CLIP | Tarbase |
| hsa-miR-195-5p | MTBP | Microarrays | Tarbase |
| hsa-miR-195-5p | MBOAT1 | Microarrays | Tarbase |
| hsa-miR-15a-3p | ID4 | HITS-CLIP | Tarbase |
| hsa-miR-3613-3p | CEBPB | HITS-CLIP | Tarbase |
| hsa-miR-3613-3p | CEBPB | PAR-CLIP | Tarbase |
| hsa-miR-195-5p | C5orf34 | Microarrays | Tarbase |
| hsa-miR-141-3p | DPAGT1 | HITS-CLIP | Tarbase |
| hsa-miR-141-3p | BSG | HITS-CLIP | Tarbase |
| hsa-miR-15a-3p | CERS6 | HITS-CLIP | Tarbase |
| hsa-miR-195-5p | STARD5 | Microarrays | Tarbase |
| hsa-miR-195-5p | RCAN2 | Microarrays | Tarbase |
| hsa-miR-15a-5p | GNB2 | HITS-CLIP | Tarbase |
| hsa-miR-195-5p | GNB2 | HITS-CLIP | Tarbase |
| hsa-miR-15a-5p | FAM195A | HITS-CLIP | Tarbase |
| hsa-miR-141-3p | GNG12 | HITS-CLIP | Tarbase |
| hsa-miR-15a-5p | GNG12 | Chimeric fragments | Tarbase |
| hsa-miR-15a-3p | SYNPO2 | HITS-CLIP | Tarbase |
| hsa-miR-15a-3p | ZNF24 | HITS-CLIP | Tarbase |
| hsa-miR-219a-1-3p | ZNF24 | PAR-CLIP | Tarbase |
| hsa-miR-15a-3p | PPP1CA | HITS-CLIP | Tarbase |
| hsa-miR-15a-5p | PPP1CA | HITS-CLIP | Tarbase |

| | | | |
|---|---|---|---|
| hsa-miR-141-5p | HCFC1 | HITS-CLIP | Tarbase |
| hsa-miR-3605-5p | HCFC1 | HITS-CLIP | Tarbase |
| hsa-miR-3613-5p | HCFC1 | HITS-CLIP | Tarbase |
| hsa-miR-15a-5p | TMEM134 | HITS-CLIP | Tarbase |
| hsa-miR-15a-5p | TMEM134 | HITS-CLIP | Tarbase |
| hsa-miR-195-5p | TMEM134 | HITS-CLIP | Tarbase |
| hsa-miR-195-5p | TMEM134 | HITS-CLIP | Tarbase |
| hsa-miR-15a-5p | ZMAT3 | PAR-CLIP | Tarbase |
| hsa-miR-15a-5p | ZMAT3 | PAR-CLIP | Tarbase |
| hsa-miR-15a-5p | ZMAT3 | PAR-CLIP | Tarbase |
| hsa-miR-15a-5p | ZMAT3 | PAR-CLIP | Tarbase |
| hsa-miR-15a-5p | ZMAT3 | PAR-CLIP | Tarbase |
| hsa-miR-15a-5p | ZMAT3 | PAR-CLIP | Tarbase |
| hsa-miR-15a-5p | ZMAT3 | HITS-CLIP | Tarbase |
| hsa-miR-195-5p | ZMAT3 | PAR-CLIP | Tarbase |
| hsa-miR-195-5p | ZMAT3 | PAR-CLIP | Tarbase |
| hsa-miR-195-5p | ZMAT3 | PAR-CLIP | Tarbase |
| hsa-miR-3613-3p | ZMAT3 | PAR-CLIP | Tarbase |
| hsa-miR-195-5p | ZMAT3 | Microarrays | Tarbase |
| hsa-miR-15a-5p | ZMAT3 | Chimeric fragments | Tarbase |
| hsa-miR-15a-3p | SLFN11 | HITS-CLIP | Tarbase |
| hsa-miR-15a-5p | SLFN11 | HITS-CLIP | Tarbase |
| hsa-miR-195-5p | SLFN11 | HITS-CLIP | Tarbase |
| hsa-miR-195-5p | SLFN11 | Microarrays | Tarbase |
| hsa-miR-15a-5p | PURG | PAR-CLIP | Tarbase |
| hsa-miR-141-5p | CFL1 | HITS-CLIP | Tarbase |
| hsa-miR-15a-5p | TMCC1 | HITS-CLIP | Tarbase |
| hsa-miR-15a-5p | TMCC1 | HITS-CLIP | Tarbase |
| hsa-miR-195-5p | TMCC1 | HITS-CLIP | Tarbase |
| hsa-miR-195-5p | TMCC1 | HITS-CLIP | Tarbase |
| hsa-miR-15a-5p | TMCC1 | HITS-CLIP | Tarbase |
| hsa-miR-195-5p | TMCC1 | HITS-CLIP | Tarbase |
| hsa-miR-15a-3p | NAA16 | HITS-CLIP | Tarbase |
| hsa-miR-195-5p | HOXC5 | Microarrays | Tarbase |
| hsa-miR-141-3p | DCP2 | PAR-CLIP | Tarbase |
| hsa-miR-141-3p | DCP2 | HITS-CLIP | Tarbase |
| hsa-miR-141-3p | DCP2 | HITS-CLIP | Tarbase |
| hsa-miR-141-3p | DCP2 | PAR-CLIP | Tarbase |
| hsa-miR-15a-5p | DCP2 | HITS-CLIP | Tarbase |
| hsa-miR-195-5p | DCP2 | HITS-CLIP | Tarbase |
| hsa-miR-15a-5p | OVOL1 | HITS-CLIP | Tarbase |
| hsa-miR-15a-5p | PDP2 | PAR-CLIP | Tarbase |
| hsa-miR-195-5p | PDP2 | PAR-CLIP | Tarbase |
| hsa-miR-141-3p | SP3 | HITS-CLIP | Tarbase |
| hsa-miR-15a-5p | SP3 | HITS-CLIP | Tarbase |
| hsa-miR-15a-5p | SP3 | HITS-CLIP | Tarbase |
| hsa-miR-195-5p | SP3 | HITS-CLIP | Tarbase |
| hsa-miR-141-5p | DMXL1 | HITS-CLIP | Tarbase |
| hsa-miR-219a-1-3p | DMXL1 | PAR-CLIP | Tarbase |
| hsa-miR-141-3p | ZNF621 | HITS-CLIP | Tarbase |
| hsa-miR-195-5p | NADSYN1 | Microarrays | Tarbase |
| hsa-miR-15a-5p | NBEA | HITS-CLIP | Tarbase |
| hsa-miR-195-5p | NBEA | HITS-CLIP | Tarbase |
| hsa-miR-15a-5p | NBEA | HITS-CLIP | Tarbase |
| hsa-miR-195-5p | NBEA | HITS-CLIP | Tarbase |
| hsa-miR-3605-5p | RNASEH2C | HITS-CLIP | Tarbase |
| hsa-miR-15a-3p | ANKRD13D | HITS-CLIP | Tarbase |
| hsa-miR-15a-5p | MYD88 | PAR-CLIP | Tarbase |
| hsa-miR-141-5p | LCLAT1 | HITS-CLIP | Tarbase |
| hsa-miR-195-5p | SH3RF3 | Microarrays | Tarbase |
| hsa-miR-15a-5p | TADA2B | HITS-CLIP | Tarbase |
| hsa-miR-195-5p | TADA2B | HITS-CLIP | Tarbase |
| hsa-miR-141-3p | ADRBK1 | HITS-CLIP | Tarbase |
| hsa-miR-15a-3p | ZNF680 | HITS-CLIP | Tarbase |
| hsa-miR-486-5p | ZNF680 | PAR-CLIP | Tarbase |
| hsa-miR-195-5p | ZNF680 | Microarrays | Tarbase |
| hsa-miR-141-3p | HECTD4 | HITS-CLIP | Tarbase |
| hsa-miR-15a-5p | HECTD4 | HITS-CLIP | Tarbase |
| hsa-miR-195-5p | HECTD4 | HITS-CLIP | Tarbase |
| hsa-miR-141-5p | FAM222B | HITS-CLIP | Tarbase |
| hsa-miR-15a-5p | BNC2 | PAR-CLIP | Tarbase |
| hsa-miR-195-5p | BNC2 | PAR-CLIP | Tarbase |
| hsa-miR-195-5p | COQ2 | Microarrays | Tarbase |
| hsa-miR-141-3p | TRMT112 | HITS-CLIP | Tarbase |
| hsa-miR-141-3p | TRMT112 | HITS-CLIP | Tarbase |
| hsa-miR-486-3p | KDM2A | HITS-CLIP | Tarbase |
| hsa-miR-486-5p | KDM2A | PAR-CLIP | Tarbase |
| hsa-miR-15a-5p | RAPH1 | PAR-CLIP | Tarbase |
| hsa-miR-3613-5p | RAPH1 | HITS-CLIP | Tarbase |
| hsa-miR-15a-3p | ADCY5 | HITS-CLIP | Tarbase |
| hsa-miR-195-5p | CYSLTR1 | Microarrays | Tarbase |
| hsa-miR-195-5p | CKS1B | Microarrays | Tarbase |
| hsa-miR-15a-3p | ABCD2 | HITS-CLIP | Tarbase |
| hsa-miR-195-5p | KIAA1919 | Microarrays | Tarbase |
| hsa-miR-141-5p | VANGL1 | HITS-CLIP | Tarbase |
| hsa-miR-195-5p | GLRX | Microarrays | Tarbase |
| hsa-miR-15a-5p | GOLGB1 | HITS-CLIP | Tarbase |
| hsa-miR-195-5p | GOLGB1 | HITS-CLIP | Tarbase |
| hsa-miR-195-5p | GOLGB1 | Microarrays | Tarbase |
| hsa-miR-195-5p | SNCG | AGO-IP | Tarbase |
| hsa-miR-15a-5p | TNKS | PAR-CLIP | Tarbase |
| hsa-miR-486-5p | TNKS | PAR-CLIP | Tarbase |
| hsa-miR-15a-5p | ZBTB21 | HITS-CLIP | Tarbase |
| hsa-miR-195-5p | ZBTB21 | HITS-CLIP | Tarbase |
| hsa-miR-195-5p | PPP1R3B | Microarrays | Tarbase |
| hsa-miR-141-5p | STOX2 | HITS-CLIP | Tarbase |
| hsa-miR-15a-5p | STOX2 | Chimeric fragments | Tarbase |
| hsa-miR-15a-5p | TRIB1 | HITS-CLIP | Tarbase |
| hsa-miR-195-5p | TRIB1 | Microarrays | Tarbase |
| hsa-miR-15a-3p | NAA20 | HITS-CLIP | Tarbase |
| hsa-miR-15a-5p | NAA20 | HITS-CLIP | Tarbase |
| hsa-miR-15a-5p | NAA20 | HITS-CLIP | Tarbase |
| hsa-miR-195-5p | NAA20 | HITS-CLIP | Tarbase |
| hsa-miR-195-5p | RNF26 | Microarrays | Tarbase |
| hsa-miR-486-3p | PPP1R14B | HITS-CLIP | Tarbase |
| hsa-miR-3613-5p | SMARCC1 | HITS-CLIP | Tarbase |
| hsa-miR-15a-3p | FKBP2 | HITS-CLIP | Tarbase |
| hsa-miR-486-5p | GMPPB | PAR-CLIP | Tarbase |
| hsa-miR-3613-3p | MOB1B | PAR-CLIP | Tarbase |
| hsa-miR-195-5p | MOB1B | Microarrays | Tarbase |
| hsa-miR-15a-5p | ZNF622 | HITS-CLIP | Tarbase |
| hsa-miR-15a-5p | ZNF622 | PAR-CLIP | Tarbase |
| hsa-miR-15a-5p | ZNF622 | HITS-CLIP | Tarbase |
| hsa-miR-195-5p | ZNF622 | HITS-CLIP | Tarbase |
| hsa-miR-195-5p | ZNF622 | HITS-CLIP | Tarbase |
| hsa-miR-15a-5p | ZNF622 | Chimeric fragments | Tarbase |
| hsa-miR-195-5p | NABP1 | Microarrays | Tarbase |
| hsa-miR-141-3p | CHD2 | HITS-CLIP | Tarbase |
| hsa-miR-15a-3p | CHD2 | HITS-CLIP | Tarbase |
| hsa-miR-15a-5p | CHD2 | HITS-CLIP | Tarbase |
| hsa-miR-195-5p | CHD2 | HITS-CLIP | Tarbase |
| hsa-miR-15a-5p | CEP83 | HITS-CLIP | Tarbase |
| hsa-miR-195-5p | CEP83 | HITS-CLIP | Tarbase |
| hsa-miR-195-5p | SULT1B1 | Microarrays | Tarbase |
| hsa-miR-195-5p | PC | Microarrays | Tarbase |
| hsa-miR-15a-5p | SCAI | PAR-CLIP | Tarbase |
| hsa-miR-15a-5p | LRFN4 | HITS-CLIP | Tarbase |
| hsa-miR-141-3p | SLC19A1 | HITS-CLIP | Tarbase |
| hsa-miR-15a-3p | SLC19A1 | HITS-CLIP | Tarbase |
| hsa-miR-15a-5p | RCE1 | PAR-CLIP | Tarbase |
| hsa-miR-15a-5p | RCE1 | HITS-CLIP | Tarbase |
| hsa-miR-15a-5p | UQCRH | HITS-CLIP | Tarbase |
| hsa-miR-195-5p | UQCRH | HITS-CLIP | Tarbase |
| hsa-miR-15a-3p | PSMD1 | PAR-CLIP | Tarbase |
| hsa-miR-15a-5p | PSMD1 | HITS-CLIP | Tarbase |
| hsa-miR-195-5p | GPR64 | Microarrays | Tarbase |
| hsa-miR-195-5p | HEG1 | Microarrays | Tarbase |
| hsa-miR-15a-5p | TOMM20 | HITS-CLIP | Tarbase |
| hsa-miR-195-5p | TOMM20 | HITS-CLIP | Tarbase |
| hsa-miR-15a-5p | JUP | PAR-CLIP | Tarbase |
| hsa-miR-195-5p | JUP | Microarrays | Tarbase |
| hsa-miR-15a-3p | ENDOV | HITS-CLIP | Tarbase |
| hsa-miR-15a-5p | ENDOV | HITS-CLIP | Tarbase |
| hsa-miR-195-5p | ENDOV | HITS-CLIP | Tarbase |
| hsa-miR-141-3p | RNF213 | HITS-CLIP | Tarbase |
| hsa-miR-15a-3p | RNF213 | HITS-CLIP | Tarbase |
| hsa-miR-15a-5p | RNF213 | HITS-CLIP | Tarbase |
| hsa-miR-15a-5p | RNF213 | HITS-CLIP | Tarbase |
| hsa-miR-15a-5p | RNF213 | HITS-CLIP | Tarbase |

| | | | |
|---|---|---|---|
| hsa-miR-15a-5p | RNF213 | HITS-CLIP | Tarbase |
| hsa-miR-15a-5p | RNF213 | HITS-CLIP | Tarbase |
| hsa-miR-15a-5p | RNF213 | HITS-CLIP | Tarbase |
| hsa-miR-195-5p | RNF213 | HITS-CLIP | Tarbase |
| hsa-miR-195-5p | RNF213 | HITS-CLIP | Tarbase |
| hsa-miR-195-5p | RNF213 | HITS-CLIP | Tarbase |
| hsa-miR-195-5p | RNF213 | HITS-CLIP | Tarbase |
| hsa-miR-195-5p | RNF213 | HITS-CLIP | Tarbase |
| hsa-miR-195-5p | PLK3 | Microarrays | Tarbase |
| hsa-miR-15a-5p | NET1 | HITS-CLIP | Tarbase |
| hsa-miR-195-5p | NET1 | HITS-CLIP | Tarbase |
| hsa-miR-195-5p | NET1 | Microarrays | Tarbase |
| hsa-miR-141-5p | DPY19L1 | HITS-CLIP | Tarbase |
| hsa-miR-141-3p | ZNF791 | HITS-CLIP | Tarbase |
| hsa-miR-486-5p | ZNF791 | HITS-CLIP | Tarbase |
| hsa-miR-15a-5p | PHC3 | PAR-CLIP | Tarbase |
| hsa-miR-15a-5p | PHC3 | PAR-CLIP | Tarbase |
| hsa-miR-15a-5p | PHC3 | HITS-CLIP | Tarbase |
| hsa-miR-15a-5p | PHC3 | HITS-CLIP | Tarbase |
| hsa-miR-15a-5p | PHC3 | HITS-CLIP | Tarbase |
| hsa-miR-195-5p | PHC3 | HITS-CLIP | Tarbase |
| hsa-miR-195-5p | PHC3 | HITS-CLIP | Tarbase |
| hsa-miR-195-5p | PHC3 | HITS-CLIP | Tarbase |
| hsa-miR-15a-5p | CBX2 | PAR-CLIP | Tarbase |
| hsa-miR-15a-5p | SPTBN2 | PAR-CLIP | Tarbase |
| hsa-miR-15a-5p | GOLIM4 | PAR-CLIP | Tarbase |
| hsa-miR-486-5p | GOLIM4 | HITS-CLIP | Tarbase |
| hsa-miR-15a-5p | RBM4B | HITS-CLIP | Tarbase |
| hsa-miR-195-5p | RBM4B | HITS-CLIP | Tarbase |
| hsa-miR-15a-5p | RBM4B | HITS-CLIP | Tarbase |
| hsa-miR-195-5p | RBM4B | HITS-CLIP | Tarbase |
| hsa-miR-195-5p | RBM4B | Microarrays | Tarbase |
| hsa-miR-15a-5p | RBM4 | HITS-CLIP | Tarbase |
| hsa-miR-15a-5p | RBM4 | HITS-CLIP | Tarbase |
| hsa-miR-195-5p | RBM4 | HITS-CLIP | Tarbase |
| hsa-miR-195-5p | RBM4 | HITS-CLIP | Tarbase |
| hsa-miR-15a-5p | RBM4 | HITS-CLIP | Tarbase |
| hsa-miR-195-5p | RBM4 | HITS-CLIP | Tarbase |
| hsa-miR-3605-5p | NRROS | PAR-CLIP | Tarbase |
| hsa-miR-141-3p | KLHL15 | HITS-CLIP | Tarbase |
| hsa-miR-15a-5p | KLHL15 | PAR-CLIP | Tarbase |
| hsa-miR-15a-5p | KLHL15 | PAR-CLIP | Tarbase |
| hsa-miR-15a-5p | KLHL15 | Chimeric fragments | Tarbase |
| hsa-miR-141-3p | LEMD3 | HITS-CLIP | Tarbase |
| hsa-miR-141-5p | LEMD3 | HITS-CLIP | Tarbase |
| hsa-miR-15a-5p | LEMD3 | PAR-CLIP | Tarbase |
| hsa-miR-195-5p | TLR1 | Microarrays | Tarbase |
| hsa-miR-141-5p | RGMB | HITS-CLIP | Tarbase |
| hsa-miR-15a-3p | RGMB | HITS-CLIP | Tarbase |
| hsa-miR-486-3p | RGMB | HITS-CLIP | Tarbase |
| hsa-miR-195-5p | CYB561D1 | Microarrays | Tarbase |
| hsa-miR-141-5p | TRMT10C | HITS-CLIP | Tarbase |
| hsa-miR-15a-5p | MGA | HITS-CLIP | Tarbase |
| hsa-miR-195-5p | MGA | HITS-CLIP | Tarbase |
| hsa-miR-15a-5p | C12orf66 | PAR-CLIP | Tarbase |
| hsa-miR-15a-5p | C12orf66 | Chimeric fragments | Tarbase |
| hsa-miR-141-5p | PRPF8 | HITS-CLIP | Tarbase |
| hsa-miR-141-5p | PRPF8 | HITS-CLIP | Tarbase |
| hsa-miR-15a-3p | PRPF8 | PAR-CLIP | Tarbase |
| hsa-miR-15a-5p | PRPF8 | PAR-CLIP | Tarbase |
| hsa-miR-3613-3p | PRPF8 | HITS-CLIP | Tarbase |
| hsa-miR-3613-5p | PRPF8 | HITS-CLIP | Tarbase |
| hsa-miR-3613-5p | PRPF8 | HITS-CLIP | Tarbase |
| hsa-miR-15a-5p | ADCY6 | HITS-CLIP | Tarbase |
| hsa-miR-195-5p | ADCY6 | HITS-CLIP | Tarbase |
| hsa-miR-15a-5p | ADCY6 | Chimeric fragments | Tarbase |
| hsa-miR-141-5p | PITPNA | HITS-CLIP | Tarbase |
| hsa-miR-141-5p | PITPNA | HITS-CLIP | Tarbase |
| hsa-miR-486-5p | PITPNA | HITS-CLIP | Tarbase |
| hsa-miR-195-5p | PITPNA | Microarrays | Tarbase |
| hsa-miR-195-5p | ZBTB4 | Microarrays | Tarbase |
| hsa-miR-141-3p | ZHX3 | HITS-CLIP | Tarbase |
| hsa-miR-15a-3p | PHLDA3 | HITS-CLIP | Tarbase |
| hsa-miR-15a-5p | PHLDA3 | HITS-CLIP | Tarbase |
| hsa-miR-15a-5p | PHLDA3 | PAR-CLIP | Tarbase |
| hsa-miR-15a-5p | PHLDA3 | PAR-CLIP | Tarbase |
| hsa-miR-195-5p | PHLDA3 | HITS-CLIP | Tarbase |
| hsa-miR-195-5p | PHLDA3 | PAR-CLIP | Tarbase |
| hsa-miR-195-5p | PHLDA3 | PAR-CLIP | Tarbase |
| hsa-miR-195-5p | EXO1 | Microarrays | Tarbase |
| hsa-miR-15a-5p | LIG4 | PAR-CLIP | Tarbase |
| hsa-miR-3613-3p | LIG4 | PAR-CLIP | Tarbase |
| hsa-miR-15a-3p | ATP2A2 | HITS-CLIP | Tarbase |
| hsa-miR-15a-3p | ATP2A2 | HITS-CLIP | Tarbase |
| hsa-miR-15a-5p | ATP2A2 | HITS-CLIP | Tarbase |
| hsa-miR-15a-5p | ATP2A2 | HITS-CLIP | Tarbase |
| hsa-miR-15a-5p | ATP2A2 | HITS-CLIP | Tarbase |
| hsa-miR-15a-5p | ATP2A2 | HITS-CLIP | Tarbase |
| hsa-miR-195-5p | ATP2A2 | HITS-CLIP | Tarbase |
| hsa-miR-195-5p | ATP2A2 | HITS-CLIP | Tarbase |
| hsa-miR-195-5p | ATP2A2 | HITS-CLIP | Tarbase |
| hsa-miR-195-5p | ATP2A2 | HITS-CLIP | Tarbase |
| hsa-miR-195-5p | ZWILCH | Microarrays | Tarbase |
| hsa-miR-195-5p | VWC2L | Microarrays | Tarbase |
| hsa-miR-15a-3p | C12orf76 | HITS-CLIP | Tarbase |
| hsa-miR-15a-5p | C12orf76 | PAR-CLIP | Tarbase |
| hsa-miR-15a-5p | C12orf76 | HITS-CLIP | Tarbase |
| hsa-miR-15a-5p | C12orf76 | HITS-CLIP | Tarbase |
| hsa-miR-195-5p | C12orf76 | HITS-CLIP | Tarbase |
| hsa-miR-141-5p | ZCCHC12 | HITS-CLIP | Tarbase |
| hsa-miR-15a-5p | ZCCHC12 | HITS-CLIP | Tarbase |
| hsa-miR-195-5p | ZCCHC12 | HITS-CLIP | Tarbase |
| hsa-miR-3613-3p | DENND4A | PAR-CLIP | Tarbase |
| hsa-miR-486-5p | ANKRD36C | HITS-CLIP | Tarbase |
| hsa-miR-141-5p | PELI3 | HITS-CLIP | Tarbase |
| hsa-miR-141-3p | AKIRIN1 | PAR-CLIP | Tarbase |
| hsa-miR-141-3p | AKIRIN1 | PAR-CLIP | Tarbase |
| hsa-miR-141-3p | AKIRIN1 | PAR-CLIP | Tarbase |
| hsa-miR-486-5p | AKIRIN1 | PAR-CLIP | Tarbase |
| hsa-miR-141-3p | MSL2 | HITS-CLIP | Tarbase |
| hsa-miR-141-3p | MSL2 | HITS-CLIP | Tarbase |
| hsa-miR-486-5p | MSL2 | HITS-CLIP | Tarbase |
| hsa-miR-15a-5p | UGT8 | PAR-CLIP | Tarbase |
| hsa-miR-15a-5p | SLC29A2 | HITS-CLIP | Tarbase |
| hsa-miR-195-5p | SLC29A2 | HITS-CLIP | Tarbase |
| hsa-miR-15a-5p | B3GNT1 | HITS-CLIP | Tarbase |
| hsa-miR-195-5p | B3GNT1 | HITS-CLIP | Tarbase |
| hsa-miR-195-5p | SH3PXD2B | Microarrays | Tarbase |
| hsa-miR-15a-5p | KIAA1551 | PAR-CLIP | Tarbase |
| hsa-miR-195-5p | KIAA1551 | PAR-CLIP | Tarbase |
| hsa-miR-3613-3p | KIAA1551 | PAR-CLIP | Tarbase |
| hsa-miR-141-5p | LARP7 | HITS-CLIP | Tarbase |
| hsa-miR-15a-5p | NR1D2 | HITS-CLIP | Tarbase |
| hsa-miR-195-3p | NR1D2 | HITS-CLIP | Tarbase |
| hsa-miR-195-5p | NR1D2 | Microarrays | Tarbase |
| hsa-miR-486-5p | THAP6 | PAR-CLIP | Tarbase |
| hsa-miR-195-5p | FZD4 | Microarrays | Tarbase |
| hsa-miR-141-5p | DENND6A | HITS-CLIP | Tarbase |
| hsa-miR-141-3p | DNAH12 | HITS-CLIP | Tarbase |
| hsa-miR-195-5p | YIF1A | Microarrays | Tarbase |
| hsa-miR-15a-3p | NDUFA11 | HITS-CLIP | Tarbase |
| hsa-miR-15a-5p | RAB1B | PAR-CLIP | Tarbase |
| hsa-miR-15a-5p | RAB1B | HITS-CLIP | Tarbase |
| hsa-miR-195-5p | PTDSS2 | Microarrays | Tarbase |
| hsa-miR-195-5p | ASPHD1 | Microarrays | Tarbase |
| hsa-miR-15a-5p | DHX36 | PAR-CLIP | Tarbase |
| hsa-miR-15a-5p | DHX36 | HITS-CLIP | Tarbase |
| hsa-miR-195-5p | DHX36 | HITS-CLIP | Tarbase |
| hsa-miR-15a-5p | DHX36 | Chimeric fragments | Tarbase |
| hsa-miR-15a-3p | CTBP2 | HITS-CLIP | Tarbase |
| hsa-miR-141-3p | CTBP2 | Luciferase Reporter Assay | Tarbase |
| hsa-miR-195-5p | CHST2 | Microarrays | Tarbase |
| hsa-miR-195-5p | UBE2C | Microarrays | Tarbase |
| hsa-miR-141-3p | VCPIP1 | HITS-CLIP | Tarbase |
| hsa-miR-15a-5p | VCPIP1 | HITS-CLIP | Tarbase |
| hsa-miR-15a-5p | VCPIP1 | PAR-CLIP | Tarbase |
| hsa-miR-15a-5p | VCPIP1 | HITS-CLIP | Tarbase |
| hsa-miR-195-5p | VCPIP1 | HITS-CLIP | Tarbase |
| hsa-miR-195-5p | VCPIP1 | HITS-CLIP | Tarbase |

| | | | |
|---|---|---|---|
| hsa-miR-141-3p | PDIK1L | HITS-CLIP | Tarbase |
| hsa-miR-141-3p | PDIK1L | HITS-CLIP | Tarbase |
| hsa-miR-141-3p | PDIK1L | HITS-CLIP | Tarbase |
| hsa-miR-15a-5p | SPSB4 | HITS-CLIP | Tarbase |
| hsa-miR-195-5p | SPSB4 | HITS-CLIP | Tarbase |
| hsa-miR-141-5p | SH3BP5L | HITS-CLIP | Tarbase |
| hsa-miR-15a-5p | SH3BP5L | HITS-CLIP | Tarbase |
| hsa-miR-195-5p | SH3BP5L | HITS-CLIP | Tarbase |
| hsa-miR-195-5p | YPEL2 | Microarrays | Tarbase |
| hsa-miR-195-5p | CADM2 | HITS-CLIP | Tarbase |
| hsa-miR-15a-5p | PSMD2 | HITS-CLIP | Tarbase |
| hsa-miR-141-3p | PPM1E | HITS-CLIP | Tarbase |
| hsa-miR-15a-5p | PPM1E | HITS-CLIP | Tarbase |
| hsa-miR-15a-5p | CTDSP2 | HITS-CLIP | Tarbase |
| hsa-miR-195-5p | CTDSP2 | HITS-CLIP | Tarbase |
| hsa-miR-195-5p | CKAP5 | Microarrays | Tarbase |
| hsa-miR-15a-5p | ARHGAP1 | HITS-CLIP | Tarbase |
| hsa-miR-3605-5p | ARHGAP1 | HITS-CLIP | Tarbase |
| hsa-miR-15a-5p | ATG13 | HITS-CLIP | Tarbase |
| hsa-miR-195-5p | ATG13 | HITS-CLIP | Tarbase |
| hsa-miR-141-3p | DOLK | HITS-CLIP | Tarbase |
| hsa-miR-141-3p | CCNE2 | Western Blot | Tarbase |
| hsa-miR-141-5p | CCNE2 | HITS-CLIP | Tarbase |
| hsa-miR-141-5p | CCNE2 | HITS-CLIP | Tarbase |
| hsa-miR-15a-5p | CCNE2 | PAR-CLIP | Tarbase |
| hsa-miR-195-5p | CCNE2 | PAR-CLIP | Tarbase |
| hsa-miR-15a-5p | GRAMD2 | HITS-CLIP | Tarbase |
| hsa-miR-195-5p | GRAMD2 | HITS-CLIP | Tarbase |
| hsa-miR-15a-5p | SMAD2 | HITS-CLIP | Tarbase |
| hsa-miR-15a-5p | SMAD2 | HITS-CLIP | Tarbase |
| hsa-miR-15a-5p | SMAD2 | PAR-CLIP | Tarbase |
| hsa-miR-195-5p | SMAD2 | HITS-CLIP | Tarbase |
| hsa-miR-195-5p | SMAD2 | HITS-CLIP | Tarbase |
| hsa-miR-15a-5p | SMAD2 | HITS-CLIP | Tarbase |
| hsa-miR-195-5p | SMAD2 | HITS-CLIP | Tarbase |
| hsa-miR-141-3p | EIF3F | HITS-CLIP | Tarbase |
| hsa-miR-141-5p | PPP2R2D | HITS-CLIP | Tarbase |
| hsa-miR-195-5p | PPP2R2D | Microarrays | Tarbase |
| hsa-miR-195-5p | MCTP1 | Microarrays | Tarbase |
| hsa-miR-3613-3p | ALG10B | HITS-CLIP | Tarbase |
| hsa-miR-195-5p | LONRF3 | Microarrays | Tarbase |
| hsa-miR-15a-5p | C11orf68 | PAR-CLIP | Tarbase |
| hsa-miR-15a-3p | FOSL1 | HITS-CLIP | Tarbase |
| hsa-miR-195-5p | FOSL1 | Microarrays | Tarbase |
| hsa-miR-195-5p | SUGCT | Microarrays | Tarbase |
| hsa-miR-15a-3p | CCDC85B | HITS-CLIP | Tarbase |
| hsa-miR-141-3p | TMEM70 | HITS-CLIP | Tarbase |
| hsa-miR-15a-5p | TMEM70 | PAR-CLIP | Tarbase |
| hsa-miR-486-3p | RMI2 | HITS-CLIP | Tarbase |
| hsa-miR-195-5p | RMI2 | Microarrays | Tarbase |
| hsa-miR-15a-3p | MLXIP | HITS-CLIP | Tarbase |
| hsa-miR-3613-3p | MLXIP | HITS-CLIP | Tarbase |
| hsa-miR-486-5p | MLXIP | HITS-CLIP | Tarbase |
| hsa-miR-15a-3p | NR2F1 | HITS-CLIP | Tarbase |
| hsa-miR-486-5p | NR2F1 | HITS-CLIP | Tarbase |
| hsa-miR-15a-5p | NR2F1 | HITS-CLIP | Tarbase |
| hsa-miR-195-5p | NR2F1 | HITS-CLIP | Tarbase |
| hsa-miR-141-5p | AURKAIP1 | HITS-CLIP | Tarbase |
| hsa-miR-486-5p | RUVBL1 | PAR-CLIP | Tarbase |
| hsa-miR-195-5p | SFN | Microarrays | Tarbase |
| hsa-miR-15a-5p | ZDHHC21 | PAR-CLIP | Tarbase |
| hsa-miR-15a-5p | ZDHHC21 | HITS-CLIP | Tarbase |
| hsa-miR-195-5p | ZDHHC21 | HITS-CLIP | Tarbase |
| hsa-miR-15a-5p | PLEKHF2 | HITS-CLIP | Tarbase |
| hsa-miR-15a-5p | PLEKHF2 | HITS-CLIP | Tarbase |
| hsa-miR-486-5p | PLEKHF2 | PAR-CLIP | Tarbase |
| hsa-miR-486-5p | PLEKHF2 | PAR-CLIP | Tarbase |
| hsa-miR-141-3p | LRRN1 | HITS-CLIP | Tarbase |
| hsa-miR-195-5p | KLHL38 | Microarrays | Tarbase |
| hsa-miR-15a-5p | B3GALT6 | PAR-CLIP | Tarbase |
| hsa-miR-141-5p | SLC35A4 | HITS-CLIP | Tarbase |
| hsa-miR-15a-5p | SLC35A4 | PAR-CLIP | Tarbase |
| hsa-miR-195-5p | SLC35A4 | Microarrays | Tarbase |
| hsa-miR-15a-5p | IP6K1 | HITS-CLIP | Tarbase |
| hsa-miR-15a-5p | IP6K1 | HITS-CLIP | Tarbase |
| hsa-miR-195-5p | IP6K1 | HITS-CLIP | Tarbase |
| hsa-miR-3605-3p | CSTF3 | HITS-CLIP | Tarbase |
| hsa-miR-141-5p | YES1 | HITS-CLIP | Tarbase |
| hsa-miR-15a-5p | YES1 | PAR-CLIP | Tarbase |
| hsa-miR-15a-5p | YES1 | HITS-CLIP | Tarbase |
| hsa-miR-195-5p | YES1 | PAR-CLIP | Tarbase |
| hsa-miR-219a-1-3p | YES1 | PAR-CLIP | Tarbase |
| hsa-miR-195-5p | YES1 | Microarrays | Tarbase |
| hsa-miR-195-5p | DLEU1 | Microarrays | Tarbase |
| hsa-miR-141-5p | CCDC57 | HITS-CLIP | Tarbase |
| hsa-miR-141-5p | CCDC57 | HITS-CLIP | Tarbase |
| hsa-miR-486-5p | CCDC57 | HITS-CLIP | Tarbase |
| hsa-miR-195-5p | SPHK1 | Microarrays | Tarbase |
| hsa-miR-195-5p | BNIP3 | Microarrays | Tarbase |
| hsa-miR-486-5p | MYPOP | HITS-CLIP | Tarbase |
| hsa-miR-195-5p | LRRTM4 | Microarrays | Tarbase |
| hsa-miR-15a-5p | ATAD5 | PAR-CLIP | Tarbase |
| hsa-miR-15a-5p | ATAD5 | HITS-CLIP | Tarbase |
| hsa-miR-195-5p | ATAD5 | Microarrays | Tarbase |
| hsa-miR-15a-5p | ATAD5 | Chimeric fragments | Tarbase |
| hsa-miR-195-5p | RTTN | Microarrays | Tarbase |
| hsa-miR-195-5p | ACBD7 | Microarrays | Tarbase |
| hsa-miR-15a-5p | SLC35G1 | PAR-CLIP | Tarbase |
| hsa-miR-15a-5p | SLC35G1 | HITS-CLIP | Tarbase |
| hsa-miR-15a-5p | CRLF3 | HITS-CLIP | Tarbase |
| hsa-miR-15a-5p | KCMF1 | HITS-CLIP | Tarbase |
| hsa-miR-195-5p | KCMF1 | HITS-CLIP | Tarbase |
| hsa-miR-3605-5p | CLK2 | HITS-CLIP | Tarbase |
| hsa-miR-3613-5p | LMNB2 | PAR-CLIP | Tarbase |
| hsa-miR-195-5p | LMNB2 | Microarrays | Tarbase |
| hsa-miR-141-5p | MEX3C | HITS-CLIP | Tarbase |
| hsa-miR-3613-3p | MEX3C | HITS-CLIP | Tarbase |
| hsa-miR-195-5p | RNF152 | Microarrays | Tarbase |
| hsa-miR-15a-5p | RNF152 | Chimeric fragments | Tarbase |
| hsa-miR-195-5p | BDNF | Microarrays | Tarbase |
| hsa-miR-15a-5p | CDK5R1 | PAR-CLIP | Tarbase |
| hsa-miR-15a-5p | CDK5R1 | HITS-CLIP | Tarbase |
| hsa-miR-195-5p | CDK5R1 | HITS-CLIP | Tarbase |
| hsa-miR-15a-5p | CDK5R1 | HITS-CLIP | Tarbase |
| hsa-miR-195-5p | CDK5R1 | HITS-CLIP | Tarbase |
| hsa-miR-15a-5p | FAM91A1 | HITS-CLIP | Tarbase |
| hsa-miR-3613-3p | FAM91A1 | HITS-CLIP | Tarbase |
| hsa-miR-141-3p | SOX11 | HITS-CLIP | Tarbase |
| hsa-miR-141-5p | SOX11 | HITS-CLIP | Tarbase |
| hsa-miR-141-5p | SOX11 | HITS-CLIP | Tarbase |
| hsa-miR-141-5p | SOX11 | HITS-CLIP | Tarbase |
| hsa-miR-195-5p | TYMS | Microarrays | Tarbase |
| hsa-miR-195-5p | C8orf4 | Microarrays | Tarbase |
| hsa-miR-3605-5p | NFATC2IP | PAR-CLIP | Tarbase |
| hsa-miR-15a-5p | SEC24C | PAR-CLIP | Tarbase |
| hsa-miR-141-3p | SMCR8 | HITS-CLIP | Tarbase |
| hsa-miR-15a-5p | SMCR8 | HITS-CLIP | Tarbase |
| hsa-miR-195-5p | SMCR8 | HITS-CLIP | Tarbase |
| hsa-miR-15a-5p | MTHFR | PAR-CLIP | Tarbase |
| hsa-miR-15a-5p | SLC38A9 | PAR-CLIP | Tarbase |
| hsa-miR-15a-5p | SLC38A9 | PAR-CLIP | Tarbase |
| hsa-miR-15a-5p | SLC38A9 | PAR-CLIP | Tarbase |
| hsa-miR-195-5p | SLC38A9 | PAR-CLIP | Tarbase |
| hsa-miR-195-5p | SLC38A9 | PAR-CLIP | Tarbase |
| hsa-miR-195-5p | POLE | Microarrays | Tarbase |
| hsa-miR-195-5p | SCN4B | Microarrays | Tarbase |
| hsa-miR-141-3p | ANO6 | HITS-CLIP | Tarbase |
| hsa-miR-141-3p | ZBTB34 | HITS-CLIP | Tarbase |
| hsa-miR-141-3p | ZBTB34 | HITS-CLIP | Tarbase |
| hsa-miR-141-3p | ZBTB34 | HITS-CLIP | Tarbase |
| hsa-miR-15a-5p | ZBTB34 | HITS-CLIP | Tarbase |
| hsa-miR-15a-5p | ZBTB34 | HITS-CLIP | Tarbase |
| hsa-miR-195-5p | ZBTB34 | HITS-CLIP | Tarbase |
| hsa-miR-15a-5p | RIMKLA | PAR-CLIP | Tarbase |
| hsa-miR-141-5p | RPS6KA3 | HITS-CLIP | Tarbase |
| hsa-miR-15a-5p | RPS6KA3 | HITS-CLIP | Tarbase |
| hsa-miR-15a-5p | RPS6KA3 | HITS-CLIP | Tarbase |
| hsa-miR-15a-5p | RPS6KA3 | PAR-CLIP | Tarbase |
| hsa-miR-15a-5p | RPS6KA3 | HITS-CLIP | Tarbase |

| | | | |
|---|---|---|---|
| hsa-miR-15a-5p | RPS6KA3 | HITS-CLIP | Tarbase |
| hsa-miR-15a-5p | RPS6KA3 | HITS-CLIP | Tarbase |
| hsa-miR-195-5p | RPS6KA3 | HITS-CLIP | Tarbase |
| hsa-miR-195-5p | RPS6KA3 | HITS-CLIP | Tarbase |
| hsa-miR-195-5p | RPS6KA3 | HITS-CLIP | Tarbase |
| hsa-miR-141-3p | CHD9 | HITS-CLIP | Tarbase |
| hsa-miR-141-5p | CHD9 | HITS-CLIP | Tarbase |
| hsa-miR-15a-5p | CHD9 | HITS-CLIP | Tarbase |
| hsa-miR-15a-5p | CHD9 | PAR-CLIP | Tarbase |
| hsa-miR-195-5p | CHD9 | HITS-CLIP | Tarbase |
| hsa-miR-3613-3p | CHD9 | PAR-CLIP | Tarbase |
| hsa-miR-15a-3p | PPFIA3 | HITS-CLIP | Tarbase |
| hsa-miR-486-5p | SAMD9L | PAR-CLIP | Tarbase |
| hsa-miR-15a-5p | TGIF1 | HITS-CLIP | Tarbase |
| hsa-miR-195-5p | TGIF1 | HITS-CLIP | Tarbase |
| hsa-miR-15a-5p | NR2C2 | HITS-CLIP | Tarbase |
| hsa-miR-195-5p | NR2C2 | HITS-CLIP | Tarbase |
| hsa-miR-486-5p | NR2C2 | HITS-CLIP | Tarbase |
| hsa-miR-15a-5p | ZBTB33 | HITS-CLIP | Tarbase |
| hsa-miR-15a-5p | ZBTB33 | HITS-CLIP | Tarbase |
| hsa-miR-15a-5p | ZBTB33 | HITS-CLIP | Tarbase |
| hsa-miR-195-5p | ZBTB33 | HITS-CLIP | Tarbase |
| hsa-miR-195-5p | ZBTB33 | HITS-CLIP | Tarbase |
| hsa-miR-15a-5p | ST8SIA3 | HITS-CLIP | Tarbase |
| hsa-miR-195-5p | ST8SIA3 | HITS-CLIP | Tarbase |
| hsa-miR-15a-5p | ST8SIA3 | HITS-CLIP | Tarbase |
| hsa-miR-195-5p | ST8SIA3 | HITS-CLIP | Tarbase |
| hsa-miR-15a-5p | SLC25A22 | HITS-CLIP | Tarbase |
| hsa-miR-15a-5p | SLC25A22 | HITS-CLIP | Tarbase |
| hsa-miR-15a-5p | SLC25A22 | HITS-CLIP | Tarbase |
| hsa-miR-195-5p | SLC25A22 | HITS-CLIP | Tarbase |
| hsa-miR-195-5p | SLC25A22 | HITS-CLIP | Tarbase |
| hsa-miR-195-5p | SLC25A22 | Microarrays | Tarbase |
| hsa-miR-15a-5p | ATOX1 | PAR-CLIP | Tarbase |
| hsa-miR-15a-5p | ATOX1 | HITS-CLIP | Tarbase |
| hsa-miR-195-5p | ATOX1 | HITS-CLIP | Tarbase |
| hsa-miR-141-3p | TBL1XR1 | HITS-CLIP | Tarbase |
| hsa-miR-15a-5p | TBL1XR1 | HITS-CLIP | Tarbase |
| hsa-miR-15a-5p | TBL1XR1 | PAR-CLIP | Tarbase |
| hsa-miR-15a-5p | TBL1XR1 | HITS-CLIP | Tarbase |
| hsa-miR-15a-5p | TBL1XR1 | HITS-CLIP | Tarbase |
| hsa-miR-15a-5p | TBL1XR1 | PAR-CLIP | Tarbase |
| hsa-miR-15a-5p | TBL1XR1 | PAR-CLIP | Tarbase |
| hsa-miR-15a-5p | TBL1XR1 | HITS-CLIP | Tarbase |
| hsa-miR-15a-5p | TBL1XR1 | HITS-CLIP | Tarbase |
| hsa-miR-15a-5p | TBL1XR1 | PAR-CLIP | Tarbase |
| hsa-miR-195-5p | TBL1XR1 | HITS-CLIP | Tarbase |
| hsa-miR-195-5p | TBL1XR1 | HITS-CLIP | Tarbase |
| hsa-miR-195-5p | TBL1XR1 | HITS-CLIP | Tarbase |
| hsa-miR-195-5p | TBL1XR1 | HITS-CLIP | Tarbase |
| hsa-miR-195-5p | TBL1XR1 | PAR-CLIP | Tarbase |
| hsa-miR-15a-5p | PIDD1 | HITS-CLIP | Tarbase |
| hsa-miR-15a-5p | PIDD1 | PAR-CLIP | Tarbase |
| hsa-miR-195-5p | PIDD1 | HITS-CLIP | Tarbase |
| hsa-miR-195-5p | GSG2 | Microarrays | Tarbase |
| hsa-miR-15a-3p | JUN | HITS-CLIP | Tarbase |
| hsa-miR-195-5p | JUN | Microarrays | Tarbase |
| hsa-miR-15a-5p | CD151 | HITS-CLIP | Tarbase |
| hsa-miR-15a-5p | CD151 | HITS-CLIP | Tarbase |
| hsa-miR-15a-5p | CD151 | HITS-CLIP | Tarbase |
| hsa-miR-15a-5p | CD151 | PAR-CLIP | Tarbase |
| hsa-miR-15a-5p | CD151 | PAR-CLIP | Tarbase |
| hsa-miR-195-5p | CD151 | HITS-CLIP | Tarbase |
| hsa-miR-195-5p | CD151 | HITS-CLIP | Tarbase |
| hsa-miR-195-5p | CD151 | HITS-CLIP | Tarbase |
| hsa-miR-195-5p | CD151 | PAR-CLIP | Tarbase |
| hsa-miR-195-5p | POLR2L | HITS-CLIP | Tarbase |
| hsa-miR-141-5p | FAM20C | HITS-CLIP | Tarbase |
| hsa-miR-15a-3p | FAM20C | HITS-CLIP | Tarbase |
| hsa-miR-3613-5p | FAM20C | HITS-CLIP | Tarbase |
| hsa-miR-195-5p | FAM20C | Microarrays | Tarbase |
| hsa-miR-15a-5p | PVRL3 | HITS-CLIP | Tarbase |
| hsa-miR-195-5p | PVRL3 | HITS-CLIP | Tarbase |
| hsa-miR-195-5p | PVRL3 | Microarrays | Tarbase |
| hsa-miR-141-3p | HNRNPA0 | HITS-CLIP | Tarbase |
| hsa-miR-141-3p | HNRNPA0 | PAR-CLIP | Tarbase |
| hsa-miR-15a-3p | CHID1 | HITS-CLIP | Tarbase |
| hsa-miR-141-3p | ZNF518A | HITS-CLIP | Tarbase |
| hsa-miR-195-5p | CCDC184 | Microarrays | Tarbase |
| hsa-miR-141-3p | GRB2 | PAR-CLIP | Tarbase |
| hsa-miR-141-3p | GRB2 | PAR-CLIP | Tarbase |
| hsa-miR-15a-5p | GRB2 | PAR-CLIP | Tarbase |
| hsa-miR-15a-5p | GRB2 | PAR-CLIP | Tarbase |
| hsa-miR-195-5p | GRB2 | PAR-CLIP | Tarbase |
| hsa-miR-195-5p | GRB2 | PAR-CLIP | Tarbase |
| hsa-miR-3613-3p | ZBTB41 | Chimeric fragments | Tarbase |
| hsa-miR-486-5p | ARL6IP6 | PAR-CLIP | Tarbase |
| hsa-miR-15a-5p | BET1L | PAR-CLIP | Tarbase |
| hsa-miR-195-5p | BET1L | PAR-CLIP | Tarbase |
| hsa-miR-15a-5p | RIC8A | HITS-CLIP | Tarbase |
| hsa-miR-15a-5p | RIC8A | HITS-CLIP | Tarbase |
| hsa-miR-195-5p | RIC8A | HITS-CLIP | Tarbase |
| hsa-miR-195-5p | FAM26E | Microarrays | Tarbase |
| hsa-miR-15a-5p | NDUFAF3 | HITS-CLIP | Tarbase |
| hsa-miR-195-5p | NDUFAF3 | HITS-CLIP | Tarbase |
| hsa-miR-15a-5p | C2orf69 | HITS-CLIP | Tarbase |
| hsa-miR-15a-5p | C2orf69 | HITS-CLIP | Tarbase |
| hsa-miR-15a-5p | C2orf69 | HITS-CLIP | Tarbase |
| hsa-miR-15a-5p | C2orf69 | HITS-CLIP | Tarbase |
| hsa-miR-15a-5p | C2orf69 | HITS-CLIP | Tarbase |
| hsa-miR-195-5p | C2orf69 | HITS-CLIP | Tarbase |
| hsa-miR-195-5p | C2orf69 | HITS-CLIP | Tarbase |
| hsa-miR-195-5p | C2orf69 | HITS-CLIP | Tarbase |
| hsa-miR-195-5p | C2orf69 | HITS-CLIP | Tarbase |
| hsa-miR-15a-3p | PDE4DIP | HITS-CLIP | Tarbase |
| hsa-miR-15a-5p | PDE4DIP | HITS-CLIP | Tarbase |
| hsa-miR-15a-5p | PDE4DIP | HITS-CLIP | Tarbase |
| hsa-miR-195-5p | PDE4DIP | HITS-CLIP | Tarbase |
| hsa-miR-15a-5p | PDE4DIP | HITS-CLIP | Tarbase |
| hsa-miR-195-5p | PDE4DIP | HITS-CLIP | Tarbase |
| hsa-miR-141-5p | SH2B1 | HITS-CLIP | Tarbase |
| hsa-miR-15a-5p | KDELC2 | PAR-CLIP | Tarbase |
| hsa-miR-15a-5p | PLEC | HITS-CLIP | Tarbase |
| hsa-miR-15a-3p | WDR6 | HITS-CLIP | Tarbase |
| hsa-miR-15a-3p | WDR6 | HITS-CLIP | Tarbase |
| hsa-miR-15a-5p | WDR6 | HITS-CLIP | Tarbase |
| hsa-miR-15a-5p | WDR6 | HITS-CLIP | Tarbase |
| hsa-miR-486-5p | WDR6 | HITS-CLIP | Tarbase |
| hsa-miR-141-3p | FAM220A | HITS-CLIP | Tarbase |
| hsa-miR-141-3p | FAM220A | HITS-CLIP | Tarbase |
| hsa-miR-141-3p | BEND3 | HITS-CLIP | Tarbase |
| hsa-miR-195-5p | KLHL11 | Microarrays | Tarbase |
| hsa-miR-141-3p | CTXN1 | HITS-CLIP | Tarbase |
| hsa-miR-195-5p | CTXN1 | Microarrays | Tarbase |
| hsa-miR-15a-5p | CA8 | PAR-CLIP | Tarbase |
| hsa-miR-195-5p | CA8 | Microarrays | Tarbase |
| hsa-miR-141-3p | EPM2AIP1 | HITS-CLIP | Tarbase |
| hsa-miR-15a-3p | EPM2AIP1 | HITS-CLIP | Tarbase |
| hsa-miR-15a-5p | CTNNBIP1 | HITS-CLIP | Tarbase |
| hsa-miR-195-5p | CTNNBIP1 | HITS-CLIP | Tarbase |
| hsa-miR-195-5p | CTNNBIP1 | Microarrays | Tarbase |
| hsa-miR-141-5p | SUZ12 | HITS-CLIP | Tarbase |
| hsa-miR-141-3p | KCTD12 | HITS-CLIP | Tarbase |
| hsa-miR-141-3p | KCTD12 | HITS-CLIP | Tarbase |
| hsa-miR-141-5p | KCTD12 | HITS-CLIP | Tarbase |
| hsa-miR-195-5p | FAM219B | Microarrays | Tarbase |
| hsa-miR-195-5p | APOLD1 | Microarrays | Tarbase |
| hsa-miR-195-5p | DPY19L3 | Microarrays | Tarbase |
| hsa-miR-141-3p | TAF7 | HITS-CLIP | Tarbase |
| hsa-miR-15a-5p | TAF7 | HITS-CLIP | Tarbase |
| hsa-miR-195-5p | TAF7 | HITS-CLIP | Tarbase |
| hsa-miR-3605-3p | TAF7 | HITS-CLIP | Tarbase |
| hsa-miR-15a-5p | TAF7 | Chimeric fragments | Tarbase |
| hsa-miR-3613-5p | RMI1 | HITS-CLIP | Tarbase |
| hsa-miR-141-3p | FBXO34 | HITS-CLIP | Tarbase |
| hsa-miR-3613-5p | FBXO34 | HITS-CLIP | Tarbase |
| hsa-miR-15a-5p | FBXO34 | HITS-CLIP | Tarbase |
| hsa-miR-195-5p | FBXO34 | HITS-CLIP | Tarbase |
| hsa-miR-195-5p | AURKB | Microarrays | Tarbase |

| | | | |
|---|---|---|---|
| hsa-miR-141-5p | MRFAP1 | HITS-CLIP | Tarbase |
| hsa-miR-15a-5p | MRFAP1 | PAR-CLIP | Tarbase |
| hsa-miR-195-5p | TMEM107 | Microarrays | Tarbase |
| hsa-miR-141-5p | RCC2 | HITS-CLIP | Tarbase |
| hsa-miR-15a-5p | PER1 | PAR-CLIP | Tarbase |
| hsa-miR-15a-5p | SPTY2D1 | HITS-CLIP | Tarbase |
| hsa-miR-195-5p | SPTY2D1 | HITS-CLIP | Tarbase |
| hsa-miR-15a-5p | EDC3 | HITS-CLIP | Tarbase |
| hsa-miR-195-5p | EDC3 | HITS-CLIP | Tarbase |
| hsa-miR-15a-5p | TCAIM | Chimeric fragments | Tarbase |
| hsa-miR-195-5p | FUCA1 | Microarrays | Tarbase |
| hsa-miR-15a-5p | ZNF664 | PAR-CLIP | Tarbase |
| hsa-miR-3605-3p | ZNF664 | HITS-CLIP | Tarbase |
| hsa-miR-3613-3p | ZNF664 | PAR-CLIP | Tarbase |
| hsa-miR-141-5p | CALR | HITS-CLIP | Tarbase |
| hsa-miR-15a-3p | CALR | HITS-CLIP | Tarbase |
| hsa-miR-15a-3p | CALR | HITS-CLIP | Tarbase |
| hsa-miR-15a-3p | CALR | HITS-CLIP | Tarbase |
| hsa-miR-15a-5p | CALR | HITS-CLIP | Tarbase |
| hsa-miR-15a-5p | CALR | HITS-CLIP | Tarbase |
| hsa-miR-15a-5p | CALR | HITS-CLIP | Tarbase |
| hsa-miR-15a-5p | CALR | HITS-CLIP | Tarbase |
| hsa-miR-15a-5p | CALR | PAR-CLIP | Tarbase |
| hsa-miR-15a-5p | CALR | HITS-CLIP | Tarbase |
| hsa-miR-15a-5p | CALR | HITS-CLIP | Tarbase |
| hsa-miR-15a-5p | CALR | HITS-CLIP | Tarbase |
| hsa-miR-195-5p | CALR | HITS-CLIP | Tarbase |
| hsa-miR-195-5p | CALR | HITS-CLIP | Tarbase |
| hsa-miR-195-5p | CALR | HITS-CLIP | Tarbase |
| hsa-miR-195-5p | CALR | HITS-CLIP | Tarbase |
| hsa-miR-195-5p | CALR | HITS-CLIP | Tarbase |
| hsa-miR-15a-3p | MAGED1 | HITS-CLIP | Tarbase |
| hsa-miR-15a-5p | MAGED1 | HITS-CLIP | Tarbase |
| hsa-miR-15a-5p | MAGED1 | HITS-CLIP | Tarbase |
| hsa-miR-15a-5p | MAGED1 | HITS-CLIP | Tarbase |
| hsa-miR-15a-5p | MAGED1 | HITS-CLIP | Tarbase |
| hsa-miR-195-5p | MAGED1 | HITS-CLIP | Tarbase |
| hsa-miR-141-3p | CLK3 | HITS-CLIP | Tarbase |
| hsa-miR-15a-3p | GATA2 | HITS-CLIP | Tarbase |
| hsa-miR-15a-5p | GEMIN4 | PAR-CLIP | Tarbase |
| hsa-miR-15a-5p | GEMIN4 | HITS-CLIP | Tarbase |
| hsa-miR-195-5p | FJX1 | Microarrays | Tarbase |
| hsa-miR-15a-5p | KLHL28 | PAR-CLIP | Tarbase |
| hsa-miR-15a-5p | KLHL28 | PAR-CLIP | Tarbase |
| hsa-miR-15a-5p | KLHL28 | HITS-CLIP | Tarbase |
| hsa-miR-141-3p | ZBTB18 | HITS-CLIP | Tarbase |
| hsa-miR-15a-3p | ZBTB18 | PAR-CLIP | Tarbase |
| hsa-miR-3605-3p | GCC1 | HITS-CLIP | Tarbase |
| hsa-miR-3605-3p | GCC1 | HITS-CLIP | Tarbase |
| hsa-miR-15a-5p | GCC1 | Chimeric fragments | Tarbase |
| hsa-miR-3605-3p | PLD6 | HITS-CLIP | Tarbase |
| hsa-miR-195-5p | CDC42EP4 | Microarrays | Tarbase |
| hsa-miR-195-5p | LACC1 | Microarrays | Tarbase |
| hsa-miR-486-5p | MAF1 | PAR-CLIP | Tarbase |
| hsa-miR-195-5p | APOBEC3B | Microarrays | Tarbase |
| hsa-miR-15a-5p | MYADM | HITS-CLIP | Tarbase |
| hsa-miR-15a-5p | MYADM | HITS-CLIP | Tarbase |
| hsa-miR-15a-5p | MYADM | HITS-CLIP | Tarbase |
| hsa-miR-15a-5p | MYADM | PAR-CLIP | Tarbase |
| hsa-miR-15a-5p | MYADM | HITS-CLIP | Tarbase |
| hsa-miR-195-5p | MYADM | HITS-CLIP | Tarbase |
| hsa-miR-15a-5p | MYADM | HITS-CLIP | Tarbase |
| hsa-miR-195-5p | MYADM | HITS-CLIP | Tarbase |
| hsa-miR-195-5p | MYADM | Microarrays | Tarbase |
| hsa-miR-15a-3p | SERTAD2 | HITS-CLIP | Tarbase |
| hsa-miR-486-5p | SERTAD2 | HITS-CLIP | Tarbase |
| hsa-miR-15a-5p | AKAP5 | PAR-CLIP | Tarbase |
| hsa-miR-141-3p | ABCA13 | HITS-CLIP | Tarbase |
| hsa-miR-15a-5p | ABCA13 | HITS-CLIP | Tarbase |
| hsa-miR-195-5p | ABCA13 | HITS-CLIP | Tarbase |
| hsa-miR-15a-5p | PDXDC1 | PAR-CLIP | Tarbase |
| hsa-miR-486-5p | PDXDC1 | HITS-CLIP | Tarbase |
| hsa-miR-15a-5p | PDXDC1 | HITS-CLIP | Tarbase |
| hsa-miR-195-5p | PDXDC1 | HITS-CLIP | Tarbase |
| hsa-miR-15a-5p | B3GNT3 | PAR-CLIP | Tarbase |
| hsa-miR-15a-5p | C14orf119 | HITS-CLIP | Tarbase |
| hsa-miR-195-5p | C14orf119 | HITS-CLIP | Tarbase |
| hsa-miR-141-3p | BBS10 | HITS-CLIP | Tarbase |
| hsa-miR-15a-5p | CRIPAK | HITS-CLIP | Tarbase |
| hsa-miR-195-5p | CRIPAK | HITS-CLIP | Tarbase |
| hsa-miR-195-5p | CRIPAK | Microarrays | Tarbase |
| hsa-miR-195-5p | TSHZ1 | Microarrays | Tarbase |
| hsa-miR-3605-5p | SOCS4 | PAR-CLIP | Tarbase |
| hsa-miR-15a-3p | TMEM86B | HITS-CLIP | Tarbase |
| hsa-miR-486-5p | RCC1 | HITS-CLIP | Tarbase |
| hsa-miR-486-5p | RCC1 | HITS-CLIP | Tarbase |
| hsa-miR-195-5p | PRKRA | HITS-CLIP | Tarbase |
| hsa-miR-15a-5p | ZNRF2 | PAR-CLIP | Tarbase |
| hsa-miR-15a-5p | ZNRF2 | HITS-CLIP | Tarbase |
| hsa-miR-141-3p | OAZ2 | HITS-CLIP | Tarbase |
| hsa-miR-15a-3p | OAZ2 | HITS-CLIP | Tarbase |
| hsa-miR-15a-5p | MTURN | HITS-CLIP | Tarbase |
| hsa-miR-195-5p | MTURN | HITS-CLIP | Tarbase |
| hsa-miR-141-3p | ZNF609 | HITS-CLIP | Tarbase |
| hsa-miR-15a-5p | PAK2 | PAR-CLIP | Tarbase |
| hsa-miR-15a-3p | MCFD2 | HITS-CLIP | Tarbase |
| hsa-miR-195-5p | GAS1 | Microarrays | Tarbase |
| hsa-miR-15a-3p | FAM73A | HITS-CLIP | Tarbase |
| hsa-miR-15a-3p | FAM73A | HITS-CLIP | Tarbase |
| hsa-miR-15a-5p | FAM73A | HITS-CLIP | Tarbase |
| hsa-miR-15a-5p | FAM73A | HITS-CLIP | Tarbase |
| hsa-miR-15a-5p | FAM73A | HITS-CLIP | Tarbase |
| hsa-miR-15a-5p | FAM73A | HITS-CLIP | Tarbase |
| hsa-miR-15a-5p | FAM73A | PAR-CLIP | Tarbase |
| hsa-miR-195-5p | FAM73A | HITS-CLIP | Tarbase |
| hsa-miR-195-5p | FAM73A | HITS-CLIP | Tarbase |
| hsa-miR-195-5p | FAM73A | HITS-CLIP | Tarbase |
| hsa-miR-195-5p | FAM73A | HITS-CLIP | Tarbase |
| hsa-miR-15a-5p | FAM73A | HITS-CLIP | Tarbase |
| hsa-miR-195-5p | FAM73A | HITS-CLIP | Tarbase |
| hsa-miR-15a-5p | NRIP1 | HITS-CLIP | Tarbase |
| hsa-miR-195-5p | NRIP1 | HITS-CLIP | Tarbase |
| hsa-miR-195-5p | HIST1H2AC | Microarrays | Tarbase |
| hsa-miR-195-5p | ZNF594 | Microarrays | Tarbase |
| hsa-miR-141-3p | PCGF5 | HITS-CLIP | Tarbase |
| hsa-miR-15a-5p | PCGF5 | HITS-CLIP | Tarbase |
| hsa-miR-195-5p | PCGF5 | HITS-CLIP | Tarbase |
| hsa-miR-141-3p | YOD1 | HITS-CLIP | Tarbase |
| hsa-miR-15a-3p | YOD1 | HITS-CLIP | Tarbase |
| hsa-miR-15a-3p | TMEM64 | HITS-CLIP | Tarbase |
| hsa-miR-15a-3p | TMEM64 | PAR-CLIP | Tarbase |
| hsa-miR-15a-3p | TMEM64 | PAR-CLIP | Tarbase |
| hsa-miR-15a-3p | TMEM64 | PAR-CLIP | Tarbase |
| hsa-miR-195-5p | TMEM64 | Microarrays | Tarbase |
| hsa-miR-15a-5p | GPR157 | HITS-CLIP | Tarbase |
| hsa-miR-15a-5p | GPR157 | HITS-CLIP | Tarbase |
| hsa-miR-195-5p | GPR157 | HITS-CLIP | Tarbase |
| hsa-miR-195-5p | GPR157 | HITS-CLIP | Tarbase |
| hsa-miR-486-5p | ZDHHC20 | HITS-CLIP | Tarbase |
| hsa-miR-486-5p | ZDHHC20 | HITS-CLIP | Tarbase |
| hsa-miR-141-3p | HOXC10 | HITS-CLIP | Tarbase |
| hsa-miR-486-3p | ZNF792 | HITS-CLIP | Tarbase |
| hsa-miR-195-5p | ZNF792 | Microarrays | Tarbase |
| hsa-miR-15a-5p | GPR137C | PAR-CLIP | Tarbase |
| hsa-miR-15a-3p | AEN | HITS-CLIP | Tarbase |
| hsa-miR-141-5p | SLC26A11 | HITS-CLIP | Tarbase |
| hsa-miR-15a-5p | EHMT1 | HITS-CLIP | Tarbase |
| hsa-miR-195-5p | EHMT1 | HITS-CLIP | Tarbase |
| hsa-miR-195-5p | F2R | Microarrays | Tarbase |
| hsa-miR-15a-5p | NPM1 | HITS-CLIP | Tarbase |
| hsa-miR-15a-5p | NPM1 | HITS-CLIP | Tarbase |
| hsa-miR-15a-5p | NPM1 | PAR-CLIP | Tarbase |
| hsa-miR-195-5p | NPM1 | HITS-CLIP | Tarbase |
| hsa-miR-195-5p | NPM1 | HITS-CLIP | Tarbase |
| hsa-miR-15a-5p | PJA1 | HITS-CLIP | Tarbase |
| hsa-miR-195-5p | PJA1 | HITS-CLIP | Tarbase |
| hsa-miR-15a-5p | HIST3H2A | PAR-CLIP | Tarbase |
| hsa-miR-15a-5p | POLR2A | HITS-CLIP | Tarbase |
| hsa-miR-195-5p | POLR2A | HITS-CLIP | Tarbase |

| | | | |
|---|---|---|---|
| hsa-miR-486-3p | POLR2A | HITS-CLIP | Tarbase |
| hsa-miR-141-5p | FRAT2 | HITS-CLIP | Tarbase |
| hsa-miR-15a-3p | HEPHL1 | HITS-CLIP | Tarbase |
| hsa-miR-15a-5p | AATK | HITS-CLIP | Tarbase |
| hsa-miR-141-5p | ZNF467 | HITS-CLIP | Tarbase |
| hsa-miR-15a-5p | ZNF678 | HITS-CLIP | Tarbase |
| hsa-miR-3613-5p | ZNF678 | HITS-CLIP | Tarbase |
| hsa-miR-3613-5p | ZNF678 | PAR-CLIP | Tarbase |
| hsa-miR-195-5p | TMEM45A | Microarrays | Tarbase |
| hsa-miR-15a-5p | RAP2B | HITS-CLIP | Tarbase |
| hsa-miR-15a-5p | MRPS23 | PAR-CLIP | Tarbase |
| hsa-miR-195-5p | TNFSF15 | Microarrays | Tarbase |
| hsa-miR-15a-5p | GPR88 | HITS-CLIP | Tarbase |
| hsa-miR-195-5p | GPR88 | HITS-CLIP | Tarbase |
| hsa-miR-141-3p | PLAG1 | PAR-CLIP | Tarbase |
| hsa-miR-15a-5p | PLAG1 | PAR-CLIP | Tarbase |
| hsa-miR-15a-5p | PLAG1 | HITS-CLIP | Tarbase |
| hsa-miR-15a-5p | PLAG1 | PAR-CLIP | Tarbase |
| hsa-miR-195-5p | PLAG1 | PAR-CLIP | Tarbase |
| hsa-miR-195-5p | PLAG1 | HITS-CLIP | Tarbase |
| hsa-miR-15a-3p | ZBTB20 | HITS-CLIP | Tarbase |
| hsa-miR-195-5p | ZBTB20 | Microarrays | Tarbase |
| hsa-miR-15a-5p | C3orf58 | PAR-CLIP | Tarbase |
| hsa-miR-195-5p | C3orf58 | Microarrays | Tarbase |
| hsa-miR-15a-5p | C5orf30 | HITS-CLIP | Tarbase |
| hsa-miR-15a-5p | C5orf30 | HITS-CLIP | Tarbase |
| hsa-miR-195-5p | C5orf30 | HITS-CLIP | Tarbase |
| hsa-miR-141-5p | AMIGO1 | HITS-CLIP | Tarbase |
| hsa-miR-15a-5p | AMIGO1 | PAR-CLIP | Tarbase |
| hsa-miR-219a-1-3p | COPG1 | HITS-CLIP | Tarbase |
| hsa-miR-195-5p | RELL1 | Microarrays | Tarbase |
| hsa-miR-3605-5p | SLC35C1 | HITS-CLIP | Tarbase |
| hsa-miR-141-3p | IBA57 | HITS-CLIP | Tarbase |
| hsa-miR-141-3p | C5orf24 | HITS-CLIP | Tarbase |
| hsa-miR-15a-3p | C5orf24 | HITS-CLIP | Tarbase |
| hsa-miR-15a-5p | C5orf24 | PAR-CLIP | Tarbase |
| hsa-miR-15a-5p | C5orf24 | PAR-CLIP | Tarbase |
| hsa-miR-3613-3p | C5orf24 | PAR-CLIP | Tarbase |
| hsa-miR-15a-5p | PRKAG1 | PAR-CLIP | Tarbase |
| hsa-miR-15a-5p | PRKAG1 | HITS-CLIP | Tarbase |
| hsa-miR-195-5p | GINS3 | Microarrays | Tarbase |
| hsa-miR-195-5p | OR4A16 | Microarrays | Tarbase |
| hsa-miR-141-5p | CHST15 | HITS-CLIP | Tarbase |
| hsa-miR-195-5p | MGAT4C | Microarrays | Tarbase |
| hsa-miR-141-5p | IDH2 | HITS-CLIP | Tarbase |
| hsa-miR-141-5p | TMEM259 | HITS-CLIP | Tarbase |
| hsa-miR-3605-3p | TNRC18 | HITS-CLIP | Tarbase |
| hsa-miR-15a-5p | FAM181B | PAR-CLIP | Tarbase |
| hsa-miR-141-3p | TDRKH | HITS-CLIP | Tarbase |
| hsa-miR-15a-5p | IST1 | HITS-CLIP | Tarbase |
| hsa-miR-195-5p | IST1 | HITS-CLIP | Tarbase |
| hsa-miR-195-5p | CREB3L2 | Microarrays | Tarbase |
| hsa-miR-15a-5p | TSEN54 | HITS-CLIP | Tarbase |
| hsa-miR-195-5p | RAD51B | Microarrays | Tarbase |
| hsa-miR-141-3p | EXT1 | HITS-CLIP | Tarbase |
| hsa-miR-15a-3p | EXT1 | PAR-CLIP | Tarbase |
| hsa-miR-3613-5p | SHMT2 | PAR-CLIP | Tarbase |
| hsa-miR-3613-5p | SHMT2 | PAR-CLIP | Tarbase |
| hsa-miR-15a-3p | B4GALNT4 | HITS-CLIP | Tarbase |
| hsa-miR-3605-3p | B4GALNT4 | HITS-CLIP | Tarbase |
| hsa-miR-195-5p | ZNF804B | Microarrays | Tarbase |
| hsa-miR-141-5p | NPLOC4 | HITS-CLIP | Tarbase |
| hsa-miR-15a-5p | NPLOC4 | PAR-CLIP | Tarbase |
| hsa-miR-3613-3p | NPLOC4 | HITS-CLIP | Tarbase |
| hsa-miR-15a-3p | TEX19 | HITS-CLIP | Tarbase |
| hsa-miR-141-5p | TSHZ2 | HITS-CLIP | Tarbase |
| hsa-miR-195-5p | TSHZ2 | Microarrays | Tarbase |
| hsa-miR-15a-5p | EXOC7 | HITS-CLIP | Tarbase |
| hsa-miR-141-3p | KPNA2 | HITS-CLIP | Tarbase |
| hsa-miR-195-5p | KPNA2 | Microarrays | Tarbase |
| hsa-miR-3613-5p | BGN | HITS-CLIP | Tarbase |
| hsa-miR-15a-3p | MFSD5 | HITS-CLIP | Tarbase |
| hsa-miR-15a-5p | MFSD5 | PAR-CLIP | Tarbase |
| hsa-miR-15a-5p | MFSD5 | PAR-CLIP | Tarbase |
| hsa-miR-15a-5p | MFSD5 | PAR-CLIP | Tarbase |
| hsa-miR-15a-5p | MFSD5 | HITS-CLIP | Tarbase |
| hsa-miR-15a-5p | MFSD5 | HITS-CLIP | Tarbase |
| hsa-miR-15a-5p | MFSD5 | PAR-CLIP | Tarbase |
| hsa-miR-195-5p | MFSD5 | HITS-CLIP | Tarbase |
| hsa-miR-195-5p | MFSD5 | PAR-CLIP | Tarbase |
| hsa-miR-15a-5p | MFSD5 | Chimeric fragments | Tarbase |
| hsa-miR-15a-3p | ADI1 | HITS-CLIP | Tarbase |
| hsa-miR-15a-3p | ACTL10 | HITS-CLIP | Tarbase |
| hsa-miR-195-5p | EPGN | Microarrays | Tarbase |
| hsa-miR-141-3p | TTC3 | HITS-CLIP | Tarbase |
| hsa-miR-141-3p | TTC3 | HITS-CLIP | Tarbase |
| hsa-miR-15a-5p | TTC3 | HITS-CLIP | Tarbase |
| hsa-miR-15a-5p | TTC3 | HITS-CLIP | Tarbase |
| hsa-miR-15a-5p | TTC3 | HITS-CLIP | Tarbase |
| hsa-miR-195-5p | TTC3 | HITS-CLIP | Tarbase |
| hsa-miR-15a-5p | CMC4 | PAR-CLIP | Tarbase |
| hsa-miR-195-5p | RGS6 | Microarrays | Tarbase |
| hsa-miR-141-5p | HOXB4 | HITS-CLIP | Tarbase |
| hsa-miR-15a-5p | PAPPA | HITS-CLIP | Tarbase |
| hsa-miR-195-5p | PAPPA | HITS-CLIP | Tarbase |
| hsa-miR-3613-5p | NGRN | HITS-CLIP | Tarbase |
| hsa-miR-15a-5p | DDX28 | HITS-CLIP | Tarbase |
| hsa-miR-195-5p | DDX28 | HITS-CLIP | Tarbase |
| hsa-miR-486-5p | ACBD3 | PAR-CLIP | Tarbase |
| hsa-miR-195-5p | ACBD3 | Microarrays | Tarbase |
| hsa-miR-141-3p | C16orf72 | HITS-CLIP | Tarbase |
| hsa-miR-141-3p | C16orf72 | HITS-CLIP | Tarbase |
| hsa-miR-141-5p | C16orf72 | HITS-CLIP | Tarbase |
| hsa-miR-15a-5p | C16orf72 | PAR-CLIP | Tarbase |
| hsa-miR-15a-5p | C16orf72 | PAR-CLIP | Tarbase |
| hsa-miR-15a-5p | C16orf72 | PAR-CLIP | Tarbase |
| hsa-miR-15a-5p | C16orf72 | PAR-CLIP | Tarbase |
| hsa-miR-15a-5p | C16orf72 | PAR-CLIP | Tarbase |
| hsa-miR-15a-5p | C16orf72 | PAR-CLIP | Tarbase |
| hsa-miR-195-5p | C16orf72 | PAR-CLIP | Tarbase |
| hsa-miR-195-5p | C16orf72 | PAR-CLIP | Tarbase |
| hsa-miR-3605-5p | C16orf72 | HITS-CLIP | Tarbase |
| hsa-miR-3605-5p | ZNF721 | HITS-CLIP | Tarbase |
| hsa-miR-141-5p | SRPR | HITS-CLIP | Tarbase |
| hsa-miR-15a-5p | SRPR | HITS-CLIP | Tarbase |
| hsa-miR-15a-5p | SRPR | HITS-CLIP | Tarbase |
| hsa-miR-15a-5p | SRPR | HITS-CLIP | Tarbase |
| hsa-miR-15a-5p | SRPR | HITS-CLIP | Tarbase |
| hsa-miR-15a-5p | SRPR | HITS-CLIP | Tarbase |
| hsa-miR-15a-5p | SRPR | HITS-CLIP | Tarbase |
| hsa-miR-15a-5p | SRPR | HITS-CLIP | Tarbase |
| hsa-miR-195-5p | SRPR | HITS-CLIP | Tarbase |
| hsa-miR-195-5p | SRPR | HITS-CLIP | Tarbase |
| hsa-miR-195-5p | SRPR | HITS-CLIP | Tarbase |
| hsa-miR-195-5p | SRPR | HITS-CLIP | Tarbase |
| hsa-miR-195-5p | SRPR | HITS-CLIP | Tarbase |
| hsa-miR-195-5p | SPATA13 | Microarrays | Tarbase |
| hsa-miR-15a-3p | MTA1 | HITS-CLIP | Tarbase |
| hsa-miR-195-5p | CADM1 | Microarrays | Tarbase |
| hsa-miR-15a-5p | NAA38 | PAR-CLIP | Tarbase |
| hsa-miR-195-5p | PCP4 | Microarrays | Tarbase |
| hsa-miR-195-5p | ABAT | Microarrays | Tarbase |
| hsa-miR-195-5p | CAMK1D | Microarrays | Tarbase |
| hsa-miR-195-5p | GAS6 | Microarrays | Tarbase |
| hsa-miR-141-3p | CEP57L1 | HITS-CLIP | Tarbase |
| hsa-miR-15a-3p | FANCF | HITS-CLIP | Tarbase |
| hsa-miR-15a-5p | SMDT1 | HITS-CLIP | Tarbase |
| hsa-miR-15a-5p | SMDT1 | HITS-CLIP | Tarbase |
| hsa-miR-195-5p | SMDT1 | HITS-CLIP | Tarbase |
| hsa-miR-15a-5p | PTTG1IP | HITS-CLIP | Tarbase |
| hsa-miR-15a-5p | PTTG1IP | HITS-CLIP | Tarbase |
| hsa-miR-195-5p | PTTG1IP | HITS-CLIP | Tarbase |
| hsa-miR-141-3p | BCOR | HITS-CLIP | Tarbase |
| hsa-miR-15a-3p | BCOR | HITS-CLIP | Tarbase |
| hsa-miR-195-3p | BCOR | HITS-CLIP | Tarbase |
| hsa-miR-3613-5p | KIAA2026 | HITS-CLIP | Tarbase |
| hsa-miR-141-5p | RIPK4 | HITS-CLIP | Tarbase |
| hsa-miR-15a-3p | RIPK4 | HITS-CLIP | Tarbase |
| hsa-miR-3605-5p | SF3A3 | PAR-CLIP | Tarbase |

| | | | |
|---|---|---|---|
| hsa-miR-195-5p | MX2 | Microarrays | Tarbase |
| hsa-miR-141-3p | FAM46C | HITS-CLIP | Tarbase |
| hsa-miR-3605-5p | FAM46C | HITS-CLIP | Tarbase |
| hsa-miR-3605-5p | FAM46C | HITS-CLIP | Tarbase |
| hsa-miR-195-5p | FAM46C | Microarrays | Tarbase |
| hsa-miR-141-3p | PRR14L | HITS-CLIP | Tarbase |
| hsa-miR-15a-3p | PRR14L | HITS-CLIP | Tarbase |
| hsa-miR-3613-3p | PRR14L | HITS-CLIP | Tarbase |
| hsa-miR-486-5p | PRR14L | PAR-CLIP | Tarbase |
| hsa-miR-195-5p | SETD3 | Microarrays | Tarbase |
| hsa-miR-15a-5p | SETD3 | Chimeric fragments | Tarbase |
| hsa-miR-15a-5p | ZNRF3 | PAR-CLIP | Tarbase |
| hsa-miR-15a-5p | ZNRF3 | HITS-CLIP | Tarbase |
| hsa-miR-195-5p | ZNRF3 | HITS-CLIP | Tarbase |
| hsa-miR-195-5p | FBXL7 | Microarrays | Tarbase |
| hsa-miR-195-5p | FAM167B | Microarrays | Tarbase |
| hsa-miR-195-5p | PSG9 | Microarrays | Tarbase |
| hsa-miR-3613-5p | ALYREF | HITS-CLIP | Tarbase |
| hsa-miR-195-5p | FAM101B | Microarrays | Tarbase |
| hsa-miR-195-5p | NOG | Microarrays | Tarbase |
| hsa-miR-141-3p | LHFP | HITS-CLIP | Tarbase |
| hsa-miR-15a-3p | CMTM4 | HITS-CLIP | Tarbase |
| hsa-miR-15a-5p | CMTM4 | HITS-CLIP | Tarbase |
| hsa-miR-195-5p | CMTM4 | HITS-CLIP | Tarbase |
| hsa-miR-141-5p | TBK1 | HITS-CLIP | Tarbase |
| hsa-miR-141-5p | TBK1 | HITS-CLIP | Tarbase |
| hsa-miR-15a-3p | CBX6 | HITS-CLIP | Tarbase |
| hsa-miR-15a-3p | CBX6 | HITS-CLIP | Tarbase |
| hsa-miR-15a-5p | CBX6 | HITS-CLIP | Tarbase |
| hsa-miR-15a-5p | CBX6 | PAR-CLIP | Tarbase |
| hsa-miR-195-5p | CBX6 | HITS-CLIP | Tarbase |
| hsa-miR-3605-3p | CBX6 | HITS-CLIP | Tarbase |
| hsa-miR-195-5p | CBX6 | Microarrays | Tarbase |
| hsa-miR-141-3p | MACC1 | PAR-CLIP | Tarbase |
| hsa-miR-15a-5p | TBL3 | PAR-CLIP | Tarbase |
| hsa-miR-195-5p | TBL3 | PAR-CLIP | Tarbase |
| hsa-miR-3605-5p | KREMEN1 | HITS-CLIP | Tarbase |
| hsa-miR-195-5p | TRAIP | Microarrays | Tarbase |
| hsa-miR-15a-5p | KCTD16 | HITS-CLIP | Tarbase |
| hsa-miR-195-5p | KCTD16 | HITS-CLIP | Tarbase |
| hsa-miR-141-5p | ZNF703 | HITS-CLIP | Tarbase |
| hsa-miR-195-5p | ZNF703 | Microarrays | Tarbase |
| hsa-miR-219a-1-3p | ZNF703 | Chimeric fragments | Tarbase |
| hsa-miR-15a-5p | RBM12B | HITS-CLIP | Tarbase |
| hsa-miR-195-5p | RBM12B | HITS-CLIP | Tarbase |
| hsa-miR-486-5p | RBM12B | HITS-CLIP | Tarbase |
| hsa-miR-486-5p | RBM12B | HITS-CLIP | Tarbase |
| hsa-miR-195-5p | LIN9 | Microarrays | Tarbase |
| hsa-miR-15a-5p | BTBD9 | HITS-CLIP | Tarbase |
| hsa-miR-195-5p | BTBD9 | HITS-CLIP | Tarbase |
| hsa-miR-195-5p | ZNF730 | Microarrays | Tarbase |
| hsa-miR-15a-3p | KIRREL | HITS-CLIP | Tarbase |
| hsa-miR-195-5p | IQGAP3 | Microarrays | Tarbase |
| hsa-miR-15a-3p | TOB2 | HITS-CLIP | Tarbase |
| hsa-miR-3613-5p | TOB2 | PAR-CLIP | Tarbase |
| hsa-miR-3613-5p | TOB2 | PAR-CLIP | Tarbase |
| hsa-miR-3613-5p | TOB2 | PAR-CLIP | Tarbase |
| hsa-miR-141-5p | PRKX | HITS-CLIP | Tarbase |
| hsa-miR-15a-5p | SMTN | HITS-CLIP | Tarbase |
| hsa-miR-195-5p | SMTN | HITS-CLIP | Tarbase |
| hsa-miR-141-5p | ACTG1 | HITS-CLIP | Tarbase |
| hsa-miR-15a-5p | ACTG1 | HITS-CLIP | Tarbase |
| hsa-miR-15a-5p | ACTG1 | HITS-CLIP | Tarbase |
| hsa-miR-15a-5p | ACTG1 | HITS-CLIP | Tarbase |
| hsa-miR-15a-5p | ACTG1 | HITS-CLIP | Tarbase |
| hsa-miR-195-5p | ACTG1 | HITS-CLIP | Tarbase |
| hsa-miR-195-5p | ACTG1 | HITS-CLIP | Tarbase |
| hsa-miR-195-5p | ACTG1 | HITS-CLIP | Tarbase |
| hsa-miR-219a-1-3p | ACTG1 | HITS-CLIP | Tarbase |
| hsa-miR-15a-5p | VPS33B | HITS-CLIP | Tarbase |
| hsa-miR-195-5p | VPS33B | Microarrays | Tarbase |
| hsa-miR-15a-3p | FAM120C | HITS-CLIP | Tarbase |
| hsa-miR-195-5p | LRTOMT | Microarrays | Tarbase |
| hsa-miR-195-5p | KCNQ3 | Microarrays | Tarbase |
| hsa-miR-15a-5p | SCFD2 | HITS-CLIP | Tarbase |
| hsa-miR-15a-5p | UBE2F | Chimeric fragments | Tarbase |
| hsa-miR-15a-5p | PPP1R2 | HITS-CLIP | Tarbase |
| hsa-miR-195-5p | PPP1R2 | HITS-CLIP | Tarbase |
| hsa-miR-195-5p | TSPYL2 | HITS-CLIP | Tarbase |
| hsa-miR-15a-5p | TSPYL2 | Luciferase Reporter Assay | Tarbase |
| hsa-miR-15a-5p | PGP | HITS-CLIP | Tarbase |
| hsa-miR-195-5p | CMSS1 | Microarrays | Tarbase |
| hsa-miR-15a-5p | PCDH9 | HITS-CLIP | Tarbase |
| hsa-miR-195-5p | PCDH9 | HITS-CLIP | Tarbase |
| hsa-miR-15a-5p | PCDH9 | HITS-CLIP | Tarbase |
| hsa-miR-195-5p | PCDH9 | HITS-CLIP | Tarbase |
| hsa-miR-195-5p | PCDH9 | Microarrays | Tarbase |
| hsa-miR-15a-5p | ALDH1A3 | PAR-CLIP | Tarbase |
| hsa-miR-195-5p | HIST2H2AC | Microarrays | Tarbase |
| hsa-miR-195-5p | HIST2H2AB | Microarrays | Tarbase |
| hsa-miR-15a-5p | TACSTD2 | HITS-CLIP | Tarbase |
| hsa-miR-195-5p | TACSTD2 | HITS-CLIP | Tarbase |
| hsa-miR-15a-5p | CCSER1 | HITS-CLIP | Tarbase |
| hsa-miR-195-5p | CCSER1 | HITS-CLIP | Tarbase |
| hsa-miR-15a-5p | CCSER1 | HITS-CLIP | Tarbase |
| hsa-miR-195-5p | CCSER1 | HITS-CLIP | Tarbase |
| hsa-miR-15a-5p | ZDHHC23 | HITS-CLIP | Tarbase |
| hsa-miR-195-5p | ZDHHC23 | HITS-CLIP | Tarbase |
| hsa-miR-15a-3p | HIST1H1B | HITS-CLIP | Tarbase |
| hsa-miR-15a-5p | HIST1H1B | HITS-CLIP | Tarbase |
| hsa-miR-195-5p | HIST1H1B | Microarrays | Tarbase |
| hsa-miR-15a-5p | PKP3 | HITS-CLIP | Tarbase |
| hsa-miR-195-5p | CSF1 | Microarrays | Tarbase |
| hsa-miR-195-5p | COLEC10 | Microarrays | Tarbase |
| hsa-miR-141-5p | COPB2 | HITS-CLIP | Tarbase |
| hsa-miR-486-5p | COPB2 | HITS-CLIP | Tarbase |
| hsa-miR-141-3p | KNTC1 | HITS-CLIP | Tarbase |
| hsa-miR-195-5p | KNTC1 | Microarrays | Tarbase |
| hsa-miR-15a-5p | CCR10 | PAR-CLIP | Tarbase |
| hsa-miR-195-5p | WDR27 | Microarrays | Tarbase |
| hsa-miR-15a-5p | ZFP1 | PAR-CLIP | Tarbase |
| hsa-miR-195-5p | SLITRK6 | Microarrays | Tarbase |
| hsa-miR-195-5p | PDE4B | Microarrays | Tarbase |
| hsa-miR-3605-3p | MED12 | HITS-CLIP | Tarbase |
| hsa-miR-3605-3p | SEPT9 | HITS-CLIP | Tarbase |
| hsa-miR-141-3p | CDCA2 | HITS-CLIP | Tarbase |
| hsa-miR-195-5p | CDCA2 | Microarrays | Tarbase |
| hsa-miR-141-5p | AMER1 | HITS-CLIP | Tarbase |
| hsa-miR-195-5p | AMER1 | Microarrays | Tarbase |
| hsa-miR-195-5p | HIST2H2BE | Microarrays | Tarbase |
| hsa-miR-15a-5p | SEPT5 | PAR-CLIP | Tarbase |
| hsa-miR-15a-5p | FAM110C | HITS-CLIP | Tarbase |
| hsa-miR-195-5p | FAM110C | HITS-CLIP | Tarbase |
| hsa-miR-141-3p | ATL3 | HITS-CLIP | Tarbase |
| hsa-miR-15a-5p | ATL3 | HITS-CLIP | Tarbase |
| hsa-miR-15a-5p | ATL3 | HITS-CLIP | Tarbase |
| hsa-miR-486-5p | ATL3 | PAR-CLIP | Tarbase |
| hsa-miR-195-5p | ATL3 | Microarrays | Tarbase |
| hsa-miR-15a-5p | NDUFA12 | PAR-CLIP | Tarbase |
| hsa-miR-15a-3p | SMIM10 | PAR-CLIP | Tarbase |
| hsa-miR-486-5p | OSBP2 | HITS-CLIP | Tarbase |
| hsa-miR-3613-5p | APOO | PAR-CLIP | Tarbase |
| hsa-miR-195-5p | PRR16 | Microarrays | Tarbase |
| hsa-miR-141-3p | TMED9 | HITS-CLIP | Tarbase |
| hsa-miR-141-3p | RBM33 | HITS-CLIP | Tarbase |
| hsa-miR-141-5p | RBM33 | HITS-CLIP | Tarbase |
| hsa-miR-141-5p | RBM33 | HITS-CLIP | Tarbase |
| hsa-miR-15a-5p | ARMCX2 | HITS-CLIP | Tarbase |
| hsa-miR-195-5p | ARMCX2 | HITS-CLIP | Tarbase |
| hsa-miR-15a-5p | BTBD6 | PAR-CLIP | Tarbase |
| hsa-miR-15a-5p | SUMO3 | PAR-CLIP | Tarbase |
| hsa-miR-15a-5p | SUMO3 | HITS-CLIP | Tarbase |
| hsa-miR-15a-5p | SUMO3 | HITS-CLIP | Tarbase |
| hsa-miR-195-5p | SUMO3 | HITS-CLIP | Tarbase |
| hsa-miR-3605-3p | IMMP2L | HITS-CLIP | Tarbase |
| hsa-miR-195-5p | NDUFA6 | Microarrays | Tarbase |
| hsa-miR-15a-5p | TMEM121 | HITS-CLIP | Tarbase |
| hsa-miR-195-5p | TMEM121 | HITS-CLIP | Tarbase |
| hsa-miR-141-3p | BRI3BP | HITS-CLIP | Tarbase |

| | | | |
|---|---|---|---|
| hsa-miR-141-5p | BRI3BP | HITS-CLIP | Tarbase |
| hsa-miR-15a-5p | BRI3BP | HITS-CLIP | Tarbase |
| hsa-miR-195-5p | BRI3BP | HITS-CLIP | Tarbase |
| hsa-miR-195-5p | ROBO2 | Microarrays | Tarbase |
| hsa-miR-195-5p | MAFF | Microarrays | Tarbase |
| hsa-miR-15a-5p | INTS5 | HITS-CLIP | Tarbase |
| hsa-miR-486-3p | RPS27L | PAR-CLIP | Tarbase |
| hsa-miR-486-3p | RPS27L | PAR-CLIP | Tarbase |
| hsa-miR-195-5p | MANEAL | Microarrays | Tarbase |
| hsa-miR-141-5p | FAF1 | HITS-CLIP | Tarbase |
| hsa-miR-15a-5p | NDNL2 | PAR-CLIP | Tarbase |
| hsa-miR-15a-5p | NDNL2 | PAR-CLIP | Tarbase |
| hsa-miR-195-5p | NDNL2 | PAR-CLIP | Tarbase |
| hsa-miR-141-3p | PURA | HITS-CLIP | Tarbase |
| hsa-miR-15a-3p | PURA | HITS-CLIP | Tarbase |
| hsa-miR-15a-5p | PURA | PAR-CLIP | Tarbase |
| hsa-miR-15a-5p | PURA | PAR-CLIP | Tarbase |
| hsa-miR-15a-5p | PURA | HITS-CLIP | Tarbase |
| hsa-miR-15a-5p | PURA | Luciferase Reporter Assay | Tarbase |
| hsa-miR-15a-5p | PURA | PAR-CLIP | Tarbase |
| hsa-miR-195-5p | PURA | PAR-CLIP | Tarbase |
| hsa-miR-3613-3p | PURA | HITS-CLIP | Tarbase |
| hsa-miR-3613-3p | PURA | PAR-CLIP | Tarbase |
| hsa-miR-15a-5p | PURA | HITS-CLIP | Tarbase |
| hsa-miR-195-5p | PURA | HITS-CLIP | Tarbase |
| hsa-miR-15a-3p | NOMO2 | HITS-CLIP | Tarbase |
| hsa-miR-141-5p | WBP5 | HITS-CLIP | Tarbase |
| hsa-miR-141-5p | PRPF39 | HITS-CLIP | Tarbase |
| hsa-miR-141-5p | PRPF39 | HITS-CLIP | Tarbase |
| hsa-miR-141-3p | SMYD3 | HITS-CLIP | Tarbase |
| hsa-miR-141-3p | METTL7A | HITS-CLIP | Tarbase |
| hsa-miR-3613-3p | METTL7A | HITS-CLIP | Tarbase |
| hsa-miR-195-5p | METTL7A | Microarrays | Tarbase |
| hsa-miR-15a-5p | FAM174B | HITS-CLIP | Tarbase |
| hsa-miR-195-5p | PARPBP | Microarrays | Tarbase |
| hsa-miR-3613-5p | BRCC3 | PAR-CLIP | Tarbase |
| hsa-miR-3613-3p | NR2F2 | Chimeric fragments | Tarbase |
| hsa-miR-15a-5p | DLK1 | Other | Tarbase |
| hsa-miR-15a-5p | DLK1 | qPCR | Tarbase |
| hsa-miR-15a-5p | SP1 | PAR-CLIP | Tarbase |
| hsa-miR-15a-5p | SP1 | HITS-CLIP | Tarbase |
| hsa-miR-15a-5p | SP1 | HITS-CLIP | Tarbase |
| hsa-miR-15a-5p | SP1 | PAR-CLIP | Tarbase |
| hsa-miR-15a-5p | SP1 | HITS-CLIP | Tarbase |
| hsa-miR-195-5p | SP1 | HITS-CLIP | Tarbase |
| hsa-miR-195-5p | SP1 | HITS-CLIP | Tarbase |
| hsa-miR-195-5p | SP1 | HITS-CLIP | Tarbase |
| hsa-miR-219a-1-3p | SP1 | HITS-CLIP | Tarbase |
| hsa-miR-15a-5p | SP1 | HITS-CLIP | Tarbase |
| hsa-miR-195-5p | SP1 | HITS-CLIP | Tarbase |
| hsa-miR-15a-5p | SP1 | Chimeric fragments | Tarbase |
| hsa-miR-3613-5p | PCGF3 | PAR-CLIP | Tarbase |
| hsa-miR-3613-5p | PCGF3 | PAR-CLIP | Tarbase |
| hsa-miR-3613-5p | PCGF3 | PAR-CLIP | Tarbase |
| hsa-miR-141-3p | LMLN | HITS-CLIP | Tarbase |
| hsa-miR-141-5p | P4HB | HITS-CLIP | Tarbase |
| hsa-miR-15a-3p | P4HB | HITS-CLIP | Tarbase |
| hsa-miR-3605-3p | P4HB | HITS-CLIP | Tarbase |
| hsa-miR-195-5p | PBX1 | Microarrays | Tarbase |
| hsa-miR-15a-5p | PBX1 | Chimeric fragments | Tarbase |
| hsa-miR-3613-5p | KRT79 | HITS-CLIP | Tarbase |
| hsa-miR-141-5p | ZFP36L1 | HITS-CLIP | Tarbase |
| hsa-miR-3613-3p | ZFP36L1 | PAR-CLIP | Tarbase |
| hsa-miR-15a-3p | UBE2L3 | HITS-CLIP | Tarbase |
| hsa-miR-195-5p | NTF3 | Microarrays | Tarbase |
| hsa-miR-141-3p | BRWD1 | HITS-CLIP | Tarbase |
| hsa-miR-15a-5p | BRWD1 | HITS-CLIP | Tarbase |
| hsa-miR-15a-5p | BRWD1 | HITS-CLIP | Tarbase |
| hsa-miR-195-5p | BRWD1 | HITS-CLIP | Tarbase |
| hsa-miR-195-5p | BRWD1 | HITS-CLIP | Tarbase |
| hsa-miR-486-5p | BRWD1 | HITS-CLIP | Tarbase |
| hsa-miR-486-5p | BRWD1 | HITS-CLIP | Tarbase |
| hsa-miR-15a-5p | BRWD1 | HITS-CLIP | Tarbase |
| hsa-miR-195-5p | BRWD1 | HITS-CLIP | Tarbase |
| hsa-miR-15a-5p | MYBL1 | PAR-CLIP | Tarbase |
| hsa-miR-141-5p | DRG1 | HITS-CLIP | Tarbase |
| hsa-miR-141-3p | ANKFY1 | HITS-CLIP | Tarbase |
| hsa-miR-15a-5p | ANKFY1 | HITS-CLIP | Tarbase |
| hsa-miR-3613-3p | ANKFY1 | PAR-CLIP | Tarbase |
| hsa-miR-141-3p | YTHDF3 | HITS-CLIP | Tarbase |
| hsa-miR-3613-5p | YTHDF3 | HITS-CLIP | Tarbase |
| hsa-miR-3613-3p | MORF4L1 | HITS-CLIP | Tarbase |
| hsa-miR-141-3p | DMWD | HITS-CLIP | Tarbase |
| hsa-miR-15a-3p | NAT8L | HITS-CLIP | Tarbase |
| hsa-miR-15a-3p | NAT8L | HITS-CLIP | Tarbase |
| hsa-miR-15a-5p | NAT8L | HITS-CLIP | Tarbase |
| hsa-miR-195-5p | NAT8L | HITS-CLIP | Tarbase |
| hsa-miR-15a-5p | DNAH14 | HITS-CLIP | Tarbase |
| hsa-miR-195-5p | DNAH14 | HITS-CLIP | Tarbase |
| hsa-miR-15a-5p | POMK | HITS-CLIP | Tarbase |
| hsa-miR-195-5p | POMK | HITS-CLIP | Tarbase |
| hsa-miR-195-5p | KLHDC8B | Microarrays | Tarbase |
| hsa-miR-141-3p | PTCH1 | HITS-CLIP | Tarbase |
| hsa-miR-141-3p | PTCH1 | HITS-CLIP | Tarbase |
| hsa-miR-3605-5p | PTCH1 | HITS-CLIP | Tarbase |
| hsa-miR-195-5p | PTCH1 | Microarrays | Tarbase |
| hsa-miR-15a-5p | ZNF267 | PAR-CLIP | Tarbase |
| hsa-miR-141-5p | IRS2 | HITS-CLIP | Tarbase |
| hsa-miR-141-3p | BICD2 | PAR-CLIP | Tarbase |
| hsa-miR-141-3p | BICD2 | HITS-CLIP | Tarbase |
| hsa-miR-195-5p | RASA3 | Microarrays | Tarbase |
| hsa-miR-141-3p | LRCH3 | HITS-CLIP | Tarbase |
| hsa-miR-141-3p | LRCH3 | HITS-CLIP | Tarbase |
| hsa-miR-195-5p | C15orf41 | Microarrays | Tarbase |
| hsa-miR-15a-5p | ANKRD46 | HITS-CLIP | Tarbase |
| hsa-miR-195-5p | ANKRD46 | HITS-CLIP | Tarbase |
| hsa-miR-141-5p | POLR1D | HITS-CLIP | Tarbase |
| hsa-miR-195-5p | KIF18B | Microarrays | Tarbase |
| hsa-miR-15a-5p | EDARADD | HITS-CLIP | Tarbase |
| hsa-miR-195-5p | EDARADD | HITS-CLIP | Tarbase |
| hsa-miR-195-5p | CYP4F12 | Microarrays | Tarbase |
| hsa-miR-15a-5p | MKL2 | HITS-CLIP | Tarbase |
| hsa-miR-195-5p | MKL2 | HITS-CLIP | Tarbase |
| hsa-miR-195-5p | TOR3A | Microarrays | Tarbase |
| hsa-miR-15a-3p | PPP1CC | HITS-CLIP | Tarbase |
| hsa-miR-15a-5p | PPP1CC | HITS-CLIP | Tarbase |
| hsa-miR-15a-5p | PPP1CC | HITS-CLIP | Tarbase |
| hsa-miR-195-5p | PPP1CC | HITS-CLIP | Tarbase |
| hsa-miR-195-5p | PPP1CC | HITS-CLIP | Tarbase |
| hsa-miR-195-5p | PPP1CC | HITS-CLIP | Tarbase |
| hsa-miR-15a-5p | BACE1 | Luciferase Reporter Assay | Tarbase |
| hsa-miR-195-5p | BACE1 | Microarrays | Tarbase |
| hsa-miR-195-5p | RXRA | Microarrays | Tarbase |
| hsa-miR-195-5p | GLDN | Microarrays | Tarbase |
| hsa-miR-486-5p | PCLO | HITS-CLIP | Tarbase |
| hsa-miR-15a-5p | RGS7BP | HITS-CLIP | Tarbase |
| hsa-miR-195-5p | RGS7BP | HITS-CLIP | Tarbase |
| hsa-miR-15a-5p | INSIG1 | Chimeric fragments | Tarbase |
| hsa-miR-141-5p | ZNF396 | HITS-CLIP | Tarbase |
| hsa-miR-486-5p | TMEM222 | PAR-CLIP | Tarbase |
| hsa-miR-195-5p | CYP4F3 | Microarrays | Tarbase |
| hsa-miR-15a-5p | GPATCH8 | HITS-CLIP | Tarbase |
| hsa-miR-141-3p | NF2 | HITS-CLIP | Tarbase |
| hsa-miR-15a-5p | NF2 | HITS-CLIP | Tarbase |
| hsa-miR-486-5p | NF2 | PAR-CLIP | Tarbase |
| hsa-miR-195-5p | KIF24 | Microarrays | Tarbase |
| hsa-miR-3613-3p | ZFP91 | Chimeric fragments | Tarbase |
| hsa-miR-141-3p | LYRM7 | HITS-CLIP | Tarbase |
| hsa-miR-141-3p | SPIN4 | HITS-CLIP | Tarbase |
| hsa-miR-141-3p | SPIN4 | HITS-CLIP | Tarbase |
| hsa-miR-15a-5p | SPIN4 | HITS-CLIP | Tarbase |
| hsa-miR-195-5p | SPIN4 | Microarrays | Tarbase |
| hsa-miR-141-3p | ZNF397 | HITS-CLIP | Tarbase |
| hsa-miR-141-3p | ZSCAN30 | HITS-CLIP | Tarbase |
| hsa-miR-15a-5p | TPCN1 | HITS-CLIP | Tarbase |
| hsa-miR-195-5p | TPCN1 | HITS-CLIP | Tarbase |
| hsa-miR-195-5p | TPCN1 | Microarrays | Tarbase |
| hsa-miR-15a-5p | HEXIM1 | HITS-CLIP | Tarbase |
| hsa-miR-195-5p | HEXIM1 | HITS-CLIP | Tarbase |

| | | | |
|---|---|---|---|
| hsa-miR-3605-3p | HEXIM1 | HITS-CLIP | Tarbase |
| hsa-miR-195-5p | ERCC6L | Microarrays | Tarbase |
| hsa-miR-141-3p | ZDHHC17 | HITS-CLIP | Tarbase |
| hsa-miR-141-3p | ZDHHC17 | HITS-CLIP | Tarbase |
| hsa-miR-141-3p | ZDHHC17 | HITS-CLIP | Tarbase |
| hsa-miR-15a-5p | ZDHHC17 | HITS-CLIP | Tarbase |
| hsa-miR-195-5p | ZDHHC17 | HITS-CLIP | Tarbase |
| hsa-miR-141-3p | PPARA | HITS-CLIP | Tarbase |
| hsa-miR-15a-3p | TEAD1 | HITS-CLIP | Tarbase |
| hsa-miR-15a-5p | TEAD1 | HITS-CLIP | Tarbase |
| hsa-miR-219a-1-3p | TEAD1 | HITS-CLIP | Tarbase |
| hsa-miR-486-5p | TEAD1 | HITS-CLIP | Tarbase |
| hsa-miR-195-5p | TEAD1 | Microarrays | Tarbase |
| hsa-miR-141-5p | ENTPD5 | HITS-CLIP | Tarbase |
| hsa-miR-15a-5p | ENTPD5 | PAR-CLIP | Tarbase |
| hsa-miR-141-3p | NAP1L1 | HITS-CLIP | Tarbase |
| hsa-miR-141-3p | NAP1L1 | HITS-CLIP | Tarbase |
| hsa-miR-141-3p | NAP1L1 | PAR-CLIP | Tarbase |
| hsa-miR-195-5p | AKR1C1 | Microarrays | Tarbase |
| hsa-miR-195-5p | VSTM2B | Microarrays | Tarbase |
| hsa-miR-141-5p | RNF220 | HITS-CLIP | Tarbase |
| hsa-miR-141-5p | RNF220 | HITS-CLIP | Tarbase |
| hsa-miR-141-5p | ANGPTL5 | HITS-CLIP | Tarbase |
| hsa-miR-141-3p | KIAA1598 | HITS-CLIP | Tarbase |
| hsa-miR-141-3p | KIAA1598 | HITS-CLIP | Tarbase |
| hsa-miR-15a-5p | KIAA1598 | HITS-CLIP | Tarbase |
| hsa-miR-141-5p | TSPYL4 | HITS-CLIP | Tarbase |
| hsa-miR-15a-5p | MT1X | HITS-CLIP | Tarbase |
| hsa-miR-195-5p | GCNT1 | Microarrays | Tarbase |
| hsa-miR-15a-5p | SESTD1 | PAR-CLIP | Tarbase |
| hsa-miR-195-5p | SESTD1 | PAR-CLIP | Tarbase |
| hsa-miR-195-5p | SESTD1 | Microarrays | Tarbase |
| hsa-miR-15a-5p | TAF9B | HITS-CLIP | Tarbase |
| hsa-miR-3613-3p | KCNJ11 | HITS-CLIP | Tarbase |
| hsa-miR-195-5p | COL4A1 | Microarrays | Tarbase |
| hsa-miR-195-5p | AGMO | Microarrays | Tarbase |
| hsa-miR-141-3p | USP7 | HITS-CLIP | Tarbase |
| hsa-miR-3613-3p | USP7 | PAR-CLIP | Tarbase |
| hsa-miR-141-3p | TET3 | HITS-CLIP | Tarbase |
| hsa-miR-15a-5p | EXD3 | PAR-CLIP | Tarbase |
| hsa-miR-15a-5p | EXD3 | PAR-CLIP | Tarbase |
| hsa-miR-195-5p | C17orf97 | Microarrays | Tarbase |
| hsa-miR-15a-3p | SAMD11 | HITS-CLIP | Tarbase |
| hsa-miR-3605-3p | SAMD11 | HITS-CLIP | Tarbase |
| hsa-miR-15a-5p | SPRY4 | PAR-CLIP | Tarbase |
| hsa-miR-195-5p | AMTN | Microarrays | Tarbase |
| hsa-miR-15a-5p | TMEM203 | PAR-CLIP | Tarbase |
| hsa-miR-486-3p | THSD4 | HITS-CLIP | Tarbase |
| hsa-miR-3605-5p | AMY1C | HITS-CLIP | Tarbase |
| hsa-miR-141-5p | NHEJ1 | HITS-CLIP | Tarbase |
| hsa-miR-15a-5p | FANCA | HITS-CLIP | Tarbase |
| hsa-miR-195-5p | FANCA | Microarrays | Tarbase |
| hsa-miR-15a-5p | SEMA4D | HITS-CLIP | Tarbase |
| hsa-miR-15a-5p | FANCM | Chimeric fragments | Tarbase |
| hsa-miR-195-5p | ZFP69B | Microarrays | Tarbase |
| hsa-miR-15a-5p | HIST1H1C | HITS-CLIP | Tarbase |
| hsa-miR-15a-5p | HIST1H1C | HITS-CLIP | Tarbase |
| hsa-miR-15a-5p | HIST1H1C | HITS-CLIP | Tarbase |
| hsa-miR-15a-5p | HIST1H1C | HITS-CLIP | Tarbase |
| hsa-miR-15a-5p | HIST1H1C | HITS-CLIP | Tarbase |
| hsa-miR-15a-5p | HIST1H1C | HITS-CLIP | Tarbase |
| hsa-miR-15a-5p | HIST1H1C | PAR-CLIP | Tarbase |
| hsa-miR-15a-5p | HIST1H1C | HITS-CLIP | Tarbase |
| hsa-miR-15a-5p | HIST1H1C | HITS-CLIP | Tarbase |
| hsa-miR-15a-5p | HIST1H1C | HITS-CLIP | Tarbase |
| hsa-miR-195-5p | HIST1H1C | HITS-CLIP | Tarbase |
| hsa-miR-195-5p | HIST1H1C | HITS-CLIP | Tarbase |
| hsa-miR-195-5p | HIST1H1C | HITS-CLIP | Tarbase |
| hsa-miR-195-5p | HIST1H1C | HITS-CLIP | Tarbase |
| hsa-miR-195-5p | HIST1H1C | HITS-CLIP | Tarbase |
| hsa-miR-195-5p | HIST1H1C | HITS-CLIP | Tarbase |
| hsa-miR-195-5p | HIST1H1C | HITS-CLIP | Tarbase |
| hsa-miR-15a-5p | HIST1H1C | HITS-CLIP | Tarbase |
| hsa-miR-195-5p | HIST1H1C | HITS-CLIP | Tarbase |
| hsa-miR-195-5p | HIST1H1C | Microarrays | Tarbase |
| hsa-miR-15a-5p | HIST1H1C | Chimeric fragments | Tarbase |
| hsa-miR-486-3p | DMBT1 | PAR-CLIP | Tarbase |
| hsa-miR-195-5p | COL14A1 | Microarrays | Tarbase |
| hsa-miR-15a-5p | DNER | HITS-CLIP | Tarbase |
| hsa-miR-195-5p | DNER | HITS-CLIP | Tarbase |
| hsa-miR-195-5p | TPRG1 | Microarrays | Tarbase |
| hsa-miR-195-5p | ARL4C | Microarrays | Tarbase |
| hsa-miR-195-5p | WNT7B | Microarrays | Tarbase |
| hsa-miR-15a-5p | COL4A5 | HITS-CLIP | Tarbase |
| hsa-miR-195-5p | COL4A5 | HITS-CLIP | Tarbase |
| hsa-miR-15a-5p | AGRN | HITS-CLIP | Tarbase |
| hsa-miR-195-5p | AGRN | HITS-CLIP | Tarbase |
| hsa-miR-195-5p | AGRN | HITS-CLIP | Tarbase |
| hsa-miR-141-5p | ZC3H6 | HITS-CLIP | Tarbase |
| hsa-miR-15a-5p | ZC3H6 | PAR-CLIP | Tarbase |
| hsa-miR-15a-5p | ZC3H6 | PAR-CLIP | Tarbase |
| hsa-miR-15a-5p | ZC3H6 | PAR-CLIP | Tarbase |
| hsa-miR-195-5p | ZC3H6 | PAR-CLIP | Tarbase |
| hsa-miR-141-3p | NCR3LG1 | HITS-CLIP | Tarbase |
| hsa-miR-141-3p | ZNF793 | HITS-CLIP | Tarbase |
| hsa-miR-15a-5p | C3orf62 | HITS-CLIP | Tarbase |
| hsa-miR-195-5p | C3orf62 | HITS-CLIP | Tarbase |
| hsa-miR-195-5p | FAM92A1 | Microarrays | Tarbase |
| hsa-miR-141-5p | FOCAD | HITS-CLIP | Tarbase |
| hsa-miR-141-5p | PPP3R2 | HITS-CLIP | Tarbase |
| hsa-miR-15a-5p | H2AFX | HITS-CLIP | Tarbase |
| hsa-miR-195-5p | H2AFX | Microarrays | Tarbase |
| hsa-miR-195-5p | SRSF10 | HITS-CLIP | Tarbase |
| hsa-miR-15a-3p | C15orf52 | HITS-CLIP | Tarbase |
| hsa-miR-195-5p | C15orf52 | Microarrays | Tarbase |
| hsa-miR-15a-5p | NBR1 | HITS-CLIP | Tarbase |
| hsa-miR-486-3p | NBR1 | HITS-CLIP | Tarbase |
| hsa-miR-195-5p | NKAIN2 | Microarrays | Tarbase |
| hsa-miR-195-5p | NANOS1 | Microarrays | Tarbase |
| hsa-miR-15a-5p | LDOC1L | HITS-CLIP | Tarbase |
| hsa-miR-15a-5p | LDOC1L | HITS-CLIP | Tarbase |
| hsa-miR-195-5p | LDOC1L | HITS-CLIP | Tarbase |
| hsa-miR-195-5p | S100A16 | Microarrays | Tarbase |
| hsa-miR-15a-3p | PTAR1 | HITS-CLIP | Tarbase |
| hsa-miR-486-5p | UROS | HITS-CLIP | Tarbase |
| hsa-miR-15a-3p | NOXA1 | HITS-CLIP | Tarbase |
| hsa-miR-15a-3p | NOXA1 | HITS-CLIP | Tarbase |
| hsa-miR-15a-3p | TMEM198 | HITS-CLIP | Tarbase |
| hsa-miR-195-5p | SPRED3 | Microarrays | Tarbase |
| hsa-miR-15a-5p | MTF1 | PAR-CLIP | Tarbase |
| hsa-miR-3613-5p | MTF1 | PAR-CLIP | Tarbase |
| hsa-miR-3613-5p | MTF1 | PAR-CLIP | Tarbase |
| hsa-miR-3613-5p | MTF1 | PAR-CLIP | Tarbase |
| hsa-miR-15a-5p | RPL14 | PAR-CLIP | Tarbase |
| hsa-miR-195-5p | RPL14 | Microarrays | Tarbase |
| hsa-miR-15a-3p | ASTL | HITS-CLIP | Tarbase |
| hsa-miR-195-5p | MSL1 | HITS-CLIP | Tarbase |
| hsa-miR-3605-3p | MSL1 | HITS-CLIP | Tarbase |
| hsa-miR-15a-5p | MSL1 | HITS-CLIP | Tarbase |
| hsa-miR-195-5p | MSL1 | HITS-CLIP | Tarbase |
| hsa-miR-15a-5p | NOC2L | HITS-CLIP | Tarbase |
| hsa-miR-195-5p | NOC2L | HITS-CLIP | Tarbase |
| hsa-miR-15a-3p | NELFB | HITS-CLIP | Tarbase |
| hsa-miR-15a-5p | LITAF | HITS-CLIP | Tarbase |
| hsa-miR-15a-5p | LITAF | HITS-CLIP | Tarbase |
| hsa-miR-195-5p | LITAF | Microarrays | Tarbase |
| hsa-miR-15a-5p | ARID2 | HITS-CLIP | Tarbase |
| hsa-miR-195-5p | ARID2 | HITS-CLIP | Tarbase |
| hsa-miR-141-3p | SF3B3 | HITS-CLIP | Tarbase |
| hsa-miR-141-3p | SF3B3 | HITS-CLIP | Tarbase |
| hsa-miR-141-3p | SF3B3 | HITS-CLIP | Tarbase |
| hsa-miR-15a-5p | SF3B3 | PAR-CLIP | Tarbase |
| hsa-miR-15a-5p | CLDN4 | HITS-CLIP | Tarbase |
| hsa-miR-195-5p | CLDN4 | HITS-CLIP | Tarbase |
| hsa-miR-195-3p | ZNF33A | HITS-CLIP | Tarbase |
| hsa-miR-486-5p | ZNF600 | PAR-CLIP | Tarbase |
| hsa-miR-486-5p | ZNF600 | PAR-CLIP | Tarbase |
| hsa-miR-195-5p | TSPYL1 | Microarrays | Tarbase |
| hsa-miR-15a-3p | PNRC2 | HITS-CLIP | Tarbase |

| | | | |
|---|---|---|---|
| hsa-miR-195-5p | FAM180A | Microarrays | Tarbase |
| hsa-miR-15a-5p | FAM179A | HITS-CLIP | Tarbase |
| hsa-miR-195-5p | FAM179A | HITS-CLIP | Tarbase |
| hsa-miR-15a-5p | PTPRT | HITS-CLIP | Tarbase |
| hsa-miR-195-5p | PTPRT | HITS-CLIP | Tarbase |
| hsa-miR-195-5p | AKR1C3 | Microarrays | Tarbase |
| hsa-miR-15a-5p | WDSUB1 | PAR-CLIP | Tarbase |
| hsa-miR-15a-5p | WDSUB1 | HITS-CLIP | Tarbase |
| hsa-miR-195-5p | WDSUB1 | HITS-CLIP | Tarbase |
| hsa-miR-141-3p | ACADSB | HITS-CLIP | Tarbase |
| hsa-miR-15a-5p | ACADSB | PAR-CLIP | Tarbase |
| hsa-miR-15a-5p | GREB1 | HITS-CLIP | Tarbase |
| hsa-miR-195-5p | GREB1 | HITS-CLIP | Tarbase |
| hsa-miR-15a-5p | SRGAP3 | HITS-CLIP | Tarbase |
| hsa-miR-195-5p | SRGAP3 | HITS-CLIP | Tarbase |
| hsa-miR-15a-5p | FAM217B | HITS-CLIP | Tarbase |
| hsa-miR-195-5p | FAM217B | HITS-CLIP | Tarbase |
| hsa-miR-141-3p | TUBB | HITS-CLIP | Tarbase |
| hsa-miR-15a-5p | TUBB | PAR-CLIP | Tarbase |
| hsa-miR-15a-5p | TUBB | PAR-CLIP | Tarbase |
| hsa-miR-15a-5p | TUBB | HITS-CLIP | Tarbase |
| hsa-miR-486-3p | TUBB | PAR-CLIP | Tarbase |
| hsa-miR-141-5p | LCOR | HITS-CLIP | Tarbase |
| hsa-miR-15a-5p | LCOR | HITS-CLIP | Tarbase |
| hsa-miR-15a-5p | LCOR | CLEAR-CLIP | Tarbase |
| hsa-miR-15a-5p | XPNPEP3 | HITS-CLIP | Tarbase |
| hsa-miR-15a-5p | XPNPEP3 | HITS-CLIP | Tarbase |
| hsa-miR-195-5p | XPNPEP3 | HITS-CLIP | Tarbase |
| hsa-miR-195-5p | XPNPEP3 | HITS-CLIP | Tarbase |
| hsa-miR-219a-1-3p | ZNF107 | HITS-CLIP | Tarbase |
| hsa-miR-219a-1-3p | ZNF107 | HITS-CLIP | Tarbase |
| hsa-miR-141-5p | IARS | HITS-CLIP | Tarbase |
| hsa-miR-141-5p | IARS | HITS-CLIP | Tarbase |
| hsa-miR-15a-5p | IARS | HITS-CLIP | Tarbase |
| hsa-miR-15a-5p | IARS | HITS-CLIP | Tarbase |
| hsa-miR-141-3p | ZBTB44 | HITS-CLIP | Tarbase |
| hsa-miR-141-5p | CD55 | HITS-CLIP | Tarbase |
| hsa-miR-15a-5p | TRRAP | PAR-CLIP | Tarbase |
| hsa-miR-15a-5p | TRRAP | PAR-CLIP | Tarbase |
| hsa-miR-15a-5p | TRRAP | PAR-CLIP | Tarbase |
| hsa-miR-15a-5p | TRRAP | PAR-CLIP | Tarbase |
| hsa-miR-15a-5p | TRRAP | PAR-CLIP | Tarbase |
| hsa-miR-195-5p | TRRAP | PAR-CLIP | Tarbase |
| hsa-miR-195-5p | TRRAP | PAR-CLIP | Tarbase |
| hsa-miR-15a-5p | TRRAP | Chimeric fragments | Tarbase |
| hsa-miR-15a-5p | FUT4 | HITS-CLIP | Tarbase |
| hsa-miR-141-3p | ASB13 | HITS-CLIP | Tarbase |
| hsa-miR-15a-3p | NBPF4 | HITS-CLIP | Tarbase |
| hsa-miR-15a-5p | TSC22D2 | PAR-CLIP | Tarbase |
| hsa-miR-15a-5p | TSC22D2 | HITS-CLIP | Tarbase |
| hsa-miR-15a-5p | TSC22D2 | PAR-CLIP | Tarbase |
| hsa-miR-195-5p | TSC22D2 | PAR-CLIP | Tarbase |
| hsa-miR-486-5p | TSC22D2 | HITS-CLIP | Tarbase |
| hsa-miR-141-5p | ARMCX4 | HITS-CLIP | Tarbase |
| hsa-miR-15a-5p | YRDC | PAR-CLIP | Tarbase |
| hsa-miR-15a-5p | YRDC | HITS-CLIP | Tarbase |
| hsa-miR-15a-5p | YRDC | PAR-CLIP | Tarbase |
| hsa-miR-15a-5p | YRDC | PAR-CLIP | Tarbase |
| hsa-miR-15a-5p | YRDC | HITS-CLIP | Tarbase |
| hsa-miR-195-5p | YRDC | HITS-CLIP | Tarbase |
| hsa-miR-195-5p | RFX8 | Microarrays | Tarbase |
| hsa-miR-141-3p | SIAH1 | HITS-CLIP | Tarbase |
| hsa-miR-15a-5p | SIAH1 | HITS-CLIP | Tarbase |
| hsa-miR-195-5p | SIAH1 | HITS-CLIP | Tarbase |
| hsa-miR-141-3p | ESRRG | HITS-CLIP | Tarbase |
| hsa-miR-15a-5p | NCOR2 | HITS-CLIP | Tarbase |
| hsa-miR-15a-5p | NCOR2 | HITS-CLIP | Tarbase |
| hsa-miR-195-5p | NCOR2 | HITS-CLIP | Tarbase |
| hsa-miR-3605-5p | NCOR2 | HITS-CLIP | Tarbase |
| hsa-miR-3613-3p | NCOR2 | HITS-CLIP | Tarbase |
| hsa-miR-15a-5p | PRPF40A | HITS-CLIP | Tarbase |
| hsa-miR-195-5p | PRPF40A | HITS-CLIP | Tarbase |
| hsa-miR-15a-5p | GDAP2 | HITS-CLIP | Tarbase |
| hsa-miR-141-3p | ANAPC7 | HITS-CLIP | Tarbase |
| hsa-miR-141-3p | SLC6A9 | HITS-CLIP | Tarbase |
| hsa-miR-15a-5p | AFAP1 | PAR-CLIP | Tarbase |
| hsa-miR-195-5p | AFAP1 | Microarrays | Tarbase |
| hsa-miR-141-3p | SPTSSB | HITS-CLIP | Tarbase |
| hsa-miR-15a-5p | SPTSSB | PAR-CLIP | Tarbase |
| hsa-miR-15a-5p | MME | HITS-CLIP | Tarbase |
| hsa-miR-195-5p | MME | HITS-CLIP | Tarbase |
| hsa-miR-141-3p | CACNA1H | HITS-CLIP | Tarbase |
| hsa-miR-15a-5p | PLXNB2 | HITS-CLIP | Tarbase |
| hsa-miR-195-5p | PLXNB2 | HITS-CLIP | Tarbase |
| hsa-miR-195-5p | XRCC2 | Microarrays | Tarbase |
| hsa-miR-141-5p | HDAC2 | HITS-CLIP | Tarbase |
| hsa-miR-195-5p | ADH1B | Microarrays | Tarbase |
| hsa-miR-141-5p | HRH1 | HITS-CLIP | Tarbase |
| hsa-miR-195-5p | HRH1 | Microarrays | Tarbase |
| hsa-miR-15a-3p | RABL6 | HITS-CLIP | Tarbase |
| hsa-miR-15a-5p | RABL6 | HITS-CLIP | Tarbase |
| hsa-miR-15a-5p | RABL6 | HITS-CLIP | Tarbase |
| hsa-miR-195-5p | RABL6 | HITS-CLIP | Tarbase |
| hsa-miR-195-5p | RABL6 | HITS-CLIP | Tarbase |
| hsa-miR-15a-5p | TTC30B | PAR-CLIP | Tarbase |
| hsa-miR-195-5p | TTC30B | PAR-CLIP | Tarbase |
| hsa-miR-141-5p | TOMM7 | HITS-CLIP | Tarbase |
| hsa-miR-15a-5p | ZNF33B | PAR-CLIP | Tarbase |
| hsa-miR-141-3p | ZNF512B | HITS-CLIP | Tarbase |
| hsa-miR-141-5p | ZNF512B | HITS-CLIP | Tarbase |
| hsa-miR-15a-3p | ZNF512B | HITS-CLIP | Tarbase |
| hsa-miR-219a-1-3p | ZNF431 | HITS-CLIP | Tarbase |
| hsa-miR-3605-5p | ZNF431 | HITS-CLIP | Tarbase |
| hsa-miR-15a-5p | NF1 | HITS-CLIP | Tarbase |
| hsa-miR-15a-5p | NF1 | HITS-CLIP | Tarbase |
| hsa-miR-15a-5p | NF1 | PAR-CLIP | Tarbase |
| hsa-miR-15a-5p | NF1 | HITS-CLIP | Tarbase |
| hsa-miR-15a-5p | NF1 | HITS-CLIP | Tarbase |
| hsa-miR-15a-5p | NF1 | HITS-CLIP | Tarbase |
| hsa-miR-195-5p | NF1 | HITS-CLIP | Tarbase |
| hsa-miR-195-5p | NF1 | HITS-CLIP | Tarbase |
| hsa-miR-195-5p | NF1 | HITS-CLIP | Tarbase |
| hsa-miR-486-5p | NF1 | HITS-CLIP | Tarbase |
| hsa-miR-486-5p | NF1 | HITS-CLIP | Tarbase |
| hsa-miR-15a-5p | NF1 | HITS-CLIP | Tarbase |
| hsa-miR-195-5p | NF1 | HITS-CLIP | Tarbase |
| hsa-miR-141-3p | VKORC1L1 | HITS-CLIP | Tarbase |
| hsa-miR-141-3p | VKORC1L1 | HITS-CLIP | Tarbase |
| hsa-miR-15a-5p | HLA-DQA1 | PAR-CLIP | Tarbase |
| hsa-miR-195-5p | HLA-DQA1 | PAR-CLIP | Tarbase |
| hsa-miR-3605-5p | GM2A | HITS-CLIP | Tarbase |
| hsa-miR-141-3p | MVB12B | HITS-CLIP | Tarbase |
| hsa-miR-15a-3p | C6orf106 | HITS-CLIP | Tarbase |
| hsa-miR-141-3p | NHLRC2 | HITS-CLIP | Tarbase |
| hsa-miR-141-3p | NHLRC2 | HITS-CLIP | Tarbase |
| hsa-miR-15a-5p | NHLRC2 | HITS-CLIP | Tarbase |
| hsa-miR-195-5p | NHLRC2 | HITS-CLIP | Tarbase |
| hsa-miR-195-5p | HIST3H2BB | Microarrays | Tarbase |
| hsa-miR-15a-5p | KPNA5 | HITS-CLIP | Tarbase |
| hsa-miR-195-5p | KPNA5 | HITS-CLIP | Tarbase |
| hsa-miR-3605-5p | ARHGEF12 | HITS-CLIP | Tarbase |
| hsa-miR-15a-5p | ARHGEF12 | HITS-CLIP | Tarbase |
| hsa-miR-195-5p | ARHGEF12 | HITS-CLIP | Tarbase |
| hsa-miR-3605-3p | PDLIM7 | HITS-CLIP | Tarbase |
| hsa-miR-195-5p | PDLIM7 | Microarrays | Tarbase |
| hsa-miR-15a-3p | FLNA | HITS-CLIP | Tarbase |
| hsa-miR-3613-5p | FLNA | HITS-CLIP | Tarbase |
| hsa-miR-141-3p | SRGAP1 | HITS-CLIP | Tarbase |
| hsa-miR-141-3p | NOP9 | HITS-CLIP | Tarbase |
| hsa-miR-486-5p | NOP9 | HITS-CLIP | Tarbase |
| hsa-miR-15a-5p | SLC39A10 | PAR-CLIP | Tarbase |
| hsa-miR-219a-1-3p | SLC39A10 | HITS-CLIP | Tarbase |
| hsa-miR-15a-5p | SLC39A10 | HITS-CLIP | Tarbase |
| hsa-miR-195-5p | SLC39A10 | HITS-CLIP | Tarbase |
| hsa-miR-15a-5p | AP2A1 | HITS-CLIP | Tarbase |
| hsa-miR-195-5p | AP2A1 | Microarrays | Tarbase |
| hsa-miR-15a-5p | ZNF585A | PAR-CLIP | Tarbase |
| hsa-miR-15a-5p | ANXA4 | PAR-CLIP | Tarbase |
| hsa-miR-15a-5p | ANXA4 | PAR-CLIP | Tarbase |

| | | | |
|---|---|---|---|
| hsa-miR-15a-5p | ANXA4 | PAR-CLIP | Tarbase |
| hsa-miR-195-5p | ANXA4 | PAR-CLIP | Tarbase |
| hsa-miR-195-5p | ANXA4 | PAR-CLIP | Tarbase |
| hsa-miR-219a-1-3p | ANXA4 | PAR-CLIP | Tarbase |
| hsa-miR-141-3p | METTL9 | HITS-CLIP | Tarbase |
| hsa-miR-141-3p | METTL9 | HITS-CLIP | Tarbase |
| hsa-miR-15a-5p | METTL9 | HITS-CLIP | Tarbase |
| hsa-miR-15a-5p | METTL9 | HITS-CLIP | Tarbase |
| hsa-miR-195-5p | METTL9 | HITS-CLIP | Tarbase |
| hsa-miR-486-5p | ZNF100 | PAR-CLIP | Tarbase |
| hsa-miR-15a-5p | GMFB | PAR-CLIP | Tarbase |
| hsa-miR-15a-5p | HIST1H4C | HITS-CLIP | Tarbase |
| hsa-miR-195-5p | HIST1H4C | HITS-CLIP | Tarbase |
| hsa-miR-15a-3p | MAFG | HITS-CLIP | Tarbase |
| hsa-miR-15a-5p | MAFG | HITS-CLIP | Tarbase |
| hsa-miR-195-5p | MAFG | HITS-CLIP | Tarbase |
| hsa-miR-141-3p | DYNC1H1 | HITS-CLIP | Tarbase |
| hsa-miR-141-5p | DYNC1H1 | HITS-CLIP | Tarbase |
| hsa-miR-15a-5p | DYNC1H1 | HITS-CLIP | Tarbase |
| hsa-miR-15a-5p | DYNC1H1 | HITS-CLIP | Tarbase |
| hsa-miR-195-5p | DYNC1H1 | HITS-CLIP | Tarbase |
| hsa-miR-15a-5p | DYNC1H1 | HITS-CLIP | Tarbase |
| hsa-miR-195-5p | DYNC1H1 | HITS-CLIP | Tarbase |
| hsa-miR-141-5p | PCBP2 | HITS-CLIP | Tarbase |
| hsa-miR-195-5p | PGAP1 | Microarrays | Tarbase |
| hsa-miR-141-5p | ZNF257 | HITS-CLIP | Tarbase |
| hsa-miR-141-5p | LRRC8B | HITS-CLIP | Tarbase |
| hsa-miR-141-5p | LRRC8B | HITS-CLIP | Tarbase |
| hsa-miR-141-5p | LRRC8B | HITS-CLIP | Tarbase |
| hsa-miR-15a-5p | ABCB8 | HITS-CLIP | Tarbase |
| hsa-miR-15a-3p | SND1 | HITS-CLIP | Tarbase |
| hsa-miR-15a-5p | MIRLET7BHG | PAR-CLIP | Tarbase |
| hsa-miR-15a-5p | NOL4L | PAR-CLIP | Tarbase |
| hsa-miR-15a-5p | NOL4L | HITS-CLIP | Tarbase |
| hsa-miR-15a-5p | NOL4L | PAR-CLIP | Tarbase |
| hsa-miR-195-5p | NOL4L | PAR-CLIP | Tarbase |
| hsa-miR-195-5p | NOL4L | HITS-CLIP | Tarbase |
| hsa-miR-195-5p | NOL4L | PAR-CLIP | Tarbase |
| hsa-miR-195-5p | NOL4L | Microarrays | Tarbase |
| hsa-miR-141-3p | ENTPD4 | HITS-CLIP | Tarbase |
| hsa-miR-15a-5p | ENTPD4 | HITS-CLIP | Tarbase |
| hsa-miR-15a-5p | ENTPD4 | HITS-CLIP | Tarbase |
| hsa-miR-15a-5p | ENTPD4 | PAR-CLIP | Tarbase |
| hsa-miR-15a-5p | ENTPD4 | HITS-CLIP | Tarbase |
| hsa-miR-195-5p | ENTPD4 | HITS-CLIP | Tarbase |
| hsa-miR-195-5p | ENTPD4 | HITS-CLIP | Tarbase |
| hsa-miR-195-5p | ENTPD4 | HITS-CLIP | Tarbase |
| hsa-miR-15a-5p | SERPINA1 | HITS-CLIP | Tarbase |
| hsa-miR-195-5p | SERPINA1 | HITS-CLIP | Tarbase |
| hsa-miR-15a-5p | KANK2 | HITS-CLIP | Tarbase |
| hsa-miR-195-5p | KANK2 | HITS-CLIP | Tarbase |
| hsa-miR-15a-5p | SYNGAP1 | PAR-CLIP | Tarbase |
| hsa-miR-195-5p | FITM2 | HITS-CLIP | Tarbase |
| hsa-miR-195-5p | BLM | Microarrays | Tarbase |
| hsa-miR-141-5p | DDI2 | HITS-CLIP | Tarbase |
| hsa-miR-15a-5p | DDI2 | HITS-CLIP | Tarbase |
| hsa-miR-15a-5p | DDI2 | HITS-CLIP | Tarbase |
| hsa-miR-195-5p | DDI2 | HITS-CLIP | Tarbase |
| hsa-miR-15a-5p | SVIL | HITS-CLIP | Tarbase |
| hsa-miR-195-5p | SVIL | HITS-CLIP | Tarbase |
| hsa-miR-15a-5p | C17orf102 | PAR-CLIP | Tarbase |
| hsa-miR-141-3p | TRIM33 | HITS-CLIP | Tarbase |
| hsa-miR-15a-5p | TRIM33 | HITS-CLIP | Tarbase |
| hsa-miR-195-5p | TRIM33 | HITS-CLIP | Tarbase |
| hsa-miR-3613-5p | TRIM33 | PAR-CLIP | Tarbase |
| hsa-miR-141-5p | PELI1 | HITS-CLIP | Tarbase |
| hsa-miR-3613-3p | PELI1 | PAR-CLIP | Tarbase |
| hsa-miR-3613-3p | PELI1 | PAR-CLIP | Tarbase |
| hsa-miR-195-5p | PELI1 | Microarrays | Tarbase |
| hsa-miR-195-5p | ADARB1 | Microarrays | Tarbase |
| hsa-miR-141-3p | HTT | HITS-CLIP | Tarbase |
| hsa-miR-15a-5p | HTT | HITS-CLIP | Tarbase |
| hsa-miR-15a-5p | HTT | HITS-CLIP | Tarbase |
| hsa-miR-195-5p | HTT | HITS-CLIP | Tarbase |
| hsa-miR-3613-3p | HTT | HITS-CLIP | Tarbase |
| hsa-miR-15a-5p | HTT | HITS-CLIP | Tarbase |
| hsa-miR-195-5p | HTT | HITS-CLIP | Tarbase |
| hsa-miR-15a-5p | HIST1H3D | HITS-CLIP | Tarbase |
| hsa-miR-195-5p | MAP3K5 | Microarrays | Tarbase |
| hsa-miR-141-3p | HNRNPAB | HITS-CLIP | Tarbase |
| hsa-miR-15a-5p | PDGFA | HITS-CLIP | Tarbase |
| hsa-miR-195-5p | PDGFA | HITS-CLIP | Tarbase |
| hsa-miR-15a-5p | MYO5A | HITS-CLIP | Tarbase |
| hsa-miR-15a-5p | MYO5A | PAR-CLIP | Tarbase |
| hsa-miR-195-5p | MYO5A | HITS-CLIP | Tarbase |
| hsa-miR-15a-5p | MYO5A | HITS-CLIP | Tarbase |
| hsa-miR-195-5p | MYO5A | HITS-CLIP | Tarbase |
| hsa-miR-141-3p | SIPA1L1 | HITS-CLIP | Tarbase |
| hsa-miR-15a-5p | SIPA1L1 | HITS-CLIP | Tarbase |
| hsa-miR-195-5p | SIPA1L1 | Microarrays | Tarbase |
| hsa-miR-141-3p | TTC30A | HITS-CLIP | Tarbase |
| hsa-miR-195-5p | RAB40C | HITS-CLIP | Tarbase |
| hsa-miR-15a-5p | ENTPD6 | HITS-CLIP | Tarbase |
| hsa-miR-195-5p | ENTPD6 | HITS-CLIP | Tarbase |
| hsa-miR-195-5p | ENTPD6 | Microarrays | Tarbase |
| hsa-miR-195-5p | ENPP1 | Microarrays | Tarbase |
| hsa-miR-141-5p | FAR1 | HITS-CLIP | Tarbase |
| hsa-miR-195-5p | CXorf40A | AGO-IP | Tarbase |
| hsa-miR-141-3p | CDC42SE1 | PAR-CLIP | Tarbase |
| hsa-miR-15a-5p | CDC42SE1 | PAR-CLIP | Tarbase |
| hsa-miR-219a-1-3p | PDCD1LG2 | HITS-CLIP | Tarbase |
| hsa-miR-195-5p | PDCD1LG2 | Microarrays | Tarbase |
| hsa-miR-141-5p | SPTAN1 | HITS-CLIP | Tarbase |
| hsa-miR-195-3p | FAM114A1 | HITS-CLIP | Tarbase |
| hsa-miR-141-3p | PHF2 | HITS-CLIP | Tarbase |
| hsa-miR-141-5p | PSAP | HITS-CLIP | Tarbase |
| hsa-miR-15a-5p | PSAP | HITS-CLIP | Tarbase |
| hsa-miR-195-5p | PSAP | HITS-CLIP | Tarbase |
| hsa-miR-486-3p | S100A10 | HITS-CLIP | Tarbase |
| hsa-miR-15a-3p | RPL37A | HITS-CLIP | Tarbase |
| hsa-miR-15a-3p | CFD | HITS-CLIP | Tarbase |
| hsa-miR-141-5p | EME2 | HITS-CLIP | Tarbase |
| hsa-miR-195-3p | KLHDC1 | HITS-CLIP | Tarbase |
| hsa-miR-141-3p | OCLN | HITS-CLIP | Tarbase |
| hsa-miR-15a-5p | ZNF181 | HITS-CLIP | Tarbase |
| hsa-miR-195-5p | ZNF181 | HITS-CLIP | Tarbase |
| hsa-miR-15a-5p | GPAA1 | HITS-CLIP | Tarbase |
| hsa-miR-15a-5p | GPAA1 | HITS-CLIP | Tarbase |
| hsa-miR-195-5p | GPAA1 | HITS-CLIP | Tarbase |
| hsa-miR-195-5p | GPAA1 | Microarrays | Tarbase |
| hsa-miR-141-3p | ADH5 | HITS-CLIP | Tarbase |
| hsa-miR-141-3p | ERO1L | HITS-CLIP | Tarbase |
| hsa-miR-141-3p | ERO1L | HITS-CLIP | Tarbase |
| hsa-miR-3605-5p | ZNF71 | PAR-CLIP | Tarbase |
| hsa-miR-141-3p | ZNF121 | HITS-CLIP | Tarbase |
| hsa-miR-3613-3p | MPZL1 | PAR-CLIP | Tarbase |
| hsa-miR-15a-5p | VPS13A | HITS-CLIP | Tarbase |
| hsa-miR-195-5p | VPS13A | HITS-CLIP | Tarbase |
| hsa-miR-15a-5p | MBP | HITS-CLIP | Tarbase |
| hsa-miR-195-5p | MBP | HITS-CLIP | Tarbase |
| hsa-miR-15a-3p | C1orf122 | HITS-CLIP | Tarbase |
| hsa-miR-15a-5p | NOL8 | PAR-CLIP | Tarbase |
| hsa-miR-219a-1-3p | ENTPD7 | HITS-CLIP | Tarbase |
| hsa-miR-141-3p | ZNF335 | HITS-CLIP | Tarbase |
| hsa-miR-15a-3p | RPS4X | HITS-CLIP | Tarbase |
| hsa-miR-141-5p | SIRPA | HITS-CLIP | Tarbase |
| hsa-miR-195-5p | DSCR8 | Microarrays | Tarbase |
| hsa-miR-141-5p | GRK6 | HITS-CLIP | Tarbase |
| hsa-miR-195-5p | PRIM1 | Microarrays | Tarbase |
| hsa-miR-15a-5p | MARCH5 | HITS-CLIP | Tarbase |
| hsa-miR-195-5p | MARCH5 | HITS-CLIP | Tarbase |
| hsa-miR-195-5p | AKR1B10 | Microarrays | Tarbase |
| hsa-miR-15a-5p | ZBTB14 | HITS-CLIP | Tarbase |
| hsa-miR-195-5p | ZBTB14 | HITS-CLIP | Tarbase |
| hsa-miR-3613-3p | ZBTB14 | PAR-CLIP | Tarbase |
| hsa-miR-141-3p | CD2AP | HITS-CLIP | Tarbase |
| hsa-miR-15a-5p | CD2AP | PAR-CLIP | Tarbase |
| hsa-miR-15a-5p | CD2AP | PAR-CLIP | Tarbase |
| hsa-miR-15a-5p | CD2AP | PAR-CLIP | Tarbase |

| | | | |
|---|---|---|---|
| hsa-miR-15a-5p | CD2AP | PAR-CLIP | Tarbase |
| hsa-miR-15a-5p | CD2AP | PAR-CLIP | Tarbase |
| hsa-miR-15a-5p | CD2AP | HITS-CLIP | Tarbase |
| hsa-miR-15a-5p | CD2AP | PAR-CLIP | Tarbase |
| hsa-miR-195-5p | CD2AP | PAR-CLIP | Tarbase |
| hsa-miR-195-5p | CD2AP | HITS-CLIP | Tarbase |
| hsa-miR-195-5p | CD2AP | PAR-CLIP | Tarbase |
| hsa-miR-15a-5p | CD2AP | Chimeric fragments | Tarbase |
| hsa-miR-15a-3p | SFI1 | HITS-CLIP | Tarbase |
| hsa-miR-141-3p | ZNF248 | HITS-CLIP | Tarbase |
| hsa-miR-195-5p | HMGN5 | Microarrays | Tarbase |
| hsa-miR-141-5p | MAN1A2 | HITS-CLIP | Tarbase |
| hsa-miR-15a-5p | MAN1A2 | HITS-CLIP | Tarbase |
| hsa-miR-195-5p | MAN1A2 | HITS-CLIP | Tarbase |
| hsa-miR-15a-5p | SVIP | PAR-CLIP | Tarbase |
| hsa-miR-15a-5p | SVIP | PAR-CLIP | Tarbase |
| hsa-miR-15a-5p | SVIP | PAR-CLIP | Tarbase |
| hsa-miR-15a-5p | SVIP | PAR-CLIP | Tarbase |
| hsa-miR-195-5p | SVIP | PAR-CLIP | Tarbase |
| hsa-miR-195-5p | SVIP | PAR-CLIP | Tarbase |
| hsa-miR-15a-5p | SVIP | HITS-CLIP | Tarbase |
| hsa-miR-195-5p | SVIP | HITS-CLIP | Tarbase |
| hsa-miR-141-5p | DDRGK1 | HITS-CLIP | Tarbase |
| hsa-miR-195-5p | HSD17B11 | Microarrays | Tarbase |
| hsa-miR-486-5p | DDX42 | HITS-CLIP | Tarbase |
| hsa-miR-15a-3p | SLC29A3 | HITS-CLIP | Tarbase |
| hsa-miR-15a-5p | STYX | PAR-CLIP | Tarbase |
| hsa-miR-15a-5p | STYX | HITS-CLIP | Tarbase |
| hsa-miR-195-5p | STYX | HITS-CLIP | Tarbase |
| hsa-miR-3605-5p | UBL5 | HITS-CLIP | Tarbase |
| hsa-miR-15a-5p | UCKL1 | HITS-CLIP | Tarbase |
| hsa-miR-195-5p | UCKL1 | HITS-CLIP | Tarbase |
| hsa-miR-3613-3p | PEG3 | HITS-CLIP | Tarbase |
| hsa-miR-141-3p | ZKSCAN8 | PAR-CLIP | Tarbase |
| hsa-miR-141-3p | ZNF442 | HITS-CLIP | Tarbase |
| hsa-miR-3605-3p | PIM3 | HITS-CLIP | Tarbase |
| hsa-miR-3613-3p | PIM3 | HITS-CLIP | Tarbase |
| hsa-miR-141-5p | ASPH | HITS-CLIP | Tarbase |
| hsa-miR-3613-5p | ASPH | HITS-CLIP | Tarbase |
| hsa-miR-195-5p | WWP2 | Microarrays | Tarbase |
| hsa-miR-141-5p | GFPT1 | HITS-CLIP | Tarbase |
| hsa-miR-141-5p | GFPT1 | HITS-CLIP | Tarbase |
| hsa-miR-15a-5p | GFPT1 | PAR-CLIP | Tarbase |
| hsa-miR-15a-5p | GFPT1 | PAR-CLIP | Tarbase |
| hsa-miR-15a-5p | GFPT1 | PAR-CLIP | Tarbase |
| hsa-miR-15a-5p | GFPT1 | HITS-CLIP | Tarbase |
| hsa-miR-15a-5p | GFPT1 | PAR-CLIP | Tarbase |
| hsa-miR-15a-5p | GFPT1 | PAR-CLIP | Tarbase |
| hsa-miR-195-5p | GFPT1 | PAR-CLIP | Tarbase |
| hsa-miR-195-5p | GFPT1 | HITS-CLIP | Tarbase |
| hsa-miR-195-5p | GFPT1 | PAR-CLIP | Tarbase |
| hsa-miR-3613-3p | FAM115A | HITS-CLIP | Tarbase |
| hsa-miR-15a-5p | FAM115A | Chimeric fragments | Tarbase |
| hsa-miR-141-3p | TXNRD1 | PAR-CLIP | Tarbase |
| hsa-miR-15a-5p | SH3BGRL2 | HITS-CLIP | Tarbase |
| hsa-miR-195-5p | SH3BGRL2 | CLASH | Tarbase |
| hsa-miR-195-5p | SH3BGRL2 | HITS-CLIP | Tarbase |
| hsa-miR-15a-5p | SH3BGRL2 | HITS-CLIP | Tarbase |
| hsa-miR-195-5p | SH3BGRL2 | HITS-CLIP | Tarbase |
| hsa-miR-15a-5p | ATL1 | HITS-CLIP | Tarbase |
| hsa-miR-195-5p | ATL1 | HITS-CLIP | Tarbase |
| hsa-miR-15a-5p | MAFK | PAR-CLIP | Tarbase |
| hsa-miR-15a-5p | MAFK | PAR-CLIP | Tarbase |
| hsa-miR-15a-5p | MAFK | PAR-CLIP | Tarbase |
| hsa-miR-15a-5p | MAFK | PAR-CLIP | Tarbase |
| hsa-miR-195-5p | MAFK | PAR-CLIP | Tarbase |
| hsa-miR-195-5p | MAFK | PAR-CLIP | Tarbase |
| hsa-miR-15a-5p | ZNF43 | PAR-CLIP | Tarbase |
| hsa-miR-15a-5p | ZNF43 | PAR-CLIP | Tarbase |
| hsa-miR-195-5p | ZNF43 | PAR-CLIP | Tarbase |
| hsa-miR-15a-3p | ZNF511 | HITS-CLIP | Tarbase |
| hsa-miR-15a-5p | ZNF627 | HITS-CLIP | Tarbase |
| hsa-miR-195-5p | ZNF627 | HITS-CLIP | Tarbase |
| hsa-miR-15a-5p | WDHD1 | PAR-CLIP | Tarbase |
| hsa-miR-195-5p | WDHD1 | Microarrays | Tarbase |
| hsa-miR-15a-3p | CTNND1 | HITS-CLIP | Tarbase |
| hsa-miR-141-5p | DDX39B | HITS-CLIP | Tarbase |
| hsa-miR-141-5p | DDX39B | HITS-CLIP | Tarbase |
| hsa-miR-195-5p | SH2D1B | Microarrays | Tarbase |
| hsa-miR-15a-5p | TLK1 | PAR-CLIP | Tarbase |
| hsa-miR-15a-5p | TLK1 | PAR-CLIP | Tarbase |
| hsa-miR-15a-5p | TLK1 | PAR-CLIP | Tarbase |
| hsa-miR-15a-5p | TLK1 | HITS-CLIP | Tarbase |
| hsa-miR-15a-5p | TLK1 | PAR-CLIP | Tarbase |
| hsa-miR-195-5p | TLK1 | PAR-CLIP | Tarbase |
| hsa-miR-195-5p | TLK1 | HITS-CLIP | Tarbase |
| hsa-miR-15a-5p | TLK1 | HITS-CLIP | Tarbase |
| hsa-miR-195-5p | TLK1 | HITS-CLIP | Tarbase |
| hsa-miR-15a-5p | LRBA | HITS-CLIP | Tarbase |
| hsa-miR-195-5p | LRBA | HITS-CLIP | Tarbase |
| hsa-miR-141-5p | BAZ1A | HITS-CLIP | Tarbase |
| hsa-miR-15a-5p | KLHL9 | PAR-CLIP | Tarbase |
| hsa-miR-15a-3p | NCOA6 | HITS-CLIP | Tarbase |
| hsa-miR-141-3p | C6orf89 | HITS-CLIP | Tarbase |
| hsa-miR-15a-5p | CALM1 | HITS-CLIP | Tarbase |
| hsa-miR-15a-5p | CALM1 | HITS-CLIP | Tarbase |
| hsa-miR-15a-5p | CALM1 | HITS-CLIP | Tarbase |
| hsa-miR-15a-5p | CALM1 | HITS-CLIP | Tarbase |
| hsa-miR-195-5p | CALM1 | HITS-CLIP | Tarbase |
| hsa-miR-195-5p | CALM1 | HITS-CLIP | Tarbase |
| hsa-miR-195-5p | CALM1 | HITS-CLIP | Tarbase |
| hsa-miR-195-5p | FAM19A2 | Microarrays | Tarbase |
| hsa-miR-141-3p | TTC37 | HITS-CLIP | Tarbase |
| hsa-miR-195-5p | SLC9A6 | Microarrays | Tarbase |
| hsa-miR-15a-3p | IPO9 | HITS-CLIP | Tarbase |
| hsa-miR-15a-5p | IPO9 | HITS-CLIP | Tarbase |
| hsa-miR-15a-5p | IPO9 | HITS-CLIP | Tarbase |
| hsa-miR-195-5p | IPO9 | HITS-CLIP | Tarbase |
| hsa-miR-195-5p | IPO9 | HITS-CLIP | Tarbase |
| hsa-miR-3613-5p | IPO9 | HITS-CLIP | Tarbase |
| hsa-miR-3613-5p | IPO9 | PAR-CLIP | Tarbase |
| hsa-miR-15a-5p | C1orf85 | HITS-CLIP | Tarbase |
| hsa-miR-15a-5p | C1orf85 | HITS-CLIP | Tarbase |
| hsa-miR-195-5p | C1orf85 | HITS-CLIP | Tarbase |
| hsa-miR-15a-5p | DLL1 | HITS-CLIP | Tarbase |
| hsa-miR-15a-5p | DLL1 | HITS-CLIP | Tarbase |
| hsa-miR-15a-5p | DLL1 | PAR-CLIP | Tarbase |
| hsa-miR-195-5p | DLL1 | HITS-CLIP | Tarbase |
| hsa-miR-195-5p | DLL1 | HITS-CLIP | Tarbase |
| hsa-miR-15a-3p | ANKRD13B | HITS-CLIP | Tarbase |
| hsa-miR-15a-5p | PPP1R14C | HITS-CLIP | Tarbase |
| hsa-miR-195-5p | PPP1R14C | HITS-CLIP | Tarbase |
| hsa-miR-141-5p | CTR9 | HITS-CLIP | Tarbase |
| hsa-miR-15a-5p | SMOC1 | HITS-CLIP | Tarbase |
| hsa-miR-195-5p | LRRTM3 | Microarrays | Tarbase |
| hsa-miR-15a-5p | ZNF652 | HITS-CLIP | Tarbase |
| hsa-miR-3613-5p | ZNF652 | PAR-CLIP | Tarbase |
| hsa-miR-3613-5p | ZNF652 | PAR-CLIP | Tarbase |
| hsa-miR-15a-5p | SMURF1 | PAR-CLIP | Tarbase |
| hsa-miR-15a-3p | SLC5A3 | HITS-CLIP | Tarbase |
| hsa-miR-15a-5p | SLC5A3 | PAR-CLIP | Tarbase |
| hsa-miR-15a-5p | MT-ND5 | CLEAR-CLIP | Tarbase |
| hsa-miR-15a-5p | TMEM184B | HITS-CLIP | Tarbase |
| hsa-miR-15a-5p | SCAMP5 | HITS-CLIP | Tarbase |
| hsa-miR-195-5p | SCAMP5 | HITS-CLIP | Tarbase |
| hsa-miR-15a-5p | LRIG2 | PAR-CLIP | Tarbase |
| hsa-miR-15a-5p | PNP | PAR-CLIP | Tarbase |
| hsa-miR-195-5p | PAX9 | Microarrays | Tarbase |
| hsa-miR-15a-5p | CHAMP1 | PAR-CLIP | Tarbase |
| hsa-miR-15a-5p | ARHGAP11A | PAR-CLIP | Tarbase |
| hsa-miR-195-5p | ARHGAP11A | Microarrays | Tarbase |
| hsa-miR-15a-5p | SELM | HITS-CLIP | Tarbase |
| hsa-miR-195-5p | SELM | HITS-CLIP | Tarbase |
| hsa-miR-15a-5p | UBE2J1 | Chimeric fragments | Tarbase |
| hsa-miR-15a-5p | DENND4B | PAR-CLIP | Tarbase |
| hsa-miR-195-5p | DENND4B | Microarrays | Tarbase |
| hsa-miR-15a-5p | ZNF277 | PAR-CLIP | Tarbase |
| hsa-miR-195-5p | TOX | Microarrays | Tarbase |
| hsa-miR-219a-1-3p | FICD | PAR-CLIP | Tarbase |

| | | | |
|---|---|---|---|
| hsa-miR-195-5p | TSEN15 | Microarrays | Tarbase |
| hsa-miR-195-5p | GRK5 | Microarrays | Tarbase |
| hsa-miR-15a-5p | SFMBT2 | HITS-CLIP | Tarbase |
| hsa-miR-15a-3p | MT-ND4 | CLEAR-CLIP | Tarbase |
| hsa-miR-15a-5p | MT-ND4 | CLEAR-CLIP | Tarbase |
| hsa-miR-486-3p | PRMT6 | HITS-CLIP | Tarbase |
| hsa-miR-15a-5p | CAPZA2 | PAR-CLIP | Tarbase |
| hsa-miR-15a-5p | CAPZA2 | HITS-CLIP | Tarbase |
| hsa-miR-195-5p | CAPZA2 | HITS-CLIP | Tarbase |
| hsa-miR-141-3p | TOP1 | HITS-CLIP | Tarbase |
| hsa-miR-15a-5p | TOP1 | HITS-CLIP | Tarbase |
| hsa-miR-195-5p | TOP1 | HITS-CLIP | Tarbase |
| hsa-miR-15a-5p | PRC1 | HITS-CLIP | Tarbase |
| hsa-miR-15a-5p | PRC1 | PAR-CLIP | Tarbase |
| hsa-miR-195-5p | PRC1 | Microarrays | Tarbase |
| hsa-miR-195-5p | MAP3K3 | Microarrays | Tarbase |
| hsa-miR-195-5p | L1CAM | Microarrays | Tarbase |
| hsa-miR-141-5p | POU3F3 | HITS-CLIP | Tarbase |
| hsa-miR-15a-5p | ATG9A | PAR-CLIP | Tarbase |
| hsa-miR-15a-5p | ATG9A | PAR-CLIP | Tarbase |
| hsa-miR-15a-5p | ATG9A | PAR-CLIP | Tarbase |
| hsa-miR-15a-5p | ATG9A | HITS-CLIP | Tarbase |
| hsa-miR-15a-5p | ATG9A | PAR-CLIP | Tarbase |
| hsa-miR-15a-5p | ATG9A | PAR-CLIP | Tarbase |
| hsa-miR-15a-5p | ATG9A | PAR-CLIP | Tarbase |
| hsa-miR-15a-5p | ATG9A | HITS-CLIP | Tarbase |
| hsa-miR-195-5p | ATG9A | PAR-CLIP | Tarbase |
| hsa-miR-195-5p | ATG9A | HITS-CLIP | Tarbase |
| hsa-miR-195-5p | ATG9A | PAR-CLIP | Tarbase |
| hsa-miR-195-5p | ATG9A | PAR-CLIP | Tarbase |
| hsa-miR-195-5p | ATG9A | Microarrays | Tarbase |
| hsa-miR-141-5p | TBKBP1 | HITS-CLIP | Tarbase |
| hsa-miR-3605-3p | TBKBP1 | HITS-CLIP | Tarbase |
| hsa-miR-195-5p | CCDC167 | Microarrays | Tarbase |
| hsa-miR-15a-5p | MT-CO3 | CLEAR-CLIP | Tarbase |
| hsa-miR-195-5p | TGM2 | Microarrays | Tarbase |
| hsa-miR-195-5p | PJA2 | Microarrays | Tarbase |
| hsa-miR-195-5p | RORB | Microarrays | Tarbase |
| hsa-miR-486-5p | SGMS1 | HITS-CLIP | Tarbase |
| hsa-miR-195-5p | COX20 | qPCR | Tarbase |
| hsa-miR-195-5p | COX20 | Western Blot | Tarbase |
| hsa-miR-486-5p | C6orf99 | HITS-CLIP | Tarbase |
| hsa-miR-195-5p | CENPW | Microarrays | Tarbase |
| hsa-miR-195-5p | FANK1 | Microarrays | Tarbase |
| hsa-miR-195-5p | PPAPDC1A | Microarrays | Tarbase |
| hsa-miR-195-5p | GDI1 | Microarrays | Tarbase |
| hsa-miR-15a-5p | OOEP | PAR-CLIP | Tarbase |
| hsa-miR-195-5p | OOEP | PAR-CLIP | Tarbase |
| hsa-miR-15a-5p | SYS1 | PAR-CLIP | Tarbase |
| hsa-miR-3613-5p | INPP5B | HITS-CLIP | Tarbase |
| hsa-miR-141-5p | MAFB | HITS-CLIP | Tarbase |
| hsa-miR-141-5p | CHIC1 | HITS-CLIP | Tarbase |
| hsa-miR-15a-5p | PHACTR4 | HITS-CLIP | Tarbase |
| hsa-miR-141-3p | TMEM57 | HITS-CLIP | Tarbase |
| hsa-miR-195-5p | ZDBF2 | Microarrays | Tarbase |
| hsa-miR-141-5p | RING1 | HITS-CLIP | Tarbase |
| hsa-miR-141-5p | BRD2 | HITS-CLIP | Tarbase |
| hsa-miR-141-5p | BRD2 | HITS-CLIP | Tarbase |
| hsa-miR-15a-3p | BRD2 | HITS-CLIP | Tarbase |
| hsa-miR-15a-3p | BRD2 | HITS-CLIP | Tarbase |
| hsa-miR-15a-5p | BRD2 | PAR-CLIP | Tarbase |
| hsa-miR-15a-5p | BRD2 | HITS-CLIP | Tarbase |
| hsa-miR-195-5p | HLA-DMA | Microarrays | Tarbase |
| hsa-miR-15a-3p | COL5A2 | HITS-CLIP | Tarbase |
| hsa-miR-195-5p | COL15A1 | Microarrays | Tarbase |
| hsa-miR-15a-5p | AGPAT1 | HITS-CLIP | Tarbase |
| hsa-miR-15a-5p | AGPAT1 | HITS-CLIP | Tarbase |
| hsa-miR-195-5p | AGPAT1 | HITS-CLIP | Tarbase |
| hsa-miR-15a-5p | AGPAT1 | HITS-CLIP | Tarbase |
| hsa-miR-195-5p | AGPAT1 | HITS-CLIP | Tarbase |
| hsa-miR-141-5p | ERICH2 | HITS-CLIP | Tarbase |
| hsa-miR-195-5p | LAYN | Microarrays | Tarbase |
| hsa-miR-3613-5p | C6orf48 | HITS-CLIP | Tarbase |
| hsa-miR-15a-5p | HSPA1B | PAR-CLIP | Tarbase |
| hsa-miR-15a-5p | HSPA1B | PAR-CLIP | Tarbase |
| hsa-miR-15a-5p | HSPA1B | HITS-CLIP | Tarbase |
| hsa-miR-15a-5p | HSPA1B | HITS-CLIP | Tarbase |
| hsa-miR-195-5p | HSPA1B | HITS-CLIP | Tarbase |
| hsa-miR-195-5p | HSPA1B | HITS-CLIP | Tarbase |
| hsa-miR-195-5p | HSPA1B | HITS-CLIP | Tarbase |
| hsa-miR-3613-3p | HSPA1B | HITS-CLIP | Tarbase |
| hsa-miR-3613-5p | HSPA1B | HITS-CLIP | Tarbase |
| hsa-miR-3613-5p | HSPA1B | HITS-CLIP | Tarbase |
| hsa-miR-15a-5p | HSPA1B | HITS-CLIP | Tarbase |
| hsa-miR-195-5p | HSPA1B | HITS-CLIP | Tarbase |
| hsa-miR-141-3p | HSPA1A | HITS-CLIP | Tarbase |
| hsa-miR-15a-5p | BAG6 | HITS-CLIP | Tarbase |
| hsa-miR-195-5p | BAG6 | HITS-CLIP | Tarbase |
| hsa-miR-141-5p | PRRC2A | HITS-CLIP | Tarbase |
| hsa-miR-486-3p | PRRC2A | HITS-CLIP | Tarbase |
| hsa-miR-15a-5p | NFKBIL1 | PAR-CLIP | Tarbase |
| hsa-miR-141-5p | MICA | HITS-CLIP | Tarbase |
| hsa-miR-15a-5p | MICA | HITS-CLIP | Tarbase |
| hsa-miR-15a-5p | MICA | PAR-CLIP | Tarbase |
| hsa-miR-195-5p | MICA | HITS-CLIP | Tarbase |
| hsa-miR-141-3p | ZNF805 | HITS-CLIP | Tarbase |
| hsa-miR-15a-3p | CCHCR1 | HITS-CLIP | Tarbase |
| hsa-miR-195-5p | C6orf136 | Microarrays | Tarbase |
| hsa-miR-141-3p | ABCF1 | HITS-CLIP | Tarbase |
| hsa-miR-141-3p | ABCF1 | HITS-CLIP | Tarbase |
| hsa-miR-15a-3p | ABCF1 | HITS-CLIP | Tarbase |
| hsa-miR-15a-5p | ABCF1 | HITS-CLIP | Tarbase |
| hsa-miR-195-5p | ABCF1 | HITS-CLIP | Tarbase |
| hsa-miR-195-5p | ABCF1 | Microarrays | Tarbase |
| hsa-miR-15a-5p | PRR3 | PAR-CLIP | Tarbase |
| hsa-miR-15a-5p | PRR3 | PAR-CLIP | Tarbase |
| hsa-miR-15a-5p | PRR3 | PAR-CLIP | Tarbase |
| hsa-miR-15a-5p | PRR3 | PAR-CLIP | Tarbase |
| hsa-miR-15a-5p | PRR3 | PAR-CLIP | Tarbase |
| hsa-miR-15a-5p | PRR3 | PAR-CLIP | Tarbase |
| hsa-miR-195-5p | PRR3 | PAR-CLIP | Tarbase |
| hsa-miR-195-5p | PRR3 | PAR-CLIP | Tarbase |
| hsa-miR-15a-5p | DDR1 | HITS-CLIP | Tarbase |
| hsa-miR-15a-5p | DDR1 | HITS-CLIP | Tarbase |
| hsa-miR-195-5p | DDR1 | HITS-CLIP | Tarbase |
| hsa-miR-3605-5p | DDR1 | HITS-CLIP | Tarbase |
| hsa-miR-15a-5p | PPP1R11 | PAR-CLIP | Tarbase |
| hsa-miR-15a-5p | PPP1R11 | HITS-CLIP | Tarbase |
| hsa-miR-15a-5p | PPP1R11 | HITS-CLIP | Tarbase |
| hsa-miR-15a-5p | PPP1R11 | PAR-CLIP | Tarbase |
| hsa-miR-15a-5p | PPP1R11 | HITS-CLIP | Tarbase |
| hsa-miR-195-5p | PPP1R11 | AGO-IP | Tarbase |
| hsa-miR-195-5p | PPP1R11 | PAR-CLIP | Tarbase |
| hsa-miR-195-5p | PPP1R11 | HITS-CLIP | Tarbase |
| hsa-miR-195-5p | PPP1R11 | HITS-CLIP | Tarbase |
| hsa-miR-195-5p | PPP1R11 | Microarrays | Tarbase |
| hsa-miR-3605-5p | GNB2L1 | HITS-CLIP | Tarbase |
| hsa-miR-3605-3p | AKT1S1 | HITS-CLIP | Tarbase |
| hsa-miR-141-5p | GABBR1 | HITS-CLIP | Tarbase |
| hsa-miR-15a-3p | GABBR1 | HITS-CLIP | Tarbase |
| hsa-miR-15a-5p | GABBR1 | HITS-CLIP | Tarbase |
| hsa-miR-486-3p | TRIM27 | HITS-CLIP | Tarbase |
| hsa-miR-195-5p | TCTN1 | HITS-CLIP | Tarbase |
| hsa-miR-195-5p | FAM216A | Microarrays | Tarbase |
| hsa-miR-141-3p | PSG5 | HITS-CLIP | Tarbase |
| hsa-miR-15a-5p | PCDHA9 | HITS-CLIP | Tarbase |
| hsa-miR-195-5p | PCDHA9 | HITS-CLIP | Tarbase |
| hsa-miR-15a-5p | SPIRE2 | HITS-CLIP | Tarbase |
| hsa-miR-195-5p | SPIRE2 | HITS-CLIP | Tarbase |
| hsa-miR-141-3p | ZBTB10 | HITS-CLIP | Tarbase |
| hsa-miR-141-5p | ZBTB10 | HITS-CLIP | Tarbase |
| hsa-miR-15a-5p | ZBTB10 | HITS-CLIP | Tarbase |
| hsa-miR-195-5p | ZBTB10 | HITS-CLIP | Tarbase |
| hsa-miR-141-3p | LGR4 | HITS-CLIP | Tarbase |
| hsa-miR-141-3p | LGR4 | HITS-CLIP | Tarbase |
| hsa-miR-141-3p | LGR4 | HITS-CLIP | Tarbase |
| hsa-miR-195-5p | VIT | Microarrays | Tarbase |
| hsa-miR-15a-5p | PDE7A | HITS-CLIP | Tarbase |
| hsa-miR-195-5p | PDE7A | HITS-CLIP | Tarbase |

| | | | |
|---|---|---|---|
| hsa-miR-141-3p | TMEM170B | HITS-CLIP | Tarbase |
| hsa-miR-3613-3p | TMEM170B | PAR-CLIP | Tarbase |
| hsa-miR-15a-5p | IPO7 | HITS-CLIP | Tarbase |
| hsa-miR-15a-5p | IPO7 | HITS-CLIP | Tarbase |
| hsa-miR-15a-5p | IPO7 | PAR-CLIP | Tarbase |
| hsa-miR-15a-5p | IPO7 | HITS-CLIP | Tarbase |
| hsa-miR-195-5p | IPO7 | HITS-CLIP | Tarbase |
| hsa-miR-195-5p | IPO7 | HITS-CLIP | Tarbase |
| hsa-miR-195-5p | IPO7 | HITS-CLIP | Tarbase |
| hsa-miR-219a-1-3p | IPO7 | HITS-CLIP | Tarbase |
| hsa-miR-15a-5p | IPO7 | HITS-CLIP | Tarbase |
| hsa-miR-195-5p | IPO7 | HITS-CLIP | Tarbase |
| hsa-miR-3613-5p | TECPR1 | HITS-CLIP | Tarbase |
| hsa-miR-195-5p | KRT81 | Microarrays | Tarbase |
| hsa-miR-15a-5p | CCDC85C | PAR-CLIP | Tarbase |
| hsa-miR-141-3p | NAP1L4 | HITS-CLIP | Tarbase |
| hsa-miR-141-5p | NAP1L4 | HITS-CLIP | Tarbase |
| hsa-miR-15a-3p | NAP1L4 | HITS-CLIP | Tarbase |
| hsa-miR-15a-3p | SMIM11 | HITS-CLIP | Tarbase |
| hsa-miR-3613-3p | DPF3 | HITS-CLIP | Tarbase |
| hsa-miR-15a-5p | ITSN1 | HITS-CLIP | Tarbase |
| hsa-miR-195-5p | ITSN1 | HITS-CLIP | Tarbase |
| hsa-miR-15a-5p | C5orf51 | PAR-CLIP | Tarbase |
| hsa-miR-15a-5p | C5orf51 | HITS-CLIP | Tarbase |
| hsa-miR-15a-5p | C5orf51 | HITS-CLIP | Tarbase |
| hsa-miR-195-5p | C5orf51 | HITS-CLIP | Tarbase |
| hsa-miR-15a-5p | PPAPDC2 | HITS-CLIP | Tarbase |
| hsa-miR-195-5p | PPAPDC2 | HITS-CLIP | Tarbase |
| hsa-miR-195-5p | KLRC3 | Microarrays | Tarbase |
| hsa-miR-15a-3p | DEFB134 | HITS-CLIP | Tarbase |
| hsa-miR-141-5p | HN1L | HITS-CLIP | Tarbase |
| hsa-miR-15a-5p | HN1L | HITS-CLIP | Tarbase |
| hsa-miR-195-5p | HCG27 | Microarrays | Tarbase |
| hsa-miR-141-5p | RAB12 | HITS-CLIP | Tarbase |
| hsa-miR-15a-5p | ANKRD28 | HITS-CLIP | Tarbase |
| hsa-miR-195-5p | ANKRD28 | HITS-CLIP | Tarbase |
| hsa-miR-141-3p | STK38L | HITS-CLIP | Tarbase |
| hsa-miR-15a-5p | SACM1L | PAR-CLIP | Tarbase |
| hsa-miR-15a-5p | SACM1L | PAR-CLIP | Tarbase |
| hsa-miR-195-5p | SACM1L | PAR-CLIP | Tarbase |
| hsa-miR-15a-5p | SACM1L | HITS-CLIP | Tarbase |
| hsa-miR-195-5p | SACM1L | HITS-CLIP | Tarbase |
| hsa-miR-15a-5p | TSN | HITS-CLIP | Tarbase |
| hsa-miR-195-5p | TSN | HITS-CLIP | Tarbase |
| hsa-miR-15a-5p | FAM127C | HITS-CLIP | Tarbase |
| hsa-miR-141-5p | ZNF580 | HITS-CLIP | Tarbase |
| hsa-miR-15a-5p | ZNF580 | HITS-CLIP | Tarbase |
| hsa-miR-195-5p | ZNF580 | HITS-CLIP | Tarbase |
| hsa-miR-3613-5p | ZNF611 | HITS-CLIP | Tarbase |
| hsa-miR-141-5p | SFT2D2 | HITS-CLIP | Tarbase |
| hsa-miR-15a-5p | SFT2D2 | HITS-CLIP | Tarbase |
| hsa-miR-3613-3p | SFT2D2 | HITS-CLIP | Tarbase |
| hsa-miR-486-5p | FGFR1OP | HITS-CLIP | Tarbase |
| hsa-miR-15a-5p | SCAF8 | PAR-CLIP | Tarbase |
| hsa-miR-15a-5p | SCAF8 | HITS-CLIP | Tarbase |
| hsa-miR-3613-5p | SCAF8 | PAR-CLIP | Tarbase |
| hsa-miR-3613-5p | SCAF8 | PAR-CLIP | Tarbase |
| hsa-miR-486-5p | NRAS | PAR-CLIP | Tarbase |
| hsa-miR-15a-5p | CHUK | Luciferase Reporter Assay | Tarbase |
| hsa-miR-15a-5p | CHUK | PAR-CLIP | Tarbase |
| hsa-miR-15a-5p | CHUK | qPCR | Tarbase |
| hsa-miR-15a-5p | CHUK | Western Blot | Tarbase |
| hsa-miR-15a-5p | CHUK | PAR-CLIP | Tarbase |
| hsa-miR-15a-5p | CHUK | HITS-CLIP | Tarbase |
| hsa-miR-15a-5p | CHUK | HITS-CLIP | Tarbase |
| hsa-miR-195-5p | CHUK | qPCR | Tarbase |
| hsa-miR-195-5p | CHUK | Western Blot | Tarbase |
| hsa-miR-195-5p | CHUK | HITS-CLIP | Tarbase |
| hsa-miR-195-5p | CHUK | Microarrays | Tarbase |
| hsa-miR-15a-5p | CHUK | Luciferase Reporter Assay | Tarbase |
| hsa-miR-195-5p | MXD3 | Microarrays | Tarbase |
| hsa-miR-15a-5p | ARL2 | HITS-CLIP | Tarbase |
| hsa-miR-15a-5p | ARL2 | HITS-CLIP | Tarbase |
| hsa-miR-15a-5p | ARL2 | HITS-CLIP | Tarbase |
| hsa-miR-15a-5p | ARL2 | HITS-CLIP | Tarbase |
| hsa-miR-15a-5p | ARL2 | HITS-CLIP | Tarbase |
| hsa-miR-15a-5p | ARL2 | HITS-CLIP | Tarbase |
| hsa-miR-15a-5p | ARL2 | HITS-CLIP | Tarbase |
| hsa-miR-15a-5p | ARL2 | HITS-CLIP | Tarbase |
| hsa-miR-15a-5p | ARL2 | HITS-CLIP | Tarbase |
| hsa-miR-15a-5p | ARL2 | HITS-CLIP | Tarbase |
| hsa-miR-195-5p | ARL2 | HITS-CLIP | Tarbase |
| hsa-miR-195-5p | ARL2 | HITS-CLIP | Tarbase |
| hsa-miR-195-5p | ARL2 | HITS-CLIP | Tarbase |
| hsa-miR-195-5p | ARL2 | AGO-IP | Tarbase |
| hsa-miR-195-5p | ARL2 | HITS-CLIP | Tarbase |
| hsa-miR-195-5p | ARL2 | HITS-CLIP | Tarbase |
| hsa-miR-195-5p | ARL2 | HITS-CLIP | Tarbase |
| hsa-miR-195-5p | ARL2 | HITS-CLIP | Tarbase |
| hsa-miR-195-5p | ARL2 | HITS-CLIP | Tarbase |
| hsa-miR-141-5p | VDAC1 | HITS-CLIP | Tarbase |
| hsa-miR-195-3p | VDAC1 | HITS-CLIP | Tarbase |
| hsa-miR-486-5p | ZBTB9 | PAR-CLIP | Tarbase |
| hsa-miR-486-5p | ZBTB9 | PAR-CLIP | Tarbase |
| hsa-miR-141-3p | HEXA | HITS-CLIP | Tarbase |
| hsa-miR-195-5p | LBH | Microarrays | Tarbase |
| hsa-miR-141-5p | PPP1CB | HITS-CLIP | Tarbase |
| hsa-miR-15a-3p | NCKIPSD | HITS-CLIP | Tarbase |
| hsa-miR-15a-3p | NCKIPSD | HITS-CLIP | Tarbase |
| hsa-miR-141-3p | SLC35F6 | PAR-CLIP | Tarbase |
| hsa-miR-15a-5p | DDAH2 | HITS-CLIP | Tarbase |
| hsa-miR-195-5p | DDAH2 | HITS-CLIP | Tarbase |
| hsa-miR-195-5p | DDAH2 | Microarrays | Tarbase |
| hsa-miR-141-5p | RPS29 | HITS-CLIP | Tarbase |
| hsa-miR-141-5p | RPS29 | HITS-CLIP | Tarbase |
| hsa-miR-15a-5p | ZNF134 | Chimeric fragments | Tarbase |
| hsa-miR-141-3p | EMP2 | HITS-CLIP | Tarbase |
| hsa-miR-15a-5p | LTB4R2 | HITS-CLIP | Tarbase |
| hsa-miR-195-5p | LTB4R2 | HITS-CLIP | Tarbase |
| hsa-miR-195-5p | DNASE1 | Microarrays | Tarbase |
| hsa-miR-3613-5p | CSNK1E | HITS-CLIP | Tarbase |
| hsa-miR-3613-5p | CSNK1E | HITS-CLIP | Tarbase |
| hsa-miR-195-5p | ZNF726 | Microarrays | Tarbase |
| hsa-miR-15a-5p | REPIN1 | HITS-CLIP | Tarbase |
| hsa-miR-15a-5p | REPIN1 | HITS-CLIP | Tarbase |
| hsa-miR-195-5p | REPIN1 | HITS-CLIP | Tarbase |
| hsa-miR-141-3p | ZNF891 | HITS-CLIP | Tarbase |
| hsa-miR-3613-3p | SMIM7 | HITS-CLIP | Tarbase |
| hsa-miR-219a-1-3p | ALG3 | HITS-CLIP | Tarbase |
| hsa-miR-15a-3p | FIS1 | HITS-CLIP | Tarbase |
| hsa-miR-195-5p | CPEB1 | Microarrays | Tarbase |
| hsa-miR-141-5p | HNRNPUL2 | HITS-CLIP | Tarbase |
| hsa-miR-15a-3p | HNRNPUL2 | HITS-CLIP | Tarbase |
| hsa-miR-15a-5p | HNRNPUL2 | HITS-CLIP | Tarbase |
| hsa-miR-195-5p | HNRNPUL2 | HITS-CLIP | Tarbase |
| hsa-miR-195-5p | ARHGEF28 | Microarrays | Tarbase |
| hsa-miR-141-3p | PHB2 | HITS-CLIP | Tarbase |
| hsa-miR-141-3p | PHB2 | HITS-CLIP | Tarbase |
| hsa-miR-141-5p | NEURL4 | HITS-CLIP | Tarbase |
| hsa-miR-15a-5p | UBE2QL1 | HITS-CLIP | Tarbase |
| hsa-miR-195-5p | UBE2QL1 | HITS-CLIP | Tarbase |
| hsa-miR-15a-5p | UBE2QL1 | HITS-CLIP | Tarbase |
| hsa-miR-141-5p | DDX3X | HITS-CLIP | Tarbase |
| hsa-miR-15a-5p | DDX3X | PAR-CLIP | Tarbase |
| hsa-miR-486-5p | DDX3X | HITS-CLIP | Tarbase |
| hsa-miR-141-5p | VPS16 | HITS-CLIP | Tarbase |
| hsa-miR-141-5p | ZNF407 | HITS-CLIP | Tarbase |
| hsa-miR-15a-5p | TMEM167B | PAR-CLIP | Tarbase |
| hsa-miR-15a-5p | IFI30 | HITS-CLIP | Tarbase |
| hsa-miR-195-5p | IFI30 | HITS-CLIP | Tarbase |
| hsa-miR-15a-5p | FNIP1 | HITS-CLIP | Tarbase |
| hsa-miR-15a-5p | NBPF1 | HITS-CLIP | Tarbase |
| hsa-miR-195-5p | NBPF1 | HITS-CLIP | Tarbase |
| hsa-miR-195-5p | EBF2 | Microarrays | Tarbase |
| hsa-miR-141-3p | PPP3R1 | HITS-CLIP | Tarbase |
| hsa-miR-141-3p | PPP3R1 | HITS-CLIP | Tarbase |
| hsa-miR-141-3p | PPP3R1 | HITS-CLIP | Tarbase |
| hsa-miR-141-3p | PPP3R1 | HITS-CLIP | Tarbase |
| hsa-miR-15a-5p | PPP3R1 | HITS-CLIP | Tarbase |

| | | | |
|---|---|---|---|
| hsa-miR-195-5p | FANCG | Microarrays | Tarbase |
| hsa-miR-195-5p | CEBPD | Microarrays | Tarbase |
| hsa-miR-141-3p | PPP2R2A | HITS-CLIP | Tarbase |
| hsa-miR-15a-3p | PPP2R2A | HITS-CLIP | Tarbase |
| hsa-miR-15a-5p | ZNF880 | PAR-CLIP | Tarbase |
| hsa-miR-15a-5p | TIGD1 | HITS-CLIP | Tarbase |
| hsa-miR-195-5p | SLC12A8 | Microarrays | Tarbase |
| hsa-miR-195-5p | FADS3 | Microarrays | Tarbase |
| hsa-miR-141-3p | CCNL2 | HITS-CLIP | Tarbase |
| hsa-miR-15a-3p | CCNL2 | HITS-CLIP | Tarbase |
| hsa-miR-141-3p | PPT2 | HITS-CLIP | Tarbase |
| hsa-miR-15a-5p | PPT2 | HITS-CLIP | Tarbase |
| hsa-miR-195-5p | C10orf55 | Microarrays | Tarbase |
| hsa-miR-141-5p | CERS1 | HITS-CLIP | Tarbase |
| hsa-miR-486-5p | CERS1 | HITS-CLIP | Tarbase |
| hsa-miR-3613-5p | ADM5 | HITS-CLIP | Tarbase |
| hsa-miR-15a-5p | ATXN1L | HITS-CLIP | Tarbase |
| hsa-miR-15a-5p | ATXN1L | PAR-CLIP | Tarbase |
| hsa-miR-15a-5p | ATXN1L | HITS-CLIP | Tarbase |
| hsa-miR-195-5p | ATXN1L | HITS-CLIP | Tarbase |
| hsa-miR-195-5p | ATXN1L | HITS-CLIP | Tarbase |
| hsa-miR-3613-3p | ATXN1L | PAR-CLIP | Tarbase |
| hsa-miR-141-3p | SMIM13 | HITS-CLIP | Tarbase |
| hsa-miR-15a-5p | SMIM13 | PAR-CLIP | Tarbase |
| hsa-miR-15a-5p | SMIM13 | HITS-CLIP | Tarbase |
| hsa-miR-15a-5p | SMIM13 | HITS-CLIP | Tarbase |
| hsa-miR-15a-5p | SMIM13 | HITS-CLIP | Tarbase |
| hsa-miR-195-5p | SMIM13 | HITS-CLIP | Tarbase |
| hsa-miR-195-5p | SMIM13 | HITS-CLIP | Tarbase |
| hsa-miR-195-5p | SMIM13 | HITS-CLIP | Tarbase |
| hsa-miR-3613-5p | SMIM13 | PAR-CLIP | Tarbase |
| hsa-miR-3613-5p | SMIM13 | PAR-CLIP | Tarbase |
| hsa-miR-3613-5p | SMIM13 | PAR-CLIP | Tarbase |
| hsa-miR-15a-5p | SMIM13 | Chimeric fragments | Tarbase |
| hsa-miR-141-5p | FAM195B | HITS-CLIP | Tarbase |
| hsa-miR-141-5p | FAM195B | HITS-CLIP | Tarbase |
| hsa-miR-486-3p | SLC26A6 | HITS-CLIP | Tarbase |
| hsa-miR-15a-5p | ERCC6 | Chimeric fragments | Tarbase |
| hsa-miR-15a-5p | SCAMP4 | PAR-CLIP | Tarbase |
| hsa-miR-195-5p | DHFR | Microarrays | Tarbase |
| hsa-miR-15a-3p | NAMPTL | HITS-CLIP | Tarbase |
| hsa-miR-141-3p | RP11-159G9.5 | HITS-CLIP | Tarbase |
| hsa-miR-15a-3p | TAPBP | HITS-CLIP | Tarbase |
| hsa-miR-3613-5p | HIST1H2BN | PAR-CLIP | Tarbase |
| hsa-miR-3613-3p | RPS28 | HITS-CLIP | Tarbase |
| hsa-miR-141-5p | TRIM26 | HITS-CLIP | Tarbase |
| hsa-miR-141-3p | ZNF736 | HITS-CLIP | Tarbase |
| hsa-miR-141-5p | ZNF736 | HITS-CLIP | Tarbase |
| hsa-miR-15a-3p | JRK | HITS-CLIP | Tarbase |
| hsa-miR-3613-5p | HLA-B | HITS-CLIP | Tarbase |
| hsa-miR-195-5p | ZSCAN31 | Microarrays | Tarbase |
| hsa-miR-15a-5p | ZBED5 | PAR-CLIP | Tarbase |
| hsa-miR-15a-5p | ZBED5 | PAR-CLIP | Tarbase |
| hsa-miR-15a-5p | ZBED5 | PAR-CLIP | Tarbase |
| hsa-miR-15a-5p | ZBED5 | PAR-CLIP | Tarbase |
| hsa-miR-15a-5p | ZBED5 | PAR-CLIP | Tarbase |
| hsa-miR-15a-5p | ZBED5 | PAR-CLIP | Tarbase |
| hsa-miR-15a-5p | ZBED5 | PAR-CLIP | Tarbase |
| hsa-miR-15a-5p | ZBED5 | PAR-CLIP | Tarbase |
| hsa-miR-15a-5p | ZBED5 | PAR-CLIP | Tarbase |
| hsa-miR-195-5p | ZBED5 | PAR-CLIP | Tarbase |
| hsa-miR-195-5p | ZBED5 | PAR-CLIP | Tarbase |
| hsa-miR-195-5p | ZBED5 | PAR-CLIP | Tarbase |
| hsa-miR-195-5p | ZBED5 | PAR-CLIP | Tarbase |
| hsa-miR-195-5p | ZBED5 | PAR-CLIP | Tarbase |
| hsa-miR-195-5p | KIFC1 | Microarrays | Tarbase |
| hsa-miR-15a-5p | C9orf69 | HITS-CLIP | Tarbase |
| hsa-miR-195-5p | C9orf69 | HITS-CLIP | Tarbase |
| hsa-miR-3605-3p | C9orf69 | HITS-CLIP | Tarbase |
| hsa-miR-141-3p | RBM14 | HITS-CLIP | Tarbase |
| hsa-miR-15a-5p | RBM14 | HITS-CLIP | Tarbase |
| hsa-miR-15a-5p | RBM14 | HITS-CLIP | Tarbase |
| hsa-miR-15a-5p | RBM14 | HITS-CLIP | Tarbase |
| hsa-miR-195-5p | RBM14 | HITS-CLIP | Tarbase |
| hsa-miR-195-5p | RBM14 | HITS-CLIP | Tarbase |
| hsa-miR-3613-3p | RBM14 | HITS-CLIP | Tarbase |
| hsa-miR-141-3p | NME1 | HITS-CLIP | Tarbase |
| hsa-miR-15a-5p | WBP1 | HITS-CLIP | Tarbase |
| hsa-miR-15a-5p | WBP1 | HITS-CLIP | Tarbase |
| hsa-miR-195-5p | WBP1 | HITS-CLIP | Tarbase |
| hsa-miR-15a-3p | GET4 | HITS-CLIP | Tarbase |
| hsa-miR-15a-5p | GET4 | HITS-CLIP | Tarbase |
| hsa-miR-195-5p | GET4 | HITS-CLIP | Tarbase |
| hsa-miR-15a-5p | LY6G5B | HITS-CLIP | Tarbase |
| hsa-miR-195-5p | LY6G5B | HITS-CLIP | Tarbase |
| hsa-miR-195-5p | AQP1 | Microarrays | Tarbase |
| hsa-miR-15a-5p | PCDHGC5 | HITS-CLIP | Tarbase |
| hsa-miR-195-5p | PCDHGC5 | HITS-CLIP | Tarbase |
| hsa-miR-3613-5p | CRCP | HITS-CLIP | Tarbase |
| hsa-miR-141-5p | ATP5O | HITS-CLIP | Tarbase |
| hsa-miR-15a-5p | PISD | PAR-CLIP | Tarbase |
| hsa-miR-15a-5p | PI4KA | HITS-CLIP | Tarbase |
| hsa-miR-15a-5p | PI4KA | HITS-CLIP | Tarbase |
| hsa-miR-195-5p | PI4KA | HITS-CLIP | Tarbase |
| hsa-miR-3613-3p | HYPK | PAR-CLIP | Tarbase |
| hsa-miR-15a-5p | ARFGAP3 | HITS-CLIP | Tarbase |
| hsa-miR-15a-5p | ARFGAP3 | HITS-CLIP | Tarbase |
| hsa-miR-195-5p | ARFGAP3 | HITS-CLIP | Tarbase |
| hsa-miR-195-5p | ARFGAP3 | HITS-CLIP | Tarbase |
| hsa-miR-15a-5p | PEG10 | PAR-CLIP | Tarbase |
| hsa-miR-15a-5p | PEG10 | HITS-CLIP | Tarbase |
| hsa-miR-195-5p | PEG10 | HITS-CLIP | Tarbase |
| hsa-miR-195-5p | PEG10 | Microarrays | Tarbase |
| hsa-miR-15a-5p | MRPL20 | PAR-CLIP | Tarbase |
| hsa-miR-195-5p | HLA-DMB | Microarrays | Tarbase |
| hsa-miR-195-5p | PSG4 | Microarrays | Tarbase |
| hsa-miR-15a-5p | C4orf48 | HITS-CLIP | Tarbase |
| hsa-miR-195-5p | C4orf48 | HITS-CLIP | Tarbase |
| hsa-miR-15a-5p | C4orf48 | HITS-CLIP | Tarbase |
| hsa-miR-195-5p | C4orf48 | HITS-CLIP | Tarbase |
| hsa-miR-195-5p | NFS1 | Microarrays | Tarbase |
| hsa-miR-15a-5p | DDOST | HITS-CLIP | Tarbase |
| hsa-miR-15a-5p | DDOST | HITS-CLIP | Tarbase |
| hsa-miR-15a-5p | DDOST | PAR-CLIP | Tarbase |
| hsa-miR-15a-5p | DDOST | HITS-CLIP | Tarbase |
| hsa-miR-195-5p | DDOST | HITS-CLIP | Tarbase |
| hsa-miR-195-5p | DDOST | HITS-CLIP | Tarbase |
| hsa-miR-195-5p | DDOST | HITS-CLIP | Tarbase |
| hsa-miR-15a-5p | TMEM199 | HITS-CLIP | Tarbase |
| hsa-miR-195-5p | TMEM199 | HITS-CLIP | Tarbase |
| hsa-miR-195-5p | IFITM10 | Microarrays | Tarbase |
| hsa-miR-15a-5p | DBNDD2 | PAR-CLIP | Tarbase |
| hsa-miR-486-5p | ENSG00000244291 | HITS-CLIP | Tarbase |
| hsa-miR-15a-3p | RBM12 | HITS-CLIP | Tarbase |
| hsa-miR-15a-5p | RBM12 | HITS-CLIP | Tarbase |
| hsa-miR-15a-5p | RBM12 | HITS-CLIP | Tarbase |
| hsa-miR-195-5p | RBM12 | HITS-CLIP | Tarbase |
| hsa-miR-486-5p | RBM12 | PAR-CLIP | Tarbase |
| hsa-miR-15a-5p | RBM12 | Chimeric fragments | Tarbase |
| hsa-miR-15a-5p | SCARF2 | HITS-CLIP | Tarbase |
| hsa-miR-195-5p | SCARF2 | HITS-CLIP | Tarbase |
| hsa-miR-195-5p | UBE2V1 | Microarrays | Tarbase |
| hsa-miR-15a-5p | H2AFJ | HITS-CLIP | Tarbase |
| hsa-miR-15a-3p | PGAM5 | HITS-CLIP | Tarbase |
| hsa-miR-15a-5p | PGAM5 | HITS-CLIP | Tarbase |
| hsa-miR-141-3p | ZCCHC3 | HITS-CLIP | Tarbase |
| hsa-miR-141-3p | ZCCHC3 | HITS-CLIP | Tarbase |
| hsa-miR-141-3p | ZCCHC3 | PAR-CLIP | Tarbase |
| hsa-miR-141-3p | ZCCHC3 | PAR-CLIP | Tarbase |
| hsa-miR-15a-5p | ZCCHC3 | PAR-CLIP | Tarbase |
| hsa-miR-15a-5p | ZCCHC3 | PAR-CLIP | Tarbase |
| hsa-miR-15a-5p | ZCCHC3 | PAR-CLIP | Tarbase |
| hsa-miR-15a-5p | ZCCHC3 | PAR-CLIP | Tarbase |
| hsa-miR-15a-5p | ZCCHC3 | PAR-CLIP | Tarbase |
| hsa-miR-15a-5p | ZCCHC3 | HITS-CLIP | Tarbase |
| hsa-miR-15a-5p | ZCCHC3 | PAR-CLIP | Tarbase |
| hsa-miR-15a-5p | ZCCHC3 | PAR-CLIP | Tarbase |
| hsa-miR-15a-5p | ZCCHC3 | PAR-CLIP | Tarbase |
| hsa-miR-15a-5p | ZCCHC3 | PAR-CLIP | Tarbase |

| | | | |
|---|---|---|---|
| hsa-miR-15a-5p | ZCCHC3 | PAR-CLIP | Tarbase |
| hsa-miR-195-5p | ZCCHC3 | PAR-CLIP | Tarbase |
| hsa-miR-195-5p | ZCCHC3 | HITS-CLIP | Tarbase |
| hsa-miR-195-5p | ZCCHC3 | PAR-CLIP | Tarbase |
| hsa-miR-195-5p | ZCCHC3 | PAR-CLIP | Tarbase |
| hsa-miR-195-5p | ZCCHC3 | PAR-CLIP | Tarbase |
| hsa-miR-195-5p | ZCCHC3 | PAR-CLIP | Tarbase |
| hsa-miR-195-5p | ZCCHC3 | PAR-CLIP | Tarbase |
| hsa-miR-3613-5p | ZCCHC3 | PAR-CLIP | Tarbase |
| hsa-miR-3613-5p | ZCCHC3 | PAR-CLIP | Tarbase |
| hsa-miR-3613-5p | ZCCHC3 | PAR-CLIP | Tarbase |
| hsa-miR-195-5p | ADH1C | Microarrays | Tarbase |
| hsa-miR-141-3p | FMN1 | HITS-CLIP | Tarbase |
| hsa-miR-195-5p | HAUS5 | Microarrays | Tarbase |
| hsa-miR-195-5p | TMEM158 | Microarrays | Tarbase |
| hsa-miR-195-5p | CHCHD10 | Microarrays | Tarbase |
| hsa-miR-15a-3p | GLI4 | HITS-CLIP | Tarbase |
| hsa-miR-195-5p | FOXD1 | Microarrays | Tarbase |
| hsa-miR-15a-5p | HOXA10 | PAR-CLIP | Tarbase |
| hsa-miR-195-3p | ALG11 | HITS-CLIP | Tarbase |
| hsa-miR-15a-5p | ATXN7L3B | PAR-CLIP | Tarbase |
| hsa-miR-15a-5p | ATXN7L3B | HITS-CLIP | Tarbase |
| hsa-miR-15a-5p | ATXN7L3B | HITS-CLIP | Tarbase |
| hsa-miR-15a-5p | ATXN7L3B | HITS-CLIP | Tarbase |
| hsa-miR-195-5p | ATXN7L3B | HITS-CLIP | Tarbase |
| hsa-miR-486-5p | ATXN7L3B | HITS-CLIP | Tarbase |
| hsa-miR-195-5p | ATXN7L3B | Microarrays | Tarbase |
| hsa-miR-141-5p | PRKDC | HITS-CLIP | Tarbase |
| hsa-miR-15a-5p | PRKDC | HITS-CLIP | Tarbase |
| hsa-miR-15a-5p | PRKDC | HITS-CLIP | Tarbase |
| hsa-miR-15a-5p | PRKDC | HITS-CLIP | Tarbase |
| hsa-miR-195-5p | PRKDC | HITS-CLIP | Tarbase |
| hsa-miR-195-5p | PRKDC | HITS-CLIP | Tarbase |
| hsa-miR-3613-5p | PRKDC | HITS-CLIP | Tarbase |
| hsa-miR-15a-5p | UTP14C | PAR-CLIP | Tarbase |
| hsa-miR-219a-1-3p | UTP14C | PAR-CLIP | Tarbase |
| hsa-miR-486-5p | RP11-108K14.8 | PAR-CLIP | Tarbase |
| hsa-miR-15a-5p | TM9SF1 | HITS-CLIP | Tarbase |
| hsa-miR-195-5p | TM9SF1 | HITS-CLIP | Tarbase |
| hsa-miR-15a-5p | MEX3A | HITS-CLIP | Tarbase |
| hsa-miR-15a-5p | CKLF-CMTM1 | HITS-CLIP | Tarbase |
| hsa-miR-195-5p | CKLF-CMTM1 | HITS-CLIP | Tarbase |
| hsa-miR-141-5p | ANKHD1-EIF4EBP3 | HITS-CLIP | Tarbase |
| hsa-miR-15a-5p | ANKHD1-EIF4EBP3 | PAR-CLIP | Tarbase |
| hsa-miR-15a-5p | ANKHD1-EIF4EBP3 | PAR-CLIP | Tarbase |
| hsa-miR-15a-5p | ANKHD1-EIF4EBP3 | HITS-CLIP | Tarbase |
| hsa-miR-15a-5p | EID1 | PAR-CLIP | Tarbase |
| hsa-miR-15a-5p | EID1 | HITS-CLIP | Tarbase |
| hsa-miR-15a-5p | EID1 | HITS-CLIP | Tarbase |
| hsa-miR-195-5p | EID1 | HITS-CLIP | Tarbase |
| hsa-miR-3613-3p | POLR2M | HITS-CLIP | Tarbase |
| hsa-miR-195-5p | AP000721.4 | HITS-CLIP | Tarbase |
| hsa-miR-195-5p | AP000721.4 | HITS-CLIP | Tarbase |
| hsa-miR-195-5p | ZNF486 | Microarrays | Tarbase |
| hsa-miR-3605-5p | ZNF253 | HITS-CLIP | Tarbase |
| hsa-miR-141-3p | KIAA1147 | HITS-CLIP | Tarbase |
| hsa-miR-3613-3p | KIAA1147 | PAR-CLIP | Tarbase |
| hsa-miR-15a-5p | GATC | PAR-CLIP | Tarbase |
| hsa-miR-15a-5p | GATC | PAR-CLIP | Tarbase |
| hsa-miR-15a-5p | GATC | PAR-CLIP | Tarbase |
| hsa-miR-15a-5p | GATC | PAR-CLIP | Tarbase |
| hsa-miR-195-5p | GATC | PAR-CLIP | Tarbase |
| hsa-miR-3605-5p | GATC | PAR-CLIP | Tarbase |
| hsa-miR-15a-5p | ZBED6 | HITS-CLIP | Tarbase |
| hsa-miR-15a-5p | FNTB | HITS-CLIP | Tarbase |
| hsa-miR-195-5p | FNTB | HITS-CLIP | Tarbase |
| hsa-miR-141-3p | CUX1 | PAR-CLIP | Tarbase |
| hsa-miR-141-3p | CUX1 | HITS-CLIP | Tarbase |
| hsa-miR-15a-5p | CUX1 | HITS-CLIP | Tarbase |
| hsa-miR-15a-5p | CUX1 | PAR-CLIP | Tarbase |
| hsa-miR-15a-5p | CUX1 | HITS-CLIP | Tarbase |
| hsa-miR-195-5p | CUX1 | HITS-CLIP | Tarbase |
| hsa-miR-15a-5p | ALG9 | HITS-CLIP | Tarbase |
| hsa-miR-15a-5p | THTPA | HITS-CLIP | Tarbase |
| hsa-miR-195-5p | THTPA | HITS-CLIP | Tarbase |
| hsa-miR-141-3p | RBM15B | HITS-CLIP | Tarbase |
| hsa-miR-15a-5p | FRRS1L | HITS-CLIP | Tarbase |
| hsa-miR-195-5p | FRRS1L | HITS-CLIP | Tarbase |
| hsa-miR-15a-5p | RP6-24A23.6 | PAR-CLIP | Tarbase |
| hsa-miR-15a-5p | RP6-24A23.6 | HITS-CLIP | Tarbase |
| hsa-miR-195-5p | RP6-24A23.6 | HITS-CLIP | Tarbase |
| hsa-miR-15a-5p | RP6-24A23.6 | Chimeric fragments | Tarbase |
| hsa-miR-15a-5p | TMEM178B | HITS-CLIP | Tarbase |
| hsa-miR-195-5p | TMEM178B | HITS-CLIP | Tarbase |
| hsa-miR-141-3p | GAN | HITS-CLIP | Tarbase |
| hsa-miR-15a-5p | GAN | PAR-CLIP | Tarbase |
| hsa-miR-15a-5p | GAN | PAR-CLIP | Tarbase |
| hsa-miR-15a-5p | GAN | PAR-CLIP | Tarbase |
| hsa-miR-15a-5p | GAN | HITS-CLIP | Tarbase |
| hsa-miR-15a-5p | GAN | PAR-CLIP | Tarbase |
| hsa-miR-15a-5p | GAN | HITS-CLIP | Tarbase |
| hsa-miR-195-5p | GAN | PAR-CLIP | Tarbase |
| hsa-miR-195-5p | GAN | HITS-CLIP | Tarbase |
| hsa-miR-486-5p | GAN | HITS-CLIP | Tarbase |
| hsa-miR-15a-5p | GAN | Chimeric fragments | Tarbase |
| hsa-miR-141-3p | TCF24 | HITS-CLIP | Tarbase |
| hsa-miR-486-3p | LY6G5B | HITS-CLIP | Tarbase |
| hsa-miR-141-5p | SRSF8 | HITS-CLIP | Tarbase |
| hsa-miR-141-5p | SRSF8 | HITS-CLIP | Tarbase |
| hsa-miR-486-5p | GDF2 | HITS-CLIP | Tarbase |
| hsa-miR-486-5p | GDF2 | HITS-CLIP | Tarbase |
| hsa-miR-486-5p | GDF2 | HITS-CLIP | Tarbase |
| hsa-miR-141-3p | DYNLL2 | HITS-CLIP | Tarbase |
| hsa-miR-141-3p | DYNLL2 | HITS-CLIP | Tarbase |
| hsa-miR-15a-5p | DYNLL2 | HITS-CLIP | Tarbase |
| hsa-miR-15a-5p | DYNLL2 | HITS-CLIP | Tarbase |
| hsa-miR-15a-5p | DYNLL2 | HITS-CLIP | Tarbase |
| hsa-miR-15a-5p | DYNLL2 | HITS-CLIP | Tarbase |
| hsa-miR-15a-5p | DYNLL2 | HITS-CLIP | Tarbase |
| hsa-miR-195-5p | DYNLL2 | HITS-CLIP | Tarbase |
| hsa-miR-195-5p | DYNLL2 | HITS-CLIP | Tarbase |
| hsa-miR-195-5p | DYNLL2 | HITS-CLIP | Tarbase |
| hsa-miR-195-5p | DYNLL2 | HITS-CLIP | Tarbase |
| hsa-miR-195-5p | DYNLL2 | Microarrays | Tarbase |
| hsa-miR-15a-5p | OTUD7B | HITS-CLIP | Tarbase |
| hsa-miR-195-5p | OTUD7B | HITS-CLIP | Tarbase |
| hsa-miR-141-5p | TXNIP | HITS-CLIP | Tarbase |
| hsa-miR-141-5p | TXNIP | HITS-CLIP | Tarbase |
| hsa-miR-141-5p | TXNIP | HITS-CLIP | Tarbase |
| hsa-miR-141-5p | TXNIP | HITS-CLIP | Tarbase |
| hsa-miR-15a-3p | TXNIP | HITS-CLIP | Tarbase |
| hsa-miR-3605-5p | TXNIP | HITS-CLIP | Tarbase |
| hsa-miR-3613-3p | TXNIP | HITS-CLIP | Tarbase |
| hsa-miR-486-3p | TXNIP | HITS-CLIP | Tarbase |
| hsa-miR-15a-3p | BAHCC1 | HITS-CLIP | Tarbase |
| hsa-miR-195-5p | RASSF5 | Microarrays | Tarbase |
| hsa-miR-3613-5p | MRPS21 | HITS-CLIP | Tarbase |
| hsa-miR-15a-5p | ZNF224 | HITS-CLIP | Tarbase |
| hsa-miR-15a-5p | ZNF224 | PAR-CLIP | Tarbase |
| hsa-miR-195-5p | ZNF224 | HITS-CLIP | Tarbase |
| hsa-miR-141-3p | SLC6A14 | HITS-CLIP | Tarbase |
| hsa-miR-486-5p | MTRNR2L12 | HITS-CLIP | Tarbase |
| hsa-miR-15a-5p | TMEM185A | HITS-CLIP | Tarbase |
| hsa-miR-141-5p | NBPF9 | HITS-CLIP | Tarbase |
| hsa-miR-195-5p | SLC25A53 | AGO-IP | Tarbase |
| hsa-miR-15a-3p | TFDP2 | HITS-CLIP | Tarbase |
| hsa-miR-195-5p | HIST2H4B | Microarrays | Tarbase |
| hsa-miR-15a-5p | RP5-850E9.3 | HITS-CLIP | Tarbase |
| hsa-miR-141-3p | TAF15 | HITS-CLIP | Tarbase |
| hsa-miR-3605-3p | TAF15 | HITS-CLIP | Tarbase |
| hsa-miR-195-5p | HIST2H4A | Microarrays | Tarbase |
| hsa-miR-195-5p | SRXN1 | Microarrays | Tarbase |
| hsa-miR-141-3p | RP11-139J15.7 | HITS-CLIP | Tarbase |
| hsa-miR-141-5p | NUDT3 | HITS-CLIP | Tarbase |
| hsa-miR-141-5p | NUDT3 | HITS-CLIP | Tarbase |
| hsa-miR-141-5p | KMT2B | HITS-CLIP | Tarbase |
| hsa-miR-15a-5p | KMT2B | PAR-CLIP | Tarbase |
| hsa-miR-141-3p | POM121C | HITS-CLIP | Tarbase |
| hsa-miR-15a-5p | POM121C | PAR-CLIP | Tarbase |

| | | | |
|---|---|---|---|
| hsa-miR-15a-5p | POM121C | PAR-CLIP | Tarbase |
| hsa-miR-15a-5p | POM121C | HITS-CLIP | Tarbase |
| hsa-miR-195-5p | CD24 | Microarrays | Tarbase |
| hsa-miR-15a-5p | DCP1A | HITS-CLIP | Tarbase |
| hsa-miR-15a-5p | DCP1A | HITS-CLIP | Tarbase |
| hsa-miR-15a-5p | DCP1A | PAR-CLIP | Tarbase |
| hsa-miR-15a-5p | DCP1A | HITS-CLIP | Tarbase |
| hsa-miR-195-5p | DCP1A | HITS-CLIP | Tarbase |
| hsa-miR-195-5p | DCP1A | HITS-CLIP | Tarbase |
| hsa-miR-195-5p | DCP1A | HITS-CLIP | Tarbase |
| hsa-miR-15a-5p | DCP1A | Chimeric fragments | Tarbase |
| hsa-miR-15a-5p | GRIN2B | HITS-CLIP | Tarbase |
| hsa-miR-195-5p | GRIN2B | HITS-CLIP | Tarbase |
| hsa-miR-15a-5p | ZNHIT3 | HITS-CLIP | Tarbase |
| hsa-miR-195-5p | ZNHIT3 | HITS-CLIP | Tarbase |
| hsa-miR-15a-5p | HIST1H2BG | PAR-CLIP | Tarbase |
| hsa-miR-15a-5p | HIST1H2BG | HITS-CLIP | Tarbase |
| hsa-miR-195-5p | HIST1H2BG | HITS-CLIP | Tarbase |
| hsa-miR-15a-5p | HIST1H3G | HITS-CLIP | Tarbase |
| hsa-miR-195-5p | HIST1H3G | Microarrays | Tarbase |
| hsa-miR-15a-5p | NATD1 | HITS-CLIP | Tarbase |
| hsa-miR-195-5p | NATD1 | HITS-CLIP | Tarbase |
| hsa-miR-141-3p | SOCS7 | HITS-CLIP | Tarbase |
| hsa-miR-15a-5p | SOCS7 | HITS-CLIP | Tarbase |
| hsa-miR-15a-5p | HIST1H3B | PAR-CLIP | Tarbase |
| hsa-miR-195-5p | HIST1H3B | Microarrays | Tarbase |
| hsa-miR-15a-5p | HIST1H2BE | HITS-CLIP | Tarbase |
| hsa-miR-195-5p | HIST1H2BE | HITS-CLIP | Tarbase |
| hsa-miR-15a-5p | HIST1H2BO | PAR-CLIP | Tarbase |
| hsa-miR-195-5p | HIST1H2BO | Microarrays | Tarbase |
| hsa-miR-15a-5p | HIST1H2AH | HITS-CLIP | Tarbase |
| hsa-miR-15a-5p | HIST1H2AH | HITS-CLIP | Tarbase |
| hsa-miR-15a-5p | HIST1H2AH | PAR-CLIP | Tarbase |
| hsa-miR-15a-5p | HIST1H2AH | HITS-CLIP | Tarbase |
| hsa-miR-195-5p | HIST1H2AH | HITS-CLIP | Tarbase |
| hsa-miR-195-5p | HIST1H2AH | HITS-CLIP | Tarbase |
| hsa-miR-195-5p | HIST1H2AH | HITS-CLIP | Tarbase |
| hsa-miR-195-5p | HIST1H2AH | Microarrays | Tarbase |
| hsa-miR-15a-3p | ZNF280B | HITS-CLIP | Tarbase |
| hsa-miR-15a-3p | MLLT6 | HITS-CLIP | Tarbase |
| hsa-miR-15a-5p | MLLT6 | HITS-CLIP | Tarbase |
| hsa-miR-195-5p | MLLT6 | HITS-CLIP | Tarbase |
| hsa-miR-141-5p | SMEK2 | HITS-CLIP | Tarbase |
| hsa-miR-15a-5p | SMEK2 | HITS-CLIP | Tarbase |
| hsa-miR-15a-5p | SMEK2 | PAR-CLIP | Tarbase |
| hsa-miR-195-5p | SMEK2 | HITS-CLIP | Tarbase |
| hsa-miR-195-5p | HIST1H4L | Microarrays | Tarbase |
| hsa-miR-195-5p | HIST1H2AK | Microarrays | Tarbase |
| hsa-miR-195-5p | AATF | HITS-CLIP | Tarbase |
| hsa-miR-15a-5p | HIST1H2BH | HITS-CLIP | Tarbase |
| hsa-miR-195-5p | HIST1H2BH | HITS-CLIP | Tarbase |
| hsa-miR-486-3p | HIST1H2BH | HITS-CLIP | Tarbase |
| hsa-miR-195-5p | HIST1H2BH | Microarrays | Tarbase |
| hsa-miR-15a-5p | HIST1H2AJ | HITS-CLIP | Tarbase |
| hsa-miR-15a-5p | HIST1H2AJ | HITS-CLIP | Tarbase |
| hsa-miR-15a-5p | HIST1H2AJ | HITS-CLIP | Tarbase |
| hsa-miR-15a-5p | HIST1H2AJ | HITS-CLIP | Tarbase |
| hsa-miR-195-5p | HIST1H2AJ | HITS-CLIP | Tarbase |
| hsa-miR-195-5p | HIST1H2AJ | HITS-CLIP | Tarbase |
| hsa-miR-195-5p | HIST1H2AJ | HITS-CLIP | Tarbase |
| hsa-miR-3605-5p | HIST1H2AJ | HITS-CLIP | Tarbase |
| hsa-miR-195-5p | HIST1H2AJ | Microarrays | Tarbase |
| hsa-miR-195-5p | HIST1H4D | Microarrays | Tarbase |
| hsa-miR-15a-5p | HIST1H2BF | HITS-CLIP | Tarbase |
| hsa-miR-195-5p | HIST1H2BF | Microarrays | Tarbase |
| hsa-miR-15a-5p | PCGF2 | Chimeric fragments | Tarbase |
| hsa-miR-15a-5p | MARCKS | HITS-CLIP | Tarbase |
| hsa-miR-15a-5p | MARCKS | HITS-CLIP | Tarbase |
| hsa-miR-15a-5p | MARCKS | HITS-CLIP | Tarbase |
| hsa-miR-195-5p | MARCKS | HITS-CLIP | Tarbase |
| hsa-miR-195-5p | MARCKS | HITS-CLIP | Tarbase |
| hsa-miR-195-5p | MARCKS | HITS-CLIP | Tarbase |
| hsa-miR-195-5p | HIST1H3F | Microarrays | Tarbase |
| hsa-miR-15a-5p | PSMB3 | Chimeric fragments | Tarbase |
| hsa-miR-15a-5p | ZNF8 | PAR-CLIP | Tarbase |
| hsa-miR-15a-5p | MYO19 | PAR-CLIP | Tarbase |
| hsa-miR-195-5p | MYO19 | PAR-CLIP | Tarbase |
| hsa-miR-195-5p | MYO19 | Microarrays | Tarbase |
| hsa-miR-15a-5p | HIST1H3C | HITS-CLIP | Tarbase |
| hsa-miR-195-5p | HIST1H3C | HITS-CLIP | Tarbase |
| hsa-miR-15a-5p | HIST1H2AB | PAR-CLIP | Tarbase |
| hsa-miR-195-5p | HIST1H2AB | Microarrays | Tarbase |
| hsa-miR-15a-5p | ACACA | HITS-CLIP | Tarbase |
| hsa-miR-195-5p | ACACA | HITS-CLIP | Tarbase |
| hsa-miR-219a-1-3p | ACACA | PAR-CLIP | Tarbase |
| hsa-miR-486-5p | ACACA | PAR-CLIP | Tarbase |
| hsa-miR-195-5p | HIST1H2BI | Microarrays | Tarbase |
| hsa-miR-195-5p | HIST1H4A | Microarrays | Tarbase |
| hsa-miR-195-5p | HIST1H2AM | Microarrays | Tarbase |
| hsa-miR-141-3p | HuD(hsa) | qPCR | Tarbase |
| hsa-miR-141-3p | JNK2(hsa) | Western Blot | Tarbase |
| hsa-miR-141-3p | cyclinD1(hsa) | Western Blot | Tarbase |
| hsa-miR-195-5p | cyclinD1(hsa) | qPCR | Tarbase |
| hsa-miR-195-5p | cyclinD1(hsa) | Western Blot | Tarbase |
