## Supplementary Table S5 for "Methylome-based cell-of-origin modeling (Methyl-COOM) identifies aberrant expression of immune regulatory molecules in CLL"

**Supplementary Table S5: List of epigenetic regulators**

| Gene.ID | Group | Subgroup | CMM_for.NGR.Epigenetic_Function | Data.browser | Function |
|---|---|---|---|---|---|
| ACTL6A | Chromatin remodeling | nucleosome remodeling factors | | IDA | |
| ACTL6B | Chromatin remodeling | chromatin remodeling factors | chromatin remodelling | TAS | |
| AICDA | DNA modifications | editors | DNA demethylation | HEMD | |
| ALKBH1 | DNA modifications | editors | DNA demethylation | HEMD | |
| ALKBH3 | DNA modifications | editors | DNA demethylation DNA repair | HEMD | Direct reversal of damage |
| ANKRD28 | Histone modification | Editors | Histone dephosphorylation | HEMD | |
| ANKRD44 | Histone modification | Editors | Histone dephosphorylation | HEMD | |
| ANKRD52 | Histone modification | Editors | Histone dephosphorylation | HEMD | |
| APOBEC1 | DNA modifications | editors | DNA demethylation | HEMD | |
| ARID1A | Not assigned | Not assigned | reg. of gene expression, epigenetic | IDA | |
| ASXL1 | Not assigned | Not assigned | methylation, chromatin | UCSC | |
| ARIP4 | Chromatin remodeling | Chromatin remodeling helicase | Chromatin remodeling helicase | HEMD | |
| ASH1L | Histone modification | Writers | Histone methylation | HEMD | |
| ASH2L | Histone modification | Readers | Readers | IDA | |
| ASZ1 | DNA modifications | writers | DNA methylation | IEA | |
| ATM | Histone modification | Writers | Histone phosphorylation DNA repair | HEMD | Genes defective in diseases associated with sensitivity to DNA damaging agents |
| ATR | Histone modification | Writers | Histone phosphorylation DNA repair | HEMD | Other conserved DNA damage response genes |
| ATRX | Chromatin remodeling | Chromatin remodeling helicase | Chromatin remodeling helicase | HEMD | |
| ATXN7 | Histone modification | Readers | Readers | IDA | |
| ATXN7L3 | Histone modification | Readers | Readers | UCSC | |
| BAHD1 | Chromatin remodeling | chromatin remodeling factors | chromatin remodelling | IMP | |
| BAP1 | Histone modification | Editors | Histone deubiquitination | HEMD | |
| BAP18 | Chromatin remodeling | nucleosome remodeling factors | | IEA | |
| BARD1 | Histone modification | Writers | Histone ubiquitination | HEMD | |
| BAZ1B | Histone modification | Writers | Histone phosphorylation | HEMD | |
| BAZ2A | Chromatin remodeling | chromatin remodeling factors | chromatin remodelling | TAS | |
| BNIP3 | Chromatin remodeling | chromatin remodeling factors | chromatin remodelling | IDA | |
| BPTF | Chromatin remodeling | nucleosome remodeling factors | | IDA | |
| BRAC1 | Histone modification | Writers | Histone ubiquitination | HEMD | |
| BRCA2 | Histone modification | Writers | Histone ubiquitination DNA repair | HEMD | Fanconi anemia |
| BRCC36 | Histone modification | Editors | Histone deubiquitination | HEMD | |
| BRD1 | Histone modification | Readers | Readers | IDA | |
| BRDT | Chromatin remodeling | chromatin remodeling factors | chromatin remodelling | NAS | |
| BRE1A | Histone modification | Writers | Histone ubiquitination | HEMD | |
| BRE1B | Histone modification | Writers | Histone ubiquitination | HEMD | |
| BRG1 | Chromatin remodeling | Chromatin remodeling helicase | Chromatin remodeling helicase | HEMD | |
| BRPF3 | Histone modification | Readers | Readers | IEA | |
| BTAF1 | Chromatin remodeling | Chromatin remodeling helicase | Chromatin remodeling helicase | HEMD | |
| BTD | Histone modification | Writers | Histone biotinylation | HEMD | |
| BUB1 | Histone modification | Writers | Histone phosphorylation | HEMD | |
| CARM1 | Histone modification | Writers | Histone methylation | HEMD | |
| CASC5 | Chromatin remodeling | chromatin remodeling factors | chromatin remodelling | NAS | |
| CBX3 | Chromatin remodeling | chromatin remodeling factors | chromatin remodelling | IMP | |
| CBX4 | Histone modification | Readers | Readers | UCSC | |
| CDK17 | Histone modification | Writers | Histone phosphorylation | HEMD | |
| CDK2 | Histone modification | Writers | Histone phosphorylation | HEMD | |
| CDK3 | Histone modification | Writers | Histone phosphorylation | HEMD | |
| CDK5 | Histone modification | Writers | Histone phosphorylation | HEMD | |
| CDK8 | Histone modification | Writers | Histone phosphorylation | HEMD | |
| CDKN2A | Chromatin remodeling | chromatin remodeling factors | chromatin remodelling | IMP | |
| CDY1 | Histone modification | Writers | Histone acetylation | HEMD | |
| CDY2 | Histone modification | Writers | Histone acetylation | HEMD | |
| CDYL | Histone modification | Writers | Histone acetylation | HEMD | |
| CENPA | Chromatin remodeling | chromatin remodeling factors | chromatin remodelling | NAS | |
| CENPH | Chromatin remodeling | chromatin remodeling factors | chromatin remodelling | TAS | |
| CENPI | Chromatin remodeling | chromatin remodeling factors | chromatin remodelling | ISS | |
| CENPK | Chromatin remodeling | chromatin remodeling factors | chromatin remodelling | IEA | |
| CENPN | Chromatin remodeling | chromatin remodeling factors | chromatin remodelling | IEA | |
| CENPO | Chromatin remodeling | chromatin remodeling factors | chromatin remodelling | TAS | |
| CENPP | Chromatin remodeling | chromatin remodeling factors | chromatin remodelling | TAS | |
| CENPQ | Chromatin remodeling | chromatin remodeling factors | chromatin remodelling | TAS | |
| CENPV | Chromatin remodeling | chromatin remodeling factors | chromatin remodelling | TAS | |
| CHD1 | Histone modification | Readers | Readers | HEMD | |
| CHD1L | Chromatin remodeling | Chromatin remodeling helicase | Chromatin remodeling helicase | HEMD | |
| CHD2 | Chromatin remodeling | Chromatin remodeling helicase | Chromatin remodeling helicase | HEMD | |
| CHD3 | Histone modification | Readers | Readers | HEMD | |
| CHD4 | Histone modification | Readers | Readers | HEMD | |
| CHD5 | Histone modification | Readers | Readers | HEMD | |
| CHD6 | Chromatin remodeling | Chromatin remodeling helicase | Chromatin remodeling helicase | HEMD | |
| CHD7 | Chromatin remodeling | Chromatin remodeling helicase | Chromatin remodeling helicase | HEMD | |
| CHD8 | Histone modification | Readers | Readers | HEMD | |
| CHD9 | Chromatin remodeling | Chromatin remodeling helicase | Chromatin remodeling helicase | HEMD | |
| CHEK1 | Chromatin remodeling | chromatin remodeling factors | chromatin remodelling DNA repair | TAS | Other conserved DNA damage response genes |
| Chk1 | Histone modification | Writers | Histone phosphorylation | HEMD | |
| CHRAC1 | Chromatin remodeling | chromatin remodeling factors | chromatin remodelling | TAS | |
| CHUK | Histone modification | Writers | Histone phosphorylation | HEMD | |
| CLOCK | Histone modification | Writers | Histone acetylation | HEMD | |
| CREBBP | Histone modification | Writers | Histone acetylation | HEMD | |
| CSRP2BP | Histone modification | Readers | Readers | TAS | |
| CXXC1 | Histone modification | Readers | Readers | IDA | |
| DAPK3 | Histone modification | Writers | Histone phosphorylation | HEMD | |
| DDB1 | Histone modification | Writers | Histone ubiquitination DNA repair | HEMD | Nucleotide excision repair (NER) |
| DDB2 | Histone modification | Writers | Histone ubiquitination DNA repair | HEMD | Nucleotide excision repair (NER) |
| DMAP1 | Histone modification | Readers | Readers | NAS | |
| DNMT1 | DNA modifications | writers | DNA Methyltransferase (DNMTs) | HEMD | |
| DNMT3A | DNA modifications | writers | DNA Methyltransferase (DNMTs) | HEMD | |
| DNMT3B | DNA modifications | writers | DNA Methyltransferase (DNMTs) | HEMD | |
| DNMT3L | DNA modifications | writers | DNA methylation | HEMD | |
| DOT1L | Histone modification | Writers | Histone methylation | HEMD | |
| DPF1 | Chromatin remodeling | nucleosome remodeling factors | histone binding | NAS | |
| DPF2 | Chromatin remodeling | nucleosome remodeling factors | histone binding | NAS | |
| DTX3L | Histone modification | Writers | Histone ubiquitination | HEMD | |
| DUSP1 | Histone modification | Editors | Histone dephosphorylation | HEMD | |
| DZIP3 | Histone modification | Writers | Histone ubiquitination | HEMD | |
| EED | Histone modification | Readers | Readers | HEMD | |
| EHMT1 | Histone modification | Writers | Histone methylation | HEMD | |
| EHMT2 | Histone modification | Writers | Histone methylation | HEMD | |
| ELP3 | Histone modification | Writers | Histone acetylation | HEMD | |
| ELP4 | Chromatin remodeling | chromatin remodeling factors | histone organization | IDA | |
| EP300 | Histone modification | Writers | Histone acetylation | HEMD | |
| EP400 | Histone modification | Readers | Readers | HEMD | |
| EPC1 | Histone modification | Readers | Readers | IDA | |
| EPC2 | Histone modification | Readers | Readers | UCSC | |
| ERCC6 | Chromatin remodeling | Chromatin remodeling helicase | Chromatin remodeling helicase DNA repair | HEMD | Nucleotide excision repair (NER) |
| EVI1 | Histone modification | Writers | Histone methylation | HEMD | |
| EYA1 | Histone modification | Editors | Histone dephosphorylation | HEMD | |
| EYA2 | Histone modification | Editors | Histone dephosphorylation | HEMD | |
| EYA3 | Histone modification | Editors | Histone dephosphorylation | HEMD | |
| EYA4 | Histone modification | Editors | Histone dephosphorylation | HEMD | |
| EZH1 | Histone modification | Writers | Histone methylation | HEMD | |
| EZH2 | Histone modification | Writers | Histone methylation | HEMD | |
| FOS | DNA modifications | writers | DNA methylation | ISS | |
| FOXA1 | Chromatin remodeling | chromatin remodeling factors | chromatin remodelling | TAS | |
| FSHR | Chromatin remodeling | chromatin remodeling factors | chromatin remodelling | NAS | |
| FTO | DNA modifications | editors | DNA demethylation | HEMD | |
| FYN | Histone modification | Writers | Histone phosphorylation | HEMD | |
| GADD45A | DNA modifications | editors | DNA demethylation | HEMD | |
| GADD45B | DNA modifications | editors | DNA demethylation | HEMD | |
| GATAD2A | Histone modification | Readers | Readers | TAS | |
| GATAD2B | Histone modification | Readers | Readers | IDA | |
| GNAS | DNA modifications | writers | DNA methylation | TAS | |
| GSG2 | Histone modification | Writers | Histone phosphorylation | HEMD | |
| GSK3B | Histone modification | Writers | Histone phosphorylation | HEMD | |
| GTF3C4 | Histone modification | Writers | Histone acetylation | HEMD | |
| H1F0 | Histone modification | Histones | histones | UCSC | |
| H1FNT | Histone modification | Histones | histones | UCSC | |
| H1FOO | Histone modification | Histones | histones | UCSC | |
| H1FX | Histone modification | Histones | histones | UCSC | |
| H2AFB2 | Histone modification | Histones | histones | UCSC | |
| H2AFB3 | Histone modification | Histones | histones | UCSC | |
| H2AFJ | Histone modification | Histones | histones | UCSC | |
| H2AFV | Histone modification | Histones | histones | UCSC | |
| H2AFX | Histone modification | Histones | Histones | UCSC | Chromatin Structure and Modification |
| H2AFY | Histone modification | Histones | Histones | IDA | |
| H2AFY2 | Histone modification | Histones | Histones | IDA | |
| H2AFZ | Histone modification | Histones | histones | UCSC | |
| H2BFWT | Histone modification | Histones | histones | UCSC | |
| H3F3B | Histone modification | Histones | histones | UCSC | |
| H3F3C | Histone modification | Histones | histones | UCSC | |
| HAT1 | Histone modification | Writers | Histone acetylation | HEMD | |
| HCFC1 | Histone modification | Readers | Readers | IDA | |
| HDAC1 | Histone modification | Editors | Histone Deacetylase (HDACs) | HEMD | |

| | | | | | |
|---|---|---|---|---|---|
| HDAC10 | Histone modification | Editors | Histone Deacetylase (HDACs) | HEMD | |
| HDAC11 | Histone modification | Editors | Histone Deacetylase (HDACs) | HEMD | |
| HDAC2 | Histone modification | Editors | Histone Deacetylase (HDACs) | HEMD | |
| HDAC3 | Histone modification | Editors | Histone Deacetylase (HDACs) | HEMD | |
| HDAC4 | Histone modification | Editors | Histone Deacetylase (HDACs) | HEMD | |
| HDAC5 | Histone modification | Editors | Histone Deacetylase (HDACs) | HEMD | |
| HDAC6 | Histone modification | Editors | Histone Deacetylase (HDACs) | HEMD | |
| HDAC7 | Histone modification | Editors | Histone Deacetylase (HDACs) | HEMD | |
| HDAC8 | Histone modification | Editors | Histone Deacetylase (HDACs) | HEMD | |
| HDAC9 | Histone modification | Editors | Histone Deacetylase (HDACs) | HEMD | |
| HELLS | Chromatin remodeling | Chromatin remodeling helicase | Chromatin remodeling helicase | HEMD | |
| HILS1 | Chromatin remodeling | chromatin remodeling factors | chromatin remodelling | IEA | |
| HIST1H2AA | Histone modification | Histones | Histones | UCSC | |
| HIST1H2AB | Histone modification | Histones | histones | UCSC | |
| HIST1H2AC | Histone modification | Histones | histones | UCSC | |
| HIST1H2AD | Histone modification | Histones | histones | UCSC | |
| HIST1H2AE | Histone modification | Histones | histones | UCSC | |
| HIST1H2AG | Histone modification | Histones | histones | UCSC | |
| HIST1H2AH | Histone modification | Histones | histones | UCSC | |
| HIST1H2AJ | Histone modification | Histones | histones | UCSC | |
| HIST1H2AK | Histone modification | Histones | histones | UCSC | |
| HIST1H2AL | Histone modification | Histones | Histones | UCSC | |
| HIST1H2AM | Histone modification | Histones | histones | UCSC | |
| HIST1H2BA | Histone modification | Histones | Histones | UCSC | |
| HIST1H2BB | Histone modification | Histones | histones | UCSC | |
| HIST1H2BC | Histone modification | Histones | histones | UCSC | |
| HIST1H2BD | Histone modification | Histones | histones | UCSC | |
| HIST1H2BE | Histone modification | Histones | histones | UCSC | |
| HIST1H2BF | Histone modification | Histones | histones | UCSC | |
| HIST1H2BG | Histone modification | Histones | histones | UCSC | |
| HIST1H2BH | Histone modification | Histones | histones | UCSC | |
| HIST1H2BI | Histone modification | Histones | histones | UCSC | |
| HIST1H2BJ | Histone modification | Histones | Histones | UCSC | |
| HIST1H2BK | Histone modification | Histones | histones | UCSC | |
| HIST1H2BL | Histone modification | Histones | histones | UCSC | |
| HIST1H2BM | Histone modification | Histones | histones | UCSC | |
| HIST1H2BN | Histone modification | Histones | histones | UCSC | |
| HIST1H2BO | Histone modification | Histones | histones | UCSC | |
| HIST1H3A | Histone modification | Histones | Histones | UCSC | |
| HIST1H3B | Histone modification | Histones | histones | UCSC | |
| HIST1H3C | Histone modification | Histones | histones | UCSC | |
| HIST1H3D | Histone modification | Histones | histones | UCSC | |
| HIST1H3E | Histone modification | Histones | histones | UCSC | |
| HIST1H3F | Histone modification | Histones | histones | UCSC | |
| HIST1H3G | Histone modification | Histones | histones | UCSC | |
| HIST1H3H | Histone modification | Histones | histones | UCSC | |
| HIST1H3I | Histone modification | Histones | histones | UCSC | |
| HIST1H3J | Histone modification | Histones | histones | UCSC | |
| HIST1H4A | Histone modification | Histones | Histones | UCSC | |
| HIST1H4B | Histone modification | Histones | histones | UCSC | |
| HIST1H4C | Histone modification | Histones | histones | UCSC | |
| HIST1H4D | Histone modification | Histones | histones | UCSC | |
| HIST1H4E | Histone modification | Histones | histones | UCSC | |
| HIST1H4F | Histone modification | Histones | histones | UCSC | |
| HIST1H4G | Histone modification | Histones | histones | UCSC | |
| HIST1H4H | Histone modification | Histones | histones | UCSC | |
| HIST1H4I | Histone modification | Histones | histones | UCSC | |
| HIST1H4J | Histone modification | Histones | histones | UCSC | |
| HIST1H4K | Histone modification | Histones | Histones | UCSC | |
| HIST1H4L | Histone modification | Histones | histones | UCSC | |
| HIST2H2AA3 | Histone modification | Histones | Histones | UCSC | |
| HIST2H2AB | Histone modification | Histones | histones | UCSC | |
| HIST2H2AC | Histone modification | Histones | histones | UCSC | |
| HIST2H2BE | Histone modification | Histones | Histones | UCSC | |
| HIST2H2BF | Histone modification | Histones | histones | UCSC | |
| HIST2H3C | Histone modification | Histones | Histones | UCSC | |
| HIST2H3D | Histone modification | Histones | histones | UCSC | |
| HIST2H4A | Chromatin remodeling | chromatin remodeling factors | chromatin remodelling | NAS | |
| HIST3H2A | Histone modification | Histones | histones | UCSC | |
| HIST3H2BB | Histone modification | Histones | histones | UCSC | |
| HIST3H3 | Histone modification | Histones | Histones | UCSC | |
| HIST4H4 | Histone modification | Histones | Histones | UCSC | |
| HJURP | Chromatin remodeling | chromatin remodeling factors | chromatin remodelling | IC | |
| HLCS | Histone modification | Writers | Histone biotinylation | HEMD | |
| HLTF | Chromatin remodeling | nucleosome remodeling factors | | HEMD | Ubiquitination and modification |
| HMGA1 | Chromatin remodeling | chromatin remodeling factors | chromatin remodelling | IDA | |
| HNF1A | Chromatin remodeling | chromatin remodeling factors | chromatin remodelling | IDA | |
| HUWE1 | Histone modification | Writers | Histone ubiquitination | HEMD | |
| IDH1 | DNA modifications | writers | DNA methylation | TAS | |
| IDH2 | DNA modifications | writers | DNA methylation | ISS | |
| ING2 | Histone modification | Readers | Readers | UCSC | |
| ING3 | Histone modification | Readers | Readers | IDA | |
| INO80 | Chromatin remodeling | Chromatin remodeling helicase | Chromatin remodeling helicase | HEMD | |
| ITGB3BP | Chromatin remodeling | chromatin remodeling factors | chromatin remodelling | IDA | |
| JAK2 | Histone modification | Writers | Histone phosphorylation | HEMD | |
| JMJD1C | Histone modification | Editors | Histone demethylation | HEMD | |
| JMJD6 | Histone modification | Editors | Histone demethylation | HEMD | |
| KANSL1 | Chromatin remodeling | chromatin remodeling factors | histone organization | IDA | |
| KANSL2 | Chromatin remodeling | chromatin remodeling factors | histone organization | IDA | |
| KANSL3 | Chromatin remodeling | chromatin remodeling factors | histone organization | IDA | |
| KAT2A | Histone modification | Writers | Histone acetylation | HEMD | |
| KAT2B | Histone modification | Writers | Histone acetylation | HEMD | |
| KAT5 | Histone modification | Writers | Histone acetylation | HEMD | |
| KAT8 | Chromatin remodeling | chromatin remodeling factors | histone organization | IDA | |
| KDM1A | Histone modification | Editors | Histone demethylation | HEMD | |
| KDM1B | Histone modification | Editors | Histone demethylation | HEMD | |
| KDM2A | Histone modification | Editors | Histone demethylation | HEMD | |
| KDM2B | Histone modification | Editors | Histone demethylation | HEMD | |
| KDM3A | Histone modification | Editors | Histone demethylation | HEMD | |
| KDM3B | Histone modification | Editors | Histone demethylation | HEMD | |
| KDM4A | Histone modification | Editors | Histone demethylation | HEMD | |
| KDM4B | Histone modification | Editors | Histone demethylation | HEMD | |
| KDM4C | Histone modification | Editors | Histone demethylation | HEMD | |
| KDM4D | Histone modification | Editors | Histone demethylation | HEMD | |
| KDM5A | Histone modification | Editors | Histone demethylation | HEMD | |
| KDM5B | Histone modification | Editors | Histone demethylation | HEMD | |
| KDM5C | Histone modification | Editors | Histone demethylation | HEMD | |
| KDM5D | Histone modification | Editors | Histone demethylation | HEMD | |
| KDM6A | Histone modification | Editors | Histone demethylation | HEMD | |
| KDM6B | Histone modification | Editors | Histone demethylation | HEMD | |
| KDM7 | Histone modification | Editors | Histone demethylation | HEMD | |
| KDM8 | Histone modification | Editors | Histone demethylation | HEMD | |
| KLF1 | Chromatin remodeling | chromatin remodeling factors | chromatin remodelling | IDA | |
| L3MBTL2 | Histone modification | Readers | Readers | UCSC | |
| LIMK2 | Histone modification | Writers | Histone phosphorylation | HEMD | |
| MAEL | DNA modifications | writers | DNA methylation | | |
| MAP3K12 | Histone modification | Writers | Histone phosphorylation | HEMD | |
| MAP3K8 | Histone modification | Writers | Histone phosphorylation | HEMD | |
| MASTL | Histone modification | Writers | Histone phosphorylation | HEMD | |
| MBD1 | DNA modifications | readers | DNA methylation | UCSC | |
| MBD2 | Histone modification | Readers | Readers | HEMD | |
| MBD3 | DNA modifications | readers | DNA methylation | HEMD | |
| MBD4 | DNA modifications | readers | DNA methylation | HEMD | Base excision repair (BER) |
| MBIP | Histone modification | Readers | Readers | IDA | |
| MCRS1 | Chromatin remodeling | nucleosome remodeling factors | | IDA | |
| Mdm2 | Histone modification | Writers | Histone ubiquitination | HEMD | |
| MEAF6 | Histone modification | Readers | Readers | IDA | |
| MECP2 | DNA modifications | readers | DNA methylation | UCSC | |
| MEN1 | Histone modification | Readers | Readers | IEA | |
| MGMT | DNA modifications | writers | DNA methylation DNA repair | IMP | Direct reversal of damage |
| MIS18A | Chromatin remodeling | chromatin remodeling factors | chromatin remodelling | IDA | |
| MIS18BP1 | Chromatin remodeling | chromatin remodeling factors | chromatin remodelling | IBA | |
| MIZF | DNA modifications | writers | DNA methylation | NAS | |
| MLF1IP | Chromatin remodeling | chromatin remodeling factors | chromatin remodelling | TAS | |
| MLL | Histone modification | Writers | Histone methylation | HEMD | |
| MLL2 | Histone modification | Writers | Histone methylation | HEMD | |
| MLL3 | Histone modification | Writers | Histone methylation | HEMD | |
| MLL4 | Histone modification | Writers | Histone methylation | HEMD | |
| MLL5 | Histone modification | Writers | Histone methylation | HEMD | |
| MLT | Histone modification | Writers | Histone phosphorylation | HEMD | |
| MORF4L1 | Histone modification | Readers | Readers | IDA | |
| MORF4L2 | Histone modification | Readers | Readers | UCSC | |
| MTA1 | Histone modification | Readers | Readers | NAS | |
| MTA2 | Histone modification | Readers | Readers | UCSC | |
| MTA3 | Histone modification | Readers | Readers | NAS | |

| | | | | | |
|---|---|---|---|---|---|
| MYB | Chromatin remodeling | chromatin remodeling factors | chromatin remodelling | IMP | |
| MYC | Chromatin remodeling | chromatin remodeling factors | chromatin remodelling | NAS | |
| MYSM1 | Histone modification | Editors | Histone deubiquitination | HEMD | |
| MYST1 | Histone modification | Writers | Histone acetylation | HEMD | |
| MYST2 | Histone modification | Writers | Histone acetylation | HEMD | |
| MYST3 | Histone modification | Writers | Histone acetylation | HEMD | |
| MYST4 | Histone modification | Writers | Histone acetylation | HEMD | |
| NAP1L3 | Chromatin remodeling | nucleosome remodeling factors | nucleosome assembly proteins | NAS | |
| NAP1L4 | Chromatin remodeling | nucleosome remodeling factors | nucleosome assembly proteins | NAS | |
| NASP | Chromatin remodeling | chromatin remodeling factors | chromatin remodelling | NAS | |
| NAT10 | Histone modification | Writers | Histone acetylation | HEMD | |
| NCOA1 | Histone modification | Writers | Histone acetylation | HEMD | |
| NCOA2 | Histone modification | Writers | Histone acetylation | HEMD | |
| NCOA3 | Histone modification | Writers | Histone acetylation | HEMD | |
| NCOAT | Histone modification | Editors | Histone de-O-GlcNAcylation | HEMD | |
| Nek6 | Histone modification | Writers | Histone phosphorylation | HEMD | |
| Nek9 | Histone modification | Writers | Histone phosphorylation | HEMD | |
| NFRKB | Chromatin remodeling | nucleosome remodeling factors | | UCSC | |
| NIPBL | Histone modification | Readers | Readers | IEA | |
| NO66 | Histone modification | Editors | Histone demethylation | HEMD | |
| NPM1 | Chromatin remodeling | chromatin remodeling factors | chromatin remodelling | IEA | |
| NPM2 | Chromatin remodeling | chromatin remodeling factors | chromatin remodelling | IEA | |
| NR3C1 | Chromatin remodeling | chromatin remodeling factors | chromatin remodelling | IBA | |
| NSD1 | Histone modification | Writers | Histone methylation | HEMD | |
| NSD2 | Histone modification | Writers | Histone methylation | HEMD | |
| NSD3 | Histone modification | Writers | Histone methylation | HEMD | |
| OGT | Histone modification | Writers | Histone O-GlcNAcylation | HEMD | |
| OIP5 | Chromatin remodeling | chromatin remodeling factors | chromatin remodelling | NAS | |
| PADI4 | Histone modification | Writers | Histone citrullination | HEMD | |
| PAK1 | Histone modification | Writers | Histone phosphorylation | HEMD | |
| PAK2 | Histone modification | Writers | Histone phosphorylation | HEMD | |
| PARG | Histone modification | Editors | Histone de-ADP-ribosylation | HEMD | |
| PARP1 | Histone modification | Writers | Histone ADP-ribosylation DNA repair | HEMD | Poly(ADP-ribose) polymerase (PARP) enzymes that bind to DNA |
| PARP2 | Histone modification | Writers | Histone ADP-ribosylation DNA repair | HEMD | Poly(ADP-ribose) polymerase (PARP) enzymes that bind to DNA |
| PARP3 | Histone modification | Writers | Histone ADP-ribosylation DNA repair | HEMD | Poly(ADP-ribose) polymerase (PARP) enzymes that bind to DNA |
| PBK | Histone modification | Writers | Histone phosphorylation | HEMD | |
| PBRM1 | Chromatin remodeling | chromatin remodeling factors | chromatin remodelling | TAS | |
| PCGF6 | Histone modification | Readers | Readers | IEA | |
| PCNA | DNA modifications | readers | DNA methylation | | FANCA |
| PHF10 | Chromatin remodeling | chromatin remodeling factors | chromatin-remodeling complex | NAS | |
| PHF15 | Histone modification | Readers | Readers | IDA | |
| PHF16 | Histone modification | Readers | Readers | IEA | |
| PHF17 | Histone modification | Readers | Readers | IMP | |
| PHF20 | Chromatin remodeling | chromatin remodeling factors | histone organization | IDA | |
| PHF8 | Histone modification | Editors | Histone demethylation | HEMD | |
| PKN1 | Histone modification | Writers | Histone phosphorylation | HEMD | |
| PPM1D | Histone modification | Editors | Histone dephosphorylation | HEMD | |
| PPP1CA | Histone modification | Editors | Histone dephosphorylation | HEMD | |
| PPP1CB | Histone modification | Editors | Histone dephosphorylation | HEMD | |
| PPP1CC | Histone modification | Editors | Histone dephosphorylation | HEMD | |
| PPP2CA | Histone modification | Editors | Histone dephosphorylation | HEMD | |
| PPP2CB | Histone modification | Editors | Histone dephosphorylation | HEMD | |
| PPP2R1A | Histone modification | Editors | Histone dephosphorylation | HEMD | |
| PPP2R1B | Histone modification | Editors | Histone dephosphorylation | HEMD | |
| PPP2R2A | Histone modification | Editors | Histone dephosphorylation | HEMD | |
| PPP2R2B | Histone modification | Editors | Histone dephosphorylation | HEMD | |
| PPP2R2C | Histone modification | Editors | Histone dephosphorylation | HEMD | |
| PPP2R2D | Histone modification | Editors | Histone dephosphorylation | HEMD | |
| PPP2R3A | Histone modification | Editors | Histone dephosphorylation | HEMD | |
| PPP2R3B | Histone modification | Editors | Histone dephosphorylation | HEMD | |
| PPP2R3C | Histone modification | Editors | Histone dephosphorylation | HEMD | |
| PPP2R4 | Histone modification | Editors | Histone dephosphorylation | HEMD | |
| PPP2R5A | Histone modification | Editors | Histone dephosphorylation | HEMD | |
| PPP2R5B | Histone modification | Editors | Histone dephosphorylation | HEMD | |
| PPP2R5C | Histone modification | Editors | Histone dephosphorylation | HEMD | |
| PPP2R5D | Histone modification | Editors | Histone dephosphorylation | HEMD | |
| PPP2R5E | Histone modification | Editors | Histone dephosphorylation | HEMD | |
| PPP3CA | Histone modification | Editors | Histone dephosphorylation | HEMD | |
| PPP3CB | Histone modification | Editors | Histone dephosphorylation | HEMD | |
| PPP3CC | Histone modification | Editors | Histone dephosphorylation | HEMD | |
| PPP4C | Histone modification | Editors | Histone dephosphorylation | HEMD | |
| PPP4R1 | Histone modification | Editors | Histone dephosphorylation | HEMD | |
| PPP4R2 | Histone modification | Editors | Histone dephosphorylation | HEMD | |
| PPP4R4 | Histone modification | Editors | Histone dephosphorylation | HEMD | |
| PPP5C | Histone modification | Editors | Histone dephosphorylation | HEMD | |
| PPP6C | Histone modification | Editors | Histone dephosphorylation | HEMD | |
| PPP6R1 | Histone modification | Editors | Histone dephosphorylation | HEMD | |
| PPP6R2 | Histone modification | Editors | Histone dephosphorylation | HEMD | |
| PPP6R3 | Histone modification | Editors | Histone dephosphorylation | HEMD | |
| PRDM2 | Histone modification | Writers | Histone methylation | HEMD | |
| PRDM6 | Histone modification | Writers | Histone methylation | HEMD | |
| PRDM7 | Histone modification | Writers | Histone methylation | HEMD | |
| PRDM9 | Histone modification | Writers | Histone methylation | HEMD | |
| PRKAA1 | Histone modification | Writers | Histone phosphorylation | HEMD | |
| PRKAA2 | Histone modification | Writers | Histone phosphorylation | HEMD | |
| PRKAB1 | Histone modification | Writers | Histone phosphorylation | HEMD | |
| PRKAB2 | Histone modification | Writers | Histone phosphorylation | HEMD | |
| PRKAG1 | Histone modification | Writers | Histone phosphorylation | HEMD | |
| PRKAG2 | Histone modification | Writers | Histone phosphorylation | HEMD | |
| PRKAG3 | Histone modification | Writers | Histone phosphorylation | HEMD | |
| PRKCA | Histone modification | Writers | Histone phosphorylation | HEMD | |
| PRKCB | Histone modification | Writers | Histone phosphorylation | HEMD | |
| PRKDC | Histone modification | Writers | Histone phosphorylation DNA repair | HEMD | Non-homologous end-joining |
| PRMT1 | Histone modification | Writers | Histone methylation | HEMD | |
| PRMT10 | Histone modification | Writers | Histone methylation | HEMD | |
| PRMT11 | Histone modification | Writers | Histone methylation | HEMD | |
| PRMT12 | Histone modification | Writers | Histone methylation | HEMD | |
| PRMT13 | Histone modification | Writers | Histone methylation | HEMD | |
| PRMT14 | Histone modification | Writers | Histone methylation | HEMD | |
| PRMT15 | Histone modification | Writers | Histone methylation | HEMD | |
| PRMT16 | Histone modification | Writers | Histone methylation | HEMD | |
| PRMT2 | Histone modification | Writers | Histone methylation | HEMD | |
| PRMT3 | Histone modification | Writers | Histone methylation | HEMD | |
| PRMT4 | Histone modification | Writers | Histone methylation | HEMD | |
| PRMT5 | Histone modification | Writers | Histone methylation | HEMD | |
| PRMT6 | Histone modification | Writers | Histone methylation | HEMD | |
| PRMT7 | Histone modification | Writers | Histone methylation | HEMD | |
| PRMT8 | Histone modification | Writers | Histone methylation | HEMD | |
| RAD54B | Chromatin remodeling | Chromatin remodeling helicase | Chromatin remodeling helicase DNA repair | HEMD | Homologous recombination |
| RAD54L | Chromatin remodeling | Chromatin remodeling helicase | Chromatin remodeling helicase DNA repair | HEMD | Homologous recombination |
| RAG1 | Histone modification | Writers | Histone ubiquitination | HEMD | |
| RB1 | Chromatin remodeling | chromatin remodeling factors | chromatin remodelling | IEA | |
| RBBP4 | Chromatin remodeling | chromatin remodeling factors | chromatin remodelling | IDA | |
| RBBP5 | Histone modification | Readers | Readers | IDA | |
| RBBP7 | Chromatin remodeling | chromatin remodeling factors | chromatin remodelling | TAS | |
| RBX1 | Histone modification | Writers | Histone ubiquitination | HEMD | |
| RERE | Chromatin remodeling | chromatin remodeling factors | chromatin remodelling | ISS | |
| RING1 | Histone modification | Writers | Histone ubiquitination | HEMD | |
| RING2 | Histone modification | Writers | Histone ubiquitination | HEMD | |
| RNF168 | Histone modification | Writers | Histone ubiquitination DNA repair | HEMD | Ubiquitination and modification |
| RNF20 | Histone modification | Writers | Histone ubiquitination | IDA | |
| RNF8 | Histone modification | Writers | Histone ubiquitination DNA repair | HEMD | Ubiquitination and modification |
| RPS6KA3 | Histone modification | Writers | Histone phosphorylation | HEMD | |
| RPS6KA4 | Histone modification | Writers | Histone phosphorylation | HEMD | |
| RPS6KA5 | Histone modification | Writers | Histone phosphorylation | HEMD | |
| RSF1 | Chromatin remodeling | chromatin remodeling factors | chromatin remodelling | TAS | |
| RUVBL1 | Histone modification | Readers | Readers | TAS | |
| RUVBL2 | Histone modification | Readers | Readers | IMP | |
| SAE1 | Histone modification | Writers | Histone sumoylation | HEMD | |
| SAE2 | Histone modification | Writers | Histone sumoylation | HEMD | |
| SALL1 | Chromatin remodeling | chromatin remodeling factors | histone organization | IDA | |
| SAP130 | Histone modification | Readers | Readers | IDA | |
| SATB2 | Chromatin remodeling | chromatin remodeling factors | chromatin remodelling | TAS | |
| SETD1A | Histone modification | Readers | Readers | HEMD | |
| SETD1B | Histone modification | Readers | Readers | HEMD | |
| SETD2 | Histone modification | Writers | Histone methylation | HEMD | |
| SETD3 | Histone modification | Writers | Histone methylation | HEMD | |
| SETD5 | Histone modification | Writers | Histone methylation | HEMD | |
| SETD7 | Histone modification | Writers | Histone methylation | HEMD | |
| SETD8 | Histone modification | Writers | Histone methylation | HEMD | |
| SETDB1 | Histone modification | Writers | Histone methylation | HEMD | |
| SETDB2 | Histone modification | Writers | Histone methylation | HEMD | |
| SETMAR | Histone modification | Writers | Histone methylation DNA repair | HEMD | Chromatin Structure and Modification |
| SIRT1 | Histone modification | Editors | Histone Deacetylase (HDACs) | HEMD | |
| SIRT2 | Histone modification | Editors | Histone Deacetylase (HDACs) | HEMD | |

| | | | | | |
|---|---|---|---|---|---|
| SIRT3 | Histone modification | Editors | Histone Deacetylase (HDACs) | HEMD | |
| SIRT4 | Histone modification | Editors | Histone Deacetylase (HDACs) | HEMD | |
| SIRT5 | Histone modification | Editors | Histone Deacetylase (HDACs) | HEMD | |
| SIRT6 | Histone modification | Editors | Histone Deacetylase (HDACs) | HEMD | |
| SIRT7 | Histone modification | Editors | Histone Deacetylase (HDACs) | HEMD | |
| SMARCA1 | Chromatin remodeling | Chromatin remodeling helicase | Chromatin remodeling helicase | HEMD | |
| SMARCA2 | Chromatin remodeling | Chromatin remodeling helicase | Chromatin remodeling helicase | HEMD | |
| SMARCA5 | Histone modification | Readers | Readers | HEMD | |
| SMARCAD1 | Chromatin remodeling | Chromatin remodeling helicase | Chromatin remodeling helicase | HEMD | |
| SMARCAL1 | Chromatin remodeling | Chromatin remodeling helicase | Chromatin remodeling helicase | HEMD | |
| SMARCC1 | Chromatin remodeling | chromatin remodeling factors | chromatin remodelling | TAS | |
| SMARCC2 | Chromatin remodeling | chromatin remodeling factors | chromatin remodelling | IEA | |
| SMARCD1 | Chromatin remodeling | chromatin remodeling factors | chromatin remodelling | IEA | |
| SMARCD2 | Chromatin remodeling | chromatin remodeling factors | chromatin remodelling | TAS | |
| SMARCD3 | Chromatin remodeling | chromatin remodeling factors | chromatin remodelling | IDA | |
| SMARCE1 | Chromatin remodeling | chromatin remodeling factors | chromatin remodelling | IEA | |
| SMCHD1 | Histone modification | Readers | Readers | IMP | |
| SMEK1 | Histone modification | Editors | Histone dephosphorylation | HEMD | |
| SMEK2 | Histone modification | Editors | Histone dephosphorylation | HEMD | |
| SMYD1 | Histone modification | Writers | Histone methylation | HEMD | |
| SMYD3 | Histone modification | Writers | Histone methylation | HEMD | |
| SMYD4 | Histone modification | Writers | Histone methylation | HEMD | |
| SMYD5 | Histone modification | Writers | Histone methylation | HEMD | |
| SOX9 | Chromatin remodeling | chromatin remodeling factors | chromatin remodelling | IDA | |
| SPI1 | DNA modifications | writers | DNA methylation, negative regulation of histone H4 acetylation | IDA | |
| SRCAP | Chromatin remodeling | Chromatin remodeling helicase | Chromatin remodeling helicase | HEMD | |
| STK10 | Histone modification | Writers | Histone phosphorylation | HEMD | |
| STK12 | Histone modification | Writers | Histone phosphorylation | HEMD | |
| STK13 | Histone modification | Writers | Histone phosphorylation | HEMD | |
| STK4 | Histone modification | Writers | Histone phosphorylation | HEMD | |
| STK6 | Histone modification | Writers | Histone phosphorylation | HEMD | |
| SUDS3 | Histone modification | Readers | Readers | UCSC | |
| SUPT3H | Histone modification | Readers | Readers | IDA | |
| SUPT4H1 | Chromatin remodeling | chromatin remodeling factors | chromatin remodelling | IMP | |
| SUPT5H | Chromatin remodeling | chromatin remodeling factors | chromatin remodelling | TAS | |
| SUPT6H | Chromatin remodeling | chromatin remodeling factors | chromatin remodelling | TAS | |
| SUPT7L | Histone modification | Readers | Readers | IDA | |
| SUV39H1 | Histone modification | Writers | Histone methylation | HEMD | |
| SUV39H2 | Histone modification | Writers | Histone methylation | HEMD | |
| SUV420H1 | Histone modification | Writers | Histone methylation | HEMD | |
| SUV420H2 | Histone modification | Writers | Histone methylation | HEMD | |
| SUZ12 | Histone modification | Readers | Readers | HEMD | |
| SYCP3 | Chromatin remodeling | chromatin remodeling factors | chromatin remodelling | TAS | |
| TADA1 | Chromatin remodeling | chromatin remodeling factors | histone organization | IDA | |
| TADA2A | Chromatin remodeling | chromatin remodeling factors | histone organization | IDA | |
| TADA3 | Chromatin remodeling | chromatin remodeling factors | histone organization | IDA | |
| TAF1 | Histone modification | Readers | Readers | HEMD | |
| TAF10 | Histone modification | Readers | Readers | IDA | |
| TAF12 | Histone modification | Readers | Readers | IDA | |
| TAF5 | Histone modification | Readers | Readers | UCSC | |
| TAF5L | Histone modification | Readers | Readers | IDA | |
| TAF6L | Histone modification | Readers | Readers | IDA | |
| TAF7 | Histone modification | Readers | Readers | IDA | |
| TAF9 | Histone modification | Readers | Readers | IDA | |
| TDG | DNA modifications | editors | DNA demethylation DNA repair | HEMD | Base excision repair (BER) |
| TET1 | DNA modifications | editors | DNA demethylation | HEMD | |
| TET2 | DNA modifications | editors | DNA demethylation | HEMD | |
| TET3 | DNA modifications | editors | DNA demethylation | HEMD | |
| TLK1 | Histone modification | Writers | Histone phosphorylation | HEMD | |
| TNP1 | Chromatin remodeling | chromatin remodeling factors | chromatin remodelling | IEA | |
| TOP1 | Chromatin remodeling | chromatin remodeling factors | chromatin remodelling | TAS | |
| TOP1MT | Chromatin remodeling | chromatin remodeling factors | chromatin remodelling | IEA | |
| TP63 | Chromatin remodeling | chromatin remodeling factors | chromatin remodelling | IDA | |
| TRDMT1 | DNA modifications | writers | DNA methylation | HEMD | |
| TRIM24 | Histone modification | Readers | Readers | UCSC | |
| TRIM28 | Chromatin remodeling | nucleosome remodeling factors | | UCSC | |
| TRRAP | Histone modification | Readers | Readers | ISS | |
| TTF1 | Chromatin remodeling | chromatin remodeling factors | chromatin remodelling | TAS | |
| TTF2 | Chromatin remodeling | Chromatin remodeling helicase | Chromatin remodeling helicase | HEMD | |
| UBA1 | Histone modification | Writers | Histone ubiquitination | HEMD | |
| UBC9 | Histone modification | Writers | Histone sumoylation | HEMD | |
| UBE2A | Histone modification | Writers | Histone ubiquitination DNA repair | HEMD | Ubiquitination and modification |
| UBE2B | Histone modification | Writers | Histone ubiquitination DNA repair | HEMD | Ubiquitination and modification |
| UBE2E1 | Histone modification | Writers | Histone ubiquitination | HEMD | |
| UBE2H | Histone modification | Writers | Histone ubiquitination | HEMD | |
| UBR2 | Histone modification | Writers | Histone ubiquitination | HEMD | |
| UHRF1 | DNA modifications | readers | DNA methylation | HEMD | |
| USP16 | Histone modification | Editors | Histone deubiquitination | HEMD | |
| USP21 | Histone modification | Editors | Histone deubiquitination | HEMD | |
| USP22 | Histone modification | Readers | Readers | HEMD | |
| USP3 | Histone modification | Editors | Histone deubiquitination | HEMD | |
| USP36 | Histone modification | Editors | Histone deubiquitination | HEMD | |
| USP7 | Histone modification | Editors | Histone deubiquitination | HEMD | |
| UTY | Histone modification | Editors | Histone demethylation | HEMD | |
| VRK1 | Histone modification | Writers | Histone phosphorylation | HEMD | |
| WDR82 | Chromatin remodeling | chromatin remodeling factors | histone organization | ISS | |
| YEATS2 | Histone modification | Readers | Readers | IDA | |
| YEATS4 | Histone modification | Readers | Readers | IDA | |
| YY1 | Chromatin remodeling | nucleosome remodeling factors | | UCSC | |
| ZFP57 | DNA modifications | writers | DNA methylation | | |
| ARID1A | Not assigned | Not assigned | reg. of gene expression, epigenetic | IDA | |
| ASXL1 | Not assigned | Not assigned | methylation, chromatin | UCSC | |
| DNMT1 | DNA modifications | writers | DNA Methyltransferase (DNMTs) | HEMD | |
