## Supplementary Table S6 for "Methylome-based cell-of-origin modeling (Methyl-COOM) identifies aberrant expression of immune regulatory molecules in CLL"

**Supplementary Table S6: List of epigenetic regulators being targeted by CLL-specific microRNAs**

| miRNA | Target.Gene | Experiments | Database |
|---|---|---|---|
| hsa-miR-486-5p | H3F3B | PAR-CLIP | miRTarbase |
| hsa-miR-486-5p | H3F3B | PAR-CLIP | miRTarbase |
| hsa-miR-486-5p | H3F3B | qRT-PCR | miRTarbase |
| hsa-miR-486-5p | H3F3B | PAR-CLIP | miRTarbase |
| hsa-miR-486-5p | H3F3B | PAR-CLIP | miRTarbase |
| hsa-miR-486-5p | MBD4 | PAR-CLIP | miRTarbase |
| hsa-miR-486-5p | CENPN | HITS-CLIP | miRTarbase |
| hsa-miR-486-5p | CDK17 | HITS-CLIP | Tarbase |
| hsa-miR-486-5p | ATXN7L3 | HITS-CLIP | Tarbase |
| hsa-miR-486-5p | BTAF1 | PAR-CLIP | Tarbase |
| hsa-miR-486-5p | SIRT1 | HITS-CLIP | Tarbase |
| hsa-miR-486-5p | SETD1A | PAR-CLIP | Tarbase |
| hsa-miR-486-5p | BNIP3L | PAR-CLIP | Tarbase |
| hsa-miR-486-5p | NR3C1 | HITS-CLIP | Tarbase |
| hsa-miR-486-5p | ARID1A | HITS-CLIP | Tarbase |
| hsa-miR-486-5p | TNFRSF10B | HITS-CLIP | Tarbase |
| hsa-miR-486-5p | H3F3B | HITS-CLIP | Tarbase |
| hsa-miR-486-5p | H3F3B | HITS-CLIP | Tarbase |
| hsa-miR-486-5p | H3F3B | HITS-CLIP | Tarbase |
| hsa-miR-486-5p | NCOA2 | HITS-CLIP | Tarbase |
| hsa-miR-486-5p | TAF1 | HITS-CLIP | Tarbase |
| hsa-miR-486-5p | CHD1 | PAR-CLIP | Tarbase |
| hsa-miR-486-5p | HLCS | Chimeric fragments | Tarbase |
| hsa-miR-486-5p | ATXN7L2 | PAR-CLIP | Tarbase |
| hsa-miR-486-5p | JMJD1C | HITS-CLIP | Tarbase |
| hsa-miR-486-5p | KDM2A | PAR-CLIP | Tarbase |
| hsa-miR-486-5p | RUVBL1 | PAR-CLIP | Tarbase |
| hsa-miR-486-5p | ATXN7L3B | HITS-CLIP | Tarbase |
| hsa-miR-486-3p | OGT | PAR-CLIP | miRTarbase |
| hsa-miR-486-3p | OGT | PAR-CLIP | miRTarbase |
| hsa-miR-486-3p | OGT | PAR-CLIP | miRTarbase |
| hsa-miR-486-3p | MEN1 | PAR-CLIP | miRTarbase |
| hsa-miR-486-3p | MBD4 | PAR-CLIP | miRTarbase |
| hsa-miR-486-3p | MBD4 | PAR-CLIP | miRTarbase |
| hsa-miR-486-3p | MBD4 | PAR-CLIP | miRTarbase |
| hsa-miR-486-3p | BAZ2A | PAR-CLIP | miRTarbase |
| hsa-miR-486-3p | ATXN7L3B | PAR-CLIP | miRTarbase |
| hsa-miR-486-3p | ATXN7L3B | PAR-CLIP | miRTarbase |
| hsa-miR-486-3p | ATXN7L3 | PAR-CLIP | miRTarbase |
| hsa-miR-486-3p | CASZ1 | PAR-CLIP | miRTarbase |
| hsa-miR-486-3p | CASZ1 | PAR-CLIP | miRTarbase |
| hsa-miR-486-3p | NSD1 | PAR-CLIP | miRTarbase |
| hsa-miR-486-3p | MTA1 | HITS-CLIP | miRTarbase |
| hsa-miR-486-3p | ATXN7L3 | HITS-CLIP | Tarbase |
| hsa-miR-486-3p | SUV39H1 | HITS-CLIP | Tarbase |
| hsa-miR-486-3p | CHD6 | HITS-CLIP | Tarbase |
| hsa-miR-486-3p | HIST1H2BD | HITS-CLIP | Tarbase |
| hsa-miR-486-3p | TUBA1C | HITS-CLIP | Tarbase |
| hsa-miR-486-3p | TUBA1C | HITS-CLIP | Tarbase |
| hsa-miR-486-3p | KDM2A | HITS-CLIP | Tarbase |
| hsa-miR-486-3p | PRMT6 | HITS-CLIP | Tarbase |
| hsa-miR-486-3p | HIST1H2BH | HITS-CLIP | Tarbase |
| hsa-miR-3613-5p | PRKAA1 | PAR-CLIP | miRTarbase |
| hsa-miR-3613-5p | H3F3C | PAR-CLIP | miRTarbase |
| hsa-miR-3613-5p | H3F3C | PAR-CLIP | miRTarbase |
| hsa-miR-3613-5p | ARID1A | PAR-CLIP | miRTarbase |
| hsa-miR-3613-5p | YY1 | PAR-CLIP | miRTarbase |
| hsa-miR-3613-5p | TNFRSF13C | HITS-CLIP | miRTarbase |
| hsa-miR-3613-5p | SMARCE1 | HITS-CLIP | Tarbase |
| hsa-miR-3613-5p | MLLT10 | PAR-CLIP | Tarbase |
| hsa-miR-3613-5p | EYA1 | HITS-CLIP | Tarbase |
| hsa-miR-3613-5p | CHD4 | HITS-CLIP | Tarbase |
| hsa-miR-3613-5p | NR3C1 | PAR-CLIP | Tarbase |
| hsa-miR-3613-5p | NR3C1 | PAR-CLIP | Tarbase |
| hsa-miR-3613-5p | NR3C1 | PAR-CLIP | Tarbase |
| hsa-miR-3613-5p | HIST1H2BJ | HITS-CLIP | Tarbase |
| hsa-miR-3613-5p | KDM5C | PAR-CLIP | Tarbase |
| hsa-miR-3613-5p | KDM4B | HITS-CLIP | Tarbase |
| hsa-miR-3613-5p | CLOCK | HITS-CLIP | Tarbase |
| hsa-miR-3613-5p | EPC2 | PAR-CLIP | Tarbase |
| hsa-miR-3613-5p | CARM1 | HITS-CLIP | Tarbase |
| hsa-miR-3613-5p | RERE | PAR-CLIP | Tarbase |
| hsa-miR-3613-5p | NIPBL | PAR-CLIP | Tarbase |
| hsa-miR-3613-5p | SETD5 | HITS-CLIP | Tarbase |
| hsa-miR-3613-5p | HCFC1 | HITS-CLIP | Tarbase |
| hsa-miR-3613-5p | SMARCC1 | HITS-CLIP | Tarbase |
| hsa-miR-3613-5p | HIST1H2BN | PAR-CLIP | Tarbase |
| hsa-miR-3613-5p | PRKDC | HITS-CLIP | Tarbase |
| hsa-miR-3613-3p | CBX3 | PAR-CLIP | miRTarbase |
| hsa-miR-3613-3p | CBX3 | HITS-CLIP | miRTarbase |
| hsa-miR-3613-3p | TET3 | PAR-CLIP | miRTarbase |
| hsa-miR-3613-3p | GADD45A | PAR-CLIP | miRTarbase |
| hsa-miR-3613-3p | UHRF1BP1L | PAR-CLIP | miRTarbase |
| hsa-miR-3613-3p | PTCHD1 | PAR-CLIP | miRTarbase |
| hsa-miR-3613-3p | PTCHD1 | PAR-CLIP | miRTarbase |
| hsa-miR-3613-3p | HAT1 | PAR-CLIP | miRTarbase |
| hsa-miR-3613-3p | SMEK1 | PAR-CLIP | miRTarbase |
| hsa-miR-3613-3p | GRB10 | PAR-CLIP | miRTarbase |
| hsa-miR-3613-3p | GRB10 | PAR-CLIP | miRTarbase |
| hsa-miR-3613-3p | GRB10 | PAR-CLIP | miRTarbase |
| hsa-miR-3613-3p | ATRNL1 | PAR-CLIP | miRTarbase |
| hsa-miR-3613-3p | RNF20 | PAR-CLIP | miRTarbase |
| hsa-miR-3613-3p | CHD1 | PAR-CLIP | miRTarbase |
| hsa-miR-3613-3p | CHD1 | PAR-CLIP | miRTarbase |
| hsa-miR-3613-3p | YY1 | PAR-CLIP | miRTarbase |
| hsa-miR-3613-3p | HUWE1 | PAR-CLIP | miRTarbase |
| hsa-miR-3613-3p | MYBL1 | PAR-CLIP | miRTarbase |

| | | | |
|---|---|---|---|
| hsa-miR-3613-3p | PARP15 | HITS-CLIP | miRTarbase |
| hsa-miR-3613-3p | RNF207 | HITS-CLIP | miRTarbase |
| hsa-miR-3613-3p | GSG2 | HITS-CLIP | miRTarbase |
| hsa-miR-3613-3p | TNFRSF13C | HITS-CLIP | miRTarbase |
| hsa-miR-3613-3p | CENPH | HITS-CLIP | miRTarbase |
| hsa-miR-3613-3p | USP36 | HITS-CLIP | Tarbase |
| hsa-miR-3613-3p | EED | HITS-CLIP | Tarbase |
| hsa-miR-3613-3p | NCOA1 | PAR-CLIP | Tarbase |
| hsa-miR-3613-3p | EP300 | HITS-CLIP | Tarbase |
| hsa-miR-3613-3p | DUSP1 | HITS-CLIP | Tarbase |
| hsa-miR-3613-3p | TNFRSF10B | PAR-CLIP | Tarbase |
| hsa-miR-3613-3p | FOSB | HITS-CLIP | Tarbase |
| hsa-miR-3613-3p | RERE | HITS-CLIP | Tarbase |
| hsa-miR-3613-3p | SMARCA5 | HITS-CLIP | Tarbase |
| hsa-miR-3613-3p | KDM1B | PAR-CLIP | Tarbase |
| hsa-miR-3613-3p | TET2 | PAR-CLIP | Tarbase |
| hsa-miR-3613-3p | UBE2E1 | Chimeric fragments | Tarbase |
| hsa-miR-3613-3p | CHD9 | PAR-CLIP | Tarbase |
| hsa-miR-3613-3p | MORF4L1 | HITS-CLIP | Tarbase |
| hsa-miR-3613-3p | USP7 | PAR-CLIP | Tarbase |
| hsa-miR-3605-5p | KDM6A | PAR-CLIP | miRTarbase |
| hsa-miR-3605-5p | TNFRSF10D | PAR-CLIP | miRTarbase |
| hsa-miR-3605-5p | TNFRSF10B | PAR-CLIP | miRTarbase |
| hsa-miR-3605-5p | TNFRSF10B | PAR-CLIP | miRTarbase |
| hsa-miR-3605-5p | PPP2R5E | PAR-CLIP | miRTarbase |
| hsa-miR-3605-5p | MIS18BP1 | PAR-CLIP//HITS-CLIP | miRTarbase |
| hsa-miR-3605-5p | PPP2R2A | PAR-CLIP | miRTarbase |
| hsa-miR-3605-5p | RSF1 | HITS-CLIP | Tarbase |
| hsa-miR-3605-5p | KDM3A | HITS-CLIP | Tarbase |
| hsa-miR-3605-5p | MORF4L2 | HITS-CLIP | Tarbase |
| hsa-miR-3605-5p | PRMT1 | HITS-CLIP | Tarbase |
| hsa-miR-3605-5p | H3F3B | HITS-CLIP | Tarbase |
| hsa-miR-3605-5p | ANKRD52 | HITS-CLIP | Tarbase |
| hsa-miR-3605-5p | ANKRD52 | HITS-CLIP | Tarbase |
| hsa-miR-3605-5p | YEATS2 | HITS-CLIP | Tarbase |
| hsa-miR-3605-5p | HCFC1 | HITS-CLIP | Tarbase |
| hsa-miR-3605-5p | KREMEN1 | HITS-CLIP | Tarbase |
| hsa-miR-3605-5p | HIST1H2AJ | HITS-CLIP | Tarbase |
| hsa-miR-3605-3p | KDM2B | HITS-CLIP | miRTarbase |
| hsa-miR-3605-3p | TRIM28 | HITS-CLIP | Tarbase |
| hsa-miR-3605-3p | MYC | HITS-CLIP | Tarbase |
| hsa-miR-3605-3p | ANKRD52 | HITS-CLIP | Tarbase |
| hsa-miR-3605-3p | PRKCA | HITS-CLIP | Tarbase |
| hsa-miR-3605-3p | CHD3 | HITS-CLIP | Tarbase |
| hsa-miR-3605-3p | TAF7 | HITS-CLIP | Tarbase |
| hsa-miR-3605-3p | TAF15 | HITS-CLIP | Tarbase |
| hsa-miR-219a-1-3p | NCOA3 | PAR-CLIP | miRTarbase |
| hsa-miR-219a-1-3p | H2AFX | PAR-CLIP | miRTarbase |
| hsa-miR-219a-1-3p | H2AFX | PAR-CLIP | miRTarbase |
| hsa-miR-219a-1-3p | CREBBP | PAR-CLIP | miRTarbase |
| hsa-miR-219a-1-3p | YY1 | PAR-CLIP | miRTarbase |
| hsa-miR-219a-1-3p | GRB10 | HITS-CLIP | Tarbase |
| hsa-miR-219a-1-3p | GTF3C4 | HITS-CLIP | Tarbase |
| hsa-miR-219a-1-3p | OGT | HITS-CLIP | Tarbase |
| hsa-miR-219a-1-3p | CHEK1 | HITS-CLIP | Tarbase |
| hsa-miR-219a-1-3p | TNFRSF11B | HITS-CLIP | Tarbase |
| hsa-miR-219a-1-3p | NFRKB | HITS-CLIP | Tarbase |
| hsa-miR-195-5p | MECP2 | Microarray//Northern blot | miRTarbase |
| hsa-miR-195-5p | CHUK | Luciferase reporter assay | miRTarbase |
| hsa-miR-195-5p | MBD1 | Luciferase reporter assay | miRTarbase |
| hsa-miR-195-5p | JAK2 | CLASH | miRTarbase |
| hsa-miR-195-5p | MYB | Luciferase reporter assay//qRT-PCR//Western blot | miRTarbase |
| hsa-miR-195-5p | OGT | PAR-CLIP | miRTarbase |
| hsa-miR-195-5p | OGT | PAR-CLIP | miRTarbase |
| hsa-miR-195-5p | OGT | HITS-CLIP | miRTarbase |
| hsa-miR-195-5p | OGT | PAR-CLIP | miRTarbase |
| hsa-miR-195-5p | PAK2 | PAR-CLIP | miRTarbase |
| hsa-miR-195-5p | EZH1 | HITS-CLIP | miRTarbase |
| hsa-miR-195-5p | TET3 | HITS-CLIP | miRTarbase |
| hsa-miR-195-5p | HMGA1 | HITS-CLIP | miRTarbase |
| hsa-miR-195-5p | ATXN7L3B | PAR-CLIP | miRTarbase |
| hsa-miR-195-5p | ATXN7L3B | PAR-CLIP | miRTarbase |
| hsa-miR-195-5p | TBRG1 | PAR-CLIP | miRTarbase |
| hsa-miR-195-5p | CHEK1 | HITS-CLIP | miRTarbase |
| hsa-miR-195-5p | CHEK1 | HITS-CLIP | miRTarbase |
| hsa-miR-195-5p | SIRT4 | HITS-CLIP | miRTarbase |
| hsa-miR-195-5p | SETD1B | PAR-CLIP | miRTarbase |
| hsa-miR-195-5p | PPP2R5C | PAR-CLIP | miRTarbase |
| hsa-miR-195-5p | PPP2R5C | PAR-CLIP | miRTarbase |
| hsa-miR-195-5p | MLLT6 | HITS-CLIP | miRTarbase |
| hsa-miR-195-5p | RNF168 | PAR-CLIP | miRTarbase |
| hsa-miR-195-5p | SRSF1 | PAR-CLIP | miRTarbase |
| hsa-miR-195-5p | SRSF1 | PAR-CLIP | miRTarbase |
| hsa-miR-195-5p | SRSF1 | PAR-CLIP | miRTarbase |
| hsa-miR-195-5p | CBX4 | PAR-CLIP | miRTarbase |
| hsa-miR-195-5p | CBX4 | HITS-CLIP | miRTarbase |
| hsa-miR-195-5p | BAZ2A | PAR-CLIP | miRTarbase |
| hsa-miR-195-5p | SALL1 | PAR-CLIP | miRTarbase |
| hsa-miR-195-5p | SALL1 | PAR-CLIP | miRTarbase |
| hsa-miR-195-5p | MBD4 | PAR-CLIP | miRTarbase |
| hsa-miR-195-5p | CDK17 | PAR-CLIP | miRTarbase |
| hsa-miR-195-5p | CDK17 | HITS-CLIP | miRTarbase |
| hsa-miR-195-5p | ASH1L | PAR-CLIP | miRTarbase |
| hsa-miR-195-5p | ASH1L | HITS-CLIP | miRTarbase |
| hsa-miR-195-5p | TAF13 | PAR-CLIP | miRTarbase |
| hsa-miR-195-5p | TAF13 | PAR-CLIP | miRTarbase |
| hsa-miR-195-5p | PRKAA1 | PAR-CLIP | miRTarbase |
| hsa-miR-195-5p | PRKAA1 | PAR-CLIP | miRTarbase |
| hsa-miR-195-5p | HIST2H2BE | PAR-CLIP | miRTarbase |
| hsa-miR-195-5p | TLK1 | PAR-CLIP | miRTarbase |

| | | | |
|---|---|---|---|
| hsa-miR-195-5p | TLK1 | HITS-CLIP | miRTarbase |
| hsa-miR-195-5p | UBE2H | PAR-CLIP | miRTarbase |
| hsa-miR-195-5p | USP3 | HITS-CLIP | miRTarbase |
| hsa-miR-195-5p | USP31 | HITS-CLIP | miRTarbase |
| hsa-miR-195-5p | RPS6KA3 | HITS-CLIP | miRTarbase |
| hsa-miR-195-5p | PPP6C | HITS-CLIP | miRTarbase |
| hsa-miR-195-5p | GSK3B | HITS-CLIP | miRTarbase |
| hsa-miR-195-5p | CDKN2AIPNL | HITS-CLIP | miRTarbase |
| hsa-miR-195-5p | ASXL1 | HITS-CLIP | miRTarbase |
| hsa-miR-195-5p | CREBBP | HITS-CLIP | Tarbase |
| hsa-miR-195-5p | TNFRSF12A | HITS-CLIP | Tarbase |
| hsa-miR-195-5p | TNFRSF12A | Microarrays | Tarbase |
| hsa-miR-195-5p | MATR3 | HITS-CLIP | Tarbase |
| hsa-miR-195-5p | MATR3 | HITS-CLIP | Tarbase |
| hsa-miR-195-5p | MATR3 | HITS-CLIP | Tarbase |
| hsa-miR-195-5p | CENPQ | Microarrays | Tarbase |
| hsa-miR-195-5p | CDK17 | HITS-CLIP | Tarbase |
| hsa-miR-195-5p | TAF11 | Microarrays | Tarbase |
| hsa-miR-195-5p | UHRF1BP1 | PAR-CLIP | Tarbase |
| hsa-miR-195-5p | HEATR6 | HITS-CLIP | Tarbase |
| hsa-miR-195-5p | PPP2R5B | Microarrays | Tarbase |
| hsa-miR-195-5p | JMJD6 | Microarrays | Tarbase |
| hsa-miR-195-5p | MBD3 | HITS-CLIP | Tarbase |
| hsa-miR-195-5p | CYBRD1 | Microarrays | Tarbase |
| hsa-miR-195-5p | FOSL2 | HITS-CLIP | Tarbase |
| hsa-miR-195-5p | FOSL2 | HITS-CLIP | Tarbase |
| hsa-miR-195-5p | FOSL2 | HITS-CLIP | Tarbase |
| hsa-miR-195-5p | BAZ2A | HITS-CLIP | Tarbase |
| hsa-miR-195-5p | PPP2R5C | HITS-CLIP | Tarbase |
| hsa-miR-195-5p | SRCAP | HITS-CLIP | Tarbase |
| hsa-miR-195-5p | RAD54L | Microarrays | Tarbase |
| hsa-miR-195-5p | HUWE1 | HITS-CLIP | Tarbase |
| hsa-miR-195-5p | ATXN7L3 | HITS-CLIP | Tarbase |
| hsa-miR-195-5p | ATXN7L3 | PAR-CLIP | Tarbase |
| hsa-miR-195-5p | ATXN7L3 | PAR-CLIP | Tarbase |
| hsa-miR-195-5p | ATXN7L3 | HITS-CLIP | Tarbase |
| hsa-miR-195-5p | ATXN7L3 | Microarrays | Tarbase |
| hsa-miR-195-5p | BTAF1 | HITS-CLIP | Tarbase |
| hsa-miR-195-5p | BTAF1 | HITS-CLIP | Tarbase |
| hsa-miR-195-5p | BTAF1 | HITS-CLIP | Tarbase |
| hsa-miR-195-5p | EP300 | HITS-CLIP | Tarbase |
| hsa-miR-195-5p | RPS6KA5 | HITS-CLIP | Tarbase |
| hsa-miR-195-5p | SMEK1 | HITS-CLIP | Tarbase |
| hsa-miR-195-5p | MYBL2 | Microarrays | Tarbase |
| hsa-miR-195-5p | SUV39H1 | Microarrays | Tarbase |
| hsa-miR-195-5p | CENPI | Microarrays | Tarbase |
| hsa-miR-195-5p | USP31 | PAR-CLIP | Tarbase |
| hsa-miR-195-5p | SALL1 | HITS-CLIP | Tarbase |
| hsa-miR-195-5p | SALL1 | HITS-CLIP | Tarbase |
| hsa-miR-195-5p | SALL1 | HITS-CLIP | Tarbase |
| hsa-miR-195-5p | SALL1 | Microarrays | Tarbase |
| hsa-miR-195-5p | OIP5 | Microarrays | Tarbase |
| hsa-miR-195-5p | EYA1 | Microarrays | Tarbase |
| hsa-miR-195-5p | PPP2CB | HITS-CLIP | Tarbase |
| hsa-miR-195-5p | PPP2R1A | HITS-CLIP | Tarbase |
| hsa-miR-195-5p | H2AFV | HITS-CLIP | Tarbase |
| hsa-miR-195-5p | EZH2 | Microarrays | Tarbase |
| hsa-miR-195-5p | ATRNL1 | Microarrays | Tarbase |
| hsa-miR-195-5p | CAMTA2 | HITS-CLIP | Tarbase |
| hsa-miR-195-5p | EZH1 | HITS-CLIP | Tarbase |
| hsa-miR-195-5p | PPARGC1A | Microarrays | Tarbase |
| hsa-miR-195-5p | CDK2AP1 | HITS-CLIP | Tarbase |
| hsa-miR-195-5p | CHD4 | HITS-CLIP | Tarbase |
| hsa-miR-195-5p | CHD4 | HITS-CLIP | Tarbase |
| hsa-miR-195-5p | CHD4 | HITS-CLIP | Tarbase |
| hsa-miR-195-5p | PPP2CA | HITS-CLIP | Tarbase |
| hsa-miR-195-5p | PPP2CA | HITS-CLIP | Tarbase |
| hsa-miR-195-5p | NR3C1 | HITS-CLIP | Tarbase |
| hsa-miR-195-5p | KANSL3 | Microarrays | Tarbase |
| hsa-miR-195-5p | ASH1L | PAR-CLIP | Tarbase |
| hsa-miR-195-5p | ASH1L | HITS-CLIP | Tarbase |
| hsa-miR-195-5p | ASH1L | HITS-CLIP | Tarbase |
| hsa-miR-195-5p | ASH1L | PAR-CLIP | Tarbase |
| hsa-miR-195-5p | MYB | HITS-CLIP | Tarbase |
| hsa-miR-195-5p | HEATR1 | PAR-CLIP | Tarbase |
| hsa-miR-195-5p | PPP4R4 | Microarrays | Tarbase |
| hsa-miR-195-5p | HELLS | HITS-CLIP | Tarbase |
| hsa-miR-195-5p | DUSP1 | Microarrays | Tarbase |
| hsa-miR-195-5p | MASTL | Microarrays | Tarbase |
| hsa-miR-195-5p | EPC1 | PAR-CLIP | Tarbase |
| hsa-miR-195-5p | TNFRSF10B | HITS-CLIP | Tarbase |
| hsa-miR-195-5p | CENPK | Microarrays | Tarbase |
| hsa-miR-195-5p | CDK2 | Microarrays | Tarbase |
| hsa-miR-195-5p | HJURP | Microarrays | Tarbase |
| hsa-miR-195-5p | CHD6 | HITS-CLIP | Tarbase |
| hsa-miR-195-5p | HIST1H2BJ | Microarrays | Tarbase |
| hsa-miR-195-5p | GTF3C4 | HITS-CLIP | Tarbase |
| hsa-miR-195-5p | PARP2 | Microarrays | Tarbase |
| hsa-miR-195-5p | HEATR5A | HITS-CLIP | Tarbase |
| hsa-miR-195-5p | KLF16 | Microarrays | Tarbase |
| hsa-miR-195-5p | PHF10 | Microarrays | Tarbase |
| hsa-miR-195-5p | MLLT4 | HITS-CLIP | Tarbase |
| hsa-miR-195-5p | DNMT1 | Microarrays | Tarbase |
| hsa-miR-195-5p | PRKAB2 | HITS-CLIP | Tarbase |
| hsa-miR-195-5p | GRSF1 | HITS-CLIP | Tarbase |
| hsa-miR-195-5p | NASP | Microarrays | Tarbase |
| hsa-miR-195-5p | SMYD5 | HITS-CLIP | Tarbase |
| hsa-miR-195-5p | SMYD5 | Microarrays | Tarbase |
| hsa-miR-195-5p | TAF5L | Microarrays | Tarbase |
| hsa-miR-195-5p | CDK5RAP2 | Microarrays | Tarbase |

| | | | |
|---|---|---|---|
| hsa-miR-195-5p | MYC | qPCR | Tarbase |
| hsa-miR-195-5p | MYC | Western Blot | Tarbase |
| hsa-miR-195-5p | HMGA1 | HITS-CLIP | Tarbase |
| hsa-miR-195-5p | HMGA1 | HITS-CLIP | Tarbase |
| hsa-miR-195-5p | HMGA1 | HITS-CLIP | Tarbase |
| hsa-miR-195-5p | PPP2R1B | HITS-CLIP | Tarbase |
| hsa-miR-195-5p | PPP2R1B | HITS-CLIP | Tarbase |
| hsa-miR-195-5p | PPP2R1B | HITS-CLIP | Tarbase |
| hsa-miR-195-5p | CASC5 | Microarrays | Tarbase |
| hsa-miR-195-5p | CENPO | Microarrays | Tarbase |
| hsa-miR-195-5p | BARD1 | HITS-CLIP | Tarbase |
| hsa-miR-195-5p | BARD1 | Microarrays | Tarbase |
| hsa-miR-195-5p | ASNSD1 | HITS-CLIP | Tarbase |
| hsa-miR-195-5p | PARP16 | HITS-CLIP | Tarbase |
| hsa-miR-195-5p | PARP16 | PAR-CLIP | Tarbase |
| hsa-miR-195-5p | PARP16 | Microarrays | Tarbase |
| hsa-miR-195-5p | ZCRB1 | HITS-CLIP | Tarbase |
| hsa-miR-195-5p | BRCA2 | Microarrays | Tarbase |
| hsa-miR-195-5p | KANSL2 | HITS-CLIP | Tarbase |
| hsa-miR-195-5p | ANKRD52 | HITS-CLIP | Tarbase |
| hsa-miR-195-5p | ANKRD52 | HITS-CLIP | Tarbase |
| hsa-miR-195-5p | ANKRD52 | Microarrays | Tarbase |
| hsa-miR-195-5p | SETD1B | HITS-CLIP | Tarbase |
| hsa-miR-195-5p | CBX4 | HITS-CLIP | Tarbase |
| hsa-miR-195-5p | CBX4 | HITS-CLIP | Tarbase |
| hsa-miR-195-5p | CBX4 | HITS-CLIP | Tarbase |
| hsa-miR-195-5p | CARM1 | Microarrays | Tarbase |
| hsa-miR-195-5p | DUSP10 | Microarrays | Tarbase |
| hsa-miR-195-5p | GATAD2B | HITS-CLIP | Tarbase |
| hsa-miR-195-5p | KDM6A | HITS-CLIP | Tarbase |
| hsa-miR-195-5p | OGT | HITS-CLIP | Tarbase |
| hsa-miR-195-5p | OGT | HITS-CLIP | Tarbase |
| hsa-miR-195-5p | TAF5 | Microarrays | Tarbase |
| hsa-miR-195-5p | CSRP2BP | HITS-CLIP | Tarbase |
| hsa-miR-195-5p | MBIP | HITS-CLIP | Tarbase |
| hsa-miR-195-5p | CENPH | Microarrays | Tarbase |
| hsa-miR-195-5p | CXXC1 | HITS-CLIP | Tarbase |
| hsa-miR-195-5p | BUB1B | Microarrays | Tarbase |
| hsa-miR-195-5p | HIST1H2BD | Microarrays | Tarbase |
| hsa-miR-195-5p | MIS18A | Microarrays | Tarbase |
| hsa-miR-195-5p | MYSM1 | HITS-CLIP | Tarbase |
| hsa-miR-195-5p | GABRG1 | Microarrays | Tarbase |
| hsa-miR-195-5p | GABRB1 | Microarrays | Tarbase |
| hsa-miR-195-5p | PPP4R2 | HITS-CLIP | Tarbase |
| hsa-miR-195-5p | DTX3L | Microarrays | Tarbase |
| hsa-miR-195-5p | RNF168 | PAR-CLIP | Tarbase |
| hsa-miR-195-5p | H2AFZ | Microarrays | Tarbase |
| hsa-miR-195-5p | WDR82 | HITS-CLIP | Tarbase |
| hsa-miR-195-5p | WDR82 | HITS-CLIP | Tarbase |
| hsa-miR-195-5p | HEATR2 | HITS-CLIP | Tarbase |
| hsa-miR-195-5p | CDK5 | Microarrays | Tarbase |
| hsa-miR-195-5p | UBAP1 | HITS-CLIP | Tarbase |
| hsa-miR-195-5p | PTCHD1 | Microarrays | Tarbase |
| hsa-miR-195-5p | NSD1 | PAR-CLIP | Tarbase |
| hsa-miR-195-5p | NSD1 | HITS-CLIP | Tarbase |
| hsa-miR-195-5p | NSD1 | HITS-CLIP | Tarbase |
| hsa-miR-195-5p | CENPN | HITS-CLIP | Tarbase |
| hsa-miR-195-5p | CENPN | Microarrays | Tarbase |
| hsa-miR-195-5p | GATAD2A | HITS-CLIP | Tarbase |
| hsa-miR-195-5p | TUBA1A | HITS-CLIP | Tarbase |
| hsa-miR-195-5p | TUBA1A | AGO-IP | Tarbase |
| hsa-miR-195-5p | DDB1 | HITS-CLIP | Tarbase |
| hsa-miR-195-5p | DDB1 | HITS-CLIP | Tarbase |
| hsa-miR-195-5p | DDB1 | HITS-CLIP | Tarbase |
| hsa-miR-195-5p | PBK | Microarrays | Tarbase |
| hsa-miR-195-5p | SETD5 | HITS-CLIP | Tarbase |
| hsa-miR-195-5p | HSPBAP1 | HITS-CLIP | Tarbase |
| hsa-miR-195-5p | BUB1 | Microarrays | Tarbase |
| hsa-miR-195-5p | CHD7 | HITS-CLIP | Tarbase |
| hsa-miR-195-5p | ASXL1 | PAR-CLIP | Tarbase |
| hsa-miR-195-5p | CAMTA1 | HITS-CLIP | Tarbase |
| hsa-miR-195-5p | JMJD1C | HITS-CLIP | Tarbase |
| hsa-miR-195-5p | CHD2 | HITS-CLIP | Tarbase |
| hsa-miR-195-5p | PPP2R2D | Microarrays | Tarbase |
| hsa-miR-195-5p | FOSL1 | Microarrays | Tarbase |
| hsa-miR-195-5p | BNIP3 | Microarrays | Tarbase |
| hsa-miR-195-5p | CDK5R1 | HITS-CLIP | Tarbase |
| hsa-miR-195-5p | CDK5R1 | HITS-CLIP | Tarbase |
| hsa-miR-195-5p | RPS6KA3 | HITS-CLIP | Tarbase |
| hsa-miR-195-5p | RPS6KA3 | HITS-CLIP | Tarbase |
| hsa-miR-195-5p | RPS6KA3 | HITS-CLIP | Tarbase |
| hsa-miR-195-5p | CHD9 | HITS-CLIP | Tarbase |
| hsa-miR-195-5p | GSG2 | Microarrays | Tarbase |
| hsa-miR-195-5p | TAF7 | HITS-CLIP | Tarbase |
| hsa-miR-195-5p | HIST1H2AC | Microarrays | Tarbase |
| hsa-miR-195-5p | EHMT1 | HITS-CLIP | Tarbase |
| hsa-miR-195-5p | NPM1 | HITS-CLIP | Tarbase |
| hsa-miR-195-5p | NPM1 | HITS-CLIP | Tarbase |
| hsa-miR-195-5p | SETD3 | Microarrays | Tarbase |
| hsa-miR-195-5p | HIST2H2AC | Microarrays | Tarbase |
| hsa-miR-195-5p | HIST2H2AB | Microarrays | Tarbase |
| hsa-miR-195-5p | HIST2H2BE | Microarrays | Tarbase |
| hsa-miR-195-5p | PPP1CC | HITS-CLIP | Tarbase |
| hsa-miR-195-5p | PPP1CC | HITS-CLIP | Tarbase |
| hsa-miR-195-5p | PPP1CC | HITS-CLIP | Tarbase |
| hsa-miR-195-5p | ERCC6L | Microarrays | Tarbase |
| hsa-miR-195-5p | H2AFX | Microarrays | Tarbase |
| hsa-miR-195-5p | SRSF10 | HITS-CLIP | Tarbase |
| hsa-miR-195-5p | TRRAP | PAR-CLIP | Tarbase |
| hsa-miR-195-5p | TRRAP | PAR-CLIP | Tarbase |

| | | | |
|---|---|---|---|
| hsa-miR-195-5p | HIST3H2BB | Microarrays | Tarbase |
| hsa-miR-195-5p | HIST1H4C | HITS-CLIP | Tarbase |
| hsa-miR-195-5p | ADARB1 | Microarrays | Tarbase |
| hsa-miR-195-5p | TLK1 | PAR-CLIP | Tarbase |
| hsa-miR-195-5p | TLK1 | HITS-CLIP | Tarbase |
| hsa-miR-195-5p | TLK1 | HITS-CLIP | Tarbase |
| hsa-miR-195-5p | TOP1 | HITS-CLIP | Tarbase |
| hsa-miR-195-5p | ANKRD28 | HITS-CLIP | Tarbase |
| hsa-miR-195-5p | CHUK | qPCR | Tarbase |
| hsa-miR-195-5p | CHUK | Western Blot | Tarbase |
| hsa-miR-195-5p | CHUK | HITS-CLIP | Tarbase |
| hsa-miR-195-5p | CHUK | Microarrays | Tarbase |
| hsa-miR-195-5p | CHCHD10 | Microarrays | Tarbase |
| hsa-miR-195-5p | ATXN7L3B | HITS-CLIP | Tarbase |
| hsa-miR-195-5p | ATXN7L3B | Microarrays | Tarbase |
| hsa-miR-195-5p | PRKDC | HITS-CLIP | Tarbase |
| hsa-miR-195-5p | PRKDC | HITS-CLIP | Tarbase |
| hsa-miR-195-5p | HIST2H4A | Microarrays | Tarbase |
| hsa-miR-195-5p | HIST1H2BG | HITS-CLIP | Tarbase |
| hsa-miR-195-5p | HIST1H3G | Microarrays | Tarbase |
| hsa-miR-195-5p | HIST1H3B | Microarrays | Tarbase |
| hsa-miR-195-5p | HIST1H2BE | HITS-CLIP | Tarbase |
| hsa-miR-195-5p | HIST1H2BO | Microarrays | Tarbase |
| hsa-miR-195-5p | HIST1H2AH | HITS-CLIP | Tarbase |
| hsa-miR-195-5p | HIST1H2AH | HITS-CLIP | Tarbase |
| hsa-miR-195-5p | HIST1H2AH | HITS-CLIP | Tarbase |
| hsa-miR-195-5p | HIST1H2AH | Microarrays | Tarbase |
| hsa-miR-195-5p | MLLT6 | HITS-CLIP | Tarbase |
| hsa-miR-195-5p | SMEK2 | HITS-CLIP | Tarbase |
| hsa-miR-195-5p | HIST1H4L | Microarrays | Tarbase |
| hsa-miR-195-5p | HIST1H2AK | Microarrays | Tarbase |
| hsa-miR-195-5p | HIST1H2BH | HITS-CLIP | Tarbase |
| hsa-miR-195-5p | HIST1H2BH | Microarrays | Tarbase |
| hsa-miR-195-5p | HIST1H2AJ | HITS-CLIP | Tarbase |
| hsa-miR-195-5p | HIST1H2AJ | HITS-CLIP | Tarbase |
| hsa-miR-195-5p | HIST1H2AJ | HITS-CLIP | Tarbase |
| hsa-miR-195-5p | HIST1H2AJ | Microarrays | Tarbase |
| hsa-miR-195-5p | HIST1H4D | Microarrays | Tarbase |
| hsa-miR-195-5p | HIST1H2BF | Microarrays | Tarbase |
| hsa-miR-195-5p | HIST1H3F | Microarrays | Tarbase |
| hsa-miR-195-5p | HIST1H3C | HITS-CLIP | Tarbase |
| hsa-miR-195-5p | HIST1H2AB | Microarrays | Tarbase |
| hsa-miR-195-5p | HIST1H2BI | Microarrays | Tarbase |
| hsa-miR-195-5p | HIST1H4A | Microarrays | Tarbase |
| hsa-miR-195-5p | HIST1H2AM | Microarrays | Tarbase |
| hsa-miR-195-3p | MYC | PAR-CLIP | miRTarbase |
| hsa-miR-195-3p | PRKAA1 | PAR-CLIP | miRTarbase |
| hsa-miR-195-3p | PRKAA1 | PAR-CLIP//HITS-CLIP | miRTarbase |
| hsa-miR-195-3p | PRKAA1 | PAR-CLIP | miRTarbase |
| hsa-miR-195-3p | PRKAA1 | PAR-CLIP | miRTarbase |
| hsa-miR-195-3p | RPS6KA5 | PAR-CLIP | miRTarbase |
| hsa-miR-15a-5p | MYB | Luciferase reporter assay//Western blot | miRTarbase |
| hsa-miR-15a-5p | MYB | Luciferase reporter assay//Microarray//qRT-PCR | miRTarbase |
| hsa-miR-15a-5p | H3F3B | Microarray | miRTarbase |
| hsa-miR-15a-5p | CHUK | Luciferase reporter assay//qRT-PCR//Western blot | miRTarbase |
| hsa-miR-15a-5p | HMGA1 | Luciferase reporter assay//qRT-PCR//Western blot | miRTarbase |
| hsa-miR-15a-5p | HMGA1 | HITS-CLIP | miRTarbase |
| hsa-miR-15a-5p | HIST1H2BK | CLASH | miRTarbase |
| hsa-miR-15a-5p | ASNSD1 | CLASH | miRTarbase |
| hsa-miR-15a-5p | CHD4 | CLASH | miRTarbase |
| hsa-miR-15a-5p | TRIM28 | CLASH | miRTarbase |
| hsa-miR-15a-5p | MYBL1 | CLASH | miRTarbase |
| hsa-miR-15a-5p | CARM1 | Luciferase reporter assay//qRT-PCR//Western blot | miRTarbase |
| hsa-miR-15a-5p | OGT | PAR-CLIP | miRTarbase |
| hsa-miR-15a-5p | OGT | PAR-CLIP | miRTarbase |
| hsa-miR-15a-5p | OGT | HITS-CLIP | miRTarbase |
| hsa-miR-15a-5p | OGT | PAR-CLIP | miRTarbase |
| hsa-miR-15a-5p | PAK2 | PAR-CLIP | miRTarbase |
| hsa-miR-15a-5p | EZH1 | HITS-CLIP | miRTarbase |
| hsa-miR-15a-5p | TET3 | HITS-CLIP | miRTarbase |
| hsa-miR-15a-5p | ATXN7L3B | PAR-CLIP | miRTarbase |
| hsa-miR-15a-5p | ATXN7L3B | PAR-CLIP | miRTarbase |
| hsa-miR-15a-5p | TBRG1 | PAR-CLIP | miRTarbase |
| hsa-miR-15a-5p | CHEK1 | HITS-CLIP | miRTarbase |
| hsa-miR-15a-5p | CHEK1 | HITS-CLIP | miRTarbase |
| hsa-miR-15a-5p | SIRT4 | HITS-CLIP | miRTarbase |
| hsa-miR-15a-5p | SETD1B | PAR-CLIP | miRTarbase |
| hsa-miR-15a-5p | PPP2R5C | PAR-CLIP | miRTarbase |
| hsa-miR-15a-5p | PPP2R5C | PAR-CLIP | miRTarbase |
| hsa-miR-15a-5p | MLLT6 | HITS-CLIP | miRTarbase |
| hsa-miR-15a-5p | RNF168 | PAR-CLIP | miRTarbase |
| hsa-miR-15a-5p | SRSF1 | PAR-CLIP | miRTarbase |
| hsa-miR-15a-5p | SRSF1 | PAR-CLIP | miRTarbase |
| hsa-miR-15a-5p | SRSF1 | PAR-CLIP | miRTarbase |
| hsa-miR-15a-5p | CBX4 | PAR-CLIP | miRTarbase |
| hsa-miR-15a-5p | CBX4 | HITS-CLIP | miRTarbase |
| hsa-miR-15a-5p | BAZ2A | PAR-CLIP | miRTarbase |
| hsa-miR-15a-5p | SALL1 | PAR-CLIP | miRTarbase |
| hsa-miR-15a-5p | SALL1 | PAR-CLIP | miRTarbase |
| hsa-miR-15a-5p | MBD4 | PAR-CLIP | miRTarbase |
| hsa-miR-15a-5p | CDK17 | PAR-CLIP | miRTarbase |
| hsa-miR-15a-5p | CDK17 | HITS-CLIP | miRTarbase |
| hsa-miR-15a-5p | ASH1L | PAR-CLIP | miRTarbase |
| hsa-miR-15a-5p | ASH1L | HITS-CLIP | miRTarbase |
| hsa-miR-15a-5p | TAF13 | PAR-CLIP | miRTarbase |
| hsa-miR-15a-5p | TAF13 | PAR-CLIP | miRTarbase |
| hsa-miR-15a-5p | PRKAA1 | PAR-CLIP | miRTarbase |
| hsa-miR-15a-5p | PRKAA1 | PAR-CLIP | miRTarbase |
| hsa-miR-15a-5p | HIST2H2BE | PAR-CLIP | miRTarbase |
| hsa-miR-15a-5p | TLK1 | PAR-CLIP | miRTarbase |

| | | | |
|---|---|---|---|
| hsa-miR-15a-5p | TLK1 | HITS-CLIP | miRTarbase |
| hsa-miR-15a-5p | UBE2H | PAR-CLIP | miRTarbase |
| hsa-miR-15a-5p | USP3 | HITS-CLIP | miRTarbase |
| hsa-miR-15a-5p | USP31 | HITS-CLIP | miRTarbase |
| hsa-miR-15a-5p | RPS6KA3 | HITS-CLIP | miRTarbase |
| hsa-miR-15a-5p | PPP6C | HITS-CLIP | miRTarbase |
| hsa-miR-15a-5p | GSK3B | HITS-CLIP | miRTarbase |
| hsa-miR-15a-5p | CDKN2AIPNL | HITS-CLIP | miRTarbase |
| hsa-miR-15a-5p | ASXL1 | HITS-CLIP | miRTarbase |
| hsa-miR-15a-5p | CREBBP | HITS-CLIP | Tarbase |
| hsa-miR-15a-5p | TNFRSF12A | HITS-CLIP | Tarbase |
| hsa-miR-15a-5p | MATR3 | HITS-CLIP | Tarbase |
| hsa-miR-15a-5p | MATR3 | HITS-CLIP | Tarbase |
| hsa-miR-15a-5p | MATR3 | HITS-CLIP | Tarbase |
| hsa-miR-15a-5p | MATR3 | HITS-CLIP | Tarbase |
| hsa-miR-15a-5p | MATR3 | Chimeric fragments | Tarbase |
| hsa-miR-15a-5p | CDK17 | PAR-CLIP | Tarbase |
| hsa-miR-15a-5p | CDK17 | PAR-CLIP | Tarbase |
| hsa-miR-15a-5p | CDK17 | HITS-CLIP | Tarbase |
| hsa-miR-15a-5p | UHRF1BP1 | PAR-CLIP | Tarbase |
| hsa-miR-15a-5p | UHRF1BP1 | PAR-CLIP | Tarbase |
| hsa-miR-15a-5p | HEATR6 | HITS-CLIP | Tarbase |
| hsa-miR-15a-5p | MBD3 | HITS-CLIP | Tarbase |
| hsa-miR-15a-5p | SMARCE1 | HITS-CLIP | Tarbase |
| hsa-miR-15a-5p | FOSL2 | HITS-CLIP | Tarbase |
| hsa-miR-15a-5p | FOSL2 | HITS-CLIP | Tarbase |
| hsa-miR-15a-5p | FOSL2 | PAR-CLIP | Tarbase |
| hsa-miR-15a-5p | FOSL2 | HITS-CLIP | Tarbase |
| hsa-miR-15a-5p | FOSL2 | HITS-CLIP | Tarbase |
| hsa-miR-15a-5p | FOSL2 | HITS-CLIP | Tarbase |
| hsa-miR-15a-5p | BAZ2A | HITS-CLIP | Tarbase |
| hsa-miR-15a-5p | BAZ2A | HITS-CLIP | Tarbase |
| hsa-miR-15a-5p | PPP2R5C | HITS-CLIP | Tarbase |
| hsa-miR-15a-5p | PPP2R5C | PAR-CLIP | Tarbase |
| hsa-miR-15a-5p | SRCAP | HITS-CLIP | Tarbase |
| hsa-miR-15a-5p | SRCAP | HITS-CLIP | Tarbase |
| hsa-miR-15a-5p | SRCAP | HITS-CLIP | Tarbase |
| hsa-miR-15a-5p | HUWE1 | HITS-CLIP | Tarbase |
| hsa-miR-15a-5p | HUWE1 | HITS-CLIP | Tarbase |
| hsa-miR-15a-5p | ATXN7L3 | PAR-CLIP | Tarbase |
| hsa-miR-15a-5p | ATXN7L3 | HITS-CLIP | Tarbase |
| hsa-miR-15a-5p | ATXN7L3 | PAR-CLIP | Tarbase |
| hsa-miR-15a-5p | ATXN7L3 | PAR-CLIP | Tarbase |
| hsa-miR-15a-5p | ATXN7L3 | HITS-CLIP | Tarbase |
| hsa-miR-15a-5p | BTAF1 | HITS-CLIP | Tarbase |
| hsa-miR-15a-5p | BTAF1 | HITS-CLIP | Tarbase |
| hsa-miR-15a-5p | BTAF1 | HITS-CLIP | Tarbase |
| hsa-miR-15a-5p | BTAF1 | HITS-CLIP | Tarbase |
| hsa-miR-15a-5p | BTAF1 | HITS-CLIP | Tarbase |
| hsa-miR-15a-5p | BTAF1 | Chimeric fragments | Tarbase |
| hsa-miR-15a-5p | BRPF3 | HITS-CLIP | Tarbase |
| hsa-miR-15a-5p | RBX1 | PAR-CLIP | Tarbase |
| hsa-miR-15a-5p | EP300 | HITS-CLIP | Tarbase |
| hsa-miR-15a-5p | EP300 | HITS-CLIP | Tarbase |
| hsa-miR-15a-5p | RPS6KA5 | HITS-CLIP | Tarbase |
| hsa-miR-15a-5p | SMEK1 | HITS-CLIP | Tarbase |
| hsa-miR-15a-5p | SMEK1 | PAR-CLIP | Tarbase |
| hsa-miR-15a-5p | SMEK1 | Chimeric fragments | Tarbase |
| hsa-miR-15a-5p | CHD8 | HITS-CLIP | Tarbase |
| hsa-miR-15a-5p | STK4 | HITS-CLIP | Tarbase |
| hsa-miR-15a-5p | CDK5RAP1 | HITS-CLIP | Tarbase |
| hsa-miR-15a-5p | SMARCA1 | PAR-CLIP | Tarbase |
| hsa-miR-15a-5p | SMARCA1 | PAR-CLIP | Tarbase |
| hsa-miR-15a-5p | USP31 | PAR-CLIP | Tarbase |
| hsa-miR-15a-5p | USP31 | PAR-CLIP | Tarbase |
| hsa-miR-15a-5p | USP31 | Chimeric fragments | Tarbase |
| hsa-miR-15a-5p | SALL1 | HITS-CLIP | Tarbase |
| hsa-miR-15a-5p | SALL1 | HITS-CLIP | Tarbase |
| hsa-miR-15a-5p | SALL1 | HITS-CLIP | Tarbase |
| hsa-miR-15a-5p | SALL1 | PAR-CLIP | Tarbase |
| hsa-miR-15a-5p | SALL1 | Chimeric fragments | Tarbase |
| hsa-miR-15a-5p | PPP2CB | HITS-CLIP | Tarbase |
| hsa-miR-15a-5p | PPP2R1A | HITS-CLIP | Tarbase |
| hsa-miR-15a-5p | H2AFV | HITS-CLIP | Tarbase |
| hsa-miR-15a-5p | CAMTA2 | HITS-CLIP | Tarbase |
| hsa-miR-15a-5p | EZH1 | HITS-CLIP | Tarbase |
| hsa-miR-15a-5p | SUV420H1 | HITS-CLIP | Tarbase |
| hsa-miR-15a-5p | CHD4 | HITS-CLIP | Tarbase |
| hsa-miR-15a-5p | CHD4 | HITS-CLIP | Tarbase |
| hsa-miR-15a-5p | CHD4 | HITS-CLIP | Tarbase |
| hsa-miR-15a-5p | PPP2CA | HITS-CLIP | Tarbase |
| hsa-miR-15a-5p | PPP2CA | HITS-CLIP | Tarbase |
| hsa-miR-15a-5p | NR3C1 | HITS-CLIP | Tarbase |
| hsa-miR-15a-5p | HDAC1 | Chimeric fragments | Tarbase |
| hsa-miR-15a-5p | ASH1L | PAR-CLIP | Tarbase |
| hsa-miR-15a-5p | ASH1L | PAR-CLIP | Tarbase |
| hsa-miR-15a-5p | ASH1L | HITS-CLIP | Tarbase |
| hsa-miR-15a-5p | ASH1L | PAR-CLIP | Tarbase |
| hsa-miR-15a-5p | ASH1L | HITS-CLIP | Tarbase |
| hsa-miR-15a-5p | ASH1L | HITS-CLIP | Tarbase |
| hsa-miR-15a-5p | ASH1L | PAR-CLIP | Tarbase |
| hsa-miR-15a-5p | ASH1L | HITS-CLIP | Tarbase |
| hsa-miR-15a-5p | ASH1L | PAR-CLIP | Tarbase |
| hsa-miR-15a-5p | ARID1A | HITS-CLIP | Tarbase |
| hsa-miR-15a-5p | MYB | HITS-CLIP | Tarbase |
| hsa-miR-15a-5p | MYB | HITS-CLIP | Tarbase |
| hsa-miR-15a-5p | HEATR1 | PAR-CLIP | Tarbase |
| hsa-miR-15a-5p | PPP2R4 | PAR-CLIP | Tarbase |
| hsa-miR-15a-5p | PPP6C | PAR-CLIP | Tarbase |
| hsa-miR-15a-5p | HELLS | HITS-CLIP | Tarbase |

| | | | |
|---|---|---|---|
| hsa-miR-15a-5p | EPC1 | PAR-CLIP | Tarbase |
| hsa-miR-15a-5p | EPC1 | HITS-CLIP | Tarbase |
| hsa-miR-15a-5p | EPC1 | PAR-CLIP | Tarbase |
| hsa-miR-15a-5p | TNFRSF10B | HITS-CLIP | Tarbase |
| hsa-miR-15a-5p | CBX3 | HITS-CLIP | Tarbase |
| hsa-miR-15a-5p | CBX3 | HITS-CLIP | Tarbase |
| hsa-miR-15a-5p | CHD6 | HITS-CLIP | Tarbase |
| hsa-miR-15a-5p | USP22 | HITS-CLIP | Tarbase |
| hsa-miR-15a-5p | GTF3C4 | HITS-CLIP | Tarbase |
| hsa-miR-15a-5p | HEATR5A | HITS-CLIP | Tarbase |
| hsa-miR-15a-5p | FOXA1 | HITS-CLIP | Tarbase |
| hsa-miR-15a-5p | MLLT4 | HITS-CLIP | Tarbase |
| hsa-miR-15a-5p | PRKAB2 | HITS-CLIP | Tarbase |
| hsa-miR-15a-5p | PRKAB2 | HITS-CLIP | Tarbase |
| hsa-miR-15a-5p | GRSF1 | HITS-CLIP | Tarbase |
| hsa-miR-15a-5p | GRSF1 | HITS-CLIP | Tarbase |
| hsa-miR-15a-5p | SMYD5 | HITS-CLIP | Tarbase |
| hsa-miR-15a-5p | TAF5L | HITS-CLIP | Tarbase |
| hsa-miR-15a-5p | SRSF1 | Chimeric fragments | Tarbase |
| hsa-miR-15a-5p | HMGA1 | HITS-CLIP | Tarbase |
| hsa-miR-15a-5p | HMGA1 | HITS-CLIP | Tarbase |
| hsa-miR-15a-5p | HMGA1 | PAR-CLIP | Tarbase |
| hsa-miR-15a-5p | HMGA1 | HITS-CLIP | Tarbase |
| hsa-miR-15a-5p | HMGA1 | HITS-CLIP | Tarbase |
| hsa-miR-15a-5p | HMGA1 | Luciferase Reporter Assay | Tarbase |
| hsa-miR-15a-5p | PPP2R1B | HITS-CLIP | Tarbase |
| hsa-miR-15a-5p | PPP2R1B | HITS-CLIP | Tarbase |
| hsa-miR-15a-5p | PPP2R1B | HITS-CLIP | Tarbase |
| hsa-miR-15a-5p | PPP2R1B | HITS-CLIP | Tarbase |
| hsa-miR-15a-5p | BARD1 | HITS-CLIP | Tarbase |
| hsa-miR-15a-5p | ASNSD1 | HITS-CLIP | Tarbase |
| hsa-miR-15a-5p | ASNSD1 | HITS-CLIP | Tarbase |
| hsa-miR-15a-5p | ASNSD1 | Chimeric fragments | Tarbase |
| hsa-miR-15a-5p | PARP16 | PAR-CLIP | Tarbase |
| hsa-miR-15a-5p | PARP16 | HITS-CLIP | Tarbase |
| hsa-miR-15a-5p | PARP16 | HITS-CLIP | Tarbase |
| hsa-miR-15a-5p | PARP16 | PAR-CLIP | Tarbase |
| hsa-miR-15a-5p | PARP16 | HITS-CLIP | Tarbase |
| hsa-miR-15a-5p | ZCRB1 | HITS-CLIP | Tarbase |
| hsa-miR-15a-5p | SMARCC2 | PAR-CLIP | Tarbase |
| hsa-miR-15a-5p | KANSL2 | HITS-CLIP | Tarbase |
| hsa-miR-15a-5p | ANKRD52 | HITS-CLIP | Tarbase |
| hsa-miR-15a-5p | ANKRD52 | HITS-CLIP | Tarbase |
| hsa-miR-15a-5p | ANKRD52 | HITS-CLIP | Tarbase |
| hsa-miR-15a-5p | ANKRD52 | PAR-CLIP | Tarbase |
| hsa-miR-15a-5p | SETD1B | PAR-CLIP | Tarbase |
| hsa-miR-15a-5p | SETD1B | HITS-CLIP | Tarbase |
| hsa-miR-15a-5p | SETD1B | HITS-CLIP | Tarbase |
| hsa-miR-15a-5p | NCOA2 | HITS-CLIP | Tarbase |
| hsa-miR-15a-5p | CBX4 | HITS-CLIP | Tarbase |
| hsa-miR-15a-5p | CBX4 | HITS-CLIP | Tarbase |
| hsa-miR-15a-5p | CBX4 | PAR-CLIP | Tarbase |
| hsa-miR-15a-5p | CBX4 | HITS-CLIP | Tarbase |
| hsa-miR-15a-5p | CBX4 | HITS-CLIP | Tarbase |
| hsa-miR-15a-5p | CBX4 | HITS-CLIP | Tarbase |
| hsa-miR-15a-5p | SAE1 | HITS-CLIP | Tarbase |
| hsa-miR-15a-5p | GATAD2B | HITS-CLIP | Tarbase |
| hsa-miR-15a-5p | GATAD2B | HITS-CLIP | Tarbase |
| hsa-miR-15a-5p | KDM6A | PAR-CLIP | Tarbase |
| hsa-miR-15a-5p | KDM6A | HITS-CLIP | Tarbase |
| hsa-miR-15a-5p | OGT | PAR-CLIP | Tarbase |
| hsa-miR-15a-5p | OGT | HITS-CLIP | Tarbase |
| hsa-miR-15a-5p | OGT | HITS-CLIP | Tarbase |
| hsa-miR-15a-5p | OGT | HITS-CLIP | Tarbase |
| hsa-miR-15a-5p | OGT | Chimeric fragments | Tarbase |
| hsa-miR-15a-5p | CSRP2BP | HITS-CLIP | Tarbase |
| hsa-miR-15a-5p | CHEK1 | PAR-CLIP | Tarbase |
| hsa-miR-15a-5p | PPP4C | Chimeric fragments | Tarbase |
| hsa-miR-15a-5p | MBIP | HITS-CLIP | Tarbase |
| hsa-miR-15a-5p | SMARCA5 | PAR-CLIP | Tarbase |
| hsa-miR-15a-5p | INO80C | Chimeric fragments | Tarbase |
| hsa-miR-15a-5p | CXXC1 | HITS-CLIP | Tarbase |
| hsa-miR-15a-5p | KLF10 | Chimeric fragments | Tarbase |
| hsa-miR-15a-5p | HIST1H2BD | PAR-CLIP | Tarbase |
| hsa-miR-15a-5p | MYSM1 | HITS-CLIP | Tarbase |
| hsa-miR-15a-5p | PPP4R2 | PAR-CLIP | Tarbase |
| hsa-miR-15a-5p | PPP4R2 | HITS-CLIP | Tarbase |
| hsa-miR-15a-5p | ATXN7 | HITS-CLIP | Tarbase |
| hsa-miR-15a-5p | RNF168 | PAR-CLIP | Tarbase |
| hsa-miR-15a-5p | RNF168 | Chimeric fragments | Tarbase |
| hsa-miR-15a-5p | WDR82 | HITS-CLIP | Tarbase |
| hsa-miR-15a-5p | WDR82 | PAR-CLIP | Tarbase |
| hsa-miR-15a-5p | WDR82 | HITS-CLIP | Tarbase |
| hsa-miR-15a-5p | NIPBL | PAR-CLIP | Tarbase |
| hsa-miR-15a-5p | HEATR2 | HITS-CLIP | Tarbase |
| hsa-miR-15a-5p | UBAP1 | HITS-CLIP | Tarbase |
| hsa-miR-15a-5p | UBAP1 | PAR-CLIP | Tarbase |
| hsa-miR-15a-5p | NSD1 | PAR-CLIP | Tarbase |
| hsa-miR-15a-5p | NSD1 | HITS-CLIP | Tarbase |
| hsa-miR-15a-5p | NSD1 | PAR-CLIP | Tarbase |
| hsa-miR-15a-5p | NSD1 | HITS-CLIP | Tarbase |
| hsa-miR-15a-5p | NSD1 | HITS-CLIP | Tarbase |
| hsa-miR-15a-5p | CENPN | HITS-CLIP | Tarbase |
| hsa-miR-15a-5p | GATAD2A | HITS-CLIP | Tarbase |
| hsa-miR-15a-5p | GATAD2A | Chimeric fragments | Tarbase |
| hsa-miR-15a-5p | TUBA1A | HITS-CLIP | Tarbase |
| hsa-miR-15a-5p | DDB1 | HITS-CLIP | Tarbase |
| hsa-miR-15a-5p | DDB1 | HITS-CLIP | Tarbase |
| hsa-miR-15a-5p | DDB1 | HITS-CLIP | Tarbase |
| hsa-miR-15a-5p | SETD5 | PAR-CLIP | Tarbase |

| | | | |
|---|---|---|---|
| hsa-miR-15a-5p | SETD5 | HITS-CLIP | Tarbase |
| hsa-miR-15a-5p | CDKN2AIP | PAR-CLIP | Tarbase |
| hsa-miR-15a-5p | HSPBAP1 | HITS-CLIP | Tarbase |
| hsa-miR-15a-5p | CHD3 | PAR-CLIP | Tarbase |
| hsa-miR-15a-5p | CHD3 | PAR-CLIP | Tarbase |
| hsa-miR-15a-5p | CHD3 | HITS-CLIP | Tarbase |
| hsa-miR-15a-5p | USP38 | PAR-CLIP | Tarbase |
| hsa-miR-15a-5p | USP38 | Chimeric fragments | Tarbase |
| hsa-miR-15a-5p | PPM1D | PAR-CLIP | Tarbase |
| hsa-miR-15a-5p | PPM1D | PAR-CLIP | Tarbase |
| hsa-miR-15a-5p | CHD7 | PAR-CLIP | Tarbase |
| hsa-miR-15a-5p | CHD7 | HITS-CLIP | Tarbase |
| hsa-miR-15a-5p | ASXL1 | PAR-CLIP | Tarbase |
| hsa-miR-15a-5p | ASXL1 | PAR-CLIP | Tarbase |
| hsa-miR-15a-5p | ASXL1 | Chimeric fragments | Tarbase |
| hsa-miR-15a-5p | JMJD1C | HITS-CLIP | Tarbase |
| hsa-miR-15a-5p | PPP1CA | HITS-CLIP | Tarbase |
| hsa-miR-15a-5p | CHD2 | HITS-CLIP | Tarbase |
| hsa-miR-15a-5p | CDK5R1 | PAR-CLIP | Tarbase |
| hsa-miR-15a-5p | CDK5R1 | HITS-CLIP | Tarbase |
| hsa-miR-15a-5p | CDK5R1 | HITS-CLIP | Tarbase |
| hsa-miR-15a-5p | RPS6KA3 | HITS-CLIP | Tarbase |
| hsa-miR-15a-5p | RPS6KA3 | HITS-CLIP | Tarbase |
| hsa-miR-15a-5p | RPS6KA3 | PAR-CLIP | Tarbase |
| hsa-miR-15a-5p | RPS6KA3 | HITS-CLIP | Tarbase |
| hsa-miR-15a-5p | RPS6KA3 | HITS-CLIP | Tarbase |
| hsa-miR-15a-5p | RPS6KA3 | HITS-CLIP | Tarbase |
| hsa-miR-15a-5p | CHD9 | HITS-CLIP | Tarbase |
| hsa-miR-15a-5p | CHD9 | PAR-CLIP | Tarbase |
| hsa-miR-15a-5p | TAF7 | HITS-CLIP | Tarbase |
| hsa-miR-15a-5p | TAF7 | Chimeric fragments | Tarbase |
| hsa-miR-15a-5p | PAK2 | PAR-CLIP | Tarbase |
| hsa-miR-15a-5p | EHMT1 | HITS-CLIP | Tarbase |
| hsa-miR-15a-5p | NPM1 | HITS-CLIP | Tarbase |
| hsa-miR-15a-5p | NPM1 | HITS-CLIP | Tarbase |
| hsa-miR-15a-5p | NPM1 | PAR-CLIP | Tarbase |
| hsa-miR-15a-5p | HIST3H2A | PAR-CLIP | Tarbase |
| hsa-miR-15a-5p | PRKAG1 | PAR-CLIP | Tarbase |
| hsa-miR-15a-5p | PRKAG1 | HITS-CLIP | Tarbase |
| hsa-miR-15a-5p | SETD3 | Chimeric fragments | Tarbase |
| hsa-miR-15a-5p | MYBL1 | PAR-CLIP | Tarbase |
| hsa-miR-15a-5p | PPP1CC | HITS-CLIP | Tarbase |
| hsa-miR-15a-5p | PPP1CC | HITS-CLIP | Tarbase |
| hsa-miR-15a-5p | TAF9B | HITS-CLIP | Tarbase |
| hsa-miR-15a-5p | H2AFX | HITS-CLIP | Tarbase |
| hsa-miR-15a-5p | TRRAP | PAR-CLIP | Tarbase |
| hsa-miR-15a-5p | TRRAP | PAR-CLIP | Tarbase |
| hsa-miR-15a-5p | TRRAP | PAR-CLIP | Tarbase |
| hsa-miR-15a-5p | TRRAP | PAR-CLIP | Tarbase |
| hsa-miR-15a-5p | TRRAP | PAR-CLIP | Tarbase |
| hsa-miR-15a-5p | TRRAP | Chimeric fragments | Tarbase |
| hsa-miR-15a-5p | HIST1H4C | HITS-CLIP | Tarbase |
| hsa-miR-15a-5p | HIST1H3D | HITS-CLIP | Tarbase |
| hsa-miR-15a-5p | TLK1 | PAR-CLIP | Tarbase |
| hsa-miR-15a-5p | TLK1 | PAR-CLIP | Tarbase |
| hsa-miR-15a-5p | TLK1 | PAR-CLIP | Tarbase |
| hsa-miR-15a-5p | TLK1 | HITS-CLIP | Tarbase |
| hsa-miR-15a-5p | TLK1 | PAR-CLIP | Tarbase |
| hsa-miR-15a-5p | TLK1 | HITS-CLIP | Tarbase |
| hsa-miR-15a-5p | TOP1 | HITS-CLIP | Tarbase |
| hsa-miR-15a-5p | ANKRD28 | HITS-CLIP | Tarbase |
| hsa-miR-15a-5p | CHUK | Luciferase Reporter Assay | Tarbase |
| hsa-miR-15a-5p | CHUK | PAR-CLIP | Tarbase |
| hsa-miR-15a-5p | CHUK | qPCR | Tarbase |
| hsa-miR-15a-5p | CHUK | Western Blot | Tarbase |
| hsa-miR-15a-5p | CHUK | PAR-CLIP | Tarbase |
| hsa-miR-15a-5p | CHUK | HITS-CLIP | Tarbase |
| hsa-miR-15a-5p | CHUK | HITS-CLIP | Tarbase |
| hsa-miR-15a-5p | CHUK | Luciferase Reporter Assay | Tarbase |
| hsa-miR-15a-5p | ERCC6 | Chimeric fragments | Tarbase |
| hsa-miR-15a-5p | H2AFJ | HITS-CLIP | Tarbase |
| hsa-miR-15a-5p | ATXN7L3B | PAR-CLIP | Tarbase |
| hsa-miR-15a-5p | ATXN7L3B | HITS-CLIP | Tarbase |
| hsa-miR-15a-5p | ATXN7L3B | HITS-CLIP | Tarbase |
| hsa-miR-15a-5p | ATXN7L3B | HITS-CLIP | Tarbase |
| hsa-miR-15a-5p | PRKDC | HITS-CLIP | Tarbase |
| hsa-miR-15a-5p | PRKDC | HITS-CLIP | Tarbase |
| hsa-miR-15a-5p | PRKDC | HITS-CLIP | Tarbase |
| hsa-miR-15a-5p | HIST1H2BG | PAR-CLIP | Tarbase |
| hsa-miR-15a-5p | HIST1H2BG | HITS-CLIP | Tarbase |
| hsa-miR-15a-5p | HIST1H3G | HITS-CLIP | Tarbase |
| hsa-miR-15a-5p | HIST1H3B | PAR-CLIP | Tarbase |
| hsa-miR-15a-5p | HIST1H2BE | HITS-CLIP | Tarbase |
| hsa-miR-15a-5p | HIST1H2BO | PAR-CLIP | Tarbase |
| hsa-miR-15a-5p | HIST1H2AH | HITS-CLIP | Tarbase |
| hsa-miR-15a-5p | HIST1H2AH | HITS-CLIP | Tarbase |
| hsa-miR-15a-5p | HIST1H2AH | PAR-CLIP | Tarbase |
| hsa-miR-15a-5p | HIST1H2AH | HITS-CLIP | Tarbase |
| hsa-miR-15a-5p | MLLT6 | HITS-CLIP | Tarbase |
| hsa-miR-15a-5p | SMEK2 | HITS-CLIP | Tarbase |
| hsa-miR-15a-5p | SMEK2 | PAR-CLIP | Tarbase |
| hsa-miR-15a-5p | HIST1H2BH | HITS-CLIP | Tarbase |
| hsa-miR-15a-5p | HIST1H2AJ | HITS-CLIP | Tarbase |
| hsa-miR-15a-5p | HIST1H2AJ | HITS-CLIP | Tarbase |
| hsa-miR-15a-5p | HIST1H2AJ | HITS-CLIP | Tarbase |
| hsa-miR-15a-5p | HIST1H2AJ | HITS-CLIP | Tarbase |
| hsa-miR-15a-5p | HIST1H2BF | HITS-CLIP | Tarbase |
| hsa-miR-15a-5p | HIST1H3C | HITS-CLIP | Tarbase |
| hsa-miR-15a-5p | HIST1H2AB | PAR-CLIP | Tarbase |
| hsa-miR-15a-3p | PRMT3 | PAR-CLIP | miRTarbase |

| | | | |
|---|---|---|---|
| hsa-miR-15a-3p | TNFRSF10D | PAR-CLIP | miRTarbase |
| hsa-miR-15a-3p | TAF1D | PAR-CLIP | miRTarbase |
| hsa-miR-15a-3p | BAZ1B | HITS-CLIP | Tarbase |
| hsa-miR-15a-3p | MBD3 | HITS-CLIP | Tarbase |
| hsa-miR-15a-3p | MBD3 | HITS-CLIP | Tarbase |
| hsa-miR-15a-3p | CYBRD1 | HITS-CLIP | Tarbase |
| hsa-miR-15a-3p | RAD54L | HITS-CLIP | Tarbase |
| hsa-miR-15a-3p | HUWE1 | HITS-CLIP | Tarbase |
| hsa-miR-15a-3p | RBBP7 | HITS-CLIP | Tarbase |
| hsa-miR-15a-3p | CHD4 | HITS-CLIP | Tarbase |
| hsa-miR-15a-3p | CHD4 | HITS-CLIP | Tarbase |
| hsa-miR-15a-3p | PPP2CA | HITS-CLIP | Tarbase |
| hsa-miR-15a-3p | ASH1L | HITS-CLIP | Tarbase |
| hsa-miR-15a-3p | MYB | HITS-CLIP | Tarbase |
| hsa-miR-15a-3p | PPP6C | PAR-CLIP | Tarbase |
| hsa-miR-15a-3p | PPP6C | HITS-CLIP | Tarbase |
| hsa-miR-15a-3p | PPP6C | PAR-CLIP | Tarbase |
| hsa-miR-15a-3p | CBX3 | HITS-CLIP | Tarbase |
| hsa-miR-15a-3p | MORF4L2 | HITS-CLIP | Tarbase |
| hsa-miR-15a-3p | USP22 | HITS-CLIP | Tarbase |
| hsa-miR-15a-3p | USP22 | HITS-CLIP | Tarbase |
| hsa-miR-15a-3p | TRIM28 | HITS-CLIP | Tarbase |
| hsa-miR-15a-3p | TRIM28 | HITS-CLIP | Tarbase |
| hsa-miR-15a-3p | TRIM28 | HITS-CLIP | Tarbase |
| hsa-miR-15a-3p | MYC | HITS-CLIP | Tarbase |
| hsa-miR-15a-3p | RERE | HITS-CLIP | Tarbase |
| hsa-miR-15a-3p | DUSP11 | HITS-CLIP | Tarbase |
| hsa-miR-15a-3p | HIST1H4H | HITS-CLIP | Tarbase |
| hsa-miR-15a-3p | SETD5 | HITS-CLIP | Tarbase |
| hsa-miR-15a-3p | SETD5 | HITS-CLIP | Tarbase |
| hsa-miR-15a-3p | PPM1D | HITS-CLIP | Tarbase |
| hsa-miR-15a-3p | KLF11 | HITS-CLIP | Tarbase |
| hsa-miR-15a-3p | PPP1CA | HITS-CLIP | Tarbase |
| hsa-miR-15a-3p | CHD2 | HITS-CLIP | Tarbase |
| hsa-miR-15a-3p | FOSL1 | HITS-CLIP | Tarbase |
| hsa-miR-15a-3p | MTA1 | HITS-CLIP | Tarbase |
| hsa-miR-15a-3p | PPP1CC | HITS-CLIP | Tarbase |
| hsa-miR-15a-3p | NAP1L4 | HITS-CLIP | Tarbase |
| hsa-miR-15a-3p | PPP2R2A | HITS-CLIP | Tarbase |
| hsa-miR-15a-3p | MLLT6 | HITS-CLIP | Tarbase |
| hsa-miR-141-5p | ADRB1 | PAR-CLIP | miRTarbase |
| hsa-miR-141-5p | ADRB1 | PAR-CLIP | miRTarbase |
| hsa-miR-141-5p | ADRB1 | PAR-CLIP | miRTarbase |
| hsa-miR-141-5p | ADRB1 | PAR-CLIP | miRTarbase |
| hsa-miR-141-5p | ADRB1 | PAR-CLIP | miRTarbase |
| hsa-miR-141-5p | SUZ12 | PAR-CLIP | miRTarbase |
| hsa-miR-141-5p | H3F3B | PAR-CLIP | miRTarbase |
| hsa-miR-141-5p | KANSL3 | HITS-CLIP | miRTarbase |
| hsa-miR-141-5p | HLA-DRB1 | HITS-CLIP | miRTarbase |
| hsa-miR-141-5p | MTA3 | HITS-CLIP | miRTarbase |
| hsa-miR-141-5p | INO80D | HITS-CLIP | miRTarbase |
| hsa-miR-141-5p | RNF8 | HITS-CLIP | miRTarbase |
| hsa-miR-141-5p | HEATR5A | HITS-CLIP | miRTarbase |
| hsa-miR-141-5p | CREBBP | HITS-CLIP | Tarbase |
| hsa-miR-141-5p | UBR2 | HITS-CLIP | Tarbase |
| hsa-miR-141-5p | PHF20 | HITS-CLIP | Tarbase |
| hsa-miR-141-5p | CENPQ | HITS-CLIP | Tarbase |
| hsa-miR-141-5p | CENPQ | HITS-CLIP | Tarbase |
| hsa-miR-141-5p | TAF11 | HITS-CLIP | Tarbase |
| hsa-miR-141-5p | ING3 | HITS-CLIP | Tarbase |
| hsa-miR-141-5p | BAZ2A | HITS-CLIP | Tarbase |
| hsa-miR-141-5p | BAZ2A | HITS-CLIP | Tarbase |
| hsa-miR-141-5p | MLLT10 | HITS-CLIP | Tarbase |
| hsa-miR-141-5p | NCOA1 | HITS-CLIP | Tarbase |
| hsa-miR-141-5p | H2AFY2 | HITS-CLIP | Tarbase |
| hsa-miR-141-5p | RBBP7 | HITS-CLIP | Tarbase |
| hsa-miR-141-5p | RBBP7 | HITS-CLIP | Tarbase |
| hsa-miR-141-5p | PPP2R1A | HITS-CLIP | Tarbase |
| hsa-miR-141-5p | PPP3CB | HITS-CLIP | Tarbase |
| hsa-miR-141-5p | SUPT6H | HITS-CLIP | Tarbase |
| hsa-miR-141-5p | PPP6R3 | HITS-CLIP | Tarbase |
| hsa-miR-141-5p | DUSP16 | HITS-CLIP | Tarbase |
| hsa-miR-141-5p | INO80D | HITS-CLIP | Tarbase |
| hsa-miR-141-5p | KDM5B | HITS-CLIP | Tarbase |
| hsa-miR-141-5p | HEATR1 | HITS-CLIP | Tarbase |
| hsa-miR-141-5p | TNFRSF10B | HITS-CLIP | Tarbase |
| hsa-miR-141-5p | TNFRSF10B | HITS-CLIP | Tarbase |
| hsa-miR-141-5p | MORF4L2 | HITS-CLIP | Tarbase |
| hsa-miR-141-5p | MORF4L2 | HITS-CLIP | Tarbase |
| hsa-miR-141-5p | MORF4L2 | HITS-CLIP | Tarbase |
| hsa-miR-141-5p | YEATS4 | HITS-CLIP | Tarbase |
| hsa-miR-141-5p | YEATS4 | HITS-CLIP | Tarbase |
| hsa-miR-141-5p | YEATS4 | HITS-CLIP | Tarbase |
| hsa-miR-141-5p | YEATS4 | HITS-CLIP | Tarbase |
| hsa-miR-141-5p | YEATS4 | HITS-CLIP | Tarbase |
| hsa-miR-141-5p | TRIM28 | HITS-CLIP | Tarbase |
| hsa-miR-141-5p | PRKAA1 | HITS-CLIP | Tarbase |
| hsa-miR-141-5p | PRKAA1 | HITS-CLIP | Tarbase |
| hsa-miR-141-5p | CLOCK | HITS-CLIP | Tarbase |
| hsa-miR-141-5p | EPC2 | HITS-CLIP | Tarbase |
| hsa-miR-141-5p | SRSF1 | HITS-CLIP | Tarbase |
| hsa-miR-141-5p | PPP2R1B | HITS-CLIP | Tarbase |
| hsa-miR-141-5p | IDH1 | HITS-CLIP | Tarbase |
| hsa-miR-141-5p | SETD1B | HITS-CLIP | Tarbase |
| hsa-miR-141-5p | URB1 | HITS-CLIP | Tarbase |
| hsa-miR-141-5p | GATAD2B | HITS-CLIP | Tarbase |
| hsa-miR-141-5p | PARP1 | HITS-CLIP | Tarbase |
| hsa-miR-141-5p | KDM6A | HITS-CLIP | Tarbase |
| hsa-miR-141-5p | TAF1 | HITS-CLIP | Tarbase |
| hsa-miR-141-5p | SMARCA5 | HITS-CLIP | Tarbase |

| | | | |
|---|---|---|---|
| hsa-miR-141-5p | PPP4R1 | HITS-CLIP | Tarbase |
| hsa-miR-141-5p | KLF10 | HITS-CLIP | Tarbase |
| hsa-miR-141-5p | KLF10 | HITS-CLIP | Tarbase |
| hsa-miR-141-5p | HEATR3 | HITS-CLIP | Tarbase |
| hsa-miR-141-5p | MYSM1 | HITS-CLIP | Tarbase |
| hsa-miR-141-5p | YY1AP1 | HITS-CLIP | Tarbase |
| hsa-miR-141-5p | RAD54L2 | HITS-CLIP | Tarbase |
| hsa-miR-141-5p | NIPBL | HITS-CLIP | Tarbase |
| hsa-miR-141-5p | PPP2R3B | HITS-CLIP | Tarbase |
| hsa-miR-141-5p | CHD3 | HITS-CLIP | Tarbase |
| hsa-miR-141-5p | BPTF | HITS-CLIP | Tarbase |
| hsa-miR-141-5p | CAMTA1 | HITS-CLIP | Tarbase |
| hsa-miR-141-5p | JMJD1C | HITS-CLIP | Tarbase |
| hsa-miR-141-5p | JMJD1C | HITS-CLIP | Tarbase |
| hsa-miR-141-5p | HCFC1 | HITS-CLIP | Tarbase |
| hsa-miR-141-5p | PPP2R2D | HITS-CLIP | Tarbase |
| hsa-miR-141-5p | RPS6KA3 | HITS-CLIP | Tarbase |
| hsa-miR-141-5p | CHD9 | HITS-CLIP | Tarbase |
| hsa-miR-141-5p | SUZ12 | HITS-CLIP | Tarbase |
| hsa-miR-141-5p | IDH2 | HITS-CLIP | Tarbase |
| hsa-miR-141-5p | HDAC2 | HITS-CLIP | Tarbase |
| hsa-miR-141-5p | RING1 | HITS-CLIP | Tarbase |
| hsa-miR-141-5p | NAP1L4 | HITS-CLIP | Tarbase |
| hsa-miR-141-5p | PPP1CB | HITS-CLIP | Tarbase |
| hsa-miR-141-5p | PRKDC | HITS-CLIP | Tarbase |
| hsa-miR-141-5p | SMEK2 | HITS-CLIP | Tarbase |
| hsa-miR-141-3p | BAP1 | Reporter assay | miRTarbase |
| hsa-miR-141-3p | BAP1 | Microarray | miRTarbase |
| hsa-miR-141-3p | CLOCK | Luciferase reporter assay//Reporter assay | miRTarbase |
| hsa-miR-141-3p | CLOCK | Northern blot//qRT-PCR//Western blot | miRTarbase |
| hsa-miR-141-3p | UBAP1 | Luciferase reporter assay//Western blot//Reporter assay;Western blot;Oth | miRTarbase |
| hsa-miR-141-3p | CDYL | Luciferase reporter assay//Western blot | miRTarbase |
| hsa-miR-141-3p | KLF11 | Reporter assay | miRTarbase |
| hsa-miR-141-3p | OGT | PAR-CLIP | miRTarbase |
| hsa-miR-141-3p | OGT | PAR-CLIP | miRTarbase |
| hsa-miR-141-3p | ATXN7L1 | PAR-CLIP | miRTarbase |
| hsa-miR-141-3p | H2AFZ | PAR-CLIP | miRTarbase |
| hsa-miR-141-3p | H2AFZ | PAR-CLIP | miRTarbase |
| hsa-miR-141-3p | PRKAA2 | HITS-CLIP | miRTarbase |
| hsa-miR-141-3p | MBTD1 | HITS-CLIP | Tarbase |
| hsa-miR-141-3p | MATR3 | HITS-CLIP | Tarbase |
| hsa-miR-141-3p | RB1CC1 | HITS-CLIP | Tarbase |
| hsa-miR-141-3p | RB1CC1 | HITS-CLIP | Tarbase |
| hsa-miR-141-3p | ADRB1 | HITS-CLIP | Tarbase |
| hsa-miR-141-3p | KDM5A | HITS-CLIP | Tarbase |
| hsa-miR-141-3p | FOSL2 | HITS-CLIP | Tarbase |
| hsa-miR-141-3p | USP33 | PAR-CLIP | Tarbase |
| hsa-miR-141-3p | PPP2R5C | HITS-CLIP | Tarbase |
| hsa-miR-141-3p | SRCAP | HITS-CLIP | Tarbase |
| hsa-miR-141-3p | SRCAP | HITS-CLIP | Tarbase |
| hsa-miR-141-3p | BTAF1 | HITS-CLIP | Tarbase |
| hsa-miR-141-3p | CHD8 | HITS-CLIP | Tarbase |
| hsa-miR-141-3p | STK4 | HITS-CLIP | Tarbase |
| hsa-miR-141-3p | USP31 | HITS-CLIP | Tarbase |
| hsa-miR-141-3p | H2AFV | HITS-CLIP | Tarbase |
| hsa-miR-141-3p | H2AFV | HITS-CLIP | Tarbase |
| hsa-miR-141-3p | DUSP3 | HITS-CLIP | Tarbase |
| hsa-miR-141-3p | DUSP3 | HITS-CLIP | Tarbase |
| hsa-miR-141-3p | NR3C1 | HITS-CLIP | Tarbase |
| hsa-miR-141-3p | INO80D | HITS-CLIP | Tarbase |
| hsa-miR-141-3p | TAF1B | PAR-CLIP | Tarbase |
| hsa-miR-141-3p | ASH1L | HITS-CLIP | Tarbase |
| hsa-miR-141-3p | ASH1L | PAR-CLIP | Tarbase |
| hsa-miR-141-3p | SRSF11 | HITS-CLIP | Tarbase |
| hsa-miR-141-3p | PPP6C | HITS-CLIP | Tarbase |
| hsa-miR-141-3p | CDK2 | Western Blot | Tarbase |
| hsa-miR-141-3p | MORF4L2 | HITS-CLIP | Tarbase |
| hsa-miR-141-3p | NCOA3 | HITS-CLIP | Tarbase |
| hsa-miR-141-3p | NCOA3 | HITS-CLIP | Tarbase |
| hsa-miR-141-3p | USP22 | HITS-CLIP | Tarbase |
| hsa-miR-141-3p | FOXA1 | HITS-CLIP | Tarbase |
| hsa-miR-141-3p | FOXA1 | HITS-CLIP | Tarbase |
| hsa-miR-141-3p | MLLT4 | HITS-CLIP | Tarbase |
| hsa-miR-141-3p | PRKAA1 | HITS-CLIP | Tarbase |
| hsa-miR-141-3p | H3F3B | HITS-CLIP | Tarbase |
| hsa-miR-141-3p | CLOCK | Luciferase Reporter Assay | Tarbase |
| hsa-miR-141-3p | CLOCK | Luciferase Reporter Assay | Tarbase |
| hsa-miR-141-3p | SRSF1 | HITS-CLIP | Tarbase |
| hsa-miR-141-3p | ANKRD52 | HITS-CLIP | Tarbase |
| hsa-miR-141-3p | ANKRD52 | HITS-CLIP | Tarbase |
| hsa-miR-141-3p | ANKRD52 | HITS-CLIP | Tarbase |
| hsa-miR-141-3p | SETD1B | HITS-CLIP | Tarbase |
| hsa-miR-141-3p | SETD1B | HITS-CLIP | Tarbase |
| hsa-miR-141-3p | NCOA2 | HITS-CLIP | Tarbase |
| hsa-miR-141-3p | NCOA2 | HITS-CLIP | Tarbase |
| hsa-miR-141-3p | RERE | HITS-CLIP | Tarbase |
| hsa-miR-141-3p | RERE | HITS-CLIP | Tarbase |
| hsa-miR-141-3p | SETDB1 | HITS-CLIP | Tarbase |
| hsa-miR-141-3p | GATAD2B | HITS-CLIP | Tarbase |
| hsa-miR-141-3p | SETD7 | HITS-CLIP | Tarbase |
| hsa-miR-141-3p | OGT | HITS-CLIP | Tarbase |
| hsa-miR-141-3p | PPP4R1 | HITS-CLIP | Tarbase |
| hsa-miR-141-3p | RNF20 | HITS-CLIP | Tarbase |
| hsa-miR-141-3p | HIST1H2BD | HITS-CLIP | Tarbase |
| hsa-miR-141-3p | EOGT | HITS-CLIP | Tarbase |
| hsa-miR-141-3p | BAP1 | Luciferase Reporter Assay | Tarbase |
| hsa-miR-141-3p | BAP1 | Luciferase Reporter Assay | Tarbase |
| hsa-miR-141-3p | H2AFZ | HITS-CLIP | Tarbase |
| hsa-miR-141-3p | WDR82 | HITS-CLIP | Tarbase |
| hsa-miR-141-3p | HEATR2 | HITS-CLIP | Tarbase |

| | | | |
|---|---|---|---|
| hsa-miR-141-3p | UBAP1 | Luciferase Reporter Assay | Tarbase |
| hsa-miR-141-3p | UBAP1 | Western Blot | Tarbase |
| hsa-miR-141-3p | ATMIN | HITS-CLIP | Tarbase |
| hsa-miR-141-3p | MECP2 | HITS-CLIP | Tarbase |
| hsa-miR-141-3p | NFRKB | HITS-CLIP | Tarbase |
| hsa-miR-141-3p | ASXL1 | HITS-CLIP | Tarbase |
| hsa-miR-141-3p | KLF11 | Luciferase Reporter Assay | Tarbase |
| hsa-miR-141-3p | KLF11 | Luciferase Reporter Assay | Tarbase |
| hsa-miR-141-3p | CHD2 | HITS-CLIP | Tarbase |
| hsa-miR-141-3p | CHD9 | HITS-CLIP | Tarbase |
| hsa-miR-141-3p | TAF7 | HITS-CLIP | Tarbase |
| hsa-miR-141-3p | SMYD3 | HITS-CLIP | Tarbase |
| hsa-miR-141-3p | USP7 | HITS-CLIP | Tarbase |
| hsa-miR-141-3p | TET3 | HITS-CLIP | Tarbase |
| hsa-miR-141-3p | TOP1 | HITS-CLIP | Tarbase |
| hsa-miR-141-3p | NAP1L4 | HITS-CLIP | Tarbase |
| hsa-miR-141-3p | PPP2R2A | HITS-CLIP | Tarbase |
| hsa-miR-141-3p | TAF15 | HITS-CLIP | Tarbase |
