## Supplementary Table S7 for "Methylome-based cell-of-origin modeling (Methyl-COOM) identifies aberrant expression of immune regulatory molecules in CLL"

**Supplementary Table S7: List of ENCODE TF ChIP-seq datasets**

| filename | cellType | treatment | antibody | description |
|---|---|---|---|---|
| wgEncodeAwgTfbsBroadGm12878CtcfUniPk.narrowPeak | GM12878 | None | CTCF | ChIP GM12878 CTCF |
| wgEncodeAwgTfbsBroadGm12878Ezh239875UniPk.narrowPeak | GM12878 | None | EZH2_(39875) | ChIP GM12878 EZH2_(39875) |
| wgEncodeAwgTfbsHaibGm12878Atf2sc81188V0422111UniPk.narrowPeak | GM12878 | None | ATF2_(SC-81188) | ChIP GM12878 ATF2_(SC-81188) |
| wgEncodeAwgTfbsHaibGm12878Atf3Pcr1xUniPk.narrowPeak | GM12878 | None | ATF3 | ChIP GM12878 ATF3 |
| wgEncodeAwgTfbsHaibGm12878BatfPcr1xUniPk.narrowPeak | GM12878 | None | BATF | ChIP GM12878 BATF |
| wgEncodeAwgTfbsHaibGm12878Bcl11aPcr1xUniPk.narrowPeak | GM12878 | None | BCL11A | ChIP GM12878 BCL11A |
| wgEncodeAwgTfbsHaibGm12878Bcl3V0416101UniPk.narrowPeak | GM12878 | None | BCL3 | ChIP GM12878 BCL3 |
| wgEncodeAwgTfbsHaibGm12878Bclaf101388V0416101UniPk.narrowPeak | GM12878 | None | BCLAF1_(SC-101388) | ChIP GM12878 BCLAF1_(SC-101388) |
| wgEncodeAwgTfbsHaibGm12878Cebpbsc150V0422111UniPk.narrowPeak | GM12878 | None | CEBPB_(SC-150) | ChIP GM12878 CEBPB_(SC-150) |
| wgEncodeAwgTfbsHaibGm12878Ebf1sc137065Pcr1xUniPk.narrowPeak | GM12878 | None | EBF1_(SC-137065) | ChIP GM12878 EBF1_(SC-137065) |
| wgEncodeAwgTfbsHaibGm12878Egr1Pcr2xUniPk.narrowPeak | GM12878 | None | Egr-1 | ChIP GM12878 Egr-1 |
| wgEncodeAwgTfbsHaibGm12878Elf1sc631V0416101UniPk.narrowPeak | GM12878 | None | ELF1_(SC-631) | ChIP GM12878 ELF1_(SC-631) |
| wgEncodeAwgTfbsHaibGm12878Ets1Pcr1xUniPk.narrowPeak | GM12878 | None | ETS1 | ChIP GM12878 ETS1 |
| wgEncodeAwgTfbsHaibGm12878Foxm1sc502V0422111UniPk.narrowPeak | GM12878 | None | FOXM1_(SC-502) | ChIP GM12878 FOXM1_(SC-502) |
| wgEncodeAwgTfbsHaibGm12878GabpPcr2xUniPk.narrowPeak | GM12878 | None | GABP | ChIP GM12878 GABP |
| wgEncodeAwgTfbsHaibGm12878Irf4sc6059Pcr1xUniPk.narrowPeak | GM12878 | None | IRF4_(SC-6059) | ChIP GM12878 IRF4_(SC-6059) |
| wgEncodeAwgTfbsHaibGm12878Mef2aPcr1xUniPk.narrowPeak | GM12878 | None | MEF2A | ChIP GM12878 MEF2A |
| wgEncodeAwgTfbsHaibGm12878Mef2csc13268V0416101UniPk.narrowPeak | GM12878 | None | MEF2C_(SC-13268) | ChIP GM12878 MEF2C_(SC-13268) |
| wgEncodeAwgTfbsHaibGm12878Mta3sc81325V0422111UniPk.narrowPeak | GM12878 | None | MTA3_(SC-81325) | ChIP GM12878 MTA3_(SC-81325) |
| wgEncodeAwgTfbsHaibGm12878Nfatc1sc17834V0422111UniPk.narrowPeak | GM12878 | None | NFATC1_(SC-17834) | ChIP GM12878 NFATC1_(SC-17834) |
| wgEncodeAwgTfbsHaibGm12878Nficsc81335V0422111UniPk.narrowPeak | GM12878 | None | NFIC_(SC-81335) | ChIP GM12878 NFIC_(SC-81335) |
| wgEncodeAwgTfbsHaibGm12878NrsfPcr1xUniPk.narrowPeak | GM12878 | None | NRSF | ChIP GM12878 NRSF |
| wgEncodeAwgTfbsHaibGm12878P300Pcr1xUniPk.narrowPeak | GM12878 | None | p300 | ChIP GM12878 p300 |
| wgEncodeAwgTfbsHaibGm12878Pax5c20Pcr1xUniPk.narrowPeak | GM12878 | None | PAX5-C20 | ChIP GM12878 PAX5-C20 |
| wgEncodeAwgTfbsHaibGm12878Pax5n19Pcr1xUniPk.narrowPeak | GM12878 | None | PAX5-N19 | ChIP GM12878 PAX5-N19 |
| wgEncodeAwgTfbsHaibGm12878Pbx3Pcr1xUniPk.narrowPeak | GM12878 | None | Pbx3 | ChIP GM12878 Pbx3 |
| wgEncodeAwgTfbsHaibGm12878Pmlsc71910V0422111UniPk.narrowPeak | GM12878 | None | PML_(SC-71910) | ChIP GM12878 PML_(SC-71910) |
| wgEncodeAwgTfbsHaibGm12878Pol24h8Pcr1xUniPk.narrowPeak | GM12878 | None | Pol2-4H8 | ChIP GM12878 Pol2-4H8 |
| wgEncodeAwgTfbsHaibGm12878Pol2Pcr2xUniPk.narrowPeak | GM12878 | None | Pol2 | ChIP GM12878 Pol2 |
| wgEncodeAwgTfbsHaibGm12878Pou2f2Pcr1xUniPk.narrowPeak | GM12878 | None | POU2F2 | ChIP GM12878 POU2F2 |
| wgEncodeAwgTfbsHaibGm12878Pu1Pcr1xUniPk.narrowPeak | GM12878 | None | PU.1 | ChIP GM12878 PU.1 |
| wgEncodeAwgTfbsHaibGm12878Rad21V0416101UniPk.narrowPeak | GM12878 | None | Rad21 | ChIP GM12878 Rad21 |
| wgEncodeAwgTfbsHaibGm12878Runx3sc101553V0422111UniPk.narrowPeak | GM12878 | None | RUNX3_(SC-101553) | ChIP GM12878 RUNX3_(SC-101553) |
| wgEncodeAwgTfbsHaibGm12878RxraPcr1xUniPk.narrowPeak | GM12878 | None | RXRA | ChIP GM12878 RXRA |
| wgEncodeAwgTfbsHaibGm12878Six5Pcr1xUniPk.narrowPeak | GM12878 | None | SIX5 | ChIP GM12878 SIX5 |
| wgEncodeAwgTfbsHaibGm12878Sp1Pcr1xUniPk.narrowPeak | GM12878 | None | SP1 | ChIP GM12878 SP1 |
| wgEncodeAwgTfbsHaibGm12878SrfPcr2xUniPk.narrowPeak | GM12878 | None | SRF | ChIP GM12878 SRF |
| wgEncodeAwgTfbsHaibGm12878Stat5asc74442V0422111UniPk.narrowPeak | GM12878 | None | STAT5A_(SC-74442) | ChIP GM12878 STAT5A_(SC-74442) |
| wgEncodeAwgTfbsHaibGm12878Taf1Pcr1xUniPk.narrowPeak | GM12878 | None | TAF1 | ChIP GM12878 TAF1 |
| wgEncodeAwgTfbsHaibGm12878Tcf12Pcr1xUniPk.narrowPeak | GM12878 | None | TCF12 | ChIP GM12878 TCF12 |
| wgEncodeAwgTfbsHaibGm12878Tcf3Pcr1xUniPk.narrowPeak | GM12878 | None | TCF3_(SC-349) | ChIP GM12878 TCF3_(SC-349) |
| wgEncodeAwgTfbsHaibGm12878Usf1Pcr2xUniPk.narrowPeak | GM12878 | None | USF-1 | ChIP GM12878 USF-1 |
| wgEncodeAwgTfbsHaibGm12878Yy1sc281Pcr1xUniPk.narrowPeak | GM12878 | None | YY1_(SC-281) | ChIP GM12878 YY1_(SC-281) |
| wgEncodeAwgTfbsHaibGm12878Zbtb33Pcr1xUniPk.narrowPeak | GM12878 | None | ZBTB33 | ChIP GM12878 ZBTB33 |
| wgEncodeAwgTfbsHaibGm12878Zeb1sc25388V0416102UniPk.narrowPeak | GM12878 | None | ZEB1_(SC-25388) | ChIP GM12878 ZEB1_(SC-25388) |
| wgEncodeAwgTfbsSydhGm12878Bhlhe40cIggmusUniPk.narrowPeak | GM12878 | None | BHLHE40_(NB100-1800) | ChIP GM12878 BHLHE40_(NB100-1800) |
| wgEncodeAwgTfbsSydhGm12878Brca1a300IggmusUniPk.narrowPeak | GM12878 | None | BRCA1_(A300-000A) | ChIP GM12878 BRCA1_(A300-000A) |
| wgEncodeAwgTfbsSydhGm12878CfosUniPk.narrowPeak | GM12878 | None | c-Fos | ChIP GM12878 c-Fos |
| wgEncodeAwgTfbsSydhGm12878Chd1a301218aIggmusUniPk.narrowPeak | GM12878 | None | CHD1_(A301-218A) | ChIP GM12878 CHD1_(A301-218A) |
| wgEncodeAwgTfbsSydhGm12878Chd2ab68301IggmusUniPk.narrowPeak | GM12878 | None | CHD2_(AB68301) | ChIP GM12878 CHD2_(AB68301) |
| wgEncodeAwgTfbsSydhGm12878Corestsc30189IggmusUniPk.narrowPeak | GM12878 | None | COREST_(sc-30189) | ChIP GM12878 COREST_(sc-30189) |
| wgEncodeAwgTfbsSydhGm12878Ctcfsc15914c20UniPk.narrowPeak | GM12878 | None | CTCF_(SC-15914) | ChIP GM12878 CTCF_(SC-15914) |
| wgEncodeAwgTfbsSydhGm12878E2f4IggmusUniPk.narrowPeak | GM12878 | None | E2F4 | ChIP GM12878 E2F4 |
| wgEncodeAwgTfbsSydhGm12878Ebf1sc137065UniPk.narrowPeak | GM12878 | None | EBF1_(SC-137065) | ChIP GM12878 EBF1_(SC-137065) |
| wgEncodeAwgTfbsSydhGm12878Elk112771IggmusUniPk.narrowPeak | GM12878 | None | ELK1_(1277-1) | ChIP GM12878 ELK1_(1277-1) |
| wgEncodeAwgTfbsSydhGm12878Ikzf1iknuclaUniPk.narrowPeak | GM12878 | None | IKZF1_(IkN)_(UCLA) | ChIP GM12878 IKZF1_(IkN)_(UCLA) |
| wgEncodeAwgTfbsSydhGm12878JundUniPk.narrowPeak | GM12878 | None | JunD | ChIP GM12878 JunD |
| wgEncodeAwgTfbsSydhGm12878MaxIggmusUniPk.narrowPeak | GM12878 | None | Max | ChIP GM12878 Max |
| wgEncodeAwgTfbsSydhGm12878Mazab85725IggmusUniPk.narrowPeak | GM12878 | None | MAZ_(ab85725) | ChIP GM12878 MAZ_(ab85725) |
| wgEncodeAwgTfbsSydhGm12878Mxi1IggmusUniPk.narrowPeak | GM12878 | None | Mxi1_(AF4185) | ChIP GM12878 Mxi1_(AF4185) |
| wgEncodeAwgTfbsSydhGm12878Nfe2sc22827UniPk.narrowPeak | GM12878 | None | NF-E2_(SC-22827) | ChIP GM12878 NF-E2_(SC-22827) |
| wgEncodeAwgTfbsSydhGm12878NfkbTnfaIggrabUniPk.narrowPeak | GM12878 | TNFa | NFKB | ChIP GM12878 NFKB |
| wgEncodeAwgTfbsSydhGm12878NfyaIggmusUniPk.narrowPeak | GM12878 | None | NF-YA | ChIP GM12878 NF-YA |
| wgEncodeAwgTfbsSydhGm12878NfybIggmusUniPk.narrowPeak | GM12878 | None | NF-YB | ChIP GM12878 NF-YB |
| wgEncodeAwgTfbsSydhGm12878Nrf1IggmusUniPk.narrowPeak | GM12878 | None | Nrf1 | ChIP GM12878 Nrf1 |
| wgEncodeAwgTfbsSydhGm12878P300IggmusUniPk.narrowPeak | GM12878 | None | p300 | ChIP GM12878 p300 |
| wgEncodeAwgTfbsSydhGm12878P300bUniPk.narrowPeak | GM12878 | None | p300_(SC-584) | ChIP GM12878 p300_(SC-584) |
| wgEncodeAwgTfbsSydhGm12878Pol2IggmusUniPk.narrowPeak | GM12878 | None | Pol2 | ChIP GM12878 Pol2 |
| wgEncodeAwgTfbsSydhGm12878Pol2UniPk.narrowPeak | GM12878 | None | Pol2 | ChIP GM12878 Pol2 |
| wgEncodeAwgTfbsSydhGm12878Pol2s2IggmusUniPk.narrowPeak | GM12878 | None | Pol2(phosphoS2) | ChIP GM12878 Pol2(phosphoS2) |
| wgEncodeAwgTfbsSydhGm12878Pol3UniPk.narrowPeak | GM12878 | None | Pol3 | ChIP GM12878 Pol3 |
| wgEncodeAwgTfbsSydhGm12878Rad21IggrabUniPk.narrowPeak | GM12878 | None | Rad21 | ChIP GM12878 Rad21 |
| wgEncodeAwgTfbsSydhGm12878Rfx5200401194IggmusUniPk.narrowPeak | GM12878 | None | RFX5_(200-401-194) | ChIP GM12878 RFX5_(200-401-194) |
| wgEncodeAwgTfbsSydhGm12878Sin3anb6001263IggmusUniPk.narrowPeak | GM12878 | None | SIN3A_(NB600-1263) | ChIP GM12878 SIN3A_(NB600-1263) |
| wgEncodeAwgTfbsSydhGm12878Smc3ab9263IggmusUniPk.narrowPeak | GM12878 | None | SMC3_(ab9263) | ChIP GM12878 SMC3_(ab9263) |
| wgEncodeAwgTfbsSydhGm12878Stat1UniPk.narrowPeak | GM12878 | None | STAT1 | ChIP GM12878 STAT1 |
| wgEncodeAwgTfbsSydhGm12878Stat3IggmusUniPk.narrowPeak | GM12878 | None | STAT3 | ChIP GM12878 STAT3 |
| wgEncodeAwgTfbsSydhGm12878Tblr1ab24550IggmusUniPk.narrowPeak | GM12878 | None | TBLR1_(ab24550) | ChIP GM12878 TBLR1_(ab24550) |
| wgEncodeAwgTfbsSydhGm12878TbpIggmusUniPk.narrowPeak | GM12878 | None | TBP | ChIP GM12878 TBP |
| wgEncodeAwgTfbsSydhGm12878Tr4UniPk.narrowPeak | GM12878 | None | TR4 | ChIP GM12878 TR4 |
| wgEncodeAwgTfbsSydhGm12878Usf2IggmusUniPk.narrowPeak | GM12878 | None | USF2 | ChIP GM12878 USF2 |
| wgEncodeAwgTfbsSydhGm12878WhipIggmusUniPk.narrowPeak | GM12878 | None | WHIP | ChIP GM12878 WHIP |
| wgEncodeAwgTfbsSydhGm12878Yy1UniPk.narrowPeak | GM12878 | None | YY1 | ChIP GM12878 YY1 |
| wgEncodeAwgTfbsSydhGm12878Znf143166181apUniPk.narrowPeak | GM12878 | None | Znf143_(16618-1-AP) | ChIP GM12878 Znf143_(16618-1-AP) |
| wgEncodeAwgTfbsSydhGm12878Znf274UniPk.narrowPeak | GM12878 | None | ZNF274 | ChIP GM12878 ZNF274 |

| | | | | |
|---|---|---|---|---|
| wgEncodeAwgTfbsSydhGm12878Zzz3UniPk.narrowPeak | GM12878 | None | ZZZ3 | ChIP GM12878 ZZZ3 |
| wgEncodeAwgTfbsUtaGm12878CmycUniPk.narrowPeak | GM12878 | None | c-Myc | ChIP GM12878 c-Myc |
| wgEncodeAwgTfbsUtaGm12878CtcfUniPk.narrowPeak | GM12878 | None | CTCF | ChIP GM12878 CTCF |
| wgEncodeAwgTfbsUtaGm12878Pol2UniPk.narrowPeak | GM12878 | None | Pol2 | ChIP GM12878 Pol2 |
| wgEncodeAwgTfbsUwGm12878CtcfUniPk.narrowPeak | GM12878 | None | CTCF | ChIP GM12878 CTCF |
