## Supplementary Table S8 for "Methylome-based cell-of-origin modeling (Methyl-COOM) identifies aberrant expression of immune regulatory molecules in CLL"

**Supplementary Table S8: List of linear B cell-specific CpGs**

| CpG |
|---|
| cg00001249 |
| cg00001269 |
| cg00001747 |
| cg00001809 |
| cg00001854 |
| cg00001930 |
| cg00002080 |
| cg00002145 |
| cg00002810 |
| cg00003298 |
| cg00003858 |
| cg00003965 |
| cg00003994 |
| cg00004421 |
| cg00004775 |
| cg00005847 |
| cg00006081 |
| cg00006397 |
| cg00006867 |
| cg00007076 |
| cg00007987 |
| cg00008033 |
| cg00008621 |
| cg00008629 |
| cg00008647 |
| cg00008695 |
| cg00009053 |
| cg00009293 |
| cg00009306 |
| cg00009553 |
| cg00009922 |
| cg00010742 |
| cg00011346 |
| cg00011513 |
| cg00012522 |
| cg00012576 |
| cg00013301 |
| cg00013349 |
| cg00013369 |
| cg00014260 |
| cg00014333 |
| cg00014377 |
| cg00015121 |
| cg00015343 |
| cg00015770 |
| cg00017437 |
| cg00017489 |
| cg00017546 |
| cg00017842 |

cg00018010
cg00018606
cg00018828
cg00018845
cg00019275
cg00019495
cg00019826
cg00019877
cg00020405
cg00020474
cg00020991
cg00021030
cg00021242
cg00021275
cg00021513
cg00022054
cg00022101
cg00022871
cg00022900
cg00022997
cg00023174
cg00023464
cg00023726
cg00024086
cg00024516
cg00025643
cg00026033
cg00026141
cg00026603
cg00026703
cg00026909
cg00027083
cg00027155
cg00028013
cg00028034
cg00028056
cg00028507
cg00028677
cg00028844
cg00028929
cg00028935
cg00029521
cg00030005
cg00030588
cg00030807
cg00031013
cg00031105
cg00031165
cg00031346
cg00032366
cg00032879
cg00032884

cg00032912
cg00033584
cg00033909
cg00034003
cg00034076
cg00034154
cg00035220
cg00035347
cg00036408
cg00036585
cg00036732
cg00037457
cg00038020
cg00038675
cg00038857
cg00039016
cg00039326
cg00039441
cg00039480
cg00039489
cg00040322
cg00040590
cg00040749
cg00040837
cg00040986
cg00041014
cg00041047
cg00041270
cg00041636
cg00041989
cg00042156
cg00043591
cg00043788
cg00043792
cg00043819
cg00044097
cg00044245
cg00044354
cg00044372
cg00044665
cg00045118
cg00045515
cg00045762
cg00045902
cg00046499
cg00046625
cg00049033
cg00049047
cg00049156
cg00049664
cg00049883
cg00049892

cg00050271
cg00051371
cg00052665
cg00053393
cg00053851
cg00054197
cg00054301
cg00054472
cg00054639
cg00054774
cg00055073
cg00055557
cg00055603
cg00055771
cg00055981
cg00055986
cg00056002
cg00056624
cg00057240
cg00058329
cg00059459
cg00059643
cg00059662
cg00060641
cg00060836
cg00061031
cg00061520
cg00061635
cg00061792
cg00062437
cg00063174
cg00063291
cg00063346
cg00063357
cg00063699
cg00063703
cg00064255
cg00064261
cg00064296
cg00064778
cg00065215
cg00065570
cg00065905
cg00066239
cg00066663
cg00067018
cg00067528
cg00067720
cg00068102
cg00068433
cg00069860
cg00070529

cg00070870
cg00071051
cg00071241
cg00071250
cg00071265
cg00071727
cg00072407
cg00073549
cg00073794
cg00073837
cg00074145
cg00074348
cg00075103
cg00075975
cg00076538
cg00076555
cg00076797
cg00078221
cg00078513
cg00078861
cg00079302
cg00079388
cg00079485
cg00079598
cg00080188
cg00080252
cg00080418
cg00080682
cg00081087
cg00081404
cg00081975
cg00082285
cg00082912
cg00083018
cg00083059
cg00084019
cg00084202
cg00084720
cg00085013
cg00085493
cg00086429
cg00086488
cg00086493
cg00087067
cg00087368
cg00087906
cg00088148
cg00088183
cg00089091
cg00089314
cg00089451
cg00089798

cg00089814
cg00089945
cg00090261
cg00090265
cg00090452
cg00090737
cg00090787
cg00091285
cg00091760
cg00092443
cg00092992
cg00093544
cg00093643
cg00093829
cg00093971
cg00094303
cg00094734
cg00094735
cg00094744
cg00094898
cg00095259
cg00095326
cg00095431
cg00095523
cg00095594
cg00095629
cg00095688
cg00096307
cg00096747
cg00096922
cg00097088
cg00097146
cg00097550
cg00097645
cg00097790
cg00097968
cg00098144
cg00098182
cg00098336
cg00098819
cg00099519
cg00099869
cg00099976
cg00100420
cg00100762
cg00100800
cg00101555
cg00101629
cg00101712
cg00101715
cg00103398
cg00104608

cg00105628
cg00106196
cg00106345
cg00106565
cg00106609
cg00107490
cg00107667
cg00107772
cg00108098
cg00109076
cg00109131
cg00109302
cg00109356
cg00110545
cg00110654
cg00110832
cg00111697
cg00111778
cg00112238
cg00112260
cg00113225
cg00113315
cg00113369
cg00113663
cg00114160
cg00115178
cg00115639
cg00116092
cg00118229
cg00118342
cg00118507
cg00118808
cg00118989
cg00119034
cg00119181
cg00119314
cg00120147
cg00120254
cg00120274
cg00120416
cg00120783
cg00121045
cg00121255
cg00121374
cg00122347
cg00122406
cg00123055
cg00124009
cg00125002
cg00125544
cg00126320
cg00126959

cg00127167
cg00128273
cg00128482
cg00128702
cg00129563
cg00129785
cg00130165
cg00130707
cg00130710
cg00130778
cg00130947
cg00131081
cg00131423
cg00131758
cg00131893
cg00131983
cg00132141
cg00132716
cg00132769
cg00132888
cg00133811
cg00135056
cg00135293
cg00135874
cg00136400
cg00136405
cg00136477
cg00136547
cg00139496
cg00139681
cg00141308
cg00141925
cg00142670
cg00143249
cg00144335
cg00145118
cg00145150
cg00145904
cg00146437
cg00147248
cg00147509
cg00147788
cg00148025
cg00148223
cg00149679
cg00149716
cg00150837
cg00150882
cg00151048
cg00151124
cg00151709
cg00151728

cg00152008
cg00152034
cg00152117
cg00152340
cg00152577
cg00152611
cg00152708
cg00152946
cg00152948
cg00153306
cg00153448
cg00153622
cg00153693
cg00153707
cg00154357
cg00154727
cg00154920
cg00155619
cg00156118
cg00156194
cg00156633
cg00156948
cg00156995
cg00157109
cg00157223
cg00157228
cg00157439
cg00157477
cg00157572
cg00157668
cg00157935
cg00157962
cg00158122
cg00158934
cg00159764
cg00159873
cg00160027
cg00160359
cg00160872
cg00160976
cg00161683
cg00162231
cg00162344
cg00162611
cg00163541
cg00163546
cg00163674
cg00163861
cg00164189
cg00164678
cg00164837
cg00166915

cg00167670
cg00167836
cg00167958
cg00168240
cg00169019
cg00169930
cg00170295
cg00170487
cg00171092
cg00171214
cg00171729
cg00171942
cg00172227
cg00172410
cg00172597
cg00172603
cg00172794
cg00173538
cg00173659
cg00174179
cg00174428
cg00174508
cg00174713
cg00174718
cg00174743
cg00175153
cg00175344
cg00175487
cg00175702
cg00176675
cg00176949
cg00177142
cg00177237
cg00177388
cg00177797
cg00178249
cg00178928
cg00179795
cg00179921
cg00179940
cg00180470
cg00180762
cg00181439
cg00181669
cg00181849
cg00182273
cg00182421
cg00182639
cg00183107
cg00183186
cg00183639
cg00184203

cg00184461
cg00184953
cg00186374
cg00187059
cg00187120
cg00187244
cg00187299
cg00187437
cg00187889
cg00188654
cg00188748
cg00189989
cg00191178
cg00192031
cg00192046
cg00193021
cg00193200
cg00193668
cg00193947
cg00193959
cg00194126
cg00194771
cg00195796
cg00196151
cg00196414
cg00196671
cg00197313
cg00198586
cg00199059
cg00199065
cg00199549
cg00199753
cg00199759
cg00201025
cg00201275
cg00201457
cg00201534
cg00201870
cg00202479
cg00203913
cg00204501
cg00204748
cg00205786
cg00205813
cg00206161
cg00206463
cg00207280
cg00207352
cg00207428
cg00207731
cg00207921
cg00208012

cg00208153
cg00208830
cg00208967
cg00209038
cg00209786
cg00210271
cg00210856
cg00211609
cg00211630
cg00212521
cg00212979
cg00212988
cg00213044
cg00213281
cg00213295
cg00214012
cg00214791
cg00214856
cg00214970
cg00215019
cg00215182
cg00215432
cg00215521
cg00218404
cg00218447
cg00218914
cg00219855
cg00220113
cg00220335
cg00220661
cg00221012
cg00221658
cg00221709
cg00223245
cg00223899
cg00225006
cg00225157
cg00225196
cg00225697
cg00226225
cg00226545
cg00226822
cg00226849
cg00226931
cg00227093
cg00227781
cg00228475
cg00228796
cg00229387
cg00229532
cg00229754
cg00229809

cg00229868
cg00230517
cg00230584
cg00230700
cg00230715
cg00230924
cg00231771
cg00232092
cg00232265
cg00232679
cg00232735
cg00233028
cg00233943
cg00233948
cg00234384
cg00234446
cg00234625
cg00234691
cg00234998
cg00235367
cg00235442
cg00235661
cg00236729
cg00237010
cg00237606
cg00238182
cg00238249
cg00238391
cg00238805
cg00239353
cg00239563
cg00239657
cg00239835
cg00240195
cg00240205
cg00240734
cg00241941
cg00242325
cg00242951
cg00242976
cg00243040
cg00243854
cg00244299
cg00244745
cg00244920
cg00245465
cg00245746
cg00245890
cg00245896
cg00246301
cg00247269
cg00247557

cg00247614
cg00248449
cg00248828
cg00248856
cg00249511
cg00250123
cg00250391
cg00250414
cg00251358
cg00251973
cg00252282
cg00252529
cg00252534
cg00252814
cg00253228
cg00253269
cg00253658
cg00253735
cg00253851
cg00254133
cg00254470
cg00254886
cg00255099
cg00255368
cg00255699
cg00255732
cg00255821
cg00256593
cg00256675
cg00256928
cg00257047
cg00257473
cg00257775
cg00257800
cg00259367
cg00259388
cg00259755
cg00259886
cg00260012
cg00260512
cg00260575
cg00260937
cg00261912
cg00262159
cg00263146
cg00263760
cg00264500
cg00264754
cg00264799
cg00266322
cg00266604
cg00267142

cg00267156
cg00267360
cg00268185
cg00268476
cg00268840
cg00268869
cg00269817
cg00270259
cg00270460
cg00270654
cg00270764
cg00271522
cg00272971
cg00273543
cg00273863
cg00274357
cg00275126
cg00275503
cg00275741
cg00275828
cg00276214
cg00278028
cg00278547
cg00280220
cg00280799
cg00281664
cg00282195
cg00282877
cg00283524
cg00283876
cg00283887
cg00284386
cg00284390
cg00284551
cg00285394
cg00285620
cg00286102
cg00286512
cg00286909
cg00287312
cg00287589
cg00288376
cg00288736
cg00288844
cg00289094
cg00289546
cg00290023
cg00290101
cg00290506
cg00291051
cg00291478
cg00292029

cg00292636
cg00292947
cg00292986
cg00293060
cg00293940
cg00293996
cg00294267
cg00294684
cg00294940
cg00295060
cg00295182
cg00295259
cg00295325
cg00295382
cg00295485
cg00295794
cg00296578
cg00297075
cg00297600
cg00297741
cg00297950
cg00298065
cg00298324
cg00298405
cg00298919
cg00298977
cg00299105
cg00299660
cg00299736
cg00299737
cg00300637
cg00300672
cg00300969
cg00301033
cg00301372
cg00301660
cg00301879
cg00302083
cg00302340
cg00303606
cg00303919
cg00304423
cg00305071
cg00305193
cg00305947
cg00305993
cg00306578
cg00306851
cg00306893
cg00307383
cg00308560
cg00308914

cg00309232
cg00309582
cg00309873
cg00309965
cg00310215
cg00310463
cg00310588
cg00311035
cg00311768
cg00311799
cg00311984
cg00312508
cg00313094
cg00313498
cg00314343
cg00314892
cg00314923
cg00314966
cg00314994
cg00315395
cg00315853
cg00316478
cg00316485
cg00316800
cg00316839
cg00317020
cg00317577
cg00317886
cg00318320
cg00318543
cg00318682
cg00319026
cg00319655
cg00320216
cg00320288
cg00320980
cg00321234
cg00321288
cg00321644
cg00321824
cg00322212
cg00322618
cg00323014
cg00323842
cg00324520
cg00325381
cg00325491
cg00325910
cg00326183
cg00326208
cg00326785
cg00326908

cg00327332
cg00327437
cg00327947
cg00328075
cg00328376
cg00328870
cg00329039
cg00329270
cg00329447
cg00330346
cg00330825
cg00331027
cg00331283
cg00332021
cg00332067
cg00332146
cg00332421
cg00332882
cg00332937
cg00333512
cg00333800
cg00333824
cg00334056
cg00334063
cg00334274
cg00334863
cg00334933
cg00334976
cg00335219
cg00335712
cg00335892
cg00336149
cg00336164
cg00336669
cg00336700
cg00337090
cg00337440
cg00337703
cg00338519
cg00338852
cg00339241
cg00339331
cg00339415
cg00339556
cg00339695
cg00340127
cg00341204
cg00341415
cg00342891
cg00343063
cg00343127
cg00343433

cg00343633
cg00344443
cg00345443
cg00346446
cg00346951
cg00347056
cg00347452
cg00347620
cg00347729
cg00347775
cg00347904
cg00348222
cg00349925
cg00350296
cg00350304
cg00350405
cg00350478
cg00350503
cg00351152
cg00351202
cg00352031
cg00352576
cg00353330
cg00353353
cg00353773
cg00353953
cg00354884
cg00355019
cg00355235
cg00355286
cg00355325
cg00355377
cg00355656
cg00356045
cg00356183
cg00356208
cg00356875
cg00357551
cg00357958
cg00357971
cg00358278
cg00358442
cg00358936
cg00359365
cg00359395
cg00359590
cg00360474
cg00360761
cg00361176
cg00361495
cg00361898
cg00362322

cg00362569
cg00362818
cg00363177
cg00364287
cg00364696
cg00365945
cg00366531
cg00366850
cg00366880
cg00367281
cg00367424
cg00367497
cg00368296
cg00368577
cg00368604
cg00369372
cg00370022
cg00370123
cg00370173
cg00371061
cg00371368
cg00372249
cg00372774
cg00372886
cg00373256
cg00373436
cg00373707
cg00374016
cg00374101
cg00374672
cg00374717
cg00375581
cg00375819
cg00375999
cg00376464
cg00376480
cg00376544
cg00377916
cg00377953
cg00378234
cg00378292
cg00378658
cg00379648
cg00379720
cg00380428
cg00380435
cg00380904
cg00381442
cg00381755
cg00382138
cg00382944
cg00383020

cg00383081
cg00383136
cg00383220
cg00383763
cg00384537
cg00384539
cg00384707
cg00385194
cg00386035
cg00386940
cg00387200
cg00387217
cg00387704
cg00388287
cg00388812
cg00388871
cg00390253
cg00390484
cg00390511
cg00390724
cg00391067
cg00391247
cg00391668
cg00391739
cg00392156
cg00392242
cg00392257
cg00392677
cg00393247
cg00393415
cg00393487
cg00393992
cg00394101
cg00394180
cg00394261
cg00394385
cg00394581
cg00395579
cg00395657
cg00396667
cg00396910
cg00397673
cg00397849
cg00397986
cg00398268
cg00398500
cg00398816
cg00399483
cg00400733
cg00403579
cg00403716
cg00404280

cg00405078
cg00405112
cg00405198
cg00405677
cg00405803
cg00406338
cg00406844
cg00407062
cg00407150
cg00407413
cg00407659
cg00407866
cg00407944
cg00408773
cg00409684
cg00409696
cg00409816
cg00409844
cg00410034
cg00410719
cg00411411
cg00411595
cg00411948
cg00412094
cg00412772
cg00413617
cg00415057
cg00415424
cg00415971
cg00415993
cg00416130
cg00416243
cg00416386
cg00416945
cg00417025
cg00417197
cg00417605
cg00418071
cg00418163
cg00418216
cg00418282
cg00418735
cg00420246
cg00420510
cg00420929
cg00421139
cg00421173
cg00421335
cg00421624
cg00422311
cg00422566
cg00422624

cg00422699
cg00422935
cg00423035
cg00423185
cg00423291
cg00423453
cg00423486
cg00423675
cg00423704
cg00424467
cg00424572
cg00424785
cg00425213
cg00425636
cg00426720
cg00426855
cg00426976
cg00427939
cg00427986
cg00428692
cg00429268
cg00430036
cg00431835
cg00432262
cg00432454
cg00432953
cg00433709
cg00434010
cg00434119
cg00434395
cg00434535
cg00435576
cg00435606
cg00435723
cg00436496
cg00437411
cg00439261
cg00440032
cg00441573
cg00441765
cg00441918
cg00442258
cg00442646
cg00443111
cg00443797
cg00444151
cg00444261
cg00444360
cg00444390
cg00445305
cg00445523
cg00445755

cg00446413
cg00446763
cg00447376
cg00448162
cg00448720
cg00448761
cg00449482
cg00449728
cg00449812
cg00450784
cg00450960
cg00451054
cg00451388
cg00451397
cg00452252
cg00452257
cg00452266
cg00452755
cg00452835
cg00453864
cg00454274
cg00455216
cg00455331
cg00455402
cg00455467
cg00455526
cg00456672
cg00457403
cg00457867
cg00457913
cg00458351
cg00458754
cg00458878
cg00458932
cg00459116
cg00459617
cg00459623
cg00460049
cg00460235
cg00460575
cg00460916
cg00461299
cg00462700
cg00463631
cg00463746
cg00463848
cg00463949
cg00463982
cg00464046
cg00464520
cg00464946
cg00465073

cg00465277
cg00465739
cg00466249
cg00466268
cg00466334
cg00466402
cg00466436
cg00466492
cg00466500
cg00466839
cg00467296
cg00467553
cg00468146
cg00468320
cg00468380
cg00468395
cg00469207
cg00469497
cg00469856
cg00470051
cg00470341
cg00470641
cg00471000
cg00471159
cg00472277
cg00472281
cg00472373
cg00472528
cg00472814
cg00472841
cg00473501
cg00473531
cg00474657
cg00474696
cg00474713
cg00476872
cg00476944
cg00476955
cg00477086
cg00477226
cg00478884
cg00479347
cg00479463
cg00479718
cg00480389
cg00481159
cg00481216
cg00481280
cg00481629
cg00481639
cg00482162
cg00482996

cg00483030
cg00483217
cg00484484
cg00485050
cg00485194
cg00486011
cg00486022
cg00486259
cg00486280
cg00486340
cg00486998
cg00487155
cg00487506
cg00488514
cg00488575
cg00489401
cg00489539
cg00489660
cg00490655
cg00490976
cg00491404
cg00491523
cg00491530
cg00491548
cg00492142
cg00492333
cg00494183
cg00495574
cg00495658
cg00495708
cg00495775
cg00495811
cg00495860
cg00496126
cg00496462
cg00496484
cg00497078
cg00497219
cg00497232
cg00497737
cg00498289
cg00498360
cg00498434
cg00498804
cg00499237
cg00499277
cg00499475
cg00499700
cg00500229
cg00500564
cg00501622
cg00502190

cg00502444
cg00502662
cg00502665
cg00502885
cg00503298
cg00503302
cg00503383
cg00504569
cg00504599
cg00504690
cg00504705
cg00504782
cg00505277
cg00505570
cg00506886
cg00507135
cg00508158
cg00508604
cg00508817
cg00508950
cg00509108
cg00509670
cg00509996
cg00510437
cg00510598
cg00510689
cg00510870
cg00511577
cg00511674
cg00512459
cg00512484
cg00512563
cg00513334
cg00513984
cg00514713
cg00514895
cg00514971
cg00515455
cg00515457
cg00515669
cg00516867
cg00516966
cg00517261
cg00517787
cg00518190
cg00518276
cg00518857
cg00519537
cg00519829
cg00519851
cg00519884
cg00520378

cg00520441
cg00521703
cg00521850
cg00522048
cg00522935
cg00523067
cg00523241
cg00523901
cg00524561
cg00524900
cg00525722
cg00525823
cg00526772
cg00527653
cg00528572
cg00528616
cg00529909
cg00529958
cg00530015
cg00530127
cg00530564
cg00530825
cg00530870
cg00530925
cg00531338
cg00532335
cg00532399
cg00532449
cg00532455
cg00532474
cg00532502
cg00532936
cg00533183
cg00533390
cg00534255
cg00534479
cg00534540
cg00535248
cg00535994
cg00536366
cg00536910
cg00537673
cg00537960
cg00537984
cg00538247
cg00538695
cg00538888
cg00539154
cg00539300
cg00539360
cg00539386
cg00539976

cg00540067
cg00540593
cg00540769
cg00541303
cg00542224
cg00543460
cg00543840
cg00544449
cg00544557
cg00544573
cg00544674
cg00545232
cg00545462
cg00545469
cg00545702
cg00545705
cg00545804
cg00546005
cg00546491
cg00547024
cg00547103
cg00547304
cg00547445
cg00547484
cg00548009
cg00548098
cg00548911
cg00549018
cg00549772
cg00549910
cg00550080
cg00550797
cg00551647
cg00551679
cg00552235
cg00552574
cg00553149
cg00553487
cg00554173
cg00554250
cg00554604
cg00554631
cg00555420
cg00555456
cg00555894
cg00557402
cg00557536
cg00557947
cg00558689
cg00559560
cg00560072
cg00560542

cg00561194
cg00561674
cg00561739
cg00561854
cg00562088
cg00562107
cg00562468
cg00562504
cg00563061
cg00563495
cg00563678
cg00563824
cg00563847
cg00563873
cg00564061
cg00564426
cg00564524
cg00564737
cg00564862
cg00564996
cg00565412
cg00565679
cg00565882
cg00566208
cg00566320
cg00566759
cg00566899
cg00566961
cg00569056
cg00569447
cg00570069
cg00570469
cg00571483
cg00571819
cg00572089
cg00572387
cg00573410
cg00573835
cg00573887
cg00574196
cg00574379
cg00574530
cg00574907
cg00575003
cg00575005
cg00577167
cg00577449
cg00577488
cg00577560
cg00578039
cg00578673
cg00579520

cg00579706
cg00579717
cg00579805
cg00580752
cg00580854
cg00580978
cg00581541
cg00582524
cg00582628
cg00582671
cg00582970
cg00582971
cg00583291
cg00583861
cg00584238
cg00584422
cg00584450
cg00584485
cg00585207
cg00585621
cg00586058
cg00586383
cg00586420
cg00586732
cg00586817
cg00587665
cg00587890
cg00588517
cg00588734
cg00589181
cg00589371
cg00589488
cg00589581
cg00590022
cg00590039
cg00590139
cg00591225
cg00591740
cg00591791
cg00591868
cg00591895
cg00592007
cg00592593
cg00592781
cg00593008
cg00593298
cg00593536
cg00593607
cg00594016
cg00594854
cg00595472
cg00596508

cg00597607
cg00598021
cg00598235
cg00598394
cg00598429
cg00598600
cg00598956
cg00599219
cg00599463
cg00599770
cg00601042
cg00601124
cg00602115
cg00602693
cg00603110
cg00604202
cg00604209
cg00604410
cg00604840
cg00606739
cg00607194
cg00607755
cg00607989
cg00608540
cg00608779
cg00609291
cg00609961
cg00610000
cg00610294
cg00610360
cg00610577
cg00611192
cg00612985
cg00613113
cg00613255
cg00613385
cg00613752
cg00613786
cg00614011
cg00614182
cg00614249
cg00614360
cg00615377
cg00615485
cg00615941
cg00616687
cg00617691
cg00618155
cg00618507
cg00618717
cg00619126
cg00619207

cg00619263
cg00620190
cg00620224
cg00620733
cg00621646
cg00621899
cg00621943
cg00622318
cg00622552
cg00622654
cg00622837
cg00623161
cg00623774
cg00624418
cg00625103
cg00625110
cg00625653
cg00625845
cg00626412
cg00626856
cg00627029
cg00628277
cg00628477
cg00628788
cg00629088
cg00629427
cg00629625
cg00630480
cg00630775
cg00630794
cg00630958
cg00631759
cg00632007
cg00632379
cg00632863
cg00633413
cg00634968
cg00635082
cg00635343
cg00635390
cg00636194
cg00636416
cg00636427
cg00636508
cg00637047
cg00637477
cg00637687
cg00637695
cg00638020
cg00638021
cg00638062
cg00638073

cg00638574
cg00638945
cg00639010
cg00639635
cg00640314
cg00640376
cg00640479
cg00641579
cg00642749
cg00643296
cg00643651
cg00643814
cg00643876
cg00644103
cg00644823
cg00644847
cg00645731
cg00646019
cg00646347
cg00647262
cg00647317
cg00647810
cg00648203
cg00648274
cg00648883
cg00649632
cg00650641
cg00650953
cg00651359
cg00652796
cg00653387
cg00653847
cg00654296
cg00654657
cg00654888
cg00655094
cg00656478
cg00656513
cg00656728
cg00656881
cg00657094
cg00657460
cg00657508
cg00658411
cg00658590
cg00658743
cg00658890
cg00659077
cg00659931
cg00660096
cg00661018
cg00661399

cg00661523
cg00661732
cg00661970
cg00662122
cg00662400
cg00662556
cg00663068
cg00663077
cg00663972
cg00663986
cg00664343
cg00664581
cg00664792
cg00665343
cg00665492
cg00665827
cg00667048
cg00667298
cg00667315
cg00667789
cg00668303
cg00668515
cg00669390
cg00670060
cg00670721
cg00671386
cg00671600
cg00671823
cg00672093
cg00672277
cg00672633
cg00672712
cg00672936
cg00673109
cg00673365
cg00674004
cg00674365
cg00674628
cg00674995
cg00675229
cg00675260
cg00676085
cg00676394
cg00676536
cg00676660
cg00676988
cg00678005
cg00678892
cg00679281
cg00679556
cg00679763
cg00679792

cg00679877
cg00681365
cg00682069
cg00682367
cg00683329
cg00683332
cg00683665
cg00684594
cg00685480
cg00685795
cg00685836
cg00686030
cg00686132
cg00686197
cg00686477
cg00686823
cg00686880
cg00686926
cg00687444
cg00688402
cg00688447
cg00688803
cg00688810
cg00688962
cg00689010
cg00689213
cg00689225
cg00689612
cg00689835
cg00690148
cg00690195
cg00690280
cg00690431
cg00690710
cg00690809
cg00690929
cg00691081
cg00691451
cg00691598
cg00691625
cg00691830
cg00691906
cg00692088
cg00692279
cg00692462
cg00692774
cg00692780
cg00693222
cg00694180
cg00694302
cg00695112
cg00695821

cg00696685
cg00697358
cg00697440
cg00697639
cg00697672
cg00697916
cg00697992
cg00698169
cg00698204
cg00698575
cg00698646
cg00698688
cg00699029
cg00699219
cg00699486
cg00699945
cg00699993
cg00700239
cg00701064
cg00701514
cg00701692
cg00702547
cg00703120
cg00703260
cg00703822
cg00705059
cg00706191
cg00706570
cg00706697
cg00706913
cg00707134
cg00707317
cg00707406
cg00707688
cg00707777
cg00707814
cg00708105
cg00708215
cg00709387
cg00709652
cg00709775
cg00709966
cg00710229
cg00710269
cg00710343
cg00710862
cg00711072
cg00711088
cg00711496
cg00711984
cg00712050
cg00712390

cg00712627
cg00712711
cg00713435
cg00713567
cg00714184
cg00714464
cg00714725
cg00714740
cg00715122
cg00715696
cg00716016
cg00716040
cg00716579
cg00717022
cg00718142
cg00718175
cg00718261
cg00718336
cg00718518
cg00718539
cg00719899
cg00720629
cg00721016
cg00721170
cg00721193
cg00721315
cg00721947
cg00722081
cg00722834
cg00723041
cg00723338
cg00723379
cg00723994
cg00724015
cg00724839
cg00725089
cg00725138
cg00725162
cg00725635
cg00726029
cg00726786
cg00727334
cg00727663
cg00727675
cg00728046
cg00729133
cg00729275
cg00730441
cg00731062
cg00731458
cg00731874
cg00732232

cg00732656
cg00732878
cg00733328
cg00733493
cg00733567
cg00733722
cg00733978
cg00735595
cg00735611
cg00735667
cg00735907
cg00735923
cg00736104
cg00736201
cg00736406
cg00736459
cg00736958
cg00737556
cg00737593
cg00737979
cg00738345
cg00738660
cg00738897
cg00739155
cg00739217
cg00739471
cg00739679
cg00739778
cg00740088
cg00740510
cg00741410
cg00741609
cg00741624
cg00741695
cg00741836
cg00741900
cg00741992
cg00742465
cg00742587
cg00742649
cg00742898
cg00743629
cg00743849
cg00744413
cg00744739
cg00744866
cg00745520
cg00745555
cg00745725
cg00746157
cg00746844
cg00747477

cg00747647
cg00747922
cg00747944
cg00748226
cg00748373
cg00748911
cg00750518
cg00752143
cg00752195
cg00752480
cg00752969
cg00753748
cg00754617
cg00754896
cg00755287
cg00755546
cg00755980
cg00756448
cg00756860
cg00757731
cg00757822
cg00757952
cg00758069
cg00758854
cg00759573
cg00759807
cg00759987
cg00760014
cg00760872
cg00760935
cg00760950
cg00762160
cg00762397
cg00763594
cg00763743
cg00764099
cg00764217
cg00764612
cg00765127
cg00765635
cg00765705
cg00765783
cg00765828
cg00765992
cg00766158
cg00767010
cg00767496
cg00767642
cg00768741
cg00768815
cg00768993
cg00769590

cg00770998
cg00772091
cg00772192
cg00772257
cg00772380
cg00772407
cg00773795
cg00774006
cg00774437
cg00774867
cg00775197
cg00775777
cg00775863
cg00776166
cg00777079
cg00777315
cg00777375
cg00777627
cg00777689
cg00778084
cg00778190
cg00779065
cg00780520
cg00780971
cg00781519
cg00782099
cg00782323
cg00782355
cg00782503
cg00782811
cg00783712
cg00784671
cg00785029
cg00785112
cg00786237
cg00786406
cg00786685
cg00787013
cg00787180
cg00787290
cg00787996
cg00788170
cg00789416
cg00789792
cg00790071
cg00790535
cg00790672
cg00790916
cg00791024
cg00791074
cg00791253
cg00791406

cg00791548
cg00791712
cg00791868
cg00792015
cg00792251
cg00792424
cg00792513
cg00792688
cg00792849
cg00793181
cg00793562
cg00793937
cg00794178
cg00794673
cg00795024
cg00795341
cg00795825
cg00795830
cg00796302
cg00799539
cg00800229
cg00800759
cg00801134
cg00802237
cg00802510
cg00802728
cg00803453
cg00803692
cg00804097
cg00804628
cg00804745
cg00804934
cg00805360
cg00806319
cg00806490
cg00806704
cg00806826
cg00807235
cg00807366
cg00807877
cg00808086
cg00808170
cg00808403
cg00808563
cg00808648
cg00808721
cg00809021
cg00809267
cg00809399
cg00809988
cg00810208
cg00810292

cg00810483
cg00810710
cg00810956
cg00811065
cg00812236
cg00812246
cg00812438
cg00812578
cg00812611
cg00812833
cg00812921
cg00813135
cg00813373
cg00813890
cg00815091
cg00815198
cg00815266
cg00815325
cg00816224
cg00816620
cg00816770
cg00817331
cg00818106
cg00819141
cg00819818
cg00820056
cg00820637
cg00821731
cg00822797
cg00823110
cg00823148
cg00824018
cg00824767
cg00825690
cg00826537
cg00827176
cg00827210
cg00827382
cg00827895
cg00828836
cg00829600
cg00829858
cg00829990
cg00830086
cg00830325
cg00830599
cg00830621
cg00831127
cg00831726
cg00831735
cg00831909
cg00832199

cg00832567
cg00832581
cg00832703
cg00833364
cg00834899
cg00837398
cg00838081
cg00839579
cg00840332
cg00840960
cg00842549
cg00842595
cg00844277
cg00844298
cg00845245
cg00845901
cg00846043
cg00846121
cg00846968
cg00846988
cg00847277
cg00847826
cg00848461
cg00849025
cg00849148
cg00849365
cg00850193
cg00851389
cg00851627
cg00851770
cg00851830
cg00852051
cg00852534
cg00852622
cg00853042
cg00853103
cg00853242
cg00853687
cg00854242
cg00854817
cg00855901
cg00855990
cg00857536
cg00857851
cg00858840
cg00859330
cg00860359
cg00860372
cg00860712
cg00860808
cg00861646
cg00861856

cg00861945
cg00862016
cg00862117
cg00862443
cg00862597
cg00862675
cg00863309
cg00863318
cg00863716
cg00864458
cg00864460
cg00864530
cg00864661
cg00864666
cg00864691
cg00864954
cg00865429
cg00865690
cg00865973
cg00866054
cg00866500
cg00866556
cg00866632
cg00867889
cg00868074
cg00868523
cg00868860
cg00870242
cg00870449
cg00871055
cg00871207
cg00871358
cg00871371
cg00871501
cg00872232
cg00872435
cg00872683
cg00872695
cg00872726
cg00872865
cg00873050
cg00874225
cg00874480
cg00874891
cg00875382
cg00875511
cg00875989
cg00876141
cg00876196
cg00876267
cg00876272
cg00877938

cg00880872
cg00881010
cg00882967
cg00883190
cg00883493
cg00883798
cg00883837
cg00884093
cg00884606
cg00885461
cg00886571
cg00886780
cg00887508
cg00887814
cg00887902
cg00887936
cg00887966
cg00888758
cg00888818
cg00889363
cg00889398
cg00890049
cg00890054
cg00890363
cg00893471
cg00893493
cg00893522
cg00894134
cg00895324
cg00895354
cg00895528
cg00895997
cg00896068
cg00896282
cg00896347
cg00897058
cg00898102
cg00898111
cg00900226
cg00900242
cg00901264
cg00901554
cg00902104
cg00902153
cg00902195
cg00902374
cg00902417
cg00902815
cg00903099
cg00903250
cg00903308
cg00903371

cg00903452
cg00903584
cg00903932
cg00904966
cg00905101
cg00905524
cg00905584
cg00905746
cg00906319
cg00906420
cg00906476
cg00906908
cg00907008
cg00907198
cg00907267
cg00907272
cg00907288
cg00908004
cg00908291
cg00908323
cg00908571
cg00908584
cg00908631
cg00908828
cg00909062
cg00909286
cg00909514
cg00909926
cg00910067
cg00910203
cg00910676
cg00910695
cg00910882
cg00910972
cg00911226
cg00911351
cg00911370
cg00911376
cg00911881
cg00912625
cg00912738
cg00912772
cg00913288
cg00913787
cg00914787
cg00914902
cg00915974
cg00915990
cg00916179
cg00916536
cg00916884
cg00917101

cg00917181
cg00917413
cg00917642
cg00918005
cg00918181
cg00918522
cg00918738
cg00919126
cg00919411
cg00919591
cg00919702
cg00919971
cg00920162
cg00920327
cg00920395
cg00920424
cg00920938
cg00921219
cg00922157
cg00922200
cg00922781
cg00923860
cg00923934
cg00923996
cg00924143
cg00924255
cg00924459
cg00924575
cg00925492
cg00925620
cg00925953
cg00926215
cg00927256
cg00927364
cg00927624
cg00928002
cg00928751
cg00928816
cg00929332
cg00929345
cg00929635
cg00929871
cg00930606
cg00931972
cg00933411
cg00933443
cg00933542
cg00933835
cg00933859
cg00934087
cg00934232
cg00934746

cg00934987
cg00935653
cg00935717
cg00936187
cg00936790
cg00936935
cg00937175
cg00937286
cg00937359
cg00938050
cg00938540
cg00939495
cg00939735
cg00940941
cg00941900
cg00942219
cg00943360
cg00943851
cg00944163
cg00944304
cg00944580
cg00945108
cg00945209
cg00945260
cg00945293
cg00945666
cg00946598
cg00946921
cg00947324
cg00947744
cg00948900
cg00949386
cg00949661
cg00949753
cg00950004
cg00950244
cg00950398
cg00950497
cg00950846
cg00951599
cg00951770
cg00953256
cg00954256
cg00954391
cg00954633
cg00955307
cg00955413
cg00955547
cg00955769
cg00955808
cg00955828
cg00955911

cg00956142
cg00956199
cg00957080
cg00957665
cg00957901
cg00958225
cg00958412
cg00959431
cg00959883
cg00960147
cg00960305
cg00960395
cg00961139
cg00961325
cg00961450
cg00961792
cg00961940
cg00962147
cg00962271
cg00962740
cg00962755
cg00963169
cg00963378
cg00963496
cg00963675
cg00963998
cg00964446
cg00965023
cg00965154
cg00965971
cg00966099
cg00966255
cg00966522
cg00966763
cg00967711
cg00968167
cg00968538
cg00969119
cg00969288
cg00969323
cg00969893
cg00970313
cg00970396
cg00970435
cg00971309
cg00971695
cg00972420
cg00973118
cg00973944
cg00974095
cg00974255
cg00974464

cg00974523
cg00974761
cg00974944
cg00975971
cg00976181
cg00976453
cg00976532
cg00976619
cg00977384
cg00977895
cg00978783
cg00978848
cg00979288
cg00980058
cg00980060
cg00980772
cg00980804
cg00980904
cg00981060
cg00981350
cg00982136
cg00982214
cg00982519
cg00983440
cg00983482
cg00983608
cg00983803
cg00983948
cg00984146
cg00985817
cg00985983
cg00986580
cg00986824
cg00987015
cg00987044
cg00987080
cg00987534
cg00988148
cg00988175
cg00988350
cg00988548
cg00988577
cg00988592
cg00988641
cg00989229
cg00989315
cg00989505
cg00989853
cg00989986
cg00990769
cg00991744
cg00991794

cg00991875
cg00992055
cg00992153
cg00992317
cg00992348
cg00992521
cg00992850
cg00993310
cg00994251
cg00995065
cg00995141
cg00995152
cg00995220
cg00995520
cg00995854
cg00995986
cg00996588
cg00996827
cg00996847
cg00997611
cg00998387
cg00998782
cg01001101
cg01002030
cg01002242
cg01002253
cg01002313
cg01002529
cg01003211
cg01003390
cg01003666
cg01003815
cg01003859
cg01003961
cg01004017
cg01004762
cg01004918
cg01005303
cg01006048
cg01007737
cg01007828
cg01008164
cg01008256
cg01008557
cg01008680
cg01009664
cg01010091
cg01010221
cg01010923
cg01010988
cg01011530
cg01012026

cg01012330
cg01012572
cg01012936
cg01013155
cg01013590
cg01014190
cg01014224
cg01014615
cg01015123
cg01015175
cg01015395
cg01015401
cg01015663
cg01015769
cg01015871
cg01016122
cg01016169
cg01016533
cg01017257
cg01018110
cg01018477
cg01018492
cg01019262
cg01019484
cg01019587
cg01019770
cg01019874
cg01020203
cg01020744
cg01021552
cg01021658
cg01022012
cg01022370
cg01023696
cg01023759
cg01024168
cg01024247
cg01024618
cg01025233
cg01025434
cg01026204
cg01026233
cg01026458
cg01026796
cg01027010
cg01027532
cg01028287
cg01028849
cg01029450
cg01029568
cg01029637
cg01029669

cg01030121
cg01030476
cg01030534
cg01030598
cg01031441
cg01031983
cg01031999
cg01032398
cg01032675
cg01033001
cg01033419
cg01033938
cg01034411
cg01035160
cg01035260
cg01035261
cg01035563
cg01035771
cg01036409
cg01037823
cg01038207
cg01039498
cg01039508
cg01039876
cg01040624
cg01040786
cg01040850
cg01040890
cg01041284
cg01041455
cg01042100
cg01042246
cg01042640
cg01043616
cg01043681
cg01044129
cg01044179
cg01044476
cg01044637
cg01045374
cg01046377
cg01046961
cg01047555
cg01047586
cg01047631
cg01047904
cg01048107
cg01048195
cg01048372
cg01048810
cg01049782
cg01049849

cg01049933
cg01051003
cg01051310
cg01052614
cg01052636
cg01052701
cg01053260
cg01054140
cg01054282
cg01054402
cg01054478
cg01055044
cg01055121
cg01055129
cg01055594
cg01055633
cg01055691
cg01056004
cg01056615
cg01057132
cg01057656
cg01057945
cg01058070
cg01058569
cg01059047
cg01059449
cg01059609
cg01059704
cg01059768
cg01059818
cg01059881
cg01060409
cg01060471
cg01060861
cg01060964
cg01062020
cg01062024
cg01062116
cg01062126
cg01063509
cg01063701
cg01063935
cg01064373
cg01064918
cg01064979
cg01065349
cg01065697
cg01065806
cg01066954
cg01067139
cg01068808
cg01068944

cg01069451
cg01069738
cg01069941
cg01070095
cg01070209
cg01071185
cg01071534
cg01071644
cg01071903
cg01071966
cg01072658
cg01072821
cg01074356
cg01074365
cg01075095
cg01075271
cg01075459
cg01075553
cg01076272
cg01076454
cg01076459
cg01076708
cg01077058
cg01077185
cg01077469
cg01078332
cg01078582
cg01079019
cg01079599
cg01080998
cg01081091
cg01081096
cg01081400
cg01081636
cg01082512
cg01082907
cg01084270
cg01084837
cg01085362
cg01085569
cg01087294
cg01088387
cg01088410
cg01089249
cg01089319
cg01089425
cg01089498
cg01089544
cg01089579
cg01089602
cg01089858
cg01090026

cg01090482
cg01090930
cg01091565
cg01091751
cg01092213
cg01092890
cg01093028
cg01093065
cg01093088
cg01093296
cg01094192
cg01095406
cg01096381
cg01096398
cg01097071
cg01097384
cg01097663
cg01097725
cg01097768
cg01097950
cg01097963
cg01099855
cg01100030
cg01100208
cg01100796
cg01100912
cg01101058
cg01101522
cg01101663
cg01102471
cg01103582
cg01104208
cg01104250
cg01104549
cg01104724
cg01105058
cg01105140
cg01105337
cg01105375
cg01105494
cg01105802
cg01106338
cg01106926
cg01107872
cg01108633
cg01109017
cg01109023
cg01109219
cg01109227
cg01109535
cg01109763
cg01110759

cg01111041
cg01111358
cg01111718
cg01112778
cg01112805
cg01112899
cg01113073
cg01113412
cg01114085
cg01114156
cg01115243
cg01115380
cg01115923
cg01116059
cg01116321
cg01116873
cg01117213
cg01117339
cg01118711
cg01119072
cg01119145
cg01119374
cg01119452
cg01119875
cg01120509
cg01121603
cg01121610
cg01122167
cg01122415
cg01124420
cg01124523
cg01124530
cg01125010
cg01125531
cg01126532
cg01126567
cg01127249
cg01127300
cg01127423
cg01127994
cg01128031
cg01128109
cg01128503
cg01128566
cg01128574
cg01128642
cg01129320
cg01130010
cg01132471
cg01132484
cg01132568
cg01132653

cg01134183
cg01134282
cg01134459
cg01134526
cg01135315
cg01135800
cg01138652
cg01138819
cg01138867
cg01138921
cg01139503
cg01140006
cg01140008
cg01140149
cg01140247
cg01141413
cg01141438
cg01142328
cg01142635
cg01142693
cg01143068
cg01143426
cg01143547
cg01144251
cg01145026
cg01145137
cg01145396
cg01146063
cg01146232
cg01146320
cg01147550
cg01147727
cg01148073
cg01148441
cg01149053
cg01149940
cg01150105
cg01150270
cg01150433
cg01150683
cg01150799
cg01150918
cg01151584
cg01151966
cg01152072
cg01152727
cg01153080
cg01153166
cg01153376
cg01153385
cg01153817
cg01153946

cg01154527
cg01154903
cg01155497
cg01156370
cg01156892
cg01158094
cg01158415
cg01158527
cg01158680
cg01158822
cg01159576
cg01160079
cg01160241
cg01160311
cg01160347
cg01160452
cg01161629
cg01162421
cg01162494
cg01162507
cg01163842
cg01164574
cg01164972
cg01165355
cg01165560
cg01165817
cg01165997
cg01166071
cg01166876
cg01167502
cg01167755
cg01168235
cg01168865
cg01169396
cg01170196
cg01170694
cg01172073
cg01172082
cg01172150
cg01172682
cg01172872
cg01173332
cg01173650
cg01173794
cg01174140
cg01174276
cg01174843
cg01175550
cg01175610
cg01175832
cg01176516
cg01177709

cg01178040
cg01178099
cg01178188
cg01178624
cg01178680
cg01178703
cg01178710
cg01178854
cg01178924
cg01180523
cg01181227
cg01181415
cg01181485
cg01182048
cg01182154
cg01182620
cg01182683
cg01182895
cg01184149
cg01184194
cg01184282
cg01184975
cg01185433
cg01185470
cg01185573
cg01186595
cg01186702
cg01187505
cg01187600
cg01187920
cg01188235
cg01188396
cg01188592
cg01188753
cg01188914
cg01189011
cg01190024
cg01190276
cg01190317
cg01190637
cg01190974
cg01191058
cg01192051
cg01192172
cg01192489
cg01192869
cg01194057
cg01194320
cg01195676
cg01196079
cg01196158
cg01196744

cg01196788
cg01196858
cg01197156
cg01197871
cg01198033
cg01198364
cg01198491
cg01198887
cg01198994
cg01200021
cg01200264
cg01200640
cg01200651
cg01201399
cg01201519
cg01201818
cg01202197
cg01202296
cg01202639
cg01202700
cg01203136
cg01203550
cg01203812
cg01204058
cg01204844
cg01204982
cg01205011
cg01205087
cg01205165
cg01205267
cg01205831
cg01206705
cg01207916
cg01208224
cg01208537
cg01208557
cg01209150
cg01209199
cg01209635
cg01211763
cg01212010
cg01212215
cg01212562
cg01212887
cg01213022
cg01213342
cg01213440
cg01214384
cg01214458
cg01215682
cg01215762
cg01216882

cg01217329
cg01217350
cg01217601
cg01217745
cg01218180
cg01219000
cg01219549
cg01219553
cg01220564
cg01220680
cg01220923
cg01221157
cg01222324
cg01223336
cg01223986
cg01224657
cg01224891
cg01225698
cg01226265
cg01226417
cg01227294
cg01227492
cg01227537
cg01227558
cg01228271
cg01229785
cg01229865
cg01229943
cg01230207
cg01230796
cg01231009
cg01231108
cg01231464
cg01231921
cg01232121
cg01233025
cg01233664
cg01233934
cg01234044
cg01236027
cg01236063
cg01236132
cg01236299
cg01236751
cg01237056
cg01238639
cg01239651
cg01240056
cg01240823
cg01241450
cg01242677
cg01243371

cg01244015
cg01244270
cg01244514
cg01244520
cg01244944
cg01245155
cg01245393
cg01245880
cg01246982
cg01247127
cg01247537
cg01247628
cg01247693
cg01247788
cg01248426
cg01248745
cg01248801
cg01249544
cg01250845
cg01251150
cg01251710
cg01251971
cg01252713
cg01253676
cg01254170
cg01254264
cg01254303
cg01254991
cg01256165
cg01256440
cg01256444
cg01257194
cg01257383
cg01257889
cg01257975
cg01258050
cg01258789
cg01260603
cg01261086
cg01261351
cg01261797
cg01261798
cg01262413
cg01263386
cg01263574
cg01264963
cg01265860
cg01266375
cg01266985
cg01267150
cg01267165
cg01268022

cg01268590
cg01269212
cg01269314
cg01269620
cg01269798
cg01270241
cg01270424
cg01270709
cg01271455
cg01271485
cg01271695
cg01271812
cg01272393
cg01272668
cg01273232
cg01273336
cg01273384
cg01274752
cg01274812
cg01274878
cg01275038
cg01275697
cg01276323
cg01277162
cg01277542
cg01277890
cg01277983
cg01278387
cg01278654
cg01278680
cg01279413
cg01280180
cg01281157
cg01281285
cg01281911
cg01282906
cg01283246
cg01283289
cg01283300
cg01284192
cg01284289
cg01284709
cg01285237
cg01285501
cg01285652
cg01285783
cg01285926
cg01286319
cg01286624
cg01286655
cg01287000
cg01287218

cg01287376
cg01287730
cg01287975
cg01288337
cg01288372
cg01288375
cg01288724
cg01289782
cg01290136
cg01290718
cg01291336
cg01291856
cg01291920
cg01292539
cg01292722
cg01292810
cg01292924
cg01293723
cg01294283
cg01294808
cg01295203
cg01295399
cg01295539
cg01296889
cg01296927
cg01297020
cg01297303
cg01297370
cg01297670
cg01297806
cg01297896
cg01298731
cg01299082
cg01299388
cg01299925
cg01300641
cg01300684
cg01301197
cg01301382
cg01301637
cg01302136
cg01302656
cg01303008
cg01303236
cg01303372
cg01303723
cg01304499
cg01304575
cg01305421
cg01305539
cg01305830
cg01305913

cg01305981
cg01307108
cg01307130
cg01307483
cg01309121
cg01309342
cg01309395
cg01310118
cg01310482
cg01310829
cg01311064
cg01311102
cg01311151
cg01311170
cg01311313
cg01311470
cg01311802
cg01311805
cg01311955
cg01312394
cg01312482
cg01312828
cg01313363
cg01313538
cg01313732
cg01314165
cg01314306
cg01316378
cg01319892
cg01320036
cg01320540
cg01320573
cg01320698
cg01321816
cg01323672
cg01323993
cg01324802
cg01324963
cg01325409
cg01326120
cg01328158
cg01328500
cg01329151
cg01330688
cg01330815
cg01331196
cg01331275
cg01331992
cg01332136
cg01332534
cg01333287
cg01333327

cg01333471
cg01333824
cg01335126
cg01337143
cg01337391
cg01337514
cg01337931
cg01337940
cg01340163
cg01340347
cg01342115
cg01342226
cg01342572
cg01342746
cg01343192
cg01344518
cg01344738
cg01344813
cg01345315
cg01345395
cg01346767
cg01347335
cg01347752
cg01348055
cg01348374
cg01350169
cg01350680
cg01350993
cg01351089
cg01351315
cg01352108
cg01352175
cg01352294
cg01353670
cg01353960
cg01354088
cg01354137
cg01354426
cg01354473
cg01354588
cg01354782
cg01355932
cg01356115
cg01356198
cg01357234
cg01357292
cg01357429
cg01357713
cg01357846
cg01358620
cg01359200
cg01359236

cg01359329
cg01359532
cg01359809
cg01360115
cg01360618
cg01361263
cg01361555
cg01362243
cg01363198
cg01363714
cg01364172
cg01364483
cg01364621
cg01364766
cg01364826
cg01364935
cg01365766
cg01366338
cg01366725
cg01366985
cg01367097
cg01367899
cg01367929
cg01368068
cg01368219
cg01369413
cg01369750
cg01370181
cg01370294
cg01370541
cg01370805
cg01371799
cg01372572
cg01372811
cg01372898
cg01372939
cg01373505
cg01374580
cg01375075
cg01375248
cg01375651
cg01375869
cg01376469
cg01376826
cg01376829
cg01377644
cg01378182
cg01378456
cg01378515
cg01378999
cg01379237
cg01379240

cg01379344
cg01379656
cg01380104
cg01380194
cg01381829
cg01381846
cg01381934
cg01382072
cg01382110
cg01382860
cg01383203
cg01383486
cg01383881
cg01384111
cg01384246
cg01384290
cg01384319
cg01384488
cg01386157
cg01387072
cg01387102
cg01388243
cg01388565
cg01388598
cg01389445
cg01389964
cg01391022
cg01391071
cg01391280
cg01391299
cg01392017
cg01392180
cg01392351
cg01392544
cg01393604
cg01393705
cg01393841
cg01394339
cg01395254
cg01396065
cg01396112
cg01396176
cg01396387
cg01396587
cg01397223
cg01397513
cg01400040
cg01400480
cg01400638
cg01400737
cg01400822
cg01401344

cg01401465
cg01401762
cg01402099
cg01402347
cg01402631
cg01403030
cg01403655
cg01404122
cg01404750
cg01404873
cg01405040
cg01405242
cg01405404
cg01405582
cg01406506
cg01406909
cg01407018
cg01407062
cg01407332
cg01407447
cg01407927
cg01408486
cg01408508
cg01408654
cg01408817
cg01409460
cg01409693
cg01410279
cg01410801
cg01410878
cg01410923
cg01411892
cg01412404
cg01413522
cg01413548
cg01413632
cg01414185
cg01414572
cg01417625
cg01417849
cg01418188
cg01419727
cg01420369
cg01420398
cg01420564
cg01420931
cg01421119
cg01421140
cg01421252
cg01421830
cg01422476
cg01422881

cg01424263
cg01425293
cg01425409
cg01425680
cg01425718
cg01425762
cg01426713
cg01426808
cg01426853
cg01426903
cg01427458
cg01427575
cg01428071
cg01428589
cg01428849
cg01429391
cg01429408
cg01429449
cg01429662
cg01429844
cg01429933
cg01430302
cg01430430
cg01430588
cg01431057
cg01431184
cg01431340
cg01431972
cg01432055
cg01432087
cg01432520
cg01432552
cg01432692
cg01433468
cg01433836
cg01435039
cg01435344
cg01435574
cg01436487
cg01436998
cg01437135
cg01437998
cg01438467
cg01439248
cg01439670
cg01440333
cg01440796
cg01440841
cg01441622
cg01441699
cg01441777
cg01442005

cg01442565
cg01442601
cg01443426
cg01443433
cg01443467
cg01443755
cg01443946
cg01444397
cg01444712
cg01444716
cg01445151
cg01445580
cg01446731
cg01446894
cg01447112
cg01447281
cg01447344
cg01448863
cg01448944
cg01449144
cg01449266
cg01449674
cg01450390
cg01450803
cg01450807
cg01450896
cg01452847
cg01453814
cg01454134
cg01454215
cg01454487
cg01455398
cg01455766
cg01455981
cg01456609
cg01456640
cg01457572
cg01458054
cg01458195
cg01458336
cg01458510
cg01458605
cg01458646
cg01459162
cg01461410
cg01461514
cg01461847
cg01461862
cg01462537
cg01462607
cg01462668
cg01463828

cg01464067
cg01464104
cg01464186
cg01464738
cg01464835
cg01465527
cg01467100
cg01468220
cg01468420
cg01468579
cg01471196
cg01471232
cg01471372
cg01471572
cg01472101
cg01472375
cg01473038
cg01473602
cg01473840
cg01474011
cg01474260
cg01475325
cg01475397
cg01475538
cg01475649
cg01476222
cg01476795
cg01477133
cg01477861
cg01477933
cg01477985
cg01478179
cg01480771
cg01482087
cg01482969
cg01483627
cg01484075
cg01484915
cg01485009
cg01485075
cg01486146
cg01486814
cg01487187
cg01487409
cg01487892
cg01488260
cg01489057
cg01489177
cg01489686
cg01490728
cg01491071
cg01492538

cg01492909
cg01493150
cg01493297
cg01493727
cg01494441
cg01494454
cg01495122
cg01495831
cg01496199
cg01496696
cg01496891
cg01497434
cg01497576
cg01498249
cg01498437
cg01498609
cg01499001
cg01499172
cg01499623
cg01500527
cg01501819
cg01502208
cg01502373
cg01502407
cg01502457
cg01502484
cg01502743
cg01502811
cg01502872
cg01503065
cg01503299
cg01503348
cg01503706
cg01503938
cg01504248
cg01505878
cg01506471
cg01507342
cg01507577
cg01507753
cg01508023
cg01508380
cg01509065
cg01511234
cg01511901
cg01513063
cg01513392
cg01513978
cg01514341
cg01514353
cg01514356
cg01514490

cg01515427
cg01515444
cg01515534
cg01515543
cg01515574
cg01517384
cg01517490
cg01517500
cg01517832
cg01517968
cg01518591
cg01518607
cg01518790
cg01518889
cg01519253
cg01519695
cg01519765
cg01520072
cg01520122
cg01520454
cg01520867
cg01521220
cg01521648
cg01521917
cg01522296
cg01522412
cg01522437
cg01522592
cg01523600
cg01523881
cg01524278
cg01524618
cg01524923
cg01525538
cg01526089
cg01526177
cg01526355
cg01526553
cg01526854
cg01527459
cg01528542
cg01529201
cg01530032
cg01530101
cg01530803
cg01531058
cg01531915
cg01532168
cg01532230
cg01532487
cg01532749
cg01533082

cg01533258
cg01533409
cg01534253
cg01534572
cg01535838
cg01536744
cg01536878
cg01537445
cg01537471
cg01537995
cg01538575
cg01538605
cg01538731
cg01539036
cg01540203
cg01540953
cg01543173
cg01543179
cg01543184
cg01543654
cg01544400
cg01544877
cg01544903
cg01545493
cg01546378
cg01546561
cg01546568
cg01546729
cg01546820
cg01546873
cg01547809
cg01549929
cg01549977
cg01550397
cg01550799
cg01551133
cg01551177
cg01551300
cg01551797
cg01552035
cg01552272
cg01552504
cg01553090
cg01553182
cg01553231
cg01553335
cg01553935
cg01554316
cg01554357
cg01554457
cg01555151
cg01555300

cg01556466
cg01556799
cg01556846
cg01557027
cg01557260
cg01557332
cg01557411
cg01558215
cg01558401
cg01558785
cg01558877
cg01559502
cg01559597
cg01559663
cg01560554
cg01560676
cg01560773
cg01560873
cg01561092
cg01561157
cg01561194
cg01561869
cg01562502
cg01564322
cg01564343
cg01564665
cg01564902
cg01565013
cg01565291
cg01565774
cg01565985
cg01566170
cg01566216
cg01566526
cg01566584
cg01566692
cg01566999
cg01567672
cg01567771
cg01568317
cg01568319
cg01568492
cg01568784
cg01568998
cg01569440
cg01569827
cg01569875
cg01570938
cg01571203
cg01571735
cg01571916
cg01571928

cg01572434
cg01572652
cg01573470
cg01573635
cg01574297
cg01574481
cg01574663
cg01574903
cg01575414
cg01575601
cg01576142
cg01576260
cg01577083
cg01577751
cg01578017
cg01578764
cg01579458
cg01579600
cg01579765
cg01580681
cg01581018
cg01581084
cg01581222
cg01581237
cg01581757
cg01582616
cg01582980
cg01583716
cg01584000
cg01584103
cg01584457
cg01584932
cg01585096
cg01585180
cg01585372
cg01585852
cg01586116
cg01586851
cg01586952
cg01587064
cg01587896
cg01588224
cg01588250
cg01588345
cg01588388
cg01588475
cg01588826
cg01589129
cg01589580
cg01589587
cg01590787
cg01591152

cg01591240
cg01591487
cg01591579
cg01592682
cg01592687
cg01593385
cg01593673
cg01593787
cg01593834
cg01593969
cg01594685
cg01595262
cg01595397
cg01595870
cg01596674
cg01596834
cg01597149
cg01597373
cg01597604
cg01597629
cg01598009
cg01598421
cg01598596
cg01598741
cg01599099
cg01599460
cg01600921
cg01601299
cg01601658
cg01601926
cg01601944
cg01602001
cg01602690
cg01603862
cg01603912
cg01603973
cg01604883
cg01604946
cg01605366
cg01605435
cg01605984
cg01606773
cg01607369
cg01607530
cg01608731
cg01608878
cg01609275
cg01609420
cg01610605
cg01610632
cg01610915
cg01611894

cg01612003
cg01612095
cg01612110
cg01612449
cg01612730
cg01612824
cg01612936
cg01613010
cg01613048
cg01613691
cg01614020
cg01614101
cg01614102
cg01614643
cg01614746
cg01614915
cg01615114
cg01615271
cg01615704
cg01615827
cg01616440
cg01616682
cg01616797
cg01617211
cg01617280
cg01617750
cg01617933
cg01618102
cg01618130
cg01618245
cg01618287
cg01618697
cg01618806
cg01618851
cg01619509
cg01620379
cg01620785
cg01621390
cg01621576
cg01622606
cg01622930
cg01624414
cg01625087
cg01626493
cg01626885
cg01627405
cg01627492
cg01627597
cg01627785
cg01627823
cg01628245
cg01629435

cg01629545
cg01630588
cg01630690
cg01631855
cg01632562
cg01632597
cg01632865
cg01634544
cg01634644
cg01635405
cg01637175
cg01637519
cg01637537
cg01637609
cg01637789
cg01638592
cg01638792
cg01639723
cg01639892
cg01639943
cg01640150
cg01640660
cg01640727
cg01641371
cg01642521
cg01643605
cg01643624
cg01643690
cg01643997
cg01644518
cg01645874
cg01646289
cg01647204
cg01647314
cg01647693
cg01647728
cg01649773
cg01649879
cg01651250
cg01651378
cg01652927
cg01653494
cg01654824
cg01655548
cg01657574
cg01657758
cg01657761
cg01658421
cg01658605
cg01659038
cg01659184
cg01659631

cg01660911
cg01661350
cg01661777
cg01663016
cg01663440
cg01663603
cg01663677
cg01663696
cg01664151
cg01664241
cg01664727
cg01664929
cg01665118
cg01665555
cg01666652
cg01667575
cg01668008
cg01669085
cg01669948
cg01670430
cg01670608
cg01670677
cg01670789
cg01670876
cg01671881
cg01672322
cg01672943
cg01673307
cg01673783
cg01674119
cg01674130
cg01674585
cg01675814
cg01676182
cg01677601
cg01677623
cg01678084
cg01679108
cg01680054
cg01680823
cg01681351
cg01681367
cg01682208
cg01683000
cg01684256
cg01684579
cg01684586
cg01686920
cg01686975
cg01687040
cg01687177
cg01687389

cg01687680
cg01688536
cg01689438
cg01690182
cg01691827
cg01692018
cg01692233
cg01692482
cg01692639
cg01692757
cg01693026
cg01693284
cg01693305
cg01693350
cg01693423
cg01695023
cg01695225
cg01695279
cg01695643
cg01696226
cg01698392
cg01699217
cg01699293
cg01699298
cg01700290
cg01700358
cg01700462
cg01701055
cg01701649
cg01702009
cg01702055
cg01702210
cg01702225
cg01706498
cg01706515
cg01708014
cg01708273
cg01708458
cg01709846
cg01709871
cg01710084
cg01711160
cg01711322
cg01712428
cg01713250
cg01713311
cg01713348
cg01714342
cg01714383
cg01714553
cg01714749
cg01714756

cg01714811
cg01715818
cg01715910
cg01716065
cg01716474
cg01716499
cg01716680
cg01716683
cg01716959
cg01716975
cg01717609
cg01717727
cg01718116
cg01718646
cg01718739
cg01718742
cg01719079
cg01719210
cg01719405
cg01719987
cg01720455
cg01720520
cg01721822
cg01721835
cg01722304
cg01722868
cg01723252
cg01723892
cg01724456
cg01726767
cg01726982
cg01727145
cg01727585
cg01727625
cg01727686
cg01728830
cg01729491
cg01729523
cg01729717
cg01729827
cg01730859
cg01731500
cg01731783
cg01732465
cg01733570
cg01733928
cg01734112
cg01734628
cg01735080
cg01735384
cg01736278
cg01736784

cg01737026
cg01737465
cg01737554
cg01738095
cg01738406
cg01738457
cg01739725
cg01739831
cg01739845
cg01740146
cg01740544
cg01741041
cg01741616
cg01741688
cg01741836
cg01742357
cg01743841
cg01743846
cg01744056
cg01744607
cg01745766
cg01747591
cg01748263
cg01748757
cg01748970
cg01749249
cg01749491
cg01749651
cg01749904
cg01750200
cg01750654
cg01750714
cg01750895
cg01751245
cg01751337
cg01751584
cg01751917
cg01753198
cg01753544
cg01753788
cg01754525
cg01754756
cg01754967
cg01755100
cg01755342
cg01755505
cg01756103
cg01756381
cg01756536
cg01756799
cg01757116
cg01757312

cg01758314
cg01758512
cg01758548
cg01758942
cg01759110
cg01759112
cg01759155
cg01759562
cg01759887
cg01760100
cg01760983
cg01761595
cg01761729
cg01763690
cg01763692
cg01764020
cg01764046
cg01764252
cg01765111
cg01765174
cg01765406
cg01767230
cg01767862
cg01767885
cg01768446
cg01769468
cg01769553
cg01770019
cg01770571
cg01771065
cg01771247
cg01771871
cg01772854
cg01774019
cg01774475
cg01774894
cg01775921
cg01776172
cg01777575
cg01777643
cg01777812
cg01778864
cg01778939
cg01779323
cg01779740
cg01779934
cg01780490
cg01780493
cg01780685
cg01781850
cg01782711
cg01782798

cg01783070
cg01783650
cg01783816
cg01784128
cg01784928
cg01785490
cg01785906
cg01786466
cg01786704
cg01787084
cg01787445
cg01787460
cg01787494
cg01787574
cg01788191
cg01788773
cg01789501
cg01789576
cg01790002
cg01790269
cg01790498
cg01790523
cg01791232
cg01791371
cg01791421
cg01791798
cg01791874
cg01792403
cg01792532
cg01792616
cg01792749
cg01793178
cg01793320
cg01794245
cg01794426
cg01794676
cg01794802
cg01796038
cg01796166
cg01796223
cg01796225
cg01796228
cg01796493
cg01796929
cg01797257
cg01797371
cg01797381
cg01797527
cg01797590
cg01797899
cg01798266
cg01798595

cg01799507
cg01799671
cg01800253
cg01800297
cg01800445
cg01800521
cg01800884
cg01802043
cg01802258
cg01802295
cg01802453
cg01802532
cg01802593
cg01802713
cg01802898
cg01803844
cg01804233
cg01804281
cg01805427
cg01805540
cg01806151
cg01806170
cg01806181
cg01806192
cg01806520
cg01806741
cg01806857
cg01806921
cg01806956
cg01807131
cg01807825
cg01807946
cg01808508
cg01808723
cg01808739
cg01808849
cg01808968
cg01809540
cg01809797
cg01809895
cg01810329
cg01810719
cg01810763
cg01810926
cg01811153
cg01811325
cg01811583
cg01811674
cg01812571
cg01813327
cg01814027
cg01814191

cg01815167
cg01815529
cg01815536
cg01815538
cg01815626
cg01815671
cg01816783
cg01816936
cg01817029
cg01817067
cg01817286
cg01817897
cg01817965
cg01818121
cg01818151
cg01818715
cg01818776
cg01819281
cg01819512
cg01819552
cg01819568
cg01821029
cg01821684
cg01821850
cg01822130
cg01823428
cg01823497
cg01823718
cg01824410
cg01824511
cg01824603
cg01824618
cg01824931
cg01825818
cg01826117
cg01826337
cg01827195
cg01827426
cg01827633
cg01827999
cg01828061
cg01828548
cg01828695
cg01829024
cg01829632
cg01829672
cg01829829
cg01830002
cg01830053
cg01830256
cg01830294
cg01830883

cg01832005
cg01832294
cg01832325
cg01833222
cg01833890
cg01834022
cg01835021
cg01835326
cg01835474
cg01835580
cg01835620
cg01836044
cg01836438
cg01836683
cg01836687
cg01837218
cg01837375
cg01838497
cg01838965
cg01839464
cg01839565
cg01839898
cg01840724
cg01840740
cg01841050
cg01841306
cg01841415
cg01841471
cg01841651
cg01841758
cg01842807
cg01843264
cg01843272
cg01843946
cg01844514
cg01845111
cg01847441
cg01848372
cg01849085
cg01849903
cg01850202
cg01850269
cg01851378
cg01851399
cg01851450
cg01852186
cg01852522
cg01853216
cg01853665
cg01853828
cg01854491
cg01855010

cg01855636
cg01856509
cg01857748
cg01857829
cg01858157
cg01858866
cg01858895
cg01859237
cg01859340
cg01859549
cg01860328
cg01860378
cg01860459
cg01862180
cg01862363
cg01862641
cg01862897
cg01863596
cg01863964
cg01864825
cg01865312
cg01866606
cg01866770
cg01866955
cg01867045
cg01867395
cg01868106
cg01868128
cg01868869
cg01869093
cg01869193
cg01869233
cg01869360
cg01869955
cg01870113
cg01870679
cg01870826
cg01871214
cg01871722
cg01871963
cg01871995
cg01872122
cg01872216
cg01872593
cg01872947
cg01874183
cg01874334
cg01874697
cg01874877
cg01875467
cg01876531
cg01877500

cg01878345
cg01878366
cg01878406
cg01878761
cg01878960
cg01879591
cg01879723
cg01880096
cg01880176
cg01880463
cg01881308
cg01881428
cg01881474
cg01881524
cg01881590
cg01881828
cg01882058
cg01882113
cg01882260
cg01882541
cg01882767
cg01882866
cg01882870
cg01882871
cg01882930
cg01883155
cg01883843
cg01884989
cg01885635
cg01885692
cg01886385
cg01886524
cg01886855
cg01887388
cg01888055
cg01889083
cg01889448
cg01889899
cg01889956
cg01890568
cg01890712
cg01891782
cg01891818
cg01892516
cg01893068
cg01893322
cg01893560
cg01893731
cg01894192
cg01894667
cg01895124
cg01895374

cg01896085
cg01896130
cg01896517
cg01896761
cg01897424
cg01897944
cg01898433
cg01898553
cg01898628
cg01898661
cg01898698
cg01899600
cg01899911
cg01900413
cg01900759
cg01900800
cg01901262
cg01901434
cg01901466
cg01903654
cg01904063
cg01904183
cg01904296
cg01905478
cg01905967
cg01906015
cg01906741
cg01907005
cg01907071
cg01907194
cg01908020
cg01908462
cg01910236
cg01910330
cg01910527
cg01910638
cg01910727
cg01912040
cg01912119
cg01913568
cg01913970
cg01914621
cg01915609
cg01915791
cg01916115
cg01916333
cg01916743
cg01916918
cg01917266
cg01917716
cg01917852
cg01918222

cg01919567
cg01919632
cg01919701
cg01919768
cg01919999
cg01920011
cg01920398
cg01920524
cg01920563
cg01920657
cg01921066
cg01921126
cg01921432
cg01922437
cg01923020
cg01923037
cg01923089
cg01923218
cg01923312
cg01923514
cg01923775
cg01923894
cg01924561
cg01925738
cg01926238
cg01926877
cg01927745
cg01928206
cg01928411
cg01930204
cg01930676
cg01930719
cg01932308
cg01932361
cg01933073
cg01933318
cg01933836
cg01934626
cg01934781
cg01935149
cg01935221
cg01935400
cg01935976
cg01936287
cg01936370
cg01937055
cg01937068
cg01937793
cg01937819
cg01938009
cg01938018
cg01938514

cg01938650
cg01939453
cg01939562
cg01939741
cg01939825
cg01939883
cg01940055
cg01940425
cg01940437
cg01940713
cg01941309
cg01941619
cg01941755
cg01942023
cg01942098
cg01942654
cg01942776
cg01942962
cg01943155
cg01943632
cg01944137
cg01944655
cg01946042
cg01946364
cg01947943
cg01948094
cg01948226
cg01948275
cg01948724
cg01948978
cg01949993
cg01950077
cg01951477
cg01951879
cg01952027
cg01952234
cg01952807
cg01953399
cg01953658
cg01953797
cg01953819
cg01954320
cg01954404
cg01956386
cg01956420
cg01956472
cg01956614
cg01957582
cg01958295
cg01958928
cg01959050
cg01959562

cg01959730
cg01960016
cg01960128
cg01960300
cg01961547
cg01962062
cg01962428
cg01962750
cg01962937
cg01963134
cg01963189
cg01963294
cg01963374
cg01963618
cg01963696
cg01963933
cg01964562
cg01964856
cg01964925
cg01965380
cg01965476
cg01965552
cg01965673
cg01966105
cg01966425
cg01967416
cg01967511
cg01967564
cg01967586
cg01967642
cg01968457
cg01968525
cg01969586
cg01969701
cg01970322
cg01970325
cg01970383
cg01971813
cg01972009
cg01972071
cg01972288
cg01972335
cg01972418
cg01972688
cg01972751
cg01972919
cg01973676
cg01973695
cg01973778
cg01973920
cg01974027
cg01974715

cg01974921
cg01976641
cg01976913
cg01976921
cg01978237
cg01978534
cg01978558
cg01978743
cg01979266
cg01979270
cg01979489
cg01980222
cg01980793
cg01980854
cg01981048
cg01982835
cg01983373
cg01983393
cg01984798
cg01984802
cg01985833
cg01986263
cg01986665
cg01987202
cg01987333
cg01988129
cg01988325
cg01989621
cg01990328
cg01990593
cg01990910
cg01991752
cg01992280
cg01993111
cg01993124
cg01993468
cg01993847
cg01994216
cg01995743
cg01995815
cg01996004
cg01996343
cg01997272
cg01997606
cg01997759
cg01998856
cg01999046
cg01999333
cg02000328
cg02000808
cg02000882
cg02001956

cg02002125
cg02002194
cg02002258
cg02002551
cg02002583
cg02003117
cg02003159
cg02003272
cg02003460
cg02003612
cg02003682
cg02004418
cg02004632
cg02004641
cg02004845
cg02005147
cg02005524
cg02005755
cg02005771
cg02005873
cg02006107
cg02006134
cg02006257
cg02006405
cg02006426
cg02007217
cg02007288
cg02008154
cg02008402
cg02008627
cg02008828
cg02009358
cg02009395
cg02010152
cg02010196
cg02010852
cg02010963
cg02011124
cg02011392
cg02011607
cg02011953
cg02012594
cg02012703
cg02012769
cg02013148
cg02013838
cg02013846
cg02014003
cg02014334
cg02014416
cg02014475
cg02014733

cg02014899
cg02015582
cg02018326
cg02019333
cg02019487
cg02019528
cg02020675
cg02020969
cg02021529
cg02021789
cg02022526
cg02023345
cg02023548
cg02023728
cg02024579
cg02026062
cg02026611
cg02026637
cg02027717
cg02027945
cg02028216
cg02028402
cg02028758
cg02029564
cg02029843
cg02029926
cg02030275
cg02030452
cg02031187
cg02031308
cg02032282
cg02032299
cg02032334
cg02032778
cg02033213
cg02033666
cg02034222
cg02034822
cg02036261
cg02036494
cg02037425
cg02037794
cg02038168
cg02038492
cg02039058
cg02039267
cg02039404
cg02039465
cg02039839
cg02041484
cg02041885
cg02042712

cg02042736
cg02043159
cg02044223
cg02044320
cg02044725
cg02044754
cg02045294
cg02045981
cg02046017
cg02046423
cg02046668
cg02046941
cg02047051
cg02047286
cg02047300
cg02047661
cg02047803
cg02047977
cg02048613
cg02049217
cg02049243
cg02049629
cg02049682
cg02049955
cg02050028
cg02050202
cg02050376
cg02051616
cg02051771
cg02051806
cg02051924
cg02051975
cg02051982
cg02052366
cg02052536
cg02052905
cg02052915
cg02052956
cg02053477
cg02054252
cg02054431
cg02054542
cg02054792
cg02055132
cg02055607
cg02055963
cg02056245
cg02056653
cg02056682
cg02056794
cg02056986
cg02057391

cg02057501
cg02057716
cg02057782
cg02057796
cg02057917
cg02058870
cg02059176
cg02059237
cg02059317
cg02060465
cg02060584
cg02060639
cg02060682
cg02060717
cg02061173
cg02061660
cg02062140
cg02062480
cg02063759
cg02063833
cg02064106
cg02064237
cg02064275
cg02064402
cg02065182
cg02065707
cg02065790
cg02066343
cg02066569
cg02066682
cg02067022
cg02067239
cg02067703
cg02067712
cg02067946
cg02068166
cg02068171
cg02068361
cg02068388
cg02068923
cg02069243
cg02069619
cg02069715
cg02069721
cg02069772
cg02070740
cg02071005
cg02071319
cg02071447
cg02071600
cg02072400
cg02072875

cg02072975
cg02073224
cg02073680
cg02074116
cg02075221
cg02075324
cg02075791
cg02075820
cg02076355
cg02077068
cg02077256
cg02077702
cg02077766
cg02078164
cg02078690
cg02078943
cg02079348
cg02079551
cg02079700
cg02080093
cg02081006
cg02081052
cg02081266
cg02081277
cg02081722
cg02081733
cg02081841
cg02082083
cg02082571
cg02083214
cg02083559
cg02084467
cg02084485
cg02084979
cg02085942
cg02086195
cg02086431
cg02086467
cg02086690
cg02087332
cg02088673
cg02088996
cg02089348
cg02089629
cg02090326
cg02091100
cg02092181
cg02092790
cg02093112
cg02093732
cg02093902
cg02094363

cg02095401
cg02095504
cg02095596
cg02095895
cg02096172
cg02096793
cg02097120
cg02097235
cg02097254
cg02098029
cg02098313
cg02098380
cg02098413
cg02099390
cg02099543
cg02100381
cg02100497
cg02100819
cg02100848
cg02101338
cg02101571
cg02102807
cg02102872
cg02102889
cg02103401
cg02103647
cg02103822
cg02103843
cg02103875
cg02104392
cg02104648
cg02104851
cg02104952
cg02105152
cg02105481
cg02105519
cg02105856
cg02106215
cg02106850
cg02106941
cg02107844
cg02108033
cg02108080
cg02108135
cg02108198
cg02109484
cg02109699
cg02109704
cg02110527
cg02110836
cg02111032
cg02111741

cg02111748
cg02111865
cg02113214
cg02113284
cg02113604
cg02113641
cg02114903
cg02114954
cg02115218
cg02115416
cg02115599
cg02115781
cg02115911
cg02115960
cg02116864
cg02116996
cg02117102
cg02117364
cg02117416
cg02117721
cg02119348
cg02120658
cg02121427
cg02121529
cg02121892
cg02122920
cg02123767
cg02124149
cg02124184
cg02124383
cg02124565
cg02124699
cg02124758
cg02124912
cg02125316
cg02126031
cg02126788
cg02127375
cg02127509
cg02127689
cg02127888
cg02127965
cg02128244
cg02128483
cg02128567
cg02129814
cg02130905
cg02131862
cg02132284
cg02132560
cg02132667
cg02132909

cg02132946
cg02133849
cg02134022
cg02134046
cg02134606
cg02134705
cg02135861
cg02136961
cg02137108
cg02137183
cg02137215
cg02137984
cg02138218
cg02139050
cg02139586
cg02139827
cg02139960
cg02140579
cg02141748
cg02142483
cg02142926
cg02143487
cg02144516
cg02145543
cg02145866
cg02145932
cg02146001
cg02146340
cg02146412
cg02146929
cg02147126
cg02147194
cg02147231
cg02147292
cg02147443
cg02149446
cg02149773
cg02149899
cg02149965
cg02150262
cg02150344
cg02150354
cg02150582
cg02150820
cg02151632
cg02151983
cg02152351
cg02152985
cg02153339
cg02153855
cg02154084
cg02154186

cg02154345
cg02154456
cg02155184
cg02155398
cg02155891
cg02156570
cg02156692
cg02156870
cg02156952
cg02157454
cg02158392
cg02161115
cg02161292
cg02162324
cg02162880
cg02162886
cg02164046
cg02164088
cg02164225
cg02164442
cg02164741
cg02165099
cg02165355
cg02166450
cg02166842
cg02167399
cg02167438
cg02167986
cg02168186
cg02168291
cg02168303
cg02168857
cg02169338
cg02169542
cg02169713
cg02169848
cg02170400
cg02171725
cg02172150
cg02172312
cg02172652
cg02172773
cg02173455
cg02173583
cg02174092
cg02174450
cg02174748
cg02175308
cg02176069
cg02176858
cg02177231
cg02178846

cg02178957
cg02179492
cg02179652
cg02179700
cg02180907
cg02181920
cg02182272
cg02182592
cg02183491
cg02185007
cg02185248
cg02185464
cg02185750
cg02186027
cg02186141
cg02186748
cg02186782
cg02187214
cg02187231
cg02188225
cg02188939
cg02189738
cg02189843
cg02189888
cg02190618
cg02190928
cg02191985
cg02192218
cg02192673
cg02192677
cg02192965
cg02192967
cg02193425
cg02193513
cg02194046
cg02194908
cg02195405
cg02196592
cg02196805
cg02197228
cg02197938
cg02198144
cg02198625
cg02198895
cg02198988
cg02199826
cg02200207
cg02201052
cg02201239
cg02202664
cg02202933
cg02203380

cg02203656
cg02203665
cg02203760
cg02204261
cg02204521
cg02205072
cg02205445
cg02206108
cg02206137
cg02206323
cg02207155
cg02207219
cg02208125
cg02208504
cg02208657
cg02209667
cg02210314
cg02210498
cg02210887
cg02211735
cg02211741
cg02212084
cg02212491
cg02212846
cg02214441
cg02215641
cg02215899
cg02216481
cg02216951
cg02217713
cg02217786
cg02217842
cg02218324
cg02219476
cg02220008
cg02220125
cg02220233
cg02220481
cg02220837
cg02221053
cg02221302
cg02221422
cg02221471
cg02222170
cg02223801
cg02223905
cg02224138
cg02224164
cg02224369
cg02224372
cg02224795
cg02224994

cg02225137
cg02225716
cg02226239
cg02226645
cg02227496
cg02227623
cg02228111
cg02228294
cg02228492
cg02229095
cg02229288
cg02229433
cg02229946
cg02230017
cg02230103
cg02230964
cg02232208
cg02232641
cg02232704
cg02232751
cg02233459
cg02233518
cg02233750
cg02234120
cg02234235
cg02234281
cg02234613
cg02234653
cg02234847
cg02235049
cg02235249
cg02235562
cg02235848
cg02236673
cg02236847
cg02237470
cg02237987
cg02238550
cg02240030
cg02240252
cg02240291
cg02240539
cg02240748
cg02241145
cg02241330
cg02241759
cg02241986
cg02243061
cg02243276
cg02243303
cg02243386
cg02243665

cg02243785
cg02244274
cg02244386
cg02244933
cg02245004
cg02245566
cg02246002
cg02246190
cg02246426
cg02246609
cg02246640
cg02246800
cg02246992
cg02248514
cg02248749
cg02248763
cg02249292
cg02250594
cg02251243
cg02251393
cg02251567
cg02251859
cg02252421
cg02253236
cg02253499
cg02253760
cg02255004
cg02255054
cg02255098
cg02255510
cg02255950
cg02256576
cg02258381
cg02258614
cg02258703
cg02259324
cg02259384
cg02259429
cg02259550
cg02259760
cg02261047
cg02261294
cg02261543
cg02263260
cg02263813
cg02263932
cg02264082
cg02265105
cg02265758
cg02265922
cg02266177
cg02266732

cg02266878
cg02267215
cg02267488
cg02268013
cg02268202
cg02268229
cg02268561
cg02269571
cg02271139
cg02271603
cg02271843
cg02271895
cg02272429
cg02272795
cg02272831
cg02272968
cg02273106
cg02273617
cg02273827
cg02274361
cg02274366
cg02274550
cg02275396
cg02275418
cg02276600
cg02276974
cg02278768
cg02278924
cg02279147
cg02279625
cg02279883
cg02281208
cg02281493
cg02281854
cg02281970
cg02282247
cg02283042
cg02283204
cg02283366
cg02283735
cg02284800
cg02285791
cg02285887
cg02286091
cg02286506
cg02286819
cg02287004
cg02287007
cg02287693
cg02287710
cg02287817
cg02288165

cg02288684
cg02288831
cg02288964
cg02288969
cg02289020
cg02289038
cg02289040
cg02289321
cg02290110
cg02290880
cg02291164
cg02291204
cg02292739
cg02293096
cg02293437
cg02293852
cg02294111
cg02294687
cg02295509
cg02295574
cg02296890
cg02297541
cg02297967
cg02298327
cg02298472
cg02298525
cg02299328
cg02299465
cg02299971
cg02300083
cg02300154
cg02300379
cg02301079
cg02301292
cg02301741
cg02302089
cg02303016
cg02303258
cg02303701
cg02303807
cg02304033
cg02304095
cg02304156
cg02304584
cg02304614
cg02304628
cg02305064
cg02305135
cg02305242
cg02305765
cg02305853
cg02306139

cg02306156
cg02306630
cg02306936
cg02307239
cg02307288
cg02308150
cg02308226
cg02308706
cg02308914
cg02308988
cg02311364
cg02311414
cg02312409
cg02312593
cg02312715
cg02313118
cg02313130
cg02313492
cg02313836
cg02313986
cg02315111
cg02315187
cg02315271
cg02315315
cg02316445
cg02317400
cg02317955
cg02318282
cg02318567
cg02318784
cg02318926
cg02319392
cg02320356
cg02320474
cg02320481
cg02320721
cg02320740
cg02322205
cg02322365
cg02322400
cg02322879
cg02323003
cg02323699
cg02323958
cg02324221
cg02324227
cg02324367
cg02324789
cg02324920
cg02325250
cg02325324
cg02325411

cg02326058
cg02326806
cg02327530
cg02327719
cg02327997
cg02328221
cg02328332
cg02328545
cg02328578
cg02328758
cg02329029
cg02329038
cg02329527
cg02330818
cg02331189
cg02331198
cg02331769
cg02331772
cg02331808
cg02332123
cg02332492
cg02332525
cg02332537
cg02332982
cg02334109
cg02334314
cg02334775
cg02335185
cg02335376
cg02335657
cg02335700
cg02336334
cg02336697
cg02337349
cg02337355
cg02338142
cg02338965
cg02339122
cg02339243
cg02339850
cg02340056
cg02340083
cg02340329
cg02340818
cg02341759
cg02342148
cg02343182
cg02343905
cg02344361
cg02344477
cg02344833
cg02345089

cg02345321
cg02345399
cg02346062
cg02346135
cg02346141
cg02346706
cg02346901
cg02347074
cg02347487
cg02347573
cg02347881
cg02347984
cg02348430
cg02348751
cg02349468
cg02349866
cg02350677
cg02351674
cg02352240
cg02352281
cg02352328
cg02352370
cg02352716
cg02352722
cg02353034
cg02353165
cg02354563
cg02355062
cg02355420
cg02355885
cg02356566
cg02358362
cg02358456
cg02358653
cg02358664
cg02360175
cg02360432
cg02360676
cg02360939
cg02361458
cg02361702
cg02362103
cg02362292
cg02362859
cg02363526
cg02363711
cg02363950
cg02363960
cg02364703
cg02364826
cg02364831
cg02365086

cg02365303
cg02365517
cg02366772
cg02367133
cg02367144
cg02367160
cg02367660
cg02368058
cg02368591
cg02368606
cg02369052
cg02369725
cg02369921
cg02370605
cg02370644
cg02370770
cg02371121
cg02371397
cg02371777
cg02372215
cg02373484
cg02374745
cg02374934
cg02375065
cg02375208
cg02376426
cg02377024
cg02377915
cg02378440
cg02378784
cg02378847
cg02379079
cg02379107
cg02379856
cg02380531
cg02380840
cg02380983
cg02381279
cg02381426
cg02381539
cg02382037
cg02382109
cg02382706
cg02383130
cg02383399
cg02385153
cg02385411
cg02385474
cg02385728
cg02385848
cg02385908
cg02386219

cg02386403
cg02386604
cg02387326
cg02387510
cg02387803
cg02387843
cg02388150
cg02388468
cg02388795
cg02388849
cg02388865
cg02389040
cg02389180
cg02389317
cg02389949
cg02390319
cg02390329
cg02390447
cg02390624
cg02390947
cg02391713
cg02391906
cg02392634
cg02393097
cg02393428
cg02393479
cg02393591
cg02393907
cg02394305
cg02394812
cg02394955
cg02395779
cg02396005
cg02397205
cg02397368
cg02397497
cg02397727
cg02397768
cg02398789
cg02399008
cg02399098
cg02399294
cg02399524
cg02399788
cg02399826
cg02400308
cg02400433
cg02400565
cg02401149
cg02401454
cg02401589
cg02402423

cg02402458
cg02402744
cg02402882
cg02402946
cg02403395
cg02403412
cg02404507
cg02404974
cg02405476
cg02406096
cg02406469
cg02407545
cg02408532
cg02409125
cg02409351
cg02409815
cg02411275
cg02412169
cg02412346
cg02412803
cg02412810
cg02414499
cg02415029
cg02415431
cg02415992
cg02416536
cg02416986
cg02417408
cg02417427
cg02417473
cg02418072
cg02419321
cg02419835
cg02420395
cg02420617
cg02421207
cg02421985
cg02422426
cg02422603
cg02423000
cg02423064
cg02423185
cg02423494
cg02423574
cg02423618
cg02424249
cg02424835
cg02425108
cg02426623
cg02426739
cg02428056
cg02428119

cg02428506
cg02428792
cg02430430
cg02430558
cg02430732
cg02430935
cg02431531
cg02431562
cg02431732
cg02431866
cg02432280
cg02432467
cg02433515
cg02433785
cg02433979
cg02434035
cg02434541
cg02436616
cg02438026
cg02438107
cg02438611
cg02438781
cg02440199
cg02440668
cg02440719
cg02441090
cg02441149
cg02441556
cg02441747
cg02441982
cg02442640
cg02443072
cg02444118
cg02444957
cg02444987
cg02446106
cg02447304
cg02447620
cg02447627
cg02448796
cg02449336
cg02449354
cg02449461
cg02449608
cg02449638
cg02450164
cg02451508
cg02453929
cg02454476
cg02454815
cg02455094
cg02455453

cg02455706
cg02455792
cg02455797
cg02456226
cg02456288
cg02456552
cg02457319
cg02457346
cg02458053
cg02458065
cg02458188
cg02458554
cg02458689
cg02459415
cg02459818
cg02460251
cg02461767
cg02463418
cg02464665
cg02465859
cg02466572
cg02466815
cg02467451
cg02467990
cg02468050
cg02468649
cg02469095
cg02469871
cg02469909
cg02470008
cg02471897
cg02472646
cg02473237
cg02473460
cg02473540
cg02473594
cg02474116
cg02474563
cg02474567
cg02477218
cg02477557
cg02478149
cg02478589
cg02478708
cg02479177
cg02479468
cg02480077
cg02480320
cg02480489
cg02480675
cg02480685
cg02481064

cg02481228
cg02481498
cg02481697
cg02481848
cg02482218
cg02482438
cg02482624
cg02484335
cg02484469
cg02484634
cg02484673
cg02484687
cg02485642
cg02485697
cg02485729
cg02485922
cg02486096
cg02486325
cg02486332
cg02486904
cg02487130
cg02487452
cg02487453
cg02487987
cg02488299
cg02488934
cg02488942
cg02489189
cg02489219
cg02489228
cg02489377
cg02489478
cg02489769
cg02489968
cg02490182
cg02490185
cg02490934
cg02490989
cg02491628
cg02492207
cg02492567
cg02492708
cg02493217
cg02493602
cg02493771
cg02493846
cg02493905
cg02493986
cg02494512
cg02494593
cg02494794
cg02496234

cg02496728
cg02497046
cg02497758
cg02498177
cg02498702
cg02499308
cg02499535
cg02499614
cg02499925
cg02500392
cg02500691
cg02500990
cg02501746
cg02501779
cg02502025
cg02503159
cg02503850
cg02503981
cg02504416
cg02505749
cg02506324
cg02506360
cg02507889
cg02508651
cg02508830
cg02509730
cg02509825
cg02510802
cg02511156
cg02511231
cg02511723
cg02512108
cg02512226
cg02512352
cg02512860
cg02514022
cg02514611
cg02515217
cg02516101
cg02516288
cg02516330
cg02516845
cg02517040
cg02517337
cg02517342
cg02517632
cg02517686
cg02517711
cg02518161
cg02518824
cg02519208
cg02519499

cg02519718
cg02519751
cg02520804
cg02521457
cg02522196
cg02522990
cg02523270
cg02523640
cg02523844
cg02524236
cg02524902
cg02524946
cg02524983
cg02525597
cg02525826
cg02526508
cg02527056
cg02527751
cg02528008
cg02528159
cg02529095
cg02529358
cg02530022
cg02530407
cg02530656
cg02530834
cg02531067
cg02531439
cg02531768
cg02531924
cg02532096
cg02532996
cg02533058
cg02533173
cg02533239
cg02533364
cg02533423
cg02534072
cg02534163
cg02534164
cg02537491
cg02537638
cg02538169
cg02538248
cg02539539
cg02539662
cg02539664
cg02540307
cg02540477
cg02540858
cg02541658
cg02541778

cg02542401
cg02542748
cg02543772
cg02545583
cg02546376
cg02546833
cg02547054
cg02547263
cg02547269
cg02547394
cg02547724
cg02548228
cg02549628
cg02550119
cg02550231
cg02550257
cg02550268
cg02550607
cg02550722
cg02550741
cg02551589
cg02551646
cg02551743
cg02553039
cg02553862
cg02555152
cg02555727
cg02555788
cg02555944
cg02556005
cg02556071
cg02556363
cg02556393
cg02556649
cg02556928
cg02557139
cg02557922
cg02558111
cg02558120
cg02558132
cg02558288
cg02558364
cg02558570
cg02558677
cg02558684
cg02558735
cg02560364
cg02561009
cg02561482
cg02561693
cg02562431
cg02562544

cg02563364
cg02563916
cg02564291
cg02564302
cg02564313
cg02564829
cg02565062
cg02565878
cg02565993
cg02565999
cg02566789
cg02566952
cg02567119
cg02567879
cg02568479
cg02568902
cg02569142
cg02569554
cg02570298
cg02570981
cg02571042
cg02571055
cg02571534
cg02571560
cg02571636
cg02571816
cg02572463
cg02573234
cg02573357
cg02573551
cg02574254
cg02574502
cg02574734
cg02574952
cg02575483
cg02575561
cg02575764
cg02576153
cg02576415
cg02576468
cg02577745
cg02578087
cg02578709
cg02579133
cg02579377
cg02579389
cg02580763
cg02580969
cg02581163
cg02581587
cg02581828
cg02582057

cg02582218
cg02582355
cg02582387
cg02582587
cg02582848
cg02582957
cg02583228
cg02583247
cg02583589
cg02583796
cg02584756
cg02584867
cg02584939
cg02585622
cg02585702
cg02585931
cg02586025
cg02586268
cg02587988
cg02589353
cg02589604
cg02589901
cg02591442
cg02592062
cg02592239
cg02593227
cg02593601
cg02593932
cg02594183
cg02595219
cg02595760
cg02595832
cg02595851
cg02596508
cg02596645
cg02596960
cg02596994
cg02597923
cg02598335
cg02598564
cg02599006
cg02599385
cg02599587
cg02600411
cg02600625
cg02601140
cg02601249
cg02601893
cg02602413
cg02603128
cg02603756
cg02604018

cg02604524
cg02605760
cg02606369
cg02606840
cg02607772
cg02608124
cg02608452
cg02608831
cg02609127
cg02609337
cg02610070
cg02610106
cg02610360
cg02610600
cg02611811
cg02611812
cg02611848
cg02612397
cg02612971
cg02613108
cg02613295
cg02613300
cg02613818
cg02614024
cg02614660
cg02615214
cg02615468
cg02616368
cg02616565
cg02616769
cg02617418
cg02619001
cg02619205
cg02619941
cg02620533
cg02621130
cg02621154
cg02621528
cg02621905
cg02621979
cg02622133
cg02623050
cg02623400
cg02624067
cg02624836
cg02625481
cg02625749
cg02626012
cg02626129
cg02626991
cg02627875
cg02628360

cg02628483
cg02628704
cg02628840
cg02628907
cg02629106
cg02629586
cg02629976
cg02630214
cg02630604
cg02630628
cg02630677
cg02630694
cg02630914
cg02630999
cg02631189
cg02631462
cg02631468
cg02631838
cg02632200
cg02632261
cg02632490
cg02632800
cg02633723
cg02634222
cg02634733
cg02635932
cg02636177
cg02636947
cg02637318
cg02637512
cg02637654
cg02638136
cg02638523
cg02638567
cg02638691
cg02640147
cg02640260
cg02640497
cg02640604
cg02641517
cg02641539
cg02641676
cg02641770
cg02641941
cg02642585
cg02642962
cg02642979
cg02643054
cg02643668
cg02644195
cg02644421
cg02644510

cg02644867
cg02645295
cg02646064
cg02646225
cg02646491
cg02646528
cg02646706
cg02646787
cg02647493
cg02647520
cg02647628
cg02647878
cg02647929
cg02647941
cg02649597
cg02649613
cg02650080
cg02650127
cg02650128
cg02650767
cg02650779
cg02651357
cg02651871
cg02652072
cg02652361
cg02652579
cg02652760
cg02653559
cg02655630
cg02655972
cg02656089
cg02656594
cg02656891
cg02657401
cg02657441
cg02657632
cg02657654
cg02657836
cg02658319
cg02658551
cg02658668
cg02659217
cg02659568
cg02659581
cg02660089
cg02660165
cg02660277
cg02660643
cg02661034
cg02661431
cg02661764
cg02662277

cg02662362
cg02662828
cg02663352
cg02663496
cg02664349
cg02664812
cg02666287
cg02666556
cg02666820
cg02667102
cg02667103
cg02667467
cg02668822
cg02668843
cg02668999
cg02669055
cg02669544
cg02671880
cg02672332
cg02673323
cg02674324
cg02674384
cg02674639
cg02674693
cg02674870
cg02675344
cg02675353
cg02675969
cg02676375
cg02677609
cg02677946
cg02678305
cg02678414
cg02678496
cg02678768
cg02679012
cg02679028
cg02679336
cg02679503
cg02679809
cg02680398
cg02680903
cg02680909
cg02681173
cg02681442
cg02683197
cg02683307
cg02683577
cg02683759
cg02683929
cg02684548
cg02684964

cg02685016
cg02685032
cg02685286
cg02685484
cg02687026
cg02687786
cg02687883
cg02688359
cg02689863
cg02690571
cg02690638
cg02691035
cg02691360
cg02691745
cg02691934
cg02692893
cg02693002
cg02693486
cg02693910
cg02694017
cg02694099
cg02694427
cg02694489
cg02694810
cg02696201
cg02696327
cg02696428
cg02696561
cg02696798
cg02696910
cg02697115
cg02697402
cg02697527
cg02697979
cg02699518
cg02700130
cg02700491
cg02700891
cg02700894
cg02701615
cg02702692
cg02703563
cg02703728
cg02704158
cg02704949
cg02705909
cg02705958
cg02706919
cg02707114
cg02707277
cg02707552
cg02707799

cg02707840
cg02708506
cg02709321
cg02709725
cg02709924
cg02710168
cg02710296
cg02711510
cg02711647
cg02711929
cg02712303
cg02712545
cg02713180
cg02713720
cg02713951
cg02714303
cg02714666
cg02715304
cg02715629
cg02716614
cg02716826
cg02716952
cg02717117
cg02717150
cg02717347
cg02717506
cg02718078
cg02718516
cg02718966
cg02721602
cg02721665
cg02721693
cg02721751
cg02721902
cg02722808
cg02723856
cg02724461
cg02724647
cg02724909
cg02725001
cg02725370
cg02725885
cg02725908
cg02726819
cg02726930
cg02727104
cg02727811
cg02728172
cg02730990
cg02731366
cg02732067
cg02732164

cg02732508
cg02734358
cg02734408
cg02734505
cg02735733
cg02735736
cg02735829
cg02735962
cg02736313
cg02736420
cg02736966
cg02737219
cg02737268
cg02737335
cg02737486
cg02737507
cg02738086
cg02738298
cg02739346
cg02739850
cg02740093
cg02740606
cg02741056
cg02741554
cg02741698
cg02742551
cg02742906
cg02743222
cg02743674
cg02744113
cg02744216
cg02744252
cg02744524
cg02744588
cg02744705
cg02744945
cg02745045
cg02745211
cg02745398
cg02745459
cg02745847
cg02746232
cg02746381
cg02746514
cg02746658
cg02746744
cg02746748
cg02746771
cg02746781
cg02746852
cg02747210
cg02747254

cg02748293
cg02748624
cg02749463
cg02749511
cg02749735
cg02749752
cg02750743
cg02751064
cg02751731
cg02752105
cg02752932
cg02753027
cg02753719
cg02754084
cg02754943
cg02755847
cg02756107
cg02756168
cg02756502
cg02756683
cg02756698
cg02757516
cg02757609
cg02757667
cg02758499
cg02758593
cg02758612
cg02759193
cg02759725
cg02760031
cg02760152
cg02760218
cg02760280
cg02760495
cg02761110
cg02761733
cg02763234
cg02763671
cg02763800
cg02763807
cg02763822
cg02764245
cg02764250
cg02764438
cg02764571
cg02764897
cg02765031
cg02765403
cg02765564
cg02765731
cg02765820
cg02765859

cg02765962
cg02766207
cg02766259
cg02766482
cg02766895
cg02766941
cg02767024
cg02767068
cg02767382
cg02767665
cg02767771
cg02768671
cg02769522
cg02770688
cg02770843
cg02770857
cg02770985
cg02771230
cg02771336
cg02771673
cg02773086
cg02773433
cg02773671
cg02774129
cg02774186
cg02774282
cg02774302
cg02774439
cg02774705
cg02775223
cg02775414
cg02775417
cg02776283
cg02776313
cg02777065
cg02777934
cg02778243
cg02779016
cg02779037
cg02779724
cg02779913
cg02779944
cg02780400
cg02780988
cg02783179
cg02783918
cg02784874
cg02785332
cg02786019
cg02786378
cg02786417
cg02787387

cg02788102
cg02788195
cg02788264
cg02789485
cg02789965
cg02790260
cg02792136
cg02792322
cg02792821
cg02793099
cg02793948
cg02794696
cg02794920
cg02795014
cg02796568
cg02797569
cg02797933
cg02798157
cg02798394
cg02798576
cg02798703
cg02798801
cg02799237
cg02799411
cg02799905
cg02800607
cg02800810
cg02801359
cg02801786
cg02801789
cg02802420
cg02802514
cg02802834
cg02803037
cg02803141
cg02803847
cg02804722
cg02805025
cg02805810
cg02805920
cg02806322
cg02806733
cg02807446
cg02807482
cg02807859
cg02807882
cg02809028
cg02809075
cg02809611
cg02809746
cg02810692
cg02810967

cg02811232
cg02811473
cg02812207
cg02812767
cg02813121
cg02813916
cg02814118
cg02814416
cg02814805
cg02814858
cg02815025
cg02815102
cg02815320
cg02815374
cg02816429
cg02816816
cg02818409
cg02818572
cg02819921
cg02820040
cg02821464
cg02822087
cg02823066
cg02823132
cg02824099
cg02824202
cg02824386
cg02824443
cg02824604
cg02824980
cg02824983
cg02825171
cg02825373
cg02825412
cg02825728
cg02826233
cg02826368
cg02826735
cg02826890
cg02827007
cg02827278
cg02828023
cg02828306
cg02828687
cg02828779
cg02829285
cg02829654
cg02829688
cg02829743
cg02830467
cg02830545
cg02830936

cg02831267
cg02832905
cg02832952
cg02833172
cg02833725
cg02834280
cg02835467
cg02835822
cg02835977
cg02835981
cg02836017
cg02836135
cg02836325
cg02837162
cg02837536
cg02838178
cg02838309
cg02838589
cg02838825
cg02839727
cg02840008
cg02840828
cg02840936
cg02841000
cg02841692
cg02841912
cg02842227
cg02842704
cg02843201
cg02843277
cg02843667
cg02844502
cg02844545
cg02844589
cg02845063
cg02845204
cg02845569
cg02846219
cg02846801
cg02846963
cg02848157
cg02848448
cg02848711
cg02849693
cg02850273
cg02850468
cg02851397
cg02852210
cg02852407
cg02852847
cg02853358
cg02853387

cg02853418
cg02853616
cg02853850
cg02854396
cg02854972
cg02855363
cg02855432
cg02855850
cg02856026
cg02856109
cg02856338
cg02856768
cg02857074
cg02857166
cg02857726
cg02858402
cg02858591
cg02858594
cg02858606
cg02859516
cg02859837
cg02859866
cg02859992
cg02860599
cg02860602
cg02860749
cg02861071
cg02861456
cg02861926
cg02863073
cg02863594
cg02863856
cg02864757
cg02864844
cg02865101
cg02865896
cg02866106
cg02866628
cg02867145
cg02867305
cg02867735
cg02868123
cg02868468
cg02868521
cg02869235
cg02869459
cg02869649
cg02870189
cg02870676
cg02871021
cg02871985
cg02871995

cg02872914
cg02873204
cg02873384
cg02873517
cg02874376
cg02875015
cg02876062
cg02876466
cg02878244
cg02878284
cg02878289
cg02878739
cg02878949
cg02879159
cg02879656
cg02879776
cg02880169
cg02880444
cg02881127
cg02881186
cg02881570
cg02882813
cg02883230
cg02883247
cg02884884
cg02884944
cg02885007
cg02886077
cg02886578
cg02886715
cg02887114
cg02887652
cg02888210
cg02888490
cg02888894
cg02889617
cg02889973
cg02890352
cg02890365
cg02890435
cg02891082
cg02891579
cg02891728
cg02891785
cg02892153
cg02892367
cg02892388
cg02892624
cg02893260
cg02893429
cg02893604
cg02893823

cg02894109
cg02895126
cg02895724
cg02895948
cg02896768
cg02896941
cg02897570
cg02897974
cg02898094
cg02899011
cg02899870
cg02899903
cg02900034
cg02902186
cg02902313
cg02902415
cg02902854
cg02902882
cg02902946
cg02903275
cg02903822
cg02904344
cg02904800
cg02906503
cg02906784
cg02906823
cg02907098
cg02907374
cg02907415
cg02907638
cg02908900
cg02909136
cg02909329
cg02909793
cg02910011
cg02910208
cg02910987
cg02912291
cg02912371
cg02912790
cg02912937
cg02914112
cg02914422
cg02915544
cg02915785
cg02916147
cg02916962
cg02917236
cg02918054
cg02918593
cg02918648
cg02919090

cg02919116
cg02919422
cg02919615
cg02919982
cg02920150
cg02920216
cg02920235
cg02920376
cg02920600
cg02920604
cg02920701
cg02922084
cg02922094
cg02922280
cg02923264
cg02923340
cg02923485
cg02923706
cg02923972
cg02924234
cg02924562
cg02924836
cg02925039
cg02926307
cg02926397
cg02926820
cg02927074
cg02927111
cg02927346
cg02927618
cg02927679
cg02927682
cg02927747
cg02927859
cg02928278
cg02928664
cg02928699
cg02928840
cg02928935
cg02929431
cg02929681
cg02929780
cg02929855
cg02929961
cg02930348
cg02930356
cg02930721
cg02930866
cg02931467
cg02932321
cg02932364
cg02932738

cg02933354
cg02933639
cg02933962
cg02934462
cg02935024
cg02935338
cg02935562
cg02935627
cg02936049
cg02936347
cg02936451
cg02936931
cg02937158
cg02937674
cg02937751
cg02937763
cg02938303
cg02938413
cg02938414
cg02938807
cg02939105
cg02939139
cg02939767
cg02940003
cg02940147
cg02940165
cg02940362
cg02940555
cg02941385
cg02943074
cg02943456
cg02943578
cg02943595
cg02943604
cg02944021
cg02944084
cg02944123
cg02944238
cg02944312
cg02944437
cg02944761
cg02944871
cg02946394
cg02946846
cg02946850
cg02946885
cg02946903
cg02948259
cg02948476
cg02948862
cg02949838
cg02951514

cg02951739
cg02952078
cg02952913
cg02952984
cg02953039
cg02953144
cg02953306
cg02953912
cg02953960
cg02954563
cg02955020
cg02955098
cg02955804
cg02955940
cg02955989
cg02956248
cg02956542
cg02956806
cg02957057
cg02957094
cg02957799
cg02958014
cg02958515
cg02958527
cg02959108
cg02959678
cg02960347
cg02960777
cg02960938
cg02961004
cg02961012
cg02961101
cg02961267
cg02961280
cg02961707
cg02961934
cg02962406
cg02962602
cg02962903
cg02963248
cg02963613
cg02963641
cg02963866
cg02964555
cg02965092
cg02965304
cg02965895
cg02967616
cg02968165
cg02968429
cg02969164
cg02969380

cg02969706
cg02970297
cg02971654
cg02973171
cg02974931
cg02975060
cg02975463
cg02975667
cg02976699
cg02976898
cg02977443
cg02977524
cg02977559
cg02977790
cg02978184
cg02978784
cg02979001
cg02979520
cg02981092
cg02981103
cg02981183
cg02981368
cg02981703
cg02981942
cg02982238
cg02982734
cg02983424
cg02983950
cg02984790
cg02984966
cg02985250
cg02985539
cg02985541
cg02985871
cg02986451
cg02986643
cg02987249
cg02987387
cg02987547
cg02987618
cg02987635
cg02987928
cg02988730
cg02988826
cg02988984
cg02989940
cg02990302
cg02990474
cg02990705
cg02990742
cg02990869
cg02991338

cg02991727
cg02991888
cg02992311
cg02992546
cg02992617
cg02992632
cg02993352
cg02993699
cg02993991
cg02994009
cg02994349
cg02994463
cg02994956
cg02995013
cg02995731
cg02996181
cg02996869
cg02997111
cg02997295
cg02997316
cg02997755
cg02997851
cg02998463
cg02998861
cg02999385
cg02999711
cg03000116
cg03000593
cg03001143
cg03001176
cg03001863
cg03002526
cg03002848
cg03003526
cg03003745
cg03003746
cg03005040
cg03005124
cg03005191
cg03005603
cg03007309
cg03007522
cg03007849
cg03008515
cg03008693
cg03009690
cg03009785
cg03010301
cg03010383
cg03010425
cg03010756
cg03010761

cg03010785
cg03011658
cg03011928
cg03012028
cg03012170
cg03012374
cg03012521
cg03012544
cg03012726
cg03012854
cg03013488
cg03013636
cg03013832
cg03013946
cg03014241
cg03014326
cg03015368
cg03015498
cg03015592
cg03016867
cg03016906
cg03017520
cg03018001
cg03018256
cg03018380
cg03019112
cg03019219
cg03019844
cg03020424
cg03020635
cg03021100
cg03022263
cg03022541
cg03023237
cg03024372
cg03024478
cg03024720
cg03024760
cg03025082
cg03025473
cg03025569
cg03025880
cg03026004
cg03026462
cg03027179
cg03027227
cg03027354
cg03027661
cg03028061
cg03029127
cg03029296
cg03029734

cg03029752
cg03030267
cg03031009
cg03031959
cg03031988
cg03032053
cg03032098
cg03032214
cg03032443
cg03032677
cg03032770
cg03033111
cg03033182
cg03033796
cg03034696
cg03035162
cg03036777
cg03037761
cg03037879
cg03039218
cg03039243
cg03039843
cg03040292
cg03040622
cg03040821
cg03041109
cg03041510
cg03041629
cg03041658
cg03041920
cg03042537
cg03043406
cg03043696
cg03043804
cg03044096
cg03044178
cg03044249
cg03044344
cg03044376
cg03044513
cg03044725
cg03045425
cg03045635
cg03046148
cg03046247
cg03046424
cg03046611
cg03047070
cg03047508
cg03047616
cg03048162
cg03048654

cg03049125
cg03049445
cg03049501
cg03049846
cg03049863
cg03049917
cg03050127
cg03050669
cg03051742
cg03052004
cg03052128
cg03052251
cg03053579
cg03054529
cg03055065
cg03055693
cg03057236
cg03058163
cg03058260
cg03058346
cg03058412
cg03059126
cg03059131
cg03059382
cg03059921
cg03060036
cg03060986
cg03062329
cg03062865
cg03062944
cg03063309
cg03064067
cg03064592
cg03064914
cg03064940
cg03065107
cg03065139
cg03065447
cg03065503
cg03066577
cg03066788
cg03068446
cg03068843
cg03069242
cg03069736
cg03069737
cg03070297
cg03071134
cg03071876
cg03072621
cg03072665
cg03073218

cg03073429
cg03073786
cg03074126
cg03074225
cg03074925
cg03074946
cg03075065
cg03075214
cg03075244
cg03075534
cg03076319
cg03077331
cg03078043
cg03078269
cg03078363
cg03078586
cg03078733
cg03078819
cg03080142
cg03081173
cg03081478
cg03082523
cg03084648
cg03084808
cg03084838
cg03084996
cg03085859
cg03085949
cg03086003
cg03086235
cg03086899
cg03087992
cg03088328
cg03088958
cg03090969
cg03091010
cg03091070
cg03091331
cg03091551
cg03091776
cg03092191
cg03092482
cg03093838
cg03094121
cg03095087
cg03095697
cg03095773
cg03095797
cg03095982
cg03096347
cg03096543
cg03097995

cg03098643
cg03098837
cg03099208
cg03100024
cg03100044
cg03100639
cg03100906
cg03101522
cg03102898
cg03103023
cg03103192
cg03103611
cg03103888
cg03104157
cg03104428
cg03105070
cg03106056
cg03106958
cg03107393
cg03107987
cg03109101
cg03109701
cg03109827
cg03110921
cg03111404
cg03111498
cg03112087
cg03113867
cg03114057
cg03114255
cg03114410
cg03114584
cg03115530
cg03116016
cg03116035
cg03116124
cg03116740
cg03117079
cg03117412
cg03117976
cg03120185
cg03120284
cg03120289
cg03120555
cg03121793
cg03122532
cg03123017
cg03124146
cg03124318
cg03127174
cg03127334
cg03127349

cg03127370
cg03127416
cg03128025
cg03128029
cg03128163
cg03128860
cg03129014
cg03129324
cg03129384
cg03130090
cg03130292
cg03130530
cg03130816
cg03131097
cg03131556
cg03132551
cg03133097
cg03133987
cg03134157
cg03134285
cg03134573
cg03134809
cg03134947
cg03134962
cg03135061
cg03135577
cg03135964
cg03137135
cg03137177
cg03138364
cg03139255
cg03139423
cg03139435
cg03140135
cg03140521
cg03140968
cg03141971
cg03142002
cg03142554
cg03142697
cg03142741
cg03143347
cg03143395
cg03143886
cg03144708
cg03144922
cg03145200
cg03145274
cg03145386
cg03145658
cg03145930
cg03146413

cg03146557
cg03146683
cg03146733
cg03146993
cg03147184
cg03147185
cg03147598
cg03149560
cg03150753
cg03151187
cg03151469
cg03151709
cg03152187
cg03154226
cg03155345
cg03156547
cg03157862
cg03158063
cg03158310
cg03158400
cg03159124
cg03160135
cg03160174
cg03160466
cg03161190
cg03161309
cg03161498
cg03162092
cg03162314
cg03162410
cg03162506
cg03164275
cg03165378
cg03165807
cg03167521
cg03167633
cg03167683
cg03167812
cg03167883
cg03168922
cg03169059
cg03169131
cg03169180
cg03171651
cg03171770
cg03171795
cg03172226
cg03172657
cg03172947
cg03172971
cg03173570
cg03174043

cg03174521
cg03174768
cg03175653
cg03176423
cg03176810
cg03177042
cg03179496
cg03180302
cg03180486
cg03181248
cg03181340
cg03181582
cg03182584
cg03182596
cg03182608
cg03182620
cg03182688
cg03183618
cg03184290
cg03185593
cg03185794
cg03186620
cg03186975
cg03186986
cg03187073
cg03187125
cg03187135
cg03187795
cg03188064
cg03188390
cg03188976
cg03189075
cg03189184
cg03189453
cg03189821
cg03190130
cg03190219
cg03190769
cg03190876
cg03191504
cg03192060
cg03192598
cg03192775
cg03193096
cg03193847
cg03194122
cg03194383
cg03195643
cg03196097
cg03196360
cg03196720
cg03197032

cg03197835
cg03197992
cg03198203
cg03199646
cg03200309
cg03200631
cg03200786
cg03202077
cg03202564
cg03202832
cg03203197
cg03203287
cg03204140
cg03204322
cg03204605
cg03205010
cg03205140
cg03205252
cg03205764
cg03206044
cg03207868
cg03208245
cg03209395
cg03209720
cg03209854
cg03210210
cg03211092
cg03211098
cg03211239
cg03211327
cg03212620
cg03214217
cg03214622
cg03215137
cg03215181
cg03215422
cg03215535
cg03216355
cg03216506
cg03216846
cg03217253
cg03217897
cg03217995
cg03218479
cg03219514
cg03219968
cg03220543
cg03220734
cg03221025
cg03221073
cg03222244
cg03222374

cg03222632
cg03222834
cg03224126
cg03224418
cg03224572
cg03225093
cg03225452
cg03225580
cg03225787
cg03225791
cg03225817
cg03225882
cg03226000
cg03226054
cg03226114
cg03226435
cg03226737
cg03227021
cg03228132
cg03228145
cg03228209
cg03228567
cg03228773
cg03229561
cg03229638
cg03229682
cg03229780
cg03230154
cg03230711
cg03230769
cg03230822
cg03231447
cg03231647
cg03231953
cg03232390
cg03233332
cg03233876
cg03234188
cg03234604
cg03234732
cg03234777
cg03235603
cg03236397
cg03236627
cg03237606
cg03238359
cg03238396
cg03238797
cg03238899
cg03239178
cg03239957
cg03240473

cg03241388
cg03242819
cg03242905
cg03243226
cg03243893
cg03244696
cg03244974
cg03245218
cg03245641
cg03246791
cg03247919
cg03248970
cg03249011
cg03250019
cg03250947
cg03251079
cg03252605
cg03253303
cg03253309
cg03253489
cg03253556
cg03254023
cg03254067
cg03254137
cg03254336
cg03254534
cg03254566
cg03254928
cg03255440
cg03256597
cg03257423
cg03260240
cg03260765
cg03260781
cg03261473
cg03261725
cg03262295
cg03262554
cg03262627
cg03262802
cg03263226
cg03263337
cg03263716
cg03263979
cg03264585
cg03264601
cg03264693
cg03264729
cg03265074
cg03265268
cg03266453
cg03267139

cg03267220
cg03267419
cg03267487
cg03267511
cg03270395
cg03270710
cg03270881
cg03271893
cg03272310
cg03272408
cg03272932
cg03273057
cg03273183
cg03273315
cg03273420
cg03273715
cg03274391
cg03275201
cg03275375
cg03275459
cg03275461
cg03275648
cg03275698
cg03275851
cg03276000
cg03276401
cg03276408
cg03276489
cg03276883
cg03277153
cg03277496
cg03277950
cg03278146
cg03278811
cg03279985
cg03280108
cg03280235
cg03281154
cg03281194
cg03282408
cg03282439
cg03283235
cg03283429
cg03283464
cg03283694
cg03284102
cg03284554
cg03285073
cg03285468
cg03286387
cg03286742
cg03287108

cg03287340
cg03287574
cg03287877
cg03288168
cg03289072
cg03289764
cg03289833
cg03289872
cg03290281
cg03290522
cg03290530
cg03290560
cg03290602
cg03291099
cg03291865
cg03292069
cg03292284
cg03292388
cg03292992
cg03293015
cg03293883
cg03294066
cg03294424
cg03294458
cg03294557
cg03295107
cg03296204
cg03296502
cg03296565
cg03296761
cg03296810
cg03296935
cg03297163
cg03297901
cg03298319
cg03299617
cg03300062
cg03300099
cg03300596
cg03300649
cg03300787
cg03301582
cg03302088
cg03302386
cg03302801
cg03302897
cg03303006
cg03303433
cg03303475
cg03303766
cg03303857
cg03304380

cg03304472
cg03304528
cg03304610
cg03304660
cg03304718
cg03305017
cg03305746
cg03306221
cg03306641
cg03306956
cg03307776
cg03308038
cg03308092
cg03308120
cg03308652
cg03309176
cg03309350
cg03309770
cg03309967
cg03310518
cg03310939
cg03311252
cg03311274
cg03312117
cg03312427
cg03312565
cg03312805
cg03312958
cg03313140
cg03313945
cg03314569
cg03315346
cg03315459
cg03315869
cg03316101
cg03316359
cg03316570
cg03317430
cg03317505
cg03318906
cg03319082
cg03319497
cg03319638
cg03319695
cg03320827
cg03321020
cg03322161
cg03322234
cg03322353
cg03322633
cg03322752
cg03323462

cg03323696
cg03323793
cg03325014
cg03325693
cg03325733
cg03326188
cg03326252
cg03326307
cg03327443
cg03327570
cg03328107
cg03328299
cg03328673
cg03328935
cg03328961
cg03329500
cg03329949
cg03329976
cg03330022
cg03330223
cg03330485
cg03330678
cg03331229
cg03332469
cg03333253
cg03333377
cg03333634
cg03333757
cg03334072
cg03334213
cg03335128
cg03335624
cg03336086
cg03336334
cg03337218
cg03337507
cg03337575
cg03338691
cg03338754
cg03339065
cg03339537
cg03340228
cg03340398
cg03340466
cg03340715
cg03341377
cg03342032
cg03342498
cg03342543
cg03342674
cg03343013
cg03343942

cg03344031
cg03344433
cg03344484
cg03344767
cg03345093
cg03345391
cg03345448
cg03345938
cg03346692
cg03346706
cg03346733
cg03346987
cg03347306
cg03347444
cg03348397
cg03348461
cg03349134
cg03349582
cg03349953
cg03350326
cg03350366
cg03351003
cg03352106
cg03352287
cg03352717
cg03353654
cg03355204
cg03355524
cg03355808
cg03356492
cg03356494
cg03356595
cg03357077
cg03357664
cg03357798
cg03358769
cg03361379
cg03361633
cg03362215
cg03362421
cg03362751
cg03362815
cg03362840
cg03362882
cg03363161
cg03363863
cg03364108
cg03364486
cg03364781
cg03364891
cg03365437
cg03365733

cg03366285
cg03366382
cg03367264
cg03368358
cg03368818
cg03369269
cg03369289
cg03369432
cg03369957
cg03370106
cg03370261
cg03370704
cg03371199
cg03371669
cg03372385
cg03372974
cg03373781
cg03376481
cg03378003
cg03378876
cg03379681
cg03381111
cg03382306
cg03383382
cg03384552
cg03384825
cg03386347
cg03386373
cg03386536
cg03386791
cg03386971
cg03387092
cg03387723
cg03388409
cg03388445
cg03388575
cg03388789
cg03389164
cg03389525
cg03389538
cg03389650
cg03389982
cg03390717
cg03391568
cg03391642
cg03391776
cg03391854
cg03392837
cg03392965
cg03393754
cg03395846
cg03396664

cg03397609
cg03398865
cg03399609
cg03400303
cg03400335
cg03400437
cg03400695
cg03400731
cg03400849
cg03401464
cg03401738
cg03401997
cg03402369
cg03402443
cg03402630
cg03402926
cg03403065
cg03403640
cg03403991
cg03404143
cg03404572
cg03405114
cg03405473
cg03405491
cg03405781
cg03405909
cg03407228
cg03408556
cg03409272
cg03410269
cg03410667
cg03410790
cg03411376
cg03411507
cg03411865
cg03412153
cg03412497
cg03412784
cg03412967
cg03413097
cg03413355
cg03413393
cg03414318
cg03414932
cg03416228
cg03416521
cg03416601
cg03417233
cg03417975
cg03418876
cg03419014
cg03419058

cg03419646
cg03420040
cg03420068
cg03420389
cg03420846
cg03421624
cg03421657
cg03422911
cg03423138
cg03423228
cg03424345
cg03425194
cg03425609
cg03425782
cg03426236
cg03427171
cg03427191
cg03427841
cg03427849
cg03427905
cg03428503
cg03428864
cg03428945
cg03429152
cg03429512
cg03429643
cg03430605
cg03430846
cg03431111
cg03431524
cg03431972
cg03432176
cg03432225
cg03432338
cg03433313
cg03433597
cg03433728
cg03433942
cg03433986
cg03434085
cg03434950
cg03435244
cg03435384
cg03436722
cg03437157
cg03437273
cg03437748
cg03437887
cg03438024
cg03438330
cg03438416
cg03438552

cg03438558
cg03439237
cg03439301
cg03440588
cg03440636
cg03440799
cg03440944
cg03441853
cg03442254
cg03442378
cg03442425
cg03442795
cg03443360
cg03443455
cg03444566
cg03444838
cg03445287
cg03445516
cg03445965
cg03446062
cg03447557
cg03448362
cg03448635
cg03448766
cg03450096
cg03450734
cg03450948
cg03451178
cg03451670
cg03451750
cg03453309
cg03453469
cg03453585
cg03453870
cg03453917
cg03454017
cg03454607
cg03455107
cg03455644
cg03455713
cg03456213
cg03456968
cg03457027
cg03457281
cg03457400
cg03457729
cg03458096
cg03458170
cg03458229
cg03458344
cg03458455
cg03458695

cg03458957
cg03459312
cg03459617
cg03459776
cg03459809
cg03460027
cg03460032
cg03460239
cg03460777
cg03461110
cg03461399
cg03461430
cg03462096
cg03462322
cg03462380
cg03463990
cg03464224
cg03464229
cg03464392
cg03464514
cg03464573
cg03464655
cg03464692
cg03464847
cg03464896
cg03465320
cg03465464
cg03466342
cg03466598
cg03467001
cg03467087
cg03467235
cg03467725
cg03467825
cg03468466
cg03469054
cg03469521
cg03469607
cg03469618
cg03469655
cg03469683
cg03469766
cg03469790
cg03471436
cg03471977
cg03472755
cg03473042
cg03473377
cg03473890
cg03474070
cg03474903
cg03475044

cg03475509
cg03475665
cg03476195
cg03476212
cg03476948
cg03477240
cg03477303
cg03478444
cg03478610
cg03478630
cg03478689
cg03478719
cg03479289
cg03479572
cg03480068
cg03480337
cg03481077
cg03481488
cg03481509
cg03482600
cg03482717
cg03483150
cg03483267
cg03483314
cg03483330
cg03483626
cg03485172
cg03485669
cg03486832
cg03486974
cg03486976
cg03487387
cg03489016
cg03489020
cg03489427
cg03491087
cg03491270
cg03491821
cg03492092
cg03492495
cg03492603
cg03493774
cg03494022
cg03495632
cg03495868
cg03496114
cg03496122
cg03496157
cg03496520
cg03496533
cg03498588
cg03499570

cg03499808
cg03500646
cg03502002
cg03502601
cg03502791
cg03502982
cg03503046
cg03503516
cg03503642
cg03503785
cg03505125
cg03505638
cg03506316
cg03506489
cg03506640
cg03507064
cg03507218
cg03508063
cg03508235
cg03508390
cg03509106
cg03509412
cg03509961
cg03510041
cg03510511
cg03510732
cg03511671
cg03512172
cg03512250
cg03512426
cg03512484
cg03512692
cg03513309
cg03513388
cg03513858
cg03513893
cg03514404
cg03514934
cg03515098
cg03515656
cg03515663
cg03515829
cg03515897
cg03515927
cg03516026
cg03516335
cg03516577
cg03517160
cg03517476
cg03517913
cg03517998
cg03518202

cg03519879
cg03519880
cg03519931
cg03520136
cg03520471
cg03521348
cg03521812
cg03521982
cg03523254
cg03523785
cg03524083
cg03525385
cg03526165
cg03526920
cg03527274
cg03527683
cg03527802
cg03528302
cg03528538
cg03529043
cg03529432
cg03529662
cg03530006
cg03530168
cg03530754
cg03531100
cg03531177
cg03531247
cg03531396
cg03531810
cg03532013
cg03532904
cg03532926
cg03533058
cg03533590
cg03533858
cg03534031
cg03534410
cg03534655
cg03535079
cg03535255
cg03535659
cg03536404
cg03536474
cg03537591
cg03538436
cg03538565
cg03538814
cg03539789
cg03539808
cg03539903
cg03541746

cg03541864
cg03542031
cg03542197
cg03542208
cg03542778
cg03542891
cg03542938
cg03543110
cg03543120
cg03543985
cg03544320
cg03544499
cg03544781
cg03545548
cg03545972
cg03546163
cg03548170
cg03548330
cg03548344
cg03548384
cg03548463
cg03548857
cg03550352
cg03550727
cg03552025
cg03552103
cg03552242
cg03553386
cg03553587
cg03553758
cg03553826
cg03554174
cg03554221
cg03554552
cg03555227
cg03555881
cg03557725
cg03558010
cg03559064
cg03559686
cg03560973
cg03561071
cg03561338
cg03561637
cg03561787
cg03562117
cg03562528
cg03562759
cg03562978
cg03563308
cg03564557
cg03564606

cg03564620
cg03564674
cg03564698
cg03564793
cg03565189
cg03565280
cg03565475
cg03566156
cg03566418
cg03566527
cg03566752
cg03566787
cg03567830
cg03568675
cg03569068
cg03570232
cg03570948
cg03570994
cg03571301
cg03571764
cg03572760
cg03573168
cg03573390
cg03573529
cg03573792
cg03574306
cg03574723
cg03574832
cg03575556
cg03575765
cg03576411
cg03576425
cg03576469
cg03576554
cg03576555
cg03576748
cg03577433
cg03577837
cg03578193
cg03578495
cg03578548
cg03578926
cg03579392
cg03579821
cg03580065
cg03580107
cg03580297
cg03580358
cg03580787
cg03580872
cg03581262
cg03582327

cg03582787
cg03584506
cg03584891
cg03585237
cg03585329
cg03585611
cg03585912
cg03586240
cg03586564
cg03586879
cg03587650
cg03589259
cg03589296
cg03589751
cg03590257
cg03591104
cg03591285
cg03591912
cg03592903
cg03593616
cg03594246
cg03594972
cg03595243
cg03595755
cg03596016
cg03597159
cg03597347
cg03597491
cg03598297
cg03598457
cg03599078
cg03599224
cg03599618
cg03600318
cg03601644
cg03602014
cg03602280
cg03602297
cg03602567
cg03603260
cg03603395
cg03603866
cg03603888
cg03603918
cg03604073
cg03604278
cg03604367
cg03604424
cg03604731
cg03604763
cg03604930
cg03605692

cg03606214
cg03606215
cg03606774
cg03607179
cg03607415
cg03607644
cg03607708
cg03607855
cg03608224
cg03608502
cg03608577
cg03609102
cg03609427
cg03609435
cg03609488
cg03609614
cg03609960
cg03610826
cg03611151
cg03611241
cg03611536
cg03611542
cg03611551
cg03611555
cg03611618
cg03612452
cg03612990
cg03613625
cg03613662
cg03614193
cg03614721
cg03614916
cg03615145
cg03615568
cg03615933
cg03616148
cg03616827
cg03617850
cg03619083
cg03620853
cg03622592
cg03623097
cg03623130
cg03624195
cg03625109
cg03625136
cg03625177
cg03625260
cg03625287
cg03625515
cg03625665
cg03625911

cg03626208
cg03626668
cg03626672
cg03626892
cg03627300
cg03627489
cg03627881
cg03628033
cg03628800
cg03629447
cg03629943
cg03630596
cg03631065
cg03631131
cg03631455
cg03631457
cg03631561
cg03632278
cg03632704
cg03633120
cg03634394
cg03634729
cg03634775
cg03634928
cg03635073
cg03635863
cg03636234
cg03636394
cg03637066
cg03637815
cg03638172
cg03638781
cg03638795
cg03638905
cg03639249
cg03639429
cg03639919
cg03640036
cg03640110
cg03641062
cg03641066
cg03642472
cg03642503
cg03642518
cg03644034
cg03645007
cg03645571
cg03645984
cg03646354
cg03646889
cg03648155
cg03649356

cg03649497
cg03650068
cg03650550
cg03650608
cg03650674
cg03650713
cg03651021
cg03651493
cg03651715
cg03652257
cg03652336
cg03652564
cg03653817
cg03653855
cg03653880
cg03654058
cg03654106
cg03654433
cg03654735
cg03654783
cg03655142
cg03655174
cg03655330
cg03655371
cg03655692
cg03655701
cg03656055
cg03656099
cg03657064
cg03658236
cg03659519
cg03660034
cg03660085
cg03660096
cg03660115
cg03660331
cg03660603
cg03660723
cg03661324
cg03661929
cg03662347
cg03662398
cg03663215
cg03663462
cg03663874
cg03663984
cg03664589
cg03665605
cg03666028
cg03666244
cg03666300
cg03666439

cg03667591
cg03667621
cg03668470
cg03668556
cg03668687
cg03669357
cg03669367
cg03669797
cg03670238
cg03670393
cg03670736
cg03670994
cg03671075
cg03671276
cg03671556
cg03671597
cg03671700
cg03672274
cg03672342
cg03673138
cg03673684
cg03674127
cg03674496
cg03676512
cg03676578
cg03677056
cg03677161
cg03677924
cg03678039
cg03678062
cg03678566
cg03679565
cg03679755
cg03680790
cg03680873
cg03680932
cg03682719
cg03683003
cg03683207
cg03683946
cg03684845
cg03684886
cg03685475
cg03685556
cg03685886
cg03686067
cg03686593
cg03687765
cg03688058
cg03688324
cg03689799
cg03690334

cg03691720
cg03691822
cg03692563
cg03692651
cg03692991
cg03693486
cg03694587
cg03694713
cg03695433
cg03695613
cg03696599
cg03696944
cg03697308
cg03698144
cg03698145
cg03698948
cg03699182
cg03699623
cg03700024
cg03700302
cg03700516
cg03701341
cg03701930
cg03701965
cg03701992
cg03702236
cg03702413
cg03702477
cg03702686
cg03703637
cg03703707
cg03704308
cg03704523
cg03705366
cg03705571
cg03705912
cg03705962
cg03706175
cg03706358
cg03706720
cg03706977
cg03707183
cg03707634
cg03707749
cg03707918
cg03707945
cg03707955
cg03708012
cg03708045
cg03708155
cg03708925
cg03708990

cg03709091
cg03709271
cg03709496
cg03709613
cg03711485
cg03712823
cg03713642
cg03714619
cg03715846
cg03716032
cg03716125
cg03716405
cg03717239
cg03717836
cg03717961
cg03718241
cg03718509
cg03718824
cg03718883
cg03719380
cg03719428
cg03719532
cg03719978
cg03720100
cg03720195
cg03720605
cg03721154
cg03721362
cg03721454
cg03721978
cg03721994
cg03722052
cg03723148
cg03723270
cg03723356
cg03723715
cg03723730
cg03724279
cg03724463
cg03724990
cg03725309
cg03726556
cg03726739
cg03727202
cg03728395
cg03728674
cg03728806
cg03729035
cg03729152
cg03729204
cg03729889
cg03730314

cg03730428
cg03730533
cg03731268
cg03731898
cg03731974
cg03732020
cg03732056
cg03732087
cg03732759
cg03733760
cg03734595
cg03734700
cg03734874
cg03734896
cg03735496
cg03736795
cg03737080
cg03738025
cg03738352
cg03738384
cg03739411
cg03739470
cg03740167
cg03740752
cg03741155
cg03741824
cg03742629
cg03742763
cg03743675
cg03745002
cg03745114
cg03745160
cg03746930
cg03747456
cg03747657
cg03747884
cg03748210
cg03748243
cg03748476
cg03748499
cg03748503
cg03749154
cg03750525
cg03752203
cg03752741
cg03752977
cg03753331
cg03753813
cg03753936
cg03754153
cg03754403
cg03754605

cg03754705
cg03756778
cg03758021
cg03758150
cg03758241
cg03758367
cg03758467
cg03758697
cg03759236
cg03759824
cg03759993
cg03760839
cg03760919
cg03760951
cg03761291
cg03761891
cg03762242
cg03762349
cg03762817
cg03762823
cg03763118
cg03764518
cg03766264
cg03768057
cg03769055
cg03769383
cg03769939
cg03770410
cg03770907
cg03772350
cg03773413
cg03773820
cg03774463
cg03774514
cg03775123
cg03775350
cg03776082
cg03776087
cg03776194
cg03776870
cg03776935
cg03777414
cg03777681
cg03778029
cg03778108
cg03778258
cg03778769
cg03779097
cg03780653
cg03780927
cg03780997
cg03782157

cg03782202
cg03783126
cg03783410
cg03784113
cg03784793
cg03785426
cg03785728
cg03786920
cg03787454
cg03787603
cg03788090
cg03788215
cg03788952
cg03789372
cg03789507
cg03789934
cg03792162
cg03792964
cg03793270
cg03793551
cg03794679
cg03794801
cg03794840
cg03794981
cg03796321
cg03797115
cg03799024
cg03799230
cg03799283
cg03799378
cg03799713
cg03801830
cg03802191
cg03802297
cg03803008
cg03803009
cg03804083
cg03804213
cg03804732
cg03805182
cg03805279
cg03805684
cg03806261
cg03806784
cg03807391
cg03807737
cg03807883
cg03808233
cg03810301
cg03810376
cg03811319
cg03811891

cg03812323
cg03815116
cg03815913
cg03816954
cg03817107
cg03817667
cg03817939
cg03818576
cg03820608
cg03821121
cg03821384
cg03822255
cg03823352
cg03823904
cg03824275
cg03825175
cg03825810
cg03825855
cg03827327
cg03827337
cg03828328
cg03830181
cg03830936
cg03831289
cg03831405
cg03831502
cg03832010
cg03832197
cg03833046
cg03833216
cg03833429
cg03834055
cg03834286
cg03834467
cg03834767
cg03835038
cg03835296
cg03835547
cg03835754
cg03835801
cg03836171
cg03836283
cg03836621
cg03836831
cg03837111
cg03838714
cg03839074
cg03839782
cg03840177
cg03840351
cg03840467
cg03840594

cg03840694
cg03841376
cg03841832
cg03842617
cg03843000
cg03843092
cg03843951
cg03843972
cg03846107
cg03847373
cg03847716
cg03849755
cg03850035
cg03850213
cg03850426
cg03850957
cg03850986
cg03851835
cg03852551
cg03852570
cg03854069
cg03854595
cg03854796
cg03856044
cg03857640
cg03857664
cg03858047
cg03858387
cg03858673
cg03858756
cg03859162
cg03860250
cg03860252
cg03861217
cg03861633
cg03862987
cg03863591
cg03864215
cg03864479
cg03865257
cg03865667
cg03867377
cg03867465
cg03868278
cg03869077
cg03869608
cg03870816
cg03872339
cg03872376
cg03872584
cg03873049
cg03873930

cg03874137
cg03874199
cg03874229
cg03874539
cg03874653
cg03874978
cg03876085
cg03876150
cg03876413
cg03876618
cg03877174
cg03877332
cg03877376
cg03878133
cg03879160
cg03879594
cg03880355
cg03880506
cg03880673
cg03881382
cg03881775
cg03882071
cg03882167
cg03882270
cg03882382
cg03882612
cg03882634
cg03882698
cg03882967
cg03883256
cg03883295
cg03884270
cg03885055
cg03885119
cg03885271
cg03885399
cg03885639
cg03886085
cg03886192
cg03887584
cg03887721
cg03887724
cg03888033
cg03888946
cg03889507
cg03890988
cg03891268
cg03891318
cg03892566
cg03892631
cg03892781
cg03893271

cg03893387
cg03894068
cg03894789
cg03894892
cg03895159
cg03895540
cg03895880
cg03896269
cg03896542
cg03896685
cg03896940
cg03897489
cg03898010
cg03898365
cg03898631
cg03899098
cg03899215
cg03899510
cg03899643
cg03900143
cg03900241
cg03900378
cg03900492
cg03900565
cg03900581
cg03901281
cg03901431
cg03901576
cg03901660
cg03902462
cg03902718
cg03903398
cg03903831
cg03904711
cg03905144
cg03905758
cg03905847
cg03906075
cg03906444
cg03906843
cg03908676
cg03910874
cg03911502
cg03911745
cg03914397
cg03914707
cg03914922
cg03915055
cg03915432
cg03915559
cg03915682
cg03915738

cg03915932
cg03915940
cg03916277
cg03916421
cg03916630
cg03916694
cg03917158
cg03917492
cg03918288
cg03918304
cg03918433
cg03918703
cg03919488
cg03919650
cg03920161
cg03921216
cg03921696
cg03922381
cg03922648
cg03922834
cg03923071
cg03923813
cg03924483
cg03924566
cg03925072
cg03925149
cg03926640
cg03926852
cg03926983
cg03928874
cg03928887
cg03929366
cg03930088
cg03930929
cg03930964
cg03931518
cg03932361
cg03932723
cg03932904
cg03933203
cg03934069
cg03934167
cg03934713
cg03935060
cg03935736
cg03936031
cg03936104
cg03937361
cg03937440
cg03938015
cg03938043
cg03938060

cg03938110
cg03938477
cg03938558
cg03938711
cg03939719
cg03940074
cg03940098
cg03940549
cg03941108
cg03941745
cg03941975
cg03942265
cg03942471
cg03942932
cg03943025
cg03943135
cg03943218
cg03943605
cg03944122
cg03945902
cg03946297
cg03946707
cg03947749
cg03947814
cg03947979
cg03948727
cg03948781
cg03949284
cg03949391
cg03949978
cg03950121
cg03950354
cg03950492
cg03950493
cg03950614
cg03950873
cg03951219
cg03951877
cg03952543
cg03952578
cg03953069
cg03953246
cg03953482
cg03953828
cg03954045
cg03954086
cg03954393
cg03954794
cg03954858
cg03955314
cg03955764
cg03956820

cg03957108
cg03957440
cg03958038
cg03958863
cg03958883
cg03958979
cg03959186
cg03959515
cg03959945
cg03960100
cg03960217
cg03960279
cg03960562
cg03960804
cg03961718
cg03961998
cg03962019
cg03962200
cg03962217
cg03962359
cg03962421
cg03962527
cg03962678
cg03962691
cg03962950
cg03963198
cg03963391
cg03963654
cg03963720
cg03963963
cg03964696
cg03964958
cg03965044
cg03965739
cg03965816
cg03966785
cg03967266
cg03967293
cg03967327
cg03967627
cg03968755
cg03968911
cg03969305
cg03969651
cg03969725
cg03969763
cg03970036
cg03970088
cg03970483
cg03970591
cg03970615
cg03971555

cg03972076
cg03972838
cg03974193
cg03974204
cg03974889
cg03975271
cg03975815
cg03975834
cg03976366
cg03976645
cg03976712
cg03976877
cg03978579
cg03978658
cg03978961
cg03979024
cg03979311
cg03979510
cg03979754
cg03980072
cg03982609
cg03982831
cg03982897
cg03983364
cg03983645
cg03983647
cg03983837
cg03984375
cg03984750
cg03984758
cg03985415
cg03985478
cg03985727
cg03985801
cg03986017
cg03986097
cg03986792
cg03986829
cg03987221
cg03987573
cg03987648
cg03987842
cg03987884
cg03988297
cg03988380
cg03988518
cg03988540
cg03989260
cg03989480
cg03989987
cg03990033
cg03990412

cg03990819
cg03990905
cg03991126
cg03991188
cg03991297
cg03991309
cg03991477
cg03991953
cg03992323
cg03993087
cg03993213
cg03993463
cg03994318
cg03994959
cg03995361
cg03995533
cg03995615
cg03996020
cg03997139
cg03997176
cg03997232
cg03997517
cg03998348
cg03998636
cg03998792
cg03999020
cg03999067
cg03999137
cg03999372
cg03999941
cg04000940
cg04001333
cg04001359
cg04001520
cg04001802
cg04002298
cg04002480
cg04002794
cg04003264
cg04003327
cg04003357
cg04003615
cg04003952
cg04004158
cg04004558
cg04004744
cg04004830
cg04005059
cg04005707
cg04006133
cg04006143
cg04006303

cg04006722
cg04007028
cg04007271
cg04007890
cg04008252
cg04008601
cg04008901
cg04008913
cg04009342
cg04010205
cg04010248
cg04010296
cg04011062
cg04011469
cg04011474
cg04012266
cg04012319
cg04012523
cg04012681
cg04012913
cg04013024
cg04013650
cg04014586
cg04015361
cg04016086
cg04016527
cg04017102
cg04017533
cg04018474
cg04018702
cg04020309
cg04020764
cg04020944
cg04021014
cg04021706
cg04021856
cg04022112
cg04022561
cg04023150
cg04023526
cg04024095
cg04025127
cg04025150
cg04027961
cg04028464
cg04028481
cg04029645
cg04030615
cg04030848
cg04031411
cg04031656
cg04032066

cg04032292
cg04033857
cg04034846
cg04035377
cg04035638
cg04035929
cg04036593
cg04037038
cg04037044
cg04037707
cg04038163
cg04038246
cg04038680
cg04038879
cg04040095
cg04041679
cg04042333
cg04043642
cg04044720
cg04045009
cg04045079
cg04047221
cg04048249
cg04048259
cg04048339
cg04048740
cg04048802
cg04049225
cg04050068
cg04050315
cg04050824
cg04050867
cg04050941
cg04052013
cg04052934
cg04053108
cg04053392
cg04053959
cg04055211
cg04055913
cg04056343
cg04056774
cg04057455
cg04057488
cg04057858
cg04057861
cg04058100
cg04058237
cg04059568
cg04059696
cg04059773
cg04061117

cg04061540
cg04061752
cg04062087
cg04062190
cg04062391
cg04063166
cg04063230
cg04063348
cg04063945
cg04064050
cg04064254
cg04064583
cg04064588
cg04064998
cg04065143
cg04065405
cg04066147
cg04066885
cg04067139
cg04067612
cg04068042
cg04068564
cg04068980
cg04070211
cg04070215
cg04070601
cg04070839
cg04071118
cg04071602
cg04071866
cg04071948
cg04072323
cg04072419
cg04074066
cg04074140
cg04074908
cg04075184
cg04075191
cg04075738
cg04075783
cg04075973
cg04076774
cg04076817
cg04077303
cg04077417
cg04077692
cg04077800
cg04078418
cg04079139
cg04079301
cg04079399
cg04079779

cg04080022
cg04080282
cg04080302
cg04082461
cg04083553
cg04083575
cg04084173
cg04085076
cg04085571
cg04086012
cg04086928
cg04086977
cg04087207
cg04087434
cg04087449
cg04087742
cg04088152
cg04088338
cg04088945
cg04089240
cg04089332
cg04089887
cg04090349
cg04090745
cg04091563
cg04093208
cg04093381
cg04093499
cg04093633
cg04094148
cg04094178
cg04094653
cg04094838
cg04095070
cg04095339
cg04095498
cg04095776
cg04096150
cg04096305
cg04096477
cg04097639
cg04098051
cg04098176
cg04098685
cg04098829
cg04098866
cg04099803
cg04100337
cg04100395
cg04100397
cg04100532
cg04100684

cg04100843
cg04101351
cg04101699
cg04102163
cg04102854
cg04102894
cg04102905
cg04103317
cg04103761
cg04103847
cg04104329
cg04104463
cg04104789
cg04104966
cg04105282
cg04105342
cg04105479
cg04106076
cg04106196
cg04106630
cg04106877
cg04107005
cg04107939
cg04108360
cg04108788
cg04109092
cg04109276
cg04109704
cg04109883
cg04110601
cg04110886
cg04111071
cg04111435
cg04111611
cg04111761
cg04112102
cg04112366
cg04112417
cg04113094
cg04113436
cg04113632
cg04114590
cg04114655
cg04115680
cg04115740
cg04115878
cg04117301
cg04117356
cg04117375
cg04117590
cg04118306
cg04118462

cg04119852
cg04119977
cg04120686
cg04120779
cg04121731
cg04121771
cg04121938
cg04123357
cg04124606
cg04125370
cg04125765
cg04126713
cg04126866
cg04127342
cg04128065
cg04128095
cg04128237
cg04128945
cg04128967
cg04129469
cg04130185
cg04130408
cg04130433
cg04130557
cg04130572
cg04130988
cg04131101
cg04131431
cg04131583
cg04131610
cg04131898
cg04132379
cg04132891
cg04134015
cg04134069
cg04134096
cg04134528
cg04134754
cg04135246
cg04136036
cg04136490
cg04136610
cg04137576
cg04137837
cg04138112
cg04138185
cg04138502
cg04138756
cg04138976
cg04139043
cg04139223
cg04139465

cg04139641
cg04139827
cg04139849
cg04140525
cg04140596
cg04140663
cg04140903
cg04141520
cg04141813
cg04142323
cg04142661
cg04142988
cg04144151
cg04144606
cg04144938
cg04145134
cg04145621
cg04145937
cg04147875
cg04147929
cg04147940
cg04148163
cg04148489
cg04148794
cg04149585
cg04149773
cg04149930
cg04150488
cg04150495
cg04150762
cg04151146
cg04151449
cg04152616
cg04153135
cg04153489
cg04154290
cg04154424
cg04154653
cg04154986
cg04155793
cg04156606
cg04157726
cg04158018
cg04158698
cg04159006
cg04159491
cg04159546
cg04161137
cg04161236
cg04161365
cg04162153
cg04162276

cg04162299
cg04163533
cg04164499
cg04164838
cg04165188
cg04165508
cg04165933
cg04166042
cg04166294
cg04166432
cg04167804
cg04168185
cg04168670
cg04168675
cg04168853
cg04168854
cg04169021
cg04169774
cg04169830
cg04170065
cg04170359
cg04170440
cg04170623
cg04171235
cg04171487
cg04171808
cg04172049
cg04172115
cg04173397
cg04173561
cg04173586
cg04174124
cg04174538
cg04174777
cg04175027
cg04175111
cg04176472
cg04176557
cg04176888
cg04176995
cg04177132
cg04177287
cg04177684
cg04178316
cg04178634
cg04178787
cg04179670
cg04179740
cg04180113
cg04180890
cg04181546
cg04181892

cg04182114
cg04182226
cg04184208
cg04184232
cg04184427
cg04184432
cg04184810
cg04186249
cg04187668
cg04188241
cg04188273
cg04188862
cg04188920
cg04189149
cg04189187
cg04192663
cg04193065
cg04193901
cg04193923
cg04193970
cg04194651
cg04194745
cg04194947
cg04195125
cg04195543
cg04195702
cg04195774
cg04195831
cg04195941
cg04196298
cg04196862
cg04198042
cg04198075
cg04198210
cg04198308
cg04198912
cg04199256
cg04199303
cg04199556
cg04200562
cg04201285
cg04201347
cg04201527
cg04201750
cg04203238
cg04204669
cg04205034
cg04205412
cg04205715
cg04205943
cg04206351
cg04206417

cg04206637
cg04207084
cg04207549
cg04207746
cg04208291
cg04208434
cg04208743
cg04209579
cg04209913
cg04210056
cg04210186
cg04210284
cg04210796
cg04211018
cg04211037
cg04211115
cg04212450
cg04214430
cg04214459
cg04214629
cg04214966
cg04215126
cg04215238
cg04215359
cg04215800
cg04215883
cg04215950
cg04216589
cg04217706
cg04217778
cg04219544
cg04219726
cg04221274
cg04221650
cg04222358
cg04222582
cg04223006
cg04223553
cg04223832
cg04223842
cg04224064
cg04225104
cg04225554
cg04226536
cg04226892
cg04227079
cg04227278
cg04227745
cg04227922
cg04227945
cg04228468
cg04229928

cg04230050
cg04230073
cg04230397
cg04230579
cg04231019
cg04231429
cg04231590
cg04231643
cg04231677
cg04231888
cg04232128
cg04232340
cg04233054
cg04234543
cg04234597
cg04235416
cg04236178
cg04236639
cg04237003
cg04237522
cg04238199
cg04238789
cg04239078
cg04239353
cg04239863
cg04241286
cg04241652
cg04241844
cg04242021
cg04242070
cg04242245
cg04242396
cg04242464
cg04243827
cg04243968
cg04245961
cg04246167
cg04246526
cg04246726
cg04247152
cg04247966
cg04248157
cg04248446
cg04248499
cg04248796
cg04249191
cg04249522
cg04249541
cg04249559
cg04250608
cg04250837
cg04250847

cg04250920
cg04251368
cg04251828
cg04252928
cg04254487
cg04254886
cg04255770
cg04256345
cg04256864
cg04256886
cg04257097
cg04257163
cg04257238
cg04257377
cg04257841
cg04257865
cg04258001
cg04258138
cg04258676
cg04258926
cg04259001
cg04259904
cg04260557
cg04260633
cg04261072
cg04261138
cg04261496
cg04262465
cg04262505
cg04262934
cg04264002
cg04264925
cg04265332
cg04265797
cg04265957
cg04266474
cg04266908
cg04267284
cg04267526
cg04267828
cg04267953
cg04268401
cg04268950
cg04269043
cg04269602
cg04269747
cg04270013
cg04272011
cg04272613
cg04272615
cg04273148
cg04273811

cg04274236
cg04274288
cg04274340
cg04274356
cg04275027
cg04275692
cg04275977
cg04276005
cg04276302
cg04276536
cg04276626
cg04276765
cg04277125
cg04277232
cg04277560
cg04277677
cg04277821
cg04277893
cg04278105
cg04278371
cg04280548
cg04280647
cg04281464
cg04281738
cg04282125
cg04282694
cg04282912
cg04283162
cg04283307
cg04284063
cg04284196
cg04284321
cg04285130
cg04285362
cg04285710
cg04285822
cg04285869
cg04285935
cg04286558
cg04288108
cg04289033
cg04289575
cg04289624
cg04289880
cg04290133
cg04290346
cg04290427
cg04291097
cg04291888
cg04291970
cg04293085
cg04293733

cg04293778
cg04293948
cg04294388
cg04294894
cg04296837
cg04297067
cg04297507
cg04298953
cg04299028
cg04299135
cg04299439
cg04300161
cg04303033
cg04303509
cg04303920
cg04304054
cg04304121
cg04304509
cg04304705
cg04304802
cg04304978
cg04305036
cg04305228
cg04305821
cg04305913
cg04306063
cg04306507
cg04307561
cg04307594
cg04307853
cg04308167
cg04309214
cg04309234
cg04309375
cg04309480
cg04311496
cg04311961
cg04311964
cg04313565
cg04313941
cg04314261
cg04315554
cg04316353
cg04316429
cg04316624
cg04317962
cg04318327
cg04319399
cg04319617
cg04319659
cg04320510
cg04321240

cg04321303
cg04321714
cg04322363
cg04323427
cg04323713
cg04323814
cg04324521
cg04324666
cg04324917
cg04324995
cg04326023
cg04326198
cg04326348
cg04326499
cg04326854
cg04327181
cg04327247
cg04328562
cg04329429
cg04329455
cg04329478
cg04330122
cg04330326
cg04330371
cg04330449
cg04330597
cg04331802
cg04331960
cg04332747
cg04333296
cg04333575
cg04334011
cg04334723
cg04335523
cg04335714
cg04336379
cg04336561
cg04337174
cg04337270
cg04337403
cg04337534
cg04337828
cg04338074
cg04339553
cg04339613
cg04339790
cg04340203
cg04340553
cg04340585
cg04340595
cg04340918
cg04341343

cg04342863
cg04342955
cg04344000
cg04344476
cg04344695
cg04344730
cg04346637
cg04347016
cg04347164
cg04347477
cg04347874
cg04348250
cg04348643
cg04348709
cg04349243
cg04349420
cg04349727
cg04350120
cg04350571
cg04350645
cg04351062
cg04351541
cg04351793
cg04351828
cg04352088
cg04352272
cg04353053
cg04354077
cg04355093
cg04357398
cg04357841
cg04358214
cg04359022
cg04359129
cg04360334
cg04360793
cg04361126
cg04361551
cg04361749
cg04362096
cg04362211
cg04362586
cg04362614
cg04362706
cg04362790
cg04362886
cg04363281
cg04363282
cg04363624
cg04364261
cg04364752
cg04365255

cg04365452
cg04365481
cg04365699
cg04366011
cg04366628
cg04367107
cg04367216
cg04367345
cg04368094
cg04369491
cg04369588
cg04369957
cg04370174
cg04370314
cg04370983
cg04371413
cg04371780
cg04373334
cg04373359
cg04373435
cg04373487
cg04373937
cg04373948
cg04374015
cg04374393
cg04374719
cg04374959
cg04375225
cg04375688
cg04376312
cg04376617
cg04376820
cg04377061
cg04377618
cg04378288
cg04380107
cg04380213
cg04380269
cg04380681
cg04381249
cg04381957
cg04383849
cg04384031
cg04384033
cg04384208
cg04384209
cg04384491
cg04384626
cg04384810
cg04385200
cg04386400
cg04386642

cg04387592
cg04387658
cg04388278
cg04388863
cg04388983
cg04389058
cg04389709
cg04389921
cg04389950
cg04390328
cg04390523
cg04391186
cg04391205
cg04391391
cg04391401
cg04391588
cg04391666
cg04391718
cg04394390
cg04394529
cg04394641
cg04394710
cg04395167
cg04395462
cg04395689
cg04395954
cg04396010
cg04396711
cg04396791
cg04397433
cg04398171
cg04398526
cg04398978
cg04399565
cg04399631
cg04399899
cg04400620
cg04401100
cg04402007
cg04402351
cg04403415
cg04403423
cg04403474
cg04403488
cg04404240
cg04404304
cg04404694
cg04404982
cg04405075
cg04405547
cg04406254
cg04406436

cg04406574
cg04406762
cg04407063
cg04407215
cg04408595
cg04409030
cg04409981
cg04410448
cg04411044
cg04411086
cg04411201
cg04411449
cg04412099
cg04412116
cg04412646
cg04413147
cg04413320
cg04413895
cg04414509
cg04415176
cg04415297
cg04415672
cg04415736
cg04415798
cg04416111
cg04416302
cg04416414
cg04418434
cg04419610
cg04419856
cg04419883
cg04420335
cg04421583
cg04421891
cg04421915
cg04422019
cg04422625
cg04422802
cg04423493
cg04424060
cg04424643
cg04425710
cg04425920
cg04426653
cg04427469
cg04427498
cg04428662
cg04431143
cg04431331
cg04431788
cg04432319
cg04433322

cg04433909
cg04435632
cg04436207
cg04436383
cg04436528
cg04436634
cg04436701
cg04436755
cg04437407
cg04437605
cg04438395
cg04438688
cg04438997
cg04439218
cg04439252
cg04440551
cg04440811
cg04441775
cg04442417
cg04442806
cg04442892
cg04443360
cg04444027
cg04444661
cg04444771
cg04444942
cg04445851
cg04445871
cg04447362
cg04447445
cg04449473
cg04449953
cg04450037
cg04450501
cg04450606
cg04450855
cg04451880
cg04452311
cg04452961
cg04453065
cg04453140
cg04453550
cg04453552
cg04454086
cg04454285
cg04454571
cg04454807
cg04456662
cg04456754
cg04456861
cg04456892
cg04457069

cg04457196
cg04457979
cg04458023
cg04458206
cg04458776
cg04459030
cg04459499
cg04460557
cg04460609
cg04461123
cg04461197
cg04461228
cg04461481
cg04461586
cg04461678
cg04461867
cg04462662
cg04463638
cg04464276
cg04464504
cg04464727
cg04465078
cg04465730
cg04465784
cg04466214
cg04466549
cg04466886
cg04467093
cg04467119
cg04467320
cg04467618
cg04468092
cg04469059
cg04470072
cg04471202
cg04471890
cg04472001
cg04472009
cg04472592
cg04472685
cg04473078
cg04473405
cg04473569
cg04473654
cg04473902
cg04474190
cg04474366
cg04475027
cg04475846
cg04476508
cg04476531
cg04476764

cg04476891
cg04477635
cg04477962
cg04478080
cg04478179
cg04478223
cg04478605
cg04478795
cg04478875
cg04478905
cg04478991
cg04479219
cg04479580
cg04481096
cg04481181
cg04481212
cg04482812
cg04482825
cg04483701
cg04483711
cg04484916
cg04484967
cg04484984
cg04484994
cg04486050
cg04486725
cg04486919
cg04487296
cg04487720
cg04487855
cg04487904
cg04488853
cg04489012
cg04489548
cg04489573
cg04490113
cg04490211
cg04491064
cg04491808
cg04491917
cg04492086
cg04492174
cg04492193
cg04492263
cg04492459
cg04492496
cg04492858
cg04493316
cg04494136
cg04494873
cg04495336
cg04495670

cg04495891
cg04496108
cg04496824
cg04496906
cg04497684
cg04497889
cg04498349
cg04498418
cg04499015
cg04499646
cg04499648
cg04499701
cg04500084
cg04500819
cg04502620
cg04502706
cg04503912
cg04504043
cg04504627
cg04505023
cg04505750
cg04505972
cg04506053
cg04506569
cg04507071
cg04507925
cg04508467
cg04508482
cg04508575
cg04508964
cg04509100
cg04509221
cg04510282
cg04510398
cg04510459
cg04510512
cg04510788
cg04511104
cg04511444
cg04512786
cg04512965
cg04512966
cg04513146
cg04513214
cg04513311
cg04513411
cg04514249
cg04514469
cg04514998
cg04515702
cg04515996
cg04517263

cg04517282
cg04517524
cg04517738
cg04517749
cg04518304
cg04518342
cg04519622
cg04519626
cg04519775
cg04519870
cg04519916
cg04520251
cg04520340
cg04520370
cg04521004
cg04521583
cg04521626
cg04522575
cg04522625
cg04522691
cg04522792
cg04522984
cg04523868
cg04524239
cg04524770
cg04525464
cg04525757
cg04525814
cg04525852
cg04525857
cg04525892
cg04526104
cg04526480
cg04526584
cg04526883
cg04527202
cg04527369
cg04527429
cg04527612
cg04527868
cg04527989
cg04528240
cg04528352
cg04528445
cg04528931
cg04529066
cg04529193
cg04529370
cg04529785
cg04529938
cg04530015
cg04530214

cg04530628
cg04532952
cg04533276
cg04534404
cg04534765
cg04536807
cg04536829
cg04536893
cg04536922
cg04537185
cg04538289
cg04539172
cg04539240
cg04539705
cg04539957
cg04540882
cg04541146
cg04541368
cg04541699
cg04542080
cg04543008
cg04543273
cg04543289
cg04543510
cg04544908
cg04546181
cg04546186
cg04546573
cg04547675
cg04548076
cg04548327
cg04548519
cg04549076
cg04549162
cg04549288
cg04549460
cg04550052
cg04550103
cg04550474
cg04550697
cg04550712
cg04550737
cg04551118
cg04551318
cg04551776
cg04552451
cg04552585
cg04552811
cg04552852
cg04552921
cg04553623
cg04553690

cg04553838
cg04554122
cg04554195
cg04554272
cg04554505
cg04554720
cg04555373
cg04555665
cg04556126
cg04556210
cg04556349
cg04556472
cg04556646
cg04557057
cg04557544
cg04557883
cg04558115
cg04558322
cg04558386
cg04559604
cg04559779
cg04559908
cg04559909
cg04560015
cg04560093
cg04560098
cg04560213
cg04560225
cg04560668
cg04560741
cg04560810
cg04561571
cg04561731
cg04561937
cg04562138
cg04562217
cg04562589
cg04562757
cg04563046
cg04563718
cg04564030
cg04564515
cg04565255
cg04565261
cg04565337
cg04566534
cg04567323
cg04567334
cg04567661
cg04568492
cg04568774
cg04568823

cg04569082
cg04569313
cg04570910
cg04571584
cg04572429
cg04572540
cg04572706
cg04573031
cg04573550
cg04573706
cg04573851
cg04574090
cg04575202
cg04575395
cg04575427
cg04577162
cg04578397
cg04578774
cg04578894
cg04579507
cg04579809
cg04580170
cg04580179
cg04581516
cg04582294
cg04582365
cg04582672
cg04583074
cg04583339
cg04583874
cg04584275
cg04584301
cg04585612
cg04585937
cg04586418
cg04587424
cg04587581
cg04587858
cg04588079
cg04588257
cg04588710
cg04589521
cg04589523
cg04589674
cg04590119
cg04590434
cg04591709
cg04592144
cg04592560
cg04593880
cg04594378
cg04596004

cg04596429
cg04597393
cg04597433
cg04597545
cg04597658
cg04597894
cg04597975
cg04597985
cg04598121
cg04598517
cg04599026
cg04599612
cg04600000
cg04600092
cg04600699
cg04600984
cg04601775
cg04603031
cg04603391
cg04603419
cg04603818
cg04604142
cg04604656
cg04605681
cg04605697
cg04605816
cg04606367
cg04606550
cg04606672
cg04606830
cg04606861
cg04607131
cg04607372
cg04607412
cg04607699
cg04608190
cg04608829
cg04609163
cg04609227
cg04609245
cg04609576
cg04609640
cg04609801
cg04611161
cg04611466
cg04612444
cg04613057
cg04614155
cg04614290
cg04614339
cg04615290
cg04615460

cg04615865
cg04616793
cg04617428
cg04617549
cg04617606
cg04617640
cg04617755
cg04617914
cg04617948
cg04618230
cg04618277
cg04618333
cg04618812
cg04620298
cg04620750
cg04620833
cg04621521
cg04621997
cg04622802
cg04623440
cg04623955
cg04624110
cg04624362
cg04625975
cg04627863
cg04628369
cg04628438
cg04629204
cg04629424
cg04629867
cg04631584
cg04631994
cg04632997
cg04633225
cg04633676
cg04635364
cg04635642
cg04636834
cg04637264
cg04637454
cg04638265
cg04638468
cg04638878
cg04639304
cg04639905
cg04640109
cg04640250
cg04640528
cg04641860
cg04643397
cg04643444
cg04643655

cg04643822
cg04643920
cg04645070
cg04645150
cg04645676
cg04645914
cg04647494
cg04647654
cg04647835
cg04648494
cg04649598
cg04649669
cg04650977
cg04651603
cg04651781
cg04652097
cg04652208
cg04652645
cg04653893
cg04654167
cg04654288
cg04654299
cg04654429
cg04654530
cg04654716
cg04655136
cg04655481
cg04655834
cg04656009
cg04656042
cg04656070
cg04656101
cg04656424
cg04656692
cg04656831
cg04657588
cg04657684
cg04658038
cg04658243
cg04658357
cg04658405
cg04658601
cg04658615
cg04658679
cg04658699
cg04658841
cg04658856
cg04660191
cg04661382
cg04661436
cg04661457
cg04662250

cg04662369
cg04663790
cg04663842
cg04664153
cg04664309
cg04664328
cg04665588
cg04665930
cg04670168
cg04670857
cg04671611
cg04671739
cg04671932
cg04672103
cg04672210
cg04672506
cg04672706
cg04673737
cg04673937
cg04674315
cg04674421
cg04674428
cg04674568
cg04674956
cg04677158
cg04677410
cg04677581
cg04677942
cg04678336
cg04678392
cg04679384
cg04679530
cg04681600
cg04683830
cg04684481
cg04685186
cg04685253
cg04685442
cg04685632
cg04686481
cg04686953
cg04687310
cg04688051
cg04688351
cg04688596
cg04688828
cg04688889
cg04689080
cg04689178
cg04690110
cg04690482
cg04690917

cg04691180
cg04691669
cg04692301
cg04692403
cg04693423
cg04693782
cg04693895
cg04694209
cg04694812
cg04695027
cg04695635
cg04696391
cg04696720
cg04697056
cg04697262
cg04698114
cg04699199
cg04699394
cg04700283
cg04700474
cg04700930
cg04701034
cg04702509
cg04703241
cg04703535
cg04703620
cg04703800
cg04704053
cg04704064
cg04704294
cg04704449
cg04705499
cg04707013
cg04707332
cg04707519
cg04707603
cg04707715
cg04708321
cg04708601
cg04708790
cg04709035
cg04709144
cg04709675
cg04710115
cg04710226
cg04710402
cg04710528
cg04711622
cg04711852
cg04712321
cg04712635
cg04713352

cg04713531
cg04714110
cg04714195
cg04714466
cg04715572
cg04715649
cg04716530
cg04716990
cg04718492
cg04719215
cg04719819
cg04719903
cg04720797
cg04721828
cg04722643
cg04722901
cg04723401
cg04723534
cg04724556
cg04724807
cg04725637
cg04726373
cg04726824
cg04727071
cg04727521
cg04728180
cg04728429
cg04729173
cg04729913
cg04730685
cg04731448
cg04731618
cg04731624
cg04731861
cg04732303
cg04733365
cg04733537
cg04733787
cg04733856
cg04734076
cg04734784
cg04735632
cg04735773
cg04737150
cg04737356
cg04737758
cg04737916
cg04738237
cg04738464
cg04738877
cg04738965
cg04739460

cg04739647
cg04739648
cg04739880
cg04740205
cg04740359
cg04741094
cg04741118
cg04741366
cg04742334
cg04742624
cg04742719
cg04742985
cg04743510
cg04744093
cg04744379
cg04744713
cg04744717
cg04744735
cg04745004
cg04745064
cg04747226
cg04747322
cg04747619
cg04747634
cg04747693
cg04747824
cg04748546
cg04749631
cg04749646
cg04750099
cg04750421
cg04751035
cg04751149
cg04751276
cg04751631
cg04751839
cg04752257
cg04752768
cg04752812
cg04752885
cg04752992
cg04753163
cg04753469
cg04753836
cg04755523
cg04755561
cg04755857
cg04757168
cg04757350
cg04757411
cg04758026
cg04759227

cg04759439
cg04759648
cg04759651
cg04760448
cg04760708
cg04761768
cg04762351
cg04762815
cg04763554
cg04763994
cg04764012
cg04764902
cg04765277
cg04765303
cg04765410
cg04765848
cg04765904
cg04766365
cg04766834
cg04768606
cg04768697
cg04768939
cg04770577
cg04771100
cg04771199
cg04771285
cg04771774
cg04772683
cg04772947
cg04772948
cg04773818
cg04774103
cg04774364
cg04774696
cg04774711
cg04775627
cg04775824
cg04776489
cg04777091
cg04777348
cg04777352
cg04778012
cg04778125
cg04779796
cg04780380
cg04780434
cg04780454
cg04780572
cg04780726
cg04781095
cg04781638
cg04783231

cg04783450
cg04784316
cg04784410
cg04784635
cg04784672
cg04785187
cg04785900
cg04785902
cg04786287
cg04787211
cg04787888
cg04787916
cg04788390
cg04788743
cg04789663
cg04789818
cg04790084
cg04790154
cg04790800
cg04790887
cg04791129
cg04791256
cg04791477
cg04792330
cg04792378
cg04792712
cg04793118
cg04794141
cg04794268
cg04794430
cg04795387
cg04796162
cg04796525
cg04796828
cg04797323
cg04797496
cg04797725
cg04800569
cg04801098
cg04801430
cg04801617
cg04801767
cg04802221
cg04802386
cg04802696
cg04804052
cg04804094
cg04804542
cg04804618
cg04804726
cg04804770
cg04805181

cg04806942
cg04806979
cg04807594
cg04809380
cg04809948
cg04810745
cg04810997
cg04811540
cg04811706
cg04812347
cg04812351
cg04812475
cg04812556
cg04812644
cg04813159
cg04813787
cg04813876
cg04814317
cg04814352
cg04815334
cg04815371
cg04815973
cg04816311
cg04816394
cg04816803
cg04818637
cg04818710
cg04818943
cg04819337
cg04819351
cg04819714
cg04820362
cg04820679
cg04820773
cg04821248
cg04821993
cg04822456
cg04822495
cg04822710
cg04823311
cg04823455
cg04826040
cg04829509
cg04829760
cg04830191
cg04830355
cg04830471
cg04830629
cg04830757
cg04831076
cg04831174
cg04831795

cg04832466
cg04833533
cg04833713
cg04833845
cg04833885
cg04833898
cg04834204
cg04834436
cg04834770
cg04835091
cg04835811
cg04836442
cg04836530
cg04836851
cg04837025
cg04837494
cg04837597
cg04837616
cg04837832
cg04837847
cg04838191
cg04839289
cg04839677
cg04839683
cg04840203
cg04840813
cg04841358
cg04841389
cg04841806
cg04842146
cg04842426
cg04842880
cg04843111
cg04843425
cg04843813
cg04843968
cg04844207
cg04844495
cg04844534
cg04844693
cg04845060
cg04845176
cg04846036
cg04846156
cg04847110
cg04847610
cg04847638
cg04847649
cg04848660
cg04848858
cg04849227
cg04849541

cg04849850
cg04850284
cg04850286
cg04850366
cg04850423
cg04850479
cg04850765
cg04850842
cg04850976
cg04850999
cg04851702
cg04851712
cg04852339
cg04852977
cg04853995
cg04854282
cg04854328
cg04855433
cg04856022
cg04856117
cg04857033
cg04857172
cg04857420
cg04857792
cg04858155
cg04858597
cg04858987
cg04859117
cg04859466
cg04859706
cg04859929
cg04861380
cg04862321
cg04863679
cg04864441
cg04864453
cg04864600
cg04864807
cg04865113
cg04865180
cg04865290
cg04865319
cg04865491
cg04865691
cg04865998
cg04866639
cg04867052
cg04867749
cg04868219
cg04868540
cg04869532
cg04869770

cg04870120
cg04870732
cg04871364
cg04871807
cg04872123
cg04872593
cg04873188
cg04874562
cg04874795
cg04875020
cg04875212
cg04875514
cg04875821
cg04876500
cg04876582
cg04877287
cg04877737
cg04877959
cg04878146
cg04878152
cg04878385
cg04878685
cg04878813
cg04878955
cg04879235
cg04879838
cg04880052
cg04880091
cg04880253
cg04880804
cg04880990
cg04881095
cg04881903
cg04882213
cg04882273
cg04883553
cg04883903
cg04884011
cg04884025
cg04884118
cg04884798
cg04885549
cg04885642
cg04885678
cg04885825
cg04885881
cg04886849
cg04886857
cg04887030
cg04887082
cg04887675
cg04888174

cg04888241
cg04888360
cg04889790
cg04890851
cg04891040
cg04891053
cg04891688
cg04891836
cg04892170
cg04892724
cg04892766
cg04893786
cg04894537
cg04894619
cg04895001
cg04895202
cg04895233
cg04895345
cg04895616
cg04896048
cg04896191
cg04896437
cg04896596
cg04897713
cg04898035
cg04898695
cg04899153
cg04899175
cg04899273
cg04899656
cg04900080
cg04900877
cg04900882
cg04901044
cg04901179
cg04902529
cg04902729
cg04902838
cg04902968
cg04903146
cg04903900
cg04904030
cg04904276
cg04904468
cg04904531
cg04904784
cg04905434
cg04905557
cg04906462
cg04906886
cg04907173
cg04907218

cg04907257
cg04907571
cg04907849
cg04908088
cg04908380
cg04908668
cg04908905
cg04908961
cg04909432
cg04909834
cg04910179
cg04910228
cg04910677
cg04910877
cg04910921
cg04910970
cg04911669
cg04911739
cg04912542
cg04912843
cg04913057
cg04913114
cg04913790
cg04913979
cg04914126
cg04914198
cg04914221
cg04914283
cg04914574
cg04914930
cg04915788
cg04915974
cg04917181
cg04917511
cg04918831
cg04919274
cg04919489
cg04919563
cg04920630
cg04921646
cg04923174
cg04923194
cg04923840
cg04925032
cg04925261
cg04926244
cg04926280
cg04926361
cg04926384
cg04926461
cg04926732
cg04926767

cg04926904
cg04927066
cg04927695
cg04927725
cg04927931
cg04928293
cg04928338
cg04929543
cg04929612
cg04930211
cg04930601
cg04930630
cg04930816
cg04930840
cg04930848
cg04931032
cg04931184
cg04931256
cg04931539
cg04931913
cg04932551
cg04933990
cg04935434
cg04935449
cg04935792
cg04936178
cg04936619
cg04938317
cg04938738
cg04938830
cg04938966
cg04939302
cg04940570
cg04940764
cg04941018
cg04941520
cg04942107
cg04943065
cg04943586
cg04943797
cg04943879
cg04943986
cg04944300
cg04944537
cg04944659
cg04945153
cg04945331
cg04945434
cg04945608
cg04945735
cg04945753
cg04946274

cg04946561
cg04946749
cg04947801
cg04947907
cg04948438
cg04948892
cg04948906
cg04949206
cg04949225
cg04950301
cg04950627
cg04950824
cg04950931
cg04951104
cg04951293
cg04951476
cg04952627
cg04953061
cg04953935
cg04954559
cg04954680
cg04955246
cg04955511
cg04957757
cg04958055
cg04958496
cg04958528
cg04958794
cg04959480
cg04959572
cg04960007
cg04960169
cg04960602
cg04961235
cg04961448
cg04961466
cg04961582
cg04961911
cg04962005
cg04962480
cg04964669
cg04965050
cg04965934
cg04966159
cg04967578
cg04967698
cg04967734
cg04968426
cg04968473
cg04970064
cg04970570
cg04971534

cg04972065
cg04972384
cg04972459
cg04972529
cg04972745
cg04972770
cg04973183
cg04973669
cg04973915
cg04973925
cg04974062
cg04974290
cg04974647
cg04975856
cg04975920
cg04976154
cg04976330
cg04977343
cg04977528
cg04977618
cg04977834
cg04978007
cg04978078
cg04978831
cg04979933
cg04980478
cg04980849
cg04980991
cg04981007
cg04981088
cg04981338
cg04981940
cg04982308
cg04982401
cg04982781
cg04982951
cg04983276
cg04983821
cg04984852
cg04986276
cg04987362
cg04988367
cg04988476
cg04988487
cg04988978
cg04990472
cg04990605
cg04990850
cg04991036
cg04991214
cg04991805
cg04992127

cg04992191
cg04992521
cg04992757
cg04993020
cg04993082
cg04993112
cg04993267
cg04993290
cg04994217
cg04994447
cg04995095
cg04995291
cg04996020
cg04996089
cg04996219
cg04997273
cg04997558
cg04998375
cg04998536
cg04999435
cg05000039
cg05000598
cg05000994
cg05001334
cg05001691
cg05001964
cg05003322
cg05004506
cg05004518
cg05004818
cg05005073
cg05005217
cg05005332
cg05006067
cg05006092
cg05006211
cg05006231
cg05006302
cg05006304
cg05007097
cg05007163
cg05007697
cg05008139
cg05008898
cg05008975
cg05009601
cg05009934
cg05011265
cg05011467
cg05011526
cg05012356
cg05012825

cg05012902
cg05012972
cg05013064
cg05013661
cg05014568
cg05017199
cg05017639
cg05018034
cg05018460
cg05018513
cg05018648
cg05019016
cg05019826
cg05020759
cg05021029
cg05021636
cg05021643
cg05021842
cg05023114
cg05024627
cg05024715
cg05024961
cg05026014
cg05026798
cg05027052
cg05027597
cg05027713
cg05028087
cg05028750
cg05029312
cg05029983
cg05030251
cg05030450
cg05030680
cg05031448
cg05032348
cg05032957
cg05034656
cg05034702
cg05034973
cg05035143
cg05035315
cg05035456
cg05036130
cg05036153
cg05036173
cg05036212
cg05036656
cg05036937
cg05036991
cg05037544
cg05037662

cg05037806
cg05038436
cg05040348
cg05040472
cg05040725
cg05041265
cg05041351
cg05042706
cg05043349
cg05043389
cg05043733
cg05043910
cg05044185
cg05044778
cg05044809
cg05045027
cg05045598
cg05045981
cg05046306
cg05046377
cg05046589
cg05047401
cg05047521
cg05047908
cg05048426
cg05048523
cg05048927
cg05049003
cg05049361
cg05049825
cg05051043
cg05051571
cg05052324
cg05052407
cg05052450
cg05052463
cg05053562
cg05055001
cg05055490
cg05055766
cg05056497
cg05056653
cg05057515
cg05057534
cg05057720
cg05057932
cg05058167
cg05058313
cg05058354
cg05058758
cg05059571
cg05059605

cg05059843
cg05060440
cg05060607
cg05061107
cg05061754
cg05062413
cg05063856
cg05063999
cg05064645
cg05065507
cg05066621
cg05068156
cg05068951
cg05069426
cg05070265
cg05070372
cg05071046
cg05072559
cg05073901
cg05074946
cg05075118
cg05075562
cg05076755
cg05076820
cg05076914
cg05077452
cg05078019
cg05078644
cg05079049
cg05079544
cg05080154
cg05080371
cg05080409
cg05080651
cg05080746
cg05080966
cg05081498
cg05082102
cg05082311
cg05083033
cg05083067
cg05083273
cg05084700
cg05085500
cg05085636
cg05086314
cg05087459
cg05087577
cg05087982
cg05089225
cg05090785
cg05091734

cg05092353
cg05093169
cg05093429
cg05093576
cg05093591
cg05093811
cg05095158
cg05095428
cg05095591
cg05096637
cg05096777
cg05096800
cg05098343
cg05098382
cg05098512
cg05098551
cg05099196
cg05099221
cg05099387
cg05099596
cg05100419
cg05100867
cg05102394
cg05104080
cg05104639
cg05104897
cg05106502
cg05106699
cg05106785
cg05109931
cg05110225
cg05110943
cg05111779
cg05112070
cg05112254
cg05113958
cg05114676
cg05114959
cg05116378
cg05116777
cg05117208
cg05117324
cg05117693
cg05119113
cg05119290
cg05119754
cg05119988
cg05120042
cg05120428
cg05121182
cg05121202
cg05121480

cg05122026
cg05122040
cg05122082
cg05122213
cg05124129
cg05125795
cg05126095
cg05126289
cg05126581
cg05127036
cg05127077
cg05127369
cg05128656
cg05128922
cg05129325
cg05129489
cg05129572
cg05129858
cg05130075
cg05130830
cg05131034
cg05131142
cg05131202
cg05131696
cg05132150
cg05132306
cg05132568
cg05132851
cg05133179
cg05133814
cg05133853
cg05134015
cg05134325
cg05134796
cg05134926
cg05134945
cg05135252
cg05135549
cg05136956
cg05137103
cg05137450
cg05138040
cg05138082
cg05138776
cg05138957
cg05139028
cg05139061
cg05139728
cg05139788
cg05139861
cg05140937
cg05141384

cg05141432
cg05141602
cg05141924
cg05142148
cg05142211
cg05142272
cg05142677
cg05142765
cg05142982
cg05143386
cg05143420
cg05143887
cg05144620
cg05144814
cg05145297
cg05146536
cg05146658
cg05146852
cg05147143
cg05147273
cg05147578
cg05147692
cg05150077
cg05151154
cg05152116
cg05152300
cg05152589
cg05153380
cg05154390
cg05155047
cg05155704
cg05155965
cg05156532
cg05157098
cg05157140
cg05157541
cg05157791
cg05159438
cg05159685
cg05160554
cg05161682
cg05162166
cg05162320
cg05162358
cg05162750
cg05163933
cg05164666
cg05164933
cg05165087
cg05165250
cg05165750
cg05165940

cg05166490
cg05166820
cg05166871
cg05167251
cg05167857
cg05168033
cg05168229
cg05168368
cg05168609
cg05168977
cg05169988
cg05170183
cg05170205
cg05170342
cg05170353
cg05170445
cg05170867
cg05171336
cg05171937
cg05172061
cg05172122
cg05172259
cg05172912
cg05173889
cg05174942
cg05174975
cg05175762
cg05175803
cg05175964
cg05176551
cg05176667
cg05176958
cg05177060
cg05177287
cg05177629
cg05177685
cg05177932
cg05178683
cg05178839
cg05179846
cg05180182
cg05180856
cg05180985
cg05181041
cg05181212
cg05181846
cg05183671
cg05183931
cg05184372
cg05184456
cg05184519
cg05184607

cg05184843
cg05185364
cg05185584
cg05185784
cg05186311
cg05186455
cg05187154
cg05187193
cg05187700
cg05188268
cg05188959
cg05189281
cg05189570
cg05190718
cg05190790
cg05191076
cg05192402
cg05192533
cg05192639
cg05193538
cg05194412
cg05194808
cg05195452
cg05196610
cg05196905
cg05198938
cg05198963
cg05200037
cg05200057
cg05200313
cg05201185
cg05201300
cg05201789
cg05201970
cg05202042
cg05202654
cg05203875
cg05204315
cg05204540
cg05205299
cg05205351
cg05205647
cg05205932
cg05206276
cg05206884
cg05207067
cg05207107
cg05207465
cg05207637
cg05207943
cg05208050
cg05208249

cg05208747
cg05209330
cg05209584
cg05210258
cg05210266
cg05210373
cg05210412
cg05210464
cg05210497
cg05210501
cg05210558
cg05210615
cg05211057
cg05211356
cg05212570
cg05213414
cg05214235
cg05214460
cg05214511
cg05215056
cg05215277
cg05215304
cg05215575
cg05215806
cg05215925
cg05215966
cg05216542
cg05217213
cg05217312
cg05217983
cg05218035
cg05218678
cg05218696
cg05220114
cg05220280
cg05220510
cg05220630
cg05220857
cg05221057
cg05221167
cg05221264
cg05221326
cg05222243
cg05222671
cg05222995
cg05223158
cg05223210
cg05223396
cg05223441
cg05223459
cg05223638
cg05223720

cg05224498
cg05224741
cg05225089
cg05225461
cg05225967
cg05226125
cg05226558
cg05226740
cg05227556
cg05228186
cg05228361
cg05229229
cg05229416
cg05230054
cg05231226
cg05231542
cg05232118
cg05232215
cg05232708
cg05232857
cg05233576
cg05233774
cg05233816
cg05235171
cg05235610
cg05237641
cg05238769
cg05238976
cg05239225
cg05239311
cg05239967
cg05240350
cg05240479
cg05240760
cg05241277
cg05241308
cg05241461
cg05241536
cg05241571
cg05241586
cg05241830
cg05242592
cg05242653
cg05243629
cg05244013
cg05245379
cg05245794
cg05246100
cg05246110
cg05246900
cg05247096
cg05247640

cg05248004
cg05248618
cg05248804
cg05249074
cg05249836
cg05249937
cg05250768
cg05251676
cg05252143
cg05253165
cg05253759
cg05254221
cg05254373
cg05254518
cg05255522
cg05255580
cg05255811
cg05256043
cg05256269
cg05256719
cg05256765
cg05256999
cg05257772
cg05258689
cg05258914
cg05259891
cg05260236
cg05260372
cg05260373
cg05260469
cg05261824
cg05261893
cg05262369
cg05262436
cg05262755
cg05262829
cg05264870
cg05264881
cg05265071
cg05265081
cg05265143
cg05265526
cg05265574
cg05265596
cg05265703
cg05266212
cg05266564
cg05266781
cg05266786
cg05266836
cg05267394
cg05267735

cg05267756
cg05267955
cg05269040
cg05269080
cg05269218
cg05269258
cg05269534
cg05270344
cg05270938
cg05271255
cg05272157
cg05272921
cg05273205
cg05273533
cg05274242
cg05274680
cg05275576
cg05275752
cg05276226
cg05277146
cg05278186
cg05278307
cg05279092
cg05279172
cg05279387
cg05280255
cg05280527
cg05281126
cg05281206
cg05281965
cg05283379
cg05283542
cg05283597
cg05283685
cg05283716
cg05283992
cg05284631
cg05284713
cg05284727
cg05285582
cg05287409
cg05287437
cg05289320
cg05289983
cg05290056
cg05290058
cg05290658
cg05290820
cg05291966
cg05292150
cg05292310
cg05292376

cg05292404
cg05292954
cg05293299
cg05293738
cg05295006
cg05295038
cg05295286
cg05295388
cg05295557
cg05295584
cg05296940
cg05298088
cg05298628
cg05299793
cg05299836
cg05300029
cg05300538
cg05301500
cg05301866
cg05302035
cg05302095
cg05302218
cg05302338
cg05302350
cg05302386
cg05302441
cg05302701
cg05302802
cg05303057
cg05303132
cg05304005
cg05304177
cg05304366
cg05304461
cg05305140
cg05306070
cg05306472
cg05306745
cg05307477
cg05307752
cg05308233
cg05308643
cg05309280
cg05309948
cg05310240
cg05310990
cg05311180
cg05311383
cg05311410
cg05311412
cg05312094
cg05312104

cg05312353
cg05313263
cg05313910
cg05314024
cg05315047
cg05315334
cg05316065
cg05316650
cg05316785
cg05317275
cg05317396
cg05317851
cg05318142
cg05318186
cg05318210
cg05318546
cg05319305
cg05320842
cg05321361
cg05322019
cg05322332
cg05322931
cg05323324
cg05323533
cg05324194
cg05324519
cg05325193
cg05325480
cg05325887
cg05326036
cg05327835
cg05327864
cg05328547
cg05328761
cg05329692
cg05329720
cg05329893
cg05329976
cg05330291
cg05331214
cg05331290
cg05331334
cg05331472
cg05331807
cg05331856
cg05332483
cg05333174
cg05334119
cg05334414
cg05334416
cg05334655
cg05335186

cg05335388
cg05335953
cg05336115
cg05336292
cg05336395
cg05336997
cg05337177
cg05337387
cg05337636
cg05338187
cg05338397
cg05338785
cg05338976
cg05339037
cg05339390
cg05339403
cg05339415
cg05339605
cg05339727
cg05340085
cg05342515
cg05342682
cg05343021
cg05343105
cg05343159
cg05343512
cg05343811
cg05344066
cg05344293
cg05344430
cg05344949
cg05345154
cg05345160
cg05346287
cg05346491
cg05346527
cg05347479
cg05347845
cg05347927
cg05348084
cg05348216
cg05348888
cg05349042
cg05349089
cg05349159
cg05349349
cg05350798
cg05350850
cg05351257
cg05351942
cg05352161
cg05352500

cg05352668
cg05352688
cg05353869
cg05353872
cg05354921
cg05355301
cg05355381
cg05355411
cg05355757
cg05357209
cg05357229
cg05357287
cg05357347
cg05357717
cg05357750
cg05358291
cg05358575
cg05358758
cg05358946
cg05359853
cg05361096
cg05361290
cg05361366
cg05361459
cg05361489
cg05361530
cg05363382
cg05363616
cg05363871
cg05363938
cg05363955
cg05364179
cg05365033
cg05365199
cg05365385
cg05365670
cg05365685
cg05366024
cg05366909
cg05367173
cg05368158
cg05368595
cg05368971
cg05369857
cg05369926
cg05370101
cg05370136
cg05370555
cg05371438
cg05371498
cg05371552
cg05371578

cg05372113
cg05372679
cg05373251
cg05373655
cg05374044
cg05374129
cg05374238
cg05374412
cg05374756
cg05374783
cg05374982
cg05376228
cg05376362
cg05376617
cg05376754
cg05376869
cg05377120
cg05377587
cg05377733
cg05377753
cg05378861
cg05378938
cg05379509
cg05380019
cg05380130
cg05380734
cg05380759
cg05380885
cg05380982
cg05381746
cg05381960
cg05382282
cg05383332
cg05384102
cg05385282
cg05385331
cg05385434
cg05385513
cg05385882
cg05386077
cg05386493
cg05386977
cg05387269
cg05387399
cg05387519
cg05387565
cg05389310
cg05389731
cg05389935
cg05390077
cg05390279
cg05390685

cg05391338
cg05391998
cg05392244
cg05392263
cg05392949
cg05393578
cg05393726
cg05394187
cg05394210
cg05395321
cg05395523
cg05395704
cg05396761
cg05397202
cg05397490
cg05398036
cg05398769
cg05398903
cg05399214
cg05399225
cg05399244
cg05399330
cg05399607
cg05400075
cg05400498
cg05401312
cg05401965
cg05402319
cg05402891
cg05403316
cg05403433
cg05403555
cg05404091
cg05404912
cg05405094
cg05405275
cg05405389
cg05405550
cg05405572
cg05405742
cg05406334
cg05406635
cg05406868
cg05407714
cg05407757
cg05408403
cg05408649
cg05409218
cg05409693
cg05409945
cg05411171
cg05411186

cg05412396
cg05413061
cg05413285
cg05413305
cg05413694
cg05415020
cg05415285
cg05415606
cg05416055
cg05416223
cg05417035
cg05417127
cg05418042
cg05418105
cg05418495
cg05419096
cg05419385
cg05419752
cg05420867
cg05420994
cg05421564
cg05421677
cg05422352
cg05422920
cg05423018
cg05423393
cg05423418
cg05424333
cg05424422
cg05426299
cg05426485
cg05426956
cg05427381
cg05427488
cg05427907
cg05427966
cg05429117
cg05429448
cg05431171
cg05431532
cg05431842
cg05431942
cg05432034
cg05432322
cg05433152
cg05434363
cg05434641
cg05435295
cg05436231
cg05437823
cg05438320
cg05438719

cg05438727
cg05440435
cg05442408
cg05442599
cg05442602
cg05444083
cg05444514
cg05445221
cg05445490
cg05446322
cg05446414
cg05446424
cg05446629
cg05448271
cg05448383
cg05448715
cg05448721
cg05449164
cg05449190
cg05450193
cg05450477
cg05450629
cg05452406
cg05452524
cg05452692
cg05452899
cg05453378
cg05453631
cg05454389
cg05454957
cg05455036
cg05455409
cg05455720
cg05455729
cg05455971
cg05456713
cg05457480
cg05457768
cg05457923
cg05458079
cg05458609
cg05459280
cg05459750
cg05461143
cg05461153
cg05461169
cg05461386
cg05462446
cg05462543
cg05462922
cg05463589
cg05464506

cg05465059
cg05465572
cg05467160
cg05467458
cg05467890
cg05468458
cg05468584
cg05468843
cg05468917
cg05469591
cg05470166
cg05471495
cg05471616
cg05471845
cg05471881
cg05472059
cg05472874
cg05472998
cg05474726
cg05474943
cg05475107
cg05475524
cg05475702
cg05477514
cg05477834
cg05478330
cg05479130
cg05480370
cg05480377
cg05481452
cg05481579
cg05481881
cg05482119
cg05483125
cg05483406
cg05483505
cg05484231
cg05484376
cg05484788
cg05484949
cg05485462
cg05485639
cg05486138
cg05486931
cg05487134
cg05488160
cg05488168
cg05488681
cg05489054
cg05489904
cg05490366
cg05490712

cg05490757
cg05490864
cg05491001
cg05492113
cg05492657
cg05493001
cg05493509
cg05494014
cg05494604
cg05495949
cg05496514
cg05497175
cg05497216
cg05497273
cg05499148
cg05500734
cg05500783
cg05500840
cg05501060
cg05501419
cg05501498
cg05501617
cg05501996
cg05502349
cg05502701
cg05503062
cg05504025
cg05505545
cg05505763
cg05505803
cg05505872
cg05505961
cg05506011
cg05506365
cg05506480
cg05506600
cg05507514
cg05507908
cg05508084
cg05509179
cg05509359
cg05509820
cg05510902
cg05511148
cg05511722
cg05511924
cg05512111
cg05512756
cg05512869
cg05513300
cg05513806
cg05514072

cg05514256
cg05514531
cg05514883
cg05514971
cg05515077
cg05515143
cg05516271
cg05516617
cg05517218
cg05517527
cg05517824
cg05519002
cg05519187
cg05520050
cg05521659
cg05522498
cg05522774
cg05523603
cg05523882
cg05523897
cg05524387
cg05525368
cg05525416
cg05526099
cg05526364
cg05526832
cg05528492
cg05529123
cg05530403
cg05530484
cg05530592
cg05530612
cg05531317
cg05531502
cg05531689
cg05532481
cg05533037
cg05533210
cg05533615
cg05534186
cg05534710
cg05534973
cg05536156
cg05536286
cg05536800
cg05536998
cg05537380
cg05537607
cg05538980
cg05539265
cg05539745
cg05541311

cg05542262
cg05542338
cg05543049
cg05543492
cg05543593
cg05544500
cg05544885
cg05545441
cg05545832
cg05545882
cg05546296
cg05546763
cg05546863
cg05547200
cg05547260
cg05547777
cg05547853
cg05548130
cg05548349
cg05548402
cg05548672
cg05549854
cg05550276
cg05550995
cg05551792
cg05552183
cg05552569
cg05552814
cg05552874
cg05553502
cg05553560
cg05553591
cg05554192
cg05554906
cg05555111
cg05555928
cg05556038
cg05556361
cg05556461
cg05556535
cg05557110
cg05557266
cg05557449
cg05557991
cg05558046
cg05558259
cg05560435
cg05563672
cg05563813
cg05565697
cg05567236
cg05567269

cg05567408
cg05567440
cg05568930
cg05569086
cg05569220
cg05569767
cg05569987
cg05571215
cg05571581
cg05571619
cg05571912
cg05573262
cg05573378
cg05573412
cg05573626
cg05573844
cg05574272
cg05575505
cg05575733
cg05576619
cg05576992
cg05577810
cg05578802
cg05579030
cg05579549
cg05580001
cg05580073
cg05580560
cg05580671
cg05581297
cg05581701
cg05581878
cg05582310
cg05582719
cg05583087
cg05583171
cg05584235
cg05585303
cg05585475
cg05585556
cg05585991
cg05586304
cg05586667
cg05586863
cg05587232
cg05587627
cg05588271
cg05589246
cg05589784
cg05590982
cg05591728
cg05591769

cg05592150
cg05592245
cg05592769
cg05593122
cg05593139
cg05593162
cg05593336
cg05593641
cg05593707
cg05593759
cg05594525
cg05595142
cg05595345
cg05596294
cg05596552
cg05596841
cg05597504
cg05597908
cg05598010
cg05599759
cg05600350
cg05600421
cg05600609
cg05601096
cg05601456
cg05602439
cg05602642
cg05602715
cg05602862
cg05603527
cg05603610
cg05604079
cg05605603
cg05606878
cg05607053
cg05608777
cg05609160
cg05609218
cg05611565
cg05612673
cg05612966
cg05614013
cg05615464
cg05615487
cg05616355
cg05616858
cg05616999
cg05617469
cg05617678
cg05617732
cg05618683
cg05618807

cg05618880
cg05618934
cg05619449
cg05619785
cg05619892
cg05620165
cg05620261
cg05620710
cg05621131
cg05622324
cg05622422
cg05622577
cg05623392
cg05623727
cg05624623
cg05624853
cg05625526
cg05625559
cg05626226
cg05626536
cg05626664
cg05626927
cg05627639
cg05627733
cg05627826
cg05628049
cg05628160
cg05629015
cg05629278
cg05629317
cg05630016
cg05631737
cg05632319
cg05632420
cg05633391
cg05633523
cg05633605
cg05634149
cg05634376
cg05634428
cg05634879
cg05635169
cg05635807
cg05636843
cg05638174
cg05638299
cg05638426
cg05638787
cg05639405
cg05640128
cg05640322
cg05640361

cg05641278
cg05641348
cg05641529
cg05641656
cg05641843
cg05641882
cg05641913
cg05642143
cg05643339
cg05643453
cg05644049
cg05644143
cg05645398
cg05646686
cg05646765
cg05647720
cg05647859
cg05648002
cg05648010
cg05648510
cg05648582
cg05648614
cg05648914
cg05649009
cg05649108
cg05649126
cg05650171
cg05650628
cg05650911
cg05652480
cg05652528
cg05652533
cg05652547
cg05652569
cg05652817
cg05653707
cg05653714
cg05654304
cg05655534
cg05655555
cg05656180
cg05656442
cg05656486
cg05656688
cg05656812
cg05657090
cg05657096
cg05657427
cg05657717
cg05658173
cg05658236
cg05658487

cg05658512
cg05658811
cg05659097
cg05659486
cg05660179
cg05660197
cg05660355
cg05660670
cg05661149
cg05662884
cg05663031
cg05663262
cg05663341
cg05663558
cg05663573
cg05664421
cg05664833
cg05666607
cg05666713
cg05667197
cg05667348
cg05667379
cg05668422
cg05668674
cg05669244
cg05670004
cg05670193
cg05670290
cg05670885
cg05671018
cg05671350
cg05671702
cg05672554
cg05673079
cg05673293
cg05673756
cg05673966
cg05674036
cg05674150
cg05674444
cg05675249
cg05676400
cg05677062
cg05678332
cg05678658
cg05678749
cg05678758
cg05678976
cg05680111
cg05681072
cg05681076
cg05681827

cg05681981
cg05683808
cg05684266
cg05684371
cg05685056
cg05685435
cg05686323
cg05686497
cg05686990
cg05687174
cg05687660
cg05688523
cg05688618
cg05689413
cg05689582
cg05691054
cg05691158
cg05691871
cg05692585
cg05692746
cg05692819
cg05693204
cg05694245
cg05694250
cg05694290
cg05694621
cg05694921
cg05695327
cg05695925
cg05695937
cg05696136
cg05696969
cg05697785
cg05697835
cg05697849
cg05697866
cg05697976
cg05698082
cg05698763
cg05699921
cg05700079
cg05700339
cg05700789
cg05701765
cg05702161
cg05703009
cg05703230
cg05703659
cg05703996
cg05704093
cg05704496
cg05705492

cg05705496
cg05705813
cg05706117
cg05706652
cg05707236
cg05707655
cg05707788
cg05708497
cg05709437
cg05709458
cg05709468
cg05709770
cg05710060
cg05710142
cg05710997
cg05711047
cg05711235
cg05711284
cg05712073
cg05712125
cg05714155
cg05714673
cg05714729
cg05714732
cg05715138
cg05715532
cg05716166
cg05716671
cg05717794
cg05719164
cg05719181
cg05719392
cg05719830
cg05720403
cg05721485
cg05722439
cg05722504
cg05722931
cg05722981
cg05723179
cg05723552
cg05724110
cg05724871
cg05724902
cg05725666
cg05726239
cg05726274
cg05726661
cg05726758
cg05728166
cg05728754
cg05729490

cg05729745
cg05729836
cg05730269
cg05730975
cg05732750
cg05733104
cg05733135
cg05733181
cg05733463
cg05733497
cg05733703
cg05735180
cg05735384
cg05735547
cg05735782
cg05736424
cg05736768
cg05736847
cg05737004
cg05738152
cg05738403
cg05739379
cg05740157
cg05740325
cg05740582
cg05740775
cg05741102
cg05743346
cg05743885
cg05744064
cg05745134
cg05746500
cg05747390
cg05747555
cg05748163
cg05748238
cg05748937
cg05749559
cg05749642
cg05749717
cg05749728
cg05750047
cg05750542
cg05750788
cg05753486
cg05754035
cg05754435
cg05755017
cg05755408
cg05755497
cg05756029
cg05756220

cg05756489
cg05756492
cg05756622
cg05756754
cg05756933
cg05757448
cg05757757
cg05759718
cg05759948
cg05760402
cg05760765
cg05761882
cg05762488
cg05762671
cg05763228
cg05764238
cg05765440
cg05765605
cg05765761
cg05766107
cg05767015
cg05767306
cg05767421
cg05768044
cg05768427
cg05768620
cg05769086
cg05769169
cg05769349
cg05771369
cg05772104
cg05772125
cg05772917
cg05773108
cg05773194
cg05773312
cg05773958
cg05775492
cg05775517
cg05775542
cg05775746
cg05775980
cg05776053
cg05776138
cg05776861
cg05777583
cg05778080
cg05778278
cg05778424
cg05779088
cg05779441
cg05780139

cg05780228
cg05780294
cg05781649
cg05781785
cg05782292
cg05782888
cg05783449
cg05783585
cg05783854
cg05783915
cg05785102
cg05785510
cg05785598
cg05785839
cg05785932
cg05785947
cg05785988
cg05786341
cg05787035
cg05787038
cg05787063
cg05787556
cg05787951
cg05788138
cg05788176
cg05788927
cg05789250
cg05790038
cg05790960
cg05791136
cg05791142
cg05791173
cg05791506
cg05791946
cg05792650
cg05793094
cg05793288
cg05795849
cg05796178
cg05797224
cg05797623
cg05797732
cg05798059
cg05798429
cg05798720
cg05799169
cg05799811
cg05800321
cg05801329
cg05801639
cg05802990
cg05803237

cg05803631
cg05803790
cg05804037
cg05804220
cg05807420
cg05807849
cg05807885
cg05807991
cg05808004
cg05808226
cg05809481
cg05809668
cg05810177
cg05810536
cg05810883
cg05810886
cg05811904
cg05812804
cg05813532
cg05813673
cg05814312
cg05815255
cg05815275
cg05816398
cg05817709
cg05818382
cg05818394
cg05818501
cg05818605
cg05818824
cg05819223
cg05819416
cg05820066
cg05821464
cg05821789
cg05822335
cg05822532
cg05822633
cg05823493
cg05823605
cg05824050
cg05824218
cg05824809
cg05824973
cg05825121
cg05825420
cg05825459
cg05825596
cg05826295
cg05826823
cg05826944
cg05827190

cg05828605
cg05828925
cg05829548
cg05830220
cg05831242
cg05831315
cg05831517
cg05833067
cg05833610
cg05835105
cg05835265
cg05836943
cg05836951
cg05836965
cg05837855
cg05838101
cg05839235
cg05839533
cg05839818
cg05839875
cg05840707
cg05841659
cg05842175
cg05842815
cg05842855
cg05843312
cg05844827
cg05845217
cg05845592
cg05846337
cg05846798
cg05846851
cg05846868
cg05847835
cg05847960
cg05848579
cg05848785
cg05848884
cg05848894
cg05849612
cg05849676
cg05849857
cg05851730
cg05852008
cg05852231
cg05852523
cg05852625
cg05852824
cg05853503
cg05854601
cg05855039
cg05855208

cg05855490
cg05855618
cg05855917
cg05856677
cg05857999
cg05858136
cg05858290
cg05858607
cg05859777
cg05860195
cg05860220
cg05860371
cg05860566
cg05860723
cg05860978
cg05861255
cg05861879
cg05862007
cg05863502
cg05863862
cg05864011
cg05864261
cg05864326
cg05865548
cg05865615
cg05865660
cg05866737
cg05866854
cg05867975
cg05868469
cg05868547
cg05869126
cg05869438
cg05869491
cg05869699
cg05870072
cg05870398
cg05870586
cg05871694
cg05871756
cg05872306
cg05872506
cg05873604
cg05875410
cg05876623
cg05876738
cg05878103
cg05878558
cg05878630
cg05878786
cg05879075
cg05879419

cg05879810
cg05880330
cg05880829
cg05881566
cg05882699
cg05882745
cg05883269
cg05883690
cg05884023
cg05884032
cg05884053
cg05884214
cg05884394
cg05885462
cg05885688
cg05885936
cg05886087
cg05886272
cg05886537
cg05886546
cg05886789
cg05887270
cg05887421
cg05887821
cg05887869
cg05888037
cg05888433
cg05889171
cg05890019
cg05890243
cg05890785
cg05891548
cg05892024
cg05892146
cg05892213
cg05892419
cg05893029
cg05893353
cg05893614
cg05893709
cg05894566
cg05895353
cg05895403
cg05896295
cg05897210
cg05897809
cg05898092
cg05898320
cg05900530
cg05900538
cg05900648
cg05901279

cg05901447
cg05901613
cg05903444
cg05903736
cg05904135
cg05904411
cg05904716
cg05904795
cg05904901
cg05904966
cg05905568
cg05905844
cg05906769
cg05906777
cg05907034
cg05908241
cg05910407
cg05910415
cg05910443
cg05910850
cg05913238
cg05914582
cg05914894
cg05914981
cg05916059
cg05917460
cg05917537
cg05918312
cg05919105
cg05919202
cg05919360
cg05919685
cg05919907
cg05920182
cg05921260
cg05921316
cg05921324
cg05921699
cg05921889
cg05922429
cg05922437
cg05922563
cg05922911
cg05923101
cg05923105
cg05923519
cg05924485
cg05924489
cg05924543
cg05924652
cg05925497
cg05925894

cg05926722
cg05927190
cg05927274
cg05927742
cg05928053
cg05928342
cg05928873
cg05929019
cg05929069
cg05929192
cg05929272
cg05929882
cg05930173
cg05930881
cg05931231
cg05931439
cg05933037
cg05933789
cg05933904
cg05933997
cg05934012
cg05934125
cg05934276
cg05934984
cg05935075
cg05935086
cg05935961
cg05936441
cg05936471
cg05936516
cg05937687
cg05937737
cg05937818
cg05938624
cg05938633
cg05938958
cg05939089
cg05940452
cg05940966
cg05940984
cg05941025
cg05941778
cg05941840
cg05941904
cg05942022
cg05942111
cg05942128
cg05942459
cg05943077
cg05943254
cg05943554
cg05943836

cg05943860
cg05943996
cg05944004
cg05944137
cg05944260
cg05944477
cg05944661
cg05945059
cg05945290
cg05945665
cg05946513
cg05946604
cg05947505
cg05947761
cg05948101
cg05948411
cg05948940
cg05948962
cg05949397
cg05949660
cg05949889
cg05949913
cg05950509
cg05950570
cg05951275
cg05951296
cg05951817
cg05951860
cg05953243
cg05953316
cg05954780
cg05955251
cg05955385
cg05955436
cg05956058
cg05956076
cg05956518
cg05956679
cg05957546
cg05957567
cg05958351
cg05958816
cg05959464
cg05960932
cg05961294
cg05961809
cg05962106
cg05962710
cg05962781
cg05963085
cg05963179
cg05964600

cg05965863
cg05965892
cg05966371
cg05967403
cg05968145
cg05968268
cg05968822
cg05969150
cg05969306
cg05970021
cg05970116
cg05970311
cg05970874
cg05971789
cg05971934
cg05972303
cg05973132
cg05973424
cg05973555
cg05974067
cg05975881
cg05976166
cg05976231
cg05976247
cg05976481
cg05976571
cg05977002
cg05977053
cg05977669
cg05977671
cg05977776
cg05977845
cg05977992
cg05978254
cg05978306
cg05978707
cg05979118
cg05979170
cg05979264
cg05979400
cg05981033
cg05981335
cg05981785
cg05981862
cg05983315
cg05983363
cg05985066
cg05985135
cg05985767
cg05986538
cg05986623
cg05987823

cg05988548
cg05988605
cg05988698
cg05988970
cg05989248
cg05989533
cg05989693
cg05989862
cg05990805
cg05990988
cg05991092
cg05991685
cg05991902
cg05992400
cg05992726
cg05993643
cg05993778
cg05994094
cg05994320
cg05994690
cg05994974
cg05995361
cg05996512
cg05997414
cg05997730
cg05999010
cg05999364
cg05999604
cg05999628
cg06000883
cg06000951
cg06000994
cg06001519
cg06001734
cg06002197
cg06003388
cg06003656
cg06003986
cg06004033
cg06005098
cg06005891
cg06006071
cg06006623
cg06006907
cg06007282
cg06007645
cg06007761
cg06007966
cg06008356
cg06008724
cg06008803
cg06008912

cg06009031
cg06009422
cg06009497
cg06010020
cg06010032
cg06010588
cg06011441
cg06011964
cg06012215
cg06012483
cg06013494
cg06013769
cg06013833
cg06015142
cg06016823
cg06019726
cg06019763
cg06020207
cg06020529
cg06021552
cg06021880
cg06021990
cg06022561
cg06023841
cg06023994
cg06024039
cg06024215
cg06024411
cg06025456
cg06027691
cg06028605
cg06029095
cg06029700
cg06029905
cg06030549
cg06030689
cg06030846
cg06032349
cg06033321
cg06033488
cg06033721
cg06033789
cg06034133
cg06034527
cg06034548
cg06035708
cg06037693
cg06038342
cg06039392
cg06039417
cg06040206
cg06040998

cg06041466
cg06042148
cg06042156
cg06042608
cg06044004
cg06044468
cg06045968
cg06046580
cg06046739
cg06047587
cg06048102
cg06048153
cg06048750
cg06049781
cg06050495
cg06050964
cg06051581
cg06052090
cg06053098
cg06053347
cg06053959
cg06056415
cg06057528
cg06058073
cg06059483
cg06060135
cg06060191
cg06061002
cg06061238
cg06061257
cg06061492
cg06062984
cg06062998
cg06063171
cg06064098
cg06064525
cg06065125
cg06065800
cg06066180
cg06066676
cg06066697
cg06066700
cg06067884
cg06068392
cg06068991
cg06069154
cg06069616
cg06070002
cg06070324
cg06071334
cg06071476
cg06071479

cg06071746
cg06071824
cg06072717
cg06072743
cg06072965
cg06073041
cg06073141
cg06073348
cg06073417
cg06074801
cg06076277
cg06076323
cg06076610
cg06076901
cg06077003
cg06077224
cg06078469
cg06078747
cg06079495
cg06079509
cg06080026
cg06080169
cg06080250
cg06080300
cg06081147
cg06081869
cg06081951
cg06082318
cg06082596
cg06082804
cg06082883
cg06082897
cg06082906
cg06083001
cg06083330
cg06083483
cg06084312
cg06085038
cg06085743
cg06085994
cg06086198
cg06086389
cg06086990
cg06087554
cg06087826
cg06088427
cg06088619
cg06088858
cg06089468
cg06090362
cg06090660
cg06090684

cg06091013
cg06091804
cg06091934
cg06092084
cg06092310
cg06092815
cg06093152
cg06093525
cg06094113
cg06094150
cg06094515
cg06094588
cg06094615
cg06095065
cg06095752
cg06095802
cg06096184
cg06096677
cg06096805
cg06096913
cg06097077
cg06097370
cg06097580
cg06097994
cg06098474
cg06098530
cg06098963
cg06099303
cg06099431
cg06099459
cg06099697
cg06099971
cg06100056
cg06100161
cg06100618
cg06100684
cg06101180
cg06101324
cg06101576
cg06101661
cg06102242
cg06103377
cg06103412
cg06103654
cg06103657
cg06103746
cg06104877
cg06105555
cg06106063
cg06106126
cg06106653
cg06108323

cg06108360
cg06108383
cg06108459
cg06109293
cg06109939
cg06110701
cg06110802
cg06111486
cg06112171
cg06112560
cg06112654
cg06113001
cg06113354
cg06114363
cg06116009
cg06116236
cg06116704
cg06118122
cg06118546
cg06118609
cg06119575
cg06120122
cg06120359
cg06120945
cg06121192
cg06121193
cg06121808
cg06121816
cg06122635
cg06122697
cg06122825
cg06122871
cg06123346
cg06123584
cg06124285
cg06124390
cg06124793
cg06125080
cg06125461
cg06126713
cg06126721
cg06127313
cg06127316
cg06127541
cg06128123
cg06128161
cg06128198
cg06128608
cg06129712
cg06130072
cg06130599
cg06130714

cg06131211
cg06131879
cg06131936
cg06132283
cg06132502
cg06132505
cg06132727
cg06133061
cg06133145
cg06133771
cg06133876
cg06135008
cg06135139
cg06135285
cg06135425
cg06136185
cg06136199
cg06137032
cg06138526
cg06138825
cg06139239
cg06139341
cg06139513
cg06139908
cg06140051
cg06140698
cg06140900
cg06141025
cg06141561
cg06142537
cg06142621
cg06143087
cg06143522
cg06144311
cg06144361
cg06144469
cg06145336
cg06145862
cg06146331
cg06147361
cg06148330
cg06149159
cg06150430
cg06150772
cg06151165
cg06151243
cg06151964
cg06152215
cg06152845
cg06152945
cg06154313
cg06154531

cg06155229
cg06155341
cg06155641
cg06155710
cg06157334
cg06158627
cg06158650
cg06158985
cg06159181
cg06159296
cg06159340
cg06159486
cg06159548
cg06159562
cg06160682
cg06160973
cg06161299
cg06162185
cg06162589
cg06163274
cg06163371
cg06163425
cg06164440
cg06164660
cg06164804
cg06165395
cg06165706
cg06165761
cg06166785
cg06167664
cg06167719
cg06167937
cg06168204
cg06169597
cg06171830
cg06171971
cg06172732
cg06173461
cg06173470
cg06173481
cg06173509
cg06173663
cg06174704
cg06174858
cg06174882
cg06175008
cg06175907
cg06176471
cg06176608
cg06176750
cg06177278
cg06178310

cg06178383
cg06178563
cg06179179
cg06179765
cg06179971
cg06180389
cg06180630
cg06180846
cg06181187
cg06181703
cg06182018
cg06182099
cg06182121
cg06182261
cg06182338
cg06183244
cg06183287
cg06183338
cg06184396
cg06184647
cg06185246
cg06186332
cg06187080
cg06187770
cg06187947
cg06188545
cg06189133
cg06189266
cg06189394
cg06190082
cg06190461
cg06190732
cg06191091
cg06191348
cg06191536
cg06191898
cg06192497
cg06192592
cg06192895
cg06192961
cg06193328
cg06193359
cg06193578
cg06193628
cg06193838
cg06194808
cg06194960
cg06195525
cg06196379
cg06196689
cg06196755
cg06196801

cg06196828
cg06197482
cg06197751
cg06198312
cg06198384
cg06199346
cg06200092
cg06200607
cg06200634
cg06201672
cg06201717
cg06201861
cg06202228
cg06203649
cg06204040
cg06204665
cg06204735
cg06204745
cg06206174
cg06207120
cg06208288
cg06210070
cg06210548
cg06210630
cg06210989
cg06211243
cg06211724
cg06211837
cg06211872
cg06212175
cg06212745
cg06212759
cg06213004
cg06213047
cg06213060
cg06213360
cg06214253
cg06215457
cg06215908
cg06216103
cg06216428
cg06217071
cg06217769
cg06218344
cg06218523
cg06218932
cg06219204
cg06219337
cg06219408
cg06219695
cg06220112
cg06220604

cg06220618
cg06221087
cg06221222
cg06221449
cg06221570
cg06221609
cg06222206
cg06222397
cg06222937
cg06223120
cg06224510
cg06226384
cg06226386
cg06226567
cg06226703
cg06226724
cg06227019
cg06228507
cg06228648
cg06228763
cg06228828
cg06230736
cg06231962
cg06232205
cg06232807
cg06233018
cg06233202
cg06233376
cg06233731
cg06233930
cg06234584
cg06235224
cg06235642
cg06236140
cg06236276
cg06237220
cg06237487
cg06237697
cg06237774
cg06237826
cg06238661
cg06238921
cg06239616
cg06240200
cg06240209
cg06240450
cg06240652
cg06241300
cg06241689
cg06244306
cg06244927
cg06245037

cg06245338
cg06245427
cg06246950
cg06247406
cg06247666
cg06248560
cg06249753
cg06250288
cg06250720
cg06251590
cg06252160
cg06252290
cg06254214
cg06254356
cg06254768
cg06255004
cg06255609
cg06256007
cg06256856
cg06256858
cg06256880
cg06257371
cg06257798
cg06258261
cg06258549
cg06259125
cg06259881
cg06261937
cg06262098
cg06262436
cg06263495
cg06263939
cg06264450
cg06264653
cg06264743
cg06265809
cg06266613
cg06266973
cg06267084
cg06268320
cg06268398
cg06268478
cg06268632
cg06268694
cg06269115
cg06270661
cg06271087
cg06271165
cg06271561
cg06271720
cg06271850
cg06271970

cg06272038
cg06272089
cg06273376
cg06273514
cg06274159
cg06275635
cg06275813
cg06275853
cg06275859
cg06277277
cg06277657
cg06278714
cg06279535
cg06280807
cg06281016
cg06281297
cg06281850
cg06282446
cg06282802
cg06283052
cg06283478
cg06284816
cg06285307
cg06285333
cg06285340
cg06286962
cg06287140
cg06287303
cg06288198
cg06288251
cg06288570
cg06288696
cg06288788
cg06288900
cg06291448
cg06291594
cg06292709
cg06292891
cg06293108
cg06293787
cg06294279
cg06294416
cg06294766
cg06294803
cg06294976
cg06295928
cg06296169
cg06296331
cg06296503
cg06296759
cg06297194
cg06298205

cg06298480
cg06299427
cg06300137
cg06300880
cg06300961
cg06301178
cg06303168
cg06303936
cg06304097
cg06304190
cg06304259
cg06304401
cg06305891
cg06306006
cg06306198
cg06307567
cg06308022
cg06308109
cg06308323
cg06309053
cg06309788
cg06311422
cg06311804
cg06312072
cg06312789
cg06312813
cg06313507
cg06313718
cg06314761
cg06315208
cg06315606
cg06315974
cg06316886
cg06318013
cg06318062
cg06318478
cg06318624
cg06319120
cg06319762
cg06320418
cg06321045
cg06321954
cg06321962
cg06322518
cg06323290
cg06323332
cg06324192
cg06324373
cg06324507
cg06325540
cg06325565
cg06326092

cg06326425
cg06326690
cg06327732
cg06328750
cg06329574
cg06330289
cg06330431
cg06330558
cg06330587
cg06330621
cg06330901
cg06331300
cg06332304
cg06332964
cg06333800
cg06334737
cg06335597
cg06335867
cg06336590
cg06336926
cg06337342
cg06338421
cg06338550
cg06339090
cg06339171
cg06339629
cg06339924
cg06340987
cg06341397
cg06341701
cg06342072
cg06342954
cg06343745
cg06344474
cg06345856
cg06346138
cg06346954
cg06346974
cg06347896
cg06347943
cg06348377
cg06348773
cg06348826
cg06350196
cg06351425
cg06351682
cg06351981
cg06352249
cg06352807
cg06352873
cg06353345
cg06353486

cg06353655
cg06354193
cg06354392
cg06354398
cg06354543
cg06355870
cg06357743
cg06357908
cg06358108
cg06358171
cg06358191
cg06358218
cg06359056
cg06359132
cg06359632
cg06359703
cg06360427
cg06360465
cg06360543
cg06361405
cg06361667
cg06362052
cg06362313
cg06362543
cg06362985
cg06363129
cg06363417
cg06363968
cg06365303
cg06365535
cg06365984
cg06366789
cg06367354
cg06367655
cg06368093
cg06368184
cg06369118
cg06369322
cg06369407
cg06369573
cg06369992
cg06370528
cg06371014
cg06371583
cg06372223
cg06372539
cg06372607
cg06372967
cg06373721
cg06373817
cg06374165
cg06374325

cg06374384
cg06374472
cg06374610
cg06374814
cg06374962
cg06375062
cg06375628
cg06376520
cg06376558
cg06377177
cg06377825
cg06378491
cg06378617
cg06378990
cg06379368
cg06379532
cg06379754
cg06380199
cg06381350
cg06381803
cg06381864
cg06382344
cg06382546
cg06383124
cg06383163
cg06383401
cg06385000
cg06385079
cg06385202
cg06386634
cg06388418
cg06389252
cg06389697
cg06390363
cg06390651
cg06390790
cg06391932
cg06393558
cg06394247
cg06395634
cg06395652
cg06396527
cg06398181
cg06398251
cg06398335
cg06398556
cg06398636
cg06398881
cg06399302
cg06400065
cg06400334
cg06400428

cg06400857
cg06401021
cg06401414
cg06401999
cg06402315
cg06402330
cg06404526
cg06404572
cg06404695
cg06404832
cg06405510
cg06405563
cg06405820
cg06407371
cg06408053
cg06409741
cg06410104
cg06410273
cg06410510
cg06410537
cg06411137
cg06411724
cg06411879
cg06412323
cg06413087
cg06413398
cg06415906
cg06416034
cg06417385
cg06417460
cg06417570
cg06418867
cg06419750
cg06419793
cg06420129
cg06421526
cg06422309
cg06422351
cg06422467
cg06422694
cg06423822
cg06425919
cg06426416
cg06427176
cg06427571
cg06427759
cg06428196
cg06428797
cg06429466
cg06430061
cg06430772
cg06431347

cg06431864
cg06432875
cg06433023
cg06433350
cg06433694
cg06433879
cg06434451
cg06434874
cg06436032
cg06436658
cg06436905
cg06437052
cg06437100
cg06437799
cg06438559
cg06438797
cg06442148
cg06444294
cg06444515
cg06444755
cg06444891
cg06446432
cg06446668
cg06446824
cg06447424
cg06448332
cg06448705
cg06449084
cg06450373
cg06450634
cg06451479
cg06451745
cg06451822
cg06451957
cg06452558
cg06452779
cg06453241
cg06453916
cg06454226
cg06454760
cg06454901
cg06454982
cg06455149
cg06455180
cg06456255
cg06456365
cg06457317
cg06457408
cg06457534
cg06458171
cg06458258
cg06458358

cg06459070
cg06459926
cg06460568
cg06460587
cg06460691
cg06462220
cg06462543
cg06462966
cg06463958
cg06464239
cg06464775
cg06465011
cg06465175
cg06465371
cg06466348
cg06467040
cg06467504
cg06468347
cg06468626
cg06468695
cg06469345
cg06469455
cg06469726
cg06469895
cg06470558
cg06471073
cg06471596
cg06473181
cg06473288
cg06473675
cg06474179
cg06474661
cg06475633
cg06476438
cg06476685
cg06476693
cg06476964
cg06477444
cg06478085
cg06478699
cg06479604
cg06479607
cg06479628
cg06480353
cg06480998
cg06481122
cg06481517
cg06482428
cg06482498
cg06483505
cg06483559
cg06483698

cg06484100
cg06484146
cg06484232
cg06484432
cg06485032
cg06485101
cg06485139
cg06485308
cg06485447
cg06485603
cg06485787
cg06485892
cg06486622
cg06488187
cg06488256
cg06488388
cg06488443
cg06488536
cg06488655
cg06488678
cg06489077
cg06489289
cg06490225
cg06491451
cg06492076
cg06492706
cg06492708
cg06492796
cg06493289
cg06494356
cg06494872
cg06495233
cg06495310
cg06495961
cg06496261
cg06496272
cg06496344
cg06496484
cg06496546
cg06496666
cg06497051
cg06497408
cg06497736
cg06497848
cg06498267
cg06498400
cg06498964
cg06499647
cg06499778
cg06500090
cg06500337
cg06500476

cg06500714
cg06501716
cg06501721
cg06501790
cg06503824
cg06504162
cg06504636
cg06504820
cg06505273
cg06505619
cg06506007
cg06506470
cg06506523
cg06506868
cg06506966
cg06507579
cg06507590
cg06507678
cg06508005
cg06508552
cg06508623
cg06509828
cg06510438
cg06511049
cg06511330
cg06513974
cg06515144
cg06515562
cg06516124
cg06516769
cg06516820
cg06517984
cg06518708
cg06519434
cg06520081
cg06520300
cg06520331
cg06520508
cg06521145
cg06521247
cg06521646
cg06521827
cg06521852
cg06522679
cg06522681
cg06522772
cg06523022
cg06523151
cg06523224
cg06523516
cg06524479
cg06524846

cg06525019
cg06525127
cg06525651
cg06525987
cg06525993
cg06526693
cg06526872
cg06526997
cg06527050
cg06528306
cg06528575
cg06528737
cg06528773
cg06529149
cg06529756
cg06530319
cg06530443
cg06530558
cg06530903
cg06531034
cg06531870
cg06532574
cg06532856
cg06533042
cg06533196
cg06533244
cg06533314
cg06533798
cg06534427
cg06534464
cg06534553
cg06535308
cg06535498
cg06536378
cg06536469
cg06536806
cg06537372
cg06537575
cg06537894
cg06538141
cg06538709
cg06539006
cg06539091
cg06539116
cg06539391
cg06540811
cg06541093
cg06541733
cg06541857
cg06543101
cg06544405
cg06544882

cg06545033
cg06545268
cg06545761
cg06546193
cg06546469
cg06546806
cg06546834
cg06547490
cg06547715
cg06547849
cg06549863
cg06550462
cg06550704
cg06550984
cg06551464
cg06551540
cg06551905
cg06552160
cg06552356
cg06552810
cg06553312
cg06553736
cg06554120
cg06554342
cg06554535
cg06555227
cg06556290
cg06556929
cg06557618
cg06557644
cg06558014
cg06559575
cg06559864
cg06560494
cg06560754
cg06560955
cg06561329
cg06562014
cg06562246
cg06562283
cg06562406
cg06563066
cg06563089
cg06563441
cg06563471
cg06564388
cg06564837
cg06564900
cg06565615
cg06566239
cg06566615
cg06568438

cg06569229
cg06569861
cg06570476
cg06571075
cg06571387
cg06572031
cg06572420
cg06573378
cg06574769
cg06575065
cg06575425
cg06575723
cg06576318
cg06576557
cg06576765
cg06576873
cg06576920
cg06576940
cg06577205
cg06577223
cg06577270
cg06577876
cg06578111
cg06578276
cg06578342
cg06579354
cg06580419
cg06580689
cg06580966
cg06581175
cg06582411
cg06584224
cg06584494
cg06585141
cg06585566
cg06585690
cg06585708
cg06585820
cg06586713
cg06587435
cg06587521
cg06588645
cg06588841
cg06589239
cg06589885
cg06590513
cg06592027
cg06593118
cg06593258
cg06594404
cg06595154
cg06595670

cg06596221
cg06596503
cg06596654
cg06597413
cg06597431
cg06597895
cg06598091
cg06598836
cg06598987
cg06599169
cg06599543
cg06599677
cg06599949
cg06600429
cg06601266
cg06601666
cg06601774
cg06602545
cg06602847
cg06602871
cg06603074
cg06604568
cg06606539
cg06606753
cg06607594
cg06607679
cg06607764
cg06607919
cg06608359
cg06608414
cg06609094
cg06609220
cg06609489
cg06609496
cg06609866
cg06610125
cg06610259
cg06610484
cg06610705
cg06610849
cg06610876
cg06611426
cg06612196
cg06613095
cg06613263
cg06614002
cg06614284
cg06615699
cg06615816
cg06615840
cg06616178
cg06616710

cg06616729
cg06617217
cg06617456
cg06618097
cg06618134
cg06618377
cg06618666
cg06619066
cg06619428
cg06619621
cg06619906
cg06619939
cg06620117
cg06620200
cg06620972
cg06621094
cg06621425
cg06621620
cg06621799
cg06622108
cg06622151
cg06622892
cg06622994
cg06622999
cg06623630
cg06623778
cg06624146
cg06624154
cg06624527
cg06624951
cg06625077
cg06625244
cg06626232
cg06626591
cg06626750
cg06626786
cg06627009
cg06627334
cg06627916
cg06628473
cg06629130
cg06629170
cg06629644
cg06629999
cg06630204
cg06630241
cg06630413
cg06630455
cg06630988
cg06631531
cg06631625
cg06633061

cg06633739
cg06634179
cg06634596
cg06634716
cg06634914
cg06635952
cg06636427
cg06636678
cg06637017
cg06637330
cg06637812
cg06638723
cg06638795
cg06638874
cg06639267
cg06639544
cg06639585
cg06639701
cg06640364
cg06640822
cg06640997
cg06641153
cg06641342
cg06641415
cg06641537
cg06642397
cg06643622
cg06643951
cg06644428
cg06644457
cg06644638
cg06644719
cg06645921
cg06645987
cg06646548
cg06646780
cg06647782
cg06647928
cg06648156
cg06648277
cg06648455
cg06648812
cg06649201
cg06649410
cg06649540
cg06649856
cg06650250
cg06651311
cg06651452
cg06651752
cg06652085
cg06652329

cg06652392
cg06652849
cg06653632
cg06653699
cg06654079
cg06654510
cg06654537
cg06655097
cg06655623
cg06655861
cg06656924
cg06656958
cg06658129
cg06658147
cg06658615
cg06659338
cg06659727
cg06660057
cg06660160
cg06660395
cg06660523
cg06661266
cg06661456
cg06662812
cg06663251
cg06663317
cg06664085
cg06664254
cg06664357
cg06664465
cg06664486
cg06665109
cg06666025
cg06666466
cg06667761
cg06668194
cg06668555
cg06669056
cg06669752
cg06669759
cg06671450
cg06671962
cg06672160
cg06672272
cg06672560
cg06674117
cg06674731
cg06675190
cg06675348
cg06675483
cg06675705
cg06676119

cg06676637
cg06678468
cg06678594
cg06679087
cg06679209
cg06679262
cg06679347
cg06679878
cg06679990
cg06680397
cg06680481
cg06680863
cg06681768
cg06681845
cg06682185
cg06683078
cg06683094
cg06683148
cg06683602
cg06684321
cg06684850
cg06685737
cg06685949
cg06686679
cg06687489
cg06687772
cg06688910
cg06688989
cg06689619
cg06689659
cg06689816
cg06690085
cg06690689
cg06690831
cg06691250
cg06691616
cg06691963
cg06694239
cg06694734
cg06695115
cg06695264
cg06695691
cg06696905
cg06697267
cg06697448
cg06697536
cg06697600
cg06698638
cg06698843
cg06698844
cg06698926
cg06699422

cg06699564
cg06700060
cg06700209
cg06700275
cg06700606
cg06701529
cg06702216
cg06702600
cg06703343
cg06704093
cg06704122
cg06704567
cg06704573
cg06704631
cg06704884
cg06705122
cg06705237
cg06705548
cg06705767
cg06705879
cg06705925
cg06705986
cg06705997
cg06706068
cg06706170
cg06707356
cg06707663
cg06708133
cg06708634
cg06708731
cg06710195
cg06710785
cg06711831
cg06712340
cg06712541
cg06712651
cg06712886
cg06713076
cg06713098
cg06713550
cg06713632
cg06714088
cg06714480
cg06714655
cg06715135
cg06716508
cg06716698
cg06719900
cg06719949
cg06720467
cg06720722
cg06721231

cg06721255
cg06721538
cg06722121
cg06722212
cg06722630
cg06724078
cg06724305
cg06724588
cg06724937
cg06725798
cg06726231
cg06727065
cg06727242
cg06728135
cg06728232
cg06728793
cg06729381
cg06729555
cg06730161
cg06730183
cg06730250
cg06730285
cg06731886
cg06732684
cg06732687
cg06732873
cg06733602
cg06734406
cg06734812
cg06734985
cg06735127
cg06735218
cg06735243
cg06736148
cg06737561
cg06737912
cg06737981
cg06738040
cg06738169
cg06738242
cg06738356
cg06738548
cg06738602
cg06738786
cg06739029
cg06739256
cg06739303
cg06740209
cg06740442
cg06740578
cg06740950
cg06740985

cg06741096
cg06741594
cg06741996
cg06742678
cg06742778
cg06743703
cg06745001
cg06745684
cg06745740
cg06745941
cg06746118
cg06746365
cg06746829
cg06747288
cg06747919
cg06748060
cg06748147
cg06748440
cg06748470
cg06748831
cg06748978
cg06749715
cg06749872
cg06750076
cg06751243
cg06751398
cg06751920
cg06752040
cg06752260
cg06752698
cg06753273
cg06753367
cg06753398
cg06753513
cg06753787
cg06754496
cg06754606
cg06754764
cg06754987
cg06755263
cg06755296
cg06755427
cg06755438
cg06755825
cg06756164
cg06757356
cg06757810
cg06758211
cg06758255
cg06758605
cg06758644
cg06759255

cg06759309
cg06759314
cg06759518
cg06759632
cg06759890
cg06760280
cg06760467
cg06760710
cg06760830
cg06763054
cg06764092
cg06764302
cg06764842
cg06765022
cg06765172
cg06765217
cg06765324
cg06765475
cg06766427
cg06767314
cg06767655
cg06768399
cg06768939
cg06769296
cg06769546
cg06770082
cg06770683
cg06770735
cg06770743
cg06771056
cg06771126
cg06771839
cg06772063
cg06772202
cg06772309
cg06772654
cg06772824
cg06773306
cg06773813
cg06773873
cg06774445
cg06774893
cg06776146
cg06776159
cg06776422
cg06776548
cg06776709
cg06776999
cg06777202
cg06777750
cg06777844
cg06779113

cg06779253
cg06779847
cg06780726
cg06781135
cg06781712
cg06782035
cg06782204
cg06782419
cg06783197
cg06784466
cg06785293
cg06785999
cg06786238
cg06786349
cg06786860
cg06787206
cg06787316
cg06787422
cg06787519
cg06787622
cg06787724
cg06788355
cg06788631
cg06789840
cg06790044
cg06791107
cg06791592
cg06792417
cg06792538
cg06793062
cg06793562
cg06793849
cg06794038
cg06794141
cg06794244
cg06794355
cg06796220
cg06796271
cg06796611
cg06797253
cg06797644
cg06798288
cg06798483
cg06798934
cg06799321
cg06799735
cg06800231
cg06800942
cg06801483
cg06801852
cg06801943
cg06802658

cg06803853
cg06804312
cg06804344
cg06804564
cg06804625
cg06804823
cg06805280
cg06805615
cg06805925
cg06806210
cg06806862
cg06806958
cg06807030
cg06807168
cg06807568
cg06807650
cg06807905
cg06807915
cg06807926
cg06808751
cg06809153
cg06809252
cg06809544
cg06810647
cg06811094
cg06811295
cg06811571
cg06811607
cg06811800
cg06812574
cg06812760
cg06812795
cg06812836
cg06812844
cg06812861
cg06812991
cg06813100
cg06813515
cg06813862
cg06814287
cg06814792
cg06814892
cg06815003
cg06815411
cg06815715
cg06815826
cg06815910
cg06815950
cg06816235
cg06816239
cg06817126
cg06817264

cg06817368
cg06817454
cg06817455
cg06817516
cg06817864
cg06818159
cg06819298
cg06819763
cg06820388
cg06820545
cg06820905
cg06821009
cg06821079
cg06821999
cg06822067
cg06822193
cg06822816
cg06822943
cg06823681
cg06824294
cg06824308
cg06824423
cg06824581
cg06824795
cg06825039
cg06825166
cg06825415
cg06825448
cg06825878
cg06826289
cg06826760
cg06827562
cg06828043
cg06829299
cg06829830
cg06829852
cg06830006
cg06830148
cg06831576
cg06832406
cg06832898
cg06833110
cg06833564
cg06833778
cg06833823
cg06834005
cg06834689
cg06834912
cg06834916
cg06836725
cg06837040
cg06837325

cg06837799
cg06838038
cg06838584
cg06838985
cg06839417
cg06839631
cg06839854
cg06841648
cg06841902
cg06842181
cg06842913
cg06842954
cg06844159
cg06844661
cg06845268
cg06846214
cg06846495
cg06846898
cg06847670
cg06847828
cg06848060
cg06848148
cg06848199
cg06848711
cg06849501
cg06849504
cg06850005
cg06850158
cg06850241
cg06850283
cg06850467
cg06850597
cg06850651
cg06850996
cg06851000
cg06851240
cg06851817
cg06852058
cg06852240
cg06852309
cg06852395
cg06852461
cg06853339
cg06853836
cg06854015
cg06854264
cg06854772
cg06854842
cg06855137
cg06855337
cg06855422
cg06856275

cg06856378
cg06856720
cg06856984
cg06857614
cg06857674
cg06857900
cg06858263
cg06858559
cg06858835
cg06859135
cg06859970
cg06860600
cg06861011
cg06861037
cg06861044
cg06861519
cg06861560
cg06861736
cg06863239
cg06864047
cg06865036
cg06865119
cg06865363
cg06866232
cg06866451
cg06866553
cg06866597
cg06866756
cg06867285
cg06867377
cg06869505
cg06869641
cg06870073
cg06870213
cg06871582
cg06872010
cg06872172
cg06872185
cg06872313
cg06872548
cg06872705
cg06873024
cg06873316
cg06873343
cg06873452
cg06873880
cg06874144
cg06874172
cg06874656
cg06875131
cg06875227
cg06875305

cg06875863
cg06875985
cg06876892
cg06877024
cg06877247
cg06878108
cg06878663
cg06878709
cg06879746
cg06879786
cg06880420
cg06880612
cg06880735
cg06880930
cg06880963
cg06881239
cg06881421
cg06882533
cg06882571
cg06883126
cg06883496
cg06883956
cg06884314
cg06884401
cg06884875
cg06885910
cg06886827
cg06887407
cg06887580
cg06888387
cg06888746
cg06888836
cg06889756
cg06890522
cg06890553
cg06891043
cg06893379
cg06893389
cg06893491
cg06893560
cg06894134
cg06894334
cg06894469
cg06894710
cg06894891
cg06895675
cg06895831
cg06895849
cg06895913
cg06895946
cg06896987
cg06897125

cg06897927
cg06898168
cg06898293
cg06899329
cg06899483
cg06899985
cg06900253
cg06900494
cg06900514
cg06900843
cg06901238
cg06901893
cg06902099
cg06902367
cg06902368
cg06902607
cg06902665
cg06903103
cg06903656
cg06903917
cg06904667
cg06905511
cg06906472
cg06906965
cg06907391
cg06907770
cg06908450
cg06908550
cg06909503
cg06909630
cg06909773
cg06911121
cg06911974
cg06912252
cg06912272
cg06912403
cg06912601
cg06912814
cg06912936
cg06913501
cg06914270
cg06915193
cg06915343
cg06915667
cg06915773
cg06916059
cg06916740
cg06917325
cg06917727
cg06917763
cg06917772
cg06918474

cg06920037
cg06923861
cg06924459
cg06924602
cg06924873
cg06924878
cg06925502
cg06925855
cg06926782
cg06926934
cg06927305
cg06927807
cg06928209
cg06928695
cg06928741
cg06928952
cg06929324
cg06930433
cg06930722
cg06931523
cg06931933
cg06933777
cg06934265
cg06934572
cg06934747
cg06934774
cg06934867
cg06935336
cg06935438
cg06935979
cg06936155
cg06937409
cg06937617
cg06937927
cg06938264
cg06938699
cg06939447
cg06939451
cg06939970
cg06940127
cg06940482
cg06940505
cg06940688
cg06940792
cg06941230
cg06941630
cg06941638
cg06942183
cg06942450
cg06942701
cg06943309
cg06943420

cg06943561
cg06943627
cg06943845
cg06944351
cg06944559
cg06944758
cg06945563
cg06945807
cg06946543
cg06946837
cg06946850
cg06947608
cg06948658
cg06948937
cg06949673
cg06949778
cg06950346
cg06951565
cg06951626
cg06951647
cg06952310
cg06952412
cg06952681
cg06952873
cg06953454
cg06953773
cg06954368
cg06954481
cg06954520
cg06955484
cg06955716
cg06955749
cg06956052
cg06956295
cg06956444
cg06956786
cg06957003
cg06957290
cg06957447
cg06958184
cg06958563
cg06959251
cg06959664
cg06960461
cg06960700
cg06960784
cg06961354
cg06961429
cg06961439
cg06962177
cg06962748
cg06962867

cg06964353
cg06964738
cg06964899
cg06966113
cg06966245
cg06966613
cg06966811
cg06967304
cg06967998
cg06968241
cg06968674
cg06968788
cg06968859
cg06968926
cg06969622
cg06970472
cg06970744
cg06970772
cg06971092
cg06971129
cg06971773
cg06971903
cg06972043
cg06972213
cg06972565
cg06972578
cg06973225
cg06974428
cg06974662
cg06974672
cg06975048
cg06975166
cg06975633
cg06975914
cg06976250
cg06977689
cg06977872
cg06978036
cg06978514
cg06978604
cg06979108
cg06979726
cg06979771
cg06981309
cg06981439
cg06981541
cg06983322
cg06984025
cg06984201
cg06985259
cg06987369
cg06987468

cg06988336
cg06988547
cg06989930
cg06990194
cg06991300
cg06992846
cg06993099
cg06993118
cg06993703
cg06994420
cg06994502
cg06994652
cg06994787
cg06994813
cg06997226
cg06997259
cg06997707
cg06997767
cg06998224
cg06998765
cg06999127
cg06999346
cg06999858
cg07000643
cg07000686
cg07001072
cg07001421
cg07001508
cg07001576
cg07002061
cg07002201
cg07002403
cg07002498
cg07003030
cg07003542
cg07003900
cg07004925
cg07004950
cg07005180
cg07005444
cg07005506
cg07005654
cg07005778
cg07006091
cg07006818
cg07006935
cg07007165
cg07007400
cg07008213
cg07008478
cg07008926
cg07009376

cg07010373
cg07010486
cg07010687
cg07011093
cg07011304
cg07012128
cg07012178
cg07012926
cg07013734
cg07014126
cg07014174
cg07014673
cg07015221
cg07015663
cg07015784
cg07015886
cg07016184
cg07016730
cg07017114
cg07017135
cg07017153
cg07017597
cg07018580
cg07018648
cg07019438
cg07019638
cg07019697
cg07019857
cg07020001
cg07020596
cg07020690
cg07021181
cg07021275
cg07022343
cg07022477
cg07022764
cg07022788
cg07024439
cg07025157
cg07025989
cg07026107
cg07026490
cg07026904
cg07027430
cg07027613
cg07027713
cg07028533
cg07028914
cg07028950
cg07029084
cg07029873
cg07030277

cg07030704
cg07032359
cg07032439
cg07032866
cg07033372
cg07033395
cg07033513
cg07033624
cg07034362
cg07035165
cg07035173
cg07035320
cg07035442
cg07035503
cg07035659
cg07035704
cg07035875
cg07035961
cg07036530
cg07036991
cg07037251
cg07037371
cg07037509
cg07038423
cg07039113
cg07039130
cg07039362
cg07040894
cg07041214
cg07042014
cg07042729
cg07043247
cg07044016
cg07044458
cg07045241
cg07045460
cg07045816
cg07045857
cg07046101
cg07047068
cg07047471
cg07047731
cg07047963
cg07048415
cg07048516
cg07048693
cg07048909
cg07049421
cg07050011
cg07050869
cg07051721
cg07051786

cg07051905
cg07052004
cg07052737
cg07053699
cg07053841
cg07054804
cg07055616
cg07055769
cg07055879
cg07057235
cg07057320
cg07057532
cg07057579
cg07058918
cg07058988
cg07059469
cg07059629
cg07060383
cg07060551
cg07061260
cg07061298
cg07061566
cg07062043
cg07062658
cg07062764
cg07063068
cg07063171
cg07063351
cg07063357
cg07063912
cg07064495
cg07064537
cg07064720
cg07065111
cg07065666
cg07067171
cg07067952
cg07068327
cg07068570
cg07068655
cg07068674
cg07068735
cg07068761
cg07068768
cg07069342
cg07069742
cg07069759
cg07070138
cg07070518
cg07071449
cg07071764
cg07071839

cg07072366
cg07072405
cg07072463
cg07072722
cg07073964
cg07074571
cg07075085
cg07075268
cg07076175
cg07076850
cg07076915
cg07077013
cg07077735
cg07078225
cg07078479
cg07079663
cg07080050
cg07080867
cg07081245
cg07081759
cg07082447
cg07082452
cg07083101
cg07084627
cg07084941
cg07085827
cg07086179
cg07086295
cg07086945
cg07087457
cg07087779
cg07088414
cg07088771
cg07089399
cg07090083
cg07090356
cg07090539
cg07090569
cg07090653
cg07090701
cg07090714
cg07090813
cg07091016
cg07091102
cg07091220
cg07091412
cg07091481
cg07092029
cg07093155
cg07093324
cg07093570
cg07094075

cg07094915
cg07095637
cg07095908
cg07095945
cg07095995
cg07096110
cg07096238
cg07097098
cg07097184
cg07098391
cg07099073
cg07099331
cg07099991
cg07100000
cg07100448
cg07102368
cg07102705
cg07103129
cg07103201
cg07103493
cg07103673
cg07104135
cg07104652
cg07104706
cg07104860
cg07105285
cg07105499
cg07106761
cg07106766
cg07107453
cg07107756
cg07108581
cg07109115
cg07109438
cg07109743
cg07109801
cg07110103
cg07110129
cg07110232
cg07110949
cg07111415
cg07112337
cg07112604
cg07113006
cg07113199
cg07114886
cg07115542
cg07115820
cg07116712
cg07116919
cg07116972
cg07117311

cg07117751
cg07118491
cg07119001
cg07119076
cg07119829
cg07119973
cg07120346
cg07120544
cg07121421
cg07121517
cg07121795
cg07121856
cg07122006
cg07122123
cg07122529
cg07122905
cg07122986
cg07123038
cg07123069
cg07123548
cg07124183
cg07124366
cg07125166
cg07125243
cg07125269
cg07125967
cg07125976
cg07126216
cg07126399
cg07126525
cg07126783
cg07127357
cg07128021
cg07128624
cg07128660
cg07129067
cg07129392
cg07129523
cg07129879
cg07129924
cg07131336
cg07132005
cg07132038
cg07132068
cg07132650
cg07133099
cg07133760
cg07133994
cg07134252
cg07134316
cg07134930
cg07135032

cg07135303
cg07135446
cg07135614
cg07135664
cg07135732
cg07136029
cg07136325
cg07136920
cg07136998
cg07137429
cg07138070
cg07138305
cg07139165
cg07139284
cg07139495
cg07139509
cg07139514
cg07139762
cg07141002
cg07141504
cg07141605
cg07141875
cg07142009
cg07142010
cg07142723
cg07143083
cg07143418
cg07143479
cg07143532
cg07143559
cg07143635
cg07143733
cg07143805
cg07143863
cg07144328
cg07145284
cg07145531
cg07146073
cg07146912
cg07147166
cg07147449
cg07147846
cg07147958
cg07148145
cg07148167
cg07148197
cg07148207
cg07148296
cg07148697
cg07148716
cg07148968
cg07149296

cg07149609
cg07150038
cg07150045
cg07150062
cg07151155
cg07151572
cg07152216
cg07152355
cg07152520
cg07152755
cg07152925
cg07154254
cg07156484
cg07156733
cg07157117
cg07157169
cg07157248
cg07157333
cg07157830
cg07158040
cg07158816
cg07159023
cg07159228
cg07159230
cg07159673
cg07160746
cg07160783
cg07160871
cg07161369
cg07161429
cg07161721
cg07162000
cg07162198
cg07162571
cg07162608
cg07162700
cg07163480
cg07164556
cg07164791
cg07165029
cg07165260
cg07166084
cg07166450
cg07166550
cg07166654
cg07167114
cg07167168
cg07167396
cg07167933
cg07168060
cg07168810
cg07168939

cg07169046
cg07169637
cg07169826
cg07170183
cg07171496
cg07171504
cg07172334
cg07172701
cg07173275
cg07173635
cg07173760
cg07174453
cg07175433
cg07175582
cg07175797
cg07176264
cg07177379
cg07177394
cg07177437
cg07177867
cg07177917
cg07178066
cg07178143
cg07178206
cg07178462
cg07178864
cg07178896
cg07179016
cg07179068
cg07179267
cg07179329
cg07179816
cg07180212
cg07180307
cg07180988
cg07181565
cg07181702
cg07182163
cg07182756
cg07183218
cg07183372
cg07184228
cg07184465
cg07184578
cg07184986
cg07185131
cg07185287
cg07185587
cg07185664
cg07185695
cg07186154
cg07186199

cg07187268
cg07187521
cg07188009
cg07188665
cg07189782
cg07190535
cg07190914
cg07190947
cg07191524
cg07191791
cg07192734
cg07193553
cg07193766
cg07194093
cg07194551
cg07194993
cg07195011
cg07196505
cg07196761
cg07197493
cg07197785
cg07197831
cg07198150
cg07198552
cg07200386
cg07200490
cg07200877
cg07200939
cg07201456
cg07201620
cg07201717
cg07202413
cg07202610
cg07203024
cg07203151
cg07203423
cg07203561
cg07203767
cg07203893
cg07204280
cg07204353
cg07204550
cg07204800
cg07205154
cg07205589
cg07205823
cg07206053
cg07206208
cg07206442
cg07206630
cg07207220
cg07208057

cg07208691
cg07209298
cg07209550
cg07209634
cg07209977
cg07210454
cg07210840
cg07211472
cg07211768
cg07212778
cg07212852
cg07213145
cg07213685
cg07213830
cg07214954
cg07215697
cg07215975
cg07216036
cg07216436
cg07217653
cg07217710
cg07217935
cg07219425
cg07219649
cg07219769
cg07220152
cg07220448
cg07220650
cg07220782
cg07220903
cg07221438
cg07221695
cg07222133
cg07222243
cg07222575
cg07222721
cg07222863
cg07223266
cg07223990
cg07224067
cg07224114
cg07225829
cg07226484
cg07226755
cg07226964
cg07228702
cg07228905
cg07229186
cg07230107
cg07230786
cg07231911
cg07232063

cg07232577
cg07232618
cg07232823
cg07234097
cg07235577
cg07235935
cg07236001
cg07236127
cg07236594
cg07236781
cg07236958
cg07237397
cg07237939
cg07238926
cg07239452
cg07239716
cg07240413
cg07240557
cg07241084
cg07241090
cg07241908
cg07242280
cg07243138
cg07243202
cg07244076
cg07244716
cg07245558
cg07246273
cg07247405
cg07247778
cg07247810
cg07249459
cg07249730
cg07249888
cg07249939
cg07250222
cg07250462
cg07251046
cg07251128
cg07252363
cg07252539
cg07252933
cg07253384
cg07253500
cg07253712
cg07253829
cg07254032
cg07254054
cg07254706
cg07256797
cg07257571
cg07258695

cg07259382
cg07259660
cg07259733
cg07259811
cg07260017
cg07260541
cg07260927
cg07261186
cg07262519
cg07262873
cg07263322
cg07263475
cg07263759
cg07264329
cg07264679
cg07265310
cg07265444
cg07265976
cg07266046
cg07267600
cg07267845
cg07268734
cg07269374
cg07270078
cg07270259
cg07270423
cg07271264
cg07271383
cg07271580
cg07272353
cg07273222
cg07273342
cg07273992
cg07274137
cg07274590
cg07274618
cg07274716
cg07275218
cg07275552
cg07276118
cg07276388
cg07276708
cg07276831
cg07277549
cg07277864
cg07277892
cg07277923
cg07278047
cg07278425
cg07279070
cg07279157
cg07279321

cg07279752
cg07279955
cg07280097
cg07280182
cg07280985
cg07281879
cg07283011
cg07283114
cg07283441
cg07285276
cg07285760
cg07287078
cg07287698
cg07288031
cg07289618
cg07290257
cg07290522
cg07291100
cg07291439
cg07292140
cg07293406
cg07294406
cg07294734
cg07295362
cg07295669
cg07295678
cg07295904
cg07295964
cg07296256
cg07296597
cg07296772
cg07297777
cg07298347
cg07298363
cg07298431
cg07298473
cg07300060
cg07300178
cg07300384
cg07300559
cg07301242
cg07301829
cg07303244
cg07303577
cg07303829
cg07303912
cg07304173
cg07304923
cg07305514
cg07305933
cg07306737
cg07307332

cg07307830
cg07308381
cg07308384
cg07309102
cg07310916
cg07311566
cg07311759
cg07312601
cg07313162
cg07313835
cg07315745
cg07316313
cg07316730
cg07316846
cg07316961
cg07317449
cg07317551
cg07318313
cg07318627
cg07318808
cg07318983
cg07319315
cg07319363
cg07319614
cg07319712
cg07322064
cg07322117
cg07322898
cg07322909
cg07323399
cg07323648
cg07323825
cg07323919
cg07323996
cg07324172
cg07325168
cg07325820
cg07326149
cg07326586
cg07326765
cg07327299
cg07327347
cg07327435
cg07328519
cg07328664
cg07328796
cg07329360
cg07329987
cg07330114
cg07330438
cg07330489
cg07331204

cg07332757
cg07333608
cg07333715
cg07333792
cg07336123
cg07336350
cg07336617
cg07337250
cg07337385
cg07337434
cg07337480
cg07337796
cg07339662
cg07339783
cg07339864
cg07340025
cg07340651
cg07341624
cg07342322
cg07342559
cg07342783
cg07342901
cg07343607
cg07343821
cg07344086
cg07344338
cg07345016
cg07345205
cg07345515
cg07345521
cg07345734
cg07345999
cg07346187
cg07347431
cg07348263
cg07348300
cg07348418
cg07348535
cg07349208
cg07349217
cg07349899
cg07350016
cg07350076
cg07350501
cg07352127
cg07352416
cg07352438
cg07352834
cg07353489
cg07355763
cg07355841
cg07356486

cg07356667
cg07356745
cg07356876
cg07357300
cg07358634
cg07359966
cg07360028
cg07360304
cg07360792
cg07361154
cg07361858
cg07362512
cg07362800
cg07362969
cg07363855
cg07364433
cg07364646
cg07364657
cg07365127
cg07366082
cg07366967
cg07367469
cg07368061
cg07369190
cg07369374
cg07369950
cg07369988
cg07370352
cg07370588
cg07370745
cg07371413
cg07371628
cg07371747
cg07371757
cg07372450
cg07372602
cg07373578
cg07373723
cg07373798
cg07374109
cg07375851
cg07376029
cg07376067
cg07376374
cg07376535
cg07376547
cg07376916
cg07376932
cg07377098
cg07377172
cg07377422
cg07377446

cg07377519
cg07377994
cg07378067
cg07378577
cg07379055
cg07379703
cg07380026
cg07380200
cg07380705
cg07381283
cg07382039
cg07382454
cg07382554
cg07382687
cg07382920
cg07382998
cg07383092
cg07383941
cg07385257
cg07385573
cg07385778
cg07386008
cg07386299
cg07386859
cg07390459
cg07390598
cg07392307
cg07392324
cg07392385
cg07392838
cg07393352
cg07393734
cg07393736
cg07394719
cg07394914
cg07395354
cg07395804
cg07397493
cg07397956
cg07398352
cg07398661
cg07398791
cg07398952
cg07399395
cg07399928
cg07400592
cg07401516
cg07402310
cg07402669
cg07403152
cg07403610
cg07403899

cg07404485
cg07404527
cg07404961
cg07405330
cg07408232
cg07408552
cg07408835
cg07408989
cg07409153
cg07409200
cg07410173
cg07410872
cg07411432
cg07411620
cg07412281
cg07413251
cg07414574
cg07415359
cg07416656
cg07416733
cg07416844
cg07417020
cg07417146
cg07418387
cg07418452
cg07418892
cg07419011
cg07419795
cg07419921
cg07420362
cg07420416
cg07420827
cg07421287
cg07421515
cg07422329
cg07422412
cg07422735
cg07422880
cg07423050
cg07423765
cg07423997
cg07424400
cg07425204
cg07425597
cg07426472
cg07426634
cg07427606
cg07427642
cg07428004
cg07428782
cg07428959
cg07429087

cg07429192
cg07429284
cg07429804
cg07430136
cg07430472
cg07431522
cg07431944
cg07431973
cg07432626
cg07433178
cg07433663
cg07433769
cg07434253
cg07434271
cg07434402
cg07435275
cg07435331
cg07435894
cg07436074
cg07436208
cg07436418
cg07436562
cg07436701
cg07436820
cg07436991
cg07437292
cg07438099
cg07438246
cg07438401
cg07439975
cg07440264
cg07440398
cg07440877
cg07440958
cg07441143
cg07441263
cg07441278
cg07441319
cg07441518
cg07441659
cg07441854
cg07442409
cg07442479
cg07443710
cg07443717
cg07443748
cg07444323
cg07444408
cg07444963
cg07445258
cg07445838
cg07446767

cg07447769
cg07447870
cg07447922
cg07448060
cg07448928
cg07448997
cg07449450
cg07450030
cg07450086
cg07450292
cg07450805
cg07451261
cg07452879
cg07453440
cg07453451
cg07453773
cg07454154
cg07454558
cg07455504
cg07455789
cg07456081
cg07456114
cg07456376
cg07456480
cg07456815
cg07457213
cg07457252
cg07457727
cg07457967
cg07458272
cg07458308
cg07458482
cg07458509
cg07459181
cg07460294
cg07460524
cg07462112
cg07462292
cg07462540
cg07462756
cg07462808
cg07462863
cg07463197
cg07463541
cg07464094
cg07464408
cg07464423
cg07464450
cg07464578
cg07465028
cg07465344
cg07466576

cg07466622
cg07466705
cg07467338
cg07467618
cg07467861
cg07467919
cg07468397
cg07468585
cg07468710
cg07468782
cg07469215
cg07469402
cg07469647
cg07469700
cg07469887
cg07470598
cg07470639
cg07471156
cg07471572
cg07471943
cg07471948
cg07472097
cg07472708
cg07472953
cg07473138
cg07473985
cg07474266
cg07474269
cg07474682
cg07474797
cg07476030
cg07476963
cg07477792
cg07478107
cg07478467
cg07479253
cg07479337
cg07479670
cg07479711
cg07480567
cg07480608
cg07480640
cg07480647
cg07481928
cg07483304
cg07483638
cg07483709
cg07484827
cg07485423
cg07486017
cg07486085
cg07486732

cg07486821
cg07488417
cg07488634
cg07488872
cg07489048
cg07489447
cg07489502
cg07489890
cg07490719
cg07491676
cg07492590
cg07492757
cg07493097
cg07493197
cg07493760
cg07493858
cg07494259
cg07494300
cg07495363
cg07495470
cg07495574
cg07496078
cg07496457
cg07498415
cg07499142
cg07499553
cg07500242
cg07500957
cg07501748
cg07501827
cg07502050
cg07502358
cg07502829
cg07503211
cg07504598
cg07505018
cg07505478
cg07506573
cg07507085
cg07507251
cg07507493
cg07507518
cg07507755
cg07508480
cg07509323
cg07510323
cg07510333
cg07510346
cg07510423
cg07510510
cg07511098
cg07512206

cg07512258
cg07512295
cg07512517
cg07512997
cg07513323
cg07513791
cg07513814
cg07513950
cg07514372
cg07515989
cg07516450
cg07516556
cg07516956
cg07517739
cg07517893
cg07518851
cg07519049
cg07519229
cg07519485
cg07519873
cg07520506
cg07520673
cg07520810
cg07521929
cg07522372
cg07522416
cg07522661
cg07523189
cg07523741
cg07523753
cg07524000
cg07524456
cg07524827
cg07524873
cg07525003
cg07525077
cg07525202
cg07525420
cg07525782
cg07525886
cg07526408
cg07526904
cg07527126
cg07527685
cg07529087
cg07529210
cg07529575
cg07529750
cg07529754
cg07531056
cg07531228
cg07531663

cg07532354
cg07532405
cg07533148
cg07534823
cg07535316
cg07535366
cg07535928
cg07536847
cg07536910
cg07537443
cg07537692
cg07537734
cg07537917
cg07538204
cg07539244
cg07539443
cg07539709
cg07539983
cg07540103
cg07540326
cg07541020
cg07541160
cg07541559
cg07542000
cg07542463
cg07542476
cg07543138
cg07543199
cg07543626
cg07543967
cg07544796
cg07545037
cg07545052
cg07545278
cg07545427
cg07545457
cg07545858
cg07546334
cg07546433
cg07546508
cg07546654
cg07547000
cg07547054
cg07547549
cg07547788
cg07548367
cg07549278
cg07549297
cg07549406
cg07551230
cg07552803
cg07554053

cg07554626
cg07554775
cg07554831
cg07554986
cg07555272
cg07555731
cg07556599
cg07556911
cg07557173
cg07557260
cg07557560
cg07558455
cg07558704
cg07559273
cg07559992
cg07560948
cg07561338
cg07562039
cg07562451
cg07562926
cg07563050
cg07564837
cg07565499
cg07565505
cg07565551
cg07565956
cg07566050
cg07566080
cg07566169
cg07566286
cg07566463
cg07567256
cg07568203
cg07568644
cg07569216
cg07569288
cg07570113
cg07570257
cg07570298
cg07570394
cg07570537
cg07570573
cg07570723
cg07570898
cg07571344
cg07571451
cg07571509
cg07572028
cg07572341
cg07572773
cg07573342
cg07573366

cg07573727
cg07573872
cg07575193
cg07575519
cg07576135
cg07576142
cg07576376
cg07576525
cg07577799
cg07578108
cg07579123
cg07579946
cg07580831
cg07581070
cg07581226
cg07581395
cg07581732
cg07582923
cg07583848
cg07583894
cg07583933
cg07584031
cg07584494
cg07585502
cg07585562
cg07586285
cg07586760
cg07586956
cg07587117
cg07587786
cg07587893
cg07588216
cg07588253
cg07588439
cg07588561
cg07589202
cg07589691
cg07589972
cg07590294
cg07591403
cg07592012
cg07592136
cg07592216
cg07592513
cg07592963
cg07593852
cg07594007
cg07594472
cg07594531
cg07594658
cg07595776
cg07596106

cg07596645
cg07596668
cg07597892
cg07598199
cg07598331
cg07598370
cg07598464
cg07598549
cg07599389
cg07599713
cg07599819
cg07600499
cg07600998
cg07601645
cg07601804
cg07602072
cg07602195
cg07602659
cg07603330
cg07603357
cg07603449
cg07604651
cg07605211
cg07605523
cg07608826
cg07609200
cg07609206
cg07609844
cg07609862
cg07610765
cg07611933
cg07612562
cg07612827
cg07612978
cg07613047
cg07613153
cg07614623
cg07614747
cg07615067
cg07615127
cg07615497
cg07616094
cg07616471
cg07618409
cg07620167
cg07621046
cg07621227
cg07622079
cg07622096
cg07622222
cg07622493
cg07622533

cg07622996
cg07623058
cg07623116
cg07624008
cg07624812
cg07624918
cg07624977
cg07625131
cg07625774
cg07625783
cg07627317
cg07627491
cg07628073
cg07628416
cg07628580
cg07628717
cg07629125
cg07630229
cg07630255
cg07630274
cg07630787
cg07630816
cg07631309
cg07631533
cg07632615
cg07632618
cg07632946
cg07633435
cg07633702
cg07633853
cg07635132
cg07636117
cg07636295
cg07636338
cg07636761
cg07637239
cg07637375
cg07637554
cg07638462
cg07638500
cg07638689
cg07638884
cg07639489
cg07639520
cg07639783
cg07640117
cg07640391
cg07641160
cg07641563
cg07643225
cg07643355
cg07643696

cg07643942
cg07644939
cg07645844
cg07645980
cg07646083
cg07646362
cg07646665
cg07647771
cg07648238
cg07648740
cg07649045
cg07651189
cg07653395
cg07653946
cg07654012
cg07654762
cg07654982
cg07655487
cg07656173
cg07658646
cg07658702
cg07659054
cg07659529
cg07660473
cg07661106
cg07661564
cg07661660
cg07661904
cg07662121
cg07663572
cg07663732
cg07663765
cg07664198
cg07664856
cg07664990
cg07664999
cg07665387
cg07665466
cg07666840
cg07668558
cg07668671
cg07669969
cg07671011
cg07671776
cg07671805
cg07672688
cg07673578
cg07673740
cg07674238
cg07674459
cg07675656
cg07675700

cg07676145
cg07676329
cg07676361
cg07676849
cg07676859
cg07677446
cg07677598
cg07678449
cg07679581
cg07679679
cg07680129
cg07680167
cg07680533
cg07681748
cg07681935
cg07681938
cg07682515
cg07682600
cg07684019
cg07684063
cg07684519
cg07685034
cg07685977
cg07687070
cg07687119
cg07687354
cg07687398
cg07687899
cg07688490
cg07690181
cg07690768
cg07691684
cg07691705
cg07692529
cg07693270
cg07693617
cg07694319
cg07694404
cg07695272
cg07695566
cg07695835
cg07695909
cg07696099
cg07696699
cg07696842
cg07697597
cg07697895
cg07698102
cg07698783
cg07699300
cg07699454
cg07699546

cg07699767
cg07699978
cg07700491
cg07700644
cg07700925
cg07701579
cg07702235
cg07702888
cg07703119
cg07703401
cg07704331
cg07704934
cg07705604
cg07705739
cg07705908
cg07705913
cg07707586
cg07707641
cg07707891
cg07708487
cg07709205
cg07709210
cg07709358
cg07709903
cg07710073
cg07710211
cg07710328
cg07710335
cg07710481
cg07711566
cg07712478
cg07713318
cg07713411
cg07713929
cg07714085
cg07714420
cg07715909
cg07716052
cg07716910
cg07717083
cg07717429
cg07718650
cg07719096
cg07719492
cg07719695
cg07719898
cg07721042
cg07721547
cg07721901
cg07722360
cg07722678
cg07723130

cg07723659
cg07724670
cg07724816
cg07724971
cg07725033
cg07725167
cg07725527
cg07725579
cg07725829
cg07727884
cg07729059
cg07729527
cg07729537
cg07731047
cg07731535
cg07732143
cg07732472
cg07733446
cg07733920
cg07733993
cg07734975
cg07735577
cg07735790
cg07736012
cg07736716
cg07737135
cg07737560
cg07737644
cg07737781
cg07739205
cg07739369
cg07740199
cg07741624
cg07742370
cg07742404
cg07743730
cg07744430
cg07744502
cg07744841
cg07744912
cg07745624
cg07745905
cg07746882
cg07746943
cg07748767
cg07748854
cg07750560
cg07750741
cg07751104
cg07753755
cg07754880
cg07755752

cg07756788
cg07757183
cg07757281
cg07757577
cg07757887
cg07759223
cg07759394
cg07759478
cg07759903
cg07760161
cg07760622
cg07762764
cg07764113
cg07764334
cg07764916
cg07766263
cg07766612
cg07766875
cg07768103
cg07768501
cg07770968
cg07772400
cg07772781
cg07773337
cg07773769
cg07775417
cg07775468
cg07776049
cg07776698
cg07777008
cg07777344
cg07777349
cg07777790
cg07777793
cg07778180
cg07778900
cg07778983
cg07779120
cg07779313
cg07779444
cg07779777
cg07780095
cg07780199
cg07780543
cg07781445
cg07782221
cg07782241
cg07782285
cg07782620
cg07782744
cg07782795
cg07783094

cg07783480
cg07783491
cg07784042
cg07784084
cg07785447
cg07785556
cg07787240
cg07787666
cg07787851
cg07787977
cg07788092
cg07788348
cg07788369
cg07789083
cg07789132
cg07789792
cg07789940
cg07790615
cg07790752
cg07790947
cg07791011
cg07791418
cg07791490
cg07791803
cg07791987
cg07792478
cg07792978
cg07793638
cg07793808
cg07794981
cg07795307
cg07795452
cg07795614
cg07795916
cg07795928
cg07795949
cg07797833
cg07798143
cg07798605
cg07798731
cg07799395
cg07799572
cg07799947
cg07800208
cg07800524
cg07800544
cg07800670
cg07800890
cg07802612
cg07802625
cg07802909
cg07803859

cg07804076
cg07804260
cg07804289
cg07804749
cg07804888
cg07805146
cg07805500
cg07805542
cg07807026
cg07807757
cg07808023
cg07808712
cg07809027
cg07810039
cg07810190
cg07811054
cg07811110
cg07811198
cg07811262
cg07812589
cg07812715
cg07812849
cg07813142
cg07813249
cg07813322
cg07813377
cg07813445
cg07813622
cg07813643
cg07813695
cg07814541
cg07814565
cg07814763
cg07814776
cg07814886
cg07815238
cg07815694
cg07816047
cg07817055
cg07817505
cg07817608
cg07817648
cg07817739
cg07817928
cg07819010
cg07819944
cg07820280
cg07820868
cg07821417
cg07821629
cg07821739
cg07822928

cg07824114
cg07824422
cg07826255
cg07826516
cg07827717
cg07828741
cg07829379
cg07829804
cg07830156
cg07830855
cg07831164
cg07831273
cg07831277
cg07832674
cg07832772
cg07833068
cg07833104
cg07833951
cg07834418
cg07834743
cg07834841
cg07835051
cg07835424
cg07835443
cg07835482
cg07836225
cg07837260
cg07837433
cg07837556
cg07837852
cg07838002
cg07838106
cg07838270
cg07838483
cg07838993
cg07839533
cg07840120
cg07841132
cg07841353
cg07841601
cg07841831
cg07841866
cg07842062
cg07842289
cg07842386
cg07842521
cg07842635
cg07846061
cg07846220
cg07846855
cg07847863
cg07848601

cg07848636
cg07849384
cg07849616
cg07849944
cg07850246
cg07850316
cg07850418
cg07850488
cg07850527
cg07850987
cg07851675
cg07851738
cg07853104
cg07853489
cg07853571
cg07854547
cg07854954
cg07855337
cg07855572
cg07855575
cg07855639
cg07856521
cg07856707
cg07857278
cg07858973
cg07859439
cg07859753
cg07859880
cg07860213
cg07860673
cg07861373
cg07861448
cg07861564
cg07861968
cg07862129
cg07862488
cg07865091
cg07865216
cg07865418
cg07866632
cg07866762
cg07866909
cg07867517
cg07868561
cg07869023
cg07869434
cg07869830
cg07870237
cg07870307
cg07870378
cg07870982
cg07870999

cg07872300
cg07872854
cg07873154
cg07873848
cg07874386
cg07874755
cg07875068
cg07875818
cg07877629
cg07877987
cg07878085
cg07879681
cg07880109
cg07880384
cg07880636
cg07880943
cg07881065
cg07881365
cg07881405
cg07882671
cg07883457
cg07883696
cg07883823
cg07884251
cg07885141
cg07887168
cg07887978
cg07887985
cg07888347
cg07889827
cg07889938
cg07890233
cg07890839
cg07891044
cg07891120
cg07891533
cg07892412
cg07892422
cg07892597
cg07893474
cg07893544
cg07893738
cg07894822
cg07895132
cg07895576
cg07896009
cg07896448
cg07896855
cg07897248
cg07897690
cg07899551
cg07899681

cg07900344
cg07901138
cg07901571
cg07902192
cg07903001
cg07903144
cg07903370
cg07904879
cg07905055
cg07905696
cg07907386
cg07907670
cg07908679
cg07908718
cg07908868
cg07908870
cg07909418
cg07910488
cg07910726
cg07910813
cg07910945
cg07911353
cg07911961
cg07912766
cg07912789
cg07913016
cg07913294
cg07913362
cg07914457
cg07914621
cg07914866
cg07915117
cg07915206
cg07915635
cg07915921
cg07917062
cg07917177
cg07917478
cg07917609
cg07917901
cg07918545
cg07919443
cg07919760
cg07920414
cg07921625
cg07921712
cg07921759
cg07922443
cg07922521
cg07922933
cg07923686
cg07925549

cg07925587
cg07926691
cg07927098
cg07928065
cg07928083
cg07928255
cg07928363
cg07929642
cg07930940
cg07931391
cg07931785
cg07931844
cg07932102
cg07932199
cg07934434
cg07934561
cg07934818
cg07935012
cg07935357
cg07935784
cg07936323
cg07936470
cg07936950
cg07937786
cg07937803
cg07938480
cg07939567
cg07939743
cg07939836
cg07940072
cg07940593
cg07941301
cg07942990
cg07943461
cg07944221
cg07944862
cg07945221
cg07945323
cg07945335
cg07946260
cg07947266
cg07947705
cg07948085
cg07948875
cg07948968
cg07949060
cg07949103
cg07949863
cg07949989
cg07950000
cg07950002
cg07950296

cg07950786
cg07951728
cg07951749
cg07951931
cg07951949
cg07952381
cg07953890
cg07954324
cg07954572
cg07955064
cg07955741
cg07956247
cg07956897
cg07958502
cg07958575
cg07959138
cg07959477
cg07959771
cg07959956
cg07959986
cg07960691
cg07960922
cg07961512
cg07961715
cg07961994
cg07962143
cg07962303
cg07962338
cg07962392
cg07963234
cg07964219
cg07964754
cg07964848
cg07964975
cg07965331
cg07965335
cg07965566
cg07965633
cg07966218
cg07967630
cg07967679
cg07967717
cg07967723
cg07968613
cg07968760
cg07969475
cg07969619
cg07969622
cg07969668
cg07969676
cg07969684
cg07969984

cg07970007
cg07970146
cg07970874
cg07971007
cg07971716
cg07972135
cg07972309
cg07972716
cg07972805
cg07973435
cg07974025
cg07974511
cg07974833
cg07976064
cg07976328
cg07976787
cg07977620
cg07977927
cg07977933
cg07978472
cg07978996
cg07979652
cg07981013
cg07981266
cg07982825
cg07983888
cg07984133
cg07984358
cg07984754
cg07985589
cg07985933
cg07986943
cg07987449
cg07987890
cg07988206
cg07988820
cg07988979
cg07989293
cg07989851
cg07990556
cg07990681
cg07991139
cg07991229
cg07991288
cg07992385
cg07993003
cg07993350
cg07993586
cg07993743
cg07995179
cg07997434
cg07997737

cg07997936
cg07999090
cg07999201
cg07999668
cg07999988
cg08000065
cg08000301
cg08001668
cg08003353
cg08003402
cg08003628
cg08003789
cg08004073
cg08004278
cg08004425
cg08005208
cg08005460
cg08005774
cg08006672
cg08006727
cg08007197
cg08007665
cg08008088
cg08008352
cg08009184
cg08009487
cg08009568
cg08009697
cg08010546
cg08010619
cg08010865
cg08012844
cg08013274
cg08013557
cg08014182
cg08014661
cg08014964
cg08015107
cg08015447
cg08015496
cg08016092
cg08017450
cg08018143
cg08018572
cg08018585
cg08019736
cg08020808
cg08022436
cg08022819
cg08023265
cg08023751
cg08023852

cg08024038
cg08026930
cg08027123
cg08027810
cg08028452
cg08029014
cg08029287
cg08031024
cg08032135
cg08032476
cg08032619
cg08032665
cg08032924
cg08033165
cg08034070
cg08034077
cg08034379
cg08034797
cg08034816
cg08035195
cg08035351
cg08035694
cg08035981
cg08036487
cg08036668
cg08037115
cg08037235
cg08037774
cg08038033
cg08038054
cg08038178
cg08038219
cg08038353
cg08039084
cg08039116
cg08039560
cg08041140
cg08041350
cg08042055
cg08042258
cg08042975
cg08043020
cg08043242
cg08043565
cg08043781
cg08043868
cg08044056
cg08044097
cg08044713
cg08044789
cg08045489
cg08047086

cg08047376
cg08047969
cg08048687
cg08049614
cg08049853
cg08050656
cg08051575
cg08051964
cg08053057
cg08053129
cg08053904
cg08055131
cg08055377
cg08056229
cg08056629
cg08056957
cg08057037
cg08058191
cg08058544
cg08058560
cg08058988
cg08059112
cg08060270
cg08060454
cg08060578
cg08060987
cg08061463
cg08062049
cg08062087
cg08062812
cg08062858
cg08063051
cg08063125
cg08063317
cg08065063
cg08065231
cg08065241
cg08065657
cg08066129
cg08066631
cg08066943
cg08067721
cg08067983
cg08068240
cg08068547
cg08069247
cg08070725
cg08070771
cg08071130
cg08071904
cg08073312
cg08073591

cg08074477
cg08074555
cg08074805
cg08076125
cg08076158
cg08076437
cg08076830
cg08077155
cg08077337
cg08077617
cg08077671
cg08077736
cg08079330
cg08079596
cg08079763
cg08080029
cg08080038
cg08080395
cg08080489
cg08080669
cg08080848
cg08081424
cg08082351
cg08082810
cg08084339
cg08084616
cg08084860
cg08084967
cg08085199
cg08086720
cg08086724
cg08087868
cg08088511
cg08089153
cg08089513
cg08090128
cg08090384
cg08090385
cg08090621
cg08090835
cg08091192
cg08092111
cg08092837
cg08093323
cg08094206
cg08094614
cg08094784
cg08095183
cg08095196
cg08095654
cg08096367
cg08096854

cg08097086
cg08097157
cg08097290
cg08097359
cg08097581
cg08097631
cg08098111
cg08098128
cg08098256
cg08098420
cg08098868
cg08099570
cg08099797
cg08100069
cg08100122
cg08100284
cg08100421
cg08100687
cg08101375
cg08103144
cg08104310
cg08104568
cg08105352
cg08105471
cg08106190
cg08106377
cg08106764
cg08106973
cg08108868
cg08109646
cg08109815
cg08110337
cg08110455
cg08110488
cg08110692
cg08110874
cg08111167
cg08111284
cg08111446
cg08111661
cg08111960
cg08112313
cg08112616
cg08113895
cg08113916
cg08114180
cg08114317
cg08114547
cg08114696
cg08116113
cg08116238
cg08116408

cg08116462
cg08116915
cg08117309
cg08117728
cg08118042
cg08118140
cg08118354
cg08119452
cg08119884
cg08120023
cg08120226
cg08120263
cg08121117
cg08121164
cg08121408
cg08121652
cg08121984
cg08122140
cg08122369
cg08123074
cg08123106
cg08123256
cg08123266
cg08123444
cg08123701
cg08123816
cg08124156
cg08124372
cg08124634
cg08124986
cg08125181
cg08125397
cg08125726
cg08125821
cg08125960
cg08126118
cg08126560
cg08126807
cg08127080
cg08128264
cg08128361
cg08128418
cg08128650
cg08129017
cg08129092
cg08129331
cg08129366
cg08129490
cg08130144
cg08130222
cg08130546
cg08130572

cg08130912
cg08130968
cg08130988
cg08131184
cg08131499
cg08132525
cg08133058
cg08133755
cg08134070
cg08134671
cg08134678
cg08134818
cg08134863
cg08135904
cg08136772
cg08136806
cg08136809
cg08137040
cg08137888
cg08137948
cg08139247
cg08139512
cg08140114
cg08140412
cg08140712
cg08141192
cg08141424
cg08141862
cg08141873
cg08141956
cg08141989
cg08142096
cg08143343
cg08144943
cg08145625
cg08145949
cg08146256
cg08146323
cg08146483
cg08146841
cg08148379
cg08148553
cg08149193
cg08149682
cg08150250
cg08151370
cg08151442
cg08151470
cg08151596
cg08151612
cg08151857
cg08152065

cg08152198
cg08152263
cg08152546
cg08153098
cg08153621
cg08154107
cg08154887
cg08155249
cg08155439
cg08155994
cg08156066
cg08156074
cg08156531
cg08156844
cg08157292
cg08157310
cg08157459
cg08157575
cg08158233
cg08158408
cg08158662
cg08158952
cg08159033
cg08159291
cg08161163
cg08161323
cg08161666
cg08162372
cg08162389
cg08162976
cg08163163
cg08163199
cg08163855
cg08165487
cg08165971
cg08166431
cg08166587
cg08166729
cg08166767
cg08167085
cg08168141
cg08168529
cg08169394
cg08169778
cg08169794
cg08169864
cg08170375
cg08170519
cg08170583
cg08170787
cg08170911
cg08172575

cg08172927
cg08172947
cg08173606
cg08174792
cg08174890
cg08175536
cg08175640
cg08176410
cg08176812
cg08177015
cg08177070
cg08177332
cg08177882
cg08179353
cg08179808
cg08180187
cg08180831
cg08180884
cg08181657
cg08181850
cg08182446
cg08182501
cg08182597
cg08183141
cg08183425
cg08184159
cg08184652
cg08185241
cg08186575
cg08186790
cg08186915
cg08186924
cg08187110
cg08188271
cg08188471
cg08189843
cg08189989
cg08190125
cg08190291
cg08190545
cg08190828
cg08190858
cg08190913
cg08191530
cg08194009
cg08194313
cg08194879
cg08196032
cg08196138
cg08196553
cg08196667
cg08196758

cg08198071
cg08198187
cg08198540
cg08198983
cg08199352
cg08199758
cg08199953
cg08200621
cg08200952
cg08202165
cg08202226
cg08202494
cg08202743
cg08204050
cg08204280
cg08204702
cg08204783
cg08206959
cg08207154
cg08209104
cg08210399
cg08210681
cg08210975
cg08211028
cg08211837
cg08211895
cg08212128
cg08212266
cg08212438
cg08212545
cg08213337
cg08213590
cg08213689
cg08213909
cg08214029
cg08214192
cg08214267
cg08214423
cg08214455
cg08214518
cg08214995
cg08215169
cg08215318
cg08215532
cg08216808
cg08217024
cg08217526
cg08217589
cg08218734
cg08218944
cg08220149
cg08220614

cg08220696
cg08220872
cg08221136
cg08221350
cg08221741
cg08221830
cg08222293
cg08223924
cg08224185
cg08224208
cg08224773
cg08225389
cg08225642
cg08225986
cg08226300
cg08227141
cg08227353
cg08227526
cg08227583
cg08228249
cg08229058
cg08229127
cg08229358
cg08229740
cg08230105
cg08230215
cg08231096
cg08231493
cg08231710
cg08231870
cg08232654
cg08233235
cg08234308
cg08234354
cg08234372
cg08235161
cg08235864
cg08236022
cg08236285
cg08236537
cg08236766
cg08236767
cg08237220
cg08237521
cg08237695
cg08237898
cg08238215
cg08239103
cg08239858
cg08239893
cg08239906
cg08240778

cg08241528
cg08241694
cg08242194
cg08243401
cg08243626
cg08243931
cg08244028
cg08244581
cg08244848
cg08245317
cg08245578
cg08245702
cg08245989
cg08246090
cg08246316
cg08246318
cg08246428
cg08246800
cg08247450
cg08247806
cg08247866
cg08248577
cg08248600
cg08249698
cg08249988
cg08250081
cg08250108
cg08250187
cg08250738
cg08250921
cg08251362
cg08251449
cg08251598
cg08251765
cg08252303
cg08252384
cg08252579
cg08253296
cg08253374
cg08254089
cg08254359
cg08255105
cg08255475
cg08255964
cg08256691
cg08256701
cg08256781
cg08257009
cg08257133
cg08257293
cg08258000
cg08258224

cg08258526
cg08258667
cg08258719
cg08258922
cg08259168
cg08259462
cg08259514
cg08260025
cg08260790
cg08261094
cg08262002
cg08263071
cg08263140
cg08263589
cg08263647
cg08263991
cg08264266
cg08264338
cg08264376
cg08264835
cg08265915
cg08265926
cg08266095
cg08266286
cg08266366
cg08267038
cg08267489
cg08268099
cg08269188
cg08271153
cg08271622
cg08271804
cg08271907
cg08272268
cg08272368
cg08272731
cg08272783
cg08272996
cg08273233
cg08273672
cg08273750
cg08273957
cg08274011
cg08274097
cg08274234
cg08275953
cg08276171
cg08276221
cg08276289
cg08276295
cg08277259
cg08277291

cg08277369
cg08277836
cg08278002
cg08278108
cg08278435
cg08279035
cg08279075
cg08279194
cg08280383
cg08281777
cg08282145
cg08283882
cg08284151
cg08284233
cg08284310
cg08284443
cg08284733
cg08284926
cg08284967
cg08285489
cg08285677
cg08286181
cg08286196
cg08288345
cg08288406
cg08288703
cg08288811
cg08288818
cg08288942
cg08289177
cg08289323
cg08289346
cg08289937
cg08290013
cg08290240
cg08290628
cg08291098
cg08291506
cg08291907
cg08292058
cg08292485
cg08292919
cg08293483
cg08293528
cg08293777
cg08294136
cg08294662
cg08294772
cg08295410
cg08295661
cg08296288
cg08296665

cg08297751
cg08298644
cg08298827
cg08299063
cg08299783
cg08300570
cg08300860
cg08301612
cg08301779
cg08302433
cg08302480
cg08304103
cg08304190
cg08305378
cg08305436
cg08305533
cg08305808
cg08305879
cg08305942
cg08306084
cg08306614
cg08307030
cg08309069
cg08309106
cg08309183
cg08309251
cg08309687
cg08310842
cg08311885
cg08312355
cg08312626
cg08313382
cg08313406
cg08313939
cg08314176
cg08314300
cg08314660
cg08314996
cg08315174
cg08315202
cg08315483
cg08315770
cg08316074
cg08316083
cg08316422
cg08316825
cg08317412
cg08318212
cg08318726
cg08318732
cg08320122
cg08320225

cg08320989
cg08321111
cg08321366
cg08321942
cg08322102
cg08322194
cg08323075
cg08323277
cg08323651
cg08324157
cg08324862
cg08325845
cg08326300
cg08326515
cg08326879
cg08327277
cg08327428
cg08327884
cg08328483
cg08328671
cg08328750
cg08328777
cg08329931
cg08330217
cg08330404
cg08330690
cg08330778
cg08331163
cg08331313
cg08331398
cg08332074
cg08332212
cg08333333
cg08333931
cg08334278
cg08334726
cg08334899
cg08335199
cg08336604
cg08337166
cg08337366
cg08337420
cg08338281
cg08339915
cg08340023
cg08340737
cg08341079
cg08343600
cg08343834
cg08344485
cg08345189
cg08345719

cg08346159
cg08346735
cg08347183
cg08347373
cg08347473
cg08347500
cg08348422
cg08349093
cg08349497
cg08349804
cg08351163
cg08351710
cg08352148
cg08352920
cg08353444
cg08353938
cg08353973
cg08354053
cg08354093
cg08354614
cg08354889
cg08355045
cg08355099
cg08355260
cg08356028
cg08356243
cg08357012
cg08357230
cg08357895
cg08358983
cg08359336
cg08359464
cg08360457
cg08362273
cg08362785
cg08362880
cg08363067
cg08363114
cg08363339
cg08363345
cg08363794
cg08364561
cg08364772
cg08364970
cg08365638
cg08365738
cg08366446
cg08367223
cg08368055
cg08368617
cg08368654
cg08369014

cg08369810
cg08369943
cg08370077
cg08370506
cg08370996
cg08371140
cg08371391
cg08371556
cg08371852
cg08371976
cg08373187
cg08373250
cg08373989
cg08375658
cg08376020
cg08376141
cg08378286
cg08378342
cg08379828
cg08380164
cg08380252
cg08380262
cg08380391
cg08381046
cg08381562
cg08382220
cg08382332
cg08382774
cg08382963
cg08383399
cg08384637
cg08386009
cg08386692
cg08386696
cg08387141
cg08387293
cg08387780
cg08388507
cg08389266
cg08389277
cg08389588
cg08390209
cg08390254
cg08390877
cg08391020
cg08391415
cg08391852
cg08392125
cg08392199
cg08392212
cg08392325
cg08393041

cg08393317
cg08393424
cg08393788
cg08393937
cg08394753
cg08397459
cg08397758
cg08397978
cg08400159
cg08400962
cg08401456
cg08401916
cg08402365
cg08402433
cg08402734
cg08402850
cg08402963
cg08403064
cg08404009
cg08404028
cg08404225
cg08406486
cg08406698
cg08407620
cg08408971
cg08408994
cg08409113
cg08409642
cg08410227
cg08411049
cg08411435
cg08412584
cg08413157
cg08413381
cg08414087
cg08414570
cg08414647
cg08415046
cg08415156
cg08415391
cg08415849
cg08416720
cg08417313
cg08418076
cg08419450
cg08420370
cg08420572
cg08421351
cg08421587
cg08421685
cg08422054
cg08422599

cg08422645
cg08423142
cg08423378
cg08423507
cg08424167
cg08424423
cg08425203
cg08425479
cg08425796
cg08426066
cg08426369
cg08426962
cg08427928
cg08428266
cg08428486
cg08430009
cg08430329
cg08430395
cg08430972
cg08431567
cg08431704
cg08431709
cg08431718
cg08431899
cg08432013
cg08432509
cg08432727
cg08432971
cg08435240
cg08435674
cg08435780
cg08437132
cg08438435
cg08438529
cg08438837
cg08439149
cg08439518
cg08439728
cg08440077
cg08440266
cg08441029
cg08441170
cg08441806
cg08442052
cg08442064
cg08442088
cg08442713
cg08445263
cg08445323
cg08445550
cg08445740
cg08445802

cg08446121
cg08446539
cg08446900
cg08447044
cg08447387
cg08447525
cg08447739
cg08447842
cg08448074
cg08448084
cg08448407
cg08448589
cg08448665
cg08448701
cg08448812
cg08449144
cg08449164
cg08449389
cg08451113
cg08451469
cg08452095
cg08452671
cg08452802
cg08452870
cg08453096
cg08453903
cg08453928
cg08454015
cg08454053
cg08454061
cg08454125
cg08454455
cg08454563
cg08454922
cg08455089
cg08455099
cg08455778
cg08456140
cg08456722
cg08456993
cg08457011
cg08457906
cg08458292
cg08458711
cg08459368
cg08460041
cg08460464
cg08460502
cg08460621
cg08460732
cg08461107
cg08461425

cg08462030
cg08462108
cg08462122
cg08462367
cg08462497
cg08462941
cg08463231
cg08463297
cg08463485
cg08463758
cg08464093
cg08464177
cg08464190
cg08464402
cg08464498
cg08465307
cg08465346
cg08465774
cg08466030
cg08466256
cg08466792
cg08467103
cg08467887
cg08468082
cg08468741
cg08468772
cg08468965
cg08469305
cg08469346
cg08469419
cg08469491
cg08469845
cg08469939
cg08470031
cg08470180
cg08470742
cg08471173
cg08471739
cg08471972
cg08472142
cg08473693
cg08473725
cg08474748
cg08474859
cg08474929
cg08475042
cg08475096
cg08475244
cg08475953
cg08476157
cg08476244
cg08476595

cg08477485
cg08477768
cg08478006
cg08478046
cg08478623
cg08479073
cg08479590
cg08479629
cg08480098
cg08480223
cg08480566
cg08480609
cg08480826
cg08480901
cg08481075
cg08481543
cg08482436
cg08482501
cg08482531
cg08482837
cg08482979
cg08484506
cg08486507
cg08486875
cg08486903
cg08487909
cg08488756
cg08488841
cg08489883
cg08490107
cg08490115
cg08490220
cg08491437
cg08491919
cg08492173
cg08492178
cg08492568
cg08492619
cg08493463
cg08493581
cg08493590
cg08493762
cg08494843
cg08495245
cg08495374
cg08495770
cg08495813
cg08495878
cg08496033
cg08496123
cg08496162
cg08496664

cg08496737
cg08497530
cg08498254
cg08498475
cg08498533
cg08498585
cg08498647
cg08498747
cg08498836
cg08499046
cg08499840
cg08500172
cg08500704
cg08500908
cg08502533
cg08503522
cg08503681
cg08505243
cg08505647
cg08506113
cg08506654
cg08506869
cg08507270
cg08507393
cg08507422
cg08507571
cg08507975
cg08508455
cg08508576
cg08508724
cg08508988
cg08509270
cg08509586
cg08509991
cg08510264
cg08511886
cg08512142
cg08512462
cg08513123
cg08513547
cg08514385
cg08514436
cg08514558
cg08514594
cg08514779
cg08515427
cg08515646
cg08515811
cg08515891
cg08516083
cg08516792
cg08517126

cg08517286
cg08519134
cg08519191
cg08520743
cg08520746
cg08521987
cg08521995
cg08522176
cg08522831
cg08523075
cg08523384
cg08523865
cg08524128
cg08525461
cg08525713
cg08525845
cg08526074
cg08526209
cg08526991
cg08527161
cg08527195
cg08527435
cg08527448
cg08528417
cg08528626
cg08529049
cg08529179
cg08529399
cg08529481
cg08529744
cg08529882
cg08530008
cg08530237
cg08530610
cg08530632
cg08531365
cg08531623
cg08531687
cg08534016
cg08534147
cg08534628
cg08535112
cg08535361
cg08535779
cg08535928
cg08535938
cg08536228
cg08536358
cg08536617
cg08536911
cg08536961
cg08537660

cg08537804
cg08538032
cg08538882
cg08540829
cg08540953
cg08541862
cg08542715
cg08543452
cg08543849
cg08544192
cg08545287
cg08548284
cg08548498
cg08548500
cg08548718
cg08549889
cg08550101
cg08550317
cg08550353
cg08550490
cg08550748
cg08552214
cg08552233
cg08553158
cg08553437
cg08553630
cg08554440
cg08554603
cg08555653
cg08555912
cg08556772
cg08557246
cg08558733
cg08560373
cg08560874
cg08561856
cg08563054
cg08563299
cg08564375
cg08564601
cg08564661
cg08564874
cg08565003
cg08565197
cg08566437
cg08566455
cg08566882
cg08567760
cg08568649
cg08568767
cg08569322
cg08569799

cg08570226
cg08570691
cg08571680
cg08572278
cg08572513
cg08572611
cg08574629
cg08574978
cg08575049
cg08575537
cg08575860
cg08575894
cg08577046
cg08577727
cg08578776
cg08579468
cg08579540
cg08579858
cg08580057
cg08580836
cg08581004
cg08581018
cg08582701
cg08583570
cg08583780
cg08583859
cg08584061
cg08584947
cg08585489
cg08587297
cg08587313
cg08587797
cg08587845
cg08588156
cg08589354
cg08591265
cg08591388
cg08592104
cg08592234
cg08593261
cg08594043
cg08594218
cg08594281
cg08594379
cg08594554
cg08595501
cg08595592
cg08595989
cg08596532
cg08597761
cg08597802
cg08598221

cg08598654
cg08601899
cg08602008
cg08602190
cg08602500
cg08602861
cg08603449
cg08603768
cg08603829
cg08604097
cg08604416
cg08604523
cg08604596
cg08605016
cg08605326
cg08605965
cg08606911
cg08607018
cg08608126
cg08608974
cg08610076
cg08610862
cg08610901
cg08610968
cg08611714
cg08612137
cg08612871
cg08613144
cg08614290
cg08614370
cg08614481
cg08614562
cg08616702
cg08616943
cg08616951
cg08617020
cg08617135
cg08617683
cg08619893
cg08619954
cg08620044
cg08620154
cg08620751
cg08620986
cg08621406
cg08621441
cg08621473
cg08621574
cg08621610
cg08621624
cg08621843
cg08621957

cg08622198
cg08622369
cg08623256
cg08623535
cg08623556
cg08623810
cg08623843
cg08623979
cg08624648
cg08625590
cg08625822
cg08626831
cg08627720
cg08628006
cg08628428
cg08628911
cg08629043
cg08629379
cg08630923
cg08631650
cg08631819
cg08632164
cg08632767
cg08632810
cg08633249
cg08635765
cg08636115
cg08636573
cg08636638
cg08637618
cg08638097
cg08638184
cg08638320
cg08638960
cg08639379
cg08639389
cg08639586
cg08639799
cg08640046
cg08640498
cg08640634
cg08641579
cg08642111
cg08642285
cg08642292
cg08642342
cg08642707
cg08642716
cg08642771
cg08643003
cg08643128
cg08643994

cg08644276
cg08644356
cg08644365
cg08644897
cg08645907
cg08646355
cg08647091
cg08647730
cg08648047
cg08648317
cg08649013
cg08650784
cg08650910
cg08651894
cg08652441
cg08652487
cg08652499
cg08652722
cg08652808
cg08653369
cg08653692
cg08653747
cg08653945
cg08655330
cg08655800
cg08656316
cg08657341
cg08657671
cg08657705
cg08657851
cg08658124
cg08658634
cg08658787
cg08658895
cg08659179
cg08659183
cg08659421
cg08660876
cg08661164
cg08661470
cg08662168
cg08662226
cg08663298
cg08663552
cg08665093
cg08665251
cg08666707
cg08666915
cg08667024
cg08667569
cg08667809
cg08668281

cg08668316
cg08668551
cg08669096
cg08669954
cg08670181
cg08670288
cg08670691
cg08671425
cg08671647
cg08672916
cg08675521
cg08675717
cg08675869
cg08676173
cg08676755
cg08677140
cg08678055
cg08678314
cg08678851
cg08679011
cg08679180
cg08680415
cg08681473
cg08681497
cg08681689
cg08681855
cg08681924
cg08681966
cg08682535
cg08683147
cg08683263
cg08683297
cg08683644
cg08683938
cg08684511
cg08684630
cg08684893
cg08685714
cg08685955
cg08686462
cg08687005
cg08687753
cg08688043
cg08689929
cg08690412
cg08690634
cg08690859
cg08690876
cg08690999
cg08691328
cg08691422
cg08691577

cg08692277
cg08692430
cg08692733
cg08693490
cg08693832
cg08694680
cg08695337
cg08695416
cg08695418
cg08695558
cg08695941
cg08696107
cg08696165
cg08696316
cg08696727
cg08696931
cg08697297
cg08697732
cg08697797
cg08698026
cg08698135
cg08698159
cg08698477
cg08698796
cg08698997
cg08699512
cg08699794
cg08699935
cg08700083
cg08700546
cg08700932
cg08701047
cg08701543
cg08701621
cg08701675
cg08701816
cg08701998
cg08702413
cg08702805
cg08702879
cg08702915
cg08702941
cg08703808
cg08703857
cg08703872
cg08704302
cg08705406
cg08705942
cg08707112
cg08707456
cg08707471
cg08707536

cg08708212
cg08708229
cg08708629
cg08708684
cg08708747
cg08709116
cg08711059
cg08711113
cg08711175
cg08711320
cg08711858
cg08712008
cg08712190
cg08712336
cg08712407
cg08712866
cg08713073
cg08713858
cg08715231
cg08715720
cg08716684
cg08717402
cg08717751
cg08718490
cg08719095
cg08719486
cg08719869
cg08720250
cg08720786
cg08721026
cg08721187
cg08721324
cg08722115
cg08722675
cg08722774
cg08723571
cg08724310
cg08724753
cg08726900
cg08727316
cg08728448
cg08729031
cg08729059
cg08729318
cg08729447
cg08729600
cg08729686
cg08729755
cg08729927
cg08730434
cg08730747
cg08732352

cg08732456
cg08732466
cg08733482
cg08733496
cg08733933
cg08734090
cg08734600
cg08734647
cg08734740
cg08735710
cg08735716
cg08736072
cg08736813
cg08736883
cg08737236
cg08738722
cg08739188
cg08739696
cg08739738
cg08740008
cg08740223
cg08740698
cg08741842
cg08741843
cg08743017
cg08743199
cg08743303
cg08744177
cg08744264
cg08744726
cg08745334
cg08745451
cg08746653
cg08746785
cg08746853
cg08747377
cg08747557
cg08747676
cg08747777
cg08748033
cg08748184
cg08748308
cg08748671
cg08748948
cg08749576
cg08749599
cg08750439
cg08750504
cg08750510
cg08750534
cg08750547
cg08751352

cg08751451
cg08751472
cg08751508
cg08751840
cg08751971
cg08752433
cg08753227
cg08753373
cg08753391
cg08753560
cg08753706
cg08754088
cg08754149
cg08754967
cg08755224
cg08756205
cg08756614
cg08756898
cg08757628
cg08759276
cg08762819
cg08763351
cg08763626
cg08763846
cg08764465
cg08765100
cg08765597
cg08765811
cg08766256
cg08766770
cg08767182
cg08767286
cg08767637
cg08768395
cg08768621
cg08768852
cg08769451
cg08770034
cg08770757
cg08770935
cg08771694
cg08772106
cg08774443
cg08774694
cg08775629
cg08776619
cg08776641
cg08778316
cg08778805
cg08778851
cg08780887
cg08781140

cg08782158
cg08782337
cg08782899
cg08783253
cg08783317
cg08783468
cg08783934
cg08784911
cg08785985
cg08786079
cg08786674
cg08787039
cg08787698
cg08787769
cg08788055
cg08789933
cg08789970
cg08790440
cg08790676
cg08792272
cg08792423
cg08792630
cg08792703
cg08792812
cg08792976
cg08793877
cg08794157
cg08794265
cg08794544
cg08794696
cg08794763
cg08795515
cg08795557
cg08795964
cg08796391
cg08797194
cg08797423
cg08797471
cg08797704
cg08797864
cg08798307
cg08799922
cg08799967
cg08800414
cg08800461
cg08800670
cg08800869
cg08801446
cg08801479
cg08802358
cg08802513
cg08802607

cg08802944
cg08803023
cg08803263
cg08803791
cg08805041
cg08805386
cg08805500
cg08806153
cg08806184
cg08806408
cg08807496
cg08808205
cg08808817
cg08809361
cg08809919
cg08810842
cg08811210
cg08811227
cg08811259
cg08812189
cg08812555
cg08812889
cg08813062
cg08813888
cg08814749
cg08814800
cg08815081
cg08815110
cg08815340
cg08815352
cg08815403
cg08815672
cg08816023
cg08816098
cg08816988
cg08817836
cg08817867
cg08818337
cg08818632
cg08821738
cg08821909
cg08821937
cg08822523
cg08822635
cg08823837
cg08824344
cg08824384
cg08824840
cg08825075
cg08825645
cg08825817
cg08826184

cg08826251
cg08826345
cg08826743
cg08826840
cg08826919
cg08827110
cg08827307
cg08827454
cg08827584
cg08828723
cg08828819
cg08829117
cg08829841
cg08830588
cg08830641
cg08830672
cg08830758
cg08831002
cg08831034
cg08831401
cg08832227
cg08832234
cg08832603
cg08832745
cg08832842
cg08832906
cg08833577
cg08833675
cg08834018
cg08834128
cg08834436
cg08834573
cg08834602
cg08835403
cg08835847
cg08835910
cg08836009
cg08836024
cg08836729
cg08836825
cg08836904
cg08837344
cg08837355
cg08837627
cg08837744
cg08837866
cg08838158
cg08838779
cg08839053
cg08839272
cg08840017
cg08841511

cg08841552
cg08843064
cg08843513
cg08843769
cg08843809
cg08843859
cg08844770
cg08844913
cg08845025
cg08845123
cg08845685
cg08845721
cg08846959
cg08847944
cg08849294
cg08849514
cg08850461
cg08850729
cg08851289
cg08851784
cg08852641
cg08854572
cg08854834
cg08855395
cg08855551
cg08855630
cg08855671
cg08856224
cg08856318
cg08856946
cg08857713
cg08857729
cg08857797
cg08857994
cg08858210
cg08858220
cg08858272
cg08858295
cg08858345
cg08858649
cg08859095
cg08859215
cg08860083
cg08860136
cg08860143
cg08860296
cg08860639
cg08861115
cg08861404
cg08862033
cg08862366
cg08862394

cg08864105
cg08864904
cg08864944
cg08865099
cg08866497
cg08867241
cg08867825
cg08867933
cg08868213
cg08868915
cg08869244
cg08869700
cg08869722
cg08870281
cg08870587
cg08870743
cg08870893
cg08871052
cg08871239
cg08871474
cg08871608
cg08871964
cg08872425
cg08872493
cg08872521
cg08874384
cg08874609
cg08874747
cg08874974
cg08875948
cg08876039
cg08876434
cg08876441
cg08876932
cg08877148
cg08877257
cg08877463
cg08878177
cg08878323
cg08878368
cg08878567
cg08878713
cg08878744
cg08879111
cg08879390
cg08879684
cg08879910
cg08880369
cg08881175
cg08881785
cg08882027
cg08882278

cg08882904
cg08884074
cg08884363
cg08884395
cg08884410
cg08884611
cg08884752
cg08884928
cg08885076
cg08885265
cg08885854
cg08886020
cg08886154
cg08886973
cg08887138
cg08888905
cg08889440
cg08889797
cg08889930
cg08890418
cg08891110
cg08892078
cg08892386
cg08893036
cg08893575
cg08893692
cg08893817
cg08893853
cg08894020
cg08894131
cg08894480
cg08895260
cg08895307
cg08897388
cg08898155
cg08898349
cg08899215
cg08899442
cg08901662
cg08901867
cg08902243
cg08902698
cg08902977
cg08903019
cg08903069
cg08903181
cg08903261
cg08903389
cg08903465
cg08904151
cg08905399
cg08906309

cg08907282
cg08907525
cg08907926
cg08908089
cg08908482
cg08908643
cg08909961
cg08911275
cg08913410
cg08913523
cg08914740
cg08914932
cg08915603
cg08916646
cg08917117
cg08917121
cg08917478
cg08917933
cg08918120
cg08918576
cg08918658
cg08918709
cg08918749
cg08918845
cg08920068
cg08920174
cg08920210
cg08920610
cg08922361
cg08923494
cg08924321
cg08924566
cg08924696
cg08926253
cg08927738
cg08927754
cg08928038
cg08928408
cg08928753
cg08928882
cg08928958
cg08928966
cg08929103
cg08929791
cg08929806
cg08930112
cg08930147
cg08930413
cg08931515
cg08931647
cg08931835
cg08931849

cg08932277
cg08933237
cg08933284
cg08933505
cg08933615
cg08934208
cg08934785
cg08935125
cg08935418
cg08935526
cg08936549
cg08937380
cg08937407
cg08938062
cg08938584
cg08938793
cg08939095
cg08939394
cg08939481
cg08939521
cg08939850
cg08939974
cg08940252
cg08941515
cg08942191
cg08943045
cg08943116
cg08943494
cg08943714
cg08944026
cg08944236
cg08944723
cg08945756
cg08946157
cg08946713
cg08947084
cg08947167
cg08947774
cg08948808
cg08948841
cg08948964
cg08949101
cg08949329
cg08950859
cg08951005
cg08951078
cg08954025
cg08954761
cg08954776
cg08955099
cg08955461
cg08955941

cg08956008
cg08956010
cg08956101
cg08956226
cg08956463
cg08956474
cg08957279
cg08957564
cg08957652
cg08958168
cg08958294
cg08958747
cg08959645
cg08959677
cg08960830
cg08961621
cg08962020
cg08962305
cg08962452
cg08962888
cg08964780
cg08965235
cg08965655
cg08965685
cg08965888
cg08967456
cg08968329
cg08969344
cg08969532
cg08969925
cg08970648
cg08970655
cg08971281
cg08971562
cg08971771
cg08972170
cg08972588
cg08973672
cg08973688
cg08974450
cg08974852
cg08975197
cg08975803
cg08975917
cg08975922
cg08976342
cg08976995
cg08977536
cg08977611
cg08977627
cg08977702
cg08978665

cg08979319
cg08979573
cg08979831
cg08979895
cg08980304
cg08980355
cg08980382
cg08981228
cg08981648
cg08981777
cg08981980
cg08982827
cg08982864
cg08982961
cg08983097
cg08983253
cg08985029
cg08985063
cg08985282
cg08985530
cg08985570
cg08985669
cg08987251
cg08989212
cg08989942
cg08991615
cg08991927
cg08992872
cg08992911
cg08993690
cg08993922
cg08994045
cg08994982
cg08996483
cg08996521
cg08998189
cg08998192
cg08998657
cg08999895
cg09000302
cg09000832
cg09001226
cg09001527
cg09001549
cg09003917
cg09005125
cg09005159
cg09005679
cg09006487
cg09008705
cg09009130
cg09009410

cg09009466
cg09009536
cg09010499
cg09010707
cg09011883
cg09011891
cg09012001
cg09012633
cg09012858
cg09013267
cg09014354
cg09016242
cg09016324
cg09016806
cg09016968
cg09017139
cg09017235
cg09017434
cg09018824
cg09019347
cg09020665
cg09021712
cg09021817
cg09022422
cg09022430
cg09022808
cg09022943
cg09023136
cg09023820
cg09023965
cg09024124
cg09024126
cg09024606
cg09025210
cg09025327
cg09026448
cg09026568
cg09026722
cg09027255
cg09027493
cg09027725
cg09028445
cg09028855
cg09029409
cg09030187
cg09030848
cg09032137
cg09032310
cg09032314
cg09032499
cg09032973
cg09033472

cg09034664
cg09034695
cg09034753
cg09035127
cg09038794
cg09039506
cg09041337
cg09041678
cg09042277
cg09042448
cg09042966
cg09043104
cg09043518
cg09044174
cg09044294
cg09044868
cg09045196
cg09045262
cg09045552
cg09046370
cg09046550
cg09047361
cg09047474
cg09047513
cg09048205
cg09048251
cg09048500
cg09048530
cg09050670
cg09050775
cg09051342
cg09051393
cg09051775
cg09051837
cg09053680
cg09054633
cg09054949
cg09054951
cg09055205
cg09055236
cg09055750
cg09056845
cg09057412
cg09057885
cg09058099
cg09058554
cg09059156
cg09061207
cg09062640
cg09063173
cg09063663
cg09063683

cg09064061
cg09064304
cg09064482
cg09067029
cg09068492
cg09068528
cg09069138
cg09069150
cg09069203
cg09069900
cg09070201
cg09070371
cg09070415
cg09070885
cg09071093
cg09071155
cg09071407
cg09072162
cg09072859
cg09073398
cg09073443
cg09073539
cg09074338
cg09074468
cg09075515
cg09076010
cg09076077
cg09076337
cg09076877
cg09077126
cg09077752
cg09077862
cg09078517
cg09079818
cg09080874
cg09081266
cg09081544
cg09082487
cg09082518
cg09082823
cg09082903
cg09083279
cg09083627
cg09083945
cg09083947
cg09084885
cg09085201
cg09086442
cg09086964
cg09087110
cg09087901
cg09087961

cg09088107
cg09088396
cg09088437
cg09088576
cg09088720
cg09088988
cg09090376
cg09090724
cg09092280
cg09094037
cg09094290
cg09094473
cg09094952
cg09095364
cg09095551
cg09095645
cg09096950
cg09098195
cg09098522
cg09098545
cg09098707
cg09098720
cg09099500
cg09099830
cg09100013
cg09100373
cg09101258
cg09101894
cg09102260
cg09102329
cg09102610
cg09102690
cg09103598
cg09104284
cg09105442
cg09105683
cg09105802
cg09106510
cg09106984
cg09107007
cg09107240
cg09109383
cg09110241
cg09111223
cg09111831
cg09113070
cg09113505
cg09113939
cg09114833
cg09115116
cg09115216
cg09115646

cg09117206
cg09117442
cg09117643
cg09117673
cg09118247
cg09118258
cg09118749
cg09119776
cg09120267
cg09120609
cg09120873
cg09121152
cg09121339
cg09121378
cg09121415
cg09121772
cg09121994
cg09122035
cg09122423
cg09122442
cg09122593
cg09122655
cg09122999
cg09123155
cg09123188
cg09123431
cg09123951
cg09124223
cg09124230
cg09125320
cg09125550
cg09127107
cg09127371
cg09127472
cg09127909
cg09128377
cg09128529
cg09128944
cg09129100
cg09130091
cg09130676
cg09131339
cg09132215
cg09132388
cg09132634
cg09132909
cg09133026
cg09134497
cg09134760
cg09135529
cg09135913
cg09136129

cg09136346
cg09137718
cg09138133
cg09139128
cg09140665
cg09140851
cg09141205
cg09141623
cg09141835
cg09141953
cg09141965
cg09142166
cg09142313
cg09142405
cg09142858
cg09143012
cg09143318
cg09143655
cg09143673
cg09143850
cg09144520
cg09144608
cg09145147
cg09145546
cg09145629
cg09146813
cg09147372
cg09149648
cg09149974
cg09150232
cg09150807
cg09150980
cg09151131
cg09151187
cg09151613
cg09151742
cg09151754
cg09151801
cg09152136
cg09154062
cg09154166
cg09154414
cg09154591
cg09154880
cg09155692
cg09155881
cg09156233
cg09157127
cg09157727
cg09158651
cg09159022
cg09160020

cg09160231
cg09160359
cg09161043
cg09161579
cg09162537
cg09164898
cg09165004
cg09165441
cg09166085
cg09166410
cg09166898
cg09167291
cg09167553
cg09167779
cg09167828
cg09167976
cg09168808
cg09168997
cg09169117
cg09169132
cg09169255
cg09169320
cg09169356
cg09169739
cg09169764
cg09169779
cg09169923
cg09171565
cg09171692
cg09172244
cg09173768
cg09174188
cg09174555
cg09174601
cg09175338
cg09175485
cg09176233
cg09176539
cg09178384
cg09178678
cg09179211
cg09180702
cg09180848
cg09181103
cg09182435
cg09182750
cg09183207
cg09184675
cg09185231
cg09185884
cg09185990
cg09186006

cg09186051
cg09186818
cg09187338
cg09187413
cg09187549
cg09187933
cg09188600
cg09188697
cg09189601
cg09189889
cg09191036
cg09191173
cg09191335
cg09192086
cg09193479
cg09193791
cg09195868
cg09196534
cg09197332
cg09198782
cg09199250
cg09199562
cg09199598
cg09201151
cg09201238
cg09201499
cg09203312
cg09204187
cg09205438
cg09208010
cg09208942
cg09209257
cg09211256
cg09211265
cg09212014
cg09212058
cg09212118
cg09213175
cg09214112
cg09214254
cg09214300
cg09216442
cg09217157
cg09217219
cg09218250
cg09219547
cg09219575
cg09220167
cg09220361
cg09220563
cg09221478
cg09221867

cg09222023
cg09222269
cg09222367
cg09222884
cg09222892
cg09222982
cg09224821
cg09225287
cg09225840
cg09226262
cg09226421
cg09226596
cg09227064
cg09227138
cg09229620
cg09230250
cg09230667
cg09230870
cg09230996
cg09231170
cg09231675
cg09231741
cg09231843
cg09232225
cg09232269
cg09232514
cg09232555
cg09232727
cg09232805
cg09232937
cg09233392
cg09233651
cg09233658
cg09233980
cg09234378
cg09234518
cg09234616
cg09234936
cg09235983
cg09236284
cg09236445
cg09237790
cg09237846
cg09237849
cg09238000
cg09238180
cg09238801
cg09238992
cg09240515
cg09241714
cg09241928
cg09242541

cg09243507
cg09243576
cg09243811
cg09244100
cg09244635
cg09244872
cg09245073
cg09245192
cg09245302
cg09245698
cg09246570
cg09246878
cg09247392
cg09247619
cg09248345
cg09248429
cg09248613
cg09249588
cg09249803
cg09250072
cg09250199
cg09250367
cg09251400
cg09251429
cg09251508
cg09251910
cg09252495
cg09253179
cg09254904
cg09255061
cg09256683
cg09256941
cg09256964
cg09257101
cg09257796
cg09257824
cg09258439
cg09258813
cg09259037
cg09259081
cg09259595
cg09260089
cg09261816
cg09262100
cg09262245
cg09262442
cg09263516
cg09265762
cg09265940
cg09267046
cg09267051
cg09267056

cg09268729
cg09269454
cg09269480
cg09269866
cg09270028
cg09270255
cg09271097
cg09271157
cg09271279
cg09271709
cg09272217
cg09273641
cg09275693
cg09275704
cg09275956
cg09276072
cg09276486
cg09276655
cg09277376
cg09277467
cg09277749
cg09278107
cg09278610
cg09279389
cg09279458
cg09279512
cg09279646
cg09279949
cg09280095
cg09281220
cg09281580
cg09281979
cg09282085
cg09282388
cg09282500
cg09282946
cg09284102
cg09284708
cg09285543
cg09286253
cg09287086
cg09287295
cg09287629
cg09287933
cg09289005
cg09290120
cg09290655
cg09290705
cg09290893
cg09291834
cg09292069
cg09292354

cg09292377
cg09293391
cg09293773
cg09293786
cg09293816
cg09294246
cg09294492
cg09294937
cg09295021
cg09295081
cg09295202
cg09295573
cg09296536
cg09297225
cg09297288
cg09297456
cg09298313
cg09299082
cg09299388
cg09299592
cg09301143
cg09301294
cg09301559
cg09302895
cg09302922
cg09303645
cg09303881
cg09304047
cg09304293
cg09305680
cg09305815
cg09305883
cg09306214
cg09307431
cg09307530
cg09307689
cg09307788
cg09308336
cg09309112
cg09310185
cg09310186
cg09311630
cg09312781
cg09312809
cg09312904
cg09313053
cg09313140
cg09313632
cg09313705
cg09313917
cg09313938
cg09314155

cg09314421
cg09314434
cg09315464
cg09315586
cg09315775
cg09316607
cg09317128
cg09317774
cg09317884
cg09318158
cg09318283
cg09319475
cg09320652
cg09321109
cg09321307
cg09321363
cg09321400
cg09322349
cg09323434
cg09323727
cg09323806
cg09324001
cg09324304
cg09325360
cg09325404
cg09326204
cg09326832
cg09327253
cg09327378
cg09329079
cg09329097
cg09329936
cg09330145
cg09330551
cg09330596
cg09331106
cg09331144
cg09331835
cg09331986
cg09332604
cg09333109
cg09333392
cg09333785
cg09334277
cg09334514
cg09334629
cg09334915
cg09335647
cg09336320
cg09336671
cg09336922
cg09337049

cg09337121
cg09337391
cg09337943
cg09338032
cg09338297
cg09339194
cg09339527
cg09340198
cg09340205
cg09340615
cg09341020
cg09341580
cg09341730
cg09341793
cg09341796
cg09342225
cg09342940
cg09344028
cg09344041
cg09344631
cg09346831
cg09347160
cg09347824
cg09349613
cg09349723
cg09350341
cg09352647
cg09353052
cg09353289
cg09353345
cg09353931
cg09353985
cg09354241
cg09354294
cg09354331
cg09354381
cg09354518
cg09355027
cg09355528
cg09356124
cg09356916
cg09357350
cg09357684
cg09358376
cg09359006
cg09359106
cg09359114
cg09360041
cg09360044
cg09360253
cg09360770
cg09361493

cg09361598
cg09361748
cg09362335
cg09362969
cg09363443
cg09363587
cg09363604
cg09363891
cg09365645
cg09366342
cg09367174
cg09367817
cg09367879
cg09367967
cg09368836
cg09369190
cg09369221
cg09371084
cg09371351
cg09372525
cg09372664
cg09372728
cg09373037
cg09373136
cg09373727
cg09375033
cg09375205
cg09375907
cg09378756
cg09379485
cg09380067
cg09380069
cg09381155
cg09381582
cg09382127
cg09382328
cg09383106
cg09383389
cg09384035
cg09384610
cg09385093
cg09385371
cg09386054
cg09386139
cg09387914
cg09389370
cg09389826
cg09390350
cg09390594
cg09391066
cg09393254
cg09394306

cg09394608
cg09395612
cg09395732
cg09395844
cg09396032
cg09396892
cg09397246
cg09398200
cg09398550
cg09399225
cg09399878
cg09400566
cg09401661
cg09402276
cg09403666
cg09403738
cg09403933
cg09404376
cg09404516
cg09404981
cg09405076
cg09406887
cg09407122
cg09407319
cg09407967
cg09408098
cg09408143
cg09408366
cg09408917
cg09408937
cg09409457
cg09409539
cg09409845
cg09409956
cg09410380
cg09410512
cg09410607
cg09411597
cg09411682
cg09411874
cg09411999
cg09412882
cg09412975
cg09413450
cg09413950
cg09414264
cg09414426
cg09416096
cg09416203
cg09416521
cg09416543
cg09416903

cg09416934
cg09417011
cg09417176
cg09417355
cg09418063
cg09418321
cg09418852
cg09418984
cg09419102
cg09419192
cg09420637
cg09420907
cg09421562
cg09422450
cg09423283
cg09423703
cg09423875
cg09424308
cg09424348
cg09424526
cg09424759
cg09425481
cg09425611
cg09425926
cg09426383
cg09426834
cg09427016
cg09427311
cg09427644
cg09429735
cg09430158
cg09430445
cg09430647
cg09430664
cg09430827
cg09431058
cg09431318
cg09432154
cg09434085
cg09434193
cg09434667
cg09434995
cg09435170
cg09436502
cg09436713
cg09436767
cg09436907
cg09437283
cg09437783
cg09438113
cg09438147
cg09438605

cg09439010
cg09439722
cg09439920
cg09440340
cg09440866
cg09441030
cg09441363
cg09442613
cg09442654
cg09442828
cg09443039
cg09443153
cg09443388
cg09443484
cg09444206
cg09444358
cg09444405
cg09444685
cg09445803
cg09445939
cg09446136
cg09446806
cg09449104
cg09449232
cg09449905
cg09449947
cg09450197
cg09450260
cg09451434
cg09451457
cg09451574
cg09452082
cg09452312
cg09453636
cg09453760
cg09454925
cg09455208
cg09455336
cg09455513
cg09455805
cg09456493
cg09457131
cg09457527
cg09458237
cg09458595
cg09458931
cg09459548
cg09459581
cg09460231
cg09461507
cg09461792
cg09462445

cg09462808
cg09463917
cg09464466
cg09465142
cg09465746
cg09467487
cg09467508
cg09467675
cg09467825
cg09467902
cg09468836
cg09469079
cg09469111
cg09469554
cg09469566
cg09469812
cg09470754
cg09470983
cg09471343
cg09472467
cg09472600
cg09473510
cg09473613
cg09473725
cg09473826
cg09473969
cg09474173
cg09474331
cg09475635
cg09475796
cg09476306
cg09476963
cg09477453
cg09477895
cg09478002
cg09478095
cg09479286
cg09480047
cg09480264
cg09480446
cg09481404
cg09483763
cg09484032
cg09484214
cg09486367
cg09486764
cg09488184
cg09488200
cg09488203
cg09488352
cg09489128
cg09489306

cg09489330
cg09490124
cg09491068
cg09491410
cg09491536
cg09491542
cg09491901
cg09491991
cg09492451
cg09492640
cg09492832
cg09493290
cg09494876
cg09495282
cg09495482
cg09496161
cg09496303
cg09497201
cg09497644
cg09498078
cg09498134
cg09498572
cg09498652
cg09499256
cg09499421
cg09499787
cg09499849
cg09499856
cg09500437
cg09500443
cg09500738
cg09500815
cg09501025
cg09501047
cg09501282
cg09501762
cg09502069
cg09502149
cg09502177
cg09502207
cg09503608
cg09504320
cg09504322
cg09505918
cg09506797
cg09507184
cg09507386
cg09507968
cg09508022
cg09508275
cg09508556
cg09508934

cg09509365
cg09509592
cg09510559
cg09510921
cg09510954
cg09511177
cg09511427
cg09511963
cg09512973
cg09513026
cg09513647
cg09513990
cg09514310
cg09514366
cg09514545
cg09515947
cg09515953
cg09516210
cg09516237
cg09516965
cg09517450
cg09518195
cg09518670
cg09520414
cg09521647
cg09521703
cg09521872
cg09522479
cg09524455
cg09524686
cg09524907
cg09526469
cg09526712
cg09526758
cg09526912
cg09527126
cg09528275
cg09528496
cg09528501
cg09529437
cg09530108
cg09530156
cg09530163
cg09530438
cg09531892
cg09533063
cg09533145
cg09533168
cg09533305
cg09533783
cg09535027
cg09535605

cg09535924
cg09535963
cg09536516
cg09537568
cg09538587
cg09539739
cg09539908
cg09540102
cg09540961
cg09541333
cg09542154
cg09543255
cg09543627
cg09544149
cg09546258
cg09546272
cg09546755
cg09546907
cg09547427
cg09547692
cg09548051
cg09549240
cg09549304
cg09549591
cg09550024
cg09550083
cg09550127
cg09550293
cg09551544
cg09551916
cg09553232
cg09553242
cg09553753
cg09554443
cg09554509
cg09554951
cg09554952
cg09555124
cg09555323
cg09555364
cg09556515
cg09556527
cg09556971
cg09557341
cg09557462
cg09557747
cg09558000
cg09558195
cg09558281
cg09558425
cg09558463
cg09558502

cg09558547
cg09559189
cg09559278
cg09559950
cg09560078
cg09560082
cg09560096
cg09561138
cg09562045
cg09562322
cg09562331
cg09562426
cg09563216
cg09563805
cg09564253
cg09564361
cg09564802
cg09565226
cg09565597
cg09566023
cg09566131
cg09566302
cg09566316
cg09567048
cg09567915
cg09568216
cg09568464
cg09568483
cg09568774
cg09569760
cg09570614
cg09570655
cg09571082
cg09571713
cg09571848
cg09571950
cg09572125
cg09573585
cg09574088
cg09574757
cg09575189
cg09576124
cg09576143
cg09577511
cg09577907
cg09577986
cg09578028
cg09578475
cg09578568
cg09579677
cg09579932
cg09579989

cg09582482
cg09582952
cg09583939
cg09584051
cg09584118
cg09584188
cg09584785
cg09584978
cg09585333
cg09586743
cg09586924
cg09588020
cg09588074
cg09588271
cg09588653
cg09589308
cg09589565
cg09590733
cg09591072
cg09591519
cg09591830
cg09594075
cg09595067
cg09595185
cg09595202
cg09595912
cg09596614
cg09596674
cg09596975
cg09599062
cg09599228
cg09599971
cg09600088
cg09600520
cg09601175
cg09602288
cg09603398
cg09603795
cg09603919
cg09605212
cg09605494
cg09606034
cg09606208
cg09606925
cg09606941
cg09607276
cg09607679
cg09608533
cg09609051
cg09610891
cg09611690
cg09612202

cg09612463
cg09613161
cg09614300
cg09614301
cg09614415
cg09614626
cg09614810
cg09614876
cg09615224
cg09615276
cg09615505
cg09615688
cg09615874
cg09616053
cg09616721
cg09617441
cg09617579
cg09618904
cg09619146
cg09619271
cg09619598
cg09620688
cg09622122
cg09622715
cg09623085
cg09623279
cg09623400
cg09623623
cg09624466
cg09624551
cg09624858
cg09624870
cg09624923
cg09624942
cg09625781
cg09626054
cg09626634
cg09626894
cg09627247
cg09627757
cg09628195
cg09628359
cg09628488
cg09628962
cg09629675
cg09631059
cg09631175
cg09631192
cg09631415
cg09631769
cg09631880
cg09631909

cg09633240
cg09633579
cg09634202
cg09635053
cg09635533
cg09635667
cg09636202
cg09636941
cg09637172
cg09637885
cg09637963
cg09638704
cg09639220
cg09639725
cg09640070
cg09640202
cg09640300
cg09640555
cg09640571
cg09640621
cg09641689
cg09642090
cg09642347
cg09642925
cg09643136
cg09643312
cg09643340
cg09643350
cg09643544
cg09644153
cg09644332
cg09644707
cg09644974
cg09646173
cg09646392
cg09646760
cg09647108
cg09648239
cg09648454
cg09649473
cg09649486
cg09649586
cg09650495
cg09650989
cg09651145
cg09651648
cg09651708
cg09652472
cg09652746
cg09652964
cg09654242
cg09654796

cg09655253
cg09655341
cg09655666
cg09656274
cg09656389
cg09657435
cg09658183
cg09659163
cg09659227
cg09659400
cg09660145
cg09660171
cg09660365
cg09660867
cg09661370
cg09661721
cg09661865
cg09661951
cg09662034
cg09663081
cg09663193
cg09663343
cg09663608
cg09664596
cg09665406
cg09665980
cg09666444
cg09667394
cg09667650
cg09667753
cg09668964
cg09669354
cg09669874
cg09670128
cg09670535
cg09670601
cg09671258
cg09672032
cg09672383
cg09673245
cg09673570
cg09673602
cg09673807
cg09674170
cg09674867
cg09675069
cg09675824
cg09676390
cg09676484
cg09676794
cg09676860
cg09677082

cg09677243
cg09677252
cg09677330
cg09678971
cg09679014
cg09679234
cg09679746
cg09679862
cg09680007
cg09680149
cg09681043
cg09681089
cg09681167
cg09682051
cg09682361
cg09682727
cg09682848
cg09683350
cg09683413
cg09683824
cg09684079
cg09684112
cg09684233
cg09684588
cg09685182
cg09685250
cg09685763
cg09686317
cg09686443
cg09686672
cg09687005
cg09687417
cg09687907
cg09688546
cg09689137
cg09689895
cg09690040
cg09690103
cg09690215
cg09690632
cg09692368
cg09692428
cg09693432
cg09694381
cg09694782
cg09695734
cg09695996
cg09696091
cg09696115
cg09696421
cg09696535
cg09696747

cg09696932
cg09697033
cg09698471
cg09699830
cg09701700
cg09702010
cg09702315
cg09702434
cg09703136
cg09703679
cg09703727
cg09704544
cg09705784
cg09705798
cg09706133
cg09706245
cg09707038
cg09707814
cg09707903
cg09708044
cg09709457
cg09709472
cg09709565
cg09709641
cg09709951
cg09710363
cg09710586
cg09711214
cg09713234
cg09713684
cg09714114
cg09714184
cg09714778
cg09715353
cg09715672
cg09716921
cg09717526
cg09717923
cg09718071
cg09718739
cg09719956
cg09721469
cg09721612
cg09721961
cg09724492
cg09725286
cg09726279
cg09726623
cg09726866
cg09727046
cg09728120
cg09728393

cg09729012
cg09729387
cg09729866
cg09730170
cg09730719
cg09732036
cg09732255
cg09733312
cg09734837
cg09734932
cg09735579
cg09735627
cg09736112
cg09736922
cg09739536
cg09739904
cg09740313
cg09740450
cg09740671
cg09741713
cg09742177
cg09743437
cg09744374
cg09744766
cg09745307
cg09746086
cg09746494
cg09747445
cg09747633
cg09747891
cg09748975
cg09750084
cg09750210
cg09750599
cg09752223
cg09752779
cg09753006
cg09755594
cg09755784
cg09755872
cg09756014
cg09757109
cg09757430
cg09757595
cg09757644
cg09758462
cg09758796
cg09758896
cg09759432
cg09759588
cg09759719
cg09760081

cg09760210
cg09761121
cg09761503
cg09762474
cg09765089
cg09765297
cg09765797
cg09766556
cg09766710
cg09767550
cg09767587
cg09767602
cg09767822
cg09769104
cg09769218
cg09769755
cg09770570
cg09770579
cg09772121
cg09772154
cg09772661
cg09772792
cg09772857
cg09773238
cg09774787
cg09774917
cg09775169
cg09775263
cg09775648
cg09775972
cg09776277
cg09778596
cg09778828
cg09779576
cg09780241
cg09780767
cg09780768
cg09781028
cg09781448
cg09781650
cg09781944
cg09783270
cg09783457
cg09783969
cg09784886
cg09784932
cg09785394
cg09786701
cg09786978
cg09787236
cg09788003
cg09788008

cg09788416
cg09789650
cg09790325
cg09790470
cg09790617
cg09790780
cg09791504
cg09792881
cg09793121
cg09793172
cg09793796
cg09793866
cg09794131
cg09794680
cg09795081
cg09795087
cg09795137
cg09795588
cg09795809
cg09797337
cg09797340
cg09797390
cg09797446
cg09797577
cg09798366
cg09798801
cg09798938
cg09799980
cg09799983
cg09800500
cg09801065
cg09801824
cg09801837
cg09802627
cg09802818
cg09802893
cg09803002
cg09803177
cg09803262
cg09803321
cg09803758
cg09803951
cg09803959
cg09804858
cg09805172
cg09805692
cg09805925
cg09806671
cg09806683
cg09807356
cg09807524
cg09807643

cg09807856
cg09807875
cg09808075
cg09808501
cg09809402
cg09809416
cg09809618
cg09809922
cg09810059
cg09811127
cg09811510
cg09811599
cg09812070
cg09812397
cg09813226
cg09813525
cg09814448
cg09816471
cg09816507
cg09816756
cg09816821
cg09816889
cg09817016
cg09817427
cg09817985
cg09818632
cg09819642
cg09820150
cg09820716
cg09821218
cg09821328
cg09821400
cg09822423
cg09822697
cg09822726
cg09823543
cg09824563
cg09824603
cg09824615
cg09824900
cg09825309
cg09826895
cg09826923
cg09826973
cg09827071
cg09828012
cg09828346
cg09828381
cg09828632
cg09828714
cg09829319
cg09829382

cg09830866
cg09831105
cg09831491
cg09831620
cg09831923
cg09832243
cg09832613
cg09833158
cg09833400
cg09833475
cg09833701
cg09834172
cg09834794
cg09835220
cg09835225
cg09835408
cg09836827
cg09836979
cg09837116
cg09837245
cg09837935
cg09838262
cg09838823
cg09839177
cg09839635
cg09840122
cg09840645
cg09841488
cg09841889
cg09842892
cg09843272
cg09843907
cg09844474
cg09844805
cg09845093
cg09845489
cg09845914
cg09846581
cg09847250
cg09847820
cg09847882
cg09848096
cg09848445
cg09848749
cg09849688
cg09850101
cg09850105
cg09850925
cg09851245
cg09851620
cg09851765
cg09851951

cg09851989
cg09852127
cg09852209
cg09852439
cg09852493
cg09852871
cg09853388
cg09853792
cg09853936
cg09854003
cg09854011
cg09854088
cg09854709
cg09854734
cg09855212
cg09855706
cg09856467
cg09856590
cg09856709
cg09857247
cg09857273
cg09857577
cg09857617
cg09858188
cg09858281
cg09858767
cg09858862
cg09858951
cg09859492
cg09860353
cg09860486
cg09860601
cg09861057
cg09861240
cg09862193
cg09862200
cg09862420
cg09862509
cg09862711
cg09863066
cg09863136
cg09863950
cg09864227
cg09864245
cg09865698
cg09866565
cg09866983
cg09868003
cg09868111
cg09868336
cg09868598
cg09868640

cg09868768
cg09868882
cg09869729
cg09869858
cg09871181
cg09871315
cg09871424
cg09871666
cg09871685
cg09871837
cg09872278
cg09873164
cg09873215
cg09873544
cg09873584
cg09873772
cg09873933
cg09874822
cg09874873
cg09875213
cg09876519
cg09876651
cg09877744
cg09878888
cg09879458
cg09880551
cg09881510
cg09883394
cg09883650
cg09883745
cg09884107
cg09884146
cg09884225
cg09884341
cg09885505
cg09885657
cg09887059
cg09887220
cg09887251
cg09887284
cg09887953
cg09888283
cg09889479
cg09889987
cg09890077
cg09890374
cg09890699
cg09890775
cg09891125
cg09892262
cg09892671
cg09893588

cg09893947
cg09894136
cg09895514
cg09895590
cg09895653
cg09896093
cg09896412
cg09896900
cg09897131
cg09897374
cg09898217
cg09898548
cg09900242
cg09902062
cg09902641
cg09904296
cg09904358
cg09905973
cg09906066
cg09906297
cg09906382
cg09906857
cg09906991
cg09906995
cg09907509
cg09907570
cg09907720
cg09907936
cg09909740
cg09911083
cg09911316
cg09911401
cg09911747
cg09911755
cg09912079
cg09912350
cg09913413
cg09913813
cg09913882
cg09914304
cg09914444
cg09915984
cg09916599
cg09916651
cg09916692
cg09917591
cg09917979
cg09918077
cg09918091
cg09918296
cg09918510
cg09919570

cg09920043
cg09920804
cg09921821
cg09922103
cg09922481
cg09923953
cg09923978
cg09925057
cg09926364
cg09926747
cg09929831
cg09930046
cg09930325
cg09930328
cg09931134
cg09931783
cg09931909
cg09932774
cg09933516
cg09934381
cg09934692
cg09935282
cg09935440
cg09935792
cg09936218
cg09936426
cg09936561
cg09936762
cg09936799
cg09936839
cg09937438
cg09938164
cg09938213
cg09938443
cg09938511
cg09939079
cg09939462
cg09939823
cg09939997
cg09940311
cg09940675
cg09940861
cg09941406
cg09942424
cg09942743
cg09943401
cg09944398
cg09946142
cg09946623
cg09949789
cg09950076
cg09950208

cg09950309
cg09950688
cg09950812
cg09951136
cg09951201
cg09951338
cg09953239
cg09954189
cg09954385
cg09954820
cg09955152
cg09955705
cg09956302
cg09956615
cg09956782
cg09957942
cg09958402
cg09959684
cg09959790
cg09959931
cg09960109
cg09960202
cg09960641
cg09961397
cg09962809
cg09963024
cg09963123
cg09963856
cg09964012
cg09964402
cg09965363
cg09967329
cg09968723
cg09969797
cg09971102
cg09971227
cg09971314
cg09971540
cg09971549
cg09971562
cg09971591
cg09971653
cg09971881
cg09972052
cg09972454
cg09974774
cg09975576
cg09975620
cg09975804
cg09975867
cg09978023
cg09978259

cg09978533
cg09978583
cg09979478
cg09979924
cg09980058
cg09980771
cg09981464
cg09981533
cg09981830
cg09981841
cg09982347
cg09983546
cg09984096
cg09984316
cg09985078
cg09985192
cg09985260
cg09985560
cg09985615
cg09985793
cg09985832
cg09987240
cg09987740
cg09988118
cg09988671
cg09989037
cg09990169
cg09990194
cg09990962
cg09991173
cg09991267
cg09991306
cg09991633
cg09991710
cg09991769
cg09992376
cg09993814
cg09994356
cg09994418
cg09994435
cg09994553
cg09994756
cg09995507
cg09996325
cg09996777
cg09996789
cg09997062
cg09997082
cg09997356
cg09997530
cg09998189
cg09998251

cg09999109
cg09999114
cg09999194
cg09999977
cg09999984
cg10000148
cg10001186
cg10001715
cg10002371
cg10002668
cg10003048
cg10004481
cg10004882
cg10004897
cg10005224
cg10005270
cg10005565
cg10005947
cg10005978
cg10006252
cg10006419
cg10006515
cg10006817
cg10007051
cg10007927
cg10008947
cg10009056
cg10009207
cg10009497
cg10010182
cg10010393
cg10010898
cg10011091
cg10011495
cg10011744
cg10011799
cg10012448
cg10012512
cg10012920
cg10015668
cg10016122
cg10016380
cg10016647
cg10016717
cg10017167
cg10017653
cg10018167
cg10020787
cg10020883
cg10020890
cg10021122
cg10021428

cg10022165
cg10022914
cg10023178
cg10023454
cg10023530
cg10024583
cg10025432
cg10025514
cg10026203
cg10026317
cg10027091
cg10027159
cg10027639
cg10027752
cg10027934
cg10028666
cg10028983
cg10029213
cg10030250
cg10030684
cg10031203
cg10031769
cg10031793
cg10032375
cg10034176
cg10034890
cg10035737
cg10036001
cg10036892
cg10037072
cg10037494
cg10037818
cg10037894
cg10037913
cg10038907
cg10039041
cg10039047
cg10039160
cg10039188
cg10039444
cg10040329
cg10040836
cg10041417
cg10042799
cg10042846
cg10043058
cg10043253
cg10043709
cg10045881
cg10046101
cg10046892
cg10047041

cg10047845
cg10049789
cg10050816
cg10051615
cg10052164
cg10052561
cg10054857
cg10055231
cg10055235
cg10055501
cg10055566
cg10055593
cg10055771
cg10056132
cg10056627
cg10056706
cg10056728
cg10057033
cg10057065
cg10057879
cg10058766
cg10059324
cg10059536
cg10059660
cg10059924
cg10060562
cg10061361
cg10061384
cg10061532
cg10061770
cg10061906
cg10061927
cg10063169
cg10063179
cg10063575
cg10063845
cg10064322
cg10064525
cg10064892
cg10065823
cg10065825
cg10067062
cg10067723
cg10068989
cg10069323
cg10069394
cg10069644
cg10069716
cg10070328
cg10070864
cg10071824
cg10074498

cg10074727
cg10074775
cg10076479
cg10076902
cg10076951
cg10077765
cg10079740
cg10080553
cg10080966
cg10081086
cg10081681
cg10081941
cg10081994
cg10082354
cg10082515
cg10082589
cg10083417
cg10083824
cg10083889
cg10083898
cg10084554
cg10084644
cg10085741
cg10088622
cg10088685
cg10088715
cg10088985
cg10089081
cg10089552
cg10089626
cg10089801
cg10090568
cg10090792
cg10090836
cg10091408
cg10091458
cg10091866
cg10092108
cg10092198
cg10092231
cg10092394
cg10092518
cg10092779
cg10092878
cg10094358
cg10094616
cg10094994
cg10095226
cg10095242
cg10096066
cg10096145
cg10096893

cg10096955
cg10097215
cg10097316
cg10097906
cg10099518
cg10099900
cg10099917
cg10100287
cg10100424
cg10100437
cg10100887
cg10102736
cg10103000
cg10103266
cg10103373
cg10103528
cg10103635
cg10104542
cg10104674
cg10105237
cg10105884
cg10105917
cg10106935
cg10107241
cg10107671
cg10108237
cg10108591
cg10109351
cg10109500
cg10110027
cg10110335
cg10110604
cg10110957
cg10111115
cg10111290
cg10111292
cg10111328
cg10111335
cg10111599
cg10111764
cg10111816
cg10112265
cg10112407
cg10112943
cg10113107
cg10114877
cg10115368
cg10116178
cg10116209
cg10116432
cg10116443
cg10116675

cg10116864
cg10117562
cg10118167
cg10118210
cg10118557
cg10119075
cg10119342
cg10120185
cg10120555
cg10120652
cg10120698
cg10121601
cg10122434
cg10122558
cg10123201
cg10123247
cg10123642
cg10123654
cg10124440
cg10124445
cg10124710
cg10124993
cg10125195
cg10125399
cg10125451
cg10125882
cg10125901
cg10126022
cg10126181
cg10126205
cg10127275
cg10127490
cg10127921
cg10129201
cg10129391
cg10129768
cg10129884
cg10130497
cg10131026
cg10131286
cg10131465
cg10131657
cg10131810
cg10132208
cg10133100
cg10133170
cg10133177
cg10134307
cg10134425
cg10134833
cg10134939
cg10134963

cg10135474
cg10135483
cg10135640
cg10136216
cg10136263
cg10136354
cg10136841
cg10137010
cg10137253
cg10137597
cg10138313
cg10138494
cg10138623
cg10139151
cg10139889
cg10140638
cg10141715
cg10142237
cg10142436
cg10142452
cg10142566
cg10142837
cg10143146
cg10143731
cg10143811
cg10143911
cg10144104
cg10144474
cg10144597
cg10145196
cg10146186
cg10146199
cg10146277
cg10146692
cg10146880
cg10147329
cg10147666
cg10147974
cg10148067
cg10148473
cg10148694
cg10148764
cg10149054
cg10150592
cg10151196
cg10151367
cg10151685
cg10151953
cg10154597
cg10154812
cg10154971
cg10155026

cg10156217
cg10156366
cg10157251
cg10157715
cg10158369
cg10158715
cg10159215
cg10160397
cg10160612
cg10161121
cg10161194
cg10162522
cg10163377
cg10163483
cg10164264
cg10164592
cg10165167
cg10165801
cg10165847
cg10166283
cg10166314
cg10166609
cg10166664
cg10166735
cg10166825
cg10167316
cg10167512
cg10168009
cg10168086
cg10168149
cg10168494
cg10169241
cg10169327
cg10169364
cg10169462
cg10169812
cg10170944
cg10171448
cg10172675
cg10172979
cg10173774
cg10173787
cg10173814
cg10174170
cg10174867
cg10175583
cg10175753
cg10176011
cg10176110
cg10176514
cg10176692
cg10177059

cg10177154
cg10177394
cg10177465
cg10178130
cg10178628
cg10179330
cg10179693
cg10180058
cg10180281
cg10180406
cg10180697
cg10181192
cg10181414
cg10181820
cg10182317
cg10182421
cg10182697
cg10185424
cg10185478
cg10185489
cg10185984
cg10186131
cg10186254
cg10186366
cg10187713
cg10187894
cg10188823
cg10188909
cg10189029
cg10189135
cg10189395
cg10189695
cg10189763
cg10190509
cg10190898
cg10190990
cg10191005
cg10192198
cg10192893
cg10192949
cg10194319
cg10194884
cg10196026
cg10196720
cg10197336
cg10198932
cg10200391
cg10201533
cg10201712
cg10201923
cg10202727
cg10203223

cg10204692
cg10204725
cg10204868
cg10204884
cg10205430
cg10205548
cg10205928
cg10206073
cg10206380
cg10206588
cg10206627
cg10206931
cg10206933
cg10207745
cg10207928
cg10208370
cg10208610
cg10209670
cg10210076
cg10210094
cg10210690
cg10211058
cg10211193
cg10211414
cg10211744
cg10211873
cg10211891
cg10212847
cg10213658
cg10213812
cg10213821
cg10213873
cg10214421
cg10214991
cg10216615
cg10216717
cg10216872
cg10217292
cg10217445
cg10217528
cg10217713
cg10219789
cg10219816
cg10219957
cg10219995
cg10221596
cg10221896
cg10222027
cg10222254
cg10222734
cg10222925
cg10223061

cg10223225
cg10223989
cg10224098
cg10224476
cg10225149
cg10225865
cg10226952
cg10227629
cg10227678
cg10228162
cg10228456
cg10230190
cg10230519
cg10231126
cg10231746
cg10231902
cg10232265
cg10233104
cg10233454
cg10233825
cg10233830
cg10233904
cg10234007
cg10234069
cg10234991
cg10234998
cg10235181
cg10235817
cg10236239
cg10236596
cg10236631
cg10236857
cg10237088
cg10237148
cg10237419
cg10237442
cg10238051
cg10239098
cg10239563
cg10239638
cg10240139
cg10240950
cg10241039
cg10241097
cg10241484
cg10241675
cg10241871
cg10242602
cg10244770
cg10244994
cg10245879
cg10245967

cg10245988
cg10246121
cg10246903
cg10247061
cg10247405
cg10248617
cg10248985
cg10249542
cg10250660
cg10250663
cg10251896
cg10255904
cg10256045
cg10256839
cg10257049
cg10257302
cg10259521
cg10260192
cg10260267
cg10260711
cg10261416
cg10262032
cg10262891
cg10263370
cg10263682
cg10264012
cg10264274
cg10264791
cg10266211
cg10267724
cg10268548
cg10268811
cg10269365
cg10269431
cg10269476
cg10269548
cg10270150
cg10271058
cg10271112
cg10271192
cg10272492
cg10272601
cg10275766
cg10275917
cg10276227
cg10276869
cg10277651
cg10277927
cg10278102
cg10278318
cg10278520
cg10278552

cg10279294
cg10279872
cg10280007
cg10280342
cg10281002
cg10281768
cg10281977
cg10282162
cg10282330
cg10282491
cg10283879
cg10284006
cg10284406
cg10285337
cg10287321
cg10287814
cg10287970
cg10288307
cg10288437
cg10288541
cg10289305
cg10289926
cg10290200
cg10290276
cg10290728
cg10290764
cg10290814
cg10292376
cg10292436
cg10293359
cg10293925
cg10294240
cg10294253
cg10294363
cg10294820
cg10294822
cg10296062
cg10296205
cg10297193
cg10297473
cg10297792
cg10298052
cg10298567
cg10299128
cg10299383
cg10300154
cg10300188
cg10300252
cg10300684
cg10300864
cg10300962
cg10301236

cg10303487
cg10303653
cg10303698
cg10303842
cg10303967
cg10304534
cg10304637
cg10305797
cg10305969
cg10307548
cg10307632
cg10308396
cg10308810
cg10309377
cg10309454
cg10310168
cg10310237
cg10310310
cg10312186
cg10312376
cg10313109
cg10315366
cg10315800
cg10315903
cg10315962
cg10316141
cg10316270
cg10316764
cg10316899
cg10317164
cg10317815
cg10318148
cg10318258
cg10318359
cg10318458
cg10318510
cg10318528
cg10319505
cg10319640
cg10319663
cg10319870
cg10319905
cg10320160
cg10320659
cg10320997
cg10321026
cg10322443
cg10322876
cg10323697
cg10323841
cg10323962
cg10324703

cg10324998
cg10325478
cg10325509
cg10325659
cg10326031
cg10326182
cg10327270
cg10327756
cg10328297
cg10328538
cg10328583
cg10328844
cg10329173
cg10329418
cg10329468
cg10330056
cg10330057
cg10330169
cg10330371
cg10330545
cg10332715
cg10333023
cg10333400
cg10333402
cg10334053
cg10334127
cg10334354
cg10334509
cg10335452
cg10335743
cg10335834
cg10336309
cg10336406
cg10336792
cg10336935
cg10339098
cg10339152
cg10340048
cg10340117
cg10340622
cg10341154
cg10341242
cg10341719
cg10342016
cg10343391
cg10343742
cg10343758
cg10344832
cg10345326
cg10345920
cg10345936
cg10347270

cg10348376
cg10351287
cg10351914
cg10353705
cg10354811
cg10354926
cg10355965
cg10356210
cg10356613
cg10356759
cg10358533
cg10358623
cg10359291
cg10359332
cg10359807
cg10361005
cg10361659
cg10362591
cg10363018
cg10363310
cg10363926
cg10364040
cg10364182
cg10364374
cg10364905
cg10365562
cg10365900
cg10366093
cg10366786
cg10366797
cg10366878
cg10368284
cg10368641
cg10369125
cg10369319
cg10369653
cg10371155
cg10371414
cg10371615
cg10372047
cg10372057
cg10372921
cg10373147
cg10373196
cg10373524
cg10373607
cg10374009
cg10374084
cg10374202
cg10374248
cg10374813
cg10375110

cg10375710
cg10375890
cg10376286
cg10376814
cg10377690
cg10378032
cg10378459
cg10378904
cg10379687
cg10380328
cg10380348
cg10381153
cg10381354
cg10382206
cg10382945
cg10383019
cg10383428
cg10383724
cg10384245
cg10384570
cg10384725
cg10385089
cg10385101
cg10385208
cg10385390
cg10386016
cg10386483
cg10387551
cg10387579
cg10389114
cg10389138
cg10389148
cg10390058
cg10390435
cg10390696
cg10390979
cg10391102
cg10391522
cg10392572
cg10393369
cg10393508
cg10393875
cg10395806
cg10395868
cg10396171
cg10396673
cg10396701
cg10398418
cg10398455
cg10398682
cg10399099
cg10399269

cg10399929
cg10399946
cg10400390
cg10400944
cg10401733
cg10401741
cg10401879
cg10402715
cg10403394
cg10403441
cg10403518
cg10405075
cg10406124
cg10406295
cg10406538
cg10407092
cg10407113
cg10408430
cg10408439
cg10409443
cg10409560
cg10409753
cg10410142
cg10410146
cg10410483
cg10412312
cg10413136
cg10413169
cg10413688
cg10413945
cg10414881
cg10414946
cg10414975
cg10414985
cg10415314
cg10416593
cg10416958
cg10417386
cg10417567
cg10420075
cg10420150
cg10420161
cg10420838
cg10421002
cg10421435
cg10421979
cg10422455
cg10422777
cg10423607
cg10423842
cg10424581
cg10424974

cg10426017
cg10426062
cg10426422
cg10426658
cg10427430
cg10427452
cg10427868
cg10428345
cg10429091
cg10429523
cg10429608
cg10430077
cg10430444
cg10430941
cg10430963
cg10431265
cg10431340
cg10431747
cg10431802
cg10432569
cg10432626
cg10433327
cg10433463
cg10433805
cg10434152
cg10434728
cg10435235
cg10435838
cg10435849
cg10436631
cg10436792
cg10437245
cg10437265
cg10437931
cg10438391
cg10438671
cg10439725
cg10439765
cg10440524
cg10440578
cg10440583
cg10440696
cg10440835
cg10441070
cg10441379
cg10441661
cg10442101
cg10442325
cg10442445
cg10442735
cg10443049
cg10443146

cg10443237
cg10444058
cg10445453
cg10446043
cg10446869
cg10447095
cg10447404
cg10447615
cg10447977
cg10448719
cg10448808
cg10449091
cg10450005
cg10450692
cg10451116
cg10452060
cg10453230
cg10453420
cg10453602
cg10453719
cg10453758
cg10453977
cg10454127
cg10454246
cg10455321
cg10455412
cg10455940
cg10456035
cg10456541
cg10457446
cg10458216
cg10458473
cg10459018
cg10459187
cg10460036
cg10460350
cg10460643
cg10460657
cg10461878
cg10462961
cg10463150
cg10463626
cg10464130
cg10464312
cg10464467
cg10465202
cg10465949
cg10466011
cg10466728
cg10468107
cg10468951
cg10469329

cg10469717
cg10469791
cg10470588
cg10470891
cg10471574
cg10471825
cg10471944
cg10472064
cg10472263
cg10472711
cg10472919
cg10473157
cg10473158
cg10475153
cg10476112
cg10476585
cg10476689
cg10476773
cg10477621
cg10478221
cg10479082
cg10479493
cg10479629
cg10479672
cg10481617
cg10482596
cg10483660
cg10483828
cg10484085
cg10484673
cg10484693
cg10485411
cg10485638
cg10485664
cg10486610
cg10486998
cg10487067
cg10487221
cg10488012
cg10488050
cg10488267
cg10489463
cg10491120
cg10491452
cg10491546
cg10491556
cg10492791
cg10492953
cg10493185
cg10493259
cg10493454
cg10493855

cg10495160
cg10495227
cg10495683
cg10497754
cg10497884
cg10498502
cg10498524
cg10498921
cg10499176
cg10499493
cg10499651
cg10500084
cg10500219
cg10500702
cg10501454
cg10501704
cg10502121
cg10502149
cg10503138
cg10503199
cg10503208
cg10503542
cg10503827
cg10503840
cg10504066
cg10504805
cg10506318
cg10506742
cg10506882
cg10507304
cg10507390
cg10507482
cg10508193
cg10508271
cg10508295
cg10508416
cg10508468
cg10508993
cg10509254
cg10509407
cg10509692
cg10510259
cg10511255
cg10511357
cg10511390
cg10511985
cg10511988
cg10512089
cg10512194
cg10512203
cg10512278
cg10512376

cg10512535
cg10512745
cg10512769
cg10512875
cg10513118
cg10513330
cg10513595
cg10514097
cg10514207
cg10514714
cg10517230
cg10517281
cg10517312
cg10518572
cg10518816
cg10519004
cg10519368
cg10519389
cg10519465
cg10519903
cg10519996
cg10520405
cg10520543
cg10520672
cg10520740
cg10520780
cg10520887
cg10521638
cg10521706
cg10521767
cg10521851
cg10522115
cg10522770
cg10522845
cg10523105
cg10523352
cg10524033
cg10524152
cg10524576
cg10524582
cg10524701
cg10525642
cg10525720
cg10526223
cg10526277
cg10526374
cg10526598
cg10526841
cg10527010
cg10528576
cg10528989
cg10529401

cg10529470
cg10529845
cg10530055
cg10530169
cg10530281
cg10530680
cg10530793
cg10530851
cg10530883
cg10531104
cg10531637
cg10531711
cg10532833
cg10533069
cg10534216
cg10534479
cg10534490
cg10534507
cg10534873
cg10535845
cg10536067
cg10536076
cg10536276
cg10536327
cg10536786
cg10536916
cg10537176
cg10537555
cg10538440
cg10538486
cg10539507
cg10539808
cg10540110
cg10540573
cg10540721
cg10541517
cg10541767
cg10541864
cg10542562
cg10543503
cg10543953
cg10546065
cg10546487
cg10546603
cg10548805
cg10548968
cg10549088
cg10549227
cg10549331
cg10549986
cg10550033
cg10550166

cg10550416
cg10550506
cg10550816
cg10551134
cg10551329
cg10552126
cg10552275
cg10553596
cg10556036
cg10557312
cg10557827
cg10558233
cg10560038
cg10560420
cg10564800
cg10566012
cg10566121
cg10566578
cg10566776
cg10567810
cg10568634
cg10569414
cg10570241
cg10572355
cg10572807
cg10573018
cg10573143
cg10574283
cg10574431
cg10574566
cg10574843
cg10575089
cg10575219
cg10575261
cg10575547
cg10576132
cg10576245
cg10576280
cg10576992
cg10577257
cg10577534
cg10577586
cg10578482
cg10578701
cg10578996
cg10579624
cg10579941
cg10580056
cg10580513
cg10581650
cg10582048
cg10582941

cg10583000
cg10583297
cg10583322
cg10583664
cg10583876
cg10583942
cg10584904
cg10584914
cg10585962
cg10586087
cg10586147
cg10586317
cg10586423
cg10586546
cg10586619
cg10587047
cg10587082
cg10588023
cg10588135
cg10588696
cg10588705
cg10588962
cg10589034
cg10589330
cg10589539
cg10590220
cg10590857
cg10591322
cg10591607
cg10591801
cg10592245
cg10592710
cg10592734
cg10592766
cg10593165
cg10594075
cg10595031
cg10595547
cg10596434
cg10596768
cg10597379
cg10598034
cg10599444
cg10601624
cg10601821
cg10601940
cg10602537
cg10602543
cg10603004
cg10603136
cg10603275
cg10604040

cg10606730
cg10606834
cg10606923
cg10608064
cg10608166
cg10608596
cg10610244
cg10610428
cg10610788
cg10611331
cg10611444
cg10612259
cg10612282
cg10612692
cg10613332
cg10613706
cg10614045
cg10614188
cg10615414
cg10615519
cg10615591
cg10616216
cg10617909
cg10619379
cg10619398
cg10620395
cg10620457
cg10620680
cg10621825
cg10621902
cg10623219
cg10623221
cg10624445
cg10625096
cg10625247
cg10625575
cg10625666
cg10625737
cg10626107
cg10626169
cg10626305
cg10626515
cg10627913
cg10628035
cg10628465
cg10628785
cg10628818
cg10631546
cg10631854
cg10631947
cg10632093
cg10632395

cg10632525
cg10632600
cg10632837
cg10632955
cg10632971
cg10633040
cg10633463
cg10633609
cg10633648
cg10633827
cg10633906
cg10634403
cg10634568
cg10634728
cg10635122
cg10635188
cg10635722
cg10636246
cg10636896
cg10637995
cg10638780
cg10639029
cg10640333
cg10643236
cg10643271
cg10643489
cg10643691
cg10644072
cg10644148
cg10644361
cg10645317
cg10645426
cg10645640
cg10646368
cg10646402
cg10647513
cg10647739
cg10648542
cg10649130
cg10649903
cg10650127
cg10650398
cg10651657
cg10652393
cg10652558
cg10653465
cg10655021
cg10655144
cg10655236
cg10656204
cg10657141
cg10657228

cg10658542
cg10658666
cg10658853
cg10659138
cg10659805
cg10659886
cg10660253
cg10660256
cg10660367
cg10660848
cg10660909
cg10661191
cg10662314
cg10662643
cg10663078
cg10663180
cg10663557
cg10664162
cg10664768
cg10665065
cg10665263
cg10666081
cg10666909
cg10667810
cg10668195
cg10668614
cg10669129
cg10669386
cg10671710
cg10671757
cg10672136
cg10672164
cg10672416
cg10673246
cg10673328
cg10675058
cg10675138
cg10675515
cg10675797
cg10676084
cg10677214
cg10678215
cg10679156
cg10679301
cg10679688
cg10680051
cg10680488
cg10681391
cg10681525
cg10682566
cg10683615
cg10683775

cg10684297
cg10684627
cg10688297
cg10688799
cg10689404
cg10691282
cg10691387
cg10691866
cg10692571
cg10692693
cg10693058
cg10693390
cg10694030
cg10694152
cg10695105
cg10695180
cg10695490
cg10696969
cg10698097
cg10698928
cg10700019
cg10700424
cg10702283
cg10702366
cg10702418
cg10702457
cg10702770
cg10703453
cg10703619
cg10703692
cg10703826
cg10704491
cg10704680
cg10705085
cg10705306
cg10705541
cg10706438
cg10707964
cg10708221
cg10708271
cg10708400
cg10708898
cg10709246
cg10711303
cg10711778
cg10711871
cg10712934
cg10713839
cg10715092
cg10715223
cg10716120
cg10716356

cg10716612
cg10717463
cg10718130
cg10719144
cg10719346
cg10719604
cg10720154
cg10720545
cg10721030
cg10721149
cg10721755
cg10721812
cg10721834
cg10721942
cg10722217
cg10722426
cg10723617
cg10724054
cg10724086
cg10724130
cg10724690
cg10725720
cg10725768
cg10726214
cg10726725
cg10726931
cg10728473
cg10729111
cg10729196
cg10729854
cg10730174
cg10730576
cg10730712
cg10731020
cg10731281
cg10732094
cg10732611
cg10733317
cg10734448
cg10734581
cg10734665
cg10735599
cg10736208
cg10736337
cg10736454
cg10737718
cg10738025
cg10738041
cg10740895
cg10740902
cg10741025
cg10741153

cg10741308
cg10741333
cg10741422
cg10742225
cg10742917
cg10743390
cg10743697
cg10745498
cg10745499
cg10746159
cg10746278
cg10746936
cg10748355
cg10748359
cg10749607
cg10749994
cg10750010
cg10750182
cg10750934
cg10750959
cg10751726
cg10751811
cg10751914
cg10752406
cg10752604
cg10753283
cg10753398
cg10754002
cg10754215
cg10754341
cg10755058
cg10755235
cg10755723
cg10755973
cg10756457
cg10756475
cg10757951
cg10758022
cg10758369
cg10758466
cg10758824
cg10758875
cg10759648
cg10760715
cg10761141
cg10761315
cg10761558
cg10761708
cg10761807
cg10761978
cg10762626
cg10763234

cg10763374
cg10763891
cg10764891
cg10765397
cg10765655
cg10766282
cg10766475
cg10766866
cg10767045
cg10767141
cg10767192
cg10767216
cg10767350
cg10767554
cg10767662
cg10767665
cg10768063
cg10768206
cg10769618
cg10770187
cg10770742
cg10772185
cg10772304
cg10772991
cg10773224
cg10773249
cg10773266
cg10773526
cg10773805
cg10774440
cg10775000
cg10776244
cg10776919
cg10777000
cg10777347
cg10777461
cg10778072
cg10778337
cg10778636
cg10778841
cg10779317
cg10779492
cg10780367
cg10780718
cg10780778
cg10781632
cg10781814
cg10781870
cg10782277
cg10782703
cg10784258
cg10784386

cg10784511
cg10784519
cg10784776
cg10784813
cg10785263
cg10785340
cg10785385
cg10785622
cg10787197
cg10787409
cg10788618
cg10788735
cg10789050
cg10789281
cg10789300
cg10789370
cg10789749
cg10789956
cg10790429
cg10790672
cg10790685
cg10791278
cg10791406
cg10791422
cg10791580
cg10792120
cg10792130
cg10792302
cg10793234
cg10793669
cg10795509
cg10795659
cg10796022
cg10796805
cg10796899
cg10798225
cg10798745
cg10798815
cg10799608
cg10799791
cg10800620
cg10801143
cg10801328
cg10802371
cg10802509
cg10802977
cg10803034
cg10804254
cg10805039
cg10805440
cg10805970
cg10806146

cg10806236
cg10806318
cg10806586
cg10806639
cg10807309
cg10807643
cg10807894
cg10808710
cg10808934
cg10809134
cg10809166
cg10809485
cg10810290
cg10810752
cg10810769
cg10810799
cg10810847
cg10811880
cg10812186
cg10812684
cg10812889
cg10814005
cg10814646
cg10816760
cg10817441
cg10817916
cg10818566
cg10819050
cg10819244
cg10819495
cg10819981
cg10820158
cg10820186
cg10820587
cg10820926
cg10821127
cg10821320
cg10821845
cg10822718
cg10824063
cg10824677
cg10824915
cg10825068
cg10825847
cg10827485
cg10827488
cg10827650
cg10828316
cg10828337
cg10828561
cg10828577
cg10829004

cg10829024
cg10830841
cg10831212
cg10831324
cg10831642
cg10832005
cg10832076
cg10832281
cg10833685
cg10835083
cg10835986
cg10836101
cg10836110
cg10836392
cg10836431
cg10836509
cg10836540
cg10837116
cg10837383
cg10837653
cg10838091
cg10838260
cg10838380
cg10838500
cg10838844
cg10839475
cg10840135
cg10840704
cg10841124
cg10841248
cg10842557
cg10843046
cg10843343
cg10843707
cg10845124
cg10846615
cg10848140
cg10848367
cg10848640
cg10848828
cg10849289
cg10849854
cg10850791
cg10850823
cg10851010
cg10851168
cg10851251
cg10851698
cg10851715
cg10852154
cg10852177
cg10852698

cg10853086
cg10853431
cg10854423
cg10854441
cg10854528
cg10855531
cg10855746
cg10855978
cg10857167
cg10857558
cg10857884
cg10857905
cg10858103
cg10858752
cg10858945
cg10859755
cg10859789
cg10859930
cg10860578
cg10861722
cg10862848
cg10862951
cg10863145
cg10863207
cg10863741
cg10864319
cg10864878
cg10864952
cg10864979
cg10868277
cg10868668
cg10868817
cg10869669
cg10870917
cg10872774
cg10872844
cg10874314
cg10876003
cg10876207
cg10876928
cg10877049
cg10877315
cg10878114
cg10879116
cg10879486
cg10879712
cg10879854
cg10880163
cg10880928
cg10881225
cg10881749
cg10881937

cg10882179
cg10882599
cg10883303
cg10884142
cg10884288
cg10884614
cg10887280
cg10887386
cg10887937
cg10888031
cg10888348
cg10888790
cg10889556
cg10889792
cg10890302
cg10890403
cg10891482
cg10892950
cg10893095
cg10893667
cg10893908
cg10894063
cg10894483
cg10894991
cg10896594
cg10897459
cg10898024
cg10898127
cg10898182
cg10899099
cg10899301
cg10899904
cg10900663
cg10901569
cg10901806
cg10902340
cg10902514
cg10902738
cg10903451
cg10903903
cg10904109
cg10904867
cg10905495
cg10905898
cg10906284
cg10907025
cg10908460
cg10908953
cg10909080
cg10909185
cg10909501
cg10910525

cg10910862
cg10911054
cg10912240
cg10912292
cg10912515
cg10912634
cg10914115
cg10916459
cg10916515
cg10916569
cg10916651
cg10917619
cg10917910
cg10917952
cg10919098
cg10919344
cg10919889
cg10920378
cg10920632
cg10921168
cg10921599
cg10921709
cg10921890
cg10922935
cg10923036
cg10923662
cg10925364
cg10925640
cg10925729
cg10925991
cg10926083
cg10926413
cg10926554
cg10928294
cg10928466
cg10929628
cg10930289
cg10930290
cg10932018
cg10932247
cg10933774
cg10934129
cg10935662
cg10936999
cg10937205
cg10937802
cg10940093
cg10940462
cg10940789
cg10941254
cg10943359
cg10943524

cg10943877
cg10943921
cg10944063
cg10944401
cg10944799
cg10945695
cg10946573
cg10947001
cg10947408
cg10948359
cg10948615
cg10948671
cg10948790
cg10948929
cg10949378
cg10949430
cg10949499
cg10949576
cg10949632
cg10950593
cg10950723
cg10951591
cg10952171
cg10952234
cg10952596
cg10953158
cg10954469
cg10955164
cg10955243
cg10955669
cg10955995
cg10956096
cg10956811
cg10956860
cg10958964
cg10959198
cg10959353
cg10959408
cg10959440
cg10959668
cg10959672
cg10959896
cg10959907
cg10959950
cg10959999
cg10960118
cg10961249
cg10961486
cg10961604
cg10961733
cg10961753
cg10962754

cg10962805
cg10963033
cg10963664
cg10963793
cg10963949
cg10964330
cg10964367
cg10964627
cg10965508
cg10966440
cg10966580
cg10966582
cg10966830
cg10967350
cg10967990
cg10968027
cg10968055
cg10968631
cg10969006
cg10969919
cg10970045
cg10970124
cg10970143
cg10972392
cg10972932
cg10973762
cg10974219
cg10974479
cg10974980
cg10975001
cg10975049
cg10975391
cg10975798
cg10976311
cg10976908
cg10976968
cg10977352
cg10977795
cg10977989
cg10978355
cg10978719
cg10978753
cg10979209
cg10979364
cg10979602
cg10980293
cg10980436
cg10981111
cg10981651
cg10981909
cg10982364
cg10982460

cg10982692
cg10983752
cg10985065
cg10985145
cg10985970
cg10987682
cg10987960
cg10988513
cg10988531
cg10989138
cg10989175
cg10989763
cg10989897
cg10990993
cg10991175
cg10991184
cg10991475
cg10991578
cg10991579
cg10991681
cg10992219
cg10992267
cg10993074
cg10993442
cg10994126
cg10994249
cg10994339
cg10994511
cg10994564
cg10994693
cg10994944
cg10995381
cg10996058
cg10996368
cg10997479
cg10997540
cg10997627
cg10997785
cg10997860
cg10998570
cg10999312
cg10999598
cg10999780
cg10999932
cg11000381
cg11001536
cg11001581
cg11002208
cg11002698
cg11002709
cg11002986
cg11003573

cg11004117
cg11005628
cg11005651
cg11006453
cg11006995
cg11007015
cg11007362
cg11007759
cg11007850
cg11009001
cg11009100
cg11009245
cg11009736
cg11010395
cg11010432
cg11010561
cg11010909
cg11010976
cg11011640
cg11011938
cg11012061
cg11013714
cg11014079
cg11014380
cg11014740
cg11015348
cg11015497
cg11016563
cg11017804
cg11018337
cg11019088
cg11019771
cg11020821
cg11021744
cg11022060
cg11022432
cg11022926
cg11023668
cg11024522
cg11024682
cg11025743
cg11026604
cg11027005
cg11027058
cg11027717
cg11028052
cg11028075
cg11028167
cg11028201
cg11030420
cg11030446
cg11031567

cg11032189
cg11032623
cg11032640
cg11032707
cg11033835
cg11034592
cg11035122
cg11036041
cg11036831
cg11037420
cg11037477
cg11037979
cg11038054
cg11038231
cg11038322
cg11039614
cg11040363
cg11040394
cg11041835
cg11042505
cg11043251
cg11043993
cg11044273
cg11046638
cg11049961
cg11050045
cg11051295
cg11051843
cg11052202
cg11052259
cg11052780
cg11053466
cg11053663
cg11054192
cg11054631
cg11055449
cg11056129
cg11056766
cg11058154
cg11058942
cg11058951
cg11059246
cg11059760
cg11061434
cg11062672
cg11063277
cg11063495
cg11063729
cg11063811
cg11063877
cg11064034
cg11064744

cg11065425
cg11065458
cg11065621
cg11066209
cg11067412
cg11067714
cg11067829
cg11068217
cg11068289
cg11068930
cg11068938
cg11068946
cg11069071
cg11069824
cg11070176
cg11070419
cg11070420
cg11070657
cg11071146
cg11071202
cg11072637
cg11072723
cg11073381
cg11073382
cg11073558
cg11073773
cg11073796
cg11074933
cg11076160
cg11076487
cg11076902
cg11076994
cg11077392
cg11077631
cg11078090
cg11078221
cg11078773
cg11078875
cg11079936
cg11080456
cg11080540
cg11080915
cg11081251
cg11081752
cg11083416
cg11083848
cg11084086
cg11084474
cg11085294
cg11085344
cg11085771
cg11086410

cg11087324
cg11087723
cg11087819
cg11088051
cg11088672
cg11088682
cg11088968
cg11089117
cg11089489
cg11089595
cg11090409
cg11090458
cg11090582
cg11091262
cg11091914
cg11092157
cg11092836
cg11093373
cg11093474
cg11094938
cg11095099
cg11095383
cg11096993
cg11097433
cg11097541
cg11097675
cg11098259
cg11100386
cg11100465
cg11100581
cg11100933
cg11101926
cg11103296
cg11104088
cg11104510
cg11105029
cg11105358
cg11106262
cg11106468
cg11106652
cg11107212
cg11107612
cg11107873
cg11108269
cg11108702
cg11109621
cg11112104
cg11112161
cg11112729
cg11113534
cg11113536
cg11113931

cg11114141
cg11114503
cg11114757
cg11115235
cg11115622
cg11116300
cg11117177
cg11117305
cg11117364
cg11118171
cg11118413
cg11118422
cg11118440
cg11118566
cg11119033
cg11120194
cg11120927
cg11121623
cg11122256
cg11122968
cg11123440
cg11123644
cg11123720
cg11123751
cg11124142
cg11124364
cg11126205
cg11126769
cg11128216
cg11128345
cg11128457
cg11129341
cg11129609
cg11129772
cg11130317
cg11130692
cg11130955
cg11131384
cg11131532
cg11131599
cg11132336
cg11132469
cg11133264
cg11134334
cg11136111
cg11136386
cg11136592
cg11137145
cg11138192
cg11138227
cg11138837
cg11139090

cg11139646
cg11139864
cg11141340
cg11141380
cg11141696
cg11142389
cg11142617
cg11142705
cg11143063
cg11143092
cg11143438
cg11143485
cg11143876
cg11144753
cg11145108
cg11145160
cg11145282
cg11145296
cg11145399
cg11145412
cg11145776
cg11146023
cg11147154
cg11147155
cg11147442
cg11147866
cg11147878
cg11148130
cg11148581
cg11148677
cg11149277
cg11149930
cg11150585
cg11152302
cg11153239
cg11153912
cg11154070
cg11154542
cg11155073
cg11155222
cg11155489
cg11155924
cg11156132
cg11156702
cg11156873
cg11157253
cg11157691
cg11158057
cg11158430
cg11158441
cg11158979
cg11159234

cg11159295
cg11160908
cg11161417
cg11161550
cg11162104
cg11162839
cg11163179
cg11163620
cg11163975
cg11164162
cg11165158
cg11165619
cg11165752
cg11165765
cg11168293
cg11169102
cg11170041
cg11170796
cg11170998
cg11171221
cg11171549
cg11171719
cg11173076
cg11173131
cg11173246
cg11173941
cg11174088
cg11174855
cg11175473
cg11176472
cg11176493
cg11177552
cg11177980
cg11178293
cg11178863
cg11181726
cg11181985
cg11183072
cg11183144
cg11183632
cg11184109
cg11185991
cg11186405
cg11187807
cg11188549
cg11189107
cg11189134
cg11189837
cg11189854
cg11190240
cg11190248
cg11191337

cg11191390
cg11192191
cg11192895
cg11193865
cg11195360
cg11195363
cg11195933
cg11196175
cg11196544
cg11197916
cg11199011
cg11199046
cg11199506
cg11199713
cg11199770
cg11200635
cg11201256
cg11201398
cg11201710
cg11201869
cg11202023
cg11202720
cg11203211
cg11203215
cg11203293
cg11203512
cg11203616
cg11203990
cg11204152
cg11204212
cg11205173
cg11205296
cg11206811
cg11207307
cg11207507
cg11208165
cg11208915
cg11208967
cg11209289
cg11209447
cg11209507
cg11209631
cg11210138
cg11210357
cg11210422
cg11210703
cg11211384
cg11211965
cg11212234
cg11212329
cg11213248
cg11213369

cg11213520
cg11213690
cg11213983
cg11214001
cg11214532
cg11214576
cg11214757
cg11215486
cg11215644
cg11215976
cg11216777
cg11217218
cg11217666
cg11217953
cg11217960
cg11218176
cg11218434
cg11218954
cg11219178
cg11219614
cg11219691
cg11220245
cg11220950
cg11221656
cg11222933
cg11223315
cg11223361
cg11223735
cg11224946
cg11226100
cg11226403
cg11226913
cg11226956
cg11227141
cg11227598
cg11228052
cg11228126
cg11228136
cg11228250
cg11228624
cg11228744
cg11228758
cg11228785
cg11229005
cg11229101
cg11229267
cg11229334
cg11229543
cg11229611
cg11229862
cg11229975
cg11230002

cg11231249
cg11231631
cg11231701
cg11231913
cg11233468
cg11234281
cg11234429
cg11235015
cg11235297
cg11235498
cg11235583
cg11235594
cg11235790
cg11236429
cg11236515
cg11236526
cg11236641
cg11236755
cg11236795
cg11237406
cg11238048
cg11239633
cg11239720
cg11240043
cg11241413
cg11241436
cg11242764
cg11242907
cg11243283
cg11245681
cg11245989
cg11246007
cg11249066
cg11249486
cg11251498
cg11252555
cg11253737
cg11253886
cg11254192
cg11255265
cg11256935
cg11257689
cg11257888
cg11258089
cg11258404
cg11260097
cg11260904
cg11261412
cg11263617
cg11264206
cg11264878
cg11264947

cg11265874
cg11266407
cg11267410
cg11267546
cg11267630
cg11267802
cg11267879
cg11268077
cg11268385
cg11268546
cg11268590
cg11268739
cg11269159
cg11270633
cg11270798
cg11271171
cg11271546
cg11271593
cg11272244
cg11273176
cg11273478
cg11273947
cg11274172
cg11275049
cg11275630
cg11277090
cg11277355
cg11278080
cg11278749
cg11280082
cg11280858
cg11281110
cg11281320
cg11281641
cg11282433
cg11283289
cg11283404
cg11283429
cg11284316
cg11284797
cg11284842
cg11284959
cg11285998
cg11286063
cg11286405
cg11287400
cg11287885
cg11287987
cg11290265
cg11290998
cg11291313
cg11291464

cg11291531
cg11291745
cg11291773
cg11291798
cg11292721
cg11292836
cg11293190
cg11293275
cg11293312
cg11293699
cg11293800
cg11294364
cg11295178
cg11295724
cg11296498
cg11296767
cg11297046
cg11297236
cg11298144
cg11298343
cg11299537
cg11299964
cg11300147
cg11300208
cg11301281
cg11301670
cg11302943
cg11303605
cg11304283
cg11304383
cg11304682
cg11305134
cg11306701
cg11306735
cg11307565
cg11308170
cg11308396
cg11308643
cg11309237
cg11309704
cg11310087
cg11310191
cg11310341
cg11310639
cg11311280
cg11312108
cg11312408
cg11312896
cg11313708
cg11313862
cg11314668
cg11314787

cg11320144
cg11320271
cg11320910
cg11321190
cg11322252
cg11322432
cg11322464
cg11323198
cg11323255
cg11323506
cg11323650
cg11324259
cg11324620
cg11324660
cg11324712
cg11324957
cg11324999
cg11325267
cg11325716
cg11326166
cg11326415
cg11326429
cg11326574
cg11328127
cg11328303
cg11328436
cg11328541
cg11328661
cg11329678
cg11330139
cg11330282
cg11330307
cg11331078
cg11332505
cg11332951
cg11334406
cg11334709
cg11334730
cg11334771
cg11334822
cg11335171
cg11335323
cg11335536
cg11336061
cg11336311
cg11336860
cg11337388
cg11337915
cg11337945
cg11338128
cg11338178
cg11339839

cg11340537
cg11340603
cg11341028
cg11341199
cg11341428
cg11341440
cg11341610
cg11341981
cg11342106
cg11342198
cg11342331
cg11342437
cg11342452
cg11342789
cg11343072
cg11344013
cg11344164
cg11344566
cg11344950
cg11345352
cg11345703
cg11346837
cg11346901
cg11347165
cg11347955
cg11348165
cg11348188
cg11348372
cg11348701
cg11349187
cg11349631
cg11351140
cg11351527
cg11352430
cg11352681
cg11353250
cg11353598
cg11353792
cg11354629
cg11355135
cg11356451
cg11357746
cg11358713
cg11358777
cg11358945
cg11359133
cg11360416
cg11360522
cg11360718
cg11361387
cg11361636
cg11361895

cg11362010
cg11362049
cg11362074
cg11362310
cg11362622
cg11363229
cg11363234
cg11364256
cg11364621
cg11365072
cg11365913
cg11368643
cg11369071
cg11370011
cg11371231
cg11373323
cg11374355
cg11374596
cg11374681
cg11375072
cg11375458
cg11375547
cg11375652
cg11375831
cg11376198
cg11376339
cg11377054
cg11377136
cg11377213
cg11377336
cg11377730
cg11378052
cg11379081
cg11379605
cg11379756
cg11380108
cg11380483
cg11380624
cg11381539
cg11382055
cg11382203
cg11382394
cg11383291
cg11384014
cg11384744
cg11385094
cg11385338
cg11385536
cg11385616
cg11386011
cg11386866
cg11387248

cg11387340
cg11388547
cg11389172
cg11389889
cg11390151
cg11390378
cg11391828
cg11392848
cg11393173
cg11394278
cg11394611
cg11394785
cg11395890
cg11396585
cg11397476
cg11397854
cg11397878
cg11398452
cg11398511
cg11399319
cg11399409
cg11399640
cg11400650
cg11400729
cg11400953
cg11401784
cg11401957
cg11402363
cg11403598
cg11404489
cg11404751
cg11406667
cg11407507
cg11407801
cg11408632
cg11408933
cg11409006
cg11410023
cg11411564
cg11411705
cg11411838
cg11411884
cg11412466
cg11412468
cg11413039
cg11413778
cg11414202
cg11414560
cg11416276
cg11416338
cg11416384
cg11416433

cg11416518
cg11416552
cg11416811
cg11417025
cg11417323
cg11417612
cg11418477
cg11418681
cg11418725
cg11419304
cg11419506
cg11419575
cg11420780
cg11421724
cg11421728
cg11422851
cg11423517
cg11424376
cg11425517
cg11425656
cg11426080
cg11426899
cg11427510
cg11428724
cg11428886
cg11429271
cg11429283
cg11429969
cg11430353
cg11430783
cg11431261
cg11431723
cg11433081
cg11433319
cg11433659
cg11433672
cg11434527
cg11435441
cg11435506
cg11435943
cg11436143
cg11436767
cg11437140
cg11437328
cg11437784
cg11438011
cg11438150
cg11438428
cg11438552
cg11438560
cg11439390
cg11439596

cg11439821
cg11441704
cg11442326
cg11443170
cg11444278
cg11444704
cg11445094
cg11445542
cg11445634
cg11446126
cg11448557
cg11448807
cg11449860
cg11451477
cg11452391
cg11452596
cg11453343
cg11453608
cg11453719
cg11453837
cg11454300
cg11454936
cg11457640
cg11458498
cg11459714
cg11460558
cg11460714
cg11460727
cg11461298
cg11461875
cg11462252
cg11462865
cg11462939
cg11464053
cg11464297
cg11464895
cg11465363
cg11465442
cg11465466
cg11465939
cg11465943
cg11466504
cg11466837
cg11467291
cg11467440
cg11468089
cg11468233
cg11468635
cg11469051
cg11469098
cg11469391
cg11470063

cg11470337
cg11471772
cg11472422
cg11472725
cg11473001
cg11473240
cg11473417
cg11474135
cg11474182
cg11474811
cg11474864
cg11475555
cg11476241
cg11476306
cg11478000
cg11479165
cg11480800
cg11481451
cg11481968
cg11482794
cg11483884
cg11484413
cg11484936
cg11485655
cg11485823
cg11486133
cg11486285
cg11486694
cg11489765
cg11490961
cg11492102
cg11492723
cg11494699
cg11494841
cg11495006
cg11496014
cg11497458
cg11497864
cg11497924
cg11499681
cg11499970
cg11500797
cg11501236
cg11501438
cg11503210
cg11503274
cg11503710
cg11503833
cg11505037
cg11505338
cg11505404
cg11505533

cg11506658
cg11506843
cg11507178
cg11507895
cg11508429
cg11510523
cg11511795
cg11511842
cg11512365
cg11514462
cg11517269
cg11517728
cg11518408
cg11518509
cg11518570
cg11519366
cg11519930
cg11521079
cg11521212
cg11521325
cg11521470
cg11521563
cg11521797
cg11521965
cg11522012
cg11522069
cg11522612
cg11523071
cg11523905
cg11524454
cg11524773
cg11525020
cg11525643
cg11525980
cg11526020
cg11527153
cg11527913
cg11527988
cg11528069
cg11528849
cg11529236
cg11529819
cg11529924
cg11530564
cg11530995
cg11531239
cg11531339
cg11531997
cg11532852
cg11534215
cg11534240
cg11535638

cg11535869
cg11536474
cg11536851
cg11537479
cg11537828
cg11538641
cg11538998
cg11539780
cg11539857
cg11540476
cg11540705
cg11540815
cg11541382
cg11543937
cg11544138
cg11544692
cg11545457
cg11545720
cg11545923
cg11546137
cg11547122
cg11547544
cg11547613
cg11547950
cg11548303
cg11549265
cg11549874
cg11549972
cg11550064
cg11551464
cg11552025
cg11552694
cg11552853
cg11552868
cg11553308
cg11553721
cg11554153
cg11554650
cg11554728
cg11554975
cg11555257
cg11556489
cg11557382
cg11557636
cg11558474
cg11558705
cg11558731
cg11560197
cg11560431
cg11560554
cg11561301
cg11561737

cg11563383
cg11563680
cg11563844
cg11564711
cg11565377
cg11565512
cg11566152
cg11566172
cg11566244
cg11566267
cg11566394
cg11566561
cg11568314
cg11568791
cg11569159
cg11569329
cg11570011
cg11571124
cg11571585
cg11571942
cg11572655
cg11574469
cg11574620
cg11575131
cg11575234
cg11576166
cg11576349
cg11576405
cg11577329
cg11577461
cg11579116
cg11579222
cg11579466
cg11579646
cg11579905
cg11580026
cg11580212
cg11580493
cg11581706
cg11582014
cg11582226
cg11582369
cg11582579
cg11583041
cg11583211
cg11583287
cg11583351
cg11583762
cg11583802
cg11584284
cg11584519
cg11584690

cg11585280
cg11585670
cg11586034
cg11586330
cg11586570
cg11587069
cg11587191
cg11587224
cg11587466
cg11587584
cg11587799
cg11588317
cg11588566
cg11589287
cg11589932
cg11590213
cg11590974
cg11591267
cg11591485
cg11591520
cg11591636
cg11591719
cg11592230
cg11592503
cg11592640
cg11592852
cg11593908
cg11594299
cg11594420
cg11594811
cg11595048
cg11595270
cg11596239
cg11596947
cg11597131
cg11597475
cg11597515
cg11598054
cg11599269
cg11599539
cg11599680
cg11599718
cg11599721
cg11599793
cg11600285
cg11600511
cg11600596
cg11600734
cg11601252
cg11601398
cg11601663
cg11603406

cg11605835
cg11605903
cg11606570
cg11608147
cg11609780
cg11610350
cg11610626
cg11610831
cg11610925
cg11611097
cg11611127
cg11611505
cg11611676
cg11612266
cg11612345
cg11612786
cg11613164
cg11613415
cg11613450
cg11613741
cg11615509
cg11617816
cg11617821
cg11617843
cg11617938
cg11618057
cg11619050
cg11619602
cg11619879
cg11620238
cg11620672
cg11621559
cg11624233
cg11624993
cg11625488
cg11625933
cg11626496
cg11628880
cg11629220
cg11629812
cg11629830
cg11630103
cg11630154
cg11631775
cg11632679
cg11633080
cg11633102
cg11633264
cg11633310
cg11633461
cg11633770
cg11634297

cg11634875
cg11637718
cg11637721
cg11637968
cg11638298
cg11638320
cg11639708
cg11639849
cg11639950
cg11640185
cg11642106
cg11642676
cg11642681
cg11643099
cg11643442
cg11644596
cg11644627
cg11644891
cg11645306
cg11645532
cg11646078
cg11647690
cg11647719
cg11648289
cg11648730
cg11648740
cg11648981
cg11649182
cg11649679
cg11649981
cg11650135
cg11650479
cg11650648
cg11650925
cg11650987
cg11651237
cg11651446
cg11651538
cg11651615
cg11651936
cg11652182
cg11652252
cg11653466
cg11654011
cg11654037
cg11654118
cg11654321
cg11654813
cg11654904
cg11655017
cg11655410
cg11655490

cg11656175
cg11657203
cg11658054
cg11658060
cg11658909
cg11659475
cg11659501
cg11660146
cg11660726
cg11661512
cg11661914
cg11662745
cg11663667
cg11664500
cg11664903
cg11666040
cg11666098
cg11666343
cg11667086
cg11667258
cg11667754
cg11668053
cg11668419
cg11668561
cg11668749
cg11668923
cg11669284
cg11669824
cg11669998
cg11670651
cg11671115
cg11671335
cg11671400
cg11672037
cg11672757
cg11674448
cg11674933
cg11675098
cg11675165
cg11675225
cg11677222
cg11677744
cg11677857
cg11678118
cg11678864
cg11679014
cg11679124
cg11679268
cg11680604
cg11680605
cg11680908
cg11682083

cg11682319
cg11682605
cg11682706
cg11683191
cg11683482
cg11683511
cg11684541
cg11684549
cg11684826
cg11685186
cg11685391
cg11685898
cg11685960
cg11686154
cg11686427
cg11687036
cg11687330
cg11687773
cg11688378
cg11688469
cg11688633
cg11688949
cg11688990
cg11689753
cg11689813
cg11690724
cg11691504
cg11691844
cg11692021
cg11692070
cg11692124
cg11692403
cg11692852
cg11694026
cg11694120
cg11696165
cg11696388
cg11697244
cg11697427
cg11697660
cg11697978
cg11698647
cg11698888
cg11699208
cg11699435
cg11699517
cg11699845
cg11700490
cg11701312
cg11701471
cg11701604
cg11701615

cg11702639
cg11703632
cg11703722
cg11704513
cg11704631
cg11705167
cg11706111
cg11706163
cg11706780
cg11706983
cg11707084
cg11708116
cg11708136
cg11708454
cg11708784
cg11709405
cg11710851
cg11711057
cg11711420
cg11712386
cg11712991
cg11713619
cg11715092
cg11715202
cg11715474
cg11715828
cg11717054
cg11717189
cg11717507
cg11717564
cg11717880
cg11719049
cg11719362
cg11720054
cg11720325
cg11720896
cg11721178
cg11721450
cg11721491
cg11721554
cg11721566
cg11721591
cg11721610
cg11721913
cg11722376
cg11722699
cg11723713
cg11723801
cg11723848
cg11724055
cg11724085
cg11724135

cg11724461
cg11724942
cg11725400
cg11725718
cg11725972
cg11726074
cg11726288
cg11726648
cg11727383
cg11728546
cg11728738
cg11728809
cg11728821
cg11729251
cg11729430
cg11729934
cg11731114
cg11731238
cg11731596
cg11731606
cg11731671
cg11732063
cg11732197
cg11732306
cg11732619
cg11733057
cg11733675
cg11734329
cg11735997
cg11736104
cg11736167
cg11736544
cg11736739
cg11736940
cg11737318
cg11738198
cg11738782
cg11739399
cg11739612
cg11740087
cg11740570
cg11741004
cg11741189
cg11741335
cg11742472
cg11743349
cg11744116
cg11745166
cg11746359
cg11746464
cg11747183
cg11747279

cg11747499
cg11747820
cg11749902
cg11750592
cg11750706
cg11750883
cg11751117
cg11751344
cg11751810
cg11752100
cg11752250
cg11752275
cg11752894
cg11754318
cg11754382
cg11754857
cg11754914
cg11755251
cg11755405
cg11755796
cg11756387
cg11756734
cg11756777
cg11756873
cg11757167
cg11757357
cg11759749
cg11760593
cg11761176
cg11761200
cg11761535
cg11762513
cg11762839
cg11762968
cg11763509
cg11763800
cg11763830
cg11764900
cg11765312
cg11765913
cg11766283
cg11766468
cg11767711
cg11770664
cg11771529
cg11772020
cg11772193
cg11772799
cg11772804
cg11775291
cg11775367
cg11775521

cg11776038
cg11776534
cg11778714
cg11778763
cg11779053
cg11779204
cg11779273
cg11780042
cg11780768
cg11781306
cg11781518
cg11781718
cg11781854
cg11782387
cg11782635
cg11782684
cg11783356
cg11783364
cg11783901
cg11784124
cg11784564
cg11784578
cg11785699
cg11785933
cg11786623
cg11787176
cg11787186
cg11787694
cg11788486
cg11788598
cg11788945
cg11789321
cg11789534
cg11789681
cg11789804
cg11790580
cg11791144
cg11791155
cg11791452
cg11791670
cg11793106
cg11793269
cg11793631
cg11793699
cg11794356
cg11795222
cg11795488
cg11795753
cg11795854
cg11796219
cg11796455
cg11796602

cg11796827
cg11796996
cg11797099
cg11797430
cg11797700
cg11797949
cg11800608
cg11800620
cg11800635
cg11800802
cg11800832
cg11801110
cg11803589
cg11803871
cg11804721
cg11804725
cg11804833
cg11804928
cg11805188
cg11805311
cg11805414
cg11805552
cg11807188
cg11807356
cg11807539
cg11808228
cg11808677
cg11809107
cg11810370
cg11810743
cg11811610
cg11811840
cg11812662
cg11813198
cg11813265
cg11813489
cg11814235
cg11814882
cg11815072
cg11816081
cg11817899
cg11818127
cg11818376
cg11818544
cg11819201
cg11819637
cg11820154
cg11820840
cg11820913
cg11820931
cg11821068
cg11821200

cg11821439
cg11821795
cg11822876
cg11823034
cg11823511
cg11823597
cg11823603
cg11823729
cg11824121
cg11824956
cg11825436
cg11825621
cg11826008
cg11826324
cg11826726
cg11826961
cg11827629
cg11827910
cg11828251
cg11828741
cg11829416
cg11830061
cg11830605
cg11831116
cg11831238
cg11831981
cg11832000
cg11832210
cg11832722
cg11835020
cg11835068
cg11835347
cg11835442
cg11835657
cg11836212
cg11837181
cg11837471
cg11837749
cg11838237
cg11838299
cg11838475
cg11838898
cg11839291
cg11839566
cg11839863
cg11841394
cg11841741
cg11841771
cg11842073
cg11842354
cg11842610
cg11843304

cg11843333
cg11844042
cg11844087
cg11844530
cg11844856
cg11844927
cg11844965
cg11845292
cg11846064
cg11846580
cg11846956
cg11847702
cg11848788
cg11849281
cg11849703
cg11850773
cg11851129
cg11852218
cg11852796
cg11853320
cg11853788
cg11853830
cg11853970
cg11854806
cg11855615
cg11855664
cg11855705
cg11855741
cg11855842
cg11855895
cg11855943
cg11856822
cg11856897
cg11856976
cg11857363
cg11857433
cg11857514
cg11857548
cg11857626
cg11857704
cg11857997
cg11858029
cg11858226
cg11859607
cg11861579
cg11861654
cg11863406
cg11863717
cg11863854
cg11863863
cg11864327
cg11864402

cg11866756
cg11867016
cg11867257
cg11867546
cg11868485
cg11869499
cg11869628
cg11869770
cg11870426
cg11870561
cg11870956
cg11871064
cg11871421
cg11871991
cg11872672
cg11872776
cg11873462
cg11873482
cg11874627
cg11874976
cg11875459
cg11875926
cg11875995
cg11876048
cg11876705
cg11876912
cg11877129
cg11877251
cg11878331
cg11878452
cg11879776
cg11880010
cg11881599
cg11882563
cg11882607
cg11884243
cg11884611
cg11884778
cg11884933
cg11886405
cg11886639
cg11886884
cg11887214
cg11887833
cg11887996
cg11888511
cg11888982
cg11889808
cg11891377
cg11891393
cg11891431
cg11891925

cg11892307
cg11893763
cg11893951
cg11894687
cg11898486
cg11898695
cg11899535
cg11900210
cg11902408
cg11902909
cg11904429
cg11905850
cg11906285
cg11906444
cg11906607
cg11906781
cg11907323
cg11907477
cg11907729
cg11907797
cg11908124
cg11908751
cg11909748
cg11909912
cg11910782
cg11911769
cg11912215
cg11912272
cg11912315
cg11912330
cg11913346
cg11913951
cg11914962
cg11915001
cg11915444
cg11915635
cg11915650
cg11915672
cg11915901
cg11916054
cg11916195
cg11916729
cg11918124
cg11920592
cg11920765
cg11924683
cg11925263
cg11925729
cg11925872
cg11926425
cg11926456
cg11926473

cg11926610
cg11927233
cg11927345
cg11929154
cg11929284
cg11930197
cg11930390
cg11930477
cg11930554
cg11931116
cg11931501
cg11932158
cg11932205
cg11932891
cg11933267
cg11933583
cg11933693
cg11933951
cg11934394
cg11934695
cg11934771
cg11934819
cg11935021
cg11935147
cg11935896
cg11936556
cg11937797
cg11938367
cg11938646
cg11938718
cg11939450
cg11940177
cg11940960
cg11941428
cg11941920
cg11942059
cg11942503
cg11942910
cg11942956
cg11943753
cg11943883
cg11944110
cg11944462
cg11944518
cg11945297
cg11945824
cg11946449
cg11946503
cg11946618
cg11947084
cg11947262
cg11947857

cg11948865
cg11949874
cg11950292
cg11950383
cg11950642
cg11950778
cg11950860
cg11952236
cg11952344
cg11953088
cg11953516
cg11953775
cg11954030
cg11954355
cg11955385
cg11955474
cg11955491
cg11955768
cg11956761
cg11956804
cg11957261
cg11957350
cg11957481
cg11957848
cg11958221
cg11958234
cg11958666
cg11960229
cg11960677
cg11960719
cg11961138
cg11961237
cg11961834
cg11961918
cg11962566
cg11963400
cg11964006
cg11964099
cg11964364
cg11964564
cg11965311
cg11965370
cg11965409
cg11965671
cg11965976
cg11966284
cg11967108
cg11967211
cg11967480
cg11967765
cg11969053
cg11969108

cg11969213
cg11969493
cg11969556
cg11970349
cg11970703
cg11970797
cg11970804
cg11971918
cg11972062
cg11972721
cg11975608
cg11976222
cg11976592
cg11976671
cg11978120
cg11978441
cg11978784
cg11979129
cg11979303
cg11979420
cg11979589
cg11980016
cg11980129
cg11980800
cg11981599
cg11982293
cg11982577
cg11983245
cg11983385
cg11983939
cg11983942
cg11984636
cg11985116
cg11985146
cg11985220
cg11986385
cg11986760
cg11987068
cg11987780
cg11987959
cg11988591
cg11988722
cg11989421
cg11990296
cg11990630
cg11991058
cg11991566
cg11991735
cg11992599
cg11992940
cg11993173
cg11993754

cg11994229
cg11994674
cg11995374
cg11996209
cg11996817
cg11997536
cg11997679
cg11998307
cg11999044
cg12000297
cg12000587
cg12001304
cg12001887
cg12002139
cg12002303
cg12002820
cg12003024
cg12004641
cg12004787
cg12005098
cg12006072
cg12006425
cg12007399
cg12008034
cg12008054
cg12008118
cg12008350
cg12009023
cg12009697
cg12009759
cg12009803
cg12010198
cg12011136
cg12012932
cg12013443
cg12014052
cg12014646
cg12014818
cg12016405
cg12017558
cg12018140
cg12018581
cg12018969
cg12019650
cg12019893
cg12020396
cg12020676
cg12020778
cg12021641
cg12021814
cg12024530
cg12024564

cg12024811
cg12025690
cg12026529
cg12026749
cg12026891
cg12027068
cg12027135
cg12028548
cg12028936
cg12030298
cg12030523
cg12030550
cg12030667
cg12030788
cg12031863
cg12032027
cg12032370
cg12032655
cg12032915
cg12033075
cg12033216
cg12033297
cg12033966
cg12034383
cg12034610
cg12036544
cg12036735
cg12036877
cg12037162
cg12037593
cg12037994
cg12038641
cg12040750
cg12040830
cg12041459
cg12042060
cg12042264
cg12042587
cg12042714
cg12043818
cg12044338
cg12044685
cg12044919
cg12045030
cg12045337
cg12045715
cg12045881
cg12045999
cg12046837
cg12047860
cg12049236
cg12050358

cg12050706
cg12051819
cg12052114
cg12053009
cg12053891
cg12054981
cg12055258
cg12055422
cg12056138
cg12057318
cg12057368
cg12058501
cg12058544
cg12059112
cg12059540
cg12059866
cg12060422
cg12061127
cg12061357
cg12062220
cg12062672
cg12062900
cg12063490
cg12063580
cg12063688
cg12064134
cg12064202
cg12064372
cg12064459
cg12065138
cg12065366
cg12065442
cg12065579
cg12066473
cg12066624
cg12066770
cg12068346
cg12068816
cg12068916
cg12069267
cg12070159
cg12070911
cg12071008
cg12071073
cg12071328
cg12071536
cg12071888
cg12072560
cg12072611
cg12072690
cg12072789
cg12072833

cg12072964
cg12074182
cg12075445
cg12075586
cg12075720
cg12075928
cg12076463
cg12076823
cg12076931
cg12077460
cg12078033
cg12078211
cg12078605
cg12078913
cg12079115
cg12079322
cg12079975
cg12080687
cg12080717
cg12081643
cg12082015
cg12082048
cg12082516
cg12082609
cg12083893
cg12084436
cg12084541
cg12085101
cg12086028
cg12087123
cg12087412
cg12087627
cg12088946
cg12089570
cg12091396
cg12091642
cg12093043
cg12093060
cg12093371
cg12093419
cg12094104
cg12094863
cg12095505
cg12095547
cg12095807
cg12096759
cg12097191
cg12097222
cg12097243
cg12097441
cg12097550
cg12098156

cg12098183
cg12098441
cg12098569
cg12098949
cg12099423
cg12100721
cg12101474
cg12102973
cg12103152
cg12104108
cg12104246
cg12104698
cg12105920
cg12105941
cg12106976
cg12107189
cg12107247
cg12109747
cg12110537
cg12111714
cg12112021
cg12112529
cg12112719
cg12113251
cg12113316
cg12113981
cg12114524
cg12114584
cg12115190
cg12116288
cg12117141
cg12118082
cg12120360
cg12120750
cg12121535
cg12121704
cg12122146
cg12122597
cg12123671
cg12124386
cg12124767
cg12124927
cg12125173
cg12126243
cg12126706
cg12126990
cg12127159
cg12127290
cg12127472
cg12127706
cg12128017
cg12128507

cg12128589
cg12129480
cg12130211
cg12130328
cg12130607
cg12130725
cg12131620
cg12131894
cg12132508
cg12133962
cg12134633
cg12135856
cg12136731
cg12136772
cg12137439
cg12138102
cg12138124
cg12138450
cg12138483
cg12140543
cg12140665
cg12140851
cg12141056
cg12141135
cg12141659
cg12141979
cg12142424
cg12142521
cg12143110
cg12143789
cg12144100
cg12144446
cg12144497
cg12144728
cg12145018
cg12145085
cg12145394
cg12146386
cg12148262
cg12148774
cg12149189
cg12149606
cg12149911
cg12150022
cg12150401
cg12151150
cg12152352
cg12152540
cg12153668
cg12154418
cg12154538
cg12154976

cg12157387
cg12157444
cg12157646
cg12157785
cg12158488
cg12159189
cg12159215
cg12159314
cg12160233
cg12160258
cg12160586
cg12161192
cg12161228
cg12161893
cg12161982
cg12162424
cg12162759
cg12163490
cg12163781
cg12164282
cg12164321
cg12164612
cg12165943
cg12166018
cg12166610
cg12166662
cg12166899
cg12167489
cg12167564
cg12168275
cg12168357
cg12170314
cg12170787
cg12171567
cg12172441
cg12172531
cg12173114
cg12173558
cg12174409
cg12174804
cg12175729
cg12175905
cg12176191
cg12176463
cg12176793
cg12177113
cg12177314
cg12177909
cg12178431
cg12178465
cg12178669
cg12179154

cg12179219
cg12179288
cg12179658
cg12179661
cg12180070
cg12180703
cg12181417
cg12182580
cg12183032
cg12183741
cg12184038
cg12184157
cg12184445
cg12184660
cg12185975
cg12186816
cg12187358
cg12187586
cg12187999
cg12188197
cg12188268
cg12188416
cg12188650
cg12188986
cg12189551
cg12190044
cg12190521
cg12190994
cg12191488
cg12192282
cg12192749
cg12193153
cg12193365
cg12193649
cg12193731
cg12193817
cg12194055
cg12194336
cg12196692
cg12197557
cg12197573
cg12197579
cg12198813
cg12198934
cg12200038
cg12200270
cg12200716
cg12201043
cg12201167
cg12201625
cg12201779
cg12202847

cg12202913
cg12203198
cg12203957
cg12204395
cg12204732
cg12205591
cg12205669
cg12206423
cg12206846
cg12208353
cg12208782
cg12209937
cg12210293
cg12210378
cg12210890
cg12211161
cg12211856
cg12212103
cg12212198
cg12212311
cg12213062
cg12213910
cg12214647
cg12214908
cg12215340
cg12216688
cg12217560
cg12217971
cg12218389
cg12219082
cg12219155
cg12219390
cg12219570
cg12219638
cg12219715
cg12220426
cg12220788
cg12221475
cg12223582
cg12223734
cg12225226
cg12226135
cg12226524
cg12226639
cg12226717
cg12226735
cg12227411
cg12228123
cg12228704
cg12229082
cg12229364
cg12229979

cg12231298
cg12231713
cg12231969
cg12232222
cg12232463
cg12233379
cg12233463
cg12234272
cg12236164
cg12236880
cg12237140
cg12237269
cg12239580
cg12240237
cg12241333
cg12241599
cg12241733
cg12241963
cg12242338
cg12242345
cg12242450
cg12243034
cg12243582
cg12243597
cg12244013
cg12244325
cg12246530
cg12246645
cg12247101
cg12248614
cg12248807
cg12248883
cg12249516
cg12250496
cg12250896
cg12251111
cg12251145
cg12251828
cg12251939
cg12252069
cg12253845
cg12253859
cg12255501
cg12255505
cg12255774
cg12255897
cg12256469
cg12256803
cg12257692
cg12257894
cg12258042
cg12258161

cg12258344
cg12259140
cg12259537
cg12259792
cg12259903
cg12261451
cg12261582
cg12262372
cg12263130
cg12263377
cg12265623
cg12266167
cg12267236
cg12267283
cg12268637
cg12269138
cg12269535
cg12271022
cg12271079
cg12271415
cg12271587
cg12271759
cg12271981
cg12272160
cg12272198
cg12272837
cg12274082
cg12274663
cg12274898
cg12275289
cg12276123
cg12277666
cg12278474
cg12279294
cg12279419
cg12279518
cg12279877
cg12280113
cg12280471
cg12280901
cg12281620
cg12282777
cg12284854
cg12285381
cg12286159
cg12286910
cg12287785
cg12288551
cg12289476
cg12289801
cg12290450
cg12291038

cg12294121
cg12296269
cg12296605
cg12296772
cg12298268
cg12299478
cg12299554
cg12300018
cg12300084
cg12300353
cg12300724
cg12300939
cg12300957
cg12301814
cg12301823
cg12302070
cg12302119
cg12302182
cg12302402
cg12302647
cg12303413
cg12303582
cg12303610
cg12304663
cg12304930
cg12305074
cg12305265
cg12305798
cg12306195
cg12306333
cg12307237
cg12307484
cg12307787
cg12307840
cg12308369
cg12308746
cg12308965
cg12309348
cg12309668
cg12310370
cg12310850
cg12311002
cg12311094
cg12311132
cg12311175
cg12311636
cg12312107
cg12312338
cg12312863
cg12313149
cg12314012
cg12315311

cg12315353
cg12315713
cg12316010
cg12316290
cg12316887
cg12317217
cg12317414
cg12317456
cg12317601
cg12317815
cg12320221
cg12320750
cg12321193
cg12322146
cg12323265
cg12323361
cg12324353
cg12324708
cg12325536
cg12326923
cg12326965
cg12328429
cg12329318
cg12329404
cg12331389
cg12332524
cg12332902
cg12333043
cg12334759
cg12334793
cg12334842
cg12335056
cg12336777
cg12337053
cg12338137
cg12338417
cg12338552
cg12338657
cg12339133
cg12339531
cg12339834
cg12340144
cg12340651
cg12341314
cg12341429
cg12342451
cg12342742
cg12343182
cg12343591
cg12343881
cg12343929
cg12344546

cg12346269
cg12346967
cg12348511
cg12348588
cg12349422
cg12349874
cg12350863
cg12351073
cg12351310
cg12351433
cg12352210
cg12352960
cg12353688
cg12354382
cg12355110
cg12356107
cg12356951
cg12357514
cg12357606
cg12358000
cg12358050
cg12359158
cg12360084
cg12360753
cg12362478
cg12362517
cg12363682
cg12363722
cg12364136
cg12364324
cg12364755
cg12365402
cg12366423
cg12366518
cg12366631
cg12367058
cg12367543
cg12367786
cg12368280
cg12368541
cg12368612
cg12368694
cg12369078
cg12369353
cg12371991
cg12373084
cg12373951
cg12374353
cg12374721
cg12375586
cg12375903
cg12376422

cg12376916
cg12377111
cg12377113
cg12377139
cg12377401
cg12377816
cg12378187
cg12378321
cg12379383
cg12380441
cg12382398
cg12382415
cg12384802
cg12384932
cg12385165
cg12385467
cg12385599
cg12385849
cg12386285
cg12386297
cg12386808
cg12387464
cg12387713
cg12388309
cg12389770
cg12390344
cg12390524
cg12391690
cg12392473
cg12392519
cg12392857
cg12393108
cg12393318
cg12393552
cg12394014
cg12394201
cg12394289
cg12395205
cg12395420
cg12395929
cg12396057
cg12396325
cg12396368
cg12397274
cg12398124
cg12398448
cg12398645
cg12398921
cg12399197
cg12399350
cg12399411
cg12399700

cg12400336
cg12400350
cg12400519
cg12400790
cg12401798
cg12402183
cg12403253
cg12404168
cg12404294
cg12404831
cg12405098
cg12405639
cg12406124
cg12407882
cg12408250
cg12409149
cg12409525
cg12409547
cg12409982
cg12411858
cg12412030
cg12412079
cg12412255
cg12412384
cg12412390
cg12413242
cg12413421
cg12413566
cg12414557
cg12415455
cg12415479
cg12415917
cg12416080
cg12417362
cg12417685
cg12417689
cg12417744
cg12417775
cg12417992
cg12419052
cg12419135
cg12419482
cg12421282
cg12421755
cg12421814
cg12421834
cg12422154
cg12423553
cg12424785
cg12425180
cg12427039
cg12428416

cg12431087
cg12431517
cg12431879
cg12432010
cg12432155
cg12432161
cg12432236
cg12432526
cg12433479
cg12433575
cg12433713
cg12433782
cg12434085
cg12434257
cg12434735
cg12434889
cg12435134
cg12435725
cg12435792
cg12436196
cg12436377
cg12436427
cg12436612
cg12436631
cg12436738
cg12436772
cg12437012
cg12437115
cg12437481
cg12438037
cg12438819
cg12439157
cg12440258
cg12440350
cg12440549
cg12441126
cg12441694
cg12441919
cg12442108
cg12442369
cg12444076
cg12444166
cg12444328
cg12445309
cg12445422
cg12445565
cg12445786
cg12446045
cg12446613
cg12447154
cg12447928
cg12448265

cg12448724
cg12448933
cg12449404
cg12449813
cg12449852
cg12450391
cg12451280
cg12452298
cg12453014
cg12453228
cg12453378
cg12453623
cg12454167
cg12454229
cg12456269
cg12456379
cg12456432
cg12456510
cg12456714
cg12456825
cg12457040
cg12457238
cg12457611
cg12457773
cg12458280
cg12458587
cg12458966
cg12459000
cg12459028
cg12459932
cg12460175
cg12461092
cg12461099
cg12461141
cg12461287
cg12461551
cg12462224
cg12463600
cg12463722
cg12464638
cg12465704
cg12465710
cg12466324
cg12466599
cg12467223
cg12467264
cg12467435
cg12467749
cg12467852
cg12468119
cg12468478
cg12468786

cg12469755
cg12469902
cg12470092
cg12472416
cg12472483
cg12473406
cg12473767
cg12473912
cg12474695
cg12475759
cg12476044
cg12476455
cg12476487
cg12476754
cg12476968
cg12478331
cg12478370
cg12478384
cg12479444
cg12479507
cg12479878
cg12480375
cg12480390
cg12480580
cg12481212
cg12482993
cg12483357
cg12483476
cg12483947
cg12485185
cg12485386
cg12485652
cg12485727
cg12485744
cg12485793
cg12486537
cg12486614
cg12486762
cg12486993
cg12487088
cg12487540
cg12487655
cg12488585
cg12488919
cg12490907
cg12491966
cg12492087
cg12492273
cg12492380
cg12492496
cg12492885
cg12492906

cg12493058
cg12493761
cg12494301
cg12494603
cg12495559
cg12496156
cg12496211
cg12496307
cg12496694
cg12497142
cg12497564
cg12497581
cg12498094
cg12498395
cg12498887
cg12499130
cg12500811
cg12500857
cg12500879
cg12502751
cg12503742
cg12503846
cg12504148
cg12505170
cg12505991
cg12506121
cg12506971
cg12507030
cg12507788
cg12508451
cg12508624
cg12510035
cg12510221
cg12510512
cg12510535
cg12510717
cg12511214
cg12511310
cg12512013
cg12513379
cg12513880
cg12514506
cg12514933
cg12515517
cg12515635
cg12515638
cg12515942
cg12516059
cg12516359
cg12516686
cg12516767
cg12516954

cg12517027
cg12517576
cg12517843
cg12518442
cg12520276
cg12520549
cg12520982
cg12521267
cg12521353
cg12521736
cg12522599
cg12523924
cg12524061
cg12524168
cg12524428
cg12524698
cg12524865
cg12525474
cg12525584
cg12526197
cg12527478
cg12528408
cg12528835
cg12530050
cg12530503
cg12530864
cg12533496
cg12533853
cg12534424
cg12535280
cg12535476
cg12535715
cg12536891
cg12537394
cg12537728
cg12539124
cg12541454
cg12541721
cg12542080
cg12542255
cg12542584
cg12542656
cg12543219
cg12543245
cg12543836
cg12543918
cg12544191
cg12544243
cg12544392
cg12545687
cg12546695
cg12547085

cg12547166
cg12547839
cg12547939
cg12548634
cg12548638
cg12549906
cg12549908
cg12550344
cg12550767
cg12552104
cg12552402
cg12553321
cg12554268
cg12554573
cg12555086
cg12555380
cg12555810
cg12555816
cg12556325
cg12556823
cg12558090
cg12558835
cg12559471
cg12559827
cg12560421
cg12560772
cg12560987
cg12562132
cg12562403
cg12562479
cg12562660
cg12563178
cg12563372
cg12565335
cg12565585
cg12565637
cg12565985
cg12566756
cg12566890
cg12568669
cg12569007
cg12569119
cg12569835
cg12570134
cg12570712
cg12570716
cg12571570
cg12571629
cg12573849
cg12573871
cg12574576
cg12575674

cg12577320
cg12579684
cg12580580
cg12580783
cg12581549
cg12581627
cg12581691
cg12581769
cg12582728
cg12582777
cg12583087
cg12583369
cg12583550
cg12583572
cg12584785
cg12586347
cg12586707
cg12587615
cg12587960
cg12588200
cg12588299
cg12588917
cg12589098
cg12589188
cg12589431
cg12589456
cg12589538
cg12590535
cg12590600
cg12590902
cg12591622
cg12591668
cg12591770
cg12592276
cg12592365
cg12592691
cg12593634
cg12593764
cg12593776
cg12594140
cg12594253
cg12594635
cg12594648
cg12594803
cg12595013
cg12595306
cg12595401
cg12595742
cg12596243
cg12596692
cg12597389
cg12597472

cg12597839
cg12598054
cg12598235
cg12598524
cg12598581
cg12598606
cg12598635
cg12598865
cg12599168
cg12599390
cg12599598
cg12599641
cg12599765
cg12600174
cg12600534
cg12600749
cg12600784
cg12600843
cg12601945
cg12602633
cg12603369
cg12604192
cg12604950
cg12605157
cg12605662
cg12605747
cg12606396
cg12606891
cg12607235
cg12607287
cg12608132
cg12608507
cg12608633
cg12608881
cg12609140
cg12610078
cg12610207
cg12610471
cg12610859
cg12611423
cg12612296
cg12612393
cg12612410
cg12613618
cg12613632
cg12613839
cg12614148
cg12614667
cg12615165
cg12615535
cg12615982
cg12616487

cg12616592
cg12616766
cg12617066
cg12617080
cg12617225
cg12617456
cg12617684
cg12618806
cg12619015
cg12619162
cg12619347
cg12620005
cg12621044
cg12621315
cg12621745
cg12623625
cg12623982
cg12624155
cg12624721
cg12626242
cg12626882
cg12627071
cg12627074
cg12627726
cg12628075
cg12628145
cg12628659
cg12629008
cg12629325
cg12629564
cg12629698
cg12629796
cg12629991
cg12630082
cg12630243
cg12630953
cg12631351
cg12631819
cg12632309
cg12632832
cg12633680
cg12634208
cg12635120
cg12635314
cg12635937
cg12636169
cg12636369
cg12636421
cg12636543
cg12636785
cg12636835
cg12637676

cg12638731
cg12639234
cg12639558
cg12639990
cg12640049
cg12640482
cg12642207
cg12642651
cg12643406
cg12643502
cg12645075
cg12645236
cg12646029
cg12646385
cg12646386
cg12646452
cg12646649
cg12646786
cg12646800
cg12647283
cg12647496
cg12647574
cg12647920
cg12648520
cg12648691
cg12649051
cg12649175
cg12650490
cg12650635
cg12651352
cg12651712
cg12651953
cg12652634
cg12653570
cg12654032
cg12655190
cg12655921
cg12656424
cg12656475
cg12656815
cg12658374
cg12658387
cg12659128
cg12659158
cg12659291
cg12659446
cg12659883
cg12659981
cg12660445
cg12660813
cg12661206
cg12662072

cg12663490
cg12663535
cg12663550
cg12664056
cg12664209
cg12664291
cg12664464
cg12664649
cg12664795
cg12664951
cg12665059
cg12665460
cg12666279
cg12666874
cg12666899
cg12667732
cg12668122
cg12668551
cg12669088
cg12669354
cg12670347
cg12672540
cg12674840
cg12676322
cg12677330
cg12679790
cg12680205
cg12680719
cg12680822
cg12681402
cg12681478
cg12681771
cg12681796
cg12681948
cg12682367
cg12682458
cg12682607
cg12682684
cg12682872
cg12683120
cg12684175
cg12684999
cg12685846
cg12686055
cg12687463
cg12689055
cg12689128
cg12689670
cg12689689
cg12689806
cg12690119
cg12690996

cg12691739
cg12692069
cg12693436
cg12693641
cg12696488
cg12698487
cg12698626
cg12699286
cg12699321
cg12699371
cg12699728
cg12699756
cg12699865
cg12700449
cg12700788
cg12700904
cg12701188
cg12702615
cg12703333
cg12705156
cg12705735
cg12706947
cg12707346
cg12708124
cg12708634
cg12710538
cg12711059
cg12712659
cg12712908
cg12713060
cg12713909
cg12714665
cg12714746
cg12715235
cg12715421
cg12715811
cg12716367
cg12717584
cg12718339
cg12718346
cg12718579
cg12719622
cg12719844
cg12719904
cg12720905
cg12721695
cg12723251
cg12723425
cg12723492
cg12724826
cg12726298
cg12726839

cg12726927
cg12727060
cg12728032
cg12728097
cg12728578
cg12729166
cg12730047
cg12730087
cg12731568
cg12731799
cg12732548
cg12732791
cg12734688
cg12734852
cg12735333
cg12735838
cg12735966
cg12736773
cg12737392
cg12737660
cg12737833
cg12738765
cg12738913
cg12739454
cg12739664
cg12739914
cg12740037
cg12740337
cg12741441
cg12742178
cg12743165
cg12743248
cg12743359
cg12743398
cg12743419
cg12743470
cg12743621
cg12743978
cg12744311
cg12744758
cg12744820
cg12744859
cg12745203
cg12745632
cg12746059
cg12746356
cg12747467
cg12748639
cg12750151
cg12750431
cg12750902
cg12751005

cg12751432
cg12751948
cg12752325
cg12752647
cg12753165
cg12754421
cg12756396
cg12757155
cg12758039
cg12758082
cg12758147
cg12758687
cg12758867
cg12759166
cg12760368
cg12760887
cg12760968
cg12761282
cg12761421
cg12761788
cg12762413
cg12762600
cg12762799
cg12763546
cg12764034
cg12764051
cg12764221
cg12764314
cg12764441
cg12765028
cg12765064
cg12765567
cg12767196
cg12768605
cg12768681
cg12769506
cg12769519
cg12769599
cg12769732
cg12771178
cg12771852
cg12773030
cg12773197
cg12776033
cg12776756
cg12777730
cg12778215
cg12778938
cg12779301
cg12781700
cg12781915
cg12782180

cg12782294
cg12783693
cg12784386
cg12784990
cg12785573
cg12786680
cg12787209
cg12787323
cg12788866
cg12789448
cg12789472
cg12789884
cg12789935
cg12790424
cg12790698
cg12790874
cg12791245
cg12791327
cg12792952
cg12793525
cg12793753
cg12794992
cg12795893
cg12796135
cg12796332
cg12797522
cg12797746
cg12797828
cg12798118
cg12798157
cg12798259
cg12798411
cg12798970
cg12799029
cg12799689
cg12799777
cg12800047
cg12800781
cg12801067
cg12801082
cg12801917
cg12802329
cg12802844
cg12802876
cg12802900
cg12804677
cg12804755
cg12805865
cg12806353
cg12806613
cg12806663
cg12806853

cg12807794
cg12808290
cg12808347
cg12810096
cg12810503
cg12810800
cg12811084
cg12811157
cg12811284
cg12811617
cg12812233
cg12812394
cg12812583
cg12812834
cg12813192
cg12814353
cg12816961
cg12816979
cg12817494
cg12818557
cg12819144
cg12819826
cg12820310
cg12821278
cg12821804
cg12823072
cg12823233
cg12824060
cg12824796
cg12825070
cg12827188
cg12827405
cg12827530
cg12828075
cg12828294
cg12829045
cg12829360
cg12829747
cg12831208
cg12832313
cg12832565
cg12832726
cg12833018
cg12833048
cg12833207
cg12833765
cg12835048
cg12835908
cg12836011
cg12836959
cg12837919
cg12838007

cg12838061
cg12838303
cg12839556
cg12840024
cg12840502
cg12843033
cg12843448
cg12843528
cg12843874
cg12844277
cg12844366
cg12844913
cg12846203
cg12846583
cg12847240
cg12848405
cg12848457
cg12849791
cg12850078
cg12852210
cg12852499
cg12853633
cg12853742
cg12854483
cg12854979
cg12856237
cg12856708
cg12856788
cg12857945
cg12858577
cg12859101
cg12859211
cg12859429
cg12860156
cg12860686
cg12862977
cg12864185
cg12865025
cg12865818
cg12865837
cg12866551
cg12866859
cg12867156
cg12868052
cg12868067
cg12868494
cg12869058
cg12869240
cg12870014
cg12870705
cg12871285
cg12871376

cg12871652
cg12873903
cg12873919
cg12874035
cg12875548
cg12875720
cg12876594
cg12877169
cg12877223
cg12877378
cg12877723
cg12878228
cg12878400
cg12878796
cg12878824
cg12881453
cg12881901
cg12882697
cg12883523
cg12883980
cg12884814
cg12886406
cg12886494
cg12888712
cg12889133
cg12889420
cg12890346
cg12891678
cg12892004
cg12892243
cg12892506
cg12892638
cg12892747
cg12893697
cg12894449
cg12894649
cg12895370
cg12895857
cg12896170
cg12896922
cg12898019
cg12898868
cg12900082
cg12901291
cg12902405
cg12902896
cg12903024
cg12903530
cg12903574
cg12903638
cg12904891
cg12905176

cg12906963
cg12907295
cg12907379
cg12907959
cg12909188
cg12910851
cg12910906
cg12911732
cg12912426
cg12912809
cg12915695
cg12915834
cg12915892
cg12916055
cg12916580
cg12917718
cg12918457
cg12918473
cg12919006
cg12919520
cg12919829
cg12920361
cg12921799
cg12922751
cg12923745
cg12924224
cg12924956
cg12925881
cg12926300
cg12926938
cg12927617
cg12927730
cg12927990
cg12928173
cg12928346
cg12929027
cg12929160
cg12929487
cg12929755
cg12929930
cg12930392
cg12930753
cg12930930
cg12931554
cg12931620
cg12932306
cg12933521
cg12934046
cg12934430
cg12934788
cg12935178
cg12935478

cg12936671
cg12937391
cg12938488
cg12939085
cg12939270
cg12939390
cg12939797
cg12940420
cg12940822
cg12940991
cg12942161
cg12942878
cg12944572
cg12945707
cg12946660
cg12947103
cg12947138
cg12947833
cg12950030
cg12950911
cg12951488
cg12951849
cg12952132
cg12954267
cg12954334
cg12954718
cg12955127
cg12955441
cg12955539
cg12956304
cg12957986
cg12958020
cg12958309
cg12958813
cg12959558
cg12960696
cg12960729
cg12961080
cg12963656
cg12965599
cg12966875
cg12966876
cg12967723
cg12967902
cg12969193
cg12969644
cg12970155
cg12970171
cg12970512
cg12970662
cg12973930
cg12973941

cg12974138
cg12974388
cg12974832
cg12975230
cg12975470
cg12975597
cg12976145
cg12977328
cg12977942
cg12978214
cg12978232
cg12978275
cg12978582
cg12978820
cg12979789
cg12980830
cg12981272
cg12981595
cg12982531
cg12983089
cg12983741
cg12984580
cg12984636
cg12984877
cg12984905
cg12985065
cg12986110
cg12986453
cg12986639
cg12987138
cg12987403
cg12988534
cg12988813
cg12989498
cg12989574
cg12990035
cg12990308
cg12990575
cg12992268
cg12993663
cg12993838
cg12994591
cg12995800
cg12997579
cg12997720
cg12998503
cg12999084
cg12999579
cg12999636
cg13000912
cg13001012
cg13001142

cg13001543
cg13001878
cg13001963
cg13002189
cg13002222
cg13002809
cg13004587
cg13004927
cg13005002
cg13005228
cg13008174
cg13008492
cg13009111
cg13010161
cg13010373
cg13010621
cg13011388
cg13011453
cg13011749
cg13012115
cg13012746
cg13012916
cg13015710
cg13016048
cg13016524
cg13016528
cg13016773
cg13017318
cg13017929
cg13019143
cg13019306
cg13019491
cg13019512
cg13019868
cg13020760
cg13020868
cg13021841
cg13021890
cg13022679
cg13022977
cg13022993
cg13023619
cg13023623
cg13023870
cg13024368
cg13024709
cg13026629
cg13026772
cg13027142
cg13027184
cg13027727
cg13027851

cg13028481
cg13028630
cg13028819
cg13030582
cg13031097
cg13031551
cg13031611
cg13033338
cg13033858
cg13034073
cg13035743
cg13035910
cg13036546
cg13036549
cg13036702
cg13037118
cg13037201
cg13038544
cg13038685
cg13039082
cg13039199
cg13039454
cg13040392
cg13041244
cg13041597
cg13042605
cg13043815
cg13044052
cg13046524
cg13046688
cg13046832
cg13047196
cg13047306
cg13047336
cg13048967
cg13049182
cg13049398
cg13049483
cg13049624
cg13051450
cg13051970
cg13052638
cg13052680
cg13052796
cg13053150
cg13054007
cg13054224
cg13054321
cg13054358
cg13054613
cg13055199
cg13056222

cg13056505
cg13057898
cg13058453
cg13059335
cg13059495
cg13060157
cg13060236
cg13060500
cg13060704
cg13061249
cg13061403
cg13061469
cg13061910
cg13062406
cg13062516
cg13062913
cg13063344
cg13064046
cg13064571
cg13064679
cg13065169
cg13065683
cg13065735
cg13066104
cg13066983
cg13067221
cg13067649
cg13067816
cg13067908
cg13069100
cg13069985
cg13070573
cg13071729
cg13072098
cg13072186
cg13072508
cg13072890
cg13073186
cg13073773
cg13074173
cg13074470
cg13074634
cg13074682
cg13075322
cg13075357
cg13076441
cg13076757
cg13076829
cg13077031
cg13077519
cg13078140
cg13078894

cg13079123
cg13079247
cg13079429
cg13079528
cg13080379
cg13080565
cg13080606
cg13081720
cg13083057
cg13083436
cg13084345
cg13084372
cg13084458
cg13085403
cg13088556
cg13088902
cg13089299
cg13091456
cg13091702
cg13092082
cg13092433
cg13092806
cg13093023
cg13093110
cg13093285
cg13093389
cg13093774
cg13095704
cg13096135
cg13096208
cg13097102
cg13097800
cg13097816
cg13098124
cg13098139
cg13098960
cg13099330
cg13099374
cg13099890
cg13099892
cg13100137
cg13101087
cg13101652
cg13101705
cg13102658
cg13102803
cg13103303
cg13103815
cg13104274
cg13106156
cg13106389
cg13106430

cg13106758
cg13107191
cg13107669
cg13107768
cg13107839
cg13109410
cg13109494
cg13110749
cg13113737
cg13117948
cg13119609
cg13120108
cg13120258
cg13120534
cg13120548
cg13121377
cg13121732
cg13121938
cg13122532
cg13123165
cg13123578
cg13123826
cg13123962
cg13124342
cg13125157
cg13125488
cg13125510
cg13126530
cg13126790
cg13126871
cg13127574
cg13128443
cg13130031
cg13130338
cg13131015
cg13131329
cg13131455
cg13132404
cg13132775
cg13133387
cg13133560
cg13133883
cg13134399
cg13134497
cg13134650
cg13134802
cg13134916
cg13134950
cg13135437
cg13136556
cg13137032
cg13139361

cg13139843
cg13139972
cg13139986
cg13141009
cg13142612
cg13142700
cg13145229
cg13145504
cg13145528
cg13146484
cg13146553
cg13146674
cg13148478
cg13148650
cg13150306
cg13151006
cg13151269
cg13151527
cg13151664
cg13153190
cg13154012
cg13154614
cg13154622
cg13155001
cg13156856
cg13157757
cg13157980
cg13159318
cg13160386
cg13160891
cg13160967
cg13161658
cg13163770
cg13163930
cg13164309
cg13165070
cg13166171
cg13166213
cg13166577
cg13167143
cg13167443
cg13168012
cg13168617
cg13168820
cg13169011
cg13169515
cg13170640
cg13170891
cg13171004
cg13171406
cg13171679
cg13171718

cg13171904
cg13173286
cg13173392
cg13174735
cg13175739
cg13175830
cg13176012
cg13177275
cg13177747
cg13178198
cg13178597
cg13179074
cg13179915
cg13180098
cg13180566
cg13180844
cg13182816
cg13183651
cg13184823
cg13184872
cg13186327
cg13186597
cg13186648
cg13187990
cg13188263
cg13189271
cg13189434
cg13191049
cg13191195
cg13191951
cg13192013
cg13192714
cg13193179
cg13193239
cg13194045
cg13194260
cg13194425
cg13195520
cg13195526
cg13195895
cg13197216
cg13197450
cg13198297
cg13198321
cg13200838
cg13201172
cg13201297
cg13201587
cg13201808
cg13202213
cg13202415
cg13202845

cg13203394
cg13205613
cg13205848
cg13206191
cg13206220
cg13207326
cg13208333
cg13208922
cg13209113
cg13209192
cg13209481
cg13209567
cg13209739
cg13209907
cg13210467
cg13210534
cg13210595
cg13210618
cg13210763
cg13212186
cg13212525
cg13213987
cg13214162
cg13216240
cg13218831
cg13219205
cg13219709
cg13219896
cg13219922
cg13220569
cg13220723
cg13221681
cg13221899
cg13222752
cg13222915
cg13223043
cg13223209
cg13223721
cg13224161
cg13224420
cg13226290
cg13226457
cg13226597
cg13227806
cg13228636
cg13228642
cg13229078
cg13231131
cg13231322
cg13233461
cg13233545
cg13234062

cg13234063
cg13234863
cg13235059
cg13235717
cg13235825
cg13236257
cg13236649
cg13237740
cg13238909
cg13238990
cg13239348
cg13239420
cg13240206
cg13241144
cg13241663
cg13242249
cg13242871
cg13242944
cg13244293
cg13244315
cg13244522
cg13244799
cg13245228
cg13246235
cg13248574
cg13249774
cg13250203
cg13250324
cg13250679
cg13252206
cg13252307
cg13252730
cg13253847
cg13254553
cg13255398
cg13255629
cg13256912
cg13257129
cg13257436
cg13257690
cg13258655
cg13258774
cg13258831
cg13259506
cg13259539
cg13259925
cg13260474
cg13260758
cg13261390
cg13261713
cg13261725
cg13261938

cg13261971
cg13262467
cg13263512
cg13264399
cg13265003
cg13265583
cg13265740
cg13265789
cg13266018
cg13266242
cg13266292
cg13266327
cg13266435
cg13266773
cg13267264
cg13267931
cg13269555
cg13271206
cg13272108
cg13272644
cg13272701
cg13273236
cg13273396
cg13274077
cg13274254
cg13275679
cg13275724
cg13276570
cg13276580
cg13276631
cg13277254
cg13277939
cg13278052
cg13278105
cg13279009
cg13279083
cg13279673
cg13280803
cg13280914
cg13281139
cg13282691
cg13283094
cg13283896
cg13283952
cg13284426
cg13285077
cg13285174
cg13285846
cg13287523
cg13288164
cg13289202
cg13289755

cg13290001
cg13290254
cg13290371
cg13290476
cg13290719
cg13291611
cg13291704
cg13292082
cg13292636
cg13292703
cg13293266
cg13295314
cg13296809
cg13297960
cg13298147
cg13298167
cg13298466
cg13298841
cg13298963
cg13298997
cg13299325
cg13299707
cg13299984
cg13300202
cg13300246
cg13300670
cg13300744
cg13300756
cg13301454
cg13301676
cg13301733
cg13302823
cg13302994
cg13305336
cg13306335
cg13306338
cg13306522
cg13307058
cg13307384
cg13307880
cg13308187
cg13309429
cg13309461
cg13310292
cg13310971
cg13311096
cg13311890
cg13312458
cg13312668
cg13313616
cg13314145
cg13314694

cg13315728
cg13315923
cg13316148
cg13316295
cg13316404
cg13316854
cg13318129
cg13318771
cg13319042
cg13319506
cg13320138
cg13320146
cg13320211
cg13320291
cg13320436
cg13320538
cg13320585
cg13320842
cg13321077
cg13321967
cg13322175
cg13323489
cg13324311
cg13324546
cg13325417
cg13326175
cg13326844
cg13327228
cg13327664
cg13328151
cg13328400
cg13328713
cg13329242
cg13329789
cg13330005
cg13330050
cg13330363
cg13330559
cg13331200
cg13332832
cg13332953
cg13333800
cg13333954
cg13334518
cg13335054
cg13335567
cg13335974
cg13336112
cg13336662
cg13337064
cg13337413
cg13338454

cg13338734
cg13338798
cg13339454
cg13339930
cg13340705
cg13341470
cg13341995
cg13342435
cg13342710
cg13343392
cg13343486
cg13343818
cg13344206
cg13344587
cg13344740
cg13344886
cg13345726
cg13346411
cg13347257
cg13347910
cg13348062
cg13348519
cg13348733
cg13348943
cg13348944
cg13349425
cg13350173
cg13350783
cg13351406
cg13351552
cg13351621
cg13351860
cg13352750
cg13353152
cg13353679
cg13354679
cg13354872
cg13354893
cg13355129
cg13355542
cg13356792
cg13356896
cg13357341
cg13357408
cg13357714
cg13357922
cg13358321
cg13358636
cg13358873
cg13359415
cg13359998
cg13360215

cg13361665
cg13361910
cg13362028
cg13362546
cg13362627
cg13364903
cg13365524
cg13365620
cg13365972
cg13366738
cg13366863
cg13367406
cg13368756
cg13368923
cg13368983
cg13369099
cg13369817
cg13370427
cg13370485
cg13371777
cg13371788
cg13371951
cg13372055
cg13372231
cg13372879
cg13373080
cg13373343
cg13373589
cg13374432
cg13375187
cg13375629
cg13376199
cg13376598
cg13378013
cg13378643
cg13378649
cg13379195
cg13379558
cg13379822
cg13381844
cg13381984
cg13382072
cg13382322
cg13382516
cg13382700
cg13382769
cg13383814
cg13384150
cg13384816
cg13384850
cg13385016
cg13385568

cg13385794
cg13386070
cg13387113
cg13387972
cg13388277
cg13388692
cg13388830
cg13389146
cg13389508
cg13390441
cg13390484
cg13390570
cg13391028
cg13391235
cg13391381
cg13391455
cg13392022
cg13392029
cg13392512
cg13392687
cg13392885
cg13393433
cg13393505
cg13393785
cg13393978
cg13394110
cg13394491
cg13395854
cg13396044
cg13396060
cg13396068
cg13396436
cg13396607
cg13396943
cg13397057
cg13397166
cg13397260
cg13397282
cg13397359
cg13397818
cg13398192
cg13398346
cg13399261
cg13400077
cg13400168
cg13400248
cg13400591
cg13401079
cg13401235
cg13401703
cg13403695
cg13403967

cg13404054
cg13404472
cg13404951
cg13405332
cg13405608
cg13405636
cg13405678
cg13405782
cg13405783
cg13406243
cg13406339
cg13406797
cg13407456
cg13407658
cg13408636
cg13408655
cg13408712
cg13408752
cg13408966
cg13409409
cg13409449
cg13409645
cg13410614
cg13410764
cg13411311
cg13411373
cg13411522
cg13411554
cg13412213
cg13412283
cg13413719
cg13413982
cg13414212
cg13414654
cg13415073
cg13415773
cg13416129
cg13416381
cg13418632
cg13418710
cg13419240
cg13419792
cg13419884
cg13420089
cg13420413
cg13420975
cg13421247
cg13421345
cg13421412
cg13422676
cg13422804
cg13423098

cg13423759
cg13423770
cg13424673
cg13425135
cg13425146
cg13425279
cg13425335
cg13426307
cg13426642
cg13427189
cg13427748
cg13427789
cg13427827
cg13427828
cg13427880
cg13428066
cg13428477
cg13428567
cg13429145
cg13429555
cg13430053
cg13430376
cg13430464
cg13430755
cg13430960
cg13431678
cg13432253
cg13433278
cg13433302
cg13433729
cg13433951
cg13434142
cg13434308
cg13434312
cg13434411
cg13434489
cg13434764
cg13434842
cg13434989
cg13435795
cg13436155
cg13436968
cg13437164
cg13437581
cg13438095
cg13438337
cg13438549
cg13438893
cg13439315
cg13439595
cg13440692
cg13441058

cg13441112
cg13441730
cg13442016
cg13442749
cg13442960
cg13443165
cg13443950
cg13444151
cg13444966
cg13445067
cg13445177
cg13445230
cg13445251
cg13445796
cg13446070
cg13446351
cg13446474
cg13446622
cg13446689
cg13446918
cg13447033
cg13447566
cg13447933
cg13448605
cg13448704
cg13448753
cg13448775
cg13449257
cg13449332
cg13449535
cg13449778
cg13450005
cg13450196
cg13450437
cg13451000
cg13451093
cg13451416
cg13451497
cg13451637
cg13451651
cg13452923
cg13453157
cg13453374
cg13453847
cg13453904
cg13454846
cg13455326
cg13455444
cg13455704
cg13455829
cg13455960
cg13456138

cg13456504
cg13457172
cg13457427
cg13457462
cg13457515
cg13457549
cg13457741
cg13457790
cg13458211
cg13458212
cg13458407
cg13458645
cg13458781
cg13458803
cg13459291
cg13459303
cg13460168
cg13460378
cg13460643
cg13460850
cg13460934
cg13461241
cg13461554
cg13461622
cg13462129
cg13462544
cg13462622
cg13463245
cg13463516
cg13463690
cg13464004
cg13464117
cg13464337
cg13464436
cg13464738
cg13464915
cg13464995
cg13465477
cg13465587
cg13466180
cg13466456
cg13466725
cg13466809
cg13466917
cg13466988
cg13467292
cg13467765
cg13467939
cg13468767
cg13469034
cg13469268
cg13470155

cg13471188
cg13471209
cg13471932
cg13472192
cg13472594
cg13473196
cg13473274
cg13473336
cg13473356
cg13473691
cg13474262
cg13474414
cg13474719
cg13474918
cg13475638
cg13476044
cg13476200
cg13477178
cg13478228
cg13478289
cg13478588
cg13478737
cg13479204
cg13480357
cg13480465
cg13480902
cg13481126
cg13481132
cg13481239
cg13481638
cg13482073
cg13482327
cg13482432
cg13482911
cg13483727
cg13483882
cg13484901
cg13485564
cg13485718
cg13485874
cg13485980
cg13486532
cg13486600
cg13486641
cg13486759
cg13486805
cg13487312
cg13487474
cg13487726
cg13487918
cg13487983
cg13487992

cg13488201
cg13488851
cg13490235
cg13490368
cg13490422
cg13490979
cg13491139
cg13491144
cg13491317
cg13491400
cg13491462
cg13491490
cg13491662
cg13491731
cg13491907
cg13492223
cg13492364
cg13492692
cg13492826
cg13493001
cg13493620
cg13494037
cg13494087
cg13494587
cg13494826
cg13495205
cg13495373
cg13496244
cg13496323
cg13497315
cg13497525
cg13498800
cg13499300
cg13499324
cg13500720
cg13500877
cg13501185
cg13501527
cg13501538
cg13502128
cg13502395
cg13502540
cg13502716
cg13502741
cg13503179
cg13503564
cg13504993
cg13505279
cg13505631
cg13506288
cg13506496
cg13506600

cg13507207
cg13507224
cg13508672
cg13508800
cg13508904
cg13509195
cg13509702
cg13509934
cg13510418
cg13510813
cg13511000
cg13511324
cg13511576
cg13511936
cg13512032
cg13512830
cg13512859
cg13512948
cg13512951
cg13513349
cg13514042
cg13515064
cg13515269
cg13515808
cg13516479
cg13517124
cg13518265
cg13518529
cg13518604
cg13518856
cg13518968
cg13519035
cg13519184
cg13519191
cg13519373
cg13520525
cg13521045
cg13521077
cg13521124
cg13521678
cg13521941
cg13522187
cg13522575
cg13522882
cg13522891
cg13524302
cg13524919
cg13525428
cg13525804
cg13526007
cg13526040
cg13526221

cg13526476
cg13526488
cg13526492
cg13526518
cg13526915
cg13527203
cg13527631
cg13528513
cg13528935
cg13529623
cg13530039
cg13530377
cg13531315
cg13531387
cg13531460
cg13532004
cg13533061
cg13533069
cg13533070
cg13533151
cg13533340
cg13534095
cg13536051
cg13537008
cg13537240
cg13538299
cg13539545
cg13540060
cg13540341
cg13540411
cg13540981
cg13541578
cg13541788
cg13541970
cg13542368
cg13543106
cg13543216
cg13543254
cg13543473
cg13543915
cg13545212
cg13545297
cg13546223
cg13546308
cg13546414
cg13546736
cg13547559
cg13547644
cg13547694
cg13547889
cg13548452
cg13548543

cg13548554
cg13549174
cg13549345
cg13549719
cg13549845
cg13549940
cg13550311
cg13550401
cg13550410
cg13550608
cg13550694
cg13551330
cg13551505
cg13551706
cg13552026
cg13552201
cg13552712
cg13553204
cg13553455
cg13553498
cg13554136
cg13554242
cg13555116
cg13555225
cg13555415
cg13555464
cg13555662
cg13555928
cg13556624
cg13556767
cg13556934
cg13557668
cg13557752
cg13558322
cg13558568
cg13558754
cg13559409
cg13560058
cg13560612
cg13561390
cg13561592
cg13561879
cg13562102
cg13562266
cg13562353
cg13562542
cg13563074
cg13563725
cg13563869
cg13563959
cg13564428
cg13564460

cg13564742
cg13565138
cg13565300
cg13566088
cg13566141
cg13566610
cg13566648
cg13567205
cg13567360
cg13567378
cg13567403
cg13568213
cg13568659
cg13569051
cg13569101
cg13569342
cg13569868
cg13570101
cg13570284
cg13570487
cg13570585
cg13570831
cg13570972
cg13571224
cg13571319
cg13571358
cg13572071
cg13572369
cg13573210
cg13573346
cg13573371
cg13573582
cg13573745
cg13574150
cg13574488
cg13576330
cg13576991
cg13577389
cg13577778
cg13579205
cg13579932
cg13580107
cg13580477
cg13580783
cg13580827
cg13581422
cg13581779
cg13581832
cg13582028
cg13582399
cg13582816
cg13583162

cg13583414
cg13583523
cg13584258
cg13585175
cg13585240
cg13585586
cg13585675
cg13585927
cg13585930
cg13586457
cg13586794
cg13587552
cg13588086
cg13588127
cg13588355
cg13588599
cg13589218
cg13590228
cg13590277
cg13590456
cg13590979
cg13591352
cg13591408
cg13592599
cg13592931
cg13593192
cg13593617
cg13594465
cg13594869
cg13595161
cg13595308
cg13596036
cg13596370
cg13596497
cg13597317
cg13597554
cg13598193
cg13598358
cg13599020
cg13599062
cg13600201
cg13600314
cg13600489
cg13600632
cg13600716
cg13601435
cg13601565
cg13601977
cg13601997
cg13603508
cg13603599
cg13605325

cg13605674
cg13606255
cg13606495
cg13606569
cg13607179
cg13607226
cg13607323
cg13608414
cg13608733
cg13609544
cg13609694
cg13609861
cg13610014
cg13610455
cg13611121
cg13611347
cg13611432
cg13612583
cg13613044
cg13613891
cg13614060
cg13614372
cg13614477
cg13614617
cg13615402
cg13616033
cg13617087
cg13617419
cg13618111
cg13619277
cg13619772
cg13620439
cg13621396
cg13621612
cg13623495
cg13624385
cg13624833
cg13624860
cg13626140
cg13626633
cg13626881
cg13628057
cg13628514
cg13629388
cg13630095
cg13630131
cg13630739
cg13632630
cg13632816
cg13632983
cg13633262
cg13633270

cg13633964
cg13635578
cg13636014
cg13637731
cg13638028
cg13638257
cg13638983
cg13639536
cg13639901
cg13640769
cg13640972
cg13641043
cg13641567
cg13641591
cg13641690
cg13641993
cg13642872
cg13643452
cg13643465
cg13643764
cg13643914
cg13644239
cg13644300
cg13644317
cg13644626
cg13645049
cg13645954
cg13646020
cg13648192
cg13648344
cg13648764
cg13649519
cg13649600
cg13649864
cg13650974
cg13651224
cg13652336
cg13652373
cg13652536
cg13652841
cg13653144
cg13654085
cg13654103
cg13654588
cg13654836
cg13655242
cg13655303
cg13655385
cg13655635
cg13656408
cg13656504
cg13656556

cg13657884
cg13658542
cg13658899
cg13659051
cg13659813
cg13659970
cg13660443
cg13660679
cg13661740
cg13661968
cg13662121
cg13662225
cg13662246
cg13662778
cg13663218
cg13663300
cg13663303
cg13663667
cg13663911
cg13664067
cg13664588
cg13665008
cg13665850
cg13666041
cg13666232
cg13666267
cg13666417
cg13667638
cg13668495
cg13669036
cg13669432
cg13669740
cg13670091
cg13670316
cg13670898
cg13671374
cg13671601
cg13671919
cg13672078
cg13672106
cg13672300
cg13672525
cg13672547
cg13672747
cg13673164
cg13673833
cg13673837
cg13674249
cg13675015
cg13675278
cg13675837
cg13676088

cg13676312
cg13676366
cg13676583
cg13676800
cg13677043
cg13677149
cg13677415
cg13677571
cg13677785
cg13677859
cg13678009
cg13678973
cg13679817
cg13679842
cg13681106
cg13681468
cg13681655
cg13681935
cg13683194
cg13683280
cg13683550
cg13683990
cg13685151
cg13685294
cg13685506
cg13685727
cg13686795
cg13686807
cg13687497
cg13687784
cg13687885
cg13688162
cg13688312
cg13688769
cg13688966
cg13689309
cg13689591
cg13689996
cg13690525
cg13690564
cg13691060
cg13691192
cg13691209
cg13691599
cg13692112
cg13692416
cg13692446
cg13692517
cg13692649
cg13693941
cg13694576
cg13694680

cg13694749
cg13694867
cg13694876
cg13695362
cg13695383
cg13695933
cg13696135
cg13696490
cg13696609
cg13696942
cg13697223
cg13698212
cg13698224
cg13698613
cg13698778
cg13698865
cg13698883
cg13699355
cg13699963
cg13700094
cg13700250
cg13701310
cg13702053
cg13702222
cg13702421
cg13702953
cg13702996
cg13703049
cg13703089
cg13703871
cg13705216
cg13705810
cg13705830
cg13706769
cg13707560
cg13707729
cg13707760
cg13707951
cg13708759
cg13709612
cg13710297
cg13710542
cg13711063
cg13711394
cg13713066
cg13713830
cg13713922
cg13714026
cg13715502
cg13715827
cg13716167
cg13716409

cg13717394
cg13718100
cg13718141
cg13718539
cg13718820
cg13719243
cg13719287
cg13719712
cg13720108
cg13720705
cg13720868
cg13721397
cg13721404
cg13721429
cg13721907
cg13722619
cg13723005
cg13723825
cg13724123
cg13724170
cg13724182
cg13724238
cg13724496
cg13724550
cg13724706
cg13724732
cg13724788
cg13725826
cg13726218
cg13726563
cg13726814
cg13726828
cg13727122
cg13727618
cg13727946
cg13728287
cg13728308
cg13728394
cg13729210
cg13730600
cg13731311
cg13732164
cg13732302
cg13732865
cg13733266
cg13733545
cg13733708
cg13733737
cg13734043
cg13734535
cg13735018
cg13735282

cg13735759
cg13736279
cg13736376
cg13737042
cg13737408
cg13737493
cg13739659
cg13740185
cg13740339
cg13740709
cg13740929
cg13740985
cg13741263
cg13741304
cg13741668
cg13741861
cg13741876
cg13742513
cg13742752
cg13743091
cg13744194
cg13744411
cg13744481
cg13745402
cg13745489
cg13745653
cg13746740
cg13746891
cg13748845
cg13749063
cg13749420
cg13749670
cg13752239
cg13752749
cg13753460
cg13753488
cg13754665
cg13754877
cg13754948
cg13755776
cg13755795
cg13756014
cg13757194
cg13758186
cg13758221
cg13758331
cg13759822
cg13759905
cg13761421
cg13761594
cg13761998
cg13763061

cg13763289
cg13763339
cg13763482
cg13763633
cg13764897
cg13764979
cg13765061
cg13770361
cg13771161
cg13771219
cg13771227
cg13771254
cg13771376
cg13772414
cg13772515
cg13773356
cg13773570
cg13773633
cg13773741
cg13774825
cg13775224
cg13775593
cg13776620
cg13776718
cg13777681
cg13777717
cg13777798
cg13777844
cg13778208
cg13778274
cg13778567
cg13780065
cg13781121
cg13781414
cg13781758
cg13781956
cg13781989
cg13782090
cg13782092
cg13782213
cg13782301
cg13782937
cg13783444
cg13783786
cg13783957
cg13784017
cg13784578
cg13785196
cg13785779
cg13786023
cg13786089
cg13786567

cg13786722
cg13787135
cg13787671
cg13788117
cg13788819
cg13790719
cg13790810
cg13790837
cg13791131
cg13791183
cg13791254
cg13791379
cg13792444
cg13792641
cg13792913
cg13793580
cg13794530
cg13794807
cg13794970
cg13794993
cg13795321
cg13795840
cg13795883
cg13795948
cg13796804
cg13797402
cg13797580
cg13798146
cg13798270
cg13798289
cg13798558
cg13798745
cg13799504
cg13799772
cg13799838
cg13800209
cg13800240
cg13800496
cg13800802
cg13801193
cg13802013
cg13802354
cg13802527
cg13802595
cg13802605
cg13802981
cg13803266
cg13803503
cg13803688
cg13803727
cg13804058
cg13804467

cg13804575
cg13804738
cg13805518
cg13805537
cg13805730
cg13805793
cg13806095
cg13806741
cg13806939
cg13807386
cg13807549
cg13807871
cg13808223
cg13808842
cg13808923
cg13810457
cg13810489
cg13810673
cg13810707
cg13810843
cg13811417
cg13811423
cg13811879
cg13811955
cg13812230
cg13812452
cg13813366
cg13813710
cg13816321
cg13816419
cg13816428
cg13816734
cg13816999
cg13818075
cg13818276
cg13818302
cg13818451
cg13819687
cg13820205
cg13820281
cg13820726
cg13821571
cg13821978
cg13823120
cg13823169
cg13823701
cg13823919
cg13824009
cg13826452
cg13826459
cg13826890
cg13827179

cg13827677
cg13828579
cg13828808
cg13829550
cg13829638
cg13829849
cg13830799
cg13832670
cg13832955
cg13832988
cg13833484
cg13833923
cg13834623
cg13834844
cg13835436
cg13835894
cg13836183
cg13836380
cg13836518
cg13838276
cg13838599
cg13839457
cg13839563
cg13839957
cg13840174
cg13840239
cg13840526
cg13841742
cg13841979
cg13842222
cg13842240
cg13842293
cg13842648
cg13844402
cg13844463
cg13844767
cg13844779
cg13845094
cg13845105
cg13845149
cg13845351
cg13845439
cg13845692
cg13845982
cg13846998
cg13847070
cg13847137
cg13847226
cg13847437
cg13847701
cg13847853
cg13848598

cg13848711
cg13848818
cg13849066
cg13849253
cg13850625
cg13850695
cg13850927
cg13851621
cg13851641
cg13851843
cg13851904
cg13852116
cg13852561
cg13852784
cg13852948
cg13853141
cg13853198
cg13854960
cg13854962
cg13855568
cg13855924
cg13856126
cg13856340
cg13856405
cg13856674
cg13856825
cg13856867
cg13857169
cg13858407
cg13858747
cg13858909
cg13858974
cg13859757
cg13859847
cg13860283
cg13860447
cg13860849
cg13861180
cg13861294
cg13862848
cg13862927
cg13864354
cg13864749
cg13865360
cg13866214
cg13867255
cg13867370
cg13867630
cg13867865
cg13868356
cg13869366
cg13870275

cg13870866
cg13870990
cg13872278
cg13872688
cg13872780
cg13873263
cg13873655
cg13873684
cg13875518
cg13876844
cg13876878
cg13878136
cg13879177
cg13879776
cg13879937
cg13879954
cg13880113
cg13880193
cg13880379
cg13880458
cg13880467
cg13881341
cg13881790
cg13882284
cg13882285
cg13882486
cg13882988
cg13883438
cg13883681
cg13884879
cg13884903
cg13885120
cg13885504
cg13885609
cg13886135
cg13886180
cg13886354
cg13887180
cg13887261
cg13887532
cg13888748
cg13889415
cg13889750
cg13889934
cg13890428
cg13890706
cg13890969
cg13891021
cg13891768
cg13892257
cg13892828
cg13894719

cg13895046
cg13895322
cg13895765
cg13895830
cg13895867
cg13896106
cg13896650
cg13897222
cg13897627
cg13897833
cg13898106
cg13898318
cg13898430
cg13899694
cg13900719
cg13901000
cg13901319
cg13904214
cg13904667
cg13904877
cg13905053
cg13905899
cg13906689
cg13906792
cg13906811
cg13908299
cg13908635
cg13908827
cg13908968
cg13909076
cg13909533
cg13909895
cg13910681
cg13910746
cg13910860
cg13911697
cg13911707
cg13911785
cg13911801
cg13911959
cg13912117
cg13912573
cg13912858
cg13914083
cg13914738
cg13914984
cg13915501
cg13915538
cg13915803
cg13916080
cg13916421
cg13916459

cg13916895
cg13916989
cg13917002
cg13917020
cg13917127
cg13917287
cg13917578
cg13917614
cg13917863
cg13917918
cg13918472
cg13918640
cg13919285
cg13919466
cg13920435
cg13920934
cg13921352
cg13921420
cg13921444
cg13922021
cg13922121
cg13922494
cg13922669
cg13922800
cg13923018
cg13923669
cg13924432
cg13924998
cg13925011
cg13926338
cg13926347
cg13927504
cg13927566
cg13927756
cg13928144
cg13928709
cg13929106
cg13929328
cg13930388
cg13930766
cg13930892
cg13931134
cg13931250
cg13931531
cg13931845
cg13932916
cg13933070
cg13933080
cg13933293
cg13933734
cg13933838
cg13934236

cg13934406
cg13934629
cg13935437
cg13935548
cg13935928
cg13935962
cg13936863
cg13936972
cg13937012
cg13937155
cg13937208
cg13937986
cg13939166
cg13939271
cg13939859
cg13940029
cg13940218
cg13940444
cg13940693
cg13940715
cg13941235
cg13941801
cg13941826
cg13942157
cg13942627
cg13942695
cg13942782
cg13943025
cg13943217
cg13944018
cg13944084
cg13944105
cg13944175
cg13944389
cg13944685
cg13944805
cg13945667
cg13946520
cg13946538
cg13946806
cg13947735
cg13948082
cg13948772
cg13949137
cg13949713
cg13950362
cg13950558
cg13950603
cg13951074
cg13951450
cg13951490
cg13952688

cg13953532
cg13954022
cg13954127
cg13954213
cg13954457
cg13954754
cg13955423
cg13955487
cg13957043
cg13957175
cg13959014
cg13959031
cg13959241
cg13959523
cg13960641
cg13960660
cg13960999
cg13961527
cg13961576
cg13962186
cg13962664
cg13962866
cg13963053
cg13963210
cg13964092
cg13964105
cg13964726
cg13965707
cg13966313
cg13966547
cg13966575
cg13966771
cg13966878
cg13967424
cg13967908
cg13967936
cg13969001
cg13970540
cg13971140
cg13971232
cg13971504
cg13971546
cg13971892
cg13972124
cg13972661
cg13973180
cg13973641
cg13973820
cg13973849
cg13974192
cg13974330
cg13974632

cg13974790
cg13975075
cg13975106
cg13975172
cg13976324
cg13976502
cg13976988
cg13977620
cg13978347
cg13978352
cg13978367
cg13978601
cg13978856
cg13979274
cg13979407
cg13979887
cg13980410
cg13980944
cg13981001
cg13981026
cg13981356
cg13981838
cg13982529
cg13982695
cg13982739
cg13982823
cg13983319
cg13983692
cg13984701
cg13985445
cg13985446
cg13985485
cg13985784
cg13986130
cg13986272
cg13986355
cg13986762
cg13987192
cg13988052
cg13988440
cg13988455
cg13989854
cg13989999
cg13990743
cg13992678
cg13993218
cg13994241
cg13994599
cg13994680
cg13995140
cg13995516
cg13996155

cg13996430
cg13996453
cg13996593
cg13996610
cg13996999
cg13997469
cg13997553
cg13997645
cg13997759
cg13998003
cg13998293
cg13998871
cg13998988
cg13999086
cg13999458
cg14000125
cg14000361
cg14001023
cg14001035
cg14001157
cg14001664
cg14001914
cg14002345
cg14002960
cg14003035
cg14003039
cg14003090
cg14003143
cg14003265
cg14004073
cg14004552
cg14005139
cg14005217
cg14005246
cg14005531
cg14006181
cg14006266
cg14006649
cg14007067
cg14007559
cg14008030
cg14008071
cg14008506
cg14008593
cg14009098
cg14009440
cg14009588
cg14010394
cg14010720
cg14011229
cg14011244
cg14011416

cg14011422
cg14011789
cg14012546
cg14013195
cg14014720
cg14015441
cg14016055
cg14016177
cg14016236
cg14016363
cg14018141
cg14018153
cg14018414
cg14018471
cg14018698
cg14020285
cg14021153
cg14021478
cg14021572
cg14022105
cg14022313
cg14023329
cg14023357
cg14024243
cg14024579
cg14024653
cg14024728
cg14025053
cg14025831
cg14025883
cg14026106
cg14026202
cg14026451
cg14026459
cg14028400
cg14028653
cg14029061
cg14029856
cg14030836
cg14030940
cg14031054
cg14031246
cg14031402
cg14031473
cg14032371
cg14032465
cg14032856
cg14032950
cg14033193
cg14033237
cg14033273
cg14034325

cg14034778
cg14034870
cg14035553
cg14035807
cg14036069
cg14036079
cg14036128
cg14036226
cg14036479
cg14036856
cg14037114
cg14037463
cg14037546
cg14037553
cg14037837
cg14038009
cg14038391
cg14038731
cg14039138
cg14039228
cg14039779
cg14040131
cg14040871
cg14040931
cg14041701
cg14041812
cg14041921
cg14041976
cg14042099
cg14042137
cg14042230
cg14042601
cg14042780
cg14042799
cg14042851
cg14042931
cg14043298
cg14043702
cg14043774
cg14043803
cg14044216
cg14044461
cg14044578
cg14044785
cg14045072
cg14046074
cg14046640
cg14047091
cg14047094
cg14047361
cg14047387
cg14048837

cg14049470
cg14049990
cg14051236
cg14051336
cg14051368
cg14051544
cg14051639
cg14051721
cg14051770
cg14053590
cg14053997
cg14054275
cg14054883
cg14054928
cg14054990
cg14055004
cg14055195
cg14055212
cg14055655
cg14055835
cg14055896
cg14056110
cg14056357
cg14056644
cg14057560
cg14057952
cg14058476
cg14058848
cg14059768
cg14059959
cg14060111
cg14060496
cg14060944
cg14062050
cg14062119
cg14065115
cg14065127
cg14065446
cg14065857
cg14066993
cg14067244
cg14067873
cg14068057
cg14068328
cg14068788
cg14069049
cg14069088
cg14069111
cg14069306
cg14069333
cg14069949
cg14070275

cg14070647
cg14071014
cg14071307
cg14072046
cg14072219
cg14072515
cg14073571
cg14073722
cg14073922
cg14074251
cg14074284
cg14074626
cg14075413
cg14075934
cg14076161
cg14076417
cg14077923
cg14078662
cg14078730
cg14079359
cg14080048
cg14080967
cg14080982
cg14081465
cg14081744
cg14082061
cg14082127
cg14082422
cg14082645
cg14083356
cg14083720
cg14084623
cg14084896
cg14085358
cg14085952
cg14086822
cg14087319
cg14087401
cg14087574
cg14087796
cg14088161
cg14088282
cg14088357
cg14089007
cg14089067
cg14089692
cg14089714
cg14090556
cg14090995
cg14091154
cg14091631
cg14091840

cg14092045
cg14092626
cg14093720
cg14093792
cg14093970
cg14094027
cg14094498
cg14094960
cg14094983
cg14095303
cg14096482
cg14096553
cg14096744
cg14096767
cg14096855
cg14096889
cg14097046
cg14097117
cg14098223
cg14098681
cg14098737
cg14098951
cg14099121
cg14099345
cg14099514
cg14099540
cg14101494
cg14101500
cg14101976
cg14102128
cg14102186
cg14102585
cg14102988
cg14103872
cg14104327
cg14106345
cg14106660
cg14106710
cg14107328
cg14107382
cg14107457
cg14108532
cg14109388
cg14109456
cg14110073
cg14110444
cg14110548
cg14111697
cg14112449
cg14113046
cg14113203
cg14113958

cg14113960
cg14114282
cg14114377
cg14115756
cg14115884
cg14116122
cg14116153
cg14116509
cg14116792
cg14117138
cg14118187
cg14118515
cg14118583
cg14118666
cg14119158
cg14119706
cg14119792
cg14119827
cg14119936
cg14119999
cg14120010
cg14120129
cg14120436
cg14120583
cg14120984
cg14121282
cg14121665
cg14122138
cg14122599
cg14122749
cg14123750
cg14123942
cg14124066
cg14124132
cg14124261
cg14124818
cg14124894
cg14125013
cg14125403
cg14125530
cg14125604
cg14126093
cg14126408
cg14126466
cg14127046
cg14127659
cg14127859
cg14127947
cg14128411
cg14129328
cg14129430
cg14129735

cg14129855
cg14130985
cg14131273
cg14131289
cg14132236
cg14134003
cg14134128
cg14134497
cg14135060
cg14135809
cg14135814
cg14136186
cg14136260
cg14136372
cg14137162
cg14137548
cg14137698
cg14138047
cg14138235
cg14139311
cg14139652
cg14140118
cg14140375
cg14140691
cg14140717
cg14141061
cg14141238
cg14141741
cg14141843
cg14142007
cg14143304
cg14143580
cg14143723
cg14144038
cg14144728
cg14145185
cg14146100
cg14146669
cg14146740
cg14147420
cg14148016
cg14148055
cg14148981
cg14150071
cg14150186
cg14150378
cg14150727
cg14150774
cg14151372
cg14151459
cg14152854
cg14153637

cg14153740
cg14154381
cg14154456
cg14154784
cg14155058
cg14155193
cg14155327
cg14155416
cg14155831
cg14156381
cg14156446
cg14156751
cg14157107
cg14158968
cg14159747
cg14160151
cg14161165
cg14161269
cg14161359
cg14161400
cg14161676
cg14161821
cg14162076
cg14162144
cg14162627
cg14163049
cg14163311
cg14163494
cg14164099
cg14165051
cg14165096
cg14165308
cg14165911
cg14166053
cg14166209
cg14166212
cg14166377
cg14166453
cg14166701
cg14167858
cg14168316
cg14168530
cg14168573
cg14168713
cg14168923
cg14169886
cg14170172
cg14170358
cg14170392
cg14170976
cg14171414
cg14171486

cg14171516
cg14171986
cg14172375
cg14173523
cg14173736
cg14174320
cg14174502
cg14174526
cg14175438
cg14175690
cg14176638
cg14176966
cg14177925
cg14178013
cg14178077
cg14178343
cg14178364
cg14178541
cg14178794
cg14178895
cg14179148
cg14179908
cg14179950
cg14180039
cg14180511
cg14181271
cg14181345
cg14181409
cg14181777
cg14182461
cg14182690
cg14182890
cg14183540
cg14183922
cg14184078
cg14184369
cg14184400
cg14184524
cg14184614
cg14184873
cg14185201
cg14185229
cg14186336
cg14186697
cg14187724
cg14187877
cg14188111
cg14188401
cg14188639
cg14188856
cg14189141
cg14189441

cg14189571
cg14189605
cg14190151
cg14191040
cg14191343
cg14191476
cg14191493
cg14191688
cg14192029
cg14192037
cg14192291
cg14192782
cg14192813
cg14192935
cg14192957
cg14193129
cg14193156
cg14193653
cg14193806
cg14194397
cg14194715
cg14195059
cg14195115
cg14195117
cg14195216
cg14196840
cg14197113
cg14197156
cg14197182
cg14198860
cg14199144
cg14199187
cg14199384
cg14199723
cg14200170
cg14200470
cg14200631
cg14201467
cg14201597
cg14202099
cg14203417
cg14203721
cg14203738
cg14204166
cg14204241
cg14204266
cg14205169
cg14205519
cg14205632
cg14205791
cg14206056
cg14206523

cg14206717
cg14207539
cg14207613
cg14208041
cg14208875
cg14209244
cg14209299
cg14210678
cg14211055
cg14211385
cg14211735
cg14211837
cg14211930
cg14212012
cg14212038
cg14212447
cg14213439
cg14213590
cg14214262
cg14215666
cg14215702
cg14216940
cg14217211
cg14218629
cg14218962
cg14219262
cg14220170
cg14220181
cg14220329
cg14221048
cg14221644
cg14222033
cg14222229
cg14222432
cg14223293
cg14224203
cg14224444
cg14225031
cg14225090
cg14225168
cg14225195
cg14225450
cg14225485
cg14225665
cg14225678
cg14226212
cg14226755
cg14227056
cg14227773
cg14227911
cg14228146
cg14228830

cg14229655
cg14230666
cg14230696
cg14230803
cg14231297
cg14232082
cg14232128
cg14232851
cg14233720
cg14234083
cg14234104
cg14234302
cg14234656
cg14235273
cg14235768
cg14236413
cg14236430
cg14236443
cg14236662
cg14236735
cg14236955
cg14236957
cg14236976
cg14237025
cg14237458
cg14238120
cg14238625
cg14238959
cg14238986
cg14239307
cg14239811
cg14240790
cg14240993
cg14241045
cg14242696
cg14242936
cg14243209
cg14243481
cg14243623
cg14243778
cg14244013
cg14244493
cg14244636
cg14245804
cg14246703
cg14246940
cg14247325
cg14247635
cg14248598
cg14248715
cg14248717
cg14249078

cg14249348
cg14250833
cg14251964
cg14252279
cg14252395
cg14252486
cg14252519
cg14253035
cg14253096
cg14253327
cg14253388
cg14254422
cg14254480
cg14254534
cg14254686
cg14254720
cg14254748
cg14254767
cg14254999
cg14255103
cg14255256
cg14255644
cg14255655
cg14256511
cg14256699
cg14257107
cg14257429
cg14257435
cg14257656
cg14258031
cg14258623
cg14258980
cg14259707
cg14259755
cg14260169
cg14260458
cg14261401
cg14261875
cg14262681
cg14262740
cg14262937
cg14263070
cg14263176
cg14263222
cg14263391
cg14263518
cg14263779
cg14264316
cg14264793
cg14264795
cg14265823
cg14266348

cg14266611
cg14266837
cg14266859
cg14267151
cg14267525
cg14267645
cg14267754
cg14267780
cg14269191
cg14269477
cg14269525
cg14269716
cg14269813
cg14269973
cg14270293
cg14270348
cg14270434
cg14270687
cg14270857
cg14271150
cg14271531
cg14271862
cg14273450
cg14274542
cg14274989
cg14275340
cg14275511
cg14275593
cg14275836
cg14276006
cg14276133
cg14276258
cg14276379
cg14276584
cg14277392
cg14277403
cg14278148
cg14279205
cg14279650
cg14280533
cg14280580
cg14281646
cg14282304
cg14282531
cg14282904
cg14283447
cg14283454
cg14283480
cg14283802
cg14283944
cg14283978
cg14285150

cg14285447
cg14285533
cg14285539
cg14286256
cg14286320
cg14286660
cg14286682
cg14286732
cg14287235
cg14287443
cg14287724
cg14287869
cg14288066
cg14288424
cg14288594
cg14289319
cg14289511
cg14289542
cg14289826
cg14289985
cg14290040
cg14290291
cg14290590
cg14290904
cg14291693
cg14291751
cg14291919
cg14292280
cg14292522
cg14292552
cg14292572
cg14292834
cg14293892
cg14294077
cg14294240
cg14294321
cg14294646
cg14294758
cg14295306
cg14295437
cg14295526
cg14295960
cg14296561
cg14297444
cg14298020
cg14298200
cg14298689
cg14298777
cg14299508
cg14299675
cg14299883
cg14299940

cg14300161
cg14300531
cg14301348
cg14301932
cg14302428
cg14303187
cg14303478
cg14303526
cg14304236
cg14304674
cg14304702
cg14304817
cg14305278
cg14305313
cg14305401
cg14305641
cg14306161
cg14306330
cg14306709
cg14306734
cg14307157
cg14307300
cg14307853
cg14308313
cg14308612
cg14310958
cg14311250
cg14311597
cg14311688
cg14312063
cg14312323
cg14312538
cg14312649
cg14312921
cg14313373
cg14314029
cg14314071
cg14314674
cg14314744
cg14315430
cg14315805
cg14315891
cg14317056
cg14317285
cg14317513
cg14317884
cg14318068
cg14320519
cg14321245
cg14321623
cg14321742
cg14323204

cg14323675
cg14324358
cg14324502
cg14325112
cg14325436
cg14325674
cg14325779
cg14326009
cg14326196
cg14326354
cg14326909
cg14326912
cg14326989
cg14327394
cg14327531
cg14328408
cg14328477
cg14328641
cg14328676
cg14329026
cg14329680
cg14330189
cg14331206
cg14331281
cg14331619
cg14331623
cg14331720
cg14332049
cg14333454
cg14333579
cg14333800
cg14334258
cg14334298
cg14334310
cg14334441
cg14335275
cg14335579
cg14336665
cg14337134
cg14337844
cg14338073
cg14338345
cg14338425
cg14338590
cg14338887
cg14339048
cg14339068
cg14339343
cg14339410
cg14340089
cg14340481
cg14341289

cg14341315
cg14341551
cg14342532
cg14342707
cg14342850
cg14343017
cg14343210
cg14343283
cg14344095
cg14344261
cg14344864
cg14344955
cg14345012
cg14345128
cg14346202
cg14346243
cg14346873
cg14346895
cg14347199
cg14347300
cg14347914
cg14348397
cg14348706
cg14349078
cg14349667
cg14349763
cg14350197
cg14350337
cg14350382
cg14350557
cg14350701
cg14350800
cg14352658
cg14352715
cg14352983
cg14353001
cg14353137
cg14353201
cg14353508
cg14353958
cg14354327
cg14354418
cg14354663
cg14354820
cg14355482
cg14355899
cg14356190
cg14356396
cg14356530
cg14356608
cg14356919
cg14359052

cg14359798
cg14360405
cg14360663
cg14361001
cg14361409
cg14361458
cg14361654
cg14361804
cg14363954
cg14365420
cg14366256
cg14366490
cg14366598
cg14367020
cg14368473
cg14369648
cg14369938
cg14370314
cg14370520
cg14370653
cg14371123
cg14372265
cg14372452
cg14372996
cg14373068
cg14373396
cg14373702
cg14373873
cg14373988
cg14374923
cg14375111
cg14375345
cg14375531
cg14376041
cg14376106
cg14376467
cg14377171
cg14377594
cg14377923
cg14378039
cg14378231
cg14378924
cg14378925
cg14379288
cg14379854
cg14380094
cg14380939
cg14382056
cg14382888
cg14383549
cg14384014
cg14384283

cg14385566
cg14386061
cg14386126
cg14386312
cg14386496
cg14386624
cg14386668
cg14386870
cg14386999
cg14387909
cg14388601
cg14389122
cg14389547
cg14391419
cg14391568
cg14391832
cg14392725
cg14393693
cg14394550
cg14394692
cg14394939
cg14396421
cg14396800
cg14397171
cg14397690
cg14397813
cg14398214
cg14398557
cg14398691
cg14399183
cg14399440
cg14399930
cg14400090
cg14400118
cg14400354
cg14400498
cg14400528
cg14400672
cg14400768
cg14400886
cg14401466
cg14402224
cg14403169
cg14403327
cg14404298
cg14404418
cg14404842
cg14405625
cg14405768
cg14405825
cg14405924
cg14406134

cg14406401
cg14406727
cg14406814
cg14406878
cg14407179
cg14407697
cg14409919
cg14409940
cg14410016
cg14410733
cg14411221
cg14411873
cg14412111
cg14412322
cg14413262
cg14413378
cg14413533
cg14414100
cg14414124
cg14414406
cg14414464
cg14415456
cg14415800
cg14416311
cg14416456
cg14417498
cg14417896
cg14417917
cg14418582
cg14418857
cg14419102
cg14419187
cg14419194
cg14419435
cg14419740
cg14419975
cg14420000
cg14420245
cg14421288
cg14421700
cg14421980
cg14422093
cg14422315
cg14423881
cg14425045
cg14425564
cg14425869
cg14425914
cg14426371
cg14427009
cg14427243
cg14428146

cg14429209
cg14429847
cg14429979
cg14430698
cg14430943
cg14431361
cg14431528
cg14431546
cg14431577
cg14432251
cg14432910
cg14433497
cg14433823
cg14433990
cg14434062
cg14434293
cg14434398
cg14434412
cg14434601
cg14434870
cg14435098
cg14435720
cg14437002
cg14437045
cg14438695
cg14438812
cg14439102
cg14439753
cg14439761
cg14439952
cg14440205
cg14441262
cg14441491
cg14441976
cg14442421
cg14442492
cg14443472
cg14443606
cg14444650
cg14444710
cg14445047
cg14445205
cg14445389
cg14445814
cg14446138
cg14446658
cg14446951
cg14448116
cg14448169
cg14448195
cg14448248
cg14448919

cg14449051
cg14449069
cg14449958
cg14451382
cg14451791
cg14451882
cg14452619
cg14453012
cg14453187
cg14453403
cg14454127
cg14454798
cg14454867
cg14454907
cg14455170
cg14455319
cg14455421
cg14456490
cg14456672
cg14456683
cg14457875
cg14458292
cg14458615
cg14458619
cg14458731
cg14459032
cg14459130
cg14461434
cg14462017
cg14463164
cg14463669
cg14463790
cg14464133
cg14464244
cg14464401
cg14465237
cg14465628
cg14465747
cg14465801
cg14465992
cg14466269
cg14466661
cg14466759
cg14467690
cg14467816
cg14467840
cg14468932
cg14469693
cg14471064
cg14471455
cg14471645
cg14472026

cg14472291
cg14472366
cg14472551
cg14472888
cg14473102
cg14473145
cg14473327
cg14473568
cg14473838
cg14473924
cg14474499
cg14475478
cg14475702
cg14475840
cg14475859
cg14476095
cg14476213
cg14476364
cg14476479
cg14476562
cg14477514
cg14477581
cg14478255
cg14478368
cg14478422
cg14478460
cg14478768
cg14479136
cg14479139
cg14479344
cg14479709
cg14480035
cg14480222
cg14480343
cg14481079
cg14481124
cg14482313
cg14482712
cg14483244
cg14483391
cg14483831
cg14484274
cg14484356
cg14485633
cg14485901
cg14486338
cg14486346
cg14486632
cg14486995
cg14487111
cg14487131
cg14487188

cg14487665
cg14487854
cg14487944
cg14488952
cg14489049
cg14489474
cg14490025
cg14490447
cg14490686
cg14491284
cg14491667
cg14491707
cg14491761
cg14491820
cg14492800
cg14494313
cg14494365
cg14494620
cg14495241
cg14495732
cg14495953
cg14497203
cg14498227
cg14498745
cg14499058
cg14499385
cg14499413
cg14499928
cg14500070
cg14500776
cg14501638
cg14502484
cg14502847
cg14503168
cg14503291
cg14503489
cg14503537
cg14503689
cg14503815
cg14504259
cg14504497
cg14504768
cg14504826
cg14505256
cg14505292
cg14505299
cg14505585
cg14505667
cg14506195
cg14507337
cg14507482
cg14507560

cg14507845
cg14507868
cg14508508
cg14508832
cg14509761
cg14510299
cg14510322
cg14510359
cg14510812
cg14511038
cg14511652
cg14511698
cg14511946
cg14512563
cg14513071
cg14513298
cg14513432
cg14513448
cg14513570
cg14513778
cg14514130
cg14514716
cg14514813
cg14514991
cg14515453
cg14515614
cg14515791
cg14515801
cg14516100
cg14516213
cg14516287
cg14516632
cg14517004
cg14517359
cg14517454
cg14517863
cg14518227
cg14518276
cg14518847
cg14518988
cg14519150
cg14519621
cg14520424
cg14521115
cg14521134
cg14521724
cg14522047
cg14522302
cg14522990
cg14523238
cg14523259
cg14523574

cg14523706
cg14523847
cg14524476
cg14524579
cg14524622
cg14524704
cg14524754
cg14524955
cg14524956
cg14526021
cg14526375
cg14526671
cg14526718
cg14526825
cg14526874
cg14527610
cg14527645
cg14528046
cg14529201
cg14529394
cg14529643
cg14529707
cg14530143
cg14530764
cg14530834
cg14531437
cg14531564
cg14531576
cg14532188
cg14532484
cg14533401
cg14533753
cg14534803
cg14535518
cg14535531
cg14535980
cg14537090
cg14537247
cg14537362
cg14537533
cg14537713
cg14537825
cg14538146
cg14538532
cg14538537
cg14538944
cg14539062
cg14539466
cg14539658
cg14539856
cg14540913
cg14541582

cg14542178
cg14542334
cg14542367
cg14542554
cg14543152
cg14543226
cg14544087
cg14544180
cg14544225
cg14544684
cg14545418
cg14546619
cg14547335
cg14547801
cg14547895
cg14548047
cg14548527
cg14548542
cg14548777
cg14548858
cg14549051
cg14549300
cg14549973
cg14550066
cg14550153
cg14550611
cg14550760
cg14550985
cg14551002
cg14551991
cg14553243
cg14553377
cg14553444
cg14553740
cg14554017
cg14554491
cg14555253
cg14555733
cg14555736
cg14556303
cg14556762
cg14556868
cg14557216
cg14557554
cg14557699
cg14557714
cg14558431
cg14558529
cg14558716
cg14558880
cg14559176
cg14559259

cg14559356
cg14560093
cg14560435
cg14560670
cg14561229
cg14561282
cg14562060
cg14562086
cg14562327
cg14562712
cg14562851
cg14563260
cg14563485
cg14563949
cg14564076
cg14564144
cg14564956
cg14565014
cg14565651
cg14565725
cg14565903
cg14566081
cg14566901
cg14566959
cg14567043
cg14567214
cg14567877
cg14567963
cg14568552
cg14568830
cg14569837
cg14570210
cg14570291
cg14571092
cg14571539
cg14572252
cg14574066
cg14574843
cg14575983
cg14576062
cg14576319
cg14578363
cg14578500
cg14578677
cg14578879
cg14579118
cg14579184
cg14579240
cg14579769
cg14579864
cg14579898
cg14580228

cg14580737
cg14580853
cg14580859
cg14581419
cg14581803
cg14582049
cg14582080
cg14582108
cg14582191
cg14582400
cg14582452
cg14582691
cg14582763
cg14582913
cg14582917
cg14582957
cg14583103
cg14583225
cg14583675
cg14583871
cg14584177
cg14584448
cg14584535
cg14584926
cg14585353
cg14585371
cg14585797
cg14586373
cg14588998
cg14589553
cg14590214
cg14591419
cg14591786
cg14591995
cg14592066
cg14592399
cg14592468
cg14592474
cg14592673
cg14592928
cg14594063
cg14594187
cg14595013
cg14595786
cg14596108
cg14596967
cg14597586
cg14597598
cg14597637
cg14597804
cg14598387
cg14598976

cg14600420
cg14600824
cg14600885
cg14601416
cg14601770
cg14602438
cg14602530
cg14603031
cg14603098
cg14603227
cg14603345
cg14603539
cg14604568
cg14605274
cg14605590
cg14605877
cg14605999
cg14606382
cg14606586
cg14607011
cg14607876
cg14609025
cg14609139
cg14609568
cg14610403
cg14611628
cg14611816
cg14612303
cg14612544
cg14612733
cg14612870
cg14613164
cg14613271
cg14614211
cg14614490
cg14614643
cg14614689
cg14614881
cg14614917
cg14615152
cg14615268
cg14615336
cg14615807
cg14615971
cg14616345
cg14616514
cg14617634
cg14618638
cg14620234
cg14620572
cg14620768
cg14621053

cg14621161
cg14621336
cg14622879
cg14622967
cg14622996
cg14623306
cg14623910
cg14624546
cg14624773
cg14625040
cg14625095
cg14625113
cg14625154
cg14625175
cg14626309
cg14626759
cg14626793
cg14627348
cg14627404
cg14627862
cg14627921
cg14628260
cg14628274
cg14628982
cg14629075
cg14629665
cg14630106
cg14630206
cg14630357
cg14630601
cg14630905
cg14630993
cg14631053
cg14631386
cg14631834
cg14632180
cg14632899
cg14632902
cg14633329
cg14633455
cg14634534
cg14634760
cg14634850
cg14635132
cg14635199
cg14635303
cg14635629
cg14636077
cg14636261
cg14636268
cg14636534
cg14637787

cg14638272
cg14638315
cg14638729
cg14638883
cg14639904
cg14640525
cg14640592
cg14640659
cg14641333
cg14641625
cg14642381
cg14642392
cg14642432
cg14642773
cg14644001
cg14644204
cg14644752
cg14646983
cg14647190
cg14647253
cg14647287
cg14647451
cg14647640
cg14647989
cg14648471
cg14648916
cg14649095
cg14649424
cg14649449
cg14649507
cg14649589
cg14650167
cg14650610
cg14651082
cg14651210
cg14651616
cg14651650
cg14651936
cg14652717
cg14653250
cg14653281
cg14653390
cg14653988
cg14654533
cg14654613
cg14654773
cg14654886
cg14655122
cg14655242
cg14655316
cg14655855
cg14656093

cg14656180
cg14656575
cg14656657
cg14657517
cg14657725
cg14658067
cg14658149
cg14658804
cg14659346
cg14659547
cg14659606
cg14659978
cg14660328
cg14660733
cg14662009
cg14662181
cg14662210
cg14662355
cg14662522
cg14662750
cg14663264
cg14663451
cg14664242
cg14664316
cg14664621
cg14664759
cg14665203
cg14665366
cg14666119
cg14666397
cg14667175
cg14667769
cg14667799
cg14667871
cg14669274
cg14669361
cg14669362
cg14669607
cg14669847
cg14670629
cg14671011
cg14671222
cg14671809
cg14671982
cg14673247
cg14673256
cg14673387
cg14673618
cg14673936
cg14674720
cg14674802
cg14675357

cg14676407
cg14676529
cg14677615
cg14677698
cg14678088
cg14679444
cg14679587
cg14679780
cg14680147
cg14681171
cg14681767
cg14682005
cg14682080
cg14682277
cg14682419
cg14683065
cg14683125
cg14684068
cg14684166
cg14684925
cg14685006
cg14685796
cg14686207
cg14686297
cg14686437
cg14687145
cg14687949
cg14688104
cg14688451
cg14688579
cg14688879
cg14688905
cg14689623
cg14689979
cg14691529
cg14691596
cg14691671
cg14691921
cg14692768
cg14693194
cg14693375
cg14694744
cg14695378
cg14695492
cg14695751
cg14696396
cg14696820
cg14696926
cg14697268
cg14697657
cg14698448
cg14698665

cg14698932
cg14699728
cg14699839
cg14699903
cg14701491
cg14701643
cg14701867
cg14701962
cg14702787
cg14703224
cg14703454
cg14703605
cg14704758
cg14704968
cg14705079
cg14705695
cg14706475
cg14707834
cg14708514
cg14708905
cg14709239
cg14709292
cg14709481
cg14711743
cg14711869
cg14711997
cg14712186
cg14712575
cg14712657
cg14712784
cg14713139
cg14713217
cg14714046
cg14714364
cg14714629
cg14714797
cg14714922
cg14715696
cg14716968
cg14717116
cg14717170
cg14717557
cg14717666
cg14717906
cg14718250
cg14718995
cg14719352
cg14719951
cg14720274
cg14720773
cg14721139
cg14721506

cg14722082
cg14722140
cg14722162
cg14722693
cg14723373
cg14724471
cg14724925
cg14725082
cg14725719
cg14728380
cg14729344
cg14730445
cg14730449
cg14730815
cg14731225
cg14731400
cg14731407
cg14731422
cg14732248
cg14732789
cg14732969
cg14733048
cg14733103
cg14733637
cg14733725
cg14734340
cg14735741
cg14736058
cg14736458
cg14737125
cg14737994
cg14738806
cg14741236
cg14741399
cg14741726
cg14742148
cg14742186
cg14742211
cg14742716
cg14743206
cg14743346
cg14743594
cg14743710
cg14744050
cg14744537
cg14744743
cg14745270
cg14746717
cg14747132
cg14747550
cg14747563
cg14748795

cg14748856
cg14749396
cg14750066
cg14750475
cg14750543
cg14750746
cg14750836
cg14750948
cg14752069
cg14752089
cg14752525
cg14752877
cg14753385
cg14753599
cg14754187
cg14754555
cg14754713
cg14754787
cg14755066
cg14756307
cg14756999
cg14757107
cg14757228
cg14757492
cg14758065
cg14758525
cg14759549
cg14760020
cg14761227
cg14761417
cg14761514
cg14761773
cg14762973
cg14763548
cg14763706
cg14764792
cg14765646
cg14768785
cg14769207
cg14769962
cg14770389
cg14771240
cg14771301
cg14771889
cg14772458
cg14772925
cg14772935
cg14773619
cg14773694
cg14774685
cg14775207
cg14775286

cg14775423
cg14776195
cg14776894
cg14777634
cg14777768
cg14778531
cg14778922
cg14779327
cg14779404
cg14779588
cg14779951
cg14780068
cg14780070
cg14780276
cg14780632
cg14780990
cg14781189
cg14781281
cg14781374
cg14782638
cg14782970
cg14783581
cg14783993
cg14784242
cg14784253
cg14784335
cg14784653
cg14784922
cg14786024
cg14788046
cg14788242
cg14788655
cg14789121
cg14789175
cg14789259
cg14789529
cg14789659
cg14789828
cg14790000
cg14790609
cg14790876
cg14791280
cg14791413
cg14791530
cg14792008
cg14792912
cg14793406
cg14794027
cg14794076
cg14794189
cg14794480
cg14794741

cg14794799
cg14795968
cg14796318
cg14797004
cg14797402
cg14798032
cg14799277
cg14799446
cg14799457
cg14800532
cg14800603
cg14801864
cg14804384
cg14805347
cg14805996
cg14806636
cg14806752
cg14807090
cg14808040
cg14808689
cg14809166
cg14810013
cg14810242
cg14810329
cg14812376
cg14813258
cg14814167
cg14815361
cg14815950
cg14816421
cg14816917
cg14817490
cg14817655
cg14817867
cg14818812
cg14819088
cg14819229
cg14819838
cg14820573
cg14820653
cg14821245
cg14821322
cg14822490
cg14822754
cg14823363
cg14823740
cg14823825
cg14823851
cg14824543
cg14824848
cg14824901
cg14825372

cg14825688
cg14825736
cg14826031
cg14826417
cg14826425
cg14826624
cg14826683
cg14826891
cg14826909
cg14827481
cg14827924
cg14828411
cg14829063
cg14829066
cg14830367
cg14830601
cg14830791
cg14830915
cg14832378
cg14832432
cg14832661
cg14832998
cg14833747
cg14833896
cg14834267
cg14834881
cg14834893
cg14834925
cg14834938
cg14835443
cg14835517
cg14835956
cg14835981
cg14836018
cg14836602
cg14837148
cg14837955
cg14838433
cg14839256
cg14839501
cg14839558
cg14839837
cg14839898
cg14841011
cg14842777
cg14842820
cg14843129
cg14843888
cg14843920
cg14844236
cg14844401
cg14844559

cg14845689
cg14845812
cg14847045
cg14847917
cg14848077
cg14849065
cg14849237
cg14849262
cg14850156
cg14851271
cg14851471
cg14851485
cg14851608
cg14851685
cg14851701
cg14852134
cg14852384
cg14852456
cg14852603
cg14852736
cg14853282
cg14853341
cg14854112
cg14854474
cg14854517
cg14855150
cg14855378
cg14855519
cg14855529
cg14856914
cg14858469
cg14859204
cg14859350
cg14859381
cg14859460
cg14859724
cg14861020
cg14861089
cg14861570
cg14862787
cg14862864
cg14863186
cg14864167
cg14864369
cg14865407
cg14865435
cg14866069
cg14866167
cg14866595
cg14866863
cg14867085
cg14868222

cg14868466
cg14868574
cg14871544
cg14871736
cg14871932
cg14872454
cg14872952
cg14873515
cg14874351
cg14874750
cg14875171
cg14876761
cg14878852
cg14879418
cg14881393
cg14881470
cg14883291
cg14883993
cg14884183
cg14884902
cg14884929
cg14884931
cg14885742
cg14885748
cg14886059
cg14886530
cg14887142
cg14887903
cg14889069
cg14889682
cg14890343
cg14890509
cg14891209
cg14891617
cg14893163
cg14894097
cg14894245
cg14894848
cg14896948
cg14897096
cg14897188
cg14897263
cg14897920
cg14898127
cg14898177
cg14898416
cg14898611
cg14899046
cg14899303
cg14901243
cg14901284
cg14903245

cg14903707
cg14904305
cg14904662
cg14904814
cg14905320
cg14905466
cg14905514
cg14905577
cg14906355
cg14906909
cg14907737
cg14908255
cg14908307
cg14908994
cg14909213
cg14909325
cg14909424
cg14909464
cg14909730
cg14910227
cg14910524
cg14910534
cg14911393
cg14912034
cg14912644
cg14912728
cg14913969
cg14914440
cg14914532
cg14914822
cg14916108
cg14916983
cg14917100
cg14917846
cg14918359
cg14918442
cg14918927
cg14921271
cg14921326
cg14921884
cg14923124
cg14923640
cg14924041
cg14924131
cg14924160
cg14924278
cg14924671
cg14925328
cg14926715
cg14927712
cg14928241
cg14928319

cg14928452
cg14928902
cg14930075
cg14930884
cg14931230
cg14931669
cg14931884
cg14932468
cg14934413
cg14934522
cg14934821
cg14935206
cg14936188
cg14936266
cg14936269
cg14936778
cg14936938
cg14938561
cg14938587
cg14939082
cg14941291
cg14942536
cg14944269
cg14947010
cg14947263
cg14947411
cg14947547
cg14947846
cg14948804
cg14948822
cg14950039
cg14950072
cg14950134
cg14950193
cg14950237
cg14950829
cg14951864
cg14953379
cg14953666
cg14954582
cg14955103
cg14955235
cg14955238
cg14957089
cg14957240
cg14958635
cg14959260
cg14959580
cg14959729
cg14960373
cg14960489
cg14960827

cg14961117
cg14961476
cg14962327
cg14962616
cg14963724
cg14964336
cg14964512
cg14965220
cg14965639
cg14966613
cg14966782
cg14967393
cg14968375
cg14969185
cg14969261
cg14969646
cg14969899
cg14969965
cg14970311
cg14970991
cg14971718
cg14972228
cg14972271
cg14972576
cg14972743
cg14972827
cg14974644
cg14974938
cg14975547
cg14975881
cg14976660
cg14977433
cg14979674
cg14980234
cg14980260
cg14980983
cg14981637
cg14981819
cg14982043
cg14982276
cg14984066
cg14986447
cg14986671
cg14987354
cg14988869
cg14989243
cg14989661
cg14990076
cg14990478
cg14991487
cg14991769
cg14991984

cg14993439
cg14993464
cg14993491
cg14994183
cg14994247
cg14994944
cg14995235
cg14997413
cg14997707
cg14998335
cg14998460
cg14998613
cg14999716
cg15000071
cg15000279
cg15000813
cg15001056
cg15001406
cg15001747
cg15001843
cg15002294
cg15002691
cg15002713
cg15003194
cg15003465
cg15003468
cg15004136
cg15004938
cg15005421
cg15005526
cg15006828
cg15007156
cg15007959
cg15008287
cg15009198
cg15009575
cg15010616
cg15011260
cg15011775
cg15012468
cg15012625
cg15013544
cg15014006
cg15014369
cg15014975
cg15015340
cg15015416
cg15015426
cg15015920
cg15016079
cg15016120
cg15017119

cg15017226
cg15017496
cg15017828
cg15018193
cg15019001
cg15020132
cg15020425
cg15020568
cg15020894
cg15021031
cg15021280
cg15022420
cg15023513
cg15024955
cg15025103
cg15025240
cg15025379
cg15025485
cg15026456
cg15026569
cg15026634
cg15026767
cg15027266
cg15027907
cg15028339
cg15029037
cg15029475
cg15029631
cg15029735
cg15029747
cg15029935
cg15030849
cg15031579
cg15031661
cg15032142
cg15032239
cg15032490
cg15032615
cg15032909
cg15033269
cg15033552
cg15034345
cg15034393
cg15034857
cg15034962
cg15035033
cg15035973
cg15036475
cg15036867
cg15036894
cg15039036
cg15039102

cg15042150
cg15042811
cg15043113
cg15044767
cg15044957
cg15045153
cg15045846
cg15046062
cg15047026
cg15047134
cg15047502
cg15048991
cg15049101
cg15049402
cg15049778
cg15050865
cg15052447
cg15053253
cg15054281
cg15054288
cg15054725
cg15055101
cg15055773
cg15055782
cg15056751
cg15056773
cg15056794
cg15056966
cg15057323
cg15057670
cg15057735
cg15058036
cg15058210
cg15058253
cg15058464
cg15058557
cg15058901
cg15059176
cg15059225
cg15059367
cg15059743
cg15059804
cg15059851
cg15060438
cg15060599
cg15061231
cg15061530
cg15061662
cg15061954
cg15062059
cg15062535
cg15063366

cg15063443
cg15063687
cg15064086
cg15065340
cg15065877
cg15066337
cg15066503
cg15066837
cg15067015
cg15067386
cg15067687
cg15067806
cg15068318
cg15068641
cg15069471
cg15072405
cg15073708
cg15074033
cg15075226
cg15075241
cg15075709
cg15076145
cg15077792
cg15078211
cg15078284
cg15080351
cg15080430
cg15080939
cg15081128
cg15081351
cg15081720
cg15081740
cg15081886
cg15082028
cg15082498
cg15082875
cg15082983
cg15083775
cg15083851
cg15084269
cg15084823
cg15084902
cg15085044
cg15085086
cg15085791
cg15085899
cg15088574
cg15089219
cg15089846
cg15090005
cg15090509
cg15090889

cg15091407
cg15092168
cg15092219
cg15092343
cg15093248
cg15093766
cg15094119
cg15094353
cg15094668
cg15094819
cg15096085
cg15097640
cg15098643
cg15099245
cg15100491
cg15101483
cg15102387
cg15102567
cg15102655
cg15102821
cg15104815
cg15104839
cg15105359
cg15106134
cg15107117
cg15107132
cg15107202
cg15107203
cg15107641
cg15107670
cg15107869
cg15108047
cg15108334
cg15108553
cg15108641
cg15108645
cg15108705
cg15108727
cg15110243
cg15111360
cg15111638
cg15111708
cg15112032
cg15112747
cg15112803
cg15112910
cg15113009
cg15113168
cg15113766
cg15114076
cg15114474
cg15114651

cg15115594
cg15116095
cg15118447
cg15118665
cg15119027
cg15119221
cg15120333
cg15121267
cg15121304
cg15122498
cg15123499
cg15123519
cg15123662
cg15123984
cg15125684
cg15125762
cg15126975
cg15127879
cg15128365
cg15128876
cg15128905
cg15129608
cg15130054
cg15130459
cg15130928
cg15131001
cg15131883
cg15132565
cg15133041
cg15133201
cg15133351
cg15133564
cg15133719
cg15134583
cg15135241
cg15135773
cg15136410
cg15136953
cg15137369
cg15137838
cg15138339
cg15138614
cg15138702
cg15139008
cg15139644
cg15140703
cg15140798
cg15142042
cg15142214
cg15143702
cg15143788
cg15145341

cg15145672
cg15145767
cg15146859
cg15147534
cg15147833
cg15148371
cg15148394
cg15148513
cg15148573
cg15148933
cg15149600
cg15150017
cg15150064
cg15150310
cg15150336
cg15150472
cg15150970
cg15151052
cg15151492
cg15152331
cg15152824
cg15153887
cg15153957
cg15154635
cg15155441
cg15157241
cg15157974
cg15158106
cg15159060
cg15159104
cg15159605
cg15160198
cg15160524
cg15162869
cg15164180
cg15164733
cg15165122
cg15166089
cg15166311
cg15167956
cg15167959
cg15168577
cg15168615
cg15168727
cg15168892
cg15168995
cg15169121
cg15169286
cg15169900
cg15173079
cg15173134
cg15173443

cg15174396
cg15174623
cg15176213
cg15177169
cg15177195
cg15179976
cg15180867
cg15181151
cg15181153
cg15181441
cg15182360
cg15182790
cg15183083
cg15184781
cg15186181
cg15186616
cg15186969
cg15187550
cg15188010
cg15188252
cg15188623
cg15189210
cg15190834
cg15191648
cg15191781
cg15191798
cg15192750
cg15193228
cg15194251
cg15194943
cg15195276
cg15195910
cg15196529
cg15197050
cg15197330
cg15197583
cg15197657
cg15197893
cg15198101
cg15198193
cg15198344
cg15199181
cg15199678
cg15201458
cg15202104
cg15202954
cg15203028
cg15203735
cg15204874
cg15205559
cg15206473
cg15207742

cg15208375
cg15208832
cg15209440
cg15209604
cg15210427
cg15210817
cg15211996
cg15212295
cg15212369
cg15213210
cg15213488
cg15214506
cg15215033
cg15217308
cg15219145
cg15221604
cg15221739
cg15222651
cg15222899
cg15224955
cg15225364
cg15225594
cg15225844
cg15226275
cg15226808
cg15227168
cg15227610
cg15227982
cg15228441
cg15228639
cg15228928
cg15229052
cg15229496
cg15229943
cg15230460
cg15230622
cg15231462
cg15232654
cg15233131
cg15233380
cg15233892
cg15234271
cg15234492
cg15235798
cg15235805
cg15236063
cg15236196
cg15236484
cg15236881
cg15237152
cg15237494
cg15237644

cg15238224
cg15238831
cg15239627
cg15240033
cg15240476
cg15240876
cg15241236
cg15241582
cg15242360
cg15242449
cg15242570
cg15243478
cg15243542
cg15244049
cg15244101
cg15245051
cg15246085
cg15246238
cg15246693
cg15246805
cg15246989
cg15247039
cg15247329
cg15247974
cg15248242
cg15250073
cg15251311
cg15251385
cg15252509
cg15252768
cg15253304
cg15254238
cg15254559
cg15255403
cg15255797
cg15256053
cg15257539
cg15257565
cg15257719
cg15257836
cg15259904
cg15259986
cg15260428
cg15260437
cg15260721
cg15260951
cg15260978
cg15261665
cg15262325
cg15262515
cg15263682
cg15264806

cg15264991
cg15266705
cg15267232
cg15270654
cg15270892
cg15271461
cg15272173
cg15272362
cg15274171
cg15275393
cg15275965
cg15276331
cg15277677
cg15277906
cg15278109
cg15278374
cg15279364
cg15279413
cg15279476
cg15279832
cg15280185
cg15280358
cg15282632
cg15282973
cg15283194
cg15283967
cg15284082
cg15284457
cg15285250
cg15285286
cg15285975
cg15286016
cg15286037
cg15286044
cg15286206
cg15287410
cg15288405
cg15288618
cg15288779
cg15288800
cg15288874
cg15289466
cg15293363
cg15294263
cg15295273
cg15296767
cg15297628
cg15297752
cg15299835
cg15301316
cg15302350
cg15303062

cg15303841
cg15304464
cg15304679
cg15304699
cg15305732
cg15306060
cg15309004
cg15309006
cg15309361
cg15309879
cg15310492
cg15310568
cg15311814
cg15312203
cg15312264
cg15315056
cg15316206
cg15316843
cg15317049
cg15317267
cg15318176
cg15318546
cg15318627
cg15318957
cg15319126
cg15319470
cg15320980
cg15321363
cg15321908
cg15323840
cg15324331
cg15324978
cg15325658
cg15325704
cg15326144
cg15327641
cg15328072
cg15328703
cg15329866
cg15330117
cg15330654
cg15331332
cg15331500
cg15331603
cg15331633
cg15332132
cg15332173
cg15332978
cg15333426
cg15333932
cg15334372
cg15334391

cg15334916
cg15335104
cg15335297
cg15335334
cg15336893
cg15337006
cg15337897
cg15338327
cg15338449
cg15339026
cg15339164
cg15341350
cg15341601
cg15341751
cg15341833
cg15342452
cg15343119
cg15343224
cg15344192
cg15345393
cg15345437
cg15346781
cg15346842
cg15347348
cg15348319
cg15349139
cg15349696
cg15350036
cg15350661
cg15351741
cg15352186
cg15352367
cg15352829
cg15353533
cg15353603
cg15355967
cg15356529
cg15357821
cg15357945
cg15358723
cg15359093
cg15359390
cg15360478
cg15361750
cg15362155
cg15362455
cg15362894
cg15362957
cg15364504
cg15367150
cg15367287
cg15367892

cg15368872
cg15368905
cg15371566
cg15371815
cg15371881
cg15372721
cg15373239
cg15373767
cg15374053
cg15374582
cg15375424
cg15375469
cg15375488
cg15376155
cg15376677
cg15376996
cg15377044
cg15377283
cg15377518
cg15378486
cg15379170
cg15379185
cg15379354
cg15379412
cg15381475
cg15382538
cg15382568
cg15382659
cg15382743
cg15382933
cg15383087
cg15383322
cg15383705
cg15384108
cg15384383
cg15384598
cg15384668
cg15386069
cg15386103
cg15386293
cg15389191
cg15389749
cg15389765
cg15391555
cg15391590
cg15391601
cg15392054
cg15392124
cg15392364
cg15392413
cg15393700
cg15393733

cg15393816
cg15393937
cg15394297
cg15395441
cg15395560
cg15396204
cg15396686
cg15397374
cg15398497
cg15399131
cg15399466
cg15399926
cg15400117
cg15400401
cg15401332
cg15401418
cg15401477
cg15401862
cg15402095
cg15402156
cg15402529
cg15403283
cg15404043
cg15404785
cg15404858
cg15405084
cg15405439
cg15406010
cg15407058
cg15407063
cg15407213
cg15407257
cg15407570
cg15407638
cg15408237
cg15408889
cg15409013
cg15409712
cg15409939
cg15410418
cg15410441
cg15411034
cg15411303
cg15411840
cg15412087
cg15412721
cg15412759
cg15412772
cg15412815
cg15415452
cg15415945
cg15416329

cg15416442
cg15416735
cg15416877
cg15417798
cg15418419
cg15418499
cg15420468
cg15420678
cg15421321
cg15421520
cg15422005
cg15422147
cg15422307
cg15422382
cg15423092
cg15423315
cg15423764
cg15423781
cg15423872
cg15425280
cg15425444
cg15425498
cg15425541
cg15426006
cg15427886
cg15427906
cg15429180
cg15429214
cg15431544
cg15431934
cg15432532
cg15434292
cg15435391
cg15436606
cg15437528
cg15438314
cg15438548
cg15439078
cg15439294
cg15439862
cg15440026
cg15440157
cg15440158
cg15440688
cg15441392
cg15442678
cg15442737
cg15442888
cg15443020
cg15444358
cg15444478
cg15445319

cg15445722
cg15445958
cg15446373
cg15447231
cg15447787
cg15448372
cg15448721
cg15448975
cg15449023
cg15449049
cg15449323
cg15449516
cg15449956
cg15449979
cg15451642
cg15451662
cg15451729
cg15452381
cg15452970
cg15453278
cg15455864
cg15456082
cg15456502
cg15456821
cg15457187
cg15457481
cg15457934
cg15459537
cg15460584
cg15460872
cg15461413
cg15462411
cg15462501
cg15462970
cg15463387
cg15463580
cg15464481
cg15465121
cg15465535
cg15466753
cg15466862
cg15467785
cg15468243
cg15468521
cg15468668
cg15468739
cg15469001
cg15469171
cg15469321
cg15469655
cg15471073
cg15471388

cg15471402
cg15471519
cg15471953
cg15472092
cg15473329
cg15474711
cg15474755
cg15476396
cg15476523
cg15476790
cg15478045
cg15478401
cg15480305
cg15481294
cg15481472
cg15482082
cg15482122
cg15482717
cg15482872
cg15483228
cg15483381
cg15484033
cg15484384
cg15484532
cg15484808
cg15484988
cg15485378
cg15486183
cg15487620
cg15488722
cg15488986
cg15489196
cg15489301
cg15490335
cg15490880
cg15491247
cg15491385
cg15492834
cg15493035
cg15493970
cg15496388
cg15496866
cg15497761
cg15498563
cg15499275
cg15500259
cg15501219
cg15503084
cg15503752
cg15504461
cg15504662
cg15505276

cg15505737
cg15505860
cg15506609
cg15507321
cg15507449
cg15507486
cg15508262
cg15508462
cg15508776
cg15508809
cg15509286
cg15509687
cg15511928
cg15512289
cg15512715
cg15513743
cg15514918
cg15515320
cg15516314
cg15516835
cg15519508
cg15519535
cg15520279
cg15521745
cg15521790
cg15522517
cg15523060
cg15524101
cg15524283
cg15526469
cg15526941
cg15527308
cg15528411
cg15529542
cg15529692
cg15530112
cg15531369
cg15531386
cg15531450
cg15532088
cg15532640
cg15534052
cg15534561
cg15534855
cg15538103
cg15538264
cg15538427
cg15538607
cg15538820
cg15539420
cg15539944
cg15540138

cg15540820
cg15541062
cg15541630
cg15542670
cg15542741
cg15542880
cg15543736
cg15544721
cg15545247
cg15545316
cg15545772
cg15545842
cg15545878
cg15545942
cg15547344
cg15549100
cg15549231
cg15549435
cg15549637
cg15549927
cg15549973
cg15551881
cg15552091
cg15552370
cg15552491
cg15552607
cg15552843
cg15552853
cg15553770
cg15553958
cg15554087
cg15554678
cg15555712
cg15555985
cg15556672
cg15556990
cg15557459
cg15557758
cg15557878
cg15558717
cg15558904
cg15559162
cg15559714
cg15560801
cg15561305
cg15562220
cg15562346
cg15562947
cg15563070
cg15564098
cg15564444
cg15564619

cg15565004
cg15567237
cg15567428
cg15567853
cg15568107
cg15570293
cg15570323
cg15571353
cg15571556
cg15572396
cg15572489
cg15572656
cg15573040
cg15574442
cg15574642
cg15575356
cg15575967
cg15576082
cg15576900
cg15576918
cg15577373
cg15577927
cg15578140
cg15579817
cg15580052
cg15580417
cg15582376
cg15582707
cg15582789
cg15583263
cg15583349
cg15583517
cg15583738
cg15584070
cg15584221
cg15584445
cg15584486
cg15585794
cg15586327
cg15586393
cg15586563
cg15586654
cg15588737
cg15588941
cg15589354
cg15590350
cg15590780
cg15591276
cg15593298
cg15594471
cg15594967
cg15595918

cg15596493
cg15596647
cg15597522
cg15598244
cg15599182
cg15600279
cg15600488
cg15600751
cg15601071
cg15601164
cg15601452
cg15601682
cg15602074
cg15602217
cg15602241
cg15602329
cg15603130
cg15603548
cg15603568
cg15604182
cg15605096
cg15605307
cg15606663
cg15606914
cg15607142
cg15607292
cg15607664
cg15607672
cg15608892
cg15609035
cg15609678
cg15609734
cg15610437
cg15611112
cg15611150
cg15611413
cg15611912
cg15612077
cg15612085
cg15613048
cg15613567
cg15614155
cg15616998
cg15617023
cg15617155
cg15617591
cg15617640
cg15617814
cg15617853
cg15618210
cg15618336
cg15618347

cg15618525
cg15618533
cg15619377
cg15621260
cg15621697
cg15621731
cg15622352
cg15623175
cg15623552
cg15624311
cg15624813
cg15625636
cg15626105
cg15626193
cg15626405
cg15626421
cg15627089
cg15627180
cg15628518
cg15630743
cg15632659
cg15632901
cg15632936
cg15633056
cg15633664
cg15634398
cg15634877
cg15635483
cg15635524
cg15636792
cg15637465
cg15638123
cg15638709
cg15639556
cg15640140
cg15641340
cg15641657
cg15642035
cg15642468
cg15642666
cg15642933
cg15644174
cg15644324
cg15644626
cg15645674
cg15645728
cg15646411
cg15646543
cg15646741
cg15646778
cg15647584
cg15647725

cg15649236
cg15649384
cg15649710
cg15650161
cg15650745
cg15652169
cg15652280
cg15652303
cg15652683
cg15653111
cg15653173
cg15653478
cg15653569
cg15653807
cg15653828
cg15654892
cg15657225
cg15657888
cg15658426
cg15658793
cg15659052
cg15659537
cg15659599
cg15660427
cg15662251
cg15662768
cg15663948
cg15664323
cg15665276
cg15666143
cg15666275
cg15666789
cg15667931
cg15668533
cg15669183
cg15669228
cg15670475
cg15672065
cg15672437
cg15672498
cg15672768
cg15672853
cg15673001
cg15674825
cg15675643
cg15676719
cg15676988
cg15677985
cg15678662
cg15678825
cg15678861
cg15679064

cg15679139
cg15679829
cg15680288
cg15680624
cg15680798
cg15681255
cg15681376
cg15681466
cg15681537
cg15683295
cg15683497
cg15684231
cg15684661
cg15684702
cg15684772
cg15685144
cg15685223
cg15685834
cg15686202
cg15686949
cg15687000
cg15687147
cg15687300
cg15687653
cg15688396
cg15688713
cg15688918
cg15689036
cg15689121
cg15689795
cg15689969
cg15690342
cg15690347
cg15690511
cg15690542
cg15690598
cg15690760
cg15691325
cg15691862
cg15692052
cg15692837
cg15692992
cg15693299
cg15693572
cg15694715
cg15694752
cg15695045
cg15696460
cg15696906
cg15697440
cg15697849
cg15698107

cg15698244
cg15698545
cg15698691
cg15698842
cg15699099
cg15699226
cg15699524
cg15700022
cg15700197
cg15700437
cg15700587
cg15700661
cg15700989
cg15701047
cg15701170
cg15701210
cg15701412
cg15702185
cg15702277
cg15702677
cg15703632
cg15703690
cg15703886
cg15704225
cg15704872
cg15705175
cg15705203
cg15705428
cg15705971
cg15706312
cg15706657
cg15706973
cg15707093
cg15707110
cg15707869
cg15708618
cg15708776
cg15709235
cg15709446
cg15709989
cg15710554
cg15711268
cg15711987
cg15712267
cg15712304
cg15712559
cg15712777
cg15713707
cg15713734
cg15713898
cg15714328
cg15714682

cg15715477
cg15717183
cg15717840
cg15717844
cg15718287
cg15718555
cg15718581
cg15718932
cg15719339
cg15719903
cg15720500
cg15720535
cg15720995
cg15721020
cg15721238
cg15721243
cg15722533
cg15723195
cg15723222
cg15723468
cg15723784
cg15724256
cg15726123
cg15726260
cg15726557
cg15727249
cg15727287
cg15729307
cg15729649
cg15731056
cg15731772
cg15732451
cg15732768
cg15733309
cg15733729
cg15733932
cg15734454
cg15735129
cg15735721
cg15736127
cg15736165
cg15736524
cg15737302
cg15738800
cg15739881
cg15740507
cg15741930
cg15741935
cg15742777
cg15743325
cg15743799
cg15743907

cg15745900
cg15746253
cg15746445
cg15746719
cg15747712
cg15748507
cg15749661
cg15749870
cg15750696
cg15751117
cg15753546
cg15753746
cg15754084
cg15754901
cg15755240
cg15755348
cg15756055
cg15756359
cg15756878
cg15757437
cg15758700
cg15759721
cg15759873
cg15763121
cg15765486
cg15765724
cg15765889
cg15766687
cg15767218
cg15768138
cg15769388
cg15769756
cg15770125
cg15770515
cg15770599
cg15771092
cg15771200
cg15773338
cg15773539
cg15773794
cg15773890
cg15773925
cg15774153
cg15774489
cg15774495
cg15775674
cg15775779
cg15776494
cg15776647
cg15777028
cg15777261
cg15777760

cg15778089
cg15778232
cg15778434
cg15778437
cg15778745
cg15779716
cg15780078
cg15781504
cg15785860
cg15785868
cg15786180
cg15786205
cg15786900
cg15787039
cg15787636
cg15787712
cg15789106
cg15790767
cg15790820
cg15791180
cg15791815
cg15792338
cg15793181
cg15795305
cg15796941
cg15797422
cg15798385
cg15798502
cg15799353
cg15800776
cg15800975
cg15802639
cg15803869
cg15803972
cg15805104
cg15805567
cg15806588
cg15806780
cg15807455
cg15807655
cg15808426
cg15808472
cg15808943
cg15810386
cg15810758
cg15810996
cg15811427
cg15811435
cg15811515
cg15812139
cg15812322
cg15812599

cg15812753
cg15813427
cg15814750
cg15815318
cg15815435
cg15815827
cg15816712
cg15817440
cg15817482
cg15817705
cg15818005
cg15818307
cg15818577
cg15818632
cg15818671
cg15819188
cg15819471
cg15819739
cg15820062
cg15820432
cg15820574
cg15820868
cg15820955
cg15821095
cg15821546
cg15822656
cg15823845
cg15824312
cg15824435
cg15825065
cg15825786
cg15826426
cg15827779
cg15828353
cg15828364
cg15830544
cg15830591
cg15831285
cg15831722
cg15832300
cg15832329
cg15832662
cg15832847
cg15833286
cg15833353
cg15834072
cg15834592
cg15835058
cg15835232
cg15835713
cg15835747
cg15835852

cg15836317
cg15837212
cg15837233
cg15837235
cg15837531
cg15838320
cg15839088
cg15839219
cg15839448
cg15839752
cg15839913
cg15840110
cg15840151
cg15840438
cg15840949
cg15841063
cg15841533
cg15841728
cg15841865
cg15842216
cg15842276
cg15842722
cg15843262
cg15843278
cg15843496
cg15843567
cg15844711
cg15845071
cg15845118
cg15846556
cg15846590
cg15847565
cg15847600
cg15848031
cg15848714
cg15849445
cg15849465
cg15849872
cg15850739
cg15850918
cg15851882
cg15852125
cg15852312
cg15852490
cg15853028
cg15853169
cg15854022
cg15854847
cg15855924
cg15855970
cg15856626
cg15856829

cg15857470
cg15857635
cg15858239
cg15859995
cg15861156
cg15861357
cg15861585
cg15861806
cg15862380
cg15863841
cg15864733
cg15865026
cg15865243
cg15866542
cg15867307
cg15867829
cg15868105
cg15868183
cg15868208
cg15868692
cg15869022
cg15869649
cg15871766
cg15871828
cg15872079
cg15873149
cg15873301
cg15873320
cg15873474
cg15874092
cg15874877
cg15876264
cg15876676
cg15877233
cg15877785
cg15880016
cg15882566
cg15883417
cg15884185
cg15884430
cg15884713
cg15884992
cg15885043
cg15885337
cg15885430
cg15886294
cg15886307
cg15887283
cg15887459
cg15887709
cg15888097
cg15888522

cg15889231
cg15889793
cg15889797
cg15889913
cg15890009
cg15890075
cg15890143
cg15890794
cg15891546
cg15892115
cg15892176
cg15892497
cg15892617
cg15893204
cg15894129
cg15894365
cg15895121
cg15895132
cg15896593
cg15896696
cg15896978
cg15897102
cg15897645
cg15897840
cg15898113
cg15898536
cg15898840
cg15899043
cg15899743
cg15900034
cg15900052
cg15900054
cg15900221
cg15900248
cg15901722
cg15901963
cg15902297
cg15902830
cg15902864
cg15903032
cg15903571
cg15904283
cg15905124
cg15905435
cg15906055
cg15906260
cg15906761
cg15907819
cg15908324
cg15908921
cg15909006
cg15909056

cg15909319
cg15909737
cg15910070
cg15910981
cg15911165
cg15912800
cg15913157
cg15913375
cg15913671
cg15914908
cg15915658
cg15916004
cg15916061
cg15916399
cg15916646
cg15917438
cg15918284
cg15920335
cg15920379
cg15920472
cg15920561
cg15920654
cg15921239
cg15921887
cg15922513
cg15922611
cg15923814
cg15924285
cg15924332
cg15924691
cg15924693
cg15924738
cg15925365
cg15927196
cg15927455
cg15927699
cg15927720
cg15928093
cg15929078
cg15929395
cg15929797
cg15930219
cg15930777
cg15931673
cg15931949
cg15932610
cg15932886
cg15933120
cg15934095
cg15935096
cg15935247
cg15935791

cg15935860
cg15937784
cg15937861
cg15937925
cg15941948
cg15943335
cg15943602
cg15944766
cg15944856
cg15945475
cg15945999
cg15946224
cg15946590
cg15946856
cg15947487
cg15947866
cg15948536
cg15948871
cg15949553
cg15949830
cg15950068
cg15950203
cg15950251
cg15950504
cg15951469
cg15952370
cg15952840
cg15953187
cg15953461
cg15953602
cg15954031
cg15954792
cg15956189
cg15956565
cg15957341
cg15957797
cg15958035
cg15958289
cg15958792
cg15959080
cg15961007
cg15961211
cg15961219
cg15961455
cg15961744
cg15962267
cg15962313
cg15962547
cg15963095
cg15963499
cg15964159
cg15964309

cg15965134
cg15965301
cg15965716
cg15966954
cg15967525
cg15968932
cg15969149
cg15969431
cg15970074
cg15970897
cg15971761
cg15973075
cg15974487
cg15974867
cg15975443
cg15975574
cg15975598
cg15975980
cg15976416
cg15976539
cg15976541
cg15977002
cg15977096
cg15977716
cg15977933
cg15978017
cg15978039
cg15978561
cg15979173
cg15981092
cg15981982
cg15982299
cg15983005
cg15983072
cg15983605
cg15984596
cg15984718
cg15985132
cg15985157
cg15985184
cg15985191
cg15985470
cg15985891
cg15986555
cg15986841
cg15987885
cg15987934
cg15988204
cg15988287
cg15988350
cg15988937
cg15989068

cg15989091
cg15989810
cg15989892
cg15990090
cg15990632
cg15990724
cg15990972
cg15991207
cg15991405
cg15991503
cg15992284
cg15993258
cg15993345
cg15993383
cg15994731
cg15994861
cg15995265
cg15996480
cg15996534
cg15997429
cg15997518
cg15997770
cg15997906
cg15998518
cg15998686
cg15998779
cg15999077
cg15999099
cg15999609
cg16000017
cg16000182
cg16000683
cg16001135
cg16001165
cg16001323
cg16001495
cg16001865
cg16001913
cg16002305
cg16002660
cg16002799
cg16002810
cg16002818
cg16002887
cg16002929
cg16003913
cg16004056
cg16004593
cg16005494
cg16006349
cg16006415
cg16007742

cg16008327
cg16008993
cg16009167
cg16009633
cg16010207
cg16011217
cg16012162
cg16012294
cg16015276
cg16015423
cg16015816
cg16016431
cg16017904
cg16020747
cg16021018
cg16021678
cg16022081
cg16023304
cg16023415
cg16023545
cg16023896
cg16024274
cg16024318
cg16024377
cg16024523
cg16024891
cg16025120
cg16026035
cg16026902
cg16027122
cg16027147
cg16027761
cg16028201
cg16029623
cg16030011
cg16031270
cg16031528
cg16032841
cg16033053
cg16033780
cg16034562
cg16034652
cg16035633
cg16035777
cg16036142
cg16036528
cg16036584
cg16037062
cg16037137
cg16037981
cg16038120
cg16039374

cg16039867
cg16040614
cg16041090
cg16042149
cg16042259
cg16043345
cg16043357
cg16043523
cg16043635
cg16043888
cg16044603
cg16045223
cg16046833
cg16047117
cg16047567
cg16048006
cg16048568
cg16048771
cg16048803
cg16049600
cg16049690
cg16050028
cg16050357
cg16050974
cg16051195
cg16051245
cg16052198
cg16052237
cg16052317
cg16052325
cg16052975
cg16053580
cg16054651
cg16055026
cg16055052
cg16055410
cg16055869
cg16056272
cg16057408
cg16057587
cg16058105
cg16058274
cg16059167
cg16060189
cg16060382
cg16061354
cg16061498
cg16061528
cg16061656
cg16061668
cg16063112
cg16063656

cg16063716
cg16063747
cg16063925
cg16064793
cg16065021
cg16065050
cg16065899
cg16066544
cg16067045
cg16067536
cg16067628
cg16068263
cg16069359
cg16069910
cg16069986
cg16070279
cg16071219
cg16071713
cg16072535
cg16072754
cg16073378
cg16073467
cg16076569
cg16076587
cg16077055
cg16078269
cg16080125
cg16080781
cg16080876
cg16081281
cg16081499
cg16083439
cg16084363
cg16085171
cg16085531
cg16086559
cg16086570
cg16086620
cg16086730
cg16087093
cg16087263
cg16087432
cg16088427
cg16093580
cg16095155
cg16096063
cg16096432
cg16096631
cg16098227
cg16098304
cg16098726
cg16099023

cg16099599
cg16099687
cg16099804
cg16099871
cg16100120
cg16100244
cg16100446
cg16100530
cg16101077
cg16101213
cg16101346
cg16101601
cg16101739
cg16102102
cg16102739
cg16102971
cg16103159
cg16103203
cg16103275
cg16103421
cg16103604
cg16103712
cg16103959
cg16104371
cg16104584
cg16105558
cg16105591
cg16105687
cg16107105
cg16107470
cg16107907
cg16108230
cg16108710
cg16108823
cg16108835
cg16109680
cg16110087
cg16110455
cg16110541
cg16110645
cg16110704
cg16111448
cg16112237
cg16112276
cg16113172
cg16113347
cg16113681
cg16114380
cg16116512
cg16117513
cg16118212
cg16118839

cg16119835
cg16120196
cg16120788
cg16121136
cg16122370
cg16122628
cg16122716
cg16123202
cg16123211
cg16123760
cg16124188
cg16124434
cg16124934
cg16124935
cg16126058
cg16126286
cg16126816
cg16126900
cg16127349
cg16127594
cg16128892
cg16129200
cg16129526
cg16130238
cg16130857
cg16132208
cg16132337
cg16132851
cg16133703
cg16134070
cg16134262
cg16134515
cg16134678
cg16134717
cg16134718
cg16135698
cg16136800
cg16137928
cg16138752
cg16139316
cg16139380
cg16139785
cg16139912
cg16140179
cg16140253
cg16140558
cg16140794
cg16141752
cg16141784
cg16142824
cg16142855
cg16143325

cg16143997
cg16144181
cg16144317
cg16144864
cg16145113
cg16145209
cg16146177
cg16146806
cg16147544
cg16148891
cg16149138
cg16149164
cg16149574
cg16149930
cg16150047
cg16150435
cg16150752
cg16151261
cg16151924
cg16152136
cg16152753
cg16155081
cg16155588
cg16155702
cg16156617
cg16157435
cg16157615
cg16158220
cg16158874
cg16160637
cg16161359
cg16161415
cg16161418
cg16162122
cg16162970
cg16164144
cg16164607
cg16164622
cg16165258
cg16165450
cg16165748
cg16166230
cg16166796
cg16166826
cg16167050
cg16167240
cg16167524
cg16167877
cg16168509
cg16170380
cg16170767
cg16171105

cg16171182
cg16171213
cg16171281
cg16171392
cg16171771
cg16172329
cg16172619
cg16172814
cg16174029
cg16174106
cg16174779
cg16175792
cg16176378
cg16176600
cg16176834
cg16177163
cg16177187
cg16177633
cg16177732
cg16177922
cg16178141
cg16178353
cg16178907
cg16179521
cg16179720
cg16179952
cg16180258
cg16180796
cg16180848
cg16180920
cg16181383
cg16181396
cg16181903
cg16183361
cg16183813
cg16184803
cg16185419
cg16185666
cg16186435
cg16186918
cg16189995
cg16190266
cg16190732
cg16190888
cg16192052
cg16192707
cg16193295
cg16194233
cg16194746
cg16195332
cg16196021
cg16196211

cg16196728
cg16196812
cg16196857
cg16197188
cg16197388
cg16197643
cg16197925
cg16198013
cg16198103
cg16198315
cg16199280
cg16199460
cg16199881
cg16200736
cg16200799
cg16201038
cg16201146
cg16201674
cg16202340
cg16202564
cg16202755
cg16203125
cg16203203
cg16204509
cg16204717
cg16204818
cg16205058
cg16205375
cg16206138
cg16206191
cg16206344
cg16206504
cg16206564
cg16206659
cg16206744
cg16208657
cg16208959
cg16209664
cg16209873
cg16210078
cg16210355
cg16210461
cg16211161
cg16211174
cg16212145
cg16212341
cg16213028
cg16215817
cg16216212
cg16216269
cg16216407
cg16217794

cg16218715
cg16218721
cg16218738
cg16219266
cg16219322
cg16219860
cg16219879
cg16221969
cg16222629
cg16223546
cg16224417
cg16225491
cg16225703
cg16225773
cg16225947
cg16226642
cg16226962
cg16227623
cg16227748
cg16228077
cg16228359
cg16229671
cg16230352
cg16231066
cg16231757
cg16231923
cg16232126
cg16232234
cg16233724
cg16233797
cg16233920
cg16234068
cg16234557
cg16234966
cg16234978
cg16235748
cg16236328
cg16237031
cg16237191
cg16237204
cg16238815
cg16238918
cg16239184
cg16239218
cg16239536
cg16239736
cg16240162
cg16240368
cg16241218
cg16241649
cg16242615
cg16242629

cg16243019
cg16243197
cg16243652
cg16243756
cg16244299
cg16244648
cg16244770
cg16244889
cg16246240
cg16246489
cg16247183
cg16247184
cg16247269
cg16247912
cg16248329
cg16248926
cg16251016
cg16251647
cg16251902
cg16252322
cg16252905
cg16253580
cg16253629
cg16253834
cg16254309
cg16254344
cg16254962
cg16255914
cg16256390
cg16256414
cg16257051
cg16257181
cg16257215
cg16258657
cg16259571
cg16259591
cg16259931
cg16260298
cg16261857
cg16262215
cg16262614
cg16263152
cg16263180
cg16263801
cg16263887
cg16264004
cg16264526
cg16264705
cg16265542
cg16266667
cg16267162
cg16269010

cg16269184
cg16269431
cg16269532
cg16269733
cg16269775
cg16269943
cg16270399
cg16270990
cg16271359
cg16271663
cg16271833
cg16272420
cg16272981
cg16273051
cg16273509
cg16273724
cg16273979
cg16274476
cg16275295
cg16276063
cg16276225
cg16276850
cg16278512
cg16278661
cg16278996
cg16279158
cg16279237
cg16279529
cg16280141
cg16280363
cg16280499
cg16281322
cg16281600
cg16282618
cg16283362
cg16284000
cg16284238
cg16284507
cg16285941
cg16286925
cg16287451
cg16287679
cg16287734
cg16288089
cg16288630
cg16289029
cg16289417
cg16289421
cg16289618
cg16289848
cg16290101
cg16290522

cg16291069
cg16291750
cg16291880
cg16292016
cg16292181
cg16292586
cg16293569
cg16293732
cg16293983
cg16294013
cg16294045
cg16294566
cg16294641
cg16294838
cg16295030
cg16295880
cg16296417
cg16296679
cg16296797
cg16297011
cg16297171
cg16298612
cg16299813
cg16300300
cg16300565
cg16301004
cg16301036
cg16301728
cg16301890
cg16302916
cg16303129
cg16303742
cg16304215
cg16304950
cg16306654
cg16306706
cg16306870
cg16306900
cg16307674
cg16307705
cg16308270
cg16309224
cg16309319
cg16309325
cg16310415
cg16310491
cg16311302
cg16312002
cg16312496
cg16312514
cg16312552
cg16312870

cg16312954
cg16313026
cg16313577
cg16313807
cg16314543
cg16314639
cg16315740
cg16316058
cg16316344
cg16316649
cg16316755
cg16316993
cg16317318
cg16317923
cg16318036
cg16318053
cg16318271
cg16318311
cg16318412
cg16318728
cg16318994
cg16319947
cg16320427
cg16320577
cg16321280
cg16321846
cg16322478
cg16322565
cg16323609
cg16325174
cg16325421
cg16326073
cg16326358
cg16326674
cg16327228
cg16327489
cg16327839
cg16328023
cg16328319
cg16329650
cg16330147
cg16330195
cg16330247
cg16330450
cg16330535
cg16330965
cg16331632
cg16331929
cg16332256
cg16332300
cg16332493
cg16332920

cg16334314
cg16334795
cg16334869
cg16335098
cg16335919
cg16336556
cg16336586
cg16336752
cg16338011
cg16338313
cg16339155
cg16339238
cg16339286
cg16339560
cg16339915
cg16340767
cg16340963
cg16341266
cg16341269
cg16343616
cg16343924
cg16344081
cg16344234
cg16345213
cg16345521
cg16345559
cg16345720
cg16346032
cg16346422
cg16346813
cg16347377
cg16347629
cg16348470
cg16348668
cg16349724
cg16349774
cg16351002
cg16351569
cg16353318
cg16353615
cg16353632
cg16353957
cg16354117
cg16354452
cg16354502
cg16354520
cg16354522
cg16355785
cg16356045
cg16356456
cg16356737
cg16356837

cg16356956
cg16357179
cg16357224
cg16357388
cg16357582
cg16357719
cg16357921
cg16357969
cg16358044
cg16358099
cg16358696
cg16358738
cg16359091
cg16360432
cg16360836
cg16360861
cg16360868
cg16361249
cg16361253
cg16361302
cg16361876
cg16361890
cg16362014
cg16362181
cg16364121
cg16364709
cg16365445
cg16365690
cg16365807
cg16366604
cg16366843
cg16367606
cg16367786
cg16368147
cg16368670
cg16369987
cg16370398
cg16370556
cg16370642
cg16370778
cg16371230
cg16373202
cg16373456
cg16373526
cg16374093
cg16375265
cg16375290
cg16375358
cg16375541
cg16376108
cg16376379
cg16377162

cg16377679
cg16377790
cg16378015
cg16378133
cg16378215
cg16379205
cg16379716
cg16380876
cg16380885
cg16381321
cg16383005
cg16384355
cg16384588
cg16384626
cg16384849
cg16385335
cg16385533
cg16385684
cg16386719
cg16386945
cg16387141
cg16387581
cg16389209
cg16389407
cg16390049
cg16390060
cg16390380
cg16391182
cg16391678
cg16391955
cg16392193
cg16392213
cg16392391
cg16393182
cg16393877
cg16393899
cg16393905
cg16394551
cg16394952
cg16395183
cg16395336
cg16396309
cg16396866
cg16397028
cg16397401
cg16397794
cg16398362
cg16398761
cg16398832
cg16399049
cg16399368
cg16399624

cg16399632
cg16399745
cg16400492
cg16400631
cg16400794
cg16401207
cg16401465
cg16402387
cg16402875
cg16403344
cg16404040
cg16404105
cg16404157
cg16404259
cg16404371
cg16404784
cg16404903
cg16405019
cg16405308
cg16405876
cg16406892
cg16406960
cg16407998
cg16408555
cg16408784
cg16408815
cg16408820
cg16408856
cg16408865
cg16409261
cg16409650
cg16409955
cg16410171
cg16410413
cg16410579
cg16412573
cg16413675
cg16413763
cg16414149
cg16414691
cg16414796
cg16415831
cg16415870
cg16416987
cg16417028
cg16417118
cg16417374
cg16417409
cg16418183
cg16418228
cg16418810
cg16419009

cg16419354
cg16419421
cg16419441
cg16419629
cg16420199
cg16420200
cg16420340
cg16420756
cg16420798
cg16420801
cg16420866
cg16421526
cg16422098
cg16422816
cg16423738
cg16424196
cg16424311
cg16425489
cg16425829
cg16426366
cg16426537
cg16427107
cg16427387
cg16427899
cg16428251
cg16428612
cg16429499
cg16429725
cg16429735
cg16430279
cg16430963
cg16431352
cg16431978
cg16432441
cg16433258
cg16433493
cg16434510
cg16434897
cg16435571
cg16435627
cg16435923
cg16436164
cg16436421
cg16436711
cg16436762
cg16437021
cg16437385
cg16437728
cg16438655
cg16438841
cg16439608
cg16440058

cg16441238
cg16441259
cg16442298
cg16442402
cg16443050
cg16443866
cg16444103
cg16444629
cg16445268
cg16445423
cg16446484
cg16446577
cg16446783
cg16448636
cg16448847
cg16449687
cg16450249
cg16450677
cg16451371
cg16451455
cg16451527
cg16451661
cg16451803
cg16452067
cg16452248
cg16453224
cg16453474
cg16454426
cg16455748
cg16456172
cg16456252
cg16456980
cg16457786
cg16458494
cg16458596
cg16458866
cg16459265
cg16459364
cg16459519
cg16459890
cg16460727
cg16460816
cg16461139
cg16461251
cg16461485
cg16462075
cg16462183
cg16462462
cg16462648
cg16463150
cg16463460
cg16463697

cg16464007
cg16465407
cg16465430
cg16465643
cg16466065
cg16466270
cg16466750
cg16466870
cg16468174
cg16468187
cg16468417
cg16468914
cg16469223
cg16469419
cg16469492
cg16469740
cg16470423
cg16470760
cg16470816
cg16471362
cg16472050
cg16472356
cg16473125
cg16473797
cg16474170
cg16474184
cg16474701
cg16475721
cg16475891
cg16476048
cg16476432
cg16476940
cg16477611
cg16477744
cg16477774
cg16478536
cg16479131
cg16479539
cg16480759
cg16481280
cg16481618
cg16481969
cg16483326
cg16483466
cg16484020
cg16484526
cg16484575
cg16485048
cg16485558
cg16485797
cg16485800
cg16485975

cg16486109
cg16488065
cg16488382
cg16488737
cg16489259
cg16489468
cg16490906
cg16491362
cg16491547
cg16491617
cg16491668
cg16491865
cg16491960
cg16492211
cg16492377
cg16493584
cg16493813
cg16494108
cg16494192
cg16494747
cg16496217
cg16496753
cg16497661
cg16498194
cg16498504
cg16498514
cg16498741
cg16498913
cg16499834
cg16499923
cg16499947
cg16500987
cg16501235
cg16501237
cg16501560
cg16501892
cg16502931
cg16503006
cg16503259
cg16503628
cg16503924
cg16504670
cg16505731
cg16506185
cg16506910
cg16507522
cg16507827
cg16508480
cg16509531
cg16509592
cg16509777
cg16509851

cg16510070
cg16510099
cg16510350
cg16510548
cg16511086
cg16511454
cg16512144
cg16512311
cg16512458
cg16512615
cg16512661
cg16512667
cg16512727
cg16513326
cg16514214
cg16514473
cg16515477
cg16516691
cg16517298
cg16517702
cg16518348
cg16518590
cg16518729
cg16519217
cg16519487
cg16519587
cg16519923
cg16520046
cg16520191
cg16520712
cg16520796
cg16520965
cg16521454
cg16521773
cg16521917
cg16522557
cg16523380
cg16523868
cg16524232
cg16524284
cg16524818
cg16524928
cg16525435
cg16525470
cg16526416
cg16526673
cg16527112
cg16527552
cg16527877
cg16528511
cg16528585
cg16528678

cg16528895
cg16529229
cg16529323
cg16529477
cg16529592
cg16530130
cg16530587
cg16530596
cg16530881
cg16530970
cg16531181
cg16532016
cg16532185
cg16532262
cg16532438
cg16532606
cg16533767
cg16534315
cg16535035
cg16535080
cg16535493
cg16535752
cg16536329
cg16536739
cg16536824
cg16537371
cg16537483
cg16538359
cg16538606
cg16538674
cg16538749
cg16539957
cg16540590
cg16541026
cg16541931
cg16542646
cg16543027
cg16543094
cg16544010
cg16544169
cg16544737
cg16544956
cg16546376
cg16546442
cg16546658
cg16547529
cg16547629
cg16548262
cg16548780
cg16548911
cg16548961
cg16549308

cg16549596
cg16549644
cg16550010
cg16550446
cg16551669
cg16551670
cg16553297
cg16553500
cg16553757
cg16554153
cg16554164
cg16554447
cg16555556
cg16555818
cg16555866
cg16555926
cg16556078
cg16556145
cg16557964
cg16558925
cg16560209
cg16560711
cg16560774
cg16560847
cg16561487
cg16561893
cg16562486
cg16562776
cg16563134
cg16563171
cg16563938
cg16564180
cg16564302
cg16565002
cg16565294
cg16565394
cg16565562
cg16565901
cg16566191
cg16566391
cg16566400
cg16566627
cg16567216
cg16568681
cg16569273
cg16569650
cg16570223
cg16570256
cg16570414
cg16570507
cg16570669
cg16571983

cg16572224
cg16572375
cg16572957
cg16573755
cg16573782
cg16573863
cg16574310
cg16575291
cg16575694
cg16576160
cg16576569
cg16576811
cg16577081
cg16577789
cg16577947
cg16578085
cg16578267
cg16578549
cg16579158
cg16579908
cg16580197
cg16580391
cg16580935
cg16580937
cg16580982
cg16581017
cg16581179
cg16581199
cg16581308
cg16581347
cg16581455
cg16581536
cg16581884
cg16582174
cg16582732
cg16583536
cg16584392
cg16584393
cg16584624
cg16586104
cg16586200
cg16586394
cg16586538
cg16586807
cg16587265
cg16588163
cg16588731
cg16589299
cg16590190
cg16590237
cg16590466
cg16590794

cg16590910
cg16591182
cg16591404
cg16592121
cg16592326
cg16592378
cg16593060
cg16593921
cg16594273
cg16594468
cg16594913
cg16595458
cg16595750
cg16596440
cg16596691
cg16596889
cg16596957
cg16597079
cg16598244
cg16598292
cg16598668
cg16599136
cg16599143
cg16599974
cg16599983
cg16601094
cg16601385
cg16601415
cg16601569
cg16602097
cg16602747
cg16602816
cg16603916
cg16604126
cg16604566
cg16604975
cg16605327
cg16606195
cg16607685
cg16607898
cg16608087
cg16608498
cg16608656
cg16609260
cg16609297
cg16609872
cg16609966
cg16609995
cg16611352
cg16611911
cg16612562
cg16615040

cg16615154
cg16615508
cg16616340
cg16617817
cg16617910
cg16618218
cg16618493
cg16618789
cg16619395
cg16619425
cg16619764
cg16619991
cg16620382
cg16620392
cg16620526
cg16620809
cg16621364
cg16621560
cg16621833
cg16622190
cg16622495
cg16622651
cg16622652
cg16622875
cg16623941
cg16624210
cg16624994
cg16625119
cg16625218
cg16626012
cg16626063
cg16626067
cg16626196
cg16626809
cg16626884
cg16626954
cg16627086
cg16627854
cg16627949
cg16628039
cg16628093
cg16628205
cg16628918
cg16629076
cg16629262
cg16629452
cg16629523
cg16629695
cg16629702
cg16629749
cg16630482
cg16630572

cg16631083
cg16631326
cg16632159
cg16632715
cg16633951
cg16634544
cg16635196
cg16635314
cg16635352
cg16635578
cg16635735
cg16635777
cg16636682
cg16637224
cg16637458
cg16637813
cg16638092
cg16638248
cg16638540
cg16638834
cg16638920
cg16639540
cg16639691
cg16639860
cg16640358
cg16640855
cg16641876
cg16642063
cg16642284
cg16642545
cg16642695
cg16642791
cg16643152
cg16643603
cg16643851
cg16644366
cg16645423
cg16645705
cg16646054
cg16646298
cg16646726
cg16646743
cg16647921
cg16648632
cg16648841
cg16649034
cg16649298
cg16650096
cg16650717
cg16651262
cg16651314
cg16651768

cg16651895
cg16652063
cg16652264
cg16652739
cg16652741
cg16652855
cg16652874
cg16653534
cg16654801
cg16654911
cg16655166
cg16655240
cg16656895
cg16657244
cg16657538
cg16657724
cg16657886
cg16658020
cg16658473
cg16658582
cg16658618
cg16659728
cg16659773
cg16659899
cg16660041
cg16660220
cg16660494
cg16660547
cg16660971
cg16661123
cg16661164
cg16662408
cg16662645
cg16663033
cg16663497
cg16664405
cg16664732
cg16666329
cg16667189
cg16667251
cg16668359
cg16668520
cg16668896
cg16668903
cg16669602
cg16670554
cg16670736
cg16671365
cg16671674
cg16671899
cg16672223
cg16672637

cg16673052
cg16673412
cg16673508
cg16673904
cg16674248
cg16675507
cg16676200
cg16676655
cg16677448
cg16677947
cg16677955
cg16678446
cg16678925
cg16679297
cg16679378
cg16679452
cg16680125
cg16680438
cg16680677
cg16681421
cg16681873
cg16682454
cg16682903
cg16682935
cg16683741
cg16684691
cg16684817
cg16685313
cg16685586
cg16686185
cg16686900
cg16686960
cg16689193
cg16689232
cg16689937
cg16691158
cg16691821
cg16692090
cg16692653
cg16693012
cg16693595
cg16694365
cg16694480
cg16694785
cg16695380
cg16695576
cg16696002
cg16696035
cg16696648
cg16696774
cg16697268
cg16697433

cg16697438
cg16697850
cg16698575
cg16699074
cg16700282
cg16700364
cg16700392
cg16701059
cg16701456
cg16701890
cg16701939
cg16702313
cg16703008
cg16703135
cg16703390
cg16703762
cg16703956
cg16704560
cg16704834
cg16705086
cg16705395
cg16705627
cg16705697
cg16706099
cg16706220
cg16706324
cg16706690
cg16706813
cg16707954
cg16708012
cg16708451
cg16708465
cg16709232
cg16709433
cg16709836
cg16710042
cg16710099
cg16710517
cg16710604
cg16710894
cg16711450
cg16711612
cg16712438
cg16712730
cg16713551
cg16716750
cg16717008
cg16717192
cg16717431
cg16717532
cg16717546
cg16717713

cg16717781
cg16718263
cg16718624
cg16718710
cg16718760
cg16720005
cg16720379
cg16720807
cg16721191
cg16721418
cg16721489
cg16722118
cg16723982
cg16724332
cg16724588
cg16724800
cg16725552
cg16725642
cg16726869
cg16727006
cg16727201
cg16727923
cg16728651
cg16728846
cg16728934
cg16729415
cg16729555
cg16730559
cg16730908
cg16732616
cg16733117
cg16733267
cg16733705
cg16734845
cg16735190
cg16735682
cg16735875
cg16735881
cg16736055
cg16736630
cg16736826
cg16736989
cg16737749
cg16739178
cg16739441
cg16739463
cg16739580
cg16739679
cg16740744
cg16740893
cg16741041
cg16742703

cg16743289
cg16743754
cg16744438
cg16745033
cg16745596
cg16746901
cg16747247
cg16747928
cg16748088
cg16749235
cg16750440
cg16750801
cg16752583
cg16752876
cg16753939
cg16754613
cg16755067
cg16755214
cg16755833
cg16755863
cg16756527
cg16757384
cg16757984
cg16758813
cg16759783
cg16759787
cg16760130
cg16760223
cg16760587
cg16760981
cg16762308
cg16762386
cg16762735
cg16762794
cg16764147
cg16764236
cg16764362
cg16764778
cg16766249
cg16766325
cg16766382
cg16767801
cg16768018
cg16768937
cg16769152
cg16769381
cg16769649
cg16770238
cg16770368
cg16770400
cg16770549
cg16771578

cg16772039
cg16773677
cg16773768
cg16773803
cg16773814
cg16773899
cg16774375
cg16774528
cg16774755
cg16776035
cg16776065
cg16776421
cg16776981
cg16777090
cg16777106
cg16778734
cg16779839
cg16780562
cg16780603
cg16780771
cg16780784
cg16780860
cg16781205
cg16781264
cg16782084
cg16782184
cg16783279
cg16783744
cg16785520
cg16786297
cg16786581
cg16786808
cg16786971
cg16787600
cg16787652
cg16788014
cg16790794
cg16790843
cg16791210
cg16791424
cg16791508
cg16791797
cg16792071
cg16792144
cg16792302
cg16792312
cg16792613
cg16792800
cg16792842
cg16793866
cg16794471
cg16794961

cg16795184
cg16795280
cg16795307
cg16796272
cg16797421
cg16797473
cg16797818
cg16797901
cg16798252
cg16798537
cg16799191
cg16800165
cg16801887
cg16802508
cg16803064
cg16803083
cg16803596
cg16803846
cg16803967
cg16804165
cg16804284
cg16804882
cg16804948
cg16805553
cg16805884
cg16807297
cg16808120
cg16808156
cg16808587
cg16809304
cg16810272
cg16810559
cg16811455
cg16811558
cg16812893
cg16813187
cg16814513
cg16814808
cg16815157
cg16816623
cg16817426
cg16818395
cg16818414
cg16818712
cg16818830
cg16819051
cg16820973
cg16821446
cg16821558
cg16822095
cg16822143
cg16822174

cg16822939
cg16823430
cg16823945
cg16824282
cg16825000
cg16826286
cg16827215
cg16827475
cg16831331
cg16831361
cg16831623
cg16831889
cg16832407
cg16833444
cg16833797
cg16834726
cg16836040
cg16836853
cg16838723
cg16839083
cg16840358
cg16840744
cg16840978
cg16841748
cg16844661
cg16844941
cg16845126
cg16845394
cg16845880
cg16846645
cg16847695
cg16847696
cg16848524
cg16848937
cg16849024
cg16849041
cg16849201
cg16850150
cg16850173
cg16851417
cg16851989
cg16853712
cg16853860
cg16854826
cg16856049
cg16856286
cg16856734
cg16856833
cg16857113
cg16857134
cg16857801
cg16857858

cg16858125
cg16858472
cg16858587
cg16861031
cg16861076
cg16861968
cg16863522
cg16864658
cg16865396
cg16865618
cg16865953
cg16866136
cg16866373
cg16866941
cg16868145
cg16868253
cg16868298
cg16869647
cg16869862
cg16870867
cg16871519
cg16871828
cg16872357
cg16873271
cg16874347
cg16874583
cg16875235
cg16875864
cg16876315
cg16877004
cg16877924
cg16878027
cg16878214
cg16878549
cg16879596
cg16879608
cg16879936
cg16880856
cg16881730
cg16883332
cg16883475
cg16883726
cg16884902
cg16885235
cg16885557
cg16885861
cg16885982
cg16886175
cg16886329
cg16886882
cg16887070
cg16887170

cg16887179
cg16887183
cg16887264
cg16888547
cg16888559
cg16889054
cg16890051
cg16890739
cg16891431
cg16891892
cg16891965
cg16891968
cg16892812
cg16894819
cg16895216
cg16895672
cg16895814
cg16896079
cg16896539
cg16896911
cg16898239
cg16898498
cg16900102
cg16900671
cg16901014
cg16901123
cg16901840
cg16902122
cg16903016
cg16903225
cg16903269
cg16903589
cg16904666
cg16904859
cg16905434
cg16906187
cg16907101
cg16907558
cg16908208
cg16908944
cg16909408
cg16909579
cg16909733
cg16910830
cg16910908
cg16911214
cg16912910
cg16915653
cg16917058
cg16918050
cg16918327
cg16918846

cg16919517
cg16919569
cg16919680
cg16921285
cg16921789
cg16922279
cg16922523
cg16922869
cg16922884
cg16923982
cg16924010
cg16924337
cg16924702
cg16924753
cg16926302
cg16926445
cg16927040
cg16927372
cg16928487
cg16928869
cg16929850
cg16930728
cg16931068
cg16931664
cg16932285
cg16932472
cg16932560
cg16933181
cg16933386
cg16934969
cg16935065
cg16936094
cg16936846
cg16937268
cg16938363
cg16938805
cg16940399
cg16941122
cg16942931
cg16943185
cg16943529
cg16943635
cg16944958
cg16944977
cg16945415
cg16946445
cg16946687
cg16947583
cg16948320
cg16949103
cg16949120
cg16949584

cg16950627
cg16950726
cg16953006
cg16953297
cg16953612
cg16954727
cg16956132
cg16956426
cg16956804
cg16957224
cg16957569
cg16957993
cg16958148
cg16958351
cg16958402
cg16958939
cg16959747
cg16960046
cg16960112
cg16960402
cg16960674
cg16961623
cg16961648
cg16962045
cg16963144
cg16964025
cg16964389
cg16964728
cg16966139
cg16966243
cg16966261
cg16966315
cg16966340
cg16967068
cg16967578
cg16967583
cg16968623
cg16969207
cg16969368
cg16971128
cg16971668
cg16973098
cg16973527
cg16973715
cg16974075
cg16974190
cg16974832
cg16974873
cg16975053
cg16975095
cg16975587
cg16976370

cg16976499
cg16976857
cg16976870
cg16977270
cg16978004
cg16978168
cg16981024
cg16981685
cg16983084
cg16983159
cg16983282
cg16983723
cg16985023
cg16985333
cg16985652
cg16986499
cg16987305
cg16987552
cg16987735
cg16987754
cg16987900
cg16988616
cg16988989
cg16989132
cg16989248
cg16989380
cg16989719
cg16989784
cg16989867
cg16990168
cg16990174
cg16990474
cg16993015
cg16993043
cg16993957
cg16994534
cg16996281
cg16996574
cg16996682
cg16997364
cg16997486
cg16998656
cg16998950
cg16999370
cg16999602
cg17000593
cg17001194
cg17001566
cg17002156
cg17002328
cg17002510
cg17003221

cg17003607
cg17003736
cg17004025
cg17006136
cg17006281
cg17006506
cg17007164
cg17007640
cg17008031
cg17008160
cg17008318
cg17009522
cg17009616
cg17010112
cg17010522
cg17010827
cg17011276
cg17011453
cg17012160
cg17012181
cg17013303
cg17014035
cg17014186
cg17014214
cg17015994
cg17016394
cg17016847
cg17016922
cg17017189
cg17017814
cg17018096
cg17018661
cg17018767
cg17018879
cg17018946
cg17019053
cg17020464
cg17020895
cg17021487
cg17021648
cg17021880
cg17022577
cg17023149
cg17023856
cg17024482
cg17025232
cg17025835
cg17026542
cg17028067
cg17028301
cg17028595
cg17028814

cg17029168
cg17029721
cg17030173
cg17030233
cg17030981
cg17031773
cg17031926
cg17032081
cg17032602
cg17033086
cg17033359
cg17033391
cg17033546
cg17033684
cg17033891
cg17034079
cg17034088
cg17036164
cg17037101
cg17037282
cg17037931
cg17038049
cg17038577
cg17038667
cg17039236
cg17039391
cg17040303
cg17040471
cg17040807
cg17041296
cg17043624
cg17043651
cg17043722
cg17046141
cg17046322
cg17046484
cg17046577
cg17047829
cg17048000
cg17048169
cg17048610
cg17048700
cg17049320
cg17050597
cg17050756
cg17052041
cg17052466
cg17052783
cg17053060
cg17054060
cg17054640
cg17054708

cg17055207
cg17055794
cg17055993
cg17056676
cg17057086
cg17057501
cg17058616
cg17059658
cg17061244
cg17061340
cg17062268
cg17062665
cg17062829
cg17063632
cg17064754
cg17066452
cg17066531
cg17067190
cg17067226
cg17071063
cg17071446
cg17071639
cg17071957
cg17072368
cg17072503
cg17072805
cg17073169
cg17073273
cg17073306
cg17073708
cg17074396
cg17075459
cg17076890
cg17078116
cg17078219
cg17078253
cg17078629
cg17078686
cg17079431
cg17079497
cg17079556
cg17080163
cg17080371
cg17080740
cg17080987
cg17081489
cg17081867
cg17081998
cg17082803
cg17082938
cg17083209
cg17083429

cg17084044
cg17084151
cg17084937
cg17085280
cg17085352
cg17086204
cg17087331
cg17087479
cg17087499
cg17089511
cg17090237
cg17090401
cg17090754
cg17091020
cg17091301
cg17091361
cg17091967
cg17092523
cg17093478
cg17093684
cg17093795
cg17095171
cg17095731
cg17095910
cg17095929
cg17096807
cg17097481
cg17097923
cg17098147
cg17098387
cg17098965
cg17098979
cg17099072
cg17099569
cg17100390
cg17101296
cg17101703
cg17102325
cg17102495
cg17104149
cg17104151
cg17104202
cg17104258
cg17104596
cg17104700
cg17105834
cg17107540
cg17107691
cg17107840
cg17107893
cg17108388
cg17108629

cg17112089
cg17112108
cg17112382
cg17112426
cg17112707
cg17112828
cg17112958
cg17112975
cg17113158
cg17113476
cg17113968
cg17114584
cg17115419
cg17116120
cg17116178
cg17118095
cg17118478
cg17119170
cg17120402
cg17121178
cg17121747
cg17122311
cg17122577
cg17123334
cg17123690
cg17124583
cg17124829
cg17124844
cg17125585
cg17126250
cg17126435
cg17126622
cg17126947
cg17128767
cg17128799
cg17129262
cg17129388
cg17130063
cg17130251
cg17130457
cg17131377
cg17131837
cg17131905
cg17132421
cg17133833
cg17135392
cg17135722
cg17136719
cg17137063
cg17137182
cg17137218
cg17137424

cg17138769
cg17138852
cg17139392
cg17139666
cg17140103
cg17140222
cg17140307
cg17140992
cg17142404
cg17142432
cg17142722
cg17142931
cg17143179
cg17143291
cg17143462
cg17143696
cg17144184
cg17144548
cg17145190
cg17146110
cg17146526
cg17146570
cg17146738
cg17147045
cg17147440
cg17147638
cg17147917
cg17148976
cg17149019
cg17149030
cg17149262
cg17149350
cg17150789
cg17150809
cg17150898
cg17151560
cg17151621
cg17152266
cg17152757
cg17153045
cg17153078
cg17153775
cg17153794
cg17153947
cg17154315
cg17156011
cg17156053
cg17156402
cg17156432
cg17156565
cg17156633
cg17156862

cg17157798
cg17158941
cg17160327
cg17160329
cg17160525
cg17160984
cg17161180
cg17161250
cg17161266
cg17162024
cg17162031
cg17162319
cg17162355
cg17162576
cg17163404
cg17163760
cg17164016
cg17164063
cg17164513
cg17165759
cg17165841
cg17167278
cg17167536
cg17167852
cg17168519
cg17168630
cg17168771
cg17170463
cg17171127
cg17171215
cg17172331
cg17172569
cg17173369
cg17173423
cg17173498
cg17173767
cg17173896
cg17174275
cg17174372
cg17175521
cg17176573
cg17176609
cg17177455
cg17177632
cg17178291
cg17178336
cg17178468
cg17178761
cg17179568
cg17179881
cg17179923
cg17181043

cg17181653
cg17182240
cg17182837
cg17183432
cg17183752
cg17185267
cg17186103
cg17186132
cg17186163
cg17186168
cg17186328
cg17186750
cg17187521
cg17187764
cg17188567
cg17189465
cg17190117
cg17190363
cg17191109
cg17191443
cg17191567
cg17191715
cg17192247
cg17192315
cg17192377
cg17192862
cg17192954
cg17193040
cg17193436
cg17195230
cg17197051
cg17197065
cg17198929
cg17199113
cg17199247
cg17199320
cg17199367
cg17199481
cg17199800
cg17200577
cg17200850
cg17200999
cg17202013
cg17202313
cg17203063
cg17203290
cg17203447
cg17204129
cg17204510
cg17205313
cg17205386
cg17206001

cg17206420
cg17208360
cg17208530
cg17209280
cg17210014
cg17210535
cg17210546
cg17210837
cg17210933
cg17210999
cg17211349
cg17212019
cg17212184
cg17212277
cg17212791
cg17213048
cg17213402
cg17213713
cg17214381
cg17215278
cg17215361
cg17215863
cg17216266
cg17216583
cg17216807
cg17217834
cg17217996
cg17218270
cg17218713
cg17218813
cg17219740
cg17220574
cg17220584
cg17220933
cg17221083
cg17221226
cg17221813
cg17222000
cg17222500
cg17222682
cg17222755
cg17222950
cg17223748
cg17223809
cg17226286
cg17226446
cg17227156
cg17227967
cg17228942
cg17229929
cg17230874
cg17231293

cg17231373
cg17231999
cg17232583
cg17235499
cg17235833
cg17236709
cg17237086
cg17238065
cg17238830
cg17239057
cg17239236
cg17239558
cg17240133
cg17240157
cg17240815
cg17241289
cg17241310
cg17242596
cg17245726
cg17246765
cg17247122
cg17250160
cg17250615
cg17251609
cg17251713
cg17252489
cg17252528
cg17252645
cg17252780
cg17252960
cg17253407
cg17253709
cg17253785
cg17254386
cg17255125
cg17255397
cg17256609
cg17257469
cg17257636
cg17258387
cg17258460
cg17258607
cg17258741
cg17259183
cg17260180
cg17260660
cg17261120
cg17262492
cg17262810
cg17264240
cg17264541
cg17264609

cg17264670
cg17264941
cg17265120
cg17265358
cg17265829
cg17266805
cg17267239
cg17267681
cg17267737
cg17270257
cg17271504
cg17272549
cg17273416
cg17273683
cg17274064
cg17274215
cg17274827
cg17275781
cg17275894
cg17276450
cg17276590
cg17277001
cg17277248
cg17277529
cg17277892
cg17277933
cg17278072
cg17278355
cg17278595
cg17279887
cg17280106
cg17280134
cg17280284
cg17280346
cg17280398
cg17280514
cg17281600
cg17282004
cg17282466
cg17282592
cg17282688
cg17282836
cg17282904
cg17283111
cg17283169
cg17283453
cg17284116
cg17284326
cg17284334
cg17284609
cg17284804
cg17284854

cg17285605
cg17286157
cg17286695
cg17286983
cg17287122
cg17287235
cg17287605
cg17288256
cg17288397
cg17288892
cg17290213
cg17291001
cg17291157
cg17291251
cg17291906
cg17292583
cg17293195
cg17293838
cg17293936
cg17293973
cg17294725
cg17295225
cg17295389
cg17295733
cg17295739
cg17296166
cg17296447
cg17296707
cg17296990
cg17297475
cg17297613
cg17298403
cg17298516
cg17298751
cg17299007
cg17299616
cg17299636
cg17300051
cg17300275
cg17301902
cg17302155
cg17302896
cg17303756
cg17304025
cg17304276
cg17305181
cg17305266
cg17305436
cg17306401
cg17306686
cg17306747
cg17306848

cg17307280
cg17307442
cg17308758
cg17308840
cg17310600
cg17310882
cg17311132
cg17311865
cg17312004
cg17312536
cg17314180
cg17314478
cg17314538
cg17314888
cg17315844
cg17315964
cg17316300
cg17316316
cg17316862
cg17317859
cg17317869
cg17317895
cg17319007
cg17319198
cg17319486
cg17319889
cg17319956
cg17320282
cg17320378
cg17320698
cg17320707
cg17321385
cg17321888
cg17322443
cg17322655
cg17322774
cg17323336
cg17324804
cg17325595
cg17325866
cg17325911
cg17327171
cg17327331
cg17328514
cg17328716
cg17329499
cg17329648
cg17330097
cg17330672
cg17331296
cg17331494
cg17333109

cg17333973
cg17333990
cg17334018
cg17335741
cg17336354
cg17336455
cg17337269
cg17337668
cg17337791
cg17338299
cg17338430
cg17339147
cg17339327
cg17340373
cg17340391
cg17340519
cg17340729
cg17341174
cg17341366
cg17342132
cg17342709
cg17342759
cg17342973
cg17343107
cg17343253
cg17343873
cg17344048
cg17344080
cg17344321
cg17344598
cg17345277
cg17345569
cg17345741
cg17346145
cg17348201
cg17349151
cg17349389
cg17349486
cg17350544
cg17351439
cg17351862
cg17352276
cg17352462
cg17352540
cg17352891
cg17354303
cg17355153
cg17355329
cg17355865
cg17356056
cg17357062
cg17357548

cg17358124
cg17358127
cg17358530
cg17359265
cg17360680
cg17361203
cg17362483
cg17362661
cg17362861
cg17364114
cg17366309
cg17367077
cg17367078
cg17367472
cg17367616
cg17368760
cg17369115
cg17369527
cg17370163
cg17370357
cg17370412
cg17370981
cg17371070
cg17371081
cg17372245
cg17372657
cg17372758
cg17372841
cg17373345
cg17373759
cg17375585
cg17376813
cg17377066
cg17378342
cg17378535
cg17378803
cg17379932
cg17380661
cg17380770
cg17381377
cg17382198
cg17382459
cg17382939
cg17383186
cg17383203
cg17383272
cg17383727
cg17383842
cg17383853
cg17384380
cg17384457
cg17385936

cg17386057
cg17386093
cg17386185
cg17386213
cg17386911
cg17387577
cg17387670
cg17389519
cg17389949
cg17389956
cg17390106
cg17390350
cg17390890
cg17391194
cg17391928
cg17392047
cg17392573
cg17394306
cg17395184
cg17396707
cg17396797
cg17397069
cg17397150
cg17397548
cg17397592
cg17397631
cg17397836
cg17400113
cg17400559
cg17401106
cg17402103
cg17403609
cg17403699
cg17404532
cg17405031
cg17405586
cg17405840
cg17406204
cg17408087
cg17408185
cg17408527
cg17410236
cg17410313
cg17410650
cg17410660
cg17411031
cg17412005
cg17412043
cg17412102
cg17412258
cg17412351
cg17412464

cg17412560
cg17412886
cg17412901
cg17413943
cg17415650
cg17415859
cg17416793
cg17417049
cg17418387
cg17418579
cg17419731
cg17420058
cg17420585
cg17421062
cg17421241
cg17421805
cg17422403
cg17422774
cg17423624
cg17425351
cg17425484
cg17425590
cg17425818
cg17426273
cg17426766
cg17427615
cg17427926
cg17428593
cg17429000
cg17429236
cg17429686
cg17430268
cg17430805
cg17430979
cg17431952
cg17432154
cg17434309
cg17434643
cg17434901
cg17435445
cg17435965
cg17436025
cg17437419
cg17437939
cg17438030
cg17439967
cg17442194
cg17442269
cg17442683
cg17443080
cg17443359
cg17443388

cg17443753
cg17444479
cg17444521
cg17444582
cg17444738
cg17444747
cg17444859
cg17445157
cg17445213
cg17448335
cg17449235
cg17449759
cg17449954
cg17450585
cg17450733
cg17450815
cg17451015
cg17451609
cg17451760
cg17452249
cg17452615
cg17453374
cg17453460
cg17453603
cg17454086
cg17454094
cg17454298
cg17454397
cg17454857
cg17455088
cg17455348
cg17456675
cg17457560
cg17457986
cg17458693
cg17458967
cg17459182
cg17459204
cg17459660
cg17460183
cg17460441
cg17460447
cg17460855
cg17461336
cg17461600
cg17462107
cg17463145
cg17463149
cg17463527
cg17463698
cg17463745
cg17465112

cg17466082
cg17466658
cg17466857
cg17467879
cg17467898
cg17468185
cg17469019
cg17470067
cg17470095
cg17470327
cg17470942
cg17471425
cg17472832
cg17473580
cg17473656
cg17473727
cg17473765
cg17474920
cg17475136
cg17475172
cg17475190
cg17475304
cg17475733
cg17475857
cg17476101
cg17477162
cg17477445
cg17477818
cg17477930
cg17478016
cg17478203
cg17478228
cg17478282
cg17478979
cg17478992
cg17479119
cg17479716
cg17479763
cg17480438
cg17480554
cg17480641
cg17481117
cg17481912
cg17482089
cg17482360
cg17482801
cg17482935
cg17484331
cg17484671
cg17484874
cg17484944
cg17485530

cg17485838
cg17486097
cg17487029
cg17487050
cg17488628
cg17489748
cg17491228
cg17491449
cg17491456
cg17491850
cg17493815
cg17494034
cg17494199
cg17494334
cg17494927
cg17495466
cg17495555
cg17495715
cg17496059
cg17496869
cg17496990
cg17497172
cg17497320
cg17497536
cg17497824
cg17498101
cg17498321
cg17499251
cg17500742
cg17500962
cg17501569
cg17502213
cg17502434
cg17503085
cg17503200
cg17503456
cg17504164
cg17504641
cg17504737
cg17504999
cg17505463
cg17505819
cg17506460
cg17507751
cg17507897
cg17508039
cg17508167
cg17508300
cg17508591
cg17508816
cg17509220
cg17509561

cg17509967
cg17510786
cg17512799
cg17513592
cg17514199
cg17514558
cg17515110
cg17515145
cg17515760
cg17515837
cg17516156
cg17516469
cg17516571
cg17516825
cg17516837
cg17516997
cg17517296
cg17517458
cg17517997
cg17518215
cg17518949
cg17519101
cg17519156
cg17519280
cg17519477
cg17519645
cg17519949
cg17520909
cg17522953
cg17525102
cg17525357
cg17526573
cg17526890
cg17526952
cg17527764
cg17527766
cg17529058
cg17529152
cg17529832
cg17529965
cg17530031
cg17533522
cg17533563
cg17534009
cg17534654
cg17534916
cg17535314
cg17535550
cg17535595
cg17535647
cg17535702
cg17536114

cg17536586
cg17536595
cg17537007
cg17537177
cg17537466
cg17538572
cg17541002
cg17541528
cg17542330
cg17542385
cg17543112
cg17544952
cg17545399
cg17545652
cg17545853
cg17546247
cg17546422
cg17547157
cg17547864
cg17547897
cg17547898
cg17548844
cg17550175
cg17551002
cg17552116
cg17552333
cg17552471
cg17554182
cg17555373
cg17555806
cg17555825
cg17556527
cg17557464
cg17559786
cg17560340
cg17560832
cg17563229
cg17564775
cg17565078
cg17565113
cg17566002
cg17566118
cg17566541
cg17567838
cg17568266
cg17569842
cg17573292
cg17573933
cg17574050
cg17574812
cg17576603
cg17576736

cg17577636
cg17578131
cg17578235
cg17578322
cg17578341
cg17578548
cg17579446
cg17579667
cg17580416
cg17580935
cg17581104
cg17581925
cg17583413
cg17584512
cg17584530
cg17585615
cg17585889
cg17587327
cg17587385
cg17588294
cg17588578
cg17589341
cg17589449
cg17589866
cg17590162
cg17590499
cg17590523
cg17591640
cg17592231
cg17592875
cg17593342
cg17593625
cg17594632
cg17595735
cg17598358
cg17598466
cg17598552
cg17598574
cg17599298
cg17599748
cg17600393
cg17601191
cg17602126
cg17602444
cg17603206
cg17603689
cg17603763
cg17604985
cg17605476
cg17607298
cg17608082
cg17608587

cg17609931
cg17610800
cg17610889
cg17611262
cg17611674
cg17611951
cg17612147
cg17612412
cg17612535
cg17613707
cg17613911
cg17615629
cg17616099
cg17616305
cg17616453
cg17616554
cg17616677
cg17617491
cg17617843
cg17619311
cg17619823
cg17619876
cg17620300
cg17621507
cg17621580
cg17621718
cg17622131
cg17622246
cg17624196
cg17626022
cg17626301
cg17626734
cg17627328
cg17627629
cg17627654
cg17628060
cg17628540
cg17628717
cg17628867
cg17629148
cg17629322
cg17629870
cg17630392
cg17631198
cg17631972
cg17632372
cg17632579
cg17632711
cg17633074
cg17633170
cg17633576
cg17634401

cg17634572
cg17635777
cg17636883
cg17636986
cg17637045
cg17638160
cg17638507
cg17639111
cg17639959
cg17640485
cg17640807
cg17641443
cg17641857
cg17641876
cg17642201
cg17642774
cg17643106
cg17643564
cg17643796
cg17644227
cg17645598
cg17645664
cg17645677
cg17647537
cg17648213
cg17650028
cg17650057
cg17650274
cg17651451
cg17651959
cg17651972
cg17652174
cg17652291
cg17652600
cg17654050
cg17654107
cg17654419
cg17654567
cg17654749
cg17655253
cg17655320
cg17655415
cg17655609
cg17656260
cg17656890
cg17657268
cg17657394
cg17657459
cg17657814
cg17657898
cg17658266
cg17658568

cg17658874
cg17661135
cg17661642
cg17661686
cg17662369
cg17664182
cg17664233
cg17664577
cg17665497
cg17665505
cg17668731
cg17669334
cg17670263
cg17670496
cg17671005
cg17672101
cg17672850
cg17673237
cg17673897
cg17675301
cg17675852
cg17676574
cg17676618
cg17677030
cg17677397
cg17678139
cg17678645
cg17678847
cg17679427
cg17679470
cg17679987
cg17680092
cg17680611
cg17681054
cg17681447
cg17681750
cg17683775
cg17685019
cg17685628
cg17687282
cg17688525
cg17689490
cg17689601
cg17689721
cg17689899
cg17690322
cg17691233
cg17692125
cg17692971
cg17693865
cg17694593
cg17694848

cg17695462
cg17696006
cg17696566
cg17697633
cg17697873
cg17700915
cg17701146
cg17701942
cg17702736
cg17704189
cg17704343
cg17704864
cg17705041
cg17705615
cg17705631
cg17705731
cg17706173
cg17707126
cg17710804
cg17711567
cg17711754
cg17712694
cg17712818
cg17713376
cg17713910
cg17714025
cg17714703
cg17714799
cg17714832
cg17715521
cg17715641
cg17716661
cg17716664
cg17716724
cg17716986
cg17717866
cg17718074
cg17718182
cg17718401
cg17718576
cg17719126
cg17719202
cg17719543
cg17719818
cg17720087
cg17720153
cg17721715
cg17722354
cg17722435
cg17723122
cg17723653
cg17723710

cg17724175
cg17724366
cg17724789
cg17725129
cg17725851
cg17727071
cg17727579
cg17728065
cg17729072
cg17729253
cg17729668
cg17729694
cg17729941
cg17731547
cg17732038
cg17732877
cg17733447
cg17735531
cg17735539
cg17735631
cg17736336
cg17737681
cg17737835
cg17738194
cg17739152
cg17739643
cg17740140
cg17740322
cg17740401
cg17740645
cg17740722
cg17740752
cg17740822
cg17741501
cg17741839
cg17741986
cg17742051
cg17742416
cg17743381
cg17744293
cg17744604
cg17746395
cg17747005
cg17747551
cg17747635
cg17747879
cg17748092
cg17748504
cg17748645
cg17749454
cg17750109
cg17751591

cg17753162
cg17753347
cg17754042
cg17754510
cg17754680
cg17754914
cg17754943
cg17755082
cg17755554
cg17755610
cg17756545
cg17756730
cg17757079
cg17757778
cg17758048
cg17758905
cg17759274
cg17759443
cg17760412
cg17762073
cg17762554
cg17765324
cg17765387
cg17765751
cg17765885
cg17766305
cg17766560
cg17767099
cg17767185
cg17767285
cg17768491
cg17768768
cg17769771
cg17771605
cg17772649
cg17772789
cg17773654
cg17773660
cg17774559
cg17774672
cg17774851
cg17775003
cg17775490
cg17776207
cg17776688
cg17777432
cg17777676
cg17777683
cg17777731
cg17778120
cg17778211
cg17778288

cg17780246
cg17781313
cg17781716
cg17781866
cg17782848
cg17782950
cg17783174
cg17784403
cg17784808
cg17784811
cg17786516
cg17786697
cg17786845
cg17786864
cg17786959
cg17787502
cg17787876
cg17788013
cg17789102
cg17789713
cg17790159
cg17790333
cg17790852
cg17793608
cg17794299
cg17794905
cg17795252
cg17796010
cg17798659
cg17798981
cg17799760
cg17799946
cg17800654
cg17801546
cg17801864
cg17802364
cg17802464
cg17802552
cg17803175
cg17803868
cg17803965
cg17804071
cg17804112
cg17804850
cg17805547
cg17806162
cg17806224
cg17806717
cg17807001
cg17807172
cg17807458
cg17807479

cg17807777
cg17807889
cg17808746
cg17809945
cg17810211
cg17810609
cg17810878
cg17811434
cg17811658
cg17811778
cg17811857
cg17811894
cg17813225
cg17813874
cg17814175
cg17814615
cg17814680
cg17814758
cg17815187
cg17815538
cg17815567
cg17815933
cg17816038
cg17816394
cg17816908
cg17817047
cg17817203
cg17817709
cg17819085
cg17819732
cg17822502
cg17822911
cg17822978
cg17823751
cg17825770
cg17826753
cg17827208
cg17827477
cg17827767
cg17829936
cg17830209
cg17830420
cg17830985
cg17831277
cg17831694
cg17831826
cg17832380
cg17832639
cg17833071
cg17833265
cg17833419
cg17833476

cg17833578
cg17833703
cg17833746
cg17835072
cg17835356
cg17835455
cg17836014
cg17836487
cg17837492
cg17838368
cg17838420
cg17838697
cg17839166
cg17839226
cg17839366
cg17839611
cg17840047
cg17840367
cg17840501
cg17840719
cg17841440
cg17842130
cg17842997
cg17843722
cg17847044
cg17847723
cg17848003
cg17848389
cg17848515
cg17848546
cg17849128
cg17850055
cg17850367
cg17850642
cg17851113
cg17851795
cg17851868
cg17852588
cg17852876
cg17853691
cg17854066
cg17854297
cg17854454
cg17855264
cg17856559
cg17857070
cg17857791
cg17858643
cg17858880
cg17859110
cg17859772
cg17861161

cg17861277
cg17861618
cg17862113
cg17862146
cg17862558
cg17863076
cg17863912
cg17864046
cg17864646
cg17864650
cg17865718
cg17866257
cg17866650
cg17866692
cg17867420
cg17868893
cg17869161
cg17869514
cg17869960
cg17869988
cg17870569
cg17870688
cg17870909
cg17871403
cg17872799
cg17872838
cg17874670
cg17875028
cg17875135
cg17875171
cg17875320
cg17875483
cg17875657
cg17875845
cg17875988
cg17876645
cg17877071
cg17877704
cg17877898
cg17878878
cg17879133
cg17881200
cg17881513
cg17882322
cg17882660
cg17884327
cg17884766
cg17884843
cg17885226
cg17885791
cg17885841
cg17886303

cg17886420
cg17886678
cg17886959
cg17887685
cg17888259
cg17888390
cg17889623
cg17890044
cg17890983
cg17891251
cg17891820
cg17892159
cg17892178
cg17892736
cg17894318
cg17894755
cg17894779
cg17895446
cg17895730
cg17896463
cg17896776
cg17896808
cg17897518
cg17898124
cg17898329
cg17899761
cg17900199
cg17900854
cg17901382
cg17901584
cg17903019
cg17903229
cg17903548
cg17903776
cg17903782
cg17904105
cg17904367
cg17904575
cg17904797
cg17905462
cg17906105
cg17906351
cg17909890
cg17910121
cg17910470
cg17911788
cg17912370
cg17912835
cg17913347
cg17914107
cg17914485
cg17915961

cg17916177
cg17916418
cg17916490
cg17917837
cg17918131
cg17918158
cg17918314
cg17918556
cg17919137
cg17919667
cg17920076
cg17921080
cg17921439
cg17921815
cg17922226
cg17922283
cg17922359
cg17922695
cg17923377
cg17924225
cg17924227
cg17924243
cg17924936
cg17925174
cg17925308
cg17926016
cg17926756
cg17926940
cg17927236
cg17927313
cg17927847
cg17928268
cg17928286
cg17928584
cg17929065
cg17929687
cg17929770
cg17929920
cg17930201
cg17930361
cg17930365
cg17930583
cg17931180
cg17931641
cg17931661
cg17932849
cg17932911
cg17933008
cg17933206
cg17933224
cg17933300
cg17933764

cg17934130
cg17934470
cg17934871
cg17936783
cg17939295
cg17939805
cg17940113
cg17941049
cg17941572
cg17942219
cg17942617
cg17942750
cg17942785
cg17944161
cg17944561
cg17944891
cg17945053
cg17945447
cg17947172
cg17948136
cg17949147
cg17949440
cg17949969
cg17950169
cg17950410
cg17952647
cg17952824
cg17952826
cg17952980
cg17953300
cg17953577
cg17953646
cg17955761
cg17956485
cg17956747
cg17956775
cg17957235
cg17958516
cg17958577
cg17959327
cg17959416
cg17959631
cg17959970
cg17960629
cg17960823
cg17961381
cg17962384
cg17962547
cg17963840
cg17964510
cg17964955
cg17965737

cg17966227
cg17966560
cg17967260
cg17967577
cg17968668
cg17968946
cg17969662
cg17969789
cg17970078
cg17971003
cg17971015
cg17971129
cg17971251
cg17971747
cg17971837
cg17971947
cg17972058
cg17972600
cg17973115
cg17973164
cg17973773
cg17974166
cg17974515
cg17974863
cg17975258
cg17975976
cg17976839
cg17977304
cg17978727
cg17978764
cg17978995
cg17979970
cg17980119
cg17980283
cg17980786
cg17982038
cg17983307
cg17985029
cg17985300
cg17985646
cg17985972
cg17986880
cg17987458
cg17987968
cg17988021
cg17988310
cg17988460
cg17988798
cg17988830
cg17989833
cg17990569
cg17991436

cg17991695
cg17991823
cg17992670
cg17993419
cg17993504
cg17994788
cg17996401
cg17996619
cg17996830
cg17996991
cg17997332
cg17997463
cg17998822
cg17999319
cg17999358
cg18000445
cg18001180
cg18001869
cg18002116
cg18002447
cg18002909
cg18003362
cg18004235
cg18004701
cg18004756
cg18006637
cg18007341
cg18007983
cg18008019
cg18008451
cg18008487
cg18009380
cg18009574
cg18010014
cg18010302
cg18011188
cg18011364
cg18011672
cg18011691
cg18011916
cg18013789
cg18014109
cg18014186
cg18014842
cg18014983
cg18015393
cg18015534
cg18016194
cg18016288
cg18016365
cg18017383
cg18017926

cg18018027
cg18018313
cg18018828
cg18018990
cg18019372
cg18019572
cg18020480
cg18020785
cg18021623
cg18022224
cg18022357
cg18023080
cg18023096
cg18023263
cg18023283
cg18023455
cg18023504
cg18023771
cg18024479
cg18025831
cg18026235
cg18026588
cg18026820
cg18027592
cg18028470
cg18029039
cg18029119
cg18029761
cg18030453
cg18031239
cg18031663
cg18031850
cg18032014
cg18032190
cg18032217
cg18032812
cg18032891
cg18032969
cg18033416
cg18033443
cg18034295
cg18034299
cg18034343
cg18034577
cg18034737
cg18035399
cg18036653
cg18037376
cg18037834
cg18039296
cg18039642
cg18040354

cg18040974
cg18042043
cg18042806
cg18044383
cg18044622
cg18044663
cg18045100
cg18045152
cg18045155
cg18045515
cg18046365
cg18047172
cg18047961
cg18048071
cg18048370
cg18049474
cg18050903
cg18051798
cg18052528
cg18053715
cg18053846
cg18053948
cg18053973
cg18054172
cg18054620
cg18057548
cg18058048
cg18058230
cg18058689
cg18058747
cg18060030
cg18060185
cg18060999
cg18061433
cg18062860
cg18063312
cg18063733
cg18064754
cg18065177
cg18065599
cg18065811
cg18066403
cg18067840
cg18068637
cg18068664
cg18070593
cg18070676
cg18072388
cg18073115
cg18073390
cg18073688
cg18073788

cg18074189
cg18074834
cg18074954
cg18075315
cg18075666
cg18076399
cg18076430
cg18076767
cg18076842
cg18077068
cg18077971
cg18078658
cg18079355
cg18079499
cg18080694
cg18080733
cg18080819
cg18081104
cg18081375
cg18081515
cg18081863
cg18082337
cg18082362
cg18082885
cg18082979
cg18083415
cg18083817
cg18084114
cg18084554
cg18085435
cg18085862
cg18086386
cg18087306
cg18087694
cg18087760
cg18087953
cg18088931
cg18089047
cg18089426
cg18090004
cg18090331
cg18090390
cg18091269
cg18092028
cg18092363
cg18093372
cg18093635
cg18094261
cg18094401
cg18094404
cg18094781
cg18095506

cg18096388
cg18096962
cg18096987
cg18097189
cg18097246
cg18097503
cg18097532
cg18098498
cg18098806
cg18099058
cg18099070
cg18099096
cg18099749
cg18100008
cg18100079
cg18100237
cg18100753
cg18101022
cg18102072
cg18104012
cg18104148
cg18104206
cg18104919
cg18105139
cg18105361
cg18105842
cg18106366
cg18106507
cg18107232
cg18108329
cg18108753
cg18109186
cg18109289
cg18109320
cg18109389
cg18109530
cg18109798
cg18110104
cg18110483
cg18110661
cg18112681
cg18112794
cg18113453
cg18113826
cg18114040
cg18114129
cg18114811
cg18114913
cg18115040
cg18115132
cg18115215
cg18115481

cg18115507
cg18115766
cg18116971
cg18117367
cg18117403
cg18117669
cg18117847
cg18118198
cg18118685
cg18118897
cg18119407
cg18119485
cg18120323
cg18121003
cg18121220
cg18121713
cg18121914
cg18121962
cg18122392
cg18122493
cg18122696
cg18122743
cg18122767
cg18123760
cg18124162
cg18124782
cg18126247
cg18126442
cg18126504
cg18127395
cg18128212
cg18128298
cg18128495
cg18128550
cg18128969
cg18129046
cg18129621
cg18129628
cg18129755
cg18129971
cg18130044
cg18132256
cg18133076
cg18133966
cg18134262
cg18135489
cg18136932
cg18137308
cg18137450
cg18138671
cg18139806
cg18140857

cg18140889
cg18141622
cg18141828
cg18142730
cg18142856
cg18142906
cg18143317
cg18143535
cg18144247
cg18144373
cg18144404
cg18144593
cg18144692
cg18145558
cg18145877
cg18147048
cg18147147
cg18147278
cg18147296
cg18147485
cg18147543
cg18147676
cg18148156
cg18148255
cg18148349
cg18148375
cg18148498
cg18148530
cg18149207
cg18149307
cg18150247
cg18150439
cg18150584
cg18150958
cg18151487
cg18151635
cg18152535
cg18152887
cg18154283
cg18154417
cg18155888
cg18156204
cg18156471
cg18156963
cg18157275
cg18158670
cg18158990
cg18159289
cg18159860
cg18159934
cg18159937
cg18160072

cg18160402
cg18160691
cg18161211
cg18161670
cg18162059
cg18162091
cg18162838
cg18164281
cg18165251
cg18165852
cg18166144
cg18166150
cg18166655
cg18166900
cg18166990
cg18167585
cg18167921
cg18168101
cg18168448
cg18168672
cg18168740
cg18169013
cg18169095
cg18169940
cg18170076
cg18170297
cg18170680
cg18171009
cg18171097
cg18171240
cg18171890
cg18172358
cg18172877
cg18173058
cg18173184
cg18173235
cg18173859
cg18174258
cg18174402
cg18174530
cg18175606
cg18175747
cg18175779
cg18175809
cg18177414
cg18177533
cg18177814
cg18178189
cg18178197
cg18180230
cg18181034
cg18181196

cg18181492
cg18181607
cg18181703
cg18182399
cg18182438
cg18182844
cg18183163
cg18183242
cg18183701
cg18183892
cg18184218
cg18185008
cg18187200
cg18187581
cg18187680
cg18187729
cg18188150
cg18188326
cg18188653
cg18188717
cg18189288
cg18189591
cg18190187
cg18190861
cg18190866
cg18191138
cg18191209
cg18192252
cg18192491
cg18193094
cg18193148
cg18193817
cg18194535
cg18194945
cg18195416
cg18196063
cg18196453
cg18197377
cg18197392
cg18198306
cg18198461
cg18199581
cg18199664
cg18200261
cg18200741
cg18200783
cg18201671
cg18202080
cg18202127
cg18202205
cg18202336
cg18202449

cg18202521
cg18202765
cg18202928
cg18203203
cg18203974
cg18204145
cg18204684
cg18204964
cg18205883
cg18206017
cg18207741
cg18208707
cg18208842
cg18209212
cg18209489
cg18210226
cg18210732
cg18212762
cg18213443
cg18213472
cg18213545
cg18214661
cg18215449
cg18215705
cg18216249
cg18216587
cg18217048
cg18217252
cg18218772
cg18219226
cg18220087
cg18220632
cg18221016
cg18221168
cg18221341
cg18221897
cg18222759
cg18222913
cg18223430
cg18223579
cg18224038
cg18224468
cg18224793
cg18226002
cg18226169
cg18226539
cg18226915
cg18227237
cg18227471
cg18229049
cg18229071
cg18229092

cg18229396
cg18229521
cg18231054
cg18231898
cg18232497
cg18232871
cg18232879
cg18233746
cg18234111
cg18234224
cg18234606
cg18234865
cg18235050
cg18235192
cg18235635
cg18235662
cg18235682
cg18235734
cg18235799
cg18235937
cg18236421
cg18236862
cg18237047
cg18237203
cg18237330
cg18237551
cg18237594
cg18238531
cg18238570
cg18239431
cg18239481
cg18241337
cg18241351
cg18241387
cg18241469
cg18241647
cg18241829
cg18242103
cg18242368
cg18243760
cg18244286
cg18244362
cg18244487
cg18244508
cg18244845
cg18245230
cg18245316
cg18246107
cg18246584
cg18246964
cg18247055
cg18247179

cg18249173
cg18249580
cg18249634
cg18250028
cg18250474
cg18250846
cg18251191
cg18251449
cg18252037
cg18252039
cg18252309
cg18252424
cg18252903
cg18252924
cg18252964
cg18253101
cg18253591
cg18254183
cg18255166
cg18255240
cg18256205
cg18256498
cg18257485
cg18258397
cg18258482
cg18258620
cg18259189
cg18259253
cg18260397
cg18261434
cg18261909
cg18262616
cg18263166
cg18263365
cg18263572
cg18264092
cg18264173
cg18264486
cg18264562
cg18265845
cg18266458
cg18266594
cg18267047
cg18267049
cg18267381
cg18268547
cg18268562
cg18269382
cg18270378
cg18270629
cg18270687
cg18271789

cg18271897
cg18272307
cg18272461
cg18273840
cg18274480
cg18275316
cg18275519
cg18275733
cg18275958
cg18276135
cg18276155
cg18276638
cg18277979
cg18278424
cg18278638
cg18279094
cg18279625
cg18279742
cg18280362
cg18280758
cg18280830
cg18281939
cg18282393
cg18282455
cg18283037
cg18283321
cg18283361
cg18284590
cg18285819
cg18286501
cg18286850
cg18286995
cg18286997
cg18287066
cg18287310
cg18288119
cg18288161
cg18289306
cg18289581
cg18290233
cg18290624
cg18290848
cg18290981
cg18291014
cg18291422
cg18291664
cg18291941
cg18292202
cg18292394
cg18292664
cg18293662
cg18294158

cg18294257
cg18294691
cg18294707
cg18295183
cg18295744
cg18296189
cg18297044
cg18297196
cg18297349
cg18297437
cg18297705
cg18297736
cg18297960
cg18300589
cg18301410
cg18301583
cg18301598
cg18301891
cg18302605
cg18303215
cg18303410
cg18303896
cg18304305
cg18305790
cg18305912
cg18306516
cg18307303
cg18307783
cg18307978
cg18308184
cg18308516
cg18309255
cg18309817
cg18310495
cg18311537
cg18312989
cg18313013
cg18313132
cg18313416
cg18313661
cg18314424
cg18315125
cg18315380
cg18315831
cg18315960
cg18315963
cg18316256
cg18317135
cg18317506
cg18317694
cg18317788
cg18318649

cg18318941
cg18319029
cg18319381
cg18319612
cg18319790
cg18319799
cg18320149
cg18320737
cg18321495
cg18322510
cg18322569
cg18323376
cg18324583
cg18324707
cg18324798
cg18324824
cg18325389
cg18325622
cg18325866
cg18326021
cg18327695
cg18328265
cg18328894
cg18328998
cg18329753
cg18330370
cg18331348
cg18331412
cg18333041
cg18333566
cg18334345
cg18334734
cg18334819
cg18334977
cg18335068
cg18335607
cg18336674
cg18336854
cg18337170
cg18338775
cg18339718
cg18339788
cg18340416
cg18340535
cg18341512
cg18342673
cg18342703
cg18343437
cg18343480
cg18343556
cg18344652
cg18345406

cg18346707
cg18347010
cg18347153
cg18347442
cg18347602
cg18347642
cg18347762
cg18348086
cg18348318
cg18348566
cg18348690
cg18348830
cg18349138
cg18349835
cg18350391
cg18350895
cg18351711
cg18351939
cg18352710
cg18352721
cg18352916
cg18352935
cg18354462
cg18354905
cg18355311
cg18355902
cg18357048
cg18358845
cg18358869
cg18358894
cg18359129
cg18359618
cg18360873
cg18361024
cg18361098
cg18361418
cg18361425
cg18362003
cg18362281
cg18364707
cg18365536
cg18366169
cg18366748
cg18367268
cg18369288
cg18369629
cg18369654
cg18369680
cg18369866
cg18369913
cg18369972
cg18370747

cg18370764
cg18371475
cg18372548
cg18372607
cg18373318
cg18374755
cg18374914
cg18375236
cg18375610
cg18375860
cg18375902
cg18376306
cg18377014
cg18377941
cg18378453
cg18380550
cg18381051
cg18381455
cg18381490
cg18382294
cg18382353
cg18382926
cg18383538
cg18383660
cg18384226
cg18384277
cg18384402
cg18384712
cg18384778
cg18385299
cg18385371
cg18386103
cg18387659
cg18387985
cg18388084
cg18388380
cg18388910
cg18389810
cg18389931
cg18390472
cg18390991
cg18391127
cg18391203
cg18391467
cg18391899
cg18391972
cg18392016
cg18392139
cg18392187
cg18392943
cg18393044
cg18393172

cg18393511
cg18394533
cg18394552
cg18394557
cg18395150
cg18395231
cg18395327
cg18395354
cg18395355
cg18395466
cg18395636
cg18395809
cg18395917
cg18396137
cg18396173
cg18396626
cg18396675
cg18396811
cg18396984
cg18396987
cg18397314
cg18397405
cg18397653
cg18398175
cg18399834
cg18399935
cg18400281
cg18400845
cg18401111
cg18401367
cg18401376
cg18402104
cg18402615
cg18403182
cg18403193
cg18403613
cg18404374
cg18404652
cg18405341
cg18405544
cg18406033
cg18406792
cg18406801
cg18407864
cg18407955
cg18408209
cg18408326
cg18409732
cg18410078
cg18411043
cg18411150
cg18412546

cg18412834
cg18412889
cg18413131
cg18413427
cg18413830
cg18414033
cg18414579
cg18415147
cg18416576
cg18417061
cg18417954
cg18418242
cg18418457
cg18419070
cg18419276
cg18419358
cg18420965
cg18421167
cg18421538
cg18421840
cg18422055
cg18422587
cg18422632
cg18422654
cg18422853
cg18423469
cg18423591
cg18423626
cg18423852
cg18423935
cg18424470
cg18424634
cg18424850
cg18425434
cg18425570
cg18426477
cg18426908
cg18427091
cg18427465
cg18427696
cg18428006
cg18428104
cg18428234
cg18428542
cg18428688
cg18428825
cg18429742
cg18430128
cg18430222
cg18430255
cg18430895
cg18431123

cg18431127
cg18431489
cg18431640
cg18431856
cg18432099
cg18432451
cg18432806
cg18432895
cg18433009
cg18433086
cg18434354
cg18434367
cg18436008
cg18436272
cg18436444
cg18436788
cg18437033
cg18437039
cg18437365
cg18437792
cg18437808
cg18438300
cg18438793
cg18438853
cg18438894
cg18438944
cg18439195
cg18439339
cg18440069
cg18440123
cg18440523
cg18440774
cg18440839
cg18440897
cg18442362
cg18442992
cg18443253
cg18443359
cg18443392
cg18444835
cg18445088
cg18445438
cg18446140
cg18446744
cg18446916
cg18447402
cg18447727
cg18447785
cg18447876
cg18448581
cg18448614
cg18448977

cg18449120
cg18449135
cg18450168
cg18450254
cg18450420
cg18450500
cg18450930
cg18451814
cg18451817
cg18452695
cg18452703
cg18453551
cg18454288
cg18454510
cg18454685
cg18454916
cg18455405
cg18455650
cg18456003
cg18456331
cg18456512
cg18456640
cg18456689
cg18456755
cg18456942
cg18457597
cg18457737
cg18458017
cg18458352
cg18459339
cg18460375
cg18460830
cg18461021
cg18461347
cg18461584
cg18462653
cg18462898
cg18463876
cg18464296
cg18464364
cg18464402
cg18464991
cg18465133
cg18465747
cg18465962
cg18466162
cg18468354
cg18468511
cg18469270
cg18469292
cg18469813
cg18470038

cg18470839
cg18471993
cg18472549
cg18472699
cg18472912
cg18473137
cg18473544
cg18473686
cg18474081
cg18475775
cg18476604
cg18476778
cg18477072
cg18477587
cg18477666
cg18477771
cg18477928
cg18477949
cg18478319
cg18478850
cg18479255
cg18479593
cg18480005
cg18480127
cg18480432
cg18480654
cg18480906
cg18482268
cg18482960
cg18483549
cg18483908
cg18484189
cg18484299
cg18484309
cg18484958
cg18485844
cg18486258
cg18486644
cg18487309
cg18487598
cg18488855
cg18489675
cg18489755
cg18490614
cg18490846
cg18491039
cg18491080
cg18492126
cg18493030
cg18493182
cg18493502
cg18496247

cg18496318
cg18497238
cg18498115
cg18498263
cg18498593
cg18498871
cg18499396
cg18499750
cg18500177
cg18501029
cg18501230
cg18501555
cg18502099
cg18502142
cg18503260
cg18506018
cg18506124
cg18506390
cg18506744
cg18507053
cg18507379
cg18508525
cg18509490
cg18510311
cg18511011
cg18511526
cg18511546
cg18512205
cg18512262
cg18512515
cg18512930
cg18513023
cg18513062
cg18515510
cg18516150
cg18516557
cg18516946
cg18517540
cg18518074
cg18519834
cg18519902
cg18519905
cg18520696
cg18520925
cg18522239
cg18522715
cg18523886
cg18524114
cg18524739
cg18525316
cg18525352
cg18525616

cg18525873
cg18527241
cg18527574
cg18527971
cg18528091
cg18528591
cg18529010
cg18529845
cg18530299
cg18530551
cg18530956
cg18531405
cg18532076
cg18533201
cg18533365
cg18533392
cg18533473
cg18534176
cg18534345
cg18535167
cg18535456
cg18536994
cg18537027
cg18537383
cg18537464
cg18537482
cg18538812
cg18538958
cg18539122
cg18540492
cg18541087
cg18541254
cg18542304
cg18542613
cg18543142
cg18543270
cg18543428
cg18543991
cg18544329
cg18544740
cg18544809
cg18545596
cg18545695
cg18546236
cg18546674
cg18547838
cg18548950
cg18550003
cg18550212
cg18550262
cg18550632
cg18551048

cg18551353
cg18552334
cg18552413
cg18552425
cg18552520
cg18552939
cg18553677
cg18553840
cg18554639
cg18554868
cg18555142
cg18556676
cg18556788
cg18557131
cg18558968
cg18560152
cg18560328
cg18560638
cg18560915
cg18562121
cg18563630
cg18563987
cg18564167
cg18564587
cg18565264
cg18565783
cg18566883
cg18567682
cg18567882
cg18567924
cg18568116
cg18568589
cg18569623
cg18569734
cg18569988
cg18571094
cg18571640
cg18572214
cg18573443
cg18573745
cg18575809
cg18576057
cg18576686
cg18577167
cg18578080
cg18578578
cg18580339
cg18580385
cg18581657
cg18581663
cg18582824
cg18583378

cg18584067
cg18584265
cg18584639
cg18586212
cg18586780
cg18586851
cg18586867
cg18587340
cg18587465
cg18587499
cg18587504
cg18587984
cg18588000
cg18588768
cg18588811
cg18589016
cg18589526
cg18589858
cg18590502
cg18591228
cg18591235
cg18591496
cg18593081
cg18593210
cg18593236
cg18593303
cg18593374
cg18593660
cg18594467
cg18595065
cg18595174
cg18595196
cg18595867
cg18596362
cg18596616
cg18596794
cg18596947
cg18597677
cg18599081
cg18599155
cg18599400
cg18599533
cg18601057
cg18602114
cg18602350
cg18603028
cg18603228
cg18603483
cg18604395
cg18604823
cg18604842
cg18607127

cg18607533
cg18608206
cg18609078
cg18609306
cg18609779
cg18610071
cg18610894
cg18613721
cg18614139
cg18614381
cg18614510
cg18614702
cg18614734
cg18615369
cg18616558
cg18617860
cg18621538
cg18622075
cg18623836
cg18623980
cg18624059
cg18624900
cg18625289
cg18625916
cg18626035
cg18626954
cg18627235
cg18627816
cg18628255
cg18629427
cg18629527
cg18629535
cg18630363
cg18630580
cg18631996
cg18632189
cg18632637
cg18633154
cg18633340
cg18633471
cg18634175
cg18635670
cg18635723
cg18636509
cg18637380
cg18637901
cg18638581
cg18639233
cg18639238
cg18639495
cg18640098
cg18640444

cg18641486
cg18641937
cg18642179
cg18642459
cg18643029
cg18643421
cg18644653
cg18645316
cg18646207
cg18647259
cg18647989
cg18648031
cg18649464
cg18649854
cg18650307
cg18650670
cg18651026
cg18651347
cg18651870
cg18651932
cg18652043
cg18652327
cg18652346
cg18653195
cg18653350
cg18653372
cg18653451
cg18653534
cg18654464
cg18655025
cg18655441
cg18656708
cg18656829
cg18657094
cg18657700
cg18657751
cg18659248
cg18659450
cg18660278
cg18660987
cg18661205
cg18665334
cg18666379
cg18666454
cg18666630
cg18666748
cg18667511
cg18669022
cg18669186
cg18669209
cg18669381
cg18669388

cg18669406
cg18669588
cg18670076
cg18670767
cg18671554
cg18672030
cg18672053
cg18672939
cg18673409
cg18674267
cg18674828
cg18675128
cg18675600
cg18675847
cg18677603
cg18678353
cg18679410
cg18680587
cg18680788
cg18680834
cg18681853
cg18682226
cg18683606
cg18683691
cg18685071
cg18685330
cg18685561
cg18687085
cg18687314
cg18687675
cg18688062
cg18688293
cg18689332
cg18689614
cg18690019
cg18691800
cg18692507
cg18692678
cg18693051
cg18693346
cg18693395
cg18693673
cg18696495
cg18696900
cg18697636
cg18698431
cg18698788
cg18698884
cg18699025
cg18699226
cg18699804
cg18701343

cg18702948
cg18702971
cg18703227
cg18703511
cg18703515
cg18703983
cg18704047
cg18704201
cg18705773
cg18705776
cg18707191
cg18707417
cg18708252
cg18708504
cg18708986
cg18709313
cg18710278
cg18710784
cg18711535
cg18711921
cg18712349
cg18712809
cg18713028
cg18713766
cg18713806
cg18714086
cg18714358
cg18714560
cg18714586
cg18716210
cg18717067
cg18717689
cg18718613
cg18718692
cg18719158
cg18719217
cg18719787
cg18720617
cg18721420
cg18721838
cg18721888
cg18722504
cg18722841
cg18722847
cg18723409
cg18723937
cg18724037
cg18724430
cg18724565
cg18725544
cg18725599
cg18725867

cg18728403
cg18729001
cg18729664
cg18729681
cg18729973
cg18730174
cg18730865
cg18731327
cg18733967
cg18734304
cg18734472
cg18734706
cg18734877
cg18735451
cg18736186
cg18736711
cg18736739
cg18738058
cg18738220
cg18738367
cg18738804
cg18738985
cg18739675
cg18739950
cg18740893
cg18741044
cg18742346
cg18743793
cg18744273
cg18744822
cg18745131
cg18745424
cg18745507
cg18745782
cg18747039
cg18748062
cg18748085
cg18749015
cg18749097
cg18750833
cg18750960
cg18751141
cg18752412
cg18753424
cg18753619
cg18754629
cg18754857
cg18755296
cg18755643
cg18755783
cg18756545
cg18757228

cg18758339
cg18758622
cg18758987
cg18759102
cg18759960
cg18760496
cg18761349
cg18761958
cg18761976
cg18762500
cg18763165
cg18763223
cg18763854
cg18763911
cg18764031
cg18764358
cg18764575
cg18764577
cg18764804
cg18765127
cg18765366
cg18765405
cg18765542
cg18765906
cg18766475
cg18766586
cg18766685
cg18766691
cg18766795
cg18766900
cg18767287
cg18768338
cg18768347
cg18768582
cg18770052
cg18770739
cg18770763
cg18771173
cg18771357
cg18772531
cg18772838
cg18773591
cg18775149
cg18777311
cg18777600
cg18777792
cg18778254
cg18778433
cg18780021
cg18780257
cg18780288
cg18781031

cg18781835
cg18782201
cg18782604
cg18783033
cg18784875
cg18785787
cg18786643
cg18787245
cg18787420
cg18787802
cg18787846
cg18788890
cg18789177
cg18789620
cg18789663
cg18789918
cg18789958
cg18790057
cg18790325
cg18790382
cg18790597
cg18790787
cg18792022
cg18792381
cg18793404
cg18793661
cg18794577
cg18794839
cg18794976
cg18795022
cg18795133
cg18795277
cg18795786
cg18795959
cg18797590
cg18798111
cg18798445
cg18799069
cg18799217
cg18800739
cg18801045
cg18801691
cg18801806
cg18801825
cg18802754
cg18802818
cg18802998
cg18803303
cg18803306
cg18804206
cg18804615
cg18804680

cg18804777
cg18804920
cg18804963
cg18804985
cg18805734
cg18806140
cg18806193
cg18806561
cg18806660
cg18806980
cg18808054
cg18808261
cg18808506
cg18808692
cg18808904
cg18808929
cg18809102
cg18810022
cg18810244
cg18810664
cg18810737
cg18811269
cg18811318
cg18811550
cg18812079
cg18813020
cg18813322
cg18813527
cg18813545
cg18813778
cg18813881
cg18815565
cg18816595
cg18816826
cg18817158
cg18817318
cg18817392
cg18817396
cg18817487
cg18817529
cg18817654
cg18817970
cg18818210
cg18818498
cg18820209
cg18821742
cg18822414
cg18822719
cg18823007
cg18823647
cg18823849
cg18824193

cg18824330
cg18824585
cg18824990
cg18825419
cg18825427
cg18827107
cg18827221
cg18827332
cg18827787
cg18827954
cg18828137
cg18828927
cg18830042
cg18830373
cg18831409
cg18832619
cg18832655
cg18833228
cg18833788
cg18833907
cg18835300
cg18835865
cg18836287
cg18836626
cg18836661
cg18838431
cg18839416
cg18839504
cg18839637
cg18841159
cg18841452
cg18841634
cg18841653
cg18841796
cg18841952
cg18842363
cg18842563
cg18843062
cg18844382
cg18845375
cg18845433
cg18845718
cg18845797
cg18846954
cg18848446
cg18853490
cg18854169
cg18854419
cg18854765
cg18854872
cg18855429
cg18855780

cg18856043
cg18856198
cg18856581
cg18856632
cg18857467
cg18857618
cg18860329
cg18861119
cg18861231
cg18861547
cg18862005
cg18862481
cg18862813
cg18863333
cg18863595
cg18863922
cg18865990
cg18866015
cg18866212
cg18866358
cg18866586
cg18867200
cg18869404
cg18869840
cg18872497
cg18873686
cg18873909
cg18874332
cg18875371
cg18875435
cg18875453
cg18875496
cg18877028
cg18877699
cg18878654
cg18879198
cg18879828
cg18879950
cg18880384
cg18882449
cg18884555
cg18884741
cg18884942
cg18885372
cg18885766
cg18887230
cg18887961
cg18888137
cg18888403
cg18888464
cg18888883
cg18889092

cg18890556
cg18891864
cg18894648
cg18894781
cg18895088
cg18895336
cg18896032
cg18896184
cg18897026
cg18897632
cg18898125
cg18898209
cg18898325
cg18899117
cg18899355
cg18899516
cg18899569
cg18899999
cg18901644
cg18902109
cg18902579
cg18903398
cg18905668
cg18906360
cg18907363
cg18907610
cg18908185
cg18909389
cg18909638
cg18911129
cg18911395
cg18911817
cg18911972
cg18912103
cg18912209
cg18912768
cg18912847
cg18913890
cg18913924
cg18914000
cg18914211
cg18914258
cg18914751
cg18916055
cg18917544
cg18918116
cg18918349
cg18918538
cg18919478
cg18920423
cg18920858
cg18921025

cg18921475
cg18922620
cg18922629
cg18924324
cg18924562
cg18927604
cg18928221
cg18928866
cg18929251
cg18929335
cg18929842
cg18930012
cg18930354
cg18930994
cg18931850
cg18932726
cg18932740
cg18933943
cg18935014
cg18935108
cg18935231
cg18935453
cg18935813
cg18936283
cg18936620
cg18937395
cg18938313
cg18939241
cg18939666
cg18940047
cg18940674
cg18941322
cg18941818
cg18942451
cg18942631
cg18942985
cg18943289
cg18943588
cg18943957
cg18944010
cg18944653
cg18944702
cg18944984
cg18945039
cg18946437
cg18946533
cg18947276
cg18947550
cg18947801
cg18948125
cg18948646
cg18949794

cg18950108
cg18951135
cg18952506
cg18952546
cg18952796
cg18952945
cg18954541
cg18954700
cg18954865
cg18956825
cg18956933
cg18958859
cg18958907
cg18959422
cg18960324
cg18961501
cg18961546
cg18961589
cg18961681
cg18961788
cg18963171
cg18963763
cg18963953
cg18964706
cg18964775
cg18965391
cg18966401
cg18966597
cg18967021
cg18967533
cg18969232
cg18970338
cg18970929
cg18971054
cg18972123
cg18972811
cg18972849
cg18973515
cg18973735
cg18973863
cg18974117
cg18975659
cg18976212
cg18977423
cg18977541
cg18977891
cg18979491
cg18979589
cg18980148
cg18982384
cg18983132
cg18983604

cg18984165
cg18984200
cg18984452
cg18984724
cg18985738
cg18986589
cg18986967
cg18987410
cg18987491
cg18988110
cg18988170
cg18988498
cg18988861
cg18989252
cg18989477
cg18989499
cg18989709
cg18989909
cg18990157
cg18990313
cg18990407
cg18990588
cg18990899
cg18991321
cg18991549
cg18993918
cg18993949
cg18994049
cg18994063
cg18996274
cg18997137
cg18998138
cg18998172
cg18998321
cg18998543
cg18999641
cg19000647
cg19001226
cg19001290
cg19002591
cg19002763
cg19003337
cg19003397
cg19003797
cg19003815
cg19003958
cg19004594
cg19004771
cg19005210
cg19005390
cg19005446
cg19005707

cg19006211
cg19007031
cg19007141
cg19007167
cg19007269
cg19008646
cg19009132
cg19009417
cg19009644
cg19009764
cg19010139
cg19011911
cg19012958
cg19013399
cg19013611
cg19014036
cg19014368
cg19014419
cg19014749
cg19015086
cg19015264
cg19015654
cg19015708
cg19015997
cg19016332
cg19016652
cg19017142
cg19017553
cg19019345
cg19019537
cg19021738
cg19021988
cg19022446
cg19022525
cg19022717
cg19024161
cg19024344
cg19025272
cg19025430
cg19025573
cg19027361
cg19027636
cg19027710
cg19029181
cg19029214
cg19029422
cg19029904
cg19030579
cg19030607
cg19031047
cg19031658
cg19031844

cg19032066
cg19032797
cg19033555
cg19033906
cg19034797
cg19035031
cg19035107
cg19035908
cg19037884
cg19038027
cg19038490
cg19039583
cg19040012
cg19040483
cg19042062
cg19042459
cg19042947
cg19043574
cg19043798
cg19044550
cg19045427
cg19046167
cg19046695
cg19046697
cg19046725
cg19046826
cg19047265
cg19047660
cg19047707
cg19047868
cg19048251
cg19050061
cg19050940
cg19050990
cg19051105
cg19051213
cg19052164
cg19052355
cg19052547
cg19053889
cg19054096
cg19054524
cg19055323
cg19056573
cg19056664
cg19056772
cg19057459
cg19059861
cg19061112
cg19062099
cg19062489
cg19063061

cg19063856
cg19064523
cg19064846
cg19066682
cg19067730
cg19068510
cg19069360
cg19070139
cg19070159
cg19070856
cg19071002
cg19071490
cg19071879
cg19071976
cg19072817
cg19073035
cg19074292
cg19074340
cg19074393
cg19075787
cg19076265
cg19076895
cg19077794
cg19077806
cg19079194
cg19079845
cg19081021
cg19082230
cg19082416
cg19082560
cg19083007
cg19083977
cg19084053
cg19084269
cg19084726
cg19085352
cg19086001
cg19086905
cg19088912
cg19088962
cg19089079
cg19089328
cg19089383
cg19089617
cg19089701
cg19090128
cg19090522
cg19091036
cg19091779
cg19092045
cg19092096
cg19092322

cg19092343
cg19092396
cg19092735
cg19093295
cg19094243
cg19094366
cg19094530
cg19095187
cg19095600
cg19096460
cg19096475
cg19096825
cg19097280
cg19098619
cg19098763
cg19098785
cg19098796
cg19098932
cg19099050
cg19099213
cg19101232
cg19101326
cg19101417
cg19102740
cg19102955
cg19103169
cg19103609
cg19103704
cg19104015
cg19104032
cg19106326
cg19106489
cg19107296
cg19107469
cg19107973
cg19108747
cg19109439
cg19109538
cg19110758
cg19111238
cg19113081
cg19113599
cg19113920
cg19115017
cg19115204
cg19115344
cg19115393
cg19115490
cg19115882
cg19116956
cg19116959
cg19117047

cg19117063
cg19117322
cg19118569
cg19118904
cg19118951
cg19118972
cg19119016
cg19119032
cg19119071
cg19119072
cg19120225
cg19121056
cg19122206
cg19123345
cg19123356
cg19124225
cg19125370
cg19125767
cg19126110
cg19126300
cg19126954
cg19128082
cg19129325
cg19129369
cg19129687
cg19130180
cg19130188
cg19131272
cg19133221
cg19133948
cg19134130
cg19134568
cg19134945
cg19136075
cg19136371
cg19136704
cg19137348
cg19139343
cg19139618
cg19139832
cg19140262
cg19140548
cg19141489
cg19142341
cg19142553
cg19143020
cg19143211
cg19145710
cg19145730
cg19145858
cg19145995
cg19146067

cg19146119
cg19147483
cg19147534
cg19148201
cg19151121
cg19152508
cg19153095
cg19154467
cg19156483
cg19156552
cg19156875
cg19157819
cg19158553
cg19159365
cg19159964
cg19160053
cg19160624
cg19161124
cg19161251
cg19161424
cg19161559
cg19162158
cg19162970
cg19164621
cg19164987
cg19165216
cg19165390
cg19165640
cg19166660
cg19168556
cg19169154
cg19170015
cg19170021
cg19170156
cg19170807
cg19171596
cg19171919
cg19174658
cg19175742
cg19176553
cg19178081
cg19178479
cg19178561
cg19180555
cg19181162
cg19182267
cg19182986
cg19183252
cg19183395
cg19183742
cg19183743
cg19184160

cg19184688
cg19184771
cg19185579
cg19185846
cg19186089
cg19186105
cg19186145
cg19187290
cg19187410
cg19188182
cg19188306
cg19188359
cg19188632
cg19188855
cg19189176
cg19189864
cg19190163
cg19190699
cg19190762
cg19191888
cg19192065
cg19192120
cg19193136
cg19193403
cg19193595
cg19193826
cg19194098
cg19194167
cg19194373
cg19195712
cg19196684
cg19198178
cg19199266
cg19200814
cg19201068
cg19202014
cg19203115
cg19204131
cg19204834
cg19204958
cg19205064
cg19205850
cg19206040
cg19206078
cg19206146
cg19206437
cg19208681
cg19208749
cg19209066
cg19209355
cg19210276
cg19210770

cg19212453
cg19212465
cg19212949
cg19214184
cg19215199
cg19215671
cg19216731
cg19216792
cg19216817
cg19216994
cg19218067
cg19219068
cg19219255
cg19219423
cg19220282
cg19220607
cg19221489
cg19222156
cg19223119
cg19223299
cg19223782
cg19224201
cg19224793
cg19225602
cg19226099
cg19226358
cg19227130
cg19230025
cg19231602
cg19233156
cg19233268
cg19233472
cg19233663
cg19234495
cg19234705
cg19234736
cg19234738
cg19234894
cg19234983
cg19235071
cg19236454
cg19237294
cg19238058
cg19238325
cg19238380
cg19238521
cg19238840
cg19240938
cg19241311
cg19241619
cg19241831
cg19241878

cg19242851
cg19242984
cg19243585
cg19243721
cg19243842
cg19244064
cg19244562
cg19244928
cg19247032
cg19247726
cg19248162
cg19248395
cg19248564
cg19248676
cg19249107
cg19249516
cg19250304
cg19250417
cg19250571
cg19250709
cg19251161
cg19251719
cg19251850
cg19252333
cg19252584
cg19253577
cg19253850
cg19254119
cg19254152
cg19255053
cg19255136
cg19256906
cg19257089
cg19257102
cg19257402
cg19257550
cg19257832
cg19258062
cg19258177
cg19259125
cg19259179
cg19260084
cg19260189
cg19260606
cg19261426
cg19261516
cg19261810
cg19262315
cg19262476
cg19262563
cg19263469
cg19264170

cg19264203
cg19265546
cg19266387
cg19267163
cg19267252
cg19267253
cg19267254
cg19267554
cg19267861
cg19268652
cg19269426
cg19269530
cg19269666
cg19270197
cg19270277
cg19270409
cg19271111
cg19272150
cg19272720
cg19272754
cg19272832
cg19273074
cg19273378
cg19273495
cg19273668
cg19274270
cg19274368
cg19274612
cg19274618
cg19274680
cg19275133
cg19275261
cg19275632
cg19275653
cg19276186
cg19277516
cg19278414
cg19279257
cg19279310
cg19279464
cg19281083
cg19282250
cg19282443
cg19282452
cg19283298
cg19284277
cg19284368
cg19285774
cg19286672
cg19286687
cg19286989
cg19287501

cg19287817
cg19288904
cg19288921
cg19290091
cg19290410
cg19290938
cg19290962
cg19291576
cg19291696
cg19292625
cg19292760
cg19293468
cg19293784
cg19295951
cg19296509
cg19296740
cg19297427
cg19297868
cg19298225
cg19299755
cg19300195
cg19300307
cg19300568
cg19301114
cg19301827
cg19301960
cg19303305
cg19303865
cg19303898
cg19304003
cg19304093
cg19304150
cg19304273
cg19304797
cg19305488
cg19305718
cg19306640
cg19306847
cg19306866
cg19308222
cg19308397
cg19308769
cg19309079
cg19309499
cg19309616
cg19310162
cg19310167
cg19310322
cg19310458
cg19310515
cg19311055
cg19312305

cg19312404
cg19313043
cg19313221
cg19313311
cg19313373
cg19313408
cg19314136
cg19314352
cg19314777
cg19314992
cg19315202
cg19315854
cg19316647
cg19317211
cg19317258
cg19317715
cg19318537
cg19319037
cg19319808
cg19320261
cg19320476
cg19320820
cg19322065
cg19323059
cg19323958
cg19324023
cg19325791
cg19327615
cg19328051
cg19328560
cg19329160
cg19329999
cg19330334
cg19331215
cg19334176
cg19334657
cg19335742
cg19338056
cg19338524
cg19341630
cg19342102
cg19343464
cg19343611
cg19344545
cg19345417
cg19345602
cg19345662
cg19346202
cg19346371
cg19346632
cg19346645
cg19346786

cg19348206
cg19349283
cg19350024
cg19350115
cg19350360
cg19350767
cg19350870
cg19351143
cg19352038
cg19352193
cg19352507
cg19352830
cg19353578
cg19353949
cg19354539
cg19355087
cg19356191
cg19358307
cg19358493
cg19358520
cg19358877
cg19359071
cg19360308
cg19360794
cg19360796
cg19360907
cg19361032
cg19363607
cg19363785
cg19364068
cg19364538
cg19365176
cg19365673
cg19366753
cg19366770
cg19366877
cg19367176
cg19368582
cg19368911
cg19369955
cg19370858
cg19371121
cg19371339
cg19371916
cg19372178
cg19372710
cg19373099
cg19373760
cg19373874
cg19374182
cg19374752
cg19375053

cg19375140
cg19375196
cg19375403
cg19375479
cg19375492
cg19375539
cg19375583
cg19375810
cg19376664
cg19377014
cg19377230
cg19377421
cg19378216
cg19378389
cg19378729
cg19379569
cg19379721
cg19380156
cg19381811
cg19383853
cg19384289
cg19384448
cg19384704
cg19385628
cg19387862
cg19388739
cg19389884
cg19390271
cg19390582
cg19390934
cg19391276
cg19391527
cg19392551
cg19392831
cg19393755
cg19393762
cg19394293
cg19395074
cg19396878
cg19397320
cg19397885
cg19397991
cg19399370
cg19399514
cg19400179
cg19400238
cg19400926
cg19401111
cg19402179
cg19402236
cg19403005
cg19403014

cg19403029
cg19403323
cg19404207
cg19404354
cg19404723
cg19404835
cg19404979
cg19405162
cg19405329
cg19405799
cg19406182
cg19406878
cg19407095
cg19407435
cg19407717
cg19409133
cg19409546
cg19410046
cg19410532
cg19410609
cg19411699
cg19411729
cg19411922
cg19412467
cg19412669
cg19412743
cg19412808
cg19414227
cg19414591
cg19414741
cg19415202
cg19415383
cg19416590
cg19417024
cg19417168
cg19417526
cg19418648
cg19418958
cg19420930
cg19420968
cg19421125
cg19421218
cg19421752
cg19423543
cg19423625
cg19423854
cg19424531
cg19425513
cg19425603
cg19425969
cg19426266
cg19426594

cg19426625
cg19427188
cg19427325
cg19427399
cg19428336
cg19429051
cg19429281
cg19429405
cg19430553
cg19430694
cg19430967
cg19431235
cg19431346
cg19433118
cg19433175
cg19433694
cg19433697
cg19433807
cg19435264
cg19435526
cg19436270
cg19437258
cg19438750
cg19439043
cg19439071
cg19439651
cg19440035
cg19441717
cg19441872
cg19442201
cg19442264
cg19442889
cg19443642
cg19443786
cg19445044
cg19445097
cg19445269
cg19445335
cg19445684
cg19445690
cg19445758
cg19446165
cg19446264
cg19446990
cg19448144
cg19448837
cg19449594
cg19450056
cg19450119
cg19450840
cg19451167
cg19451848

cg19452248
cg19453093
cg19454081
cg19455189
cg19455965
cg19456540
cg19456938
cg19458697
cg19459056
cg19459094
cg19459470
cg19460792
cg19461621
cg19462523
cg19462635
cg19462712
cg19463531
cg19464252
cg19464876
cg19465002
cg19465033
cg19465320
cg19466037
cg19466563
cg19468528
cg19469722
cg19469742
cg19470652
cg19470832
cg19470841
cg19471040
cg19472098
cg19474571
cg19474725
cg19474865
cg19474888
cg19474983
cg19475270
cg19475988
cg19476426
cg19476529
cg19476647
cg19477293
cg19477674
cg19477689
cg19478111
cg19478298
cg19478500
cg19478820
cg19479744
cg19480724
cg19481531

cg19481686
cg19482540
cg19483007
cg19483465
cg19483679
cg19485748
cg19485804
cg19485911
cg19486949
cg19488206
cg19488922
cg19489512
cg19489885
cg19490609
cg19490761
cg19490899
cg19491035
cg19492047
cg19495351
cg19495444
cg19495650
cg19496364
cg19496479
cg19496772
cg19496978
cg19497451
cg19497767
cg19498794
cg19499581
cg19499614
cg19499884
cg19500415
cg19501108
cg19502050
cg19502744
cg19502867
cg19503341
cg19503826
cg19503977
cg19504702
cg19505136
cg19505194
cg19506253
cg19506435
cg19507032
cg19507068
cg19507267
cg19508285
cg19508458
cg19509125
cg19509303
cg19509393

cg19509675
cg19510409
cg19512268
cg19513004
cg19513232
cg19513727
cg19513903
cg19514381
cg19514854
cg19516340
cg19516404
cg19516457
cg19517476
cg19517718
cg19518666
cg19519310
cg19519747
cg19520280
cg19520372
cg19520710
cg19520763
cg19521511
cg19521610
cg19521646
cg19521693
cg19522511
cg19523085
cg19524362
cg19524723
cg19525872
cg19526568
cg19526659
cg19528338
cg19528654
cg19529273
cg19529583
cg19530885
cg19530924
cg19531032
cg19531604
cg19532402
cg19534753
cg19535000
cg19536281
cg19536407
cg19536810
cg19537184
cg19537490
cg19537558
cg19537969
cg19538089
cg19538655

cg19539568
cg19540425
cg19541479
cg19542238
cg19542816
cg19543017
cg19544065
cg19545560
cg19546583
cg19547124
cg19547319
cg19548181
cg19549758
cg19549902
cg19550034
cg19551055
cg19551912
cg19553721
cg19553910
cg19554281
cg19554457
cg19554461
cg19554569
cg19554797
cg19554939
cg19555116
cg19555331
cg19555988
cg19556343
cg19556613
cg19556774
cg19556854
cg19558802
cg19559587
cg19559891
cg19560370
cg19560588
cg19560971
cg19561607
cg19561972
cg19562321
cg19563365
cg19563510
cg19563932
cg19564450
cg19564648
cg19565171
cg19565396
cg19565423
cg19567740
cg19568115
cg19570244

cg19570558
cg19571015
cg19571715
cg19572157
cg19572637
cg19574327
cg19576099
cg19576738
cg19577074
cg19577080
cg19577377
cg19577395
cg19577779
cg19578175
cg19578183
cg19578305
cg19578660
cg19578835
cg19579005
cg19579167
cg19579176
cg19580752
cg19580930
cg19580937
cg19581683
cg19581963
cg19582128
cg19582222
cg19582822
cg19583265
cg19583934
cg19585646
cg19586027
cg19586288
cg19586709
cg19587237
cg19587777
cg19588016
cg19588649
cg19589060
cg19589277
cg19589283
cg19589396
cg19589728
cg19589800
cg19589939
cg19590900
cg19591696
cg19592235
cg19592336
cg19592439
cg19592472

cg19592720
cg19592732
cg19592898
cg19594305
cg19594433
cg19595277
cg19595295
cg19596110
cg19596509
cg19596655
cg19596804
cg19597382
cg19598094
cg19598567
cg19599386
cg19599803
cg19599972
cg19600023
cg19600081
cg19600368
cg19600583
cg19602315
cg19602728
cg19602938
cg19603100
cg19604323
cg19604896
cg19605623
cg19605644
cg19606153
cg19606290
cg19607021
cg19607474
cg19608357
cg19609242
cg19609334
cg19609644
cg19610529
cg19611002
cg19611612
cg19611809
cg19612191
cg19613205
cg19613432
cg19614216
cg19614643
cg19615017
cg19615651
cg19616339
cg19617373
cg19618279
cg19618601

cg19618897
cg19619028
cg19619405
cg19619576
cg19619909
cg19620086
cg19620333
cg19620724
cg19621065
cg19621076
cg19621338
cg19621427
cg19622415
cg19622474
cg19622569
cg19622623
cg19622662
cg19622675
cg19622874
cg19623438
cg19623717
cg19623835
cg19624318
cg19625388
cg19625689
cg19626213
cg19627213
cg19628148
cg19628600
cg19628739
cg19628874
cg19630039
cg19630629
cg19631576
cg19631648
cg19631876
cg19632772
cg19633205
cg19633236
cg19634252
cg19634693
cg19635571
cg19635644
cg19635695
cg19636240
cg19636302
cg19639427
cg19639490
cg19640007
cg19640166
cg19640424
cg19641315

cg19641924
cg19642007
cg19642168
cg19642408
cg19642727
cg19643467
cg19645221
cg19646165
cg19647114
cg19650157
cg19650520
cg19650573
cg19650920
cg19651003
cg19651578
cg19651694
cg19652416
cg19653211
cg19653783
cg19653907
cg19653967
cg19654195
cg19654437
cg19656282
cg19656577
cg19656950
cg19656974
cg19657174
cg19657198
cg19657351
cg19658665
cg19659215
cg19660744
cg19660806
cg19662554
cg19664013
cg19666600
cg19666687
cg19667811
cg19669199
cg19670883
cg19670885
cg19671120
cg19671533
cg19672145
cg19672546
cg19672751
cg19672997
cg19673125
cg19673176
cg19673350
cg19673376

cg19674032
cg19674091
cg19675599
cg19675778
cg19676181
cg19676227
cg19677607
cg19677733
cg19677739
cg19678111
cg19678828
cg19678959
cg19678968
cg19679327
cg19679543
cg19679633
cg19679989
cg19681610
cg19681956
cg19682537
cg19682666
cg19682835
cg19683073
cg19684151
cg19684887
cg19685066
cg19685479
cg19685521
cg19685532
cg19686637
cg19687529
cg19687985
cg19688403
cg19688761
cg19689211
cg19689368
cg19689427
cg19690214
cg19690404
cg19691505
cg19691563
cg19692710
cg19692782
cg19693151
cg19694010
cg19694453
cg19694460
cg19695386
cg19696317
cg19696701
cg19697232
cg19697382

cg19697530
cg19697725
cg19698294
cg19698789
cg19699140
cg19699170
cg19699200
cg19699815
cg19700035
cg19700380
cg19700682
cg19701087
cg19701540
cg19701879
cg19702176
cg19702194
cg19702271
cg19702771
cg19703240
cg19704451
cg19704630
cg19704805
cg19704853
cg19705386
cg19705853
cg19706446
cg19706534
cg19707040
cg19707503
cg19707677
cg19708201
cg19708340
cg19708432
cg19710451
cg19710672
cg19710916
cg19711530
cg19711783
cg19712198
cg19712291
cg19712603
cg19712659
cg19712979
cg19713038
cg19713567
cg19714132
cg19714279
cg19714489
cg19714737
cg19714985
cg19715094
cg19715167

cg19715410
cg19716018
cg19716073
cg19716238
cg19716930
cg19717150
cg19717216
cg19717347
cg19717586
cg19718662
cg19718771
cg19718882
cg19719207
cg19719391
cg19719478
cg19719987
cg19720260
cg19720753
cg19721221
cg19721889
cg19722471
cg19722698
cg19722706
cg19723596
cg19723952
cg19724187
cg19724555
cg19725903
cg19725935
cg19727439
cg19727641
cg19727827
cg19728223
cg19728577
cg19728718
cg19729536
cg19729693
cg19730691
cg19730699
cg19730924
cg19732144
cg19732340
cg19732886
cg19733463
cg19734013
cg19734045
cg19734190
cg19734301
cg19735514
cg19736047
cg19736128
cg19736179

cg19736334
cg19736503
cg19737787
cg19738281
cg19738283
cg19739082
cg19739126
cg19739482
cg19740240
cg19740964
cg19741868
cg19743545
cg19743686
cg19744173
cg19744908
cg19745415
cg19745637
cg19746343
cg19747271
cg19747945
cg19747968
cg19748027
cg19748436
cg19749001
cg19749445
cg19749916
cg19752602
cg19752855
cg19753174
cg19753864
cg19754282
cg19754462
cg19754554
cg19754707
cg19754709
cg19755016
cg19755069
cg19755435
cg19755767
cg19756068
cg19757261
cg19757478
cg19757535
cg19757573
cg19757631
cg19758873
cg19759135
cg19759671
cg19760241
cg19760651
cg19760823
cg19760898

cg19760965
cg19761640
cg19761848
cg19762427
cg19763108
cg19764048
cg19764066
cg19764096
cg19764540
cg19764555
cg19765175
cg19766441
cg19766624
cg19767020
cg19767622
cg19767800
cg19767857
cg19768329
cg19769182
cg19769613
cg19770281
cg19770426
cg19771626
cg19771648
cg19771748
cg19772011
cg19773311
cg19773474
cg19774846
cg19774973
cg19775250
cg19776080
cg19776282
cg19776366
cg19776515
cg19776793
cg19776833
cg19776877
cg19777067
cg19777664
cg19777783
cg19778647
cg19778822
cg19779057
cg19779211
cg19779886
cg19779893
cg19779965
cg19780993
cg19781376
cg19782190
cg19782271

cg19782446
cg19782686
cg19784545
cg19786648
cg19786808
cg19786983
cg19786988
cg19787125
cg19787448
cg19787891
cg19788272
cg19788831
cg19790323
cg19791379
cg19791606
cg19791637
cg19792581
cg19792599
cg19793640
cg19793809
cg19793904
cg19794507
cg19795074
cg19795150
cg19795151
cg19795292
cg19795338
cg19795722
cg19795898
cg19796532
cg19796981
cg19797376
cg19797700
cg19798224
cg19799702
cg19800032
cg19800728
cg19801062
cg19801256
cg19801646
cg19801921
cg19802138
cg19802243
cg19802368
cg19802378
cg19802564
cg19802865
cg19802929
cg19802981
cg19803131
cg19803721
cg19804605

cg19805033
cg19805748
cg19806310
cg19806642
cg19806995
cg19807257
cg19807286
cg19807317
cg19808205
cg19809039
cg19809499
cg19809667
cg19809988
cg19810038
cg19810433
cg19810598
cg19810954
cg19811144
cg19811148
cg19811761
cg19811934
cg19811994
cg19812283
cg19813025
cg19814280
cg19814981
cg19815017
cg19815042
cg19815271
cg19815904
cg19818271
cg19819280
cg19819285
cg19819320
cg19820058
cg19820287
cg19820705
cg19821128
cg19821297
cg19821536
cg19821682
cg19822251
cg19822309
cg19822343
cg19822502
cg19823494
cg19823512
cg19823649
cg19823847
cg19824006
cg19824055
cg19824396

cg19824833
cg19824894
cg19824907
cg19825277
cg19825898
cg19825974
cg19825988
cg19827089
cg19827346
cg19827555
cg19827923
cg19828869
cg19830128
cg19830270
cg19831575
cg19831962
cg19832521
cg19832529
cg19834045
cg19834071
cg19834134
cg19834585
cg19835128
cg19835595
cg19836431
cg19838087
cg19838963
cg19839325
cg19839798
cg19839801
cg19841369
cg19841740
cg19843036
cg19843457
cg19843540
cg19844101
cg19844232
cg19844283
cg19846096
cg19846295
cg19846389
cg19846491
cg19846500
cg19846991
cg19847130
cg19847244
cg19847945
cg19848082
cg19848140
cg19848629
cg19849428
cg19850275

cg19850370
cg19850728
cg19851416
cg19851560
cg19851594
cg19851909
cg19852958
cg19853760
cg19854376
cg19854521
cg19854901
cg19855215
cg19855559
cg19855777
cg19856845
cg19857925
cg19858413
cg19859270
cg19859290
cg19859445
cg19859729
cg19861312
cg19861741
cg19861824
cg19862235
cg19862242
cg19863224
cg19863426
cg19863592
cg19863655
cg19864060
cg19864758
cg19865425
cg19865472
cg19865770
cg19866195
cg19866811
cg19866866
cg19867250
cg19867579
cg19867649
cg19867991
cg19868125
cg19868631
cg19868730
cg19868751
cg19868934
cg19869971
cg19871173
cg19871588
cg19871597
cg19871903

cg19871940
cg19873297
cg19873701
cg19875547
cg19876302
cg19876388
cg19876649
cg19876657
cg19876814
cg19877064
cg19879483
cg19879906
cg19880077
cg19880942
cg19880947
cg19882784
cg19882857
cg19884262
cg19884492
cg19884950
cg19885306
cg19885587
cg19885761
cg19886365
cg19886448
cg19886655
cg19888509
cg19888870
cg19889666
cg19889795
cg19889856
cg19890774
cg19893585
cg19893664
cg19893751
cg19894640
cg19896824
cg19897071
cg19897113
cg19899385
cg19901801
cg19902173
cg19902593
cg19903071
cg19903805
cg19904122
cg19904905
cg19905414
cg19906534
cg19907380
cg19907479
cg19907483

cg19907520
cg19907915
cg19908028
cg19908151
cg19908485
cg19908556
cg19908768
cg19908812
cg19908856
cg19909063
cg19909117
cg19909239
cg19909349
cg19910327
cg19912141
cg19912142
cg19912309
cg19912436
cg19913118
cg19913291
cg19913551
cg19913563
cg19913582
cg19915413
cg19915738
cg19917300
cg19917530
cg19917989
cg19919303
cg19919383
cg19920134
cg19922137
cg19923650
cg19924120
cg19924619
cg19925186
cg19925849
cg19926144
cg19926250
cg19926385
cg19926444
cg19926480
cg19926902
cg19927100
cg19927816
cg19928084
cg19928195
cg19928377
cg19929126
cg19929194
cg19930116
cg19930296

cg19931733
cg19932079
cg19932515
cg19935065
cg19935326
cg19935531
cg19935845
cg19936757
cg19937056
cg19937331
cg19937513
cg19937938
cg19938199
cg19939130
cg19939178
cg19939250
cg19939793
cg19940482
cg19940537
cg19942213
cg19942256
cg19942495
cg19942731
cg19944217
cg19944367
cg19944705
cg19945554
cg19946742
cg19947104
cg19947463
cg19948807
cg19949550
cg19949591
cg19949932
cg19950606
cg19951107
cg19952303
cg19952454
cg19952821
cg19953290
cg19953427
cg19953917
cg19954017
cg19955019
cg19955173
cg19955500
cg19955823
cg19956449
cg19956836
cg19956889
cg19956914
cg19957411

cg19959477
cg19960114
cg19960616
cg19960778
cg19961720
cg19962116
cg19962424
cg19963178
cg19963231
cg19963589
cg19964320
cg19965221
cg19965300
cg19965312
cg19965589
cg19967668
cg19968790
cg19968840
cg19968890
cg19969104
cg19969606
cg19969694
cg19971102
cg19971388
cg19971716
cg19972193
cg19972558
cg19972648
cg19972696
cg19973085
cg19973758
cg19974227
cg19974448
cg19975800
cg19977179
cg19977966
cg19978105
cg19978859
cg19979108
cg19979590
cg19980718
cg19980929
cg19981409
cg19981828
cg19982668
cg19982684
cg19983801
cg19984355
cg19987060
cg19987122
cg19987768
cg19988235

cg19988426
cg19988449
cg19988492
cg19989018
cg19989277
cg19989498
cg19991046
cg19991723
cg19991948
cg19992474
cg19994023
cg19994779
cg19995117
cg19995130
cg19995899
cg19998073
cg19998328
cg19998368
cg19998432
cg19998533
cg19999705
cg20000602
cg20001791
cg20001810
cg20002843
cg20003179
cg20003377
cg20004315
cg20005760
cg20007130
cg20007894
cg20008148
cg20008164
cg20008332
cg20009101
cg20010506
cg20011562
cg20012308
cg20012366
cg20012537
cg20013688
cg20013942
cg20014049
cg20014398
cg20014412
cg20015401
cg20015412
cg20015689
cg20015855
cg20015986
cg20016416
cg20017381

cg20017995
cg20019163
cg20019356
cg20019985
cg20021336
cg20021570
cg20021790
cg20023231
cg20023788
cg20025238
cg20025677
cg20025970
cg20026346
cg20026613
cg20026633
cg20026737
cg20026939
cg20027133
cg20027296
cg20027684
cg20029745
cg20029881
cg20031717
cg20032789
cg20033070
cg20033619
cg20034100
cg20034792
cg20035127
cg20036377
cg20038070
cg20039073
cg20039211
cg20039443
cg20040364
cg20040743
cg20040872
cg20041105
cg20042291
cg20042662
cg20043649
cg20043679
cg20043699
cg20045394
cg20045492
cg20047055
cg20047561
cg20049415
cg20049422
cg20049427
cg20050237
cg20050877

cg20051292
cg20051314
cg20051875
cg20051949
cg20052865
cg20052933
cg20053885
cg20054111
cg20054157
cg20054248
cg20054309
cg20054385
cg20055426
cg20055581
cg20057782
cg20058043
cg20058148
cg20058187
cg20059140
cg20059377
cg20060206
cg20060685
cg20060759
cg20061155
cg20061873
cg20062492
cg20062681
cg20062806
cg20063450
cg20063640
cg20064008
cg20064558
cg20065005
cg20065520
cg20065905
cg20067049
cg20067418
cg20067612
cg20067719
cg20067801
cg20068209
cg20069688
cg20070618
cg20071670
cg20072162
cg20072171
cg20072442
cg20073007
cg20073553
cg20073893
cg20074340
cg20075114

cg20075156
cg20076442
cg20076659
cg20078074
cg20080245
cg20080282
cg20080624
cg20080983
cg20082196
cg20082641
cg20082779
cg20083186
cg20083442
cg20084504
cg20085268
cg20085588
cg20086208
cg20086597
cg20086660
cg20086916
cg20090108
cg20090143
cg20090162
cg20090194
cg20090283
cg20090301
cg20090482
cg20090497
cg20090970
cg20091384
cg20091795
cg20092728
cg20092736
cg20094279
cg20094784
cg20097219
cg20097440
cg20098015
cg20098420
cg20098471
cg20098887
cg20099104
cg20099449
cg20099511
cg20099830
cg20100006
cg20100768
cg20100987
cg20101015
cg20101352
cg20102086
cg20102280

cg20102639
cg20102955
cg20103018
cg20103093
cg20103150
cg20103311
cg20103758
cg20104055
cg20104432
cg20105087
cg20105257
cg20106459
cg20106776
cg20107346
cg20107395
cg20107987
cg20109058
cg20111269
cg20112984
cg20113802
cg20114528
cg20114732
cg20115827
cg20116199
cg20116804
cg20116869
cg20116935
cg20117799
cg20118364
cg20118717
cg20118823
cg20119101
cg20119106
cg20119308
cg20120165
cg20120939
cg20121012
cg20122043
cg20122518
cg20122925
cg20122943
cg20124376
cg20124551
cg20125516
cg20126656
cg20126775
cg20127733
cg20128319
cg20128928
cg20129082
cg20129242
cg20129853

cg20130098
cg20130213
cg20130789
cg20130989
cg20132590
cg20136855
cg20136951
cg20137237
cg20137312
cg20138067
cg20139706
cg20140054
cg20140110
cg20140333
cg20140377
cg20140940
cg20141733
cg20143773
cg20144359
cg20146536
cg20146541
cg20148796
cg20149170
cg20150017
cg20150640
cg20151187
cg20151301
cg20151452
cg20151576
cg20152088
cg20152300
cg20152382
cg20152413
cg20152871
cg20154003
cg20154938
cg20156165
cg20156353
cg20156659
cg20157281
cg20157577
cg20158826
cg20159095
cg20159415
cg20159490
cg20162146
cg20162381
cg20163959
cg20163980
cg20164505
cg20165520
cg20165545

cg20165746
cg20167074
cg20168230
cg20168495
cg20168964
cg20169015
cg20169101
cg20169185
cg20169823
cg20169952
cg20169988
cg20170147
cg20170777
cg20170943
cg20171011
cg20171396
cg20171429
cg20172500
cg20174636
cg20174730
cg20175390
cg20176648
cg20176971
cg20177507
cg20177522
cg20178603
cg20180050
cg20180364
cg20180513
cg20181079
cg20181209
cg20181887
cg20183519
cg20183619
cg20185017
cg20185461
cg20185615
cg20185718
cg20185945
cg20186445
cg20187049
cg20188017
cg20188739
cg20189244
cg20191453
cg20191740
cg20192747
cg20193057
cg20193203
cg20193324
cg20194314
cg20194364

cg20195005
cg20195049
cg20195319
cg20197622
cg20198768
cg20198923
cg20199836
cg20200460
cg20201128
cg20203210
cg20203352
cg20203768
cg20204921
cg20205704
cg20207829
cg20207911
cg20207973
cg20208009
cg20210051
cg20210482
cg20210689
cg20211134
cg20211604
cg20212364
cg20214477
cg20214973
cg20215212
cg20215257
cg20216139
cg20216655
cg20217449
cg20217649
cg20217702
cg20217978
cg20218614
cg20219379
cg20219457
cg20220242
cg20220678
cg20220737
cg20220755
cg20222568
cg20222926
cg20224285
cg20224984
cg20224991
cg20225915
cg20226253
cg20226764
cg20227056
cg20227165
cg20227326

cg20229931
cg20231084
cg20231299
cg20231883
cg20232102
cg20232986
cg20234910
cg20235871
cg20236424
cg20236995
cg20237264
cg20237595
cg20237888
cg20238105
cg20238197
cg20238368
cg20239174
cg20239912
cg20240154
cg20240715
cg20241270
cg20242070
cg20243424
cg20244649
cg20245568
cg20246113
cg20246273
cg20246277
cg20247486
cg20248093
cg20248163
cg20248516
cg20249252
cg20249877
cg20249919
cg20250981
cg20252837
cg20253105
cg20253268
cg20253561
cg20253872
cg20253994
cg20254658
cg20254790
cg20255272
cg20255713
cg20256009
cg20257553
cg20257866
cg20259814
cg20260619
cg20260841

cg20261431
cg20262722
cg20263045
cg20263392
cg20263853
cg20264481
cg20264543
cg20265661
cg20265733
cg20265951
cg20266154
cg20266316
cg20266320
cg20266715
cg20267441
cg20267590
cg20267775
cg20268039
cg20268352
cg20268992
cg20269359
cg20269544
cg20269876
cg20271029
cg20271057
cg20271865
cg20272028
cg20272619
cg20272884
cg20272962
cg20274006
cg20274462
cg20274673
cg20274950
cg20275558
cg20276202
cg20277504
cg20278269
cg20279493
cg20281375
cg20281554
cg20282550
cg20283498
cg20284891
cg20285128
cg20286347
cg20286882
cg20287234
cg20288445
cg20288916
cg20289688
cg20289854

cg20289913
cg20290367
cg20290672
cg20291049
cg20291363
cg20291636
cg20294962
cg20295442
cg20295537
cg20297976
cg20298273
cg20299399
cg20299572
cg20299573
cg20300093
cg20300343
cg20300500
cg20300541
cg20301308
cg20301780
cg20302414
cg20303033
cg20303331
cg20303399
cg20303724
cg20305489
cg20306062
cg20307598
cg20308351
cg20309353
cg20309891
cg20310100
cg20310894
cg20311002
cg20311601
cg20311730
cg20311863
cg20312026
cg20312179
cg20312205
cg20312390
cg20312687
cg20313362
cg20313628
cg20315057
cg20315124
cg20315739
cg20316538
cg20316883
cg20317521
cg20317748
cg20318166

cg20318580
cg20318608
cg20318845
cg20319253
cg20319654
cg20319765
cg20320200
cg20320283
cg20320574
cg20320946
cg20321270
cg20321319
cg20322193
cg20322837
cg20323571
cg20323874
cg20324206
cg20324687
cg20324771
cg20324858
cg20325280
cg20326410
cg20326674
cg20326936
cg20327444
cg20327845
cg20328399
cg20329153
cg20329958
cg20330521
cg20332645
cg20333014
cg20334115
cg20334313
cg20334849
cg20335152
cg20335384
cg20335735
cg20336172
cg20336388
cg20338410
cg20339650
cg20339692
cg20339720
cg20339859
cg20340302
cg20340508
cg20340596
cg20340716
cg20340866
cg20341089
cg20342079

cg20342458
cg20343063
cg20344040
cg20344925
cg20344956
cg20345556
cg20345562
cg20345923
cg20346388
cg20346912
cg20347666
cg20348189
cg20348703
cg20348858
cg20350518
cg20351495
cg20351875
cg20352178
cg20352728
cg20353489
cg20353653
cg20357628
cg20357862
cg20358275
cg20358313
cg20359285
cg20359360
cg20359445
cg20359565
cg20359793
cg20359888
cg20360285
cg20361205
cg20361924
cg20362308
cg20362817
cg20363022
cg20363192
cg20364183
cg20364222
cg20364776
cg20365074
cg20365451
cg20366239
cg20366831
cg20367258
cg20367923
cg20370281
cg20370678
cg20371142
cg20372666
cg20373012

cg20373894
cg20374054
cg20374118
cg20375122
cg20376009
cg20376082
cg20376953
cg20377305
cg20378687
cg20379919
cg20380768
cg20382146
cg20382154
cg20383007
cg20383794
cg20384702
cg20385001
cg20385229
cg20385368
cg20385395
cg20385461
cg20385708
cg20386303
cg20386316
cg20387256
cg20387258
cg20387391
cg20387775
cg20388294
cg20388428
cg20388729
cg20388823
cg20388916
cg20388967
cg20389653
cg20389678
cg20390015
cg20390119
cg20391432
cg20391793
cg20391948
cg20392088
cg20393324
cg20393695
cg20394609
cg20395040
cg20395881
cg20395967
cg20398418
cg20398486
cg20398571
cg20399534

cg20399616
cg20399720
cg20399871
cg20399965
cg20400904
cg20401130
cg20401252
cg20401521
cg20401567
cg20402653
cg20402826
cg20403528
cg20403541
cg20403938
cg20405017
cg20405697
cg20406033
cg20406460
cg20407365
cg20407861
cg20411340
cg20411756
cg20413331
cg20413415
cg20413441
cg20414015
cg20414364
cg20414618
cg20416129
cg20417024
cg20417128
cg20417424
cg20417527
cg20419307
cg20420651
cg20422099
cg20422212
cg20422722
cg20424036
cg20424973
cg20425154
cg20426042
cg20426415
cg20426768
cg20426959
cg20427144
cg20427378
cg20428196
cg20428720
cg20428976
cg20429277
cg20430631

cg20431250
cg20433382
cg20433386
cg20433455
cg20433822
cg20433952
cg20434422
cg20435370
cg20435387
cg20435474
cg20435535
cg20435725
cg20436086
cg20436458
cg20437132
cg20438286
cg20438306
cg20439030
cg20439656
cg20440093
cg20442232
cg20442289
cg20442379
cg20443278
cg20443357
cg20443778
cg20444370
cg20445197
cg20445727
cg20445806
cg20446404
cg20446641
cg20447508
cg20447966
cg20448089
cg20448594
cg20449074
cg20449134
cg20449382
cg20449404
cg20449685
cg20449692
cg20450058
cg20450198
cg20450336
cg20450507
cg20450689
cg20451455
cg20451566
cg20452657
cg20452739
cg20452961

cg20453394
cg20454002
cg20454336
cg20454887
cg20455687
cg20456243
cg20457275
cg20457796
cg20459138
cg20460227
cg20461188
cg20461912
cg20462100
cg20462795
cg20463298
cg20463526
cg20464246
cg20464719
cg20465661
cg20467957
cg20468163
cg20469625
cg20470223
cg20471413
cg20471736
cg20472643
cg20474425
cg20475082
cg20475114
cg20475322
cg20475718
cg20477147
cg20477574
cg20478080
cg20478129
cg20479672
cg20479688
cg20479774
cg20480334
cg20481032
cg20481287
cg20481340
cg20481655
cg20481720
cg20482145
cg20482390
cg20483324
cg20483374
cg20483857
cg20484653
cg20484832
cg20485058

cg20485165
cg20485191
cg20485758
cg20486417
cg20489453
cg20490088
cg20490724
cg20492807
cg20493526
cg20494374
cg20494635
cg20495052
cg20495183
cg20495370
cg20495476
cg20495903
cg20497212
cg20497304
cg20497968
cg20498859
cg20498872
cg20500126
cg20501434
cg20502536
cg20504856
cg20504993
cg20505223
cg20505457
cg20506347
cg20506659
cg20507066
cg20507963
cg20508508
cg20509513
cg20510033
cg20510207
cg20510869
cg20511815
cg20512711
cg20513206
cg20513418
cg20513777
cg20514083
cg20514239
cg20514290
cg20514520
cg20515679
cg20516032
cg20516364
cg20516712
cg20518155
cg20518362

cg20518446
cg20519121
cg20519577
cg20519670
cg20521198
cg20522006
cg20522104
cg20522387
cg20522398
cg20522675
cg20523466
cg20523861
cg20524040
cg20524636
cg20526800
cg20530843
cg20530855
cg20531169
cg20532925
cg20532999
cg20533952
cg20534694
cg20534702
cg20535067
cg20535254
cg20536049
cg20536794
cg20536983
cg20537629
cg20537640
cg20537886
cg20538211
cg20539752
cg20540566
cg20540714
cg20541233
cg20541752
cg20541785
cg20543426
cg20543478
cg20543651
cg20544216
cg20546518
cg20547131
cg20547407
cg20548182
cg20550154
cg20551181
cg20552220
cg20552747
cg20552903
cg20553982

cg20554142
cg20554882
cg20555305
cg20555682
cg20555772
cg20555922
cg20556503
cg20556803
cg20556936
cg20556951
cg20557595
cg20557687
cg20557778
cg20557801
cg20559385
cg20559422
cg20559736
cg20560115
cg20560169
cg20560600
cg20560862
cg20562446
cg20563468
cg20563910
cg20563971
cg20566484
cg20566766
cg20566970
cg20567368
cg20567768
cg20568497
cg20569369
cg20570313
cg20570458
cg20570611
cg20571462
cg20571592
cg20574436
cg20574490
cg20574695
cg20575761
cg20575856
cg20576162
cg20576393
cg20576726
cg20576955
cg20578258
cg20578612
cg20579480
cg20579564
cg20582941
cg20583430

cg20583930
cg20584157
cg20584905
cg20585530
cg20585869
cg20587394
cg20587874
cg20587945
cg20588045
cg20588069
cg20588285
cg20588868
cg20589078
cg20590797
cg20591472
cg20592390
cg20592447
cg20592734
cg20592995
cg20593059
cg20593611
cg20593774
cg20594126
cg20594465
cg20595818
cg20595862
cg20596162
cg20596329
cg20596493
cg20596724
cg20597172
cg20597215
cg20597264
cg20597403
cg20597619
cg20599086
cg20599405
cg20599967
cg20600210
cg20600916
cg20601276
cg20601919
cg20605134
cg20605413
cg20605886
cg20607169
cg20607287
cg20607293
cg20608100
cg20608192
cg20608329
cg20608338

cg20608699
cg20609302
cg20609883
cg20610452
cg20611334
cg20611371
cg20611882
cg20611911
cg20612299
cg20612962
cg20614854
cg20615910
cg20616108
cg20617669
cg20617917
cg20618109
cg20618433
cg20618622
cg20618695
cg20619751
cg20621200
cg20621765
cg20622131
cg20623350
cg20624923
cg20624941
cg20625334
cg20626249
cg20627572
cg20628376
cg20628718
cg20628734
cg20629551
cg20630887
cg20631014
cg20631351
cg20632573
cg20632887
cg20633213
cg20634266
cg20634442
cg20634514
cg20634967
cg20635370
cg20635409
cg20636248
cg20636526
cg20637438
cg20638140
cg20638525
cg20638582
cg20639396

cg20640266
cg20640433
cg20640774
cg20641026
cg20642630
cg20643012
cg20643675
cg20644554
cg20645058
cg20645878
cg20646280
cg20646533
cg20646846
cg20648847
cg20649017
cg20651616
cg20651995
cg20652371
cg20653075
cg20653144
cg20653375
cg20654462
cg20654619
cg20655761
cg20656525
cg20656566
cg20658779
cg20659435
cg20659752
cg20659941
cg20660989
cg20661303
cg20661965
cg20662501
cg20663432
cg20664201
cg20664238
cg20664247
cg20665007
cg20665259
cg20665341
cg20665774
cg20666307
cg20667664
cg20667964
cg20668321
cg20668718
cg20668765
cg20670075
cg20670939
cg20671415
cg20672264

cg20672430
cg20672477
cg20672549
cg20672708
cg20672981
cg20673075
cg20673829
cg20674212
cg20674577
cg20674658
cg20675439
cg20675440
cg20676047
cg20676413
cg20676696
cg20676716
cg20676788
cg20677058
cg20678043
cg20679157
cg20680045
cg20680265
cg20681321
cg20681948
cg20682222
cg20682573
cg20682639
cg20683057
cg20683656
cg20684174
cg20684197
cg20685419
cg20685736
cg20686125
cg20686403
cg20687098
cg20687242
cg20687553
cg20688114
cg20689224
cg20689476
cg20689730
cg20690028
cg20690782
cg20691580
cg20693334
cg20693607
cg20693690
cg20694933
cg20695203
cg20695534
cg20695936

cg20696173
cg20696451
cg20696605
cg20696698
cg20697044
cg20698113
cg20698421
cg20698781
cg20698862
cg20698924
cg20698942
cg20699076
cg20699079
cg20699549
cg20699711
cg20699736
cg20700731
cg20701183
cg20701486
cg20701531
cg20702913
cg20703137
cg20703581
cg20703614
cg20703928
cg20704530
cg20705236
cg20705321
cg20705390
cg20705812
cg20706196
cg20706468
cg20707222
cg20707323
cg20707774
cg20707780
cg20707921
cg20708282
cg20709275
cg20709280
cg20710258
cg20710394
cg20710519
cg20710667
cg20710697
cg20710709
cg20710824
cg20710902
cg20711030
cg20711047
cg20712054
cg20712631

cg20714110
cg20714487
cg20715295
cg20715438
cg20716812
cg20717123
cg20720059
cg20721320
cg20722987
cg20723355
cg20723412
cg20723436
cg20723935
cg20723941
cg20724168
cg20724476
cg20725021
cg20725135
cg20725239
cg20725716
cg20726070
cg20727434
cg20727623
cg20728105
cg20728419
cg20728791
cg20729460
cg20730333
cg20730595
cg20731384
cg20731623
cg20731798
cg20732076
cg20732137
cg20732478
cg20733108
cg20733250
cg20734219
cg20734569
cg20734996
cg20736321
cg20737500
cg20737708
cg20737812
cg20737995
cg20738274
cg20738500
cg20738665
cg20738719
cg20739526
cg20739984
cg20740024

cg20740328
cg20740434
cg20740485
cg20741040
cg20741605
cg20741620
cg20742548
cg20742696
cg20742981
cg20743662
cg20743708
cg20743744
cg20743881
cg20744108
cg20744362
cg20744943
cg20746459
cg20746552
cg20746795
cg20747012
cg20747266
cg20747462
cg20747577
cg20748065
cg20748397
cg20748559
cg20750319
cg20750528
cg20750832
cg20751706
cg20751926
cg20752233
cg20752683
cg20752695
cg20752831
cg20754034
cg20754261
cg20754324
cg20754844
cg20755170
cg20756702
cg20758789
cg20758834
cg20759047
cg20759084
cg20759281
cg20759486
cg20759600
cg20760920
cg20761187
cg20762480
cg20762586

cg20762805
cg20764887
cg20765100
cg20765441
cg20766179
cg20767111
cg20767344
cg20768043
cg20768399
cg20768506
cg20768743
cg20769177
cg20769487
cg20769856
cg20770006
cg20770409
cg20770435
cg20772037
cg20774676
cg20775162
cg20776829
cg20776947
cg20776977
cg20777390
cg20778071
cg20778199
cg20778451
cg20779964
cg20780180
cg20780302
cg20780811
cg20781516
cg20781967
cg20781981
cg20782816
cg20783464
cg20784259
cg20785098
cg20785796
cg20786876
cg20787173
cg20788020
cg20788149
cg20789595
cg20790648
cg20790998
cg20791593
cg20792062
cg20792294
cg20792590
cg20792735
cg20792744

cg20793071
cg20793341
cg20793665
cg20794208
cg20794351
cg20794828
cg20795680
cg20795889
cg20796968
cg20798604
cg20800199
cg20800391
cg20800892
cg20801110
cg20801476
cg20801645
cg20801711
cg20801892
cg20802214
cg20802942
cg20803211
cg20803857
cg20804050
cg20804191
cg20805334
cg20805920
cg20806172
cg20806175
cg20806180
cg20806345
cg20808234
cg20808363
cg20808613
cg20808693
cg20809087
cg20809224
cg20809378
cg20810198
cg20810222
cg20810396
cg20810478
cg20810675
cg20810975
cg20810997
cg20811755
cg20811788
cg20812354
cg20813518
cg20813798
cg20813827
cg20813996
cg20814718

cg20814774
cg20814813
cg20815339
cg20815571
cg20815643
cg20815992
cg20817073
cg20817257
cg20820063
cg20820107
cg20821042
cg20821170
cg20821314
cg20821544
cg20821800
cg20822540
cg20822693
cg20823091
cg20823348
cg20823695
cg20824189
cg20824200
cg20824761
cg20825022
cg20825110
cg20826024
cg20826151
cg20826165
cg20829347
cg20829834
cg20830514
cg20832020
cg20832320
cg20832559
cg20833090
cg20833786
cg20834073
cg20837557
cg20838323
cg20839205
cg20840069
cg20840783
cg20840795
cg20841588
cg20841748
cg20842326
cg20842396
cg20844427
cg20844619
cg20844851
cg20845494
cg20847018

cg20847586
cg20847701
cg20847766
cg20848988
cg20849150
cg20849286
cg20850016
cg20852846
cg20853889
cg20854539
cg20856497
cg20856834
cg20857280
cg20857416
cg20858526
cg20860124
cg20861489
cg20861822
cg20862283
cg20865082
cg20866075
cg20866121
cg20866810
cg20867746
cg20869310
cg20870000
cg20870512
cg20871721
cg20871964
cg20872072
cg20872252
cg20872937
cg20873136
cg20873416
cg20874827
cg20875807
cg20876760
cg20876937
cg20878317
cg20879768
cg20880499
cg20881175
cg20881910
cg20882260
cg20883194
cg20883831
cg20884298
cg20884362
cg20884887
cg20885615
cg20886721
cg20886791

cg20887396
cg20888142
cg20889394
cg20889395
cg20889479
cg20889841
cg20892260
cg20892840
cg20892990
cg20893714
cg20893838
cg20895054
cg20895785
cg20896728
cg20897479
cg20899321
cg20899354
cg20899718
cg20900103
cg20900540
cg20900927
cg20901032
cg20902353
cg20902817
cg20904010
cg20905460
cg20905984
cg20906291
cg20907409
cg20907456
cg20908297
cg20909656
cg20910424
cg20910490
cg20910746
cg20910807
cg20912169
cg20912770
cg20912874
cg20914464
cg20916066
cg20916535
cg20916578
cg20917348
cg20917891
cg20919730
cg20919781
cg20920419
cg20921479
cg20924428
cg20924825
cg20925233

cg20926035
cg20926651
cg20926939
cg20927567
cg20927649
cg20927656
cg20927661
cg20928974
cg20929609
cg20930320
cg20930366
cg20931033
cg20931134
cg20931474
cg20931541
cg20932397
cg20932440
cg20932535
cg20933297
cg20934765
cg20934839
cg20935165
cg20935293
cg20935542
cg20936920
cg20937028
cg20937139
cg20937981
cg20938170
cg20938346
cg20938699
cg20938708
cg20938802
cg20939084
cg20939114
cg20939319
cg20939662
cg20940024
cg20940398
cg20941110
cg20941855
cg20942223
cg20942429
cg20943032
cg20943353
cg20944195
cg20944278
cg20944283
cg20944315
cg20944701
cg20945085
cg20945221

cg20945261
cg20946805
cg20946991
cg20947082
cg20948431
cg20948740
cg20948923
cg20949644
cg20950167
cg20951266
cg20951444
cg20951539
cg20951642
cg20952257
cg20952286
cg20953187
cg20953257
cg20954179
cg20954180
cg20954219
cg20954460
cg20954777
cg20954893
cg20955839
cg20956114
cg20956815
cg20957316
cg20958486
cg20959189
cg20959403
cg20959460
cg20959480
cg20959866
cg20960383
cg20961940
cg20962117
cg20962215
cg20962532
cg20963030
cg20963227
cg20964635
cg20964856
cg20965527
cg20965743
cg20966089
cg20966148
cg20966659
cg20967441
cg20967585
cg20968678
cg20968717
cg20968963

cg20969194
cg20969424
cg20970205
cg20970845
cg20970886
cg20971158
cg20971571
cg20972294
cg20974724
cg20974826
cg20975182
cg20975414
cg20975835
cg20976831
cg20976833
cg20979799
cg20979979
cg20980271
cg20980653
cg20981000
cg20981106
cg20981163
cg20981451
cg20981919
cg20983647
cg20983854
cg20983876
cg20984065
cg20984924
cg20984972
cg20985014
cg20985269
cg20985399
cg20985450
cg20985479
cg20985521
cg20986076
cg20986370
cg20986665
cg20986935
cg20987067
cg20987283
cg20987924
cg20988565
cg20989454
cg20989855
cg20989926
cg20990620
cg20991421
cg20991483
cg20991819
cg20992002

cg20992180
cg20992439
cg20992708
cg20993253
cg20993909
cg20995172
cg20995304
cg20995564
cg20995928
cg20995977
cg20996561
cg20996682
cg20997661
cg20997993
cg20998127
cg20998200
cg20998511
cg20998539
cg20998791
cg21000999
cg21001184
cg21001273
cg21001441
cg21001487
cg21002264
cg21003377
cg21003581
cg21003878
cg21003902
cg21004104
cg21004129
cg21004604
cg21005218
cg21005438
cg21005457
cg21006031
cg21007645
cg21007852
cg21008040
cg21008145
cg21008399
cg21008969
cg21009400
cg21009824
cg21010569
cg21010859
cg21011133
cg21012139
cg21012296
cg21013206
cg21013866
cg21015669

cg21015805
cg21022395
cg21022868
cg21022949
cg21023770
cg21025142
cg21025494
cg21026460
cg21026566
cg21026830
cg21027535
cg21027599
cg21028182
cg21028981
cg21029631
cg21030160
cg21030300
cg21031035
cg21031128
cg21031917
cg21033494
cg21033632
cg21033709
cg21033861
cg21034784
cg21035374
cg21035669
cg21036840
cg21037180
cg21038156
cg21038461
cg21038740
cg21039708
cg21041701
cg21041775
cg21042276
cg21042456
cg21043558
cg21044104
cg21044933
cg21045464
cg21045575
cg21045770
cg21045828
cg21046160
cg21046951
cg21047322
cg21047761
cg21048162
cg21048260
cg21048812
cg21049302

cg21049501
cg21049809
cg21049958
cg21051473
cg21051580
cg21052415
cg21052677
cg21052682
cg21052687
cg21053529
cg21053831
cg21054521
cg21055546
cg21057473
cg21057494
cg21058391
cg21058511
cg21059786
cg21063500
cg21063722
cg21063758
cg21064451
cg21064916
cg21066636
cg21066748
cg21067750
cg21069005
cg21069434
cg21069965
cg21070172
cg21070222
cg21070527
cg21071075
cg21071237
cg21071793
cg21073650
cg21074015
cg21074260
cg21074797
cg21075643
cg21075784
cg21077957
cg21078330
cg21078963
cg21079003
cg21079591
cg21079805
cg21079946
cg21080533
cg21080562
cg21081803
cg21081971

cg21082557
cg21082678
cg21083556
cg21084702
cg21086326
cg21086551
cg21087481
cg21088119
cg21088438
cg21088983
cg21089930
cg21090182
cg21090344
cg21091227
cg21092790
cg21092808
cg21093166
cg21093192
cg21094405
cg21095561
cg21095673
cg21096141
cg21096457
cg21096898
cg21096915
cg21097788
cg21100077
cg21100167
cg21100712
cg21101356
cg21101465
cg21101995
cg21102121
cg21103227
cg21104449
cg21104971
cg21105620
cg21105928
cg21106407
cg21106719
cg21107210
cg21107581
cg21107767
cg21108519
cg21108826
cg21109025
cg21109045
cg21110505
cg21110769
cg21111471
cg21111500
cg21112391

cg21112452
cg21113592
cg21113740
cg21113768
cg21113776
cg21114004
cg21114773
cg21115391
cg21115977
cg21116812
cg21117476
cg21117734
cg21118780
cg21118819
cg21120249
cg21121136
cg21121843
cg21122366
cg21122474
cg21122605
cg21122774
cg21123160
cg21123417
cg21124375
cg21125190
cg21126020
cg21126453
cg21126487
cg21126506
cg21126583
cg21126692
cg21126943
cg21127898
cg21128568
cg21128953
cg21129006
cg21129253
cg21129864
cg21130124
cg21130236
cg21130255
cg21131024
cg21131402
cg21132564
cg21133868
cg21134096
cg21134182
cg21134737
cg21134771
cg21135135
cg21135560
cg21136443

cg21137557
cg21138935
cg21139496
cg21139587
cg21142398
cg21142743
cg21143428
cg21143441
cg21143587
cg21144009
cg21144063
cg21144270
cg21144871
cg21144928
cg21144988
cg21145136
cg21145248
cg21145284
cg21145416
cg21145449
cg21145524
cg21146170
cg21146184
cg21146209
cg21146652
cg21147063
cg21147203
cg21147708
cg21148025
cg21148923
cg21149466
cg21150026
cg21150053
cg21150108
cg21150537
cg21151061
cg21151122
cg21151960
cg21151963
cg21152177
cg21152628
cg21152690
cg21153040
cg21153898
cg21154627
cg21154876
cg21155834
cg21156006
cg21156200
cg21156276
cg21156970
cg21157065

cg21157094
cg21158075
cg21158087
cg21158561
cg21158631
cg21158633
cg21158795
cg21159128
cg21159370
cg21160105
cg21161126
cg21161138
cg21161526
cg21161568
cg21161891
cg21162344
cg21162356
cg21162528
cg21162779
cg21162828
cg21162938
cg21163444
cg21163617
cg21163717
cg21163972
cg21164019
cg21164050
cg21164131
cg21164877
cg21165051
cg21165221
cg21165812
cg21165909
cg21166330
cg21166380
cg21166775
cg21166852
cg21168134
cg21169762
cg21170041
cg21170304
cg21170583
cg21170658
cg21170752
cg21170820
cg21170920
cg21171314
cg21171625
cg21172322
cg21172377
cg21172497
cg21172944

cg21173192
cg21173239
cg21173447
cg21174533
cg21174565
cg21174686
cg21174903
cg21175685
cg21177112
cg21177426
cg21177494
cg21177778
cg21177940
cg21178333
cg21178618
cg21179088
cg21179618
cg21179654
cg21179705
cg21180080
cg21180201
cg21180286
cg21180599
cg21181989
cg21182263
cg21183256
cg21184216
cg21185255
cg21185662
cg21186235
cg21186560
cg21186708
cg21186966
cg21187195
cg21187352
cg21187554
cg21187669
cg21187769
cg21187920
cg21188242
cg21188533
cg21188652
cg21189438
cg21190895
cg21191088
cg21193143
cg21193744
cg21194726
cg21194807
cg21195231
cg21195235
cg21195256

cg21195462
cg21196265
cg21196930
cg21197224
cg21197233
cg21197796
cg21198021
cg21198219
cg21198586
cg21199093
cg21199495
cg21200129
cg21200539
cg21201109
cg21201357
cg21202406
cg21203902
cg21204310
cg21204395
cg21204764
cg21205222
cg21206147
cg21206652
cg21206770
cg21207403
cg21207436
cg21208539
cg21208682
cg21208686
cg21208744
cg21208943
cg21209214
cg21209315
cg21209510
cg21210220
cg21210232
cg21210529
cg21210962
cg21211213
cg21211478
cg21211645
cg21211780
cg21211873
cg21211882
cg21214232
cg21214293
cg21214521
cg21214566
cg21214743
cg21215409
cg21215550
cg21215899

cg21216258
cg21216268
cg21216471
cg21216477
cg21218687
cg21219157
cg21220403
cg21220731
cg21220873
cg21220965
cg21221516
cg21222350
cg21222426
cg21222703
cg21222888
cg21223423
cg21223577
cg21223983
cg21225563
cg21226024
cg21226253
cg21226516
cg21226735
cg21227253
cg21227867
cg21228005
cg21228792
cg21229179
cg21229268
cg21229570
cg21230317
cg21230400
cg21230493
cg21230503
cg21230535
cg21230547
cg21230578
cg21230745
cg21230793
cg21231400
cg21231715
cg21231739
cg21232620
cg21232935
cg21234265
cg21234506
cg21234955
cg21235151
cg21236500
cg21236569
cg21236860
cg21236864

cg21237418
cg21237586
cg21238061
cg21238234
cg21239317
cg21240420
cg21240736
cg21241410
cg21241560
cg21242002
cg21242009
cg21242079
cg21242212
cg21242356
cg21242918
cg21243459
cg21243589
cg21243631
cg21243848
cg21243889
cg21243919
cg21244086
cg21244135
cg21244461
cg21244580
cg21245300
cg21245718
cg21246665
cg21247874
cg21248196
cg21248287
cg21249371
cg21249640
cg21249659
cg21250356
cg21250433
cg21251140
cg21252105
cg21252523
cg21253043
cg21253590
cg21254753
cg21254883
cg21254912
cg21254939
cg21255356
cg21255942
cg21263208
cg21263710
cg21265770
cg21266502
cg21267232

cg21267936
cg21268144
cg21268650
cg21268658
cg21271015
cg21271420
cg21271945
cg21272713
cg21272716
cg21272774
cg21272919
cg21272996
cg21274857
cg21275801
cg21276022
cg21276056
cg21276413
cg21277243
cg21279996
cg21280719
cg21281009
cg21282131
cg21282630
cg21282907
cg21284015
cg21284680
cg21284779
cg21285056
cg21285198
cg21285773
cg21286967
cg21287322
cg21287356
cg21287936
cg21288496
cg21289015
cg21289191
cg21289461
cg21291134
cg21291385
cg21292008
cg21292068
cg21292473
cg21293216
cg21293902
cg21293934
cg21295295
cg21295678
cg21295811
cg21296230
cg21296320
cg21297493

cg21297772
cg21298082
cg21298174
cg21298978
cg21299458
cg21299542
cg21300373
cg21300460
cg21300993
cg21301148
cg21301240
cg21301505
cg21302133
cg21302796
cg21302951
cg21304453
cg21304488
cg21305834
cg21306240
cg21307155
cg21307484
cg21309351
cg21312148
cg21312482
cg21312554
cg21312929
cg21314318
cg21314480
cg21314603
cg21316923
cg21318213
cg21319323
cg21319932
cg21320123
cg21320573
cg21320793
cg21323630
cg21324884
cg21324936
cg21325154
cg21325715
cg21327694
cg21328033
cg21328389
cg21330553
cg21330577
cg21330949
cg21331088
cg21331386
cg21331510
cg21331791
cg21331821

cg21332500
cg21334045
cg21334198
cg21334510
cg21335600
cg21336434
cg21337120
cg21337909
cg21340621
cg21340815
cg21340845
cg21340944
cg21341134
cg21341391
cg21341520
cg21341822
cg21342994
cg21343895
cg21344159
cg21345145
cg21346031
cg21346966
cg21347444
cg21347874
cg21349061
cg21349625
cg21349637
cg21350575
cg21350697
cg21350860
cg21351241
cg21351647
cg21351852
cg21352006
cg21352720
cg21353911
cg21355793
cg21355828
cg21356998
cg21357291
cg21363348
cg21365298
cg21367256
cg21367907
cg21368063
cg21368143
cg21370924
cg21372914
cg21373806
cg21374754
cg21375105
cg21375204

cg21376136
cg21376148
cg21376526
cg21376774
cg21377489
cg21379733
cg21379953
cg21380024
cg21381511
cg21381845
cg21382567
cg21383534
cg21384645
cg21384691
cg21385666
cg21385821
cg21386611
cg21386863
cg21386904
cg21386933
cg21387009
cg21387604
cg21388753
cg21389723
cg21390574
cg21390682
cg21391133
cg21391452
cg21392204
cg21392341
cg21392385
cg21393387
cg21394039
cg21395077
cg21396949
cg21397848
cg21398386
cg21398469
cg21399492
cg21399832
cg21400296
cg21401879
cg21402035
cg21402096
cg21402748
cg21402921
cg21403759
cg21404700
cg21404980
cg21405799
cg21406409
cg21406518

cg21406786
cg21407055
cg21407117
cg21408061
cg21408294
cg21408581
cg21408624
cg21408915
cg21410954
cg21410991
cg21411084
cg21411702
cg21411705
cg21412973
cg21413557
cg21414251
cg21414505
cg21415035
cg21415292
cg21415530
cg21418052
cg21418854
cg21419003
cg21419383
cg21421984
cg21422623
cg21422871
cg21423404
cg21424090
cg21425099
cg21426003
cg21427902
cg21428001
cg21428990
cg21429780
cg21430539
cg21430752
cg21431617
cg21431929
cg21432954
cg21433231
cg21433239
cg21434376
cg21434530
cg21435473
cg21436938
cg21438252
cg21439129
cg21439672
cg21439763
cg21440776
cg21441674

cg21442730
cg21443133
cg21443659
cg21444002
cg21444086
cg21444421
cg21444500
cg21446772
cg21446955
cg21447117
cg21447167
cg21447170
cg21447717
cg21448057
cg21448870
cg21448927
cg21449397
cg21449463
cg21449897
cg21450228
cg21451906
cg21452188
cg21452281
cg21454161
cg21454965
cg21455927
cg21456320
cg21457856
cg21458721
cg21459229
cg21460582
cg21461649
cg21461720
cg21461856
cg21461981
cg21462097
cg21462477
cg21462934
cg21464115
cg21464289
cg21464903
cg21466107
cg21466315
cg21466736
cg21468929
cg21468949
cg21469370
cg21469707
cg21471828
cg21472165
cg21472198
cg21472468

cg21472639
cg21472642
cg21473545
cg21474062
cg21474257
cg21475771
cg21475781
cg21476666
cg21477176
cg21479226
cg21480165
cg21481141
cg21481950
cg21481966
cg21482659
cg21483415
cg21483922
cg21484315
cg21484749
cg21484834
cg21484910
cg21485303
cg21485555
cg21486233
cg21486694
cg21487856
cg21487950
cg21488279
cg21489565
cg21489991
cg21490635
cg21491107
cg21492127
cg21493516
cg21493633
cg21494075
cg21494093
cg21494105
cg21494367
cg21494379
cg21494953
cg21495956
cg21497082
cg21497480
cg21497594
cg21497990
cg21498326
cg21499006
cg21499078
cg21499175
cg21499949
cg21500064

cg21500092
cg21500166
cg21500300
cg21500895
cg21501167
cg21501175
cg21501525
cg21501724
cg21502551
cg21503476
cg21504345
cg21504505
cg21505231
cg21505886
cg21506439
cg21508639
cg21508713
cg21509889
cg21510348
cg21510549
cg21510841
cg21511321
cg21511948
cg21512179
cg21512502
cg21512644
cg21512950
cg21513316
cg21513803
cg21513991
cg21514164
cg21514529
cg21514871
cg21515305
cg21515956
cg21516291
cg21517683
cg21517739
cg21517904
cg21517947
cg21518208
cg21518947
cg21519058
cg21519781
cg21519980
cg21520299
cg21520772
cg21521758
cg21522988
cg21523496
cg21523564
cg21523613

cg21524899
cg21525092
cg21525369
cg21526205
cg21527621
cg21528112
cg21528399
cg21529807
cg21531126
cg21531389
cg21531416
cg21532325
cg21534264
cg21534423
cg21534578
cg21535670
cg21536074
cg21537049
cg21537214
cg21537235
cg21537297
cg21537915
cg21538013
cg21538208
cg21538450
cg21539678
cg21540437
cg21540530
cg21540588
cg21541833
cg21542078
cg21542643
cg21542842
cg21543102
cg21543589
cg21543670
cg21544415
cg21544416
cg21544585
cg21545720
cg21545862
cg21545952
cg21546184
cg21546211
cg21546821
cg21546950
cg21547095
cg21547283
cg21547628
cg21547690
cg21547982
cg21548021

cg21548152
cg21548155
cg21548340
cg21549161
cg21549299
cg21550107
cg21550219
cg21550372
cg21550580
cg21550612
cg21550785
cg21551549
cg21551985
cg21552290
cg21553182
cg21553596
cg21554233
cg21554399
cg21554569
cg21554588
cg21554616
cg21554723
cg21555085
cg21555177
cg21555940
cg21556527
cg21556967
cg21559386
cg21559452
cg21561953
cg21561970
cg21562740
cg21563683
cg21563912
cg21564128
cg21565196
cg21565823
cg21565914
cg21565960
cg21566086
cg21567022
cg21568106
cg21569398
cg21569854
cg21570843
cg21570988
cg21571060
cg21571339
cg21572688
cg21572719
cg21573200
cg21574244

cg21574853
cg21575923
cg21576003
cg21576886
cg21577365
cg21578219
cg21578351
cg21579209
cg21579239
cg21579294
cg21579556
cg21579975
cg21582240
cg21582611
cg21583016
cg21584518
cg21584691
cg21584862
cg21585162
cg21585306
cg21585438
cg21587375
cg21587469
cg21588838
cg21589375
cg21589858
cg21590021
cg21590108
cg21590264
cg21591742
cg21591938
cg21592003
cg21592158
cg21592613
cg21592954
cg21593015
cg21593149
cg21593409
cg21593554
cg21593952
cg21594383
cg21595092
cg21596290
cg21597025
cg21597152
cg21597692
cg21598389
cg21599081
cg21599230
cg21599324
cg21600346
cg21601493

cg21601845
cg21602160
cg21602518
cg21604136
cg21604244
cg21604856
cg21605178
cg21605238
cg21605271
cg21605283
cg21605341
cg21605369
cg21605772
cg21606042
cg21606405
cg21606928
cg21606953
cg21607236
cg21607248
cg21607965
cg21608519
cg21609106
cg21609584
cg21609624
cg21609640
cg21609684
cg21609706
cg21610436
cg21610554
cg21610556
cg21611093
cg21611262
cg21611722
cg21614201
cg21614211
cg21614682
cg21614690
cg21615831
cg21615841
cg21616031
cg21616089
cg21616142
cg21616425
cg21616720
cg21618273
cg21618455
cg21618549
cg21618620
cg21618730
cg21619594
cg21620078
cg21620246

cg21620524
cg21621204
cg21621248
cg21622202
cg21622709
cg21623566
cg21623722
cg21624165
cg21624311
cg21624833
cg21625097
cg21625568
cg21625570
cg21625817
cg21627466
cg21627775
cg21628539
cg21629500
cg21629640
cg21629821
cg21629895
cg21630608
cg21631156
cg21631427
cg21631443
cg21633026
cg21633105
cg21634628
cg21634818
cg21635854
cg21635870
cg21637670
cg21637761
cg21637843
cg21638161
cg21638625
cg21639114
cg21642224
cg21643095
cg21643295
cg21643547
cg21643649
cg21644130
cg21644489
cg21644856
cg21644914
cg21645554
cg21645630
cg21645973
cg21646032
cg21646199
cg21646392

cg21646598
cg21647227
cg21647278
cg21648069
cg21648202
cg21649013
cg21649258
cg21649668
cg21650436
cg21650866
cg21651795
cg21651857
cg21653184
cg21653474
cg21655830
cg21655915
cg21655969
cg21656246
cg21656937
cg21657043
cg21657059
cg21657087
cg21657577
cg21657704
cg21659346
cg21659714
cg21659949
cg21661983
cg21663916
cg21664636
cg21664866
cg21665448
cg21665905
cg21666857
cg21667061
cg21667069
cg21667215
cg21667878
cg21667998
cg21669679
cg21670003
cg21671052
cg21671806
cg21672450
cg21672829
cg21672843
cg21674813
cg21675115
cg21678312
cg21678377
cg21678445
cg21678461

cg21679730
cg21681109
cg21681320
cg21681377
cg21683105
cg21683390
cg21684179
cg21685808
cg21686287
cg21686944
cg21686987
cg21687740
cg21688152
cg21688264
cg21688288
cg21688533
cg21690050
cg21690064
cg21691893
cg21692159
cg21692246
cg21692434
cg21694161
cg21694941
cg21695830
cg21696256
cg21696975
cg21696987
cg21697512
cg21697851
cg21697944
cg21698185
cg21699044
cg21699124
cg21699506
cg21700214
cg21700332
cg21700772
cg21700774
cg21702431
cg21702971
cg21703255
cg21703322
cg21704448
cg21704793
cg21705324
cg21705897
cg21707172
cg21707187
cg21707633
cg21710255
cg21710324

cg21711132
cg21711827
cg21712322
cg21712377
cg21713614
cg21714266
cg21714557
cg21716061
cg21717508
cg21718267
cg21719418
cg21720373
cg21720385
cg21720802
cg21721022
cg21722170
cg21722242
cg21722785
cg21722795
cg21723707
cg21724798
cg21725716
cg21726079
cg21727141
cg21727145
cg21728840
cg21729798
cg21730118
cg21731304
cg21732056
cg21733013
cg21734996
cg21737020
cg21737421
cg21737444
cg21738443
cg21739530
cg21739719
cg21741284
cg21741562
cg21742048
cg21742790
cg21742836
cg21743826
cg21743867
cg21745595
cg21745612
cg21746275
cg21746991
cg21746997
cg21747070
cg21748201

cg21748223
cg21748418
cg21750428
cg21750887
cg21750986
cg21751023
cg21751684
cg21752270
cg21752413
cg21752525
cg21752578
cg21753226
cg21753682
cg21753973
cg21754194
cg21754201
cg21754388
cg21754400
cg21755087
cg21755755
cg21756208
cg21756647
cg21757872
cg21758126
cg21758171
cg21758446
cg21758672
cg21758962
cg21759331
cg21759685
cg21760128
cg21762695
cg21762788
cg21763211
cg21764190
cg21764358
cg21765235
cg21767537
cg21767657
cg21768418
cg21768539
cg21768702
cg21769093
cg21769117
cg21769820
cg21770617
cg21771125
cg21771682
cg21772178
cg21772773
cg21773162
cg21773633

cg21773665
cg21773967
cg21774338
cg21775122
cg21775463
cg21775570
cg21775619
cg21775668
cg21776417
cg21776813
cg21777095
cg21777364
cg21777553
cg21779088
cg21779611
cg21780393
cg21782409
cg21783183
cg21784036
cg21784274
cg21784383
cg21784716
cg21784917
cg21786289
cg21786381
cg21786601
cg21787567
cg21787848
cg21787965
cg21789008
cg21789085
cg21789545
cg21790570
cg21790851
cg21791023
cg21791662
cg21794994
cg21797001
cg21798630
cg21798815
cg21799346
cg21799469
cg21799473
cg21799607
cg21800400
cg21800989
cg21801176
cg21802055
cg21802703
cg21803927
cg21805731
cg21805788

cg21805880
cg21806015
cg21806100
cg21806139
cg21806242
cg21807215
cg21807262
cg21807594
cg21808053
cg21808287
cg21808937
cg21809054
cg21809913
cg21810108
cg21810793
cg21811143
cg21811896
cg21813377
cg21813905
cg21814238
cg21814290
cg21815220
cg21815667
cg21815807
cg21816308
cg21816330
cg21816849
cg21817858
cg21818274
cg21818713
cg21820652
cg21820677
cg21821214
cg21821833
cg21822146
cg21822187
cg21823425
cg21823426
cg21824620
cg21824985
cg21826978
cg21828173
cg21828319
cg21828432
cg21828517
cg21828629
cg21829336
cg21829923
cg21831174
cg21831463
cg21831467
cg21831485

cg21831649
cg21832142
cg21832537
cg21832546
cg21833076
cg21834207
cg21834684
cg21834754
cg21836117
cg21836358
cg21836370
cg21837308
cg21837323
cg21837358
cg21837569
cg21837826
cg21838554
cg21838640
cg21838979
cg21839588
cg21841963
cg21844184
cg21844291
cg21845052
cg21845182
cg21845300
cg21846009
cg21846159
cg21846461
cg21846488
cg21846790
cg21846949
cg21846970
cg21848407
cg21849844
cg21850518
cg21850722
cg21851351
cg21851395
cg21851672
cg21851956
cg21852117
cg21852342
cg21852408
cg21852439
cg21852513
cg21853510
cg21853568
cg21853587
cg21854348
cg21854408
cg21855205

cg21855316
cg21855392
cg21855816
cg21856067
cg21856603
cg21856670
cg21857190
cg21858108
cg21858485
cg21858961
cg21859019
cg21859290
cg21859562
cg21859781
cg21860285
cg21860968
cg21861192
cg21862081
cg21863546
cg21863721
cg21863949
cg21864016
cg21864732
cg21865150
cg21865751
cg21865845
cg21868699
cg21868779
cg21869027
cg21869586
cg21870229
cg21870274
cg21870545
cg21871091
cg21872009
cg21872204
cg21873967
cg21874404
cg21874706
cg21875330
cg21875401
cg21875839
cg21876925
cg21877680
cg21877907
cg21878275
cg21878598
cg21878650
cg21878759
cg21878934
cg21879716
cg21880020

cg21880128
cg21881184
cg21881330
cg21883291
cg21883598
cg21884064
cg21884122
cg21884231
cg21884421
cg21884509
cg21885046
cg21885107
cg21885251
cg21886089
cg21886186
cg21887097
cg21887193
cg21887430
cg21887591
cg21888330
cg21889448
cg21889703
cg21890423
cg21892295
cg21892967
cg21893764
cg21894914
cg21895324
cg21895387
cg21895526
cg21896766
cg21896800
cg21898884
cg21898944
cg21899310
cg21899461
cg21899473
cg21899542
cg21899939
cg21900069
cg21900073
cg21901643
cg21901718
cg21902327
cg21902817
cg21902964
cg21903221
cg21903684
cg21904329
cg21904807
cg21906947
cg21907107

cg21907579
cg21908129
cg21908235
cg21908420
cg21908886
cg21908960
cg21908972
cg21909390
cg21910196
cg21910355
cg21910673
cg21910709
cg21911343
cg21913159
cg21913301
cg21913319
cg21913528
cg21913652
cg21913681
cg21913787
cg21913888
cg21913981
cg21915313
cg21915962
cg21916108
cg21917090
cg21917728
cg21918126
cg21918969
cg21919257
cg21919719
cg21919820
cg21919900
cg21919968
cg21920859
cg21921361
cg21921384
cg21922574
cg21924447
cg21924449
cg21924472
cg21925310
cg21925465
cg21926038
cg21926708
cg21926782
cg21928701
cg21929183
cg21929472
cg21929473
cg21929784
cg21929943

cg21930668
cg21930712
cg21930938
cg21931174
cg21931713
cg21932542
cg21932672
cg21932934
cg21933495
cg21934273
cg21934504
cg21935402
cg21936261
cg21936906
cg21937128
cg21937395
cg21937969
cg21938148
cg21938261
cg21938436
cg21938718
cg21938894
cg21940220
cg21941611
cg21942150
cg21942184
cg21942490
cg21942572
cg21942734
cg21943117
cg21943416
cg21944224
cg21945260
cg21945989
cg21946006
cg21946631
cg21946698
cg21946824
cg21949333
cg21949781
cg21949830
cg21950037
cg21950720
cg21951478
cg21951729
cg21953058
cg21953905
cg21956318
cg21957191
cg21958090
cg21958743
cg21960303

cg21960364
cg21961487
cg21961694
cg21961766
cg21962204
cg21962316
cg21962423
cg21962760
cg21963436
cg21963768
cg21963854
cg21963920
cg21964481
cg21964551
cg21964798
cg21965995
cg21966764
cg21966975
cg21967668
cg21967789
cg21967790
cg21969395
cg21969477
cg21970438
cg21970554
cg21971621
cg21971800
cg21973914
cg21974506
cg21974580
cg21975390
cg21977057
cg21977075
cg21977377
cg21977522
cg21978128
cg21979032
cg21979559
cg21981096
cg21981144
cg21982455
cg21983192
cg21983520
cg21984645
cg21985251
cg21987785
cg21988244
cg21990700
cg21991633
cg21993547
cg21994818
cg21995304

cg21995800
cg21996227
cg21997401
cg21997465
cg21997510
cg21997672
cg21997746
cg21998983
cg21999471
cg21999591
cg21999726
cg21999939
cg22001496
cg22001533
cg22001557
cg22004069
cg22004422
cg22004830
cg22005145
cg22006708
cg22007666
cg22008551
cg22008851
cg22009923
cg22010052
cg22010446
cg22011804
cg22012156
cg22012248
cg22012281
cg22012476
cg22012759
cg22013019
cg22014112
cg22014430
cg22014491
cg22014887
cg22014891
cg22015486
cg22015527
cg22017213
cg22019247
cg22020299
cg22021597
cg22021934
cg22021975
cg22022067
cg22022379
cg22022798
cg22022957
cg22025849
cg22027075

cg22028075
cg22028624
cg22029284
cg22029396
cg22029774
cg22030072
cg22030890
cg22032137
cg22032364
cg22032385
cg22032900
cg22033586
cg22033604
cg22033759
cg22034269
cg22034968
cg22035229
cg22035259
cg22035738
cg22035959
cg22036879
cg22036966
cg22037030
cg22038124
cg22038738
cg22039909
cg22039960
cg22040469
cg22040631
cg22042348
cg22042908
cg22043168
cg22044002
cg22044342
cg22044630
cg22044951
cg22045977
cg22046201
cg22047013
cg22047387
cg22048348
cg22048474
cg22048922
cg22048936
cg22049569
cg22050495
cg22052659
cg22052720
cg22052948
cg22053720
cg22054362
cg22054793

cg22054912
cg22055397
cg22056336
cg22056544
cg22057680
cg22059073
cg22059469
cg22061561
cg22061628
cg22061831
cg22061832
cg22062984
cg22063369
cg22063654
cg22065513
cg22065514
cg22065614
cg22065851
cg22065923
cg22066599
cg22067069
cg22068464
cg22068629
cg22068687
cg22068747
cg22070406
cg22070991
cg22071015
cg22072375
cg22073152
cg22073332
cg22073468
cg22073750
cg22074114
cg22074467
cg22074950
cg22075222
cg22075362
cg22075918
cg22076123
cg22076676
cg22076769
cg22077197
cg22077553
cg22079008
cg22079077
cg22079098
cg22079161
cg22079685
cg22080436
cg22081879
cg22081905

cg22082015
cg22082046
cg22082176
cg22082350
cg22082438
cg22082462
cg22082709
cg22083166
cg22083298
cg22085702
cg22086834
cg22087053
cg22089701
cg22091053
cg22091236
cg22091421
cg22092292
cg22092397
cg22094045
cg22094551
cg22094781
cg22095604
cg22096465
cg22096687
cg22097425
cg22098115
cg22098660
cg22101141
cg22101701
cg22101977
cg22104103
cg22105145
cg22105162
cg22105545
cg22106280
cg22106810
cg22107147
cg22108469
cg22108646
cg22108980
cg22109530
cg22110267
cg22110922
cg22111078
cg22112267
cg22112360
cg22112587
cg22115076
cg22117438
cg22117498
cg22117666
cg22117819

cg22117918
cg22118082
cg22119466
cg22120095
cg22120825
cg22121495
cg22122013
cg22122321
cg22122808
cg22122862
cg22123464
cg22123865
cg22124590
cg22124859
cg22125176
cg22125717
cg22125805
cg22126171
cg22126211
cg22126505
cg22127570
cg22127773
cg22128954
cg22129111
cg22129180
cg22129183
cg22129220
cg22129241
cg22129276
cg22129809
cg22131234
cg22131691
cg22132982
cg22134162
cg22134325
cg22134923
cg22135064
cg22135102
cg22135105
cg22135561
cg22136363
cg22136365
cg22136749
cg22137236
cg22137572
cg22137815
cg22138564
cg22138627
cg22139363
cg22141456
cg22142137
cg22143064

cg22143387
cg22143766
cg22144328
cg22144875
cg22145445
cg22146612
cg22147084
cg22148827
cg22150051
cg22150335
cg22151644
cg22151773
cg22152407
cg22152446
cg22152605
cg22152728
cg22153181
cg22153994
cg22154447
cg22154659
cg22154859
cg22156126
cg22156700
cg22156899
cg22157629
cg22158462
cg22158650
cg22159815
cg22159835
cg22160263
cg22160463
cg22161476
cg22162097
cg22162435
cg22162638
cg22162757
cg22163631
cg22165175
cg22166883
cg22167515
cg22167763
cg22169427
cg22169791
cg22169874
cg22169990
cg22170177
cg22170457
cg22170801
cg22172610
cg22173632
cg22175494
cg22175873

cg22176913
cg22178613
cg22178798
cg22179082
cg22179905
cg22180580
cg22180738
cg22181664
cg22182666
cg22183016
cg22183706
cg22183731
cg22184096
cg22184136
cg22184210
cg22186216
cg22186503
cg22187125
cg22187630
cg22187733
cg22188918
cg22189286
cg22190361
cg22190437
cg22191665
cg22192374
cg22193385
cg22193912
cg22195627
cg22196977
cg22197050
cg22197830
cg22199118
cg22202689
cg22202692
cg22203237
cg22203628
cg22203776
cg22205607
cg22206370
cg22207135
cg22207405
cg22207584
cg22207732
cg22208022
cg22208536
cg22209010
cg22209573
cg22210145
cg22210627
cg22211507
cg22211672

cg22212218
cg22212250
cg22212691
cg22213022
cg22214349
cg22214800
cg22215508
cg22215942
cg22216015
cg22216413
cg22218806
cg22219089
cg22219379
cg22219450
cg22219913
cg22220502
cg22220722
cg22220806
cg22220986
cg22221870
cg22222663
cg22222999
cg22223402
cg22224043
cg22224704
cg22226131
cg22226775
cg22227306
cg22228748
cg22228988
cg22229016
cg22229114
cg22229594
cg22229906
cg22231424
cg22231602
cg22231902
cg22232500
cg22233189
cg22233974
cg22234119
cg22234345
cg22235080
cg22235258
cg22235407
cg22236681
cg22236711
cg22237425
cg22237937
cg22238777
cg22239727
cg22239963

cg22240472
cg22240515
cg22241124
cg22241820
cg22242539
cg22242614
cg22244122
cg22244192
cg22245446
cg22245494
cg22245615
cg22246344
cg22247041
cg22248052
cg22249291
cg22249754
cg22250390
cg22250531
cg22250556
cg22250605
cg22253945
cg22254135
cg22254313
cg22254464
cg22254606
cg22255095
cg22256354
cg22259097
cg22259126
cg22259765
cg22259831
cg22260508
cg22260625
cg22261207
cg22261467
cg22262704
cg22263131
cg22263591
cg22264930
cg22265644
cg22266009
cg22266072
cg22266693
cg22268164
cg22268341
cg22272803
cg22272840
cg22274395
cg22274414
cg22274539
cg22274813
cg22275276

cg22277936
cg22278782
cg22279027
cg22280038
cg22280333
cg22280475
cg22280671
cg22281380
cg22281494
cg22282089
cg22282405
cg22282410
cg22283058
cg22283643
cg22283754
cg22283925
cg22283959
cg22284311
cg22284398
cg22284860
cg22285914
cg22286382
cg22286936
cg22287731
cg22287800
cg22289115
cg22289815
cg22290648
cg22291153
cg22291974
cg22293823
cg22294181
cg22294577
cg22294740
cg22295787
cg22295864
cg22295925
cg22298437
cg22298487
cg22298860
cg22299454
cg22299782
cg22300282
cg22301128
cg22302772
cg22302985
cg22303227
cg22304262
cg22304522
cg22305167
cg22305209
cg22305574

cg22306038
cg22306408
cg22306928
cg22307297
cg22307444
cg22307471
cg22307508
cg22308101
cg22309632
cg22310279
cg22310648
cg22310770
cg22311200
cg22311615
cg22314684
cg22315844
cg22315928
cg22317989
cg22318514
cg22318991
cg22319784
cg22320059
cg22320365
cg22320962
cg22321318
cg22322343
cg22322465
cg22322562
cg22322679
cg22324487
cg22324567
cg22325673
cg22325764
cg22325828
cg22326681
cg22327543
cg22327646
cg22328396
cg22329515
cg22329579
cg22329776
cg22330021
cg22330467
cg22330625
cg22330638
cg22330841
cg22330875
cg22330924
cg22331862
cg22332076
cg22332257
cg22332722

cg22333583
cg22333663
cg22333960
cg22334059
cg22335261
cg22335340
cg22335847
cg22335872
cg22335876
cg22337128
cg22337407
cg22337624
cg22337920
cg22338446
cg22338567
cg22338798
cg22339219
cg22340475
cg22340508
cg22340811
cg22341256
cg22341471
cg22341513
cg22341788
cg22341848
cg22342100
cg22342329
cg22343083
cg22344188
cg22344581
cg22344611
cg22344793
cg22345214
cg22346124
cg22346461
cg22346476
cg22346765
cg22346959
cg22347967
cg22349442
cg22350070
cg22350438
cg22352772
cg22353329
cg22353551
cg22354595
cg22354646
cg22354782
cg22355224
cg22355506
cg22355517
cg22357679

cg22358066
cg22358121
cg22360016
cg22360318
cg22361106
cg22361859
cg22363368
cg22363783
cg22365313
cg22366799
cg22366938
cg22367191
cg22367250
cg22367549
cg22367556
cg22368204
cg22368262
cg22368281
cg22368664
cg22369420
cg22369590
cg22369786
cg22370006
cg22370716
cg22371492
cg22371972
cg22372194
cg22372285
cg22372668
cg22372815
cg22373221
cg22374057
cg22374165
cg22374233
cg22374742
cg22374940
cg22375192
cg22375763
cg22376214
cg22376262
cg22376706
cg22376805
cg22377043
cg22378095
cg22378459
cg22379436
cg22379463
cg22379708
cg22379915
cg22380139
cg22380663
cg22380921

cg22381154
cg22381590
cg22383292
cg22383874
cg22385297
cg22385463
cg22385702
cg22386008
cg22386073
cg22386311
cg22386774
cg22387369
cg22387575
cg22388232
cg22388347
cg22388608
cg22388948
cg22389209
cg22389375
cg22389950
cg22391205
cg22392178
cg22395811
cg22396119
cg22396498
cg22396555
cg22398226
cg22399133
cg22399646
cg22399790
cg22399984
cg22400015
cg22400454
cg22401262
cg22401679
cg22402753
cg22402850
cg22403811
cg22405226
cg22405653
cg22408108
cg22408430
cg22409100
cg22410057
cg22411165
cg22412536
cg22412680
cg22413603
cg22414525
cg22415472
cg22416002
cg22416376

cg22417613
cg22418307
cg22418337
cg22418565
cg22418620
cg22418829
cg22418909
cg22420044
cg22420514
cg22421579
cg22421804
cg22422090
cg22423207
cg22424213
cg22424444
cg22424746
cg22425186
cg22425466
cg22426139
cg22427208
cg22427313
cg22428027
cg22428147
cg22428541
cg22429121
cg22430565
cg22430700
cg22430861
cg22430985
cg22431753
cg22431967
cg22433418
cg22434409
cg22434594
cg22434923
cg22435313
cg22436103
cg22436527
cg22437074
cg22437276
cg22437987
cg22438763
cg22438810
cg22439792
cg22439922
cg22441157
cg22441882
cg22442090
cg22442730
cg22443102
cg22443330
cg22443940

cg22446206
cg22449085
cg22449745
cg22450138
cg22450342
cg22450968
cg22451233
cg22451412
cg22451632
cg22452520
cg22453435
cg22454022
cg22454313
cg22454371
cg22454564
cg22454686
cg22455082
cg22455694
cg22455914
cg22457238
cg22459630
cg22459739
cg22460173
cg22460204
cg22460896
cg22461018
cg22461324
cg22462109
cg22462240
cg22465362
cg22466350
cg22466678
cg22467594
cg22468258
cg22471129
cg22471401
cg22472023
cg22472304
cg22472488
cg22473637
cg22474464
cg22476252
cg22476268
cg22476504
cg22476960
cg22477971
cg22478258
cg22478317
cg22480742
cg22481162
cg22481632
cg22481960

cg22482700
cg22484737
cg22485363
cg22486834
cg22486928
cg22488081
cg22488256
cg22488278
cg22488719
cg22488970
cg22489321
cg22489498
cg22489759
cg22489883
cg22490454
cg22490780
cg22490845
cg22490861
cg22490991
cg22491141
cg22491548
cg22491927
cg22494078
cg22494876
cg22494907
cg22495008
cg22495310
cg22495388
cg22495968
cg22496729
cg22496809
cg22497336
cg22497847
cg22497856
cg22497907
cg22498676
cg22498784
cg22499381
cg22499797
cg22499809
cg22499893
cg22500738
cg22501143
cg22501604
cg22504024
cg22504204
cg22504523
cg22504898
cg22505977
cg22507023
cg22507069
cg22507154

cg22507406
cg22507727
cg22508338
cg22508831
cg22508969
cg22509057
cg22509138
cg22509164
cg22509504
cg22509845
cg22510662
cg22511368
cg22511889
cg22512663
cg22512830
cg22512888
cg22515475
cg22515654
cg22516643
cg22516975
cg22517356
cg22518208
cg22519158
cg22519356
cg22519420
cg22519422
cg22522920
cg22523050
cg22523074
cg22523679
cg22523852
cg22524061
cg22524065
cg22524373
cg22524657
cg22526555
cg22528044
cg22528172
cg22529645
cg22529822
cg22529990
cg22530462
cg22530519
cg22530668
cg22531266
cg22532234
cg22532370
cg22533099
cg22533517
cg22533606
cg22533689
cg22534145

cg22534374
cg22534509
cg22534585
cg22534724
cg22535349
cg22537167
cg22537474
cg22537597
cg22538355
cg22538780
cg22539420
cg22539670
cg22539738
cg22539744
cg22541254
cg22541400
cg22541735
cg22542222
cg22542373
cg22542803
cg22543374
cg22543892
cg22544881
cg22545168
cg22545294
cg22545356
cg22545385
cg22546130
cg22546168
cg22546374
cg22547540
cg22548438
cg22548485
cg22548714
cg22548735
cg22549086
cg22549408
cg22549610
cg22549870
cg22550847
cg22551546
cg22551578
cg22552966
cg22553301
cg22555517
cg22556143
cg22557091
cg22557662
cg22559013
cg22561203
cg22561329
cg22561727

cg22561757
cg22562494
cg22562853
cg22562942
cg22563937
cg22565248
cg22567315
cg22568703
cg22569359
cg22570805
cg22570970
cg22571654
cg22571664
cg22572159
cg22573118
cg22573731
cg22574202
cg22574825
cg22575043
cg22575261
cg22575603
cg22576976
cg22578373
cg22578425
cg22578926
cg22579075
cg22579828
cg22583147
cg22583148
cg22583305
cg22583981
cg22584138
cg22585927
cg22586221
cg22586603
cg22586653
cg22588144
cg22588275
cg22588850
cg22589161
cg22589628
cg22589728
cg22589778
cg22590902
cg22591875
cg22593432
cg22593953
cg22594416
cg22595235
cg22595920
cg22596016
cg22596049

cg22596861
cg22597040
cg22597733
cg22598247
cg22598669
cg22599122
cg22599150
cg22599740
cg22600043
cg22601348
cg22602592
cg22605303
cg22606205
cg22606224
cg22608160
cg22609474
cg22609497
cg22609511
cg22610106
cg22611308
cg22611523
cg22612642
cg22614239
cg22614521
cg22616255
cg22616430
cg22616525
cg22617682
cg22618337
cg22619018
cg22619376
cg22620027
cg22620090
cg22620111
cg22620221
cg22621663
cg22622676
cg22623033
cg22623223
cg22623319
cg22624022
cg22624147
cg22624391
cg22624697
cg22625098
cg22626522
cg22626985
cg22627406
cg22627427
cg22627570
cg22627753
cg22627841

cg22628512
cg22628608
cg22628926
cg22629065
cg22629266
cg22629375
cg22630169
cg22630875
cg22633024
cg22633590
cg22633936
cg22634633
cg22634891
cg22635088
cg22635402
cg22635491
cg22636722
cg22637867
cg22638461
cg22639011
cg22639242
cg22640233
cg22641280
cg22642777
cg22642909
cg22643811
cg22645427
cg22645961
cg22646210
cg22646408
cg22646995
cg22647031
cg22647713
cg22647738
cg22648135
cg22648996
cg22649181
cg22649187
cg22650099
cg22650458
cg22650617
cg22650942
cg22653878
cg22653976
cg22654142
cg22654517
cg22655436
cg22655696
cg22656940
cg22657044
cg22657059
cg22657457

cg22657632
cg22657682
cg22658311
cg22658316
cg22659564
cg22659676
cg22660033
cg22660196
cg22660578
cg22661893
cg22663859
cg22663933
cg22664258
cg22664296
cg22664425
cg22664614
cg22665096
cg22665383
cg22665692
cg22666015
cg22667294
cg22668218
cg22668740
cg22668995
cg22669120
cg22669623
cg22669656
cg22670675
cg22670868
cg22671611
cg22671986
cg22672060
cg22672790
cg22674699
cg22675598
cg22675604
cg22675767
cg22675937
cg22677100
cg22677240
cg22677650
cg22677715
cg22678044
cg22678136
cg22679014
cg22679740
cg22680204
cg22680536
cg22681190
cg22681279
cg22681344
cg22681721

cg22682172
cg22682200
cg22682646
cg22682739
cg22683277
cg22684316
cg22685215
cg22685409
cg22685502
cg22685822
cg22686240
cg22686253
cg22686718
cg22686722
cg22687807
cg22688535
cg22689597
cg22689833
cg22689904
cg22690720
cg22691587
cg22691776
cg22691824
cg22691926
cg22692545
cg22693055
cg22693994
cg22694818
cg22695532
cg22695729
cg22695986
cg22696426
cg22696438
cg22696652
cg22697325
cg22697365
cg22697821
cg22697825
cg22698115
cg22698177
cg22698272
cg22699026
cg22699361
cg22699462
cg22699620
cg22700148
cg22700220
cg22700848
cg22701672
cg22702328
cg22702443
cg22702857

cg22703124
cg22704057
cg22705225
cg22706106
cg22706610
cg22706883
cg22707016
cg22707207
cg22707381
cg22707446
cg22707857
cg22708087
cg22708150
cg22708818
cg22708897
cg22709144
cg22711111
cg22711676
cg22712403
cg22713752
cg22714668
cg22716262
cg22716738
cg22716949
cg22716950
cg22717014
cg22717080
cg22717825
cg22719004
cg22719902
cg22719992
cg22720041
cg22720431
cg22721334
cg22722340
cg22722777
cg22724153
cg22724228
cg22724921
cg22725222
cg22725685
cg22725830
cg22726155
cg22726349
cg22727810
cg22728782
cg22728904
cg22729681
cg22729726
cg22729973
cg22730007
cg22730636

cg22731271
cg22731637
cg22731763
cg22732253
cg22732731
cg22732749
cg22733133
cg22734058
cg22734236
cg22735331
cg22735650
cg22736144
cg22736483
cg22736787
cg22736807
cg22736956
cg22738219
cg22738289
cg22738642
cg22739115
cg22739858
cg22740783
cg22741264
cg22742965
cg22743617
cg22743884
cg22744818
cg22745102
cg22745143
cg22746058
cg22746681
cg22746865
cg22747092
cg22747142
cg22747380
cg22747507
cg22747746
cg22748407
cg22748722
cg22749589
cg22749642
cg22749855
cg22751954
cg22752023
cg22754866
cg22755445
cg22756866
cg22757362
cg22757399
cg22757447
cg22757824
cg22758104

cg22758454
cg22758806
cg22758846
cg22758916
cg22759096
cg22759823
cg22760037
cg22760338
cg22760420
cg22762091
cg22762492
cg22763649
cg22763718
cg22764289
cg22765178
cg22765299
cg22765612
cg22768222
cg22769941
cg22770135
cg22771332
cg22772440
cg22772698
cg22772930
cg22773522
cg22773899
cg22775138
cg22775600
cg22776451
cg22776856
cg22777186
cg22777724
cg22777952
cg22778014
cg22778435
cg22778788
cg22779896
cg22779991
cg22780174
cg22780193
cg22780242
cg22780731
cg22782271
cg22783180
cg22784836
cg22784954
cg22786472
cg22786748
cg22788065
cg22789065
cg22789545
cg22789982

cg22791030
cg22791045
cg22792674
cg22794304
cg22795331
cg22795586
cg22796355
cg22796507
cg22796572
cg22797031
cg22797461
cg22797735
cg22797773
cg22798121
cg22798201
cg22798214
cg22798707
cg22798737
cg22799141
cg22799691
cg22799963
cg22801561
cg22801690
cg22801949
cg22802102
cg22802439
cg22802461
cg22803222
cg22803223
cg22803285
cg22803642
cg22803868
cg22805381
cg22807449
cg22807576
cg22807585
cg22807822
cg22809653
cg22809683
cg22809798
cg22809826
cg22809871
cg22810403
cg22810423
cg22810434
cg22811478
cg22812275
cg22812280
cg22812955
cg22813711
cg22813950
cg22814801

cg22815110
cg22816909
cg22817258
cg22817993
cg22818319
cg22820108
cg22821300
cg22821355
cg22821554
cg22822599
cg22822630
cg22823146
cg22823273
cg22823644
cg22824738
cg22825230
cg22825871
cg22826333
cg22827827
cg22827864
cg22828110
cg22828383
cg22828602
cg22829706
cg22829917
cg22830507
cg22830638
cg22830694
cg22830707
cg22831029
cg22832061
cg22832254
cg22833175
cg22833603
cg22834052
cg22834443
cg22836229
cg22836400
cg22836769
cg22836787
cg22837015
cg22837321
cg22837763
cg22837767
cg22838079
cg22838399
cg22838685
cg22838882
cg22839180
cg22840361
cg22841810
cg22842189

cg22843146
cg22843424
cg22843538
cg22843950
cg22844862
cg22845912
cg22846067
cg22846582
cg22847391
cg22847608
cg22848598
cg22848646
cg22848704
cg22849635
cg22850505
cg22850587
cg22851295
cg22851420
cg22851557
cg22851694
cg22852013
cg22852353
cg22852503
cg22852840
cg22853427
cg22853542
cg22855020
cg22855237
cg22855933
cg22855968
cg22855988
cg22856324
cg22856496
cg22857085
cg22858169
cg22859061
cg22860740
cg22860848
cg22861587
cg22862094
cg22862112
cg22862319
cg22862420
cg22862635
cg22864077
cg22864340
cg22864462
cg22864519
cg22865470
cg22865713
cg22865769
cg22865908

cg22866015
cg22866121
cg22866754
cg22866875
cg22867063
cg22867719
cg22867816
cg22868282
cg22868943
cg22869030
cg22869726
cg22869944
cg22871098
cg22871318
cg22871653
cg22871668
cg22872002
cg22872478
cg22872549
cg22873167
cg22874305
cg22874407
cg22874893
cg22875410
cg22875527
cg22875823
cg22875984
cg22876356
cg22876812
cg22876908
cg22877024
cg22877504
cg22878054
cg22878148
cg22878441
cg22878489
cg22879667
cg22880677
cg22881033
cg22881174
cg22881266
cg22881325
cg22881914
cg22882121
cg22882665
cg22882902
cg22883332
cg22883472
cg22884082
cg22884714
cg22885558
cg22886575

cg22888055
cg22888418
cg22888848
cg22889163
cg22889448
cg22889573
cg22889887
cg22890545
cg22891099
cg22891191
cg22891500
cg22891707
cg22891862
cg22892860
cg22892904
cg22893605
cg22894276
cg22894301
cg22894983
cg22895969
cg22897419
cg22898797
cg22899392
cg22900075
cg22900217
cg22902532
cg22902550
cg22904815
cg22906524
cg22906692
cg22907034
cg22907258
cg22907952
cg22909759
cg22910382
cg22910449
cg22911184
cg22911200
cg22911422
cg22911462
cg22912610
cg22912701
cg22914595
cg22914729
cg22916722
cg22917314
cg22919728
cg22919908
cg22920258
cg22920300
cg22920586
cg22920603

cg22923409
cg22923895
cg22924250
cg22924840
cg22924958
cg22925840
cg22927076
cg22927302
cg22927788
cg22928016
cg22928082
cg22928599
cg22930445
cg22930650
cg22930773
cg22931085
cg22931182
cg22931604
cg22931622
cg22931795
cg22931822
cg22932649
cg22933107
cg22933549
cg22934308
cg22935609
cg22936990
cg22937462
cg22937649
cg22938061
cg22938901
cg22939709
cg22940372
cg22941178
cg22941646
cg22941974
cg22942576
cg22942804
cg22944499
cg22946066
cg22946150
cg22946562
cg22946774
cg22947000
cg22947965
cg22948288
cg22948791
cg22948959
cg22949077
cg22949256
cg22950223
cg22950831

cg22950899
cg22950904
cg22950982
cg22951411
cg22952459
cg22952787
cg22953017
cg22953267
cg22953865
cg22954171
cg22954449
cg22954810
cg22954818
cg22955555
cg22955595
cg22956023
cg22957898
cg22958047
cg22958090
cg22958248
cg22959894
cg22961335
cg22962923
cg22963382
cg22963554
cg22963629
cg22963843
cg22963979
cg22965752
cg22966302
cg22967008
cg22967080
cg22967140
cg22968936
cg22969666
cg22971038
cg22971153
cg22971157
cg22971191
cg22971594
cg22971600
cg22971939
cg22972318
cg22972989
cg22974210
cg22974463
cg22975521
cg22975568
cg22975913
cg22976218
cg22976224
cg22976366

cg22977317
cg22978087
cg22979783
cg22979807
cg22980483
cg22980771
cg22980833
cg22981158
cg22982570
cg22983083
cg22984256
cg22984631
cg22984992
cg22986077
cg22987447
cg22987487
cg22988346
cg22988566
cg22988581
cg22989103
cg22989463
cg22989958
cg22990178
cg22991101
cg22991512
cg22992266
cg22992966
cg22994720
cg22995042
cg22995692
cg22996170
cg22996783
cg22997262
cg22998020
cg22998959
cg22999502
cg22999786
cg23001131
cg23001918
cg23002238
cg23002708
cg23002761
cg23003720
cg23003783
cg23003832
cg23003926
cg23004758
cg23004985
cg23006040
cg23006227
cg23007325
cg23008718

cg23009042
cg23009441
cg23009818
cg23010077
cg23010538
cg23010585
cg23010900
cg23010913
cg23012054
cg23012579
cg23012591
cg23012700
cg23013137
cg23013402
cg23013473
cg23013664
cg23013850
cg23014549
cg23015341
cg23015361
cg23015605
cg23016129
cg23016600
cg23016632
cg23016726
cg23016734
cg23016776
cg23017728
cg23018092
cg23018181
cg23018448
cg23018755
cg23019565
cg23020514
cg23020600
cg23020891
cg23021268
cg23021771
cg23024136
cg23024158
cg23024326
cg23024444
cg23024967
cg23025244
cg23025416
cg23025804
cg23026024
cg23026106
cg23026675
cg23026754
cg23026864
cg23027296

cg23027574
cg23028209
cg23028436
cg23028721
cg23029159
cg23029597
cg23029655
cg23030069
cg23030200
cg23030863
cg23031141
cg23031444
cg23031493
cg23032032
cg23032129
cg23032364
cg23032421
cg23033158
cg23034362
cg23034373
cg23034536
cg23034757
cg23036385
cg23037386
cg23037403
cg23037411
cg23037871
cg23040011
cg23041447
cg23041822
cg23042076
cg23042387
cg23042510
cg23042540
cg23042706
cg23042775
cg23043139
cg23043779
cg23044087
cg23044270
cg23044884
cg23045972
cg23045991
cg23047671
cg23049226
cg23049291
cg23049458
cg23049823
cg23051091
cg23052830
cg23053525
cg23054188

cg23054189
cg23054394
cg23054840
cg23054883
cg23055682
cg23055772
cg23056157
cg23056923
cg23057567
cg23057597
cg23057648
cg23058185
cg23058719
cg23058745
cg23059452
cg23059709
cg23059797
cg23060047
cg23060801
cg23061725
cg23061795
cg23062096
cg23062112
cg23062198
cg23062810
cg23063047
cg23063070
cg23064082
cg23064481
cg23064601
cg23065100
cg23065185
cg23065934
cg23066159
cg23067228
cg23067856
cg23068499
cg23068621
cg23068623
cg23068913
cg23068939
cg23068989
cg23071186
cg23071791
cg23072051
cg23072408
cg23072559
cg23073467
cg23074732
cg23074881
cg23074932
cg23074996

cg23075139
cg23075337
cg23075364
cg23075555
cg23075771
cg23076913
cg23077606
cg23078228
cg23079808
cg23080908
cg23081213
cg23081357
cg23081527
cg23081542
cg23082003
cg23082281
cg23082387
cg23082454
cg23082862
cg23083984
cg23084016
cg23085281
cg23085662
cg23086176
cg23086243
cg23087149
cg23087992
cg23088551
cg23089272
cg23089445
cg23089825
cg23089838
cg23089840
cg23090046
cg23090529
cg23090824
cg23091122
cg23091159
cg23091549
cg23092040
cg23092086
cg23092820
cg23093217
cg23093404
cg23093589
cg23093609
cg23093754
cg23093846
cg23093870
cg23094903
cg23095192
cg23095451

cg23095556
cg23095584
cg23095729
cg23095743
cg23096144
cg23096806
cg23097296
cg23097402
cg23097534
cg23097961
cg23098068
cg23098069
cg23098305
cg23098371
cg23098529
cg23098803
cg23099740
cg23100626
cg23102197
cg23102514
cg23104248
cg23104436
cg23104539
cg23105839
cg23106115
cg23106912
cg23106940
cg23107301
cg23107916
cg23109206
cg23109714
cg23109897
cg23110019
cg23110407
cg23110422
cg23110514
cg23110957
cg23111106
cg23111196
cg23111830
cg23112326
cg23112425
cg23112950
cg23114183
cg23114435
cg23115083
cg23117999
cg23119026
cg23119463
cg23119487
cg23119977
cg23120541

cg23120725
cg23120936
cg23121156
cg23121170
cg23121993
cg23122109
cg23122650
cg23125492
cg23128056
cg23128495
cg23128949
cg23129024
cg23129374
cg23129573
cg23130097
cg23130131
cg23130165
cg23130254
cg23130369
cg23130849
cg23131355
cg23135908
cg23138161
cg23138250
cg23138373
cg23138413
cg23141902
cg23142048
cg23142394
cg23142799
cg23143090
cg23143104
cg23143313
cg23144445
cg23144681
cg23146663
cg23146741
cg23146998
cg23147597
cg23148701
cg23149072
cg23149454
cg23152742
cg23153661
cg23153991
cg23154021
cg23154270
cg23154495
cg23154497
cg23154781
cg23154849
cg23155911

cg23156226
cg23156711
cg23156916
cg23157239
cg23158500
cg23158731
cg23159250
cg23159678
cg23160016
cg23162479
cg23162647
cg23163573
cg23163653
cg23164076
cg23164535
cg23164982
cg23165166
cg23165899
cg23166252
cg23166765
cg23166773
cg23166781
cg23166857
cg23166865
cg23167287
cg23167425
cg23168531
cg23169051
cg23169059
cg23169762
cg23169957
cg23170057
cg23170410
cg23170439
cg23170495
cg23170988
cg23171103
cg23172057
cg23172853
cg23172892
cg23172995
cg23173817
cg23174201
cg23174481
cg23175023
cg23175533
cg23176577
cg23176632
cg23177095
cg23179265
cg23179976
cg23180841

cg23181035
cg23181133
cg23181170
cg23181282
cg23181580
cg23183422
cg23183568
cg23184556
cg23184690
cg23186823
cg23187159
cg23187669
cg23187727
cg23188704
cg23188950
cg23189044
cg23189175
cg23189410
cg23190190
cg23190719
cg23191354
cg23191801
cg23191956
cg23192749
cg23192824
cg23193059
cg23193365
cg23193616
cg23193639
cg23194295
cg23194364
cg23195687
cg23195753
cg23196720
cg23196831
cg23198262
cg23198334
cg23198558
cg23198793
cg23198815
cg23200020
cg23200506
cg23201199
cg23201265
cg23201575
cg23201817
cg23202058
cg23202887
cg23202913
cg23203234
cg23203302
cg23204096

cg23204113
cg23204707
cg23205034
cg23205387
cg23205454
cg23206209
cg23206630
cg23206745
cg23206822
cg23206873
cg23207054
cg23207304
cg23207412
cg23207527
cg23207542
cg23207990
cg23208271
cg23208430
cg23208513
cg23208590
cg23209976
cg23209990
cg23210075
cg23210303
cg23211240
cg23212916
cg23213876
cg23214764
cg23216612
cg23217126
cg23217512
cg23217527
cg23217575
cg23217946
cg23217965
cg23219748
cg23220343
cg23220435
cg23220439
cg23221737
cg23222057
cg23224042
cg23225050
cg23225103
cg23225476
cg23225508
cg23227824
cg23227851
cg23228019
cg23228178
cg23228279
cg23229770

cg23230362
cg23230564
cg23230584
cg23230830
cg23230904
cg23230929
cg23231163
cg23232188
cg23232719
cg23232755
cg23234726
cg23234811
cg23234847
cg23235989
cg23236165
cg23236554
cg23237183
cg23237733
cg23238293
cg23238677
cg23238790
cg23238834
cg23238918
cg23239150
cg23240171
cg23240927
cg23242052
cg23242898
cg23243267
cg23243343
cg23243408
cg23243463
cg23243867
cg23243982
cg23244226
cg23244253
cg23244289
cg23244488
cg23244540
cg23244559
cg23245835
cg23246885
cg23247191
cg23247945
cg23247968
cg23248351
cg23248357
cg23248887
cg23249140
cg23250593
cg23250750
cg23251170

cg23251200
cg23251746
cg23252243
cg23253053
cg23253569
cg23253752
cg23254393
cg23254528
cg23254538
cg23255934
cg23256150
cg23256802
cg23257289
cg23257461
cg23257697
cg23257859
cg23257935
cg23258585
cg23258611
cg23259289
cg23259681
cg23260502
cg23260554
cg23260573
cg23260578
cg23260625
cg23261375
cg23261443
cg23261640
cg23261743
cg23262218
cg23262312
cg23262325
cg23262897
cg23262973
cg23263802
cg23263937
cg23264682
cg23265210
cg23266063
cg23266594
cg23266743
cg23267550
cg23267665
cg23267759
cg23267987
cg23268102
cg23268630
cg23270523
cg23270841
cg23272978
cg23273418

cg23273758
cg23273809
cg23273843
cg23274824
cg23274984
cg23275064
cg23275972
cg23277328
cg23277830
cg23278137
cg23278235
cg23278885
cg23279261
cg23279503
cg23279522
cg23279756
cg23280305
cg23281529
cg23281555
cg23283187
cg23284720
cg23286903
cg23287435
cg23287485
cg23288059
cg23288099
cg23288563
cg23288838
cg23289779
cg23290204
cg23290344
cg23291145
cg23291161
cg23291280
cg23291301
cg23291305
cg23291376
cg23291534
cg23291682
cg23291854
cg23291886
cg23293467
cg23294090
cg23295623
cg23296678
cg23298047
cg23298114
cg23299254
cg23299807
cg23300372
cg23300732
cg23301462

cg23302649
cg23302945
cg23302998
cg23304023
cg23304298
cg23304647
cg23305408
cg23305899
cg23307798
cg23309843
cg23310345
cg23310835
cg23312431
cg23313266
cg23313445
cg23313963
cg23314545
cg23315637
cg23316253
cg23317501
cg23317604
cg23318063
cg23318703
cg23318764
cg23318812
cg23318967
cg23319477
cg23320693
cg23321038
cg23321397
cg23322812
cg23322933
cg23323158
cg23323297
cg23323879
cg23324101
cg23325313
cg23325364
cg23325384
cg23325413
cg23325963
cg23326197
cg23327334
cg23328194
cg23328208
cg23328308
cg23328335
cg23328622
cg23328707
cg23329411
cg23330006
cg23330212

cg23330281
cg23330788
cg23331421
cg23331778
cg23331961
cg23332610
cg23332813
cg23333820
cg23333915
cg23334264
cg23335460
cg23335916
cg23336321
cg23336905
cg23337361
cg23337382
cg23337647
cg23338195
cg23338408
cg23338993
cg23339482
cg23339656
cg23340017
cg23340034
cg23340505
cg23341085
cg23342105
cg23342148
cg23342625
cg23342718
cg23343066
cg23343291
cg23343486
cg23344395
cg23345292
cg23345298
cg23345340
cg23345977
cg23346544
cg23346945
cg23347094
cg23347718
cg23347790
cg23347850
cg23348010
cg23348270
cg23348723
cg23348743
cg23349242
cg23349517
cg23349790
cg23349982

cg23350273
cg23350580
cg23350754
cg23350904
cg23351290
cg23351479
cg23351583
cg23351724
cg23351738
cg23352424
cg23353281
cg23353952
cg23355492
cg23356017
cg23356302
cg23356505
cg23357130
cg23357150
cg23357250
cg23357805
cg23357981
cg23358612
cg23359363
cg23359665
cg23360019
cg23361092
cg23361764
cg23361809
cg23362032
cg23362884
cg23363039
cg23363971
cg23364373
cg23365071
cg23365637
cg23366050
cg23367119
cg23367341
cg23367478
cg23368458
cg23369107
cg23369632
cg23369670
cg23370512
cg23370548
cg23370820
cg23370883
cg23371208
cg23371746
cg23372297
cg23372422
cg23372684

cg23372795
cg23373401
cg23375068
cg23375593
cg23375617
cg23375781
cg23375948
cg23377395
cg23378033
cg23378108
cg23378142
cg23378565
cg23380039
cg23380496
cg23380558
cg23381232
cg23381749
cg23383138
cg23383871
cg23384017
cg23384960
cg23385208
cg23388392
cg23389465
cg23390871
cg23392162
cg23392389
cg23392804
cg23393637
cg23393800
cg23394462
cg23394916
cg23395341
cg23395449
cg23395692
cg23395715
cg23396177
cg23396732
cg23397013
cg23398109
cg23398508
cg23399011
cg23399286
cg23400222
cg23400451
cg23400512
cg23400715
cg23401251
cg23401684
cg23402144
cg23402541
cg23402821

cg23403099
cg23403895
cg23403980
cg23404248
cg23404476
cg23405575
cg23406159
cg23408285
cg23408558
cg23409403
cg23411043
cg23411426
cg23413307
cg23413924
cg23413954
cg23414387
cg23414759
cg23414861
cg23415398
cg23415756
cg23415916
cg23416002
cg23416081
cg23416909
cg23416994
cg23418465
cg23419606
cg23419635
cg23420260
cg23420541
cg23421166
cg23421560
cg23422763
cg23423933
cg23424003
cg23424145
cg23424933
cg23425242
cg23425316
cg23425533
cg23426156
cg23426958
cg23429036
cg23429306
cg23429510
cg23430388
cg23432932
cg23433285
cg23433370
cg23433532
cg23434074
cg23434592

cg23435118
cg23435594
cg23435671
cg23436058
cg23436081
cg23436703
cg23437216
cg23438309
cg23438841
cg23438893
cg23440155
cg23440413
cg23441441
cg23442217
cg23442637
cg23444251
cg23444264
cg23447202
cg23448139
cg23448348
cg23448390
cg23448729
cg23449588
cg23449659
cg23449696
cg23449764
cg23450509
cg23451129
cg23452119
cg23452220
cg23453353
cg23453372
cg23454517
cg23456212
cg23456595
cg23456739
cg23458887
cg23459212
cg23459363
cg23460943
cg23461824
cg23461926
cg23462514
cg23462788
cg23463186
cg23463269
cg23463700
cg23464269
cg23465427
cg23465891
cg23466059
cg23466749

cg23468130
cg23468144
cg23468878
cg23469117
cg23470520
cg23471482
cg23472215
cg23472708
cg23473114
cg23473443
cg23473819
cg23474211
cg23474501
cg23474723
cg23475018
cg23477348
cg23477774
cg23479922
cg23479949
cg23480479
cg23480730
cg23481246
cg23482259
cg23482427
cg23482746
cg23482898
cg23483799
cg23483840
cg23484454
cg23484899
cg23485307
cg23485591
cg23486561
cg23487201
cg23487759
cg23487859
cg23488055
cg23488414
cg23489236
cg23489273
cg23489883
cg23490246
cg23490822
cg23491599
cg23491807
cg23491841
cg23493016
cg23493018
cg23493119
cg23493260
cg23493497
cg23493787

cg23493867
cg23494863
cg23495279
cg23495748
cg23495837
cg23495995
cg23497016
cg23497383
cg23497644
cg23498494
cg23498925
cg23499133
cg23499632
cg23499956
cg23500122
cg23501406
cg23501610
cg23502492
cg23502639
cg23502778
cg23502883
cg23502888
cg23504211
cg23504463
cg23504522
cg23505303
cg23506979
cg23507802
cg23507819
cg23507834
cg23508908
cg23510131
cg23511531
cg23512231
cg23513211
cg23514016
cg23514936
cg23515702
cg23516370
cg23516463
cg23516824
cg23517013
cg23518037
cg23518039
cg23518497
cg23518532
cg23518655
cg23519155
cg23519626
cg23520574
cg23521507
cg23521666

cg23521835
cg23521919
cg23522601
cg23522611
cg23522981
cg23523093
cg23523364
cg23523535
cg23524195
cg23524735
cg23525051
cg23525438
cg23525708
cg23526071
cg23526504
cg23526960
cg23527468
cg23527920
cg23528247
cg23528705
cg23530268
cg23530330
cg23530473
cg23530553
cg23530586
cg23530596
cg23531468
cg23531734
cg23532138
cg23532322
cg23532454
cg23533000
cg23534142
cg23534216
cg23534301
cg23534766
cg23536473
cg23536629
cg23537305
cg23537419
cg23538064
cg23538344
cg23539633
cg23541921
cg23541926
cg23542356
cg23542948
cg23542968
cg23543795
cg23544107
cg23545250
cg23545458

cg23545793
cg23546343
cg23546474
cg23547017
cg23547157
cg23548374
cg23548885
cg23549117
cg23549972
cg23550429
cg23550826
cg23551605
cg23552820
cg23553762
cg23554286
cg23554497
cg23554932
cg23555120
cg23555395
cg23556443
cg23556715
cg23558601
cg23558842
cg23559636
cg23559689
cg23562011
cg23562479
cg23563014
cg23564178
cg23564243
cg23564432
cg23564535
cg23564700
cg23565821
cg23565826
cg23565942
cg23566335
cg23566793
cg23568538
cg23568643
cg23568737
cg23569968
cg23572315
cg23572908
cg23573114
cg23573526
cg23574243
cg23574427
cg23574519
cg23574526
cg23574897
cg23574899

cg23575275
cg23575732
cg23578346
cg23578733
cg23580038
cg23580945
cg23581793
cg23582982
cg23584144
cg23584308
cg23584647
cg23584743
cg23585168
cg23586138
cg23586322
cg23586992
cg23588217
cg23588462
cg23591039
cg23592467
cg23592953
cg23593232
cg23593694
cg23595413
cg23595451
cg23595927
cg23596233
cg23596244
cg23596291
cg23596471
cg23597288
cg23597629
cg23597650
cg23598089
cg23598604
cg23599820
cg23599850
cg23599973
cg23600701
cg23601416
cg23601468
cg23602752
cg23603057
cg23603782
cg23603891
cg23604876
cg23606396
cg23606421
cg23609450
cg23610154
cg23611925
cg23612570

cg23613162
cg23613219
cg23613882
cg23614000
cg23615483
cg23615741
cg23616527
cg23618440
cg23618477
cg23619217
cg23619357
cg23619365
cg23619399
cg23620184
cg23621013
cg23622047
cg23622129
cg23622353
cg23623060
cg23623251
cg23623851
cg23624929
cg23625466
cg23625628
cg23625660
cg23626122
cg23626733
cg23627335
cg23629166
cg23629393
cg23629775
cg23629998
cg23630430
cg23631133
cg23631578
cg23632393
cg23632656
cg23632875
cg23632985
cg23633028
cg23633568
cg23633856
cg23633905
cg23634124
cg23634134
cg23634395
cg23634413
cg23634554
cg23636055
cg23636571
cg23636606
cg23636802

cg23636833
cg23637354
cg23637385
cg23637396
cg23637494
cg23638252
cg23639120
cg23639134
cg23639642
cg23640411
cg23640925
cg23641108
cg23641267
cg23641283
cg23641319
cg23641852
cg23641899
cg23642130
cg23642250
cg23642392
cg23642827
cg23642929
cg23643343
cg23643435
cg23644960
cg23645091
cg23645210
cg23645476
cg23647640
cg23648082
cg23649280
cg23649435
cg23649708
cg23650244
cg23650423
cg23650765
cg23651495
cg23651708
cg23651812
cg23651826
cg23652093
cg23652987
cg23654219
cg23654401
cg23655160
cg23655242
cg23656213
cg23656300
cg23656322
cg23657299
cg23657694
cg23658462

cg23658477
cg23658503
cg23658684
cg23658739
cg23659056
cg23660197
cg23660611
cg23660927
cg23661013
cg23662331
cg23662734
cg23663547
cg23663774
cg23664186
cg23664361
cg23664512
cg23665568
cg23665710
cg23665778
cg23665824
cg23666300
cg23666362
cg23667432
cg23667734
cg23667759
cg23667868
cg23668028
cg23668057
cg23668184
cg23668195
cg23668581
cg23669440
cg23669876
cg23670766
cg23673919
cg23674882
cg23675084
cg23675524
cg23676151
cg23676302
cg23676551
cg23676588
cg23676682
cg23677103
cg23677352
cg23677657
cg23678274
cg23678364
cg23678722
cg23679018
cg23679344
cg23679471

cg23679992
cg23680099
cg23680787
cg23682310
cg23682822
cg23683012
cg23683254
cg23683497
cg23683501
cg23683588
cg23684198
cg23684316
cg23684410
cg23685712
cg23685965
cg23686352
cg23687000
cg23688350
cg23688713
cg23689428
cg23689630
cg23689712
cg23690350
cg23690488
cg23690808
cg23690893
cg23691006
cg23691410
cg23691781
cg23693169
cg23693245
cg23693485
cg23693487
cg23693705
cg23693749
cg23694492
cg23694605
cg23695037
cg23695133
cg23695504
cg23696193
cg23696212
cg23696618
cg23697410
cg23697417
cg23697796
cg23697855
cg23697997
cg23699196
cg23699918
cg23701643
cg23702615

cg23702644
cg23704362
cg23704517
cg23704770
cg23705989
cg23706782
cg23708209
cg23708361
cg23709079
cg23709231
cg23709528
cg23709902
cg23710226
cg23710594
cg23710611
cg23710751
cg23710945
cg23711059
cg23711422
cg23711760
cg23712342
cg23712458
cg23712594
cg23713079
cg23713724
cg23713772
cg23713900
cg23713998
cg23714019
cg23714454
cg23714481
cg23715603
cg23715667
cg23715715
cg23715830
cg23716309
cg23716800
cg23716866
cg23717441
cg23717593
cg23717725
cg23719069
cg23719209
cg23719342
cg23719393
cg23719877
cg23720384
cg23722204
cg23722778
cg23723793
cg23724641
cg23724935

cg23725924
cg23726408
cg23726427
cg23726483
cg23726537
cg23726566
cg23726831
cg23727983
cg23729107
cg23729200
cg23729763
cg23729777
cg23730027
cg23730109
cg23730181
cg23730558
cg23730649
cg23731089
cg23731595
cg23731742
cg23732483
cg23732608
cg23732729
cg23732845
cg23733568
cg23733739
cg23733845
cg23734234
cg23734418
cg23734973
cg23735290
cg23735682
cg23735745
cg23737061
cg23737768
cg23738308
cg23738443
cg23739862
cg23740474
cg23740491
cg23741520
cg23742217
cg23743114
cg23744372
cg23745947
cg23746497
cg23746574
cg23747675
cg23748098
cg23748340
cg23748514
cg23748737

cg23749046
cg23750365
cg23750514
cg23751110
cg23751407
cg23752086
cg23753006
cg23754918
cg23755033
cg23756005
cg23756143
cg23756264
cg23756768
cg23756872
cg23757446
cg23758305
cg23758354
cg23759189
cg23759393
cg23760103
cg23760687
cg23761616
cg23762579
cg23762632
cg23762657
cg23763043
cg23763647
cg23765633
cg23765993
cg23766360
cg23766591
cg23766918
cg23766996
cg23767994
cg23768534
cg23768816
cg23770904
cg23771185
cg23771366
cg23771585
cg23771603
cg23771985
cg23772226
cg23772463
cg23772843
cg23772928
cg23773532
cg23774016
cg23774356
cg23774975
cg23774988
cg23778149

cg23779993
cg23780146
cg23780488
cg23780947
cg23781075
cg23781495
cg23782083
cg23782758
cg23782833
cg23784046
cg23784413
cg23785790
cg23786152
cg23786205
cg23786209
cg23786545
cg23786806
cg23787937
cg23789148
cg23790387
cg23791308
cg23791568
cg23791592
cg23791600
cg23792592
cg23793172
cg23793500
cg23793599
cg23793965
cg23795217
cg23796578
cg23796713
cg23797252
cg23798065
cg23799393
cg23799901
cg23801028
cg23801338
cg23801782
cg23803022
cg23803223
cg23803225
cg23803337
cg23803431
cg23804607
cg23805357
cg23806621
cg23806894
cg23807890
cg23808421
cg23808946
cg23809424

cg23809645
cg23810114
cg23810463
cg23811117
cg23811289
cg23813179
cg23813257
cg23814172
cg23814714
cg23815000
cg23816247
cg23816347
cg23817096
cg23818246
cg23818621
cg23818827
cg23819145
cg23819330
cg23820816
cg23820828
cg23820933
cg23822732
cg23824376
cg23824801
cg23826980
cg23827572
cg23828170
cg23828467
cg23829256
cg23830540
cg23831143
cg23831864
cg23831898
cg23832237
cg23832814
cg23833033
cg23833452
cg23834593
cg23834919
cg23835453
cg23835646
cg23835677
cg23835783
cg23836594
cg23836704
cg23837282
cg23839136
cg23839680
cg23840027
cg23840053
cg23840988
cg23841538

cg23841763
cg23842170
cg23842255
cg23842572
cg23843022
cg23843090
cg23843407
cg23843953
cg23844013
cg23844085
cg23844527
cg23844904
cg23845249
cg23845250
cg23845310
cg23845889
cg23846712
cg23847017
cg23847069
cg23847381
cg23847712
cg23848052
cg23848624
cg23850212
cg23850277
cg23850283
cg23850899
cg23851515
cg23851932
cg23853026
cg23853165
cg23854008
cg23854087
cg23854488
cg23854728
cg23855715
cg23855989
cg23856611
cg23857078
cg23857700
cg23858197
cg23860325
cg23861668
cg23861841
cg23862011
cg23863157
cg23865980
cg23867441
cg23867494
cg23867638
cg23868250
cg23869158

cg23871273
cg23871276
cg23873669
cg23874437
cg23875391
cg23875663
cg23876056
cg23876072
cg23877734
cg23878206
cg23878264
cg23878404
cg23878494
cg23879460
cg23879743
cg23881278
cg23883265
cg23883632
cg23883696
cg23883802
cg23884217
cg23885005
cg23885415
cg23885809
cg23885889
cg23886009
cg23886165
cg23888215
cg23889730
cg23889967
cg23890008
cg23891087
cg23891273
cg23891360
cg23891487
cg23892568
cg23892836
cg23893332
cg23893346
cg23893897
cg23893898
cg23894439
cg23894587
cg23894854
cg23894980
cg23895055
cg23895340
cg23895558
cg23895613
cg23896164
cg23896695
cg23897083

cg23897913
cg23898497
cg23898998
cg23899057
cg23899424
cg23900141
cg23901212
cg23901852
cg23901920
cg23902076
cg23902105
cg23902471
cg23902994
cg23903129
cg23903244
cg23903301
cg23903774
cg23905216
cg23906060
cg23906261
cg23906612
cg23906641
cg23906738
cg23906876
cg23907860
cg23908015
cg23908269
cg23912186
cg23912266
cg23912509
cg23913077
cg23913350
cg23913421
cg23913679
cg23913995
cg23914328
cg23915699
cg23915792
cg23916689
cg23916878
cg23917006
cg23917513
cg23917744
cg23917844
cg23918490
cg23918821
cg23919479
cg23919865
cg23919924
cg23921338
cg23921838
cg23922259

cg23922724
cg23922755
cg23922819
cg23924526
cg23924737
cg23925173
cg23925201
cg23926543
cg23928118
cg23928126
cg23928713
cg23929194
cg23929344
cg23930711
cg23930945
cg23931049
cg23931819
cg23932261
cg23932378
cg23932491
cg23933241
cg23933289
cg23934071
cg23934072
cg23934404
cg23936070
cg23936215
cg23936402
cg23936766
cg23937078
cg23937381
cg23938300
cg23939182
cg23941014
cg23941354
cg23941527
cg23942508
cg23943136
cg23943495
cg23943829
cg23944102
cg23944251
cg23944408
cg23944572
cg23944804
cg23944920
cg23945333
cg23945382
cg23945481
cg23946286
cg23947376
cg23947928

cg23948080
cg23948240
cg23948391
cg23948876
cg23949542
cg23949611
cg23949667
cg23950233
cg23950454
cg23950728
cg23951526
cg23953052
cg23953174
cg23953396
cg23953697
cg23954153
cg23954268
cg23954465
cg23954787
cg23954968
cg23955593
cg23956680
cg23956737
cg23956760
cg23956771
cg23957742
cg23958319
cg23959474
cg23960022
cg23960324
cg23960736
cg23961466
cg23962048
cg23962250
cg23962528
cg23962746
cg23963517
cg23963591
cg23964682
cg23964820
cg23967544
cg23968154
cg23968213
cg23968552
cg23968893
cg23969200
cg23969274
cg23969342
cg23970546
cg23971170
cg23972298
cg23972738

cg23972785
cg23973371
cg23973429
cg23973801
cg23973885
cg23974418
cg23976652
cg23976846
cg23976934
cg23976981
cg23977888
cg23978062
cg23978358
cg23978449
cg23979405
cg23979832
cg23980070
cg23980760
cg23983710
cg23985388
cg23985420
cg23985641
cg23985818
cg23986143
cg23986185
cg23986671
cg23987938
cg23989821
cg23989963
cg23990012
cg23990303
cg23990624
cg23990814
cg23991358
cg23991413
cg23993511
cg23994043
cg23994443
cg23994714
cg23994802
cg23995459
cg23995769
cg23996071
cg23996800
cg23996846
cg23997185
cg23997263
cg23997339
cg23997911
cg23998391
cg23998645
cg23998903

cg23999080
cg23999833
cg24000223
cg24000241
cg24000285
cg24000437
cg24000873
cg24000937
cg24001345
cg24001710
cg24001728
cg24002071
cg24002887
cg24003028
cg24003850
cg24004483
cg24004665
cg24005158
cg24005685
cg24005701
cg24005950
cg24006126
cg24006305
cg24006547
cg24006715
cg24007254
cg24008634
cg24009030
cg24009250
cg24009650
cg24011260
cg24011261
cg24011511
cg24013351
cg24013589
cg24013954
cg24015522
cg24017555
cg24017800
cg24018190
cg24018321
cg24019054
cg24019564
cg24020157
cg24020826
cg24021846
cg24021956
cg24022301
cg24022375
cg24024528
cg24025567
cg24025721

cg24026619
cg24026690
cg24027068
cg24028218
cg24028627
cg24028798
cg24028809
cg24029426
cg24030232
cg24030277
cg24032056
cg24032252
cg24032304
cg24032445
cg24032666
cg24032691
cg24033330
cg24033633
cg24034357
cg24034715
cg24035035
cg24035245
cg24035447
cg24035745
cg24035928
cg24035931
cg24037651
cg24037746
cg24038105
cg24039149
cg24039310
cg24039697
cg24039816
cg24040115
cg24040450
cg24041118
cg24041541
cg24042242
cg24042923
cg24043916
cg24044501
cg24044560
cg24044707
cg24044957
cg24045498
cg24046110
cg24047400
cg24047736
cg24047802
cg24047810
cg24047926
cg24048008

cg24048338
cg24048921
cg24049468
cg24050775
cg24051103
cg24051242
cg24051481
cg24052039
cg24052097
cg24052359
cg24052475
cg24052772
cg24053664
cg24054180
cg24054871
cg24055029
cg24055703
cg24055836
cg24055855
cg24057107
cg24058132
cg24058145
cg24058732
cg24059623
cg24060173
cg24060317
cg24060475
cg24061141
cg24061208
cg24062909
cg24063043
cg24063912
cg24064041
cg24064224
cg24064884
cg24065504
cg24065807
cg24066316
cg24068468
cg24068972
cg24069188
cg24069602
cg24071177
cg24071532
cg24071582
cg24071719
cg24072093
cg24072224
cg24072451
cg24072947
cg24073777
cg24074033

cg24075412
cg24076399
cg24076537
cg24076884
cg24077277
cg24078043
cg24079051
cg24079251
cg24079381
cg24079811
cg24080008
cg24080247
cg24080571
cg24082174
cg24082368
cg24082983
cg24085232
cg24085502
cg24085570
cg24085713
cg24085809
cg24085885
cg24086068
cg24087515
cg24087681
cg24087736
cg24087743
cg24087887
cg24088250
cg24088496
cg24089118
cg24089171
cg24089600
cg24089932
cg24089936
cg24090990
cg24091195
cg24091315
cg24091474
cg24091995
cg24092179
cg24092282
cg24092295
cg24092914
cg24093182
cg24093763
cg24094029
cg24094450
cg24094775
cg24095592
cg24095864
cg24096120

cg24098008
cg24099067
cg24099605
cg24100168
cg24100636
cg24100671
cg24101337
cg24101359
cg24101492
cg24101979
cg24102242
cg24102420
cg24103475
cg24104268
cg24104938
cg24105457
cg24106636
cg24107848
cg24109739
cg24109744
cg24109980
cg24110052
cg24110063
cg24110184
cg24110635
cg24111025
cg24112010
cg24112454
cg24112562
cg24112628
cg24112692
cg24113409
cg24114556
cg24114587
cg24114651
cg24115019
cg24115681
cg24116870
cg24119500
cg24119674
cg24119869
cg24119990
cg24120041
cg24120696
cg24121503
cg24121765
cg24122498
cg24123116
cg24124079
cg24124145
cg24124443
cg24125282

cg24125710
cg24126847
cg24127239
cg24127719
cg24128434
cg24128943
cg24128979
cg24129198
cg24129390
cg24130568
cg24131595
cg24131671
cg24132325
cg24133115
cg24133547
cg24133784
cg24135300
cg24136205
cg24136314
cg24136340
cg24136431
cg24136690
cg24136700
cg24136932
cg24137567
cg24138234
cg24138857
cg24138964
cg24139093
cg24139443
cg24139720
cg24139737
cg24139843
cg24139874
cg24139912
cg24141516
cg24141724
cg24141911
cg24142603
cg24143221
cg24143287
cg24144083
cg24144440
cg24145020
cg24146685
cg24147596
cg24147838
cg24148085
cg24148361
cg24149502
cg24149590
cg24150172

cg24150244
cg24150325
cg24150623
cg24150662
cg24150986
cg24151352
cg24151755
cg24152251
cg24152264
cg24152605
cg24153574
cg24153714
cg24153874
cg24154722
cg24154839
cg24154951
cg24155427
cg24155961
cg24156393
cg24156983
cg24157349
cg24157814
cg24157892
cg24158141
cg24158157
cg24158205
cg24158599
cg24159247
cg24159436
cg24159514
cg24159636
cg24160312
cg24162030
cg24162603
cg24163575
cg24164296
cg24164564
cg24165056
cg24165638
cg24166018
cg24166100
cg24166457
cg24166463
cg24166814
cg24168061
cg24168914
cg24169315
cg24169417
cg24169669
cg24171152
cg24172416
cg24172675

cg24173753
cg24174232
cg24174608
cg24174959
cg24174996
cg24178440
cg24178897
cg24179719
cg24180759
cg24181062
cg24181216
cg24182329
cg24183098
cg24183909
cg24185357
cg24186774
cg24186915
cg24187101
cg24187645
cg24187758
cg24188017
cg24188415
cg24189303
cg24189734
cg24189904
cg24190139
cg24190223
cg24190603
cg24191287
cg24192786
cg24193456
cg24194279
cg24195359
cg24195524
cg24195878
cg24195971
cg24196784
cg24197051
cg24197567
cg24198558
cg24198944
cg24198990
cg24200753
cg24201941
cg24202123
cg24202131
cg24202485
cg24202916
cg24203741
cg24204847
cg24205387
cg24205914

cg24206537
cg24207068
cg24208886
cg24209126
cg24211006
cg24211214
cg24211504
cg24212267
cg24213248
cg24213507
cg24213719
cg24214471
cg24215459
cg24215812
cg24216555
cg24217005
cg24217729
cg24218109
cg24219058
cg24219966
cg24221541
cg24221648
cg24221738
cg24222546
cg24222995
cg24224020
cg24225517
cg24225668
cg24226115
cg24226238
cg24226371
cg24228819
cg24229691
cg24229707
cg24229750
cg24231148
cg24231854
cg24232444
cg24233194
cg24233594
cg24234270
cg24234899
cg24235518
cg24236409
cg24237081
cg24237609
cg24237862
cg24238345
cg24239148
cg24239329
cg24239961
cg24240626

cg24240870
cg24241192
cg24242280
cg24243314
cg24243790
cg24243914
cg24244374
cg24244695
cg24244747
cg24244752
cg24245418
cg24245726
cg24247370
cg24248367
cg24248680
cg24248713
cg24249648
cg24249778
cg24250684
cg24250902
cg24251135
cg24251218
cg24251439
cg24252148
cg24252536
cg24253500
cg24254206
cg24254317
cg24255032
cg24255327
cg24256211
cg24256786
cg24259244
cg24260156
cg24260882
cg24262066
cg24262325
cg24262691
cg24262870
cg24263283
cg24264285
cg24264382
cg24266670
cg24267699
cg24269074
cg24269377
cg24269617
cg24269863
cg24270629
cg24270710
cg24271203
cg24274579

cg24275315
cg24275769
cg24276445
cg24276787
cg24277007
cg24277586
cg24277715
cg24277791
cg24278076
cg24278087
cg24279243
cg24279418
cg24280540
cg24280925
cg24284973
cg24285903
cg24286765
cg24286789
cg24287460
cg24288751
cg24290756
cg24291087
cg24291490
cg24291747
cg24291991
cg24292761
cg24293507
cg24293614
cg24294544
cg24294950
cg24295381
cg24296452
cg24296920
cg24297509
cg24298280
cg24298421
cg24298684
cg24299136
cg24299177
cg24301288
cg24301290
cg24301620
cg24302235
cg24302683
cg24303806
cg24304309
cg24304338
cg24304425
cg24304714
cg24304919
cg24305861
cg24306397

cg24306422
cg24306962
cg24306982
cg24307292
cg24307341
cg24307499
cg24307601
cg24308389
cg24309428
cg24309962
cg24310246
cg24310500
cg24310780
cg24310924
cg24310959
cg24311485
cg24311643
cg24311693
cg24311704
cg24311922
cg24312063
cg24312105
cg24312283
cg24312520
cg24312680
cg24312874
cg24312985
cg24313473
cg24313572
cg24314434
cg24314474
cg24315209
cg24315421
cg24315508
cg24315586
cg24315710
cg24316949
cg24317069
cg24317310
cg24317406
cg24317647
cg24318076
cg24318763
cg24319171
cg24319902
cg24320225
cg24320612
cg24321467
cg24322366
cg24323597
cg24324984
cg24325215

cg24325551
cg24326030
cg24328095
cg24328944
cg24330514
cg24331282
cg24332570
cg24332770
cg24332783
cg24333189
cg24333629
cg24333798
cg24334833
cg24335310
cg24335895
cg24336089
cg24336674
cg24337215
cg24337235
cg24337318
cg24338411
cg24338941
cg24339273
cg24339574
cg24339597
cg24340333
cg24340655
cg24340935
cg24341611
cg24341770
cg24343594
cg24343720
cg24343823
cg24343835
cg24344074
cg24345062
cg24345261
cg24345551
cg24345856
cg24346223
cg24346464
cg24346629
cg24346637
cg24346868
cg24347098
cg24347611
cg24347994
cg24349665
cg24350771
cg24351076
cg24351452
cg24352349

cg24352757
cg24352878
cg24353443
cg24353740
cg24354612
cg24355048
cg24355907
cg24357161
cg24357447
cg24357619
cg24357935
cg24358047
cg24358467
cg24358603
cg24359048
cg24359109
cg24359323
cg24359868
cg24360034
cg24360137
cg24360745
cg24360747
cg24360909
cg24360993
cg24361079
cg24361391
cg24362726
cg24362812
cg24362813
cg24362824
cg24363955
cg24364282
cg24365215
cg24365464
cg24365626
cg24365984
cg24366314
cg24366493
cg24366557
cg24366665
cg24367957
cg24368031
cg24368848
cg24369185
cg24369728
cg24370357
cg24370613
cg24370881
cg24371033
cg24371534
cg24373369
cg24373608

cg24374013
cg24374738
cg24374861
cg24375364
cg24375399
cg24375470
cg24375626
cg24375646
cg24375938
cg24376266
cg24376295
cg24376339
cg24376776
cg24376955
cg24378250
cg24378692
cg24378940
cg24379252
cg24379915
cg24381587
cg24382063
cg24382276
cg24382417
cg24384051
cg24384057
cg24384120
cg24384244
cg24384437
cg24384676
cg24385450
cg24385652
cg24387367
cg24387380
cg24387472
cg24387800
cg24390428
cg24390519
cg24390913
cg24392299
cg24392646
cg24394172
cg24394340
cg24394624
cg24394631
cg24395452
cg24395801
cg24396686
cg24396745
cg24396875
cg24397545
cg24399122
cg24399337

cg24399349
cg24399830
cg24399959
cg24401026
cg24401262
cg24401487
cg24402070
cg24402151
cg24402667
cg24402683
cg24403338
cg24403845
cg24404533
cg24404823
cg24404878
cg24405248
cg24405951
cg24406162
cg24406619
cg24407086
cg24407596
cg24408469
cg24408656
cg24410255
cg24410535
cg24410546
cg24410945
cg24411312
cg24411817
cg24411929
cg24411961
cg24412079
cg24412659
cg24412714
cg24412895
cg24413662
cg24413879
cg24414627
cg24414788
cg24415208
cg24415941
cg24416513
cg24417135
cg24417845
cg24418420
cg24418941
cg24419040
cg24419324
cg24420742
cg24421216
cg24421397
cg24421870

cg24421906
cg24422797
cg24423088
cg24425149
cg24425175
cg24425705
cg24425807
cg24425838
cg24425972
cg24425984
cg24426483
cg24426822
cg24427089
cg24427243
cg24427376
cg24427724
cg24427915
cg24428218
cg24428877
cg24429533
cg24429622
cg24430140
cg24430703
cg24431693
cg24432464
cg24432632
cg24432977
cg24433265
cg24433287
cg24433738
cg24434056
cg24435669
cg24435939
cg24436446
cg24436715
cg24437104
cg24437715
cg24438217
cg24438644
cg24439256
cg24439334
cg24439401
cg24439686
cg24440898
cg24442572
cg24442766
cg24444059
cg24445116
cg24445167
cg24445674
cg24446242
cg24446326

cg24446586
cg24446932
cg24447775
cg24448171
cg24449091
cg24449706
cg24449777
cg24449914
cg24450239
cg24450653
cg24451141
cg24452128
cg24452260
cg24452347
cg24452602
cg24452765
cg24453118
cg24453580
cg24453600
cg24453664
cg24453699
cg24454144
cg24455365
cg24455634
cg24455969
cg24456002
cg24456365
cg24457403
cg24457611
cg24459227
cg24459498
cg24459556
cg24459561
cg24460268
cg24461171
cg24461242
cg24461524
cg24462934
cg24463476
cg24464397
cg24464789
cg24465429
cg24465469
cg24465685
cg24465935
cg24465943
cg24467387
cg24468070
cg24468736
cg24468890
cg24468934
cg24468955

cg24469013
cg24469464
cg24469742
cg24469849
cg24469980
cg24470332
cg24471254
cg24472231
cg24472495
cg24473832
cg24475272
cg24475782
cg24476103
cg24476497
cg24476584
cg24477315
cg24477886
cg24478420
cg24478779
cg24480012
cg24480695
cg24480735
cg24480810
cg24481801
cg24481841
cg24482350
cg24483247
cg24483785
cg24484138
cg24484296
cg24485354
cg24485927
cg24486958
cg24487856
cg24488391
cg24488891
cg24490256
cg24490338
cg24490529
cg24490633
cg24491776
cg24491784
cg24492749
cg24492778
cg24492886
cg24494102
cg24495062
cg24495350
cg24495585
cg24496475
cg24497361
cg24497686

cg24498885
cg24499516
cg24500711
cg24500832
cg24500959
cg24501230
cg24501758
cg24501831
cg24502330
cg24502478
cg24503003
cg24503056
cg24503796
cg24504289
cg24504361
cg24504854
cg24505167
cg24505341
cg24505618
cg24505713
cg24505892
cg24506604
cg24507551
cg24507762
cg24509810
cg24512093
cg24512138
cg24512216
cg24512400
cg24512581
cg24513381
cg24513522
cg24513684
cg24513791
cg24514391
cg24514678
cg24515861
cg24516799
cg24516901
cg24517323
cg24519147
cg24519867
cg24520479
cg24521811
cg24522478
cg24522654
cg24523948
cg24524352
cg24524414
cg24524485
cg24525138
cg24525286

cg24525400
cg24525988
cg24526596
cg24527098
cg24527434
cg24527466
cg24527591
cg24527776
cg24528164
cg24528307
cg24528350
cg24528829
cg24530250
cg24530432
cg24530460
cg24530728
cg24531009
cg24531396
cg24531401
cg24531494
cg24532610
cg24533564
cg24533720
cg24533864
cg24533913
cg24533917
cg24534566
cg24534767
cg24535823
cg24535910
cg24536244
cg24536691
cg24537237
cg24537836
cg24537942
cg24537973
cg24537993
cg24538292
cg24538331
cg24539090
cg24539517
cg24540569
cg24540810
cg24541193
cg24541426
cg24542929
cg24543552
cg24543596
cg24543696
cg24543759
cg24543937
cg24544177

cg24544490
cg24544656
cg24545125
cg24545728
cg24546155
cg24546683
cg24547910
cg24549161
cg24549289
cg24549659
cg24550456
cg24550525
cg24550933
cg24551324
cg24551418
cg24553184
cg24553332
cg24554789
cg24554839
cg24554944
cg24555071
cg24555204
cg24555286
cg24555906
cg24556441
cg24557027
cg24558112
cg24559044
cg24561305
cg24561411
cg24561419
cg24561661
cg24562018
cg24562081
cg24562110
cg24562814
cg24562819
cg24563501
cg24563570
cg24563967
cg24565369
cg24566400
cg24566595
cg24566824
cg24567424
cg24569447
cg24569492
cg24569831
cg24571875
cg24571899
cg24573261
cg24574793

cg24575234
cg24575275
cg24575624
cg24576425
cg24576804
cg24576995
cg24577131
cg24577933
cg24578937
cg24579534
cg24579604
cg24579887
cg24579970
cg24580066
cg24580724
cg24581183
cg24581200
cg24582743
cg24583987
cg24584161
cg24584721
cg24585203
cg24585210
cg24585422
cg24585690
cg24586039
cg24586444
cg24586528
cg24587080
cg24587185
cg24587208
cg24587312
cg24587349
cg24588278
cg24589834
cg24590430
cg24590788
cg24591082
cg24591861
cg24592194
cg24592546
cg24592621
cg24593372
cg24593464
cg24594224
cg24594454
cg24594459
cg24594504
cg24595152
cg24595510
cg24595580
cg24596116

cg24597073
cg24598316
cg24598495
cg24598948
cg24599329
cg24599434
cg24601327
cg24601424
cg24601835
cg24603464
cg24604013
cg24605023
cg24605046
cg24605325
cg24605948
cg24607283
cg24607398
cg24608717
cg24609101
cg24609819
cg24610044
cg24610231
cg24610349
cg24610790
cg24611351
cg24611446
cg24611608
cg24612667
cg24613080
cg24614671
cg24615945
cg24616382
cg24616385
cg24617008
cg24617352
cg24617647
cg24617696
cg24617723
cg24617992
cg24618169
cg24618387
cg24618467
cg24619276
cg24619998
cg24620617
cg24620761
cg24620797
cg24621042
cg24621437
cg24621790
cg24621972
cg24622007

cg24622209
cg24623760
cg24624090
cg24624586
cg24624873
cg24624901
cg24627789
cg24627900
cg24628076
cg24628394
cg24628676
cg24629356
cg24629392
cg24629443
cg24629972
cg24631428
cg24631735
cg24632139
cg24632241
cg24632873
cg24633312
cg24633390
cg24633978
cg24634333
cg24634420
cg24634526
cg24635109
cg24635581
cg24635722
cg24636336
cg24636477
cg24636944
cg24636999
cg24637035
cg24637252
cg24637261
cg24637364
cg24637974
cg24638326
cg24639100
cg24639117
cg24639703
cg24640561
cg24640637
cg24640676
cg24641186
cg24642065
cg24642372
cg24642825
cg24642933
cg24643102
cg24643158

cg24643381
cg24644026
cg24644605
cg24645149
cg24645611
cg24645671
cg24646414
cg24646841
cg24647015
cg24647428
cg24647724
cg24648119
cg24649332
cg24650501
cg24651828
cg24652919
cg24652956
cg24653263
cg24654264
cg24655262
cg24655669
cg24655701
cg24655706
cg24657535
cg24659054
cg24659696
cg24660120
cg24660640
cg24660670
cg24662231
cg24662614
cg24662698
cg24662789
cg24662867
cg24662961
cg24663256
cg24664347
cg24664351
cg24664855
cg24665290
cg24666702
cg24667407
cg24667437
cg24667758
cg24668440
cg24669401
cg24669528
cg24669602
cg24670000
cg24670151
cg24670219
cg24670241

cg24670445
cg24670453
cg24670927
cg24671330
cg24671344
cg24671666
cg24672685
cg24673043
cg24673385
cg24673605
cg24673742
cg24673765
cg24673769
cg24673771
cg24674635
cg24674703
cg24674754
cg24674887
cg24675150
cg24675673
cg24675730
cg24675998
cg24676071
cg24676244
cg24676514
cg24676755
cg24677222
cg24677278
cg24678137
cg24678581
cg24678767
cg24679007
cg24679043
cg24679124
cg24680320
cg24680570
cg24680632
cg24680758
cg24681086
cg24681307
cg24681438
cg24681499
cg24681573
cg24681963
cg24682036
cg24682512
cg24683222
cg24683623
cg24685755
cg24686153
cg24686187
cg24686610

cg24686679
cg24686845
cg24686918
cg24686983
cg24687432
cg24687841
cg24687970
cg24688563
cg24688690
cg24688837
cg24690054
cg24690071
cg24690588
cg24691461
cg24691792
cg24691891
cg24691965
cg24692170
cg24692595
cg24693053
cg24693189
cg24693341
cg24693551
cg24694329
cg24694665
cg24694675
cg24694691
cg24694702
cg24695849
cg24696151
cg24696967
cg24697648
cg24698488
cg24698655
cg24699146
cg24699685
cg24699871
cg24699895
cg24700316
cg24700823
cg24700929
cg24700959
cg24701725
cg24701780
cg24702091
cg24702253
cg24702341
cg24702826
cg24703336
cg24703468
cg24704177
cg24704730

cg24704759
cg24705960
cg24706674
cg24706793
cg24706839
cg24706992
cg24707311
cg24707432
cg24708471
cg24708674
cg24709033
cg24710073
cg24710179
cg24710631
cg24710671
cg24711070
cg24711336
cg24711649
cg24712482
cg24712501
cg24713204
cg24713259
cg24714905
cg24715445
cg24715735
cg24715767
cg24716152
cg24717078
cg24717325
cg24717898
cg24718015
cg24718197
cg24718617
cg24718903
cg24719005
cg24719086
cg24719321
cg24719489
cg24720242
cg24720355
cg24721899
cg24721964
cg24722055
cg24722073
cg24722099
cg24722886
cg24723373
cg24724428
cg24724567
cg24724596
cg24724618
cg24724630

cg24724664
cg24727130
cg24727311
cg24727399
cg24728099
cg24728456
cg24728698
cg24728949
cg24729221
cg24730157
cg24730705
cg24731597
cg24732137
cg24732702
cg24733450
cg24735226
cg24735489
cg24736734
cg24736935
cg24736989
cg24737374
cg24737570
cg24737612
cg24737857
cg24738611
cg24738924
cg24739407
cg24739950
cg24740218
cg24740320
cg24740868
cg24741666
cg24741888
cg24742349
cg24743122
cg24743155
cg24743156
cg24743290
cg24744014
cg24744172
cg24744430
cg24744697
cg24747537
cg24748769
cg24748786
cg24749089
cg24749265
cg24749470
cg24749947
cg24750539
cg24750674
cg24750887

cg24751129
cg24753662
cg24753760
cg24753988
cg24754199
cg24754223
cg24754633
cg24755189
cg24755931
cg24756698
cg24756865
cg24757159
cg24757755
cg24757926
cg24757978
cg24758181
cg24758452
cg24758816
cg24759237
cg24759599
cg24759654
cg24759795
cg24761467
cg24761507
cg24761597
cg24761668
cg24762231
cg24762932
cg24762962
cg24764372
cg24764810
cg24765360
cg24765446
cg24765648
cg24766030
cg24766398
cg24766821
cg24767002
cg24767148
cg24767540
cg24768113
cg24768167
cg24768463
cg24769348
cg24769846
cg24770677
cg24771111
cg24772753
cg24773412
cg24773418
cg24773770
cg24774306

cg24774357
cg24774396
cg24775496
cg24776142
cg24777101
cg24777351
cg24778004
cg24778016
cg24778683
cg24778776
cg24779170
cg24779941
cg24780236
cg24780759
cg24781100
cg24781699
cg24783211
cg24783356
cg24783642
cg24784730
cg24784794
cg24786705
cg24786875
cg24787130
cg24787823
cg24788028
cg24788034
cg24789424
cg24789447
cg24790676
cg24791025
cg24791428
cg24791666
cg24792113
cg24792289
cg24792365
cg24792658
cg24793584
cg24794992
cg24795274
cg24795627
cg24795903
cg24795958
cg24796852
cg24797276
cg24797431
cg24797508
cg24798540
cg24798826
cg24799175
cg24800630
cg24800754

cg24800887
cg24801023
cg24801835
cg24803614
cg24803687
cg24804060
cg24804707
cg24805985
cg24806326
cg24806371
cg24807294
cg24807547
cg24807850
cg24808105
cg24808162
cg24808891
cg24809011
cg24809529
cg24809973
cg24810305
cg24811685
cg24811864
cg24812167
cg24812837
cg24813038
cg24813206
cg24813736
cg24813857
cg24813995
cg24814257
cg24815792
cg24815934
cg24817257
cg24817358
cg24818200
cg24819686
cg24821307
cg24822696
cg24823222
cg24823764
cg24824371
cg24825210
cg24825693
cg24825722
cg24825966
cg24826270
cg24826696
cg24827357
cg24827381
cg24827660
cg24827839
cg24828167

cg24828208
cg24828257
cg24828648
cg24828746
cg24828864
cg24829430
cg24830241
cg24832498
cg24833674
cg24834394
cg24834436
cg24834590
cg24834659
cg24835153
cg24836281
cg24836946
cg24837075
cg24837149
cg24837219
cg24837277
cg24837370
cg24837653
cg24838349
cg24838764
cg24838825
cg24842142
cg24842733
cg24842859
cg24843003
cg24843380
cg24843701
cg24844971
cg24845329
cg24845998
cg24846199
cg24847035
cg24847169
cg24847230
cg24847427
cg24847685
cg24847829
cg24848035
cg24848351
cg24850296
cg24850711
cg24851249
cg24851854
cg24852713
cg24855045
cg24856658
cg24857040
cg24859352

cg24859393
cg24859546
cg24859986
cg24860169
cg24860429
cg24861399
cg24862000
cg24862510
cg24863175
cg24863254
cg24863335
cg24863544
cg24863552
cg24864189
cg24864663
cg24865270
cg24866700
cg24866717
cg24866764
cg24866792
cg24867468
cg24868292
cg24868305
cg24868359
cg24868960
cg24869601
cg24869815
cg24869834
cg24870273
cg24870340
cg24870391
cg24870676
cg24871414
cg24872117
cg24872692
cg24872815
cg24873202
cg24873280
cg24874191
cg24874254
cg24874350
cg24874602
cg24874783
cg24874998
cg24875288
cg24875595
cg24875808
cg24876164
cg24876187
cg24876368
cg24876404
cg24876837

cg24876960
cg24876998
cg24877616
cg24878374
cg24878483
cg24879273
cg24879335
cg24880056
cg24880701
cg24880818
cg24881205
cg24881420
cg24881834
cg24881993
cg24882332
cg24882551
cg24883413
cg24883441
cg24883742
cg24884084
cg24884142
cg24884217
cg24884343
cg24884961
cg24885937
cg24886257
cg24886267
cg24886439
cg24886708
cg24886710
cg24886788
cg24886831
cg24887211
cg24889512
cg24891312
cg24891434
cg24891520
cg24892091
cg24892966
cg24893837
cg24894353
cg24894783
cg24895173
cg24895497
cg24896053
cg24896649
cg24898236
cg24898914
cg24899294
cg24899767
cg24900164
cg24900654

cg24901488
cg24901811
cg24902174
cg24902196
cg24902278
cg24903497
cg24903887
cg24904788
cg24905248
cg24905254
cg24906316
cg24907075
cg24908814
cg24909309
cg24909466
cg24909660
cg24910161
cg24911113
cg24911827
cg24913349
cg24913355
cg24913825
cg24913868
cg24914077
cg24914219
cg24914244
cg24914884
cg24915503
cg24916013
cg24919348
cg24919468
cg24919476
cg24919806
cg24919884
cg24921221
cg24921240
cg24921794
cg24921808
cg24921858
cg24921943
cg24922143
cg24923516
cg24923694
cg24923953
cg24924505
cg24924779
cg24924936
cg24925945
cg24926042
cg24926276
cg24926465
cg24926589

cg24927800
cg24928053
cg24928333
cg24928391
cg24928546
cg24929264
cg24929514
cg24929538
cg24929847
cg24930218
cg24931242
cg24931346
cg24931355
cg24931520
cg24931632
cg24931713
cg24932585
cg24933856
cg24934063
cg24934400
cg24935208
cg24935896
cg24936204
cg24936695
cg24937747
cg24938761
cg24938830
cg24939819
cg24939838
cg24940230
cg24940583
cg24940601
cg24942458
cg24943066
cg24943198
cg24944017
cg24944328
cg24945231
cg24945701
cg24946597
cg24946729
cg24947694
cg24947764
cg24948189
cg24948743
cg24949579
cg24951100
cg24951137
cg24951989
cg24952240
cg24952640
cg24952936

cg24953703
cg24954121
cg24954350
cg24954809
cg24956176
cg24956452
cg24957489
cg24957628
cg24957964
cg24960565
cg24960947
cg24961523
cg24961583
cg24962576
cg24964061
cg24964130
cg24964368
cg24965187
cg24965322
cg24965356
cg24965791
cg24966363
cg24966575
cg24966702
cg24966958
cg24968721
cg24968730
cg24968931
cg24969222
cg24970918
cg24971162
cg24971209
cg24971846
cg24973864
cg24974217
cg24975222
cg24975342
cg24975564
cg24975642
cg24975986
cg24977541
cg24978630
cg24979093
cg24979203
cg24980572
cg24980653
cg24980709
cg24980856
cg24980995
cg24981058
cg24981097
cg24981833

cg24982697
cg24983189
cg24983539
cg24984147
cg24984506
cg24984653
cg24985151
cg24985442
cg24987622
cg24987667
cg24987745
cg24988345
cg24988625
cg24989962
cg24990212
cg24990400
cg24992471
cg24993400
cg24993711
cg24994611
cg24995674
cg24995976
cg24996161
cg24996179
cg24996282
cg24996482
cg24996491
cg24996758
cg24997585
cg24997703
cg24997896
cg24998078
cg24998197
cg25000696
cg25002426
cg25002781
cg25003924
cg25004193
cg25004427
cg25004471
cg25004981
cg25006194
cg25006829
cg25007422
cg25007511
cg25007705
cg25008147
cg25008504
cg25009504
cg25009842
cg25009965
cg25010691

cg25011395
cg25011475
cg25012919
cg25013011
cg25013052
cg25013095
cg25013303
cg25013443
cg25014026
cg25014229
cg25015416
cg25015613
cg25017369
cg25017584
cg25017774
cg25017777
cg25018403
cg25018801
cg25019378
cg25019503
cg25019526
cg25019648
cg25020073
cg25020204
cg25020286
cg25020293
cg25020550
cg25020678
cg25020850
cg25020969
cg25021259
cg25021970
cg25022311
cg25023165
cg25023275
cg25023298
cg25023666
cg25024321
cg25024717
cg25024734
cg25025437
cg25025966
cg25025992
cg25026125
cg25026529
cg25026755
cg25026898
cg25026904
cg25026992
cg25027125
cg25027699
cg25027798

cg25027832
cg25027863
cg25028293
cg25028404
cg25029561
cg25029797
cg25030049
cg25030832
cg25030862
cg25030888
cg25031824
cg25032321
cg25032595
cg25032722
cg25032756
cg25032793
cg25032963
cg25032968
cg25033990
cg25034395
cg25035200
cg25035376
cg25037394
cg25037677
cg25038082
cg25038311
cg25039722
cg25040937
cg25043986
cg25044186
cg25044701
cg25044765
cg25044876
cg25045013
cg25045219
cg25045228
cg25045972
cg25046651
cg25046772
cg25047056
cg25048100
cg25048344
cg25049733
cg25049844
cg25050447
cg25050565
cg25051529
cg25051805
cg25052274
cg25052664
cg25053018
cg25053142

cg25054816
cg25055341
cg25056510
cg25057461
cg25060339
cg25061289
cg25061451
cg25061610
cg25061755
cg25063189
cg25064395
cg25064551
cg25064552
cg25065131
cg25065191
cg25066134
cg25066857
cg25066972
cg25067242
cg25067686
cg25067940
cg25068253
cg25070253
cg25070340
cg25071602
cg25072666
cg25072962
cg25073089
cg25073137
cg25073563
cg25074066
cg25074426
cg25075504
cg25075794
cg25076324
cg25076408
cg25076881
cg25077137
cg25078026
cg25078061
cg25078137
cg25078150
cg25078444
cg25078908
cg25079102
cg25079628
cg25079847
cg25079865
cg25080501
cg25080924
cg25081106
cg25081201

cg25082710
cg25083732
cg25083922
cg25084101
cg25084878
cg25085245
cg25085755
cg25087341
cg25087589
cg25088758
cg25092681
cg25092764
cg25092838
cg25092915
cg25092989
cg25093045
cg25093492
cg25093813
cg25094569
cg25095722
cg25096107
cg25096264
cg25096566
cg25096615
cg25097308
cg25097705
cg25097946
cg25098174
cg25099065
cg25099389
cg25099516
cg25099892
cg25100880
cg25101184
cg25101291
cg25101380
cg25101405
cg25102370
cg25103103
cg25103160
cg25103275
cg25103417
cg25103492
cg25103772
cg25103895
cg25104105
cg25104234
cg25104397
cg25105522
cg25105633
cg25105917
cg25106322

cg25106645
cg25106913
cg25107254
cg25107417
cg25107522
cg25109019
cg25109238
cg25110782
cg25110832
cg25111926
cg25112002
cg25112284
cg25112414
cg25113237
cg25114523
cg25114611
cg25114913
cg25116216
cg25116388
cg25116430
cg25116503
cg25117523
cg25117600
cg25117725
cg25119077
cg25119155
cg25119261
cg25119946
cg25120290
cg25121292
cg25121513
cg25122395
cg25122699
cg25123521
cg25123670
cg25124030
cg25124045
cg25124241
cg25124276
cg25124402
cg25124956
cg25125060
cg25125340
cg25128453
cg25130385
cg25130728
cg25130939
cg25131620
cg25132510
cg25132772
cg25133376
cg25133753

cg25134283
cg25134701
cg25135018
cg25135436
cg25135457
cg25135654
cg25136495
cg25137314
cg25137372
cg25137841
cg25138327
cg25139636
cg25140347
cg25140385
cg25141642
cg25142390
cg25142500
cg25142614
cg25143359
cg25143990
cg25144574
cg25145360
cg25145680
cg25145687
cg25145765
cg25145840
cg25146017
cg25146557
cg25147026
cg25147632
cg25147941
cg25148572
cg25148632
cg25149122
cg25149218
cg25149238
cg25149888
cg25150520
cg25151498
cg25151559
cg25151919
cg25153233
cg25153726
cg25154038
cg25154236
cg25154699
cg25156198
cg25156960
cg25157472
cg25157850
cg25157999
cg25159668

cg25160034
cg25160605
cg25161029
cg25161310
cg25162117
cg25163611
cg25165358
cg25165880
cg25165932
cg25165975
cg25166437
cg25167643
cg25168494
cg25170017
cg25170822
cg25172185
cg25174412
cg25174438
cg25176746
cg25177452
cg25178500
cg25178645
cg25179685
cg25179853
cg25179876
cg25180759
cg25181170
cg25181651
cg25182165
cg25182523
cg25183989
cg25184088
cg25185173
cg25185454
cg25186802
cg25188006
cg25188032
cg25188395
cg25188398
cg25188760
cg25189564
cg25189873
cg25190513
cg25190534
cg25191628
cg25191850
cg25192855
cg25193276
cg25193494
cg25194322
cg25194470
cg25195210

cg25195309
cg25195987
cg25196088
cg25196158
cg25198275
cg25198661
cg25200616
cg25201047
cg25201101
cg25201255
cg25201511
cg25201707
cg25201811
cg25201881
cg25201998
cg25202070
cg25202141
cg25203007
cg25203085
cg25203286
cg25203351
cg25203481
cg25203916
cg25204186
cg25205321
cg25205489
cg25205699
cg25206073
cg25206709
cg25207248
cg25207366
cg25207627
cg25208258
cg25209111
cg25209672
cg25210796
cg25211267
cg25211705
cg25212453
cg25213547
cg25213720
cg25213968
cg25214305
cg25214900
cg25214914
cg25215028
cg25215890
cg25216196
cg25216580
cg25216634
cg25216669
cg25217710

cg25217765
cg25217842
cg25218152
cg25218351
cg25218716
cg25218793
cg25220751
cg25220781
cg25220990
cg25221235
cg25221239
cg25223567
cg25223664
cg25224090
cg25224232
cg25225483
cg25225607
cg25227672
cg25228625
cg25230074
cg25231776
cg25232725
cg25233024
cg25233208
cg25233271
cg25234117
cg25234732
cg25235766
cg25236971
cg25237113
cg25237894
cg25238382
cg25240964
cg25242423
cg25242557
cg25242729
cg25242988
cg25243475
cg25244722
cg25244921
cg25244980
cg25245161
cg25245192
cg25246082
cg25246431
cg25247083
cg25247122
cg25247177
cg25247290
cg25247402
cg25247558
cg25247888

cg25247976
cg25248278
cg25248415
cg25249068
cg25249539
cg25249681
cg25249713
cg25249728
cg25250013
cg25250358
cg25250650
cg25252708
cg25253677
cg25255850
cg25255988
cg25256067
cg25256924
cg25257748
cg25259697
cg25260137
cg25260543
cg25262147
cg25263189
cg25263454
cg25264101
cg25264393
cg25264630
cg25266521
cg25266629
cg25267808
cg25268100
cg25268678
cg25268681
cg25269388
cg25269404
cg25269756
cg25270178
cg25270844
cg25271063
cg25271525
cg25273160
cg25273233
cg25274468
cg25274975
cg25275372
cg25276223
cg25276521
cg25276892
cg25278144
cg25278347
cg25278941
cg25279613

cg25280526
cg25281029
cg25281820
cg25282285
cg25282354
cg25283486
cg25283916
cg25284213
cg25285440
cg25285720
cg25285904
cg25286012
cg25286482
cg25286579
cg25286679
cg25287257
cg25287372
cg25287708
cg25288068
cg25289028
cg25289569
cg25290175
cg25290482
cg25291404
cg25291978
cg25292309
cg25294504
cg25294906
cg25295727
cg25296222
cg25296374
cg25296923
cg25297790
cg25298189
cg25301649
cg25301756
cg25302284
cg25302646
cg25302957
cg25303150
cg25303831
cg25304107
cg25304136
cg25304405
cg25304439
cg25306087
cg25306838
cg25307074
cg25307168
cg25307332
cg25308381
cg25308508

cg25308562
cg25308742
cg25308878
cg25308885
cg25309041
cg25309493
cg25309759
cg25310233
cg25310567
cg25310676
cg25311623
cg25311666
cg25311963
cg25312229
cg25312594
cg25312636
cg25312939
cg25312970
cg25313172
cg25313930
cg25314245
cg25314624
cg25314733
cg25316305
cg25316663
cg25316749
cg25317435
cg25317585
cg25318189
cg25318211
cg25318296
cg25318809
cg25319067
cg25319237
cg25320328
cg25320443
cg25320665
cg25321885
cg25322847
cg25323444
cg25323497
cg25323554
cg25324047
cg25324068
cg25324786
cg25325322
cg25325454
cg25325932
cg25326048
cg25326378
cg25330843
cg25331309

cg25331880
cg25332499
cg25333529
cg25334575
cg25336198
cg25336892
cg25336900
cg25337124
cg25337965
cg25338000
cg25339073
cg25339112
cg25339566
cg25339815
cg25340098
cg25340403
cg25340973
cg25341925
cg25342059
cg25342568
cg25342642
cg25343227
cg25343331
cg25343618
cg25343734
cg25344265
cg25344503
cg25345102
cg25345178
cg25345603
cg25345625
cg25345746
cg25346709
cg25346915
cg25346936
cg25347615
cg25348841
cg25349154
cg25349373
cg25349900
cg25349939
cg25350252
cg25350703
cg25350825
cg25351550
cg25351606
cg25351763
cg25352328
cg25352781
cg25352940
cg25352980
cg25353171

cg25353287
cg25353308
cg25353384
cg25354587
cg25354657
cg25355006
cg25355599
cg25355803
cg25356715
cg25356719
cg25357359
cg25357706
cg25357729
cg25357825
cg25357830
cg25360059
cg25360385
cg25360518
cg25360861
cg25361651
cg25361694
cg25362648
cg25362652
cg25363080
cg25363337
cg25363387
cg25363564
cg25364343
cg25366404
cg25366586
cg25367332
cg25367912
cg25368103
cg25368482
cg25368647
cg25368824
cg25369185
cg25370412
cg25371634
cg25372195
cg25372290
cg25373401
cg25373430
cg25373553
cg25374161
cg25374494
cg25374854
cg25375162
cg25375656
cg25375757
cg25375916
cg25376875

cg25377206
cg25377289
cg25377802
cg25377865
cg25378181
cg25378920
cg25379026
cg25379762
cg25380207
cg25380351
cg25381017
cg25381737
cg25382032
cg25382687
cg25384000
cg25384556
cg25385819
cg25385940
cg25386234
cg25386579
cg25386707
cg25386845
cg25388349
cg25388527
cg25388528
cg25388738
cg25388823
cg25388861
cg25388882
cg25389393
cg25389804
cg25390063
cg25390243
cg25391092
cg25391162
cg25391681
cg25391820
cg25392995
cg25393429
cg25393613
cg25394452
cg25395087
cg25395619
cg25397111
cg25397597
cg25397973
cg25398315
cg25398727
cg25400358
cg25400746
cg25401219
cg25402049

cg25402083
cg25402228
cg25403205
cg25403721
cg25403895
cg25404339
cg25405138
cg25405922
cg25405962
cg25406011
cg25406665
cg25407540
cg25407979
cg25408329
cg25408617
cg25408620
cg25410117
cg25410668
cg25411239
cg25412230
cg25412387
cg25414209
cg25415384
cg25415966
cg25415989
cg25416319
cg25417529
cg25417675
cg25417766
cg25418001
cg25420253
cg25420254
cg25420502
cg25420747
cg25420945
cg25420952
cg25421209
cg25421530
cg25421908
cg25422678
cg25422880
cg25422992
cg25423174
cg25423647
cg25424029
cg25424194
cg25424525
cg25425074
cg25425089
cg25425390
cg25425667
cg25425705

cg25426302
cg25426892
cg25427625
cg25429137
cg25429446
cg25430442
cg25430713
cg25430878
cg25432696
cg25432792
cg25436126
cg25436157
cg25436486
cg25436634
cg25436766
cg25437359
cg25438801
cg25439479
cg25439496
cg25440035
cg25441207
cg25441879
cg25442600
cg25443512
cg25443943
cg25444339
cg25444386
cg25444821
cg25445707
cg25446061
cg25446460
cg25446612
cg25446789
cg25448281
cg25449484
cg25449821
cg25451456
cg25452231
cg25453154
cg25453841
cg25454931
cg25455408
cg25455811
cg25455865
cg25456020
cg25456144
cg25456468
cg25457064
cg25457832
cg25459301
cg25459558
cg25459800

cg25460031
cg25461801
cg25462858
cg25463470
cg25463688
cg25464921
cg25465322
cg25465483
cg25466245
cg25467833
cg25469314
cg25469406
cg25469414
cg25469474
cg25471639
cg25471644
cg25471802
cg25471987
cg25474372
cg25475742
cg25477456
cg25477843
cg25478113
cg25478600
cg25479708
cg25479788
cg25481253
cg25481454
cg25481630
cg25482132
cg25482454
cg25482786
cg25482900
cg25482983
cg25483596
cg25484139
cg25484904
cg25485734
cg25485875
cg25485994
cg25486618
cg25486738
cg25486757
cg25486957
cg25487404
cg25488160
cg25488288
cg25488990
cg25490551
cg25491558
cg25493276
cg25493335

cg25493589
cg25494395
cg25496678
cg25497250
cg25497399
cg25498489
cg25499099
cg25499650
cg25500001
cg25500080
cg25500196
cg25500948
cg25502399
cg25503381
cg25503410
cg25503804
cg25504884
cg25505069
cg25507121
cg25507767
cg25507845
cg25508181
cg25508633
cg25508830
cg25509184
cg25509872
cg25510090
cg25510201
cg25510614
cg25511813
cg25512683
cg25513321
cg25513740
cg25513776
cg25513853
cg25514273
cg25515091
cg25515758
cg25517810
cg25518132
cg25518365
cg25518868
cg25520440
cg25520910
cg25522046
cg25523332
cg25523477
cg25523509
cg25523538
cg25524052
cg25524464
cg25525825

cg25526039
cg25526519
cg25526960
cg25527090
cg25527547
cg25527770
cg25528121
cg25528298
cg25528449
cg25528806
cg25529535
cg25530299
cg25530583
cg25533387
cg25535316
cg25535587
cg25536014
cg25536171
cg25537119
cg25537669
cg25537993
cg25538881
cg25538883
cg25539045
cg25540871
cg25541209
cg25541259
cg25541621
cg25542878
cg25545500
cg25546118
cg25546253
cg25547580
cg25549141
cg25549459
cg25549720
cg25550425
cg25550629
cg25551622
cg25552018
cg25552843
cg25553916
cg25554316
cg25554340
cg25554740
cg25554943
cg25555059
cg25555068
cg25556035
cg25556051
cg25557432
cg25557835

cg25558027
cg25558103
cg25558288
cg25558617
cg25560518
cg25561401
cg25561792
cg25562031
cg25562059
cg25562958
cg25563256
cg25563456
cg25565082
cg25565203
cg25566568
cg25567054
cg25567280
cg25567448
cg25567594
cg25567938
cg25568066
cg25569300
cg25569624
cg25569880
cg25570278
cg25570453
cg25570522
cg25570913
cg25570929
cg25571189
cg25571414
cg25573386
cg25573480
cg25573915
cg25574024
cg25574111
cg25574227
cg25574691
cg25575937
cg25575961
cg25576011
cg25576093
cg25576788
cg25577367
cg25577463
cg25577489
cg25577670
cg25578225
cg25578773
cg25579854
cg25580497
cg25581222

cg25581822
cg25584474
cg25584668
cg25585043
cg25585185
cg25585712
cg25586011
cg25587069
cg25587091
cg25588844
cg25589070
cg25589371
cg25591260
cg25591301
cg25591451
cg25591888
cg25592353
cg25592910
cg25593476
cg25593565
cg25594106
cg25594736
cg25595388
cg25596287
cg25596289
cg25596584
cg25597390
cg25597623
cg25597784
cg25597797
cg25597894
cg25598488
cg25598685
cg25599065
cg25599521
cg25599538
cg25599567
cg25599573
cg25599761
cg25600410
cg25600766
cg25600951
cg25601886
cg25602242
cg25602520
cg25602684
cg25602692
cg25603108
cg25603636
cg25603742
cg25603927
cg25604731

cg25605174
cg25605243
cg25605731
cg25606046
cg25606097
cg25606842
cg25606969
cg25607920
cg25609434
cg25610200
cg25610428
cg25611057
cg25611736
cg25613180
cg25614094
cg25614364
cg25614935
cg25615327
cg25616160
cg25616535
cg25617092
cg25617157
cg25617340
cg25617725
cg25619312
cg25619551
cg25619586
cg25621286
cg25622154
cg25623768
cg25623833
cg25624359
cg25624690
cg25625127
cg25625146
cg25625296
cg25625670
cg25626166
cg25626375
cg25626611
cg25627098
cg25627226
cg25627242
cg25628433
cg25628898
cg25628989
cg25629905
cg25630910
cg25631489
cg25632105
cg25632577
cg25633067

cg25633678
cg25634141
cg25634742
cg25635916
cg25636159
cg25636481
cg25636933
cg25637966
cg25638443
cg25638611
cg25638626
cg25639095
cg25639531
cg25640096
cg25640519
cg25640822
cg25641439
cg25643208
cg25644095
cg25644224
cg25644243
cg25644556
cg25644649
cg25645491
cg25646708
cg25648203
cg25649000
cg25649188
cg25649641
cg25649765
cg25650360
cg25650956
cg25650964
cg25652701
cg25652821
cg25653204
cg25653368
cg25653449
cg25653802
cg25653816
cg25654653
cg25654677
cg25655234
cg25655854
cg25656098
cg25657713
cg25658010
cg25658276
cg25662041
cg25662428
cg25662463
cg25663326

cg25663892
cg25663970
cg25664021
cg25664054
cg25665528
cg25665622
cg25667676
cg25668093
cg25670330
cg25670451
cg25670685
cg25671242
cg25673112
cg25673241
cg25678587
cg25678815
cg25680014
cg25681339
cg25681469
cg25684640
cg25684999
cg25685262
cg25685359
cg25685455
cg25685510
cg25685675
cg25687179
cg25687507
cg25687642
cg25689193
cg25691167
cg25691899
cg25692507
cg25695568
cg25696972
cg25697881
cg25698842
cg25699727
cg25699851
cg25700513
cg25700897
cg25702335
cg25703338
cg25703534
cg25704278
cg25705508
cg25706281
cg25706683
cg25706990
cg25707238
cg25707298
cg25707767

cg25708328
cg25710298
cg25711239
cg25711246
cg25711375
cg25711575
cg25711963
cg25713185
cg25713625
cg25714096
cg25714115
cg25714306
cg25716102
cg25716149
cg25717032
cg25717406
cg25717464
cg25718132
cg25718467
cg25719786
cg25720804
cg25720815
cg25720825
cg25722465
cg25722880
cg25722983
cg25723050
cg25723866
cg25724283
cg25724842
cg25725843
cg25726847
cg25728328
cg25729094
cg25729166
cg25729432
cg25729445
cg25730685
cg25731261
cg25731428
cg25731959
cg25732028
cg25732462
cg25734089
cg25734198
cg25734539
cg25734624
cg25734726
cg25734928
cg25735305
cg25735572
cg25735823

cg25736183
cg25736198
cg25737836
cg25738340
cg25738714
cg25739043
cg25739875
cg25739882
cg25740457
cg25740565
cg25740602
cg25741116
cg25741118
cg25741452
cg25741487
cg25741562
cg25741810
cg25741837
cg25743622
cg25744476
cg25744613
cg25744767
cg25745642
cg25746778
cg25746890
cg25747190
cg25747427
cg25747655
cg25747670
cg25748521
cg25748524
cg25748958
cg25749107
cg25749254
cg25749341
cg25749512
cg25750004
cg25751961
cg25752514
cg25754143
cg25754485
cg25756406
cg25756617
cg25756780
cg25757115
cg25757598
cg25757869
cg25757877
cg25758195
cg25759507
cg25761793
cg25762078

cg25762654
cg25762679
cg25762974
cg25763036
cg25763393
cg25763716
cg25764120
cg25764899
cg25764996
cg25765192
cg25765264
cg25766271
cg25766774
cg25767154
cg25767345
cg25767433
cg25768734
cg25769127
cg25769417
cg25770225
cg25770267
cg25770340
cg25771113
cg25771271
cg25771451
cg25772185
cg25773223
cg25773259
cg25774643
cg25775322
cg25775386
cg25775770
cg25778446
cg25778497
cg25779483
cg25780109
cg25780607
cg25781210
cg25781755
cg25781936
cg25781963
cg25782440
cg25782665
cg25783241
cg25783987
cg25784136
cg25784219
cg25784220
cg25784280
cg25784308
cg25785303
cg25785397

cg25785495
cg25786366
cg25786651
cg25787300
cg25787499
cg25788088
cg25788103
cg25788549
cg25789277
cg25789861
cg25790298
cg25791330
cg25792473
cg25792581
cg25793377
cg25793597
cg25795774
cg25796439
cg25798460
cg25798600
cg25798782
cg25798792
cg25800765
cg25802093
cg25805521
cg25806591
cg25806701
cg25806808
cg25806932
cg25808809
cg25809290
cg25809861
cg25811526
cg25811633
cg25812683
cg25813290
cg25813942
cg25814096
cg25814649
cg25814752
cg25815024
cg25817111
cg25817165
cg25817351
cg25818109
cg25818292
cg25818605
cg25818697
cg25818763
cg25820699
cg25820728
cg25820791

cg25821168
cg25821245
cg25821399
cg25823128
cg25824218
cg25824236
cg25824509
cg25825488
cg25825506
cg25826173
cg25827670
cg25829531
cg25830182
cg25830696
cg25831075
cg25831752
cg25832084
cg25832305
cg25832867
cg25833566
cg25834692
cg25836827
cg25837803
cg25837984
cg25838150
cg25838818
cg25838968
cg25839663
cg25839724
cg25840094
cg25840173
cg25840237
cg25840608
cg25840685
cg25840780
cg25841309
cg25841608
cg25842285
cg25843174
cg25844133
cg25845326
cg25846723
cg25850629
cg25850808
cg25851597
cg25852019
cg25852472
cg25853097
cg25853612
cg25855592
cg25856811
cg25859099

cg25859468
cg25860737
cg25861340
cg25862140
cg25862345
cg25862444
cg25863223
cg25864024
cg25865120
cg25865368
cg25869196
cg25869950
cg25872816
cg25872852
cg25874108
cg25874148
cg25874953
cg25875049
cg25876227
cg25876586
cg25878149
cg25879062
cg25879561
cg25879836
cg25880347
cg25880537
cg25880834
cg25880980
cg25882366
cg25883066
cg25883149
cg25883348
cg25883762
cg25884399
cg25888561
cg25888914
cg25889504
cg25889581
cg25889711
cg25890391
cg25890838
cg25891355
cg25891433
cg25891647
cg25892286
cg25892296
cg25894334
cg25894551
cg25895695
cg25895913
cg25895948
cg25897251

cg25897951
cg25898192
cg25898520
cg25898550
cg25900085
cg25901381
cg25903158
cg25903783
cg25904166
cg25905215
cg25906705
cg25906711
cg25907132
cg25907215
cg25908283
cg25908985
cg25911468
cg25911968
cg25912037
cg25912428
cg25912611
cg25912699
cg25912911
cg25913882
cg25915512
cg25915838
cg25916135
cg25916172
cg25916711
cg25918821
cg25918827
cg25918833
cg25919687
cg25920406
cg25920512
cg25920535
cg25920792
cg25920857
cg25921357
cg25921502
cg25921518
cg25921544
cg25922329
cg25922610
cg25923056
cg25923609
cg25923629
cg25923788
cg25925006
cg25926515
cg25927708
cg25928199

cg25928603
cg25928819
cg25929046
cg25929731
cg25929955
cg25930639
cg25930644
cg25931371
cg25931491
cg25932761
cg25933341
cg25934064
cg25934728
cg25934824
cg25935985
cg25936054
cg25936358
cg25936381
cg25936902
cg25937230
cg25937451
cg25937790
cg25938010
cg25938530
cg25938646
cg25938891
cg25939350
cg25939647
cg25939793
cg25939869
cg25940008
cg25940485
cg25940517
cg25941151
cg25941751
cg25942450
cg25942761
cg25943276
cg25943503
cg25943588
cg25943727
cg25944168
cg25944419
cg25944717
cg25947121
cg25947408
cg25947945
cg25947970
cg25948075
cg25948180
cg25948591
cg25948982

cg25950100
cg25950112
cg25950235
cg25950278
cg25950625
cg25951075
cg25951210
cg25951981
cg25952037
cg25952192
cg25952964
cg25953239
cg25953417
cg25953672
cg25953788
cg25954134
cg25954269
cg25954705
cg25955126
cg25955692
cg25955817
cg25956106
cg25956483
cg25956985
cg25957301
cg25957599
cg25957967
cg25958283
cg25958582
cg25959131
cg25960038
cg25960272
cg25960479
cg25960785
cg25961292
cg25961618
cg25962657
cg25962699
cg25962829
cg25963041
cg25964032
cg25964728
cg25965576
cg25965776
cg25966599
cg25966682
cg25967418
cg25967459
cg25968835
cg25968937
cg25969460
cg25969932

cg25970726
cg25970832
cg25971844
cg25972997
cg25973534
cg25973580
cg25974617
cg25974888
cg25974922
cg25975712
cg25975805
cg25975856
cg25976242
cg25977206
cg25977900
cg25977938
cg25978208
cg25979244
cg25979559
cg25979644
cg25980042
cg25980168
cg25980489
cg25982880
cg25982983
cg25983553
cg25983901
cg25984068
cg25984344
cg25984671
cg25984996
cg25985347
cg25985659
cg25986240
cg25987452
cg25987744
cg25987842
cg25987923
cg25988034
cg25988118
cg25988603
cg25988830
cg25989526
cg25991631
cg25992016
cg25993718
cg25994968
cg25995825
cg25996077
cg25996119
cg25996388
cg25996663

cg25996688
cg25997110
cg25997187
cg25998402
cg25999442
cg25999578
cg26000011
cg26000663
cg26001333
cg26001902
cg26002370
cg26002512
cg26002568
cg26002634
cg26003056
cg26003119
cg26004203
cg26004221
cg26004710
cg26005232
cg26005329
cg26006870
cg26008369
cg26008785
cg26009944
cg26010412
cg26010751
cg26012927
cg26014036
cg26014391
cg26014409
cg26014538
cg26014689
cg26014994
cg26015134
cg26017043
cg26017484
cg26017754
cg26018073
cg26019112
cg26019295
cg26020069
cg26020982
cg26021810
cg26023228
cg26023709
cg26024424
cg26024682
cg26024843
cg26025224
cg26025487
cg26026598

cg26026951
cg26027576
cg26028371
cg26028440
cg26028571
cg26029221
cg26029547
cg26031046
cg26031192
cg26031678
cg26032325
cg26032419
cg26033349
cg26033526
cg26033681
cg26034629
cg26034658
cg26034769
cg26034811
cg26034934
cg26035366
cg26035542
cg26037026
cg26037848
cg26037936
cg26038589
cg26038697
cg26039217
cg26039298
cg26039623
cg26040244
cg26040761
cg26040903
cg26042300
cg26043322
cg26043460
cg26044566
cg26046136
cg26047127
cg26048334
cg26049998
cg26050159
cg26050864
cg26050906
cg26053480
cg26053864
cg26053881
cg26054057
cg26055120
cg26055370
cg26055770
cg26055899

cg26056166
cg26056703
cg26057616
cg26057784
cg26058016
cg26058884
cg26059153
cg26059468
cg26059632
cg26060835
cg26061177
cg26062204
cg26062581
cg26062856
cg26063719
cg26064142
cg26064774
cg26065057
cg26065799
cg26066361
cg26067203
cg26067319
cg26068079
cg26068180
cg26068215
cg26068551
cg26068654
cg26068911
cg26069068
cg26069230
cg26069291
cg26069539
cg26069919
cg26070020
cg26070379
cg26070487
cg26070540
cg26071135
cg26071414
cg26071526
cg26073060
cg26073101
cg26073885
cg26074326
cg26074603
cg26076793
cg26077454
cg26079428
cg26079753
cg26079857
cg26080567
cg26080826

cg26081162
cg26081584
cg26082257
cg26082814
cg26083180
cg26083458
cg26084511
cg26084529
cg26085533
cg26085987
cg26086358
cg26086468
cg26086785
cg26086902
cg26087411
cg26089160
cg26090045
cg26090619
cg26091021
cg26091232
cg26091304
cg26092233
cg26093145
cg26094232
cg26094484
cg26094554
cg26094789
cg26096752
cg26097549
cg26098117
cg26098816
cg26099018
cg26099134
cg26099316
cg26100099
cg26101714
cg26101767
cg26101942
cg26102295
cg26102819
cg26103306
cg26103845
cg26104541
cg26104640
cg26104690
cg26105283
cg26106316
cg26106778
cg26106948
cg26107597
cg26108690
cg26109145

cg26109510
cg26110064
cg26110281
cg26110605
cg26111030
cg26111757
cg26111881
cg26112170
cg26112574
cg26112639
cg26112909
cg26112999
cg26113233
cg26113278
cg26114043
cg26114324
cg26114352
cg26115633
cg26115667
cg26116485
cg26116542
cg26116669
cg26116950
cg26117369
cg26117398
cg26118045
cg26118131
cg26118326
cg26118553
cg26118698
cg26118906
cg26118943
cg26119806
cg26120226
cg26120279
cg26120636
cg26120813
cg26120958
cg26121234
cg26121591
cg26121639
cg26122129
cg26122710
cg26122980
cg26123532
cg26123605
cg26123647
cg26123763
cg26123824
cg26124016
cg26124318
cg26124980

cg26125956
cg26126417
cg26127114
cg26127479
cg26127662
cg26127778
cg26127925
cg26128079
cg26129108
cg26129110
cg26130053
cg26130242
cg26130488
cg26131286
cg26131315
cg26132320
cg26132511
cg26132774
cg26133091
cg26133399
cg26133769
cg26134090
cg26134692
cg26134895
cg26136794
cg26137350
cg26137940
cg26138637
cg26140366
cg26141385
cg26142132
cg26142412
cg26143577
cg26144658
cg26144999
cg26145228
cg26145268
cg26145511
cg26147279
cg26147480
cg26148020
cg26148774
cg26149244
cg26149275
cg26149567
cg26149658
cg26149682
cg26150300
cg26150514
cg26151467
cg26151761
cg26151910

cg26152427
cg26153096
cg26153353
cg26153410
cg26153688
cg26153885
cg26154831
cg26155170
cg26155492
cg26155674
cg26155802
cg26156256
cg26157385
cg26157386
cg26158133
cg26158180
cg26158270
cg26158406
cg26158528
cg26159090
cg26159205
cg26159860
cg26160008
cg26160189
cg26161104
cg26161885
cg26162295
cg26162554
cg26162564
cg26162616
cg26163057
cg26163101
cg26163234
cg26163319
cg26163458
cg26163537
cg26163578
cg26163699
cg26163880
cg26164310
cg26164488
cg26164577
cg26164773
cg26165108
cg26165146
cg26167249
cg26169081
cg26169213
cg26170569
cg26170604
cg26170805
cg26172195

cg26172504
cg26173447
cg26173630
cg26174018
cg26176717
cg26177096
cg26177760
cg26177968
cg26178251
cg26178657
cg26181334
cg26181818
cg26181894
cg26182253
cg26183021
cg26183708
cg26184391
cg26185056
cg26185071
cg26185079
cg26185508
cg26186331
cg26186546
cg26186727
cg26187031
cg26187205
cg26187724
cg26187962
cg26188131
cg26188212
cg26188226
cg26189021
cg26190363
cg26190386
cg26190835
cg26190966
cg26191586
cg26191882
cg26191923
cg26192309
cg26192612
cg26193268
cg26193418
cg26194447
cg26194477
cg26194775
cg26195356
cg26196087
cg26196124
cg26196424
cg26197699
cg26198638

cg26199241
cg26199552
cg26199631
cg26200118
cg26201596
cg26202964
cg26205890
cg26206196
cg26206396
cg26207568
cg26207909
cg26208159
cg26210364
cg26210445
cg26211060
cg26213224
cg26213438
cg26213458
cg26215982
cg26216243
cg26216433
cg26216876
cg26217441
cg26217588
cg26217832
cg26218277
cg26218380
cg26218698
cg26218809
cg26219069
cg26219233
cg26219540
cg26219843
cg26220018
cg26220336
cg26220673
cg26220985
cg26221111
cg26221801
cg26222731
cg26223968
cg26224018
cg26224314
cg26224781
cg26225417
cg26225478
cg26225694
cg26225893
cg26226988
cg26227044
cg26227119
cg26227465

cg26227592
cg26228024
cg26228464
cg26229018
cg26229087
cg26229201
cg26229224
cg26230338
cg26230851
cg26231094
cg26231267
cg26231272
cg26231761
cg26232282
cg26232359
cg26232451
cg26233331
cg26234223
cg26234233
cg26234721
cg26234900
cg26235215
cg26235369
cg26235748
cg26236177
cg26236235
cg26237512
cg26237628
cg26237651
cg26238041
cg26238771
cg26239297
cg26239725
cg26239984
cg26240939
cg26241863
cg26241893
cg26242304
cg26242531
cg26242583
cg26243740
cg26244013
cg26246074
cg26246342
cg26246572
cg26247093
cg26247309
cg26247508
cg26248075
cg26249106
cg26249135
cg26249245

cg26249834
cg26249852
cg26249873
cg26250086
cg26251098
cg26251585
cg26251680
cg26251865
cg26252045
cg26252077
cg26252167
cg26252281
cg26252592
cg26253134
cg26253577
cg26253663
cg26253898
cg26254193
cg26254678
cg26255089
cg26255314
cg26255996
cg26256630
cg26257822
cg26258274
cg26258396
cg26259363
cg26259381
cg26259537
cg26259818
cg26261291
cg26262055
cg26262988
cg26263234
cg26263239
cg26263263
cg26263664
cg26263674
cg26264032
cg26264232
cg26264896
cg26266099
cg26266429
cg26266708
cg26266876
cg26267340
cg26267388
cg26267678
cg26267852
cg26268145
cg26268565
cg26268742

cg26269286
cg26269966
cg26270038
cg26270261
cg26270879
cg26271435
cg26271776
cg26271970
cg26272069
cg26272214
cg26272220
cg26272623
cg26272651
cg26274662
cg26274891
cg26274946
cg26274995
cg26275110
cg26275373
cg26275430
cg26275647
cg26275799
cg26276014
cg26276249
cg26276301
cg26276327
cg26277556
cg26277754
cg26278103
cg26278151
cg26278454
cg26279183
cg26279372
cg26279668
cg26279783
cg26280179
cg26280326
cg26280713
cg26280998
cg26281107
cg26281310
cg26282347
cg26282384
cg26282731
cg26282761
cg26283220
cg26283240
cg26283480
cg26283496
cg26283874
cg26283921
cg26284264

cg26284735
cg26284782
cg26286082
cg26286198
cg26286365
cg26287609
cg26288715
cg26288892
cg26289049
cg26289295
cg26289788
cg26290632
cg26291655
cg26291848
cg26292150
cg26292521
cg26292918
cg26293019
cg26293512
cg26295057
cg26295774
cg26295786
cg26296364
cg26296488
cg26296752
cg26297547
cg26297819
cg26298192
cg26298273
cg26298599
cg26298855
cg26299169
cg26300500
cg26300652
cg26300672
cg26300785
cg26301215
cg26301562
cg26301690
cg26301788
cg26302157
cg26303222
cg26304088
cg26305174
cg26305683
cg26305756
cg26305827
cg26306056
cg26306530
cg26306909
cg26306976
cg26307359

cg26307701
cg26307814
cg26308330
cg26308512
cg26309194
cg26309311
cg26309498
cg26310719
cg26311677
cg26311734
cg26311804
cg26311995
cg26312149
cg26312191
cg26312888
cg26312920
cg26313421
cg26313647
cg26315277
cg26315862
cg26316423
cg26316702
cg26316946
cg26317237
cg26317318
cg26317549
cg26317834
cg26318872
cg26320696
cg26321840
cg26321999
cg26322137
cg26322248
cg26322711
cg26323752
cg26325174
cg26325286
cg26326621
cg26327040
cg26327548
cg26327666
cg26328058
cg26328626
cg26328757
cg26328951
cg26328954
cg26329192
cg26329274
cg26329618
cg26330116
cg26330479
cg26330518

cg26330809
cg26331243
cg26331343
cg26332052
cg26332191
cg26332253
cg26332285
cg26332310
cg26332560
cg26333450
cg26333652
cg26333822
cg26334154
cg26334166
cg26334507
cg26334804
cg26335377
cg26335577
cg26335620
cg26335898
cg26336935
cg26336955
cg26337259
cg26337277
cg26337312
cg26337390
cg26337841
cg26338386
cg26338428
cg26338735
cg26339990
cg26340725
cg26341848
cg26342535
cg26342676
cg26343001
cg26343116
cg26343154
cg26343532
cg26343889
cg26344227
cg26344776
cg26344818
cg26345142
cg26345203
cg26346167
cg26346796
cg26347408
cg26347632
cg26347851
cg26348614
cg26348902

cg26349048
cg26349957
cg26350007
cg26350784
cg26350921
cg26351229
cg26351951
cg26353448
cg26353598
cg26353764
cg26353903
cg26354092
cg26354128
cg26354367
cg26354398
cg26354908
cg26355573
cg26355577
cg26355755
cg26355894
cg26356694
cg26357344
cg26357403
cg26357587
cg26359335
cg26359389
cg26360026
cg26360616
cg26360924
cg26360930
cg26361041
cg26361999
cg26362105
cg26362368
cg26362453
cg26363579
cg26363695
cg26363935
cg26364091
cg26364205
cg26365147
cg26365254
cg26365534
cg26365545
cg26365813
cg26366091
cg26366958
cg26367719
cg26368431
cg26368842
cg26369269
cg26369594

cg26370729
cg26371320
cg26371327
cg26371501
cg26371913
cg26372498
cg26373051
cg26373071
cg26373874
cg26374826
cg26375057
cg26376241
cg26376245
cg26378179
cg26378984
cg26379258
cg26379475
cg26379859
cg26380116
cg26380692
cg26381210
cg26381919
cg26382173
cg26383193
cg26384465
cg26384697
cg26385172
cg26385283
cg26385743
cg26386189
cg26386624
cg26386653
cg26386826
cg26387064
cg26387689
cg26388204
cg26388707
cg26388816
cg26389500
cg26389888
cg26389913
cg26389955
cg26390214
cg26390660
cg26391219
cg26391902
cg26392708
cg26392737
cg26393534
cg26393630
cg26394055
cg26394191

cg26394257
cg26394380
cg26394624
cg26395382
cg26395507
cg26395694
cg26398044
cg26398341
cg26398656
cg26398921
cg26400341
cg26400546
cg26401796
cg26402410
cg26402417
cg26402435
cg26402555
cg26402630
cg26403171
cg26403416
cg26403580
cg26403608
cg26403843
cg26404511
cg26405020
cg26405841
cg26406394
cg26406441
cg26406891
cg26406924
cg26407161
cg26407917
cg26408612
cg26409237
cg26409494
cg26409712
cg26410450
cg26410540
cg26411015
cg26411222
cg26413192
cg26413528
cg26413691
cg26413942
cg26416887
cg26416905
cg26417361
cg26417554
cg26417713
cg26417912
cg26418147
cg26418671

cg26419265
cg26419676
cg26420013
cg26422161
cg26422458
cg26422762
cg26422784
cg26423145
cg26424147
cg26424300
cg26425256
cg26425305
cg26425711
cg26426334
cg26426470
cg26426632
cg26426690
cg26426938
cg26427498
cg26428069
cg26429140
cg26429381
cg26429636
cg26429925
cg26430027
cg26430081
cg26430560
cg26431018
cg26432088
cg26432256
cg26432539
cg26433322
cg26433494
cg26433554
cg26433816
cg26433955
cg26434048
cg26435109
cg26435172
cg26435575
cg26437203
cg26437826
cg26438846
cg26439139
cg26441291
cg26441627
cg26442107
cg26442903
cg26443093
cg26444528
cg26446071
cg26446133

cg26446499
cg26446593
cg26446803
cg26447413
cg26448394
cg26449119
cg26449294
cg26449680
cg26450230
cg26450574
cg26450717
cg26450750
cg26451198
cg26451345
cg26451930
cg26452056
cg26452081
cg26452771
cg26452868
cg26454666
cg26456223
cg26456304
cg26456399
cg26457075
cg26457160
cg26457700
cg26457809
cg26458286
cg26458557
cg26458748
cg26459819
cg26460366
cg26460471
cg26460483
cg26460530
cg26460678
cg26461267
cg26461477
cg26461695
cg26462586
cg26462595
cg26462924
cg26463800
cg26464446
cg26464701
cg26464821
cg26464882
cg26465405
cg26465837
cg26466145
cg26466258
cg26466375

cg26466587
cg26466781
cg26467528
cg26467536
cg26467999
cg26468007
cg26468693
cg26468696
cg26468878
cg26469011
cg26469099
cg26469387
cg26469608
cg26470479
cg26470501
cg26470696
cg26470798
cg26470926
cg26470937
cg26472133
cg26472802
cg26472910
cg26472922
cg26474014
cg26474549
cg26474814
cg26475500
cg26475742
cg26475911
cg26476852
cg26476925
cg26477387
cg26477488
cg26478079
cg26478205
cg26478215
cg26479022
cg26479374
cg26479630
cg26479868
cg26480039
cg26480662
cg26481302
cg26482052
cg26482626
cg26482939
cg26483075
cg26483229
cg26484108
cg26484464
cg26485825
cg26485937

cg26486070
cg26487582
cg26488544
cg26488999
cg26489750
cg26490054
cg26490274
cg26490626
cg26492368
cg26492446
cg26492480
cg26492514
cg26492686
cg26493188
cg26494044
cg26494929
cg26495092
cg26496795
cg26498132
cg26498574
cg26499286
cg26500142
cg26500398
cg26500588
cg26500804
cg26501046
cg26501154
cg26501671
cg26502018
cg26502127
cg26502489
cg26502666
cg26503038
cg26503073
cg26503792
cg26504021
cg26504110
cg26504263
cg26504421
cg26504467
cg26504906
cg26505158
cg26506007
cg26506680
cg26506790
cg26506932
cg26507629
cg26508164
cg26508170
cg26508928
cg26509305
cg26509318

cg26509870
cg26510017
cg26510178
cg26511075
cg26511108
cg26511321
cg26511585
cg26512226
cg26512635
cg26512948
cg26513304
cg26513654
cg26513689
cg26513818
cg26515460
cg26515755
cg26516287
cg26516935
cg26517171
cg26517663
cg26517714
cg26520739
cg26521139
cg26521207
cg26521404
cg26521766
cg26522157
cg26522239
cg26522684
cg26522708
cg26523175
cg26523338
cg26523478
cg26524638
cg26525162
cg26525313
cg26525592
cg26526374
cg26526397
cg26527263
cg26527984
cg26528311
cg26529516
cg26529687
cg26530275
cg26530713
cg26530942
cg26531286
cg26533107
cg26533193
cg26533368
cg26534460

cg26534493
cg26534677
cg26534938
cg26535871
cg26537098
cg26537209
cg26537335
cg26537478
cg26538164
cg26538442
cg26538691
cg26538782
cg26539042
cg26539631
cg26540004
cg26540410
cg26540441
cg26540515
cg26541867
cg26542218
cg26542254
cg26542567
cg26543333
cg26543539
cg26543908
cg26544247
cg26544499
cg26544719
cg26544722
cg26544981
cg26545040
cg26545162
cg26545519
cg26546155
cg26546373
cg26546687
cg26546912
cg26547506
cg26547898
cg26548095
cg26548251
cg26548682
cg26548834
cg26549601
cg26549701
cg26551092
cg26551913
cg26552621
cg26552733
cg26553513
cg26553763
cg26554141

cg26554265
cg26554326
cg26554592
cg26555966
cg26556336
cg26556385
cg26556489
cg26556719
cg26557540
cg26557617
cg26557693
cg26558111
cg26558485
cg26559209
cg26559315
cg26559829
cg26560222
cg26560367
cg26560769
cg26561254
cg26561358
cg26561681
cg26562566
cg26562772
cg26563242
cg26563889
cg26564172
cg26564768
cg26565021
cg26565544
cg26566660
cg26566927
cg26567423
cg26567496
cg26568372
cg26568762
cg26569309
cg26569349
cg26569800
cg26570389
cg26570574
cg26570830
cg26571002
cg26571105
cg26571889
cg26572548
cg26572608
cg26572770
cg26572865
cg26572896
cg26572905
cg26572992

cg26574265
cg26574680
cg26575173
cg26575914
cg26576047
cg26576454
cg26577252
cg26577529
cg26577836
cg26578156
cg26578274
cg26578682
cg26578726
cg26578923
cg26579145
cg26579556
cg26579578
cg26579986
cg26580869
cg26581200
cg26581449
cg26582706
cg26582733
cg26583474
cg26584120
cg26584545
cg26584747
cg26585899
cg26586508
cg26586719
cg26586917
cg26586952
cg26587228
cg26588076
cg26588688
cg26589669
cg26590003
cg26591679
cg26592076
cg26593267
cg26593946
cg26594179
cg26595278
cg26595643
cg26596350
cg26596719
cg26597242
cg26600461
cg26601073
cg26601317
cg26601574
cg26602558

cg26603047
cg26603050
cg26604359
cg26604799
cg26604895
cg26605164
cg26605476
cg26607031
cg26607228
cg26607897
cg26608883
cg26609631
cg26611328
cg26611809
cg26612418
cg26612735
cg26612794
cg26613778
cg26614154
cg26614264
cg26615188
cg26615224
cg26616258
cg26616780
cg26617787
cg26618441
cg26618703
cg26618903
cg26619624
cg26619725
cg26620157
cg26620314
cg26620655
cg26621046
cg26621491
cg26621780
cg26622174
cg26622320
cg26623444
cg26623753
cg26624489
cg26625897
cg26626251
cg26628069
cg26628847
cg26629749
cg26630231
cg26630859
cg26631039
cg26631979
cg26631984
cg26632171

cg26632700
cg26632831
cg26633899
cg26635219
cg26635451
cg26636045
cg26636193
cg26636312
cg26636379
cg26638069
cg26638266
cg26638505
cg26639596
cg26639747
cg26639809
cg26639820
cg26640601
cg26640901
cg26641996
cg26642224
cg26642655
cg26643047
cg26643914
cg26644395
cg26644674
cg26645237
cg26645468
cg26645655
cg26646411
cg26647200
cg26647242
cg26647453
cg26647566
cg26647771
cg26648579
cg26648725
cg26648818
cg26649384
cg26649851
cg26650064
cg26650480
cg26650494
cg26650638
cg26651233
cg26651775
cg26651837
cg26652266
cg26652771
cg26653777
cg26653824
cg26653867
cg26653879

cg26654798
cg26654934
cg26655294
cg26655340
cg26655542
cg26655955
cg26656751
cg26657045
cg26657090
cg26657648
cg26657675
cg26657725
cg26658927
cg26659079
cg26659582
cg26660002
cg26660631
cg26661034
cg26661623
cg26661640
cg26661718
cg26662201
cg26662656
cg26662732
cg26663651
cg26663772
cg26664161
cg26664528
cg26665507
cg26665701
cg26665764
cg26665785
cg26666538
cg26666835
cg26667946
cg26668042
cg26668906
cg26669393
cg26669717
cg26669806
cg26670249
cg26670875
cg26671281
cg26671784
cg26671848
cg26671966
cg26672452
cg26672573
cg26673012
cg26673377
cg26674445
cg26674546

cg26675200
cg26675289
cg26675485
cg26675654
cg26675682
cg26676090
cg26677406
cg26678245
cg26678605
cg26678609
cg26678978
cg26679004
cg26679047
cg26679855
cg26680350
cg26680644
cg26680995
cg26681097
cg26681123
cg26681139
cg26681383
cg26681458
cg26681770
cg26682048
cg26682094
cg26682517
cg26682580
cg26682661
cg26683252
cg26683361
cg26683398
cg26683881
cg26684131
cg26684191
cg26684679
cg26684946
cg26685596
cg26685743
cg26686415
cg26687318
cg26687842
cg26688574
cg26689186
cg26689666
cg26690138
cg26690758
cg26690949
cg26692791
cg26693467
cg26693953
cg26694290
cg26694437

cg26697065
cg26697334
cg26697605
cg26697929
cg26698693
cg26698819
cg26699292
cg26700320
cg26700407
cg26700702
cg26700919
cg26701043
cg26701815
cg26702929
cg26703245
cg26703507
cg26705266
cg26705408
cg26705829
cg26705953
cg26705960
cg26706520
cg26706614
cg26706759
cg26706803
cg26706896
cg26707013
cg26707887
cg26708220
cg26708319
cg26708427
cg26709285
cg26709720
cg26710371
cg26711508
cg26712821
cg26716753
cg26717554
cg26717935
cg26718433
cg26718878
cg26721193
cg26721264
cg26721868
cg26721877
cg26722577
cg26723045
cg26723228
cg26723237
cg26723355
cg26723635
cg26723928

cg26724967
cg26725665
cg26727231
cg26727372
cg26727488
cg26727693
cg26729956
cg26730048
cg26730050
cg26730369
cg26730416
cg26731575
cg26732544
cg26732808
cg26733897
cg26735353
cg26735478
cg26735503
cg26735846
cg26736440
cg26736645
cg26736812
cg26738437
cg26738987
cg26739230
cg26739849
cg26740109
cg26740230
cg26740639
cg26740937
cg26742770
cg26745143
cg26745402
cg26746224
cg26746469
cg26746560
cg26746573
cg26746878
cg26748297
cg26748898
cg26748945
cg26749543
cg26749978
cg26750747
cg26750789
cg26750927
cg26751356
cg26752663
cg26753393
cg26754262
cg26754664
cg26755141

cg26755793
cg26756193
cg26756782
cg26757229
cg26757673
cg26757793
cg26757820
cg26758986
cg26759223
cg26761110
cg26761572
cg26762224
cg26763486
cg26763542
cg26763877
cg26765567
cg26765743
cg26766005
cg26766064
cg26766373
cg26766885
cg26767081
cg26767299
cg26767922
cg26767974
cg26769700
cg26770890
cg26770907
cg26771097
cg26771557
cg26771582
cg26771969
cg26772172
cg26773791
cg26775558
cg26775658
cg26776656
cg26777475
cg26777729
cg26777883
cg26779147
cg26779306
cg26779608
cg26779676
cg26780125
cg26780915
cg26781129
cg26781464
cg26784011
cg26784929
cg26785157
cg26785220

cg26785370
cg26785702
cg26785954
cg26786407
cg26786974
cg26787020
cg26787787
cg26788216
cg26788672
cg26788812
cg26789779
cg26790091
cg26790247
cg26791148
cg26791242
cg26791266
cg26792089
cg26792948
cg26793720
cg26794346
cg26794830
cg26794885
cg26794938
cg26795248
cg26795312
cg26795540
cg26795848
cg26796283
cg26796518
cg26796642
cg26797159
cg26798452
cg26798702
cg26798861
cg26799474
cg26799877
cg26800551
cg26800884
cg26800915
cg26801037
cg26801278
cg26802289
cg26802786
cg26803268
cg26803289
cg26803305
cg26804381
cg26804595
cg26804848
cg26805749
cg26805779
cg26806779

cg26806835
cg26807367
cg26807432
cg26807939
cg26808154
cg26808583
cg26808618
cg26808687
cg26808747
cg26809635
cg26810146
cg26810230
cg26810908
cg26810970
cg26811385
cg26811904
cg26812481
cg26813137
cg26813646
cg26813718
cg26814276
cg26815395
cg26816432
cg26816636
cg26817217
cg26818573
cg26818645
cg26819695
cg26820000
cg26820055
cg26821433
cg26821462
cg26821634
cg26821681
cg26821745
cg26821962
cg26822097
cg26822728
cg26823666
cg26823762
cg26824006
cg26824947
cg26826729
cg26827876
cg26827987
cg26828601
cg26828839
cg26829101
cg26829418
cg26829529
cg26829913
cg26830291

cg26830958
cg26831087
cg26831119
cg26831566
cg26832294
cg26833120
cg26833652
cg26834627
cg26835324
cg26835419
cg26835506
cg26836233
cg26836696
cg26836743
cg26837399
cg26837477
cg26837708
cg26837730
cg26837853
cg26840082
cg26841013
cg26841040
cg26841048
cg26841157
cg26841647
cg26841894
cg26842310
cg26842423
cg26843074
cg26843783
cg26843807
cg26843872
cg26844246
cg26845278
cg26845817
cg26847503
cg26847642
cg26847866
cg26848086
cg26848524
cg26848718
cg26849111
cg26849637
cg26850624
cg26851500
cg26851803
cg26851984
cg26852712
cg26852723
cg26852879
cg26853368
cg26853415

cg26854096
cg26854588
cg26855148
cg26855918
cg26856475
cg26856578
cg26857192
cg26857408
cg26858065
cg26858238
cg26858414
cg26859042
cg26859365
cg26859878
cg26860935
cg26861457
cg26861703
cg26861706
cg26861946
cg26862964
cg26863013
cg26863600
cg26863750
cg26863821
cg26863842
cg26864297
cg26864766
cg26865084
cg26865478
cg26865494
cg26866174
cg26867987
cg26869723
cg26870536
cg26870570
cg26870584
cg26870746
cg26871347
cg26871372
cg26872137
cg26872482
cg26874367
cg26874464
cg26874542
cg26875135
cg26875233
cg26875384
cg26875655
cg26875958
cg26876077
cg26876101
cg26876444

cg26876914
cg26877508
cg26877678
cg26877720
cg26878190
cg26878209
cg26878734
cg26878836
cg26878949
cg26879581
cg26880549
cg26880708
cg26881677
cg26881761
cg26882204
cg26883033
cg26883171
cg26883251
cg26884581
cg26885220
cg26886462
cg26886875
cg26888012
cg26888422
cg26890181
cg26890480
cg26890621
cg26891150
cg26892635
cg26893451
cg26894278
cg26894288
cg26894354
cg26894523
cg26894807
cg26894841
cg26895804
cg26895931
cg26896572
cg26896762
cg26896771
cg26896979
cg26898336
cg26898567
cg26898869
cg26899156
cg26902216
cg26903216
cg26903766
cg26904017
cg26905112
cg26905258

cg26905261
cg26905720
cg26906258
cg26907313
cg26910384
cg26910870
cg26912105
cg26912131
cg26912251
cg26912314
cg26912426
cg26912602
cg26912890
cg26913155
cg26913248
cg26913344
cg26913563
cg26913669
cg26914205
cg26914267
cg26914334
cg26914931
cg26915329
cg26915370
cg26915704
cg26916166
cg26916780
cg26917745
cg26918041
cg26918305
cg26918577
cg26918808
cg26918954
cg26919100
cg26919347
cg26919618
cg26919735
cg26919780
cg26920627
cg26920864
cg26921482
cg26921612
cg26922451
cg26922483
cg26923490
cg26923754
cg26924223
cg26924440
cg26925025
cg26925231
cg26925688
cg26925980

cg26926221
cg26926521
cg26926612
cg26926765
cg26927516
cg26927939
cg26928418
cg26929163
cg26929247
cg26929272
cg26929806
cg26931050
cg26931063
cg26931101
cg26931526
cg26931862
cg26932614
cg26932779
cg26932790
cg26932873
cg26933021
cg26934362
cg26935168
cg26935219
cg26935304
cg26935391
cg26936171
cg26936219
cg26938014
cg26938272
cg26938314
cg26938597
cg26938599
cg26939283
cg26939503
cg26940847
cg26943731
cg26945241
cg26945780
cg26945996
cg26946015
cg26946218
cg26946259
cg26946288
cg26946311
cg26946769
cg26947626
cg26948823
cg26948907
cg26949694
cg26950557
cg26950652

cg26950756
cg26951410
cg26951440
cg26952618
cg26952928
cg26953137
cg26953453
cg26953640
cg26953954
cg26954101
cg26954174
cg26954197
cg26954519
cg26954968
cg26955835
cg26955850
cg26956007
cg26956135
cg26957988
cg26958124
cg26958741
cg26959303
cg26959358
cg26959392
cg26959827
cg26960562
cg26960719
cg26962618
cg26963271
cg26963356
cg26963632
cg26963797
cg26963892
cg26964875
cg26966083
cg26966828
cg26967100
cg26967596
cg26967723
cg26968039
cg26968748
cg26969179
cg26970983
cg26972479
cg26972614
cg26973018
cg26974383
cg26975787
cg26976108
cg26976908
cg26977256
cg26977737

cg26977759
cg26977944
cg26978698
cg26979518
cg26980244
cg26980529
cg26980729
cg26980954
cg26980978
cg26980987
cg26981268
cg26981966
cg26982279
cg26982447
cg26982601
cg26982892
cg26983177
cg26983198
cg26983710
cg26983991
cg26985447
cg26985674
cg26985686
cg26985711
cg26985818
cg26986937
cg26986989
cg26987521
cg26987597
cg26988213
cg26988283
cg26989103
cg26990102
cg26990587
cg26991025
cg26991946
cg26992245
cg26992949
cg26993132
cg26993729
cg26993947
cg26994225
cg26995244
cg26995942
cg26996626
cg26996656
cg26996818
cg26997028
cg26997090
cg26998274
cg26998481
cg26999345

cg27000659
cg27000831
cg27001250
cg27001715
cg27002303
cg27003682
cg27003782
cg27004195
cg27004523
cg27005118
cg27005487
cg27005749
cg27005794
cg27005847
cg27005946
cg27006650
cg27006914
cg27007937
cg27009439
cg27009475
cg27009703
cg27010159
cg27011060
cg27012021
cg27012108
cg27012421
cg27012952
cg27013696
cg27013931
cg27014135
cg27014354
cg27014380
cg27015057
cg27016307
cg27016988
cg27017414
cg27017580
cg27019059
cg27019712
cg27019717
cg27020216
cg27020362
cg27020668
cg27020966
cg27021406
cg27021492
cg27021666
cg27022324
cg27022420
cg27022827
cg27022956
cg27023597

cg27023727
cg27024161
cg27024844
cg27025137
cg27025320
cg27025384
cg27026405
cg27026615
cg27027230
cg27028923
cg27028938
cg27029518
cg27029744
cg27030352
cg27031099
cg27032146
cg27032184
cg27032232
cg27032352
cg27032760
cg27032905
cg27034935
cg27035514
cg27035708
cg27035734
cg27035999
cg27036111
cg27037013
cg27037018
cg27037551
cg27038382
cg27040525
cg27040644
cg27040709
cg27041424
cg27042065
cg27042584
cg27044591
cg27044841
cg27044861
cg27045264
cg27045447
cg27046936
cg27047353
cg27047406
cg27049076
cg27049539
cg27050031
cg27050111
cg27050153
cg27051089
cg27051260

cg27051315
cg27051683
cg27051931
cg27052709
cg27053040
cg27053542
cg27054144
cg27054847
cg27055156
cg27055365
cg27055685
cg27056742
cg27056759
cg27057329
cg27057408
cg27057524
cg27058077
cg27058486
cg27058889
cg27058931
cg27059354
cg27059379
cg27060294
cg27060355
cg27060793
cg27061485
cg27061889
cg27062243
cg27062514
cg27062927
cg27063969
cg27064708
cg27065478
cg27066256
cg27067035
cg27067618
cg27067781
cg27068095
cg27069616
cg27069753
cg27070951
cg27070962
cg27071405
cg27072813
cg27072996
cg27073068
cg27073113
cg27073349
cg27073361
cg27073431
cg27074041
cg27074174

cg27074221
cg27075917
cg27076454
cg27077070
cg27077673
cg27078786
cg27078927
cg27079680
cg27082426
cg27082881
cg27083614
cg27084232
cg27084654
cg27085741
cg27087243
cg27087553
cg27087809
cg27088072
cg27090062
cg27090073
cg27092191
cg27092605
cg27093376
cg27093379
cg27093637
cg27094229
cg27094244
cg27094376
cg27094856
cg27095222
cg27095388
cg27095915
cg27096144
cg27096172
cg27096779
cg27096807
cg27097034
cg27097386
cg27097762
cg27097923
cg27099088
cg27099262
cg27100236
cg27100370
cg27100436
cg27100990
cg27101034
cg27101095
cg27102350
cg27103585
cg27104671
cg27104822

cg27105094
cg27105446
cg27105634
cg27106673
cg27107094
cg27107171
cg27107685
cg27109129
cg27109238
cg27109461
cg27109482
cg27109600
cg27109672
cg27110177
cg27110886
cg27110994
cg27111678
cg27112565
cg27112585
cg27112666
cg27112809
cg27112972
cg27113015
cg27113059
cg27114644
cg27115863
cg27116114
cg27116510
cg27116912
cg27117439
cg27117476
cg27118533
cg27118825
cg27119456
cg27119877
cg27119919
cg27120405
cg27120816
cg27120999
cg27121538
cg27121661
cg27122213
cg27122441
cg27122536
cg27123256
cg27123351
cg27125364
cg27125605
cg27125631
cg27125849
cg27126743
cg27128001

cg27128883
cg27129725
cg27130075
cg27130665
cg27131703
cg27132732
cg27132814
cg27132966
cg27133493
cg27133711
cg27133959
cg27134910
cg27135915
cg27135989
cg27136719
cg27136844
cg27137084
cg27137887
cg27138204
cg27138600
cg27138949
cg27139419
cg27139805
cg27139836
cg27140295
cg27141176
cg27142093
cg27142530
cg27142536
cg27142680
cg27143070
cg27143605
cg27143664
cg27143688
cg27143842
cg27144368
cg27144592
cg27144670
cg27144719
cg27145048
cg27145458
cg27147114
cg27147858
cg27148193
cg27148452
cg27149179
cg27149471
cg27150412
cg27150460
cg27151135
cg27151677
cg27152190

cg27152299
cg27153759
cg27153942
cg27154343
cg27154651
cg27154725
cg27158519
cg27158573
cg27158862
cg27160524
cg27161340
cg27161702
cg27162122
cg27164770
cg27165039
cg27165794
cg27166033
cg27166177
cg27166718
cg27166993
cg27167562
cg27168033
cg27168114
cg27168493
cg27168844
cg27169020
cg27169073
cg27170427
cg27170720
cg27170782
cg27171201
cg27171509
cg27171837
cg27172140
cg27172331
cg27172866
cg27173965
cg27174108
cg27174438
cg27175311
cg27175788
cg27176828
cg27177158
cg27177296
cg27177709
cg27178045
cg27178677
cg27179002
cg27179101
cg27180153
cg27180443
cg27180671

cg27181013
cg27181295
cg27182150
cg27182159
cg27182172
cg27182527
cg27182923
cg27183435
cg27183726
cg27184438
cg27184903
cg27185501
cg27187541
cg27188243
cg27188358
cg27189087
cg27189986
cg27190239
cg27190398
cg27190654
cg27190946
cg27191906
cg27191921
cg27192708
cg27193519
cg27194211
cg27195421
cg27195956
cg27196158
cg27196745
cg27196880
cg27198931
cg27199384
cg27199811
cg27199820
cg27199827
cg27200236
cg27200630
cg27201767
cg27202075
cg27202622
cg27202779
cg27203351
cg27203372
cg27203895
cg27203924
cg27204266
cg27205798
cg27205928
cg27206367
cg27207776
cg27208079

cg27209072
cg27209571
cg27209993
cg27210136
cg27211576
cg27212428
cg27212729
cg27213386
cg27213509
cg27213533
cg27216807
cg27217214
cg27218227
cg27218500
cg27218796
cg27219182
cg27219221
cg27219691
cg27219748
cg27219955
cg27219973
cg27220062
cg27220070
cg27220117
cg27220999
cg27221538
cg27222147
cg27222157
cg27223047
cg27223158
cg27223882
cg27224809
cg27225130
cg27225220
cg27225309
cg27226153
cg27227317
cg27227804
cg27228136
cg27228168
cg27228272
cg27228578
cg27228660
cg27228712
cg27228986
cg27230067
cg27230118
cg27230510
cg27230705
cg27231068
cg27231441
cg27232145

cg27232482
cg27232494
cg27232569
cg27233071
cg27233151
cg27234067
cg27234864
cg27235662
cg27235955
cg27236146
cg27236154
cg27237300
cg27237537
cg27237541
cg27237814
cg27238547
cg27238903
cg27239243
cg27239280
cg27241808
cg27241889
cg27243312
cg27243405
cg27244242
cg27245185
cg27247252
cg27247689
cg27247934
cg27248029
cg27248148
cg27248887
cg27250032
cg27250236
cg27250607
cg27252019
cg27252325
cg27252696
cg27253807
cg27254338
cg27254468
cg27254482
cg27254661
cg27254852
cg27255329
cg27255672
cg27257408
cg27257452
cg27257453
cg27257955
cg27257987
cg27258399
cg27258696

cg27260174
cg27260772
cg27261597
cg27261921
cg27262041
cg27262298
cg27262398
cg27262412
cg27262821
cg27262850
cg27262870
cg27263998
cg27264049
cg27264249
cg27264437
cg27264993
cg27265170
cg27267617
cg27268486
cg27268501
cg27269149
cg27269279
cg27269492
cg27269790
cg27269917
cg27269921
cg27271937
cg27272220
cg27272487
cg27272677
cg27272679
cg27273054
cg27273643
cg27274382
cg27274706
cg27274989
cg27277081
cg27277212
cg27277463
cg27278500
cg27278787
cg27278843
cg27278953
cg27279244
cg27279545
cg27280332
cg27280535
cg27281191
cg27281882
cg27282206
cg27282264
cg27282530

cg27283345
cg27284032
cg27287167
cg27287305
cg27287357
cg27288226
cg27288968
cg27290215
cg27291181
cg27291258
cg27291468
cg27291602
cg27292099
cg27292417
cg27292667
cg27292977
cg27293090
cg27294168
cg27294268
cg27294678
cg27294813
cg27295342
cg27295373
cg27295412
cg27295434
cg27295595
cg27295832
cg27296293
cg27296944
cg27297033
cg27297088
cg27297993
cg27298262
cg27298506
cg27299033
cg27299047
cg27299776
cg27300829
cg27300950
cg27300957
cg27302539
cg27303211
cg27304110
cg27304332
cg27304729
cg27305009
cg27305060
cg27305460
cg27305510
cg27305939
cg27306191
cg27307183

cg27307810
cg27308021
cg27308576
cg27308654
cg27308754
cg27308982
cg27309038
cg27309276
cg27309631
cg27309905
cg27310092
cg27310311
cg27310319
cg27310594
cg27312388
cg27312652
cg27312775
cg27312916
cg27312979
cg27313012
cg27313021
cg27313201
cg27313577
cg27313642
cg27314422
cg27314569
cg27314580
cg27314798
cg27314998
cg27315109
cg27315191
cg27315497
cg27315556
cg27316453
cg27316567
cg27316626
cg27316886
cg27317017
cg27317356
cg27317439
cg27317909
cg27318539
cg27319123
cg27319445
cg27319530
cg27320518
cg27321175
cg27321439
cg27321750
cg27321942
cg27322397
cg27323784

cg27324128
cg27324797
cg27326882
cg27328080
cg27328179
cg27328797
cg27329559
cg27329908
cg27330502
cg27331233
cg27331241
cg27331316
cg27331524
cg27332575
cg27332583
cg27333271
cg27333283
cg27333706
cg27335999
cg27336026
cg27336379
cg27337194
cg27337902
cg27339290
cg27340506
cg27340584
cg27341128
cg27341472
cg27341926
cg27342314
cg27342720
cg27343150
cg27343325
cg27343836
cg27343917
cg27344776
cg27345534
cg27345757
cg27345946
cg27346707
cg27346806
cg27347269
cg27348223
cg27348266
cg27348420
cg27348779
cg27349302
cg27350104
cg27350402
cg27350428
cg27351783
cg27352639

cg27353143
cg27353346
cg27353629
cg27353824
cg27353891
cg27354094
cg27354586
cg27354635
cg27355501
cg27356296
cg27356513
cg27357306
cg27357571
cg27358936
cg27359279
cg27359472
cg27360098
cg27360300
cg27360326
cg27360727
cg27361577
cg27361914
cg27362255
cg27362525
cg27363021
cg27363617
cg27363745
cg27363829
cg27364242
cg27364741
cg27365499
cg27365535
cg27365701
cg27365991
cg27366072
cg27366280
cg27366532
cg27366888
cg27367045
cg27367170
cg27368515
cg27368709
cg27368718
cg27369423
cg27370104
cg27371466
cg27372422
cg27372452
cg27373015
cg27373604
cg27374199
cg27374508

cg27374881
cg27375286
cg27376136
cg27376182
cg27376707
cg27376870
cg27376872
cg27377632
cg27379109
cg27380227
cg27380499
cg27380599
cg27380904
cg27382409
cg27382568
cg27382910
cg27383124
cg27383203
cg27383216
cg27383859
cg27384342
cg27384918
cg27386326
cg27386614
cg27388966
cg27389562
cg27389684
cg27390627
cg27391093
cg27391267
cg27391334
cg27391934
cg27391982
cg27392286
cg27392792
cg27392804
cg27393010
cg27393726
cg27394460
cg27394566
cg27394794
cg27395200
cg27395342
cg27396284
cg27396403
cg27396824
cg27398640
cg27399031
cg27400216
cg27401891
cg27402883
cg27403265

cg27403635
cg27404351
cg27404824
cg27405706
cg27405791
cg27406091
cg27406975
cg27407147
cg27408184
cg27408285
cg27408683
cg27408897
cg27409771
cg27409802
cg27409991
cg27410595
cg27410679
cg27412093
cg27412902
cg27413462
cg27413508
cg27413563
cg27414087
cg27414481
cg27414655
cg27415425
cg27415552
cg27416171
cg27416337
cg27416372
cg27417606
cg27417609
cg27418204
cg27418851
cg27420049
cg27420123
cg27420236
cg27420415
cg27420819
cg27421679
cg27422141
cg27423027
cg27423323
cg27423627
cg27423692
cg27423760
cg27423838
cg27424244
cg27425169
cg27425996
cg27426287
cg27426835

cg27427054
cg27428051
cg27428414
cg27428520
cg27430561
cg27431274
cg27431545
cg27432085
cg27432242
cg27434086
cg27434509
cg27434984
cg27435498
cg27438798
cg27438832
cg27438854
cg27439480
cg27440592
cg27440715
cg27442308
cg27443071
cg27443224
cg27443867
cg27444062
cg27444437
cg27444913
cg27444994
cg27446185
cg27446442
cg27447477
cg27447874
cg27448110
cg27448426
cg27449507
cg27449793
cg27450668
cg27451587
cg27454298
cg27454412
cg27454698
cg27455211
cg27455540
cg27455890
cg27456382
cg27456561
cg27457457
cg27457631
cg27458177
cg27458265
cg27458987
cg27461196
cg27461824

cg27462475
cg27463141
cg27464065
cg27464184
cg27465569
cg27465849
cg27466709
cg27467929
cg27468830
cg27470047
cg27470087
cg27470486
cg27470827
cg27472176
cg27472937
cg27473406
cg27473871
cg27473895
cg27474382
cg27475088
cg27475132
cg27475652
cg27477067
cg27477251
cg27477419
cg27478485
cg27478659
cg27478707
cg27480427
cg27480444
cg27480727
cg27480819
cg27480913
cg27481499
cg27483571
cg27483694
cg27484456
cg27485269
cg27485596
cg27485790
cg27485810
cg27486427
cg27486786
cg27487860
cg27487931
cg27488095
cg27488648
cg27488680
cg27490380
cg27490391
cg27490568
cg27490875

cg27491025
cg27491398
cg27492103
cg27492286
cg27492467
cg27492725
cg27493010
cg27493411
cg27493772
cg27494470
cg27495223
cg27495903
cg27495908
cg27498197
cg27498387
cg27499105
cg27499185
cg27500292
cg27501293
cg27501362
cg27501458
cg27501612
cg27501748
cg27501878
cg27502088
cg27502580
cg27503360
cg27503920
cg27504292
cg27504555
cg27504802
cg27504861
cg27505273
cg27506210
cg27507254
cg27508055
cg27508071
cg27508314
cg27508586
cg27508904
cg27509105
cg27509213
cg27509347
cg27509381
cg27509823
cg27510041
cg27510354
cg27511169
cg27511824
cg27512316
cg27512716
cg27513308

cg27513573
cg27513574
cg27513684
cg27513979
cg27514017
cg27514080
cg27514538
cg27514730
cg27514777
cg27514986
cg27515201
cg27516159
cg27517345
cg27517681
cg27518240
cg27518483
cg27518631
cg27518892
cg27518976
cg27519140
cg27519910
cg27521109
cg27522078
cg27522154
cg27523087
cg27523417
cg27523779
cg27526649
cg27527018
cg27527642
cg27527798
cg27527874
cg27528032
cg27528222
cg27528351
cg27528510
cg27528660
cg27529911
cg27530576
cg27531336
cg27531366
cg27531490
cg27531496
cg27531553
cg27531571
cg27531731
cg27532621
cg27533288
cg27533767
cg27533999
cg27534396
cg27534534

cg27535502
cg27535687
cg27535690
cg27536700
cg27537692
cg27537740
cg27537795
cg27537972
cg27538859
cg27540865
cg27542729
cg27543320
cg27543334
cg27543385
cg27543409
cg27544405
cg27544673
cg27545296
cg27545615
cg27545630
cg27546012
cg27546066
cg27546237
cg27546911
cg27547281
cg27547291
cg27547375
cg27548256
cg27549186
cg27549720
cg27549952
cg27550998
cg27552493
cg27552679
cg27552857
cg27553626
cg27553667
cg27553971
cg27554565
cg27555582
cg27555761
cg27556492
cg27557378
cg27558283
cg27558693
cg27559224
cg27560132
cg27560250
cg27560796
cg27561819
cg27562005
cg27562259

cg27563328
cg27563985
cg27564715
cg27565899
cg27566805
cg27566947
cg27567206
cg27567401
cg27567416
cg27567430
cg27567922
cg27570249
cg27570256
cg27570354
cg27570873
cg27571769
cg27572072
cg27572927
cg27573017
cg27573344
cg27574595
cg27574973
cg27576241
cg27576694
cg27577972
cg27578439
cg27578734
cg27578977
cg27579131
cg27579313
cg27579501
cg27579805
cg27579953
cg27581493
cg27581871
cg27582527
cg27582546
cg27582585
cg27582790
cg27583010
cg27583220
cg27583740
cg27584135
cg27584546
cg27584713
cg27585641
cg27586207
cg27586885
cg27587307
cg27587952
cg27588076
cg27588083

cg27588093
cg27588653
cg27589088
cg27589368
cg27589654
cg27589809
cg27590049
cg27590397
cg27591349
cg27591407
cg27591554
cg27592410
cg27593384
cg27595576
cg27595860
cg27596495
cg27596649
cg27596828
cg27596890
cg27597459
cg27597681
cg27597956
cg27599211
cg27600205
cg27600804
cg27601695
cg27603000
cg27603283
cg27603796
cg27604416
cg27605228
cg27605300
cg27605701
cg27606397
cg27606473
cg27606499
cg27607438
cg27608102
cg27609986
cg27610486
cg27612129
cg27612389
cg27612447
cg27614723
cg27615388
cg27615938
cg27616543
cg27616666
cg27616833
cg27617086
cg27617109
cg27617159

cg27618216
cg27618240
cg27618777
cg27618792
cg27619006
cg27619457
cg27619796
cg27620077
cg27621514
cg27621745
cg27622610
cg27622701
cg27623769
cg27623948
cg27624313
cg27624334
cg27625055
cg27625491
cg27627246
cg27627381
cg27628839
cg27629589
cg27629673
cg27629782
cg27629977
cg27629986
cg27630311
cg27631199
cg27631389
cg27632402
cg27632959
cg27633342
cg27634020
cg27634164
cg27634195
cg27635069
cg27635135
cg27635271
cg27635714
cg27635859
cg27636058
cg27637382
cg27637706
cg27637730
cg27638035
cg27638126
cg27638196
cg27638288
cg27638458
cg27638727
cg27639046
cg27640064

cg27640528
cg27640903
cg27641136
cg27641532
cg27641961
cg27642416
cg27642554
cg27642618
cg27643147
cg27643890
cg27648056
cg27648075
cg27648224
cg27648556
cg27648738
cg27649094
cg27649239
cg27651070
cg27651090
cg27653595
cg27654142
cg27655158
cg27655168
cg27655206
cg27655954
cg27656362
cg27656459
cg27656949
cg27656973
cg27657603
cg27657983
cg27658068
cg27658330
cg27659049
cg27659505
cg27659557
cg27659787
cg27660388
cg27662505
cg27662639
cg27663200
cg27663249
cg27663476
cg27663938
cg27664687
cg27665181
cg27665580
cg27666046
cg27666123
