## Supplementary Table S9 for "Methylome-based cell-of-origin modeling (Methyl-COOM) identifies aberrant expression of immune regulatory molecules in CLL"

**Supplementary Table S9: List of CLL-specific CpGs**

**ClassA**

| chromosome | start | end | CpG |
|---|---|---|---|
| chr1 | 879958 | 879959 | cg23731742 |
| chr1 | 944783 | 944784 | cg24004483 |
| chr1 | 967561 | 967562 | cg02288964 |
| chr1 | 988623 | 988624 | cg22627753 |
| chr1 | 1062237 | 1062238 | cg27582546 |
| chr1 | 1084473 | 1084474 | cg20254790 |
| chr1 | 1101348 | 1101349 | cg13449535 |
| chr1 | 1101462 | 1101463 | cg14144728 |
| chr1 | 1149501 | 1149502 | cg23166857 |
| chr1 | 1150322 | 1150323 | cg23867494 |
| chr1 | 1150410 | 1150411 | cg21815220 |
| chr1 | 1150416 | 1150417 | cg17084044 |
| chr1 | 1150936 | 1150937 | cg16252905 |
| chr1 | 1261797 | 1261798 | cg05598010 |
| chr1 | 1294881 | 1294882 | cg23973885 |
| chr1 | 1610660 | 1610661 | cg01189072 |
| chr1 | 1695585 | 1695586 | cg00992048 |
| chr1 | 1731415 | 1731416 | cg03335128 |
| chr1 | 1820656 | 1820657 | cg03043696 |
| chr1 | 1833078 | 1833079 | cg24706992 |
| chr1 | 2003650 | 2003651 | cg11227141 |
| chr1 | 2003864 | 2003865 | cg00095688 |
| chr1 | 2004057 | 2004058 | cg24842354 |
| chr1 | 2004163 | 2004164 | cg08439149 |
| chr1 | 2019526 | 2019527 | cg06176608 |
| chr1 | 2037476 | 2037477 | cg00409816 |
| chr1 | 2038940 | 2038941 | cg12713909 |
| chr1 | 2043761 | 2043762 | cg13918804 |
| chr1 | 2043853 | 2043854 | cg19257562 |
| chr1 | 2046928 | 2046929 | cg26910870 |
| chr1 | 2065263 | 2065264 | cg16048006 |
| chr1 | 2090814 | 2090815 | cg24578937 |
| chr1 | 2119531 | 2119532 | cg17412102 |
| chr1 | 2163472 | 2163473 | cg17942763 |
| chr1 | 2165579 | 2165580 | cg25139649 |
| chr1 | 2171834 | 2171835 | cg21775570 |
| chr1 | 2180697 | 2180698 | cg21414505 |
| chr1 | 2182126 | 2182127 | cg08468599 |
| chr1 | 2182373 | 2182374 | cg21442419 |
| chr1 | 2188767 | 2188768 | cg23645210 |
| chr1 | 2190850 | 2190851 | cg24481594 |
| chr1 | 2203589 | 2203590 | cg01949002 |
| chr1 | 2206662 | 2206663 | cg19025786 |
| chr1 | 2371057 | 2371058 | cg01309121 |
| chr1 | 2435087 | 2435088 | cg06350196 |
| chr1 | 2456436 | 2456437 | cg00295182 |
| chr1 | 2515692 | 2515693 | cg02393107 |
| chr1 | 3163710 | 3163711 | cg16694785 |
| chr1 | 3382237 | 3382238 | cg11587191 |
| chr1 | 3427479 | 3427480 | cg06353486 |
| chr1 | 3447348 | 3447349 | cg17163527 |

**ClassB**

| chromosome | start | end | CpG |
|---|---|---|---|
| chr1 | 1021299 | 1021300 | cg26338735 |
| chr1 | 1133215 | 1133216 | cg09021712 |
| chr1 | 3136528 | 3136529 | cg26053881 |
| chr1 | 9795934 | 9795935 | cg03541746 |
| chr1 | 18606374 | 18606375 | cg15081740 |
| chr1 | 36866445 | 36866446 | cg26653824 |
| chr1 | 47489282 | 47489283 | cg26558485 |
| chr1 | 59043255 | 59043256 | cg16699148 |
| chr1 | 63790044 | 63790045 | cg18279094 |
| chr1 | 91183051 | 91183052 | cg25026529 |
| chr1 | 91190620 | 91190621 | cg15452970 |
| chr1 | 92329399 | 92329400 | cg00308438 |
| chr1 | 110008990 | 110008991 | cg20591472 |
| chr1 | 211308349 | 211308350 | cg10968055 |
| chr1 | 228400217 | 228400218 | cg03054277 |
| chr1 | 234764068 | 234764069 | cg24339574 |
| chr2 | 5836713 | 5836714 | cg23668184 |
| chr2 | 28114447 | 28114448 | cg16651262 |
| chr2 | 28114462 | 28114463 | cg05517527 |
| chr2 | 47916139 | 47916140 | cg07464408 |
| chr2 | 54483498 | 54483499 | cg04098176 |
| chr2 | 65571891 | 65571892 | cg19672145 |
| chr2 | 65571948 | 65571949 | cg07588875 |
| chr2 | 68654146 | 68654147 | cg08464093 |
| chr2 | 72371838 | 72371839 | cg06049781 |
| chr2 | 172945698 | 172945699 | cg16472050 |
| chr2 | 176948759 | 176948760 | cg07411620 |
| chr2 | 176972068 | 176972069 | cg05500840 |
| chr2 | 176981328 | 176981329 | cg18115040 |
| chr2 | 176987605 | 176987606 | cg02885007 |
| chr2 | 176994764 | 176994765 | cg24416513 |
| chr2 | 234111145 | 234111146 | cg01591579 |
| chr2 | 237076811 | 237076812 | cg21426003 |
| chr2 | 237076815 | 237076816 | cg19761848 |
| chr2 | 237123218 | 237123219 | cg21300993 |
| chr2 | 240196769 | 240196770 | cg05870586 |
| chr2 | 241386846 | 241386847 | cg13393505 |
| chr3 | 9788958 | 9788959 | cg13312668 |
| chr3 | 12045662 | 12045663 | cg20888142 |
| chr3 | 107148372 | 107148373 | cg19401033 |
| chr3 | 124653659 | 124653660 | cg22907034 |
| chr3 | 147127012 | 147127013 | cg01227537 |
| chr3 | 149753446 | 149753447 | cg15440157 |
| chr4 | 2321957 | 2321958 | cg22488278 |
| chr4 | 4670703 | 4670704 | cg05870398 |
| chr4 | 7940579 | 7940580 | cg22079161 |
| chr4 | 53614727 | 53614728 | cg05077452 |
| chr4 | 102737794 | 102737795 | cg14779735 |
| chr4 | 188953126 | 188953127 | cg10380328 |
| chr5 | 1807733 | 1807734 | cg00561739 |
| chr5 | 59189120 | 59189121 | cg03323696 |

**ClassC**

| chromosome | start | end | CpG |
|---|---|---|---|
| chr1 | 991708 | 991709 | cg13439946 |
| chr1 | 1003529 | 1003530 | cg15899727 |
| chr1 | 1268243 | 1268244 | cg06395167 |
| chr1 | 1268469 | 1268470 | cg16174225 |
| chr1 | 1502606 | 1502607 | cg20490900 |
| chr1 | 1549699 | 1549700 | cg13897348 |
| chr1 | 1549799 | 1549800 | cg19246761 |
| chr1 | 1549818 | 1549819 | cg01229567 |
| chr1 | 1564920 | 1564921 | cg17122213 |
| chr1 | 1713676 | 1713677 | cg02769781 |
| chr1 | 1713944 | 1713945 | cg24163210 |
| chr1 | 1982791 | 1982792 | cg07797073 |
| chr1 | 1988681 | 1988682 | cg26059839 |
| chr1 | 2004222 | 2004223 | cg13077150 |
| chr1 | 2004346 | 2004347 | cg00359465 |
| chr1 | 2043799 | 2043800 | cg04315214 |
| chr1 | 2065242 | 2065243 | cg09560161 |
| chr1 | 2066006 | 2066007 | cg22366001 |
| chr1 | 2066167 | 2066168 | cg07836663 |
| chr1 | 2066430 | 2066431 | cg20975074 |
| chr1 | 2069012 | 2069013 | cg24577455 |
| chr1 | 2106222 | 2106223 | cg15468976 |
| chr1 | 2106365 | 2106366 | cg27264462 |
| chr1 | 2106400 | 2106401 | cg23648809 |
| chr1 | 2106620 | 2106621 | cg02393699 |
| chr1 | 2121039 | 2121040 | cg26227225 |
| chr1 | 2179933 | 2179934 | cg24377150 |
| chr1 | 2180027 | 2180028 | cg02360367 |
| chr1 | 2183438 | 2183439 | cg07114422 |
| chr1 | 2191299 | 2191300 | cg26880297 |
| chr1 | 2191354 | 2191355 | cg25314600 |
| chr1 | 2191402 | 2191403 | cg18934822 |
| chr1 | 2191558 | 2191559 | cg01881525 |
| chr1 | 2191658 | 2191659 | cg12222277 |
| chr1 | 2230923 | 2230924 | cg23022809 |
| chr1 | 2244585 | 2244586 | cg06952577 |
| chr1 | 2245988 | 2245989 | cg06784691 |
| chr1 | 2426035 | 2426036 | cg11692307 |
| chr1 | 2615690 | 2615691 | cg18854424 |
| chr1 | 3569899 | 3569900 | cg11504517 |
| chr1 | 3806715 | 3806716 | cg17879856 |
| chr1 | 3806817 | 3806818 | cg12594933 |
| chr1 | 3807168 | 3807169 | cg16349667 |
| chr1 | 3809431 | 3809432 | cg03921357 |
| chr1 | 3809532 | 3809533 | cg27446570 |
| chr1 | 4619207 | 4619208 | cg16264595 |
| chr1 | 6085995 | 6085996 | cg07015749 |
| chr1 | 6645463 | 6645464 | cg23642826 |
| chr1 | 6664103 | 6664104 | cg27539986 |
| chr1 | 6664134 | 6664135 | cg09159285 |
| chr1 | 6664268 | 6664269 | cg19884658 |

**ClassD**

| chromosome | start | end | CpG |
|---|---|---|---|
| chr1 | 976707 | 976708 | cg20594982 |
| chr1 | 21818863 | 21818864 | cg13170600 |
| chr1 | 55462274 | 55462275 | cg02392553 |
| chr1 | 63786271 | 63786272 | cg18175808 |
| chr1 | 64014643 | 64014644 | cg18803079 |
| chr1 | 94703098 | 94703099 | cg26229607 |
| chr1 | 120435245 | 120435246 | cg22167148 |
| chr1 | 150335507 | 150335508 | cg25066665 |
| chr1 | 162039224 | 162039225 | cg17096191 |
| chr1 | 180886483 | 180886484 | cg00864916 |
| chr1 | 220863513 | 220863514 | cg01491219 |
| chr1 | 227635322 | 227635323 | cg04117972 |
| chr2 | 3607301 | 3607302 | cg19540797 |
| chr2 | 7005733 | 7005734 | cg11044724 |
| chr2 | 7057402 | 7057403 | cg25705900 |
| chr2 | 27616206 | 27616207 | cg17424134 |
| chr2 | 40679582 | 40679583 | cg14411266 |
| chr2 | 47798396 | 47798397 | cg27320127 |
| chr2 | 88650618 | 88650619 | cg15581269 |
| chr2 | 118617499 | 118617500 | cg22545206 |
| chr2 | 133426418 | 133426419 | cg17674725 |
| chr2 | 154335250 | 154335251 | cg04011228 |
| chr2 | 183902928 | 183902929 | cg09997497 |
| chr2 | 183903098 | 183903099 | cg22224337 |
| chr2 | 183903302 | 183903303 | cg15171982 |
| chr2 | 200524012 | 200524013 | cg27211284 |
| chr2 | 222436947 | 222436948 | cg03776464 |
| chr2 | 241771687 | 241771688 | cg19584530 |
| chr2 | 242904738 | 242904739 | cg18948877 |
| chr3 | 13692202 | 13692203 | cg03553786 |
| chr3 | 13692212 | 13692213 | cg23620822 |
| chr3 | 98620518 | 98620519 | cg01754155 |
| chr3 | 147096130 | 147096131 | cg25645064 |
| chr3 | 157812475 | 157812476 | cg06570224 |
| chr3 | 160167977 | 160167978 | cg07553761 |
| chr3 | 181445514 | 181445515 | cg25382400 |
| chr3 | 192958832 | 192958833 | cg24710320 |
| chr3 | 195490309 | 195490310 | cg16034541 |
| chr4 | 206874 | 206875 | cg11836119 |
| chr4 | 1513236 | 1513237 | cg22244039 |
| chr4 | 3683268 | 3683269 | cg03965172 |
| chr4 | 4109441 | 4109442 | cg03844506 |
| chr4 | 6021538 | 6021539 | cg14480507 |
| chr4 | 54960259 | 54960260 | cg18099189 |
| chr4 | 113442251 | 113442252 | cg05024939 |
| chr4 | 125380468 | 125380469 | cg12454925 |
| chr4 | 172735924 | 172735925 | cg05106058 |
| chr4 | 187549458 | 187549459 | cg01483139 |
| chr5 | 676441 | 676442 | cg05871207 |
| chr5 | 958299 | 958300 | cg02514143 |
| chr5 | 2038743 | 2038744 | cg25090514 |

| | | | | | | | | | | | | | | | |
|---|---|---|---|---|---|---|---|---|---|---|---|---|---|---|---|
| chr1 | 3508884 | 3508885 | cg17183752 | chr5 | 140457585 | 140457586 | cg06883126 | chr1 | 6676673 | 6676674 | cg12907598 | chr5 | 50258893 | 50258894 | cg11606459 |
| chr1 | 3540478 | 3540479 | cg13059495 | chr5 | 140725147 | 140725148 | cg01561869 | chr1 | 6696166 | 6696167 | cg12218771 | chr5 | 75473201 | 75473202 | cg02515553 |
| chr1 | 3547523 | 3547524 | cg26354191 | chr5 | 153577937 | 153577938 | cg13579752 | chr1 | 8427379 | 8427380 | cg26915618 | chr5 | 77843030 | 77843031 | cg26658897 |
| chr1 | 3555442 | 3555443 | cg16563938 | chr5 | 156817610 | 156817611 | cg10606923 | chr1 | 8427556 | 8427557 | cg01771201 | chr5 | 87437400 | 87437401 | cg26246127 |
| chr1 | 3575674 | 3575675 | cg04137576 | chr5 | 176101928 | 176101929 | cg01603073 | chr1 | 8456466 | 8456467 | cg09246103 | chr5 | 134364968 | 134364969 | cg04101060 |
| chr1 | 3585424 | 3585425 | cg09248714 | chr5 | 179075765 | 179075766 | cg02586025 | chr1 | 8518145 | 8518146 | cg15148088 | chr5 | 172110569 | 172110570 | cg07485775 |
| chr1 | 3695198 | 3695199 | cg07572028 | chr6 | 3850738 | 3850739 | cg14275511 | chr1 | 8536454 | 8536455 | cg15701149 | chr5 | 172110579 | 172110580 | cg06933824 |
| chr1 | 3807709 | 3807710 | cg19375403 | chr6 | 5133475 | 5133476 | cg01392180 | chr1 | 8623473 | 8623474 | cg25722041 | chr5 | 172144013 | 172144014 | cg22118147 |
| chr1 | 3819354 | 3819355 | cg09275693 | chr6 | 10415597 | 10415598 | cg16871519 | chr1 | 8692312 | 8692313 | cg07145988 | chr6 | 26988150 | 26988151 | cg04020984 |
| chr1 | 3823918 | 3823919 | cg14264496 | chr6 | 10415667 | 10415668 | cg14909730 | chr1 | 8729713 | 8729714 | cg00154554 | chr6 | 28411287 | 28411288 | cg09682330 |
| chr1 | 3827065 | 3827066 | cg09362969 | chr6 | 11044888 | 11044889 | cg24724428 | chr1 | 8748155 | 8748156 | cg14154131 | chr6 | 28956312 | 28956313 | cg16622920 |
| chr1 | 6062462 | 6062463 | cg04088152 | chr6 | 19838211 | 19838212 | cg00468146 | chr1 | 8759920 | 8759921 | cg19872136 | chr6 | 28956320 | 28956321 | cg00667751 |
| chr1 | 6085872 | 6085873 | cg13614083 | chr6 | 26988141 | 26988142 | cg06781135 | chr1 | 9127501 | 9127502 | cg04901835 | chr6 | 28956324 | 28956325 | cg18803104 |
| chr1 | 6419767 | 6419768 | cg16306654 | chr6 | 26988148 | 26988149 | cg11369071 | chr1 | 9686058 | 9686059 | cg02787675 | chr6 | 29829406 | 29829407 | cg08460078 |
| chr1 | 6534951 | 6534952 | cg25142390 | chr6 | 26988177 | 26988178 | cg11549265 | chr1 | 9751353 | 9751354 | cg16125214 | chr6 | 56819429 | 56819430 | cg05022673 |
| chr1 | 6649337 | 6649338 | cg18263572 | chr6 | 50818268 | 50818269 | cg10679301 | chr1 | 9880771 | 9880772 | cg23261073 | chr6 | 80657428 | 80657429 | cg25643118 |
| chr1 | 6672620 | 6672621 | cg06778853 | chr6 | 78172691 | 78172692 | cg00181439 | chr1 | 9895799 | 9895800 | cg14037769 | chr6 | 80657430 | 80657431 | cg05492839 |
| chr1 | 8000314 | 8000315 | cg08840010 | chr6 | 85482612 | 85482613 | cg08445802 | chr1 | 10314029 | 10314030 | cg23716690 | chr6 | 80657436 | 80657437 | cg18417423 |
| chr1 | 8388411 | 8388412 | cg08644356 | chr6 | 100896055 | 100896056 | cg08979319 | chr1 | 11042201 | 11042202 | cg02454025 | chr6 | 144329672 | 144329673 | cg21526238 |
| chr1 | 8570225 | 8570226 | cg14286456 | chr6 | 156428912 | 156428913 | cg26739230 | chr1 | 11711141 | 11711142 | cg19095000 | chr7 | 1452425 | 1452426 | cg02381192 |
| chr1 | 8662670 | 8662671 | cg12764441 | chr6 | 156429014 | 156429015 | cg27070962 | chr1 | 11862907 | 11862908 | cg17514528 | chr7 | 2883003 | 2883004 | cg01139696 |
| chr1 | 8720599 | 8720600 | cg21268984 | chr7 | 1120247 | 1120248 | cg21126453 | chr1 | 11867308 | 11867309 | cg14472778 | chr7 | 27239763 | 27239764 | cg26512254 |
| chr1 | 8824177 | 8824178 | cg20263853 | chr7 | 27225058 | 27225059 | cg12997720 | chr1 | 12185435 | 12185436 | cg07707505 | chr7 | 101005846 | 101005847 | cg07883117 |
| chr1 | 8875053 | 8875054 | cg10211414 | chr7 | 45961569 | 45961570 | cg26434048 | chr1 | 12185845 | 12185846 | cg22123711 | chr7 | 135662562 | 135662563 | cg21537187 |
| chr1 | 8963950 | 8963951 | cg10207928 | chr7 | 98592094 | 98592095 | cg24310924 | chr1 | 12186307 | 12186308 | cg10425291 | chr7 | 150901654 | 150901655 | cg01572810 |
| chr1 | 9086585 | 9086586 | cg04599612 | chr7 | 100463508 | 100463509 | cg19279257 | chr1 | 12221410 | 12221411 | cg20300911 | chr7 | 150901733 | 150901734 | cg04795774 |
| chr1 | 9143538 | 9143539 | cg25602242 | chr7 | 127658301 | 127658302 | cg15067687 | chr1 | 12225569 | 12225570 | cg23097499 | chr8 | 1592736 | 1592737 | cg14485877 |
| chr1 | 9145034 | 9145035 | cg14565651 | chr7 | 128845217 | 128845218 | cg20489453 | chr1 | 12230509 | 12230510 | cg18124517 | chr8 | 24857567 | 24857568 | cg03533519 |
| chr1 | 9182171 | 9182172 | cg20284891 | chr7 | 139577589 | 139577590 | cg24710631 | chr1 | 12238390 | 12238391 | cg22677556 | chr8 | 40753096 | 40753097 | cg05614657 |
| chr1 | 9375627 | 9375628 | cg06779253 | chr7 | 157204364 | 157204365 | cg18362281 | chr1 | 12251820 | 12251821 | cg15622556 | chr8 | 54790372 | 54790373 | cg27106513 |
| chr1 | 9405823 | 9405824 | cg00890054 | chr7 | 158936739 | 158936740 | cg21038156 | chr1 | 12251869 | 12251870 | cg06740893 | chr8 | 67872678 | 67872679 | cg00374056 |
| chr1 | 9651205 | 9651206 | cg16582732 | chr8 | 23564717 | 23564718 | cg16544169 | chr1 | 12260067 | 12260068 | cg13391664 | chr8 | 130877076 | 130877077 | cg02711724 |
| chr1 | 9687880 | 9687881 | cg25593631 | chr8 | 24772513 | 24772514 | cg26980244 | chr1 | 12268883 | 12268884 | cg11665613 | chr8 | 136469585 | 136469586 | cg27186774 |
| chr1 | 9788715 | 9788716 | cg07499142 | chr8 | 28974428 | 28974429 | cg06043201 | chr1 | 12559783 | 12559784 | cg10987347 | chr8 | 141540813 | 141540814 | cg08610326 |
| chr1 | 9788767 | 9788768 | cg12033075 | chr8 | 100559503 | 100559504 | cg22564448 | chr1 | 13912116 | 13912117 | cg17826518 | chr9 | 998547 | 998548 | cg13952963 |
| chr1 | 9788769 | 9788770 | cg03971555 | chr9 | 34370894 | 34370895 | cg13746854 | chr1 | 14031383 | 14031384 | cg25450806 | chr9 | 123854335 | 123854336 | cg13817287 |
| chr1 | 9844683 | 9844684 | cg15678052 | chr9 | 90340652 | 90340653 | cg11154542 | chr1 | 14033779 | 14033780 | cg10375300 | chr10 | 8096991 | 8096992 | cg18599069 |
| chr1 | 10494322 | 10494323 | cg26896979 | chr9 | 93523028 | 93523029 | cg14125403 | chr1 | 14105607 | 14105608 | cg12909925 | chr10 | 65186953 | 65186954 | cg17983571 |
| chr1 | 11985115 | 11985116 | cg01005303 | chr9 | 122132052 | 122132053 | cg14445814 | chr1 | 15208255 | 15208256 | cg05046721 | chr10 | 88149324 | 88149325 | cg06647751 |
| chr1 | 12186079 | 12186080 | cg10058766 | chr9 | 127215669 | 127215670 | cg14620572 | chr1 | 16264490 | 16264491 | cg07180355 | chr10 | 100993556 | 100993557 | cg08927006 |
| chr1 | 12213045 | 12213046 | cg24941292 | chr9 | 139565151 | 139565152 | cg14197156 | chr1 | 16273971 | 16273972 | cg14091149 | chr10 | 123368181 | 123368182 | cg14157844 |
| chr1 | 12218933 | 12218934 | cg16549957 | chr9 | 139565154 | 139565155 | cg04074066 | chr1 | 16285251 | 16285252 | cg00529422 | chr10 | 123731471 | 123731472 | cg19360212 |
| chr1 | 12233852 | 12233853 | cg20136855 | chr9 | 140400693 | 140400694 | cg13706769 | chr1 | 16335452 | 16335453 | cg14529268 | chr11 | 2930322 | 2930323 | cg22858288 |
| chr1 | 12238546 | 12238547 | cg15526535 | chr9 | 140711975 | 140711976 | cg15150472 | chr1 | 16340746 | 16340747 | cg06159980 | chr11 | 35644854 | 35644855 | cg13916633 |
| chr1 | 12261884 | 12261885 | cg16513984 | chr10 | 1084866 | 1084867 | cg25614935 | chr1 | 16472728 | 16472729 | cg27582323 | chr11 | 75140865 | 75140866 | cg08160331 |
| chr1 | 12538678 | 12538679 | cg05654304 | chr10 | 1084935 | 1084936 | cg14453012 | chr1 | 17287501 | 17287502 | cg14973240 | chr11 | 77907656 | 77907657 | cg15243034 |
| chr1 | 13992541 | 13992542 | cg23799025 | chr10 | 3215966 | 3215967 | cg11586382 | chr1 | 17528238 | 17528239 | cg19414383 | chr11 | 78614522 | 78614523 | cg13228701 |
| chr1 | 14029875 | 14029876 | cg25402049 | chr10 | 4697267 | 4697268 | cg12359895 | chr1 | 17766995 | 17766996 | cg01713950 | chr11 | 132952915 | 132952916 | cg12658947 |
| chr1 | 14030010 | 14030011 | cg19359071 | chr10 | 30336565 | 30336566 | cg20032789 | chr1 | 17865713 | 17865714 | cg10123662 | chr12 | 214224 | 214225 | cg01239735 |
| chr1 | 14030141 | 14030142 | cg23267665 | chr10 | 43601862 | 43601863 | cg09848445 | chr1 | 19232926 | 19232927 | cg09181054 | chr12 | 1675846 | 1675847 | cg06880438 |
| chr1 | 14077143 | 14077144 | cg11750706 | chr10 | 50820278 | 50820279 | cg01295399 | chr1 | 19752812 | 19752813 | cg12424867 | chr12 | 33205862 | 33205863 | cg12582896 |
| chr1 | 14134260 | 14134261 | cg16652739 | chr10 | 50887472 | 50887473 | cg09402276 | chr1 | 19756350 | 19756351 | cg06788764 | chr12 | 64080144 | 64080145 | cg00087746 |
| chr1 | 14847711 | 14847712 | cg13107839 | chr10 | 50887488 | 50887489 | cg09233392 | chr1 | 19774989 | 19774990 | cg01832549 | chr12 | 120868468 | 120868469 | cg04880546 |
| chr1 | 15954209 | 15954210 | cg21004104 | chr10 | 79422304 | 79422305 | cg23888215 | chr1 | 20142497 | 20142498 | cg09195271 | chr12 | 133659856 | 133659857 | cg14774371 |
| chr1 | 16273212 | 16273213 | cg06640695 | chr10 | 135113191 | 135113192 | cg20090482 | chr1 | 20737787 | 20737788 | cg16357353 | chr13 | 27131676 | 27131677 | cg18781240 |
| chr1 | 16275539 | 16275540 | cg26334507 | chr10 | 135113239 | 135113240 | cg11685186 | chr1 | 21607409 | 21607410 | cg22077962 | chr13 | 27131683 | 27131684 | cg08453926 |

| | | | |
|---|---|---|---|
| chr1 | 16393791 | 16393792 | cg18800143 |
| chr1 | 16473143 | 16473144 | cg26371327 |
| chr1 | 16689696 | 16689697 | cg09234164 |
| chr1 | 16763671 | 16763672 | cg26845278 |
| chr1 | 17746597 | 17746598 | cg15995125 |
| chr1 | 17877451 | 17877452 | cg08947790 |
| chr1 | 17951330 | 17951331 | cg14780070 |
| chr1 | 18010807 | 18010808 | cg11742472 |
| chr1 | 18693462 | 18693463 | cg13840174 |
| chr1 | 19230460 | 19230461 | cg01366338 |
| chr1 | 19250279 | 19250280 | cg09012001 |
| chr1 | 19255712 | 19255713 | cg10992634 |
| chr1 | 19996240 | 19996241 | cg01791421 |
| chr1 | 20141781 | 20141782 | cg05393025 |
| chr1 | 20306641 | 20306642 | cg22220722 |
| chr1 | 20501558 | 20501559 | cg04605617 |
| chr1 | 20807386 | 20807387 | cg02951739 |
| chr1 | 20985249 | 20985250 | cg14519150 |
| chr1 | 21615863 | 21615864 | cg16603916 |
| chr1 | 21645238 | 21645239 | cg05954918 |
| chr1 | 21650610 | 21650611 | cg16514473 |
| chr1 | 21652298 | 21652299 | cg16416158 |
| chr1 | 22290005 | 22290006 | cg27522078 |
| chr1 | 23878540 | 23878541 | cg11078090 |
| chr1 | 23880481 | 23880482 | cg15102567 |
| chr1 | 23884373 | 23884374 | cg15357156 |
| chr1 | 23886730 | 23886731 | cg27518976 |
| chr1 | 23887324 | 23887325 | cg11336311 |
| chr1 | 23887514 | 23887515 | cg09858862 |
| chr1 | 23887816 | 23887817 | cg18150584 |
| chr1 | 23902856 | 23902857 | cg12646386 |
| chr1 | 23945686 | 23945687 | cg16685586 |
| chr1 | 24528471 | 24528472 | cg17172066 |
| chr1 | 24528833 | 24528834 | cg15571353 |
| chr1 | 24862381 | 24862382 | cg00716040 |
| chr1 | 25062925 | 25062926 | cg27425146 |
| chr1 | 25069086 | 25069087 | cg16521555 |
| chr1 | 25070672 | 25070673 | cg10546487 |
| chr1 | 25292215 | 25292216 | cg16529592 |
| chr1 | 25868969 | 25868970 | cg06395634 |
| chr1 | 26098310 | 26098311 | cg16417374 |
| chr1 | 26219198 | 26219199 | cg24809011 |
| chr1 | 26643109 | 26643110 | cg19777853 |
| chr1 | 27060511 | 27060512 | cg09849688 |
| chr1 | 27087961 | 27087962 | cg21454760 |
| chr1 | 27109220 | 27109221 | cg00759987 |
| chr1 | 27152694 | 27152695 | cg04427651 |
| chr1 | 27155559 | 27155560 | cg06216065 |
| chr1 | 27189349 | 27189350 | cg21950166 |
| chr1 | 27240669 | 27240670 | cg16177693 |
| chr1 | 27527276 | 27527277 | cg02799237 |
| chr1 | 27627058 | 27627059 | cg27531366 |
| chr1 | 27755803 | 27755804 | cg09588020 |
| chr1 | 27819060 | 27819061 | cg22226904 |
| chr1 | 27901805 | 27901806 | cg22429121 |
| chr1 | 28385657 | 28385658 | cg27509347 |
| chr11 | 1981264 | 1981265 | cg06498964 |
| chr11 | 18476131 | 18476132 | cg27316453 |
| chr11 | 27740813 | 27740814 | cg13974632 |
| chr11 | 43604758 | 43604759 | cg17592231 |
| chr11 | 69707303 | 69707304 | cg26579578 |
| chr11 | 70261373 | 70261374 | cg16709342 |
| chr11 | 108570019 | 108570020 | cg04967734 |
| chr11 | 113945432 | 113945433 | cg03597491 |
| chr11 | 125774406 | 125774407 | cg19792599 |
| chr11 | 133837711 | 133837712 | cg19522930 |
| chr12 | 39539422 | 39539423 | cg03468072 |
| chr12 | 56364837 | 56364838 | cg09596975 |
| chr12 | 113916664 | 113916665 | cg09887059 |
| chr12 | 122651762 | 122651763 | cg04820362 |
| chr12 | 129298759 | 129298760 | cg15324978 |
| chr12 | 132839125 | 132839126 | cg14058027 |
| chr12 | 132936807 | 132936808 | cg18053948 |
| chr13 | 43702940 | 43702941 | cg01403655 |
| chr13 | 43703025 | 43703026 | cg16760130 |
| chr13 | 53775108 | 53775109 | cg17555373 |
| chr13 | 78492678 | 78492679 | cg15699226 |
| chr13 | 96204978 | 96204979 | cg25032595 |
| chr13 | 112721950 | 112721951 | cg16705627 |
| chr13 | 113671965 | 113671966 | cg22186503 |
| chr13 | 113974478 | 113974479 | cg11451477 |
| chr14 | 34420415 | 34420416 | cg08122140 |
| chr14 | 91742453 | 91742454 | cg18729681 |
| chr14 | 100069535 | 100069536 | cg18414021 |
| chr14 | 100069657 | 100069658 | cg16717713 |
| chr14 | 100069666 | 100069667 | cg05406635 |
| chr14 | 101193397 | 101193398 | cg02874376 |
| chr14 | 106331803 | 106331804 | cg14387626 |
| chr15 | 68121563 | 68121564 | cg24880701 |
| chr16 | 335830 | 335831 | cg05703996 |
| chr16 | 459792 | 459793 | cg26946311 |
| chr16 | 541243 | 541244 | cg16776421 |
| chr16 | 542535 | 542536 | cg27581871 |
| chr16 | 1183462 | 1183463 | cg00701514 |
| chr16 | 2140870 | 2140871 | cg01965047 |
| chr16 | 2801730 | 2801731 | cg04484916 |
| chr16 | 51188372 | 51188373 | cg02352240 |
| chr16 | 54962963 | 54962964 | cg09293786 |
| chr16 | 57394387 | 57394388 | cg04453552 |
| chr16 | 85427738 | 85427739 | cg08006672 |
| chr16 | 85427847 | 85427848 | cg05824809 |
| chr16 | 87440505 | 87440506 | cg07178066 |
| chr17 | 88246 | 88247 | cg10016717 |
| chr17 | 88286 | 88287 | cg11751810 |
| chr17 | 723242 | 723243 | cg17842130 |
| chr17 | 14201063 | 14201064 | cg15909319 |
| chr17 | 47113489 | 47113490 | cg18129628 |
| chr17 | 70680294 | 70680295 | cg27219182 |
| chr17 | 72740524 | 72740525 | cg02797539 |
| chr17 | 75316383 | 75316384 | cg15401418 |
| chr17 | 77987482 | 77987483 | cg26350921 |
| chr17 | 79652244 | 79652245 | cg22209573 |
| chr1 | 21611224 | 21611225 | cg11834658 |
| chr1 | 21649784 | 21649785 | cg18116670 |
| chr1 | 22142491 | 22142492 | cg06830981 |
| chr1 | 23449368 | 23449369 | cg11825603 |
| chr1 | 23496648 | 23496649 | cg01435109 |
| chr1 | 23505138 | 23505139 | cg24437797 |
| chr1 | 23879671 | 23879672 | cg11580897 |
| chr1 | 23880210 | 23880211 | cg02690117 |
| chr1 | 23880736 | 23880737 | cg02978140 |
| chr1 | 23883338 | 23883339 | cg13977835 |
| chr1 | 23894436 | 23894437 | cg18027654 |
| chr1 | 23895366 | 23895367 | cg23664396 |
| chr1 | 23925874 | 23925875 | cg06933796 |
| chr1 | 23942134 | 23942135 | cg00458949 |
| chr1 | 23953480 | 23953481 | cg00171783 |
| chr1 | 24511042 | 24511043 | cg16117757 |
| chr1 | 25061987 | 25061988 | cg07191152 |
| chr1 | 25105430 | 25105431 | cg19857379 |
| chr1 | 25246854 | 25246855 | cg15498134 |
| chr1 | 25246922 | 25246923 | cg08544331 |
| chr1 | 25258679 | 25258680 | cg20670361 |
| chr1 | 25344342 | 25344343 | cg22169866 |
| chr1 | 25354366 | 25354367 | cg04281159 |
| chr1 | 26096954 | 26096955 | cg00978203 |
| chr1 | 26126233 | 26126234 | cg07352544 |
| chr1 | 26198360 | 26198361 | cg23799313 |
| chr1 | 26250290 | 26250291 | cg18143792 |
| chr1 | 26603970 | 26603971 | cg21646082 |
| chr1 | 26617780 | 26617781 | cg10402321 |
| chr1 | 26646801 | 26646802 | cg27430637 |
| chr1 | 26648467 | 26648468 | cg04893145 |
| chr1 | 26857774 | 26857775 | cg24585377 |
| chr1 | 27113219 | 27113220 | cg24313303 |
| chr1 | 27240694 | 27240695 | cg01942646 |
| chr1 | 27287181 | 27287182 | cg21921474 |
| chr1 | 27462916 | 27462917 | cg20816789 |
| chr1 | 27468772 | 27468773 | cg26313337 |
| chr1 | 27686328 | 27686329 | cg13449606 |
| chr1 | 27899025 | 27899026 | cg13375463 |
| chr1 | 27940619 | 27940620 | cg03018771 |
| chr1 | 27953336 | 27953337 | cg12465678 |
| chr1 | 27962021 | 27962022 | cg09845000 |
| chr1 | 27962412 | 27962413 | cg16897005 |
| chr1 | 27977270 | 27977271 | cg04089743 |
| chr1 | 28288853 | 28288854 | cg25107788 |
| chr1 | 28504285 | 28504286 | cg07821160 |
| chr1 | 28848173 | 28848174 | cg16691863 |
| chr1 | 31206669 | 31206670 | cg10709350 |
| chr1 | 31231214 | 31231215 | cg12654790 |
| chr1 | 31231363 | 31231364 | cg10001720 |
| chr1 | 31231488 | 31231489 | cg08463932 |
| chr1 | 31492493 | 31492494 | cg20790367 |
| chr1 | 32236337 | 32236338 | cg02238760 |
| chr1 | 32716212 | 32716213 | cg01525376 |
| chr1 | 32716557 | 32716558 | cg11683242 |
| chr1 | 32717620 | 32717621 | cg06517940 |
| chr13 | 47214871 | 47214872 | cg16503053 |
| chr13 | 92050720 | 92050721 | cg03036557 |
| chr13 | 92050726 | 92050727 | cg12678562 |
| chr13 | 114064536 | 114064537 | cg10072464 |
| chr14 | 100150802 | 100150803 | cg04276484 |
| chr14 | 100437639 | 100437640 | cg02268225 |
| chr14 | 101317622 | 101317623 | cg03039990 |
| chr14 | 104172109 | 104172110 | cg23702848 |
| chr15 | 23035493 | 23035494 | cg27514038 |
| chr15 | 31775895 | 31775896 | cg04875128 |
| chr15 | 32162729 | 32162730 | cg03256884 |
| chr15 | 39872186 | 39872187 | cg23849826 |
| chr15 | 62516405 | 62516406 | cg03455316 |
| chr15 | 93615039 | 93615040 | cg08334034 |
| chr15 | 98964982 | 98964983 | cg00718470 |
| chr15 | 99192520 | 99192521 | cg19620752 |
| chr15 | 100882162 | 100882163 | cg04822808 |
| chr15 | 100882165 | 100882166 | cg05027458 |
| chr15 | 100882334 | 100882335 | cg20941820 |
| chr16 | 970802 | 970803 | cg08227260 |
| chr16 | 970818 | 970819 | cg01207650 |
| chr16 | 8619943 | 8619944 | cg04066637 |
| chr16 | 57562740 | 57562741 | cg05004142 |
| chr16 | 57562841 | 57562842 | cg02102533 |
| chr16 | 84690554 | 84690555 | cg07831918 |
| chr16 | 89119002 | 89119003 | cg05350293 |
| chr17 | 925669 | 925670 | cg24439623 |
| chr17 | 14201938 | 14201939 | cg23280506 |
| chr17 | 26443366 | 26443367 | cg11792281 |
| chr17 | 42980188 | 42980189 | cg19026811 |
| chr17 | 59572908 | 59572909 | cg21018997 |
| chr17 | 64961440 | 64961441 | cg10355837 |
| chr17 | 70117530 | 70117531 | cg06391468 |
| chr17 | 79011140 | 79011141 | cg11620135 |
| chr17 | 80043349 | 80043350 | cg03407524 |
| chr18 | 60383672 | 60383673 | cg20317272 |
| chr19 | 613111 | 613112 | cg06657917 |
| chr19 | 613818 | 613819 | cg12159023 |
| chr19 | 5827811 | 5827812 | cg18517266 |
| chr19 | 17942411 | 17942412 | cg11624708 |
| chr19 | 19216451 | 19216452 | cg25612391 |
| chr19 | 34269211 | 34269212 | cg03148562 |
| chr19 | 51227943 | 51227944 | cg02759846 |
| chr20 | 3644151 | 3644152 | cg06919203 |
| chr20 | 44803411 | 44803412 | cg17412248 |
| chr21 | 39047855 | 39047856 | cg10848373 |
| chr21 | 44088823 | 44088824 | cg27168131 |
| chr21 | 44898090 | 44898091 | cg23732182 |
| chr22 | 46860105 | 46860106 | cg11747897 |
| chr22 | 49765229 | 49765230 | cg22471075 |

| | | | |
|---|---|---|---|
| chr1 | 28571667 | 28571668 | cg23394462 |
| chr1 | 28572299 | 28572300 | cg02251754 |
| chr1 | 28576552 | 28576553 | cg11844530 |
| chr1 | 28765031 | 28765032 | cg00956142 |
| chr1 | 28971003 | 28971004 | cg03287108 |
| chr1 | 29098920 | 29098921 | cg11654813 |
| chr1 | 29225533 | 29225534 | cg23840027 |
| chr1 | 29240554 | 29240555 | cg04844693 |
| chr1 | 31221135 | 31221136 | cg18411043 |
| chr1 | 31234437 | 31234438 | cg25310233 |
| chr1 | 31247617 | 31247618 | cg06747288 |
| chr1 | 31459731 | 31459732 | cg08136313 |
| chr1 | 31709364 | 31709365 | cg11793106 |
| chr1 | 32039682 | 32039683 | cg03211098 |
| chr1 | 32321494 | 32321495 | cg00988548 |
| chr1 | 32405083 | 32405084 | cg21545720 |
| chr1 | 32405360 | 32405361 | cg09373727 |
| chr1 | 32572611 | 32572612 | cg20633875 |
| chr1 | 32577363 | 32577364 | cg00958944 |
| chr1 | 32708485 | 32708486 | cg02830936 |
| chr1 | 32739493 | 32739494 | cg20239639 |
| chr1 | 32787489 | 32787490 | cg18857467 |
| chr1 | 32799069 | 32799070 | cg25893831 |
| chr1 | 33150185 | 33150186 | cg22350438 |
| chr1 | 34329348 | 34329349 | cg27559224 |
| chr1 | 35836009 | 35836010 | cg21199093 |
| chr1 | 36351841 | 36351842 | cg16356622 |
| chr1 | 36771781 | 36771782 | cg21418854 |
| chr1 | 36825645 | 36825646 | cg05798664 |
| chr1 | 36840028 | 36840029 | cg25722983 |
| chr1 | 36856866 | 36856867 | cg20272463 |
| chr1 | 36948570 | 36948571 | cg14523574 |
| chr1 | 37784177 | 37784178 | cg05846868 |
| chr1 | 38100467 | 38100468 | cg13448605 |
| chr1 | 38276835 | 38276836 | cg04496824 |
| chr1 | 38356176 | 38356177 | cg23025416 |
| chr1 | 38942403 | 38942404 | cg00725089 |
| chr1 | 38942581 | 38942582 | cg03216506 |
| chr1 | 38942644 | 38942645 | cg11959697 |
| chr1 | 39041399 | 39041400 | cg11848788 |
| chr1 | 39456103 | 39456104 | cg03130816 |
| chr1 | 39658128 | 39658129 | cg10316899 |
| chr1 | 39873077 | 39873078 | cg15158106 |
| chr1 | 39875238 | 39875239 | cg07570723 |
| chr1 | 40307257 | 40307258 | cg00255699 |
| chr1 | 40435396 | 40435397 | cg15616998 |
| chr1 | 40780905 | 40780906 | cg10512203 |
| chr1 | 40843940 | 40843941 | cg27264437 |
| chr1 | 40887988 | 40887989 | cg26042300 |
| chr1 | 41128874 | 41128875 | cg13935127 |
| chr1 | 41807865 | 41807866 | cg03962019 |
| chr1 | 41892144 | 41892145 | cg27521109 |
| chr1 | 41984243 | 41984244 | cg08139729 |
| chr1 | 42271216 | 42271217 | cg20149359 |
| chr1 | 43304307 | 43304308 | cg09856274 |
| chr1 | 43351860 | 43351861 | cg06372967 |
| chr17 | 80928561 | 80928562 | cg19046167 |
| chr18 | 76739409 | 76739410 | cg05080154 |
| chr18 | 77545223 | 77545224 | cg23579185 |
| chr19 | 643457 | 643458 | cg02775417 |
| chr19 | 2428677 | 2428678 | cg18147048 |
| chr19 | 17213063 | 17213064 | cg20091114 |
| chr19 | 19729048 | 19729049 | cg00591868 |
| chr19 | 39051817 | 39051818 | cg16865475 |
| chr19 | 45174681 | 45174682 | cg23147597 |
| chr19 | 49657000 | 49657001 | cg11297046 |
| chr19 | 52222446 | 52222447 | cg01024444 |
| chr20 | 11871375 | 11871376 | cg14603345 |
| chr20 | 11871384 | 11871385 | cg23944804 |
| chr20 | 11871396 | 11871397 | cg12894649 |
| chr20 | 13976093 | 13976094 | cg04624110 |
| chr20 | 22565228 | 22565229 | cg25218716 |
| chr20 | 37054900 | 37054901 | cg13610455 |
| chr20 | 48099146 | 48099147 | cg26709285 |
| chr21 | 34396506 | 34396507 | cg13524919 |
| chr22 | 24796628 | 24796629 | cg27501362 |
| chr1 | 32731477 | 32731478 | cg16348358 |
| chr1 | 32738251 | 32738252 | cg14781242 |
| chr1 | 32739049 | 32739050 | cg10325497 |
| chr1 | 32739114 | 32739115 | cg17223520 |
| chr1 | 33813321 | 33813322 | cg07570498 |
| chr1 | 35585672 | 35585673 | cg13254203 |
| chr1 | 35908609 | 35908610 | cg16815882 |
| chr1 | 36108206 | 36108207 | cg12413579 |
| chr1 | 36275039 | 36275040 | cg02639366 |
| chr1 | 36351470 | 36351471 | cg00712762 |
| chr1 | 36589944 | 36589945 | cg06501835 |
| chr1 | 36771021 | 36771022 | cg10622536 |
| chr1 | 36771714 | 36771715 | cg02865595 |
| chr1 | 36771793 | 36771794 | cg19373545 |
| chr1 | 36785859 | 36785860 | cg00426377 |
| chr1 | 36788416 | 36788417 | cg16394290 |
| chr1 | 36789786 | 36789787 | cg15095906 |
| chr1 | 36852956 | 36852957 | cg00913954 |
| chr1 | 38474788 | 38474789 | cg06444374 |
| chr1 | 38493031 | 38493032 | cg24033742 |
| chr1 | 39500070 | 39500071 | cg14656441 |
| chr1 | 39873697 | 39873698 | cg03518138 |
| chr1 | 40311142 | 40311143 | cg22016731 |
| chr1 | 40557841 | 40557842 | cg04560534 |
| chr1 | 40626599 | 40626600 | cg18646240 |
| chr1 | 41828130 | 41828131 | cg19471466 |
| chr1 | 42129921 | 42129922 | cg25202370 |
| chr1 | 42208040 | 42208041 | cg10054398 |
| chr1 | 42288825 | 42288826 | cg11256186 |
| chr1 | 42385662 | 42385663 | cg16145216 |
| chr1 | 43281794 | 43281795 | cg23839159 |
| chr1 | 45193615 | 45193616 | cg07780762 |
| chr1 | 45250440 | 45250441 | cg15502465 |
| chr1 | 45250840 | 45250841 | cg00280609 |
| chr1 | 45265337 | 45265338 | cg07368443 |
| chr1 | 45268975 | 45268976 | cg08259083 |
| chr1 | 45274048 | 45274049 | cg03434029 |
| chr1 | 45274099 | 45274100 | cg11448683 |
| chr1 | 45274513 | 45274514 | cg14565781 |
| chr1 | 45274600 | 45274601 | cg00043599 |
| chr1 | 45276537 | 45276538 | cg17093466 |
| chr1 | 45279329 | 45279330 | cg12275060 |
| chr1 | 45292204 | 45292205 | cg26297462 |
| chr1 | 45350531 | 45350532 | cg17298989 |
| chr1 | 46633153 | 46633154 | cg06974930 |
| chr1 | 52348865 | 52348866 | cg17403995 |
| chr1 | 53014805 | 53014806 | cg11690826 |
| chr1 | 53016396 | 53016397 | cg04682580 |
| chr1 | 53019814 | 53019815 | cg03950476 |
| chr1 | 53040491 | 53040492 | cg00063982 |
| chr1 | 53054907 | 53054908 | cg05749906 |
| chr1 | 53067637 | 53067638 | cg18087326 |
| chr1 | 53094893 | 53094894 | cg17438457 |
| chr1 | 53125850 | 53125851 | cg04568981 |
| chr1 | 53145415 | 53145416 | cg04341526 |
| chr1 | 53217358 | 53217359 | cg00692173 |

| | | | |
|---|---|---|---|
| chr1 | 43402591 | 43402592 | cg05942022 |
| chr1 | 43416103 | 43416104 | cg01924561 |
| chr1 | 43422337 | 43422338 | cg00737593 |
| chr1 | 43429380 | 43429381 | cg00568910 |
| chr1 | 43695264 | 43695265 | cg06907770 |
| chr1 | 43703989 | 43703990 | cg21267936 |
| chr1 | 43748663 | 43748664 | cg19234495 |
| chr1 | 44402353 | 44402354 | cg23954153 |
| chr1 | 44444305 | 44444306 | cg17331566 |
| chr1 | 44446600 | 44446601 | cg09666532 |
| chr1 | 44903330 | 44903331 | cg07348418 |
| chr1 | 45138784 | 45138785 | cg14009440 |
| chr1 | 45249541 | 45249542 | cg02072975 |
| chr1 | 45266782 | 45266783 | cg17233819 |
| chr1 | 45267345 | 45267346 | cg27583604 |
| chr1 | 45277354 | 45277355 | cg02923972 |
| chr1 | 46053650 | 46053651 | cg26068180 |
| chr1 | 46646158 | 46646159 | cg19321695 |
| chr1 | 46646224 | 46646225 | cg04975920 |
| chr1 | 46664598 | 46664599 | cg04036898 |
| chr1 | 47079672 | 47079673 | cg06760830 |
| chr1 | 47081524 | 47081525 | cg03345668 |
| chr1 | 48458049 | 48458050 | cg04084173 |
| chr1 | 48458463 | 48458464 | cg27522154 |
| chr1 | 48508839 | 48508840 | cg00055603 |
| chr1 | 48688453 | 48688454 | cg21297772 |
| chr1 | 48703955 | 48703956 | cg05042706 |
| chr1 | 48704831 | 48704832 | cg25469314 |
| chr1 | 48960137 | 48960138 | cg04865113 |
| chr1 | 51730449 | 51730450 | cg09941363 |
| chr1 | 52498044 | 52498045 | cg00041014 |
| chr1 | 53019727 | 53019728 | cg00754357 |
| chr1 | 53067365 | 53067366 | cg01400822 |
| chr1 | 53068834 | 53068835 | cg09161043 |
| chr1 | 53072410 | 53072411 | cg03330022 |
| chr1 | 53098299 | 53098300 | cg04882349 |
| chr1 | 53878650 | 53878651 | cg24699914 |
| chr1 | 54359880 | 54359881 | cg17453374 |
| chr1 | 54822031 | 54822032 | cg00172603 |
| chr1 | 55682652 | 55682653 | cg17004733 |
| chr1 | 56069436 | 56069437 | cg25286579 |
| chr1 | 56397447 | 56397448 | cg08118140 |
| chr1 | 56636154 | 56636155 | cg03724279 |
| chr1 | 56756750 | 56756751 | cg02086690 |
| chr1 | 56842338 | 56842339 | cg26837800 |
| chr1 | 57040807 | 57040808 | cg17902947 |
| chr1 | 58098908 | 58098909 | cg23710611 |
| chr1 | 58333307 | 58333308 | cg00944580 |
| chr1 | 60158474 | 60158475 | cg19746186 |
| chr1 | 60216631 | 60216632 | cg19773937 |
| chr1 | 61636722 | 61636723 | cg16001865 |
| chr1 | 62123940 | 62123941 | cg20013688 |
| chr1 | 64238366 | 64238367 | cg24502478 |
| chr1 | 64578227 | 64578228 | cg00327437 |
| chr1 | 64660725 | 64660726 | cg19248395 |
| chr1 | 64961535 | 64961536 | cg17181043 |

| | | | |
|---|---|---|---|
| chr1 | 53782077 | 53782078 | cg11950754 |
| chr1 | 54358862 | 54358863 | cg17734802 |
| chr1 | 54739176 | 54739177 | cg09787089 |
| chr1 | 54822503 | 54822504 | cg08641990 |
| chr1 | 56901693 | 56901694 | cg26248114 |
| chr1 | 63142359 | 63142360 | cg16991569 |
| chr1 | 63150905 | 63150906 | cg10583485 |
| chr1 | 63151692 | 63151693 | cg01097102 |
| chr1 | 63901727 | 63901728 | cg15832574 |
| chr1 | 64169293 | 64169294 | cg18973238 |
| chr1 | 64242960 | 64242961 | cg17298776 |
| chr1 | 64293886 | 64293887 | cg06743782 |
| chr1 | 64301568 | 64301569 | cg03140289 |
| chr1 | 64379958 | 64379959 | cg03950655 |
| chr1 | 64602654 | 64602655 | cg07797372 |
| chr1 | 65528997 | 65528998 | cg17940858 |
| chr1 | 65531864 | 65531865 | cg13390284 |
| chr1 | 65535397 | 65535398 | cg04924028 |
| chr1 | 67399983 | 67399984 | cg15930380 |
| chr1 | 67900465 | 67900466 | cg00176043 |
| chr1 | 68215783 | 68215784 | cg04849985 |
| chr1 | 84755291 | 84755292 | cg12743388 |
| chr1 | 85313079 | 85313080 | cg07110659 |
| chr1 | 85412601 | 85412602 | cg13227206 |
| chr1 | 85872804 | 85872805 | cg15991082 |
| chr1 | 87596934 | 87596935 | cg01445399 |
| chr1 | 88231424 | 88231425 | cg06712674 |
| chr1 | 89333529 | 89333530 | cg13109106 |
| chr1 | 89592433 | 89592434 | cg11068071 |
| chr1 | 90229631 | 90229632 | cg10576151 |
| chr1 | 90307339 | 90307340 | cg05578673 |
| chr1 | 92056542 | 92056543 | cg24006770 |
| chr1 | 92083173 | 92083174 | cg07922843 |
| chr1 | 92350112 | 92350113 | cg00342358 |
| chr1 | 92350133 | 92350134 | cg01336162 |
| chr1 | 92350430 | 92350431 | cg17074704 |
| chr1 | 92545650 | 92545651 | cg26580095 |
| chr1 | 93415910 | 93415911 | cg05036103 |
| chr1 | 95238920 | 95238921 | cg12719836 |
| chr1 | 97418099 | 97418100 | cg01566312 |
| chr1 | 101493828 | 101493829 | cg05376756 |
| chr1 | 108039344 | 108039345 | cg08950364 |
| chr1 | 108337960 | 108337961 | cg02978220 |
| chr1 | 109265387 | 109265388 | cg11481351 |
| chr1 | 109291831 | 109291832 | cg04124962 |
| chr1 | 109503393 | 109503394 | cg05048348 |
| chr1 | 109850837 | 109850838 | cg09332784 |
| chr1 | 110449308 | 110449309 | cg09167608 |
| chr1 | 110452482 | 110452483 | cg07568430 |
| chr1 | 110452616 | 110452617 | cg02361903 |
| chr1 | 110564742 | 110564743 | cg16698212 |
| chr1 | 110752047 | 110752048 | cg20966270 |
| chr1 | 110756235 | 110756236 | cg09898793 |
| chr1 | 110933345 | 110933346 | cg25320780 |
| chr1 | 111320416 | 111320417 | cg04140834 |
| chr1 | 111729823 | 111729824 | cg07019187 |

| | | | |
|---|---|---|---|
| chr1 | 65358393 | 65358394 | cg07798602 |
| chr1 | 65421982 | 65421983 | cg15082498 |
| chr1 | 65433360 | 65433361 | cg07453451 |
| chr1 | 65889855 | 65889856 | cg15466952 |
| chr1 | 66708864 | 66708865 | cg09320652 |
| chr1 | 67135552 | 67135553 | cg22949256 |
| chr1 | 67807960 | 67807961 | cg14849855 |
| chr1 | 68166689 | 68166690 | cg13784017 |
| chr1 | 68225713 | 68225714 | cg00147943 |
| chr1 | 68594186 | 68594187 | cg22482700 |
| chr1 | 68841715 | 68841716 | cg27173965 |
| chr1 | 69552291 | 69552292 | cg12658930 |
| chr1 | 77307268 | 77307269 | cg06446432 |
| chr1 | 77411902 | 77411903 | cg16926302 |
| chr1 | 79085765 | 79085766 | cg00855901 |
| chr1 | 79627622 | 79627623 | cg17175521 |
| chr1 | 81974159 | 81974160 | cg01425718 |
| chr1 | 82165326 | 82165327 | cg13471209 |
| chr1 | 84767878 | 84767879 | cg03204322 |
| chr1 | 85159003 | 85159004 | cg10096893 |
| chr1 | 85458682 | 85458683 | cg12277336 |
| chr1 | 87621160 | 87621161 | cg03434085 |
| chr1 | 87799914 | 87799915 | cg20701183 |
| chr1 | 89989507 | 89989508 | cg01128109 |
| chr1 | 90205170 | 90205171 | cg03899643 |
| chr1 | 91989267 | 91989268 | cg01782024 |
| chr1 | 92179761 | 92179762 | cg15326069 |
| chr1 | 92216881 | 92216882 | cg21116284 |
| chr1 | 92952702 | 92952703 | cg24517501 |
| chr1 | 92952897 | 92952898 | cg04777348 |
| chr1 | 92953037 | 92953038 | cg25320328 |
| chr1 | 93301669 | 93301670 | cg08661164 |
| chr1 | 93425802 | 93425803 | cg22128645 |
| chr1 | 94074034 | 94074035 | cg05941840 |
| chr1 | 94145223 | 94145224 | cg02342148 |
| chr1 | 94147831 | 94147832 | cg06221222 |
| chr1 | 94155657 | 94155658 | cg21937969 |
| chr1 | 94167631 | 94167632 | cg04165508 |
| chr1 | 94585452 | 94585453 | cg04491808 |
| chr1 | 95508854 | 95508855 | cg09795087 |
| chr1 | 95561502 | 95561503 | cg21263566 |
| chr1 | 96828042 | 96828043 | cg04095498 |
| chr1 | 101272300 | 101272301 | cg01789501 |
| chr1 | 101830939 | 101830940 | cg27057524 |
| chr1 | 107680714 | 107680715 | cg01939741 |
| chr1 | 107682552 | 107682553 | cg00758069 |
| chr1 | 108409977 | 108409978 | cg12047260 |
| chr1 | 109099375 | 109099376 | cg08707476 |
| chr1 | 109204325 | 109204326 | cg00328227 |
| chr1 | 109242775 | 109242776 | cg20706196 |
| chr1 | 109399734 | 109399735 | cg05295584 |
| chr1 | 109914235 | 109914236 | cg09131339 |
| chr1 | 110036141 | 110036142 | cg12597839 |
| chr1 | 110306591 | 110306592 | cg00491404 |
| chr1 | 110548562 | 110548563 | cg16269943 |
| chr1 | 111175082 | 111175083 | cg05656442 |
| chr1 | 111742515 | 111742516 | cg26518580 |
| chr1 | 111743200 | 111743201 | cg19269039 |
| chr1 | 111743202 | 111743203 | cg20317872 |
| chr1 | 111743537 | 111743538 | cg23631538 |
| chr1 | 111768837 | 111768838 | cg03916225 |
| chr1 | 111934770 | 111934771 | cg26681414 |
| chr1 | 112046026 | 112046027 | cg15673896 |
| chr1 | 112046915 | 112046916 | cg25674286 |
| chr1 | 112161864 | 112161865 | cg24160066 |
| chr1 | 112280233 | 112280234 | cg16136290 |
| chr1 | 113216382 | 113216383 | cg03417473 |
| chr1 | 113242047 | 113242048 | cg15095622 |
| chr1 | 113252666 | 113252667 | cg20247781 |
| chr1 | 113255818 | 113255819 | cg06740440 |
| chr1 | 113337251 | 113337252 | cg03208044 |
| chr1 | 114298184 | 114298185 | cg22495801 |
| chr1 | 114300298 | 114300299 | cg12587818 |
| chr1 | 114423726 | 114423727 | cg03325407 |
| chr1 | 114468988 | 114468989 | cg18342866 |
| chr1 | 116193783 | 116193784 | cg00819310 |
| chr1 | 116236487 | 116236488 | cg17465013 |
| chr1 | 116521539 | 116521540 | cg22353097 |
| chr1 | 116925613 | 116925614 | cg17107017 |
| chr1 | 116925753 | 116925754 | cg20309677 |
| chr1 | 117114916 | 117114917 | cg21167563 |
| chr1 | 117208496 | 117208497 | cg04140633 |
| chr1 | 117398001 | 117398002 | cg04878826 |
| chr1 | 144539026 | 144539027 | cg17745262 |
| chr1 | 145466550 | 145466551 | cg17877786 |
| chr1 | 145471727 | 145471728 | cg08916374 |
| chr1 | 146515238 | 146515239 | cg26810966 |
| chr1 | 146516011 | 146516012 | cg09441331 |
| chr1 | 149396788 | 149396789 | cg03525266 |
| chr1 | 149782411 | 149782412 | cg00721080 |
| chr1 | 150808973 | 150808974 | cg19750321 |
| chr1 | 150983722 | 150983723 | cg10832186 |
| chr1 | 151132257 | 151132258 | cg25306932 |
| chr1 | 151799073 | 151799074 | cg21784096 |
| chr1 | 151805241 | 151805242 | cg03005293 |
| chr1 | 152132853 | 152132854 | cg05765254 |
| chr1 | 153517384 | 153517385 | cg23048494 |
| chr1 | 153588927 | 153588928 | cg13098855 |
| chr1 | 154164994 | 154164995 | cg16586406 |
| chr1 | 154243781 | 154243782 | cg00718681 |
| chr1 | 154402922 | 154402923 | cg17427986 |
| chr1 | 154415062 | 154415063 | cg20688791 |
| chr1 | 154968781 | 154968782 | cg23998749 |
| chr1 | 154980743 | 154980744 | cg17329534 |
| chr1 | 155152367 | 155152368 | cg01412907 |
| chr1 | 155177237 | 155177238 | cg01428953 |
| chr1 | 155291345 | 155291346 | cg26876647 |
| chr1 | 156074135 | 156074136 | cg12593793 |
| chr1 | 156098772 | 156098773 | cg26667550 |
| chr1 | 156187819 | 156187820 | cg11084266 |
| chr1 | 156309258 | 156309259 | cg05532239 |
| chr1 | 156408887 | 156408888 | cg08548089 |

| | | | | | | | |
|---|---|---|---|---|---|---|---|
| chr1 | 111213827 | 111213828 | cg17516825 | chr1 | 156551787 | 156551788 | cg09319828 |
| chr1 | 111216000 | 111216001 | cg14200574 | chr1 | 156551873 | 156551874 | cg01010328 |
| chr1 | 111743020 | 111743021 | cg08864105 | chr1 | 157523253 | 157523254 | cg03329572 |
| chr1 | 111743271 | 111743272 | cg24641737 | chr1 | 157672074 | 157672075 | cg24881910 |
| chr1 | 111743368 | 111743369 | cg00619207 | chr1 | 157737828 | 157737829 | cg12282588 |
| chr1 | 111743411 | 111743412 | cg19268695 | chr1 | 157811790 | 157811791 | cg11139878 |
| chr1 | 111744108 | 111744109 | cg20970369 | chr1 | 158984321 | 158984322 | cg07196514 |
| chr1 | 111744309 | 111744310 | cg09826895 | chr1 | 159772162 | 159772163 | cg25698205 |
| chr1 | 111770111 | 111770112 | cg01440333 | chr1 | 159772194 | 159772195 | cg00936728 |
| chr1 | 111770274 | 111770275 | cg26366091 | chr1 | 159795589 | 159795590 | cg10179949 |
| chr1 | 111770291 | 111770292 | cg10045881 | chr1 | 159868242 | 159868243 | cg04258124 |
| chr1 | 112048205 | 112048206 | cg21419003 | chr1 | 160050381 | 160050382 | cg27315635 |
| chr1 | 112171207 | 112171208 | cg17598574 | chr1 | 160368823 | 160368824 | cg12662929 |
| chr1 | 113047945 | 113047946 | cg04571584 | chr1 | 160371455 | 160371456 | cg24482544 |
| chr1 | 113218414 | 113218415 | cg24344074 | chr1 | 160393253 | 160393254 | cg21213571 |
| chr1 | 113239645 | 113239646 | cg00719568 | chr1 | 160546243 | 160546244 | cg21454485 |
| chr1 | 113263764 | 113263765 | cg00342891 | chr1 | 160550488 | 160550489 | cg02945019 |
| chr1 | 116310712 | 116310713 | cg05190718 | chr1 | 160696468 | 160696469 | cg09248276 |
| chr1 | 116917619 | 116917620 | cg04403423 | chr1 | 161008127 | 161008128 | cg11578532 |
| chr1 | 116925411 | 116925412 | cg17469276 | chr1 | 161011634 | 161011635 | cg18646186 |
| chr1 | 117109623 | 117109624 | cg11379605 | chr1 | 161091980 | 161091981 | cg15516558 |
| chr1 | 117114263 | 117114264 | cg17215278 | chr1 | 161179735 | 161179736 | cg05043311 |
| chr1 | 117117195 | 117117196 | cg17032184 | chr1 | 161184305 | 161184306 | cg20609803 |
| chr1 | 117129532 | 117129533 | cg18786643 | chr1 | 161332601 | 161332602 | cg08161491 |
| chr1 | 117280833 | 117280834 | cg07570898 | chr1 | 161476548 | 161476549 | cg12643083 |
| chr1 | 117281442 | 117281443 | cg09312904 | chr1 | 161602002 | 161602003 | cg26561570 |
| chr1 | 117296242 | 117296243 | cg16719404 | chr1 | 161693987 | 161693988 | cg18889147 |
| chr1 | 117610620 | 117610621 | cg10021122 | chr1 | 163136667 | 163136668 | cg04353106 |
| chr1 | 118151821 | 118151822 | cg09992184 | chr1 | 165174748 | 165174749 | cg03428399 |
| chr1 | 118157624 | 118157625 | cg20306265 | chr1 | 166823514 | 166823515 | cg23694811 |
| chr1 | 118198739 | 118198740 | cg10523000 | chr1 | 167602860 | 167602861 | cg01809639 |
| chr1 | 119502537 | 119502538 | cg17868079 | chr1 | 169081894 | 169081895 | cg26009832 |
| chr1 | 120530251 | 120530252 | cg26196087 | chr1 | 169556022 | 169556023 | cg16054275 |
| chr1 | 144998869 | 144998870 | cg08708629 | chr1 | 171226702 | 171226703 | cg11192688 |
| chr1 | 145471701 | 145471702 | cg19913118 | chr1 | 171226993 | 171226994 | cg16034517 |
| chr1 | 145715636 | 145715637 | cg10798745 | chr1 | 171283893 | 171283894 | cg02694395 |
| chr1 | 145715672 | 145715673 | cg12832565 | chr1 | 172351357 | 172351358 | cg25722073 |
| chr1 | 145715719 | 145715720 | cg15892497 | chr1 | 172628065 | 172628066 | cg06983746 |
| chr1 | 147233887 | 147233888 | cg12740910 | chr1 | 173443209 | 173443210 | cg10881451 |
| chr1 | 150436947 | 150436948 | cg08856268 | chr1 | 174769548 | 174769549 | cg02958263 |
| chr1 | 150533123 | 150533124 | cg27630771 | chr1 | 174844490 | 174844491 | cg07285983 |
| chr1 | 150537574 | 150537575 | cg24674754 | chr1 | 175162550 | 175162551 | cg17845761 |
| chr1 | 150552817 | 150552818 | cg17724175 | chr1 | 175162553 | 175162554 | cg00958927 |
| chr1 | 150595206 | 150595207 | cg11147155 | chr1 | 175163045 | 175163046 | cg07916615 |
| chr1 | 150944634 | 150944635 | cg02459569 | chr1 | 175533566 | 175533567 | cg04923875 |
| chr1 | 150954826 | 150954827 | cg13320146 | chr1 | 178514897 | 178514898 | cg07396407 |
| chr1 | 151021364 | 151021365 | cg03603260 | chr1 | 178693595 | 178693596 | cg25384333 |
| chr1 | 151172905 | 151172906 | cg15690542 | chr1 | 178693891 | 178693892 | cg06788278 |
| chr1 | 151798432 | 151798433 | cg23114183 | chr1 | 178693897 | 178693898 | cg10559803 |
| chr1 | 151798824 | 151798825 | cg09171692 | chr1 | 179945973 | 179945974 | cg18278694 |
| chr1 | 151805634 | 151805635 | cg18149207 | chr1 | 180494132 | 180494133 | cg05334419 |
| chr1 | 151809534 | 151809535 | cg02493846 | chr1 | 181087920 | 181087921 | cg16273734 |
| chr1 | 151812435 | 151812436 | cg24425838 | chr1 | 181449699 | 181449700 | cg08943293 |
| chr1 | 151831144 | 151831145 | cg06872185 | chr1 | 183957982 | 183957983 | cg10312498 |
| chr1 | 151970131 | 151970132 | cg09809416 | chr1 | 184504284 | 184504285 | cg24162801 |
| chr1 | 152783489 | 152783490 | cg19250709 | chr1 | 192542576 | 192542577 | cg24886076 |

| | | | |
|---|---|---|---|
| chr1 | 152783674 | 152783675 | cg08878744 |
| chr1 | 152815843 | 152815844 | cg13366738 |
| chr1 | 152880324 | 152880325 | cg15059851 |
| chr1 | 153014310 | 153014311 | cg10812889 |
| chr1 | 153174759 | 153174760 | cg24091195 |
| chr1 | 153329013 | 153329014 | cg07039113 |
| chr1 | 153509284 | 153509285 | cg24375627 |
| chr1 | 153746211 | 153746212 | cg08129092 |
| chr1 | 153777721 | 153777722 | cg07567724 |
| chr1 | 153919638 | 153919639 | cg24678869 |
| chr1 | 153920347 | 153920348 | cg15831722 |
| chr1 | 153950843 | 153950844 | cg03655142 |
| chr1 | 153950867 | 153950868 | cg01588224 |
| chr1 | 153960103 | 153960104 | cg24543596 |
| chr1 | 154127629 | 154127630 | cg17383186 |
| chr1 | 154299179 | 154299180 | cg06811467 |
| chr1 | 154374951 | 154374952 | cg26538164 |
| chr1 | 154376344 | 154376345 | cg06980173 |
| chr1 | 154392070 | 154392071 | cg00791854 |
| chr1 | 154492947 | 154492948 | cg09861057 |
| chr1 | 154492952 | 154492953 | cg02097120 |
| chr1 | 154577895 | 154577896 | cg24304309 |
| chr1 | 154978980 | 154978981 | cg16618493 |
| chr1 | 154989197 | 154989198 | cg25604536 |
| chr1 | 155048476 | 155048477 | cg23217965 |
| chr1 | 155176341 | 155176342 | cg10017653 |
| chr1 | 155223726 | 155223727 | cg01417714 |
| chr1 | 155530423 | 155530424 | cg09851245 |
| chr1 | 155881846 | 155881847 | cg08256691 |
| chr1 | 155911238 | 155911239 | cg08403419 |
| chr1 | 156025757 | 156025758 | cg13216331 |
| chr1 | 156122709 | 156122710 | cg14847134 |
| chr1 | 156122730 | 156122731 | cg17971328 |
| chr1 | 156422580 | 156422581 | cg02488934 |
| chr1 | 157085163 | 157085164 | cg07398352 |
| chr1 | 157669111 | 157669112 | cg07184316 |
| chr1 | 157716415 | 157716416 | cg05777583 |
| chr1 | 157802305 | 157802306 | cg00995854 |
| chr1 | 158174027 | 158174028 | cg17945429 |
| chr1 | 158324471 | 158324472 | cg26014409 |
| chr1 | 158799935 | 158799936 | cg25119415 |
| chr1 | 158900644 | 158900645 | cg15036529 |
| chr1 | 158978468 | 158978469 | cg19939130 |
| chr1 | 158978504 | 158978505 | cg24539090 |
| chr1 | 159138217 | 159138218 | cg11518938 |
| chr1 | 160314061 | 160314062 | cg08015496 |
| chr1 | 160336712 | 160336713 | cg00881010 |
| chr1 | 160391109 | 160391110 | cg02660165 |
| chr1 | 160429714 | 160429715 | cg22602513 |
| chr1 | 160520346 | 160520347 | cg12034938 |
| chr1 | 160520374 | 160520375 | cg24691964 |
| chr1 | 161017007 | 161017008 | cg10701110 |
| chr1 | 161065862 | 161065863 | cg06974388 |
| chr1 | 161068781 | 161068782 | cg02957185 |
| chr1 | 161069839 | 161069840 | cg27582585 |
| chr1 | 161185061 | 161185062 | cg24465943 |
| chr1 | 198131619 | 198131620 | cg11521274 |
| chr1 | 198136518 | 198136519 | cg09372617 |
| chr1 | 198204074 | 198204075 | cg12750917 |
| chr1 | 198828157 | 198828158 | cg25349643 |
| chr1 | 198859579 | 198859580 | cg10096122 |
| chr1 | 202130344 | 202130345 | cg01437204 |
| chr1 | 202130692 | 202130693 | cg25754958 |
| chr1 | 202130728 | 202130729 | cg12436568 |
| chr1 | 202172912 | 202172913 | cg06825163 |
| chr1 | 202584465 | 202584466 | cg11071448 |
| chr1 | 203244158 | 203244159 | cg16311306 |
| chr1 | 203246482 | 203246483 | cg12067421 |
| chr1 | 203320190 | 203320191 | cg04704856 |
| chr1 | 203320223 | 203320224 | cg03764585 |
| chr1 | 203320386 | 203320387 | cg26987645 |
| chr1 | 203320396 | 203320397 | cg16289210 |
| chr1 | 203372112 | 203372113 | cg20464155 |
| chr1 | 203449947 | 203449948 | cg25143508 |
| chr1 | 203490804 | 203490805 | cg00577847 |
| chr1 | 203561580 | 203561581 | cg13235761 |
| chr1 | 203578680 | 203578681 | cg14600368 |
| chr1 | 203670963 | 203670964 | cg13115222 |
| chr1 | 203733042 | 203733043 | cg25625457 |
| chr1 | 203787773 | 203787774 | cg12736206 |
| chr1 | 204373163 | 204373164 | cg19692604 |
| chr1 | 204466432 | 204466433 | cg10950251 |
| chr1 | 205173544 | 205173545 | cg00889227 |
| chr1 | 205242518 | 205242519 | cg26891661 |
| chr1 | 205271231 | 205271232 | cg15512534 |
| chr1 | 205313913 | 205313914 | cg23638226 |
| chr1 | 205326059 | 205326060 | cg21996351 |
| chr1 | 205618102 | 205618103 | cg23228450 |
| chr1 | 206300911 | 206300912 | cg24700388 |
| chr1 | 206856073 | 206856074 | cg15015980 |
| chr1 | 206857579 | 206857580 | cg08055786 |
| chr1 | 206943566 | 206943567 | cg15096505 |
| chr1 | 207070539 | 207070540 | cg23909633 |
| chr1 | 207081344 | 207081345 | cg13981287 |
| chr1 | 207082479 | 207082480 | cg21607172 |
| chr1 | 207083336 | 207083337 | cg05721773 |
| chr1 | 207143870 | 207143871 | cg19204946 |
| chr1 | 207262706 | 207262707 | cg26407558 |
| chr1 | 208137626 | 208137627 | cg12277870 |
| chr1 | 209877614 | 209877615 | cg13781869 |
| chr1 | 209917237 | 209917238 | cg08865695 |
| chr1 | 210013268 | 210013269 | cg24661168 |
| chr1 | 210612391 | 210612392 | cg16166061 |
| chr1 | 211430780 | 211430781 | cg06123700 |
| chr1 | 211498962 | 211498963 | cg27433482 |
| chr1 | 212587188 | 212587189 | cg08270613 |
| chr1 | 212604918 | 212604919 | cg10487521 |
| chr1 | 212769641 | 212769642 | cg10985914 |
| chr1 | 213080934 | 213080935 | cg25880211 |
| chr1 | 214510205 | 214510206 | cg11957699 |
| chr1 | 215147891 | 215147892 | cg27589988 |
| chr1 | 216269240 | 216269241 | cg13376246 |

| | | | |
|---|---|---|---|
| chr1 | 161185092 | 161185093 | cg13853198 |
| chr1 | 161186839 | 161186840 | cg20806175 |
| chr1 | 161678601 | 161678602 | cg02305590 |
| chr1 | 161706293 | 161706294 | cg18200075 |
| chr1 | 162500634 | 162500635 | cg08538581 |
| chr1 | 163395211 | 163395212 | cg08005208 |
| chr1 | 163437170 | 163437171 | cg23261375 |
| chr1 | 164627790 | 164627791 | cg12629991 |
| chr1 | 164717625 | 164717626 | cg22729681 |
| chr1 | 164725227 | 164725228 | cg06909503 |
| chr1 | 166124419 | 166124420 | cg24428232 |
| chr1 | 166139996 | 166139997 | cg11979420 |
| chr1 | 167217689 | 167217690 | cg11132582 |
| chr1 | 167471264 | 167471265 | cg13210595 |
| chr1 | 167508884 | 167508885 | cg12396057 |
| chr1 | 167904284 | 167904285 | cg12749559 |
| chr1 | 168197198 | 168197199 | cg08738280 |
| chr1 | 169078316 | 169078317 | cg13104274 |
| chr1 | 169679199 | 169679200 | cg06816239 |
| chr1 | 172114419 | 172114420 | cg03322057 |
| chr1 | 172515025 | 172515026 | cg00452835 |
| chr1 | 172608644 | 172608645 | cg17756730 |
| chr1 | 172628020 | 172628021 | cg10161121 |
| chr1 | 172628263 | 172628264 | cg00071250 |
| chr1 | 172863499 | 172863500 | cg07393570 |
| chr1 | 173159746 | 173159747 | cg24633390 |
| chr1 | 173176366 | 173176367 | cg21439763 |
| chr1 | 173381511 | 173381512 | cg23091549 |
| chr1 | 174844397 | 174844398 | cg13130398 |
| chr1 | 174933574 | 174933575 | cg17516247 |
| chr1 | 175511259 | 175511260 | cg22941315 |
| chr1 | 175671657 | 175671658 | cg15067015 |
| chr1 | 176172985 | 176172986 | cg16372976 |
| chr1 | 178832759 | 178832760 | cg19807716 |
| chr1 | 178992788 | 178992789 | cg03393754 |
| chr1 | 178998656 | 178998657 | cg23933289 |
| chr1 | 180922636 | 180922637 | cg17237951 |
| chr1 | 180981015 | 180981016 | cg16122793 |
| chr1 | 181002669 | 181002670 | cg01040850 |
| chr1 | 181075209 | 181075210 | cg21220403 |
| chr1 | 181104570 | 181104571 | cg25604316 |
| chr1 | 182024689 | 182024690 | cg06361127 |
| chr1 | 182586153 | 182586154 | cg14651210 |
| chr1 | 182807962 | 182807963 | cg16243197 |
| chr1 | 183256588 | 183256589 | cg05262549 |
| chr1 | 183274638 | 183274639 | cg16373880 |
| chr1 | 183274661 | 183274662 | cg05380885 |
| chr1 | 183537770 | 183537771 | cg09472600 |
| chr1 | 184403226 | 184403227 | cg11201256 |
| chr1 | 186954259 | 186954260 | cg09067029 |
| chr1 | 192509700 | 192509701 | cg09822423 |
| chr1 | 192548609 | 192548610 | cg06357908 |
| chr1 | 192602520 | 192602521 | cg13092082 |
| chr1 | 192778890 | 192778891 | cg05374271 |
| chr1 | 196312876 | 196312877 | cg14518227 |
| chr1 | 196945530 | 196945531 | cg25840094 |
| chr1 | 218472324 | 218472325 | cg04210779 |
| chr1 | 219558325 | 219558326 | cg06516956 |
| chr1 | 220917572 | 220917573 | cg14777702 |
| chr1 | 221818304 | 221818305 | cg02919721 |
| chr1 | 222759637 | 222759638 | cg05595429 |
| chr1 | 223897463 | 223897464 | cg13574099 |
| chr1 | 224896204 | 224896205 | cg11526665 |
| chr1 | 226012507 | 226012508 | cg03337430 |
| chr1 | 226068334 | 226068335 | cg20240791 |
| chr1 | 226270724 | 226270725 | cg01205584 |
| chr1 | 226296868 | 226296869 | cg10483871 |
| chr1 | 226296871 | 226296872 | cg00672802 |
| chr1 | 226308547 | 226308548 | cg05704832 |
| chr1 | 226308960 | 226308961 | cg15260921 |
| chr1 | 226833134 | 226833135 | cg23038277 |
| chr1 | 226878419 | 226878420 | cg25287071 |
| chr1 | 226885709 | 226885710 | cg17933936 |
| chr1 | 226905595 | 226905596 | cg04213163 |
| chr1 | 226911669 | 226911670 | cg16265553 |
| chr1 | 226918736 | 226918737 | cg23676767 |
| chr1 | 226922948 | 226922949 | cg03199014 |
| chr1 | 226927988 | 226927989 | cg19030737 |
| chr1 | 227003060 | 227003061 | cg03547355 |
| chr1 | 227015418 | 227015419 | cg08654091 |
| chr1 | 227015456 | 227015457 | cg16985288 |
| chr1 | 227954102 | 227954103 | cg22356061 |
| chr1 | 228275828 | 228275829 | cg18882687 |
| chr1 | 228278432 | 228278433 | cg06625004 |
| chr1 | 228661675 | 228661676 | cg09791366 |
| chr1 | 229294380 | 229294381 | cg25349990 |
| chr1 | 229460525 | 229460526 | cg12133078 |
| chr1 | 229480001 | 229480002 | cg16672421 |
| chr1 | 229480105 | 229480106 | cg04724659 |
| chr1 | 230201582 | 230201583 | cg08141553 |
| chr1 | 230204526 | 230204527 | cg00923744 |
| chr1 | 230249292 | 230249293 | cg17220749 |
| chr1 | 230260837 | 230260838 | cg23230158 |
| chr1 | 230279135 | 230279136 | cg20703242 |
| chr1 | 230284198 | 230284199 | cg05784532 |
| chr1 | 230367682 | 230367683 | cg06801857 |
| chr1 | 230883300 | 230883301 | cg03704061 |
| chr1 | 230883552 | 230883553 | cg06585893 |
| chr1 | 231553510 | 231553511 | cg21875980 |
| chr1 | 231561414 | 231561415 | cg16855929 |
| chr1 | 231747906 | 231747907 | cg08458637 |
| chr1 | 233426334 | 233426335 | cg17894435 |
| chr1 | 234623575 | 234623576 | cg20848218 |
| chr1 | 234793300 | 234793301 | cg13777730 |
| chr1 | 234905772 | 234905773 | cg00851028 |
| chr1 | 235107837 | 235107838 | cg22628240 |
| chr1 | 235423736 | 235423737 | cg24710309 |
| chr1 | 235468148 | 235468149 | cg02985240 |
| chr1 | 235587038 | 235587039 | cg00773060 |
| chr1 | 236559646 | 236559647 | cg04311403 |
| chr1 | 241804421 | 241804422 | cg08162224 |
| chr1 | 241804939 | 241804940 | cg26687670 |

| | | | |
|---|---|---|---|
| chr1 | 197745025 | 197745026 | cg24461171 |
| chr1 | 197745159 | 197745160 | cg21534578 |
| chr1 | 197778303 | 197778304 | cg24302127 |
| chr1 | 198669986 | 198669987 | cg12390585 |
| chr1 | 200358418 | 200358419 | cg10540721 |
| chr1 | 200707086 | 200707087 | cg15027266 |
| chr1 | 200847096 | 200847097 | cg03040292 |
| chr1 | 200864307 | 200864308 | cg00971747 |
| chr1 | 202130920 | 202130921 | cg23506842 |
| chr1 | 202171723 | 202171724 | cg12761421 |
| chr1 | 203257946 | 203257947 | cg18319612 |
| chr1 | 203273571 | 203273572 | cg20138067 |
| chr1 | 203278044 | 203278045 | cg06373167 |
| chr1 | 203525604 | 203525605 | cg10720154 |
| chr1 | 203626524 | 203626525 | cg20312390 |
| chr1 | 203711066 | 203711067 | cg01953850 |
| chr1 | 203733971 | 203733972 | cg23893629 |
| chr1 | 203734256 | 203734257 | cg10117369 |
| chr1 | 203734478 | 203734479 | cg06958535 |
| chr1 | 203734559 | 203734560 | cg00942920 |
| chr1 | 204034928 | 204034929 | cg00100762 |
| chr1 | 204071770 | 204071771 | cg08620986 |
| chr1 | 204224211 | 204224212 | cg11131599 |
| chr1 | 204452127 | 204452128 | cg20828284 |
| chr1 | 204535000 | 204535001 | cg07972458 |
| chr1 | 204796491 | 204796492 | cg11253737 |
| chr1 | 204993424 | 204993425 | cg26271435 |
| chr1 | 205254523 | 205254524 | cg10529555 |
| chr1 | 205263738 | 205263739 | cg09030187 |
| chr1 | 205286927 | 205286928 | cg05699921 |
| chr1 | 205289384 | 205289385 | cg01012936 |
| chr1 | 205313833 | 205313834 | cg21538450 |
| chr1 | 205317032 | 205317033 | cg09319656 |
| chr1 | 205326201 | 205326202 | cg07429623 |
| chr1 | 205470209 | 205470210 | cg27511824 |
| chr1 | 206135024 | 206135025 | cg01279219 |
| chr1 | 206289463 | 206289464 | cg13685151 |
| chr1 | 206289775 | 206289776 | cg24056232 |
| chr1 | 206603444 | 206603445 | cg00796611 |
| chr1 | 206729040 | 206729041 | cg01442843 |
| chr1 | 206729099 | 206729100 | cg02035751 |
| chr1 | 206729553 | 206729554 | cg14163311 |
| chr1 | 206729613 | 206729614 | cg17220933 |
| chr1 | 206729685 | 206729686 | cg05756492 |
| chr1 | 206848389 | 206848390 | cg10336193 |
| chr1 | 206876996 | 206876997 | cg24141516 |
| chr1 | 206879950 | 206879951 | cg06419268 |
| chr1 | 206941464 | 206941465 | cg14789529 |
| chr1 | 206947477 | 206947478 | cg14284394 |
| chr1 | 206972113 | 206972114 | cg20998511 |
| chr1 | 206972130 | 206972131 | cg11582579 |
| chr1 | 206976689 | 206976690 | cg15392364 |
| chr1 | 207070825 | 207070826 | cg16417028 |
| chr1 | 207070834 | 207070835 | cg06796611 |
| chr1 | 207082900 | 207082901 | cg00008647 |
| chr1 | 207143533 | 207143534 | cg16729555 |
| chr1 | 242002464 | 242002465 | cg24361198 |
| chr1 | 243641454 | 243641455 | cg17216385 |
| chr1 | 243904118 | 243904119 | cg15962538 |
| chr1 | 244214593 | 244214594 | cg01105418 |
| chr1 | 244214679 | 244214680 | cg23829949 |
| chr1 | 244216231 | 244216232 | cg22812005 |
| chr1 | 244480459 | 244480460 | cg12039595 |
| chr1 | 245836835 | 245836836 | cg03327643 |
| chr1 | 245855541 | 245855542 | cg16362305 |
| chr1 | 247496922 | 247496923 | cg22036872 |
| chr1 | 247578953 | 247578954 | cg21806273 |
| chr1 | 247581417 | 247581418 | cg21991396 |
| chr1 | 247616686 | 247616687 | cg14830002 |
| chr1 | 248903300 | 248903301 | cg04024799 |
| chr1 | 249137995 | 249137996 | cg12657014 |
| chr2 | 3200012 | 3200013 | cg17494897 |
| chr2 | 7016509 | 7016510 | cg23213327 |
| chr2 | 7029602 | 7029603 | cg00186143 |
| chr2 | 7569878 | 7569879 | cg05187797 |
| chr2 | 7910535 | 7910536 | cg04653800 |
| chr2 | 8634827 | 8634828 | cg08212913 |
| chr2 | 9705768 | 9705769 | cg20899781 |
| chr2 | 9788946 | 9788947 | cg22406334 |
| chr2 | 10184650 | 10184651 | cg18825594 |
| chr2 | 10470465 | 10470466 | cg04618171 |
| chr2 | 10470961 | 10470962 | cg23728189 |
| chr2 | 10487314 | 10487315 | cg10283844 |
| chr2 | 10517323 | 10517324 | cg18913843 |
| chr2 | 10517352 | 10517353 | cg00141688 |
| chr2 | 10517361 | 10517362 | cg22614203 |
| chr2 | 10720198 | 10720199 | cg21626540 |
| chr2 | 11968394 | 11968395 | cg24826252 |
| chr2 | 17699243 | 17699244 | cg07709606 |
| chr2 | 17997660 | 17997661 | cg11046593 |
| chr2 | 20817916 | 20817917 | cg02458141 |
| chr2 | 20868568 | 20868569 | cg07789367 |
| chr2 | 20876220 | 20876221 | cg00011239 |
| chr2 | 20996747 | 20996748 | cg20324777 |
| chr2 | 23751324 | 23751325 | cg00450990 |
| chr2 | 23755343 | 23755344 | cg01837846 |
| chr2 | 25498444 | 25498445 | cg20669908 |
| chr2 | 25524877 | 25524878 | cg00050692 |
| chr2 | 25563141 | 25563142 | cg05896193 |
| chr2 | 25600451 | 25600452 | cg02068690 |
| chr2 | 25704759 | 25704760 | cg05847790 |
| chr2 | 25744763 | 25744764 | cg26870999 |
| chr2 | 30450574 | 30450575 | cg13401212 |
| chr2 | 30495331 | 30495332 | cg14993140 |
| chr2 | 30496325 | 30496326 | cg21335012 |
| chr2 | 30516374 | 30516375 | cg16783186 |
| chr2 | 30557562 | 30557563 | cg07066814 |
| chr2 | 30568530 | 30568531 | cg24721179 |
| chr2 | 30687453 | 30687454 | cg23457506 |
| chr2 | 31439687 | 31439688 | cg15941014 |
| chr2 | 32678846 | 32678847 | cg11971852 |
| chr2 | 37381088 | 37381089 | cg24523650 |

| | | | |
|---|---|---|---|
| chr1 | 207144812 | 207144813 | cg23912186 |
| chr1 | 207152909 | 207152910 | cg22032961 |
| chr1 | 207490952 | 207490953 | cg22687766 |
| chr1 | 207928706 | 207928707 | cg22500738 |
| chr1 | 208005598 | 208005599 | cg17877600 |
| chr1 | 208021736 | 208021737 | cg15646741 |
| chr1 | 208040203 | 208040204 | cg00085493 |
| chr1 | 208040253 | 208040254 | cg24315421 |
| chr1 | 208132787 | 208132788 | cg10210594 |
| chr1 | 208281523 | 208281524 | cg21144009 |
| chr1 | 208290896 | 208290897 | cg07908868 |
| chr1 | 208405868 | 208405869 | cg01876531 |
| chr1 | 208444590 | 208444591 | cg23843022 |
| chr1 | 209105888 | 209105889 | cg23962746 |
| chr1 | 209549585 | 209549586 | cg04239078 |
| chr1 | 209928895 | 209928896 | cg08655071 |
| chr1 | 210858075 | 210858076 | cg12706474 |
| chr1 | 211499638 | 211499639 | cg10177528 |
| chr1 | 212004782 | 212004783 | cg21790344 |
| chr1 | 212004804 | 212004805 | cg08892386 |
| chr1 | 212108591 | 212108592 | cg13990743 |
| chr1 | 212463238 | 212463239 | cg12417775 |
| chr1 | 212588848 | 212588849 | cg21400549 |
| chr1 | 212607111 | 212607112 | cg24511404 |
| chr1 | 212626474 | 212626475 | cg06817455 |
| chr1 | 212662017 | 212662018 | cg06093152 |
| chr1 | 212671907 | 212671908 | cg00426089 |
| chr1 | 212966209 | 212966210 | cg03040821 |
| chr1 | 214006138 | 214006139 | cg00852051 |
| chr1 | 215196452 | 215196453 | cg24464500 |
| chr1 | 215228588 | 215228589 | cg04923840 |
| chr1 | 215804508 | 215804509 | cg14524956 |
| chr1 | 216144819 | 216144820 | cg05126581 |
| chr1 | 219261864 | 219261865 | cg17216266 |
| chr1 | 219958132 | 219958133 | cg11789804 |
| chr1 | 221049931 | 221049932 | cg15469321 |
| chr1 | 221054825 | 221054826 | cg12479878 |
| chr1 | 221913576 | 221913577 | cg05941778 |
| chr1 | 222340714 | 222340715 | cg16651768 |
| chr1 | 223903862 | 223903863 | cg10100437 |
| chr1 | 224075899 | 224075900 | cg02582848 |
| chr1 | 224620806 | 224620807 | cg18137450 |
| chr1 | 225648715 | 225648716 | cg19867250 |
| chr1 | 225662350 | 225662351 | cg09694381 |
| chr1 | 226069189 | 226069190 | cg21710324 |
| chr1 | 226129824 | 226129825 | cg20742696 |
| chr1 | 226272763 | 226272764 | cg26612648 |
| chr1 | 226300601 | 226300602 | cg14522302 |
| chr1 | 226306890 | 226306891 | cg22489204 |
| chr1 | 226373204 | 226373205 | cg18385299 |
| chr1 | 226375174 | 226375175 | cg02583796 |
| chr1 | 226739902 | 226739903 | cg11211263 |
| chr1 | 226825606 | 226825607 | cg16177187 |
| chr1 | 226842455 | 226842456 | cg01910727 |
| chr1 | 226844154 | 226844155 | cg05773194 |
| chr1 | 226884832 | 226884833 | cg12221970 |

| | | | |
|---|---|---|---|
| chr2 | 37553621 | 37553622 | cg19190900 |
| chr2 | 38297759 | 38297760 | cg11656478 |
| chr2 | 38358147 | 38358148 | cg11130441 |
| chr2 | 38358225 | 38358226 | cg27141474 |
| chr2 | 42410249 | 42410250 | cg02593043 |
| chr2 | 42725387 | 42725388 | cg12447236 |
| chr2 | 42991343 | 42991344 | cg13820899 |
| chr2 | 43027940 | 43027941 | cg10588617 |
| chr2 | 43364053 | 43364054 | cg09489567 |
| chr2 | 43397515 | 43397516 | cg00259518 |
| chr2 | 43398171 | 43398172 | cg01243072 |
| chr2 | 43405947 | 43405948 | cg09308580 |
| chr2 | 43540749 | 43540750 | cg14807446 |
| chr2 | 44843165 | 44843166 | cg10979368 |
| chr2 | 44996068 | 44996069 | cg05037556 |
| chr2 | 45880618 | 45880619 | cg08203192 |
| chr2 | 46004187 | 46004188 | cg24788291 |
| chr2 | 46163986 | 46163987 | cg00793946 |
| chr2 | 46930327 | 46930328 | cg23900606 |
| chr2 | 47290974 | 47290975 | cg17647823 |
| chr2 | 47298563 | 47298564 | cg22360765 |
| chr2 | 48545621 | 48545622 | cg18064411 |
| chr2 | 54751507 | 54751508 | cg24544082 |
| chr2 | 54783773 | 54783774 | cg01286930 |
| chr2 | 54789491 | 54789492 | cg23486701 |
| chr2 | 54797285 | 54797286 | cg03017850 |
| chr2 | 54857270 | 54857271 | cg10929758 |
| chr2 | 54861563 | 54861564 | cg08556541 |
| chr2 | 55200201 | 55200202 | cg02986924 |
| chr2 | 55512437 | 55512438 | cg21384729 |
| chr2 | 55636555 | 55636556 | cg13359689 |
| chr2 | 55642894 | 55642895 | cg01953317 |
| chr2 | 60102800 | 60102801 | cg23598480 |
| chr2 | 60229772 | 60229773 | cg26040101 |
| chr2 | 60687392 | 60687393 | cg02334333 |
| chr2 | 60722802 | 60722803 | cg09670127 |
| chr2 | 60735530 | 60735531 | cg03606511 |
| chr2 | 60737387 | 60737388 | cg26149739 |
| chr2 | 60755359 | 60755360 | cg16951074 |
| chr2 | 60761027 | 60761028 | cg01616628 |
| chr2 | 60766554 | 60766555 | cg03981213 |
| chr2 | 61645604 | 61645605 | cg01723929 |
| chr2 | 62520912 | 62520913 | cg01406456 |
| chr2 | 64291800 | 64291801 | cg26099045 |
| chr2 | 64320678 | 64320679 | cg13101088 |
| chr2 | 64330510 | 64330511 | cg12974773 |
| chr2 | 64488961 | 64488962 | cg15132282 |
| chr2 | 64754308 | 64754309 | cg01663407 |
| chr2 | 64868515 | 64868516 | cg17206596 |
| chr2 | 64877006 | 64877007 | cg00063909 |
| chr2 | 65282029 | 65282030 | cg01447702 |
| chr2 | 65352897 | 65352898 | cg09154256 |
| chr2 | 68483298 | 68483299 | cg08898515 |
| chr2 | 68646210 | 68646211 | cg23809597 |
| chr2 | 68663346 | 68663347 | cg07128759 |
| chr2 | 69970294 | 69970295 | cg12518775 |

| | | | |
|---|---|---|---|
| chr1 | 226887044 | 226887045 | cg02896872 |
| chr1 | 226900896 | 226900897 | cg07795928 |
| chr1 | 226921576 | 226921577 | cg05262829 |
| chr1 | 228273674 | 228273675 | cg06779113 |
| chr1 | 228289747 | 228289748 | cg14255644 |
| chr1 | 229275061 | 229275062 | cg25023165 |
| chr1 | 229692519 | 229692520 | cg00355656 |
| chr1 | 229997842 | 229997843 | cg19871173 |
| chr1 | 230241764 | 230241765 | cg13359998 |
| chr1 | 230248121 | 230248122 | cg23735745 |
| chr1 | 230410329 | 230410330 | cg11527913 |
| chr1 | 230413174 | 230413175 | cg16517298 |
| chr1 | 231504339 | 231504340 | cg06096015 |
| chr1 | 231842206 | 231842207 | cg23172853 |
| chr1 | 233248709 | 233248710 | cg00980058 |
| chr1 | 234616222 | 234616223 | cg10780718 |
| chr1 | 234635574 | 234635575 | cg06511330 |
| chr1 | 234651180 | 234651181 | cg15095427 |
| chr1 | 234734475 | 234734476 | cg09691134 |
| chr1 | 234977572 | 234977573 | cg02889973 |
| chr1 | 235122803 | 235122804 | cg17819085 |
| chr1 | 235477322 | 235477323 | cg19314777 |
| chr1 | 235500468 | 235500469 | cg19269426 |
| chr1 | 235665822 | 235665823 | cg07836225 |
| chr1 | 236047034 | 236047035 | cg12167564 |
| chr1 | 236273181 | 236273182 | cg19811148 |
| chr1 | 236651140 | 236651141 | cg27631057 |
| chr1 | 238054569 | 238054570 | cg26810908 |
| chr1 | 239450936 | 239450937 | cg09334915 |
| chr1 | 240375464 | 240375465 | cg07418387 |
| chr1 | 241795170 | 241795171 | cg04918831 |
| chr1 | 241799063 | 241799064 | cg17461214 |
| chr1 | 241801700 | 241801701 | cg08634082 |
| chr1 | 242163388 | 242163389 | cg14382888 |
| chr1 | 244208721 | 244208722 | cg17965737 |
| chr1 | 244617756 | 244617757 | cg10029259 |
| chr1 | 245002565 | 245002566 | cg17718182 |
| chr1 | 245812674 | 245812675 | cg01565291 |
| chr1 | 245855322 | 245855323 | cg11587466 |
| chr1 | 248903100 | 248903101 | cg23230830 |
| chr1 | 249116587 | 249116588 | cg11681597 |
| chr2 | 444632 | 444633 | cg09757158 |
| chr2 | 1051107 | 1051108 | cg17600393 |
| chr2 | 3659466 | 3659467 | cg19183742 |
| chr2 | 3741732 | 3741733 | cg05977776 |
| chr2 | 5850120 | 5850121 | cg26584747 |
| chr2 | 6110966 | 6110967 | cg24478779 |
| chr2 | 6305776 | 6305777 | cg07900344 |
| chr2 | 6614169 | 6614170 | cg01407018 |
| chr2 | 6644914 | 6644915 | cg19618601 |
| chr2 | 7056075 | 7056076 | cg22070232 |
| chr2 | 7713711 | 7713712 | cg22891707 |
| chr2 | 8011432 | 8011433 | cg08700932 |
| chr2 | 8397810 | 8397811 | cg19256423 |
| chr2 | 8714417 | 8714418 | cg14093127 |
| chr2 | 8746519 | 8746520 | cg01099150 |

| | | | |
|---|---|---|---|
| chr2 | 70366299 | 70366300 | cg01613454 |
| chr2 | 70915210 | 70915211 | cg18067420 |
| chr2 | 70960959 | 70960960 | cg06915226 |
| chr2 | 71212157 | 71212158 | cg04076766 |
| chr2 | 73201477 | 73201478 | cg24803346 |
| chr2 | 74603226 | 74603227 | cg10785537 |
| chr2 | 74607536 | 74607537 | cg14601444 |
| chr2 | 74645769 | 74645770 | cg09569569 |
| chr2 | 74647813 | 74647814 | cg21655923 |
| chr2 | 74729502 | 74729503 | cg06768010 |
| chr2 | 77235218 | 77235219 | cg08963013 |
| chr2 | 85034766 | 85034767 | cg04361556 |
| chr2 | 85060011 | 85060012 | cg05242065 |
| chr2 | 85622094 | 85622095 | cg27090078 |
| chr2 | 85637834 | 85637835 | cg07215695 |
| chr2 | 85638053 | 85638054 | cg10664272 |
| chr2 | 85640001 | 85640002 | cg03718845 |
| chr2 | 85764282 | 85764283 | cg02196422 |
| chr2 | 85895048 | 85895049 | cg03380645 |
| chr2 | 85895360 | 85895361 | cg07313956 |
| chr2 | 85933069 | 85933070 | cg19805943 |
| chr2 | 86215764 | 86215765 | cg05021291 |
| chr2 | 86225602 | 86225603 | cg14017402 |
| chr2 | 87566763 | 87566764 | cg26324057 |
| chr2 | 88313351 | 88313352 | cg08952844 |
| chr2 | 88352949 | 88352950 | cg12691004 |
| chr2 | 88390376 | 88390377 | cg18115724 |
| chr2 | 96930546 | 96930547 | cg13672342 |
| chr2 | 97746622 | 97746623 | cg19708848 |
| chr2 | 98329691 | 98329692 | cg08859278 |
| chr2 | 98329811 | 98329812 | cg09006159 |
| chr2 | 98329830 | 98329831 | cg15933451 |
| chr2 | 98552472 | 98552473 | cg04223924 |
| chr2 | 99124870 | 99124871 | cg18455616 |
| chr2 | 100498600 | 100498601 | cg21959618 |
| chr2 | 100863359 | 100863360 | cg04611493 |
| chr2 | 100863380 | 100863381 | cg19364276 |
| chr2 | 100863404 | 100863405 | cg04369903 |
| chr2 | 101978981 | 101978982 | cg16529548 |
| chr2 | 102359027 | 102359028 | cg01611777 |
| chr2 | 102425416 | 102425417 | cg21698385 |
| chr2 | 102608155 | 102608156 | cg14325025 |
| chr2 | 102644247 | 102644248 | cg21674927 |
| chr2 | 103039542 | 103039543 | cg13897122 |
| chr2 | 103090077 | 103090078 | cg02567476 |
| chr2 | 103352296 | 103352297 | cg13607230 |
| chr2 | 105484823 | 105484824 | cg27560561 |
| chr2 | 105759884 | 105759885 | cg00612633 |
| chr2 | 105851295 | 105851296 | cg08323766 |
| chr2 | 105877824 | 105877825 | cg23371476 |
| chr2 | 105911437 | 105911438 | cg27273675 |
| chr2 | 106360709 | 106360710 | cg14131038 |
| chr2 | 106512578 | 106512579 | cg21661768 |
| chr2 | 106713059 | 106713060 | cg20995188 |
| chr2 | 106885926 | 106885927 | cg05468156 |
| chr2 | 107192564 | 107192565 | cg24751773 |

| | | | |
|---|---|---|---|
| chr2 | 9027671 | 9027672 | cg08430813 |
| chr2 | 9518324 | 9518325 | cg19426625 |
| chr2 | 9761025 | 9761026 | cg13724182 |
| chr2 | 10950238 | 10950239 | cg20725704 |
| chr2 | 11097873 | 11097874 | cg21844291 |
| chr2 | 11485561 | 11485562 | cg13266327 |
| chr2 | 11513527 | 11513528 | cg22107147 |
| chr2 | 11515164 | 11515165 | cg11972534 |
| chr2 | 11518493 | 11518494 | cg27375286 |
| chr2 | 11607488 | 11607489 | cg02819921 |
| chr2 | 11968342 | 11968343 | cg03238522 |
| chr2 | 11968388 | 11968389 | cg09346503 |
| chr2 | 16105823 | 16105824 | cg19893929 |
| chr2 | 16832247 | 16832248 | cg04163638 |
| chr2 | 17700224 | 17700225 | cg20313628 |
| chr2 | 18150633 | 18150634 | cg05417035 |
| chr2 | 18621149 | 18621150 | cg24903497 |
| chr2 | 23749087 | 23749088 | cg21206147 |
| chr2 | 24387250 | 24387251 | cg09769104 |
| chr2 | 25095040 | 25095041 | cg11023668 |
| chr2 | 25141901 | 25141902 | cg08266286 |
| chr2 | 25142016 | 25142017 | cg26752663 |
| chr2 | 25424387 | 25424388 | cg02642979 |
| chr2 | 25473529 | 25473530 | cg15150970 |
| chr2 | 25562512 | 25562513 | cg06748978 |
| chr2 | 25603640 | 25603641 | cg04256886 |
| chr2 | 26048283 | 26048284 | cg27492103 |
| chr2 | 26266360 | 26266361 | cg20628718 |
| chr2 | 26470049 | 26470050 | cg04375688 |
| chr2 | 27068081 | 27068082 | cg08806184 |
| chr2 | 27375563 | 27375564 | cg17495302 |
| chr2 | 27545006 | 27545007 | cg14948822 |
| chr2 | 27629738 | 27629739 | cg06369118 |
| chr2 | 27649143 | 27649144 | cg23571126 |
| chr2 | 29121607 | 29121608 | cg20319654 |
| chr2 | 30372795 | 30372796 | cg02766770 |
| chr2 | 30373187 | 30373188 | cg26855302 |
| chr2 | 30441244 | 30441245 | cg16444826 |
| chr2 | 30458365 | 30458366 | cg18793512 |
| chr2 | 30524251 | 30524252 | cg14113353 |
| chr2 | 30633168 | 30633169 | cg19791637 |
| chr2 | 30967383 | 30967384 | cg19304003 |
| chr2 | 31497155 | 31497156 | cg03065107 |
| chr2 | 31638511 | 31638512 | cg09239591 |
| chr2 | 33152037 | 33152038 | cg21314603 |
| chr2 | 33720546 | 33720547 | cg06971903 |
| chr2 | 33737453 | 33737454 | cg21244038 |
| chr2 | 36586710 | 36586711 | cg09530108 |
| chr2 | 36598278 | 36598279 | cg01958295 |
| chr2 | 36667594 | 36667595 | cg20391432 |
| chr2 | 36677570 | 36677571 | cg17516469 |
| chr2 | 36716635 | 36716636 | cg00296578 |
| chr2 | 36964593 | 36964594 | cg20385708 |
| chr2 | 37310396 | 37310397 | cg02255510 |
| chr2 | 37545307 | 37545308 | cg21247948 |
| chr2 | 37604081 | 37604082 | cg26986629 |

| | | | |
|---|---|---|---|
| chr2 | 109190922 | 109190923 | cg08055994 |
| chr2 | 109257429 | 109257430 | cg05310309 |
| chr2 | 111979319 | 111979320 | cg02116283 |
| chr2 | 111979398 | 111979399 | cg13551117 |
| chr2 | 113544232 | 113544233 | cg23407561 |
| chr2 | 113593785 | 113593786 | cg15836722 |
| chr2 | 113914442 | 113914443 | cg06961278 |
| chr2 | 120107745 | 120107746 | cg23562675 |
| chr2 | 120979401 | 120979402 | cg01109047 |
| chr2 | 121020189 | 121020190 | cg06876354 |
| chr2 | 122402890 | 122402891 | cg22346073 |
| chr2 | 134882048 | 134882049 | cg23640664 |
| chr2 | 135017168 | 135017169 | cg23806727 |
| chr2 | 135018453 | 135018454 | cg11878712 |
| chr2 | 135050894 | 135050895 | cg14356440 |
| chr2 | 135125439 | 135125440 | cg22763593 |
| chr2 | 136873409 | 136873410 | cg07784959 |
| chr2 | 139272578 | 139272579 | cg10033694 |
| chr2 | 145090814 | 145090815 | cg07780074 |
| chr2 | 145779560 | 145779561 | cg12683813 |
| chr2 | 146359434 | 146359435 | cg24536827 |
| chr2 | 149481321 | 149481322 | cg02072515 |
| chr2 | 152117363 | 152117364 | cg05960677 |
| chr2 | 152906698 | 152906699 | cg05105477 |
| chr2 | 153347526 | 153347527 | cg26207245 |
| chr2 | 153413442 | 153413443 | cg12505929 |
| chr2 | 153419601 | 153419602 | cg11173609 |
| chr2 | 157291462 | 157291463 | cg03230175 |
| chr2 | 157395835 | 157395836 | cg23077884 |
| chr2 | 158184535 | 158184536 | cg21563471 |
| chr2 | 159327413 | 159327414 | cg24519780 |
| chr2 | 159386608 | 159386609 | cg17648451 |
| chr2 | 161228737 | 161228738 | cg01256139 |
| chr2 | 161826714 | 161826715 | cg24964215 |
| chr2 | 162047158 | 162047159 | cg27447753 |
| chr2 | 162209962 | 162209963 | cg08135437 |
| chr2 | 166634454 | 166634455 | cg15343463 |
| chr2 | 166649614 | 166649615 | cg02345886 |
| chr2 | 169025892 | 169025893 | cg06754557 |
| chr2 | 169311665 | 169311666 | cg00185988 |
| chr2 | 172014013 | 172014014 | cg01970904 |
| chr2 | 172014861 | 172014862 | cg24772525 |
| chr2 | 172015098 | 172015099 | cg10033675 |
| chr2 | 174058660 | 174058661 | cg02904488 |
| chr2 | 174123660 | 174123661 | cg13297097 |
| chr2 | 174989267 | 174989268 | cg17452006 |
| chr2 | 175355057 | 175355058 | cg10907868 |
| chr2 | 179568746 | 179568747 | cg24739899 |
| chr2 | 179916184 | 179916185 | cg25889918 |
| chr2 | 181388538 | 181388539 | cg09613192 |
| chr2 | 189315917 | 189315918 | cg04322162 |
| chr2 | 190628945 | 190628946 | cg22332666 |
| chr2 | 190701251 | 190701252 | cg20546098 |
| chr2 | 190984844 | 190984845 | cg25301173 |
| chr2 | 191276183 | 191276184 | cg04144521 |
| chr2 | 191395606 | 191395607 | cg18008221 |

| | | | |
|---|---|---|---|
| chr2 | 37816339 | 37816340 | cg08478623 |
| chr2 | 37897716 | 37897717 | cg15293422 |
| chr2 | 38155843 | 38155844 | cg15657390 |
| chr2 | 38894198 | 38894199 | cg20799917 |
| chr2 | 38976288 | 38976289 | cg04441847 |
| chr2 | 39236214 | 39236215 | cg01361555 |
| chr2 | 39696979 | 39696980 | cg06900514 |
| chr2 | 39740720 | 39740721 | cg27509105 |
| chr2 | 40392494 | 40392495 | cg09296536 |
| chr2 | 40472772 | 40472773 | cg15146122 |
| chr2 | 40508306 | 40508307 | cg00870449 |
| chr2 | 40545033 | 40545034 | cg15491134 |
| chr2 | 40599006 | 40599007 | cg24503369 |
| chr2 | 40622490 | 40622491 | cg00415424 |
| chr2 | 42326070 | 42326071 | cg26672098 |
| chr2 | 42479385 | 42479386 | cg00162001 |
| chr2 | 42793517 | 42793518 | cg08009902 |
| chr2 | 43188851 | 43188852 | cg23008718 |
| chr2 | 43188940 | 43188941 | cg21514164 |
| chr2 | 43334941 | 43334942 | cg26685998 |
| chr2 | 43398079 | 43398080 | cg00360761 |
| chr2 | 43398118 | 43398119 | cg00055073 |
| chr2 | 43398154 | 43398155 | cg02871985 |
| chr2 | 43398271 | 43398272 | cg09446583 |
| chr2 | 43398339 | 43398340 | cg07502661 |
| chr2 | 46119607 | 46119608 | cg22284398 |
| chr2 | 46768111 | 46768112 | cg22307974 |
| chr2 | 47879734 | 47879735 | cg10809134 |
| chr2 | 48540671 | 48540672 | cg00032884 |
| chr2 | 49084769 | 49084770 | cg06481517 |
| chr2 | 50234533 | 50234534 | cg09044294 |
| chr2 | 50718584 | 50718585 | cg12884814 |
| chr2 | 52695221 | 52695222 | cg20342458 |
| chr2 | 53731598 | 53731599 | cg13724732 |
| chr2 | 54828502 | 54828503 | cg21487856 |
| chr2 | 54832147 | 54832148 | cg09404516 |
| chr2 | 55391421 | 55391422 | cg25249728 |
| chr2 | 60580559 | 60580560 | cg06629713 |
| chr2 | 60585866 | 60585867 | cg16665442 |
| chr2 | 60698937 | 60698938 | cg04588436 |
| chr2 | 60763626 | 60763627 | cg10853543 |
| chr2 | 61406491 | 61406492 | cg06949933 |
| chr2 | 62533140 | 62533141 | cg25918827 |
| chr2 | 62680940 | 62680941 | cg00752480 |
| chr2 | 64440295 | 64440296 | cg26164488 |
| chr2 | 64501134 | 64501135 | cg06233202 |
| chr2 | 64680425 | 64680426 | cg06645778 |
| chr2 | 64868710 | 64868711 | cg05783449 |
| chr2 | 65623541 | 65623542 | cg24872815 |
| chr2 | 67094729 | 67094730 | cg21711827 |
| chr2 | 68149422 | 68149423 | cg20436458 |
| chr2 | 68557960 | 68557961 | cg09360044 |
| chr2 | 69046817 | 69046818 | cg17754943 |
| chr2 | 69271042 | 69271043 | cg13358873 |
| chr2 | 69994225 | 69994226 | cg25099065 |
| chr2 | 70025869 | 70025870 | cg06635952 |
| chr2 | 191876673 | 191876674 | cg00676801 |
| chr2 | 192554623 | 192554624 | cg25981329 |
| chr2 | 198001840 | 198001841 | cg19592231 |
| chr2 | 201182777 | 201182778 | cg14361895 |
| chr2 | 201332878 | 201332879 | cg24151841 |
| chr2 | 201568315 | 201568316 | cg09396865 |
| chr2 | 201675739 | 201675740 | cg23892690 |
| chr2 | 201675829 | 201675830 | cg10637955 |
| chr2 | 202023415 | 202023416 | cg08267148 |
| chr2 | 202506468 | 202506469 | cg00656820 |
| chr2 | 204732687 | 204732688 | cg05074138 |
| chr2 | 204734181 | 204734182 | cg26091609 |
| chr2 | 204773790 | 204773791 | cg18914852 |
| chr2 | 206946028 | 206946029 | cg06470936 |
| chr2 | 208494443 | 208494444 | cg24758280 |
| chr2 | 208629716 | 208629717 | cg01383128 |
| chr2 | 208631081 | 208631082 | cg19744498 |
| chr2 | 208631259 | 208631260 | cg19957162 |
| chr2 | 208678058 | 208678059 | cg12008501 |
| chr2 | 213777611 | 213777612 | cg18303608 |
| chr2 | 216947149 | 216947150 | cg13500113 |
| chr2 | 218487226 | 218487227 | cg08401872 |
| chr2 | 218734017 | 218734018 | cg07492051 |
| chr2 | 219235515 | 219235516 | cg19701828 |
| chr2 | 219747459 | 219747460 | cg03630015 |
| chr2 | 219751342 | 219751343 | cg12001120 |
| chr2 | 219756890 | 219756891 | cg04067066 |
| chr2 | 220035927 | 220035928 | cg00552892 |
| chr2 | 220036660 | 220036661 | cg26320219 |
| chr2 | 220040571 | 220040572 | cg04955914 |
| chr2 | 220072753 | 220072754 | cg17109049 |
| chr2 | 221795858 | 221795859 | cg23689579 |
| chr2 | 222053559 | 222053560 | cg25146261 |
| chr2 | 222064243 | 222064244 | cg01415058 |
| chr2 | 222333289 | 222333290 | cg13364311 |
| chr2 | 222355484 | 222355485 | cg00335735 |
| chr2 | 223290515 | 223290516 | cg16171484 |
| chr2 | 223291007 | 223291008 | cg14280181 |
| chr2 | 223771010 | 223771011 | cg25947773 |
| chr2 | 223778280 | 223778281 | cg03040554 |
| chr2 | 224663259 | 224663260 | cg06751322 |
| chr2 | 224703232 | 224703233 | cg07813214 |
| chr2 | 224751989 | 224751990 | cg23363263 |
| chr2 | 224773702 | 224773703 | cg17778556 |
| chr2 | 224780409 | 224780410 | cg15457217 |
| chr2 | 224824500 | 224824501 | cg11268834 |
| chr2 | 225410509 | 225410510 | cg25502818 |
| chr2 | 225774214 | 225774215 | cg17075808 |
| chr2 | 225775142 | 225775143 | cg15930703 |
| chr2 | 227655319 | 227655320 | cg00305996 |
| chr2 | 227773531 | 227773532 | cg16852955 |
| chr2 | 227834762 | 227834763 | cg10439859 |
| chr2 | 227850069 | 227850070 | cg09157320 |
| chr2 | 231190414 | 231190415 | cg05217331 |
| chr2 | 231740398 | 231740399 | cg14396306 |
| chr2 | 231743193 | 231743194 | cg17674726 |

| | | | |
|---|---|---|---|
| chr2 | 70368784 | 70368785 | cg02039404 |
| chr2 | 71463632 | 71463633 | cg06614944 |
| chr2 | 73297338 | 73297339 | cg17639959 |
| chr2 | 73297389 | 73297390 | cg17329648 |
| chr2 | 74210857 | 74210858 | cg02251243 |
| chr2 | 74210890 | 74210891 | cg19445690 |
| chr2 | 74210960 | 74210961 | cg05981785 |
| chr2 | 74228640 | 74228641 | cg04648494 |
| chr2 | 74612759 | 74612760 | cg13312337 |
| chr2 | 74641453 | 74641454 | cg02466544 |
| chr2 | 74708767 | 74708768 | cg01584932 |
| chr2 | 74728623 | 74728624 | cg08973382 |
| chr2 | 74729429 | 74729430 | cg08355260 |
| chr2 | 74783039 | 74783040 | cg22989958 |
| chr2 | 74811430 | 74811431 | cg15025240 |
| chr2 | 79252794 | 79252795 | cg01357846 |
| chr2 | 79312664 | 79312665 | cg17863076 |
| chr2 | 79365892 | 79365893 | cg15359390 |
| chr2 | 80551812 | 80551813 | cg25237113 |
| chr2 | 81427138 | 81427139 | cg15704225 |
| chr2 | 82864753 | 82864754 | cg04011469 |
| chr2 | 85551850 | 85551851 | cg01054402 |
| chr2 | 85555865 | 85555866 | cg03728395 |
| chr2 | 85637116 | 85637117 | cg04881903 |
| chr2 | 85637739 | 85637740 | cg27139457 |
| chr2 | 85637851 | 85637852 | cg26476820 |
| chr2 | 85667381 | 85667382 | cg11651615 |
| chr2 | 85971548 | 85971549 | cg21512179 |
| chr2 | 86038423 | 86038424 | cg08482837 |
| chr2 | 86092937 | 86092938 | cg06909254 |
| chr2 | 86114036 | 86114037 | cg09113070 |
| chr2 | 88327530 | 88327531 | cg04195125 |
| chr2 | 88390167 | 88390168 | cg19851560 |
| chr2 | 88390175 | 88390176 | cg13172001 |
| chr2 | 96805411 | 96805412 | cg02377024 |
| chr2 | 96807942 | 96807943 | cg18145558 |
| chr2 | 96808802 | 96808803 | cg13389146 |
| chr2 | 96809174 | 96809175 | cg13725590 |
| chr2 | 96809970 | 96809971 | cg02431562 |
| chr2 | 96824799 | 96824800 | cg03604731 |
| chr2 | 96868790 | 96868791 | cg17526890 |
| chr2 | 96869976 | 96869977 | cg21616425 |
| chr2 | 96907142 | 96907143 | cg11107560 |
| chr2 | 97171137 | 97171138 | cg23824762 |
| chr2 | 97561669 | 97561670 | cg05368724 |
| chr2 | 98329942 | 98329943 | cg13853141 |
| chr2 | 98330020 | 98330021 | cg12332902 |
| chr2 | 98330087 | 98330088 | cg21773162 |
| chr2 | 98344873 | 98344874 | cg14537825 |
| chr2 | 98365143 | 98365144 | cg20820876 |
| chr2 | 98409290 | 98409291 | cg02489228 |
| chr2 | 98607810 | 98607811 | cg17836487 |
| chr2 | 98609870 | 98609871 | cg16445423 |
| chr2 | 98615797 | 98615798 | cg19124151 |
| chr2 | 98703888 | 98703889 | cg08026735 |
| chr2 | 99081350 | 99081351 | cg11555067 |
| chr2 | 231770489 | 231770490 | cg05940122 |
| chr2 | 233561679 | 233561680 | cg14046043 |
| chr2 | 233741790 | 233741791 | cg14345131 |
| chr2 | 233765616 | 233765617 | cg10259518 |
| chr2 | 234620463 | 234620464 | cg15299997 |
| chr2 | 235401950 | 235401951 | cg24441922 |
| chr2 | 235403141 | 235403142 | cg05204104 |
| chr2 | 235403218 | 235403219 | cg15016771 |
| chr2 | 235406291 | 235406292 | cg05308656 |
| chr2 | 238362492 | 238362493 | cg19839281 |
| chr2 | 238609807 | 238609808 | cg19497511 |
| chr2 | 238805257 | 238805258 | cg10846682 |
| chr2 | 238877903 | 238877904 | cg24302344 |
| chr2 | 239195537 | 239195538 | cg23072629 |
| chr2 | 240172612 | 240172613 | cg04216051 |
| chr2 | 240172645 | 240172646 | cg20415945 |
| chr2 | 241076208 | 241076209 | cg07628769 |
| chr2 | 241936844 | 241936845 | cg16937168 |
| chr2 | 241976244 | 241976245 | cg10717312 |
| chr2 | 242034995 | 242034996 | cg01796333 |
| chr2 | 242306625 | 242306626 | cg16844818 |
| chr2 | 242308655 | 242308656 | cg14684937 |
| chr2 | 242787716 | 242787717 | cg07728865 |
| chr2 | 242793078 | 242793079 | cg10057601 |
| chr3 | 3081722 | 3081723 | cg04427437 |
| chr3 | 8560168 | 8560169 | cg26083045 |
| chr3 | 8561593 | 8561594 | cg16784366 |
| chr3 | 8564151 | 8564152 | cg06266100 |
| chr3 | 8577752 | 8577753 | cg23180235 |
| chr3 | 8615444 | 8615445 | cg03557445 |
| chr3 | 9291739 | 9291740 | cg26184741 |
| chr3 | 9859360 | 9859361 | cg03375833 |
| chr3 | 9956506 | 9956507 | cg06619959 |
| chr3 | 10137658 | 10137659 | cg19476342 |
| chr3 | 10146330 | 10146331 | cg13053408 |
| chr3 | 10192855 | 10192856 | cg16373217 |
| chr3 | 10284280 | 10284281 | cg18527651 |
| chr3 | 11178064 | 11178065 | cg26864526 |
| chr3 | 12701776 | 12701777 | cg09856107 |
| chr3 | 13045896 | 13045897 | cg18644782 |
| chr3 | 13082887 | 13082888 | cg08096381 |
| chr3 | 13462465 | 13462466 | cg24642820 |
| chr3 | 14494726 | 14494727 | cg13496041 |
| chr3 | 15366862 | 15366863 | cg18444702 |
| chr3 | 15382723 | 15382724 | cg01788682 |
| chr3 | 15383022 | 15383023 | cg05046996 |
| chr3 | 15384005 | 15384006 | cg21283680 |
| chr3 | 15397117 | 15397118 | cg27375330 |
| chr3 | 16343197 | 16343198 | cg16456087 |
| chr3 | 16738417 | 16738418 | cg24264042 |
| chr3 | 16843952 | 16843953 | cg14521077 |
| chr3 | 16852000 | 16852001 | cg21497139 |
| chr3 | 17069926 | 17069927 | cg15605530 |
| chr3 | 18772063 | 18772064 | cg21454600 |
| chr3 | 19549017 | 19549018 | cg02863493 |
| chr3 | 19576826 | 19576827 | cg15149374 |

| | | | |
|---|---|---|---|
| chr2 | 99434688 | 99434689 | cg02787615 |
| chr2 | 99759406 | 99759407 | cg09298484 |
| chr2 | 99760272 | 99760273 | cg18482960 |
| chr2 | 100720179 | 100720180 | cg13353679 |
| chr2 | 100722155 | 100722156 | cg19071879 |
| chr2 | 100759623 | 100759624 | cg21988972 |
| chr2 | 101678574 | 101678575 | cg27587307 |
| chr2 | 101922427 | 101922428 | cg03191504 |
| chr2 | 101922922 | 101922923 | cg11962007 |
| chr2 | 102313242 | 102313243 | cg06095752 |
| chr2 | 102350276 | 102350277 | cg05874806 |
| chr2 | 102369368 | 102369369 | cg03876150 |
| chr2 | 102615524 | 102615525 | cg12910851 |
| chr2 | 105757726 | 105757727 | cg13149147 |
| chr2 | 106363603 | 106363604 | cg25961567 |
| chr2 | 106364229 | 106364230 | cg20703928 |
| chr2 | 106373553 | 106373554 | cg21101995 |
| chr2 | 106417701 | 106417702 | cg17501982 |
| chr2 | 106567892 | 106567893 | cg00160872 |
| chr2 | 106776775 | 106776776 | cg15341833 |
| chr2 | 106777038 | 106777039 | cg00118342 |
| chr2 | 108885152 | 108885153 | cg07795916 |
| chr2 | 109813176 | 109813177 | cg09500437 |
| chr2 | 111788355 | 111788356 | cg18282393 |
| chr2 | 111857973 | 111857974 | cg08886790 |
| chr2 | 111914486 | 111914487 | cg06363163 |
| chr2 | 112293267 | 112293268 | cg01439754 |
| chr2 | 112659978 | 112659979 | cg07748767 |
| chr2 | 112913178 | 112913179 | cg19744173 |
| chr2 | 112945506 | 112945507 | cg24743301 |
| chr2 | 113669912 | 113669913 | cg02229946 |
| chr2 | 113821936 | 113821937 | cg21854348 |
| chr2 | 113885116 | 113885117 | cg02543462 |
| chr2 | 113941371 | 113941372 | cg13224420 |
| chr2 | 113960266 | 113960267 | cg00499539 |
| chr2 | 114430284 | 114430285 | cg19143020 |
| chr2 | 115425220 | 115425221 | cg05868547 |
| chr2 | 118419342 | 118419343 | cg09860353 |
| chr2 | 119066603 | 119066604 | cg21428001 |
| chr2 | 120097181 | 120097182 | cg07051721 |
| chr2 | 122286796 | 122286797 | cg02176858 |
| chr2 | 122290763 | 122290764 | cg04075184 |
| chr2 | 125672110 | 125672111 | cg10962805 |
| chr2 | 127533123 | 127533124 | cg01919567 |
| chr2 | 127734952 | 127734953 | cg06006907 |
| chr2 | 128142155 | 128142156 | cg00021275 |
| chr2 | 128404486 | 128404487 | cg15869022 |
| chr2 | 128458248 | 128458249 | cg24925163 |
| chr2 | 129397288 | 129397289 | cg06974672 |
| chr2 | 129548156 | 129548157 | cg17588294 |
| chr2 | 129636330 | 129636331 | cg23274984 |
| chr2 | 129658241 | 129658242 | cg02054542 |
| chr2 | 130344243 | 130344244 | cg01787445 |
| chr2 | 133888263 | 133888264 | cg24072451 |
| chr2 | 134083495 | 134083496 | cg07695909 |
| chr2 | 134477298 | 134477299 | cg09446806 |

| | | | |
|---|---|---|---|
| chr3 | 23478304 | 23478305 | cg18560178 |
| chr3 | 23653462 | 23653463 | cg23340218 |
| chr3 | 23770435 | 23770436 | cg21772953 |
| chr3 | 23846810 | 23846811 | cg09560953 |
| chr3 | 23849114 | 23849115 | cg19719475 |
| chr3 | 23853015 | 23853016 | cg03668982 |
| chr3 | 23861312 | 23861313 | cg21610839 |
| chr3 | 23867300 | 23867301 | cg18805870 |
| chr3 | 27498161 | 27498162 | cg00907274 |
| chr3 | 27684452 | 27684453 | cg05499736 |
| chr3 | 30667873 | 30667874 | cg07285675 |
| chr3 | 30691493 | 30691494 | cg24719910 |
| chr3 | 31935279 | 31935280 | cg26337497 |
| chr3 | 32367045 | 32367046 | cg12823512 |
| chr3 | 37291314 | 37291315 | cg12435059 |
| chr3 | 38040073 | 38040074 | cg05620558 |
| chr3 | 38065265 | 38065266 | cg03084350 |
| chr3 | 38080203 | 38080204 | cg15681239 |
| chr3 | 38209834 | 38209835 | cg07293188 |
| chr3 | 39054994 | 39054995 | cg25123427 |
| chr3 | 39095348 | 39095349 | cg27140914 |
| chr3 | 39193251 | 39193252 | cg00501876 |
| chr3 | 42002230 | 42002231 | cg02108620 |
| chr3 | 42073868 | 42073869 | cg25651608 |
| chr3 | 42093455 | 42093456 | cg25209153 |
| chr3 | 42094749 | 42094750 | cg20372886 |
| chr3 | 42103367 | 42103368 | cg19230867 |
| chr3 | 42121737 | 42121738 | cg19704102 |
| chr3 | 42131702 | 42131703 | cg07886398 |
| chr3 | 42132870 | 42132871 | cg01486558 |
| chr3 | 42132991 | 42132992 | cg09654300 |
| chr3 | 42201898 | 42201899 | cg24877842 |
| chr3 | 43332555 | 43332556 | cg10288995 |
| chr3 | 45053351 | 45053352 | cg09626867 |
| chr3 | 45595231 | 45595232 | cg26135172 |
| chr3 | 45601455 | 45601456 | cg15765353 |
| chr3 | 45641120 | 45641121 | cg08062273 |
| chr3 | 45646427 | 45646428 | cg23825522 |
| chr3 | 45674281 | 45674282 | cg27588384 |
| chr3 | 45708680 | 45708681 | cg06808089 |
| chr3 | 45984838 | 45984839 | cg08450017 |
| chr3 | 45984942 | 45984943 | cg25226014 |
| chr3 | 45989744 | 45989745 | cg26466027 |
| chr3 | 46283902 | 46283903 | cg14312439 |
| chr3 | 46394550 | 46394551 | cg07743747 |
| chr3 | 46394845 | 46394846 | cg06280059 |
| chr3 | 46395191 | 46395192 | cg03928384 |
| chr3 | 46586857 | 46586858 | cg04522866 |
| chr3 | 48232722 | 48232723 | cg07088513 |
| chr3 | 48310544 | 48310545 | cg01458495 |
| chr3 | 48310616 | 48310617 | cg11307417 |
| chr3 | 48310897 | 48310898 | cg05070273 |
| chr3 | 48507165 | 48507166 | cg14208102 |
| chr3 | 49384054 | 49384055 | cg05383531 |
| chr3 | 49394622 | 49394623 | cg18642234 |
| chr3 | 49569379 | 49569380 | cg21553656 |

| | | | |
|---|---|---|---|
| chr2 | 134781671 | 134781672 | cg04876407 |
| chr2 | 134880302 | 134880303 | cg19185146 |
| chr2 | 134927417 | 134927418 | cg10820587 |
| chr2 | 135041380 | 135041381 | cg18482303 |
| chr2 | 135464256 | 135464257 | cg26681139 |
| chr2 | 136763629 | 136763630 | cg12313149 |
| chr2 | 136884116 | 136884117 | cg03776082 |
| chr2 | 142890700 | 142890701 | cg21340845 |
| chr2 | 143044057 | 143044058 | cg23753006 |
| chr2 | 143632190 | 143632191 | cg12192282 |
| chr2 | 145182344 | 145182345 | cg10502206 |
| chr2 | 145267984 | 145267985 | cg12571570 |
| chr2 | 145269644 | 145269645 | cg19353947 |
| chr2 | 145271307 | 145271308 | cg23095192 |
| chr2 | 145439424 | 145439425 | cg11974162 |
| chr2 | 145795797 | 145795798 | cg07679370 |
| chr2 | 146798769 | 146798770 | cg06541218 |
| chr2 | 147807867 | 147807868 | cg18448977 |
| chr2 | 150932623 | 150932624 | cg13430053 |
| chr2 | 151477988 | 151477989 | cg14472888 |
| chr2 | 151484439 | 151484440 | cg05543492 |
| chr2 | 152153974 | 152153975 | cg07972309 |
| chr2 | 152155003 | 152155004 | cg09862721 |
| chr2 | 152991620 | 152991621 | cg20625334 |
| chr2 | 153268704 | 153268705 | cg14261271 |
| chr2 | 153271840 | 153271841 | cg16794471 |
| chr2 | 153310258 | 153310259 | cg11994229 |
| chr2 | 153381623 | 153381624 | cg04215359 |
| chr2 | 155719622 | 155719623 | cg14660733 |
| chr2 | 157291759 | 157291760 | cg09290120 |
| chr2 | 157292018 | 157292019 | cg14754555 |
| chr2 | 157292113 | 157292114 | cg22033586 |
| chr2 | 157292127 | 157292128 | cg15736127 |
| chr2 | 157393600 | 157393601 | cg23836163 |
| chr2 | 160958810 | 160958811 | cg15152331 |
| chr2 | 161233779 | 161233780 | cg26055370 |
| chr2 | 162915902 | 162915903 | cg15353603 |
| chr2 | 165760725 | 165760726 | cg16710042 |
| chr2 | 166061891 | 166061892 | cg17754914 |
| chr2 | 168189841 | 168189842 | cg06702600 |
| chr2 | 169058521 | 169058522 | cg07992385 |
| chr2 | 169326407 | 169326408 | cg23490396 |
| chr2 | 169887819 | 169887820 | cg18912768 |
| chr2 | 171943243 | 171943244 | cg05590818 |
| chr2 | 173150101 | 173150102 | cg21335378 |
| chr2 | 173461819 | 173461820 | cg04033559 |
| chr2 | 174821978 | 174821979 | cg02927202 |
| chr2 | 175463829 | 175463830 | cg02558288 |
| chr2 | 175645413 | 175645414 | cg10142566 |
| chr2 | 175652652 | 175652653 | cg08579540 |
| chr2 | 175660591 | 175660592 | cg19705386 |
| chr2 | 176464572 | 176464573 | cg03483330 |
| chr2 | 177518121 | 177518122 | cg00370173 |
| chr2 | 177682959 | 177682960 | cg21118939 |
| chr2 | 178116991 | 178116992 | cg27507284 |
| chr2 | 179148245 | 179148246 | cg00829858 |

| | | | |
|---|---|---|---|
| chr3 | 50279038 | 50279039 | cg17641710 |
| chr3 | 50337305 | 50337306 | cg20912205 |
| chr3 | 50355416 | 50355417 | cg06721473 |
| chr3 | 50360097 | 50360098 | cg00840516 |
| chr3 | 50360690 | 50360691 | cg27091787 |
| chr3 | 52087583 | 52087584 | cg05504759 |
| chr3 | 52231691 | 52231692 | cg11012543 |
| chr3 | 52233019 | 52233020 | cg15698299 |
| chr3 | 52255721 | 52255722 | cg00774149 |
| chr3 | 52274561 | 52274562 | cg07757252 |
| chr3 | 52302252 | 52302253 | cg25236791 |
| chr3 | 52334181 | 52334182 | cg22626659 |
| chr3 | 52348951 | 52348952 | cg15439202 |
| chr3 | 52487733 | 52487734 | cg26738080 |
| chr3 | 52926322 | 52926323 | cg02331649 |
| chr3 | 53051694 | 53051695 | cg26850469 |
| chr3 | 53078218 | 53078219 | cg04503182 |
| chr3 | 53764604 | 53764605 | cg03562531 |
| chr3 | 53839251 | 53839252 | cg19128261 |
| chr3 | 53878642 | 53878643 | cg04071104 |
| chr3 | 56809796 | 56809797 | cg26319282 |
| chr3 | 56810151 | 56810152 | cg12325605 |
| chr3 | 56810320 | 56810321 | cg15012214 |
| chr3 | 56810749 | 56810750 | cg14633892 |
| chr3 | 56833426 | 56833427 | cg18482892 |
| chr3 | 57134628 | 57134629 | cg00770158 |
| chr3 | 57263543 | 57263544 | cg05963604 |
| chr3 | 57999988 | 57999989 | cg24204017 |
| chr3 | 58158888 | 58158889 | cg19600372 |
| chr3 | 58200326 | 58200327 | cg10235741 |
| chr3 | 59352714 | 59352715 | cg24852972 |
| chr3 | 59393503 | 59393504 | cg19411557 |
| chr3 | 61889812 | 61889813 | cg27179041 |
| chr3 | 63922542 | 63922543 | cg07139503 |
| chr3 | 63929152 | 63929153 | cg22679474 |
| chr3 | 63978545 | 63978546 | cg18405880 |
| chr3 | 64636717 | 64636718 | cg16526133 |
| chr3 | 66306234 | 66306235 | cg23247879 |
| chr3 | 67060217 | 67060218 | cg26376025 |
| chr3 | 69928232 | 69928233 | cg04535746 |
| chr3 | 70231495 | 70231496 | cg24317217 |
| chr3 | 71050832 | 71050833 | cg16697896 |
| chr3 | 71056608 | 71056609 | cg15472885 |
| chr3 | 71099193 | 71099194 | cg21423741 |
| chr3 | 71103443 | 71103444 | cg08173709 |
| chr3 | 71106252 | 71106253 | cg00052246 |
| chr3 | 71111489 | 71111490 | cg25481160 |
| chr3 | 71119415 | 71119416 | cg19402876 |
| chr3 | 71127516 | 71127517 | cg23830205 |
| chr3 | 71128840 | 71128841 | cg24864984 |
| chr3 | 71201264 | 71201265 | cg13726313 |
| chr3 | 71205277 | 71205278 | cg22798400 |
| chr3 | 71217277 | 71217278 | cg22828884 |
| chr3 | 71274624 | 71274625 | cg15871371 |
| chr3 | 71293767 | 71293768 | cg07278181 |
| chr3 | 71294341 | 71294342 | cg05384123 |

| | | | |
|---|---|---|---|
| chr2 | 179344725 | 179344726 | cg05098551 |
| chr2 | 179395885 | 179395886 | cg15609237 |
| chr2 | 179395935 | 179395936 | cg08691235 |
| chr2 | 179396214 | 179396215 | cg08736813 |
| chr2 | 179897289 | 179897290 | cg27228986 |
| chr2 | 180722610 | 180722611 | cg10671757 |
| chr2 | 181843717 | 181843718 | cg04166432 |
| chr2 | 182523146 | 182523147 | cg01494441 |
| chr2 | 182850059 | 182850060 | cg19742470 |
| chr2 | 183086277 | 183086278 | cg21471828 |
| chr2 | 183123847 | 183123848 | cg07496624 |
| chr2 | 188391145 | 188391146 | cg15957797 |
| chr2 | 188483530 | 188483531 | cg04144365 |
| chr2 | 189193728 | 189193729 | cg07791987 |
| chr2 | 189247165 | 189247166 | cg21218363 |
| chr2 | 189298330 | 189298331 | cg19475988 |
| chr2 | 190309186 | 190309187 | cg09775648 |
| chr2 | 190706964 | 190706965 | cg24396875 |
| chr2 | 191044481 | 191044482 | cg04391205 |
| chr2 | 191207820 | 191207821 | cg17516156 |
| chr2 | 191543902 | 191543903 | cg17118478 |
| chr2 | 191754464 | 191754465 | cg03962451 |
| chr2 | 191844998 | 191844999 | cg08946713 |
| chr2 | 192114017 | 192114018 | cg12404831 |
| chr2 | 192155833 | 192155834 | cg24376339 |
| chr2 | 192169585 | 192169586 | cg06844526 |
| chr2 | 192229245 | 192229246 | cg02523585 |
| chr2 | 192476634 | 192476635 | cg03022510 |
| chr2 | 192712920 | 192712921 | cg10082589 |
| chr2 | 192712975 | 192712976 | cg17809945 |
| chr2 | 196525457 | 196525458 | cg27314761 |
| chr2 | 198062750 | 198062751 | cg03771282 |
| chr2 | 198077081 | 198077082 | cg20402826 |
| chr2 | 198127245 | 198127246 | cg02641917 |
| chr2 | 199639143 | 199639144 | cg05392844 |
| chr2 | 201194453 | 201194454 | cg14106933 |
| chr2 | 201242945 | 201242946 | cg06226703 |
| chr2 | 201245077 | 201245078 | cg13144059 |
| chr2 | 201312367 | 201312368 | cg06240652 |
| chr2 | 201320027 | 201320028 | cg04945753 |
| chr2 | 201829669 | 201829670 | cg25714096 |
| chr2 | 201993795 | 201993796 | cg25567938 |
| chr2 | 201995729 | 201995730 | cg13368983 |
| chr2 | 202313532 | 202313533 | cg18786374 |
| chr2 | 203244123 | 203244124 | cg14787880 |
| chr2 | 203504161 | 203504162 | cg01745766 |
| chr2 | 204973770 | 204973771 | cg16134718 |
| chr2 | 205385002 | 205385003 | cg18990899 |
| chr2 | 206231192 | 206231193 | cg05260373 |
| chr2 | 206544411 | 206544412 | cg23833033 |
| chr2 | 206789816 | 206789817 | cg01491071 |
| chr2 | 207854583 | 207854584 | cg25923788 |
| chr2 | 208338079 | 208338080 | cg22543026 |
| chr2 | 208485495 | 208485496 | cg15035973 |
| chr2 | 209028478 | 209028479 | cg11729430 |
| chr2 | 210292579 | 210292580 | cg01441622 |

| | | | |
|---|---|---|---|
| chr3 | 71301786 | 71301787 | cg05098381 |
| chr3 | 71481486 | 71481487 | cg02336104 |
| chr3 | 72433837 | 72433838 | cg10805254 |
| chr3 | 72440793 | 72440794 | cg26108120 |
| chr3 | 73150125 | 73150126 | cg10639411 |
| chr3 | 93789237 | 93789238 | cg05431656 |
| chr3 | 99586990 | 99586991 | cg13612275 |
| chr3 | 99664011 | 99664012 | cg17241937 |
| chr3 | 99721850 | 99721851 | cg24156261 |
| chr3 | 99790878 | 99790879 | cg19533955 |
| chr3 | 99979355 | 99979356 | cg04730047 |
| chr3 | 100053096 | 100053097 | cg13016775 |
| chr3 | 100322493 | 100322494 | cg13709487 |
| chr3 | 100322629 | 100322630 | cg26645827 |
| chr3 | 107805635 | 107805636 | cg09227934 |
| chr3 | 107823424 | 107823425 | cg17892917 |
| chr3 | 108020451 | 108020452 | cg02124498 |
| chr3 | 108041508 | 108041509 | cg24769830 |
| chr3 | 108180898 | 108180899 | cg21480966 |
| chr3 | 112012867 | 112012868 | cg01411366 |
| chr3 | 112021179 | 112021180 | cg00813264 |
| chr3 | 112054506 | 112054507 | cg26652155 |
| chr3 | 112067412 | 112067413 | cg25086874 |
| chr3 | 112143970 | 112143971 | cg13537646 |
| chr3 | 112360952 | 112360953 | cg21307628 |
| chr3 | 112695305 | 112695306 | cg02155111 |
| chr3 | 112932345 | 112932346 | cg24572745 |
| chr3 | 113213220 | 113213221 | cg10572188 |
| chr3 | 113249814 | 113249815 | cg06273125 |
| chr3 | 114012316 | 114012317 | cg22577252 |
| chr3 | 114018592 | 114018593 | cg09246203 |
| chr3 | 114037617 | 114037618 | cg23637607 |
| chr3 | 114419469 | 114419470 | cg24146776 |
| chr3 | 114807036 | 114807037 | cg17875356 |
| chr3 | 119011925 | 119011926 | cg09081994 |
| chr3 | 119499190 | 119499191 | cg02863947 |
| chr3 | 121150715 | 121150716 | cg12200866 |
| chr3 | 121312164 | 121312165 | cg07378309 |
| chr3 | 121380533 | 121380534 | cg15458222 |
| chr3 | 121972621 | 121972622 | cg09447811 |
| chr3 | 122128730 | 122128731 | cg18797132 |
| chr3 | 122286677 | 122286678 | cg02519879 |
| chr3 | 122334330 | 122334331 | cg19889855 |
| chr3 | 122509111 | 122509112 | cg09359351 |
| chr3 | 124303389 | 124303390 | cg20925263 |
| chr3 | 124303404 | 124303405 | cg04516112 |
| chr3 | 124304960 | 124304961 | cg04915618 |
| chr3 | 125077434 | 125077435 | cg01289769 |
| chr3 | 126679199 | 126679200 | cg11629955 |
| chr3 | 127393517 | 127393518 | cg03596635 |
| chr3 | 127535191 | 127535192 | cg14476212 |
| chr3 | 127935096 | 127935097 | cg14972564 |
| chr3 | 128746334 | 128746335 | cg14511273 |
| chr3 | 128838350 | 128838351 | cg05859076 |
| chr3 | 128845626 | 128845627 | cg07730360 |
| chr3 | 128994810 | 128994811 | cg19497709 |

| | | | |
|---|---|---|---|
| chr2 | 210443311 | 210443312 | cg15310568 |
| chr2 | 212232130 | 212232131 | cg25817111 |
| chr2 | 213933386 | 213933387 | cg16345559 |
| chr2 | 214857742 | 214857743 | cg25824236 |
| chr2 | 216660598 | 216660599 | cg05232118 |
| chr2 | 218466307 | 218466308 | cg26501046 |
| chr2 | 218531045 | 218531046 | cg12820310 |
| chr2 | 219148914 | 219148915 | cg13835894 |
| chr2 | 219435282 | 219435283 | cg04726824 |
| chr2 | 219610103 | 219610104 | cg01827633 |
| chr2 | 219697159 | 219697160 | cg09244100 |
| chr2 | 220222765 | 220222766 | cg07434457 |
| chr2 | 220862582 | 220862583 | cg10977352 |
| chr2 | 220908336 | 220908337 | cg22573731 |
| chr2 | 221864241 | 221864242 | cg05318546 |
| chr2 | 221940988 | 221940989 | cg19406878 |
| chr2 | 222147678 | 222147679 | cg02550257 |
| chr2 | 222367110 | 222367111 | cg03850035 |
| chr2 | 223901462 | 223901463 | cg11754382 |
| chr2 | 224808252 | 224808253 | cg02225786 |
| chr2 | 224821113 | 224821114 | cg00474696 |
| chr2 | 225266580 | 225266581 | cg18890556 |
| chr2 | 225435131 | 225435132 | cg01474011 |
| chr2 | 225792082 | 225792083 | cg19411699 |
| chr2 | 225874509 | 225874510 | cg05629278 |
| chr2 | 225907875 | 225907876 | cg02441556 |
| chr2 | 227657192 | 227657193 | cg18018313 |
| chr2 | 231731972 | 231731973 | cg02806733 |
| chr2 | 231732249 | 231732250 | cg18346707 |
| chr2 | 231735434 | 231735435 | cg18665334 |
| chr2 | 231773530 | 231773531 | cg24416609 |
| chr2 | 231849693 | 231849694 | cg04519775 |
| chr2 | 232321649 | 232321650 | cg08338359 |
| chr2 | 232396241 | 232396242 | cg24430754 |
| chr2 | 232455630 | 232455631 | cg27232569 |
| chr2 | 232549224 | 232549225 | cg20793665 |
| chr2 | 233734115 | 233734116 | cg25237894 |
| chr2 | 233739114 | 233739115 | cg17611093 |
| chr2 | 233741920 | 233741921 | cg13759905 |
| chr2 | 233981885 | 233981886 | cg22666015 |
| chr2 | 235178208 | 235178209 | cg06644719 |
| chr2 | 235319934 | 235319935 | cg17342709 |
| chr2 | 237468416 | 237468417 | cg23138373 |
| chr2 | 237874990 | 237874991 | cg10104542 |
| chr2 | 238525524 | 238525525 | cg18141622 |
| chr2 | 238598832 | 238598833 | cg07063912 |
| chr2 | 239134945 | 239134946 | cg12047086 |
| chr2 | 240143979 | 240143980 | cg26767974 |
| chr2 | 240172742 | 240172743 | cg17532753 |
| chr2 | 240176050 | 240176051 | cg07134930 |
| chr2 | 240207606 | 240207607 | cg16360836 |
| chr2 | 241854588 | 241854589 | cg08440077 |
| chr2 | 242039676 | 242039677 | cg00726786 |
| chr2 | 242087883 | 242087884 | cg21399778 |
| chr2 | 242169936 | 242169937 | cg21663667 |
| chr2 | 242296992 | 242296993 | cg15832329 |
| chr3 | 128998952 | 128998953 | cg08166362 |
| chr3 | 129148047 | 129148048 | cg10474404 |
| chr3 | 129148362 | 129148363 | cg03221715 |
| chr3 | 129323778 | 129323779 | cg02544002 |
| chr3 | 129369724 | 129369725 | cg00986598 |
| chr3 | 130469275 | 130469276 | cg17586154 |
| chr3 | 130568122 | 130568123 | cg07892374 |
| chr3 | 132139298 | 132139299 | cg03396594 |
| chr3 | 132176123 | 132176124 | cg13011713 |
| chr3 | 133282378 | 133282379 | cg22932385 |
| chr3 | 133956516 | 133956517 | cg16943019 |
| chr3 | 136649333 | 136649334 | cg00382999 |
| chr3 | 138552648 | 138552649 | cg00738869 |
| chr3 | 141456305 | 141456306 | cg11626153 |
| chr3 | 142844036 | 142844037 | cg15171155 |
| chr3 | 142888303 | 142888304 | cg26442397 |
| chr3 | 143148893 | 143148894 | cg18151422 |
| chr3 | 143255785 | 143255786 | cg02501410 |
| chr3 | 146213648 | 146213649 | cg03075662 |
| chr3 | 149167242 | 149167243 | cg00214395 |
| chr3 | 149212391 | 149212392 | cg19422253 |
| chr3 | 150996563 | 150996564 | cg09571369 |
| chr3 | 150997160 | 150997161 | cg21793358 |
| chr3 | 151985869 | 151985870 | cg12360736 |
| chr3 | 152046751 | 152046752 | cg02061820 |
| chr3 | 152092595 | 152092596 | cg08535978 |
| chr3 | 155594778 | 155594779 | cg14694205 |
| chr3 | 156394398 | 156394399 | cg07574216 |
| chr3 | 156395056 | 156395057 | cg09379146 |
| chr3 | 156423325 | 156423326 | cg13065203 |
| chr3 | 156784781 | 156784782 | cg15697575 |
| chr3 | 156793711 | 156793712 | cg16699540 |
| chr3 | 156806144 | 156806145 | cg11979011 |
| chr3 | 156809115 | 156809116 | cg02551745 |
| chr3 | 156827696 | 156827697 | cg01535205 |
| chr3 | 156966620 | 156966621 | cg27558387 |
| chr3 | 160546500 | 160546501 | cg04747996 |
| chr3 | 160586959 | 160586960 | cg09007331 |
| chr3 | 160587737 | 160587738 | cg25203361 |
| chr3 | 160611917 | 160611918 | cg15171488 |
| chr3 | 167454339 | 167454340 | cg18123677 |
| chr3 | 169629136 | 169629137 | cg05721199 |
| chr3 | 170910597 | 170910598 | cg14924866 |
| chr3 | 170924582 | 170924583 | cg11095839 |
| chr3 | 171146337 | 171146338 | cg02499139 |
| chr3 | 171844005 | 171844006 | cg00589541 |
| chr3 | 171858472 | 171858473 | cg19980260 |
| chr3 | 176773597 | 176773598 | cg09226275 |
| chr3 | 176868896 | 176868897 | cg03930153 |
| chr3 | 177002388 | 177002389 | cg03889263 |
| chr3 | 177039618 | 177039619 | cg11270278 |
| chr3 | 177066294 | 177066295 | cg00554341 |
| chr3 | 177239931 | 177239932 | cg09247235 |
| chr3 | 177370839 | 177370840 | cg20187173 |
| chr3 | 177514999 | 177515000 | cg22694983 |
| chr3 | 179061771 | 179061772 | cg12250561 |

| | | | |
|---|---|---|---|
| chr2 | 242810748 | 242810749 | cg09034753 |
| chr2 | 242810927 | 242810928 | cg11654325 |
| chr2 | 242811149 | 242811150 | cg23491841 |
| chr3 | 1811915 | 1811916 | cg10466011 |
| chr3 | 4549756 | 4549757 | cg24131262 |
| chr3 | 5071083 | 5071084 | cg10125901 |
| chr3 | 5095248 | 5095249 | cg00874032 |
| chr3 | 5246986 | 5246987 | cg22250390 |
| chr3 | 5259831 | 5259832 | cg21887591 |
| chr3 | 8308781 | 8308782 | cg13744481 |
| chr3 | 8363442 | 8363443 | cg06367354 |
| chr3 | 8499933 | 8499934 | cg07162700 |
| chr3 | 9273803 | 9273804 | cg26389316 |
| chr3 | 9291773 | 9291774 | cg06858559 |
| chr3 | 9406960 | 9406961 | cg24817003 |
| chr3 | 9921760 | 9921761 | cg16777090 |
| chr3 | 9943884 | 9943885 | cg07832674 |
| chr3 | 9950984 | 9950985 | cg10314459 |
| chr3 | 9958686 | 9958687 | cg12487088 |
| chr3 | 10379281 | 10379282 | cg09477124 |
| chr3 | 10437269 | 10437270 | cg12115190 |
| chr3 | 11623800 | 11623801 | cg03370106 |
| chr3 | 11624023 | 11624024 | cg07330114 |
| chr3 | 12197630 | 12197631 | cg10110335 |
| chr3 | 12202995 | 12202996 | cg16858253 |
| chr3 | 12713226 | 12713227 | cg13724170 |
| chr3 | 12996362 | 12996363 | cg25157472 |
| chr3 | 13008096 | 13008097 | cg09437283 |
| chr3 | 13008273 | 13008274 | cg25484139 |
| chr3 | 13082751 | 13082752 | cg01044637 |
| chr3 | 13117754 | 13117755 | cg18496318 |
| chr3 | 13457267 | 13457268 | cg04992150 |
| chr3 | 14567529 | 14567530 | cg12890346 |
| chr3 | 14699141 | 14699142 | cg03788239 |
| chr3 | 14825290 | 14825291 | cg16922884 |
| chr3 | 15032403 | 15032404 | cg22691776 |
| chr3 | 15085146 | 15085147 | cg19730887 |
| chr3 | 15244698 | 15244699 | cg26712821 |
| chr3 | 15352493 | 15352494 | cg07480608 |
| chr3 | 15359338 | 15359339 | cg04858987 |
| chr3 | 15374534 | 15374535 | cg15145672 |
| chr3 | 15374586 | 15374587 | cg08359336 |
| chr3 | 15850965 | 15850966 | cg23458145 |
| chr3 | 16766180 | 16766181 | cg22626825 |
| chr3 | 16974217 | 16974218 | cg19448816 |
| chr3 | 16974681 | 16974682 | cg24159436 |
| chr3 | 19055533 | 19055534 | cg24200753 |
| chr3 | 19752610 | 19752611 | cg25650360 |
| chr3 | 23243300 | 23243301 | cg02794920 |
| chr3 | 23727711 | 23727712 | cg21450228 |
| chr3 | 24565992 | 24565993 | cg07762764 |
| chr3 | 27353034 | 27353035 | cg07047471 |
| chr3 | 28982939 | 28982940 | cg20187049 |
| chr3 | 30638618 | 30638619 | cg14656948 |
| chr3 | 30651317 | 30651318 | cg04916416 |
| chr3 | 30670335 | 30670336 | cg06270049 |

| | | | |
|---|---|---|---|
| chr3 | 179067057 | 179067058 | cg08058131 |
| chr3 | 179110334 | 179110335 | cg17582444 |
| chr3 | 180629523 | 180629524 | cg01816191 |
| chr3 | 181568620 | 181568621 | cg27024196 |
| chr3 | 182506874 | 182506875 | cg18548400 |
| chr3 | 183236384 | 183236385 | cg18772205 |
| chr3 | 183702886 | 183702887 | cg11320499 |
| chr3 | 184054901 | 184054902 | cg00205703 |
| chr3 | 186083922 | 186083923 | cg18800785 |
| chr3 | 186651116 | 186651117 | cg11386711 |
| chr3 | 187452863 | 187452864 | cg21739584 |
| chr3 | 187467427 | 187467428 | cg23664385 |
| chr3 | 187664092 | 187664093 | cg01388693 |
| chr3 | 187770324 | 187770325 | cg12626957 |
| chr3 | 187870713 | 187870714 | cg00119073 |
| chr3 | 187870929 | 187870930 | cg14091103 |
| chr3 | 187871024 | 187871025 | cg19599666 |
| chr3 | 187871034 | 187871035 | cg18127680 |
| chr3 | 187872975 | 187872976 | cg21427213 |
| chr3 | 187930704 | 187930705 | cg06547766 |
| chr3 | 187960622 | 187960623 | cg02579706 |
| chr3 | 188021823 | 188021824 | cg23181899 |
| chr3 | 188291633 | 188291634 | cg04387984 |
| chr3 | 188664747 | 188664748 | cg04252044 |
| chr3 | 188664887 | 188664888 | cg03176993 |
| chr3 | 191046547 | 191046548 | cg19066520 |
| chr3 | 191100959 | 191100960 | cg07393854 |
| chr3 | 191715823 | 191715824 | cg06204618 |
| chr3 | 193495351 | 193495352 | cg06583921 |
| chr3 | 193545238 | 193545239 | cg20186588 |
| chr3 | 193803934 | 193803935 | cg24939710 |
| chr3 | 193815787 | 193815788 | cg26345916 |
| chr3 | 194646995 | 194646996 | cg18762125 |
| chr3 | 194754642 | 194754643 | cg20074526 |
| chr3 | 194797099 | 194797100 | cg06704969 |
| chr3 | 194797138 | 194797139 | cg25921813 |
| chr3 | 194866144 | 194866145 | cg02041946 |
| chr3 | 194866280 | 194866281 | cg07463963 |
| chr3 | 194903259 | 194903260 | cg03916038 |
| chr3 | 194924602 | 194924603 | cg15722952 |
| chr3 | 194979832 | 194979833 | cg26062560 |
| chr3 | 195310883 | 195310884 | cg05624196 |
| chr3 | 195310887 | 195310888 | cg23720929 |
| chr3 | 195600311 | 195600312 | cg20092936 |
| chr3 | 195600608 | 195600609 | cg08767936 |
| chr3 | 195678176 | 195678177 | cg03732638 |
| chr3 | 195682326 | 195682327 | cg23245007 |
| chr3 | 195714100 | 195714101 | cg20120631 |
| chr3 | 195863402 | 195863403 | cg10699852 |
| chr3 | 195869691 | 195869692 | cg15932961 |
| chr3 | 196749317 | 196749318 | cg02470858 |
| chr3 | 197350982 | 197350983 | cg25351687 |
| chr3 | 197387652 | 197387653 | cg27537046 |
| chr3 | 197401399 | 197401400 | cg15435900 |
| chr3 | 197401451 | 197401452 | cg10579986 |
| chr3 | 197401572 | 197401573 | cg06336230 |

| | | | |
|---|---|---|---|
| chr3 | 30673459 | 30673460 | cg19615017 |
| chr3 | 30689835 | 30689836 | cg13724812 |
| chr3 | 30706645 | 30706646 | cg23485307 |
| chr3 | 30708746 | 30708747 | cg07613391 |
| chr3 | 30893353 | 30893354 | cg10336406 |
| chr3 | 31432787 | 31432788 | cg09627302 |
| chr3 | 32023722 | 32023723 | cg25681469 |
| chr3 | 32023996 | 32023997 | cg25692785 |
| chr3 | 32442229 | 32442230 | cg15872079 |
| chr3 | 32941716 | 32941717 | cg15235805 |
| chr3 | 33096255 | 33096256 | cg11236746 |
| chr3 | 33701209 | 33701210 | cg00744866 |
| chr3 | 34187750 | 34187751 | cg16171771 |
| chr3 | 34429403 | 34429404 | cg09619751 |
| chr3 | 36218374 | 36218375 | cg07047731 |
| chr3 | 37998565 | 37998566 | cg06027691 |
| chr3 | 38216354 | 38216355 | cg14989661 |
| chr3 | 38529102 | 38529103 | cg06120492 |
| chr3 | 39235366 | 39235367 | cg12413566 |
| chr3 | 39509081 | 39509082 | cg04442806 |
| chr3 | 40150853 | 40150854 | cg23024967 |
| chr3 | 41265374 | 41265375 | cg05726118 |
| chr3 | 42113624 | 42113625 | cg05628049 |
| chr3 | 42132310 | 42132311 | cg26334154 |
| chr3 | 42158765 | 42158766 | cg19028706 |
| chr3 | 42452862 | 42452863 | cg21374754 |
| chr3 | 42701690 | 42701691 | cg25824509 |
| chr3 | 45166560 | 45166561 | cg20849286 |
| chr3 | 45505334 | 45505335 | cg02489202 |
| chr3 | 45579354 | 45579355 | cg05380759 |
| chr3 | 45734030 | 45734031 | cg17207780 |
| chr3 | 46033800 | 46033801 | cg11211486 |
| chr3 | 46068932 | 46068933 | cg15888522 |
| chr3 | 46282766 | 46282767 | cg04111761 |
| chr3 | 46411447 | 46411448 | cg04131610 |
| chr3 | 46411474 | 46411475 | cg00803692 |
| chr3 | 46448084 | 46448085 | cg18599081 |
| chr3 | 46448134 | 46448135 | cg23350385 |
| chr3 | 46974965 | 46974966 | cg06240200 |
| chr3 | 47040292 | 47040293 | cg01892547 |
| chr3 | 47040357 | 47040358 | cg26432350 |
| chr3 | 47269695 | 47269696 | cg17579089 |
| chr3 | 48507087 | 48507088 | cg01870865 |
| chr3 | 49531973 | 49531974 | cg02522485 |
| chr3 | 49573805 | 49573806 | cg09251508 |
| chr3 | 50337494 | 50337495 | cg03655330 |
| chr3 | 50360667 | 50360668 | cg26460678 |
| chr3 | 50376216 | 50376217 | cg19854901 |
| chr3 | 50404127 | 50404128 | cg03625136 |
| chr3 | 50686177 | 50686178 | cg22950982 |
| chr3 | 51745321 | 51745322 | cg20984065 |
| chr3 | 52280329 | 52280330 | cg07773769 |
| chr3 | 52407081 | 52407082 | cg14092988 |
| chr3 | 52492874 | 52492875 | cg07386061 |
| chr3 | 52724044 | 52724045 | cg21199854 |
| chr3 | 52813652 | 52813653 | cg20219911 |
| chr3 | 197401931 | 197401932 | cg12023911 |
| chr3 | 197459812 | 197459813 | cg07255038 |
| chr4 | 979183 | 979184 | cg09359391 |
| chr4 | 979331 | 979332 | cg24226891 |
| chr4 | 1148524 | 1148525 | cg20084184 |
| chr4 | 1148569 | 1148570 | cg16118592 |
| chr4 | 1160842 | 1160843 | cg17456304 |
| chr4 | 1201881 | 1201882 | cg23045716 |
| chr4 | 1202780 | 1202781 | cg14505741 |
| chr4 | 1828603 | 1828604 | cg09371344 |
| chr4 | 1828697 | 1828698 | cg03008204 |
| chr4 | 1855934 | 1855935 | cg04814467 |
| chr4 | 2475103 | 2475104 | cg07799157 |
| chr4 | 2572546 | 2572547 | cg12833118 |
| chr4 | 2763026 | 2763027 | cg25585279 |
| chr4 | 2792582 | 2792583 | cg08320703 |
| chr4 | 2797201 | 2797202 | cg17749374 |
| chr4 | 2802612 | 2802613 | cg24492202 |
| chr4 | 2813458 | 2813459 | cg03443467 |
| chr4 | 2844730 | 2844731 | cg15457390 |
| chr4 | 2932447 | 2932448 | cg04498801 |
| chr4 | 3418911 | 3418912 | cg04203824 |
| chr4 | 4335184 | 4335185 | cg08179817 |
| chr4 | 4587217 | 4587218 | cg07652189 |
| chr4 | 6611128 | 6611129 | cg25892537 |
| chr4 | 6929045 | 6929046 | cg26562462 |
| chr4 | 6940915 | 6940916 | cg06496803 |
| chr4 | 6964527 | 6964528 | cg10681992 |
| chr4 | 6992752 | 6992753 | cg10117400 |
| chr4 | 7052788 | 7052789 | cg24709285 |
| chr4 | 8386188 | 8386189 | cg08206079 |
| chr4 | 8386198 | 8386199 | cg18999185 |
| chr4 | 8412369 | 8412370 | cg25963412 |
| chr4 | 11368007 | 11368008 | cg17748061 |
| chr4 | 11369349 | 11369350 | cg04203702 |
| chr4 | 11408659 | 11408660 | cg21300019 |
| chr4 | 11427700 | 11427701 | cg14047130 |
| chr4 | 11652441 | 11652442 | cg15634747 |
| chr4 | 11652483 | 11652484 | cg14006800 |
| chr4 | 11652505 | 11652506 | cg23971987 |
| chr4 | 14891007 | 14891008 | cg07230621 |
| chr4 | 16261642 | 16261643 | cg19604703 |
| chr4 | 24583375 | 24583376 | cg12857541 |
| chr4 | 25913666 | 25913667 | cg06129544 |
| chr4 | 26095438 | 26095439 | cg16847428 |
| chr4 | 26152723 | 26152724 | cg03002688 |
| chr4 | 26332763 | 26332764 | cg26924822 |
| chr4 | 26414901 | 26414902 | cg22992279 |
| chr4 | 26832099 | 26832100 | cg04873627 |
| chr4 | 26856624 | 26856625 | cg17470198 |
| chr4 | 37604497 | 37604498 | cg19029127 |
| chr4 | 37891376 | 37891377 | cg04710179 |
| chr4 | 38867304 | 38867305 | cg16030869 |
| chr4 | 38869779 | 38869780 | cg20484352 |
| chr4 | 38980960 | 38980961 | cg14209186 |
| chr4 | 39364570 | 39364571 | cg17550495 |

| | | | |
|---|---|---|---|
| chr3 | 52909345 | 52909346 | cg11485012 |
| chr3 | 52927548 | 52927549 | cg04865290 |
| chr3 | 53198666 | 53198667 | cg07687398 |
| chr3 | 53807593 | 53807594 | cg13757263 |
| chr3 | 53855929 | 53855930 | cg17184165 |
| chr3 | 53877436 | 53877437 | cg27567495 |
| chr3 | 54226648 | 54226649 | cg23487721 |
| chr3 | 54934591 | 54934592 | cg23516560 |
| chr3 | 55005522 | 55005523 | cg19631651 |
| chr3 | 55340399 | 55340400 | cg03046611 |
| chr3 | 55546118 | 55546119 | cg17578322 |
| chr3 | 55693182 | 55693183 | cg19551485 |
| chr3 | 55721067 | 55721068 | cg16718760 |
| chr3 | 56822059 | 56822060 | cg20888995 |
| chr3 | 56889795 | 56889796 | cg16451527 |
| chr3 | 57774266 | 57774267 | cg06672696 |
| chr3 | 57945695 | 57945696 | cg24033633 |
| chr3 | 57995180 | 57995181 | cg23730027 |
| chr3 | 58059919 | 58059920 | cg05093811 |
| chr3 | 58103186 | 58103187 | cg07557173 |
| chr3 | 58389654 | 58389655 | cg23549972 |
| chr3 | 59703569 | 59703570 | cg05709770 |
| chr3 | 60066405 | 60066406 | cg17894779 |
| chr3 | 62535030 | 62535031 | cg23003926 |
| chr3 | 62571255 | 62571256 | cg16915733 |
| chr3 | 63263076 | 63263077 | cg04013592 |
| chr3 | 64711384 | 64711385 | cg23078228 |
| chr3 | 65887859 | 65887860 | cg02761110 |
| chr3 | 66519998 | 66519999 | cg09716921 |
| chr3 | 69812125 | 69812126 | cg15449049 |
| chr3 | 69819855 | 69819856 | cg03489020 |
| chr3 | 69915189 | 69915190 | cg13151171 |
| chr3 | 69924342 | 69924343 | cg08652499 |
| chr3 | 70800230 | 70800231 | cg03275461 |
| chr3 | 70853598 | 70853599 | cg21444500 |
| chr3 | 71013485 | 71013486 | cg02220284 |
| chr3 | 71032216 | 71032217 | cg19628600 |
| chr3 | 71057560 | 71057561 | cg06033488 |
| chr3 | 71057616 | 71057617 | cg19776833 |
| chr3 | 71116326 | 71116327 | cg06175008 |
| chr3 | 71136296 | 71136297 | cg21379733 |
| chr3 | 71149599 | 71149600 | cg06691963 |
| chr3 | 71354056 | 71354057 | cg08038033 |
| chr3 | 71355122 | 71355123 | cg12431087 |
| chr3 | 71439885 | 71439886 | cg22589728 |
| chr3 | 71457237 | 71457238 | cg09459774 |
| chr3 | 71598380 | 71598381 | cg15958828 |
| chr3 | 71626654 | 71626655 | cg25811633 |
| chr3 | 71629054 | 71629055 | cg02520804 |
| chr3 | 72424982 | 72424983 | cg09006487 |
| chr3 | 73596205 | 73596206 | cg17804071 |
| chr3 | 73789574 | 73789575 | cg22379436 |
| chr3 | 73923792 | 73923793 | cg23012591 |
| chr3 | 74337905 | 74337906 | cg09614932 |
| chr3 | 74451852 | 74451853 | cg13336112 |
| chr3 | 76076212 | 76076213 | cg20197622 |

| | | | |
|---|---|---|---|
| chr4 | 40623291 | 40623292 | cg09141635 |
| chr4 | 40628704 | 40628705 | cg10459087 |
| chr4 | 40632788 | 40632789 | cg10019467 |
| chr4 | 40633269 | 40633270 | cg21151541 |
| chr4 | 40675171 | 40675172 | cg09126279 |
| chr4 | 41262458 | 41262459 | cg09270434 |
| chr4 | 47423779 | 47423780 | cg19935756 |
| chr4 | 47840095 | 47840096 | cg02553486 |
| chr4 | 48501509 | 48501510 | cg07193667 |
| chr4 | 48777471 | 48777472 | cg19212779 |
| chr4 | 53523482 | 53523483 | cg10098221 |
| chr4 | 56411138 | 56411139 | cg04264638 |
| chr4 | 71506651 | 71506652 | cg05147262 |
| chr4 | 71511557 | 71511558 | cg01088658 |
| chr4 | 76640579 | 76640580 | cg08528970 |
| chr4 | 76772075 | 76772076 | cg19499591 |
| chr4 | 77131705 | 77131706 | cg13870520 |
| chr4 | 78098654 | 78098655 | cg07196992 |
| chr4 | 80991289 | 80991290 | cg27529018 |
| chr4 | 81407975 | 81407976 | cg15870832 |
| chr4 | 86673958 | 86673959 | cg17425014 |
| chr4 | 86874205 | 86874206 | cg25310097 |
| chr4 | 87849250 | 87849251 | cg02262553 |
| chr4 | 87852752 | 87852753 | cg11634531 |
| chr4 | 87886892 | 87886893 | cg14209344 |
| chr4 | 88008667 | 88008668 | cg21988461 |
| chr4 | 90172038 | 90172039 | cg10497345 |
| chr4 | 95429383 | 95429384 | cg07236912 |
| chr4 | 99186446 | 99186447 | cg18936471 |
| chr4 | 99402843 | 99402844 | cg22404125 |
| chr4 | 99450728 | 99450729 | cg02015219 |
| chr4 | 99583945 | 99583946 | cg02304751 |
| chr4 | 100996832 | 100996833 | cg01354411 |
| chr4 | 101110248 | 101110249 | cg00360769 |
| chr4 | 101904283 | 101904284 | cg01949353 |
| chr4 | 102097226 | 102097227 | cg09444818 |
| chr4 | 102227740 | 102227741 | cg05571448 |
| chr4 | 103686232 | 103686233 | cg04605316 |
| chr4 | 103998497 | 103998498 | cg14459011 |
| chr4 | 105513274 | 105513275 | cg08824227 |
| chr4 | 108637177 | 108637178 | cg09720012 |
| chr4 | 109030040 | 109030041 | cg19296305 |
| chr4 | 109087312 | 109087313 | cg24023289 |
| chr4 | 109087419 | 109087420 | cg08460812 |
| chr4 | 113329985 | 113329986 | cg02266725 |
| chr4 | 114367529 | 114367530 | cg10154973 |
| chr4 | 114375605 | 114375606 | cg23670529 |
| chr4 | 114561619 | 114561620 | cg05257260 |
| chr4 | 114621848 | 114621849 | cg06633243 |
| chr4 | 117520432 | 117520433 | cg15256972 |
| chr4 | 122137696 | 122137697 | cg05559643 |
| chr4 | 122184241 | 122184242 | cg09730689 |
| chr4 | 124225943 | 124225944 | cg05805445 |
| chr4 | 137199590 | 137199591 | cg25277462 |
| chr4 | 139107328 | 139107329 | cg21877274 |
| chr4 | 139110589 | 139110590 | cg15441006 |

| | | | |
|---|---|---|---|
| chr3 | 78917133 | 78917134 | cg00906420 |
| chr3 | 79028263 | 79028264 | cg08147563 |
| chr3 | 86224327 | 86224328 | cg15484033 |
| chr3 | 96844879 | 96844880 | cg01556799 |
| chr3 | 97151261 | 97151262 | cg19963231 |
| chr3 | 99463106 | 99463107 | cg17918158 |
| chr3 | 99566949 | 99566950 | cg21589858 |
| chr3 | 99591722 | 99591723 | cg08350549 |
| chr3 | 99742090 | 99742091 | cg00718336 |
| chr3 | 100213794 | 100213795 | cg12137665 |
| chr3 | 101284166 | 101284167 | cg12422199 |
| chr3 | 101567360 | 101567361 | cg01771416 |
| chr3 | 101659630 | 101659631 | cg05429448 |
| chr3 | 101659687 | 101659688 | cg12963656 |
| chr3 | 101798142 | 101798143 | cg07772781 |
| chr3 | 107240645 | 107240646 | cg23518214 |
| chr3 | 107552866 | 107552867 | cg14029061 |
| chr3 | 107921136 | 107921137 | cg23169051 |
| chr3 | 108019540 | 108019541 | cg11326415 |
| chr3 | 108021106 | 108021107 | cg02059214 |
| chr3 | 108021214 | 108021215 | cg10431989 |
| chr3 | 108065259 | 108065260 | cg14703454 |
| chr3 | 108204398 | 108204399 | cg16467929 |
| chr3 | 110027072 | 110027073 | cg03517913 |
| chr3 | 110110880 | 110110881 | cg23471798 |
| chr3 | 110250679 | 110250680 | cg21977522 |
| chr3 | 110754331 | 110754332 | cg19855559 |
| chr3 | 111721343 | 111721344 | cg00373436 |
| chr3 | 111842345 | 111842346 | cg00714564 |
| chr3 | 112053517 | 112053518 | cg07681748 |
| chr3 | 112096801 | 112096802 | cg13671536 |
| chr3 | 112109797 | 112109798 | cg02919122 |
| chr3 | 112184271 | 112184272 | cg00477226 |
| chr3 | 112486072 | 112486073 | cg17086773 |
| chr3 | 112494836 | 112494837 | cg24431372 |
| chr3 | 112814660 | 112814661 | cg07034145 |
| chr3 | 114012659 | 114012660 | cg19421218 |
| chr3 | 114012806 | 114012807 | cg19456938 |
| chr3 | 114012833 | 114012834 | cg13669740 |
| chr3 | 114012912 | 114012913 | cg20832020 |
| chr3 | 114052269 | 114052270 | cg02556155 |
| chr3 | 114057297 | 114057298 | cg21179654 |
| chr3 | 114238429 | 114238430 | cg13666648 |
| chr3 | 114478256 | 114478257 | cg19981243 |
| chr3 | 115378386 | 115378387 | cg09463650 |
| chr3 | 115524031 | 115524032 | cg23075139 |
| chr3 | 116428610 | 116428611 | cg09237790 |
| chr3 | 116432221 | 116432222 | cg14476095 |
| chr3 | 117075024 | 117075025 | cg12837919 |
| chr3 | 117158482 | 117158483 | cg12699321 |
| chr3 | 117516743 | 117516744 | cg00205813 |
| chr3 | 117520653 | 117520654 | cg16634544 |
| chr3 | 117999270 | 117999271 | cg23717441 |
| chr3 | 119316673 | 119316674 | cg06073417 |
| chr3 | 119361670 | 119361671 | cg13008492 |
| chr3 | 119963930 | 119963931 | cg08414647 |

| | | | |
|---|---|---|---|
| chr4 | 140003444 | 140003445 | cg07002832 |
| chr4 | 140076137 | 140076138 | cg22027435 |
| chr4 | 140245762 | 140245763 | cg14286594 |
| chr4 | 144104893 | 144104894 | cg01719995 |
| chr4 | 144275126 | 144275127 | cg12710519 |
| chr4 | 144300134 | 144300135 | cg14142817 |
| chr4 | 146446271 | 146446272 | cg02884773 |
| chr4 | 146467791 | 146467792 | cg08618463 |
| chr4 | 146807291 | 146807292 | cg14384960 |
| chr4 | 146861608 | 146861609 | cg03279633 |
| chr4 | 150999042 | 150999043 | cg05561193 |
| chr4 | 151001971 | 151001972 | cg03779973 |
| chr4 | 151038391 | 151038392 | cg12115081 |
| chr4 | 151287731 | 151287732 | cg12035325 |
| chr4 | 153611560 | 153611561 | cg17385088 |
| chr4 | 153699460 | 153699461 | cg04676766 |
| chr4 | 154005381 | 154005382 | cg06270779 |
| chr4 | 154144800 | 154144801 | cg15410675 |
| chr4 | 154269418 | 154269419 | cg12614529 |
| chr4 | 154386136 | 154386137 | cg21908828 |
| chr4 | 160514445 | 160514446 | cg18127012 |
| chr4 | 162298751 | 162298752 | cg12225651 |
| chr4 | 164440990 | 164440991 | cg15265116 |
| chr4 | 165628435 | 165628436 | cg03466428 |
| chr4 | 169242366 | 169242367 | cg17384323 |
| chr4 | 170187451 | 170187452 | cg18949666 |
| chr4 | 176892927 | 176892928 | cg10523370 |
| chr4 | 178364890 | 178364891 | cg07155684 |
| chr4 | 185238627 | 185238628 | cg12654155 |
| chr4 | 185309524 | 185309525 | cg21533544 |
| chr4 | 185345707 | 185345708 | cg11802666 |
| chr4 | 185362534 | 185362535 | cg02354388 |
| chr4 | 185468777 | 185468778 | cg22905866 |
| chr4 | 185684040 | 185684041 | cg03977443 |
| chr4 | 185726782 | 185726783 | cg11168687 |
| chr4 | 186123008 | 186123009 | cg02148270 |
| chr4 | 186253454 | 186253455 | cg20461310 |
| chr5 | 471171 | 471172 | cg10902033 |
| chr5 | 1414806 | 1414807 | cg16614020 |
| chr5 | 10708717 | 10708718 | cg16362140 |
| chr5 | 14283950 | 14283951 | cg05649906 |
| chr5 | 14286263 | 14286264 | cg21531038 |
| chr5 | 14342307 | 14342308 | cg07611121 |
| chr5 | 14707001 | 14707002 | cg07573020 |
| chr5 | 14870083 | 14870084 | cg11792786 |
| chr5 | 17488879 | 17488880 | cg16487297 |
| chr5 | 34535730 | 34535731 | cg18833994 |
| chr5 | 36302349 | 36302350 | cg20384132 |
| chr5 | 36686601 | 36686602 | cg17839232 |
| chr5 | 36744026 | 36744027 | cg21558053 |
| chr5 | 39203755 | 39203756 | cg07669182 |
| chr5 | 40227327 | 40227328 | cg21879236 |
| chr5 | 40693569 | 40693570 | cg00988227 |
| chr5 | 41748833 | 41748834 | cg24851100 |
| chr5 | 42991657 | 42991658 | cg10615765 |
| chr5 | 43017351 | 43017352 | cg27628536 |

| | | | |
|---|---|---|---|
| chr3 | 121312127 | 121312128 | cg12456825 |
| chr3 | 121312241 | 121312242 | cg02845063 |
| chr3 | 121953249 | 121953250 | cg09326204 |
| chr3 | 122281939 | 122281940 | cg08122652 |
| chr3 | 122296027 | 122296028 | cg22503489 |
| chr3 | 122296029 | 122296030 | cg09384183 |
| chr3 | 122514814 | 122514815 | cg10627428 |
| chr3 | 122830175 | 122830176 | cg27383859 |
| chr3 | 123950018 | 123950019 | cg21618455 |
| chr3 | 123987591 | 123987592 | cg04816013 |
| chr3 | 124232190 | 124232191 | cg06690085 |
| chr3 | 124284219 | 124284220 | cg06218523 |
| chr3 | 124303035 | 124303036 | cg14688451 |
| chr3 | 124505038 | 124505039 | cg18634175 |
| chr3 | 124719009 | 124719010 | cg18008247 |
| chr3 | 124837734 | 124837735 | cg06713373 |
| chr3 | 124866646 | 124866647 | cg00316080 |
| chr3 | 124873768 | 124873769 | cg17478992 |
| chr3 | 124873848 | 124873849 | cg01432055 |
| chr3 | 125235415 | 125235416 | cg17252780 |
| chr3 | 126074344 | 126074345 | cg00540067 |
| chr3 | 126191181 | 126191182 | cg01357892 |
| chr3 | 126671985 | 126671986 | cg24673769 |
| chr3 | 127348828 | 127348829 | cg02782895 |
| chr3 | 127790462 | 127790463 | cg01693830 |
| chr3 | 127831673 | 127831674 | cg26861995 |
| chr3 | 128079636 | 128079637 | cg27292264 |
| chr3 | 128226565 | 128226566 | cg18453551 |
| chr3 | 128420513 | 128420514 | cg15607142 |
| chr3 | 128444572 | 128444573 | cg27297137 |
| chr3 | 128501650 | 128501651 | cg07156336 |
| chr3 | 128771903 | 128771904 | cg06285307 |
| chr3 | 129349316 | 129349317 | cg09463220 |
| chr3 | 129420294 | 129420295 | cg26175815 |
| chr3 | 129513427 | 129513428 | cg04208434 |
| chr3 | 130232371 | 130232372 | cg25663326 |
| chr3 | 130300740 | 130300741 | cg13067649 |
| chr3 | 131430364 | 131430365 | cg18512963 |
| chr3 | 132261993 | 132261994 | cg19770281 |
| chr3 | 133193741 | 133193742 | cg11642298 |
| chr3 | 133394743 | 133394744 | cg00828377 |
| chr3 | 133464047 | 133464048 | cg02621528 |
| chr3 | 133777381 | 133777382 | cg16782084 |
| chr3 | 134371285 | 134371286 | cg00310463 |
| chr3 | 135688971 | 135688972 | cg11162839 |
| chr3 | 137301727 | 137301728 | cg16338011 |
| chr3 | 137321245 | 137321246 | cg20566970 |
| chr3 | 137844436 | 137844437 | cg09866565 |
| chr3 | 137938348 | 137938349 | cg04554122 |
| chr3 | 138101415 | 138101416 | cg11209289 |
| chr3 | 138581702 | 138581703 | cg21742790 |
| chr3 | 139400319 | 139400320 | cg08526209 |
| chr3 | 141118615 | 141118616 | cg27084654 |
| chr3 | 141119262 | 141119263 | cg01879723 |
| chr3 | 141119541 | 141119542 | cg14500070 |
| chr3 | 141122632 | 141122633 | cg20959189 |
| chr5 | 52306165 | 52306166 | cg26722342 |
| chr5 | 53411773 | 53411774 | cg03978682 |
| chr5 | 53502584 | 53502585 | cg26814089 |
| chr5 | 53765871 | 53765872 | cg01606885 |
| chr5 | 53814625 | 53814626 | cg08544436 |
| chr5 | 53814891 | 53814892 | cg21995347 |
| chr5 | 53815303 | 53815304 | cg16358034 |
| chr5 | 53817073 | 53817074 | cg23001370 |
| chr5 | 53817963 | 53817964 | cg01582396 |
| chr5 | 53821365 | 53821366 | cg14952488 |
| chr5 | 53829957 | 53829958 | cg14684434 |
| chr5 | 53966914 | 53966915 | cg19698472 |
| chr5 | 54089309 | 54089310 | cg25133422 |
| chr5 | 54128677 | 54128678 | cg03861379 |
| chr5 | 54140068 | 54140069 | cg19481513 |
| chr5 | 55645530 | 55645531 | cg06651783 |
| chr5 | 55761274 | 55761275 | cg23506143 |
| chr5 | 55772865 | 55772866 | cg09078527 |
| chr5 | 55977468 | 55977469 | cg01164141 |
| chr5 | 56107058 | 56107059 | cg10597362 |
| chr5 | 56128454 | 56128455 | cg06478987 |
| chr5 | 56133370 | 56133371 | cg04452534 |
| chr5 | 58479549 | 58479550 | cg07348982 |
| chr5 | 58824741 | 58824742 | cg06974135 |
| chr5 | 59783035 | 59783036 | cg21023952 |
| chr5 | 60631028 | 60631029 | cg14742809 |
| chr5 | 64334489 | 64334490 | cg11190987 |
| chr5 | 64399755 | 64399756 | cg21123686 |
| chr5 | 71613149 | 71613150 | cg26293423 |
| chr5 | 71895997 | 71895998 | cg01941278 |
| chr5 | 75903700 | 75903701 | cg08604223 |
| chr5 | 75938513 | 75938514 | cg26072352 |
| chr5 | 76654783 | 76654784 | cg12559197 |
| chr5 | 77255129 | 77255130 | cg27629992 |
| chr5 | 77949077 | 77949078 | cg26964651 |
| chr5 | 78758952 | 78758953 | cg14468634 |
| chr5 | 78813058 | 78813059 | cg02125365 |
| chr5 | 79489249 | 79489250 | cg10531355 |
| chr5 | 79490511 | 79490512 | cg06677021 |
| chr5 | 79501131 | 79501132 | cg14978242 |
| chr5 | 80542999 | 80543000 | cg07727547 |
| chr5 | 89823130 | 89823131 | cg01310768 |
| chr5 | 90459841 | 90459842 | cg07706111 |
| chr5 | 95514729 | 95514730 | cg00283187 |
| chr5 | 96078678 | 96078679 | cg24982541 |
| chr5 | 96298493 | 96298494 | cg22566142 |
| chr5 | 102364835 | 102364836 | cg17259761 |
| chr5 | 112022460 | 112022461 | cg07710833 |
| chr5 | 112047824 | 112047825 | cg23497707 |
| chr5 | 112074043 | 112074044 | cg01240931 |
| chr5 | 115178810 | 115178811 | cg09523691 |
| chr5 | 115181403 | 115181404 | cg09504737 |
| chr5 | 115194332 | 115194333 | cg24770230 |
| chr5 | 115696151 | 115696152 | cg01419670 |
| chr5 | 118608672 | 118608673 | cg26305729 |
| chr5 | 118658343 | 118658344 | cg27621513 |

| | | | |
|---|---|---|---|
| chr3 | 141122887 | 141122888 | cg27242726 |
| chr3 | 141132146 | 141132147 | cg09025210 |
| chr3 | 141514782 | 141514783 | cg00686926 |
| chr3 | 144899243 | 144899244 | cg17052041 |
| chr3 | 145734655 | 145734656 | cg13425146 |
| chr3 | 148916713 | 148916714 | cg24876069 |
| chr3 | 149192377 | 149192378 | cg13688966 |
| chr3 | 149238478 | 149238479 | cg02013148 |
| chr3 | 149327651 | 149327652 | cg19483007 |
| chr3 | 149797095 | 149797096 | cg10729196 |
| chr3 | 149825195 | 149825196 | cg05684266 |
| chr3 | 150456069 | 150456070 | cg23214895 |
| chr3 | 150477925 | 150477926 | cg13590228 |
| chr3 | 150500902 | 150500903 | cg15711288 |
| chr3 | 150644434 | 150644435 | cg08653369 |
| chr3 | 150663357 | 150663358 | cg10625737 |
| chr3 | 150692080 | 150692081 | cg27322397 |
| chr3 | 150948274 | 150948275 | cg25117092 |
| chr3 | 150997688 | 150997689 | cg09410045 |
| chr3 | 151048406 | 151048407 | cg00091349 |
| chr3 | 151177940 | 151177941 | cg25499537 |
| chr3 | 151619923 | 151619924 | cg20790648 |
| chr3 | 152017240 | 152017241 | cg13702222 |
| chr3 | 152121157 | 152121158 | cg25342059 |
| chr3 | 152557119 | 152557120 | cg07546334 |
| chr3 | 153202275 | 153202276 | cg00731062 |
| chr3 | 154741095 | 154741096 | cg13233545 |
| chr3 | 154848194 | 154848195 | cg27235955 |
| chr3 | 155031033 | 155031034 | cg17630138 |
| chr3 | 155422971 | 155422972 | cg02344477 |
| chr3 | 155431380 | 155431381 | cg20943723 |
| chr3 | 156332967 | 156332968 | cg08237220 |
| chr3 | 156807520 | 156807521 | cg26331343 |
| chr3 | 156873744 | 156873745 | cg19824894 |
| chr3 | 159733028 | 159733029 | cg19465002 |
| chr3 | 160544991 | 160544992 | cg16233724 |
| chr3 | 161214690 | 161214691 | cg01015123 |
| chr3 | 162210236 | 162210237 | cg25713736 |
| chr3 | 167293814 | 167293815 | cg18142906 |
| chr3 | 168268421 | 168268422 | cg21490751 |
| chr3 | 168891416 | 168891417 | cg03804083 |
| chr3 | 168967238 | 168967239 | cg18060030 |
| chr3 | 169247421 | 169247422 | cg27502580 |
| chr3 | 169351419 | 169351420 | cg04414509 |
| chr3 | 169656700 | 169656701 | cg25345722 |
| chr3 | 169687379 | 169687380 | cg10936995 |
| chr3 | 169975155 | 169975156 | cg21140290 |
| chr3 | 170073454 | 170073455 | cg20085953 |
| chr3 | 170332444 | 170332445 | cg25962699 |
| chr3 | 170484840 | 170484841 | cg10799327 |
| chr3 | 170975382 | 170975383 | cg18052528 |
| chr3 | 171088451 | 171088452 | cg14865094 |
| chr3 | 171241974 | 171241975 | cg21086551 |
| chr3 | 172635656 | 172635657 | cg04461197 |
| chr3 | 173119212 | 173119213 | cg10623221 |
| chr3 | 173992754 | 173992755 | cg17482360 |

| | | | |
|---|---|---|---|
| chr5 | 118683890 | 118683891 | cg01057573 |
| chr5 | 123775157 | 123775158 | cg04226232 |
| chr5 | 124077966 | 124077967 | cg06055229 |
| chr5 | 124126863 | 124126864 | cg01876548 |
| chr5 | 124536183 | 124536184 | cg05599160 |
| chr5 | 125800778 | 125800779 | cg13049992 |
| chr5 | 125800797 | 125800798 | cg09168797 |
| chr5 | 130596889 | 130596890 | cg08861957 |
| chr5 | 131092116 | 131092117 | cg24629111 |
| chr5 | 131596602 | 131596603 | cg06947286 |
| chr5 | 131723239 | 131723240 | cg19040266 |
| chr5 | 131763756 | 131763757 | cg13948857 |
| chr5 | 131827918 | 131827919 | cg00255919 |
| chr5 | 131829935 | 131829936 | cg24749385 |
| chr5 | 131992430 | 131992431 | cg04303330 |
| chr5 | 132009630 | 132009631 | cg12377972 |
| chr5 | 132010740 | 132010741 | cg18145937 |
| chr5 | 132155450 | 132155451 | cg11322797 |
| chr5 | 132577108 | 132577109 | cg12828656 |
| chr5 | 133446318 | 133446319 | cg09701062 |
| chr5 | 133510077 | 133510078 | cg19957803 |
| chr5 | 133710015 | 133710016 | cg14711145 |
| chr5 | 133775258 | 133775259 | cg09101941 |
| chr5 | 133866097 | 133866098 | cg08295111 |
| chr5 | 134294026 | 134294027 | cg16525762 |
| chr5 | 134780343 | 134780344 | cg24708145 |
| chr5 | 135290288 | 135290289 | cg21783004 |
| chr5 | 135361707 | 135361708 | cg18665526 |
| chr5 | 135365401 | 135365402 | cg21583694 |
| chr5 | 135365890 | 135365891 | cg09096234 |
| chr5 | 135367668 | 135367669 | cg05838308 |
| chr5 | 137071788 | 137071789 | cg08748913 |
| chr5 | 137609681 | 137609682 | cg26721890 |
| chr5 | 137797039 | 137797040 | cg22509422 |
| chr5 | 138088442 | 138088443 | cg02276665 |
| chr5 | 138311077 | 138311078 | cg05462339 |
| chr5 | 138495127 | 138495128 | cg07536018 |
| chr5 | 138605898 | 138605899 | cg10795552 |
| chr5 | 138704493 | 138704494 | cg17263916 |
| chr5 | 138720490 | 138720491 | cg18115658 |
| chr5 | 139039026 | 139039027 | cg01008405 |
| chr5 | 139045301 | 139045302 | cg13707793 |
| chr5 | 139056794 | 139056795 | cg23573900 |
| chr5 | 139064216 | 139064217 | cg07879474 |
| chr5 | 139136991 | 139136992 | cg18695746 |
| chr5 | 139142898 | 139142899 | cg00271204 |
| chr5 | 139726689 | 139726690 | cg00521255 |
| chr5 | 141295517 | 141295518 | cg26513854 |
| chr5 | 141338604 | 141338605 | cg12845808 |
| chr5 | 141338826 | 141338827 | cg18849725 |
| chr5 | 141391533 | 141391534 | cg25940447 |
| chr5 | 141477782 | 141477783 | cg08928100 |
| chr5 | 142431272 | 142431273 | cg27579771 |
| chr5 | 142432538 | 142432539 | cg10670376 |
| chr5 | 142499847 | 142499848 | cg07618462 |
| chr5 | 149157216 | 149157217 | cg09814231 |

| | | | |
|---|---|---|---|
| chr3 | 175139160 | 175139161 | cg01150105 |
| chr3 | 176295392 | 176295393 | cg00075103 |
| chr3 | 176850227 | 176850228 | cg13273540 |
| chr3 | 176904010 | 176904011 | cg13072186 |
| chr3 | 176916496 | 176916497 | cg01132484 |
| chr3 | 177308016 | 177308017 | cg00840990 |
| chr3 | 179324813 | 179324814 | cg07583756 |
| chr3 | 181577820 | 181577821 | cg07753755 |
| chr3 | 182840603 | 182840604 | cg04119852 |
| chr3 | 183245952 | 183245953 | cg03805566 |
| chr3 | 183533873 | 183533874 | cg18489755 |
| chr3 | 184034132 | 184034133 | cg08681689 |
| chr3 | 185270308 | 185270309 | cg08099797 |
| chr3 | 185928164 | 185928165 | cg10498502 |
| chr3 | 186080868 | 186080869 | cg00434119 |
| chr3 | 186194046 | 186194047 | cg02346442 |
| chr3 | 186682634 | 186682635 | cg12632309 |
| chr3 | 186739029 | 186739030 | cg23965109 |
| chr3 | 187635183 | 187635184 | cg10168494 |
| chr3 | 187929592 | 187929593 | cg00869941 |
| chr3 | 187959138 | 187959139 | cg15417654 |
| chr3 | 188002729 | 188002730 | cg16464007 |
| chr3 | 188034983 | 188034984 | cg04535966 |
| chr3 | 188506715 | 188506716 | cg00902153 |
| chr3 | 188664928 | 188664929 | cg27409802 |
| chr3 | 188868788 | 188868789 | cg04474190 |
| chr3 | 190024765 | 190024766 | cg10475153 |
| chr3 | 191046380 | 191046381 | cg25342125 |
| chr3 | 191094694 | 191094695 | cg25939630 |
| chr3 | 192576964 | 192576965 | cg10755723 |
| chr3 | 193096626 | 193096627 | cg06318062 |
| chr3 | 193848133 | 193848134 | cg14643001 |
| chr3 | 193859889 | 193859890 | cg22176709 |
| chr3 | 194403234 | 194403235 | cg20668765 |
| chr3 | 194696866 | 194696867 | cg04252203 |
| chr3 | 194786218 | 194786219 | cg10348826 |
| chr3 | 194786989 | 194786990 | cg19775763 |
| chr3 | 194815259 | 194815260 | cg12085101 |
| chr3 | 194815339 | 194815340 | cg27134832 |
| chr3 | 194815434 | 194815435 | cg02504211 |
| chr3 | 194868843 | 194868844 | cg19760965 |
| chr3 | 194893148 | 194893149 | cg21501175 |
| chr3 | 194979565 | 194979566 | cg05486924 |
| chr3 | 195167302 | 195167303 | cg16834320 |
| chr3 | 195267087 | 195267088 | cg27230534 |
| chr3 | 195436154 | 195436155 | cg27349302 |
| chr3 | 195620147 | 195620148 | cg17640485 |
| chr3 | 195623792 | 195623793 | cg23548201 |
| chr3 | 195868640 | 195868641 | cg13403369 |
| chr3 | 195897833 | 195897834 | cg09393619 |
| chr3 | 195897904 | 195897905 | cg06305891 |
| chr3 | 195897912 | 195897913 | cg10959668 |
| chr3 | 195920019 | 195920020 | cg03891268 |
| chr3 | 196358703 | 196358704 | cg17657502 |
| chr3 | 196363641 | 196363642 | cg04840813 |
| chr3 | 196388687 | 196388688 | cg07364240 |
| chr5 | 149867577 | 149867578 | cg11100481 |
| chr5 | 149867965 | 149867966 | cg12636435 |
| chr5 | 149868855 | 149868856 | cg07038243 |
| chr5 | 149870395 | 149870396 | cg15538767 |
| chr5 | 150002231 | 150002232 | cg06732545 |
| chr5 | 150003037 | 150003038 | cg00089453 |
| chr5 | 151148420 | 151148421 | cg00601432 |
| chr5 | 153787169 | 153787170 | cg20540306 |
| chr5 | 155822151 | 155822152 | cg17739626 |
| chr5 | 156083022 | 156083023 | cg06882854 |
| chr5 | 156696185 | 156696186 | cg23462242 |
| chr5 | 156696189 | 156696190 | cg08224785 |
| chr5 | 156722426 | 156722427 | cg05195318 |
| chr5 | 156944020 | 156944021 | cg07257747 |
| chr5 | 157003983 | 157003984 | cg23214435 |
| chr5 | 157340439 | 157340440 | cg20462493 |
| chr5 | 158602264 | 158602265 | cg19479086 |
| chr5 | 158937447 | 158937448 | cg07289335 |
| chr5 | 159519152 | 159519153 | cg19535980 |
| chr5 | 162936453 | 162936454 | cg22017733 |
| chr5 | 169725927 | 169725928 | cg02676052 |
| chr5 | 171234429 | 171234430 | cg15717448 |
| chr5 | 171384415 | 171384416 | cg25959472 |
| chr5 | 171410890 | 171410891 | cg23985408 |
| chr5 | 172216846 | 172216847 | cg27652869 |
| chr5 | 172305986 | 172305987 | cg24699021 |
| chr5 | 172306099 | 172306100 | cg16119483 |
| chr5 | 172306112 | 172306113 | cg16345114 |
| chr5 | 172306136 | 172306137 | cg21204530 |
| chr5 | 172430563 | 172430564 | cg22907189 |
| chr5 | 172447063 | 172447064 | cg09199609 |
| chr5 | 172447213 | 172447214 | cg19870567 |
| chr5 | 173068163 | 173068164 | cg19945535 |
| chr5 | 173171737 | 173171738 | cg04483863 |
| chr5 | 173251865 | 173251866 | cg11119275 |
| chr5 | 173317332 | 173317333 | cg18757087 |
| chr5 | 173317342 | 173317343 | cg06889108 |
| chr5 | 173318755 | 173318756 | cg12665414 |
| chr5 | 173342997 | 173342998 | cg16290431 |
| chr5 | 173351052 | 173351053 | cg18693985 |
| chr5 | 175799756 | 175799757 | cg10845391 |
| chr5 | 175936021 | 175936022 | cg27349687 |
| chr5 | 175955925 | 175955926 | cg20748242 |
| chr5 | 176080248 | 176080249 | cg07347294 |
| chr5 | 176861823 | 176861824 | cg02523893 |
| chr5 | 176862098 | 176862099 | cg22017187 |
| chr5 | 176917390 | 176917391 | cg17037976 |
| chr5 | 176930735 | 176930736 | cg13370819 |
| chr5 | 176930754 | 176930755 | cg18649283 |
| chr5 | 176938522 | 176938523 | cg07265015 |
| chr5 | 177036766 | 177036767 | cg18036734 |
| chr5 | 179181715 | 179181716 | cg01971149 |
| chr5 | 179218853 | 179218854 | cg17614703 |
| chr5 | 179230002 | 179230003 | cg23011886 |
| chr5 | 179232533 | 179232534 | cg12619504 |
| chr5 | 180244832 | 180244833 | cg12950829 |

| | | | |
|---|---|---|---|
| chr3 | 196594273 | 196594274 | cg10772991 |
| chr3 | 197382627 | 197382628 | cg17473580 |
| chr3 | 197383170 | 197383171 | cg10631854 |
| chr3 | 197639960 | 197639961 | cg11938367 |
| chr4 | 671791 | 671792 | cg17073306 |
| chr4 | 671851 | 671852 | cg23595927 |
| chr4 | 683772 | 683773 | cg25510614 |
| chr4 | 732749 | 732750 | cg13917589 |
| chr4 | 903575 | 903576 | cg24620617 |
| chr4 | 909785 | 909786 | cg20085268 |
| chr4 | 979307 | 979308 | cg09153458 |
| chr4 | 1148456 | 1148457 | cg24262654 |
| chr4 | 1208960 | 1208961 | cg00917642 |
| chr4 | 1223170 | 1223171 | cg06166341 |
| chr4 | 1234993 | 1234994 | cg15877785 |
| chr4 | 1295078 | 1295079 | cg21167046 |
| chr4 | 1656414 | 1656415 | cg11721566 |
| chr4 | 1724428 | 1724429 | cg21505886 |
| chr4 | 1724449 | 1724450 | cg02089348 |
| chr4 | 1754069 | 1754070 | cg08745334 |
| chr4 | 1853351 | 1853352 | cg06854772 |
| chr4 | 2262318 | 2262319 | cg03840289 |
| chr4 | 2839781 | 2839782 | cg23746574 |
| chr4 | 2944550 | 2944551 | cg24015522 |
| chr4 | 2944692 | 2944693 | cg18912103 |
| chr4 | 3390332 | 3390333 | cg15436123 |
| chr4 | 6623589 | 6623590 | cg16025611 |
| chr4 | 6857986 | 6857987 | cg16051718 |
| chr4 | 6918868 | 6918869 | cg19906672 |
| chr4 | 6965057 | 6965058 | cg06109284 |
| chr4 | 6993556 | 6993557 | cg19734190 |
| chr4 | 8359499 | 8359500 | cg09979924 |
| chr4 | 9371043 | 9371044 | cg27201767 |
| chr4 | 9558077 | 9558078 | cg08174890 |
| chr4 | 9892887 | 9892888 | cg02387843 |
| chr4 | 9990048 | 9990049 | cg13841979 |
| chr4 | 10686300 | 10686301 | cg03485217 |
| chr4 | 11369730 | 11369731 | cg06606539 |
| chr4 | 11398983 | 11398984 | cg17366225 |
| chr4 | 11471730 | 11471731 | cg11021317 |
| chr4 | 13977635 | 13977636 | cg16631326 |
| chr4 | 14362239 | 14362240 | cg05626927 |
| chr4 | 14746310 | 14746311 | cg04285362 |
| chr4 | 14918405 | 14918406 | cg13797402 |
| chr4 | 15200268 | 15200269 | cg06257371 |
| chr4 | 15374950 | 15374951 | cg06742778 |
| chr4 | 16708336 | 16708337 | cg25889581 |
| chr4 | 16846297 | 16846298 | cg23452220 |
| chr4 | 17310680 | 17310681 | cg14599380 |
| chr4 | 20574555 | 20574556 | cg08415391 |
| chr4 | 22476976 | 22476977 | cg05029312 |
| chr4 | 22913868 | 22913869 | cg02378063 |
| chr4 | 23776092 | 23776093 | cg06861395 |
| chr4 | 23791473 | 23791474 | cg09356916 |
| chr4 | 24915774 | 24915775 | cg02289038 |
| chr4 | 25845210 | 25845211 | cg09461545 |

| | | | |
|---|---|---|---|
| chr6 | 226092 | 226093 | cg20359147 |
| chr6 | 299762 | 299763 | cg26822870 |
| chr6 | 312058 | 312059 | cg18134411 |
| chr6 | 312105 | 312106 | cg11787218 |
| chr6 | 348772 | 348773 | cg05698911 |
| chr6 | 373609 | 373610 | cg18114053 |
| chr6 | 401311 | 401312 | cg11417701 |
| chr6 | 474500 | 474501 | cg17645074 |
| chr6 | 474629 | 474630 | cg11682724 |
| chr6 | 709725 | 709726 | cg02409090 |
| chr6 | 1464075 | 1464076 | cg10136452 |
| chr6 | 1663171 | 1663172 | cg23120243 |
| chr6 | 1994056 | 1994057 | cg09841915 |
| chr6 | 2375895 | 2375896 | cg05612654 |
| chr6 | 2768616 | 2768617 | cg09393448 |
| chr6 | 2874553 | 2874554 | cg01703162 |
| chr6 | 2944352 | 2944353 | cg13254586 |
| chr6 | 3722818 | 3722819 | cg05919456 |
| chr6 | 4134485 | 4134486 | cg11647493 |
| chr6 | 4909143 | 4909144 | cg17283752 |
| chr6 | 5764816 | 5764817 | cg14029759 |
| chr6 | 7113738 | 7113739 | cg09754197 |
| chr6 | 7140931 | 7140932 | cg26991433 |
| chr6 | 7242625 | 7242626 | cg22335826 |
| chr6 | 7907918 | 7907919 | cg22986947 |
| chr6 | 9711044 | 9711045 | cg26226306 |
| chr6 | 10520874 | 10520875 | cg14112601 |
| chr6 | 10521142 | 10521143 | cg24690479 |
| chr6 | 10521452 | 10521453 | cg22378252 |
| chr6 | 11233592 | 11233593 | cg26844213 |
| chr6 | 11810356 | 11810357 | cg05235344 |
| chr6 | 11972963 | 11972964 | cg12594290 |
| chr6 | 12050371 | 12050372 | cg21041472 |
| chr6 | 12279872 | 12279873 | cg17617182 |
| chr6 | 13248958 | 13248959 | cg03891302 |
| chr6 | 13372573 | 13372574 | cg24320816 |
| chr6 | 13463084 | 13463085 | cg19084822 |
| chr6 | 13466329 | 13466330 | cg10946816 |
| chr6 | 13469700 | 13469701 | cg11007195 |
| chr6 | 13483832 | 13483833 | cg00728848 |
| chr6 | 13767628 | 13767629 | cg00698916 |
| chr6 | 14211985 | 14211986 | cg27629753 |
| chr6 | 14321559 | 14321560 | cg01061480 |
| chr6 | 14369697 | 14369698 | cg20218040 |
| chr6 | 14720183 | 14720184 | cg05797097 |
| chr6 | 14729030 | 14729031 | cg22164009 |
| chr6 | 14729722 | 14729723 | cg24219974 |
| chr6 | 15090163 | 15090164 | cg26076724 |
| chr6 | 15304885 | 15304886 | cg05985988 |
| chr6 | 15358299 | 15358300 | cg16248435 |
| chr6 | 15385378 | 15385379 | cg21089584 |
| chr6 | 15428301 | 15428302 | cg26405097 |
| chr6 | 15551489 | 15551490 | cg12577411 |
| chr6 | 15575773 | 15575774 | cg02830023 |
| chr6 | 15616502 | 15616503 | cg11203401 |
| chr6 | 15691258 | 15691259 | cg18061140 |

| | | | |
|---|---|---|---|
| chr4 | 26032635 | 26032636 | cg18470101 |
| chr4 | 26143392 | 26143393 | cg19274680 |
| chr4 | 26177098 | 26177099 | cg10937802 |
| chr4 | 26362792 | 26362793 | cg13002809 |
| chr4 | 26789915 | 26789916 | cg19719391 |
| chr4 | 26861261 | 26861262 | cg13621396 |
| chr4 | 26865310 | 26865311 | cg27025079 |
| chr4 | 26995889 | 26995890 | cg08639389 |
| chr4 | 27004697 | 27004698 | cg19621160 |
| chr4 | 33151488 | 33151489 | cg14749573 |
| chr4 | 37059651 | 37059652 | cg23651495 |
| chr4 | 38667430 | 38667431 | cg22453435 |
| chr4 | 38673130 | 38673131 | cg05205842 |
| chr4 | 39641752 | 39641753 | cg08821747 |
| chr4 | 40193114 | 40193115 | cg26137103 |
| chr4 | 40517938 | 40517939 | cg15967525 |
| chr4 | 40518041 | 40518042 | cg00511464 |
| chr4 | 40518065 | 40518066 | cg14160419 |
| chr4 | 40518143 | 40518144 | cg25117091 |
| chr4 | 40518663 | 40518664 | cg06332621 |
| chr4 | 40519094 | 40519095 | cg18183961 |
| chr4 | 40704257 | 40704258 | cg22990388 |
| chr4 | 42296896 | 42296897 | cg04135540 |
| chr4 | 46874750 | 46874751 | cg21472835 |
| chr4 | 47841314 | 47841315 | cg02955940 |
| chr4 | 48272513 | 48272514 | cg07026107 |
| chr4 | 48779921 | 48779922 | cg02886077 |
| chr4 | 53968201 | 53968202 | cg21253943 |
| chr4 | 54518744 | 54518745 | cg12114584 |
| chr4 | 56504657 | 56504658 | cg04528240 |
| chr4 | 57485847 | 57485848 | cg22911184 |
| chr4 | 57937220 | 57937221 | cg23287435 |
| chr4 | 62111221 | 62111222 | cg16368147 |
| chr4 | 62606570 | 62606571 | cg14974938 |
| chr4 | 67950111 | 67950112 | cg05825121 |
| chr4 | 69829409 | 69829410 | cg19045700 |
| chr4 | 71262957 | 71262958 | cg06585690 |
| chr4 | 71575507 | 71575508 | cg09215582 |
| chr4 | 71587640 | 71587641 | cg03173722 |
| chr4 | 72996777 | 72996778 | cg24923953 |
| chr4 | 73437692 | 73437693 | cg11202023 |
| chr4 | 74921425 | 74921426 | cg12684958 |
| chr4 | 75366983 | 75366984 | cg11521212 |
| chr4 | 75722231 | 75722232 | cg14231845 |
| chr4 | 76554724 | 76554725 | cg10131286 |
| chr4 | 76596784 | 76596785 | cg03862987 |
| chr4 | 77011716 | 77011717 | cg22301128 |
| chr4 | 77119206 | 77119207 | cg01624414 |
| chr4 | 77815872 | 77815873 | cg04028481 |
| chr4 | 77994145 | 77994146 | cg17080371 |
| chr4 | 79338837 | 79338838 | cg02102872 |
| chr4 | 79693986 | 79693987 | cg01333327 |
| chr4 | 81047158 | 81047159 | cg10713839 |
| chr4 | 81103968 | 81103969 | cg05007549 |
| chr4 | 82127274 | 82127275 | cg10060562 |
| chr4 | 83675306 | 83675307 | cg09363604 |
| chr6 | 16323285 | 16323286 | cg00151370 |
| chr6 | 16699028 | 16699029 | cg14791639 |
| chr6 | 16742123 | 16742124 | cg10262747 |
| chr6 | 18215340 | 18215341 | cg02903589 |
| chr6 | 20118318 | 20118319 | cg07057444 |
| chr6 | 24499591 | 24499592 | cg00346970 |
| chr6 | 24859863 | 24859864 | cg16313640 |
| chr6 | 24877462 | 24877463 | cg12213680 |
| chr6 | 25088999 | 25089000 | cg12449049 |
| chr6 | 25297487 | 25297488 | cg17031967 |
| chr6 | 25369563 | 25369564 | cg11784243 |
| chr6 | 25407628 | 25407629 | cg13089568 |
| chr6 | 26235699 | 26235700 | cg05554718 |
| chr6 | 26276143 | 26276144 | cg08578491 |
| chr6 | 26520793 | 26520794 | cg27490387 |
| chr6 | 26602900 | 26602901 | cg09364178 |
| chr6 | 28226885 | 28226886 | cg21251018 |
| chr6 | 28226905 | 28226906 | cg00765128 |
| chr6 | 29602034 | 29602035 | cg04324598 |
| chr6 | 29633673 | 29633674 | cg18488651 |
| chr6 | 29717656 | 29717657 | cg17514431 |
| chr6 | 29893168 | 29893169 | cg09470907 |
| chr6 | 29913343 | 29913344 | cg19585676 |
| chr6 | 29933511 | 29933512 | cg04007531 |
| chr6 | 29987762 | 29987763 | cg15874113 |
| chr6 | 30169768 | 30169769 | cg08850243 |
| chr6 | 30175299 | 30175300 | cg07523043 |
| chr6 | 30175792 | 30175793 | cg12663302 |
| chr6 | 30176076 | 30176077 | cg19593182 |
| chr6 | 30291314 | 30291315 | cg13818708 |
| chr6 | 30291349 | 30291350 | cg18286080 |
| chr6 | 30454518 | 30454519 | cg25393388 |
| chr6 | 30581820 | 30581821 | cg13522406 |
| chr6 | 30582333 | 30582334 | cg23583454 |
| chr6 | 30594037 | 30594038 | cg01245721 |
| chr6 | 30619137 | 30619138 | cg18184941 |
| chr6 | 30619173 | 30619174 | cg01152616 |
| chr6 | 30619190 | 30619191 | cg13584784 |
| chr6 | 30737436 | 30737437 | cg25063165 |
| chr6 | 30737455 | 30737456 | cg03813377 |
| chr6 | 30782080 | 30782081 | cg20802826 |
| chr6 | 30853569 | 30853570 | cg25607383 |
| chr6 | 31090929 | 31090930 | cg19456997 |
| chr6 | 31105872 | 31105873 | cg03034365 |
| chr6 | 31133910 | 31133911 | cg07764829 |
| chr6 | 31437927 | 31437928 | cg20757966 |
| chr6 | 31438205 | 31438206 | cg08859411 |
| chr6 | 31496949 | 31496950 | cg12259379 |
| chr6 | 31582285 | 31582286 | cg15343510 |
| chr6 | 31582837 | 31582838 | cg24211388 |
| chr6 | 31680228 | 31680229 | cg14258501 |
| chr6 | 31681882 | 31681883 | cg13123009 |
| chr6 | 31695027 | 31695028 | cg00175838 |
| chr6 | 31695061 | 31695062 | cg13318914 |
| chr6 | 31695064 | 31695065 | cg16515974 |
| chr6 | 31744391 | 31744392 | cg16672203 |

| | | | |
|---|---|---|---|
| chr4 | 85604328 | 85604329 | cg18033333 |
| chr4 | 86851225 | 86851226 | cg22017213 |
| chr4 | 86851411 | 86851412 | cg14376467 |
| chr4 | 86851425 | 86851426 | cg21446955 |
| chr4 | 87230737 | 87230738 | cg05181212 |
| chr4 | 87235054 | 87235055 | cg14155327 |
| chr4 | 87287889 | 87287890 | cg00864661 |
| chr4 | 87308061 | 87308062 | cg16737749 |
| chr4 | 88775480 | 88775481 | cg27060793 |
| chr4 | 88812922 | 88812923 | cg17127329 |
| chr4 | 88876673 | 88876674 | cg26678245 |
| chr4 | 89207307 | 89207308 | cg05850997 |
| chr4 | 89534723 | 89534724 | cg24750887 |
| chr4 | 89829015 | 89829016 | cg22073750 |
| chr4 | 89978566 | 89978567 | cg04536922 |
| chr4 | 90360897 | 90360898 | cg02054697 |
| chr4 | 91570600 | 91570601 | cg15687147 |
| chr4 | 91689972 | 91689973 | cg24055836 |
| chr4 | 92879355 | 92879356 | cg11947262 |
| chr4 | 94607780 | 94607781 | cg23985818 |
| chr4 | 95328682 | 95328683 | cg11291712 |
| chr4 | 96025493 | 96025494 | cg12007466 |
| chr4 | 96080181 | 96080182 | cg14866069 |
| chr4 | 96445830 | 96445831 | cg04281738 |
| chr4 | 97948716 | 97948717 | cg16652264 |
| chr4 | 99427937 | 99427938 | cg03298704 |
| chr4 | 99502614 | 99502615 | cg11693424 |
| chr4 | 99534176 | 99534177 | cg25841008 |
| chr4 | 99552952 | 99552953 | cg23246886 |
| chr4 | 99830489 | 99830490 | cg17031926 |
| chr4 | 101715708 | 101715709 | cg00353330 |
| chr4 | 102258631 | 102258632 | cg11820929 |
| chr4 | 103740913 | 103740914 | cg15289466 |
| chr4 | 104021583 | 104021584 | cg03276883 |
| chr4 | 108069982 | 108069983 | cg05132150 |
| chr4 | 108672936 | 108672937 | cg05885462 |
| chr4 | 108913443 | 108913444 | cg08746653 |
| chr4 | 110107068 | 110107069 | cg26442903 |
| chr4 | 110128090 | 110128091 | cg13279083 |
| chr4 | 110625080 | 110625081 | cg17714799 |
| chr4 | 111758936 | 111758937 | cg10142837 |
| chr4 | 113933301 | 113933302 | cg11474135 |
| chr4 | 114505779 | 114505780 | cg02379856 |
| chr4 | 115527443 | 115527444 | cg12931707 |
| chr4 | 115576003 | 115576004 | cg01991861 |
| chr4 | 117624749 | 117624750 | cg01238639 |
| chr4 | 119888794 | 119888795 | cg21614201 |
| chr4 | 120056829 | 120056830 | cg07598199 |
| chr4 | 121613795 | 121613796 | cg01963189 |
| chr4 | 124791563 | 124791564 | cg09839177 |
| chr4 | 124917187 | 124917188 | cg15624719 |
| chr4 | 128778193 | 128778194 | cg09542154 |
| chr4 | 128778381 | 128778382 | cg24501758 |
| chr4 | 129582864 | 129582865 | cg06567920 |
| chr4 | 129643491 | 129643492 | cg01266375 |
| chr4 | 137718563 | 137718564 | cg15713734 |

| | | | |
|---|---|---|---|
| chr6 | 31759806 | 31759807 | cg05797367 |
| chr6 | 31760047 | 31760048 | cg04959196 |
| chr6 | 31760233 | 31760234 | cg23781467 |
| chr6 | 31760345 | 31760346 | cg15864153 |
| chr6 | 31760796 | 31760797 | cg02597894 |
| chr6 | 31833072 | 31833073 | cg26434400 |
| chr6 | 31833104 | 31833105 | cg07705820 |
| chr6 | 31833291 | 31833292 | cg20498033 |
| chr6 | 31833386 | 31833387 | cg18263014 |
| chr6 | 31833452 | 31833453 | cg24732991 |
| chr6 | 31937731 | 31937732 | cg05828520 |
| chr6 | 32087190 | 32087191 | cg23782145 |
| chr6 | 32154715 | 32154716 | cg20765522 |
| chr6 | 32155141 | 32155142 | cg04363228 |
| chr6 | 32182002 | 32182003 | cg22155039 |
| chr6 | 32182174 | 32182175 | cg21462693 |
| chr6 | 32189610 | 32189611 | cg16809040 |
| chr6 | 32192024 | 32192025 | cg25317746 |
| chr6 | 32294260 | 32294261 | cg03282991 |
| chr6 | 32303790 | 32303791 | cg19988162 |
| chr6 | 32304143 | 32304144 | cg02808240 |
| chr6 | 32304276 | 32304277 | cg16564946 |
| chr6 | 32339208 | 32339209 | cg21195277 |
| chr6 | 32373620 | 32373621 | cg09514561 |
| chr6 | 32407289 | 32407290 | cg25764570 |
| chr6 | 32849783 | 32849784 | cg20666438 |
| chr6 | 32903598 | 32903599 | cg24523259 |
| chr6 | 32904317 | 32904318 | cg10411221 |
| chr6 | 32904621 | 32904622 | cg26964592 |
| chr6 | 32905012 | 32905013 | cg01026383 |
| chr6 | 32905114 | 32905115 | cg19268453 |
| chr6 | 32905127 | 32905128 | cg01374870 |
| chr6 | 32905190 | 32905191 | cg04903089 |
| chr6 | 32905320 | 32905321 | cg10453850 |
| chr6 | 32921791 | 32921792 | cg09713850 |
| chr6 | 32921805 | 32921806 | cg24290947 |
| chr6 | 32966587 | 32966588 | cg21066720 |
| chr6 | 32966622 | 32966623 | cg17945766 |
| chr6 | 32972093 | 32972094 | cg11527772 |
| chr6 | 32974843 | 32974844 | cg00540941 |
| chr6 | 32974897 | 32974898 | cg15737554 |
| chr6 | 32974906 | 32974907 | cg18043773 |
| chr6 | 32974924 | 32974925 | cg26690103 |
| chr6 | 33037690 | 33037691 | cg22037249 |
| chr6 | 33079692 | 33079693 | cg25438492 |
| chr6 | 33085565 | 33085566 | cg07735952 |
| chr6 | 33092802 | 33092803 | cg09578643 |
| chr6 | 33136859 | 33136860 | cg10397710 |
| chr6 | 33161438 | 33161439 | cg16634404 |
| chr6 | 33161589 | 33161590 | cg17560929 |
| chr6 | 33163760 | 33163761 | cg01359676 |
| chr6 | 33219335 | 33219336 | cg01101354 |
| chr6 | 33219392 | 33219393 | cg20595453 |
| chr6 | 33219410 | 33219411 | cg26987613 |
| chr6 | 33219654 | 33219655 | cg21571162 |
| chr6 | 33270914 | 33270915 | cg05105110 |

| | | | |
|---|---|---|---|
| chr4 | 138435350 | 138435351 | cg08053129 |
| chr4 | 139146513 | 139146514 | cg24869834 |
| chr4 | 139238668 | 139238669 | cg14800532 |
| chr4 | 139480684 | 139480685 | cg03732759 |
| chr4 | 140432613 | 140432614 | cg24475272 |
| chr4 | 140782938 | 140782939 | cg11868485 |
| chr4 | 140787316 | 140787317 | cg26131315 |
| chr4 | 140985719 | 140985720 | cg06456255 |
| chr4 | 141034510 | 141034511 | cg01342226 |
| chr4 | 141217605 | 141217606 | cg00321234 |
| chr4 | 141648570 | 141648571 | cg10685128 |
| chr4 | 144266241 | 144266242 | cg25911551 |
| chr4 | 144277796 | 144277797 | cg17453767 |
| chr4 | 144303008 | 144303009 | cg06855485 |
| chr4 | 146462169 | 146462170 | cg24827839 |
| chr4 | 146729164 | 146729165 | cg15029037 |
| chr4 | 148283066 | 148283067 | cg20168495 |
| chr4 | 151163526 | 151163527 | cg10190898 |
| chr4 | 152017137 | 152017138 | cg03479572 |
| chr4 | 153443405 | 153443406 | cg11089117 |
| chr4 | 153561975 | 153561976 | cg25351755 |
| chr4 | 153586440 | 153586441 | cg06536988 |
| chr4 | 153601466 | 153601467 | cg06371582 |
| chr4 | 153602251 | 153602252 | cg11844042 |
| chr4 | 153771421 | 153771422 | cg06712886 |
| chr4 | 153871345 | 153871346 | cg00170487 |
| chr4 | 154391944 | 154391945 | cg20367304 |
| chr4 | 154419554 | 154419555 | cg24020152 |
| chr4 | 154439980 | 154439981 | cg03464224 |
| chr4 | 154443833 | 154443834 | cg25570522 |
| chr4 | 154451068 | 154451069 | cg02830749 |
| chr4 | 154515617 | 154515618 | cg26103306 |
| chr4 | 157227668 | 157227669 | cg06669631 |
| chr4 | 157311747 | 157311748 | cg17170463 |
| chr4 | 160026608 | 160026609 | cg00861646 |
| chr4 | 165108732 | 165108733 | cg02570981 |
| chr4 | 166323062 | 166323063 | cg06549863 |
| chr4 | 168865149 | 168865150 | cg13078894 |
| chr4 | 169481028 | 169481029 | cg17914107 |
| chr4 | 177455600 | 177455601 | cg06655097 |
| chr4 | 178364342 | 178364343 | cg00398048 |
| chr4 | 181688500 | 181688501 | cg06690831 |
| chr4 | 184578976 | 184578977 | cg03427849 |
| chr4 | 184723252 | 184723253 | cg09155692 |
| chr4 | 184829966 | 184829967 | cg08346735 |
| chr4 | 185205197 | 185205198 | cg18317788 |
| chr4 | 185306187 | 185306188 | cg25556051 |
| chr4 | 185339075 | 185339076 | cg05501617 |
| chr4 | 185382704 | 185382705 | cg12453504 |
| chr4 | 185398007 | 185398008 | cg21960364 |
| chr4 | 185668393 | 185668394 | cg24839529 |
| chr4 | 186132412 | 186132413 | cg12836958 |
| chr4 | 186393468 | 186393469 | cg12347792 |
| chr4 | 187509603 | 187509604 | cg25124956 |
| chr5 | 195609 | 195610 | cg25621286 |
| chr5 | 195698 | 195699 | cg00099519 |

| | | | |
|---|---|---|---|
| chr6 | 33396610 | 33396611 | cg08371951 |
| chr6 | 33397122 | 33397123 | cg26257411 |
| chr6 | 33397233 | 33397234 | cg01340991 |
| chr6 | 33558900 | 33558901 | cg08255724 |
| chr6 | 33559603 | 33559604 | cg02484633 |
| chr6 | 33587496 | 33587497 | cg08355863 |
| chr6 | 33595160 | 33595161 | cg06376233 |
| chr6 | 33601177 | 33601178 | cg17865955 |
| chr6 | 34284953 | 34284954 | cg16767886 |
| chr6 | 34352082 | 34352083 | cg20159193 |
| chr6 | 34563243 | 34563244 | cg10519906 |
| chr6 | 34633133 | 34633134 | cg10163122 |
| chr6 | 34663155 | 34663156 | cg14725580 |
| chr6 | 34802849 | 34802850 | cg18118738 |
| chr6 | 35027147 | 35027148 | cg14601709 |
| chr6 | 35206618 | 35206619 | cg11562411 |
| chr6 | 35311534 | 35311535 | cg26668919 |
| chr6 | 35461236 | 35461237 | cg10893014 |
| chr6 | 35631736 | 35631737 | cg07061368 |
| chr6 | 35694245 | 35694246 | cg00052684 |
| chr6 | 35697759 | 35697760 | cg08915438 |
| chr6 | 35700382 | 35700383 | cg13719443 |
| chr6 | 36305071 | 36305072 | cg04434994 |
| chr6 | 36645100 | 36645101 | cg05460965 |
| chr6 | 36645608 | 36645609 | cg11920449 |
| chr6 | 36645648 | 36645649 | cg24425727 |
| chr6 | 36756830 | 36756831 | cg23309670 |
| chr6 | 36935591 | 36935592 | cg04633721 |
| chr6 | 37533112 | 37533113 | cg12612465 |
| chr6 | 37660403 | 37660404 | cg18821972 |
| chr6 | 37913121 | 37913122 | cg22995064 |
| chr6 | 38121432 | 38121433 | cg06300222 |
| chr6 | 38207741 | 38207742 | cg27655671 |
| chr6 | 38673615 | 38673616 | cg14607286 |
| chr6 | 41020847 | 41020848 | cg22682201 |
| chr6 | 41411455 | 41411456 | cg11894474 |
| chr6 | 41419370 | 41419371 | cg14732815 |
| chr6 | 41471014 | 41471015 | cg26242687 |
| chr6 | 41517062 | 41517063 | cg26432961 |
| chr6 | 41552580 | 41552581 | cg22791678 |
| chr6 | 41752769 | 41752770 | cg00975876 |
| chr6 | 41998500 | 41998501 | cg14029001 |
| chr6 | 42231202 | 42231203 | cg17291767 |
| chr6 | 42540883 | 42540884 | cg14282280 |
| chr6 | 42719001 | 42719002 | cg01323212 |
| chr6 | 42757805 | 42757806 | cg19352726 |
| chr6 | 42893750 | 42893751 | cg19638876 |
| chr6 | 43228649 | 43228650 | cg05606072 |
| chr6 | 43244304 | 43244305 | cg06241901 |
| chr6 | 43334003 | 43334004 | cg01751549 |
| chr6 | 43445012 | 43445013 | cg14009632 |
| chr6 | 43663647 | 43663648 | cg10024501 |
| chr6 | 43692678 | 43692679 | cg10794973 |
| chr6 | 44189984 | 44189985 | cg00920254 |
| chr6 | 44235210 | 44235211 | cg08855022 |
| chr6 | 44396701 | 44396702 | cg05585911 |

| | | | |
|---|---|---|---|
| chr5 | 195790 | 195791 | cg14465992 |
| chr5 | 314642 | 314643 | cg14982043 |
| chr5 | 527640 | 527641 | cg00658890 |
| chr5 | 607516 | 607517 | cg21194115 |
| chr5 | 1491600 | 1491601 | cg10217292 |
| chr5 | 1509723 | 1509724 | cg09809922 |
| chr5 | 1510088 | 1510089 | cg11468635 |
| chr5 | 1510512 | 1510513 | cg06984201 |
| chr5 | 1510544 | 1510545 | cg24093182 |
| chr5 | 1510684 | 1510685 | cg11960719 |
| chr5 | 1511473 | 1511474 | cg11576486 |
| chr5 | 1515837 | 1515838 | cg20962974 |
| chr5 | 1526929 | 1526930 | cg06940720 |
| chr5 | 1608614 | 1608615 | cg20414364 |
| chr5 | 3965271 | 3965272 | cg08907926 |
| chr5 | 6491550 | 6491551 | cg09156073 |
| chr5 | 6722123 | 6722124 | cg10857441 |
| chr5 | 7870577 | 7870578 | cg10300154 |
| chr5 | 9782142 | 9782143 | cg16086730 |
| chr5 | 10663596 | 10663597 | cg01184194 |
| chr5 | 11851713 | 11851714 | cg13685506 |
| chr5 | 14744206 | 14744207 | cg00975971 |
| chr5 | 15865364 | 15865365 | cg01572460 |
| chr5 | 17306335 | 17306336 | cg25421209 |
| chr5 | 17395952 | 17395953 | cg12995004 |
| chr5 | 17436715 | 17436716 | cg00308914 |
| chr5 | 19886600 | 19886601 | cg15050865 |
| chr5 | 30390070 | 30390071 | cg12642207 |
| chr5 | 30558470 | 30558471 | cg15704315 |
| chr5 | 32532223 | 32532224 | cg15686949 |
| chr5 | 32768395 | 32768396 | cg26647771 |
| chr5 | 33855241 | 33855242 | cg21569398 |
| chr5 | 33889461 | 33889462 | cg08768395 |
| chr5 | 34617800 | 34617801 | cg12184038 |
| chr5 | 34667887 | 34667888 | cg06622892 |
| chr5 | 34673551 | 34673552 | cg05661149 |
| chr5 | 34732408 | 34732409 | cg14854112 |
| chr5 | 35146528 | 35146529 | cg19433694 |
| chr5 | 35860337 | 35860338 | cg01027405 |
| chr5 | 35902250 | 35902251 | cg00275126 |
| chr5 | 36239673 | 36239674 | cg21586453 |
| chr5 | 36725464 | 36725465 | cg12681478 |
| chr5 | 36744219 | 36744220 | cg03995615 |
| chr5 | 38534081 | 38534082 | cg03864479 |
| chr5 | 38535574 | 38535575 | cg10959950 |
| chr5 | 38554778 | 38554779 | cg06182018 |
| chr5 | 38671111 | 38671112 | cg06495233 |
| chr5 | 39415351 | 39415352 | cg26186331 |
| chr5 | 40570344 | 40570345 | cg23192749 |
| chr5 | 40685989 | 40685990 | cg11821200 |
| chr5 | 41219519 | 41219520 | cg24883742 |
| chr5 | 52334107 | 52334108 | cg17480554 |
| chr5 | 52657514 | 52657515 | cg26341848 |
| chr5 | 53816563 | 53816564 | cg26118943 |
| chr5 | 54100067 | 54100068 | cg22421804 |
| chr5 | 54467378 | 54467379 | cg01988325 |
| chr6 | 45517025 | 45517026 | cg11630605 |
| chr6 | 46258880 | 46258881 | cg26677394 |
| chr6 | 46326932 | 46326933 | cg00588393 |
| chr6 | 47444224 | 47444225 | cg23250715 |
| chr6 | 52438690 | 52438691 | cg15807046 |
| chr6 | 52439185 | 52439186 | cg18397730 |
| chr6 | 52440429 | 52440430 | cg08966889 |
| chr6 | 52441068 | 52441069 | cg11998049 |
| chr6 | 52840469 | 52840470 | cg15448894 |
| chr6 | 53058587 | 53058588 | cg09752583 |
| chr6 | 53147641 | 53147642 | cg08597067 |
| chr6 | 53222886 | 53222887 | cg01461368 |
| chr6 | 53305580 | 53305581 | cg03282312 |
| chr6 | 53396138 | 53396139 | cg08349400 |
| chr6 | 53407186 | 53407187 | cg02731193 |
| chr6 | 53513754 | 53513755 | cg06555281 |
| chr6 | 53661064 | 53661065 | cg15726169 |
| chr6 | 53678839 | 53678840 | cg27022531 |
| chr6 | 70507230 | 70507231 | cg03001484 |
| chr6 | 70573798 | 70573799 | cg04092581 |
| chr6 | 70579618 | 70579619 | cg02400725 |
| chr6 | 72199637 | 72199638 | cg00206507 |
| chr6 | 73329475 | 73329476 | cg02351713 |
| chr6 | 79911416 | 79911417 | cg22032366 |
| chr6 | 83904319 | 83904320 | cg18674899 |
| chr6 | 87861261 | 87861262 | cg05316627 |
| chr6 | 87884827 | 87884828 | cg19372952 |
| chr6 | 88040249 | 88040250 | cg06087457 |
| chr6 | 89340909 | 89340910 | cg15692086 |
| chr6 | 90807675 | 90807676 | cg25670076 |
| chr6 | 90951134 | 90951135 | cg18477569 |
| chr6 | 90997723 | 90997724 | cg20768122 |
| chr6 | 91007365 | 91007366 | cg09790829 |
| chr6 | 91007563 | 91007564 | cg15380603 |
| chr6 | 91007625 | 91007626 | cg10913050 |
| chr6 | 97369501 | 97369502 | cg24500972 |
| chr6 | 101448078 | 101448079 | cg01349655 |
| chr6 | 106041074 | 106041075 | cg23316191 |
| chr6 | 106551368 | 106551369 | cg01146808 |
| chr6 | 107423930 | 107423931 | cg23796329 |
| chr6 | 107731096 | 107731097 | cg19411084 |
| chr6 | 108167599 | 108167600 | cg01063280 |
| chr6 | 109193762 | 109193763 | cg15608714 |
| chr6 | 109802446 | 109802447 | cg02803604 |
| chr6 | 110964408 | 110964409 | cg25202367 |
| chr6 | 111887243 | 111887244 | cg01520105 |
| chr6 | 112081490 | 112081491 | cg02816367 |
| chr6 | 112081538 | 112081539 | cg08061598 |
| chr6 | 113198826 | 113198827 | cg18446641 |
| chr6 | 116338847 | 116338848 | cg06127202 |
| chr6 | 116381904 | 116381905 | cg26893134 |
| chr6 | 119027122 | 119027123 | cg01113488 |
| chr6 | 119031922 | 119031923 | cg12761472 |
| chr6 | 119043982 | 119043983 | cg17323739 |
| chr6 | 119674313 | 119674314 | cg04408448 |
| chr6 | 126036146 | 126036147 | cg23965144 |

| | | | |
|---|---|---|---|
| chr5 | 54737053 | 54737054 | cg02008511 |
| chr5 | 54987767 | 54987768 | cg07262873 |
| chr5 | 55903504 | 55903505 | cg09790325 |
| chr5 | 55994653 | 55994654 | cg23977888 |
| chr5 | 56109540 | 56109541 | cg25148456 |
| chr5 | 56115499 | 56115500 | cg20439101 |
| chr5 | 56268061 | 56268062 | cg16534315 |
| chr5 | 56620429 | 56620430 | cg02402882 |
| chr5 | 56629471 | 56629472 | cg07362800 |
| chr5 | 56972268 | 56972269 | cg11818544 |
| chr5 | 57702195 | 57702196 | cg25026755 |
| chr5 | 58388744 | 58388745 | cg16377679 |
| chr5 | 58893722 | 58893723 | cg07267294 |
| chr5 | 59126518 | 59126519 | cg27583655 |
| chr5 | 59783668 | 59783669 | cg18704047 |
| chr5 | 59784544 | 59784545 | cg24844211 |
| chr5 | 60603263 | 60603264 | cg23111196 |
| chr5 | 60624229 | 60624230 | cg10166664 |
| chr5 | 64330880 | 64330881 | cg01061714 |
| chr5 | 64330915 | 64330916 | cg25366639 |
| chr5 | 64330935 | 64330936 | cg09571848 |
| chr5 | 64330959 | 64330960 | cg06294700 |
| chr5 | 65443428 | 65443429 | cg21977218 |
| chr5 | 66296978 | 66296979 | cg09005125 |
| chr5 | 66486419 | 66486420 | cg18670483 |
| chr5 | 66488838 | 66488839 | cg04813904 |
| chr5 | 66493682 | 66493683 | cg14137162 |
| chr5 | 67535553 | 67535554 | cg01251150 |
| chr5 | 67583627 | 67583628 | cg09101894 |
| chr5 | 68473620 | 68473621 | cg13647309 |
| chr5 | 71724363 | 71724364 | cg25351763 |
| chr5 | 71883710 | 71883711 | cg11664417 |
| chr5 | 72660351 | 72660352 | cg12942161 |
| chr5 | 73690902 | 73690903 | cg20969424 |
| chr5 | 74571706 | 74571707 | cg08623843 |
| chr5 | 75794475 | 75794476 | cg21806100 |
| chr5 | 76132547 | 76132548 | cg22816131 |
| chr5 | 76247679 | 76247680 | cg26196496 |
| chr5 | 76372556 | 76372557 | cg02141543 |
| chr5 | 77256916 | 77256917 | cg26485844 |
| chr5 | 78054648 | 78054649 | cg12189389 |
| chr5 | 78282099 | 78282100 | cg24893837 |
| chr5 | 79438093 | 79438094 | cg07172701 |
| chr5 | 79479211 | 79479212 | cg02417427 |
| chr5 | 79514577 | 79514578 | cg14711743 |
| chr5 | 79542470 | 79542471 | cg01267797 |
| chr5 | 81452987 | 81452988 | cg25363080 |
| chr5 | 82858676 | 82858677 | cg17599241 |
| chr5 | 86042549 | 86042550 | cg26899012 |
| chr5 | 88841819 | 88841820 | cg24652205 |
| chr5 | 95060853 | 95060854 | cg24834165 |
| chr5 | 95206387 | 95206388 | cg16133703 |
| chr5 | 95299519 | 95299520 | cg10841248 |
| chr5 | 95583220 | 95583221 | cg05612966 |
| chr5 | 95666291 | 95666292 | cg16168509 |
| chr5 | 95996644 | 95996645 | cg01741836 |

| | | | |
|---|---|---|---|
| chr6 | 126080132 | 126080133 | cg00066750 |
| chr6 | 127781873 | 127781874 | cg06003184 |
| chr6 | 130007252 | 130007253 | cg25347356 |
| chr6 | 130017522 | 130017523 | cg20760057 |
| chr6 | 130787470 | 130787471 | cg04529078 |
| chr6 | 131690266 | 131690267 | cg16736964 |
| chr6 | 132825654 | 132825655 | cg01140783 |
| chr6 | 135271056 | 135271057 | cg02396253 |
| chr6 | 135271222 | 135271223 | cg12491555 |
| chr6 | 135271333 | 135271334 | cg20352402 |
| chr6 | 136174534 | 136174535 | cg14623715 |
| chr6 | 136914796 | 136914797 | cg18914962 |
| chr6 | 137421461 | 137421462 | cg22980427 |
| chr6 | 137493017 | 137493018 | cg00415333 |
| chr6 | 137518909 | 137518910 | cg02480602 |
| chr6 | 139307985 | 139307986 | cg00631221 |
| chr6 | 139614104 | 139614105 | cg10484756 |
| chr6 | 140204424 | 140204425 | cg15224626 |
| chr6 | 143132934 | 143132935 | cg10295609 |
| chr6 | 143167279 | 143167280 | cg13829419 |
| chr6 | 143206075 | 143206076 | cg06770429 |
| chr6 | 143269881 | 143269882 | cg25685456 |
| chr6 | 143861183 | 143861184 | cg13363748 |
| chr6 | 144217883 | 144217884 | cg24525744 |
| chr6 | 144259320 | 144259321 | cg15350652 |
| chr6 | 144281452 | 144281453 | cg21776667 |
| chr6 | 144329922 | 144329923 | cg03562868 |
| chr6 | 144329997 | 144329998 | cg04840930 |
| chr6 | 144330162 | 144330163 | cg18593039 |
| chr6 | 144330233 | 144330234 | cg23840797 |
| chr6 | 144330345 | 144330346 | cg00981250 |
| chr6 | 144654512 | 144654513 | cg05360911 |
| chr6 | 145050022 | 145050023 | cg09878408 |
| chr6 | 146057943 | 146057944 | cg05220968 |
| chr6 | 148878402 | 148878403 | cg11241455 |
| chr6 | 149336332 | 149336333 | cg17384769 |
| chr6 | 149401166 | 149401167 | cg17072494 |
| chr6 | 149658548 | 149658549 | cg16439130 |
| chr6 | 150068017 | 150068018 | cg17341370 |
| chr6 | 150390894 | 150390895 | cg00432049 |
| chr6 | 151087821 | 151087822 | cg00994250 |
| chr6 | 151188259 | 151188260 | cg16336494 |
| chr6 | 151288153 | 151288154 | cg14081861 |
| chr6 | 151670448 | 151670449 | cg26390965 |
| chr6 | 152011399 | 152011400 | cg18745416 |
| chr6 | 152126180 | 152126181 | cg07584093 |
| chr6 | 152379044 | 152379045 | cg07455133 |
| chr6 | 152620697 | 152620698 | cg17979719 |
| chr6 | 152953820 | 152953821 | cg05750323 |
| chr6 | 152960531 | 152960532 | cg25326491 |
| chr6 | 154477894 | 154477895 | cg26415516 |
| chr6 | 154535120 | 154535121 | cg10276116 |
| chr6 | 155442993 | 155442994 | cg00931843 |
| chr6 | 155537930 | 155537931 | cg04228709 |
| chr6 | 156488510 | 156488511 | cg18393175 |
| chr6 | 156711764 | 156711765 | cg20394376 |

| | | | |
|---|---|---|---|
| chr5 | 96078430 | 96078431 | cg11214001 |
| chr5 | 97703334 | 97703335 | cg03849299 |
| chr5 | 97949352 | 97949353 | cg10659138 |
| chr5 | 102598034 | 102598035 | cg21184216 |
| chr5 | 103820591 | 103820592 | cg24905254 |
| chr5 | 109029868 | 109029869 | cg22931822 |
| chr5 | 110886711 | 110886712 | cg01301660 |
| chr5 | 111090142 | 111090143 | cg06838283 |
| chr5 | 111191530 | 111191531 | cg18054620 |
| chr5 | 111869581 | 111869582 | cg03620123 |
| chr5 | 112315632 | 112315633 | cg06817368 |
| chr5 | 112721426 | 112721427 | cg02021529 |
| chr5 | 112882744 | 112882745 | cg16210979 |
| chr5 | 113388886 | 113388887 | cg19965300 |
| chr5 | 113744106 | 113744107 | cg19062099 |
| chr5 | 114887230 | 114887231 | cg02637031 |
| chr5 | 117822219 | 117822220 | cg08442713 |
| chr5 | 118609965 | 118609966 | cg00524900 |
| chr5 | 120179610 | 120179611 | cg01214384 |
| chr5 | 121022314 | 121022315 | cg11619879 |
| chr5 | 124043109 | 124043110 | cg01154355 |
| chr5 | 124044098 | 124044099 | cg00323842 |
| chr5 | 124081751 | 124081752 | cg26413942 |
| chr5 | 124256096 | 124256097 | cg17604985 |
| chr5 | 124526339 | 124526340 | cg13607323 |
| chr5 | 124745818 | 124745819 | cg04469998 |
| chr5 | 125800764 | 125800765 | cg14800014 |
| chr5 | 126205081 | 126205082 | cg06918474 |
| chr5 | 126424797 | 126424798 | cg04134528 |
| chr5 | 126988029 | 126988030 | cg27666123 |
| chr5 | 127805611 | 127805612 | cg14790000 |
| chr5 | 127868982 | 127868983 | cg07355763 |
| chr5 | 130510020 | 130510021 | cg24610854 |
| chr5 | 131343261 | 131343262 | cg25309041 |
| chr5 | 131408382 | 131408383 | cg13259290 |
| chr5 | 131430859 | 131430860 | cg11225089 |
| chr5 | 131760037 | 131760038 | cg20781981 |
| chr5 | 131879579 | 131879580 | cg23208285 |
| chr5 | 131999237 | 131999238 | cg06451479 |
| chr5 | 132009352 | 132009353 | cg25368824 |
| chr5 | 132111426 | 132111427 | cg18090331 |
| chr5 | 132232586 | 132232587 | cg22212250 |
| chr5 | 133404710 | 133404711 | cg19882512 |
| chr5 | 133451383 | 133451384 | cg25947408 |
| chr5 | 133452188 | 133452189 | cg18338046 |
| chr5 | 133904674 | 133904675 | cg27024654 |
| chr5 | 134120499 | 134120500 | cg12499311 |
| chr5 | 134987654 | 134987655 | cg13005228 |
| chr5 | 136314223 | 136314224 | cg22612642 |
| chr5 | 136374727 | 136374728 | cg17221083 |
| chr5 | 137071523 | 137071524 | cg13847070 |
| chr5 | 137090964 | 137090965 | cg19101232 |
| chr5 | 137143726 | 137143727 | cg00813373 |
| chr5 | 137365576 | 137365577 | cg24566595 |
| chr5 | 137419242 | 137419243 | cg09513026 |
| chr5 | 137588237 | 137588238 | cg21210529 |
| chr6 | 156715760 | 156715761 | cg12199954 |
| chr6 | 156718177 | 156718178 | cg07292773 |
| chr6 | 156718262 | 156718263 | cg21795221 |
| chr6 | 156718288 | 156718289 | cg05225634 |
| chr6 | 156718398 | 156718399 | cg09056876 |
| chr6 | 156718546 | 156718547 | cg20021506 |
| chr6 | 156720309 | 156720310 | cg01608493 |
| chr6 | 156885985 | 156885986 | cg06010163 |
| chr6 | 157137791 | 157137792 | cg07212702 |
| chr6 | 157168834 | 157168835 | cg15618758 |
| chr6 | 157184114 | 157184115 | cg16017800 |
| chr6 | 157254584 | 157254585 | cg03561416 |
| chr6 | 157264715 | 157264716 | cg16455339 |
| chr6 | 157345266 | 157345267 | cg17164954 |
| chr6 | 158178235 | 158178236 | cg17071432 |
| chr6 | 158194541 | 158194542 | cg11924238 |
| chr6 | 158234655 | 158234656 | cg25252017 |
| chr6 | 158243386 | 158243387 | cg04064963 |
| chr6 | 158245646 | 158245647 | cg24492119 |
| chr6 | 158260990 | 158260991 | cg15264757 |
| chr6 | 158424174 | 158424175 | cg15142819 |
| chr6 | 158744008 | 158744009 | cg06096446 |
| chr6 | 159359940 | 159359941 | cg01207473 |
| chr6 | 159462964 | 159462965 | cg27084622 |
| chr6 | 159592134 | 159592135 | cg18426860 |
| chr6 | 160219791 | 160219792 | cg00585742 |
| chr6 | 160401562 | 160401563 | cg17154159 |
| chr6 | 160512657 | 160512658 | cg03634777 |
| chr6 | 166673846 | 166673847 | cg10922521 |
| chr6 | 166722773 | 166722774 | cg05089968 |
| chr6 | 166887879 | 166887880 | cg00424286 |
| chr6 | 167171394 | 167171395 | cg05573767 |
| chr6 | 167263401 | 167263402 | cg16100486 |
| chr6 | 167490870 | 167490871 | cg06454157 |
| chr6 | 167490980 | 167490981 | cg14705155 |
| chr6 | 167530185 | 167530186 | cg05540913 |
| chr7 | 553423 | 553424 | cg20063728 |
| chr7 | 556429 | 556430 | cg15454385 |
| chr7 | 650249 | 650250 | cg16190265 |
| chr7 | 872293 | 872294 | cg22972055 |
| chr7 | 872735 | 872736 | cg10770230 |
| chr7 | 914964 | 914965 | cg01259220 |
| chr7 | 955497 | 955498 | cg01017397 |
| chr7 | 955551 | 955552 | cg24586376 |
| chr7 | 960246 | 960247 | cg13687274 |
| chr7 | 960528 | 960529 | cg23667602 |
| chr7 | 1004748 | 1004749 | cg09363068 |
| chr7 | 1007557 | 1007558 | cg21728650 |
| chr7 | 1066396 | 1066397 | cg27287438 |
| chr7 | 1545385 | 1545386 | cg06942110 |
| chr7 | 1584061 | 1584062 | cg00715835 |
| chr7 | 1632620 | 1632621 | cg18560936 |
| chr7 | 1855709 | 1855710 | cg20396385 |
| chr7 | 1858440 | 1858441 | cg21639553 |
| chr7 | 1889727 | 1889728 | cg00860321 |
| chr7 | 1961968 | 1961969 | cg04342202 |

| | | | |
|---|---|---|---|
| chr5 | 137610645 | 137610646 | cg19908207 |
| chr5 | 137793707 | 137793708 | cg08519191 |
| chr5 | 137896757 | 137896758 | cg26829184 |
| chr5 | 138608519 | 138608520 | cg25391820 |
| chr5 | 138612986 | 138612987 | cg15962267 |
| chr5 | 138613030 | 138613031 | cg20385395 |
| chr5 | 138614407 | 138614408 | cg05585556 |
| chr5 | 138641864 | 138641865 | cg14717906 |
| chr5 | 138723816 | 138723817 | cg20566766 |
| chr5 | 138743684 | 138743685 | cg17264541 |
| chr5 | 138861832 | 138861833 | cg03317505 |
| chr5 | 138861847 | 138861848 | cg04560810 |
| chr5 | 138861855 | 138861856 | cg14655316 |
| chr5 | 138862441 | 138862442 | cg16983159 |
| chr5 | 139024407 | 139024408 | cg13626633 |
| chr5 | 139027024 | 139027025 | cg01408486 |
| chr5 | 139082468 | 139082469 | cg02478589 |
| chr5 | 139088316 | 139088317 | cg07708521 |
| chr5 | 139140550 | 139140551 | cg16691158 |
| chr5 | 139141138 | 139141139 | cg06629170 |
| chr5 | 139334561 | 139334562 | cg06658615 |
| chr5 | 139362307 | 139362308 | cg24724630 |
| chr5 | 139929838 | 139929839 | cg10912634 |
| chr5 | 140895708 | 140895709 | cg26217846 |
| chr5 | 141001788 | 141001789 | cg13294753 |
| chr5 | 141346577 | 141346578 | cg06073041 |
| chr5 | 141488016 | 141488017 | cg23435118 |
| chr5 | 141488047 | 141488048 | cg08523384 |
| chr5 | 142435797 | 142435798 | cg24559044 |
| chr5 | 142586466 | 142586467 | cg03615933 |
| chr5 | 142692961 | 142692962 | cg19457823 |
| chr5 | 143191565 | 143191566 | cg10678215 |
| chr5 | 143883482 | 143883483 | cg19452248 |
| chr5 | 146184364 | 146184365 | cg02154084 |
| chr5 | 146268595 | 146268596 | cg12524428 |
| chr5 | 146333034 | 146333035 | cg15021031 |
| chr5 | 147257456 | 147257457 | cg04257865 |
| chr5 | 147286378 | 147286379 | cg18818210 |
| chr5 | 148208041 | 148208042 | cg08370787 |
| chr5 | 148654912 | 148654913 | cg07127357 |
| chr5 | 148908014 | 148908015 | cg03569876 |
| chr5 | 149185704 | 149185705 | cg17645677 |
| chr5 | 149202020 | 149202021 | cg07226964 |
| chr5 | 149462180 | 149462181 | cg20754844 |
| chr5 | 149778964 | 149778965 | cg04242245 |
| chr5 | 149793436 | 149793437 | cg22975568 |
| chr5 | 149868359 | 149868360 | cg17781669 |
| chr5 | 149962501 | 149962502 | cg08818337 |
| chr5 | 149996740 | 149996741 | cg08413646 |
| chr5 | 150524635 | 150524636 | cg15892650 |
| chr5 | 150538414 | 150538415 | cg00407659 |
| chr5 | 150727812 | 150727813 | cg10345936 |
| chr5 | 151076195 | 151076196 | cg12550344 |
| chr5 | 151119245 | 151119246 | cg19939793 |
| chr5 | 151120070 | 151120071 | cg17574050 |
| chr5 | 151137430 | 151137431 | cg01201215 |

| | | | |
|---|---|---|---|
| chr7 | 2007588 | 2007589 | cg22500280 |
| chr7 | 2019608 | 2019609 | cg05533001 |
| chr7 | 2057835 | 2057836 | cg15225688 |
| chr7 | 2059796 | 2059797 | cg07027305 |
| chr7 | 2205745 | 2205746 | cg22493042 |
| chr7 | 2289585 | 2289586 | cg02326253 |
| chr7 | 2291960 | 2291961 | cg08027265 |
| chr7 | 2319604 | 2319605 | cg13641082 |
| chr7 | 2444534 | 2444535 | cg10152449 |
| chr7 | 2445331 | 2445332 | cg21159568 |
| chr7 | 2549030 | 2549031 | cg09125127 |
| chr7 | 2551598 | 2551599 | cg21531604 |
| chr7 | 2551978 | 2551979 | cg21136499 |
| chr7 | 2556962 | 2556963 | cg07041720 |
| chr7 | 2557446 | 2557447 | cg05829046 |
| chr7 | 2557466 | 2557467 | cg22902289 |
| chr7 | 2563670 | 2563671 | cg20297638 |
| chr7 | 2563816 | 2563817 | cg03676485 |
| chr7 | 2563909 | 2563910 | cg09723161 |
| chr7 | 2565944 | 2565945 | cg06160397 |
| chr7 | 2566027 | 2566028 | cg11623476 |
| chr7 | 2566113 | 2566114 | cg19071989 |
| chr7 | 2566213 | 2566214 | cg20711703 |
| chr7 | 2566534 | 2566535 | cg17226348 |
| chr7 | 2600161 | 2600162 | cg27189465 |
| chr7 | 2608671 | 2608672 | cg20870668 |
| chr7 | 2674181 | 2674182 | cg06398114 |
| chr7 | 2678208 | 2678209 | cg02154914 |
| chr7 | 2679067 | 2679068 | cg01561864 |
| chr7 | 2679148 | 2679149 | cg22788953 |
| chr7 | 2679546 | 2679547 | cg14893935 |
| chr7 | 2680532 | 2680533 | cg27229251 |
| chr7 | 2681369 | 2681370 | cg26551200 |
| chr7 | 2758116 | 2758117 | cg05123755 |
| chr7 | 2968406 | 2968407 | cg10939949 |
| chr7 | 2968559 | 2968560 | cg23770265 |
| chr7 | 2968595 | 2968596 | cg22989995 |
| chr7 | 2968627 | 2968628 | cg00578304 |
| chr7 | 5269245 | 5269246 | cg27152128 |
| chr7 | 5270512 | 5270513 | cg21585409 |
| chr7 | 5272846 | 5272847 | cg08891989 |
| chr7 | 5293333 | 5293334 | cg06595212 |
| chr7 | 5422234 | 5422235 | cg23058673 |
| chr7 | 7224869 | 7224870 | cg19473623 |
| chr7 | 7291514 | 7291515 | cg16101574 |
| chr7 | 8011838 | 8011839 | cg21380380 |
| chr7 | 20447707 | 20447708 | cg14959425 |
| chr7 | 21410396 | 21410397 | cg00572966 |
| chr7 | 21452797 | 21452798 | cg01686177 |
| chr7 | 21469772 | 21469773 | cg10365563 |
| chr7 | 21470054 | 21470055 | cg24101039 |
| chr7 | 22485718 | 22485719 | cg17972105 |
| chr7 | 23050722 | 23050723 | cg11357953 |
| chr7 | 26327799 | 26327800 | cg04924454 |
| chr7 | 26437192 | 26437193 | cg00749672 |
| chr7 | 27704065 | 27704066 | cg01662334 |

| | | | |
|---|---|---|---|
| chr5 | 151771606 | 151771607 | cg07429087 |
| chr5 | 151908977 | 151908978 | cg25116430 |
| chr5 | 152591026 | 152591027 | cg10976311 |
| chr5 | 152869009 | 152869010 | cg07633435 |
| chr5 | 152869882 | 152869883 | cg17987968 |
| chr5 | 152896564 | 152896565 | cg22798121 |
| chr5 | 154146853 | 154146854 | cg15111708 |
| chr5 | 154230141 | 154230142 | cg26655340 |
| chr5 | 154230308 | 154230309 | cg10257049 |
| chr5 | 155134244 | 155134245 | cg09586743 |
| chr5 | 156340946 | 156340947 | cg27144719 |
| chr5 | 156692395 | 156692396 | cg07750741 |
| chr5 | 156696520 | 156696521 | cg20541456 |
| chr5 | 156934938 | 156934939 | cg03647778 |
| chr5 | 157733288 | 157733289 | cg26645237 |
| chr5 | 158757456 | 158757457 | cg18307303 |
| chr5 | 159072916 | 159072917 | cg12446613 |
| chr5 | 159191650 | 159191651 | cg19013399 |
| chr5 | 159246470 | 159246471 | cg17397548 |
| chr5 | 159247072 | 159247073 | cg18144404 |
| chr5 | 159848299 | 159848300 | cg23024444 |
| chr5 | 160048394 | 160048395 | cg09705798 |
| chr5 | 168254759 | 168254760 | cg03726739 |
| chr5 | 169008622 | 169008623 | cg05722931 |
| chr5 | 169815771 | 169815772 | cg27044591 |
| chr5 | 169817247 | 169817248 | cg17425484 |
| chr5 | 169929560 | 169929561 | cg23479949 |
| chr5 | 170845610 | 170845611 | cg15699524 |
| chr5 | 170850143 | 170850144 | cg26874542 |
| chr5 | 171333801 | 171333802 | cg13277254 |
| chr5 | 171354691 | 171354692 | cg04973915 |
| chr5 | 171485294 | 171485295 | cg14641625 |
| chr5 | 171569243 | 171569244 | cg26941823 |
| chr5 | 171616253 | 171616254 | cg19782271 |
| chr5 | 171616388 | 171616389 | cg19873297 |
| chr5 | 172099681 | 172099682 | cg21742048 |
| chr5 | 172203954 | 172203955 | cg20745620 |
| chr5 | 172263112 | 172263113 | cg24581650 |
| chr5 | 172443740 | 172443741 | cg05201300 |
| chr5 | 172740954 | 172740955 | cg26371913 |
| chr5 | 172892099 | 172892100 | cg25304439 |
| chr5 | 173125575 | 173125576 | cg18256498 |
| chr5 | 173141761 | 173141762 | cg15618908 |
| chr5 | 173216171 | 173216172 | cg05871756 |
| chr5 | 173314531 | 173314532 | cg25122699 |
| chr5 | 173314559 | 173314560 | cg14454798 |
| chr5 | 174944792 | 174944793 | cg17747551 |
| chr5 | 174962968 | 174962969 | cg00290023 |
| chr5 | 175955524 | 175955525 | cg05517697 |
| chr5 | 175991129 | 175991130 | cg02006915 |
| chr5 | 176316988 | 176316989 | cg15357821 |
| chr5 | 176855304 | 176855305 | cg23327334 |
| chr5 | 176856845 | 176856846 | cg19459094 |
| chr5 | 176875120 | 176875121 | cg14098951 |
| chr5 | 176898673 | 176898674 | cg21091766 |
| chr5 | 176916434 | 176916435 | cg15197657 |

| | | | |
|---|---|---|---|
| chr7 | 28070030 | 28070031 | cg12528349 |
| chr7 | 28189139 | 28189140 | cg22892036 |
| chr7 | 28214983 | 28214984 | cg15170557 |
| chr7 | 28216851 | 28216852 | cg16286055 |
| chr7 | 28574260 | 28574261 | cg03226887 |
| chr7 | 28747836 | 28747837 | cg06647007 |
| chr7 | 29237766 | 29237767 | cg20304988 |
| chr7 | 29304984 | 29304985 | cg15030712 |
| chr7 | 30185315 | 30185316 | cg01541443 |
| chr7 | 30202350 | 30202351 | cg13619044 |
| chr7 | 30287790 | 30287791 | cg11994390 |
| chr7 | 30919932 | 30919933 | cg03475190 |
| chr7 | 32552212 | 32552213 | cg06133097 |
| chr7 | 35836570 | 35836571 | cg05903223 |
| chr7 | 35844045 | 35844046 | cg17168428 |
| chr7 | 35948145 | 35948146 | cg06334484 |
| chr7 | 36916137 | 36916138 | cg05998950 |
| chr7 | 38880852 | 38880853 | cg04957903 |
| chr7 | 41875113 | 41875114 | cg02998075 |
| chr7 | 41931493 | 41931494 | cg23796130 |
| chr7 | 43151168 | 43151169 | cg04442388 |
| chr7 | 44077761 | 44077762 | cg20608215 |
| chr7 | 44128218 | 44128219 | cg16073408 |
| chr7 | 44155564 | 44155565 | cg23671792 |
| chr7 | 47518657 | 47518658 | cg20782724 |
| chr7 | 47764246 | 47764247 | cg20430104 |
| chr7 | 47778880 | 47778881 | cg19195077 |
| chr7 | 47797675 | 47797676 | cg26203572 |
| chr7 | 47803072 | 47803073 | cg17614506 |
| chr7 | 50343298 | 50343299 | cg01139861 |
| chr7 | 50445456 | 50445457 | cg07849052 |
| chr7 | 55540725 | 55540726 | cg12015615 |
| chr7 | 55545188 | 55545189 | cg15599795 |
| chr7 | 55552130 | 55552131 | cg09286421 |
| chr7 | 55553375 | 55553376 | cg05391282 |
| chr7 | 55553462 | 55553463 | cg14876996 |
| chr7 | 55553545 | 55553546 | cg16182302 |
| chr7 | 55589803 | 55589804 | cg07459478 |
| chr7 | 55641049 | 55641050 | cg26106590 |
| chr7 | 56147772 | 56147773 | cg26790897 |
| chr7 | 64847121 | 64847122 | cg20049309 |
| chr7 | 72719729 | 72719730 | cg08192872 |
| chr7 | 74202486 | 74202487 | cg00668927 |
| chr7 | 74378824 | 74378825 | cg06796482 |
| chr7 | 74378838 | 74378839 | cg11322879 |
| chr7 | 74573718 | 74573719 | cg20111234 |
| chr7 | 75112219 | 75112220 | cg05817295 |
| chr7 | 75956931 | 75956932 | cg03707599 |
| chr7 | 75957040 | 75957041 | cg10738119 |
| chr7 | 75957061 | 75957062 | cg26162326 |
| chr7 | 75957123 | 75957124 | cg13616508 |
| chr7 | 75994082 | 75994083 | cg11549025 |
| chr7 | 76967860 | 76967861 | cg26205859 |
| chr7 | 77179087 | 77179088 | cg24401957 |
| chr7 | 84671582 | 84671583 | cg02607810 |
| chr7 | 86574444 | 86574445 | cg22719826 |

| | | | |
|---|---|---|---|
| chr5 | 176921820 | 176921821 | cg15225325 |
| chr5 | 176928448 | 176928449 | cg16548961 |
| chr5 | 176931422 | 176931423 | cg17200320 |
| chr5 | 177849197 | 177849198 | cg16181383 |
| chr5 | 177980755 | 177980756 | cg07841353 |
| chr5 | 178645682 | 178645683 | cg25550425 |
| chr5 | 179280345 | 179280346 | cg01213455 |
| chr5 | 179284731 | 179284732 | cg05432322 |
| chr5 | 179372900 | 179372901 | cg01169396 |
| chr5 | 179372946 | 179372947 | cg11810370 |
| chr5 | 179402240 | 179402241 | cg04265672 |
| chr6 | 236559 | 236560 | cg25019526 |
| chr6 | 290800 | 290801 | cg16103275 |
| chr6 | 312026 | 312027 | cg24319143 |
| chr6 | 317607 | 317608 | cg03624871 |
| chr6 | 348323 | 348324 | cg09779827 |
| chr6 | 394966 | 394967 | cg24319508 |
| chr6 | 397469 | 397470 | cg07817739 |
| chr6 | 447226 | 447227 | cg23001918 |
| chr6 | 447497 | 447498 | cg09757644 |
| chr6 | 447777 | 447778 | cg13299325 |
| chr6 | 466935 | 466936 | cg05444514 |
| chr6 | 503773 | 503774 | cg14101659 |
| chr6 | 503807 | 503808 | cg20949502 |
| chr6 | 525032 | 525033 | cg27562005 |
| chr6 | 611523 | 611524 | cg09806625 |
| chr6 | 691319 | 691320 | cg03107817 |
| chr6 | 1374513 | 1374514 | cg25640096 |
| chr6 | 1624442 | 1624443 | cg27095915 |
| chr6 | 2842550 | 2842551 | cg22977317 |
| chr6 | 2891973 | 2891974 | cg09046168 |
| chr6 | 2892050 | 2892051 | cg22376758 |
| chr6 | 2892148 | 2892149 | cg20726195 |
| chr6 | 2892150 | 2892151 | cg10863922 |
| chr6 | 2892152 | 2892153 | cg01345354 |
| chr6 | 2894011 | 2894012 | cg11296767 |
| chr6 | 2901712 | 2901713 | cg23963071 |
| chr6 | 3798615 | 3798616 | cg11611676 |
| chr6 | 4772890 | 4772891 | cg14913512 |
| chr6 | 4820670 | 4820671 | cg25938891 |
| chr6 | 7053734 | 7053735 | cg16554099 |
| chr6 | 7110773 | 7110774 | cg04418434 |
| chr6 | 7129714 | 7129715 | cg15815726 |
| chr6 | 7145478 | 7145479 | cg20893838 |
| chr6 | 7225223 | 7225224 | cg16958939 |
| chr6 | 7234581 | 7234582 | cg05295286 |
| chr6 | 7243643 | 7243644 | cg09326832 |
| chr6 | 8066098 | 8066099 | cg14267151 |
| chr6 | 8181746 | 8181747 | cg23164076 |
| chr6 | 11272495 | 11272496 | cg11406667 |
| chr6 | 11278005 | 11278006 | cg02029843 |
| chr6 | 11656844 | 11656845 | cg01413548 |
| chr6 | 11710597 | 11710598 | cg16533359 |
| chr6 | 11711971 | 11711972 | cg06193328 |
| chr6 | 11768095 | 11768096 | cg25589070 |
| chr6 | 11806115 | 11806116 | cg01699293 |

| | | | |
|---|---|---|---|
| chr7 | 86819455 | 86819456 | cg00221736 |
| chr7 | 89809444 | 89809445 | cg04070188 |
| chr7 | 90335803 | 90335804 | cg27403658 |
| chr7 | 90346445 | 90346446 | cg08583610 |
| chr7 | 91874085 | 91874086 | cg00035937 |
| chr7 | 92382963 | 92382964 | cg17959129 |
| chr7 | 92397161 | 92397162 | cg23705979 |
| chr7 | 92414346 | 92414347 | cg13788918 |
| chr7 | 92676158 | 92676159 | cg11416650 |
| chr7 | 93190462 | 93190463 | cg13916255 |
| chr7 | 95099917 | 95099918 | cg02819992 |
| chr7 | 95865631 | 95865632 | cg20782117 |
| chr7 | 97912641 | 97912642 | cg11366934 |
| chr7 | 100073281 | 100073282 | cg00376553 |
| chr7 | 100082322 | 100082323 | cg08626653 |
| chr7 | 100461644 | 100461645 | cg26426488 |
| chr7 | 101518619 | 101518620 | cg24420432 |
| chr7 | 101585659 | 101585660 | cg02131870 |
| chr7 | 103449893 | 103449894 | cg17713912 |
| chr7 | 104567387 | 104567388 | cg16314851 |
| chr7 | 104619961 | 104619962 | cg03997145 |
| chr7 | 104681396 | 104681397 | cg20770516 |
| chr7 | 104681440 | 104681441 | cg25076197 |
| chr7 | 104752143 | 104752144 | cg05400200 |
| chr7 | 104881282 | 104881283 | cg00571519 |
| chr7 | 104887042 | 104887043 | cg21261709 |
| chr7 | 105079663 | 105079664 | cg02813710 |
| chr7 | 105399252 | 105399253 | cg16963093 |
| chr7 | 105705154 | 105705155 | cg01083148 |
| chr7 | 112063079 | 112063080 | cg19724567 |
| chr7 | 115608503 | 115608504 | cg06493963 |
| chr7 | 115608801 | 115608802 | cg00927226 |
| chr7 | 115609683 | 115609684 | cg11479380 |
| chr7 | 115969424 | 115969425 | cg24249542 |
| chr7 | 115977777 | 115977778 | cg20301677 |
| chr7 | 116377467 | 116377468 | cg11319403 |
| chr7 | 116511339 | 116511340 | cg07485754 |
| chr7 | 120968288 | 120968289 | cg12073479 |
| chr7 | 121033957 | 121033958 | cg03485909 |
| chr7 | 124538660 | 124538661 | cg21792432 |
| chr7 | 127031314 | 127031315 | cg21841620 |
| chr7 | 127231698 | 127231699 | cg14938677 |
| chr7 | 127439670 | 127439671 | cg22282131 |
| chr7 | 127544716 | 127544717 | cg05967129 |
| chr7 | 127547490 | 127547491 | cg08625996 |
| chr7 | 127750971 | 127750972 | cg19414360 |
| chr7 | 128551672 | 128551673 | cg17150663 |
| chr7 | 128924101 | 128924102 | cg15488143 |
| chr7 | 129275471 | 129275472 | cg14735364 |
| chr7 | 129516344 | 129516345 | cg22145318 |
| chr7 | 129996313 | 129996314 | cg20917077 |
| chr7 | 130134488 | 130134489 | cg20876852 |
| chr7 | 130673538 | 130673539 | cg02695677 |
| chr7 | 130679521 | 130679522 | cg17001464 |
| chr7 | 130698005 | 130698006 | cg27337540 |
| chr7 | 133531244 | 133531245 | cg18217874 |

| | | | |
|---|---|---|---|
| chr6 | 11878092 | 11878093 | cg00955089 |
| chr6 | 12045381 | 12045382 | cg20583432 |
| chr6 | 12234001 | 12234002 | cg14622879 |
| chr6 | 13192742 | 13192743 | cg02220008 |
| chr6 | 13455261 | 13455262 | cg24594504 |
| chr6 | 13471187 | 13471188 | cg18342964 |
| chr6 | 13715593 | 13715594 | cg16278996 |
| chr6 | 14129370 | 14129371 | cg26525121 |
| chr6 | 14209372 | 14209373 | cg05168368 |
| chr6 | 14549941 | 14549942 | cg09571082 |
| chr6 | 14716193 | 14716194 | cg01559597 |
| chr6 | 14755316 | 14755317 | cg05368595 |
| chr6 | 14755320 | 14755321 | cg25162117 |
| chr6 | 14755359 | 14755360 | cg00530015 |
| chr6 | 14925093 | 14925094 | cg19917989 |
| chr6 | 15266782 | 15266783 | cg13077031 |
| chr6 | 15280024 | 15280025 | cg14662750 |
| chr6 | 15906515 | 15906516 | cg04541146 |
| chr6 | 16305180 | 16305181 | cg05567408 |
| chr6 | 16472305 | 16472306 | cg26794346 |
| chr6 | 16483997 | 16483998 | cg05968188 |
| chr6 | 16513829 | 16513830 | cg24776142 |
| chr6 | 16636157 | 16636158 | cg26843612 |
| chr6 | 16758889 | 16758890 | cg00464814 |
| chr6 | 18053826 | 18053827 | cg04918770 |
| chr6 | 18261541 | 18261542 | cg20895054 |
| chr6 | 18262523 | 18262524 | cg08451113 |
| chr6 | 18390303 | 18390304 | cg02059055 |
| chr6 | 19155350 | 19155351 | cg07523189 |
| chr6 | 19834248 | 19834249 | cg21904807 |
| chr6 | 19835100 | 19835101 | cg00909062 |
| chr6 | 20209127 | 20209128 | cg11639708 |
| chr6 | 20211373 | 20211374 | cg04369491 |
| chr6 | 20583779 | 20583780 | cg03477240 |
| chr6 | 20920015 | 20920016 | cg06008803 |
| chr6 | 21157865 | 21157866 | cg21498062 |
| chr6 | 21598595 | 21598596 | cg13995516 |
| chr6 | 21711700 | 21711701 | cg11728738 |
| chr6 | 22009091 | 22009092 | cg23902105 |
| chr6 | 22011878 | 22011879 | cg03146683 |
| chr6 | 22079302 | 22079303 | cg01639892 |
| chr6 | 22136363 | 22136364 | cg20414618 |
| chr6 | 22574143 | 22574144 | cg10459018 |
| chr6 | 22658507 | 22658508 | cg19077794 |
| chr6 | 23214308 | 23214309 | cg21196930 |
| chr6 | 24494462 | 24494463 | cg03967327 |
| chr6 | 24900051 | 24900052 | cg13708298 |
| chr6 | 24950870 | 24950871 | cg23876056 |
| chr6 | 25341389 | 25341390 | cg19938535 |
| chr6 | 26401691 | 26401692 | cg14004232 |
| chr6 | 26539327 | 26539328 | cg18635670 |
| chr6 | 26600049 | 26600050 | cg21579726 |
| chr6 | 27106564 | 27106565 | cg21777095 |
| chr6 | 28552582 | 28552583 | cg27333426 |
| chr6 | 28963785 | 28963786 | cg08811210 |
| chr6 | 28964909 | 28964910 | cg06527050 |
| chr7 | 134845376 | 134845377 | cg22121647 |
| chr7 | 134853645 | 134853646 | cg20830848 |
| chr7 | 134889789 | 134889790 | cg23081855 |
| chr7 | 134893191 | 134893192 | cg26988523 |
| chr7 | 134916876 | 134916877 | cg26502514 |
| chr7 | 134917144 | 134917145 | cg11649846 |
| chr7 | 134922321 | 134922322 | cg04839520 |
| chr7 | 137609343 | 137609344 | cg23697157 |
| chr7 | 137660322 | 137660323 | cg13016916 |
| chr7 | 137671105 | 137671106 | cg05041511 |
| chr7 | 137675881 | 137675882 | cg18091615 |
| chr7 | 138719784 | 138719785 | cg05576245 |
| chr7 | 138745709 | 138745710 | cg00303210 |
| chr7 | 139332087 | 139332088 | cg15535471 |
| chr7 | 139333352 | 139333353 | cg23999318 |
| chr7 | 139468310 | 139468311 | cg06079564 |
| chr7 | 139637446 | 139637447 | cg06750564 |
| chr7 | 139759577 | 139759578 | cg19798832 |
| chr7 | 140026277 | 140026278 | cg14385298 |
| chr7 | 140043078 | 140043079 | cg01012242 |
| chr7 | 140174476 | 140174477 | cg19494591 |
| chr7 | 140556696 | 140556697 | cg12750675 |
| chr7 | 141398961 | 141398962 | cg02574033 |
| chr7 | 141811005 | 141811006 | cg26445785 |
| chr7 | 142937561 | 142937562 | cg06081238 |
| chr7 | 143013207 | 143013208 | cg04039957 |
| chr7 | 147987450 | 147987451 | cg15208525 |
| chr7 | 148420058 | 148420059 | cg21828951 |
| chr7 | 150067896 | 150067897 | cg09378425 |
| chr7 | 150125199 | 150125200 | cg04301738 |
| chr7 | 150130069 | 150130070 | cg26837768 |
| chr7 | 150669058 | 150669059 | cg15510325 |
| chr7 | 150936167 | 150936168 | cg05878101 |
| chr7 | 151077901 | 151077902 | cg02468770 |
| chr7 | 151109656 | 151109657 | cg06263887 |
| chr7 | 151328116 | 151328117 | cg09449988 |
| chr7 | 155790771 | 155790772 | cg14635659 |
| chr7 | 155790956 | 155790957 | cg09016797 |
| chr7 | 155791233 | 155791234 | cg04591648 |
| chr7 | 155796902 | 155796903 | cg15078841 |
| chr7 | 155796939 | 155796940 | cg05325633 |
| chr7 | 156409058 | 156409059 | cg16771407 |
| chr7 | 158578750 | 158578751 | cg27159987 |
| chr8 | 615351 | 615352 | cg08885114 |
| chr8 | 1726903 | 1726904 | cg16317553 |
| chr8 | 1734990 | 1734991 | cg02236941 |
| chr8 | 6287904 | 6287905 | cg11043092 |
| chr8 | 6565084 | 6565085 | cg24470692 |
| chr8 | 6584320 | 6584321 | cg26814335 |
| chr8 | 6590103 | 6590104 | cg25004868 |
| chr8 | 8176714 | 8176715 | cg23015041 |
| chr8 | 8182003 | 8182004 | cg27000590 |
| chr8 | 8188483 | 8188484 | cg25539628 |
| chr8 | 8256332 | 8256333 | cg10597661 |
| chr8 | 8420735 | 8420736 | cg18226700 |
| chr8 | 8681198 | 8681199 | cg22680252 |

| | | | |
|---|---|---|---|
| chr6 | 28966423 | 28966424 | cg09088396 |
| chr6 | 29528119 | 29528120 | cg07326586 |
| chr6 | 29601489 | 29601490 | cg17099072 |
| chr6 | 29601491 | 29601492 | cg10234998 |
| chr6 | 29601498 | 29601499 | cg21481950 |
| chr6 | 29601556 | 29601557 | cg12061917 |
| chr6 | 29601705 | 29601706 | cg19022254 |
| chr6 | 29705939 | 29705940 | cg16289618 |
| chr6 | 29717651 | 29717652 | cg06097213 |
| chr6 | 29717917 | 29717918 | cg23606396 |
| chr6 | 29833169 | 29833170 | cg27632493 |
| chr6 | 30010457 | 30010458 | cg26354367 |
| chr6 | 30033850 | 30033851 | cg06083200 |
| chr6 | 30038791 | 30038792 | cg17322683 |
| chr6 | 30136790 | 30136791 | cg24592546 |
| chr6 | 30167065 | 30167066 | cg19988490 |
| chr6 | 30169120 | 30169121 | cg03782662 |
| chr6 | 30174845 | 30174846 | cg05850884 |
| chr6 | 30175074 | 30175075 | cg14799927 |
| chr6 | 30175092 | 30175093 | cg19714347 |
| chr6 | 30175305 | 30175306 | cg20345556 |
| chr6 | 30175327 | 30175328 | cg14971718 |
| chr6 | 30176093 | 30176094 | cg02749752 |
| chr6 | 30179435 | 30179436 | cg00206463 |
| chr6 | 30227001 | 30227002 | cg14296767 |
| chr6 | 30290443 | 30290444 | cg26193268 |
| chr6 | 30565190 | 30565191 | cg22366909 |
| chr6 | 30569105 | 30569106 | cg16586538 |
| chr6 | 30580581 | 30580582 | cg26503792 |
| chr6 | 30582352 | 30582353 | cg04694022 |
| chr6 | 30619080 | 30619081 | cg26088561 |
| chr6 | 30619113 | 30619114 | cg18048542 |
| chr6 | 30619167 | 30619168 | cg01676996 |
| chr6 | 30619209 | 30619210 | cg05143092 |
| chr6 | 30619221 | 30619222 | cg22752886 |
| chr6 | 30704955 | 30704956 | cg09179723 |
| chr6 | 30704998 | 30704999 | cg13534892 |
| chr6 | 30709012 | 30709013 | cg11285284 |
| chr6 | 30720491 | 30720492 | cg16137928 |
| chr6 | 30737007 | 30737008 | cg14129735 |
| chr6 | 30737068 | 30737069 | cg20873416 |
| chr6 | 30850543 | 30850544 | cg08951271 |
| chr6 | 30851638 | 30851639 | cg03270204 |
| chr6 | 31038713 | 31038714 | cg16519217 |
| chr6 | 31091345 | 31091346 | cg22402850 |
| chr6 | 31091383 | 31091384 | cg26888012 |
| chr6 | 31094068 | 31094069 | cg17598552 |
| chr6 | 31105927 | 31105928 | cg27314422 |
| chr6 | 31106045 | 31106046 | cg13671374 |
| chr6 | 31127863 | 31127864 | cg16095155 |
| chr6 | 31132637 | 31132638 | cg15316118 |
| chr6 | 31238388 | 31238389 | cg18511546 |
| chr6 | 31322914 | 31322915 | cg23923934 |
| chr6 | 31507879 | 31507880 | cg26408612 |
| chr6 | 31522510 | 31522511 | cg03077969 |
| chr6 | 31528915 | 31528916 | cg15852826 |
| chr8 | 8860050 | 8860051 | cg18237548 |
| chr8 | 8997420 | 8997421 | cg01887881 |
| chr8 | 9416736 | 9416737 | cg17969579 |
| chr8 | 9522664 | 9522665 | cg13088432 |
| chr8 | 10823238 | 10823239 | cg20858107 |
| chr8 | 11413043 | 11413044 | cg15685006 |
| chr8 | 11413186 | 11413187 | cg26161004 |
| chr8 | 11413234 | 11413235 | cg14702231 |
| chr8 | 11417517 | 11417518 | cg04335815 |
| chr8 | 11471395 | 11471396 | cg09817024 |
| chr8 | 11717853 | 11717854 | cg16624891 |
| chr8 | 17822162 | 17822163 | cg04443233 |
| chr8 | 17943293 | 17943294 | cg04912297 |
| chr8 | 19310134 | 19310135 | cg18741439 |
| chr8 | 19369752 | 19369753 | cg17487784 |
| chr8 | 19555033 | 19555034 | cg00002190 |
| chr8 | 19796335 | 19796336 | cg16125291 |
| chr8 | 20080666 | 20080667 | cg16479633 |
| chr8 | 21771540 | 21771541 | cg08288130 |
| chr8 | 21771668 | 21771669 | cg16476235 |
| chr8 | 21780820 | 21780821 | cg22272713 |
| chr8 | 21961713 | 21961714 | cg03520938 |
| chr8 | 22224272 | 22224273 | cg14348540 |
| chr8 | 22421029 | 22421030 | cg03478916 |
| chr8 | 22485721 | 22485722 | cg25764534 |
| chr8 | 22561445 | 22561446 | cg03720745 |
| chr8 | 22561893 | 22561894 | cg23302740 |
| chr8 | 22839080 | 22839081 | cg12082089 |
| chr8 | 22847209 | 22847210 | cg03437605 |
| chr8 | 22987034 | 22987035 | cg09938634 |
| chr8 | 23418389 | 23418390 | cg24917065 |
| chr8 | 26171697 | 26171698 | cg08201604 |
| chr8 | 27306846 | 27306847 | cg08498406 |
| chr8 | 28174455 | 28174456 | cg13077562 |
| chr8 | 28206278 | 28206279 | cg17199018 |
| chr8 | 28241186 | 28241187 | cg23475021 |
| chr8 | 28244769 | 28244770 | cg08606497 |
| chr8 | 28260790 | 28260791 | cg26734167 |
| chr8 | 28727543 | 28727544 | cg26890195 |
| chr8 | 28745680 | 28745681 | cg11385906 |
| chr8 | 28908912 | 28908913 | cg20212339 |
| chr8 | 28912915 | 28912916 | cg16847760 |
| chr8 | 28918821 | 28918822 | cg11893955 |
| chr8 | 28961315 | 28961316 | cg07023317 |
| chr8 | 28961333 | 28961334 | cg25596405 |
| chr8 | 28961356 | 28961357 | cg09322573 |
| chr8 | 29033680 | 29033681 | cg00328219 |
| chr8 | 29082964 | 29082965 | cg20106866 |
| chr8 | 29605992 | 29605993 | cg20521277 |
| chr8 | 29630611 | 29630612 | cg23223330 |
| chr8 | 29937472 | 29937473 | cg18813599 |
| chr8 | 29944442 | 29944443 | cg19993591 |
| chr8 | 29957344 | 29957345 | cg16536330 |
| chr8 | 30254923 | 30254924 | cg13456241 |
| chr8 | 32985289 | 32985290 | cg06106484 |
| chr8 | 33341954 | 33341955 | cg23663332 |

| | | | |
|---|---|---|---|
| chr6 | 31529154 | 31529155 | cg20826024 |
| chr6 | 31529897 | 31529898 | cg19367859 |
| chr6 | 31550612 | 31550613 | cg02249390 |
| chr6 | 31550699 | 31550700 | cg06887580 |
| chr6 | 31550842 | 31550843 | cg18384277 |
| chr6 | 31560441 | 31560442 | cg27490128 |
| chr6 | 31596847 | 31596848 | cg15449023 |
| chr6 | 31617472 | 31617473 | cg08899215 |
| chr6 | 31617517 | 31617518 | cg26804733 |
| chr6 | 31617530 | 31617531 | cg27000659 |
| chr6 | 31625701 | 31625702 | cg00812578 |
| chr6 | 31627653 | 31627654 | cg26886875 |
| chr6 | 31627678 | 31627679 | cg15415945 |
| chr6 | 31629900 | 31629901 | cg26969933 |
| chr6 | 31629909 | 31629910 | cg23666117 |
| chr6 | 31701110 | 31701111 | cg18909389 |
| chr6 | 31733619 | 31733620 | cg00686197 |
| chr6 | 31780526 | 31780527 | cg24143221 |
| chr6 | 31801032 | 31801033 | cg17552909 |
| chr6 | 31801042 | 31801043 | cg24375209 |
| chr6 | 31832962 | 31832963 | cg02775414 |
| chr6 | 31833153 | 31833154 | cg23701033 |
| chr6 | 31833162 | 31833163 | cg05686323 |
| chr6 | 31833199 | 31833200 | cg07546508 |
| chr6 | 31864106 | 31864107 | cg14955103 |
| chr6 | 32038177 | 32038178 | cg27073361 |
| chr6 | 32041191 | 32041192 | cg02248749 |
| chr6 | 32054441 | 32054442 | cg23636802 |
| chr6 | 32055007 | 32055008 | cg22547540 |
| chr6 | 32134865 | 32134866 | cg16468631 |
| chr6 | 32140758 | 32140759 | cg04528352 |
| chr6 | 32155082 | 32155083 | cg09001226 |
| chr6 | 32158953 | 32158954 | cg03280235 |
| chr6 | 32165321 | 32165322 | cg13185046 |
| chr6 | 32167840 | 32167841 | cg14004552 |
| chr6 | 32181764 | 32181765 | cg15318627 |
| chr6 | 32182013 | 32182014 | cg16602097 |
| chr6 | 32185171 | 32185172 | cg06535498 |
| chr6 | 32187918 | 32187919 | cg08939974 |
| chr6 | 32294470 | 32294471 | cg06192961 |
| chr6 | 32294503 | 32294504 | cg17039645 |
| chr6 | 32299617 | 32299618 | cg17201409 |
| chr6 | 32299649 | 32299650 | cg06545464 |
| chr6 | 32310527 | 32310528 | cg09966430 |
| chr6 | 32311517 | 32311518 | cg00861856 |
| chr6 | 32373089 | 32373090 | cg26657086 |
| chr6 | 32410873 | 32410874 | cg05500783 |
| chr6 | 32411067 | 32411068 | cg25645491 |
| chr6 | 32411144 | 32411145 | cg06968241 |
| chr6 | 32609783 | 32609784 | cg13778567 |
| chr6 | 32714334 | 32714335 | cg17524162 |
| chr6 | 32796216 | 32796217 | cg08327277 |
| chr6 | 32800591 | 32800592 | cg09187413 |
| chr6 | 32810833 | 32810834 | cg16186435 |
| chr6 | 32819858 | 32819859 | cg26033526 |
| chr6 | 32819911 | 32819912 | cg01673307 |
| chr8 | 33356290 | 33356291 | cg26426774 |
| chr8 | 37556386 | 37556387 | cg14602087 |
| chr8 | 37700802 | 37700803 | cg03171419 |
| chr8 | 38090208 | 38090209 | cg04500377 |
| chr8 | 38125449 | 38125450 | cg25405452 |
| chr8 | 38125865 | 38125866 | cg19272266 |
| chr8 | 38133060 | 38133061 | cg20170882 |
| chr8 | 38236714 | 38236715 | cg24258108 |
| chr8 | 38770414 | 38770415 | cg13514772 |
| chr8 | 38782090 | 38782091 | cg15717617 |
| chr8 | 41140500 | 41140501 | cg17486234 |
| chr8 | 41500461 | 41500462 | cg23100365 |
| chr8 | 41504993 | 41504994 | cg25883020 |
| chr8 | 41523480 | 41523481 | cg13432087 |
| chr8 | 41523968 | 41523969 | cg12003463 |
| chr8 | 41873091 | 41873092 | cg21706771 |
| chr8 | 42188597 | 42188598 | cg07696514 |
| chr8 | 42234798 | 42234799 | cg09297903 |
| chr8 | 42234803 | 42234804 | cg02571142 |
| chr8 | 42659977 | 42659978 | cg06815361 |
| chr8 | 47761906 | 47761907 | cg12861622 |
| chr8 | 52801982 | 52801983 | cg00836007 |
| chr8 | 52852168 | 52852169 | cg26275360 |
| chr8 | 54757132 | 54757133 | cg12047375 |
| chr8 | 56737276 | 56737277 | cg14591655 |
| chr8 | 56869208 | 56869209 | cg03359337 |
| chr8 | 58168222 | 58168223 | cg23826993 |
| chr8 | 59170968 | 59170969 | cg00028299 |
| chr8 | 59650783 | 59650784 | cg11413576 |
| chr8 | 60020513 | 60020514 | cg06779553 |
| chr8 | 61563866 | 61563867 | cg21205855 |
| chr8 | 61593696 | 61593697 | cg05575639 |
| chr8 | 61777711 | 61777712 | cg14719959 |
| chr8 | 61789893 | 61789894 | cg26460578 |
| chr8 | 61823027 | 61823028 | cg20904475 |
| chr8 | 61861713 | 61861714 | cg24800616 |
| chr8 | 61910949 | 61910950 | cg12171761 |
| chr8 | 65496126 | 65496127 | cg07205627 |
| chr8 | 66711941 | 66711942 | cg27410091 |
| chr8 | 67023811 | 67023812 | cg15882353 |
| chr8 | 67417740 | 67417741 | cg15798873 |
| chr8 | 67417790 | 67417791 | cg09622557 |
| chr8 | 67417875 | 67417876 | cg15070801 |
| chr8 | 67434522 | 67434523 | cg27131563 |
| chr8 | 67434868 | 67434869 | cg08781403 |
| chr8 | 67446912 | 67446913 | cg09610332 |
| chr8 | 67467992 | 67467993 | cg03656863 |
| chr8 | 67591714 | 67591715 | cg08633239 |
| chr8 | 67592861 | 67592862 | cg17724184 |
| chr8 | 68659328 | 68659329 | cg19250907 |
| chr8 | 68879947 | 68879948 | cg10419957 |
| chr8 | 70584832 | 70584833 | cg07998216 |
| chr8 | 70845447 | 70845448 | cg08976334 |
| chr8 | 71060638 | 71060639 | cg20699548 |
| chr8 | 71063694 | 71063695 | cg18227776 |
| chr8 | 71527496 | 71527497 | cg26228558 |

| | | | |
|---|---|---|---|
| chr6 | 32819921 | 32819922 | cg24111025 |
| chr6 | 32820102 | 32820103 | cg06473288 |
| chr6 | 32820201 | 32820202 | cg17626301 |
| chr6 | 32820214 | 32820215 | cg26234900 |
| chr6 | 32820249 | 32820250 | cg10666909 |
| chr6 | 32823292 | 32823293 | cg06791592 |
| chr6 | 32823728 | 32823729 | cg10817441 |
| chr6 | 32828996 | 32828997 | cg23636571 |
| chr6 | 32904074 | 32904075 | cg13524037 |
| chr6 | 32904402 | 32904403 | cg05525374 |
| chr6 | 32904419 | 32904420 | cg09017117 |
| chr6 | 32906857 | 32906858 | cg23527920 |
| chr6 | 32916606 | 32916607 | cg15418419 |
| chr6 | 32917065 | 32917066 | cg26905720 |
| chr6 | 32917093 | 32917094 | cg01910638 |
| chr6 | 32917165 | 32917166 | cg07627491 |
| chr6 | 32917240 | 32917241 | cg24713259 |
| chr6 | 32918123 | 32918124 | cg21992044 |
| chr6 | 32918306 | 32918307 | cg01782709 |
| chr6 | 32918419 | 32918420 | cg18031239 |
| chr6 | 32918427 | 32918428 | cg06006623 |
| chr6 | 32918479 | 32918480 | cg02015582 |
| chr6 | 32918659 | 32918660 | cg04111611 |
| chr6 | 32919427 | 32919428 | cg12682872 |
| chr6 | 32919431 | 32919432 | cg14851271 |
| chr6 | 32919433 | 32919434 | cg25285720 |
| chr6 | 32942714 | 32942715 | cg07275218 |
| chr6 | 32945906 | 32945907 | cg02707277 |
| chr6 | 32975145 | 32975146 | cg14490686 |
| chr6 | 32975336 | 32975337 | cg06954660 |
| chr6 | 33043072 | 33043073 | cg23750365 |
| chr6 | 33043220 | 33043221 | cg21151963 |
| chr6 | 33046582 | 33046583 | cg10136841 |
| chr6 | 33049983 | 33049984 | cg20981163 |
| chr6 | 33050741 | 33050742 | cg23333490 |
| chr6 | 33081024 | 33081025 | cg10735599 |
| chr6 | 33081351 | 33081352 | cg25119261 |
| chr6 | 33083457 | 33083458 | cg17763566 |
| chr6 | 33084608 | 33084609 | cg27264993 |
| chr6 | 33084741 | 33084742 | cg08088295 |
| chr6 | 33084825 | 33084826 | cg24465429 |
| chr6 | 33084888 | 33084889 | cg24185832 |
| chr6 | 33084922 | 33084923 | cg04315646 |
| chr6 | 33084928 | 33084929 | cg05337177 |
| chr6 | 33085063 | 33085064 | cg06210070 |
| chr6 | 33130918 | 33130919 | cg02890435 |
| chr6 | 33161290 | 33161291 | cg25459558 |
| chr6 | 33180402 | 33180403 | cg16529229 |
| chr6 | 33180415 | 33180416 | cg12721695 |
| chr6 | 33243775 | 33243776 | cg26912426 |
| chr6 | 33246105 | 33246106 | cg09080120 |
| chr6 | 33246149 | 33246150 | cg16580935 |
| chr6 | 33246157 | 33246158 | cg14023774 |
| chr6 | 33247509 | 33247510 | cg02299465 |
| chr6 | 33248616 | 33248617 | cg16428857 |
| chr6 | 33249075 | 33249076 | cg21334198 |

| | | | |
|---|---|---|---|
| chr8 | 72761320 | 72761321 | cg04910834 |
| chr8 | 74416786 | 74416787 | cg21585977 |
| chr8 | 74750772 | 74750773 | cg25597976 |
| chr8 | 81077480 | 81077481 | cg23342367 |
| chr8 | 81408357 | 81408358 | cg20176142 |
| chr8 | 81963379 | 81963380 | cg10609868 |
| chr8 | 82028084 | 82028085 | cg09291217 |
| chr8 | 86090773 | 86090774 | cg00849990 |
| chr8 | 91017761 | 91017762 | cg27040700 |
| chr8 | 91650159 | 91650160 | cg05458554 |
| chr8 | 91659356 | 91659357 | cg01528492 |
| chr8 | 94898169 | 94898170 | cg00630761 |
| chr8 | 96142671 | 96142672 | cg07556057 |
| chr8 | 97658325 | 97658326 | cg03613618 |
| chr8 | 101662246 | 101662247 | cg04410446 |
| chr8 | 101662628 | 101662629 | cg17808631 |
| chr8 | 102898712 | 102898713 | cg01971590 |
| chr8 | 103135208 | 103135209 | cg27008565 |
| chr8 | 103780068 | 103780069 | cg23312086 |
| chr8 | 103794922 | 103794923 | cg20596543 |
| chr8 | 103816049 | 103816050 | cg21629325 |
| chr8 | 103817950 | 103817951 | cg11313780 |
| chr8 | 103818062 | 103818063 | cg06892907 |
| chr8 | 103821171 | 103821172 | cg19611886 |
| chr8 | 103943308 | 103943309 | cg20627646 |
| chr8 | 104362929 | 104362930 | cg12726906 |
| chr8 | 105341338 | 105341339 | cg00206779 |
| chr8 | 105351661 | 105351662 | cg08657449 |
| chr8 | 107672853 | 107672854 | cg08924256 |
| chr8 | 115301887 | 115301888 | cg13732895 |
| chr8 | 116575902 | 116575903 | cg27039118 |
| chr8 | 117650383 | 117650384 | cg24004040 |
| chr8 | 119121281 | 119121282 | cg05783185 |
| chr8 | 119152014 | 119152015 | cg07463445 |
| chr8 | 123798823 | 123798824 | cg03314365 |
| chr8 | 123981935 | 123981936 | cg23822407 |
| chr8 | 124284176 | 124284177 | cg10451262 |
| chr8 | 124284393 | 124284394 | cg20987431 |
| chr8 | 126302396 | 126302397 | cg02006142 |
| chr8 | 128191730 | 128191731 | cg20197756 |
| chr8 | 128264282 | 128264283 | cg17095489 |
| chr8 | 128902684 | 128902685 | cg06986734 |
| chr8 | 129526327 | 129526328 | cg18720393 |
| chr8 | 129604293 | 129604294 | cg05584990 |
| chr8 | 131414614 | 131414615 | cg00659129 |
| chr8 | 133699791 | 133699792 | cg04445427 |
| chr8 | 133773093 | 133773094 | cg10054641 |
| chr8 | 133773099 | 133773100 | cg05598678 |
| chr8 | 133773115 | 133773116 | cg23313885 |
| chr8 | 134039099 | 134039100 | cg17500680 |
| chr8 | 134062192 | 134062193 | cg06772221 |
| chr8 | 134066590 | 134066591 | cg17088014 |
| chr8 | 134081000 | 134081001 | cg11883141 |
| chr8 | 134307597 | 134307598 | cg17129188 |
| chr8 | 134585815 | 134585816 | cg23336702 |
| chr8 | 135634128 | 135634129 | cg06293872 |

| | | | |
|---|---|---|---|
| chr6 | 33273477 | 33273478 | cg11396585 |
| chr6 | 33273716 | 33273717 | cg02191909 |
| chr6 | 33273965 | 33273966 | cg00988641 |
| chr6 | 33278333 | 33278334 | cg10036001 |
| chr6 | 33286244 | 33286245 | cg26887212 |
| chr6 | 33289660 | 33289661 | cg18113453 |
| chr6 | 33386985 | 33386986 | cg25767433 |
| chr6 | 33387003 | 33387004 | cg01320573 |
| chr6 | 34002114 | 34002115 | cg01140008 |
| chr6 | 34437227 | 34437228 | cg27000690 |
| chr6 | 34660294 | 34660295 | cg06810293 |
| chr6 | 35228319 | 35228320 | cg06656924 |
| chr6 | 35466777 | 35466778 | cg22548485 |
| chr6 | 35654363 | 35654364 | cg03546163 |
| chr6 | 35696870 | 35696871 | cg25114611 |
| chr6 | 36088380 | 36088381 | cg11075561 |
| chr6 | 36094917 | 36094918 | cg04449473 |
| chr6 | 36304683 | 36304684 | cg14673936 |
| chr6 | 36305155 | 36305156 | cg18520925 |
| chr6 | 36761843 | 36761844 | cg23024326 |
| chr6 | 37102603 | 37102604 | cg00377681 |
| chr6 | 37458535 | 37458536 | cg18391972 |
| chr6 | 37465801 | 37465802 | cg00885461 |
| chr6 | 37504610 | 37504611 | cg26579986 |
| chr6 | 38123312 | 38123313 | cg04316353 |
| chr6 | 38945383 | 38945384 | cg14464563 |
| chr6 | 39683259 | 39683260 | cg03139423 |
| chr6 | 40291048 | 40291049 | cg07581070 |
| chr6 | 40324498 | 40324499 | cg06112171 |
| chr6 | 40347475 | 40347476 | cg12797522 |
| chr6 | 40698276 | 40698277 | cg22653878 |
| chr6 | 41512853 | 41512854 | cg04617914 |
| chr6 | 41665888 | 41665889 | cg15783349 |
| chr6 | 41885417 | 41885418 | cg05689413 |
| chr6 | 41903172 | 41903173 | cg17159550 |
| chr6 | 41907198 | 41907199 | cg05568549 |
| chr6 | 42209899 | 42209900 | cg04291097 |
| chr6 | 42397121 | 42397122 | cg09804858 |
| chr6 | 42716310 | 42716311 | cg26069291 |
| chr6 | 42900123 | 42900124 | cg07710211 |
| chr6 | 42900231 | 42900232 | cg13387050 |
| chr6 | 42931581 | 42931582 | cg11336178 |
| chr6 | 43197675 | 43197676 | cg00741333 |
| chr6 | 43231063 | 43231064 | cg10965508 |
| chr6 | 43279607 | 43279608 | cg06407843 |
| chr6 | 43604584 | 43604585 | cg02447620 |
| chr6 | 43744388 | 43744389 | cg16907527 |
| chr6 | 44088275 | 44088276 | cg22020299 |
| chr6 | 44101190 | 44101191 | cg04074908 |
| chr6 | 44222845 | 44222846 | cg00071051 |
| chr6 | 44229440 | 44229441 | cg19707503 |
| chr6 | 44229550 | 44229551 | cg11913951 |
| chr6 | 44234511 | 44234512 | cg19109431 |
| chr6 | 44698962 | 44698963 | cg23155089 |
| chr6 | 44796705 | 44796706 | cg08493590 |
| chr6 | 46720370 | 46720371 | cg00170295 |

| | | | |
|---|---|---|---|
| chr8 | 135842613 | 135842614 | cg11466449 |
| chr8 | 141588750 | 141588751 | cg06895124 |
| chr8 | 142159992 | 142159993 | cg12807790 |
| chr8 | 142200140 | 142200141 | cg16494567 |
| chr8 | 142217653 | 142217654 | cg08205396 |
| chr8 | 142262761 | 142262762 | cg08958952 |
| chr8 | 142263053 | 142263054 | cg21539907 |
| chr8 | 142288508 | 142288509 | cg11547696 |
| chr8 | 142288517 | 142288518 | cg25877009 |
| chr8 | 142297519 | 142297520 | cg25489732 |
| chr8 | 144099482 | 144099483 | cg17052170 |
| chr8 | 144105210 | 144105211 | cg16400320 |
| chr8 | 144416327 | 144416328 | cg08597832 |
| chr8 | 144416485 | 144416486 | cg12188860 |
| chr8 | 144543486 | 144543487 | cg11848483 |
| chr8 | 144543568 | 144543569 | cg04180114 |
| chr8 | 144543750 | 144543751 | cg20156237 |
| chr8 | 144577094 | 144577095 | cg11996592 |
| chr8 | 144577185 | 144577186 | cg11303301 |
| chr8 | 144577238 | 144577239 | cg24974704 |
| chr8 | 144597448 | 144597449 | cg23318523 |
| chr8 | 144650595 | 144650596 | cg04022537 |
| chr8 | 145014989 | 145014990 | cg03958308 |
| chr8 | 145015122 | 145015123 | cg02623114 |
| chr8 | 145025760 | 145025761 | cg17327067 |
| chr8 | 145045235 | 145045236 | cg03792042 |
| chr8 | 145061234 | 145061235 | cg10787442 |
| chr8 | 145061291 | 145061292 | cg16796959 |
| chr8 | 145068773 | 145068774 | cg07212837 |
| chr8 | 145161551 | 145161552 | cg22481882 |
| chr8 | 145239316 | 145239317 | cg02389682 |
| chr8 | 145578911 | 145578912 | cg04843168 |
| chr8 | 145812770 | 145812771 | cg03657589 |
| chr9 | 176893 | 176894 | cg14436491 |
| chr9 | 211689 | 211690 | cg14500300 |
| chr9 | 857563 | 857564 | cg13619074 |
| chr9 | 4984206 | 4984207 | cg20394284 |
| chr9 | 5760346 | 5760347 | cg14419441 |
| chr9 | 22203744 | 22203745 | cg13819609 |
| chr9 | 33140269 | 33140270 | cg13858803 |
| chr9 | 33168018 | 33168019 | cg05779272 |
| chr9 | 34098451 | 34098452 | cg14412639 |
| chr9 | 35610380 | 35610381 | cg13918628 |
| chr9 | 35689475 | 35689476 | cg13969788 |
| chr9 | 36985761 | 36985762 | cg03261800 |
| chr9 | 37002685 | 37002686 | cg13482308 |
| chr9 | 37040454 | 37040455 | cg13753527 |
| chr9 | 37574338 | 37574339 | cg13893555 |
| chr9 | 82278827 | 82278828 | cg13598109 |
| chr9 | 82308107 | 82308108 | cg14048465 |
| chr9 | 88419213 | 88419214 | cg12472181 |
| chr9 | 91995929 | 91995930 | cg14541848 |
| chr9 | 91996229 | 91996230 | cg02702041 |
| chr9 | 92050313 | 92050314 | cg13619387 |
| chr9 | 94184131 | 94184132 | cg04521449 |
| chr9 | 95475751 | 95475752 | cg14184780 |

| | | | |
|---|---|---|---|
| chr6 | 47234372 | 47234373 | cg07989740 |
| chr6 | 47326353 | 47326354 | cg16769649 |
| chr6 | 47749714 | 47749715 | cg22852840 |
| chr6 | 50947261 | 50947262 | cg14529201 |
| chr6 | 51481908 | 51481909 | cg01776172 |
| chr6 | 51814354 | 51814355 | cg27255329 |
| chr6 | 52009154 | 52009155 | cg25284213 |
| chr6 | 52147973 | 52147974 | cg02243303 |
| chr6 | 52225658 | 52225659 | cg06052888 |
| chr6 | 52284109 | 52284110 | cg07654982 |
| chr6 | 52363701 | 52363702 | cg22388471 |
| chr6 | 52382441 | 52382442 | cg26020069 |
| chr6 | 53181687 | 53181688 | cg16946445 |
| chr6 | 53181709 | 53181710 | cg04658615 |
| chr6 | 53187374 | 53187375 | cg09387914 |
| chr6 | 53208623 | 53208624 | cg12297970 |
| chr6 | 54172183 | 54172184 | cg12397274 |
| chr6 | 55763658 | 55763659 | cg06210548 |
| chr6 | 56387378 | 56387379 | cg01033419 |
| chr6 | 56400181 | 56400182 | cg05537607 |
| chr6 | 62976042 | 62976043 | cg26556196 |
| chr6 | 63038652 | 63038653 | cg06551177 |
| chr6 | 64226341 | 64226342 | cg01986648 |
| chr6 | 66493769 | 66493770 | cg20159490 |
| chr6 | 70501068 | 70501069 | cg01735991 |
| chr6 | 70501081 | 70501082 | cg01386735 |
| chr6 | 72113416 | 72113417 | cg20815778 |
| chr6 | 72114693 | 72114694 | cg16506910 |
| chr6 | 72455356 | 72455357 | cg01546561 |
| chr6 | 74268449 | 74268450 | cg19399370 |
| chr6 | 74289980 | 74289981 | cg21282907 |
| chr6 | 76636725 | 76636726 | cg22803222 |
| chr6 | 79940953 | 79940954 | cg16343616 |
| chr6 | 80121727 | 80121728 | cg20393695 |
| chr6 | 80411369 | 80411370 | cg14515801 |
| chr6 | 81252663 | 81252664 | cg26031192 |
| chr6 | 82460558 | 82460559 | cg22388948 |
| chr6 | 86171768 | 86171769 | cg24702826 |
| chr6 | 88414961 | 88414962 | cg01767230 |
| chr6 | 90598826 | 90598827 | cg24360137 |
| chr6 | 90878637 | 90878638 | cg09818385 |
| chr6 | 91194267 | 91194268 | cg03718509 |
| chr6 | 94509563 | 94509564 | cg09103598 |
| chr6 | 98373013 | 98373014 | cg26342141 |
| chr6 | 100581833 | 100581834 | cg08881610 |
| chr6 | 100624512 | 100624513 | cg21496187 |
| chr6 | 100680152 | 100680153 | cg18446744 |
| chr6 | 106535425 | 106535426 | cg02108623 |
| chr6 | 106957404 | 106957405 | cg27632402 |
| chr6 | 107057773 | 107057774 | cg27007060 |
| chr6 | 107078648 | 107078649 | cg07476030 |
| chr6 | 107235623 | 107235624 | cg14326909 |
| chr6 | 107816677 | 107816678 | cg05726239 |
| chr6 | 108026281 | 108026282 | cg02258846 |
| chr6 | 109127593 | 109127594 | cg01659184 |
| chr6 | 109266451 | 109266452 | cg12271487 |

| | | | |
|---|---|---|---|
| chr9 | 95737070 | 95737071 | cg14263098 |
| chr9 | 95819930 | 95819931 | cg05675356 |
| chr9 | 95889381 | 95889382 | cg14474777 |
| chr9 | 99615182 | 99615183 | cg13457522 |
| chr9 | 100502742 | 100502743 | cg14132620 |
| chr9 | 100952629 | 100952630 | cg12800915 |
| chr9 | 101759749 | 101759750 | cg14259208 |
| chr9 | 107687761 | 107687762 | cg14564724 |
| chr9 | 111880884 | 111880885 | cg05043794 |
| chr9 | 112887903 | 112887904 | cg21171115 |
| chr9 | 112889386 | 112889387 | cg14170787 |
| chr9 | 112914149 | 112914150 | cg13430173 |
| chr9 | 114545810 | 114545811 | cg01822570 |
| chr9 | 114546247 | 114546248 | cg07751669 |
| chr9 | 114658195 | 114658196 | cg01472113 |
| chr9 | 114706971 | 114706972 | cg13633625 |
| chr9 | 114836121 | 114836122 | cg14618635 |
| chr9 | 115631083 | 115631084 | cg14350114 |
| chr9 | 116114796 | 116114797 | cg14245185 |
| chr9 | 117228122 | 117228123 | cg14623754 |
| chr9 | 117265998 | 117265999 | cg00305629 |
| chr9 | 117266029 | 117266030 | cg13657200 |
| chr9 | 117993982 | 117993983 | cg13965272 |
| chr9 | 125873915 | 125873916 | cg01729066 |
| chr9 | 126031747 | 126031748 | cg14186304 |
| chr9 | 127018815 | 127018816 | cg13958199 |
| chr9 | 127054804 | 127054805 | cg14536906 |
| chr9 | 128412220 | 128412221 | cg14146657 |
| chr9 | 129643082 | 129643083 | cg14360444 |
| chr9 | 129678298 | 129678299 | cg13931285 |
| chr9 | 130640220 | 130640221 | cg14005294 |
| chr9 | 130648201 | 130648202 | cg14533412 |
| chr9 | 130672636 | 130672637 | cg00677730 |
| chr9 | 130675438 | 130675439 | cg14081251 |
| chr9 | 130723444 | 130723445 | cg13558912 |
| chr9 | 130877638 | 130877639 | cg21245729 |
| chr9 | 131313189 | 131313190 | cg01182455 |
| chr9 | 131681980 | 131681981 | cg14520423 |
| chr9 | 131682162 | 131682163 | cg13707645 |
| chr9 | 134109022 | 134109023 | cg14548264 |
| chr9 | 134400392 | 134400393 | cg13989834 |
| chr9 | 135820767 | 135820768 | cg19393006 |
| chr9 | 135989682 | 135989683 | cg14541011 |
| chr9 | 135995491 | 135995492 | cg10585321 |
| chr9 | 136003437 | 136003438 | cg14468055 |
| chr9 | 136006118 | 136006119 | cg13520893 |
| chr9 | 136287409 | 136287410 | cg14802951 |
| chr9 | 136655966 | 136655967 | cg14415079 |
| chr9 | 136721183 | 136721184 | cg14266770 |
| chr9 | 136921209 | 136921210 | cg14472181 |
| chr9 | 137277819 | 137277820 | cg13931640 |
| chr9 | 137335080 | 137335081 | cg14022501 |
| chr9 | 139133513 | 139133514 | cg14599362 |
| chr9 | 139300299 | 139300300 | cg05145428 |
| chr9 | 139512327 | 139512328 | cg14360448 |
| chr9 | 139538535 | 139538536 | cg14276557 |

| | | | |
|---|---|---|---|
| chr6 | 109615419 | 109615420 | cg02774282 |
| chr6 | 109778433 | 109778434 | cg25318211 |
| chr6 | 109800627 | 109800628 | cg24153874 |
| chr6 | 110015052 | 110015053 | cg03492495 |
| chr6 | 110732115 | 110732116 | cg11645539 |
| chr6 | 110841946 | 110841947 | cg04700474 |
| chr6 | 111888567 | 111888568 | cg05423393 |
| chr6 | 112081102 | 112081103 | cg08130572 |
| chr6 | 112084071 | 112084072 | cg20706496 |
| chr6 | 112165062 | 112165063 | cg26846943 |
| chr6 | 112306329 | 112306330 | cg21391373 |
| chr6 | 113580203 | 113580204 | cg27158519 |
| chr6 | 114126759 | 114126760 | cg24364282 |
| chr6 | 114384871 | 114384872 | cg18192491 |
| chr6 | 114515500 | 114515501 | cg08893817 |
| chr6 | 114551190 | 114551191 | cg06678468 |
| chr6 | 116381609 | 116381610 | cg16176600 |
| chr6 | 116409276 | 116409277 | cg12673978 |
| chr6 | 117001170 | 117001171 | cg00564061 |
| chr6 | 118709682 | 118709683 | cg08169794 |
| chr6 | 118886798 | 118886799 | cg12311636 |
| chr6 | 119032202 | 119032203 | cg10164393 |
| chr6 | 119069624 | 119069625 | cg11883418 |
| chr6 | 119221279 | 119221280 | cg04183425 |
| chr6 | 121769777 | 121769778 | cg02580095 |
| chr6 | 123033536 | 123033537 | cg08198075 |
| chr6 | 123839720 | 123839721 | cg19143211 |
| chr6 | 124236421 | 124236422 | cg14458292 |
| chr6 | 124982296 | 124982297 | cg05308643 |
| chr6 | 125121219 | 125121220 | cg02285725 |
| chr6 | 125534847 | 125534848 | cg02807882 |
| chr6 | 126072671 | 126072672 | cg13799242 |
| chr6 | 126110790 | 126110791 | cg02027561 |
| chr6 | 128554002 | 128554003 | cg27618252 |
| chr6 | 128602411 | 128602412 | cg11155222 |
| chr6 | 129203783 | 129203784 | cg15801019 |
| chr6 | 130288563 | 130288564 | cg24011511 |
| chr6 | 130690629 | 130690630 | cg11584284 |
| chr6 | 130757762 | 130757763 | cg26931050 |
| chr6 | 130788615 | 130788616 | cg27216807 |
| chr6 | 131147216 | 131147217 | cg21775668 |
| chr6 | 132468790 | 132468791 | cg15985157 |
| chr6 | 132557557 | 132557558 | cg05949913 |
| chr6 | 132834914 | 132834915 | cg10097215 |
| chr6 | 133084805 | 133084806 | cg18493502 |
| chr6 | 134304883 | 134304884 | cg04254487 |
| chr6 | 134538515 | 134538516 | cg21834463 |
| chr6 | 134856977 | 134856978 | cg02936931 |
| chr6 | 135051046 | 135051047 | cg25704278 |
| chr6 | 135217576 | 135217577 | cg15010109 |
| chr6 | 135516936 | 135516937 | cg02127509 |
| chr6 | 136358605 | 136358606 | cg24777101 |
| chr6 | 137094444 | 137094445 | cg24451117 |
| chr6 | 137416146 | 137416147 | cg04598035 |
| chr6 | 137538584 | 137538585 | cg03065308 |
| chr6 | 138247491 | 138247492 | cg09340403 |

| | | | |
|---|---|---|---|
| chr9 | 139608277 | 139608278 | cg14172126 |
| chr9 | 139637747 | 139637748 | cg21157923 |
| chr9 | 139737793 | 139737794 | cg00755588 |
| chr9 | 140611316 | 140611317 | cg14156842 |
| chr9 | 140657381 | 140657382 | cg14470976 |
| chr9 | 140815010 | 140815011 | cg14353506 |
| chr10 | 237794 | 237795 | cg15287806 |
| chr10 | 1981672 | 1981673 | cg16158484 |
| chr10 | 2971684 | 2971685 | cg26254345 |
| chr10 | 3594801 | 3594802 | cg26852331 |
| chr10 | 3809095 | 3809096 | cg02813265 |
| chr10 | 3818357 | 3818358 | cg03216580 |
| chr10 | 4174368 | 4174369 | cg13701910 |
| chr10 | 4693210 | 4693211 | cg25681379 |
| chr10 | 5936166 | 5936167 | cg18764192 |
| chr10 | 5936216 | 5936217 | cg09395021 |
| chr10 | 6014983 | 6014984 | cg12128445 |
| chr10 | 6105656 | 6105657 | cg26105232 |
| chr10 | 6130578 | 6130579 | cg00242839 |
| chr10 | 6130580 | 6130581 | cg15953921 |
| chr10 | 7311500 | 7311501 | cg05103064 |
| chr10 | 7564401 | 7564402 | cg10983206 |
| chr10 | 11755103 | 11755104 | cg00594191 |
| chr10 | 13830045 | 13830046 | cg03775372 |
| chr10 | 18240316 | 18240317 | cg10368935 |
| chr10 | 18241116 | 18241117 | cg19856444 |
| chr10 | 18430412 | 18430413 | cg03634479 |
| chr10 | 18629006 | 18629007 | cg22858500 |
| chr10 | 18629497 | 18629498 | cg19998294 |
| chr10 | 18629545 | 18629546 | cg05719612 |
| chr10 | 18629674 | 18629675 | cg03103919 |
| chr10 | 18629677 | 18629678 | cg15169374 |
| chr10 | 24614999 | 24615000 | cg20054872 |
| chr10 | 24995583 | 24995584 | cg11122255 |
| chr10 | 25240660 | 25240661 | cg23418510 |
| chr10 | 27389908 | 27389909 | cg08856118 |
| chr10 | 27541626 | 27541627 | cg03848129 |
| chr10 | 31016591 | 31016592 | cg00941203 |
| chr10 | 31747241 | 31747242 | cg18516609 |
| chr10 | 32213946 | 32213947 | cg17158149 |
| chr10 | 32216031 | 32216032 | cg04359828 |
| chr10 | 32216066 | 32216067 | cg03015672 |
| chr10 | 32216101 | 32216102 | cg09642378 |
| chr10 | 32216311 | 32216312 | cg18675610 |
| chr10 | 32216359 | 32216360 | cg22706007 |
| chr10 | 32229887 | 32229888 | cg03781748 |
| chr10 | 32627605 | 32627606 | cg15986326 |
| chr10 | 35703322 | 35703323 | cg16832648 |
| chr10 | 35740216 | 35740217 | cg00201133 |
| chr10 | 44318634 | 44318635 | cg08478125 |
| chr10 | 45872735 | 45872736 | cg06127294 |
| chr10 | 45966142 | 45966143 | cg13709271 |
| chr10 | 50724286 | 50724287 | cg27383873 |
| chr10 | 52340628 | 52340629 | cg26665667 |
| chr10 | 52379467 | 52379468 | cg03727333 |
| chr10 | 60041363 | 60041364 | cg04776120 |

| | | | |
|---|---|---|---|
| chr6 | 138528677 | 138528678 | cg17480438 |
| chr6 | 138539706 | 138539707 | cg19823847 |
| chr6 | 139293014 | 139293015 | cg14814167 |
| chr6 | 139349120 | 139349121 | cg13265740 |
| chr6 | 139540827 | 139540828 | cg27020966 |
| chr6 | 143023932 | 143023933 | cg22751954 |
| chr6 | 143176015 | 143176016 | cg05519606 |
| chr6 | 143234184 | 143234185 | cg17717866 |
| chr6 | 143857467 | 143857468 | cg00416243 |
| chr6 | 143876139 | 143876140 | cg16198013 |
| chr6 | 143891455 | 143891456 | cg20298778 |
| chr6 | 143891515 | 143891516 | cg20435387 |
| chr6 | 143932803 | 143932804 | cg22660196 |
| chr6 | 144386422 | 144386423 | cg25291978 |
| chr6 | 144386474 | 144386475 | cg25535418 |
| chr6 | 144386848 | 144386849 | cg21113768 |
| chr6 | 144901284 | 144901285 | cg13800496 |
| chr6 | 147225846 | 147225847 | cg06488536 |
| chr6 | 148829044 | 148829045 | cg27113015 |
| chr6 | 149099724 | 149099725 | cg23828170 |
| chr6 | 149657288 | 149657289 | cg16237031 |
| chr6 | 150068006 | 150068007 | cg20335735 |
| chr6 | 150918915 | 150918916 | cg01013955 |
| chr6 | 151322717 | 151322718 | cg03708590 |
| chr6 | 152126250 | 152126251 | cg10441070 |
| chr6 | 154535553 | 154535554 | cg02864954 |
| chr6 | 154640892 | 154640893 | cg24635109 |
| chr6 | 154651158 | 154651159 | cg24873280 |
| chr6 | 154651280 | 154651281 | cg14295437 |
| chr6 | 154678326 | 154678327 | cg19690214 |
| chr6 | 154691940 | 154691941 | cg25296374 |
| chr6 | 155686868 | 155686869 | cg01113073 |
| chr6 | 157110510 | 157110511 | cg10574283 |
| chr6 | 157129618 | 157129619 | cg04606397 |
| chr6 | 157131429 | 157131430 | cg08268892 |
| chr6 | 157289645 | 157289646 | cg06019273 |
| chr6 | 157316222 | 157316223 | cg13602547 |
| chr6 | 157746584 | 157746585 | cg00647262 |
| chr6 | 158404061 | 158404062 | cg15323840 |
| chr6 | 158404082 | 158404083 | cg22242614 |
| chr6 | 158404121 | 158404122 | cg13735018 |
| chr6 | 158749367 | 158749368 | cg25019378 |
| chr6 | 159220343 | 159220344 | cg15118447 |
| chr6 | 159227784 | 159227785 | cg16978168 |
| chr6 | 159230857 | 159230858 | cg23496730 |
| chr6 | 159231891 | 159231892 | cg13433729 |
| chr6 | 159463103 | 159463104 | cg00187889 |
| chr6 | 159515345 | 159515346 | cg01046905 |
| chr6 | 159547894 | 159547895 | cg01832757 |
| chr6 | 160219090 | 160219091 | cg22844862 |
| chr6 | 161592221 | 161592222 | cg08424166 |
| chr6 | 163776782 | 163776783 | cg24192505 |
| chr6 | 165933115 | 165933116 | cg11175473 |
| chr6 | 166100605 | 166100606 | cg05787951 |
| chr6 | 166100693 | 166100694 | cg11759749 |
| chr6 | 166908574 | 166908575 | cg11599721 |
| chr10 | 62423377 | 62423378 | cg14413904 |
| chr10 | 63753550 | 63753551 | cg25953130 |
| chr10 | 63764155 | 63764156 | cg18980410 |
| chr10 | 63829033 | 63829034 | cg22863838 |
| chr10 | 64279749 | 64279750 | cg00236005 |
| chr10 | 64373395 | 64373396 | cg10223233 |
| chr10 | 64451393 | 64451394 | cg22633096 |
| chr10 | 64566678 | 64566679 | cg16863332 |
| chr10 | 64568066 | 64568067 | cg19580792 |
| chr10 | 65224441 | 65224442 | cg17775079 |
| chr10 | 69992004 | 69992005 | cg20156072 |
| chr10 | 70845585 | 70845586 | cg06486994 |
| chr10 | 71087364 | 71087365 | cg23177739 |
| chr10 | 72237473 | 72237474 | cg00391031 |
| chr10 | 73303234 | 73303235 | cg04457572 |
| chr10 | 73471634 | 73471635 | cg18235465 |
| chr10 | 73472315 | 73472316 | cg07571519 |
| chr10 | 73844130 | 73844131 | cg14614581 |
| chr10 | 73848817 | 73848818 | cg12623536 |
| chr10 | 73848823 | 73848824 | cg06970090 |
| chr10 | 73848825 | 73848826 | cg00661320 |
| chr10 | 73848830 | 73848831 | cg15331996 |
| chr10 | 73856465 | 73856466 | cg20571967 |
| chr10 | 73869442 | 73869443 | cg23723835 |
| chr10 | 73988469 | 73988470 | cg18515031 |
| chr10 | 74004071 | 74004072 | cg26033520 |
| chr10 | 74082489 | 74082490 | cg05988358 |
| chr10 | 74842425 | 74842426 | cg08695113 |
| chr10 | 74857663 | 74857664 | cg18195276 |
| chr10 | 75486391 | 75486392 | cg19500311 |
| chr10 | 75639879 | 75639880 | cg25725418 |
| chr10 | 75683980 | 75683981 | cg16353350 |
| chr10 | 75800004 | 75800005 | cg18908419 |
| chr10 | 75935758 | 75935759 | cg18786171 |
| chr10 | 76589109 | 76589110 | cg00432328 |
| chr10 | 76974078 | 76974079 | cg15075988 |
| chr10 | 78871238 | 78871239 | cg05907577 |
| chr10 | 78923827 | 78923828 | cg02636348 |
| chr10 | 79133935 | 79133936 | cg12614113 |
| chr10 | 79150517 | 79150518 | cg16055185 |
| chr10 | 80842848 | 80842849 | cg15522953 |
| chr10 | 81165685 | 81165686 | cg07491858 |
| chr10 | 81583959 | 81583960 | cg01423845 |
| chr10 | 81946545 | 81946546 | cg11900509 |
| chr10 | 82225771 | 82225772 | cg18923906 |
| chr10 | 85898700 | 85898701 | cg12706330 |
| chr10 | 88559053 | 88559054 | cg25261694 |
| chr10 | 90253615 | 90253616 | cg02977388 |
| chr10 | 90656978 | 90656979 | cg14785527 |
| chr10 | 91405677 | 91405678 | cg08661112 |
| chr10 | 91405685 | 91405686 | cg25770176 |
| chr10 | 91598023 | 91598024 | cg09509179 |
| chr10 | 92689909 | 92689910 | cg07620928 |
| chr10 | 92950766 | 92950767 | cg09107180 |
| chr10 | 92971269 | 92971270 | cg04092437 |
| chr10 | 93150018 | 93150019 | cg04595479 |

| | | | |
|---|---|---|---|
| chr6 | 167363345 | 167363346 | cg11301670 |
| chr6 | 167366683 | 167366684 | cg17991206 |
| chr6 | 167530343 | 167530344 | cg20143758 |
| chr6 | 167535171 | 167535172 | cg16523158 |
| chr6 | 167571324 | 167571325 | cg15706657 |
| chr6 | 167571328 | 167571329 | cg01859669 |
| chr6 | 167571803 | 167571804 | cg12588299 |
| chr6 | 167632873 | 167632874 | cg19445758 |
| chr6 | 170200938 | 170200939 | cg27348420 |
| chr6 | 170784959 | 170784960 | cg02245832 |
| chr7 | 553693 | 553694 | cg00639010 |
| chr7 | 625028 | 625029 | cg24028627 |
| chr7 | 625089 | 625090 | cg20750319 |
| chr7 | 625136 | 625137 | cg21769820 |
| chr7 | 642127 | 642128 | cg21381845 |
| chr7 | 643155 | 643156 | cg02217713 |
| chr7 | 691506 | 691507 | cg00665343 |
| chr7 | 691761 | 691762 | cg00479718 |
| chr7 | 872208 | 872209 | cg05357209 |
| chr7 | 873449 | 873450 | cg11634705 |
| chr7 | 947151 | 947152 | cg18664596 |
| chr7 | 960350 | 960351 | cg19445335 |
| chr7 | 961322 | 961323 | cg27287951 |
| chr7 | 961384 | 961385 | cg19471433 |
| chr7 | 961457 | 961458 | cg06589239 |
| chr7 | 1066650 | 1066651 | cg04816311 |
| chr7 | 1095416 | 1095417 | cg10009497 |
| chr7 | 1095595 | 1095596 | cg18377014 |
| chr7 | 1196370 | 1196371 | cg26608667 |
| chr7 | 1584275 | 1584276 | cg03664589 |
| chr7 | 1632867 | 1632868 | cg14343017 |
| chr7 | 1855682 | 1855683 | cg12163781 |
| chr7 | 1858916 | 1858917 | cg22963979 |
| chr7 | 1883251 | 1883252 | cg25814293 |
| chr7 | 1966361 | 1966362 | cg03586240 |
| chr7 | 2059921 | 2059922 | cg15896696 |
| chr7 | 2140321 | 2140322 | cg18319104 |
| chr7 | 2247473 | 2247474 | cg25314733 |
| chr7 | 2298727 | 2298728 | cg10745498 |
| chr7 | 2328746 | 2328747 | cg19497702 |
| chr7 | 2339170 | 2339171 | cg05516271 |
| chr7 | 2340959 | 2340960 | cg22319052 |
| chr7 | 2557152 | 2557153 | cg12497581 |
| chr7 | 2563184 | 2563185 | cg14603031 |
| chr7 | 2564256 | 2564257 | cg02613295 |
| chr7 | 2566750 | 2566751 | cg27186851 |
| chr7 | 2567893 | 2567894 | cg23510807 |
| chr7 | 2582200 | 2582201 | cg01269212 |
| chr7 | 2606796 | 2606797 | cg02072875 |
| chr7 | 2606824 | 2606825 | cg10688297 |
| chr7 | 2677986 | 2677987 | cg10648960 |
| chr7 | 2681436 | 2681437 | cg17550566 |
| chr7 | 2755000 | 2755001 | cg24375364 |
| chr7 | 2757342 | 2757343 | cg19799702 |
| chr7 | 2774414 | 2774415 | cg11884933 |
| chr7 | 2774543 | 2774544 | cg01492538 |

| | | | |
|---|---|---|---|
| chr10 | 97049193 | 97049194 | cg08799394 |
| chr10 | 97049610 | 97049611 | cg18389639 |
| chr10 | 97470786 | 97470787 | cg16300094 |
| chr10 | 97514275 | 97514276 | cg03257107 |
| chr10 | 97982467 | 97982468 | cg23715056 |
| chr10 | 98062747 | 98062748 | cg03930857 |
| chr10 | 99223744 | 99223745 | cg12182708 |
| chr10 | 99255236 | 99255237 | cg07052063 |
| chr10 | 99894078 | 99894079 | cg23108673 |
| chr10 | 101732382 | 101732383 | cg02732134 |
| chr10 | 101736260 | 101736261 | cg26002091 |
| chr10 | 101759257 | 101759258 | cg17993335 |
| chr10 | 101767352 | 101767353 | cg27319563 |
| chr10 | 101767908 | 101767909 | cg19212550 |
| chr10 | 101768580 | 101768581 | cg18522740 |
| chr10 | 101768701 | 101768702 | cg14405603 |
| chr10 | 101779926 | 101779927 | cg01099300 |
| chr10 | 102028800 | 102028801 | cg10743378 |
| chr10 | 102102351 | 102102352 | cg12714759 |
| chr10 | 102131012 | 102131013 | cg07504977 |
| chr10 | 102730130 | 102730131 | cg05428706 |
| chr10 | 103114734 | 103114735 | cg24537986 |
| chr10 | 104436518 | 104436519 | cg27316828 |
| chr10 | 104536695 | 104536696 | cg25223634 |
| chr10 | 104562697 | 104562698 | cg16054018 |
| chr10 | 105218971 | 105218972 | cg03485533 |
| chr10 | 105451331 | 105451332 | cg24911198 |
| chr10 | 105454768 | 105454769 | cg00277384 |
| chr10 | 105589064 | 105589065 | cg11014810 |
| chr10 | 106078025 | 106078026 | cg23523388 |
| chr10 | 106078922 | 106078923 | cg24168991 |
| chr10 | 112256517 | 112256518 | cg12323347 |
| chr10 | 112256640 | 112256641 | cg20925954 |
| chr10 | 112256825 | 112256826 | cg06110816 |
| chr10 | 112259695 | 112259696 | cg25417042 |
| chr10 | 112261994 | 112261995 | cg18756771 |
| chr10 | 112582591 | 112582592 | cg04840051 |
| chr10 | 114135092 | 114135093 | cg02402091 |
| chr10 | 115612802 | 115612803 | cg04281204 |
| chr10 | 116282534 | 116282535 | cg12649038 |
| chr10 | 120489658 | 120489659 | cg12880685 |
| chr10 | 121013116 | 121013117 | cg08650639 |
| chr10 | 121155124 | 121155125 | cg15580458 |
| chr10 | 121259610 | 121259611 | cg04182483 |
| chr10 | 121260443 | 121260444 | cg17675386 |
| chr10 | 121482923 | 121482924 | cg11788307 |
| chr10 | 121487556 | 121487557 | cg07003611 |
| chr10 | 121493706 | 121493707 | cg20704028 |
| chr10 | 121497344 | 121497345 | cg10053779 |
| chr10 | 121503164 | 121503165 | cg07679322 |
| chr10 | 121578639 | 121578640 | cg25045785 |
| chr10 | 121609179 | 121609180 | cg12002455 |
| chr10 | 125805493 | 125805494 | cg08082692 |
| chr10 | 125806390 | 125806391 | cg17606491 |
| chr10 | 125825166 | 125825167 | cg00864684 |
| chr10 | 125847615 | 125847616 | cg14267222 |

| | | | |
|---|---|---|---|
| chr7 | 2864192 | 2864193 | cg17705631 |
| chr7 | 2948081 | 2948082 | cg16586807 |
| chr7 | 3025634 | 3025635 | cg15025536 |
| chr7 | 3025689 | 3025690 | cg23352157 |
| chr7 | 3025736 | 3025737 | cg15638239 |
| chr7 | 3067980 | 3067981 | cg20308351 |
| chr7 | 3082168 | 3082169 | cg24404878 |
| chr7 | 3082233 | 3082234 | cg03534655 |
| chr7 | 3084077 | 3084078 | cg06173509 |
| chr7 | 3613164 | 3613165 | cg17756105 |
| chr7 | 4762236 | 4762237 | cg10106935 |
| chr7 | 4762371 | 4762372 | cg06660057 |
| chr7 | 5257805 | 5257806 | cg22663933 |
| chr7 | 5269290 | 5269291 | cg23122899 |
| chr7 | 5270553 | 5270554 | cg21764632 |
| chr7 | 5270567 | 5270568 | cg02667577 |
| chr7 | 5272208 | 5272209 | cg15277378 |
| chr7 | 5272275 | 5272276 | cg20099458 |
| chr7 | 5272697 | 5272698 | cg27021327 |
| chr7 | 5272900 | 5272901 | cg23562222 |
| chr7 | 5276991 | 5276992 | cg18237047 |
| chr7 | 5413872 | 5413873 | cg06262098 |
| chr7 | 5535641 | 5535642 | cg25343388 |
| chr7 | 5709854 | 5709855 | cg24534583 |
| chr7 | 5816628 | 5816629 | cg26724841 |
| chr7 | 6680085 | 6680086 | cg24862131 |
| chr7 | 7261594 | 7261595 | cg17994788 |
| chr7 | 7711168 | 7711169 | cg19739774 |
| chr7 | 14192484 | 14192485 | cg15549435 |
| chr7 | 16570042 | 16570043 | cg07088414 |
| chr7 | 16794623 | 16794624 | cg11384744 |
| chr7 | 17585471 | 17585472 | cg08993283 |
| chr7 | 18079924 | 18079925 | cg15698691 |
| chr7 | 18329602 | 18329603 | cg24350771 |
| chr7 | 18575633 | 18575634 | cg26465837 |
| chr7 | 19961027 | 19961028 | cg03729152 |
| chr7 | 20269648 | 20269649 | cg19086110 |
| chr7 | 20346927 | 20346928 | cg18396789 |
| chr7 | 21065099 | 21065100 | cg08889440 |
| chr7 | 21488742 | 21488743 | cg19644145 |
| chr7 | 21986672 | 21986673 | cg20983531 |
| chr7 | 22602133 | 22602134 | cg16110788 |
| chr7 | 22817223 | 22817224 | cg02463988 |
| chr7 | 23401203 | 23401204 | cg02302089 |
| chr7 | 23412236 | 23412237 | cg06632790 |
| chr7 | 23502744 | 23502745 | cg21025494 |
| chr7 | 24474486 | 24474487 | cg16717546 |
| chr7 | 25601773 | 25601774 | cg23485591 |
| chr7 | 26193607 | 26193608 | cg08822075 |
| chr7 | 26193805 | 26193806 | cg12510708 |
| chr7 | 26206579 | 26206580 | cg07945582 |
| chr7 | 26236069 | 26236070 | cg19142553 |
| chr7 | 26897597 | 26897598 | cg12252069 |
| chr7 | 26897685 | 26897686 | cg03730533 |
| chr7 | 26897714 | 26897715 | cg10180281 |
| chr7 | 26958919 | 26958920 | cg15700661 |

| | | | |
|---|---|---|---|
| chr10 | 126211704 | 126211705 | cg03499943 |
| chr10 | 126211892 | 126211893 | cg16426764 |
| chr10 | 126234119 | 126234120 | cg09985802 |
| chr10 | 126234138 | 126234139 | cg00932063 |
| chr10 | 126274650 | 126274651 | cg14985591 |
| chr10 | 126303677 | 126303678 | cg06139794 |
| chr10 | 126311927 | 126311928 | cg15986847 |
| chr10 | 126390317 | 126390318 | cg09519218 |
| chr10 | 126425471 | 126425472 | cg02293775 |
| chr10 | 126428199 | 126428200 | cg23088510 |
| chr10 | 126428818 | 126428819 | cg26517376 |
| chr10 | 133743048 | 133743049 | cg26789365 |
| chr10 | 134038395 | 134038396 | cg25917732 |
| chr10 | 134045514 | 134045515 | cg01458105 |
| chr10 | 134211857 | 134211858 | cg11579421 |
| chr10 | 134236975 | 134236976 | cg24643922 |
| chr10 | 134322334 | 134322335 | cg26118208 |
| chr10 | 134358535 | 134358536 | cg09421083 |
| chr10 | 134451193 | 134451194 | cg10808367 |
| chr10 | 134451258 | 134451259 | cg08277036 |
| chr10 | 134748215 | 134748216 | cg26896918 |
| chr11 | 250989 | 250990 | cg05600342 |
| chr11 | 288301 | 288302 | cg18727742 |
| chr11 | 288305 | 288306 | cg08829299 |
| chr11 | 312518 | 312519 | cg05432003 |
| chr11 | 312560 | 312561 | cg01886988 |
| chr11 | 312841 | 312842 | cg27032101 |
| chr11 | 313120 | 313121 | cg04582010 |
| chr11 | 353567 | 353568 | cg14293323 |
| chr11 | 413594 | 413595 | cg08869273 |
| chr11 | 421691 | 421692 | cg03365403 |
| chr11 | 491086 | 491087 | cg16150393 |
| chr11 | 493125 | 493126 | cg23552637 |
| chr11 | 504784 | 504785 | cg14468692 |
| chr11 | 542109 | 542110 | cg19171175 |
| chr11 | 542226 | 542227 | cg02837194 |
| chr11 | 705692 | 705693 | cg11329129 |
| chr11 | 705748 | 705749 | cg13555772 |
| chr11 | 832357 | 832358 | cg03885332 |
| chr11 | 928439 | 928440 | cg16730825 |
| chr11 | 955120 | 955121 | cg20877443 |
| chr11 | 955328 | 955329 | cg12914733 |
| chr11 | 971527 | 971528 | cg05626073 |
| chr11 | 1328479 | 1328480 | cg10379476 |
| chr11 | 1328486 | 1328487 | cg16472011 |
| chr11 | 1778547 | 1778548 | cg22680424 |
| chr11 | 1890901 | 1890902 | cg04650653 |
| chr11 | 2321955 | 2321956 | cg11525052 |
| chr11 | 2322050 | 2322051 | cg25961099 |
| chr11 | 2705153 | 2705154 | cg01693193 |
| chr11 | 2709914 | 2709915 | cg08160246 |
| chr11 | 2710486 | 2710487 | cg02180189 |
| chr11 | 3887906 | 3887907 | cg17696660 |
| chr11 | 3899234 | 3899235 | cg26296653 |
| chr11 | 6418966 | 6418967 | cg07552374 |
| chr11 | 7532359 | 7532360 | cg19673705 |

| | | | |
|---|---|---|---|
| chr7 | 28201106 | 28201107 | cg22938901 |
| chr7 | 28208557 | 28208558 | cg25941154 |
| chr7 | 28221667 | 28221668 | cg04266607 |
| chr7 | 28469901 | 28469902 | cg02124565 |
| chr7 | 28713246 | 28713247 | cg22681186 |
| chr7 | 28848902 | 28848903 | cg19588408 |
| chr7 | 28848933 | 28848934 | cg19559984 |
| chr7 | 29271780 | 29271781 | cg18529010 |
| chr7 | 29446751 | 29446752 | cg19410046 |
| chr7 | 30321030 | 30321031 | cg15822656 |
| chr7 | 30473983 | 30473984 | cg14784253 |
| chr7 | 30590918 | 30590919 | cg27107685 |
| chr7 | 30891837 | 30891838 | cg14370520 |
| chr7 | 31725748 | 31725749 | cg23348723 |
| chr7 | 33146808 | 33146809 | cg01016102 |
| chr7 | 35615527 | 35615528 | cg08753560 |
| chr7 | 38580353 | 38580354 | cg00911226 |
| chr7 | 38710069 | 38710070 | cg19162970 |
| chr7 | 39994617 | 39994618 | cg13791719 |
| chr7 | 40755270 | 40755271 | cg12494603 |
| chr7 | 40855639 | 40855640 | cg00423291 |
| chr7 | 40961163 | 40961164 | cg07737644 |
| chr7 | 41611383 | 41611384 | cg26649851 |
| chr7 | 41744327 | 41744328 | cg09498134 |
| chr7 | 41817771 | 41817772 | cg02088996 |
| chr7 | 42656871 | 42656872 | cg24645671 |
| chr7 | 43562162 | 43562163 | cg01213645 |
| chr7 | 44134432 | 44134433 | cg12399411 |
| chr7 | 44189684 | 44189685 | cg17959631 |
| chr7 | 44253807 | 44253808 | cg18446472 |
| chr7 | 44645306 | 44645307 | cg07408740 |
| chr7 | 44662296 | 44662297 | cg23929194 |
| chr7 | 44791211 | 44791212 | cg11425656 |
| chr7 | 45075791 | 45075792 | cg12573289 |
| chr7 | 45617892 | 45617893 | cg25322847 |
| chr7 | 46185114 | 46185115 | cg09824603 |
| chr7 | 46495783 | 46495784 | cg15956565 |
| chr7 | 46972700 | 46972701 | cg25983901 |
| chr7 | 47539765 | 47539766 | cg27275374 |
| chr7 | 47539837 | 47539838 | cg23895495 |
| chr7 | 47611235 | 47611236 | cg17996401 |
| chr7 | 50343131 | 50343132 | cg14216940 |
| chr7 | 50443901 | 50443902 | cg18117847 |
| chr7 | 50934682 | 50934683 | cg04384491 |
| chr7 | 51303122 | 51303123 | cg07152355 |
| chr7 | 54768889 | 54768890 | cg10668195 |
| chr7 | 55304314 | 55304315 | cg18461584 |
| chr7 | 55584387 | 55584388 | cg05170183 |
| chr7 | 55606442 | 55606443 | cg23654401 |
| chr7 | 55614512 | 55614513 | cg15902297 |
| chr7 | 55620641 | 55620642 | cg07006935 |
| chr7 | 55621681 | 55621682 | cg24430703 |
| chr7 | 55621712 | 55621713 | cg06750076 |
| chr7 | 56066988 | 56066989 | cg04023936 |
| chr7 | 56301065 | 56301066 | cg16559630 |
| chr7 | 64126140 | 64126141 | cg12856392 |

| | | | |
|---|---|---|---|
| chr11 | 8215201 | 8215202 | cg05261592 |
| chr11 | 8662007 | 8662008 | cg25942880 |
| chr11 | 8983288 | 8983289 | cg16733419 |
| chr11 | 9638619 | 9638620 | cg03169242 |
| chr11 | 9775858 | 9775859 | cg05804846 |
| chr11 | 9992187 | 9992188 | cg01053763 |
| chr11 | 10476023 | 10476024 | cg18593305 |
| chr11 | 10534506 | 10534507 | cg25040934 |
| chr11 | 11893602 | 11893603 | cg24383467 |
| chr11 | 13461802 | 13461803 | cg08114852 |
| chr11 | 18611549 | 18611550 | cg25976340 |
| chr11 | 18630052 | 18630053 | cg21107103 |
| chr11 | 20010227 | 20010228 | cg01896424 |
| chr11 | 20465288 | 20465289 | cg13544027 |
| chr11 | 26298241 | 26298242 | cg19182008 |
| chr11 | 27250166 | 27250167 | cg07535191 |
| chr11 | 27492378 | 27492379 | cg22715531 |
| chr11 | 33084877 | 33084878 | cg02813542 |
| chr11 | 33809547 | 33809548 | cg07164065 |
| chr11 | 33914088 | 33914089 | cg09358725 |
| chr11 | 34167855 | 34167856 | cg19045191 |
| chr11 | 34457835 | 34457836 | cg04464122 |
| chr11 | 35086165 | 35086166 | cg14378420 |
| chr11 | 35374336 | 35374337 | cg14746605 |
| chr11 | 35808688 | 35808689 | cg04237436 |
| chr11 | 36383991 | 36383992 | cg16829924 |
| chr11 | 43941594 | 43941595 | cg01928068 |
| chr11 | 44578801 | 44578802 | cg14780449 |
| chr11 | 44617598 | 44617599 | cg09664479 |
| chr11 | 44626708 | 44626709 | cg22697239 |
| chr11 | 44626750 | 44626751 | cg01199327 |
| chr11 | 44785768 | 44785769 | cg15576669 |
| chr11 | 44846667 | 44846668 | cg11972564 |
| chr11 | 44927976 | 44927977 | cg06401614 |
| chr11 | 44973110 | 44973111 | cg04745100 |
| chr11 | 45102239 | 45102240 | cg23299618 |
| chr11 | 45825356 | 45825357 | cg19527995 |
| chr11 | 45844932 | 45844933 | cg13702370 |
| chr11 | 45942027 | 45942028 | cg16335858 |
| chr11 | 45950696 | 45950697 | cg06508085 |
| chr11 | 46079554 | 46079555 | cg19076587 |
| chr11 | 46121221 | 46121222 | cg14394264 |
| chr11 | 46368045 | 46368046 | cg00530720 |
| chr11 | 46382544 | 46382545 | cg07092212 |
| chr11 | 47024513 | 47024514 | cg18033092 |
| chr11 | 47554407 | 47554408 | cg12979590 |
| chr11 | 47571658 | 47571659 | cg14232165 |
| chr11 | 47873225 | 47873226 | cg26858581 |
| chr11 | 47959926 | 47959927 | cg07160014 |
| chr11 | 48036852 | 48036853 | cg07401324 |
| chr11 | 48038449 | 48038450 | cg19268720 |
| chr11 | 48040772 | 48040773 | cg22976120 |
| chr11 | 57279398 | 57279399 | cg10964211 |
| chr11 | 58291656 | 58291657 | cg02375166 |
| chr11 | 58385947 | 58385948 | cg12027899 |
| chr11 | 58389290 | 58389291 | cg00271311 |

| | | | |
|---|---|---|---|
| chr7 | 64730944 | 64730945 | cg15082875 |
| chr7 | 66144066 | 66144067 | cg13204977 |
| chr7 | 69304751 | 69304752 | cg17184370 |
| chr7 | 69427388 | 69427389 | cg05608790 |
| chr7 | 70163455 | 70163456 | cg07470570 |
| chr7 | 70322401 | 70322402 | cg20866121 |
| chr7 | 71578347 | 71578348 | cg01584103 |
| chr7 | 73181157 | 73181158 | cg24311485 |
| chr7 | 73207968 | 73207969 | cg00987534 |
| chr7 | 73770020 | 73770021 | cg07814763 |
| chr7 | 73940352 | 73940353 | cg20448594 |
| chr7 | 74058031 | 74058032 | cg19313311 |
| chr7 | 74604867 | 74604868 | cg12494301 |
| chr7 | 75043671 | 75043672 | cg14113960 |
| chr7 | 76032544 | 76032545 | cg00208012 |
| chr7 | 76109050 | 76109051 | cg26558111 |
| chr7 | 76956562 | 76956563 | cg07265444 |
| chr7 | 77468418 | 77468419 | cg19818271 |
| chr7 | 79848290 | 79848291 | cg17986880 |
| chr7 | 80544479 | 80544480 | cg15556990 |
| chr7 | 81420133 | 81420134 | cg17578341 |
| chr7 | 81805241 | 81805242 | cg26613778 |
| chr7 | 81860423 | 81860424 | cg25585043 |
| chr7 | 82222420 | 82222421 | cg08951078 |
| chr7 | 86413438 | 86413439 | cg04062190 |
| chr7 | 90355905 | 90355906 | cg09781028 |
| chr7 | 90406191 | 90406192 | cg21700774 |
| chr7 | 90794392 | 90794393 | cg03834767 |
| chr7 | 92459517 | 92459518 | cg10481072 |
| chr7 | 92667406 | 92667407 | cg07325692 |
| chr7 | 93114192 | 93114193 | cg04650977 |
| chr7 | 93763461 | 93763462 | cg16906641 |
| chr7 | 93977625 | 93977626 | cg17933009 |
| chr7 | 94278830 | 94278831 | cg22086834 |
| chr7 | 94955348 | 94955349 | cg15887283 |
| chr7 | 95433546 | 95433547 | cg18127395 |
| chr7 | 95435309 | 95435310 | cg03716590 |
| chr7 | 95696973 | 95696974 | cg23381232 |
| chr7 | 95811907 | 95811908 | cg27185510 |
| chr7 | 97833154 | 97833155 | cg13427789 |
| chr7 | 98989843 | 98989844 | cg02257708 |
| chr7 | 99054834 | 99054835 | cg23957742 |
| chr7 | 99767035 | 99767036 | cg24616382 |
| chr7 | 99816988 | 99816989 | cg25225693 |
| chr7 | 99817027 | 99817028 | cg23221013 |
| chr7 | 99970448 | 99970449 | cg03121834 |
| chr7 | 100139917 | 100139918 | cg13695383 |
| chr7 | 100142441 | 100142442 | cg03431524 |
| chr7 | 100144475 | 100144476 | cg18991321 |
| chr7 | 100285650 | 100285651 | cg05922723 |
| chr7 | 100798809 | 100798810 | cg16914953 |
| chr7 | 100899765 | 100899766 | cg17920076 |
| chr7 | 101257675 | 101257676 | cg24830241 |
| chr7 | 101596404 | 101596405 | cg01973676 |
| chr7 | 101932902 | 101932903 | cg00100114 |
| chr7 | 101936318 | 101936319 | cg12315284 |
| chr11 | 58389438 | 58389439 | cg09514875 |
| chr11 | 59439293 | 59439294 | cg06022671 |
| chr11 | 59950405 | 59950406 | cg03055440 |
| chr11 | 59950745 | 59950746 | cg26250585 |
| chr11 | 59951557 | 59951558 | cg04353769 |
| chr11 | 59951663 | 59951664 | cg06881914 |
| chr11 | 59952134 | 59952135 | cg24026212 |
| chr11 | 59952153 | 59952154 | cg20284999 |
| chr11 | 60679546 | 60679547 | cg05778528 |
| chr11 | 60716250 | 60716251 | cg02939292 |
| chr11 | 60722107 | 60722108 | cg10704729 |
| chr11 | 60858720 | 60858721 | cg20418308 |
| chr11 | 60869601 | 60869602 | cg16001613 |
| chr11 | 60869634 | 60869635 | cg27301655 |
| chr11 | 60870121 | 60870122 | cg22202088 |
| chr11 | 61110186 | 61110187 | cg04468281 |
| chr11 | 61115736 | 61115737 | cg20242427 |
| chr11 | 61900852 | 61900853 | cg13883397 |
| chr11 | 62315320 | 62315321 | cg23447569 |
| chr11 | 62321136 | 62321137 | cg18825531 |
| chr11 | 62481055 | 62481056 | cg07094785 |
| chr11 | 64322994 | 64322995 | cg18158438 |
| chr11 | 64514658 | 64514659 | cg03663788 |
| chr11 | 64631357 | 64631358 | cg10495164 |
| chr11 | 64638190 | 64638191 | cg11321083 |
| chr11 | 64765098 | 64765099 | cg23175322 |
| chr11 | 64936920 | 64936921 | cg23079866 |
| chr11 | 65150416 | 65150417 | cg26615176 |
| chr11 | 65150871 | 65150872 | cg04261607 |
| chr11 | 65196622 | 65196623 | cg20889322 |
| chr11 | 65196696 | 65196697 | cg13119578 |
| chr11 | 65201759 | 65201760 | cg24772412 |
| chr11 | 65201834 | 65201835 | cg10416861 |
| chr11 | 65210600 | 65210601 | cg04238983 |
| chr11 | 65211709 | 65211710 | cg13167340 |
| chr11 | 65256983 | 65256984 | cg03390187 |
| chr11 | 65314913 | 65314914 | cg21584461 |
| chr11 | 65352232 | 65352233 | cg26667091 |
| chr11 | 65360449 | 65360450 | cg23343073 |
| chr11 | 65378622 | 65378623 | cg05436845 |
| chr11 | 65557185 | 65557186 | cg11569703 |
| chr11 | 65687892 | 65687893 | cg10467098 |
| chr11 | 65839402 | 65839403 | cg24127061 |
| chr11 | 66049756 | 66049757 | cg10336131 |
| chr11 | 66092918 | 66092919 | cg18259535 |
| chr11 | 66311334 | 66311335 | cg05516272 |
| chr11 | 66311348 | 66311349 | cg07977202 |
| chr11 | 67032472 | 67032473 | cg16061947 |
| chr11 | 67089352 | 67089353 | cg01464484 |
| chr11 | 67182672 | 67182673 | cg25414605 |
| chr11 | 67183498 | 67183499 | cg13708910 |
| chr11 | 67183546 | 67183547 | cg20234060 |
| chr11 | 67221691 | 67221692 | cg03686027 |
| chr11 | 67353032 | 67353033 | cg10552496 |
| chr11 | 67775972 | 67775973 | cg09982224 |
| chr11 | 67776576 | 67776577 | cg18229767 |

| | | | |
|---|---|---|---|
| chr7 | 101947698 | 101947699 | cg09863136 |
| chr7 | 102073120 | 102073121 | cg21253590 |
| chr7 | 104530865 | 104530866 | cg26268565 |
| chr7 | 104621597 | 104621598 | cg06579481 |
| chr7 | 104623427 | 104623428 | cg05816398 |
| chr7 | 104650933 | 104650934 | cg07995893 |
| chr7 | 105128488 | 105128489 | cg09490124 |
| chr7 | 105313992 | 105313993 | cg10575219 |
| chr7 | 105478993 | 105478994 | cg21095673 |
| chr7 | 105621846 | 105621847 | cg04215126 |
| chr7 | 105817781 | 105817782 | cg15107117 |
| chr7 | 105923321 | 105923322 | cg05004518 |
| chr7 | 106125168 | 106125169 | cg00139681 |
| chr7 | 106407970 | 106407971 | cg09735627 |
| chr7 | 106708700 | 106708701 | cg26104690 |
| chr7 | 106842512 | 106842513 | cg10840361 |
| chr7 | 107587365 | 107587366 | cg02052536 |
| chr7 | 107744567 | 107744568 | cg18929251 |
| chr7 | 110753971 | 110753972 | cg10769531 |
| chr7 | 110838461 | 110838462 | cg07373723 |
| chr7 | 111200944 | 111200945 | cg02787120 |
| chr7 | 111571767 | 111571768 | cg03705962 |
| chr7 | 114053988 | 114053989 | cg02647628 |
| chr7 | 114559768 | 114559769 | cg14847045 |
| chr7 | 114561804 | 114561805 | cg01550799 |
| chr7 | 114561850 | 114561851 | cg05338976 |
| chr7 | 114648973 | 114648974 | cg26275647 |
| chr7 | 115456904 | 115456905 | cg22578425 |
| chr7 | 116222436 | 116222437 | cg21786601 |
| chr7 | 116449640 | 116449641 | cg23451876 |
| chr7 | 120918492 | 120918493 | cg15239627 |
| chr7 | 121006948 | 121006949 | cg04873577 |
| chr7 | 122840109 | 122840110 | cg00895997 |
| chr7 | 123080165 | 123080166 | cg15083851 |
| chr7 | 123459295 | 123459296 | cg09630417 |
| chr7 | 126752911 | 126752912 | cg24294950 |
| chr7 | 126990845 | 126990846 | cg23362032 |
| chr7 | 127295153 | 127295154 | cg25747190 |
| chr7 | 127566935 | 127566936 | cg04359022 |
| chr7 | 127747345 | 127747346 | cg27179002 |
| chr7 | 128559623 | 128559624 | cg03603395 |
| chr7 | 128574410 | 128574411 | cg09387224 |
| chr7 | 128587827 | 128587828 | cg08751971 |
| chr7 | 129266958 | 129266959 | cg02041484 |
| chr7 | 129464288 | 129464289 | cg01822759 |
| chr7 | 129781681 | 129781682 | cg19145858 |
| chr7 | 129991187 | 129991188 | cg22664614 |
| chr7 | 130124971 | 130124972 | cg09425926 |
| chr7 | 130630462 | 130630463 | cg16852756 |
| chr7 | 130712346 | 130712347 | cg22544881 |
| chr7 | 130904147 | 130904148 | cg12724086 |
| chr7 | 134345856 | 134345857 | cg03950253 |
| chr7 | 134443864 | 134443865 | cg03188976 |
| chr7 | 134585278 | 134585279 | cg22369318 |
| chr7 | 134832221 | 134832222 | cg21363348 |
| chr7 | 134854551 | 134854552 | cg00464046 |
| chr11 | 67949358 | 67949359 | cg20252769 |
| chr11 | 67978654 | 67978655 | cg23796243 |
| chr11 | 67979188 | 67979189 | cg20100049 |
| chr11 | 68142198 | 68142199 | cg14624207 |
| chr11 | 68142234 | 68142235 | cg21746120 |
| chr11 | 68603379 | 68603380 | cg22911054 |
| chr11 | 68924956 | 68924957 | cg15234197 |
| chr11 | 69257736 | 69257737 | cg00211115 |
| chr11 | 71165839 | 71165840 | cg04774822 |
| chr11 | 71737549 | 71737550 | cg11196788 |
| chr11 | 71749760 | 71749761 | cg04462378 |
| chr11 | 72437871 | 72437872 | cg23249399 |
| chr11 | 72447419 | 72447420 | cg00242341 |
| chr11 | 72467029 | 72467030 | cg02470043 |
| chr11 | 72880846 | 72880847 | cg05426662 |
| chr11 | 73681281 | 73681282 | cg13953458 |
| chr11 | 75230135 | 75230136 | cg05521767 |
| chr11 | 75242705 | 75242706 | cg05962522 |
| chr11 | 75295186 | 75295187 | cg04463386 |
| chr11 | 77125157 | 77125158 | cg20232503 |
| chr11 | 77776607 | 77776608 | cg03493668 |
| chr11 | 77852880 | 77852881 | cg09298680 |
| chr11 | 77898785 | 77898786 | cg00808969 |
| chr11 | 77903617 | 77903618 | cg21167761 |
| chr11 | 82556674 | 82556675 | cg17098857 |
| chr11 | 82609924 | 82609925 | cg19733740 |
| chr11 | 82689648 | 82689649 | cg23271915 |
| chr11 | 82718527 | 82718528 | cg24227371 |
| chr11 | 82865069 | 82865070 | cg15129876 |
| chr11 | 85378422 | 85378423 | cg11527436 |
| chr11 | 85519653 | 85519654 | cg09405463 |
| chr11 | 88059527 | 88059528 | cg25636441 |
| chr11 | 92080444 | 92080445 | cg09250365 |
| chr11 | 93254589 | 93254590 | cg12895631 |
| chr11 | 94824192 | 94824193 | cg27242134 |
| chr11 | 94824747 | 94824748 | cg12675417 |
| chr11 | 95071108 | 95071109 | cg10658307 |
| chr11 | 95408124 | 95408125 | cg07563725 |
| chr11 | 95828805 | 95828806 | cg26986443 |
| chr11 | 95974543 | 95974544 | cg03371275 |
| chr11 | 95990968 | 95990969 | cg08426823 |
| chr11 | 102150686 | 102150687 | cg21616785 |
| chr11 | 102300650 | 102300651 | cg16100721 |
| chr11 | 103480630 | 103480631 | cg02730303 |
| chr11 | 103830847 | 103830848 | cg06560282 |
| chr11 | 103905296 | 103905297 | cg04142010 |
| chr11 | 104769411 | 104769412 | cg18471635 |
| chr11 | 111648010 | 111648011 | cg17089665 |
| chr11 | 113775900 | 113775901 | cg06961323 |
| chr11 | 114043903 | 114043904 | cg07631435 |
| chr11 | 116723030 | 116723031 | cg04087571 |
| chr11 | 116948652 | 116948653 | cg06646174 |
| chr11 | 116966699 | 116966700 | cg06928797 |
| chr11 | 116969870 | 116969871 | cg05256304 |
| chr11 | 117068115 | 117068116 | cg11029270 |
| chr11 | 117698911 | 117698912 | cg22335223 |

| | | | |
|---|---|---|---|
| chr7 | 134924584 | 134924585 | cg12086281 |
| chr7 | 136558341 | 136558342 | cg07116919 |
| chr7 | 136641740 | 136641741 | cg26253500 |
| chr7 | 136768089 | 136768090 | cg06902367 |
| chr7 | 136815227 | 136815228 | cg22207672 |
| chr7 | 137620397 | 137620398 | cg05811904 |
| chr7 | 137620868 | 137620869 | cg09271097 |
| chr7 | 137665118 | 137665119 | cg15975598 |
| chr7 | 137671793 | 137671794 | cg05723179 |
| chr7 | 137683602 | 137683603 | cg04472009 |
| chr7 | 137760128 | 137760129 | cg25271841 |
| chr7 | 137854552 | 137854553 | cg15114607 |
| chr7 | 138816336 | 138816337 | cg01686975 |
| chr7 | 139180385 | 139180386 | cg24624586 |
| chr7 | 139760375 | 139760376 | cg03873220 |
| chr7 | 140206867 | 140206868 | cg07958575 |
| chr7 | 140452281 | 140452282 | cg08428486 |
| chr7 | 140567399 | 140567400 | cg07843733 |
| chr7 | 141285961 | 141285962 | cg14953379 |
| chr7 | 142494244 | 142494245 | cg16767506 |
| chr7 | 142828903 | 142828904 | cg05667197 |
| chr7 | 142829304 | 142829305 | cg26628847 |
| chr7 | 143013161 | 143013162 | cg04182207 |
| chr7 | 143895753 | 143895754 | cg23231902 |
| chr7 | 144049427 | 144049428 | cg01046377 |
| chr7 | 144081684 | 144081685 | cg19242984 |
| chr7 | 145395986 | 145395987 | cg27635135 |
| chr7 | 148035996 | 148035997 | cg24429310 |
| chr7 | 148394542 | 148394543 | cg18597411 |
| chr7 | 149319535 | 149319536 | cg02826890 |
| chr7 | 149557121 | 149557122 | cg01844514 |
| chr7 | 150067807 | 150067808 | cg21784940 |
| chr7 | 150072968 | 150072969 | cg06101576 |
| chr7 | 150074779 | 150074780 | cg07385974 |
| chr7 | 150105713 | 150105714 | cg17640893 |
| chr7 | 150109704 | 150109705 | cg10658768 |
| chr7 | 150130551 | 150130552 | cg00066351 |
| chr7 | 150786044 | 150786045 | cg25788549 |
| chr7 | 150786051 | 150786052 | cg22169990 |
| chr7 | 150786082 | 150786083 | cg21887193 |
| chr7 | 151502322 | 151502323 | cg22615330 |
| chr7 | 151512686 | 151512687 | cg20534694 |
| chr7 | 151512698 | 151512699 | cg26482626 |
| chr7 | 151522573 | 151522574 | cg13686795 |
| chr7 | 154557556 | 154557557 | cg22255773 |
| chr7 | 155091523 | 155091524 | cg21341645 |
| chr7 | 155438919 | 155438920 | cg27637730 |
| chr7 | 155556590 | 155556591 | cg13639901 |
| chr7 | 155556652 | 155556653 | cg14430943 |
| chr7 | 155794752 | 155794753 | cg11314668 |
| chr7 | 155794827 | 155794828 | cg15507449 |
| chr7 | 156412301 | 156412302 | cg16975053 |
| chr7 | 158352181 | 158352182 | cg17748769 |
| chr7 | 158494734 | 158494735 | cg05703659 |
| chr8 | 615409 | 615410 | cg16779811 |
| chr8 | 840398 | 840399 | cg07881065 |

| | | | |
|---|---|---|---|
| chr11 | 117701909 | 117701910 | cg26502105 |
| chr11 | 117800571 | 117800572 | cg00971332 |
| chr11 | 117801332 | 117801333 | cg05382782 |
| chr11 | 117855870 | 117855871 | cg26661481 |
| chr11 | 117947555 | 117947556 | cg05775918 |
| chr11 | 118086770 | 118086771 | cg05471842 |
| chr11 | 118753476 | 118753477 | cg01257799 |
| chr11 | 118767238 | 118767239 | cg06394082 |
| chr11 | 118774009 | 118774010 | cg08983011 |
| chr11 | 118781813 | 118781814 | cg20029201 |
| chr11 | 118900111 | 118900112 | cg06567290 |
| chr11 | 118958046 | 118958047 | cg20929545 |
| chr11 | 118958129 | 118958130 | cg21638712 |
| chr11 | 119190850 | 119190851 | cg27475169 |
| chr11 | 121477737 | 121477738 | cg07356488 |
| chr11 | 122543428 | 122543429 | cg26684673 |
| chr11 | 123229577 | 123229578 | cg03374729 |
| chr11 | 123309227 | 123309228 | cg18159802 |
| chr11 | 123322182 | 123322183 | cg25243766 |
| chr11 | 124481614 | 124481615 | cg08191915 |
| chr11 | 124933814 | 124933815 | cg03800939 |
| chr11 | 124974175 | 124974176 | cg00578690 |
| chr11 | 126180465 | 126180466 | cg02052726 |
| chr11 | 126329507 | 126329508 | cg11252310 |
| chr11 | 128180077 | 128180078 | cg22668906 |
| chr11 | 128276612 | 128276613 | cg05448032 |
| chr11 | 128318549 | 128318550 | cg21121082 |
| chr11 | 128566958 | 128566959 | cg03798942 |
| chr11 | 129900945 | 129900946 | cg26054130 |
| chr11 | 129992368 | 129992369 | cg27437304 |
| chr11 | 130076148 | 130076149 | cg04103418 |
| chr11 | 132184930 | 132184931 | cg10677017 |
| chr11 | 134095863 | 134095864 | cg09084200 |
| chr12 | 731864 | 731865 | cg26416981 |
| chr12 | 741507 | 741508 | cg23971638 |
| chr12 | 1428039 | 1428040 | cg12165841 |
| chr12 | 1598243 | 1598244 | cg03507641 |
| chr12 | 1656337 | 1656338 | cg21864961 |
| chr12 | 1691867 | 1691868 | cg17856495 |
| chr12 | 1700011 | 1700012 | cg17824906 |
| chr12 | 1726028 | 1726029 | cg10696062 |
| chr12 | 3862810 | 3862811 | cg22221074 |
| chr12 | 4224915 | 4224916 | cg12977447 |
| chr12 | 4226577 | 4226578 | cg15988261 |
| chr12 | 4397947 | 4397948 | cg17558623 |
| chr12 | 4922070 | 4922071 | cg02722637 |
| chr12 | 6572482 | 6572483 | cg22129323 |
| chr12 | 6747405 | 6747406 | cg19345219 |
| chr12 | 6881303 | 6881304 | cg22777668 |
| chr12 | 6881997 | 6881998 | cg04671742 |
| chr12 | 6882083 | 6882084 | cg01820374 |
| chr12 | 6883826 | 6883827 | cg20652042 |
| chr12 | 7023262 | 7023263 | cg19056004 |
| chr12 | 7023346 | 7023347 | cg09692396 |
| chr12 | 7023352 | 7023353 | cg04604946 |
| chr12 | 7038866 | 7038867 | cg10162209 |

| | | | |
|---|---|---|---|
| chr8 | 3253958 | 3253959 | cg23564432 |
| chr8 | 6405105 | 6405106 | cg21193744 |
| chr8 | 6433775 | 6433776 | cg18537383 |
| chr8 | 7671497 | 7671498 | cg10673328 |
| chr8 | 8182454 | 8182455 | cg07890238 |
| chr8 | 8182618 | 8182619 | cg09292244 |
| chr8 | 8182747 | 8182748 | cg18717967 |
| chr8 | 8241752 | 8241753 | cg02239258 |
| chr8 | 8702053 | 8702054 | cg12077460 |
| chr8 | 8746679 | 8746680 | cg27280535 |
| chr8 | 8870185 | 8870186 | cg20165746 |
| chr8 | 8966732 | 8966733 | cg01527562 |
| chr8 | 8993102 | 8993103 | cg02326058 |
| chr8 | 10573804 | 10573805 | cg21547283 |
| chr8 | 10982836 | 10982837 | cg03438558 |
| chr8 | 11311868 | 11311869 | cg08224773 |
| chr8 | 11323474 | 11323475 | cg01383082 |
| chr8 | 11366745 | 11366746 | cg21497594 |
| chr8 | 11402219 | 11402220 | cg07608611 |
| chr8 | 11421594 | 11421595 | cg16861076 |
| chr8 | 11628346 | 11628347 | cg09768257 |
| chr8 | 11664728 | 11664729 | cg02711665 |
| chr8 | 11763828 | 11763829 | cg01359236 |
| chr8 | 12584192 | 12584193 | cg24921221 |
| chr8 | 12610812 | 12610813 | cg00598600 |
| chr8 | 12623485 | 12623486 | cg19931644 |
| chr8 | 13191645 | 13191646 | cg02049217 |
| chr8 | 13434991 | 13434992 | cg26965236 |
| chr8 | 19419126 | 19419127 | cg17509561 |
| chr8 | 19444419 | 19444420 | cg12059540 |
| chr8 | 19588125 | 19588126 | cg19859445 |
| chr8 | 21771059 | 21771060 | cg03732056 |
| chr8 | 21771278 | 21771279 | cg21031128 |
| chr8 | 21771446 | 21771447 | cg14386061 |
| chr8 | 21911015 | 21911016 | cg27601695 |
| chr8 | 22145101 | 22145102 | cg05079448 |
| chr8 | 22288224 | 22288225 | cg18042586 |
| chr8 | 22422657 | 22422658 | cg19675778 |
| chr8 | 22555243 | 22555244 | cg22631350 |
| chr8 | 22846006 | 22846007 | cg20642630 |
| chr8 | 22853281 | 22853282 | cg10951691 |
| chr8 | 22860884 | 22860885 | cg20091591 |
| chr8 | 23102091 | 23102092 | cg17048073 |
| chr8 | 24242109 | 24242110 | cg13059335 |
| chr8 | 25227128 | 25227129 | cg20227326 |
| chr8 | 26298067 | 26298068 | cg03280108 |
| chr8 | 26431736 | 26431737 | cg00122406 |
| chr8 | 26457789 | 26457790 | cg26187031 |
| chr8 | 28175003 | 28175004 | cg03642518 |
| chr8 | 28227321 | 28227322 | cg19507160 |
| chr8 | 28232176 | 28232177 | cg26101942 |
| chr8 | 29661687 | 29661688 | cg08887138 |
| chr8 | 30402570 | 30402571 | cg07575193 |
| chr8 | 30413794 | 30413795 | cg06945634 |
| chr8 | 30505290 | 30505291 | cg03655174 |
| chr8 | 30670732 | 30670733 | cg04927725 |

| | | | |
|---|---|---|---|
| chr12 | 7071383 | 7071384 | cg16642299 |
| chr12 | 7902721 | 7902722 | cg01050005 |
| chr12 | 9106586 | 9106587 | cg13955347 |
| chr12 | 9269671 | 9269672 | cg00146928 |
| chr12 | 10455788 | 10455789 | cg00059699 |
| chr12 | 11184005 | 11184006 | cg23146713 |
| chr12 | 11215083 | 11215084 | cg20031632 |
| chr12 | 11244899 | 11244900 | cg25106783 |
| chr12 | 11339792 | 11339793 | cg25023291 |
| chr12 | 11812062 | 11812063 | cg18763536 |
| chr12 | 11821908 | 11821909 | cg14830166 |
| chr12 | 11905390 | 11905391 | cg00001793 |
| chr12 | 11905431 | 11905432 | cg08064997 |
| chr12 | 11909593 | 11909594 | cg05161923 |
| chr12 | 12031304 | 12031305 | cg26373351 |
| chr12 | 12225262 | 12225263 | cg20141578 |
| chr12 | 12620523 | 12620524 | cg08764599 |
| chr12 | 13062947 | 13062948 | cg24765748 |
| chr12 | 13359372 | 13359373 | cg16022049 |
| chr12 | 14514673 | 14514674 | cg19679081 |
| chr12 | 15698215 | 15698216 | cg13554714 |
| chr12 | 15700393 | 15700394 | cg01574823 |
| chr12 | 22218253 | 22218254 | cg02702107 |
| chr12 | 22688743 | 22688744 | cg21005240 |
| chr12 | 23714914 | 23714915 | cg04575699 |
| chr12 | 24104007 | 24104008 | cg21862372 |
| chr12 | 24228975 | 24228976 | cg16095464 |
| chr12 | 24549226 | 24549227 | cg13588483 |
| chr12 | 24608519 | 24608520 | cg01766743 |
| chr12 | 25147218 | 25147219 | cg21156602 |
| chr12 | 25216962 | 25216963 | cg19199865 |
| chr12 | 25436262 | 25436263 | cg11258277 |
| chr12 | 26987065 | 26987066 | cg07645296 |
| chr12 | 27973808 | 27973809 | cg10855773 |
| chr12 | 31255279 | 31255280 | cg00218628 |
| chr12 | 32287236 | 32287237 | cg00832457 |
| chr12 | 39981136 | 39981137 | cg13406085 |
| chr12 | 40496213 | 40496214 | cg14061497 |
| chr12 | 42299446 | 42299447 | cg10802176 |
| chr12 | 42865986 | 42865987 | cg02372037 |
| chr12 | 44410498 | 44410499 | cg07020967 |
| chr12 | 46101447 | 46101448 | cg15251256 |
| chr12 | 46169000 | 46169001 | cg22526226 |
| chr12 | 46581199 | 46581200 | cg12494373 |
| chr12 | 46637975 | 46637976 | cg26069615 |
| chr12 | 46654629 | 46654630 | cg10091798 |
| chr12 | 46941377 | 46941378 | cg02352391 |
| chr12 | 47469754 | 47469755 | cg05838358 |
| chr12 | 47472810 | 47472811 | cg18893793 |
| chr12 | 48195549 | 48195550 | cg04381888 |
| chr12 | 48199277 | 48199278 | cg23522915 |
| chr12 | 48743861 | 48743862 | cg25479963 |
| chr12 | 49583496 | 49583497 | cg22064451 |
| chr12 | 50091449 | 50091450 | cg12905592 |
| chr12 | 50366280 | 50366281 | cg05502439 |
| chr12 | 50366542 | 50366543 | cg07034276 |

| | | | | | | | |
|---|---|---|---|---|---|---|---|
| chr8 | 32715985 | 32715986 | cg01544400 | chr12 | 50400225 | 50400226 | cg16614527 |
| chr8 | 33866594 | 33866595 | cg02943456 | chr12 | 50901878 | 50901879 | cg05512157 |
| chr8 | 36894241 | 36894242 | cg22523679 | chr12 | 50962271 | 50962272 | cg23925540 |
| chr8 | 37743949 | 37743950 | cg17339440 | chr12 | 51029261 | 51029262 | cg13294447 |
| chr8 | 37755254 | 37755255 | cg03318906 | chr12 | 51154575 | 51154576 | cg01352472 |
| chr8 | 37761565 | 37761566 | cg03978658 | chr12 | 51421225 | 51421226 | cg25493658 |
| chr8 | 38387291 | 38387292 | cg24909309 | chr12 | 52305332 | 52305333 | cg20131897 |
| chr8 | 38423639 | 38423640 | cg17508816 | chr12 | 52444020 | 52444021 | cg12541478 |
| chr8 | 38790375 | 38790376 | cg13971849 | chr12 | 52476217 | 52476218 | cg19107511 |
| chr8 | 38832267 | 38832268 | cg27258399 | chr12 | 53591490 | 53591491 | cg11510999 |
| chr8 | 39792769 | 39792770 | cg11251498 | chr12 | 53592259 | 53592260 | cg20135711 |
| chr8 | 39806701 | 39806702 | cg25981315 | chr12 | 53612734 | 53612735 | cg09107344 |
| chr8 | 41464795 | 41464796 | cg12859101 | chr12 | 53627106 | 53627107 | cg17436656 |
| chr8 | 41523177 | 41523178 | cg17753169 | chr12 | 54784180 | 54784181 | cg17465423 |
| chr8 | 41557540 | 41557541 | cg11696388 | chr12 | 54805345 | 54805346 | cg20909017 |
| chr8 | 41814854 | 41814855 | cg02707799 | chr12 | 54806218 | 54806219 | cg01213231 |
| chr8 | 42009622 | 42009623 | cg18404652 | chr12 | 54809239 | 54809240 | cg09583374 |
| chr8 | 42038135 | 42038136 | cg01419713 | chr12 | 54942853 | 54942854 | cg06793728 |
| chr8 | 42163265 | 42163266 | cg11283404 | chr12 | 54942934 | 54942935 | cg21370074 |
| chr8 | 42266931 | 42266932 | cg01012595 | chr12 | 55376897 | 55376898 | cg25486399 |
| chr8 | 42608060 | 42608061 | cg20148796 | chr12 | 56082002 | 56082003 | cg17294610 |
| chr8 | 43046482 | 43046483 | cg24048338 | chr12 | 56111944 | 56111945 | cg12926596 |
| chr8 | 48174649 | 48174650 | cg03714676 | chr12 | 56327217 | 56327218 | cg15354065 |
| chr8 | 49780701 | 49780702 | cg05631737 | chr12 | 56587526 | 56587527 | cg06850687 |
| chr8 | 49801972 | 49801973 | cg13457462 | chr12 | 56624552 | 56624553 | cg00668685 |
| chr8 | 50729333 | 50729334 | cg16883726 | chr12 | 56652945 | 56652946 | cg03847895 |
| chr8 | 52807648 | 52807649 | cg04544196 | chr12 | 56771656 | 56771657 | cg04554217 |
| chr8 | 53319176 | 53319177 | cg10923561 | chr12 | 57421088 | 57421089 | cg01395351 |
| chr8 | 54757048 | 54757049 | cg18835815 | chr12 | 57422435 | 57422436 | cg21582956 |
| chr8 | 56805218 | 56805219 | cg21441674 | chr12 | 57655696 | 57655697 | cg26247373 |
| chr8 | 56813677 | 56813678 | cg21301224 | chr12 | 58152793 | 58152794 | cg05220083 |
| chr8 | 57103638 | 57103639 | cg21448513 | chr12 | 58210661 | 58210662 | cg04046364 |
| chr8 | 57212793 | 57212794 | cg16790374 | chr12 | 58210716 | 58210717 | cg26620147 |
| chr8 | 61613721 | 61613722 | cg10232395 | chr12 | 65008851 | 65008852 | cg00771490 |
| chr8 | 61785026 | 61785027 | cg25014229 | chr12 | 65067877 | 65067878 | cg19992375 |
| chr8 | 61789834 | 61789835 | cg10420756 | chr12 | 68024459 | 68024460 | cg16011250 |
| chr8 | 61825469 | 61825470 | cg18808929 | chr12 | 68551655 | 68551656 | cg01940810 |
| chr8 | 62624872 | 62624873 | cg07118376 | chr12 | 68592548 | 68592549 | cg20596773 |
| chr8 | 65603587 | 65603588 | cg00313094 | chr12 | 69632231 | 69632232 | cg09446329 |
| chr8 | 66699054 | 66699055 | cg08961450 | chr12 | 77159365 | 77159366 | cg18837713 |
| chr8 | 66743912 | 66743913 | cg06222774 | chr12 | 77457099 | 77457100 | cg16743048 |
| chr8 | 67343633 | 67343634 | cg16642695 | chr12 | 90125911 | 90125912 | cg25148705 |
| chr8 | 67344936 | 67344937 | cg01988129 | chr12 | 90234041 | 90234042 | cg02352252 |
| chr8 | 67452722 | 67452723 | cg00522276 | chr12 | 90268539 | 90268540 | cg02347249 |
| chr8 | 67457376 | 67457377 | cg26852879 | chr12 | 90341955 | 90341956 | cg01871427 |
| chr8 | 67834312 | 67834313 | cg00060836 | chr12 | 92456539 | 92456540 | cg19979853 |
| chr8 | 68110902 | 68110903 | cg10311161 | chr12 | 92815336 | 92815337 | cg04845867 |
| chr8 | 69355417 | 69355418 | cg15241236 | chr12 | 92880258 | 92880259 | cg04397652 |
| chr8 | 71073335 | 71073336 | cg25676074 | chr12 | 93400542 | 93400543 | cg02727832 |
| chr8 | 71279864 | 71279865 | cg10456591 | chr12 | 94536509 | 94536510 | cg15256743 |
| chr8 | 72824512 | 72824513 | cg10842557 | chr12 | 95266710 | 95266711 | cg09893576 |
| chr8 | 72893465 | 72893466 | cg01384246 | chr12 | 95941571 | 95941572 | cg03162823 |
| chr8 | 77602161 | 77602162 | cg16591182 | chr12 | 95945277 | 95945278 | cg27100916 |
| chr8 | 77712148 | 77712149 | cg16877004 | chr12 | 96762309 | 96762310 | cg12149299 |
| chr8 | 80706829 | 80706830 | cg24728099 | chr12 | 100557738 | 100557739 | cg17316564 |
| chr8 | 81947741 | 81947742 | cg00476430 | chr12 | 102137391 | 102137392 | cg08412215 |

| | | | |
|---|---|---|---|
| chr8 | 86099504 | 86099505 | cg08155591 |
| chr8 | 88712003 | 88712004 | cg00064296 |
| chr8 | 89336217 | 89336218 | cg11630103 |
| chr8 | 89969235 | 89969236 | cg11367259 |
| chr8 | 90916395 | 90916396 | cg01421994 |
| chr8 | 91637884 | 91637885 | cg24030920 |
| chr8 | 93005307 | 93005308 | cg04563046 |
| chr8 | 93075561 | 93075562 | cg22334059 |
| chr8 | 93103361 | 93103362 | cg03760919 |
| chr8 | 94567610 | 94567611 | cg10975798 |
| chr8 | 94892613 | 94892614 | cg10558233 |
| chr8 | 96163530 | 96163531 | cg02480685 |
| chr8 | 96167327 | 96167328 | cg12930753 |
| chr8 | 97376970 | 97376971 | cg07364657 |
| chr8 | 97658774 | 97658775 | cg15133564 |
| chr8 | 98361938 | 98361939 | cg05313910 |
| chr8 | 98446695 | 98446696 | cg08245578 |
| chr8 | 99838488 | 99838489 | cg20673075 |
| chr8 | 101678439 | 101678440 | cg03629107 |
| chr8 | 101736708 | 101736709 | cg18817529 |
| chr8 | 102063914 | 102063915 | cg07546433 |
| chr8 | 102083111 | 102083112 | cg03483314 |
| chr8 | 102110139 | 102110140 | cg00734471 |
| chr8 | 102167694 | 102167695 | cg20162696 |
| chr8 | 102509885 | 102509886 | cg15679829 |
| chr8 | 102944342 | 102944343 | cg26064870 |
| chr8 | 102984290 | 102984291 | cg01666490 |
| chr8 | 103407566 | 103407567 | cg03575122 |
| chr8 | 103548145 | 103548146 | cg20611272 |
| chr8 | 103549622 | 103549623 | cg09916692 |
| chr8 | 103568792 | 103568793 | cg01553616 |
| chr8 | 103662317 | 103662318 | cg07171504 |
| chr8 | 103671986 | 103671987 | cg17594632 |
| chr8 | 103751623 | 103751624 | cg23203234 |
| chr8 | 103872978 | 103872979 | cg02243276 |
| chr8 | 103932076 | 103932077 | cg01142735 |
| chr8 | 103937374 | 103937375 | cg19589396 |
| chr8 | 103952396 | 103952397 | cg07549297 |
| chr8 | 103973799 | 103973800 | cg08236537 |
| chr8 | 104117866 | 104117867 | cg03090969 |
| chr8 | 104132683 | 104132684 | cg10172675 |
| chr8 | 105899330 | 105899331 | cg06033789 |
| chr8 | 106511081 | 106511082 | cg11399319 |
| chr8 | 107870405 | 107870406 | cg02102791 |
| chr8 | 110099750 | 110099751 | cg22268164 |
| chr8 | 110657877 | 110657878 | cg16891965 |
| chr8 | 110990365 | 110990366 | cg17164016 |
| chr8 | 112612368 | 112612369 | cg21187920 |
| chr8 | 116822442 | 116822443 | cg07147958 |
| chr8 | 118683119 | 118683120 | cg02902186 |
| chr8 | 118993090 | 118993091 | cg22747380 |
| chr8 | 119586905 | 119586906 | cg04034290 |
| chr8 | 119890170 | 119890171 | cg03489427 |
| chr8 | 120106427 | 120106428 | cg03073218 |
| chr8 | 120615567 | 120615568 | cg09444531 |
| chr8 | 121140957 | 121140958 | cg00288736 |

| | | | |
|---|---|---|---|
| chr12 | 102218837 | 102218838 | cg03756485 |
| chr12 | 102270723 | 102270724 | cg27425327 |
| chr12 | 102873312 | 102873313 | cg14240353 |
| chr12 | 103828459 | 103828460 | cg08027643 |
| chr12 | 104230671 | 104230672 | cg01164094 |
| chr12 | 104302786 | 104302787 | cg10884544 |
| chr12 | 104500900 | 104500901 | cg21050282 |
| chr12 | 104776085 | 104776086 | cg10163220 |
| chr12 | 104849615 | 104849616 | cg02908942 |
| chr12 | 104975516 | 104975517 | cg03318667 |
| chr12 | 105065028 | 105065029 | cg00608661 |
| chr12 | 105130282 | 105130283 | cg15404019 |
| chr12 | 105143826 | 105143827 | cg20883723 |
| chr12 | 105323396 | 105323397 | cg23855818 |
| chr12 | 105323744 | 105323745 | cg20495009 |
| chr12 | 105486126 | 105486127 | cg18166987 |
| chr12 | 105744023 | 105744024 | cg26459229 |
| chr12 | 106100228 | 106100229 | cg23088810 |
| chr12 | 106639919 | 106639920 | cg13249833 |
| chr12 | 108078821 | 108078822 | cg00632811 |
| chr12 | 109031953 | 109031954 | cg09907439 |
| chr12 | 109035219 | 109035220 | cg23816140 |
| chr12 | 109040983 | 109040984 | cg23429240 |
| chr12 | 109046825 | 109046826 | cg06251181 |
| chr12 | 109064006 | 109064007 | cg05679686 |
| chr12 | 109107559 | 109107560 | cg10155304 |
| chr12 | 109899089 | 109899090 | cg11130630 |
| chr12 | 110427408 | 110427409 | cg27069285 |
| chr12 | 110488701 | 110488702 | cg08726598 |
| chr12 | 110811045 | 110811046 | cg06703803 |
| chr12 | 111048024 | 111048025 | cg23953290 |
| chr12 | 111086880 | 111086881 | cg10453481 |
| chr12 | 111284228 | 111284229 | cg24389359 |
| chr12 | 111345314 | 111345315 | cg17348388 |
| chr12 | 111521899 | 111521900 | cg03884572 |
| chr12 | 111889200 | 111889201 | cg20595271 |
| chr12 | 112284205 | 112284206 | cg19563158 |
| chr12 | 112822378 | 112822379 | cg27180555 |
| chr12 | 113515332 | 113515333 | cg11787160 |
| chr12 | 113515969 | 113515970 | cg01313387 |
| chr12 | 113587581 | 113587582 | cg15033653 |
| chr12 | 116571240 | 116571241 | cg03621504 |
| chr12 | 116586898 | 116586899 | cg09230679 |
| chr12 | 116946758 | 116946759 | cg08153404 |
| chr12 | 117036895 | 117036896 | cg24901447 |
| chr12 | 117036972 | 117036973 | cg23126203 |
| chr12 | 117037220 | 117037221 | cg03432814 |
| chr12 | 117523153 | 117523154 | cg10856812 |
| chr12 | 117579332 | 117579333 | cg21441555 |
| chr12 | 117579501 | 117579502 | cg25725899 |
| chr12 | 117579690 | 117579691 | cg01613406 |
| chr12 | 117588951 | 117588952 | cg18860868 |
| chr12 | 117613954 | 117613955 | cg04402948 |
| chr12 | 118313657 | 118313658 | cg06711448 |
| chr12 | 118404706 | 118404707 | cg10084383 |
| chr12 | 118405988 | 118405989 | cg16300531 |

| | | | | | | | |
|---|---|---|---|---|---|---|---|
| chr8 | 121791273 | 121791274 | cg01443285 | chr12 | 118407487 | 118407488 | cg01386425 |
| chr8 | 122118171 | 122118172 | cg14936938 | chr12 | 118471960 | 118471961 | cg10617224 |
| chr8 | 122273909 | 122273910 | cg11476306 | chr12 | 118500931 | 118500932 | cg14474811 |
| chr8 | 122302722 | 122302723 | cg10864603 | chr12 | 118501232 | 118501233 | cg01932247 |
| chr8 | 124055146 | 124055147 | cg21139496 | chr12 | 118583229 | 118583230 | cg27065869 |
| chr8 | 124215033 | 124215034 | cg00831735 | chr12 | 118583242 | 118583243 | cg16572268 |
| chr8 | 124284379 | 124284380 | cg00078299 | chr12 | 118782453 | 118782454 | cg17627898 |
| chr8 | 124666123 | 124666124 | cg22342100 | chr12 | 120578192 | 120578193 | cg14667104 |
| chr8 | 125631251 | 125631252 | cg05013438 | chr12 | 121150686 | 121150687 | cg06302566 |
| chr8 | 125640145 | 125640146 | cg24630782 | chr12 | 121164194 | 121164195 | cg09813248 |
| chr8 | 126154086 | 126154087 | cg20390587 | chr12 | 121569969 | 121569970 | cg23801057 |
| chr8 | 126619805 | 126619806 | cg08843064 | chr12 | 121570571 | 121570572 | cg15965241 |
| chr8 | 128265902 | 128265903 | cg08599249 | chr12 | 121570616 | 121570617 | cg10852165 |
| chr8 | 128342705 | 128342706 | cg14437045 | chr12 | 121688317 | 121688318 | cg23383531 |
| chr8 | 128428869 | 128428870 | cg14289643 | chr12 | 121726240 | 121726241 | cg16942632 |
| chr8 | 128808554 | 128808555 | cg00780520 | chr12 | 122015596 | 122015597 | cg17169037 |
| chr8 | 128922521 | 128922522 | cg03458957 | chr12 | 122021013 | 122021014 | cg22246785 |
| chr8 | 128931535 | 128931536 | cg23896695 | chr12 | 122235560 | 122235561 | cg08421051 |
| chr8 | 129008054 | 129008055 | cg03566001 | chr12 | 122611269 | 122611270 | cg22501243 |
| chr8 | 129336547 | 129336548 | cg16489468 | chr12 | 122712212 | 122712213 | cg09123760 |
| chr8 | 130215373 | 130215374 | cg08587313 | chr12 | 123468780 | 123468781 | cg18637761 |
| chr8 | 130389960 | 130389961 | cg05084525 | chr12 | 123548027 | 123548028 | cg05384139 |
| chr8 | 130472165 | 130472166 | cg09631175 | chr12 | 123707536 | 123707537 | cg10169515 |
| chr8 | 130898833 | 130898834 | cg19123356 | chr12 | 124014409 | 124014410 | cg23825057 |
| chr8 | 131268256 | 131268257 | cg00162859 | chr12 | 124248044 | 124248045 | cg20752847 |
| chr8 | 131289180 | 131289181 | cg02771336 | chr12 | 124492549 | 124492550 | cg25459280 |
| chr8 | 132929147 | 132929148 | cg15420926 | chr12 | 124497105 | 124497106 | cg09436681 |
| chr8 | 134059874 | 134059875 | cg22845912 | chr12 | 124971111 | 124971112 | cg13982220 |
| chr8 | 134571813 | 134571814 | cg18977891 | chr12 | 124971775 | 124971776 | cg24461063 |
| chr8 | 134576933 | 134576934 | cg03965649 | chr12 | 124972783 | 124972784 | cg10929873 |
| chr8 | 134734839 | 134734840 | cg11745166 | chr12 | 124979813 | 124979814 | cg26524350 |
| chr8 | 134895500 | 134895501 | cg10902738 | chr12 | 124979893 | 124979894 | cg22611259 |
| chr8 | 135697767 | 135697768 | cg03616827 | chr12 | 124982002 | 124982003 | cg08192337 |
| chr8 | 136245114 | 136245115 | cg00937286 | chr12 | 125023878 | 125023879 | cg04822278 |
| chr8 | 138493473 | 138493474 | cg20316883 | chr12 | 125223540 | 125223541 | cg19194645 |
| chr8 | 138493933 | 138493934 | cg08832234 | chr12 | 125345167 | 125345168 | cg03319070 |
| chr8 | 141594473 | 141594474 | cg14835938 | chr12 | 127226034 | 127226035 | cg10948698 |
| chr8 | 141599185 | 141599186 | cg11006453 | chr12 | 130084035 | 130084036 | cg16048915 |
| chr8 | 141599208 | 141599209 | cg23731089 | chr12 | 130311289 | 130311290 | cg26614129 |
| chr8 | 141608316 | 141608317 | cg26291848 | chr12 | 130961453 | 130961454 | cg13802583 |
| chr8 | 141608401 | 141608402 | cg01980793 | chr12 | 131303247 | 131303248 | cg11011512 |
| chr8 | 141608434 | 141608435 | cg21208682 | chr12 | 132259591 | 132259592 | cg15817163 |
| chr8 | 142217405 | 142217406 | cg04560093 | chr12 | 132259908 | 132259909 | cg22325958 |
| chr8 | 143554875 | 143554876 | cg04436528 | chr12 | 132282367 | 132282368 | cg24209874 |
| chr8 | 144103587 | 144103588 | cg14392283 | chr12 | 132537711 | 132537712 | cg24545687 |
| chr8 | 144396827 | 144396828 | cg12843874 | chr12 | 132537749 | 132537750 | cg19865523 |
| chr8 | 144410815 | 144410816 | cg08170427 | chr12 | 132537775 | 132537776 | cg11075854 |
| chr8 | 144410972 | 144410973 | cg17117981 | chr12 | 132549292 | 132549293 | cg14622549 |
| chr8 | 144460468 | 144460469 | cg06330431 | chr12 | 132549404 | 132549405 | cg21703988 |
| chr8 | 144543951 | 144543952 | cg11210357 | chr12 | 132551398 | 132551399 | cg14248360 |
| chr8 | 144544041 | 144544042 | cg24010658 | chr12 | 132551483 | 132551484 | cg19172447 |
| chr8 | 144599347 | 144599348 | cg03345925 | chr12 | 132551592 | 132551593 | cg24980032 |
| chr8 | 144656997 | 144656998 | cg18047920 | chr12 | 132562425 | 132562426 | cg01101020 |
| chr8 | 145003653 | 145003654 | cg24891660 | chr12 | 132562495 | 132562496 | cg07359518 |
| chr8 | 145012644 | 145012645 | cg13389508 | chr12 | 132639139 | 132639140 | cg22561460 |
| chr8 | 145022842 | 145022843 | cg16001422 | chr12 | 133028853 | 133028854 | cg23393615 |

| | | | |
|---|---|---|---|
| chr8 | 145023013 | 145023014 | cg03280622 |
| chr8 | 145025128 | 145025129 | cg21672292 |
| chr8 | 145061318 | 145061319 | cg25477843 |
| chr8 | 145067245 | 145067246 | cg21813905 |
| chr8 | 145494342 | 145494343 | cg10146692 |
| chr8 | 145495665 | 145495666 | cg25958911 |
| chr8 | 145539893 | 145539894 | cg10718130 |
| chr8 | 145543014 | 145543015 | cg09565397 |
| chr9 | 1871778 | 1871779 | cg21160429 |
| chr9 | 4742079 | 4742080 | cg13993274 |
| chr9 | 5184060 | 5184061 | cg01003815 |
| chr9 | 5831999 | 5832000 | cg13979887 |
| chr9 | 8234760 | 8234761 | cg13556767 |
| chr9 | 10232876 | 10232877 | cg14406814 |
| chr9 | 14082328 | 14082329 | cg14431257 |
| chr9 | 15147652 | 15147653 | cg14405768 |
| chr9 | 19228799 | 19228800 | cg01364172 |
| chr9 | 21996207 | 21996208 | cg14069088 |
| chr9 | 27408242 | 27408243 | cg14643264 |
| chr9 | 33857698 | 33857699 | cg13525804 |
| chr9 | 34316383 | 34316384 | cg01062470 |
| chr9 | 35564052 | 35564053 | cg21212076 |
| chr9 | 35616244 | 35616245 | cg03487430 |
| chr9 | 35649906 | 35649907 | cg13832290 |
| chr9 | 35814283 | 35814284 | cg00193668 |
| chr9 | 36999934 | 36999935 | cg13695362 |
| chr9 | 37001111 | 37001112 | cg00318543 |
| chr9 | 37529128 | 37529129 | cg14178364 |
| chr9 | 37576958 | 37576959 | cg04153536 |
| chr9 | 37985583 | 37985584 | cg13599062 |
| chr9 | 38091405 | 38091406 | cg13415773 |
| chr9 | 72735830 | 72735831 | cg13829849 |
| chr9 | 74524266 | 74524267 | cg07194321 |
| chr9 | 75449747 | 75449748 | cg14184524 |
| chr9 | 87449508 | 87449509 | cg14273545 |
| chr9 | 89422493 | 89422494 | cg13445230 |
| chr9 | 89722432 | 89722433 | cg13676800 |
| chr9 | 90208854 | 90208855 | cg14286732 |
| chr9 | 91924749 | 91924750 | cg14307853 |
| chr9 | 92040726 | 92040727 | cg13584258 |
| chr9 | 92216472 | 92216473 | cg13737493 |
| chr9 | 92224930 | 92224931 | cg14567214 |
| chr9 | 93619467 | 93619468 | cg14054883 |
| chr9 | 95525261 | 95525262 | cg01028796 |
| chr9 | 95766607 | 95766608 | cg13863007 |
| chr9 | 96940848 | 96940849 | cg03275648 |
| chr9 | 97405553 | 97405554 | cg13573745 |
| chr9 | 97767954 | 97767955 | cg03456213 |
| chr9 | 98245640 | 98245641 | cg13563132 |
| chr9 | 99145525 | 99145526 | cg15582789 |
| chr9 | 99146400 | 99146401 | cg13771161 |
| chr9 | 99262208 | 99262209 | cg14585797 |
| chr9 | 99318213 | 99318214 | cg14276584 |
| chr9 | 99416629 | 99416630 | cg14591340 |
| chr9 | 99773358 | 99773359 | cg03776935 |
| chr9 | 100068361 | 100068362 | cg13739659 |

| | | | |
|---|---|---|---|
| chr12 | 133028905 | 133028906 | cg07562175 |
| chr12 | 133100051 | 133100052 | cg02656049 |
| chr12 | 133383619 | 133383620 | cg01230386 |
| chr13 | 23898864 | 23898865 | cg04591230 |
| chr13 | 23919407 | 23919408 | cg00893424 |
| chr13 | 24041251 | 24041252 | cg09323835 |
| chr13 | 24417943 | 24417944 | cg18057306 |
| chr13 | 24793488 | 24793489 | cg02537941 |
| chr13 | 24837937 | 24837938 | cg04462561 |
| chr13 | 24847935 | 24847936 | cg08472727 |
| chr13 | 27829014 | 27829015 | cg09363424 |
| chr13 | 28578045 | 28578046 | cg15126273 |
| chr13 | 28752067 | 28752068 | cg17987059 |
| chr13 | 29291701 | 29291702 | cg02097152 |
| chr13 | 31245234 | 31245235 | cg26756921 |
| chr13 | 31309032 | 31309033 | cg21054703 |
| chr13 | 32611483 | 32611484 | cg25274185 |
| chr13 | 32614209 | 32614210 | cg16995086 |
| chr13 | 32835962 | 32835963 | cg23705155 |
| chr13 | 33403686 | 33403687 | cg17994441 |
| chr13 | 34063815 | 34063816 | cg13583316 |
| chr13 | 40670783 | 40670784 | cg09331166 |
| chr13 | 41055185 | 41055186 | cg13750802 |
| chr13 | 41055261 | 41055262 | cg26226408 |
| chr13 | 41111493 | 41111494 | cg22615203 |
| chr13 | 41154103 | 41154104 | cg24858665 |
| chr13 | 41223175 | 41223176 | cg23413567 |
| chr13 | 41594904 | 41594905 | cg12691679 |
| chr13 | 42035731 | 42035732 | cg12421110 |
| chr13 | 42070600 | 42070601 | cg19266276 |
| chr13 | 42803600 | 42803601 | cg19307180 |
| chr13 | 42840965 | 42840966 | cg07583851 |
| chr13 | 44690989 | 44690990 | cg11650298 |
| chr13 | 45545983 | 45545984 | cg02951237 |
| chr13 | 47240378 | 47240379 | cg04092803 |
| chr13 | 47371635 | 47371636 | cg27021512 |
| chr13 | 49975418 | 49975419 | cg17777592 |
| chr13 | 50802085 | 50802086 | cg00843117 |
| chr13 | 51103373 | 51103374 | cg05091135 |
| chr13 | 51946399 | 51946400 | cg21028785 |
| chr13 | 52375776 | 52375777 | cg04786592 |
| chr13 | 60648567 | 60648568 | cg05170711 |
| chr13 | 62245229 | 62245230 | cg10453777 |
| chr13 | 74713082 | 74713083 | cg19526076 |
| chr13 | 74732386 | 74732387 | cg19731286 |
| chr13 | 77283725 | 77283726 | cg13711538 |
| chr13 | 77289721 | 77289722 | cg14287311 |
| chr13 | 77364735 | 77364736 | cg00901878 |
| chr13 | 97873877 | 97873878 | cg02838877 |
| chr13 | 97877417 | 97877418 | cg06340704 |
| chr13 | 99103124 | 99103125 | cg07340007 |
| chr13 | 99148667 | 99148668 | cg10914815 |
| chr13 | 99174875 | 99174876 | cg09504131 |
| chr13 | 99209304 | 99209305 | cg09110502 |
| chr13 | 99218287 | 99218288 | cg01185345 |
| chr13 | 99218657 | 99218658 | cg17176619 |

| | | | |
|---|---|---|---|
| chr9 | 101570703 | 101570704 | cg14400354 |
| chr9 | 102208423 | 102208424 | cg13630114 |
| chr9 | 102588232 | 102588233 | cg13655635 |
| chr9 | 102589165 | 102589166 | cg06145669 |
| chr9 | 104237697 | 104237698 | cg13683194 |
| chr9 | 107525704 | 107525705 | cg13888748 |
| chr9 | 110797033 | 110797034 | cg14206056 |
| chr9 | 112191008 | 112191009 | cg13510418 |
| chr9 | 114663626 | 114663627 | cg14278148 |
| chr9 | 114936583 | 114936584 | cg14195117 |
| chr9 | 115512530 | 115512531 | cg09619598 |
| chr9 | 116262291 | 116262292 | cg14124261 |
| chr9 | 117157614 | 117157615 | cg13910813 |
| chr9 | 117159457 | 117159458 | cg13967908 |
| chr9 | 117657237 | 117657238 | cg14113958 |
| chr9 | 117694183 | 117694184 | cg14361804 |
| chr9 | 123637816 | 123637817 | cg14614490 |
| chr9 | 123640474 | 123640475 | cg13501527 |
| chr9 | 123687270 | 123687271 | cg14518276 |
| chr9 | 123688715 | 123688716 | cg15551881 |
| chr9 | 123689193 | 123689194 | cg04517263 |
| chr9 | 123691850 | 123691851 | cg14580859 |
| chr9 | 123694717 | 123694718 | cg14368473 |
| chr9 | 123745713 | 123745714 | cg13567360 |
| chr9 | 124029926 | 124029927 | cg17071957 |
| chr9 | 125281227 | 125281228 | cg24137660 |
| chr9 | 125796809 | 125796810 | cg04655481 |
| chr9 | 126118355 | 126118356 | cg02010852 |
| chr9 | 126943462 | 126943463 | cg13669432 |
| chr9 | 127018840 | 127018841 | cg13505631 |
| chr9 | 127032953 | 127032954 | cg14285539 |
| chr9 | 127040562 | 127040563 | cg14109456 |
| chr9 | 127284971 | 127284972 | cg14156446 |
| chr9 | 127888818 | 127888819 | cg13856340 |
| chr9 | 129624994 | 129624995 | cg13916080 |
| chr9 | 130546644 | 130546645 | cg14286292 |
| chr9 | 130564509 | 130564510 | cg00188748 |
| chr9 | 130662451 | 130662452 | cg05517711 |
| chr9 | 130677549 | 130677550 | cg13687735 |
| chr9 | 130739838 | 130739839 | cg13992008 |
| chr9 | 130743943 | 130743944 | cg06096805 |
| chr9 | 130888547 | 130888548 | cg01311102 |
| chr9 | 130910995 | 130910996 | cg01871963 |
| chr9 | 130964374 | 130964375 | cg00407062 |
| chr9 | 130964918 | 130964919 | cg14532894 |
| chr9 | 131447995 | 131447996 | cg14097046 |
| chr9 | 131545865 | 131545866 | cg13583162 |
| chr9 | 133807443 | 133807444 | cg13478737 |
| chr9 | 134293793 | 134293794 | cg13543915 |
| chr9 | 134497566 | 134497567 | cg13781989 |
| chr9 | 134612644 | 134612645 | cg04086928 |
| chr9 | 136018794 | 136018795 | cg14099540 |
| chr9 | 136390861 | 136390862 | cg01263574 |
| chr9 | 136547862 | 136547863 | cg13824009 |
| chr9 | 136654631 | 136654632 | cg13754665 |
| chr9 | 136809652 | 136809653 | cg21157725 |
| chr13 | 99960957 | 99960958 | cg10736330 |
| chr13 | 100019647 | 100019648 | cg10898310 |
| chr13 | 100149311 | 100149312 | cg00344422 |
| chr13 | 100252695 | 100252696 | cg12127863 |
| chr13 | 101185965 | 101185966 | cg13776095 |
| chr13 | 101186172 | 101186173 | cg07218357 |
| chr13 | 101186247 | 101186248 | cg02923162 |
| chr13 | 101186274 | 101186275 | cg00336946 |
| chr13 | 109568637 | 109568638 | cg11300777 |
| chr13 | 110431691 | 110431692 | cg12085119 |
| chr13 | 110874267 | 110874268 | cg21478902 |
| chr13 | 111305805 | 111305806 | cg18358662 |
| chr13 | 111305822 | 111305823 | cg26801538 |
| chr13 | 111305863 | 111305864 | cg27510257 |
| chr13 | 111306273 | 111306274 | cg02402141 |
| chr13 | 111306312 | 111306313 | cg18116097 |
| chr13 | 111317935 | 111317936 | cg21067963 |
| chr13 | 111318160 | 111318161 | cg17394623 |
| chr13 | 111318490 | 111318491 | cg08400494 |
| chr13 | 111318555 | 111318556 | cg12423658 |
| chr13 | 111318640 | 111318641 | cg08289839 |
| chr13 | 113343376 | 113343377 | cg03992114 |
| chr13 | 113348391 | 113348392 | cg09507215 |
| chr13 | 113423096 | 113423097 | cg12917253 |
| chr13 | 113423178 | 113423179 | cg14864184 |
| chr13 | 113425199 | 113425200 | cg27096043 |
| chr13 | 113425621 | 113425622 | cg21513610 |
| chr13 | 113425756 | 113425757 | cg00902895 |
| chr13 | 113428161 | 113428162 | cg21897676 |
| chr13 | 113567732 | 113567733 | cg24722198 |
| chr13 | 114825082 | 114825083 | cg19100884 |
| chr13 | 114834683 | 114834684 | cg05296619 |
| chr13 | 114834922 | 114834923 | cg11997468 |
| chr13 | 114867544 | 114867545 | cg16521443 |
| chr13 | 114867701 | 114867702 | cg17672262 |
| chr13 | 114868344 | 114868345 | cg27378358 |
| chr13 | 114875412 | 114875413 | cg21261158 |
| chr13 | 114892556 | 114892557 | cg17061183 |
| chr13 | 114896192 | 114896193 | cg10002066 |
| chr13 | 114908876 | 114908877 | cg12701674 |
| chr13 | 114908901 | 114908902 | cg24635112 |
| chr14 | 20945319 | 20945320 | cg23173586 |
| chr14 | 22938675 | 22938676 | cg23062357 |
| chr14 | 22945181 | 22945182 | cg18887769 |
| chr14 | 22965472 | 22965473 | cg15849446 |
| chr14 | 22966633 | 22966634 | cg20507742 |
| chr14 | 22968031 | 22968032 | cg02903983 |
| chr14 | 22974144 | 22974145 | cg09953583 |
| chr14 | 23015657 | 23015658 | cg09232358 |
| chr14 | 23284518 | 23284519 | cg19872681 |
| chr14 | 23285365 | 23285366 | cg18960218 |
| chr14 | 23308294 | 23308295 | cg10418289 |
| chr14 | 23351487 | 23351488 | cg04539476 |
| chr14 | 23352488 | 23352489 | cg24129115 |
| chr14 | 23367828 | 23367829 | cg14183602 |
| chr14 | 23530542 | 23530543 | cg04854749 |

| | | | |
|---|---|---|---|
| chr9 | 136844768 | 136844769 | cg13755204 |
| chr9 | 136845816 | 136845817 | cg14069928 |
| chr9 | 137215364 | 137215365 | cg04329455 |
| chr9 | 137711757 | 137711758 | cg21208686 |
| chr9 | 138742730 | 138742731 | cg14479377 |
| chr9 | 138904715 | 138904716 | cg13959241 |
| chr9 | 138904964 | 138904965 | cg13549345 |
| chr9 | 139131620 | 139131621 | cg14094347 |
| chr9 | 139247374 | 139247375 | cg14040131 |
| chr9 | 139299357 | 139299358 | cg14108380 |
| chr9 | 139300582 | 139300583 | cg14115884 |
| chr9 | 139302769 | 139302770 | cg01080407 |
| chr9 | 139427171 | 139427172 | cg12076931 |
| chr9 | 139437479 | 139437480 | cg14363954 |
| chr9 | 139496844 | 139496845 | cg14085952 |
| chr9 | 139618608 | 139618609 | cg14149774 |
| chr9 | 139627019 | 139627020 | cg13491139 |
| chr9 | 139635449 | 139635450 | cg13754355 |
| chr9 | 139642875 | 139642876 | cg11873854 |
| chr9 | 139643351 | 139643352 | cg14611112 |
| chr9 | 139776893 | 139776894 | cg13823169 |
| chr9 | 139859471 | 139859472 | cg13917020 |
| chr9 | 140408634 | 140408635 | cg14487665 |
| chr9 | 140651049 | 140651050 | cg13628057 |
| chr10 | 264920 | 264921 | cg14301030 |
| chr10 | 346227 | 346228 | cg13351028 |
| chr10 | 370756 | 370757 | cg23983710 |
| chr10 | 468386 | 468387 | cg02594677 |
| chr10 | 468412 | 468413 | cg24461627 |
| chr10 | 980765 | 980766 | cg01503450 |
| chr10 | 2247630 | 2247631 | cg12763430 |
| chr10 | 2812567 | 2812568 | cg00806826 |
| chr10 | 3269930 | 3269931 | cg18602350 |
| chr10 | 3494422 | 3494423 | cg11721591 |
| chr10 | 3627796 | 3627797 | cg14043803 |
| chr10 | 3666755 | 3666756 | cg05180182 |
| chr10 | 3789788 | 3789789 | cg10593713 |
| chr10 | 3817124 | 3817125 | cg09076960 |
| chr10 | 3822379 | 3822380 | cg12570716 |
| chr10 | 3823790 | 3823791 | cg23680451 |
| chr10 | 3823844 | 3823845 | cg06048750 |
| chr10 | 3823907 | 3823908 | cg25749254 |
| chr10 | 3824555 | 3824556 | cg08614871 |
| chr10 | 3935314 | 3935315 | cg24432632 |
| chr10 | 4782836 | 4782837 | cg12279397 |
| chr10 | 5544140 | 5544141 | cg27250032 |
| chr10 | 5593926 | 5593927 | cg17115419 |
| chr10 | 5593949 | 5593950 | cg06250720 |
| chr10 | 5617743 | 5617744 | cg10410412 |
| chr10 | 5735349 | 5735350 | cg09411597 |
| chr10 | 5938186 | 5938187 | cg07633835 |
| chr10 | 6183528 | 6183529 | cg04189187 |
| chr10 | 6183575 | 6183576 | cg18264092 |
| chr10 | 6392160 | 6392161 | cg15084823 |
| chr10 | 6665001 | 6665002 | cg09017139 |
| chr10 | 6691515 | 6691516 | cg25812683 |

| | | | |
|---|---|---|---|
| chr14 | 23623935 | 23623936 | cg01790464 |
| chr14 | 24439157 | 24439158 | cg14214739 |
| chr14 | 24458208 | 24458209 | cg01597387 |
| chr14 | 24458299 | 24458300 | cg10199763 |
| chr14 | 24475906 | 24475907 | cg22926554 |
| chr14 | 24549943 | 24549944 | cg04774505 |
| chr14 | 24601808 | 24601809 | cg13167816 |
| chr14 | 24780167 | 24780168 | cg07509935 |
| chr14 | 24780540 | 24780541 | cg26310551 |
| chr14 | 31484468 | 31484469 | cg08274139 |
| chr14 | 31484496 | 31484497 | cg10195619 |
| chr14 | 39579505 | 39579506 | cg14289929 |
| chr14 | 39739199 | 39739200 | cg08267072 |
| chr14 | 40142517 | 40142518 | cg11835955 |
| chr14 | 50559312 | 50559313 | cg25716415 |
| chr14 | 50689394 | 50689395 | cg18976542 |
| chr14 | 51137626 | 51137627 | cg18406010 |
| chr14 | 51606299 | 51606300 | cg23942311 |
| chr14 | 52347968 | 52347969 | cg09261791 |
| chr14 | 53576363 | 53576364 | cg13654573 |
| chr14 | 53616150 | 53616151 | cg18866817 |
| chr14 | 53658573 | 53658574 | cg13507452 |
| chr14 | 54980058 | 54980059 | cg23020857 |
| chr14 | 55228808 | 55228809 | cg07449447 |
| chr14 | 55794598 | 55794599 | cg04971161 |
| chr14 | 56264572 | 56264573 | cg14442746 |
| chr14 | 56587633 | 56587634 | cg01208566 |
| chr14 | 59007136 | 59007137 | cg15913815 |
| chr14 | 59102696 | 59102697 | cg16715687 |
| chr14 | 62011349 | 62011350 | cg03050508 |
| chr14 | 64191548 | 64191549 | cg26013396 |
| chr14 | 64364050 | 64364051 | cg16778211 |
| chr14 | 64676792 | 64676793 | cg09975367 |
| chr14 | 64676831 | 64676832 | cg05050858 |
| chr14 | 64935374 | 64935375 | cg19792099 |
| chr14 | 65652792 | 65652793 | cg24017995 |
| chr14 | 65696480 | 65696481 | cg23373640 |
| chr14 | 67656029 | 67656030 | cg02809404 |
| chr14 | 67656082 | 67656083 | cg19566774 |
| chr14 | 68745571 | 68745572 | cg02893348 |
| chr14 | 68825875 | 68825876 | cg18842157 |
| chr14 | 68830704 | 68830705 | cg10975863 |
| chr14 | 68839222 | 68839223 | cg11724735 |
| chr14 | 68871428 | 68871429 | cg13572771 |
| chr14 | 68944441 | 68944442 | cg21503275 |
| chr14 | 69274280 | 69274281 | cg07502892 |
| chr14 | 69286344 | 69286345 | cg14491362 |
| chr14 | 70077191 | 70077192 | cg03505866 |
| chr14 | 70097358 | 70097359 | cg26947583 |
| chr14 | 71377000 | 71377001 | cg07137751 |
| chr14 | 71831132 | 71831133 | cg10134583 |
| chr14 | 71953890 | 71953891 | cg27053629 |
| chr14 | 72061746 | 72061747 | cg13720639 |
| chr14 | 74957673 | 74957674 | cg21915100 |
| chr14 | 75303022 | 75303023 | cg11165667 |
| chr14 | 75897841 | 75897842 | cg22143352 |

| | | | |
|---|---|---|---|
| chr10 | 10436568 | 10436569 | cg03478630 |
| chr10 | 10837818 | 10837819 | cg00982519 |
| chr10 | 11616142 | 11616143 | cg05265574 |
| chr10 | 12241348 | 12241349 | cg12089187 |
| chr10 | 12390599 | 12390600 | cg03575602 |
| chr10 | 13749382 | 13749383 | cg13707951 |
| chr10 | 14215314 | 14215315 | cg04941706 |
| chr10 | 14628301 | 14628302 | cg04638265 |
| chr10 | 14695349 | 14695350 | cg18450420 |
| chr10 | 14861832 | 14861833 | cg05335277 |
| chr10 | 17689470 | 17689471 | cg15609734 |
| chr10 | 18334434 | 18334435 | cg16284507 |
| chr10 | 18629568 | 18629569 | cg07731871 |
| chr10 | 22608858 | 22608859 | cg10763374 |
| chr10 | 23100186 | 23100187 | cg19835128 |
| chr10 | 23732088 | 23732089 | cg16152753 |
| chr10 | 24370705 | 24370706 | cg11009100 |
| chr10 | 25183825 | 25183826 | cg20633213 |
| chr10 | 25303159 | 25303160 | cg26938599 |
| chr10 | 27032461 | 27032462 | cg01401971 |
| chr10 | 27070113 | 27070114 | cg26142490 |
| chr10 | 27150407 | 27150408 | cg18825419 |
| chr10 | 31610754 | 31610755 | cg13303654 |
| chr10 | 32345864 | 32345865 | cg13612317 |
| chr10 | 32345866 | 32345867 | cg14776578 |
| chr10 | 32438418 | 32438419 | cg16489259 |
| chr10 | 33244887 | 33244888 | cg06538549 |
| chr10 | 33299147 | 33299148 | cg00146027 |
| chr10 | 33321730 | 33321731 | cg00453735 |
| chr10 | 33426312 | 33426313 | cg07661660 |
| chr10 | 33432514 | 33432515 | cg20286882 |
| chr10 | 33483109 | 33483110 | cg25179876 |
| chr10 | 34189023 | 34189024 | cg24919806 |
| chr10 | 35676874 | 35676875 | cg17262810 |
| chr10 | 43637359 | 43637360 | cg24376955 |
| chr10 | 43881239 | 43881240 | cg12766287 |
| chr10 | 44287023 | 44287024 | cg17952824 |
| chr10 | 44426803 | 44426804 | cg04816803 |
| chr10 | 45395734 | 45395735 | cg09623279 |
| chr10 | 45458834 | 45458835 | cg01484915 |
| chr10 | 45474317 | 45474318 | cg17186163 |
| chr10 | 45474339 | 45474340 | cg08856318 |
| chr10 | 45916904 | 45916905 | cg23084016 |
| chr10 | 46634749 | 46634750 | cg14266837 |
| chr10 | 46996357 | 46996358 | cg05694250 |
| chr10 | 48391548 | 48391549 | cg01042100 |
| chr10 | 49378921 | 49378922 | cg15015416 |
| chr10 | 49459949 | 49459950 | cg23085281 |
| chr10 | 49815243 | 49815244 | cg11095743 |
| chr10 | 50010529 | 50010530 | cg19685285 |
| chr10 | 50329828 | 50329829 | cg24385652 |
| chr10 | 51489980 | 51489981 | cg20129853 |
| chr10 | 51542259 | 51542260 | cg00061792 |
| chr10 | 51575702 | 51575703 | cg06098215 |
| chr10 | 51575763 | 51575764 | cg16814786 |
| chr10 | 52262623 | 52262624 | cg13799263 |

| | | | |
|---|---|---|---|
| chr14 | 75936789 | 75936790 | cg09929567 |
| chr14 | 76450658 | 76450659 | cg00780666 |
| chr14 | 77504156 | 77504157 | cg03017135 |
| chr14 | 77507461 | 77507462 | cg23925650 |
| chr14 | 77588860 | 77588861 | cg05937046 |
| chr14 | 77593161 | 77593162 | cg10361494 |
| chr14 | 77767650 | 77767651 | cg22825202 |
| chr14 | 78079686 | 78079687 | cg24171453 |
| chr14 | 89866110 | 89866111 | cg05347794 |
| chr14 | 89884608 | 89884609 | cg25008263 |
| chr14 | 89976587 | 89976588 | cg24725246 |
| chr14 | 90301560 | 90301561 | cg20582976 |
| chr14 | 90862689 | 90862690 | cg20250935 |
| chr14 | 91579859 | 91579860 | cg06607384 |
| chr14 | 91580129 | 91580130 | cg06907680 |
| chr14 | 91719356 | 91719357 | cg15390894 |
| chr14 | 91755524 | 91755525 | cg05245430 |
| chr14 | 91766170 | 91766171 | cg18821098 |
| chr14 | 91841955 | 91841956 | cg26120073 |
| chr14 | 91842149 | 91842150 | cg10564112 |
| chr14 | 91842949 | 91842950 | cg10511902 |
| chr14 | 91859797 | 91859798 | cg16505891 |
| chr14 | 91880061 | 91880062 | cg23165913 |
| chr14 | 91881497 | 91881498 | cg20303561 |
| chr14 | 92979305 | 92979306 | cg07669002 |
| chr14 | 92981121 | 92981122 | cg03345232 |
| chr14 | 92994936 | 92994937 | cg09317403 |
| chr14 | 93005917 | 93005918 | cg10139508 |
| chr14 | 93151082 | 93151083 | cg06762891 |
| chr14 | 93156816 | 93156817 | cg20072528 |
| chr14 | 93507882 | 93507883 | cg16426321 |
| chr14 | 93556554 | 93556555 | cg10040594 |
| chr14 | 93570854 | 93570855 | cg02039485 |
| chr14 | 93707435 | 93707436 | cg00846498 |
| chr14 | 94759767 | 94759768 | cg19937039 |
| chr14 | 95760031 | 95760032 | cg06753163 |
| chr14 | 95956325 | 95956326 | cg06634140 |
| chr14 | 100631798 | 100631799 | cg16800939 |
| chr14 | 100631897 | 100631898 | cg19741660 |
| chr14 | 102230231 | 102230232 | cg04726250 |
| chr14 | 102274741 | 102274742 | cg05865327 |
| chr14 | 102487020 | 102487021 | cg12127149 |
| chr14 | 102650356 | 102650357 | cg04387969 |
| chr14 | 102653321 | 102653322 | cg01755336 |
| chr14 | 102685619 | 102685620 | cg04313071 |
| chr14 | 103377010 | 103377011 | cg26434328 |
| chr14 | 103992856 | 103992857 | cg01118348 |
| chr14 | 104158878 | 104158879 | cg01024069 |
| chr14 | 104160076 | 104160077 | cg10028434 |
| chr14 | 105113991 | 105113992 | cg27289402 |
| chr14 | 105130703 | 105130704 | cg21562656 |
| chr14 | 105147461 | 105147462 | cg23592421 |
| chr14 | 105147537 | 105147538 | cg26256521 |
| chr14 | 105155145 | 105155146 | cg05873568 |
| chr14 | 105370983 | 105370984 | cg05071383 |
| chr14 | 105812713 | 105812714 | cg21170836 |

| | | | |
|---|---|---|---|
| chr10 | 52271899 | 52271900 | cg00936935 |
| chr10 | 53182118 | 53182119 | cg16401207 |
| chr10 | 53367157 | 53367158 | cg11486694 |
| chr10 | 53852689 | 53852690 | cg11964006 |
| chr10 | 54114224 | 54114225 | cg23931150 |
| chr10 | 54223605 | 54223606 | cg04972770 |
| chr10 | 54532059 | 54532060 | cg18972123 |
| chr10 | 57394513 | 57394514 | cg10271112 |
| chr10 | 60966072 | 60966073 | cg02788195 |
| chr10 | 61147668 | 61147669 | cg00722081 |
| chr10 | 61596106 | 61596107 | cg09082518 |
| chr10 | 62148959 | 62148960 | cg22150335 |
| chr10 | 63683346 | 63683347 | cg24495062 |
| chr10 | 63746909 | 63746910 | cg10502149 |
| chr10 | 63800880 | 63800881 | cg18547838 |
| chr10 | 63809073 | 63809074 | cg14789659 |
| chr10 | 63809098 | 63809099 | cg00928816 |
| chr10 | 63809108 | 63809109 | cg20746552 |
| chr10 | 63809121 | 63809122 | cg16389209 |
| chr10 | 63809149 | 63809150 | cg07520810 |
| chr10 | 63809170 | 63809171 | cg16401465 |
| chr10 | 64572581 | 64572582 | cg00963675 |
| chr10 | 65846243 | 65846244 | cg12044919 |
| chr10 | 69455804 | 69455805 | cg19996026 |
| chr10 | 69611755 | 69611756 | cg04100124 |
| chr10 | 70096482 | 70096483 | cg16896539 |
| chr10 | 70816008 | 70816009 | cg24417845 |
| chr10 | 71077221 | 71077222 | cg09273059 |
| chr10 | 72221844 | 72221845 | cg05240350 |
| chr10 | 72235645 | 72235646 | cg18776616 |
| chr10 | 72344881 | 72344882 | cg14260169 |
| chr10 | 72432573 | 72432574 | cg14612544 |
| chr10 | 72459629 | 72459630 | cg00618155 |
| chr10 | 73408185 | 73408186 | cg04567334 |
| chr10 | 73497609 | 73497610 | cg19429405 |
| chr10 | 73497629 | 73497630 | cg23056923 |
| chr10 | 73516760 | 73516761 | cg04179740 |
| chr10 | 73534338 | 73534339 | cg11633461 |
| chr10 | 73849106 | 73849107 | cg12752647 |
| chr10 | 73849124 | 73849125 | cg02909446 |
| chr10 | 73849167 | 73849168 | cg13665981 |
| chr10 | 73976790 | 73976791 | cg15013801 |
| chr10 | 73976798 | 73976799 | cg17372758 |
| chr10 | 73976821 | 73976822 | cg23685969 |
| chr10 | 74544059 | 74544060 | cg12580580 |
| chr10 | 75336622 | 75336623 | cg15402162 |
| chr10 | 75351076 | 75351077 | cg05970116 |
| chr10 | 75416760 | 75416761 | cg05697785 |
| chr10 | 75677011 | 75677012 | cg04084348 |
| chr10 | 76179907 | 76179908 | cg23198334 |
| chr10 | 76349278 | 76349279 | cg26757820 |
| chr10 | 76742856 | 76742857 | cg02976898 |
| chr10 | 76806873 | 76806874 | cg01354426 |
| chr10 | 77021847 | 77021848 | cg17814175 |
| chr10 | 77878257 | 77878258 | cg22990178 |
| chr10 | 78635553 | 78635554 | cg23533270 |
| chr14 | 106028568 | 106028569 | cg19700328 |
| chr14 | 106030981 | 106030982 | cg16296017 |
| chr14 | 106033512 | 106033513 | cg19952651 |
| chr14 | 106133799 | 106133800 | cg26308506 |
| chr14 | 106153356 | 106153357 | cg22070177 |
| chr14 | 106165316 | 106165317 | cg15989433 |
| chr14 | 106235984 | 106235985 | cg18674446 |
| chr14 | 106239286 | 106239287 | cg22865700 |
| chr14 | 106319730 | 106319731 | cg10256210 |
| chr15 | 22922392 | 22922393 | cg24851866 |
| chr15 | 23062853 | 23062854 | cg09873524 |
| chr15 | 23082284 | 23082285 | cg20587188 |
| chr15 | 27189789 | 27189790 | cg27310090 |
| chr15 | 28448838 | 28448839 | cg07726456 |
| chr15 | 31393845 | 31393846 | cg14323928 |
| chr15 | 31624584 | 31624585 | cg07814318 |
| chr15 | 40353216 | 40353217 | cg05951095 |
| chr15 | 40389891 | 40389892 | cg07712904 |
| chr15 | 40674438 | 40674439 | cg13984173 |
| chr15 | 40697419 | 40697420 | cg26886850 |
| chr15 | 40745035 | 40745036 | cg02726739 |
| chr15 | 40762864 | 40762865 | cg26767214 |
| chr15 | 41231049 | 41231050 | cg14157570 |
| chr15 | 41806234 | 41806235 | cg03321553 |
| chr15 | 41983773 | 41983774 | cg22894805 |
| chr15 | 42258950 | 42258951 | cg17244218 |
| chr15 | 42261154 | 42261155 | cg12655112 |
| chr15 | 42749748 | 42749749 | cg06794543 |
| chr15 | 44096850 | 44096851 | cg22038409 |
| chr15 | 44969129 | 44969130 | cg18059012 |
| chr15 | 44969244 | 44969245 | cg01647795 |
| chr15 | 45669010 | 45669011 | cg10760299 |
| chr15 | 46589259 | 46589260 | cg27090409 |
| chr15 | 49675495 | 49675496 | cg24789138 |
| chr15 | 50150583 | 50150584 | cg20023184 |
| chr15 | 50642820 | 50642821 | cg14821257 |
| chr15 | 51694383 | 51694384 | cg17811589 |
| chr15 | 51762495 | 51762496 | cg17080902 |
| chr15 | 52130305 | 52130306 | cg21078654 |
| chr15 | 52408529 | 52408530 | cg19382136 |
| chr15 | 52584919 | 52584920 | cg15605746 |
| chr15 | 55788778 | 55788779 | cg12417176 |
| chr15 | 57510460 | 57510461 | cg25688583 |
| chr15 | 57510575 | 57510576 | cg00027650 |
| chr15 | 57590030 | 57590031 | cg02402565 |
| chr15 | 59024033 | 59024034 | cg02816525 |
| chr15 | 59154450 | 59154451 | cg00126173 |
| chr15 | 59700360 | 59700361 | cg05061769 |
| chr15 | 59701812 | 59701813 | cg02454802 |
| chr15 | 59733552 | 59733553 | cg12556586 |
| chr15 | 61114065 | 61114066 | cg09984602 |
| chr15 | 63845726 | 63845727 | cg11426726 |
| chr15 | 63889093 | 63889094 | cg01767116 |
| chr15 | 64197566 | 64197567 | cg21838952 |
| chr15 | 64442561 | 64442562 | cg14004161 |
| chr15 | 64442578 | 64442579 | cg07196571 |

| | | | |
|---|---|---|---|
| chr10 | 78946146 | 78946147 | cg19273378 |
| chr10 | 78954388 | 78954389 | cg26091990 |
| chr10 | 78956948 | 78956949 | cg04912778 |
| chr10 | 79079736 | 79079737 | cg19730814 |
| chr10 | 79238225 | 79238226 | cg06828335 |
| chr10 | 79285311 | 79285312 | cg15200789 |
| chr10 | 79289545 | 79289546 | cg26442890 |
| chr10 | 79393414 | 79393415 | cg26611809 |
| chr10 | 79446426 | 79446427 | cg26465532 |
| chr10 | 79605162 | 79605163 | cg11778714 |
| chr10 | 79977519 | 79977520 | cg25817351 |
| chr10 | 80052820 | 80052821 | cg03455990 |
| chr10 | 80063568 | 80063569 | cg08185255 |
| chr10 | 80063791 | 80063792 | cg24711336 |
| chr10 | 80270722 | 80270723 | cg15619377 |
| chr10 | 80959688 | 80959689 | cg09936762 |
| chr10 | 80963128 | 80963129 | cg13843840 |
| chr10 | 81955971 | 81955972 | cg04329351 |
| chr10 | 81961468 | 81961469 | cg23258615 |
| chr10 | 81963408 | 81963409 | cg02291020 |
| chr10 | 82263737 | 82263738 | cg23463186 |
| chr10 | 82687542 | 82687543 | cg21367907 |
| chr10 | 88658528 | 88658529 | cg19166799 |
| chr10 | 88724584 | 88724585 | cg09218924 |
| chr10 | 89418206 | 89418207 | cg24243914 |
| chr10 | 90147030 | 90147031 | cg17766305 |
| chr10 | 90428384 | 90428385 | cg18148156 |
| chr10 | 90749920 | 90749921 | cg26478401 |
| chr10 | 90847168 | 90847169 | cg23195687 |
| chr10 | 91002590 | 91002591 | cg12555086 |
| chr10 | 91111929 | 91111930 | cg12083893 |
| chr10 | 91133941 | 91133942 | cg00460983 |
| chr10 | 91408027 | 91408028 | cg23204113 |
| chr10 | 91717815 | 91717816 | cg25278353 |
| chr10 | 92987457 | 92987458 | cg16578636 |
| chr10 | 93100919 | 93100920 | cg25065131 |
| chr10 | 93171002 | 93171003 | cg06520331 |
| chr10 | 93174498 | 93174499 | cg02075791 |
| chr10 | 93916288 | 93916289 | cg10754794 |
| chr10 | 95536738 | 95536739 | cg02300379 |
| chr10 | 96997377 | 96997378 | cg20714347 |
| chr10 | 97457068 | 97457069 | cg04276302 |
| chr10 | 97515372 | 97515373 | cg04451770 |
| chr10 | 97515398 | 97515399 | cg10381771 |
| chr10 | 97624891 | 97624892 | cg09217157 |
| chr10 | 97785444 | 97785445 | cg19715167 |
| chr10 | 98745012 | 98745013 | cg13383814 |
| chr10 | 99185264 | 99185265 | cg25603927 |
| chr10 | 99443455 | 99443456 | cg08181251 |
| chr10 | 99648780 | 99648781 | cg04888360 |
| chr10 | 99894087 | 99894088 | cg03970483 |
| chr10 | 100638900 | 100638901 | cg25970726 |
| chr10 | 102100212 | 102100213 | cg00653847 |
| chr10 | 102107666 | 102107667 | cg06400428 |
| chr10 | 102494320 | 102494321 | cg10022914 |
| chr10 | 102593679 | 102593680 | cg02308150 |

| | | | |
|---|---|---|---|
| chr15 | 64649128 | 64649129 | cg05933189 |
| chr15 | 64649139 | 64649140 | cg03840511 |
| chr15 | 65115218 | 65115219 | cg05869754 |
| chr15 | 66062176 | 66062177 | cg27331863 |
| chr15 | 66116727 | 66116728 | cg27665985 |
| chr15 | 66678782 | 66678783 | cg14155397 |
| chr15 | 66709285 | 66709286 | cg21070009 |
| chr15 | 67411858 | 67411859 | cg27130493 |
| chr15 | 67538446 | 67538447 | cg22940988 |
| chr15 | 68133972 | 68133973 | cg23793686 |
| chr15 | 69107085 | 69107086 | cg11797485 |
| chr15 | 69613382 | 69613383 | cg08437576 |
| chr15 | 69843549 | 69843550 | cg00758816 |
| chr15 | 69867297 | 69867298 | cg23529249 |
| chr15 | 69883460 | 69883461 | cg15554863 |
| chr15 | 70541438 | 70541439 | cg20648004 |
| chr15 | 73044064 | 73044065 | cg02259723 |
| chr15 | 73921385 | 73921386 | cg02329430 |
| chr15 | 73992274 | 73992275 | cg06478102 |
| chr15 | 74679906 | 74679907 | cg08051804 |
| chr15 | 74729482 | 74729483 | cg19251514 |
| chr15 | 75071042 | 75071043 | cg20743868 |
| chr15 | 75095006 | 75095007 | cg10601181 |
| chr15 | 75336436 | 75336437 | cg18176723 |
| chr15 | 75490372 | 75490373 | cg14832223 |
| chr15 | 75915192 | 75915193 | cg19013753 |
| chr15 | 77287243 | 77287244 | cg26227523 |
| chr15 | 77289047 | 77289048 | cg21322248 |
| chr15 | 77519170 | 77519171 | cg08130265 |
| chr15 | 77734286 | 77734287 | cg21298249 |
| chr15 | 77800637 | 77800638 | cg05908442 |
| chr15 | 79298209 | 79298210 | cg24231513 |
| chr15 | 80430079 | 80430080 | cg24312321 |
| chr15 | 83515335 | 83515336 | cg01959287 |
| chr15 | 83736925 | 83736926 | cg16239296 |
| chr15 | 85215021 | 85215022 | cg19606131 |
| chr15 | 85945991 | 85945992 | cg25950769 |
| chr15 | 85949116 | 85949117 | cg17178777 |
| chr15 | 86042851 | 86042852 | cg17133734 |
| chr15 | 89764836 | 89764837 | cg10173075 |
| chr15 | 89764895 | 89764896 | cg15808008 |
| chr15 | 90068229 | 90068230 | cg00502017 |
| chr15 | 90547692 | 90547693 | cg18443571 |
| chr15 | 90548037 | 90548038 | cg16558770 |
| chr15 | 90606494 | 90606495 | cg18568872 |
| chr15 | 90611340 | 90611341 | cg08001256 |
| chr15 | 90611414 | 90611415 | cg07315755 |
| chr15 | 90717994 | 90717995 | cg21152753 |
| chr15 | 91476398 | 91476399 | cg05272099 |
| chr15 | 91497412 | 91497413 | cg01641174 |
| chr15 | 91503707 | 91503708 | cg13170728 |
| chr15 | 91531151 | 91531152 | cg04775232 |
| chr15 | 93361889 | 93361890 | cg14588003 |
| chr15 | 93362247 | 93362248 | cg00103299 |
| chr15 | 94841334 | 94841335 | cg21467108 |
| chr15 | 97322504 | 97322505 | cg00114784 |

| | | | |
|---|---|---|---|
| chr10 | 102732224 | 102732225 | cg26991946 |
| chr10 | 102760970 | 102760971 | cg03271893 |
| chr10 | 102792835 | 102792836 | cg15428620 |
| chr10 | 102821565 | 102821566 | cg26605164 |
| chr10 | 102823812 | 102823813 | cg02891579 |
| chr10 | 103594334 | 103594335 | cg11685391 |
| chr10 | 103601188 | 103601189 | cg17962384 |
| chr10 | 103889899 | 103889900 | cg16167639 |
| chr10 | 103910870 | 103910871 | cg13388830 |
| chr10 | 104194843 | 104194844 | cg09701700 |
| chr10 | 104236741 | 104236742 | cg00122347 |
| chr10 | 104406990 | 104406991 | cg17782974 |
| chr10 | 104434010 | 104434011 | cg06361405 |
| chr10 | 104535792 | 104535793 | cg03161190 |
| chr10 | 104535794 | 104535795 | cg17894755 |
| chr10 | 104535854 | 104535855 | cg15227982 |
| chr10 | 104535920 | 104535921 | cg25104397 |
| chr10 | 104535990 | 104535991 | cg14939082 |
| chr10 | 104597435 | 104597436 | cg04471890 |
| chr10 | 104611004 | 104611005 | cg02418072 |
| chr10 | 104674367 | 104674368 | cg09279458 |
| chr10 | 104953608 | 104953609 | cg00035347 |
| chr10 | 105205887 | 105205888 | cg07240689 |
| chr10 | 105209452 | 105209453 | cg02167201 |
| chr10 | 105219068 | 105219069 | cg10081681 |
| chr10 | 106082537 | 106082538 | cg16301004 |
| chr10 | 106088711 | 106088712 | cg19640166 |
| chr10 | 106089203 | 106089204 | cg24199006 |
| chr10 | 106094976 | 106094977 | cg18149009 |
| chr10 | 107301935 | 107301936 | cg02696798 |
| chr10 | 108690391 | 108690392 | cg19040385 |
| chr10 | 109641026 | 109641027 | cg08327428 |
| chr10 | 110084660 | 110084661 | cg05673079 |
| chr10 | 111659903 | 111659904 | cg18780288 |
| chr10 | 111766049 | 111766050 | cg25341032 |
| chr10 | 112113375 | 112113376 | cg06601203 |
| chr10 | 112118688 | 112118689 | cg16947025 |
| chr10 | 112258641 | 112258642 | cg03652339 |
| chr10 | 112259305 | 112259306 | cg09357462 |
| chr10 | 112375475 | 112375476 | cg26878209 |
| chr10 | 112427911 | 112427912 | cg22623319 |
| chr10 | 112546110 | 112546111 | cg15880704 |
| chr10 | 112553035 | 112553036 | cg10955669 |
| chr10 | 112722440 | 112722441 | cg21022395 |
| chr10 | 112834861 | 112834862 | cg12121535 |
| chr10 | 115442231 | 115442232 | cg25763292 |
| chr10 | 116109825 | 116109826 | cg05120428 |
| chr10 | 116394557 | 116394558 | cg12403162 |
| chr10 | 116585424 | 116585425 | cg18325866 |
| chr10 | 117754806 | 117754807 | cg07760622 |
| chr10 | 118561572 | 118561573 | cg07109438 |
| chr10 | 120001266 | 120001267 | cg22682200 |
| chr10 | 120971516 | 120971517 | cg09337049 |
| chr10 | 121075316 | 121075317 | cg07551060 |
| chr10 | 121298236 | 121298237 | cg05241536 |
| chr10 | 121300857 | 121300858 | cg15611912 |

| | | | |
|---|---|---|---|
| chr15 | 99143542 | 99143543 | cg20891771 |
| chr15 | 99978986 | 99978987 | cg22700246 |
| chr15 | 99982524 | 99982525 | cg02378208 |
| chr15 | 100146819 | 100146820 | cg09494293 |
| chr15 | 100154002 | 100154003 | cg20240243 |
| chr15 | 100270048 | 100270049 | cg18742785 |
| chr15 | 100350984 | 100350985 | cg16484450 |
| chr15 | 102282101 | 102282102 | cg06486467 |
| chr16 | 279746 | 279747 | cg27362010 |
| chr16 | 357249 | 357250 | cg10354730 |
| chr16 | 358248 | 358249 | cg16412914 |
| chr16 | 358490 | 358491 | cg01517185 |
| chr16 | 434356 | 434357 | cg00354542 |
| chr16 | 617420 | 617421 | cg03815552 |
| chr16 | 690468 | 690469 | cg07284546 |
| chr16 | 1361806 | 1361807 | cg06161540 |
| chr16 | 1391501 | 1391502 | cg04663017 |
| chr16 | 1413159 | 1413160 | cg00122851 |
| chr16 | 1414422 | 1414423 | cg02791338 |
| chr16 | 1414480 | 1414481 | cg27297104 |
| chr16 | 1673702 | 1673703 | cg27624753 |
| chr16 | 1782905 | 1782906 | cg02994291 |
| chr16 | 2084192 | 2084193 | cg04751761 |
| chr16 | 2094602 | 2094603 | cg27370991 |
| chr16 | 2388350 | 2388351 | cg08900781 |
| chr16 | 2549625 | 2549626 | cg04659439 |
| chr16 | 2567316 | 2567317 | cg27576259 |
| chr16 | 2652696 | 2652697 | cg01244128 |
| chr16 | 2760410 | 2760411 | cg09372449 |
| chr16 | 3021802 | 3021803 | cg05737682 |
| chr16 | 3629620 | 3629621 | cg02127172 |
| chr16 | 3781749 | 3781750 | cg27318635 |
| chr16 | 3781770 | 3781771 | cg04432137 |
| chr16 | 3784851 | 3784852 | cg04979551 |
| chr16 | 3847222 | 3847223 | cg09519954 |
| chr16 | 3906833 | 3906834 | cg08364730 |
| chr16 | 4131584 | 4131585 | cg07906046 |
| chr16 | 4397291 | 4397292 | cg01406317 |
| chr16 | 4579402 | 4579403 | cg02642565 |
| chr16 | 8815355 | 8815356 | cg09410453 |
| chr16 | 9030093 | 9030094 | cg00607630 |
| chr16 | 9058365 | 9058366 | cg12914014 |
| chr16 | 10843654 | 10843655 | cg03287527 |
| chr16 | 11187192 | 11187193 | cg26765295 |
| chr16 | 11220376 | 11220377 | cg10364862 |
| chr16 | 11406995 | 11406996 | cg07312240 |
| chr16 | 11715746 | 11715747 | cg09417209 |
| chr16 | 11734383 | 11734384 | cg09481483 |
| chr16 | 11777108 | 11777109 | cg05036502 |
| chr16 | 11794641 | 11794642 | cg03382501 |
| chr16 | 11852095 | 11852096 | cg09095525 |
| chr16 | 11886668 | 11886669 | cg08422172 |
| chr16 | 12088037 | 12088038 | cg02498072 |
| chr16 | 12156525 | 12156526 | cg01858712 |
| chr16 | 12359297 | 12359298 | cg16712866 |
| chr16 | 12411139 | 12411140 | cg05849013 |

| | | | | | | | |
|---|---|---|---|---|---|---|---|
| chr10 | 121301041 | 121301042 | cg00291478 | chr16 | 12554036 | 12554037 | cg04724177 |
| chr10 | 121447367 | 121447368 | cg20341089 | chr16 | 14219324 | 14219325 | cg00808545 |
| chr10 | 121620547 | 121620548 | cg06601266 | chr16 | 14552306 | 14552307 | cg07181528 |
| chr10 | 122614164 | 122614165 | cg13789938 | chr16 | 16082095 | 16082096 | cg10111352 |
| chr10 | 123408913 | 123408914 | cg05162358 | chr16 | 16190393 | 16190394 | cg01332683 |
| chr10 | 124138853 | 124138854 | cg19863426 | chr16 | 17398605 | 17398606 | cg09548718 |
| chr10 | 124380473 | 124380474 | cg27230705 | chr16 | 17472479 | 17472480 | cg07248440 |
| chr10 | 125805783 | 125805784 | cg26306435 | chr16 | 17477485 | 17477486 | cg05634915 |
| chr10 | 126301842 | 126301843 | cg21390852 | chr16 | 17562960 | 17562961 | cg06321596 |
| chr10 | 126302546 | 126302547 | cg18593374 | chr16 | 17563300 | 17563301 | cg04359840 |
| chr10 | 126303795 | 126303796 | cg24428218 | chr16 | 17565129 | 17565130 | cg06129210 |
| chr10 | 126329354 | 126329355 | cg17879101 | chr16 | 19180934 | 19180935 | cg02157872 |
| chr10 | 126330905 | 126330906 | cg07081759 | chr16 | 19185373 | 19185374 | cg08781257 |
| chr10 | 126782397 | 126782398 | cg06332304 | chr16 | 19185556 | 19185557 | cg05712435 |
| chr10 | 128190000 | 128190001 | cg20544216 | chr16 | 20420222 | 20420223 | cg06686156 |
| chr10 | 129239566 | 129239567 | cg25247976 | chr16 | 20420716 | 20420717 | cg10938486 |
| chr10 | 129844711 | 129844712 | cg06585141 | chr16 | 20686073 | 20686074 | cg02675179 |
| chr10 | 129845375 | 129845376 | cg18166144 | chr16 | 20702872 | 20702873 | cg09620718 |
| chr10 | 131819853 | 131819854 | cg12736773 | chr16 | 20910405 | 20910406 | cg06126593 |
| chr10 | 131905654 | 131905655 | cg00057517 | chr16 | 21390421 | 21390422 | cg26909864 |
| chr10 | 134211874 | 134211875 | cg25303150 | chr16 | 21514953 | 21514954 | cg07813747 |
| chr10 | 134221503 | 134221504 | cg00818106 | chr16 | 21623651 | 21623652 | cg01574390 |
| chr10 | 134300809 | 134300810 | cg24495044 | chr16 | 22020630 | 22020631 | cg07571928 |
| chr10 | 134300922 | 134300923 | cg05170353 | chr16 | 22023163 | 22023164 | cg08262496 |
| chr10 | 134360977 | 134360978 | cg07402396 | chr16 | 22217246 | 22217247 | cg11059800 |
| chr10 | 134364082 | 134364083 | cg17352462 | chr16 | 23522771 | 23522772 | cg01522095 |
| chr10 | 135205162 | 135205163 | cg07562039 | chr16 | 23957272 | 23957273 | cg26685404 |
| chr11 | 194498 | 194499 | cg18276135 | chr16 | 24663848 | 24663849 | cg03012370 |
| chr11 | 315118 | 315119 | cg21686213 | chr16 | 24748339 | 24748340 | cg09018299 |
| chr11 | 315262 | 315263 | cg03038262 | chr16 | 24997394 | 24997395 | cg06999243 |
| chr11 | 492997 | 492998 | cg11503833 | chr16 | 25011481 | 25011482 | cg08067346 |
| chr11 | 503807 | 503808 | cg19223782 | chr16 | 25023604 | 25023605 | cg00396427 |
| chr11 | 628677 | 628678 | cg04150762 | chr16 | 25027571 | 25027572 | cg14045374 |
| chr11 | 646580 | 646581 | cg21531416 | chr16 | 25029018 | 25029019 | cg27313271 |
| chr11 | 698201 | 698202 | cg07161179 | chr16 | 25043992 | 25043993 | cg02829279 |
| chr11 | 698760 | 698761 | cg21191255 | chr16 | 25100355 | 25100356 | cg01658444 |
| chr11 | 724525 | 724526 | cg18770763 | chr16 | 27237122 | 27237123 | cg04094640 |
| chr11 | 811791 | 811792 | cg20550458 | chr16 | 27281055 | 27281056 | cg03368690 |
| chr11 | 844686 | 844687 | cg07047068 | chr16 | 27367172 | 27367173 | cg01165142 |
| chr11 | 970389 | 970390 | cg12893697 | chr16 | 27378255 | 27378256 | cg05013912 |
| chr11 | 993372 | 993373 | cg02128483 | chr16 | 28915234 | 28915235 | cg27473355 |
| chr11 | 995425 | 995426 | cg14804384 | chr16 | 28915665 | 28915666 | cg27347003 |
| chr11 | 1406719 | 1406720 | cg22366799 | chr16 | 28939378 | 28939379 | cg02007534 |
| chr11 | 1506936 | 1506937 | cg13534999 | chr16 | 28995882 | 28995883 | cg08331345 |
| chr11 | 1507119 | 1507120 | cg15470102 | chr16 | 29296693 | 29296694 | cg01719179 |
| chr11 | 1507263 | 1507264 | cg11611244 | chr16 | 29296704 | 29296705 | cg23515125 |
| chr11 | 1507321 | 1507322 | cg03565674 | chr16 | 29296797 | 29296798 | cg02033694 |
| chr11 | 1606664 | 1606665 | cg25145687 | chr16 | 29800596 | 29800597 | cg02876297 |
| chr11 | 1674770 | 1674771 | cg14909325 | chr16 | 29837267 | 29837268 | cg09309024 |
| chr11 | 1862020 | 1862021 | cg13133304 | chr16 | 30106682 | 30106683 | cg26709300 |
| chr11 | 1862187 | 1862188 | cg06753398 | chr16 | 30131641 | 30131642 | cg00192773 |
| chr11 | 1862477 | 1862478 | cg05258834 | chr16 | 30563514 | 30563515 | cg05194689 |
| chr11 | 1945564 | 1945565 | cg02556649 | chr16 | 31188401 | 31188402 | cg09425279 |
| chr11 | 1945607 | 1945608 | cg02821464 | chr16 | 31213045 | 31213046 | cg27615603 |
| chr11 | 1945648 | 1945649 | cg11654118 | chr16 | 31214880 | 31214881 | cg07461837 |
| chr11 | 2222076 | 2222077 | cg14271862 | chr16 | 31884740 | 31884741 | cg03405560 |

| | | | |
|---|---|---|---|
| chr11 | 2239813 | 2239814 | cg15194251 |
| chr11 | 2323801 | 2323802 | cg03983837 |
| chr11 | 2433468 | 2433469 | cg02743674 |
| chr11 | 2482284 | 2482285 | cg17291157 |
| chr11 | 2482303 | 2482304 | cg07210454 |
| chr11 | 2703128 | 2703129 | cg20406758 |
| chr11 | 2722539 | 2722540 | cg03654058 |
| chr11 | 2800047 | 2800048 | cg05438727 |
| chr11 | 3858493 | 3858494 | cg24497361 |
| chr11 | 3901552 | 3901553 | cg07109745 |
| chr11 | 4118978 | 4118979 | cg10113107 |
| chr11 | 4686966 | 4686967 | cg08426891 |
| chr11 | 4716867 | 4716868 | cg20201128 |
| chr11 | 4726105 | 4726106 | cg20743708 |
| chr11 | 4843021 | 4843022 | cg18352935 |
| chr11 | 4927915 | 4927916 | cg05046377 |
| chr11 | 5270556 | 5270557 | cg18768582 |
| chr11 | 5645703 | 5645704 | cg03306641 |
| chr11 | 6805810 | 6805811 | cg02634507 |
| chr11 | 7092719 | 7092720 | cg07103673 |
| chr11 | 7597814 | 7597815 | cg04363470 |
| chr11 | 7694524 | 7694525 | cg00859655 |
| chr11 | 8228147 | 8228148 | cg16311302 |
| chr11 | 8816428 | 8816429 | cg01902758 |
| chr11 | 9333358 | 9333359 | cg08757628 |
| chr11 | 9587743 | 9587744 | cg25225070 |
| chr11 | 9769192 | 9769193 | cg08010619 |
| chr11 | 9860910 | 9860911 | cg20072162 |
| chr11 | 10332052 | 10332053 | cg14359798 |
| chr11 | 10339086 | 10339087 | cg13054358 |
| chr11 | 10590282 | 10590283 | cg03908676 |
| chr11 | 11258914 | 11258915 | cg22176913 |
| chr11 | 11371451 | 11371452 | cg20438286 |
| chr11 | 11590474 | 11590475 | cg14158968 |
| chr11 | 11778177 | 11778178 | cg15032941 |
| chr11 | 11866554 | 11866555 | cg20781421 |
| chr11 | 11887055 | 11887056 | cg20728419 |
| chr11 | 12562492 | 12562493 | cg27185501 |
| chr11 | 13124294 | 13124295 | cg25068253 |
| chr11 | 13388349 | 13388350 | cg01055121 |
| chr11 | 14335070 | 14335071 | cg17509893 |
| chr11 | 14340592 | 14340593 | cg20007622 |
| chr11 | 14376795 | 14376796 | cg08923349 |
| chr11 | 15136505 | 15136506 | cg12170314 |
| chr11 | 15217886 | 15217887 | cg13208333 |
| chr11 | 15229502 | 15229503 | cg19115017 |
| chr11 | 15796715 | 15796716 | cg12804755 |
| chr11 | 15960510 | 15960511 | cg06359632 |
| chr11 | 16424673 | 16424674 | cg17284609 |
| chr11 | 16763554 | 16763555 | cg06617449 |
| chr11 | 17944560 | 17944561 | cg25608555 |
| chr11 | 18128177 | 18128178 | cg00345366 |
| chr11 | 18287231 | 18287232 | cg06522679 |
| chr11 | 18334871 | 18334872 | cg26089997 |
| chr11 | 19362896 | 19362897 | cg09032137 |
| chr11 | 19388224 | 19388225 | cg01388979 |

| | | | |
|---|---|---|---|
| chr16 | 46919021 | 46919022 | cg00622166 |
| chr16 | 46919075 | 46919076 | cg00270789 |
| chr16 | 46919112 | 46919113 | cg03533472 |
| chr16 | 46919194 | 46919195 | cg23684449 |
| chr16 | 46920836 | 46920837 | cg07710971 |
| chr16 | 46921663 | 46921664 | cg05380921 |
| chr16 | 48398409 | 48398410 | cg03524700 |
| chr16 | 48402112 | 48402113 | cg00041311 |
| chr16 | 48406857 | 48406858 | cg06183001 |
| chr16 | 50244074 | 50244075 | cg01719268 |
| chr16 | 50730385 | 50730386 | cg16771652 |
| chr16 | 50730737 | 50730738 | cg08554257 |
| chr16 | 53114571 | 53114572 | cg04770195 |
| chr16 | 56229180 | 56229181 | cg04507426 |
| chr16 | 56963627 | 56963628 | cg16343503 |
| chr16 | 56979519 | 56979520 | cg04170948 |
| chr16 | 57099057 | 57099058 | cg00589550 |
| chr16 | 57294356 | 57294357 | cg04495354 |
| chr16 | 57406955 | 57406956 | cg15195412 |
| chr16 | 62988282 | 62988283 | cg07700524 |
| chr16 | 66442374 | 66442375 | cg27326306 |
| chr16 | 66582592 | 66582593 | cg07512993 |
| chr16 | 66583078 | 66583079 | cg03448915 |
| chr16 | 66584358 | 66584359 | cg09238666 |
| chr16 | 66585210 | 66585211 | cg02295216 |
| chr16 | 66596317 | 66596318 | cg04701716 |
| chr16 | 66651242 | 66651243 | cg07589273 |
| chr16 | 66756614 | 66756615 | cg04860008 |
| chr16 | 67139348 | 67139349 | cg06886153 |
| chr16 | 67212681 | 67212682 | cg04550079 |
| chr16 | 67550511 | 67550512 | cg03737424 |
| chr16 | 67973788 | 67973789 | cg04582263 |
| chr16 | 68003649 | 68003650 | cg08571821 |
| chr16 | 68117460 | 68117461 | cg08281590 |
| chr16 | 68262681 | 68262682 | cg01380533 |
| chr16 | 68265587 | 68265588 | cg00806259 |
| chr16 | 68268252 | 68268253 | cg05510337 |
| chr16 | 68291720 | 68291721 | cg00346883 |
| chr16 | 68881254 | 68881255 | cg00818680 |
| chr16 | 69098945 | 69098946 | cg06285909 |
| chr16 | 69555896 | 69555897 | cg00449930 |
| chr16 | 70441937 | 70441938 | cg05246530 |
| chr16 | 70453185 | 70453186 | cg09426937 |
| chr16 | 70469236 | 70469237 | cg08641155 |
| chr16 | 70471055 | 70471056 | cg01389386 |
| chr16 | 70473841 | 70473842 | cg05261349 |
| chr16 | 70771142 | 70771143 | cg08329113 |
| chr16 | 71657362 | 71657363 | cg08769634 |
| chr16 | 71763859 | 71763860 | cg05493952 |
| chr16 | 73080909 | 73080910 | cg10483525 |
| chr16 | 75604552 | 75604553 | cg00294149 |
| chr16 | 76521159 | 76521160 | cg05630943 |
| chr16 | 81066575 | 81066576 | cg26789859 |
| chr16 | 81347551 | 81347552 | cg04543865 |
| chr16 | 81602667 | 81602668 | cg02575319 |
| chr16 | 81898406 | 81898407 | cg00773142 |

| | | | |
|---|---|---|---|
| chr11 | 19411473 | 19411474 | cg04231888 |
| chr11 | 19744280 | 19744281 | cg18009574 |
| chr11 | 19779691 | 19779692 | cg19252584 |
| chr11 | 19870761 | 19870762 | cg11129341 |
| chr11 | 20499293 | 20499294 | cg03635073 |
| chr11 | 20987359 | 20987360 | cg27258696 |
| chr11 | 23811425 | 23811426 | cg24599329 |
| chr11 | 23980861 | 23980862 | cg04538990 |
| chr11 | 23994937 | 23994938 | cg17104596 |
| chr11 | 24015048 | 24015049 | cg01823718 |
| chr11 | 24532473 | 24532474 | cg05788927 |
| chr11 | 24849257 | 24849258 | cg08227141 |
| chr11 | 26523296 | 26523297 | cg05146852 |
| chr11 | 27105814 | 27105815 | cg24585203 |
| chr11 | 27504612 | 27504613 | cg15933729 |
| chr11 | 30444984 | 30444985 | cg05798059 |
| chr11 | 30771240 | 30771241 | cg07082447 |
| chr11 | 30929224 | 30929225 | cg21758962 |
| chr11 | 33758505 | 33758506 | cg16568681 |
| chr11 | 33758529 | 33758530 | cg02224369 |
| chr11 | 33759043 | 33759044 | cg03292992 |
| chr11 | 33913538 | 33913539 | cg20147326 |
| chr11 | 33913586 | 33913587 | cg12004276 |
| chr11 | 34244798 | 34244799 | cg22219450 |
| chr11 | 34350440 | 34350441 | cg08663298 |
| chr11 | 34535357 | 34535358 | cg12728032 |
| chr11 | 34641925 | 34641926 | cg05503887 |
| chr11 | 34642885 | 34642886 | cg18414381 |
| chr11 | 35138084 | 35138085 | cg26025505 |
| chr11 | 35195513 | 35195514 | cg25096745 |
| chr11 | 35201133 | 35201134 | cg14593191 |
| chr11 | 39689583 | 39689584 | cg00544674 |
| chr11 | 40310738 | 40310739 | cg26553513 |
| chr11 | 44597119 | 44597120 | cg05175803 |
| chr11 | 44638520 | 44638521 | cg21957813 |
| chr11 | 44786104 | 44786105 | cg03236627 |
| chr11 | 44855796 | 44855797 | cg18395231 |
| chr11 | 44881716 | 44881717 | cg01571916 |
| chr11 | 44970771 | 44970772 | cg01859616 |
| chr11 | 44994687 | 44994688 | cg02828306 |
| chr11 | 45109122 | 45109123 | cg02674693 |
| chr11 | 45115390 | 45115391 | cg26821540 |
| chr11 | 45115819 | 45115820 | cg20227165 |
| chr11 | 45231545 | 45231546 | cg22731637 |
| chr11 | 45725227 | 45725228 | cg10519389 |
| chr11 | 45924191 | 45924192 | cg07566286 |
| chr11 | 46261352 | 46261353 | cg14487944 |
| chr11 | 46383924 | 46383925 | cg02211741 |
| chr11 | 46940613 | 46940614 | cg15161667 |
| chr11 | 47175143 | 47175144 | cg07438103 |
| chr11 | 47239549 | 47239550 | cg15500466 |
| chr11 | 47430812 | 47430813 | cg19668951 |
| chr11 | 47510670 | 47510671 | cg15755476 |
| chr11 | 47517381 | 47517382 | cg03566787 |
| chr11 | 47529924 | 47529925 | cg11987802 |
| chr11 | 47608487 | 47608488 | cg13999086 |
| chr16 | 84154549 | 84154550 | cg06929843 |
| chr16 | 84578173 | 84578174 | cg04102161 |
| chr16 | 84676746 | 84676747 | cg05128520 |
| chr16 | 84687549 | 84687550 | cg03054598 |
| chr16 | 85160569 | 85160570 | cg02380595 |
| chr16 | 85600639 | 85600640 | cg16544989 |
| chr16 | 85780385 | 85780386 | cg04497015 |
| chr16 | 85783575 | 85783576 | cg08774871 |
| chr16 | 85860163 | 85860164 | cg07272042 |
| chr16 | 85931840 | 85931841 | cg06020352 |
| chr16 | 85966197 | 85966198 | cg07508446 |
| chr16 | 85966425 | 85966426 | cg06951664 |
| chr16 | 85966435 | 85966436 | cg04211501 |
| chr16 | 85984070 | 85984071 | cg27632179 |
| chr16 | 85990019 | 85990020 | cg00206468 |
| chr16 | 86018811 | 86018812 | cg26947831 |
| chr16 | 87415168 | 87415169 | cg04255044 |
| chr16 | 87751605 | 87751606 | cg05613083 |
| chr16 | 87866696 | 87866697 | cg07558761 |
| chr16 | 87979202 | 87979203 | cg03240511 |
| chr16 | 87988631 | 87988632 | cg02331262 |
| chr16 | 88038949 | 88038950 | cg02145701 |
| chr16 | 88039257 | 88039258 | cg01946510 |
| chr16 | 88100413 | 88100414 | cg07930510 |
| chr16 | 88292973 | 88292974 | cg07099640 |
| chr16 | 88513513 | 88513514 | cg02197669 |
| chr16 | 88525221 | 88525222 | cg01278564 |
| chr16 | 88528734 | 88528735 | cg04940214 |
| chr16 | 88532245 | 88532246 | cg09789939 |
| chr16 | 88534398 | 88534399 | cg02383666 |
| chr16 | 88535609 | 88535610 | cg06126698 |
| chr16 | 88536950 | 88536951 | cg05183538 |
| chr16 | 88537187 | 88537188 | cg04387347 |
| chr16 | 88538355 | 88538356 | cg10195726 |
| chr16 | 88543627 | 88543628 | cg06212670 |
| chr16 | 88545042 | 88545043 | cg03310838 |
| chr16 | 88545571 | 88545572 | cg08478121 |
| chr16 | 88547861 | 88547862 | cg05446615 |
| chr16 | 88747591 | 88747592 | cg04848686 |
| chr16 | 88806610 | 88806611 | cg08497311 |
| chr16 | 88845204 | 88845205 | cg04914290 |
| chr16 | 88927403 | 88927404 | cg01379825 |
| chr16 | 88950197 | 88950198 | cg01168962 |
| chr16 | 88969224 | 88969225 | cg08229905 |
| chr16 | 88977278 | 88977279 | cg08484992 |
| chr16 | 88977887 | 88977888 | cg26655856 |
| chr16 | 88978082 | 88978083 | cg08345246 |
| chr16 | 88980433 | 88980434 | cg05531174 |
| chr16 | 88982220 | 88982221 | cg06821871 |
| chr16 | 88984067 | 88984068 | cg27635976 |
| chr16 | 88984237 | 88984238 | cg05540133 |
| chr16 | 88991238 | 88991239 | cg27248474 |
| chr16 | 88991343 | 88991344 | cg00121135 |
| chr16 | 88995416 | 88995417 | cg09910271 |
| chr16 | 89003833 | 89003834 | cg08940410 |
| chr16 | 89004908 | 89004909 | cg26824780 |

| | | | |
|---|---|---|---|
| chr11 | 48034953 | 48034954 | cg11468953 |
| chr11 | 56956373 | 56956374 | cg09667650 |
| chr11 | 57252371 | 57252372 | cg09888840 |
| chr11 | 57529614 | 57529615 | cg09878888 |
| chr11 | 58870494 | 58870495 | cg20302957 |
| chr11 | 59270409 | 59270410 | cg20181079 |
| chr11 | 59435407 | 59435408 | cg15844711 |
| chr11 | 59807594 | 59807595 | cg08505647 |
| chr11 | 59950068 | 59950069 | cg07170641 |
| chr11 | 60553755 | 60553756 | cg11231631 |
| chr11 | 60684332 | 60684333 | cg27119919 |
| chr11 | 60869760 | 60869761 | cg06417460 |
| chr11 | 60869960 | 60869961 | cg24674703 |
| chr11 | 61349648 | 61349649 | cg14851471 |
| chr11 | 61598025 | 61598026 | cg01400685 |
| chr11 | 62309707 | 62309708 | cg05427381 |
| chr11 | 62315034 | 62315035 | cg20518446 |
| chr11 | 62361794 | 62361795 | cg16469419 |
| chr11 | 62477624 | 62477625 | cg03433986 |
| chr11 | 62553869 | 62553870 | cg14712784 |
| chr11 | 62574403 | 62574404 | cg21702188 |
| chr11 | 63634673 | 63634674 | cg26904017 |
| chr11 | 63679532 | 63679533 | cg11797099 |
| chr11 | 64163340 | 64163341 | cg15009090 |
| chr11 | 64528569 | 64528570 | cg10296062 |
| chr11 | 64635728 | 64635729 | cg14378925 |
| chr11 | 64635774 | 64635775 | cg07019638 |
| chr11 | 64635846 | 64635847 | cg04038246 |
| chr11 | 64642516 | 64642517 | cg14043527 |
| chr11 | 64642558 | 64642559 | cg06154903 |
| chr11 | 64898766 | 64898767 | cg05300538 |
| chr11 | 64918439 | 64918440 | cg20714487 |
| chr11 | 65102891 | 65102892 | cg09609051 |
| chr11 | 65152051 | 65152052 | cg24215776 |
| chr11 | 65195039 | 65195040 | cg02345399 |
| chr11 | 65195995 | 65195996 | cg03249011 |
| chr11 | 65196227 | 65196228 | cg11573318 |
| chr11 | 65255266 | 65255267 | cg12589431 |
| chr11 | 65262358 | 65262359 | cg15915658 |
| chr11 | 65263917 | 65263918 | cg02652361 |
| chr11 | 65311181 | 65311182 | cg23495995 |
| chr11 | 65315030 | 65315031 | cg23272978 |
| chr11 | 65315284 | 65315285 | cg16327477 |
| chr11 | 65315466 | 65315467 | cg20210376 |
| chr11 | 65315625 | 65315626 | cg09801334 |
| chr11 | 65319844 | 65319845 | cg03101763 |
| chr11 | 65325158 | 65325159 | cg08965235 |
| chr11 | 65325249 | 65325250 | cg16477774 |
| chr11 | 65327156 | 65327157 | cg17451760 |
| chr11 | 65354231 | 65354232 | cg12758867 |
| chr11 | 65550444 | 65550445 | cg09419102 |
| chr11 | 65586170 | 65586171 | cg02230254 |
| chr11 | 65778210 | 65778211 | cg19088912 |
| chr11 | 65842323 | 65842324 | cg20938379 |
| chr11 | 65904369 | 65904370 | cg24278087 |
| chr11 | 66048759 | 66048760 | cg06155341 |
| chr16 | 89009557 | 89009558 | cg07630078 |
| chr16 | 89009691 | 89009692 | cg04540712 |
| chr16 | 89009929 | 89009930 | cg08843248 |
| chr16 | 89034099 | 89034100 | cg22023435 |
| chr16 | 89044681 | 89044682 | cg27292079 |
| chr16 | 89130931 | 89130932 | cg01937601 |
| chr16 | 89131017 | 89131018 | cg03060214 |
| chr16 | 89183643 | 89183644 | cg10512427 |
| chr16 | 89183678 | 89183679 | cg10202714 |
| chr16 | 89384721 | 89384722 | cg09164168 |
| chr16 | 89390609 | 89390610 | cg27150870 |
| chr16 | 89394864 | 89394865 | cg16386046 |
| chr16 | 89439933 | 89439934 | cg08583277 |
| chr16 | 89455108 | 89455109 | cg06270206 |
| chr16 | 89462661 | 89462662 | cg05900234 |
| chr16 | 89471650 | 89471651 | cg09293925 |
| chr16 | 89475959 | 89475960 | cg06669446 |
| chr16 | 89498634 | 89498635 | cg09907628 |
| chr16 | 89509760 | 89509761 | cg26536240 |
| chr16 | 89539230 | 89539231 | cg09300089 |
| chr16 | 89541815 | 89541816 | cg05320533 |
| chr16 | 89638519 | 89638520 | cg00562180 |
| chr16 | 89826330 | 89826331 | cg27582168 |
| chr16 | 90050128 | 90050129 | cg02760445 |
| chr17 | 585596 | 585597 | cg15137213 |
| chr17 | 1057343 | 1057344 | cg00025496 |
| chr17 | 1123634 | 1123635 | cg22697034 |
| chr17 | 1123812 | 1123813 | cg20964216 |
| chr17 | 1123836 | 1123837 | cg12102682 |
| chr17 | 1124131 | 1124132 | cg12889817 |
| chr17 | 1124191 | 1124192 | cg18700329 |
| chr17 | 1130376 | 1130377 | cg12630479 |
| chr17 | 1366221 | 1366222 | cg09236126 |
| chr17 | 1396953 | 1396954 | cg00946187 |
| chr17 | 1553484 | 1553485 | cg24096415 |
| chr17 | 1553661 | 1553662 | cg04049033 |
| chr17 | 1640472 | 1640473 | cg10426084 |
| chr17 | 1686627 | 1686628 | cg16616918 |
| chr17 | 1686737 | 1686738 | cg23679982 |
| chr17 | 1731852 | 1731853 | cg09901733 |
| chr17 | 2041773 | 2041774 | cg07058377 |
| chr17 | 2300911 | 2300912 | cg15416352 |
| chr17 | 2302144 | 2302145 | cg12685560 |
| chr17 | 2394399 | 2394400 | cg24035107 |
| chr17 | 2908068 | 2908069 | cg15030415 |
| chr17 | 2908369 | 2908370 | cg14999189 |
| chr17 | 4108148 | 4108149 | cg21653149 |
| chr17 | 4235082 | 4235083 | cg27019093 |
| chr17 | 4495735 | 4495736 | cg16082620 |
| chr17 | 4539072 | 4539073 | cg23873735 |
| chr17 | 4674220 | 4674221 | cg01487803 |
| chr17 | 4800583 | 4800584 | cg22110517 |
| chr17 | 4806344 | 4806345 | cg01739167 |
| chr17 | 4872603 | 4872604 | cg12815142 |
| chr17 | 6348200 | 6348201 | cg20124188 |
| chr17 | 6353762 | 6353763 | cg07890785 |

| | | | |
|---|---|---|---|
| chr11 | 66631469 | 66631470 | cg19272754 |
| chr11 | 66631536 | 66631537 | cg06244306 |
| chr11 | 66632197 | 66632198 | cg25483596 |
| chr11 | 67033594 | 67033595 | cg14519515 |
| chr11 | 67179310 | 67179311 | cg02610064 |
| chr11 | 67183616 | 67183617 | cg10422334 |
| chr11 | 67202122 | 67202123 | cg12128589 |
| chr11 | 67205739 | 67205740 | cg25256924 |
| chr11 | 67205869 | 67205870 | cg16408081 |
| chr11 | 67206263 | 67206264 | cg23468927 |
| chr11 | 67206308 | 67206309 | cg12044599 |
| chr11 | 67206418 | 67206419 | cg02740606 |
| chr11 | 67206434 | 67206435 | cg17690322 |
| chr11 | 67210033 | 67210034 | cg16334425 |
| chr11 | 67210192 | 67210193 | cg23010585 |
| chr11 | 67210225 | 67210226 | cg05750788 |
| chr11 | 67210282 | 67210283 | cg15336893 |
| chr11 | 67259452 | 67259453 | cg06754606 |
| chr11 | 67340709 | 67340710 | cg16132851 |
| chr11 | 67340713 | 67340714 | cg02539539 |
| chr11 | 67351786 | 67351787 | cg11566244 |
| chr11 | 67352041 | 67352042 | cg22224704 |
| chr11 | 67417958 | 67417959 | cg11096993 |
| chr11 | 67982703 | 67982704 | cg20092736 |
| chr11 | 68037299 | 68037300 | cg25633067 |
| chr11 | 68038406 | 68038407 | cg15332978 |
| chr11 | 68065710 | 68065711 | cg16228359 |
| chr11 | 68081207 | 68081208 | cg20310100 |
| chr11 | 68095662 | 68095663 | cg02793948 |
| chr11 | 68139350 | 68139351 | cg08648997 |
| chr11 | 68238943 | 68238944 | cg05246110 |
| chr11 | 69061911 | 69061912 | cg22779330 |
| chr11 | 69240805 | 69240806 | cg13988440 |
| chr11 | 69240844 | 69240845 | cg15297628 |
| chr11 | 69241075 | 69241076 | cg22203628 |
| chr11 | 69339188 | 69339189 | cg12597472 |
| chr11 | 69486584 | 69486585 | cg07545052 |
| chr11 | 71710614 | 71710615 | cg16385335 |
| chr11 | 71710643 | 71710644 | cg13944685 |
| chr11 | 71710885 | 71710886 | cg26972614 |
| chr11 | 71710982 | 71710983 | cg05446414 |
| chr11 | 71752919 | 71752920 | cg03350366 |
| chr11 | 72280409 | 72280410 | cg21659346 |
| chr11 | 72380070 | 72380071 | cg23149454 |
| chr11 | 72397418 | 72397419 | cg06838038 |
| chr11 | 72452482 | 72452483 | cg01380194 |
| chr11 | 72860376 | 72860377 | cg06429138 |
| chr11 | 73061896 | 73061897 | cg02391615 |
| chr11 | 73087016 | 73087017 | cg02855207 |
| chr11 | 73668757 | 73668758 | cg04044120 |
| chr11 | 73697917 | 73697918 | cg13719084 |
| chr11 | 73711382 | 73711383 | cg14136260 |
| chr11 | 73716367 | 73716368 | cg10859442 |
| chr11 | 74440516 | 74440517 | cg22902532 |
| chr11 | 74654369 | 74654370 | cg15389191 |
| chr11 | 75052065 | 75052066 | cg22110267 |

| | | | |
|---|---|---|---|
| chr17 | 6478486 | 6478487 | cg10608948 |
| chr17 | 6921400 | 6921401 | cg02329670 |
| chr17 | 6921465 | 6921466 | cg21040575 |
| chr17 | 6943341 | 6943342 | cg05174446 |
| chr17 | 6945159 | 6945160 | cg05031424 |
| chr17 | 6950669 | 6950670 | cg23148631 |
| chr17 | 7253189 | 7253190 | cg08324090 |
| chr17 | 7341436 | 7341437 | cg13492227 |
| chr17 | 7383488 | 7383489 | cg01781446 |
| chr17 | 7453377 | 7453378 | cg00031162 |
| chr17 | 7529693 | 7529694 | cg01479396 |
| chr17 | 7791882 | 7791883 | cg12870750 |
| chr17 | 7960092 | 7960093 | cg07840332 |
| chr17 | 8056985 | 8056986 | cg13759852 |
| chr17 | 8311635 | 8311636 | cg14191955 |
| chr17 | 8535468 | 8535469 | cg22897522 |
| chr17 | 8798062 | 8798063 | cg01394255 |
| chr17 | 8844506 | 8844507 | cg03366884 |
| chr17 | 9940227 | 9940228 | cg16768966 |
| chr17 | 10059923 | 10059924 | cg01929340 |
| chr17 | 11953402 | 11953403 | cg00947023 |
| chr17 | 12704978 | 12704979 | cg07029411 |
| chr17 | 16176891 | 16176892 | cg19135352 |
| chr17 | 17109239 | 17109240 | cg01412970 |
| chr17 | 17715520 | 17715521 | cg23875758 |
| chr17 | 17744282 | 17744283 | cg16460860 |
| chr17 | 17753516 | 17753517 | cg03845363 |
| chr17 | 17817462 | 17817463 | cg04324276 |
| chr17 | 17824756 | 17824757 | cg17514485 |
| chr17 | 18905779 | 18905780 | cg25708364 |
| chr17 | 20026062 | 20026063 | cg22675702 |
| chr17 | 21186789 | 21186790 | cg25538809 |
| chr17 | 21194718 | 21194719 | cg03858720 |
| chr17 | 21451571 | 21451572 | cg17529235 |
| chr17 | 25872931 | 25872932 | cg04561005 |
| chr17 | 26380792 | 26380793 | cg09120171 |
| chr17 | 26990202 | 26990203 | cg07042338 |
| chr17 | 27034448 | 27034449 | cg15734651 |
| chr17 | 27263598 | 27263599 | cg17445097 |
| chr17 | 27482619 | 27482620 | cg18748374 |
| chr17 | 27505257 | 27505258 | cg20029153 |
| chr17 | 27507678 | 27507679 | cg27385272 |
| chr17 | 27901067 | 27901068 | cg13872326 |
| chr17 | 27913793 | 27913794 | cg22961402 |
| chr17 | 27934880 | 27934881 | cg20649847 |
| chr17 | 28907176 | 28907177 | cg13915843 |
| chr17 | 29235812 | 29235813 | cg27606337 |
| chr17 | 29817279 | 29817280 | cg25709050 |
| chr17 | 32561989 | 32561990 | cg07431106 |
| chr17 | 33306307 | 33306308 | cg00762550 |
| chr17 | 33759512 | 33759513 | cg03251655 |
| chr17 | 33759573 | 33759574 | cg19963856 |
| chr17 | 33760419 | 33760420 | cg18587557 |
| chr17 | 34434265 | 34434266 | cg24656005 |
| chr17 | 36885965 | 36885966 | cg27492942 |
| chr17 | 36889069 | 36889070 | cg07095330 |

| | | | |
|---|---|---|---|
| chr11 | 75061838 | 75061839 | cg02449461 |
| chr11 | 75428910 | 75428911 | cg20938170 |
| chr11 | 75522097 | 75522098 | cg16253834 |
| chr11 | 75736215 | 75736216 | cg11274172 |
| chr11 | 76340765 | 76340766 | cg20430631 |
| chr11 | 76614479 | 76614480 | cg19314992 |
| chr11 | 77123331 | 77123332 | cg13039082 |
| chr11 | 77787848 | 77787849 | cg20267732 |
| chr11 | 78136642 | 78136643 | cg25486618 |
| chr11 | 78215445 | 78215446 | cg01088387 |
| chr11 | 78542948 | 78542949 | cg23361764 |
| chr11 | 82708076 | 82708077 | cg01250845 |
| chr11 | 82735870 | 82735871 | cg15468739 |
| chr11 | 82745712 | 82745713 | cg12453687 |
| chr11 | 88059873 | 88059874 | cg03596479 |
| chr11 | 88178390 | 88178391 | cg07702100 |
| chr11 | 92001577 | 92001578 | cg05283685 |
| chr11 | 92670490 | 92670491 | cg08001668 |
| chr11 | 93277841 | 93277842 | cg03333377 |
| chr11 | 93280125 | 93280126 | cg13411373 |
| chr11 | 94307971 | 94307972 | cg16436762 |
| chr11 | 94536536 | 94536537 | cg23265210 |
| chr11 | 94841384 | 94841385 | cg26402417 |
| chr11 | 94975774 | 94975775 | cg18829068 |
| chr11 | 95413642 | 95413643 | cg06893560 |
| chr11 | 95785126 | 95785127 | cg18925664 |
| chr11 | 95815452 | 95815453 | cg01806722 |
| chr11 | 95846117 | 95846118 | cg04677410 |
| chr11 | 95870414 | 95870415 | cg18219522 |
| chr11 | 95889454 | 95889455 | cg15521790 |
| chr11 | 95980186 | 95980187 | cg02322400 |
| chr11 | 96466206 | 96466207 | cg19574327 |
| chr11 | 100211450 | 100211451 | cg10748359 |
| chr11 | 100734094 | 100734095 | cg05925497 |
| chr11 | 101323578 | 101323579 | cg04554505 |
| chr11 | 102583226 | 102583227 | cg08747557 |
| chr11 | 102745934 | 102745935 | cg09002726 |
| chr11 | 102746365 | 102746366 | cg02368591 |
| chr11 | 103791252 | 103791253 | cg19992785 |
| chr11 | 103964219 | 103964220 | cg04813876 |
| chr11 | 104030857 | 104030858 | cg23852348 |
| chr11 | 106892666 | 106892667 | cg03615568 |
| chr11 | 108991744 | 108991745 | cg14533401 |
| chr11 | 108994504 | 108994505 | cg25007511 |
| chr11 | 109172798 | 109172799 | cg07680368 |
| chr11 | 112035945 | 112035946 | cg05687149 |
| chr11 | 113135432 | 113135433 | cg02323699 |
| chr11 | 113285594 | 113285595 | cg08948964 |
| chr11 | 113403184 | 113403185 | cg24868960 |
| chr11 | 114031120 | 114031121 | cg25009965 |
| chr11 | 115238331 | 115238332 | cg01850954 |
| chr11 | 115790529 | 115790530 | cg11113536 |
| chr11 | 116663103 | 116663104 | cg25682080 |
| chr11 | 116891433 | 116891434 | cg22657772 |
| chr11 | 117355376 | 117355377 | cg06923861 |
| chr11 | 117740722 | 117740723 | cg09193791 |

| | | | |
|---|---|---|---|
| chr17 | 37024169 | 37024170 | cg14624145 |
| chr17 | 37720009 | 37720010 | cg17288560 |
| chr17 | 37782685 | 37782686 | cg15070513 |
| chr17 | 37808339 | 37808340 | cg03572260 |
| chr17 | 37894027 | 37894028 | cg08284496 |
| chr17 | 37896175 | 37896176 | cg14575854 |
| chr17 | 37897076 | 37897077 | cg00697095 |
| chr17 | 37907944 | 37907945 | cg04587617 |
| chr17 | 38220694 | 38220695 | cg11512009 |
| chr17 | 38254714 | 38254715 | cg01346501 |
| chr17 | 38263416 | 38263417 | cg08324801 |
| chr17 | 38297856 | 38297857 | cg15597662 |
| chr17 | 38337188 | 38337189 | cg12041100 |
| chr17 | 38511592 | 38511593 | cg04956244 |
| chr17 | 38511950 | 38511951 | cg05442717 |
| chr17 | 38599056 | 38599057 | cg02627216 |
| chr17 | 38601676 | 38601677 | cg17980404 |
| chr17 | 38707719 | 38707720 | cg27109912 |
| chr17 | 38708809 | 38708810 | cg23375169 |
| chr17 | 38764244 | 38764245 | cg03917473 |
| chr17 | 38784596 | 38784597 | cg19712600 |
| chr17 | 39405817 | 39405818 | cg00550617 |
| chr17 | 39662206 | 39662207 | cg16023337 |
| chr17 | 40117578 | 40117579 | cg05182416 |
| chr17 | 40123227 | 40123228 | cg20212624 |
| chr17 | 40438511 | 40438512 | cg00690082 |
| chr17 | 40465546 | 40465547 | cg07780782 |
| chr17 | 40554519 | 40554520 | cg04726374 |
| chr17 | 40799848 | 40799849 | cg13250566 |
| chr17 | 40810559 | 40810560 | cg10180165 |
| chr17 | 40995546 | 40995547 | cg11961813 |
| chr17 | 41121153 | 41121154 | cg22396850 |
| chr17 | 41159289 | 41159290 | cg08090640 |
| chr17 | 41365094 | 41365095 | cg25918947 |
| chr17 | 42189331 | 42189332 | cg08531489 |
| chr17 | 42199063 | 42199064 | cg11049075 |
| chr17 | 42427639 | 42427640 | cg16500497 |
| chr17 | 42427647 | 42427648 | cg06800040 |
| chr17 | 43099144 | 43099145 | cg19407385 |
| chr17 | 43298345 | 43298346 | cg00933182 |
| chr17 | 43657317 | 43657318 | cg22687902 |
| chr17 | 43715162 | 43715163 | cg03954353 |
| chr17 | 43976002 | 43976003 | cg10955972 |
| chr17 | 44135007 | 44135008 | cg22192454 |
| chr17 | 44859215 | 44859216 | cg00140914 |
| chr17 | 44898510 | 44898511 | cg13550419 |
| chr17 | 45785206 | 45785207 | cg23673974 |
| chr17 | 45808345 | 45808346 | cg09529165 |
| chr17 | 45813433 | 45813434 | cg21821643 |
| chr17 | 46507895 | 46507896 | cg24322968 |
| chr17 | 47436584 | 47436585 | cg07164631 |
| chr17 | 47443155 | 47443156 | cg23104592 |
| chr17 | 47651453 | 47651454 | cg08250742 |
| chr17 | 47723651 | 47723652 | cg23876355 |
| chr17 | 48211226 | 48211227 | cg13686739 |
| chr17 | 48242070 | 48242071 | cg04582295 |

| | | | |
|---|---|---|---|
| chr11 | 117856007 | 117856008 | cg01697865 |
| chr11 | 117947459 | 117947460 | cg13159318 |
| chr11 | 117947611 | 117947612 | cg03634928 |
| chr11 | 117947628 | 117947629 | cg27300950 |
| chr11 | 117947657 | 117947658 | cg25116503 |
| chr11 | 118402280 | 118402281 | cg15694715 |
| chr11 | 118585993 | 118585994 | cg26062581 |
| chr11 | 118657363 | 118657364 | cg05587627 |
| chr11 | 118763859 | 118763860 | cg04537602 |
| chr11 | 118763863 | 118763864 | cg13298528 |
| chr11 | 118764003 | 118764004 | cg03523129 |
| chr11 | 118773983 | 118773984 | cg16130802 |
| chr11 | 118790354 | 118790355 | cg18445438 |
| chr11 | 119248136 | 119248137 | cg15648239 |
| chr11 | 119291765 | 119291766 | cg02632800 |
| chr11 | 119458741 | 119458742 | cg22722340 |
| chr11 | 119893672 | 119893673 | cg07507085 |
| chr11 | 120106598 | 120106599 | cg21886186 |
| chr11 | 120210293 | 120210294 | cg00395063 |
| chr11 | 120217718 | 120217719 | cg15690696 |
| chr11 | 120283690 | 120283691 | cg04919489 |
| chr11 | 120386165 | 120386166 | cg12562132 |
| chr11 | 121323934 | 121323935 | cg22327175 |
| chr11 | 122073211 | 122073212 | cg18730174 |
| chr11 | 122562201 | 122562202 | cg05229759 |
| chr11 | 123236871 | 123236872 | cg09077443 |
| chr11 | 123242457 | 123242458 | cg07270423 |
| chr11 | 123388890 | 123388891 | cg24402151 |
| chr11 | 123389435 | 123389436 | cg26678144 |
| chr11 | 123885967 | 123885968 | cg01264963 |
| chr11 | 124043067 | 124043068 | cg19550524 |
| chr11 | 124043158 | 124043159 | cg09991267 |
| chr11 | 124134914 | 124134915 | cg22476960 |
| chr11 | 124439945 | 124439946 | cg23941354 |
| chr11 | 124471596 | 124471597 | cg02589353 |
| chr11 | 124761914 | 124761915 | cg02150262 |
| chr11 | 125288301 | 125288302 | cg09642090 |
| chr11 | 125361336 | 125361337 | cg02493544 |
| chr11 | 126009830 | 126009831 | cg18226915 |
| chr11 | 126188994 | 126188995 | cg15958289 |
| chr11 | 128169380 | 128169381 | cg15339435 |
| chr11 | 128321662 | 128321663 | cg10616216 |
| chr11 | 128321749 | 128321750 | cg10334741 |
| chr11 | 128321880 | 128321881 | cg08750439 |
| chr11 | 128420863 | 128420864 | cg14718848 |
| chr11 | 128446070 | 128446071 | cg25578781 |
| chr11 | 128673534 | 128673535 | cg16024530 |
| chr11 | 128712280 | 128712281 | cg07168810 |
| chr11 | 129920411 | 129920412 | cg03113867 |
| chr11 | 129991544 | 129991545 | cg23729763 |
| chr11 | 130026698 | 130026699 | cg08210681 |
| chr11 | 130026798 | 130026799 | cg01668099 |
| chr11 | 130076257 | 130076258 | cg16358741 |
| chr11 | 130631528 | 130631529 | cg16924753 |
| chr11 | 131122594 | 131122595 | cg08084339 |
| chr11 | 132080258 | 132080259 | cg05184519 |

| | | | |
|---|---|---|---|
| chr17 | 48502312 | 48502313 | cg03919657 |
| chr17 | 48502923 | 48502924 | cg22002216 |
| chr17 | 48581622 | 48581623 | cg03661110 |
| chr17 | 49295249 | 49295250 | cg25670376 |
| chr17 | 49336082 | 49336083 | cg19213808 |
| chr17 | 55503603 | 55503604 | cg16323330 |
| chr17 | 55517831 | 55517832 | cg05251593 |
| chr17 | 55543388 | 55543389 | cg22233512 |
| chr17 | 55554341 | 55554342 | cg06247967 |
| chr17 | 55601697 | 55601698 | cg13916877 |
| chr17 | 55630218 | 55630219 | cg10249814 |
| chr17 | 55683689 | 55683690 | cg25373624 |
| chr17 | 55981642 | 55981643 | cg06723357 |
| chr17 | 56071745 | 56071746 | cg01821751 |
| chr17 | 57407818 | 57407819 | cg24865110 |
| chr17 | 57444078 | 57444079 | cg01653532 |
| chr17 | 57449015 | 57449016 | cg13855435 |
| chr17 | 57465550 | 57465551 | cg10708739 |
| chr17 | 58165865 | 58165866 | cg03120558 |
| chr17 | 58166202 | 58166203 | cg12320164 |
| chr17 | 58166296 | 58166297 | cg10325253 |
| chr17 | 58499700 | 58499701 | cg12131208 |
| chr17 | 58499706 | 58499707 | cg04413853 |
| chr17 | 58499720 | 58499721 | cg21122199 |
| chr17 | 58499816 | 58499817 | cg06752482 |
| chr17 | 58499854 | 58499855 | cg17628249 |
| chr17 | 58499911 | 58499912 | cg02172058 |
| chr17 | 62010607 | 62010608 | cg25729716 |
| chr17 | 62011014 | 62011015 | cg23251296 |
| chr17 | 62062565 | 62062566 | cg18041642 |
| chr17 | 62067215 | 62067216 | cg08036285 |
| chr17 | 62206650 | 62206651 | cg14603886 |
| chr17 | 62208434 | 62208435 | cg12779575 |
| chr17 | 62400642 | 62400643 | cg00879592 |
| chr17 | 62401855 | 62401856 | cg02459519 |
| chr17 | 62401956 | 62401957 | cg08699174 |
| chr17 | 62656813 | 62656814 | cg02960948 |
| chr17 | 66254884 | 66254885 | cg04495944 |
| chr17 | 66308577 | 66308578 | cg13093111 |
| chr17 | 67497879 | 67497880 | cg12323063 |
| chr17 | 67603831 | 67603832 | cg06677321 |
| chr17 | 71638531 | 71638532 | cg22895083 |
| chr17 | 71639029 | 71639030 | cg00290483 |
| chr17 | 72212478 | 72212479 | cg25060035 |
| chr17 | 72779428 | 72779429 | cg22312249 |
| chr17 | 72833123 | 72833124 | cg11763337 |
| chr17 | 72999242 | 72999243 | cg08757721 |
| chr17 | 73061848 | 73061849 | cg07392740 |
| chr17 | 73077622 | 73077623 | cg20519525 |
| chr17 | 73266070 | 73266071 | cg07104495 |
| chr17 | 73314451 | 73314452 | cg01040259 |
| chr17 | 73339287 | 73339288 | cg24244854 |
| chr17 | 73520759 | 73520760 | cg06665453 |
| chr17 | 73821508 | 73821509 | cg13298379 |
| chr17 | 74184866 | 74184867 | cg21265072 |
| chr17 | 74270190 | 74270191 | cg09812376 |

| | | | |
|---|---|---|---|
| chr11 | 132816803 | 132816804 | cg15618210 |
| chr11 | 133286631 | 133286632 | cg04745004 |
| chr11 | 133706356 | 133706357 | cg19415202 |
| chr11 | 133899912 | 133899913 | cg03961718 |
| chr11 | 133900322 | 133900323 | cg23564535 |
| chr11 | 133910729 | 133910730 | cg15003194 |
| chr11 | 134095652 | 134095653 | cg26723847 |
| chr11 | 134603744 | 134603745 | cg13107191 |
| chr11 | 134877533 | 134877534 | cg23219748 |
| chr12 | 566229 | 566230 | cg18436709 |
| chr12 | 618772 | 618773 | cg15580052 |
| chr12 | 671027 | 671028 | cg27143842 |
| chr12 | 772688 | 772689 | cg20781967 |
| chr12 | 776351 | 776352 | cg12013258 |
| chr12 | 858759 | 858760 | cg22609908 |
| chr12 | 977110 | 977111 | cg00321644 |
| chr12 | 1641924 | 1641925 | cg05170445 |
| chr12 | 1726033 | 1726034 | cg24854010 |
| chr12 | 1726076 | 1726077 | cg07896667 |
| chr12 | 1759064 | 1759065 | cg10147974 |
| chr12 | 2660504 | 2660505 | cg04006520 |
| chr12 | 3446781 | 3446782 | cg10047041 |
| chr12 | 4398032 | 4398033 | cg14834893 |
| chr12 | 4398508 | 4398509 | cg27317813 |
| chr12 | 4398618 | 4398619 | cg09799980 |
| chr12 | 4479889 | 4479890 | cg08956010 |
| chr12 | 6470176 | 6470177 | cg26244013 |
| chr12 | 6543527 | 6543528 | cg17458693 |
| chr12 | 6570167 | 6570168 | cg15719903 |
| chr12 | 6571600 | 6571601 | cg17092523 |
| chr12 | 6578523 | 6578524 | cg16770549 |
| chr12 | 6649349 | 6649350 | cg02410245 |
| chr12 | 6679571 | 6679572 | cg17820065 |
| chr12 | 6720114 | 6720115 | cg17781716 |
| chr12 | 6881118 | 6881119 | cg04153135 |
| chr12 | 6882856 | 6882857 | cg19421125 |
| chr12 | 6939048 | 6939049 | cg14538537 |
| chr12 | 6996791 | 6996792 | cg26538140 |
| chr12 | 7032793 | 7032794 | cg10201141 |
| chr12 | 7032991 | 7032992 | cg25316512 |
| chr12 | 7055153 | 7055154 | cg22335340 |
| chr12 | 7066614 | 7066615 | cg24437859 |
| chr12 | 7066825 | 7066826 | cg26004710 |
| chr12 | 7070317 | 7070318 | cg22260508 |
| chr12 | 7260546 | 7260547 | cg21439672 |
| chr12 | 7940546 | 7940547 | cg10660253 |
| chr12 | 8191646 | 8191647 | cg17815933 |
| chr12 | 8765336 | 8765337 | cg07999668 |
| chr12 | 9162872 | 9162873 | cg04548204 |
| chr12 | 9799858 | 9799859 | cg03289764 |
| chr12 | 9914965 | 9914966 | cg13918228 |
| chr12 | 10163532 | 10163533 | cg20645058 |
| chr12 | 10516240 | 10516241 | cg26485376 |
| chr12 | 10870175 | 10870176 | cg18636739 |
| chr12 | 11733759 | 11733760 | cg05931231 |
| chr12 | 11877740 | 11877741 | cg24279017 |

| | | | |
|---|---|---|---|
| chr17 | 74308979 | 74308980 | cg10318678 |
| chr17 | 74709662 | 74709663 | cg20832875 |
| chr17 | 75207115 | 75207116 | cg09013599 |
| chr17 | 75207259 | 75207260 | cg04308797 |
| chr17 | 75207380 | 75207381 | cg17940464 |
| chr17 | 75208129 | 75208130 | cg20242937 |
| chr17 | 75209460 | 75209461 | cg00915372 |
| chr17 | 75209578 | 75209579 | cg13817725 |
| chr17 | 75209644 | 75209645 | cg07166159 |
| chr17 | 75238948 | 75238949 | cg20959920 |
| chr17 | 75398014 | 75398015 | cg14608550 |
| chr17 | 75399899 | 75399900 | cg06380320 |
| chr17 | 75404563 | 75404564 | cg19422205 |
| chr17 | 75422925 | 75422926 | cg06622199 |
| chr17 | 75425180 | 75425181 | cg26079706 |
| chr17 | 75445261 | 75445262 | cg20557159 |
| chr17 | 75445905 | 75445906 | cg07324245 |
| chr17 | 75462916 | 75462917 | cg02397064 |
| chr17 | 75463984 | 75463985 | cg21808045 |
| chr17 | 75471491 | 75471492 | cg10997810 |
| chr17 | 75473610 | 75473611 | cg20923885 |
| chr17 | 75473895 | 75473896 | cg05626616 |
| chr17 | 75881093 | 75881094 | cg09889477 |
| chr17 | 75883546 | 75883547 | cg24093538 |
| chr17 | 76129475 | 76129476 | cg05637296 |
| chr17 | 76138514 | 76138515 | cg03596178 |
| chr17 | 76141038 | 76141039 | cg14771877 |
| chr17 | 76142461 | 76142462 | cg27606002 |
| chr17 | 76171191 | 76171192 | cg25069807 |
| chr17 | 76171208 | 76171209 | cg06098276 |
| chr17 | 76743501 | 76743502 | cg07823562 |
| chr17 | 76886742 | 76886743 | cg24211657 |
| chr17 | 76886835 | 76886836 | cg12125614 |
| chr17 | 76886912 | 76886913 | cg04397137 |
| chr17 | 76887989 | 76887990 | cg26963090 |
| chr17 | 76888675 | 76888676 | cg21576055 |
| chr17 | 78116968 | 78116969 | cg19010410 |
| chr17 | 78357934 | 78357935 | cg17530782 |
| chr17 | 78800574 | 78800575 | cg16636468 |
| chr17 | 78800631 | 78800632 | cg25337513 |
| chr17 | 78963221 | 78963222 | cg09164913 |
| chr17 | 79267785 | 79267786 | cg14486098 |
| chr17 | 79600681 | 79600682 | cg14399228 |
| chr17 | 80006775 | 80006776 | cg04294971 |
| chr17 | 80197412 | 80197413 | cg17749033 |
| chr17 | 80199608 | 80199609 | cg15296856 |
| chr17 | 80215375 | 80215376 | cg05385188 |
| chr17 | 80278914 | 80278915 | cg09237133 |
| chr17 | 80278947 | 80278948 | cg09340279 |
| chr17 | 80328714 | 80328715 | cg17842879 |
| chr17 | 80329175 | 80329176 | cg09787311 |
| chr17 | 80358850 | 80358851 | cg25490145 |
| chr17 | 80390068 | 80390069 | cg09921719 |
| chr17 | 80741558 | 80741559 | cg02948444 |
| chr17 | 80766090 | 80766091 | cg09086312 |
| chr17 | 80818599 | 80818600 | cg17477946 |

| | | | |
|---|---|---|---|
| chr12 | 12222913 | 12222914 | cg27366072 |
| chr12 | 12223841 | 12223842 | cg18223430 |
| chr12 | 12619103 | 12619104 | cg15429134 |
| chr12 | 12620789 | 12620790 | cg24834394 |
| chr12 | 13525681 | 13525682 | cg26547898 |
| chr12 | 14118880 | 14118881 | cg00791406 |
| chr12 | 14475272 | 14475273 | cg05485976 |
| chr12 | 14721288 | 14721289 | cg04473902 |
| chr12 | 14765994 | 14765995 | cg27486692 |
| chr12 | 15938047 | 15938048 | cg22677100 |
| chr12 | 19635946 | 19635947 | cg16822174 |
| chr12 | 20263199 | 20263200 | cg27099262 |
| chr12 | 25055381 | 25055382 | cg20342079 |
| chr12 | 25103643 | 25103644 | cg07259733 |
| chr12 | 25150626 | 25150627 | cg21368143 |
| chr12 | 25156888 | 25156889 | cg04915788 |
| chr12 | 25177892 | 25177893 | cg21232844 |
| chr12 | 25204113 | 25204114 | cg04009342 |
| chr12 | 25246439 | 25246440 | cg17965159 |
| chr12 | 25535797 | 25535798 | cg26462457 |
| chr12 | 26150045 | 26150046 | cg01234248 |
| chr12 | 26876374 | 26876375 | cg09640070 |
| chr12 | 26881038 | 26881039 | cg00813135 |
| chr12 | 26951217 | 26951218 | cg13203394 |
| chr12 | 26989568 | 26989569 | cg10837653 |
| chr12 | 27010516 | 27010517 | cg18784875 |
| chr12 | 27587175 | 27587176 | cg11307865 |
| chr12 | 29879189 | 29879190 | cg16833797 |
| chr12 | 31406441 | 31406442 | cg14306366 |
| chr12 | 31926899 | 31926900 | cg15417287 |
| chr12 | 32120103 | 32120104 | cg01871722 |
| chr12 | 32299085 | 32299086 | cg15061954 |
| chr12 | 32414514 | 32414515 | cg23791308 |
| chr12 | 40622688 | 40622689 | cg12664938 |
| chr12 | 40938465 | 40938466 | cg25890391 |
| chr12 | 40938571 | 40938572 | cg26081584 |
| chr12 | 42873202 | 42873203 | cg16931707 |
| chr12 | 43253257 | 43253258 | cg12493761 |
| chr12 | 45182192 | 45182193 | cg24686187 |
| chr12 | 45608691 | 45608692 | cg02278646 |
| chr12 | 46657338 | 46657339 | cg15777028 |
| chr12 | 46782837 | 46782838 | cg01656976 |
| chr12 | 48149200 | 48149201 | cg13353550 |
| chr12 | 48152204 | 48152205 | cg21166775 |
| chr12 | 48152428 | 48152429 | cg14854517 |
| chr12 | 48192939 | 48192940 | cg15820955 |
| chr12 | 48196167 | 48196168 | cg20995304 |
| chr12 | 48205446 | 48205447 | cg27220401 |
| chr12 | 48214523 | 48214524 | cg01088404 |
| chr12 | 48228889 | 48228890 | cg23539261 |
| chr12 | 49362475 | 49362476 | cg12580783 |
| chr12 | 49413435 | 49413436 | cg05581469 |
| chr12 | 49763288 | 49763289 | cg00562262 |
| chr12 | 50067887 | 50067888 | cg04070601 |
| chr12 | 50303785 | 50303786 | cg01057945 |
| chr12 | 50306886 | 50306887 | cg17960823 |
| chr17 | 80819208 | 80819209 | cg04099036 |
| chr17 | 80823310 | 80823311 | cg16538568 |
| chr17 | 80825520 | 80825521 | cg25754888 |
| chr17 | 80873656 | 80873657 | cg02092514 |
| chr17 | 80941542 | 80941543 | cg09575521 |
| chr18 | 2909789 | 2909790 | cg06769774 |
| chr18 | 2916738 | 2916739 | cg18061327 |
| chr18 | 3015372 | 3015373 | cg25461300 |
| chr18 | 3063385 | 3063386 | cg21518709 |
| chr18 | 3067532 | 3067533 | cg00522579 |
| chr18 | 3266232 | 3266233 | cg06860460 |
| chr18 | 3453798 | 3453799 | cg18204321 |
| chr18 | 3495888 | 3495889 | cg21013431 |
| chr18 | 10474524 | 10474525 | cg03396005 |
| chr18 | 10524509 | 10524510 | cg05199682 |
| chr18 | 10955104 | 10955105 | cg14570051 |
| chr18 | 11884680 | 11884681 | cg25033076 |
| chr18 | 11909634 | 11909635 | cg14599440 |
| chr18 | 12322027 | 12322028 | cg18348731 |
| chr18 | 20713759 | 20713760 | cg22158648 |
| chr18 | 20916335 | 20916336 | cg16206090 |
| chr18 | 21593174 | 21593175 | cg00287012 |
| chr18 | 29677086 | 29677087 | cg20338754 |
| chr18 | 42281763 | 42281764 | cg00037763 |
| chr18 | 43845381 | 43845382 | cg05520141 |
| chr18 | 47863988 | 47863989 | cg21292912 |
| chr18 | 48721148 | 48721149 | cg27511678 |
| chr18 | 51747223 | 51747224 | cg16200116 |
| chr18 | 56701518 | 56701519 | cg25730020 |
| chr18 | 60988465 | 60988466 | cg24408313 |
| chr18 | 72912760 | 72912761 | cg20936529 |
| chr18 | 76763078 | 76763079 | cg14299739 |
| chr18 | 77157760 | 77157761 | cg00151768 |
| chr18 | 77158471 | 77158472 | cg01277438 |
| chr18 | 77172633 | 77172634 | cg15908877 |
| chr18 | 77181423 | 77181424 | cg05264214 |
| chr18 | 77193464 | 77193465 | cg04516518 |
| chr18 | 77193759 | 77193760 | cg06937978 |
| chr18 | 77194663 | 77194664 | cg25332265 |
| chr18 | 77196899 | 77196900 | cg27100149 |
| chr18 | 77199432 | 77199433 | cg05753993 |
| chr18 | 77204561 | 77204562 | cg22279865 |
| chr18 | 77205869 | 77205870 | cg25546405 |
| chr18 | 77214960 | 77214961 | cg13207250 |
| chr18 | 77218480 | 77218481 | cg25951288 |
| chr18 | 77280586 | 77280587 | cg03511628 |
| chr18 | 77392535 | 77392536 | cg06976213 |
| chr18 | 77400225 | 77400226 | cg24425316 |
| chr18 | 77560089 | 77560090 | cg22095490 |
| chr18 | 77733105 | 77733106 | cg26501139 |
| chr19 | 933707 | 933708 | cg03763874 |
| chr19 | 941126 | 941127 | cg18598117 |
| chr19 | 941243 | 941244 | cg08894487 |
| chr19 | 941571 | 941572 | cg02466798 |
| chr19 | 1033606 | 1033607 | cg05220090 |
| chr19 | 1035093 | 1035094 | cg10730425 |

| | | | |
|---|---|---|---|
| chr12 | 50366624 | 50366625 | cg00604736 |
| chr12 | 51015475 | 51015476 | cg06489912 |
| chr12 | 51423248 | 51423249 | cg18480005 |
| chr12 | 51636921 | 51636922 | cg24637417 |
| chr12 | 51782159 | 51782160 | cg24537836 |
| chr12 | 51783393 | 51783394 | cg21253043 |
| chr12 | 52245356 | 52245357 | cg27042523 |
| chr12 | 52305328 | 52305329 | cg21415035 |
| chr12 | 52579609 | 52579610 | cg23243343 |
| chr12 | 52586127 | 52586128 | cg03747456 |
| chr12 | 52642706 | 52642707 | cg14645027 |
| chr12 | 52689098 | 52689099 | cg18190861 |
| chr12 | 52692031 | 52692032 | cg04950824 |
| chr12 | 53579063 | 53579064 | cg00078245 |
| chr12 | 53602179 | 53602180 | cg15889057 |
| chr12 | 53608721 | 53608722 | cg27036638 |
| chr12 | 53693054 | 53693055 | cg03624316 |
| chr12 | 54655894 | 54655895 | cg08269188 |
| chr12 | 54746067 | 54746068 | cg20474425 |
| chr12 | 54763081 | 54763082 | cg02734505 |
| chr12 | 54811762 | 54811763 | cg02419321 |
| chr12 | 55002180 | 55002181 | cg06113001 |
| chr12 | 55375859 | 55375860 | cg01874152 |
| chr12 | 55379228 | 55379229 | cg07880109 |
| chr12 | 56078391 | 56078392 | cg26979518 |
| chr12 | 56118524 | 56118525 | cg11317254 |
| chr12 | 56120816 | 56120817 | cg21008399 |
| chr12 | 56121015 | 56121016 | cg11848173 |
| chr12 | 56229821 | 56229822 | cg22810434 |
| chr12 | 56322667 | 56322668 | cg08338281 |
| chr12 | 56323816 | 56323817 | cg21587469 |
| chr12 | 56414508 | 56414509 | cg20054248 |
| chr12 | 56438419 | 56438420 | cg07355346 |
| chr12 | 56554264 | 56554265 | cg01526553 |
| chr12 | 56690194 | 56690195 | cg16268734 |
| chr12 | 56754826 | 56754827 | cg26031046 |
| chr12 | 56989385 | 56989386 | cg25880980 |
| chr12 | 57143114 | 57143115 | cg11592377 |
| chr12 | 57405251 | 57405252 | cg12178465 |
| chr12 | 57452111 | 57452112 | cg12649051 |
| chr12 | 57488961 | 57488962 | cg19466605 |
| chr12 | 57581972 | 57581973 | cg04688889 |
| chr12 | 57624371 | 57624372 | cg25110782 |
| chr12 | 57624530 | 57624531 | cg11851129 |
| chr12 | 57872430 | 57872431 | cg08134678 |
| chr12 | 58160843 | 58160844 | cg24722099 |
| chr12 | 59287786 | 59287787 | cg06587521 |
| chr12 | 59319720 | 59319721 | cg10283871 |
| chr12 | 59325286 | 59325287 | cg07180988 |
| chr12 | 59508147 | 59508148 | cg16792144 |
| chr12 | 61806201 | 61806202 | cg05150077 |
| chr12 | 62650767 | 62650768 | cg03726817 |
| chr12 | 62652662 | 62652663 | cg27552493 |
| chr12 | 63039927 | 63039928 | cg17742617 |
| chr12 | 63144886 | 63144887 | cg21058511 |
| chr12 | 63547716 | 63547717 | cg23335356 |

| | | | |
|---|---|---|---|
| chr19 | 1065688 | 1065689 | cg11714200 |
| chr19 | 1415757 | 1415758 | cg07825294 |
| chr19 | 1851995 | 1851996 | cg11434705 |
| chr19 | 1852127 | 1852128 | cg07355926 |
| chr19 | 1854819 | 1854820 | cg08525314 |
| chr19 | 1921199 | 1921200 | cg26526953 |
| chr19 | 1955395 | 1955396 | cg02968914 |
| chr19 | 2073190 | 2073191 | cg20152841 |
| chr19 | 2140048 | 2140049 | cg09261902 |
| chr19 | 2216482 | 2216483 | cg06426736 |
| chr19 | 2242181 | 2242182 | cg25431916 |
| chr19 | 2275015 | 2275016 | cg23245997 |
| chr19 | 2546877 | 2546878 | cg06996175 |
| chr19 | 2547067 | 2547068 | cg19032532 |
| chr19 | 2550027 | 2550028 | cg04603130 |
| chr19 | 2579221 | 2579222 | cg22746789 |
| chr19 | 2753457 | 2753458 | cg23683620 |
| chr19 | 3364892 | 3364893 | cg09773897 |
| chr19 | 4102601 | 4102602 | cg08644106 |
| chr19 | 4338769 | 4338770 | cg05517572 |
| chr19 | 4338932 | 4338933 | cg15797834 |
| chr19 | 5217431 | 5217432 | cg23325999 |
| chr19 | 7535526 | 7535527 | cg16618417 |
| chr19 | 7539710 | 7539711 | cg03260991 |
| chr19 | 7712095 | 7712096 | cg15460739 |
| chr19 | 7712195 | 7712196 | cg24433207 |
| chr19 | 7767584 | 7767585 | cg05641903 |
| chr19 | 7795192 | 7795193 | cg05842480 |
| chr19 | 9253668 | 9253669 | cg12976832 |
| chr19 | 10195914 | 10195915 | cg13480898 |
| chr19 | 10332165 | 10332166 | cg06655349 |
| chr19 | 10509596 | 10509597 | cg26734579 |
| chr19 | 10533016 | 10533017 | cg12186909 |
| chr19 | 10627841 | 10627842 | cg25946304 |
| chr19 | 10748778 | 10748779 | cg26847100 |
| chr19 | 10823761 | 10823762 | cg02796790 |
| chr19 | 10855405 | 10855406 | cg09530594 |
| chr19 | 11093828 | 11093829 | cg08151828 |
| chr19 | 11517079 | 11517080 | cg24937727 |
| chr19 | 11649571 | 11649572 | cg01045132 |
| chr19 | 12036981 | 12036982 | cg15570966 |
| chr19 | 12145947 | 12145948 | cg27344791 |
| chr19 | 13317326 | 13317327 | cg03242834 |
| chr19 | 13947212 | 13947213 | cg08198483 |
| chr19 | 13947229 | 13947230 | cg16572540 |
| chr19 | 13947262 | 13947263 | cg19357865 |
| chr19 | 13993318 | 13993319 | cg07606495 |
| chr19 | 14064872 | 14064873 | cg10609371 |
| chr19 | 14089284 | 14089285 | cg06905453 |
| chr19 | 14115611 | 14115612 | cg18109874 |
| chr19 | 14142047 | 14142048 | cg04755674 |
| chr19 | 14142201 | 14142202 | cg09538921 |
| chr19 | 14593766 | 14593767 | cg16905207 |
| chr19 | 15367059 | 15367060 | cg14548611 |
| chr19 | 15375319 | 15375320 | cg04152610 |
| chr19 | 16284256 | 16284257 | cg03169836 |

| | | | |
|---|---|---|---|
| chr12 | 64355151 | 64355152 | cg14122749 |
| chr12 | 64415282 | 64415283 | cg05508355 |
| chr12 | 65044027 | 65044028 | cg05942111 |
| chr12 | 65060814 | 65060815 | cg15345743 |
| chr12 | 65676662 | 65676663 | cg20315554 |
| chr12 | 66251661 | 66251662 | cg17131837 |
| chr12 | 67073266 | 67073267 | cg22912610 |
| chr12 | 68150177 | 68150178 | cg02496507 |
| chr12 | 68751981 | 68751982 | cg08031211 |
| chr12 | 69002178 | 69002179 | cg04697624 |
| chr12 | 69357333 | 69357334 | cg02266731 |
| chr12 | 69485428 | 69485429 | cg09307431 |
| chr12 | 69637167 | 69637168 | cg26873338 |
| chr12 | 69902497 | 69902498 | cg07295520 |
| chr12 | 69909397 | 69909398 | cg19563525 |
| chr12 | 70329240 | 70329241 | cg14125530 |
| chr12 | 70463309 | 70463310 | cg16797818 |
| chr12 | 70645360 | 70645361 | cg19390271 |
| chr12 | 71837175 | 71837176 | cg15909056 |
| chr12 | 72083379 | 72083380 | cg12552402 |
| chr12 | 73057369 | 73057370 | cg14905634 |
| chr12 | 76644561 | 76644562 | cg14808689 |
| chr12 | 76927207 | 76927208 | cg16165258 |
| chr12 | 77456902 | 77456903 | cg18031663 |
| chr12 | 78635247 | 78635248 | cg14847917 |
| chr12 | 79693302 | 79693303 | cg25214900 |
| chr12 | 79812901 | 79812902 | cg13320585 |
| chr12 | 81189688 | 81189689 | cg09145629 |
| chr12 | 86240461 | 86240462 | cg03707749 |
| chr12 | 88186400 | 88186401 | cg07725829 |
| chr12 | 88784937 | 88784938 | cg10516080 |
| chr12 | 89619329 | 89619330 | cg06655216 |
| chr12 | 89744150 | 89744151 | cg27365701 |
| chr12 | 89749080 | 89749081 | cg01093028 |
| chr12 | 89923541 | 89923542 | cg18474081 |
| chr12 | 89968675 | 89968676 | cg16962463 |
| chr12 | 90177012 | 90177013 | cg15282731 |
| chr12 | 90393807 | 90393808 | cg21444421 |
| chr12 | 92464328 | 92464329 | cg14995202 |
| chr12 | 92524968 | 92524969 | cg05976481 |
| chr12 | 92543184 | 92543185 | cg02587993 |
| chr12 | 92814084 | 92814085 | cg11881599 |
| chr12 | 92955405 | 92955406 | cg20451050 |
| chr12 | 93454991 | 93454992 | cg12254517 |
| chr12 | 93809819 | 93809820 | cg08336604 |
| chr12 | 93817346 | 93817347 | cg23815787 |
| chr12 | 94699141 | 94699142 | cg07200939 |
| chr12 | 95541112 | 95541113 | cg10917910 |
| chr12 | 96720813 | 96720814 | cg04741118 |
| chr12 | 96895446 | 96895447 | cg06102602 |
| chr12 | 97951712 | 97951713 | cg27109238 |
| chr12 | 98577192 | 98577193 | cg02416536 |
| chr12 | 98926723 | 98926724 | cg09840122 |
| chr12 | 99029458 | 99029459 | cg06353075 |
| chr12 | 99090758 | 99090759 | cg07278047 |
| chr12 | 100579233 | 100579234 | cg16217794 |

| | | | |
|---|---|---|---|
| chr19 | 16284421 | 16284422 | cg02900183 |
| chr19 | 16466636 | 16466637 | cg11013793 |
| chr19 | 16528319 | 16528320 | cg11414610 |
| chr19 | 16985068 | 16985069 | cg07826747 |
| chr19 | 17219073 | 17219074 | cg19418318 |
| chr19 | 17350726 | 17350727 | cg01303222 |
| chr19 | 17407198 | 17407199 | cg17941109 |
| chr19 | 17626408 | 17626409 | cg22546818 |
| chr19 | 17831720 | 17831721 | cg06649165 |
| chr19 | 17861017 | 17861018 | cg26307105 |
| chr19 | 17881640 | 17881641 | cg00395894 |
| chr19 | 18404194 | 18404195 | cg06442294 |
| chr19 | 18545441 | 18545442 | cg22161383 |
| chr19 | 18549723 | 18549724 | cg06953865 |
| chr19 | 18795207 | 18795208 | cg25561382 |
| chr19 | 19312540 | 19312541 | cg05302915 |
| chr19 | 19644143 | 19644144 | cg16640973 |
| chr19 | 30100831 | 30100832 | cg26963285 |
| chr19 | 30155590 | 30155591 | cg08445687 |
| chr19 | 30315074 | 30315075 | cg05236705 |
| chr19 | 33470852 | 33470853 | cg22062383 |
| chr19 | 35740509 | 35740510 | cg14747903 |
| chr19 | 35772917 | 35772918 | cg02131995 |
| chr19 | 36399346 | 36399347 | cg21495704 |
| chr19 | 36450896 | 36450897 | cg01519784 |
| chr19 | 38571163 | 38571164 | cg25995870 |
| chr19 | 38681615 | 38681616 | cg02268510 |
| chr19 | 38803369 | 38803370 | cg02993606 |
| chr19 | 39217721 | 39217722 | cg25383568 |
| chr19 | 39217856 | 39217857 | cg04517274 |
| chr19 | 39361447 | 39361448 | cg02812809 |
| chr19 | 39368345 | 39368346 | cg23022999 |
| chr19 | 39369353 | 39369354 | cg15581429 |
| chr19 | 39827384 | 39827385 | cg25599012 |
| chr19 | 40195028 | 40195029 | cg26274090 |
| chr19 | 40410325 | 40410326 | cg05010309 |
| chr19 | 40422963 | 40422964 | cg17499324 |
| chr19 | 40873973 | 40873974 | cg20690695 |
| chr19 | 41225005 | 41225006 | cg21869046 |
| chr19 | 41281080 | 41281081 | cg25152942 |
| chr19 | 41815095 | 41815096 | cg27530093 |
| chr19 | 41873930 | 41873931 | cg13736811 |
| chr19 | 42392337 | 42392338 | cg07796002 |
| chr19 | 42409117 | 42409118 | cg05503433 |
| chr19 | 42900540 | 42900541 | cg03731797 |
| chr19 | 44009353 | 44009354 | cg17121205 |
| chr19 | 45445486 | 45445487 | cg04347059 |
| chr19 | 45461093 | 45461094 | cg22565251 |
| chr19 | 45512122 | 45512123 | cg18788725 |
| chr19 | 46147548 | 46147549 | cg00859314 |
| chr19 | 47123947 | 47123948 | cg19043207 |
| chr19 | 47125648 | 47125649 | cg15430294 |
| chr19 | 47151769 | 47151770 | cg20120512 |
| chr19 | 49135710 | 49135711 | cg15439033 |
| chr19 | 49564932 | 49564933 | cg07470708 |
| chr19 | 49689673 | 49689674 | cg01997599 |

| | | | |
|---|---|---|---|
| chr12 | 102874286 | 102874287 | cg01305421 |
| chr12 | 102875301 | 102875302 | cg25163611 |
| chr12 | 104213449 | 104213450 | cg19312404 |
| chr12 | 104693930 | 104693931 | cg19722698 |
| chr12 | 105114162 | 105114163 | cg07161429 |
| chr12 | 105114399 | 105114400 | cg09237846 |
| chr12 | 106630712 | 106630713 | cg01595262 |
| chr12 | 107259568 | 107259569 | cg17138415 |
| chr12 | 107836508 | 107836509 | cg11362622 |
| chr12 | 108078835 | 108078836 | cg25345738 |
| chr12 | 109025846 | 109025847 | cg25165932 |
| chr12 | 109072293 | 109072294 | cg09679862 |
| chr12 | 109186856 | 109186857 | cg26663772 |
| chr12 | 109250438 | 109250439 | cg25632883 |
| chr12 | 109250515 | 109250516 | cg24698536 |
| chr12 | 109250545 | 109250546 | cg12492906 |
| chr12 | 109491989 | 109491990 | cg03124318 |
| chr12 | 110034952 | 110034953 | cg12495807 |
| chr12 | 110440527 | 110440528 | cg02927618 |
| chr12 | 110885245 | 110885246 | cg09600411 |
| chr12 | 110986864 | 110986865 | cg15698107 |
| chr12 | 110993883 | 110993884 | cg05202616 |
| chr12 | 111016566 | 111016567 | cg08752433 |
| chr12 | 111128005 | 111128006 | cg20713761 |
| chr12 | 111128247 | 111128248 | cg07414574 |
| chr12 | 111176852 | 111176853 | cg13598358 |
| chr12 | 111805149 | 111805150 | cg03874539 |
| chr12 | 111873555 | 111873556 | cg26501657 |
| chr12 | 111892971 | 111892972 | cg19122402 |
| chr12 | 112550460 | 112550461 | cg04853995 |
| chr12 | 112860691 | 112860692 | cg24032304 |
| chr12 | 113343409 | 113343410 | cg09151598 |
| chr12 | 113343413 | 113343414 | cg17445535 |
| chr12 | 113343603 | 113343604 | cg25668626 |
| chr12 | 113370524 | 113370525 | cg17415650 |
| chr12 | 113375880 | 113375881 | cg22260958 |
| chr12 | 113415930 | 113415931 | cg11601443 |
| chr12 | 113519288 | 113519289 | cg00199753 |
| chr12 | 113527336 | 113527337 | cg10877430 |
| chr12 | 113542981 | 113542982 | cg21110505 |
| chr12 | 113547430 | 113547431 | cg26621780 |
| chr12 | 113569046 | 113569047 | cg06787724 |
| chr12 | 113599272 | 113599273 | cg22252931 |
| chr12 | 113645509 | 113645510 | cg25940517 |
| chr12 | 114985224 | 114985225 | cg02596960 |
| chr12 | 116757896 | 116757897 | cg08693490 |
| chr12 | 116881705 | 116881706 | cg18790325 |
| chr12 | 117539595 | 117539596 | cg02867305 |
| chr12 | 117580007 | 117580008 | cg06597431 |
| chr12 | 117627223 | 117627224 | cg01817067 |
| chr12 | 117630719 | 117630720 | cg12656611 |
| chr12 | 118142003 | 118142004 | cg06296759 |
| chr12 | 118405665 | 118405666 | cg00791253 |
| chr12 | 118583290 | 118583291 | cg08946014 |
| chr12 | 120543182 | 120543183 | cg06139513 |
| chr12 | 120975110 | 120975111 | cg07704181 |

| | | | |
|---|---|---|---|
| chr19 | 49828833 | 49828834 | cg10186564 |
| chr19 | 49927244 | 49927245 | cg08340482 |
| chr19 | 50005045 | 50005046 | cg03737367 |
| chr19 | 50056081 | 50056082 | cg05262463 |
| chr19 | 50062005 | 50062006 | cg18734095 |
| chr19 | 50190825 | 50190826 | cg23668335 |
| chr19 | 50192100 | 50192101 | cg00569418 |
| chr19 | 50412996 | 50412997 | cg22949004 |
| chr19 | 51104400 | 51104401 | cg04308443 |
| chr19 | 51382591 | 51382592 | cg00996764 |
| chr19 | 51400998 | 51400999 | cg03547958 |
| chr19 | 54604187 | 54604188 | cg18447740 |
| chr19 | 54692102 | 54692103 | cg00864768 |
| chr19 | 54849738 | 54849739 | cg01204985 |
| chr19 | 55174624 | 55174625 | cg05922591 |
| chr19 | 55995271 | 55995272 | cg12155450 |
| chr19 | 56047884 | 56047885 | cg00149708 |
| chr19 | 56200391 | 56200392 | cg19648552 |
| chr19 | 56612525 | 56612526 | cg14704483 |
| chr19 | 56914764 | 56914765 | cg08342619 |
| chr19 | 57989282 | 57989283 | cg23391173 |
| chr20 | 207803 | 207804 | cg26281051 |
| chr20 | 310001 | 310002 | cg15090834 |
| chr20 | 310635 | 310636 | cg12675996 |
| chr20 | 825634 | 825635 | cg16406078 |
| chr20 | 2780114 | 2780115 | cg26445608 |
| chr20 | 2852292 | 2852293 | cg23904377 |
| chr20 | 3024204 | 3024205 | cg07549715 |
| chr20 | 3232711 | 3232712 | cg04969203 |
| chr20 | 3746315 | 3746316 | cg05753799 |
| chr20 | 4804581 | 4804582 | cg19614321 |
| chr20 | 4990973 | 4990974 | cg10150176 |
| chr20 | 17552881 | 17552882 | cg02245875 |
| chr20 | 24930099 | 24930100 | cg09971811 |
| chr20 | 25057444 | 25057445 | cg00699986 |
| chr20 | 30060884 | 30060885 | cg02612927 |
| chr20 | 30073537 | 30073538 | cg02991085 |
| chr20 | 30073546 | 30073547 | cg25502144 |
| chr20 | 32254216 | 32254217 | cg12710480 |
| chr20 | 32266794 | 32266795 | cg14030719 |
| chr20 | 32579442 | 32579443 | cg00782275 |
| chr20 | 32856747 | 32856748 | cg19218509 |
| chr20 | 33128552 | 33128553 | cg20980879 |
| chr20 | 33299610 | 33299611 | cg16352321 |
| chr20 | 34605662 | 34605663 | cg09760574 |
| chr20 | 35089174 | 35089175 | cg12492027 |
| chr20 | 35092466 | 35092467 | cg15730121 |
| chr20 | 35220415 | 35220416 | cg10566581 |
| chr20 | 35274281 | 35274282 | cg21565575 |
| chr20 | 35493378 | 35493379 | cg06477069 |
| chr20 | 35494835 | 35494836 | cg09273716 |
| chr20 | 35504064 | 35504065 | cg19448292 |
| chr20 | 35504371 | 35504372 | cg09241885 |
| chr20 | 35504553 | 35504554 | cg18062064 |
| chr20 | 36153865 | 36153866 | cg03226872 |
| chr20 | 39765163 | 39765164 | cg18347630 |

| | | | |
|---|---|---|---|
| chr12 | 121570752 | 121570753 | cg00364287 |
| chr12 | 121678720 | 121678721 | cg04615865 |
| chr12 | 121893668 | 121893669 | cg00707814 |
| chr12 | 121971790 | 121971791 | cg08461425 |
| chr12 | 121976833 | 121976834 | cg21423404 |
| chr12 | 122019006 | 122019007 | cg07793808 |
| chr12 | 122019031 | 122019032 | cg07020001 |
| chr12 | 122019760 | 122019761 | cg26509318 |
| chr12 | 122020166 | 122020167 | cg21340621 |
| chr12 | 122716170 | 122716171 | cg12954334 |
| chr12 | 123007555 | 123007556 | cg08407408 |
| chr12 | 123009494 | 123009495 | cg23029597 |
| chr12 | 123215010 | 123215011 | cg00357958 |
| chr12 | 123215119 | 123215120 | cg22534509 |
| chr12 | 123355562 | 123355563 | cg16853770 |
| chr12 | 123357128 | 123357129 | cg11599718 |
| chr12 | 123379308 | 123379309 | cg22608507 |
| chr12 | 123469669 | 123469670 | cg09931783 |
| chr12 | 123591123 | 123591124 | cg01973483 |
| chr12 | 123599873 | 123599874 | cg19515108 |
| chr12 | 123707825 | 123707826 | cg21805179 |
| chr12 | 123707827 | 123707828 | cg07732037 |
| chr12 | 123713553 | 123713554 | cg11143193 |
| chr12 | 123753212 | 123753213 | cg14844236 |
| chr12 | 123753272 | 123753273 | cg20181887 |
| chr12 | 123892724 | 123892725 | cg06160572 |
| chr12 | 123918498 | 123918499 | cg09694527 |
| chr12 | 123921688 | 123921689 | cg05861255 |
| chr12 | 124948068 | 124948069 | cg03406367 |
| chr12 | 124971861 | 124971862 | cg02859042 |
| chr12 | 124990897 | 124990898 | cg09825309 |
| chr12 | 124990942 | 124990943 | cg22700848 |
| chr12 | 124991087 | 124991088 | cg04403415 |
| chr12 | 124991139 | 124991140 | cg27179101 |
| chr12 | 125048224 | 125048225 | cg19660243 |
| chr12 | 125223650 | 125223651 | cg09740450 |
| chr12 | 125223679 | 125223680 | cg03117611 |
| chr12 | 126105704 | 126105705 | cg10319331 |
| chr12 | 129283486 | 129283487 | cg21830849 |
| chr12 | 129300720 | 129300721 | cg17288256 |
| chr12 | 129304956 | 129304957 | cg19717347 |
| chr12 | 130961479 | 130961480 | cg06134790 |
| chr12 | 132259779 | 132259780 | cg06555661 |
| chr12 | 132259869 | 132259870 | cg13621113 |
| chr12 | 132549361 | 132549362 | cg25708790 |
| chr12 | 132637860 | 132637861 | cg02578709 |
| chr12 | 132637988 | 132637989 | cg13031551 |
| chr12 | 132639041 | 132639042 | cg16720971 |
| chr12 | 133028201 | 133028202 | cg25455773 |
| chr12 | 133055588 | 133055589 | cg06624951 |
| chr12 | 133101587 | 133101588 | cg19510206 |
| chr12 | 133172465 | 133172466 | cg16946687 |
| chr12 | 133342347 | 133342348 | cg06839896 |
| chr13 | 20420028 | 20420029 | cg26262576 |
| chr13 | 20754025 | 20754026 | cg08583570 |
| chr13 | 20994066 | 20994067 | cg11970797 |

| | | | |
|---|---|---|---|
| chr20 | 39767358 | 39767359 | cg07893801 |
| chr20 | 39767490 | 39767491 | cg06457011 |
| chr20 | 42344913 | 42344914 | cg15099490 |
| chr20 | 42350482 | 42350483 | cg20774413 |
| chr20 | 42350667 | 42350668 | cg21426218 |
| chr20 | 42354839 | 42354840 | cg24118151 |
| chr20 | 42539946 | 42539947 | cg21293933 |
| chr20 | 42543034 | 42543035 | cg00789528 |
| chr20 | 42574624 | 42574625 | cg24403644 |
| chr20 | 42697667 | 42697668 | cg16970323 |
| chr20 | 43919338 | 43919339 | cg07095818 |
| chr20 | 44030646 | 44030647 | cg05072995 |
| chr20 | 44330621 | 44330622 | cg17890298 |
| chr20 | 45947025 | 45947026 | cg03082779 |
| chr20 | 45947123 | 45947124 | cg02341503 |
| chr20 | 46415744 | 46415745 | cg12238291 |
| chr20 | 47862976 | 47862977 | cg14468280 |
| chr20 | 48549978 | 48549979 | cg15310506 |
| chr20 | 48626621 | 48626622 | cg06260709 |
| chr20 | 50110833 | 50110834 | cg18740175 |
| chr20 | 50719777 | 50719778 | cg12097791 |
| chr20 | 52198164 | 52198165 | cg07617814 |
| chr20 | 52277572 | 52277573 | cg00722350 |
| chr20 | 54986793 | 54986794 | cg00387658 |
| chr20 | 54986888 | 54986889 | cg26712809 |
| chr20 | 54987020 | 54987021 | cg23676369 |
| chr20 | 54987662 | 54987663 | cg24519157 |
| chr20 | 55959405 | 55959406 | cg25225756 |
| chr20 | 55982844 | 55982845 | cg15636859 |
| chr20 | 55982978 | 55982979 | cg12255554 |
| chr20 | 57486076 | 57486077 | cg26452915 |
| chr20 | 57578329 | 57578330 | cg27146036 |
| chr20 | 61290162 | 61290163 | cg10601171 |
| chr20 | 61468123 | 61468124 | cg08021508 |
| chr20 | 61491699 | 61491700 | cg01412764 |
| chr20 | 61536649 | 61536650 | cg19615731 |
| chr20 | 61825707 | 61825708 | cg01082229 |
| chr20 | 61848099 | 61848100 | cg24092253 |
| chr20 | 62204908 | 62204909 | cg01190666 |
| chr20 | 62282182 | 62282183 | cg00121366 |
| chr20 | 62369956 | 62369957 | cg06226150 |
| chr20 | 62370310 | 62370311 | cg22961457 |
| chr20 | 62437033 | 62437034 | cg23831891 |
| chr20 | 62437035 | 62437036 | cg21291985 |
| chr20 | 62574023 | 62574024 | cg01073332 |
| chr20 | 62574092 | 62574093 | cg21221377 |
| chr20 | 62583946 | 62583947 | cg03196440 |
| chr20 | 62738880 | 62738881 | cg03102788 |
| chr21 | 30675125 | 30675126 | cg07083941 |
| chr21 | 33034963 | 33034964 | cg19949530 |
| chr21 | 34774882 | 34774883 | cg22669060 |
| chr21 | 35746798 | 35746799 | cg15125868 |
| chr21 | 35898717 | 35898718 | cg21301258 |
| chr21 | 35898975 | 35898976 | cg05156137 |
| chr21 | 36412119 | 36412120 | cg05644223 |
| chr21 | 37695738 | 37695739 | cg26473301 |

| | | | | | | | |
|---|---|---|---|---|---|---|---|
| chr13 | 21276233 | 21276234 | cg10236247 | chr21 | 40557788 | 40557789 | cg13686331 |
| chr13 | 22560659 | 22560660 | cg23976981 | chr21 | 42679822 | 42679823 | cg14179031 |
| chr13 | 24614176 | 24614177 | cg18442992 | chr21 | 42791867 | 42791868 | cg16785077 |
| chr13 | 24650209 | 24650210 | cg25199372 | chr21 | 43347995 | 43347996 | cg19390317 |
| chr13 | 24798133 | 24798134 | cg27026786 | chr21 | 43348008 | 43348009 | cg09148268 |
| chr13 | 24807902 | 24807903 | cg08010865 | chr21 | 43619310 | 43619311 | cg20214535 |
| chr13 | 28597227 | 28597228 | cg16639691 | chr21 | 43619510 | 43619511 | cg05046272 |
| chr13 | 28630662 | 28630663 | cg16936846 | chr21 | 44832350 | 44832351 | cg16708938 |
| chr13 | 28975690 | 28975691 | cg22341788 | chr21 | 45199094 | 45199095 | cg09468832 |
| chr13 | 29177556 | 29177557 | cg15806588 | chr21 | 45224678 | 45224679 | cg10377299 |
| chr13 | 30421866 | 30421867 | cg08751352 | chr21 | 45399400 | 45399401 | cg04312999 |
| chr13 | 31004719 | 31004720 | cg01511901 | chr21 | 45558301 | 45558302 | cg21705505 |
| chr13 | 31256790 | 31256791 | cg21030160 | chr21 | 45581008 | 45581009 | cg03738656 |
| chr13 | 31506270 | 31506271 | cg05888037 | chr21 | 45658683 | 45658684 | cg06033443 |
| chr13 | 31949086 | 31949087 | cg08543015 | chr21 | 46505130 | 46505131 | cg21913632 |
| chr13 | 32524761 | 32524762 | cg00648883 | chr21 | 46523729 | 46523730 | cg21024676 |
| chr13 | 32575905 | 32575906 | cg22593432 | chr21 | 47561781 | 47561782 | cg10063047 |
| chr13 | 32596585 | 32596586 | cg08514385 | chr21 | 48056967 | 48056968 | cg22517527 |
| chr13 | 33105476 | 33105477 | cg05397490 | chr22 | 17569018 | 17569019 | cg16389078 |
| chr13 | 34129205 | 34129206 | cg15391555 | chr22 | 18258640 | 18258641 | cg05481243 |
| chr13 | 37064804 | 37064805 | cg24598495 | chr22 | 18258664 | 18258665 | cg05647602 |
| chr13 | 41055209 | 41055210 | cg04064327 | chr22 | 18266804 | 18266805 | cg03291548 |
| chr13 | 41055491 | 41055492 | cg26687842 | chr22 | 18506180 | 18506181 | cg00040575 |
| chr13 | 41055588 | 41055589 | cg00156995 | chr22 | 19845442 | 19845443 | cg19535685 |
| chr13 | 41064545 | 41064546 | cg01830053 | chr22 | 19898335 | 19898336 | cg09149541 |
| chr13 | 41161439 | 41161440 | cg10733317 | chr22 | 20306749 | 20306750 | cg08515285 |
| chr13 | 43563347 | 43563348 | cg09843907 | chr22 | 21210457 | 21210458 | cg08615820 |
| chr13 | 44706965 | 44706966 | cg14643399 | chr22 | 21318995 | 21318996 | cg05156805 |
| chr13 | 44786667 | 44786668 | cg13048723 | chr22 | 21985496 | 21985497 | cg14201946 |
| chr13 | 44953908 | 44953909 | cg01154353 | chr22 | 24089744 | 24089745 | cg08088970 |
| chr13 | 45912168 | 45912169 | cg04766834 | chr22 | 24185121 | 24185122 | cg22510462 |
| chr13 | 45970273 | 45970274 | cg18865990 | chr22 | 24188134 | 24188135 | cg21829328 |
| chr13 | 46276342 | 46276343 | cg05014103 | chr22 | 24823141 | 24823142 | cg04250930 |
| chr13 | 46277286 | 46277287 | cg23101680 | chr22 | 24824362 | 24824363 | cg04990420 |
| chr13 | 46743535 | 46743536 | cg02696428 | chr22 | 24824483 | 24824484 | cg27381549 |
| chr13 | 46956959 | 46956960 | cg19077761 | chr22 | 25115120 | 25115121 | cg02596486 |
| chr13 | 46962963 | 46962964 | cg08879684 | chr22 | 26822032 | 26822033 | cg17084653 |
| chr13 | 47130714 | 47130715 | cg06473578 | chr22 | 27014116 | 27014117 | cg02455383 |
| chr13 | 49168127 | 49168128 | cg22355224 | chr22 | 30003603 | 30003604 | cg17505852 |
| chr13 | 49384181 | 49384182 | cg10514207 | chr22 | 31479561 | 31479562 | cg00341732 |
| chr13 | 50653048 | 50653049 | cg06228828 | chr22 | 31492692 | 31492693 | cg04242821 |
| chr13 | 50950265 | 50950266 | cg10393508 | chr22 | 31643997 | 31643998 | cg00520380 |
| chr13 | 50961428 | 50961429 | cg14651504 | chr22 | 31675144 | 31675145 | cg07713946 |
| chr13 | 51375420 | 51375421 | cg27126505 | chr22 | 31682909 | 31682910 | cg07969918 |
| chr13 | 51995368 | 51995369 | cg24460268 | chr22 | 31736976 | 31736977 | cg11163555 |
| chr13 | 53454712 | 53454713 | cg08544712 | chr22 | 32024802 | 32024803 | cg00611469 |
| chr13 | 53602385 | 53602386 | cg08119452 | chr22 | 33670007 | 33670008 | cg10097977 |
| chr13 | 55475951 | 55475952 | cg16021678 | chr22 | 35695234 | 35695235 | cg14983728 |
| chr13 | 55576092 | 55576093 | cg20845494 | chr22 | 35695301 | 35695302 | cg21146537 |
| chr13 | 55794927 | 55794928 | cg27363745 | chr22 | 37495019 | 37495020 | cg18617091 |
| chr13 | 58299015 | 58299016 | cg07629125 | chr22 | 37499649 | 37499650 | cg17745234 |
| chr13 | 59983940 | 59983941 | cg10550033 | chr22 | 37579522 | 37579523 | cg14264901 |
| chr13 | 61753592 | 61753593 | cg15946856 | chr22 | 37608611 | 37608612 | cg03389133 |
| chr13 | 73211779 | 73211780 | cg09511815 | chr22 | 37608626 | 37608627 | cg04682967 |
| chr13 | 73715706 | 73715707 | cg01450807 | chr22 | 37654641 | 37654642 | cg00364281 |
| chr13 | 76445108 | 76445109 | cg19050940 | chr22 | 38615368 | 38615369 | cg18396403 |

| | | | |
|---|---|---|---|
| chr13 | 76957970 | 76957971 | cg27391982 |
| chr13 | 77387018 | 77387019 | cg25992016 |
| chr13 | 77567402 | 77567403 | cg22952787 |
| chr13 | 78467552 | 78467553 | cg23648516 |
| chr13 | 78551078 | 78551079 | cg13937208 |
| chr13 | 79360870 | 79360871 | cg26856578 |
| chr13 | 79977499 | 79977500 | cg04261072 |
| chr13 | 80315399 | 80315400 | cg06568438 |
| chr13 | 80612156 | 80612157 | cg05317600 |
| chr13 | 81018323 | 81018324 | cg13054007 |
| chr13 | 86298441 | 86298442 | cg04814317 |
| chr13 | 89037260 | 89037261 | cg06531870 |
| chr13 | 95132211 | 95132212 | cg09189889 |
| chr13 | 95933097 | 95933098 | cg16708465 |
| chr13 | 98154591 | 98154592 | cg26298599 |
| chr13 | 99101247 | 99101248 | cg25448281 |
| chr13 | 99101389 | 99101390 | cg09809402 |
| chr13 | 99136580 | 99136581 | cg19300307 |
| chr13 | 99195819 | 99195820 | cg13222768 |
| chr13 | 99736304 | 99736305 | cg27462475 |
| chr13 | 99960369 | 99960370 | cg09626634 |
| chr13 | 99963609 | 99963610 | cg11667400 |
| chr13 | 100008200 | 100008201 | cg18560638 |
| chr13 | 100544747 | 100544748 | cg18530956 |
| chr13 | 100886412 | 100886413 | cg25084760 |
| chr13 | 101241225 | 101241226 | cg27248148 |
| chr13 | 103720291 | 103720292 | cg09125550 |
| chr13 | 107159723 | 107159724 | cg02441149 |
| chr13 | 108817948 | 108817949 | cg07743730 |
| chr13 | 108921052 | 108921053 | cg09646392 |
| chr13 | 110412178 | 110412179 | cg21601493 |
| chr13 | 111127505 | 111127506 | cg24995674 |
| chr13 | 111295728 | 111295729 | cg13491737 |
| chr13 | 111318022 | 111318023 | cg14677983 |
| chr13 | 111565111 | 111565112 | cg15855924 |
| chr13 | 111699131 | 111699132 | cg09588074 |
| chr13 | 112324767 | 112324768 | cg02530656 |
| chr13 | 112633175 | 112633176 | cg25384556 |
| chr13 | 113000032 | 113000033 | cg08296665 |
| chr13 | 113328979 | 113328980 | cg10125399 |
| chr13 | 113348722 | 113348723 | cg15980707 |
| chr13 | 113422497 | 113422498 | cg09714789 |
| chr13 | 113423268 | 113423269 | cg18748064 |
| chr13 | 113425536 | 113425537 | cg17411031 |
| chr13 | 113428094 | 113428095 | cg08162124 |
| chr13 | 113428172 | 113428173 | cg01831771 |
| chr13 | 114786825 | 114786826 | cg02588809 |
| chr13 | 114786850 | 114786851 | cg21017641 |
| chr13 | 114786887 | 114786888 | cg23013564 |
| chr13 | 114786904 | 114786905 | cg10318725 |
| chr13 | 114790112 | 114790113 | cg21236457 |
| chr13 | 114825323 | 114825324 | cg12474090 |
| chr13 | 114829733 | 114829734 | cg18020065 |
| chr13 | 114866221 | 114866222 | cg25324786 |
| chr13 | 114880831 | 114880832 | cg11288831 |
| chr13 | 114883769 | 114883770 | cg16087432 |

| | | | |
|---|---|---|---|
| chr22 | 38710725 | 38710726 | cg03228516 |
| chr22 | 39154591 | 39154592 | cg05949181 |
| chr22 | 39436183 | 39436184 | cg10016348 |
| chr22 | 39471675 | 39471676 | cg26082280 |
| chr22 | 39472544 | 39472545 | cg13256546 |
| chr22 | 39477953 | 39477954 | cg06398754 |
| chr22 | 39492189 | 39492190 | cg06229674 |
| chr22 | 39494175 | 39494176 | cg24424889 |
| chr22 | 39547181 | 39547182 | cg12193277 |
| chr22 | 39548131 | 39548132 | cg23124451 |
| chr22 | 40139626 | 40139627 | cg11757326 |
| chr22 | 40589892 | 40589893 | cg04136484 |
| chr22 | 41463770 | 41463771 | cg22822910 |
| chr22 | 41845659 | 41845660 | cg26643870 |
| chr22 | 42612320 | 42612321 | cg20051177 |
| chr22 | 42947198 | 42947199 | cg21425232 |
| chr22 | 43036734 | 43036735 | cg02935298 |
| chr22 | 43044071 | 43044072 | cg18535283 |
| chr22 | 44568913 | 44568914 | cg03903451 |
| chr22 | 44576268 | 44576269 | cg14942952 |
| chr22 | 45124555 | 45124556 | cg22018086 |
| chr22 | 46443395 | 46443396 | cg00209037 |
| chr22 | 46934981 | 46934982 | cg11533712 |
| chr22 | 47052970 | 47052971 | cg00319157 |
| chr22 | 47057371 | 47057372 | cg05002837 |
| chr22 | 47069632 | 47069633 | cg13788891 |
| chr22 | 47135258 | 47135259 | cg16154810 |
| chr22 | 47191940 | 47191941 | cg13554549 |
| chr22 | 47423986 | 47423987 | cg07397612 |
| chr22 | 47559425 | 47559426 | cg14550778 |
| chr22 | 50171827 | 50171828 | cg26073959 |
| chr22 | 50360628 | 50360629 | cg23762465 |
| chr22 | 50451054 | 50451055 | cg14278345 |
| chr22 | 50451108 | 50451109 | cg15565576 |
| chr22 | 50451151 | 50451152 | cg26206185 |
| chr22 | 50457208 | 50457209 | cg20792408 |
| chr22 | 50603005 | 50603006 | cg17837500 |
| chr22 | 50741674 | 50741675 | cg03149749 |
| chr22 | 50883366 | 50883367 | cg05152764 |
| chr22 | 50986813 | 50986814 | cg18533225 |
| chr22 | 51017067 | 51017068 | cg00983520 |
| chr22 | 51171263 | 51171264 | cg09635994 |
| chr22 | 51223343 | 51223344 | cg07660283 |

| | | | |
|---|---|---|---|
| chr13 | 114890566 | 114890567 | cg02422603 |
| chr13 | 114905640 | 114905641 | cg04388548 |
| chr13 | 114909333 | 114909334 | cg05799596 |
| chr13 | 114912269 | 114912270 | cg10753283 |
| chr13 | 114917199 | 114917200 | cg25808809 |
| chr13 | 114917223 | 114917224 | cg16206504 |
| chr14 | 21212718 | 21212719 | cg10239638 |
| chr14 | 21216325 | 21216326 | cg19906534 |
| chr14 | 21270407 | 21270408 | cg13787135 |
| chr14 | 21288715 | 21288716 | cg13061910 |
| chr14 | 21535394 | 21535395 | cg09295202 |
| chr14 | 21738473 | 21738474 | cg13673563 |
| chr14 | 22292347 | 22292348 | cg01010091 |
| chr14 | 22433693 | 22433694 | cg12072611 |
| chr14 | 22945457 | 22945458 | cg20073893 |
| chr14 | 22948498 | 22948499 | cg07532405 |
| chr14 | 22951241 | 22951242 | cg00986580 |
| chr14 | 22953113 | 22953114 | cg20976833 |
| chr14 | 22958399 | 22958400 | cg05011265 |
| chr14 | 22958402 | 22958403 | cg16130238 |
| chr14 | 22963867 | 22963868 | cg08198540 |
| chr14 | 22969695 | 22969696 | cg16302916 |
| chr14 | 22973366 | 22973367 | cg01536878 |
| chr14 | 22975248 | 22975249 | cg04284196 |
| chr14 | 22975384 | 22975385 | cg02867735 |
| chr14 | 22975486 | 22975487 | cg13252730 |
| chr14 | 22975521 | 22975522 | cg17311865 |
| chr14 | 22978073 | 22978074 | cg23198262 |
| chr14 | 22978195 | 22978196 | cg01797899 |
| chr14 | 22978729 | 22978730 | cg24065807 |
| chr14 | 22979162 | 22979163 | cg20481655 |
| chr14 | 22985112 | 22985113 | cg15899743 |
| chr14 | 22989081 | 22989082 | cg18222759 |
| chr14 | 22989580 | 22989581 | cg26271776 |
| chr14 | 22989583 | 22989584 | cg00009306 |
| chr14 | 22991735 | 22991736 | cg26053876 |
| chr14 | 22993036 | 22993037 | cg10414881 |
| chr14 | 23005866 | 23005867 | cg26345203 |
| chr14 | 23018807 | 23018808 | cg14175932 |
| chr14 | 23284559 | 23284560 | cg02999224 |
| chr14 | 23284968 | 23284969 | cg15070897 |
| chr14 | 23289114 | 23289115 | cg16465430 |
| chr14 | 23298774 | 23298775 | cg07436208 |
| chr14 | 23305548 | 23305549 | cg13094752 |
| chr14 | 23305780 | 23305781 | cg09208010 |
| chr14 | 23305835 | 23305836 | cg08321366 |
| chr14 | 23305941 | 23305942 | cg10599444 |
| chr14 | 23305957 | 23305958 | cg05931439 |
| chr14 | 23525040 | 23525041 | cg03770548 |
| chr14 | 23772548 | 23772549 | cg13408636 |
| chr14 | 23772616 | 23772617 | cg23620184 |
| chr14 | 23772695 | 23772696 | cg23604876 |
| chr14 | 24044442 | 24044443 | cg19477689 |
| chr14 | 25079357 | 25079358 | cg02150910 |
| chr14 | 30142862 | 30142863 | cg07133994 |
| chr14 | 33832387 | 33832388 | cg24686983 |

| | | | |
|---|---|---|---|
| chr14 | 33949227 | 33949228 | cg10415314 |
| chr14 | 34320864 | 34320865 | cg12138450 |
| chr14 | 35810887 | 35810888 | cg18331348 |
| chr14 | 35871954 | 35871955 | cg00689225 |
| chr14 | 39734964 | 39734965 | cg24743156 |
| chr14 | 39735211 | 39735212 | cg13277939 |
| chr14 | 45133024 | 45133025 | cg10373147 |
| chr14 | 47908980 | 47908981 | cg23629393 |
| chr14 | 49360792 | 49360793 | cg15471388 |
| chr14 | 50506110 | 50506111 | cg14007559 |
| chr14 | 50809588 | 50809589 | cg19930116 |
| chr14 | 51256954 | 51256955 | cg08124634 |
| chr14 | 51263199 | 51263200 | cg02324789 |
| chr14 | 51291467 | 51291468 | cg16435627 |
| chr14 | 51299002 | 51299003 | cg01708014 |
| chr14 | 52171626 | 52171627 | cg07507518 |
| chr14 | 52333365 | 52333366 | cg22967080 |
| chr14 | 52351055 | 52351056 | cg27019717 |
| chr14 | 52487779 | 52487780 | cg20550154 |
| chr14 | 55203969 | 55203970 | cg07921759 |
| chr14 | 55222988 | 55222989 | cg13994599 |
| chr14 | 55226755 | 55226756 | cg21554616 |
| chr14 | 55333357 | 55333358 | cg16387581 |
| chr14 | 55569959 | 55569960 | cg09298313 |
| chr14 | 56225464 | 56225465 | cg09909740 |
| chr14 | 56231604 | 56231605 | cg25101184 |
| chr14 | 56289611 | 56289612 | cg26578726 |
| chr14 | 58543900 | 58543901 | cg05939089 |
| chr14 | 59038939 | 59038940 | cg14832378 |
| chr14 | 59742285 | 59742286 | cg12126243 |
| chr14 | 60179825 | 60179826 | cg10829004 |
| chr14 | 60195269 | 60195270 | cg15815318 |
| chr14 | 60633047 | 60633048 | cg18906360 |
| chr14 | 60770205 | 60770206 | cg07125412 |
| chr14 | 61712019 | 61712020 | cg03482600 |
| chr14 | 61972660 | 61972661 | cg18417061 |
| chr14 | 62396305 | 62396306 | cg04724807 |
| chr14 | 62414451 | 62414452 | cg04137405 |
| chr14 | 64677007 | 64677008 | cg00755751 |
| chr14 | 64687388 | 64687389 | cg08130912 |
| chr14 | 64926333 | 64926334 | cg05143420 |
| chr14 | 64974907 | 64974908 | cg01944137 |
| chr14 | 65020060 | 65020061 | cg04213522 |
| chr14 | 65623632 | 65623633 | cg08164191 |
| chr14 | 65881787 | 65881788 | cg21217221 |
| chr14 | 67653661 | 67653662 | cg02903275 |
| chr14 | 68648563 | 68648564 | cg14518185 |
| chr14 | 68714746 | 68714747 | cg25603108 |
| chr14 | 69263504 | 69263505 | cg10395868 |
| chr14 | 69264333 | 69264334 | cg00320765 |
| chr14 | 69280381 | 69280382 | cg20056212 |
| chr14 | 69284017 | 69284018 | cg02941385 |
| chr14 | 69472652 | 69472653 | cg17345741 |
| chr14 | 69604488 | 69604489 | cg04947533 |
| chr14 | 69730478 | 69730479 | cg04192663 |
| chr14 | 70086897 | 70086898 | cg12838007 |

| | | | |
|---|---|---|---|
| chr14 | 70104457 | 70104458 | cg14369329 |
| chr14 | 70116951 | 70116952 | cg00842095 |
| chr14 | 70152683 | 70152684 | cg23525708 |
| chr14 | 71107039 | 71107040 | cg03762817 |
| chr14 | 71128147 | 71128148 | cg01634544 |
| chr14 | 71131956 | 71131957 | cg04176452 |
| chr14 | 71712645 | 71712646 | cg05859308 |
| chr14 | 72990359 | 72990360 | cg05111779 |
| chr14 | 73358241 | 73358242 | cg22924250 |
| chr14 | 73756251 | 73756252 | cg08147389 |
| chr14 | 73934847 | 73934848 | cg07965566 |
| chr14 | 74178976 | 74178977 | cg10523105 |
| chr14 | 74179101 | 74179102 | cg09238801 |
| chr14 | 74179674 | 74179675 | cg27165794 |
| chr14 | 74220238 | 74220239 | cg16398761 |
| chr14 | 74227499 | 74227500 | cg03519879 |
| chr14 | 74230413 | 74230414 | cg23367119 |
| chr14 | 74251926 | 74251927 | cg16702313 |
| chr14 | 74766220 | 74766221 | cg05878073 |
| chr14 | 75478802 | 75478803 | cg18459040 |
| chr14 | 75489610 | 75489611 | cg15467112 |
| chr14 | 75886161 | 75886162 | cg22875410 |
| chr14 | 75936582 | 75936583 | cg04407490 |
| chr14 | 75936668 | 75936669 | cg01919011 |
| chr14 | 76002440 | 76002441 | cg01589587 |
| chr14 | 76818832 | 76818833 | cg10024505 |
| chr14 | 77489763 | 77489764 | cg15953461 |
| chr14 | 77493278 | 77493279 | cg24570624 |
| chr14 | 77495672 | 77495673 | cg01553004 |
| chr14 | 77504364 | 77504365 | cg24774357 |
| chr14 | 77504561 | 77504562 | cg13248574 |
| chr14 | 77961897 | 77961898 | cg21822146 |
| chr14 | 78907020 | 78907021 | cg09209257 |
| chr14 | 81426577 | 81426578 | cg12196847 |
| chr14 | 81938575 | 81938576 | cg10042132 |
| chr14 | 84014672 | 84014673 | cg24136340 |
| chr14 | 89804447 | 89804448 | cg18657988 |
| chr14 | 89895742 | 89895743 | cg01210589 |
| chr14 | 89910404 | 89910405 | cg21571793 |
| chr14 | 90087792 | 90087793 | cg02916283 |
| chr14 | 90851046 | 90851047 | cg16598679 |
| chr14 | 91720527 | 91720528 | cg27042278 |
| chr14 | 91751773 | 91751774 | cg17002328 |
| chr14 | 91836354 | 91836355 | cg07112604 |
| chr14 | 92960827 | 92960828 | cg05200313 |
| chr14 | 92981666 | 92981667 | cg02268192 |
| chr14 | 92991035 | 92991036 | cg01288337 |
| chr14 | 93206959 | 93206960 | cg09963856 |
| chr14 | 93500118 | 93500119 | cg01619562 |
| chr14 | 94423704 | 94423705 | cg19949550 |
| chr14 | 94423872 | 94423873 | cg03389164 |
| chr14 | 94423943 | 94423944 | cg03562528 |
| chr14 | 94857275 | 94857276 | cg24621042 |
| chr14 | 94944009 | 94944010 | cg27266824 |
| chr14 | 95722106 | 95722107 | cg05068951 |
| chr14 | 95826997 | 95826998 | cg14013080 |

| | | | |
|---|---|---|---|
| chr14 | 95830175 | 95830176 | cg04380681 |
| chr14 | 95983383 | 95983384 | cg05546763 |
| chr14 | 98536125 | 98536126 | cg11092157 |
| chr14 | 99183876 | 99183877 | cg04914574 |
| chr14 | 100068637 | 100068638 | cg04242464 |
| chr14 | 100540291 | 100540292 | cg14245199 |
| chr14 | 100611694 | 100611695 | cg16378015 |
| chr14 | 100614973 | 100614974 | cg01513978 |
| chr14 | 102310830 | 102310831 | cg19478371 |
| chr14 | 102312533 | 102312534 | cg02220617 |
| chr14 | 102446939 | 102446940 | cg21919445 |
| chr14 | 102647731 | 102647732 | cg05100846 |
| chr14 | 103088924 | 103088925 | cg16139785 |
| chr14 | 103265614 | 103265615 | cg00876272 |
| chr14 | 103272938 | 103272939 | cg27118298 |
| chr14 | 103294656 | 103294657 | cg26115667 |
| chr14 | 103416122 | 103416123 | cg04715649 |
| chr14 | 103416268 | 103416269 | cg09262100 |
| chr14 | 103468473 | 103468474 | cg13164814 |
| chr14 | 103911341 | 103911342 | cg23010693 |
| chr14 | 104094619 | 104094620 | cg23090046 |
| chr14 | 104228343 | 104228344 | cg22973515 |
| chr14 | 105147585 | 105147586 | cg01854676 |
| chr14 | 105173980 | 105173981 | cg05018513 |
| chr14 | 105184057 | 105184058 | cg23956771 |
| chr14 | 105184176 | 105184177 | cg23343291 |
| chr14 | 105236538 | 105236539 | cg13986355 |
| chr14 | 105263191 | 105263192 | cg13774825 |
| chr14 | 105263522 | 105263523 | cg05726935 |
| chr14 | 105371541 | 105371542 | cg16793370 |
| chr14 | 105779952 | 105779953 | cg16162970 |
| chr14 | 105796964 | 105796965 | cg06783197 |
| chr14 | 106003576 | 106003577 | cg21899542 |
| chr14 | 106037781 | 106037782 | cg25096107 |
| chr14 | 106039191 | 106039192 | cg06844496 |
| chr14 | 106040464 | 106040465 | cg10859844 |
| chr14 | 106090302 | 106090303 | cg11190240 |
| chr14 | 106110146 | 106110147 | cg19684990 |
| chr14 | 106215515 | 106215516 | cg13579030 |
| chr14 | 106240524 | 106240525 | cg02129962 |
| chr14 | 106306830 | 106306831 | cg13106215 |
| chr15 | 22936259 | 22936260 | cg17872861 |
| chr15 | 26047499 | 26047500 | cg15531450 |
| chr15 | 30106104 | 30106105 | cg06525993 |
| chr15 | 30216986 | 30216987 | cg12217971 |
| chr15 | 34332433 | 34332434 | cg10772974 |
| chr15 | 34628731 | 34628732 | cg10550166 |
| chr15 | 34628769 | 34628770 | cg22999786 |
| chr15 | 34628815 | 34628816 | cg27566947 |
| chr15 | 35999901 | 35999902 | cg00658704 |
| chr15 | 37338491 | 37338492 | cg05334416 |
| chr15 | 38192897 | 38192898 | cg16967068 |
| chr15 | 39568837 | 39568838 | cg26841157 |
| chr15 | 39796985 | 39796986 | cg13497315 |
| chr15 | 39886363 | 39886364 | cg13375187 |
| chr15 | 40080369 | 40080370 | cg14208041 |

| | | | |
|---|---|---|---|
| chr15 | 40330108 | 40330109 | cg20752233 |
| chr15 | 40348303 | 40348304 | cg14482712 |
| chr15 | 40398402 | 40398403 | cg07887985 |
| chr15 | 40509518 | 40509519 | cg18039797 |
| chr15 | 40538101 | 40538102 | cg04658699 |
| chr15 | 40599680 | 40599681 | cg16543027 |
| chr15 | 40631573 | 40631574 | cg12732548 |
| chr15 | 40759208 | 40759209 | cg15431934 |
| chr15 | 40803244 | 40803245 | cg27057329 |
| chr15 | 40846280 | 40846281 | cg17140476 |
| chr15 | 41058059 | 41058060 | cg19433807 |
| chr15 | 41185800 | 41185801 | cg19969694 |
| chr15 | 41775282 | 41775283 | cg11942439 |
| chr15 | 41850612 | 41850613 | cg24147582 |
| chr15 | 42102381 | 42102382 | cg17847044 |
| chr15 | 42193477 | 42193478 | cg17750114 |
| chr15 | 42209178 | 42209179 | cg10142436 |
| chr15 | 42411135 | 42411136 | cg01878366 |
| chr15 | 42568411 | 42568412 | cg21574244 |
| chr15 | 42749885 | 42749886 | cg12492087 |
| chr15 | 42843303 | 42843304 | cg10327976 |
| chr15 | 42888179 | 42888180 | cg01143068 |
| chr15 | 43387679 | 43387680 | cg23701314 |
| chr15 | 44120341 | 44120342 | cg00074771 |
| chr15 | 44579510 | 44579511 | cg00093643 |
| chr15 | 44956192 | 44956193 | cg25008614 |
| chr15 | 44968935 | 44968936 | cg02341571 |
| chr15 | 45027253 | 45027254 | cg10517312 |
| chr15 | 45028083 | 45028084 | cg22107533 |
| chr15 | 47734755 | 47734756 | cg05877048 |
| chr15 | 47967175 | 47967176 | cg03953246 |
| chr15 | 49462118 | 49462119 | cg18736186 |
| chr15 | 49462125 | 49462126 | cg25860737 |
| chr15 | 49462157 | 49462158 | cg08036668 |
| chr15 | 50180387 | 50180388 | cg19599368 |
| chr15 | 51632611 | 51632612 | cg05303280 |
| chr15 | 52394679 | 52394680 | cg05495005 |
| chr15 | 52470919 | 52470920 | cg03211327 |
| chr15 | 52587077 | 52587078 | cg06687489 |
| chr15 | 53001117 | 53001118 | cg26653392 |
| chr15 | 53422311 | 53422312 | cg27356115 |
| chr15 | 56537579 | 56537580 | cg03413355 |
| chr15 | 57414740 | 57414741 | cg24492778 |
| chr15 | 58758468 | 58758469 | cg20968717 |
| chr15 | 59514661 | 59514662 | cg25970356 |
| chr15 | 59595945 | 59595946 | cg15851882 |
| chr15 | 59659880 | 59659881 | cg02794990 |
| chr15 | 59662087 | 59662088 | cg25980168 |
| chr15 | 59730852 | 59730853 | cg05841929 |
| chr15 | 60688948 | 60688949 | cg14397690 |
| chr15 | 61491665 | 61491666 | cg05956150 |
| chr15 | 63807633 | 63807634 | cg11938386 |
| chr15 | 65151592 | 65151593 | cg17927313 |
| chr15 | 65661080 | 65661081 | cg24741685 |
| chr15 | 66097959 | 66097960 | cg03382306 |
| chr15 | 66125463 | 66125464 | cg25587091 |

| | | | |
|---|---|---|---|
| chr15 | 66350184 | 66350185 | cg10583322 |
| chr15 | 66652179 | 66652180 | cg04792910 |
| chr15 | 66790390 | 66790391 | cg16330965 |
| chr15 | 67007131 | 67007132 | cg16553757 |
| chr15 | 67022087 | 67022088 | cg01678084 |
| chr15 | 67078890 | 67078891 | cg09434667 |
| chr15 | 67814880 | 67814881 | cg22706883 |
| chr15 | 67904302 | 67904303 | cg24247231 |
| chr15 | 68865587 | 68865588 | cg24179719 |
| chr15 | 69084888 | 69084889 | cg25139636 |
| chr15 | 69099262 | 69099263 | cg20352178 |
| chr15 | 69860750 | 69860751 | cg14560670 |
| chr15 | 70087153 | 70087154 | cg14417896 |
| chr15 | 70317022 | 70317023 | cg24241192 |
| chr15 | 70317124 | 70317125 | cg02628483 |
| chr15 | 70340064 | 70340065 | cg02961472 |
| chr15 | 70387268 | 70387269 | cg09747445 |
| chr15 | 70528789 | 70528790 | cg11551560 |
| chr15 | 70690061 | 70690062 | cg04787775 |
| chr15 | 72564179 | 72564180 | cg23331961 |
| chr15 | 72651198 | 72651199 | cg24495350 |
| chr15 | 73975176 | 73975177 | cg04289575 |
| chr15 | 73979250 | 73979251 | cg10586317 |
| chr15 | 74699653 | 74699654 | cg18915456 |
| chr15 | 74945851 | 74945852 | cg14389122 |
| chr15 | 75091082 | 75091083 | cg08273635 |
| chr15 | 75091516 | 75091517 | cg00516515 |
| chr15 | 75093165 | 75093166 | cg27211576 |
| chr15 | 75136278 | 75136279 | cg13055199 |
| chr15 | 75322058 | 75322059 | cg06070324 |
| chr15 | 76016335 | 76016336 | cg00622552 |
| chr15 | 76355522 | 76355523 | cg17515837 |
| chr15 | 76630962 | 76630963 | cg26266708 |
| chr15 | 77359477 | 77359478 | cg05914442 |
| chr15 | 77411982 | 77411983 | cg05673287 |
| chr15 | 77695188 | 77695189 | cg26438942 |
| chr15 | 77711092 | 77711093 | cg17467968 |
| chr15 | 78463330 | 78463331 | cg14517390 |
| chr15 | 78526836 | 78526837 | cg27257408 |
| chr15 | 78527113 | 78527114 | cg13642872 |
| chr15 | 79048467 | 79048468 | cg22726373 |
| chr15 | 79134331 | 79134332 | cg25957301 |
| chr15 | 79238723 | 79238724 | cg18738367 |
| chr15 | 79243526 | 79243527 | cg17506061 |
| chr15 | 79296446 | 79296447 | cg02587988 |
| chr15 | 79296585 | 79296586 | cg22821300 |
| chr15 | 79297210 | 79297211 | cg21840434 |
| chr15 | 80325923 | 80325924 | cg04384912 |
| chr15 | 81596332 | 81596333 | cg19620333 |
| chr15 | 83619037 | 83619038 | cg19256368 |
| chr15 | 85143219 | 85143220 | cg20353653 |
| chr15 | 85294586 | 85294587 | cg18231690 |
| chr15 | 85431307 | 85431308 | cg22294181 |
| chr15 | 85534863 | 85534864 | cg07964527 |
| chr15 | 85552114 | 85552115 | cg09425164 |
| chr15 | 85973746 | 85973747 | cg02810291 |

| | | | |
|---|---|---|---|
| chr15 | 85987419 | 85987420 | cg14508021 |
| chr15 | 86167192 | 86167193 | cg12412575 |
| chr15 | 86192745 | 86192746 | cg13374432 |
| chr15 | 86336631 | 86336632 | cg20737812 |
| chr15 | 89168305 | 89168306 | cg20808613 |
| chr15 | 89180799 | 89180800 | cg00854314 |
| chr15 | 89181425 | 89181426 | cg12017722 |
| chr15 | 89181615 | 89181616 | cg17459635 |
| chr15 | 89181732 | 89181733 | cg19009305 |
| chr15 | 89879515 | 89879516 | cg01192051 |
| chr15 | 90066806 | 90066807 | cg00006867 |
| chr15 | 90539856 | 90539857 | cg08305808 |
| chr15 | 90548043 | 90548044 | cg09939948 |
| chr15 | 90569017 | 90569018 | cg26330304 |
| chr15 | 90604217 | 90604218 | cg16785520 |
| chr15 | 90605618 | 90605619 | cg00624799 |
| chr15 | 90608429 | 90608430 | cg15188623 |
| chr15 | 90703444 | 90703445 | cg17186132 |
| chr15 | 90727783 | 90727784 | cg26312071 |
| chr15 | 91395896 | 91395897 | cg27535502 |
| chr15 | 91426667 | 91426668 | cg24888049 |
| chr15 | 91428056 | 91428057 | cg05180856 |
| chr15 | 91429916 | 91429917 | cg20992439 |
| chr15 | 91477073 | 91477074 | cg11145826 |
| chr15 | 91536180 | 91536181 | cg14398957 |
| chr15 | 92428182 | 92428183 | cg00314343 |
| chr15 | 92459857 | 92459858 | cg19621065 |
| chr15 | 92580100 | 92580101 | cg26677406 |
| chr15 | 92667819 | 92667820 | cg01240823 |
| chr15 | 92673594 | 92673595 | cg12667732 |
| chr15 | 93115885 | 93115886 | cg01626493 |
| chr15 | 93424933 | 93424934 | cg06605053 |
| chr15 | 93450671 | 93450672 | cg08899105 |
| chr15 | 93457211 | 93457212 | cg26694290 |
| chr15 | 93814737 | 93814738 | cg19550034 |
| chr15 | 94803071 | 94803072 | cg18513919 |
| chr15 | 94840953 | 94840954 | cg12299554 |
| chr15 | 95889928 | 95889929 | cg03275459 |
| chr15 | 97316699 | 97316700 | cg25271525 |
| chr15 | 98199925 | 98199926 | cg06755296 |
| chr15 | 98507208 | 98507209 | cg18010014 |
| chr15 | 99227735 | 99227736 | cg09824900 |
| chr15 | 99789777 | 99789778 | cg13401703 |
| chr15 | 100886002 | 100886003 | cg09783969 |
| chr15 | 101591436 | 101591437 | cg07217653 |
| chr15 | 101708575 | 101708576 | cg13106758 |
| chr15 | 101715747 | 101715748 | cg12042714 |
| chr15 | 102216116 | 102216117 | cg11292721 |
| chr16 | 126397 | 126398 | cg03462322 |
| chr16 | 358591 | 358592 | cg26616258 |
| chr16 | 575420 | 575421 | cg02143778 |
| chr16 | 671171 | 671172 | cg06714655 |
| chr16 | 728084 | 728085 | cg06983322 |
| chr16 | 728470 | 728471 | cg26510178 |
| chr16 | 945795 | 945796 | cg03962950 |
| chr16 | 1308103 | 1308104 | cg05848579 |

| | | | |
|---|---|---|---|
| chr16 | 1353627 | 1353628 | cg06908857 |
| chr16 | 1389301 | 1389302 | cg01881308 |
| chr16 | 1394089 | 1394090 | cg09526712 |
| chr16 | 1477870 | 1477871 | cg04138881 |
| chr16 | 1507265 | 1507266 | cg05948101 |
| chr16 | 1665900 | 1665901 | cg02127888 |
| chr16 | 1670473 | 1670474 | cg08776660 |
| chr16 | 1988798 | 1988799 | cg00905101 |
| chr16 | 2210434 | 2210435 | cg00812236 |
| chr16 | 2254692 | 2254693 | cg08737236 |
| chr16 | 2302519 | 2302520 | cg09970941 |
| chr16 | 2345275 | 2345276 | cg09263316 |
| chr16 | 2559709 | 2559710 | cg02878544 |
| chr16 | 2581414 | 2581415 | cg17023419 |
| chr16 | 2823300 | 2823301 | cg06216428 |
| chr16 | 3021794 | 3021795 | cg04696391 |
| chr16 | 3117862 | 3117863 | cg08659421 |
| chr16 | 3597151 | 3597152 | cg16429499 |
| chr16 | 3740774 | 3740775 | cg06170683 |
| chr16 | 3782990 | 3782991 | cg04850976 |
| chr16 | 3873444 | 3873445 | cg05194552 |
| chr16 | 4300186 | 4300187 | cg10522979 |
| chr16 | 4700737 | 4700738 | cg04071866 |
| chr16 | 5092012 | 5092013 | cg05393086 |
| chr16 | 8767706 | 8767707 | cg02564313 |
| chr16 | 8814755 | 8814756 | cg06604249 |
| chr16 | 8941747 | 8941748 | cg08676173 |
| chr16 | 8986308 | 8986309 | cg01601926 |
| chr16 | 9048985 | 9048986 | cg03935096 |
| chr16 | 9051317 | 9051318 | cg07636761 |
| chr16 | 9055526 | 9055527 | cg10359807 |
| chr16 | 9190392 | 9190393 | cg08534055 |
| chr16 | 10868974 | 10868975 | cg03777414 |
| chr16 | 10989530 | 10989531 | cg05371498 |
| chr16 | 11012292 | 11012293 | cg07371747 |
| chr16 | 11181986 | 11181987 | cg04616529 |
| chr16 | 11194243 | 11194244 | cg03995533 |
| chr16 | 11480327 | 11480328 | cg26789698 |
| chr16 | 11591651 | 11591652 | cg02301079 |
| chr16 | 11681487 | 11681488 | cg07143635 |
| chr16 | 11764847 | 11764848 | cg00022235 |
| chr16 | 11812041 | 11812042 | cg04450037 |
| chr16 | 11834960 | 11834961 | cg06994787 |
| chr16 | 11876111 | 11876112 | cg08160706 |
| chr16 | 12071814 | 12071815 | cg08957564 |
| chr16 | 12072256 | 12072257 | cg06865119 |
| chr16 | 12637499 | 12637500 | cg00548009 |
| chr16 | 15583606 | 15583607 | cg03116016 |
| chr16 | 15740113 | 15740114 | cg09714184 |
| chr16 | 16083164 | 16083165 | cg09109383 |
| chr16 | 17458534 | 17458535 | cg05152300 |
| chr16 | 17553784 | 17553785 | cg04986899 |
| chr16 | 17561865 | 17561866 | cg04313565 |
| chr16 | 17562419 | 17562420 | cg16794579 |
| chr16 | 19182453 | 19182454 | cg02395637 |
| chr16 | 19239848 | 19239849 | cg09758796 |

| | | | |
|---|---|---|---|
| chr16 | 21608593 | 21608594 | cg27662505 |
| chr16 | 21650166 | 21650167 | cg07917978 |
| chr16 | 22039309 | 22039310 | cg02146383 |
| chr16 | 23144103 | 23144104 | cg06271652 |
| chr16 | 23504930 | 23504931 | cg00736406 |
| chr16 | 23691827 | 23691828 | cg06112560 |
| chr16 | 23962572 | 23962573 | cg03503785 |
| chr16 | 24087793 | 24087794 | cg09327847 |
| chr16 | 24999103 | 24999104 | cg01902704 |
| chr16 | 25005762 | 25005763 | cg05583636 |
| chr16 | 27237190 | 27237191 | cg06388418 |
| chr16 | 27246970 | 27246971 | cg10037913 |
| chr16 | 27322356 | 27322357 | cg08008352 |
| chr16 | 27412496 | 27412497 | cg02656594 |
| chr16 | 28193063 | 28193064 | cg00468395 |
| chr16 | 28886581 | 28886582 | cg03328388 |
| chr16 | 28913517 | 28913518 | cg02304156 |
| chr16 | 28995843 | 28995844 | cg16462073 |
| chr16 | 28995994 | 28995995 | cg00607627 |
| chr16 | 29296186 | 29296187 | cg08027123 |
| chr16 | 29604068 | 29604069 | cg02596508 |
| chr16 | 29673933 | 29673934 | cg09946623 |
| chr16 | 29674618 | 29674619 | cg03549705 |
| chr16 | 29674903 | 29674904 | cg16644366 |
| chr16 | 29674959 | 29674960 | cg08480068 |
| chr16 | 29674972 | 29674973 | cg26769927 |
| chr16 | 29675160 | 29675161 | cg26691434 |
| chr16 | 29675214 | 29675215 | cg10126903 |
| chr16 | 29676071 | 29676072 | cg01681367 |
| chr16 | 29690271 | 29690272 | cg02640604 |
| chr16 | 30126595 | 30126596 | cg05329317 |
| chr16 | 30756300 | 30756301 | cg08119498 |
| chr16 | 30770794 | 30770795 | cg01688936 |
| chr16 | 30906273 | 30906274 | cg00283876 |
| chr16 | 31119067 | 31119068 | cg05299836 |
| chr16 | 31119196 | 31119197 | cg08328483 |
| chr16 | 31123892 | 31123893 | cg01016533 |
| chr16 | 31216531 | 31216532 | cg08211301 |
| chr16 | 31451097 | 31451098 | cg03925821 |
| chr16 | 31539169 | 31539170 | cg05772125 |
| chr16 | 46893929 | 46893930 | cg08998657 |
| chr16 | 48109635 | 48109636 | cg05216542 |
| chr16 | 48351958 | 48351959 | cg06363417 |
| chr16 | 48413428 | 48413429 | cg08088511 |
| chr16 | 48468609 | 48468610 | cg03977385 |
| chr16 | 48533930 | 48533931 | cg01014262 |
| chr16 | 48559770 | 48559771 | cg09971653 |
| chr16 | 49869755 | 49869756 | cg01502811 |
| chr16 | 50218944 | 50218945 | cg10459187 |
| chr16 | 50278918 | 50278919 | cg06861037 |
| chr16 | 50715529 | 50715530 | cg27081230 |
| chr16 | 50730813 | 50730814 | cg26954174 |
| chr16 | 50730881 | 50730882 | cg18177814 |
| chr16 | 50743027 | 50743028 | cg04172533 |
| chr16 | 53145386 | 53145387 | cg04495891 |
| chr16 | 56456934 | 56456935 | cg06097580 |

| | | | |
|---|---|---|---|
| chr16 | 56650508 | 56650509 | cg04578397 |
| chr16 | 56946503 | 56946504 | cg02751693 |
| chr16 | 57017595 | 57017596 | cg27612129 |
| chr16 | 57022486 | 57022487 | cg08159663 |
| chr16 | 57023022 | 57023023 | cg07839457 |
| chr16 | 57080116 | 57080117 | cg09624807 |
| chr16 | 57147202 | 57147203 | cg09100343 |
| chr16 | 57294639 | 57294640 | cg03573390 |
| chr16 | 57391655 | 57391656 | cg04250732 |
| chr16 | 57415838 | 57415839 | cg06166915 |
| chr16 | 57918670 | 57918671 | cg02358664 |
| chr16 | 58155189 | 58155190 | cg02990302 |
| chr16 | 58768565 | 58768566 | cg13883681 |
| chr16 | 59363443 | 59363444 | cg06003388 |
| chr16 | 59790180 | 59790181 | cg07570113 |
| chr16 | 61854084 | 61854085 | cg00009553 |
| chr16 | 63219709 | 63219710 | cg02560364 |
| chr16 | 66285054 | 66285055 | cg09781650 |
| chr16 | 66360304 | 66360305 | cg08237521 |
| chr16 | 66585799 | 66585800 | cg09728392 |
| chr16 | 66876872 | 66876873 | cg06438797 |
| chr16 | 67345737 | 67345738 | cg05762671 |
| chr16 | 67580610 | 67580611 | cg01872947 |
| chr16 | 68001415 | 68001416 | cg08958168 |
| chr16 | 68027839 | 68027840 | cg10512278 |
| chr16 | 68123248 | 68123249 | cg02570644 |
| chr16 | 69105306 | 69105307 | cg27133493 |
| chr16 | 69200251 | 69200252 | cg08327038 |
| chr16 | 69499075 | 69499076 | cg04406620 |
| chr16 | 70463218 | 70463219 | cg04957757 |
| chr16 | 71914820 | 71914821 | cg26585899 |
| chr16 | 73067525 | 73067526 | cg27262054 |
| chr16 | 73078071 | 73078072 | cg03805182 |
| chr16 | 75654252 | 75654253 | cg07861456 |
| chr16 | 77709272 | 77709273 | cg02295341 |
| chr16 | 80265530 | 80265531 | cg03666439 |
| chr16 | 81512025 | 81512026 | cg01567051 |
| chr16 | 81682484 | 81682485 | cg07574924 |
| chr16 | 81743881 | 81743882 | cg00050271 |
| chr16 | 81818852 | 81818853 | cg06031301 |
| chr16 | 82474489 | 82474490 | cg07128660 |
| chr16 | 82671520 | 82671521 | cg02168291 |
| chr16 | 82735595 | 82735596 | cg02495250 |
| chr16 | 83882901 | 83882902 | cg02228294 |
| chr16 | 84118667 | 84118668 | cg26868097 |
| chr16 | 84166874 | 84166875 | cg00049380 |
| chr16 | 84172431 | 84172432 | cg04006133 |
| chr16 | 84627507 | 84627508 | cg05344293 |
| chr16 | 84628969 | 84628970 | cg03738384 |
| chr16 | 84629008 | 84629009 | cg06363968 |
| chr16 | 84637476 | 84637477 | cg03100906 |
| chr16 | 84745615 | 84745616 | cg05962793 |
| chr16 | 85019868 | 85019869 | cg09938443 |
| chr16 | 85062881 | 85062882 | cg07165260 |
| chr16 | 85315212 | 85315213 | cg01824618 |
| chr16 | 85682440 | 85682441 | cg02758499 |

| | | | |
|---|---|---|---|
| chr16 | 85682651 | 85682652 | cg08315483 |
| chr16 | 85787024 | 85787025 | cg07897180 |
| chr16 | 85787176 | 85787177 | cg06144110 |
| chr16 | 85940654 | 85940655 | cg08277259 |
| chr16 | 85947219 | 85947220 | cg02549595 |
| chr16 | 85964200 | 85964201 | cg02178957 |
| chr16 | 85966009 | 85966010 | cg05677184 |
| chr16 | 85966031 | 85966032 | cg01810713 |
| chr16 | 85966237 | 85966238 | cg03127886 |
| chr16 | 85970529 | 85970530 | cg01372811 |
| chr16 | 85989730 | 85989731 | cg08705942 |
| chr16 | 85989874 | 85989875 | cg00989229 |
| chr16 | 86016387 | 86016388 | cg08215318 |
| chr16 | 86018680 | 86018681 | cg10520672 |
| chr16 | 86018936 | 86018937 | cg09395612 |
| chr16 | 86552258 | 86552259 | cg05260236 |
| chr16 | 87415899 | 87415900 | cg26771582 |
| chr16 | 87757018 | 87757019 | cg01538731 |
| chr16 | 87757033 | 87757034 | cg05830220 |
| chr16 | 87908672 | 87908673 | cg09610891 |
| chr16 | 87990693 | 87990694 | cg04337828 |
| chr16 | 88101013 | 88101014 | cg04252938 |
| chr16 | 88103629 | 88103630 | cg05166473 |
| chr16 | 88111009 | 88111010 | cg08250921 |
| chr16 | 88121781 | 88121782 | cg16609966 |
| chr16 | 88482824 | 88482825 | cg27250607 |
| chr16 | 88545340 | 88545341 | cg06928887 |
| chr16 | 88547533 | 88547534 | cg00352652 |
| chr16 | 88547579 | 88547580 | cg00216061 |
| chr16 | 88547743 | 88547744 | cg04160201 |
| chr16 | 88549074 | 88549075 | cg27362718 |
| chr16 | 88551191 | 88551192 | cg02232529 |
| chr16 | 88551223 | 88551224 | cg26899496 |
| chr16 | 88551264 | 88551265 | cg03080043 |
| chr16 | 88551422 | 88551423 | cg08466082 |
| chr16 | 88591451 | 88591452 | cg05067156 |
| chr16 | 88700005 | 88700006 | cg03918510 |
| chr16 | 88747752 | 88747753 | cg06719532 |
| chr16 | 88747903 | 88747904 | cg23802887 |
| chr16 | 88747919 | 88747920 | cg10016175 |
| chr16 | 88810993 | 88810994 | cg04063936 |
| chr16 | 88866627 | 88866628 | cg10491452 |
| chr16 | 88884315 | 88884316 | cg02669544 |
| chr16 | 88933996 | 88933997 | cg00056002 |
| chr16 | 88934475 | 88934476 | cg01502457 |
| chr16 | 88967911 | 88967912 | cg03353765 |
| chr16 | 88978463 | 88978464 | cg06326865 |
| chr16 | 88988174 | 88988175 | cg09141835 |
| chr16 | 88988235 | 88988236 | cg04910877 |
| chr16 | 88989069 | 88989070 | cg05579598 |
| chr16 | 88989212 | 88989213 | cg08698997 |
| chr16 | 88989387 | 88989388 | cg27061485 |
| chr16 | 88991518 | 88991519 | cg26776806 |
| chr16 | 89003641 | 89003642 | cg05557991 |
| chr16 | 89024511 | 89024512 | cg02059317 |
| chr16 | 89024557 | 89024558 | cg01534572 |

| | | | |
|---|---|---|---|
| chr16 | 89035228 | 89035229 | cg03797660 |
| chr16 | 89035532 | 89035533 | cg02948862 |
| chr16 | 89036057 | 89036058 | cg07625774 |
| chr16 | 89037481 | 89037482 | cg06204040 |
| chr16 | 89050731 | 89050732 | cg03339704 |
| chr16 | 89129633 | 89129634 | cg03406609 |
| chr16 | 89139048 | 89139049 | cg10217183 |
| chr16 | 89163418 | 89163419 | cg16754613 |
| chr16 | 89167018 | 89167019 | cg09427016 |
| chr16 | 89171772 | 89171773 | cg05231226 |
| chr16 | 89189383 | 89189384 | cg04033022 |
| chr16 | 89375113 | 89375114 | cg02930721 |
| chr16 | 89390685 | 89390686 | cg07929642 |
| chr16 | 89390968 | 89390969 | cg08134863 |
| chr16 | 89405478 | 89405479 | cg07054502 |
| chr16 | 89424793 | 89424794 | cg06159562 |
| chr16 | 89510331 | 89510332 | cg16437021 |
| chr16 | 89550474 | 89550475 | cg08726900 |
| chr16 | 89552259 | 89552260 | cg05288253 |
| chr16 | 89777763 | 89777764 | cg00261912 |
| chr16 | 89922397 | 89922398 | cg06988336 |
| chr16 | 89922539 | 89922540 | cg01739831 |
| chr16 | 89966879 | 89966880 | cg00206161 |
| chr16 | 90019897 | 90019898 | cg07329987 |
| chr17 | 179611 | 179612 | cg05399244 |
| chr17 | 632478 | 632479 | cg02084467 |
| chr17 | 998504 | 998505 | cg20497635 |
| chr17 | 1009219 | 1009220 | cg04724319 |
| chr17 | 1069505 | 1069506 | cg14335816 |
| chr17 | 1366164 | 1366165 | cg05981034 |
| chr17 | 1397079 | 1397080 | cg12741184 |
| chr17 | 1412950 | 1412951 | cg07185664 |
| chr17 | 1480710 | 1480711 | cg04196298 |
| chr17 | 1509883 | 1509884 | cg02188939 |
| chr17 | 1509953 | 1509954 | cg25212453 |
| chr17 | 1510000 | 1510001 | cg19880947 |
| chr17 | 1510041 | 1510042 | cg21949830 |
| chr17 | 1510080 | 1510081 | cg01438467 |
| chr17 | 1549108 | 1549109 | cg08677655 |
| chr17 | 1554577 | 1554578 | cg00717678 |
| chr17 | 1616635 | 1616636 | cg02308706 |
| chr17 | 1665303 | 1665304 | cg06940127 |
| chr17 | 1666607 | 1666608 | cg00777079 |
| chr17 | 1799727 | 1799728 | cg09021817 |
| chr17 | 1973400 | 1973401 | cg19293468 |
| chr17 | 1979495 | 1979496 | cg12789448 |
| chr17 | 2060623 | 2060624 | cg23403980 |
| chr17 | 2213196 | 2213197 | cg20205704 |
| chr17 | 2300514 | 2300515 | cg09635667 |
| chr17 | 2310278 | 2310279 | cg16002660 |
| chr17 | 2498642 | 2498643 | cg12364755 |
| chr17 | 2908023 | 2908024 | cg00739778 |
| chr17 | 3442281 | 3442282 | cg19300422 |
| chr17 | 3462255 | 3462256 | cg22092292 |
| chr17 | 3561358 | 3561359 | cg06470149 |
| chr17 | 3561497 | 3561498 | cg05719052 |

| | | | |
|---|---|---|---|
| chr17 | 3600461 | 3600462 | cg13246592 |
| chr17 | 3623472 | 3623473 | cg02225720 |
| chr17 | 3794828 | 3794829 | cg03439237 |
| chr17 | 3819339 | 3819340 | cg22044342 |
| chr17 | 3819363 | 3819364 | cg25031824 |
| chr17 | 3819390 | 3819391 | cg01526089 |
| chr17 | 3819429 | 3819430 | cg16611352 |
| chr17 | 3827633 | 3827634 | cg19821988 |
| chr17 | 3863976 | 3863977 | cg23181573 |
| chr17 | 3977385 | 3977386 | cg06753513 |
| chr17 | 4151473 | 4151474 | cg18906596 |
| chr17 | 4272875 | 4272876 | cg09034695 |
| chr17 | 4546450 | 4546451 | cg11334406 |
| chr17 | 4674371 | 4674372 | cg06928209 |
| chr17 | 4675193 | 4675194 | cg27488095 |
| chr17 | 4675292 | 4675293 | cg21066636 |
| chr17 | 4795550 | 4795551 | cg13332193 |
| chr17 | 4795672 | 4795673 | cg00663068 |
| chr17 | 4923126 | 4923127 | cg03877174 |
| chr17 | 4936114 | 4936115 | cg25535257 |
| chr17 | 5113924 | 5113925 | cg21846970 |
| chr17 | 5185005 | 5185006 | cg20802942 |
| chr17 | 5185073 | 5185074 | cg27093376 |
| chr17 | 5189032 | 5189033 | cg07967207 |
| chr17 | 5820294 | 5820295 | cg04960007 |
| chr17 | 6473106 | 6473107 | cg21737020 |
| chr17 | 6555742 | 6555743 | cg15288800 |
| chr17 | 6658898 | 6658899 | cg23571857 |
| chr17 | 6913521 | 6913522 | cg20216752 |
| chr17 | 7144367 | 7144368 | cg19646165 |
| chr17 | 7186718 | 7186719 | cg06891043 |
| chr17 | 7253720 | 7253721 | cg02676175 |
| chr17 | 7324093 | 7324094 | cg06606753 |
| chr17 | 7341936 | 7341937 | cg13194425 |
| chr17 | 7381288 | 7381289 | cg07565956 |
| chr17 | 7742126 | 7742127 | cg24893073 |
| chr17 | 7791630 | 7791631 | cg22695339 |
| chr17 | 7792674 | 7792675 | cg00066663 |
| chr17 | 7965013 | 7965014 | cg00253269 |
| chr17 | 8012841 | 8012842 | cg10466728 |
| chr17 | 8013202 | 8013203 | cg18700133 |
| chr17 | 8067759 | 8067760 | cg25479708 |
| chr17 | 8092054 | 8092055 | cg24655262 |
| chr17 | 8198295 | 8198296 | cg19697725 |
| chr17 | 8525165 | 8525166 | cg20482145 |
| chr17 | 9861712 | 9861713 | cg12642651 |
| chr17 | 9940121 | 9940122 | cg18369972 |
| chr17 | 9941095 | 9941096 | cg00052665 |
| chr17 | 12332606 | 12332607 | cg06013895 |
| chr17 | 12541750 | 12541751 | cg21875401 |
| chr17 | 15586170 | 15586171 | cg13649519 |
| chr17 | 16397992 | 16397993 | cg25325454 |
| chr17 | 17011443 | 17011444 | cg09171562 |
| chr17 | 17267288 | 17267289 | cg16533363 |
| chr17 | 17616235 | 17616236 | cg08024038 |
| chr17 | 17642247 | 17642248 | cg12581691 |

| | | | |
|---|---|---|---|
| chr17 | 17669973 | 17669974 | cg06685437 |
| chr17 | 17696044 | 17696045 | cg24136700 |
| chr17 | 17696392 | 17696393 | cg24931355 |
| chr17 | 17718525 | 17718526 | cg19619576 |
| chr17 | 17929033 | 17929034 | cg21187669 |
| chr17 | 18139506 | 18139507 | cg09658183 |
| chr17 | 19152434 | 19152435 | cg25132536 |
| chr17 | 19174122 | 19174123 | cg23767840 |
| chr17 | 19220813 | 19220814 | cg12137220 |
| chr17 | 19651190 | 19651191 | cg15796819 |
| chr17 | 19656554 | 19656555 | cg08817867 |
| chr17 | 20105626 | 20105627 | cg00018010 |
| chr17 | 20107320 | 20107321 | cg21637761 |
| chr17 | 20839291 | 20839292 | cg07470639 |
| chr17 | 25819112 | 25819113 | cg08041350 |
| chr17 | 26728914 | 26728915 | cg10762038 |
| chr17 | 26800820 | 26800821 | cg00085732 |
| chr17 | 26850322 | 26850323 | cg10213812 |
| chr17 | 26990458 | 26990459 | cg01859360 |
| chr17 | 27051499 | 27051500 | cg03702413 |
| chr17 | 27073614 | 27073615 | cg10881616 |
| chr17 | 27892866 | 27892867 | cg00574412 |
| chr17 | 27899465 | 27899466 | cg22141459 |
| chr17 | 27972099 | 27972100 | cg14426167 |
| chr17 | 28098852 | 28098853 | cg23534301 |
| chr17 | 28562142 | 28562143 | cg05951817 |
| chr17 | 28562220 | 28562221 | cg22584138 |
| chr17 | 29549474 | 29549475 | cg19856701 |
| chr17 | 30179118 | 30179119 | cg26421997 |
| chr17 | 30184997 | 30184998 | cg26284483 |
| chr17 | 30592033 | 30592034 | cg18033416 |
| chr17 | 30842946 | 30842947 | cg18483549 |
| chr17 | 30844705 | 30844706 | cg12561776 |
| chr17 | 32690255 | 32690256 | cg01387036 |
| chr17 | 32690412 | 32690413 | cg26101277 |
| chr17 | 32690569 | 32690570 | cg17118262 |
| chr17 | 33672713 | 33672714 | cg26039662 |
| chr17 | 34207663 | 34207664 | cg19411729 |
| chr17 | 34271705 | 34271706 | cg25997187 |
| chr17 | 34304462 | 34304463 | cg22647738 |
| chr17 | 34325375 | 34325376 | cg24929264 |
| chr17 | 35211226 | 35211227 | cg06569861 |
| chr17 | 35847200 | 35847201 | cg21694161 |
| chr17 | 36811232 | 36811233 | cg25918821 |
| chr17 | 36832423 | 36832424 | cg00739155 |
| chr17 | 36890321 | 36890322 | cg23901967 |
| chr17 | 37028977 | 37028978 | cg08553913 |
| chr17 | 37063649 | 37063650 | cg24882332 |
| chr17 | 37521150 | 37521151 | cg25661397 |
| chr17 | 37688455 | 37688456 | cg20936107 |
| chr17 | 37894014 | 37894015 | cg14263391 |
| chr17 | 37895886 | 37895887 | cg00735611 |
| chr17 | 38119198 | 38119199 | cg24910161 |
| chr17 | 38119207 | 38119208 | cg26162295 |
| chr17 | 38252123 | 38252124 | cg22082015 |
| chr17 | 38261988 | 38261989 | cg10493185 |

| | | | |
|---|---|---|---|
| chr17 | 38350695 | 38350696 | cg19063856 |
| chr17 | 38498914 | 38498915 | cg16276850 |
| chr17 | 38501519 | 38501520 | cg08446900 |
| chr17 | 38501773 | 38501774 | cg11522012 |
| chr17 | 38511770 | 38511771 | cg24073994 |
| chr17 | 38599366 | 38599367 | cg11961138 |
| chr17 | 38646740 | 38646741 | cg00979803 |
| chr17 | 38696507 | 38696508 | cg25248278 |
| chr17 | 38698893 | 38698894 | cg26611328 |
| chr17 | 38705895 | 38705896 | cg18960324 |
| chr17 | 38708399 | 38708400 | cg16530881 |
| chr17 | 38956837 | 38956838 | cg16048803 |
| chr17 | 39067081 | 39067082 | cg11809014 |
| chr17 | 39143567 | 39143568 | cg06821079 |
| chr17 | 39151857 | 39151858 | cg13183651 |
| chr17 | 39221567 | 39221568 | cg12188197 |
| chr17 | 39254439 | 39254440 | cg20663846 |
| chr17 | 39323742 | 39323743 | cg24872384 |
| chr17 | 39662723 | 39662724 | cg13430376 |
| chr17 | 39914156 | 39914157 | cg15386103 |
| chr17 | 39993002 | 39993003 | cg17228560 |
| chr17 | 40073688 | 40073689 | cg27470486 |
| chr17 | 40250727 | 40250728 | cg01597398 |
| chr17 | 40268381 | 40268382 | cg16550651 |
| chr17 | 40270638 | 40270639 | cg21016267 |
| chr17 | 40438312 | 40438313 | cg05117208 |
| chr17 | 40442580 | 40442581 | cg12259792 |
| chr17 | 40489785 | 40489786 | cg17833746 |
| chr17 | 40516068 | 40516069 | cg06270615 |
| chr17 | 40570233 | 40570234 | cg27196880 |
| chr17 | 40910438 | 40910439 | cg17933008 |
| chr17 | 40954091 | 40954092 | cg14343283 |
| chr17 | 41053112 | 41053113 | cg10596768 |
| chr17 | 41186713 | 41186714 | cg10424581 |
| chr17 | 41837048 | 41837049 | cg11287987 |
| chr17 | 42426483 | 42426484 | cg12837296 |
| chr17 | 43340243 | 43340244 | cg10327769 |
| chr17 | 43376882 | 43376883 | cg16826777 |
| chr17 | 43388897 | 43388898 | cg08823240 |
| chr17 | 43442340 | 43442341 | cg12856708 |
| chr17 | 43464525 | 43464526 | cg26980978 |
| chr17 | 43510478 | 43510479 | cg16281322 |
| chr17 | 43510575 | 43510576 | cg04487659 |
| chr17 | 44105544 | 44105545 | cg09764761 |
| chr17 | 44222207 | 44222208 | cg13732302 |
| chr17 | 44342336 | 44342337 | cg14438695 |
| chr17 | 45286354 | 45286355 | cg09299082 |
| chr17 | 45335202 | 45335203 | cg10753610 |
| chr17 | 45767653 | 45767654 | cg05363382 |
| chr17 | 45809727 | 45809728 | cg06927323 |
| chr17 | 45812243 | 45812244 | cg16928046 |
| chr17 | 46026612 | 46026613 | cg05446659 |
| chr17 | 46035183 | 46035184 | cg03496533 |
| chr17 | 46506947 | 46506948 | cg05106502 |
| chr17 | 46871794 | 46871795 | cg19893820 |
| chr17 | 46982592 | 46982593 | cg09931134 |

| | | | |
|---|---|---|---|
| chr17 | 47100007 | 47100008 | cg07017153 |
| chr17 | 47467221 | 47467222 | cg11906021 |
| chr17 | 47661136 | 47661137 | cg20925233 |
| chr17 | 47838533 | 47838534 | cg14503291 |
| chr17 | 48231729 | 48231730 | cg12212103 |
| chr17 | 48239271 | 48239272 | cg06474661 |
| chr17 | 48242743 | 48242744 | cg19286672 |
| chr17 | 48243707 | 48243708 | cg07826255 |
| chr17 | 48266268 | 48266269 | cg25026926 |
| chr17 | 48275324 | 48275325 | cg18618815 |
| chr17 | 48425014 | 48425015 | cg24804060 |
| chr17 | 48552276 | 48552277 | cg09151131 |
| chr17 | 48768755 | 48768756 | cg14592673 |
| chr17 | 49340654 | 49340655 | cg24948782 |
| chr17 | 53502986 | 53502987 | cg11438150 |
| chr17 | 55335955 | 55335956 | cg06078469 |
| chr17 | 55337654 | 55337655 | cg11876048 |
| chr17 | 55338456 | 55338457 | cg18875496 |
| chr17 | 55371727 | 55371728 | cg26871347 |
| chr17 | 55446979 | 55446980 | cg12430029 |
| chr17 | 55540412 | 55540413 | cg13790810 |
| chr17 | 55562964 | 55562965 | cg22801561 |
| chr17 | 55993519 | 55993520 | cg07373578 |
| chr17 | 56401734 | 56401735 | cg17525495 |
| chr17 | 56565286 | 56565287 | cg27366766 |
| chr17 | 56590729 | 56590730 | cg16264807 |
| chr17 | 57700024 | 57700025 | cg12037593 |
| chr17 | 57861641 | 57861642 | cg02766539 |
| chr17 | 59222476 | 59222477 | cg15723222 |
| chr17 | 59545890 | 59545891 | cg03969651 |
| chr17 | 61216274 | 61216275 | cg09230870 |
| chr17 | 61520112 | 61520113 | cg15773539 |
| chr17 | 62010456 | 62010457 | cg24091162 |
| chr17 | 62051709 | 62051710 | cg05818824 |
| chr17 | 62051767 | 62051768 | cg25507121 |
| chr17 | 62153284 | 62153285 | cg17463083 |
| chr17 | 62208374 | 62208375 | cg20998539 |
| chr17 | 62401212 | 62401213 | cg00427260 |
| chr17 | 62495003 | 62495004 | cg14548777 |
| chr17 | 62500377 | 62500378 | cg23225508 |
| chr17 | 62961609 | 62961610 | cg20869310 |
| chr17 | 63100218 | 63100219 | cg25027367 |
| chr17 | 63823864 | 63823865 | cg25363387 |
| chr17 | 64973939 | 64973940 | cg23968154 |
| chr17 | 66255048 | 66255049 | cg25456728 |
| chr17 | 66290505 | 66290506 | cg15642209 |
| chr17 | 66291422 | 66291423 | cg08815110 |
| chr17 | 66511183 | 66511184 | cg07425204 |
| chr17 | 67057775 | 67057776 | cg01517431 |
| chr17 | 67137953 | 67137954 | cg22081096 |
| chr17 | 70405164 | 70405165 | cg05033295 |
| chr17 | 72028714 | 72028715 | cg16736055 |
| chr17 | 72203503 | 72203504 | cg01406909 |
| chr17 | 72231318 | 72231319 | cg08594554 |
| chr17 | 72480644 | 72480645 | cg14018141 |
| chr17 | 72830151 | 72830152 | cg19134130 |

| | | | |
|---|---|---|---|
| chr17 | 73056145 | 73056146 | cg09185884 |
| chr17 | 73056187 | 73056188 | cg26475911 |
| chr17 | 73073329 | 73073330 | cg00874073 |
| chr17 | 73074561 | 73074562 | cg20767344 |
| chr17 | 73084494 | 73084495 | cg04529938 |
| chr17 | 73234288 | 73234289 | cg18531405 |
| chr17 | 73238441 | 73238442 | cg07309864 |
| chr17 | 73238513 | 73238514 | cg16554534 |
| chr17 | 73256558 | 73256559 | cg24670151 |
| chr17 | 73316721 | 73316722 | cg07316730 |
| chr17 | 73316889 | 73316890 | cg11818853 |
| chr17 | 73316961 | 73316962 | cg14852456 |
| chr17 | 73347837 | 73347838 | cg25879514 |
| chr17 | 73399189 | 73399190 | cg25589929 |
| chr17 | 73500739 | 73500740 | cg10008947 |
| chr17 | 73782472 | 73782473 | cg25814096 |
| chr17 | 73823769 | 73823770 | cg02627403 |
| chr17 | 73955839 | 73955840 | cg26205890 |
| chr17 | 74131752 | 74131753 | cg17581104 |
| chr17 | 74266324 | 74266325 | cg04912316 |
| chr17 | 74375745 | 74375746 | cg01779934 |
| chr17 | 74416551 | 74416552 | cg12547839 |
| chr17 | 74557104 | 74557105 | cg05727573 |
| chr17 | 74992942 | 74992943 | cg20813798 |
| chr17 | 75105832 | 75105833 | cg15965301 |
| chr17 | 75186962 | 75186963 | cg11872645 |
| chr17 | 75208309 | 75208310 | cg12840847 |
| chr17 | 75210996 | 75210997 | cg06024215 |
| chr17 | 75371476 | 75371477 | cg00871371 |
| chr17 | 75431407 | 75431408 | cg17112975 |
| chr17 | 75451779 | 75451780 | cg02442640 |
| chr17 | 75451809 | 75451810 | cg17922695 |
| chr17 | 75451932 | 75451933 | cg14843920 |
| chr17 | 75470567 | 75470568 | cg01513063 |
| chr17 | 75475973 | 75475974 | cg26415789 |
| chr17 | 75880308 | 75880309 | cg05264101 |
| chr17 | 75880542 | 75880543 | cg04837616 |
| chr17 | 75959756 | 75959757 | cg00854273 |
| chr17 | 76121276 | 76121277 | cg19950606 |
| chr17 | 76129099 | 76129100 | cg16935597 |
| chr17 | 76129402 | 76129403 | cg04945945 |
| chr17 | 76134099 | 76134100 | cg01125010 |
| chr17 | 76166303 | 76166304 | cg05006092 |
| chr17 | 76169212 | 76169213 | cg10456035 |
| chr17 | 76170799 | 76170800 | cg20740903 |
| chr17 | 76774832 | 76774833 | cg14134128 |
| chr17 | 76795278 | 76795279 | cg17910470 |
| chr17 | 76899627 | 76899628 | cg00689612 |
| chr17 | 77041801 | 77041802 | cg06700275 |
| chr17 | 77704175 | 77704176 | cg21462477 |
| chr17 | 78073854 | 78073855 | cg04028464 |
| chr17 | 78472159 | 78472160 | cg00066239 |
| chr17 | 78623601 | 78623602 | cg18758433 |
| chr17 | 78638554 | 78638555 | cg01432609 |
| chr17 | 78716983 | 78716984 | cg06872548 |
| chr17 | 78773453 | 78773454 | cg10585621 |

| | | | |
|---|---|---|---|
| chr17 | 78773498 | 78773499 | cg15946337 |
| chr17 | 78773661 | 78773662 | cg15815120 |
| chr17 | 78800774 | 78800775 | cg16638092 |
| chr17 | 78800806 | 78800807 | cg08939850 |
| chr17 | 79024087 | 79024088 | cg09043456 |
| chr17 | 79024447 | 79024448 | cg06768939 |
| chr17 | 79127372 | 79127373 | cg07971007 |
| chr17 | 79235968 | 79235969 | cg13871900 |
| chr17 | 79258732 | 79258733 | cg15215033 |
| chr17 | 79485529 | 79485530 | cg14316944 |
| chr17 | 79533843 | 79533844 | cg25072666 |
| chr17 | 79632335 | 79632336 | cg03764274 |
| chr17 | 79632380 | 79632381 | cg16299813 |
| chr17 | 79827882 | 79827883 | cg06457408 |
| chr17 | 80005395 | 80005396 | cg03636200 |
| chr17 | 80024177 | 80024178 | cg19576556 |
| chr17 | 80279077 | 80279078 | cg27666046 |
| chr17 | 80280330 | 80280331 | cg14728380 |
| chr17 | 80282538 | 80282539 | cg22156899 |
| chr17 | 80361098 | 80361099 | cg06601666 |
| chr17 | 80648875 | 80648876 | cg08795515 |
| chr17 | 80649886 | 80649887 | cg23117250 |
| chr17 | 80662407 | 80662408 | cg01540203 |
| chr17 | 80818684 | 80818685 | cg27586410 |
| chr17 | 80818772 | 80818773 | cg22210779 |
| chr17 | 80819020 | 80819021 | cg00830755 |
| chr17 | 80819028 | 80819029 | cg11872040 |
| chr17 | 80819108 | 80819109 | cg08811443 |
| chr17 | 80829718 | 80829719 | cg22432760 |
| chr17 | 80873120 | 80873121 | cg17184161 |
| chr17 | 80941472 | 80941473 | cg03585013 |
| chr18 | 615496 | 615497 | cg23713900 |
| chr18 | 2572965 | 2572966 | cg06454226 |
| chr18 | 3445098 | 3445099 | cg25385819 |
| chr18 | 3446566 | 3446567 | cg26701815 |
| chr18 | 3447016 | 3447017 | cg07529654 |
| chr18 | 5195201 | 5195202 | cg19718771 |
| chr18 | 9707031 | 9707032 | cg27298007 |
| chr18 | 9712461 | 9712462 | cg13049483 |
| chr18 | 10524910 | 10524911 | cg07392482 |
| chr18 | 12896164 | 12896165 | cg21149466 |
| chr18 | 13501701 | 13501702 | cg19455642 |
| chr18 | 18687151 | 18687152 | cg05835414 |
| chr18 | 20717411 | 20717412 | cg26052586 |
| chr18 | 21018562 | 21018563 | cg25436157 |
| chr18 | 21596915 | 21596916 | cg24331475 |
| chr18 | 29171878 | 29171879 | cg03180302 |
| chr18 | 29268737 | 29268738 | cg23888440 |
| chr18 | 30347555 | 30347556 | cg03477049 |
| chr18 | 30349111 | 30349112 | cg05134926 |
| chr18 | 32446543 | 32446544 | cg18517540 |
| chr18 | 33048493 | 33048494 | cg18767920 |
| chr18 | 42261980 | 42261981 | cg24217948 |
| chr18 | 42262885 | 42262886 | cg16705273 |
| chr18 | 47019224 | 47019225 | cg11857433 |
| chr18 | 48719384 | 48719385 | cg02895724 |

| | | | |
|---|---|---|---|
| chr18 | 55469021 | 55469022 | cg12517027 |
| chr18 | 55714670 | 55714671 | cg26034769 |
| chr18 | 60385273 | 60385274 | cg05875463 |
| chr18 | 61222687 | 61222688 | cg11435943 |
| chr18 | 67624393 | 67624394 | cg12929678 |
| chr18 | 72509555 | 72509556 | cg12573871 |
| chr18 | 74536226 | 74536227 | cg06398181 |
| chr18 | 74633150 | 74633151 | cg06161299 |
| chr18 | 74692145 | 74692146 | cg17061340 |
| chr18 | 74832261 | 74832262 | cg16990945 |
| chr18 | 74836070 | 74836071 | cg19310162 |
| chr18 | 74845829 | 74845830 | cg16604566 |
| chr18 | 76828521 | 76828522 | cg06556497 |
| chr18 | 77135101 | 77135102 | cg14071758 |
| chr18 | 77157986 | 77157987 | cg13608166 |
| chr18 | 77158180 | 77158181 | cg16848717 |
| chr18 | 77158676 | 77158677 | cg03586564 |
| chr18 | 77161214 | 77161215 | cg15363134 |
| chr18 | 77162457 | 77162458 | cg06751398 |
| chr18 | 77165150 | 77165151 | cg15260951 |
| chr18 | 77182405 | 77182406 | cg16997486 |
| chr18 | 77185451 | 77185452 | cg13580107 |
| chr18 | 77196320 | 77196321 | cg05302701 |
| chr18 | 77196450 | 77196451 | cg27308982 |
| chr18 | 77211963 | 77211964 | cg26915704 |
| chr18 | 77213901 | 77213902 | cg02736255 |
| chr18 | 77266975 | 77266976 | cg15678825 |
| chr18 | 77386828 | 77386829 | cg14940871 |
| chr18 | 77386989 | 77386990 | cg18476557 |
| chr18 | 77560205 | 77560206 | cg07345205 |
| chr18 | 77865873 | 77865874 | cg26914205 |
| chr19 | 575412 | 575413 | cg03233876 |
| chr19 | 617133 | 617134 | cg19865472 |
| chr19 | 617195 | 617196 | cg12111149 |
| chr19 | 617267 | 617268 | cg15726557 |
| chr19 | 619447 | 619448 | cg06169597 |
| chr19 | 711001 | 711002 | cg20631351 |
| chr19 | 827715 | 827716 | cg02147126 |
| chr19 | 827739 | 827740 | cg14663914 |
| chr19 | 891782 | 891783 | cg03721994 |
| chr19 | 891986 | 891987 | cg04340203 |
| chr19 | 935259 | 935260 | cg25298189 |
| chr19 | 940967 | 940968 | cg02001279 |
| chr19 | 946148 | 946149 | cg22809826 |
| chr19 | 951681 | 951682 | cg22035959 |
| chr19 | 993855 | 993856 | cg21046951 |
| chr19 | 1084367 | 1084368 | cg11629220 |
| chr19 | 1117109 | 1117110 | cg17284854 |
| chr19 | 1280967 | 1280968 | cg03256597 |
| chr19 | 1576714 | 1576715 | cg14305641 |
| chr19 | 1851750 | 1851751 | cg08620751 |
| chr19 | 1851882 | 1851883 | cg13382072 |
| chr19 | 1852165 | 1852166 | cg05006231 |
| chr19 | 1861356 | 1861357 | cg17841440 |
| chr19 | 1923307 | 1923308 | cg07662734 |
| chr19 | 1955419 | 1955420 | cg02990598 |

| | | | |
|---|---|---|---|
| chr19 | 2000559 | 2000560 | cg04246726 |
| chr19 | 2001580 | 2001581 | cg18431970 |
| chr19 | 2044540 | 2044541 | cg07723431 |
| chr19 | 2189267 | 2189268 | cg11776115 |
| chr19 | 2215718 | 2215719 | cg05345285 |
| chr19 | 2546598 | 2546599 | cg10181414 |
| chr19 | 2581343 | 2581344 | cg06371583 |
| chr19 | 2607726 | 2607727 | cg17617491 |
| chr19 | 2700927 | 2700928 | cg20091384 |
| chr19 | 3122304 | 3122305 | cg07628073 |
| chr19 | 3177310 | 3177311 | cg20656868 |
| chr19 | 3193804 | 3193805 | cg26794830 |
| chr19 | 3556718 | 3556719 | cg14034476 |
| chr19 | 3617614 | 3617615 | cg18057548 |
| chr19 | 3638778 | 3638779 | cg03540494 |
| chr19 | 3672411 | 3672412 | cg08301518 |
| chr19 | 3966842 | 3966843 | cg18748062 |
| chr19 | 3979758 | 3979759 | cg18789620 |
| chr19 | 4059988 | 4059989 | cg16242615 |
| chr19 | 4089526 | 4089527 | cg27198931 |
| chr19 | 4094324 | 4094325 | cg19423175 |
| chr19 | 4221628 | 4221629 | cg25440353 |
| chr19 | 4338852 | 4338853 | cg12229841 |
| chr19 | 4443084 | 4443085 | cg19682666 |
| chr19 | 4444770 | 4444771 | cg03362751 |
| chr19 | 4654722 | 4654723 | cg26756949 |
| chr19 | 5253997 | 5253998 | cg22950899 |
| chr19 | 5288631 | 5288632 | cg08462941 |
| chr19 | 5699060 | 5699061 | cg16694293 |
| chr19 | 5901268 | 5901269 | cg23477380 |
| chr19 | 6670865 | 6670866 | cg16043888 |
| chr19 | 7460671 | 7460672 | cg11734329 |
| chr19 | 7764345 | 7764346 | cg12261095 |
| chr19 | 7794952 | 7794953 | cg09772661 |
| chr19 | 8204480 | 8204481 | cg07638689 |
| chr19 | 10213817 | 10213818 | cg12161893 |
| chr19 | 10216651 | 10216652 | cg23098529 |
| chr19 | 10216676 | 10216677 | cg12819826 |
| chr19 | 11074303 | 11074304 | cg08315613 |
| chr19 | 11545491 | 11545492 | cg27646205 |
| chr19 | 11664023 | 11664024 | cg11691571 |
| chr19 | 11688711 | 11688712 | cg21207418 |
| chr19 | 12445173 | 12445174 | cg26924223 |
| chr19 | 12895288 | 12895289 | cg14294321 |
| chr19 | 12895529 | 12895530 | cg22996170 |
| chr19 | 13010694 | 13010695 | cg21044833 |
| chr19 | 13127873 | 13127874 | cg25556035 |
| chr19 | 13904774 | 13904775 | cg04391391 |
| chr19 | 13946770 | 13946771 | cg20281375 |
| chr19 | 13951168 | 13951169 | cg05339037 |
| chr19 | 14073064 | 14073065 | cg17977474 |
| chr19 | 14692759 | 14692760 | cg25770495 |
| chr19 | 14692863 | 14692864 | cg25945273 |
| chr19 | 15375283 | 15375284 | cg02681400 |
| chr19 | 15391124 | 15391125 | cg02533173 |
| chr19 | 15391339 | 15391340 | cg19287817 |

| | | | |
|---|---|---|---|
| chr19 | 15488902 | 15488903 | cg22765626 |
| chr19 | 15491808 | 15491809 | cg10225149 |
| chr19 | 15751949 | 15751950 | cg27067618 |
| chr19 | 16186244 | 16186245 | cg13978367 |
| chr19 | 16284242 | 16284243 | cg09686308 |
| chr19 | 16284351 | 16284352 | cg09424896 |
| chr19 | 16394366 | 16394367 | cg02381279 |
| chr19 | 16466421 | 16466422 | cg23304156 |
| chr19 | 16466480 | 16466481 | cg17702388 |
| chr19 | 17190259 | 17190260 | cg12229979 |
| chr19 | 17347791 | 17347792 | cg12301814 |
| chr19 | 17613063 | 17613064 | cg08220872 |
| chr19 | 18044745 | 18044746 | cg07930940 |
| chr19 | 18108895 | 18108896 | cg00764668 |
| chr19 | 18451112 | 18451113 | cg00420929 |
| chr19 | 18704214 | 18704215 | cg24425705 |
| chr19 | 18790693 | 18790694 | cg27493220 |
| chr19 | 18830892 | 18830893 | cg07780199 |
| chr19 | 18876246 | 18876247 | cg26572992 |
| chr19 | 19106904 | 19106905 | cg12289801 |
| chr19 | 19278512 | 19278513 | cg22470465 |
| chr19 | 19515518 | 19515519 | cg12488919 |
| chr19 | 19737143 | 19737144 | cg19379721 |
| chr19 | 19780767 | 19780768 | cg17692971 |
| chr19 | 30102004 | 30102005 | cg06738169 |
| chr19 | 33876779 | 33876780 | cg06478085 |
| chr19 | 34989280 | 34989281 | cg02014733 |
| chr19 | 35598494 | 35598495 | cg09960641 |
| chr19 | 35606534 | 35606535 | cg01408817 |
| chr19 | 35772657 | 35772658 | cg14554776 |
| chr19 | 35940174 | 35940175 | cg03548857 |
| chr19 | 36205201 | 36205202 | cg18593194 |
| chr19 | 38570495 | 38570496 | cg14972228 |
| chr19 | 38756382 | 38756383 | cg17344048 |
| chr19 | 39217070 | 39217071 | cg27548256 |
| chr19 | 39224370 | 39224371 | cg26623753 |
| chr19 | 39303775 | 39303776 | cg06394229 |
| chr19 | 39444822 | 39444823 | cg12955539 |
| chr19 | 39523626 | 39523627 | cg02992311 |
| chr19 | 39889222 | 39889223 | cg03301498 |
| chr19 | 39896713 | 39896714 | cg10753398 |
| chr19 | 40379680 | 40379681 | cg24811685 |
| chr19 | 40788787 | 40788788 | cg06055845 |
| chr19 | 40788926 | 40788927 | cg26620021 |
| chr19 | 40939429 | 40939430 | cg14085840 |
| chr19 | 41197903 | 41197904 | cg21956507 |
| chr19 | 41260294 | 41260295 | cg16596691 |
| chr19 | 41619952 | 41619953 | cg11267879 |
| chr19 | 41805850 | 41805851 | cg00781658 |
| chr19 | 42380725 | 42380726 | cg05921699 |
| chr19 | 42411405 | 42411406 | cg12168357 |
| chr19 | 42445462 | 42445463 | cg27267617 |
| chr19 | 42787174 | 42787175 | cg26163284 |
| chr19 | 42882838 | 42882839 | cg16696774 |
| chr19 | 44285594 | 44285595 | cg24900654 |
| chr19 | 44285940 | 44285941 | cg26890181 |

| | | | |
|---|---|---|---|
| chr19 | 44285974 | 44285975 | cg18506018 |
| chr19 | 44286076 | 44286077 | cg04833845 |
| chr19 | 45445437 | 45445438 | cg17769836 |
| chr19 | 45445449 | 45445450 | cg04401876 |
| chr19 | 45445491 | 45445492 | cg02912790 |
| chr19 | 45445521 | 45445522 | cg27353824 |
| chr19 | 45445693 | 45445694 | cg25017250 |
| chr19 | 45574210 | 45574211 | cg08692295 |
| chr19 | 45943865 | 45943866 | cg14890509 |
| chr19 | 45957466 | 45957467 | cg10732094 |
| chr19 | 46170946 | 46170947 | cg09997082 |
| chr19 | 46214108 | 46214109 | cg07636338 |
| chr19 | 47223398 | 47223399 | cg26482609 |
| chr19 | 47253604 | 47253605 | cg00838415 |
| chr19 | 47736018 | 47736019 | cg25731261 |
| chr19 | 47991237 | 47991238 | cg17705615 |
| chr19 | 49116870 | 49116871 | cg27530053 |
| chr19 | 49184628 | 49184629 | cg27262821 |
| chr19 | 49252020 | 49252021 | cg11293312 |
| chr19 | 49471563 | 49471564 | cg06745001 |
| chr19 | 49659299 | 49659300 | cg10695490 |
| chr19 | 49828629 | 49828630 | cg00041047 |
| chr19 | 49828641 | 49828642 | cg01453814 |
| chr19 | 49828658 | 49828659 | cg24188017 |
| chr19 | 49828667 | 49828668 | cg06237487 |
| chr19 | 49840176 | 49840177 | cg19753641 |
| chr19 | 49863290 | 49863291 | cg13487726 |
| chr19 | 50002551 | 50002552 | cg07569288 |
| chr19 | 50064054 | 50064055 | cg08097359 |
| chr19 | 50161702 | 50161703 | cg23960022 |
| chr19 | 50391354 | 50391355 | cg14234302 |
| chr19 | 50868879 | 50868880 | cg27285056 |
| chr19 | 50869007 | 50869008 | cg19901005 |
| chr19 | 50869332 | 50869333 | cg14894368 |
| chr19 | 50931515 | 50931516 | cg15690347 |
| chr19 | 50931622 | 50931623 | cg04508467 |
| chr19 | 50931875 | 50931876 | cg19387862 |
| chr19 | 51508927 | 51508928 | cg07798605 |
| chr19 | 51847299 | 51847300 | cg10125636 |
| chr19 | 51891339 | 51891340 | cg08861091 |
| chr19 | 51922139 | 51922140 | cg03582327 |
| chr19 | 52004630 | 52004631 | cg12530021 |
| chr19 | 53192577 | 53192578 | cg08318732 |
| chr19 | 54848878 | 54848879 | cg02434051 |
| chr19 | 54850582 | 54850583 | cg23899409 |
| chr19 | 54851571 | 54851572 | cg03575041 |
| chr19 | 55141979 | 55141980 | cg02340056 |
| chr19 | 55173997 | 55173998 | cg24140775 |
| chr19 | 55174156 | 55174157 | cg05329879 |
| chr19 | 55625184 | 55625185 | cg02474116 |
| chr19 | 55887905 | 55887906 | cg25921357 |
| chr19 | 55987415 | 55987416 | cg01290153 |
| chr19 | 55998490 | 55998491 | cg03645984 |
| chr19 | 55998931 | 55998932 | cg19820609 |
| chr19 | 56156430 | 56156431 | cg00007987 |
| chr19 | 56182200 | 56182201 | cg03734391 |

| | | | |
|---|---|---|---|
| chr20 | 400361 | 400362 | cg02185402 |
| chr20 | 400412 | 400413 | cg09713225 |
| chr20 | 816362 | 816363 | cg20289911 |
| chr20 | 825798 | 825799 | cg08137888 |
| chr20 | 826424 | 826425 | cg14170976 |
| chr20 | 1377284 | 1377285 | cg00424467 |
| chr20 | 1472419 | 1472420 | cg07725579 |
| chr20 | 1610358 | 1610359 | cg10131232 |
| chr20 | 1825918 | 1825919 | cg01674585 |
| chr20 | 2276571 | 2276572 | cg12761282 |
| chr20 | 2855312 | 2855313 | cg24983189 |
| chr20 | 2963112 | 2963113 | cg27423323 |
| chr20 | 3644719 | 3644720 | cg21597152 |
| chr20 | 3868713 | 3868714 | cg23908269 |
| chr20 | 3896885 | 3896886 | cg24825299 |
| chr20 | 3897762 | 3897763 | cg12617066 |
| chr20 | 4801487 | 4801488 | cg04132379 |
| chr20 | 4802684 | 4802685 | cg22042908 |
| chr20 | 4991028 | 4991029 | cg17978727 |
| chr20 | 5170968 | 5170969 | cg17697105 |
| chr20 | 5589042 | 5589043 | cg08938062 |
| chr20 | 9491559 | 9491560 | cg25294504 |
| chr20 | 10157028 | 10157029 | cg15449979 |
| chr20 | 17660838 | 17660839 | cg05955436 |
| chr20 | 19815086 | 19815087 | cg06786349 |
| chr20 | 19869979 | 19869980 | cg08200952 |
| chr20 | 19916087 | 19916088 | cg18952506 |
| chr20 | 19982870 | 19982871 | cg12201988 |
| chr20 | 20180469 | 20180470 | cg23732608 |
| chr20 | 21368840 | 21368841 | cg06051581 |
| chr20 | 24446527 | 24446528 | cg22775600 |
| chr20 | 25177389 | 25177390 | cg23656213 |
| chr20 | 25277813 | 25277814 | cg06082906 |
| chr20 | 25583652 | 25583653 | cg21878598 |
| chr20 | 25988587 | 25988588 | cg12659446 |
| chr20 | 30200497 | 30200498 | cg12889133 |
| chr20 | 30644728 | 30644729 | cg23496571 |
| chr20 | 30696827 | 30696828 | cg20672708 |
| chr20 | 30720375 | 30720376 | cg11360718 |
| chr20 | 31025866 | 31025867 | cg08217589 |
| chr20 | 31070190 | 31070191 | cg24484138 |
| chr20 | 31169126 | 31169127 | cg02030275 |
| chr20 | 31595115 | 31595116 | cg02191985 |
| chr20 | 32662344 | 32662345 | cg19110982 |
| chr20 | 32856846 | 32856847 | cg16440058 |
| chr20 | 32856898 | 32856899 | cg05267394 |
| chr20 | 32954013 | 32954014 | cg21001667 |
| chr20 | 33293445 | 33293446 | cg18133477 |
| chr20 | 33300430 | 33300431 | cg15636879 |
| chr20 | 33589992 | 33589993 | cg06244927 |
| chr20 | 34118924 | 34118925 | cg26287609 |
| chr20 | 34127524 | 34127525 | cg17454857 |
| chr20 | 34195484 | 34195485 | cg06575425 |
| chr20 | 34200395 | 34200396 | cg19701649 |
| chr20 | 34202721 | 34202722 | cg23713742 |
| chr20 | 34995278 | 34995279 | cg01982835 |

| | | | |
|---|---|---|---|
| chr20 | 35091947 | 35091948 | cg24216770 |
| chr20 | 35094679 | 35094680 | cg17566560 |
| chr20 | 35241157 | 35241158 | cg09191335 |
| chr20 | 35274941 | 35274942 | cg14329026 |
| chr20 | 35399961 | 35399962 | cg06313775 |
| chr20 | 35504403 | 35504404 | cg09227119 |
| chr20 | 35724900 | 35724901 | cg17113158 |
| chr20 | 35914908 | 35914909 | cg02298220 |
| chr20 | 36011999 | 36012000 | cg24956391 |
| chr20 | 36012008 | 36012009 | cg23410129 |
| chr20 | 36012016 | 36012017 | cg08553950 |
| chr20 | 36037325 | 36037326 | cg03001863 |
| chr20 | 36153479 | 36153480 | cg11421182 |
| chr20 | 39762753 | 39762754 | cg12485727 |
| chr20 | 39769764 | 39769765 | cg24961795 |
| chr20 | 39998953 | 39998954 | cg14816917 |
| chr20 | 40040268 | 40040269 | cg14276730 |
| chr20 | 42955196 | 42955197 | cg23526960 |
| chr20 | 43377579 | 43377580 | cg05658173 |
| chr20 | 43593364 | 43593365 | cg00386940 |
| chr20 | 45179226 | 45179227 | cg11173131 |
| chr20 | 45179345 | 45179346 | cg17801864 |
| chr20 | 45179354 | 45179355 | cg17775490 |
| chr20 | 45179413 | 45179414 | cg01880463 |
| chr20 | 45187423 | 45187424 | cg07727594 |
| chr20 | 46415859 | 46415860 | cg21500064 |
| chr20 | 46988400 | 46988401 | cg10167508 |
| chr20 | 47536468 | 47536469 | cg07090569 |
| chr20 | 47801453 | 47801454 | cg02733918 |
| chr20 | 48428318 | 48428319 | cg02003117 |
| chr20 | 48530654 | 48530655 | cg19221489 |
| chr20 | 49345687 | 49345688 | cg10187713 |
| chr20 | 49415761 | 49415762 | cg15549231 |
| chr20 | 49416093 | 49416094 | cg02990474 |
| chr20 | 49454326 | 49454327 | cg18551353 |
| chr20 | 49455783 | 49455784 | cg05331472 |
| chr20 | 49509062 | 49509063 | cg22920603 |
| chr20 | 50214496 | 50214497 | cg00949661 |
| chr20 | 52198225 | 52198226 | cg12032027 |
| chr20 | 52198378 | 52198379 | cg01692482 |
| chr20 | 52199520 | 52199521 | cg17098965 |
| chr20 | 52274268 | 52274269 | cg19152255 |
| chr20 | 55835020 | 55835021 | cg13236649 |
| chr20 | 55968730 | 55968731 | cg17799760 |
| chr20 | 55969282 | 55969283 | cg10336935 |
| chr20 | 56100021 | 56100022 | cg25397426 |
| chr20 | 56247302 | 56247303 | cg26681770 |
| chr20 | 57265991 | 57265992 | cg20477147 |
| chr20 | 57583520 | 57583521 | cg06942450 |
| chr20 | 57583709 | 57583710 | cg23907860 |
| chr20 | 60757239 | 60757240 | cg06914270 |
| chr20 | 61562955 | 61562956 | cg25957332 |
| chr20 | 62055252 | 62055253 | cg22368204 |
| chr20 | 62174758 | 62174759 | cg22298487 |
| chr20 | 62179485 | 62179486 | cg00369372 |
| chr20 | 62183964 | 62183965 | cg15332132 |

| | | | |
|---|---|---|---|
| chr20 | 62251227 | 62251228 | cg20732137 |
| chr20 | 62574095 | 62574096 | cg07507493 |
| chr20 | 62574114 | 62574115 | cg20252837 |
| chr20 | 62660605 | 62660606 | cg02888894 |
| chr20 | 62665676 | 62665677 | cg11945905 |
| chr20 | 62745242 | 62745243 | cg06373721 |
| chr21 | 22175001 | 22175002 | cg09614626 |
| chr21 | 26733563 | 26733564 | cg11575131 |
| chr21 | 26735730 | 26735731 | cg21498326 |
| chr21 | 26933554 | 26933555 | cg00565412 |
| chr21 | 30676184 | 30676185 | cg03732014 |
| chr21 | 30677172 | 30677173 | cg22076474 |
| chr21 | 31655654 | 31655655 | cg09892262 |
| chr21 | 32128800 | 32128801 | cg22405653 |
| chr21 | 32412191 | 32412192 | cg06790305 |
| chr21 | 33100923 | 33100924 | cg12921799 |
| chr21 | 33105288 | 33105289 | cg01066046 |
| chr21 | 34918578 | 34918579 | cg17369406 |
| chr21 | 35755964 | 35755965 | cg15274171 |
| chr21 | 35899448 | 35899449 | cg05056497 |
| chr21 | 36395663 | 36395664 | cg11704631 |
| chr21 | 38349937 | 38349938 | cg21871091 |
| chr21 | 38350069 | 38350070 | cg24573261 |
| chr21 | 40182073 | 40182074 | cg23110422 |
| chr21 | 42559652 | 42559653 | cg16365807 |
| chr21 | 43197205 | 43197206 | cg00152948 |
| chr21 | 43619559 | 43619560 | cg02316713 |
| chr21 | 43658316 | 43658317 | cg00177237 |
| chr21 | 44062183 | 44062184 | cg11852796 |
| chr21 | 44585506 | 44585507 | cg26741373 |
| chr21 | 45379721 | 45379722 | cg19057779 |
| chr21 | 45573410 | 45573411 | cg06486129 |
| chr21 | 45575832 | 45575833 | cg08039560 |
| chr21 | 45722068 | 45722069 | cg12444076 |
| chr21 | 45729326 | 45729327 | cg04643920 |
| chr21 | 46235052 | 46235053 | cg10871721 |
| chr21 | 46349059 | 46349060 | cg21006727 |
| chr21 | 46496510 | 46496511 | cg19810954 |
| chr21 | 46890849 | 46890850 | cg04760448 |
| chr21 | 47269627 | 47269628 | cg07685034 |
| chr21 | 47647656 | 47647657 | cg20229931 |
| chr21 | 47710409 | 47710410 | cg26722225 |
| chr22 | 17587971 | 17587972 | cg01760983 |
| chr22 | 18257720 | 18257721 | cg01618287 |
| chr22 | 18325505 | 18325506 | cg21103992 |
| chr22 | 18511180 | 18511181 | cg11961401 |
| chr22 | 18512081 | 18512082 | cg12870518 |
| chr22 | 18592555 | 18592556 | cg19678959 |
| chr22 | 19436948 | 19436949 | cg06501716 |
| chr22 | 19838287 | 19838288 | cg09072576 |
| chr22 | 19973978 | 19973979 | cg01138652 |
| chr22 | 20114683 | 20114684 | cg17561788 |
| chr22 | 20123223 | 20123224 | cg11732063 |
| chr22 | 20134463 | 20134464 | cg02521457 |
| chr22 | 20134578 | 20134579 | cg23798065 |
| chr22 | 20134676 | 20134677 | cg17814680 |

| | | | |
|---|---|---|---|
| chr22 | 21275401 | 21275402 | cg02210498 |
| chr22 | 21318106 | 21318107 | cg17470095 |
| chr22 | 21321253 | 21321254 | cg05279387 |
| chr22 | 21321273 | 21321274 | cg12360084 |
| chr22 | 21981360 | 21981361 | cg01220923 |
| chr22 | 21985434 | 21985435 | cg20703614 |
| chr22 | 22086335 | 22086336 | cg09476306 |
| chr22 | 22293118 | 22293119 | cg27653384 |
| chr22 | 22597965 | 22597966 | cg04776489 |
| chr22 | 22843648 | 22843649 | cg04971769 |
| chr22 | 23487048 | 23487049 | cg14497203 |
| chr22 | 23521267 | 23521268 | cg00675878 |
| chr22 | 23602765 | 23602766 | cg23225050 |
| chr22 | 24032426 | 24032427 | cg13253729 |
| chr22 | 24115568 | 24115569 | cg27532722 |
| chr22 | 24190340 | 24190341 | cg18132256 |
| chr22 | 24190721 | 24190722 | cg02612971 |
| chr22 | 24671137 | 24671138 | cg08808258 |
| chr22 | 25799160 | 25799161 | cg26636045 |
| chr22 | 25799504 | 25799505 | cg26253663 |
| chr22 | 26826684 | 26826685 | cg04360334 |
| chr22 | 26875806 | 26875807 | cg21711214 |
| chr22 | 26876075 | 26876076 | cg13345299 |
| chr22 | 26877690 | 26877691 | cg00488514 |
| chr22 | 26938937 | 26938938 | cg09856467 |
| chr22 | 26962071 | 26962072 | cg18895476 |
| chr22 | 29430271 | 29430272 | cg20080983 |
| chr22 | 29705157 | 29705158 | cg01259126 |
| chr22 | 29706704 | 29706705 | cg24844449 |
| chr22 | 29706754 | 29706755 | cg02241145 |
| chr22 | 30637461 | 30637462 | cg26581200 |
| chr22 | 30817820 | 30817821 | cg19015086 |
| chr22 | 31475122 | 31475123 | cg10237442 |
| chr22 | 31644160 | 31644161 | cg15504662 |
| chr22 | 31644190 | 31644191 | cg20742981 |
| chr22 | 31889592 | 31889593 | cg02481923 |
| chr22 | 32150934 | 32150935 | cg16160637 |
| chr22 | 33402686 | 33402687 | cg23855715 |
| chr22 | 36560801 | 36560802 | cg00965578 |
| chr22 | 36563027 | 36563028 | cg08390994 |
| chr22 | 36563048 | 36563049 | cg06703343 |
| chr22 | 37256352 | 37256353 | cg22305209 |
| chr22 | 37256356 | 37256357 | cg25863223 |
| chr22 | 37309487 | 37309488 | cg01064979 |
| chr22 | 37467720 | 37467721 | cg21188731 |
| chr22 | 37491595 | 37491596 | cg17956747 |
| chr22 | 37496929 | 37496930 | cg19979738 |
| chr22 | 37499859 | 37499860 | cg16971128 |
| chr22 | 37523151 | 37523152 | cg24509815 |
| chr22 | 37545423 | 37545424 | cg26757673 |
| chr22 | 37621745 | 37621746 | cg10603757 |
| chr22 | 37655614 | 37655615 | cg11789534 |
| chr22 | 38431726 | 38431727 | cg19603202 |
| chr22 | 38609534 | 38609535 | cg01403010 |
| chr22 | 38611369 | 38611370 | cg18266458 |
| chr22 | 38711891 | 38711892 | cg02516101 |

| | | | |
|---|---|---|---|
| chr22 | 38790207 | 38790208 | cg00248449 |
| chr22 | 38792966 | 38792967 | cg21472198 |
| chr22 | 39418130 | 39418131 | cg08665930 |
| chr22 | 39491975 | 39491976 | cg18696237 |
| chr22 | 39778604 | 39778605 | cg17537935 |
| chr22 | 39966711 | 39966712 | cg18001869 |
| chr22 | 40079626 | 40079627 | cg02206108 |
| chr22 | 40289840 | 40289841 | cg20623350 |
| chr22 | 40322629 | 40322630 | cg18545596 |
| chr22 | 40573054 | 40573055 | cg01381829 |
| chr22 | 40573076 | 40573077 | cg25276892 |
| chr22 | 40814966 | 40814967 | cg17237086 |
| chr22 | 40815101 | 40815102 | cg06803995 |
| chr22 | 41633219 | 41633220 | cg26364091 |
| chr22 | 42324281 | 42324282 | cg00253346 |
| chr22 | 42325444 | 42325445 | cg00400964 |
| chr22 | 42335021 | 42335022 | cg27450867 |
| chr22 | 42548868 | 42548869 | cg13677785 |
| chr22 | 42907591 | 42907592 | cg02431732 |
| chr22 | 42948520 | 42948521 | cg06575723 |
| chr22 | 42950376 | 42950377 | cg17573270 |
| chr22 | 43040721 | 43040722 | cg08690876 |
| chr22 | 43044387 | 43044388 | cg20950843 |
| chr22 | 43523016 | 43523017 | cg02601893 |
| chr22 | 43609331 | 43609332 | cg04352088 |
| chr22 | 44576381 | 44576382 | cg07968613 |
| chr22 | 45310578 | 45310579 | cg04556646 |
| chr22 | 46402185 | 46402186 | cg25591418 |
| chr22 | 46650375 | 46650376 | cg14686297 |
| chr22 | 46770063 | 46770064 | cg04332442 |
| chr22 | 46770195 | 46770196 | cg12974599 |
| chr22 | 46770440 | 46770441 | cg24408469 |
| chr22 | 46833806 | 46833807 | cg00253398 |
| chr22 | 47057502 | 47057503 | cg05821464 |
| chr22 | 47069799 | 47069800 | cg09233980 |
| chr22 | 48900624 | 48900625 | cg23400715 |
| chr22 | 49498376 | 49498377 | cg27166177 |
| chr22 | 50013649 | 50013650 | cg06355422 |
| chr22 | 50219754 | 50219755 | cg06057569 |
| chr22 | 50223573 | 50223574 | cg00791190 |
| chr22 | 50356763 | 50356764 | cg11801110 |
| chr22 | 50357214 | 50357215 | cg20698113 |
| chr22 | 50451975 | 50451976 | cg27068297 |
| chr22 | 50702460 | 50702461 | cg26790091 |
| chr22 | 50747014 | 50747015 | cg19553402 |
| chr22 | 50847567 | 50847568 | cg21757633 |
| chr22 | 50884509 | 50884510 | cg01781850 |
| chr22 | 50950506 | 50950507 | cg13022461 |
| chr22 | 50986511 | 50986512 | cg07833467 |
| chr22 | 51175515 | 51175516 | cg25232725 |
